# Wnt signaling restores evolutionary loss of regenerative potential in *Hydra*

**DOI:** 10.1101/2025.03.18.643955

**Authors:** Sergio E. Campos, Sahar Naziri, Jackson Crane, Jennifer Tsverov, Ben D. Cox, Craig Ciampa, Celina E. Juliano

**Author notes:** **Correspondence to:** Celina E. Juliano Department of Molecular and Cellular Biology University of California, Davis One Shields Avenue, Davis, CA 95616.

## Abstract

The regenerative potential of animals varies widely, even among closely-related species. In a comparative study of regeneration across the *Hydra* genus, we found that while most species exhibit robust whole-body regeneration, *Hydra oligactis* and other members of the Oligactis clade consistently fail to regenerate their feet. To investigate the mechanisms underlying this deficiency, we analyzed transcriptional responses during head and foot regeneration in *H. oligactis*. Our analysis revealed that the general injury response in *H. oligactis* lacks activation of Wnt signaling, a pathway essential for *Hydra vulgaris* foot regeneration. Notably, transient treatment with a Wnt agonist in *H. oligactis* triggered a foot-specific transcriptional program, successfully rescuing foot regeneration. Our transcriptional profiling also revealed *dlx2* as a likely high-level regulator of foot regeneration, dependent on Wnt signaling activation. Our study establishes a comparative framework for understanding the molecular basis of regeneration and its evolutionary loss in closely-related species.

## Introduction

Regeneration is defined as the restoration of lost or injured tissues, appendages or organs ^1^. While most animals have some capacity to regenerate, this potential varies widely across animals. Thus, a fundamental question is: how are some animals capable of regenerating large portions of their body after injury, while others exhibit only a fraction of this capacity? Mapping animal regenerative abilities on the phylogenetic tree suggests a complex evolutionary history, with numerous gains and losses. Regardless, it is likely that regenerative abilities were present at the dawn of animals and have been largely lost by vertebrates, especially mammals ^2^. Many studies have already advanced our understanding of the genes and pathways that control regeneration in highly regenerative animals ^3,4^. However, we have little understanding of the mechanisms that drive loss of regenerative potential. Comparative approaches between closely related organisms with contrasting regeneration abilities offer a promising avenue to both reveal the mechanisms of regeneration, as well as uncover how regeneration has been shaped throughout evolution.

Cnidarians, a group that includes sea anemones, jellyfish, and corals, exhibit high regenerative potential. Furthermore, given their phylogenetic relationship as sister group to bilaterians ^5^, research in cnidarians is critical to understand the evolutionary history of regeneration. The cnidarian polyp *Hydra vulgaris* is capable of whole-body regeneration and is a well-established research organism for studying the molecular mechanisms of regeneration. The body plan of *Hydra* is arranged around a single body axis, with a head, composed of a tentacle ring around a hypostome or mouth at the oral end; and the peduncle and an adhesive basal disk (foot) at the aboral end. These structures are connected by a cylindrical body column. When bisected, both halves of *H. vulgaris* are capable of regeneration: the top half regenerates a foot, and the bottom half regenerates a head. However, a few studies on other *Hydra* species suggest that regenerative potential varies in the genus, a topic that remains largely unexplored ^6,7^.

The Wnt signaling pathway plays a key role in regeneration across a diverse range of species ^8,9^. In *H. vulgaris*, the maintenance of tissue patterning in the uninjured animals relies on a Wnt signaling organizing center located at the oral end. Upon bisection perpendicular to the oral-aboral axis in *H. vulgaris*, the process of regeneration includes the formation of a new Wnt organizer on the oral-facing wound which will direct the regeneration of a new head ^9–11^. However, injury-induced transcriptional activation of Wnt ligands occurs at both sides of the wound, and treatment with a Wnt inhibitor impairs or delays both head and foot regeneration ^11–13^. In planarians, a similar regeneration pattern is observed after bisection, with Wnt activation required at both sides of the wound during the early injury response, followed by restriction to tail regeneration where it directs posterior development ^8,14^. However, in acoels, early Wnt activation after bisection is restricted to aboral wound sites where it similarly directs posterior development, ^15^ exemplifying variation in the role of Wnt signaling during whole body regeneration. In vertebrates, Wnt signaling is rapidly upregulated early in regeneration and Wnt inhibition results in impaired regeneration of mouse digits, demonstrating its importance even in animals with more limited regeneration abilities ^16,17^.

In *H. vulgaris*, injury-induced bZIP transcription factors (TFs) likely drive the transcription of Wnt ligands, which in turn are maintained by a self-reinforcing Wnt signaling amplification loop at the oral wound, creating the head organizer ^12,18–20^. In contrast, the molecular mechanisms responsible for foot specification at the aboral wound are less understood. While injury-induced Wnt signaling is necessary for this process, the Wnt signal is quickly down-regulated, giving rise to a foot-specific program through mechanisms that remain unclear ^12,13^. In this study, we discovered that foot regeneration ability has been lost in the Oligactis clade of the *Hydra* genus, including the laboratory species *Hydra oligactis* (Fig. 1A). We therefore reasoned that a comparative study of *H. vulgaris* and *H. oligactis* foot regeneration could uncover the regulatory mechanisms required to initiate foot regeneration in *Hydra*. Towards this goal, we performed RNA-seq over a time course of *H. oligactis* regeneration and compared the results to existing data sets from *H. vulgaris*. Our analysis revealed that injury-induced Wnt signaling activation is significantly attenuated in *H. oligactis*, resulting in slower head regeneration and a foot regeneration block. Using pharmacological manipulation, we demonstrated that activation of the Wnt pathway rescues *H. oligactis* foot regeneration. This approach also allowed us to identify key transcriptional regulators involved in foot formation, providing new insights into the regulatory control of foot regeneration in *Hydra*.

**Figure 1.**
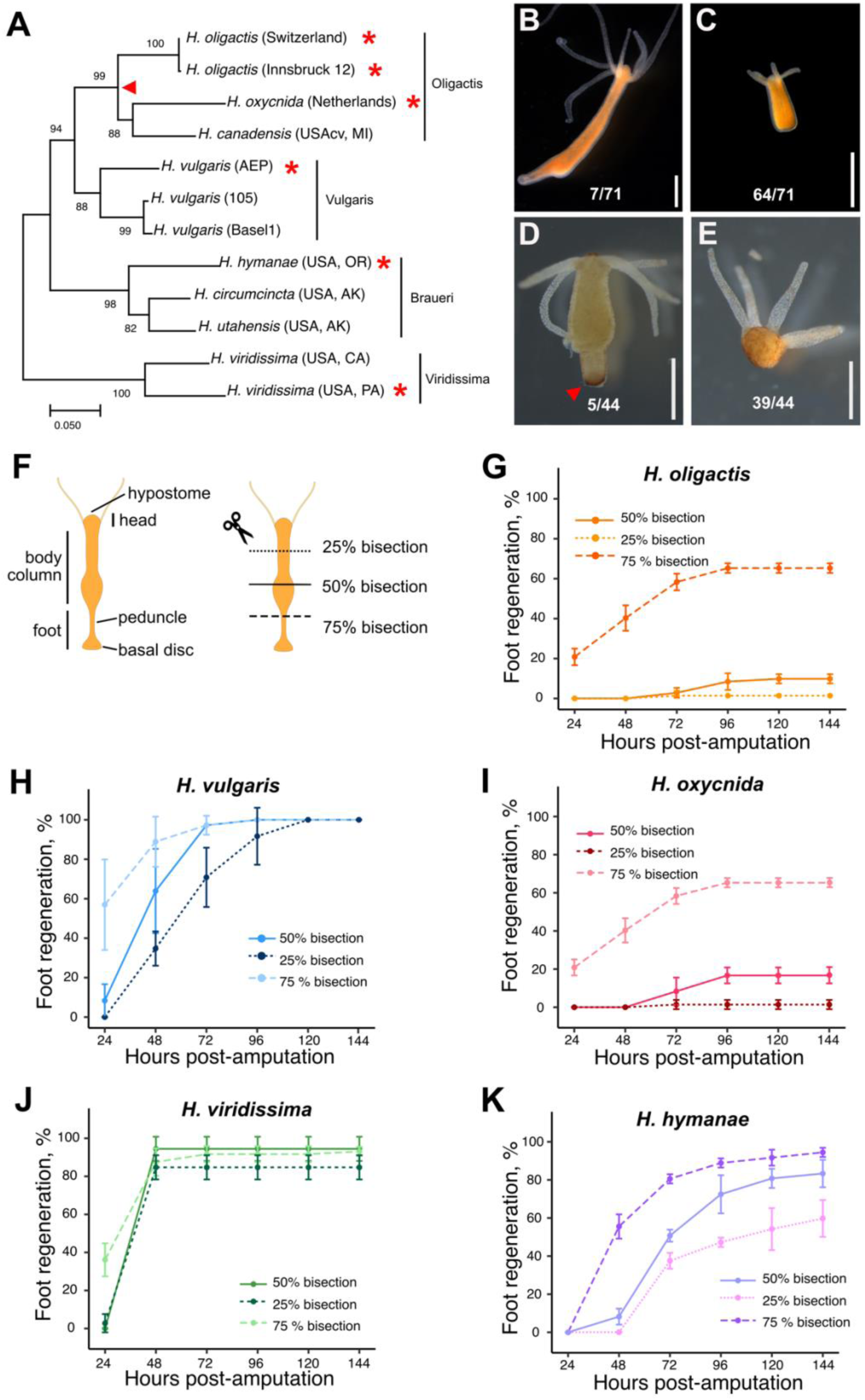
Foot regeneration ability was lost in the Oligactis clade. (A) Maximum Likelihood phylogenetic tree of the *Hydra* genus based on mitochondrial cytochrome oxidase 1 sequences. The genus is divided into four clades: Oligactis, Vulgaris, Braueri, and Viridissima ^27^. A red triangle marks the likely point of foot regeneration loss in the Oligactis clade. Asterisks denote strains used in amputation experiments. (B-C) Following 50% bisection, 90% of *H*. oligactis fail to regenerate their feet, resulting in a stably footless phenotype. (D-E) Peroxidase staining correlates with foot morphology in *H. oligactis*. The assay was performed on 44 *H. oligactis* polyps 9 days after 50% bisection (5 with feet, 39 footless), confirming the presence or absence of a foot in all cases. Red triangle indicates peroxidase staining in the foot. Scale bars: 1 mm. (F) Amputation strategy to evaluate foot regeneration abilities across the *Hydra* genus. The top half of each bisected animal was visually monitored for foot regeneration. (G-K) Foot regeneration kinetics following amputations at various positions along the oral-aboral axis for (G) *H. oligactis*, (H) *H. vulgaris*, (I) *H. oxycnida*, (J) *H. viridissima* and (K) *H. hymanae*. **Source data file 1.** Text file containing the accession numbers and sequences for CO1 used to build the phylogenetic tree. **Source data file 2.** Excel workbook containing head and foot regeneration data for the experiments shown in Figure 1 G-K.

## Results

### Aboral regenerative potential varies across the *Hydra* genus

The common laboratory *Hydra* species, *H. vulgaris*, can regenerate a complete head or foot from body column tissue. By contrast, it was previously reported that *H. oligactis* has a foot regeneration deficiency ^6^. To investigate this further, we first characterized the foot regeneration abilities of our *H. oligactis* laboratory strain (Innsbruck 12) for which we recently assembled a high-quality genome ^12^. We bisected *H. oligactis* midway between the head and foot (50% bisection), perpendicular to the oral-aboral axis, and monitored the rate of foot regeneration for the top half (Fig. 1 B-G). We found that after 6 days, only ∼10% of the animals regenerated their feet (Fig. 1B, C, G), while the remaining animals became stably footless, characterized by a flat aboral end (Fig. 1C) and an inability to stick to surfaces. We also evaluated the absence or presence of a foot using a standard colorimetric assay that reveals the activity of a foot-specific peroxidase at seven days post bisection ^21^. We confirmed that morphologically footless *H. oligactis* also lacked foot peroxidase staining (Figure 1D-E). In contrast to *H. oligactis*, we confirmed that in the same conditions, *H. vulgaris* shows complete foot regeneration after bisection, consistent with the published literature (Fig. 1H).

We next asked if the rate of successful foot regeneration in *H. oligactis* depends on the location of the cut along the oral-aboral axis. To test this, we performed bisections at three different positions along the body column. In addition to mid-body bisection (50% of body length), we also bisected at 25% body length from the head, just below the tentacle ring, and at 75% body length from the head, just above the peduncle and foot (Fig. 1F). We performed the same bisections in *H. vulgaris* as a positive control for foot regeneration. Consistent with previous literature, *H. vulgaris* exhibited nearly 100% successful foot regeneration after all three amputations, but with different kinetics depending on the level of the amputation (Fig 1H) ^6,22–24^. Also consistent with previous literature, the success rate of *H. oligactis* foot regeneration depended on the location of the cut; after 75% bisection, ∼65% of animals successfully regenerated their feet, while foot regeneration did not occur after 25% bisection (Fig. 1G) (Hoffmeister, 1991).

We next tested foot regenerative abilities across the *Hydra* genus to better understand the evolutionary history of this trait. While a handful of regeneration studies have been done in different *Hydra* species ^6,7,25,26^, no systematic study of foot regeneration ability has been done across the genus on validated species. We therefore performed foot amputation assays (Fig. 1 F) on three additional *Hydra* species to cover each phylogenetic clade (Fig. 1A, I-K) ^27^, as well as on the Swiss laboratory strain of *H. oligactis* ^28^ (Fig. S1) to ensure that our observations were not specific to the Innsbruck 12 strain of *H. oligactis*. Our experiments revealed that both *H. oligactis* and *H. oxycnida*, which are in the Oligactis clade, show similar foot regeneration defect profiles (Fig. 1G, I, Fig. S1). By contrast, *H. viridissima* had robust foot regeneration ability, with faster kinetics than *H. vulgaris* (Fig. 1J). *H. hymanae,* a member of another distinct group within the *Hydra* genus phylogeny, the Braueri clade, exhibited slightly reduced foot regenerative ability, which varied depending on the bisection location. While 75% bisections resulted in nearly 100% successful foot regeneration, 25% bisections led to only ∼60% regeneration (Fig. 1K). Overall, our survey of foot regeneration abilities across the *Hydra* genus revealed variability that enables comparative studies. Furthermore, given the relatively high rate of foot regeneration in all species, except those in the Oligactis clade, we conclude that the common ancestor of the *Hydra* genus likely had robust foot regeneration ability, which was lost by the ancestor of the Oligactis clade.

### The foot transcriptional program largely fails to be activated during *H. oligactis* regeneration

To identify key aspects of the transcriptional response that result in limited foot regeneration in *H. oligactis*, we generated RNA-seq libraries from the regenerating tips of bisected *H. oligactis* polyps during a time course of both head regeneration and failed foot regeneration at 0, 3, 12, 24, and 48 hours post amputation (hpa) (Fig. 2A). For the foot regeneration time course, it was not possible to determine which samples would successfully regenerate their feet *a priori*, therefore these samples contained a mixture of both successful and unsuccessful regeneration. However, given the low rate of successful foot regeneration after 50% bisection (∼10%, Fig. 1G), most of the signal is predicted to come from animals that will ultimately fail to regenerate their feet. To define the transcriptional end point of successful regeneration, we also collected RNA-seq data for head and foot tissue in uninjured animals. We used these data to identify head- and foot-specific genes in *H. oligactis* by determining the differentially expressed genes in uninjured animals between the foot, head, and body column tissue. The regenerating head and foot tissue at 0 hpa were used as uninjured body column for this analysis, given that these tissues were taken just above and below the plane of a 50% bisection right after injury (Fig. 2B). This approach uncovered 404 foot-specific genes and 987 head-specific genes in *H. oligactis* (Figure 2C, D; Supplementary data file 1).

**Figure 2.**
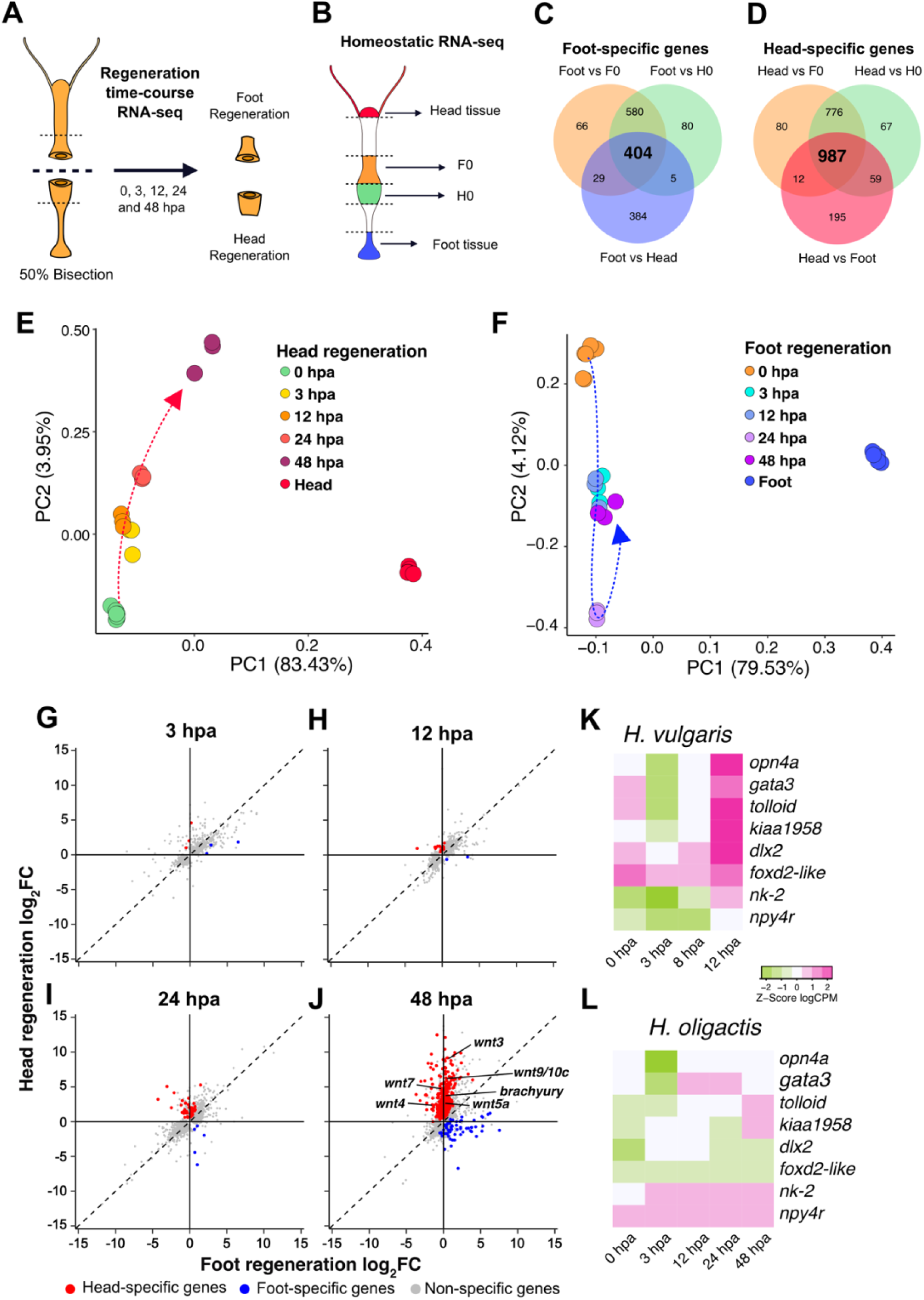
Foot-specific gene transcription is largely absent during *H. oligactis* regeneration. (A) Strategy for RNA-seq library preparation over a regeneration time course: Regenerating animals were incubated in 0.05% DMSO, allowing these samples to serve as controls for the data shown in Figure 5B. (B) RNA-seq strategy for different regions of homeostatic *H. oligactis*. F0 and H0 correspond to the 0 hpa foot and head regeneration samples in panel A. (C-D) Venn diagrams showing the differential gene expression strategy used to identify 404 foot-specific genes (C) and 987 head-specific genes (D) in *H. oligactis*. (E) PCA plot of *H. oligactis* oral-facing wound gene expression showing movement along the PC1 axis toward the homeostatic head profile during regeneration. (F) PCA plot of *H. oligactis* aboral-facing wound gene expression showing no progression along the PC1 axis toward the homeostatic foot profile during regeneration. (G-J) Comparison of log_2_ fold change (log_2_FC) transcript abundance between *H. oligactis* head regeneration and foot regeneration at 3 hpa (G), 12 hpa (H), 24 hpa (I) and 48 hpa (J). Red dots indicate differentially expressed head-specific genes at the oral-facing wound and blue dots indicate differentially expressed foot-specific genes at the aboral-facing wound. (K) Heatmap of foot-specific gene expression in *H. vulgaris* (data from Cazet et al., 2021), showing upregulation by 12 hpa during foot regeneration. Values represent Z-scores calculated from log₂ counts per million (logCPM). (L) Heatmap of *H. oligactis* expression for the same 8 genes shown in panel K. These genes do not exhibit rapid upregulation during regeneration. **Source data file 3.** Excel workbook containing differentially expressed gene tables for the conditions compared in Figure 2 C-D. **Source data file 4.** CSV file containing normalized log2CPM used to plot Figure 2 E-F **Source data file 5.** Excel workbook containing log2 Fold Change values from comparisons in Figure 2 G-J. **Source data file 6.** Excel workbook containing log2 CPM for selected foot-specific genes in *H. vulgaris* and *H. oligactis* shown in Figure 2 K-L.

To analyze the transcriptional changes that occur over the course of *H. oligactis* head regeneration and failed foot regeneration, we used Principal Component Analysis (PCA) (Fig. 2E, F). In these analyses, we included homeostatic head and foot RNA-seq libraries which represent a successful regeneration end point. Notably, we found that *H. oligactis* exhibits nearly 100% head regeneration after bisection, but the process takes up to 72 hpa to complete. This is approximately 24 hours longer than in *H. vulgaris*, as determined morphologically by the presence of tentacles (Fig. S2). Our RNA-seq time course of *H. oligactis* head regeneration ended at 48 hours and our PCA analysis confirms that head regeneration is not transcriptionally complete at this point in *H. oligactis*. Although differences between homeostatic head tissue and regenerating head tissue explain 83.34% of the variation among samples (PC1), regenerating head tissue exhibited a progressive upregulation of key PC1-associated genes, such as *wnt3* (Supplementary Data File 2), suggesting a shift toward head identity (Fig. 2E).

For failed foot regeneration, we did not observe a progressive shift of regenerating tissue toward the homeostatic foot transcriptional signature. Like head regeneration, the most variation (79.53%) for failed foot regeneration was explained by PC1, which denotes the difference in the expression profiles between homeostatic foot tissue and the regenerating samples. Importantly, 93 out of 100 of the topmost genes explaining variation within PC1 correspond to foot-specific genes (Supplementary data file 3). Whereas only 4.12% of the variation is explained by PC2, which shows changes in the wounded tissue occurring during the time course of regeneration, but the regeneration time points did not change along the PC1 axis (Fig. 2F). This suggests that the gene expression profile of the failed foot regeneration samples do not resemble that of the homeostatic foot at any timepoint.

We next examined the expression dynamics of head-specific and foot-specific transcripts over the time course of *H. oligactis* head regeneration and failed foot regeneration respectively. To identify differentially expressed head and foot-specific genes we compared gene expression on each side of the wound at each time point. We found that in *H. oligactis* head regeneration, only a small number of head-specific genes showed tissue-specific upregulation at 12 hpa, with larger transcriptional changes becoming evident only after 24 hpa (Fig. *2*G-J; Supplementary data file 4 containing lists of tissue-specific genes at each timepoint). This contrasts with *H. vulgaris*, which shows significant structure-specific gene expression by 8 hpa for both head and foot regeneration ^12^. For example, in *H. vulgaris*, several canonical Wnt pathway genes, including Wnt ligands become specific to the oral wound by 8 hpa ^12^. In *H. oligactis,* the expression of head-associated Wnt genes became specific to the oral wound by 48 hpa and overall, 472 out of the 987 (47.8%) head-specific genes were expressed at this timepoint (Fig. 2 J). In contrast, we identified only 63 out of 404 (15.6%) foot-specific genes as up-regulated in failed foot regenerating tissue at 48 hpa (Fig. 2G-J). Therefore, this analysis showed that by 48 hpa, head regeneration in *H. oligactis* although incomplete, is progressing appropriately, whereas foot regeneration is largely failing at the transcriptional level.

To gain a deeper insight into the differences in foot regeneration potential between *H. vulgaris* and *H. oligactis,* we compared the injury-induced transcriptional activation of seven foot-specific genes that were identified in our previous transcriptional profiling of *H. vulgaris*. These genes were upregulated specifically at the aboral wound site by 12 hpa and were enriched in the foot as determined using the single cell expression atlas (Cazet et al., 2021; Siebert et al., 2019; the transcript IDs for genes discussed in this study can be found in Table S1). This analysis showed that transcriptional activation of these seven genes is weak in *H. oligactis* as compared to *H. vulgaris* (Fig. 2K-L). In addition, we compared the expression of foot-specific genes from *H. oligactis* to the expression of their orthologs in *H. vulgaris* (154 genes) and found lower expression in *H. oligactis* at every time point, including 0 hpa (Fig. S3). The same comparison performed with a random gene set did not show expression differences (Fig. S3). These data suggested that the initial foot competency of the *H. oligactis* tissue is lower (Fig. S3).

### Injury-induced transcriptional activation of Wnt pathway genes is reduced in *H. oligactis* as compared to *H. vulgaris*

We next compared the early injury-induced transcriptional response between *H. vulgaris* and *H. oligactis* to identify differences that could lead to downstream failure of foot regeneration in *H. oligactis*. To accomplish this, we compared the *H. oligactis* regeneration RNA-seq data collected in this study to our previously published *H. vulgaris* regeneration RNA-seq data set, collected for both head and foot regeneration at 0, 3, 8, and 12 hpa ^12^. To establish a framework to compare the gene expression patterns of *H. oligactis* and *H. vulgaris* we used OrthoClust, a computational method that groups gene expression patterns based in gene orthology across multiple species into discrete clusters ^30^. We first looked for orthologous genes by conducting a reciprocal BLAST analysis of the RNA-seq datasets for each species, obtaining 9,371 transcripts for *H. oligactis* with a direct ortholog in *H. vulgaris.* We next used OrthoClust to compare the expression data of these transcripts over the course of regeneration to identify conserved co-expression modules across the two *Hydra* species. With this method we identified nine gene co-expression clusters containing 566 transcripts from *H. vulgaris* and 489 transcripts from *H. oligactis* (Supplementary data file 5).

While most of the OrthoClust modules contained transcripts that decreased in expression during the regeneration time course in both species, transcripts in clusters 5 and 6 showed an up regulation in head regenerating tissue in both *Hydra* species (Fig. S4, Fig. 3A, B). Cluster 6 contains several genes involved in canonical Wnt signaling. Notably, in *H. oligactis* head regeneration, cluster 6 genes such as *wnt3, wnt7,* and *brachyury* were delayed in their transcriptional activation, which correlates with the extended time frame of head regeneration in this species (Fig. 3A-H Fig. S2).

**Figure 3.**
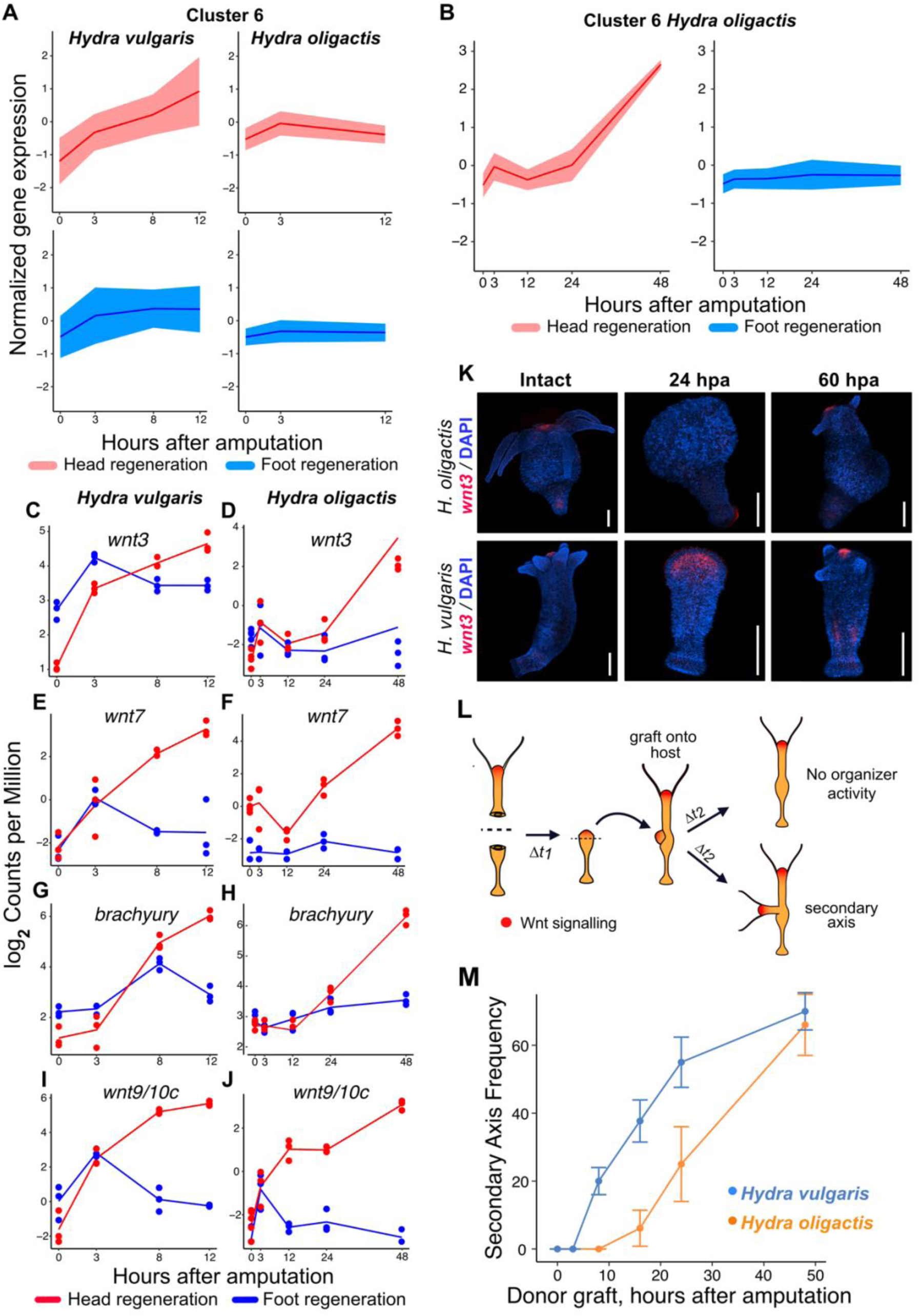
Delayed transcriptional activation of Wnt signaling genes during regeneration in *H. oligactis* as compared to *H. vulgaris*. (A) Ribbon plots showing normalized expression patterns for genes in cluster 6, identified with OrthoClust, in both *H. vulgaris* (left panels) and *H. oligactis* (right panels) over the first 12 hours of regeneration. Expression patterns at the oral-facing wound are shown in red and expression patterns at the aboral-facing wound are shown in blue. (B) Ribbon plots showing normalized expression patterns for cluster 6 genes in the *H. oligactis* regeneration time course extended to 48h shows that cluster 6 genes are upregulated between 24 and 48 hours. (C-J) RNA-seq expression profiles for *wnt3* (C, D), *wnt7* (E, F), *brachyury* (G, H) and *wnt9/10C* (I, J), during head (red) and foot (blue) regeneration in *H. vulgaris* (left panels; data from Cazet et al., 2021) and *H. oligactis* (right panels; this study, Figure 2). Note the extended time course for *H. oligactis*. (K) Confocal images of fluorescent RNA in situ hybridizations for *wnt3* in homeostatic (left) and regenerating tissue in *H. vulgaris* (top) and *H. vulgaris* (bottom) at 24 hpa (middle) and 60 hpa (right). Scale bars: 500 µm. (L) Experimental strategy used to evaluate head organizer activity acquisition during head regeneration. (M) Kinetics of secondary axis induction following grafting of injured oral tissue onto host animals at indicated times post-amputation (0, 3, 8, 16, 24 and 48 hpa). The blue line represents *H. vulgaris* and the orange line represents *H. oligactis*. **Source data file 9.** Excel workbook containing secondary axis formation frequency from grafting in *H. vulgaris* and *H. oligactis* Figure 3M.

We found that some cluster 6 genes also behaved differently during foot regeneration when comparing the two species. In our previous study, we found that Wnt pathway genes are transiently upregulated from 0-3 hpa during *H. vulgaris* foot regeneration. This included the Wnt ligands *wnt3, wnt7*, and *wnt9/10C* as well as the conserved Wnt signaling target *brachyury* ^11–13^ (Fig. 3C-I). By contrast, *wnt3, wnt7*, and *brachyury* were not significantly upregulated at 3pha in *H. oligactis* foot regeneration (Fig. 3D,F,H). Noteworthy, *wnt9/10c* is significantly upregulated in both *Hydra* species at 3 hpa during foot regeneration, although absolute levels of transcript are lower in *H. oligactis* (Fig. I,J). Altogether, these results revealed that injury-induction of Wnt signaling is either absent or highly reduced in *H. oligactis* as compared to *H. vulgaris*.

### The formation of the head organizer is delayed in *H. oligactis*

The hypostome in *Hydra* species has a conserved function as an oral organizer ^31^. While it is almost certain that Wnt signaling plays a conserved role in directing axial patterning across the *Hydra* genus, the expression of Wnt ligands specifically in the hypostome has not been formally tested in *H. oligactis*. To address this, we performed fluorescent RNA *in situ* hybridization (FISH) to analyze the expression of *wnt3* in *H. oligactis*. We found that like *H. vulgaris*, *wnt3* was expressed at the hypostome of *H. oligactis*, similar to *H. vulgaris* (Fig. 3K). Next, we used FISH to investigate the temporal and spatial expression of *wnt3* over the course of head regeneration. In contrast to *H. vulgaris*, the expression of *wnt3* was not detected by FISH in *H. oligactis* at 24 hpa. However, by 60 hpa, the expression of *wnt3* became apparent in *H. oligactis* (Fig. 3K). These results are consistent with slower head regeneration kinetics in *H. oligactis* (Fig. S2) and suggest that delayed activation of Wnt signaling may contribute to differences in head regeneration timing between these species.

Given the morphological delay of head regeneration we observed in *H. oligactis* (Fig. S2), along with the reduced levels of Wnt gene transcription (Fig. 3D, F, H, J, K), we hypothesized that formation of the oral organizer is delayed in *H. oligactis* as compared to *H vulgaris*. Tissue grafting is a classical method for determining the timing of head organizer formation: regenerating head tissue is grafted onto the body column of a host animal to test if the donor tissue can induce a secondary axis. Using this approach, previous studies demonstrated that regenerating *H. vulgaris* head tissue acquires organizing ability as early as 8 hpa, which also correlates with high levels of Wnt signaling pathway gene transcripts at this timepoint ^12,32^. To test the timing of head organizer formation in *H. oligactis,* we conducted classic grafting experiments. We also performed these experiments in *H. vulgaris* as a positive control. We bisected *Hydra* polyps and grafted the tissue from the oral injury site onto a host polyp at 0, 3, 8, 16, 24 and 48 hpa (Fig. 3L). We then quantified the frequency of secondary axis formation at 5 days post grafting (Fig. 3M). While we observed organizer ability in regenerating *H. vulgaris* head tissue as early as 8 hpa as expected, regenerating *H. oligactis* did not begin showing organizer activity until 16 hpa. *H. oligactis* head organizing ability was reduced as compared to *H. vulgaris* until 48 hpa (two-way ANOVA, *p_species_* = 2.55 X 10^-5^), at which time a similar capacity to direct secondary axis formation in both species was observed. Overall, these results support the conclusion that injury fails to strongly activate Wnt signaling in early stages of regeneration in *H. oligactis*, which may contribute to slower head regeneration kinetics.

### Transient pharmacological activation of Wnt signaling during the injury response rescues foot regeneration in *H. oligactis*

We next asked how injury-induced Wnt expression impacts foot regeneration abilities in *Hydra*. Given a previous study showing that treatment with a Wnt inhibitor negatively impacts *H. vulgaris* foot regeneration ^13^, we hypothesized that the lack of injury-induced Wnt expression contributes to *H. oligactis* foot regeneration failure. To test this, we pharmacologically activated Wnt signaling during the injury phase of *H. oligactis* foot regeneration using alsterpaullone (ALP), a Wnt signaling agonist that stabilizes β-catenin and induces oral patterning in *H. vulgaris* ^10^. We transiently treated the upper halves of 50% bisected *H. oligactis* with different concentrations of ALP (2.5, 5, and 10 µM) for 3 hours after injury, to mimic the short window of Wnt signaling activation in *H. vulgaris* foot regeneration. We then monitored the animals over 6 days for the presence or absence of foot regeneration (Fig. 4A). Treatment of regenerating *H. oligactis* with ALP, regardless of the concentration, resulted in a partial but significant rescue of foot regeneration, with the concentration of 5 µM having the largest effect (Fig. 4B-D). Notably, at 5 µM, ∼7% of animals regenerated a second head at the aboral wound site instead of a foot, with this percentage increasing to 20% at 10 µM ALP (Fig. S5). The appearance of the two-headed phenotype suggests that a precise balance in Wnt activation is required to determine whether foot or head regeneration occurs at the aboral wound site. In addition, the higher occurrence of two-headed *Hydra* in the 10 µM ALP treatment likely explains the reduced number of animals that regenerated a foot under this concentration. Overall, this result supports the conclusion that a transient burst of Wnt signaling activation during the generic injury response is essential for foot regeneration in *Hydra*.

**Figure 4.**
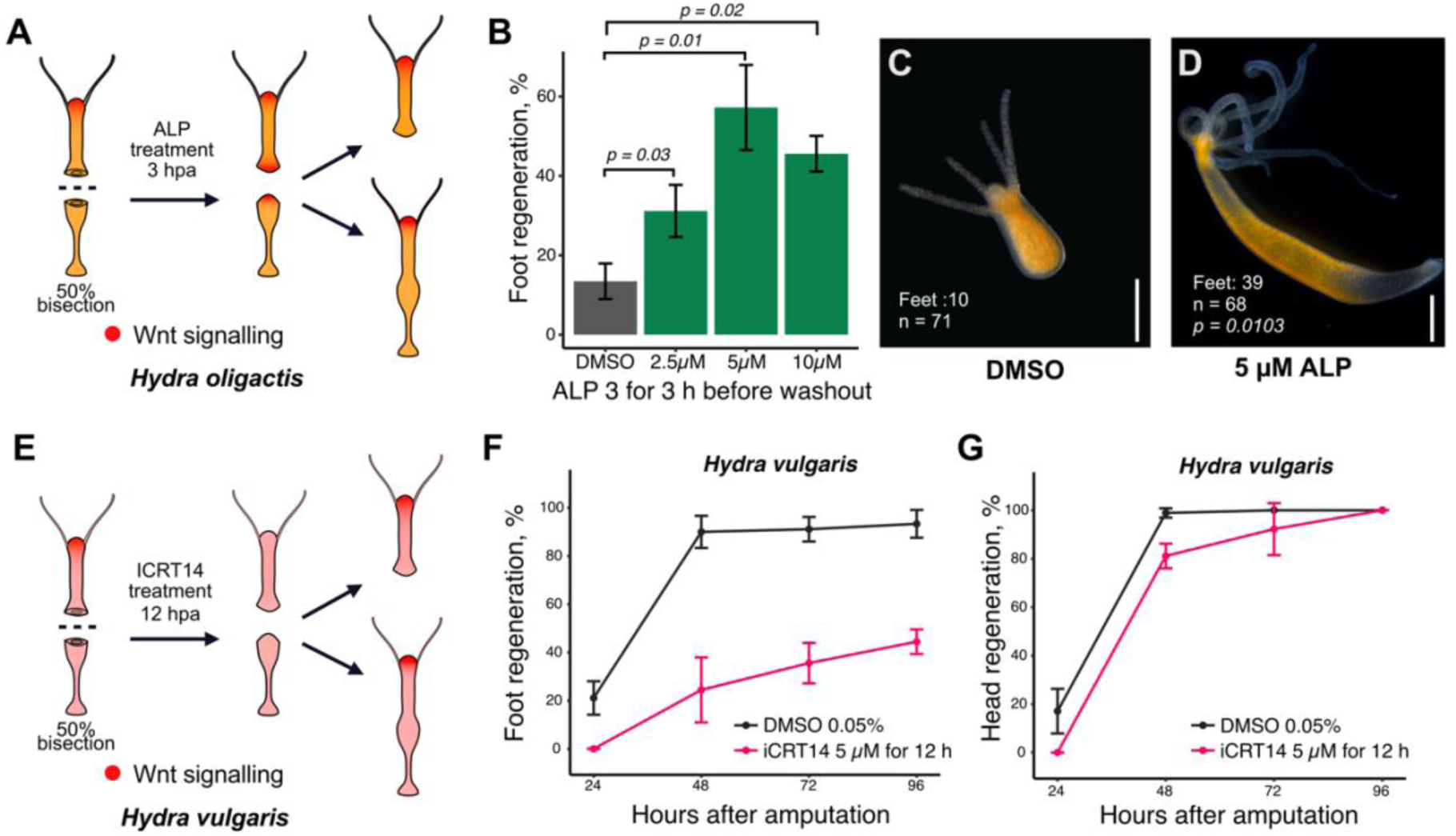
Transient activation of Wnt signaling rescues foot regeneration in *H. oligactis*. (A) Experimental strategy for treating the top halves of 50% bisected *H. oligactis* with the Wnt agonist Alsterpaullone (ALP). (B) Bar graph showing the percentage of successful foot regeneration in *H. oligactis* following a 3-hour post-bisection treatment with DMSO (grey bar) or ALP at 2.5 µM, 5 µM, and 10 µM (green bars). Statistical comparisons of DMSO versus ALP treatments were conducted using a two-tailed t-test (p-values: 2.5 µM vs DMSO = 0.02935, 5 µM vs DMSO = 0.01026, and 10 µM vs DMSO = 0.02302). (C) Representative image of *H. oligactis* showing failed foot regeneration in DMSO-treated control group. (D) Representative image of *H. oligactis* showing successful foot regeneration after treatment with 5 µM ALP. Scale bars: 500 µm. (E) Experimental strategy to assess foot and head regeneration in *H. vulgaris* after treatment with the Wnt inhibitor iCRT14. (F) Line graph showing foot regeneration kinetics in *H. vulgaris* treated with iCRT14 (pink) and DMSO controls (black). (G) Line graph showing head regeneration kinetics in *H. vulgaris* treated with iCRT14 (pink) and DMSO controls (black). **Source data file 10.** Excel workbook containing foot regeneration percentages for *H. oligactis* treated with the different ALP concentrations shown in Figure 4B. **Source data file 11.** Excel workbook containing foot regeneration percentages for *H. vulgaris* after 12 hours of iCRT14 treatment in Figure 4 F-G.

### Transient inhibition of Wnt signaling in *H. vulgaris* mimics the regeneration phenotype of *H. oligactis*

Inhibition of Wnt/beta-catenin signaling by iCRT14 in *H. vulgaris* inhibits both head and foot regeneration upon continuous treatment over the course of regeneration (Gufler et al., 2018). Given our findings that in *H. oligactis* the injury response does not elicit a strong Wnt signaling response, here we sought to mimic that effect in *H. vulgaris* by transiently treating animals with iCRT14 for 12 hours after bisection (Fig. 4E). This treatment caused *H. vulgaris* regeneration to more closely resemble *H. oligactis* regeneration. Specifically, we found that transient inhibition of Wnt signaling in *H. vulgaris* resulted in reduced foot regeneration (two-way ANOVA, *p_treatment_* = 0.002824) and an apparent delay in head regeneration (Fig. 4F, G, Fig. S6). Notably, this treatment produced stably footless *H. vulgaris* mimicking the *H. oligactis* phenotype (Fig. S6). Our results support the conclusion that a lack of injury-induced Wnt signaling significantly contributes to foot regeneration deficiency in *H. oligactis*.

### Transient activation of Wnt signaling triggers the transcription of foot-specific genes in *H. oligactis*

Our finding that transient activation of Wnt signaling rescues foot regeneration in *H. oligactis* is consistent with published literature showing that injury-induced Wnt activation is necessary for *Hydra vulgaris* foot regeneration ^12,13,20^. However, the mechanisms by which Wnt signaling promotes foot regeneration is unknown. To shed light on this, we generated RNA-seq libraries from a time course of *H. oligactis* foot regenerating fragments after a 3-hour treatment with 5 µM ALP (Fig. 5A). These data revealed that ALP treatment elicited the transient upregulation of Wnt pathway genes such as *tcf*, *wnt3, axin1, wntless and sFRP3*, demonstrating that ALP treatment transcriptionally activated Wnt signaling as expected (Fig. S7). Furthermore, PCA of ALP-treated (rescued) and DMSO-treated (control) samples revealed that by 48 hpa, transient activation of Wnt promoted transcriptional changes in the direction of the homeostatic foot along the PC1 axis (Fig. 5B).

**Figure 5.**
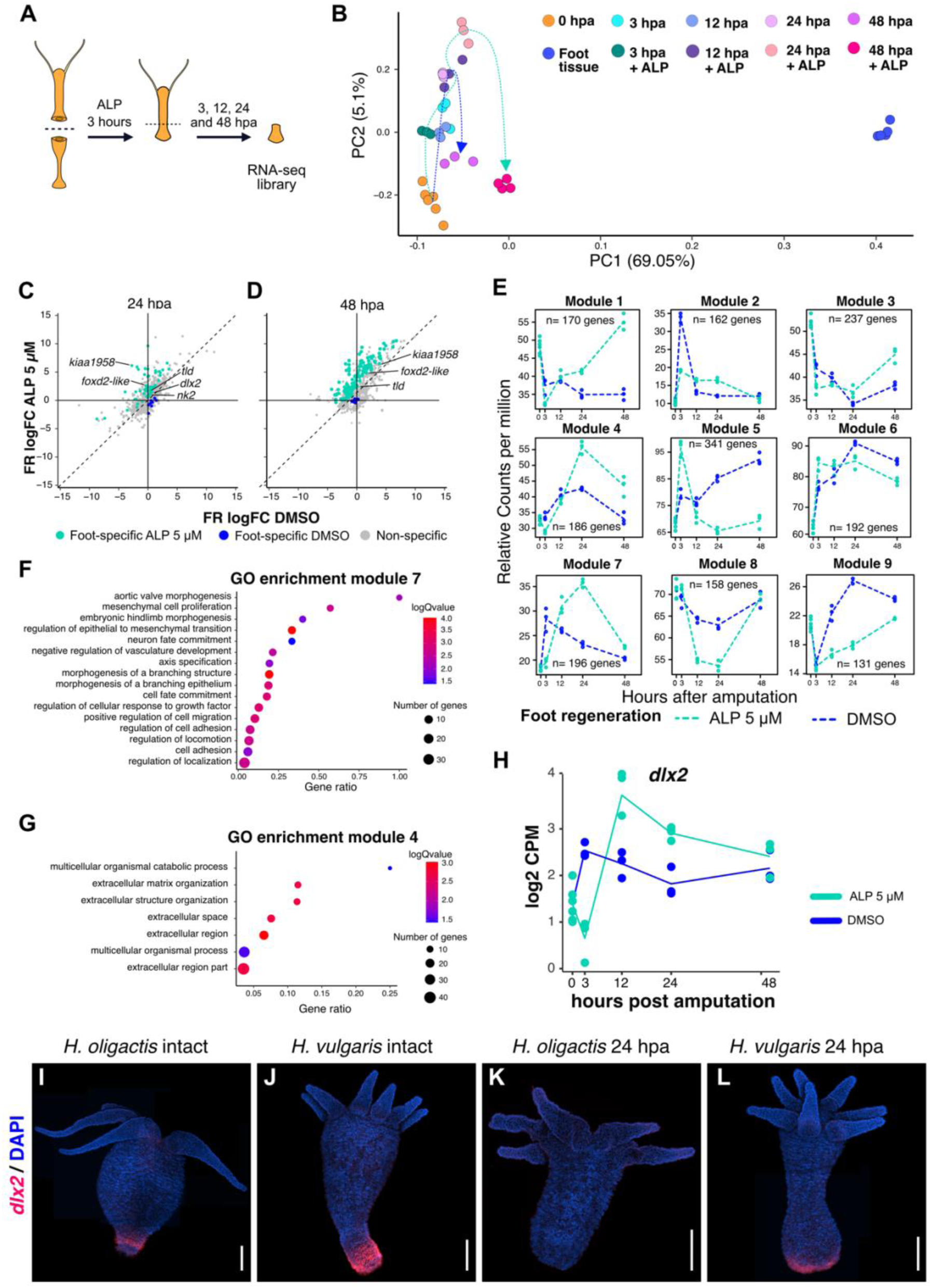
Wnt-dependent activation of the foot regeneration transcription factor *dlx2*. (A) Experimental strategy for producing RNA-seq libraries from ALP-treated aboral-facing wound tissue. (B) PCA plot showing transcriptional trajectories during regeneration in DMSO-treated (blue dotted line; data from Figure 2) and ALP-treated samples (green dotted line). (C, D) Comparison of average log_2_ fold change (log_2_FC) in transcript abundance between failed foot regeneration (DMSO-treated control animals) and rescued foot regeneration (ALP-treated animals) at 24 hpa (C) and 48 hpa (D). Blue dots represent foot-specific genes enriched in failed foot regeneration tissue; green dots represent foot-specific genes enriched in ALP-treated rescued foot regenerating tissue. (E) Gene expression modules for failed foot regeneration (DMSO-treated control animals, blue dashed line) and rescued foot regeneration (ALP-treated animals, green dashed line) identified using maSigPro. (F) Gene Ontology enrichment analysis for genes in module 7. (G) Gene Ontology enrichment analysis for genes in module 4. (H) RNA expression plot showing *dlx2* expression in log_2_ counts per million (log_2_ CPM) over time. The blue line represents DMSO-treated failed foot regenerating tissue; the green line represents ALP-treated foot regenerating tissue. (I-L) Confocal images of RNA in situ hybridizations for *dlx2* in untreated *H. oligactis* and *H. vulgaris*, showing intact polyps and aboral regenerating tissue at 24 hpa. Scales bars: 500 µm. **Source data file 12.** CSV file containing normalized log2CPM used to plot data shown in Figure 5 B and H. **Source data file 13.** Excel workbook containing fold change gene tables for the timepoints comparing foot regeneration in DMSO and ALP in Figure 5C and D. **Source data file 14.** Excel workbook containing Gene Ontology enriched terms for maSigPro modules shown in Figure 5L and M.

To characterize the transcriptional changes elicited by ALP treatment, we performed differential gene expression analysis comparing ALP-treated and control samples. We found that Wnt pathway activation induced the transcriptional activation of foot-specific genes, with 52 and 134 foot-specific genes being upregulated in ALP treatment conditions as compared to control conditions by 24 hpa and 48 hpa respectively (Fig. 5C-D). In contrast, just a few foot-specific genes were upregulated in control conditions as compared to ALP treatment conditions at these same timepoints (Fig. 5C-D).

To improve our understanding of the mechanisms which transient ALP treatment promotes foot regeneration, we used maSigPro, an R package that analyses time course expression data in different conditions through a regression approach to identify differential gene expression patterns ^33^. This analysis uncovered 1,773 genes with differential expression patterns grouped into 9 modules (Fig. 5E; Supplementary data file 6). The expression pattern of module 5 showed upregulation at 3 hpa only in ALP-treated tissue, followed by downregulation at later timepoints, and included Wnt signaling pathway genes (*tcf, axin1, lgr5, sp5,* and *nphp3*), confirming the validity of this method. In addition, several TFs whose orthologs are involved in cell differentiation and development were included in module 5, such as *foxl1*, *dmrta2, 14-3-3zeta, aatf, rfx1* and *tle3*. These TFs are also expressed in the homeostatic *Hydra*, but are not foot specific. To better understand the functions related to each module we performed Gene Ontology (GO) term enrichment analysis for all modules (Supplementary data file 7). Module 5 was enriched for gene expression regulatory functions, including transcription factors and coactivators. Contrastingly, module 2 showed downregulation in response to ALP at 3 hpa of transcriptional regulators involved in the generic injury response, such as bZIP TFs *creb, jun* and *fos* ^12,34^ (Fig. 5E, S8).

We also identified modules with foot-specific genes upregulated in the ALP-treated samples at 12 hpa (module 7) and 24 hpa (module 4) (Fig. 5E-G). GO analysis of module 7 showed enrichment for morphogenesis-related functions (Fig. 5F), including β-catenin and genes in the BMP and Notch pathways. Moreover, the foot-specific TF *dlx2*, also part of module 7, showed upregulation at 12 hpa in response to ALP, making it the earliest foot-specific TF to be transcriptionally activated upon Wnt activation (Fig. 5H). A previous study showed that *dlx2* is essential for foot regeneration in *H. vulgaris*, highlighting its likely role as a key regulator of foot regeneration (Ferenc et al., 2021). Since *dlx2* was the earliest foot-specific transcription factor activated by ALP, we used FISH to examine its spatial expression in *H. oligactis* and *H. vulgaris*. In uninjured *H. oligactis*, *dlx2* expression was restricted to a small peduncle region, whereas in *H. vulgaris*, it extended to the basal disk (Fig. 5I-L), suggesting differences in *dlx2* regulation under homeostatic conditions. At 24 hpa, *dlx2* was clearly expressed at the aboral injury site in *H. vulgaris* but was undetectable in *H. oligactis* (Fig. 5K-L), reinforcing its correlation with foot regeneration potential.

GO analysis of module 4 revealed enrichment for terms related to extracellular matrix organization (Fig. 5G). Notably, Wnt signaling has been found to induce extracellular matrix remodeling as part of the tissue patterning that occurs during *H. vulgaris* regeneration (Veschgini et al., 2023). In addition, this module included TFs *gata3* and *nk2,* previously identified as foot-specific in *H. vulgaris,* as well as other TFs that are upregulated temporarily at 24 hpa with ALP treatment (Fig. S9). Overall, these findings highlight the hierarchical gene regulatory network underlying foot regeneration in *Hydra*, where transient Wnt activation triggers a transcriptional cascade that sequentially activates non-tissue specific transcription factors, followed by foot-specific factors—starting with *dlx2*—and ultimately genes involved in morphological changes such as extracellular matrix remodeling, leading to successful tissue regeneration.

## Discussion

While previous work identified a role for injury-induced Wnt signaling in *H. vulgaris* foot regeneration, the specific contribution of early Wnt signaling and the mechanisms driving foot regeneration have remained unclear ^13^. To investigate this, we used *H. oligactis*, a species with limited foot regeneration capacity, and performed comparative transcriptomics to identify key regulatory mechanisms. Our findings revealed that weak injury-induced Wnt signaling activation in *H. oligactis* contributes to its reduced foot regeneration potential. Short-term treatment with the Wnt signaling agonist ALP partially rescued foot regeneration in *H. oligactis*, establishing Wnt signaling as a key regulator of *Hydra* foot regeneration. Furthermore, differential gene expression analysis of ALP-treated *H. oligactis* indicated that increased Wnt signaling promoted the expression of several TFs associated with early foot regeneration in *H. vulgaris*, including *nk-2, foxd2-like* and *dlx2*. In *H. vulgaris,* these TFs exhibit foot-specific expression between 8 and 12 hpa ^12^. Notably, *dlx2* was the earliest upregulated foot-specific TF in *H. oligactis* following ALP-treatment, suggesting this TF as a key regulator of foot regeneration.

The molecular basis for weaker injury-induced Wnt signaling in *H. oligactis* compared to *H. vulgaris* remains unclear. However, one possibility is the presence of a stronger Wnt inhibitory environment in *H. oligactis*. In *Hydra,* it is hypothesized that the head organizer establishes a morphogenetic gradient through a locally self-reinforcing Wnt signal and a secreted long-range inhibitory signal, which determines cell fate along the oral-aboral axis ^9,35^. Previous studies suggest that this head inhibitory signal is a secreted Wnt antagonist ^19,36^, though its precise identity remains unknown. Notably, a prior study demonstrated that *H. oligactis* has higher levels of head inhibitor compared to *H. vulgaris*. If this inhibitor is indeed a Wnt antagonist, this would be consistent with our findings that injury-induced Wnt signaling is attenuated in *H. oligactis*. In addition, we found that foot regeneration potential in *H. oligactis* increases with greater distance from the head, further supporting the hypothesis that a Wnt-inhibitory gradient emanating from the head influences the strength of injury-induced Wnt induction and, consequently, foot regenerative capacity.

In addition to a strong Wnt inhibitory environment in *H. oligactis*, reduced transcriptional activation of injury-induced Wnt signaling genes may also contribute to its impaired foot regeneration. In *H. vulgaris*, bZIP TFs are implicated in activating the expression of Wnt signaling genes during the general injury response ^12,20^. Although *H. oligactis* demonstrates injury-induced transcription of bZIP TFs comparable to *H. vulgaris* (Fig. S8), it is possible that in *H. oligactis*, bZIP TFs fail to activate the transcription of Wnt genes due to reduced protein stability or weaker binding affinity to Wnt gene regulatory regions. Supporting this, previous research revealed stronger CREB-binding activity in nuclear extracts from *H. oligactis* injured oral tissue compared to injured aboral tissue, whereas *H. vulgaris* showed equal CREB binding activity at both wound sides ^37^. Moreover, our data indicate regulatory feedback between Wnt signaling and bZIP TFs. ALP treatment causes down regulation of bZIP TFs in *H. oligactis*, consistent with the extended bZIP transcriptional activation observed in *H vulgaris* when inhibiting Wnt signaling with iCRT14 ^12^. This suggests a bidirectional regulatory relationship between bZIP TFs and Wnt signaling. Future research is necessary to investigate if differences in bZIP TF-mediated transcriptional activation of Wnt signaling genes contribute to the *H. oligactis* foot regeneration defect and how this process may be influenced by the stronger Wnt inhibitory gradient proposed for this species.

In our previous study, we found that prolonged injury-induced Wnt signaling is sufficient to induce ectopic head formation ^12^. However, our current findings reveal that injury-induced Wnt signaling activation is not strictly required for head regeneration. Transient inhibition of Wnt signaling activation during the injury phase of regeneration delays, but does not block *H. vulgaris* head regeneration, mirroring the extended timeline we observed for *H. oligactis* head regeneration.

Our findings suggest two potential, non-mutually exclusive mechanisms for establishing the Wnt organizer during *H. oligactis* head regeneration. First, the weak injury-induced expression of *Wnt9/10C* in *H. oligactis* suggests that while injury-triggered Wnt activation may contribute to head organizer formation, the associated feedforward loop could take longer to establish due to the low initial signal. Second, an unidentified mechanism may activate *Wnt* gene transcription later during head regeneration, either as the sole mechanism in *H. oligactis* or in addition to weak injury-induced Wnt activation. A similar late-acting mechanism may also function in *H. vulgaris*, enabling head regeneration even when injury-induced Wnt activation is inhibited (Fig. 4G).

In contrast to head regeneration, *Hydra* foot regeneration *requires* injury-induced Wnt signaling. Our findings indicate that a key event in this process is the Wnt-dependent expression of the TF *dlx2*, consistent with a prior study showing that *dlx2* knockdown inhibits foot regeneration in *H. vulgaris* ^38^. The TF *gata3* has also been identified as a positive regular of basal disk fate ^39^, and may be downstream of *dlx2*, based on its expression dynamics in *H. oligactis* ALP-rescued foot regeneration. However, *Hydra* possess two epithelial layers, the endoderm and ectoderm, which must interact to specify the foot ^40^. Interestingly, *dlx2* and *gata3* are both expressed in the ectoderm ^29^, whereas *nk2*, another foot-specific TF, is expressed in the endoderm ^41^. Although *nk2* has been proposed as a regulator of foot specification ^41^, functional evidence remains lacking. Our study shows that *dlx2* is upregulated by 12 hpa in ALP-treated foot regenerating tissue, whereas *nk2* upregulation occurs at 24 hpa, suggesting that ectodermal specification may occur before endodermal specification. Future studies should investigate how Wnt signaling activates *dlx2* in the ectoderm and how ectodermal and endodermal regulatory programs interact during foot regeneration. Constructing tissue-specific gene regulatory networks will be essential to understanding how Wnt signaling coordinates regeneration across both epithelial layers.

Our findings suggest that foot regeneration ability is ancestral in the *Hydra* genus but was lost in the Oligactis group. In contrast to most *Hydra* species which can generate gametes continuously throughout life, species within the Oligactis group exhibit semelparity, a reproductive strategy where organisms produce many gametes before dying ^42^. In *H. oligactis,* low temperatures induce gametogenesis, leading to somatic senescence through stem cell exhaustion ^28,43^. One possibility is that the foot regeneration defect in *H. oligactis* evolved as a trade-off associated with semelparous reproduction. To explore this possibility, future research should explore the potential function of Wnt signaling in the induction of gametogenesis in *H. oligactis*. Similar trade-offs involving Wnt signaling may occur in other animals. For example, the semelparous planarian worm *Procotyla fluviatilis* cannot regenerate its head when cut near the tail due to excessive Wnt signaling activation upon injury; pharmacological downregulation of Wnt signaling rescues regeneration ^44^. In addition, sexually mature male zebrafish show impaired regeneration of amputated pectoral fins due to androgen-driven inhibition of Wnt signaling, which can be reversed by pharmacological activation of Wnt signaling ^45^. These examples suggest that adaptations modulating reproduction may compromise regenerative capacity, highlighting the need for further research on how Wnt signaling mediates the trade-off between reproductive strategies and regeneration in *Hydra* and other animals. While Wnt signaling activation during regeneration is a shared feature among distantly related animals, the gene regulatory networks linking early injury response to Wnt signaling activation are not entirely conserved across taxa ^1^. Such variation limits our ability to draw broad conclusions about the evolutionary gain or loss of regenerative abilities, underscoring the need for more comparative studies. However, comparisons across long evolutionary distances can be difficult to interpret due to the distinct biological contexts in which these molecular pathways function. For this reason, studies of closely related animals with varying regenerative abilities, such as the one presented here, provide valuable insights into specific evolutionary rewiring events in the gene regulatory networks underlying regeneration. Such research can reveal the molecular changes driving the diversity of regenerative capacity observed across animals. By identifying these evolutionary changes, we may uncover the molecular requirements necessary to induce regeneration in animals with limited regenerative potential.

## Methods

### *Hydra* strains and culture

Here we used two strains of *H. oligactis*. The Cold Resistant Swiss strain provided by Brigitte Galliot, was used only in regeneration experiments shown in Figure S1 ^43^. The second strain used for all remaining *H. oligactis* experiments, was collected in Innsbruck, Austria by Bert Hobmayer ^12^. *H. vulgaris* AEP strain was used for comparative approaches throughout this work. The remaining *Hydra* species used for the regeneration experiments shown in Figure 1: *H. oxycnida, H. hymanae* and *H. viridissima,* were provided by Robert E. Steele. These strains form part of a larger collection of *Hydra* species validated through sequencing of cytochrome c oxidase I (CO1), their collection and taxonomy have already been reported in detail ^27^. All *Hydra* species were kept in *Hydra* medium (0.38mM CaCl_2_, 0.32 mM MgSO_4_ X 7H2O, 0.5 mM NaHCO_3_, 0.08 mM K_2_CO_3_), and fed three days per week with Brine Shrimp during experimentation.

### Phylogenetic analysis

Representative validated species from each major group in the of the *Hydra* genus phylogeny were selected and the sequence for CO1 was used to build a maximum likelihood tree with a GTR +G +I model using MEGA ^46^. Sequences of CO1 used are provided in the Source_data_1.txt file (See **Source data** section).

### Regeneration assays and foot peroxidase staining

Amputations were performed under a stereoscopic microscope using a millimetric grid to precisely measure the distance from the head. Head regeneration was measured by assessing the presence of the first two tentacle buds. Foot regeneration was primarily assessed visually by identifying basal disk morphology under a stereoscopic microscope. Foot regeneration data for all species shown on Figure 1 is provided in the Source_data_2.xlsx file (See **Source data** section). To confirm the presence of a regenerated basal disk, foot peroxidase assays were performed in *H. oligactis* polyps that appeared to have regenerated their feet nine days after amputation. Animals were relaxed in 2% urethane for 5 minutes, then fixed in 4% formaldehyde at room temperature for 1 hour. Animals were then rinsed three times in phosphate buffer saline solution (PBS) + 0.25% Triton X-100 for 5 minutes each. Foot peroxidase staining was performed by incubating animals for 15 minutes in a solution of 0.02% diaminobenzidine, 0.25 % Triton X-100 and 0.003% hydrogen peroxide in PBS. Following staining, animals were rinsed in PBS + 0.25% Triton X-100 for 30 minutes, then placed in 25% glycerol for 5 minutes, followed by 50% glycerol for mounting.

### Lateral head organizer grafting

For each time point tested, three batches of at least 10 *H. vulgaris* and 10 *H. oligactis*, were bisected. A small portion of tissue from the oral wound site was excised and set aside for grafting. Host animals were punctured at the mid-section of the body using a scalpel. The regenerating tissue from the donor oral wound site was picked up with entomological tweezers and inserted into the puncture site of the host animal. The grafted tissue was held in place against the host’s body column using tweezers and a dissection needle for 5 minutes, allowing it to adhere. Lateral head formation in the host animals was evaluated every 24 hours for 6 days, after which total number of animals with lateral heads was counted. Lateral head formation data is provided in the Source_data_9.xlsx file (See **Source data** section)

### Alsterpaullone (ALP) and iCRT14 treatments

For ALP treatments, three batches of at least 20 top halves of bisected *H. oligactis* were incubated in either 0.05% DMSO or ALP at 2.5 µM, 5 µM or 10 µM in *Hydra* medium for 3 hours. Following treatment, animals were immediately washed, and foot regeneration was assessed 144 hours post-amputation. During this period, animals were fed 3 times per week. For iCRT14 treatments, 4 batches of at least 29 *H. vulgaris* were incubated in 5 µM iCRT14 or 0.05% DMSO for 12 hours post-bisection. After treatment, both halves of the animals were washed and monitored for head and foot regeneration every 24 hours for 96 hours. Animals were fed every 48 hours following amputation. Foot regeneration data for ALP-treated *H. oligactis* is provided in the Source_data_10.xlsx file (See **Source data** section)

### RNA-seq library preparation

For each regeneration sample, at least 3 replicates of 30 *H. oligactis* were starved for 48 hours prior to bisection. Following 50% bisection, animals were incubated in either 0.05% DMSO or 5 µM ALP for 3 hours, then washed with *Hydra* medium. For the 0 hours post-amputation samples, animals were exposed to DMSO or 5 µM ALP for 30 seconds, then washed before RNA extraction. Regenerating animals were allowed to regenerate for 3, 12 or 48 hours, after which they were rinsed with *Hydra* medium and the tissue adjacent to the oral and aboral injury sites was excised for RNA extraction. The amount of tissue collected for each sample was approximately one sixth of the total body length. For ALP-treated samples, only the aboral injury site was collected. In addition, 3 replicates of 30 animals were used to collect homeostatic head tissue by decapitating just below the tentacle ring, and 3 replicates of 30 feet (peduncle + basal disk) were obtained as homeostatic foot samples. A second batch included 3 replicates of 24 hpa in DMSO and ALP, as well as 3 replicates of 0 hpa samples in DMSO and 2 replicates of homeostatic head and foot following the same conditions. Total RNA was extracted from the excised tissue fragments using Trizol^TM^ (Thermo Fisher Scientific), followed by DNA decontamination with DNAse I (R1013, Zymo Research). A final RNA purification step was performed using the Zymo RNA Clean and Concentrator kit (R1017, Zymo Research) according to the manufacturer’s instructions. Poly(A)-enriched mRNA libraries were prepared, and 150 bp paired-end sequencing was performed on a NovaSeq 6000 by Novogene Co.

### RNA-seq and computational analysis

Low-quality base calls and sequencing adapters were filtered out using Trimmomatic V.0.36. The reference transcriptome for mapping the trimmed reads was obtained by downloading the *H. oligactis* Cold Resistant (Swiss Strain) transcriptome from https://hydratlas.unige.ch/ ^47^. BUSCO analysis ^48^ revealed a high degree of redundancy (26.3% duplicates) in this transcriptome. To reduce the redundancy, Evidentialgene tr2aacds.pl (v2017.12.21) was used to decrease redundancy ^49^. BUSCO stats for both the original transcriptome and reduced transcriptome are shown in Supplementary Table 2. The resulting reduced reference (Supplementary data file 8) was then used for mapping the trimmed reads and calculating transcript counts using RSEM^50^. For annotation, predicted coding sequences were analyzed using IntersProScan^51^. Batch effects were corrected using ComBat_seq R-package ^52^. Normalization of transcript counts and differential expression analysis were performed using Edge R ^53^, with gene expression differences assessed using glmTreat and a false discovery rate of 0.01. Results are provided in the Source_data_5.xlsx and Source_ data_11.xlsx files. Additionally, *H. vulgaris* counts from *Cazet et al., 2021* were reanalyzed using glmTreat for comparison. Principal Component Analysis, heatmaps, and gene expression plots were generated using *in house* scripts.

For Orthoclust analysis, FPKMs for *H. oligactis* and *H. vulgaris* mapped reads were calculated with RSEM, and reciprocal orthology was determined using ReciprocalBlastHit.py ^54^. OrthoClust was run with a p-value of 0.001, correlation threshold of 0.975, and a Kappa value of 2.

Differences in gene expression patterns over time between DMSO and ALP treatments were analyzed using maSigPro, with Q set at 0.05 and R-square at 0.8. Gene Ontology enrichment analysis of maSigPro clusters was performed using FuncAssociate 3.0.

### RNA fluorescent in situ hybridization (FISH)

FISH was performed following a previously published protocol for *H. vulgaris* ^29^, with the only modification being the use of 250 ng of probe per sample for hybridization. Probe detection was carried out using Alexa Fluor 594 tyramide reagent (ThermoFisher). Confocal images were acquired using the Zeiss 980 LSM with Airyscan 2 confocal microscope. Z-stack images were analyzed with Fiji ^55^.

## Statistical information

Results for secondary axis formation from a regenerating graft are shown in average percentage of three different batches for each time point and species. Error bars represent the standard deviation. Three batches of at least 10 grafted animals were recorded for each time point and species of *Hydra.* Detailed numbers for each batch and timepoint can be found in Source_data_9.xlsx within the Source_data folder (See **Source data** section) Significance of the difference in secondary axis formation between species was evaluated through a two-way ANOVA, using the ‘aov’ function from the ‘stats’ package in R. This analysis showed that there is a significant difference (*p_species_* = 2.55 X 10^-5^) in secondary axis formation between *H. oligactis* and *H. vulgaris*.

Results for foot regeneration after ALP treatment with different concentrations was shown as the mean foot regeneration percentage of three batches for each tested concentration. Error bars represent standard deviation. The difference in foot regeneration percentage between tested ALP concentrations and DMSO treated control was evaluated with a two-tailed t-test using the ‘t.test’ function from the ‘stats’ package in R. The 2.5 µM ALP showed a significant difference when compared against DMSO controls (*p* = 0.03). The 5 µM ALP showed a significant difference when testes against DMSO controls (*p* = 0.01). Finally, 10 µM ALP showed a significant difference when compared against DMSO controls (*p* = 0.02). Detailed number for these batches of treated and control animals can be found in the Source_data_10.xlsx file within the Source data folder (See **Source data** section).

Results for head and foot regeneration for iCRT14 treated *H. vulgaris* are shown in average percentage of three different batches of 30 animals for each time point and four batches of at least 29 animals in the case of DMSO treated controls. Error bars represent the standard deviation. Detailed numbers for each batch and timepoint can be found in Source_data_11.xlsx within the Source_data folder (See **Source data** section). Significance of the difference in foot regeneration and head regeneration percentage across time between species was evaluated through a two-way ANOVA using the ‘aov’ function from the ‘stats’ package in R. This analysis showed that there is a significant difference in foot regeneration percentage (*p_treatment_* = 0.002824) between DMSO-treated and iCRT14-treated *H. vulgaris*. No statistical difference was found in head regeneration percentage between DMSO-treated and iCRT14 treated *H. vulgaris*.

## Data availability

All scripts used in this study are available both as a git repository at https://github.com/cejuliano/oligactis_foot_regeneration. FASTQ files of raw RNA-seq reads and raw counts for RNA-seq are available through the Gene Expression Omnibus under the Bio Project accession number: PRJNA1231128.

## Source data

A comprehensive list of the raw data used to produce the graphs in this study is provided here.

## Supporting information

Read Me for Data and Source Files

## Acknowledgements

We thank Bert Hobmayer and Jack F. Cazet for their constructive comments on the manuscript and all members of Juliano Lab for discussion and suggestions during the development of this work. We also thank Bert Hobmayer for providing the *Hydra oligactis* Innsbruck 12 strain. This work was supported by a National Institutes of Health (NIH) grant R35 GM133689 (to C.E.J.), and a Human Frontiers Science Program Postdoctoral Fellowship (Reference number: LT000496/2020-L; to S.E.C).

## Author information

### Contributions

C.E.J and S.E.C conceptualized and designed the study. S.E.C, S.N., J.C., J.T., B.D.C and C.C. performed experiments. S.E.C performed data analysis. C.E.J. oversaw all the experiments. S.E.C and C.E.J. contributed to writing of the original draft and the remaining authors read and critically revised the manuscript.

## Ethics declarations

Competing interests

The authors declare no competing interests.

---

**Supplementary Figure 1.**
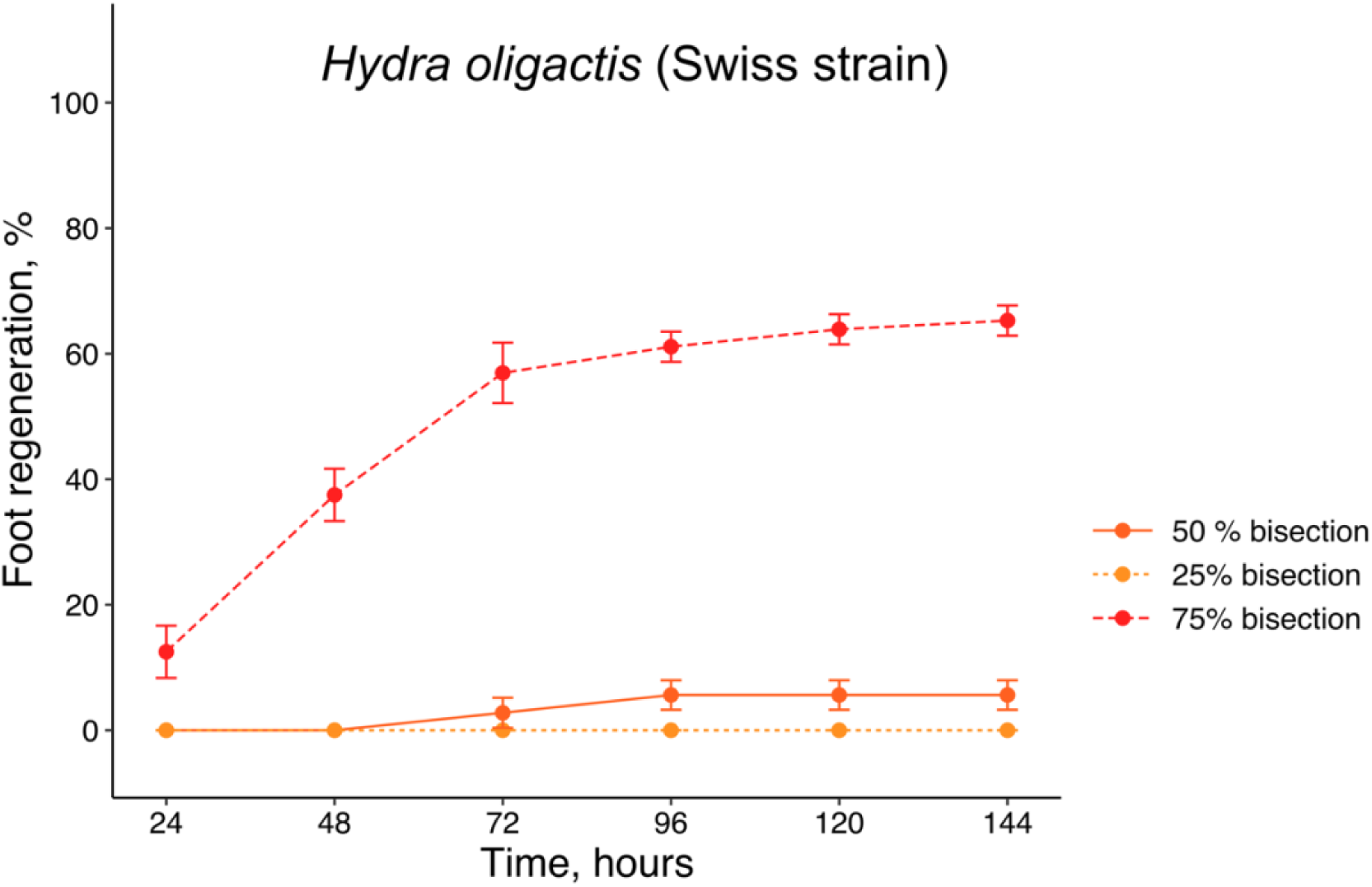
The *Hydra oligactis* Swiss strain displays a foot regeneration defect like the Innsbruck strain. Approximately 5% of *H. oligactis* Swiss strain polyps bisected at 50% body length successfully regenerated their feet (solid orange line), while the remaining ∼95% remained footless. In polyps bisected at 25% body length, 100% remained footless (light orange dotted line). Polyps bisected at 75% body length exhibited a higher foot regeneration potential, with over 60% successfully regenerating feet. See Figure 1F for diagram of amputation sites.

**Supplementary Figure 2.**
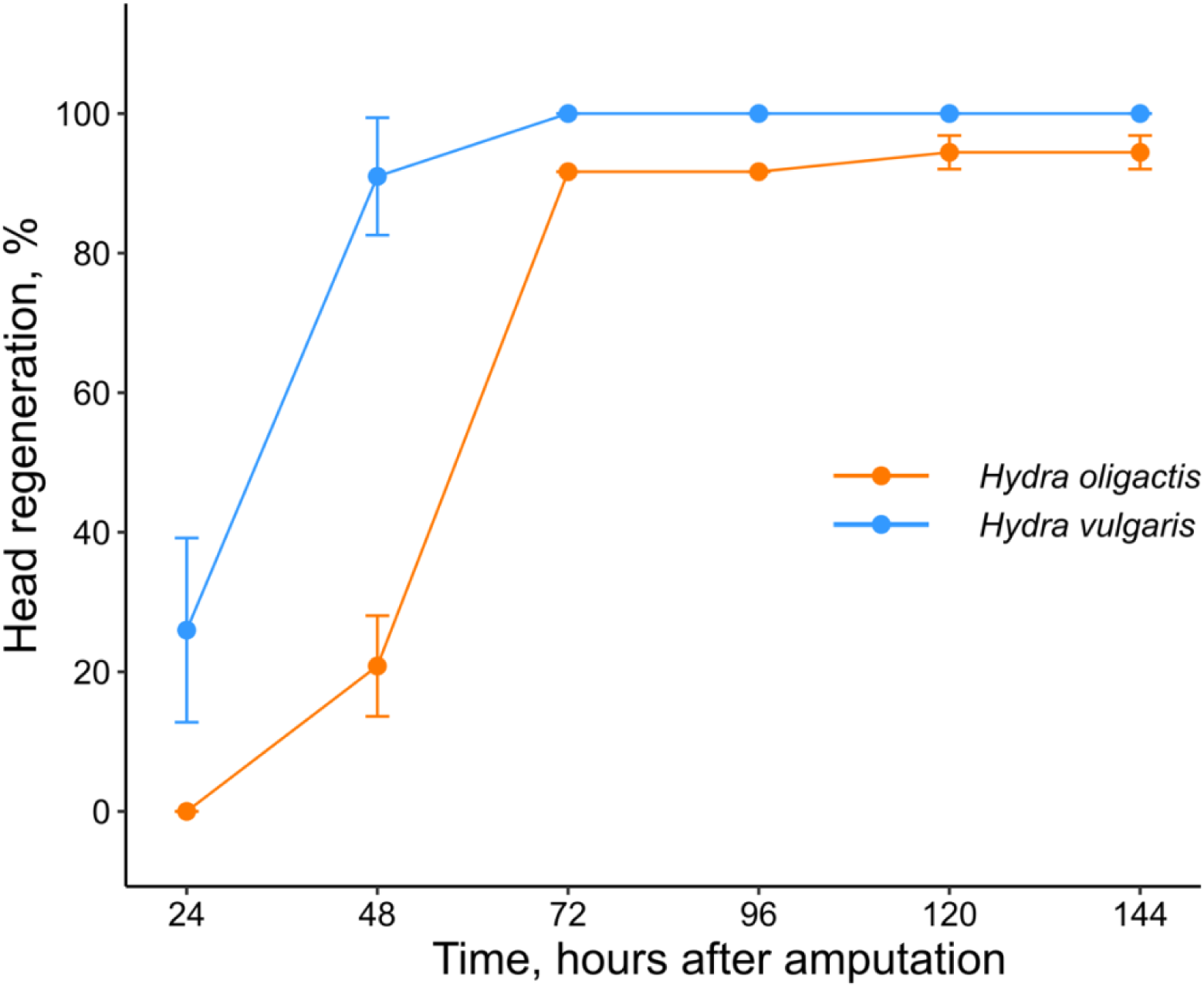
Head regeneration occurs at a slower rate in *H. oligactis* as compared to *H. vulgaris.* By 48 hpa, approximately 90% of bisected *H. vulgaris* regenerate their heads as determined by the appearance of tentacle buds (blue line plot), whereas only about 20% of *H. oligactis* achieve head regeneration within the same time frame (orange line plot).

**Supplementary Figure 3.**
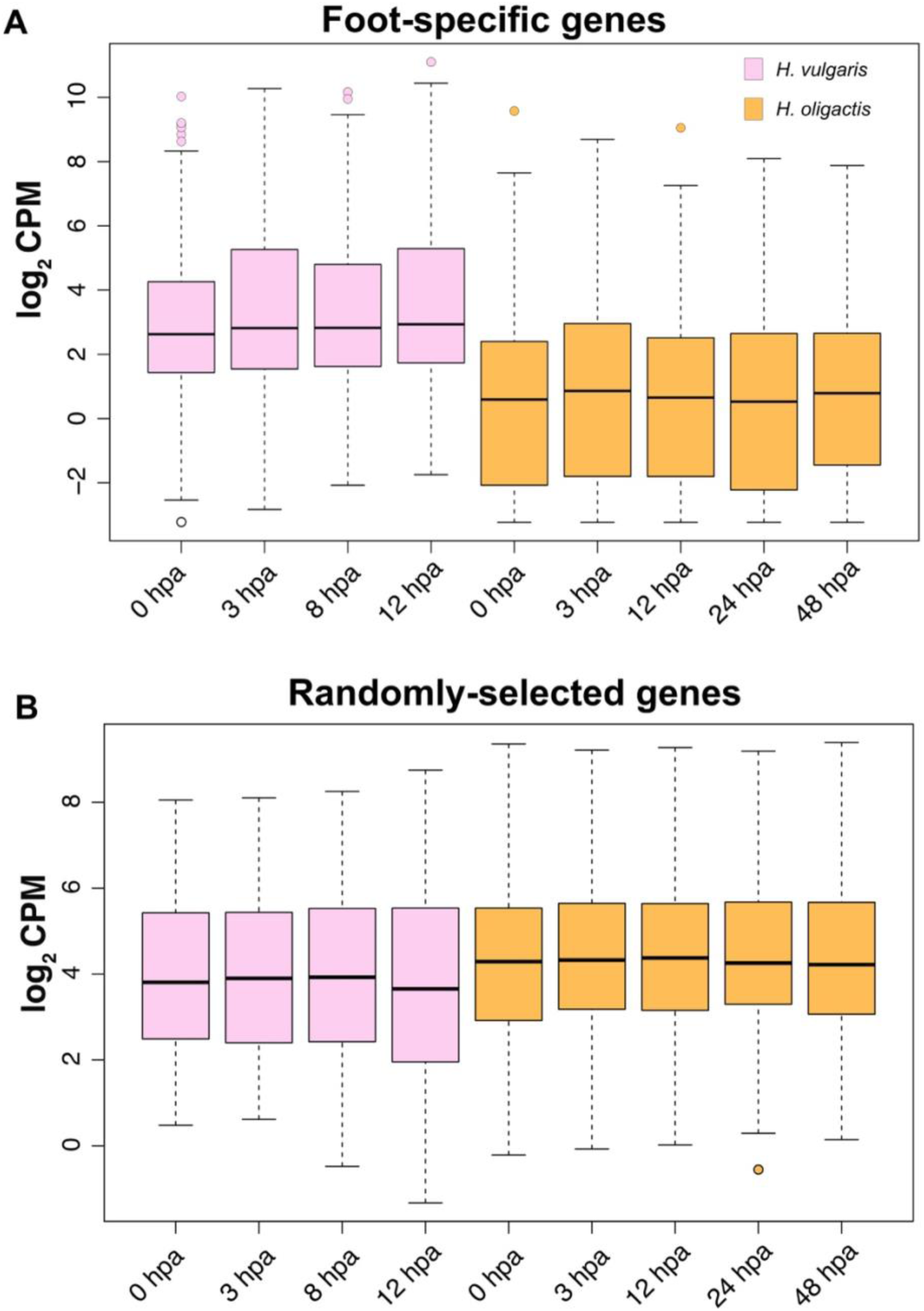
Foot regeneration genes are expressed at lower levels in *H. oligactis* compared to *H. vulgaris.* (A) Boxplots showing expression levels of 154 orthologs from the foot-specific gene list identified between *H. oligactis* and *H. vulgaris* reference transcriptomes. (B) Boxplots of 65 randomly selected orthologs between *H. oligactis* and *H. vulgaris* plotted as controls. The log2 counts per million (CPM) for these transcripts were plotted over the course of foot regeneration in *H. vulgaris* (pink boxplots) and *H. oligactis* (orange boxplots). Foot-specific genes were consistently expressed at lower levels in *H. oligactis* as compared to *H. vulgaris*, including at 0hpa.

**Supplementary Figure 4.**
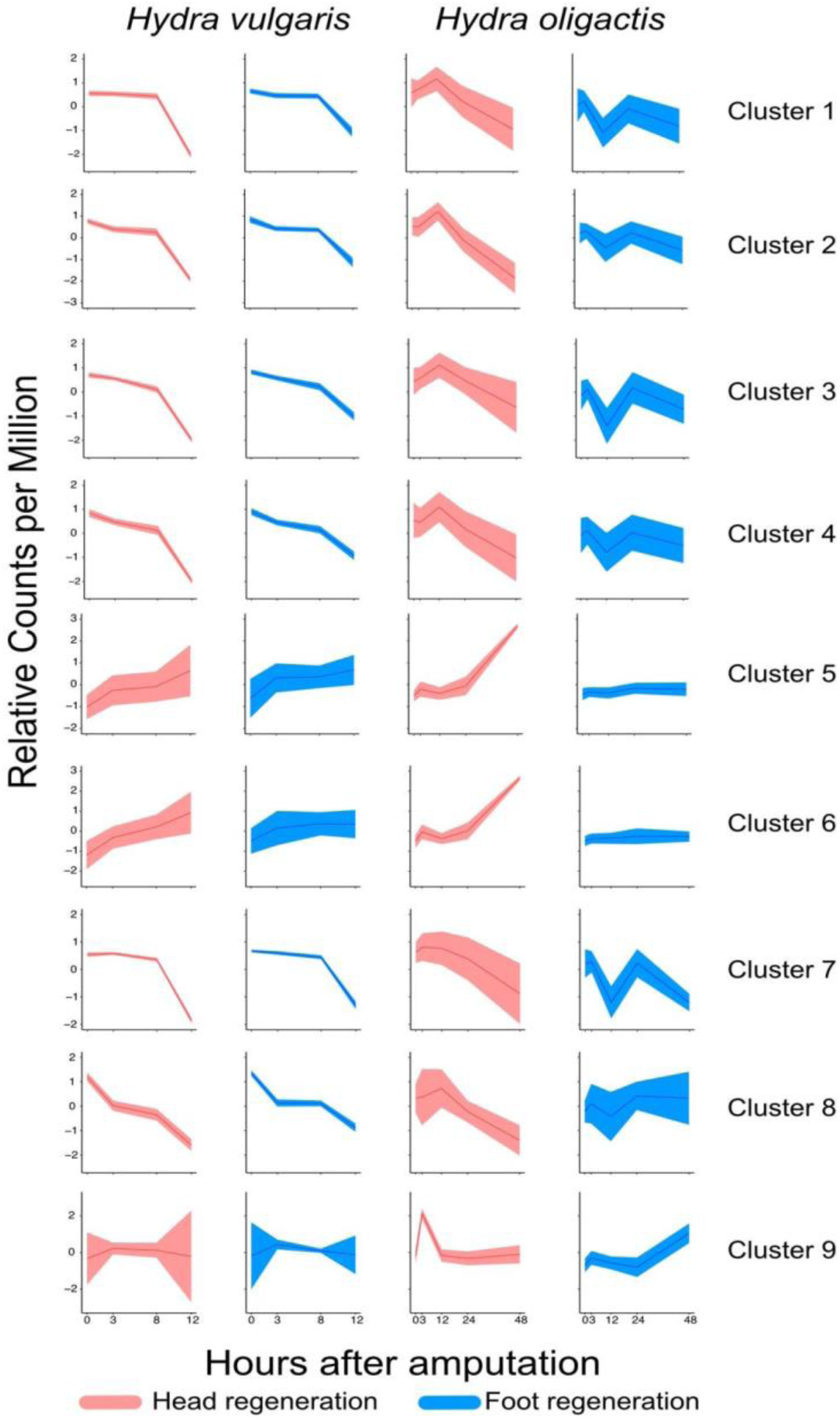
Gene co-expression analysis using the OrthoClust pipeline identified nine distinct clusters of orthologous genes with varying expression patterns between *H. oligactis* and *H. vulgaris.* Red ribbon plots depict the expression patterns of clustered genes in oral regenerating tissue, while blue ribbon plots represent expression patterns in aboral injured tissue. Expression profiles for *H. vulgaris* clusters are shown on the left, corresponding profiles for *H. oligactis* are shown on the right. Note that the *H. oligactis* RNA-seq data spans a longer time frame (0-48 hpa) as compared to the *H. vulgaris* data (0-12 hpa).

**Supplementary Figure 5.**
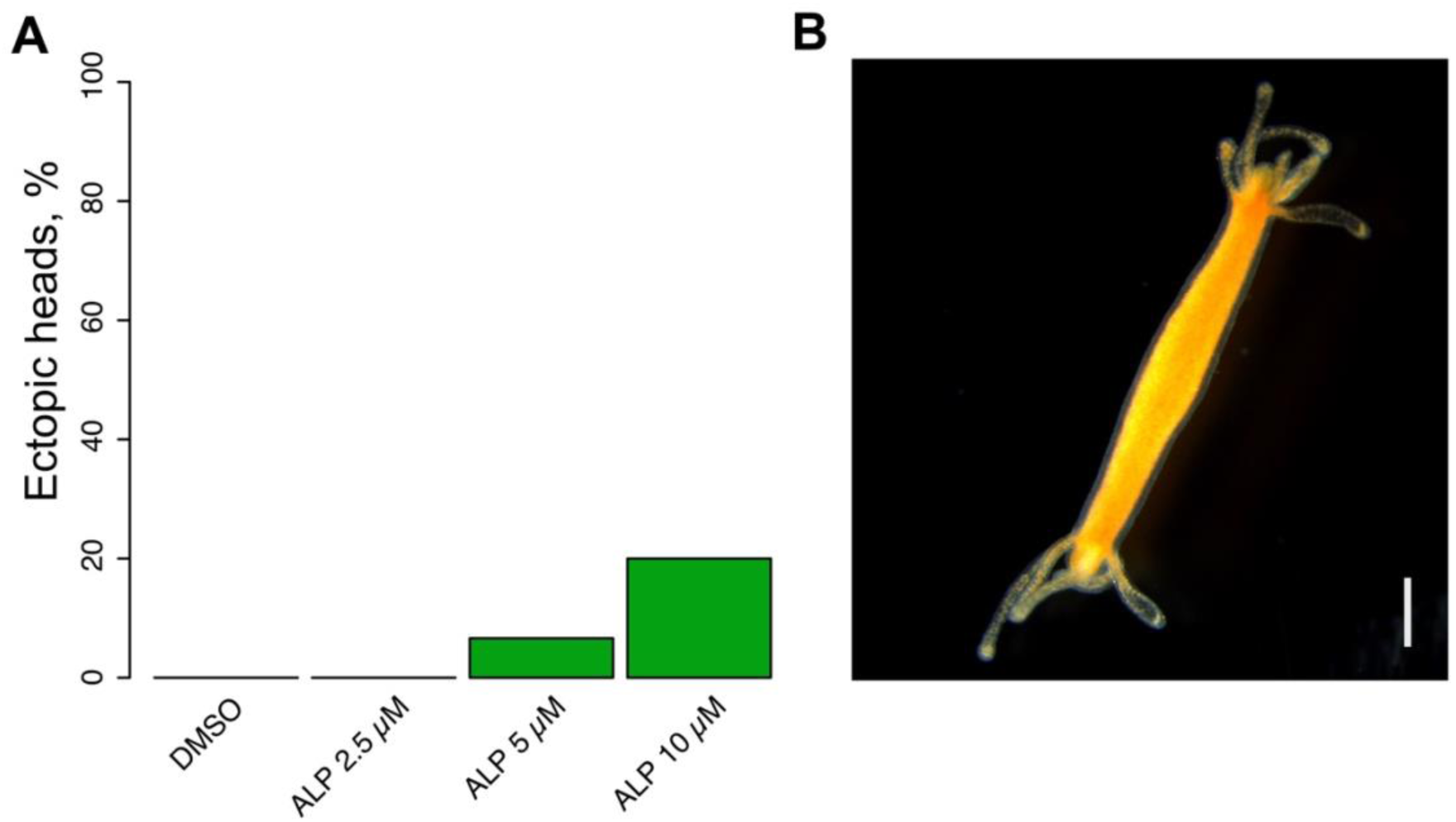
High Wnt signaling activation induces ectopic head regeneration at the aboral wound site. (A) Bar plot showing the percentage of ectopic head regeneration in *H. oligactis* upper halves treated with ALP at different concentrations for 3 hours. Ectopic head formation at the aboral wound site was assessed 6 days after injury. (B) Representative image of ectopic head regenerated at the aboral wound site in *H. oligactis* treated with 10 µM ALP. The image was taken 14 days post-injury. Scale bar: 1 mm.

**Supplementary Figure 6.**
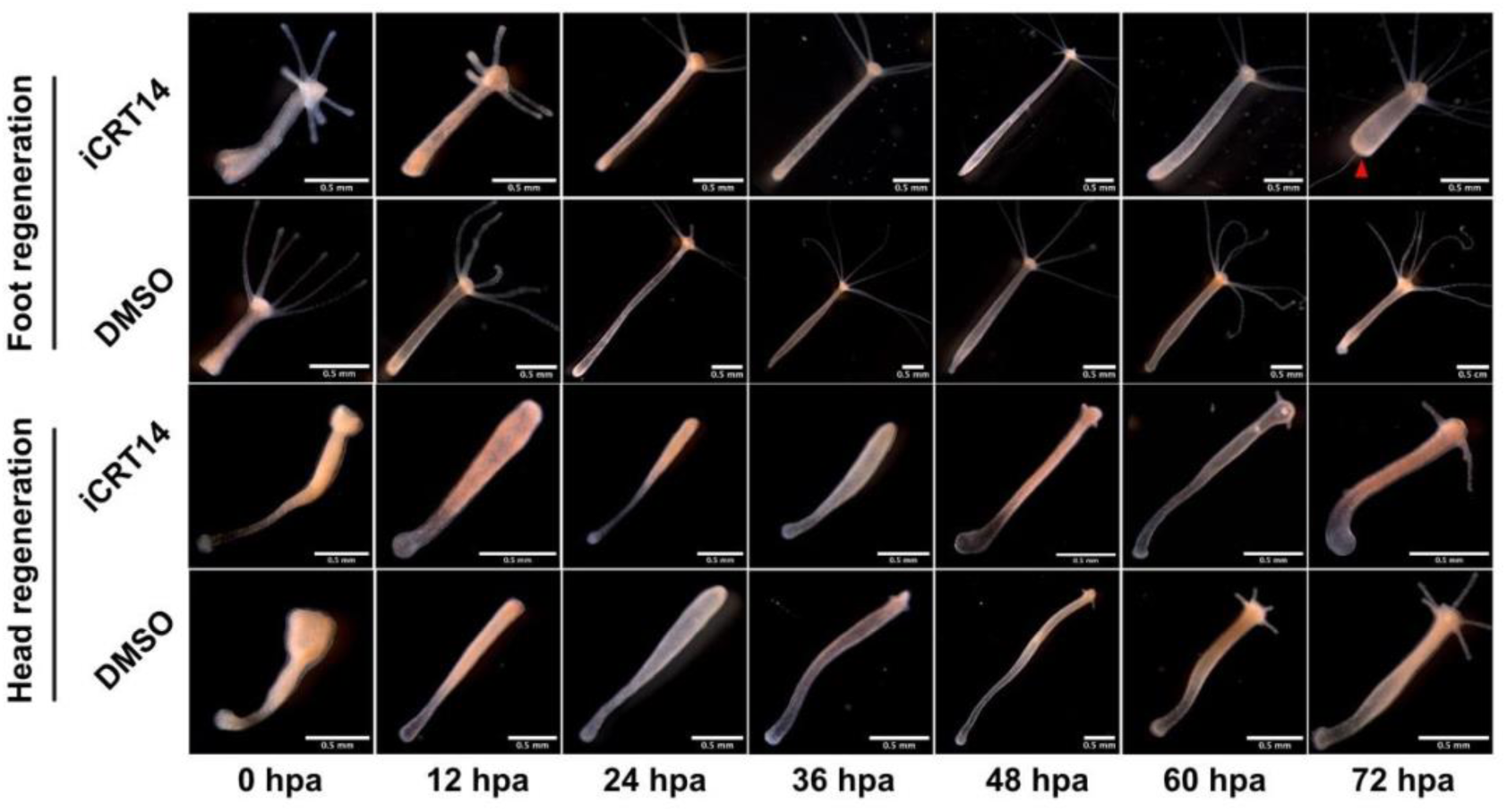
Wnt inhibition blocks foot regeneration and delays head regeneration in *H. vulgaris*. *H. vulgaris* polyps were pre-treated with 5 µM iCRT14 for 2 hours prior to 50% bisection and maintained in iCRT14 for 12 hours after injury. Representative images illustrate foot regeneration (top) and head regeneration (bottom) over the time course. Foot regeneration is inhibited by iCRT14 compared to DMSO treated animals (See Figure 4F), with inhibited animals becoming stably footless (red triangle points at a stably footless aboral end in treated *Hydra* at 72 hpa). Head regeneration showed a delay in the first appearance of tentacles in iCRT14-treated samples as compared to DMSO treated animals (See Figure 4G).

**Supplementary Figure 7.**
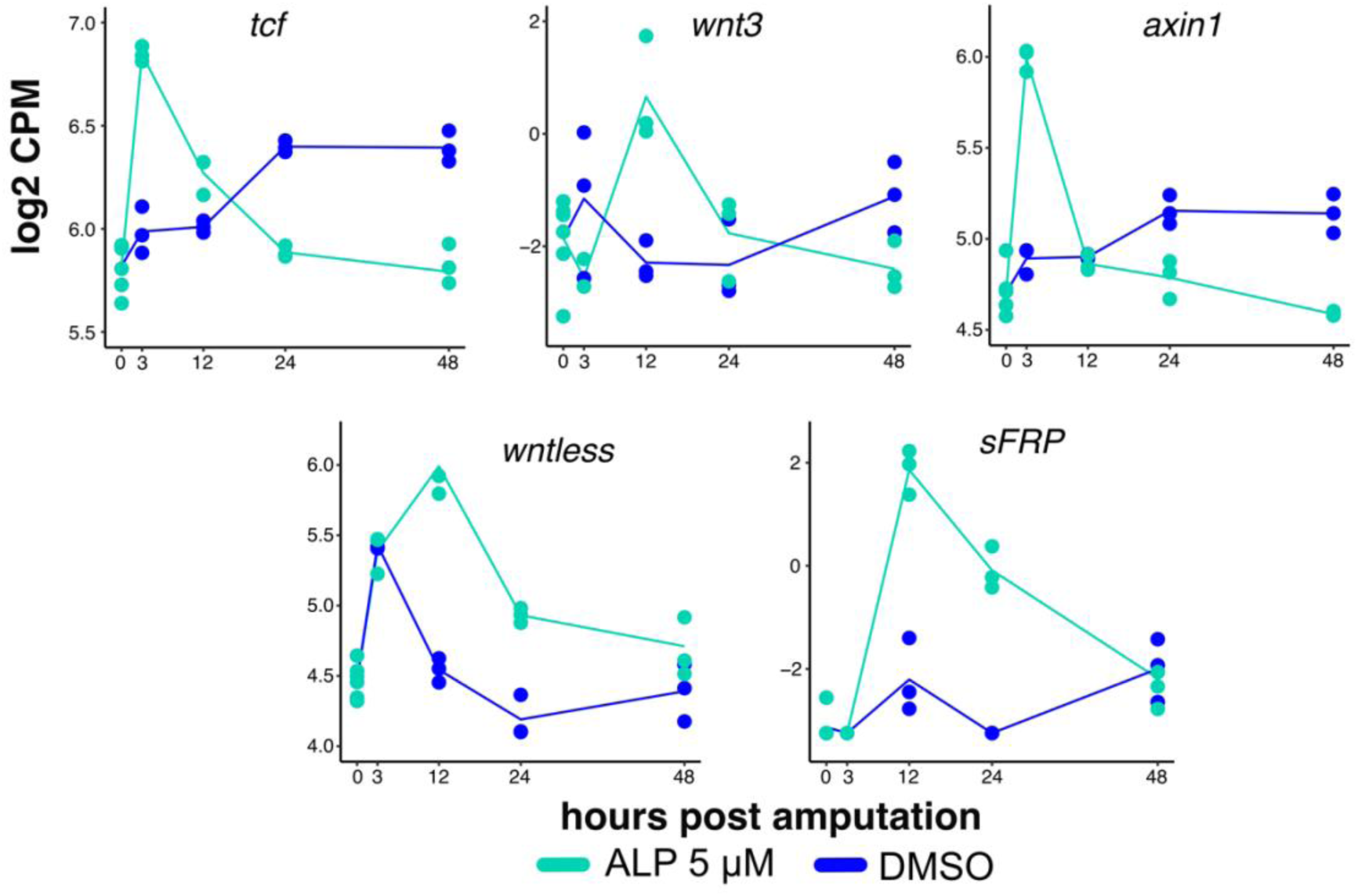
ALP treatment induces the transcriptional activation of Wnt pathway genes. RNA expression plots show log_2_ counts per million (log_2_ CPM) for Wnt pathway genes *tcf, wnt3, axin1, wntless* and *sFRP*. The blue line represents expression in DMSO-treated failed foot regenerating tissue; the green line represents expression in ALP-treated foot regenerating tissue.

**Supplementary Figure 8.**
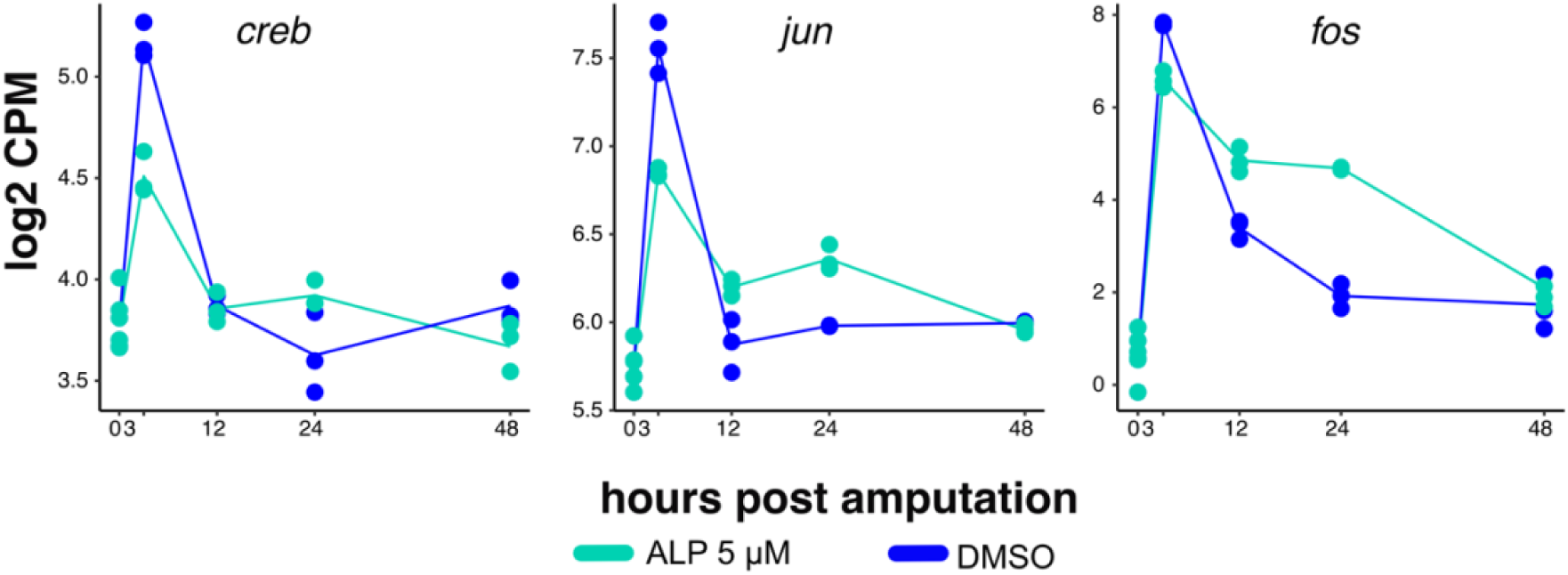
bZIP transcription factors are downregulated with ALP treatment at 3hpa. RNA expression plots show log_2_ counts per million (log_2_ CPM) for bZIP transcription factor genes *creb, jun* and *fos*. The blue line represents expression in DMSO-treated failed foot regenerating tissue; the green line represents expression in ALP-treated foot regenerating tissue.

**Supplementary Figure 9.**
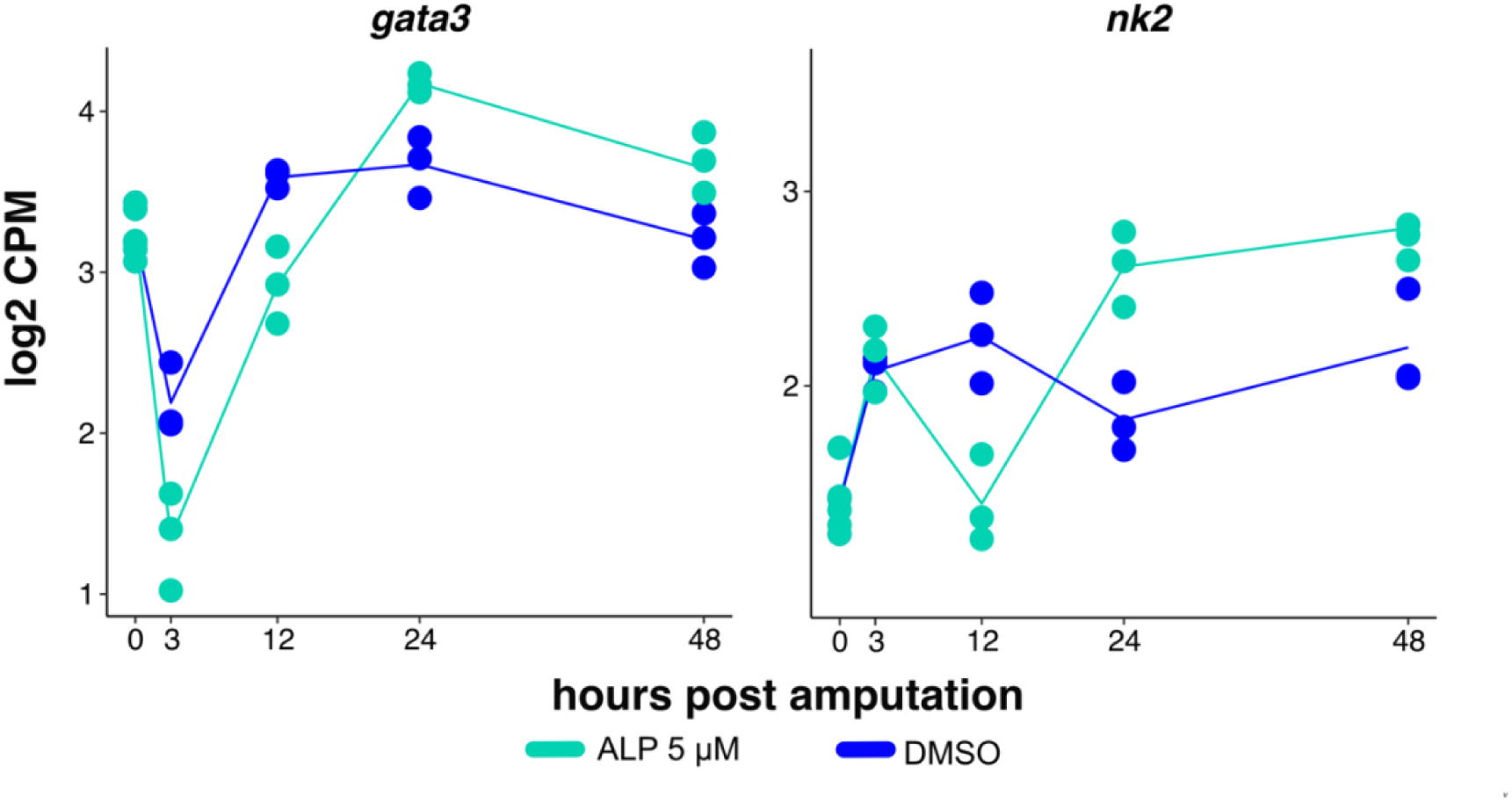
ALP promotes the upregulation of foot-specific TFs at 24 hours post amputation. RNA expression plots show log_2_ counts per million (log_2_ CPM) for foot-specific transcription factors *gata3* and *nk2*. The blue line represents expression in DMSO-treated failed foot regenerating tissue; the green line represents expression in ALP-treated foot regenerating tissue.

