## Supplementary material for "Wnt signaling restores evolutionary loss of regenerative potential in *Hydra*": Read Me for Data and Source Files

This folder contains the data used to plot all the figures in this work as described below:

Supplementary data files 
Supplementary data files are references within the manuscript
Supplementary Data 1: Spreadsheet file containing Foot-specific and Head-specific genes.
Supplementary Data 2: Csv containing the top 100 genes driving Principal Component 1 in the foot regeneration timecourse PCA analysis.
Supplementary Data 3: Csv containing the top 100 genes driving Principal Component 1 in the head regeneration timecourse PCA analysis.
Supplementary Data 4: Spreadsheet file containing foot-specific and head-specific genes shown on Figure 2G-J.
Supplementary Data 5: Spreadsheet file showing clustered genes and their expression values from OrthoClust analysis.
Supplementary Data 6: CSV file containing a list of modules and the genes contained in each module from maSigPro analysis.
Supplementary Data 7: Spreadsheet file containing Gene Ontology enrichment analysis for each gene module from maSigPro analysis.
Supplementary Data 8: Text file containing sequences for transcripts in the Reduced reference transcriptome used in this study. The orginal fasta file will be made available through Dryad upon publication. 
�

Supplementary source data files

Source data 1. Text file containing the accession numbers and sequences for CO1 used to build the phylogenetic tree.
Source data 2. Excel workbook containing head and foot regeneration data for the experiments shown in Figure 1 G-K.
Source data 3. Excel workbook containing differentially expressed gene tables for the conditions compared in Figure 2 C-D.
Source data 4. CSV file containing normalized log2CPM used to plot Figure 2 E-F
Source data 5. Excel workbook containing log2 Fold Change values from comparisons in Figure 2 G-J.
Source data 6. Excel workbook containing log2 CPM for selected foot-specific genes in H. vulgaris and H. oligactis shown in Figure 2 K-L.
Source data 7: FPKM for H. oligactis transcripts used for Orthoclust analysis shown in Figure 3A-B.
Source data 8: FPKM for H. vulgaris transcripts used for Orthoclust analysis shown in Figure 3A-B.
Source data 9. Excel workbook containing secondary axis formation frequency from grafting in H. vulgaris and H. oligactis Figure 3M.
Source data 10. Excel workbook containing foot regeneration percentages for H. oligactis treated with the different ALP concentrations shown in Figure 4B.
Source data 11. Excel workbook containing foot regeneration percentages for H. vulgaris after 12 hours of iCRT14 treatment in Figure 4 F-G.
Source data 12. CSV file containing normalized log2CPM used to plot data shown in Figure 5 B and H.
Source data 13. Excel workbook containing fold change gene tables for the timepoints comparing foot regeneration in DMSO and ALP in Figure 5C and D.
Source data 14. Excel workbook containing Gene Ontology enriched terms for maSigPro modules shown in Figure 5L and M.
