## Supplementary Data 8 for "Wnt signaling restores evolutionary loss of regenerative potential in *Hydra*"

>R000007c0g1_i01|Hydra
AAAATAGCCAAGATGTTCAAGCCTCTCCAAGTATTAGTGACATTAGAGAGAAAAGAAGAAAACTCAGAGCTGAGAGAAGAATTGACAGAGAAGTGAGAAGAACAAAAGAGAAGGTTGATGGAAAAGTCGGGCAAGCTGCATCTGAAAACCAAGTCAGAAGAGAAAAATTGAGATTGGCCAGAGAAAAGAAAAGACTTGCAAGGGAAAAGGAAAGACTTGTTAGAGAAGGTGCAGAACCCGTAACTGAATCAGTGATTCAAGATATGGTAAAAAAGATTAAGGTCAAGAAGAGCAAAAAGAAGAAACAATGCGGAATCCCAAAGTCTTCCAAAAAGCCAAAAAGTAGAA
>evgR000018c0g1_i01_Hydra
TCCAGCCACCCATGTTTGTAAACAATGCAACTTACACTATTGTGACAATCACATTAAGATTCACAACAATTCCAACAAGACCAAGTCTCACATTGGTGACGTTTCCGAAGTTAGTGGAGGTAAATCTTCTGGTGGTTCTTCTGCAAGTGGTGGATTTGTTGAATTCAGTGGTAATTATACTCATATTGCCACATTGGGTGAAAATGAAGAATTCAACTTTAACTTTGCTGGTCCAAATGCAATTGCTATTGACCCAGTTAATCGCAGTCTCTTTGTTGCCGATCGTTGGACTTTCTTTGAATTTGATTTAAA
>evgR000064c0g1_i01_Hydra
CTAGCTTATCAACTTAAAAAAATAAAAAATGGTTTCAAAAACCGATTCAAAACTTTGTTTGTTTACTTTTGGTATTGTACCACGTTGAAATATCCAAAATGTTTGGTTCAATGAATTTTATTGAAAGTATTTTGCCTTTGTCAAAAGTTGTTATGTGTCCTGCAAATTGCAGTGGCAATGGTGAATGTGTCAGTGGAATATGTTCCTGTGAGGTTAGGTTTTCCGGTTGGTCTTGTAATGATCGAAACTATAGCTATTTTTTATCTTTCGGGATCTTTTTCTACATTGTTTGTTTCATGGCTATTACACAGCTGGTTATGTGTATCTATTCCGACTTTGTGCAACTTAAAAGGCGAAATATTTTTTATGCATTTCGATTTACTATTCAAAAACTTATTTATGTAGCTACTATTTTTGCTTCTGGATCACGTGCAATATATTTCACACTTGATTTGCATAATCTGATTCCTTCTTCATTTAAAGAAAACTTGTTTTTAACTTATTATGCCTTTGTGATGACAGGACTATCTTTGATTATTTGTTTTTGGGCAGAGTCATTCCATCTCTCTGGATTGTCCTCTGATAAGCCACAGTTTTTGACTAAATCTATCAGTGGGTTCGTCATTTTTAATATTATTATTTATACAATGCTGGTGGGACAGTTAGTTTCTAACAATGTTCTGGCTCAGCCTATAAAATCAAAGATAAATATTATTATTAACAGTGTCTACGCTTTCCTCATGTTCTTTGTTCTCGTGTTCTTCCTAATTTATGGTGTTGAAATATTTTGCAAGGTTCAGGGGGCATTTAAATCAGAAAGAAGTTCTAAAGAAAGTATTAATATACATCAACTGTTTCAGTCAAGGTTAGGACTGATTGCTCAGGCATCTTTGCAATTAGCGATTACAATGTTTTTAACAAGTGAAGTTCTCGGTGAACTTTGGAAACGAAAGGCAAGAATAACCGATCGCAACATTGTCAACATTGTGTTTCATATTGCCGAATTAGGTTGTGCTTTATGGTTTCCATGTTGTTTGTGGAATGTTTTTAGACCAGATGAGTTATGGATATTAAATCCTCGTAAACTATTACTGCTCCGTAAAAACTTTGAAAATCAACCAATTGCATCTTTGCGAAAATCTTATGAGAGTTTAGTAGAAATATCTCAAGTGGAAGATGAAAGCAAAGGAAAACAAAATCTTTGTTGGATATGTTACGATGATGATAATAAAGTTGAAATCATTGAGCCATGCAACTGCAAAGGTGGAATGAAGTCAGTTCATCACGACTGTCTAAAGAAATGGCTTCAAGAACGTCCAGAAAATTCTGATTCATCAACTTTATGTTGTAGCGTTTGTAAAGTACAGTACGATGTGACAAGTGATCACTCGTTATTCTGGAATCCTGACAAACTTCAATTTAGAGCCTGGGCACAAACATTTTTTGTTGTGCTAGTAATGGCTTCAATGCCAGTACTTTTGTTTGCTCTTTGGCAAAAAATTAAAATTGCTCCGTTAAAAGTTGGACTTGTGATGCTGATGGTCATTGCTGAGTACTTTATGCTGAGAATGCTTGGTTTCAACTACATCAAAGCGTATAAGTACACTAAGACCCGTTCTTTAAAAATTTTAAGCAGAGGGGCTAAAATTGAAACTTCTGCATTGAATATCACCAGCTAGATCTTGGTTTTTAAAAAGATTTTAAGTTATAATTAATTATTAGTAGTAAAATATTATTGTATAAATTATTGTTTGTTTCAGATATACTTTATTTATGTAAATGTAGATATTTTTGTTCGATTTTTATCTTAGCGTATTAAAGTTTAACGTAATGAAGTACAACTAATAGTTGTAGTAATTGGTGATTTTTTGTTTGTAATATATTAATTTTGTGTTAACTAAATGTAACGTAGAAGTAAACAGAAAATTATTTGAAAGTATGTAAATTTTTGGAAAAAAAAAA
>evgR000076c0g1_i02_Hydra
TCAGCTCACCGCTCAGCTCACCGCTCAGCTCACCGCTCAGCTCACCGCTCAGCTCACCGCTCAGCTTACCGCTCAGCTCACCACTCAGCTCACTGATCAGCTCACCGCTCAGCTCACCAAAAGTGTACCAAAAAAGAGTTCCCCGTCGCTCTAGCGAGCCATTAAAACCATTAAATAAGATTTATAATTATGTAAGAAAGTTTCAAAGCTTTGACGCAGGTTCAAATGTTAAAAAATGCAAAGTATTTGAGGAAAAACCTGGTGAAGCCAATTTAAAAGAACTTTCAAATAAGTTGAAAAAACTTTCTTTAAAAAATCCGCTGCAACATGTATTAAATCAGGAGGTTATACTTCCTAGCAAAAAACATAATAAAAGTTCGACATCTGTTTTTAATGTTTTTCCGCCTATACCGAAATGGAGTCCTAATCCTAGCCCAAATTTTGAAACTTTCTGTACAGGGGTAATTAATTCAGAAAAAAAAGATTTTCCCCCAACACTTTCACCTAAAGAACTTGTATATAATGTACTAGACAACGGCAGTATTGAAGTCATTGGTCAATCAAATCAAATAAAAGAAACGCGGTATTTCAATGATTTTTCATCAGACTATCTAAATTCAATGCAAATGACAGATACCCAGTTTTTCAAAGATTTATCGTCAGATCATGTAAATTCTATTCAAATAAGCAATACACAGTATTGCAAAGATTCGTCACCAGATCTTGCAAATTTTAACAATTTTTTATCCGCTCATGCAAGCTTTGACGATTTGTCATTAAGTCAAGCAACCTTTGAGAATAAGTGCTATGATCAAGAAGTATCTATACAGTCAACCAGTTATATGTATCAACAAAGAAATGATGAACCAAATATTGGTGTAGATTATCAAGCAAATATTCAACAAAATAATTAATCAATAAGTTCGCCAACAACCACTTTTTGTTTGATACGTCCTACAACGTTCCTACAACGTTAGCTGTTAACACAGAATATTTAGATAATGTATTTGCTTCTCAGAATGATACCAAAATCTTAATGCAATTCAACAAAGAGTGTAACAACATACTGGATGTTAACAAAATGATGGATTAAAATGAAACGCCTTCAATAAAACATTCGCAGAGTATTGCTGGTTCGAACACCTTCACACCTTCTTCTTTACCAAAACTATTTGAGGAGATTTCAACTCTATTCCCTTAACCCCAAAATCTTTTTATTTAAATCCATCGTGTGAGTATGAAAAAAATCATTTCCCTAAATTTCAATTACGATTACTACCTAATAACAATATTTATAGATAATAATTGAATACAGAAAATAATATATTGTTAATATATTATATATTATTTAAATATAATTTATAAAGAATAAAAAAAAA
>evgR000082c0g1_i01_Hydra
TTGACTTCTATTACACGATCTTAACAATACACAGCTGTACTTTTGACTTCTATTACACGATCTTAACAATACGCTGTACTTTTGACTTCTATTACACGATCTTAACAATGCACAGCTGTACTTTTGACTTCTATTACATAATCTTAGCATTTCATAGCTGTACTTTTGACGTCTATTACATGATTTGTACATTTCATAACTGTACTTTTGACTTCTATTACACAATCTTAGCATTTCATAGCTTTTCATTTGACTTTTATTACATGATTTGTACATTTCACAGCTGTACTTTTGACTTTTATTACATGATTTGTA
>evgR000093c0g2_i01_Hydra
GTTAGCAGTACAAAAGATAAAAAGATTGTATCACCCGAAGAAGTGGTTATTAAAGAACCTAGAAAGTTTGGATCACGAGAAGAAGTGGCTTTTAAAGAAAAGACAAAGATTGAATCACGAGAAAAAGTGGATTTGAAAGAACTGAGTAATTGTTAAGAAGTGAATAACAAGTAGGAGCGTTTAAATTACTATTTATTGTTTGTCTTCAAAATGACGAAAACCTGTTATGTTCAAAATTGTTCAAACAACAACAAAAAAGATAAAAATATAAAGTTCTTTTATCTTCCAAAAGATAAAACTTTACGAAAGAAATGGTTGCTAGCTATTAATCGTATAGATATTAAAAATGTCACAAAAATATGGTGTCCCAGCTCTAAGAATGTTTGTATATGTTCTGCACATTTTATTACTGGAAAACGAGATTTTGCACAAGTTCTACAGAATATGTACCCTCTATCTTTCCACAGAGCACCTCTGGGAAAATTCCATTTTCACCTCATCGTCAATCAAGTATAAAACATTTTGAACGATCAGCTAATCGCTTAACAAAGAATACAAATAAAAAAGTTCAAAAGAAGTTACTTTTGGCAACTATACCAGAAGTGAATGAATTTGATATTGACTTAAACGAGAGTGTTAAAGGTCGATGCTTTGAAAAAATTGTGAATCTTCAATCACCACTTTGTGCTACCGAAAGAGAGCTTGTTTATAATGAAATGGATAGCCTAAGAAGTGATAAAAAAAAAGCAATGGAGTAACTTGAACAACAAAGTTTTAATCCATTGAAGTTTTCTTCTTTGGATAAACATCCTGAAAAATCTAATATGCTTACAGGTTTAAAATTTGATGTGTTGTGTAGTGTTATCAAATATATTTCGATGGAGAAAAAAATAAATCCTCAAACTCAACCGCATAAGATGGTCCTTGAAGATCAAATTATATTTACTCTTGTTAAGTTAAAACACAACGTGAATTTTGAATTTTTAGCATGTTCTTATGGAATATCAAAGACTACTGGAATTAAATATTTCTGGACTTGGGTAGACATTATGTTTTCAAAGTTATCTCAAGAATCGGTATTTGATTTTAAAAGATATTCTTCCTCTTCGAACAATTAAAAATATAGCAGATGAATCATTTGATTCACACTTATCTAACTGCGATAAGATTGTTACTGTTTGTGCAGCTTTAGTAAATTTAGGAGAAAGCATTGTCTATAGGCAATGATGTAAAATTATTATTAAAGTATGATTTAAG
>evgR000105c0g1_i01_Hydra
AGAAAAAATAAGAAAAACCAGTTTTAGGAGTTTACACCTAATTCTGATTGTCGCAGTTTTACTTCCGGTTATCACATTTTGTGCTTGTTACATAGTTGGGTTTAAACTACAAAAAATTAGTCCTTCTTTTGTTCCTTTTGTTAGTGACACTGGAGACACGAGACCTAATTCAAGTATATTTACATTTGGATTGTCGCTTAGTGGAGTTTTAACTTTGGTTATTGTTACAATAAGATATTACCAAGTAGAATATATTTTGTTAGAAACAGAAAAACAAAGTTGGATAAACTTGATTTCGTTAACTTGTGGTTTATTACTTGTATTTGGGGAATTTACAGTAGCAGCTTTTCAAATTTCAAATATAAAAGCTGTGCACTTTGTTGGCGCTGGTTTGCATTTTGGCGGCGCAGCATTATTTTGCGCATGTCAATGTTACTTTACTTATAAAAATTCAAAGAATAAATGTCAAATAACTTTTCTTGTTCGAGTTGCATCTACATCAATAATGACTATTTCGTCAATATTATTCAGTGTTTTTATGATACCTTCATTGAACAAGTACAATCGCAAAGGCGGAAGTGTTGGGCAGACCTTTGAATGGCTTCTAGTTACCAGCCAACTCATATTTATGCTCACATTTTTCAACGAATTTCGCCAAATAGAAATTGAGTTCAGAATAAAGAAGAAAGATACTGGCGAAGTTATTAAAGAATTAGAAAACGTTGAGTCTTTTAGATTTTGGAAGACATATCAGAGAATAGACTAGACAATTAACAATTCTAAAGTGAAATATTTCAAATATTAATGCATTATTACTCTTTGATTTTATTATACATTTTTTTCGTCGGGAC
>evgR000107c0g1_i01_Hydra
CATCGTCACTCGCTATTGTTGTTGTGTCCATTATCACTTTGGTCATTATGGCCACAATTGCTGAAGGCTTGCCATTGAAAAAGAGTCCAATTTCCTTCTCTATCAAGAAGATTGATGAAAATCCTAATGTAAAGATTATTGCGAATGGTGAATTGTGCCCAACATGTATTGATTTGTTGGATTTTGGTATGGCTGATTTGATGGCTGCTGTTGACAGTGGCGTTGTCTTTTCATGCTTGAGTTTGTGTAACTATGTCAGTCCATTGGGTCAATACGCTGTTGCAGGATGTGATGTTGTCTGTACTGCCATTGGTGTTAATGCATTCATTGCTGTTCTTAAGAAGGCTGATTTGGATCCAATTTATGGATGTCAATTGGCTAATATTTGTCCAATTCATGATTGTACTGCCAAGACATGTGCTGTCATTGATAATGCTGGTGTTGTTCCATCCAATAGTCCAGCTGGTGGTTCTTTATCATTCTACTCCCATTTGAGAGTTTTGAACGAAAGTGGTCCTGGTGAAGTCACTTTGGAAGTTTTCGGTGCAAGCATTGCCAAGAATCAAGAAGTCCAACGTTACTACCAAGCTGCTGGTTTCAAACCAGGT
>evgR000108c0g1_i01_Hydra
TGTTTTGACGTTATTATTTTTTCTTTACGTTAATTTTCCGGAGAAACAACAGCATGTGTGGTGTTTATAGTATACCGTAATATACCGGTATGTTACTGAGCATGGATAGAAAATATTATGCATCTTTTTTGTGTGTGTTTGGGGCATTCACCTTTTTCTTAAAGTGGCTATACAACCATTCTATTTGCACAACAACAACAAGATTAGATGGAAAAGTAATTATTATAACAGGTTCAAATACAGGAATTGGAAAAGAAACCGCTGTTGAAATAGCAAGGCGAGGGGGAACTGTAGTTATGGCTTGTCGAGATTTAAAAAGAGGACAAGCTGCTCTAGAAGATATAAAAAGACAGAGTAATAGTCACAGGATTTTTTTAAAAAGACTTGATCTTGCATCTCTTAGTTCCGTACGCAAATTTACCTCTGAATTTATTAAAGAATTTGATTGCCTACATATTTTGATAAACAATGCAGGAATAATGATGTGTCCTTACTGGAAAACTGAGGATGGATTTGAAATGCATTTTGGTGTTAACCATCTTGGACATTTTGCTTTAACAAACTTGCTTTTGAAACATTTTTCAGTTCATGGTCGAATTATAAATGTTAGTTCAGTTGCTCACAAATATGCAACAATTAATTTCGAAGATATTAACTCTGAAAAAAATTATTGTAGAAGAAAAGCCTACTGCCAGAGTAAACTAGCAAATGTTTTATTTACCCGTGAGCTTTACCGAAAACTTATTGGTTCAAATATTTCTGTATATTCTCTACATCCTGGAATAATAAACACTGAGTTAGGAAGGCACTCTTTTTTTAAATATTTGCTTCCACTTCCAGGTTTTAAATCCCCTATGCAAGGGGCTCAAACATCTATTTATTGTGCAACAAAAGAAGGATTAGAAGACCAGTCAGGAAACTATTTTTCAGAATGTAAACTTGAAAAAAGTACTAATAAAGCTTTCTTCGATGAAGGACAAGCCAAGAAATTGTGGGAGTTGAGTGAATATTTAACTAAAACAACTTACCCTTTATAAAGTTAATTTTTTTACTTATAACCTTTTTCTAAAATTGATAATAATATAAAATAATTTATGATAGAAATAAAATATTTTTTATAAAGTTTCTAAAAGTTGTCTGTATTATGTTGTCTGTATTATGTTGTATTAGTTTATGTTGTATTAGCCTCC
>evgR000113c0g1_i01_Hydra
TTTTCCAGTTTTGATCCAACAATCCCAGAAGATTTGCTCACTCAACAACCTGATGTCTCATGGGAATACCTTTTCCAACCAGATCAATGGTATATTACACCAGAAGTAATCTCTGAAAACTTCTTTGCTCTCTTCTTGAGTTTGGCTGCTGTTGCTCTCGCTGCTGGCATCTTCTGGTTCGCCAGAAATAAGGGTATCAAGACTAAATGGTATAGAGATTTGGAAAAAGGAAAATATTACTTCTCAGAATCTCTTATTTTCAACATTTGGATTGCTTTGTACATTCCATTGGCCATTGGTTCTTGGTTGGCATATGTTCATGGTGGCCAAACTTGGAACAGAGCCTTGACTGTTTATGCCTTGCACTTGGTTGTTAATGTCCTCTTTTCTGTTTCTTTATGGTGGGTTCAAGATTTGAGCTTGGCTTTGTTGAATCTTATCACTTTGATTGGTGTTTCCATGTTCACAACAAGTCAATTTAATAGTATTTTGAAATTCTCAGGCTACATTAACACTCCATATATGTTGTGGTTGTTGATTTTCACTGCCCAATATGCCTATTTCTGGTACTTGAATGAAGGAAAAGAATTAATGGAAGTTGCCAATCTTGCCAAGGGTGGTAGTGCTAAGAAGGCTCTTAAGAAAAAGAAGGGTCTCCCAACCGATGTCAAGAAGAAGCTTCAACAACAAGTCCAAGAACAACAAAAGACTATGACACCTTCCTCTGATAAGGATGAATAATTTTTGAGAAGAATTTTTCAATATTAATGACGCGACTAGTCAATTGTTG
>evgR000121c0g1_i01_Hydra
TTTGGACAGTACCAAAAAGTCACATAAAGTGTTTTATCTAAACACATAATAATATCTAAGTGTTCTCGGTTTATATTATCCAACTTAGATTCTAAGCAAGATCAATTTATATCAATCAAAATCGTGGATACGTTATGGCAGATAATTCTTATATAGCCACACGCAAAAGATCAAACCCTAGTACAAGCCCGATCAAAAGACTTTACTCAGTTGTAACTCTTAATGAAGATAATTTTCATAAAAAGTGTATGGTTCCATTTCATTGGGTAGAGGAAGATGAAAAATGTGTCTACTGGCCTCCAGTTGGAGAACATCCCCTACACTACTACATATCAAAATGGCTTTCTCCTTGTCAAAATTGGCAACAATTTCCTCTAATCAAAATCTCCTTGAGTAAAGGCACCAAAGAAATGTGCAATGTATGCTTGAATATTGACTCAGAATTATCAGAGTGTCCTCCTGCAGGTGTTGTACTAGAAACATCTGAAATGATCCGCGAGCCTAGTCCGGATCTTGTAATTAAAAGAATACCAAAGTTTAGTGGTAGTATTAAAAATGTTAAATCCGAACAATGCAAAAGAAGGATTAAAGCTCATTCTAATGAAAGCGAAAGCTGTTCTTCTGATTCCCATTCCAGCCCAATAAAAGTTGCAATCCAACAAGCAAAAAAATCAAAAAATTCTCACACTCATAATACTGTACCCAAACACAACTATACTACCTCTACTTCATTTTTTCATACTTCAGCTTCTAACAATAACAATAATCAATCCCCAAGGAGATCTCCATTAAAAGAAAATACTTTATCTCATTCCATGCTCAATTTAAGTCAATCTAATTATACTCCCTGTAACAAGGATGCATCACTGTCACTCTGCCCACCAAAATCCACTCAAGCATGTCGTTTTGGAAATTACGGTGACCCACCTTCTTCGTTTATGTTAATGGATCAGAGGCGAATGCAGTATGGGTTGTTTATGGAGTTGGCTGCTATTAAAGCCAGTCAGGCTGATATTGTTACCAAATTGGAACGCTTAGGTAGGAGAATAGAGCCAGCAGATTCAAGTTTCCATCTTACAAAGTTTGACACCGTGGAAACATTTGAGATGTTTGATAAAAAGTTGGTTGATGAAAATGAGTATACTCTTCTGGTTAGTCGTTTAAAGCAACTGGGCGGACAGTCACCTTCGCAACTAGTAAGTAAATGTATAAATGGGTTGATGGAGAAACATGTACAGGCGCAGTTCAACAAGATGGGTAACAACGGGAAAAAATCGTTTAAGATTACTTTGTCATATAAAGCAATACTAGCTGCACTTGTATCTCAAACTAACACAAGGGAAGCGATAGATAAATCCATTGGAGAACATTTAAAGCGTGCTCCTCGTGGATTGACTGCAAATTATGAATAACAAGTTATTTTATATAAAAGATATTTACA
>evgR000129c0g2_i01_Hydra
AAAAAGTTCAATTACAGATTACTGAACTTATGTCATCAATTGAAGAAGTTTTATTGGTAGAAGTTGTTTCATTAGAATCGGATTTTAATGATAGAGTTCGCTGCACGTTTAATGTATACCTATTGTTTCAAAACGAAAATGTTATCAATCAGATAACCAATCTCAAGAACAGTGATATAACAAATGTTGTTTCAGACAACGATGCAGCATATATGAGAGATGATATTCTTGAATATTATTCAAAGAGTGCTACTTTTGACGTTCTTCAATCTCAAAAGACGACTGTTGTACTTTCGTTAGAGATTTCTGAAGATGTTAGGTACCCTGTTAGTAAAATATTGGTTCAGTACGGACCTTTTGTTAATGAAAGTTTTAAAAAACTATTGGTTAACAATGCAAAGTTCATTTTCATTGATGGACTGGAGCTGTTTAAAGAATATAAAGTTTGTTTTTCATTTCAAACTATAAGAAAAAGATTTTTTTCTTCTTGTGTGCAACAAAATAAAACGTTTACAACTTTTCAAGGAGAACCTACAGAATCCCCTGTGTTTAACTGTTCACTTAATGAAGAAATAAAGACAATTTATTGTAATATGATGCAAATCAGTAACCAATCGTGGAATGGTGTCCCTTATAGTTACCAGCTAGTTTATTGGTTGAAAGGTTTGACAGACTCTTGTAAAAATTTATATATTTCTGGTCCATATAAATACTCAATCATGATAAAATACAATACCACAAGCAATTTAAAATTTCAAATAAAAACAGCTGATTTTCATTTTGTGGAGTTATGTATTGTCGGCTATGCAACAACAAATGGTGGGAAAGGTATTGGCGTCTGGAAAGAAACTATAGTTCGTACCCAAAGAGAAGCACCAAACACTGCACCTGAAAATTTTACTGCCATAATAAATCGTAACTCTATTCAGTTTTATTGGAATTTCCTTGAAAACACTATAGACGCTTGGAATGACTATAAAATACCTGGATTTTATCAACTAAGATATGGAATTGTGGCAAGTTCTTTTTTTAATGATATTGAGTTCATCACAAAAAATGTAACAAGTAATGAATTTGTACTAACAGATTTTAATGCTTGCTACTTATATATAGCTAGTGTAAACGCATTTTCTGTTGCTAGTGGTCCTAAAAGCTACTTATGTCTTTTAGATTATCTATCAAAGCCTGTCTGCATGACACCAGCCATTGAAAATTTCTATGTTGTCAGCTCTAGAAGTGCTTCGTTTACTACCAGGAGTTTTCTCCCAACTTTTTTTCGTGGCACAAATATTTATTGGATGTTTCATGTAGATGTTATAGATGAAAATGTTAACCCTGAACTCCCCTCTGACAAATTTCTTCATAAACCGTTTTATCGTACAAACTCATTAGAGAAATATTCTGGAAGGTCAAGTCAATGCACACAAAACGATTTGGCAGAATACTTCAACCAAAACAATGATTTTGTTTTAGATTTGATTGGCTTACATCCATATACCCAATACCGGGTATCAGTTCAACCTTGCAATGAAATAGGTTGTGGTCAAATTTCAAACAG
>evgR000136c0g1_i01_Hydra
ACAATCGTTTATCATCATTATTTAAATATAGAATGGATTGCGACGTTTTATACAAAGTTGTTGTTATTGGAGATTCTGGTGTCGGAAAATCAAACTTAATCACCAGATTTACTCAAAACAAGTTTACCAAGGACACAAAACCAACTATTGGTGTTGAATTTGGTGCTAAGTCAATTGAATATGATGGTGTAACTATCAAGGGTCAAATCTGGGATACTGCCGGTCAAGAACGATTCAGAGCCATTTCTGCCGCTTATTATAGAGGTGCCAACGGTGCTCTCATCGTTTATGATATTACAAACCAAGAAAGCTTTGATAATTTGGAAAAATGGTTCAAGGAAATTGAAAATCAAGGAGAAAATGGTTGTATCAATATTTTAGTTGGTAACAAGTGTGACTTGAACCATTTGAGACAAGTTGAGACATCAACTGGAAGAGCTTTTGCTGAAAAACATAACGTTTCATTCATGGAAACATCTGCTTTAGATAGCACTAACGTAGATGAAGCTTTCACAACACTTTTGAAGGAAATTTACAAGACCCAAAGCCGAAAGAGTAAGAAACCAGAAGAAGAACGTAGAGATGAACCAATAGGTCCAAGTGACAAGGTTATCATTGATGATGGTGAAGGTGAAAAGCCAAAACCAAAGCCAG
>evgR000142c0g1_i01_Hydra
ACCACGATCAATCGAGGAGATGTTTATCTATTTAAAGTGTGATATTTATACTTTTGAAGAAAATTTTTTGAAAAATATTGAGGAAATTTGTTCTCATCCAAAAGACTTAGTTATCACAAATATGAACCCAATCTCAGAAAACTCAATTCGAAACCTTCGAGAGCGATTATTTGCTGAAATTGTTGAGACATTTTGTGCAGATGAATTCAGCGAGGTAAACATAACGCTAGACCTCACAAATCCTGTTGAAAAGAATCTTCGTAAAAGATATAAAACAAGTTTATGTTTAGAAGATATTTATATACTCTCAATAACATTATGTGAAAAGCAATTTCACAAAGATATTGTTAAGGTAATTGTCTCAACAAAGAATTTGGACACTCTGCCCAGCGATAAACTTTTAATTAAATCAGTCAAAGAGTTG
>evgR000143c0g1_i01_Hydra
ATAAATCAAGACTCAAGATGCAAGGCTCAAAAATAAAGAAGTCAATATATTATGAAAAGTCTGGCCTAAAAATATGAAATATAACAATGGAGAAAACATTTTTACAATTAATAATGAAAACATTTCACCTAGTTTTGATGAAGTCATGGTTGCAATGAAACCTTCAATAAATACATCTACACTGGCAGCTGCTAGAACAAATATACCTTCTTCAATCCATACAAAATTAACAGACTCTAATCTTAAAATTGGATTGTCTCTGAAGCAAACTTTTAGTTTTGAAAATGAACTTGTTGTTGGACCAGACAGTCCTGTTTCATTAAGCAGTATTTATTCATCTCCTTCTATTAGTCCTATTAGATCTTTGAGTCAGTCAGCAGGATGCCATAAACTTCATTCAGCAATAAAAAAAGCACAAATGAATCTTAGTCAAAGTTGTTTTGCGGATATAGGTGGTAGTAAAGCTGGAGATATTGACAGTGAACTTACATGTTTAAATTGGCTTCAAAATGGTAATTTAATAAATGGTTTTACACCTGGTACGGCTAAAAAAGAGAGTCTACCAGAGGAGGACGATAGCTTACAAGAGTTACGAACTACATCTGATGCTACTGAGACTATTAATGATGTAGACCAATCATTAAGGAAACCTGCTTATTCATTTTCAACCCTTATTTTTATGGCTATTGAAAATGCAAAAGACAAACGTTTACCTGTTAAAGATATATATAGGTGGATACAAGAAACTTTTCCTTATTTTCAAAAAGCTCCTGTAGGCTGGAAAAATTCAGTACGTCATAATCTTTCCTTAAACAAATCATTTAAAAAAGTTGACAAGGAGAAATGCATTGGCAAAGGTTCTCTTTGGACCATTGACCATGAAGCAAGAGCAGGGCTTGTGCAGCAGTTACGAAAAACGACATACCATTCTTATCCATACGCTCGCCCTTATCCACAAAATTATACAAAATATTTTTCAAAAGATAGGGCAATGTGCAAAAGAGATTCTGCTGGAAGATTTGTTGCTAATAGAGATGATGCTGAGTTTGATGCAGCTGCTATTATGTGTTCACTCAGTGCACCATCTCTTTTTCGTTACATGGAACAGCGCTCAAAGCTAAGCTTAGATAAACAATCAAGTGCAGAACTATTACAAATATGGAGAACATTTATTGAGCCTGGTGATTGGCAAGAGGATGTCAATTGGAAAGACAACAATAGCTATTCAGTAGATAACTTTCTTGACGAAGTTCATAAAAAGTTGTATGGAAAACGGAAAAATAAAATCCATGATCGTGAAAGAGATAACGATGATGGCATAGACTCTAGTCTTGATGAAGAATACGAGTTTTGCAATTCAGATGACGATTTATCAGAGGATGATGAGATGACAAAGAATATGAAAAGACTTGTTGATAAAGAGATTGACTATGATCTTGGAGATTCTGGTTATTCAGAAAATTTATTTACTTTAGATCACTCTTACTCAAAGCCAATAAACAGTCTCATTTTACCATGTTTTAAGCGTGAATCTACTGGAGTTGCTGCATTATTACACTTAGCAAACGCTGCAGCGATGCAGCTTAAGCATCAAAGTTGTGGTCCAAAACTCACTAAAGAACAAGCAACCCTTACAGAGTTTAACGATAACAATCAAATGATAGATGCCGAATTGGACTTTACAATAGAATCGAAATTTACTGGTCTAGTTGAACTTGTCAGTTGAATACTCTAGCTCATTTCTTTGTGTTTTCTTTCTAGTTTTGACGATGATGCTGCTATTTTTGTATAATAAATTAATATTTTTTGCTGACGTAGTTTTGAATTGACATATTTTATAAAATTTATTATATAACGAATTCGTACACCTAACACATTATATATTCATTTAATAAAAGATTTGTCATACGTGTTATGTATTTATTCTTGTATTTTAAATGAGCTTGATGCATTTTATTATAAAATTACTTTTAGTGATACTTACAAAAACTTTATGTTCGCATTCGGTTTTTTAGAGTTGCGTTTCATAATTTTTTTTTATCACTCTAGGTTTTGGAAACGAAACGATATTGTAAATCGTTTTTAATGTTTTATTATTATTATTATTTTTTTTAGTACTTATATTTTAAGAATTTTATTTGAATTTTGCTTGGTT
>evgR000176c0g1_i01_Hydra
TTATATCTTTTTACTTTAATTTTATGCGATGAATTGAATTTGTCTCACTTTCGGTATGCTTGTTGCAAAGATTTATTTTCAAATGAAGCTAAAACATTATTAAGGTACATCTGTAATATTGATTTATTTGAATAAACTGAAATCAAAGTAAGAAAAAAAAGAAGAATTCTAGAACTATTGTAAAGTTGGGGAAAACAAGCGATTTCTTCATTTAATCAAAGTCGATCAATCCGTTTTCTTTTTGGTAAATATATGGATGTTGCACGCTTAAAAGTCTTTGTCGATGTTACATGGGAGCAGTACTGCAGACCGCTGGTGCGGCTGAATTAGCAGCAGAACTTACTTGTTGTGCATGTTCTTCTGTTTGTGGTCTTGTCACAAGGGACAGTGCTACTCGTGCAAAGTTTCGCTATTGTTCTTTTTTGCTGTTTGCAACAGTGACCTGCATCATTTTGCTTATACCGGGATTACGGTTTAAACTTGACAAAATTCCTGGGTTTTGTTCAAAGGTTGTATCTTCGGACACTTGCGATAAATTTGTTGGCTTTGCTGCTGTCTACCGCATACTAACAGCATTGGCTTTGTTTCATTTTGTCCTTTCAATTCTTACAATTAGAGTTACTCGAATAAAAAGTTTCAGAGCAAAGTTTAACAACGGGCTTTGGTTTTTTAAAATGGGATTAATATTTGCTTTAACAATTCTATTATTTTCGATTCCGAAAAAAGCAGGGATCTTTAGAATGTGGATGTACGTAAGTATGACAGCTGGTTTTGTGTTTATAATGTTTCAGATTATGCTTATCATCGACTTTGGAAATTCTTGGTCGTTATCGTGGGCAGAAAAGTTAGAAAGTGGTAATGCAAAATTTTGGTACGCAGCGATGGCTTTTGTCACATTATTTATGTTTTCACTGTCATGCGGAGCGATAATAACGTTTTATGTTTATTTTACTCATGCTCCAGACATTCGAAAATGCCATGCCAACGTTTTTTACATCTCGTTTGTAGGACTACAATGTTTTCTAGCGGTTCTTATATCAGTTACACCATCCGTACAACAGGAGCTTACTGGAGCAGGACTCCTTCAAAGTTCTGTTGTTGTATTGTATACAATGTATTTGACATGGAATACTTTATCATCTGAACCAGACTCCACCTGCAATCCACTGGGTAGTATCATACTTGAATACAATTCGCTTACCGGAGTAAGTGGCGAGGCTATTTTTGGATGTGCCATTACACTTATTCTGTTGGTTTTTGCATGCTCTGTCAGAGCCAACACTTCGCATTTAGGAAAATATGGATTATCATTGGCTGAAAGTGAAGATTTTGCAATGGCTACTTACAAAGAAGATAAATCTCGTGCAAATTTTGAAAATAGTCTTGAGACCGGAGGAAAAGAAGTATTTTTACACGAATACGTTGGATATAATTATTCATTTTTTCATATGGTTATGTCTATAGGATCTTTGTACGTATTAATGATACTGACCAATTGGCATTCTCCAGAAGAAAACTCTGACCTGCAAAGATTAGTAAAAAACTGGGCCTCTGTTTGGGTCCAAATGGCCACAAGTTTTGTTTGTATTCTTATTTACGTTTGGTTTTTAGTAACACCATTAGTAAAACGAGTTTGGGGACCAACGTTTGGTATACTCTTAGAAGATGAATACTATAAAGACGATCAAATTAAAGACAAAACAACCATTTTACTAAAATCTCATCAAACTAATTTAAAAAAAGATTCTGTTACATCATTTAACTATAAACCTGATTCTGATGATAAAATTCATTGCATTGACGCGCAACCAAGAAAACTTTCTCAATCAAAAAATACTTCTCCTGATCGACTAGTTAAAGAAGATATAAAAATTAATAGAAACACCTTTAACGCAGTCGCCATTAATGGTAAAAAATCCCGTTTGAGAAGAGTTGATTCGGAAGAACTGATTGATTCCGAAACTAGATCACAAATAACATCGCAATCAAACAGATATCAAGAAAACACAAAAGGTGTTGAACAACGGCGTAAAAACTCAGGGTCCTTAATAAAAGGAGTCGCAGAAAAATGCAGCGAAAACTTACAAGATAACAACGAAAGTGAAAAAGAAAGTATCACAAGCTATGCATCCCGCAACAAAAAGACAATTAAAGGAAAATTAAAAAAAGAGCAATTATTAAGTTTACCAGACACAAACCTAGCAGATGAACCACACAGAGCTAAATTAAAAAATTCATTTAAATCAAGTTCAGAAAGTTTATATAAAAAGTTTGATAAGAAAAAAGTAACTAAATCTAACTTAAAACCAACTAAATGCTTAGTTGACAAACACACGGAAACTAGCATTCAAGATGAACTCTCTCTTGTACCAAATGTATTTAAAGTAGTAGAACAAGACCCTGAAACGGCTCAAGAAATACTCAAACTTCAATGGAAAATTTTAAGGACTCAAGCTAAAGTAGTGAAAGTTCAAGAACGAATAATCAAACTACAAGAGTCAAATAAAACGACTGGATCAATTTCTCCACAGACTAATCATAAGCAGATAAAAAAAGTCAATAAAAGTGAACACGGCTGAACTTCCTC
>evgR000185c0g1_i01_Hydra
GAAAAGATACTAGAAAAAGTAGTTGGTTGGTTTCAAACTAGTAACAACTGGTAAATTGATCTAAAATTGACTCTCAAATATTATAGTTAATCAAACAATCATGACCGAATTAGTCCCAAGTTTTGTCCAAATGAACGTCACTCGAATAAATATAAGAAAAGTACTCGAAGTTCAATCAATACAAAACCAAGAACTTATGCATTCTTCCAAAGGAAAAGACATTGTTGTATTTGTAGGCAGAACTGGGGCTGGCAAAAGCACTCTAATAAATTATTTAGTCGAAAAAGAATTAACGGTTGACCAAAATGATAACATTGTCTTGAAAAACTTAAACGATCCAAGTGCTATGGTCGTTGGAGTTAGCGACGTTTCGGAAACTTTACTTCCAAAATTTATCGAATGTAATCAGTTTTTACTCTACGATTTACCAGGATTTGGTGATACTAGAGGAACCGCAGTAAATCTGGTTAATGCAGCATTCATAAAAAACATAATAGAAAATGCTGAAACAGTCAGAATTGTTTTTGTGACTGGTCAGGACGAAATCACTGCAGGTAGAGGCGAATTAATTAAGCAACTTATGTTAAGTTTGAAAACACTTATACCCAATGAATCAATCGAAGAAATAAGTGCCTTGGTAGTAACAAAAAGTTCAGCTAACCGAAATATTGAGCAGCTCGTAGAAAAACTTAAAGAAAAAACGGAAGGGGATACTGTTACTAAGTGGTTTAAAGTCGGGCGCCTTGCAAAAATGGAATCGGCTTACAATAACAGGATCAACGAAGATGACAGAAAATGTATTTTGAACACAATTATCAATATTAAAAAAAAACGCATAACAACTGTCAACATTGGCGCTATTATTAGCTACAGTGATCAAAACACTATTAAAAGCATATATATTGACGAGATGCAAGAAATATTTGACCGCTTTCTTTTAAGCATAATTGATACAACAGCAATTGAGGCTCTTTCGTTACCATATTTAGAAGATAAAATAAATCAACTAAACAGATCAATCGACGAGTGTCAATTTAATGGTTCTACTCTTATTCCACTTTTAAGAACAATTTCAAGTGACGTATACGATTTAGCTTTAATTGAATTCAAAAAAATAACCAAGTCTAACGTTGAAAAAATAGTTTCTTCAATTGTCGTTGAAATAAAAGAAAGACAACGAATAAAAGCTGAAAAAGATCTGAAAGAAGAAGCTAATAAAAGACTTTTGGCAGAGCAACAACACCAACAACATATCCAACATCTACAAAATGTTCAAAACCAACTTATGCAAACGTTGCGTGAACAAGGAAATGGGAGTGATTGTGTTATTCTGTGAATGTTTTGAAACTTTATTTAATTGATGCAAACTTAATAAGTAATTAATTAGATTTTATCAAATTAAAAAATACATATAATATATGTATTTAAGAGACATTTTTTATAAGAATCATGCAGTGAGTGTATGGAATCCACCTCCAAATTGATTAAGATTTTATATGTTTTCCAGAATATAGAGAAAATCGGTTAGGAAAAAAAAATTCTTAAAGTGTTTCGTTTGCTCAGAAAAAAAA
>evgR000186c0g2_i01_Hydra
GTTTTTATAATCACAAAAATATTCAGAAAAAATTTAATTAAGTAACCTCATGTATTTTAGTGGGTTTACAGGTAGGACTGTTAATCAAACAGATTTAATTATTCAAGATGACTCAATTTTACGATCAATACAATCTCAGTCAGCGCCAGAATTGATTTCTGATTTGTTAGGAGAGCTTTTTCTTCCAACTACAAGCCATATATATTTCACCAAAAATCACTTATTACATCAGTATGAGTTAAATTCTAGGATTGTCAATATAATATGCAGGGAGCCTGGATTATCCTATAATGTATTAGAATGGTTTAGAGATAATTTATTTATAGTTTGTGATCTTCATGGAAAACAAGTTATTAAAGTTTACTCCGAATGCGATGAAACCTGTGTTACTATGGATACCTGCTTCAATATTAGTAATCTATCCATTGGCAAAACAATTGGTCTGTTACTTACTCACGATGGAAAGGTTTACAAGTGCATAAATATGAACTGTGATGCTAAGGTCTTACTTGCTGAAGAAAATCTTTTATTGCTTGAGTCTTTTTTTACAATTAAAAAAATATGTTGTGGGCTAGAACATGGCATTGCTTTAACTAACGATGGCATTGTTTATACAATAGGTGGTGGGTGGAGAGGCCAACTTGGTCTTGGTGTTATTGATTCAGCTGATAGTTTAACTAAAGTTGATCTTTTAGAACCTCTTAAAATGAAAGATGTATCAGCTGGTGGTTGGCATTGTTTGGTTATTTCAGATTCTGGTGATTTATATGGCTGGGGATGGAATGAAAGTGGACAAGCTGGTGTTGTTAAAGAGGAATTAGATGAATCTGAGGAAGAACTTTTGAGCTATCCAAAACTTATATTTTCTCCGTCATTGGTTGATACACCTATTGAGGAAACATTTATCTCAGTATCAGCTGGGTCAAGGCATTCGTCTGCTGTATCTAGTAATGGTTTTTTATTTTCTTGGGGTTGGAATGGATATGGGCAGTTAATGCATGGTGATTTTATTAATAAATTGAAACCTAAACTTGTTGAAAGTTTGTGTAAAAAATCTTTATCTAAAGTTGTTTGTGGAAAATGGTCTACAGTTGTTGTGTGATACATGTTCTCCAATTTAATGGTTGCAGCTTCAGAAAATAAAGCTCTTTTCAGGAGACAATTTATACACGATACAAAACACAAAAGATTTTAAATTGTTTATGTTTTTAAAATGGTGAAAATGTATTTTGTCTGTAACTTTTTGACTTTTGTGAATCGTTTTATTTTTCAAATAGGTTTTTATGTGCTTTCTTCGTTTTTCTTATATGTTATATTAGTTGCTATGGGTTTTTTGTTTAAGTAAAGTATTTGGATTCTAGATAGTCC
>evgR000197c0g1_i01_Hydra
ACATGTTATAATGGGGTAAACCTTATATTTACATGTTATAATGGGGTAAACCTTATATTTACATGTTATAATGGGGTAAACCTTATATTTACATGTTATAATGGGGTAAACCTTATATTTTTGTGTAAGAATAGTAAATGCATTGAGACAGTTATAAATTTTACTCCTGCTTTGCATTCTTTACGTGACCGGATTTTAACTGTCAGCTTTTACCAATTTTCTTTTGTTTTAAGCCTGTTTGCAATGAGTCCTTCCATTTCCCATAAATTGTTATGTTATATTACCATTAGGTTAATGAGGTTTCGGCATTTTCCAGCTGATGAATACAAAAATATTGTTATCTTTATGTTTTTTGAATGATCCACAATTTCCTTGTTTTGCACGATAGCTCCGTTTATTTC
>evgR000220c0g1_i01_Hydra
TTAGACATGGTGAATCAGAATGGAACAAGGAAAATAAATTTACTGGTTGGTATGATTGTGGCTTGAGTGAAACTGGATTGAAGGAAGCTAAGGAAGCTGGTGAAATTTTGACCAAGGAAGGTTATCAATTCGATGTTTGTTATACAAGTTTATTAAAACGTGCCATCAAGACTTTGTGGATTGTTTTGGAGGGTATGGACATGATGTATTTGCCTGTTCATAAGAGTTGGAGATTAAACGAAAGACACTATGGTGCTCTTCAAGGATTAAACAAGGCTCAAACTGCTGAAAAGCACGGAGAAGAACAAGTAAAGGTTTGGAGAAGATCCTACGATATTCCACCACCAGCTCTTGAAGTTAGTGATGAAAGATATCCAG
>evgR000236c0g1_i01_Hydra
CTTCGGTACTTTTTTTTGTGTAGCTATGCTAATTATATTTATAACAGTTAAAGTTCTTTGTTTTCACATTTTTCTTCCTAAAGTTAGCAAAACTTCTCCAAGTAAATTCCCCACGTTTTTCTTGAAATAATTTTATTGTTTCCTTATTTCTATTTATTAAACTTCAACATTATAATTTAAAGTAGATATAATCACATGACAGTTCTATGTAAATTGCTAATAAAAGGGTTGGTTCCTGTTATAGAATGACGCACACTAATGATAAATTTTTATCCTTTTTACAAAGTATGTAATAGTATATTTTGGCTCCATCAGTTTGGAGTATAAACTTAATTGTAGGGCCTGTAAAGTTGTTTCCCTTGCTACATCAATTCCTAATCTTTCCAATGACATGACCTTTTCCCAGCAAACTCTACATGCTCGGTCGATGCATGGGATGGACCAAGCAGTATGAATGATGACCAGTACCACTGCGCTGGTGATGCTGTTCAAATTGCAAGCTGTTCAAATTGCAATTGTTGAACGTGTTCTTTTACGATTGGTCGACTGTGCCGATCATGCTGACCTTGTGCCTAACTTCAACCATGTTTGTAATAAAATAAAGGTTTTGAATAGGCACATGCGTTATTTATATAAAGACTCGCCATCTTCATATAAATATGTTCCACTAGTGATTAGACTTCATTGTTTTAGTTGCAAACATTATGTACCAGTATTTATCATTATGTACCAGTATTTAACATTATGTACCAGTAAAGCGTTAGCGCATGATGACATAAAATTGTAATGGTGGTTGAGCTTTATTCCTTTGCAAGTTAAACGTGTTTAATGAAATCTTTTATTGAAATACTGAATATTTTTAACTAAGTTTGTGTATTGCTAAGATGACAATGCGTTATTATTTTTAACATGATTGAAG
>evgR000238c0g1_i01_Hydra
GTACAAACAGCATGTCTCTTATTACAGGGAGTATTAAATCTATTTTGTCAATTAAAATTAAACATAGACATGACAGCTACAGTGATCAAATATCAAGAATCTTTGTTGCAAAATCCTTTATGGTGGCAAGTTTAGTAATTGGTGTCGATTGGTTCCGCGATAATGTTTTCTGCACTCTCCCACACAACTCAATCATAAATGAAGATTTCGTACACAAAGCATGTTGGATCCAGGGTTTTTATATATATCCACAAATGGAAAACTACGCTGATCGTTCTGCATATTATGGGATACCACAGTATGTTGAAATAGATGGAATTAGGAAAGACACTATGGACTTGTGTAAAACAAGAAAAAAGCTTTATGAATTAGACAAGATGTGCACACCAATGGAAAAATACTACTTTTCTCACTATCAATGGATGCCATTCTACATGGCTAGTTTATCAACTATATTTTATCTACCTTATGTTTTATACAGACTAATTAATGCTGATATAATTAGTTTGAGACAACATTTAAAAAACTCCAAGACTGAAGTCAATGATATTGTTGATAACTACTTCAATCGAGAGTTGAACACAAAGCGTAAAATGAGATACAGAATTGGTGCAAACATTTTAATAAAATGCGCTTACGTTGGTATTAATGTATTTACGTTTAATCTTCTCAACCATATTTTAAACCATCGATATCGAGATTACGGCGTAAATTGGTTAAAGTGGCTTCAATATAATTCATCAATGGCTTTAAACTATAATTTACGAAAAATACCTACTCCTGGTAATAAAATGCTTCCAACTTTTGGATATTGTGATGTACACGAGCTTGCAATGGATGGTGAAACAGCTACTGACAACAAGAATAAAGTTTTATGCGAAATCTCATCAAATATATTGTATCAATATGTGCTGATGGTGATATGGTTTGTGTTGATCCTCTCAATTATTGCATCAATTATCGGTGTTTTTGTATACATTATAACACATTTGATTAATATCTTTTACCTCGTTAGAAACGAAAAAAATGCAAAATTTATTTACCGTTTTTTGACATTTAGAGAATGCGAGTATTTAGAATTTATTCGTCAAGGAAATCTTGCTCTTTATGGTAATGTGATTAGAGAAATAAATTCTAAAATGGTTCTTAATTCAAACACAAACAATCAACAATCAATTCAAATACAAACTAATAATCATATAATAAATAATAAGCAATTAGCAAGTGATACAATAAAAAACAATATAAAAGAAAACGACCTTAGGTTTATTATGGCAAAAAAAGAAGGAAAAACACAAACACTCAATAGAATATTGGGGCAAATAAATTCTTATGAAACAAATGCATAAAACTCGCAAATTTATTTTTTAATTCAGTTTAAACATTTACTCACCGAATAGTAACGAAGTTATGTTATACACAATGTCTACTATGCTTTGAATGTCTATGGTTTATACATATTACATTTAATGATAGTAATATTTTTATTTATTTAAATTGTATTAACTTAGAAAATGGTCAGTTAATTATTGCCAGCTATTAGATTGC
>evgR000288c0g1_i03_Hydra
TAAAGTTATAAATACATAATACATTTGTATACTATTTAAAAAAATATGGAAGAGCAGAACTCACATAAGATCTTAATCAAGTTACGAGATCTGAATCCTTATTTGGTTTGTATGCTTTGTGCTGGATATTTTGTTGATGCAACAACCATTATAGAATGCTTACACACATTTTGTAAAAGTTGCATAGTACGACATCTTCAATCTAGTAAACGGTGCCCCACTTGTAATATTCAAGTTCATGAGACTGAACCCATCTCAAAACTTATGCTTGATCGAACCATGCAAGACGTTATTCAGAAGTTGATACCATGGGTTTTTATAGATGAAACTCGTCGTGTACAAGACTTTAAATGTAAAATAAACAAACAAGCAGTAAAAGATTTACCTAAAAATGATGTTCAAAAGTATGAATTGCTAGAACTAAAAAATAGTTATATAAGCGACGAACAAATAAATTTGTGTTTGTGTTTGGAAAGATGTCCAGAATTCTTCAATAACATGGAATTTGAGGAAATTCAACATAAATACGTGCGCTGTTCAGTACGATCACAAATACATCATATTCGCTATTTATTGAGTAAACTATACAGCGTATCTCTCAAAGAATACACGATACAAATCTGGTGCAGTGGAAAAATACTTTCCGATTTAAATAATCTAAAATTGATATATTTTGTACATTGGAAATGCAAGCCTCAGCCAATGGTTTTGAATTATACTTTGGAAGAAAAAGACGAAAATATAGTAGTTGATGTTGTGGAAAGATTACTTCAACATGTTAATGCAATAGATTCAAAATAGTTTTATAAGATAAACTTAAGATTTTTTTTACCAGCATTGAATATGGTTTGGTTTAGTGATCCAGGTGGAATTTTTAAGGAAATTTTTTTCTTTAATATAATTAAAAAAAGCAATTTTGGTTTGAAAAGGAAATAGATAATGATTTATTATTACAAAATTAA
>evgR000292c0g1_i01_Hydra
TACATCTCAACCAGTACTCCTGACACAAGTCAATAATTTTGATTGTTTAGATATATCAATTATTTCATTTTTTCGTGTTTGGTTTTTAGGCTTATCTGGGTGTATTTTTATGATTATTTCTGTTGAGCGATATTTTACCGTGTTTTATGATAATAAGATTTGTGGAGTCTTCATTAAAGATTCGTATTTCGTTGTATACATTATTTGTATTATATTTTTTACATTTGGTACGGCTATTTTATATGGAGTTGTCTATTCTACATCAAGTATGTATTTACAGTTTATTCTTTTCATTGGAACTGGGTGTTTAACACTAGCGACACTCTTTGTAATTCTGGCTGCAAATATTCTAATGCTAGTAAAAACTAAAGAAAAATTAAGAAGTACGTCGATTAGTATCCAGAAAAACGCTGAAATTGAGAATTATATCACAAAAACCTTGGTGATAATATCGCTCTCTCACGTAATGTTTTACACTCCGTGCCTCACAGCACAATATCAT
>evgR000325c0g1_i01_Hydra
AAGTAATTAAGTTTTATGTTCTAGTCAACAAACAACATGCCCGAAATACACAAATAATACAAATACGTCATAATAAACAAGCTATTATCCGTAATGAAATTATGGCATTTATTTTGCTAATGCAGTTTGGAAACTTTCTCCCACAAAATTTTCTAAGTGTCACCGAATATTCGGCTATTCGGCGGAGAGTCAAGCCGAATATTCAGTATTTTGTATTTGGCAAAATTACTATTCGTGGCATCTCTAATAATAGTGTTTGTTTTCAAAAGGTTAAAATATATATTAAAATAATGGAAGAGAATGTTTCCAAACAGTGTGTACTGTGTAGAAGACCAACCAAGACTGTCAAAATCAATTTTGTCA
>evgR000329c0g1_i01_Hydra
AAAAAAACATTTCGAAGTCGCAGTTTTAATTTATGTTGCACACTGTATCTTACTAATATTCGCCACCATCTTGAGTTTGCTGTGCAAGCCTGGTCACCGCTTTTAATGTCAGACATTGCCATTCTAGAGCAGGTCCAACACCGAGCAACAAAAACAATATCAACTATTAAATACCAGCTATACAAAGATCGACTTCAACGGCTTGGTTTAACAACACTTGAAGAACTCTGCAAAAGAGCAAATCTCATCGAGCAGTTCAAAATCACTCGTCAACTTGAACATGTCAGTTTTTTTTTTAAAAACTTTAAAAAAAGTAAAAACAAGTGCATTCTAAGAGGCCATAACTATAAACTAGAACGTCAGCTGATAAAAAACTGTGACGAGCGATATAAC
>evgR000338c0g1_i01_Hydra
GAATTGGACAACAAAATGGAAACTTCGGATGAAACTCGACAAATTAACTTAAAATGTAAAAAAAATATATGTCTATGTCGACAGCAAGAAAGACCATGCGAAGAATGCTATAGCGTTAATTATTCTATTGCTTGCTATGGAATCAATGATCCCTTTGTGGAGGAAAATAAATGCAACTATTGCATTATGGAAAAGATTTTTTCTAAGAATCAAGCCTTAAAACTAGATGATATATTTTATAAAGATACAAATGGATTTGCAACCATTTGCGCCATCACTCAAGCCGATCAAGATTTAAAGAGAAAATCGTTTTGGTCAAAAATTCATAAAACAATCCAAGAGGAAGTCCAAAGCATGGAAGAATGCGCATTCACAATTTGTG
>evgR000366c0g1_i02_Hydra
TTTAGTGCAAAAAGTTTGCTTATAGAAATATTTGTGAGTGCAATCCAGTTAAATAAAATGAAATACTAAAAAGTTTACAAAATGTCAGTGAAGAATCGACACTCCAAAAATTATGACTATTTGTTCAAGATTATATTAATTGGTGACTCTGGGGTTGGAAAAACATCTTTATTGGCAAAGTTTGTAGATGAAGAAGTTGTTCAATCTCATATATCAACCATTGGAATTGATTTTAAGATGAGATGTTTGGTCATAAGTGGAAAACAAGTCAAAGTGCAAATATGGGATACTGCAGGACAAGAGAGATACGAAACAATAACGACTCAGTATTACAGAAGAGCCCATGGAATTATTTTAACTTACGATGTAACAAGACACGAATCATTTGTGAATGTTCGAAAATGGCTTCGATATGTAGAGGAGTTTGCTGATGGAAATGTTAAACTAGTACTTCTTGGAAATAAAAGTGATTTGATAGATGCAAGAAAAGTAACAAAAGTTGAAGCTCAAGCTCTTGCTGAAGAGTTCAAAATACAATGGTTTGAAACAAGTGCCTACACAGGTGAAAATGTTGAAGATGCTTTTTTAATGATTACTCGTCAAATATATGTCGATGTTTTGCGCTCTGAAGAAGATTCGAAAATTACGGAAGAATTAAAAGGAAACCTCACTTCGTTTCCGACTCCGCTCGAAAATGATGCTCCAAAATGTACCTGTTAATTTTGATTTTTTAAACTTTTTTACCATTATTTAAATGTATTTATTGTTTTTATTACCGCCTGGTAAATTTATATTATATCTTTAATATTTAAAAAGTTTTCATAGCATTTCATTGCTGTTTTTTTCAATATTTATTTGCATATTCCAATGTATTTTTTTATCTTTTTTTTTTTTTTAAAGTTATGTTCCATTTTAATAAGTTTTTATGAGATTTTGTACTTCAAAGTGATTTTGTGTGCGTAGGTCAACAAAAGAAATTTAAGTAACTGTTTTTAAGCCTTTTTATGTGGATATTAAGTATTTCTTTTTTGTGTGTGGTTTTTA
>evgR000375c0g2_i01_Hydra
GTAACGAACGTACTCTGAAAAAGGCTATGGATACCTCTGAAAATGGTGATAACCTGCCCAGTGACGACATACCTTTGTCAAGTGAGGAATTGGTTTCGTCAACTTCTATACTTAAAAAAATCATAAAATTTGACCGCTTTGATTCATTTGCTGATTTTGAATCGAAGTTACTAGAGTACTGTGAAGAAAATAGTCACCCTATACACTATAGAAGTAGTCACAAGCTTAAGGACTTAATACCAGCTCACAAAGATAAACCAGTCAACAGAATTTGCATTGTTTTGCAAGCATGGGGGGTTAATCATCTGCTTGGACTGTTTGTCTTGATACTACTGCAACCAATACTGGAATGATTGGTGGTTCAATATACAGAAATAGTTCTTAAACATTTTTGGATGGCTGTAACTATTGAACTAACAAGTGGTCCTGAAAATGTTCTAATCAAAGGTCTAAAGGAAAAAATGGCATGACTTGAATATTGTAAACTGTGTCACCGCCCACTTCAAGTGGCAATATTACCGTGGATCTTGGTTGGAAAAGCAGGCATTTTTTGCAAAGAATCAGCTTAGGAAGATTGTTGAAACTAAAGTTTTTAAAAATTATCAGTCAAAAGAAAAGATTATCATGAACTAGCTGAGTTGGCCCTCATGTTCTTGGATCCAGAAATTAAACCTACCCACATTTACAAGCCTGGTGCAGTTCATCGTCCACGGTTTATGTGTGTAATATCCAACATAGTTGGTGGTAATATTAAGGGTCAACTGTATTAATAATATTAGTTATTATTTTTGGATAGTATATTGTTGTATAGTATAGTATATAGGGTCTGATTTGCATATATACTGAAAAAAATTAATTTCATTTCTAAATAAAATGCCTAATGTAATTGTATTGCT
>evgR000376c0g1_i01_Hydra
TCGACAATGTTGACGATGAAAGTGTTAGAAATATAAGAATGTACATTTCAAGACAATCAAGTTCATTTAACTTGATCACCTCTCAATGGAAATCATGGTCAGATAATGTATATCAGATGTTAATTGATGTTTTTTCTCCTGAAGAGGCTTTTAGTTACATAAAAAATAACATTGAAACAAACAACGAAGTACAACAAAAAGAGTTAATTGAAGAGCTTAGCTTTCATCCATTTGCCATCACACAGGCGATAAAGTATATCAACATACAAAACGTATCAATTGAAACTTATCTAACACTTTATAGATTGCATCCTACAGAACTTCTTGACAGCGAACACTGTCAAATCGAAAAACAATCAAAGTCCGCAATAAAGTCAATCAACTTATTTTTACTAAAATTAGAAAAAACCGAAACTACTCCATTTAATATATTAAATATTTTGTCTCA
>evgR000378c0g1_i01_Hydra
GATAAAATAATGAATTTAACTGAAGGATTTTATTACAGTCTTTCTTTGGTTCAAATAGTTCAATCAGACATTTTAAGTTTCTATCATGCATCCATTCAACCTGACTGGCAGTTGCAAGGGTGTTGCATACTATGCGTTTTTTTGGTTATACTCATAGTTATTATTAAATTAGAATGGGAGTGGTATGGATCAACTATAAGCAATCCTAATTCATTTGGAACTTCAAAAAATAACTACATTGACGAACAAGATGGTGAAATTGAAGCTTTTCCTGATGAAGATCCTTTAACATTTTTAAAGAAAACAAAATAATAATTTTGTAGTTAAAATTATATTTTAAAAAAAAA
>evgR000380c0g1_i01_Hydra
ATTATTAGCATAGGTATTACTAATGCAGTGAATAAAAGTGAGTTGTCGGCCATTGCTGGTGGAGAAAAAAATGTTTATATTGCTGATACCTTTGAAAAGCTTAAAGACAATAATTTTTTAAATGAAATCAAAACTAAGTTGTGTAAAACAGCATCAACCAGCTTGTCATTATCAGTGGTTTCAACTACAACTGCAAAACCTGCAGTTACTACACAAGCTACGGAGTCTGCCTCAACATCATCCATTAAGACAACACAGTTTGAATCCATAGAGACAACACAGTTTGCAGGAACAGAACTAACTGAGGGCAGGTCATTTGAGATGGATGTTGTATTTGCAATGGATT
>evgR000403c0g1_i01_Hydra
GTTTTATATTTATTATATGTTTTGCTCTGACCCAAGGTTGGTATTCAGCTGACTATACATTGCAACACATCATACATTGCTACATGCAATGCCATATATTGATACACACAATGCCACCTTATAAACATTTTATCTTTTTATCAGTAAGTCAAGAAATCATAGGAATATTGTATGCTAGAAAGCGAATTAAATACATCAAACTTAGGAAAAGATTTGCTATTATTGCTAAAGGCCCAATTACAAAATTATTTTTAAAAAGAATATTTAAAAAATGTGTGTGTGTGTGTGTGTGTAGTGACATTATCTGCAGGC
>evgR000410c0g2_i01_Hydra
TAATGATTCTGAATCTGATGGTAATTTTTTTATTATTTTGTATCAAAAAGATATATACTTACAACTATAAAAAATTGTTCTTACGTCAGTCAATAACAGGGTGTCTGCCAAATCAGGGAAAAGTCAGGAAATTCAGTTTTCTGCAAAAAAAGTCAGGGCATAATCAGGGAGATTTAGTTGAATTTAAAAAAAATCAGGGAAAATGGCTGTGGCAATCTCACATAACCAGTAAGTTGTCAGTTGTATATCATACATGTTACACAAATTTGTTTAGCTTTTTTTATTATGCCATTTTATTAATACTTTAAATGTAG
>evgR000420c0g1_i01_Hydra
CATTCTTCAAAGAAAATTGAAACAAACAGATAAGATGATGTTGTTCAAGTTTTATGGAGAATTTGATATAAAGTTGTATGATGAAGTAATTGTGTCGGAGTTAAACAAATATTTTTTTACAACAGTTGATGCACTTATAAATGAGGTTAATTTTTTAAATTTGCTACCATCAATCCAACAAAACTACTCTACCTCACTACTTACAAAGTATTTTACAAACAAATGGCGTAATCTTAATAGTGCAGAAGATAAATTGAGGTTTTGCTTAGATTGGGAGCCAGCCAAAGTATTTTTGCAGATTATAAAAATGGTAACCAATGATGATGCAGCTGAAATTATTAATGAGTATATATCTGTGATTCAAGATTTGTTTAAAAACATTAAGGACGGAAATATTCAAATTTCACTATTTACTTACTTATCTAAGAAAAAAATAACATTGTTTGAGCTAAGGAAAATTTTAGGTTTTGATATATGTGATAACATTGAAACTTGTTTTGCTGTAAGAACAAC
>evgR000427c0g1_i01_Hydra
CAATTATATCTCCTTTGCCACAAACTCCATTACCTGAGTCTGTATCACCACTTCCTATTTCTCCAACTATATTATTTCCACTTTTAACTAAAAGTTCTGCGTTGTTACAATTTGAAGATTCATCTGTATTTGCTCTAAAACCCCAAATAATACAAACAGCACATAAAATCATTACTGGTTTAAATTTTGAAGAATCTACCTTTCAATGTAATATTGTGAAACCAAAAGTAAAATGTGTGTCATTTGCTGCAAAGCGTGCACTATTGCTCAATGATTTAAATAATTTAAAACACAAAAATGCAAAAACTGAAAATCAAAATTGCCGATCCTCAACCATAGTTACTACAGCTTCGACCATAGTTGCTACATCTGTAACCATAGTTACTACATCTTCAACCATAGTTCCTATCGCTTCAAACATAGTTACTACAACTTCACCCATAATTACTACAGCTTCAACCATAGTTATTAAAGATTCTAGTACTGCCACAATTAATTTAGTAAAAACAAACTTTAATAATGTATCTCATGATAAAGAAGTTGTTGATGAAAATGATCTAAAGAACGTAAAAACTATTATGACTTTTGAGAAGAAAAATATGAAACGAAAACATTCACTTGATGAAAACTTATGCAAAAATGCAAAAATTAGTAAAAATAGTTTAACTGATACTTTTAAACATCCTTTAGAGAATTTAAATGATGTATTGCAATTTTTAAAACAATTTTCAGAAAAGTCTTTAAATAAACAACATCTTGTTGCAGCATTGAGTAAGAAAGTAAATACCAGTGCAGCCTCTCTTTCAAAAGCTATAGTAAAACTTGTTGTCTCTTGTGAAATATTTTTGTTGCCAGTTGC
>evgR000431c0g2_i01_Hydra
TTAGTGCAAAAAGGTTTTTTTGGAAGGGAAATGTTTTTGTTTATAAATATTTTATTATGCTTTTGTCGTGTTGTTGAACTACTAAACTGCGATAGTCTACTGCCTGGTCAATATTTGTGTGATCCTGCTGAAATTGACTTTGATACTCAAGCTGAAAAAGAGTGTCAACCTAACAAAACGGTCCAGGTATGGTGTAAAATTGCTCCTGGAATAATGTGCGATGGAATGTCATTCTCTAATACAAAATCCAATTCAACTGGGTTTTATAAGAAAAAATCATGTCGATATGTTAAAGGAAAATCATTTAAGACAGCTATGCTGTTATCAATATTTCTTGGTGTATTTGGAATTGATCGTATTTATTTAGGGTATCCAGCAATTGGACTTTTCAAGTTTTGTACACTTGGTTTTTTCATGATATTTCAGTTTGTTGATGTTATTTTGTTAGCTACACAAATTCTTAAGCCAGCTGATGGTTCTGACTTTTATATGGATTATTTTGGCCATAGATTAATTCATATTGCAACAGATAATGAAACCATATGGTACTAATAAATACCTAATATCAAAGAAGAAAGTATATGTACTTTTAAGTTTATGAATACCACTATTTATTGATAAATTTATGTACTTTAGTGGATGCCATTATTTATTGCTGGTATTTTCTGTGTTCATATGATGGTTTTATAATATGTCAGCAAATCCAAACAAAGAAGAAGAGAATCATTAATGTTAATATTGTTAATAATAATGTTATTATAATAAAAAAA
>evgR000433c0g1_i01_Hydra
AATGGAGGTTGCATCAGAAGAGTCAAAAAAATTAAGCACTGCGTTTTCAGAAGATTCTGTAATATCTAATACTTTAAAAAATGTTACATCTAAAACCGAAAATGAGAGAGTTTCCAAAATATTAGACGATTCATCAGACAATTCTTTAAAATCTCAAAAATGTAGAAACTCAACAAATTCATTAGAGAACATTATTTCATATCCATCTTCGATTACATTAAGTTCTTCAAGTAATATTTATATATCACTTGTTGATCTACTTCAAAAAAAAAGCCGAAATAACAGCACAAATAAAAGCACAGTATTGGAGACTATTGTTCCTCTAAAAGAACGTTTGAATTCAGAGGCAAATAACATCAATAAGCAAAATGACGTAACAACAGAAAATGAAGATATCATTATGGAGGAGAGTATTTACAAAGAATATTCAGATGAAAATCTCTTAAGACATGGTCTTGCTGACTGTTCTGTTGAAAAAATTATGCCTATACCTCCTCACCATGCTGTTGCTAGTGGAAAACTTGACTCTGCAGCTGCAGTACTTGAGTATAAA
>evgR000434c0g1_i01_Hydra
GTTTACACAGAAAAAAAAATTGAATTATTACAGAGATGCCGCATGGATTTGTCGAGAACGAGAAAAAATATATTTTTGTGAGCTCTATAAAAAACGTATTCAACACGTTAAAAAAAACAACCAGATTCAATACTCACAGGAGTTATGGAAAAAAGTAAAAAATCATCCTTGTGTGAAAAAGAGTAAACAATTATGGATCAATCATGTCATTCTGGGGAATTTATGTTATGGATCAATCTGGATCAACAATTATGGATCAATCTTGGGAAATGGTTTTTCACTACCAAGCAAATATAAAAAAGATTTTCTATAG
>evgR000436c0g1_i01_Hydra
ACAGAATCAGTTTGTCAATTAATTCATCAACTTGCATGAGGTCTAAAATCAGTTTGATATTGCCAAATGTTTCAGGGGTATTTTCAGAGATTGCTACAATTAGTTGCTTTTTGACACCTGTGCTTTTAAAATGTTCACTTGAAGCCGACTTAATTGACTTTTTTATTGGACTTTGTTGTTCTGTTTGTTATTTAGAAATTAGACTCAAACTAGCTTTGAAAAAGAATCCTCCACCATCAAGGCCCAGTTTCACAGTCTCCGAGCTCAACAAATCTTTTGAAACTGTGATGTTTTCCTTCAGGGTTTTTAACACAAATCTCATTATAATTTG
>evgR000437c0g1_i01_Hydra
AACAATCTTAATTATTCTTTGTTGCAGCATATTGAATTTAAAGATGTTTTTACGAAGAACACGTTCCTTCCAACCTTTTAATCAGAAATCAATCGAAAAAGAAGAGAGCTTAATCGAAAAATTGCGTATTGCTACATTTGGCTACGGTGGAGTGGGTAAAACTTCGCTAATAAAGAGATTTTTATACGACGAGTTTAGAGAAGAGTATTGCGAAACGATTGAAGACGACTACAGACAGGTTCTTGAATACAACGACATAACATGCGATGTAACGATTCTAGATACGTCTGGAAACCACCAGTTTCCAGCCATGAGGAAATTAGCGATTGAAAGTTGCCATGGCTTTATTTTGGTATACTCAATTGATTGTCAAAAATCATTTGATGAAGTGAAGCGTTTATACAATATAATTATCGATATAAAAAAAACTCCAAACGTTCCAATTATATTAGTTGGAAATAAATCCGATAGTCATTGCCGGGAAGTTCCAAACGACGACGCCTACATCTTGATAAACGCTATGGGAAACAAATGCGAGTTTATTGAAGCATCAGCAAAATTTAATTTAAATACTAAATTAGTTTTTTATGACTTAATGCGTTTAATTTCTGAAGAGAAAAATGAAATTCTAGAATTAGAACTTATAAACAAAAACCGATCTAGATCCCGCATATCGAGTTTACGAGCCTCGTTTAGCCACGTTATTAAACTTTAAAGCCCAAAACAAAAGATTTACTTTAAAAAAAAAATGAAGTTTTAATAAAGTTATTAAACAGTAACATGTCTGGACATCTTTTACTCGTTGAAATTTATGAGTATGGAATGCAATTTGTAATTTTA
>evgR000467c0g1_i01_Hydra
TCAAAAAATATATGAATGGTCAGATGGTACACCTTTTTCATACAACAATTGGGATAGTGGTGAACCTTCTGGTGGAAACGAGAACTGTGTAATCATATGGAAGTTGGGGGTTTGGAATGATGTTTTTTGTGATAATTCATATGCATATATTTGTAAAAAGAAAGCTTCTCCAACTACACCTCCACCTGGACCTACACCTGTACCTGAACCAGGACCTTGTGTGATTGCGGCTTTTCAAGCAAAACAAAATGCTTTGGGTAAAGTTGGAGTCTTTATACCAGATTGTGACCAAAACGGAAACTATAAACCTTTGCAAGTTAATCCATCTACTGGTCAAAAATGGTGTGTTGATGTGAAAACAGGT
>evgR000491c0g1_i01_Hydra
CAATGTCCCTTACCAAGAACGCTAAGAGTAACATCTTGGACTTGACTCCAACTGATGTCCAACAATTGTTGGCTTGTGAATGCCACAAGGGTACCAAGAATTTGGAAACCGACATGCAAAAGTATGTTCACGGAAGAAGCCAAAAGGACGGTACTCATGTCATTAACGTTGCTAAGACTCACGAAAAGTTGATGCTCGCTGCTAGAGTCCTTGTTACCATTGAAAACCCAAAGGATATCTGTGCTATTTCCACCAAGCAATTTGGTCATAGAGCTGTCCTCAAGTTCGGTTCTTACACTGGTGCTAACTATTTGGCTGGTAAATTCACCCCTGGTACATTCACCAATCAAGTTCAATCCAAATTCATGGAACCAAGAGTTTTGATTGTTTCTGATCCAAGAAGCGATTATCAAGCTATCAAGGAAGCTTCTAAGGCTAACATCCCAGTCATTGCTTTCGCTGATACTGATTCTCCACTCAAGTTTGTTGATATTGCTATCCCAGTTAACAACAAGAATAAGAATTCTATTGCTTTGATGTACTGGATGTTGGCTAGAGAAGTCCTCAGATTGAAGGGTGTCGTCTCCAGAAAGAAGGAATGGGACGTCAAGCCAGACTTGTTCTTGCACAGAGATGTCGATGAAAAGAAGGAAGAAAGAGAAACCAAGGAAAGTCAAGAAACCAAGGTTCAAGCTGTTGTTGCTACTGAAGGTGCTGAAGAAGCTGAAGAATACGAAGGTGAAGGTGGTGCTACTGCTGAATGGGGAAATGATCAATAAGTTCTCAACTCTTAATTATATAAA
>evgR000500c0g1_i01_Hydra
AATTTCTTTTGTTGAAAAATGTTGCCTTTTCAATTTTATACACCAATTAATGTAATAATTTTTATCATGGTTATTATGTTAGTGAAATCGAAGTTCCGTGAAACAACTGCAGAAATAATAATTACTAGTCAAGAATGGGCATCAACGCCATTTTTAAATGTAACTGTAGAAACATCTGCAACTGTAAAGCAATCATCATTTGTAGAAACTTTACAAGTATTTAGCTCAACGGTTAAGCAAAGTTCTTACGTAACATCACCAACTAAAACAGAAACTCTTGATTCATTATTAAAATGGTTTCTCAAATTTGACTTATCAGCTAAAATAGAAAAACCGTTCAACGAAAATCTTAGAGACTCAAAATCAAAGGATTTTTTTAATTTATCTACACAAATAATTAATGACTTTTCTGAAAATCTTCAGCCAAATCCACCTGTTGAGATTATAATTCACAGTTTCAGTAATGG
>evgR000509c0g1_i01_Hydra
TTCAAATACAAAAATAATACATAAATAAAGCATTCCTATACGACACAGAAAAGTAAAATAACATGACATCCATCACCTTGGTAATATCATTTGCTGTTTTGGGAATCATTACGTCAGTCCAATCGCTTTGCAAAACATCATACACTGAATGGAACTCAGAAGGCGGAGGTAACATATTCTATCTTGATCGTCATAACCTAGCTTGCGATCGAGGTTTTGCTCTTAACCAATTTACAGTCCAAAGAGATGAAACAAAAGGAAACTTTCGTTACGCTGGCGAGTGTTGTTCTCCCCCAAAATCATTCGTCAACCGCTTCTACACAAATCCTTATACAATTGTAGTAAATGACCAACCAATTTATCTTGATCGACAAATTGTAGCATGCTATTCAAATTATATTACCAGTTTCGGACTAATAAGATCAAATAGTCTAATATCGTATCATTTCTCCTGCGGTGAATCCTCGGAACCAAAATCATGCTACGAGGGTAGTACCAATTGGGAGCCTGATGGTACGGCATACCCTTTCGGTATACCTTATTCTTTGAATAACTTAAAGGTCGAATGCAGAAAAGATTACTTTATTACTAAATTCCAATTAATCCGAAGCCCGAATCAAATAGAATGGAAATATGTTATTAGTTGTTGCAGTTGAATATTGAAGTAAGTGTTACATTCATATATGAATATGAAAAAACACAGAGGCAAAAAACAAACACTTGAAGAACAGTC
>evgR000565c0g1_i01_Hydra
TGGATGCTTTGAAAATATTTTAAACTACTGCCGTACTGGCAAGCTTCATTGTCCAAATGATGTATGCGGACCGCTGTTTGAAGAGGAACTTGCATTCTGGGGAATTGAAGAGTTAAACATGCAGCCATGCTGCTGGGCAACGTACAGCCAATATCGCGATGTAAGCAAAAACTTAAAATTTATACATGAAGGAAGAAAAAATGTAACCGAAAAAGTGCAATTCAAAAGCAAAGAAGAAAATATTTGTCACAAGGGTTTTCTAAAAAACATTTTTAATACAATATGGAAGTTTTTGGAAGTTCCACAAAAAAACTTTTTGACAAAG
>evgR000575c0g2_i01_Hydra
TATACTGATAAATAACTTCAGTTTGAATTCACTTAAAAGTTACTCTGTACCATGCCTTAGTTATATTGAGAATGAAAGAAAAGAGTTGTTTCATATTAAGAAACAGCAATTCAAAAATCTTGAACTTTTTTTTACAATTACTGATTATAATAAAGTCAACGATGGCGCCGAATTTCTTTTAGATGTTACGAAAGAACATGAAAGAGATGAAAGTATTTCAATGGAAGAAAAATGCCATTGCAAATAAGTGATAACACTGAAAATTTCAAAAAGAGAACTATTAGAAGAAACGTGTACAGCTGATAGATCAATATGCTACTGTTCAAGTCATATGGATCGAGTTAATTGCAATCAGTTTGAGAAATTAAAGATATTCAATGTAGATTACAATGAGTTTGACAATTTAAAGATTGAAAAACTACCTAGTGATGTAATAAACTCGCTTGCACGTGACTTACCTAAAGATCGGAGAGAACTTGCTTGTTTCTTGGGAATAAACAATACAGAAATAGAAACAAACCAAGAACAAGCTTTTAAAATTTTAAATAGTTGGTGTCAAAATAATCCCTTTAAAGAATGGAAAGATATAAAATCTAAATTGATTTTTTTTAAACGAAGAGATGTTATCAGTAAATGCAAAAAAAAAAAA
>evgR000577c0g1_i01_Hydra
AATTTTTACTGCTACAGAGATGTTTTTACTGCTATAGATTGTTTAACTTTAACTACTTTAATGGTTTCAAGCCGAAAGTCTTCACTTGCTACAAGAAGAGGAACAGTTTTTTCTGAAGATTTTTATCCGACTAATCATTTATTTGTTTTAAAGCCTCGCAACCCTCTTCGCAGAGCATGTGTTAAAATTTGTTCATCTAGAATTTTTGAATCTCTTATAATGCTTGTCATAGCTGGAAACTGTGTTATTATGGCTCTAAATGCTCCTTTGCCTAACGATGATAAGTCAAACCTTAACATTAAGCTGGAGTCACTCGACAAGTATCTGCTCGGAATATATATAACAGAAGCTCTTCTAAAGATTGTCTCTCAAGGTTTTATTTTACACCCATATTCATATATGCGGATTCCTTGGAACATTCTTGATTTTGTCGTCATTATTACGGGTATTCTACCTTACGTTGGTTTGGGAAACAGCGGACCAACTAAAGTTATTAAGGCAGCG
>evgR000586c0g1_i01_Hydra
TTTGTTGCTGAATCTGAAACTGATACAGTTGACAATAAGGAAGGTGGTGGTATTCAATTGGTTGAATTTAAGAGATGGTTCATGGATTATTTAAATAGAGATCCAACTGATTTGTGTTGCCTCAGAAGATTGAAAATTCCATCCATTGAAAATAGACCAAAAGATGAAGTCATCTCTGAAACATTCCCAAATCTCGTTGACTATCTCAAAGTTGCCCACACTAAAAAGTATGAAACCTCAAAATCTGAACTCGTAAAGTGTGTCATTAAGAGAAATAAGAAATCAGACGATTCCAATGTTTTCTGTTCTGAATTTGTTGCTGAATGTTATGTCAAGATGAATTTGCTTCCATCCACAACAATTACTTGTAATTATGGTCCTCGTGATTTCAGTACCGAGAGTAATTTGGTCAACACAGTACTATTGAAGGGAGCTAGTTTGACTCACGAGAAGAGAGTGAGAGTTAGAGAATGGGATGAAGAACACAAATAGCC
>evgR000600c0g1_i02_Hydra
TAAATTTTTTATAGCATTTAAAATTTTGTCATGGCTCGTTTAGACTTAAGCATGGCGGAAAACTGTCGAGATCCGTCTACACATGTCTCGAATACTGATCCAGAGCAAATGGCTTCTGATGCCGAGGGAGTTTGGAGTCCAGACATTGAGCAGAGTTTTCAAGAGGCTTTAGCTATATATCCTCCATGTGGAAGAAGAAAAATAATTCTTTCTGATGAAGGAAAAATGTATGGAAGGAATGAACTTATAGCAAGATATATAAAATTACGTACGGGAAAAACACGTTCCAGAAAACAGGTATCTAGTCATATTCAGGTTTTGGCTAGAAGAAAGGATCCTCATGTTAAAGATAAAGCTGGTATTCATCCTATGAATTCAATGACATCATCTCAGATTGCATCTGTTAGTGCTATACACTCAAAGCTTGGTCTTCAAGGTGTTGATTCAGTTGCAGTTAAACGAGAAATGAATTTAAGTCCACCTCTTGGGTATTGGGCACCTGGTTTGCCACAACCTGGACTTCCAATTCCTCCACCACCTATTGGTTTTCCATATACTTCTGCAGGGGCTTATTATGCAAATGCTATTACACATGCAACATCTATTTCACAAGCTGGAAATTTAATTCGATCTTCTAATGCTCCGCCTGGCAACTCTTCTTCGCCAACTACTCATCACATGACTAGTTCTTTACCACATGGTTCTTTACCACATGGTTCTTCCAACCTTCATGTTAATCACAGCTCTTCGTGGCAATCTCGAGTATCTACGACACAAATGCCTCTGCGCTTAATAGATTTTTCTGCTTATATTGACCAACAACGACTTGAACAACCAGAAACGTACCACAAACATCTGTTTGTTCACATTGGTTCACAAAGAAGTTTTGAAGACCCAAACTTAGAAGCCATTGACATTCGACAAATATATGATAAATTCCCAGAAAAAAAAGGGGGACTAAAAGAATTTTACGATAAAGGCCCACAACATCTGTTTTTTCTTGTAAAATTTTGGGCGGATTTAAATACTAGCATTCCTGACGATGCTGGCTCATTTTATGGTGTATCTACAACATATGAATCATCTGAAAATATGACAGTCACTTGCTCAACAAAAGTTTGTTCGTTTGGTAAGCAAGTTGTTGAGAAAGTTGAAACTGAATATCCTCGCTACGAAAATGGACGCTTTGTTTATCGCATACAGAGATCCCCTATGTGTGAATACATGATAAACTTTATTAGAAAACTCAAACATCTTCCTGAAAAGTTTATGATGAATTCAGTTCTCGAAAATTTCACTATTCTTCAAGTTGTAAGTAATCGAGATACTCAAGAAACACTTCTTTGTCTTGCTTTTGTATTTGAAATATCATCCAGCGAACATGGTGCGCAGCATCACATATTTAAATTAGTCCGGGACTGATATAGTATGCTGAATGTTTGAAACTTTGTAAACACGTGACCGAACTTTCAGCCAATCAAACTTGTAAATATTATTCATCAAATGTTTTTTAATCCTATTTCTTATCAGATTTTTAGCTTTTTATTTTTGTAAAACTAGAATTATTTGTCAAAAGATTTAACTTCATTATTATATTTTTACCAGGTAAAAAAAGAATTCTTTCTTTAATTTTTTCATAGTTTTATGAAAGGTCTGTTATTTTTGTAGAGCACGTATTTTTCAAAATGTTTGTGTTTATTTATCTCTTTTTAAAGAGTTCCAACTTTGTTTGTAAAGTTGAAAAAAT
>evgR000606c0g1_i01_Hydra
AATTTTATCCATTGCTTTCCTTTCATTGCTTTCTTTTACTACAATAGCACTAAACGGTGTAAGCCTAGTTGTTATAAAAAAATCAAAGAAAATGTTAAGCAGACCATCAATACATTTCATTATCAATCTATTGATAGTAGATTTGTTACAAGGGTTGTTTGTAATTCCAATCTATGCCCTTAAAAAACAAAGCATTCAAAGCAAATTTTGGGATAGTTTTATATGTAATTCTTTTCGATTTTTATATATGTTAACTTATTATATGTCAATATTTTGTGTTCTACTTATTGCGTCCGATCGATACATTGC
>evgR000609c0g1_i01_Hydra
ATTAATAATACCAAATCTAAACGTTTTTTCAATTCTTTTCGCCATTCCTACTGTTGCTGCAGTTGTTTGTAACTTTTTTGTGGTTATTATTATCATCAAGATGCAAAAAAGACCGGTATCTAAGTATCTTTTGCTGTCTCTATCTATTTCTGATTTGCTTGTTGGCGTGATACTTGGCCCACTTACCTTGACACAAGTCTTAGATACTAGCCTGCTATCAAACTGCACACTGCATTTCATTCGTGGTTATGTAATAGTTTTACTTGTTGGATCATCTTTGTCAACATTAGCAGTGATAGCATACGATAGATATTTATTACTGACTAAGCTTGCTAATTATAATCGACATATGAGTAAAAAAAAAAGTTTTTTACTGATAGGATTTTCCTGGATTTTTCCAGGATTGGTTCCTATAGTCAGGACATATAATATGACACTTTACGTCGTTCTGC
>evgR000626c0g1_i01_Hydra
CCACATGGCCAATAACTAATCTCAACAGACTTATGAAAATTGTTTTGACTTCATGGATTGTGGTAAGAAGTGGTTTGTATACTGAATCTTTTCACCCAATTAGATCTGTCAAATTATCAGCATTATTGTTTGTGCATAGGTTTTTTGAATCTTCTTCAAAATGGGATTTCATTCATTCAAATCAAAATTACCATCTCCTTTCCCTCTTATCTCTAGCATTTTTTAGACTCCAAACTGGGGCACAAAAGAGCCTGAATAACGTATCAAGTAAATGCAAACAATGCAGATCTCTTTACAGCTATCATTACTATTGTAACAATCCCTTTTTTATATTTTAGTTGTTTTCATTAAAGCAAAATATCCCGAGATCCATCAAACCAGGAGTTTTTAGCTTTGAAACTTGCAAGGAAATTGTACATCAACTATAACTTAAATAGTTTAACCATGTTGGCTATAGGCGTAGGTTTGTCCAGTCGAGTTCATTCAGC
>evgR000627c0g1_i01_Hydra
TGGTCCTAGTCAATTAAATATAAATGAAGAAAACGAAACGTTGAAGTTGAGTTGGAAGGAACCTGATTCTCCAGTTTCAATTCTAAGATATAAAGTAAGCTTCCAATCACTAAAGCCATACAACAATTCATTTCTTTTTGCGGGAACAATTGGAACAGTTACTTCTAAAATTGCTATCGAAGTTGACTTTCATGCAGCTATCTTCATCATATCTGTAGATGTTGTGTACGATGAAAATTATTTTGTCAGTCAAAGTATACAGTATTATTCCAACCCATATTCACCTCCAGTACCAGTTTATAAATTGTTAAAATTGTTTGACAACTACACTGAAGTTTATGAATTCCAATTTTATAGTGTATCCGATATTAATGGACCTGTTAGTTTTTATGAGATCATTATTGCATATGAGAACATATCCTCAATGAATATGGAAACGCT
>evgR000652c0g1_i01_Hydra
ACTTTATATAAGGCTATGGACATTGGTGCCATTCCAAGATTTATTCTCTTTTTCATTGGTTATGGTGTTGCCATTTTGTACTGTGCTCTCGGTGTTGCTGGTTATTTGACATATGGTTTCAGTGGTATTGTCACTGCTATTTATTATTTCCCAACCACTACTACTGGTAGTATTCCTGGATTTGTCTGTAACTTGGTTATGGCTGTCATTTGGTTTGCTTTGATGTGTAACTTTATTGTTATTTTCATTTTGGTTATCAAGGTCTTCAGAGCTCTCAAGGGATCATTCGGCAAGATTAAGGAATATGGTGCTGGTTTAGTTCAAAGTGGTGCTGTATCTGCCATAGGTTCAGCTGTTAAAAGCTCATTGACAGGTGGTAACAAGAGCGAGCAAGTTTAAGTTATCAAAAAATAACAAATGTATTTATCATGACGGCAAATACATCTCGTATATAATTATCTATTTT
>evgR000654c0g1_i01_Hydra
GGTCGTCGTACACGGCCATCTTTCTTTAGCATCCTCGAGTGGAAGGCCGCAGGATGAAGTGTCTCGGCGAATTAAAGGAATACCGAATCATTGGAAGGCGTATACCAAATGCCAAGAATCCTCAACCACCTCTTTACCGTATGAGGATATTCTGTTCAGATGTTGTATCTGCTAAATCAAGATATTGGTATTTTATGAAAAAATTGAAGAAACTTAAAAAAGCTGTTGGTGAAATAGTTCAATGCTCTGAAATCTTTGATAAAAGCCCAACAACAGTTAAAAATTTTGGTATATGGCTGCGTTACAATTCTCGAAGTGGTACACATAATATGTATAGAGAATATCGTGATGTTACCACAACTGGTGCCGTAACACAGTGTTACCAAGATATGGCTGCCCGTCATCGTGCACGTGCTAGTTCCATACAAATATTAAAAGTTGAAACTGTTGAATCAAAAAATTGTAAACGACCTCATGTGAAGCAAATGCATAACTCAAAGATGCGTTTCCCTCTTCCTCACAGAGTATTGAAGTCAAGTTTCCGTACTACCTTCCAAGCTCGCAGACCTCATACTTTCCTGTAATTTTTATGAAAGTAAATTAACAGACAATATTTTTGTTTTTACTTGTTATGTCGGAGTAAATGTGGAATCCCGTTAATTTGGACTCTGAAGGGAAATATATAAATTATTAGTCTTACTGAAAAAAAAAAA
>evgR000657c0g2_i01_Hydra
CTTTTAATTTCATAGTCCTAAGTTGAAAAGTTTACCGCATACGTATATATACTTTTCCATTTATTACTTATCAGTTTCAGTCATATATAGAAGTATACTTTTCTGTTGAGTATACTTTTGTAGTTGCCATCAAATTTAAGAAAACCGCGGCTATAAACACATGCTTTCAAAATTAGATACCAGTCTTTAAAAACTCAATTGAATAACGCGTTTCCCTGATAAACACCAAAACATTCGGAAACAATCGAGCGATGCAATACTAAAACCGGCGTAGGTATTACAAGCATGTGTGTCAAGGCATATAGTTTTATAATTATAATATTAACAATAAAAATAACAATTCTTCTTCAACTTGTTCCAAGTATTTATGGGAAATGTGGATGTCATAAAGCTTCAGGACAAGGATTCTTTGCACAAGAGCGTCAAGTTATAAATAATTTTGAGCCCTTATCAAAGAACATTCATGATGGAAGTCCTAATGCTGCGTCTTTTCAATACTCAGCACCAAATTTTAAAATACTTCTTTTTGAGCCAACATTTCTTGATTTTAAAGATCAACCTGTTGGAATGCCTGATGTGCGATCAATTATACTCTCTAATCCAAGTGATAAAAATGATATTGAGATTGCGGCTGTCATTGTGCATCCTCCATTTCATTCATCAAAGTTTCAAAAAATAGTTCTCGCCCCTTTCAAAAATGTAACTATTGATTTGGTATTCTTAGCTACAATTTTAGGGAATGTAGAAAACACTGTTTATATATATACTTCAAAAGGAAATTTTCCTTATCAGGTATTTGGTGTTGGGATAAAGAATCCATACAGATTACGCTCTTATGTATCAGCACGGATACCAGTCAATGCTGTATTTTCTCCTTATATAAATATGTATAATCCATATAGTAATTCTATTCAAATTAATGAAATGTATTCAAGTGGGAATGACCTGCATTTGGATCCACTATCAAATAGCAACTACAGTAGTGTAAGCAAATTATGGGAAATACAGCCTTATGAAACAAAATCTATTATGAAAGCCACTTACCATTCAAAAGAAGAAGGACATTCTTCTCGCTTCATTCGTATTGTTGTAGGTCCACCGGCTCCTTCTCATATAATTATATTGCCTTTCGAAATTAGTGTATCAAGCAAACATGGATTATATACAAGTATCGATATGCTAGATTTTGGAACTCTGAGAACAATGGATGAACCAAAGTCTCTTAATCTCAATGTTATTAATTATGGCAACAAATCTTTGCAAATTACGTCTATACAAGTGGTTCCTCCGAATAGTGCTGTTAACATACAATTTACTCCTCTTGTACTGAAGCCTACGAAGAAGTATACAAAAATAGGAATAGTTAGTTTTTTAGCTTTAAATTCAACACATAGAAGACAGAATAGTGGAAAAATAGTCGTATCTACTGCAGATGAATCTGTAAAATTAGAACTCCAGTACCATGCTGATGTTTTATATGGGACATTGGCATATTCTTTTGGTAATACATCATTTTTTACTGGAAAATCCCCCTCTTCTCAGCCATTAGCAATCACAAATACGTTTAATGTATCTCTTGTCATATATAATGCAACACTTCCTCCTGAAATTCAAGATATATTTTATATTTCTAATTTTGAGCCACCAGTGCTAATACCACCTGGAATACTGCTCTCTCCACTTACGATAAATTTTGTAGCAAACCATTCATTGCTATCAATTTCTTCTGTTTTACGACTATATACAAATGCTTCTATATTCACTATACCAGTCAGCTGCTATAATGGTAAACTTGAGTACCACGTTCAAAGTACAAGCTCAAACAAAAGTATACTTGATTATGGCACTGTTGGCATCTATGAAAATAGAAGTTTACTCTTTACTATTGAAAACAATAATCCTATAGAGGTATTGGTTAAAGAATATAGAACTGATTTTGAATTTGGTCGTTTAAGACTTCTTAATGTCAAAAGTTCCACAAAGATAAACAATAAAATAAATATTGAAAATAATACAGTGTATTCTGAACAGAAATGGGATATGCACGACTTTGTTTTAAAACCAAGTTTTGCTGCTACTTTTACGCTTGATACAACTGCTACTCAAAAAGAAGGAATCTTTCATGGAGAAGTAGTAATTGAAACAACATACGAGAACATCACAATACCAGTGTATGCAAAAATGGTTAGAGGTGAGCTTTTGGCAAGACCCAAGAAAATAAAGATAGACAAAGCTTTTCCGGGGCGTTCTGAAAATTATCCCTTATCCGTAGAATCAACCTTTGATTATCCAATAAGGTTAAAAAATATAACCATATATCCTCCAGATTTAAGGTTTGAACTGGTGCTTCCAGATTTATTACCTGAAATTTTTCCAAAAGAAAAGACTCATCTTGGTTGGTTGAAATTTGATCCATTTTATGGTGATATAAAGCAGTGTTACACTTGCATTTGTGAAATTGATCCTGCGTGTGAGCAATGGCATCATAGTATTCAATTACCTTTAAATGCTTGGGAGATGGACTTTAATTTATTAGAAAAGCTTAGAAGAAAATGGGATAATTTTGAAAATATTCACAAATCAAGCTTTAATACATCCATAGTACTTGATTCCTCAATAGTTAAAAGTTACACTGTTCCGATCCAGGTATCATTGGCATGGCCTTCTTTAGTTGATCAATCAGTTAAATTTCCATGTACTCATGTTGGAAACTATTCTTTTGAAGATATCTTTATTACAAATCCATCAAATCATCCTTTAGTTGTGCAGGTCATTACTTTGTTAAATTACCCTCAACCTGATGGGGGATTGGATCTGCTATCTGACAGATTAATTATGGATTCGTTTTCACTAGATTTAAATGGACAATCAAACTTTTTTATACCAGAAATTAAGGATGGAAATCAAGACCATAAAGCAATTCATTCTGTGCTAGGTGTTTACCCATCACCAAATAGTTTAACAATGTTGCTTGAGCCCAGGCAAAAAAAATCATTTAAAGTGGGTTTTATTCCAGTTGATGAAACATTGAAAACTTCAATCTTAGTTATTCGAAATAATTTAACAGTTCTTGACTTGGTAATAGTTAGTGGCCAAGGTGCACAAACGTACTTTAGCATTAATGGACATTCACCAGGAGATAAGAATTCTATTTTGCATTTTGAAATAAAGCCTGTTCACCTAGAAGACTGTAAAAAGTCAACTCCTAAAGCTCGTTTATATCCTTCATTTACTGTGAAACGATCATTTACAGCTACTAACAACGGTCAACTTCCAGTTCATATTGTTTTTATGAATATTAACAGTTATGAATGTGAAGGCTTTGGATTTCGTATTATAAATTGTGAACCTTTTGTTCTTACACCAAACTCTTCTCATAAGATTGACATATCTTTCTCTCCTGACTTTACAACTTCTCGTGTTGCAAGAACATTGAAAATAGTTACTTCTATTGGTCAAACAATGGATTTTGCTCTTCTAGCGACTCTTCCTCATCATTTACTGCCTTTATGTGCTGATGCACTTCCTCGTCCATTTTGGGAGCCATATTTGCATGTTGTGGCAACTGTTGTTATGTCTGCTATATTCATTTTCATAATTTTCTTTGCTTATTTTGATGCTCAGAAGTATGTAGTTCAGTGTACTTTTACTGCTGGTCGTAGTTCTACTATGGAAAAAAATATTGATCTATATACTCCAGGAAAAGTATTTGATTTAAATGCTATAGCTGGAGTTAAAGTCAGAATAACTGATAAAAAAGATAAATATAATGAAAGTAAGTTACATCAGCGAGCTTCTAAATCAAACAGTTGCCCTGTGTCTAAAGTTAATAACCAACCAGTTTTAAAGCGCAGTGATTCTATAACATCTATATCATCTTGTACTCGAGAAAGTTCTACTCAATTTTCGGACTCTAGTAATTCTTCATCTCGTAAAAATTCAAATACGAACAGTCCACAGCCATATATGTTGGAATTAAAGCCAGAATCAAAGATCGAAAATAAACGTACTAAAAGAAAAATAGTTGATTTAAATAAAAGCCAAACTAAGTGTGAGTTTATACCCAGTGATCCTGTTATTATTGCAGAAGAAATAAATGGCTCCTGGGATCAGAATCTAGTTGACAAAACAATAGTTGACCAAAAAAAAAA
>evgR000693c0g1_i01_Hydra
ATTTACCAGACTCTTTAAAAGATTCTTTACCAGACTCTTTAAAGCTAGAAATAGATTCCAATTTAACAGGTTCTTCGAATAAAGAACCAATTTACATTTTAACAGACTCTTTGAAACATGGAGTATCTTTATCAGATACATTATCAGATAATAATAAAAGTTTTGATGTTGTAACTTTGCCAGAACTAAATTTGACTAATGTTCATATACCATCGCATACACAAACAGTTGTTTCTAGACATTGTTTTTTAGATGGTTTAAAATTAGATATAGATAAGTCAGATCTAATAGACTGTTTAAAAAAGCATCAGGATCAATATAGTGATCTCCATACAAAGAATGAAACCAATATAGATAACAATCCTAATGTAGATACAAATACTGAAATTGTGAGTGACTTGATTAAGTTTTTACTAGATAAAGTATGTCAAAATCATTTTTACAAGAAAAATATTTTTTGTATGGCTTCCAACAAAGCTTGCAAG
>evgR000697c0g2_i01_Hydra
GTGCAGCTGGTGGAGGTATAGCTGGTGGAGGTGCAGCTGGTGGTGAAGGAGGTTCAGCTGGTGAAGGTGGTGCAGCGGGTGGTGAAGGAGGTGTAGCAGGTGAAGGAGGTGTAACAGGTGGAGGTGGAGGTACAGCTAGTGGAGGTGTTGCTGGGGCAGGTTCAGCTGAAGAAGGAGGTGTAGCTGGAGAAGGAGGTACTGGTGGAGGAGGTGTATCTGGTGGAGGAGGTGTAGCTGTTGGAGGTGCAACTGGTGGAATTGTTGGAGGTGCAACTGGTGGAACAGCTGGAGAAGGAGGTGCAACTAGTGGAGGTGGAGCCGAAGGAGTAGGTACCAATGGCGGTGCAGTTTCACCTCAATCTGGAGGTAATAACGGAGGTTGCAATAAAGAATTATTGACATGTACTTGCTCTCCGAAATGTTCTTTATGTTGTCAACCAATTATATGTAATCCTAAAGCTCGAAACTGTATAACACGAGTGGAACCATTAATAAAATCATTTAGTAATTCAGGAAAACATGGATATAGTGTAGGCAATGGCCAGACATCTAACGATGAAAAAGAAATTAACACTATTCAGCAGTTGTTAGAAGACTTCTATGGCGAAGATACTGTTAAGAAAAGAAGCTTTCATCCACGATTATCTTCCGGTAGATTTGGAAATAAATGACCAACTAACGGGATCTTGAGGAAACATTGCATCTCTTACGACAAGCATAATACTAAGCCAAATAAAGTGTTTTCTGAAGAAAGGCGTTGTGCAAAATAGTTTGTTGTGACGTATTTTTGCAAACTTTTGCAATGCAACAACTCAACGCTCTACTTTAAAGACTATATGAGTTTTTAAGAAAGATGTTGCTAATATACACTTGGAATAACGTAACCAACACGTTGCAAACGTGTTTGCCGTTGCGTAGTTAAAGGCCATTGGTTAAAAAAGCGAACTTTGTTGCTATGTAGATATATTTGTGTTTTATAAATAGATTTTTCAATGTATATAAAAAGTAAAGGAAAATACTTATTGTATTTAATATATTTACTGTAAAATTATTGTCTAAGATTGTTTTTATTGACAATTTTATTAAATATTGTTTGGAATTTTG
>evgR000703c0g1_i01_Hydra
TCGATAATTTTTGTAATATATGGTTTTTGTTTACTTTTAGTTAAAATAACCAATAACTTTCTAATGTTCTTTTATTATGAGTGCACCAATAGCAGCTGCCCGATTATGTAGGGTTGGAAGAAATCTTTTAAAGAAAGGAAAACCACGCAAGTATCCTGAATGGAAGATTAATTCAGGCAAACAAATAGTCAAAGTTTTCGATACAAACGAGAAAGTATTTATTTTCCAATACAATTCATTAACTGCTCTCGAATGGAATGATTTTCGATACGATATGGGTAAAGAACAAATATCCGTTAAAGTTTTCCCAAATGAAATTTCAAGAGTGTATCTCAATGATTCTCCTTATAAAAACATAGCACCACTTTTTTACACTCAAACCGCTATTGCTTTTGGTAACGATGTTAATATGAAATCGGTATTGAAGTTATTCAATCGTAATCCGAGAATTCACCTCTTAGGCGCTGTAATTGAAAAAACAATAATGAACAAAGACAATATCGTCAGTTATGCTAATTTGCCATCACTGAAGGATATGCATAGCGAACTGCTTAGCATGCTATCGCAGCCGGTTCGAGAGCTAAGTCGTTTGTTGAAAACGAATCAAAGTAACCTTAGTGTTTCTTTAGAGCAACACTTAAAGCAAATTCCATAAATGATTTAAATACAAAATTTATTTTACAAAATACTTTTTTTAAGTATGTATAAATTTACGGGTATAAATTTATTTTTAATTTCTAAAAAAAAAAAA
>evgR000707c0g1_i01_Hydra
AACACTTTCATCATCATAATGCTGTCTGTTTTGAGATTCAAATATGTTCAAGGTGATTACGTAACAAACTGTACTGCTGGTTGGTTAGCATACAAAAACGCTTGCTATTTTTTTCAAAAAGAAAACAAAACTTGGTTGGATTCCAATTCGGAATGTACAATAAACGGAGGAAATTTGTCAAGTGTGGCTGATCTAGGAGAATCTAATTTTATAGAATCCAACTTGAATGGCAATACTTATTATATTGGCCTTAAACGTTTTAAAAAAAATGCAGAGTTTTTATGGTCTGACGGATCACCTAATACCTTCAGTAATTGGGGAAATGGTCAACCTGATATGTATAATGAAGAAATTATGTGTGTTGCAGTTTATCTTAAACGATGGAATGATTTTAAATGTGATCGACTTTTTGGGCATATCTGTAAAGTTGTTCAGATCTCCTGTTTAAATGCTTCGTCTTTGCTATCCATATTTGCGTCTCCTTCGGTTTTCACAGTTTTGACAACTTCGATATCCATCGTTGCGTCACCTTCGGTTTTCACAGTTGCGCCTCCATCGACTTCCATCGTTGCTCCTGCATCAATTGAATGGATTGGAAACTGGGC
>evgR000715c0g1_i01_Hydra
TTTATATGTTTTAGATATCAGCTTTCATTTAAACTATAATAATTCAGAAGAATCTGCAGTTAACGTAACTTGGGAATATTTTATTCCACCGTTCATAACGCTTCAGTCAGAAAATTCATCAGTGGATGTTGTAAAAATGGATTCTAAAAACCTAAAATATAAAATTTTTAATTTACCATCCAATGGAGTCAATCAGACTATAACTGTTGTAATAAATAGTACTCGTTGTTCATTTGGTGGAATTTATAATATTGATATACCAATAAAACTACATTTTGAAAATGCAGAGGGGAGTTCTTGGAAAACATTTAAAACATTTACAAAGCAATTAAAGCAAAATAGCGCAAACCACATTTTACCAATTAGCCCGAATCACGTGTCTGAATATTATGGTCGAGGCATATACTGGGATGACACTAGTTCTCACATATATCTATGTATGAACC
>evgR000722c0g1_i01_Hydra
ATAGCTTTTTTACCGTTATAAAAAAGCGACCGCCATTTGCGATGCCATTGCAAAGAGTTCATCAAAACTTCAACAACCACGTAAATCTAATACTTTCTTGTCAAAAACAACAAAAATGGAAGCCTGTCTTTTGCAAGCGAACTGTTTACTTCAAGATGAAGACTGCTCAAAGAATTTGCATAACATCTAAAACCTCAAATTCAAAAACTTCAACCTCAACTTGGAAAACTTTAACAAAAATTAAACCTCAAAACTTATACCTCAACCTTAAAAACTTCAACAACAACATCAATAACAACATCAACAACAACATCAATAACATAACCATTATTGCCAAACTTGTTTATGATATTATAGAACATTATGAATTTACGGATTGATTGATATGTATAACCATGAATTTACG
>evgR000725c0g1_i01_Hydra
ATGCAATGGTTTTAAGCTATTTACTTCAATCTGAAAATAACGCTTATTTTCATTTTAATTCTAGTACAGAAACTTTGGATGAATGTATTTTAAACAAAATGATAGAAACTGGTGCACGTGTTTTAATTGATGTTGGGGCACTAATGTTACAAATGGACAATAAGAATGTTGCTCTTGAATTGCTTAGAAAGTTTTCTAATCTTTCTTTAGTTTCAGCAGCGGTGTATTTTGATAAAGACGACAGATTGATGGTAAGCGATAGCAAAGGTTGTGTATGTCCTCTTTCAATTTCTCCGTATATGCATCAACTTGATCGGTGTGTATTTTATATTGATGATATTCATACCCGAGGGACTGATTTAAGATTTCCTTTTAATACTCTTGCTTGTGTTACGCTGGGCAAAGGACTTACAAAAGATCGCCTTGTTCAAGCTTGTATGCGAATGCGATTGTTGGGCCATGGCCATCGTGTGTTCTTCTTTGCCGCAAGTGACGTTCACTTATCTATTACAAAACATATGAAATCCTGTACTAATATACCCGAAACTGCCGATGTCCTTCGATGGGCTATTAATAATAGTTGTACCACTATTCGAGATGGGCTTACACATTGGTCTTCGCAAGGACTTCGCCGTTCAATAAAGGAAGCAGCAGAATCATTTATGAATAACAACGAAAGTAAAAGTAAAATAACACTATATGACTTAAAGGAACTAGGCACTAGGTGTACAGAAGACGAAGTATTTGCTTTACGCGAATTTTAT
>evgR000728c0g2_i01_Hydra
GATGAAAACATTTGTTCTTCTGTTGATGGAATGGAAATTTCTGATGGTATTTCTGTTCCTGATCTAAAATCTAGTTTGAAAGTCATCGATCCTAAATCGGATGAAGTTAAGAGTGAAGACAGCGTAAGCCTAATGAAAAATGACAATCAAAAGAACGATGAAAACATTATTCATTTTGTTGATAGGATGGAAAATTCTGATGTTGTCCCAGAACATAATTCTAGTTTGGAAATCCACAAACCTAAATCGCATGTAAATAAAAGTGAACGCAATAAAACTTCAATTGAAAATGACAATCAAATGAACAAATATGTTTGTTCTTCTGTTGGTGGAACGGAAAGCTCTGATATTTTGTCAAGTACAAATGAATCGAAAACAAATCAATCTCAAAATTTTTGTTCAGTTATACCTATTCACACTAAGATGACATATTCAAGTGTAGTTCAAATGGGGGTTTTAACCCAAGATGAGAATAATAATAAAATCTTAAATCAACTCCAGTGTGAAGTATCAAATCCTGCTAATAACAGGTGTTTTGGAAACGGAAAAAATAATGAACTAGCACAACAAAATCTCTCATTTGGAAATAAAGTAGCTGAAAGCAAATGTTTTGAAAATAGTTCTTCATCTTTTTCTATTAAAGAGGACGATGATGCTAATCAATCTTGTAACACATCCAACAATCCTACTAAAACTAGAGAACTAAGGTCAGCAAATCAACAAAATTCAGATGATGCAAATCTAAAATACAACAATAGAGAGAACAGAGCAAAAAATGGACAGTCTAATCAAGGTGGAGTAACCAATTCCACTGAACATAGATTCATTCATATACATTTCTTTTATATTTCACACAAGTTTGTTGAAAATCTTTGTATTGTATTTGGGCATAATTATTTATATGGATGGGATGGTATTGGACTTCAGATGAGACCATTTAGGCGTGACTCAGAAGAAGTAATTTACATCGGAGATATGACCATACGCAAAGATTTATTTCCTTTAGAATATAAATATGTTATACAAGATAGAGAAAAATCTTGGGAGTTTCTTCATTCTTCTTTCCCACAGAACTCGAACAGAGTTTTGATAATATCAAATGATGCAATAGAACTTGGTCGGTTTTTTCAAGTGGATGACTGTTTTATGTATGAAAATGTGGAAATCAATTTAG
>evgR000738c0g1_i01_Hydra
GGCGCTTGGAACTCACAGATATAAACTATTTCATCTCTTATAGACCGGGCCGCAAAAACCTTGCTGCTGACGCCCTCTCCTGATGTTGTTCTATACTAACTTCTTCAAAATTATTTTACAATATCTATTCCCAGCTATGTCATCCGGGAATTAAAAGGATGCTGCATTATTGTAAAACTAGAAACTTACCTTATTCGACCACAGAGATCAAGCAATTTACAAACCAGTGTACAACTTGTAAAGAGCTTAAACCACGTTTCTACAAACCTCCACCAGGTCGCCTTATTTAATCAACCAGACCATGGGAACGCCTGTCTATGGACTTTATTGGTCCTTTGTATTTTTCCATGTAACGATATAACTTCAGATACAGTTATACGTCACCTCCTTAATCTATTCTTGTTATTTGACTCACCCTCATCAATACACACCGATCGGGGAACGCAGTTTGAAAGCGTCAAACTCCAGAAATTTCTTGAAAGAAATGGAGTTATCAAAACCAACACCTTTCCACCCACAAGGCAATGGGCAGTGCAAAAAGACAAATAGAACTATACTTAAAGCTGTCAGTCTCGCCTTAAAAACGCTAAACCTAGGAAAAGAAAATTGGGAAGTAGTTCTACAAATGGCTTTATCCTCAGTAAGAGGATTACTGTGTACGGCCACAATTGAAACACCAATTCAGAGGATGATGAGTTTTCCACGATCATCCATAAAAGGTACCGTACTACCAGATTTTCTAAATGACAAGGATTCAACAATCCTTTACACGAGGCAAAACCGACAAAAGGGAGACCCACTTGTTAATAAAGTTCATCTGATAGAGACGATATCCCCTTATTTTGCACGTGCTAGATTTCAAAGTGCAAAGATTGATACAGTGTCTACCAAAGATCTTGCTCCACCTCAAGAGGTAGACCTATCTGCAGAATTGAGCGGTTATGTAAGGCCGGTCCAACCGCAATCCAACGAACAAAGTACTGACTAAATTCATATTACTACCAAAACATCTAACCAGGCAAACGGAGAGCGTTATGTTTTGACAGAAACCAGCAAAATAGTTTCACCCCCACCAAGCATATTTGATAGGTCGCTAATAAATGACACCACAAACATCAAAGCACCAGAAAAAGACGTATATCGCCAGCTTGCTACCCATAGTCCTGCTACTGTAGAAACACAGAATAGTTTATTTTCAAGATTTGGAGGGAAAGTTGAAAAGCCGAATTACTTAAACGACTATATTACAT
>evgR000745c0g1_i01_Hydra
TACAACAATTGCATCTGAAATATTAACAAAATTAACTGAACCTACAAAAACCACACTTGAAACTACAACAACTACACCTACAACAACTAAAACTACAACAACCACACCTATAACTACAACAACCACACGTACAGTTAAAACAACCACGCCTGTAACTACAACAACAACATCTGAAACTGCAACAACCACGCCTGAAACTACAACAACCACACCTGAAACTACTAAAACTACACCTGAAACTACAACAATTGCATCTGAAATATTAACAAACAGATTTGAAACTACAACAACCACACCTGAATCTACATCAAGAATCGCTGAATCTGTATCAACTCAATCTTCAAGCACAATTCCTGGATCTTCTACTTCTGAACCTGAAAATTATTCTTGTCCATGTTTGATAACATTTTGTGATCACCACTATAAAGAAGAGTTTTGTTCAGCTGTATATCAATCATGCGGAAACACTTCAAAATGTTCTTCAACAACAACAACTTCAAAAGCCTAAGCTACCAAAACTAGAACTGTACTTAAAAC
>evgR000749c0g1_i01_Hydra
GCACATAGTGCGTTCTAGCAATTAAAAAACAAAAAAACAAGAAAATGAAAAATGTTTAACAAAATAAAAATATTGGTTATTGGACCAAAGCAGAGTGGGAAAACATTTTTATCAAATTTTATTGCTGATGCTACTGAATCTACAAGTGGAGAGTATAGATCAACAAAAGGTGTCAGGATATTAGAATTTGAGGTCGAAGGTGTAGATGTTGGTGGAAATCACACAGTCAATTTAGAGGTGGAGTTGTGGGATTGCAGTGGCGATCAAAGATATGAAGGATGTTGGCCAGCATTGCAAAAGGGTGCAAATGGATTGATATTTGTTTACAATCCTGATGTAACCGCTCATGAAAAAGAATTAGAAAGTTGGTATCTTCATTTTTCAAAGCAACAGGGTATAAAAGATCAGCAGTGCATGGTTTTTGCTCACTCAAAGTCTGGTAATTCAGTGCCATCTGTTAAGCTGAACGGAAACTTAGGACGAGTTTCTTTAACAAATACGAACTTTGAAGAAGATCCTGATGGCATTAAAAGTGCTTTTAAAAGATATTTAAGTTCTGTTTTGTCGCACGTAAATGATAGCAGAGATCAAGAAGAACTTAGCATTATTGAGTAATCAAGAAGAAGCATTGTTGAGTAATCAAGAAGAACTTAGCATTATTGTGTAATCAAGGAGAACCTTTATTGAGTGACTTATATGTTTATAAAAATGATAATAATATTTATGTAAATAATAAATAAAATAGAGAAATCCACAATAAAAAAAAA
>evgR000752c0g1_i01_Hydra
TTATCTTGACATTAATAAACCAATGGGAAACGGATTAACATGCTTACACATGGCCGTGATGTTTGGAAATGTGAACATTATTCAAATGCTACTAGGTTGCAAACCATTCAAAAAAGAGTCAGAGGAAGAATTGTTTCCAGGATGTTTGGTTATTGATAAAGTATGTGATACAAAAACTGCATTTCATTTTTCATTAGTTGCAGGCAGTAAAAACAGCTTAAAATGTGCCGAGTTATTAATTCAAAATGGTGCTGATGTATACAGAAAGTTTGATGACGAGGAAATGTTTCAAGAAGGATATGTGTTATGTACTCCTTTACAGTATGTTTGTAATAATGAAAAAATTGAATTTATAAAGTTGCTTAAAAAATATAATGTTGAGGATATTAATGGGGTAGCTTTAAAGAAATCAATGAAAAAAAAAAATGAAAGAGCTGAAATGGAGCTCCTAATAGATGGTGTTTTCTCTAATAAAGATAGTAAGCTTGTTCTTGCTGCAAGTGAAGTAGACTATAAATTAAATCCCACATCTATATCATGGATGAAGCGTCATTTAAAATTAATAAAAGAAAAGTGGATCTTAAATTCTGCATTAGTTATGAATTCACATACAGATTATATAAAAAATAACGAATCTCTTCAAATGAAAGTTATCACTCACATTAATTTATCAGAAAATCTTTTGAAATCAATTCCTATTATCATTTTTCAAATGCCTTCATTAAAAAAGCTAAATCTTTCACATAACAAGCTGGTAAATTTGCCTGTTCCCATTAATGATGATAAAATGGATCCATTTATAAACAGCATCAATGATGATGAGAATAACAAACTTTCAATGAACTGGAATTGCCCCAGTTTGGATGAATTAGAGCTTCAACATAATTTACTAAAAACAGTTCCAATAAATGTTTTTAAAATGCCAGCATTGCAATCTTTAAACTGCGGAGACAATAAAATAGCTAGTTTACCATTTGAAATATGGACGGCTAAATCTTTAGCAACACTAAATCTAGATAACAATTGCTTAACTGATTTGCCGATTTTTGATGAAAGTGCCAAATCTAGATTGGAATCCCAAAATGAAACTAATAGCAGAGTTAAAGAAAAAGAACATTTAGATGTTCGTATTAAGAAAAGTGCATTTGCAACTTCTAAAGAATTTCTTTTCAAGAAAAGACCTGCATGGATAAAAGAATCTGATGATTCTGATGAGGAGGAAATTTCTAAAAAAATTGGTTTGAAAAAACTAAGTTTATCAAATAACAAATTGACAGATGTTCCAATAGGACTAGCATGCCTTGCACCTAATTTATTAAAAATAAACTTGAACAATAATAAAATAACTGAAATATCTAACTGTGTTTATTTATTTCCGGAACATTTACAATATTTAGAGCTTGCTGGGAACAACATTAAAACATTTAATTTAACTGAACCAAAAATTTTAGGATGCTATGCTTTTAAAATAAGCCAGATTAGTAAACGTGCATCTCTTTTAAATGAAAGTTGTCAGCATCGTAAGCACAGGCGTTTAAATGAATTATACCTCCTTAATCTCTCAAATAACTATCTCGATCATATCAATTTGTTTGTTTCAAATACTCAATCTATGACTCCAAACACGCCCTCAAATGAAAAAGTGTTTTGTAAGAACTTAAAAGTGTTAAATATTTCAAGCAACAAACTGAAAAGTTTACCTAATGGAATTGAAAAATTTAAAAAATTAGAGTCATTAAGTGCAAATCAAAATGCAATTAGCACTCTTCCATCTGAGTTTGGACTTTGTTCTGAGATTTATGAATTGAAACTAGATAATAATCTTTTGAAAAATTTTCCTCAGCATGTTATTGACAATTATGAGAAAAAAAAGATGAAACCATTGATACGTCACCTAAAGTCACTGCATGATAAAGCACTTTCCTTTCCAAATATAAAGTTGATGTTTGTTGGTAATCATGGAAAGGGGAAAACTACCTTGCTTACATCCTTGAGGAAAAAAGGAAATGGTAACTATCAACATCCAGAATATAAATTTCATTTTCTTGAACGAATTAAAGATAAAGGTAGTGTTAGAGCAAAATCTAATTTGAGCACTGTTGGTGTGGATATTTGTGATTGGAAATATGGCAAAATACGGGAGAGGTCCATTAACTTCAGTACTTGGGATTTTGGTGGCCAGAGTGAATACTATACAACTCATCAATGTTTTTTGTCTCAACGTGCCATTTATTTGGCAATATGTCG
>evgR000760c0g1_i01_Hydra
ATAAATTTTTCCTACCTACCTAAGATGCACGCGTTGCCTGTGAGGAACTTAAGTTGAAAATAAAAAGTCCTTTTCTTAATGACAACGAGAAAAAATACGCAATAGCAGAGCTTATGGTCCGCAATAGGAGAGCTAAAAAGTTTTACAACAAAATAAAAGATACCACAAAGAAATGCATAAAAGACCCTAAACAAGTTGGATTATTTGTAGACTGTATGTCTAATATGACATTACCGTGTATACCAGTACAAGATACTTATTATTTCAAAAAACTCACAGTAAACGTGTTTAGAATTCACGATATTGCTTCCCGAAAAATTACTGTTTTTATTTATCATGAAGGCGAAGGAGGTAAAGGTTCAAACGAAGTGTGCTCTATGCTTATGTGGTACGTAAAAAACAAAATTTCAGATGAAGTTAAAATTTTTCATCTTTTTGGTGACAACTGTGCTGGTCAAAACAAGAACCACACCCTAGTTAGACTTATGATGGCTCTTTGTGAAACTACGATGTTTGAAAAAATAAATTAGGTGTTTCCTATTAGGGGACATTCTTTATGCATAATGACCGTGATTTTGGAATTATAAGGAGAAAACTAAAAAATGTAGAAAGATTTTATGAAACTG
>evgR000775c0g1_i01_Hydra
AACCACTTTGCTTTTGTGTACGCAGTGGACGCCTCATTGTGATGACTGTTATACTTGCCATCAGGTAAAAAATCTGAGTCAAGGTCTCAACATTTCAAAACCTCAAATAAGAGATAAAACAAAGTTTATTGGCCATCCTAGAATTGGTGAAAAAGTTTGGACTTTTAAAATGTTAAGCATTTTAAAAGAAAATATTTTCACAATACACAATATGGAAATAGATAATAGCGATTTAAACAATGAATTCAATCCTTATTTACAGCTATGCATATGTAATTTATGTGGTAAAATTTTAAAGCAACCGGTGTCTTTAAAAAAAT
>evgR000781c0g1_i01_Hydra
AACAAAAAATGATGTTCAAATTAAATTTGAGGATGATCTGTTTGAGATGCGTACTGTGCATAGCTACAAAACGGATCTGATTAATTTAGTAGCAAGTAGGGCAAGTAATAAGACCAGTTCTAATAAATTGAAAAATCACTACAGAGGTGTAAAGACTGAATGCCCACTCAATAAAATAAAAGGTTATCATGTGACAGATAATTGGTGTTTAGATATCATGCACACTCTCTTGGAAGGTGTTGTGCTTGTAGAGCTAGGGTGTATTTTTCATGGTTTGTGCATATTAGATAAGTGCATAACTCTATCAGATATAAATAAAGCTATTTCTTTATTATGGGGAAAAATAACTATTGAAAAAACTCACAAACCCGCAGAAATTTTCAAAATACAAGAACCAGGGCATGTACTTACACCCTCTATGAAAGCAGTACAATGGTGGGCTCTTCTCAAGTATTTACCAATGGCTGTAGGAAAAATGGTCCCTCTAAAGAATAAAAACTGGAAATTTCTACTTCACTTGTCTCATTTAATTGATTTAATTTTTGCACCATGTTTTACACATGAAATGGTTGTTTATATGAAGCATGTTATTAGTGATCATTTATCTATGTTTAGCAAGTTATACTGCAGCGATGAAGTAAGGCTACGCCCAAAGCATCATTTTCTCGTGCATTTGCCCAATATTATAATCAAGTCTGGTCCACTTGTTGGCATGAGTTGTATGCCATACGAACTGAAAAATTCATTTTTTAAAAGGTGTGCACATATCGTATGCAACTTTACCAATATTTGTCACACTCTAGCACATAGACATCAGCAACAGGCTTTAGAGTCACAGCTCTCTAACAAACACATTCGCAAAATTGTCAC
>evgR000782c0g1_i01_Hydra
GAATATATTGTTTATATTGACTATGACGTGCAATTTGAACTTGGGAACGAGTTTGATTTATTTCCAACCCATTATAATATTACTCTAAAACCAGATCAGCTTGCCAAAGATTTTAAAATATTGCGTATAATCCTTAATGAGGTTTTTAAACAAAAACACGTAAAGTTACTCGGCCCTGACGTTGCAACGTTAAATCGCTATGAGTTTTTTGAAACCTTTTTAAAATCCGTTGAAAAAGATATTTTAGACGGAATAACATTTCATCATTATTATTCATCGAGCGATGACATAAATCCTAAAAACTTTACAAATATAAAATATCTTGACAGTTTTCTTGACTATGGGTTAAATGCCGT
>evgR000790c0g2_i01_Hydra
ACGATATTTTTAATATATTCTTTAATCGTATTTGTTCTCAAGAAACTATATTAGATCAAATGGAAAATGGCTCAAGAAAACAAGGAGGAGGCAACATCGCTGATATCAATAGGTTACGATACAAAATATCAAATTACATGGGAAACCAAATTGGAGAAAGAGATCTCCGAGAACAATTGTACTTACTTACCGACAACGTTAGAGACCTTAAGTTCGTGTACTACTGGAATGATTCCAAAGACTCCGCCGGCATGGGAACACTCCAAAAGAACCCTTCAGTTACAAGACCAAATAAACAAAGCAATTCCTAAGTTTAAGATTAAGAAGAATTATTCGGCTCTGAATCAAACTACTCATGTCCAGCATCACCATCAATCCTTGATGGACAGTTTCATCCTGCTAGATTTGAATCTAAAACTCCTTGCAAACAACTGCAATGAAAAATCAACTTCGTGTATTAACGAGAAACCCAATGTAACAAATAGAAAAAGAAAAATTTATCCGATTGTCGAAATCCCCGATGACAACAACCTCGAGTTCATAATGGATGAATCGATAAGAATTAAACCTTATTATTTTAAAGATTGGGATGGAGATTTTAAACTTGCA
>evgR000805c0g1_i01_Hydra
GGAAACCAACTTTTATGTTTTGTGGTGACTCAATCCAAAGATATTTTTACGCGTTTTTAGTAAAAAATTCCAAATTGTGTGAAATTTTTGTTTGTGAACTTACTTACACACATTTATATGTAATAAAGCCGGAGAATGACTGTGTTAACGGAAAAGATTTCAACCGCACTAAATTTGTTGGTGAAATTAGAGATGTAATTATATCAGAACCAATGAGAAATAAGAACAGCGTTTTTCTAATAAATTTTGGACTTCATCACTTAAGAAGCATTAATATGAGTGAATGTAAAATGTTATTTGACGAACTACTAACAATGCTTCAATTATTAAAATCTACATCAACAAGCGATTTACCTAGGATTATTTGGAAAACAACTACGCCTACATTTAATACAAACAGCAGAAACAGAAATTATATTCGTAAACAGAGAACAATTGAGTGGAATAATTATTGCTTGGAAAAAGCATGCAAAGTAAACTTACCAGTGCTTGTCGTATTTTGGATGGCTGGTTCTTATCCAGGCGGGACATTTGATACTGTTCATTATGCTGATAATGTGTTTATAGAAGCAGAAAAATTTTTGTTGCATATTCTAACATAATAGATTTTCTAATTATACAGTTATGTTTTTTATTAAATAAAGGTTTAGTTAAGATTTTTACTGAAATG
>evgR000806c0g1_i01_Hydra
TATTCAAACTAAATTGATTTCCTTCTGGGGAACCAAGTTGGTTGGTCTTGAACCATGGTTGGGTGTCATTCTTGAAGATAAGGTTGGTAACTCACAATTGGGTGAATTGGGTACATTGTTGCTCAAGACACAATTCGAATCTTTTGTAAAGGGAGATCCATGTTTCTATTCACACTCTCAAAAGTGGGCAACTACTCTCTTCAGAAAGGAAATTCAAAAGGTTACCCTCGCTGATATCATTGATGTCAACACTGGAGTCAAGAAGAATGAAAAGGCAATGGGAGAATCTCCATTCACTGCCAAGTATGGTCCATTGATTTAATTCGTGGAATTCCATTTGTAC
>evgR000812c0g1_i01_Hydra
TCAAAAATTGTTTTTGAAAACAGATCGTTTCTTTATAATATATGAGAATTGCAAAGATTTTGTTGTTCGTGGAGCTAAAAAGGATTTTGACCAATTTAACGAAAACTCACAGTTCAACATTTTTCATAAAAACAACTCAATTTATAAAACTATGGCGATTGATCTGTCTAAACAAGCAAGAACAAGAATTTTAAAGTTTAGGAGAGCATCTGTTGCATTAAAAGATTTCTATAAACAAACCTATAAGAATATTGGTTTTCTGGATCCACCACTGAATGCATCTGCTAATGCTGGTTTATTAAATAAATTTATTAACTTGTGTATTGCTGAAGGAATTTATGTTGAG
>evgR000813c0g1_i01_Hydra
CAGTCAACAATAATGTTGGTGATTTCTTTGGTGGTGGTGTTAATATTGCTGCTCAAGCTGCCAATTTGGGTAAAGCTAAGAAGGAACAAAGAGATACTACAGGATTCAACTTTGATGATGTTGTTGTTGCTAACAGAGATAATGTTCTTGAACAAAACGAACAAGTTCATCAAGACGCTGCTGTTAAACATCAAGTTTTACTCAAGTCCAATGATGGTGTTGCCATTGAAGATGAACATTTACAAGTCAAGATCAAGACTGAATATCACGGATGTAAGGGTAGAATTATTCTTCTCTACGGAAATAAGACTACTAATCCATTAACCCAATTCAAGGTTCACATCTTGGTTCCAAAGGAATTGGAAGCTCAAATATCTAAGGTTCCACCAATGGTTTCACCAAAGACTCAAGTCCAACAATATGTGGATTTGGTTGCTAAGGGTCCATTCAGTCAATTGACTGATTTGGACTTCTACTTTGAAGCTAACCGTATGATACGATAGATTTAACATCAAGTTTCCAATTGTTGTTCACAAGTTC
>evgR000814c0g1_i01_Hydra
TATTAAAAAATAAAAGATGATCACATATATTATTTTGTTAATCGCCTGCGCTGTTTCAAATAAAGCGCATATAGATATTACAGACGACCTTGACTATGCAGGAGTTAGTGGAATTTTTGTTCCTATTAATGCGAATCTTTACTATGGTACTTCTGAGGAGTTATCAGCTCGTCAAATTCCTTATAAAAGACTAATAGATCAAAATTTTTTGAACCACATAGAAGATGCTTTCAACTTGTTTCAAAAGATTAAAAATGCGTATGATACAGGAGAAGTTGATATTTTATTATACAATGATTGCATTGTTGATATATCGTTTAAAGAACCAAACTATTTTATAACAGCCGAAGACTCAAAATGTAAACTAGGAATTAATCTTGGCAGTGTTATTGAAAAGATATTTAAAG
>evgR000819c0g1_i01_Hydra
TTTTTTTTTTTTTGAAACTGTTAATTATAACATTAATACTTTAAAATGCAACAGCTTACATTAGTTTGCATTTTTTTGTTGGGGCTAGTTTATAGTGAACGAGTTCATTACAATGGCGATAAAGTTTTTCGAATAATTCCTAAAAACGCTGAAGAATCTGATTGGCTGCATTCATGGATCGAATCAACTTTAACAAAGCTAGACGTTTGGCGCAATTTAAAAGGAGTCGGCCGGATAACTGACATTCATGTCCGAGCAACTGATGTTGAAAGTGTGACAGAAGTCTTAAAACAAAAATCTATAAACTTTACAATAACAATTAAAGATTTAGAAAAGGTGATAAATCAAGAACACTTTTATAACCAGCTCTTCTCGTATAATAACGGTGTATATTCTTTCGACAGGTTTAACAGATATACTGAAATTGATGCACAACTAACACAGTTTGCAAACACAAACTATAATCATGCTAAAGTCACCAAATT
>evgR000822c0g1_i01_Hydra
AGCAGAGTTTATGTCTAAAACTATCGCTATAATAAACCAAAAGGGTGGTGTAGGCAAATCAACGGTAGCTGTCAATTTAGCTTATGAATTAGCAGAGCAAAAAAAGAAAACACTATTAATTGATTTAGACCCTCAAGCCCATTCAAGCTGTATTTACTGTGAAGATACAAATAAAAATAAAGCTATAAATATTGTTTTTGAAAAAAATGATTACGAAATTGCAGAAATATTACATTCAGCTACAGTAAAAGGTCAAAAACTAGATTACCTAGACATAATACCATCAACTATTCACTTAGCAATGGTTGCAGAAACTGCTAATCTAAAAATATTTAGAGAAAAAATAATAAAACGTCTAATAGAGAAAACAAATAAATCATACGATTATATTATATTAGATTGCCAGCCTACATTGGG
>evgR000825c0g1_i01_Hydra
CAGCTCGTCTGAATTTCGACAGCGCATTGCTAATACCATGCGCTTCCCTTTTAACGGTAAATATTGTTTCTTGGTTTTTTAATAGTTTGTTAGTTTCTTTGAAAAAATATTTTTTCAGTTCTTCGAGTTTTGACGTTATAAGTCGTTCAATATTTATCATTGTATCTTCCATTATTTTTTACAGAAAAATAAGTTTTGCTCTACAAACCAATGCGGCAAAAAAAATCATACCAATTACTTCCCAGCAAGGACAAGACGTATTTAACACGTATTATTCTAACATTATTTACGTCTTTTTTAGAGTACACAATAAAAATACGCATTAAAAACGAATTTTTAAGATGAGTCTCAACGGAATAAAAAACTTATCCTCAAA
>evgR000839c0g1_i01_Hydra
TAATGCAGTTGAAGATATTTTTGAAACACTTAGTAGAGACAAATGTACTACAGTAAAATGTGAGTACTATGCAGTTTGTATCCAAGATGGTAGAGATGTGACAAAATGTGTATGCAAAACAGACTGTTCTATAGAGTATAAACCAGTCTGTGGAAGTGACGGTGTTAGTTATCCAAATAAATGTGCAATGAAAGCTGCAAGTTGCAAGAACCAACACGAAATTTTAGTTATCAAGGATGAACCCTGTGGCATTGAAGAAAACTCAAATCTTTTTATTGCGATAGAAAACTCTTTTGAAGTTGATGAGGG
>evgR000843c0g2_i01_Hydra
CTAAAACTAATCTGCCATTGTCTAAATAAGGTACAAAGTTAAATCTGAAATATTTTTATTATAATTGTATCACCGAGCACCTGGGATAAATATTTTACTAGCAAAAATTTATATTATAATTTTAAATGGAGGCTGCATTAATAACAGATGCTACTGGTGAGCAAAGTTGTTTATCTTTTTGGAATCTTACTACTGGTGTAATTTTAAAGAGCTATAAAAATGGAGCATGTGCTAGAAACAGTTTTGGTTTGATAGGAAATGATTATATCATATCAGGACAACATGAGCAACAGCTGTTGCATATTTATGATGCGCATAATGAGCAAATTGTCAAAAAGATTGTCTGTCCTGGAAAAATATCTTCTCTGGCTATTTCACCTGATGGTCATTATTGTATTGTTGCATTAGTAGAAAAAATATATGTTTGGCAGATAAACACTGGTAACTTAATAAAGATTCTTTCAAAGCATTATCAAAATATCAACGCCATTCATTTTTCACGTGATGGTAGCTTTTTTCTATCATGTGGAGATGATAACCTTTTGCTAATTTGGAATTTTTCAAATATTTGTCAGCCAAAAGATGATTTCAATGATGATTCTGAGAAGCCTGCATTTATTTTATCCCATCACTCTCTACCGATCAAAGATTTTTATGTTGGAATTGGAGGGATGCGATGTCACGTTGCAACATGCTCCTTAGATCAAACATGCAAGATATACGATATGAGCTCAGGAACATTGGTGTGTTCTTTTCTGTTTGAAAATAGTTGTTCATCGGTTGCTATGGATATTGATGAAAAACGACTTTTTGCTGGTACAACAAATGGGAAGATACATTTTGTAAATTTGTATTCAAAAGAAAAAGTAAGAGAAGTCCATGTTGCGACTAATGAATCTACACTTCAAGCACATACAAAGGAAGTTAGTTCATTATCAGTTAGCATGGATGGTACTAAACTGTTGTCTGGTTCATTTGATTGTAATGTAAAGCTTTGGCATATATCAAGTAAGCAATGTCTCAGAACATTCCAGTTTAAAGGTGCAATAACAAATGCTGTTATTATTTCAATTCCAACCTTTTTACAAGAAAATCATAAATCAATAAATGTAAAAATTCCTATTCCGAATTTTAACAGAACAGTCTACACACCTTTTATGAAAAAACCAGACTTTGATTCATTAGAATGGGACGATAAGATTCCAGTCATTATAAAACCATTGGATACGAGCTTCCATGAGCAATTTGTTCTTCCAGTTGATGAAATAAATTGTGATACTCAATTTGATGATATTATACAGCAGTGCTCAACAGTTCATTCTCCAGAAGAATTTTCATTAGAAAATGCAACTCGAGATCAGTTAATTGAACGAATAAGCAACCTTGAAGCTATTTCAAAAGAGTTTTATTCATTGGTATGCCAAACTATCTTCAAAGAAACATAAAATATTTTCAATAATTTCAATATAAAAAATTATCTAGATAACTTTATTATTTTAAAAAAAAA
>evgR000855c0g1_i01_Hydra
CTTATTTGCCTCATCAGTTGATAAATCGATAATGACTTTTACTCTGATAGTGCAAAATTTGTTGGGCATCATTTTTATGATAAAATCGCCTCCACGTGCTATACCAATTTTAAATTACGACAAAAAGACCGTAACATTTCAATGTACGGGAGATCCTCAATTAAGTCCTCAAATCTTTTATCTTGGTTTTATTCTATTAGTTTGCTCCATTCAAAGTTTTAGTGTTCGAAATCTTCCTTCTTACTATAATGAGACACTCTCTATTACATACAGTACTGTAGTATCATTTATATTTCTCATTAGTTATTTTCCTCTCTACTACGGTAATAATGACATCAAATTAAGAACCACTACTGTCGC
>evgR000855c0g2_i01_Hydra
GTTGATAAATCTATAATGAGTTTTACTTTGATAATACAAAATTTGTTGGGCATCATTTTTATAATAAAATCACCTCCACGTGCTATACCAATTTTAAACTACGACAAAAAGACCGTAACATTTCAATGTACGGGAGATCCTCAATTAAGTCCTCAAATCTTTTATCTTGGTTTTATTCTATTAGTTTGCTCCATTCAAAGTTTTAGTGTTCGAAATCTTCCTTCTTACTATAATGAGACACTCTCTATTACATACAGTACTGTAGTATCATTTATATTTCTCATTAGTTATTTTCCTCTCTACTACGGTAATAATGACATCAAATTAAGAACCACTACTGTCGC
>evgR000857c0g1_i01_Hydra
CCTTTAGTAAATTATAACTTATCGTTATTTATTTGGAAATAACATTCGATGGAGTGGTTCAGTAATTTGCCTCGTCCATTACAGTATTGTTTCATTTTTGGTGCTTTTTTTATTTGCACTACTTCTGTTGCGTTTATATCACAATATTATCCTGTGCATAAAGTTAAAAAAAAGAAGCTTTTTGTTGTTGACAGTTCTTTGGAGAAAAATCATGGTTTAGAATTTAATATAGAGAACAAACCTATTAAAGCAGAAGAAGATGACTCTGATCTTAAGACCGAAAAGTATTTTAATTTTTTCTCTACTGAAGATGAATTTGAATATTTAATAAACAAAAATGTTTCATCTTTAAAAGTAGCTGTAGAAAATGAAGATGACGGTTTTTCAGAGCAAAAGATTGTAAGAAAAATTGTTATTCTTGGTTTGGAAAATTCAGGTAAATCTACACTTCTAGCTGAACTTTCAAAAAAAGGAATTCACTACATGGAATATAAACCTACCAAAGAGTTTAATGTTAGATGCATTTCATTTGATTGCTGTGATCTCAGCATATGGGAAATAGGAGGAATGGAAGATTATCGCATTTATTGGCGTAACTTTTGTGATACCACAGACATGATTATATTTGTTATTGATTCTTCTGATAGAAATAAATTTAAATTGGCAAAAACATTCTTCTCTGAAATATTTGAGAATTTACACAATACAGTTATGCATATAGTTGCTACAAAGGTTGATAAAAATCAATCGGCGTCACTTGAAGAGTTATATGCAGCTCTTGAATTGGAACAATCAGTTTGGAAAATACATAGAGTGGCAGTGGTTACAGGAAGTGTAACTGAAAATATAGGTATCGAGTGGCTGCAACAGCTTTGTTTGGATAATGGTGAAAAGTAATTATTTTTAAATCAACAAAAAATGTAATTATTTACTTAATTATTTTTGTCTGTTTCTTTTTTGTGAAATCAATTTTCTTAGTTTATTAAATAAAATTAAAAAATCAAAGTAAAAAAAAA
>evgR000863c0g1_i01_Hydra
CAGATAAAAAGTTATATTAAAAAAGTCATTTAGATCAATCAGCTAAGATGACCACATATTTTGAAAATATGTGCAACAGTGTGGTTTTCTACGACTTTAATATCAAGGTGCTAGTGGCTGTTTGTGGAATCAATGCAATACTTACAGCAGCTGCAAATTCTTCTTTAATTATTGGAATACATAAAACTAAAAAAAATATGAAATTTTCACGAAACGAGAAGTTAATGCTTCTTTTGTTTTCAACGGACCTGGTTGTAGCAATAGTACAGCTTTTAATACCGTTCATAAAAACAACCAACTTTATCGATTGCATGGAGATATCGATATTAGGTTTCTGCTTTGTGTTTATTGTTTCTTTTACTGGATCAATTTTCTTCCTTATTTCAACGGAGCGACTGATAAAGATTTTTTTTAAAAACTCATTATGGGGTGTCGTTTTTGAAGATAACCACATGATTCCTGGTGTTGTTATTTTAACTTTAACAAACATTGCATTAGGAGTGTGGTTTGCATTAGTTTCAAACTCACAAAGTATTTTCCACCAATTTATTTTGTACATAAGCATCGGAACCTATTCATTTGTTCTTCTGCTGATTATATCAGCAATAAACATTTCATTGATGTTGGGAACAAGAAAAAAAATTGAAAGTTGTCAACTACCCATTCCAAAATGTAAAAATAATGAAATTCAACTGACCGTAACTACAATTTATACTTCTTGGATAATAGTAGTATTGTATGTTCCATTCGTTACAGCTGTAATTTACTTTGGAAACTTAATAAATAGCTCCTCGTATTCATATCTAAATAAACAGCATTTGCACTGGGTACTAGCTTGCACTCGTCTTGTTCCAATATTAAACTCATTCATTTATATAACAAGAAGCAAAAAAATTAAAAAAATGTATAGGTTCATCTTTGAAAAGTTTATAAATAAGATGAGATTTTTACACCATGCTATTGACGAACACTGAAATAACAAAAAACTATATAAATTTCTGCTTGTGGTTCAAAATAATGACAAATAAATAGTTATTAACGATCTGTTGGTTAAATGAAG
>evgR000864c0g1_i01_Hydra
TTTAGTGCAAAAAGATTTTTATAAAAAAAGTTGATGGATAGAAAAGTATTGTTTCTAGTATTTATTGTTCTACTAACTATCCTGACACCAGAAGTTGAAAGCTTTAGTAAGCATTATTTACAAAAAAAGTTATACAAAGCATTACTTACAAACAATTGGCCAAAAATATATGGGAGATGCTCAAAAATACCTGTTTGGAACTGGAATAATACATACCAGCCTGTTTTAGATGCAGCAGCAAAAGGTCGTGTGAGTGTATTATTTTTAACAAAAGGCTCCAGTAAATATTGCACAGAGCTATTGCCTCCGCTTCATGATTTAGCTGCACATTACAGGAATGGGATAGGAAATGTCACATTTATTGCTCTCAGCTATCGATTAAAGAAAGTTCCTGACAAAATCAAATCATTATATCCTCTTATTAATTTTTATGAAGAACCTGTTGAATCTAGAATATTTTCCCAGCTTAAAGCTAATACAGGACATTTTTTAGTCTATGACAAATGCGGACGCCAACAGTACCATATTGGATATCCGTTTGGTTCATTTTATTATAATTTGTTACCACATGCAATTGCAAATTCTCTTTTTCATCATATAACCTTGTGCGGTAAATGCAATGAAGATGAAGTTACGTTCGGAAATTCGACTCGTCTGCACAGTACAAGTTTTGCATTTTCAGGTACTCCGCCGTCCATAACTGAAGTCTAAAATTCAGTTCATGCTGTTCGTTACTTTTTATTTATTTTTAGTAAGAAAGAAAGAAGAAAAAAGAATTATTACAAATAATAATCACTTTTTTTTTAAACAATAAATTTATTTTTAAAAATATTCATTTAATAATAAGTTTAAGTTTGAATTACATTTTTTATTTATTTTAGACTTTTATATATTTTTTCATATTAAAAATGTTTTATATAGTTTTTTAATCACAAAC
>evgR000865c0g1_i01_Hydra
TATTTGATCATATTTCTAAATCAGTGATACATTTAAAGGAGTGCAGATTTATATTTATTTAAAAGTTTGTAGTAGTTTAATTGTCATTCTAATTTTTAACCTCATTGGTTTTATTTTATGGCAAACATAGATATTAAATATGAACTATCTTCAAGCGACGAAAATAAATCTGATAACGATGATAAAGACAGGCAGTTTTCAAACGAGGAATGCTTCGACATTAAAGAACTACATAAAAAAATAATAACTCAAGCGGAATCTTATTTTAGTGATGAAAATTTAATAAAGGATAAATTTTTATTAAAGCACATAAAACGAAATAAAGAAGGTTATGTAAATATTAGGCTAATTGCTTCTTTTAAACGAATGCGATCTTTAACTAAAGATTGCAATTTAATTGTTGACGCTCTTAAAACTTCTTCAAAACTTGTAGTAGACGAAAATAAGCTTAAATTGAAAAGGCTTGAACCTCTTCCCAAAGAACTTTTAGAGCAAGCACATGTACAACATCTTGTATTAAGTGATATACCATCTGAAAATCCTTCTGTTGATTTCATCAAAAATGCCTTCATTTTAATGAAGAATGACATTTTGTCTGTTAAAATTGTAAAACCTTGCAAAAGTTTTCCTAATGATCTTCAGTCACATTACTCTAAACATCCAGAACTAAAAGAAAAGTTGGTTGCTTTAGTTGAATTTACAAATACAGATTCTGCAAAAGAAGCGTTAAAAACTCAGTTTACTGATGCTTATACAGGTTTAAAAGTATCTTTTCTTGAACTAGGGCCAAAACAATTAAAAAATATTGCAAGCGGTGAAACCCACTCAGATGCGGATTCAGACACAAGTTGCGGGAAAAAGAAGAAGAAAAAAAAGAAAAATGTTGTTAAGTATCTTTCACCTAGAGAAGATAGCTATAGCAGTTGTGCAAGTTCTTCTGATAATGAGTTTAACTCTTTTACAAATTGCAGTAAGAGGTATTCTCGCTCGAAAAAACATCCTGGGGACAATATTTCTCAGCATTATTATAAAAATGAAAGGTATAACTCCTCACAAAATTCTTCGCCTTTATCAAGCCCTTCATTAAGTCCAGTTTATCCTCGGAAAAATAAATCTGGTTACCAATCTCCTAAAGAATACAAATTATCTCCATTAATAAGCAATTTGAGTCCATCGTCCAGTCCAGATATAAGTCGCAGAGAAATTCCTAAATTTCATGAGCCATATGAAAGCTCCCATAGTGCGTGGATGTTAAAAAGAATGGAATTGAATAAGAATAATAATGTTGAGTCTGTATCAACAAATGTCAATAGTTTAAATAAATTAGAGATAGTTAGACAACCAAAAGGTCCAGATGGATCAAATGGTTTTAATTCACTTCCCTTAATTAAGTATGTATTGATTCACCCTGTTTAGCATTTTTGCATTTAATTGCTTAAATTTTTTTTCTATAACGTTTCATTAAAATATTCAAATTATATTTAAAAAAATTTATGTATTTTTGAAATGCAATTTTTTTAAAATTATTATTTTATGCTTAATTTGATGTAATTTGAAATGCTCATTTTAAAAGTTTTTGCATTGAAATAATAATAAAAAGTATTTATACATTCTGTAAATATGCACAAAACTATATTTAAGCTGTAAATAAATAAACTTTAATATAAACTTGTAAC
>evgR000867c0g2_i01_Hydra
CTTTTATTTTCTAAATAAGATTAATATCAAAATGTCTATAAAGCAGATTCCTTTGACCTGCCCTGGTCACACGAGACCTGTAGTGGACTTGGCTTTTTCTGATGTAACCGATTATGGTTATTTTTTAATAAGTGCTTGTAAAGATGGCAAGCCAATGTTACGCCATGGTCACACTGGTGACTGGATTGGAACGTTTAATGGTCACAAAGGAGCAGTCTGGGGTGCAACTCTCAATAAAAAAGCAACATTGGCCGCGACAGCAGCAGCTGATTATACAGCAAAGATTTGGGATGCGTTAAAAGGAACAGAACTCTCTACTTTCACACACAATCACATTGTAAAGACAGTTGACTTCTCAAAGGATGAAGATAGACTATTAACTGGAAGCAATAATAAAACTATGAGGTTATTTGATCTCAGCTGTCCTGATAAAGAGCCTTTCATTATAGAGGGTCTAGGAGGTAATCCTAAGGCATGCATATGGTTAAATAATGATACAAACATATTATCAGGGTGTGATGAAAAAACAGTGAGATTATGGGATGTTCGTTCATTTAAACAAACAAATACCATTGTGCTGGACAAACCATTAACTGGCTTAGAATTATCTAAAGACCAATCAACGCTTTCTGTTGTATGTGGTAACCAAGCCCTTTTTTATGACCCTTTAAGTTTAACACTACTGAAAAGCTATGAATTAAAGACTGTTACTAATTCTATTTCATTGAGTCCCAAAAAAAATGTTTTTGTGACTGGTGGAGAGGATTTCTTGCTTTATAAGTTTGATTATAACACTGGAACGCAACTAGAAGCATATCACGGACATTTTGGTCCAGTCCATTGTGTAAGATTTTCCCCTGATGGTGAATTATATGCCTCAGGATCTGAAGACGGTACTCTAAGATTATGGCAGACAACTGTTGGAAAAACATATGGCTTATGGAAAGCGCAAGGCGATGAAGAAGATTCAAAACTCTGCGTCAATGAACAAAATTTATGTAGTGAAGTTGCAGTAGAATCATAGTTATTTTATATTGACAACAATGGGATGTGAATTGACTAGATATTTAAAAGACAATAAGAAATTTATTATTGGTTTTAGAAAGATTGATACCTATTTATATATATTAAAAAAAAAACTAGAATTCAATTTTTAATTGTTTACTTTTTTTTTTATGTTGCAATTTTTTAAATTTAAAAGTTTCATGATGTTTAGAATTTACTTTCTGTGTAAATTTTCA
>evgR000868c0g1_i01_Hydra
GATACAATGATGACAGAAGATCCTACGATAGAAGACCACCACCAAGTGAAACCAACAATGCTGCTCTCCAAGAAGAAAAGTCAAAGGCAACATCTATTTTCCTGGGTAGTTTAGATAGCACAACTACTGAACATGATTTACGTGATGCCTTCTCCAAGTGTGGTGAAATTTCTAGTATTAAGCTCGGTTTCACTAGACAAGGACATTTCTTGAGATATGCCTTCATTTCATTCGTCAACAGAGAAGATACTGAAAAGGCCTTCAATGCTGTTAAGAGTGGTGAAATTTCCGGTCAAAACAGATGGGATGTTGACTATGATGTCGGTAGAGACAAGAAGAGTCAACTCGGAATTCCTCCAGCAGGTCCACGTGGAGGTGGTAGAGGTGGAAGATTTGGTGGTAGAGGTGGTTCTCGTAGATATGACGATGATCGTTACAGTGGTAGTTATAGAAGAGAACAAGGAGGTAGTAGATATCATCCTTATTCCAGAAGATCCCCACCATTCAGAGGTGATAGAGGAGGTGAC
>evgR000870c0g1_i01_Hydra
AACAATACATCTAAAATGTCCAAGTTATCTAGTGACGATCTCAGAGAAAGCGTTCAAAAGGTTTTGAGTGAAAAGGAAAAGAGAAAATTCGTTCAATCCGTCGATCTCCAAATCAAGTTGAAGAACTACGATCCATCTAAGGATAAGAGATTCAACGGTTCTTTGAAGTTGCCAAACTACGTCAAGAAGAGTTTGAAGGTCTGTGTTTTGGGTGACCACAACCATTGCGAAAAGGCAAAGGAATTGGGTATCCCATTCAAGACTGTTGATGATTTGAAGAACACCAACAAGAATAAGAAGATTGTCAAGAAATTGGCTGATTCTTATGATGCTTTCTTGGCTTCTGACTCTTTGATCAAGAGAATTCCAAGATTGTTGGGACCTGGTTTGTCTAAGGCCGGTAAATTCCCTCTCGTCTTGGGTCAAAATGATGACATCACTTCTAAGATTGAAGAATTGAAGAAGACTGTTAAATTCCAATTGAAGAAGGAAATTAACTTGGCTACCGCCATTGGTAACGTCCAAATGCCAGAAGAAGAAGTCCAACAAAACATCACCATTGCTTTGAACTTTTTGGTTTCTTTGTTGAAGAAGAATTGGCAAAACGTTGGTGCCGTCAACATTAAGGCCACCATGGGTAAGCCAAACAAGATCTATCCATAAATCTTTTAACCAGCTCTCCACCACACCAATACTCCGAGTTTAAACACGCAAGTGTACTTGGATGCTGTTGGAAGTCACACAATAAAATCAGATGTAAACAGATCAATCCGAAAATTTAATATCTTGCGG
>evgR000872c0g1_i01_Hydra
ATTTGTATAAAAAAAAAATTAGACATTAGCATTATTAGTGATATTACCAACAAAATAATTTTCATGTATCAAAATGGAGGAAAACTTTTAATTGCAGGCAATGGTGGTTCCGCTTCGGATGCTCAACACTTTTCAGGAGAACTTATGTCCAGATTTTCCAAAGAGCGCAAACCTTTGCCAGCAATTGCACTTCATTGCGATACATCAAGTTTAACAGCGTGGTCAAATGACTATGATTTCCACTCGTTTTATGAAAGAATAGTGGATGGGCTAGGAAAGATTGGCGACATTTTTTTTGCAATATCCACTTCTGGTAATTCAAAAAACATGATTAACGCAGCAATAATGAGCAAAAAAAAAGGTTTGTATGTTATCGGTCTTCTTGGAAAAGATGGCGGAAAATTAAAAAACTTTTGCGATTTGTTTCTAATTGTTCCGTCCAATCAAACACCAAGAATTCAAGAATCTCATATCCTTGTGATACACATAATTTGTCAACTAATAGACAACACGTTTAATAATTAAAAAAAATTTCTCTAAAACTCATTGTACATTATTTACGTATGTATTACATTATACTACACTAATTATCTGACGTTTTTATTATGTTAAACTAATTGTCTGACG
>evgR000877c0g1_i01_Hydra
GTCAAAAAGCGAGTCTGTAGACCAAAGAACATCTGATACAATAATAGATATACCATGAAGACTGTTAAGCATGGGGCGTCTGCTATGGTTTGGGATGTTTCTCAGCACTAGGGACTGGTCTTATTTGCAAAATCAATGGAATTAGGGACTGCTTTGTGTACAAAGATATAATGGAAGATGTTATGTTACCACATGCTGAATTGGAGATGCCCCTAAAATGGATTTTTCAACAGGACAATGACCCCAAACATACATCAAAAATTGTAAAACAATGGTTCAAGAACAAGAATATTACCGTAATGGAGTGGCCTGCACAAAGCCCTGACTTGAATCCCATAGAAAATTTCTGGGAAATAATTGACAATAAGATTGAGCATGGAAACATAAAGAGTGAGGACTTATTTGAACAAATCGAGAAGGCCTGGCACGCGATTGACATGAGAATTGTTAACTCATTAACTGAGCCTATGCCTCGAAGAATGAAAGCAGTAATTGAATCTAAAGGAGGCC
>evgR000883c0g1_i01_Hydra
ATATATTTGCCTTTTTTTACTTTATATACATACATTGTATAACATTTAATCTACTGATCTTTAAAATAGTTACTTTTGGTTTTCCTTTTCTCGTTTATATCAATTTAATCACAATTACATTCGTCACTAACTTTGATAGACTGAATCAATTCTCTCTTAAGGTGGTTTTACACTCATCCTTTTTTCTACGGGAAAAGGAAAGGGACGCGAAAAACAATCACGTGAGCATGCACAGAATTGTCGTCCAAACACTTAATTACAAAAGATTGTTTTTCCCGTCCCTTTCCCTTTCCCGTAGAAAAAAGGATGAGTGGAACCACCTTTAGACTAGACGGACTCAACTTTCGTATCAGATAAAAAGTCTTAAAATATATAATCTTCAAGTTCATCTCTGAGAGAATAGCAGCCGTGGCACAGTGGTTAGAGCATTGGACTCGTAACTCAAGGGTTGCGGGTTCGATCACGCTCAAAGCACTTTAATGTTATCGGTGAAGTTGGAAACAAAAATCAACTTAATAAATGGTATTCTTTATGTAGCGGTGCTCTGTGACCATCAGGACTTCTTAGAACACCTAAATAACTGTTTAATTATTTATTAAATATACTATTTCGAGGGAGATTGACTTGATCTTGGAGATTTTCTTGGGAGATTGACTTGAGCCAATGACTTAATACCAAGGGTATCGATACTCCTAGTAATATTATATTTTATAAATAATAAATTAAAGATTTTATCCATTAAAGATATT
>evgR000888c0g2_i01_Hydra
AGTGCAAAAAGAGATATAGATATAGATATATAAATACAGATATGTAGATATATTTAGCTCAAATCAAGTATACAAACAGATGGTTGTTGAAATGGATAGCCCTTTTCAATGTTTTCCGAAAGAATGTCATAGGAAAATAGTTCTTCAACAAAATCCATCAGATCTTATTACATTAAAACAGAAGACTGGTGCTAGCTTGTGTTTAAATCAATCAGAAAAAATCGGAATAAACTTTAATGATATTTATTCAAAAAGTAATGATCTCTCAGATGAATTAAGTCCTAAACAGGTTTCTATTCAATGCGGAGAAGAAGAAATTAAAGAACGCCCATTCGTTTGCCATCTTTGTGGAAGAAAGTTTCGTCAACGCTGTCATGTTGACCAACATTTAAGAACGCATACTAATGTGCGTCCATATCACTGCAGTTATTGTGCAAAATCTTTCAAACAAAAAAGTCAGATAAACCAACACGAACGAATTCACACAGGCGTAAAACCTTACAAGTGTGAAATGTGTGCTCAAGCCTACCCTCAAGCAACGCAACTTCGTTATCACATGAAAAGTCACGTGGATAATGTCTGTGATAAGCCTTTAACTAACTTTTCGGATACTATTAAATCAAAAAGAGGACGTCCACCGAAATCATTAGGAGATTTACTTTTAGTTTGAAACGTAAAATCCGCTTTCATAGAATCGTCAATGGGTTCCAAATCGTTAATCATTAGGGATTTAATAAAATGCGTTAAATCTTATTTCTGAAATAATTTTTTTTATCAAAAATAAAATAATGCTCGTTAAACGCTTACTTTATTAGCATATTTACGAATAGAGAATAGTGAAACTATGGAAACAATTTATATTAAAAAGATATAAGAGAATTGTTTTATAGAATGTTGCAAAATATTTTATTAAATTTTGAATTGAATTTGTGTAGTTTTTATTAATTTAAAATATTTTTTGCTCGCCAAAAAAAAA
>evgR000893c0g1_i01_Hydra
AGTGCAAAAAGGTATAATTAAATTAATAAATATGTCTGAAATAGCCAGCTTAAATGATATAAGTTATCTTATGACTACATTGGGCCAGGAAGCTGTTGCCGCCGATGATCCAACGACAAATTTAGGTATGGTCGATTCGCTCCCTCCAATCTATGAACAGCCCTCCGCTCCTCCATTATCAATTCCAAACTCACCTGCTTCTAGTCTGAAGAGAAGAAAGAGGAAAGAAGCAAGACGATCCCGATCGAGCAGCACTAGCAGTAATGATTCAATATCAACAATATTGTCATTTCAAGATAGCTATCTGGGGCCTAAGTCTGTTGGAGAAGCAGTTAAGACCGCAATTCTTGATTCCGGTATCGCAAAAGACATATCCAATGTTCTGGTAGCTGTTGATCAAGGTATATCCACGTTAAACAATACGTTGGTTTTGTCAAAATGCCCCAGAATTCCAGATTTTAGGCCTCTTATAACTGAAAAAATAGGTGAGTCAATTACAGGATTAAATTTGTCAGGCGTAATTGGTGTAGCAATGGCCAAATCAACAAGCGACTCTATTCCCAAAATAGTACAGGCTGTCAAGGATAGTCTCGCCCATGTTACTCTACAAGTACCTCCTTTTGATCCGACCCCGATCATTGATCTCATTGATAAGCGTCTTATATTAATTAGAAATCAAGAAGACCAAAGATTGGAGAATCTACTTAAGGATGTAAACTCAAAATATGTAGCCGTGAGCAATGAGTTAGCGGATATTAGGCTAACACTCAAAGATGCCATGAATAATTTATTAAGCGCTGCTAAACAATCTCAGCAATTACCAATTGATAGTAGTAACCCAAGGAAGAGGGTTCTAACCAATATTAAGGTTATTGCCAACACTGAGGAAACAATATTTATTCACGAAACGATTAAGATCTGGCAGGATCTAAGAACAAAAATAATGCTCCTTGTTAAGCATGTCACAAATTGGTTAAACCGTCATCATCCTTCGAAGAACAGTCATATATCTGAGGCCGAAAATGCGTTTCTTTTATTCCTTCAAGGGAAAATTTGTGTTGGCTCGCTTAATGATTTAATTAATCCTAGTATGGACATAGACTTAAATCTTAATGAGACATTTATTAATTAGTTAATTCATTATGTAATTTTATTACGAAGATTAAATACTAAAGTTACTTAAAAAAAAAA
>evgR000897c0g1_i01_Hydra
GATTTGGTCTCTAGTGTTATCCAAAAAATCGTTAGTGCATCTTTACCTTCGCCAAAGGTAACTGATCGTGTTATTTCAATTGTTGATACTCTGCTAAGCCTCGATGACAAGGTAATCAAGTCGGCAAATGAGAAGTTTAATTCTTCTAAAATAATGTTATCTCAATTAGATGATTTGGGTAAACAGCAGACAGAAAACATTACAATTTCAAAGAAAAACTTAGCGTTTGCATCGTACTCCATTAAAAACAATCAAAACCCAATCTATATTTATTCGACCAAACAAAAAAATGCAAGTGTTGATATTTCCAGTGAAGATCGTATAAATTCGTCTGAAAATATTAAAGCTTACATAATTTTACCCTCTGATCTGTTCACTGGGCGTAATATATCACAAATTTACTCTTTCGTGTTTTTAGAAAAAACATTTTTTTTACCAAGTATACAAAATATAGATATTTTATCAAAAAAAGATA
>evgR000898c0g1_i02_Hydra
CAAATAAAGATTTCTTGCTTTATATTTACAGGAAATATTGTTTATATTGATTATTAATGAAAATTTAATAATGTCAAATATTGGAGATCTTGTAAGTGAACTAAGACAATATTTTCCTGCAGATACACCTGAGGATGAAATGATTGAATTATTCAGCAATCGTGAGATGTTCAATGAATATTTAGCACAGCTTGAACTACATTACTCAGATTTACAACAGAAGTACAATGTTTATGAGGAATGTTTTCAGCCAATTTTATCTCAAACGTTTGAATCTATATGGATGATTTTGCTTTTTAGTTTAATATTGCGATTCTTTTCTCTTGTGAGACCGCCACCAATTTTAATTCACAGTCTTTCTGCGTGTTTAGGCTGTTGTTTATTATTTAAGTTCTTTGATTATGATATGATTTATATTATTGCTCCATGTTTATCTTTATTTCTTCTTGTACAATTGGATAAAAATGGAGTTCTTGTTATTGTGTACGCTGCAGCATACCTTGTTATATGCGAATTTTACCTTCTTGACTCAGTTGTATGGACCAAGATAAAAGGATCAGTAATGATATCATTTATGAATGCCATTTCAATTACTCTTCAAATGGGCAAGAGTGGTATCAGTAAAGTATCGTTTCTAGAATTTTATGGATATTTACTTAATCCTTCTACAGTTATATTTGGACCTTTTCTTACCTATGATAATTATCAGCAAATAATGAGTGCTCACCCTTTGAGTCTCAGCTGGTTATTAAATAGCTCGAAATGTTTCATTATGTCAATATTTTGTTTATTATGGTCAAGCTGTGGGACAAGTTACTATTTACCATTGTTAGAAAATTATATACCTTCTCTAAGAACTAGCAGTTGGTGGGATGCGTACAAAACAGCACAGTCATTTCGCTTTAGTCATTATTTTGTATGTTATTTATCAGAATCAGCTTGTTTGTTTGCTGGTATATCAATGATAATCACTGATGAACACAATTTTGAAGTTAAAAAAGTAACAAAAAAAGATAATAAACACAGTAATGAACACAATCTTGCTTTTAAAAAACAAACTAAAAAAGACATAAAAATCATTGATGAAAAAAACACTGATAAAAACACAATAAAAGTGATTTGGTCTCATTCTGTTGCTAGAATGTTTCAAATTGAACTTCCTCGATCACTGGTAGATGTAGTCACAAATTGGAATCTCCCTCATCATTATTGGTTAAAACAATATATTTTTAAACCTGTACGAAGATTTGGAGTGTTACCAGCCATTTTGATTACTTATATAGCTAGTTCATTCTTGCATAGTTTAAACTTTCAATTGTCTGCTGTGCTGCTCTCAATTGGTCTGTTTGCATATATAGAAAGTGTTTTTCGCAACTTTATTGCGGTTTTACTTAGTGCTTGTGTAGAGGCTCGCAAGTGCAGACCTGGATGTAACCATTTGCATAAAAAAAACACTTTATGGGTGATGATGCTTAATGTGTTTTTCACTATTACTGCGTTATGGAATCTTTCTTATCTTGGTTGTTTATTTGACAATGGAGGGGAAGAAGAAAAGGGGTACTCATTCCAACACACATTAAAGAAGTGGTCAAGTTATGGATTTTTAAACCATTATGTCATGTTATTAATATTTGGTATTAATTGGTTTTCCAAAAAATTTCATTTATTTTAGCCATGTTTTTATTGCTCTTACTTTTTTTTAAGTTTTTCTTTTACAAATTTTTTATTGCTTTTTTAAAAAAAACTACTTTTTAGTTCTTTTTTACACAACTATTTTTAGTGCTTTTTAAC
>evgR000916c0g2_i01_Hydra
AATTAAGTATAATAAATTAAGCCTCCAGTTATATAATATGTCAGAAAACCATTCCATTCAACCATTAGCTGGTATAAAAAAACTAAAAATTAACTTGGCTGAGAAAATGAAACTCTACATGTTTGCAATATTTATTTTGCATCTTTCGCAAGGATCAACTATAACTTCCTGTATTATAAATAACTGCGTGGTTGATCCCTGCTTAAATCATGGTTGTCCTGTTGGTTTTCAATGCGTTCAAAACTACTGTAATGGTTGCTACGCAGAGTGCAAACCTTGCGAACCAACCTACTGCAAAGTTGACCCCTGCTTAAAATATATTTGTTTGCCTGGATACAAATGTTTCTCAAATTACTGTAACGGCTGCAATGCTGAATGCAAAAACGAAGTCACCGAATAAAGAAGAAATTAACAATGCTGACAGAATTTTTATTATTTCAAAATTGTTCAACAAATAAAATGCAGAAACAAATATTTAATTTTACTTGTATCCAATGGCTTAGAAATGAGCCCGAATTCACTAGAGAGGGGGAGGGAAGGATATTTTTGGAGCCAGGTATAAATCCAATCGTAGAGCGCAAGAAGAATCAATGAATAATTGATATCGTACGACAATAATTTTAAAGTTAAACTTATTGAATCTTACAAAGAAAATCATACACCGTCTTTTTA
>evgR000920c0g2_i01_Hydra
CAAAAAATAGTAATATTTCATTGTTCAGACATGAAGTCGTACCAATGTGGGAATCTTTCCCTCAAGGTGGATGTTGGATGGTTAAAATCAGAAAGGGAGAAAATGGACAAACATCTTCATTCTGTGATAGATTGTGGGAACAACTCATCTTCTCTACTATTTGTGAACAATTCGAAACTGGTAACGTTGTTGGTTGTGTTTTGAGTTTGAGAAGCAAGGAAGATGTCTTACAAGTCTGGATTAAGAATTCAAACCATCATGCTGACAGAATTCACATTTGTGACCGTTTCAAGTGGATTTTGCACTTTGGAGAAGGTGTTATTGTCCAATACAAACACAACAAGTTGTCCATCAAGGATGGTAGTACTTTCCGTAATGCTCGTACCTTCTTGGTATCTAGCTCTGAAACATCAAATGCCTCCAATTCAATTGATGATGAAGCTGAAAAGACACCTGCCAAACCATAAAAGGATA
>evgR000922c1g1_i01_Hydra
TATTTGCACTCCAGGGGCTTACCATCGTGCAAGACGGATGGCTAAAGGAATATACTGTTTAAAAAGTTTTGGTTTTTGTCATCAATTAAATGTTACAGATAAAGAAATCGGGATTTTAAGAAGAATTTGTCTCTTCATAATCGCAATTTATATATGCTTCTGGTTTTCAGCTCCACTCACAACAGAAGCACCAAAAAATGATCTCCTTCTTTTAAAAGTTATCGAGAACTATTGTCAAGTTGACAGCAAAGTTGCAAATGTTGCAAAAAATAGAATTAAACTTCAATTATGGTATTTAAGTGAAGATCTAGCTGCTTTCTCATTATTCAGTGAAAAATTTTCAGTTAAAGAGAAAATGGATATTGTAAATGCCTTACAAAAA
>evgR000924c0g1_i01_Hydra
TAACTTTACTAGATGTCAAAGGTTGTCTGTTGCAGTTTCTTTGTTAATGACTTCTATGGTAGCAAGCACAATGTTCTATGGAAAGATGCCACCAGCAAATCCATCAACCGAAAATAAAGTGGCCAGCTTTTCATTCACATGGGGACAGGTTTTTATAGCAATTGTTAGTAGCTGTATAACGATACCTGTAAATGTCATACTTGTTGGAGTTTTTCGTTCAATAAAACCATTTGAAAATATAAAAGACAGTTCAGACGAAAACTTTCATTGGAATGATGAAGAGGGATCATCTTCTCCATCGCCGTTGATCAGAAAAAATTCAAATAAGAAGACATCAACTATATATATTGAAAAAAGTACACCATCCCTGATAAAGAAACAGAAAATATTTTACCTTCCTCGTTGGTGTTTGTACATGGCGTGGTTTATCTGTGTTTGCTGTATTTTTGGTTGCGGTGTTATTGTTTTGTTATATGGAATAGCTTTTGGAAATAATAAATCTTTGAATTGGCTATCTAGTGTGTCT
>evgR000926c0g1_i01_Hydra
AATTAATTATTAAAAGTATGAATATTTTAAAATAATTGATTTTTGTAAATTGACTACACATGGCAAGTATGCATTCTGCATGTTCTCAGTGTCAATCGCAAAATCATGAAATACAAGATTGTCCATTTTTTAAAGTTACTTCAAAATTTGCCAACTCTGATATTGAATCCATAAATAATAAGAGTTACTGCGATATCCCTATCTATGATACCCATTGTCATATTGATTATGTATTTAAAAGATTAAAGCACCATAATCTATGCTTTAAGGACTTTAAACAATCTTATTTTTTCCCATCGAACTTTAAAGGCTGTATAGGGTCATTTAGTGATGTTTCTGTTTATAACAACTGTAGTGTATGGTTAGATCTACTCTGTCAAGATGACATATGGGGGACATTTGGTTTACATCCACACCAAGCAAAGTATTTTACAGACCATGTTCAAACAGCTATGCTTCAATGTCTAAAGCATCCAAAAGCTCTTGCGGTTGGTGAATGTGGTTTAGATTTGTCACGTTGTAATCCATTATCAACTTATTCTGAGCAGAAATCAGCTTTTATAAAACAACTTCAAATAGCAAAGCAAATTAATAAACCTGTTGTTATTCATTCAAGAGAAGCTGAAGACCTTGTGTACGAAATTTTGTGTGATGAAGATATGAGGGAATGGCCAATTCATATACATTGCTTTACAGGGAACTGTACACATCTTCATCGTTTTGTAAATGAATTCCCGAACATGTTTTTTGGTTTTACAAATCTTATTTCTTATTCTACAGCAAAAGTTACACATGAGGTTGTTAAAAGTGTTCCGTTGGATAGAATACTACTTGAAACTGATGCTCCTTATTTTGTTCCAAAAAAACTTAAAAATAAATGTAAATTTAGTCATCCAGGTAATATTTTATTTGTAGCAGAGGAAGTTGCACTTATTAAAAACAAAGATATTCATGTAATCTTGGATAATTGCTTCAAAAATGTTGAAAAGCTTTTCAAAATATAAGTACCAAAAACAACCATATCATGGTAAAATAATTAAAAACACTTTTTTTTTAATTGCTTATATTAAGTATATATACTTATTTTTTACATACTATTAAAACAAAGTGCAATAAACTTGATTTCTTTTATGTTATGTAAAAAAATTAATATTTTATTTGAATAGTGAATTATAAATTATAAATGTAAAATAATGAAATTATAGATTGATACTTGTTATTAGAAATTGTATATATCCTTTTATTTTTAT
>evgR000927c0g1_i01_Hydra
GTAATATTTAGAATGATGACTATGGAGAATCGAGCTGAAAGGTTCAATATTCAAGAAGATATTCGTTTAAACATTCTTGTGAATTCCTATGAAAAACAAAAGCACAACTTTTCAAGAAACATGGAAAATTCTCAAAGACATTTGGCTTTACAATTTCAGCAAAGGTTTTCGAAAGAAACGAGTAAAAAAAACGAACACATTCTTTTACAAAGAGAAAAAATGAATTCTGTTATAAGTTTACCAGTATTATCAAGTAGCTATGAAAACATTTCCCACCTGAAAACAAAACACAATTCTGTCAAGACAAATCTTGTATCTCCTTTTAGCCTTAGAAGATATAGTTCAACACAATTATTAATACCACAATTGAATAATAATGGTAAAAAATTATCTCCACTAATCAGTGCCAGCAGTTTGCCGTCCAGTAACAGCCTTATTAATAAACAAACACAACAGATTTATAACAGAATCACAGAAAATAAAGATGGCGAAAAAACACTAATAAAAAGCAAAAGTGACAGTTCACTATCAAAATCAAACAATCTGACCAAAATTTTAAAGAAAGATCTTACTGAAAATAATTTCAATAAAAAAAATACTTCAACAACATCAAGAAAAAGTTCATATAAAATAAGTTATGAAGACAGAGACTACTTAAATAAAGAAGAGATTCAGAAAGAAAAAATGGAGCATTTTTTTAAGAGCATTGAAAAACTAAAATTAAAGAAAAAATCAAATAATGAAAATGTTTTTATAAATAGAAACATTTTGACTTATGACCATTTATTTGATTAACATTTACAAGTTTTTGATTTATGACTATATATTTGATTAATATTTACACATTTTTGTATTATGGCTGTATATTTGATTAATATTAAATTAAAAAAATTAATTTAAAAAAGTTTTTGACGTAAGAATATATTTAATTTACACTGACAAGTTTTTGACTCATGACTATATATTTTATTAACATTAACAAGTTTTTGACTCATGACTATATATTTAATTAACATTAACTAGTTTTTTACTTATGTTGACCATATATTTGATTAATTCATCAATTAGTTAGTAATTTATTTATTCAATAGACATTAATTATTACTACAAACCCCCTAACATCAATCACCAATCAAAAAGTGAAAAATGACAAAGAAATATGACTAATTAATTAATTGAAATGACAAAAGGACTACTCCTGAATTTTCAACATAAATTTAC
>evgR000933c0g1_i01_Hydra
TGTCAACCGTATGAAGTCTCAAGATAAGAATATTATTTTGGATAAGCTTCGTTTGGTTCTTCGTTGGATGAAGGCCAATCAAGAAAAGGTTACTGTCAAGAAGGAACACTTTATTGCTCAATATAAGAAGTTGAGAAGTACTGCTGGTCCAATTTTGGCTAAGCTCTATGGTGGTGCTGATAGCTTGACTGATGAAGAAATTTTGGGTGAAGAATTCGAGGATGATGAATTCGGTGATGATGCTCCATTGGATGATGATGAAATGGGTTTGAAGAATGATGATGATAAGCCAATCGATGATTCTCCAAAGAAGGATGAATTGTAAATTAACAATTTGCTAAGCTC
>evgR000951c0g2_i01_Hydra
GTAAACATCTTGACCAAGCAACATAAAAATTATCATTAGTAAATAAATCTCAATTTTTGTATTAGTCCTTATTAAAGATGTCATTGTGTTTGGTATGCGTCAAAAGTGGTATAATTGGTAATAGAAATAAAAGCAACCTTACAACAATAGAACAGAAGCTCAAGTTGGAAGCTACAGCAGGTCCTATGTTTACAATATGCTTTTGGACCTTGTCTGCAATAAAAAAAAAAACCATTGGAAAAACCAGCCCATACGTAACAACAATATTCAGTAAAGGGACAAACAAGAGATTTATGAAGGTAGAAAATGGCAAGCACAATAAAGATAAAGTATTTAAAAATATAAAAGACGAAAAACCATTGTCACCTCCTATTTCATTATTTTCCAGATATTCATGCCAAAAAAGTTGGTTTTAT
>evgR000953c0g1_i01_Hydra
GTGACAGTACCAATTTAAATACTGGATGGGCAGGAGGAGTGATACAGTTGGTTGAAAAAAAACTTAGTAGGAAACTAGTATGGATTGTATCTGACCTACACACTGGAGAACTAGGGTTACGACGTCTAATTAAGTTTCTCTATGGTCACACTAAATCCAGCAATCAATGGTCAGGACCACTTGGTAAGATGTTGAATAATGCTACTAACCTTGAAATTAATCCACAGTTTATAAAAATTTGTAAAGCTTCTCCACTAATCCATCTTGATCCTAAAATTATTAAAGATCTGAGTACCGATCAGAGCTATGCATACC
>evgR000961c0g1_i01_Hydra
GTTGCCATATAAAAAATTTATGTATCTAAAATTTCTGAACTTTCTATTAACTTCTGACGCTTGGGCATCTAAACGGAACGCTCGTAATTGATCGTGGAAGAAAAACTAAAAACTTGATTGTTAATAGTTTTCATCGACTGCATTTTTCAAAAACGTTATATCTCTAAAACGTTTCTTCAATGTTGATAGTAATACTAACGGATCATGCAAAAGTGTAAGATTACTTTCTATATCGTCATCATCCGACAAGTCTGCATTCTGATAATTGTTATTAAATGCTTTATCGGGTACTTTTTTCTCTCGATTTGTTCAAGTACTTTCATCAATTTGCGTAATTATGTAACGTTATTGTCCTCGTTGAATATCTCCAAATTCAATATCCATTTCTTTGCCGAAATATATATTTATGTTTTCATTAAAATAATTTTTTATTACTTAATCAATATCTGTACAACATTTTCATTAAGCATATCAGTTGGAATTTTCCATGCACTACCAGCATACATTCTGTTGTGCGAACCTAAATGATTTACACCAAATAACTCAAGTGTGGTGTGACAAAACGCGCAACATAATTTTGATATTTCAATATAATCAATAGTTTTAATATCTTTTATTTTTATTCAATGTTAAGTGTTTAACTAATGCTATTTATGAATGATAATTTGATAGATTATCAGCAAATTCAGATTTAGGATGTAGCGCATAGTTTTCTTTTTTAATGACATTTTTTCGACGTCTTGATTTACTCAAACCCAACTTTCAATGAGCGCATCAAATTTAAGCTCAAACGGTTTGCTAAACAATATCTCTTTTATAGAATAAAATACTATATCAGCATCTTTTAATACTTCTGTTTTTTTTATAAATTTCGTAAAATTGTAG
>evgR000968c0g1_i01_Hydra
CAGTTTCTTTTCAAATTAAACCAAACTCTTATTCAAAAGAATTCCAAAGTGTTATTCATTTGACTATTGATTTCGATATTGGACAATACGGAGAAAGAACTCCAGGTGTATGGTTTTCAAAAGATGGATCAGGAAAGTTATCTATAGCGTCTGCAGTCAATGATGATGTTAACTATTCTTTTCTAACAGAACCACTACCGTTGGCTAAGTGGACTAGTATAAAAATTTCTCAAGTTGAGCTCAAAGGAGTGTATAGTTTCAATTGCTATGTAAATGGTGAGCCTGTTCATAGTATTGAAAATAAAAATCCACAATTGTTTGAACATGTCAAAGTGTACGTAGCTGATCCGTGGTATAATGCACAGGATGGTTCCATT
>evgR000969c0g1_i01_Hydra
TTTTAATTTTGTCAAATTTGAAACAATACAATTCTCAAAATCAAATTCGAAGAAGTAAAGTCAGGCCCTGCATCAGGTCAGCATCTACTCTGTCAGTGCTGCAAAACTCAACTAACAGCTTCACAACTCAGCTCGCTAAACCTAGTTCTTCAACTAGCGTTGCAATTATAAACGCCTTGACCAAAAACACCAAACTTCAGCAAACCAAAGCAAAACGAGTAATTATCGCTGGGCTCCCAGATTTATACCAAATAACTGAAGGAGCCGTTGACAAAATAGAAAAAATAAAAAATGTATTAGACTCCTTGAAGTTCCCAAATGCTTTCAAAGTTGTAGGTCCTATCCGAGTTAAATACTTGTCTGCAAACAAAACTCCTCTTTCCGCACCACTCATCGTCAAACTAGAATCGACAGAATCTTGTAACAACCTACTGTTTATATCCAAAATCTTTCATTGCTTGAAAACTTCAAAACCATCTATATTTGCCCCGACCTCACCCCAGATGAACAAGAAAATTTTACAAAACTTAATAAAGTTAATGCTAACAACTTTTAACAAACTTAAAAAACTTAGAGCCTCTCTCAACCAGCCCTTTCGATTTGTCATCCGTGACGACCAGATCCGCTGTATTAACACCGCTGCAAAAAAACTCTTTATCAATGGTACAGAGAAATGCCCATTCGGCAACGGGAAAGCTGCTTGCAATGCCGCCAAGCCTAAAATTCTGGTCTAATTCTTCAATCCAGAAAAACAGTTACCTTCAATATTTCACCTTTTCTTAAACTACTAAAATGCCTCTACACCAATCCAACTCAACTTTCTACTTGTTCAGAAAATTTATCTAAATGTACTTTGTTTCTTTGAGAAATGGTTCGACCTAGAAAAACCCCTTTTTGAATT
>evgR000978c0g2_i01_Hydra
AGGCTACTTTCCGAGATAGATAACAATTATTTTTGCGTATGTAAACAGAAATTTGATACGTTAGTAAGTTTTTTTAAATAAAGCAAAATGTTGAAATTAGACACAACAATATCTATAGAAAAGGAAAAAGAAGTAAAAATGACTAGTGAAACCTCAAGTTTGAAATATATCAGCTTAGCAGTGTTAATTTTTCAAAATGCTGGATTAATTCTTGCCATACGCTATTCCAGAACACTTGAAGGTGATATGTACATTTCTAGTACAGCTGTGTTTTTGGCTGAGATTCTTAAACTGTTGGCTTGTTTGATTTTAACTTTTACAGAAGTTAAAAACTTTAGTTCTTGGATCAGGTACCTGTACAATAACATAATCGCAGACCCGTTATCAACTTTTAAAGTTGCAATACCTTCATTTATTTATGTTTTACAAAATAATTTACAGTTTATTGCTATATCCAATCTTGATGCTGCTACATTTCAAGTAACGTATCAGTTAAAGATTTTGACAACAGCTTTATTTTCAGTATTCATGTTAAATAAAACTTTATCGAAATTTCAATGGTTTTCATTAATACTGTTGTTTGTTGGGGTAGCACTAGTACAGTTTCAACCAAATCAAGCTAAAAGTTCTCTCACAACTCAAAACCCTATTGTTGGTTTTACTGCTGTGGTTGTTTCAAGTTTGTGCTCAGGATTTGCAGGTGTATACTTTGAAAAGATTCTAAAGGGATCTGGAAATGTCTCAATATGGTTAAGAAATATTCAACTGGGTGTTTTTGGTGCTTTAATTGGTGCTGTTGGAATGGCGGTAAATGATGGCACTAAAATAAAACAAAACGGATTACTTTTTGGTTATTCAACAATTGTTTGGTTTGTAATTCTTATGCAAGCGTTTGGTGGTTTATTAGTGGCTGTTGTAGTGAAATATGCAGATAATATATTAAAGGGTTTTGCAACTTCGTTTGCTATTATTGTTTCATGTATTATATCTATTTATGCATTTAACTTTGTATTAAGTTTAGAGTTTGTTGGCGGTTCAGTTTTAGTAATGGTTGCTATTTATATTTATGGTCTTCCACAAAAATAAGTAAAAATGGTAAATAAATTAGTTGGAAGTAATTTTTT
>evgR000994c0g2_i01_Hydra
CGAGAAACCAATATACGAAAAATGTTAAGTTTATCATTGATTTTGATGATGGTTACATCTTTGTTTGCAGATAAAAACTGCGATTGGGCATGTCTGCAAGTATACGAACCAATCTGTGGTAGCGATGGTATAACACATGGCAATGAATGTGAAATGTCAGTTGCAGCATGTACTAAAAAAAAGACCATTACAAAAGTCAACGACGGTGAATGCTCAAGAAATGATCAAGATTTAGGCTGTAATATGGCATGTATTGAAGCGACTCAATATGTATGTGCTAGTGATGGTATGACATATGACAATGAATGTAAAATGTTATTTGCATCATGCATGCAGAAAAAAACCATTACTAAAATTCACAACGGTGTTTGCCATGTTAAAGACTGTAAGAAGGTATGTCCAAGAATATTTCTACCTGTGTGCGGTAGTGACGGTAATACATACAGTAGTAAATGTGAATTATCTGTTACATCTTGTAAAAAAAATCAGACCATCACTAAGCTCTACGACGGTCGATGCACTCTTCAAGAAGTAAACTGCAAAACAGCATGTCCTAAAATAATGGAACGAATGTGTGGTAGCGATGGTAAAACTTACAGTAGTTACTGTGAATTATTAGTTGCGTCATGCTTAAAACAAAAAGCAATTATTAAAGTTTATGATGGAGAGTGCAACAATGAAGGGAAATGTAATATTCCTTGTAATCGAATGCTGGCACCTGTGTGCGGAAGTGATGGCATTGTATACTCAAACGAGTGTATTATGAGAACGGCGTCGTGCAAACAAAAAAAGGCTATCACATTAATACGAAGCACAGATGATGAAAGTTGTTCATGTCTCCTTGAATGCACAAAAGAATATAAGCCAGTGTGCGGATCAAACAGAATAACGTATTCTTCAGAATGTGTAATGCGTCGATACAGCTGTCTAACTAAAAAGCCAATTATTGTTATTAGAAAAGGAATATGTAAACTTCCAAGAACAACAGACGACCAGGACTACTTTTCTAGTGAAACAGAAATATTATGAAAC
>evgR001011c0g1_i01_Hydra
ATAATGGAGTTTTGTGCATCTGTATATGCATATATTAGCATTAGTGGTAAAGATATTATGGCCAACATTAAGCTTAGAAACTGGTTACCAATGATCTGTGGATTATTAAATAAAAATAATAAAAGTTTTTTAAAGTCTCTTGCAAACCTGGATGATACAAACCTAGACAAAATAACATTATGTCATATTCGTGATGTTATTCCGAAAGGATATCAGATATCAGAACAATTGAAGATCAGAAATTATCACAGATTTTCACAAAGTTTTAGGAGAGTTGTTCACAATGATGGCCAGAGGTTATTCATTGAATGCTTTTATGAATCTCAATCACTATTCACCGATGAGATGAAAACATCTATTGCCGAAATAGAATGGTTCATTTCAATTGAAGATGGAAAACCTTCTTATGATATTTCTTGTGTGACATATTTTGTCGATCTCTTAAACTCTGATAAAAAGGTATTGCATTTATTTGTTAATAAAAAAATATTTTCTGAAGAAGAAAAGAATCTGATTAAGAAATGTTCGACAAACGCTTTAAGTGTAAGGTTTTTTTATCCTTGTAAAATCGATGAATGGAAACCGACTAACAAAGTTAAACTGTTGCTGATAGACTGTCAGATATATTGTATACCAAAAGAAGACTTTCAATATTTTGTTCCATGGATCAAAATGTGTGAAGAGTTAGTTATTTTGCTTCATGATGAATGCAATGAAAGCTATCTGAGGAGTATTTGGAAATGGACTGGAAAGTTAGAACTCCGTAGTTTTATGATATTCTTTAAAGGACAGAAATTTACGAATAGAAAATCTTTTAAAAAAATAATAATTCATAAAAAGTAAGTATATAATTATATATTCAGAGTGGCCATTAATTCAGAGTGGCCATTCATAAAGAGTGGCCATTCATTAAGAGTGGACTTTCATTCAGAATGGATATTCATTCAGAGTGGCCATTCATTCAGAGTGGCCATTCATTCAGAGTGGACTTTCATTCAGAGTGGTCAGCAACTTAGAAAACATGGAAAAGTCAAG
>evgR001032c0g1_i01_Hydra
ATTTTTAGAAATCGAGTACCTTTTTTTTTTCTAAGAAACAAAAATGTCATACAAAATACAAAAAAATTCAACATCCCTCCCCCCAAAAAAAAAATTCCTGGATCCGTCACTGAAGGCACTGTATATATATGTTATTCTTTTTTACAGAGTTTATTAATTACCGTCTTTCCGATAACAGCGTTTGCAAAGTTAGAGGGACTTATTTCCCTTATTTTGCACGAGGGCGGCAAAACGACTAGGGAAAAATTAGGAAAGTATCCACTAACTCTAAATATAATAAGTTTTTTATTAAAAAGTGACTTAAGCGAGTTTTGAAAATTTTGGAAATTATTGTTTTTCTTTTACCAACAAAATACCTTTAAATGAAAGGGCCCTATGTTTTTGACACTTTTTGACAAAAGGTCCAAACTCCAGATGACCCTGCAACAAAAGGCCCTACAAACTGTTCAAAAGTTACCTAACTATCATACTGATACCGCCTGAGTTAATGCGTGTATTTGTGGTATCAATTTTATTTTAAAATACGTATGTGCTTATGTTTAATTAAATATTTTGTGTTTGGTTTTAGTCGGCG
>evgR001034c0g1_i01_Hydra
TGGAGTGTTCGTTTGTATCACGACGGTAAAATGTTGAATAAAGATTTATTAAGGATTCTTAAAGATAAATATGCAAATTTGTTCTTTTATGACGTCAGTTCTTTATCAGAATACGGAAACGTATCTTATATTAACGGAAGAGCATGGCGGTTTCTTGCTACTGGCGATAGTAATGTAGAAGTAATGTGCAGCAGAGATTTGGATTCTCCTTTATTAGAGAGAGAGGAAGATGCTGTCAAGGAGTGGTTAAATAGCGGTAAAACATTTCATGTGATGAGAGATAACCCAATGCACGGAGTTCAAATACTTGGTGGAACGTGGTGTTTTCGAAACAAAAAAGATTCATCTTTTGGTATAACTATTGCCAAGTTAGTTATCGAAAAAAGTAAAACTCGTAGATTTGACAAAGAAGCCCCCACTGACGATGACCAGTCAGTTTTAGAAACATATATATGGCCGCTAATAAAAAATAATGTCCTACAACACGACTCATTTCTATGTTCTAGATATAACGGTAGTAAGCCGTTCCCATCAAAGAGAAGTGACAGGAAAATATTTATCGGAATGATAGATCAATTAAACGATCTTCCTCCTTGTCCAATTGAATGTCGACCTGAACAACACAAAGATTGGGAATATTGTTAGTATCGTTAAAACTATATGAAACATAAATG
>evgR001039c0g1_i01_Hydra
GAAGTTGCTCCAGAAAATATTGTTTATGACTTTGCTTGCAGCTTGTCAGAGTTTTGTCACAACCGTGAATCGGGATATTTTAAAAATACTTCTTTTTTTCATGATGTGTTTCATGGCTATACACATAAATGTGCAGATGTTTTCCGATGCAATCGTTTGGGGAATTTTTATCCAATTAATTCATCTATTTGCGAACAGTTTAATAGCTTTTTACAAAATATAAAATCTTCTGCTAAATTAATGACGCAAACTCATTTTTGTTTCTATCTACAATTTTTTATTCATATTTGGAATCAGCGGAAAAAGCATTTATTTGAAAAAAGAATTATTGTATCAGAATACACTAGAGATTAGCATATATAATATTTTAAAGGAGAAAATAATCAATTATGTTATAATTTATTTTATGTTTGAATGATAC
>evgR001061c0g1_i01_Hydra
CCATAGATTTAATGCATCTAATTTATTATTGTTGCACAATCCTCAAGGTAGAAGAATTACCACAGATAACATAGGAAAGTTATTTTGTGAATCCTATATGAATACTGCAACACCTTCTATTGCTGCTGCAGGCTTTAAAAATGCATGTATTTGGCCTCTAAATGCTAATACGATTCATGAGAGTGATCTTATTGCGTGCAAAAATGAACTATTTACGAGATCAAAAGTAAATTCAGAAAATCATAATAATGTAGATATCGATAATAGCTATGTTCCTGGATGCAATATTACAGTAAAAGTTACCATAGTTCTTAATGTCTCATTTAATGATATTGTCTCCATACCCAAGTTAAAACCAACAAAAAGAAACAGCAAAGAGTCTAGAAACATCACAAGTTCTCCTAACATGATAAGAACTGAAGAATCTAATAAAAAAGCTGGTCAAAAAAATAACTTAAAAATAAGTAAAAAGATCTGGCCCTCTAGTCCCTTGCACACAAAAAAAGGAGCAGTTAAAAGACCACCCACTCACGATGAGGATATCTTACCA
>evgR001064c0g1_i01_Hydra
TTCAAAATTAATCTTAACAAAATTAGTCTTAAAAAATAAACAATGATGACTTAAAAAAGTCAGTTTGTTAAGTAATAGTCTAGCTTAAAATATGACTGAATCTTCGCCTGTGGAAAGTGCCAGAGCAAGAGGTAAAACTGTTGTGAAACAAATTGTGTTTGGAAATGAATCAAAATATTTTGGGAAGAAAAGAGAGAATGACGGCCATACTCACTCATGGACTGTGTACGTGAAGCCTTTTATTGAAACTGAAGATTTATCTTCATTTATTAAAAAGGTTCAATTCAAGTTACATGAAAGTTACACAGATGCCCTTCGCACTGTTTCTAAACCTCCATATCAAGTCACAGAGACAGGTTGGGGAGAGTTTGAAGTAATCATAAAGATATATTTTATTGATGCAGCTGAAAGACCTGTAACACTTTATCATCTTTTAAAGTTATTTTCAAATCCACTCCTAGCCAATCAACCCCATGTTGGTACACAAGGTCAGCTTGTTTCAGAATATTATGATGAGATTATTTTTCATGATCCTAGCATATTTCTTTATAAACAATTAACTACTCCAAAAGTGTCAACAGCTGTTCGTAAGCATGAAACTGACTGGAAGACTAAAGGCAATAAAACATTAGCATCAATAAAATCAGCCAAGAAAAGGGTTGCAGAGGAAATTGCGCAGATGAATGACAGATTGAAGAAAAGAAAAGTTGCTATTCAGAAATTAAAGGAAGAAATACAAAAGATCGAAAAGCAAAACCAAAAGCAATAAACTATTGAAATATGTATTAGAAATGTCTTTGTAAAATCTATATTTATTTTTGAATATTTTTTAATTGAAACAAAAATGTTTTTTTTAAATATTTAGTGCTTTATATGTGTTCTTAATTAGATAGTACCGATTGGATGTATTAGTTTTTAAGAGTAATACACATGTAATTAGGCTTATGTAGTGGAGTTGTAAGACTTTTTCACTTCTTGGCTCTGGCTTTGATGAAATATACAGCCTCAGCTGACGGCTTCCAAATTTTATGTGCAAAAATTACAATATAGATTTTGTTAAAAACATTAATTAAGCAGTAAATTTAGCTTACTGTGAGAAAAATTGTGATTGCTCGGAGCCACTGGCATTCTGCGGCTCCAACAATAAAAATACAGCAACTTTATGGCTCCACAAGCCTGCATGTAAATAAATTTTGTTTTTGTCAGTAATATTATTTTTTTATTTAACTGGTAAAATATGAGTATATGAATATAAAAAAAAAA
>evgR001070c0g1_i01_Hydra
AGTTTACCAAAATAAGTAAGTCAAAATGGCTGAATACAAATTTTTTTTTGTTAAAGCGAAAAAGATTTTTAGAAGTGGTTTGTGACGAACAAACAACGAGTGACGTAGTTGACAAAAGAATATCATTTTTTTATTAAAATTGTGCTTGCATGACCAGAGAAGGAATAGCGACAAATATGTTAATTAGTTTAATCTGAAAAAAGATACTATGTCCCTGGTTGCCGAACAAGACAAAATCTATGATAAATATAAAAAGCAAGAAAGCAAATATCTCATTAAAAATGAAAATGTCGATCACAATAACAAAGAAGAAATCAATGACTTCAAAATGTTAGATGTCGATGAAATACTCCTTCTGATCGGAGAGTTTGGGAAAAGTCAGCTATTGTTATTGGTTATGCTGAGTTTGCTTATGATCCCAGTTGCTTACCAGTCCCTTTCCATCACATTTATTGGAATGAATCCGTCTTGGCGTTGTACAAATACTAGCGAGGAATGTAACAGGCTGGGCGAGTTTTCAATAAATGACGAGTTTT
>evgR001073c0g1_i01_Hydra
ATTTGCGTATGGCTCTGAAAACTGTATCCATGGTTTTTTCAATTAATTGAAATGCTGAAAATTCAAAAGCTTATTTTATTTTTTTAATTTGATGATCAATTCCTAAAAACAGTTCCCACTTAAAAAAATGCATCTTGTGTTCTTTTTATTTTTTATATTTATGTATTTTGGATGTGCAAAGACGGGTAAATCGTTTTTTGAATCTCCTGCAGCACTTCCACGAACTAAAGCATCTACTGTTGTATTTCTACCAAAGATAAAAGTGTCAGATTCAGGAACAATTGACACTTTGTTAGATTCGCCTAGTTCAACCCATTGCGTAGAGAGCTGCTTGAAACCTTGTGAAAACCCAAAACTTACATGTAGATCCGTCAGTTTTAATTTGCCCAGCACAATGTGCATATTGTCAGCTAATGAGGTTTGGAATGTCACAACGCAAAGAAGAAATTTTTTGCCGAACGATGATTTTGACTCGTATGCTCTTTAAAAAAAATATTTTGTGAGTCCTTAAACTTTTTAGAGTCTTAAATTAAACAAAGTTGTGCGTGTAAACGCGTGTAACTGCAACTCCAATTTTTTGAATTAGTTAACTCTTGCGATTTGAATTATTTTTGTTTATTGAGGAAATCTAGTAACTTGTAACATTTTCTTTTTTTTCTTTCTAATTTTTTTATTTGCCGTACATATATATATTTAAAAGTTTCGTAAATTAAAATAGTACGGGCCGGACTACAAGAAGTCCGTACTAATCTTATCACAAGCCTTTACATTTTTTACATTTTGATCTTATCACAAGCCCTTACAATTTCTATTTCTTGATGAAAGTTTATGGGAACATTTTCACGTGTTGATGAAATCTTATGGGAACATTTTTATTTATAAAGATTTATGGTAATACTTATATTACTATTAATATATATTACGAGTATTATTAAAAGCTGCGTTATTATAAAAAAAAA
>evgR001078c0g2_i01_Hydra
CTTTAATTAACCTACTTTAGTGCAAAAAGTCAAAAATGATTTTTGTTTAACAATGGCATATAGTCAAAGAGTTAAAAAATCAAGCACAGTTCAAACATATTCAAAAGAAGATATTCTTGATGTCTTAGACAAATTTGGTGGAGACAAAAATTTAATATCTACTGATGATACTGGGAATAATGAGAAATTGTTTGATACAAAATTTGTTGATGACCATGAAGAAAGTTTACAAGAAGAATGGGATCAACTATTCCAAGATGGATTTTTAGGAAAGTTAAAAGCTGCCGCTCTAAATGGTAACCTTGGTACACAACACTTCCGCAGCATTTGTTGGAGGGTATTTTGGAGATGTCTATCTGAAAATTCAGTGCATTGGGTAAATGAAGTCAAAGAGCAAAGGAATATGTATACCTCCATAAGAAATAAATATGTGATTAACCCTTATGAACAGGAGAAAGAGTTAGATTTAGAAAAACATCATCCACTTTCGCAATCAGAAGATAGTGTTTGGCATAAGTATTTTCAAGATAACGAGCTGCGATCATTAATTATAATTGATTTAGATCGACTATCGCCTGAGCTAGAATTTTTTCAAAATGATAATATAAGGTCTATGATTCTTCATATTTTGTTTTGCTATGCTAGAGAAAAACCCAAAGTTTCATACAAACAGGGAATGCATGAAATAGTTTCGGTCATCATATATGTAATTATGAAGAACACTACGGCTGTTAAAGATTCAAATGAAGTTTTAAGCAATGAGTTGAAGACTATTATGGATGTTCAATTTGTTGAACACGATTCATACATGTTGTTTGCCAAGATAATGGAACCCATTCAATCTTGGTTCACAAATGTTCCGCAACCGCAGAGTCTAAAACAAGTATCTGCTGATCAGCGAATATTCAAAAATCCTGAAAATGAAATAGTAACTGCTATTTCAGAGAAACTTAATCGGATACAAGGTTCTATGTTAAAGATATACGACAATCCCCTATGCCTACATCTTCAAAAAATTGATATAAGTCCACAGCTGTATGGAATTCGATGGATGAGGCTTCTTTTTGGACGAGAATTTCCTTTGAATAAATTATTAATCCTTTGGGATGCACTATTTGCTTATGGACTTGATTTAGTTGATTATGTTTTCCTAGCCATGCTTATAAATCTTCGAGCAGTTTTACTCCAGAATGACTTCAACAAATGTTTGTCTTATTTGATGAAGTACCCAGCAAACACTGATTCAAAATATATACTCAAGCAAGCTCTTTTACTTAAGAATTATTCTCAAAGAAAGGTGAGCAATGATCAAACAACAACCAAACATCCACAACAACATAAAGTCCATCCTGTACGAAACAGATTAGGGAAAACAAAGGAACAGTTGTTTTCACAACCTATAAATTCAGGGGGTCAAATTGAAAAAACAGAAGAATTAGTAGATAAAAAGTATTTTGGTGAAAATCTTGAACCCTCGCCGTCACCTTTATTTAATGAAAATCGAGCTGATGCAGGTGTGCGTGTCGATGCCAGTGTGCGTGTTGATGCTAGTGTGCGCTTTGCATCACAGTTATCACAATTGAAAACCAAAGTTAATTCTATTCAGAATCCTTTAAAAGGTATGGGTACAAGTAAGCAGACAAAAGAAATTGTTGAGGATAATGAAAACTTGCAAGAACAGTTATATGACATGCAAACTGAGTTTTACAAGCTACAGTCACTTTGTTGTTATTGCGGAGATAAAATGGACTCATATATAAATCTTTTGCAACATCGTCTACTTAAAGAAGAAATGTTTAACGATGAAATTGCAATTTTAAGCATTGCTGGTTTAAAACAGATTCGAGAAATTTTCAAAGGAGGAGTAGATATAAATTTTCTTGATCATGAGGTAGATTTTAAAGACATTGACAGTCATATTGCTTCAGCTTCCGATAATAATAAAAACTTTGCTGCAGCTATTGACCCTTTGAATTCATTAAATGGAACAAACTTATTATTAATAGAGAAAAATTCTAGGGAAGATAAACATCATGGACACTCCATTGAGAAAACAGCTTACGTTTCTTGCTCTATAATTCCAGGTAATCTGAATGTCTAAGTAAATCACATTTAAAGAATCATTCGGTATCCTACTCAAGTTGCGGATATTATTTCGATAATTTTGGTTTGATATGTTTTTATGACTCGAAACTATTTAATGTTTATATACAAGCGTTTCTATGTTACCACAAAATTGTTGCCATGTTACCACAAAATTTAAACAAAGAACACGTGGAATAAGATCTATAGAAAATTTCTTCGATTGATTGTCAATGGTTATCAGTGTCGATGAAAGTTTTATCTCAAAATATATTTATTTGCTCTAAAACGATTGAGGAGGTATAATGCCCATTTTATCGATTGTTTGAATACCGATGTTTATCTTTGTCGATATATGTTAACGGATACGGTTGAAGTGGTAAAGATAAAGGAATGTTTAAGATTTATGCAACATTTTTGTAAATTTTTAGTTAAAACTTAAAATACTGATAAAAAGAGATTTTTATTTTAAAAATCTATAATTACGGTAATGCTTTGCGAAACATTAATCTGTTCATGTTATAGTTACAGTAATGCTTTGCGAAATATTTTTATGTTCATGTTATAATTACAGTAATGCTTTGCGAAAC
>evgR001080c0g1_i01_Hydra
AGATATTACCACTTGGTCGCCACAAGGCAGATTGCACCAAGTTGAATATGCTATGGAAGCCGTTCAACAAGGAAGTGCTGCTGTTGGATTAAAATCAAAGGATTTTGTAGTATTGGCAACATTGAAGAGGGCTCCTCATGCCGAACTTTCTAGCTATCAAAAGAAGCTTTTCAAGATTGATGATCATATTGGTATTGCAATTGCTGGATTGGCTGCTGATGGACGTGTATTGAGTAAATATATGAGAAATGAATGTATGAGCCATCAATATGTATATGAAACACCAGTTCCTGTACAAAGATTAGCATCAAAGATTGCTGATAAATCCCAAAAGAGCACACAAAGCTCATCTGGTCGTCCATTCGGTGTCGGATTATTAATCGGTGGTTATGATGAAACTGGAGCACACTTGTATCAAACTTGTCCAAGTGGTAATTTGTATAGTTATAAGGCAATTGCCATTGGTGCAAGATCACAAAGTGCCAAGACTTACTTTGAAAAACATTACGATACTTTCACAAATGATGTACAGGATCTTATTGTTCATGCTTTGAAGGCCCTTAAAGGAACAACCGGAGACAATGTAGAATTGACAACAAAGAACTGTTCTGTTGCTGTTGTTGGAAAGGATTTCGCATTCACTTTATATGAGGATGATGCTATTGAAAATTATTTGAATTTATTGGACACATCTTCAAAGAAGATTGATGAAATGGTCG
>evgR001083c0g1_i01_Hydra
GGTTTAGATATGGACCCAACCGTTAAGAAACAACCATCTGAAGATATCAAGATTGGTATTCAATCAGCTGGTGGTGCTGCTGCAGGTGGAGATGAAGAAGATGAAGAAGGTGAAGACGAAGAGGGTGATGAAATGATGAGCTTTGAAGACGAAGATGCTCCAAAGACAACTGAAGAACTTCTCAAACTTTTGGAACAAGAAGATTACAAGGAAAAGGTTTGGTATGTGACCAACAATGAACATTACGCAAAGAAATTGCAAGAAATGAATGCCACTGTTGCAATTGAAGTTGCTTGTACAGGTATTAGTGGTGGTATCCTTGCTGAAGCCCCATTCAGAGGTGTTCCAGTTTTGAGCTTGTTCGCTCCACTCAATCGTTTCCTCAAGATTGGTACTAGAGCCAGTATTGCTCTAATTCATGCTCTTAATAAGCTCATCAGTGAAGACCAAACCAG
>evgR001089c0g1_i01_Hydra
GTTTGTGATGACGTTAGACGATTTGGCACTCTAGATCAATACAGTGCTTTTCCTTTTGAAAACGCATTAGGCATGTTTAAAAAAATGATTAGAACTGGCAACCTCCCTTTAAGACAAATATGCAGGCGCTTATCTGAATTTGATGAAAAGCCATTTGCGTTTTGCAGGCCAATTTCTCGACAGCCAAAACTGTCTAAACAACATTCTAATGGACCAACGTTAGGTTATCAAGGCACACAGTATATGCAACTTTGTTATGCAGGTTTTACTTTTTGTACATTTAACTGCAACAATTGTGCACTACTCACAGAAGGAAAAGTTATTGTTATTGAAAACATAATTGACACTAACGAGAATTATGTTATTGTATGTCGTCAATTTTGCAAACAAAGAAGTTTTTATTCCTATCCTTGCAAATCTTCATTGCTTCAAGTGTATCGTG
>evgR001103c0g1_i01_Hydra
TGACAATCATAAAAAAAGAAAATTTTACAAAATTTTTAGGAGTCATAATAGATAAAAACTTATCTTGGAGTCAACAAATTGATAATATTTCAATGAAAGTTGCTAAAAACATTGAAGTTCTATATAAAGCTCGATGTATACTAAATAAACACCAACTAACTCAACTATATTACTCATTTATTCATTGCCATTTAAACTATGCTAATATAATATGGGGAAGCACTCAAAAACCAAACTTTAATCTCTTTATCGGCATCATCTTTATCGGCATCAAAAACACGCAGCTTGCACAATTCATTTTGCAAATCGTTTTTCAAGCTTAACACCACTTCTAATAGAAATGAAAGCTTTAAATATATACGAACTAAATATTTTTAATGTTTTATGT
>evgR001106c0g1_i01_Hydra
GGAGACAGTGAATGTTTTTCTTGTCCTCATGGAATGTTTAGTGTTGCTGGATCTGATTCTTGTCAAAAATGTACATCTAGACAACTTTTCACTGCTCCATCAGAATTCATAAAACCATCAAGTGAATGCCCTTTTGGGTATTATTCAAGTGAAAATGGTTCTTGTTTGTGGTGCCCACATGGTTCAAGAATAGTATTAAATGGTTCAGGATGTATTCCTTGTGATCCTGGATACTATTGCCCAACAAGTTACCTGATGTTGGCATGTTCAGCAGGAATGTATTCCCTTGGCAATGCTCATCTAAATTGCACAATTTGTCCTCCAGGCTTTGCATGTCAGGTTTCTAATGCTGTTCCTGTACCTTGTTCTGGAGCTACCTATTCATTGGGATTTGCTGTATCTTGCTATCCATGCTTAGCTGGTTACCAATGTCCTAGAAATTCAGCTATTGAACCTTGC
>evgR001107c0g1_i01_Hydra
GAACGTATTCAAGGTAAACAATTCTTGCAAAAGGTTTTATTCGATAAGATTTTGGAACCAATCATTGAAGAAAAGGATTTGAACTTGAATTTAAATGCTGTCACAATCTTGAAGGAAAGAATTTCCCAACAAGAAGTTGAAACTGGTGTCAAGAGTGATATCAACATTAAGGATATGACCTATGAAAAGGCTATGCAAGATGAATATGTTAGCAAGCTTGTTCAACAACGTACCAAGAAGATGATTGACATTTGTCAAGGTGTTTTGGATACTTTGAATGATAATATTAATGAATTGCCATATGGTTTGAGATACATTTG
>evgR001113c0g2_i01_Hydra
TTATTATTTCATGGAGTCTCACAATTTATCCAGGTTGAATTTGCCCCAAAGTGAAACATATGTTCAAGAACGTTCCTCAGATTCTTTTCTATATATAACAGATCTGCAGAATGTAAAACCAGCTTCTTTAACAGATTTGAAGAATGTACAACAATCATCTCTGACATTTAAAGATTTGATAAATATACAAACAACAATGCCATCTTTAACAGAACTTATGAATGTAAAACAAATAGCTCTGCCATCTTTAACAGACTTTATGCATGAGCAACCAACATCTCTTCCATTTTTAACAGACCTTATGAATGTTCAATCAACAACTCTACCATCTTTAAAAGACCTTATGAGTTTGCAATCAACAACTCAACCATCTTTAACAGACCTTGTGAATGTGCAATCAACAACTCAACAAACGTTAACAAACCTTATGAATGTGCAATCAGTAAATCAACCATCTCTAACAAACCTTATGAATGTGCAATCAACAACTCAACCATCTTTAAAAGACCTTATGAATGTGCAATCAGCAACTCTGCCACCTTTATCAGATTTATTGAATGTGCAGTCAAGAAATCTGCCATCTTTATCAAATTTTATAAATATCAATCCTACATCTGTAACATCTTCAGATTTTATTAGTGTACAGTCAATGTCTCAATTAACTTTAGTTAATGCAGTAAATAACCATGAGAAGTCATCAATTAAAAAAATAGATGAAAAGTTATCTGATGTTCCTTTACTTAAAAATGAACCAATGTCTGAAAATTATGATAGTTATGCAAAAGTAAATATGAACATACCATCGTTACCATCTTTAGCATTGTTACCATCGTTAGCAAATTTACTTGCCTCACAAATGGCTATTAAGAATTCAGAGCTTTATTCATTAAATAACGTTAACAATATTGCACAATCATCAGACAGTAATTTAACAAGTATTTCACAATTATCAACTGAATTCAATACTCTGTTTTTAGAAAAGAAAAACTTTTTTAATGTTTTATCTGATAACCGAGTGCCATTTCAAAATATTTCAAAAACAAGTAGTTCATTTCAGAAAAGGCCGTCACAATTCGGAAGAATTGTGGGTGCTCAACTTAAAAAAAAACAAAAAAAAATTCCTTTGTGTGTTCATCCTTATTCGCTACCTGTCGAAGTGTTTTTTTATGAAAATGTTTGCCCACAAAACAATCAGATCTTTAGTTCTCCTTCACCAGATGACATTGTTTTAAGTTCACAAATGAAAGCATTTGAAAGAGCTGGCTTCACTGACCTGACTTGAATGTTTTTATATATTCTCTAAATTTTAATGTTTTAATCC
>evgR001142c0g1_i01_Hydra
CTGATAACCATTTATGGTTTTCAGTTTTTGCTCGCCCACCTAAAAGTACCTTTACCAGGTGTCAAAGATTGTCTGTTGCTGTTTCATTAATTATGACTTCCATGATGGTAAGCGCGATGTACTATTTGCTAAAAGTTAAGGAAAGCCCTGCTGAAGAAAATAGAGCTATTTTTGCAGTTACAACTGCACAAGTTTTTGTGGCTTTGGTTTGCACAGCTATAAAGTTTCCTATCCATTTTATATTAATGAAACTTTTTCATTTAATAAAACCATTAAATCAAAGTCTTAATAAAGTTTCGCCTGAGAGCAGCAATACCCAAAGCAAAACACATACTGTTATTGTAGAAGAAAAACCAAACTCAATAAGAAAAAGAAGAGTATTACAAAGTTGGTGTTTGTACTTAGCATGGTTTATTTGTGTTAGCAATATTTTGGTTTGCGGTTTTGTTGTCCTATGGTATGGAATGTTCTTCGGAAATAAAAAGTCGTTAAATTGGTTAGCAAGCATAGTAATAGATTTTACAAAAGACATTTTTTTAACTGGACCAATAAAGATATTTATTCTGGCAATTTTTTTTGCTCTCTTTGTCAAGAAGATTAAGGAAGAAGAAAGTGAAGTTGAAAAGAAAGATATGCTGTTACTTAAAGATGAATCTTGGCTATACAGACTAAAAGATGTAGCAACTATTTTTGAAAATGTTAGTTTGCAACCCCTTGACTCTGTTAAAGTAAATGATATGCGTGAAAAAAAGATTAAACAAATAAAAATGGGTGCTTTAACAAATGAGATAATAGTATTTACAATCTATGCAGTTTTAGTGATGTTCATTGGTTATTCAACACGTAGTAATGTTGCTTTTAAACAAACTCAAAATGTTCAAGACCTTTTCAACTTGACACTGAGAGGAGTTCCATGGCCAAGAGATTACAGTAAAATTTATGGACAGATTGTTTCAGCACAAGATTTTTGGTTTTGGATAAAGGAATTTTTTCTCTTGCAAGTTTATCCATTTCAATGGGATAGTTTTAGTGATTTCTCTGCTAATACAAAAAAACAAGATTTACCTGGAAAATTGTTTTTGAATGATCTTAACTCTAAGATTGTGAATGGAATAAGAATTCGACAAATTCGTGTTCAACCAAATTCATGTCTTAAAGCTTACGCAGTTGCTAAG
>evgR001145c0g1_i01_Hydra
CTTTCTTTCATTCTCAATATAACTAAGGCATGGTACAGAGTAACTCTTGGTACAAAATATTTTATCGCAGTTATAATGTACCAAATGGGAAAGTATAATGAGGCTTTAGATATTTTTGATACTGTTATTAAAATACAAACTAAAGTTTTAGGTTGTAACCATCCGTCAACAATGGATACAAAGATTCAAATCGCATACTGCTTGCAGCAAATTGGAAGGTATGACGAAGCTTTAGAAATTAATAATGCTGTCGATAAAATACAAACTGAATTATTAGGTATTAACCATCCGTTAAACATTAGTGCAAAAACTAATAAAGCAAATTGTTTGCTTTTAATGGAAAAATTTAACGATGCTTTGGAAATTAGTTATTCTGTTCATCAAATATTAGCTGAAACTTTTGGTATTAACCATCATTTAACACTGCAACACAAAATCAGTATCATAACCTGTTTGATTGTAATGCAAAAATATAGCGAAGCCTTTACAATTTGTTGTTCTGTTGAAAAAATACAAATTGAAATTTTAGGTATCAACCATCC
>evgR001180c0g1_i01_Hydra
GCTTACTTAAAAGTTAGTAATTAAAAAAAAAGATGAAAGTCTTAGCTTTTTTTTTCTTATTTTCTTTACAACGTACATTGTGTAACGTTAGAAAAACTAGAGAGGCATGCGTTAACAGAGTTGCTCATGAAATAGATATTGTCCAAATTATAGCGGAAACAAGTAAAGTTCTTATCTGCAACAATGGGCTCGAAGAATGGCTATTGAAAACTGATTATACCATAGACTCTCCTTGTGCAAAAGATGTTGTTCCATCTGCCGAATATCGTTCAATATATGCGGTTTGCGTTCAAAATACTCCTACTATTTGCTATAAAGGGGACGGCTTAACACTAATCAACTGGCTTCACCATCCACATAATCCAACAACTGTTCGAGCTTTGTTTCCTCATTTGTATAATAAATTGAACAAGGCAGAAAAAACCATTGATGATAAAATATAAAATAAAGCAAACTATTATAACAAAGAGTTATAAGTGTGTATAGTGGAGCATAATAAACCCCTCGTTTTGAATTAGAAAGAACAAACACGATGTAAGAAGTTTTGCGTTTCAGAAACAATACTTACTTTGTAAAGGTTTATGCTTAATTTTTCTAAAGTTAACAATGATTAAAGTATTCCAATTGAAAAATAAATATTGTGTAAAGTATAATACAGAAGTGTTTAAAAGAAATTCCATTGTCATTAGTCGAAATAAAT
>evgR001183c0g1_i01_Hydra
TGGCGTGCAAAGTGTCTATATTACTGATACCACCTTTGCACTGCGATAATGCAAGTTTTGAAGCAATTAGATGTTTTACTTTGTTAGTGAAGCATGCACCTCAATTAACAAGGGAGATAGTTGAGCTAATATCAACTACATGTATAAAAGAGATACACAAACATATTGAAAATTGTTATGCTAATGAAGCACTTATAGAAATAGTTAAAACGGTTCCTCAGTTTACAGAATATGTGGTAAAACAAATAATTCTACTTTGCTCTCATTCTGATGCTTTCGTCAGAGTTAATGCTTCCAATACACTTTGCAGTATAACTGTTGCAGTTCCTCAGTTTGCAAGTGATGTTTTAAAACTAATAATCACACTTTTGTCTGATTCTGACGTTTCCGTCAGAGCTAATGCTGCCCATATACTTGGAAATATAATTATAGAAGCTCCTCAGTTTGCCGATGATGTGTTAAAACTTTTAACTCAAATAGAGTCCTAAAGCTAACTAAATAGAGTCCTAAAGCTAACTATATAGAGTCCTAAAGGAAGCCCACAAGTTACGGTAGATGAAAATGTTTTTTTATTAGTATATTATGATTAACGGTATCGAAAGCTTTTGACAGATCAATGAAAACTCTTAAGGTAAAGTAGTTACTACTAAATGCAC
>evgR001191c0g1_i01_Hydra
TTGATATATACTTCTATCAAAATTTACTATGTTTTTTGTTTATTGGTTGTTTCAGTGAAAATCAATTGAAACTAAATGTTTGGTTAAACATAGAAGTGAGTCGAATATTTAAAAACATGGTTTTTGTTTACAGAATTAAAATAAACGGAAATAGTTTTCAGTCAATAATTACACAGTCCTTTAGTTTTACGGAAGTTAAAGTTTATGCGTCAGATTCATTTATACTTGATGAGGACACAGATGCTTCAATAAAGAATTTGTTAATTATCAATGGCATTTCAAATAGAATCATGCCATCCACTGTTGTTTTTTCTGTTGATTTTGCAGATTATGTCTACTATAAAAAAGCGTTTCCTATTAGAAAAGGAAAACTTGCTAATCTTAAAACATTGATGAAAAGTTATGCTGTTTCTTTGTGCATAAGACCAACCAGTTATGCTAACGGTATATATAATGTTTTTAACCTAACTTTGAATGATGATATTGAAGATTATGGTGACAGTAATCTAGGTATTTGGTTTAAAAATGGGAAACTTTTTATTTTTGTTGCAGTAAATAGAAATGCTTCCTACAACATTGAAACACCTCCTTTAATAATTGATGAATTTTCTACTTTTCATATTTATCAAAGTTTTTTAAATGGTAGTTTTTTGCTATCAATTCGCCTGAACAAAAAAGTTATTCACAGCGTAATAAACCATGATGCTCGAGACTTTAATCACATTAAGGTGTATGCTTCTGATCCTCGGAACAATGCTCAAGATGGTTATATTGATCATCTCTTGGTAATCAATGGTAAGCTATCAGAATATATTGTTAGTACCACTACTGCAATAGAAAGAGCAAAGAAGATTGCTGTAATACCAATGCTTGAAACGGAATATTTTCTTTCTTTTGATATTATGCTTACAAAATTAGAATCCACATATGATAATAAAAGTGTTATTTCTTTCTCAAATGGTTCAAGTAGTATTTTGCATGTTGTTTTAATGGGGGAGAAAATTAAATTTAAATTTTTTAACAATGAAGTTTATGTAAACATGGAACTCAATATTAAATGGCATAATGTTGTAATTACTCAATTTTTAAAAAATACATTTTACATAAATACAATTAAAATTGATGATAAAATTGTCTCTTGTGAAATAAACAATTATGTCAATAGGTTTGAAAATACCATTGTATATATGTCTCATCCTGATTTCAAACATGTAGATGGTTCTATAGATAATCTGTTTATTGTCAATGGTATTACAAATAGTAGTTTGCAAAAAATTGTTTATCCTTCTAGCAATGTAGCAAATAAAAGCACAGAAAGTTTACAAAAAATTGTTCATACTTCTAGCAATGTAGAAAATAAAAGCACAGATAGTAGTTTGCAAAATAATGTTTATCCTTCTAGCAATGTAGAAAATAAAAGCACAGACAAATCATCTATTGTCATTGGTGTAACAGTACCTCTATTTGTTGTTCTTGTTATGATAACTGCTGTCG
>evgR001194c0g1_i01_Hydra
ATTTCCACTAATAGCGAAGATGATGATTAATCTATGGATCATTTTGATAATATTTTCAAGTAGAAGTTTGGAAGAAGCCGTATTAAATCTAGCAGTTCTCATACCATTGGATGAAAGTTGCACTAAGAATGCTATTGATATGGCAATTGAAGATATAGAAAAAAAACCTGACATGATTTTCAGAAATCTTACGCGTTATAAGCTTACTGCATATTTCATGAAAACAGATAAAAGTTTAGGGAACGCAATCGAAGCGCAAAATCGTTTGTATAAAACAAATCCAACAAAAATTGTTACTATTGGTCCGCCGTACACTGAGCAAAACGATTATTGTACTGAATATGCGACTGCACAAGGAAACTTACAAATATCATACACAGAAGCAAATGAACCTTTTGAGCCATATTTTGTCATGTATTTCCAAACGCCACCTTCAATAGTTTCAAGTTTTATAGCAGTTTGCTATTTTCTTAACAAATATGAATGGTCTAGAGTTGCAATTGTATTCGACTATTCCCAAGTGAGATATCGTAACAACCCAGAAAGTTTGAAACATATTTTAGAAAACTGCTTAAATAAAGTAAACTTATTAGTTGTTCAAGGCGTACGGAGTTATCCAGTAAGTTATGGAGTTTCTAATGTAATAGGAGAACTGCAAAATAAAGGTGTTCGGATAATATTGGCTCTCATTTCAGTTAGTGCTGCTCGTAAAGTCTTTTGCGAAGCCTACAAAAGAAAAATGACAAATCCTAAAGTACTATGGATACTGTTTGAGATTCTTCCAAATGGATGGGCATCAGAAATCTATAACGCTGAACCTATACCAGGTGGCTTTAGGAGAGAAATAAATTGCTCAGAAGAACAACTTCTTGAGGCAGCACAAGGATATATTTCGTTTACAAAACAAAGTCTTAGAAGAGACAATAAGACAACTTTGAGTAATGTTACTTCGGAATCATTTCTCAATAGACTAAATAACAACATAGGTAAGCAATGTGATGAAAATTTAGCATATGCTTATGACGCAATCTGGGTTGCCGCAATGGCTTTTAATTTAACAGACAGGACTGCTAACCTTTCTGAATATTCATATAACTCGTTCGATGTTGTATTCAAACTAGCCAACAGCTTTCAAAACGTTACATTTGAAGGAATTACTGGGCCGATTTCATATCAAACACAATACTCCGGAGGTTTTCCCTCGCGAATCGGTCAAATTTCTATACATACATTTGTTAAAGAATACGGAATGAAACTAATAGGAACACATGACACGGCAAGTGAAGTTTTGCTATTTGAACCTGACGCGGAATTAATTTTGTTTCCTAATAACGTTATACCGAATGACCAATCAACGTATAATTTTTACTTTTCTTCTTTCAGTAATGCTCTACTCATCAC
>evgR001199c0g1_i01_Hydra
TGATACACCTGTTACAACAACTGGAAACCAATTCAACAATAATGGTTTTGGAAATAACAACGGTTTTGGAAATAATGCTGGATTCAATACTGGTTTCTCTCAACCTCAACCAGTCTCTAATAACACTGGTTTCAACCCAAGAGGAAATACTCAACCAACTAACTCTGGATTCAATCCAAGAGGTAACACATCATCTGGATTCAGCACTGGTGGTAATGAATTCGAAGCTAATTTTGATAACTCTGGTTTTGATGATGCTTTCGGATCAAATGATGCCTTCAACACTTCCTCAACACAAGAAAAGAAGGATTCAACATTGTTCGACATTTCAAGTGGTTTGGTTAACTTGGATCTCAATGAAACTAAGCAACAATCAACCCAACAAACCAACACAACACAACCTTCTTCTAATATGTCATTGAAGGAATTGAAGAGATTGCAAGGTGGCAATACTCCAACAAAGCAACCTGTCATGAATAATACTCCATCCTTCACAAATCAACCAAACTTCCCACCACAAAATAATAACTTTGCCGGGTATAATAATTATCCACCACAAAACAATTTCCAAGGAGGTTATCCACCACAAAACAATATGCAACCAAGAGGAGGCTATCCACCAAATAACAATAATAATATGCCAAACTCTTTCGGTGGAGGTTATCCATCTCAACCTAGTGGATGGGGAAATCAACCACCAGTCAATAATAACCAAAGACAACCACCACAAAACAATAACAATGGAG
>evgR001201c0g1_i01_Hydra
CTTGCTCTAAGATTCTTCTTGTCTCTTCCCCGACAACTGCTGCTCTTCGTTTTTGCACAAATAGCTCTTAATTGTTCCGAATTATCCTTATTATTATCTTCTGTTAACACGTCAATATTTTTTGCTAAACTACTAACTTCCAAAGGAATTAATTTTTGTAGAGGCCTAACAGTTTTTGTTTGTCTATTCTCCTTAGAAATAACCGCTAACTCTGCTAACTCTGCTCCTCGTACAACACCATCTCGACCAGTAATTAACTTTTTAGTTTTTCCGAATCCTCCATTGTCCACGTGGAGTTTTGTTTTCATCTTTAATTAATACAAG
>evgR001210c0g1_i01_Hydra
CAGAAAGGTTTGTACAAGGCAAGTAGCACACTGAAAATGTTTTATCTTAAGACTTATGAAACAATTGATGAAATCCGACCTCCACTTAACAAAAGCACTGAAATTAACTTTTTGGAAAAATTTGTTGAATTGTCTGTTGTCGATGCTGATAAGGTTCAAACAGCTGTAATTAACATTACAGAGCGAGGTCAATTACTAAAGCAACAAATGAGTACATCAATTCTATTTAAAAAGCGTTTGAAAGATCGTTCTTTATTTAAAAGATCTCTTCTAATATCTGGTATTGCCGGTATTGGCAAAACATGGTTACTTAGAAAGTGTTTACTTGATTGGTCA
>evgR001223c0g1_i01_Hydra
CGGAAATTTTGCACGAGAAGACAGAACTGCTGCTCAGCAAGAAGCGTTCAACGATACTCGAGCTAGACTTAAAGCGAAAAAAGAAGAACTGAAAGATGCTGGAATTCACGACAAACCGTTCCGTGTCATCCACAAAAGAACCGGTCAATTATGCTGTATTAGTATCACAAAGTCAAACGAGCAAACGAAATATGTGTTCGCCTCAGCAACGACAGCTCTCAACGAATATCGACAAAGCAATACAACAGTTTCAACAAGCACCACTCAAGAGAACTCGCCCACCAATAAAAAAACCACCCAGCATTCCTGTAATAAATAAAAATAATCTGTCTTCTGTAAATCTCCGCTTTTGAGCCAACAACCCATGCTCGTTAAATAAAAAGAAAAGCGACGATGCCATAGCCCGGTTACACAACTTAGACGTTAAAAATACTCCACATTTTATTCTTCACTGAAACTTGGTACAACAGAACGAGTAATATAGTTATACCAGGCTATCAAATTCATCGATGAGACAGACGGAAAAGGAGGTGGCGTGGCCATTTATATACACGAATAAATAACTACATTAGAAACCAATTTTGCACAACTGAACTCAGACTCCATTGAAAAAA
>evgR001234c0g2_i02_Hydra
AAACACTTTAAAAAGCGCGTTGTACTCACTTGCAGCGTTTAAGCAAGAGTTTAAAAAAGTATAATGTTTGTTGCGTTAGACACGTTAGCGAAGAATAGATCCCATTTACAATTCCTCTGAAGCAATGCATTTATATTTTAAAAGAGATTTTAGAGATGAAAATAAGTTTGCTTTTGGGGCGGCATATTTTATTGTTAGACATATTTTTTTATCTTTTACGCCTCAGCAGTACCACTGGTGATTTGGGAAGAGTTTACGATGAACTTGGTGGGCGTTCATATCATTTGAAAAGAAAAACAAAACAACACGTGCACGGCAACCATCATGTTAAAGGAAGTAATGTTTCAACAATTCCGCGTGAAACAAAAAGCCGAGACCAGATAAATGAGAATGACCACAGTTTTTCATTAAGAAGTACATCGTACGCTTACATAATGGAAATGGAATTAGGAACACCTGGACAGAAACTAGAATTTTTAATTGATACTGGTTCAGCTAACATGGCAATTGCAGGTCCCGAATGCCGAGATGAACACGGAGTACGTTGCAAAATTGAAAACTTTTACTTTCCGGACAAATCATCGACATCTCTACGAATCGGTCATCCTGTTGAAACAGAATACGGAAAAGGTTCTTGGTCAGGAGATGTTTATAGAGACGTTGTAAGTTTTCCTGGTGGACCAAGCATTGTTGCAGAATTTGCTGTAATTAAAAATGAAAAAGAATTTTTTATAAAAAACTCAGTAAATGAAGGAATATTAGGTTTAGCATATAAATCGTTGTCTTCGAGTCAAATAGAACCGCTATACGATCAACTAGTATTCGAAAAAAAAGCTCCCGACGTATTTTCATTATCTTTATGCAATGGAAACGGTCGTTTTTGGTTGGATTACCCTAAACCAAATGAGTACCATCACCCGATATATTGGACTAGAGTTATTCACAGAGCTTGGTATACAGTGCGCATGACCGGAATTGATGTAGCCGGAGAAAGTTTGGGGCTATCTCCATATAGTTATCCTCCTGCTATTGTCGATAGTGGAACTACTGATATATTATTAGAACCTGAAATTTATATAGCAGTCATTGCCATGCTTAAAAAAAAAGGACCTCTGGTTGATCAGAGATTTTGGAACAATTACTGTGTTATTACAGATCCTTTTAAATGGCCTTATATAACTATTTATTTAAAAGACATGCGTGGAGGTTCTTTTGCATTAACTGTACTTGGAAAGCATTATGTGCGCAAACAAGACCAATTTTATTGCTTGAGCATCGGAGCAAAGCAGTTTGGAGCTGTTTTAGGTGAAGTAACTTTAGAAGGAAATGTTGTTATTTTTGATCGGGCAAACCATCGTGTTGGGTTTGCTCCAAGCAATTTAACCGATATTGAAAGCGGTGTGGTTTGCGGAAACCCTATCAATATAAACACTACAGCTTCAGGTACATTGCAAGACTATTGCACAGCACATTTTATGCCTCGTCTTAGTTGTCACCGAACTTGTGATAAGTGCATTGCACAAAATGGCGCATGGTGTCCGAAAGGCAGTATATACTTGTGGATTAATGGAGTTAAATCTTCTTTAGATGGATCGCTAGGATTTTGCTGGTTAGGCAACTTATATAGTTTAGAATCAACACGTTACCAAACGTATTTAGACGGTCATGGAAGTGTTGAATTTGAGCTAGAATGTTTTTCAATGTTGCCAATGTACAAACAATGCGTTTTAAGAGGAGAGTGGCTGTTTGCTATTGTTATTGTAAGCATTTTTACATCGTCAATTTTATGTTTATGTCTTTGCTGCAGGAGAGTTTATGAAGTTGACGCAAGAGATCAATATCCTGTTTATAATAATGTCAATCAAAGAGAACAGTTTCCGGTTTACAATTCGTTGCGAGATCGTACTTTTCGTTAGCTCGTATGCAACCAACATCACATATGTCCTGCGACAAAGCATTTTTTTAAATGTTACTGTCGTAGTTGGAACTTGTAAACATGTGTTAATTCACCAAAAACGTTTTAATTATTATGATCACGACTAACCACTTAACCTCTTTATAAAATTTATAAAATGATTCAAAAGTAAAAAAATGAATGAATAAAAATGAATGAAATGTACACAAGCATAACACAAGTGTACTTTCTGTACATAAAAGTAACTGAAGAAACATCTTGTTCAGTAGTCTAGCCAGGGGCGAGGGAAATGGCCGAGGGCGCAAAGCTGGAACGCCTAAAAAGAAAAACCATTATATTATAAAAATTAATACTCTTTAAATTTTACTGTAAATAAGTTTAATATTTCTTATTTTTGTCCAGTGCGCAAAAATCCTTGCTACGCCACTGTTTCTGTTACTCTACCTGTTTTATAGTATTAATACGGCATAATACATCTGGAAGATCATGGCCGTTGCCAACAAAGGGGGCGGGGAGGGATGCTATGACTCCCCTCTTCCCCCGCACACATTTTTTTTTAAGGAGCG
>evgR001246c0g1_i01_Hydra
TCAAAAATGATGCTTCTTTTATAATACGGGAAAACTATTATTACCCTGGTTATATTTCGTTTGAGTCAACAAACTACCCTGGTTTTTTCCTCAGACATCAAGGCTATACTCTAAAATTACAAGTAGAAGAACCTCTTGTTGAACTTTACAGAAAAGATGCAAGTTTTGGGCTCGTTGATATAACAAAAACTACAAATGGTCTTACTCGATCACCAGCCCTCGCGCAACATATTGGAAAAATCTATTCATTTGAATCAATCAATTATCCACAATATCGAATTGGTGTTAAAGATGATTCAACTGCCGCCATTCTTTTATTTGGTGTTGACAAATTTACTGTTGTTAAAGCACTTAATGGACGAG
>evgR001253c0g2_i01_Hydra
AATCTTTTATTTTCTAAATAAGTTTAATTTACAATGTGGTTCTATTTTGTTCCTCATGTAGCCTATCATTTCACTAGATGCTTTCCTAAAGATGGTCGTTGTGGTGCCAGAATTTTTGTGATGCTTATTGCACTTCTTTTTTTGCTACCTCAATTTATAATTTTGACACGTCGAACAACTGAAAATTACTGCGAACAGCCGATTCTTAATATGTTGGTAGTATCCATCTGCATGTCATTTGCAATGTTAGCGTTTACTTTTCTTTTTACTATGATGGAGCCTGTTCCATGGAAATTAAAAATTGGTTTTCATTTTTTTGGATTCTTTTCTCTTATTATTGGATTGGTTCAGTTTGGAATAACTGTTGATTCAAATTGTTCTGAAGCAATACCTGAACTTTATTTGTTATCTTTGTCATTTGGTATTTTTTCTCTAGTGTCAGCAATATTTTTACTTCTATTGGTTCCATTTTGGTTTGTGAACTTTTTTTGGCCTGGATCCGTTTTAAGCTTGAAAGAGAGAAGAGGTGTCTGCTATGAGCCAGTGAAATCATTTTCTTGCGTTTGGCACGTTTAAAATAAACCTTTTTATTTGAAAACTAATTCTGAAAATAACTCAGTTATTGGATTTCTTGTATATTTTGTGTATATTTTACATAAATATTTTTTGTGTATGTTTTACATAAAATTGAATGAGATTAGGAAGAATAAAATATGTTTTAATATACAGTAAACTATTGGTGATAATGGAGTTTTAAACGAAATCGCTCTAGTTAGTTTTACACTTCATTAAAAAATAGAAATATTGAATCATTTAGTAATATTGAGTCATAA
>evgR001255c0g2_i01_Hydra
AAAAAAAAAAAAAAAAAAAAAAAAAAAAAAGCTATTAGAAGTACTAGTGCAAAAGACTTAGTTATTGAAAGAGTTGATGAGGAGATGTATATTTTTCCAGATAGTCAAGGAAACCAAAATAACTCTTTACGTACTCCCCATAGACTAAGCTCTCCAGTTCTCTCTGACACAGAGCAAATATCGGATAATCATTCACAATGGTCAATTAGCCTAAAAAGAAGTTTAAACTTTTTTCGAGATCGTACAAAATCTGGTGGACCTCTCCAAAGAAAAAAGAATATTAAAAAAGATAATGAAATGTCGAATTTACCTGAAAATGTTGAGTTGTATACTGGTTTAATTGACGACTCTGAATCATTATACGAAAGGCTTCCTACAGATATGGCTTATGAGTCTCTTGATCTCATTAAACATATTCAACAAACTTTAGTCCCTTCAATTCCATCTGTTGAAACTAATGGTGATAGTGGCTTTGATGAAATAGATCAGGCTAAAATCAAAGATTGGAGAGCTAAAATGATGCCTCCTCCACTTCCAGGCAATCACCCTCTTAAAAACCGTACTTCAGTAGCAAATGAGGATATTTATGAAGAGGCAATAGATAATAAAAAAACTGACACAAACCATGCTCAACAGTCATGGACACAATTCTATAGTATTGTTGAACAGAGTGCAAAAGAAATTTCAACATGGAGTATGCAAGATATTTCTGATTGTCTGAAAGATTTAAATTTAGCTAAACACACAAATGTGTTTAACGACTCAATGGTTGATGGTCAACTTTTACTTGATTTAGATGAGAGCGTATTAAGAGAATTGGGTTTAAGTTTATTTGAAGCTCGTAAATTGCGTAAGTTTGTATTTGGTTGGAGACCAGATATTGTAAGACCAGTAAACTATCCTGTTTTAAAAGGTTTCACCAGTGATGACCCTTCCGAGTGGAGTGAAACTGATGTTATAAATCATTTGAAAGTTATTGAAATAAATGACTTTGCTGAGTTTTGTCATCATAACCAAGTGAACGGTGATTTGCTGAAAGATATTTGCATAGATGACTTATTAATGCAATCTATTCCTAATTTAAAAGATAAAAAATTTAAAATTTTACAAATAAAAAACTATGTTATTGACAAATGGAGACCTAAAAAAACTGGAGAAGGCAACTATGTTTCATCAACCGTGCACAAAAATAATCCAAATAATCAATCATTTTCCTTGGAAGAAACCATTTGCTCATCAACAAGTAATTATCCTAAAATAGAATCACCATATTCTAGTCAAAGGTCTACTACTCTTGGACTTGCTAAAAAACCTGTTGCTGATGCACCTTTAGTAGCAAAGATGAAAATGCAGTTAGAAGAAAAGCAAAATGATTGGCAAAATCGAAAGAAAGTTGAATCAATTTCTTAACATAAGCTATAATTATGACAAACTATTGAAATAAATTTGTATATATATATATGTGTGTGTGTGTG
>evgR001267c0g1_i01_Hydra
AATCAGCTCAAAATATAGTTATACAAATAAAAAATGATGTTTTTTTAAGTGCAGATTCTTATTTTGTTGTTGAACTAGATAGCATTGGAATAACAAAGCCAACCCTTGTTCCACCATCATCTCCACAATTAACTACACAACAATCTAGGAGAATTAATATTACCAATAACCTTGCGAATGGGGAGATAGGGTTTTATAATATTCCAGTGTACTTTTCCATTTCTGAACCAAGCGAAAATTACTCTGTTGTCAATTTAACACTAATGCGTGATGGGACAGCTTCTCATACAAGCGTATTGTGGAATATTTC
>evgR001278c0g1_i01_Hydra
GTCAAATTCTGAGTTTGAAAGGCAACTCCCTTATGGATTTATCTTTACTCATGCAAGAAAATGGTGTGTTTTCCGCCAACATTACACCACCTGATTTGCCGTTTAAAATAAAATGTTATGGTCTTACCAAACATGAATCGAATTTTGAACGTCTATCTCTTGGAGTCTCGACAGCTAAAACTGCTTTTATGGTTCCCAGATATGCCGGAAATATGTTTACTGCAAGTTCATCGAATACTGTTGATGCTCAAGTAATGGTTTTCAACTTTGGTTACGAAGAGTCTATTACTTTCACTATAAAAGCAAGTCAAGGAAGTATTTCACCAGCAACACAAACTTTGACTATTAGAAATTTTGGCTTTATACAATTCAGATATAGCGCACCCAGTGATTCAACCCTTGTTGGAACTACAACCCGC
>evgR001285c0g1_i01_Hydra
TATAATCATAATGCCACTCCCAACATTTGATCCTGAAGATGCTTCTAATGAAGATCCATCAAACAAGAAAATGAATGTTGATTATACCGAACAAACCGAGGCAACACTCCAAAGTGCTTCTTTGGCTTTGGATCAAGGTGGAATTAACAAATTAAAGGAAATCCTTGAACCTGTTTTAGCATTGGAAAAGATTGTTCGTCTTGCTGAAGATTTTGAAAATCTCTCTAAAATCCTTGTTCAAATAATTGCCTTCTGTGCTAAAGTTCAAGCTTGGGATGAATTAAATGATCAATTAATAAGTCTTAGTAAGAAACGTGCTCAATCTAGAGGTGGTACCCAAAAGATGATCCAAGAAGCAATGAAATATATTGATGTTACACCTTCTTTGGAAAGAAAGATTGAATTAATTAATACAATTCGTAAAATTACAGAAGGTAAAATATTTGTTGAATTAGAAGGAGCTAGAGTTATTAGAATGTTGGCAAAGATTAGAGAAGAAGAAGGAACTGTTGCTGAAGCTGCTTCATTGTTAC
>evgR001293c0g2_i01_Hydra
ACACGACAGCAGCTGCACACAACCCAACTTGTACATTTTTGAATAGCTAATAGCCAAATTGTTGTGAGCCTATCCACATACCATATTGATAAAATTATTTGAGAGAGAATCAAGGCTTTGAAAATGACAATCACCATTCTTAAGGTTAAATATGTTTCAGAGGATATCCTCTTCGAGCATGGAACTCTTCCATTGGAATTTGATATTTGTTGCTCCAAAACCTCGTCACCAACCAAATCTAAAACTAGTGCAAAGAGAAAACAACAAGTCAAGTGGTCTGAAGATGATGATAGAATTCTCAGAATGTATGCCGAGAAGGAATACTCTTGGAAGGAAATTGCTACTAACTTCCTCCACGACAAGTTTACTCCTCAACAACTCCATCAACGTTGGAAGAGAGTTCTCAATGATAACATCAATAGAGAACCATGGTCAGAAG
>evgR001297c0g1_i01_Hydra
AAAAAAACTCTGGAACCTTCAACTACAACAGCAGAAACAGTGTCCTATGAATCTGCAACAAATATTATAAACACTACCACAACAGAAAAAACCATTACAACTACAGCAGATTCTACAACAGCATTTAACAATGATATAACACCAATATCAACCACTCAAATTACAGGCACAACAACAGCAGAAGAACCTCAATACACAGCTTTGGTATCTTCAACTGTTACAGAAAGCAAAACAGTAAAACCTACAACAAGTTTAAATTTCGAAATAAAAACAGATTCAACAGAATCTACAAATACAAATACAAAGACTATAACTAAAGATTCAACAACAATCAACTCTACACCAACAACTTTAAAATCAACAAGTACAACATCCGAATATACAACCACAACCAAATCTGAATTTTCAACTGAACCTACGACTACTAAACATAAATATTCAACAAATGAAAAAATTATATCTGCAGCTACAACAAACCAACCTTCAAATCAAACCAAAAATACAACAAACGAAACTGAGAACACAATAACAGCAATATTTGAGCCTACAACTACT
>evgR001302c0g1_i01_Hydra
TTATGCCAAGAATACTGTTACCGTCGGTAAAACATGTACCACAACAAACAATGCTACCAAGACTTATGATACTTGTGAAGATTTCTCAAGAATGTTCTGTAACTCAACTCTTGGTCTCTGTGTTGAAAAGTTGGCTGATGGTTCTTTTTGTAATACTCACTCTCAATGTAAGAGTGGTGACTGTGTCAGATCCGATCTCACTGATCCATGGACTGCCACTTGTAAGACTTTGAAGACTGCCGGTGCCACTTGTCAATCTGCTCTCGAATGTACCAACTCCTTGACTTGTCGTTCAACTGTTGCAAAGGCTTCCCGTGTTTGTACTACTCCTGCAGCTTTAGGTGATTATTGTGCTGCTGACTCGGATTGTTTCTTCAATAGTGGTTTCACTTTTGGAGGTAATGTAATCTCAACAGAATTGGTTGTTTGTGCAAACCAAAGATGTGTTCGTCAAAATGGTGTTATCAATGGTCAAAAATGTGAAAAGGATGCCAATTGTCACCCTG
>evgR001328c0g1_i01_Hydra
CTATTTACCAAAGATTTTTATTTTTTTAAACAAGAAGCGATGAGTACACCAAACTATATACTTATCATAGTTGGATTTGGAATGCTACCAATTGAAATACTTGGAGCATATTTTTTGTTTAAAGTAAGAAGAAAACTTGATCAAATTCAGTTTTTGTTGTTGCTTAGTTATTGTTTTGCAAATACTAAGATAAATATTTTGATTCCTACTAATATTATATGCATAACTTTAAACATAAAAATATCTAAAAATGTTTCAATTTACCCAATAGCCGAAATGTTTATATCCTTGGAATACTTCTGCATTATTTTTATTATAACGGCTAATCGTTTCTCTCAAGTATATCTTAACTTACGTTATCAACAAACTTGGACTTATTGTCGTGTAAAAATAGCTTTAAGTATTATTTATATAACATTAACTATTCTATCAGCGTGTGTCATTATTTTTGATAAGAATTTTAAGAGTGTAG
>evgR001330c0g1_i01_Hydra
TACGTTGTCATTTGTTCATTAATTAGGTATGCTAATAGTCAAGTTCCAATTAATTTGGCACAGCTGTCTGAAGGAGCAAATAGACTTTGGTTGATTATAGACTATCACATTAATAAAATGACAAATTACAAAAAGGTGGACATTGAGTTCAAGAAAATATCTGATAGTATTACGATCAAAAAATTAAATGCTTCAAATACTAAAGATACATTTATTTTGGGTGTTCAAGATATTTTATTGAACAAAAAAACAGCGCTACAAAACTTGAAAGAAAACGTGACTGCTATTGAAAATACGTACATTTATGAACCTAATATCGAGGAGTTTTCTTACCCGACTTCAAGAAACATGTACGATTCCTTCAAAAAAAATATTAGTTTCGTGAGAGTTGAAAAAGAGTTTCCAACAGATGTGGCCATAAATAATACAGAAAGCTTTTTGCACGTCCCTACAAACATTTATGAAAAGAACAAAAACGTACTAAATGTTGTAAGGTGGACTAAAAATCTTGTTGATGTGTTTAACGAAAATCAGCATCGTGACCAAACGATTATTCTTCAATATTATTGCGACAAATCAGGAGTTTTACGAATGTTTCCAGGGAGTTTAAATGGATTGAACGAAACTAATGTTGATTTGTATGACTGTAGACGACGGTTGTGGTATCAGCAAAGTTCTGCTAGTCCAAAAGATGTTGTTATTGTTATTGACAGATCAGGTTCAATGACAGGAAACAATATAGTGATAGCAAAGATAGCTGCAAAATCAATTATCGATAGTTT
>evgR001346c0g1_i01_Hydra
GGAAGGCGGGGTTTTTATGTATGTTTATCAAGAATTCATTGTTATTTAAGTTTAATTCAAATTTAAGTTCAATCAGTAATGATTATGAGTCATTATGCGTGGAAATCATTAATAAAGCCTCAAAAAACATTGTCGTGCATGCATTATACAGACCACCCTCCGGAAAAATTAATGCGTTTGAAGATTACATTAAAAATATAATTGAAAATAAATCGAGTTTAAATAAAAGTGTTTTTTTTGTGGGTGACATTAACATAAATATTTTAGACTATGACGTAAATAAACAAACAAGTAAATTCTTTAACGCTATTTTTCAAAGTGGATACATTCCATTAATAAATAAACCTACAAGTTAGACTAACGATACT
>evgR001353c0g1_i01_Hydra
GATATTCCACTTATGGGTTCTACAACAAGAATAGACAAAGCCCTTCGCCTTGCGCAAAAAGAGTTATTTCTGGAAAAAAATGGAGGTCGACCAGGGGTACCAAAATTACTTTTTTTACTTACTGATGGTTCCCAAACTTTAGATCCTGGCTCTGAAGATCCTGACGTTATAGCAGATGAAATGCGCTTGATGGGAGTTAACATTATTGGCATAGGTATTGGTAAAGGAATTAATGTTACAGAGCTTGCTCACATAGCTGGAGGTAAAAATAACACATACAGTGCAAATTTTGATGAGCTAGAAGAAAGCAATTTTTTACATGACATCAGAACAAGTTCATGCAAAACAGTTTCTGTTCAACAAAGTTGCGATATTGCGGTTGATGTTGCATTTATTTTGGATACTTC
>evgR001357c0g1_i01_Hydra
ATTTAAAACTAATCTTCTATTGTCTAAATAAGATACAAATCAAATAATTTGCTAAACATGGATCATGAAGATGAAGAACTCAAGCAAAGAATGCGCAATCAACTCATGCAAAGTCAGAGTGGTTCATTTTATCCGTCTAATAGTCCGATGCCAGTTATCTCTTCCATTAATGCTTATTATCCATCTAATCCATCAATAGCACCAAATACTCCTATGGTTGGGCCTCCAATGACTCCATTAACCCCTATGATTGCTCCAATGACACCGATGTCTGCAGAAGCTTCTGGTATTATTCCACAACTTCAGAATATAGTTTCAACGGTTAATCTTAGCTGCAAGCTTGACTTGAAAAAAATAGCATTGCATGCACGAAATGCTGAATATAATCCCAAGCGGTTCGCTGCAGTTATTATGAGAATACGTGAACCGAGGACAACTGCTCTCATTTTTAGCTCAGGGAAAATGGTATGCACTGGAGCAAAAAGTGAAGAACAATCAAGACTTGCCGCTCGAAAATATGCAAGAGTTATTCAAAAGCTGGGATTTCCTGCTAAATTTGTGGACTTTAAAATTCAAAATATTGTGGGAAGTTGCGATGTTAAGTTTCCGATCAGATTGGAAGGATTGGTTTTAGCTCATAGTCAGTTTTGCAGTTATGAACCCGAGCTATTTCCTGGATTGATTTACCGCATGGTAAAGCCTCGGATTGTACTACTAATTTTTGTTTCTGGGAAAGTTGTACTTACCGGAGCAAAAGTACGCTCAGAGATATATGAAGCATTTGAAAGCATCTATCCAATTCTACGAGGTTTCAGAAAAACATAGTGGCAGCTTTTTATGAGCAAAATATTCTTATACAAAAGACTTCGTAACGTTAACAAGTTTACAGCTTCATGTAGAAGTTTTTTAAAGTATAACATTTTGTTAAATTCTTTAAAAAAATGTAGTTACTGATGACAGTCAAATTATAATGATATATAATAAACATAGTATTGTAAGGAATATTAGGTTGAAATATTTATTATGTCTTTAAAAAAAAAAAA
>evgR001361c0g1_i01_Hydra
TACAGAAGAAAGGCTCTATTGCTCCTGTTTACTGAGTTTTTGTAAATATAATTCTGGGGATGAATACTGTTCACAAATATATCAATCATGTATGTTTACAACAACTGAAAATTCTACTGCAGTTTTATCCACAGCTACAACTGAATATAAACAATCTACTACTGTATCAACTGAAGTTTTAACTTCTGCAGTAAGTAACAATGTTAAATGTCCTGATAGTTGGTTACAAAATGCTGGTAATTGCTACTTTTACCAGGAAAGTCAAAAAACTTATCAAGACGCCTTGTTATCATGCGAGAGTTATGGAGGAAACTTATTAAGCATCGAAGATAGTGCAGAAAATGATTTTATTATAAACTATGTTAATACCAATAACTTACAAAACACTGATATCTTTATTGGTCTTAATACTAATCAAAATAAAGCTGAAAGAATATTTGCTTGGTCAGATGGTAC
>evgR001368c0g1_i01_Hydra
ACATATTTCTTAAGTAATCAACAAGTAATTGAAAAATTAAATATTTCTAAAACTTTACTTTTGTTGACATATAGCATTAATGATTCAGCTTCTTATGCAGGGTATTCTTGTCAATTTTGTGATTTTTTTTCTTGATGAATCTTCAGCTGAGTTCTTTAACAATCTACCGACACTTGAAAAAAGTATTACTTTACAACCAAGATCTTAAATTAGTCGACAGAGGTGGATTGAAAATTCTCTTTGATAATAATATTCAGTTTGTTTTTGTCTCAATTGTAATCTTTACTTCTATTAAAGAAAAAGTTTATCGGACATCCATGTGCAACATTTTCATGTTAATATCAGAGCATTATTCTTTTTATATGGAAAGATATTATGGTAATATAAGGTCCAATATTTTTATTAAAAACTATTCCAAGCTATCAACACCAAAATCAAACAAAAAAC
>evgR001378c0g1_i01_Hydra
GTTTTATATGTATGGCCCAAATGATTGGCACATCTTATTTAGCACGTATTTATTTAAGGATTATGTCACATTAAGTCGAATAGTTATATGGATCGCATTATGTGTTCCACATGTCGGCCGCGCTGATCTTCGTCATATACGTTATTTATCAAATGTAATATATTTTCGCACTTTTAATCCAAAAGAAAGCAGTATGGAAACATGTGTGTCACTGATAGCATTGTTGGCGGATATGAACTTAGATGCAAACAATTCATTATCAATCATAGTTTGTAGATGCAAGCAAAGGGATATGATTCAACAATATAAAAAATCACATAACATAAATTATGTACAATTGATCAAACTACAAATAAATATTCCAGATCGTGCTAATCAAAATTAAATCACCATTAATACAGGAAACAATATTAAATTACCAGCCTAGATATTGTTTTTTTTTTACAAATGGTATTGTCTCTAAATGAAATGAACTCCCCAAGCAGCCATCAAATTCCGAAGCTATAAAATCGTTCAAGATATTTCTCAAAACTTTGGACAAAAAACCAGCAAAAGTGATACAGGGCTTCATCAACAGATGACACGTTCCGCACTATTTAACCCTTGTTTAAATGTGCAGCAGATTATTATACTATTGCTACTACTATTACTATAATCCCTCTCACAATTTTCCAACATGCCCTCATGTTTCTAAACAGTCTAGCGTATAACTGGGAAACACAAATTTTGTAAAGTCTAATTTTTTAAAATGTTTTTAATATAAATAATTACAGCAGAAAATTGGTTAAATGGTTACAGCAGAAAATTATCACACAAAATGACAGTAGTTG
>evgR001402c0g1_i01_Hydra
AACATTTAGGTTCTAATAAAATGTAAGTTTTACAGTTTTAGTGACTATGTAGTGACTATGTATCTGGTTATGCAGAAATGAAGCTTATCAAATGAAATACTTATCAACTATAATCATCAAGAACTTAAGGCATTCTTTTGTTAAATAAATACTAAGTCAACTTTTTTGTTGACTTAAAACTGAAAAACCAACTTCTATAAAGCTATGGAGAATTTTACTAGAACTATGTGGTAACCAACACTTTCATTCGTTCATCCATTTGAAACCTTGATAATCTTAGCACTTTTATAAAATGGCTCAAGTTTTAATGGCAAATAAAATTAACGCAATAGAAACTTTGAAACAAGTTGACGAGATAGCAATTATGAAACAAATTGAAACAATATCAGCATTGGTATCGACAATCAGCTCAACAATCAGCACAACGACAAATTTTAATGATTCGTGTAAAACTAATTTTTTAGTTACCCTAAAAGAGGCGAAGTACTTTTTATTTGTTTATTGGTTTAATTGCGCGTTTGTAACACTCCCTAACATGCTATTAATAGTTGGAATCTTCCTTTATAAAAAACCAAAAAAAATTAGCAGAAACGACAGGCATGTTTTAATTCTATCCACCGTTGATTTAATAATTGGTTTAGTTCAAATTCCGTTAGAGATTCACTTTTTACAGTCGTCAAATTTTCGGTGCAATGAATATTTTATTCGGAGTTTCTTCAAAATGTTTCTCGGTAAAGCCTCGGTATGTTGTATAGTTTTAATTGCTGTTGATCGATTAATGACGGTATTGTACGAAGATCAAAATTTTATTGAATTTCACCCGAGATTAGGCGTTGCTACCGTTTTTGTAACAAGTTTTATTTTTGGAGTTTGGTATGCGTTCGTTGCAAGTGTCAAATCGCGCATCAAAATATTTTTTTTCCTCTTTAACGCAAGTTTTGATTTTTTGATTTTCTTAGTTGTCAGTATAGTGAATATCGGCTTACTTTATAATACCAAAAAAAGATTAAAATTATTGTCTGGTTTTAGATCGTGCAATTTAAATTCTGACCAGCGTCTAACTCGTACAGTAGGATTAATATCAACAACGCTCGCAATCATTTACTTGCCTTTTATAATAATGCAGTTTGTTTACGCAGTGGAGTATAAAGATGAAAAGGTTAAATTTTTGTTTTTTGGAACCACTACGTTAGTTCAATTCAACTCGGGAATCAATGCTTTGATATATATTTCAAGAAATAAAGAACTGTTGAAAAAAAGTAAAACAAATTTCCGTATTGCAATAAAAAAAATTCGTGCAATAATGAGG
>evgR001403c0g1_i01_Hydra
TTGTTGTCGTAACATTTTCTAATGGTAGAGTTTTATCTTTCCCTGATTGGTTCATGGTTGATATTAATGGTTCATATCACAACATGAAAACATCTAATAAAAGTTCACCCACTCATCCTTTTGAAAAAATTATTAAACTTTTTCCATCAAGGAACTTATTATTTGAATATCCAATCCATTCCATGGCTATTGTGTCTTCTAATTTTGGTTTGAAACAAAACTCTACTAATGGTCTAAAACTGCCCACAACCAGCTATCCAACTGAAGGGCTTGAGTTATGGTTTGGGCAAAAAAATGGACAAGTTACTATTTTAAATGCAAAAACAATGACTCCGATTGAAACCATTTCAGTAAAAAGTAATGATATAGACAATGCAGTGTTTAATGATTTGACTGTTACTTTTTTGGAAACCCCTCGATCTTACAATTTATTAACTGAAACTATAAATCAAAATTGTCATATTTGGATTGTTACTTCTCCTGGAACTCAAGTTGAAAAGTGGAATGTGGATACTCGACAAATTGAAAATATATTTTTTGC
>evgR001428c0g1_i01_Hydra
GATAATGAGGGAAATGGAAAATCAGTTCCAAAACATCAATCAAAATGTGTTCAAAGCGTTACCATCGTCCTTTAAAGGGGAGACTGCAGTTCCAGTAATTAGTAGTAAGGGAGATGACAATATGTACAGGCTTGTCCTTGATTTAAGTGGGTTTAAGCCCGAGGACGTCAAGATTGATCTAATGGACAGGAACCTGAGAGTAACTGGAAAGTGCGAGCAGAAAACATCTGATGGCTGTAGAATGTACCATGAAACTCAAAGAGAATACCTTCTCCCCGAAAATGTAAACCTTAATGAGTTAAAATCGGCCTTCACTGACAGTGGTTACCTGACTATTGAAGCACCCATGCCAGAAGGAATGAAACCTAAAGAAATCCCTATCAATCGCGGGGCGCAGCAAATTGAAAGTGAATCAAAAGAAAGCAAACGAGAGGACTGATATTCTTAGACTAGACAAACATGGATTAACTGATTTCTTAATTTAGAATACCTG
>evgR001440c0g1_i01_Hydra
CAGAGTTGCTAGCGCCAGATGATGTTGTTGATTCCTTTGTAATTAGAAAAAATCTTACATCTTCTGGAATATTTTTTAGCTTTGCTTCAGTTACAAATTGGGCGTACGTTTATAACCAAAAAGATATCCATCGCCTTTATGTTATTTTGACAGATGATGCATGGGCATATAAAAATATGGTGGATGACGATGAAGTGAAAATTTATCTTTTTAATTATGAATTATTAAGTTTAGAGGCTTACGGAGTTTATATACCTAATCAGTCTCACAGACCACTTTACTATTACTATGTAGAGTTTACATTTTCTTCATTCAACACACAAAACTTAAATATTTGGGCTTTTCATATGAACTATTTTCTTAACAGTTTGATTGATGTACTTGGTAGGAGAACTTATGGTTCAAAGATTTATTTAAGATGGTTTTACTTAAACGGGGGCGAGTTTGCAACTCAGAAGAAAGACACATCCGGTTATCTTGCTTGGGGGATTGATTTCAGAGAAGTGTGTGACAATGCTCTTATTCGAGCTACGCAAAAAATTATGTTTTTAAATGCAAGTTCATTGATTATAAATCCATTATTTCTGAAAAACATGAGTAATGTACTAAATTTGACAAGGATAAAAGAATCATTTAATGGTATATGCAATTTTAACCCACCGGTTTTAAAAAATGTTATTCCCACACTGCAATTTTCGTATTGTGGAATTAATGCGTTCCAAATCCCACAAAATGCTTTTATGGATATTGAGGACGGTGATACAAGAAATTTAAAATTGACACTGTTGGAAGAAAACAGTCAAGTATTGACTCTTAACTCATGGATTCAGTTTAATGTTAGTACACAAACAATTTTTGGGTTAGCGAGTGCCAGTATTGTTAGCAAAAGTTACAAGTTTTTATTGATAGTTACAGATAGTTCTGATTTATCAATTCAATCTTCTATAAATGTTTTTCTTAATGGAACTGTGGCCGAAGAAACATTCTTTATTACAATGAAAGGTTTGTTTAACCCTACAAATTTAAATGTTGCATATACTGTTTGGAGTTACATAAATATATGGCTTTATAACCGTGGATATCTTTCAAAAGATTTTCGTCTACTTCAAAATACAGTTACTTCAGTTAATGAGAATTTTCTCTCATTCACTAGCTGCTCTTTTCGAATCCAGCCGTGTGATGCTCTGAAAATAAAATTTTATAAAGATTTTTTTTTTGAAAGCGCTACATCAGTTAAACAAGACTTTTTAGCATATTTATCTCCCTTATTTTCAAATCTTAAAATTAGTTCAGATGTGGCCGGTGCTTGTCAAACTAACAATCCACCTGTACTGGTTAAATCTTGGGGTCCCTTGTATGTGTCAACATGTGAAACATTCGCAGTGGCTGTTCCCGTGGAAACTTTTAACGATATTGAACAAGGCAATACTCGCTTCTTATATCTTACCTTAACTACTTTTGATGGAACGATGGTTCCAGAATGGATTCTATTTTCTAGTGAGAACCAAACTCTCACAATTGTGACTTATCAACTTTTGAAAGATACTAGTTTTAATTATCAGTTAAAAGCAACGGACCTTAGTGGACAAAGTGTTATTCAAATTTTAAATATTGTTATCATTAAATCATTAAAAGAGCCGTCCTTATTGGTAAGTATGAATTTTTCTTCAAAATTTCAACAAGCCAGTGACTTTGTTACAGTTTATAAAACTTTACGAAACATAATTCTATCCTATTTTGGAGATAAGAGATCGACAGTGGAGTATATTGGTTTGAGTTC
>evgR001469c0g1_i01_Hydra
GGTTAGAATATTAAATTTCTTCAAATCCTTGTTGTCAATATTGATCAAGAATTTGTTACACATTTAAAGTTTAATGTTATTCTCTTGATGTCAAATCAACTGGCCTTTGATGTCAAATTAACTGGCCTTTGGTGCTTGCTTTATCGATACTCTTTTTTTCTTTTGGTTTTAAAAATTGTTGCTTTAATTTTGGGAACAAATGAAAGATTGCAGAGTTGAACTTTTTTTGTGACTGATCTTTGCAGCACTAAATAAATGTCCCATTAGCCAAAATCATATGCAGATTTTAAAATAAAATATTTATTACATTCCTATAAAACTTACTTATTAATCTAATATATTTATTGAAACAGATGTTAGCAATACTTTAGTTCCGATCCACGTAAAGCCACCTGCAGTTACACAGAAAACAGAAAATAATCTTTATTACTCTAACACTGAAATTGAATCTTCTAATATAAAAATTAATAAAAAAAGTATTAAATCTCAAGAAGGTTGTAGTAAAGATAGTTTTGGAAATATTTCTAAAAATCGTTTTGTGGTTCCCATGTCTCAAGAAAAATCAATTGTTACAGATTCTCAATCCAATTCTGTTGTTACCACTTTTTTAGATGATCTTAATGTTCCTCCTATTTCTGTGTTTACAACTGCACAAGATTGCTTTGCCATTGAAAATGCTCCAAATAATGGAGATGATACTCCAGATAATGAAGATGATGCTCCAAATCAATCTACTTATTCTACTGCTCTTTTTGAATGTTTGGAAAAACATCCTAAAAGTTTAAAAGAAATCAAAAAGATTAGAGTAAAATATAGAGAACATAAATCTAAGAGAAGTAGTATAGTTTCACGATTGCGTGAAAAGAGTTCGGAAGATTCAGATATTCAGCAAACTCAGAAAAAACAAAAGTGCTTCATGACTACCAATAAGGAAATAAAGAGCCTACCTGCCTCTTACTTTAAAATGAATCAGAACTATTTCACAAACAGTGTGTTAGAATGTGAAGTTCATTTAAAACATCAGTTCTGTGAAACACAAAAGCATTCTCCCTCATCAAACAATTCTAGTAAACACAAGATACTTGAAGAAACAGCTCAAATTATAAATTCCATCTTACCTAAAGAAGCCAGTGAAACAATTGAAAATATTTGTAATAAAACAACCATTCCACAAAATAAAGAAAGAAAAAAAAATTCTGAAATGATGAATAAAATTAAAAAAGCCGATAGCCTTAAAAATTCCATTGATTTTGTAATAGCTAAAAAAGATTTGACGCGTAGAGTTAAAAAAGGTATCTCGTTTAAGACAACATCTTCAAGCGGTAATTTTTGTGTAGGCGATATATTAAAAGACCATATGAGTAGGAAACGATCTTTGTCTAGTAATGCAAGATCTTTGTCTAGTAATGCAGGGAAAACCACTTTATCAACTATATGCTGTGACATAATAAATAGTGCAGCTGATAAAAAATTAAGTTATTTAAAGCAGCAAGATTCATGTACATCAATAGATACCACGGATACCGTGAATAGCAAACAACATAGTGTTAACCCAAGTGGAACAGCGACTTCCTCGAAAACTTTGCCTTTTCAATCTTTATCATCTAAAGTAAATTTACCTTTATTATTTTTAACAGCTAGCAATATTTCAACATCGTTGACATCTTCAACAGTTACCAAATCTGTCATGTTATCTTCAATATCATTAGCAACTACTAATACGGTCATGTCATATGTAATATCACCAAAAGCCACTGAGACTACCATTTCACCTTTGATATTTTCTCCAGCTAGCAAGAAAATTACACTACCTTTGATATCTACTGTTACTTGTGACAATGCCATGCCACCTTTAATTTTACTAAGATCAAATGAGTTTATGGCTTTACCTTTAAATTCATCATTGTCTGCAAATATAACATTTAACAGTAACTTTTATATGCAGCCTCGCCTGACAATCTCATCAGCTAATGCTGAAACTTTCGTTTTACCCAAAAACTCTCATGTTGGATTATTTAACCCTAATCCAATCACAATGAAAAGTTTCAACACTGATCCAATCATAATTGAAAGTTTTAACACTGATGCAGCAAACAGCAATAAAATGGCAATTTCTGCTTCGTATTGGAACATTCAAAATAATGCAATTAATAAAAGTAAAAATTTAGTTCTTAAAAATGATTGCACAACATCAGAAAATACTTGCACTACCTCAGAAAATAATTGGACAACATCAGAAAATACTTTCACAACATCAGAAAATACTTGCACAACATCAGAAAATACTTGCACAACATCAGAAAATACTTGCACAACATCAGAAAATACTTGCACAACATC
>evgR001482c0g1_i01_Hydra
TGCAAAAAGCGAAGAGACAATAATAATAAAAGGTCAATTTCTTACTAATTTAACAATTCCAATAGAGTATGGAGTGTCCGTTGAAGTAAACTTAGATTCATTTACAGGCTTTAGCAATTTAATTCATTTATCAAAGGGAATTGCAGACACATTTTGCTATGGTTGCAGAATTGCTACAATTTTTACTCAAAATCAACAAGCATTTATCTGCGGCCCAATCAGAGGAGAGGGCAATTATTACATAACAACAAACTTATTAAAACTTAATACTTGGGTCAGAGTCACAGTTAGGCAATACTTGAAAAAACATGAATATATTTATACTATTGAAATAAATGGAAAAATAATTTTCTCAATCACAAATACAATAGTAACAGTCTTCGAAAAC
>evgR001493c0g1_i01_Hydra
TTATTAACTCCACATTTTACTTTCATACGTTTTGGCTTGTTTTGGATAGGTACCTTCATTTAAAACTAAATATAAAATACTTAACCTGCTGGTCCAAACGAAAAACATTAATTAGTGCAATAGTTTTATGGATTTTTGGAGCATCTATTGGAGTATTTGGACATTTTTTTTCACAACAACTTTATTTTGTATATGCAACTTTTGATTTAGCTATAACTACTTTCAGCATGTACGTGTACACATATGCTATAGTGCTTCTCAAAAAGCAAAAAACAAGGATTTATTCCAATCAAAGCAACCGTGGTATATTTAAAGGCTTATTTTTTTCTTTCCTTATCATAAGCATTTATGTTGTTGTAGTTGCTATACCAGATATCACATGGTTGTTTATGTTTGTTTACAATAATCCTAACTTTGAAAACAATAACACCG
>evgR001508c0g1_i01_Hydra
TTTTATTTTATATAACAAATACTACATACTGATAAACCCGATTTTGCAATAAAGATGTCTGATCTGATATCTTGAAGTTCTGCAACATTTTTACATATGTAAGATTAAACATTTTTGAGGCTAAAAGTTAAAAAGAATTAATAATGAGAAAAATATATACTTATATTAGAAAGACAATGAAGTTTATGCTACCAGTTGCGATGATTGTGTCCATTTTATTCTTAAGCAATAACATAATAAATGTTTCAATGGATAAAATCATTTTTACAAACTATTTTAGTTTAAGTTTAGAAGAAAAAAACGAAATAAATTGTACATCTACCGAGAATGCAGATTACTTGCAAAGAACATTAATTCAAACAACGACAAGCACAAATGTGAAAAGAACCACAACGACAAGCACAAACCCAATTAAAAACCTTGAAAAAGAAAAGGAAGTACTTTCTCATATAATATTTGACTCAGTTGAAAAAGACAATACACT
>evgR001517c0g1_i01_Hydra
TGGTTTGGAAGCCCTTGATGGCTCCGAAGGAAGATGTGTTCAATCACTTATGGAAAGCCCAAACGTTTTCTTTTCAGTTGTTGCTCACTTTTTAGTGTGGATCCAACCAGTTGTCCACAATTATTACATTTTACATAACACCAAAAAAGGAAAGCTTGCTTTCAAAATGACATTGACTTTGTCTTGGATGACACTCATCTTGGCTACAGTTTCTTTGTATATGGGTTATAACAATTTGATGGGAATGAATTCTGATTTACCTGCCAATCATATGGAAGGAAATATCACACCAGTTCAACCAGTCTATCAATTCGGAGTCCAAATCCACAATGCTCACACC
>evgR001524c0g2_i01_Hydra
AGAAGTATAAACTACTTTACCAAAAACATAAGCTGCTTCACGGGAAAACAGCTGATATTCATGGTATAACAACGATATGGTTATAGAATAACATTTCTACTCAGAGTTTTGTGTGTATTTGCTTTGTATAACAATACATTTTTTTGTAGCAATACAACTAGAAAATAGAATTTTGTAATTCTTGTCCTGTTTCTTGAGCAGTATCTTTGTAAGAAGTTTTATTATCTTTAACGAGCTTGGTCATGAATGAATTTTCGAACAGGAGTTTTAAAACTCAATCAAATCAAAAACAATATTTAATTTACGGAAATGAAAACGAAAACGAAAGCGAAGAACTTTCGATAGAGCTGAAAGAAATATCTGAAAAATTACCACTAATGAGACGTTATGGTCAAAGTAATGACCAGAATAATAGTGAAAGGCCAGCTACTAACTATGTACCAGTTACGATGGTTACAGAGTCCAACTTTGACAGGGTTACTATTGAAGAAAACGACGGTCAAACACAAAATCTAAGCACAATGAAACTTTATGAACAAACAGCTCCAATCGACCCACAACATAACTTATTTTCCAGCAATGGTTGTGAAAAAAGTAATAACAATTATACATTAATAGATTTAATTACTGCTAAGAAGAATTTGTCTAAAGAAAAAATCGTTTTAAGAGATTGCAATATTAACATTGATGGTAACATGCCATTAGTCATGCAACCGGTTTTTCAGAGTTTTCAAAAACGAACGCTTGTCAAGCAAAACACATCCAAGTTGTCAATGCTCCGTAAATTTTTTGGCGCAGAACAAAAAGATTCTAAAGAGAGTGCAGAACTACTGCCTAAGCAAGAACTTTCGTTACTAACCAGTCCTTTCCAAGTATATTTTCCAAGATTGCTTAAAACTCAGGAAGCCCCAAAAGAAAACAATTTGCCTTGCGATTTACCCGGTTTATCAAACTTGTCGCAATTAACAAGTGTTTCCGACAAGAAGAAAAAAAAATCGCTTGCCATTGTGTTACTAGCTTTAGCAATCGATGCAGAGTTAGACGAATCATTTTTTCATGCAATCGAAAAATCGGAACAGTTAATAAAGTTGCTGAAAACGATAGCAAAAACAAACTTATCCCGAGATGCTTTGCTGCTATTAAAACGGTTGTCAACTGATATCTAATATTTACTAATTTTATAAATTACAAACTTGTTTAATAAAAATATTTCGTTAATTATGGGAAAATGATTTTTTTTTATTTGTACTTTATACATTATTTATATTACATATATTATATATATTAC
>evgR001538c0g1_i01_Hydra
TTTTCTAAATAAGATATTTATATACATAATTAAAAATGCCTACATTTTCAAGGACCATGATGATTGCAGAAGGACAGTTTGTGTTGTTTGTATAAAGAAAAGTGACAAAAAAAGGAATTTGACAAAATTGAATGATGGAGGTAATGCCTTAATACAACTCTACAGTTTTGATTCCATTGTTGTGAAGCCATTAACAAGACTTACAACCTGTTGTGATTGCATTATTTGTCAAAATGCAAAGGTGAAAGGCAAAGAAAAGCATCCATTTGAAAAACGTTTTGCATCTTTGCAAGCAGAAAGAAATTCAGTTGTAAAACAATGCATAAAATGCTTGTCTATCATTGCTCGTAGAATTTCTCACAATTGCAGTGATACAACTCGCCGTGCAAAACTTATATCAACTTGCATTGAAAGATTATGTGGGCGCAAAACAAGTTGCTTCATCTGTCATTGCGAAAGATGTCTTTCCGGGTGGAACAATTTTTTTTGGTCTAGACTTCCTCTCGCACTCGCAAAATTCAGTAACTTGTTGTGAAGCCCCCTGCCGTGTTCCTCACAACAAGTTACTGAAATTTTACATATAATATATTATTTTTATAAAAAAAA
>evgR001545c0g1_i01_Hydra
TTTTCAAATGCTGATTATTGGTGTCTTGGTAACAGTTCTGAAGTAATAAAATGCATTGAATCCTTGATATTGGAAAATGAGAATATGATATTAGGAGGAGATTTAGTTAATACAATTACTTTACTTCGTAAAGAATACTCAGTATCTTTTGAAATAAAACCAACAACTTTTTTGGATGATTGGGCTAATATCATTCATTTAACTATTGGAGAAGCAAATACAAATTATGGAGATCGTATTCCAGCTGTGTTTTTTCGTCCTAATAAAACAGGAATTTTGCATATTTGTTCAGCTATAAATAGCAGCAAAAGTTATAATTTTAATACAGTTCCATTGGAGCTTGGAACATGGTCTTCAGTAGGAATATCCCAGCAGTTTATTGATGGTAAATATATTTATTCAATTGATCTTAACTGCACAAATATCTTGAAAAAAGAAAACATCGATGCAAGAGACTTTTTTAACGTTAAAGTTTATATTTCCAATCCTTGGACTGCAAC
>evgR001546c0g1_i01_Hydra
CCAAGACAACATGTCGTAGTTATAGCACTTTGAAGCAACGTGGTGCTCAAATGGACCAAGAAATGTCCATCACCAAGACACCTCTTGAACGTATGTTGCTCGGTAAACAAACAAAGGCTGATTTCGCTGGCGTTGATATCAAGGATGTAAAGACTGTTGGTAAATGGATTAACAAGTCTGCCAGAGAAGGACCAAAAATCATGACCAAGGATGTCGATCAATTGGCTCCTCCAGTAACCGTCACTGTTTCCGGTGCTGCTGGTCAAATTGCTTATAGTTTGTTGTTCAGACTTGCTACTGGTGAAATGTTGGGTGAAAATCAACCAGTCAGATTGAGATTGTTAGACGTCCCACAATCCATGAAGATTTTGGAAGGTGTTAAAATGGAATTGGAAGATGGTGCTTTCCCACTCATCGAAGAAATTTTCATTACTGATAGTGAATCTGAAGCTTTCAGAGGAACCGACTATGCTGTCTTGTTGGGAGGTAAGCCAAGAGGCCCAGGTATGGAAAGAGCTGATGTCATGAAGGATAATGCTGCTATTTTCAAGAAGCAAGGTGAAGCTTTGAATGCCAATGCTAACGGTGATGTTTTGGTCTTAATTGTTGCCAACCCAGCCAATACAAATGCCATGATCACATCCCTGAACGCTCCAGATATTCCACAAGAAAATATTACTGCTTTGACTAGACTTGACCATGATAGAGCTTTGGCCCAAGTTGCCAACAAGCTTGGATGTAACATTTCCGATATTTCCCGATTTGTTATCTGGGGTAATCACTCTGCCACTCAATATCCAGATCTTTCCTTCACTACTGTTAAGGGACAATGGGCTCTCAACATGATCAATGACGAAAAGTGGATAACTGAAGATTTCATTCCAAAGGTTCAACAAAGAGGTGCTGCCATTATTGCTTCTAGAGGTAAATCATCTGCCGCTTCTGCTGCTGATGCTGCTGTTAAGCACATGAGAGATTGGGTTTTGGGTAACAACGAATGGGTTTCCATGGCTATTCCAAGCAAAGGTCAATATGGTATTCCAAGAGGTATTTGGTCTTCCATGCCAGTGCAATGTTTCGGTGCTGGTCAATACGGTGTCGTTGAAGGTTTGCCAATTAACAATTTCTCTGCTGACAGAATCAATGCTAGTGTTAAGGAACTCATTGAAGAAAAGAAGATTGTTGAAGGTTTATTACCAAACCCAGTCTACAGATTAGTCCCTGTCGATAAGGTTAAGGTTATCAACAAGTCTTTCATCTCACAATAATTTATATCATTCCTCAACTTTTTGTTACAACACAGGGCCTTCCCACCTCCATTAGGGTCATTCAATCAAAAAATTAAAAACCTTTATGTATAAATATAATACTTGC
>evgR001547c0g1_i01_Hydra
ATTCGTTCTGTTTTGTGTGAGGTTTTAGATAATTAATAAATTTTGTTTTATGCAACAAATCTCAGTCATATTGAATTCCATAAATTCGCGAGTGATAAATATAAAGTTACGGTTGTTGCAGTAATTATTATGCATTTCAAAAGTCTTAAAAGATTCCATCATTAAACTTTTAAAGATCTACTGATTTTCTAACGTATAAATAAAATCGTGAAGCATACTTGTTCTTTAGATTGTAAAACCTAAAAATAAAAATAAATTAAAAATGAAAAGCTCGCCCATTGTTGTGGATCATACTAAATCATTAAATACTGTTGAAGAATTAGGCATTTGTACAACATCTCTTAGCAATGACGAGAAATCAACCGATCCTCTTTCTTGTACCATTATGAGTATAATAGAACCTAGTGTTTCAATTGATTTCAATTCCAGCACTTTGAAAGAGACAAAAAAAAAGGAACAAAACGATAGTTTTCCAGCTAATACTGTCTTGTGTGTGTCTGAGAAAATACCCCAATACACTGAAAGAGCTTTGATAACAAAAAAATTAAACGCGAATTCAATTGTTAATTCAAAAAATAAATCTATAAGAAAAAATGATTTAGATTTGCTTACTTTTAGCAATGATGGGTGTCAAGTAAAACTATCTCAGAGTGCACAATCTTTTAATTTAGTCGCACATAGTTTTAGCGCAGATAATTCAAAAACGACGAAAACCTCATTGTGTACGAGTGTTAGTTGTAAACCTTCTATTTCTGTAATTTCAAATAATAACTCTTCTACAGTTTCTTTATTAACTAATAAGGCCGATTGTTTTTCAGTTAGCTCAAATTTAGTAAAAAATAAAATATCTTTTCCAAATAAGTGTACAGAAACCTCAGTAAATGTTATAAAAAATTTGACGACTAAAGTGACTACAAAAAGCCTATGTCTGAATTTGCCACCTTTAAAAGCAACTGTTGTGATTAATCCGAGTTCTTTATCTACTTCCAATTTTTCGGAAAAACATTCAGAAATAGATCATTCCTACAATTTAAAAAATTATGACGGTAAAGGAACGACTAAAGCTTTTTTAGAACAGAAAAAAATTGTTTATGGCAAACTTAGTGTTATAACTGCGCCAAATAGCGAAGGGGTTAATAAAAAGAAACTGGTTTTCCTACCTTCTAAATCTACCGAGTTTAATTTTTCTAAGGAACTTCCTCTGAATTTATTTTCAAACTCCATTAACCCTCAAACGTCACAAATCGTTCTTAAGTCGCTAACAGTTCCTCAAAGAACAACGATAGGTACTACAGCAACACCACAAAAATCGGTGATATACAAAATTCCACCTGTTCATCAAAAATCGCCGATATGCAAGACTTCATCTGCTTCCCAAAAATTACCTATAATGGTTTCAAATCTTTCTAAAAATTTATCCGACTTGAATGTGATCGAACGTAATAAATTTTCTTCTCAAGTAACGCATTCAAATAATATCAAGCTATCGGCTAGTCGCGTTTCAAATTCTATAGAACGTTCTCTTGTTAACGAATTAAAGGAAGGAGTTGCTACTAGTGACAGTAATTCCAACATTTTGCAAACTTTTATTTTAAACATGCCGAGATCTAAGTTTGGTTTAAAGTTTGGTGTACACTGTTTACCGCCTACCACAGTTATTACTACTACGTTTTTAAATAGCGCAAAAGTTCCAGTCTCAAATGAAAAATCTAATATTAGCAACCTAACCGTTTCACATTCAAATGAAAAATCATATATTTGCAACCCAACCGTTTCACAATCAAATGAAAAATTAAATATTTGCAATTCAACCGTTTCACATTTAAATGAGAAATTGAATATTTGCAACACAACCGTTTCACATTCAAATGAGAAATTGAATATTTGCAACCCAACCGTTTCACATTCAAATGAGAAATTGAATATTTGCAACCCAACCGTTTCACATTCAAATGAGAAATTGA
>evgR001548c0g1_i01_Hydra
TAGTTTCACTAAGAACTCTTATAATCTGGTTCGCCGTTTACGGCTTTTTAATTTTTCAGGTTGGCGGATATGATTACTTTGAAGGGAAATGCATGCCGGGGGTAATCGAGAGACATGTTGGCGATATTAATGTAGCAGTTATATTTAGTTTTCTAGAGCAAAATTTAACTAAACTGTACGTCAGCTGGCTTAAGTATATTATCGAGAGTTATGAGTCGTGGTATCATAAGCCATTTACGATTGGATACGATTTCTACAAATCTTGTGGTAACAGTAACACAACAAACTTGAAAGAGCTGTTTAATTTCAGCTTATGTTCTACGGAAGAAGTTTCATTAAGTCCTTGTAAAAACCGCTCGAAGATCAATTCGGTTTATTATTTTGGAATAAATTTTGATGTGGCGAGTATAAATTTAATAGCTCAAGAACATGGCGTGCCTTTTTATAATATGGTTAATTTTACATCTAGTTTTTATAATTATAATATTAAAGATTTAAGTAAGGTGGTTCTATTTGAGAGAGCATTGCGCCAATTTAAACAGAACGAAATAGCACTGTTGAATCCTTTCTCATCTGATATCGGTAAAATATTATGGGAAAGAAATTTTAAAAATCGTTCTTGCTTGACGGTGTTTACGTACGAAAAAAGCAACTTTAGTATATTTGAG
>evgR001552c0g1_i01_Hydra
TTATTTCATCAGTCGCTGATATAACAAATACTTTGGTTGTTTCAACATCTCAAGTTTTATTACTTAAACGAGTTGATAATTTTGGCTGCTTAGAAATCTCGATAATATTATTTTTTCGCGTTTGGTTTCTAGGCGTTACCTGCTCAATTTTTGTAATGATATCCTGGGAAAGATTTTTTACTGTTTTATATAACAACAAGATTTGTGGTATTACTTTAAATGATGCACATTCAATAGGATACTTTGTCTTAGTTGTTTTTTTCTCTTTTGTAGCTGGTATTTGTTACGGATTTGTTTACGCTTCTTCAGACTTGTATCTACAGTTTTTGCTGTACATAGGAACAGGTTGCTTTACGTTGGTTTTACTTTTAATGATTGTTGTTGCGAATGTCTCAACATTAACTGGAATCAAAAAAAAATTAAAGCATAGCTCTCTAGAAGTCCAGAGAAATGCGGCGCTTGAAAGTCGTATCACAAAAACGGTTGTTATTACTTCTCTCGCACATGTTTTGTTTTTCTGCCCTTGTATAGCGGCGCAGTACTATATGTCATTTATATTTTCAAAAGATGATGTTTACGTTATTCCTCCTGCTTTGAAGATACTTATATGGACACTAGTAATTAGGAAAACAAGCTCTTGGATCAACGCAATTATTTATACTCTGAGGAACAGAAAACTTTCAAGATTCTATATTGCAAGATTAAACTCTATACTTGGAAACAGAGACACTAAAACCAGTTCTACCAAATACAAACGTTGAAATCAAATAATGATATTTGCAAGATTAAACTCTATACTTGGAAA
>evgR001554c0g1_i01_Hydra
TGTATTTCTCGTTTGCCAATAATTGCTTCTATTATTTATAGAAAGACTTATCACAATGGACAATATATTAAGAGTGATTCATCATTGGATTGGGCTGCCGATTATGCCTCCATGATGGGTTTCGGTGGAAACAAGCAAATTGCTGAATTGTTCCGTTTGTATTTGTCTATTCACACTGACCATGAAGGTGGTAACGTTTCTGCTCACGCAACTCACCTTGTTGGTAGTGCTCTCTCTGATCCATACTTGTCACTTTCTGCTGGTATGAATGGTTTGGCTGGTCCATTGCACGGTTTGGCCAATCAAGAAGTTTTGAAATGGTTGAAGGAAGTTCAAAAGGACTTGAACGGAAAGGAACCAACTAAGGAAGTTTTAAGACAACTCATTTGGGATACTTTGAATAATGGACGTGTTGTTCCAGGTTATGGTCACGCTGTTTTGAGAAAGACTGATCCAAGATATACTTGTCAAAGAGAATTTGCC
>evgR001558c0g1_i01_Hydra
CGACAAAAAGATCTTATAACTTTCTTATTTTGCTTCAAAATGTGGATTTCGTGGTGGACTCTTGTTTTTTTTTCCCCGTGTGTCTTGAGTTTAACATGCTGGTATCAAGATCCAAATGAATTAAAAAAGCAACGTTGTATTCCACTTTTACAAAACACATGCGCAACCCTGACTTTAGTCACTGTTTACAACGGGACAAGTTCTAAAATGATATTAAAAATGTGTTCAAATGATTATTCGGGTTGCAATTCGTTATGTGATGAAACTTTAAAATCAATGGGAAAATATTTTTCTTCAATTGATGTCACTTCATGCGATACACAGTGCTTCAATGGGGACTTAAGCAACACATCTTGGTCAGCAAAAGCTAGTTGGATCGTGATTATTGCATCACTTTGCTTGTTAACAGTTGCTCGAACATTTACGTTATAAATTTCTTTATATAAGAATTATTTTTAAAGTTTTTTTATTTTTTTTCGTGAATTTATTTCGTGAAGTTTTTCATTATTTGCCAAAGAAAAGTTTTTTT
>evgR001560c0g1_i01_Hydra
TTTTGTTTATGAAAATTAACCGGTTGAAATAAATAATGATTGGTTGAAGTGATTATTGATCGGTTTGGTGATTACTTATCAACAAAGATGATTAATAACAGGATGAGAACACTTTACAAAGTTTATCTCGCTATCCGCGTAACTGAGTTCATTTAAAGGGTTGCTACATTTATTTTGTAATTGCTGGATTTACTAAAAGTGTTGTTCTGGTTGGATATCACCTGATCAATGGCATTGGTTTTTATTATTGTATTGTTATCGTTTTTATGTGGATTTTCTGTGATATTGAACGTTACTGTAGTCTTTACAATACTAGCAAAAGGAAATTCTAAAAACATTCGCGATGTTATCTTGATGAGTCTTGCAATATGCGATGGCATACAATGCACTTTAGGTTATCCGGTTGAACTTTTTGGTTATGCTAATAACGATAACCCATCACGAACTGAAAAGTTCTGCAAAACTAGTGGCTTTATTGTAATGTATCTTGCTTTAACTGCAATCACACATTTGGTTTGTTTATGCGTATATCGTTACTTGACAATTGTCTATCCGCTAAAATCACAAAGGTTTTCCACAAACTCTAACTGGAGTTCATTCTCTCTTATTACCTTTTGCTGGATTTATGGCTTGTTCTGGTCGATAAGTCCTCTGCTTGGCTGGAATGAGATAGTTCGAGAAAAACAAGATACGTATAGGTGCTCAATAAATTTATATCCTGATGATGAGACAAAAGGTAGTTATTTATATGCGCTGGCAATATTTTGTTACCTTATTCCTGTGATAATTATTATTTATTGTAGTTTAAAAGTACATATAGAACTTCGCAACATGTTAAAATTGTGCAAACAGGTCTCTGGTGTTGAAGCAAATATTACGAAAGTTACATATCGAATAGAAAAACAAGATTTCATATCCATAAGCCTAATTATAGTATCATTTTTTACAGTTTGGACACCATACGCCGTGTGTGTTGTTTTTTTAACAATTGGTAAAAAATTACCAACAAGTTTTTTAACTTACTGTGCTTTGTTTGCAAAATCATCAACGATACTTAATCCTATTATTTACTGTCTCATGTATAAAGAGTTTCGTCAAACAATGCAACGTAAACTTCGAAGATTTTTCAGAAGTGCAGTGGTTGCACCAGCTGCATAGAAAGAAGCTCAAGCAAGCTTGACATTTGAACAACTAGTTACGGGTTATCCATAAAGTAAGTCACGCTAAATTTTAATTTAAAATAGAAGAAGGATTGTAAAGTTATTCTTGCCCAACGATTTTTAAAGTATTCATCACATTATTTTGAAAAATAAATCTTTAAGCAGCAGGGTAAACGATCTCTTACTTCCGTTTTAAATTTTACTCAGCGTGAC
>evgR001561c0g1_i01_Hydra
CTAAAAAGAATGAAATTATCATTAAAAAGAAGGAAGAAGCCATTTACAGCCTTGGTAAAATACATGCTGCCAAAGGTGATGCCCAAGCAATCATGTTATTGAATAAATCAATTCGACCATTCTTCCAAGACCTTCCAAAAGCTAAGACTGCCAAAATTGTTAGAACACTTATTGAATTAGTTGGTACAATCAAAGGAAGTGAGGCTATTCAAATCGACATTTGTAGAGAATCAATTGAATGGGCAACCAATGAAAAGAGAAGTTTCTTACGTCAACGTATCGAATCAAGATTAGCAAACTTGTTGTTCAACAAGAAGGAATTTACTGAATCATTGGAAATACTTACCAGATTATTGAGAGAAGTTAGAACATTG
>evgR001564c0g2_i01_Hydra
TTAGATCACATACAACTTTGCTTAGAACTAAATAATTTTTCTAAAATGTATCATAAGTTAAAAATGTCTACAACAGAAAGAACTAAAAAAATATACCGCCAATTTGATACTGATTCAGATGCAAACACAGATAACGATGAAGAAAATATGAATTATGATGCAGCATTTTCTTGTAACAACAAAAACAGCATACATGATACAAACAATTGTTTACAATGTGTATACAAACTTGTATACAATTTAAACATGCATGTCTCGGCTTATACTCAACTATGTGCTTCATATGAGTTTTTTCTTACTTTGTCAGTCACCGAAGTCAACTGCGAAAGGACATTCAGCAAACTTAAGCTTGGTAAAACAAGACTCAGAGCCAATATAAGTCAAGATAATTTAGAAGCACTTTTAGTTACAAGTGTTGAAAAACAATTACTCAATGAAATTGAAATTTCAAATGTCATTGAATATTTAAAAGCTAGTTCTACC
>evgR001567c0g2_i01_Hydra
ATGTTTCTGATTCTACAGAATATTTGTTTGTGTTCCTCCAAGGATATAAGATAAGTGTTGATTGTCAGGAAGAAATAAGAAATACAATTTCTCTAGAAAGAAACTGAACCTTTAAAATATTGTATAAATGGATCATTGCAAATTATATTTAAAGAAACAGCTTCACAACTAAGATGGATGAGCTTTCACCAAGAAGGGAAATAAAGCAGGGGAGAAGGGCTGGAAATGTGTTTCTGTCTTCATCAACTTCTCTTGTATCAGAACCATCAACAACGTCTGTAACACAACCACAGAATTTAAATAGCTCATCTCCAAAACTAGCTCGAAGACAAAGTGGTTGGTCTGATTCTGTTGATAAAAAAAAAGGAAAAAATATTCGAGAGGATGATGAAGCAGATGAAAGATTAAAAAATGATTCAACAGTTTTGTTAGACACTAATATCGATGACATTCCAGTCATTCCAGATCTTGAGGAAATACAAAAAGATGATACTACACAACATGTGTCTGCTCCTCCCAGTGTGCAAGTAACAAGAGTGGCGACTTATAAAGAGCTGGATCACGATTTTGAAAAGCATGCAGGTTTATTAACCTTGGACAGAGATATCGATCTTAAATTGTTAGGATGTGTATTATCTTCAGAATCAGATGTTTGCGAGGAAGACAAAGTATGGGAATGGGACCGAGTGTTTGCTGAAGTATCTTCAGAGTTACGAAATGATTGGGAAAGTATCAAAATTATAAGTGATAAAAGCGAATAAAAACTTGTGTGCGTTTGTGTTTATATGTTTTTAATTATTATTAAATATTTATATGAAAATTTAATGTCATACGGTATAAATTATGTATTTTGTTATAAAGATACGCCGATTGTAGATTAAAGATTTTTTTTAGTCTCGATTAATACAGATTAATTTTTGAAAAAAAAAA
>evgR001568c0g1_i01_Hydra
TTCATATTTTATAGTTTTACAACACTGTGCTATATTTTTGTCGCAAAGAAAACTAAGTTTTCTTTTTCAAGTATTGGTAATCATTTTCGCTTATTAAAAGTTCATGGGAAACAAAGTGTATGTGAGTGTTTCTACAATTTCCAACAATTTCGCTCGCAAATTAACTTTGAAAACGATGACTACAAAGAAAAGCTGTAAAGCAATACTCCTTGGAGACTCGAATTGCGGAAAAACAAGTTTATGGAAAAAACATATAGAAAGAGACACTAACTATGATCATATTCCTACTGTATTCGAGTCATCGGAAACAATGATATTTGTTGATAAAAACATGGAGATAAAACTTACAATATGGGATACACCAGGTTCGTCAGAGTTTGACCACGTGCGGTTGCTATCTTTTAATGGAGCTGACGTTTTTGTACTATGTTTTGATTTAAACCAACTAGAATCACTAGAAAATATCCAAACCAAGTGGGCAAAAGAGGCTAAAGCATATTCTCCTAATACGCCCATCGTCCTTTGTGGTTGCAAATGCGATTTACAAGATGTCAACAAATGTGATAAAGATAAGATCGACAAAGTTATACAACAAATAAACTCGTCTGTGTATATAGAGTGCTCGAGTCAAAATAATACAAACGTTGAAGAGATCTTTAAAATTATTTCTTTACTCGCTATTGGTGACGTTAACAAAATAAAAAACGAATTTAAATGCAAAACTAAGAAAAAACATGTTAACAGGTGTTGCGTCACATGATATTGCTTTTGTCAGTGAGTATCCTCGTATCAAGGCTGCATGAATTTTCCACACACAAAAATCTTATATTTATGTATTTATTTTAAAATATTGAGCTTTAATTTTATATATTTTATACAATTTTTATTGTAAATTAGACAGCATATGTCTTAACCCGTAATAAAGTTACGTAATAGTATAATATAA
>evgR001569c0g1_i01_Hydra
GCTCCGTTGGTTTAGACATTATACAAAAAGTGTCAGATGCTGCAGTCGAATCTGTTCGAGAAGCACAGAGAAAAGCAAATGCAGTATTTGAGCAGGCAAAAAGGACTTTAGAGAAGGCTCAAGAAGATATCCGTAAAGTAAAACAAAAAATTGAATCTTGGAAAAGTGGAATTTCGAGTTATAAAATAAGATTAAAAGAAAGAAGTGATCAAATTCAAAAAGAGAAGATAGCAATGGAAAAATTGTGTGTTCCTGAATGTGGGACAGTTTGTGTTCCATTCTTTGGGTGGAAATCTCATTGTACTACTTTTTGGGGTAGGCCAGTTGGTTGTCTGACATGGGATCATTGTAAGTGGGAAGTTCCTAATCTTATATGCATTGCTGCTTGTGAAATACAAAAGGGTTTTGCTAAATTTGCAGCTTGGGCTGAACAAATAGGAATAAAGATTATGCAGGGTTTAGCAAGCATTGGTGATTTTTTCCTGGATGCTGCTTCAGGATTAACGAAGATTGGTGATGGTATTCTTGCAATGGCTGAGGAATTAATTGATTTTGCGAAAGCTACAACAGCAGCTGTGCTGAAAATGGCTGAAAGTCTAGTAAGTTTAAGAATCGATCACATTTGCTTAAATGCTAATTTTGACCCAGCATCCAAAACTTGCGTTGGATATTTTTTAAATGGATCCTACATTGGAGTGGATTTCCGTTTTTCTGGATCACTCTGCATAGACTTGTCTACATTTAAGACTCTGGGTTCTAAATCAGCTAAAACAGCGGAACCTGCGCTTGAGAAATTGGATGAAAATGCCGCAAGTATTGATGCCAAATCAAAAGAAGTGGAACGAGCATGTGATGAGATGGCTAAAAATACAACACACCTTGAAAGTGAAGTCAAATCTAAGACTTCAGCTAAAGTAGCGAAACGGTCACATGTTCCCACAGAAGCTGAAATGCGCATCCACAAATTAATTTATGATCCTTTACCATTAGTTAACTTGAGAAGCAAAGAAACAAACAGTGCTATGAGAAGTGCTGTGCCCTGGGAATATCTTAATATTGATGACATTGGCACTAAAGATGGGATTGTGCGGTTTGTAAACAACACTTCTTTGCGTCATCAAATTCCTAATGAAGCGTCTCATGACTTTTGTCATCAACACCAAAACGTTCTTTCAAGGTACAGCACAATAGCTGATGGATTTTCTTTTGCTATCAAACATGTCAATGAAGCTAAAAAAGGCTATATTAATCAAAAAAGAACATATTTAAATCAAATCGACTATTTAAGTTTTATGGTTCGTTCAGGGCCATTAAACAACATGACCTTTAGCGAACAGGAGGATATTACGCACTGGAGGGACTATGCTGACAACCATATTAAAAGCTACTTGGATAAAGCCACAAGAGCATTTCACGGAAAGAAACGTGATAGCATTAATCATGTTCGTCAGGCTATTAATGATGTTATTGTAAGTAAGCGAGGTATGAGGCTGCCTGAATATATCAAATACTTGCATTCTCTAGCAGTTAAAGCTGATAAAAAATCTCAAGTTAACAAAAAGCCATTTAAAGATAATTCTGTTTCTTGGCATCATATAAAGGAGATTCTATTAAACGTTGCAAGCGATGAGCAACTACCAACGTTAGAGTTAGCTAAGGAAATCGGTAAAGTTCAAAAGATGCTATCGACTATGAAAAGATCAAGTATCCCTTGTGCAGCATAGTTTTTATTATTATTTATGTAAAGTATATTTATGTAAAATATTTATATATACG
>evgR001571c0g1_i01_Hydra
TCGGTTTTATAAAACCACGTGATTGAAACGCAAGTTTTTGTTTATTTCACTTGCTTTACAAGGTGAGAAATATAAAAAAATTAAGAATGGATGTTCTTACATCATATTTATCTGATGAGGAGGAGCATACTAATTGTGATGAAGAATCTCCAAAAGTTGAGAGCAATTCTTTAGAAAAGTCTAATTCAGATCAATTTTCAATAGAAAATTTAAAAACAAAATATCAACTAAATATTATACCTCCCGTTATAGATCGGAAGTTTGAACATGATGAAAAAAGATACTTAGTTGACTCCTCAACAAAAGAGCTTAAGCTGAATATCAAATATGACGATCTCTTCAAACCTGAGATTGGTCCTGTTCATCCATACAAAACACAGCAACAGTTAGCTAAAAGAAACACATTAGCTGGTTTTGTAGAAACAGCCAATGTGAGTGATTTTCAGTTTGACAATCAACATAAGACATTTATTAGCTATGGATATGCATTGGATCCAAGCACTGAAGGGGAAACTTACGTTGGAGCTGTTGCAAAAATGGTTGAAAATGGAGGAAAGACTGTATTTGAGTCAACTAAAAAGCGTCTTGGTGATAAAAGAAAAAGATTAAGTCCAGGGAATCCGGAAGATATTGATGGATATAAAGGTCCGTGGGCTCCATTTGTTGATCAAAAGGTTTCATCTAAACCAAACGAGGAGCAACAAACAATTCTTACTGAATATGAAGAAAAGAAAGAAAAGAAAAGCAAAAAAGATGATGAGAAAATTGAAGAAAAATCACGTGCACATATTGACGACTTGTACGATTATCAAGGTCGATCTTTCTTACATATTCCTGTTGATGTGGATGTTGATTTAAAAAACGACGAACCACCAGAAAAGTGTTTTCTTCCCAAGAAGCTTATTCATACATGGTCCGGTCATTCAAAAGGGGTTACAGCAATTCGTCTTTTTCCAAAATCCGGTCACTTGCTCTTGTCTGCGAGCATGGATTGTAAAATTAAGATTTGGGAAGTATATAACAAAAGAAGAAATATCCGCACTTATATAGGTCACACAAAATCAGTACGTGATATTTGCTTCAATAATGATGGTACAAAATTTATAAGCTGTGGTTACGACAGGTGGATTAAGTTGTGGGACACTGAGACAGGGGAGTGTTTAGGGCGTTATTCAAATAAGAAAATACCATACTGTATAAAGTTTAATCCGGACGAGGACAAGCAGCATCTTTTCATAGCTGGTATGTCAGATAACAAAATGATTACTTGGGACACAAGGGAAAACGAAATTGTTCAAGAATACGACAGGCATCTTGGTTCAGTTAATACCATTACATTTGTAGACAAAAATCAGAGAATTGTAACTACTTCTGACGATAAAAGTTTGCGCATATGGGAATGGGATATACCAGTTGATGCAAAGCTTATTCAAGAACCATCAATGCATTCAATGCCAGCTGCTACTTTATCACCAAATGGAAAATGGCTTGCCGCTCAGTCAATGGACAATCAAATTTTAATATATAGTGTACTTGGTCGCTTTCGACAAAACCGTAAAAAGGTTTTTAAAGGGCACATGAATGCTGGATACGCATGTCAAGTGAACTTTTCTCCAGATATGAGTTATCTTGTTTCTGGCGATGCTGATGGCAAACTAAATATATGGGACTGGAAGACAACTAAACTATACTGCAAATTTAAAGCACACGATCAAGTTTGCATCGGTTGCGAATGGCTCCCTCATGAAACTTCGAAAATAGCTACTTGCGGTTGGGATGGCTTAATAAAACTATGGGACTAAGATTATAGAAGAATTATAATAAAGCTAATTAATAAAGTGCTACGATAATAATTATAAGATATGGAAATTATTCCACCAGACCAGGAAATTATTTAAACAACTATTCGAACAATCCTCGTTATTTAAATTCGTCGACAATGCGGCGCATTGTAATATTTGTTAAAATCCGACGGTAATACTACGCATTGTAATATTTAGTTTATTTAAATAAATACTTTATGCGAAAAAAAAAAA
>evgR001575c0g1_i01_Hydra
AAAAAGAATGTTCTCTATAAAACATTCTCTCGTGTTGTATCAGTTTGTTCAACAACTACTTATGCTAACTTTGAAAAATTGTATTCAATAATGCTTTCCTGGGTTATTGCCGCGTTGGTTATTGACACATTGGTTATTATTTTAAACGCTTTGATCATACTTTTAACAGTTAGAAAAGGCAACACTTCTAACCGATTTGAACAGTTTATTTTAAACCTAGCGGTCGCGGATGTTTCATCAGGAATCGTTATGGCAATACAAGATTTTTTATTGTTAAAGTTTTTATTGTATAGAGTTTTTCCGAAAGACACACTGGGGATCGATATTTTAGAAGGTTTTTTCCTCTTTTTTCTATTTGCTTCTCATGTCAATGTTTTGTGCATGGCGATTGAGAGAATTATCGGTATGTTATTTCCAACTTACTTTTACAAATTTAGTCTGTATAAATACAAAGTAGCACTTATTTTACTACCGTGGGGTGTAGCTTTAATATCCGCTACTGTAACAAAGTATTTGATGAACGAGGAGTTTGGGTATAAATTTTATGGCTATATTTTACTTGTTGGGTCCTCTCTGATTTTATTTTTGTATATGATATTGGTTGGACTGCTTATAATCAGAAAAATAACAACATGTGATATGCTTTTCAACCAGAATCAAGACTTAAAAAATAGCGGCATATGTATTTTAATTGGACTTTCTTTTTTTACCTGCTTCTTCCCTACCGCCTACGGTCAAACTTTCAAAACATTTGCACACCCTTTTTCTACTTTGCTTCTAAATTTATATCATCTAGTGAATCCCTTGCTTTACTTTTTCCGCCAATTTTATTGCTCGACAGTACAGCCACGAGTGCCTAGCAACCATCGTCAAAGTAACATAAATCGAAATGAAGAAAACATAGACCTACATACGTACAGCGATTGAAACCATTTTTTAATAGAGTTTTAGTTAAGGACTTTGGTGAAGATATATAGTTTGTAAAGATAATTTTATATTGATTTTTTGTTTTGTTTTGATATTATATTTAAAGATAACGATAGCAATGGCTTAGAAACAGGGTGCGTGGGTGGAATTTCGATTTTTTTAATTCTGGCTGCCTTAGTCTCCCTTAATTTCCGTCTCACCTCTCCCACAACTTTATTGACTTTCCTACGCCACTGAATGATCGTTATGAATAATCTAAGCTATCGTAAGTTATTCTAAACTCTAAGTTATAAAAAAAACGGATTTTTTGTTTGTTATAATTTTAGTGGATATGAAGTTTCTGGCTTAGCAATTACTCCAATAAGCTTCTTTGCATCGTATCAACTTTTTTTGATTTTTGGTTTTTGTTCTATTTCGACTGGTC
>evgR001577c0g1_i01_Hydra
TGGCGACTATCGTTTCAAGAATAACAAATCTGAATTCTATAATCACTTGTATTCTATTGAAGAAGGTTTGAAGTGTTTGGTTTGGCCAACAACTCCTGCACCTGTTTCACACGTTAAAGAAATGGTTAATGCTGCTCAATTTTACAACAATAAGATCTTGATGACCTTCAAGAATACCGACAAGGTTGAATTGCATAGAGGTTTTGTCAGTCAATTCAAGACTGTCATTGAAGAATTGGCCGTCTACGTTAAGGAATTCCACATGAGTGGTTTGTCATGGAATGCTAAGGCAACTACTGTTGCTACTGCTGCTGCTTTGAATGGTTCAGCAACCACCACAACTCCAAGTGTTCCTTCCACTCCTAGTGTTGGAGGAGGTGTTCCAC
>evgR001585c0g1_i01_Hydra
GGTGAAAGTTTTAGTTAGAAATATATATTTATGTATATAAATATTCATAAATATGGCTTTAAATAATATCAATTGGTATAAAACAGAAACAGTATGTGACGCTATAGAAAACTTTCATAAATCAAAAGCCGCTGTGCCAAACATACAAAAGGAAAAAGAAGATCTTAAAACGATAATGAAATTACTTAAAGTATTTTTAAAGGAAAATTACAATTTATGTGATCGAGTTTGGTACTCAGGTAGCGCATTTGAATCTTTAAATATACCCGAAGAAGTACCCGAGTTTGACGTCATGTTTGTTGTTCAAAAAAGCGATTATCTTGATTTGATTGACGAATCGAATGGATTTTGTACTATCAAATGCAATAGTAAGATGATATGCTATCCTGTTGATAATAATAGCAATCTTGTTGCAGATAAATTTCGAAATGAGTTTAAAGGACTTGTTCAAAAATGGGTAAACATTATGACTAAACAAATGAAAAAAGAAATTTGTGTCGTTGAACATGGGGTTGCAACACAGATTGATGTTTCACTAAATCGCAAGTTGTGGTATCATGTTGATTTAGTTCCATGTTTTGTAATCAAAGATTCAAG
>evgR001589c0g1_i01_Hydra
TATATTTTATTTCCTTTGAATTTTTTTTATTTCTTTAAGACAATACAATACAATTCTCTAACGCATGGGCATTCTCTTAAGAAATGTCAGGGACCGTGCTCAAAATTAATTGAGACAGTTTTGCAGTCATTGAAAGTTCAAAGACAGGCTTACCATCGAAAAAGTTTTGTAGGCAATCCTGTGCATAAAATGCTTAAGAAAACATCGATACTTAAGTTATGTAACTCGATTCCAAAGTTTGTGTACGATCAAGGGTCAGGTATCAGGCACAAATGTGCATCAACGAGCAATTGAGATCAGCACAAAATATAAAAAACTTTTTGATAAGTTTGCTCAGTGCTACAATATATTTTCTTCTAAAAGTACTATGACATCAGATACATTAAGGCTTCTGGAAAAAGCGATTGATGATTTAATGATTTTTTTTCGTGCTACTTGGCCAGATGCTTCAGTTACACCAAAACTCCATATGCTAGAGTCCCACGCTGTTCCATTTTTGCAAAAATGGGGGGCAGGATTTAGTTATTATGGTGAGCAAGGTGGGGAATCTATTCATATGGAATTTAACAAATTGAAAACAGCTTATCAATCAATTCCATGCCCAAGTATGAAACTAAAAAGTATTCTGAAAAGTCATCATCAAAAGACAGATCATACCAAAAAGATAAGAACATACAACTTAGACCTCGTTTATTTAAAAG
>evgR001592c0g1_i01_Hydra
TTGGTTTTTATTTATTGGAATGATAGTATTATACTATTTATTTGAAAACGTTACTTATATAGCAAGTTTGCAAAACCAAAAATATGCTCCCGAAATTTATTATACTCTGAATGATAGCTTGATATACATAGCTGGAGTTACTTTACAAAAAGATATGGATCCAGTTAATCCAGTAAAAACAGGCGCAAGGTTTACAGCAATAGTATATACGTTTGGAATGGTTATTCTGGTTAGCACTTACACCGCAAAGCTTGCAGAATATAATATTCAAACGCAGGAGAAAAATCCGTTTCTTGGATCAAAAGATGAAAGGATCCGTAACCCAACAACGGGTTTTAAGTTTGCAACAGCAAAAAGCACGAGCTTTGTATCGTTTTTTAGATCTTCTGATATACCCGAATACCAGAGAATTAGTAAGTTCATGGATACGTACAACGTAGACACGTTGAAAGAAGGAGTAGATTGTGTTAAAAATGGTTCATTAGGAGCATTCATAAATGATTATATCTTTATGTTTTGGTATGCATCAAATGACCACGATTGTAAACTTCGCTTGTTCAAAGATTCCATTGCTAAATCTAGCAACTCATTTGCTTTAAAAAAAAACTCGCCGCGATTGGAAGATATGAACAACGCAATTCAACGCCTAATAGATGAAGATAAAATTAATGATATGCAGAATAAATGGATGTCTAAATTTTGCCAACCCGTTTTCTTTGTTCAAACTAAAACTCCCCTTGGTATTGATAATTTTGGCGGTCTGATAATGGTTCTTGTTGTAACTACGATTTTATGTTTTCCACTCTTGGTTGCAGAGAACTTGTATTACAAATACCATATTCCTCAAACACTGGGAATACTTTTAAAAAAAAAACTCGGTCAAACTAATAACTTTAAAAAAGAGACTATAGAAGAAATATGTGGCGAGGATTTAAATTCGTCCGTTATAGTTAACCGCTCAGGCGTTGTTAAGTTTAAGGATGCAAAGAAATTAGAGTTCTTAAGTTATTGAGAATACTATTGATACAGATCTTTTAAATTTTAACTAATCGGTAAAAGGTTTTGAGAAACGTTTTTGCTTTGTTAAAGCTCTTGCTATTCATTAGTGTACCGGTTGCGTTGCGTTTCAGTAATAAATTTCCA
>evgR001594c0g2_i01_Hydra
AGCTTCAAATAATTGTAATTGCATTATTGTGAGTGATAGTCGTAACAATAGAATTCAATTCTTTGATGCCAAGACTAAGGAATATAAGGCAACACACAAGACACCTAAGAATCCAAGATATTTGGCAATTGAACCAAACTTTGATGGACAAAATAACGATGCTTTGATTTATTCATGTGATAATCATTGTGTTTACAAGTATAATTTGAAACAACTTGTGGCAGGTGAAAAGGTTGAACCAATTTGGACTTCTGGTAAGAAGAAGGGAAGTGATATCTCATTGTTCCATGAACCAGCTGGTGTCATTGTTTGTACATCAAGAAAGATTAATCAAATGTTCCCAGAATTGTCACAAACTGGTGGAAACACTGTTTTCGTCTGTGATGGTGAAAATCATCGTGTCAAGGTTTTGAGAGCTTCTGATGGTAAACCACTTAAGGTATTCGGATCCAAGAAAGATGATAAAGATGGTATTGTTTTCAATCAACCAGAAGGTATTTACTTGACAGAAAACAATCAATTATTTGTCACTGAATACGGTAGCTCTAGAATTCATGTCATTGGTAAGGACCAAACTGAATTCGTTTCTTTGAGAACATTCGGTAAGGAAGGTCAAGGAAGTATGGAATTCAACAATCCACGTGGTCTTGCTGTTGATGAAACACTCAAGAATGTCATGGTTTGCGATGGTGGTAACAATAGAGTTACCTTCTTCAATCAAGATTCAGGTGAACACAGAGCCACTTTCAATGGTGGTAAGGATCAATTCGGTCGTATTGTCGGTTCATACTTTAATGACTTGTCTGGTGAATTGTATCTTGCTGATTATACCAATG
>evgR001596c0g2_i01_Hydra
CTTCTATTGTCTAAATAAGTGGAATAATTAAATGAACGATAACAAGGGATGAAACTAACTATATACTCAGCAACTTTTTTAACGTTTGTGGCTTGTGCAACAAATGTCATCTTTTTGGAACATCTTGTTAAATTTTCTCCGGGTTGTGGTGAACTTGTAACTTTTGCACAGTTTTTATTCATTGCTTCTTATGGTTTCATCACAGTGGCAAAGTTTGGAACTGAGCGACCCAAAGTTCCTATTAGGGAATATCTGTTTGCTGTTGTTCTATTTTATGGAAGCAGTATATCTGGAAACCTTGCTTTTGACTGTCATATATCTATGCCAATACAGATGATTTTTAAATCAGGATCAGTTATGGCTAGTATGGCATTGGGTGTCTTGTTGCTAAAAAGAAGTTACTCTCTCACTAAGTACATATCCGTTATCATGATAACAACTGGTATTGGAATGTGTTTATTATTCTCAACAAAAGGAAAAAAGGATGAAAGTGAACCTGAAGTAAACTTTTTCACTTGGCTATGGGGTGTGTTTCTATTGACATGTTCTTTGTTTATGGGTGCAAGATTAGGGGTTTGCCAGGAAGAGATATCATTAAAGTATGGAAAGTATCCAGAAGAGAGTTCTTTTTATTTACATGCTTTAGCACTTCCAGGATTTTTATTCTTCTCAAAAAAGATATACTCCCAAGCATCTGTTTTTACAACCTCAGAAAACTTTGAGTTACCTATTATTCAAACTGGCTTTCCAATAATTTGGTTGTACTTAATTGTAAATGTAGCTACTCAATTTGTTTGTATACGAAGTGTTTACCAACTGTCAACTGAGATGTCGTCGTTGTCAGTGACGGTTGTCTTGACTTTAAGGAAATTTCTTAGTCTTTTGATTTCAATATATTATTTTCAAAACCCGTTTACAATTTATCATTGGATTGGCTCAGCGTTGGTTTTTACTGGCACGCTGCTTTTTACGGGTGTTCTTCAAAGTGCTTTGTCTTTTTCAAAAGACACTGCAAAGAAGACGAACTAAAATATTAAAAAATAATCCACAGATATAAATATGTGTGGGCAAACGCAAATCTTTGTATATTCATAATATATGTTAGTGTGTGTGGCCACGCGCTAAAACTATCTACGTTTTTCAAAAGGTAAAACCGGATGGCCCGATGCAAACAAGATATTAAAAAAAATATTTAATTTTAATATTATTTATATTAATAAAGTAATAATAAAAGTTATTAAATACTATTTGTATTTTTTATTTAAAGACATTGTTATCAAAAACTTATGTTATATAGTTATTCCGAGGAATTTCTGGTATAGTGCAGCTTCGTTATGTAGGTGAAATGTTTTAATATATCATACATATCATATCATT
>evgR001602c0g2_i01_Hydra
TTTATCACACTCAACTCTATAAAAACTACAATTAGGTTTAAATATAAATAAAAAAAGCAATTTTAAATCATAACAAAGTTTAAAATGGGTAAATGGGGAGAACGATTTGTAATGCTTAAGAAGGTTACTGGCAAATGTTTCTATAGCGATCATCGCAATGGTTACCAAGAAACAACTTGCTGTAAATCTTTAAGGGTGCTTGTTCTTGGTGCTCCAGGAGTTGGTAAAACAAGTGTACTAGAACAGTTTTTATATGGAAAGTTTTGCAATGATTACTTGCCTACCGTATCAGAAGTTTATAAATCCAGAGTATTTATCAAAAAAGACAAAACTAAACGAGAAATAAATTTAGAACTTGTTGACTTCAATGGAAAAATGAATCGAATGTTCACAAAACTTTATCAAGAGGAAGTGATCAAAGCAGACGCCTTTATCATTGTTTACAGCGAAAATAGTGAGCGAAGCTTTGCTAAAGTTGTAGAAACAGTTGCTGATATCAACAATATTCGAAACGGAAGATCTCCGATTCTTATATTACAAAATAAAATAGACTTAAAAAGTTATGAAGAACTTTCCAAGTATTTTCATATGATAAACACTAAGCATGCAGAGGTATCCGCTAAAGACAACAAAAACATTCATGACGCTATTGAAGATTTGATTTTACATCTTGAATCTGATTTTTAATCAATAATAAAATAACTCTATTTATA
>evgR001608c0g1_i01_Hydra
CTTTTATTTTCTAAATAAGTTGTTTAGATATCGACACAAAATATTTGCATGTTGAAATGGATGATTCTGAATCTTTTGAGGTAGAGATGGCTGGTTCAGCAAGTAATCTTAAAGTAGACACCATTTCTACGTTGAATCGACTGCATGACACATCAACAACATCAGAGGAATCAAATTCTTCTTTAAGTGATCTTGCTTCTCCTGAAATGTCACATGAACCTAAATCATTGCGATTCCAGCTGAAAACTGATTTTGGTAACGGCTTAGAAGAATCGTGTAAACAAATGACAGAATCAGTGGTAGCCTATCTCAATGGCGAATTGACAGCAACAGTTTCAGATTACCAACTGTTAACTAAGATGAATACTTTAGCTGCTGACAAGTACCAGGAAATGTCATCATCAACAAACACTTTAATATCGAAAATGGTCGAGCTAAATGAATCGTATAAGCTGCTTCAACCTTACTTGATACAGATTGATTTATTGGAGGAGAGTGTTAACACATTAGAGCAAACAGCTTATAAGTTAGACTCATATTCGAAGCAGCTAGAAACAAAGTTCAAGAAAATGGAATGGAAATAAGTTGAAATGAAAGTTTTCTTAAAATGAGAAGCAGGTTCAGTTTTCAGTAAAATGAAAAAGAATGCTACAAACTGAAGAAAATGTTTGTTTCACTTGTAAAAAAATTTTAAAGCTATTTTGAGAAATCTTTTAGTATTGATTAAAAAAAAAAAA
>evgR001609c0g1_i01_Hydra
GTTAAAAAGTGATGTCATAGCCTAGCTAAAAGTATTTGGATACAAAATTTTTGCTGTTTATAAGTTAATTTTTCAAAATTTAAGGTTACCTAACGTAAAGAGAGTTCAAAGACCAATTTATAACTGCGGTGATGGTGGAATCAATTTTTAACTATGTTCTGGAATGCATCAAAAGAGTTATCAGAAGTTGCCGACCAATGGATCCAACTTCCATACACAATCATTCAGTACATTGTTCTCATCCTAAGTATATTAGGGAACTCACTTCTTATTGTTACACTCGTTCGCATAAGAAAAATGAAAAAATTTGTTGATAAGTTGATTTTTAACTTAGGGGCGTGCAATCTTCTTATGGTAGTTGTTTCAATACCAGCTGATATTTTGCTTCGAGCATCTTACAATTTTCCATTTGGCTTGTTTGGATGTAAAACAATTAATCCTGTTTCCACGTACGCAGTTAATACAGGAGTTTTTACATTATTGGTAATAGCTTTTGAACGTTGGAAAGTAGTGGCCCAACCTTTTGGATTAGTCAACAAAGGAATTAGAGCAGCACTCATTATAGTGACAATTCATGGTTGTGGTCTTGGCACTATCATACCTTATATGTTTACTTTACGCCTTAAACTCAAAGAAGGAAAAATAATTTGCGAAGAAGCATGGTCTAATGATTCTGCGCGTGTATACACAGTTGTCCTATTTGCTGTGCAGTATGCAGTGCCAGTTCCTATTATGTTAGGATTTTATACAAGAGCGTGGTTAATAGTGCAACATATGAACATAAAAATTGTCAAGCCTCATTCCACACCCTCAGATACGTCTATACGAAATGAGAGAAGCTCGATTAGTATATTTTCAAAATGCGATCTTAAACAAGCTAGCGTCTTGTACAAGATCCATGAAAATTTAATGAATAAGAAAATGGGAAAGTTATATCATCTTCCTCTTCCTCCACCACAAAAGGGAAATTCGGTTACTTTAAGCAGATCAAATAACTCAAACTTGTTTGTAAACGACTTAACTTCTTTAAGAAACAGCTTCCAATCAAATAAAGATTTCATGCTACAATCTAATATAATAGAACAAAATAAAACTATATATCTTCCTAATCGATTTGAACGAAAATTTTCTACTAAAATACTTGATGGCGATAATGAAGAAAAAATTCGACGATTTAGTTTAGGCTATCCGCGATTTGCCAAATTTTCTCCAAGAAGGTTTTCTTTAAATGACAAAGACTTTTTCAAAGAAAAAAAAAGTCCTTTCAAAACTCAAAATGGCTGCTCAAACTACAAGCAAAATACGAAAACACGGTCAAATCAGATGCTCTCAAGCTATATCTTCCCAAACATTCTAATAGAACCCACTGCTTTACAACACAAAGCGCTACAACGAAGATTTCGTCAGAGTCGAGACTTATTAAGAACATTTACTGTTGTAGTTACAGTTTTTGTGATATTCATGCTTCCTAATCAAATCTATTGGATGACCTTGTATTTTAGTCATATATCGCAGAATGAACGTGCTGTTAATATTGTCTATATACTTACGTACGCTAATTGTGTTTTTAACCCTTGGATATATGGGGGGTTAAATCGAACATTTCGAAAAGATTTCAAATATTTGC
>evgR001617c0g2_i01_Hydra
TCACGTCATTCAAAGATAATCTGTTATTTAACTCAATGAAATGGCTGGATCTACAATATTGGACCAACGAGCAAGAAAATGGTTCAAACGATATTACCAAATGGCTGGATCCTCAATATTGGACCAACAAGCAAGAAAATGGTTCAAACGATATAAAATACATTATGCAACATTTTAACCAATCGCTTGCAGAAGCCAATTTCCAGGAAGCATATGATAACGAAGATAATGTTTTCCAAGAGTGGAGTTTGGTACAAATAATTGTAAAAACACGCTATTCAAAGTTACAGTGTCTCAAAGTTTGGGAGAAGTTATGGGAGTTTGAAAAACCTTAATTTCCAAACATTTTTCTGTTAGCTGAGTTGTGCTTCTGTTTGTCGTCATCAAACACCGCAGTTGAAAGATGCTTCAGCACACTGACGAC
>evgR001619c0g1_i01_Hydra
AGTGCAAAAAGGTTTAAACTTTACATATAGCCTCATAATAAAAACATGAGTATAAAAAATTTTGGTGCACTTTTTAATGGTATCACAAGGCAAATCTTTTATGAAGATGCAGAGATTACAGATGAATTTCTAAAAGAGCAGTTGTACCCTGAGATAACTCAAGAAGAGTTTCAGATTCTTTTGTCAAAATGTAAGTCATGGTTAGAAAAAATGTCGTACTTTAACATGGACTTGAAACAGCTTGAAGCTTTTTTGAAATCTCAAATGACGAGGAGCAAAGACAATTTAACTGAGGACCAGATGGCAGCGATTATCAAGTATTGGAAAAGTAACAAAGATAAGATCCATGCTACATTATTGTCAAGAACGAATTGGGAAGATAAACTTAGAACGGTTAAATGGCAAATAGATCTTGGTTATAAAAAAAATAAAGAGATAGACAATTCAAAAGCAGTTTTTCAATTAGGAATAACAAAATGGAATGGTACGGATGGAGAATTAGAATTTGAAGTAGACAGTGAATCTTTAAACAGTATGATATCACAAGTTGAAACAATTCAATCAAAAATAGACTGTTTATTAAAGTAGTATCCACATAGCGTACTGAAGATTATGAATTAGTATTGAAGTATTATAGTTGTTTTAAATTATTGCCGTACTCAAAAATCTTTATCGAAATTATGAGTATTATTTTGTTTTTCAAATTTCTTCAATATTCTAGACAAAAATTGTTACTGTTCTCAGAATTTGTATACTTAATATGTATCCTAATATAATTCTATAAATTAAGACTTTTAAAGTGGACCAATTTTATATTCATATCTTTTTTTAATTATTTTTATTATATAGAAACATAATAAGTTGTAG
>evgR001628c0g1_i01_Hydra
CTTGCTTTCTCAATTGGCTTCTCAATACATTCGTGTTATTTTGTCAGGTGAAGGAAGTGATGAAATGTTTGGAGGTTATGTTTACTTTAGAGATGCCAAGGACTCTCAACAATTGCAAGATGAATTGGTTACCAAGGTTAGAAACTTGCATTTGGCTGACGTTATGAGATGTGACAGAGCAACTATGGCTCATGGTCTTGAAGCAAGAGTTCCATTCTTGGACAGAGACTTTTTGGACCTTGTCATGAACATCCATCCTGACCACAAATTGCATTCTTATCATGCTGATGATGGCGAGGAACACAACATGGAAAAGTATATTCTCAGAAAGGCATTTGATGGATTGGACTTGATGCCTGATGAAATTTTGTGGAGAACTAAGGTTCAATTTGGTGATGGTGTTGGTCATGGTTGGATTGACTCATTGAAGGAATTCTGTGATGCTCAAGTCTCTGATGATGACTTTGCAAAGGCCGCTGAAAGATTCCCTTACAATACCCCAATCTCTAAGGAAG
>evgR001629c0g1_i01_Hydra
TTATTAAACTTGAGGACGTGGATTTTCAACATGCTATTGATAAACAATTTTATATTCATGAAAAGGATTCATATTTTCTAATAGAATCACTATATAAAAATAATGGGAATTCACTTTATATTGGAAAAGAAGGTGATCATTTTAAAATAAAGCAAGATGCAAGTATACTAGCTTTTACTCTTGTTGTGATTGGTTTGACAATTGCAAAGATAGAAAGCGTCGATAGATATTCAGTCAAAATTACAATAAATTCTAATTATATATGGAATATAACAACTAATTATATAATAAAATGCAAAGATCTTAATAATTTACACAGTTCAAGTTATGCAGTCACTAATAATGAAGCTTTTATTTCTTCGTTGCAAAGCTACACTTGGTACAACTTTTCTGCTGGTATGAAAAATGAGTTTGGTTTTTATGTGTATGGGCGAATGTCTTCCTTTCAAACTGATGAAGATGTACCTGCTGGGTATCCAAATAATTTTACAGTTAAAAAAGTAAATTTTACTACTCTGTTTCTATCATGGGATAATGTTAAAAGTTCGATGAGAAAAGGAATCATATTAAATTATACATATTTGTGTAACAGCATTGAAGATAGTGCTTCAAGCATAGTAGACAGCAGTACATTTAATGTAACAGTTTACAATCTTAATCCAGATGTTATTTATAACTGTTCGGTTGCGGCATGCACCAAGGTTGGTTGTGGTTTGCCGGCATTTGTGTATGGTTTTACATTAGCTGGAATACCT
>evgR001631c0g1_i01_Hydra
TTTGTAAAATGAATTCAATAAATTAAGTTAGGACAACAGAAATTTTTACTAAATCGTACACAAGACTCTTATTTATTAGGAAGAGATGAGTACATTAAACGTCATAGTTATCTGGATTGGATATGTAATATTACCCATTGAAATACTTGGAACATATTTATTGTTTAAAGCAAGAAGAAAACTCGATAAAGTTCAGTTTTTATTACTGCTGAGCAACTCTGTTCTTGATATCAAAATGACCTTTTTTTACCCTACCGCTCAGTTATGCCGCATCAAATTTCTTTCAATTTATGCTGATGCAACTGTTGCGTTGGAATTTTACTGCATTGTTTTCATTATAACGGCTAATCGCTTTTCTGAAGTATATCTAAACTTACGTTATCATCAAACATGGACATATCATCGTGTAAAAGCAGCTTTAACTATTATTTTCATAATATCACATTTTTTATTCGCATGCATTATGAGCTTTGATAAAAGTTTTAAAACCGAAGCATTTTTTTACAGGTATTTATTTCCTATTCTGAATATTCTTATACTGTTAACTGTCGTGGTCTTTTATGCTTATATATGTTTTAAAAATTGTGACAATAAAATGGAAAATTTAAACAAGAACAAAAATCATGCAACCTTATTTATCGGTCAACCATTTTCTAAAGAAATAATAAAAAAG
>evgR001632c0g1_i01_Hydra
AGCACTTTCAGAGTTTTTGTGTTTTATAAAAAATATGAAGTGAGTTTGTGACTTATGTATGATTTATAGCTTTTCATAATGAGTGAATCAAACAGATTTTCTTGCGGTACGTTAAAAACATTTAGTACGCCTTCGTTCTTGAAAGATCGCTTAAACGCCATACCTAAAGTAACTTACACTTCAGTTGGTTGCATTCAAAAAAACGATCGGTATATTCGTTTAAAAACGTTAGTTCATTATGCCAGCTTGAAACTTTATTCCAATTTGTTTCACAGCCCACAATATGGTGGTTTTCCGACCAAAAAAAAGGAGCTATATAAATCGCTAAAACAATTTGAAAGTAAACTTCGCCATTCTCGGGATACTACTATTACTGAGAGTGACTGGATAAAGTTATTCCCTGATAATAAAAAATCAAATTCACTAGAATTTTCTGTTGAACTATTTCAATTTCTAGTTAAATGCTGCGTGTCACGATTTCCTGAGACTGCAAAAGATGAGAGCGATGTCAAATTTGAAACAGATACTTATATGCCATTAATAGAAAAACTTCACGATACAGTATATAATCCTAAATTTGCTCAAGCTGTAAATAACTCACACGATTATTTTATTTCTTCGTGGAATTTTCTTCAAGGAATTCTTCTGAAATTAAATTTCGATGTTGATCTCATAAAAGATCTTGAAAGTGGAGATTTAGATAAATTAACAACTTGTCGCTACTCTCTTGTCACAATACAACTGCAGGTACTTTTAGAAGAATGCAAGGAGCTTGAAAAATCAACAAGTATGAATGGCCTGGCTGTCACAAAACTAATTTCAGATTATAATGAAATGATATCCAAACAGGAGAAAGGCTTCTATGATGAAGAGTTCCCTAAAAAAGCAAAATCGCTTCAAAATAATATAGCTTTAGTGATGCAGCTTTTAACATTGATAAAGTTGAGCATCTCGGAATTAACATTAATCGTTCCACATTGGAAGGGAAAAAATATAGAAGGTGATATTATTATTTATGAAGAAAAACTGTCTAAGTTAAGAAACCAAGTAAATGAACAAACTGGAGAGCTCGGGGAAATTTTCCATTCAACATCTTTAGATATTGTAAAAATTAAGGAAAGGCTTCACTTTCAAGAGGATAATATCCACGTAATTCACCGGCAACGGTCAACTAGTTCTCAATCGTTAAAGGAAACAGTCAACTCTTCAATAAACTAATAAACCTTGTAGTTAAGTGCATTAGTTACCTGCAATGGTTCCTTTAAGCAAATAATAAAATAAAGTAAAGATTGCAATGTTAAAAAATCCCCCGCTCCTTGATCCGCCACTGTTGCAATGTTAAAAAATCCCCCGCTCCTTGATCCGCCACTGTTGCAATGTTAAAAAA
>evgR001637c0g2_i01_Hydra
ATTTTAATAGAGTAGATGCCAATGTCATTGAAAATATTCTGGATTACGTTTTTTCTTCCAAAACAACAAAAGCTTTTTCACCATTACTAGATTATGCACTAGATCCAACCCCTGTTATTCAATCATTTTTGCTCCAGCAACTTCTTCATAGGAATTACAAGGTTCAAGTTCAAACATACCTCAGCCAATATTTAGCTACGCTATATTCACATACAGATCAAAGTCAGCTTCTTGAAGTTGCAAAGCTTTTCATATATTGCATTGAGGACAAAAAGGAATTAGAAACTTCTATGCTAATAAAGGAGACAGTTGTTAACTCAACTTTGCTTGAGAACATATACGAGAAATTGCATGCCAGAAATAATTTTGTACAATCAGTCGAGCCTGGAGAGTCTATAACAAAAATTTTATTACTTAAAATAGTTGCTGAATTGAGGGCAGTTTTAAAGATATCAGCCCAAGTGTTGTGTACCTGGTTTTTTAAAGAAAGTTATGAAGAAAGACAGATTAAAGAACACTTTGATACTTTTTTAGCTGAATTAACAAAGTTTTTTATCAAAAATAAAGACAGTCAACCTTACTTTTATTTTATGAAGCAAATTGTTCGAAACTATGGGATTCAAGGTTTAAAAGAGCTTGCAATAAAATCTAAACTAGATTGGCTTCTTAATCAAG
>evgR001645c0g1_i01_Hydra
GATGGAATTTTTATTTTGTTGAAAGCAGCTATTTTTAGGACAATACAATAAAAAAAATGATGCGAGTAATACAAAAACAGTACAAATTTTTAAAATCCAACACAGTGGGGATCATAGGAGTCTCTGTAAAAGAAGGTCAAAATTTACCAGGTGTTGAACTTGCCCCTGAAAAGATTCGAAGAGGAGGACTCTTTGAAGTTATAAAGAGTTTAAACTGGGACTACTCAGACAAAGGAGATATCAAAACCGAAAACATTCAAAGACAAGCACCCTCTTCAAAGATATACAAATATAATGGAATTAAGAATGCAGATATTATTGGCGCAGTTAACGAAAAGCTGCACCACACAACTTCAGAGATTGCAGAATTAGGACAATTTTGTCTTACCTTGGGAGGTGATCATGGAGTTGCTTCTGGATCAATTTCAGGACTAAAGCAGGTTTATAAAGATTTAAAGGTAATATGGATAGATGCACACTCCGATTGCAATACTCCCGAAACATCTCCAAGTGGAAACTATCACGGAATGCCTGTTGCTCATTTACTTGGTTGGATTTCTGATGGAAGTTGTCCTGGTTTTGATTGGTTCAAAAACTGTTTGAAAAAAGAGGATCTTGTTATAATTGGTTTACGAGATGTTGACAGCGAAGAAAAAGCTTTACTTAAAAAACATGGAATTAAATGTTTTTCAATGCATGATGTGGTCAAATACGGAATCTCAGGGGTAATGGAAAAAACATTCGAGTATTTTAGAAAAGACGGAGTTGATCACCCAATTCATATTAGCTATGATGTAGACGGTATTGATCCAACAGTAGCCTATGGAACAGGAACAAAAGCTAGAGGTGGGTTATTGTATAGAGAAGCTCACTATATAGTAAGAGAGGTCGTAAGTACTGGATTATTTGTCAGTATGGATGTTGTTGAAGTAAACCCTTTACTTGATAAAGAAAAGGAGTATTTTCATGGAGACAATAAGTTAATTAAAGGAACCGAAACTGTTTGCCTGGCTATTGAACTTTGTGCTTCAGCACTGGGTGATACATTGCTATAAATAGCTTAATTTCATATAAATAAACATAATCAGATATTATAAAAAAAGTTGGATTAGAAAAAATAATTAGGAATCAAATAAAGTTTTATTATTCAAAAAAAAAA
>evgR001653c0g1_i01_Hydra
TAGAAATCTGGCTTCTTTGATCGCCGCGAAGCTGATTTATGGCTTTGGTGAACACAAAAAATAACAGTAAAAAAAAAAACGTTTGAAAAAATTCTTAATGAACCACTATGCCAAAACAATTACCATACTTAAATTAGAATAAATAATTGGTCTTTAGCTCAGCTTTTAAAAAAAATCATCGGATTGATTTTTTTTTTGTTTTAAAGAAATATATTGATTGAATAGCTGAGGAATAAGTTTTCAGAAAGGTTTTGTAAAAAGGGTTAAGCTTAAGAAGTTAAGCGCGGAGTTATTTATATGGCTAGCCTTGGAACTGCATCTCACACTGTTCACGAAGTTTGCTCTCAAGTAAAGGTTGAACCTGATATTCTTCCAGATGCGCCTCAACCACAATTTATAAAACCAGTATCAATTTCTGCTACCCGAAGTTTATCGGAAGGAACGACGATTGCTGCTTTAGTATCAAATCTCTCCACTTTAGTAGGAAACTCATCTTCAAATAGAGGTGCTGTAACAAGCTTTATAACATCCCGGCCTTCAATAAATATTTCTTCTGCGTCAAACATTGCTCATATTACCAAACCTTTAGTAGTTACCAAACCTTTAGTAGTTACTCAAAATTCACAACCATTGTTACTACCAAAGCCGAGTATCCAACAAAAACAGAATGTTCAAAGTACACCTCCAAAGTTTGCCTTTCCTCCAAATCCATCAGTATTGTCTCCACATCCTGCAATTCCAAGCTATTCATGGCCAGGATCATCACCTTCCCAAAATAAAAAAAGACCCAATGATGAAGTTGAGGATCAATTTGACAGTAAGCGGCGCCGTATACCAAATCGATGTGGAGTGGAAAAAGGTGGAAAAGGTCTGCGTCACTTTTCAATGAAAGTTTGTGAAAAGGTACAAAAGAAAGGAATTACATCTTATAATGAAGTTGCAGATGAATTAGTTCAAGAATTTAGTGAACAGTCGTCGCGCAATCTTCTCTCAAACATTGATCAGACTTATGATCAAAAGAATATTCGTCGTAGAGTTTATGATGCACTCAATGTTTTAATGGCGATGAACATTATTTCTAAAGAAAAGAAAGAAATTAAATGGGTTGGTTTGCCTACTAATTCTGCTCAAGAGTATAAACATTTAGAGGACCAAAAACGTGAACGCATCGAAAGGATCAAGCAGAAACAGCAACAGTTGAAAGAACTAATTGTCCAGCAAATTGCTTTTAAAAATCTTGTTGAACGAAATAAACATTTGGAAAAGTTGAATGGTCCACCACAACCCAATGCATGTATTCATTTGCCATTTATAATTGTTAATACAAGCAAAAAAACTACAATCGACTGCAGCATCTCTAGTGACAAATCAGAATACTTGTTCAATTTTAATGAAGCTTTTGAAATTCGTGACGATATAGAAGTGTTGAAGCGAATGGGACTCTCGTTTGGGTTGGATAAAGGGGAATGTAGTGAGAAGAATTTAAACACAGCTAAAAGTATGGTTCCAAAAGCTATTGCACCTTACGTTAATCAAATGGCAAATGGAAGCTATGGTAACATTGCTCTTATTGATGCATTCCGAACGGGACTTAAATCGCCCTTTGCGACACCTGTAACGAACACTAAGGAATCTCC
>evgR001655c0g1_i01_Hydra
ATGGATGAAAATAATGTACTAAAACTTTTATCAAATTTAGATTTCTACAAAGCAAGTAATACACTAAAAATGTTTTATCGTAAGACTTATGAAACACTTGATGAAGTTGGAATACAACAGAACAAAATACTTAAAGTTAACTTTTTGGAAAAATTCGTTGAACTGTCTGTTGTTGATGCTTATTATGGCCAATTAGATGTAATTAACATTACAAAGCGAGATCAACTTTTAAAGCAACAAATGAAAACGTCAATTCTATTTAAAAATTATTTCGAAGATCCCTCTTTATTTGAAAAATCTTTTCTAATATCTGGTATTGCCGGTATTGGCAAAACATGGTTGCTTAGAAAGTGTTTACTCGAATG
>evgR001661c0g1_i01_Hydra
GGCCACCCTGTGCTTATAATCGAATTACTGTAAATGTGGATTATCAGATAGTTAGTACCTTTATTTGTGAAGGAGAGTCAGCGGAAAATATTCGAAATGCTCTTGCTAAAATAAAAGAGTGGAATCCTGATTGGAATCCTTTATATTTTATGGTTGATTGTTGTGCTGAAGAGATGAATGCCATTGAATCTTTACATCCAGAGTGCAGAGTTCTCATTTGCGACTTTCACCGGGAGCAAGCCTGGGAACACTGGTTCAATAAAGTAAGCAATGGTGCTAGTGACATTAAAACAGAAATGCTAGGTAAACTTAGAAGAATTACTAGAGCTCAAAAAGCAGTGGTTGCCATACATCGAGAGATGGGCATACTTTTACCGAAAGGACAGATTAATGGTGAGTATAAATACCAACAATGGTGTTGAGAGGCAAAATAAAACTTTCAAACATGCATATCTAAAAA
>evgR001663c0g1_i01_Hydra
TTCATTTCATTTTCTATTTCATTGTCTAATATTATTCAAACTGTTATTATGTCAATAATGCGAGGAATTGCAATAAAGTATCCTTTCTTCTACATGATGCATTGTAAAACAATGAAGTTAAGAATAATGCTTATGTTGTTTTCTTATTTATACGGGTTTTTGTGGCCTTCTTTTCCGTTACTTGGATGGTCAAAATATGACGTCGATTTAGATAAAAAGCGATGTTCGTTAGATTGGAATTTGACGCATCCGGGTTCCTTATCGTATCTCGTAGTTGTGATTTTATTTTGTTATATTCTAC
>evgR001669c0g1_i01_Hydra
CTGAGCTTTTAGCAATATGACGGCAGCATTTTTAAAACTTTGCCCATTATTAGCGGTAATTATAATTGAAAATTGGGTCCTTGCAGCTAATCACTTAGAGGAAAAATGGGAAGAACAGTATAAAAAGGAAATTGGATCATTAATAATGAACAAATTAGATATGACTGAAAGACCAAAAAACTTTAGTAAATTATATTCAAAATCGTTCATAGATAGATTAACGGAAGAACAAGAGGAAAAGACTCAAATTATTATAACAGCAGAACAAGATTGGAATTCATTTGGTGAAATTAACTTTCTTTCAAAAAACACTGTTGAAGCAGATTCTGTAATAGCATCGGCTATATTTTCGGTTTATATTAAGTCTTTTACCACAGAAACAGATCTTATGATAGCTATTCTAAAGGAAAATAACGTTATAATTTCAAGCAGACCAATATTAAAAGACTATTTTGGCTGGATTGATTTTGATATTACTTTAGCATTTCAAAAGTGGACTAAGTCAAATAACAATAGTTTGAGTTCTCTCTCTTTATCTTGCAACAAACTTGGTAAATGCCTTCAAATATCAACTGAAAAACAACATACTCCGTATTTGCATGTAAGTTTTTTTACAAAAAAAGGTCCACCACGCCAGCGACGAAAAACATGGAAGTGCAATGAAAGCGTCAGTTGCTGCTCCCGTCCAGTTTTTATTAGTTTTGAGGATATAAGTTGGAGATGGGTGTTGTCTCCTAAAGGTTTTTGGACTAATAAGTGTATGGGAAGCTGCCATAATGAGAATCCAGATAAAACTACCTGTTGTGCTCCACTAACACTTGGATCAATTACAATGTACTATCACAATGGTGACTACATTGTATATGGCAAAAAGCTAGAAAGTATTGTCATAAAAGATTGCGGTTGCCAATAGTGTAAGTTGCACGGACAGAAAAAAAAATGTCACTACTTCACTTTAAAGCTGCTCCCAGACTATTAATTCAACTCAGAATATAAAGTTATCAAGTGAAACTCAAAGATGTTGCTGGCTGCCATACTTTTAAATAAATTAAAAAGAAAAAACAATAAATAAGTTAATAAGTTTCTTTGTTATTTATTTTAAGTGTTATTTTTTTGTTTACATGACTATTTTGTTAAAAAAAA
>evgR001670c0g1_i01_Hydra
TCAAAAAACAAAATGGGTAAGAGAATTAGAGCTCAAAGAAAGGGTAACGGTGCTGGTCCATACAAGACTAGATCTATCTACAAGATCGCCAAGCCACAATATAGACATTTGGACTACTCTGAAAGAGAAGGTTTCATGAAGGGTGTCGTTAGACAAATCCTCCACGAACCAGGTCGTGGTGCTCCATTGGCTGAAGTTGCTTTCAGACATCCATTGAAGCCAAGAAAGGATATCCAATATTTCCTTGCTCCAGAAGGTTTGTACACTGGTCAATTCGTTTTCTGTGGACAAAAGGCTGAATTGAAGATCGGTAACGTTCTCCCAATTGGTAACTTGCCAGAAGGTACCGTCTGTTGTAACATCGAAAGAAAGACCGGTGATAGAGGTTCTTTGGCTAGATCCGCTGGTGAATCCGCTGTTATTATTGCCCACTCTGAAGATGGTAAGACTAGAGTTAAGCTCCCATCCGGTGTCAAGAAGACCCTCCCATCTGATAACAGAGCTACCGTTGGTGTTATTGCCTGTGGTGGTAAGACTGAAAGACCAGTTTTGAAGGCTTCAACCAACTACTTCAGATTCAAGGCTAAGGGTAGAAAGCAATGGCCAAGAGTTAGAGGTGTTGCCATGAATCCTGTTGAACATCCACACGGTGGTGGTAACCATCAACACATTGGTAAGCCATCTACTGTTTCCAGATTGACACCTGCCGGTAGAAAGGTCGGTTTGATCGGTGCCAGAAGAACTGGTAGAATTAGAGGTGGTAAGGCTAACGTCAACCAAGTTGAAAAGGATTAAATTTATTACATCTTCAAACCCAATAAATCATAAAAAAATTATCAAAAATTTATTAAAAAAAAA
>evgR001674c0g1_i01_Hydra
ATGTACTCCTTCGCTACTCTCTACAAGCCAGTCAAGATCATCCTCTCCACTGGTGCTTCTGATAAAGTTTCTCTCTCTGTTAATACCAAGGCTAACCCAGATTTCAGCGTTGGTGTTAGTGGTACTAGATCAAATGTCGATGGAAAGGTTGATGCTTCTCTCTCATGGAGCCAAAAGCTCAAGCAAGGAGGTGTTGCTTACACTGTTAACGGATCTTTGGACCAAGATGCCACTGTTGGTGTTACCGTCGTTGCTAACGGTGTTGCCAAGGGTTTGAATGCTGATCTCTCATCAAAGATGAAGACTGCTGCTTGTGATAACAAGGTTACTGGTACTTTGGATTACACTGCTGATAAGTTCCATTTAACTGGTAACTTTGACCACTCTTGTGGTAAGGACTTGCTCTTGACCTCAAGCGTTACTGGACAAGTTGTTGAAGGTTTGGTTGTTGGTGCTGAAGCCCAAATCAAGGCCTATCCAAAGCCAGAAAAGGACCAATTGTTGAAGAAGTACACTGTTGGTGCTAGATACTCCATGAAGAATATCCATGTTGCCACTCAATTGGAAGAATCTTTGAAGAAGATCAGAGTTGGTTATGCTCAACAAATTGACGAATCTGTCTCCGTTGCTGGTGAATTCGTTCACGAATTGAAGAGTGACAAGGCTCCAGTTTTCACTTTGGGTACCTCATACACTATTGACAAGGAATCATCAGTCAAGGCTAACATTAATACTAAGGGTGTTGCTAATGCCACTTACACTTTGACTGTTAACCCACGTTTGACAACTGCCATTTCTGCT
>evgR001675c0g1_i01_Hydra
TTGATCATCCAAAAAAGAAAAACAATTCAATAAATCCTGATTTTAAAGAAAAGTTTCAAGTTTCAAACTCATTAAACTTGAAAAAAGAAATCAATTACGGAAATAAAGGGAAAAAAGAAAATGCTCAAAGTAATTACATTAAAATAAATAGTTTGCAAAAAAACTCAATAAATGACGATTACTTAATAGAGGGACCTAATTCACTAAAAGATGTTCAATCAAATAAAGTGATAAACCAATCAAATGAATTGTTATCTAACCTTTCAAAGAATCAGTCCTCAAATGACAAGCTTTCAAAGAAGGAAAGTCGTAAGTTAAAAAGAAAAATTTCATTTAAATTAAAACTGAAAAAAAGGCGAACTAAGTTGAAGGAAATAAAACGTAAACAAAAAGCAAGTGATGCTGATACTGAGAATGATATTAACCAAAATAGTGTTAAAAATGGTGTAATGGAAGAAATTAGCAATGTTGTGGGGAATTTATTTTATGTATCCCTGAGACACAAAAT
>evgR001678c0g1_i01_Hydra
ATGCAACTGTGCTGCTCGCTGGACCAATTTAATGTTGCAAAACAGTTTTAAAGTTATAAAACTTAAAAAAAAATGTATATATATGCCAATAACTCTAATAGTACAAATATTTACATGAAAATGAGTTCTTATGTGCTAATCTTTGTTTCCGTTCTGTTGATACCATTAACCACATCTACTCAACGCGACGAATTTTCTCGATATACAAGATGTACTAAATCCGAGTGTGATTCAACTGGAGCTTTGTCGTATTGTGGACCAATCGAAAAACCATGTAATCCCAACCACGAAGACGCACCTGCAAGTAGTTGCATAGACTCTAATAAAAGATGTTGCCGCTCTATATGTAACGGACCCAAGTCTCATTGTCTTCATAACAGTCGAGCATGTTTACCAGGATACGAAGAAGCCGAAGGAAAATGTTTAGTAGATGGACATAAATGTTGTAAACCCGTTTCTTAAAATAAATAAAACTGTTTAAATAATTCAACAAAAACAATTAGTTAATAATAAGTATGTTAATTAGTAACAAGTAGTAATTAATAATAGATTTTAGTAAC
>evgR001681c0g1_i01_Hydra
CAGTCGAATAAGGAATAATAAATAACGTTAATATAATTTTTTTAGGAGTTATGGAAGCCACAATTTAAGAATTGAGATGGACATTCAAAGTAAAGGAAACATTTTGTTGGAGTTAATGCAGGAGGCCATAGCACATGCTGACACAAACTGTAAAAATGCAGCGAAATCAGCATACTATGGAATCGAACAAACAGTTGAAGCTTTATATAATCGTAAAAAAAATTTAGAAGAGGCGTGGAAAGGTAGAAAGAAAATCTTAGAGCAATCAGTGGACTCATGTAAACTAGATAATGAAATTAAAGTGTGTGCCCAATGGTTGTTAACTGATGGAGAAAAGTATTTAGCAAGTTTAAAAAATTGTTCGTCGATAGAAGAGTTGAGAATCATGCTACAAAAACATCACGATTTTCAAGGTCTTTTTGAGATAAAGAATAAAGATGCAAAAAAACTTATTGCTGATGGTCAGAGGTTGATTCAGCGAGGTCATTTTGATGTTGCTGCAGTAAAGTATATGATAAGTAAACTAGATCGATTGTTTATTTCTTTCCAGATAAAGCTTGATCACTTAAGAAATCTTATGGAACAATCTTTAAATTTTTTTGAATCAGTTAAATCGGTAACTGAATCATTAGAAAAGATAACTGCCAAACTTGATAACAAAACGAGTGATCCAGAAAAAGCAAAATATCGTAAAGAAGCTATTCAAATTGCTTCATTAGTTTTGCGGGAAGGTCAGTTGCTTCTAAAAAAAACAAATGATATAAAGGTTGCTGAGCTCTTAAACAAATTGAATAACGTTCACAAAAGTATTATGGCTAAGACTGATGGAACACAATTTACTGGAGATGCTGACCTTAGTGCATTAAACTCAGAAATTGAAATAATATTTAATTGGATTACTAATATTGGGGAGAACTTTTTAGACACTCACACTGTTGTTGGGGATCGACAGCAGATGGCGGCATCATTTCTTAATGAACACCAACAATTATTTGCTGAACTTGCGAACTACGAAAAGGAGTTACTTCGAATAAAATCAAGAAATCAAGAAATACAGAAAAAGAAGAAGAATGCCATTAGTGATAAAATAGACAATCTTCAAGTAAAATCTCTATCATTATCAGAAAGAATAAAAGACAGAATGAATATTGTTACTATGATGATTGCTTTTATTGACCATACAGATGAAATTTTGAAAAGTATCTTTCTTTTGGAAGAGGAACTCACCAGCAATCAATCCAATGAAAATGTAGATGCTTCTATTACCACAATCAACAGACAAAAGCAAGAAAAAGAAGAGATCATAAACAGAGCAGAAGTTTTGCAGGAAGAAGGTCAGAGAATATTGGATGACTTAATTAAATCTGAGAAATCTGACAAAGCTCTGGATCAAGCATCAATTAAATTGAAAATTGGAAAAGAACTTAAAAGTATTGTTGTGCATTCTATGCGGCTAGAAGAACTTTGGGATCAACGTGAAAGCCAATTAGACTCTCACAAAAGACTCAGTACACAGATTTTTGATTTTGAAAAGGAAATTCGGAAGGTGATAAATGGAGTTCACAATGAAAAGGATGATATCATTCAAACATTTGTAAATGACAATCAAAATGGTTTTAGATCAATTCAATTAGAAGAAAAGTTACAACAATTCAAGAGTGTTGCAAAAATTGCAAATGGAGAGATTGAAAATATTATGAAAGAAGAATTAAAGTTTCAAAATGAAAAGTTGAAAGAAAACAATGAGACATTCAATTTAACACTAACTCATCAGAGACTGAAAACCAAGTTAAGCAAATATGAAGTTCTGGATGCCATGTATTCTACATTCTATTTATCTACCGAGAAGTTGGTTACTTTAATGAACGAATTATTTCAAGATTGTTTGTACGGTGTTCTACCAAGAACAGTCGCATCTGCACAATATAAACTACAAGATCACTCTGTGACTTTTAGTGCTTTTGAACAACTTTTTACTCAAACTATCCAAAGTAGTCATTTTATCTTAGAGTTTTTTGAACCTACCGAGATAACCCCCTGTGTTATTCCTTTGGATGGAGTTAAACTGTCTGAGGAAAAGAATACAGATGATAACGATTTGCAATTTTTGATATTTCTGCAAGAGTTGTTTGAAGTGATGTTGGCAACTGAAGAACATATTCGCCAGGTCCTGAACTTTAGTAAGAAATTTAGCGATTCCCTAATATCTGCAGTAACACTTTGTCAAGAATCAGATCAGTTTCATGATGAAATAAAAAAAACAGAGCAGCGAATCAACATTTTACGTACGAATGCGTTCATAAGCTCTACTCATACTGAATATGGTCAAATGGAAAAAAAAAAAATACTGTTAGAAAAACTTTGGGGGAAACTTGGTCAAATGATGGAGAACCATCGAACACTTTTAAACACTTCTGTAAAATTTTATCAAGTTTGTGACCAGGTGGAAACTTGGC
>evgR001682c0g1_i01_Hydra
ACTCTCTAAAAACCAATAATAGCGTTACTACTAGCAACATTGAAATCGCCGAAGTGTTTAAAAAACACTTTGGAAGTGTTTTTACGGTAGATAACGGAACTTTACCTAATACGACTGACTTTGTTGGGGAGTGCATTAAGATGGATTTTGTTGATTTTTCAGAAAATATAGTTTATGAAAAACTAAAAAATCTAAAGCTTAGTACGTCGTTTGGACCTGACGGTATACCAAACATACTTTTAAAACAACTAGCACATGTAATATGTATTTCACTTTCATATATCTTTGAATCAAGTTTCACTTCAAGCTCGTTAC
>evgR001684c0g1_i01_Hydra
TCATAGTGTTAAGAATAACAGTCCAAAATCCTTTAAACACGTCAAAGTGTACGTAGCTGATCCATGGTATGAACCACAGGATGGCTCCATTAAGGAAATCATAATTCAAAATGGCAAAGAGGAAAGTCCTAAAGGAAAGTGTGCCAATGGTTTGTTAGAATATAAAAATCATTGCTACTTTTTCCACAAAGTTACTCCAATAATACCTGGGAGAACTTGGAAAGAATCACATGATTCATGTAAGAAGATGGGAGGGCATTTATTAAGTCTTACTGAGCAGTCAGAAGTTGCGTTTGTTATGAACCAGCTCAAGGATGAAGGCGTTAAAACTAAGAAGCACTGGATAGGTCTCAATGCAATCCAGAATAATAGAACATTTGTGTGGTCTGATAAATCACCATTGCTGTTCTTAAATTGGAAAACCCAGGAACCTAATAACTTTAACAACTTGAATGAAGACTGTGTGGAAATGACCAATGATGGTTGGAATGACAACAACTGTGATCGCCAATTTGGATTCATTTGCAAAATTAAAAATGATAATGGATTGGATAAACTCGAAGAATTTGATTTGGAAGATGATAACTTGATAACAACATTGTCATCATTAGAAAAAGTATATAGAGTTTCGTTCATGTTAAAACCTAGTTCATATTCGCCAGGATGGAAAAATGTTATTCATTTAACTATCAATGGAGATCTTGGGAAATATGGAGAAAGAAGTCCTGCTGTATGGTTCCTTGATGATGGGTCAGGTAGATTAGCTATTTCATCGGCTGTTAATGGAAATGCCAATTACTATTTCATTACGCAGCCGTTACCGCTTAATGAGTGGTCTAGTATACAAATAAGTCAAGTTAAGCTTAAAGGAGTTTATTTTTTCACTTGTTATTTAAATGGAGTACTCATTCACAGTGTGGAAAATACTAAACCGCGATCATTTTTTAATGTAAAAGTGTACGCCTCAGATCCATGGTATGCACCACAAAATGGTTCTGTTAAAGATCTTTTGGTTATTAATGGAAATGTTGAGCAAATGATTGATAGAGAAACAAGAATGAACCAAACAAACGTGGTGGCAGTTATACCAGTGTTAGAAAAATCATACAGAATATATTTTGATGTAAAGCCAAATTCATTTTCTAAAGGATTTCACAGTGTTCTTCATTTAACTATTGGCTCTGATTTGAGTAAATATGGTGATCGAGCTCCGGGGGTATGGTTTCATGAAAATGGTGATGGCACTCTTTTTATTGCTGCACCTATAAATGGTAATCTTAATAGAGTCATTAAAACAACACCTGTTCCTCTTAAAAAATGGTCACGCGTTGAAATCACTCAACAACTTGTTTCGAATGTTTATATGTTCTCAATAAAGTTAAATGGAAAAATTGAATTATCAGAACAAAACAACCAACCTCAAATTTTTGAAAACGTTAACGTATTTGCTTCTGATCCGTGGTATCCGCCTCAGGATGGCTTAATCAAAAATCTTTTTATCATTAATGGTGAATTAGATACAAATGTTCCTAACATTAAACGTTTGACATATCAATGTCCGGATGAACTGAAAACTGGAAAATCGACAAGAAAAGACGGTCAGTTAAAGAAGTCGTTTATTTCTAAGTTTTCTGGACATAGTTTTGACTAATTAAAACTAATACACTATTTAAATTGTAATTTTTTAAACTATAGAAATATTAATTGAAAAAAAAAAAAAAAA
>evgR001686c0g1_i01_Hydra
CTATTTTAGTGCAAAAAGCATGACATAATATAATATTACAAGCTGGACCCAGGAAAGACAGAATCTGGTTTAGCAACTACTTTAGAAGATGTCTATGGTTGCAAATAAACAAGTATTGTATTCCATATACAAAAATATATCAAGAGCTTTGGTTTCACAGGAAAAACATTTTGGATCGTTTTGTAGTTCTACCTTAATAACTCGCCAATTTTTCAAACCGAATAAATATCTCTCAATATTTAACTATGGTAACAAGAATGTATATCGTATGAAAAGTTTTTTTAGTATTTCTCAGAACAGTTTACAGAAAAATATTTTACCCCCGCCATTGCTACCTTCTTATAATCGCTATGAGTCAATAACAATTAAAAGTCGTCTTGGAAAAATCAAAAGTTGCAAAGCTGTTTTAAAGCGTTTTCGTAGGGTTCGTGGTGGATTCCTAAAAAGGTGGAGACCAGGAAAGCATCATAAAATGTTAAAAAAATCCAGCTGGCGAAAATTCAGATTAAGAGGTAGCATTTTAGTGAAGCGAAAATCTCACCTGAGAGTTCTTAACAAAATGATGTATAAACATTAAAATATTGCAGAGTTTTAAATGTTGTTTGGTTATATATTTTGCTGAATTTTTAATGGTGTTTTTGAATTTTAAATGGTGTTTTGCCATATTTTTTGCAGAACTTCAAATATATATATATATA
>evgR001689c0g1_i01_Hydra
ATGTTGTTACTCACTGTAAACCGATTTTTAGAGATATATCTGAACATTCATTACAGTAGATTAATTACAAAACAAGTGATTTGTAAAATCTTATTCGTATGTTGGGGAATCGGTGCTGTATGTGGAGCATCTATAGTTAAATTTTCGAATGAAAACATTGTGATATTCATTTTTCATTATGTACTGCCATCGCTAGAAGGGATTTTATTTTTAATAGCACTTTGCACATATTTGTACATATGGAAAGTTTTTAGGAAAATAAGGCGTAGCGAAGAAAGAAAAAATTGTTTTAAAGGAAAATGTTTCACTCCTTTTCTTATTATCATTTCCTTCATTGCTTTTTTTATTGCACCGGATATAGCAAGTTTTGTATTAAATTTTATATTTTTGTTGGAAAATCCAAATGTTTCGAATTCTTTGCTTATGTTACACGGGATTGGTTTTATATGTGATGCCCTTATTTATACATTTTTGCAACACCATCTGAGAGTAAGACTTAAAAAACAAGTGAAGGCCTTAACTGAAACAATAACTTTGTTTTTCCGAATGTTATGTTTTTTTAAATTAAATGAAATACCTCAAGAATACGCCATCACCATTGCCAATACGCCATCACCATGTACGCCATCACCAAGTACGCTATCACCATGTATAAACTAAGGGAATGAAAAGCATCAAAATATCAATGTAAAAATCTTCATCAATAATTCTCTCTATATGTTAAACAATATAGCAAATATATTCTACTAAAAAAAATATGTTAT
>evgR001690c0g1_i01_Hydra
AATTAACAGCGATCCACAATCCACATGGAAGGCTGGAGTTAACAGAAATTTGGCTGGTAAGACCATTGCTGAAATTAAGAGCTTGTTGGGTTTCCAAAAGAAGGAAGGTCCAGTTAGATACTCTGCTGAAGAACTCGCCACTATTAAACACTACAATCAAGCCAAGATGAACGCTGTCAAGGCTGTTGGTGCTGAAGAAGCCGCCAAGCAATTCAAGGGTCTCGGATTGCCTGCTAACTTTGATTCTAGACAACAATGGGGTAAATGCATTCACCCAATCAGAAATCAAGAACAATGTGGTTCATGTTGGGCTTTCTCTTCTTCAGAAGTCTTGTCTGACAGATTCTGTATTGCTTCTGCTGGTAAGGTTAACGTTGTCTTGTCACCTCAAGATTTGGTCAGTTGTAACTGGTACAATAACGGTTGTGATGGAGGTATTTTGTGGGCTGCTTGGTTGTATCTTAAACATACTGGTATTGTTCCAGATAGCTGCATGCCATATCAATCTGGTAACGGTGTTGCTCCATCTTGTCCAAAGTATTGTAATGGTACTTCCACTCCAATTGAATCTGTCAAGTATAAGGCTAAGGACTGGTATGAAGTTGGTTCTGTTGGTGGTTTCTTCATCAAGGAAGAAAAGATCATGAATGAAATTGTTCAAAATGGTCCAGTCCAAACTGGTTTCTCTGTCTATCAAGATTTCATTTCATACAAGGGTGGTGTCTACTCTCATCAAACTGGTAGCTTCTTGGGCGGTCACGCTGTCAAGATTGTTGGTTATGGTGTTGATAATGGTGTCAAGTACTGGTTAGTTGCTAACTCTTGGGGTCCAACTTGGGGAGAACAAGGTTTTTTCAAGATTAAGAGAGGTGACAATGAATGTGGTATTGAAGGAGATGTTTACACTGGCTTGGC
>evgR001694c0g1_i01_Hydra
TTTCTATAAATGCCTAATAAGCATATATAGGCATTATAAATTATATAAGTACTGCTTATATAAAAGTTGTACAAATCAATTAAGAAAAAAACTATTAAATTTGTAGAAATGTGGTTTATCAAATGCTCTATATTTAGCACGATTGCTCTACTTTCCACTGCTCAGTTAGTGGATCACCAACAGTGTGTAACTAAACTGACGAATGGATGCAGCATACCTCTAAATCTACCTTTTCCTTACAAATTTGTTTTTTTACCAGCTTGTACATATCATGATGTTTGTTACTATTGCGGGAGAGTTTACAATTGGACAAGAACTCAGTGTGATCTTGCTTTTCACACAAACATGATGAACTTATGTCAGTTAAAAGAAGAACTTCAGCTTGTAAAGTTTAACTATAAGGTGTCTCCTTGGGACAGGTTAAGAAGTATTTTTCATATTGTTGCGCAACTGTTAAAATGGATTGGTATTAAATCAGGTTCTCTCGAGCATTGTAATCATGGAGCAAATGTGTATTACTCATCTGTGGACAAATTTGGCCATACTAAATATGGCACAGGTGACCAGACGTTATGTAAAATGCCGTGTGCCCGTATACTGGGTAATCCGCACAATTTAATATAAAATTTTTTTCAATGTATATTTAAATGATGTTAAGAACAATGACGTAAAAAAAAAAA
>evgR001697c0g1_i01_Hydra
CATTTGGTTTAAATGATGATTTTATGAGCTTTTTTGATAAGAATGATTTGTGCAATCTAATGATTAGTGATGAAATGAATAATGAAATAGTAAAAGAAAAGGAAGGTGCTCAATTTGATGCCACATTTCAAATTCAAAATAATGCTGCACAACAAATACAAAATGTTTCTTATAAGAATGATCTGCTGAGGACCCGGAAAATGATTAATGATGAAACAATTGATGAAGCGTTGATAGAAGAACAAAATTGTGAAAACATTCAATCTAATACCACATTCCATATTCAAAATAATAGTGCACAACAAATACAAAATGATTCTTATAAGAATGATCTACTGAGGACTCAGCAAATGATTAATGATGAAACAATCAATGAAGCAGTGATAGAAGAAGAAAATTGTGAAAACATTCAATTTGATGCTATTTTCCAAACTGATGGTACACTCCAATTTCAAAGTGATCATAATGAAGTTATTGTTAATAATATTTATCAATATTCTATAGACAATACTATTTCTGAGGATGTTCCAAATAAACTTGATTCAACAAATACTAAATCAGATCAGGAGGCTGAAGAGATAAAATCTCTTATACAGAAAAATGAAGATCTTATACAGCTTTTGAAATCCGAGAAGAGGAATCTTAAAGAGAAAATAAAACATTCTGAGCAGAAGATAAAAGAATACGAGTCACTAGTTATCAAGTTTGCTCAAGAAACTGATAACTCAAAAGCGGATCTTTCAAACATGAAATATCATTATGATATCCTAAATAATGACTTTTATCAGCAGCAACAGCAGAGTATGTGGCGGGAGAATGAACTTATTGAAAAGCTAGTTTCTTGTCAAGCAGAACTTGCTCAACTTAAAGGAAGATTGACAATTGAAAATGAAGAATTTCAAAAACTAAATCAGATTCTTATTTTCAAAGAATCTGAAATTGCACACTTATCAGAAGTGGTTCAGGAACAAGAAGCTTGCATCCAAAATATTAGTGATCAAATATCTCATAAAAATATTGAAATCAATAAACTAAAAGAAACTGTTCATGGACTTGAACTTCAAGAGATAACACTAAAAAATGAAATAATTCAAATCAACTGTTTATATAACGAAAAAAGTAATTTATGTGTCTCTGTTTCTGACGAATTGGCTTCAACAACTAAAAAATTGCAACAAGTGACTGAAGAGTTTGATGAGCACAGACACATTTACAATGATAATATAGGTTCTCTGCGGCAATGTTTTGAAGAAATGACTAACAATAGTGCTGTTATTGAACAGCGTTTGACTGAAGTTCAACATTTTCATAATGAAAAATCAATCGAGTGTGAACAACTTAAAAAGCTACTTACAGATGAACAAC
>evgR001699c0g1_i01_Hydra
CAGGTAAATAATAATGAATTTATATTCAAAGAAATTATTGGATATTTAATGAAAAATAAATTGGATGATTTTTGTTATAATGGCACCAGCTCAATCAATGGCTGTTCAAGACAAAGTTGCAGAATTTGCTCATGCAATGAAAGAACGGTTTTCACATAATTATCTTAAATTTTCATCTGTTAATTTAATGTCTTTGATATCATGGGCGGCATCTGGTGCTATGATGTTTGGTGGAGTTGTACCTTATATTCCACAGTATTATGACATCTACTCTACGTCAAACACTGATGGTTTCTCTATATATGTTTGCTTTGCATTACTTGCTGCAAATATTTTAAGGATTTTATATTGGTTTGGACATCCTTTTGAGCTACCTTTATTTCTGCAAAGTGTTATTATGATCATTGCAATGCTTGTTATGTTAGAACTTTGTGTACGTATTAGGAGAGGCAATGAATTTACTGCTAAAAGAAGAAAATTTTCAGATTTTGACTGGAATTATTTTTGGAATTGGACACACTTTGCAGATTATATGCAGTTTCTAATGGTGTTCAGCTTGCTTGGTTCTATTTTAACTTTTACATTAAGTAACATACCTATATATATGGAAACCCTGGGTTTTATATCAGTCTTTACTGAAGCAATGTTAGGTTTACCGCAGTTTTACAGAAATTATAAAAATAGGTCTACAGTTGGTATGAGTTTAAAAATGGTTGCAATGTGGTTTTGCGGTGATACCTTTAAAACTGGCTATTTTATTATCAAAGAGGCTCCTATACAGTTTAGCATATGTGGATCTTTGCAGGTGTCAATTGATATAGCGATTCTGTATCAAGCTCATTTTTATAATCGCCCCAAATCTCTGAGTAAAGATTAGCCATTAGGATTTAGTGACTGCAAGTATTTGCGAGTTTTAAACAGTTTACGAGTAAATTTGATGCCACAGACCTATCAGAGCACATTAAAGAATTTTCGTATGTTGCATCATGCCATTCGTGCCGTACAGATGTTGTTGCTGGCATATTTCACGAAATTAAATAGCAATTCTTTATAGTTTTTTAACATATTTATATATCAATTAAAAGATGTATATAAATAGTATTTTTTTATTTATTTGTGAATTTGGCATATAAATTTATATTATTGAAAAATAATTATATAAATATTTCTTAAAGATGTACAAAAAAAAAA
>evgR001701c0g1_i01_Hydra
ACTATAACTACTATATTTATTACATAAATACAAGTAACTACTATATATTGGTACTATATTTTATTCAAATACCCACGGCCTACAACTAGTAACAAATTATATATGGCTGAAAAGATGACTGTTATTGGATTGCAAGGTTTTGTAATGATTTTGATGTTTACAATGAAAAGAAAGGTAACAAATGCCGACGTTACCTGTAACCGACTCATAAGTATGAATATATCGAATGAAAAGCAAACATCTTTGTCCACGTATACCTGTATTGCTGCTCAATATTGTTGCAACGGTGGATGTTGTCCGGTGTACAACCTCTGGTACTTTTGGGCAATTATTGCATTTTGTTTGTTATCATGCTGCACTGGGTCTTTCTACTACGGATATCGCAAAAAGAAATCTTTAAAGATATGGCAAGCACCAATGTTTATAGACATGAGCTTGAAAGAACAACGAGATATATACACAAACGCTGTAAAGTCCCACGGCCATCATACTTAGAATACTTTTTAATTACAGATAAAAATTTCCAATGGAAAGAAAGTTTACTTTAAAAAGCTGCAAGGTATGGTATAGCTGGCGCAATGTTGTTTTGTTTGGCTGGCGCAATGTGGCGTGATTCAAATTATATTTATCGCCGTTCGACATGTTTTGTATTTTTTGACGACAGGCAGTTAAAAAATTTTAACTCCTTTAAAAAACACAAAATTATCAGTTAGTTCTCAACTCCCGAAATCAAGAATTTGTGTTCAATCAACTTTGATTGGTAACTAACTTTATGAATTGTTCTAATTTAAATTAACCTATAGTATTAAATAATGTACATTTTAAATTATAAAAAATTAATAATTTTTAAATATTTTGCATTTGATCAAATAATTATATAAATTTGAGCAATTTATACAAAGTTGTTGTTTGTGTTTATGTGTGACTGGACTGTGT
>evgR001716c0g1_i01_Hydra
TAGAGATTCTTCTCAGATATCTAAGCATAATTCTGTTTCGTTTGCACCTACAGATCTTGAAGAATATCGTGTTAATTCTTTAGCCTTAACTTTGGAAGAGTTAGTTAAGTCTGCTCTTTATAGAAGAGGTATTAGCACTACGTTTGATTTCTATGATCAGCATTACATTAGAAAGCTGAGTAAATATATAATTAATGATTTTAAAAATGAAAGACAATCACCAAGACCAATACAACCACCACAACCATCACAACCACCACAACAAGCATTAATAGACAGAAATGAAGCTGTTAACAACGCTCAGTTGATTTCTCAGAATTTCACTCACACCATTAAAAATGAACCACAACCACCACAACCA
>evgR001737c0g1_i01_Hydra
AGCTTCAATCAACCTGAAGGACCTATCAGTGCCTTTTTCTGACTGTGAAAAACCAGTTGATTACTTTTTAAAATATTTTGATAACGAAATAATGGATTCCATCTTGTTCCAAAGTAATTTGTATATCAATCAAACACAAAAAAGAGTCACTACATTATCAAAAGCTGAGTTATTTTCATTTGTGGGGATAAACATGGGCATGGGATATCACAAACTACCATCATGGAAACATTATTGGAATTGCGAACCAGATTTTTCTGTACCATTTATTTCAAATGTATTGTCTCAAAATAGATCTTTACAGATACTGTCGAATATACATATTAATGAC
>evgR001743c0g1_i01_Hydra
TTTTACGTGATTAAAAGAATTTTCTTAGAAGGATATTATTGTTTCACATTTTGGTTGGTATTCGACAGATACTTACATATAAAACTAAGCATTAAATACGTTATATACTGGACAAGAAAGAAAAGAATAATTTCAGCAACAGTTCTTTGGTTTATTTCTGTTTTAAGTGGGTGCTTAATTGAATTTTTTCTTAATCGTTACGACGCGTTTATTGATGTCGGTTGTGATCTGATGATAATTAGCGTTACGGCATTTCTCTATAGTTACGCTTTGATTATTTTTCACAAGCAGAAATCAAGTGTTCGTTCCAACAATGACAAAGGTATATTTAAAGGATTGTTACTAGCAAGTCTCATAATAGCCATGCTTATCATATTTATTGCAATACCTGATATATTGTGGGCTTTACAGGTTTTTGGATTTTATCGATATTCCAGATTGAACTCACAATTTGGCATTGTTTTTATCATATCTTTATGGATGGATTCACTGTTATACATATTTGTTTCGCCTCGAGTGAGATTTGTACTGAAAAAAAAAGTAACTTTTTTACTCAACATCCGAAGAAAAAACGAGAGAAAAAAATCTTCCCTCTCAAAATCATCTTTTACAGTAGTAAATGAGAATACGCGTTTCTAGAAAGTTTTAAAATTTAAATTTTTTCCAATTAATTTTTGAAAAAATAATTTATTCCTAAATCTTGGATTAATTATAAGACGAATTATTACTTATTATTATTTTTTTATTTAAACTTTTTTT
>evgR001748c1g1_i01_Hydra
TTATCAACGTGTAATTAGACTTTAATTTTTGAAAATAATTGTACAACTGAGAGGAAGATGAAAACAATTTTCTTGGTTTGTATACTTCCCTTGATAGTGGCTGGTGGTACTATTAGAGGTAGCGACAATGCATCAGAGAAAAAAAGTTTGAAATGGGTATTGTTAAAACCGAAAGCTTGTTTAGAAGGAAAAGATAGTAAACCTGCAATCGTTCCTGTGCTTACAAAAGGTCTTTTAGTTGCAGTTAAGCTGAAATATAACAGATACAAAGGTCAGACTGATGCTATCCAATGCGATCCATCTTATAAATCAACAATTTTCGGTTGCTCAAGAGACGAATCTTATTTCAATATATATATTACTGATGACAAAAACATGATATTGTTCCCGTCTCCCAGGCTCTCACTAGCTGTCGACCATACGTACAGTTATTACGAGTATAAATACCCTGGTTATAGTCCAAAGGACGAGGAAGTAATTTTTGCCGACTTTCCTCATCCTACTGCCTTTGATGACGAAGAATATGAAAATAAAGTAATCAAAATCTGGAACAGTGAAGATCTTTATAAATCGGCCGAGGGGAATAACATTGGTAGACTTTGTGTAGATGTATATGGCGCTTTTAAGGAGGAAAGCTCTTTGGCGTAAAAACCGAAGTATGCTTTCAATTTAATAATTTTTCATGGTTAATTTTTAAAAAAAAATTTTCATATTCAAAAAATTGTAATTCATGTTATAGATCAAACGTTTAAAATTAAAAAAAAAAAA
>evgR001749c0g1_i01_Hydra
GCTTACAATGTAATACGTAAGATTCGTGGCTTTAGCTAAATAAAAGGAGCAAAAAAAGTCTTTAGTAATCGAATGACTATATTTTAAAAAACAAATGTTTTAAAAACGACGTTTAAAGTGAACAATATGAGGAGAATTAGGCTGTTAATTGGACTGAACGAATTAGAGGAATTATGATGTAATGAACAAAATATGGATTGCTTACATTTAATTTAAAAAAGAATTCGCAGCGTAAGAAGGGAAAAGGTTTCAATATTCCATCAGTTTTAAAACTAACGCATAGTTCAGATTTGCCTAACAATGAAGACTGTCATTGTTTTCCTTGCTTTTATTTCGTTATCTTTACTTCAAGAAAAAATGCCTTCTAGTAACGTAAACCAAGATACTGACACTTTAAAACAGATTAACCGCAGTTTATCTGTTCAATGTGCCACGGACACTCTCACAATTGCTGCCGGTATGGTTGCCCAGTGCAGCAGTGGTATAGGAAATGATGCACCATGCGTAACTCCTGCATGCCCAGACAGTTCTCCTGCCACACCTACTCAGTGTATTTATTTTTACAATGTTTTTAACTATCCTACCGGATCATATTGTTTGGCCATTGCTAACTTTTTATTAAATATCAACAATGTCGTTGTATTTTTTAATTGTTGTCAATGATCATTTGCTAATATTATCGTTTATTGATCTTATCGTTCATTGATATTATCGTTCATTGATATTATCGTTCATTGATATTATCGTTCATTGATTTAATCGTTTATTGATCTTATCGTTAACTGACATTTTTTTATTATGGAAAAATGGAAGTTGAAAACTTAAGTTTTAGTTTATGTATATGGATTTAAAAAAAGCAAGATTATTGAGCTTTGAAGAAAAAAGATTGGATTTTAAAGAACCACAAATAAAAAGAAGTGAAGAGTCCATGTACCAACAAACTATCTTACAAATAAATTGTGTATGTGAACAAAAATTCGCTATATATACAAATAATTTGCAGTTTTTTGGGAATAAAATTAGTTACATTTTTATAATAGCGATTATTATTTCATTAGTGTTAATAGCGAT
>evgR001751c0g1_i01_Hydra
TTAACATACTTTAGTGCAAAAAGGAAATAAATAAGATCCGGTTGTAACTGCATCAAATGTGTATGTTCGGAGGGTGAAAAATGTCGACTATAAATTTACTTCAAACATCGCAACAATATGATAACCAAATCAACAACATAGGGAAAATGCTAAACAGCACATCAACATCAATCCCTATGCTTGATAACAGCAATCTTTTAGTAACAGATCTGTCATATGTTTTAGTTATATGTTGTCCAATCGGAATACTTTTACATGTCATAGGTATATATTTGATAACAAAAAATAATAACTTGAGAATGCATAAAATTGATCGGCTTATAATTTTAAACTTATCACTCGCTGAAATGCTAACCAACGCTGTTCTTTTCTCGACCGGAGTTTTAGACTTTGCTGAAAAAAAGATGCCTATTTGGCTAAAATTTTGTATTCGATCTTTGGTATTTGCTAACTATTTAAAAACCTTTTTGCTAACGTTGAACCGATTTCTAATATTTTATTTAAACGTCAAATACTACTGGGTCTTGTCAATGAAAAAAGCAATTTGCTGTTTAGTTGCATCTTGGGTGTGTAGCATAGTTTACGGTTTCACAGTATCTTTAAACAGCAGGACATTGACATTTTCAACAAAGAAAGGTCAGGTCAGTAATGCTATTACCGACCTCACTTTCGATGTATTAATAACTTTATTATTTCTTATTGTATACGGTTATGTGCTTTGGAAACAAAATCCTTCTAAAAATGATATTCTTTTACGAGATAGACACCGTGTTTTTAATGCGTTAATATTACCAACGCTAATTGTATCAAGCTATTTTATATTTTCCGTTTTGCCCCACATAGTTTTTGCACTTTATAAAATGTTTAATGTTGCTATTAACAACAAACTTGCACTTAGTTTAGATGTTTTCTTGGCGTTATCCTATTGGAATGATGCAGTTTTGTATATATACATTAGAAAATTTCGAAAGTTTGTATTTAAACGTTCTAGAAATCGAAGGAAGACCTCTTATATTCTCAAAAGGAAATCGGTTGTTGATACTTTAAAACCAAACTTTTTTGGAAGAGCGCGTCTTGCAAAAAATAACGGTAGTAGTATGTTTCCGGTCAGAAATTATTAGCTAACTTTGTTTGCAACAGGTTCACAGTTATAGGAAATTAATAAACAAGTCAACGAAAATATTGCAAAGTATAATTTTCAAAGTATAAATTTTAGTATAGTAAATTAATTAGTGAAGTATGAATTCATTGTTAATAGTTATAACACCCTAGTCCGCATTAGTTAATAATAATAAAAAAATTTATCTAGCTATAAATACATGAACGTGTCATACACAAATATTTGTAGATCCACAAGTGTTTTCATTGTAACAAATTGTACCATAAAATTTTA
>evgR001752c0g1_i01_Hydra
TAATCTTTTATTTTCTAAATAAGGGATTTTTGCACCGTGTGTATTCCACAATGTAAATATACCCAGGTTTGTGTGTTAGCGAACATATAATCAGAGTAGTATGATGGTAACAATGTGTGGCCAGATAAAGTAGATATGTGGTAAAAAGTGATTTTTTGCAAATTTTATTTTCAGAATAGTATTAGTACAAACCTACAACTGAATACAAAAAATGTTTTTATCAATCCATGTTGCATTTAAAATAATTATCAATACATGTTGCACTTAAAACTTACAAATTTGATTCAGCTTGCTGCTGTATCAAGGTTGATACTTCTTATCGTGCAATTCGTTTTCGATGTTTGCTTCGAAGATTTTGATACATCAACGAATCAGATTACTTTGTTCGAAGCAAATATTAGTAACAGCCAATCGAATATCAGTAGAAAAATTTTCAACGGTGTAAATAAATGGGATTCAGTTTATTTTACTTATATTTCTATAAATGGTTATGAATATGAACAAATGCTTGCTTTCTACCCTGGGTTTCCGCTGTTGCTGAATTTATTTTCAAGATTAACCGCTTGTTTATTTGCACATGAGTATTTAATAACTTGTATTACCAGTTTTTGTATGAATTCTTTTTTTTGTGTTGTATCCACTTTCTATCTTTATCAATTAACAGACAGATTATTTGGTGAAAAAATGGCTGTTGTTTCGACGTTTTTGTTTATTATCAATCCTGCAAATGTTTTTATGATTTCATCGTACAGCGAGTCACTATATTCAAGTTTGCAATTTATTTTACTGTTTTACCTTGAAAAGGATTGTTTTTTAAGTGCGTGTACAGTCATATGTCTATCGATTATAACTCGATCAAACGGTATTTTAAATATAGCATTTTTGATGCATTTCTTTTTGAAAAGAAACGCTAAGACTCTGATACGTCAGTTTTTTAATAAACCAGCTTCACTTATAATTTTTTTACATCCGAATAGTTTTTTACCAAATCTTCTCGGTTATTTAAATTTTGCTTTACCACTGTGCTTTTATTTGAGTGCGCCATGTGTCGTTTGTGGCGTACATTTATTTTGGGTGTATGAAACTTATTGTCATCATAATCATGTGGTAAATTCTTCGTCTCATTGGTGTTATAATATGATTCCGTCGTCATATTATTACGTTCAATACAAGTATTGGAATGTCGGCTTACTGAAGTATTTTGATACAAAGCAAATACCAAATTTTCTATTAGCTTTTCCAGTAATTTTCATTGTAATAATAGGATTGAAAAAGTTCGCATCGCTCCAACATTCATATCTCGCTATAAAATCACTTGGGTTGATAGACACAGAGAACTCACGTATAGGAGAAGATTCTTTAAAAAATGCATCCGCCATCTTTGTTTATTTATCACATTGTTTATTTTTAACTACTCTTGGATTGGTTGGTTTTCATGTTCAAATATTAACAAGAATGTTGTTTTCATCGTGTCCGTTTTTATATTGGACGTTAGCTTCTACTTTCATAGATAATAAAAGAAATGAAGTTATGATATGTATAGTAACATACTTTGTTATATATAATATTACAGGTGTTTTATTACATTGTAATTTCTATCCTTGGACATGATTTTAAATAAAACTGTAATTCATAACAATTTTTCCTATATTTAAAGGGTCTATAAAGTACCACAAAAAAA
>evgR001755c0g1_i01_Hydra
ACAATATTTTGATTGTCTATTATATTGCGATATTGTCAATAGCTTGTCTTATAAGAAGATATCGCCATTGTTACGTTTCTGCGTGTTGGAATGCAATGACGTCATCATGCACAAATAATGATTGTTAAACGATTTTATATTAAATAGTTAACACGGAATTGTAGAAACTTATTCAATATGGCGGATGAAATAGAAACTCTGTTTCACGAAGTTGACTCGAGTGGAGACGGAAAAATTTCATTAAAAGAGCTGGGGGTAATGTTCAAAAAGCTAGGCATAAAAGTAACTGTAAACGATGTAAAAAAAATGATATACGAATTTGATAACGATGGAAACAGGCAACTTAGTTTACCTGAGTTTCGAGAACTAATATCAAATGTAACATCATTTGACAAGTCTTACAACGAAGCCTATGAAGTTTTTAAAGTCTTTGACAAAGATGGAAGTAATACTCTTTCCATAGATGAAGTTAAGGAAGCTTGTAAATGTTTGCCAAATAAGCTATCTGAAGAAGAAATTAATGAGTTTGTAAAGAAATTAGACGCTGATGGAAACGGCATTATTGATTTCAAAGAGTTTGCTAAAGCTTATGCTTCAGGGGTTTGATTTATAAAGTGTATTGGATTTTGGACGTACTTGAACTTTACTATGTTCGTGAAGTTATTTCTCGAAGTTTTAATTTATGTTAAATATTATTATAACTAATAAAAATTTAATTTATTTTCTTATTGACAACAATATTTTGCTAAAAGGTTTAAAGTAACATAAAAATCTTTTTTATTATTTTTAATTATGCGATATTTATTATTTTTAATTATGCGATATTTATTATTTTTAATTATGCGATATTTATT
>evgR001764c0g1_i01_Hydra
TGTTTATAAAAAAAAGTATAAAATAATATTCATAATGTATTAGAAATTATAAAATTATATTTTTATAAACTATGGCAATCGTAGAAGATTTCTTAAAAACTCAAGGCTGTATAAATGTGTTTGGTTACGGTTCTCTAATATGGAATCCAGATTTTGAATATGACTGCTCTTATCTTGGCTATATAAGTGGTTACGAAAGAAGATTTTGGCAAGGAAGCACTTTTCATAGAGGAACTTTAGAACAACCTGGGCGAGTTCTTACAATGACCAAAGTAAACGAGGGCTGTTGCTGGGGCAGAGTTTTTGTAATAAAAGGCGCCGATAACGTTCGAAAAGCGCTGGATTCGTTGCAAACGCGTGAGCAGCAACTAGGACATTATGATATTAGAATTGTGCCAGTAAATGTATTAGAAACGAACGAAGTTTTATATTCAGTAGTGTATTATGCAACGCCAGAAAATCCGTTGTTTTTAGGGGAATCTTCTTTAGAAGAAATAGCAGAAAATATTGTACAATCTTGCGGTGTCATCGGTCACAATATTGAATATTTGTTTCGATTAACAGACTTTATGCGCGAAAATTTACCAGATGAAAAAGACGATCATCTTTATGCTCTAGACAAACTAGTTCGCTCAAGAATTGGTTTGCATGCTAAAAACGTGTTGCCATGGCATAAACTTTTAGAATGCGATAGTTTTAAAAAACAAATTTTTGATATAAAAGAAAGTAATTATAACGACAAAGTTGGTTTAAAAACTATTGTTGTTCAATCATAGATAATATTTAAATTATAATATTTTTTAGGTTTAGGAATAGTTTATACTTTAATTTCTGTTACCAATTTTTATTAATTGTTATTTTTATAATATGTATATTGGCGTTTATTTAATTTTTCTAATTTATAAAAAAAGCGTTTATTTTAGTACTAAAATAAAAAAATACATAAAAAAAAAAAA
>evgR001768c0g1_i01_Hydra
TCCAATGTATACTCCATTCTTCATGACTAAGGATCAAATGAAGAGAGTTGCTCAATTATCTCAATTTGATGACGAATTATACACAGTCTCTGGTGAAGGAGAAGACAAGTACTTGATTGCCACTTCTGAACAACCAATTGCTGCTTTCCATTTGGACAAGAGATTTGACGAAGTTGAACTTCCATTGAAGTACTGTGGTATCTCCACTTGTTTCAGAAAGGAAGTTGGTGCCCACGGTAAGGATACTTTGGGTATTTTCCGTGTCCATCAATTTGAAAAGATTGAACAATTTGTTGTCACTTCTCCAAAGGATAACAAGAGTTGGGAAATGTTTGAAGAAATGATTGGTAATTCTGAACTCTTTTATCAAAGTCTTGGCCTCCCATACAGAGTTGTCAATATTGTCAGTGGTGCTTTGAATAATGCTGCTGCTAAGAAGCTTGACTTGGAAGCTTGGTTCCCAGGTGCTGAAGAAGGTAACGAATATCGTGAATTAGTTTCTTGCTCTAATTGTACTGATTATCAAGCTAGAAGATTGAATGTCAGATATGGTAAGGCTGCCACTAAGGGTGATGCTGAATTCTGTCACATGTTGAACTCTACATTGAGTGCCACTTCTAGAACTATCTGTTGTATTGTTGAAAACTACCAAACCCCAGAAGGTATCAAGATTCCAGAAGTTTTGCAACCATTCATGGGTGGTACCA
>evgR001771c0g2_i01_Hydra
GTTTTTTATCGCTTCCTTATTTTCGAAAACTGTTACGGACATTAAAGATAAAAAGGAACGCGTGGAAGTAGCTTTAATATTATATTTGTTACCTAAATTTACTATTTAATTGACTTTTCAAAAATGTGTTTACTGATGTTTTATACGTTATGGTTCTACTTAAAACAATGTTATAATTTCGTAAATCTTGGATATTCACCCATATGGAACACACTGCCTCAGTTGCCTTTCTTTTTTGGTAGTTTAAAAATTATTTGAGACAAAAAAAACATAATTGTATTGATTTTCACATTACGCATGTACATAATTTTGGAGCAAGAAATTTGGAATTAATAAGCTAAGGAAATCCAAATGTCGAAACCAACGTTGGAAATAATAGAAGCAATTGTCTTAATTTTAATTGCAGCATGTGCACTTTTGGGGAATTGCATGATTATTCTCGCTTTTATCATTGGTCCTAGAGCAATAAAAACTTTTACAAACTACTTTGTCGTTAACTTAGCTGTTGCGGACTTAATGGTAGTATGCTTATCGTTACCATTTTGGATTGCAAGTCGTTTAGACTACGAACTAATCGATGACAGAATTCGCAAGTTTTTTGATGCTCTCGACATACTTTGTGGAACAACGTCAATATTAAACTTAGCGGCAATAAGTTTAGAACGCATGTACGCTGTGAAATTTCCCACACGTCACTTTAACTTATCGCCCCATCCTGTTGTTGGAGTGATTGCGTTAACTTGGACGACTGGATTATTTTTCACTATAACGCAATATCCGGCTACAACTGGGAAAAGCATTAGACTATACACTTGTTTTATTTTTACGATGGCATTTCTTATTCCGTTACTTATAATTGTTGCTTCTTACTGCATTACTTTTCATTGCGCAATATCTACCTTAAAAATTAATAGTAAACGTGAACTGAAAGTTGCTAAAACAATTTCTGTGATAATTGCTTTATTCTTTGTTTGCTGGAGTCCATTTTTTATTGTGAACATGATATTTGCATTTTGCGAAGGTACAAATTGTTCAAAGCTTCCACGTTGGTTGGTTGATGTTACTAAAATTATGCACTATAGTAACAGTATGATGAACTTCTTTGTTTACGGACACCGAAGTCCAGATTTTAGACGCTCTTTTAAAGCTTTTTTTCACTGTAATTTTAGACCTTTGCAACGAAGAGTGCATCGGTAATATTTATAAAAGATAAGTTAAAAATACAATATAAAGTGTAAATAAATAAAAACTAAGTAAGGAA
>evgR001772c0g2_i01_Hydra
CAATGTCTGGAAGTTAACTATTTCAAACACCAAAAATACGCCCTTGACACGGGAGGGATTAAGTATAAATTTAAGAGTCAAGACCATCTAAAAATGCAGCATCAAATAAATCTAACCTTGTCAGACGAGGAATGGTTAAATGAAGTCTGTTCCAAATTTATGCCTGGCTACTATGCATTTAAACTGAAAGATTTTAATTGTTTCCCAACATATTCATTTAATGATAGTGTAAAAAGAAAAACATCAGCAACCAATTGGTGGAGAATTATAGAAATGAAGGAGTCTTCCAAATTGCCTGCAAATTTTGCACAATTTTTACAACAATTGCAAAAGCTGTACAAGTCAGCTAGTACTGGATCCATTGAAAGATTTTTCTCATTCTTTGGCATTATCTGGAGCAACATCAAATCTTGAGTTTTAATGGAAGTTATTTTATGTATTGATTATATGTAATTACTATAAACAAAGTTTGATACCAAAAAC
>evgR001773c0g1_i01_Hydra
CCATGCTCGATCAAGTAGAGTCATTGGAAAATGATCTGTTTTTAAGTAAAAAAATATAGAAAATCTTTTGCTATCTATTAGTGAAAAAGACTTGTTTTCAGAATCATTAAAAATATCTATTGCAGAACAAGCTGTTGTTATTGAGCGACTGAAGGAAGAAATTACCAACACTGAAATATCCTATCAAAATGATCTTAGAGAAGCTAATGACATTTATAATGATCTTCAAAAACAGTTTGAAAGTGCTTCAATGTCTGAAGCAGCATTACAAACAATACTTAACAATAAAGAAGTCGATTTTGAAAATAGATTAGCTCAGATGAATAGCATAACCAATGAGTATGAAGAAAACCTTTCTATTTTGAATAGTACCTTGATATCAAAGAATGAAGCACTGAAACTACTACAGTCAAAACTTCAACATACTAATGACCATGACCATATGTTAAAACAAAATAATGAAGAAATTAATGGACTATACAATTGTTTAACAAATATGGAAAACACAGTGAGTGTATTAAACAAGGAAAAAGAAGATATTGTTGTTTTGTTAGATAATAAAGAGTTGGAGTTAGCAGACACACTAAAGCAGATGCATCAGTTAAGGGAAAATTTTGAAGATTTACAATCCAAAAGTGTTGAAAATAATTTAATCAGAGAAAAACTCAATAATGCTTTACTAAATACAGAAGAAGAAAATAAAAGCTTGAAAATATCACTTGATTCAAAAATAGAAGAATTTGGTTCACTTGACGCTGAAATCGGATCGTTGCTAGAAAATCAAAAAGAATTAATGAATCTGCTTTCTTTTGAAAAAGAAAATAATGCAGTGAAAGATAACCAAATAAAAAATCTGGCGGAACAAATTGAAGGTCTAAAAGTTGAAGTTAATTTCAAATGCGACTTGGAGAAACTTGTGGAAACCAGAGACAATGAAATAGTTAGACTTGAATTATTACTAAGTGAATTAAAGATTAAAAATGAGGAAGAACACAATGAATTGTTATTAAATCTATCAAAGTATGAGACTTGTAGTTTAAATCTAAATATGGAAGTGCAAAGATTCAATGTTGATAAAGAAGCTTGGAAACAACAACAAGAACATTTGTATTCCATTATTTCATCAAAAGAAAATGACTGTTACTGTATCACTGAAGAATTAAAAAAACTTAGGGAAACATTCAATTATCTTGAAAATGAGTACAACCAACTAAATTTAGTTTTAAGTGATGCACAGAACACTGTTAACAACAATCAAGAGTTAGTTGCTGTATTAAAATCAGAGTTAAATGAGAAGTTGCATCAAATTGCAGTTTTAGAAGACATCAGCAGAGAACAGTTGAATAAATTTGAAAATGAACTCCAAAAAAAAGAGGAACTTTATGAAGAAGCTGAAAAGTGCCTTAAAGTCAGTGAAGATCGTCTTTTAAAAATTGAAAAGGATCTTATTACTAGCTTAGAGTCTTACTCTAAATCTCAAAATGATTTGAAAGTAAAGGAGGAAGAGTGTTCTAATATCATGAAAGAACTTTCAATAAGTAAGGAGTGTTTCTTGAAAGTTGAAAATGAACTTATAGTTAAAGAGAACGAATGTTCTAATATTATGAAAGATTTTCAATTAAATAAGGAATATTGCTCTAAAATAGAAATTGATTTCAAAGTAAAAGAAAAGGAATGCTCTGATGTTATAAACGAGCTTATGTTTAGCAAAGAGTATTGCTCTAAACTTGAAAATGAGCTTAAAACTAAAGACGAAGAATACATTACTATTGCAAGGAAATTTAAAGATAAAAGTGAACTTCTTTTAAAAGTTGAAAACGAGCTTAAAACTAAACAAGAGTACTGTTGTTGTATTGAAAATGATCTTGAGATTTACAAAGAAAATTGCTCTAAGATTGAATTTGAATTGAAACATTTGAAAGAGTATAGCAATAATGTTGAAAGAGGATTACAGGAATCTTCGAAAGATTTATCAGCTTTGGCTCAGGAAGAGATTTTAGGTTTTGAAAGAATAATTTATAGTAAATCAGCAGAGCTTTTATATCTTAACCAAGTAAATGATGATTTGACATCAAAGTTGCAGCAGGCTCAATTTGAAAAAGATTCAAGCAATCAAATTATATCGGAGCTTCAAAGTGCTGTTAGAGTTAGAACCACTGAATATGAAGCTTTATCTAACTCTTTAAAAAATCTTGCAGCTTATTGTGATGAACAATTGCCTAATCAAATACCCGGTGAAAACATACCTGGTGTAGACATATTGAAAACAGATAAAATTAAAACATTTTTGGATTTAAAGGAAACAGAGTTAAAACAGACCATGGAGATGCTTAACAAAAAAGAAGAAAATATTTCCCTCAAGGACACCATGATTAACGATCTAAATCATGAACTGATATCAATCAAAAACATATTAAAATTAAAAGAAGATGAACTAAACACAACTAAAGAAGTTAATAGAAGTTCAACAGAAGATTACAACTCTTTAAAAGCAAAAACCGCACGAATTTTAAAACAATCTCAGACTTTAAAGAAAGATTTTGCTGAAAAACAAAATGAATTTGAAAAGGTTTTAAAATCTTACACTGCTTCTGAAAATAGTCTTGTCAGTTTGAAGAAATGTTACAGTGAGTTGCAGACAGAGCAAGAAACAAACCAAATTACAATAATAACTTTAAAAGATGAAATAAAAAAATTAAAAGAAAGAGCAGAGGATAATCAGTTGGCTACATCAGATAAAATTATTTTAAATGATAAATCTTTGTTAAATGAAGTTTGTATGTCAGAAACACTTTTGCAGTCTTCAGATCTTACATTACAACAACTAGAAAATAAAACATGCAGTGTTGAGTTGCATCATGATAACGAAGAAAATATAACAAAGCTTGACAATTTGAAATTTACCCATGATAAAGAAGACTTCTCAAATTTAGTCAAATCTAAAAATGATGAAATTTCAGAGTTGAAGGTAATAAATGAAGATCAAAAAACAGTTATTGTAAAGATGAAGAAAAAATTGTTACTAATGAAGAAGGCTGAAATACTGCTTGCCAAAGAAAACGAAAAACTAAATAGTGAAATAGCAAAGCTAAAACTTCTTTTGGAATCATCAAATATAGAAAAATCTTTGCTTGCTTCAAAAAATGAAAAGCTATTAAGTTTAATGAAAATTCTTGGTGAGGAAAGTGAAGACTTTGATAGTATAGGAGAGTTTATTAACAATTTTAAGTCTTTAAAAT
>evgR001776c0g2_i01_Hydra
TTTTTTTTTTAAATGTCATGTGGGAATATTGCAGCAACAACTTATTGCGTTATGACTTTGTTTAGAATAAATTTCTTAGTTTAGTTTTAAGAAAAAGCAGATATTAAAGTTTTATATTTTATTTTGAAACTTTAATACATATAATATTTACAATGAGAGTATGCTAGTGATGGCTAATCAAAATACACGTGTTACTAAAAAGCGTAAGTGGATGGGAAGAGAAGAAAAGTTTCCTAACGACGACGAGATAGTAGTATCTGGAAATGAAATGAATCTGTTTTCAACGCTAAAAACCCTTATATGGCCTGCTATTGGTGAAAAGAAAAAAAGAATAGACGAAAATGGAAATGATGAGCAAGTTCATTGTATGTCATCATATGTAGGTAGATCTTGTGATGCTAATAATCAATCAATAAATAATACTTTTCAGAATGACGATAAAAAATTTAAACCTGATTGTGATAAAAATAAAGGAGGTTATGTACATAAAGGAATTATTGTAACTACAGAATGTAACAATTATATTAGTGACGAAAGTGAACATGAATCTATTAGTGATGAGTTTGATTATGAAAATTATGAAAATGATAAATCAAAAACAACTAACCAAACAATATTAGGATCTATTTTTTCTCCAGTTCTCTCTTTCTTTAATAAACCATGTAGCAAAACAGTTTACAGCGTACATGATTCAACTGTTTCACAAAACCTTGATGGAAATAACTTTTCATACACTGAAATAAAACCTTTTCATTCAATTTTACACCAAGAGGAAAAAACAGATGAGTTTGATCCTTTTTATTTCATAAAACATATCCCACCGCCTCCATTAGAAGCCTATTTAAGGAAAACTGTACTTCCATTACCAACCCGAAGAACTCCCCAAATGACTTTAGTTTTAGATTTAGATGAAACTCTGGTTCATTGTAGTCTTAGTAAACTTGAAGCCTACAACATGACTTTTAATGTAGTTTTTGATAATGTGACTTACCAACTTTTTGTAAAATTACGACCTCATTTACTTGAATTTTTGGAGAGAGTTTCAAAGTTATATGAAGTTATTTTATTTACTGCTTCTCGCAGGGTTTATGCAGACAAACTTTTAAACATTATTGATCCACGTAGGCAGTTTTTTCGCCATCGTTTATTCAGAGAGCATTGTTTACATGTCCAGGGAAATTATATTAAAGATCTGAATATTCTAGGACGTGATTTAGAGCGTACTATGATTGTTGATAACTCTCCACAAGCATTTGCATATCAAATGGCAAATGGTATTCCGATTGAATCGTGGTTTATGGATGAAAACGATTGTGAGTTATTGGAGTTGTTACCTTTTCTTGAGAAGCTAGCGGAAGTAAATGAAGATGTAAGGCCACACATTAGAGATAGGTATCGCTTACATGAGCTTCTACCTCCTTGAGGGTTCAACATTTTGTTATAAAAAAGTTATAGCAAAATTGATATCATTTATTTTTTTCTATTTTGTATATAAATTATTATTTTATTTTGCGTTAAAATGGGTTGTATTAAATGTACTAATTAGTTCTGTACTAATTTTACAAAAATTAGGAAAATATGAATATTTTTCATTTTTATTATAAACTATATTTTTTGCTATATACTTTTGCGCTTGTTTTATTTATTTTTTGTTCATTGCAAACATTTCTGCAATCATGATTTTTGTAAATATT
>evgR001779c0g1_i02_Hydra
TTTTTTTTTTTTTTTAAAAGATGGCGGAATCAGAGGGCATGTGTGAAACACATAACACTAATACTGTATTACGTACTACAATTTCAGATCGTATTTATAATTTTCTCAAAAAAAGCCTAGAATATCTTTTTAAATTTTGGTTTATTGTATTTTTCATGATAGTTTGGTTTCTTAGAGAGCCTATAAAACTAGCAGATGCAAGTAATTCAGTAATAATGTTTCTGTCCACTCTTACTCCGAAGTTTTATGTGGCATTAACTGCAACATCGTCTTTTTTGTCTGGTTTAATACTAATTTTTGATTGGTGGTACTTTTGCAAATACGGAAGTTCTTTTATAGAATATTTTTCAGTCTCTTATTTTGGTCCATTTCTAGGGCTAATACCAAATCAACAAAATACAAAGCAATTAATTGCTGAGTGCAAAGTTTGGAGAAATCCAATGAGTCTTCTTCGAGGAGCTGAGTACCAAAGATTAAAATGGGCAATGAATTTGCAGCCACTGACAGAGTATGATTTGTTATTGTCAGAAAAAGATCATCTGTCATTTTTTACTTGCTCAACTGATAAAATCAATGGAAGGCCTGGAGACTATAAAATGTTAAAAGCGTTTAAAGAAAAAGAAAAAGACAAAAGGATATCCATGGCTCATGAAGCACTTACTATTTCACAATCCAAGTGTGCAACTGCGTTTGTGTTACTTGCAGAAGAAGAAGCTACTACAATTGAAGAGGCTGAAAACTTACTCAAGCAAGCATTAAAAGCTAGTGATAATAACCTCAGACAGTATAATAGTCTCTCTGCTGATGAAGAAGCTGAGATCTGCCATAGGCGAGATACAAATGTGCATGTATATGTGAAACGAAGGCTCGCAATGTGTGCACGAAAATTAGGAAGGACAAAGGATGCTGTTAAAACAATGAAAGATTTACTTAAAGAACATCCATTCTTAAGCGTATTAAATATTCATGAAAATCTTATTGAAGCATTGTTAGAAATGCAAGCATATTCAGAAGTTCAGCAAGTACTTACAAAATATGATGATATTAGTCTTCCAAAATCTGCAACAATTTGTTATACTGCTGCTCTCTTGAAAGCAAGGGCTGTGGCAGATAAATTTTCTCCAGATGTAGCTGCAAAACGTGGCTTAAATCCTGCAGAAATAAGTGCTGTTGAAGCAATTCACAGAGCTGTAGAGTTTAATCCACATGTTCCAAAATATTTGCTGGAAGAAAAAGCACTCATTTTACCCCCTGAACATATATTAAAGCGAGGAGATAGTGAAGCTGTTGCATACGCCTTTTTTCATCTTCAATATTGGAAAAGAATAGAAGGAGCTCTTACTTTGCTTGATTGTACTTGGAAGGGAACTTTTAGTCTTGTACCTTATCCTCTTGAAAAAGGACACTTATTCCATCCATATCCATCTTGTACTGAAGCAGCTGACAAGGAGCTTATGCCGGATTTTCACAAAGTTTCAGTATATCCTCAAAGAGAGCTTCCATTTTTTATATTATTTACTGCTGCTATGTGTTCCATATCTGCTGTTCTTGCATTACTTACCTCACAGTTTCCAGAAATTATGATATGTGCAGTCAACATGAATTTGGAAGCGTGTTATACATTAAAGAAGATTACTTATTATTATTTTGTAAATTTTATATTTATATTACGATGGAAAAAAAGTTTTTTTTTTATAAAAAAAA
>evgR001782c0g1_i01_Hydra
GAGTCTTGCAATATGCGATGGATTGCAGAGCAGTGTTGGATTTCCAAATGAAATTTATGGATATGCCAACAAAAATAGTCCTTTTAAAAATGAAAGTTTGTGCAAAACCAGTGGGTTTGTTGTAATGTACTTAGCTCTCACTTCCATTTCTCATTTGGTAGCTTTATGTATATACCGATACGCCGTTATTGTATATCCTATGAAAACTCGGGCTTTTGTTACAAACTATATAAAAATTGCTTTATATTTCATCACACCAAGTTGGATTTACGGATTGTTTTGGTCAGTTACTCCTCTTTTAGGATGGAATGAAATTGTGCGTGAAAAACAAGATAACTACCGGTGCTCAATTAATTTACATACTAATGATCATTTAAAACAAAGTTATCTGTATGCACTTTCTGTTTTCTGCTACTTTGTACCATTACTAATTATTATTTACTGCATATTGAAAGTGCATCTTGAACTTCGGAACATGCTAAAGTTGTGCAAACAGATTTCAGGGGTGGATGCAGACATTACAAAGGCAACATACAAAATGGAAAGAGAACACTTTATATCCGTTTTCCTAATTGTAGCTTCTTTTTTCACTGTGTGGACTCCTTACTTTATTTGCGTCTGTTACTTAGTTTACGGTCGAAATCTACCTACTGGTTTTTTAACCTACTGTGCTTTGTTTGCAAAATCATCAACAATTATCAATCCAATTATTTATTGCATAATGTACAAGGAGTTTCGTCAAACCGCTTTAAATAAATTTCGACAACTTTTTTTAAGGTCAACTGTGGCACCTTTGGAGGTTCCTAAACTCTTAACAACACAATTTGCTTTAAATGAGTTACCTAGATTATAAGTCAATTTAACTAACTAGCTAACTTAGTAAATGTTTGGCTATTATATATTGTTAGAATATTTATAAAGATGGATGTATTTTCC
>evgR001784c0g1_i01_Hydra
CTATAACCAACTATTTTGGCGTATTAGTATTTGGCGGCAAAGAGAGACAAAGATCGAAAATATTTTAAGCATTCGAATTCGAAATATTTTCTTAGTTTGAAACCTTTTTTCTTAAATAAATAAAGATTTACGCGGTTTGAATTTATCTAATTTGAAATAGCAACAGAATGGCTGGAAGTCCAAAGAAAGGCAGTCCTAGAAAAGCAAGTCCGAAAAGAGGCGGTAGTCCGAAAAGAGGTGGTAGTCCTAAAAGAGGCGGTAGTCCTAAAAGAGGAGGCAGTCCTAAAAAGACCAGTCCTAGAAAGACAAGTCCTCATCGATCAAGACCTACTAGCCCGAAAAAAGCTAACCGAGGTAAAAAGGCAATCAAAACTACTCATAAAAAGCACAAGAAGAGTCATCACCGACGAAAGGGAAAGCAAAGTTATGGCATTTATATATACCGAGTGCTTAAGCAAGTTCATCCTGATGTTGGAATCTCAAGCAAGGCTATGAGTATAATGAATTCATTTGTCAACGATATTTTTGAGCGAATTGCGGGAGAGTCATCAAAACTTGCAATGCACAATAAGAAAATGACCATTTCTTCACGCGAGGTTCAAACATCAGTACGCCTTATCCTTCCTGGAGAGCTTGCAAAACATGCTGTAAGTGAAGGAACAAAAGCCGTTACAAAGTATTCGAGTTCAAGATAACATTAAAATAGGATAGCGGCTTTTTTTTAAGCCATGTATTTATAAACGCAGGCGATAAAAGGATAACTTATTGTAAT
>evgR001785c0g1_i01_Hydra
ATGCCGAAGGTGAAAGAGTTAAGATTAATGAATGGGTTTCCTCCAAGACCAATCAATTGATTAAGGACTTGATTCAACCTGGACTTATTAATGTTCTCACCAGATTGGTTCTCGTCAATGCAGTTCACTTCCTTGGTCAATGGAAAAATTCCTTTGATAAGACAGAAACCTATCCATATTCATTCTATGGAGTTGATGGTGAAACAAAGGCTCCAACCATGTCAAAGAGTGAAACAAAGGAACAATATGGCGCTGAAGGAAATTACCATTGGGTTACCATGCCATTTTCTGATACCAAGTACAAGATGACAATTGTTGCTGTTAAGGATCATGAACATCAGGCTACTCTTGAAAGCTTTGTTGCTCAAAAATTGA
>evgR001786c0g2_i01_Hydra
TTCAAGATAGTCTAATAAAAGATCTAAGAATCATTAACGGATTATCTGATGATGTTTCTTTGGCGAATATTGAGGTTATCGCAAAAGATTTTAATTTTCCTGCTGAAGTGGAATTAAAAATGAATAATTTAATTGGAAAACTGCCATTACTAAAAGCATCATATACAATTTCTTTTAAGATTAAGCTAACTGCGTTGCCGGTTGGTTGGAGAAGTGTCATTCACTTAACTACCAGTGGTGATTATTATAGAATACCTGCTATATTTTTTTATGATAAAACCGGAATGTATATCTGGAATAGTTTTGCATCTAGTCTTTTTTTTAAAACACCTGATTTGCCGTTAAATACCTGGTCAAATGTTCAAATATCTCAATATCAAACTAATGGAATTTATTCATTTCGTGTGTATATAAATGGAACTATTACAAATAGCATGTCGAATATAATTGCGCTATCGTTTGAAAATGTCATGGTATATGCTTCGAGTCCATACTTTTATCCACAGTTAGGTTTTATTAAGGATCTAAGAATTGTAAATGGAAATCAATGATCGCACTCTAATTGAACTAAAATCTGTAAAATATTGAAATAAATCATATTAAATATTATCCTATTACATATTACACCTGCGTTACATTAGAGCTTTATTTTCATTTTGAGACTATACTGCGAGAATATTATTAAAACATAAATCTATTATATGAAGGTGAATACACTAAAATGAATTTTTATTATATTTTTATCACTGTATATGTTTATATAACATTTCCATCTTAAAATAATTTATACACATAGGAATCAGTGTTTTGTAGGGTAAAGGGTAAATTAAACATCTCGGCG
>evgR001790c0g1_i01_Hydra
AAAATATATAACCATCAGCAAAATATTATAAAGTATTTAAAGTTAATTCAATTTAATGAGGTAACTAGAGAACTGCTTGAGAATTTTATATTGAGGGAGGCACTAAGATTAGAGTCAATTAGCCTACTTAAAACGGAGGCTATACAATTCTTGAAAGGAAAAGGCATACTATCTCCCATAGAAGAGACATTAAACCGCTTAATCTCGACTCAAAGAAAAAAAGCCAGACATTATATATTTGATACAGTTCAGTCAAAGCTTTCAACGACTTCCATTGCTAAATTGGCTGAACTTATATCTAGCGTAGGTAAATACTCGAAACTAGATAAATTAAAAGAACAGCCCGCTAGG
>evgR001792c0g1_i01_Hydra
TTAATGGAAGTTTGATAAGACATTATTTTATAAGAAGCTTTGTTTTAACTGCATCACACTTCACATTATAATATTTTTTGAAATGGCCGATGTTGTAAAGTACGGTATTCAGATTGCCAATCGAGTTTTTGTTGGAGGAGTTGCCTTTGATACAAAGGAAGAGGAACTAAAAGATTTTTTTTCACAGTATGGTAAAGTGAAAGAATGCAAAATTATTAAAGACAATCAAAGCATCAGTCGTGGTTTTGCGTTTGTCACATTTGAATGTAAACAAGATGCAGTCAAAACCCAACAACTGGGAACAGTATTTTTCAATCAAAAGAAATTAAATTTGGGTCCTGCAATTCGACAAAAGGGTGTAGTGTTTAATGGAACTAAAGAAGTTGAGAACGACCCAGTTTTGTCAACTGGTGTTTATATTCACCCAAGTGGTTATTCATATACAGTAGCACCAAATGGTGTCTGGTATTTTCACAATTCTGATGGTAAGCCACCAACACGAGTACCTGGCTCACCTCCAATGGCATATGCACCAAATATTTTGCAGCGTCGAAGGAGCTCTAATAGTCAAACTGAACAAACTATTTTGCTGAGAGAACGCCAAACACCAGGTGAGTGTTGCTCATCTCCTTCTTGTGAATCATTATCTCCTACATCTGTTAACTCTACTGATGTAAACTTGCCTATTTTCCATGGTGTTCCACCAGCAGAAGTTCACCCTATAATGCATAACCGTCAAGTCATTCAACAAACTGTTCTTCCTCCTCAACATGAATTTATTCAGCATTCACCGTTTGTAAATCCTGGTCTTCAATTTGCATTCCAAAGTATGTCTATCACATCTCCACCATCTCGTGCTCAAGGATTAACAGGGAATGACGCAGTTCATCAAATACAACACCCTCCTTTGACTGCTTTTCATGGATTTAATGCAAATTCATCAACTCAGATTAGTGAAATGTATTTTCCATATTACCAGAATAAGATAATGCCTGGAAACATTCCAGTAAATAACATTAGTCCACAGCAGATTCACCCGTTACATTACCACTATAATTTTCCAGTACAGAAAACAGAGTGCTATATGGTGCCTCCTATATGCATAAATCCCATCAAAGATACGTTTTTATACCCTATGTGTACAGACCAATCACTACCTTTTTTTTTTGCGCCTAATGAGATGAATTCATACAATAGTAATGGGTGTTATAAGCGAAAGTCTTATAAAGAATCTTTTGACACTGGAAACAATTTAAATCGTATAGCGGTTTAATAAATTAATTTTAAGCAATTTTTCAGACTATTTTTCCAGCGAGTGAAAGATTTATGTTTCGTATTTTAATTTATACCAAATTGTTTAAGTTATTTTTCATCTTTATTTTCTTTTAAATATGATTTTAAAATGAAAAATTAGCATACTATAGTTTTCATCTTATCTTTAAGTGACATGTGACTCACTGTTAGATTTTTTTAAAATATATTGCTGTTTAAAAATGTTTTAGTATCCTTTGTTGATTGTTACGTTCCTAAAGAATGCTGTCATAAGTATAAAC
>evgR001794c0g1_i01_Hydra
CAATAAAAAAATAATCAATTTTAGCGACACAAAACACCAACAATGAGATTCCATATACGCTTTTTGTTTTGTATTCTGACATTGATTACCTGGATATTAATTTATAGCTTCAAAGTTCAATTTTACGATTTCGTCATGACAACCTCACATGAAACAATTTTGCATGACGCAAGCTTACATAAAACACTTTTAAACAAAAGACCCTCGCATGAACTATTTTTAAATGAACATTTGCATGAGATATTGCAGGAAAGCATAAAAAAAAAAACTAACGATGTTCTCCGTGTTGCTGTTGTTTGTTACATACCTTCGTCACCTGAAGAAGTACACATTTTTTTTAATATGCTGTATGGTTCCTGGAGGTACATAAGTAATAAAAAAAATGAGCTTAATGAAGCAAATTATTTAGACAATGTAAATCTTTTAACATTTACGCATGCTGATAATATAGTTGTACTAAAAACAGTTTGTCAAGAATATAATGCAAGATTGGATAGGGAAAACAATGACGAGCGTTGCTGGTATATTAGACAAGATTATGAG
>evgR001795c0g1_i02_Hydra
TTTTTAAAGTACTTTTTTTTTGCCGGTGCTAGAAGCTATGTCTGGAATTCTTTTCTTAAACAGATCTGTTCATACTGTTAAATATAAATCTATTTATTTTATATGGACCCATTTATGATATGAGTGGTACCACAAAATCGGGGATTAATTGCTTGACATGTGATTTTTCTAACATGATTGAATGAAACATGAAAATTTATATTATTGCTTTGAAAGTAATTAAGTAATGTTTTTTGGTATATATCTGGCCATCTTTCTTTCGTTATTTGGCACTTTTTGTAATGCAAAAAATAATCCTGAAATAGATTTAAATGTGCCCGAAATTATATCCTTCTATGGATATCCAGCTGAATCACATTTGGTTACAACAGATGATGGTTATATTTTAACTCTACATCGAATCTCTCATGGGAGTAAAACACTACCATTTACCAGACCTGTGATATTTCTTCAGCATGGGTTGCTGTGCTCCAGCGCAGATTGGGTTATGAATAGACCAGACGGAAGTTTAGCTTTCTTATTAGCTGATTCTGAATTTGATGTGTGGCTAGGCAATAGTAGAGGAAATAAATACTCCACAATGCATAAAGTACTCGACATAAATTCGGATGAGTACTGGAAGTTTAGTTGGGATGAAATGGCAAGCAAAGATTTGCCGGCATTCATAAATTATATAACAAATATTACAGAGCAACAACAAATATATTATGTTGGTCATTCACAGGGTACCATGATAGCATTTGCTGAGTTTTCACGCAACAAACAATTAGCTAGCAAAATCAAGAGATTCTATGCTCTAGCACCAGTAGCTTTTGTAGGTTCTATGACGAGTCCTTTAAAATATTTAGCTCCTTACGTTCCTGAAATTGAGTGGCTGCTCAAAGTAATCGGAGTACGCGATTTTCTGCCTCAGTCATGGGTAATAAAATGGCTAGGTTCACACATGTGTAGCCAAACTTTGATTGATGAAGTTTGTGCTAATATTGTTTTTATAATTTGTGGCTTTGATGAACCACAAATGAATAAGACGCGCCTCGATGTTTACATAACGCATACTCCAGCTGGCACTTCAGTGCAAAATGTTGTTCACTTTGAACAGTTGTACATGTCAAAGAAATTCCAAATGTATGACTGGGGTAATAAAGAAAACATGAAGAGGTACAATCAGTCAACACCGCCTATATACAATATCTCTAACTTTCATATACCAACTGTAATGTTTTCTGGCGGTAATGATTGGCTAGCGGATTCTAATGATGTCGACAAGCTGTTACGGCTTTTACCTAAAGAAATATTAGTATCTCATAAAGTGATTAAAACTTGGATGCACCTCGATTTTATATGGGGGATGGATGCACCAGAAGAAGTCTATAATGATTTGATAGCTGATGCTTTAAGGCAAGAGACTTTAAATAAACATTCTGATTAGAGTGAAATGGTTGTTTTTGGATATTCATTACTGATTTTGTAAAGTAATTGTTTTTGTTTAGATATTATAAATTAATAGTTAATTAAATAAAATATTTTTAAAGATTAAAAATAAATTTTGTAAACATTTTGTTTTTTGATTAAACGGATAAAAGATTTTCTCTATAAATATGAAAAAAAAAA
>evgR001796c0g2_i01_Hydra
TATTAAAATATATAAATAAAAAGAAATGAGTTTCTATAAACTTGATGATTGTATACATAAGCTTCATGAATTCGATTCAGTTATTCAAGATGGTATTGAAATAAGCTGCAAAGTTGCCCTTGATTGCGATGAAGATGAATCTGGAAAAGAGGTTGTAGATAAATTGCATGAAGTTATTATTCAATATGCAAAATTGAAACGCGATCTTAAACAATGTTGTGAGGCAGCAAATGATTCCAAAAAACATTTTCAGAGTATTTTGAACAGAAAAAATGGAAGTCAAGAAATTGGAAATTTTTGTGATGTTTACGAAAAAAATCTTGAAGCACTGCAGTGTCGAAACACAGATGTTGAATTACTTGAATTTAGCCATGTTGTCGATTTTAAGAATACTGTGTGGAATCAGCATCATAAAATTAGTGAACCTTTAGATGTGGAAGTAGATTGTGAATTAGAGATGACCCAGGATAACACAAATCAATACATTTGTCCCATTACAAAGACCAAGTTTGTTGATCCAATGCTAAATACTGAGTGCGGTCATTCATACAGCAAAGAAGCTATACTTTCTCATATTCAACTAAGTAAAAGAAAATGTCGATGTCCAGTAGGTGGTTGCATTCAACTCGTAAGATTAGAATGTCTAAAACCAAATGTTGTTATTTCTTTTCAGATGAAAAATTTGTGCAAAAGCTAGTTTTTTAAACATTGTTGTTTACATTGGTTTTTAATGTTAGGGGACATCCACAAAGTACGTACGCAGTAAAACGACTGATTTAGAACCCCCCTGTACGCTTTTCAGCGATCCCCTTCCCCCATCCTGCGTACGTACGCATTATTTTACAAAATCAAACCCTCTCTCCAACGCATAAATATTTCAAAACGTTATAAACAAAAACATTTTCTTAATTCAACGTTTCTCTAATTAAAACTTAAAATGTAACCGAATATACTTATTTAAATCTTAAAATGTTAC
>evgR001801c0g1_i01_Hydra
AACCTATTTTAGTGCAAAAAGTTAACTAACAATGGATGTCTTGGGAAGATTCTTAGTTACAATTGGTATTGTTTCTTTAATACATTGTGGTTACTCAGCTATTCAATATCGAACATATTTGAAACTTATTGAAGAAGAGTTTACATCTGTTCCAATTGATATAACACTTCAGGTATTGATGAGTGTTTTGATATCAATATTTGGTATAGTTCGTGTTGCTGGGGAACTAAAGGACATTCACGCTGCTGCAGAATTAGCTAGCAAGTCTTGGGAAACACTTGGAAACCGAGCTTCGTTTTTTACATTTACTCATAGAGGTCAGCAACTATTCCAAGACAAGTAAGAAGAATATATTCAGTGAAATTTCGGCGAGAGCATATCCAGTGAACATTTCTATTATAGTTATTTTTCACAAATTACTAAAGTTGCTTCAACTAATCATTTTGTTGTACTAAACTTTTAGACATTTTATATTTGTTTTATTTTATATTTGTTTAATTTAATTTATTCATCAAAAGTTTAATGATTTTTCTTAAAGTTATAATTATGACGAAATAATTTATTTTTCAATGTGTTTGCAAAAAAAAAA
>evgR001844c0g1_i01_Hydra
AACAAAATAATTTCTGATGACCATGCATACGCTCTAAATAACATAAGTTATTCAAGAAAATTATCGTTATACGTGTCTGATGTTGTTGAATATATTGCAGGATTTGTTGTAAAAAAAGTTAAAAAGGTTTTAGATTGTGACATTTGTTTAGCCTTAATTCAATCAAAGACCCCTACTTCAACTTTAATAAAAAATAAAAATTACAGCAGTGTTAAGTTCTGTTTGGTTCAACCATCCAAAGACGTAATCCACATATGTAAACTTGCAGAAAGTGAAATTAGAATTTTTCAGGAAGAAAATAGGTTGCTTCAAAAAAATATAATGACAAAATTGATTATTAATATTCATAACAGGGTTAGCAGTTCTATTTTTATTGATAATAGAGATCATATGTTTGACAATGATGTTCAATCTAATCATTTGGTCAATTTAGTAAAAATGATTAGCTTCTATTATTGTAACCTTAGGTTATACCATCAAGCAGATACCTTAAAT
>evgR001889c0g1_i01_Hydra
TATAGAAATTTTTTGAAATACTTAAAATATTTGTTTTTTCAATCAGACATGGCTTCCTCACCAGTAAATATATCATGCAATGGCAACCTAACGTATTCAACAATGATGCCTTTTAAAATGGCAATGACATGGTTATATGCTACAATTTTTGTGATTGGTTTGCCCGGTAACATATTGGTAATCTTAACGATCATTAGAGTAAAGCGCTTACGAAATGTAAGAAATGGGTTTATTGCAAACTTAGCAGTAGGTGATTTAATAAATATTCTTTGGTGTTTACCAACTTCACTTGTTTCACTTTATGTGCCGTGGCCTTATGGAAGGTTTGTTTGCAAGTATATTTTTCCGATTTCTGATGTCGTTATTGCAAATACAATTTTTACTATGATGCAAATCACCATAGATCGTTATCGAGCCATTTGTTATCCATTTTCCAGAAAAGTTTCCTTTAAAACTACTTTGTATATAATCACTGTGACGTGGATTTTTTCTTATCTTTGTTTTGGACTTCCATTAGTTGGATCTTTTGAAATTTCGTCTGGGTATTGGGTCAAAAAAAGTTGCAATTTGAATTTGAAAAATGCAGAAAAAATTGCTCATCGGATTCTACTTTTTGTGATTTTGTTCATTCTTCCATTTATAGTGGTATTATTTTGTATATTGCACATGCGCCGTGGTTTGAGAGATATGTTACAAATGACAAATGGTTCGTTAGGTAAAATGCAGTTGTTTTATCGAAC
>evgR001894c0g1_i01_Hydra
TCACTTTGCTGATAATATCTTCAAGGCTGCCCAAGAAAGAGGAGCCAAGACCAAGAGTGAACACGATGATGAACAAAGAAGAAATAAGCCAACATTTGAAGGAAGTGGAAGAAGATTAGGTAATACTGAACAATCCAGCTCACTTATCCCATCTGAACAAAAGAGAAAGGAAAAGGAAGTGACAATTACATTCTGGCATGATGGTTTCACTGTTGACGATGGACCATTGAGAGGATATGATGATCCACAAAATCAACAATTTATGGATGCTATCAACAAGGGATTTGCTCCAAGAGAATTGGCTGAACCAGGTGTTGACATTGCTGTAAATTTAATCAACAGGAAAACTGACAAGTACACTGAACAACCTAAAAATTTCAAGGCCTTTGTTGGCTCAGGTAGATCTTTGGGAGCTCAATCTAGCTCTTCTG
>evgR001904c0g1_i01_Hydra
GGCATATGGCGTACCTACAATTATAATGGTATCAACATATTTCCTAATTATACGTGAACTTTGGAAATCCAATGCTCAAATTAATCCCAACGACAGTAATTCACCTCGAACTCATTTTTTATTGACTATTAAACTTGTAAAGTTAATTGCTACTTTCGTCATATGTTGGATACCTTTTTTTGTTTGTTTTATGTTGCAAATTAAAACTGTAAACACTGCGTTAATTATTGCAACTAATTGGACAAAGTTATTGGCTTATTGTCACTGCTTTCTTGATCCTATTCTATATTCTATTTCGCACCATGAAATCAAATATGAAATTGGAAGAACTATTTTATCTGTAACGGGAACCGATGAACATTTAAAAAAAAAA
>evgR001915c0g1_i01_Hydra
AATTCTTTTTTGAGTATACAAGGCAAAAAGGTTGCTGCATTGAATAATTCTCTCGAGTTTTTTATTGCTCATAAAAATTATAACACATGGTATAAAAAAGGTCTAACTCTATACCTTATGCCAATCGTTTTTTCAAACCAAGTTTCGGAAAACCAAGCATACAACTCATACGAAGAGGTACTTGAAGAAGTAGAGTCAAATAATGTTGAATATGGAGTTATAAACATCGATGTCTTACTAAATATGGACTATAAAACAAAATATAAAAATGTTGTTCTTGTGAAGTTACTAGACACCGCTTCACCGGTCATACTAGCTTTTGGTAATGAAACCAACTTGATGGCAAATCTATATCGATTCAATATTGACCAAAGCGATACTGTTTTGGCCTGTTTAAAAAATTTAAATTTCAAAGATATTTTACAAAATGTACAAGAGCTGTTTAAAAAAATTCCAAACATAGAAACAGAATCTTCACAAACCCTCGATGACTTTATGAAAAC
>evgR001926c0g1_i01_Hydra
TAAACTTTAAAATTAAATATAAAAGTAAATTAAAATGTTTTAGTTTTCATTGAGTAAAAACTTTTAAAATGATAGTAAACATTCATTCGGTTGATGAAAGTAGAAATTTTATGAAGTTTAATAACCACATCGTTAATGGGTACAGAGTTAGTTTTTCAAAAATTCAATGCTTAAAAAGTGTATTTATGATAACAAATGAAACTATCAATATATGGACTCACTTGCTTGGTTCATTATATTTTTTTTACACAACAGTTAATATTAACGTTTCTGTTTTACCAAGTTTCTCTGGCTCAACAACAATCGACTATTTTATCATGACAGCTTTCTGTTTGTCCTATTCTTTGTGTTTAATGCTTTCCACACTTTACCATGTGTTTAATTGTCACTCAGAAAAATATTACATCGTGTGGTTAAATAATGATCGTCTGGGTATTACGCTAGGATTATTAGGCAGCTACATTCCTTCAATTTACTATGGATTTTACCATGAATATTTGCTCCAAGTTTTTTACACAAGCTCTGTACTATTGCTAATTGTGTTTGTTATTATATACCAGCTAACCCGAACTCGATATAGTCTACACGATAGTGTTTTAGTAATTTTTTATATAGTAGTGGCAATATTTGGTATTGTCCCAACAATCCATTTTGTATTGCTGCATGGATTTATGAATCCAATTGTTTTACAATTTCTCCCTAGAATTTGTATTATGTATCTTTTAATTCTGACTGGTTATTTTTTTTATCAAAGCAAATTACCAGAGTGTTTAATTATAGGGCTATTTGATTATGTAGGCGCCAGTCATCAGATCTGGCATATATTTGTGTTTCTCTCGCTGCTTTGGTGGTATAACACATCCGTCACTCTCATTCGCTTGCGGTTGGAATCCAAAATTGAATAAGTTTTTTAATAATTGAAATTGTTTCTTTTTTATTATAAATAAATAAATTTTTACAGTTTGTTACACATAGAAATTGTACATTTGATAATTATTATTAATTGTTATTTTTTAAATAATCAGTATTTCAACTATATATGTAAAAATATATTCAAATACAAAAT
>evgR001927c0g1_i01_Hydra
ATTGTTTATGACCATTTCTTCATTTTTACTAATAAAGACAAAGCTAATATTGGAACTCCTCGAGCAAAAATCTAAATTTCAATTGTTTGCTTCACAAATGTTGATAGTTGCCTTTTTATTGCTCGTGCAAATATGTACAGCAGCAACACATTTAATCATTGATTCAGGACAAGCTCCTATTGCAAATCAGTCAGTTAATAACGATTCTTTATTAGAAAAATTTTGCAATGTAGAATTGACTCCAGCAAAACTTGTTTCTATTTTATTTCCAATCATTTTACTCACAATCTCGACAATTTTTGCTTTTCGTGACCGAAATAGTAAACATACATTTTACGAGCCTAAATTTTTATGCTTTTCCTGCGCTTCTTTGAGCATCATAATGGCAGCTTTTTTATCGAC
>evgR001944c0g1_i01_Hydra
CTTTACAGTAGTTGCAACGACATTTGGTATTGCCAGCGCTTTAGGGTATTCATTACCAGCATATGGACTAATTCCACTTACTACATGGTTTGGGTATTACGGAATTTGGGTACTATTCGCACCAGCAATCATTGGCTTTATATGGGCATTGAGTTATTTACTGAAGCTAGAGAAGTCAAGCGGCGCTTACGATAGTTATCCAGACGAACCAAAAATTAAAGACACTGCCTTAGAAGAAGGTGATTTTAAATATGAGCTGGATGGCGAATATGAACAATTTAGCAGCAAATGCGAATATAGCGTAGACCTGCTGAATAAACTGGAGGCACTAAGTAAAGAGCAAGATATAAAGCTAAACATGAAAGTGATAGAAAAGGCAATAACCTTTACTAAAAAGTGGCATACCAAACAGATGCGAAAAACAGGGGAGCATCCATTTTATTGGCATCCTCTTAGAGTTGCTGAAATGGTTGCTACACATTATTGCAAAACGGAT
>evgR001947c0g1_i01_Hydra
TAAAAATCAGCTCGGAGAACAGTGGAGATATTAAAAAAAGTGTTTTGAACGCGAAAACAAGTCTGTCATAAGTTGTTGTTAAAAGTGTGCAATGCAGTACAAATACATCATCAACCATGCGTTGCTATTTATGCAAAGTCGCTTCGATAGATTTAAACAAACTTTCAAAAATATTGAAACTCATAATAGATTTCAAAAACATATAGTTTGGATTATCAATTTTACATGCCAGAATCCAATCTTTCGAAACTTTACTGGACATTTCGTACAAGATAACTTTAGAAAAATGCCAAGCGAGAACACCAACTGAAAAAAACATTGTTAAAGAACGAAAAGCAAAAATTCAAGAACAATTTAAAAATGAGCTCGGATTGATTGTCCATGTACCAAAACCATATTATGGGAACAGCAATAATGGGAACTCCAGTAGAAAATTTTTCAGTAATCCTGAAATATCCGCAAATATCACCGGAATCGACAAAAAAATGATAGATAGGTTTGCTATAATATAGAAGTCATGTCAAGTGAGCACAAAATTGACATAAACAAATTTCGTATTTATTGTTTAGATACCAAATATATTACAAATACAAATACAAATACAAATATTTACATTGAACTCTATTCGTGGTATCCTTTGTCAACCACTGTACATAAAATACTTATGCACGGACCTACAGTCATAGAACATGCAATCTTACCTATGGGAGCCTTGTCTGAAGAAGCAGCAGAGGCCAGAAATAAAAATTCCAGACAATATAGGGAACGTTACAGCAGAAAATTAAGCAGAGTTGCCTGTAATATAGATGTCATTAACAGACTTTTGCTAACATATCCATATCTGAGTAGTATAAGGCCCAAAGCCTCGAAAAAAAGAAAATCTTATAAACCTGAAACTAAAGAAATGGTATAAAATGGCCGAATGAAGGTATTACTGCTAATAAAAATGACGATGACAATGTCGACGAATCAGATGAAGATGAGTGGATTGCGTCATTAGATTGAACTTTAACATTTAAATTTATTTAAAATAATTTTGTAGAATTGAAAATAAATAACTGAAATAACATTTTATAGACTACCAGTAACCCTAATTGCATAATTAAATGCATTATAAGCATGCTTTGTTTTTTTTCGACATTGAAATGATT
>evgR001952c0g1_i01_Hydra
GTTTATTGTTTTAGTATGGTTTATTATTAACAATATTTTCCATACACAAATAAAAATAAAGATGCAATTTGGCTCTCTTATTGATTACAACATATTTTCATCCAGCTCAGTAAAAGAATTTCTTAGTCCATTGATAAAGAAAGGAAAGCGAAAATATTTTGGGTTATTGGAGTCACCATTAAATTATGGTGCAGCTGAAAAATTATATGCCACTGAGCTTCAGTATAAAATTTTCGTTTGTGGTAATGGGGCTATTGGTAAAACGAGTTCAATACTAAAACTTTCAGGGCAAAAGATTCCAGATAATGTACCTGAAACACTTGGCATACAAACAACAATTATTTTTTGGCCTGCTCGAATACTTGACACCAATCAATATGTTTTGTTTAAGCTTATATTTTTTGATACAAGTGAACAAACTTTAAGCACATACAATCATGTTCTTCCAAGTTGTACTGATAATATTGATTGTATTGTATATATGTTTTCTTTCTCAAGCTATACTACATGGGAAGATATACCTAAGTTAATTTCCAGAGTAGAAGTTGGTGAGGATGTACTTAAGGTATGCACAGGAGTACAAAATGCAAATCTTTTTTCGAAAAACATTGAAGTAACAAATAAAATGATTGAAGACTTTGAGCAAGTTTGGAGATTTCCTGTTTTAGCTATTAATAATGAAAATAAAGAAAACAGCACTATTTTTGAATTATTTGAAGAAATAAGTATATTCATGAACAAACTATGTGAGCTATTATGGTACAGAGATCAAGTTAGAGCCAATTTGATTCCTAAGAATCAAGTGAACTTTTGTTTTAAAAATAACACAAATATCTCTGATAATCAGAGTTATTCTGATAAAGCTTTTTGTTGAAATTAATAAAATTAAATTAGTAAATCCTTTATTTTTCCTGCAAATATAGTGATGTATATATTAATTTTATAATTATAATACATATATTTATTGTTAC
>evgR001953c0g1_i01_Hydra
CTTCAAATGCCAGCTGAAAACGTTGAAACTGTGGAACAAACTGTCGTCGATGAAACCATCGATAACCCAGAAGTCATCAAGAAATACAAACTTGCTGCCGAAATTGCTAATGAAACCATGAAGCACATTCAAGGTTTGATGAAGCCTGGTGTCAAGGTTATTGAATTGTGTTCTGCTGGTGATGCTTTCATGGAAGAAAAGATTAAACACGCTTATAAGAAGTCAAAGGGTTTGAAGAAGGGTATTGCTTTCCCAACTTGCGTTAACGTTAACGAAATTGTTTGCCACTTTTCTCCACTCGAAAATACACCAGAAGCTGCTGCTACATTGAAGGAAAATGATGTTGTTAGAGTTGATTTGGGTGTTCAAATTGATGGATACATTTCCAATATTGCTCAAACTTATATTTTGACACCAAACGTCGTCACCGGAAGAGAAGCTGACTGTCTTGCTGCTGTCCAACAAGGTATGGAAGTTGCTCAACGTTTGATGAAGCCAGGTAAGACCAACACTGAACTCATTCAAGCCCTCCAAAAGGTTGCTCAAACCTTCAACGTTAAATGGGTTAACGGTGTTTTGTCACACGAAATCAAGAGAGATGTTGTTGACGGAAAGAACACTATTTTGTTGCACGAAGATGCTGACCAAACTGTTGATGAATTCAAGTTTGAAACTAACCAAGTTTTCTGTTTGGATTTGGTTGCTACAACTGGTGATGAAGGAAAGTTGCGTGAAGGTACTTCTAGAACTACTGTCTATAAGAGAGAACCATCCGTTCTTGTTGATTTGAAGGTTAACGCTTCTCGTAAGATTTATGAAGAAATTAAAGAAAAATTCGGTGCTGTTCCATTCACAGTTAAGCACTTAGATCCTAAATTGGGTAGAATGGGTGTTTTGGAATTGTTAGGTGCTAGAATGATTGATCCATATCCAGTTTTGGCTGCCAAGAAGGGTGAATTTGTTGCTCAACAAAAGTCAACAATCCTCATTGCTAAGAAGCAAGTCCAAAAGATCACTGGTTTGACAACCCAATCTTTCCAATCTGAAAAGAGCATTCAAGATGAACAAATCCTCTCTATCCTCAAGTCTGATTTGAAGATTGTTCCAGAAGCCAAGATGGACGAAACCGATT
>evgR001972c0g1_i01_Hydra
AGAGCAAACAATAGAAACACTGGAAACTGAAAACGAGTGGTTGCTGAGCATAAAACATAAAGTCGTCACAGAACTTACAAACTTATTATCTTGCAAAGATTCAACAACGTTTCCAAGAGATGATTACTGTGACACTCTAATGCTCTTGGGTGTAGTTCCCCCACGTAGGGCCCACTTTCTCAAACCAGGAGCAATGCATCAGGCTCATTGGATGGCCAACAACATTTATGCTAGAAAAATGTATATGTTTTGCAAACAGATGCAGTATGATGATGACATGGTTACTAAACTCATGCAAATGAATGGATTCTTGGCTCATTTCTACACTCCTGCGTGGATGAAAGCATCATGTGGAGCAGATGCACCTATAAACAACCTCCAGTTTATTCATGACATGATTGATTTTAGAGTTGTTGACAAAGATATTGCTGACATTGTGATTAACAAGTTCCGTAATCATCATTGGTATTTATCTGAAGAAGTCATACCATTTGCCTTGTTTAGTAAACATCCTCTGATGACACAAGAAATAAAAAAAGAAATGGCAGATCAATTGCTTTTGACTCCAATACCAGAAAACTTCGGCCTTGGAAAGCCAGTCTTCAAATTGTCAAATTGCCAAAGCCAGTCAAATTGTTTGTGATACAACTTTGACAAGCTTGATTGGCCCTGAATCTCATTCGTTGTTCCACATTTTGAAAGTGAGCACTGACTGGCTTTCCAAAAATTT
>evgR001976c0g2_i01_Hydra
TAAAACTAATCTTTTATTGTCTAAATAAGATAAAATTAAGGGAAAAGATGAAGTCGCAAATTTTTCATATCCTTATTGGAGCTGGGTTGTTTATAATCATCGAAGGATTAAAAAATATCGACAGCAACGGAATACCTTTGAAATACGGTTGCCATTTGGATGAAAACCATTGCATTAAGATTATTAATAAAAAGAAGGAGTGGTGTTACTCAAATGAAAACGGATTAGACATTAAAGTGCCTTGCAAAAGCAATGACGACTGCATAAACTCAATTGAATGCGTAACCTATCCAGAAAAATCGGTAAACCTTCTTTTTATCGATCAACCTTCAACGTCAGATGTTCCTCCGTCGGTTTCCGGATCTCCTCCGTCATTAGAAATTAATGTCACAATCGATTCAAATCAAGCAAACAAAAGACCTGTATCTCTTCTTTCTCATTATTCGTATGACACCGAATCTGGAGTTGGACCTCAAAACTGGTACAAAGCATTTCGTAAATGCTTAAGTAATATTAATAAAAATCAATCTCCAATAGACATTAAAACCTATGAGGCTAGAAGAATGGAAGATTGGAGTCCTGTCATGGTCATTTTCCCGCCAACCTCCACGCCTTTTAAAGATGCTGTTTTAGAAAACAATGGGATTGCTCTAAAACTGTCAATTTTAAATGACCCACCAATTAAATTAAAGGCTGCTGGTCTTAAAAATGACTATCTTTTACAAGAGGTGTATTTTCACTTTGGATGCGAAGCAGATAGAGGATCAGAGCATCAGATTGATGGCAAAAAATATTCATTAGAAGTACAGTTTGTGTTCTGGAATAATGTGGAATATAATTCTTTTAATGATGCCAAACTTTCCGATACCGTAAAACTTCCAACTGGAGAACGAAAGGATATCCAGAATCCTGTTGACAATGGTCTTGCTATTGTTGCAGTTCTATTTGATGAAGATTATAACGTTGATATGAAGAATTTTGATGTCTTAAATAATGCAATAAACGAAATCGGTGACAAACCAAAAGGAAAATCAGTAATTACAGAAAATTTAGGAGAGTTATTACAAGGACTCTTACCAGAGGATATCAAAATTGGTCATCATGATTTTTTTAAATATAAAGGATCTTTAACAATGCCTCCGTGCTCTGAGTCTGTAGATTGGTTTGTTTTGGCTGACTGTTGCAAGCTTAAAGTGCCTAATAAATTTTTACAAACGTTACAAAAACTTCGAGGCAATCCAAGTAATCGATTTGCCGAACTATGTGATAACTTTAGACCAATCCAGAAGAGAAATTCTCGGTTTAGAACGGATATTTTATGGTAACATTAAATAACTAGAAATAAAATCAAAAAAAGAATTGATAGTAAAGTTGTACTTAAAAAAGATGTTGTATTTATCTACTTGTATGTATTTAGCAATAAATGTTACATTTTTTGAAAAAAAAAAAAA
>evgR001979c0g1_i01_Hydra
ACCACAAATCATATCAACTAAATTGCAACAATTAAAATATAAGAAGCAACCAAAAGATAGTTTTACTCAAAAGTTAATAGAACCAACATTAAAAAAGATATAAAGGAATGATCTGTTGGAGAATAAAGTTGCATCATTTATCCAATTTAACAATTGTAGGGATCCTTTTTTGTATGGCAACAGTTGTAGCTGCAGTCAGTAACTTGTTTGTATTGATCATTATAGTTCGTATGAAAAAAATATCTACATCTTATCGTATATTGATTTCACTTGCATCTGCTGACCTTCTTGTTGGGCTAATATTAGGTCCAATAACTATAGCACAAGTTTTTGATGTCGTACTGTTAGATAATTGTTTAATACATTCAATTCGAATTTCTTTTCTAATTTTGCTGGTTGGTTCTTCATTATCTAGTTTAGCATTAGTAGCATACGATAGATATCTGCTTCTGACAAAGC
>evgR001986c0g1_i01_Hydra
AACTGGTTATTTTGACGATATCATGTATCTTCAATTTGGAAGTGTTAATGATGTTACAAACAACTGGAATTCCTTATACATAAGATATTGTCAAGATAATTCACCAACTGGAATTACTAAACATACTGCAATTAATTGTAACTGCAAAGGAAGGTCCTATTATACATTTAATTATCCCATTCAACAAAGCTTTTTGTATGGGGTTATTTATGAAGAATTTGTAATAGCATTTGATATGTACAAAATTGATGAAAGTGAAAGAAATATTTTGAGTTTTTTTGAAAACGGTGACCTCTGTGAAGATTTGCCGCAAGTTAGTTATTTT
>evgR001989c0g1_i01_Hydra
AAACAATCAATTGCCACCATTCTATTGATGGCTTCCTTGATGGTCGTAGCCTGCTTTTTGTTGGTTGGTTTTGGAGACGCTCAAGTGAGTTCTGCCGTTGAAACTGCTTTGGAAAAGACTGATAGTGGAATTAACAAGTTACCTGCATCTGCAACCACCAAGGATCAACTTTCTGAACTTCATACTTTTATTCTCTCTCCAATTGCAAAGAATGTTGCAAATAAGATTGAAAAAGTAGCCAAGGAACACCCAAACGAAGTCCAACAAATCAAAAATGATGTCAATGCTTTGAGAGGAAAGTTTGAGAATAAAGTCAAGGCTGAGGCACAAAAACGTAAAGATATCAAGAAGTTCATGAAAGAAAAGGAAGCTGAATTGCTCCAAAAGCTCAGAC
>evgR002008c0g1_i01_Hydra
GTTGTACGATTGTGGGATATGGCTACTGGTGATTTAATTGAAGCTTTTAGTGGTAATGGTGGTCATGATGACTCTGTATATAGTGTTGCTTTTTCTCCTGATGGTAAAACACTTGCTTCTGGTTCTTTGGATAGAACCTTAAAGATCTGGGATATCAAATCTGCTTCATGTATTGCTTCACTTTCTGGTCACAGAGATTTTGTTCTCTCTGTTGCTTATTCTCCTGATGGTAAATGGTTGGTGAGTGGTTCAAAGGATAGAAGTGTTCAATTCTGGGATCCTCGTAGTAATGTTTTACATTTGATGCTCCAAGGTCATAAGAATTCTGTAATTAGTGTTTCTTTGAATCCAAAACGTCCACTTTTTGCAACTGGTAGTGGTGATTTCCGAGCAAGAGTTTGGAGG
>evgR002012c0g1_i01_Hydra
ATATCCACATCCACCACAATCAGGCAGTGTTCCTCCTCCAGGTTATGCTTATCCACCACAACCAGGTTATTCCTATCCAGCCCATTCTGAACAATATCCATATCCTTACCCACCTACTTCAATGGTTCAACATCCTTATCCTCACCAAATGACTCCTTATCCTCCACAACCAGTTCAAGTTCCAAAACCACAACAATCAGTAACTCCTGTCATTATTTCGACATCTGCTGCTACTGAATCTCAACCTCTTTCTGGAACACCTTGCAGAACTGTCCCAAGTGGCGAACAAACACCAACAACACCTGTTGGTTCTCCACAAATGAAGATGGTCTACCAAAGGGAAACCAAGGTTATCCATCCAGTTAGATGTTCAAAGGTTAGTCCAACTGAAACCACTTTTGAAATTGCTACCAAGATCAAGAGAGAAAGAACTGACGAAGTTGAAACACCAACCAAGAAGATGATTGTTCCTGGGTGTGCAACAGCTTCAATCATGGTGACACCAACAACAGTTCAAAACGGAGGTATGGTAACACCAATCAAAGTAGCTGACAAATAATTTATAAAAAACCAAC
>evgR002022c0g1_i01_Hydra
AATGTAGGAATTCCTTTGGGAAAGGTGAATTTAGACTCATGTAGCCAAACATTATTGGCTCTTAATGAAAGTTACAATCAATGGTTTAGTTGCTATGCGAGAGTAGAAAGTGGAGTAGGAATGGTTTTTGAATGGGATTTCAGTGAAGAAAAAAATGTTGTTTATAAAGTTGCCTCATCCCAGATTTGCCAGAGCTCACCTTGTTTTTACAATTGTGGAAATAATACACTTAGCATATTGGCGATGATATCTCCGATTTTATCAATTGCTGGCGTGTTTTTGTTTTCTCAAACAATTTTTTATTACGATTGTTACTTGATTGGTATTGAGTTATATGCAAATAAGAATAGTTCATTGATTTTACAGATATATGAAAAGAACTCCCAGGTTTATGTTGCTACACAAAGCTTCAATTTAAGCTTAACTCAAGGTTATAATGTTATAAAGATTCAAAGTCAAATAGTTGTTTTAAAAGGCTCTTTTATTGGTTTTTGGACGTCTGACATCAAAGCTCTTCAACTAGTTAATTCTGAGAGTGATTCAATTGAATTTTATAATGCGAGTGCTTATGACACTCTATTTACAAGAACTTATAAATTTGAAAATGTTACCTTTCAAGCAAGAGTTTTTGCTTCTGCTTTATCATACGTTTATGTTGATTACCAATTCAAACTAGCGGGCTATCATACTATATCATTTAATGTGTCGGACATGTTTGGAAATTCAAAGGTTTCAAATCAATTGGTTGTGGCTGTCATATCTCATAATGTTTCAATACTGGTTGACAACAGTATAAAGTATGCTGTTGCTACAAAAGAATATTCT
>evgR002040c0g1_i01_Hydra
TTTTTTTTTTTTTTATTGTTTTAAAAACAACATTTTTGTAAATGCTTTATTACGTTTTTAACTTTTAATATTTTATAAAAAAATGTTCGAAATACACAAACAAATAATGGCTTAGAATTGGTAGTACCTTCGCCGAATTTACAAACAGTGAAATGATTGAACCTTCGCAAGTAGACGTTTTTGTTTTACCAAGACAACAAAAGAAAGTATTACGTACAATATTAGCAGTTTTAGTTGCATCAGTATTATATGTACCTTTTGGCAATGGTTTGAGATTTGCAAACGAAAGTGACGATCCAGGAATTCAATATTCACGATTTTCTAGTGCATTTATAGGCAGTTTGCTTTCTGGAATTTTTATCGACAAGATTGGAAGAAAATTTACTATTATTATGGCAATAATATTTTATCTTCTTGGTTATATATTTGCGCTGCTTAGTTGGCCAACATTAAATAGTTCTAAACAAATGTCTACAATTTGCAGTTTTGTTGGAGGCATTTACTATGGAGTTGGCGTTGGAGTGACTTCTTTGGTGGTCCCCATATATATTGCTGAGATATCATCTACATGTTTACGCGGACCAATGGGAGTAACATGCAACAACTCTTTGCTATTTGGTGCATTTTTACTGTTATTTGTTGAGCTCAAACTTTCAAACGATTTATATTTTAGTAATATTATTGGAATTATAATAATTGTATTAAGCTTTTTAATGATTCTTATGCCAGAAACACCAAGATGGCTTTTAGCGCAAAACCGAAGAGATGAAGCTCTTAAAAGCTTATTATGGCTTCGTGGAAAACTACCAGATGCAGAAGATGAATGCAATGACATTCAAATAAGTGTAGAATATCAGCAAGTCGCATCTGTAAACGATTATCGCTCACTGGGGTTGTTTAGACCACTTTTTATAGGAAGTTATTTGTGTTTTATTCAGCAATTTATTTATCATCAATACTTTTTTAATCTTCTAAATTTTTTAAAATCTCAAAACAAAGTACTTGTTTTGCCTATTTGTTTGCTGTTGCAGACCCTACTTGCATTACTTGGATGCATCTTTGTGCATAAATTCCGTCGAAGACATCTTCTTATGTCTGGAAGTTGTTTATTGTTTTTGTATAGCATTGTTATTGCCATTTTTTATCAGGTAAAGCAAGATTTTAATGAAAGAGTTATTTCTATATACACTACAGTATTTTATGGTATTGTTTTCACTTTATCATGGGGTCCGTTGCCTTGGATAATTTTTTCGGAAATATTTCCCCCACGCGCTAGAGGATTTTTTGGTAGCATTGCTATATCTGCCTATTGGTTGTTGAATATCGCAAGGCAGTTTTTCTCAAATTGGCTGATAGAAAATGAAAAGTTTTCATTTTGGCTCTTATCTGGCATATATTTAATAAGTATTCCGTTTATTTACTTTATTGTACCTGAAACGAAAGGCAAAACGTTAGAAGAAATTGAACATTATTATGTTATGTACCGAACGTTCCGTAATTATTTGTTTTAAGTTATTATTAGTGTTTTCCACTCACTTAAATCGACTAAAAGTCTAAATTTGAAAAACTAGTCAAAATTTGAAATTAGTGGATTTCAAAAAATTTAATAGTAAATTTAATCGACTTTGTATTTTTTATTAATTGACCTAATGTCGATTTAGTCGGAC
>evgR002061c0g1_i01_Hydra
CTTGAGTATTTGTGGTGTTGAAGGTGAAGAAGCCGAAAAGATTATCTATGCAAACTGGCTCCTCATGGTTAGAGCTGGTCTATTGGGTCTCCAATTCTACAATCCTGAGACCAAGACCTTCTCACAAGCTCACATGCAAGCTAGATACAGCATTCTCAGAGTTTTACTCGAAGCTGGTTTGGTTAAGATCAACAAGTTGGAGAATGATTTACAAATTGTTTTGGATGGTAGCAAGATTGAAACTGTTGGTAAGGAAGCTATTGGAAAGTACTTGCTTCATATCAATGTTTCTAAGGCAACTGCTGACAAGGTTGGTGCAACTGAACTTTTTGGCAAGTATACAACTGTTGAGGATGAATTTGTCCAAATGAGAAAGATTGTAATGGAAAAGAAGCAACCAAGAAGAGTCTTTGCTCAACCTCACACTGTTCTCAACGAAGAAGGTAAG
>evgR002081c0g1_i01_Hydra
TATCGACGATCACAAACTCAGTATTATCGCCAACTAGCATATGAAAAAATTCGAATGAGCAAACAACAAAAATTAAATAGTCCTTGTTATGATGAAAATGAGCCCGAGTTGAGCAAAGCTGCTCATAATGTTCTTGAAAGACAGAGAAGAAATGAACTGAAATTACGATTTAATATTTTGAGAGACGAAATTCCAGATTTAGCAATGAACGATAAAGCGCCAAAAATACAAATTCTAAAAAGAGGCCGAGAAATGCTGAAGGACTTAAAAGCGCAAGAGCAAAAACTGATTGCTGACCATGAACTAGAAAAGCAACGCCAAATAATTTTATTAAATCGCATCAAACATTTGCAGTTGGGACTAATATAAATATTAGGCGGGAAAATTATAAATAATCGGG
>evgR002086c0g1_i01_Hydra
TAATCAATCAAACTCTACAGACGACTTGATGTGGCAAGCATTAATAACTGATATGACAAACAATAAAAATATTGGATCAATATGTGAAAATAACTCCCCTTGTGGTCCGTTATTTTGCAGGGATGTATGTTTAAACATTAACAACGGTTTAGTGCAGAAAAACGCTTGCTCTGGAACAACGGATATAACCAAAACTGCAATAACTGCTATTTCATCAGTGTGGGGCCCAACTTTCTTAGCAAGTAATGCGATTGATGGTAAACTATCAACTAGTGCAGTAACAGATGATATCTCTGGTTTACCCTGGTTTAAACTTGATTTAATTTTCATATACCAATTAACTCAAATTGTTGTCTACAATCGGGATTATTTACCGGAAAGAATGATTGGTAATAAACTACTCATAGGTATAACTGACAATGCGGCTGGTTTACATGTCATTGCCACGCTAAAGATTGATGCACAACAAACTTTCATAGGGTTGTATATTACCAGATACATTTCAATTACTAAAGTTCCTAAGAGTAATGGAGATATATTACAGATAGCAGAAATTAATGTTTTTGTATGATTTCAACATTATATACAATATTTGCTTTAATACACCTAAATGTTTCCTTAATAAGTGACGGGTAAGCCGGAATGATAAAATAGGACTCTGGGGGAAGG
>evgR002088c0g1_i01_Hydra
TACAATAACCTAATCACAAAAGAGCAGCATGGTTTTATTCGTAAACGCAGTACCTGTACAAATATTCTAGAAAGTCTTCATGACTGGACTTTAAATCTTCAGCACCACTTTCCTACAGATATTATTTACTTTGATTTTAAGAAAGCTTTCGATTCAGTTTCCAATCACAAGTTACTGATAAAATTACCCGCTTATGGAATAGTTGGCAATCTTCTGAATTGGTTTTTGCACAAGAGATTACAGGTAGTAAGTTTAAACAATGTTACATCCCACACTATATTAGTAACAAGTGGTGTTCCACAGGGGAACGTTCTAG
>evgR002115c0g1_i01_Hydra
TGTAGCTCAAACACCAACAAATGACACATTTAAGGTAAACTTTTGGGATATTGGAGGTCAAAAATCGATTAGAAGCTATTGGAGAAATTACTTTGAAAATACAGATGGTTTGATTTGGGTGATTGATAGTGCAGATACCGACAGATTAGAAGATTGTAAGAGAGAACTTAATCTGTTATTGGGAGAAGAAAAGTTGGCAGGTGCCAGTCTATTGATTCTGGCCAACAAGTCCGATTTAGATAACGCCCTTAGTGTAAATGATATTGCTCAATTCCTTGGCTTGGAACAATTATCTCAATCAACAGAAACAACAATCATTGCATCAGGC
>evgR002137c0g1_i02_Hydra
CCTAAGACATTCAGTAAAGAATCATTTTTTGCTGAGTATGCAATATTTTTACCCAATCTGGTTATTACTATTGGCTTTGTAATTGGAATTGTTTGGGGGTTGTTTGTTATTCCTAAGGGTATACGTGCTCCCAAACATGAGAAGATTTGTCTAGTTACAGAGGAAAAGTCAGAGCTTGATGATGTGAGAGAAAGACACGACTATTTTATGAATCTTTCTCTAACAGTAGATGATCACAATACATCCGAAACATCTTCATTGCTTATTAAAAGTAAAAATACAGAGTCATTGTTTTGCAGATTTGTTTATATGTTGAAGTATTCATCTATTGGTAGAGTGTTAAGATCAAAAGTAAGCTTGTCATGTGTCATTCTTTATGGCGTGTACGGAATTATTGGCATTGGATTTAGAGAAATTTCACCTGTATTTTTAGCAACTTCTATAAAATATGGTGGAGATGGAATGACAGTATTTGACATCAGCATTTTAATGTTAATATATGATGTTATCCTTGTTGTTGTACAGCTTGGTGCTTCAAAGTTGCAATTTTGGTTTGGAGCCAAAAAGACATTTATTGGATGTATTCTGGTATTTGCTTTTATGACTCCCCTCTTACCACTTTCTGCATTAATTAAAAACAGTGCATTGCGATGGATCTTTCTAATGATTGTACAAGTTTCTGTTAATGCAGCAAATAATGTATGTTTCATCTGTATAAATATTTTTTTGGCTAACAGTGTTGGACCTGATCTTGTTGGTACGGTCAATGGTTTGGGAATGTCTGTGTCTAGTGTTGGGCGGGCACTTGGACCGATGATCTTTGGATTTTCCTACAGTTGGTCATTGTCAAATACTGAGTATCACAGACTTGGATTTCCATTTAATCAATATTTTGCGTTTTTTCTAATATCCATTTTCTGTTTTCTAAACCTTGTTTACGTTTACTTGCTTATTCCAACTTCGTTGAATAAAAGAAGCATACAACAAGATAAAACTGAAATTGTTATTAGGC
>evgR002159c0g2_i01_Hydra
GCCTTTGACAAAGTTCCACATAAATTTCTTATTTCTAAACTTAAAGCTTATGATATAAATGGTTCAATACTATATTGGATAACAAACTTTTTGGAAAACCGTAGACAAAGAGTTGTTATAAACGGTAATTATTCCGACTGGGTGTCTGTTAAAAGTGGAGTTCCCCAAGGCTCTGTCCTTTCAGCGTTACTTTTTATTATATATGTCAACGATATCCCTGAATTAATCAAACCAAAAGCTGCTCTTTTTGCTGATGATACCAAAATCTTTCAATCTATTGAAAGTACTCAATCTTTGAAAAATTACAAAACGACAT
>evgR002160c0g1_i01_Hydra
ATTATTGCTTCATCTTGTACAAATGAAGCTTTTAAATTGGCTTCTTTCTGTACACCATTCTTGGATGATTACATGATGTACAACGGAAGTGAAGGTATTTACACATTCACTTACAAGAATGAAAGAAAGGAAAATTGTTTGCAATGTGGTCCTGCTGGTGTTTCAAAGACAATTAATTGTGCATCAGATATTACTTTGGATGAATTTAGAGATTTGTTGAGAAGTGATAAGTCAATTCAATTTAACAATGCTTCATTGCGTGATTTGACTAATGATCAAACTTTGTACTTGACAAAACCAGCTGCTGTCAGACAAATGACTGAACCAAATTTGAAGAAAAAGCTCAAGGATTTATTCCAAACTGGAGCTGAACTCTATGCTACCGATACTGAAGTTATTGCTGATCGTGGTGTAAACATTGTTGTTCAATTCACCGACTAAAC
>evgR002161c0g1_i01_Hydra
ATTTCTCTCCAAAATTTTCGAAAATAGGGACAACAAAGGCTGATTTCCAGCTGGATGCAAAACAAGACTCTAATAAGCACTTGTTAAATAGTTTTGAAAATATAGACGACAGCTCGGGACAACATGTTTGCAAGACTATAACAGATATGTTGTCCGGACCACAAGCTGTAGAAGAATCTAAGCTTTTAGATACCGAAGCTGGTGTAATGCAAATGTTAAGCAATTTATCAACCTGTTTGTCAGCTACATCAGGTTTGGTGAGTTTTGCAAAAGATAAGACTCAACAGGAAAAAAGTTACTTTGAAG
>evgR002206c0g1_i01_Hydra
TTAGCAACTACCGACCCATTTCTATCTTACCTATTTTTTCAAAAATATTAGAAAGAATTATGTATAATAGAATCTATGATTATTTTCAATGCAATAATCTACTTTATAACAACCAGTTTGGCTTTAAAAAACACAACTCCACGGAGCATGCAATCATTCAATTTGTACGCGAAATTTCAAATTCCTTTGAAAATTCGCAATACACACTAGGTATCTTTATTGACTTATCTAAAGCTTTTGATACAGTTGATCACACTATCTTACTCCATAAGCTAAAATACTATGGTATAAACGATAAAACTCTAAAGTGGTTTGAAAGCTACCTATCAAACCGTAAACAGTTTGTTTCTTGTAATGGTGACTATCATAAAGATTTATTAGATATATCATGCGGTGTTCCTCAAGGCTCTATATTGGGTCCCCTCTTATTCTTAATTTACATTAATGATTTAAATAAAGCTTCAAACCTAATTAGTATTATGTTTGCTGATGATACTAACCTATTCCTTTCTAACACTGATATCTGCAAACTGTTTTGTGATATGAATGACGAACTTAAAAAAATATCTAATTGGTTTAACTATAACAAATTAACTCTTAATGTTAATAAAACAAAATGGATAATTTTCCATCCGCTTCAAAAAAGACGTTATATACCCGAAAATTTGCCTCAAATATTTATTAATCAAGTTGAGATTAAG
>evgR002208c0g1_i01_Hydra
ATGTGAAAAGGATGCCAATTGTCACTCTGGATATTGTGATATTTCTGGTGGTGTCTGTAAAGTTGCTCCTGCTCTTACATGTGATGGTAAGAATACTGATGGAAAGTGTCCAAGTTGTGCCTGCACTTCAGGTGAAAACAGTGCTGGTGTCTGTCTCCACAATGTTGCCACAAATTGCCCAGGTTATCTTTTGGATCTCCAAGTTTGTGTCTACAATAACTTTGTTAACACATTAACTGACCCACTCATTTCAAGTTATTTCTTATTGGCTAGATCCAATGTTGATTCTCCATTCCTTGATACCGATAGCAC
>evgR002210c0g1_i01_Hydra
ATGAAATTGGAACTTTTTTATTATAAAAGAGGTTATTATTGTTTAAAAAAAAAGGCTATTATTACAAAGGTTCAATATAGTGTTTGTATATTTGCGGCTCCTTTGACTATACCCAAGACCGTCTTAAGGGGGCAATCAATAACTGTTGATTTTGTCAAAATAATGAACAGTAGTTGGATGCTCCCTCGTTTAAGACTTTGGAATGGCCCAAGGGGGTGGGCGATTTTCGTGAATTTTTATTTTAAATTCTTGTTAAGGTGCTTTAAGAAGTTCAACCGGTCTTATTACAGAGAACCAGCACATGAAGCATTTAAAAGATTAGCTCTACGTTTCTAACCTTACAAAAAATGTTTTTGTGCTAAGCCGAATCCACTAAAT
>evgR002247c0g1_i01_Hydra
AATTTTTGAACTTAAAAAGCTACCGCATATGAAAAAACTTTTTATTTTTCTTAAACAACAAAGTCGCCATGCTGTATTTGAAGGACTAGGTAAAAGGTTGTCGTGTGAAGTTTTAGCATTGTGTCGATTGTATGCTGTTGTAAGTCGAATAACTAACAAACTCAATGAGCTTATCTATTTTTTATATATCCTTGCATATGATCAAGTGCCTCACACTAATTTTGCAATGTTTTTTATTACCTGTTATTCGATATGGCCATCTCTCTTATTTCAATCAAATCTTATTTCTCAAAATAAAGACAATAAACTTGAATATCACAATGTACAACTTTCACCTATGTCTGCAGCCTTTCAAATATTAGCATTTTATGAAGCTGCTTTGAAAGATGATGCTAACTCTTTAAAAATTCTGGAATATTTGTTTTCAATAAATGTGAGAGTAAC
>evgR002265c0g1_i01_Hydra
AAAACTTGGGTGTTGGAAAGACAGCTCAAATCGAACATTAACTACTCTTGAAGGCATCTCACCATTCACCATTCTTGATGGAAGTTATACAAATCGAATTGATCCTTTTGGTAAATGTTACAAAACTGCATCAGAAAGAGGCTATAGATATTTTGCTTTGCAAAATGGTGGTTATTGTTTAGCTGGAAATGATGAATCCTACAAACAGTTAGGAAGGTCAACTGATTGTGAATCAGGTGGAAAAGGAGGTGTGTGGGCAAATGAAGTCTACAAAGTAAACAATAACTGTTTTGAAGTGCATGGTATGCTAGAATTTAGAACTTTCTTGTATATAAGCTATTGGAGTTTGGATGTACTAGCTGTTAACGCAACTTGGGAATACTTTTTTCCATCTTTTATAATGCTTCAAAATATTAATACATCAGTAGATGTTGATCGTTCAAACATACTTAAATTTAAGACAATCAATTTCCCTGATAATGGAATTATTCAGAGTATTATTGCC
>evgR002312c0g1_i01_Hydra
CATGACAGCAATTGCTCTAGACGACACCAATGCGATCAATGAGGTTTCTGTACCAGTCACCTATGGAACTGCACTCAGAATTGTCCATGGTAATTCTGGTTTTCGTTTACATTCCCACAATGTTAGTATTGACGAGACTGGTAGAGAGGTGGTGTGTTACCAAGCAGTCGATGAAAATGACCTTTGGATTTTCAAAAGTGAAAAGGAGAACATGGATGGAAAGAACATCAAACAAGGAGATATTATTACTCTCCAACACAAGTCAAGTGGTTATTTCTTATGCTCTTTGAAGGGATCTAAAGCACCAAAATCACCAA
>evgR002325c1g1_i01_Hydra
GAGTTGTTTCCATTAGATTATATGCATCTTGTATGTTTAGGCACTGTTCGTAGACTACTAAATTATATAAAAAATGGTCCGTGTGGAAAAATATCGGCATGCCAAATAAGTAAAATATCAACTTTACTATTAACATTTAATACACACATGCCCAGCGAATTTGCTTATCGCCCACGCAGTCTTCAAGATATGGACCACTGGAAAGCAACAGAGTTTGAATATGCTCATGAACTTATCAATTGTTTTGTTTATAACTGCAAATTTATGGGCTTCACTGTTTATAACATTCATAATATGCTGCATTTACCATAATTTGTTGTTAACTACGTTAACAAAATCTGTTAGAAGTTCTTCTAATCCAATTGCTCAGGTGAAAAAGCGACAAGCTGAATTCCAAAATGCTTTCACATCATTATCCAAGAAAACATTTTATTAAGGCCATCAAAGATAAATATTTTATCTGCT
>evgR002379c0g2_i01_Hydra
GTTGATAAACAACTTTAATTGGTACATATAAATAACAATAAAGAAGGAAACAAAAAGCTGATTAAATTAAAAAATGAAATTCAAATTAGGTTATCTGCAGAGATTGCAATATTTCTTCGTATTGGCAACTATGATATCGATTGAGGGAATTCTTTCAACAGAGTGCGTTAATAGTAGGCTATGTCGAGTAGAAACACTAAGTTCGTCGTTAAAAACAAAACCAAGTTATTCGAGCTTAAAACGCTTCTCGTCAACGTTGTTAAACAAAACCGCTCTCGGAAAATCAATTGTAAAAAACGAAGAACAATGCATTAAAAGTTGCTTGATCTCTTTAACCTGTTTCTCAATAAATATAATTACAAACTCAACATCAATTCCAGCTGTTATGGAGTGTCAGTTACTTACACAGCAAGCATACACCAATAAAGTTGATTTAGTACACCAAGAAAACTCTACACACCTAACCGTTTTGAATCTAGGTTGCGAAAAGAATCTCTGCGCAGGCAATGCAACTTGTGCTCCAAATTATGTTGACGATTCATCATCGTGTGTTTGTCCCTCTTCCATTTTTAGCGGAACGTATTGTGAAACAGAAACAAAAGTGAGCTCAGTTTGGTATTCCAGAGCAACTGATGCAACTGATAATATGAATATAGTGACAAGTGATCCAAACTTCAAACACGTAATACAAAGATCAAATTGTTTACCAAATTTCATTTATAGTGCACAAGAGGGCATAAGAGCTCACGTGTTTGCTTACTTTATGGTTCCTGATACTGGATTGTTCTACTTTCAATTATATTGCAGTTCTGAATGCAGATTATATATTTCTACATACAGAAAACAAGATAGGAGAGTAATTTTATGGGGACCTGGAACAAAATCAAAGGATGTTTATATGGTGAAAGAACAAAAAATAGCAATTGAAATAGTAGCAGGTTGCCCTCCAGGTACCACTAATAGTTATACCTTGGGTGTAACTTTACCTAACGGAACAATATATGCAAGCATTCCTGACAGTTTTTTGGTTTCTTTGTTCTCTAATCAAATATTAAAATAACATTTAAATAATAC
>evgR002434c0g1_i01_Hydra
CAAAGTGATGAAAAGGAAAATAAGGAACCAGAACAAGAAATTACCGAATCAGATGAACAAGAAATTAAGATCTTGTCATCCAAGGTTCCAAAGCCATCCGAATTGGCTGGTTATCGTTTGAATGTCAGTGAATTTGAAAAGGATGATGACTCAAACTTCCATATTGATTTTATTGCTGCCACAAGTAACTTGCGTGCTAGAAACTACAAGATCCCAGAAGCTGATAGACACAGAACTAAGGGTATTGCTGGTAAGATTATTCCTGCCATGGTTACCACTACTGCTCTCGTGACTGGTCTTGCTTGTTTGGAATTCTACAAGTTAATGCATGGTGCTCAAAAGATTGCC
>evgR002465c0g1_i01_Hydra
CGCGGTTTATATCATTATTATTTTGGATTCATGAAAACATTTAGCAAAAGTTATACGGCGGCTTTGTCAGTGATAAGTGGTCAACTTGCTTTAAATACAGACATCAATATTGTAAAGGCATCTACGTTTATAATTAATTTGATTCCTTTTTTTGGTAAGGTTTTAGCAACTTCTATGAAGGAAATAATGGAATTTTTGAAATCCGCAGAAATTCGAAATGCTGCCGTTCATTTATCGTCGCTTTCTCCTCGTGCGAATGAATTCGAGGATATGATACAAGATATAATTGTCGAAATAATTTTTGATAATATGGAAAAATTTCTAACAAAG
>evgR002499c0g1_i01_Hydra
CTAATCTTTTATTTTCTAAATAAGATTTAGATATGAATGCTCTTATATTGGTTGGTGGGTATGGGACAAGATTAAGACCACTCACTTTATCAAGACCAAAGCCTTTAATTGAATTTTGTAATAAACCAATGCTTTTGCACCAAGTCGAAGCATTAGCAAAGGCTGGTGTTAAGCATGTAATCCTTGCTGTCAGCTATCTTTCAGATATGCTGGAAGAAGAACTTAAGAAGGAGGAAGAAAAGCTTGGAATTAAAATAAGTATGTCACACGAACGTGAACCCTTGGGGACAGCTGGTCCGCTAGCACTTGCTAAACATTGGCTTGAAGAATCAACAGAACCTTTTTTTGTGTTAAATAGTGATATAACATGCGAATACCCATTTGAAGAGATGATTCTTTTTCATAAAAAGCATGGAAAAGAAGGAACATTAGTTGTCACAAAAGTTGATGAACCTTCAAAATATGGTGTTGTTGTGTTTGATCCGGCAACTGGAAAGATTGAAAACTTTGTTGAAAAACCAGCTGAATTTGTATCCAACAAAATTAATGCTGGAATGTATATTTTCAATCCCAGTATACTTAAAAGAATTTTGCCAAAGCCAATGTCAATTGAAAAAGAAGTTTTCCCTTTTATGGCTAAAGATGGTGAATTGTATTGCTTTGAATTAAAAGGTTACTGGATGGATATAGGGCAACCTAAAGATTTTATAACTGGAACATGCTTATATTTAAATGCATTGAAAAAAAATTGTTCAGATGAGTTGTACAAAGGCCCAGGAAATATTGGAAATGTTTTAGTACATTCTACTGCAAAAATTGACGATGATGTTCAGATAGGTCCAAATGTTGTTATTGGCCCTGATGTTATTGTCGAAAAAGGAGCTTGTCTCTCAAAATGTGTAATAATGAAAGGCACTTTGATTAAATCTCACTCATGGATTAATAACTCAATTGTAGGCTGGAAATCCTCTGTAGGAAAGTGGGTTCGTATGGAAGGTGTCTGTGTTTTAGGTGAAGATGTTCATATTCAAGATGAAATTTATTTAAATGGGGCAAAAGTTCTTCCTCATAAATCATTAGCAGCAAGCATTCCAGAGCCAAATATTGTTATGTAGTTTATTAACATTGTTGAATGTTATGTAAATTAAGAATAATTTTTATGATATTTATTTAGGCGTTATTCATAATTTTTGTATAGATTTTGATAATGAAGTTTTATTATGAAAAAAAAAAAAAAA
>evgR002520c0g1_i01_Hydra
GTCACTGTTCGTTCTGCAGTCAATGGTGATATTCACATGTATTTTGAACGTGATTTGGTTTCCACTGGTAAAACCAATTCCAAATATTATTATGATTTCGACAGTACTAGAGCTGATTCTAATTTAATGTTTGCCACTTCACCTGCTGCTGCCATTAATACCTATGATGATTTCTCCAATTCTCAACACACTGCTGCAAAGATTTTGAGATATAATCTTGTTGGTACTTCTGGTAATGAAGATGCTGGACAATTGGCAACTTGTACTTCTGCTGAAATGACTGCTGCCATGCCAGCTTATCGTACTCCAAAGGAAATTGCTCAAGCTTCAACTGTTCCTCCAAAGAGTGGATCTCCG
>evgR002523c0g1_i01_Hydra
TGTAGTGAAAAAAAAGTTTGTATAAACACTAAAGACAATTTAGTTTTAAAAGAACATTGTGACGATGAAACACCTAAATTTTCCTTTACAAAAGAAAGATCGCTTCAACATATAAAAGGTAAAAAATGTGTTGGATGTGTTTCAGATGGTGGTTGGCCAGTTGATGGATGTATTGTTGGTATAGATGATGAATGTGATCAATCAGAGGATCGAAGAGTTATAATCATGAATGCTTTAGATTGTGAGGTACCATTGTTAAAGGCATTACCCGATTCTGCATATTCAGCTAGTAGTACAAATGAAAGTAGTAAAGCTTCAAATGTAGTAAACAGTGATGGTTGGTGTCCCGATGATGCAACTAATACTTGGTTCCTAATTGATTTTGGGCAGAGTGTCAAGATATCAGCTTTGGCAATGGGAAAAACGAAGTTATATACCACTTGGTTTACTTTAGAATATTCGGACGATGGAGATTCGTGGAATAAATATTTAGAAAGTGACGTACTTGAGTTAAAGAAATTGAAAAGTGTCTGCTTTTCTGGTTCATGTAAACAAACATTAACTGAG
>evgR002524c0g1_i01_Hydra
TGATTTTGATATTAACAATCACACAGTAATATTTATAATTGATGGATTAGATGAGTTTAAGTATTTGAATGAATTTTTAAATCCAAGTTTGAGTTGTAACGATCCAATTGTTGATACTTTAGCAGAAATTCTAAAATACAATTATGTAGTAGCTGGTAGAGTTCATGCAATACATCGATTTCAAAATGCATGCAAATGCAGAATATTGAAAAATTGCAAATGCATTACAGAGAATTGCAAGCTTACCATTCAAATAATGGGTTTTAATGAAAATGGAATAAATAATTATATTGATACTAATGTTGTTGGTGAAAATAAAGAAATTGTAAAAACAATTTTAAAAGAATCTTCTATTGCAAAATCCATGGCATCTGTTCCGTTTTATTTATCTTGTATGTGTAAAATTATTAGTGAGTCAAAGCAAAGAGACTTTTTATTCTTAACTATGACTG
>evgR002535c0g1_i02_Hydra
ATCGTTATCAGATAACACATTGAACTCTACCGGTCAACCGATAACAAAATCAAATGGAAATCAATCATCGCTATCAGATAATACAAATAACGAAACCGGTCAACCAATGACAACATCAAACACATCAAATGGAAACTCTTCATCGTTATCAGATAACACAATAAATTCAACCAGTCAACCAATGACAACATCAAATACATCAAATGGATACCCAACATCGTTATCAGATAACACATTGAACTCTACCGGTCAACCGATAACAAAATCAAATGGAAATCAATCATCGCTATCAGATAATACAAATAACGAAACCGGTCAACCAATGACAACATCAAACACATCAAATG
>evgR002554c0g1_i01_Hydra
TGTCCCCACTGGTGCTGGGCCTATGGATTTGAATTCAGCCCTTCAAGAGGTCCTTAAACATGCTCTTATCAATGATGGCCTCACAAGGGGCTTACACGAGGCTGCTAAAGCTCTGGATAAGCGTCAGGCTTTGCTCTGCGTTATTGCTGACAGCTGTGACGAACCACTCTACAAGAAACTGGTTACAGCATTGTGTACAGAGCACGAAATCCCTCTGATCAAGGTTGACAGCAACAAAAAGCTGGGTGAATGGGCTGGACTCTGTAAAATTGATAAAGAAGGAAAAGCCAGGAAAGTGGTTGGATGCTCCTGCGTTGTTGTTAAGGATTGGGGTGAAGAGACACAAGCTACACAATTTCTTCTCGACTACGTCAAGAAGCAGTAAAACCGCTGCTATATTGTATATCTGATACTAGATGTACGATTTTAAGATACAATTTACAAATTTT
>evgR002566c0g2_i01_Hydra
TCCAGGACCAGTTACACCACCAGTTCCAGGACCAGTTACACCACCAGTTCCAGGACCAGTTACACCACCAGTTCCAGGGTCAGTTACACCACCAGTTCCAGGACGAGTTACACCAACAGTTCCATCACCAGTTACACTACCAGTTACACCAACTGCAGTGAGTAACTGTCCTGGTGGTTGGTTACAGAATGCAGGCAATTGCTACCTTTACCAGAGTATTCAAAAAAATCGTCAAGATGCATTGTCATCTTGTCAGAGTTATGGAGGAAGTTTATTAAGTATAGAGAATAAAGCAGAAAATGATTTTGTTATAGACTATATTAAAACCAATAAGGCACAAAACACTGGTATCTACATTGGTCTAAACAATATAAAAAATAAAC
>evgR002578c0g1_i01_Hydra
ATATAGAATACAAGAAGTGCCAAGAAGTCATAATAGTACATGCAGAAACAGAGAATGAGTTCTCAATTCTTTTTTACTCTAAAAGTCTTCTATTTACTGCAATATTCTATCTCTTGGGTATTGTGACTGCTGGAGTTATTTTTCTTGTAGGGTACTGGTATCCTATTTGCCGTTTAAAATTCACACATAACAGATGTGCCGTAAAAGAAGCATCATCGGTACAAATAAAAAAAACTGAGGGGAAAAATGCTTATGTATCTAAAATAAAAGAGATTTGTTTAGGTGATATGCAATATGACGAAAAAGACTTTACTGAGTTTGAAAAGATCATTGTTTCAAAATCAAGTGTACGATATTTTGAATTTATGTTTTTACGTTCCATAATAAAAG
>evgR002594c0g1_i02_Hydra
TTCAATTTATTTACATCCAGAATCTTTAATTATATTTAAAGGATTCTTATGGAATGATTTAAGTGTAAATGAATCATGTGTTAAAGAACATGGTGATTACTTATGGGTTCAGGTAGACACATTGTCAAAATATTCATATGCCTCGTTCTCAGCGATTTTTGTTTATGGTAAAAAAGTTCCTGTTGGAAAAAAATATATTTCTGTGGAATGTTATTTAACATGGACTTATTCAAATGGAAAAACTTTGGTATATAATGTTAATCGACAAATAATAGAGTTGCCTGTTGAAAGAATGTTTCTAAAAGTCTTTAAAATGGAGCCAACATGGAAAGACAAACATCCGATTGGAACATCATTTGAAGCAGAAGCTTATCATTTTGTGTGTTCAACCATGCAGAAAAGACAGCCAAGTCCTTGCTATCAGCGCGAACTATCAACAGGAGTAATTATATTTCTGCCAATTCAGG
>evgR002596c0g2_i01_Hydra
CAGGTCATTCAATACGGTTACAAGGTTGAGAAAGAATTGTTTAATGGACAGAAACTTGACTCTGTATCAATCGTAGCTAAAGTCAATAGTGACTTACAACAGCTTTGGTGTAGGGTTAATCCAAATCTACCTCTGATAAAAGATAAACCATTGAAAAATAAAATATCAAGAACATTTCTAAAGGTGCTGTCTGCAGAACGAACTCAATCTAAAAACCATAAGTCCAGTATAAGCTGGCTAGAAAAGAAACTGGAAACATTGTTTGATGTGTCTGCTTGCCGGTGCGATCTTCTAATTATTGATGATTGTCATGATAGACGAATTGGATGTCGGAAAAGTAATTGCACAATGAAACACATAGTTTGTCTCTGTGAGGAAAGTAGAAAGGTCCCTGTTCTGGAAAGAGAATATTTAAAGGACCAAAGATGCAAGATAGGACCAAAAGGAAGTTGGCAGTTAGGCAATGTGGACCATGTGGAGGCAACAAGGAACGCCAAAAAGTATATTAGAGAGGAAAACGAGGCAACTAAAAAGCTAATTAGACAAAAAAAATTCAAGAAACATCAGAA
>evgR002611c0g1_i01_Hydra
CTGCGTTGTCTGAAACATTACGGTTATCTTCAGTAACTCCATTAGGAGTACCTCACAAAGCCATGGATGATACAACTCTCTTAAATAACTTAAATATTCCAAAAGGTACCACAATTCTAACGAACCTTTGGCAATTACATCACAATGAAAACAGCTGGGAAAATCCACGAGAATTTAATCCATATAGATGGTTAGCCACGGAGGACGGTCAAACACTTGACTCGATGAAATCGCCAAATTTTTTACCTTTTTCGGCCGGCACTAGAGTTTGCTTAGGAAAGGGTATTGCAGAAGTAGAACTGTTTCTATTTTAT
>evgR002614c0g1_i01_Hydra
TTAGTGCAAAAAGATCTGATAAAAATGGTAAGATAATCCAAATAATAAGAGTGAATTGAATTTTATAACAAATGCAATGTTTCTGTGATGGTTTTATCAAGTGAGACAAGGAAATGTATTGTTCACTTATTGGATGGTCAAAAGCTAGATATTCCACTTCAGCCTAAGCTTTTAACAAGCGAGCTTTTGGATATTGTTGGTTCTCATTTTAATCTGAAAGAAAAAGAATATTTTGGATTTTACTATTATGACACTAAAGAAAACAGGTGTTGGATGAGGGAGAATAAAAAAGCATTAGATCATGAGTTTCAAAAAACGAATCCAATTGTTTTATACTTCTCTGTAAGAAATTATGTTCCTAGTGTAGCAACTCTAAAAGAAAGCTACACAGTTGAATTATTCTTCCTTCAACTTCAAAAATTGATATTCCAGGGAGAATTGGAATGCAACAGTGAAACTATTTTTAAACTTGCAGCCTATGTATTACAGACCACGCATGGAGATTTTGTAAGTAATAATATAGCTTCAAACGATTTGAAAACTCTAAATGTAATCCCAATAAAGACTCAAAATGAGTACCTGACACCCTCATATTGTGATTTGAAAATAACAGAACACTATCGTAATTTTGATGGCATGACAAGGGGTGAAGCCATTGTTAATTACCTTGGAGATGTAGAAAGATTACCTATGTACGGGAATCATTGTTACAATGTTAAGGACAAAGATCAAAATTCATGGATACTTGGTTTAAGTCATCGTGGTGTATCAGTGTTTGATAATCAAAATAAAACTTTACCTCAGAAGATATTTCGTTGGAGTCACTTAGAAAATGTTTTTTATAAAGATAAAAAATTTTCCATTGAATATCGTGAAGGACCAAGGCAACAACAATCAATGTTAAAATTGAACACTTTTCGGAGGAGTAATTGTCGAGTAACTCTACAAGCATGGTATTCCTCTACAACAACTCAATGCCGGGAGATTTGGTTAATGGCTGTCTCGCAGCATCAATTCTATTTGGATAAAAAACTAAATCAGCATGCTCCAAAATCGTTAAGAGATGTTGCTAAAAGCTTGTGCAAATCTTCAACTTCATTATCCTCAATGTATTCTGGTAGCATTGGCGGTTATAGTAGTAGCCTCAGTGATACGGCTTCCTTTTACTTTTCTTTGGATGGTGAGAGCATTACTTCTTCTCAAATGGTTGTAGCTGAAAAAGACATGTACTATGCATTGAAAGCTCGAAAAGAATTTCTCCAAGAACTTTTAATTGAAAAGAGAGAGCTATTAAATGAGCTGTGCTTGCGTGAAGCAGATTTAACTGGGCAGCTGCCAACTGAACTTCCAGAAAGTCCTACCCATCCTGGAATGCCAATGCGCAGAAAGATGGGAACTAGCTTTCAGTTTGACGAATCTTTATTTTCAAACCAACATAGCAGTGATGAGCAAGAATATGACCGGTTACTGCGTGATTACCAGATTCAAAAGCAAATTACGAATGCTGCTCAGCGTATAGCTATGGATACATCGGCTCAAAAGAAAGTTCGTAAGGCCAGGAAGATTACTTATCAAAAGTCTTTAGTCAAGTTAAAAACCATGGAGCAACAGTTGGAATATTTAAAACGTGATGCTGCTAAAGGAACCAGTGAAGCAATTTCTCGTTTAATTAAGCAAGATGAAAATGAAAGTAATGGCGATAGTTTAGGGCCAATTGATGGAGAATCCTTGGATTCAGATAGAAATTATCATTCTCTTAATCGACCGTCTCGTTACTCCATGTTTCTCGCCACTCCGGTTGTTAATTCTGCAAACAAAGGATTAGATCAAATGAAACGTCGTCCGTCTCCTGTTATGTCTCTTGTATCTCCGCGTTATGATGCTGTTAAAGACTGTTACCGTAAAAATGATTTTTTTGGTGAAGGCGCAACACGTAACGGTTCGCATTTACCTTTAAGAAAATACTCTGACGATTGTCTAACAAAAAAATGTTCTCGTTCATCTTCTGATATAACAGCAATAAGACACGATAGTGCTGATAAGCTTAACATGTCCCAATCTCATATAGATGATAGTTTTGTTTGTGGAGAATTTGAAAATACCGGAATGTACAGCATAGGTGCAGTAAAGAAAATTGACTTTAACTCCACTTATTCGATACCTCTATCAAAAAGCAATCCAGGGATTGCTCAAACCCGCATGCACGCTTCGCCAAGTACACTTAGACGATTACAAATGCTAGCACTATCGCCTTCTCATAAAAAGGTAGGATCTCTAGGGTATATGTATTCCCAGTCATCGCTAGCAAGTAGCAGCGAATATTCTGAGACGTCGTGGTCAGATGTTGTGTCAGAGCAAACTTTGGTTTAAAAATAAAAAAATGAAAGAAATTTTAAAATAACTTATAGAAATATTCGGCGCGTTGAATCTTTTTACCCATGTTTATTATTATATTAACTGCTAACTTCATAATAATACAACATAATTTTAACATTACGATAAACTGGTTTATTCGATTTTTAATTTTTGCTTGCGCTAGGACAAATATTTTTTAAAGGCATCGCGGTATTTATCAAAGCTGATTTTCTATTGGTTTTCGCATCGCGGTATTTACCAAAGCTG
>evgR002636c0g1_i01_Hydra
ATGCTACAATAGCAGGAAGCACTTGCTCTCCATATTATCATTTAAAATGTTACCAAAAGTTCACTGACAAAGTCCTTATTGCAAGAATAGAAAAAAAAATTAATGAATCTAAAGCTAAAAATAAAAGAAGATCTAATTCACCGACAACACTTTGCACTAGAGGTAGAAAGTCCACTAGAAAATGCAACAGCTCTAACAGTAATCGTGATAGAAAAAACACTCACATTTTACCCGAAATTTGTTTGGTTTGTGAAAAGGTGGAGAAGTACTTTATGGAAAAGGTGCACCTAAAGCGATCTGTGCAATACCTAACTCTGGCAGAAACAAAAGATGCAGGATTACTTCGTGAAGCTGCAGAGCAAAAAAATGATGAGCGTATACTCTTGCAGATAAAAGAAAAAGACTGTGTTGCTGTAGGGGTACGATACCACCAAAATCTGCTACA
>evgR002638c0g1_i01_Hydra
CAATACACTAGGCTTTTTATTGGATATGTAATGACAATCATCTTTGTCATGTCACATGCTGCCCTCACTGATCGTAAAAGATCTCAACCTTCATTCCAACAAGATGATTCAAATGGAGCTAGTAGAATTTTCGACCCTGAAGAAATCATTCGTAGAGACGAAATGCAAAGAAGAAGTTTCCAACCAACAATGTCTGGTGGCTACTCATCAAGTAATAGATTTATGACTCCTGGTACCATCCAACCAACTTCTTATGAACCATCATCATCAACTGTATCCGAGTCATCTGATGATGCTGATTCAAAGCGTTTCAAGAAGGAAAAGATTGAACAAACTAGAAAACGTATTAAGG
>evgR002640c0g1_i01_Hydra
TACCAAATGTTCAATTTCTCTAGCGCTTCAATTCAATTTTAAATTGTCTAAAGCGTTTTATATTCCAAAGATAGAGCCTATAAGTATTCTAATTCCGTGGGCCCTTCGAAAAACTATTAAAAAGGTGGAATCTAAAAAATTCAACTATTTTTTATACCGTTCCGCTGTCAAAGATGATGTAGCAGATAAATCCTTCAATGTGACAAATGGTGAAAGACATCATGACACTCTTTTTTGCGTACAAAAAAAAGGTGCTATTAATAGAAACTATTACAATGATTTGATTCCTCACTGCGTTTCAATTCA
>evgR002641c0g1_i01_Hydra
ACAGGGATGTGATGGAGGTTTTGCTTACTTGATCAGCAAGTGGGGTGAAGATTTTGGTATCATTGCTGAACAATGTGATCCATACACTGCTGCTCCACACAAGTGTAATTTGAATCAAGCTTGTTCCACCAGACAATACTGGACCAATTACAAATATGTTGGAGGTTATTACGGAGCTAGTACTGTTGAAAACATGCAATTGGACATTTTAAAGTATGGTTCCTTGTCTGTTTCTATGGAAGTTTACAATGACTTGTTCCAATATCATAGTGGTGTTTATAGACATGTTCCATCCTCAAAGCTCTCTTCTCCAGTTCCAAATCCATTCGAACTTAC
>evgR002651c0g1_i01_Hydra
TCCAAACGATCAAGGTAACAGAACAACTCCATCTTATGTTGCTTTCACTGATGAAGAAAGATTAATCGGTGATGCTGCTAAGAATCAAGTTGCCATGAACCCACACAATACTGTTTTTGATGCTAAGCGTTTGATTGGTAGAAAGTTTACCGATGCTTCAGTTCAATCAGACATGAAGCACTGGCCTTTCAAGGTTATCACCAAGGGTGATGATAAGCCATACATTCAAGTTGAATATAAGAATGAATCACACACTTTCACTCCTGAACAAGTTTCATCAATGGTTTTGACCAAGATGAGAGATATCGCCGAACAATATTTGGGTAAACAAGTTAAGAAGGCTGTTGTTACTGTCCCAGCTTATTTCAACGATTCACAAAGACAAGCTACCAAGGATGCCGGTGCTATTGCTGGTTTGGAAGTTTTGAGAATTATCAACGAACCAACCGCTGCTGCTATTGCTTACGGTTTGAACAAGAAGGGCGAAAGAAACGTCTTGATCTTCGATTTGGGTGGTGGTACTTTCGATGTCTCCATCTTGAACATTGAAGATGGTGTCTTTGAAGTCAAGGCTACTGCTGGTGATACTCACTTGGGTGGTGAAGATTTCGATAACAGAATGGTTGATTTCTGTTGCCAAGAATTCAAGAGAAAGCACAAGAAGGACTTGCAAGAAAATCCAAGAGCTTTGAGAAGATTGAGAACTTCTTGCGAAAGAGCAAAGAGAAATTTATCTTCAGCTGCTAATGCTACTATCAATGTTGATTCTATTATGGATGGTATTGATTTGGATATCACAATGACTCGTGCTAAATTCGAACAAATGAATATTGATATGTTCAGAAATTGTTTCGAACCAGTTAAGAAGGTTCTCCAAGATTCTGGTTTGGATAAGGCTCAAATTGATGATGTTGTTTTGGTTGGTGGTTCTACTCGTATTCC
>evgR002669c0g1_i01_Hydra
GTCCTCTATTAGAATATGTCAAAATTATTTTGATAGAAAATATTGGTTGTCTGTACATATTAATGGAGTAGTTATTAAAATTATAAATAACTCGGATGCTAGAAATTTTACCAATATAAAGGTTTATGCCTCTTATCTTCAATATGATGCTCAAATTGGTTCTATCGCTGATCTTTTGATTATCAATGGTAAGGCTGAAAATCTTGTTGGTAGTGTAATAACCCCATTAGTAAAAGGAAAAGTTACTGCTGAGATACCAAATCTTAGCATTGAATATTTAATTTCGTTTGATTTTATTCCTTATGAGTATGCTGTTGGTTTGCGT
>evgR002678c0g1_i01_Hydra
TTTTATCATGGCTTCAACAAATGATGAAGATTTAAAAAAAGGTTTTATCACAGATGTGACTGATTACTCAAAATGTATGATCGATATATATAAAGAAATCGATTGTTTATTAGTAAATCCAAACCGTTTTATGTCGATAACTGAACGAGAAGAAAAAAGCGTTTTTGATATTGAACTTATCAAAAAAGTATCGAAAGGAAAATATACAACTCGATGGCGAGGGATTAGCATGATGAAGGATACATTGGATATGACCATTTATGAACAACTCCTATGGGAAATAAAACCTACAACCATATTTGAAACTGGTGCTTACACAGGAGCATGTGCACTATGGATGGCTGATGTTATGAAAATGTACAACCTTCCTGTTCAAATATACTCGATTGACATCAGTTTGGACCTCATCGACCCGCAAATAAAGTCATGCAAAAACATCAGTTTACACCAAATAGATGTTAAAAAGATCAATCAACTATTCTCAGGAGAATTTTTAAAAACCTGCCCACATCCGTGGATAATATCTGAAGACTGTCACGTTGATGTAGTTGGTGTCATGGAGCATTTTCACCGTTATATGTGCAAAGGGGATTATTTTATAATAGAAGACACTTCACCAGATACACCCTTGACAAGTGGCCAAGGTCTACTTCAACAACAATATGAACAGTGGGGAACCAGAAAATTGACATTAATGAAAGAATTTTTGAAGAAATACAATGAATTTTACAGAGTGGACACATTTTACACTGATTATTTTGGTTATAATGGCACCTTTAATTGGGACGGCTACATCAAAAGAATAAAAGATTAATTACAAGATTATGGTGTAATAATAACAATAACAATAAATAATGTTTATTAAAACGAAAAATAAGATTAAAAGTGGAAAATAAGTAAAAAATTATTGAATATAATTGATTATATAAAGTAAGAAATGTTAAAAAATATTTTTTTTAATTAAATATTATTAGAGGCGACAACA
>evgR002679c0g1_i01_Hydra
CTGAAGTGAAAGACGAAGTAGTTATTATAAGCAATGAATATGATGAGAGAGAGGGTATAGATTTAATAAACTTAGAAGATATTGCAGAATATTTAAAAGATAAAAACAAAGTAGTTATAAGGCAGATAAATAATTTAGATGCAACAAAACTTATAAAAGAGAAAGTGGATAAATTAATCAATAAAGGAGGAGTGCTAAATTATTTAAAAGAGTATGTAGGGTATGCAAGTGTTGAAATAAGTGAAACAAAGAGAGTTGAGGCTAGTGAGAGTATTAAGTTAGAAATTATGCAGCTACTTGGTAAAAAAGATGTAAATTACGATGAAGTGGAGGTAATAATTAAAAATGCAATAGATATTATAAAATCAAAATACTTACCAAGTTTATATAATAGCATAATAAATAATGAAGTATTAAAAATTTTAAAAGAAATAGAAGAGGAAGTATTAGTTGTAAAAGATAAAGACGGAGGAAGAGATAAAGGAAGTAAGGAATATTATACTGAAGAAGTAAGTAGGGTGATAGTAGGGAAAGATGGAGAAGGAATAACAATTATCGAAGTATCGCCATATAAGCAACTAACAAGTCAAACTAGTGAAGAACAATTATATTGTAATAGAATGAGTAAAGAAAATGAGGAAAATAAAATATATGAAAGTATAAAATGTATAAGGAAAGATTTAGATATTGCAGATATTACAAATAACAAATTACATTTACCTATGATGATGGGATTAGATGAAGCAGTAGCAGATATATTAATTGATTCTAAAAACAATAATTCTAAAGATAATAACAAAGATGTTCTTTATTACTGGAATAAAGAACATAAAACAAAGTTAGAGAAAGCATTGATAAATTTATCTAATAATGAAGAGCTGATTAATATCATAACTAAGGAAATAGAAGAATCAAAGAGCTGGATAATACCGCAATCTGTAAAACAAGAAATTATAACGGAAGAGTATATAAAAGAAAAAACAGAAAAGGAGTTAGGTATTGTTGGGAGTGTTATGAATTATGGTAAGTCGTTCTTTGTATCAGATAATAGGAGTTTAAATGATAGATTGGAAGAGTACCTAGTAAGTAAAGTAACTGATAAGAGTAAAGAATCTGATGTAGCAACAATGATAAAAATAATGAGTGTTAATAAAGAAGGATGCTATATGCATGGGTATATATTAGATAGCTATGCTGAGATAATAGATCATGAAACAGTAAAGTCTAATAACAGTGAGTTATGTAATAATGAAATATCTAATTATATTAATAAAGGAGATATTGATGCGTAAAACTATGCTTTGCAAAAATCCTATTTTAGGTCTACAATTGGTATTAATGGCATTCGGTTAAACCATAAGTAAGGGAAACAGCGGATTATAAGAATATTAAACTCCATAAAAAACATTTAAATAATACTTAATGACAATTATTTTTTTAAGAGATTAGTATCAGTATATCTGGATTTATAATCCAGATACAAGAGATTATGGAAGATACACTTAACCGAATGCCATTACCCTCAGCGCAGCTACCTTAACACTAGTCCTAATAAATTTTTTAAGTTAGGTTCATATATTTGACATACTCCTTCCGATAAAGCTTTGATTTTTAACTTGTCGTTGTCAACACATACTACCATATGTCCTAGTTCTGCTAAACAAACCCCTGTCACTAATCCAACATAACCTACTCCAACTATTGTAATTTTCATATTTTTAATTATTACTTGAAAATAATTATATTTAAAAA
>evgR002690c0g1_i01_Hydra
AACAACTAATTATGAAAATTTATTAATAATAAAAATTCCTTATAATATTCAACTTTGATTATTTATAGGATTTTTCATTGGATTAGCTGTGAAAGTTCCTATGATCCCTTTTCATATATGATTACCGCAAGCTCACGTAGAAGCTCCAATAGCAGGCTCAATTCTATTAGCAGGAATTCTTTTAAAATTAGGGGGTTATGGTTTAATTAGATTTTGCTATCCGATTTTTCCTTTAGCATCTCAATTTTATAACCCTTTAATTATTATTATAAGCATAATTGCTATAGTGTATGGTGCTTTAACTACATGTCGTCAAATAGATATAAAACGTTTAATAGCTTATTCTTCAGTTTCTCATATGGGTTTAGTTACTATAGGAATTTTTTCCCATTCTATTGAAGGTATAACAGGATCTATATTAATGATGATAGCACATGGTCTATCTAGTTCAGGATTATTTATTATTTCTGCTATTATATATTTTAGATTTCATTCTAGAATTATAAAATATTTTAGAGGCTTAACAATTACTATGCCTATTCTAAGTGTGATTACATTTATTTTAATCTTAGCTAATATTTCATTTCCTTTAACATTAAATTTTATAGCTGAATTGTTTTTAATAATTTCTGCTTTCAAATTTCCTTTATTTATTATTCTATTAATTCTAATAGGGGTATTTATTAATTTAATTTATTCATTATGGGTTTACAATAGAATATTTTTTGGTTCTAATTCATCTCATTTAAAATTTTCCAGAGATATAATTAATTTCGAATATCAAAGTTTAATTCCTTTAATCATATTAATTATAATATTAGGCGTTTATCCTGATATAATTATAAACTCAATTATATTTAGC
>evgR002695c0g1_i01_Hydra
ATTTTAGTGCAAAAAGTTTTTTATTAATTAAAGTAAAAAATGGGTCGAGCGGAAGCCGAACTCTTTGTTAAAGAACAAATTGATTCAAATTTCATTGTTGTCTTTTCAAAAACGTATTGTCCATATTGTACTATGGCAAAGAAAGCACTTGATGATGTTAATGCTTCTTATACAGTTTTGGAGCTAGAAAACAGAGATGATTGTCAAGATATTCAGGATGTACTTATGGATATGACTGGCGCTCGTACTGTTCCTAGAGTGTTTATAAATCGCAAATTTATTGGAGGAGGAACTGATTTAAAAATGCTGCAGGAAAACGGAGAGTTAGAGAAACTTGTAAAATCAAACTAAAACCGAATTATAAACATTGTATGTTGTAATTTTTTTTCCTTATCTATATTATACAATTGTATCGATAATTGTTCAAAAAAAAAAAA
>evgR002700c0g1_i01_Hydra
CTATGCTGATGTTCCTGCCGGTGATGTTTTCTATCAGGGTCAAGTTATTGATCACTGGTCAATGATTCCATTCCAATTATTCTGTTTCAATTTTGGTGCTACTCATATTATTCATCACTTTGTTCCAAATCAACCATTCTACATTCGTGAAATGATCTCCAGAAAGGCCAATGCTGAAATGATTAGACAAGGTGTTAGAAATAATGATTGGGGTATTGTTGCTCGTGCAAATAGATTCTTTGAAAGTTCCAAGACTTTGACTGAATTTGACAAGAATCAAATTATTCAACCAAATAACTGGGCAACTCTCTTGAAGGAAAAATCAGATTAAATTCCAATTAGTTTGATTGAATTGCCACCTTAGGTCTCCAGACAAG
>evgR002707c0g1_i01_Hydra
ATGATCACAAACATTCTTTTATTCATCGCCTTCGTTGTGTACAATGAAGCACATATAGATTTTACAGACGACCTTGACTATGCAGGAGTTAGTGGCATGTTTGTTCCTATTAATGCAAACCTTAACTATGAAGCTTTTGACGAGTTATCAGCTCGTCAATTTTCTAGTCAAAGAATAGACGACTCTGGAAGTTTTTTGAACAAAATTATAGATGATGCCGACTTGTTTGATAAGATTGATAAAGCATATAATACATATAAAAAAAGAAAATTTGAAATGAAGTTATTCGACAAATGCGTTGTTGATATATCGTTTACGAATCCAAACTATTTTATAAAAGCTGAAGACTCAAAGTGTAGCAGTGGAGTTGACTTTGGCAGAGTAATTTAAAAGATATTTAAAAGTATAAACATAGATACTCTATCTTCAGACGACTCTGGAAGTTTTTTGAACAAAAT
>evgR002711c0g1_i01_Hydra
GTTACTGACAATATTGTGTATCCACGCTCGACTGCATTTACAAGTAGAGTTGGTTAGGAAATCAATATCATCTCTAACTCCAAACCCAATTAAATCTAATCAAGTGCAATCAAGTAAAATTAAATACAATCAAAATATAAGTAGGAAAATAATTACGAGGATTCGTGATGCAGCAGCTGAGAAAAAAGTCCAAAAAGTGAACAAGTTTACAATAATGGTGTACGTTACCACATGTGTTGCGTATTTACCATTTGTTGCAATTTTTTATTTGTGGGGGTTTAAAAAGATTTCGCTGAGTTGGAAATGGACTCGTAATTTTTATTTGTGGGCACACGCACTAATGATAACAAAATCTTTAGCAAATCCAACAATTTATTTTTATCGATCAAGAGTATTGCGCCGAAGTCTAAGGCGTGTTAGGCAATCACACTTTAAATCAAATCACAGTATAAAGGAATAGCTTCTAATGATGATCTATATGGAGATTCTGCGTAGCGTTCTGGATTTTAAGAGTGATGGAGTTATAAATGGTGCATTATAAATTTAAACGAGATTTACAAACGTTTCAAACAAGAAGAGGCCTAAG
>evgR002714c0g1_i01_Hydra
TTGACGAAGCAAGAAAACTAGAAAGAGAAAAATCAAATTACAAACCCGTTGAAAATTTAGTCATTATTGAAGAACCATTTACGGAAGTGAAAAGTAAAACACAAGATAAGAAAGAACGATTAACATCGAAGAAGAAAGAAGATGGCGAAGTATCCAGCCATAACGAGAGCAGTAACAACAGGGAAGTCACGGTGGTGGAAAACGTAGGAAAATCGAAAACACTACCCAGTAAGGACAAGACGCATTCAGTACGTATTATTTTCATACTATTTACGTCTAAGATTCTTTCTGAAGTAGACAGCGGAATATAAAACTTATCTTTACAAGACGTATTAAATACGTATTTTTTTTATATGTGAAAACCCTCTTTTGTTTACGAATATAATACGTGTTTAAAATGTTTTTAAAAGACTAACGTCATCTTTAAAAGATGTTTTAAATACTTATTTTTGTAGTTTGTTTTTAGACGACGAAGGCATGTG
>evgR002717c0g1_i01_Hydra
ACATTGAAATAACTTAAAATAATTGAAATACATTGAAATAAATTGAAAATGTTTTTGACAATAAAATTATTAAATGCTGTTTGTCAACGGTCGAATCCAGCAAAGTTTTTTTACGTAAAAACTTCAGAACATAACTTTTTAAACAGTGTGTTTCGAATTGCAAAAAATGAACTGCGTTTTTCATCTTTTGCCAGTGGAAATCTCAAAGGAAACTTAATGACTAATGAAAATAGAAGATTATCAGAGTTAAATATTAAATGGGATGTATTATATACGTTTCCACATATAAAGATTTTAGGAATAATATCTCGATTCAAAATGTACCAGGCAGTCATAATGGTTGGAATATTTTATCCTATGTATAATATGCATTTGGAAGGGAACCTATCTAATGTTATGTTAGTATATGCTGCTGCAAGCTCTATTGGTACAGGGTTGTTATTTATTATTTATAGTTATTTTGCTACTAGAGTTGTGGGTCAACTTGCTGTTAATCTGCATGACAATGTCATTAGAATTTCTCATTTATCGTTTTACGGAAGAAGAATAGAGGAATATGTTGAAGTAACGAAAATCATTCCAATGAGCGATTTTATGGAAGATTATGATCTTGAAAATGTATTTCAAAGACTTTATATAGATAAAGGAAATAAAAAGTTAAATAAATATGTATACTCGTTAAGATATGGAAAGGTATTTGATCGTGAACAGTTTAATAGATTATTAGGAATTCCATCTTTTTGAAAAAAAAGATGGAACCACATGATGATTTGAAAAAAAAAATCTTTATTTATTTGAATTACTTTGTAAAAATTAGTCTGTTAATACTTCTTCTTTTGGTCTTTTAATACTGCTACTTTTGGTATGTTAATACTGCTCATTTTGGTCTGTTAATACTGCTACTTTTGGTCTGTTAATACTGCTACTTTTGGTATGTAAAAAGATTATAATAATAGTTTATTTGATGATTTTATAATTATAATGATAGTTTATTTTTGGTTCATCTTTGTAATATCCATAATGCTACATATATTTATTAAAACAGTTTGCAACATTGCATCCATAAAGGAAGTCATCCAAAATAAAGAGGAAGAGATTGAGATAAAAATGACCAAAAAAGGATAAAGGGGAGGGATGGGGTTGTGAGAAAAGTACGTCACTTAATAATTTTTGTTTTTTTTTGTTTACATGTTTAACAATTTAAACTATATTTACATGGGAAACTATGTAAGCAATTGTTTTTATGAAGGTATTGATAGAGAGTATTTTTTTGTCAAAAGTTGGATAAAGATAGTTATATATATATGTTCTTAAAGTCTTAATATAGTAAAT
>evgR002726c0g1_i01_Hydra
AACATGAAACCAGGAGAAATTTCTCAATTTTTGGAATATCTCAGCAATCATTTTGGTGATCTAAATGAAATCAACCTGAAAGACTATGCACTTTCTTTTACTTCACTCAAAGACGTGTTTTTGAGAGTCACCCATGGTGGTACCCAAGATTTTGAGGATGAAGCTGCTGAAGTTGTCGAAACAACACATGATGATGAGGAAGAGGAAATGGAAGAACATTTGACAACGGTTAACAAACGATTCTTTAGTGTCCTTTCCCATGTTCCTACCGACAAGATTAAATTGATTATGGATAAGAACTATTCTATCGCCGATTTGAATAGAAA
>evgR002734c0g1_i01_Hydra
AAAATATTTAATGAAAGGTCCAGTTGGAAAAATATCAGCATCACAGGTAGCTGAAATTTCCAATAGCTTAACTTGTTTAAATGGGTATCTGCCAAGTGAGTTTGCTCGTGAACCAAGAAGTCTTCAATTTTTAGATCGTTGGATAGCTACTGAGTTTCAACAGTTTTTGATATATTTAGGTCCTGTTGTATTAAAAGGTGTTGTATCTGAACAAATATATGAGCACTTTCTTCTGTTAAGCACTTCTATTGGTATTATGTTGACATCTAACGACCAGTTTCGCAACAATCAGCTAGATTTAGCTCGCAGTATGATAGCACATTTTGTTTATTCATGTAAGTATATTTATGGAGAAACATTTACTGTATATAATGTACATAACCTACTTCATTTAGCTGATGATTGTATCAATCATTGTTGCTCACTTAACAGTATTTCTTGTTTTCCGTTTGAAAACTTTCTTCAAAGAGTAAAGCAGTCAGTTAACAACTCTAGTAGCCCTATTGTGCAAGTGTGTAAAAGACAAGTAGAATTTCAAAAAAATTTTAAACATGTTC
>evgR002749c0g2_i01_Hydra
CTCCAAAAATCACCATTGTTTATGCATTTGGTAGCTTTACTTGAGGTTTATAACTGGCCGTTAAATGTGAATGACGGTACTTTTGGCGATAAAGCTATCGGTGAAGTTGTTGAACATTTTTCTGAATTTTTAATTGGTGTAGGATGTGATCTGAATAATATACTTCATGAATGGATCATTTTAAAAGCATATATGTTTCCCATCATAAGTAACAATCAAAATGAGTATTATTTAAAAATTTGGGAAATTGTTTTTTCCAAATCAAGTTTGGTCAAAGAATGCCGAAATATTCTTGATATTTTTGAACTATTTTTATTTTGTCCTTTTACAAATGCGAATGTTTTGATGTATGAATTGCATAAAAAATGAC
>evgR002757c0g1_i01_Hydra
GTCACAGCTTTTGTATCAAATTTTATTGTGAGTAGTACTGGAGTTTTCAATTCAACATTTACTGACTCTACCAATGTTACAGTGTTTTGCTTGTTATCCCCACTAAATATCATAACTGTACCTTCTGTTTCACTAAATGGATACATTGAATTATCCCTCACCAAAAGTTTAATGTTGATTATTAATCAAGATTCTGGATCACACAGGAATTACTCAATTGATTGGGGTGATTCTACTTATTCATATAATAATCAATCAGTTATCAATCTAAACGTATATTCTCCTGTTTCTTTTGTAGTTCAGCATTTTTATATAATAGAGAACAAATATAATGTTTC
>evgR002766c0g1_i01_Hydra
AACCGCAGTTAAAGTCTTTGACCATACCCAACTCAATGATACCCATACTCCTCAACAAGTTATTGTTGAATTGATGAAGCTCTACTATACTCAAGGATGGGTGAGTGGTACCGGTGGTGGTATTTCAGTTAAGAAGGATGGTAAAATTTATATTGCTCCTTCCGGAATCCACAAGGAAAGAATTAAAGAAGATCAAATCTTTGTCATGGAAGAAGATCCAACCGTTGAAGTTGTTGATATTCAATCATGTCAACACTTGAAGACTGTCTATACTCCTAATGATAATGGTTATAATTTCAAGATTTCAGCCTGTCAACCACTCTTCCAATTGTGTTATGATGTTAGAGGTGCTGGTGCTGTTATCCACTCTCATGCCTTGTCAGCAATGTTAGC
>evgR002769c0g1_i01_Hydra
CTTAGTAATTTAGAGAAATCTTTTGAGTACTACGTAGTAATTTACGTAGAAATTTATGTGTTCTTGAAGTTAACATTTGCTTTAATCATTTGGAAGTTGCAAGAACATCGTCTGTGGAGTGTTTTATCATGAGTCCCATCACAGTTTTTTCATTTGACAGATACAATGTGTTCATTTCAGCTCTTGTCACGATTGGTATTCAGTTTTCTTTCTTTATCGTGGCTGCTACATGCAAATTTGACAAAGTAACAGATTTTGCTGGAGGGAGCAATTTTATAGTTTTGAATGTGCTAACTTTTTTTCTATCCAATACTTATTATCAGCGACAGATTGCAATCTTTGTTTTGGTTTTGCTTTGGGGATTTCGGCTAACTGGTTATTTGTTTTATCGAATTTTAAAGATTGGCGAAGATAAAAGATTTGATGATAAAAGACAGGATCCTGTCAGATTTGCTGTTTTTTGGATATTGCAAGCTATTTGGGTGTTTACTGTTAGTCTACCTGTCATATATGTTAATGCTCCTGTTAATGAATCAGCCAAACAACTTCAAGTTACAGATTATGTTGGTTTTTCAATTTTTTCATTTGGTTTATTAATTGAGGCTATATCTGATCAACAGAAGTTTAATTTTAGAAATAATCCTAGTAACAAAGGTCTTTGGTGCAATGCTGGTTTGTGGTCTTGGTCCCGTCATCCCAATTATTTTGGTGAAATTCTTGTTTGGTGGGGTTCATTCATTATAAGCATAGCTGTTATTAAAGAAAATAAACTTTGGACATCAATCATGAGTCCATTAGTGACCATGCTGCTGTTGCTTTTTGTTAGTGGAATACCTTTACTAGAAAAAACTGCAGATGAGAAGTTTCATTTAAATACGAAATACGTCGAGTACAAGCTTCAAACAAGTCCCCTAATATTATGTCCACCGTTTCTTTATGCAAAAATGAACAAATTTCTCAAGCTTATTTTCTGCTTTGAATTTCCTTTTTACAGTAAATTAGATAGTGAAGACTACAAGCGAATTGATCCTTGAGATGCATGCGATGAAAAAATCAGTTAAAATTAGAAAGATTACGAAGTTTGAAAAGTATTCGCAATGGTTTAAAAAAAAAAAAAAATCTCCGACAACAGAAAAACATGGTTTCAAAGTTTGGAGCATAAACTCTTTCTTTATACTCATTTTTAATATTGTTGTTTTCCGAACTTTTGTTAATAGAATTAAATTTAGGAGTCATCTTTAAATTGTTCAGAGTAGATTTTTACCATTTTAACATTAAAAAGTTTTGAATGTCAGCACGTTCAAATAGTATTTCTGTTTCGAATTTTCTAGTTATGC
>evgR002776c0g1_i02_Hydra
AGAGATTTGATTTATTTAGAAGAAACCAACAAGACGATCTTGAATACCCTAACATACCAACGAATTGCATACACTTTCTTATTGGGAACGATTCTGTTAAGTTTGACGAAAGAGATTTGATTTATTTAGAAGAAACCAACGCGTGTAATGCTTCTACTACAACTATTAGCACATCAACCAGCGCATCAATTACTACATCAATTACTACATCAATTATCACATCAACCAGCACTACTCTACCAACCCCTAAACCGCTACCATTATTTTCTGATATGCCAGCGTTAATAAAGAAGAAAGCTATAAAAAAGAAAAAAAACAAAAATATAAAAGAAAAAAGGAGACTTAAGCGACGAGAAGCGATGCTATACTTGGAAAACGAAAGACGCTCGTTAGCGAAGAAAAACCTGAAACGTCAAAATAATATGAACTAAAAAAT
>evgR002788c0g1_i01_Hydra
CCTTTAACTAAAACTACAACAACTGCAACATCTACATACCCATCTACAACTACTACATCTCCAAATACAACAACTAAATCAACAAATACAGTAAATATAAAGCCTCTAACTACAGTAACAACAACTACATACCCATCTATAACTACTACATCTACAACTTCTGAAGCATCGATTACAGCAAATGAGGCACCAACAATTCAATCATCGACAACAACAACTCAAACTCTGATATTTGAAGCAATTCAAAATTTGAATTTAATTTTAACTGCTGTGACATCTTCATCTTTAACAGTAGTTATTCCAGATTTTTTGTCGTTTTTGAAAATAATTTACATCATTTTGGTTAAGTTGAAATCAAAATCTTATCCTAAAAGAAATCCAATCACATATTCATGGAATGAAATAAGCGATCATACAGATGGTTTATTTTTAGTAGGAATTATTGATGGCAATAGCATGGTTAAAAGTTTTAAAGTTGGAGGAGGATTAAACCGTAAACGAAGAGATA
>evgR002831c0g1_i01_Hydra
TTTGGCTTTGACTATTTATGCTGATTTTTTGAAGAAAAATGCTATGCTATCGTTCCTAAAGTTTGCAACCATGAATTTGTTATATTTCTGTTTCATCGCTATTCTTATTGAACTCAGCGGAATTAAAATGCAAGTGTTTCAAGGTGATTTTGAATATGTTCCACGTGATAATTTCTATTATGAGATTGATGGTTTCAGGACAGCTTGGGCTAATTGCCGTCCACCAAAGTCACAACCTCTAGTTATTCCTATTTGGATTAATGGTTCCAGTAATTTAATTGTAAACGAATCAGATCCTCGCATAAAAGTCACTGTTGTTGCTAGTATTGGTATCTTCTCCCAAATGATTATCAACCCTGCAACTAAATGGGATTCAGGAAAATGGGGATGCAGAGCAATGAACAGCACAACTAATTTGACACATTGGTTTAATATCACTCTTTATGCTTCCCCCCTTGCTCCCCAAAATATGCATGTGAAACAGTATCGAC
>evgR002845c0g1_i01_Hydra
ATTCTTTGGACTACACTGTATCAAAAGATCTAAAAGTTTTGCTCCAGATGGTTGGCAAACAGACAGCAACAGCTAAACATCCTTACCCATTCTGTAAAACCTCATCTCCCAAAATGATGAAATCTGAACATTATACGTTGTCTTCTCTTTGCAAACTGTACAATGATTGGATGGTGGACGGTGCAAAATTAGAAAATGCAAAAGAGTTTAGTAATGTTGTACATCCACCTCTGTTAACTGGGGACCCAACAAAGAAGATAATTGAAATAGTTAATATACCAGGTCTACATATACTGCTAGGTGTAGTGGACAAGTAAATCTCAGAAATGGAGTCTATTCATCAAGACTTCAAGCAGTTTTGGGAAAGTCGAAAAGTTGGGCAGCATAGCCCAAATTATCTTACAGCACTAAAAAAAGCAACTGTTGCTTATAACTTTAACCATTTTTTATCTAATAAATTTAATTTTTTGCAAAAAAAAAA
>evgR002847c0g1_i01_Hydra
GAAAACCTTGGTGTGGAGATGACGAGCACACATTTTTGTTATTTGATTTTGTTCATGTTATGAAGTGTGTTCGTAATAATTGGTTAACTGAAGAAAACGGAGAATTAGTTTTTGAGTTTGATGGTAATAAACATATAGCAAAATGGGCTGATTTAAAGAATCTTTATGAGTTAGAGTCTGGCAATCTTTTAAAGCTTAGCAAATTGAATGATATTTCCATCTCACCAAAACCAATTGAGAGGCAGAAAGTTAGTACTTGTTTGCGAGTTTTCTGTGATGAAACAATTGCAGCATTAAAAAATCATCCATCCATTAAAAATAATGATGTTATTGGCAC
>evgR002848c0g1_i01_Hydra
TTAAAAAAATTGTTATCAAAATAATTTAAAAATAGAAAAACAAGTTTTTTGTTGTATTCTCTAAATATGGATTTTGCAGCCATTGTTTTATTTTACTTACAATATGAATGGATAATACTGTATTCATTCACTATTTCAATGTTTATTATTGGACTCATTGTATTTATCGTGTTACTATGGTTACCTGCACCCTATGGACGATATTCACAAACTGGAAAGTTATTAAGTTTGGGGATATCTCCTCGCTTAGGTTGGTTTTTGCAAGAGTTGCCAGCATTTCTTGTGCCTGTTCTGTGTATTGCAAGTTCTTACTTTGGTATCAGCCAATATAAGTTTTCTAACACACAGCTTGTTGTTCTTGGGTGCTTTATTCTTCATTACTTTATTAGAACGTTTTACTACTCATATAACATAGTTGGAGGTAAAGAAACGCCACTCTATACTATGTTATTAGCTTTTTCGTTTTGTTGCGTAAACGGGTATATTCAATCACGTGGCATGATGTTGTCCGAAAATGACTATGAAGATAACTTTATGCGAATTTTTTGTGGAATTTTAATCTTTTCTTCTGGTTTGGTTATAAACCTTCATAGCGATTCAATATTAAGAAATTTGCGCAAACCTGGCGAAACGGGATACAAAATACCAAAAGGTGGTTTGTTTAATATTGTTTCATGTGCAAACTACTCAGGTGAAATATTGGAATGGACTGGTTTTGCCATAGCTGCATGGAATTGGTATTCATTTGCTTTTGCTTATTTTACTTTAGCAAATCTTGTACCCAGGGCTCTGCATCACCACAGGTTTTATCTGGAAAAGTTTGACGATTACCCAAAGCACAGACGAGCAGTTCTTCCATTTTTAGAAGTAAAATCTGCAACCTTGATATCCACAAATGCA
>evgR002852c0g1_i01_Hydra
TTTTTGGAATCCATACCTGTTACTATTTTTCAATTGCCTTCATTAAAAAGGCTAAATCTTTCACACAATAATCTAGTAACTTTGCCTGTTTTGCTTGAGGACGAAGAAATAAATTCTCTATTAAACAGCATCAATGGTGAGAACAGTAATCTTTCAATGAACTGGAATTGTCCCAGTCTGGAAGAATTAGAGCTTCAACATAATTTATTAAAAACACTCCCAAAAAATGTTTTTGATATGCCAGCATTGCAAGCTTTAAACTGTAGAAAAAATGAAATATCTAGTTTGCCATTTAATATGTGGACTGCTAAATCGTTGAAAACATTTATTCTAGATGACAATTGTTTAACTGGTTTGCCAATTCTTGATAAAACTGCCAAATCCCAACTAGAATCCCGACATGAAACTATCAGTGAGTCCCAAGTTGGAAACGAAAGAAGAGTTAAAGAAAGAGAACATATTGATGTTCGTATTATGAAAAATGCATTTGCAACTTCTAAAGAATTTCTTTTCAAAAAAAGACCTACATGGGTAGAAGAATCTGATGATTCTAATGAGGAGGAAATAGATTTTTACCAAAAAATTGGTTTGAACAACTTGTTATTGTCAAATAACAAATTGACAGAGGTTCCAATAGGGCTAGCATGTCTTGCACCTAATTTAATAAAGTTGAACTTGAGCGG
>evgR002856c0g1_i01_Hydra
TTTATTTTCTAAATAAGAATTAAAGCTAAAATATAATGTCATGGAAAGGTGTGCTTGTAATTTTCTGCTTGAGAAACATTCTATCGGACATCCAAGTTCAAGGTGATGTTACACAAACCACAGACATAGATGAATGCCAGATGAAAACTACGGTTTGCCCAAGTTTTTCGACTTGTGAAAATACTATCGGCAGCTATATTTGCAAATGTCGACCAGGATACTCTCATATTGGTTCATTGAACACAGAAACTGTTTGTAAAAATTTATTAACTGAACATAACAGAAACGTCCTCACAGTTGTAGATTATTTAGTTCTGAAAAATGCTCAAAGAGGCTGTCGTTCTCTTCATAAAAAGTGTCTTCCTAATAAAAATTGAGGATATTACGTGTTTAAATTATCAACATTATAAATAAACTTGTTAAATATTTATTATTTAATTTATTGTAATTTTACTTTAAAATTATTATTATTTAATAAAATGAGACTAATTGTAATTATATATTTTTTTAATTATTTAATTTTTCGAAAAAAAAAA
>evgR002865c0g1_i01_Hydra
CTGGTTTAGTTCCAAAAGGATCAGTTATTGTTCAGTTGACTAAAAATAAGGGTGTTGTCATTAAATCAAGCCCATCTCTCACAACAGCTACTCCAACAGATGATGAATCAGATGATGACGATCAAGGACCAACAACTCTTCCAGTTATTGGAGGTCGTACTTTGGGAGGATTTACAAAGCCAATTGATGAAGAACCAGCATTGAATGCAATTTTGAAACAGACTTCAAATTTCAACAAGGAAGCCTATACTTTACTCTTATGTGAAAAACAGTGGATTTTATCAAAGTCTGAAGGAAATGAACAATCTGGAGTTACAGTTTATAGACATTTAGGAGATTCTAACTTGGGTTCAAGCTCAATTAATGAATGTATTTTATATTCTGCACATGGCCATGTCAGTATGTCAAGTCATGGAGACACAATGGTTGGCACATGGAGATTTGTTGATCAAAATGGAG
>evgR002872c0g1_i04_Hydra
ATGTTTAATCTGTTTCTCCATCAATGTTTATTGTACAACCCATGTTTACTCTGTTTCTTCATCAATGTTTATTGTACAACCCATGTTTAATCTGTTTCTCCATCAATGTTTATTGTACAACCCATATTTAATCTGTTTCTGCATCAATGTTTATTGTACAACCCATTTTGACTCTGTTTATCCAACAATGTTTATTGTACAACCCATGTTTAATCTGTTTATCCATCAATGTTTATTGTACAACCAATGTTTAATCTGTTTCTCCATCAATGTTTATTGTACAACCCATTTTTACTCTGTTTCTCCATCAATCTTTATTGTACAACCTATGTTTAATCTGTTTCTTCATCAATCTTTATTGTACAACCCATTTTTACTCTGTTT
>evgR002874c0g1_i01_Hydra
AAATAACAAACGCTCGCCAGAAAACATGAAGCTCAAAATTTCTGATCCATCCACTGGTATGACCAAGATGGTCGTCATCGAAGAAGACGCCGATATCAAATTGAGAGCTTTCTTTGATAGAAAGTTGTCTGAAGAAGTTGAAGGTGATAACATTGGTCCAGAATACAAGGGATACGTCTTCAAGATTGTTGGTGGTCAAGATAAGTCTGGTTTCGCCATGAAGCAAGGTGTTTTGGCTAACACAAGAGTTAGACTCCTCCTCGAAAGAGGTTCCCTCGGTTTCCAAAAGTGGAGAGGTAGACAAGGTGAGAGAAGAAGAAAGTCTGTTAGAGGCTGTATTGTTGGTCCTGATCTCTCAGTTTTGTGTTTGACTGTTGCCAAGAAGGGTGATGCTGAATTGCCAGGTGTTACTGATACTGTCAAGCCAAGAACCCACGGTCCAAAGAGAGCCTCAAAGATTAGAAAGCTCTTCAATTTGACCAAGTCTGATGATGTCAGAAAGTACGTTATTAAACGCGAAAGAAAGGTCGCCAAGAATAAGAAGGTTCTCAAGGGTGTTAAGGTCCAAAGATTGGTTACCCCAGAAAGACTCAGCAGAAAGGCTAAGGAAATCAAGAGAAGCATTAAGAGAGTTGCTGCCACAAAGAGTGAAAAGAAGGCTTATGCTAAACTTTTGGCCAAGAGAAAGTCCGAATCCCAAAAGGCTACTGAAACAAAGACCAACTAAACTTATTTAACCAAACACATAAACCAAAAAAATTCTAAAAAAAA
>evgR002888c0g1_i01_Hydra
TATTTTAGTGCAAAAAGGTAGAAAGTAATTCAGTTATGGATGAAGAGTCATTTGATAATGATTTCAGATTAAAAGATAGTGAACTTTATAAAATAAAATTGGCTTTATCTTCTCTAACAAACACTGTCATAGTATGCCCAACAGTTGCTACTTTATATACTCAAGCTCTAAACGATGATGAAACATGGAATTTCTTAAGAAAAGGTGTTCCTGGTGTAATTTTAATTTACAATAAAAACAAGGAAGTTACTGCAACCGAACTTTGTTTAGCAGATCCTGCATCAGGGTTTGCTGTTTGGAGAGAAATTTTAAACGAGAATTCTGAATATAGAGCTGTTCAAAAAAACTTTCATAGTTTTAAGCTTTATTCAAGCGGACAATGTTCAATGGGAGGTTTAAAGTTTCCAATCACAGAAGTTGCAAATAATTTTCTGCAAGATGTTTTAAATAGTATACCAAAGTTTACTGTACATGAACTTCTTGAAAAGCACAACGAAAAAAGAACTTTTTCATTTCGTAGACTTCACAAAGAAGAAATTTCACTGCCGTGTATGTTTTTACATGTATCTGGTATAAAAAATAAACAAAACGAGAAAAAAAAATCAATAACCAAAAATCAGCATCTTTCATAGATATTTGTATAAATTCAAATGTTAGTTAAATGAGTAATGATGTTTGTTCAAGCAACAAAAAACATCTATATGCATGCATCTATATTTGGAAATAATGGCAATTACTGGCATTTACTATCTGCAGCAGATTCTGATTTTATGAACTGGAATGAAATGAAGACATTTCACTCTTATTTTGTGTTGGAATATGATAAACCGATTTCATAGTTTTGTGTGCTGGAATATGATAGAATAGATTTCATAGTTTTGTGTGCTGGAAT
>evgR002904c0g1_i01_Hydra
CTCATAGTGAATCATACCCTTTCCTTGTGGATCAGCCTCCTTCAACATATCATCAACTTCAACATCAGACAATTTCTCACCCAACGAAGTCATTATATGCTTGAGTTCATCCACTGGAATGAAACCATCCTGATCACGATCAAACAAACGGAAACTCTCCAAATATTCCTCCATATTTCCAACTTCCGAAGATTTACCATATCGATTACGCATTTCATTCACAAAATTATTAACGTTAATGTTACCAGATTGGCCAGGATCGAGAATGGAGGTAATTTCGTGAAGTTCGCGGTCAGATGGAGCGAAACCCAAGGCACGAATGGCATTGTTGAGTTTGTCGTGTGGAATGGAACCGGAGCCACCATCGTAGAGGGAGAAGGCTTCACGAAAT
>evgR002915c0g1_i01_Hydra
TCCGGATCGCTAGTTATTTGTTTTGCTGTTAATAATGATCTTAATTGCTGCATTAACACAAATCCACTTACTTTGGATGAATGGAATGATATTAAAATATATCAGCGTCTCAATGATATGAAATATATTTTATCTGTTTATCTGAACAAAAAGAATATTTGTTCAGTGGAAAATTCTGCACCTAATGATTTTTTTAACATTAAAGTATACGCTTCAAATCTTTGGGACAATGCACCGAATGCTTCTATTTCTAAACTTTTTATTGAAAATAAAGTCAATGGAACAGCTGATTTCATAAATCACAGAGAGGAGTTTCTTCTCACTCGTGGTTTAGTTCTTGGAACACTGACTGATTTAAAAAAGGAGTACATTGTGTTCTTCAATTTAAAACCTACTAGTTACTCTAAGGGTTTGAAGAATGTTCTACTTTTATCATTAAG
>evgR002932c0g1_i01_Hydra
GTTTCAAACTTTAGTATCTTTAATAGAACTATAAATAATATTGGACTAGAACTAAGAACGATTGGATCATCACGGGTCATGAAACGAATTATATTGCAGTTGCTGGTTGTTTTTTTATTCTTCAATTTTATCGAATTGAATATTGAGTTGAACTCTAATGCGTTTCGTGTTTTTCAACAAAATCAAGTAACAACAGAAAAAAACTATATCAAAATTCGATTTAAAACAACTGAAGCAACAGGTTTAATAATGTTAGCTCAAGGAGCTGATAAAGAACGGTTTTTATCAATGGAGTTGTTCAGAGGAAAAATAAGAATTGCGGCTAACTTCGGAAATGGACCGTTAAATCAACTACGTCCAGTAAATGGGTTTTATGAAATGTTTCTTGGAGTAGATCTGGCTGATAACAAATGGCATACTGTTGAACTCATACGAAACGTGAGAGACAATATTATTTATATTGATCGTGGTCGAGGAAAAATGGAAAAGTATTCTTTCGTTCTTTCACCGCCTACTTATATTGAGCTACCTGTATCAATGGTAATATTTAGTGGGTATTACTCATTCCTGGTTTCAGAAATAAGCACTCAAAAATCGCTATCACGTAAAGGAATCAAAGCTTGCTTTAGCGAAGCTTTGTTTAGTCAATATCTACCAGATACTAATGGTGAAACTTTCAACTTTTTATCAATGAATTCTGCTCATTTTAAGCATATAGGGGAACCTGCTACACAATGTACTGAACTTGAATATTCTCCGTTATATTTCCCTTCATCTGCTGTTCATATTTCTTACACACAAAACTATACAATACAGTCGATGAAAATTGATTTAAAATTTCGTACTGTTATTAAC
>evgR002943c0g2_i01_Hydra
AATGGTACCGTTTCACACGAAACCTACGAACAGTATATGAAGAATGAAATTGAGAAAGTTGTTCGTTTACAAGAAGAAATCGGGTTGGATGTACTTGTTCACGGAGAACCAGAAAGAAATGATATGGTAGAATATTTTGGTGAGCTCATGAATGGTTATTTGTTCACTGAAAATGGTTGGACTCAATCATTTGGTAGTCGATGTGTTAAACCACCTGTTATCTATGGAGATATTTCACTTGATAAACCAATGACAGTTGAATGGTCTTCCTATGCACAATCTTTAACTCAAAAGGCAATGAAGGGAATGTTGACTGGTCCAGTTACAATTTTGAATTGGAGTTTCGTCAGAGATGATCAACCAAGAAGTGAAACTTGTAAACAATTGTCACTTGTATTGAGAGATGAAGTTCTCAATTTGGAAAAAGCAGGTATCAAGGTTATCCAAATTGATGAACCAGCTATCAGAGAAGGATTGCCATTGAGAAAATCTGATTGGAATACCTATCTTAAATGGGCAGTTGACCTCTTTCAAATTAACAACATCAGGTGTTGCCATCGATACTCAAATCCA
>evgR002973c0g1_i01_Hydra
GTCAATGTGGTCTTCTTTGTCATTGTGCAGCCAACCATGTGGAGATGGTTTACAATCAAAATATCGCAGCTGCTCAAACCCTGCACCTAGTTTTGGTGGTCAGGATTGCTATGGAAACTTTGAGCAGTTTTTCAATTGTTCAATGCAAATTTGCAAAAGTGTGAAATTGAATTTGAATATGAATTTCATTGATGAAAAATATGACTCGTACTCGATGGAAGCGACTACAGATATACTAAGTGATAAAATCCAAGAGGCTATAGCAAAACTTTACAGTAATCGCAAAGTGGGCTTCCATGTTGTTATTCATTCTATAGAAAATGGACTTTAAGCATTTTTACCGATGTTTTTGTCTTATTTTAAAACTGCAAAGAAGATCAAATGAGACTTGATAATAAAGAATGGTGAAGAAGTTTATGCGAGTTTATTCGAAAACGAAC
>evgR002977c0g1_i01_Hydra
ACATGAAAATAATGTAGCTAATGAAGATTCGTATGATGAATATGAACATGAAAATAATGTAGCTAATGAAGATTCATATGATGAATATGAACATGAAAATAATGTATCTAATGAAGATTTGTATGACGAATATGAACATGAAAATAATGTAGCTAATGAAGATTCATATGATGAATATGAACATGAAAATAATGTATCTAATGAAGATTTGTATGATGAATGTGAACATGAAAATAATGTAGCTAATGAAGATTTGTATGACGAATATGAACATGAAAATAATGTAGCTAATGAAGATTTGTATGACGAATATGAAC
>evgR002985c0g1_i01_Hydra
GATATTTGCACACTTCACACATGCTTTATCAGTCAATATGACTGCAACTATGGCATAATTTAATTCTAACCAAGGTAACACTAGCATAATTTGTGCAGCATATTAAATATAATTATTTTTAAGCTATAAATAAAGGCCTCGAGCCTTCCGAGTCATAATTTTATTTGCCAAAGAATTGTTGGGTTAAACATGGAGCAAAATTCAACACTTTTTAGTTTTAAAAATAATTATGATTTTGGGGCCATTTTAACTCTCTTTGTAAGTCATGAGTTACAAATTTTAAACTCAATAACGACGTCGATGTTTACGGGTGGGTTTATTGTTTTTATAACAACACTTTGGCTTCCTGCGCCTTATGGACGTTACACATATACCGGGAAACTACTTTTGTTACCAATATCTCCTCGTTTGGGATGGCTTGTACAAGAATTTCCAGCTTTTTGCATACCTTGTATTGTAATTGGCAGCTCTTTCTTTGATATATCGCAAAATAAGTTTTCTAGTTTACAGTTGACCGTTCTCGGATGTTTTGTGCTTCATTATACAATGAGGACGTTTTATTATTCATACAATATCAACGGGGGTAAAGCGTCACCGCTTATTGGCTACTTATTGGCTGGTTCATTTTGTTTTATAAATGGATACGCTCAATCACGTGGCATGCTGTTGTCTAAAAACACTGTGGATGATAATATTGCACGAGTTATATGTGGAGTTTTTATTTTTTTAGTTGGTTTGGTAATAAATGTTCACAGCGACTATCTTTTAAGCACTTTACGCAAACCTGGTGAGACAGACTACAAGATACCAAAAGGTGGTTTGTTTAATGTTGTTTCGTGTGCGAATTACACCGGAGAAATTACAGAATGGATTGGTTTTGCCATTGCGGCTTGGAATGTGTATTCGTTTGCATTTGCGTATTTTACAATTGCATTTTTAGTACCAAGGGCCATACGACATCACCGGTTCTACCAAGAAAAGTTCCAAGACTATCCAAAAGATAGATATGCTGTTCTGCCGTTTTTATTATGAAAGGAAGCATTCCATAGTTCAAGTTCAAGTCAAATTCCTTTTATTATTTCTTTTATAAAGATTTTTTATATTTATTTTTTATTTTAACGCAGAAAGTTAATTGTTTGTGTAATTGGAACAAAGATTGGACTTTTTAATTATAATATTATCTAGGAAATAAGTTAACACAAAAACGTAAAAAAAAAAA
>evgR002994c0g1_i01_Hydra
GTTACTGGACGATGTCTGTCATTTGAATTTAAGTTTGGTGGAGGTGGATATGTGAGTCTAGCTGTATTAGGCTATACTAATAATGTTTCTAAAGAACTGGCATTCTATTACCGAAGAGAGGGTCTAGATAAATGGGTTAAGAGTTTTATTGATTTGGATCAAAATAGCAGAAATGTTTCAATGGTTAGTTTTGTGGCTTTAAAAAATGGCGTTGGCACAGGTTACATTGCACTTGGTGCCATTCGGTTTTTGGTTAGCTGTAATGATCTTTTCACCAATGAACTGAACACATATGGTATAACTGACATATTAAATGATTGGAAACTTTGGAATGATTCTTCTTTATTTTTAAGTCCAGTCATCTTTATTAGAG
>evgR003000c0g1_i01_Hydra
TTCTATTGTCTAAATAAGCTTAGATATGGAAAGAAACAAAAAAAAATTAGCTGATTCAAAATTTAATTTAAAAAAAGAAGATATTCCACCGGAATTCCACGATCCATTTATCATCAGAGGATATCGAAAGCCATATTTGTCATTTAAAGACTGCCTCAAAAGTTCTGTAGTATTTTCGTGCAATGAGTCCTTTAACATTTTGTCTCATTTCTTATCCTTCATATTTTTTATAATACTATTTATTTTCTATTTTAAAACAGAATTGGATATAAGTGACGTTAAAACATGGCCTCTAGCCCTCAACTTTATTGGAAATATTGGATTTAGTCTGATGAGTACAATAGCACATACTTTGAACTCTATGTCAATTAAAGCGAGACACAAATGTTTTTATCTCGACTACGCAGCAATATCAATATACACGTATTGTTCAGGACAAGCGACATTTTTCTACACAAGAGCTTTATTTTCAGAAAAAGAGTTTTTCAAAAACTCGTACTATTTTAATTTTGGTGCAATGTTTATATCGATCATGTCAACCTTATTAAATTGTTTATCGAGAAAAAAATGGCATTCTGCAAAATACATTATACGGACAATGGCATATGTAACAGCTTATATATGGAACTGTGTTCCATATGCTTATCGACTAATTATATGTAACAATCATATAGACTGCGATAAAAGTGGGTTGTTTTTATCTCTGCTCAGCGCACTGACATTATTAGGAGCTGGAATATTAAATATTACAAAGTTTCCTGAACGTTTTTATCCTGGTAAATTTGACTGTTGCGGACAAAGCCATAACTTAATGCATATTTTAGTGACGTTGGGATGCTTTTTTCAGTTAGAATTTGCTAAGAACGAAATGTTGCGCGGAATTTCACAAGGATTCAATTACGAAAGTTCTTTAAACTTGACAATTTTTACCGTTTTCGTTAGTTTATTGATAGCTGCGGTTGTTCCAGCCGATGATGACAAAGATAAAAACTATCAGAAAAGTAATTGATCATTGCTTTATCTTCCTTTAATTTTAGAATTATTAAAACAATGCTCAACCAATATTAATATTAATAGCATATAGAATTGATCAGAGAAGATTTAGTAATAAATCAAAGATAAAAATAAACTCAAATGATGTTATAAATGAAATATGTGTATAAGTATAAATATGAAAAATAGATATGAAGTATAAATATGTAGTATGAAGTATAAATATGTAGTATGAAAAATAGATATGAAG
>evgR003007c0g1_i01_Hydra
CTGTCAATTCGAAAAGATTGTACAACTAAGGGGAAAAAATAGTGGAAAATATTATAAAATTTATTCAAACAATTTTCTTCCTGAATTTAGTCTAATAAGCAATACTGAACAATTTTTTAATGCTTGTAGTTACCTATCAGCTGACCCACCAACATTTGGAGCACAACAATTTCAAAATCTTGAGGTAGATTCTGGAAGTCTATTTTCTGAAAGTATTCCCAATGATCTTTTTAGACCTCCATTAAGTGGAAGTACAATGATACTTACTATAGAACTTAGAAACCAAAATGGCGAACCTTTATTAATACCATGGATTGCATTAAACGGGAATATATTTCAATATGGCTTAAGAATTGAAGGATGTATTCCTTCTGATATTCCTTCAGCTTCTGGGAATACAAGCTTTCTCCTCCAGCTTGTTGCAATCACTCCGTTATTTGAAGAAGTTGTTAATGTTTTTAATATCATTATATCAGATCAAAAATCTTTACAGCTGAACTTTGTTGTAACTTTTTCTTCTCAATCATATGCAATCGGTAACTATACTTTTTTTTCTAATTTCATATCACACCTATCAAGTTACTTGTCAGTTCGAAAGAATTCTATTATTATTATTAAGTTTTTTACTACACCGTCATCGACTGATATACAGTGGACTCTATTATCACTATCTGATTCTTGTTGTGATAATGACTTGGTTCAGTCTCTTACTTCAAAAATTAAGAGCAAAGGAGCATCAACTCCAAGTATAGACCATTTGCTAAACAAAATGAACTTTGTAAACACAAGTATAACTTATAAATTAGGGTTTGCATGCAATAATGATAACACTCCTCCTGTACCTGATGTAAACGCAGAAGATATAACTATTCAAAAAATTGTTTATCGGTTTAGGCTCACATTACCATTAACGACATTTGTTGATAAAAACGATGGGAACATGAGCATGCT
>evgR003012c0g1_i01_Hydra
TTAATTTTAGCCGTGCTCGCTGTTTTCGTAAATATCATTTTTCTTTTGTCTCACATAATATTGATTCGAATAAAGTCATTTACAGTTCAGTGAACAAGCAACAAGATTGACTTTGTAAAATGTTAGATTGGGTTGATTTTAAAATGCGATCAACGATTTTGTTATTGGCGTTTATTTCGGTTTCTTTTACCAATGTTGTTAAAGAAAAAGACTCAGGTATTAGCCAGAAGATTATCGCCCAACCTCAAAGTAAGGACGTCAAGATTCAAAAATGCGCATCTGATACTTTTAAATTTGCTTCCGCCATGATGAAAAACTGCAGTGCAGTACTCAGTGCACCCGTTGGTAAAGATTCAGTATGTGCAATTCCTCCATGTAATAATCAAGCGTTTGCCTCACCAACCCAATGCAATTATTTTTTAGATGTGTTCAACTATCCTACTGGAACGTATTGTCAATCGGTTGCTATGTTCTTAGCTGCAATCAACAACGCTGTTAATTACAGCAATTGTTGCTCGTAACTTGTTTCATTTGTCAGGTCAAAACAAATTTTGCAAATCTTTTAGTTTTTATGGAGTACTTTTTTGATGAACAACCTTAAAATTTTCTTAAAATTGTTGCATATGTAAAGAGTTTTTTTTCTAACGGAATCTAAGATTATTTTCCAGAAATTTTGTTTAACTTTTAATTTTATTTTTATAGCCTTCGATTTTTTTTGTGTAAATTTCAATATTATGTATATTATGTCTTCCAACGTCAAAAGTATGCCCCAAACCTTGATAATCTCAAAAATGGATTTTATTTGCAGAATTTAGTTAAAAAACAATTTTTTTTTATTGAAAATAAAAGAACA
>evgR003015c0g2_i01_Hydra
CAGGTCTGCTTCTCGGACAAGTCAACATTTCAAATTTTGATGGACAAAAGCTCCTTGGTTTGACAACGCCCAGGAGAAAATTTCACCATCACTGCCTTATAGAAAAGGTTAAACATCCTGTGAGTGTAATGGTGTGGTCTGTGATCTCAGCCAAGGGTATGGGCTGCCTATACATTGTGAAGGAGACCATGAGTCGGAACCAGTACAAGCGAGTACTGGAAAACAGTCTCATACCAGGCATTAGAATACACGAAAAGATACCCCAGGTCCAAAAATGGCTCCCTGGTAATAAAGAGTATATGTTCATGTATGACTCGGCACCG
>evgR003024c0g1_i01_Hydra
GAAGGTAACATAGAGTTTCTTTTAAACGATTTTATAATTGCTGGATCGGAAACATCTGCAAACACTATTCTTTGGTTTATTGTTTATATTTTGCATTGGCCAGAGTATCAAGATAAGCTTTATGATGAAATAGTAAAAGTAACGAAAGGTAGTCGATATCCATCCATAAATGATCGCCCTATGCTTCATTTAATGCAAGCTGTAATTCACGAAACTTTAAGATTGTCATCGGTAGGACCACTTGGTTTATGTCATAAGGCAATGGAGAACAGTAGCATTTGTGGAAAATACGTTCCAAAAGGAGCTCTAATACTGACTAATTTATGGAGTATACATCACGATGAAAAGTATTGGAACAATGCCATGTCTTTTTATCCAGAACGTTGGCTTGAAAAAACTGGAGAGTTTAATTCTAAATTAGGATTTGCATATTTTCCATTTTCGAGTGGACCTCGTAGTTGCATAGGAGAAACATTGGCAAAAACGGAATTGTTTGTGTTTTTTTCGCGATTAGTCACAGATTGTCTATTCGAAAAACCGATCGGAAAAGAGTTACCTAGATTAGATGGTCAATCTGGTGTTACTTGCCCTCCCTTTGAATTCGATGTTGTAATAACCCCTAGGGATTGAATTAAAAAAAAAAAAA
>evgR003039c0g1_i01_Hydra
CTAATGTTATCCTTAAATTTGTTCATTTCAACAACAATAATGAACTCATTCATGATCTCAATAATTCTAAGAGAGACGGAGTGCACGTGCTCAGACCATTCCTCAACATAACCATAGAGCAAAAGTGTCGAAAAGTGTATAATCCAGTACAACATTTTCGCCGATATATTAAAACTAAAAAAACAGGATTTGGAATAAAGTTGGATGAGTTAACTACGTCTGATGGTATTACCATCGAATTTTTAGTGTACAGTGGCACTGGGATGTATGATGATGACAAAAAATCGGCACTGCTTTCATGTCAGCGCATACCTGTAGACCTGATGGATCCACATCAAAACTTTTTTTTTTCGAAAAAATTTATTTTCAGTTGAGTTTATACAATAGTTTTAAAACATAAATTGGCTTTGCATTGGAAAAAAGTGCCAACGATCGCGATGGTTTTACATATGTTGGCCCAATGTAAGTATGCTAGTTTGGATAGACGTCGATAGGACGTCCGCGTGTCCACTTGGCAAGGGTTACCTCTAGTTTAAGACTTCATTCGCACGATGAATTGGGCTGCTCTAAA
>evgR003044c0g1_i01_Hydra
GCGCCTATTGTATCACAGAGGGCGGAGAATACAAAAGAAAACAGATGCAGTCAGAGGTGTATTGCAATTTGAAGATGTGTGCTTCGACTACAACATCTACACCAACAACAACTCAATCTACATCTACTAAAACTTCAACTACAACAGCTACACCAACAACAACTCAACCCAAACCTCCTATAGATATTTCCTGTCCGTGTTTGCAAAGATATTGTGAAAGGTTTTTTGACGAAGAGTTTTGTTCATTAGTTAATCAATTGTGTCCAAAAACTAACTGCACACCAACATACAGGCCAATTACCACAAATTTATATACAACAACCACAACTCAGTCTACATCAACCATAACGCTACCTACAATAACGCTACCTACAACAACCACAACTCAGCCCACAACAACAACAACGCCACCTACAACAACCACAACTCAGTCTACATCAAC
>evgR003050c0g1_i01_Hydra
GAGATTGGAATTCTTGAAAATAAGACACGTAATCCTTTTCCTGATTTATAAAGTTGCTTTTGGTCGACAAATTATACAAACAATAGAACATGGTAGCGGCTTTACATGTCGAAGTTATTGCGATGGGAATTATGACTTGTGTAAGACAATTGTTAAGTCATTCCATGAAACTTTTATATGCGTCAAGACAGCAACGCTGTGTAGAGCGACCTGCGGTAAAACAATAAAAGAAGTTGAAAAAAAATTAGAAACATTTGCTAAAAAAATAGAAGCAGTCGAGAAAAAACAAGAAGTAGTTGATAGCCGTCCTCAGTTGAAACCTAACAAATCTGATCACAGCAGGAAACAAAACCAGGATCGTAACAATATTAAAATTAGTTTACCGCGAACAAAAATGTCTGTTCGGTTAGAAAGAATATGGCTTAATAATAATATATAGTATGTCAACTACCAATCAACAACTACCAATATTTCAACAACTACCAATATAATAAATGCCAATAATAAAAATACTACTGATATTAGGAAATGTGAAATTAAAGAAGATTACAAATCGTAACAACGTTTACAATCAAAAATCAGATTAAAAATAAATCAATAAACAACTATCAACAACACCAAACAGTCAACAATACCAAAAATCTGTAAAAAATCAGCACCCAAAAAAATTGCTAAAGCTAATCTATATTTATCTGTCAAAGCGTGCAACTCCTAAAGACTTGTTTTAGAAATACAAATGCC
>evgR003053c0g1_i01_Hydra
TTCAAATTTAGTACTAATTTATTTAAATTTATATTGCTAAATATATATATGACAGACATCATTAAAATGGCAATACCAAACAGAAAGTCAATTCCGGCAAGAACGCAATCATGTAACATTCAAATAAAAAGACGGCGAGTAAGAAGATCCTTTTCAAGGCCTGATAACGAACTGTTTCTAATTAATGCTGTAGTAAGGCAAGACGTGAACGATGCTCAGTTTATTCTTGAAAGAAAGAAGGTAAACATTAATTCAATGCGTTACCTTGGTTGGGCCGCTCTACACTATGCTTGCAAAAATAATCAATTAGAAATGGTGAAACTGCTGTTACACCATGGCGCAAACACTAACTTAATGACAAACGACGAAAACTTTCCGTTAAAAATTGCCGTTACTTGGGGAGCATTTGACGTTTATTCTTTTCTATTACGAAATGGCGTTTTTAATAATTTCAAAGAAAACGGTTTACAAATAAGTTACTTTTAATAACTTCTGTATTTTGATTTTATCTTTATCGTTAGATGGGTACAATAACTTTCAAC
>evgR003061c0g2_i01_Hydra
ATGTTATTATTTATTGTAAATTGTTATTTTGATTTTTTTTAAATATTATTCTGATGAGCAGAAATTTATTGAAGAATTTTTTTGTTATAGACAATTCTTGTTTTGATAAGGTTCGATGAGTTCTTCAGCAGGACATTCCGCATTGTCTTGTGGCAGTATTCACAATATTACATCTCTTTCAGCCTTTACTAGCTCCGTTTCCATTATTCTGTGTTTTACTACAACGCCTTGTAATATTGCACTTATTATTGCAATATTTGGAGATAAGAAAAATGTTTTAAAAAAGTCTATTTTTTATAAGCTAATACTGAACAGTGCAATTGCAGATCTTGCAATTGGTATTGTGGCTGACATTACATCAGTTTCATTTCATGTTAAAGAAGCCTTGAGAATTAAGATTTATCCTAAAGATATTAAACTGCTTCATTTAGAGCTTTTCGTTTTAAGTAACGCTTCAATTTTGTCCATGGCATTATTATGTATTGATCGAATATTTGCATTAATGAAGCCAATAGCTTATTATAGAAATTCTTTATCTAACCGCATGTCTTACATGGTTCTTGTTGCTACGTGGATAATTTCAGGTCTTATTGTGTTGCCTTATTTTTCTGTTGGATACATTACCTATCTCGGTATTTTTGCATTTACAACCATTACTATAGCTTGTACATCTTTGATATCGGTTGTTTATCTTTATAAAACTAAGTTTTCGGTTTATTGTAACTTTAATAAATCGTGTAATGTTGTTTACGAAGCCAATAAAGAAAATTCCACAAATAAAAATTCAAATCAGAATTCAGCTACCGTAATTTCAATAGCAAACATTGTTGAAAGCAAAAAAATCGCCACTGCTAAGAGTAAAGATATAAAACAAAAGACTGCAGAGCAAAAGGTGAACCAATCATTTATTATAATGCTAATGGTGTTTCTACTTACCTATATGCCTGCATGCTGCATGACAGTATACATGAACGTGTGCTTCGAGTGCAACTGTACTGTTGTACAGGCTTTTAGAGATTGCGTTTATCTCGTGTTACTATCAGGTTCTTTGTGGAGATCGCTGAACTTTGCATATAAAATTACAACCTTGAATAAAAAAATAAAAGAAATGCTATCTTTTGGACACAGCAGTGCTAGAAAATAAAAGCTATTTCACATATCTATTATAACTGAAATAATGTGTTTTAGGCACATTAATTTTAGAATGTTTCACGTATCTTTTAAAAACGATTAAC
>evgR003062c0g1_i01_Hydra
GAAACAGCCGCAGATGATGAAGTTAACAATAGCCAAGAAATAATATGATTTCGGAAACAAAAGAAAACATTTTGAATCAAAGCGAAAGCATAAAAAGGCGGAACACGATTAGGCATCTAGACGAATACTACGACGAGCACATAAATGAAACTTTACCTCAAGATTATCATAGAGGAGACGTATCGTCATTAAACAGAGTGCTCGCTTATTTTGACGCTGTTATGGCAGCTTGTGCAACATTTCTTGCTATCCCTTTACGAGGTTTTCATACAGATGATGAAAACAAGTCATTTTATGACCATATTAATTCTATTTCAGACAAACTTATAAATTATTTTGTGGGATTTTATATTGTTCTTTCAATTTGGGAAAATATGAACAATCGGGCCATTGTAATAAAACGCGTAGATGACTTTGTATTGTCATTTGTCATTTTTGAAATATTAGTAACTTCGTTTCTTCCATTTTCGTTGGCATTGCAAGGACATTATCCTCTTGAGAAGGTTTCATTTTTAACAACATGTGTTGTTTTAGCTATACTTCAAACGATAGATATTGTAATAATTTTGTACGCCATTCATTCTCCAAATCTAATTCACGTTGATTTAAAAAATTGGACAAAGTCGGATCTTCGAGAACTAACTCTCATGATGATTTTTAAACCTTTGGTAAGTTTGTTTTTACTTATAATTGCTGGTGCGTCTGTTTTAGTTCACTATGGCGTGTCTTGGGCTTTTATTGCACTGTTGATTATAATACACCCAATACGCAAGTTTTACTGGTTTGTTCGTAGAAAAATGAAAATGCTTAAACAAACAGAAAAAAGTATATTTTTGGAGCATTTTTCTAAGGGTAATGTTCCAAAAGAACGAGTTGAAATTATGAGCGATGCTGTTGTTGCCATCGTTGCTTGTATTCTTATACTCGAAATTACCGACTTAGGGTTTCCGCCTCCAAAAGATGTTAAGGAATATGGTCTCATCAACGTGTTGAAACATATGAATTCTGAATTTCTTACGTATGCCGCTTCTTTTTGCTTGATTTCTGCTCTATGGTATGTTAACCACGCCGTTCTACATTTAATTAAAACAGTTAGCTCAGTTATGTTATTCCTTCAAAAAGTTTTTCTTTCTTTTTGTTGTCTTTGTCCATTAGCTGGTAACATGGTATTACACTTTGCAACCAAAAAGAATAATGAATCTCATGCCGCTATCCGCTATTCATCCATGATTATTTTTCTATCAAGTATTGCAAACTTCTTTATTTTGTTGTATGGCTTGCTCACTGGAGAAAAATACTTCCACAAATGGGCTTCAATTAAACATTTTAAAAAAAACAAACGACAGCAACTATATTCTCTGACTAAAGCATTAAATTTACCTTTTTGGTCTTTCATTTGCACATTAGGAAGTTTGGGATCCCCATCTGCAGCACTTTATATATTGTATGCAATAATTGTTGCCGCACCTTGTTCTTTTTTTGTTTCAAAAATAGTACTTATGAATCACTTAGGTAAACTAGCAACATAGCTTTCGTAGCAACATAGCAACATTAGCTTTTTTTGTCAAACTTCTTTACCAAAGATAAAAAGCAGAAAAAGAATAACAATCTTTAATAAGTAATAACTTTTAAAAACATATTTTATATTAGAAACAGTCATTTAGGTGTCAAAGAGATTATTTATGTTCATTTAAAGTTACGGATGGTGAACACGAATGATACCAAGATATGGAGCTTGTTGTGTTTGATTCTCCATATCATTACTAGAACTATTCGAAGGATAATTATGCGTAAAATACATACTTAAACTTCAGTATATTAAAAAAGTTTTTTTAACAGCTAACTTTGTTAACCCCCTCCTTCACCTATATGAGACATAACACAAAAAAAAATAATCTTTGTTTACAATTTTTTTTGTAAAAATCTTACAAACCTGTGC
>evgR003091c0g1_i01_Hydra
GCAAAAAGAATAATAACTAGTTAATTGTCTGAAAGTTACTGTGATTGACTGAAAGTTACTATGAATGACTGGCAAAATGTTTGGAGAAACGTTGATGCTGTGTGTTTTGATGTTGACAGTACTTTGATAGTTTCAGAAGCAATTGATGAATTAGCAGAGTTTTGTGGAGTTGGTAAAGAAGTTGCTGAACTAACAACAAAAGCAATGGGAGAAGGAACTTCATTTCGTGAAAGCCTGTACCAGAGGCTCTCAATTATCAAGCCAAGTAAGGATGTGCTAAATTCATTTATTAATAGGAATCCATTTAAACTTAGCCCAGGAGTTGAAGATTTAATAAAAGCTTTGAGGATTAGAAATGTTGAAATCTATTTGATATCTGGAGGTTTTCGTTCGATTATAAGTCCAATTGCTGATGCTCTAAACATACCTCCAGAAAATGTGTTTGCTAATGAAATAAGGTTTAACCAAAATGGTGAGTATGAAAGTTTTGACGAAGAACAACCAACTTCAATGTCTGGTGGCAAAAAAGTAGTTGTTGGAAAGTTAATTGAGAAATACAGTTATCATAACATTGTCATGATTGGTGATGGAATGACAGATTTGGAAGCCTTTCCGCCCGCTCATCTTTTTATTGGATATGGTGGTAATAAATTGAGGGAAAATGTCCGCATGCTGGCGCCCTACTACGTGACACACTTTAATGAACTTATTGACGCTTTAAATTAATTTTATTTCCGTTTTGTTTTGAATAAAAATAGTTACAATTTTTTTTTAAATAAAGGGTTAAAATAAGTATTTATATATCGAC
>evgR003093c0g1_i01_Hydra
AGCGGCCTTCTCCAGTTGCGTAAAAACTCTCCGTTGTATGTAACCATAAAATGTAAAGAATCATTTTCAATTAGAAACACAAGTTACTTTTCAATAACGCCCATAAAAAATGAGATTTTAGCATTTTTTGGATATGTAAAGAAAACGGAAGAACTCACGTGCAAAAATCAATTAATGCCTTTAAATGGTATATATAGTCATTTTAATAATACCTTGAGCAATTTTTTGTCCAAACACAACAAGTTTATAGTGCCAACTATTGGAGTGTATTTAATATTAATTAATGCACTAGTCAAAACATCAAAAACACGGGATAGTAAAATACGTCTCGCTGTTTTTTCAAAACGCTTCAACGAGGAGAGGATGATAATAGCGGATATACAAAACTCGTTAACAACTGAGTCAAATGAACTGTGGATATCTAATATTTTTCAATTAAAATCTGGTGACGAATTAGAACTAATGATTAATTGTAGTGAGG
>evgR003115c0g1_i01_Hydra
ATGATGATCTTACTTACCGAAATGAAATCAGTCGATATAGCTCAAAAAAAAATGTTGTCTTTTTTCATGAATCTGGTATACTCTATATAAAAGCAGTGACTGATGGTAGTGGATATTTATCTGGTTTTACTCTAACATGGAACAGTGAAAGTGCTGAAAATTCACAAGTTCTCGATTTCAATTACAACTGTGGAAATAAAAGTACATTTAATAATCCTTTTGGTAAAATTCGCTTTCCATTGATTGGTAAGTATTTAGGAAACGTTGAATGTAGATGGTTGATTAATGTCTCTCCTTACATACTAAGAGTTGAAATTAATGTTGATCTGGCTTGTTGCGGTTACACCGATAACTATGTTGCC
>evgR003118c1g1_i01_Hydra
TTTCTATCTGTATTTTCAATGACGACATTTGCTCTTTATATTAAACTTCATAATTGTGAAAGATTGGTATTTTTGAAGAATAACACTTCCAAGATTTGCCATAAAAACAAAGTAGTTATGAAAATTTTGGTTACGATGGCATTGGTGTTTATAATCTCTGTTGTACCACGTGATATATTTCTTTTATATTATACAATATCGTGGCTTCCTGGTGCTGGAGGAATATATGGAGAAAGCGTAATAAACATAAGTCATTGGCTTCGATTACTTCAGGCTAGTAATGGAATATACAACGTATTTATTTATGGTAAAATGCACGATTACTTTTGGAAAAGCTTGC
>evgR003120c0g1_i01_Hydra
CTATTGATCGAGAGAGTATTAAATGTAATTGGTCAAGTTGACAAGGCCTATCAATCCATTAATCTGCGTTTAGTAGTGGTTGCCATTGATATTCAAGAAATAAACCCTTCTTTTCAGCGATCAACTGATTCTAAAGATTTAAAGCAATTTTCAAATTACGTTCGCGACGTAATTAAGAAGACTTCACCTTTTAAAGACGTGGAATTTGATGACGCCATACTTTTAAGTCATGAGGGATGGGGTAAAGCATCAGGAATGGCTTTGATAGATGTCATTTGTTTAAAAACCAAGGAACTATTTCCTGCAAGTATCAATCATTGGAGCTACACCGATATAAGCTTCTCAAC
>evgR003129c0g1_i01_Hydra
AGAAACTTTTTTATTATCAATGGGGTTTCAGAAAGTGAAGTACAATCTACAACTTCCATTCTTTCAACAGATTATATAAACTACAGAGAAGAGTTTGCACCTGCTTATGGTACAATTCTGGGAACATTGAGTGTTGTAAAAGAAGTCTACACTTTATCTTTTAATTTAAAATCAACAAGTTTCTCCAAAGGCTATAAAAGTATTCTTCATTTATACAGCAATACTCAAAGTTTTGGAGACTCCAATCCAGGTATTTGGTTTCATGAAGATGGTTCAGGAAAGCTTGTTATTTTATCTATAATAGATACAAAAGTTAACATTGATATTAAAACAGATCCACTTATGCTGGGTGAATGGTCAAATATTAAAATATGCCAGTATTTGCAGGATTTTCAATATTGGTTCACTGTTGAATTGAATGGAGCCATTGTAAGAACAGTAAAAAATGTGAATCCAAAAGTTTTTCAAAATGTAAC
>evgR003143c0g2_i01_Hydra
GTGACCCTTTTCTCCCCATCCCCCTTATTGCATGACACCCTTTATGGATGGCCCCATACTACATAAAAATATTAGTCATTATGCATGATATTTCTAAGTGGTATGGCAATAATCATGAATACTAAACAAATCGTTCACATTTTTTATAGATTGGATATAGATAAAAGTGCATCTCTTCCACAAGAAGAAAGCAACTTCATGGTTACAGGAAGAATTAAGAGACAAATTCATGAAAAGAAATATTCATTAGGAGAGCTAGTTGTACCACAGTCATTTGAAAAGTCAGTGATAAAAGAAAAAAATTTAGAAAAAGAAGTGTTTACAGTAGAAGGACGAAAACAATCATTTGATACCATAAGAAAATATATAAATGAAAAGCACAAAGAGTTCATGAGAATCAAAAGTGACAAAGAATATATTGATATGTCAGAAACCAATATTATAGATTTTCTGAAAAGAATTCACGAGTACAATGAAACTCAACAATTAACTACAGAAGAACTCAGATCAAGACTTAAAATATTTGAACGAAAACGTCATTTAATGTTCTGGCATGATGCTTCAACTGTCTGTAATCATGGGTATATTCTTATGACAGTTGCAGCTTTATATGATACCGCATTATACTACACAGATGAAGAATATAATTTAGTTAGTGGGCAAAAAATTAATATCCAGGCAATTGTTG
>evgR003159c0g1_i01_Hydra
CTTCTATTGTCTAAATAAGTAGAATGAACCAAACTGAAGTTAAAAATGAAACCATTTGCGTATACGAGCCAACTTTCAAAGTTCTAATACTGTTTCTAACTGGCTCAACTTCTGTTGTTTCAGTCATTTTAAACTTTTCGTTTTGTTTTGCAGTTTACTTCAATAAAAAGCTTCACTCAAAGTCAAACATTCTGGTTGTAAGTTTAGCATTCACAGATATAATAACAACTGCTGCATTACCAAGCCTTCAGTCTGTCGTTATCTACTTTTATCCGTCATGGCCAATTGGTAGTACTGGAACTGTTGTACTTAACTTGTTTTGGATATTTTCGTTTGTATGCCCTTTTATTACGGTAACTACAATAACTGTTGAACGATTTGTTATAATTACTGGTAAAACAAAATGGAAAGCCTTTTTAACAAAGAAAGTATATCTGGTCATTTTGATATTTATTTGGGTTTATTCAATTATCATTTGCTTGGTTACAGCTGTTACTATGAAGCCTGCGTTAAAAAATAAGTATACTTGGAATGTGAATTTTACCCTTTACTATATATTACTTAGCATTCATATTGTATTTCCTTTGCTTTTATTACCCG
>evgR003163c0g1_i01_Hydra
TAGATATTCATATTGAGGATCAGGATAGATTACAACTATTTGCTAAAAAAATAATTAAAAATTTTGCAGATAGAATATATGATACATCATATGACTTTACTGGCATTCCTCTTCAGACCAGAATGATTGCTGAAGTTTTTCAAGATCCGTGTAACATATTCTTAGAATCAGATGAGCTTATGCCTGAACCAATTGAAAAGTTTAAAATATTTGATTTGTATGAAAAAAATTTCAAAAACAAATATAATATTTACCAGAGTGAAAAAACTAAAACAGGTTTAAGTAAACCAGAAATAAATAACGATGTACCGTATGAAGACTTTATTGGTTTTTACCAGAACATGGCTTTGCGTGCAATATTCAATGAAGAGATATCAAACGATTTAATTAATGAAAACTTTTTGCAACAATTTCTTCTTAAAGAACCGCAACTATTTGCTTTAATTCCAGATAAGACACAGCTAATGCAACTATTATCTGAACAGTTTTCAAAAAAATTCAAATCTTTTATTAAAAACATTAAAAGTGGTAACCATGGTAGCATTTGTAAAATATTAGAAGGCAAGTCGCATTTCATGCACCGTACCTTTGCGGAATATTTAGTTGCAAAATTTTTGATCGATAGTATAAAACAACCAATATTGGTACCATTATGCTCAATATTAAAAGATATATTAATTGCAGATGATTATCAGATTATTAGAGTCTTCCTTAACGATCAGTTAGAAACCCCTTTGCCTGATTTATTTTTATCAAATAATCAAAATCAAAATGATTGCTCAAACATCATTGGGATTGCTCAAACATCATTGGGATTACTCAAAC
>evgR003173c0g1_i01_Hydra
GTTTATAAAAAGTCTTTTTTGGCAATTAAAAAATTTGACATTGATATTTACGTAATAAATCAGGTGTTATAGTTGTATTGAATGCGTAGTATAATATAACTTATTAGAAATAATGAACAATAATGATTAGAAATCGTCTTTTCTTTCGTCCGACCACTTAGAAAAGAAAATCCACATAAAGATTATCGAAGGCGTATTAACCACGTTTTGATCTTTTTAAAAACCAGTTTATTTTCTGAACAAAGAAATACCGGTATTTCTTTTGTTGGAAAAAGTCGTCATATTCAGAGTTTTCAACCCATATATCATTTGAAATATACTGCTCACTTGAAGGAGTATACCGAAAGTTAACCAATGTCAGAGCGAGATATTACAATGCCCAAAGCATTTCTAGTAAAAAACAACAGAAAAGGCGGAGTTCCAACAAAATCTGAAGTAGAAGAGCCGGTACTTGATCAGCAAGTTCCGTACTTTTACCATCCTTTTTACTATCCACCGACCGATTATTGCAACACATACATACCTTCGATATGGCCTAGACATTACCCCGATATAAACTCGTGGAACAGCTACCATGAAAAACCTAATAATTTTGCAAATAGTTCCGCTATTCCTGGTTTAAATCTTCCGCAGCCTTTAAATCTTGGACCACAAATTGCAGCTCCTTTTTCACACAATCCTTTTAGTCTAAACAATCAACCAATCAATCCACAGCAAACAAATGTTGCTAATTTATTCCAAAACCACCACGTTCAATCAGATCAAAACTTTCAACTCACGCAATTTATACATCAGCAACAACATCAAAATTTTGTGCAGCAGCAAGAGTATGTCAATGAACAAGAAAAAGGAAAGCGAGATTTGTATAACGAAATATTTGATCTTGACAGTAATGCAAGCAAAAAGTCTCATATGTTGTTACCAAATGCTATAGCTACTTTAGGAAATATAAAGCAGGAGATGCTAGATAACGATGACGTGTTATCTACCGACTCTTCAAATGCCTGCAGCGAAAGCAGTTCGAATGCGGTCGTTGACAATATTATAAACAGTGCAATTTCTCCTACAATGGTTAAAAATGAACAAATGAAAAACTTTACTCCCAAAAATCTAAGTGATTTATCTCCAAAAAATGAAGATGGAGGAGACAAAATATATCAATGCGAGTTCTGCAATAAAAACTTTACTAGATCTTGGAACTACCAGCGTCATGTTTTAATACATTCAGGAAGAAAACCTCACAAATGCGAAATTTGTCCAAAAGCTTTTGTTTTGGCAGCTCATTTAAAAATTCATATGAGAATTCATACAGGAGAAAAACCATACAAGTGCACCGTTTGTGGTCGCGGTTTTGCGCAATTAACAAACTTACAAAGGCGCGTGCGCCCGCATACGGGACAGAAGCCGCACAGGTGTCAGCACTGTAATAAAGGATTTGTATCATCAAGTGATTTGCGGAGACACATACGTATTCACACAGGAGAAAAACCTTACCAATGCCAAGCTTGTTCAAAAGCATTTACCACAAGTGGCAATCTTCAAAGTCACGTTCTCACCCATACAGGGGTTAAACCATATAGTTGTTCAACTTGCAAATGGAAGTTTATAAGTAGTTCGAACTTACGCACTCATATCAGAACTCATCATTGCAATGAAGACAATCAAATGGTACCAGAAAACTTGGGTGTTCCTACACCTCTTGGCCAAGAAGAAAGGGTTTATTTACCGTCTGTAGTTAAATCGGGTTAAAAACGAACAAAAAATTGTTTGACTCTTTATTACTAGCTATATAGTTTTGTGAGTTTTATCATAATGCTGATAAAGTTTTATCCTTCGTAATCTTGTTAAAGATTTTCTATGTGGACACATGTTAATAAACATTATGCACTTTGGTATATTTATTTGTTGAATATCTATATATTATGGAAATTTTGCAACGTATCGCAGTGGAAATTTCAAGAAGCAGCTTGATTTCATCGAAGCATAGAGACTTTTTTATATTTGTTGAAAAAAGAAAAGAATTTTTAAATTTTTAAAATAATATTTTAATATAAAAAAAAAAAA
>evgR003177c0g1_i01_Hydra
AGACAAAAACTTTTTAACTGTTCTAATAAATTTTTCCGCCGTGAAATTTTTTTTTTATGCGGCATGAACATTTTTTGCTCTAAAACCAAAAAAACGCAACAACAACAACAACAAAAAACATTACACGAACGATTCGAAAAAAAATTTAGTTTTTGTTCAGTTGTTTATTCTCACCATACAAGAAATAAAAAATATATGCAACATAAATCTACTAAATCAGGTAATACTGATTTTTAACATGTTGGTTGAATATCTTACAAAATGGAAGAATCAGTTGATCGTTTAATTTATGAGTCCGAATCATATTTTAGTTTAGCTTTTAGAAAAATGGCTGAATTTCGCAATCGCGAAGAGTTTACAGATATAGTTTTGGTAGCTGGAAGCAAAGAAATTCCAGCTCACAAAGTAATTATATCTTCATTATGTGACTATTTTAGTGCAATGTTCAGAAATGAACTCTCTGAAAGTCAACAACATGTTGTAACAATTAATAATGTTGACCCTGATGCTTTAGAAGCACTGATAAACTATGCTTACACAAGTAGAATAGAAATTCGGGTTGATAATGTTGAAAATATCCTAGCGTCTGCTTGCTTATTACAAGTCACTGATGTGCGCAATGCATGCTGTACATTTATGAAAAGTCAATTGCATCCATCAAATTGCTTGGGAATAAGAGCATTTGCAGATGCCCATGCTTGTGAACAATTGTTTGATATTGCCAACAATTATGCAAAAGATAATTTTGCTGATGTAGCACGAAACCAAGAGTTTCTGTTGCTAGATAGTGAGCAACTTGTGGAGATACTTCAAAACGATGAACTAAACGTGTTGAAAGAAGAAGATGTATTTCATGCTTTGTCTTTATGGGTTAATCATAATCCAGACAGTAGAAAGAATCAAATGCATAAAGTTTTACCAGCTGTCCGATTGGCATTACTTTCACCACAATTTTTAATTGATGAGATTGAACCTATGATTAAATGCGATGAAAAGTGTAAAGACCTGCTTATAGAAACTATGAAATACCATTTACTACCTGATCGCTCATTGAAAAATAGTCTGTTATCTTTTAAGGCTAGGAAAGGTACAATAGGTGTGATTTATGCTGTTGGAGGTATTGATGAAGTTAAAGGAGCCGCCACTGGGATTGAAGAGTATAATCCTCGTAAAAACATTTGGTCACAAGTTGCTTCGATGGAAACTAAAAGATTGCAATTTGGTGTTGCTGTTGTCAACAATAAATTATATGTTACTGGCGGTCGAGATGGTCTTATGACATTAAATAATGTTGAGCGCTTTGATCCAAAGTGTAATAAATGGGAGACAATGACATCAATGCTGACTCATCGGCATGGCTTGGGAGTAGCAGTTTTATGTGGGCCTTTGTATGCTGTTGGTGGACATGATGGTTGGTCATATTTAAACACTGTCGAAAGATTTGATCCTCAAACTTCTAAATGGTGTTTTGTTAAAGAAATGAACACTCCTAGAAGCACTGTAGGTGTTGCTGTTTTAGACAGTAAACTTTATGCTGTTGGAGGAAGAGATGGATCGTCATGCTTGAATTCTGTAGAAGTGTACGATCCGCATACTGACAAGTGGAAAGTAGCTGCACCAATGGTTAAAAGACGGGGAGGAGTGGGCGTTGCTGTACTGAGAGGGTATTTATATGCAGCAGGAGGTCACGACGCTCCTGCCAGTTGTGAATCTTCCAAACAGTTTTCGTCAGTTGAGCGATATGATCCTCGAAGTGATCAATGGAGTTTAATTGCATCTATGAATAATTGTAGAGATGCTGTTGGGATGACTGCATTAGGAGATCACCTTTATTCTGTTGGTGGTTACGATGGTCAAGCGTACCTCGATGCTGTTGAATCTTACGATCCAGACTCTAACAAATGGGTAGATGTAGGAAAGCTTGCTCATCCACGTGCTGGCGCATGCGTAGTAGCCATCAAGTTTAATTGATTGATACTTGAACTTTTCCTTATTTATTAAAATGTATATATTTTCTGTATTTATATGAAATATTGCTTTATGTACAATATATAAATTAAGTATGTATATGATATCTATATGATGTGGAAATATTGTGTTTTGTAACTTCAAAAAAAAAAA
>evgR003196c0g1_i01_Hydra
TATAATGAATACTGCAAATAATAAATTAAAAGAAGTAGAAGAAAGAAAATGTAATGTAACAGACATGACGGTAATTATAGTAAAAGAGAAGAAAACAGACGAAGAACTTTTATATGAGGATTGCGTTAATAAATGCAGAAGTTTAAGCAAAAAAGTTAAAATTGTTCTAGAGGATGTATCAACAGTTCTTTGTTTGGAATATTATTTGGATGGTTTTTTTGTTTCCAAAAATCTTAACATTATCGTAGAAGTGTTGAAAAAAACAAAGAAAGGCAATGAATGTGCATTGGTAAAAGAAATGATTTTGCTTCATTGGTATTTAGATGAGCTTTTGAATAAACTATTGAATCAATGGTGGATTTCTGAATTTTCTTTTTGGCAAAAGATTTCAATATATCTACACATGTGGAGAAATATTGTTCAATGCTATTTTAAAGATTATAATTATAATATTTTACAACCTGAT
>evgR003197c0g1_i01_Hydra
ATAGTGGTATATTTGATGATCGGTTGTCTTCTGCACAGCAAAGTATAATTTCTGAGCAGTTGTTAAAATTCAATGGAAGATTGCCAAGTGAGTTTGTCCGCCAACCAAGGTCATTATCAGAATTGAATTGGTGGAAAGCTACAGAGCTACGTAGTTTTTTTCTTTACACTGGTTTGGTTTCACTAAAAAGTGTGTTAAGTAAACAAAGTTACAAGTATTTTTTGAGTTTGTCAGTTGCTATTCGAATGCTATGTGAGAAGAACACTATTAAACGAAATAGTAACATTGAGTCAGCTAGGCAGTTACTTAATTACTTTGTTAACAATTCCAAAGAGCATTATGGCTCCTCTTTTTGTGTATATAATGTACATGGACTGTTACACATTGCTGACGATGTGGAATATTTTCAGTGTAGTTTAGATGAAATATCTGCTTTTCAGTTTGAAAACTACTTGTATGAGATTAAGCGATTAATTCGGGGAAAACATAATCCAGTTATGCAAATTGCTTTGAGACTAAGTGAGCTTGAAGATAATCCAACTTTGAAGGAAGAGAGTATTTTTTAAATACGAGTTGGTAAAAAAGATTCATGTTATTTGGTTCGGAATGGTGTTGTTAATGTCACTGCTATATGTCAAGATGGAGGCTTTCAATGTGAATTGTATCCAAAATCTGCACTTTAATATTTTTTTACCCATTTTGTTGACTCAAGAGACCTTGATATATATTATATTAAAGAAAATGCCTGCTCCATAAAAGGCTTTATTCAAAGAACTGATATAATTCAAAAATGTGTTTGCTTGCCATATCAAAATAGTTATGTTATAATACCTTTAATTCATGATGTATAAACAAGTTTGAGTGGTTTTAATAAATAAATGTCCTTTTAATCTTTTTTAGTGTTTTTTCAATTCATTGTTATCCAGGGTGGGTGAGTCAATGTTAATATTAAATTAGGCGGAGAAAAAAGGTATGGTTCGGCTTCAAGTAAGTAAAG
>evgR003199c0g1_i01_Hydra
ATTTAATTCTGTAAAAGATCTTTATACTCTTTTATTAATTTCATATCTTTAATAGGTATTTACAGAGATATTATAGATACATCAAAAATGACTTAAAAAGGAAAAACGACTTTATTAAGAATTTGATTTTTAATTTAAGCATGGAAAATGTGTCAACAAATTCACGACATAAAAAATGGTACAAGCCAAAGTTGTTTGTTTATAACTTACTTGACAAACCTAAAACAAAACTAGCCTATGCTTACCATGTTGTTGTTTTACTTTTAATAATTGGAAACATGATTTTAAGTTTCGCCTCAACCGTACCCGATTTCAATTTGAGTTATAAAGTTCGATTGTCTCTTTCGGTTTACGAGTTTATTTTGCTCATTATATTCGTTATGGAGTATTCGTGTCGATTAATCGTTTGCAGTTCTAATGTAATGTACCGCGGAATACATGGTAAATTAGTTTATATAAGGAGCTTTTACATGGTAATTGATATTTTAGTTATAGTTTCTTCTATTATAATTGTTTTTTCAAATGTTGATGCATCTAATGTTGGTTTTCTCCGTTACGTGAGACTTTTTCAAATACTTCGATTGTTGCGAATGGATAGGACGAGAAGGGATTTGCACACCATGTGGCATGCTATCTACAGCCATAAAAAAGAACTTTTGACATGTTACTTTACTTGTATTGTTGTCCTCTTACTGGGATCTTATGCAATTTATCTTATCGAACATGATAGAGGAGAATCAGATGGGAGAATCGATAATATGTTAACTGGATTATACTGGGGAATAGTAACGTTTACAACAATTGGTTATGGCGATTATACTCCTCATACCTTATACGGCAAGTTAATCACTGGTCTTTTATCTATGGTAGGTTGTGCATTTTTTGCGCTACCAGCTGGTATATTAGGATCAGGGTTTGCTTTACAAGTTTCTAAGCAAAAAAAAGAAAAGGGTTCAATTAAGGTCAGAAGTCCAGCTGCATTTTTAATACAATGTGCGTGGAGAAGTTATGCAGTGCGTAACTCACGTTTAACAGCAACGTGGAGTTACTTTCTCATGAAAAAAGTTGCAGCCAAAGAAAAAGCCGAAGCAGCTAACAAAAAAATGTATACTAGTAATTTTGTCCAATTAAAC
>evgR003223c0g1_i01_Hydra
TAACAAAGTGTTTCCTCAGGTCTCTAAAAAAGGTGTATCTTTTCTTTTTTTATGGTTTTGTCCTATGCATGGACATTGTTATGGCTTCCATATAATTCCGGGTTCAGAGGGAAGAAAAGATCCACATGCTTCTTTATACACTCATCTTGAAGTAGCTCCAGAAAATATTATTTATGATTTTGCATGTAGTCTATCTGAATTCTGCCACAACCGGGAATCAGGATATTATAAAAATACGTCGTTTTTCCATGATGTATTTCATCGTTATACACATAAATGTGCTGATGTTTTTCGATGCAATCGCTTAGGTTATTTTTATCCAATAAATACTTCTATATGCGAACAGTTCAATAGCTTTTTGCAGAATATTAAGTCTTCAGCTAAACTGATGACACAAACTCACTTTTGCTTTTATCTTCAGTTTTTCATACATATTTGGAATCAAAGGAAAAAAAATCTCTTTGAAAAACAGTTGATTGTGTCTGAGAACACTATTCATTAATATTGTTATAATGTTA
>evgR003263c0g1_i01_Hydra
GGAGATTTTGTGAGCGCTTGGACCAGATTTTTGATATCGCTCACCAGAATGTACAAGCACTGATCAAGATCCCCGAGGATTGGTCCTTCTTGATTGCTCAGAGAGAAGGCAGAAAGGGTTACATGGCAGGAGTAGATAAAATTATGAAGCAGAAAGTAGAATTAAAAGAAAATAAAAGGAAACAGGAGGAAGCTCATCATATGAAATCAAAGAAAGATATTGAAATTCTAACCAGCAAGGTTCCATTATTGTTGAGCAGTAGCTCTGAGACAGATAAAAATGATGCAGCGAGAATAAACCAAAACCAAGGAAGAGCCAGTTTTTTGTAAAAGATTGAAAAAGCAACCATCTGCATTGAAACCAAACAAAATCACT
>evgR003270c0g1_i01_Hydra
AAAAAATATTTTCTCAAGGTTTTCAATATTTTCTCAAAGATAATTTAATCAAGTACCAAATACTTATTTTCAATTCAAAATTGTTAACTAAATATAACTTAAATATAAACTTTTTTTTCCCCACCCTGATTGTGGGTAAAAGTACAAGGAAACCTGTAAACAGTGGCAGTCATGCTATTGGTTATGTTACATTAAAAATACTTCAGGTTAACATCAGAAAAAGCATCAAGTTTAAAAAACTTCCATTAACTCCTCTACATAATCAAATGAATGAAAACCTATCCAAAACCACACAAGTGTGGAGCCATAGATGTAAAACTTGAAAAATACGTATTTGCAGTTTGCTGGTCATTAAAGAGTCATGATGAAGTTAGCTAAGAGTCATAATTAAGTTATTAAAGATACAAAATTAGAATTGAAA
>evgR003294c0g1_i01_Hydra
GAAAAATTATTGCAGAGGTGCCAAAGCTAGATAAAGAGTACTTGGTTTCTTTTGATGTCTTTCCAACTAAACTTGTTGCTGGTTTACGTAGCGTTATTTATTTCTCTATTGGTTCAAATTCTGCTGGTAATAGTGTTCGATTTCCTGGCATCTGGTTTGATGCTGATGGAAAAGGAGGAGTACTCATTTCAACACCAATTAATGGTAAACCTAGTAAGGTTGCAAGCACAAATCAAATTCAGTTAAATATCTGGTCAAATATAGAGATATCCCAAAATTTGAAAGGCTCAGTTTATGTATACACCATCAGAATAAATGGAGAAGTTGTCTTCTCTGAAATTAACAACCAAGTTCAAAGTTTTGAGAATGTAAG
>evgR003302c0g1_i01_Hydra
ACTTTTGCAACTTGTTGAAGTTATCAGTGGAGCATCTTCTCATAACCATTACACTTTAGGTATGTTCCTAGACTTGTCTAAAGCATTTGACACCGTTAACCATCAGATTTTATTAGAAAAACTAAAATATTGTAGAATAAGCCACGAATACCATAAATGGTTTTCCAGTTACTTAACTGATCGAAAACAGTTTATTTCTCTAAACGAAGGTTTTCAAACAAACAAATACACTCAAAATATTATCTGGGGTTCCACAAGGATCAATCTTAGGTCCTCTGCTGTTCTTAATCTATGTCAGCAATCTATATAAAGCATCAAATACTATGTCTAC
>evgR003310c0g1_i01_Hydra
ATAGAAATGACCACTAACATTGCTCCAGAAATCGATGCTGCTTGGGAAAACTTAAAGTCAGAATCCATCAAATGGTTCACTTTCGTCCCAGAAAAGACTGGTTTGATTATGGATACTGAAGGTACTGGTGATTGGGATGAATTCGTTTCTGCATTCGGTCCATCTGATGTTAAGTTTGGTATCTATCGTGCTGTTGCCGTTGATGATGATTCTAGAAGAACTAAGTTTGTTTTGATTGCTTGGACTGGTAGTCAATGTAATGCCATGAAGAGATCAAAGGTTGCTGCCAATAAGACTATTGTCACCCAAAAGTACTCTGTTCACATTGATATTCAAGCTTCTGAACACTCTCAATTGACTCAAGAAGAAGTTATTGCTAAGCTCAACTCATCAACAGGAGCTCACAAGCCAAAGAATTACGAATTTTAAATTGGAAAGTTTTCTAGTTTTTGAACAATGATTTTAAATCATTAAATAGATATTTT
>evgR003312c0g1_i02_Hydra
TCGATATTAAAACTCTAGAAGAAATGGGTGTACTAATTCCGTGTAATGGGTGTTCTTTATTCCGTATAAATACACTTCCGCTAAGAAATATGGAATATAAAAAAGAAGAGGATAAAATAAAAGCAATACTCAAAAGACTAGAACAAATGGAAATGGAAATCAAAGTACAAAATAATAGACAAGAAAGAATCGATATTAAAACTCTAGAAGAAATGGGTGAACTAAATGGTAGTGAAGAACAAGCAACAATTATCTCGAGTATTGTTCGAAGAGCTCAACAAAGATACGATCAAAATCGAGCGTCCAATAAAGAAAATAACACTGAAACGTAAACTGTACTTTGAAGAATCTAATTTTAACAAGAATGGTAACATGATAGACCCTGAAAAATTTATGTTTTTTTATAATTGTGAGTTTTTTTAAGTCATTTTTACTTTTTTAGAATATAATTAAAAAAACATGGAAAATAAAAATTATATGTATGATTCTAAAAGTACATTGCAGCAAGTTATGGATGTAATTTTAAAGAATTATAAAAGCGATGTCGAATTAACAGAAGAAGAAAAGCATTTGGAGATTATGTACAAGCAGAGGGAATATATTGGTCAATTAAAAGGAGAACTTGAAGAACAATCGAACATAATCGAGCATCTACAAAAGGAAAATTTAATTTTTAGAGAAACAATTTCGAATTTGCAAGAAACAATTTCAAATTTAATCAAATTATAATACATTATTTTTTAGAATTTTGTGG
>evgR003346c0g1_i01_Hydra
GCTACATTGCTGTCACTCGCTGGGATTGTCATTTCAAGTGCTAACCACTTCCCTCACTGCAAATGGATTGTTTTGCAATCATCCAAGAAAATTTCAAAGTAGCTCAAGCATACTACAAAAGATTTAAACTGGTCAACCTTGACAATATTTCCTGGACACTCATGAAAAGGAATTTACTTTTCCTCTTCAAGAGTCTTCTTCATTGCTTTTACAACTTCTTCAAGTTGCTCACTTGTAATGTGAGAACCTGTTTCAAGTTTGGAATAGTAGCAGTTACCACTTTCATCAGTAATTTCAAAAGCACCGACACGAGCCTTGCCTTTTGTATTGCCTTCTACGGTGGAACCTATAATGTTCTTCTCGATAAATAACTTAGCCGATCTGAAGCGACTGGCATATCCTCACGCAGTGCAATATTCAATGTGATACTTCATTAGTGAAGAGAGTAGGA
>evgR003376c0g1_i01_Hydra
TTAATAAAAAATTTCTCTCATAAATCAATTAACATTTTATTAGTTTAAAATGAGTTATAGATCGACTCGCGATTCTGTCAATCGCGATTCTGTCACGTGCTTACAAATAAATCAACTTAGTTTTAAATTGCGCCATGAACTTGTGTGCGTTAATAGTCGGGATAATAATGGAAGAACATTGTTACATTTAGCTATAATCTACTCAAATATACCTGCAGTAAAGCTTTTAATGAAACATGGAATAAATCCAGACTTTCAAGACAATGAAGGTAAAACAGCCCTTCACTTAGCAATTCAACATAGCAATATACAAGCAATTAAAGTTTTGTTAGCTCATAACGTAAAGATGAATGTTAAAGATTTGTTAGGCTTAACGCCTCTCCACCTTGCATTGCTTTCAAGAAACTATTTATCAAACAAAGAAGCAGTATCTATGCTAGTTCTGAACGGGGCTGATGTAAATACAGCAAGTTCAGCTGGAAAAACGCCATTGCATATGGCAGTCAGTAGTTTTAATAGATTTTCAACGGCATTACTCTTAGAAGCTGGTACAAATGTAAATGCTAAAGACGAAACCAAATCAACTGTTTTACATTATGCGGTACATCAAAACGACCTTAGTTTTGTACAGCTTTTCTTGATAAACGGAATAAACGATTCAGTTGTAAATGCATGTGGAAAAACAGCATTAGAAATGTCACGAGGAATGTCCGATTGGCGAATTGCTGCTATCTTAAAAAGTTCCAAAGCATCTAAATTCAGTAAAAAAAGTATTATAAAGTATGATAGCTGTTACAAGTAAGACATATAGAAACCAAAGGGGATAATCATTAAAAAATATTGTAACACAGCGTTTTTTTGTTGCTGATGTATAAATTAAT
>evgR003391c0g2_i01_Hydra
GGATTTGAGTCTGTCATAAACACTGCTATTCATTTTGGTGCTACTTATCGGGGACATATTTCTGTTAAAGACATTCTTCCTTCCAAACACAAAGTTGCTGAAGCCATTAAAACTAGTGTTTCAAAGTCTAGAATCACAATCAGCCGAATACTCAAAGCGAAACTAGAAAGAAGTGGTGTAGGCATAACTACTGACATTTGGACTGATGATTTTCACAGAAACTCATATCTATCGATGACGCTGCATTGGATTGAAAATAATCAAGTACAGGAGGCGGCAATGGCCTTTTCTGTGTTACCTCCTATTTCAAAAACTGCCAATGTCGTCTCC
>evgR003405c0g1_i01_Hydra
ATAAATTTTTAGAAATGCAAAATCAAATTATGAAACTAGAATCTGAGAATCAAAACCTAAGAATTAGAGTAGAAATTCTTGAGAAGCCAATGCAAATTATGACGAAACCCAGATTATTCATTAACTTATTTAATTCCATTAAAAATCCAGATAAACTAAAAACTAATATACTTAACGCAATTAAAATTGAAAATAAGCAAGTTGAAAAAAGGGAACGAAATATACTTATTTTTGGAACAACACGTTCAGAAGGAAAAGAAGACAAAAATGTTGTAAATGATATTCTACAGGAAATTGGAGCCAATGTAAAAGTAGCTTATATTAGAAGGTTTAAAAACTCAG
>evgR003412c0g1_i01_Hydra
GAGTTTTACATTTTTAAGAATTGGTTTAGTAATCAATTCAATATGTTTCATTTGAATCCTTACCTTGACTATAATAAAACCTTTTTTTGGGAACATGTCCAACTTAAAACTCGATATAGTTTTGCTACTAACAATACGAATGTACTTTCAATATCAAAATGCGTTATAAACTGCCGACCTGGATTTGAACAAAGAATATTATCCATAACAACAAATTGGAAAAAGACGCAATATTGCACATGCGCAAAATGCAAAGAAAATTATTTTAAAAAAGATTTCGGTACAAACCAATGTGAAAAATGTTCTCTTGGTTATATTTCAGACAAAGAGTACACGACATGCAAGTTAACGTTTGCAGCTGACCTTACTTTTCATACAATTGAAGGAATCATAATTACAGTAGTTGCTGCTTTTTCTTTCTCAATTGGTTTAGTTATTACAGTAACGTTTATACTGAACAGACATACACCTGTAGTGAAATCAATACACTTTACCTTATCGCTG
>evgR003435c0g1_i01_Hydra
ATTGATCAAAATAGTTTAATTAAGTTTGGAACATGGAAATATAATCCTAACAATGTAGTTATATATGAAAACCTTATTGCGTGGAAAAATAACATAAAACCTATATCATCATGCAGTTTGAACTGCAATCCAGGAACATATAGAGTTAATTCTACGTTATCAAAATGTTGCTGGATTTGCGTACCATGTCCAATGGGTTCAATTTCGAACTCAGTAAATGAAAACTCGTGCATAAATTGCTCACTTATGTCTATAGCTAATTCAAATCAGACAAAATGCAATAACATGGTAGAAATCAGCTTAAATGTTTTTAGTATTTCCGG
>evgR003488c0g2_i01_Hydra
GTTCAATTACACATTTAGGCTTTTTTCTAGTGTCAAGATCATATTGAGTCTGAATGAACATCCAAAAATAATCAGAAATTGGATCATATTGAATATTGAAACACCAGTAAAGTTTGAGCATTCGATCTATACCTTCCGTTATAGATTTGCCTGCATTGATTATTTTTTTTTTCATATGCAATGTTTATCGATAAAATTGTTTCGCCTTCCTGCCCTTGTATTAACAAACAGGATATGGGTTCTCAAGATTTTCTGCGAAAAATGCAATCGACTCGCCGACTTCTCAAATATGCTGTATGAACATTTTCATAATGTGTTCCATTATCTCAGGGGCA
>evgR003492c0g1_i01_Hydra
TGGATTAAAAATAATTTGCAAGATTGCTTTGTCATTCGTTCGCATTAACCTTCAAATTTGAAACTAAAGTCAAAATGGTTTGATGTTTTCGTAATTTAAGGGGCCTTCGTGTTTTTTTAAACGAAACATTGCAAAAGAGAATTATGACATGACGTAAGCTACGTGGAATTTATAGATATCGTATTTCTCGTATTTTTTAACATCAAAAGCAAAATCATCAAACTTCGAAACCGATACAGAACAAGACCAAACACAAATCTGCTCAAACCAAAAAAGATGTTTTTAAAAGAGCTTCAAGTTGTTTTTGTTGTCTTTCTTTCATTCTCTGGTGCAACAACACTTGATAACGTAGATTGTGGTTCATATGTTGACTATGTGGAGACTGACGACCAAGGTCATTTTCGTTATATACCTCGAGTTGTAAAGTTAAATCGATGCCGAGGAGTGGACAATTTCTTATCCAACCCCCATAATTATAAATGCGTTCCTTCACCTAACGGAGTTAAAAAAGTAGATTTGAACGTTTTAAAAGAGCAAACGTTTCAGTTTGTTACTATTGCGGTTGAGAATAATACAGATTGTCAGTCACTATGTAGTTTAAACGATAAATCGTGTAACCCCTACCAAAAATTTAGTAACAACAACTGCGAATGCACCTGTAACTATGACAACAGCTCTCCCGTATGCAATCTCAACTTTGAATGGAGCGCAACGGATTGTAACTGTATTTGTTCCCCATCAAAATCTAGAAGCTGCCCTTCAGGTAAACAATTTAACGAAGATCAGTGTGAATGTAAATGTTTAGCTTCAACATGTCCAAATCCAAACCAGGTACTAGATCCCATCTCATGTAACTGTCTAGACAAGTCCAAATATGCAAGTATTAAAGAAGTTCAGATTTGCAAAAATGGAATCGATAAATTTAAATTTATAATCGCACTAATATCCGAAGGTTTTGTTTTTATCTTGTTGTTTACATTAGTCTACTGTTTTTACTTGAAACCTAAGTTGACTGTCTTAACTAAACAAACCGTTAAGTTGGAAAACTCAGTCAACGCTTATTGCAATGGTCCTCACGCTACAGACATTTAGCTATGGTTATAACTCAAGTAACAAATGACGTCATTAATCATATAACCAATAATTGTATTAAACAATATGTTCCTAATATTGATTAATATTGATTACAATTTGTAAAGAATGATAAACATGAAATCCTGTTGTAATGCATTGGGGAAAAAAATGTTTTTTTTCAATTTCAGTTTAGATAAAAATATAAATATATAAATATTATAAACTAAATGTATAAATAATTTAATTTTTCAAATTTGTTAACAAATTTTTTTAAATTGTTTTTATTTTTAAATCGTAGTTACGTTACGTAGTTACGTTACGTCAACTTTAACATAATAAAATATTTATTTGTAATTAATAATTCTTTTTTTTAATACTTGAAAGATAAAATGTATTTGTAATAAAAAAAAAAAAA
>evgR003520c0g1_i01_Hydra
CCGTTAGTAGTCCTTTTAAGACGATATCTCTCTAAGATTAAAAAGAAAAATCTCTCAATGTGGAACCTAATCTTTTAAGAGTTAACATAAAGAAAACAATGAAAGAGTCGATTGAAAAGACTTTTAAAAAAACTAGCGAAGAAACATTTGAAGATTCTATTGAAGATTCTATTGATGATGTTACTGAAAATATAATTGATGAAAATATTGTAGAAACCATTGAAAAAACAATTACTGAAATCGTTGAAGAGACTATTGAAGAAACTTTTGAAGATAGAGCCGAAGACATTACAGAAGAAACTATTGAAGAAACAATTGAAGAAACTACCAAAGAAACTATTGAAAGTGCACAGCGTCTTCACATTTGCATCACATCACAGAATTCACATGTTGAGCAAAATTTATAATTTGTTGTTGTTCTAGTTAAATATAAATAAAATAAAAATTCGAAAATATGTTTAAAATGAGATCAATGTTGTAAGTAAAAAGGTTTTTTTTAATTGTAAATTCAATTTTTTAATTGTATATTAAATTTTTTAATTGTAAAAAAA
>evgR003533c0g1_i01_Hydra
GCTTTCATTTCTGGAATAAATTCAAATTGTATTTATCAACGACTTCTTGAAAATTCGACATTGAATTTGCAATCAGCATTTAAACAAGATTGCTCTCTTGACATAGAACAAAGGAACAAAGTGTTTTATATGCAGAAAACTTCTTCATCTTCCTCCTATTCAAATATTGTAGCTATAAAATTACCAGAAGAAACCTCTGCAGCACCTTATGCTTTGCCTGTTAATGGACAAACAAAATGCATTTATTGTGGTAGAAAGTCTCACAAACAACTTAATCAAATCTCTCCTAGTAAATATTATTCACCTCAAAACGTAACATGTTTTAAATGTAGAAAAAATTTCATTTTTACAAAGTTTGTCTTGGCATGCCGATGAATAAAAC
>evgR003554c0g1_i01_Hydra
GTCATCTTGAACGGAATTTTCATCATTTTATTTATTTTTGCTATCACTTTTGTGTCATCCTGTGTTAAGTCATTGATGAATATTACAAAAAGAAAAGGACCAAGACCTAATCCTTGTGGTACTCCACTTTTTACAGTTTTCCATTCTGATACACAATTCCCTAGAACTACTTTTTGCTTTCTTTTATTCAAAAACGAAACTCTCCAATTAAGTAAACAAACTCTTTTATACCGTTTGACCCTAATTCAAATTTACTTTCTATGTGCAACCGACTCGAAAGCTTTTGCAAAATCTAAAAAGGTATCAGCACAGATGCTATTTTCATTAAAATCAGTAAGCATATCTAATTTTTTTTGCAGATTTGAAATTCAGCTTTTTGTTTATATAAAACCATGTTGGTCTTTGG
>evgR003556c0g1_i01_Hydra
GTTTAGTATTTGATGAGATGTCACTTAAGCAGCATATAGAATATGAAAGAGATCTCGACCACATTATTGGTATTGATGAAGATGGAAAACCATTAAATCAAGCTTTAGTTATAATGGTTCGTGGTATTGCAATAAAATGGAAGCAGCCAATAGCGTACTTTTATAGCAACAACTCAGTTCAATCAGTTAAACTATCACAAATAATATTAGAAGCCATTGACAAGCTGAATTCAATTGGACTAAATGTACGGGCTGTTGTGTGTAACCAATGTTCAACAAATATCTCCGCTTTAAAGTTACTAGGGTTTCTTAAAACAAAACCATACATATTTCCTTCGATAATCCAAACAAAAGTATACATCATTTTTGACCCTCCGCATTTGATTAAAAGTCTTCGAAATAATTTGATTAGACATGATATAACTACAAACGGAAAAACTGTATCATGGAAGTACTTTAAGTCTTTTTATAATATAGACATATAAATATAATCAGATGGTAATAGTTTTGTATAACTATGTGAGTTTTTTAACAACATGTGAG
>evgR003568c0g1_i02_Hydra
AAATTTGTATATTGTGTACTTCTTGTGATTGGAAATATAGTTTTTATTCATCACCTGAAATAAACATAGATAAAAGAGGTAAAAAAAGCATGATATAAATTATAGAGCAGTAATTGCATTCCGATAAATTGGTTAGGAACATAAAGATATGAAGGTATTTACAACTATGATGAACATGCCACCACCGATGACTAGTAATTCATATGATGCAATAAATGTAAATCTTCACTAGGCTTATACAGAAAGTGCAATTTCTAGTATGGAAAATGCTGTTAACGAAATACATACCCAATTTTCAAATGGTTTGTTGTTGAAATGCTTAGATGGTACAACCCAAAATCCTAATGAAGCTTTTAACCAAATCATATGGAAAAAATGCCAAAAGTTTTTTAAACATCTTAACTTTTTGTTTGGAAAATATAGCGTTATAGGAGCTCACGATGTAGACCATTCAAGGGTAAAAAACATGAACAAAAAAAGTCAGACACCTCAAATAACTCAGAGGAAAAAATTAAGAGCTCGTAGAAAAGGTTTCATAGACAACGAAAAAGAAACTGAAGATTTAGAGTCATATTCTGCA
>evgR003576c0g1_i01_Hydra
TTTAGTGCAAAAAGATAAGAATAACATGCGCAATAAAAATATTTGGTTCACATTTCGGAATATAGCTACTCATGCTATTTTTATTTTGTACTCTTCGCTGCTGTGGTGGAAGCTAATAGAAATCGGTTTGATATCAAAATACAAATGTTACAAGCTAATTACAGCTTGTTTTGGCTGGTGGATTGCATTTGTAGCGTCTATCTCTATTTCTGGTTGTTTGGCAAATACGGAAAAACCTCCCAAAAGTTACTCAAAGACCTGGTTAAACATACCAACGTTAATTTTTATTATGTTTATTCTTCCTCCATATGGTTTTACGTTAAACCATATTTACGGGTCGCAAGAGAACAATTATGTAAATATGTTGGAACACAGTAAAAATGTCTCGTACATGGAACATAAGTCGAGAATAGCAGATTTTATCATTCTCCAGATGGCAATGCCTACACTACTATTGACGTTTATAATTTACCCTCTACCACCAAGAGCGTATTCAGGAGCTGAAAATTTAGCAATTGCGTTGGAGTTTATGAATGCATTTGATATTATGGATATGGTATCGGA
>evgR003611c0g1_i01_Hydra
TTGATTTCTTCAATGGAAAGAAGCCAAAGCAAGGTATTAACCCTGATGAAGCTGTAGCTGCTGGTGCTGCTATTCAAGCTGGTATTTTGGGTGGTGAATTGGATGATGATATTGTTTTGGTTGATATTGCTTCCCTCTCATTGGGTATTGAAACTGTTGGTGGTGTCATGACTAAATTGATTCCAAGAGGTACTAAGATTCCAACTCAAAAGTCTCAAGTTTTCTCCACTTATCAAGATAACCAACCAGGTGTTTTGATTCAAGTTTTCCAAGGTGAACGTGCCATGACTAAGGACAATAACTTGTTGGGTAAATTCGAATTGACTGGTATCCCACCAGCTCCTCGTGGTACCCCACAAATTGAAGTTGCTTTCTCTGTTGATGCTGATGGTATTCTCCACGTTTCCGCTAAGGATACTAAGACTGGTACTGAAAATAGTATTACTATCACTAACGATAAGCTCAACATGGATAAGGATGAAATTGAAAGAATGGTCAATGAAGCTTCCAAATTTGAAGAAGAAGATAAGAAGCAAAAGGAAAGAATTGACGCTAGAAATGCCCTTGAAGGTTATGCCTATCAATTAAAGAATCAATTGGACGATGAAAAGTTCAAGGAAAAGCTCCCAGCTGAAGACAAGAAGACTATTGAAGAAGAAACTAAAAAGTCAATTTCTTGGTTGGAACAACATCAAGATGCTACCAAGGAAGAATTCGAAGAAGAAAAGAAGAGATTAGAAGAACAAATCAAGCCAATTATGACTAAGTTCTATCAATCTCAAGGTGGTGCTCAACCAGGTGGTGCTCCAAACACAGCCGATGCTCAACATGATGAATTGTAAATAAAAGTACAATCTCAC
>evgR003624c0g1_i02_Hydra
ATTTTAGTGCAAAAAGACATAGACATTGAAAAAAGCCATTTTTTATGATGGCTGTTTGTTCAGCTGACAATAAGTTTGCAATATCATGGCATGATAGCGCTTGGATGCCTATTTTAAATTCAGAAAATGTATTGGATTATTTTTGTCAACGCAGTAATACATTTTATGATAAAACTTGCAATAATGAGCACATAAAAATGCAGAGACTTGACCCATCCCAATTGCAGTTTATGTCTGGCGTAGAGTATCAGCTGATACATTCCCAGGATCCTATACTCTACATCATCAGAAAACAGATGCGATATTCACAAACACAAGCGACTCCTCTGTCATTTTATTATATCTTGGCTGGTGTGGTTTATCAAGCTCCAGACATTGGTTCTCTTATTAATTCAAAGTTGTTATCATCGGTTCACCAACTTCAGTCTGCATTTGCTGAAGCTCGTTCATATGCACGCTATAATCCCTCAAAAGGTTATTGGTGGCAGTTTACAGAAGAGTCAAAAACAAAGAGTAATGCTATCAAGAAAGCACCCAAACCAGGCTCCGTTTTTCAACGCCATAGAGTTGATTTATTGCTATCTGATCTTGTCGAAAAGTATCCTCCAAGATTGGCTAAAGTCACTGCTGTGAAAGTTGAGAGCAACAGTAATGGCAACGATATTCCGACGAGTTCGACCGAGTCGATTGACGTGAAAGGTGAACTAGTTAATACGGCAAGAGTAAAAATTGAACCTGATTTGGAAACGACATTTTCACAGAATCCAATAATCCGGAAAAAAAAATGAAGTTTAACTTTACGGAAACGAAAATGAAATAAAAAAGGAAAATTTAATCAACAACAATATTAATAAATAAAAATATAATTTTAATATTTTTCCTTCGCTGAAAAATCTTGATTATTGTTATTTATTTAGTTTTATATATGTAATATGTCTTTATAAATATATGACATTTTATTTGTACTGCGACGCAAAATTGTAAAAATCTTCCATGTCAAATGAATTTTAATTTTTTTT
>evgR003625c0g1_i01_Hydra
GTTTCTTATTCATTTCAATCCATTAGCTTGTTTTTTTAAGTATATTTTAAATAGTTCAAAATACAAATATGGTCTGACATTGGTCTAAATATGGATACTGGTAGTAGTGAACCACGTAATATGTTTGATGGTACTAACAAAATGATGTCTATCCCTGAGAGAATCGTTTTATCTGCTGGTTATGACGGGAACGATAATGAAGATGAAGCTAGATCAAAGTACATTATTACATTTGATGAGCCACCAATTATTAATCCAGATGAAGTCATGGCAGGAATGCATACTCCTCCGTCTGTCTTAACAATAGATCATAGCTCAGGAATATATAAGCGAAATTCACCTAACTGGCACCAGGAAGGTGATGGAGCAGATGTAAATGACAGCGAGAGTTCACTGTTCCAATCAACTCAAGATAAGTTTGATCCTGAGGATGATTACTCAAGTATTGTATCTCCAAAACGGTCAGTTATGATAGCTGGACAAGAAGACTACTATACAAATTCATTTGAAGACATGCAGACGATGCAGCAGTACTACGCTAGCATGCAACGGCGATTAGGAATTCTTGAAAATGAACTGGAATATCAAAATCGTCTTTCTAATGTTTGGAAATATTTACTTCTAACATTGACAATAATTAATCCTATCGTTATAAATTATTTATTTTTTAATCGCAGGCGTTAAATTTTCAAATTTTAGTAAATAAGTGTTTTTTAGCAGGGTTATTTAACTATAAAACATGATTTATTGAAATATTGAATATTATATATAAAATAGCAAAAAGATAAAATTAGATAAAAGGCAATTTCATAAACTTAAAAAAAAGAGGCG
>evgR003629c0g1_i01_Hydra
AATGTCTAGTATTTCAACCATGCACCCAATTCTCAATGATTCATCCAGAGCAAGATCAAGATCCTCCCCTGCTACATTGGTTGGATTGAGAAAATCAGAATCATCATCAGAACAAACACCATTCTCCTACAATAAGGATCTTCAAAAGACCCCATCAGTAGTCTCAGACTTAATTGGTACCAATGGTAGAGATGGAAAGAAGGAATTTGATTCACCAACTCTCTTCCTAATGAAGAAGATGAAGCAAAAGGTTGAGGAAGATGGCTTATCTCCAAGAAATAGAAGCCAAAGTTTGATCACCATTATGAAACAACAAGGAAATGTTGTCCTTTCTCCTCAACAACAAACTGATAGATTCGAATCACGATGAATAACTGTGGATTCTCTTTAAACCGTATCACCATTACAACTCGAGCCTGTCTATTAATTGCTTCTAACTGTTGAGTGTTGTCTCTGCAACTAAATGTCGGGGAAAGAGACCGCACTCATTGTACTATATTCAAAACACATCTTACAATCTTGTCTAACTTGCCAAATTATTATTAAATTGTAAAATAAACAAC
>evgR003631c0g2_i01_Hydra
CACCATTGTTTTTTTTTATCAATCCTAAACTCTTCAGAAGATCGTGTTGAGTCTTTATCAAAAATGCTAAAGTTAATAGTTAAGGCCACACTATGCAGAAGAAACATTAATAGTACACTAGATTGCGTAGACCGAGGTTGCATTAGATCATTGAGCTTTTTTTTACTTAATGAATGTAAAGAAGGTGCAATATGGGAGGAAGCATTAAAAATAGTTAACCAATTGAGGCAACTATCCCAACAACCAATAATCAACATAAATGAAGTTGCCAGCAACGTTGACGTCGACCACCAAAATATTATACGCCCCCATTGTATTTTTTTATCTTTTATCCTAAACCTAGTAGATCGTGTTGAGTCTTTATCAAAAGTGATAAAGTTAATAGTTAAGGCCACACTACGCA
>evgR003638c0g1_i01_Hydra
AAAAATAACACAACACAAAAAATGAGTTCAGGAAAGGAATCCATTGATCAAATCCTCCAAAAGGATACTGAAGATGAATCTCTCAGAAAATACAAGGAACAATTGTTGGGTGCTGCTGCTAAAGGTGTCCCAAAGACTGATGACCCACGTCGTGTTGTTGTTGAAACATTTGGATTGAAATTCCCAGATGGTGAACATCCAGATATCCTCAGAGAAGTCAACACCAAGGAAAAGGTTGCTGCTTTGGAAAACGAACCAATCAACATCAAGGAAGGTGTTAAGTATCAATTTGTTGTTTCCTACAGAGTTCAACACGATATTGTCCCTGCTTTGCACTTTGTCAACACTATTAAGAAGACCATGGTCACTGTCGATCAACAAAATACCATGATGGGTTCTTATGCTCCTCGTGCTGAACCATTTACTTACAAGTCACAACCTCAATATGCTCCATCTGGTATGTTGGCTCGTGGTACTTTCAAGGCCAATACCAAATTCGTGGATGATGACGGAAACTGTCACTTGGAAATGAACTACAGTTTGGTCATCAAGAAGGATTGGACTGAGTAAAT
>evgR003644c0g2_i01_Hydra
TGGACTTCTTGGATTTTTGTAAATTATAATACCACTAGTAGTCGGAATCGCACATGCTTACTATTTGTTGATGCAAATACAAATGATACCCAAGATTGTGCAGATGATCTCACCAATTGGACTTCATGGAGCGATTGCTTAAATGCTTCCAACCGAACTAGAAGTCGTACGTGTAAAGAGGAATATTTTGGAAATAATTGTTCAGGATCAGTAGAAGAACAAATACAATTATGTCCTGGTTTTGCGTATACATATATTGAAATTCAATTGAATAAAACTTGGAGCAATGATTATTTTGACTCATCATCAAAACTATATAAAG
>evgR003649c0g1_i01_Hydra
AATGCTCAATATGGTTCAATCACTGACCTTTTGATTATTAATGGAAAAGCTGAGAAAATTGTTAGTGACATCAGCATTCCATTGGAAAAAAAAAGGGTGGTTGCAGAGATACCAGTTCTTGAAAAAGAATATTTAGTTTCATTTGACATTTATTTTTCTGCTGGGTATTTTAATAATTTGGTTATTTATTTCACGCATTATGAGTATAACTTTATAGAAATCCGGCTAAATTTTGGAGAAATAAGTGTTATAGTCAATGAGGAAGAACGTTCTAAAACTAATCAAATCAGTTTGAATACGTGGTCAAGTATTGAAATAAGTCACATTTTAAAAGATGCAAATTATATATTCACAATTCGAATAGATGAAGAAATTTCCTATTCTACTCCAATCAAAGTACCCCAATATTTTGATAATGTTAAAGTTTATACTTCTGATTTTCCGC
>evgR003667c0g1_i01_Hydra
CTTGTACTAAGTCTTCTACTTTTTTTTTCTGGTTTGTTTCACATTGGCCTCACATGGAAACTGTATGTATAAACGATCGTTAGCATGCAAACCACAAGCATATGGTAAGGTTGCCTCTCTTTAAGATTGCTTCTTTTTATCGTGCTTGCCATTTTCTTATTCCTGATTCCATTTTCTCCATAAATCTCCATAAATCTCTTGTTTGTCCCTTTGTTGAATATTGTTGTTACATTTGGGCTGGTTTTTTCAATGGTGCTTTTTTTATTCCAGATAAGGTCCAAAAACGTATTGAAAATGTAGTTAGACCTGCTCTAGCTTCCAAACTTGAGCTTCTGTTTTATTGTTGTAAGGTTGCTTTTATTTCTCTTTTTTTACAAATACTACTATGGTTGCTTCTGAAATGAGATAACATATCTTGTTCCATTGACTAAAACTTGCCCGGCTCCTCATTCAAAAACAAGTCACATCCTTTTACCTGTTCCTTGCGCTCCAAAAACCTCTATTACAAACCTGCTCATTTTGTTTAACACAACAAAAGATGCGCAACACAACAAAAGATAGAGGTATA
>evgR003691c0g1_i01_Hydra
ATTGATGCTAATTTCTACTTACTGTATTTGTTTTAACCAGAAGGTTCTGAAGATTAGTGCATTATTACAAAAGACAAATGATACATTGCCTCAGTATTTAGTCAACGAAGACAATGCAAGGCTTAAAGTGAATTCTGATTTTTTAGAACTGAATGAAGAAAATAGAAACTCATCCGTTTTGAGAGAAAATTTAGGAAATAAGAATTTCCATAAAATTCCAGAAAAAGAATTTGAATTTGAGGAAACCAAACGAAGAAAAAGATCTCCTAAATATCCTTATAATAGTGGATCTCAATATAGTTTTTCATTGAACTCTCCTAACGACATTATCAAGAGTCCTAATGTTATATCTAATATGCGTAGAAATAGCATTTCAGATTCTTATTACACTGGTAACATTGATGATGTAATCAGTGAACATTTTATTGATTCTAGTGATCCCGAATTTAGGTCATCTCCAATTCTTGAAGAAAGAATATCCATAGAACCAGTGAGTGATAAAACAATGCAAGATCAACATTTTATTGATTTTCCAACCAATGAAAAAATTGATAAGCTTATTAGAAATGCTGACAATGAACGCAAACATTTTGAACAAATGAATTCAGATATTGCTTTAAATTCAGTATTCCAAAACCAAAATGGTTTCCTCAGTAATCCTCAAAATATTATTGGATTTCAACCTAATGCTTTTATTGCTCCTGAAAAAAAAGATAATTTATTGAAAATAAGAAATCAAAATGAACTTGTTTATCCAAATAATATTCTGCGTAAAAATATACCTTCTGTTTCTAATAATAATAATGTTTTTCCCGTTCAAACGAGCAAAGATATACAACCTATTTCTGATGATATTTTTCGAAGGAGCAAAGAGAAAAATAGTAAAAATATTCAATCATTTTTGGAGTTTGCTAGTTTAGATAAAAAAATCAGACCTTTAGTGAATAATTCTGAGAAAAGAACATTTAGTTTAAAAAAAAGTTTTCCATCAATTCCCTCCAAAAAAAGTTCAAAGCGTACTTTAGATCTTTCAATTACAGAAGAGATTTCTAGAAATCCTGGAGTTTCTAAATTTCCTTCAAATGATGCAAAATTTATTAAAATCAAGTTAAACTCAAGTTCATTAAATGATGAAATCATTGACTATGCACAGCAATACCATGGC
>evgR003693c0g1_i01_Hydra
TCAAAACAGAAGAATATTTGGTTTTTTGATCTTTTAGTGCAAGTTTTTAGATTTATTACAGTTGTGGCTCAAATGATCGGTCACAAAGCAAAAATACACCGATGAATTTAAATAGCGGCGTACAAATAGTGGGTTTTAAAAGAGTATGCGTGCAAGGATTACGCGTACGCGGATTTTATTTATATTAGAAGTAGTGTATTTATTATCTTCGGTAAGAAACGAAGATGGAGTTGATGAAAATCGACATAACAACTTTCTCTTTCACGACATTCTTCGTGCCGATACCAAAAATAAAGATCTTTTTAATAAACCGTCATACATAGAAAAGGTTAGGCGACAGTTAATTACTATCAAAAAACATTATGATCACCAAAATGAACTTGTTGAAGATACGTTTAAAAAGCTAAAGAATATTCTTCAACTGAAAAAAACAGCCATTTTGACAAACAGTTCTGTTCGTTTCAATCGTGAGCTTAAAAAGAACTTATATAAGAAAGCGGTCGAGTATATGAAAACTATAGATGGAGAACAAAATAAAAAAAGGGGGAAAGCGCAAATATTAATGAAAGAAAAAAGATTTGCAAAAAATAAACCCAACATTTTCTCTACGCAACAAAAACGAGACTTCCCTGCTCACGATAGTGAAACCGATATTGGTTTAGATGAGAAAAGTTTGCGTTTGTTTGCTGGAGAAGATGATAATAATTCAATCGATGAATCTTTTAGTTCGTCTACCGCGTCCGAAATTTTGGGTACCATGGCAAGTATTTTTGAGGTTGCAGAGAAGAGCTCAGCTACAGAGCAACACGAACTTAAAAAACTTAAAAAAAACGTTATACCCTTACGCAATAACTACAAGGACATTTTAGATGCTTACGATACTGATGGTGAACCCTATAAGGAATCAATCAATGATGACGCCTATGTAAGCCAGTTTTATAACCCTTTTGTCGACAACATTGAAAGTAGCACAATAAGAAGATCTAGAAGAAGTGTGGGTGATGATATTGAAACTATTTATAAAGCTTCAAGACTTGATAATTTAAAAACATCCATTGCTAAAGATAGTTACCCTGCACAATTGTTAAAAAATGAACTTATAAAACATGATATCATAAGTTCCGATTCTATCAATTCAAATGCAAAATATTTAAGTGAACAAGACTTAACGGATAAAGTTAATGCATTAAAAAAGGGATCGAAATATGAAGGATATATTGGAACAAGAGACGAAATTGATAACATCAATCATCGCTCATCCATTGATATTAACAATGAATCAAATGATAAAAGTTTGGAATATTTAGATGATCAAAGCTATAGTGGTAGCAGTTACAGCGGCGACCAAGCTGTGCCTGCCAAAAACTTTGAAAATGTTTTAACTGGTGATCAGAATGAAATTATTTATGGAGAAAATTTTAATGGTGAAAAAACAAATAAAATACTCAATAGACCTAAAGGTTCAAATCACATGCAAATTGTTTGGAAACACGTTCCATTAAATTTAGCACAATTAAAACAAAATAAAGATAAAGAAGGGAAAATTGAATGGAGGTGGACGCCATATTATTGGAATTGTAATAGTAAAAATACTGGTTGCAAACTACTGCCTTTAAAAACACCTGACGGTAAAGTTTTAAAATTAAAGAATAAATCAGAAAAATCCTCTTCCTCTGGAAAATCTTCCAGTAAATCTTCTAGTAAAAGTTCTGGTAAATCTTCCAATAAACCATCAAAAAGTAAATTGTCTTCTGGTAAATCTTCCAATAAACCCTCA
>evgR003701c0g1_i01_Hydra
AGAATTGCGGATTTGAAAGTAAATGAAAATGCAAATGCAAATGATTTTTTCAATCTAACCATGGTTGATTTATATGCATATAATTTTGTGTACTTTTTTAAACGACACATTTTCAATGAAAGTCAATCAGTTTTTAAGATGATGGAAAATAATTTGAATAAGCAAAATATTTTAAACATATGTAAAATGGCCTATTATTTTTTTATTGAAAATAAAGTTATTTTTTCGAGTCAAGAAATGCAATCTTTCATCAGAGATGATCAAATTGAAATTGTAGGTTTTATTGAAAAGATTGAATCAGATTACGGTGATTATTATCAATTTTGCCAACTTGCAATGATGGAGCTGTGCGTTTCTGTTTATGCATAC
>evgR003708c0g1_i01_Hydra
GAGCTCCAAAGATCCTCGTGTGGCAAGGAATTTGCTCTTGTGACCTCAATAGTGCTCCATTTGTCACTCGAAAGACTTTGACTTCCAATTTGTACATCACACAATGTCTTCAGAAACCTCTTTTGCCATTTATTAACAAGCACAGCAAGGCCGTTATCTTTTGGCCGGATTTAGCAACGATCCACTACTCCAAGAAGACACTGGAGTGTTACAACCAAAATGGCATTGAAATCCTCCCAAAAGATGCGAACCCTCCAAACTGCCCTGAGCAGTGCGTTAATGAGAGTTACTGGGAAGTTGCAAAAGCAACAAATCAGCAAAAGATGACAAAAAGTTCAAATGAAAATGGATA
>evgR003713c0g1_i01_Hydra
AATTGGGATACATATAATCAAGAGATTTTGTCTATGGTTGGAGGTCAGACAGTGGATGATACACTTGATCTACTTTATATTTGTCGTGCGCTGGTTAGGGAATCAGAACATAGGTCTAAATATAAGCCCTTTAACCCGACTGCAGGCAAAATTGAGTACTTCAAATATTGTGCTTGCGCTCAATGGTTGTTTCTAAATCACCCTGAGAAGAACGAAGAAGAACTTAGGGAACTGGCTTCCAAATTTAGATTAAGTAAGGATGATCAAATTTACCTTACTAAAAAGATTCAAGCAGCTAAAGAGGATATAGGACCCGACCACAAGAGGCTAATAGAATCTGGGGTGTTACCAGAGATTGTCAATAGGATGATGACATGTTTGAAGATTAGTAGCAAAGGAGAGAAAGATTCTAGGCTGGAAGCAAACGAGGCTTACCTAAGATATGATCCAGATAAAGGTTACGAATATTATAAGGTCACCTGGGATTGTGCTAATCATGTCTTTTCAATCTTTTATACAAATATCATCTCTATAGCAGCAGACAATATACAAGAGAGATCAAGTGATATAGAGAAAGTGATGTTGGAAAAATTTGTACCTTCTTCAAAAGCAAGAATAGAAGCAATGAATATGAAAGTTACGAGTTTGAACAAAATCTATGCCATGCCTTTTGACTTGAAAGTTTTAGACCCTGAAAGTAGAGAAGCTTTAATGTTCTTACTTATTGCTCACGTTAAAGAAGTAGTCTACTCAAAATTCCTTGTAGGTGTACAAGCTGACGTGGAATCATCATCCTTGGTAGGATCTATCTGTGCTGCTATGTACTCTAAAGAAGGACTAAGCGTAAGCTATTATGGTGAAGGATTAGAATCCTCACTAATTGTTTTCTCATATGCTTGTAAAACTGCTGAAGAAGCTC
>evgR003716c0g2_i01_Hydra
AAAATGCTTACTCAAAATGAAATTAGAGCGAAGATTATAGGTTTAGTTAATGCTGTTGCAAGTAAAGTTGCGCGAGTTCTGAGCGAGCCAAATGCAAAATTAGCTAGAGTTATAAAATCTTATGCTGAAAAACAACAAAATTTATAGGAGACTAAAATGGCTGAACTTGATAAAATAGTTGATCAATTATCAAAATTAACAGTGATGGAGGCAGCTGATCTGGCTAAGATGCTAGAAGAGGCTTGGGGTGTTAGTGCGTCTGCGCCCGTAATGGCTGCTGTGGCTGCGCAAGGAGAATCTGCGCCTAAAGCAGCGGAAGAAAAATCGGAGTATGATATTATTTTGGAATCTGCTGGAGATAAAAAAATCAATGTTATTAAAGAAGTTAAGGCAATCACTGGGCTTGGATTGAAAGAAGCGAAAGAGCTTGTTGAATCTGCGCCAAAGCCAATTAAACTGGCTGTTCCAAAAAGTGAAGCTGAAGAATTAAAGGGTAAATTAGAGGCTGCTGGAGCAATAAAAAAAA
>evgR003721c0g1_i01_Hydra
ATTCCAATCATGATTATTCAACCTCACAACAAAACCAAGAACAACGTAGTAATTTTTCAGATGATGAGGAAGAAGAAGAAGAAGTTTCAACACCAAAAAGAAAAAAACAAACAAGACAAAAAGTTACAAGGGTGGAATCTAGTTCGTCGGATTCTGAACAATCAGAACCAGAAGAACAACAACAAAAAAAACAAAAGAAAAAACAAAGTCAACCTTCTCCAAAAAAGAAACAATCACAACTGACTGAATCAGATGATGAGGAAGAAGAACAAACAAAACGTAAAAAGAAAAAAACACAAGCTTCCCCAAAAAACAAAAACAAGGGTGCTAATGTTTCGGAAGGAGAAATTGTAGAATTAGTGACATATTTTGTCCAACACAAGTCATGGAATAATATTGTCCAACACATGTCAAATAGATATCCAATTAGAAAGAAGGAAAATTATATAAGAAAATTATCTAGTAGATATACTAAATATCTCACAAATTTAAGAAAAAGTGATGATAAGATTAA
>evgR003766c0g1_i01_Hydra
CAACAATCATGCAAGCTACCAACAGCTTAAAATCAACGCTGGTTGTTCTTGAAGCAACCCTCCTCGTTTTATCATTCGTGGTTCAATTCATTGCTGGTCAAGCCTGCACTAGTCAAGCAGGTTTAAAACCAACTACAGCAGCATATTTCTGTCCAATTGGTTACGTTAATACTAAAGTAACAGATTTGGTTGGTGGAGTTACTTATGAATGTAGAAAAGTCAATGATATGATCTCACCAATTACTAAAGGAAAAGCTTGTCGTACAGCAATTGACTGTATGGTCACTACACAAGATGATCCAAATGTTTACTCAACCAAACTTTATTCCCAATGTTCTAATGGTCGTTGCGTTCGTGCAAGCACCAGACTTCCAGGTGATAATTGTGAATGGGGTTCACAATGTTTGGGCGGAAGATGTGACATGTCTACAAAGAAATGTATCTCATCAGCCTTGTACACTGCAAAACAAGGTGACCGTTGTGAAGTTG
>evgR003787c0g1_i01_Hydra
AACAAAAGTCTGGTACTCAACCAGGTGACTCAAGTATCATGATGATGATTCATGGACAAAACCAATCACAAGGAATGGGTATGGGACAAATGTCTCAAGGAGCTGGTGCAACCCAAGCATCACAATCTGCTCCACAAAGTGATTCAGCTGATTCTCCACCACAATCCGGAAGTAACTCTAAAAAGAGAGCAGGACGTAGGAAAATTAACATTGAATTTATTCAAGACAAATCACGTAGACACATTACCTTCTCAAAGAGAAAGGCTGGTATCATGAAGAAGGCTTATGAACTTTCCACTTTGACAGGTACTCAAGTTCTGCTGTTAGTTGCTTCTGAAACTGGTCACGTTTACACATTTGCCACTCCAAAACTCCAACCTCTCATTACCCAAACCGAAGGTAAAAATCTTATCCAACAATGTCTTAATGCTCCAGAAGCTTCTCAACAACAAATGATGGGAGGTGGTGATGACGAAGAAGATGAAGAAGGTGACGAAGATGACTATCAAGGTGGAATGATGCAACCAATGATGGCCAATCCACAAATGATGGGACAACAAGGTATGGGTAATCCTCAAATGATGGGACAACACGGGCAACAACAAATGATGGGACAAGGTGGTCAACAACATATGGGGGGAAACATTCCTCAAACCCAACATATGGGTCAATCTGGTCAACAACAAGTTCCTCAAAATCAAACTCAATACTATAATCAACAAACTCAACCACAACAAACTGGTATGCCACCACAACGTATGATGAATCAACAAAATGCCAACAGTGGTATGGGTCATTTCCAAGTCCCAATGAATAATCAACCTCAATCCCAATCTGGTGGTACTCACTATCAATTCTAAGCTAAATAATTATCTGTATACTACCCTCACTTTAGTATTTAAAAATAAATTATTA
>evgR003790c0g2_i01_Hydra
GAATCTATGTTATTAGCAAAAAAATGAATGACCTCATTTTTTACCTGCTTACCTTAGAACGTGAAATCAATAATCTTTTCTCCATGAGCGTGTTTCATTAGCAAGGATACAACAGTATAAGATTTATTTGGAGCTTTATCACAGAAGCTCCTCTACTATTTGCCCTCTTCACCCCATCTTCGTTAGAACGGATTAGCTAGGGATCAGTGGAGGAAACTGGTAGATTGCGTAAAACTGCATCAATCCAGCCAATTGCATCAAAAGTTATGTACGAGTTGGAGGCTTCCATTAGTCTTACCATCTCCAATGTTAAATGTAATTCAAAGCTCCAAAAGCAAATAAATTCTGAGACAACTAAATAGAGGACAATTCTTTGTGGATCTTTGAATTTGACTATTTTCTTGGCTTCTAAGAATTTGGCTTTGCTAGGTGAATGAGAAAGAAAATAGAAACCTTTATCTGTCTGGAACTAATTGGTTGCCACAATCTCTTCATTGCATCCCACTTAGAAGAGGTAAGTTGCCATAAAAAAAAGGGCTCTAGAATTAACGCCTGTTTAACAGTAAAAGGTTATCTTTTCCCGGCGTCTCAAAATGGTTCAGAGAATCTAGCTGTTTCATTTGAAGTGCGGCTTTCGAAGAAATCCAGGATTCCATCTTGTTTGCTAAATTACAGAATTTCTAGAAAATCAAAGGACGATTCTTGCGTGTCAAAGTTTTCGAAATCTGGCAAGTAGTCGATATAGCCCATCTTATCACTGATGTGATCGTTATGTTAGGTTTGGATATTTAAAACTCTGCGGTCAGGGATATGATAATTTATTCAAAATATCAGGTGCTTACAAGGGAGCTCAAGCCTTAATAAGGCAAAATTATTTGGAAGTTATATACGCGCCCTGTCCGGTGTGCCGGCAGTTGTCCAGTCATTGAAATCAAAGTATCCTTTGGAAATATCAAATCTTTGTACGTATTTTTTAGTGCAAGTCCTGCCCGCAGTAAGATTGTC
>evgR003812c0g1_i01_Hydra
TCCATCATTACCTCAATTATCTCTAACTAAAAGTTTGGAAATTAATGACCTAATTCAACCTGCAACACCGGCTCCGTATATAAATGTAATGAGTGTAAATGCCATTAATGTAACATTACCACTTAATCTATCCGTAACTTTAATGCCACCAAATTTGAAAGTTGTTGAATGGTCAGTATGGAGTGAATGCTCTTCAACATGTGGAGGAGGTAGCAGAACTCGAAGTCCTCTTAATATGCTATCAAATTCTGTTGTTGAAAACTGCATTTTGACTTTTTGTCCAGTGGATGGAAATTGGGGTAGTTGGAAC
>evgR003815c0g1_i01_Hydra
TTTTTTTTTTTTTTTTAATTTTTAAATTAAAATGGAACCTATGAAATTTATTTCCATATTTTGGATTTTACCACTTGTTACTTGCGCCTCAACGAAGTTTGCATGCAATTCCACGCAAGTTGAAGGACTTGATGGAGAACAAGTGAATAATTTTTCTACAAAGTGTCTATGTGATTTCGAACAATGTAATGAGAAAGTTGGATGGTTTGAAATTTTCGGAAATTCTTGCAATAAGGCGGCATTTGAATGTTGTTCAAAAAGAAGACAAACTTGCAGGATGCATTCATCTTCAATCGAAGAGTATGATAACTGTTCAAAAGAGTATATTACTTGCAGTTTGAAAATCAAAAGTTTAAAAGACAACAATTAAAAGATACAGTGCATATTTAATACAAACAAAGTATTCACGAAAAAATAAAAAAGTTTGTCTACTTTTTTGATACACGTATGGAACGCATTAGTACTTTTTATTGCCTATATTTCGGGTTGAGGTGGGGAGGAGGGGGGTTGCTGAAACTAAAATAAAAGTGGGTAAGGCAGGTAGAATTTTCTCTTACGCACACTTTATCTAGGCCAGTGCTTTTAAAGTTTTTTATCTTTATAATTTGAAAGGCATTATTGCTGTTATGTAAATAAATATTTATTTATAACTTTTAGAAATATGTTAATTAAAAAAAAAA
>evgR003823c0g1_i01_Hydra
CAAACCGTAGGAATATAGAACGTCATGAAGAGTCAAAAATAATTTGTACAAGGTTTTTTATTCATTTTCTAGATATGGTTCAACAATCTAATGAAAAGTCCTGTGTTAGATGCCAAGAAATTTATAGAGTTTTATATGATCACATCACAGATCAAAATCATTATTTTTCAAACAATGACGTTCTAAATAATTTGCTTCTTAATGTACTTTCTTTAATGGTTAACAAAAGTTTGCCTCTGCAATCAACATTGGCAGATCAGTATTCTGAATTTCAATCAATTCATCTACCAACTCAAGCAACATTTGTTGATAAAAATGAAGAGCCTAGTAAAATTTACTCAGAAAAGTATGAAAGTTCCTCAGATGTTTCTCGATCTTCAAAAATTGACCATGACGTAATTAACCGAATGACATTTCAGCAAAACTCATTTTATCCTTCATCTACAAAAAGCGAACTCTGTAAAAATAATGTGAACTCGTTAGGTAACATTAACATGCCACATTCTAATAAAAGTCAAAATTTCAGTGGTTCTAGTTATGATTTTCCATTTTTATTAAATACCAGCGATATCCAACCAACATTTATTTCACCTGATGGGTATCTGACTCACATGGGTTTGTCTCTTTTGTTAGGTAATCAGTCAAAACAAAGTTTATCAGGAAAATCAGATCATCTTGATATTGCAGACATACTTGTAAACAAAATGCCAATACCTAGTTTAAATAAAGCATTTGATTCATCATTACTATCAACTTTATATTTACACCAGTTAGGAGTTGCACAATCATCTGTGGGAGTAAATGACGTGTTTAACTTTCCGGACCTATCAAATGCTGATGAAAACTCTTCTAATTATGCATCCCAAAATGCAACACTTCACCAGTTATTAAATCGTCATAAGCAAATATCAGAGCTTATTGAGTCCATGAATAATCAAACCAGAAAAAAATGCCCAGCAAAAGATTACCAAGAAAAAGGGTTTTCTCATCCTGAGGAAAGTTCAAGTTCTGTTTCAAATATGTTTAATTTGGATTTAAATAGTAAACCTTTATCAGATTTTACCAAAATAAATGATAATCTGATGTCTGACCAAATACCAAAAACTCTATTTAATAGTTCTGATAGTGAAATAAGCTCATCTATACATTCTAATGTATCTCTCGGCTCCATTCCTCTCAGCTCCTTGCAAAGTTTGATTTTAGAAACTGCAAAGGAAGTGGACGAAGATGAAAGCAAAGCCTCTGAGCCTAGTTCTAAAAGTTGTTCTGTAAACTATTCTGTTATTCAAGCAATGTCACCTGTAAAAGGAGATACTGGTGAATCTCATCGACCTAAAATTCTTTATAAATGTGGTGAGTGTGGAGTAACTTTTAGCGTATTGGCGACGCTTCAACACCATGCTAAAACTCATAACTCTGCAACCGAGATATGTAATTTTTGTCGATTAGAATTTAACAATCGAAACGATTATCACCAGCATCTAATCACTCATAGAGGAGAAGATAACATTCTTGTTTGTCAATTTTGTACAAAAATGTTTACTAGTAAAGGAGAGTTCAATAAACACGTAGGTACTCATACTCAGAGGCGCCCATATGTCTGTTCTCATTGCCAAAAAGCATTTCGAGATCCCGGTTCGTTAACAAAACATGAAAGAATTCATACTGGAGAACTTCCGTTTGTATGTTCAGTCTGTGATAGAGGATTTGCGGAGAAATCTTCTTTGCGCAAACACTCTAGAACTCATTCTGGTGAAAAGCCTTATAAGTGTAATCAGTGCCCTAAAGCCTTTTCTATATCAGGTAATCTTCAACGACATATGTTTATTCATAATGGTGAAAGACCTTTTAAATGCTACCACTGTATGAAAACATTTAACAATCCTAGTCATTTACGAAGGCATATAAAGAACTTGCATCAAAAGTTAAACGATAAACAGCATGAAACTCCGGATGATACTGAGGAAAGTGAAGCATGTTAAACCATGGGTTTCGTTGTTTTATTTTGTACCGATG
>evgR003826c0g1_i01_Hydra
TAGACAATCAAAAATAATAAATATTTTGTAGAAAAAGTATTTTATAAATTTTATTTGCTTGTACGAACGCTCAAGTGAATTTAAGGAATATGAAGTACGCTGTTGGCTTATGCCTTGTTTACATACTTGGGTTTTTAAGAGATATAATCTCAGGTGGATTAGATACACCTTGCAAGAAAATCAATCCACTTGAGCGAATTGAGTGTGGGTATGCTGGAATACCAGAAACAGAATGTTACAAGCGAAACTGCTGTTGGGATAACTCTGTCCCAAATGTTCATTTTTGTTTTTCCCCAGACGAGTTGATTGTTGGCAATGTTGCCACTACATTAACGAAAGGAAAGTTGCTTGCTGAAATACCAAAGTTAGATAGAGAACATATAATATCAGTTGAAGTGTAT
>evgR003828c0g1_i01_Hydra
TATTAGATTAGGGAAATCAGATGATGTTGTTCTCAAAACTGTTTCCTTAGAACTAGAATTTGGAAGTACTAGTAACAAAATTTCTTTAAATTGTGAATTTCATATCAAATATGGTTCTGTAGAAGAGGGGACGAACGACATACTTAAAGTCAAAGGTTTTATTATGCTTGATTTAAGTGGACAGTTGAAAGTTGGCGGCAGTATGGATGGTATGATTAGAAAACCATTTCAAATAAATTGGCTTGCGTTTGGAAATCTTCGATTAAACTTTGGCCTTGATCTCAAGACAGGTCTTCCATCTATCGAAATGAGTGCAGAAATCTGGATAGGACAAATTCCATTGACAAATGTGAGTACTGGGAACACTGATGTTTTGTTTCAATTTCAAGGTGCTTTTGGGATTAATGCGGCAAAACCCACAGAATGCTTTTTTTATGCTCGTAGTTTCGGAGCAAATTTAACTATTGCAAATATTTTAAAAGTACTTGGTAGCAATACAACATTGCCTCAAATGGTTGAAGACAGTGGTTTTGTTGGTGAAATGAAAGTTTCTGTGAGTTTTTATACTGAAGATAAAAGAATAGAGGAGTTTGATTTTTCGATATCACCTGGTTTCATGTTTAAAGGAAAAATTTCTATTTTGGGTTATATTATTGAATGTGACTTAAGGTTTAATCACAATACGGGAAACTTTTTCTTGGATGCAAAGTTTGATCCAATAGATGATTGGGGAGCGGGTATGGCGTTTGAATTTATTTATTTCTATACCTAAGAGCTTATTTATACATGGCATGTATTTAAAAGTAAGTACTTTTGCAGTAGGAAAAAGTGGCATGTTAC
>evgR003842c0g1_i01_Hydra
GTTCAAACATCGTAAGATTGATGATGCTGAAGTTGTTAATTCCAACAGTGATGAATCTCACCATGTTGCCATCTTGGAAGGTACTGACGGAACTGATAACGCTCACATGCATGAAGGTCACGCTGAAATGGAACCATTCTCTGAAATGTTCATTAAGCAACTTATCCACACTATTGAATATGTCTTGGGTACCGTTTCCAACACTGCCTCTTACTTGCGTTTGTGGGCTTTGTCTCTTGCTCACGCTCAGCTTGCTGAAGTCTTTTGGCAAATGACCCTTGGTCTCATCCTTAATATGCTCGCAGAACACCCAGAAATTGAAGATGCTGTTGTTCACAGTGGTGTTGGTGTCTTCTTTGTCTTTGGTCTCTGGTTTGGTATGACTATTGGTGTTTTACTTGTCATGGAATCTTTGTCTGCTTTCTTGCACGCTTTGAGATTGACTTGGGTCGAATTCCAAAACAAGTTCTTCAAGGGTACTGGTTCTCTCTTCCGTCCATTCAGCTTTGAAACCTTCGGTAAATCTGAAGAATAA
>evgR003843c0g1_i01_Hydra
AGTTTGTAATGTAAATGCTTGAATGATTAGATTATATGATATTTATAGTTGATTTAGTATTTTTAATAGCATAAATTGATTATTAACCAACCAAATGTCTTCACGCGGTCAATTCGAAACAGAAAAGCTTCATCACAAACTAGAGACTCAACTAGATCGACTTGTTAATCAACTTAGTGATTTAGAAGAGTGCAAAGATGATATGGACAAAAATGAATACGAAGAAACTAAGCAAGATACTTTAGAGCAGTTGCGCGAATTCGAAATTTCATTGAAAAAAATGATTGAAGGCAACATAACTCTTGTTGATGAATTTAATGGGATACAAATTGCTATTCAAGCTGCTATTTCTGAGGCCTTTAAAACTCCAGAAGTTATTAAAATGTTTGCTAAAAAACAACCTGAGCAACTCAGGCAAAAGTTATCTCAGTTACAGCAAGGAATTAGTTTAGGAAAGTTATGTGGTGTATTTTATCAACAACAAATAGTTGAAATTTTAAGTGCACTGAAAAAACTTGGTGAAGTTTTAACTCATTCTGAAGAAGAATATTTAAGAAAAAATGCATCAGCCAAGTTATATCAATTTTCAAATGTTAGTGAAACTATTGGAGATGGCCTTTTGAAAGTTGTTCACGAAGAAGTAAAACAAAGTAGTGAATAGTCTGTATTATTAAATAATAAATAAAGATTTAAAAGATATCAAATAAACAAACAGTTTTTAAAAATGTTTTTCTTGATATAAAATTACACATTCCTTGCCTAAAATAACAAGC
>evgR003860c0g1_i01_Hydra
TTATGATTGGGTGGTTCAAGAAAATCAAACGTTTTGTTTAGATATTGAAAATAGTTCACATATTATACAAACAAAATGTGTTGACTATGCTATGGAAAACAACAATATCAACATTAATGGGCTTGAACCATACACAAACTACAGTGTATTGGTATATGCCAAGAACAAAGCAGGAATCGGGTTCAAGGACAATGACAAAAAGAAACTAGTTCAAACAGATGAATTTATTCCGACTGGTTTTCCACTAAATTTTACTGTTACAAAAGCCAACTATACTGCATTGGTATTATCATGGGACAATGTTGAATATTCTTTGAGAAACGGAATCATAATAAAATATTATTATTTGTGTAATAATACTAAAGATACTATTTTTAATACAGTGGACAGCAATGTTTTTAAAGTCGTAATCATCAACCTTAACCCAGATGCAATATACAATTGTTCAGTTGCAGCATGCACTAAAGTTGGTTGCGGTGTAGAG
>evgR003864c0g1_i01_Hydra
AGAAATTCATATTTAAGTTTAAATAATCGGGACACAAGTCACTTTTTGCAACCACTAGATGTAGGAGTTTTTAAAACAGTGAAAGCGGAATGGAAGCGAATTGTAGAAGTTTACTTGGTTAGAAATTCATATTTAAGTTTAAATAATCGGCAGTTTTCTGCTATGCTAAAAGACCTTATTGCAAACAAAGGCTTTAAAGCGGAAAATGCGCGTAGTGGTTTTGAAAATGCAGGTATATTTTCACCTAATCGTTCAAAAATAACTCCTAGTAAAACTGCAATCAGCGCTATTTTTCAAGATGTAACAAATGACAAAATTGGTAATCTTTTTCTGGAAAGTCTAACGGTTCACAAGTCTCACTCAATAAGACTAATTATTTAAAAAGACTCAATGAATAGACTTTCAACAAAAAAAGTATTGCTCAAAGAATTAGATAGCATCAATAATGCCGTTAAACAAGCATCTTTTGATATTAATAAATCAGTGCTAATTAAGCTTAGAGATAGATTGATGCCTCTTTCCAGTAGTAAAGAAAGAGGAACAATAGTTAAAATAACAACAGGTTACAATATGACTGAACTATCAGCAATGGTATCAGGCTCAGGATTACATGGCTCAGGATTAAGATACACAAACTGCTAAAAAAAGAAAACTTGATGAAAAAGAAGCCAGAAAAGTTATTAGAGAAAAAAATATTGGAAGCAGCAAATTTCAAAGATTTAAAAAAAAAAAAGAGATTTAAAAGTTTTAAAAAAAAAGAAGAAAAAAAGCTTTTAAAACATAAGAAAACCCGAGAACCGTCTCAGTAATGTAACGAAAAAATAAAGATATCTACTGGTCCAATTATTGACTGTCTTCCTGCCGTGAGCTGTAATGCCAACCATATCAATCGTATTTGTTATTTCTGCGGATGTCTTTGGGACGAAGTTACAAGCACCAAATGGGTTGCATGTGAAAATTGTTCAAACTGGTCGTGCAGTCCTTGTATTGCTAATTATCATGTAACTATAGAATTCGCGTACAAGGAGTGTGAACAAGGAGTGTGAATAACTTTTATTAGTTTACATTTTTGTAGCTTGATGTTTTAAAACAATACTTTTTAGGTACTAAAAAAAAA
>evgR003865c0g2_i01_Hydra
AAATATAGTGTAAATATAGTGTATAAATTAAAAAAAAGCATTGCAATAAACATAAAAATGCGTTTTTGGAAGCTTGTTACCAGGAGTAAAAGATCATCACGGCATATAGACTTCGGAAAGGAATCTTTAAACAAGATTAGAATCGCTGTTTTTGGATATGGTGGCGTCGGAAAAACCTCAATAATAAAAAGATATCTTTTTAATGTGTTCGAAGAAAGTTATAAACGCACTATCGAAGACGATTATACTCAGATTATAAAATATCGAAAACAAATATTTTGTGTTAATATACTTGATATGGCAGGTAGTTACCAGTTTCCCGCAATGAGAAAACTTGCTATTGAAACTTGTCAAGGCTTTTTGTTGGTATATTCTATCGATAATAAAAAGTCGTTTGAAGAAGCAAAAAGGCTTTTAAAAACAATTATGGAAATTAAAGGCAAAAGTAATTTTTCCTTACTTTTAATAGGAAATAAAGAAGACTTAGAAAATCGTGTTGTTTCAAAAGATGAAGCACGAGAACTAGTAGAAAATTTTGGACCAAAAGTAGAATATATTGAAACATCTTCACGATATAACACAAATATTGATTTTTCGTTTAAAAGTATAATTCGTTTAATAGAAGAAGGAAAGGTTGATGCATTGGAAAGAACTAAAAACATAACAAACACTGATACTCAGATGCATGCAACCGCGTCTATCAAAACCGATATCAAAAAAGACTGTAATATTCGAGAAAGGAAATATGGTTCAATCAACATTAGATCATTTTCACGTCATCGAGAAGAATACGTTTATCAATTGTAGAACATTACTCTTGTCAATCAATAAACAATGATAAATATACACGCAGTGACGTATTTATAGTTTTTGAGACAATAGCAGCAATTTTAAAAAGATTTTTACAATTTGTAAAATGAAACTTAAAACCTTTAAATAAACTTAAGTAAACGTAATAAAACTTCCGAATCCTCCAAGAGAATTATAAATCATATCTATATATATTTCTTATATAAAAAAATTGATAAACTTTAGTTAGGTTAGCAAAGGGGAGAGGCAATGAAATCAATTGTATTAAAAAAACGATTAGCAAAAACAAGTCTTTTACCTAAGAGATAGAATTGCTGAATGGAGAACGAAGTAAAGTTATGCCATTTTTTCACAGAACAGTCTTTATAGCATTTTTACAGAACTTTCTTTGTAGAAATTTCACAGAACAGTCTTTAAAGCAATTTTACAGAAAAGTCTTTGTAAAAAATTTTCAAAACAGTTTTGTAGCAATACCGTGTAATTATCATAGTTTAATAAGTCGC
>evgR003869c0g1_i01_Hydra
GTCATATATACATTGATTGTTGTTATGTGCTAACTATTCAAATGACATTGATTGACAGCAAATTGGCTTTATTGATAACTGGGCTATCCATTTTCCCTTTACACGTCATTGGACTTTATGTGCTATCAAAAAACAAGGCGTCACTCGATAAAATTCAAGTTGTATTTCTTTTCAACATGTGCTTAGTTGAGGCTCAGATGATATTTTTTAAAAATTTAGAAGTACTCCTTAGTTTGTTAAAATCAAATAAAGCTGAAAATTATGTAGGTATTTACCTAAACGTCGGTTGCTGCGTTCCGTATTACATCACGATGTTGCTGTATACATTGAATAGATTTTCAGAGCTCTATCTAAACCTCAGATACACCACATATTGGACAACCAAGCGTGTTAAGAGATCAATTTTTGTAAGTTGGGGGCTTTCTGTAAGCGCAAGTGCTTATTTAATGGCTAAAAATGTAGAGATAGACTTTGTTTACCGCATAAGTTATAAATATGTTTTCCCAATATTAAATTCTTTTGCAGCATTTTTGATATCTTCGACATATATTTACATTAGCTATAAATGTTATCACTTCTTCAACAAAAGTAGGATCGTAAAAACGCAGTTAAAACATACCGTATATACAGGAAAACAATCATTTTCCGTAGGAAGAAAAGCGATGCGCAGATATTTAATGCCGACCATTTTTATATTCAACTTCTTTTTCTTTATAGTAATACCAGACTCCGTATATTGTTACAAGTATTTGACTGAAGTTAAAATAGAGGAGTGGGTCCCAATATTGGTTAATTTATTTTATTTCATCGGATTTTTTACCGATGCGTTTATTTATATATTTTTATCTCCGGCTATTAGAGCTAACTTGCTTAAAAAACTTACATCAAAAAAGAAGCATTGTAATGGATATAATATAAGCATAATTACAGGATCGTAA
>evgR003885c0g1_i01_Hydra
GTAAGATTTGTAATAAAAAATACAATCAGCTAAATACTGTGCGTTTTTGTTGTTATGATTGTATTTCGGAACTTTGCAGTTGTGAATATGTGGTGCCAGATGAAGAAACGTCTATTTTCATTGGTAAATTGCCTTTTCAATGTCCCAACGAAGACTGCATACAGATTGTTGTATTGGATGAAATGAGAGATCATTTAGTAAACCATTGTATGTTTCGCCTTATTTCTTGTGTGTGTGGAATGGCTATACCTTTTTTGAATTTCAATCATCATAAAGAAAACAGTTGTGTTGCTCATATTCTTAATATAACTTGTAGACAATGC
>evgR003887c0g1_i01_Hydra
ACGAAGTCCAAAAGCCAAAAGGCATTGGTTCTCTCTCCTTCGGTTTGTCCTTCTCACAAACAGGAAAATCCCAAAAGGATCAAACGGAACAAGAACTTTTAGGTGAGGAAGTAACAATTATCTTCCAACTCGAATCCAAGGGAGAAAGTAAGAAGCATGTTGTTAATATGGGCCGTCCTGTTGAGTGGCTTAAGTTACTCGTTGAAAGAGAATACGGAATTGCATTTGACTCACAAAAGATGTTTGTTGGATCTAAGGAATTGGCTAACTTTTTGAGTTTGCAAGATATCAAGGAAATTATTCCAAAAGCTGAAAATGTTATTATTGTAAAGGATTTGTAAAGAAG
>evgR003890c0g1_i01_Hydra
CATCGATCAACATCGTTTAAGGTTTCGTCATAGTCTGATTCGCCGATTCGCAAAGATTCGCCGATGTTTCCTAAAATCAGAACTAAAAACTACGTTCCCGACGTGCAATGTAATGTTTCTCTTACGCCAAATAGAGGCGAAATTAGATATGGAAGCTTTAGCGTGAATCTGGGAATTAATCATAATAAGTAGTTAAAAATATATACACAATAGAAATATTGCTGAAAAATTTACGTTAGTTGCAAATGGCAAAGTTAATTGCTGTATGTAGAGATGATGAATACCGATTTGAACGCAGACAAATTCCACTTATGATAGAAGAGTCTTTGACGATGGTTATGGAAATTCCTGAAACTATTATTTCTGCTGAATATGTCAGCGCTGCTGATGTAGCACACTTTTTTAAAAGACACAGTTGTTTAACACAATTAGAGATGAACAAGGCATTAATGGTTGGTCAAAAAGAGTTAATTGCATTGCTGGCTATCTCTGTTCCTTGTGTTGGATGTAGAAGAAGTGTTGAAAGGCTTTACAATCAGCTTTGGAAATCGAGACAGCCTGCTATTGAGCCACTCGTAATTACAGCAACTGGAATTTTAACATTGACAGAAGGTTTTCTTGTTGATGCTAAACTAATATATGCATTATTCTACATTCATGGATCACACTTAAATGATGTTGTTGATACCATTCCAAAGAGTCGTCGTAACAGGCGGTGTATATTACATTCGTTGGAAACACATAAAACTAGAGCTTCTGGAAGCTGGATAGATGTCTGGGATTTACTTTCACAAGAGTGCCGAGATGAGGTTGTTCTTATTGATGCTGATTCTTTATTGGATACATTGGATAATTACCTTCGCAAACACAGATTCTGTTCAGAATGCAAATCGAAAGTTTTAAAAGCGTATAGTATATTATCTGGTGATATAGATGGAGATTCAGAAAAAGGCTATTGTGCTGCGTTATACGAAGGTTTAAAAAGTTGTCCAAAAGAGAGACATATTCATGTAGTATGTGATACAGATTTTATAGCACATTTAATAGGAAGAGCTGAGCCAGAATTAGCTGGTGGCAGAAGGGAACGGCATGCAAGAACAATTGATATAGCTCAAGAGGAGGTTTTGACTTGCCTTGGGATTCATTTGTGGGAAAGATTGCATAGATTGTGGCAGAAGCTGCGAGCCGAAGAGCAAACATGGCAAATGCTTTTTTACTTGGGTGTGGAAGCCTTGCGCAAAAGTTTTGAGACAGCAGTTGAAGAAAAGCAAGGTGTATCACAGTTAGAACTTGTTGTAGAAGAAATATCTGAAGCAGAGAGAGCTAAAGAACAACGAAAAGAAAATAAAAGAATGAAAAAAAAGAAGAGAAAAGAAAATAAAGCGAAATTAAACGAGGACAAAATACACGAGTTACAAGAGGAAGAAAAACAGACTCTAGATGATGATGAATTGAGTGAAAATGATGAAAAAATATGCAATGGGGTTGATGGATCTGAATCATTAGACGGAAGTGAGAAGTCAAGTTGTGGATCCCAAGTAAGCCCTCGTAAAACAGACGAAGAAGATGAAAATTTACAAGAGAAAAAAATTTCTTGCTCGCCCTGTCACATTTCAAATATAACAGATTGTACAAACTTTAATAAATCTAGTACATTTAATTCGACCCCATGCAGTCCAACAAAAGAACAATTAGGTTTCATTGAAGTTTCTAAATTACATAAATGCAATCCTAGTGATGTAAATACAACTGATAAATCAAATTATTTTTTTTCTAATTGCAATCATTCCACTTGTACAGACCGTGACCAATTTGAAAATAAATTGAGTACAGCATGTAGGTGTGATAATGAAAATTTGCCGGCAAAACGTAATGGACATTCACATAATCTAAATTATAAAGGAAATGAACACTTGTTATACAGTAAGCGTAATGGTTATACTTGCAAAGGAAGCAATCCGGATACTGGTCCACGGTTTAATAGGTGTGATAATCTAAGTCGTCATACATCAGACTCTTACAATAATGTTTATGATAAATGTTTTTCATTTAATAATCCAGTGGTTAATGGACGCGGAAGAGGAAAGCGAAAGGGTTTAAAGGACCTGTGTCTTTCTGATCACTCTAATATTTATTGTAAAAATCATCACATCAATGGAAATGATTATCATACCACCATAAAAAATGATGGATTTTCATCATTGCGCAGCCCTGATTTATTTCAAGATTCGACTGTTGACGATGAGGGAGAGCTTCGTCTTTTAAAAATGATGGGCTGGAACTCAGAGGACTCTTCTGGGATTTCCGAAAAAGATATTCAAGATTTTGTTGCAAATCGATCAGAAGTGCAAACGCGTAGACAGGAGCTACGCGATCATTTGCGTGATCAATTTCATAAACTATCAGTTAAATCTATGCACGCAATTCCTGTGTCTATTAATGGCGTTCAGCACATTATTTAAAATAATTCCTAAATATTTTTTTAGTGGGTTGCCTTGTATATGTTTTTTTATACTTCAACGTCAACGCGTGCCGATAACCAATTCTTTAACTATTGCGAAATACTTCACAACTCGTACTTAATTTCAATATTTTTTCTCTCAAAGCTAATTTAAATTAATTTAAATAGTTTTTTTTTTATAATTTTTGTGAGATAGTGAATAAGCTTTTTCTGAGAATCTGCGTCTAAACGGCATTTATTTGTGCGTAGCAAAACAGATTTTTTTACATTGCAATGAAATGATTAGATAGAATGTCAATTTGTGTATAGAATCAAATTTTAGTGGAATCTATTTCACTTTTCTAATTTAAAAAAAAAAA
>evgR003896c0g1_i01_Hydra
AGATTCAATTATAAATAAACTGAAACCCGTATTGAAAAGATCATTTAAAAATGTAAACTTTGAAGACCTCAATAAAAGAATTGAATCAATTAAACTTGAGCCATCAATGACAACAAACGAACTAGTAAGTACTTGGTATAATGCAATTAGAATAATTATAGATGAATTTGCTCCAGTTAAAAAATTTCCATACCGAACAACTAGACCTCAATGGATTGATGAAAAAATAAAAATTATGATAAAAAGAAGAGATTATTTAGCTAAAAAAAGGGGGAAATATATAGAGTTTACTAATGAAAAAGAAAAAGAACTAAAG
>evgR003911c0g1_i01_Hydra
CTTTGGTGATCAAGCTCGTGTTGTTCAAAATAGCTTAGAAAGATTTTTCTGTGTTGAAGCACTTGACGGAATGAATGATTCAGTTTCATTTCAATCAGCAACAATTATGTATAACAAGTATGGCTACATACGTACTGATGGCAAATTAAGAATACAATCTCCAATTCCACCTTCGATGTGTAAAACTTATTCAAATTACCAAGAGTGCACTTCTTTTGTTATTGTTGATAATAAGTTTTTTCCAGGCTCTGTTTCATTTCTAAACAAGTATTGTAATGGTATGGAAATGTTATTAAGAAATGATAACAATAC
>evgR003928c0g1_i02_Hydra
ACTACTGCAATAATTTAAAACGAAGCACGATGCATACTTCAAAGCATGTCGACAACAAACTTCAAAAGTCAAAAGATATTACAAAAGTTAATAAATATTCGTTACCAGATATTTCCAGCAGTAAAATTTACATTTCCAACGGTAAAACAGACATTTCTAGCGGTACCACAGACGTTTCCAACGGCAAAACAGACAAATCCAGTAGTAGTAAAACAGCTCTACTTCAGTCTGGTTTTCTTCAGAAAAAAGAAAACGCTATTCAGGACATTAACTTACAAAAGATTTTATATAATTTAAATCCTTATCATCCAAAAATTATTAATTCAGATGCTACGCATAACCTAATGTTAAATAGAGACTTGACAAACTTTGACATTTACAATATGGATATAGCTAATTTTATAAACAATCAAAAAAATGACGTTATCGGGGAAAAAAGAATATTTAATAGCCATAACTATGGCAGCGTAAAAAGTAGTAAAACTGATTACACGCAGATGAATCTTCCAAAAAGCAATGCACAAGTATACTATCTTGACAATATTAACAATAACAAAAACATTTTCAAAATAGCAACTAACAAATTATTCACAAGTAACTTAGTAAAAGACAAATTAACTAGCTTTCCGAAACCGTTTCTAAGCAAAGACATCATGGATAATAATCGTCTTAACTTTTTTAAACAACCATCACAATCTAACAGTTATTCCAACATTGCAGGTAACAATAAGAATGATTGGAATTTTTATGGAAATAATGTACCATCAACAAATGGTGTAACAGTAAAAAACAATCCCATTAGCAGCCAAGTGCTAATGGCAACCAGGCGACAGAGCCTAAATACAAACAAGCCATCATTTGATTCAAAAAATATTATAATAAAGAATGGCCAAATAAACTTAAAGAAACCTCCTCTATTACTCCAAAAACAAGGTGACTATCCTTTATTTGCACGATTAGCCAATTTTACAAATAGCATGAATAGAAACCAGCATGTGAATTTTTTAAAACAAAATGGTTTCCATCTAAACATTCCAAACCCACCAAAAAAAGAAACATTTTCAAATATGGGATACCAATCAATATATCCCACTTTTCGACAACAAAATAACAAAATGTCAAACCCGCTCTTATTGCATCCTTTTTATAAACTTAAAGCACATGTTTTCCCCAAAGTTTCTAATCAAATATTTCATGTTAGTAATGACATTATAAATCGCTTTGATAATCAAAAACCTTTAAAGCCAATATACGATTTTAATCAAAAGGATGTTCTATTTAATAAATTTTTAAATCCGTTATACGGTTCGGACCAAAAGCATGTTCCATTAAAAAGTAGTTTAGTGAAGATTTTGAAAGATCCGAAAAATAAGCATGTTCAATTCGGAAACAATTTAGTAAATGATATGAGAAAGTTAG
>evgR003944c0g1_i02_Hydra
CCTATTTTAGTGCAAAAAGTAATATTAAAACTGAAATGTTTTGAAACTAAAAAAAATGAAAGAAGATGAAAAATGGGAAACAGATGACAATGAAGATTACGATATTATTGATAGTCCGATAGCAATAAAAAAACTGAACAACTTCAGTGATACAATTTACGAAGAAAAAAAAAATCTTAACGATTTAGAATCAGAAAGAGAGAAAGAAATCGTTGAAACACTTGAAACAGAAAATGCATTTTTACTAGAAGAAACTAAAAACTCACAAGCTGAAAGTCCGGATGAAATAACAAAATTGATATTGGCATACAAAAGTGCAACCGTCCTTCAAAATACAAATCAAAATACAAAAAATGAAGATGATAAAAAACTTGGTGATGTTAAAAAACTTGATGACGTTAAAAAACTTGATGACGTTAAAAAACTAAATGTTAAAGAGCTTGATGACGTTAAAAAACTTGATGTTAAAGAGCTTAATGACGTTAAAGAGCTTGATGACGTTAAAGAG
>evgR003957c0g1_i01_Hydra
AAACACTTGTTAAATGGCTTCTGTTCAGTGGAGGTATCCTAATTTTGATTTATATCTACAAGGCTAGTGGAATTAGTAAATTGATGCGTAGGAAACCAAGAGCTATCCGTAACCCAAATATCGATTCTACTTATCGTGCAAAGCAATCATCCCTCAAAGAAAACGAATATGATGTCTGTATTGTTGGTGCTGGTCCTAGTGGAGCTACCTGTGCCTATTATCTTGCCAAGTATGGATATAAGGTTTTAGTGTTGGAACAAAAGTACTTCCCTCGTGAAAAGACTTGTGGAGATGGTTGTACTGCCAAAACACTTGCATATTTGAAGGAAATGGGTGTCTTGCAAGAATTAGAAAG
>evgR003961c0g1_i01_Hydra
TATTCTTGGTAATTCCAACATTCCATTAGTAAAAGGAAAAATTGTTGCAGAGATACCCAAGCTAGATAAAGAGTATTTGGTTTCATTTGATGTCTATCCAAATAAGTTTGTTGCTGGTCGGCATAGTGTTATTCATTTTACGATTTGTTCTGATGTTGCCAAATATGGTGATAGAGTTCCCGGTGTCTGGTTTTTTTCAGATGGATTAGGAGGTCTTCATATTTCAGCACCAATAAATGGTAATGTTGACAGAGTTTTTACCACAAAGCCAGTTGAATTGAAACAGTGGTCTAATGTAGAAATTAG
>evgR004004c0g1_i01_Hydra
TTGTATGCTGTCGGTCCACAGTTGTATTTTGCTAAGTTGTATGACAAGTTTGAATTTGCTGTTATGACTTTTTGGCTTGTGGATTTTTTTTTATTGAAATATGCTGGTTTCAGCTATGCTTTTTTTGTTTTGCATCAGCTACAACTGGTTAGACTTTTCAAGCACACAAAAGCGTGGGGATCTCTTAAGAACTTAAAGACATCAATATTTAATAGCATATCATCAGTTTTGAGTTTGATTCTGCTTTTTTTTATTGTTGCAGTCAGTTTTGCTTTACTCGGAATGAGTCTTTTTGGAGCAGTGTGGTACAATATAGTTCCTAAGCCACGAACAAATTTTGATGATTTCTTTAATTCTATGGTAACAGTTTTTCAGATTATGATGGGGGATGGTTGGAATTATGTTATGTATGATGCAATTAAAGCTTATGGTGGTGCTGAAAAACCCTTCGCCCTTCTATCAACCTTATATTTTATTGTTGTGACTATTATTGGAAACTATGTTTTGATAAACATTTTTTTGGCTATTGCTATAGACAACATTTCATTAGCTCAAAATGATAGCAAAGAATCCCTTTTTGCTAGAAAAGAAGCTCTACTGATGCTTAAAGCAAATGAAAAGTTTACAAATAAACTTAACCAAGGTAACCAAGGTGATTCTCCTACTCTTGGTATTGAAACAAGTCATGGTGACAAAGTGATACCATGTGACGAAGTTATACCATGTGACGAAGTTATAGCAAAACCAGAAGCAGTGATGATTGATGTCAACACACTATTTATTTGTTTTCCGAAAAATCCATTACGAAAGGTTGCTCATAAAATTGTGAACGCTCAATTCTTCACCAATCTGACGCTGTTTCTTATAATTATAAGCAGCGGCATGCTTTCTCTAGAGAATGCAACTGATCCAGATGCTCAAATAAATAAGATTTTGAGATATTTTGATATTATATTCACTGTCTACTTTACATTTGAAGTTCTTCTCAAGATTGTTAATTTTGGTTTTTGTTTACATAATGGAGCTTATTGTCGAGAAATATGGAATTGTTTTGATTTGTTTATAG
>evgR004026c0g1_i01_Hydra
ATTGTTGTAGTTTAAAGTTAAAGAAGAGGATTATAACAACATAATAAAAAGTCATATTAATCGTTCTATTGTTCATACACCATCGGAAGAATGTTTTCTTGGCTGAATAATTTTGTTGAAAAAGAAAAAACTCATAGAAAGACTTTTATTGACATGAGTAATCGACTGTCGCCAATACTTAGTGATAAATTTAAACCAGTTGCTTCTCTTGATTCAAGTCTATCTGAGGCTATAAGTTTAGAATATGTTCAAAAACCAGGCTGGTACCTAACTTGTAAAAATCGAGTGTGTTATATCGAAAAAAAGAAAAACACTAGGAACTACGAACGCCAAAGTTCATTCTTTCCTCTCCACAATTTATGGTTTGATGGTTTCACAGCATTTGAGTCAACAATAAAAGCTGGCTTTTATTTGCGACTGAGTAAATTTAATAAGTTTTTCTTGGACAAATATGATTGTAGCGAAACCTTCAAGCAAGAGGCTAGCTTTTGTGTACAAAAATGTTCAGTGGTTGTTGAAGGTCCGTACAATAAAGACAGCATTGCATCACGAAATGCATTGGCTGACGGGATAACAATGTTAAACTCTACTATTCAAGATGAAAATACTGGTTTTCTAGAATCTGACGAAAAGAATGAAAATTGAAAATTAACGACGATAATAAAGTTTTTTATATGAAATATTTATATTAATTTATAAAAAAAAAA
>evgR004039c0g1_i01_Hydra
TACAGTTACAAATAATGTGCGTGTATCAGGTGTCAGTACATATCCCACTTTTTTAGATGAGTTGTTTGTTGGAGATGTTACTCCTAGTATTTTATTTTGGAGACAATTAACAGGTAATCAGAATCCAAAAAGGATCAAGCAGATGATTGAGTCTTTTCAACTTGACTTCTGTTTCAATGAACAATGGAATTATGGTGATGATGTTCTTGGCGGATATCATAATGAACAACTGGCTAACAAATTAAAGTTAAAGGTTATGGAAATACTTCGTTATTCAATTAAAGAAGTTTTGTTGCTTGAAGTTATTTCACTAAGAGAAGGTTCTGG
>evgR004057c0g1_i01_Hydra
TGGTATTAAAATTGTGTAAAATGTTACTATGCTTTTATGCATTGCTCTCCTTATATCTATTACAAACGAGCATTAGTTGCGATACTCCCAACAATATAACTATAGACTCAGTGGAGATTGTGTTTTGGCCTACTAAACCTTTTATTTTTTACAATGATAATAATGAACTAGATGGAATAATACCATACGTCTTCAAAAGACTTAGTGCTTTTTGCAATTCAAACAAGAGTTTGGCAAATTTTTATTTAATAAACCAATTAGTTACTCAAGATAACGTGTACGAAACATTAAAAAACACTAAATATAATACTAAGGAACTTCAGAATGTAACTGGAAATATGACTGTATTTTTACCAGTATTTATTAAACCAAATTCTAATATTACAGATCAAGGAAAGTTATTGATAGATAATATTTTTACTGAATCAAAATTAGCGGTTATTGTTCATAAGAGCAAGCTAACTATTACAAACAAAATGATCCATGCTATTAAGAATACATATGCAATTATATTATACGCTTTCGTATTAGTTTTAATTTTCTCAAACATTGTGTGGCTAGCTGAATGCATAAAAAACAAAGATTTCAGCAAATGGTTTTGTAAAGG
>evgR004088c0g1_i01_Hydra
ATAGTCACTTTAATCATTCATCAAAGGCATGTACAGAATTTAAGTTACTACAACAACAACAAAATGAAAACACACCTCTACTTTTGGTTCAAGATGTGCCTACTTGCTGGAATAGTGCATACTTAATGTTAGAGCAACTTCAAAAACTTAAACGAGCAGTTCAAATGTATTTAGCTGATCATAATGAATTGCCAACATTTGCATCAAATGATTGGAAAATCATTGAACATTTATTATATATTTTAAAGCCTTTTTATGAACTTACTAATGTTATGAGTTCTGAACTCTGTAGTTTATCAACATTAA
>evgR004100c0g1_i01_Hydra
ACGTCCGTATCTTGAGTACCATCGACAGAGAACTTCTTAAGCCAAATGCTTCTGTTGCTCTCCACAGACACTCAAATGCTTTGGTTGAAGTTTTGCCACCAGAAGCAGATTCTAGCATACAGTTGGTCTCAGCAGATGAACGTCCAGATGTTACCTATTCCGATATTGGTGGCTTGGATATTCAAAAGCAAGAAATCAGAGAAGCAGTCGAACTTCCACTTACACACTTTGGATTATATGAACAAATTGGTATTGATCCACCAAGAGGTGTGTTACTCTATGGTCCACCAGGTACAGGTAAAACAATGTTGGCTAAGGCTGTTGCTCACCACACAACTGCTGCTTTCATTAGAGTGGTCGGCTCAGAATTTGTACAAAAGTATTTAGGTGAAGGTCCAAGAATGGTTCGTGATGTTTTCAGATTGGCTCGTGAAAATTCACCTGCTATTGTTTTCATAGACGAAGTTGACTCAATTGCCACAAAACGTTTCGATGCTCAAACTGGTGCTGATCGTGAAGTGCAACGTATTTTACTTGAATTACTTAATCAAATGGATGGTTTCGATCAAACAACCAATGTTAAAGTTATCATGGCCACAAACAGATGGGATACTTTGGACCCTGCTTTGCTTCGTCCAGGTCGTTTGGATCGTAAGATTGAATTCCCACTTCCAGATCGTAGACAAAAGCGTCTCGTTTTCCAAGTTTGTACAAGCAAGATGAATTTGGCTGAAGATGTTGATTTGGAAGATTTTGTATCTCGTCCAGACAAGATTTCATGTGCTGAAATTACAGCCATTTGTCAAGAAGCAGGTATGCAAGCTGTGAGAAAGAACAGATATGTCGTAACGTCAAAGGACTTTGAAAAGGGTTACAAGACTCACGTTAGAAAGACGGATGGTGAAC
>evgR004102c0g1_i01_Hydra
CAACAACAGACCGGAAGTTTGGTTTGTGGATTATTTGCTATAGCTAATGCTGTGCACTTGGCTTTCGGTAAACAAACTGAAAATTTATCATTTAATCAAAAATTTATGAGACAACATCTTGAAAAAATGTTTCATGCTGGACAGTTTGAACCCTTCCCATTACTTAATAAGCGAAGTAAATTGAATATTACGCGTAACAATCCCTCAAATGTAAATGTTCAACTGCATTGCAGTTGTCTAATGCCTTGCAATTATGATTTTATGGTTGCTTGCAGCAAATGCCTAAAATGGTTCCATATTTCATGTGTCCAATATAAAGAAGACAATTCAGTGTAGGAGTGGAATTGCGG
>evgR004104c0g1_i01_Hydra
CAACATGGTCACAATGGTCCTGGAATTCCCTGCTTCTGGGGGAGCAATTCCTAATTTTTCTGTACTTACTTTTAAACTGTATGCTTATACAAAACAAACAGGTGTTGCTATGTTAGGATCTCAAATATTGTTTATTTTGATGATGTTGATATTTGCTATTCGTGAGTGTCGATTTTTGTATCGAACTGGTCGGAGGTACTTTCTTGAGTTTTGGAACTTAGTAGAGGTTGCATTGATATTAATATCGGCTGTTGCTGTTGTGTTTTTTTTTTATAAAGATCACCTGGCCAAGGTATTATTACAGCGACTGCCAGCTAAGAAACCACAATCATTCATTAACTTTCAGTTTGCTTCTTACTGGGATCTAATTTTTATATACATTGTATCACTCATCGTGTTTTTTGTAACTTTGAAGTTCATTAAGGTTTTGCGATTCAATCGTCGTGTTTCAATGCTATCAACAACATTGAAAGCTGCATGGTATCCTTTATCAATG
>evgR004118c0g1_i01_Hydra
CTACAATGCCTCACTCATTCGGTTACAGAGCCAGAACTAGACATACCTTCGCCAGAGGTTTCAGAGCCCACGGTATCCTCTCCTCAAAGTTCTTCTTGACCAACTTCAAGCTCGGTGACTTTGTTACCATCAAGGTCAACAGTAACGAACATAAGGGTATGCCACACAGAATTTACCACGGTAAGACTGGTAGAATCTGGAACGTTACCCCAAGAGCTGTTGGTGTTGAAATCACTAAACAAGTCAGAGGTAGAATTATCAACAAGAGATTCCACGTCAGAATTGAACACGTCGTGAAGAGCAAGTGTAGACAAGACTTCTTGGAAAGATGTAGACTCAACAACAAGATTGCTCTCGAAAATAAGGGTAAGCCAGTTGAACAAAGAAAGAAGTTCGTTAAGAGACTCCCAGCTCAACCAAGACCATCTTTCACTTTGAAGGTTGAAAACGTTGTCGATGTCATCCCACCAGAATTCGTTGAAACTGCTTAAATTGTTCAATAAACTAAAAAATTTCATAAATGCA
>evgR004120c0g1_i01_Hydra
GTTTATGGTAATATTTAAACATAAAATTATAAAGTTTCCAAATATATGTCTTCTAGTTTCCCTAGTTATGTCAATATCTGAATTGAACTTGGAAGTCGAGAGAACTTTTAGTATTTTCACAAACATATTAGCAGATAAACGACTATCAATGAACCACGACACACTTGCTGATTGTGTCATAATTCTTGGCAACAAAAGTCTTCGGTCAAAAGATGAATGTGATAATATTATTGACCTATCGTTGTTGAAGTATAAGAAAAAACGAAGAAAATCTGATGTTGGTAATTCTAAAAAAGTTAAAATATCAAATATTCAAGATATACAAAGATGCAGCGATACTGATGATGTGGACAGCAAAGATGTTAAACTAGCAGACAATTTTAAGGAATTAAATATAAAAATTATTTTTCGTCAAATAATTTTAGAAATGCAC
>evgR004139c0g1_i01_Hydra
CAACAACAACAACATGTCTGACGAAGATACTACATTCGAAAGTGCTGGTTCTGGTGCTTCCCACACCTTCCCAATGCAAGCTGGTAACTTGAAAAAGGGCGGTCACGTCGTCATCAAGGAAAGACCAGTCAAGATTACTGAAATCACTACCTCAAAGACTGGTAAGCACGGTCACGCCAAGGCCAACATTACTGGTAACGATATTTTCACTGGTAGAAAGTATGAAGACGTCTGTCCAACTTCCCACAACATGCCTGTTCCAAACGTTTCAAGAAACGAATATCAAGTTCTTGCTATTGATGGTGACTTCATCCATCTTATGATGGAAGATGGTACTTCAAGAGAAGACATTAAGTTGCCAGTTGATACTGACGAAGATAAGTTGTTGACCGAAAAGATCAGAGCTGCTTATGACAATGGTGATGAATTCAACGTCATCCTCGTTGCTGCTATGGGCATTGAAAAGGTCATGGAAATTAAGATGCAATAAATTTTCTCCACCCG
>evgR004142c0g1_i01_Hydra
TTCACAAAATAAAGAATTAATTACAAACTGTGGAAAAAAGTATCTTTTCAACGACTTGTCTGGAAAAATTCGTTTTCCAGCAATTGGAAATATTATTTCAAATAAGGCTGAATGTACATTTTCGATTAACATTGCTCCTCAGTTCCTCAGAATTGATATCAATAACATAGGTTTAGAGAATAATTACCATTGTAAATATTACTATGTTGCCTTTTACGATGAACCTTCTTACCAAAATGAAATAACGCGAAATTGTTCAATTAGTTCATTTGTGACGATTCTTAAATCAGGTGTAGTATATATAAAAGTAGTGACTAATAATAACAATACTAAAGCTGGTTTTAATTTAACATGGAGCAAAATAGAAAATTCTACAACAAAAGTTACCAGCTACACAGATACGTTGTTAAAACATTAGAACAATGTCGCGGATTTCGTTGGCCCAACGTCGGTTGCCAACGTTGTTATCATTATGGTTACAAAAACGACATCGCCAAC
>evgR004146c0g1_i01_Hydra
TAATTACAATTTCAGTAATGTAGGAGATTATCTAACTAGTGTTTATGATCGTCACATACCAAATCTAACAAAATATGATTATACAATACTTGCAATCCACACGTTTAAATTTTCCCTAGCAGTTTCTAATTTTGTTGCGTTAATTCGAACAGGGCATTTAACAAGTTCACCGTATAAATACACATTTTTTACAGGCATTAGTATTTCAGACGATATTTTTTCTATTTATAGGCAAGTGACAAAAGTTAGTTATGGCGTGTATGATGAGGTAATTAATGTTGCAAGTTTATTAGTAACAACTGCTTCTTCACTTGCTTTAACTCACACTGTAGCTAATCAATTTGAGCGATTATTCAATGAAATTAGTAGTAGTAAAGCAGAACATCAAAAAGCTTTTGATGAAGCAAGTAAACAAAGCATAGAAGGAGTTAATGTTTTGACAACAATTATATTACCTACAGTTGTTGAAGGTTCTGAATTATCTACTAGTGATTCTGAGTTCAAATCAGCTTTCCAACCTCAAGTATATGCAAGTAAGATTCATGAGTGTGTTGAATCTGAAAATCAACTTACTAAATCACAGGCTCATATTGGCACACAAGATTATCTTTTTTTCTTTATGAATAATGCTAAATCTTTTACTCAAATTCAAGAAATAATTGAGAAAAAAGGATTTGATTATAACTTGTTAAAAC
>evgR004152c0g1_i01_Hydra
AAATGATCAAGACTGTAGCTGTTTTACATCCAAAGTTAATCGATTCCTTGAAGAAGGTTAGAGAACAAAGAGCATTCAACCCAGAAGAGTGGATAGAAAAGAAGAGCGATTTATTTAATGATTACATGAAGAGATGCGGATTGAAAGCCTGTGTTGTTTCTGTTTCTGGAGGTGTTGATAGTGCAGTTACATTTGCTTTGGCTGAAAGAGCTATGAGGAAACCAAATTCACCAATCAAACAACTTGTTGGTGTTGCACAACCAATTAAATCAACAGAAAAGATTTGGAAGAGAGCTCTCGAACTAGAAAAAGCATACGGATCAACCGAAGGTGTCAAAGTTGTCTCAGTTGATCAAAGTGAGATTCACACACAAATTCAAGAAAGAACTGAACAAGCTCTCGGAATTAAGAGTGGAGACTTTGCTAGTGGACAACTTAAATCATACATGAGAACACCTGTCGGTTTCTTTACAGCTCAATTACTGAGTCAAGCTGGCTTTCCGTGTGTTGTTTTAGGTACTGGTAACTATGATGAAGATGGTTATTTATTCTACTTCTGTAAAGCTGGTGATGGTGT
>evgR004155c0g1_i01_Hydra
TATGGACATTTTAATTATATGGGAAAAATATGGAACCCCGTGTAGATTTTTAATGAAGATTTGAAGTTGCATATACCTTAAAGTGCATATATGTTAAAGCTCAAAATATGAAAAAATCTCAAAAACAATGGATATCAGAGTAAAAGAAATCCTTTTAGACAAAGTTACTTCTGATTGGGAAAGGTTTGCTCGTTGCTGTCTTGTTGATATCAGCAAAGCAAATGCTGTGATTGACCATATAAATGGAGATGAAATTAAAGAAGATAACAATTTTAAGATGATATGTTTATTGGACATACTCAATAAAATGTATCCTGAAAATTACAAGACAAAGATTATAAGTTCATTGAGAGATATGGATAGAAATGATGTATTAAAAGATTTATCAAATTTAGATTTATCAGAAAAATATGGGGCACTTAAAAGTTTCTATCTGAAAAACTATCAGAAAAATGCTGAATGCCGACCACCACTAAAAATTGTTTCCAGTGTTGATATACTACATGAATTTGTTGAACTGTGTCTTATTGATGCAGTAGATGTTCAAATGGATGCTGCTCATAG
>evgR004167c0g1_i01_Hydra
GAAGGAAGTGATATCCCAAAAAATTGTCCAAAAGGATATTATTGTCCTGAGAGCTCAGGGACACCTTTTCCTTGTCCAATTGGAACATACAATAGCAAAGAAAAACTGCAGTATCGAGAAGACTGTACTCCCTGTACACCTGGATGGTACTGCGATGCCCAAGGTTTACCAACACCTCGTGCACCTTGTGATCCAGGGTATATTTGCATATTAGGTGCTTATAGAAGTACACCAAATGATAATAAGACTGGTAGTCTATGTCCTGCAGGTGGGTTTTGCACCTCTGGTCAGTTTAAGAGTGAACCCTGTCCTCCAGGGACATATTTTAATAGGGACATAAACACAGCAGGTGCAACAAATAATCAAGATTGTCAAACTTGCGACCCAGGCTATTTTTGTAAGAAATCAAGTAGTCTATCTCCTGATGGACCTTGTAATCCAGGTTATTACTGTTCTAAAGGAGCTGTTTCTCCGAATGAAACTGAATCTTTTCCAGGTCACTATTCATCTCAAGGATTTTCAGATCAAAAACCATGTGATGTTGGTACATATCAACCTTATAGAGTCCAGAAAAGATGCTTTCCTTGTAAACCTGGGCATTTTTGCAACGAAACTCGAATGAGCACTACCCACCCATGTCCTGTTGGTTATTTTTGTCCTGAAGGAAGTGAGGTACCTCAATTTTG
>evgR004171c0g1_i01_Hydra
CCTCAACTCCCACTCCGTCACAAGTAATGTACGCTATAAATAAACTCGTTTGAAATTTGTTTGATGACATTTTTCAAAACATACAAATTAACAAACATAGTTGTTGTGCTTCGAAACAAGTCAAGCAAGGAACAGTGCATTGGTGGAGTTCGGTGTCTGTACTTATTGCTTCAAAGACAAGCTTTTTTCACTATTCAACACACTCTACCATACTTCATTCTCACTCTACCACACTTCACTCATATTCTACCACACTTCCCTCTTACTCTAGCACACTTCACGATCACTCTACCACACTTCACTACCACACTGCACTACCACTCTACTACACTTCAGACTTACCCTACCACACTTCTCTCTAACACACTTCACTCTCACTCTGACATACTTCAATCTTAACCTAACACACTTCATTCTCTTAAACGCTTTACCAAATTGAATCTCACTTAACTATCCTTCCCTCTATACAACCTCTTTATTTTAATCCTATAAACTATAAACTTTAGATTAATCTTTTTTTTTTAGTTATTTAGGTGCTCTAAGAAGTCCTAATGGTCTTGTCACAGAGCACTG
>evgR004176c0g1_i01_Hydra
ACAAAACAATCCAGTTCCATTGAATTGGCAATATACTAATAATTGTCTTCCTCCACCACAATTCTCTCCACTCTTGGATCAAATTCACGTACAATTGTCTCAGGCTCAAGCTTCTGAAAATAATGAACAACAAGCAGTTTCTATTCTACCAAAGTACAAGACTTGTATTGAAGCCTACACTGATTCAAACTTCTTCTTCAATGAATGGGCTAAACGTGAGTTGGAACGTTTAAAGAAGGCAAAACAAAAGAGAAAGGAAGAACGTAAGAAACAACGTTCTGAACGTGCCAAGGGTACTGTTGAAAAGAAACAAGTTAAACAAATGAAAATCCATCGTAAAAAGTACAATTCTGAAGGTCAAATCATTGAATCTGGTTCAGGTGGTCAATCACTCTCTAGAGCTTCCTCTACAAGAAATATGGATATTCCAAACAATAATAGTGTTGGAAGTGGAATGGATGAATATAGTCCAAATAATA
>evgR004181c0g1_i01_Hydra
AACAAAATTCGGTAAAGCTGAACAAACTGAAGATGAAGAATTTACCATTCTCTCCACCAAATTCAAATCAGCAGAAAAGGCTTGCGAAAACTTTGTCAAGCACACCAATCAATTGATTGAAAATTTCCATGCCTTTTCAACAACATTAGCCTTCTTATCTGAAGATTTCCGTGATGTTAATACTCAAGGATCAGCTAGAGTTGAACAAGCAGGTATGGTTTTAGAAAAGGTATCCAAGACTGTTGAAAGTAATTGTGTTCTTCCATTCCAATCTAATGTAACACAAACTGTTGTTGCTCCTGCCAACGAATATTTTGTTTTATTTGAACCTGTTAAAAAGATTTTCAAGGAAAGAAAAGATGTTACACTTCAATATGATTACTATAGACAACAAGTTGAAAAGTTAACTGAAAAACAATCAAAGAATCCACTTGATT
>evgR004185c0g1_i01_Hydra
CTACTGCTAATGGTTTACCACTGTACAACAAAACTATTGAAAACTCGAAACAAAACGACAACTCGATACAAGTTCAGCCAAGTTTTTATATTTCTCAAATAACCTCAACTATTTATGAAAACGAAGCTTTTCAAGAAGCAATTTATTCATTGTATAACACAAATGCAAGTAACAAGCCAGAACAACTCGTAAGCACTTTCGTATCAAAAAATATGCCGCTTTTTCAAGTGTCAACTGTTTCTATAAATTCATTATCTTATGAAAATAGCCAAATCCATCCATCAAATATCCATCGAATCACAACCGTGCACAATTCTTCAAATACAAAGCAACACGTGTTTATCGATTTTTCTGTTTCCGAATCTCAAAAATCACTAGAAATAACAACCTCAACATTGTTTCCCTCAGCTGTAACGTCACCGGTATTATTTACGACAAAATCATTTAAAACTCTTTCTTCTGTTAACTACGGTTGGAACTTAAACAATTATTTTGTTGTTTGTCAACCGGAGGAAAACGTCTGTCTAAACAAAGGAAATTGTTATAAATCATC
>evgR004201c0g1_i01_Hydra
TCTTGTAATCTGCTGTCCAATTGGAATAATAATGCATACATTTGGTTTTTATTTGCTGAGGAAAGCTAAAAAAACTAAAATGGAATCCATTGAAAGGATTATTATAACAAACTTGTGCTTGTCAGAAATGTTTATAAACATCGTCTTGTTAGTTTTAGGGTTGCTCGACTTTGCATCTATTCCAAGTCAAATGGGTTTATATATTTGTTCAAGAATTTTTTATGATGCGTACTATTTCTCAAGTTTGTTTTTGTTAGTCAATCGTTTTCTGCGCATTCGTTTTAGTATTCAATATTGTTCAATTTTACCGATGAAAAAAGTTCGTTTTTTAATACTTGGATTGTGGATTGTTAGTGCAGTGTTTGGTATTTTAGTTTCTAAACTATTCTTGCATAGGAATGTTGTAGGCATTGTCGAACTTTTCTTTGATCTATTAATTTTTATAATGTCAATTTTCGTTTACAGTTATGCAATGAAATTATCAAGAGATATAGCAAATAATATTCATCTGCTGCAAGAGAGAAAACCATTAACTAAAGGACTTCTGTTATCAATGATGATTGCGCTCTCTTATACTATTTTTACTGCATTACCTCATACAATTTTTCTTTGTCACTATTTATCGGGAATAAAGGTTAATATGACAGTAAACTTGGTTTTAGATTTAGTATTAGTTATGGCTTTTTGGTTTGATGCACTTATCTATATTTATATTAGTAAAGAAATCCGAAAATTGCTTAAAAGCACTTTTAAAGTTTTATTTTTTAGAAAAACCGTTACAAGCAACCCTAAATATGACGGTAATATTGCCATGGTTGAATATAGTTAACAACAACTACAAAACGGAATATTTATTTTCC
>evgR004205c0g1_i02_Hydra
TTAATACAAAATATGTTGTATATAACAAAAATATTTAAATGGTTACATATAAAAATTGTTTTGTGCGAAGTATAAAATTATGAGATTTCTAAATACTTTGGCCCAATGGAATGTAAAAGGGATAATCATGTTACACAAATTGTGAAAATGTTGGCAGTTATAGGGGATGTACCAAGAAAAATTGAAAATTTGCAGTTGTTTGAAGATTGTCTTGAAGAAGTAAACAAGTTAGGTGATACATTGCAAACAAACAGTTCTGAAGTAGTGTGTGTTTCTTTAAAAGCTTTAAAGATTATTTTCTTATTTTGCAAGGATGCAAGTATTCAACTTGATGAGAATATTGAAAAACAGCGGGTTGTTGCTCTATTCAAGAATTCTTATCAAAGTTTAAAAGTTTTCATTAAAGTTGTTGAAGGATGGTCTTTTCAATTTCAAGATTCATGTAGTATTTTAACAAATGTCTTACATTCAATATTTGAGTCAATAAAAGTACTTTTGTATTTGGACGCCATGGCTGCAGCATCTATTTGTAAATTGTTTAAAAGATTATATGTCAATCAGAAAAGCTTAGTTTTTAAGCATGTTGAAGAATTCAAATGGTTGATTAATATTTGTGATGTTGTTGAGCTTATTTTCGAAAAGTATTTTCAGGATGAAGTTGTTCAACGTAATATAATGTTTTGTAAGTTTGTCATTGGTCTTGCTCTGCAAATCACTCTGGACTTTAAAGATGTGTCCGATGATGCTATAAAGAAAATTTATTCATTTTCATGTTCAGTTCAAAGAATATTTATTAAATATACTACACAAGACTTGAATTCATTACTGGAAGCTACATTAAAAACTTGTGTTAAAGAACTATTGAGTTATAAAAACGCAAACCCTCACTTTTTTAAAGTCTTGCTTTCTGAATGTGATGGTTTGGAACACCTAATTTTATTGTCGCGTTTCCTTTCTGAATTAAATGAGACATTTCAAGGTGATATTCCATTACATTGGTGTCCTTCTGAAAATAAAGATAGCTTATTAGGATTGTTTTTAAATCTCCTTTACCACAACTCGCCTTTAATATGTGAAGATAGTAAAATCTTGCTACCGAGTTCTTTTGGCATGGTTAATGTCTATGAATATCTACTTGTGAATTTGTGTTCTTATATTGCTAATCTTCCTGCCTGCAGCATCCAAAGTATTACGGATATTATTGTTTCTTCAGTGCTTATATCAAATATTAATTTTATATTTGCTTCCGACCTTCTCAGTTTTATTGCAAGATGGAGTAGTGAAGACTTGTGTTTTCAGTATGTTAACATGCTTGGAGTAATGCTTATGAAACTATGCTCAATGGGAATGGAAAGTACATCAGAATATAACAACATGACGTTGCTTTTTCAAAGATTATTTTTGTTTCTCAGTGAGGAACAACTTAATATCAAGTTGTTTCTAAACAATCACCCTTTCTCTCAAGCGATTTATCTATGGTCGCATATACCCATTGATAAAATGAAGTACGAAGCTGAAGACCTTAATAAGATTGCTGGAATGGCAATTTCTGTTATTCATTTAAATAGTGAAGAACAAGATGGTTCTGAAAGAAATATTGTTGGTGCAATTCGAGTGTTGTGCAATATCTCTAAGTATTTTTTGGATGTTTTGAACACAAGTAACCAAGATGCAATAGTGAAAGAAGTTGTAGAGTTTTGGAGTTGTTTTAATGATCTAGAAGAGAATATTGAAATGAGTGAAAGTGTAATGACTGCAATGCTGTCCCTTGGATCATCTGTTTTTCATATTTTTCAACCCGAGCAAATGTTACTGGTCCTATCTGCTGTTAAAGACATTTTATTTCATTGTGACAATCAATTATTTTGTTTACAAGTGACATATTTTCTTTCTGATCTTAAAAATATACAGTTTGATGAAACTCATCAGGTTGAGTGTCTATCAGTTGTTGCATCCTTGTTTAATTACCTTATAGATCTTAAGGATTGGATAGTTGCATTTTTTGCTATGTCTGCTCTTCGTGATTATGCTGAGGTTACCATTTATTCACATAAGTTGTCAACTTTTTTGCCACCAACCAAGAAAACTGATTTTACTAACTTTCTGCAAAAGAAATCAAAAAATCATTGGTCGGTATTACAACAAAGTACTTATAAAATCTTGCAAAACAACCATGAAAACACAAACCAAGAAATTGACTCTGAAGTTGAGCAATCGATTCGGGAATGTCTTGCTAACATTTTTTCCAATATAGAAGTTCTAAGAAAACATTCAAGTTCCATATCATCTCCTTCTTATTATCAAGTTGAAGTGGAGAAGATGATTGGTTTACTTTCATCTATTGTAAATTAAGATTTGAATACTCTGTAAATTTTGTTTCGTTAATAAAGTTATTTCAGATTTAAAAAAAAAAAA
>evgR004206c0g1_i01_Hydra
CTATTTTAGTGCAAAAAGGATAAAGATAAAAAAAAATGAATAAACCAAATATATTTAAAGGTGTTAAATATTTTTTCACAGCAGACAAAGAGGAACTCAAACAATGTTTAAAAGATGGTGGAGCGATGAGAGAGTTTTATATCAGTGATTTAGTAACACACGTTATCTCTGATAGCACTGATTTCCCTCAGTTGAACCAAGTCGTCGAAAGCAAGATACCAATTGTTACTAGCAATTGGGTGCACATGTCTGTAAAATGTGGAATAATGTTACCCACTAAACCTTTTTCACCAATGGCTGATGGTTTATTTCAAAATATGGTTTTTTGCCCTTCACAGATTCCTGTAAAGGACAGGGAACTTTTGCTTGCCATGATTGTATTTCATGGTGGAGTGTACAAGATTAACTTGACAAAAGATTGCACTCATCTTGTTGTTGGACAGCCGTATGGTAGAAAATATGACTTTGCATTAAAACATCCACAACTCAAGATAGTCACTGCTGATTGGGTTGTTGACTGTTCAAAGGAGGAGCGTATTTTGCCTGAGGAGGAGTATGCTCCAACTTTATCTCAACAATATCCAAGGCCAGCCGCACTTTTTGATGGAAACAGGGAATGTTTAGAAAAAGTAGATGAAAACTTAGAAAAAGTACCAGAACAGTTGCTGGCAGGCTGTGTCATTTTCTTTAGTGGTTATGAAGATAGATTGCAGCCTCTAGTTTACCAAGTGTGGAAAGATGTGATAAGGAAAGCATCAGGTCAAATAGATGAAACATATTCTGACAAAGTTACACATTTTATTTGTTTGCACCAACAAGTTCCATTTTTCAAGAAGGCTCTTGCAGAAAAAAAGAAGATAGCAACAGCTTATTGGTTAAATGATGTTTTAGTTGCAAAAAGGCTTTTTGCTCCTAATACTCCATTGCACCTTCCTTCACCTTTCACTGAAGTTATTGCAGATATTCAAAACAGTCTTGTCACTGTGAGTGGGTATAATGGTCATGAGCGTTTACTTGTCAAGCATATGATTAATTTCTTGGGTGCACACTATACTGGTCACATGACTCGCTCGCACACACATCTTATATGCAAAGCACCTTCTGGTGAGAAGTTCAAAAAAGCTGTAGAATGGCAAATCCCTGTTATAAGCGCTAGTTGGCTAGGAGACATGCTACAAACTGGTCAGTCATTTCCAACAAATGCCAAATCAAAATACACTGCACTTGGCCTTAGTGACGAACTTTCAATAAATCAAACATTTTCTTCATACATCACTGAAGCGTGGAATAAAACTTATCAAGAGGAAAGAACGCAGAATTTAAATCAGGCTGAAGAAAAGTTAAGAAAGTTGGACACATCATCTGGAGTCAAGCGGGTATTATTTACCGGACTAACTGGAAATATAGTTTATCGCCTTCGGCAGAATGTACAAAAAATGAATGGTGAACTGGCGCATGGTGTCAATGACTGTACACACCTCGTTGCACCAAAGATCACCCGAACAGTGAAGTTTCTTTCTGCTGTCTCTATTTGCAAATTCTTAGTTGCACCTGCCTGGGTAGATGATTCATTTGAAGCACAGAAGTTCTTAGAAGAAGCTCCTTATACATTAGTTGATCCGGAGTCAGAAGAGCTGTTTGGGTTCAAATTAAGGCGATCTTTGCAACGAGCTCAAACGAGACAAGTTTGTAAAGGTCTACACTTCCACGTTACTCCCTCTATTTTGCCGGCTCCAAACGCAATGAGAGAAATTATTGAATGTGCAGGAGGAAAGATGTCAGAAATAAATACTTTAAAAGATATCCACAGCGTTTTTGTTACAAATTCTCTTCAGGACCGTACTCATAACAGTTACTTAGTATCATGTATGGATGACAAGGAATTTTGGAGTGAGCTTTCCAACAATGGTTATGATGTTTACAATGTTGAACTTATACTTTCGGGTGTTATGAAGCAAGAACTGGAATGGAATCTCCATCACTTATAAATCTCGAAAATGGTACAATATTAGTGCATTTTTACGCTACTAAAGTTTAAAATCGATTAATCGTTTTCTGAAAAATCTTCGAATGCAAAATGATCGAAAAACAAAGAAATTGTTGTAATGAAAGAAGAAATTACTATCATGTTAAAACTATGATTGAAGTTATTGAACCTCAAAATACAGATAAATATTTGTGTTTTTTAGTTACTAGCTCAACCAGTATGGAGGTATGTTTTCTCACCACGTGTTAGGAATATAATTAATGCACCTGTATTTAAATAATCCGTCTCTTAATTGTGTCTTCTCTTAATTGTGTCTTCTCTTAATTGTGTCTTCTCTTAATTGTGT
>evgR004209c0g1_i01_Hydra
TTTTTTTTTAAATGATTTTCGGATCCATAAATAACTATTTGTAAAATTTGCTTTCGTTAAAAAAAAAATCACAGCACCATGGCAACAGCAGAGGAGGAGTTGTCTCCAACAGTTCAACATTTACTAGATCAACCTTCATTAAAATGGATCTTTGTTGGTGGTAAAGGAGGAGTTGGAAAAACTACCTGCAGTTGCAGTTTAGCATCACAGCTTTCTTTCTGTCGTAAGAGTGTTTTAATTATTTCCACGGATCCTGCTCACAATATTTCAGATGCTTTTGACCAAAAATTTGGAAAGAAACCGACATTAGTAAATGGATACTCAAATCTATTTGCCATGGAAATTGATCCTAACTTTGGAATGAGTCAACTACCAGAAGATGTCATAGAAGAAGAAGGTATTACTTCAGTTGGGAAGAAAATGATGGCGGAGCTACTTGGAGCATTTCCTGGTATTGATGAAGCTATGAGCTTTGCTGAAGTAATGAAGCTGGTTAGAAGTATGGACTACTCTGTTGTAGTGTTTGATACAGCCCCTACTGGACATACATTGCGTTTAATTTCATTTCCTTCTGTGATTGAAAAGTCGCTGGAAAAACTTATTTCGTTAAAATCTAGAATACAACCATTTTTATCGCAGATGACTGGATTATTAGGGATGGGAGATTCAAGTATTGATATGTTTACAAGTAAACTTGAAGATACTTTACCCGTGATTAAAGAAGTTTGCAACCAATTTCAAAATCCTGATCATACAACATTTGTTTGTGTGTGCATCTCAGAGTTCTTATCCCTTTATGAAACTGAGAGACTTGTTCAAGAGTTAACTAGAATGAATATTGATATATCTACAATAATTGTTAATCAACTTGTCTTTCCAAGAGCATCATCTGGTTCAAGGTGTGATCTTTGTGTTGCTAGGTCGAATATTCAAAACAAGTATTTTGAACAGATCAAAGACCTATACGAAGACTTTCATGTTATTAGGCTACCCTTATTACCAAGAGAAGTTCGTGGATATGCTCAAATCAAAGATTTTTCAAACTTTCTTGTTAAACCATACAGTCCTCCGACATGATATTCTTGGCATAATATAGGAAATTATTTTGTTGAAAAAAATTCTTAGGTGTTAAATAAATGGAAAGATGAAAACATTTCATGACAATATTTTATATTTTTTCTCTAATTTGACAAAAAATATTATTTTGGGTAGATTTAATTAGTTTTATTTGAGTGATATGAGTTTATGAATCTTTATAAGTCATAATAG
>evgR004211c0g1_i01_Hydra
TACTTTTTTTTCAACTAATCTTGCTTTCTTGTTTCTGTTATGGCATATATTTGTTATGAAGTTGGACTTGAATACAGTATGCAAAACAACTAACTGATATTACCATTTGCTGATGAATTATTTAAATTATGAAACCTCTTATCAGTACTTGGAATCAGTTTTTGTTGTCATTGGCTGATGAATTAGTAAAAGATAATGTGAGGAAAATCAAATTTATTTTGGGAGATCGCATACCTGCTGGTGTTCTTGAGAAAATAAATGATGCTCATGATCTCCTGAATGAATTGGACAAGCGAAATATAATCTCAAGTAATGACTTATCTATGTTAAGCCAGCTACTTGCTGCTATTGGAAGGAATGATCTTGTATCAAAGTTAAAATATTATACAGATGAAAAAAACAGTGCAACAACCAGCAAAAACAATGCAACAATCAGCAAAAACAATACAACAACCAGCAAAAGCAATGCAACAACCAGCATAAACAATGCAACAACCAATTCCGTTTTAAGTTCAGAAAGTTTTGTGAATTTTGTAGGTAATGCTTCAAATCAGATGGAGGTATGCACCATGCACGCAAGACCTACAAATCAGTCAGAAGTATGCAGCATGCACACAAGTCACCCAAATCAGTTGGAAGAATACAGCATGCACAATGATCCTTGTGGAATATGTATTATATTCAGCAACTATTTCGACAAAGAAGTAATCATACCTGATGAAAATAAAAGTTTAAATCATCGCTTAGGGAATGGAGCAGACGAAATGAAATTAAAGGAAACATTTAGTCTGCTGAACTTTAATGTTTTGGTGTACCGAAACAGTAGTTCAAAAGAAATAGTAAATATATTATATAATGATCTCAAAAACCAATCGATTGAAAATTGTTTTGTATGTTGTATCTTATCCCATGGATTTAGAAATGGAGTTTATGGCAGTGATGGGTCCAGGCTAACATTTGAAGAATTGTGGACCACTGTCAATGATGCATCTTCTGAAAAGCTGAAGGGTAAACCAAAGTTATTTTTTATTCAAGCTTGTCAATCCAATATAGATGTTGATGAAATGTCTTTATCAGATATTGGACTTCAAGTTGAGTTCCAAGATTGTTTGACATCTGTAGCAACTATATCAGGTAAAGTTGCGTTTCGTGATAAAACATTTGGCTCGTGGTATATTCAAAAACTCTGTGAAATTCTAGAAAAACGTGCAATTGATTCATCATTACTTGATATGTTGACAGAACTTAACTCTATTGTCGCAATAAAAGAAGATTTTTACGAGGAAGGTGGTATTAAAAAAAAAGCCACTCAAATGCCTACGATTCAAAGCTGCACCTTATCAAAAAAATTATTTTTCAGACCATTGGTCGAAATGGCTCTCTCACAGTTACATATTTATTAAAAGTTACGTTTTTATTTGAGCAAAAAAAATGTCTACTTTATATTCTTTATAATAGCTTTTTAATTATTTTGTTGCTTCGTTGGTCTTTTAACTTCTTCATTGCTCTTTAACTTTATCCTTGCTTTTTAACTTCTTCTTTGCTCTTTAATACTTTTCTTGTTTTTTAAATTCTTCTTCTTTTTTTTAACTTCTTTCTTCTGGACATCTTTCTTGCTTTTTAACTTCTTCCTTGCTTCTGGGCATGCTTCTTGCTTTTTAACTTCCTCCTTTCTCTTAAACTTCTTCCTTTGTTTTTAAATTCTTCCTTGCTCTTTAACTTCTTCTTTTGTTTTTAACTTCTTCATTGCTCTTTGATTTCTTCCTTGCTTTTTATTTTCACCGTGCTTTATTTTCTCTTAGATTTGTTTTGTTCTATCTTCTTTACTTTCTCATAACTCCCAACTTAAATAGAGAATTAAAAAAGAAAATCATTTAGAATAGATTTTTTTCAAATATTTTCAAATAATATTTACTCTATTTATACAATCTATTTTTCCATTTATTTTAGTTAATTGTTTTATTATTTAAAAAAATGTGTAATTAAATTTAGGTTAATAGTTTTTTTAAATCTCCACTTTTATAATAATCAATTTTTTTAAATCTCCACTTTTATAATAATCAATTTTTTTAAATCTCCACTTTTATAATAATC
>evgR004213c0g2_i01_Hydra
CCGTAAAACTCGCAGACTTTCGCAACATTATAAATAAATAAAGTATTAAGAATACATCCAAACAATGTTTCAACCTCAAATTTATGATACCATATCAAGATGTGCGGTCAAGCTAATAAAAGAATTTGTTTTAACGTCTCAGGCTACATTTTTGTAACATTTTCAGAAACTCTAAATCGATTTCCAGATACACTTTTAGGAGACAAAAAACGTCGAGATGAATATTTTCATTCTGTATCCAACACGTATTTTATTGATCGCTCAGCAAAAGCATTTGAAGCAGTTTTATTTTTTTATCAGTCAAGCGGTCGTTTAATAAAACCAACTGACATGTCAACTAAACTCTTTGAACAAGAATGTCTATTTTACAACTTGGGAGATTCTGTAATTGAATCAATGAAAGTACTTGACGGATTTGATATGATTCCAAATAAAACTATCCAAAGTCAAAGTAACCGTACTTTAAATAAAATATGGCACTTTATTGAAATGCCAGAGTCCTCAACGTCCGCTCAAATATATGCATATATATCATTGCTTTTAATTCTTTTTGTCGTCACAATAGATTGTGTGAGCACAGTAGATACTTATGGCAGCTCTTCACTTTTCAAGCTGTTAAACAAAATAAATTACTACTTAAATGTATTTTTTGCGGCAGAGTTTTTCATAAGGTTGTTATCTTCACCATCAAAGTGGAAATTTATGACGTCACTAGGAAACGCGCTGGAGGTTTTTTCAATATTCCCATCCTTTTTTTTAACGATTGTCGAAGAAAAACATATGGGTGGACTTATTTTTGTTCGAGTTCTTAGAACTTTAAGAATATTGAGACTAATGCGGTTATCCAAAAACTTTCGTTCACTCTCAGTCGTTTTAAGCATATTGGTAAACTGTTTCAAAGATATAATCATGTTGGTGTTCATTATGGCTATTTTGTCTGTTTTTTTCGGTAGTATTGTTTACCACGCGGAGCTGGAAAATCAAGAAAGCCCTATATCTAGTATACTTGAAGGAATGTGGTTCGCTTTACAAACGATAGTCACGTTAGGTTATGGCGATGTAGTTCCAGTTACCTTGCTTGGTAAGATTTCATCTGCTTTTACTGCGGTAATTGGTGCATTGTTAATGATTATTCCACTACTGTACCTTGGAGGGAAATACTTTTCGATATACACAAAAGCTTGTGGAAACACGGCAACGGCTGAATAGAGATTTACAAAAATAATTTAATAATTTTTTTATTTTATTTTTGTTTATCTATTTGAAGATTTAGTTAATAAATGTAGCATTATCGTGTGGAACATGACAAGACCAGGTAAAAATCTCGTCATTTAACCCCTGGAAACATCATTTATATACTTTATGAACTAGATACGATCTATACAACAAATAAAATATAATAAACTAAAAAATATATTTACATATGTTTTTTAAACACATTTTGGAAATAAAAAATATTAAAAAATTTCATAAACATTGTTTAAAAATATTTAAAAGTGAACAGAAATTGAGATACGTCATAACTGTTTAATACATCAAGTTTTGCACCAAACTAGAAAAATAAAATATA
>evgR004217c0g1_i01_Hydra
TGCTAATAGTGATAATCGGTTTTTAATTCATTACATTTTTTTTCATGCAGATGGTGGTGGAAAACTTCAAATTGCCATTCCATCTAATAGTATTTCTAATATAATACGCTCTTTTAATACAAAACCAGTTGGATTGAATCAATGGTCAAACATAGTGATAAGTCAAGCTTTAAAAGACTCAGTTTATAAATTCAAAATCCAACTAAATGGCGAAATTATGATATCTGTTGATAACAATTATTTTCAAAGTTTTGAAAATGTCAGTATATATGCATCAGATCGAAGTTATGATAATGCAGATTGTTTAATTAAAAACTTTTTTGTCATCAATAATAGTGACATGCAGCCAGTTATTCTTCGTCCAACAGTTTATATCAATATACGAAATGAAATATCTTTGACCAGAGGTTCATTGCTTGGAAC
>evgR004221c0g1_i01_Hydra
CTATTAATTTATTATTTTAAATGGCTGCAAGTAATGATATCTTATGTTTTACTTGTTGTACATGTGGTGCACAGAAAAGTTCGCTTAAAAAGTATTTACGACATTTGTATATTATGCATGAGCATACCGTGGGTTTTCTACAAGTGTGCAATATTGATGGTTGTCCTGCTAAATACCCTTCTGTCAAAAGTTTACGTCACCATATGCGAAAGAAGCATAGTTTCCTTTACAATGCCCAAATCAATGAGTCTGAAAATATTAATTGTGAAACAATGTTGTCAGATGAACCAATTTTATTAGAGTCCACTTTTGATAACCCAATTGAAGAGTTTGATTGTATCACCGACCGCAACAGTCAAGTTAGTAGTCATTTTCATTCTTATTTAGATCGCACTTCTATGTCTCCTTCTACTTCTATGTCTCCACTAGACAATCTGCTTTGTAATTTACAACAACATTTTGCAATGTTTATAGTATCAGTAGGAGAAAGGCATTCAATACCATCTGTAGTGCAAGTTAATGTAGCAAATGAAGTTCAAGCTTTGTTGACATATTTTAACCATTGTTTTTGCAATCTTGTAAAGCAGCAACTAGGTAACAACTTTATATATAGTAGTGATTTTAATGAACTTTTAAATTGCGACATGCTTTTAGATACAGTTTTATCTAAAGTTAATTCAAGCAAAAAAATTATTTCATTCAGCAAGTCACATTTAGGTCTTATTATGCCAGTAGAAATAAAATTGCACAGCATAATAGACAATTTATTTCAAGGTGATTTAACAACACCAAAAAGTGAAACTTTTCAATACATACCAGTGCTACCATTACTAAAAAAATTAGTTGAAAATGAGTTTATCTGGAATTCGATCAAACGAAAACTTGAGAAAGATAATTCTAACAATACAGATTTGTTATGTTCATATTCGGATGGTTTCATTTGTAAAAAACATGATATTTTTAAAAATGACAAGTATGCATTAAGAATTCATTTATATTGTGATGAGTTTGAGGTGTGTAATCCCATAGGTGCCCATCGTACAGTACATAAACTCTCAGCTTTTTACTTTTTCTTAGGGAACATTGAAGAACAATATATTTCACAGCTTCAAAATATAAACTTATGTATTCTTGTAAAAGAAAAGTTTATAAAGAAATATAAAACTTACAAGCAAGTTTTAAGACCCTTAATTCAGGATTTGCTGACATTGCAAAATGAAGGTATTTTAGTCATGG
>evgR004223c0g1_i01_Hydra
CTTTTATGGTACAAAGGGTCCAGAATTCGTTCTTGAAAGAAAGAAGCAAGAAACTGATTTGACATCTCAAATGATCTCACACGGTGATTGGAAGAATGCATCATTCAAAGAATACAATTTCAAAGCTGCTGGTAAGAGAAATGAAAATGGTTACCTTCACACATTGTTAAAGGTCAGAACTGAATTCCGTAAGGTTTTCCTTGAACTTGGTTTTGAAGAAATGACCACCAATAACTTTGTAGAAAGCTCTTTCTGGAACTTTGATGCCTTGTTCCAACCTCAACAACACCCTGCTCGTGATGCTCACGACACCTTCTTCTTGGAAAATCCAAAGGAATGTATCTCTGTACCAACAGATGGTTATCTTGAAAAAGTTAGAGAAAT
>evgR004225c0g2_i01_Hydra
ATTTAAAACTAATCTTTTATTTTCTAAATAAGATGTAGTTATTAGTTAAATAATGGGTCAAACATCTGGAAAGTATCGTGGTGATACAGATTTATCCTCGGCTCTTGACTTTAATCCTAAAAAAACAGTAACACATGATGAAACACATGAAATCAAGCCTGAATCATGTGAAAACTACAGTTTGCGTCATGTCATTGATGATGGCGGTGATGGTTTTGTTCCAGTTGATTTGGATGTCTCTTCGAAAAGGTCAACTGTTCCCACCATCTTTCGTTGGGAATATGGTGGAAAAGAAGTATATGTGTCAGGGACATTTAATCATTGGAAAACACGTTTTCCTTTAACATATAGTGGAGGCGAACATGCTGCCATTATTGATTTGGAAAGTGGAGAACATCATTACAAGTTTTTAGTTGATGGCAATTGGGTGCATGATCCACACCAGAAAACAGTTGATGATGAACATGGAGGACACAGTAATGTTATTTATGTCAATGAATCAGATTTTGACCCTACACAGGCACTACTTAATGATGAATGCGGTAACAATGTACCTCGTGGAGCTAGCTCTTCATCACTTAGCAGAATGTCTCCACCTGGTAATTACGGCCAGTTTGTTCCGGAAATAAGTCAAATCTCTCAAGAACCTTGGGCAAAGACTGGCAATTTAATTCCTCCTTTATTACCTCCTCATCTGCTTGATGTTATTTTAAATAATGATGTTGTTGACGAGGATGATCCAATCGTTTTACCTATTCCAAATCACGTGGCATTAAATCATCTCTACGCACTTTCAATTAAGGATAATGTTATGACTTTGAGTACAACTCATCGCTACAAGAAAAAGTATATCACGACGGTCATGTACAAACCTATAAGTGTTCATTAAAATTTATTGTTGCTTTTAAAGCGATTCTTTAATTCATTCTACGTTGTTTGTTTTTTGTTTCGATTCCGATTTTTTTTTTCTTTAAACCTAATTTTTTATCTCGAAAAAACAAGTAGCTTTTTAAAGATTTTCATCTTTCAATGACCACTATTTATTGCGCTAACAGTTCTACGTAACTGGCAACGCATTTTTTATTTTTTATCAAAATTTTGTATAAAATTAAACTTGTTCTAGAAAAATAATGTAGATATTTTATTGCTTTTTTGACATTTTTTTCCATATTTTATTAATGACCTTTATAAGCGTCAGAACTAATATAAATAAAGGTCTTAATTTTGTTCTATTCTATAATATAGAATTAAT
>evgR004230c0g1_i01_Hydra
ATAATATATAAAACAAAATATAATATATAATTGAGTAAAAAAGAAAAGTTAATATGACATTATACTCTGTAATAAATGTTGGACCATACAATGACAATATCTCACAACCATTTACATCACCTGATGTAAACACTTTGTCTCATTCATTTGATTATGAAGATGACAGTTGGACAATTGAGAAGATTACAGCATCGATAGACCGCAAAGGGGAGGGAAAATTTTACAAAGTCAAGTGGGTCCGATACATATGGCAAGCAGAAGAAACAATTCAACCACTGCAACATTTGATTGATAAGTACTGGCAAGATTATTCTGTTGATACCACAGAACTACTTACTCAAAGCATTCAACGACAACCCCCTCAATCATTTGAAAATCACATTGATCATAACCAACTTTCTGGACACACATACAGACTTCATAAAGAAATGCATGAAAATATGGAACAACTTCAAGATTTTTATGATACTAATGCACCATGCAGTGAGTTGAACGATCACAGTGTTGATCATAGTGTTCTTTGTAATAGCACTGGTGTTGGGATACCTGATAGTTTACATGCTACTCAGTCACAATCAGTTTTAGAAAACCATTCAGGCAACATTTTAAATGATGTAACGTCTCCACTTGTTAATTTTGGATTTAAAGCAGCTTTAGTGAAACACAGTCATAATAAAAGTTTAAACGAAGGAAGTGTGCACTCTGATGGTATCATACAAAACTATAGTAAAAGTTTAAACAAAGGAAGTGTGCACCCAGATGATATTATACAAAACTATAAGCAGCATGATGATAATCTGCCAGAAGCAAATTTTTTCTCATCTAAAACTGATGAAGTAAAGTTTATCAGCAGTGGGGATAAAGTGAACTCAGCAGCTGATTTTTATCAAGATCGTGTAAATTTAGAACTACCTACCGAAACAATCGTGGTTCAAAAACAGACAAAAGAATTCATTTGCGAAGTATGTAAACGAACTTTTAATCGCAATAGTTCTCTTAAAAGACATTCAATAATTCATAGCGGAGTGAAGTCTTGGCGTTGTGAATACTGTAACAAGCCGTTTACTCAGAAAGAAGTTTTGATTCGTCATTTAAAAAGTCACACCGAGAAAAAATCATACAAATGTTCGGAGTGCGACAAATGTTTTGTCGAGAACGCTGCTCTACTTCGACACCAGGCTGTTCATAAAATTGAAAAAGATTGGTTTTGTGATATATGCGGTAAAAGGTTTGTGATTAAAGAATATTTAAAAAAACATTTACAAATTCACAGTGGAGAAAAGCCATACCGTTGCTACGTGTGTAGCAAATGTTTTGTTGACAGTTCCGCGTTAAGACGACACGAAAAAACTCATAATAAATCACTTATGTGTGAAGTATGCAATAGTTCGTTTATTAACTCTTTAAGTTTTACAAAGCATAGATGCACTTAGCACCATATTGTAAGTTGTAAAATTAACAAAAAATGAAAATAAACCTAAAATAAACCAAAAGATCCTAATTTTGAATAGTTTTTAATCAACAATTCAAGAAGCACAAAATATATAAAACGTTAGTCAACTAACGGAAGGTATCCGTTAGTTGACTTCTGGCTCGCTTTGTTAAATATTTTTTTTCAATTCTAGAATACTTAAATATTCTTATATTATAATCTTATATATAGGCTTATATTTTATCATTTAAAACATATTTTTCTCGTATTTTACGTAATATATTTGTATATGTATATATATTTTTTAAATAAAGATATTGTCTTAAAAAAAAA
>evgR004231c1g1_i01_Hydra
ATTTGAGAGTTTGTTGAAGACTTTATTTACAAATAGTAAGTTCAGGATGTTATAAGTGTTTCAACTTCATTAACATATACTTAACATTTTTGCAATTATCCCCCTGGTTTCAAAAGTGAAAGTTAATATTAAGGTCAATTTTAACGCATAGCAAAAACTATAAATACAGGTGAAAAATATTTTAAAAATCATCCCATTAAAAATGAAGGCGTTTATGATTATTTGCTTATTTTTGACTATAGCATTCTCGTATGCAGAAATTGATTTTCAGGATAATTATGATGGCGCTGAATCTGAAGACATCTTTTCTAATCTAACAAACGAACTAGAAGACAACGACGAAGAAAGAGAAGAAAACGACGATGAAAATTATGAAATTCAAAATGAACTTCAATCAGATCCGGCAACTTTTATCTTACCAAAGGAATCGAAATGGTCAAAGTTAAAGAGGAAGATGAAAAATTATAACAAAAAACTTAAGAACAAGTGGCGAGAAATTTGGCTCAGAAGAAAACTCAAAAAGCACTTAAAAAAGGAACAATAAAGATCTAATTTTATTAATTAAAAAAAAATTAAAGAATGTAAAATTAAAAGTTGTAAACTGTAAACAAATTAAAAAAAAAAAAAAA
>evgR004233c0g2_i01_Hydra
TTCTATTGTCTAAATAAGGACTTAAAAAAGAGAATGAAGTTTTTTATTTTTGCTAGTGTATTTCTGATCTTGATGGTTGATGGGGCGTATAACATTGGGGCGTATAACATTGACAACAAAGGGGTAGTAGATTTAAAATGGGTGTTGTTACAAAAGAATGCTTGTTTTGAAGCAAAAAATAATGCACCCGCGATTGTTCCTATCAATAAGAAAGGTTGGTTAGTTGCAGCTAAGCTTGTATATGCTAGTGGTGCTTTGAAATGCTTTTCTGGTTCTAAAGGGAGCAACTTTGGTTGTGTTAATGATCAAACTTTAATTAACGTCTACATTTGTGATGACAAGAAAAAAGTAATATTCCCGTCGCCAAACCAAGCTATTACTTATGACAGCAGCTTTAGATCGTACCAATATCGTGGCTTTAATGCAAACGATAACGAACTAATTTTTACCGAATTTAGCTATCCAGTCCCTTACCTAAAAGAAGGACAACATATTACAGTTTGGAATGGAGAAGATTTATTTAACCTTGACGAAGAGAATAACAACGGGAAAGTGTGTGTGGATTTATATGGTTCATTTATTGAATCTTTTAATTAGATGTAATAAGTGCGATTAATTAGTGCGTTTGTTTTTTTAAATTTAAATACAATTTTGTTTTTATTATTTTA
>evgR004236c0g3_i01_Hydra
TGCAGTTCGTTTACAGTACATGGACTTACAAAGGTTTTTACAGGTAATTGGTTTGAAAAACTGTTTTGGGGGTCTACAATCCTGTTAAGCCTATGCGTAGTTATATTCATTGTTTCAACTTTGACTATTAAGTTCAACAAAAGAGAAGTTTATACAAAAGTTCATGCAAGGGAGTATTTAGCTTCAGACATGCCCGTATTGACAATTTGTCTAGCTGATACGCCACAGCTTAGCAACACAAGTTATTTAATCAATAATATCTCTATGGATATAAACAGTACTAACGAGTGGAAAAACGATCATATATATTTGTATGTATATTATTTGATGAAGGAAGGATATTATGTTGGCGATTTTATTAAGCCTTCATATTTTAACTGCGTAAGGATGGATTTAAGTTTTTTAGACACGTCATCAAAAGCTTTTATTTCTTTTGCAACAATTTATCCAAATGAAGCCATTCAAGTATATGTTCACGATGCTGAAGAACTTTATCCTTTTTTTCAAAAAAAGCCAATTGTGATTTTCCGCAATAACACCTGTGAAATGAGTATAGAAATGAAAGAATATAGAAGATTACAGTATCCTTTTCCATCAAAATGCGTTGAAAATGTTGATTATCTAATTTATCCTGGTAAATATTCAAAGCACAAATGTCAGGAATCATTAAAGTGTCTTCATG
>evgR004238c0g2_i01_Hydra
ATTTGTTGAAAGTCATGTTGATAGCGTAGAGTTTGTTGAAAGTCATGTTGATAGTGTAATATTTGTTGAAAGTCATGTTGATAGCGTAGAGTTTGTTGAAAGTCATGTTGATAGTGTAGTATTTGTTGAAAGTCATGTTGATAGCGTAGAATTTATTGAAATTCATGTTGATAGCCTAGAGTTTGTTGAAAGTCATGTTGATAGTGTAGAATTTATTGAAAGTCATGTTGATAACATAGAATTTATTGAAAGTCATGCTGATAGCGTAGAATTTTTCGAAAGTCATGTTGATAGTGCAGAATTTGGTTGAAAGTCATGTTGATACTGTAAAGTTTGTTTAAAGTCAAGTTGATACTGTAAACTTTGTTGAAAGCCTTTTTGATAGCGTAGAATTTGTTGAAAGTCATCTTTGAAGCGTATAATTTATTGAAATTCATGTTGATAGCCTAG
>evgR004249c0g2_i01_Hydra
TGGTGAAAGTGATTTTGAGGATCTTAGTGAGTCAGACTCTGATTATGAATCTAATGTTTTAACTGCAGTTTTAAACAATAATAAGGTTGTGCATGATGACAATAATGACAACATCCTCTTTCCAACATCGGAGTCTCTTACTCAGGAAACTAGTACAGATGATGATATGCTTTATTCCAACTCTTCAGCCCAACCCCCAAAAATAATCAACAAAAAAAAAAGAAAGAAAAAAAAGAGTTTACCACTTTGGAAAAAGGTTCTACTAGATAATGATTTAATTGGTGATATTACTTTTCAAGAGGTACCTATTGAGCCTGTTCCTGATGAAACTACCACTCCATACAAATATTTTTAAATATTTGTTGATGACAATCTTATCGACATTATTGCTGAACAATCTAGTTTATATAGTTGCCAGTCATCATGTACAAACATTTCTGTTACATCGAATGAAATAGAGTCTTTTATTGGTATATTTTTTCGTATGGGTCTTGTAAAACTACCATTAGTGAGATCTTATTGGGAAACCTTTATGAACTATGATGGAGTAAGTTCAGTATTATCAAGAAATAGATTTCTTTTTATTCTCCGCATTTACACTTTGTTGACAACTTAAACGTGACAGATGAAGTAAAGAAAAATGATCGAGCTTGGAAATTAAGACCTTGGTTGGAAAAACTGAGAGAAAATTTCTTAAAAGTTTCTCCTGAGGAAAATCAATCAATTGATGAGATAATGGTTGCCTTCAAAGGAAAGTCATTTTTGAAGCAATACCTTCCAAAAAAACCAAATAAGTGGGGGTTCAAACTTTGGGGTAGATGTGGTGTCAGTGGGTATTTGTATGACTTTGATTTATATCAAGGAAAGGAAATAAAAAAGAAAGATGAAGTTTCTCCTCATGGTGTTGGGGCATCTGTTGTAATCAAAATGACGTCAACTTTACCAGAAAATCATAATTTTAAAATATTTGCTGACAAATATTTTTCTTCATTGCCTTTGCTAGATGAGCTGAAAAAGTGTCGAATATGGTATGTTGGAACAGCTCGCCTTAAGCGCCCTAACAAATGCCCATTCACTTCAGAAAAAGAATTGAAAAATCAAGGACGTGGATTGTTTGACTATCTTACTGACACCAAATCTAATAACATAGCTGTTTCTTGGATAGATAACAAGGCAGTTACTTTAATTTCATCTTTTGGTAGTATTGAACCAATCAACACAGTGAGGCGTTATGATAGAAAAACCTGCAAATATATTATGGTAAAGCAACCATATATAGTCCAGTTTTATAATAAAAATATGGGCGGAGTTGACAAACTGGATATGATGTGCTCATTCTACAAACCAAATTTAAAATGCCA
>evgR004252c0g1_i01_Hydra
GTTATGTTTTCGGGCCCATTTTATTTAACCCCAGCAAAACCCCCTTTAAGCGTCGTGCTTGTTTTCAAAACAATCGACAGACATCTCACATCAAATCGAGGTCATAGGAGAGTACTCCAAAATGGAAGAAAATGGGGTATATTCAAGGTCCCAAGATAAATATGTAAATCTTCATGAAAAGTGGGTGACAAAAGGAAATATTTCTTATATAAATTCGGATAGGGAAGAGTATACTGACGGTAATGGTTTGTCGACAGACTATGATTTATCTTTGCAAATTGAAAACATAAAGCTCAAGTCTAAATTAGAGCAGGCTAACGCCAATATCATGAGGTTATTAAAAGAAAAAGATGATTTGCAAGAAGAATTGCGTTCTACCAAAACAACTTATCATCATTTGGATATCAAAGGGGTAGTCCAAAAGTTAGAGCAAGAAATATATGGCTTGGAGGAACAGTTGGATCGGCAAAGAAAACACTATAATCAAGAACACACTGTTCTAAACAATGAATTAAGAGAGGAACGCAAGAAACAAGAGGAATGGAAGAACACATATATAAAATCACAAATTGCCATAATGTCGTTCAGAAGAAAAATTGCTACCATGAAAGACAGAATTTCATTTGAAGACAGAGCAGAGCTTTCAAGATGTTTGGAAATTGTTATTGAAAGTGAATTAGATATCTCGTCAAATTTAAGAGATCCAAATGCTGATCATACTAAAAATATTTTAAATCATAAAACTACTGAACAATTAACTCATCATAAAGATGGACAGTTAAAATACAACACACAACTAAATCACCAGATGCGAGTTTCTAGCGAGCTAGTTAATAAATATTCACCTTCAGTAATTATTCTTGATGAAAAAAACTCTTTCACTGATACTAATAGTCCAACTCAATGCAATGAACTTGATAATGGTTTTAATAAAAGTTTTAATTCTGTTGCGTATGGATTTTCACATAAAATTGATATGGACTCAGGTGTTAAACAAGAAAATCAACCTGTTTATAAGCACTGGCATAATTATGATTCACACAACTATCCTCGGTTTATTGATAAAAGAAATGTTTCTCAAAAACCTAATGAGTTTAAATCTAAAGAAAATAATGTTTCTTATTTAGATATTAATGCTAACAAATTTTTAGAGCTTAAGTTTGATTCTTCTGCTGTCAATCCTGATTACATAAATAAGGAGCAGACTAAGAATACTACTGTTGAAGCTTCAAATTCTCAAAGCTCTCCCTCTAAACGAGTAATGAATGAAGGAAAAGTTCCTGCTGGTACCCCTTTGGCAGAACATCGTAAATGGTGTTTAACTTGTGATATAGAGGTAACAAAGAAGTCTTGCGGTCACCAAAATTTGATAACCCAAACAAATGTAAAAGAGTTTCCATGTCCTACCTGCTCTCGTGTTTTTAATAACAGATCTCATCTTAAAAGACATAACATGATTCATTCTGGTGAAAAGCCGTGGCCGTGTTCATACTGTGATAAACGGTTTAATAGAAAATCACACTTAACTAGACATTTACTTACTCATACTGGTGAAAAACCGTTTAAGTGTCCGATTTGCAATAAAGGTTTTGCAGATAACAGTGACTTAAAGCGTCACCGTCAAACGCATCTAATGAATATACCCTCGACTGCTACATCTCCAACAACTTTAACAGTTAGTGTGACTTCACCATCTTTTCCGGATTCAAATTATTCCTCCAGACCAAATAGTTTAGTAACTATACTTCCTAAACTTTCTAAAATAGAAAACGCAGACAGTTTATCGTCTGGTAATTTGTCAACAACACCTTCGCTTGAAAATTCTTCTTTAAATGTGTTCGACAAAAAAGAGCGTAAGAAGGGCGACCCCCGGTTTCCATGTAATAAATGTCATAAAGTGTTTACAAGAAAATCCCATTTAACTCGTCATAAGTTAACTCATGACTTAGAAAAAATTTATTGCGATTGTGGAAGAGTTTTCAAACAAAAATCTCATATGCAAGCTCACTTGCCTTCGTGCAAACGAAGACGCGATGCTATTGAAGCAAAATATAAGGCTGAACATGGTGACTGTAGCGATCTCGATCTCTCGACGTCTATGGTTATGAATTCTTTATATTCAAGCAATACCTCTATAAAGACTGAACTAGATAATGATAATTTTGGGGAGAATGATCAAATACCTAATAATATTGCGCTTAGTAGATTTGGAACTTTGCCTTCACAAATTTAGTAAAATGATTTATAATTTTTAAACCCAAATATCTATTTTTTTATATATTCATTTCTTTGAATATTTAAGTGAAATCTCTCAGACATTTCTTTTTGATATGTGCTTAGTCATTTAACAAAATTTTTGTATTTGTGTAAATTTAAACGTAATTTAATTGTAATCAAACGATTTAAATGTGAATGTGGAACGCGAAAAAAAAA
>evgR004258c1g1_i01_Hydra
CTGCTCAAACAACTCTTGCCGCAACTGCTCCAGTAACAACAACAATAATGCCAGATTCAACCGCTGCGGCTCAAACTTCTACAGCAGCACCAATAACAACAACAGCTGAACCAGCAACGACAACTTCTGCTCAAACAACTCTTGTCGCAACTGCTCCAGTAACAACAACAATAATGCCAGATTCAACAACTGCTGCTCAAACTTCTACAGCAGCACCAATAACAACAACAGCTGCACCAGTAACAACAACTACTGCTCAAACAACTCTTGCCGCAACTGCTCCAGTAACAACAACAATAATTCCAGATTCAACAACTGCTGCTCAAACCTCTACAGCAGCACCAATAACAACAACAGCTTCACCAGCAACAACAACAGATGCTCAAACAACACTTGCCGTAACTGCTCCAGTAACAACGACAATAATTCCAGATTCAACCGCTGCGGCTCAAACTTCTACAGCAGCACCAATAACAACAACAGCTGAACCAGCAACGACAACTTCTGCTCAAACAACTCTTGCCGCAACTGCTCCAGTAACAACAACAATAATGCCAGATTCAACAACTGCTGCTCAAACTTCTA
>evgR004262c0g1_i01_Hydra
TACATTACATCACCAAATTAAAAAGAGTTGCTTTCAACTTCATGAAATTTACTTGCAAAAGTTGCTAACGTTTTTACATTTTAAAGACGAACAAAAGAAGACTAAATGAAATTCAATGAAAAATGGCCGTAGCTTCTTCCTGCAATGATACACATAACATAACATCATTGTCTATGTTTTCATTTGGTTTCTCTTCGTTACTATGCGTGGGAACAGTTACTATAAATGCTTTATTAATAGTAGCTATATTAGGAGACAAAAGAAAAGTTTTTAAAAAAGCCGTTTTTTATAAACTATTACTCAACATAACAATGGCCGACCTACTTACTGGTGCGGTAAACGATGCAAGTGCCGTCTCATTTCATTTTAAAGAAGCCAACAGAAAATCTATTAGTCGTTATGAAGTTCTTTTAGTTCATTTAGGACTTTTTATGTTTAGCAATGTTTCTATAATATCGATGGCATTATTGTGCGTAGATCGCATTATAGCGCTAATAAGACCAATTGTGTACAGAAAAGGATTATCTAATCAAACTTGCAATTTAATATTAGTTTCAACTTGGTTTTTTTCATTGCTACTATTGCTGCCTTATTTTTCAATGGGCTATGTTAAATACCTTACCGTGTTTTTATTTACTTCGGTGTCCGTAACTTTTATCGCACTCCTCTTGATGATTTATCTTTATAAAACAAGATTCGTTTTATCTTGTAATTTGAACAACTTATCTTCGACATCTGACAAAAAAAAAGAAAACAAAACTTTCCAATCAAATCAAATAACTAAAGAAAATCAGTTCAAAAGAGTTTCTAAATTAAGGAGCAACGATAAAAAATTAAACGAAAAAGAAAAATATATTTTGTATAAGCTTGGTCTATTTAATCCAAATTTTATCTTTAATCAAAGACACGATAAACAAGTCACCAATCAAAGCAATTTTATATTCAACATCAATCAAGTTAGTTTGAAGTCGTACTCTGCAGAACAGCGAGTCAATCAATCTTTTATTATTATGCTATTTGTTTTTTTTATAACTTATTTTCCGGCTTGTGGGGTAACTTTGTATCTAAACTTATGTATAATATGCAACTGTACTTTAATACATGTTTTGAGAGATTTCACTTATCTAACGCTGTTATCTAGCGCTTTGTGGAGGTCGATCAACTTTGCATGTCGAATTACAACATTGAAGAAGAGAATAAAAGATATTATCACACAAGTAAAACGCATTAAAAAACAAACAAATTCTACATGACCATATAATTTGTTGCTAATAGTAACCTTACGCTTTAAAGCAAAACGTATTACTATAAGTTTCTACTAATTTGCTGTAAAATAAGTTTAACGTTGAACTAGAATCGAACATTTTGATTGGTATAATAAATAAAATTTTGATTGGTAGATCATTTTTTAAAAAAAAA
>evgR004272c0g1_i01_Hydra
GCATATAAGAAATATTTTCAGCATTCTTGTAATAAACAAAAGTTGGTTAATAATACTAAGAAAAACTATAATATTGAGAATATAAAAAAATCGATTTTACTCTGGAATGGCCGCTATGGTAGAAATAATTGGGGTTTTCCAAACGATGCCACTAAAAGTTTTAAAGTTTTAGGTTGTCAGGTACACAACTGTTTTATAACTACTAATAAAAGTTCAGTTTTAGAATCGGATGCAGTTATCTTTCACATTATTGATGATCTATCATCTAGACCGAAACAAAAGCGTGAAAACCAACGATGGGTTTTCCTCTTGCATGAACCTGCATGTGGTGACTATAATTTTTTAAATGAGTGGAATGGACTTTTCAATTGGACAATGACATATAGATTAGACTCTGATATTCCGATTCCTTATGTCCACACATGTAAAAAACGCCGTAAAAAAAGGTATTTACAAAAAGAAAAAATTATATCTAAAAAGAAACATTTAGTAGCTTGGATAGTTAGTAATTGTTATGCACAAAGCCGCAGGATAGAATATGTAAAAGAATTAATAAAGTACATGCCAGTGCACGTGTTTGGTAACTGTGCAACTCATTTTGGGCAGGCAAATAATTGCGACAGAAGACTTGAGGACAGCTGCATGAAAATGATAAGTGATCAATACAAGTTTTATCTCAGTTTCGAAAATAGCTTGACCCAGGACTATGCAACTGAAAAGTTTATAAAAATTCTCAACACGTACGCTGTTCCTGTGGTTCGTGGCGACGCTAACTACTCTCTTCTCGGTCCCCCTAAATCTTATATTGACACTAAGAACTTCGTGTCCCCTAAGCATTTAGCAAACTATCTACTACAACTTGACAAAAATGATGAAGAGTATTTGGAGTTTTTGAAATATAAAGAGGACTATAATAGTAATAATTGTGGTCCTGAAGATATGATTGAAAAAGCCTTTTGTAATATATGTAAAAAACTAAACTCCCGCAATGCTGAAAAAAAGGTTTACACAAATATTTATGATTGGTTTTTCAAATGCCGAGAATCTGATGACATAAATATTGCTATTTAAAGTGTGAAAAATTTCGCTAATCTTATCTTACTTAACTGATGCCAGAAAAAAATCATCAATAGAATCATTATTTGCAAATATTATATGACACAAAACACGCAACATATTTTTTTCTTTATTTAGTAAATCCATTTTAATTCAATAATTTTTACATTGTTTACAACCGATATCTTTATTTAAATAAATATAACAATTTATTTTATCTTTTGAATTTTGGAG
>evgR004275c0g1_i01_Hydra
TAGCGAGGTCTTACTTAGTAACTAGGCAGTGTGTTTATAAATTAATTTGTTCCAATTTTAATAACTACTTTATTGTTAGTAATTATAATGTTAACACAAGAAACCAAGGTTGCTCGATAAAGTTCCCAAAAGTCAAATTAGAGTTTGGAAGAAAAAGATTTGATTTTATGGGTGCGAAATTATACAACGACCTTCCGACTTCAATTAAATCAGTTGACAATTACACCTTGTTCGAAAAACTCAATGAAACTCAGTATAATGATGATCGCTCACAAACATTAATCAGCAACGTATGGGTGGGTCAGAAAT
>evgR004276c0g1_i01_Hydra
TATTGATAAAATACAAACTGAAATATTCGGTATCAACAATCCATCAACAATGCAAACAAAAAATAATATCGGAAACTGTTTGTTTAAAATTGGAAAATATAACGAAGCTCTAGATATTTATTATTCTATTGAAGAAATAAAAACGAAAGCGCTAGGTATCAACCATCTATCAACAATTAATACAAAAAACAATATTGGTAACTGCTTGTGCAAAATGGCAAACTATAACGAAGCTCTGGAAATTTATTATTCTGTAAATAAAATGCAAATTGAAATTATAGGTATCAAACATTCATCAACAATAAAAACAACAAATAATATTGCTAACTGTCTCTACGAAATGGGAAAATATAATGAGGCTCTGGATGTTTTTTATTCTATTGATAATATACAAACTGAAGTATTAGGTATAAACCATTCGTCAACAATGGAAACTAAAATTAATATTGCAACCTGTTTGTATGAAATTGGAAATTATAACGAGGCTTTAGAAGTTTATTATTTTGTTAAAAGAAAAAAAACTGAAATGTTAGATCTTGGCCCTTTATCAACAGTAGAAACAAAACACAATATGGATAACTGTTTTTATAAAATTGGAAAAAATAGTGAAACTTTAAATATTTGTCATACTGTCGAGAAAATAAAAACTGAAACTTCTTGTATAAACCATTCATCAGCAACTATGGAAACAAGAATGAATATTGCAAATTGTTTGTACAATATCGGAAAATATAGCGAAGCGTTAAATATTTATTATTCTGTTGATGAAATACAAAATGAAGTTTTAGATATTGACTGTAAATTAAAAATGACTTCGAAACATAACTTTGCAAACTGTTTATATGAAATGGGAAAATATAACGAAG
>evgR004277c0g1_i03_Hydra
CAAAAAGTTTAAAAAATGACGCTGATGTTTCAGAAAATAGCTTTTAGTTGTTTCATGTTAACAGTCTGTTCTGAAATTATTTCATACACACCTCCAACTTATGCAGAGCTGACTTCCAATTTGTCATCCAAACCTCTAGGTTATAGCACGGGAAATTTGTACAACCCACCAAGAGTTTTCGGTATAACAATATCAAGTCTTAACAATAACTACCTTGATTTAAAAATTCAAACAGTTAATGAAGGTGTTGCCATATGCTTTAAAACGAATCAAATCAATGGTGAAAGCACCTGTGGAAAGGATAAGCTTCCTAGTTGCTTTATTTCGCCAACTTCTTCAATAACAATTGAGTTTTACTGCTCAGAGAGTTGTTCTACACAAGATGTTGAAAGTTTTTATTACAGAGTTGTCATAAGTCCAGATGATGCCCTAGACACCTGGTGCAGCGAACGCCAGACAACAATAGAATATCCTAAAGATCTAGAGGTTGTTGAACTTATTACGATAAAACCTCGTGCAACTAAAACTTATACTCTTAACAACGCCTTTAGTTTGGGCCCCTCAGTTTTTGTTTTGTTTATTTTATTGTTTGTGATGTCATTTAATATGTTTTAATTGTTGCTATTACTTAATAGATGATAGAAAAGTATAGATATTTTTTTATAATTTTGTTAAAAATATATTTGTAAATATATTTTGTGTATACTGTTTTTTATATTTGTTATATATTTTGTACATGAATGGGC
>evgR004291c0g1_i01_Hydra
AATACAAGCTCGCCGTTGTCGGTGGTGGTGGTGTTGGTAAAAGCGCCTTGACTGTCCAATTCATTCAAAACATCTTTGTTGAAGAATACGATCCAACCATTGAAGATTCCTATAGAAAGCACGCAAAGGTTGATGACAAGCCATGTTTCCTCGAAATTTTGGATACTGCTGGTCAAGAAGAATACAAGGCTTTGAGAGATCAGTACATGAGAACTGGTGATGGTTTCTTGTTGGTTTACTCCGTAATTGATAGAAAGACCTTTGAAGAAATTAATGATTTCTATGAACAAATTCTCAGAGTTAAGGATGCTGATAAAGTTCCAATGGTTCTTGTTGGTAACAAGTGTGATTTGGAATCTGAAAGAGTCATCAGCAATGATGAAGGTAAACAATTCGCTAGACAATTGGGTATCCCAAGTTTGGAGACATCAGCCAAGCAAAGATTGAACGTTGATGAAGCTTTCATGGCTCTTGTTCGTGAAGTTAGAAAATCAAAGCCTTCTGCCGCTGTTGGTAAGGAAAAGCAAGGTGGAAAGAAGAAGGGATGTAACCTCTTGTAAATTCTCTTTCTCAAACATCAACAAAACTTGAACACTCTCCCTGTATGATGTATCATTTTACTAATTGGTAATATATAGCTGCGTTTTCTAC
>evgR004295c0g1_i02_Hydra
CAGAAAATGTGGGTTTAAAGATTCAAAACAGTCTTAATAATATTGCAGTCAATCAAGCAACTATCAAGCGATCGAAGCAGTCAAAACACTTGCTAAGTTGATGCCATCTGTTAAAGTTAATGGCGATGCAGTCCACGTAGATCCTAATGTATTATTTCAAAGACTAGTTATGCTAATTAAGAGAGTTGAAGATTTGACTAACTACTGCGAATATGAGTTAACACCGGAGCCCACATCTCTCTTCAAAGATGGATTAATGAGGAAACCAAACAAAGCAGAACTTGGTAGAGAACTTATAAAGGATTCCGCAGTTTTAACAGAGAATAATGACACAACCTATATTCTTGACGGTGGAGCTTTGCTGCATCGAGTTTTGTGGAGTCTCCCAGCTACATATTCAGACATTATTGA
>evgR004298c0g1_i01_Hydra
ATTAATTGTTAATATATATCATTGGTTCACTTATTGTCAATAAGTGAACTAATGCTGTTCGGAACTAGGGACTGTTCGGAATTAGGGAAATCTACGGTAGATATACATAAATAAAAACATAATAATATTCTGTCGTTGTTGATACATGATATATGAAAATGGCGAAATCAACTGGAATTCAATGTGCTTCTTATGGATGTTTTTCATATAGCTATGATTTTGTAGATGGAATTAGAGTTTCAACAAATATTAGTTTTTTTACATTTCCACATGAAATAGCAGAGAGAAATATTTGGTGTAAATTAATAAAAAGGCTACCTAATAAAGACAACTTCAAGATATTAAAGTCTACTCGAGTATGTCAAAAGCATTTTTTGCAAGAAATGATACATAGGACTCCAGGAGGAACACGGAAAAGACTAATAAAAGGTGCTAGGCCAATACTTCACTCATGGAATGACTTTTCAATTTCAGATGTATCCAAAACGCCACCAAGTTTCAGGCATTCTCCTCGAAAAAAATACAGACCCCAACTTGAAAATGCGGATGATTCTGCATCAACATCATCATTAATTTCACTAAAGGAAAATGACTTAACAAAACATATAATTGAAGTTGAATCAGCTGATTCAACTTTACTAGTAGAGGGTGAAAACAAACCTAAAGAATAATACAAAGTAAATTACACAAAGTATACAATAATACTCAAATTTACTTTTTTAAAACGTAAATCAATAACTTTTTTTCCAAAATTCGTTTGTCTCTTTAAAATATATTTTTTTAATTTTTGATTTTTAAAATTATGTTGAAGTTAACATCTATATATTTTAAAATGTTTTGCAAAGTAAAATACGATAAACTAGTACTAATAGTAATCAAAG
>evgR004304c0g2_i01_Hydra
CTTAATTTTCCGCTTGTAATACAAAATTACCAGGTACTTTTTCATGTTTCATGCAACATTGCAAAAGGAGGTGTTAAAGCCTATTTTAAAACTAACCTTACCGTCTATCAAGTTACAGAATCCCAGCTACCTACTGAAATTATCAAGCAAGTCTGGTGTGGTCTCAATGTAGAAAAGAATTGCATTCTTATTGGTTGTGTTTACAGGCTCCCATCATCAGACGTAGCTACAAATAACAAAATAAAAAAATCAATCAAAAAAGCGTCCTCACTTGTAGATCAAGTCAAATTTAACCATCTTTTATTAGCTGGTGATTTTAACTATCACATTATTGGTCAAGGTCTGGTAAATTATAAAGAAC
>evgR004307c0g1_i01_Hydra
TTTTTTTAAGCTTACAAATAATTAGCTAAAATGTTGTGTTACCAACTGTGCTTCATGCTAGTGACAACAACAACACTGGGAGGAGTTGATGCTTCTTTATCAAATGAAAATATCCCATCGTACTTGCTTGATCTTGACATAAAACCTGAACTTCGATGGGTAGAAATAACTAAGAAATACTCACAACTTGCTGATCCTTTCCGAAATTTAATTCAAGCAATAATTCCGAAAAGGTATGTTAAGATTGCTAAAATAGCTATAAACGTTTTTCATTTAGAGAATTACTTCCCTAAGGAATACGCATCTGAATTAGTTGGTGTAGCTGAAGGCTTAAACTTGACTTTGGCAGAAGTATTTCTAATAAATGTAATTTATGAAGCTGGGGCTTTGTGCACCAGTATCATAGCAAAAAATGAAAACGGGACGATATTTCACGGAAGAAATCTTGACTATGGCTACCCAACAGTACATCTTTTAAAGGACTTAACTGCAAATATTGAATTTACGCGCAAAGGAAAATTACTCTATAAATCAACAACATTTATCGGGCAGGTTGGATTTTTCACTGCAATGAAACCTAACAAATTTACCGTTACCTTAAATCAAAGAGAAACATCAAATTTGCTGGTAGCTATATTAGAAGCTATATTTGATAGGAAAGCAAAAACAGCTGCAATATTTATACGCGAAGTTGTTGATACCTCTTCGTCTTATACTGAAGCTGTTTCTATGTTATCTGAGGCTCACATGATTGCTCGACTTTATTTTATTGTTGGTGGAGTTGCACCTAACGAAGGCGTAATAATAACTCGTGGACAGTATTTCTTTAAAGAAATGTATTATCTAAATGAAACAGAATATGGCTGGTTTTTGGTTCAAACCAATTATGACCATTGGTTAAATCCTCCATCCTGGGATGATAGACGTCATCCTGCTATACGGCATATGATACAGATAGGTATTCACCAATTTTCGTTAGATGGCATGAATGAAGTATTGTCAACACCCCCAGTTTTTAACAATCGTACAATTTATACTGCGCTAATGTCGGCTAAACATCCAGAGTATTATTTTTCGTATAAGCACAACTAAATAT
>evgR004319c0g1_i01_Hydra
TGCTAAAGTGTGCCAAAAACATTTATACATAATGGAATCAACAGTGGAAATTCTTACAAGTAGTAAAAATCATAACATATTTACACGAGCTTTATCATTTACAGAAGAACTTGGTACTAATCTTATATTTAAAAGCAATCCGGATTCTTTTAAAAATGCACATATTACTTCTATGGAGCAATATAATAGCTTATACAAAGAGTCTATAGAAAATCCCGAAAAGTTTTGGACAAATATTGCAAATGATTTTTTCTGGAAGAATAAATGGAATGGGAAATTTATGGAGTACAACTTTGATTTGTCTAAAGGAGATATTATTGTAAAGTTTATGGAGGGAGGAAAGACAAATATATGTTATAATGTCCTTGATAGAATTGTTAATGAAAAAAAACTAGGAGACAAAGTTGCATTTTACTGGGAAGGCAACGAAGAAAGCGAGTGTAAAAAATATTCATATTCAGATTTGTTGAAAGAGGTGTGCAAGTTTGCCAATGTTTTAAAAGAAAAAGGAATAAAAAAAGGTGATCGTGTAGCTTTATATATGCCTATGATTATTGAGTTGGTTGTTGCAATGTTGGCTTGTGCAAGAATCGGAGCAATCCATTCAATTGTTTTTGCAGGTTTTTCTGCAGATTCCCTTGCCGATAGAATGATTGACGGTCAGTGTGCTATATTAATCACAGCAGATGGTGGTCGTAGAGGAACTAAAGAACTTGAAATAAAATCTGTTGCTGATGCTGCTATTGCAGAATGTGCTGAAGCTGGTCTAGTTGTTTCATTGTGCATTGTTGTCAATAATTTAAAAACAAATCTTGTTGATGAATGCCCTGCAACAAAAAGATCAAAGGACAAAACTACCAGTTGGAATTCTAACATAGATGTTTGGTGGCATGATTTAATGGAAAATGTAAGCGACGAGTGTGAATGTGAATGGATGGATGCGGAAGATCCTTTATTTATTTTATACACAAGTGGTTCGACAGGCAAACCAAAAGGAGTTGTACATACTGTTGGAGGATATATGATTTACACAGCTACTACATTTAAATATACATTTGATTATCACAATGAGGATGTTTATTGGTGTACTGCTGATATTGGTTGGATTACTGGACATTCCTATATTGTTTATGGTCCCATGGCTAATGGTGCAACAAGTATTTTATTTGAAGGTCTCCCAATTCATCCTGATGCTGGACGAATGTGGTCCATTTGTGAAAAGTATAAAGTGTCCAAATTTTATACTGCTCCGACTGCTATTCGATTGTTAATGAAGTATGGAAGAGAGTTTGTCAAAAAGTATGACCGATCAAGTTTAACTGTTCTTGGTACTGTTGGAGAACCCATCAATCCAGAAGCTTGGTTATGGTATTATAATGTGGTCGGTGAAGAAAGATGCTATATTGTTGACACATTTTGGCAAACAGAAACTGGAGGTCACATGATTACAGGATTGCCTGGAGTTACTCCTATGAAACCAGGATCTGCTAGTTTACCATTTTTTGGTATTGTACCAGCTATTGTTGATGAAGAAGGGAAAGAATATGAAAAGACAATTGTTGATCAAGGATATTTAGTTTTCAAACAACCCTGGCCTGGTATGATGCGCACATTATACAATAATCATTCAAGATTTGAAAGTGCTTATTTTGAGCGCTTTAAAGGCTTTTACTGTGCAGGAGATGGGTGCAATCGAGATGAAGACGGGTACTATTGGATTACAGGTCGAATTGATGATTTAATGAATGTATCTGGTCATTTATTAAGCACAGCCCAAATTGAAAGTGCGCTGATAGAACATTCTTCGGTCGCTGAAGCTGCAGTTGTTGGATTTACGCACAGTATTAAAGGAACAGGAATTTATTGCTTTGTAACGCCTAAAAATGGAATTCGTTTTTGTGATGCTCTCGTATCAGAGTTGAACGAAAAAGTTCGTGCTCGCATTGGACCAATTGCTACCCCAGATTTTATCCAGTGCGCCCCAGGTTTGCCTAAGACTCGATCGGGTAAAATAATGCGACGTATATTGCGCAAAATTGCGGATGGTGATCGAGACGTTGGAGATACAACAACTTTAGCTGACCCGAACGTGGTAGAGGAACTTTTTAAAAATAAACCTAATTAATTTTTTTTATCAAATAATCTTTAGGCGAACTGAGAAGTTCTTTTAGAGTAGGAAAATATGAAATGTATTTTTTTATTTTTATTTATTTCTATCGTTTTTCACAACAAAACTTTTTAGCATTTTTTTAAATTATTTTTTGGATTTGTTAATGTAAAATCTGTTTGAAATTCAGTAACTAGTAGCAACAATAACTTGGTGTGTATTTTTTCTGAAGTTTGGATTATTCTTTTTATTTGATAAATATATTTATTAGGTTTTTTTTTAAAAAAAAA
>evgR004324c0g1_i01_Hydra
GGACGTTTACGTCCAACTATGCTTAAGTACAATTATAAGTTGTAGATATCTTTTTATAAGTGACGAAGTCTGCCTTAATAGGTATATTGAACCAATCAATAATTTGCCTCGTTTTGAGATTTTAAAAATAAGTAATAATACAAATGTTGATAGTTTAACAGCAATCTTTAAAAAATACGGCGACAACACCACCGTTATTTTTTCTTTTACTAAACATGCTCCATTTATTAAAAACATGTATTTTTCTAGCTATGAACATTATTACAGTTTTGGAAGTAATCGTGCGCATTCCGAAGCACTGCAGAAGGATCTACCGTCCAGTGTTTGTTTTCGTTTCACTCAACTTTACAAAGATATTCTTAAAGTTTTTTTAAAAAGTTTGTGGATAAAAAATAACGCTGCAATAATTGAAAACAGCTACTGTTCAAAAAGGTTTTTGTATGGAATCCTTAAAGATCTTGAAACAAGCTACAACTCAAATGTCGCACTTAGTTGGACAATAATTGCTATCGACTTTAGAAAAGACAATAACTCAGTTATACTTGAAAAATTTTACCCGCTAAGAAGCGTTACTGTCAATTTAATAGTAATGTTGACGCCGGTAGAATACTTCAGTACGATAGCAAGCCAATTAAATCTCGTTAATTCTCCGAATATTTGGTTAGTTCCGGACTACGAGAAAAATGTGACGCTACTTCAAAATTTTCAGAAGGTTATTTATTTCAAATCAAACAATTTTGATTGGATGTTTTATTCGCTTACAAACCTATCTCTTCATTTACTTATTAATGCGTACGAAGAAAGTTCAAACAACTTACAGCATGACAAAGTAGATTATTGCTTGAAATGCAGACTAGCCGAAACGGTCATCAATCAAATGGAAAATCCTAATACAAAAAGGTATCCATTGAGAATATACCAAATTTCAAAAAGCGCAACTGGAAGCCTAACCTATGAAAACTCTAGTGCAGAAATTAATCTAGAGGATTTATCAATCTCTGATCAAAATGGATTATTAGCTTTTCAATCTTTAACAGATCAAACCTTTCAAGATAAATTGCGGATAGTTACTTTAATAGAACCACCTTTTATATTTATGCTTGATAACAAGAATGATATCAACGATGAAAAATGCAACGGAATTGTTTGCAGAATATACGAAGAAAAAAAATCAGAGTGGAAACAAGTTTGTTGCTATGGCTACGGTGTTTCTCTACTATTATCTATTATGAACTATATAAATATTGACGTAGATTTATACATAGTCAAGGACTCAAAGTACGGCGAATTTCTAAACGGTAGATGGAATGGACTTATAGGTGAATTAGTAGGGGGGGATGCTGACATGGTTTTGGCTATGCTGACGGTTACCGAAGAAAGATCGAAATATGTAGACTTCACAATACCTTTCATGGAACAGTACACCGGTATTTTAACAAAACCGGAAGCTTTTCAATTATCGTTTCTTAATTGGGAGTTTATGGCGCCTCTTTCTGGAACTATTCAATTAGTTATTTGGTTTTTATTTATTGGAATGATA
>evgR004325c0g1_i01_Hydra
CATTTCTATACTAACAATCATGAAACAAATTAACGAATTGGTACAGTTGAGTCATGCCTCAATCTCTTCATCATCAGATTGCCAATCTACAAATTCATCCTGGATACAAGGTTCTTCCAAACAATCATCAAATCCACTCAATATTCCTTCAAGTTTAGTTGCACCAAAGCGTCAAAGACGTTCCACTGACCATTCCAAAGTTGTTGTAAAGACTTACAAAAATGCTCCACAAGAGACTGGTGCTGTTCTCACTCATTTCAAGGCACTTTCATACAAACCACAAAAACGTTTGAGAAAGAAGAAGAAAGCTGACCCAAAATTAGTTGATTTTTTATGGCGTCAAATTGATTCTGCTGCCATTCATGCAGTTTCTGTGTTTGGTGACTGTCAAGAATCCAATGGTTGGTGCTGTGCTACCAATAAGCATCATACTGGACATCACCATTTTGCTTGTGTGAACCACAAATCATCAAAGTCGTCTAAAAAGACCCAAGAATTTGTCAATCCTTCATCCTCTTTATCCAATGAAGAAGAAGGAAAGAAGATTAGAACTGCAATTGCCATCAGAGAGTTACTTCTTTAAACTCGCAATACGATACAACTTTTACTTGTTGTTTGGCCTCACACACTACCGAATATTAGTTTATTCAACAATGAATAACTTCCAAGGAGAGTCTGCTGACACACGATGTCGTGTTGTTTATACTGGAACTAGCATTGTGCTACTTGAAACAAG
>evgR004326c0g1_i01_Hydra
ATATTTCCACCACTGAAATGAGAGAATCCACATTTACTATTGACAATGATGAAAATTGTAATTTGAATGTTATTGTATCGTGCGATGGATCATAGCAACGAAGAGAATTTGCATCTTTAAATGACGTTGTTACTGCAATAGCAAGTGACATAGGAAAGTGTTTAGCTTACGTGTACGATCCAAAGTTTATAAGTCATGTCAGTCTTGAGGAGCTAATAGGGTTCTGGTAAATATGAAGAATATATTTCAATGCATGATTGCAACATCAATCATACTGGTTCTATGGATTCCGCTAATTTAGTTGATTGTTTTGTTGCTTCAGAAGATAGTCGCAAACTCCGATATACAAAATACATAGGGGACGAAGACTCTAAGTCATACTCTGATATTGTAGCTAGAGATCCATACAAAGGGATAATTGTTCAAAAATTGGAGTGTGTGGGACATATCCAAAAGCGAGTTGGAGGACGGTTACGGAAACTCAAAGCAACAAATAAGCAGCAATTATCTGATGGAAAAACGTTAGGTGGTGCAGATCGTCTTACTGAGAAAGCTATAAATAAACTGCAAAATTATTTTGGCGTTGCAGTTCGGCAATCAACAGGTTTTACCGTTTACCAGTTAAAAAAAGCTATTGGAGTCCTTTTTGAAAAAAAGCTATTGGAGTACTTTTCCATTGTTCTGAGGCAAAAGACGTTAAATCAAGACACCAAATGTGCC
>evgR004329c0g1_i01_Hydra
TTTTTTTCAGTTTGACAAGGTTTATGCCAGATTAAATAATGAAAATAAATGACAAAAGTTTGATATCATATGCATGTTCTAAAAGTGAGTTCGATAAAAAAGGATACTTAAACAAGAAGGGAGAGTTAAACCATGGTTATAAAAGAAGGTGGTTTGTCTTAAAAGGAAACTTGTTATTTTATTTTGAACAACCAGATAAAACTCCTCTAGGTATGATTGTAATTGAAAACTGCAGTGTTGAAGTATGTGATGTCGAGCGTTTTTCATTTTGCATCCGTTATCAAACGGAAAACCTAACTAATAACACCAGGGCATATATTCTATGTGCAGACTCCGAAAGTGATATGGAACAATGGATTAGAGCGTTAGCATCAACGTCATATATGTTTATATCGATGGTTGTAGCAGAATTTCAGAAAACATTACTTGGTTTTAAAAGTAATAACGATTTTGATTCATTGTCAAAGTCAAGCGTTGCTGATAATATTGACGGTAACATTAAAAACCGCAACACAAATAGTAGCACTTACCGCTCCTCGTCGAAAGGTTTTCCTTCAAATGACGATAGTGTCTTGAGAATAAGAACTGCTACATCAAAACCACTTGGTAGAAATGATAAAAACAAGACTGACATGTTTACAAATAGTCCTAAAAACAAAACGAAATATAAAACATTAGACACTATGCGTCGTACTAAAAGTACGGAGAACTTATATAAAATTGATAATCTTCAAATTGACTGTCTGCAAAACAAGAAAACGGATTTCACCAATTCCACTAATTCCAACAACGATTTCTGCAGGGATTTTATATATAAAAATGATAGAACAACATTTGAAACTCTACATACCAGATTCGGCGCAGCTATATGGACAAGAATTGGAGAGACTTTATGAGTCATGTTGTTTGATCTCTTTAGTTATTTTTAATGTCTTCGATTTATTAAGGTTTCTATAATTTTTTGTAATTTTTTACAGGTTTACTTCAATTAAAGATGTATTCGTCTTCTCGGTCTCAGCGCCTTTTAAAATTGTCGCGTAATATTTTCAATTCAATTTATAATATTGTGCAATATAGTTATGCAATATCGTGTGATATAGCTATAGTATTTGATTTATATAGTATAATGCAGCATAGTTAAGCGGTTTACATGTAAATAAATTCAAAATCTTTTTTTTTTGATAAACG
>evgR004331c0g2_i01_Hydra
TTTGTTTTCTGAGCTATGCATACAGTGATTTAGTATATGAAGCTTGCAAAGATAATGTACCTAATAAAGCATGCTTTGATTGTACAGATCCAGTTATGAGTCAATTTTGTCAATTCACTTGCAAAACATGTGTTGATATTAAAGGTACTCAATCATCTGTTAGTTATCCCTTTGGTCAACCTAAATCTGGCATCAAATATTTAATCAAGTACAGTGATTCTATTACTGAAGATTGCTTTTTTGGTACCAAAATCAATGATACAATAGTTTACTGTAATCCCATCAATGGGACAGAGTTTTTTCTAGTTGATGTACCATCAACATTTTACCCAACAACAGAATTCCGACTATCTTCTTCTACTGTGGATTCTTTTTATACACAATTAACTATGAACTTTTTAAGCACAGACAAGATGTTATTTGGACAGTTTAATGGTTATCCAACAACAGTTATT
>evgR004335c0g2_i01_Hydra
ATACAATAACAGCAAATGCATTGAATATTCTTGATAACGATGTAACAAAAGCTGTAGCATTGTTAACTGATTCTAAAAATGTTACAGTGTTTTGCTTACTATCTCCACTAACTTTTAAAACTTCACCTTCTATTTCACTAAATGGATACATTGAATTATCCCTCACAAACACTATAATGTTATACATTAATCAAGATGCTGGATCATACAGGAATTATTCAATTGACTGGGGTGATACTATTTATTCATATAATAACCAATCAGTTACGAATTTCAATGTTTATTCTCCTGTTTCATTTCCAGTTTATCATATTTATAAAATAGAAAATAATTACAATGTCTCTGTGTGTTCTATGAATGTGTTGCAGAAGCTTTGCCATGTTTTTAAAGTTAAAGTAAAAAATTGTTCTGTGCCACAAGTTAGTTTTTACTATGGGACAACTTTAAATCCAATGACTTTTTTTATTAGTGAAGAGAAAGTTTTCACAGCTTTTATTGAAAAAGTTAGTTCATCTTGTGCAATTGAAAGTACTACTTTTGAGTGGACTCTTACATTTTCTGATTCAAAAGATCTAGTAACTCAAAAAAAATCTACACTATATCAGCAAAAAGTTAATTATACAGTTGTAAAACATTCTCTATCTGTTGGTCGTTACATGCTTTCATTGCAATATAACTATGGAGATGTTTTTACCATTTATACTGCATATTTAAATGTCATATTATCTCCGTTGTATATATATATTGAGAATGGGTTTTTCAATTCTATCGCATATAAAAAAAATAATGGAAGTTATCAAAATTTCACAATAAGTGCAATATCAAGTTATGATCCAGATGATGCAGCTGTTGGAGTTCAGAATATCACATTTATATGGCGATGCAAAGTTGCTTTAAATTTTTCAGATGCTCTAATTGCAATGGCAAAGTTTAGGTCTTTAAATTTGACTATTCATGGTGATACATGTTTTAATGAGACCTGGGAGAGGATTTCATCTACAAATCCATATATTAATTTAAGTACACATCAGTTCTTGGAAGGAATGAACTATCGTATTGAAGTCTGTGGAATAAAATATGCAGGAAATGATACTGGTTTAAATGACCAAAGCAAGACAGGCTGTTTTATTCAAGAACTGTTAATTGTTGCTGAAAGTGTCCCTACTATCTCTTTAAAGTGCATATCTAACTGTGATACCAAACTGAATTTTAAAGATCGTGTGATATATTCATTTGACTGTGA
>evgR004336c0g1_i01_Hydra
GTTATACGACATACAAGACTTAGTAAGCAATGATAAAATTGTTAAGTTTACTGTAATAGTTCTACATACTTTATTATGAAAAAATATACAATTCTATGATACTAAACTTCATATTTTTTCGATTGTGATTTAATACAATGATGAATATTTTGGCTACTTTGATGTTACTAGTAACACAAGTTCATCAATGTCAAGGTTTTGTTGAAGGATTGTATTGTGGGGAAGAGAGTTGTTATGATGTACTTCAAGTTTCTAGAGATGCCTCAAGAAGTGATTTGTCAAAAGCTTATCGTAAACTAGCAAAATTGTATCACCCTGATGTTAGTAAACTTGAAGATGCTGATATCAAGTTTCGTAAAGTTGCCACCGCTTATGAAATTTTAAGAGATGATGAACAGAGGAAAGATTATGATTATATGTTAGACAACCCTGAAGAAGCATACTACCATTACTATAAATATTATAAACGAAGATTAACACCTAAAGTTGATGTCCGTATTGTTGTTGCTGTCACCATATCAATTATATCTGTTTTGCAATATTTACAAAAGCGCCATCGCTATGAGGAAGCAATAAATTATGCAATGCAAAATCCAAAGTTTCGAAATCAAGCCATGCAGGTTATTCATCAAGAGGGACTTCTTTCTTCTAATATTAATACAAAGAAAAACAAAAACAAGAAAAGCAAGGATGAACGAAAACAAGAAGAAGAAAGAGTTTTAAGGGAGATTGTTGAAGATTCTATTGATATAAAGGGTGGATATAGTAAACCAAGTTATCAAGATGTTTTATGGATACAGTTAGTAGTTTTACCTTTTTATTTAATTGTTTATTTAAAATGGCTATTTAGATGGATATGGAAGTTTTGGTTTTTAAAATTAGAATATGGAATTGAGGAAAAGGAATACTTAACTTATAAAGGTTTAGGCTTTACTCAACAATATTGGGAAGCTCTGGATAAATATTCAAAAGAAAACTATTTAACTCGTGATCTTTGGAAGAAGGAAAATCTTGCATCATATAAGCTTGAATTAGAGCAAGAATATAGGGTAAAACTTGCAGAAAGTGGTCGTCATAAAATGTGGAGAAGATATATGAAAAAAGGTGGTCCAGGCCAAATGTCATTTGCAGAAGATTAGGAAAACACAATAACATAGTTAAATTCATTTTTATAATGAAAAAAATTGAATTATAAAATGTATAGTATTTAAATTTTCAAGTTTTTAAATTTATTTTGTTTATTTTTAATCTTAGTAAATTCTTAGTGAGCTGTTATTTTATTTATACTTATTTTTAAAATGATTTGTTTTA
>evgR004346c0g1_i01_Hydra
TTTTAGTGCAAAAAGAGTTATATTTTATGGGTAACACACCTTCACAAGTTTCTGTTGAATCTATTGATAAAAAGATAAAAAATGATTGTGTAGTATTATTTTCAACAACTATTTGTGGTTACTGTGACAAAGCCAAAGAACTTTTGAATACCATGAATATCCAATATAAATGTATTGAGTTAGACAAGATGGAACCGCCAGAAGCTGGAAAATTAACTTTTGATCTCATGAAAAAAACGAATTGTAGAACTGTTCCACAAATATTTATAAATGGTTCATTCATTGGTGGATATACTGAGCTACTTGATTTACAAAGAGTTGGATTGTTATTTGAAAAGTTGTCAAAATGTTCCAAGCCTTGTTTTTAAATTGATTATATATTGGTGTTTTTGTTTAAAATAAATAAAAAATTTAATGAAAAAAAAAAAA
>evgR004352c0g1_i01_Hydra
CTATAGATAACAAAATTGCATGACGGAGGAAGAAAATATAACAATACTAACAAGAAAAATCCGTAATTTAACTTTCAACATACGAGCTAAGTACTTCGGCATGTTATGTGCGGCTATCAACAATTCTAGTTTGGGTTTAAGCACTTCTCAAACTTCTCGAAACAAAATGTTAAGTACGGAAGATTATTATAAACAAAGCGAACAAAGAATGAAAATGGAGAATATTTTTAACTATAATAATAATAACAATACATTTTTGCCTCATGATACGCACCTGCAGGAAAGTACTTCGGCGCTTTTAAATGATTTAAACGAGTTTAAAACTTTAACCAGTCAGAATAACTCGGGTAGTTTAGACGAACACAGAGCTTTGCAACTTGCTTTAGAATTGTCTATGCTAGGTTTAACAAACGAAGATGAAATAAATTTTGCTGATATGCGAACTAAAAAAAGTATGAACACAACAGAATGTGTACCCGTACCCAGCTCCGAGCACGTCGCTGAGATTGTTGGAAGGCAAGGATGCAAAATAAAAGCTCTCCGCTCCAAAACCAACACATATATCAAGACTCCAGTTCGTGGGGAGGAGCCTGTTTTTGTTGTTACAGGCAGAAAAGAAGATGTTCATCTTGCTAAACGAGAAATTATGCAAGCTGCTGATCATTTTAGTCAGATCAGAGCTTCTCGAAGGAATAGCACTTCTTTATCTCCAATTGGTCCACCTTCGCCAACAACTCCTGGACATGTGACTAAACAAGTTCGAGTTCCTTACCGCGTTGTTGGATTAGTTGTGGGTCCTAAAGGTTCTACAATTAAAAGGATTCAACAAGCTACTAATACTTATATAGTAACTCCATCTCGTGATAAAGAACCTGTATTCGAAGTTACTGGACTTCCAGATAATGTTGATCGAGCTAAGAGAGAAGTTGAGCATCATATTGCATTAAGAACTGGAGGAATTTTTGATCCCACTCATCCAGAAGCAAGTGAACATCTGTTGAACGGATATTCATCAAGTTTGTTTTCTCCAACTGGATCAACAAATAGTGGAGGATTTTCATCACTCTTAGATCATAATCTCCCTGGACATCCTCATCAACGAGCACAGTCAATGGATTCGTATAACTATTTAAGTGGTCAATCTTCGAATTCATCAAAGTTTTCCGACCTTGGTGGTGTTTCAACTGTTAACAGCCCATTTACCAACAATTCATTTCTTTATGATGCTGTTATCCCTTCACCAACAGCTGATATTGATAACCTGGGTCAGTTTGATCCAGCACCTGGCTCACCATCACCAAGTATTTGGCCAGACATATTGAGAAAAGGTTCTCTGATGAATAGGACTGTTGGTTCTCAAGGGTACTCTCACCGCTCTATGAGCACAAGCACATATACAAATGGAGAACGCTCTAGCCCTGCACATTTACGGCGATTGATGAGTGAGGGTGGAAATGAATCTTATTCTGTTCCTTCTATTTCTTATTCACAATACAATGGTTCAAGTTCGAATATTCCATCTTCAGCACATGACATTCTTTCCAATGGAGACCTAAAAATATCCAGTGATGGAGAATCAGGGAGTTCGACTCCAGACTGGTTAATAAACAAAGGTAAACAATGCTATGTTTGTAAAGGTAAAAATGTGGTTGCTGCATTAGTACCATGTGGTCATAATTTATTTTGCATGGAATGTGCTGAACAAGTTAAAGAAGCTGATGGAGAGTGTCCTGCTTGCCATAAAAAGATTGAAGCTGTATTGCGAATCTTTAGTTAAAGAATTATAAAGCTTCGTTTATTTTTCTAAAAATTCATTGTTGCTAAGATGTTATTTTGTAATATTGTTGTGACAAGCAGAAAAATATTCCTAATTTTAAAAACAAATCTTTCAATGATACTTCTATATTTTTGCATTTGATAATTTTTTTGGAGATAAATTTTTAAGAAAGATTGTTTTCTATATATAGTTGCATAAAAAACATTTTACGTTTTGTTTCATTTTTGTATTCTTTAAGTTTGCAACATTTTATGTTTCCTTCATTTTTTTAATTTTTTTTTTTTTTGAATTATTATAGCTATAATTACTAAAGAAAAATTCTATATTATTAGTAACTGGAGAAATGGTTTATTTTTATAAGATTCTTGTTAAAATAAAAAAATAAATACTTAAGAGATTCCATTTGTCATGAATATTATAAGAGACATAATTTTTTATTCTAAAGATATTATTTGGAGGACAATTACCTATCCAATTTAAACTTAATATAATAAGAGGAAATATTTTGCAAAAACAAAT
>evgR004353c0g1_i01_Hydra
GTTAAAATTCAAGTTGAAAATATATTTTTCTAAATAAATAAAATAATAAAGATTATGTCTCAGCAATGGTACACATTAACAAAAAGTTTGGCTTCAGCAGTTGATTTAATTAATACTATTGAAGTTGGTAAGCTATCTAAGCTATGCAAAAGAGTTCTTGAAACTATACATTTAAAGAACGACAAAGCGTTTACATCAGAAGAAGAGGATAAGTTGCAACTAGCATTTGATTTATCCGGCAATGAAGTGGAATTGTTGTTAGAAATAATATCATTTATCTGGGAACAAGCTGCATACCATTTAGCAAAGTACAATATATTTGCACAACAGTTAAGCATTATTGGAATTAATGAAGATAAGATAGAAGTTTTTACAAGTTTATGGAAAGATTTAGGGCCATTAGTCATAGAGCATTTTCGAAAAAGGAGTTTGGCACCAAAGCAACTTGATACAATAAAATGGGATTTAAACGTGGAAGTTGCACAAGCTAAAAGAATCAAGCTGAAGACACCGAATGCAATTGTTGAACTTGGTTTAGATAATACTGATAATGCTGATAAAAATATCACTGAGAAGATTCAGTTACAGTTTGACCACAGCCAGTTATATTCATTTTTCAATCAACTAGAAGCTATTCAACTCCATCTAGATGCTCTTAATTGATCTTCTATTTGTAATAATCTTGTCATCTAAAGAAGTTTTTTTACTGTTATTATTATCAATATTGTTATTTTCTTTTATGATATAATTATAATTTTCAAAGATATTTTTGAATCTTTTTTAAACTTTATTATTTATCCAATACATTTGTTTAAGGTCTGCTGTTATATTAAGATCTGCTGTTTTATAAAGGTCTACTGTTTTATTTGATTGCGGTTTTAAATAGTTA
>evgR004354c0g1_i01_Hydra
AGAAATTGTTTTGTGGCGTGATATAAAAAAGTATACCATGATAGAAATTTGTCGCCATCAAAGCTAAACATCAGCACAATTAGCCGCCCATCGAAAAACACCAAGACGATAGGTCGCTCATCGCTAATCATCGGTGCGATACGCCACCCATTACTATACACAAGAACAATACACCGCTATTCTGAAATTATAGAGAACAGGTCAAGTCATCAAAAATGAGTTTTTCATATGAAAATGAAGATGAAGATAAATTACCGATGAAAACATGGAGCAGAGAATATTCACGAAAAAATCATAAAACGTCATCATCAACGTTGAGAAATGGTTTTGAATCAGATAAAGTTTTGTTTTTGAAAATGCAAGAAAATCTAGAAAAAAGAAAAGAAAATTATCAGGAAGATTTAAAATCTAACGACCTTGAGTTTGAAGCAAAGGGTATTCTGAAATTCAAATTAAATGCTGGTGGACCGCAAAACAACTACGCAAAGAAATTAATAGCCCATTGCCAACTGCCAGAAAGAGAAATTAAAAATAAACATTTTTCAAACTCCAACCGAAGAATTTTACCCTCGCTCGGTTCATTTGAAAAAAGCGAGCTCCTCAATTCCACAAAAGATTTAATAAACTTTAACACAACATTGGGAGTTGATGAAAACCTATTTGACTATGAAGGAAAAGTTATGAAGTCTATGATAAAAAAGAAGTATATACCCCCTCTTAAAGAAAGGAACTCAATCAACTTTCACACAAAAAATAATCTGCAACTCTCTCCTCTTCCTACTCAAAATACAGATAAAATAAATAGAAAACTTAAAAGAATTGGACCAAATTTAAACGAAGTTATAGATGTTGCAAAATCAGCCCCATACGCAACCGATTTAAATGCAACAAAAGCTAAAAATCTTTTATCATACGCGCCTTTTGATAAAAAGGTTTGGCAGTAACGTTGTACTAATAAAGGCAAACATTCTTATAAAGAAAAAATAAGTTAAAAGTTGCAAATAAAATATCTATGATTTGAAAAAAAAAAAAAA
>evgR004357c0g1_i01_Hydra
CACTTTTTTGAAAATATACTAAAATGGATATTTATGAAGATCTTAAGCTTGAAGAAGCTGAAATTACAGAACTTGCTGACACTGCTAAGGATTGGGTAACTCTTCATGGAATGATCATGCGAAATAACAATGACAGAAAATTGCTTAACTATGCTCCGTTTATGTTGTTTCCTTCGCCATTTCCTGAATACCTTTATCAAGAAGCTTTTTCTGTACAAGCTGATCTACAAGAGTTGGTTTACAAAGCAAGTTGTGACCATGAATTTATATATGATGCTTTAAAAAGTGTCATTGTGCATGATGAATTTACAAGAAAGCAGTTTGAAATCTATGATCAAGTCAGAAAGGAAGGAATTAAGCAGAAATACATCTTCAATATAACTCGGTCTGATTATATGATAAATGAGTGCGAGAATCCAGATGAAGACGAGCGCAAATATAATATTAAACAGATTGAGATGAACATGATAGCTGCTTCATTTGGTGGTCTTGGCTCTTATGTTGAAAAGTTACACAGGTATATGTATGATGTAATGGGTGAGAATGCTCCGTTCAGAAAAGTTCAGGTTGCACCAAATTCAGCAATAAAGAATCTTGGATTAGGAATGGCAAAGCTTTGGAAGATTTACAACAAAGAAGATTCTATTGTGATTTTTGTTGTTCAAAATGGTGAACGAAATCGGTTCGATCAAAGGCTTTTAGAGTTCAATTTTCAAGAAAATGTGCAAGAGCTAGGAGCTGAGAAACACGTTCCTGTACTATTTCGATCTCTGGAGCAAATATACAAAAACGGAAAAATACTTGATGACAGGACTTGCTTTGTCGAAGGTCATGAAGTTGCATTGTTTTATTTTCGAGCTACATATACACCAAATGACTACACAAGTCCTGAGTGTTGGGAAGCTCGTCTAATGATTGAGAGATCAACAGCAGCTAAATGTCCATCAATGGCTGAAAATCTTATTGGAACAAAAAAAGTGCAGCAAGTTTTCGCTGCACCTGGTATGGTTGAAAGATTTGTGGACAAACCAGAAGTCGTTAAGAAAATCAGAAAAACCTTCGCTGGCTTATATTCTCTAGATGAGACCTTAGAAGGCGATGCTGCAGCAGAAATGGCTCTTGCTAAACCTTCGCGTTATGTTTTAAAACCTCAAAGAGAAGGAGGAGGGAATAATATGTATGATGCTGATATGGTTGACGAGTTAAAAAGAGTTACTGCTACTGAAAGAAGTCAATATATTCTTATGGAACGCATTCAACCTCCTTCTGTAAAAAACTATATGGTACACGTAAATTTTGATCGACCGCAGTTAACAGACACAGTAACTGAACTTGGCATCTTTGGAACGTTTATTGCAGTTGACGGCGTGATTGTACATAATGAACCAGCTGGCCATTTAATGCGAACAAAAAGCAAAGATCAAAAAGACGGAGGAGTTGCTGCTGGAGTTGCTGTTCTTGACACACCCTATTTAATTTAATTTATTTTTGTTTCAAATAAATATTTATTTTGTAGAATTAAGTAAAATTATTTATACATTTTTCATTATATTTTTATTTTTCCTTTTATTCATTATTTACCTGTTGCTGCTATAGTTATCTATAGTTGCTATCTTTACTTATTATTGATACGAATACTTATTGTTGCTATCGTTACCATATGTTGCTTATGTTTGTGTGCTTTTTTTTATTTTTTGCTTTGTTTTATACTTTTTTCTCATTCTGCATTTTTCATATCCACGTTGTTATTTTACGGTTTTATAGTTATACGTTTTTATAGAATACTTTGACTTATATTGTAAATTATGTTTGTAATATTTATTATCCTCTATAAATTATGTTCGTTTTTTTTTTAAAAAAAA
>evgR004360c0g1_i01_Hydra
ATATTGGTAGTATTTCTTCTATCATATAGCATACCCTAAGTTTACACATTTTCTTTGCATCTTCCAAATTTCTTTGGTTTATTTTGAGCAGCACATCCTCTACTAGGAGCAGCACTTTTGGTTGAAATGTCTCTGGAAAGTGTGCCAAAAGAGCTGCGTCATTTACGCTCATGTTTATTGTGCTCCATGGTAAAAACATTAGAGCAGTTTGAACGAGATGGTTGTGATAATTGTGAACGGTATCTTCATCTAAAAGGTGACCATGAAAACACGCTGTCTTGTACTAGCGCAAATTTTAATGGTATCATTTCCATGATGACACCTGAAGATAGTTGGGTAGCGAAATGGCAGAGAATAGATAAACTTAGACCTGGTTGCTATGCAACTTCAGTTTATGGAAAGCTACCTTCTGATATTATTAAAGATTTAAAATCAAGGGGAATCCCTTATAGGTCACGAGATACATCTAGTGTTTGAGATTTTTTTTATAATACATTTTGTTATAATTTAAGCAACACCATTACAAAACAATTGTTTACATTTGGAGTAAATAAATTTCAAATAATTTTTCTAAACACTTTGTACACTATCTGGTCAAACTTTCTGTTATTCTCAATAAATAGAGAAGTTGATGCATTTCTAATATATATTCAATGAAATATCATATTTATGTTTTTAACTTATGTCTTGTTATTATAATACTTAATTTAAATTTATCATTGTATTTTTTTGTTTAACAGAATAAACCATGATAAGATACTGAATAAAAACTTATCTATCTCTATG
>evgR004362c0g1_i01_Hydra
CAGCTACTGTACAAGTAGCATGTACAAACATGGCAAAGAAAGGGGTAAACATATTCAGTCTTATCTTTTACCAAGTTCTCCAAGTCTTATCTCTAAGTCTTATCTCTTACAAAATTTACTAGACTAACTTTCAATTTGTCAGACTAATTTGAGTTCAGCTGTCTTATCCTCTGATCATAGTGTTAATGGTTCTTTTGGTTTGTAAAGACTCAAATAGTCACATGCTTGACCTGGGCATTTACATTTGTAGGAAATTTGCAGCCATTTGTCAGGAAACTAGATTTGAATCCTTTGACTATCCTTTTATGTGTTTTCGTTTAGCAACACTTCACTCTATGACCTTTCTCTTTATTCTATATCGCTCTCCTTCTACTCATGAATGCACTCTTTTAGATGTTAGATCTGATCAAATTGGCCAAGACCTTTTCTCTTCCTTTATTCTTCAGCAAATATTTTTGTTATTGCTCATCAAGTATTGCTCATCAGTTAATGTTCATCAAACTGAATGGCTTGGCTCTAATGTCAGTGACTGCAGGCGTTAAGGTTCACAACCTTTGCTTTTCTTAATTTCTAATACAATTAGTCAACTTACCAACTTTTTTTCTAGACAATCCGACTCGTTCACCTTTTTTTTCTACACTTTTGTCTTGTTTTTTATCCTAGTCAGTGCTCATTTTCTCAACATTTACCTTGGGGTGCTTCTGTTTAATCTTTATAAAACTCTAATCTCATTCTTCTTCATCGTCAGAATCCCTCTATAATCGTAAATCTTAGAACTACCTTAAGTCTGACTGGGACTCTTTCCATGATTTTCTTCCTGATGGCCCTTAGGTAAAAATATTTTGTTTTCTGCTAAAAAATGTGCTTCTTATGTAAGTTCTTGGATTCTGACTGGCATGGAATCTTTCATTCCTTCTTGACGATTTGAAGTCCTCTGTGTGGTTTTCTTCTCATTGTACTGGTGCAATTCTAAGTTGTAGCCATTGCTTCCATATCTATCAGTAAAACAATTCTTCAGAAAACAAACGTCTGTTTACTATTGCTAGAAACCATTGTAAAAAG
>evgR004363c0g1_i01_Hydra
TTAAATCAAAGTTTCCGAGATTGACATCAATAATTGATTGCTTTGAGATTTTTGTTGAATCTCCATGCTCTTTGATGGCAAGAGCACAGTTATATAGCCAATACAAAAAACACTGTACTATTAAAGTTTTTATTTCCTGCACTCCGCATGGTTCTGTTTCTTTTGTATCTAAATGCTGGGGAGGAAGAGCCAGTGATAACCAAATTGTTAGAGAATCGCAATTTCACTCATTGAAGTATCACATGCCTGGTGATCAAATATTAGCTGATCGGGGCTTTACTCTAAAGGATGACTTTGCTCTTGGAAGCAGTACTCAACTAATTACTCCAGCTTATACAAGAGGAAAATCACAGTTATCAGCCAAGGAAGTTGAAGATTCTCGAAAAATATCCTCAGTTAGGATACATATAGAGAGAGTTATCGGTCTTATGAAGAACAGATATACCATTTAAAAAGGAATAATACCCTTACGAATGATCAAAAGTATTAAAGATGAAGCAACTTCATCCGATTTAGCCAATTTTGACAAGATTATTACTGTATGTGCAGCATTAACAAATTTAGGCGAAAGTATTGTTTCTAAACCACATCCAAATTTGGTTGATGACACTATATAATAATTTTTTTTAATTTTGTGTTTTATTAGAAACAAACCATTAAGCAAACTGTATTGTCAG
>evgR004367c0g2_i01_Hydra
CATAAAATTAAAATAAGCCGCATACATAGCGCTGCCGCTGCTTGCAACTTACCTTTTACGGTTTTCCCACCTTTAACTTACCTTCCACCTTAAAGTCTTTCAAGTCTTGTCTTTAAGTCTTTCAAGAGTTGTATTAACAAAATGTCAGACCATGATAAAGCTGAAGATCGGAACATCAAAAATACTCATGATGAAGTCAGACATCATAAAGGAAAAAATCAAAAACATATTGAAAATAACGAGCCAATTGAAGTGACTTTTGAAGATGTGAGTGCAGCTGCATTTCGGATAAAAAATGGAATAAGAAAGACAATATGTGAGAAATCTTGTGCATTATCAGATCTACTTCAAATGGATGTCTACCTAAAGAGAGATTACATGCAGGTGACTGGCAGTTTTAAAGAAAGAGGTGCAAGAAATACTCTTTTAATGTTAACCAAGCAACAAAAAGCAGTTGGTGTTATAGCTGCATCAGCTGGTAATCATGCATTAGCTTTAGCTTATCATGGTAAGCAGTTACATATTCCAGTTACAGTATGTATGCCTGCCAATGCCCCAATCACTAAAGTCAGTAGATGTAAGAAATATGGTGCCACTGTTCACTCTGTGGGATCAGATATTATTGAGGCCAAAGAGTTTGCTTTAAAGATAGTTCAAGAAAAAGGATGTGTTTATATCAATGGATATGACCACCCTGATATATTGGCTGGTCAAGGTACCTGTGGTATAGAAATTGTTGAGGATGTTAATGGAATTGATGCAGTTGTAATACCTATTGGAGGAGGAGGGTTGATCGCTGGATGTGCAGTTGCTATTAAAAATCTTCAGCCAGATATCGAAGTCATAGGTGTAGAAAGCGAGAAGTGTGCAAGTTTTAAAGCGGCTTTAGAGGCTGGGAAACCAGTGAAAATAGATGCAGACCAATCGCTGACGTTAGCTGATGGGTTGTGCGTTTCAAAAGTTGGAGATAACGCATTCTTTACAGCTCGACCATATGTTGATAAAGTCATTTCAGTTAGCGAGTCATACATTGCTATGGCAGTCCTCCGCTTGGTTGAAAATGAAAAATGTGTAGTTGAAGGCGCTGGTGCAACAGGTTTGGCTGCTGCATTTTCTGGTCAATTATCCCATTTAAAGGGAAAAAGAGTTGTTTTTCTACTTTGTGGTGGAAATATTGATTCAACTGTTCTCGGAAGAGTTATTGAGCGAGGATTAGCTGCTGATGGCCGGTTAGTCAGATTTTCTGTAGTAATATCAGACAGACCAGGAGGTTTGGTGGAACTTACAAAGTTACTAGCTGAACATGGCGCAAGTGTTAAAGATATCTTCCACGAGAGAGCGTGGTTGCAAACAACAGTATTTAAAGTAGAAATTAAAGTGGTTTGTGAAGTGCGCGATTACGACCATGGTTGTGGATTGCGAAAGAAATTAGAAGACAAATACGACGACGTTAAATGGTTGTCTTACGCAATTCATGATGAATCCGAAATTTAACAACCCGAAAATATTTTTAAAATTGAAAACTTTTTATGGTTTTTTTGTAAATTGATTTGTGTTTATTTATATTGCTATATTTGTTGATATGATTTATTTAAGATTTTTAGAAAAAGTTTATCTTATGTTTTCAAAAACAAATTAAATTCCATAGCATGGTTACGTTTTAGCAGAATAATTTATCTGTACTTTTGAGGTATTGTTTTAAAAACTCAC
>evgR004368c0g1_i01_Hydra
GGTGTGGCATGATAACAGTGGTGGTGGATGGTATGTTAAAAATATCACAATCATTGATCTTCAAACTAAAAAAAAATATTTTTTTATTGCTCATTGCTGGATTGCTGTAGATCGTGGCTTGTGTAAGTTAGATGTTATTTTACCAGTAGCATCAGTAGAAGAATCTTCAAGTTTTATGTATGTTTTTATAAACAAAGCACAGCAAAAATTGCATGATGAACATTTATGGGTTTCAGTCATTTTTCGTTCGCCTCATAGTACATTTACTAGATGTCAAAGGCTTTCTGTGGCAATCTCTTTGATTATGACTTCAATGCTGGTAAGCGCTATGTACTATCAGCAACTACCTCCAGTAAAACCATCAGTAGAAAATAAAGTTGGTTCCTTTTCCTTTACATGGACTCAGGTTTTTGTGATTTTGATATCGACATGCATTAAGTTTCCAGTGAACTTTTTACTTCTCAAATTGTTTTGTTCAATAAGACCATTTGAAAATGAAAAAATCTCAGATGACTGCTTAACTTGTAATAAAGACAAAGACTCGCGCAAAAAATCATCATCTTGTATGCACGTTCATGAAACCAACAATCTGTCTTTAAACAAACAAAAACTACACATTTCTATTGAGCAATTGAAATGGTATATTGCATGGTTTATTTGTGTTGGAAATATTTGTGCATGCAGTTTTGTTGTTTTATGG
>evgR004385c0g1_i01_Hydra
TTTTTTTTTTTTTTTCGGATATTTTTCGGTTACGATCTCGGTTCTGATTTCATTATTTGCCTTATTTGTTATTTTCATACTGTTTTTTCCATGCTATACGAAAAATTTACAAATTGTTGATATAGTACATTGTTGATAAAGGCACAAATTGATATAGTAAAACTCACACAAACATACAAAAACTCAAAACTATTTTAAGTTTTACTTACTACGTAACATGGAAAATAAAATTCAAAATTTTAATGCAATTTGGTATCAGATATTTTTATATTGCCTTGCAACTGTTCTCTTTGTTCACATAATAGCAGCATTAGTGGCGTTTTTCTCACTCCGTAAACATCAAGTTGGAAGATATTTATGTATTGTTTTATTTTTATTTGGTTTTTTATACCCACTGACTGGAGGCTTTATAACAAGTGTACTTATTGCTTGGATTTACTCTGCAGCTGGGTATACTATGGAAAAGAATTTTGCATTTGTATATGGAACTGGTCAAGCCCTTGTATTATTTCTAGCATCGTTTATACATCTTTACGGATTGCTGTGAGTCCAACGTAAAAAATGGCCACTTCTTAGCGATTTGAAACTCAGTATATTTTAAAATCAGAGTATAAAAATATCATTCCAAGTTTCGCACCAACTTTCCATATAAAATGTTTTGAAAAAGAGGTTGTTTTTAAAGTTTGTGGAGAGTTCAAAGGTCATGGAGTTTCAAGACTTAAAAGTTGTGCTCTGATAGCACATCGACTGACTCATAATGTTGAAAGTATATTACATAAAAACAAAAATGAATATAATTGTTTGAAAGATTTAAAATGAGCGTTTATTTTGTTGTTCTAGAAATAACACAATATTGTTGCAGTGATTATTGTAATGTTATACATAAATTGTTTTAAATATATACATCAATGGTGTTATGATAAAAAAAAAAA
>evgR004386c0g1_i01_Hydra
TAATCTTCTATTGTCTAAATAAGTACCATGTCTGCTCCCAAAAAAAGATTAAATACTACAGCTTCCGCTAGATTAAAACAAGATTACATGAGAATCAAGAGTGATCCTGTTCCATATATTGTTGGAGCTGAACCTCTACCAGCTAATATTCTGGAATGGCATTACGTTGTTAAAGGACCAGAAAAATCATTATACGAAGGCGGCTTTTATCATGGAAAGTTAATTTTTCCAAAAGAATTTCCATTCAAACCACCCAAAATAATTATGGTAACACCAAATGGTAGATTCAAAACACATACAAGATTGTGTCTCTCAATAAGCGATTTTCATCCTGACACCTGGAACCCAGCATGGTCTGTATCAACAATATTGACAGGGCTTTTGAGTTTTATGCTTGAGAAAGCACCTACACTAGGTAGTATTGAAACAACAGACTATCAGAAAAAACAGTATGCGATGAATAGTCTTGATTTCAATATAAAAGATGAACTTTTCTGCGAACTCTTCCCTGATATAACAAATGAAATAAAGGATATATTAAAAAAAAGATCAGAACAAAACGAAGAAAATGAGTTTACATCGTCGACTACCCCGTTAAATGACTTTCGTTCTGTTGATAACGATAATTTTATGAACGGCAGCATCTTTGCAAATATGTGCGTTTTTCTAGGTTTTGCTATTTTTGCGTTAACTGTCCACTATGTACTAAAAGCAGTATCCCCAGATGAATTGAGTTGATTGAAGTATTTTTTTATTTATTTATAAACGTTATTTAGATTATTTTTTTATTAAATAAGATTCTTAAAAAAAGATTCTTAAAAGAAGGTTTAAAAAAAAA
>evgR004394c0g2_i01_Hydra
TATGGTATTTACAAAACAAAAAGGAGAAACAAATACTCACGAAATGAAAAAATGGTTTTATTATTATCATTTATCGATGTGCTAGTTTCTTTAATTCATACCCCCCTAGAAATAATTTTGTTTACTTACTTGAATTCTATTGGTTGTTTGACCCTCTCAGTTGTTGCCTTTTGGATAGGATTTCCGTTGATGCTCTCTGGTTCGATGGTAGCAATTATATCAATCGAACGTTTTATCGTCGTATTTAATGATAAAAAATGTTGTGGTTTGTATTTTAAAGACCAATACATGGTACCGATATTTCTTTTTTACTTATTTTCTTCTATTGGGTTAGGCGCTTGGTATGCCCATCTCCACTATTTTTTTCCAGAAACCATTTTAAATTTTCGTAACTTTTTATACTTCACGGCAACATTTACTATTATCAATCTAGTTTTTGGAATATGTATTAACGCCTTACTTCTTAGAAGAACTTTGAAAAGATTAAATAACAAAGACATTCACGTCGAAAAGCACGTTTTAATCGAGAAAAAGCTAAACAAAACAATGATGATAATTTCAGTAACGCTGTTAGTATCATATATACCAACAATTACTACCAACTATTATATGGCTGTTACGTTTTTGTTAGGAGACTTAGATTCTACTCTTTTTGCAATGAAACTATCCCTTTGGGCTTTAATTTTATGCAAATTGAACTCAGCTTTAAATGCCATTATATACATGTCAAGAAGTAAAGTTGTTTTTCAATTATACCGATCATATTTGCGTTATTTTATAATAAAACGTGTAAATCAAAGCGTTTCCCGAATCCAGAAATCTAACAACTATAAACTAGAAAGTAACAAACAATTCAAAGAAATTAGTTTTATTGAAACATTTTAAAATTTTTATATGTCTTTGTATGTTATTATTATATGTCTTTATTTAAAAAAAAA
>evgR004400c0g1_i01_Hydra
AACATAATTGCACGAGTCTTATTTAAACATTACCAATAACTGCCGACGTAATGGAATAAGCTGATTATTGATTACATCGGTGGAATATTTTTAATTATTGATTGATGTCCAAAAATAAACGTACTCAAAGATTTTTTTGGTTTGTTTTATTGGTCTAGAACAATTACTTTTGCTGGCACTTAACAAAATGCCCATTCAAATCTATGACGATAGTCAATTTGATTGTCTCCAGTGTATACCGTACATAAATAAGCCGCCATAAAAGTTACAAGTGGAACAAATGTCATAGTTCACTGATAAGACAAGATCAAGAAATTAATAAAGTAGTTATAATGGAAAAAGGTGTTCGTAATTTTTTATACTGGGTTGATAAACAAAGCAAAGATATGTTGAAGCAAAGTCGAGACATGTGTTCTGAATCATCCCGATCGTCAAAATCCGACTTAATGAAATCGAATCGTTTCCCGAAATGTGCAAGATGTCGTAACCATGGAATGGTAAATGATTTAAAAGGCCATAAATATGTATGCCGTTGGAGAAACTGCACTTGCCAAAACTGCGCCGTTGTTAAAGATAAACAACGTATTACTGCATCGAGAGTTGCCAAGTTAAGAAAAATGCGAAAAGTTTACTCAGACAAATTGGATTCAAACGCTGACTTGATAAATAGCGGAAATAATTACAACGGGTCTTATAATGAAGATCTTGACGAAAGATCAACCGGATCACCTGCATCGATTTCAACTATTTCAACAAATTCTGAAAGTCATACATCTTTTGATACAATAATCGATATCGATCACCACTATCGAGAAATATCCCCTCAGTTTCGTCCGCTAATAGAAAAATACCCTGTAAAATGCCCTTGTAATGAATGCAACGTATGCATCCAACGAGATCATGAAAATGATTATTACTCAAACTTAAGATACCTTGCAGATCTTTTCCCAATGTGCTCAATAAAAACTCTTGGAGAGGCATTGCTGAGAGCCAAAGGAGATGTTAAAGGGGCTTTAGATCTAATGAGGTAGAAACATTTCAACTGATTGTTCTAAATATTTCTTGAAAAGCTTGTTTTTATATAATTTTATCTTATAGTTTTTTTATTTTTAAAAAAGCCATGTAAATATAATGTAATATCTTTGAATTAAAATTGTAAATATGTATTTATAAGAAGAGTTTATAAAAAATCTTAAAATAATTTTTTTCTTGGCAAAAGGGGAGGC
>evgR004406c0g1_i01_Hydra
ATTTGGGAACAGTGAGTGATATTGCCACTGCTTATAATGCAGATATTTATAAGGCATTTGGTGGAATCCAATTTGTTAATGGTGTTTTGCAAATTCCAACAAAAGATGGCAAGTTGACCAATTTGAATGTGAATACATTTGATGATTTGACAGCTGCCAATACTGCACAAGTCAAATTAGGTCATGTTTATTTGGTGAAGATTGTTTATTCATCCACCAATACTGCACAAGTCACTAAATTCTTGACTACTCAAGTGGTTAACACTGCTGATGGATCAGCTGTTCAATCTGTCATGATGAGACATCAAACCATGTATATTAATAAGGAAAATTCTCAAAGAACTACTTGTGCCCATCATAATCAACAATCTAAATCAGAAGCTGACAGCGCTGTTGCATTATCAGTTTTTGCATTGTTATTCT
>evgR004415c0g1_i01_Hydra
TGACGCTTTTAAACGTCACAAAAAATTAATTTATATAGCTCAATCCTTTTTAGTTTCAATCCAAAGTTTAGCTGACTTGATATTGAATATAAATATTCTGAAAACAGTTGAAAATCAAGATAGTAATAAAGGTTTTTTTTAAAGAATTGAGTTAAATTTCTACCCCCAAAAAATGTATGAAGTTGGAGAACCCTCTACAAATAAAGCAGATAATAGCACAATTAAAAATGCGGTAAATGGACACAAAGAAAAACACTATGAAGCTGATCTATATAAGCTGTTGCCGGAAAATCATCATTGTGGTGACATTGTTTCTTCAAATCGAATGCTTGCATATTGTGACACAGTAATGGCAACCTGTGCAACGTTTTTAGTCCTTCCTTTGAGAACTGTAAAATCGATAGATGAACTTAAAAGTAATTGCAACGCAACTATCAACTCAACAATCAACTCAACATGCAACTCAACAAAAAGCGAATCAATCATTTTAAAATGCAACTCAACAATCAACTCAACAATCAACTCAACAATCAACTCAACAATCAACTCAACAATCAACTCAACAATCAACTCAACAATCA
>evgR004428c0g2_i01_Hydra
AAAAATCACAAAAAGAAAAATGGCAAAAGTAAATACCCATTATGACATGAATTGTTTAGTGAAAATAGAAGAACTTTTTCCAAACTATAATGACGAAAAAACACGTGAACGTTTTCTAAGTTTTTTTGCTGAGTCAATGTTTTTATCAAGTCAAAAAAATGGTAGTATGAAGTTTAGTTATACGGATGTAATAAAAAAAGGAGATGCTGTTTTAGAACAATTTAAAAAAATATTAAAAAATCAGGCTGAAACAAAAAATCCAAACAGTGAACTTTCTTTTATGGTTGAAAAGATTCAATTAAATCAGAAACATTATGAGGAAGCCTTTGACGTTGAATATAAAAGTCAATTTGAAGAATCATTTAATCAGTATTTAAATTTGTGTGTAAATAAATGGATGCGTAAATCGTTATATGCTCCGATTGCAACAATATGCCAGAGCTCAGGTCATGGTAAATCTAGAACAATCAAAGAATTTTCAAAAAAGAACCTGTCACTTTACATTTGTCTGAGAAAACCAATCAGCAATGGATATGGATATCCAAAACGGTCTCAATCAGCTAATGAATTTCTGAGCTCAATGGACACAATGTTACAAGGAACAAAATAT
>evgR004437c0g2_i01_Hydra
TTATTGGCTCTGGAGATTTAAAGTATCTTTTAAACACAATATATGTTCCTGTTTTCAAAAAACCAGTTATTAAAATATACCAAGGAATGAACTGGGTGGCACAACTTTCTATGCTTTTTGGAGTTCTTGCATTTGCTATAATGTCAGTAATGGGAATAACGTCAATTTCTTCCGTGTCTACACAAATGAGTTGGAGAGAATGGAGATTCATTCAAAGTCATCTTGGATTCTTATGTTTAGTACTAACTTTTGGTCATATTATATGCCAAGGCTACAAATTTTGGAATTCAAACTACATTTACGGTTGGCGTTTATGGAAAACAAACGCAAATGGAATTTATCCGTACCCAATATTTATAATGGGAATATTTCCACTTTTGGTTATATTTCTAAAACTTGTATTGATGATTCCCTGTGTATCGCGTTATCTTGAAAAAATCAGAAATGGAAGCATTGATGCACGAGTTAGAAATAAAAAAATAAAAGATATTTATAACTGCGCATAGACAAACTGCGCATAGACAAACTGCGCTTGGTACAAAATTAAAATTAATTATGTGTATACAATAAACTTTTACTTAGACGTAAAATTAACCATATATAGTTAAGATCTTAGCCAACAAACTAAAACAAATTTAGTTAGACCTTAATAAGCATTTTAATGTAAAGTATTTGAAAGCAG
>evgR004443c0g1_i01_Hydra
TTGAGAACCCTTAGGTTTCTAATAGTTAACTATGGGTGAATCGAATTTGTTAAAACTCTTCGGAAATAAAAAAGACGAATTTTGTCGCTTTCAATATTACAATGCTGCTGCTGGATGGAATGGAGAAATGCTAAAATTAATGAATGGAATTGTGGGATTGTTAATTTTTATTTCTAACGCGTTATTAGTATACGCCATTTTTAAATCAAAAAGAAGAAATAATTACTCACGAAATGAAAAAATGGTTTTAATATTATCTTTTAGCGATATTATGGTTTCAATAATACACACTCCTCTTGAAATTGTTTTACTTGCTTACTTAGATTCCATTGGGTGTCTTACCATTTCGATAATTGCGTTTTGGATAGCATTTCCACTAATGTTGTCCGGAACGACGGTTATGATAATTTCCATTGAAAGATTTATTGTTGTATTTAATAACAAGAAATGTTGTGGATTTTATTTCAAAGGGTGTTACATGGTTCCAATATTTCTATTTTATTTTATTGTATCTTTTGGATTTGGTGTTTGGTATGTGCTTCTTCATTATTATCTTCCCGACGCTGTTTTTTCTTTTTGCTATTTTTTTTACAGCACTGCATCCTTTATTATTATAAATTTAATTTCTGGATTGTCAATAAATC
>evgR004446c0g1_i01_Hydra
TTATCTAGTAATATCAATTAGTAACTGCGTAATTTTTGGAAAGCAATGTTTTCAAACAGGTTTTTGAATCGAAATGCATCTCATGATGAGAAAGAAAAAGAGAGTAGAGTCGCCGAACTGATCTCTATTCCTTTATTATATCAACAAAATGCAAAAAAGTCATCCTGGTCGTCACTACGCACTATCATGAAAGAAAAAATTTTCGTAAAAAACTTCGAAAGTTCAAAAAAAAGATGCTCTGGAAATGATCTCTGTGAAGAGGCATATGTCAACCAATCGGCTACTACTAGTTATAAATCTAAGATGCCAAATTTATCTCGAAGATACACGGCCGAAAGTTTTGAAGACTTAATTTTCATTTCGATACTCCAAAATGATTACACCACTTTAGTAAGAGCTTTAAAAACATATAATGTAAACGTAAATTACATGAGGCCCCCAGGGTTAACACCGTTGCATCAAGCATGCGTTGTTGGAAATCTTCAAATGGTGGAAACTTTAATTCGTCACGGTGCCGATACGAAAATAAAAACATGGTCTAACCTGACACCACTTTTAATAGCAAACTTGTTCGGACATTTTGACGTTGCTTTGTTTCTAATAAACCAAGGAGCGAATACCCGCGACATTCAAGACGGAGTTACTTTTGATTCAAAATTAGTATTATCTCTTTTGATGAGCTCAAAAGAAAGCGAGGAAGAAGATGTTCTAGAAGAAGGAAGCTCAAGCACCATCATTTTAAGTTTTAATAAAAATAATACCTAATAAACCTTTGAA
>evgR004457c0g2_i01_Hydra
TTATCTTTGTTATACCAGTTTGTAACGGGAATTTTGAATTTTTCCGTCAGGGCCGGCGTTTGTTTTTTTTGTGTGGGGAGAGGGATGCCATAAAAGTTTTGCCTACTATTAAAAAAAAAAGGTACTCGATTTTTAATATTTTCCGGCTAGTTCCAACCATGTCCATTTTTGCCGACTTTCCCGACTCAAAATCAAGATTTTGGCGGGGGATGGGTACACCCCATGCACCTCCTAGTGCCGGACCTTTTGAAATTTCCGTTAATTTTCAGAATACGAACGTCCTAGAATACATCCTGTTGAATGTGAACATTTTAGTCCAAAATATTTGCGTAATTTTGCGTAACTTTTACATAACATAAAATATTACATTTAG
>evgR004469c0g1_i01_Hydra
CATCCTCCATAAAGCATTCGAAAATCGAAAAATCTTACAAAAGAACAATTATGATTGGCAACCGGGTTTGTGGTTCGAAGAAACACATTGACATCCATTATCAAGGTGAAATTTCAAAAAAAGGAGAAGTTGTAGAAGATATCAGCTCGAGTCTTCGTGCCACAGACATTTGTTGTAACGAACATAATGAATGTCAAAGAGTTGTGCCAGCTAAAAGTACTAAATATGGTTTCCAAAATGACATTGAGTATGATATTATGGCTTGTTCTTGCGACGCTAAATTTCGTACATGTTTAAAAAATGCGAATTCGTACACAGCCGATGCAGTTGGCCATCTCTACTTTAATACGTTAAAAATACCATGCTTGACCTTTGAACAGGAAATCATTCCCGATGTAAAAAACTCAATGAAAAAGGAATCAAATATTGTTAGTGTTGTTCCTGATGTCGAAACTGAGCGAGAAGTAGTGGGTGGTGCTCAATTAGTTGTTTCCCCGAGAGGATTAAGTCGTATGGAAAATGATAAAGAAATGGTAAAAGGCTTATGGAATAAACTCATAGGCAGAGGTTCCTCGGTATCCAAGGATAATATTTTGGAACCCTTACTAGCGACTTCATCATTTCGAGCTAATGCACAGCAAACCGTTCTTAAAACCGAGCCAGAAGATCAACTAGTTAACATGTCAATAAAGAGAATTGATCACGAGTCTGAAGCTTTATCTAATTTACTTAATAACAAAGAAGACCAATTAACAGCTTCATTAGTCCACAAAGAAGATTCATTAGCTATGCAACATTTAATAAAAGACTCCGAACAATTAAAATTGCCTCAAAGTGACAATGAAAAACATATTTATTATGTTAAAAAGTTATCTGAAGAACCAGTAACTTTTCAAAAAGATGAAGATCAATTTCGTTATCAAACAGATCATTTTCGTATTCAAAATGAATCTGAGCAACTTCGCTCTCATAAAGAACACGAAATTCAAAATTTGCAAAAAGAAATTGAACACTTCCAGAAGCAGTCTGAGCAAAACCATATTCAAAAAGACATTTATTTAGCTAAATTTCAGAAAGAATCAGATCAACCTCACATTCAGAAAGAAATAGAGCAACCTCGTAATCAGAAAGAGGTAGAGCAACCTCGTAATCAGAAAGAGATAGAGCAAAATCATCCTGGATTAGAAATAGAGCAGCAGAGAATTCGTTTGCAAAAAGAAATTGAACAAATGCAGGAGCAAGAAAATAACTTAGAAAAAATCAAGAAAAGAAAAAGACCAAATATAATGAATCCAAATCCTAGCCCAAACAATCCAAACCCATACAGTAAGTTGAGAGGTGTGGTAACAGTCCCAGAGGCGTATTATCCACGGGTTTCTTTATATTGAATCAGTATGAGTTTCAGTATCAAAATCAATTTAGATAATCTGTATCAGTATGAGAGTGATAATAACAACATTGTTTGCAAACTGTAATTAAAAGGAATTTAAACTAAAACATTTTAAACAAAAAGTAAAAATAAAAAAATTAGTAAAAATAAAAC
>evgR004486c0g1_i01_Hydra
AAATGAAGATGACACTTTCGAATATGACGTAGACCAAGATGTTTTCTACATTTCTGGTGTTAAAGCTTCTATGGCTGTCGGTCAAAAATTCAGCTTGGGCTCAGGAGATATTCAAAGAACCAATGGTAACGCTGCCGTGGTTACTGCATCCAATGGAGTCAAGATGACCATGTCTTGGGAACGTAATTCTGCCTGGAAGACTCAATTGGGTTCTGTTGGTTACATTAGTGTTGTAATTGACGCCCCAGAAAAGGTTGAATTCTCCAAGGGAATGTGTGTTGATGTTTCTCAATCATCTCAAAAAGCCACTGGTATCATTCACAATCACAAGGAAAGAAGAGTCATCCGTACTGTTGCTCTCAAGAAGGAAACTCCAAAGAGAAGAACAGCAGCTGGAAAGATCTGTAAAAAAGCTGGTTTGAGCAAGAAGAAGAATCCAAATGCTTGGAGAGCTTGTGTTAAGGATGCCATTCAATCTGGAAAGAAAATTGGTCTTGCAATTGGTAAGAATATTAAAAAAGTCAAGAAGACAGAAGAAAAGGTCATGAAGAAAGAAATCAAGAAGAAGGGTAAGAAAATTGTCAGAAAGGTCAAGAGAGCAATCAAGAGACAAGAAGAAAAAGCAAAGAACTTGAAGAATAAGGCTGAAAAGGCTGCCAAGAAGGCTGAACAAAAGAAGGCCAAGTGTGCCAAGAGAGCTTCCAGAAAGGCAAATAGAGCTGCTAAACGTGCTGCTCGTGCTCAAAAGAAGGTTCAAAAGAGAAAGAACTTGTTGGACAAGATTAAGAGAAAGGTCAAGAATGCTGTTACTAAGGCCAAGACTGCTGTTGTTAGAGGTTGGAAGAACATCTTGAAGAAGGTTAGACAAGATAGAAAGAAGGCTCGTCAAGCTGCCAAGAAAGCAAAGAAGGCCGCCAAGAAGGCTCTCAAAGTCTCAACCAAGAAGGCCAGAAAGCTTGCTGCTCAGAAAAAGAGACAAGCTAGAGCTGCTGCCAAGAAACTCCAAAGAGATGCCAAGAAAGCTGCCAGAAAATTGAGAAAAGTTCTTGCTCTTGCTGCTAAGAAAGCCCAAATTGCTGCCAGAAAAGCCAGAAAGGCTGTTAAGAAGGCTAAAAGAGTAGCAACCCGTCGTGCTAGAGCTGCTGCTAAGGTAGCTGCAAAGAAAGCAAGAAAGTTGGCTAAACTCGCTGCCAGAAAGCAAAAGCAAATCAAGAAGGAAATTGCAAGACTTGCCAAGTTAGCTGCCATCAAGAAGGCTAGAGCTGAAAAGGCTGCCAAGAAGAG
>evgR004493c0g1_i01_Hydra
GCAGCCTCTTTTTTAAACCAAACAAAGGTCTAGGCGTTGACGATGTTTGTAGTAATGTTATATTAAAATCAATCTCTTATTTAAAATCTCCACTCATGCATATTTTAATATGTCTCTGGAACAAGGAATCTTTCCTGATAAATTTAAACTTGCAAGGGTGGTCCCAGTCTTTAAATCCGGAGATGTTACAAGCGTGTCAAATTACAGACCAATCTCAATATTATCATGCTTTTCTAAACTGTTAGAATACATTATGTATCACCGACTATTTCCCTTTTTTGAAAATAACAATATTCTCTATAATAAACAATTCGGATTCAAAAAGAATCACTCCACTGATCAAGCAATTATTCATCTTGTTCAAAATATTTTTCAGGCATTCGATGAAAATAAATATACGTTAGGTGTATTCATTGATCTTAGCAAAGCATTTGATACTGTTGATCATTATATTCTCTTAAATAAACTAGAAAGTTATGGGATTAAAAATTCAAATATTAAATGGTTTCAAAGCTATTTGTCAAATAGAAAACAATACATTTCTTTTAGCTGCGGAGAAACTGATAGTCTGAATATAAACTGCGGGGTACCCCAAGGATCAATTTTAGGACCACTCTTATTTTTAATTTATATTAATGATTTAAGTAGCTCTTGTAATTTACTAGACGCTATT
>evgR004499c0g1_i01_Hydra
CATATTTCCTACTATAAATAGCCAGATTCAAAATTTTTTTTATTGGTCTGCATGGAGTGAGTGGTCAAGTTGTAGTATATCATGTAACGCATCTTCAATTAGAACTCGTAACTGCAAGTCTCTTTTTGGGTACATCTGCAAAGGAAATAGTAGCGAAACAAAATCTTGCCCTGCAATTATAGGTAACTGGAACGAGTGGAGTAATTGGTCAATTTGTGTTTCTTGTGGATTCCAGAATAGAACGCGCTTTTGTTCGGCTGGTTTAAATTGTAGTGGGATAGACACTCAAAAAAGTAATTGCTCTTGCATACTCTCGATTTGCTGGCTTCAAAAAAAGCAATTTTTCAAATAAGCTCAAGATTCAGTTTGGATTACCGAAGATTAGCT
>evgR004505c0g2_i01_Hydra
TCATTCAGCAAATACTGAAAATACTCTAAATGATATACCTTTACGTTTCCTTAATGATGAGACTACCATTATGGAGTCTAGTTTAAAAACATACGAAAACGGTAACCAAAATAGAGTCACATATGCTAAAAAGGTTCAAATGCAACCACCGATTATTGAAAAAAGACAGGAATGGAAAAAAAATGATTGTGCCTTTATGGTAGCAAAAAAAAACAATGCAGAAAGTAAAGCTCTAACTGAAAAGCCTAAAATAAAGCCAATTTTTGGAAATAAAATTGAGTTCAACAGAAACTCAATTGCAGGTGAACGTAGGGTGCAAGAATTTTATATTTTTGTAGGAGGGGTCTGTAACACAATATCGGAAGAAGATCTGTC
>evgR004508c0g1_i02_Hydra
TGATATCGTTATTAAATCTGATGGTGGAAAGAGAGAAGTTGTAAAGAAGAGTCTTACTCTCTCTGAAAGATTCAGCCAAGTACACGAGAACAAGTCCTCACCAGTTAAAGGTATCACCAAGAAGTCTAATACAAGAGGTGGTTCAAGAGGAGGTTCTAGAGGTGGATCTAGAGGTGCCCCAAGAGGAGGATCTAGAGGTGGAAGAGGTGGTAGAGGTGGAAAAGCTCCTAGAGAAAAGAAGGAAGCAAAGACCAAGGAAGAATTAGACAAGGAAATTGATGAACACCTTGGCCACCACGATAAGCCAGCTGAAGAAGCTTCAAACAATACTGGTTCAACAGAATAAACCAACAGATGATACTAGAAGAATCTAGAGATTGTAATAATAACAGATTTGTCATTAACTTGACGATCACCTAAATAAT
>evgR004519c0g1_i01_Hydra
CGTATACGCAACTTTGGCGTGCTCGTATGCCGCACCCTCCAATTATGGGTATGCACCAGCAGCAGGACAAAGTGCGTATGCACCAGCACCCTCATACGGCGGTCCAGGCCCAATGGCAGCCACTATGACTGCCATGAAGGATACAGCAGCCATAATGGAGCAACTCCCTGTTGCTCAACCACCCCCACGTGATTACCAGTTCAACTACAACATTAACGATGAAATGTCAGGAAGTCTCCACCAGAGAGTTGAGGAACAGAAGAACGGCGTCATTACCGGATCATTTGCTTACACCGCACCAGATGGAAAGCTGATGACAGTAAAATACGTCGCAGATGAGAAAGGATACCGACCAGAAATTAGTGAAGCTCAAGCACCGATTATGAAAACAAACACTGCATCTACCTACACAATGAACCTTCCTGATGCTCCTGAACAAGTGCAGGTCTCAATCACTCAACAAGAGATGGATGCTTATCGTACCAAAGCAGGTGAGATGGAAAAGGCAGCAGCTTACGGCACATTCCAACAAAATTCACCTCAAACACATATGACATACCAGCAAAGTGCCCCACAGGCGCCAATGATGTATGGATCCTATGGTGCATCTGCACCGGCTGCACCCGCTATGATGCCAATGCAAGCACCAGTGATACAACCGGCGTCAGCACAAGCTCAAGGTCCTCAAAAGACAATGCCCATTAGATATTGATTTTCTCTCATTGCATACCATTTTTTTGTGAAAATATTTTTATATAATGCCCAAATGAATATATACTTTTGTCTAAATAAATCTTATAAATGACCATCTAAAAAAAAA
>evgR004528c0g1_i01_Hydra
AATCATCAAACAAAATGAAGGCTTCAAACAGTAGTGGAATTGCTTTTGGCAAGGAATCTGTCCTCTTACTCGAACACATCTTTTCCGATATTGATCCAAGATTTGTTGTACAGGTCAGAAAAGGTGATTTAACGATGGAAAAAGTTGATGTAATAGTCAATGCTGCCAACTGTAGATTACAACACATGTCTGGATTGGCTGGAGCCATTGTGAAGAATGGAGGTCAAATCATCCAAAAAGAAAGTAACAAATTGATCAATGAGCTCAAAAGGGAATTGGACAATGGTGAAATTGTTGAAACAACAAGTGGAGATTTGCCTTGTAAAACTCTTTATCACGCTGTTGGACCAATATGGTCAAGTAGAAAGACCAGTAATTTGCAAAATGTAGGAACTGAAC
>evgR004529c0g2_i01_Hydra
TGGATGTACATCACAAAATAGAAATGCCATTGTTTTAAACGCACTTCTGAAATTGGCAATGCAAAAAAAAAGTAGTCATCACACAAAAATTTTTAGAAAAGGGTCATACTCAAATGGAGGTAGATTCTGTTCATTCGGTAATAGAAAGAAAGTTACAACACCTTGAAATATTTTTACCTAGTCAATACTCTTTAGTTACCAAAGAAGCTAGACGCAATCCATCTCCATATGAGGTAGCAATGGTAGATCATACATTTTTTAAAGATTACAGTATTAAAAGCTACCTTATTTATGATACAATCCGTCCTGGAAGAGGTACTGGATCAAAATGTGTGGTTGATATCAGAGTTCTAAAATACAATCCCAATGGTACAATAGATTTCAAACTAAGTTACGATGAACAGTTTAAACAGTTACCACAAAGACCAAAGAAAATTCAACTAGTGATTCCACCACAACTCTTTACTGAAAGATTGCCAATCTCAAAATCAAAATATGATCATTTACAACAGTTAAAAAGTGTTATACCCAAGGATTGCCATGAATTTTATGACAACTTACCTTACAAAAATGATTAATTTTCTTTTATTCTGTTAAATTATAACTACAATGGCAGTTGTTATTTTTTGAGTAGTTGAAATA
>evgR004531c0g1_i01_Hydra
TGAACACGAGCATAAACACGAACACAAACATGAACACGGTGATTCATGTTGTTCACACGATCACCATCATGAACACGAGCATAAACACGAACATAAACATGAACATGGTGAAGAAGAAAAGTGTTGCAAAGGATTGAGCCAAGAAGAATGTTGTAAAAATAAGGAAGGAGCTGAAAAGTGTGAAAATGAAGAAGAATGGGAAGATGATGAATGTTGTGATGATGGATGTGGATGTTGTGGAGATGTTGAATTGGATTTGAATCCTAAAGTTATTACCGGTGAAACAAGTGGTGAAATTGTTCTTCCAAGATCAAGTGAAGGCCCAGCATTTGGATGGGATCCTATTTTCCAACCAACTGGTTTCACACAAACCTATGCTGAAATGTCAAAGGAACAAAAGAATGCTATCAGTCATAGATATAAG
>evgR004535c0g2_i01_Hydra
TTAAGTAAAAAATAATATAATTTTTAATTTAGAACTACATTTATTAAAATATGGCCGGAGAAAGTAAAGGGCACAATTTTGACGATGTAAAAAGTATTACAGATTTCCTTCTTTCAAAAACACAATTACGACCAACTGTTGGGGTAATTTGTGGATCAGGTCTTGGTGGTTTGGTGAATAATGTAGAGAACAAAGAAGAGTTCTCCTACGAAGATATTCCTGGATTTCCAGTCAGCACAGTGAAGGGTCATGCTGGAAAATTGGTTTTTGGAACTATAAAAGGCAAAACAATAGTGTGCTTGCAAGGGAGGTTTCATTTCTATGAAGGACACGGAATAAATAAGAGTACTTTGCCTGTAAGAGTGATGGCTTCATTAGGAATTAAAACTTTAGTTGTCACAAATGCGGCAGGGGGTATAAACAGAGGTTTTAGTGTTGGTGATATTATGATAATTAAAGACCATATTGGAATTCCTTGTTTAGCTGGAATTAGTCCATTAGTAGGGTTAAATGACGAAAGATTTGGCCCGCGTTTTCCATCTATGAATGGAGTTTATGATAATAATTATCAAAAATCTGCATTGAACATTGCTGGCGAGTTGGGTTATGGAGATTTTATTAAAAGTGGGGTTTATTGTATGGTTAGTGGACCTTGCTATGAGACTCCGGCAGAGTTAAGATTTATACAAACAATCGGTGGGGATGCTGTTGGGATGAGTACTGTACCTGAAGTAATCGTTGCTGCACACATGGGTATTAAATGTCTTGGCTTATCTCTTATCACTAACTACTGCATATTAGACTATGAAAATAAAAATGCTCCTAATCATACAGAGGTGCTTGAAGTTGCTCAACGAAGATCAGAGTGTGTTTTGGAGATAGTTACAAAATTTATTGCAAATATTGAATGATGATTTTTCAACATTTATAAATTCTTTTTCAAACTTTTTATTTTGTTATAATTTTTATTTTTTAATAATACCCAAGGGTAATCAATAAATATTTTTTCTAATTAAAAAAAAAAAA
>evgR004543c0g1_i02_Hydra
TGGTGGCGCTTTTTAGAGAGCTTCTTATCGTAAAATATTTACCTTGTCTTTAAAAATAAAAGGTAGACTTAACCTAGTTGTCTTTAAAAATAAAAAAAATGTAGATCACTTAACCTAATTTTAATTGTTATAAAAATTTCAAATGGAGTCAGTGCCTAACGAAAACGCAAGCGCTGCTTCAAAAAAGACGTTCATTTCGCATCTTGAATCTCTGAAAACCTTTGTTAAGAGTTTTTGCAAGGAGCTAAGAATGTCACCCTGGAAATCACTTATTGGTCCAGGGGAGTTACACGAGAACGACTTTTCCCTTATTCATATGTGGTATGTATTTATTTTATCGTTTATTCCTGTGTGCAGTTTATTAACTAATGTTGGCTTAATCCGCCCTTATATTGGTACAGACAGCGGATTATGCGACTGGGATGTTAATCGACGTAAGTGTGAGAGCATTATTGTTTTCTTTATTACTTACAGAATAAGTTTCGCTCTTTCTGCATTTTTTTTGCTGGCGGCAATACTTTTGACACAAGTTAAAAAAGAAAATGCGTTCGGCACATCAAGTTTGCACACAGGATTCTGGATTGAAAAAATTATAGTTGTTCTTGTTATATCCTGCGTGGCTTTTGCTTTTCCACCGAGTACATTTGATAGAGTTTGGTTATATTTTGTACTTTTTAGTGACATTGTGACATCAACTTTACAAATGTTTTTAATACGGGACGCATCAAAATATCTTCATGAAAAATACTTCGTTATGAAGAAGAAGAATAACACAAAGCATGTTCAGTTTTTTTTAGTTTTGTTTTGTGGTATGTTTTTATCCACGTTTATCACTATTTTCTTTACTTCTTTTTCAATTTACGAACAAATTATATCGCCAAGTAGCACGCAGTTCAGTATACTTGCAACAATTGTTACGGTTACAATGATATTGAGTTTTTTGATTTACACTGCAGTCCGTGAAAAACAAGAGTTTTTTTCGTCTACTCAAGTAGCTTCGTTTTGTGGATTACTCCTATACAAGTCGTCAACTTATCACGTTTCAAAAGATGAATCTAACGTAATTACAACGGATTTATCAAAAGTTATTTCCGTTGCAATATTATACATCACAATAATATATGCTTTTGTCAGAAAAGATCACGTGTTTTATTTTTTTTATTTTAATTGGACAATCTGTCGAGAACACTCAATTGTTGAGAGACGTTTAAAAACACCTTGGAATATTAATAAAAAAAAAAGCAACAAAATTAATCCGAATGAGACCTTGCTTGATCTAGACGGAGGCATTGATCCAAAGTCACTTGAATCTTTAGATTTAAAACAATCTCAGATTTTATTAATTGATGTAAAAGAAACTGGTCATCAAGTCTCATATAACGAGGCGTTGATTCATGCAGTTTTGTTTGTAGTTGCCTTAAAATCGACTGGTTTAGTAACAAACTATTATCAGCTTTCCGGGTCAACCGAAAAACTGCATTTAGAAAAAACTTTATCTGCATCAGCAGCATTGCTGTTTAGCAGTGTTTTTGTTCCCCTACTTTACTGTTGGTTTTTAGCGATTAAACATTTTATGCTTGAACCTATAAATCCACTGTCAGCAAACAAGTTGCTTGAAGAGTTTTCGAAGATTCTTTGGCACGTGATCTTTATAACCTTTATTAAAACTCCGCTATGTTTTAAAAAGTCAACTCTATCTCGATATGTTTACACTATTTTTTTTTTAACGACATTCACTTTGGCGTGCATTGGCCTGACGCCCCTTTTTAGGCGTTTTTTAGAAAAAACAACTTTTTTTTGCGACCACGTTGGCGCATTAGGCGCATGCATGTCAAAAGATCCAGCATTTATTGGACTCTATCGTATTTTTGCAACAGCTGGAATTTTTTTTTTAATTCTTTCGGTTCTTTTGATAAGAACAAAGACTATTGACAACCCCCGTGATTCAATTCATAATGGAAGTTGGTTACTTAAATTCATTCTACTAATATTAATATTTATAATACTTGTTAATTTGCCGAATAAATTCAGTCGTGTTTGGCTTTACACTAGTTTATTTGCAACGTTTGGTTTTGCGTTTGTGCAGCTATTATGTCTTCTTGATATTGTTGACATTATAAATACTTCATGGACTGAAACCACAAAATGGTCTCCGAATACTGTCTATGTTAGTTCAACTAGTTTAACTACTTTTATGTATGCTATCTCTACAGCAGCATTTGTTTGCTTTTATGTCTATTTTGCACACAATTATAACTGTCGTGTCAATCGCCTTTTTATTAGTATCAACTTGGTTATATGCATTTCAGCAAGCATCATTTCCATACATCCAATTATCAAATCCGGAGGTCTGTTACGTTCTGCACTTGTGACATCATTTTGCATGTACCTCACCTGGTCTGCACTGAACTATAATCCAAACGAAAAATGTAATCCTTTGGCGCATACTATAATAATGCTAGAGCCAAAACCAACGCGAGATGCTGTATCGATTACGGACATTTTTTTTCTCGTTATCACCCTCATTTATTTTACTACCCGTGTTGAATTTCTGTCTTCAAATATCGAAGAACTTTTTCCAAAATATTTATTTTACATTAAAATTTGCAATAAGACCGAAACAAATAATGAAAATCTTACAACAGAGCAACAAAGAGAAATTAAACAAGACAAACAATTTCAAACCAATAAAACTTTTCGAGATAGAATACTTTTTGGTGAAAATTGCTGTAGAGAACATTTGTACAACAATTTAAAATGTGAACAAGCAGAAGTACCCGTTCAAACTGCTTATAGCTACACCTGCCTTCATTTTGTATATTATCTTGCTACCTCCTATTCTTTCGCTTTACTTACGCACTGGTTGGAACCTGTTCCAGGATCTGCATTTAAAATAAATATTCACTGGGCAATTATGTCAGTAAAAATGCTTGGAAGTTCGACATGTGCGTTATTATATGTGTGGACATTAATTGCTCCAACTTTAAAAACCATGTACAAGAAACGGCAAGAAAATTAATTGTTTATTATTTGTTAATATTATTTGTTATTACCAATAGTTACCAGTATTTCAATTGTTTAATAATTATAAATAGCTAG
>evgR004546c0g1_i01_Hydra
CATTAATAAATCTTAAATGTTTATTTTCCCAGGAGTTGCTTATCTTCTTACTACAATGGAGGCATCGATGAAAGAGCATCGTGAAAATGACAAAAGGATTATTTGCGTGTTAAGTTCATTAGAAAGATCAACTCAAGAGTTATCCAATGCAGTTTTGCAGTTAAGCAAGCCCCCTATTCTTTCTTTGTCACATAGTATTCCAAAAAGTCTTACTTCTAATTCAAATCAATATTCAGATTTATTAAATAATTCAGTTGAATTAGAGAATGTATACTTTGTAGTATACAGTAGAGAATGTAGACTTTAACATGGCTCAAAGTTCTCATTCATCTTTTCAGAAGTTGAAGGCGCCCAGCAATTCTAGCATAGTAACTGCTCACACTCTAAACTCTTCAGATATGTCTGATCAGCTTAATAATTCATTTAACATAACAAGTAATGACTTTAATAATTGTCTTCTGTCTAAATCAAATTTTAACGAAACTGTTAGTTTATCCATTCCAATCTGGAAAGAACCTAATTACCCAGACGGAACAAGCAGAAAACAAATAGTGCCTGGTCACAAAATATACATTGACCATGTAAAATTCAGCAAATGCAACAAGTCGCATGGTACAAATAGATTTGTAAATGATCTTTTAGTGTCATTGGTGGGAAGCAATAGTTATCTGAGCAGTGTCAGTTTTTCTGGCAAGCAAAGTAATGCTCACTCTGACAAAGTTGCAAAACCCAGAATTTGTCAAGAACTTGTCGATGGTGTAATTGCTTGTGCCTGTGAGATACTAGAAGTGTCGTCGAAGGATGTGCGAACAGCAGTCAAGATTAAACTAAACATCGCATCGAAAGTTAAAAAGATTTAAACAATTTCGAAAGATTTTTA
>evgR004551c0g1_i01_Hydra
GCTCAAGTTTTTTTTCACCTAAGTGTTTCGATTAGTATTCCTTTGGTGAATGCAAATACTTTAACCTATGCATTATGCACCTGCCGTCATTTAGTTGTTGGGTTTTTATTTACAGCAACTTCATCAATAATTTTTGTAAGAACCCAAAAATATGTCAAGATTTTTAAATTTTTTTATCGTCTTTCAGCTCAAGAGAATAAATTTTCCACGTCGTTAGAAGTTTGTATAATAACTATTATGTTGTTAACGCAAGTTCTCTTGGGAATCATTTTTATGATTAAGTCACCGCCATCTGTTTTAATAAGCCTAAATTATGAAAAAATGACCAGAACTCGTTATTGTTCAACTGACTCTCAACTGGGTGTTCAAGTACTTTATATTGCCTTTCTTTTTTTCTTTTGTTCATTGAAAAGTTTTCAAGCAAGAAATCTTCCATCTTACTATAATGAAACATCTACAATCACGTACAGTATGGGAATATCTTGCATCTTGTTGCTATGCTACTTCCCAGTCTTCTATGGTAATGCTGATAAAATAAATCAAAACACAGCAGTTTCTATTTTATTTCTAATAATTAACTTGATAGTGATGTTGGTCATGTTTTTTCAAAAACTGTATATAATATTATGCAGACCAGAGCAAAACACTAAACGAAACCTTCAAATAGAAATACGTG
>evgR004554c0g1_i01_Hydra
GGTCAAACCTACTAAATAAAACACAAATCTTGTTAATTAAAAATAACTTGAGTAACAAAGGCATTCTGTTACTACATAAAACACCGAACATCAAGTCAAGTTTCTCTTTAAGTGATTTTTAAAAACAGTAATATCAGTTGGATAGATCTAAAAATGAAGGTAGAACTGCTAAGTGCTTCTACAAAAGCAGTCTTGATCACTTTTGACGATCTTTCGATTTATACTTCAGTTGGGGAAATAAAAGAACTTTATAGAAAGGTAAAACCACATTTGTATCCATCACGGCAAGCTTTTCGCATAGAGCCACGCGGTCCAATACTCGACGACAAGCAAACACTTCAATCATTAAAGTTTGATAGTACTGCAATATTTTATTTTAAAGATCTTGGTCCTCAAATTCAATGGAAAACAGTTTTCTTAGTTGAATATGCAGGTCCTTTATTTGTCTATTTAATATTTTATCTTCGCCCAAGTATAATTTACGGTGATGGTTCCAACACGAAGTTAACTCATCAAAATGTTCATCTTGCTTGTGGTTGTTTTGTGTTTCACTATGCTAAACGTTTGTTGGAAACTCAATTTGTTCATAGATTTTCTCATGGTACAATGCCAATAGGAAATTTGTTTAAGAATTGCACATACTACTGGGGTTTTGCAGCTCTTATCGCATACTTTATAAACCATCCACTTTATACACCGGCTACCTACGGATATTTGCAAGTATACACAAGTCTTTTCGGATTTTTGATTTGCGAGCTTGGAAATTATTCTATTCATATGGTTTTAAGAAATTTAAGACCGGAAGGATCAAAGGAGAGAAAGATTCCGAACGCAACATCAAATCCTTTTACATGGTTGTTTTATTTAGTATCTTGTCCTAATTACACTTATGAAATTGGCGCTTGGGTATTTTTTACAATAATGACCCAAACTCTTACAAGTGGCCTGTTTACCCTGGTAGGTGCAGCGCAAATGGCTGCTTGGGCAATAAAGAAACAAAAAGCATACAAACGCGAATTCAAAGAATATCCACGCCGCCGAAAATCGATTTTCCCATTTTTACTCTGAAGAATTTTAGAAATAAACTTAATTGACTCATTTATTTTTCCTCCTATATATTTAAGAATTAAATAAAGAAATTTAAGCAAAAAAAAAAA
>evgR004563c0g1_i01_Hydra
ATTTTTCTTATTTTTGATAAAAATTATGAACGCATTCAAAGTATCATTTTTTATACTACTATGTCTGGATTTGATTTGCTTTGGTAAACCTCAAGGTCCCCGTCAAGGGTCTCTAACTCAAACAAAAATTGACGATGCTAATTTTGCAAGACAGTGTGTTTTATGCAAAAAGAGTTTACAAAAATTATCTTTGAATCAAAGCTCTGAAACAGTTATCGAGAGATGTAACAAACAAACGTTGCACTCTTCTGATTGGTGCAAAGATATGGTGCAAAAAGTCTCAGACTCATTGATCAATAAAAACATTTGTGTAAATGTGAGCATTTGCGAGTGTTATCATTTTTGTTAAAGAAAGACAAGCGAAGAAGTTTTTTTGTCTTAATTATGTAACTTTTGTCTTAATTATGTAACTTTGTCTTAACTATGTAACTTTATTTCTTAATTATGTAACTTTTTTAATTTTGTAATCTAAATTTGAAAATGTCACAAAAAAAAA
>evgR004566c0g1_i01_Hydra
CCAGAAACAACAGCTGAGCAGCTTGCACAAGAATCAATTTTGGACACTCATGTGCCAATAGCTTTAAGTGCAAAAAGAGTAAAGATGGACATTTCATCGTATTGCAAGGTTGTGACAAAATATGGAACAAACGACAAGCGACAGTTAATAGCTGACTTGGATATTGTGAAATATTTGTGCATGGGAAATTTAGCTTTTCAACATGTTTCATCACCTGCTTTTAAAGACTTTTTGTTAAGCCTTCGTCCAGAAATTATTATTAGAGCACCATCAACATATTCAAGTGTCAAACTTCCAATCCTTTATTTTAATGTTGTTGAGGCAGTTCTTAATGTTTTAACACATGACTTAAAAGATGTAGATGCAGTTGGATTTACTTCAGATCTTTGGTCCAGCAAAAGCAATGAGGCATTTACAGCCTTGACGCTTCATTATATTACAACTGGCTTTGAACTAAAGACATTTTTGCTGGGCTGTAATGCTTTTCCAGACAGCCACACAGGTGACAATATTTCAATTAAAACAGACACATTAATTTCTG
>evgR004583c0g2_i01_Hydra
TATTTCTTATGACTATGGTTATTCTTGGTATTGCGGCTCGCGTTTGCCTCATTTGCATCGTTAACCGTAAATTACACTTAACGTCCCGTAAATAAATACGTCTCGTAAAATTTTTTTTTAAAGAGTTAATTTTTTTGAAAAAAATGACAAACAATTTAGAAAACATCTCAGGTCAGATTGACAATGAAATCTCTTCAAAGTTTCTCAACGAAATTGACAAAACGAAACTACAAGTAATAAAAGGATGGTTCATAGAGCGTAACTCGCAGTTACAACCATGGAAATCTTTTTGTAATATAAACAAATTTTCAAAACCCAAAAATGCAGGAGAAGTATCAAAGCGATTATTTACGAATATAAAAATTTACCAAGGAAATTATATTGCAGTTTCTGTAATGCTTATTCTGTATTGCATTTTAACATCTCCTTTGTTGCTATTTGGTTTGGTGATGTCCCTTGGAGGCTGTTATTATATCTCAACAAGAGGTCAAGGAAAGTCGATTAAGTTGCTTGGGCGCGATTTAACAATGCCTGAACAGTATGGATTGGTGTTTTTGATTTCTCTTCCACTCTTCTTTTTGGCATCCGCTGGATCAACTGTCTTTTGGATAATAGGTGCATCGGTGTTCATTATTCTTCTCCACGCATCTATGCTTGAGTTGAAATCTGTTGACGATGACCTTGAAATGCAAGAAGTCAACATTTTATAAGACCAATGAATTTTGAACCAATGAACTGTTATCTTTTGTTCATTAATGTTTTTTGCACATTATTAAGAATTGTAAAATAAAATTTTGGTGGATTTTTTAAATTATTTAAAACATTTTAAAGGTTTTATTTATTGTTTTACTTTGTATATTTCTTTTTCTTTTCTTCCCTTCTTTTTAATATCCTTTCTTTTTAGATTTTCAAAAAGGAAAAAAAAAACTAATTTTTTTTGATTTATTTCAAT
>evgR004584c0g1_i01_Hydra
AACCTGGAAAAAGAATATTTAGTTTCACTTGATTTTATTCCTTATAAGTTTGTAGAAGGTTTGCATGGTGTTATTTACTTTTCTTATCGTTCAACTAATGGAGTTCTTGGTATCTTCTGTAATGAGGATGGAACACTGCTAATTGTAACACCTCTTTATGGTTACTTTGATCTAATAAACGGAAAATATCATTCTTACACCACCTATAAAATTAAATTATTACGCTGGTTGAACATTGAGGTAGCTCAAATTTTAGAAGGCAGGAATTATGTATTCTCAATCAGATTAAATGGAGAAGTTGTTTTTTCTCAATCAAACAAGAGGTTTCAAGCCTTTGATAATGTTAAAGTATATGCTTCTTGTCTTGAATTCTGTGACGCTCAAGATGGATCAATAAAAAACCTTTTTATCATCAACGGCATATCAAACAGCAGCATGCAAGAGGTTTTAGTTCTTCCTAGAGATTTTATCAACCATGGAGAAGAAATTAATTTAACTCAAG
>evgR004586c0g1_i01_Hydra
TTTGTATAGTCATGTTTTTGCTTATTGTTTGTATAGTCATGTTTTTGCTTATGTTTATATTGGCATGGTTTTGCTTATTGTTTCTATTGTCATGGTTTTAATTATTGCTTGTATTGTCATGGTTTTGCTTATTGCTTGTATTGTCATGGTTTTTCTAATTGTTTGTATAGTCATGGTTTTGTTTATTGTTTGTATTGTCATGGTTTTGCTTATTGTTTGTATAGTTATGTTTTGGGCTATTGTTTGTATAGTCATGTTTTTGCTTATTGTTTGTATAGTCATGTTTTTGCTTATGTTTATATTGGCATGGTTTTGCTTATTGTTTCTATTGTCATGGTTTTAATTATTGCTTGTATTGTCATGGTTTTGCTTATTGCTTGTATTGTCATGGTTTTTCTAATTGTTTGTATAGTCATGGTTTTGTTTATTGTTTGTATTGTC
>evgR004594c0g2_i01_Hydra
CACAACAACCAATGATTCACCATAGACCTCCAATGCAACCAGTAATGATGACTCCACAACCAACTCTTATTGCTCAAAGAACTGTAACAACAACCTATACCCCTCCTCAAACTGTTCTCCAACCTGTTTTTGTGACATCTGGCCCAGTTGTCACCAGACAATACCCAACCGGAAACACAACCACTATTGTAACAACAACCACTCCAACAAATATTGTTCAACCTGTCAGAGTTGTCCATGGTCCACCACCTTCAATGCCTCCAGGTGCTGTAGGATACAGTGCTTACTATCCTCAACATTCTGTTCCTGTACC
>evgR004604c0g1_i01_Hydra
TGGTACTTGGTCTTTGGATATTTCTCAAGAAGCTATCAAGAAGAGACGTGAAATGGAAGGTCTCTCTGAAAGATTTAGAGAATTGACTAAAGGAATGGAAGATGAAGACGAAGAAGAAGAAGAACAAGAAGTTTCACCTGTATCTATTTTAAAGCAATTCATGGAAGTTAAACCAAAAGCTACCAATACTCAAATCTATCAAGCCACAAGAAAATTGAAGGCTGATTATTCATTGACTGATAAGGATTGCGTTTGTTTGCTTTATGAAACTTTCTTCACACCTGAAAATATTTTGAATGACATTTCAAAGAGAGCTCCTTTATTCAAG
>evgR004606c0g2_i01_Hydra
CTTTTTAACTCTGACTCAGTAGAGTGAAAAGTTTTTGTGTCAGGCAAAAAGAGGAAGAATTTATTAACCAAACGAAATTTAATTTACGTTACATTTTGAGAAAGTTTTTTGTCGAACTCCCCATATATAGTCGATAGGTATAAAATACCTCGGATAACTTATGATATCAATATTGTTTATTTAAGTGGAAAATGAATAACAGGCTACTTCTTGAAGAAGTGCCACGGCTTTATAATATTACAACTAACTCTCCAAACATTTTAATTGAAACATCTATAATTCCATTGGACAGCGAAAATTGCAAAATGGTTATCTCCCTAAAAACAATTATTTACTCCATACTAATGTTAGTAATCTTATTGGCAAGTGTCATTGGCAACGTTGTTGTTGTTACCGCTATTTTCATCTCAAAAGCGTTGCGAAGAAAAACTACAATGAATTTCGTCATCAGTTTAGCATTTTCAGATCTGTTGTTATCTTGTTTTAAAATTCCGTTAATTGTACTGACAATTATTAGAAATATGGATTTTTGTCAGAGTCTTGCGGCCTGTTACTTGTTTATTATCACGGACGTTATTGGAAACGTAGCCAGTATGTTAAACTTATTATTAATAGCCATCGATAGATTTATTGCAGTATCGTTGCCTTTTCGATATTCTGATTGGATAACGCTAAAACGTACTAAAATCTTATCGACTGGAATATGGTTATTTGCATTTTTCTGGAGCGCCACTGGTTTGTTTAAATGGGGCAATGCGAAAAAACAAGCTATTATAATTGATAAACGTATTAAATATGTTTGCAAAAACTCGAATGTTGCCTACTACGCTGTTTCATTTTATGGCATTTACATCACTGCGTTGATAGTAATGACAAG
>evgR004607c0g2_i01_Hydra
CCTATTTTAGTGCAAAAAGATATACCTTAGTAAAAGGCTCAGTAGTTCATGGCTTATAAGTTTACCTTGGAAAATAACTTACTTACTACAGAAGAAAGAGACTTCTATGAAAAAAAGGGCTTCTTAATAGTTAAAAAATTATTTTCATGCGAAGAATTAGAAGTGTATGAAAAAAGATTTTCAGATATTTGCAACCAACGTGTTCCTTCTGGTGAAATGACTGTTATGCGGGATGTGAACCAAACTTGTTCAAAAAATTCTGACCTTGTAGTTTATAAAATTCAAGAGTTTCAAACTGATAGTGTTCTATCAGGCTATTTTAAAAATCCGAATTTACTGAAATATGCCGAGTGTTTTGTTGGAGAGCACATTACTGCTATCCATACCATGTTGATCAATAAACCTCCTGATTGTGGTAAAAAATCATCGCGACATCCGATTCATCAAGATCTTTTTTACTTTCCATTACGGCCTGCAGAGCGCATAGTCTGTGCTTGGACTGCTATGGAAAATATAAACCGTAGCAATGGTGGGCTTGTTGTTAAACCTGGTAGTCATACTCATGAGCTCTTGCCACATCAATACCCAGAGTGGGAAGGAAAGGTTAACAAGGGGTTTCTTGGAATAATTGGATTTGATAAATTAAAAAAGAGTGGATTTGAATATTTAGAAATGGATACAGGGGATACCGTCTTCTTTCACCCATTACTTATGCATGGTTCTGGAAGAAATAAAACAAAGCACTTTCGTAAGGCTATATCTTGTCATTATGCTTCTGCAGACTGTGAATATATTGATGTTGCTGGTACAATTCAAGAAGTTATTAAAAACGAAGTTGATGATATGGTTAAGAAAAGATTTCCGAATTTAACTGTTAGTTTTGAGGACGTGTGGCGATTTAAGCAAGTGATCGTGCAAGGCGATGAAAAATCAAAGTTATGAACCTTGCTTATTATTTTTATTTTTTTTAATATATAATATAAATATAAATAATATAAAATCTATACTTTAATGATTTTTTAAAGATGAAAAATAAAATTATAAACCAGTATATATTAAAATATTGTGTCAATATTTTTATCAAATTATTAAATTTTTTTTTTAATTTATTAAAAAA
>evgR004609c0g1_i01_Hydra
AATAGCTTAATATAAAGAAATTGGTTCATATGAAAAAGCGATTTTTATTGGTAATGGACTTTTTGACGATATGAATTTGTGTCTCTCCGATAGTCTTCAATAGTTCGTGTTTTAAGGTGGCGAGTGATTGCGCCAATATAGCAAAAATTACATTTTGTACAAAAAGTACAGTAAATATGAATATCAAATGCAAAAAAGCCCTAAAGATTGCAACTAAAACTTCAATATGGCAATACGTTTGCCAGTTAGACGAGTACAAAGATAGTTTAGAAGAAATGTCAGTACTTACAAAATCAAGAGTATTGACAAAATTCACGGGTGAAGTGTTTGTTTGCGGTGAAAGTAAAATACATGCTAAAGTTGGAAGTATGCTAGAGATATCACTTGAGAGTCGTTTAAATGAAGAGTTTCCATCTAGCAACGGAATGATGAATTTCTTTCCTGGACCAAATGAAATCCCATATTCATACATATTTCAGTGTGGAAAATGTTTTCACAAAGAAATGGATTTCGGGATTTCAATTAAAAATAATAGCTCTAATTTCGAGTTAAAATTACCTTACGATATGTTTATAGCGGGTGAAACAGTATTTCACCACGAACGTTCGTTCAGTGAACTACTTGCAGAAAGAGTTCAGCACTTAACAAAATTTACAAAAGCTAAATTTTTTGTTGGAATTTTTCTTCATACCACCGTTCGCAATGAAGATGATTTTTATATAGAGCTTGTTATTTTAGGTCGAACTCCTAATAACATTTTCTCGCAAGAAATGTTTGATATATTAAATTCTGCACTTGAATCCAAAAACAAAGCTGAAAGAGATCAGACTTTTTGCGAAAGAAGCCTCCCTAATAAAGCTTTCAAGCTAAGTGACTCCAAGTTATTGAGTGAATTAGGGGTTGTCATAATTCAACATACTATAATGACAAGTCGTCAAATAAGCGATGGCGAAAGTTTAACTTTGAATTTTCATGTTAATGAGTTTGGAGTTGAAAAAGATCTTGTGGTTACAATTCCCAGTTACCAGTTACAATATATTAAAGCAATTTACCACCAATACTTCAGGAGTCATCATGCTGAATAAATTGTGTACCACATAACTAAAAAACTTATGTATTTCATAAATATTTTCT
>evgR004612c0g1_i01_Hydra
CTTTTATTTTAAAATGTAAGAGAACAAATATGGCTTCGAGACGAAGGTCGTCAGTAAGTACAGCAAGAGCGCAATCCGTTGTATCAGTTGATATAGAAAACTTGTATCCTCACAATCACTTATTTGTTTTAAAGCCAGACAACCCGTTTCGCAAAGCATGTGTTTCGATTTGCACTTCTAAAACATTTGAGTCCTTTATAATGTTGGTTATTGCTGGGAACTGTGTCATGATGGCTATGAATACTCCTTTACCTAATGAAGATAAATCAGATATTAACATAAAACTGGAAGCACTTGATGTGTATTTGCTTGGCATATTTATGATTGAAGCATTTTTAAAAATTGTATCTCAAGGTTTTCTTTTACATCGACACTCTTACATGCGAACCCCCTGGAATCTTCTTGACTTTGTTGTTATTGTTACAGGTATTCTGCCCTACGCAGGTCTTCAAGGTGGACCAGTTAAAGTTATAAAAGCTGCAAAGGTTCTTAGGCCTCTAAAACTTGTTTCTGGAGTACCAAGTTTGCAATTGGTCATGAGTGCAATTTTAAAATCGATGGGCTCGTTGATGGAGATTTCACTT
>evgR004618c0g1_i01_Hydra
ATTGGTTTGCTATTGATCTAAATGGAGTAAATATTCATAAAGTTGAAAATTCAAGAGTTTTAGATTTCAAAAACATCAAAGTTTATGCTTCTAACAATTGGGATGATTCACAGGATGGCTTTATATCAGACCTTCTTATTGCAAATGGAAATACTGATAATCTAGTTGGAAATATTTCTACTCCATTAGTGAGAGAAAAGATAGTTGCTGAAATACCAAAACTAGATAAAGAATATTTAGTTTCATTTGAAGTGAACCCTAATAAACTGAATGATGGCAATATTCTTCATTTTACTGATGCTTCTGATGAAGTTATTAAGGATCTTCAAATTGGCTATTTAGTTGGTGGTAAAAGACTAGTATTAAATATGTGGTCAATCATAGAGATATGTCAAGTTTCAAAAGGCTTAGTTTATTTATACACAATTAAATTAAATGGAGAAGTTGTTATTTCTAAAATTTACAAAGAAGTTCAATATTTTGACAATGTCAAGGTATATGCTTCAGGTCCAGGATTTGATGCTTATAGTGGTTCAATAAAAAACCTTACAGTTAATGGCTTTTCAAACAGCGAGATGAAGCATGCTATTATACTTCCAACAGATTATATTTTCCTCAGAGAAGAATTTACTCCATCTCCAGGTTCTTTACTTGGCACACTGAGTTTTTTAAAGAAGCAGTATACTTTGACTTTTAAAATAAAACCTTTGAACTACTCAAAAGGTTGGCAAAGTGTTCTTCATTATGTTCTGGATAAAGATTGTATTGCTTTGGGTGGAAATCCAGGAGTGGGGTTTTGTGAGTATGGTTCTGGGAAACTTGTTATTTTTGCTTCAATTGATGGTAGTGTCAAAAAATGTGTTGAAAGTGCATCACTTCCTTTGGGAAAATGGTCATCTGTTAAAATATATCAATGTTTAGTAGATGGAAAGTATTGGTTTGCTGTTGATTTAAATGGAGTCAATATTTACAAAGTCACAAATTTAGTTGTTAAAGATTTTAAAAACATAATGTTTTATACCTCTGATCTTCGATGTAATGCGTTTGATGGTTCTATTTCTGAATTTTTGATTACCAATGGAAATGTTGAATATCTTGTTGGTAATGTTGGCACTTGTATACAGCACAATAACTTGATTGCTACGATACCAATTTTAGATAAAGAATATTTAGTTTTATTTGATGTTAATATTTAGAGTTTTATTACAGATAAGGTTGGTCAAAAATATAAAGTTATTCAATTTTCAATTTATTCTACTTATATTTTTTACATTTCGTACCT
>evgR004620c0g1_i01_Hydra
CTGTGGTTTTTAATATATGGATTTGCAAGATAACGGTAGTAACGAAAATTCGTCAAAAGTAGAATTATCAGTGGATAAGCTAACAACTAACAAGTACGCTAATAAAAAGGTGTTGCTAATAGCAGCGGTATTGTTGATTGGAGCACCAGTTTTGGGTACGCTATTCTATAAATATAGCAAAATTAGGTCTATTGAACACGAAGAGATGAACGAAAGTAAAAAAGCAGAGAATCTCTATTATGATTTAGCTCCTATAGTTGTTAATTTAGATACTAACAATCAGGAGAGTAAATTTTTGAAAGTTTCTATCACACTTCAGTTTGACAATAATTCTAGCTTAGAAGCTGCCAAAAAATTAGAGCCAAGTTTGAGGGATGAGTATTTTGTTTTTTTACGTCAGTTAAGGCCTTCTGATGTTACTGGAGCGGTAGCGCTCTATAGATTAAAAGAAAGTTTGTATATGAGGACAAATAAGATTATTTATCCTGCAGAAATTAAAGATGTGTTGATTCACGATATATTAATTCAATAGTTTTTATGGTATTAAGTTAAATGGTAGGCATAAATAAGAAAGCAGAATTCCCGCAAGATTTAAATCGCAG
>evgR004621c0g1_i01_Hydra
CTGCCGGGGTCCGCAGCATCCCTCCGCGAGGTGGAGAGCCTTTGCCTGGGCTCGATGTCGCTCCTCACCCTATCACGCCGGGACCACGCCGGCTTGCTCACCTGGGCCCACGGCCTCCGACCAGAGGGATACCCCCAGGACGTGGCCGACAAAATCATGGCCTCCGCGGCCGAAGGTGGTCACATGACCTGCATGGCACTCGTCAAAAGCTGGGGCGCAATCGACTACAACAAGGCCATGGCCAACGCGGCCAAGGGTGGCCACATAGCCTGCATGGAACTTGCCAAAAGCTGGGGCGCAACTGGCTTTAACC
>evgR004646c0g1_i01_Hydra
CCAATATCAAATTGATTTAAAAAAGTATATATACTTTTTAAATCAGTCAGGGGTGTACCTATACCTATATGCTTTAGTTTACCCTCTACTGGAACTGCAGCCATATTTAATTTTCTATTTTATTTATTTTCAACCAAAATATACAGTTAGACCTTACCTAAAGATTAATCTTTATGTAAGTAAATATTCAGGTGATCCTTTTACTTTCAGCTTTCTACAGTTGATCACTACGTATTATATTACACGTAGGGATCAACTGCAAGTAAACCTAAAATATAAAGAACACCTAAAAGACTCAGAAGTAGGATCAAAAGCCCTTGAAATCAGGATCGGCGTTCTGAAATATCGACTTAACTGCATACCTAAAAGTGTCAGAGTGTCAATTTTTTATGGTGAAACCTTAATGAAAAATTATTTTTTGCTAACTTGACGCCAACCCAAAAC
>evgR004665c0g1_i01_Hydra
TTTCTGACAAAACAGCTTTTTATGTAATCATGAAAACTATTTTTACTCGTCAGGGAATACCTTCAAATCTAATTGCTGACAACAACCCCTACAACAGTGCAGAGTTTTGTAACTTCTTAAAAGAATGGGTTCAACTTTACTTCATCAAGCCCTAGTTACCCCCAATCAAATGGTCTCAGTAAAAATCGTGATAAAAATACTTAAAAAATGTGACAATCCTGAACTTGGTCTTCTGGAATATCTTAACATGCCTCTATCAGGTATGGACTACTCTCCCTCCCAACTTCTCATGAACAGAAGAACAAAAACAACCCTACCAGTTCATCGTGACTTACTTATATCATCAGTTCCTTTAAATGCTTATAGTCAGATCATTAAGTCCAGAAATCGTCAAAAACAATACTATGATCGGAACGCATCCGAACTCCCAAATCTCATAGCAAACAACACCGCCCGGATGTAAAAGGAAGGAACATAAAGAAAGTGACAGTTAAAGGAAAATTAACAGTAGCTAGATCATACAATGTGGTAGATAAATTCGGAAAAGTATATTGACGCATCGCAATCGCAAGCATCTGATTAAAATTAGCGAACAACTGCTTCTATACCAC
>evgR004671c0g1_i01_Hydra
GGAAAATGAGTTCATGTATTCAACAAAGTCAAACATTTTGGACAGCTGAAGGAGAGCCTGTTGAATCTCCCAATATAAGTTGTAATTTTAAAACTGCAAGCAAGATTATGAGTTGTGATATAAATCCAATAAATGCAAATTCTTGGAAAGGAATTCCATATAGCTACAACCTAATATACAGAACTAAATCACCAGAAGATAAATGCGACAAAGTAGAAAGCAAGCTCAACCATAATCAATCCATGAATTATAGTCTTTCTGCTACTGTGTATAAATTTAACTTTAACACTTCCGATTTGAAAAGTTACCAATTATGTGTTATTGCTTTTGCTACTACTAAGC
>evgR004676c0g1_i01_Hydra
ATATACTAGGTAGTGATTATGTAGCAAGTGATGAGACAACAGTACGAGTGTTAGATAATGAAGGGAGTACTTCATATATGTGGGTTCATCTAAGTGGAGAGAGGGATAAAAGAGCGATAGTATATGATTATCAGGACAATAGAAGGGGTGAGAATGCAGAAACATTTTTACAAGATTTCAAAGGATATCATCAAGCAGATGCGTATAGCGGCTATAATGTGGTGCATAATAGAAAAGAAATAACGTGGATAGCGTGCTGGGCACACGCCAGAAGAAAATACATTGAAATAACAAAAACAATTAAAGAGCCTGGTATAGCTCATAAAATGGTGAAATATATAGGGCTATTATATAAAATAGAAAGAGAGGC
>evgR004700c0g1_i01_Hydra
TTCACAAGGATGGATTTTGAATCAATATAAAAATGGTTGTATTGATATCAACGAGTGCATTAACTCTAATCCATGTTGGAATAACAGCTTATGTCAAAATACAGCAGGAAGTTATTTGTGCTTATGTGCAAAAGGATGGTCTTTGGGTAATGATAAAATTAGCTGCGTGGATAACAACGAATGTATAACCTCAAATCCATGTTCTTGGAAAAACAGCATATGTGAAAATACAATAGGAAGTTATTTGTGTTCATGTACGAAAGGATGGCTTTTGGGTAATGATAAAACTAGCTGTGTAGATTACAATGAATGCATTATCTTAAATTCGTG
>evgR004704c0g1_i01_Hydra
AGCAGTCTTTGCAAAATAGTTTAGAAGACAATGCGCAGTAAAATCATGGAACTTGATACACACGAAATATTATGCAGATACAGGATATCTTATTCCTCTTCGCATAGAACTACAGCGTTATTTATTGAGAAGGTTCAATGGTTTATGGTTTTTATAATATTTTTCCCAAACGTTATCCTGATCGCTGTTCTATTTGCCTCTAGAGAAAGAAACTTTTCTTTTCAAGAAAAACTAGTATTACTTTTATCAGTTGTAGACATAACTGTTGCAGTCATTCTAACACCATTTAAAATACTTATTACAACAAAATTGAGCAATTTAAATTGCTGGCATATTGGCATAGCCGCATTTTGGCATGTGTTTCCAACAATGTTCTCCTGCTGTATTATAATAGTGATATCAGTTGAACGTTACTTCACGGTTGTTCATAACAACAGGTGGCATGGCGTTAAGATCAAAAAGAG
>evgR004705c0g1_i01_Hydra
CATCAAGCAAGATTCTCTGCAACAACACCAAGAGATCATATGCCGTCGTTGCTCAAACTCCACGTGCCTATGATACTTCCAAGGAAGACTTCAAAGCTGCTTCTCCACTCCAATCCCTCTGGAATAACGGTGCTTATGAGGATAGATTAATTTCTTTGTTGACTGCCATCCGTCCATATAAAATTACCCCAGATTTTGTTGCTGCTGCTAACGAAGCTCTTAACAAACAATACAAGGCTTTTGACCTCATCCCAAAGGACAAGAAGAATCTTGATTTTGCTAGAACCATGGATGGTGTTGAAACTATTGAATTGGACGAACATACATTGACTTCCTTCTACGGTGAAGATTACAAGGAAATTTTGGATGAATTCTTTGAATTGAAGCCAGTCCTCAGCCAAAGAGCTATGGATAGATATCTTCAATTGAACTCAGAACAATATCTTTCAATCAAGAAGGAAAGAGCTGAAAAGATTTTGGCTTGGTGTCAAGAAACTGCCAAGAAGTTGTAAATTCAAAGTATCATATAAATT
>evgR004706c0g1_i01_Hydra
GATTTTTATCTTTTTTAACAGTATGTATAACATCCTTCGATCGTTATCTTGCAGTTTTTCAACCTTTTTTTTACCAAACGTACATAGATAAGAAGCCACGTTTTTACTTAAAATGTGTTTGTATTATTGCTATACTAGTAATGCTTGTTACAGGTTCGAGTTTTCTAACACAAAACAGAGTTTTATATGAGCTCTTTATGCTATCAATCTCTCCTATACTTATATCAACAGTTTTATATGTCCATACTCGATTGCATGTAAGAGTGAAACGGGTTAGAAGTGAAATATTGGCACATACTCCCAACCAATATAAGTCTAATGAGACCGATAATGTTCATGAAACAAAAAACATTGATTTGGTGACAAAGAAGCGTATCAACCATGACGTAAAAATTACCAATTTAACCATACTTGTATATTTGTCAACGTGCTTTGCATACTTTCCAATCATGACAGTTTTATGTTTGTGGCATATAAAGAAAACATCTTATCATTGGCAATGGACTCACACCTTTTATTTATGGGGCAGTACATTCTCAAAGCTAAAGTCAGTAACTAATCCAATAATATACTTCTATCGATCTAGAATATTGCGCAAGAGTCTTAGGCGTATAAGTCAATCATATTTTTAACGAATACCTATATCATTAACTTATTTTTTGCGCTCGTGCAATATTTGTGTC
>evgR004707c0g1_i01_Hydra
AGATGAATTTAATTACGTTTTCAGTTCTTTTATACGGTTATGCCAATGGCGCAGTTGAACTGTTTTCTTTAAATGGCAAAAGAGATATATCCCAAGGCCAACTTATTACTACACTTGATGTATTTCCTATTGAATATGAAATAACATTTGAAACATGTTTAACATTATTTACATCTGGTAACTCTTTTTCTTTTTATTCAAACTTATTCGATTTCACTACTGGAAACTCAAATGATTACAGTCCAACTGGCAAAAGAGTTATCGGTATTTGGTTTAATGTAAATGGCATAATATATTTTTTTGCAAATATTAATGGAACTGTATATAGCCAATTATTTACTAATGCTCTTTCACTTGGATGCAAAAAATTTGAAATAAGACAACTGCTATATCATGGTTATTACAATTTTACTATAAGTGTAGATGCAATAAGCATTTATTCAACGATAAACGTACATGCT
>evgR004708c0g1_i01_Hydra
CCCACTCCACGAAAAAGGATCTAAACTTGATGCATCAAACTACAGAAGCAAAATCCTAGAGTGTATTGTAAAAGATTGATTATCAAAAATAAAAAGTGTGAATAATCTTTTATCTCAAAATCAGAACAGGTTTGATCTAAATAAATCATGCATTGTTTATCAAATCTCACACTTAAAAGAAGGAGGCTATAGTATAAATACTTTATATCTAGATTATGAAAAAGTGTTTGACACACCTTCCCATAATCTTACGCTCCAAAAACTGGCTATGGTATCTCTAATAAGCTTCTTAACTAGATAATTTATTTTCTTTCTAACAGAACACAACGAGTTGTCATGGATTCGGCTACTTCAGATTGGTTACCAGTCACCAGTAGTGTCCAGCAAGAATCAATTTTAGGACCTCTTCTATTCATCTTACACACTAATGACATTACAGACCTAGTTAGCAATCCAATGAAGTTATATGCTGACAAAAGCAAAGTTATTTCCACTAATTCAACACCTGAGAACCCTGCTCTGCTTCAGAAGAACATAAATCATATCTTGTCTTGGACTCTAACCTTGGACTTACAACTCAACTTAAAGAAAGGAAAGCAGCTTCTCGATTCCTCACACAAGAAAAGATATCTTGGCATTTCACGCAACCTCAAATGGGATACACATGTTGAAATAATATAATCAAAAGCTAACTTGGTTCTTGGTCAACTAAAGAAATCATTTAGCAATAAAAACTAAATGAGCACAAGTTTGGAAGAAACTACACATCGCTGGTCAGACCTCACCTTGAGTATGCTGCTCCTGTGTCGAATTCTCGTTCTGCCCACAAAGCCTTAAAACATGAAAAAGTCCAAAAAGCTGCCACTAAACCAACTAAAGAATGACAAATAACTAGAGGCACTCTTAACATTTCAACTATTCAAAGAGTTCTTTAATGCTTCTGACCTCAATAACATACATGCTATTTTTACTAAACTTAATTTCTTATCTAACAGAGTAGTCAAGCAATGGAATTCACTCCCACAGACTGCAGTTGACTCTTTTATTTGAAAATTTTAGTTTTAAAAATTTGAAAAGAATTTTAAAAAAGCATCGACAAACACATTGCTGCTACAGC
>evgR004711c0g2_i01_Hydra
GTTTGTTGATCGCTCTTGTTTCGGGTGCCACTCTCGATGTTCAGGAAAGTGCTGTTGGTAATACCAATTTTGCTGAAATTTGTAACTTGGCCTCTAATAATTGGGCTGGTATTTTGGGAACTCCTTGTAATCCAAATAATTTTACCTTGTTGGTTCCTCCATCGACTAGAGTTCAATTGACTGGACCATCTTTTACTTCATCATGTGGTGTTACTTATGCTCCTGTTTATACTTTTGATATGAATGCTGGTTATATTGACGTTTCTAATTTGTGTACTATCAAATCTGTCTTTAGAGGTGGTGACAGAGTTGCTCCATGTAAATTGCCAGCTGGTATCTCTGCATTGAGCATGTATCCAACTCCAATTCTCAGTGGAAAGATTGCTGGTAGTTTCTATACTAATCAATATGGTTTGATTTTGTCTGATCCAGTTTATACTGGTCAACGTGTTTATGTCTCTAAGGTTTTGGTTGGATCTCTCATTGGATATGTTCAAAGTGATTATCTCTGTTTGGCCTAAATTGGAAATTATAAAAC
>evgR004713c0g1_i01_Hydra
TATGGTACTGTTTCCATCTTAAGGGATGATGGTGGTGAACTCCAATCTATTTATGGTGAACCAGAATTTATTTATGAAACAAGTGCTACTGAAATTTCTGGTTCTAAGATTACCAAAAAGACTGCCAACTATTCCAGTATCTATGTAAACACAATGCTTGGATATGGTAGCTCTACTGCCATTATTAGATGGAAGGTTAAAATCCTCACTTACAACTCTTGGATTGGTATCGGTGTTGCAATGAGAGATGTTGTTGAAAAGAAGAAGCTCACCTTCTCACCATACGACAAGCACGGCACATATATGCTCAGTCAAAATGGTTATGTTTGGAATCAATTCATGAAGGGATCAACCAAGTATGAATCAGTGGGATGTGCCTACACTACTGGTGACACTGTCATTCTCGAATTAGTTTGTGCAAAGAAGGAATTGAACTTCTACAAGAATGATACCACAAACAAACTTGCCTCTTTCTCAAATATCAAACTTCCAGTTTACCCAAGCTTTATGTTGTGTGGCACTGAATCCATTGAATTCAGTTGGGACTTGCAAGAAAGCGA
>evgR004714c0g1_i01_Hydra
CTTGGTAGACCAGAAATGAAGAAAAATATCCTTGAATCATCAGATGTATTATCAGTCCTTCACAAGTTACCAAATGCCGAACAATTTGTCAGATCGTTCTACAACTGTCAATACAATTTATTCTTGGTATCATTGGTCGAATTGATGGACAACCAATTGATCAAAGATCGTTACTTGTCAAAACATGTTAGATACTATTCCAGAGAAATGAGAGTTCGTGCCTACTCTCAATTCTTGCAAGCTTATAAGAGTGTAACTGTAGATGCTATGGCAGAACAATTTGGATTAAGTTCTGAATTCTTGGATATGGAACTTTCAAGATTTATCTCAGCTGATAAATTGAATGCCAAAATTGATAAGGTTTCCGGTGTTATTGAAGCTAGCCGAACAGATGAAAAGAATACATTGTATCTCCAAACTCTTAAACAAGGTGATCACCTCTTGAACAGAATTCAAAAATTGTCAAAGGTCATCGATTTGTAAAAATAAAACCAAAATAT
>evgR004719c0g1_i01_Hydra
GACCATGTTGTACAGAATACTTACATTCCATGCTTGTGTTTATAAATTAAAAATGTTCGCAACCTCTTGTGCTAATTTTTTATAGTAATTTAAGAGATTTTAAAAGATGTCTTTTCCTTATCATAAAAATAAACATTTTATTCACTTAACTTACAAAGATGCTTCCAAAATGGCTTCAAAGAGTTATTCCTGCATAGAAAGAGTAGAACATGAAAAGGTATTATCAATGTCTCCAAAGTTTATTAGTAGTATTCGTATTGGTTTGAAAGAAACTTTAAATAAAGAACTTTTTGTATATAATACTAGTTTAAATGGAGTTCCTTTTGCATTTGGAAAAATACATATACCTAAAAGTGAAATAATTGACGACCTTGAGAGGTTTAATTTAACTTTGAAAGTGACATTTATGGTTTTCAAACCGAGACTTGGAAAAATTTTACGCGGAACCATTAATAAAATTGCTGATCATCACTTAGGATGTTTGATTCATGGATGTATAAACGGATCTATTTTTCAACCTAGATTTGATTTACTATCTGCATCACAAAAAGAAGTAATTAGCAACATTAAAGTTGGCAGTGAAGTTTTATTTTCTGTTACAAAAATCGACAATCATAATGATATACTTTTTGTTCTTGGGGAAATTGCTCCACAATTTTATTTGTGTAAAAGGAAAGCTCAAAGTTCAGAAGTATTTAACACTATCAAGAAAATCAAAGAAGAAATAGAATAATGTAGTCGACGAATATTCAAAAACGCTCAACGCAATAAAAAAGTAACTTGTTCACGCCTTCTAAATATAGTTTTTAGAGATAAAAAAAAAAAA
>evgR004721c0g1_i01_Hydra
TTACAATAAAAAATAAAAACAAAAAACAATGGTGGCTGCAACGATACAAACACCAGTAAGCGAAGTAAAAGTAAATGATGAAACGGTCGAAACAGATAGTTCCACGAAAGTTGAAACAGTTAATTCCACGAAAGCAACGAATCAATCGAAAAAAAATTTCAGAAACTTGCGTATCGAGGTTCCAAAAGTACCGATTAGGGAATTAAGTAGCTTATCGGTGGGCTCCGCCAACTATTACTCGACTGATATCCAAATAGAAGTAAAAGATGTTCCCGGAAATTCCCCTACGACGGTAAAAAATGTAGACGACGAAGGAACATATGGTACTCATTACCCCAACAGGATTTTTGTTGGCCATTTACCTGGAAAAGCTAGTGCTGGAGACTTAGCAAACTTTTTTTGTTCTTACGGAACTGTTTTGGAGGGGAAAATTGTATTAGATTCATTTGGAAGGTCTAGAAGATTCGGTTTCGTGACTTTTGCTTCTCACGATGACGTCGAGAATGTATTATCAAAAAACGATCTTTATCTACATATCTAATTAACATAGCACAAAGTTGAAACGTTGCAAT
>evgR004726c0g1_i01_Hydra
AACAAAACAATAGATATTATTCCTGTTCTGAAAAAACCAAAAAGAAATTCAGCGTGAGATCTCAACAAAACAATAGATATTATTCCTGTTCTGAAAAAACCAAAAAGAAATTCAGCGTGAGATCTCAACAAAACAATAGATATTATTCCTGTTCTGTGGATGAAAATGAAGTTGAATGCACGAATGAAGAAAATGACTTTGCTGATAAATATCAATGTGAAGAATGTTCTTACTTTTCGGTGGATAAAAGAAACTTTAAAAAGCATATGAGAATCCACAAGAAGATTAAGCCCTATTACTGTAAACCTTGTAATAAGTGGTTTACCGACAAAAGTAATTGGAACAGACACAATAAATCACTACACCATTAATGAAAAATAATTTTAAATTAATTTTTTCTCTGGTTATGG
>evgR004732c0g1_i01_Hydra
ACCTCTTATAGATCTTATAGAAGAGATTTATGAATCAAAGGAATATAAAATTGATCATTTTAAAGAGATGAAAGATTATCAGATGCTTCATGGTTTTGACCCTATTTTCAATAACAGAGTAAACAGCCTTTGTTTAGCTGAAATAAGTGTTAAGAATGAAAACAAAAGCGATGAGTATAGAAGCAGTCTTTTTGAAGAAAAAAAATTGTTCTAACAACCATTAAAAAATTTAAAGAAATTGAAAATAATCGGAATTTGTATCATTATTTTTTTGGGTTTATAAAAACATTTAGCAAAAGTTATCATGCGGCTTTGTCAGTGACTAGCGGTCAACTTGCATTAGATACAGAAAGCAAAGTTTTAAAAACATCTTCGTTGATTATTGGATTGATTCCTTTGTTCGGTGAGTTTTTAAAAAATTCTATGAATGAAATAATGCATTACATTAAATCCGTAGAATTGAAAAATGCTGCAATATATTTATCATCACTTGCTGCTCGCTCAAATGAATTCGAGGACATGATACGTGATATTGTTGTTGAAATAATTTTTGATAATAAAGAAGAGTTTCTAACAAAATCATTTGACCATAAAGCGAAAAACTGGTTCGACAAAATGAATAACTTTTGTAGAAATATGCTTGAAAATATTGACGAAGCGATTTATGTCCAAGAGTTTCCAGAAGACATGCACAAACTGGGTTACAGAGACGCCGTCAAAATAATTCAAAAATGGATTTCAAACGGCAACATATATGGCGGACAACATGGCGCCTCTCTGCTTCCAAACGATAAAAAAGAACTTCTCAAAAAAGTCATATTCAATGAAGTAATAATTTGTCAACTATCAACACGAATTTGTTAGTTTTATGTTGATATAATGTTTTATATATATTTTGATAAGCAACGGAATCAAGCGAATTTAGACTTTATTAATTTGTGTTGTATTTTC
>evgR004734c0g1_i01_Hydra
ACACTCAGCAAATATCCATTCACAATAACTATGGCAACTACTGGTACAACAACATAAAAATGTTCAAAATAGCTAGAGCTGCTAAACACTTGCAAAAGCTCAATCGTAACTTTCATTTGTCTTTCACTTTCAAAAATGTTATAAGCGAGGATAATGATCATTTGCACTTAACAAACAATCCATATTTTGAAAAATATGCTGAAAAAATCAAACAAGTTAAAGATTCTGGAATATATACTGTTAAGAAAGAAAAACAGAATCTTCAAGTTACAAAAGAGACAGCAAAATGGTCTGACCAGATTCAACAATTTGAAATGTCTCTTTCAAGACAGCAACAGCAAAATGCTGAAAAAGCTGGTTCAAAATTACCAGGAGAACTAGATAAACTTATGAAGTTAAATTTATTAGAAGATAAATCTGGGGAAGAAATAGGACAAATTTGGAATGCATATTTTAAATCTACGGACAGTGTAACAGCTGTAATTCCTTGTTCAACATTTGATCAAATTGTAAAAAGAAAAAAAGAGTATCCAATGTTCTTGTATCCATTACCAAAATCAGAAGGTTACGAATTTGTATTGTCGAATTTTACAAACCAAAATCGTTGCTTTTTTACTTCATTGATAAATTATCAGGCTCACGGTGACAATGCCCCATGGCAGTTAAGTCTGGTATACTATACAGAATTTAGAGAGAGTAAAGGAATCGTGTTGATGGCTGGTGAGTTTGATGCGAAGTGTATGAATATTTTAGAGGTGCAATGCTTAGTTCAATTACAACAATTATTTTATGCTACAGATAGCCCTGCTCGCTTGAAACTCCTAATGCAATTCAATTGTGAGCCAAATTCTTTTAAGCACATGAATTTAATAAAAGAGATGGAAAGTAGTGAAATGGTTGTTAAAATGTAGTTAAGAATATGTATCTGATCAGAATAGTATGTTAGATATATAAATAGTATGTGATATTGGTAAAGTGGCAGAGCGCTTGTCTTGTAAGCCAGAAAAGAATTTTCTGTCTCACTAGAATTCATGTCTTTGGTATTTACCACGTCTTTGATAGTTACCAGTGACGTTGGTTTTTTTACTAAGACTTAAGTGGACTACCACTTTTTTTTTTAAACCGACCAAGACGAACAAGGCCTGGACAATGAAATAAATTGAGTGAGGTACTATCATAGACATGGTGAAGATTGAACTCAAAACTTCTTGTTTACAAGTTTACATGGCGAGCGTTCTACTAATACACCACTAACGTATACCACGCCACAGTGGGATGGTTACATAGAAAACGGGGGCCATAATTTTTTTTAATTTAAAAATAAATTTATATGTATGTGTATTAAAGTTTCACATAATTTTAAACTGTAGTTTGAGAGAAACCTAACCTTTTAGCATAGCTCAGTGATACAGTAAC
>evgR004738c0g1_i01_Hydra
AAATCATCTTTTCATTGTTCGTATTGCTCAAGGTTTATTATGTGCTGGTAAAGGCTTAATTACCATGAATCCATTCTACTCTGATCGTTCCATTCTTTCACAAGCACAATTGTGTGGTTTGATGGCTCTCTGTTTTACTGGTCTTGACTTTAAGAATACCATCTTGGGTAATTACCATTACTTGTTGTACACTGCCGCAACTGCAATGTATCCAAGAATGCTTATTTTGGTTGACGAAAACCTTGAACCAGTCCAAGCAAACGTTCGTGTTGGTCAATCAGTCGATGTTGTTGCCCAAGCCGGTAAACCAAGATCAATTACAGGTTTCCAAACACACAAGGCTCCAGTCATCCTGCAGTTCGAAGAAAGAGCTGAATTGGCCTCTGAAGAATTTATTCCACTCAGTTCAACACTTGACCATGTAATTATTGTAAAGAAAAACCCAGAATACGAAGCTGACCTG
>evgR004753c0g1_i01_Hydra
ATGGAAGAATCTGGACAAGGTGTGGAGAATCATCAAGTAAAATCAACAATGAAACAACCACACGGAATTCAAAAGACACACAAGATGATTTCCAAATTACCAACTTCACATGTTTCAATTTCATCCATTCCTTCTCAATCCAACAATAACAGTAATTCTTCCTCACCTGCTTCATCTTCCAATTATCATGATCCAACCACAACCAACATGCAACAACAATCAGATTCTCAAAGTGGAAGTAAATTCTCTCGTCCTCTACTCAAGAATGAGACTGTTTTGTGCGAGAAGAATCCAAAATTCAAACAAAATTACTTTGTCTTTGAGAAGAAGAAGGAAAAG
>evgR004754c0g1_i01_Hydra
TCAAGGAGAAATCACCGAAACAAAAAAATGAATTTGATTACGCAATCGAAGGTAATGGACTCCTTCCCGTAGAATGCAATGGGCAACAAAAATATACAAGATCTGGACGATTTGAAGTTGATAATAAAGACTATATTAAATCCGTAAGTGGATGCAATTTACTTGGCAAATTATATACCGATATAAATCAAGAACAAGTTACATCAATTATAAAAGACGATTTAGTGCCTATAAAATTAGATAGATCGCCTAATAAAGGAAAAGTTACTGCAAACGTAGGCATTGATGTTTATTTGGATGCTAAGGTGGTACCACTAACACCAAATACTTATGATATCAACAATGTTGCTAAAATTATGGCTAATGGAAAAATTACACCTGACATAACATCAGAGTTTGAAATAATAGATTCCTTGGGTGTTTCACATAAGGCTTTTATAGCATTTAAAAAAGTAGCGCATTATGTATATGCAGTAGAAATGTATGCTGCACCCGGTTCTATAACAACAACAAGAGCAGACCAGTTAGTGGCTGCAGGATCAATAAAGTTCGATCCGAAGACTGGTCAGGGAGAGATTATAAATTATACACCACTTCCGACGGACGTTTTAGATCTTGGTAGTGCAATAACATTTAAATGGAATCCAGCCAATCCCAAATCAGGGGATGTTGGAGCAGACACTACGGTAAATTTTAGTTGGAATACATTGTCAGAGCATAGTAATTATAATAATAATATAAAGGTTACTTCCGATGGTAAGTTAGGTGGAACTCTTACTGGGACTAATATTGATGACAAAGGAAATTTAATTGGTACATATACCAATGGCGAACAAAAAAAACTGGCAGCTATTCCAGTTGCATTATTTGATGATCC
>evgR004756c0g1_i01_Hydra
AGTTTGATAACAACTGAAGATTGCTACAAAACTGCAGACTGCTTTCTTTTTCCTGAGTACTGTACAAATAGTGATAACTGTGATGTTTTAGTCAAATATAGATTTGATTCTGTTAAAAGCATATTCTCAATTACTTTAGCAACAAACAGAATCTGGATTGGTTTTGCACAAACATACAATTTAAATATGACACAAAAAATGCAAAATATAAAAGGAGAGATTTGCTTCAACAACAATTCTATTTCATCCTTAGCTAATTTTCATGCATTGCGTAACACAAGGCCAAATTATTTTTCATTGGTAAACGGGATCAATGTTCAAGAAATGTATGTAACCCCAAGTGGCAGTGTTGTTTGTAAATACAGTAGAAGTTTGTTAACTCCAGTCGGAAGTGAAGAATATATGTTTTCCCTTGACACCCCAGTTAGCATTGTTTATGCATTTGGAACAAAATTAAGGACAGATGGTTACCCTACTTATCATGGATCCAAAAACCATAGCTATCATTCTGAAAAGATTGACATGAAGAAAATAAGACCCAGAAGTGACAGTCTGTTGGATTTTTCTAAATGTGGAAAAGAGTTTAGCTGTTTTCTAAAGCCTCCTGGTTGCATCAAAACCTGTAAAGTTGCAGCAACATTTGAGTATGATCCAAATTCAGAGAAAGTTGTTTTCCAACTTTGGGCAAGCAAATCATTGAAATATGTTGCATTTGGACAAAAGCCAAACAATAACGATAATTTTATGATTAAAATACGTGGTGAATATTGTTCTCGGTCACAGGTTGTAGGTTTTGGTCATTTTAATGGTAATAACTTAAAGGATGCTGGAACACCACAATGGGTTTCTAATGTAGACCAAGTGAAACTTCTTTTATCAGAAGTCTTACCAGATGACTCTTTTCTTTGTAGGTACGAACGCCCATTAACATTATCTAGTAAAAACATGGATTACTTATATGACATGAGTTATCCTTTGTATGCTGCAATTGCTTATGGCAATGAATTATCAGGAAGTTTCCCTGACTACCACAATGGTCATGAAATATCTAGTAGTGCGATCAATTTTAAAGTAGTTAGTGATGTAGATGTTGACACCATATCAATTAAATTAAAAAAAGCTCATGGAAGTCTAATGGTGCTGTCTTGGATTTTGTTTGTAACTTGTGGCATATTTATGTCGCGTTACATGAAACCATTTTTAACTAACAAAATAGCTGGTAAAGATGCCTGGTTTAGAATTCACCACATCTGCATGGTTATGGCATTGCTGTGCATGATTGTTGGTTTCATTGTTATTCTAGTATTTTTTAAGGGAAAGTTGTACCTGAATGATATACATCATTGGTTAGGATTCTCTGCTTTTATTCTTGGTGTGCTGCAGCCAATACTTGCAGTTTTTCGATGTGCACCAGACCATAAAAATCGCTATATGTTTAATTGGCTTCACAGATTTATCGGAATGACGGCTTGGTTAATAGCAGTTTCAGCTGTTATTTTCGGATTAAAAAAACTATCAATCGATATTGTACCAATCATTGTATGCGCATGTGTCATCCTTGTACTTTTTGTAGCTCTTGATTTCGCACAACTTTTGATAAAACGCAGTTCCGCTACAGAAAATTCGACCTATGCAAAATTTATTTCCAAGTCTGACTCTTCCAATAATTTAGAAGTTAGCGCTAAAGTAGAAAAGAAGATTTCGAAATGCCATTTCGTCTTTATGGCTATTGTTTATGCGATATCTTGTGGGGTAGCAATATATTACGTAGTTATGATAATAAAACATTAACACAAATTCACTAGATTTAACAATAAATTTATTTAACGGGTG
>evgR004758c0g1_i01_Hydra
CGGAAAAAAAAGAACATACAAATTTATGGACTTTTACACTTTCACATGGATGTAATGTTCAAAAAGTACATTTTGCAGGATTTTTCTGGCTTTTATTTTGGAAAGGCATTTGTGCACAACAATCATACTATTTGGCACCTAATAGCAACGATTATAAAATATCAGATGCAACATATTACAACCATTATTCAAGTACCTTAGAGGATATTTGTCAATTGTTGAACATCACAAGTACTTCTTATTTATCAATAAGTCCAAATGACATAGAGTTTGACTCATCACCAGTTGCGTACTTTACTGTAAAGATGAATAATGTTAAACTAGTTCACCAGTTTATTGTTAAATCTGGTACTTATAATGAATTTAACTCGGTTGGTGGAATCTGG
>evgR004761c0g1_i01_Hydra
TTGTCAATTGCTAGAATGCTCCCATACATTATCAAGAAGCTCGAAGACCACAACTCTCCAGATCAAATCTGTGGTGGTTTGCACTTGTGTTCTGCCAAGGCTGCTTTCATTGCTGCTCAATTCAAAGCTCCAGAAATGACCTCTGAACTCAAGAAGTTGTCAATCTGCCCAGCTTGTATCATGGCTATGGAATTAGTTCAACAAGAAATTAGTCAACAATCTGTTGAAAAGTTCATTGAAGATAAGTTGAACAAGGTCTGTGCTAAGTTGCCAGGTATGTTTGGCCCTGCATGTGAAAACTTGATTGATCAATATTTGCCAACCCTCATCCAAAAGTTGCTCCTTGCTGTTTCTCCAACCAAGATTTGTAAATTCGTCAAGGCTTGTCCAAGTGCTGCTCCACAAAATGTTATCAAGTCATTGAAGGCTAACATTGTCAAGCT
>evgR004761c0g2_i01_Hydra
CTCTCCACCGTTATTATTGCCGTTTTGTTGGCCATCATTGCCTCCATTTCATTCACTCGTGCTGCTCCAACCCAATTCTGTGAAGTTTGCAAGTATGCCGTTCAAACTGTTGAAGCTCTTGCCCTCAAGGAAGAACCAGCCATCGAAAAGGCCCTCCTCAATGCTTGTAGCCAATTGGGTGATAACCAATTTGGAAAGATCTGTAACACTGTTGTTCTTGTCTATGGTCATGAATTGTTGAATGCTTTGATCACCTATGAAACTCCAGTCAAGGTCTGTACTGATCAATTGCCACTCTGTCCAAAGAATCAAACTGTTGCTGCTATTGCTGCTCCAGTTGAAGAAAAGACTGAAGAAAAGGTTGGTAATGCTGAATGCGAAATCTGCAAGTATGCCATCGCCCAAATTGAAAACTTCATCAACTCAAACCACTCTCAACAAGAAATCCAAAAGAAGTTGAACCAATTGTGTGCCAACATTCCATCAATCTTCACTCAAACTTGTTTGTCAATTGCTAGAATGCTCCCATACATT
>evgR004769c0g1_i01_Hydra
AGTTAAATAATTAACAAATGAAGTGCCAAATATTGTTTACTCTTATTGGAATTGTCTTGTTTATAACAATTGAAGGCTTAAAAGATATCGACAGAAACGGAAATACTTTTAAATTCGGTTGTCATTTTGAAGAAAACCATTGTATTAAGATTATCAATGAAAAGAAGGAGTGGTGTTACACAAATGAAAACGGATTAGAAACAAAAGAACCTTGTAAAACCGACGACGATTGCATTAACATGGTTGAATGCGCGACCCTTTTGGATAATTCAACCACTTATGACAGTGATCAAAATAACAGTCAAGTTATTAAAGCACATGCGAGATCTGAGCCAATTTATAGTTGTAGTAGCTTTCGTTCTGAAAATCAATCTCCAATAAACATTAGAACTCATTTAGCAAAAAGAATGGTAGATTGGTTCCCAATTCAAGTGATTTTTCCACCCCCTTCAACACCTTTTAAAGAAGCTCATTTAATTACTGCATACAATAAACTGAGAATTCTCAAACTATCAGTTCCAAGCGACCCACCAATTGAATTTAAAGCACCTATCCTTGAAAACAATTATCTTTTAGTTGAGGTTCGTTTTTACTTTGGATGCAAGTCAGATAGAGGATCAGAACACCAGATTGATGGCAGAAAATATCCGTTAGAAGTGCAGTTTGTGTTCTGGAACAATTTAAAATACTCATCCTATATTGATGCTCGAGAATCCAAGACTAAAAATGCACTTGCTATTGTTGCAGTTCTATTTGAAGAAGATGAATTTAGCAACAAGATTAATAATTTGAACGTGTTAACTGATGCAATACACGAAATTAATAACAAATATAAAGGACAAACAACCATAAAACAGAATTTGGGACAGATCTTACAGGCTCTTCTACCAGATGATATTGACACTGATCAAACTTATCGTAAATTTTTCACGTACAGCGGGTCTTTAACAGTACCTCCGTGCAGGGAGGTTGTAGAATGGTTTGTTTTGGCAGACTGTTGTAAACTTAAAGCCCCTGCAGATTTTATATTTGCGTTACAAAATCTTTACATGATAGATCTCTATAAAAGAGATTACCCTATGTGTGATAACAATAGAGAGATCCAGGAACAAGCTAGACAGGAGGTTTCGTGGTAACAAATATTCATAATTAAAGTCGAAACTATAGTTTTGAAAGTTGTAAAATTCCCTGGTTATCTGTTTTTATTTGAATGTATTTAGTAATGAAATATTTTGGTAATAAAATACTGTGTTCTGTAAGACTGTTCTGTAAAG
>evgR004787c0g1_i02_Hydra
ACCTAATTAATAACGATATTCAAAGTGGTTTTTCATAATTAAATTTTACTGTGGAATTTTTGAAACGAAGGCAAAATCAAATAGAAATGACTGAAAGGAGGGAAAAAGCGGAGAATGTATTAAAAGAAATACAGAAAGAAAAAACTGACAATGTGATGCAACTTATACAGACGGAAGAAACTAAAAAGGAAGTTGCAGAATCTAACGACAATAAGGGTTTGTTATCTCGTAATTTGTTATGTATGGAGAGAGCATTTGGTGCAACTGGTTATTGTTTTATTGCAATAATTATTGTGATTTTGATATCTGGTTTACCTATTGCGATGATTGTCATAGGAGAGCTAAAAAGAGAGAAATGCTCTGTCGAACCAATGATTCCAATATTTTTAATAGTTGCTGGAGCAACAAGTCTTCTACTTTTGTTGCTTGCAGCAGTAAATGGTTTCCAAAAACTGTCAATAACTAAATTCTTTGTTTCACTGATTACTCTTTTTTGGTTTTGCTGGTTTATAACGGGAAATGTTTGGGTGTTTAGAAACTATAAAAAAAGATTCAACTGCGACCGCACTGCATACTACTTGGCGTTCTCTTACATTATTTTAAGTTACATTTTAACTGTTTTCAGTTTCTGTTGTTATCTCTTATAGCGGAGATCAAAAGTTTCTGTTGTTATCTCTTATAGCGGAGATCAAAAGTTTAGGTTGTTATTTCTTATAGCGGAGATTAAAAATCGAATTTAAACTTTATTTTATCACTTTTTCAGCACTTTATTGTAAAATATTTTCTTATTTAAAA
>evgR004788c0g1_i01_Hydra
ATAATAAAAAAAAAATGTTTCAATATATCATCGACATGGAAATGGTTTCCAATAAAAAATGCGAACTAAAATTTAAACGATGTATAAGAGAATTTGCTTTGTTGCGTGTAGAAAATTTTCAATTACTTCACTTTCATTGTGTATGTTGTACAAGGTTTGTAATTTGTTTTTTTTTTTATTTTTTTATGTTTGTAAATTTATTTTATAATTATATATTTTTTTTTAGTTATAGGATGTTGACAAAAAAAGAACAAAAAACGGCGTCGTACGTTACTAAAAATATTACTGGACTTTTAACCGATTGTCAGATTAAAAGAACTGTCGATTGTACTTATAATGATTATAATGTTATCCATTGTCAAAACATTAAAGAAGAAATTATAAAATTTATAACGAAATACGACAACATTGAAGACGTTGTAATAGCGCATAAAGGTGGTTTTGACGGTAAAATATTAAATAGTTTTAACCTCAGAACAATCGACTTGAATCAATTTAAAATTGAAAAGTTTAATAGTGATAAAAGTTACTTTGGTTTAATAAAATGTTTGGACCATATATACGCAAACCAAAATGCTCTGCATGCAAGCCAACCTATTCATTGTCCAATTCAAGAAATTGGATATTTTTTAAAAGAATGTGTCGAAAAACAATTAATTACGATACAACCGTATACA
>evgR004791c0g2_i01_Hydra
AACATGGAGTGAATACAAGCACCATAACACAATGAAAGTTCTTGTTTGTATAGCTCCTAATTCTTCAATAATGTTTTTGTCTCATTGTTATGGAGGATCTATCTCAGACAAAGCAATTACTAATTTAAGTGCATATCTTGACAAAGTTCCACCTCATTCTCAAATTATGGTTGACAAAGGTTTCCTTATTAAAAAAGAATGTGACGCTCGACACATTATTTTAACTATGCCACCTGGAAGAAAAGGTTCAGAGCAAATGACACCTTCTGAAATTGTTAGCACGCAAAAAGTTGCAAATCTAAGGATTCTAGTAGAACAAGTTATTCGGAGGTTAAAGACTTTTAGAATTTTGGCTGGAGAATATCCAATTAACATGATAAGATGTTTTGATGACGTAGTTACAGTATGTGCTGCATTGTCAAACTTACGGAAACCTATATACTTGTAATGAAATTATAGACTACTATTCTATTTGCGTTCGCTTTTAATAGTATAAAAAAATAATAATTTATCTGTATGCATTTGTAGTATAGGCTGTGTTAATAAAC
>evgR004796c0g1_i01_Hydra
CCAGTTTGGTTGCCGCATCATTGATTTAAAGTCGCTTGAGATAAGATTGATCTGTTATGTCATTGTCAAAAGATATTTCTTACCAAAGTTTGTATTGGTGTCTTGGTTCAAGTATTGTATCAGGATTATTTTATTATGGATATTATCGCAGAAGCTGTTTTATGAATATTTTAAAGAATGTGGAAGTTTTTCAAATAAACAAAGATTTAATAGCAGAGTTAAATAATTCAGATGACCACACCCTTTCATACGTTGCTGTAACTGGTATAGTAGAGTCTGAAGGTAAAACAATACGTGGGTGTTACAATGCCGCTGTTGATGGGGTTGTGCAAAAAATATTTCTTAAAGAACACAAGACAGAATGGAATACTTTTACAAAGGAGTGGTCAGATAAAGTTCGTATATTATCAACAAAACAGCAGGTTGAACCATTTTGTTTGGTTGATAAGTCTGATATTAAAGTCCATGTATGTGACGTATTAAAAGCTGAAAATTTGAAACTTTCTACTATCTATGATCTATATGAACCTAGTTCAACTTCGATTACACAAGGAGTTATTGACTTTGTTGCAGGAGATAGAGTGAGAGGTATACAAACTGTTGAAGAAATGCTTCTCGTTGGGACAAAACTTGTTGGAGTAGGCCAAATAGTATTAATTAATGATAAACATATATCAATCAAATGTCCTTCACAACCTGGCCTGAAATATATCTTGTCCTCAGACTCTATTAGTGATCTTATTGAGGCCGAAAAGCATGCTGGGAGTATCTATAAAGGTTTTTCTATATTTTTTGCTCTGATAAGCGTTGCTTTTATGTCTTATTGGCTATATAAGTGGTATAAGCTATACAAAATTGAAAGCGGTTATAACACATTAATGACTGAAAATTCTGTTGAAAATGCATGCTCGGTTTGTTTGTATGAACCGCGCAGTGTAGTGATATTACCATGTGGACACGTTTGTTCCTGTAGGTCATGCACTGAACAATTGTCAGCATGTCCAATCTGCAGGACAATTATAGAAAGATACGTTCCCATTTTTAATAGTTAATAAACCAACTTAATAGCTCTTTTTTTCTTATTTCTTTTGCAATTATAAACTGATTAAGAGATTTTAGACCTATATTTTTGTAACTTGTATTTATAATATATAAACATCTGCTGAATTGCGGTTTAAGTTATTTATAATAAATAAATATCTGCTGAAATTTGATTTAAGTTATTTTAATTATATAAACATCTGCTG
>evgR004802c0g1_i01_Hydra
ATTTTTATAACAGCTTTTTTTGTACCCTTGCGTTTTACTCAAAATACAAATAAAGGATTGCTTCTAGTGGAAGTATACTACGTTTTAGCTGGAAACACATCATTGTTTTCTGTTCTGCATGGTTATAAGTGCTGGGTTATCATATTTAGACCGCAGAAAAATAGAACTTCACTATTTCGTAAGAATATTACAAACCATGTTCTTAAGGGAGTTAATGGAAATTTTTTATCCCTTCAGAATTGTGAAAGTTTTAATTCAAAGGTTTCTACAAGAACAACTTTCCATGATGTGACAGATTTACACAGTCCCCAAACAAGAAATTGTGGGAGCTTATAGCCTTGGAAAATCAGTACTATATTTGGAACGTAAAATTTGAACGAATATTAAT
>evgR004802c0g2_i01_Hydra
ATACAACTTTTGAATAACAATACATATTTGATTGACGTTCTTATTAGAGCTAATCTTAAATACAATTTTAGCCTCGCAGAAGTTAGCGAGGCAATTGAACACACTCAACATTTTGTTTTCACGGTTTTGAAAGATCGGTATCAACCTCAACTGGATTTACAAAGCATTCTAGATTTATACCAGAACTTTTCAAGTGAGACACAAAACATCTCAACTCATTTCGAAAAAACGTGTTCTTTTACGCACACAACCGGTGAAATGATTTTTACAGAAAACAACAGAGTGAATTGCGATTGCATCAACAGAAAGGATGAATACAAGCGCCTTTTTGAAATTAACATCTCTCGAGTTCAGTGTAAAAAATGCGAAAATGAAATGTATGTCAGTGATAAAGAGCCAACTAAGTGTTTGCAGTTTATGAAAGACTTTCTTTTTTATAAAGAAACATTAAGCTATTTGTCGTATTTTGCTATGACTTTGGGAACTGTGTTAACAATTTTTGTCGGTATTACCTTTGTACGGTATCGGAATACTCCATTAGTAAAATCGGCTAACTTAAAGATCTTGCTTATTCAACTACTTGCCCATAGTTTATTGTACGTTGCTCCGATACTGTTTTTTGGAGAACCAACAGTAATAATATGTGCAGCTCGTCCCTCAGTTCACGGGGTTCTACTCATTTTTGCTATAGCAATGGTCATGACTAAGACTCAAAAATTAGTTTTTATTTTTCAAGCTCGCATTCGAGTTTCTCACGGAGAGTATAAGATGTCAAAAACAACGGAAACAATGTTAATAATTTTAATGCTAAGTGTTCAAGCAATGATTAGCGTGTTAACATTTAAGGTTCGACCTGCTAAGGTTCTAATATCCAACGACGAAATAACTGGATTGCAAACTTTTAAATGCGGAAATGATTACGACTTTCTTCTTCAGCTATTATATGGTTTGGTCCTCATTATAATGTGCTTAATGCAAGCCTTTCGTGCTAGAAATCTTCCTGGTAACTTTAGTGAAGTGAGAGTTATAATAATAGCAATGCTTTCAGTCATTTTTATAACAGCTTTTTTTGTACCCTTG
>evgR004803c0g1_i01_Hydra
TGAACATCCTAATTTTATTTACGCTCCAAGCGATCTTTGCAAGTTTGATTGGAAGGAGGTTAGTGGTAGCGTAAGTGAAAAGGCTGCTGAAGATTCTTTGAACTTTTGCAAAGCATTTATAGAAAGAAAGGGGATAGTAGGAGAAGATAATCGTGAAGACAGAAGAGAGCTGGCTGAATTAGTTGTGACTTATCTATCACCTTCTGGTTATAAAATCAGGAAAACTGGAGCTATCCATCATGCCCGTTTTCTTGCTAAAGCAATTTACTATCTGAAAATGCAGCTTTTGAATTCTCAACTTGACTTTGTTCGACAAGACAACGATATTAGAAAAGAGTTCAGATATATTTGTTTGAATCCAAA
>evgR004805c0g1_i01_Hydra
ACAACACAAAATTAGTCAAATAAATATTGGACGATAAAAAACTACAAAGGAGGAGTTTTTCTAAGAAATATACTATCGTCCGCCTTATAAAATAATTTATAAACTTCCTTGGTATAAGATCCTGTCTTTGGAACCGATTTTCTTATAAAATTGATTTTACTTACATAAAGCCATAGTAATGTCTAATGTTAATATTTATGGATCAGTAACTGCTGTAAATATTTCGAATAATGGTTCTCAAAAAGAACTTCCTATTTCATGCATTCCTCCAATGAAAAATTTGTTTCCTCAATCAAATGAGCAATATTTACCAAAAAGTGTTTTGAAGATTGGTAATTGTTTTACTGATGGGATACTAGAAAAAACAAAGAGTGAAAATGAGATGGCAGAAGAAGCATTCATATCAGTAGCGGAAGCATTTCACGAAAAAGATTTCTTTTACTCAATGAGTACCGGTTTTGTAATGATCATTAACAACGAGAAGTTTGAAGACGAAGAAAATAGAGATGGATCAGAATTAGATGTTCAAGCTTTAAACTCTTTTTTCAAAGACACTTTGGGATGGTATGTAAAATCCTGGAGTAATCTTAAAAGAGAAGATATTATTAACATTGTCTTTTCTCTCTCTCAAGTTACGTTTTTCCCATTCTCTGCATTATTTATTGTCATTTTGTCACACGGATCAGAATCAGGAATTCTTGGTGTTGATTCTACTGAAGAGATAAAAATTAATAGTCTGCTCGAAAATTTTAAAGCAGACAATGCTCCTGCGTTTGATAAAAAACCAAAAGTTTTTATTGTACAAGCATGCAGAGGACAAAAAGACGATATCAGGAAAGATGGAGTAAAAGATTCTAAGAGAAATTATTACTATGAACCAACGATTCCTATGTTAAGTGATTTTGTTGTTGCTTACCCATGCGCAGCTGGATATACTGCTCTTCGATCAACAGTAAAAGGTTCGTGGTTTATTCAAGAGTTGGTTAAATGTCTATCATTGTTTTATGATAAGGAAGACTTCTTGTCTATATTAAACCGGGTAAATTTCAAAGTTGCAAATGTTATGCTTCCTCAAGGTTTAAAAGAACAGCCATGTTTTGAGGTTGGTTTGCGCATGCGATGCTTTTTTCACGAATATTTCGAAAAATCATATAATGTTTTAAAGTATTACAAGTCGTTAAACTCTGATTTAAAAAATTTAATTATTAAAAGTGATTATTTTTGATAGTTTTTGTTTTCTTTTAGAATTTACAAATAGGTATCACTTTAGTTTGTTTATATATACACAGTGCTAATGTGGGTCAAAAGAGTATCCGGAACAGGTTATAAACAGGTTTTTAATGATTTTAAATACCGTCCGTGCATTATTAAAAAAGGTCGCTGAATATAATACTGCTTACTTTGAACACTG
>evgR004811c0g1_i01_Hydra
AGATGTGGAGATGACAGTATACAACCGAAATATGGACAAAGTAATTGTTCTCTTTGCTCAGAAAATAAGATGCCAAACACGATGAGAACAAAATGTGTTGTACCCAAGAAAATTTATATCACGGCGTATAGCAAAGTGGGCTTTATTTTAATAACTTTTTCTTTTATTGTTTTTATGTTAGCTTTATTTGCTACCGGTACCTTTCTGATGTACAAAGAAACCCCTATTGTAAAAGCTTCGAACATAAACTTGTCGTTAATACAGCTTCTAGGTATAATGTGTACTTTAATGCTACCAAGTTTTTGTATTGAAAAAGTGACAGCAACAAAAATATGCAATGGTCAGCTGATTTTTTTTGCCTGTTTTAATTCAATAACTGTTTCCGTG
>evgR004813c0g1_i01_Hydra
CTCCTCAATAACAATGTCCAACAACCCATACGGATCCACCGATGTTGATCCATACGATTATGATTATCAACCACCAAATTATGGAAATCCAACATCTGGATATGTTCCACCAGTAGGTGCTATGAGTCAACCTGCCCAAATTAATATTGAACCAATGGGTGCTTCTTTAGATGATGAGGCTCTCAGACTTAAAGCCATGGAACTTGCTCGTAGAGAAGCTGAATTGGAAAAAAGAGAACACGAAGTTCAAAATTTGGATAACGCCAATGCTGCAAAGAAGAACTTCCCAAAGTGTTATCCAATTGTTTACCACAACATTTCAGGAATCCCACCTGAATTCCTCCGAAGGAAGCTTGCTTACTGGGGTTTCATTTCTGCCATCGTTTTCCAAGTTGTTGTCCTTTTGAATGCCCTTGTTGCATATATTACCACATTCTATCCACCAGCTGGTTTGACCAGTGAAGTCACTGTCATGACTCTGATTCAATATTGTGTTATTTCCACAGTGATGGCTGTTATTTCTGGTGCTGGCCACTTCTTCCTCGGATATTGGACTTTATATAAGGCTATGGACATTGGTGC
>evgR004816c0g1_i01_Hydra
GGTCAGTGTTCTTCAAACTCATTCAAATACGTACCCTTGTACCTGGTGTGACGTCAAATTCAAGTGTCTTGCCAAATCTGGAAACTTGCGAACACTGGGCTCAATCAATTCAAGCCTCATGTGTTTTCAGCAATCTGGCGGGGTTGATGATAATGCAATGTTGTTCTGCAATGTCATTCAAAAGCCTATCATCTTTGGGCCTGTCGATGTTCTGATTTTGAAAATTATCTCGCCAGTGGAATTGCATCTCCTGTTAGGAGTTGTCAACCATTATTTTAAGATTCTTAAAACTGCTTGGATGTTAAGTGGCCAGCCGCCCTCCACATTTAACTGAGCTTATATCATAGAGCTCAGTTTAATGGAAAGGATTACAGGAAGCTTCTTAAAAATAAAGATTTGATTTAGCAACTTGGTGATACAAACATGGTTTCCGACATTTTGTGCTTAATTAATATTACATAATCTTTTGTGGAG
>evgR004826c0g1_i03_Hydra
TTGATGACGTCAAATGACTTCTGATTTGGCGTTATATAAATCGCCGTCATTTAAATTTAAATCAAGTTTAAAATTTCTTCACTAATATGACGAGCAAAACAACAGATAAGGGCGGACTTTTGGGATTCCTGTGCCGACAAAATGTGCGCTGTAAAAACTTACCTTTTTCTGCTAAATCACACTTTTTTCGTGAATCAAATCAGTTTAAAACGCAGAGTGTTCCAAACTACTTCTGTCCAAAGGCAACAAAACTTTACAATACGCCTAGTCACTACATTTTGCCGCATTTTCCAGTTAAACCAACAAGAAATCATAGTCTTCCTATTGAAGTTACGACAACAAATGCAAAAATACTAATAAATAACGAGGCAATCGTGGTTAAAGAAGAAAGAGAATACAAATTACATCGCTGCAAGAAAATGTATAATATTTGCATTAAAAATGACGATTTAACTCCTAAGAAGTCCGAAAGTATCCTTCCTGAAGAAAATGCACTTATTTCTTGAAAATAAAACACTAGATCTAATTAAAATGTTATTGTTCATGAAAATCCATAAAAATTGACTAAAAAAAAAA
>evgR004833c0g1_i01_Hydra
CCAACAATTAAATATTGTTGATAATTCAAGTGATGAAAAGTTTAAAGCTTTCATGAAGCAATTAGATTCTATGGGATATTTCAAGAATACAACAGCAACTGATACATATGAAATGCGTTATGAAAAGGCCAAGCAAAAATTCCAAGAAAAGATGAAGGCCCACCAATCATCATCAACAACAACAACAATCGATGTTGAAAAGGCTGAAAAGGCAAAAGAAGAAGGAAATGAAAAGTTGAAGAACAAGGACTTTGAAGGCGCTGTTGAATGTTACAAGGAAGCCATGAAACACAATCCAGAAAATCACATCTATCCTAGTAACATGGCTGCTGCCTATATTAGTTTGGGTCAATATGAAAAGGCTTTGAAGTGTTCCGAAC
>evgR004839c0g1_i01_Hydra
TTGGTATCCAGTATTATATACGTATGCACATCAATTAGTACTTTAACATTAGTTTTTCTAATTGTTGTAATCTGTATTAATATTAAATTATTGACTGGAACGCAAAATATACTCCGAGATAGTGACGTAAACATTGAAAGAAATCTTATAATAATAAAAAAACTAAACAAGACGATCCTATTGATTTCGATTGCACTTATTGTATTGTATTTACCTACCGTTGCTGGTCTGTTAAACTTTGTGGTGTCAGCCCAATATTCAATGAAAAATGATCACTATGACAACATTAGTTTAGTTTTAACTAACTGGACGGTTCTTCTTTGCCAAGCAAATTCAGCAATAAATTCCTGGATTTATATTTTAAGAAGTAGAAAAATAATTAGTTATTATCGATGTATTTGGTCTCGATATCAACTCTCGATGTTTCGATG
>evgR004849c0g1_i01_Hydra
TGGTTTCATGAAGATGGCTCTGGAAAACTTGTTATTTTTTCTGCAGTAAATGGAAATGTTAACTACTACATTGAAACCCCTCCACTTGCATTAAATCAGTGGTCTTTTGTTCAAATATATCAAAATTTTGTTGACCATAAATATTGGTTTGCTGTTGATGTAAACAGAATAAATATTTATAAAGCAGAGAATTCTGATGCTCGAGATTTTGAAAAAATTAAAGTGTATGCTTCAGATCTTTGGTATGATTCCCAAGATGGTTTAGTCTCTGATCTTTTAGTTTTCAATGGGTATCCTGAATATCTTGTTGAAAATATTCCTACGCGATTAGTCCGAGGAAAAATTATTTCTGAGATGCAAAAGTTTGATAAAGAATATGTTGTGTCACTGGACATTTTTCCTAATAAGTTTGTTTCCGATGGGCATAGTGTGATTTCTTTAATTGTAAATTCTGGTAATAATAAATATTTTGTTCTTGCCATCTGGTTTCATCAGGATGGTAGTGGAAAGCTTCAGATTATAGCCCCAGTTAATGGCGATCCTTATCGATATTATAACAGTTATCCAGTTAGATTAAATTTTTGGTCAAATGCAGTTGTAAGTCAAATTTTTAAAGATAATACTTACATATTTTCAATCACAATAAATGGGCAAGTTATATATTCCGACATTAACAGTCAAGCACAAAGTTTTGATCAGGTCAAGGTATTTGCTTCAGATCCAGATATTGACGTTCAAGACTGCTATATAAAAAACTTGTTTATTATCAATGGGATTTTAAACAGTGGAATGCAACCTATTTTACTTTCAGATTATATCTACAATAGAGAAGAGTTTATTCCAAAAACAGGCTTGTTGCTTGGCACACTAAATGTTTTGAGGAAGATGTACACTGTATCTTTTAACATAAAACCAAGGAATTATACAAAAGGGTTTAAAAATGTTCTTCATTTAACTTTGGGTAAGGATTTTGGTGGTTATGGGGATAGGAATCCAGGTGTTTGGTTTTATGAAGATAGC
>evgR004850c0g1_i01_Hydra
ATGATAAAAAAATCATTGGAATCTTTGCTAAAGCATTGGTTTTCAAAATCTGATTGTTTATCAAATGATATGATTCATGTTTGGTGTTATGCAATTGTTATGATACATATTATCTCAGATGAAAACAAGAATTGTTTAAAATTACAAGATTTTGACGAGTCTAATTGTGGGTTACCAGCGTTCTTTAAAAAAATGGGGAAGAAGAAATTGGTTTTTGAAACTTTCAATAACTTGGATAAAATCCGAATCAGCCTGAAAAAAGATGCTTTACTACAAAGAGGTCTAGCTTATTTGTTGGAACCACCATGCTTTCATAATGCATTAACTACTGGATATTTTAAGGAGCGTCTTCTAATTAATATGATAACTTTTCGTCTTTCACTATCAAGTGGAATAGAGATAAATGAATACCAATTTTTTTTGAAAGCATTGATGACAAACATTCCAAAAGAAGAAAAAGAAAGAGCGTACGTAGTTGAAGCTATGTTGAATATGCTCTTAAAAGTTATTGAAAAGGTTAAACTATCAGAAAGCAGTTTCAATTTTGTATCGATTATCTTTAATTGCATTGAAACATTTTACAAGTCTATTGATAAGAATAAAACACTACTAGCTCTACATAAAGAAGTTTCAACAACTTTGTCTGCTATAAAATCTTGTCTACCTAATTATTTTGTTGATTGGGATGAGGAAAAAACTATAAATTTGCAACTTGAGGTAATGCAAATTTTACTTCGTTTCCAAATTGATAGAAAAAGAGACTTTCAAGAAGATGTACTTTCCATTCTTCAAAGAAAATTGAAACAAAC
>evgR004861c0g1_i01_Hydra
TAATCGTGAACTTCTCCCTCTTTTGAAATCTGGTTATCGAATGGATAGACCAGAAAACTGTTCAGAAGCAATGTACAACATAATGCTACAGTGTTGGAATAAGGACCCGTTACAGCGACCAACTTTTACAATGTTACGAGATCATTTCGATGGAGTAATGTCCCAAGGTGATTGTTATTTCAGTTTCGAGTTTGACGAAAATTATGCGCCGTCGTTTCTGCCGTTTCAAACAGAAGATGATGATAATGAGGTTGCAGACGAGATCTTTCAAAATCCTGTGCGCGTAAAGGGAATTGAGGAAATGAAAAAACTTAGCAATAAAATGATTTCTTCATCGGACGGTAGATATGCAAATGTTACCTAAAACCACTCAGCCATATCAAATTTAATGTTTTTGTTTTAATGTTATTCAATAAAAATTGGTTAGGCACTAGGTAAAAAATCCGTTAACCAATTCTATCGACAGAATCCTATACAAAAC
>evgR004868c0g1_i01_Hydra
ATTAATGGATCAATTACCGGACAAGTTCAAATCCGTAGTGTCAAGACAAAGAAACAAAGTTTGTTATTTGACTCTAAGGAAGTTGAACATGCTCCTTGGTTTGTGAAGAAGATTGCTTCTCAAAAGCCAAATGAATCTCGTAGAGTTTGGCATAAGGTAACAAAGGCATTGGCTGATAGCGAATTTGATGATGCCAACATTCAAAAGAATCTCATTGAAGAAAAACAACGTGGATTGAGAAAGAAGAGAGAAGAAAATGGTCAACAATGGTCACAAGAATTATTCAAACTGAAAGAAGGAACTGAAGATGAGTACCAATTCAAATATGATCCAACAAGTGCAGAATACAAAGATAAGGCTTTAGGAGACTATATTGATTTCCTCAAGGAAGATGAAGAAGAAATCGAATTCTCCAAGAAACTTTAATCCTTTTAACCCTTTTTAATTTATTTTCCACCACCATGTCATTCATATATACCATTTTGTAAATCG
>evgR004873c0g1_i01_Hydra
ATTGATTACTTTGAGTTCTTTATCGATGGCTAAACGAACATGTAAAAATAGTCAAAGTTACAAACTAAGTGTTATGATTGGTTTTGCAAGTTATTCTGAGTTACCCACATTAAGAGATGTTATAAAGTATGGTCTGCTTCTAAGAAGAGATGAAAACTACAAAGCAAATGCAAAACTTAAAGATACATGCATGGCAACCGTGGTTTTCTCTAAAGTGAAAACACAATGGTTGAAAGCGAATCCTGAAATTCCTATCGTGACGGACAAAATGGGCGTTAACAAAATAATAAAAGTATGGAAAAAAGCTGCCGATATTATTAACTACAGGTGTAAACCTGAAATTATTTCTAAGTTTAACTCTACTCTGGACAAGCTATTTGATACAGCAAAGTGCAAATGTGACATTGTTTCTTGTGAGGAGGTTGGAAAGGAGGTATCATTCATTTTTTAGAAAAAATACTTCAAACAAAGCTTATTTGGATAGTTTGCGCTTTGCAACCAAATAAACTACCCTTGAGGCACTTAATATCAACTATTGACGGG
>evgR004873c1g1_i01_Hydra
ACCCACATTAAGAGATGTTATAAAGTATGGTTTGCTTCTAAGAAGAAATAAAACCTGCAAAGCAAATTCAAAACCTAAAGATACATGCAAAAAGGTTAACTCTAAAGTGAAAACACAATGGTTGAAAGCAAATCCTGAAATTCCTATTGTGACCGACAAAGTGGGTGTTAATAAAATAATAAAAGTATGGAAAAAAACTACCGATATTATTAACTACAGGTGTAAACTTGAAATTATTTCTAAATATAACTCTACTCTGGGCAAGCTATCTGATATAGCAAAGTCCAAGTGTGACATTGTTTCTTGGGAGGAGGCAAGTTGTAGTTCTTGTAATTTTAAAGCGCATATTTTTTGTCAATGCAATAGAGAAAACAAAAT
>evgR004883c0g1_i02_Hydra
TAAATTAAGTTTAATATTATTTTAAATTATTTGTAAAACAAAAATTTAAAATATCAAATTGTTTTCTGGAAAATGTTTATAAAAACAAATTACAAGCGATCACCACAATCGTTTTTGAAAATTTAGATGGAAAACATAAACAAAATTGGTGGTTATGATGAGCATTTTGTAGATCCCGTTGAAGATAAGCACAAATGTCCTGTGTGTTTATTGGCTCTTCGAAAACCTATTCAAACACAACCATGTGGTCATCGATTTTGTGAGTCATGCTCAGAAGAAATTAAAAAAAGTAATGATAAAAACAAGTTTGTCTGCCCATTAGACAAAAGTACTCCTACGATTTTCTATGATAATGCTTTTGAACGAGAAGTTCTCGGTTTAAAAGTTAAATGCAATAATCACGAACAAAGTTGTCAATGGAATGATGAACTAAGTAAAATTAATAAACATTTAGATTCATGTGAGTATCAAGAAGTGAGATGCATCAATGAACAGTGCACTATCTCACTTTTGCGTAAAGATCTATTAAATCACATGGAAACAAATTGCATCTATCGTTTAGTTACCTGTGAATACTGCAATATAATGATTATTTATGACAATCTAAAGTTACATCTTGCAAACTGCGATAGTTATCCACTTGATTGTGTAAATCAATGCGGAATAATAATATTGCGTAAAGAGATGACTATTCATGTTGCTGCTGATTGCGCCCTCACAATTGTTCCTTGTCAGTACTCAAACATTGGATGCAAGTTCACGGGAATGAGAAAAGAACAAGATGCTCACACAAACTTGTCAACACAAAATCATCTTTCTTTGGCTATATCAAAAATGATGTCAAACGAAATAGAAACAAATGCAATAATAAAGTGTCTTGAAAATAACGTCCTGTCCTTCAAAAAAGATATAAATGAAAAAATGACAAGTTTGGAAAATATAATTATTTCAAATGAAAATAAAATGAGAGCAACAATTACCGGTCTTGAATACAAAATTAAGTTAAACGAAACAGAAATGACAACAAAAATGACAAACCTTGAAAAAAAATTGTTAAATAAAGAAGAAATAAATTCAGAAATGGTGAATTTTTTTAATGAGAAAGATTGTCAAGTCAATAAACCGGAGTTTACAGTCAATCAACAAAAAGTAGAGTTACCAAAGGCAATACCGGACACAGAACAATTGTGTTACCAAACAGCATTTGGTAATAAAAACACCGAATCATTACCCAATCAAAATAATACAACATATAATTATTTTGCTAAGAACAACACCACAACCCCCACTCGTCGTTTTTGCGTATTCTGTGGTACTGAGACATCTGAAAGAGCTTGCATAAAATGTAAAAAATATATCAAGTGAATAATAAGAACAACACCACCATCTTAACGAAACCACATTGCCCAAATTATGGTACTTTGGACTTTAGAAGAGGTTGCAAAAGATGTGGATATATTATTATATGAATAAATATAAAATATATAAAAAAAAAAA
>evgR004895c0g1_i01_Hydra
GAATACTTCAAAAGGCCATACATTTAGCCATAGCTGCATAACAAATTGTGCTGCAAAGTTAAACACAAAAGATAAAGTTGTTTATATGTTTTCAAGTTTGAGCATAGTTGAGTTGTCTTTTATTTGGAAAGTTTTTAATGATGACGGCGAATGGCCAGAGGAATTGCTACACCAAAATTCAACAACAACTGGGTTTTTAGAATCCTATCTTGTGATTAATTCTGACATTTTATTGATTTCCAAGAACTATACTTTTATGTTAACAGTTTGGTATGTAGGATCACCATACAAAGCTTCGTTTAAAATAATTAAACAAACATCATCAGTACCTATTCCAGGAAACTGTTTTGTAAGCCCTCAACTAGGATATGCTATTATTACAAAGTTTAAACTTGTTTGCCAAGATTGGTCTGATAATGAGAATAGTGTATTAAGATATGAATTTTTTTATGACAGCGGCAAAGAGAATTACACAATGACACGCAATGGGTATATAGGTTATCCATTGTTAAATGCCAAATCTCTGAACAATCCAATGCTGTTCGATTTTGTGTTAGGCCCTGGTAACTTGGATAAAAATTTTACAGTTAGCATATATGTCAGAGTTATTGGAATTTATAAATCTTACACACAATATCCTGATCTCTTTGTTCAAGTTTTACCTTTCCCAAATAATGTTTTAAATGAACTTGCTATTGTTGGCATACCTGATACACAGATTTCTTCCAGTTTTCTATATGGTATATCCAATGTTCTAAATGATAAATCAAACTCTGTGAAAATGTTGAACGCAAAAACCAATAATAATATAACAGGATTTAATGATGCAAATGATGAAATAGTACTTGACAAGCAAAACTTGCTGGTTCAAATGCAATCTGTTCGCACTCAAGTTGTTCAATTGCTTTCTTCAGTACCCTTGAATAGTTTAACTGACGTAGAGGATGTTAGTGATGTCTTAGCAATTATTTTAACTGAGCCAGATGAGCTCACTCCACAGACTATGACCAATGCTGTAGATATCATAGCAAATATGAGTTCTTTGTTGACTGAAGAAAATATTAAAGATATTGGCCCTTATTTATATGATCAGATTACTCAACCGTTTTTAGTTTCTGTTTCACATTTGTTTATTGCGATTGCTAACTCTGATGATAATATGGAGTCAGTTAAAGGTTTAAACACAACATTACTACCAGATTACACATTACTACCTGGTTACTTACCAAAACAGGTACAAAACAAGTCGCAAATCATCCTAAAACTTTTCAAATCTATTGACGATTACTTAAATGGTTTGCAATACTATATTGCTAATAATGAAGACCCTATAATTGGTGATACACCTGCATTTAAGTTTTTTGTGAAAAAGGTTTATGGAGCTTATTCTGGGAATATTTCGTCCATTTCGTCAGATAAAAACAATGACGAACAAAGCGGATTTAGTATCTTTAATCTCACATCTCTTTTGAATGAATCAATAAAGTATTCTACAGTTGTCATTAAGAATCTTAATATTAGGAATCAAGTATTCAGTTG
>evgR004897c0g1_i01_Hydra
TTTATTTGAGAATACTTTGTTCACAGTATCTCAAAGAATCTTTGTTCGTCCATTTCGAACTTCAGAAATCAACTTAAGTTGTTTGAAATGGTTTTCAAACAAGGAACAAAACATAACCATTACTTTACTTGGTGATGATTATCTTTCAGCTGACTACCTTTGGTTAAAAACTGATTTTACTGAGTATACAGGAAAAAATTACTACATCTACTCAAGCACCAGGGGAAGACAGAAGTTGTTTGGAAACTCAAAAAATTTAAAAAAAAACGGTTTTTCAACTAGACTAATGAGTGATTATTTTGGAAAAGATTGTTTTACTCAGCCTCACTTGTGTGTCAGCGGATTTACAATTCAACTTGATGTTGATGTTTCATTTTCATTAAATTCTGATTGGTTACCACTTATTGCAACAGATTTGTTTTCTTCCAATATATTATTGAACGAGTCAAAAAATATAATCTATTTTGGTTCCAATGTAACAAATACATTTTTGCTTTTCAATCAATTTACATTTCCAGCTACAGCAAATTCAATTCAATTGCATCTCAATATTTCTGGCTTGCTGAGTGCCTTAAATTCCAGTATACAATATTCAGCAAATATTTATGCATTTTCAATTCTTTCAAACGAGACGATGACTGCACAGAAATGTTCAAACTATTATGATAATGGGTTTTTTCCTGCTAATAATATAACAATAACTTTTGATGGTTTAGATAAAGATTATGGTTGTGCTTTGACAAGTATTATAAGCATTTTAAAATCTCAGAATAACGACTTGATTAATTATTTATGTTTGATGGTCCAGCCGATTAGTC
>evgR004909c0g1_i01_Hydra
CCTGTAAACGATTTTGAAAATAATCAATGCATTGTGCCATCTTCTATAGGGTCTACATCTGCAAACACAGCAGTACCTGTTAATGAAACCTTAACTAATAATAAGACCGAATCAGGTAAAGAAACCCAAAATATTAAAGAAAGCCAACCTGCAAACAATTTTGAAAATAATAAATACACCGTACCATCTCCCATAGGGTGTACATCTGCAAACAAAGCCGTACCTATTAATAAAACTGGAACTTATAATGAAACCGAACTTGGTGTAGAATTGCAACATATTAAGGACAGCCAACCTATAAACAATTTCGCAGCTAATTATAACACTGTTTCATCTTCAAAAAAATCTGTATCTTTGAACGAACACGTACCTGTTGAAGAAATCGTAACTAATGATAAAACCGAACCTGGTGAAGAAACCCAAAATATTAAGGAAAGCCAAC
>evgR004910c0g1_i01_Hydra
TATTTTATTTATAAGGCAAAAAAGGCTAAAAAGAGACCTTGCAGGAGTAATACCTGCAATAATGATAAATCCAAATGTGGAAATTGAACTGGAACCTCGAGTTCCGGAAAACAATATAGAAGGGCATGACAATATTTCATTTGAGATTGTTAATGAAGATCCTGATAAAGATAAAGTTGATGAAAATCTTAACAAGGATAAAGTTAATGAGAATCTTCATGAAGATAAAGACGACAAGGAGCTTCATAATGATGAGGATCTTGCCAATGATAAAAAGCCACCCGAAGTTATTGATATTAATTCTGATTATCCACCAATTTGCTTAAAAGATTTCCCCAACCATTGGAATTTGTTGCGTAAAGATAGTAATTGTGCTCTTTCACAAGAATTCAAATGTTTGAATACTGGTCAACAATACACCTGCAATGATGCTCTCTTACCAGAAGTCAAAGTTAAAAATCGCTATTCCAACATAGTAACTTACGACCATTCACGAGTGCATATTGGAAACAGTCCAGAAGTTAAAAATAATTATATAAATGCTTCCTTTATTCATGGTTATACAAAGCGTAATGTCTACATAGCAACACAAGGTCCAAACATATCATCGTTAAATGATTTTTGGAAATTGATCTGGGAACAAGACGTTCCTGTAATTGTAATGTTAACAAATCTTATAGAAAAAGGAACGAAAAAATGCGATCGGTATTGGCCAACTTCTGATGCAAT
>evgR004918c0g1_i01_Hydra
AAATGCTTAAGACAAAGACTTTATTAAAAAATTTTAAAAGACACAATGGTTATCTATCAATTCCTACTCTGAAAAAATCATCAGTTTGGCTATCTCTATCAAATTGTGTTTATGAAAATATTTCATTTGAAGAATATATTTACGACAATGTTTCATTTGATGTTTTTGATAAAGTTTTATTTATTTGGAGAAACACTCCTTCTGTTGTGATAGGGCGTCATCAAAACCCTTGGAAAGAAGTTAATTTAAAAAAAATTGAACAAAGCAATATCAAACTTTGTAGGCGTAACAGCGGGGGAGGAGCTGTTTATCATGATTTAGGCAATGTAAATTTTACAGTTTTTAGTAATAAAGAGTTATACAATAGAAGAGAAAATTTAAACTTTATTGTGGAATTTTTTAAAGAAGTTTATATGATCAATCTTGAAATAAATCATAAAGATGATATTATTTTTGAACGTTTTAAAGTATCAGGCAGTGCAGCAAAAGTTGGCTTGAATAACGCATATCATCATTTTACATTATTGTGCAATACTAGTGTTAGTTCTTTAACTGAAGTCTTACAAAATTCTTTTTCACAAATACAAACTAGTGCTACCCAAAGTGTTTCCTCACCTATTAAAAATTTGTTCTATGATGGATACAATTATTCAGATTTTGTTAAAAATTTAACTGATTTTTATCACAATAAATACAGGAAGTTAAATAGCTATGCACAGGTCCACCCTATGAATTATCCTATTTCCCATAAAGTGGATTTGCTAAAAGCATGGGATTGGACATTTGGTAAAAGCCCAAAATTTATTGTTAATGGAATTCTAATAAAAGAAGGTGTGCATGTTGGAGTTATTAATATAGTGGTTCAGAAAGGAATAATTGCAGAAATACAGTCATCGTGTGAACACAATGAAAAACTGAATTTTTTTGTTGGAACTAGGTTTGACCCTTATAAAATCATAGCTATTTTAAAGCCTTACGGTGTAACAATAAAATTTTAACTTTTATAAAATGATTATTTTTTTAAAAGGGTTATTGTGCAACAATATTTTTTCAGATTTTAAAT
>evgR004923c0g1_i01_Hydra
ACTCTTTGATAATGATAGAAACAAACTTTGTCGCTTTCAATTTATTGATGCAGCAACAGGGAAAAATGGAGAAATGTTTAAATTGACAAATGGAATTGTGGGATTTTTAATTTTTATTTCGAACACGTTGTTAACATATGGTATTTACAAAGCAAAAGGAAGAAAAAAATTCTCACGAAATGAAAAATTGGTTTTATTATTATCTGTTATTGATATTATAATTTCAATAATACATACCCCCCTAGAATTTATTTTATTTGCCTACCTGAACCATATTGGATGTCTCACTATTTCGGTGATTGCGTTTTGGATAGCATTTCCCTTAATCCTGTCTGGGGCAACAGCTACAGCAATTTCAATTGAACGGTTTATTGCTGTATTTAATAACAAAACATGTTGCGGATTTTATTTCAAAGGAATTTACATGATCCCAATATTTTTGTTTTATTTTATTGGCTCTTTTGGATTTGGTGTTTGGCATGCGCTTCTTAACTATTTTATTCCCGATACTTTTTTTTTGGTTCGTTATTTTTTTTATAGTACTGCATCCCTTATCGCTACAAATCTAGTAATTGCAATGTCGATAAATATTATACTGCTGAAAAGAATTTCTAAAAGGATAAATGATAGCGAAATTCATGTTAAAAGACATATTGCTGTAGAGAAAAAATTAACCCAGAC
>evgR004930c0g1_i01_Hydra
CTGAGGGTTTTAAAAATATTCTTCATTTAACACAAAATGAAAATATTAAAGCTTATGGCTATGGAAATCCAGGTTTTTGGTTTGATGAAGATGGTTCCGGAAAGCTTGTTATATATACTGCTGTGAGTGGAAACGCTTCTTACTTTATTAAAACAGCTGCTTTAATGATAGATCAATGGTCCTCTATTAAAATATGTCAAAGTTTATTTGATAGTCATTATTGGATTTCTGTTCATCTTAATGGAACACTTATTAAAATTTTAAATAATTCTGATGCTCGAGATTTTAAAAACATAAAGGTTTATGCC
>evgR004939c0g1_i01_Hydra
CATAAGACCAAGAGCAAACTTAGTTTTTATCTCTAAAAAAATAGTAAGTTATACTAAAGATTCAATTACGCTAGAGGCTCATGGCGTTATAGATTTAACAATCGGAGAAACCGTCCAACTAATAGTGTATGGTGAAAAACAAGAGAAAATGACATTATGTACGAGAGTAGTATGGACCATGGTTGAAGTTCTCCCCCCTGTTCCACAGCAATCATTTGTTAACTCATTAAAGAATTCTTCGTTACACATTCTTGGAGACTACTTAAAATTCGATGAAGAAGATGAAAGCCATAACTTTGACGTCGTGCAGTTTCATGATGAAAACTCTGACGCAAATCTAACTAGTGAACTGACAAGTTTTTATCAGATTAGAGATAAACAAAGTTTGATATTTAGATGTTCTGAAACTCTTGATGATTTGGTGCTGAACCCATTTCAGCATGTTTACACAAAAGCTTCTAATGAAAGACAAACGTTATGGTCGCTAGTAAACGAACCAAT
>evgR004942c0g1_i01_Hydra
AATGGAATTAGTCAATACTCGTTCACACACCAAATATGTGGACATTGAGGAATCGATATTCTTGGATACTGTCCTTAAGAGCAAGTATTGTGTGTGTCACTTCTATCACAAGGATTTCGAGAGATGCAAAATTGTTGACAAACATTTAGGAACAATCTGTCAAAAGTATTGGCAAACCAAATTCTTTAAAGTTAATGCTGAGAAAGCCCCTTTCTTTGTTGGAAAGCTCCAAGTCAGAATGCTTCCTTCTGTGATTTGTTTTAAGGATGGCGTGATGGTGGATAGATTTGACGGTTTTGAAGAACTGGGTGGAGCTGATGACTTCCCAACAGTGAAAATGGAAATGCGTCTTTTGAAATCTAAATGTGTTGGTCCTGAATTGGAATTGTAAATTAA
>evgR004950c0g1_i01_Hydra
CAAAGCAGAAAACAAGAAAAAAATTAATGGAAAAGTTGTAACAGCAGAATTAGTAAATAATGGTTCACAAATTAATGATAAAACATTAAATATAAAAACAGGGGAAAGTAAATATAATGGAGGAAAAAATGTAGACAGTTTCAGAATAGAAGCTTCTGATGGATCTAAGGTTACTGCTGACAAAGGGCAGAATAAATTTTTTTCTGGGGTTAAAAATAAAGGTAAAGATGATAAACTGCCTGGTAATACAGTTGGAGTTGGAACAGGAAAAAATAAATCATTATCAAAAACCGGAGTATCTTCTGGAGCATTAATTGCAGAAACTATAGGTACAGAAAGTAATACAAATATTGCAGGAGCAAGCAGCGGAACAAATAGAAAAACAAACGGTCGAATAAATGGGAATAGAAATGTTAAATTTAACAGTGGAACTATAAATAGTTTAAATATTTTTTCCAACGGTAGAACAGGTGGAGCAGGTAGAACAGGTGGAACAGGCGGATCAAATAATGATAAAGAAAGAAACGGATCAAATGGTTTAAATATAAGTAAAAATAACAAAGTAAATGGAGGAACAGGTGGCGCCAAGAAAAAATTTTATTATAAAACTTCTAGTGTAAAGAATGGTGGAATCAACGGTGCATCACGTGCAGCAAGCAATACAGCAAGCAATGCAGCAAGCAATACAGAAAGCAATGCAGCAAGCAATGCAGC
>evgR004954c0g1_i01_Hydra
TCGCAGTCTGCAAATGCTTGGACATAAATCACTGCAATTGTGAGAAAAGTTTTAAGGTACCAGTTGCAGAAAGACCATTTTTGGAAGATCAAAGAAGTACAAGAAATATGATGATTGGAAATATAGATAAAGTTGTTACAAATCGATTGAAAGAACGTGATAATCGCAAAAGAAAACTTACCAATATTTACAAAACTCCAACAACAGCCACTACATCTAACTCCCTATTGTCTTCAAATATTGTTGAATTCAGTTCGTCTTCTGATACTGAAACTGACAACGATACTGATGTCCATTTTCCCTCAATACATAAGGTTCCAACTACTGGAACAGCAGAGAAAAAACAGATGAGAGCAAAGATTCCAAATATCTTAAAGGAATGTGATC
>evgR004961c0g1_i02_Hydra
AATTGATAGCGATGTCTTTATTATGGAATATAAATTTTGCAACTATTGCACTGGTTACGTGTCAGTACAACAATTCAATGTTTATCGACAAAATTGAGAGTGTTTTCACTTCCACAAGTTTAAATAACCTGTACGACGAGTTTTTTGTATCTTTTACATTTTTTTCAAATGGAAAAAACATCACACCTATTGTTGGTATATATGATCTAAATTTTGGTAATCCCATAGCTTTTAGTATTGAGAACAATGGTTCTTTATTTTTACAGATTTTTACATTAAATAGTGCTTATAGTTTTAAAATAACAGCTAATACTTGGAGTTCGATTTTCTTTTACCAAATTTACAACAAGATAAGTAGTACTTATGATATCTATGTAGCTGTTAATGAAACTGTTCAATATTTTTCTGTAAATACTAAACCAATGGTGTTCAACAATGTTGTTTTGTATACTTTTGGTGAATGTTTGTACACTTCTCCTTGTAATCTTGGATCAATTAAAGATTTAGTAATTGACAGTAAAACAGAAGTTTTTTGGTCCCAATGGAGCGATTGGTCACAGTACAATTCTTCCTGCAATGCAACTAGATTTAGAAGCTGTAGCAATAATAAATGGTTAAATTGCTCAGGAAATAACATTGAGAAAGAATCATGCAATATTTTGCCTAAAGGAAATGTTGAAGTACAGACCACATGCATAGGAGAAATATATTCAATTGAAACAATGGATCCACAATATAGAATTGATCTTGCTCCAGGCTTTCATGATCAAGCTCGTGTTGTTCAAAATAGCACGGAAAGATTTTTCTGTGTTGAAGGTTATTTAACATCCTATTTCTTTAAAAAACGTTTAGGGAAGACCAGGGTTAGTTAGAAAGTTTTTTTTTATTTTTGAATTTGTGTCTGG
>evgR004962c0g1_i01_Hydra
GAAACTTACCAAACCCTAATGATCGAAATCGAGATAGATTATACAAATTGAGACCAGTTATAGAAGAACTTTTTGAAAATTTTCAAATTGTTTATACCCCTAATCAATGCACTGCAGTAGACGAGTCATTGTTACTGTGGAAAGAAAGGCTGATTGTTAAACACTATCTACCACTTAAAAGAGCCAGATTTGGAATTAAACTATTTTGTCTATGCGAAGATTCTGGTTACATTTATCAATGTTGCCACTGCTTAATAAAGGTTATATTATTTGGATGGATAATTGTTATTCTTCATGTAAGCTTTATCACTTTCTGCAACACTACGGAACAACTGCATGTGCAAGACTACGGAACAACTGCATGTGCAACACTACGGAACAACTGCATGTGCAACACTACGGAACAACTGCATGTGCAA
>evgR004974c0g1_i01_Hydra
GATTAAAATGGAAAGTTATTCAAATAGAATAATAAAGACGAAGTGCTTAGTTATGCAATCTTTAAGTGAAGGAATTCGCCAGCGCAAAACAATATCTAAAGGTTTCAAAGACTTAGATGCGTTTCCTAAAATACCTGAAAGCTATCAAGAGACCTCAGCAAGTGGTGGCACTGTTTCCATTTTGGTTTTCTTGTTTATTTCTATGCTAATTATATCTGAGTTTATATATTATACTGGATCCATTTTAACTTACAAGTATGAAGTTGATAAAGATGCTGATAATAAATTTCGTATAAATATTGATTTAACTGTTGCTATGGAGTGTGACGACATTGGTGCTGATGTGTTAGATCTTTCTGGTGGTAATGTTGATACAGGTGAAAATCTTCATTTGACACCTGCTCATTTTTCAATGAGTTCAAATCAAAAGCAGTGGTGGGATGCATTTCGATCGGCTCGCAAAAGTGATGAAGGTTATCGATCTATTAATAAAGTAACACAAATTGATATGATATTTGGTGATGTTATGCCCACATATATGCCAGATGAAATGGCATCTGAATTTGAAGGAAAAGAGTTTGATAGTTGTAGAATATATGGAAATATTGAAGTAAATAAAGTTGCCGGAAATTTTCATATAACTGCAGGGAAGTCTATTCCTCATCCACGTGGTCATGCCCATTTAAGCGCTTTAGTTTCTGAACTAAATTACAATTTTTCTCATCGCATAGATTTGCTTTCTTTTGGAGAACCCCATCCTGGTATTATAAATCCTTTGGATGGAGATCTTATGATTACTAACACTCCTTACCATATGTACCAATATTATATAACGATTGTACCCACAACTATTAAAACTTTGAAAAATACAATTAAAACAAATCAATACTCAGTAACTCAAAGATCTCGTCAGTTAAACCTTAATTCTGGATCTCAAGGTGTACCAGGCATATTCTTTAAATATGATTTCCATGCTATATCAGTATCTGTTAACGAAGAGCGCGGTTCTTTTAAAGAGTTTCTTATTCGGTTGTGTGGAATTATTGGTGGAGTTTTTGCAACATCTGGAATGCTGCATTCAGCTATTGGTGCATTAGCAGATTTTATTTTATGTCAGATGAGGCAAAAAAGAGGCAAAACCTTATTTCAAACAATAAGTAATGTTAAAAAAGAACAATTAGATTGTAAAGTTCCATTTGTATTATTAAAACAGGACAACATTAACTTGCAAAATGTATCTTTAGTTCAGCAATAATGAATGTTTAATTAGCTGCTTTTTTTGTTTGTTTAATAAATGTTATTTACTTTGTTAAAAAAAAA
>evgR004980c0g1_i01_Hydra
GGAAGTTAACTGAAGTGGAGAGTGTGATAAATTTTATTAAGCAATGTTTCAACACAATGAACGGATTGGAAGCTTCATCAAAATAATAAAAAAGTAATAAACAATCGATGGAAATTCAAGCCGTAATATTTCTGTCGGTATGTGGCTTCTTCGTTGTGGCTCTTGTTGTTTTGTTTTGTATTTTAAAATATAGAGAAAAATCACGTTCAAAGTATGAAGAAATAGCCAAAGATGAAGAATTCGACGATGATTTTCCAATACCAGAATACAATAAGTCTACATTATCACTTGTAAGTAAAGACAGTACCAACAAAGAAAGCTTAAACTTAGAAGAGCAAATAACGGAGGTTATAACTGATGAAAAAATGTCAGATGTTGATTTCAACCTGCAACTAAAGTTTTCAATACTTTTTTCAGACGAGCAACATTTTTTAATGCTAAACATTATTGATCTTACCGGCAAATTAGACAATCTAGGATGCAAGCATTTTAGAATCAATCTTACTCTACTACCGGAGAAGAAGTATCGAATGAAAACGAAATTTATTAATCTCACAAACCTTACTGATGAAAATGGTATATCTGTTTCCTCGGTGAAAGAAAATTTTAAATTTTCAAACATAACGCGAGAAAAACTGTTTAGCGCAGCTTTTAGGCTTCGTTTATATGGAAAACAAAAAAATCATTTAAAAGAAAAATGTTTTGGTGAAATAATAACACAACTTGCGGATGTCGCGCAAAGAGACGGTGGATTTGAAACAGTTCGAAAATTTTTAACAAAAAAAGAATAATTAAATTTCTACTTATAATTCTCTAGATACCAATTAACATATTTTTTATTTTTATTCTTTAAAGAACAAACGAATAGGATGCAAATATGATCTGTTATAAAAAATGTTTCAGTGATGTGGAAAATTGCATGCAACAATAGATCACGTGCAACACTTCTGATTGGTTTGGATTTTGGGTAACCAAACTAGACATTTTTTTAATATATACAAAGAATTTATTTTTGAACAATTTAATTTATTCTATATAACTTTTTTTGGAGATAGGATATTCTAACGTAGCTTCATATTCTTTTTTAGTAATTGTTTTTTAATAGTTTATTTTAAAGTATTATACGTTACAGTATATATTAATAACGATAAAATATTAAAAATATATGTACATGCATCATTCAAAAAAAATATGCATCAATCAAGAATATATACATACATATA
>evgR004981c0g1_i01_Hydra
ACAAAGTTTAAGATGTCTTCATTCAATCTATATAGTTTTTCTGTTAATACATCAAACAAGGTTGCTGCATGCAAAAGACGCCGCTACAGTGTCGTTCAACGAGACTATCATAACGAGCTAGAGCGTCGTCGTCGTCAACTTATCTCCTCGAAATTTGTTCTGCTACAGAATGCACTGCCAGAAAACGCATTTCCTGTTAAAGCCAATTTGGAAAAGGTTTCTCGGTGCTCAATACTAAATGCAGCATGTAGATATATGAACAGCATGGAATCGAAGAAAAACGCTATTTCAAATGACATTTATGTACTAGAGAGCGAAAACGAAGTTCTTAAGAAAAAAATTGAAATTCTTGAAAAAAATAGTGAACTTTCTCAAGACGATATTTATTTGTTGCTAAATGGAGACCAAGCAATAACAAAGAAAATGGATGATACTTAG
>evgR004989c0g1_i01_Hydra
CGCGGAATTTATTGGGATAACGATAACTCTCATATATATGTGTGCATGAACCAACATGTACCAAGATATTGATATTAGGATTGGCTCAATTATTGGGCACCATTTGATAACAAAAGAATTATATGGTATTCATCGCAATCAAAAAGTATACATGTTGTTTCATAAAGTACAAAAGAAGTGGTTAGCCATTACAAATCAACAGTTTAGAGAATCATCGAACTTTGATAATAGAATGCGTAAAAATTTAGAAAGTTATAATGATCAAATTAAAAATCTTGGACCAAATCAATGGCTAGGAAACGCTGAAGGTTTATTTTTTCGCAAACAGGTGAATGAATCGTGGATAAAAAGAGTTAATTGGCGCTTCTATAAGAGTTAATATATCAAGCAAACTTTAGTTTTTTAGATTGTTTGCAAGTAGATGTAAATATTCATGAAAGATTATAGATTGTAATATAATATTTAATTTTTTATACATTTGTGTTCATTGTTGTCTTTTTTTTAGTAAACATTTGATTCTGGTTAAGTTTAAACCAACGTACTTTAAAGGTTATTATTGTTATTATTATATTGATTATTTTTATAATATGTATTTGTACAAGGTTTAATTGACT
>evgR004993c0g1_i01_Hydra
CTGGATTTGGTCTTCAGAGCATCATTAGTCATCCACAATACTCGTGGATTTTTTGATCTTATCGGTGGAGTTAAAGGGGGTTTTGAGGGATTCTTAAATTTCACTTGGTCTATTATCCACTCAATTATCATGAGAATATCTCTAGAATCGACAGGGGTAAGTGGTAAACTTCTTACATTTTCAGATGATGGTCTTCTACAGTTTTATACATCAAGACAAGAGACTAAGGAAACTATAAGAAACAAAGTGAATTCTATAAAGGACACTTACTCCGCTTTGGGTTTAGAGTTCAAGGTGACAAAAACTATCGTTTCAGGGAAAGTTTGGGAATATCTAGGTGATGTGTGCTACCAATCAAAACTATTGCCCTGCTTCATTAAGGAGATGTGTTCTTTAGGCAAGAGAGAAATGAATAGAGGTCTTGAACTATTCTATGATAGGATTAAATCTATTCAAGCTCAAGCTGCAGCTTTCTCTAATAGTGGTGGTTCACCTGTTTTATCATACATCCTCAAGCGATTTATGTGTTCAGTAGCAATTGAAGAACGATTTGGAGTCAAGAGTGAAAGGCTAATAGAGTGGTTGTTAATCACCCCAGCATCTGCTGGTGGGTTTAGGATAATGTCACCTGCAGAGTCTAGCCTATTAAGTGATGTATCACAGGATGCTGAGTATTTAGCGGACCTAGAACTATACAGAGTTTACTACTCTAACATAATGCCGCAGTTACTGTCCACTTATATTCATGGCTCTCAAGCATGTGCCAACCCAGTTAGAGCAATCATGGGAGCCTCAATCTTCCACCTTAAGGGGTTCTCTACAACTGGTGGTGGTATTGTAAGAGCAGCTATAGAGGAAATTAAGAAAAATCCTAAAGTTGAGCATTTGGTCTC
>evgR004995c0g1_i01_Hydra
ATTTTGTTTGCTTAAATATTTTTTAACATTAAAAAAAAAGAAGATATTAAAATAAAGAAAGTGCGCAAAAAAGGTTTTGACAAAATGGAGAGACCTGAATGGAGAATATTTTTTATTTTGTTTTGCTTTTTATTCGCGTATTTGATTTTTACACAAAATTTTGCATTACCAAATACTCTTAAAAAACCAAGCTACTACGCTAATGAAGTTTTCAATTTTACATTCAATGGAGGTGAAAACATAAAACATCCAGGTATGCCATATCAAAGCCAAAACATTCTTATTTTAACGTATACAACATTTTTTGGTGATGGATGGCCAATCGATTCGGCATCTCTTTGTAGCTTCAACACATATAAATTTAAAATAACAGATGATAAAAAACAGTTTACAAACAGTAACGTGGTAATGTTCCATTCGCGAAATATGCCATCAATACAAGAGTTAGAGCAACTTAATAACATGAGAATAAAGGAGCAGCTGTGGATTTATTTCACAATGGAATCTATAATCAATAACCCTGGTGTTAGTGAAATAGATAAGTTTTTTAATGCTACTGCCACATATGGATTAGATACTGATATACCGTTTCCTTACAGATATCATGAAAAACGCTCTGAATTAGATGAAAACATTAAC
>evgR004999c0g1_i01_Hydra
TCAAGAGTACCTTCTCTTTGGTTTCTAAACCATCAACACAATCCATGTTGTTCCATGGTAGTGCTAGATTAATGGCCGAACAACAACAAGTTGAAAGACTTGATGTTACTTTGTTGATCAAGGAACAAAAGGAAAAGCAAAAGCTTAAAGAATTGGAAAAGAAACTCCAACGAGAACACTCTCTCAAGATTAAACAACAAAAACAAAAATCCAAGGAAGCTATTCAAGCTGTTAAACAAAAACAAGCAGACAAGATTAAAGAAAAGGGTACTCGTAAAAAGAAGGATGAATCTTTACCAAAAAGAGCCTTGACACCAATGCTTTGGTTCACTACTCGTAATTTTGCAATTATTCAAGCTGAATTAGTTTCTAATGGTGTTATTAAAACAGAAACATCAAACAAATTCACTCTTGTTCAAGCTGAATTAAAGAAGAGATGGGATACTTTACCTGAATCTGAAAAGAATATCTATATTGATTTGGCAAGTCAAGATAAATCTCGTTATCAAAAAGAAATGAAACTCTACAATAAGAAGAAGGAAGCTAATAAGAGACCAATGACTTCTTATATGCGTTTCTTTATGGAAATTAGACCACAATTGGTTCAAGAAATGCCAAATGCAAAGGTTACTGAAATTACAAAGGTTGCTGCTCAACGTTGGAAGG
>evgR005002c0g1_i01_Hydra
CGTTCATGCGATAACTGTTTTATGCGGCATGCTAACATGCCAACATAAGAAACTATTTTAACAATAAAAAATATAACTGATTATTAATATTGATTTTTTATTATTAAATTTATGGTTGAGTTTATAAAAATAGTCCCCAGAGGACTATTAACAAAAATATCAAATAAAAATAATGACAGTGTAAAACAAATCAAGCCAATAAAGTCTGTAAAATTCTCTAGTTTGGAAAACGATCACAACGAGCAAAACACCGATAATTTCAATATTGCAGACGAACTGAAGAAAGTGTTCAATGACTTAACAAACGAAGACAATAAATCAAAATTTAATCGACTAAAGCGATGCTATTTACATCAACAAAATAGCTTGATAGAAGAGTTTGAAAAAGAAAAAAGACATTTGTTAGAATTGTTCAAGCAACAAAAGGTTTTAATGTGTCGTGCGTTTGAAAATGAGAAAATTGAATTATTAAAAGAAATCGAAGAAAAAGATAAGATGCTCTTTCAGTTAGCTTCACTTATGAGAGAAGAAAGCGTGGTTGAATTTAATCTAGAAAAGCAAGAAGTAATGCCAGCAGAAAATACACGAAATACACGTTTACACGAAAAAACACAAGATAATTTACTTCAAATAATAAATAATTTACATGAAATTATATGTAACAACAATACCACTTTTAAAAATTTGAGAGCTTTTGAAAACCTTGAAAATCAATTTAAAATTCTGCTAGATGCGATTAAAATAAAGGAGCACGACTCTTATTTGCGACAACACTGTTGGACAATCGACGACAACACTATTGGACTCCGACAATTGAAGCTTAATAGAAGTAAAGGATTAATTAATAATGAAAAACATGACGATAATAACGAAGTTGTAATCCAAAAAGAGTTAGGCCAAGCACATAGAAAACAAAAAACTGAACTTTTAAAACTTTTTGCAAAAGAAAAAGATGAATATATAACCCAGTTAGAACTAGAAAAGAATATATTCGAAAAAGAAACTCGCTTAGAATATGAATCTCGAATGATTCTTGAAAGAAAAGCATGGCAAGATACCATAGAAGACTGCGAAAGAGAAATTGCAATTTTAAAGTATGAAAGGGAGCAAATGGACCGGAATTATTGCATAGGTATGAACGAGTTAAAAATGGAATTTGAACTGGAAAAAAAGAACATTTACAACCAATTTAAGGTTGATCATTTCGGTTTCAATGTACTTCTTACTGACACGTATGAACCCTCACAAGGGATATATTCAAAATAGTAATTTAAAATATTTAAAAATTTGGAACTTTTTTTATCCTACAAATTGTAAATAATACAAATCAGTTTATGTGATTTACAGTTTACATTTACACAACTATATATATTTTAATTTTAAGAATTTTAATAAAGTATCTTATTTTAAAAAAAAA
>evgR005005c0g1_i01_Hydra
CTATTTTAGTGCAAAAAGTACTAGATTTATATTACAAGCATAGTAGATAAGAATAATAAAAAATGAGTATTATACAGAAATATATGGGTTATCATCCAAGGCGCTTATCTGTACTTTGTATTTATTCAGCTGCATCTGGAATTTTTTTTGGCAAGATTTATTTTAATTTAAAGAAGCATCCTGGAACCGGTGTAAAAATCATAAAACATTCTCTCGCTACAGGTTTAATTGCTGGTACATGTGGTTTAACTTATGGATTTTTAAAAAATGCACGACCTTTGAGTTTATTTTTAACACAAAATATGTATGCTACATTAGTTGCAGTAATCTTTTTGAATATTCGATCTGTGATTAACAATGCAGAAAATCATTTTCCAGCTCAGACAGATCATAAGCGTATGAATGATCTTTCTATATCAACAGTGAGTGCCGCAATCACAGGGGGATTGATAGGTTTTGTCAGAACTCAAGGGGTTGCTTTATTTACAACTGGAACTGTTATTGGTTCGCTTACAGGATTCTGCGGTCATCTGCTGTATGAGAAAGTTTGCGCTTTACGCATTAAATATTTTTTACAACAGCAATACCCAGACCTCGTGTCCAAACACAAGGAATGTGAAGAATGGATGTTAGAACAGACTATGACCAACCCTGATTATACTGGACCAAGAAAGAGTGATCCGTATATTGTCCAAATGCTGGATAGTTTATGGGGCGATTACCGTAAGACAAAGAAATCTCTGGATATTGATTTAAATTCATAATTGCCGACTACATTATCTTTTATGGCTAATCACGCAAGTTAGTTGCTCTACCGCAAACTCGGAAAAGTGTTTTTTGCTTGTTGCTTTTGTGTTTTTCGGAGATTCAAATTTAGTAACCAATGTTTTAAAATTGATTGAAAGGAAATGTTGAAATTTCAACGTTATTTTTATCTTGCCAATAAGCAAATGCAATATGAAATTATACAAAAGTAAATAACTCTCATCATAGGTGATTGGCGAGAAGAAATGCTTGATATAGCTTTATAAATGCATGAGCATGATGCATTCTTATGCAGTTTGTATATTTTCCTATTCGAGTACCGCTATTCAATCACTAGTTCTTCGATAACCGTTATTCAAGCGCCGTACCTATTGTTAATTATTGGCATCCGGTTATTACTATTTCATCACCAACGTGTCATTTTGATCACCAGCGTAATTAACAAATAAATTAACGAACAAGTAATTTAGTAATTTAGCAGGCTTTGTTGGTAATTAGCGTAATTAACAAATAATATAGCGAACGAGTAGACCTTAATTATGTTGAAGCATAAAAATAATGGATTAACTTTTAATTTGTAAACCAAAATATTTACTTCAAAGAAAAAATAAGGATGAACTGGAACCGAAATTCTGGTTCCGGTTCTGGCTAGATTTATATTTTCGGTTTTTGCCGGATTTCGGTTCAAAAATGCTACCGAATAAATATATATCATACAGCATATAATATATATACCAAAGTATATATATA
>evgR005008c0g1_i01_Hydra
TTGCAGTTTCTTAATTTGTATTCGGATCTATCCATATATAGACTGTTGCTTATTGCTGACATGGCTAGCCTCTTTGAATCTATCAAAAAAGATGATGCTGGATTAGACAAAACAATTTTGCTATTGAGTAAAAAAGAAAGAGCATCAGATGTTGAAATTCTTAAGTTTTTGTTCGACCCAATTAATAATGTCCGTCAAAGATTTTGTATCAGTGTATACCATGGTAATCCAATTAAATACCCCACAATTGTAGCTTATACTGATGTATTAAAACAAAATGAAATGCGTTTTCCTCAAGAAGTAACTTTTTGCAGTGAGACCGAATTACGTGGGTCGTGTTACAATTTAAATTTTACAACTGAAAAAAGGTTTGAAATAAAGGACTTACATGGATGGGATTCCATCGTAAAATCAATATACGTTCCAGATAACTTTGAAGCAACTGGATATGCATCTGGTGCGTTTGGAGGTACACAATATGGTTGGTATAGTGGACCAACTGTAATTGGTAAGGTATCAAGTTCTAATGGGAAGTGGTATTCATTTGAAGTGCGTCCTTCTAAATTCTTAGCACATAAAAAAGTTCGTTTTTGTAAGAATAGGAATTTAAATCAGGATGATTTGTGTCATCAGTTG
>evgR005010c0g1_i01_Hydra
GAGAACAAACATTGAGAGCAAAACTTGCATGGGCTAACGAGAAAGGCTTTCCCTGTACAATAGGAGCTATCGACTGCACACATGTTAGAATCGATAAATCCCATGGAGAATTTGGTGACGAATTTATCAATAGAAAAGGATTCGCTTCGATTAATGTACAAGCAACTGTAAATGAAAAGGCTATATTCACCAGTGTCAATGTCAGTTGGCCAGGGTCTGTTCATGACTCAAGAGTATTCAAAAACTCGAATATATATCCGCTACTAAAAGGTTTTGAAAACCAAGGTTTTTTACTGGGAGATGCAGGGTATGGCATAGCCCCATTTTTGATGACGCCATTTGAAAATGCTCAAACGGTACTTGAAAAGAATTTCAACAAAAAACACGCTAAGAATAGAGTTATTGTTGAACATGCGTTTGGTCACTTAAAGCGACGCTTCCCAGTTCTTAGATATGGCGTAAGAATCAAGTCAGATAGCAACATGTATCACTGCATGCTTTATACTGCACAATATTGCAAAAAAATTTAAACGATCCGGATTTCGAGGATGAAGATACGGGTTTTCAAGAAGGAGAAGAAGAAGAAATCGCTCCTATAATTGAAATAGGAAATGAACAACAACTTCGTTTACTTGGTCAACAACGGAGAAACGAAATTGCCTTTTCTAACTATTATTATCAAAGTTGGCTTAAAGTACCAGAAGTGAGAAAATGCAAACACTTATGATTTTTCAAAAATTAAAAACAATATGCTTTGTTTTTCA
>evgR005012c0g1_i01_Hydra
TTACGTGTTGTTGCCAAATCAGTAGCGATATTCGAAAATAATCTTTCATTTCAATGTAAACACTGCTTAAGCAATAATATTGATTTTAGATCATCAAAAAACTCTTTCATCAATTTGTGGTTTTGTATTTCATGCGGCAATAGTATCTTTCTCTTTAGTTCTATATCGGATGATAAACTACGTCTCCAATCATCAATCGACAGTATACTACATCTCAATGAACTGCCAAATGAACTTTCAATCTTTCCTAGCTCTCTAGAATCTTATATTTTCAATCAACTCGATAAGTTTTTTACTTCCCAGTTAAATAACAACGATGACGATGATAATTGAGAGCCCAATCAATTGTAAATATAT
>evgR005017c0g1_i01_Hydra
AAAATGAAGCCATCAAGAGACAAAAGGGTCTTCAAGGAAAGAAGAAGGCACCAAACCACCTCTACAGTAATGTTACTGGCGATGCTGCTGTCAAGTTGAACAAACTTTTGACTGGTAAGTCTGGTAACACTAAGGCTGTTGTTGAAAAGGTTCTCTCTGCAAGCAAGTTCATCTTGAGTCTTCCAAATGATAACTCTGTTATTACATTCTCTTTGACTGGTATTGCTATTCCATCATCTGACCGTAATAAGGAGTTGGCTGAAGAAGCTAGACAACTCACTTTGATCACTGCATCATGCAGAGATGTTGAACTCGTCCTCGAACCAAACGTTGACAAGGCTGGTTCATTTGTTGGTCAATTATTGATTGGTG
>evgR005032c0g1_i01_Hydra
ATTACTTTGGTCTATTCGTTATTTTATTTTCTACTTCCTTAAAAACCCATGGTTAATCATACCTCTACAACTTACTTTACACGGATTAAGTTTATCGCTGTTTTTAAGTGTAAAGTCTATTCACTTAAAAACTATTGCTCCAAACAGTATCCATAATACAATGTTTGGTATTGTTCAATGTTTATATAGTGGTGTATCTATGGTTATTGGAAGTTTTATAGGAGGACAACTTTATTTTAGATATGGTGCACGATTAACATTTTTGTATGTTTGCTTATTTGCTCTTGGATGGACAGCATTTTTAAATGTTTACGTAATTGCAAAAAAACTTTGGAAAATAAACGTTGGTGAAAGGTCAAAATCGACTTACCAGTTAGAAACAATTGACGTTTCAATTGATGTTAAAGACACTTCATTGTAAGTTGGAAAAGTTCATATGATGCTATAGACACTTCAATTGATGATGGCAAAACTTCAGTTGATGTTAGTGATGAAAATATTACATCATTG
>evgR005052c0g1_i01_Hydra
CTGCTATCAGAACGTCGCGTCAATTTTGTAAACAAAAAATAAATGATTACACATGTATAACTATTGTATATATTTTATAAGTATACATTATTCTTATGCGAAATTTAAAATTTCAATTTTTTAATGAACTTTTTGGTTATGAGAAACTTAAATTTACCATATTTATAATCTGTATATATACATAATTTCATGGATTACAAAGAATGCAAAAGTTCTACAACAATCAATTTTATTGCTGGAGCAATTGGAGGTACTAGTGCTGCTATTATAACATGCCCACTTGAAGTTGTTAAAACAAGATTGCAGTCTTCAGGTGAGGTGTTTAAGCCAATAACAAGACAGATTTCATGTGTGACTACTATTACACGAATTCAACCCGAATATATTAGGCCCAATGGCATTCTTGGTTGTATGAGGCATATTATTAATACTGAAGGCTTCAGATCTTTGTATAAAGGCTTGGGGCCTAATCTTATAGGTGTTGCTCCAGCTAGAGCAATCTACTTTGCAGTATATGTTAAATCTTCGAAATACATACAGCAGACAGGAAAAGTACACCAAGATAACCCAGTAACCCATATGCTGGCTGGTGCTTCTGCTGGTTTCGTCACGCATACTTTAACAGCTCCAATTTGGTTTGTTAAAACAAGACTTCAACTAAGCAGTCATAAATACGGAATGATGCGATGTATTGCGAAGGTTTACAAATCAGAAGGAATAAACGGTTTTTATAGAGGATTATCGGCATCATATTTTGGGATTATCGAGACAATTATTCATTTTGTTATATATGAGCGTTTAAAAAAGATACTACGTGAGTATCACAGGGATAATAGTTTTGATCACCTCGAAAAAAACCAGCTAACGGACTTTATGTTATCTGCAGGATTTTCAAAAACTTGTGCTTCTATTTTTGCGTACCCTCATGAAGTGGTGAGAACACGAATGCGTCAAGAGACTCGTGATAAAGAGCGAAAGTACCGTAAATTTTTTCAAACTTTAGTATTGGTTTACAAAGAAGAAGGTAGGTCTGCATTATATGGCGGTCTAGGTACTAAATTACTTTGTCAAATTCCAAACTGTGCGGTAATGTTTATGGTATACGAGTCTGTCATTTTTTATTTATGTGAAAATGATCAAAACGACTGAATTGCTAAAATTATATTGAATCATGATATTTATTTTGAAAGGCCTAAGTATTAGATTTCACAAATACGTCAGAAACCTAATTGAGCTTTATTTTCGTTTTATATTTTGTCATAAAACGGAGATTTATATATAGGTATATATTATATTTTATTGTCGTATGTATCTTAAAAGTCACACAAGGAAAGTGTTTTATTTCTGTGAAATAAATTTACTTCAAATTCGTAAAATAAAAAAAAAA
>evgR005061c0g1_i03_Hydra
TAAATGTTGCTTGTTTTATCAGCAAAGACCAAAATATTGGGTTTCAACAGTTTATATATTTTGATGGATATAGGAATATCAAGCTGTGAAAGAGATCTCAATGTGCTACTGGCTAACCGTTTAGCAGAAGTTCATTGTTTGAGAGAGCAACTGATGCGAAAAAGTAGAGATTCTTCGCAGATATCTAAGCATAATTCTGTTTCGTTTGCACCTACAGATCTTGAAGTGACAAACTTTATTACGGAGGATGGTCAAAATGGTCAAATAATTATATCAAATAATGTGGGCTACCTTAACTGTAACAATATAAATGGTGTAGACTACTCTTCAAAGTTAAATGGTATAAGTGGTGTTTCACTTGGTTCTACTGTTTCGCTGGGAGATCTAAGAAAAACAACAATGTTACTTGATGCACCAATGTTATCTGTTAAAGATTTGAACCACAAACTAGTGAAAGCTTTTGCTGAAGTTCACCGCTTGAGAGAATTAATAGAAAAAAATACAGATGTATCGCAAGCTTTTACTAAAGATCATGAATCTGTTTGTAATGCTTCATATACGCTGACAACATCAATTGTTAATTCAACTTCAGCGTTTGAAGATATCAAAACTCATTCAACTTCAATGTTTGAAGGTATCAAAACTAGCTCAATTTCAGTGTTTGAAGGTATCAAAACTCGCTCAACTTCAGTGTTTGAAGTGAAAAACGTTATCGTGGCTGGTCGACCAGGTCAAATAATTATATCAAATGATGTAAGCTTTCCCGTCAAACTTAGGGGTGTTGATGAAGAGTCTATAACAAACGATTTAAATAAATTATCTGGTGGTGATTCAGTTAGGTCAATATCAGGTAATTTTGAATTTAAAAAGGTACATATCTCCCCAAAGTCGATGGGGCCACTTTAGTCTAGGAGGTTGCG
>evgR005066c0g1_i01_Hydra
TTTGGCCAAGAGAATTACTGGTAGAATGAAGATTGCTGATGATGCAATTAAGGTTTCTATTGTTCGTTTCCACGATACTGGTACTCTTGCTTTAGGATTGACTTCTAATGGTGCACTGATCAATCAAACTTTAACTGACATGGACTATGACAGTGGATCCACTGCTCAAATTCCAGGTTTGAGAGTTGCTGTAGATGCTTTGAGACCTGAAAATGGTGGTAGAGCTGATGCTGACAAGGTCATTTATATTTTGACAGATGGTGTTGCCAACGTTCCATGTGGCTGTTGGTCTTGTGAAAATGAATGGGGTAACAAACCAAGTATTTTCAAGAAGACTGTCGAACAAGATATTATCAACTCCCTCTCTGGAAATGCCAAAAGTAATGCTTTGAAAGCCATGTGTAAATACAAATTCACTGACAACCCTGATTCTTTGGGCATGAACGGTAAAGATTGTAACTACTGTGCGTGGGAATGGTATTCCATGTGTTTACCTTGTAGTGATGCTGTTCCAGTTGCTCAAAAGATTAACAAGTGGAAACGTGATGCCACTGGTGTTGTTCCATCTGATCTTGATAATCCTTACAATGGAAATAACAATGTGCAATGGAAGATTGTTGCCATGGCTGTTGGTAATGCCATGACTGATGTTTGGGGATCTCGTCAAATTAAGGGTATGAACTACGATCCAGCAAGAGCTATGACTGTTTCTTGGAGTGACTTGTCATCAACTATGAATGAAATTGTCGATCAATCTTGTAACCAAGTTAACGTTCAAGTTCCTTAAATAAAATTTTGAACAGTCGGC
>evgR005080c0g1_i01_Hydra
AAACATGCGGTTTTACAATCCATTTTTATTCTCGTTTGTTGTTTTTTGGATATATTTGGAGGTATAATCTACTTAATTGAACCCTATGGTCTTCCGAACTGCTATTTGGATTTAATTTCTGGATCGACCCAATTTACTTACAATTGCAATGGAGGTTCAAAGTTTTGGCTTGTTAAAGGAGTAGACGGACTGGAAGGAACAGTTTCATTTCAATCTGCAGGTAACGTAAAATACCTTCGTCATTTTTGGTATTTCATCTATTTGGAGACACTAGCTGAAAGAACCAATAATCTTGGTGATCGTTCATTTCATGTGAAAAATAGTAGTTATGATTCATATAATTTGATTTATTCATCAAACTTCCCAACAAATGTTTTGTGGCGTGATAGTTCTCTTTTTAAATTTATGGTGATTCCAAACCCATGGTTAGTTCCAGAATCAAATTTTAAACTTTTAGTTTTAGACTTAAACATTCAAGAAATAATGAGTGTTGCAAAGTATTCAGTCAATGTTACTATTAATGCTTCTTTATCCATCTGGAATATCTCTCAAAGCTTTGTTATTAAGTGTAGAGATCCATACAACATTACAAGTTTATATAATTCAACTTTTGTTGCACCACTACTTAGCTACACTGTTTATGAGTTTTCTGCAGGG
>evgR005096c0g1_i01_Hydra
AGAGCTGAAATCCCAGCCATTGCTCCATATCAGTTATATCAAGGTTTGAGCACCTTATGAAAAGCGATGTTAGTTGAAATTCAAAAAAGTCATTTTTTTGGAACACCCTAATTATTAGTTTTGCGTTTTAGAGTCATTTTGAATCAAGATGTCTAAGTCTAGAACTAGGCACAACTGAACGATTAGTAGAGGAAAATCGTAGAAGTTCCTTATAGGAAGGCAATTTGAAAAATAGCGATTGATAATTCAAAATGGCTGGCAATAGCGATCTACAAAAGACTGCCAGCACTGCTAATTTACTAATGGTAGGAGCTGGAGGAATTGGTTGTGAACTATTGAAAAATTTGGTGTTAAGTGGCTTCATAAATATTCATATTATAGATTTGGACACTATAGATGTCAGTAACCTAAATAGACAGTTTCTATTTCAGCGCAAGCATGTCGGAAAATCAAAAGCTTGTATAGCTAAAGAAAGTGTTTTATCATTGAGACCAAATTTAAATATAACAGCATTACATGACACTATTATAAGTTCTGAATACAATATAGACTTTTTTAAAAAATTTGATTTTGTTTTAAATGCTCTTGATAATAAAGTTGCTAGGAACCATGTAAACAGAATGTGTCTTGCTGCTGATATTCCTCTTATTGAAAGTGGGACAGCAGGGTATTTAGGGCAAGTAACTTTAATAAAGAAAGGTTTTACTGAGTGCTATGAATGTCAGCCTAAACCATCAAATAAAACTTACCCTGGTTGCACTATAAGGAATACTCCATCTGAGCCTGTTCATTGTATAGTTTGGGCAAAACATCTATTTAATCAACTTTTTGGTGAATATGACGAAGAAGCTGAAGTTTCACCGGATACTGCTGATCCAGAGCTTTCAGGGGAAGCAGGAGAACAGGCACTAGCTTGTGATGATGTGACTGCAGCACGAATTTCTACACGTGAGTGGGCAAAAGAAATTAATTTTAATGCTGAAAAATTATTTACAAAGTTTTTCTTCAATGATATTAAGTATTTACTTTCTATGGAAAAGTTGTGGCAGAAAAGGCAACCCCCAACACCTTTATTATGGAATAATGTTGAAATAGAAGAAGAAAAAACTCAATCAAATAATGTGCTGTTAGATAAAAAAATTTGGAGTATTTATGAATGTCGAAAAATATTTCAGGAATGTGTTGAAAAATTAAGTGAAAGGTGTAAAAGCTCAGAACTAGTTTGGGATAAAGATGATACTGTTTCAATGGACTTTGTTGCTGCAGCAGCTAATATACGATCTTTTATTTTTCATATTCCACTTAAAAGCAGATTTGATATTAAAGCGATTTCTGGAAATATTATTCCAGCAATTGCAACAACCAATGCTATTATAGCAGGATTGATGGTAATAGAAGTTCTTAAACTGCTTTCAGGTCGGCTACACGATTGTCGCACCATATTTTTAAACAAACAAGTCAAGGTTAAAAAACAGTTATTGGTTCCTTGTTTGCTTGAAGCACCAAATCCAAAATGTTATATCTGTGCTAAAAAACCAGAAGTAACTATATTTTTAAATTTACAAACTGTAACAGTTAAACAGTTAGAAGACAAAATACTTAAAGATAAGTTGTGTATGGTTGCACCAGATGTTGAGATAGATGATGGAAAAGGTACTATTTTGATATCAAGTGAAGAAGGAGAAACAGAAGAGAATTGGGATAAGGTATTATTTGATTTTAAAATCTCTGATGGCACTCGATTGAAATGTGATGATTTCTTGCAAAATTTTGAAATTGCAATTACTTTACGTGATAAAAAAGACATAAATGCTGAACAATTATTCTTCTTGGAAGGTGAAATACCTGTGCCAAAAGATGAAAAATCTGAAGATAGAACACTGGAAAATGGCAGTGGGGCTAAAAAACGAAAAATTAGTTCAGAAATTGAAAATAATAGTGCAATAAAAAAATCTAAAACAGAGTTAGATATAGATTCTGAAGATGTTGTAATTTTGTAAGAACATATTGTAGGTTGTACTGTAAAATTTCTTTTATTTTGTTATGGGAAGAGAAAGGTTGAATAAAGTATTTCTTTTCTTTCTTTAATTTTTTATTTTTGTAGTGGAAAAAGGTTAAATGATAAAGTATTTATTTTGTAAATAAGGCTGATATATTTTAAAAAAAA
>evgR005099c0g1_i01_Hydra
GAGTTTTATATGAAAAGGGAAAGATTGACTCATAGAAATCTCTGTTACTATGAATCATATGTGTCTCATAGAAATTGATATGAATCATGTGTCTCAAAACAATTATGGTAGATCTAGTTTAATGGTTTTAGTTTCCCAATTGATTTTTATAAAAAGCTTTTGCTTTTTTATAGTTGAGAAACATTGTGCAAATGTTTTAGGATCTATATCGGCTAATGAGTGGTCGAAAATGGGTGCAGGTACGGCAGTGAGGGTCACCAAAAGACCTATAAAGGGAGTGACTATCTTTCTAGTAAATTTTTTGTTAATTGGTTTCTCTTTTTTGACAAAAAACAATACATGTTTTATTTACA
>evgR005107c0g1_i01_Hydra
GGAGAACTCTACCGAAGAAGATAAGTTTGTTGCAACAATGGAAAGTTTCGAACAAAGAGTGAAGGAAGATCTTGAATTTGTTGAAGGAAGGTTCAAGGAAATGAAAACCGAAATTGAATTGCTTGCCAAGTCATTCGACGAACGACCAGAAGATTTACAAAAAGAACCTGACAAATTCTTGGCCCTCATTGACCAATTCGTTCAACAATTCAAACAGGCCAATGAAAAGAACGAACAAGCTAAAAAAGCAGAAGAAAAGAAAAAGAAGGTCGAAGAAGAAAAGGTCAAGAAGGAAGCTGCCAAACAGCAACTTAAAATCAATTCAGCCAAGCAAAAGCTTGACATTCTCAAGAATAAGACCACTGTAGGTGGTGGAACAGAAGGTAGAGGAG
>evgR005109c0g1_i01_Hydra
TATCAAGTAAGTCATCATATTATTGTAAGTATTGGTCTGTCTTCAATGAGATTATTCATTACTTAAAAAACAAAGATATTTGCCATAAAAAAAATATATTACTCGAACATATGGAGTGTATGTCTCAAAAGAAAATTGGTGAATTTACTTACTCTGCTGATATAATTACTAGAGCATTTGAATATTTTGCTATATCCAGGTCACTGTACAGTCGTATTTCCAATGACTATAAATTACCAAGTATTAGAACTTTGACAAGAATCACTTCAAAAGTTGGATCACAAGATAGTATGGAATTTTTAAAAAATGTTTTGATGAGTATTGAAGATACACAGCGTAAATGTGTATTGTTGGTTGATGAAGTTTATATCAAATCTATATTAACTTATCATGGAGGTACATTGTTTGGGTATTCTGCTGATCAACCCGGTGTCCTTGCCAAAACTATGTTGGCTATAATGGTAAAATGTTTATTTGGTGGACCAGAGTTTCTTATAAAAATGATTCCAATCAGTTGCTTAGATTCTGATTTTCTTTTGAATCAATGCAAACCAATTATTGAAATGATTAACAATCAACCTAATGCAAAATTGTTATCGGTTATTGCTGATGGAAATAGAGTAAATCAAAGATTTTTCAAAATGATGAATAAAATTGAAGGAAAACCTTGGTGTGGAGATGACGAGCAC
>evgR005122c0g2_i01_Hydra
CAATATGAACAGTGCTTCTAACATTAATTGTTTTGAATTGAGTTCAAAACTTTTTGATAAAGTTGCCTATAATGTTGCAACAGATTGGTCTAGACTTGGACGCTATCTTGACTTAACAAATGAAGAAATAGCAATGATCGACTCTGACTACGGTAAAACTCATGAGAAAGCTAATCAAGTTTTAGTAAAGTGGAGGCAAAAAATTGGAAATGTGACATGGCAGCAATTAAAAAAGAACTTGGAATCAATTGAGCGTTTTGATATAATTAGACAAATTGAAAAAGATTTAATAGGTATTCTAACTCGACCAGGCACCATTGGTGGTGAAAATACTCAACTAAGTATCATTAGTGGTGAAAAAGCTCAACCAGCCAAGAGCAGAGTTGCTGATAAGAAATCATTGCTTCAAAATCCTGGTGTTGATGGTATAAGGAATTATAAAATTCCACCTTATTTTTATGTCCATATCCTAGAC
>evgR005122c1g1_i01_Hydra
AATTATAAAATTCCACCTTATTTTTATGTCCTATCCTAGACCAAAATACTAATGTTGCTCGTGTTGAAGTTGGGCCACAAAATTATATCTGTCACGATAATGAAAATTATATATTTGGACCATATCCAATGGTAACCATTCCTACACATCATTACTGCATTATTGAAAATCCAGTTATTTATTATGAAAATGTTGTTGATCTTGATTTAAATGGTCAAGCTAGAATCAAGTATGGTGAATTAGAAGTACGCTTTACACAAGATCCATTTCCTCTTTATCCTGGAGAAGTCGTTAAACAATCTCCAATTCCATTTACAGTTTTAGAAGCAAACACAGCATTGCATTTAAGAGCAGCAATGCAATTTTCTAATAATGATGGGAAAAAATATGTTGCAGGTGATGAGTGGCTCTTTGAAGGGCCTGGTACATATATACCACGTAAAGAAGTTGAC
>evgR005123c0g1_i01_Hydra
AAAAATAAAAGAAAATCAGCTGTTAAGTAATTTATCTTATTTTCCAAAAGAATACGAATTATCGTTTGATTTCAAACCAACATCATATATGAATGATTGGGCAAGCATTCTTACTTTAACTATCGGTGGAAATAAAAAAAATTGTGGCGATCGAACACCATCAGTTTTGATCAAACCTGATAACACATTATCTTTTGCATCATGTATAAATGGTATTGTCAACTCTGTTTTTGACACAAACGTTGTATACCCTTTAAATCAATGGTTAAAAATTAAAATATCACAAATTATTATCAAAACAAAATATATTTTTGCAGTTGAAGTGTCAAACAAATGCATTTACAGTAC
>evgR005125c0g1_i01_Hydra
ACAAAATTAGACATCAAAGAATGAAAGTATTAACATTATGGCTTTGATGGTTCATATATGGTTATAGTTTGATGGTACACCCGTGGTTCACACGTAGTTCGGATAGTTTAAAACAGTTTTTTACCAAAGCAATTTTTATCAAAATTTTTTTAATCGATTTTTTAAATGAATGATCAAGATGAAATATTCTACGAGCCAAAAAGTTACTTTACAGAAAAAATGACAAATGCTTCTATTTCCGATGAAAAACAATCTTCCGGTCGACGCAAATCTCACCGACTGTCGGCTCCTGTTGATCAACGTACAAATTTTAATGCAAGTTGTCAAAATATTTCTCTTAAATCACAAGATTATAAAACGAAAAGAATATCTCAAACTTTTCTCGATCATATAAGTCAGCAATTTAAAGATATTTCCATAAAGAGATCATCAAGTTTTGATAAAACAGTTTCAAAGACATCAACCAAAAGGACAAAAAGTGCTTCAACTGAAAATGCACATATAGTTAATTCAAGCTTAGAACCAATAGGAAATTCATGTTTACAACCTTTAAAAATTGATCTAAAAAGATCTAAAAGTAATGAATCAGCACTTAAGTATCAAAATGTTCATGATTCGAATATCGTCACCAAACCTTTTGAAGGATTATATTTTAAACCCAATGTTTTTTATAAAAAAGAAGAAACTTTTTCCGAAGTTCTAATATAATTGCTTTTACTTCAAGAATTATTATATCCCGGATGATTCAACCTGTCTCCGTAATATTCAATCAGTCACCATAAAAACCATGCTTTTTAAGATAATACTTTTTTAATTTAAAGAGAAC
>evgR005148c0g1_i01_Hydra
AACAACCACTGGTCAAGCAACCACTAGCGTAACTGGTATCATCAGCAGTGGTGGTGGCCGTAGCACTGGTTATCAAATTCGTGTTCAAAAAGATCTTGACGAACTTGAAAAGAAGTGGTCTGATAATTTAAAAATCGAAGATGCCAACAAACTCAACAAGATGACCCTCGTCCTTGTTCCTGATGACGGAATTTGGAAGGCTCACCCACACGAATTTGAAATTAACATTCCTAATGAATATCCAATCAAGCCACCAAAAGTTATCTGTAAAACTAAAGTATGGCACCCAAATATTGATCAAAACGGTGCTGTTTGTTTGAATATTTTGAGAGCTGACTGGAAGCCAGTCTTGAACTTGAATGCCATCTTTACTGGTCTTCACTTCCTCTTCCTCGAACCAAATCCCGATGATCCTTTGGATGTTGATGCTGCCAAGCACTACGTTGATCAATTCGAAGATTTTAAACGTAAAGCAAAGAATTACATGAAGGGTTCTTATTATTAATTTTAGAAAGACTCGAGAGAACTAAGTCCACCCACACCATACAAACATTAGAGTTTGTTTTTATTTTGAT
>evgR005166c0g1_i01_Hydra
TCATTCATCAAGTAATTCACTGCACTAAGTAGGGTTCATACAATAAATTCTGTATAAGTTGAAAATGGTTTGGAAATTAAAGTCTATCTTGGTTACTGGTTCTAACAGAGGGATTGGGTTGGAGTTAGTTAAAAGATTAACTGAAAGAGCCGAAATTGTGTTTGCATGCTGTCGTTCTGCTCAAAATGCTCAAGAGTTGAACAAACTTGCAGCAAACCATGAGAATGTCAAAGTAGTTGAGATAGACGTAACTAATATGGATTCTATAAAAAGTGCTTATGAAAAAGTGTCTAGTTTACTTAATGGTTCTGGCTTAACTTGCCTTGTAAATAATGCAGCTATAGCTTTTTGGTCAACGTTCAACGAATTCTCTGCAGATCACTGCAAAGATGTTTTTCTTACAAATTCTATTGGACCTGCTCTTGTAACACAGACTTTTCTTCCATTAATAAGAAAAGCAGCAGTGGAGAGTTCTGAAACTGAGTTGAGTGTGTCTCGAGCTGCAATCTTGAATATATCATCTACCATGGGTTCAATATCTGAAGCTACAACTACGCAAGGTATTGAATACAGAATGTCAAAGGCTGCACTTAACATGCTGACAAAAACTTTAGCTTTTGAATTGAAATCTGATAAGATTTTAGTTGCTTCTATATGTCCTGGTTGGGTTCAAACAGATATGGGCGGCGTTAATGCTACTCGTACTCTAGACCAAGCTGGCACCGATTTAATTGCTTTGTTTGAAAAGCTAAACGAAACTAATGCAGGATTTATGACGAATTGGAGTGGTAAAATTGTCGGTGCTTAGTTGTCGGTTTGTTTTAATGGCGTGATAATGATTTTTTAATCTTTTTATGATAATTATTAACATAAATTATCAACTTTTTTTCTTCTTTAGTAATATTTCTTTTTTTAAATTTTATGTAGGCTTTTATAAAGTTATTTATCGATTATTTTTAAAAAAATTTTAAGTACCACAGTTGTAAATTAGGAATCTTGGTAATAAGTTAAAATTGCTTAAACAATTATAAATAATTAATTAATTGCTTTTTTAATAATTTGGTAAATCCCTTGAAAGCCCCCTTGCC
>evgR005169c0g1_i01_Hydra
CCGAAATATTCTTGATGTTTTTGAACTATTTTTAATTTGTCCTTTTACAAATGCGAAGGTAGAACGAATGTTTTCATGTATGAATCGCGTAAAAAATGACTGGAGAAGTTGTCTTAGACGAGATCGTTTAGAATCTCTACTTCGGATAAGCGAAGAAGGTCCTGAGATTGACAAATTTGAGCCTGATACTGCTATGGACGAGTGGTATAATGACAAAGTTAGACGATTGTCCGCAAATTTTCATCGTTATCCGGAAAAAAGAAAAAGAAAGTCAGGAAAAACAGACATTGATCTAGCGACAGTTACAATATCAGATTTCGAAGAAAGTGAGAGCGAGGAAGAATATTGTGACAACTTTTAATTTTATTTTGGCGCATTGTAACCCCTTAAAATTAAAATATCAAATAACAACTTTGAATGGCGTATGATTTTTAGCTTCGTAAAAAT
>evgR005171c0g1_i01_Hydra
AGAAATTTTTGGACACAGAACTAAGAGTGGTAGCAAAAGTCGTGTTCACAAAAGCAGAACATTTAATCGCTTGATATATGAGTGCAAAGGTGGCTTAAATGAGAGTCTGAAAAAACTAATAACGGAGCTAGATAAAAATGATGTTTGTCTTTGCTACCATCTAATTGCATCTTGTCTTCATAAAGACCAAATGTGGGAGGACATTCATCAAGGAAACAAAAAGTTTTTCATTTGCCAACCTACAGAAAAAGCAGGAGAATGGGAAAGTTCTCTCATGCTTCATCGAAGTAACCCCCAGGGAGCGTCAATGTACGGAAAACTTTGGACGGCATTATTTGAAAACCCATCGCTAGAAGACCAATTCAGAATTTTTTCAGATCTTCCAAACCATCATAAATGGATGTGGGATAAACTTGGTAAAGGTGTGAAATATGAACAGAAGTTTTCAAATTCATGCCGATATTTTCCTGGTTCACCTAGAAACAGAATTCTTCAGGCCTCATTTTATCGAGTTAACTCAGAAAAAATTATTGTATCAATTTTAAAAAATGTTTTTGGTGTTAAAGAGACAGTTATATATCCATCTAGAAAAGAAAACTTATCTCTTATAGTCGCAAAAAAGAGCAGGATATCTTCTGAAGATAGTTTTCTTGATCAAGAGTTTCAATATAATGGTTTACATTTTGGAAATAAAGTGGTTGCAAACAATATTTGCATCACAGATTCTATTAATGAGGTTAACTTGGTTCAGCATATAAGCACAAGAAACAATCTCTATGAATACTTTTGTGCTGAACTTTTTAGTCATGGCGTTCTTTGTTGGCGTGTTCGAAAACCATCTCTAAGTATATGCGTTATGAACAATTACAATCATGAGAATGGGAATTTTCAAGACAAAGAATTTGTTCATGTAAAAGCTGAAACAATGTGTTCTTCTACAATATACAACTGCTCCTGTAAAACATTCTCATCGTGTGTGTTTTTTCAATCAAACAATATGACTAATGTCAATAATTGTTGTCATTGTCGTCTGTTAAACCAACTTATGCATATGCTTGATAATCCTGATTTAATCCCAGAAAATTGTTTTTTAAACAAAAAAAAAGTTATGTGTAGTTTAGTCTATCTAACATCAAATGTTTTCAAGTTAACTTCAAAATCTGGAGTTGAAAGATATTCAGTAGTAGCTGATTCGTGTGCATTTGTTACAATTTTTGAAATTTTAAAATCATCTCGCAAAGTAATAAAATG
>evgR005172c0g1_i01_Hydra
TAGTTTTTGAAACTATATATTTCTTATATTTTGTTAAAAATGATGCTACTAGAGGTTCTGGTTACGCTAACAATAGTTGGTGTGGCGAGTTTATCAGATACCACTCAACCACATACTGTACGAGCTCTAGGCAAAAGAGAAATCCCGCGTTTAGATATTTATGGTGGCGTATCATCCATACTCTTTATAGGAAAATCGGCTTATAATTTAATTAGCTCGATTGAATCTAAAAAAACAGATAAACCATTTCAATGTATAAATAATGCCAGAGTTCAAGTTGAAGCATCACCAGCAACTATAAATATGTGCACCAGAGATTGGAGTTCAGTATGGCAAAAAAGTAGTCTGGATATCAAAAAATTTAAAAATGGTTTTCAATATAAACATGATGTTTTAGAGATCAATAGCAGTCATCTCAAGTATGTATCATTTGAGTACCTCGG
>evgR005173c0g1_i01_Hydra
AGAAATTCTAGGTGTTATTAATTTCTTCATGATAATTAAAAAACTATAACAACAACAGCAACAACAAATTTAAAAAAATGGTTGAAAATTTAAACTGTACCGAGTTGTTAACGAATGTGTTTGGCAATATAACCATTGGAAGCGAAAAAATTAAAGCGGCAAAATATTTCTGTGAAAAAATAAGCACGTTCAACGGCTCATTGTCCGCTAAAGAATACTTTAACGAGTTTTGTTGTAAAAATGATTTCTTAAATCAAATGTGTTCTACGTTTTTGAAAAAACATATTAAATACGAAGGTTCTAACATAGCTTGTTCTTCATATCTAATCTATTCATATGTATTGATTGGATTGTGTGTACCACTCGGTGTTCCAGCAAACTTCTTAATTTTAATCTTAAACTTTCGTCACCATATCTTGCCTAAACAAACTAGTTATTTTTTGGGGAGTCTGTCTGTTGCAGACATCGGAGTCATAATAAACTTTATCGCATTAGCAGGGTATCAAAAAAGCCACCCCAATCTTTCAAAACAAGTTAGCGTGTTTTTGATGCCATCAGCCCACATTTTTTTTGTATCTATATCTATGTTGCAAATTTGCGTAGTTAGCATTGAGAGAGCTATTGCAGTAACTTGGCCTCTTCAGTACACAATCTTTTTTACACCAAGAAGGGCAAAGTTCGTTACACTAGTAGTATGGTTATGGGGCACTATTAATTTGACTTTATCACTACTTCGTATACACGTGCAATCAAGTTCGTACAAAAATTTTGTTATTTACAACTTTTTATTAACTTTGTTGGTTTTTCC
>evgR005185c0g1_i01_Hydra
TTTATATCTGTTATTTAATTGGTGTTTATTGGCAACATTAACAAGTGCTATTGCCAAAGATACAGAAACTGTTTCATGGTTACCGCAATGGTTAATGGAGGCGATTGTTGATCAAGACCATATTGGAAAGATTCGTATTAATGATGGGTGTGATATGGTTTATCGCTCAGATAATTGTACTATACATTTGAAACCTATTGGTTCAAGTTGTAAACAATTATTAAATTTTCGAAACGTTTTTAAAGACCAATCAAAACTAGAAGTATTTTTTAGTTTAACCGATTCAACATTTCCATCGGATAACTTTTATATCCATAAAGTTGATGTGGGATTGTGTAATAACGACATAACTTTTGTTACTAAAAATACAGTTTCAATTAACTTTATTGATCGATATATTTTACTTTCTG
>evgR005187c0g1_i01_Hydra
AAAAAAAAAAAGCATTATAATTAGCAGGACCATGATAAACCACTTCAAAATAACTCATTGAATGTATAGTGAAAAACTTTCAGGAATATACCGTTGGAACCCTTATGATTTAAATTGAGAAATCGTCCCGCAGTCTTTGCGTAAATACCTCCGGAAATACACTCATAAGCTTAACAATGAACCGACAAAAGTAAAGATAGCTCATTTTTAATCATTCTTGACGAAGAGAAAGCAAAAACCGGTCCCGCCTTTCCTACGGAAGTTCTTGTAATATAAGAGCCTTAAATATTCACATTTTGAATAAAGTGAAATCAGAAAACACTGCAAAGATGAGGCTACTGGCGATTTGTTTTTTACTTGTTCTTAGTTTTTGCATGTGCACGAGTTTAGATGTCGATAACGATGTAATTATAAGAAACGACGATCTATTCTACGTTAAGAAAAAGACTTATACAATCGAACAGATTCAAGATATTTTAAGCAAGGGACCGTGTAAAAGTGTTAAATATACTCAGTATATAAAAAAAGAAGGTTGCTTTCCAGTTTCAGTAGAAAATAAAATATGCGGAGGAACTTGTAATTTGAACAATAATAAACAAGGTGCTAAATGCGCTGCATGTGGACCTTCAAAAATAGTTGAAGAGACATTAAAAATACTCTGCAAAGAGTCGAATGCAGACGACGCTGAGCTAAAACTTGTACCAATAAAAACTAAAGTGGTAAAAGAATGCTCTTGCGAACAGTATAAATGCACGAGTCTATAACAACTGAATATTTATAATTTATTTATCACAGTATTTATTCGTCTATTATTATTTTCAATTTTATAAATATTATTAATTTTTAACATTTCTACCTAATATTTTTTATTAAAAAAAA
>evgR005193c0g1_i01_Hydra
CTATTGTCTAAATAAGTTAAAATATGGCACCAACAACTCCAAGACAAAATAAAAAGCGTCCGAAAACTACTACTCCGCAATCAGAAGTCTCAGAAAGATCAGCACATCAATCATCAGACTTGCAGCCGTTATCTGATGATGTCTCTTTACAGCCACCTTCACAACGTAGGCGTATAGATAGAGATGCACCTAGCTCACCTCTTCTTGAAGTTCCTGTTTCTCCTAATACTTCTCAACAAGACCATGGAGTACTTGATACATCAATGAACAGTCGCACAGGCATTCTAAACACAAGTGAAATTGATCCTCAAAGTTCACCATTAAATTATGGAACTCCTTCTTCACGAATGGGAAGTGTAAGAAGTACATCGAGAGGAACACCTGCTAGATTACGAAGTGATGTAAGAAGCAACAAAAGTTATAGACAAGTCAATGTGGGTGCCGCTGAACAGTCTGATGCAAATGATCCATTATCAAGTGAAGTTCAAAGTGAGCCAAAACTTGTCATTTGGGGTACAGACGTTGTTGTCACAGACACAAAAGAAAGGTTTCGTCAGTTTGTTTTAACATACATTGATGATGACTTTGACGTAAACAATGAATACGATTATGATAAACCATTGTATATGCAAAAGCTTGAAGAGATAAGCATCACTGAACATGCGTTTTTAAATATTGACTGTAAGCACTTGAAGAAATATCACAGTGATTTGTACAGACAGTTGATATGTTATCCACAAGAAGTTATCCCTACTTTTGATGTTGCTATCAATGAGTTATTTGAAGAAAAGTTTCCTGATACAACTCTTAATCATCAGATACAAGTTCGAACATTCAATGTTGATAAAACAATGAACATGCGCTCACTAAATCCAGAAGATATTGACCAGATGATTACTATAAGTGGGATGATAATTCGAACTTCCAGTATTATACCAGAGATGAGCGAAGCATTCTTTAAGTGTAATGTGTGTCACATGACTCAAACTGTGGAAATTGACAGGGGAACCATTATTGAACCAACTTCTTGCTCGAACTGTAACACACAACAAGGAATGGGGCTGATACATAATCGCTCTAGATTTACTGACAAGCAAATGGTTAAAATGCAAGAATCTCCTGATGATATGCCTGCTGGACAAACACCGCATACTGTTATTATGTATGCATACAGTGATCTTGTAGACACTGTTCAGGCTGGTGACAGAGTAACAGTGACAGGAATATATCGCGCTACTCCACTTCGTGTTAATCCAAGAATGCGAAACGTGAAAGCTGTATACAAGACTTACATTGATGTAATTCATTTTCGAAAGTCTGACAAACGAAAGTTGTATGAGAGGGACTCAGACAGTCAGGTGACGTTTACCCAAGAAAGAATCGAATATTTAAGTGAACTTTCAAAGATGCCGGATATTTATGAAAGACTTGCACAAGCATTAGCCCCGAGTATCTATGAAAATATTGATATCAAAAAAGGTATTTTGTTGCAGCTTTTTGGCGGTACAGCCAAAGATTTTACGCATGCTGGTCGGGGAAAGTTTAGATCAGAGATAAACATTCTTTTGTGCGGCGACCCTGGAACAAGTAAATCACAACTATTACAATATGTTCACAACCTTGTCCCCCGTGGTCAATATACCTCTGGAAAAGGTTCTTCAGCGGTTGGTTTGACAGCTTATGTAACTCGAGATCCAGAAACAAACCAACTAGTATTACAAACAGGAGCACTTGTTCTAAGTGACAATGGCATTTGTTGCATTGATGAATTTGATAAAATGTCTGAAAGTACACGGTCAGTTCTTCATGAAGTTATGGAGCAGCAAACACTTTCTATTGCAAAAGCTGGAATTATTTGCCAGTTGAATGCTCGTACTTCTATTTTGGCAGCAGCAAATCCACAAAAGTCACAATGGGATCCTAATCTTACAACTGTAGAAAATATTCAATTGCCACATACCCTGCTATCAAGATTTGATTTGATATTTTTGATGTTGGATCCTCAAGATGAGCATTTTGATAGAAGATTAGCATCACATTTAGTTTCACTCTATCACATGTCAGGCATAGAAGCTGATGCTGAATCAGTGGATATAAGTACACTAAAAGATTATATATCCTATGCCAAAAATAACATAGTTCCAAAACTTAGCGAGGAAGCTGGACAACTCCTGATAAGTTCTTATGTGGATATGAGAAAAGCAGGGGGAAGCAGAGGGGCAATTTCTGCATATCCTCGTCAGTTAGAAGCTTTAATACGGATGGCTGAAGCTCATGCAAAAATGAGATTCTCAAAGTTTGTCCAAGTAGTAGATGTAGAAGAAGCAAAACGATTACATCGTGAAGCATTAAAACAATCTGCAATGGATCCAAAGACTGGTATGATTGATATCAATATCCTCACTACAGGATTGAGCATTACAGACAGAAAGAGGAGAATTGATGCATCCAAAGCATTGTTAGAGTTGTTGAAAGCTAAAGGTCGGGTTCCGACAATAAATTACTACAAGCTTTTTGAAGAATTTAGAGACCAATCTGAAGCGAAAATAAGTCGTTCATTATTTGACGATGCGTTAAGAATACTAAAAGATGACGATCTGATTCTTATGCCAGATGATCGAAAAACTATACGGTTGATTTGATTGGTGGATTTAAATTTTTGGTTTAAAAATTTAACTCATATAGGCCAGTTGAGGAAATAAGTTTAGATTGGACAGTCGAGACATCAGACTATATATTATTTCAAGTAACAGTCGAGACAGTAAGCTATTTTGGATCACAATGTGAAGTTATTCGTCGGTTTTGTGAACTTTTGATAGACTAAAGTGTATATATTAAAAAAAAGTAAAAAATATTGTTATTAAAAAAAAAAA
>evgR005207c0g1_i01_Hydra
CTTCACAACCCACATGGCTTCAGGTAGCCCAAAGTCTGACATGACTGGTGTCGTCTCTGATTCTCCAATTTATAAAACTTTGAAGAATTCTTGCAAGGATATTCAATGTCAAGTTGATGAACTCAAGAAGCAAGGAAATACTTTGGAAAACCAAGCTGAAACTATTGAAACTATTTGCTCAAAGCAATTAGATTCTGGCGTTTGCAAGTCCCTCATGAGCGGAAACTTTGAAAAATTCAAGACTGGTACTTTTTCTAATAACGATTTGATGGAATATGTTGCCCGTCAAATTGCTATCGAAAGAAAGACTAACCCACATGGTCATCATGGTCACCATTAATCAATTTATTTGATTAAAAATCACTCGAGTTTGAATTCAGCTTCAGGGAAATTGCGCGTTTTGCTC
>evgR005243c0g2_i01_Hydra
AATTGAATATCTAAGAATATGTTATTTTTAGATTAATTCGGGGAAAAAATGAGTAATTTAGATCCTATGCTGCAAAGTCAAATAACTGAAATCACATTAAGAACAATTACAGAAATTGAAGAACTAGCTGAAGAAATAATAAGCGATAGAGCAGAAATAATAGATTTTGATAAAAAACGTTGTGGAAATCGAGAAGCTTTACGAGAAACAAAAGTAAATCCACAGGATAACCAATGGATTTGCATTGGAAACTTATTTCTTAAAATTACCCAAAAACATGCTATTGAAATGTTAAAAAATGATCAAAGCACAATAGAAAAAGAAATATTGAATATTCATAATGGTTTAAAGCCTAAATTAAAAAAGCTTCATGAGTTAGAAGGTTTACCTAGTACAGATGGGTTTGATTTAAAAAGTATCAAAACAGAGTCAAAATAAATATAAACTCTTCAAAATAAAATAAAGTCAAAAATCTCTCTCTCTCTCT
>evgR005261c0g1_i01_Hydra
CAATGGGACAGAAAGGATACTAAGAAGGGTGAATCTAACTCAATTATTACTTCATACAACAGAAACTTTACTGCTCGTAACGATGGTAATTCTGCTACCCATGCTTTCGTTGCTTCACCAGATATTGTCACTGCCTTGTCTATTGCTGGTTCTTTGAAGTTTAACCCAATGAAGGATTCTTTGGTTGGTTCTGACGGTAAGGAATTCAAGCTTTCCCCACCAAAGGGTATGGAATTGCCAGCTAAGGGATTCGATGCTGGTGAAAATACTTTCCAAGCCCCACCAGAAGATGGTTCTAAGATTAATGTTGATGTTGCCCCAACAAGTGACAGATTACAATTATTGTCACCATTCTCTGCATGGGATGGTAAGGACTATAATGACTGTTCTGTCTTGATTAA
>evgR005284c0g2_i01_Hydra
CAGCTTCTTTTTTATTAAAGAATACATAGGATTTGCTAAAAGGAGTGGTGACATCAAAATGCATGGATATCTTTGAAGGCCTCCTGAGTTCAGATTTAGACCCGCACAACCACATAGCTAAAACAGAATGCATTCGTAGTTTTTTTACAGAGTGGAACGAAAGCAAAGCAGGTCTCGAGACGCAATTAAATTGATGAGGTTGCATCTATATGCGTGAGAGTGTCTTTATGATTTAATTTTTTATGTGTAGAATGATAAAAAGACGTCGTTCAATTAAAAAAGTGTTCCCTGTCTACTTTGTAGCATTATATGACTATAATTACAAAATAAATAATAACACGCCAATTGTTAATGCTCCCGATGCTGCAACAGTTTTGTGTTTTAAACGAGGCGAAATTTTGTGTATCTTAAGCTCTGTGTTAGATCATTGGATATTATGTAGATCTAACGTGACTGGCGAACAAGGTTACGTCATGACATCGTTGCTTGCTCCATTGAGTGGCGCTGTAGACAACACCGTTTGCATACATAATAATGAAAATTATCCAAGAACAAGTTGGTTGGAACCAGAATTTTTGTTATTTTTTCCGAACTTTCCTTTATCAAAACCCAAGAAACGTTGCCCAGTTGATGACTCAGGAGTAATATCAAGAATTTTTTTCGGATGGGTTACAAGCTTAGTAATTAAAGCATATAAAAAACCATTAGAAGATAAAGATATTTGGCAACTTCAAGAAAATAACACCGCTACTTATCTTGGACAACAGTTTGATAAATGTTGGAATACGGAAATGAACAAATATAAAAAGAGTCTTTGGAATGAGAATGATGTAGAAAAAAGTAAAAAGCTAATTAATCTTCAAAATGTCATAGAAGAGATTGCTTATAAAAACCTGGTTAAACATCGCGAGTACAAGATTTCAATATTAAAAGTATTACTGGAAGCTTGCGGCAAGCGTTATATATTTGCAGCCCCTATGCTTCTATTACTTTATGTAGTTTTATTATTTGTGCAACCATGGTTATTAGGGAAATCAATTGAAGTGATAAAAAATAGACAGATTAATAAAAGTTGGGGGTACTTCTACGCATCTATGTTGTTTATAACAGGTTTATTAGGATCTTTTATTGTACAGCACTACTTGCAAGTTATATATATAACCAGTTTACGAGCACGATCATCATTATTAACAGTAGTTTATAAAAAGATTTTTCGAATTTCCAACACTGGTAACCTTGAATTTACTACTGGGAATATGGTCAACTTGATGACCGTGGATACACAAAAATGCTTTGATTTATTAACTTATCTTAATTTAGTGTGGTCTTGCCCTCTTCAAATTATAATTGCATTATTTTATCTTTATTTTCTAATGGGATACTCTGTTCTTGCTGGTTTGTGTG
>evgR005287c0g1_i01_Hydra
CCCTCATAAAATATTTATTTATATGGCCACTCAAAAAATATTAATTTGGTTGGCCATTCAAAAATAATCATTTAGTTGGCCATAATAAAAAATAATCACCCTTAAGGTTGGCAGTTATATTGATCTCACCATAACTCAATAAGCGCCATCTTATAAAAGTAATTGGAATTTCTGATATGCAACAAAATTAAAAAAACCCTACATAAACATAATTGTTTTATCACAAATCATATACATATTTAAATCCATAAACTTTATTCTAGGCTTACCAGGAAAACAGATTTTCAAGTTGGCAGCAAATAATTTCTGTGAAAAATTGATAACAATAAAAGATGTAAAGCTCTCCACCATGCTGAATATCTTGATATCTTTTATCTTTCTGAATCAATTTATTATCTTGTACTTTTATTATTTTAATCTTCTTTATCATATGAATCCCCTATCCACCGTATGTCTTACAACAACCTTAAAGTTGACCGGAACTCTTTCTGTAATTATCTTTCGGACAGCCCATGGATCGAAATACTTTGTCTTCCTGACAAATGTGCTTCTTATGTATACAAGCTGGCATAGAACTTTTCATTTCCTTTTGACTATTTCAAGTCAAACCTCACTATTCTTTGTGGTTTTCTTTTCATTTTGCTGCTGCAAGTTCTAATCATAACAAATTATCTAACAATAAAAAAATCGGAAAACAATTATTTACTATTGCTAGAAACCACTGTAAAAAGCTTTAGTACAACACCAAAG
>evgR005301c0g1_i01_Hydra
CCTGTATTTATCGAAGATATCAAAGAAGGTGGTGATGGTGAAATTCAATACTATCCAGCTGGTTTGGAAAATTTAGGAAATACTTGTTATATGAATTCTGCTATTGAAAGTTTTAGATTTGTCCCAGAATTGAGACAAGCTATCAATGAATTTGATGGATCATCAGTGAGTGAATTGTTGGAAAAGGATAAGGAAGTTACCACAGAATTAAAATCATTGTACAAACAATTGGATACTCGTAGTGAATCCATTAATCCAATTACATTTTTGGAAGCTTTCAGAAAGGCCTATCCACAATTTGCCGAAAAGGAACGTAGTGAATTGGGAGAATTCTATGCTCAACAAGACTCTGATGAATTTATTACAAACTTGTTAGGATCTTTGGAAAATATGCTCAGAACTAATGAAAGC
>evgR005343c0g1_i01_Hydra
AAAAAAAAAGCGATTCTAATTTTATTAACTTGCGTGGTTACATACATTTTTTACCATACTGCGATTTTTAATTTTTTGGTTTATTTAAAAAAAGAAAACTCGGGAGATTTGTACGGATTGGCAAAAGATAAAATTGTTAGTCTTTACTCGGAAATTTATGATTTAAATAAAAAAGCGTTCAGCAACTCCACTTATTTTAATAAAGTTTCGTCAATAGTCAACGTCTTGCCATTGTGTAGTAACCTAAAAATATACAGCTCTAATGTACTATCCTATGCCTATTTCCAAAGTACTCGAAACAAAACTTATAAGAACTATATTATTGACGTTGTTAGAGAGGTCAAAAGTAGTAGACTTTACCGTAATTGGACTGTAAGGATATATCATGATTTGAACGATGTGGGCAGTCATATATATGACCAGTATGACAACTTATTATTTTGTGATGTGAGAAAAACTCCGTATGTAGGAGATTTATCTAACGTAGATGGAATGATGTGGCGTTTTCTGCCAGTTGGTGATTTAACTGTTAATGTTGTATGTTCTAGAGATTTAGATTCTCCCATTTTAGAAAGAGAAGAAGATGCAGTCAGTGATTGGTTAAAAAGTGAAAAAATGTCACACGTGATGCGCGATCATTTTATGCATACTTCATCAATAATGGGTGGCTTGTGGTGTATAAAACCTTTTAAGAATAGATTACTAGCCAAAAGACTATTTGATTTGGCTCTTACTAAGTCCTCAAAAAGGATTTCTAAATATTCACCACCCTATGGAAGTGATCAAAATACGCTGAACCGATACATATGGCCTCTTATAAAAAGTGATATTATGGTTCACGATTCTTACAGGTGTAACGGTATTGCTATTCCTTTTCCAACAAAACGAAAAGAAAATCATTTGTTTGTTGGCTGTCCACGCCCATGCAACGTAAAAAAAATACAAACATGTCCTGTTAAATGCCGTAAAGAAAAGGAATGGTTATTTTGTTGATTAAATTTCTAGAAAAGTTGAAAATTTTTAAATAACTGAAGTTTTTTTTAAAAAAAGTGGCTGGGTTGTCAGTAAAATAGGCGACTTTATATTAGCTGTTAAAAAGTTTCGAGCATTTTATAAATTTTAAATCTTTTACCGTTGATTAAAAAAATATATGTTATTTCGATTCTGAATTAGTAAAAACTACAATCAGTCAGAAACAAGGGGTTTGTATGATTAAAAAGTGAC
>evgR005348c0g1_i01_Hydra
TGCTGTTTCGATATGTTTCATAAAAATTACTTTAAAAAGATTCCAGGAAAAAACTACAATATAGCAAACCATACAATAAAATCATCAAACAGTCGTTTACTTTAGGAGTTATATACAAATATTGTTCACTAAAACTAGGTTGTATAAGTCATAACTTGAATACACTTTAAAATATGTTTATTGGCTTAACGGTGTTGCAATAAAATAAAATAGCAAATAAGAACGAGATAATCCATTTATAAATTCGAATTTTTATTCAAAGTTAAAGCGTTAGTGTAAAAAGCAAGTTAAAAGATATGGATACAGTACTCATTGATTCTGCTTATTCTAATTTAGTTGAGTCGGCTGAAAACAATTCATTCATCAAAAGAATTTTGACTAACTCTGATGAAAAGAAGAAAAATAATTTAACCCGCGTACAACGTCGCACAAGAGAACGAAAAGCCTTTAAAAAACTCAAAGATACCGTTCCAACATTAAGAGATAAGCGTAAATGGACAAACCGTTTAGACATAATAAAACATACTAGTGACTACATTATAGAATTACAGAAATTGTTAGAAAGTTCACGCGAAGAAAAACGAAAGCAACAAGAGATTTTCAAAACATTCATTTTCTAACAGTCTTCATTTTTCTTTTCAAACAGTCTTCATTTTTCTTTTCAAACAGTCTTGGTTTTTTTTTTTTTAAATCCCAATTTTTATTAAAACCCATTAAAACTATTTATTTTAAATTTTATTTTCCA
>evgR005378c0g2_i01_Hydra
TAACAAACTTGATAATTATTTTAAGTAAAAGCACCTGAGTAAAAATGAACTTGTTCACCTTTTTGTTTTTCATGCCCTCTGTGACAAGTTTAACATGTTGGAATGGTACGTCAAGAATTATTTTTGGTGTTTTTTTAAATTTAACTGTTCCTGCACCATCCTGCAAACAATGCATGACTTTAAATATAACAAATTCCGTTGGCGCTATAGTTGAACAAAACTTTTATTGCAGCACGGATTCCAATTGCACCGCTCCTGTGATTGCCACGTGTTCGAAATGCTTTACGAGCAAGGCTATATGTTGTACTACTGATTACTGCAACGTGCAGAGTAGTAGTAGCTTTTCTTCGGAATACCGTTTATGGATTTATGTTTCATTTGGAATACCGTTGATTTTAACTGGGATTTTAACAGAAAACTTTTGAAAAATAAATAAATAATAATAAAAAGTTATTAAACAAAAGTTTTAAAACTGGTTTCAATTTAATTTTTTTTCATTTTTTCATTGGTTTTCAATACTAATAAATAATCCTAGGTTTTTAGTTTCTTTAATGTAAAAAGTTTTCAATGTAAATTAAAGTTAAATGTAAATTTAATGTAGTTAAATATAAATTAGCTTTCCTAATACGGTCTAACTTGCAAATATAGCATTAATATAAAAAG
>evgR005380c0g1_i01_Hydra
AAATATTAAGAAAGAATTAATAAAAATGTCTATCGTTTTGTCATTAAAAGAATATAATGATTTGAAAGAGAATTATAAATTTTTGATAGAAAGGAACGAGGAGTTGATGAGTGATTATCAAAATATGCTGCTTCGGGATATTGAACTCCTCGAACAAAATACTAAATTTCGCCGATATATAATCAACGGAGATCTTGTCGATACATGTTCGTCTTGCAGAGGCATACTTCTATTTAAAGACATTGAAAAATGCCCAATATGTGGTTTTATTGTATGCGAACGATGCGAGAAGCTACATCGTCAGACGTCAGACAAACATTAATTATGTTCATTGCACTATTTAATATTATTTATATATATTTTATATTTTTTTAATATTTTTTAACATTGAATTATTATAACACTAC
>evgR005389c0g1_i01_Hydra
ATGATTTGGTTAACTTATTCATTGGCCTTCCCATTAATGTTCTCTCATCACACAACAACTCACATTGTTATTTTATGGGCTCTTGCTTACATGGTTGCTTCATTCATCTTGGTCTTGATGTTCCAAGTCAATCACGTTACACCACTCACTGATACTTATCATGTCGAAAAGGATGGTATGGTTCATACTGACTGGGCTGTCACACAAGTAAGCGGATCAAGCAATTTTGCCTGTGGTTCATGGCTCTGGAACCACATTAGTGGTGGTTTGTGTCACCAAACTGAACACCATCTTTTCCCAACTGTCTGCCACATTCACTATCCA
>evgR005430c0g1_i03_Hydra
GAAACATGGATCAACCAAGAATCTACAATATCAGCACAAGATATTAGTTATGCATTAAGCCCAACTTCAATGTTGGATATAAATAGCACTTACTTAACTGATTTGATGCAAGTATCATCTTTAATTTCAATTCAAGCTTCTGATATTTTAGAAAGTGAAACACTATTAGATGAATCAGAATTTCTAGAGGAAACAAGGATCGACAAAGAAAAAATAACTGCAGCTCAAGATATTAGTTATACACTAAGACCAACTAAAATAATAGGTGTGAAGAGCACTTACTTAACTGATTTGATGCAAATATCATCTTTAAATTCAATTCAAGCTTCTAATATTTTAGAAAGTGAAACAGTATTAGATAAAACAGTATTTCAAGAGAAAACAAGGATCGACAAAGTAGAAATTACTGCAGCTCAAGATATTAGTTATTCTTCAAGCACAACTTCAAT
>evgR005432c0g1_i01_Hydra
GGACGGCTTAAACAAAAGTCTACTAAAACTCGGATATTGTCTGTCATGATCTGTAACTTAATTAAGACTACTGCCTAAAAGATTAAACACAATTAAAGGAAAACGACACGTTAAGACAGTTCCAGTGAAGTTGATTAGAATTGCTAATTCAGCTCGTGAAAGGCATATAGAAGCTGGATTTACATTTTCGTTCATGCAGTACATGGAGGTAATCAGTTCAATGTTTGGTCTAGATTTGTGCTTTTATTTAAGCATAGACGACAAAGCTAAAGTGCCATTAGGATTAGCAGCTGCAACAAAACAATCACCAATTTTGATGCATCTTGATTATCAAGTACGTTTGCCCGATCATGATTTTTTAGTGGCTTCCAGACATAAGCTTATACCTTCAATTTATGCAGCTTGTGTTATTGACCCAGAGAAATTTGCAGATGCAGTGTCATATTCTGGACCAACATATATTGCTATTAGATCTATGAAACACATGGGCGTGTTTTAGAGCTTCTTGTTTCAATTGAAAACTTTAAGATTCATATTTGTATTCCAGAAGGTCAGGTGAAACCTATATTTTTTGTTGCATGTGACGGAGGGCCTGATCAAAATCCACGTTTTCCCAAAACCTTGCCTTCAAAAAACAGTTGCAATTGCAAGATTTATTAAATGGGATTTAGATGCCTATATATGTGAGGTGCATAAACTCATGCACCTCACTATTCAGCATATAACAGGGTTGAACGTCGTATGGCTCTGTTGAGTCGAGAGAAAGATTAGTTTTGTCACATGATCATTTTGGGACACATCTAAATGCCTCTGGAAAGACTATTGACGTTAGAAAAAGAAATTTTTGCTACAGATGCAGAAATTCTAGCTGAGGTATGGTCAGGTATAGTAATTGATACATACTCAGTTTGTGCTGAATATGTTGAACCTTCCCAACTTCCACCAATACCAGAGCAAGGTTTAATATGGATTGCAAAGCACGTGAGACAAAGTCGATACTTTTTAATGATTGTTAAGTGTGATGTTCCGAATTGCTGTAAACCTTGGCTTCTACGTTTTTTACCAGGTCTAGTAGTATTATCTCATGCTGCTGAATGAATTTGCATTCCTGAGCCAAAGGTAGCACAAAATATTCCAAAACCTTACTTTTCTTCTCTTGCACAGCGATTAGGATTTATAGGCT
>evgR005449c0g1_i01_Hydra
ACATAAGTAACGAGACACAATTTTTAGTTCCGTCAAAATTCATTCATATTACTGGCATATACTCAATCGTGCTTTCAGTTATAGCTTTCATAGAAAACGGCTTGGTGATCCTTGCTTATTTCAAAGATCCAACAGGAAAACTGCAGACTCCGTTAAATTATTTTCTTATAAGTTTATGTTTTAGCGACATAATGATAAGTAGTATTTCATTGCCGGTATTTGCATACGACAGAATGTTTCAAACTCCGTTACCTAATGTATTGCACACGACAGTTGACGTGTTATTTTTTTTAACAGTAACTGCATCAATGTTGAATATGTTGGTGCTTGTGATTGACAGGTTTGCAGCCATCCAATGGCCTTTTTTATACAGAGTTAACATAAACATGATTCGATCACGTGGTATAACTATAGCTGTGTGGGTGGTTGCAATACTTATTAGTATTGGT
>evgR005460c0g1_i01_Hydra
TTATTTAAAATTCCAATTTTTGTTTTTAATTTTAAACTTTTTAATCACGCTCTTGAAAAATAAAGAAGAAGTTTAAAAAATGGAAATCGAAATGAAAACGACTAGTTCAGATATTTCAGACAGCAATGACACAGACATATCAAAAGATGAAACAGAAGATGTTTTAACTCTACAAGAAAGCAATACAGTTTGTATTTACAATTCTGTTGATGAAGTTGAGCGATGTCTTTATCTGTTAACGAGCCAACAAAAGTTCCAAGTAAAATTGACTCCTACATATGCAATACAGTTGAGGAAGCAAATCAATGTCTTCATAACTATGAAATATACAATACTGTGCGGTTTGCTGCTTTCTCAACTCCAAAAGACTTTGGTAGTACAGGTTGACAATCCAACACATTATCGCAAGCTTCAAGTGTC
>evgR005469c0g1_i01_Hydra
TTTTATTTTCTAAATAAGTGCTATGCAATATAAATATCACAAATATATATTAAGGTCTCGATATCAGAAAATTAGGGTCTTGTTATCAGAAATATAGATTATAGTCATCGATGCCTTCACCAGAAGAAGTTCTAGTTCTTGGTGAAATTCTTCGATTGAAAATACATAAAGCTAAGTTTTTAATAAAAACATGCAGAGATGATCTAGGAACTTTTCCATATTCATTTGTTGGGCATGAAGTTGTCGACTGGCTTGTAAAAAATGAAGAAGTTCAAAATAAAGCTGTTGGTGTCACAGCTATGAGAATTCTCCAAGAAAATGGAATTTTACATCATGTATACGACGATCATAGTTTTAACAATGATAGCTGCTTTTATCGTTTTCGAATTGACGATGGTACATTTTTTGTTGACTCCAAGTATATATTCTATCATCAAGGAATAGTAATACATGACCGAATATTAACAGATAAAGGAAAGACGTTTTTCCAAAATTACACAACTGATGATCTATGTGTTGTGGAGAACTGTTTTAAAGGGTCAATGTTGGTTCAATGGTTGATTTCAAACAATGTTGATACTGAATCGTCAGCCATCAGATTGTGTCAGGCGTGGCTGAATATTCTTATAATTAAATCTGCACTCTTGGAATCTACTGAATTTAATAATGATGGTACACTATATCAATTTGTATTTGATTTGAGATCTCCAACGAAGTTGAGAGAGTCGTTAAAGTTATATCCTGATAATAACTCAAGAAAAAGTTTTACTGATATGGACAATACTAGGAATCTTATGGCAATATGGGCAGGGTATAACAACAATAAAGTGCATATCAACAGCAAAAATGCTATAAGTTTAAGTGAGCCTAATGAACCTACGACTCCTATACAGCCAGCTGTGTTCCGTGATATAACAAGCGAAGAACTCCTTGGCAGTGACAGTACGTACTTGTGGCAAAATATTAGAATTGATTGCGACTTGTTTGGATTTGGTTTTGTAGTTAGAGGTACTGGACCTTGCTATGTACAAACTGTTGATCCTACAAGTCCAGCAGCAGCAGCTGGTCTAAAGATACGTATGTATATTTACTCAGTTAATGGCAAAGTTGTTTTAAACATGAACCATAAAGAGGTTTCGCACGAAATCGATAAATGTACATTTGTAGAACTTGTAACACTGTACAGTCTTGGAAGAAGAAAGCGAGTCTTGGGAGAAGATGAGTTTACTGAAACGACTGAAGAATAGAATGAGTATAAAAACTTTAAAAAAAGAAAAATGTTAGATTTGTAAATAAATATTTAAACGTTTCCATAAATATGAGTATATATTTATGTTTATAGATATATATATATATATGTGTGTGTGTGTGTGTG
>evgR005499c0g1_i01_Hydra
GCAGATCTTTACCACCACAAGGAAGTGATTCCAAAATCGCTCAAATGATTAATTTCATCAAGAGTGAAGCTAAAGAAAAGGCTGATGAAATTGAACTCCAAACCAGAGAAGATTTTGCCATTGAAAAACAAAACCTCGTTGAAGAAGGTAAAAAGAAAATCAGAGAAGACTATAACAAGAAGGAACAACAAGTTGTTGTCAAAAAGAAAATCGATCACTCAAACGAAATTAAATCTGCTCGTTTGGAAATTTTGAAATTGAAGGAAGAAATTTTGAAGGATATGGTTACCGATGCTTTGGAACAAATTAAGAAAGCTATCGTTAACAAGGACTTGTACAAAAAGCTTTTGCACGATTTGGTCTTGCAAGGTACACTCCGTTTCCTTGAAACCGAAGTCAATGTTTACTGCAAAGAACAAGATTATGATTTAGTTGCTTCTCAAACTGCTTCCATTCAATCTGCCTACAATTCAAAGACTGGCATGAATGTTAAAGTTACCGTTCAAAAGAAGAACTACTTGCCTGCAAACTCTGTTGGTGGTGTTATTATTGCCTCAAACAATGACTTGATTAAGATTGATAACACTTTGGAAAAGCGTGTTTATTTGTGTCAAGAACAAAAGCTGCCAACACTCCGTAAGATGCTCTATGGTGATATTGATGTCAAGC
>evgR005517c0g1_i01_Hydra
CGTTAAACAACTACCGACCAATCTCCATACTTCCTACTTTCTCTAAACTACTTGAAAGAATAATTTACAACAGATTATATGATTATCTAATTCAAAACAAATTCTTAAATAAAAAGCAATTCGGTTTTCAAACGCAACATTCGACCGAACATGCAATTCTAGATCTTGTAAATACCATAATAAATTCATTCAAAAAAAAAAAATTTGTACTAGGAACTTTTATTGACCTATCCAAAGCGTTTGACACGGTCAATCATAATATATTACTTAAAAAAATGGAAAAATATGGGATAAAAAATGTAAACCTTGATTGGTTTAAAAGTCTACCTTACCTGTCTAGCAGACAACAATGCG
>evgR005533c0g1_i01_Hydra
ATGATTAAAATAGCATTAAAAAATGGTGCAGTAATTCCAAGTAAAGCGACGATTGGATTGGTTGGGATGGATTTGTATATATTAAAAGACGCAATTCTAAATCGGTTTGAAATTTTGATATTAGATACTGGAGTTATTGTTCAAGCTCCTATTGGAAAGTATGTGGAGATAAAGTCAAGATCTAGCTCTTTTTTGGATGACATTACAGTTCATAATGGAATTATTGATAGCGATTACACAGGAACTATAAAAATTTTAATTCGTAATAATACACCAAATACCTTGTTTTTGTCTAAAGATATAGCCATGGCGCAAATGATTGTAAAAAGAAAAGAAG
>evgR005546c0g1_i01_Hydra
TCAAGAGATGACTCGCGCTAACCGTCAAAGATACTCTGCTAACCCAGAAACCTCCAAGCACTCCAAGGCTAAGGTTGAAGGTGTTAGAGTTCACTTCAAGAATACTTGTGAAACTATCAACGCCATCAAGGGTTTGTCTTTGAGAAGAGCCAAGAAGTACTTGCACAACGTCTTGAGACACACTGAAGCCATCCCATTCAAGATCTACAACGGTGGTCCAGGTAGACATGCTCACTCAAAGGAATTCGGTGTTGCTAACAGCAGATACCCAACAAAGTCAATTTTCGCTGTTTTGAAGCTTTTGAAGAATGCTGAAAACAATGCTAGAGTTAAGGGTTTGAACATCAGAGATTTGGTCCTTTCCCACACTCAAGCTAACAGAGCTCCAAAGATGAGAAGAAGAACTTTCAGAGCCCATGGTAGAATTAACCCATTCATGGCTAACCCAACTCACATGGAAATTATTTTGACTGAAAAGAAGACCTCTGTTGCTAAAGCCCAAACAGCTGTTGCTCAAGAATAAATTCATAAACTATAAAATATTAAAA
>evgR005550c0g1_i01_Hydra
TCGCCATCCGGAAAACTTGTACAAATCGAATATGCTCTTAATGCCGTACAGGCTGGTAGCACATCATTGGCAATAAAGGCAACAAATGGTGTTGTAGTAGCAACTGAAAAGAAACTTCCTTCCCCACTTGTTGATCATTCTTCTCTTCAAAAAGTTACCCCTCTCACCGATCACATTGGAATGGTTTACAGTGGAATGGGTCCAGATTCACGTGTACTTGCAAAGCAAGGTAGAAAACATGCCGAAGAATATTATAGAATTTACAAGGAACGTATTCCTGTCTCTCAATTAGTTAAACATCTTGCCTCTGTTATGCAAGAATACACACAAAGTGGTGGTGTTAGACCATTTGGAGTGAGTTTATTAATTGCTGGTTTTGATGATATTGGTCCACATTTGTATCAAGTTGATCCTAGTGGATCATACTGGGCTTGGAAAGCCTCTGCAATCGGAAAGAACATGGTCAATGCCAATACATTTTTGGAAAAGCGTTACTCTTCTGATATAGAATTGGAAGATGCTATTCACACAGCTATTTTAACCTTAAAGGAAGGTTTTGATGGACAAATCAATAGTAAGAATATTGAAATTGGTATTGTTGGAAAGGACAAACTTTTCCGTGTGTTGACACCAGCTGAAGTTAAGGATTATTTGGAAGAAGTAGAGTAAGCTTAATTTCTATAGCAAC
>evgR005572c0g1_i01_Hydra
TTTCTTTTGATCAAGATTATTATATGACTGGAGTAGAGTTGTGGAGTGAATCACCTCTAGATATAGATTTATTGTTGCAAACTAAATACATGTATATGGGTGCCAATTACTCTTCTTATGTTACAATAATCAAGATTTCTTTACATGGTGGAGAGAGTCAATACGTTGTTTTTAAAAATGTGGTTAGAGCAAAATATTTTAGAATTTTGTTAACAACTCAAATCATTATAAAGTTTAAATTTTATGGAACAGAGAATACATTTTGGACTTCTCCTATAATATCTGCTGGTTTAGGAGGATATCAAACAGCTACCCATGGAAATGGATTTCAAATTCATTTAACTGATAATGAAATTTTGGGAACAGTTGTTACAAGTTATGGCAAGTATAAAGTTTCAATGCTCAATCCTTCATCAGCTCCTTACACTCTTACTCTTGTCTGGTCAGTAACCAATGATACTTTATATTTGTATTATAATGATAAGTTGAAGGATACTGGAGGACTTATTGAAAAAAAAAATTTCTCAAACTTTAAGTATAATGAATATATTCTATTATTTGCTGATTACTATAATCTTGAATCTGACATTCGAGCAACTCTTTCGAAGATCATAATGTGGAGCATACCTTTATCAAATAGTGAATTATCTCAAGTTATTTATAAAGCACCATTTGCTGAATTTAGTTATTCAAATATTAAAGCAGCAGATTTTTCATATGGTTCAGATTGTCATTGCGTAAATGAAGAATGTCCTTGTGAAACTCTTGAAACGACTGCAACTGTGCGCTATGAGAGC
>evgR005582c0g1_i01_Hydra
CTCAAGTTCCAATTCCAAATTCTGAATTAGCCATTGATATTGCACTTGGTTCTGAACATACTCTTATCATTACAGGTAAGAAACCATGTCCAGGAGATTGTAACTATAAGGGAGTTTGTGATTCAGTGACAGGTTTCTGTACCTGTGAAGATTCATTCACTGGTTATGATTGTGATTTGTATAATTGTGTAGATCCAAAATGTAATGGTCATGGTTCATGTGATACAAGTGTTGGTATTTGTAAATGTGATGATTCATACTCTGGTGTCAGTTGTGAACGTAGAAAATGTCCAAACGATTGTAATGGTAATGGTTTGTGTTCCAGTGAAGGTGTTTGCGTTTGTTTTGCTGGTTATACTCAAAAGATTGATTGTGCTTCA
>evgR005589c0g2_i01_Hydra
AGCCAATGCCAATGATGCTGTCTCAAACATGAATGACGCAGAATATTTTGGATCAAGGATCAAGGTTGAGGTATCTGCTCCAAGACGTGGTGGAAGAGGAGGCCGTGGTGGTGGCCGTGGCTTTGGTGGAGGCCGTGGTGGTTATGACAGAGAAGGTGGTGGCCGAGGATTTAGTCGTGGTGGTCGTGGCAGTTATGGTGGTGGTAGTCGTGGCAGTTATGGTAGCCGTGATGATGACTACCGCTCGTCCCGTCCACGAGGAGGTTACAGTAGTGGTGGCGGAAGCCGCTATGGCGACTCATCAAGTTACGGTGGATCTTCTTACGGCAGTGGCGGTGGTGGCCGATCCTATGGTGGATACTAAGAACTTGACAGTATCTACAAAATACTCTTCTTCTTCTTCTTCTACAATGTGCAAAATTATCGCTGAGATGTTGTTCCTTTACACTCGACCAATTCGCAATCGGAAAGCTTCGTATACGTTCAGCGTCTTCAGCGTTCTTCGATACGGAC
>evgR005592c0g1_i01_Hydra
CATCTACAAGAGTGGTAGAATCAAAGTTGTTATTGCCAGAACAAAGAACAGTTTTATCCTTTTTTGGCTATACTAACCCAGTTGTTTATCAAATGTTAACAGACTTTGAACTGCAGACTGATGGAAACCTTCTTCATGCCGAAGTAATCAAATATGACATCGATATTTTAAAAGACTTAGTTAATCCAGACACATTTAAAAAACTTCAGGGAAATGTTCTAAATCTTAAGCAAAGTTGGAGTTGTCCGATTTGTAAAAAACCCCAGTTTCAAGACATGATTGAGTGTGAAGATTGTTTGACTTGGTATCATTGGAATTGCATTTCTCCAGACAAAGAAATAACAGATAAATGCTGGATTTGTAGTGATTGTAATGCCAACAAGTTTTATAAGATATCCAAAAAATGTTAAAACAAG
>evgR005601c0g1_i01_Hydra
AATTGTTGAAAAGAATATTCGTGTAATTGCTCAATTCTATTCAAGAATTAATTTTAACCGTTTGTGCTCATTGTTGTACTTGGGTGAAGATGAATTGGAAGGCTTTATTTCAAGAATGGTTTCCAGTAAGACAATTCTCGCAAAGATGGACAGATTAGATGGTCTTGTTGTTTTCAAACAATTACAAACTGGTGATAGCAATGAAGTTTTGAACTCTTGGTCACACGATATTGTGAATTTACTTCAATTTGTTGAAAAGTCAAACCACCTTATTAACAAAGAGTATATGATCCAAGGAGCAAGGCAAT
>evgR005619c0g1_i01_Hydra
ATTTTGTAACCAATTGTCAAAGATAGTAAAAAGTTTTTCGACTAGATTTGTGAGGTAAAAAGTTGAAAAGGAAAATTCTCTATGAAGTCTTAAGAATCTGGATAAAGATATAACTGTTGTGTTAGCTTCAAAAAACTGTTGAAAAACGCATTCGTACACCATTAAAGATAAAAGATAAACAGAGACTTTTTAGTTTAAGAAACTAAAAAGTTTGAAAAGACCATCAAGTAGAAATATTTTATCATGGCATTTAGGAAAAACACTGATTCCGATGGGAAAAAATTGTATTTTCCTGAAGGAGCTCTGCTAAATCAAATGGAAGTTGCAGTTGTTGAATCAGAACGAAGGAAAAGTGACATTTTAAAAGAACCTTTTATTGTATATCTAGTTGAGTCAAGGGCGGTAGATGGTCTGGAACCAGGATGTACATTATGGAGAAGATATAGTGAGTTTGAAACACTTCGAAATTACTTAATGGCAATCTATCCTTTTATAGTTCTCCCTCCTCTACCTGAAAAACGAATTAATGTTGCAAAGTTTCAGCTTGCGGCTGATAAGTTTGATCCCGATTTTATTGAGCGTAGGCGACAAGGATTAGAATGTTTCTTATTGCGTCTTGCATCTCATCCCGTTATATCTCAAGATGGAATATTTAGAGGATTTCTTAATCAGGAAGTAGGTTGGAAAGAAAGTGTTTCTGAGACAAATTGGCAGCAAAAGTCTGACAGTCGATTTAAAAGTATTAGTGCTTCAATGAGTATCAAGAAACCTGACCAGCGATTTGATGATATCCGCAGTTACAGTGAAAGTCTATTCTCGAATATAACATCGCTTCTAAAAATACAGCAGAAAATATCAACAAGACAGCAAGCGATTCTAAAACAACACATTGAATTTGGAAAACTTTTCAGCGAATGGAGTGCAATTGAGACTGATATTGGTGATTCATTACAACAAGCGGGTCATCAAATGGACAGACTGGGATCTCAATCAGAATCTATATACAATGAAGAACAAGTTGCATACATCGACCAGCTCAAAGAATATCTTTATTTTACAGAATCTTTAAGATCTATAGTGCGTAAACAATACTTAAAACAATTTGAATTAGAAAGGGCAGAGGAGGCCCTTATCACTAAGAACAAACAAAAAGAAGAATTAGCAAAAGAAATTGCTGCAACAGAGAATGGTACGGAAGTGCCATCTAGTGGTTTTAATTTTAAATCTGTTTCTAATATGCTGTTTGGTCAAGATAGCATGGAAGTACGCCAGGAAAAGTTGTTACTTCTAGAAGAACAAGTTAAAGAAGCTGAGGAAAATCTTAAAATGGTTACTGAAGAGGCAGAAGTTTTCATTGAGAAATCACTAAAAGATATTGATCGGTTTAAAAAACAGAAAGTTCGTGACATCAATGAGATCATGGAGAACTATGTTAAAACTCAATACAAAGTCAATAAACTGGGTTTACAACAATGGCAATCTATGAAAGAGTATTTTATGCAAATGTAATTTTGGATTTTTTCTTTAAATCACATTGTCTTCAAATCTTTGCATCCTTTTGCAACTTTTTGCTGAAACTTGAAAAGATATTTTTTTTACAACAATTATTTTTTAAATTAATGCGAAAAAAAAACATTTTTTCTATCAAAATTTTTTATTATTTTTATTAAAAAAAGGCTTTTGCACTATAAAAACTTATATTTATTAGATTTTTGTTGGTTTATAGTCTTAGTATGTATTTTTATTATCTTAGAATTTTTTTTTTTTTAATATTTCTTAAAAGAGTTAACTTTTCAATGTTTATGTGTTATTATTTTTCAATACTTTGTTATCATTTTTCTAAGTGCTTTTAGAAACATATCTCAAAATTGTTTTTTGTTGTCTTTATCTACCTTATGAAAAAGATTAACCCTATTGTCAGCCACAACTAGCCTTTTTTTTTC
>evgR005644c0g1_i01_Hydra
GTATCTATGTTTTCATAGATAAACATAGTATCTATGTTTTCATAGATAAACATAGATACCATATATATGTAGTGTCTATGTTTATCTATGAGTGTTTGAGTGCGTCTGCAGCATATGAGTGTATTTGTCATATTGAGTCATTGTCATTGAGTGTGTCTGTGGCGATCTATGGCGTAAAAATGTCATTTTCATTTCTTCTTGTGAATCATTTGTTTGTAATTGAATGCAACCTCCTTTGTATAATTTTGCATCTGTTTACGACTCCTTTGTCTACATCATCCTCTCTATGTAGTTTAAACCTGATTTAAATTATTATTTATACAAGTATTTTCAAATTACTTAGGATCGCGCTATGTGGTTAAAAATGTGTATTAGTAATGTGGCG
>evgR005647c0g1_i01_Hydra
TAAAAATGCCAGCAGCAAAGAAGACTGAAAAGAAAACAACTGAAAAGAAGGAAAAGAAAGAAGTCAAGATCAGCGAAAACCATTGGTCAAACAGAAGAACCACCGATGTTGACAAGAACAAGAGAACCGCTGTTGAAGGATCAAAGATCGACTCTGCTAACATCATTGCTACTCCAAGAAAGAGAAAGAATATTGATTATAAAGCTGTCAATATGACTGGTGATGTAAAACCTGTTAAGAGAGCCACTGCTAATACTTATGAAAAGAGAGCTTCCAAGAAGATTGAAAAGCCAAAGAAGACAAGAAAGACAACCACCAAGAAGACCAAGAAGGACGAAGCTAAGGAAGAAGAAGAAAAGAAGGAAGAAAAGACAGATTAAATCCACTCCATTCACTCCATCAATCACCAACCGAAAGGCATACTAGTCATGGAGACTGGTGTTCTTCTCACCTAAATGTAACAATTTCAAATTCTGG
>evgR005656c0g2_i01_Hydra
TTTTAGTGCAAAAAGAAACGATGACATTATCTAGAAAAAGTGCTAAAAGGTTTAGATACTTGGTTTATATCATACTATTTATTATTTCGTTCATTATTATCTTCAGATACTATTTTGGGAACAAGAAACCTTCACCGGAGCTTGTTTTAAATACAAAAAATCTTGAACCAAAGCTTATTTTAATTTACAATACTTTATGGGGAAACAAACTATGGACAGGTTTAGAAACAACCGAAAAATGGAATAATTGGGGCGGACACCCATGCAAAATTCAAAACTGTCGAGTGACTTATAACAAAAAGCTACTTGGTAAAGCTGACGTAGTTCTTTTTCACGCGTTTGGTAATGACATGCTAACACCTAGAGAGTTGCAAAAAATTCAAAAAGATCGAAATCCAAATTCCTACTGGATATACTTTCTTCACGAATGTCCTCATAATGCTAAACCGGAACCATACCTGTATGATGGACTGTTTAATTGGACTATGGGATATCGTCATGACTCAGATATTTTTGTTCCGTACAACTGGGAATGGGGCACTTGGGAGGAAAGAAGTCGAGATGAAAAACCAATACTGTTTAGAAATCATGCTGAAGACAAAGACAAGTTAATATGGAGTGGTATAAGCCATTGCGGTTGCATGAGAGAACGTTATATTCACAAGTTAAGAGAGTTAGTTAATGTAGATGTATTCGGTAATTGCGCCAATAAGTTCTATAAAGATGTATCTCCACCGGTGTGCGAACGAGGTACATCTGAATGTAAAACTCGACTAAAACGCTACAAGTTTTATTTAGCTTTTGAAAACTCATTTTGCGATGATTATGTTACTGAAAAATTTTCGGAAACAATTTTAGATGGGCATACAGTACCCATTGTGATGGGGGGCGCAAATTACAAAAAGATTGCGTTGCCTAACTCTTATATCGATGTAAACGATTTTGATACAGTAGAAGACCTTGCCAATTACATAAAATATTTAGACGAAAATGATAACGCATATAATAAACATTTTGAGTATAAAAAGTTTTGGAAACTTGGTGAACCACGTGCGTGGAGCTGCAAAATATGTGAGATGATTAACTCAATCGATTTAAAACCAAAATCTTATCAAAAGCTAGGAGAATGGTATTCAGTACAAAACACATGTGGAACTAAGCTGAAAAAACTCAGAAAAATTATGCACAAGTCGGGTGTTCCTCATCCTTATACAGATGAATATTATTTGTACGAATATGCAGTTTGAATGATTATCCAGATAATTTTTAATTAGTTACTACATAATTGCTGCGTAATTGTAACGTATTACAGACAATATTTTGTGAGATACATGTTAACAACGTTTCAGGTACATAATAGTTTAAATAAATAAATATATAGTTCATATTTAAATATATTTTACATAGATTTTGAAATGTATTTGACTATTTGAG
>evgR005664c0g1_i01_Hydra
AATATAAATGAATGTGATGAACGGATTCGCACTTGTTTCTGGAGCAAACGTGTTTGCAAATATTCAAATGGAAGCTTTTTTAGTTCGTGTAATATTGGATGGCAGATGGGTCAAGATAAAGCTAAATGTGTTGGTATTGATGAATGTATTGGAACTAATCGTGCTTGTTCTTCGGGAGATTGGGGATGTAGTAGTTCAAACGGTAGCTTAGTATGTTCATGCAATACAGGACGGAAGTTGGGTCAAGATAATGTCAGATGCGTTGGTATTGATGAATGTGTTGGAAGTAATCGCGCTTGTTCTTGGG
>evgR005671c0g1_i01_Hydra
TGTGCATATTATACAGGTGTTGGCGATTCAAACATGGCTATCATTGATATTCAGATGGTATCAGGATTTGAACCCAATAAAGAATCACTTGATAAGGCTCGTAATGATAAAGAATCAAGTATCAAAGATATTGAAGCTAAAGGGAAAAGTGTTGTATTTTATTTTGACAAGATAAGCAGTAGTGGTACTTGCATTATGTTTCAAATTGTTCAAACTACAAATGTGGAAAAAATAAAACCTGCGGCCATCAAAGTTTACGACTACTACGATAAAGGAAAATCCTCTACTACTCTTTACGAAGTTACAGAAAAACAATGCCTTTGATTTGTTCTGAGAGCAAATGAGCTTTCGAGTTTTGCTGTTCATGTCTAGTTTTACGCGTTCTGCAAATTTACTATTATTGTAATCTGTTCTACAAATTTATTATTATTGTAATCGCGTTCTGCAAATTTATTATTATTGTAATCTGTGTAGGAATTTAAACAAAAATCATTTAAAAAAAAAA
>evgR005686c0g1_i01_Hydra
CTATGCTATGTGATGCATCGTGCTTGTCATATTCTTACAGCTGACTATTATTTATCTCTACTGTTGCCATATCTGGAGCGGATCTTCCAATGATGTCTTTTTTTTTAGACAAAGTGCAAAAACGCATTGTAAACGTAGTTAGAGCTGCTCTGCAGCCAACCTCCAGCCATTGTTGCATTGTTTTAATGATGCTTCTTTTTCTCTTTTTTACAAATACTAATATGAGTGCTGCTCTAAAGAGCATGCGTCTCTTGTGTAAAAAATCACTTAAGCCCAGTATTAGGGATTATTGGTCAAAATTCAGCTTCAGTCTCTGGGAAATTCCATTCCAACCTGACTGTTACAGACATGCATATACACACAAAAAATGCACAAGTACGCTCACCTTGTAT
>evgR005694c0g2_i01_Hydra
TGAATTTAATCAATTTAAAATGGATACTGTTAAAAACATGAGACAAAAATGGAAAGAGATTTTTAAATGGAATCAAGAAATCTTGGTGAGTAGACCCTACCAATTCATAAAAGGGTCAAAAGAAAGTTCAATAGGATGGTCAAAAATTGCAGACATTTTGAACAAAAATTTATTTGGAACTTCTTTCACATCGAAGTCTGTCAGAGACCACTTTAACATTCTAGTTTTAAAACGAAAATCTCAGAAGAGCATAGAAAACAAAGAGATTGAAAAAGGAGACATGTCCCCAACGCAACAAAAACGAAGATCGGGGACAGACACGCTAAAATTTTTAACTGAAAAAAATGAAACTATGAAGGCACAAAATGCGAAAGATTTAAAACTTCGAAAAGAAGAATTACAAATTAAAAAAGAAGAAGTTGAAAACTCAAAAAATATTATGAATGGTATGATGCAAAATATGCAAATGCAACAACAACAAATGAACCAGCAGTTTCAAATGCAACAGCAACAAATGCAGCAACAAGCTGTTTTGTTTGCTGCGTTAATTGAAAAACTGAATAAATAAATTTAAATAGAAATTTCCATATAATATATTATATGAAAATAATATGTAGTATGATGCAAATGTTTAACCTTGACTTGTG
>evgR005696c0g1_i01_Hydra
ATTTTAGTGCAAAAAGCACAACAACGTTATGATACTTATACTTACAGTTGTATATTCGAGCTCTCACGGCTAATTTCCAAAACCATTATATTGCCAACCACTGTTATGCTCCGACGGCATTCAGAATCCGATCCGTCGTTTTGGGATAAGTCTTTGAACGCCTTTTATGAGGGTTTTTCGTCGGCAGATAGTATTTGGATAGGCTTGCATCAACTTGAGCGTTTGACTACAACATTTTTAATTGATTTTAAAGTAATCATGGTTGATAACCTGTTCAAAGATACATTCACAACCTATTATATGAATGTTTAAGTTAAAAGTAACAACGGATTGTTTGTAATCACAGTCGGAAGTTTTGATGCAAACAATGCAAGAACAGAAGGATTTTATGATTCTTCACAGTTCAGTTCCGGTAAAACATTCAATGATTGTAATACACTTGGGACGTGGTGGACAACTTGTTCTCCTTTATCGCTAAC
>evgR005723c0g1_i01_Hydra
CGTCAGTGGGCAACTATTTATTACAACGACGAAAATATTAAACAACTTTCCTAATTCGTTATTAGGAAACAAAGGAAATAGAGATCAGATTCAAAAAATCAACAACGGAGAATTATATTTAAACAGACATCCTAAAGTGTTTGAATGCATCTTGAACTTTTACCGATATGGTATTTTAGAGCAACCTTCTCAAGTGAATCAATTAATTTTTGTTAACGATGTCAAATATTATGGACTTTTTAATGAAGCGTTGGAAGTTGGAACTAAAGGATTAAATTACAGGGAAAAATATTTTTCGCTTAATAAAACAGATAATGAGGGGAGCAAAGTTCACCATCAGAGGAAATGGAAAACATACATATGGAGTGTTTTAGAGTATACTGATTCTTCCTTTACAGCTCGAATATTTTCATTTCTTTCTCTGTTAATAATAATTCTTTCAACTGTCGTATTCTGTTTAGAAACAATGCCGACTTATAAAGAAAATGATCATTCAAATGAAAAAGTAGTATTAAAAAATACTTTATCAAATTTGGAGAAGTTCTGTATAATATTTTTTACTTTAGAAAGTGCGACCCGTTTCTTCACCTCTCCAGATAAATGGTGCTTTGTTAAAAACATTTCCAATATTATTGACATTATCTCAGTTTTGCCGTTTTACGTGGCGTTAACTTTAAACGGTACCGAAACAGTATCGGCTTACGTCCTGCGCGCAATACGACTTACTCGGGTGTTTCGCATATTTAAGTTATCGCGTTATTCAAGAGAAATGATAGTTCTAGGCATTACAATAAAAGAGAGTTTTCTTGAACTTCGTGTGTTTTTACTTTTCATATTTGTAGGTGTGTTAATTTTTTCAAGTGCAGTTTATATTGCAGAAGATGGTGATACAGAAACACACTTTTCCAGTATTCCTGACGCTTTTTGGTGGAGTATTGTAACTTTGACAACAGTAGGGTATGGAGATGTGTATCCAAAATCGCATACTGGCCAATTTGTTGGTGCTTTATGCGCAATTACGAGTATCTTAGTGGTCGCTATCCCGGTACCTATCATTGTTAACAACTTTACCCAGAATTATCAATTTTTAAAACCAGTATCAAAGTACTGGGATGTTTTACAGCATAAAGAAAACAAAAACTGCAAAAAAACTGGTTGGAAATTATCCATATAGTGCATATATAGTCAAAATACTACCACTAGAGTTAGTTCAAATGGTTAGAAGACTATTGGTGTTGCAATATACGTAGAATATGGATACAATTGATACGTAGTTAAAGTAAACAAAGATGTGACACAAAAGCAAAGATGTGACAACGATGTAAACAAAGATGTGACACAAAAAGTCAT
>evgR005737c0g1_i02_Hydra
CTCAGGGCGATATAGATGTGGTCGATAAATATCCCAGAAATGAATTTGCCGATCATGAATTCAAGATTACTGCAAGGAAATAAAGATTGCATCTTTGATGAACGAATAATCAACAATTTAGATGAATGGTTGAGAAATAATGCTGGGTGTAGTATTAAAAACGGGGCATCAACAGGTCTTTCGGCTTGTAATAAGAGGGCAAAAATTATAAAATCCACTTCACCGATTCCTCAATTGTTAGCAGCGCAAAAAATGGTTCATGACGAGTTAAGTGCTTCAGAATACCTAAATCAACTTTTGCAAGATAAAACTACGGTATCAAAAATGCCGAAAATATTCCTCCATTCCGAAAGATTACTCGACGAAGAAATTAATCGAGTTCGAAATGATATATTTTGCCCTTCAAATTTTTTAATGTCATTACCTAAACCTCAAGGAAATATTGTAAAACTCACTGAGAAGGTTTATGCAAAAGTCAAAGAATATCCAAAGTTTAACTTTGTGGGAAGAATTATTGGACCTCGTGGATTAACTTTAAGACAAGTAGAACAAGAAACTGCTTGCAAATTACTTGTAAGAGGTAAGGGTTCTATGAAAGACAAAAAGGCTGAGGATGAGAAACGTGGTTTGCCTAACTATGAACACTTAGATGAAGATTTACATGTTTTAATTATGGTAGAAGATACAGAAGAACGTGCACATTTAAAACTTCAAAAAACAGTTGAAGAAGTTAATTTTCTTTTAACTCCTCCAAGAGATGGCGAAGATGACATTAAAAAAAAACAGCTTCAAGATTTAGCTATTCTTAATGGAACATACAGAGCTCCAGTTAATGAAGTAAAAAGTCAATTTAATGGATACAATTCAACAACTTCTTCAACAGTGCAAAATGGGCTATTAACATTTCCAAGTCCAACTCCTGTTATGAACCACAACAATACACCTTTTCCTGCTTTGCGTAGCACATCAGTAACTCCGAGTGTTAACCAACTGAAATCTCAAATAGAAAGAGTGAGGAGGATGTATCATCCTTATAACAAGTCAGCTGCTTGACAGTATTTATAATGTCGATTTCTATATGGTGATATCCCGACAAAAATTCATTAATACCACTTTACTGACGAGTTCAATTTGCAAGTCATTCTTAGTTTGAGGTTTTGAATCCAGAGTGAGTCGTTAGTGTTTTATGCTTGTGGCAATTTAGTGGTTTTTATTATCATATTTATAATCCTGTTTGTTTTGTTATGTTTTGATTTTATTTGTTTTATTGTTATGTTTATATAGTACTGATTCTTAGATTTTGTAAATACTAAAATTACATTTATATACAAATATATTTTTTACAGACAAATTATACAATTTAAATGTTTGATTTTTAACTTATTTTAAAAATTTATTGTGAATACTTCATCTCAATTGTATTAATATTCGAATTGTATTCGAATTGTATTAGTATAAAATTCGAATAAGTTTATTATTCAATAAAAAAAAAAAAA
>evgR005752c0g1_i01_Hydra
TATATATAATAATATGTATATAAATAATATGTATATATATAATAATATGTATATATATATAATATATAAGCCTGAATATGTGTATATGTTAGTACTAACATATACATATCAAGTAACTAACAAAACAGTGACAACCTACAAATATTCACTGTACTGTAGTGTAGTATTCAGTATTCAAAGTATTCATATTCACATGACAAAAACAATAACACTTAAAAACAATCTAAGAACGACACTATATAACAATATCATTACAGTAACAAGAACATCAAGGATAGCAAAATTATTAAAGAAAAAGGCAAGATTAGGTTAACTTAGATTTTAAAGGTAAAAGGGATAAGATCAAAATAACATGAAAACTATTT
>evgR005754c0g1_i01_Hydra
GAAAGACAAGAAGACCAAGAATTATCATTCTTGCATTCAATAGAACTCAATTCTTTAAGAAATTATTGAGAAATAAGAAACATCGTTTCTTTGGAAAGAGAGGATTCAGAAAGTATGCTCGTTTGTTGAGTTTGTTGAAATGGGTTAGAGCAAAGAAGGCTGCCTTGTCAAAGAAGAGATCATTCTTGGCTAGAAAGGAAAAGAAGATTGTTAAATTGTTAAGAAAATTGAAACAAAAATGGTCAATTGGAAAGAGCAAAAGAAAGTCAAACACAAAGATTACTAGAATTGTTTTGAAAAATGTTGAAAGCAAGAAGAAAGTCAAGTTGGTTATCCCAAAAAAGTTGAAGAAGAAGGTTAAGAAGATTGTTG
>evgR005764c0g2_i01_Hydra
TTTTTAAATCCATTTAAAGTTTGGACGAATTTTATGAAATGGAAGACTTTCAAACTACTACGCGCTGGTTCTTACCGACATTTCAATCTTCTAATAATATCCTATCCCCTTTGATAACATGCAGAAGCTATATTGAAAAAGAATCTTTTGCAAAAGGTGTGTCTTCAATTCAGGAATTGCAATCAATTTTGGGAGCAAATGGGTACCACGATAGATTTCCTTCATCAAAAATTAATTACTCCTTAGAATCTCTAAAATTGTTGCCGTGCTTGCATGGCGATATATCTGATGGAAAAAATAATTTTGATTTGGCAAAAACCAGAGGACGTCCTAGCACCCATACCTTGCAAAAGC
>evgR005767c0g1_i01_Hydra
TGCGTGAATTTTTGACACATGTTTCACAAGTGAGTCCGGAATTGTATCAAATCATTCAATCAAACCCACGTGAATTTTTGGAAATTATTAATGAAACTGGACCTGTTTCAACCACTGGTCAACAACTCCCACCAGTTGGAGGTGAACTTCCACCTGTTTCTGGAGAAGAAGGAGCTCAACAACAACCACCACCAGGAACAATTCTCATTTCTCAAGAAGACGAAAGAAAGATTAATGAACTAGTTGGTTTGGGATTCACAAAGAACGAAGCAATTCAAGCATATTTGGCCTGTGACAAGAATCAAGAAATGGCTGCTAATTTGCTCTTTGAAAATCGTGACCGTGGATTTGAGGATGACAGTGATGATGAACAAGGAATGCACGACGAATAAACAGTCTAAAGAGAC
>evgR005778c0g1_i01_Hydra
CCGTCATTGGTGTGAACGGTCCTCTCGTTATTTTGGACAATGTTAAGTTCCCAAAGTTTGCCGAAATCGTTAACTTAACTTTAGGTAACGGAGAAAAACGTAGAGGTAAAATCTTGGAAATTGTTGGTAATAAAGCCGTCGCTCAAATTTTTGAAGGAACATCAGGTGTTGATGCAAAGAATACTAAAGTTGAATTTACTGGTGATGTCTTGAAGATGCCAGTCTCTGAAGATATGTTGGGAAGAATCTTCAACGGATCAGGTAAACCAATTGATAACGGTCCACCAGTTTTGGCTGAAGACTTTTTGGATATTGAAGGTCAACCTATTAACCCATTTGCTCGTCTCTATCCTCAAGAAATGATCCAAACAGGTATTTCCGCTATTGATACTATGTGTTCTATTGCTAGAGGTCAAAAGATTCCAATCTTTTCTGGTTCTGGTCTTCCACACAATGAAATTGCTGCTCAAATCTGTAGAC
>evgR005786c0g1_i02_Hydra
AATAAAATAACCTCCTATGCAATAATTTATAACGTACCACACGTGCGCAGAGTGTCGCTGGGTTATAAGTTTCCTTCGGCTCATTCCGAGGTTTATTTACTATCGAAGCTGATTAGCATGCGTTATTTTAGACGCTTTACTTCTAAACAATGTCGCTGAATCAATTTGCAGAACACAAGCCTCTGTTTTCAGAACAGAAACCTCAAATAAAACACTCTGTTGCGGAAATAACTTCGCCCTTCAACGAAGATGCTTCAAGTCAAATCGACAAGGATATCAAAAACGTCTTCAAAGAAGAACAGTTAGACAACGATTTATCTTGCAATAAAAAAGTTGAGCCCGAAAAAATTGCATCTTATACTGAGATGATTGCAAAGGCGATATTTTCAGGCAAAGGAAATATGTCAACTCTTCAAGATATTTATGAATTTTTAATAGAAAATTTTCCAATTCTAAAAAGTCGAGGCAAATCTTGGAAAAACAGTGTTCGTCACACACTTTCTCTAAACGAATGGTTTGTTAAAATTCCACGCTCTGATAACGGAAAATCATGCTATTGGTCGATTCATCCTATTTATTTAAATCGATTCCGTAAAGGGGATTTTCAAAAGCAACGAAAATCCGGTATTACTCGACTTCACCAATCATCTAAAAGAACACATGTATTTCCTCCTTACTTTTACCCTGAAAGTACTCCTTCAGATTACCCACACATGGTACCAACACTTCCGTCTCCATATAGAGAACATTGTCCTTATGTAAGCAACCCTGCACTACCTAATTTATCGTTACAATGCTATCCTTCAGCGGCTATACCAAATAGATCTCACACTGATACTGACTCGCAGTCATATATAAAAACGGACAAAAATGACATGAAGTCTTATAGCTTAAAATCGCCAAGTAGTTATTTTTACTCTCCTTACCAGAAGTATCCTCAAGATCAACTTAGCCCATGTGGCGCGCCAAAATTCTACATGGAATCTTTACGCCAATCTCCTATGATACATTTCCCACAGAACTTTTCGCCTTCTTACGGAAACACATATTCTGGAGACAATAATGTAAAAAGTTCAATAGAAACGCGAAATATGTACTTTCCTACTGACTATCGTACATGTCACTCAACCGACTTTCCGTATACAACATACACACCTTATTAAATTAGATTTGTGATTAAATGTCTTGGTGTTTTAAAAAAAAAAAA
>evgR005789c0g1_i01_Hydra
TTTTATTTTCTAAATAAGAAAGTGAAATATGGTTAAAAACTATAAAAGGAAAACCGATCGTGGCAATTTTGCTGAGCAAAGCATGTTATTAGCAATCGACTCCGTAAAATTAAAGGAAATGTCACTGAGAAAAGCGGCTTTGACATTTAATGTAAATAAAGATGCTTTACATAGAAGGATTCAAGGAAAATTAAAGAATTTAAAATCTTGTAATATTCATAAAAGATCACTTGGTGCATTTAAAAACGTATTGCCAGATGATACTGAAAATCAATTAGTTAGCTATTTTAAAGAAAGGGATTCAGTTTATTATGGATTGTCCATAAGTGATGTTCAACAAATTGTTTTCCA
>evgR005791c0g2_i01_Hydra
AAAAAGAAAGAGCATGTTCAACAAATGCTTAAGTTCATATATCCTTTTGGATTTCAAATTACAGAAGATACTGATATGGATAGTTTGCTAAAATTGTCGCATGAATATCAGATTAAAAAAATTCTGAACTTGTGTGAAAATTACCTTTTGAATAAAACACCAAGTATCCAGCTGTTAATTACTGCTCAAGATTTTAATTTAGCGGTGCTTGAAAAGCATTGCATTAAATATTTATCCGAGACAGCATCATTAGTTTTACAAAACAATCTATATTACAATGAAGTTAGTGAAAGTAATAAACTTAAACTTGCTGAGGGACAATTAAAGACTTTTCAAACTTTTTGGCGAAATAATAGAGGAAGCTTTCGTGGCTCTGTACCCAATGGATTAGAAGAAAAGTTTTTTTCAAATTCCACTCTTCAACCAATAAGAATACAAAAAATCGAAGATATAGAATATATAGAGGAAGATAGTGAAGATGAAGTGGAAGATAGTAAAGATTTATTACGAAAAAATTAATTCAATCATTTACATTTGCGTTACAAAATTAATTTTGTAACGTACGAAAGTACATTATTAGAAATAGAAATCGATAAATTTTGATTTTGCACCACATAATTTTTTCAAAAATGTAATATTATTTAGACAGCCTAACAATAGTATTTCTGTTTTTAAACAACTTTTTAGCAGTT
>evgR005795c0g2_i01_Hydra
TATACGTGAAAAACGTATCAGCAATGGTAAATGTTCAGTCAACAGAATTTTATGTATTCAAAGCAGAGCAAGAAAAGCTTATTAAAGCATTAAACATGACGATCGAGGATCTAAAAAATTCAAACTTACTGTTATCAGAAAGAATAAAAATGCTCGAAAATGCCGAACCTGTCAAATCAACACCTTTAGTATCGTGGGCAAGCGTTGTGAATGGAAAAGCACCAACAAAAAAATCAGTAGAGCAGCTGCACCTAATAAATGTTGTTACAGCCGAAAAAAAGGAAAGAGAGAAACGTGAAAATAATGTAATCATATTTGGTATTAATCCATCGAATGTAATAGATAAGTCAAGTGCTA
>evgR005799c0g1_i01_Hydra
TAAAAAACAATGCAGTTGATTCTCTAGTTGTTGTACGACTTGTTCGACTGCTTCGTTTATTTAGATTTTTTAAAAATTTAAGTGGAATGCAAGTTATAGGCCAAACTTTGAAAGCAAGTTTTAAGGAACTAATGTTGTTAGCGTTAATTTTTATGATACCCATGGTTGTATTTTCCACATTGGTATATTACGCAGAAAGGGACAATTTGCCTAGCGACTTTGAAAGCATTCCAGAAGCTTTTTGGTGGGCCATTGTTTCCATGACGACGGTTGGATACGGTGATATGGTACCTCGTTCTGGACCAGGCAAAATTGTTGGCGCAATGTGTGCTTGTTGTGGTATTTTAATTGTTGCTTTGCCCGTGTCAGTAGTTGGAAGAAATTTTGCTTTGTTTTATTCATATGCTCAAGCAAGAATAAAACTGCCAAGGAAAAAAAGTCCTACTCTAATATTGGCTGATAGAAATCTGGTTCACGACTATAATCAAAAAGTGGTTGACACAAAACTGAAAGAAACAGCATCATTAATGCTCAACTTGATAGTACCTCGAATGAATGGACGTCGAAATGCATATACACCATCCATAGAACCACGACACTGTCGACCTATTACTATGAAACCGTTCACATCATCTAATAGCGTATCACAAAGTTCAATTTTTTTGAACTCTAATGAAAGCATAGAATCAATTTTTAACAAACCAAGTTCAATTACTGACAACTCATATAGCTTAACAACGGAGAGAAATCATATTATGTAGATTGGCATTTTATTATAATGAACAATTTTGTTTATAATTTATGGGAGAAATGTTTTAAGAAATTTATACAAGAGATATATATTTATAAAGTTTTATATTTAATAATATTTTTATTTAAAAAAATATCAACA
>evgR005801c0g1_i01_Hydra
TTTAAAACAATTTTGTTTTGATTTTGTTTGATATCTGTCTTCATCTTTCTGATCGATATAGCTGGTTTTAGAGTTAGCGTTTTACAGAATGCAATGACGATAACAGTTACTACATACGTTGTTGTAAGCTGCTGCCCCTTGACAATATTGCTGCATGGATATGGACTTTTCCTTTTAGCAAAGGTAAAGAGATCTGGTATAGATTCTACTCAACGCACTATTATTTTTCATCTTTGTGTATCCGAACTTTGTTTTAATATAACACTTCTCACTTCAGCGTTATTCAGTTTTACTTTCACAAAAACCCCATTATATATTTTTTGCTTGCTCAGAATATTTACGACCGCGTGTTATTGCACAAACATTTGGCTCGCCTTTGATCGTTTTTTGCATATACAATTGACCATCAAATATCCCGTGTATTGGTCAAATAGACATACTAAATTTTCCGAGGTACTTTTATGGATATTCAGTATCGGATGCGGGTTTTTAGTCTTTATTAAGTTCAACCAAGCAATAATAATTGCAGATCTTACGTTTGATACCATAGTTTTGATTTTTCCAACTTTTGTGTACACAAATGCAATTTTGCTTTCAAAAAAAACAAAAAATGCAAGCCACAAAAAAATCAATCGAAAAGGACTTTGGTTTACAGCGATAATAGTGTCGAGCAACGCAGTTTTAATTGTTATACCACATTTGATTTTGTCTACATATAGTTTCTTAAAAATGAGAAATAGTGTCCCTAAATACTTAAGATTAGTATTGGACCTTATTTTAGTGTCAGGGTATTGGAGCCACGCATTAATTTACGTATTTATGTGTCCCAATGTTCGCAATACTTTCAATCGATTTTTATACATAACAAGAACTATAATTAACAAAAAGAAAGAACTTACAGAAGCTAAAACCGAATTTTTTCAGCAGCAAGATTCTTTAAAGTTAAAACTATTTGAAAGCGAGAATCAAGATCAAGTATTTTAATGTGAACTAGCAAATTATAAATATCCAATACGTAATTTGATGAAAAAGAACAATGAAATTTTGTGAGTTATGAAAAAGGTAGTGAAGAAAATGAAAAAAAAAAC
>evgR005881c0g1_i01_Hydra
TAGTCTAAAAATACATTATCAATGAGTGAATACTTTGCATCATCTAAGTTTTGCACTTGTTCGTTATACGTATTGTTTTTTGTTTTTCAAGCTATTTGTGTTGGTAGTTGTCCGGTTAACACGTATGAGTATAATGGCTTTTGTTATTTAATCAAAAGAGTCTCTGCTAGTTTTAAAGCATTAAATTTTATGGATGCCAACTTTTTTTGTAAAACTAATGGAGGAAATTTGATAAGTCTCGTTGATTCTTCCGAATTTAATAATTTATTAAAAATTATGGCTTTTGGGTCAAGCCAGTCGTTTTATATAGGTCTCAATGATATAGATCAAGAAGGTCAATTTGTGTGGTCTGATCAAACAGTTTCATCATTTGCTAATTGGAATCCGGGTGAGCCTAACGGCGGTAGAAGTGAGAATTGTGTTATAATTTTGTCTACTACTAATCCTAAGTTTAACGATGTTTCATGTAGCACGCTTTGTGGTTTTATTTGTAAACTTAAAGATTACTGTGCTGGAAACAATTGCTCAGTAAACGCTATATGTAACGATATTGTTGGTGGATACAATTGCTCATGTAAAAATGGTTTTTCAGCTAATAATGGTGGTGTAGAGCAGCTTGGCAATCAATGCCAAGACATTAATGAATGCGCTTTACCAATGACAAATAACTGTTCAAGTAATGCTATTTGCAACAATTTTGATGGTGGGTTCAATAGCACATGTAAATGTGGCTTCAAAAACAATAACGGTGGATTAGAATTATACTCCGATAAATGTCAAGTCGAAAGTTGTAGCTTATTATTGAAGCAATCTGCAAATTACTATAATGAAGAGATAGATATTCACATCAATATGTCGACAGACTGCTTATGCGATGTTCAAAATTTAACAGTTTATCTGGAACCAGAATCTTTACTAACGTTTAAAAAATTTGTGTGGAACAATTCTAACATAAATGATTTAAATGTACGAGCAAATGGTAAATATTTGGTCATTCATGTTAAGACACTTTCTAAAACAAAATATGCATCATTTGCTTCTATTTTTGTTTTTGGTGAGAAAATTCCTCATGGTAAAATCAACATGAC
>evgR005895c0g2_i01_Hydra
TTATTAAAAAAATATAAATATAAATAAAACAAGTTTATTCAGAAAATAAATTTAATAAAAAAAAATTAATAAAAAAAATGGCCGAAGCATTAGGTAATTTACAGACTAGATATTTAAATATCGTTAACAAGGAGTTTACAAAAAGAGACGAGGCAAATGCATTACATTTACAAAATCATAATTTGTGGCAAACAAATGTTACCGCAAGAATTGCTGCAATAAATGTCGCTTTAGTTGCTGGTGGTTTTCCTGCTAATCATATTACTTTAGAGGATGCCATTTTAGCATATAGAAATGCTTTGGGTAATGCAGTTGATCCTCTTCTAGTAACTCCTTTAATTGAACCACTAAATACATATGAACCAAGCGTGTATGTAAAAGTAATTAAACAATATGGTCTCATTACTTCA
>evgR005899c0g1_i01_Hydra
AAAAAGCCCATACATCATAAATACAATGAAGGTCGTCGCAGCTTATTTGTTGGCAGTCTTAGGTGGTAACGAACAACCAGCCAAGAAGGATATTGAAAAGATTTTGAGCTCCGTTGGTGCTGAAGTCGATGGTGAACAAATTGATAGACTCTTGAAGGAATTGTCTGGTAAGAACGTTTTCGACGTTATTGCCTCAGGAAAGGAAAAGTTAGCTTCCGTCCCAACTGGTGGTGCTGCTTCATCTGCTCCTGCTGCTGGTGCTGTTGCCGCTGCTGTTGTTGAAGAAAAGAAGCCAGAACCAGAAGAAGAAGAAGAAGAAATGGGTTTCGGTGGTCTCTTCTAAATCTACTTAAAATAAAA
>evgR005902c0g1_i01_Hydra
ATATTATTACAACAGATATCCTTAATAAAAACATACAAAAAGGTATTTTGAAAACTGATATTTCCGATCACTTCCCGATATTCTTGACATTTAATTGCAGCCCCGAAACTATGATTAAGTCAACCCAAAAAATTAAAAAGCGTCTTATTAACAAGGAAAATACTAACCAGTTTAAAGAGCAGCTATCACTGCTCCATTGGAAACATATAAACTTCAACGATGATGCTAACTCTATATATGATAACTTTTTCAATACAATATATCCTATCTACGATGCAAATTTCCCATTCGTGGAAATTAATTTAAAAGACAAAAGTAATCCCTG
>evgR005907c0g1_i01_Hydra
ATTGTTATACACATCAAGACATCAGTTTAAGTGAGAATACATCAAGATTGGATCACTAGAATTTTCTGTAAATTATAATCAATATTAAACCAAAAACAAAACAATGGACGAACTAGGACTCTCCAAAAAGAAACAATCCACAGGAGGTGATAAGGATCAAATGGATGTTGATTCTGAAGGCACACAAGTTGAATTGAAGAATACAGAAGACGAATTAAGTGTCTTGCTTAAACCTTTGCAAGATATCACAGAACAAGACCTTTACAAAAAGCTTAAACACATTGAAAGTGAACTCGAATTTCACGATATCCAAGAAGAATATGTCAAATACGAACAAAAGAATCTTAAGAGAGAATTGATGAGAGCAAAGGAAGAGGTTAAACGTATCAAATCTGTTCCTCTTGTTATTGGACAATTCTTGGAAATGATCGATCAAAATAGTGGTA
>evgR005915c0g1_i01_Hydra
CACTAATACCAACTAATCTCTCCTCATTGCGGAATAGCCTCTCGTCTCTGGAAGAATATAAAGGTCAACTAGATACTGTTAAAGGTCTAATTGATAGTTACGAAGGTAATGGCAATATAATATTAGCAGGTGACTTCCATTCTTTTTCACACCAAATGTATGATTTATTTGAAAGATCAAATTCTAAAAGAAATAGTTATTCTGTGCACCTATCGGAATTTTTAAAAGCAAATCAATTAGAACTAGTAGATGTAACTAAAGGTTCTGGCCCTAATTATACATATCGACACCAGACACTACCGAATTCGTCATATATTGATCACTTCGCCG
>evgR005918c0g2_i01_Hydra
TTTAGTGCAAAAAGTTTTCTTAGTTAAATATACTGGTATTAAATTGTTGATCACAATGCCTAATAACGACCCCATGTATAGTTCATTGCAAATCAATATTCCCCCAGAATTGCCTGAAATACTTAAACAGTTTACAAAAGCTGCTATTCGTACTCAACCGAAAGACCTGTTACAGTGGTCTGCTGCTTATTTTCTTGCACTTTCTAAAGGTGAAAGACCCCCCGTTAAAGAACGACTTGATTTTCAGCTTGGTCAAGTTAGTGAAAAAACACTGACCAAAGAAGCAATTATTGTTCTACATAGGCAGTTAGGCCATAAACCTATAGTTGAACTTTCAAAGGTTGAAGAAAAATGGAATGCACTTTGTTTATCGGATGAAGATTTAAAAAATATTTTGCGAGTCGGTAGTTTTGCGGAAGAGTTTGAGTGGTTAAAATTTCTTGCATTATCATGTAGTCATGTATGTAAAACAATAACTGAGGCATTGACATTAATTTGTGAAGTCATTACAAAAGATCTGGAGGGGGGAAGAGCAAGAGTAGAGTTTGAAACATTTAAATATCTTTATAAATATCTTGCAGTTATTGACGGAGACATTCCGCTATCACATGTGACGAAAGTCTTAGAACATCTTGACTTCGATGTCAAGCAGCAAAGTGGTATGATTGCTCCTCGCAATTTTACCAGTAATTCTTGTCCTCTCCTATTCATTGAAAAGTCTTAACTTTGAGAGCATCTATTTTGGACATCTAAAGTGTATTTAATTTTGAGGGTAGCAGTCATATTGTTTTACATTTTAATCACTTTTATAATTTTAAAATGTTTTTGAAGGTTATTTATATCACTATTTATATTGAATATAAATATTTGTGAAAATTATAATATTAGTCTTGATAAAACAACTTCAATATTTGTAAGACATGATACTTTGTAATTTATGGTATAAAGCGAAAAAAAAAAAA
>evgR005930c0g1_i02_Hydra
GCGGTATTATACCACAATATATTAAAATAAAAACAAAAACAAACAAAAACTTTTCTTGATAATAAATCATGCTATACTTCAGTCTTTTTCATTGCTATACTTCGGTCTTGTTTATTGCTATACTTCAGTCTCGTTTATTGCTATTCTTCAGTCTTGTTCATTGCTATTCTTCAGTCTTGTTCATTGCTATACTTCGGTCTTGTTCATTGCTATATTTAAGTCTTGTTCATTGCTATTCTTCAGTCTTGTTCATTGCTATTCTTCAGTCTTGTTCATTGCTATACTTCAGTCTTGTTCATTGCTATACTTCGGTTTTGTTCATTGATATAAGAATAACTTGCTGCCAACATTATTTTCTGATATTTTTACATCGGCTTTTTCTCAAAACTACAATCTTCGATCAAATACCAGCCGCAATTATAACTTGAGCAGAGTAAAATCACATGTGTCAAATTTTTCAATTATAAGCCAAAATCCGTTATTATGGAATCAATTGAAAAATAACAGCATACAAAATTTAAAAAATACGCTCTCTTTTAAACGGCAAATGAAATTGGATCTCCTTAATTTCTGAGATATATAGAGCGGAGCACGTAACAGATGCTAGTTTTATCATCATGCAGGACCATATAATAAGGATTTCAAGTTATTGGGTAAAAAAAAATATACTCAATATATATTAACAAATAGTCGATACCGAAAGTACTCGTTTTTATCAATTTAACAAGTCTCTCTCCTACACACTTTAGAGCAAA
>evgR005943c0g1_i01_Hydra
AAAAATGGTAGCCTGGGTTGTAAGTAACTGCGAAACGCAAGAAAGAAAGAAACTAGTGAAAAAACTACTTTCTTATGGAATTGAAATTGAAGTTTCTGGCAAATGCGCCCAGGATTTTCCTAAAACATTTACTTTATCATGCAAAAGGTTTATCAAAAACACCCCGTGTGTGGATTTAAAAAACTATAAATTCTATTTTTCTGCTGAAAACAGTTTATGCGAAGATTATATTACAGAGAAGTATTGGGAAACTGCGTTAGATAATAACATCATACCGATAGTTTTAGGGGGGGCAAACTATTCCGATCATAGATTGGCTATCCCTGGTTCTTTTATTGACGCAATGAGTTTTAATTCGCCAAAAGATTTAGCTAATCATATTCTAGATATCAGCACTAACCAATCACTATATAACTCTTACTTCGAATGGAAAAACC
>evgR005975c0g1_i01_Hydra
GGAGAAGCTTTCGTGAAGCATTATTACAGCACTTTTGATCAAAATAGACAAAATATTGCTCCATTGTATACTGACAATAGTATGCTCACCTGGGAAAAGGACAAAGTTCAAGGAAGAGCCAATATTTTGAAGAAATTAACTGAATTGCCATTCACTTCTGTCAGACATGATATTAGCGTTTGTGATTGTCAACCAAGTGTTTCCGGAGGTATCAATATTTTGTGTACTGGTAGCGTTGTTATTGATAACGAACACCCACACCCATTCTGTGAATATTTCCACTTGACCTCTGAAAATGGTAACTTTTATTTGACTAACCACATCTTCAGATTCAACTACGGTGC
>evgR006014c0g1_i01_Hydra
GGAGAAAGTGAAACAAAACACTCCCAAATATGTTGCGTTGCATTGCGATGGTAAGCTTTTAAAAGACTTAATGGGAGACACATATGACTGGCTTGCAATTCTTCTTTCTGGGGCTCCGGAATACAAAGAAAGAAAATTATTAGGTGTTCCTTCTCTAGCTAACTCGCAAGGAGCAACTCAGGCTGATGCAACGCTTGATCTGCTAGACGTTTGGGATCCGTCGGACAGAGTTGTAGCTCTAGTGTTTGAGACAACGGCCAGCAATAGTGGTATACACAAAAGAGCTGCTAAGTTAATAGAGTCACATCTAGATAGAAAGCTGCTATACCTAGCTTATCACCACATTTTAGAACTTGTAGTTGGTGCTGTA
>evgR006021c0g2_i01_Hydra
TGAAAAAATCTGTTTTAGATAAAAAGCTTGCGTATGAAATGCACGGTATACAAAGAGTACCAAGCTTAATTTTTTTTGAACCTGAATCACTATTTAAGGATCATGGTTTACAAAACTATGAACTTCTAATGGAACCTCTCCACGACATTTCTAACCATATTAAAAACTTGTTTATAGAACTACCTTCTCATTTTTCAAAGCCAGAGTGTAAACAACTTGAAAAAGTAACTAAAGCAGCATTTGACGGAAAAGAATCTAGACGAGCAGCAGATTACCGAGCTGCAATTATAAAGACAACAATGTATTGCCAACATCACTAACCAAATAATCCTTTGACTCAAATATTTTTAATTATGTGAAAAATCCTGGAGTACATTTATAAACCTGAATCTAAACAAACAAATGAACAAATTCTTAATTTTTATTTAAAGACATTTGAACATATAATCTTATATAAAGAAAACATTGGTTTTAATCTTAAAATGTTGACTAAAAGGAAATTCTATGGAAAATATTTCTACTGCCTATTTATGCACATTTCAGACCAGCTGAGAGTTTTTTCAGGACGTGCTGCCGTGCTGAGAAAGAAGAAAGACAATTTAATTTTATTAAAAACATGTCTAATTCTACATCAAACAGATATTTTGACAACATTATTTCGAATGCTATAGTAAGAATTCAAATATATCAG
>evgR006057c0g1_i01_Hydra
TGGCGGAACGCCCTGATGCAAAACATAAACGAGCTGTTTTTCAGTATGAGTTTGATTAGGTGGCAGAAATGTCTCGTTGGTATCATGGGCGGTAGTTCAGGGGGGAATTTAATGTTGTGTAAGTGTTTGGTGATGAAGTTTTTGATTGTCCGGATGCCGTCATTATGTGGTATGTTGGTATATAGTGACGTCACGTTTGCAGTTATGAGTGTAATTGGGCTGTCTGGGGAGTCTATGTCAGATAGTATTTTCAATAGGTGTGTGGTGTCTCTGAAGTATGCAGGAAGCATTTCCGCCATGGGTTGCAGATACTTCACCACATACT
>evgR006062c0g1_i01_Hydra
TTTAGAACTGACATATCATGCAAACTTTATTTACAATCGATTGCAAAATTAGCATATGCTAAGATTGCATTTCCTTATTGTGCTTGTCATTTTCATAGTCCTGATTCCATTGTCTCTAAATCTTATACTCATTTATTCCCTTTAAGGCTGGTTATTCTAATGATTTTTTATTTCCTTCTAGACCAAAAAAATACAAGTGTAGGTAGACCTGCTCTAGCTTCCCAGCTTGAGTCTCCTATTCTCATAACTGCATCTCTTTATTTATTTTTAAATTTCTCCCATGTTTGCTTCTCAAACAAGAACCTCTTGTACAACCAAATTTTGCGAGACACATTTTTTCATCTACATGTTACCCTGAC
>evgR006079c0g1_i01_Hydra
AAATTGATGTATTTTTAGAAAGGTCAAAACTAAAAGATCCTCTGGCGGTAAAAATTGAACCAAAAGAGGGCAAAATATCACAATATAGTAAAAGCTATAAAGCTATTAAAAACTTACTTTTTAGTTGGGGATTTTTTTGGTCGAATAATAAAGCTCATCCGTTTGTTAATGGGCATGAAAGTGAAGATGTTGTAGTTTATAGAAAAACATTTGTCAATTATTTTGCAGACAACTTAAAAAACTATTGGATGTTTTTGAAAACATCAACTTCTGGATACATTTGGAACATTCCTAAAACTCAAACTGCGCGAATAGTTATGGCTCATGATGAAAGTACGGTGCGTTCGGGTGATACTCAATCATTTAAGTGGCTTCATGTTTTTTTTAGTCCAATGTTTAATAAAGGATTGTGAAAGGAAAGTGATTTTATAGTGTTCCATCCAAAATTAACGACCTTTTCTTTAAATGAATGTGAATGGAAAAAAGCTTTAAGTGAGGAACCACAACTAATTAAAAAAACTGAGTTGAATTACATTGTAACATTGCTTGGGTAAGTACTTGCTCATCAGCATTAAAGAAAATATATAAAAAACAAAAACGAACTAGTAGAGTTATATGCAAAAAGCATAAATGCGCCCACGTCAAGCCGTTACTCCGAGAAATCAAACCTTTAAACGTCTATAAGTTAAATTTGTACTAAAACTTATTATTTATGTTTAAATATCAAAGCGAC
>evgR006082c0g2_i01_Hydra
GTAACATTAAAGCAGCCTCTTACTTTCTGGAATAAAAATGAATTTGATATCTTTCCAAATGTTGATTATCACCAAATTTGTTGGCTATTTTTTATTTGAGACAAAAGCTGCTTGTTTGCACAATAATCAACCGAATCAACTGAATGAGCATCTTGTTGGAAACATCTCTACTCCATTAGTGAAGGGGAGGATTGTTACCGTGATACCAAAGCTGGATAAGGAATTTTTAGTTTCTTTTGATGTAAATCCTAATAATTTTACGTCTGGTTATTGCAGTGTTATTCATTTTTCTATTGGTTCGAATGATTGTAAGTATGGTGATAGAGTTCCTGCTGTCTGGTTCCATGATAATGGAAAACTTCACATTTCATCAGCAAT
>evgR006091c0g1_i01_Hydra
ATAAACTTGTTATGCATATTACATATAAAGATCCTACGGTAAGATCAACCTTTGAATACATTCAAAGTTTTTTAATGAATAAAATAGTCATTGTGATGCATATGGACATATCCCTTGGGAGAGGATTTCATTTAATCGAAAAACACATACTAGCGAATAAAATCATGTATGCATTAACACGTCACTCAATAACAAATGATACCCGTTGCAAATCAGCAAATTATGCAACTTGTGATGTTCATTCGCGGTATATTGGTTCCCATGATGCTTTTGTCTTCTATGTAAATAACTATTTTAGTCGTGATATG
>evgR006106c0g1_i01_Hydra
AGATTGCCATTGCTTTGGAAGAAATGGGACTTGAATATGCTGCCCATACTATTGATATTAGAAAAGGAGAACAATTTACTGCTGAATTTTTAAAGATCTCACCAAACAATAAGATTCCAGCTATTGTTGATCAAGAAAATGGGTTGGCATTGTTTGAAAGTGGTGCCATTTTGCAATATTTGGCTGATAAGAGTGGAAAGTTCATTCCTCAAAAGGGGTCAAAGGAATATTATCAATGCTTGCAATGGTTGTCTTGGCAAATTGCTGGTCTTGGACCAATGTCTGGACAAATGGGTTTCTTCCACAAGTATTGTCCATCTGATGTACCAATCGGAAAGGAAAGATACTTGAAGGAAACTCAAAGGTTATGGGGAGTTTTGGAAAAGCATTTGGCTGAAGGTGACAAGAAATATATCATTGGAGATGAATACACTATTGCTGACATGGCTGTCTTCCCATGGTTGAGAGTTTTGCCATTCTATGGTGAACACGTTGTTGAACAATTCCCAATGGAAAATTATCCACACATTAAGGCATACATTGACAGAATTAGTGAAAGACCTGCTGTTAAAAAAGGATTGACTGTAACACCATTCCTC
>evgR006129c0g2_i01_Hydra
ACTGCTGCTCCAACAACAACAACTGCTGCTCCAACAACAACTGCTGCTCCAAAAACTACTACGGCAGCTACAACAACAACTGCTGCTATAACAACTACAACTGTAGCTTCCACAACAACTGCTATCCCAACAACAACAGTTGTTTCTCTAAAAACTACGACTGCAGCTCTAACAATTACAACTGCTGCTCCAACAACAACAACTGCTGCTCCAACAACAACTGCTGCTCCAAAAACTACTACGGCAGCTACAACAACAACTGCTGCTATAACAACTACAACTGTAGCTTCCACAACAACTGCTATCCCAACAACAACAGTTGTTTCTCTAAAAACTACGACTGCAGCTCTAACAA
>evgR006152c0g1_i01_Hydra
TGAATTAAAAAATATGAAAAAAGAAATTGATACTGAAGTAATGGCTAATAGTTTTACAACAATTATTAAAATTAACATAAATATGCAAGAAAAGGCTTATAAAACAAATAAGAAAATCAACAACAACAATGCATCTATTCACAATAACTGGTTAAGGAATTCTAAATACACAGATGCAGTAAAAGAATATAATCAAGAATGGAAAACATATGTTTGGTTTAAATACCATGGTTTCAGTGAAAGCAAGTGTAAAATACAATATAGCAAATTTAAAAAAATTAGAAATAAAAAAACATCACTATTGAGAAAACTAAAAGCTGATTACTATAAAATAAAAATTGATTCAGTCAAAAATGATCCTAAAGCTAAGTGGAATTCATTGAAATATCTAACAACTATCAAACCA
>evgR006152c0g2_i01_Hydra
CTTTTGAATTAAAAAATATTAAAAAAGAAATTGATATTGAAGTAATGGCTAATAGTTATACAACAATTATTAAAAATAACATTAATATGCAAGAAAAGGCTTATAAAACAAATATGAAAATCAACAACAACAATCCATCTATTCACAATAACTGGCTAAGGAATTCTAAATACACAGATGTAGTAAAAGAATATAATCAAGAATGGAAAACATATGTTTGGTTTAAATACCATGGTTTCAGTGAAAGCAAGTGTAAAATACAATATAGCAAATTTAAAAAAATTAGAAATAAAAAAACATCACTATTGAGAAAACTAAAAGCTGATTACTATAAAATAAAAATTGATTCAGTCAAAAATGATCCTAAAGCTAAGTGGAATTCATTGAAATATCTAACAACTATCAAACCA
>evgR006153c0g1_i01_Hydra
CGCAATAGAAATTAATTTCACAATAGTTTTCCTTGTTTTGCTCCAAACAAACCTGACAAGATCGTTTTTAGAAAAATGCACATTGATTAACTTAGAATATATCGAAGATTGGTTCAGATGGGGCCAGTTATATTCTTATTGCAGTTATTTAGAGGTTGAAAAAGGTCATCTGAGTTTATCGCAGTTTGATAACGGTTGCAACACTATTTCTTGCAAGTGCATGAAAGTTTGCAATCTCTATAGTATCTTCAATCTAACATGTCATTTGTCTCTGTCATTTTGCAACAATTCAACTTTATGTACACATAGTTGTTCGTACTACCGTTATATTACAAATAGTAGTAATACTGGAGAAAATTTTCAAATTGATGAACAGAACAAACCTGACATAATTTCTTCACTTAAAATAGTTCAAAGTTTTAAAAAATCTTTTAAAGTTGTTCTTGTTAATTCAACTGTTTTTATTTGGC
>evgR006159c0g1_i01_Hydra
TATGCCAAGTCAAAAATATGGATAAAGATCCTACAAAATTGACATCTTTTCGACCTCTGAATCCTGTTTCTTCATCATCATCGTTTTCTTTTTCTCGATTTATATCATTTGGACGTAAAAAAACAAGCCAGCCACCAAGATTGCCAGTTACACCAGAACTGTTCAGGAAAGATTCATTAATTAGCAATGAAGATTCAAGTGCATCAGACGGAATTAGTAATTCAGAGCCTATTTATGAAAAGAAAGAAAGTCTATATACCTGGCTTACAAGAAGTTCTAACAACGAGAATACTATTTCCTTAAATAATTCGACAAATAATTCTGTTAAAGATAAATGGAAATATCGAAATGTACGGACTATTATGCGAAGACTGAGTGCATTAACCATTGATAGAAGGTGGCATCAGAGTAATGGTGACTTCAAGCATAACTTTAAACAGTATTGGATGCCAGATGAACATTGCAAAGAGTGTTATGAGTGCTCTGATAAGTTTAATACTTTTAGAAGGAGACACCATTGTAGAGTATGTGGTCAAATTTTCTGTAACAGATGTTGCAATCAAGAAGTTCCTGGAAAAGTTATGGGGTTTTCAGGTGATATTCGAGTCTGCAATTACTGCTGCAATATTGTTCAAGCCTACTTAAATGACGATGGTTTTGATAAAAGCATAGCAGCTCTTAATGAAGATTGGAAGATGGGAAGTGACCATAATTGTTTTTTGTATGATTATGGAGGAAGTGCAGGCTCTGTAAATTCTGCCAAAACAAATCAAATTAAAGAAGATCCATTTCGAATGAAGCGTGCGACAAGTACCAACAGCTTGGGAAGTACCATTGACCGTGTTGCAACACCTACTCAACGATCACCTTTCGATGCTTTTGGAGTTGCCCCAAGTGAAGCAAATAATATTAAGCAGGATTATCTCTTGCGCGAATTATGGCATCAAATTAGTGATCTTCGAACTGGTTTAGAATTACAAAGTCATCGTTTTAGAATGAAGACGCATCATAATTGCTTCATAGGAAGTGAGTTAATTGATTGGTTGCAAACTCACAACAAGAGTCATTCAAGATTACAAGCTGTTAAAGTCGTGCAAGCTTTGTTAGATGGTGGTTGGATTGCTTGTCTATATGCTAATGAAGAAAATGTTGTTAAAGATGAGTATATGCTATACCAACCTACTAAGATGGCTATTGCTGAAATTGAAGTGGCAACAATTAATTCTCAACTTCTTAAAGATGATAATGATCTAACCTCTTTAGAGAAAAACTTGACTGATGATGATGCAGAAGCACCCGATTGGTTGAAACAAATCACTCATTCTGCTATTGAGGATGAAAATGAATCAGCTCATCCTGTTCCTGGAGCAAAGACTATTTCAGCTCGTTTACTAGATTTGGATCAAGTTCCAAATGATACTGACCAGAATAAAACAGATTTATCGATTCAAATGAAACTAAATGCTCAAGTTGTTGATGATGACAGCAAAGAGCCATTTGATCACGTTTCACAAATTTCCTTGATAACCAACCAGCCACCTGTTCCTTTAGATCATTGGACTTTAGTTGAAGTAAATGAATTATCAAAAAAAAAAAAAATTGCTAAACAAAGTAGTTCAGTAGCTTTAAATGAAG
>evgR006173c0g1_i01_Hydra
CTGCAAGATATTATTAATATACTATATTTTTTTATATATATATTTACCAAACAGAAATAGTTGGAACCCTAAAAAATCAAGATTTTAAACTGTTTTTGCCAGCTGTCAATGTCAATGCATCAAACTAAAAAAACTCTGCCAGTTCCTTTACAGAAAAATTTCAATTTTTTCTAATTTGATTGCGTTATATATTTATAAAAATTATAAAACTTTTTAAAAACATTGTTAAATTAGAAAATTAACCAAGGATGAATGAATCGTTTTACGTCATCAATTGTTCTAAAACCCATGTTTTAGTGCTTAACGTACAAGCTAACGGTAACCTATATGCAATTATTTGTTTACTGCTGTTTTTAACCGTTCCAACTATTACCATGAACAGTTATATTATTGGAATAACCTATTACTGCCGAAATATGCGTACAGCTGCCAACATTTTGTTAGCTATTACATCGTTTTGGGATTTATTAGTTGGAATATTTTCAATTATTCCCCTGGCAATAGTTTTGATAATGTTAATTTCGCGCACTCACAATTGCAATTTGTTTTTGTTTGCAACTTTTACGAGTTACTTGTTTGGATCAATATCATTCTTAACAGTTTTCTTGAGCTCGATCGATCGTTACCTTGCAATTTTTCAACCTTTTCTATACCAAAAATACTTACACGAAAAGCCAAGCTTTTACATAAAATGTGTTTGCGTTATTGCATTATTTGTTTTGTTTATTACAAGTTTGAGTTATTTGACATCTCAAAGAAATTTGAATTTACTGTTTGTCATGTATATAACTCCAATACTGATACCGACAGTTTTATATATTCATGCTCGATTGCACATCAGAGTTGAATTGGTTCGAAATGCGATATCGTCTTTGAAAACTAATCCAAATTGGTCCGATGCAAATATAGGACCTGAAAAAAAAGTCAGCAAAGATCTAAAAATGAGTAACTTTACAATGATGATTTACGCTTCCACATGTATTGCTTATTTACCATTTGCTACAATTTTATATTTGTGGCATTTTATAAGCATTTCGAATACCTGGAAATGGACTCACAATTTTTATTTGTGGGCGTATACATTGATATTAACAAAATCATTAGCTAATCCAACTATTTATTTCTACCGATCTAAAATATTGCGCAGAAGTCTAAACCGTGTTAGGCAATCACACTCCAAATCAAATCATGATAATGATTAACAATGGTAATTGCGCATACACCTGTATTTGTACATTTGTTAAAAATGATAATTTTCACGCCTATCATAAATTAAATATTTCTTCAAGATTTCTTTAACAGTCATGATTCTCTACTACCCGACATTTTGAAAAACAAAATATCTATTTGTTGAGTTTAAATAAAAACGAGTAAGTAAGTAGAGTCGAAAACCAAACTGTACTTAAATAGCGGAGAATATAAAAAAA
>evgR006180c0g2_i01_Hydra
TGGGAAAATATAACGAAGCCTTAGAAATTTTTTATTCTGTTGATAAAATACTTACTGAAATTTTAGGTATTAACCATCCCTCAACAATGGCATCGAAACATAATATTGCAAATACTTTGCGCGAAATGGGAAAATATAAAGAAGCTTTAGAAATATTTTATTCTCTTGATAAAATGCAAGCTGAAGTATCAGGTATCAACCATCCGTCAACTATGACAACAAAACATAACATCGCACAATGTTTGCACGAAATGGGAAAATATAACGAAGCCTTAGAAATTTTTTATTCTGTTGATAAAATACTTACTGAAATTTTAGGTATTAACCATCCCTCAACAATGGCATCGAAACATAATATTGCAAATACTTTGCGCGAAATGGGAAAATATAAAGAAGCTTTAGAAATATTTTATTCTCTTGATAAAATGCAAGCTGAAGTATCAGGTATCAACCATCCGTCAACTATGACAACAAAACATAACATCGCACAATGTTTGCACGAAATGGGAAAATATAACGAAGCCTTAGAAATTT
>evgR006191c0g1_i01_Hydra
GCATATGTTATAAAAAGAAGGAGTATAAGAATTTAATAAATATTTATAACAATAATTTATTTAATATTATATTAAACAACTTTTTATCTCTCAGATCCTACAAGCTGTCTGGAGGTATCAGCATTTATTTGAAATGATTCTTAAAGAGCTGTTTGATGTGTTGAAAAATGAGGAAAATACAAAAGAACCAATCTATAGCAACTTGTTGCAAGTTATCAGCAGGGTCAAGTCATTGATCTTTAGAATACAAAATATAATACAGCCAATCTCTGAATGTATGTGCCAAGTTGCTGACAATGTTCCAGAGGTCTACATACAAAGTTATAATAATCTGAGAAAAAAGTACAGTCATAAAAAAAGCATGTTTATTATTCTTAAACAATTGAACATATTAAGAAAAACAATTAAAGAAGATATTTGGAAAAGACAGAAAAAATACTCATGTTTGCATCAATAAAAATTGTGATGTCGTATTTATAAATTGCTAGAATTTTTATAACATAAATATCTAATCTTATAACATAATTGAAAAGTAAAT
>evgR006224c0g1_i02_Hydra
AGTATCTATAAAACTTACCCTAATGCTGAAAAGGCTTTAATCGGAAAGAATCCCAAATTGAGTTTGTTATGGAATTGGCTTTGGAATTTCTCAAAATTAATTGACTTCACTACGTTAAATAGTTCTGTCTTGTTGTTTTCTAGTTCCTTGCATTGCGGTGGCGTGTTTTGTTGTTTTAAAACCAAAATTATCTTGTGTGCTTGTTTCATTGTTGTTTATTTGTGCTTAAGTGGAAATGTTATCCAGCAAATTATTTTTATAAATTCTATTTTTGAAATTAATGAAACTATTAGTTGTATGTATATACTTTCATTCTTTATGTTTTAACGATTCCAGGAAGAACACATTGGTAGTAGAATCTCACCTTACAATTTTTGCAAATAAATTCCATGCACCAGTTGTCGCACCTTTTAAGAAGGTTTCATAATATTTGATCTGCTTGTTTATCAACATAATTTTTATGGATATTATTTTTCTTTTGTCAGCATACATTTTAATTGAGTTCAGCGCAACTTCTGGCATAATATTTGTAAATAGCAGAAACAAGATGGGACCGAAAAGCAAACCCCTATGTACTACCACTGGTTACAGATAACCCATTCTGAAATTTTGAATCACTAATAGTAACTTTGTTTTTTACCTTAAGGAAACTTTTAATTGAATTATAAATTACCAGTTAAAAAAAAAA
>evgR006237c0g1_i01_Hydra
AATGTTAAAATGTATTTTTCCGATCCGTGGTATTCAGCACAACCCGGAAACATAACATATTTTCTAATGACAAAAGGATGTTCAGAAAGCAATTATAACTGCAAACCTCTTCTAAAATTAACTTACGATGGCATCTTTTTTTTAAAACTTGGATGTTGGGCAGACAGTTTAAATAGATCTTTAGTTCTTCTTGAAGACAAATTATCAATGAATACTGGATCATATAAGAAACGTCCTGATCCATTTGGTTATTGTTACACAGTTTCATCAAATCATGGCTACAGCATATTTGCTTTACAGTACGGTGGACAATGTTTAGCAGGAGCTAATTTATCTTATCAGAATTTTGGAAAATCGAATAGCTGCAATGCAGATGGGAGAGGGGGTTTTTTGGCAAATGAAGTGTATTCAGTAGAAGACGATAATTTGAATGGTGAAATAATGCATCATAATGCGGAAACAAAAGTGTAATTATATTTTGAATTTTTTATTAACTTTTAAATAAAGAAAATATGCCTAGAG
>evgR006249c0g1_i01_Hydra
TTTACTGTGGTATATTTACTGTAGAGTTCCTTATTCGATTAATTTGTTGTCCTTCGTATAAATCGTTTTTTCGATCTGCTCTAACGTATATTGATTTTCTATCGACTATTCAATTTTATATATCGTTTACTATGAAAACTAAAGATTTTGACTTTCTCTTTGTAACTCGCCTTATACGAATATTCCGCTTGTTTCGTTTTTTTAAACAACTTAGTGGAATGCAAGTTATTGCACAGACATTGATAGCAAGCATGAACGAGTTGATGTTGCTGTTGATGTTGGTAACAATTCCCATGATTATCTTTTCCACGCTAATTCATTACGCTGAAAAAACTGATAACAGTAATAACAAGAACTATGCAAGTATTCCAGAATACTTCTGGTGGTCGATTGTTACTATGACAACGGTTGGATACGGAGACGTGATCCCTAATTCGTTTTCTGCAAAAATTGTAGGAGCTTTGTGCGCATGCTCAGGTT
>evgR006256c0g1_i01_Hydra
AAACTAGTTATAAAAAAAACTCTCTAAAGTTTGTTAGTATACTTGCTAACTTTAAATTAATAGCAATAAATGGATGCAAATAAGGACCACAGATGCAGATTATCTACAGTTAAAAAGTGGGAAAAACTTTTGAATTGCGAGTTAGAATACGATTTAAATGGAAATGAAGTTATAAGATTAAGATGCAGTTTATGTAAAAAATTTGAAAAACGGATAAGTCAAACAAAATTGTTTTCTATGACCTGGATCAAACCAGGAACAATATCAATAAAAAAAGACTCACTGGCAGCGCACTTAGGTAGTGCACAACATAAAGAAGCCACGCGAATTCATCAACAAACAACATTAGGTTCAGTTTCATTATTTAACCATTAATTTATTCATTATTTATTCATTATTTAATGTTATTCATAATACTCCTAT
>evgR006260c0g2_i01_Hydra
AATTTTGTTATAACCTTATCAACACTTTCTAGAATAAATACTAATACTGTGTTTATAAACCAATTTGTGTTGACAATCGCATATTATTTGTCATTTTTGCCATCATCTATACTCATAATCAGCTATGATACAAACTTATCGCTGACAAAAATTCAATGGACACTTTTTCAGTTCACAAGCGCTAGGTGTGATAGATCTTTGGCTTCTTCGATTTCCATGAAACTTCTTGATGCAAACGGAAAAATCAATCCTGATTTGGTTAGTGTTTTAGCAAACATTAACTTTTACTTAAACTCAGTTGATGTTTACTCTTCAGACAACTGCGCTAATGATCGCAACGGACCTGTTCCAGATCAGCCATCACAGGAGTATACACTTAATATAAATGTTTACCGCTTTTCGTTTAATCTACC
>evgR006290c0g1_i01_Hydra
GGTTTGGCAGCGTTTTTTGATTTGAATCTTCTCAAGTGTATCGGTTTAACATTTTCTTTTCCTATTGATTTCAGTAAGTCCTCAATTGCTTTCTTGTCTTCTTCAATTTTATCTTTGCTCTCAGTTTTGCTCGATTCATTAATTCCATATATAATGACGTTATTTGAACGTTTCATTCGCTCCTCCTGCTCATTAATTACTGCGTTAACCACAACAAGTTGTTCTTTTGTTTTTTTTGCGTTACGCGACACCACATTAACCGAATCATTTCCAATGTTCACTGTTGAAGTTACAGCATTTGGTTGTTTTGAGCTCCACCGTTTTTACTCTTTTCTCCAATTCTACAATATGATTATCTTTCACAACAATCGCTGTTTCAAGAACAGATATTCTCTTGATTATTTCTTGATTGATTCTATTTCCGTCTTGGTTAAAATTTCCG
>evgR006291c0g1_i01_Hydra
GGATTGGGTGGTGTATTGTTAGTTGGACTTAGTGCCGCTTCATTCAAGAACGGATATGATTCTGTCAATGAAGGTGGAAAATTCACTACTCAAAAACTCACTGAAACTTTGACTGCTGTGGATACTGATATTAAAGCACAAGTTAAAAAACTCAACGAACAAAACCACGATTTCTCCAACAGAGAAGAAGTTATTGAAGTCATCTGTTCAAAGCAACTCCCTGTCCTCCAAAAGTCTTGCAAATCATTGTTCGATAGATATTATCATAGCTACAAGGCCGGAAAGGGTGCCGATATTGAAGAATATTTAAGAGTTGTCCTCAAGAAGTAAATCACCAATATTTAATTAATATAATTAATTTAAAAATTGTTTTTTCATGATGAACACATAATGGGAAATAAAAAGTAATCATGGATCTTTGTACCACCATCATTAATCAGGAAATAATAAAAAAAAC
>evgR006303c0g1_i01_Hydra
GAAAAGGAGTTCATGTATCCAACAAAATCAAACATTTTGGACAGCTGAAGGAGAGCCTGTTAAACCTCCTTATATAAATTGTAATTTTGAAAAAAGAAACATGATAATGAGTTGCAATATAAATCAAATAAATGCAGATTCTTGGATAGGAATTCCACACAGCTACAACTTGATATACAGAACTAAATTACCTGATGATAATTGCAACAATATAGAAAGCGAGCTCAACCATTATCAATCCATGAATTATAGTGTTTCTGCTACTGTATATAATTTTAACTTTAACACTTCCGATTTAAAAAGTTACCAATTATGTGTTATTGCTCTTGCTACTACTCAGCG
>evgR006308c0g1_i01_Hydra
TGTAAAAAATGTTACTCGAATAAAAAAAGTTATGAAGCACACAATTCTTAATCTAATATCACGTAAAAAAACAAACTCTTTTAGAGTGGTTTTGCTTGGCTCACCGGGTGTTGGAAAAACGGCATTAATTGTTCGATGTCTATCCAACCAGTTTTTTAATAACTATGTCAGAGGAAAGGAAGAAGCATACAGATATAAAGTTGCGGTCGGTGCAAAAAAGATTCAGCTGGAAATTATGGATTCTAATGGGAAGAACGACGCCCTTATTTCTTACGCAGACTCAGTTATTCTCGTTTATTCAGTAACTAACAAAGAAAGTTTTGAAAATGTTAAAATATTGATTCAAAAAGTTCGCGCGTTAAACTTAAATGATATACCGATAGCAGTTATTGCAAATAAAATTGATCAGTGTCAGCAACGTGAAGTGAGTGTTAAAGATCGTTTATCTTTATCTGCTGAATACGGTGTTGAGGTTTTTGAAAGATCAGCTGCTATACCCACAGAGGATTTGAAAAGTATTCTAGTTGAAATGTGTAAACAATCTCGATTTGAAAAGCGTAAAATAAGTAGGTCTTTTATGGCGGAAGATACAAAATCAAACGGATATCAGAAATATGCAAGTATTCGATCAAAGTCATATAGTGCGTTAGACTCAATCAAAGTTCTATCCAATATTTAATTTCATATTTAATATAGTATTTAGAGTATTTACAGTTTTTTTGCGTAATAATATTTTTTTTATCACATTCAGAATTTTTGGAAATATATTTATAAAGTTATTAATTCTATATTTATCGGCAATATGAC
>evgR006310c0g1_i01_Hydra
GAAACTGTCATGTTTTATAACGTCGAAGAAACAACCAACGAATCAGGTGACAAGAAAATTTCCTTCCAACCAAGAAGTGACATTGTAATTCAAAGAAAAGTTCACGCATTTGCCCTTTCTCCAAAGAGCACTTTTGCTATTTATACTTATAACACCAACAAGTCAACTGAACAAAACAGTGTAAGGTTGTACAAATTACCAAATGTTAAGCGTCCAACTGCTACAAAGCATTTTTCACATGATGTAGATTCTGTTGAAATGCGTTGGAATAAGGTTGGTACTGCTTTACTTGTTTATACCACCACAAATACTGACCAAAGTGACAAGAGTTACTATGGTGAAAATAAGCTTTACTATGTCA
>evgR006313c0g1_i01_Hydra
AATCTATTGAAATCAATGTATTTGATCAACATTAAACCTGGATCTTTTCCAGTATCAAGTGCACTCACTCTTGCTTCTGATTTTTTTTCTACATCTCAAGGAGGAAGGGATCATTCAAAAAAATCTCTAATTTTATTTTTAACAGATGACAATGATTTAAATGCAGCCAAGGAGTCTAAATTGCTAAATAGACTAGGAATCTCCGTTTTAGTCATTGGATTTGGAAACTTTACTCAAGATTATCTTACTGAAGTTGCTGGCAATAAAGATAGCGTTCTCCAAGTAAATGATATAACAAAATCTCCAAATGAAAATTTGAATTTTGGCAAATTAGTGTCGGATGCACTCATTAAAGAGTCATGTAATGCATCTTTATGTGGAGATAACAAAGTTTGTATGAACAACCTTGATGGATCTTACACATGTGTGTGTCCTGTATGTTCAGACATTCAGTCATCTTCTTTTGTTTGTGGATCAGATACGATAACTGAAGCAAATGAATGTTTGGTGAAAAAAAAATCATGTTTAA
>evgR006330c0g1_i01_Hydra
CTTTATTGGTAATATTTATTATAAAACATTTATACAGGCCCGTACTGTGGAGAAGGATGCGTCGCACTTACGTCGGAATATCAGATACATTGGACCCCCTTTTTTGCAGCAACTCTCCTCTGTTTTCTACCAATTGCACTTTGTTTTGTATAAATTTTATAACAAAAATCGAGCAATATATATTTGCCACTATAAACTGTCTCTTCTTTCAAAAACGAACCCTCCCTCCAAAAAAGTCTATTTTAACAGTCTGTTGCGTTCAAGACTCTCATAAAATTCAGTGGTTTTACTGCGTATATACTATTTTATAGAAGCCCTTTTTTAAAGTTATATAATCAATTTATATTCAATAACAAATTATAAATATCATCGATTAATTATCGGAAAGATATGACCTGGTCAAAGTTCAAACCCCG
>evgR006339c0g1_i01_Hydra
AACCTATTGCTGTCTCTGCTATCAAGACTGTTCCATTGGAGGAACAAGCTAACTTCCCAGTTAGAATTGGTGAAGATATTGGATTGAAACGTGTAGTTTGTAATCAATTGGATCAATATTACTTGTCTTGTAAAAAGGTTTCCAATATTTGTCAAAGAAACAAGGCAACCAACTTGGCTTCTTGGGAAAAATTAACACTCTGCCATGAAGATGGTAAATCAGTGACCAAGGATGAACGTAATCGTCTCTTGACTCAACTCTATACACAAACAAACTATCAAAGAGATAGAACTGTCATTGTTAGTTTTAGAAACCAATTCAAGAAGGCTGTTCCAAAGGAAAATATTCCAACAGAAGAGGAATTGAAGACTGTCATTGCAACTCAAATTGGATTCCCAGTTGAAGAAGTGACCTATTTGGATTCGATGAATGACTGGTCCAATGAACAAGATGAAGACGAAATTGATGAAGAACCAATTTCTACGCTAGGTGAAGCACAAAAGGAGTTTGAAACTCCTAAAGTTCTTCAAACAGAGGCTAAGGATAAGAAGATTTTCACAAAGTATGCCAAGGTTGTCACCTCTACTAAATTTGATAG
>evgR006346c0g1_i01_Hydra
CAAATATCAACACAATGTCAAGCGATTCTGAATTTTATACCTTCACTGTCAAGGATGCTCAACTGAACGAAGTTTCTATGGCTGACTACAAGGGAAAGGTTGTCATGATTGTAAATGTTGCATCCAAATGTGGTTTTACTAAACAATATGATGAAATTCAAGAACTCTACAACAAGTACAAGGATCAAGGTCTTGAAATATTGGCTTTCCCTTGCAATCAATTTGGATCTCAAGAGCCTGGAACCAATGAAGAAATCTGTTCATTTGCCAGAAGTAAATTCAAAACAACTTTCAAAATCTTCGACAAGATTGATGTTAATGGTTCTGGTACTATTCCATTGTACAATTATTTAAAGAAGGAGGGTGGTGGCTTTTTGATTGATGCCATCAAGTGGAATTTTACCAAATTCCTAGTCTCTAAAACTGGAAAGGTTTTGAAGAGATATTCACCAACTACCAATCCAATGG
>evgR006371c0g2_i01_Hydra
TGGCAATTTTGCCAGAAGTTGCCAAGCCTTCTAAGACTGTTAAATTCCAAGAAAAGGTCTTGTGGACTGCTGTTGCTCTCTTTATTTATTTAGTTTGTTGTCAAATCCCAGTATACGGAATTCGTAATGCTTCTTCAAGCGATCCATTGTATTGGATGAGAGTTATTTTGGCCTCTAACAGAGGTACTTTGATGGAACTCGGTATTTCTCCAGTCATCACTGCTGGTTTGATTATTCAATTGCTCAGTGGTGCTAGATTAATTGATGTCGATGAAAATAGTCAAGAAGATCAAATTTTGCTTAACGGTGCTCAAAAGTTCTTCGGTCTTATCATGACCATTGTAGAAGCTGTTGCTTATGTATCCTCAGGAATGTACGGTGATGTCAGAGATCTTGGTCTTGTTACTGCTTTGTTAATTGTCTTGCAATTGTTCGTTGCTGGTATTGTTTGTTTGCTCCTCGATGAATTGTTGCAAAAGGGTTATGGTTTGGGCAGTGGTATTTCCTTGTTCATTGCTACTAACATTTGTGAAACCATTATCTGGAAGAGTTTCTCACCATCCACCATTAACACTGGTAGAGGTACTGAATTCGAAGGTGCCATTATTGCTCTTTTCCACATGTTGATTACTCGTAGCGATAAGGTTAGAGCTTTGAAGGAAGCTTTCTATCGCCCAAATCTTCCAAACATTACCAACTTGTTGGCTACTATTTTGGTCTTCTTGATTGTCATTTACTTCCAAGGATTCCGTGTTGAGTTGCCATTGAGATCATCAAGACAATCCGGTAACGCTGGTGCTTATCCAATTCGTTTGTTCTACA
>evgR006373c0g1_i01_Hydra
TTTTTTTTTTTTTTTTTAGATCATTAAACGATCTTTAACATGAGAAATTAAAACATGATAAACATGACAATCACAATTAATGTAACACTCAATATATCTTTCAATTCACCAATAACAACAAACGCTCCACAAACGGAAGAAATACCTCAAGCCTTGCGTATTGTGACACTTATAGTTTGTTTTTTTGTAATTACTCTTGGTGTTATTGGAAACAGTTTAGTATTGATTGTGTTTGGGCAAAAATGGTCAAATTTAAATTCATGTGAAATCTTCTTGGTCAATTTAGCTATTGCTGACTTTTTAAATGCAACTGTTACTCCTATAAAAACTGTTTTGGATTTAGTGCACATCAATTTGTTTAGCATAGGAAACGATGGATGCAAAGTTATTTCATTTTTGAGTATATCAACTATAACTGTTTCCGCTTTTACTTTATTGACAGTATCTATAGATCGATTTATTATTGTGAAATGGCCTTTACACAAAAGACCATCCCTAAGTACTTTGTTGGCATTCGTAACTGTAATGTGGTTGGTAGCAGGTGTAGTTGGATGTTTTTATTTTGTTGGTGAAAATATTAAATTAGGGCTTCACTCGCACAACGTCTATTCGTGTTTTAATTATATGAATCAGAAAGATTTTGTTGCTTTCACCCTTGCTGTTTTCTGCATACAATGCGCTATACCAATTATAATAATGACTGTTATCTATATACTGGTTATACAAGAGCTGAAGAAAAATGCACAAAGTGGAATTTTTGCTCATTGTAAAAACGAAATGAAGTTGAGACTGTCACAAAACAAAAAAGTAACAAAAATGATTTTTTTGGTTGTATTAGTTTTTTATTCGTGTGTTTTACCATCGAACAGTTTTTATATGTGGTACTTGTTTAGTGACTCCAATTATTCAATTCGAATTATTAAAATTGTTTACGACATACTAATAATGCTTCAGATGTGTAACAGCGTTGTAAATCCAATAATATACAGCCGGTTACATACTTCTTTTAAAAAAGATATGCGTAAAGTGGTTTTTTCGTGTTGCTTTAACAAATCGGAGAAGTGGGAGACATTTCAAGATACAGTGCGATCGATGCTTTATCACGGTTCATTATACAGAAGACGTTCAAGCGCGTATTCAGTAAAAACACGGGCATCTATAGTTTCTTTACACCGTCATATTTCACTTAGTAAAAGAAGAAGCACATACGACGATGACTCGCGCAGACCATCTGTTGAAACATCATTTATCAGTGACAGTTGCAGCAAACTAGATCATAACAAAGAAGAAAATAGGATACCATATTACAAATCTCCCTTATTATCACCAAATGTGTAACGAAACAACTTATTTTGTTATGACAAAATATGCAACATAGCAATTTATTTTATTATGACCAAATGTGCAACGTAACAACTTATTTTATTATGAGGCAATGCAAAATAAATTTTAATATAAATAAAAGAAAAATTATTTAGTAACAAATGTAGATTTTTAGGATAAATAATGTTTTTTAACGTTAATGATCTGGC
>evgR006432c0g1_i01_Hydra
CAGAAAAAAGGTAATATAAAAACTTACGGTGAACGAATCGTCTTAAACCGCGGGCTCTTTGGGTTTACTTCGCTAAAGCTGAAAAATGACAATCAAGAGGAGAAAGATTCTTCTAACAAAACAAAATCGAGTTCAAATAGCCCACGCTTAAGTTTTTCGGATGATATTGCTTACTCTGGGTCCAATGAAAATCAAAACACAAAACCTTACTCTCCCACAGAAGAGTTTCAATTAATTGAATATTCATACCCTAAAATACATTTTTTTCAGGAATCTGTTTTTCAAGTCTCAGATGAAGTACGACAGATTTCGACAGAGCACAACAACGAAATTGGGCCAACAGCGGACCAGTTTTGAAGTTTGAATTTTAG
>evgR006461c0g1_i01_Hydra
AAACATCTTAATAACTTAAGCTGTTTTTATACAAATGCGACCTCATTAAACACAGAGAAACTTGCTGAGCTACAAAGCCTTATTCATACTTCAAAAGCACCACAGAGTGTCGATTTTATAACAGAAACATGGTTTAATGAAAGTTCTATCTTTAATCTCACTAATTATATCATATATAGACATGATCGCAAAAACAATCCTCATGGTAGAGTCTGCTTCTATATATCTGAAACACTAATAAGCCATGAAGTAACACATTATGAACTAAACAATGACTCTGAACAAGTTTGGTGTTCTGTAATCTTGAAATTGAAATAATTTTAGTTGGTTGCATATATCGT
>evgR006471c0g1_i01_Hydra
TTATATAAACTTAACAGAGTTTAAAAGTGAAGTTGAAAGTTTTATTTTATCGTTAAAAATGGATTGTGCTTACTTTGAAAAACAATACACTCCCAGCTATTGGACCTCCAAACATTCTAGCAGAGACGAATGCTTGCAGCATTATTTAAAGCTCTCTTTGCACGAGTCTGAAGTTGTCAGAAGCAAGTTTCGATGCGAATTAAACGTACCATACGGAGTTTCAGATAGAGAAAAAGTTGATTACTTCTTACCTGAAAAAGATGACGATAATAAAATACTTTTTGTATATATTCATGGTGGATTTTGGTATGAAGGCAACAAAGAAATATATTCAATGATAGCTGGTCCTTTTTTGAAAAACGGACACCCGGTTGCCGTTCTTAGTTACGACTTATGCCCTTCAGTTAGTATATTTGAAATATCAAAACAAATTTCAAAGGCATTTGATGTTATAAAATCTCGTTTTCCAGAATTTAAACTATGCGTAATTGGTCATTCGGCTGGAGCATATTTAGCATTTGCGTTTGCTAGAGAACGCCGAATGCTCATTAAATGCTTGGTTTTAATAAGCGGAGTTTATGATGTAACTCATTTAATTAAGACCAAGATGAACGAGCTATTGCAAATCAACGACGAAACTGCTGACATGTTGAACTTATACCATTGCTGTGATACTCTACAAACAGAGAGCATTTTATTTTTGGTTGGCACTGATGAGTCTTCAGAATTTCAACGACAAACTCGAGAGTTCTATGAACACGTTTTAAAGTTCAATTCAAATGTTCATTTTTGCGCTGTTAAAGACGAAGATCACTTTAGTTTAATTGAAAATTTCTCACAGAGTCGATCGGAAGTTATTGACCAAATATTTTCGACCTTAAAAAGTTTGTAATGAGGATTTTCTTATGTTTTGCATTTTTACAATAGAGTCATTCTAATAAATGTAAAATTAGTTTTAGGTAGCCATTTTTCATTTAAATAATATCCAAAGTAACATACATAGAATCCCA
>evgR006498c0g1_i01_Hydra
TCAACTGTTACAGTATTGGTTGTAAGCAGTTTTTTATTTTTTCTAAATGATACTGGACTTATTGAGATACCAGTTCAAAACAATAACAATGAAATTCAAGAATACGACATGATGTGGGTTGGTGGAAATGTCAGTTTATTTTGCAATGAAAATGGACAAGTTCTAAAAACTGTGCAACATTTTGACGAGATGGTCTCTGAAACAGTTACTTTTCTGACAAATTCTCTAGCTCCTATTGATTACTTTCTATCCATTTTTATATCAAAACCATTTGAGTACTTAATTCCTATACCTTCAAGTTATGGATTATTCAATATAAATGTTGATGCATCAAATCATGTTCCTTCTTTTAAGCAAAAAACATTTTCTTTTAAAAACCAGATTC
>evgR006525c0g1_i01_Hydra
TTCATGATGGTTATTTGCATTGAATTGCGCATTCCTTTATTTGTTGTTGGAAAACCTGGGAGTTCAAAGTCTTTAGCAAAATCCATTGTGCAAGATTGCATGCAAGGAAAAACGTCAAAATCCTTTCTTTTTAAAAAATTCAAACAGATTTTTATGCTATCGTATCAATGCAGTCCTTTGTCAACTGCAGAAAGAATAATAAATACATTCAAGCAGTGCAGTAGATTTCAGGTAGAAAAAGATTTAGACACATTTGCATCAGTTGTAGTTTTGGATGAGATTGGTCTTGCAGAAGATTCCCCAAAGATGCCTTTGAAAGCCTTACATTCTTTGTTAGAAGATGGTACTGATGGTTCTGAAGAGCTAACAGCTGATGGTTCTGCACAC
>evgR006553c0g1_i01_Hydra
AAAATCAGAGATAAAAATATTAGATTTGTCTAGACAAAAAGATTAGGTTTAAGTGGAAAAAAAGCATTCAGGTAAACAATGAATCTCCATCAATTATTTCTGATAAAAAGTATTTACATTATTGCAGTTTTCAGCTCAGTCATAAAAAAAGCAACAATAGAATATCCAGGAATATATAAAGAAAAGGTTTTTCAACTTATACAAAGACTAGAAATGGACTTTACGCAGTTTTGGATATATAAAGATATATATCCTAACCCTGAAAGCCTTGGTGTTACACCAGCGTTAATACAAAAATCGTTTTCAAACTTTGGAGATGAGAAACGTTTGAAAAACATTTTAATTAAAGCACTGTATGGATATGCAATTCAACTGCCTATCGTAGGGGGTTCAATTTCTCGAGGAGCTCCGTTTGCAGAGTATGGAGAAGGAGATCGTGTGTATTTTAACGCTATAGTAAAATGGTGGAATGACATTTTCATGTCAATAACCACTTCAACAATGCAAGCAAAGTCTATTTCAATTGGAGGAGTGGGTACAGATTACTTTTCATACTGCTTGAACTCACATTTAAGTGAAGATGCTGAACCAAAAATTATACTATGGGAGCTGGCTGCCAATGATCGTGGAAGGTATGATGATAAACCATTTCCACCAGGGCAACCTCTAGAACAATTCACAAGAAATATTTTACAAAGAGAATCAAAACCAGCTCTTTTTTTTCTAAATTTTTTTCGTGGTCATGATTATTTAGATGGTAAATGTATGAACTATGAAGATCAAGGTGGATTGCTTGTAGCTAGACATTATAGAATATCGTCAATAAGCTGGAGAAACTTTGTTTGCGATGAAATAAAACAAATGACACCAGGATTTACAATAGAAACACTGTTTTCTGCTGATAGACTGCATCCAAGTATATTAGGTCATGCACAAATGGCATTTTTAGTTATAAATCATATTAAAAACAGTTTTTTACGAATGCTTATCAACCATGCATCACGGGTTCCATCATTAGCAGATTTTCAACAGCAAACGTACAACCATGGTGACGTAGTTCTAGCAGATATTTTATATCACGAGACATCAAACAATAAACCATTGTGTTATACATATCTTAAGTACAACCAATTTGAACCAAACAATACATTAGCTATTAACATTCTCCGTCAGGATGATTTTCATTTTAATATTTTTAAAAAATTTAAAATCCGCTCAGATGAACTTAGAGGTATGCAAACATATCTTGCCGAACAGTTACTACAGATAAGTTTTTATTTAATCCAACCCTATTCAAGACTTGTAATAACTACACATAGTTCCACTGGCAGTAGTCAAACGTGGTTAGATACAGATACTCCTGTAATTATTGAAACTGATAAGTACCACATGGGTACTAAAGTTGAATTAATTACAACCAATTTAAAAACAGGAATGCATATACTAAGTATTTTATCTACACAAAATGGATTTGTAGTTTGCGCAATTAGTGTTATATAGTTTAAACATGAAATTTGAAAAAAAAAAAAA
>evgR006608c0g1_i01_Hydra
AGTAGTAGTACCTCTGGTATGAGCCCATTCGTTCGTGATTTTGCATTGGGAGGAATTGCTGGTGCCATTGATAAATCTGTTACTGCTCCGATCGAAAGAGTTAAATTGTTGATCCAAACTCAAGATGCTAATCCTTTGATCAGATCAGGACAAGTACCACGTTACACTGGTATTGGTGATTGTTTTGTTCGTGTTGCAAAGAATGAAGGTATTCCTGCTTTCTGGAGAGGTAACATGACTAATATTTACAGATATTTCCCAACACAAGCTTTTAACTTTTCTTTCAAGGATGCCTTGAAAAAGATGTTCCCAGATTATTCTCCAAAGGAAAACTTTGGTAAATTCTTCTTGGTTCAAATGGCTTCTGGTGGTTTGGCTGGTGCTGGATCATTGTGTATTGTTTATCCATTGGATTATGCTCGTACTCGTCTTGCTTCCGATGTTGGTAATCACAGAGATTTCACCGGTTTGGGTGATTGTATGAAAAAGACTTACAAGCAAGGTGGTTTCAGAGCTTTATACAATGGATTTGGAGTTTCTGTTGCTGGTATTATCCCATACAGAGGTGTTTACTTTGGTTTATATGATACTTTGAGAGAAAAGAATCCATACAAGAGTGATCTCGGAGTCGTTGGTATCCTTTCTAAATTTGCCCTTGCTCAAACTGTTTCAATTGCTGCCGGTTATGCTTCCTATCCATTCGATACTGTCAGAAGAAGATTGATGATGCAATCTGAAAAACCAAAGGAACAATGGGTTTACAGAGGAACAGTCCACTGTTTCTCAAAGATCCTCAACGAAGAAGGTACTGCCG
>evgR006644c0g1_i01_Hydra
AACTGTTTTTATTCACGTCTTTGGTAATGATACCATGCCAATTTTCAACATTTTCATGAATAGTATTTATTATTGGGGATATGCTTTCCTTGTTGCTTACTTTACAATCCATCCAAATTATCAAGCTCCATCTGATCTCCAAGTTTACATTGGTGCTGGTGGTGTTGTCTTGTCAATGATTGGTAACTTTATTTGTCACATTATTTTGAAGAATTTGAGAGCTCCAGGAAGTAGAGAACGTAAAATTCCAAGAGGTTTCCTCTTTGAATATGTTACTTGTCCAAATTATACTTTTGAAATTTTGGAATGGGT
>evgR006666c0g1_i01_Hydra
GTACTTTATGTTTTGATTAAAAAAGATTTACCTTTGCAGAAGAGAGCTCTATGTAAAGTGAAGCATAATGTTGAGTTTGAGTCTCTAATATCAAAATTTCAGCAAATACTGTGTAAAGTTGTCCAACAACGAAGAGAGTCTTATTTAATGAAAAAAGAAGAACTTATTTGTGATATATGGGATGGTAATTTGTTTGCAAAATTTTGTTTTATATACTGTAAACAGTTAAATGAAGGAGAAGTTATGTTTGACACAGGTGTTTTCTTCAATTATAAAAGTATTATTGATAAATTTGTTGTTAAGTGTGGTGGACGATTGCAATCAACACTTCCTTTTCATCTTGGATCAAAAGAATCTTGTGAGAGACATCCTGAATCATTGCAAGCTTCTAATAAGG
>evgR006680c0g1_i01_Hydra
AATTTTCTGGAAAACATCAGAAGGTACATAGCTATCCTATCTCCCAAAAACATCAGTGTCAATGACATCAACATTCAAATACAAAGCCTGTTACCTGGAGATTCCAAATGCTATAAGTCAATCGACACAGTAACAGATAGTGCTCAATCCGTCTATTATCCAACAGAGTTTCTAAATTCATTGGAGCCACCAGGTATCCCACCTCACAACTTGCTGTTGAAGATTGGTTCACCCATCATGCTGCTCAGAAATTTGGACCCTCCCAAGCTCTGCAATGGAACTCGGCTATGCATCAAATCTCTACTTCCTCATCTCATTGAAGCCACCATTATGACTGGATTTGGAAAAGGAGAAGATGTATTCATTCCCAGGATTCCTCTGATCCCAACTGACTTGCCTTTTGATTTCAAACGATTGCAGTTTCCAGTTCGACTTGCTTTTGCAATGTCAATCAACAAGGCTCAAGGACAGTCACTGAAGGTTGCTGGAATCCACTTAGAGGAACCTTGCTTTTCTCATGGACAACTTTATGTTGCTTGCTCCCGAGTCGGGACACCAAGAAATCTCTACATATTTTCACCAGACAATGGAAGAACCAGGAACATTGTC
>evgR006716c0g1_i01_Hydra
TCAGGTCAAACGTAAAAGGTTAAAACAAACTTATTTTGTAAAACAAAAAATACGGGTTATGGTTTACCTTGTAAATTTCATTAGAATAATGCATCTTTGGCTCATTTCTGAGGCTTATTCAGTAATACATAATACTATCACCGACAATGTAAATATGGTGTGTCCTCCAATTTTAAAAACCTTTCAAGGGGATTTCATTAATCGTCTTGTAGCATTATCCTTTGATATAAAAAATGGCAGCTATCAAATTGAAGGTTTTATTAGAAGTATCGATTTTTCACAAACAATTCTTCCAAACAATTCTTTTTATCCGAGTG
>evgR006725c0g1_i01_Hydra
CTAATAGTAATTAGTAATTAGATAAAAAAACTAATAAGAAAAGTTATATTAGGCACTAATAGCGCAATTAGTTCTTATTTGTATAAGATGGCATTGCTGTTGCTAATTTTTGGGGGGTTTTTCACGCTAAGTGTTTCATTAGATGATCCAGAGGTTTTTTTAAAACGCTATGGCTATTTACATTCCGGGTACAGTGACAGTAATTCTGTTACTTCAGCTATAAAGAAAATGCAAAAATTTGCGGATATACCAGTAACAGGTGTTCTTGATAAGCAAACATTAAATTTGATGTCGTCTCCGCGTTGTGGAGTTTCTGATCATAGTTATTCAGGATACTCAAGTAAATGGAACAAAAATAGATTAACTTACAAAATAATCAATCATACTCCAGATTTGGGTGCATCAGAGACTGCAAGAATTATCCAAGAAGCTTTTAACAAATGGTCCCAAGTAACGCCACTAACTTTTACACGAGCACAAGGCGCTACGGATATGATTGTGGATTTTGGAAACCTCCAGCATTCTTCATGCGACTATGATTTTGATGGAATTGGTGGGGTTTTAGCCCATGCTTTTTTCCCTCCTGATGGTCGATCGCATTTTGATGAAGACGAGAATTTTACTGATAAGACTACATCTGGAACTAATTTGCTCTGGGTTGCTGCTCATGAATTCGGTCATGCACTTGGACTTGATCATTCGGATGTAGAAGGATCCCTTATGTACCCTTACTACCAGGGATTCAAAGAACCGTTTGTGCTTCACCAAAATGATATTCAGCGTATTCAGGAATTGTACGGCGGAAGAAGTCTTCCAGCAACTTTACCTCCAAAAACAATCGTACCTCCTACAAAAACAGTTAAACCTCCTACAAAAACAGCTGTACCTCCTCAAAAAACAACTCGCGCCCCAGCTGCCTGTACTAACTCCATTGGAGACGTTGCTTGTACGAACCTAGCATACCTATGTAAAAATATTGGTATGCAAGATTTTATGACATCCAGCTGCTACAAGACTTGTTTTTGTTAAATTTAACGTTACACGATGATGGTACACGTTAATTGAATGTTACAAGATAATATTTTACGGTAATTTAATATTACAAGATAATATTTTACGGTAATTTATTATTTCAAGATAATGTTTTACGTTAATTTAATATTTCAAGATAATGCTTTACATTTATTTAATATTAAACAATTTTGTTTTAATTTAATGTTACACGATAATGTTTTACGTTAATTTAATGTTACACGGTAATGTTTTATGTTAATTTAATGCTAAACGACAAACAAAATAAAAAAATATTACTAACTAAAAAAAAA
>evgR006766c0g1_i01_Hydra
CTAGACATGATTGCTTGGGTAACATCACTGTTTAACACGAAAGCAAACCTTCACTTACATAACTTTCAATTAATTGACATCATTTTGAAATATAGTTTCGATATACCAGGTTCTTCATATATTAAATATAAAGTTGATTCCTCGACTGGTTTGAAACAAGAAAGCGGTTTAGACGAAAAAACTAATATAAAAATAATTTTCCGAGAATTTGAAATTAAAAATTATAAAAAGAGCGGCGGCGACAACGGTGACGGCAAATGTCTAATAATAAGTTGTTCTCCTGGGTTTGAACAAAAGTTAGGAATAGTTACACTCAACTATTCGCAAAAAGAGCAGAAAAGATTTTGCGTAAGTTGCTCACAAAACTATTACAAGCCGAATTTTGGTCCCGAG
>evgR006768c0g1_i01_Hydra
TGAAATTGGAGGTGAAAAGTGGCACGAACAAGGCATCCTTGACAGAAAACAAAATGTTTTTGATGACATGATCGCAGCTGCTGAATATCTTATCAAGGAAGGTTATACTTCTCCATCAAGAATAGCCTTGAACGGAGGAAGTAATGGTGGTCTTCTGGTATCAGCCGTCACAAACCAAAGACCTGATCTCTTTAGTGTCAGTATTCCAATGGTTGGTGTTCTAGATATGCTCAAGTTTCACACTTGGACTATTGGTTCAGCTTGGAAGTCTGATTATGGATGTTCCTCAGATCCAAAAGGATTTGACTATCTAATTGAGTACAGTCCACTTCACAATGTTAAGGCTCAAAAATATCCTGCTGTTTTAGTTGTAACCTCTGACCATGATGACAGAGTTGTTCCTCATCACTCTTTCAAATATACTGCTCAATTGCAACATGTTGCTGGTTCAGTTAATGATTCTCCTCTCCTTGTGCG
>evgR006786c0g1_i01_Hydra
GCCTTGTTTTAGAACAGAATCAACAGGAGCTGAAACCTGTTGCGCATTACGTTTTACGCGAATCATGTCAGCTTCTTTAAGAGATTTCTTTATCTTTTTGCGCAGATATTCTACATTTTTTACTGCCAGTTTTTCGTATAGACCTTGCCTATCAGAATGGTCCAACGTTTTTTTGATTGGGCATAATACTGTGAAATTTGGGGGCGAAAATTTTCATCATTTAGGACAAAATTCACATTTTCGGCCTCCCAGTACAAAATTACGGATTTTGCTTAATGCGCCAATGATAGGTCAGGCCAAAATATTTAGGGATTGTCTGCAATGAATGGC
>evgR006797c0g1_i01_Hydra
GTCGAGAATGTATTATCAAAAAACGATCTTTATCTAAAAGGAAAAAAAATTAACATTGGACCAGCTGTTAAAAAAATGTTTGCTGCTGAGTTCTCGCCCCCTTATGCTTCACTAGAATCTTCTGACTCCGATACTCCATCGCCTTCTGAAGAATCCTTGAGCGACGAAAAAATAATCACAGTTCCTCCTGTGAATGAAAAAATTGATGCCAGTGGTTCACCGGCTCCACGTTTAAGTAATGTTTGGCAAAAAGTAAATGTTAAGCGAACATCGGCCAAAACTCTTACGCCCCCTACGCCATGCTCTCCCGCCGTTGCTTTTACTCAGCCG
>evgR006818c0g1_i01_Hydra
GAAAATAAATCTAGAACAGCTGACAACTATTATTGAATTATTATATTTGCAGAATTATTATTATAATGGCTGGTTTGGATTTAGTATCACCTGAAGGACTTCGATTCGATGGACGAAAATGTCATGAACTGCGCAAAATAACTGCAAAAAAAGGAGTTTTTACACAAGCTGATGGCTCAGCTTATATTGAACAAGGTAATACTAAAGTTATTGCCAGTGTTTATGGTCCACATGAAGTTAGCAACAGATCAAAGACTCTGCATGACTCAACTTTAATTAATTGCCAGTTTAGTATGGCTACATTCAGTATGAGTGAGCGTAAAAATCGTCCAAGAGGTGATAGGAAATCAACAGAAATATCAATGTTGTTAGAAAAGACATTTGCAACTGCCATTATGACTGAACTTTATCCACGTTCACAGATTGACATTTATGTTCAAGTTATACAATCCGATGGTGGCGTTATTGCTGCATGCATTAATGTTGCAACGCTTGCTCTGATAGATGCTGGTGTGCCAATGAAAGATTTTGTGTGTGCATGTACTTCAAGCTATGTCCAGGAGAAAAATTTGACAGATATAAACCATTTAGAAGAGAGTTCTGGTTCTTGTCAACTGACTCTTGCAATGTTACCAAGAACAGCAAAAGTGGTTGCTTTTCAAATGAGTTCTAGATTACATTTAGACAATCTTGAAGAGTTATTAAAGATAACTAAACAGGGTTGTTTAGATATTTACAACGTTTTACAGCAAGCTGTTGAAGCAGACTCTTTGCAACAAGCCGTTACCTTAGTTGTTTGATTTCGATGATATGTCAATTTTTTTAAATTTATAGAATTATGTTATTTATATAATGTAGAAACGATTTATTAAAAATTCTGCTGGGTGCTTTTTTTTTTAATTTTTTTAAAACAGGACTCATCAACCTTACTTTTCTGTTCCACCAAACTCTCTAGTAACATATTCTTGTCAAGAGTGTGTTACACTTGGTTTTTGATAACACCCAATCTTTTTATTTTGTAGTGTTTTTTATGATATATTAAAATGCTATTTTAAAGTTAAAAAAAA
>evgR006832c0g1_i01_Hydra
TACTTGTCTATTCCCATTGGATTTGACTGGATTTAACAACAGAAATGAAAAGATGCCTATTTGGAGAAATCAAAAGTCATCATCTACAAACTATTACAGACCGATAAGATTTGCATACAAAACGGAATCCAAAAAATCTGTGCTCGAAGAAGATCAGTACATTAAAAAAGAGATTGAAAGCTTTGGTATGATTTCATTACATATTTACGGTTCTTCTATTAAAGTGAATTGTATTATTGAACTTACAATGATTAATGGGAAAGTTCAAACAAAACTATCTGGAAAAACTAACTCATACCAATGCTGTTCAGTGTGTG
>evgR006835c0g1_i01_Hydra
GATGAAAAGGTTAATGTTGATACTCGTATGAAACAAATCAGAAAGGATGTTACTGATATGAAGTACAAACTTGAACAAGAAAGACAAAATGTTGTAACTGTTGAAGCCCAAAAACGTAAGCTCGAAGAAGAACTTGCTCTCCAACATACAAAATATGATCAATTGCAACAAGAATTTTCTGCCATCAAGAGAGAAGTCAAGAAGGCTGAACGTGAAGTTCAAAAGGCTGAAAAGAGGGCTGAATCTGAAGAACAACGTTCAAATGAATTGAAGGTCCAACAAAACCAATTGCGTGATGAAATTCGTCAATGGAAGGAAAAGGTTACAAAGGCTGAATTCGAAACTGATAAGGTTAAGAGAGAAAAAGAAAGTCTTGAAAGAACC
>evgR006859c0g2_i01_Hydra
ACATTGTTAATTGAGATCAGTCAAGTTTTTAATGGGACTGTTTATATTTATACAATTAGTTGTAATAAAACCAATTTATTTTCTAAAACTTACAACCAAGCTCAAGTTTTTAATGATATCAAAGTTTATATTTCTGATCCATGGAGTGATAGTCTATATGCCTCAATAAAAAATTTAAAATTTGTCAATGGAATTACAATTGGAGGAATGCAACCTGTTGCTATTATTCCTGCAGGTATTGATATGGGAAGCACAATGATTTTATCAAATCTTAAACAACAGCTAATCCTTAACAGCAAAATCCTTAATTTCCCTAATGGAGGTTTAAATAATCAACCTTCAGGTATTTTTTGGCAGTATTCAGAAGGATATTTTGAAGGAAGATTATATGTAGTTTTATATCCATTGGTATATTATGGAGGCTTTATTAAATTTTCA
>evgR006868c0g1_i01_Hydra
AAATTTTCGAAAATGAATTGACAAAGTTAAGCGCTCCAGTTAAATTGGATGTTCATTTGACTTCTATCGAAAATGAAGGAAATGCATGGGAATTAACAGCAGAAAACAAGTTAACTCTCTCTGAATCCAAGAAGGAACAAGGTAATGACTTCTTCAAGAAGAATAGACTCGAACTCGCCAAAAAGAGGTATGAAAGAGCATTGAAATTTATTGAAGATGAACATCCAGAAGATGAATCAGAAGAACAAAAGAAGAAGAGAGCTCAAATCATCTCCTCTTGCCATAGTAATTTAGGTGCCATCTTTGTTAAGCAATCTGAATGGAAGCTTGCCATTGATGAATGCAACAAGGTTTTAAACGTTGACCAACAAAACATCAAGGCTTACTATAGAATGGCTCAATCTTATCAATCTCTTGGTGAACTCCAAGATGCCAAGCAAGCTCTCGAAGAATGTGTTGAAGTTTGTGGAGGTTCGACCGATGAAACCCAAGTTCAATTACTTGCTTCAGTTAAAACTTTATTGTCCAAGGTCTCTAAGGCTATCACTGAAGAAAAGAAGAGAGAAAAGGCAATGTTTGGAAAGATGTTCCAATAATTCAAC
>evgR006872c0g1_i01_Hydra
TTAGCAATCTACTCATAAAAAAATCTTCATCAGGAAAAAAACTTATTGAGGAGGAGTTAAGAATACTCAAAAGTATTGCTCCTGTTCAAGAAAATAGTTGCTTTAAAACAGCGTTGCTGACTGAAAATTCAAATAAAAATAGATATCCTCAATTTTTACCACGAGATGACGCATTGGTTTCTAGTGAACCCTACCTGAATGCTGTGTTTGTTGATGGTTATAAAAGAAAGAAAGCTTTTATTGCGACCCAATGTCCACTTCAAACCACTGTCACAGATTTCTGGGATGTTCTAAAGAAACTAAATGTTTCAACTATAGTATAT
>evgR006910c0g1_i01_Hydra
TTCGCCTGCTATATTAGGTTTAAAAGATGTATCAGTTTTATTCTACATTCAAGAAAAAAAACTGACTACATACTCCATAGAAAATAACCAGGGTTTTACTGATGCAAATTCTTTGGTTACAGGTAGCCTCCTTGTTGAAAGAACAGATAATAATATTACAGTAAAGTTGGTGTGGAGTTCAAACATCTTTGTATCTAACAAATGTTCTGTGGAAAAAATTACAGGCTTTAGGTCTAAATCTACTGGAACCTTCGCTTCTATATATTATCAGAGTTTTCAACCAGAGTTTATATATGTGAAAGAAAGTGAATCTTTACGTGGGCCGTGTATAGATTTGTCAACTGATCCACCTATAATTGGTC
>evgR006929c0g1_i01_Hydra
AAATTATATAATAAAATGAGTCAAAACAAAAAGATTTTATTTGTAATTTGAAACTTGAATTGATGAATTTGTTTACTTACTGAAAAATCTTGATTTTGTGCCGACCGCAAAAAGGCTTTAATAAAAGGTTGCATGAAAGAATATCATTCATAGATTCTTAATCAAACGCATTACAACATTCTTCTAATATGAACAATCATATTTATTTTAACGGTTATATTACTAAGTCCGCAAGCGATCTTGACAAACCGTTTCGCACTAGATATTGCACAGTTTCCATGGAAAACATATCTCAATCAGATCAAATGAGCAAGAAGAAAAAAAGAGCTTCTCTCCGAAAAAGATCGTCTAAGCTATTTGATCAGTCTTTTAATCATCTCGACAACATTGATCCTCCTCAACAAAGATTAGTGCTTGCTTACTACGAAAAAGAAAATAAACCAAATCTCGGAAAGTTTATACTCGACGATTGTGACTTCAGAGATGTAAGTGAAGGCATAAAAGGTTTTCCTTTTGCGGTTTGCATCAAACTTAAACATTTGAATACCCCAAGAGATTTATATTTGTGTTTCAAAATCAAGGAAGATAAAAACAAATTTTTAGAAGTTCTTCGGAAGCATACGCCAAATAAACTATTCGACGAGATAGAAGACTTTTATAAACCGATTCAAAAAGAAATTCCAAATGACGAGAATAATCCGATTTTGGAAAAGTTGATAGAAATAACTAAGCC
>evgR006931c0g1_i01_Hydra
TTTTAAAAAATGAAGGAGCATGCTAAACTGCTTTCAAGCTGTTGTAGGGTTCGTGGAAAAAAAACAAATCTTAGAGCAAAATCAGTTGGGCGTTTCAAAGAGGTTTTAATGTATGCCTACAATATCGATATTGATTTAGACAATTCACAAATCTGCCCATCTACACTTTGTATAAAATGTTACATGAAGTTGTATTATATTCGAAAACAAAAAAATAAGAAGGATTGCACACGTATGTCTTTATACCAATTTCAACAACATAATGATGAGCAATGTTGTTTATGTAAATCTAAACTGTTTAGAGTACACACATTTTCTAATCCCACAAATGATATAAAAATAGCTACTGATTATTATAATATGAAGTGTTTTAATAACAAATTTGTATCCATTTATAT
>evgR006940c0g1_i01_Hydra
AAAAAATGTTCTTCTGACGTCAACTTTTTGGAAAGCGAAAGCGACTTTTTAAGTTGATTGTGATTTCGATTCTACATATTTTTAATCGATTTGTTAAAACAACGCACGTCATGTATTTCAAACAAAGAGCCCCAACATAACCGCTGACATTGGTATTTCTATATAAATAGACAATTATTTGATGTGGCAGAAATCGAAGAGACAAAATCATTTTAAAATGAGTCCGACTAAAATACTTTCAAATCTGCTTAAAACATTTTGTTTAATAAAGAAACATAAAATAAAAAGGCAAGTATCATTTGCTGGTAAATCTAGGGAACATTCTAATTGCCAACAGCACGACAACATAATAAAATCGGACTTATGTTCAAAAGATGATGATTTAAATATGATATTTTGCGCCATTTTATACTACGACATAGAACTATTATCAAATGTTTTAAAACGATTTAACCGCATAAATGAATTAAACGAGGATGGGATTAGTGCACTACACTTTGCCTCGATAGTAGGAAAAACACCTTGCGTTAAAATTTTAATAGAAAATGGTGCTAACATTAATAATATAGATGAAAGAGGAAAAACTGCACTTCAGTATGCCATTTCTATGAACAACTTTGACACCATCAAATATATAACGAC
>evgR006955c0g1_i01_Hydra
CAAAATTGACCCAAGAACAACTTTCTGATGCTCAAGATGCCTTCAATCTCTACGATAGAGATGGTGATGGTAAGATCCAAATTTCTGATCTTGGTACTCTCCTTCGTTGTTGCGGTAAGAACCCAACCAATGCTGAAGTTGAAAATTTGAAGCCTGAAGCTGATTCATCAAACTCAGGTTCATTCAACTTTGAAACTTTGAAGACTCTCTTGGAAAAGTATACTTTCAAGGCTGATAATGAACAAGATGTTAAGGATTGTTTCCGTATTTTCGATAAGGACGGTACTGGCTTCGTTCCTGTTTCTGAATTGAGACACGTCTTGACAACATTAGGTGAAAAGTTGTCTGCTGCTGAAGTTGATGATTTACTTCGTGATGCTGAAGTTGACGACAATGGTAACCTTAACTATGCTACCTTTGTTCACATGCTCATGCCAAGCAAGAACTAAGTTATTTGTTTTAAACTTTCAAATCCCCACATTTACCTCCTTTCCGTTCATTCCCATCATCAATCCTATCTTTCCCTCCAACATCCTTTTAATTTAAAGATAATTATGAAGATCATATTAAATTAT
>evgR006980c0g1_i01_Hydra
AATGCCAATCCCACAAACCTGTGCCGAACTTAAGTCATCCACCAGTCACTCCAAGCTTGTTGTTGTTACCCCAGAAACCACATTGGACAAGTGTTTGATGGCTATGGTTGAAAATGATGTTAGACATTTAATGGTTGTTCACAGTGAAGATAGTGGAAGAAAGATTGTTGGTCTTATTTCTGAAAGAGACATTCGTCTTGCTATTGGATCACCATTGATTCACAAGGACATGGATATCAAGAAGGAAATTGATGAATTCTCCAAACAAGTAGCTTCTACAATTATGACAAAGAATGTTTTCACTGTAAAGGAGGATGATTCCGTTCTTGAAGCCGCAAAGATTATGAGAGTTTCTAGAATTGGTGCATTGCCAATTGTTAATAATGATGATAATGTTGTTGGAGTTATTACAAGATCGGATATGTTGGATCAATTGATTCGTGTTTTGGAACCTACGAATTAAG
>evgR006987c0g1_i01_Hydra
GTTCATGGCGCAACATCGAAACAAACATAAGCTATTATGTTGACTTTAAAAGAAATAGCCGAACGGAATAGAGTACAAAAGGACATGGGGAAAATTTAACAAAACTGATGACACAATATATTGATGGCACTAAAATTATTATTACTTTATAATTTTAATATAAACGACAAAATAAAAATCCAATTTACATTTGCTTTATAGTAAAACTATCCAAACTTTACATTGAAGTAAAACCATGGAAACTACGCGGATTTTACATTAAAACCATGGAAACTACGCGAATTTTACTTGCAATTTTATTAGTTTTTATTTGCGATGTTGATACAGATTATGTTAACAATTTGAAAGCTAATTTAGCAGAAGAAATATTGAATGATTTAGAAGATGACAACTATGAAAATGATGAAATCGCTGATATAATATTGTCTGATAGCCAATGGAGAGACCCTAGATTTAGAATTCGTATTAGAATTCCTACTATTAGACTGCCTACTATTAGACTGCCTACTATTAAAATTCCTACTATTAGACTGCCTACTTTTAAAATTCCTAATTTTAAAATTCCTATTATCGACATTCCTGCCTTTAGAGTCCGTGAATCTCTAATCCGCGAGTTAAGACTCCACAAAAACAGAATCTCTCAATTTAGAATACCCAGAGTTAACATACCCAGAATTAGAATTGTCCAATTTAGAACCCCTAAAATTCCTCAATTTCGCGTTCCGATGACACGGTTTCCTAATGAAGACAGTATATTTTTAAAGTTGATGAAACCTACATGCCTTGGTTGCGCTGGAATTTGTGAAAAACCCGTCCCTGGAAGATGCGCATGTCGAATAAGACCAAACTGCAAACATGAATAATAAAGGTAAAAGATGTTTGAACTCTCGATTTAAAGACTTTATTATACTTAATATATATACATTATTTGTTACATAGATATATATAAATAATTATAAATAAATAAAAAGTATTATTTTTTTATATTTACAATATATCCTGG
>evgR007004c0g2_i01_Hydra
GAGAAAAAACTTAAAGAAGAAACTTCTAAACAAAGGTTGGAAGTTTCGCAGCTCAATAAAGCTATTTTATCGATTGAGGCTGAATTAAATTCAAATCGGCAACTCCTTCAAAAAAGTTTGGAATTAGAAACTACTAAACGATTGTTACTAGAGAAGGAAATGAGTAACAAACAACAAAAACTTGAAGATATTTTGCAACAAGAAATTAGTAAGAAAAATATCGCGGAACAAAAACTAAAAGAACAACAACTGTCAATGATGAAAGAAATGAAAGAAAAAAATAGAATAATTGAAGAAGAAAGAGAGAAAATTAGAAGAGAACTTCAACAGAAGAAAGGTTCCTGTTTACTTTTGTGAAAATAAACAATAATATTTGAAAGTTTATTAATAAAGTGCTCAATAAGTTAAATTTTAAGTTAT
>evgR007012c0g1_i01_Hydra
TGTTCAAACTTATATTTATTTTTTGAACATTTTATTTGAAACGCGATACCTAAGGAACAAACGTAACGAACATTGTATGAGTATCCTTGTGTGAGTGTTTTTAGGAGCAAATCCATTTGTGTTATTGTTTTCATATCGGTCTGAAAGTATGTTTAAATTTTCTCCGTGAAGGCAAATAAAAAAAACATTATGTAACGAAAATGATATAAACACTAGTGAAGCAAATTAATGTGTATCTGCCTTTGTAAATATTAGTTTAAAATGGATAGTTTTGAATGGGTGGAAATTATTGAACCAAGCAGACAAGAACATATGTTTGCTAATGTTAAAACAGGCCAGTGTATGTGGGAAGCACCAGAAGGTGTCAGCGTAAAACCAACCCATGCTAACCAGTGGTGGGAGCTTTTCGATGTAAACACGGAAAGAAATTATTACTATAATGCAACATCTCAGAAAACAGTTTGGCGAAAACCAAAAGATGGCGATATTATACCTTTATCGAATCTTCAGATGTTAAAGGCTAAAGAAGCTTCTCAAAAGTTATTGAGCAAATCATCTCCAACTGGTAGCGTAAGCGAGGGTTGTGCTACTTCATCTGGTTGTGTAAGCGAGGATCATACTTCTAGCTCTTTGAATCATTACGAAAATTGTGATTTAATAAAACAAAAGAAATATGTTGCGCCACAAGGAGATATACCATCAAACAATAAAAAGGCTAATAATTACTATGAAAATGTTGTACCTATTGCATCTGAAAATCACCAACCAATTAAAAATCAATTAAGTAATAATCAATTGTCTGAAGTTGATAAACATTCTGAAACACAAACCACTGAAACTGTTAAATCACATGAAATTCTTGATCATTGTTCTACACAAAAAAACACTAGTTCGTTCCGTTCTAAAGCTCCTCCAAAGCCTCCACGCTGTGATAAGTCACTTTTTAATTCAAATGAAGGTTTAACTTCAAACTCGTCTGATGACAACCATGTTTCAAATCATGTTGCAGCTGATCCTAAGGTTGTTGGTGTTGCAGCTGATCCTAAGGTTGTTGGTGTTGCAGCTGATCCTAAGGTTGTTGGTGTTGCAGCTGATCCTAAGGTTGTTGGTGT
>evgR007048c0g1_i01_Hydra
TATGTTATTCAATCAACTTGCACACCTGTCAATGACAATTTTATGGAATTATGTCTCATTGTCGACGCTTTGAAGAGAGCTAGTTGCAACAGAATTACTGCTGTTGTTCCATATTATGGATATGCTAGACAAGACAGAAAGACCAAACCTAGAGTCCCAATTTCAGCTGCCCTTGTTTCTAGACTGTTAGAATCATCAGGTGTTGACCGTGTATTGTCTGTTGACTTGCACTGTGGTCAAATTCAAGGTTTCTTTAATATTCCAGTTGACAATCTCCACGCTAAGCCAGTCTTAGTTGATCAATTCTATCAAAGAATTGTTTTGGGAGAATCATTAAAGACTGAAAGCGATTATAA
>evgR007056c0g1_i01_Hydra
ACGTTTGAACAATTCTAGAAATTTGCTTGCAATTCAAATGAACAACAATACTACAAATACTTATATTGCATCTGATTTAAATGATTCATTTGGCATTCGATTGGCAGTTTTAGATAAAATGATTCAACTACCCATTTCAAAAGATAATATAATCAACCACGTGACAACCAAGTGGTATGCAGAAAAACTAAATGAAAAGCTTTTGTATCAAAATCTAAATGTATTAAAATTGAGTAAAGTCGAACAAACAAACGTAATTCTTATTGTATCCTCTGCACCATCTCGATTTGATCGCAGACTTGCAATTCGTGAAACATGGTGGAAAAAATGCAAATCTAATTCCAAAATAAACAATTTTTAAAACAGACTTTAACTAGATCGGTACACATTGGGCCAGAATTCACTTTTAATGGACACAATTCAACACTAATTTAGTTTAAAAATTAGCATGATTTGCAATAGCAACAACCTATTTTTGGTAGTCGCTCACCTATTTTTGGTAGTAACAAGTGGCGACGTTAGAAAAACGTCGCAAAGTGCTACAATAACTGATTTTAAAATAAGTATTAACATGAGTTTGCAATTCTTATAAATGCAAT
>evgR007062c0g1_i01_Hydra
AGTAAAGAAAATGAAGCTTCGAAATTTTTTTGATTAAAGAACTTGAAGCTTCGAAATTTTTTTGAGTAAAGAACTTGAAGCTTCAAAATTTTAGTACAAGTCATCGAAAATATAAAAAAAGCATAATGGTATGGAAGACATCCACAGTTGACGTAGTTTTCTCATCTCTTGGTGTTTGTATATGCGGTGTAGGATTGTATATTTTAATAAAACTTAAAAAAACGAAATTTGATAGAACGCAATGGTATATTATTACCAACCTCTGCATTTTTGATTTTTGCTCTTGCTTTGTTCTTACAATAAACTCATCTTTATATGTATCAAATGCCGAGTACTCTGCGATAGTTTCACAAGCGTTTGCTGACATTTTTATGCTTGGTTACTATGGATCAACATTATGGTTGCTATTTGACCGATACTTGCACATTAAACTAAATAATAG
>evgR007065c0g1_i01_Hydra
GTAACATTAAAATGGGCTCAGGCAAACCAAGAGGTATTAAAGCTGCCAGAAAGATCAAGGCCATCAGAAGAGACTCTAGATGGGCCGATATTGACTACAAGAAGATCGCTTTGGGTACTTTGGTCAAATCCAACCCATTCGCTGGTGCTTCCCACGCTAAGGGTATCATCCTTGAAAAGGTCGGTGTTGAAGCTAAGCAACCTAACTCTGCCATTCGTAAGTGTGTCAGAGTCCAATTGGTCAAGAATGGTAAGAAGATCACTGCTTTCGTTCCAAACGATGGTTGTTTGCTCTTCCTCGAAGAAAACGACGAAGTTTTAGTTTCTGGTTTCGGTAGAGCTGGTCACTCCGTCGGTGATATTCCGGGAGTCCGTTTCAAGGTTGTCAAAGTTGCAGGTGTCTCCCTCCTCGCTTTGTACAAGGGAAAGAAAGAAAAACCACGTAATTAAGATTTTCCTCATTCCATTATAAAACAA
>evgR007072c0g1_i01_Hydra
CAGAATCAGGAGACTATTTTTTTAATGTCTCTTGTAATGATGACTGTGAGTTACTGTATACAGAAGATTTATCTGACATAGGTTCTGCTAAAAAGATAGCTAGCTGTCCTCGTTTTACTGATCGATACATGTGGAACAAATATGAAGAACAATCGTCGTCTGCAATCAACCTAAATGTTGGCAAACTTTACTATTTGGAAATCATCTTGTTTAATAAATTTTCCCTTTCATCCGGTGCGGCTGCTGTTTTATTACCCAACGGTACTTTTGTGGCGCCAATAAGTTATGAGTACCTGGAAAGATTTAATTGATTGTCGGATGTACTACGATAATTTTCAAG
>evgR007072c0g2_i01_Hydra
ATGCAGTAATAAATTTGTTTGTCCTAAATCTGGGATAATCTTGAGTTCTACGATGGTAATTAGTTCTACTTGTATTGACTCTGTCTCAATGCAAATACAGCGAACGATTTGGGAAACTTTACAAATGACAGAATTGCTAAAAACATTTTGTTTATCTCCATCAAAAGGCAAATTAATTAGTGGTGCTAAAGTAGAACTTGCAAATTGCACTAAAAAAGAACAAAGGCTTTTGATGTCTGGACTTGATAATGGTGATGTTTTGGTTTCGTTTTATGAAAATTTTGCAGACATGGAAGATTTAAGTAAATTTCTAAACTATCCAACAACTATAGGTTTTCTTGATAACTTAGATTTCCCATCCTATAAGATTGGCACAGGTGCAATTCGAATGCAGGCATACTTCAGAGCTCAAGAATCAGGAGACTATTTTTTTAGTGTTTCTTGTAATGATGACTGTGAGTTACTGTATACAGAAGATTTATCTGACATAAGTTCTGCTAAAAAGATAGCAAGTTGTCCTCGTTTTACTGACCGATACAAGTGGGACAAATATGAAGAACAAGAGTCGTCTGCAATTAACTTAAAAGTTGGCAAACTATACTATTTGGAAATCATCTTGTTTAATAAAATATCCATTACATCCGGTGCGG
>evgR007076c0g2_i01_Hydra
GAGGTTTTTCTGATGAAAATAGAACTCTGTTTTGAGGTTTTGTCCATCCATCTGGCCCACTTTTCTAATCACCATCTGACCATTTTTGAGGTTTTGTCCACCATCTGGCCCACTGTGCAGCCCTGTAATCATTTTGAATACTTTTTACCAGTTGAAATCACTGGAATCCAGGTTTTTCTGCCGGAGGTTTTTCTGATGAAAATAGAACTCTGTTTTGAGGTTTTGTCCATCCATCTGGCCCACTTTTCTAATCACCATCTGACCATTTTTGAGGTTTTGTCCACCATCTGGCCCACTGTGCAGCCCTGTAATCATTTTGAATACTTTTTACCAGTTGAAATCACTGGAATCCAGGTTTTTCTGCCGGAGGTTTTTCTGATGAAAATAGAACTCTGT
>evgR007090c0g1_i01_Hydra
AGGTGGATTTGGTGGAGCTGCTGCTGGCGGAAAGAAGGTCATTGTCGAACCACATCGTTTCGAAGGTGTCTACGTTGTCAAGGGTAGAGATGAATCTATTGCCACATTGAATATGGTCCCAGGTGAATCTGTCTATGATGAAAAGAGAGTTACAGTTGAAAAGGGATTCGGTGATGCTGTTCAAAAAGTTGAATACAGAATTTGGAATCCTTTCCGTTCCAAGTTGGGAGCTGGTATCATTGGTGGTGTTGAAAACATTTATATCAAGCCAGGTACCAATTTATTGTATTTGGGTGCTGCCTCTGGTACTACAGTTTCTCACTGTTCTGATTTGGTTGGTGCTGAAGGTATGGTTTATGGTGTTGAATTATCCGAAAGATGTGGACGTGATTTAATTGAAATGTCAAAGAAGAGACCAAACATTGTTCCAATCATAGAAGATGCTCGTCATCCATTGAGATACAGAATGTTAGTTGGTTCTGTTGATACTATCTTTGCCGACGTTGCTCAACCTGATCAAGCTCGTATTGTTGCTTTGAATGCTAAGTACTTCTTAAAGCCAGGTGGCAAATTCGTCATTTCCATCAAGGCTAATTGTATAGATTCAACAGTTGAAGCTGAAGTTGTTTTTGCTAGAGAAGTTGAACAATTGCAACAATTCGGTTTTACTTGTGAAGAACAATTGGACTTGGAACCTTTTGAACGTGATCACTGCTTGGTCGTTGGTACATTGAACAATGGTGATTGTATACAATTTTAAAT
>evgR007107c0g2_i01_Hydra
TTTTCTCTTTCTTCGGTCAAGAAATCAAAACTGGTGCTCCACAAGCCTTCGAAATCCCATTCGGTGAAGTCGTTCTCCACTTGTCCACCGTTTCTCTCGCCAAGGATACTCCAAAGGGATCTATCACTAGAGTTTTTGTCCACACCGTTGATGAAGAAGAAAAGGATAGCAAGTATGTCATTTGTACACTTGTTGGAAAGGAAAAGGAATCAATCTCTATTGATTTGAACTTCAGTGAAGATGTTGCTCTCTCAATTGAAACCTCTGCTGGTGCTGATACCACTGTCCACGTTACAGGTTATATCAACTTGATTGAAGACGGCGAAGAAGGTGAGTATGGTGGATATTCTGTTATTGACGGTGAAAATATTACAGATGAATCTGATGAAGAAGAAAAGACTAAAATTTTGAGAAAGATGTTGGAAGAAGAAGATGATGAAGACGATGAAGAATTCAAGCCAGACTTGAACGAATCCTCTGAATCTGCAACCATCGAAGAATTATCTGATGAAGATGATTTGGGTGAAGATGAAGTTGATCTTGATGATGACCAAACTGAAGATGTTGTCAAGAGAGTTCAAGATCTTGAAACCAGATTGGGTAGAGAAGCTAATGATGACGAAATTAAGGCTATTGTTACCAGAGTTCAAGCCGGTGAACCTGAACCAGCTGCTCCAAAGAAGAAGGAACAACCAAAGAAGGAACAAAAGAAGGAACAACCAAAACAACAACAACAACAACCAAAGGAAGGTCAAAATAAGAGAAAGCAAGAAGAAGCCCTAGGAAACAACAAGAATAAGAAGAATAAGAAGAACAAG
>evgR007145c0g1_i01_Hydra
ATGAAAATTCGATATTTTATATAGATATAAAAACATTGTTGTTTTTTTACACGACAAAACAAGTAGTGATGTGAACTCGAAACATTTTATAATGGTCTGGTGTTTGAAATACGATGCTACCTATTTGCGCTCTTTTCTCGCAATTATAAAAGTTATACAAGTAATTTTTCTACTTATTGCGTATGCTATTATACAGTCGTTTATAAACCATGTTTCATTCAGTAACTCTAAGTTTTCCAAGATGCATTTTTTTGTATTAGTAACCGCTATTGGTTGGATATCATATTTATTAATGTTCTGTCTAAATTTGTTTATGGTAGCAAACCACTTTCTAATACCTTGGTATTATGTTTATTTAATTTCAGACGTTATATACGCTTTGTTTCTATCCATTGCAGTCGTAGTCATAGCAGTTCAAATTAATACTATAAACAATACACAACGTCTCTACATACCTGATATGGGAATAGGAGAAAATCTTGTTCAAGAATGTGAGCTTATAACTTTAAACTCGAGCGCTTCGTGTTCTTTATTAAACGCAGGAGTTATCTTTGGTGCACTGGGAATATGCGCCATTTTATTGGATATAATCGTTTGCATTCACAAAATTAGAAAAGGAAAAATTAATCACTATAACTCAAGATTTCTGAACGACAACTTTACGGTATCAACAGCTGATACTGTTACTAACTCATCTATACCATCTCGGCAGTCAAACCATAGACTTGATATTCAAGAATATCGAACTGAGGGGATGTTTTGAAATAATTAGCTAACTATTGGAACACCTTAAGAAGATTATCTAAATTAAAAATTGTTTTTAATAAACTCGAAGAAAAGTTTGTTCATCAGAATTTTATTTTTTTATAAAAATTATTTAATTTTTTTAAAAAAAAAAAA
>evgR007147c0g2_i01_Hydra
TGCATCCACATGGACCAGTTGAAATTGAAGAATACACTTCAAGAAAATTAGAATGTCCATTAAATGCATTAATAACAGTTTTAAATGCTGTATATAGTCCCACTATATGCGAGGGCTGTAATTATACTTCAGAAGAATTTAAATGCCCAATAAACATAACTAATTCAATTAAAAATTTGTGTGATAAAAAAAAAGATTGCAGTATTGAATCTAATAATATTAAATATGGTGAACCATGTAATGGAGAAATAAAGTACACACGTGTTAACTACAGTTGTCATCTAGAAAGTTCAATATTTATTTCTGGAATAGTGAATAATTTTACAAATAATGTTTTGAATGGTTTTTATGATGAATTTGTGAAATAACTAATGAATGGAGC
>evgR007152c0g1_i01_Hydra
TGATTTTTTAACCCATGCAGTTGCTAAGTAAGGAGGACTTTTTAACCACACAATGCAATCTTTGTTGAGTTCCTTTGTTAAAAATTCTCTTCTCTTTGTGAGAATGCAAGGGTGGGATAATCCTGGTCCAATAACAGCCTTGCATCTTTTGCGTATTTTTATTGTTCCAGGGGATGATATATTTACTCTAGCCAGCTTGATTGTTCTTCTGTTAATGGTAGTTCGAGCAATTTCACAGATGTTACAAACACAGAACGATTGTTTTCTAGTGAGTATGCTAACTTTGACTTTTGAGAAGTCATACGGTTCACTCAGAACAAAGTTTTTCTCTCCTTTTTCCGGTCTTTCTAATGTTTTTATACATTTATTACAAATACCATTAGGAAAATGTTCGTTGCTTAAATCAAATGATTTTAAAAAGAACTTAAATATTCTATTCAATGTCACAGTTCCTTCTTTAACTTCAATCATTCTTGATCCTTGTGAAAAGCAAACTACAAAACTTCAAATTGAGGCCCCAAGGTACAGTGGTAGCCAAAAGTCCTGTATAAGTTATATAAAGCAAACAAAACAAAAGTTTGATTTCCATATCAATGGTTATAAT
>evgR007167c0g1_i01_Hydra
CTCCAATAACAATAACGCACAAAACACTGATCAATATGCCAAGTTGGATGATGAACAGCAACAAGTAATGTACTCTTCGATTGATACGACCGGATCTATGAGTGGATCCTTCTATCAACAACAAGTACCTCCACACCATCAACAAGGAACAATTGTTAATTTGAACCAACCTCCAATTCAACAACCACTTCCAAATGTCGTTCATACCACAACTACTGTCAATCCACCACAGAAAAGTGTTTTTGATTCCCTTGGTGATTCATTTACGAGTATGGTCAATTCAATCTCACTCACACCAGCTACTGAGAAGAATCCAGCCTATTTACGACCACAATTCATTCCAACTACACAAAATCCAGAAAGTATTTTGGAATATACACTTGATGCAGGAATTTTAGATTGTCAATTGAATCATCCAACTCAATTGGGTCAAGTTAGTTCC
>evgR007183c0g1_i01_Hydra
AATTGATATAACAATGATATGTTTCATTTTGATTGTGCATATTTTGCTGGCAGCCTTTTCATTTGTAAAATTGCCACCAGCTGTTATATCGTCTTTAAACCACGAAAAAAAAACAATAACATTTCAATGTACTGGTGATTCTCATTTAAGTTTTCAAGTCATATATATCTGCTTCATGTTGCTAATTAGCTCCCTTCAAAGTTTTTATGCTCGTAATCTTCCTTCCTACTATAACGAGACACTCTCTATTACATACAGTACAGGGATATCATTCATATTATTGATATGTTACTTTCCAATTTATTTTGGTAACAAGGATGTCAAATTGAGAACCACTGCTGTGTCATTATTGATTCTAACAATAAACCTGATTTTAATGACAGTCTTGTTTATACCTAAAATATTTGTAATTTATTTTAAACCTGAATTGAACAGCAAACGTAGCATTCAGAGTGAAATGATGGATTACGCACGATCACTTTCTTCAGCTTCTTCATGATGATTATTTTATAAATAACTATATAGCAATTTTCTTTAATATTTATGTTTCAAAATAAAAACAAGCAT
>evgR007207c0g1_i01_Hydra
GCTCGTATAATGGATTCTATAAACTTACGCATATCTTGATTGGCTCGTGAGCACAATTCTTGCAATAATTCTGAGACCTTTTCGCAAACTCTATCTTGAGTTCGGCACTCAACCGCAAAATACGTAACTCAACGGCAAAATACGTAATTAAATAATTTTCTCTTTACCAATGCGTCATATCTTTATGATCTCAATCTTTTGGATTTCCATAGCTTTAGTTATTGCAATTCATTTGATTTGGAACGGACGAGAACTTTCCGAAATATTTGTTCAGCTAAGTTATAGGCAATTTAAAACTATTTTCAAACGAAGCAAATCAATCGATCAATTTAGTAAATCCAACAAAAATACAGTAGAAATTCCCCTATCAAGTTATAAATCAGAAAGCAATTTAGTAAACCAAGTCTTCAATAAATCAAATAAGAAAGACTTCATAAACCAATTTAACGGTCAATTTATAACTAATACTTCATCCTTATTACAAATAAAATAAATGAAAAATGAAATTGAGATTAGAGAAAAAATAATAAAAAAACTAAAAGAGCAGAATCAAAAACTCATTGATAAAGAAAAAC
>evgR007211c0g2_i01_Hydra
TTGAACGAATTCACTAATGGTAAAAGTAAGACTTTGGTTGAATTAATCTCCAAATTGCAAAAGGGTCAAAAGGTAACTGCTGTCACTTCATCAATTTTGGATGGTGCTAAGGGTGCTAAGGAAAGTATTGAAAAGTTGAAGAATTTGAAGAAGGAAGATTTGGAAAAGTTATTGACTGCTGAAGGTTTCCAATCTGCTTTGAACAATGTTGCTCAAGCATTCTCTAAGGGTGAAAACATTGCTTCACAATTGAACGAATTAACTAATGGTAGAAATAAGGATCTCGTTCGTTTGGTTACTGCTCTCAGAAATGGTCAAAAGATTGCCACCACTGGTTCCAATGTTGTTGATAAGGTTAAGGCCATCAAGTCATCAGTTGAAAGTTTGAAGGAATTGAATGGTAAGGATCTCTTGAATGAAGAAAATTTAAAGAAGGCTCT
>evgR007216c0g1_i01_Hydra
TCCATGGTATCAAGCTCAAGACGGCTTCATAAAAGATTTAGCAATTGTAAATGGTGAAGCAGATAACACAAGTTTAGCAAGCATTGAATTTGTAACAAGGGATTTTTTTGAGTTTTCCACTGAAGGGGTATTGAAAAAAGACAATTTTATTGGAATAATACCATTGCTGGAAATAACATACTCAGTTTATTTTAAATTCAAGCCTACTTCATGTTCAGTAGGGATGAGAAATGTTATTTATTTTGCTAATGCAGATGCAGGAAATAATATATTGGAGCTCTCGTTCTATGATTTGTGGTATTCTGGTAAC
>evgR007235c0g1_i01_Hydra
AGTAATTATTTTTAACCTGCAATTAAAATGATAAAAGTAGTTTTTTTTCTTCTTTTTCAAAAGTGTCTTCTATGTAAAGCATTGCTGGTCTTTGCTCAAAATAACATAGAGTCCACTCAAGCATTGGTTATGATTTTTGAAGAGCACGATTGGATTTCACCAGAGTTCTATGATGCAGTTAATCAATATTATGAGCTTCAACTTGAAAATACTCTTTTAATTTTATCTTCCCGTAATCACACCGATGCACAAGAAATAATCATAGAACTAATCCTGGGAGATAGATTTAATCAAAGTGTTAACCAGTTTACGGGTATGTTAATTTTAACAAAGCTAAGCGATAGTTCAACTGCTTTTATAATTGATCTTATAGAACCGTATAAAGCTTTCGTTCATTACGAC
>evgR007240c0g1_i01_Hydra
TTGTGTTTGGCCGCTTGTTATATTAAGAAATTCCTCCCAGCCGACACTAAAGTTTTCTTCCCAAATCCTACTTGGGTTAATCACTTTAACATTTTCCGTGCTCAAGGTTTCTCTGATGATCGTATTCCAACCTATAGATATTTTGATAAGAAGACCAATGGTATGGATGTAGCTGGTTGTTTGGAAGATATTAAGAATGCTCCTGCCAAGTCTGTCATTTTGTTGCACGCTTGTGCTCACAACCCAACTGGTGTTGATCCAACAATGGATACTTGGAAGCAAATTTCCGATATCTGTAAGGAAAAGCAACACTATGTCTTGTTCGATTCTGCATATCAAGCTTTCGCCTCTGGTGATGCCAACACTGATGCTCAATCATTCCGTATGTTTGTTAAGGAAGGTCATCAAATCATGCTCTGTCAATCATATGCCAAGAACTTTGGTTTGTACGGTCAACGTATTGGTGCTTTCAATGTTGTCACTGAAAATGCTGAAGAAAAGAAGAACGTCGAAAGTCAATTGGGTATTATTATTCGTACTCAATATTCTAATCCACCATTGCATGGTGCTAGATTGGTCACAACTGTTTTGAACACACCTGAATTGAAGGCTCAATGGGAAAGAGATGTTAAGGAATTAGCTGACAGAATCAAGTTGATGAGATCTAAATTAGTTGAAGAATTGAAGAAGGCTGGCAGCACTCGTGACTGGTCACACATTACCAAGCAAATTGGTATGTTCGCTTTCAGTGGTTTGAACGAACAACAAGTCACCAAATTGAAGGATGAATATCACATTTACATGACCAAGGATGGTAGAATTAGTATTAGTGGTTTGAACACTAACAATGTTGCTACTGTTGCTAAGGCTATGCATGAAGTCACAAAGAACTAAATTAACCCTGTAAAATACTATCTCAAATAATTTATAAAATCACATTTAAATTTAAAAA
>evgR007247c0g1_i01_Hydra
TTTTTTTCGGAATTTTGTTTGGCGGGTGTTTATGTTTTAAGCGGAGGTGTTATTAACTATGAAAAACCATGAAAACTAAACGAAATTATGAAGAATTTCCAGGAAAAAATAAATTTTATTGCAACGGCTTTATAATTATGTCTACCAGTAACTGTACATTTTTAGTAACCGCTGCTCTCATCATAGGAACCTCAACTCTTTTCTTTATATTCGACGCTCCTTTTTTGGCTAGGAAATTGTCTATAGCCTTCCCAATAGTTGGAGGTTGTTTGTTTCTCTTTGTCATGGCAATGCTATTATGGACATCTTTTAGTGATCCAGGTATTATTCCACGTGCTGAGGATGCAGAGGCAATATACATAGAGAAAACAATAGCTCCAGTTATTGTTGCACCAGGACCTCCAACTTATGTACCCCCACTAAGACAACTAAATATATCAATTAAGGGAAAACAGTTTACATTTAAGTACTGCTTCAATTGTAAAATATTTCGTCCACCAAGAACAACACATTGTAGTATATGTGATAATTGTGTTGAATGCTTTGATCATCATTGTAAAATGGTTGGAAATTGTATTGGCAAAAGAAACTATCGATATTTTTATTTGTTTGTAGTATCACTCTCATTATTGTGTGGGTATATCTTTGCTGGTGCTGTTATGCATTTAGCGTTATGTAGTAAAGAGGAGGGTTCACTTTTAGAAGGGTTAGAGAAAAGTCCAACCAGTGCAGTAGTTGTGTTAATTTGTTTTTTTGGAGGTTTGCCAGTGGTTGCACTTGCTGCATTTCATTCGTATATGATAAAGGGTACGATGGCGACACGTCGAGGTGAAAATCCTTATCATTCTGGTTCTGCATTTCACCACTGTTTTAGAATTATATGTGGTCCAACATACCCGAGTTTGATACGGCGCCGTAACATTGTAGTGACTGATCCATTAGAGTATGCTTCATATGGTGCAATACAGATCACACCTAATAATTCTTAAAAGATTTCTAGGTCCACCGACTCGGTGTTAAGAACGCCATAGTGATTCATATTGCCGTTGATCTTTGTTTATTTGTAAACTTCAAATGAGTTTTTACCAAAATCAAGTTTAGACGATTTAGGATGACGTTCCTCGTATCGTAACGCACAAATTATATTCTCTCTATGTACATTATTGAAACGCTACTAAGCACCGTAAACACAATATGTGTTTAAAATGTACACAGCAAACATTATTTAATAATGACTTATATTGTAT
>evgR007252c0g1_i01_Hydra
CAAATGATTCATGGTTGCAACAATAAAATGTTGTTTATTAAAAGTTTATTTGAATCTCAAACATTTTTTAAAGATGATTTGAAAATGACGATGATTGTGTTGGATGATTGGTATAACAATCAAATTGAAACTGACACCTTGGTTGAAACTGACACCCTGGTTGAGAAATATCTTTTAAATAAAACTATAGAATTTTACCGCCTAATAGAATTTAGCAATATAAAAACTCGTGTTATGCAAATAATGAGACAAATAAATGAAGCTAGTCAAGACAATAAAGACTTGAGTTTTAAAAAACAATTGAATGGTTCAGGAAATTCTTCAAAAGAAGTAGAACAAAGTGAAATATTAAGAGAAAATCACAATGATGCAATATCCAAAATATTAGGCATTGCAATGTGTTTATACGATGAAAAGTGTTTTAGTGATGCGAGATATTTATTTCAAGAAGCAAAAATTATGCAAAAAAAAATTTTCAAAGAGAAGCATATTGATGCAATTACTACAAAATACTTTGTTGGAATGTGCTTATATAATATAAAAGATTATGACGGCGCAGAAAAAATATTGTCTAAAGTGCACAGTATACAAAACAAAATTTTAGGAGAGAAG
>evgR007283c0g1_i01_Hydra
CAAATTTGGTTTAGATAAAAATTTGTTTATTTGTAAAGATGACTTTGAAGATGAAAGTTTTGAAGGTTGGTGGAGATCGCATATTGTTGCATATTGTGAAGAGAAAAATACTGTTCAAAGCAATAATAAAACATTATTTAAACAACTCATTGGTTTAACTATGATGTCAATGTCTAAAGTTGACCTATCTTTGCCAACAGTGGAACCTGATGCTCAAAAGCAGATCAAATCATTAATTCTAAATGCTGAACAAAGAAATGCAATTAATGACAAAGAGAGAAAAAAAGTTATTACTGGAGGATATGGTTCAGGAAAATCAATAGTTGGAAAAGAGATTATTAAAAATATTATAGCATCAATGTCAGAAAATCTGTTTACCTTGTATTACATATGTTGTAACCACTTTACATTGTTTGAATGTGAAATGAAAGAGTTTGTTAAAAACTTAGAGAAAACCTCAAATGCTAACGTTGTTTGTAATAACTTGTATCAGTTATGGAAAAACATGTGTGAGAAAGAAAATATTTTGGAAAAAAAT
>evgR007296c0g1_i01_Hydra
TGTTAAAACACATTATCAAAAACAATGAATTAGTTTATCAGATGATGGAAAATCCAAATAATAAGAAATATATTTTAAACATTTGTAAAATAGCTTATGAGTTGTTTATTCAAAATAAGTTTATTTTTTTTAAAGAGGAAGTTTTGCTATTTATTAATGACTTTAACGAAATTGAAGAAAAACTTTTGGGTTTCATTGAAAAGATTGAAACGCATCTAGGTTGTCATTACCAATTTGCACATTTAACAATAATGGAGCTCTGTGCATCACTATATGCATATAATAATTTTACTTGTAAAGAGATTATGGCAAATAAAAAGCTGG
>evgR007303c0g2_i01_Hydra
GACAAGCTAGAAATAATTTAATTCAGAAATATAAGATTACGCATGGTGTCAAGAAAATCGAATTTGTTGTGCCTCAAGTCTACGCGCAAATTTTTTTCCGGAGTGACTTTCCCGGCAATTACTTGAGAATCACTAAGCCATTAATCAAGAACTGTGTTGTAAGGAAAAAGATCTTCACAAACTGCGAAGTCAACCCATGGAATGCTTTATCTTCAAGCATTGTTAACTCTTTTTTAAACGTCGCACAGTTGGCCAGAGATCGGTATCGGCAGAGAATGCTAAAGATGAGTAATTTTTTTATTTATTTTAAAAGTAAATACCCCACAATTAAAAAATAAGTAGTTAAAATATTTTAATTTTTTTATTTGCAGTCATAGGAAGGTTTTCAAG
>evgR007324c0g1_i01_Hydra
TTTTTATCGGTGTTTCATTTGGCCCTTTCTAACTGCTACGAGTTTTATTACTCGTTTACCTTGCAATTTGCATGCAAAGTTTAACAAAGTTAATAATTCGTATATGGTCGGAAATATTATTAAATCGATAAATAACGTTAATTCTTCTGATTGCGCTGTTGCATGCGTCGCTAACACTGAATGCACATTAGTTAATTGGAAACAAGATGAGATGATATGTGAGCTAATAGCAGACTTTAATGCTCAGCAGATTACAAAGAATCTTTGGATTGTTTTACAACCCGACAACACAAATAGTCAAGTTGTTGGTCAAATTTGTG
>evgR007345c0g1_i01_Hydra
AAAACCTTGAAGTTGAGAACTATAATGACTTTTATTTTGCGTCCAATAAAGATACTTCTGGAACTGAGAGTTTCGATCTAGACTTTGAGGTACATATGAGCGATGATAATCAAGATCAAAATGGTGGAGTTCTTGTTACAGAGATTGCCAAATGGCTTTTAAAACACAGAATAACTAGAAATGCAGGTGACGAACTGTTAACTATTTTACACCGGTATGGTCATCATGTTGATTTACCGAAATGCACAAGAACAATTTTAAATACTCCTAGGAATGTTCCAAGTGAGATAAAATGTGGCGGTGAATATGTTTATCTTGGTATTTCGAACGGAATATCAAGAACTTTAGCAGTCAACCAGCATTATAGATTTGAGTCAAATACTATTTCTCTAGTTGTCAATGTAGATGGCCTTCCCTTATTCAAGTCATCAAACACAACAGCATTATTTGTTGGTGATACTAAGCCGAATAATCGTGATGATTTTTTAAATGATTTTTTGCGTGAATATGAGTTATTATTGGCAAATGGTTTTCAGTATAATGGAAAATTACTGGCTATAAAAATTTCAGCTTTTGCGTGTGATGCGCCTGCGAGGCAGTTCTTAAAGGGTATTAAGTCTCAT
>evgR007350c0g1_i01_Hydra
TGGTGGAAAATCAATAGTATTCCCTTGGTGACACTTTATTGCATTAAAGTATATTTTTTAGTCACCTTACACAGTATTTGGTCTTGTACTGTCAGGCAACCACAAGTGTACACCTCTGTTATATCACTTTTGTTTTACCAAATACTTTATAAAGTACCAGGGGCACCCAGTGACTTGGGTTTCAAACTTCAAATTTTAGGCACCCTTGCAAGATTTTCAAAACAGGCTCTTCATTCTCCAATCAAATGGGCCCCTTTTTGCCTGGGCCTCTTGCAGTTGCGGCCCCATAGTTTTGTTACTGGGCGCACCACTGGTGCAATCTTTTGTCCTAGAA
>evgR007401c0g2_i01_Hydra
GATGTAAATATTTATTTATATATAAAAAGAAGACAATGGAAGGTTTACAAAAGTAATATGTCAATGAAAAGAAAAAGAAAGATAAAAAAATTGCATAAAGTCCCTTTGCAATTTAAGGAAAAGTGTTTTGAAGGTGACATTCATTTTGCATGTTCAATTAATGTTGGCGACCCTATTAAATTACCTTTACAGAAATTAACTTACCATACTTCATCTAATTGGATTTCCCCACAAGTTCAAAATTTAGAGGTTACTCATAGCTCTTACAAATCAAAATTGAGGCGAACCAAAGCAGGTGAGGTGAGGCAACCCAAAGAGCTTAAGAAAAAGATAGTTTTACCTTCTCAAGAGTTAATTTTGAACTCAAAATTGGATTATATCCCATTAAACATTTCAGTTACATTAACACCAAAACGTTTAGAAAATTATATTCCAGAAGCAAATAATATTTCTACTCCCATTTTTAATACAACACCAGAAGAATATGCAAAATCATTAAAGAGAATCTCCTCATCTGATATGCATTTTTTATTTCAAGGTGTTACTCCTGTAAATGATAAAAGTCATCTTGTGTTGGTCGAAGACTCCCCGTTATCAGTTCAATGATTGTCTTTCACCATTTGTTATTATTATTATGTTGTTGATTTTTTTAAACTGTGAAACATGTAGTTGCATTTTCTTGGTTAGTTAATTAGTAATTAGTAATTTTATAATTTGATTATAATTACTTTTATTTAATATCTATGTGAAATATTTATATAATAGTATAACTATTTATACAATTGTTTTTGTTATACAGTTTCACTTATTATTTAGTTTCACTATAGCATATATTTAGCACCATGTCAAGTACTACCAATTGATTATTTTTAATTGAAATTTTCTTTATAAATTCCTAATGTATGATCCTTGATTATGATTTTATTTAAAAAAAA
>evgR007433c0g1_i02_Hydra
TTGCTATCAAGTTTGAGGGATTAGCAGAATATCCTATGACCGAAGGTTCAATAGAAAATGCCACAGGATTTACAACACCAGTACCTGTGACAACAACAACAACAATTACAACGACAAGCCCAAGTACGACAGTTAGCACCACCATTCCAACTACAACATCAAAGTTGGAGTCTACGACGTTAGCCTCTACAACGCCTACAACAACTACTACTAGAAATTTACCAACTAGTCCAAAATCAACTATATCTACGACAAATATGACGTTAACTTCTACGACAAAAAGTACCAGTTTACCAATTACCGCACCTACAGATCCTATGACCGAAGGTTCAATAGAAAATGCCACAGGATTTACAACACCAGTACCTGTGACAACAACAACAACAATTACAACGACAA
>evgR007439c0g1_i01_Hydra
TCAGTAATATTATATATCAATATAAGGATCGTGGTCTCTCTATGGGTACTATGATTGCTGGTTGGGATCATACTGGTCCAAACTTGTTCATGGTTGATAATGACGGAACTCGTTTGAAGGGAACTCGTTTCTCTGTTGGTTCAGGATCTTTGTATGCTTATGGTGTATTGGATCAAGGTTATCGTCCAGATTTAACTGAAGAAGAAGCTTGTGAATTAGGTCGTCGTTCCATTTATCATGCCACTCATCGTGATGCTTATTCTGGTGGTATTATTAATGTATACTTGATTCGTTCTGATGGTTGGACTAAGATTTCCAGCACCGATATGAATGAATTGCACTATGGTAAATATGCTCTTGAAAAGGGTATTTTGGATAACGTTCCATTGCTTGATGGAAATTAAAGACAACCTGTATATTCACTATGTATAACAAAC
>evgR007444c0g1_i01_Hydra
AATTCTTAATAGCAAACTGGACACTGACATATCAAATGTTCTTATCCTCAATCATTATCATTACAAATTCAAACAGTACAAAATCCACTGTAATTCGAACAGCACCAAGTTTTTTCTTTCTAACAGAATTATTAATGAATGGAGTAGTTTACCGGATAAGGTGATGTCTGCTTCGTCTGTCTTGGAGTTTAAGAACCGTTTTGCTAAAGTGCTTTGTTATAATTTATACTTTTGTTCGAACCTATTCAACAATTTATATAATTTAAAATTTCAGTACAAACTCTTTGACTGTATACTCCAGTATATTCATATATACCGGACTAT
>evgR007459c0g1_i01_Hydra
TTTTCTTGAAGATCAACTGTGTTTCTGCAGAAATGCTTCACTTCCAAACTTATATTTATCAAGAAAAAATACTTGTGTTACAAGTACATGCTTAAATGACAATAGCACATTTGATTGCACTTTGAGCAATTTTGCATTTGTCTCTCTATCATTAGGCATAATTGAATATAACATAACTTTTAATCCTAGTGTTGCGTTTGTGGATAAGGAAGTTCAAATAAATACAAAAATTTTACGAGGTAATGCTTCAAATGCATTTGTGGTACCACTATCAAATGGCTTACCATTATCTTTTCTGAATTATGGAAACAATACAACAATTAAGACACAGTTTTATTTGCCTGGTATAAAAAAAT
>evgR007474c0g1_i01_Hydra
AATCCATTCACTCCTATCAGAGATGATAAATCACTCTATGATGCCTGTACTCTTTTGGTCAAGGAAAAGTCACACCGTTGTCCAGTCCTTGATAGCCATGGTAAGATGATTAGCATTTTGACTCAAGCTCAAGTTGTTAATTTCCTTGCTCTTCACCAAAAACAAATGGGTGACATTGCCCACCAAAAGGTTGCTGCTGCTGATTTGGGTGTTACTCCAGTCATTACTCTCGAAAAACACAATAGAACTATTGATTGTTTCAAAAAGATGCAACAAATGAAAGTCAGTGGTTTGGCTGTTGTTGATGCTACAGGAATTTTGATTGGCAATTTGTCAGCAAGAGATATTAAGGCTATTAACCCATCAAACCTCTACCATTCTCTCCACCAAAGTGTTCACTCATTCGTCCAACACATCCGAGAACAATCATTCAATGAAAGTCATCCAGCCATTTCTTGCTCTGAAGAAACTGAACTAGGATTTGTTATTGGAAGATTAGCTGCCAACAGAATCCACAGAATGTATGTCTGTGACAAACACCTCCACCCAGTCAAGGTCATTTCCCTCCGAGACATTATTTCCTATGTCATGG
>evgR007484c0g1_i01_Hydra
AATGGTTAAGGAAAAAATATTTATGTTTAAGGAAATAATTATTTATTGGTAATAGATTTTCAACAATAAGGATCTACATTCATAGGTTTTCAAGGATGAGGATTCATGGTCATATATTTTTAACAATGAGGATATGTATTCATAGATTTTCAACAATGAATATGCCTGGTCATAGATTTTCAACAATGAGGATGCTTGGTCATAGATTTTCAACAATGAGGATCTCTATTCATAGGTTTTCAAGTATGAGGATTTGTGGTCATAGATTTTCAACAATGAGGATGTGTATTCATAGATTTTCAACAATGAGTATGCTTGATCAGAGATTTTCAACGATGCGGATCTGTAGTCATAGTTTTTCAACAATGAGGATTCGTGGCCATAGATTTTCAACGATAAGTATGTGTGGTTGTAGAACTTCAACGATGAAGTTTTTTTTTTATAAGAGATTTATCAAAGAGAGATCTTATAATAATGACCCCACAAAAGAGAGATCTTGAAAAATCATTATTTATTTTTCTGTGCCAGGTGGTAATTATTCATAAGTGTTCTAGAGGAGAGATTCAAACCACACTAAACACATTGTTTAACATCTAAAGATAACAGAGCTTTTAAATCAATTCACGTTGATCCATAGTGAGCAGCATGATATTCATTTTAATTGTACTTTTATTTTGTGTGCATAATCAAATGATTTATTCAACATTTAGAGAAGAAACTAGTTGAAACAAAGCTGGATTATTTTATGTTTTTTGTATTAATACGTTGCCAATTAATAATAGCTTTGAGTGGAACAACTGTAACTATGAC
>evgR007489c0g1_i01_Hydra
ATTGGTTGGACATTTATAGGGCTTGTTTTAACTTCAATTTGTGTTGGAAGTTTGATTTCAGCTCTTACAGTTAATGTTCTCACTTCATTAAACTATTTATACCCAAAGGGAGATGTTATTGCAAAGTTAGATTCACCAGAGTATATATGGGCGAGTAGGATATCATCTATAAATCTTTTAAAAGATAAAAATTATTCCAGTCATTCAGAAGTTTTGGATGCATTAAAAGCAGGTGAAGCAAAGTATGCCATCATTGATCAATATTCTGTTGAAGCATACAGAGAGGTTTTGCAACAATATGATATTAAAATTGTGAAAACTATAAATGCTGATAATTCATTTTATG
>evgR007492c0g1_i01_Hydra
TTCGACTTAACTCTAGAGGACAAAATGATTTTTATCTTTACGTCAGGCACTACTGGATTACCAAAAGCAGCTGATATTAGAGGCACTCGGTTTCTTCTTGGAGCTTTAGCTATTGGTCAAAGTATTAATGCAATTTCAGATGATAAAGTATACAACACTTTGCCGCTATATCACAGTAATGGTGGTATTGCAATGGCAGGTTTACCAATTTTTTTCAGTGCAACAATGGTCATTCGTAAAAGGTTTTCAGCAAGTAAGTTTTTTGAAGATTGTCATAAATCTGGAGCAACTGTAGTTAATTATATTGGAGAAATATGTCGATATTTATTGGCAACCCCAATTGTGCCGTTTGAGTCTCAACATAAGGTCAGAGTGGCAGTTGGTAATGGGCTTCGGGCATCTATATGGACGCAATTTTCATCTCGCTTCAACATTCCTGTTATAACTGAATTTTACGGATCTGCGGAAGGAAATGCAGGTATGATAAATACATGCAATCGTGTTGGAGCCGTTGGTTTTGGTTTTGTCTTAATGCCATGGATTTATCCAATCAAGCTTGTTTCGATTAACAAGGATACAGGAGAAATTATTCGTGATCAGAATGGCATGGCTGTTTCTCCACAATGTGGAGAGCCTGG
>evgR007505c0g1_i01_Hydra
GAATTGCAGTAACACGACGATATTCGGGAGATGAATTTAGAAAAAGTACAAATGATCAAGTACAAAATTGTAACGAAGAACGTTTAAAACTAAACTTTGGTGGAGTTGGCGTTGGTCATTCGTCTCAAAGTTGCAAAGAAATATCTTCTACAACTGAAAGTGGTTCAGTAGATGGATATGAAAGAGAAATTGTAACGACTTATGGTTCTAAACCAGAAACCGATCTTTACACATGGTCAAAACAAAAATTTAATACTCCGATGCCAATCAAAGTAGAGCTAAGTTCATTAATTAATTTATTTTCCAAAACCTATATGAAAAATTATCCTAGTATAGATTACAAACGCATTTTAAAGGGGATTGGGCCATTATATTATAATTATTGCGAAACAAAAAAATCTGACCTAAATATTGAAACTTGCAATCCTAAAGCTAAAAAAGGGTGTGGTTGGAGTGATAATTGCATCCCACGACAACAAGTGTGCAATAATAATCTTCACGATGACGGTTTTACATGCTGCACCCCAAAATGTCTCGATATTCCTTGTAAAAACGGAGGTAAATGTAAAGATATTACTGATGTTTCTTGTACATTTTCGTGTACCTGCAAAATTGGCTGGGAAGGATCAACTTGTGAAATAATGGTTGTTACACAAGATGTTTTGAAAGTAGAGATTGAAAGACCAATGGATTTTCACCACCAGCAAAATAATAAAATATTCTTAATGAAACTTGAATTATATTTAAAAAAACTTTTTCCAAAGTATTTTTTTAGTATTGTTGTTTACGACGAACTTCCCGACAATCCGACATATAATTCGGGAAGAGGTACGTACATCAGCCTTACAAAGCACAATCGACATGTTGTTGTTGGATGGTCTAAAAGTAATTCAAAGTTCCCTCCAGTTTATGAACTTTTTGCAATTGAAAGATCCATTACAATAGGAGATCTTTATAGTGCTTCTGATTGCGTGGATAAAGCATATAACGTAGCCAAATCCTTCGGTTATCCTGTTGTATTTGCTTTAGTTGTTAGAATTCCAAACGTTATTAATAGCTTGATTACCGATCAGGTTTCGCCATTTGATTTCCGAACGAAATTTTTGACGAAACGAACATTTTTGTTTAGTAAAAACACTATTTATAAAGAATTTAAACATCTTAGACTTGTCATAATGTTTGGAAATCCTTGACATTATATTGAATTTTAAAGTTTTATTTTTGATTATATTTTTATTTTTAAAAAAAAA
>evgR007509c0g1_i01_Hydra
CACAAATACATATATATATACAATTTTTATTGACTGGCTCTGTACCTGATTATTGGAAAAAATCAAACATTAATCCCATCTTTAAGAAAGACAGCAAACTTAAGGTCTACAGTTATAGACAAGTCTCTCTCACCTCAATACCGTGCAAAGTAATAGAAATAATTTACAAATATACTAGGGCGCATTGTGTGGAAAACAACCTAATTTCCAAACATCAGCATGGTTTTATATGTAAAAAGGGATGTCTAACTAACCTCTTTGAAGCTTGCGATATACTTACCGAGGCTATACACCAAGGTTACAGTGCTAATATAATCTAGTCCGATTTTGCTAAGGCATTCGATTAAGTCCCGCACAAACATCGCCTACATAAACTGAAAGCATACGGCATTTATAAAAAGCTGCTACTTTGGTTCAAAATGTGGCTAAAAGATTTTAAGCAACGTGCTGCCTTAGGAAAACAAGTTTCTGAATGGAAAACGTCACAAGTCGAGTTCCGCATGGTTCTGCCTTAGAGCCAATTATTTTTATTTTATTTATAAACAATCTTCCTGATAATATTGTTCACAAGATAATGCTATACGCTAATGACAGCAAAGTTATTGGGATCATAAAGTTGGCATCCGACATCGCGACAATTCAGACAGACATTGATAACGTGGTCACAGGGTCAAGAACTTGGCTCGTGCAGTTTAATATCGATAAAAATAAATTGATGACTGAAGGTCGTCCTAAAAAACGTTCATCAGATAACTATTCAATGAAAACTGCTTATATATGCAAAATATGCGCCACACAATAGCAACTATAACTATTGAATGTGGCCTAGGAGTTCTCATATCAAACAACCTGAAAGTCAAAGCGCAAGTTAAAAAAGCGACATCTAAAACATACACATCCAATCGCATGTTAGGAATGCTCAAAAATGCTTTCCGCAGTAGAAACTTAACCTTGTGGCGCACACTCCATTCGTATGTATCGTATGTCCAGCCACATCTAAAATTTGCCATTCAAGACATGTCCCTGTATCTCAAAACAGACATTAAGTTAATTGAGGATTTTTTAGACTTTCAGGTACTCAAACTAACCACCTTCAAAGAACGTCGTTTACGTGGAGACATAATTTAACAATTCAAGATTTTCAACAAATATGACGAAGTTCATTTTTTGGTCCCACAGTACCACAGTATAGCGTGAGGTGTACAACCTGCGCTGCCATGACAAGCAGCTTGATCGTTAATTTGTAGGAGGGTGCGAGGAGAGATCAAACTTTTTCACAAACCGTGTTGAGATCCATTGGAGCACTCTTACACAGCATACTGTAAACGCTTCAACTTTAAATGCATTTAAAGATTGTATTTCGATGCGGTAAACTGTTAAACCAGCATAGAGACTCCTCGAAAAGTCATCAAGTTAATAATTCAATATTTAAATACACACATATGTATATATTTTTAAATTATTTTTCGG
>evgR007510c0g1_i01_Hydra
AGAGCTCAATATCCATTCGAACCAATCACTTTCAACCAAGAAACCAACGTTAGAATTCCATACTCTGAAGCTATTGGTTTGCTTAATGAAAAGCACAATTTGGGTCTTGCTAACGATGAAGATATTAACTCATCTCAAGAAAGATTGTTGGGTGAAATTATCAAGGAAAAGTACAATACCGACTTCTTTATTGTTGATAAATTCCCAATGAAGTTGCGTCCATTCTACACAATGCCATGTCCAGAAGATGCTACTTTGTCTAACTCTTATGATATTTATTTGCGTGGTGAAGAAATCTCATCAGGTGCTCAACGTATTCACGATTCTGAC
>evgR007512c0g1_i01_Hydra
CTTTAATGGCGTTGTATACAAATCACTTGTAGTGAAGTCAAGAAAAAGTGGGATGGGAAAATCTTTTTTTGTAGAAAAATGCGGATCACGGCTTGCTTCTCATTTGTATGCTAACTACCAAACAAATTTGAATAACAGTCAAAATAAAAACAACATTGTTATTGTAGCTGTGCATGGAACAGCGGTTAACATTAATGCAATTGTTGAAAGATTACTTCAATTTGAAGAAAAACCTCATTCTCATTTTCCCAGAATATATCATTTTGATATCACACCAATGGTTAGAGAAGGGCTAGATTCATTTCTATTTAACTTGGTTGTTCTGGGATCTTTAAAATCAAGTACAGGAAAACTTTGGCGGCTGAA
>evgR007519c0g2_i01_Hydra
AGAAGTTGATGCAAATAAAATGTGCGATTAATAGATAAACACAAAAAATCCAACGAAAATATAAAGATGGTTCTAAAACGTGGGATTTTTCTTCAAAAAGATGACGATTTCGTAACACACACAACTGAAGAACAGAGAAAGGTGGAGAAGCTAGTGCTAAATAACTATGAAAGTGTTAAGGATAACAATGGGATCAAACTGCTGAGATCAAAGCATATATCCTATCTAAAAAAGGGTCTCCTGCAATTATCCGACTCATATGAGTGCCTTGATGCAAGTCGACCGTGGCTTTGTTATTGGATTCTTCACAGCTTAGAGCTGTTAAATGAACCTATACCTGAAGAAGTATGCCAACAAAAATTTTCAATTTTTAGAGTTGTAGGCTTCCTTGATAAATGCCAAAATCATGATACAGGGGGTTTTGGAGGAGGACCTGGACAATTATCACACTTAGCTCCAACTTATGCAGCTGTTTGTTCTTTGTGTATATTAGGAAAATATTGGCTTGCTGCTTATCATATTATTAATAGAGAAAAACTTCAAAAGTTTCTAACCAACTGTAGAACACCATCTGGTTCATTTATAATGCATCATGATGGGGAGGAAGACATAAGAGGAGTTTATTGTGCAATGGTGGCAGCTCGTTTAACTAATGTTTTTACAAAGGAAATGTTTAAAGGTACTGCAGACTGGTTAGCAAGTTGTCAAACATATGAAGGAGGATTTGGAGGTCTTCCAGGGTTAGAGGCGCACGGTGGATATACTTTTTGTGGATATGCTTCACTAGCTCTACTTGGACAGGAACAAAAAGCTCGTACCAAGCAACTACTTAAATGGGTTGCATTTCGTCAAATGAAGCTCGAGGGAGGATTTCAGGGGCGCACTAACAAATTGGTTGATGGCTGCTACTCTTTTTGGCAAGGGGGGATATTTCCTGTTATTCATAGCGTACTTCAAATGTACAGTGATGAAAACCTTAGCAACACTAACTGGATGTTTGATCAAAGTGCGTTAGAAGAGTACATATTGATTAACTGCCAGTGGCCTCAAGGTGGTTTGATTGACAAACCTGGAAAATCTCGTGATTTCTACCATACTTGTTATTGCCTAAGTGGACTCTCAGTTGCTGAGCACTTTATAGGAGATGAAGTTACTAATACACATATTGAAGGAAGACAAACAAATTTACTACAACCAATTCATCCGATTTTCAATATATGCATTGATAGTGCAAATGAAGCTTTGTTGCATTTTAATAAACTTGCTGTTCCTTCGTTTGGTTGATGGGTTTTTCGTCAGCAGTTACAGTCTAATGTTTAGCGACTGTCGTCTATTGTGTGGTTATCGTCTTTTTATTCTTCAACTACGAGCGATATTGAATTTCTAAAATGATAATGTACGATAAATTTTTGAATTAATCTCGAAAACAATTTTTTTTTCATATTTATATTTTACAGATTTTTTCTTTAATTTTATTATTAAAATGTTAAATTTTAGTTACTATATTTAATAAAATTTTATTAATTTTTATGCAAAAATTTAACGATACCCTTTTTACAAAGAAGAAAAAGAAAAAAAAAA
>evgR007532c0g1_i01_Hydra
AAATCAATAATATAATATGGAATATGCCATAGTTTATTTTGAGAAAGAAAATTCTGTAGCTATAGTACCAGTTAGCTGGATTTCTGTGGATGCCGGATTATGTTATTGGTCACCATATAGGGCAAACAGATTAGAAGCTGCCATCAAAAATAAAGAAATGCCTGAAGAAAATTCGGAATATTATACAAATACTCGAGTTTTAAGACGCTATGAAAGAACCTCAATAACGCTGAATTGGACACATCTTGATTCTGAAAGTCCAGAAATATTTGACAGAAAACGAAAAAACTCTATAGAAAATAATACTGAAGCTACTCAAATGAATTTCACTCCAACGTCATGCCCACCAGTACCATATCCTTCTATTCCGAAACACATGACAGGTTTAGTTGCTGTAACTGACAAAGCTGCTGTCCATCTTAACAAAGTCG
>evgR007537c0g1_i01_Hydra
GATGCGAGCAAAAATATTATGTTTTGCTTTACAAATTCTCGTGGAACATTTTACCGACCTGGAGACACAATGCCTTTATTGAAAAAACTAATCAGTTCAAATAAAGGTGTTACAATCCCGATTTCTTACAAAACTGTATATTGCTTTGACAGTGAAGCATACCGATTTCTAGCAGCTGTTAAGAATAAAACAAAACCAGTAAAGTTTTCAGATGATGATCATAAAAACTTTGCTGAGAGTTGGCAAAAGTCCGTCACTGAAACTAAAAGAATGATACAACATATCACTCAATTAGTACCTCATCCAATAAAAAATACCCTAAGTCTCAATGCTGCCCGAAATCTTGTTCAAAAGCTTACACGGCCTATGGCTGAAATATCTAAAAACATACAAACCAACAGAGAGGTTCTGAAAGATCGCATAAACGAAATTAAGGAGACAGAGGCCAACATTAAAAATCTTGAAAAGGAGAAGACCTTTAAAAAAGTTGAGCTGGAGGTCATTCCGATTGATTATCCAAAGACAGTGTGTACAAATATTGATTGTAAAAAGATAATCCGCCGAGGGGAATTTATACAAGTTAATTACACAACTGTTTGCCATGATCATTGTCAACTTAAAAATGTCGACCATGACACTGTTGGAGATGCGAGGCTCTTGAGTTGCACTGCTATTATAGGTGGAATATGCAAACATTGTCGCCACAGCTACAAGGAACACATGCATATTTACTATGTGAGCAATGAAGTCGAAAAAGAGTTTGTAGACCCGGAAGTGGTTAAAATGCTAAATAGCAAGTTATCGGAAGCCGAAAAAAAACAAAGGCTAATTACTTTCAAAGAAAATGAAATTATTGAACTCGACAAAGAGTTAAAAATTATAGAAGATTCCAGCGTAAAATTTGCATATTTTCTCAAGCACAACGCCATCACTCCCTACAACGATGCAATGTTAAAATACTTGAATCATTTGATTCGAGAAGAAAAAGATAAAGTAAATAACGGAGGAAGTGATAAAACATTGAAATCTTTAGAGATCTATAAAAGTTCTTACAGAAAGAATATTGAAATCTTGAGCAAAGAAATGGAGAAAGGTGATGTTTTTGGCGATATTTTGATTCCTGAAATTTCGTCTTTAGTGGAGAGTCTGTGCAAATTAAAACATTCTGGAAAACATCTTCAAACCTGTTTAAATGTTGAATCATCCACTCAAAAAGAAACTGCAATTTATCAAGAAGTCTCTGCTGCAATAAGTGTAAAAGAAAATAGAAGTCCATTGAGAAAACTTGCCGATTGGGTCATTGCTAAAGTTGTCAATTAAACAGCAAAATATTATTGTATGAAACACTTTTTTACAGATGGATTCAATATTTGACTATTTTATTTTTAATTATTTCACAAAAAAATGTAGGATAGAAATGTAAATTCTATCCTACATTTATTCTATAAAGAGAATATGTTAAATGATCTTGTTAAATTTTTTTTTTACATTTCTTGGTTGTATTACAGTTATATATTGCCAAAAATAAAAATAATAAATAATAAACTTTCACAAAGTTTTGATAAA
>evgR007561c0g1_i01_Hydra
ACAGCTGAGAGTTGGACTCTCAAATCAAAGACGTCCAAGGAATTGATGGACGATGTTGACATTGTTCTCAAAAAGTTGAAAATTGTTGAAGATAGCTGCACTAAAGGAAGTTACACTTCCAATGCTGATCTTGTCAATTCAATCTCTTATTCAGGTGCATGCAAAGCTGTTCACTTTATCAATGATGAAAAGTTTGGTGATAGAGGTGCAAGATGTTTCCTTGACTTTAAGCAAAGTGCCAAACAATTTGGTGACATGGTAGTTAATACTGCTCCTGGAATTCATTATGTTGAGGAAACATTCACCAACACTGGAGAGCACAGTTTGTTTGTCAAAATTCGTGAAACAGGAGGTTCTAGTTGTGCTTCCACTGCGACTGTGGCTGGATTTTATGGTGGATTTGTTGGTGCTGTTTCTGTTGCTATGTCCAGCTCTAGTCAAAATCACCATGCCAGCACTAATTCTGACCAGGTCATTACTAATCGTGTAATTCATGAGAAGAGTGTGGTTGTTTCCATGAATTCATTCACTTTAAGTCCTAGCAAGTTTGTACTTAATGTTGATGCCTTGAATGAGTTGAAGGCCATCTCTTCATTAGAAATGGCAAGAAACTTCCTTCACCAATACGGTTCCCATATCTTGTTGACTGACCATGTAACTGGTGGAACATTCATTCAAACAACTGACATTTCCTTCTCCGCCAATGTCAACAAGAGTAAAGCAGAGACAGCCATTGCCAAAGCCAATGAGAAATCCGGAAGCATTTCTCTATCATATGGTGCTATTGGAGGAACTGGAGGTTTGGCTACCTCAAGTCACCACGAGAATGCTAAATTTTCAGATGCTCTAACCAGTGGTGAAACCTATAATTGCAGTGTCACTTTCAAAACTATGGGTCCTCAAGTATTTGATCTGAGTACTTTCAAAGTTGCTCTTGGTTCTAGCAACATGACTCATTGCTTGATTTCAAATGGTTTCGACCCAACTATTCCCTCCACTTGGTCATATGTTCCAGTTTGGAGTGTAGCAGCCAATAATTCAGAGATTAAGAATAATCACAAGGCAGCTCCAAAATTATTGAGAGATGCGTGGTTGAGCGTTGTACTTGATAAGAAGACAAATCTTGAGAATTTCATGGTACATGTCGTTGAGAAATGTTTGAGTGAACCATTGCCTGACGATTCATTGGATTGGAATTCATCTTGGAGAACTAAACTAGAATTGGG
>evgR007573c0g1_i01_Hydra
CATCAATCGAGTCGCCATTTATTGGGTAGCTGTTAAGCGTTCTGATTTTCGCAAAATGCCAGTGAAATGTTTTTCTCTGATCCCTAGTCAAGCTCTTATCTTGAAATGTGGTTGTTTGTGAAATGTTTAGAGTTAAAAATGAGTTGTTCAGATGAAGATGCCAATCCTTTAAAGAAAAGAAAGAAAGGAGTTGTTCATTCAGACAGATATAAAAAGAGTGTAATTAAGAAGGCTAAAGTGGCAGGAAAAATACATATGAACATGGTTGGAAAAATAATACCAGCAAAGAAACCAGCATCTAATGTTTGCAGTTGCAAAAAAAACTGTTTGAAGATTTACACAACGGATGACATGGTGGAAATATTTACCAAGCTACATGATATGGAAACAAAAACTCAACAAGACATCTACTTGCAGTCACTAATGGAGGTTTCCCCAGTTAACCGTCAAAGATCTAGACCAGGTTCTTCTTCTACAAAACCTAAATCTTTTGGAGTGAAGTACTATATTATGCCCAATGGGGTTCGTACTAAGGTATGTAAAAATGGATTTATGAAATCATACGGATGTAATTCCAAACAGTGCTACAGGCTCTCACATCTGTTACAGCTAAGCAAAACTCCAGTCGATAAAAGAGGCAGAAATGTTTCTGTCAATGCCATTCCGGGTACTGTAATCACTTGTTTGAGAGCTCATATTGAACAATATCCTACAGAAACTGAACACTATACTGTAAGAGATAAAGAATATTTACCAGCTAGTTTGAACGTAAAAACCATGTATGAGATGTTTATAAACAAACATAAATATTTCCTTTTAACTACACTTGGTAGTGATAAAAGGTTGTCATATAAATTTTTCTGGCATTTTTAAGGAAAACTACTCATTAAGTTTTGGTCAACCTGCAAAAAATGCATGCGTTGCCTGTGAGGAACTTAAGTTGAAAATAAAAAGTCC
>evgR007594c0g2_i01_Hydra
CCGCGACTAGCCATATTCTATCATGTGTCGATTCTCTATTAAGTTCAAATAAAGAGTTGATTACAAGTGCAAATGATCAGTTCAATTTTTCTTCTGTGTTTTTATCTCAGTTGGATGAACTCGGAAAGCAGCAAACTGGTAATATTACAATAGCAAAACAGAATTTGGGTTTTACATCATATTCTTTCAAAGGCAATCAAAATCCTGTATTTATTTATTCTACGCTTAATGAAAATAATATCAGTGTAAGTATATCAAATGACGATCAGAGAAAAAATTCTCGAAATATTAAAGCCTACATTATTCTTCCCTCAGAACTGTTTTCTGGAGAAAATGAAACTAAAATTTTTTCGTACGTGTTTCTAGAAAAAGCATTTTTTACAACGAATTTGAATAATATAGACAGTGTAGTCCTTTCTGCAACAATAAATGGAAAAAAAGTGGAAAATTCTACAATATCAATGATGTTTTTGCGTGCCACAAATAGTACAGGTGCAAGAACATGTCGCTTCTATCAAACAACAGAACAAGTTTGGTCAACTGAAGGATGTATTACAACTAACAACAAATCAGAAGAAGTTTTTTGTCAATGTAACCACCTTACTAACTTTGGACTAATATTAGATGTTGACCAATCAGGAAACAATCCTTTGACTGTTCGCGTTATGACATGGATTCGATGCGGC
>evgR007600c0g1_i01_Hydra
AGTAAGCAATTTTAGTTAAGCAAGTTTTTATTTTATTATTCTAATTTTGTTACATAAAGAGTAAAAATTACTTGACTTTACAAAAATATTTAAGAAAAAGACTGAAAAAAAAACTAGCTTAATGCTTAAAACAGGGCCTGCATAATGGCATTTTATGGTGACTTAAATTAATCAAAATAATGAGGATTTTGAATTATGGAGATTTGATTTTGAGAGTTTTTATTGTATATTGTTTTTTACCTGTTTATAGATTTATAAATTGGTTTTTCTCAATTCTTCTTCTTTATTTTATACTAAATTTAGATCTGAATATCTTTAAAAATTATGAAAAGTTCTTCCTAAAGTATTTATTTTTAACCTTAGTTTTCAAATTTCATAATTCAAATTTCATGTCCCATTACCCATACAGTTGCAATACAAAGACCCAAAGAAAAAGATACAAAGACAATACAAATACCCATGACCCATACAGTTGCAATACATATAAAGTCTTGTTTAAAATTGCGTATTTTTTTAATCTTGTTGGTTGGGTGTTGAGTTCTGATGTATAAATATAAATAAATAAAAAGATTGATATGAT
>evgR007619c0g1_i01_Hydra
GAAGGGTATGCTTTAGAAACAAAATTTATTATTGTTTGTTATAATTGGAGTATTTCTGATGGGTTACTAAGCTACATGTTTTATTATGACAATGGTCTATCTTATCGTATGCACACGAGTAGTTCAGAATTCAGTGCCTATCCTGTGTTAAATGCAGCTACTACTTATCAGCCTAGTTTGATCAATTTTATGATGGGGCCTGGTGACCAGAACAATGATTACATGATTAGAATTATCATACAGGTTTTTGGTAAATACCAAGGATTCACAGAATATAATCTTTATATTAAGGTGTATATCAACATTAAAGCAGTCAGTATAACAGAGTTGATGAGCGGAATAAATATAAATGATACACAGAGTGTCGCATTACTAGTTCAAGCTGTTAGTTCTACTGCAAACATAAATTCGTCTAAATTTTTTAACAGTATACTACCATCAGTAAATGCGTTTGGGATGCCTGATAATAGCGCTTTTGAAGAATATAAGCAAAAGTTAACTTTGATGCAAGAACTACGTACTCAAATGATTGACCTACTCAGCAAACTTCCTATTAATGATTTGTATTCATTCAAA
>evgR007623c0g1_i01_Hydra
GTCCGACGTGGCTACTGGGGGAATAAAATTGGTAAACCACATACAGTACCATCAAAGGTGACTGGCAAATGTGGCTCAGTATTGGTGCGTCTTATCCCAGCACCCCGTGGTACCGGCATTGTATCTGCGCCAGTTCCCAAGAAACTGCTTCAGATGGCTGGAATTGAGGATTGCTATACAACAGCGGTTGGTTCCACTGGTACCCTTGGCAACTTCGCCAAAGCGACCTACACCGCCATTGGCAAGACCTATTCATACTTGACACCTGATTTGTGGACTGAGACTGTATTCCAGAAGTCGCCATATCAAGAGTTCACTGACTACTTGTCGAAGAATCATAAGCCAATTGGCGCTCAAAAACAGGAGACCAGAGTATATTAATGTTCTAAGTTGAG
>evgR007625c0g1_i01_Hydra
CTATTAAGGACTTTAGAATAATAAATGGGTTATCTGTTGGAAGAAATAGTGGACATATCAATTTTTTTGACCAAGCTAAAGATTGTCTGTTGGTGCCTAGTAATATATTAACAATATTGCCATGGTTAGAAAAAACATTTTCAGTTTCTTTCAAATTTAAACCCAGATCATATCCAAGCGAATATGGAAGTGTTGTTCATTTGACAACTGGATTAAACAGAGAAGAATATGGGAGTAGGTGTCCTGGTATATGGTATTATATTTCTGGGCTATTAACTATAAGTTTTGCAGTCAGCGGAAATAATGATTTTCAGCTACCTCAAACTCAGCAGCTACCGCTAAATGTGTGGTCAAGCATACATGTATCACAAACTCAAATTAATGAAGTTTATACATATTTTGTGCGTGTGAATGGAACAGTTATTTATAGGATTGTGAATAATAGGCCACAGTCTTTTAGAAATGTAATAGTGTATGCAGCCGATCCTTGGCATCCTGCACAAGATGGTTTTATTAAGGAACTTGGAATCATTAATGGGTATAGTGATTCAGTGGAACAACTCAATAAGCATGAGTTAAACAAAGACAACTTGATTGCAAGATTATCATCATTGGAGAAAGAATACAC
>evgR007627c0g1_i01_Hydra
GTTTAGCTCTAAAAATTTTTATTGTAAATTCTAAATGTTTTATTTTCTTACTTATTATATAGATGATTCTACGAAATTTATGCTAGAAGTTTCTTCATCTTCAAACGAACTAGAAGTTACAAAACTAAAATTTCCCTCATCAGTTGTATGTCGTGGGAGGCCAAAATTAAAAGTAAAAAAAAATGCGATTGGCCTACGCATTCCGCAAAACAATCATTGTGTTCCATTTCAAAAGAAGTCGATTCCTCAAAAAGTTGAAATTTTGCTTGGACGTTTTTTAAAAGGAGGGAGGCCAAAAGATGTCACATCTGCAACAGCTGCAGATGTGCAAGATCTTTACATTGAATCTCATGAACTGCCCTCGTCTTTTACAAACGAAAACGTTGTTATTGAAATATTGAAACCTTTCTTAACTTTTCAAGCATGGCAGAAACTTAAATCTTCTTTGAAAAAACGTCTTAAAATGGCCATTTGGAATTGCCCAACTTGTAATTTAAATGCATCTCGAGAAAGTATCGAGTGTGATAGTTGCTTGGAATGGTTTCATTTCAAATGTGTTCAAATTACTAACAAAATAAAAGGCAATTGGTTTTGCAACAAATGTAAATTGAGCGATGAAAGCCAAAAGTGTTGAGATTTTATTAGTTTATGTTTGAAAGTATTTATAAATTATAAATTAAAGTTCTTTATTTATTTCAACATTAAATGTTTTTCTTTTTCATACAACAAACACCCTTACAGTTATTGGGGTGTTTTTTTCAGCAAGC
>evgR007628c0g1_i01_Hydra
ATGTGAAGAGAAGGCATTTTCTCAACTGCAAAATATAAATTCAATGTCTGTTGAAAAAGAAGCTGCTTCACTTTTACAAAGAAAACATTCAACAGGTTTGATGCCTTCAAATTTGCCTCCAGGATATTATCAGCAATATAAAAACTACAAAGGAAAAAAGTATTTTGTTTCAACTTCTTTACGTGACAGTTGTTTTAAAATTCATGGGATATTATGTTTTGTCAAAAATATCATTCTATCAAATAATAACGAAATTCTTGTTGTTTACAATTCTGTTGCTGAATTTAGTTCCTTTTTTGATTGTCCTCTTCAGTCTTCTATTTTGGAAATATTTG
>evgR007636c0g1_i01_Hydra
ATTTTAAAGATTTTAAACAAATTAAATAATTGTAAAAGTTTCATAAAATATTAAATGGCACTCGATCAGGTGGTTGCTTGTGGCATTCAATGAAGTTGTTGCTATTGACACTCAATCAGGTGGTGGCTTGTAACACCCAATGAAGTGGTGGCTTGTGACATTTCTGTTTGCAATTGTTGCAGTTTTGATTGCTTGGAAAAATTATGATTATGGAAGGTATGAACTAGTAATTGGCATATTGTCTTCTCGAGATCATTTTGACAATAGAAGAGTTATTCGGGATGGACTTAAAGATTTCCTGTCTCCAAATGTCATGCTAAAGTTTATTGTAAGCAAAAATGCATGTGATATTCCTCCAGAAGATCGCATCGATGAATTCAGATGTGAAGCCAGCTCACCCAAAAAAATAGATTTTGAAATGCATACAACATATCATCTACAAAATATTCATGCAAATGTTCAAATTCTTAATTCTGGTCATTATGGACTTGATTTTCAGGTAAATCATCCAATTGTTATACATGAGTTGGGTGTTTACGACCATGACCAAGATGGCATTAAAGGAATGCTTAAAACTGTTTTGTATAACTATGGTAACAAGGATGTAGTCACAGAAGTTGTCTTCACATCATTTAGTGCAGGTGAAAAAGATGGTTTCTATTTATATAAACCTGCTAAAAGATTTGTATTACCAAAGGGTTTTTATGGTTCTCTAGTAGTACAAGGAGTTTCAAATGACGATCCAGTGTTGGGAAATGCAGATTGTACAGTTAATAGCGGCGGAGGTTTGATTACACATAAAAAGATTTACCGGAGTTTATCTGCAACGGACAGCATTTTTCCTGGATTCCAGAATAAATATTTCAATACTTCTTGTTTCATTGCTGGACCTTCTTTCAGATATTCTTCTTATGGAAATGATGATAGTGTTGATAGCGATTTTCAGAGAATCAGACGATACAGACTCTATTCAAATCAAGTTGAAGAGCAGGAGAAACGTTTGCAGAAAGAATATTTGAAACATAATGATATATTATTTGTTGATGTTGTTGATAGTTATCGTATGCTGCCCATGAAAATGCTGCAATTTTATTCTTGGCTTTCAACAAAAGATTTCAGTTTTGTTTTAAAGTCTGATGATGATTGCTTCATTAATGTGACTGCAATACTACAGATGTTGAAATATACTTTAAGAAATGGAAAACATAATGTTGTTTTTGGAAGTTTCAGACAAAATTTTGCTTTGGATCAACACGGCAAGTGGGCTGAGCATGATTTTGTATCGTTAACTTATCCACCATTTCCGTGCGGATCTGGTTATTTGTTATCAAAAGATATTGTTGATTGGATATTCTACAATAAATTTCATCTAAAGATTTATCAAGGAGAAGATGTATCTCTTGGTATATGGCTGTCGGCTATAAATGTTAATCTAATTAACAACAATCGTTTCAGATGCTCTCAAAATGAACATGATGGCGATTCTTTTACATCTCCAGACCACACATTAGATGATCTAATGTATCATTTTAGACCACAGAAAAATGTTTTAGTTTAATTATCTATTGTGAACTTATTTATTCAGTTTAGGTTATATAGTTGTATAAAAACTGTTTAGAATTGTTTTGAAGTTGCCCTAGAAACATTGATTAAAAAAACACTAAATATTACATTTATTTAAAAAAAAAA
>evgR007637c0g1_i02_Hydra
CACAAATATTGCAACATCACACATACGATATTCGTTTGTTGCAGTTGGCTTCATTTTTGTTTTCCAAAGTTCAAGCGAAAGAAATATTTACTGCGACCCAGCAATTAACTTTTGGGGTGTATATTGCTGAGATATCCTCTATTCGATTACGCGAATTAATGGGAGCTACAACTCATCATCTGTTCATTTATGGAGCATTTTGTCAAATTCTTCTTGGTCATAATTTACGTTCTTATGTAATTAACATTGCACTAGTTATTTTAGGCTTGCTAATTTACTTTATGCCAGAAACCCCAAGGTGGCTTTTAGCTCATAATAAACCAAATGAAGCATTTAAAAACCTATTATGGCTTCGCGGTATAAATCAAAATGCAGAAGATGAAGTCAATGATATCAAAACAAATTTAGGAAGTGCTATATTATCATCATATAGTGCCGTTATAGCAGTTTATTGTCAGAAAAATTCAATGTTTAATTTGACAATGTTTTTGAATTCCATTATTGTTGTTTATACCATGGTTTTAAATTTAACATGGGGACCGTTGTCTTGGATACTGGTTGCTGAAATCTTTCCTCAACGCGCTCGAGGACTTATTTGCGGAGTTGTTAAATCTAGTTGGTGGTTACTACTTCCTATTCAATTATTAGCTATTGGTAGTATAGATAAACTGAATTTTGTATGGTTCCGTACCTGTTTAAGCGCACTTTGTATTCCATTTGTCTACTTTTTTGTACCCGAAACAAAAGGTAAAACTTTGGAAGAAATTGAACATTATTATCTAATGTATCGTGGATTTCGTAATTACTTGATGTAATTTTATTAATTTATTTAAAATGTTTGTACTTGATAGTAAAACTTTTATAAATATAATTATAAATAAATGTGAATGTTATCATAGTAAAACCT
>evgR007639c0g1_i01_Hydra
TTAAAAACATAATTTTTTTAAATGATTTCTTCAAGTTTTATTTCTTTAACGTCAATTAAAAGGCACAAGAAAAAACGGGTAAAGAAGCATCAGAAACAAGAAGGACTAGAGGATATGAGGGCTTTGTAAAAGTATAATCTATAAAGAAAAAATATTTTAAAACGTTATACAAGTGAATGTTTAAGTGGCAGATATGATTTGTTGGCACACAAAATTATTGGAGTTACCAAATCTAAAGATTTTCGGAATCCTTCTTTTTATACCAATTATCGCCGCAATAATTTGTAACTTTTTTGTTTTCATTGTTATTATAAGGATGCAGAAAAGAGCAATATCTAATTATCTTTTATTGTCTCTTTCCATATCTGATTTACTTGTTGGTTTGATTCTTGGTCCTTTAACCTTGGCACAAGTTTTTGATATCAACCTTTTATCGAACTGCACTGCGCATTTTATTCGTAGTTATTTCCTCGTTTTACTTGGTGGATCCTCCTTGCTAACCTTAGCCGTGATAGCTTACGATCGATACTTATTGATGACTAAGCTATCAAACTACAACAAACATATGAACAAAAAGAGAGGCTTAACACTTATTGCGTTTTCTTGGATTTTTCCTGGAATGATACCTGTTGCCAAAGTAATTAATATGAAGATATATGTTGTTCTACGTATTGTAAATTTTACCCTTTCAATAGTTGCTCTAGGAACTTTTTATTTTTTAATAATTAAAGAACTTAATAAAAAACGTGAAGAGTTTCGTCGAAACAATCAAATTTATCAATGCGTGACCTGCCATAGCCACGTTGACAAAAATATAAATAGGCAAAGCAAGTTAGCGAAATCATCTGTAAACTATTTTCTAATTAAAAATTTAAAAAAACATAAGTATTACACAAGAGTAGCGAAGTCTGTCACTTTATTATTCACGTGTTATTTTGCATTTATTTTTCCACTAAACATTTGGTTGATTTTAGAGCTTTCGGATTTAAAATATGACGTCAATGCTCATCGAATCTTTTATATGTGTGCAATCTTCCTTATGCAAGCCAACTCTTGCATAAATCCTCTTGTTTACTATTTGAAGCAAAAAGACATAAAAATTGGCATGCGGCATGTTTATAAAACGGTTATACGAAAAAAAAGCATTTCCGATACATGAGACAACACCAGGCAAAAACGCATACGCAAAACTTAAACGCTTACGCAAAGCCCAAAATTGAAACAATAAATTTTCAATTTTATAAAAAGCAATACATTTTAAAATTGTAAAATTGTACATTAAAAGTGTAATAGTAATATTTGATGAAAAATATTGGTCCGTGTTGCCAGTGTTGGCATAGTAACAAATATTGATTATGAACAGAACTTGTTTACTTCACGCACGGCGTGTTTATAACAAAATCTAATAA
>evgR007643c0g1_i01_Hydra
CTTTATCTCAAATAAATTTGAGTTTATCTTTACAATTAATTGCTGAAAAAAATTCATTCCCTACTTTAACACTTGTTCTTATTGGCGAAGAGAATGATTCCACGATATTGGCCATTGAAAAAATGCAAATATTTTACTATTGGAGAAGATCGAACTTTATATCTTTTTTTTCCTTGAGTCCTTTTAACGCACAAGAAATTTTACTGCAATTAATTATAGGAAATTTTAGCAAGCTCTGGAACGACCAACGCAATTTTTATAACATTGAACCTTTGGTTATTATAACGAAGTTAGATAAAAAGTCAACAGATTTTATTATAGATCTCATTTCTCCGTATCAAAATGTTTTTCATTTTGATACAAATTGCAATAACTCTGATTTAACTTATAGTAATAACTCAATACCATATTATTATCTTCCTTTCTATAAAGAGTATTTTGAATATATTGTAGAAGATTTTAACGTGAGTAAC
>evgR007649c0g2_i01_Hydra
CTTTAATTAACCTGTTTTAGTGCAAAAAGTAAAGATGTATAAGCTATCTCTGTTGGCAAAGTTTGGCTGTATTGCTGTAATTATATTCCAAGGTTTCTTGTTAAACAAGTATTTAACTTTGTCTTATAATGTGAAGTGGTGGGCGTGGTATATTGGAGACGCGTTGGTTGTAGGTGTTTGGGCGTTCATTTTATTTTACCTACATAAAAAATTTAAGAATAAATATGTTGACGAAGTTGATGCAGAAGATCCTACAAAATTTCCAGATGAAATAAAATACGCTTTTTGCGCTTGGATGACTTATATTGTATTTTTACTACCAAGAATTGTTATTTTATTCAGCAAAAATCCTCCAACAATCAGAGAAAGTGGTTTTTTTGGACCTAACTTTTTAAAAGTTTCATTGGCGTGTACTCCACTTGTTTTTTTGTTAATGATCTTCGGTTATCACAATGCCAATACAAGTCAAGATCGCAAAATATATATTTCTTCATTAGCTTCTAGTGTAACCCTAGATCTTTTTGATAGCATGGATTTGCTTGAGTTGCTTTTTGAACCCAGGGGAGTTCCTATATTATTTTTAATACTTACTTTAGTTTTTGCAAGCATAAATCTATTCCTTCCTACTATGGCTCTTTACGAGTTGTATGTAAATAAGTTTTCAGGCAGGGTTCCTGGATTTTCGTTCAAAATTTGCTATGTTTTTGGATATATGTTTTTAGTGAATGTACCAAATTTGATTTTGCGATCATTATTATGGCATACCCACGATGCTGATGTTTCTGTTCTTATCATGAAGAATGTCATGTGCATTTTAATTGGTATTAATGAAATTGTTGAATTTATTTTTGAAGACAGACCAAAGCAATGTGATAATTGCAGGTACCATTTCAAAGTAGCAGCATTCAAAAAACATTGTGAAAAATGTGTATCAGCAACAAACAATTCGAGTAAACAAAAATCGGATATTTTATTAGTTGATGATAATTTATAAAAACTTTATATCATTTTATAGACCCTAGTTTATCGTTTTTGAATACAAATATTTTTTTAAAAAAAAA
>evgR007651c0g1_i01_Hydra
AATGCCGAACAAACATTAAGAATGCCGAAGAAGATTAAATATTCGTTTCAAGATTTTTGCATATGCATAGGGGTCCGGATTATGCACCCGAAAAGAATGAGTGAAACCAGAGGTAAAAACCAAATTTAAAAAGGAGCCATGAGGCTGAGACTTAAAATCAACCAGAATTTTGGTTCAAAACTGTTTGTTGCCTTCATATTTTGCTTTTACGTCCTGATTCGATACAATTATTTTTTGACCGAAAATGGTAATGGAAATAACTTTCTCACTAGAAAGCTTTCTGTGTTTGAATCTGATATTAGAGAAAAATGCAAGCCTGCTAAAGGAGCAGTAGACGAGTTCCCTCCAGATTTTATGACTCAGGAGCAACGATACAAAGGTGGTATAATTTTTCATGTTACTGTGACGTTTTACATGTTTGGCCTTTTAGCGGTGGTATGCGATGACTATTTTGTTCTTTCGCTGTATCATATATGCTTGCGACTTAAAATGGATAAAGATGTTGCTGGTGCAACTTTTATGGCAATCGGATCATCTGCCCCAACATTATTTATCTCTATTGTCTCCATTTTTTTTACTGATAGCGCTGGTGATGTTGGACTAGGTACTGTAGTAGGCTCAACAATATTTAATACCCTGTTCATTGTTTCAATATGTGCTATGGTTACAACTATACCAGCACGTGTTTCTCAATGGCCACTTTTAAGAGATTCGTTTGTGTATGTTCTTGGAACAATATCACTTGCAATAACAATTCGAGATAGACGCGTTTATTGGTACGAAGCATGCTTGTTTGTAGTCGTTTACATGTTTTACATTGTCATTATATATTTCAATAAAGAACTTGGAGCTGTGTTTAACAATAATGTCAAACAAGTATGCGGTGAACTTGAAGACGAAGACTCAGATATGTTGACAATTAATTCAAAAACAACTATGGATGATTTACTGAAACCTACAGGAAACGAAGGACCACCAAAACACCACGATGGCTCAAGTAACAACTCAATGATGGACGATCTTGAACAATTCGAGCATAAACAAGAATCTCCACTACAATTTCCAGAGGGATGCATTGCCAAAATTGTTTGGATATTAGGTTTTCCTGTAATGATATTATTTTATATCACAATACCACCGTGCTGCAAGAAGCGCTGGTCAGAATGGTTTTTAGTGACTTTTGCAATGTCCGTAATGTGGATGGGATTTCTCTCTTATATTCTTGTCTGGATGGTTGTTATAATTGGAGATACATTTGATATACCTGATTGTATTATGGGCATGACGTTTTTAGCTGCGGGTTCAAGTATCCCCGATGTAATGGCGTCTGTAATTGTTGCACGTCAAGGAATGGCTGATATGGCCTTGTCAAACGCTATTGGATCAAACGTGTTTGATATGCTTTGTTTAGGATTACCGTGGTTATTAAAAACAACAATCATGGAACCAGGAAGCTTTATCACAATTCAATCTCAAAGTATAATGATATCCACTCTACTTTTAATTGGCTCTATTGTATTTACAATCGTAGCAGTTCATTTGAACAATTGGAGATTAGATTTTAGATTAGGGATTATATTTCTTATCATATACATAGTATTTATATCACTTGCAACTCTGATCGAATTTTTAGCTTGTCCTTGCCAGATTGAAGATTTTCTGCCAGTCTAATTAACTTTACAATATTCTATGAATAAAAATGCAAGTAATTTAATTTTCTAGTAATCTAATTTCCCAGCAAAGGAACTAAAAAACATTTCAAAATAGTTCATTAAGAGTAATTAGAAAACTTGAATTTACAGTGGCATCTACTGAAACACGAATTTACAGCGACATCCACTGAAATATGAATTTAAAGCGGCATCTACTTATTCCAAAAGATTCAAGATAGATGCGTCCTAATAACTATAATTGACCACACAATACATAAAGAGTGATAAAAAATCAGTGATTTTGAGTTAGATACTCAACACTTGGATACTTTAGATACTCAATCAGCTTGGATACTTTAGTTACTCAATCAGTTTGGATACTTTAGGCTCAAATACTTCATCACTTTAGACTATTTATACTATTACCTTTACCTCAAATACCTGATTGTTTGAAATACGTCAAACACCTGATTGTTTGACTTAAGTTCCTAACTAATTTTGATTTTACAATTCAGACTGTATTATATTTTTCAAGTGCCTGCGAATACTTGTTACATGTTTACAAAAATACATCTTTTGTTTAAAACCTTTTGTACTTGTAGTTTTTTTGTCTTAATGTTCCTTTACACCATTTGCATTTTTTATTAATTTTTTTTTGTAATTATACTTATAATTAAG
>evgR007652c0g2_i01_Hydra
TGTATAAATATTTTTTTAAATTAAAATGAAGATTGAATGGTTTTGAAATAAACATTGGAGAATTATAGATGGATTCATTACTTCAGCAAACAAACAGGTCTCTTTTGGAGCTTCATCATCAAATGGACACTCTTGTTAAACATTCTGGAAATGGATTACATGATTTAGAGTCCAGTATTGTGAAAGCGATCAATGAATTGTATGAAAGCTGCACTCGATTAGAAGATTATTTATTCAAAGAACCTTCTCACAGAAAGCAAACATCAAAACTAAAAGTTGACCAGATCATGTACGATGTTCGACATATTGAAGCCAGCTATAATTCATATATTCGCAAACGCAAACAGAGGGAGGAGGATGAAAATAGAAGACAACAGTTGTTGAATACTAAATTCACAACTAATGCAGAAAGTAACGATACAAAAATTGCTTTGGATTATGCTTTACAGCATCACGATGCCTTGCAAAGTGCAGATAAAAATATTGGACAGATGCTCTACACTGGTAGTGCTGCTTTGGATCAAATGAGAAACCAGCGAATGTCATTAAAGGGTGTCCGCAAAAGAATTCTTGATATTGGTAATACACTTGGATTGTCTAATACAGTGATGCGGATGATTGATAAGCGCGGAATGCAGGATAAATGGATTGTTTATGGTTGCATTATTGTCACTTTGGTTATAATGTTTTTCACGTTAAAGTACCTGACATAAAATTTGTAAACATTTTTTATTTAATTGTATAATGAAATATTATTTGTAGAATTTAGAAGTAAAACTAAAATAAAAAAAAAAAA
>evgR007657c0g1_i01_Hydra
AGGAAATATATGTCACAGAAGAAAAACCGATTAAAAGTAGTAGATCATTGAAAATGACCAACGAGACTACTGCCACAATCAGAACAAGGAAGTTCCTGACCAACAGGCTGCTATGCAGAAAACAAATGGTTGTCGATGTTCTTCATCCTGGCCGTTCGAGTGTACCAAAAACTGAAATTTGTGAGAAACTCGCCAAAATGTATAAAACCACTCCTGATGTAGTCTTTTCCTTTGGATTTAGGACAAATTTTGGTGGTGGAAAATCCACTGGTTTTGCTCTGATATACGACACACTCGACTTTGCAAAGAAATTTGAACCAAAATACAGACTGCAAAGAAAGGGAATGTTTGAGAAGAAGAAGCTGTCAAGGAAACAGCGAAAGGAAAGAAAAAACAGAATGAAGAAAGTACGCGGTACCAAGAAAGCCAAGGTTGGAGCTGCTTCTGGCAAGAAGTGAGCTCGACTTGGTTTCAATTTGGCGAAAGAAGGAGAATTTTCATTTGTTTTGTATGTGG
>evgR007663c0g1_i01_Hydra
CCACCACTCAGCAATACTCTGCCACTGGTAAGACAAACGCAGCTGTATTCAATAATGGTGTTACAGGTTGTTCTTCTAATGCTTCTACCACCAATACAACACAACAATTTAACACTTCGAGTCAAAATATCACCAACCACCATTTAAATTACGAGAGTACCTTCATTAGTCAAGGTCAACAAACATGCGAGTTATCAATCCTATCTTCTGACGAGGTATCACCAACCTCTGAAGATTGTAATGCTTTCAATGACTTTTCAATGGGAGAAGAAGACGATGAAGAATACTACAGCGAAGAAGATGAGGAAATGCAAGAGTTTCAACAACCACCAGTCCAAGTTATTCTTCCTGTTGATCCAGATCTCTTCAACGCTGCTGACAAGCTTAGATTACAAGCTTATATGGATTTTAACATTCAAAATCCATTCAATCTTCTCCCAAAGAATGGTTCTCAAGCATCTTTAATTACTGGTCATCTTCCAATCGACGTTGAAGATGACATTTTGTC
>evgR007667c0g1_i01_Hydra
TTTGTACCTAAAAATTTTCACTTTGTGTTTAACCAAATACGCACTAAAAACAAATCGCTCTTGATCAAGGGTTTGCAGGCCTACACTGCAACTCCACGCTTAGCAAAGAAAATATGGCTGAAGAAATTAGACGTCGTGTAATTGAAACTGAAACAGAAGAGGAGGATTCTAAACTTGTGTTTGAAACATCTGAAGATGTTAAAGTTACCAAAACGTTTGAGCAGATGAAGCTCCGCGAAGATCTTATACGTGGAATATATGCTTATGGTTTTGAGCGTCCTTCAGCTATCCAACAGCGATCTATTATACCCATTATAAAAGGAAGAGATGTTATAGCACAAGCCCAATCAGGTACTGGTAAAACAGCAACGTTTTCCATTGCAGCTCTACAACAAATTGATACTCGTCTACGTGAAACACAAGTGTTAGTGTTATCGCCAACTAGAGAATTGGCTTGTCAAATACAAAAGGTTGTTTTAGCATTAGGTGATTACATGAGTATTCAATGTCATGCTTGCATTGGTGGAACAAATATAGGTGAAGACATAAGGAAACTTGATTATGGGCAGCATGTTGTCTCTGGTACACCTGGAAGAGTTTTTGATATGATAAAGAGAAGAAATTTGCGAACAAGGTCTATTAAACTCCTGATTTTGGATGAATCTGATGAAATGTTAAACAAAGGGTTTAAAGAACAAATCTATGATGTTTACAGATTTCTTCCTCCTGCAACCCAGGTTGTTTTAGTAAGTGCTACTTTGCCACATGAAATTCTTGAAATGACTCAAAAGTTTATGACTGATCCTATACGCATTCTTGTCAAGCGTGATGAGTTGACTCTTGAAGGAATCAAACAATTTTTTGTTGCTGTTGAACGTGAGGAGTGGAAGTTTGAAACTCTTTGTGATCTTTATGACACTCTCACGATCACTCAAGCTGTTATATTTTGCAATACAAAAAGAAAAGTAGATTGGCTCACAGAAAAGATGAGAGAATCAAATTTTACCGTTGTTTCCATGCATGGGGATATGCCTCAAAAAGAAAGAGATTCAATTATGAAGGAATTCCGATCTGGATCAAGTCGTGTGTTGATTTCAACTGATGTCTGGTCACGTGGATTAGATGTTCCTCAAGTTTCTCTTGTTATCAATTATGATTTACCGAATAATCGAGAGTTGTATATTCACAGAATCGGTCGATCAGGTAGATATGGGCGTAAGGGTGTTGCGATTAATTTTGTAAAGTCCGATGACATACGTGTTTTACGTGATATAGAGCAGTACTATTCTACACAGATCGATGAAATGCCGATGAATGTTGCGGATTTGATTTAACTTCCACGCAGAGAATATTTCACTATCATATACTCCCAGATACAGAGGATATTTCATTATAGTGTATAGCGGTTCGGTATTTTAACGTCAACATTTTACAAATGAGAATATTACGACGAAGTTTCAAAACGAACTTATGCTTGCTATGCACATTTCATATTGATATAAAGTTCTTCTTGAAGATATTTACATTTTTTAATTAGATGTAAAACATATGAGGTTAAAATAATATTATAAAACATTTTTATATATTTTTTACTTTTTTTTTAATGTAGAAAAATTTGAAGATGCGAGCAGTTTGAGTCTTCTTTCTGCGTTCCTGTTGAGGAACAGTTTTAATTACACAAAAAGCTTTAATTAACTTTAAAAACAGAAAAAGCTTTAATTAACATTAGAAATTTATACATATAAT
>evgR007668c0g1_i01_Hydra
AACGGAGCAAAAACGGTTTGATATATTTGAAAAACTTTGGTATTGCATTTATTTAAGTACTCTAGCACAAAAGTAAATATTAACTATTAGAAAAGCAATTAATGTTACTTAAGAACAATTAAAAAGTATTATGAACGAGTCAAACGGAACCTGCTTTAACGCGTCAAACGATTCTGGTAAAGTTGAATTTATATTTATAGTTATAAATACAATACTGTGTGCGTTGGGGCTTGGAACTAACCTGTTCATTGGAATTGCATTTTACAAAAATCGTGCGGCATTAAAAATTCCTTCTAATCGCTTAGTGTTTAGTCTATTGGTTAGCGACACTTTAACTGGCTTTAACCTTATGGCGCAAATCACAATATCTGTAAAGTTTAAACACGTCATAGTAGTAAGAGTATTGAATGACGTCATAACAGCGTTTTTAACTCACAATGTTGTTTTGCATTTATGTGGCATCACATTGGACAGGTTTATTGCTGTGTATTTTGCATTAAGGTACAAAAAAATTGTAACTTTCAAATTTATTACAAGATACATAAAAGTAGCTTGGCTGTTATCATTTCTCGCTTCAGGAATACAACTTCTATGGGTTTATAAGGACTTAAAAAAGGAAAAATCTTATAATGGAAGAAGTAATGAAGAACTCATAAAAACAACAAATATAGAAAAATGGTACAGTCTCATGATACTTTTGATTTTTTTACTATTACCTATGCTGTTTTTAGGGTTAGCATTAGGTGCTATGTTTTACGAAATACGCAAGATACTGGAAAAAACCAAACATCTTATGATTGAAAAACCATACGATGTAAAAAAAAAGCAACTGAGAGTTATCTGCATGTTTACTTGTATGTATGCTTGTTTTTTTATTTTTGCAATGCCGTATTTTACAATAAGAATGATAGAAGATTTTCAAATATGGTGTGCAAGTCAAATAATACTGCAAATTGTTTATACTTTGAAAAGTACACCCCACTTTGTAAATTCCATTTTGTATTCGTTCTTCAACAAAAAG
>evgR007681c0g1_i01_Hydra
TTTTTTTTGTAATTAATTAGGTGTTTAATTTAATAATCATAATTTTTATTAACAGTTGTTCATCAACAAAGTTTTATTAACAGTTGATTAAATATTGGACCATCTTCTTCTTAAATCAACCAAGTTTCCAAATCTAAAGACCTCTAATCGGGTATGAACTCTAGATATGGCAGTGATTAAAATTCATTTGTCATTGCTAATGTCACCCGACATATGTATAAAAAACAAAACCTTAGCGCATCAAACTGCTATCCATAAAAATTTAGAGATACAGTTGTTGACATACAACAACTTGACATGCATAAAATCCTTGCAAATTTTTACTTGGACCCTCTTGGTTCATTCTCACCCTTCCAGAATGTAATAATTGATGCGCAAAATATACATCATTCTCTGCGTACAAATAAATTCGGTCGTGATCAAGAATGGATGACGGCAGATGTCCTACAACGTCATAAAAAAACGTCACGTATAGACATAAAACAACGTCAAAGCTTACACAATGCTGGATTTTACGACCAACTTATTGCTTACTGGAAAGCACCATGCTATACAAATATATATAAATATGCCAATAATAAGAATAGTATGTGGTCACGTGCTTGCACACTTTCAATCTTGATCCACATTTTACCAGCAGATACTCAATTTGCTGCAAAACTAAATGTGCAACTTTCAGATATAATTTGACGAGGAAATAATCTCGAGGAAATAATCTCAAAATAATCTCCTATCCCCCAGAATTTTTGTAACTTCTCCTACGTCAGTGAACAACTCAGTCGCATCAAGGTGCCACCTGCCAGCCATGACAGCATATCCGGCAGATTACACTATGTAATCCTTGAAACTGTTGAAGCTCTAGTAATATTATTTAATCATTGTACCGAGAAATAGCTTTATGTAAACATTATATCCATTCTGAAAACCAATAACCCAAATTTACTAAGTGACTACCAACCAATTGCTGTATCATCTGCACAATCCAAATTAGTTGAACGAATCTTTGCAAAACTTATTATACCGCATGTTAAAGATATATTATTTTAATTTTGG
>evgR007686c0g1_i01_Hydra
CATATGTCCTGATGAATCATTGATAGAAATTCAAAGTGCCTTTTATGGTTGCTTAAATTCATCTGCTACAATAAATGTTACATATTTGATTCAAAGGTTGTGCTACAAATTGAACAATTGTTTTATTATTTCCAATAATTCTGCGTTTGGATATAATCCATGTCCTGAAAACTTGAAGTACACAGTTGTTAATTATACTTGTTATTTAGAAAATTCAATGTTTATTTCAATGATAGAAGATAAATTCACTAACGTAACTTTGAATGCTTTATATGATGAGTTTGTGGTATCATTTGATATCTTATCACATCCAACCACATCTATAATATGGATTGCTAAAGGAGAGGAAGAAATGATGAATATAGATATTAATGAGAATTTACTTATTTTACTTATTGCCGCATATGTTTCGTGTCATGTTGAAATAATACTTAATACATGGAATTCTATTATAATTTCTAATTATTTTATAAAC
>evgR007693c0g3_i01_Hydra
GAAAGAGAAAAAAAAAGCGCAAAAAAGTGAGAGTGTCAGACGACCATTATATCTGTTAACTAACCCAATTTCACAGCTACCAAATACACAGCTTCCAACCAATGGTAGTATTTTAAGAAATTATCAATATCTTATTTCAAGTAAGAAAAAAAGATTTATCAAAACACAAATTAACATGGCCTATAAGAAGCAAAAAGGAACAAGGAACTTGGTATGCAAAGGTAGAGATTTTTGTGACCTTGAGGCGCAATCAAACTGCTTGATTAGACAAATAGTCAACATTTGGACCAAAGCAGGATTTGAAGCATACATCATTTCAGAATATTCAATTCTCAACAAATTTCAAG
>evgR007697c0g1_i01_Hydra
ACAATCTTTTTAACTTTGGAACCGTATTCTTCAATTTTATATCATTTCTGGGTTATAATCAATTCTGGAAATCCTTCTCTGCAACTCTTTTTTAAGTTGGCTACATGTCCGCAACAAGTTAACGGATGGTTTCTTCCTGCTATGCCAGATGATAACTTATCAGCAGTGTTTCATTTAAATTTTAGTGATAAAAACAATTCTTTTCTATTTTATGCTTGTCCTCGTGATCATCCATATGTTATAACAAATTGTGGTCGTCCTAGTCAAACCTATACTTGTCCGGTTTCTGATTGTCAATCGCCAATTGGTGGAGAAATGCTTAAACTACATCCCACAAATCAAATTGTTTCTAATAGAGTTGATGCGACAAAATCTGGGCATTGTTTGGGTGCAGCTTCTGAGCGTGCAAATTTTTCTATCCCACAACGTGAATTGACACCTTTGTCTGTTACTCTTCAAAATTTATTAGTACATCTTTCATTACTTGTTGCAACATTGACTG
>evgR007703c0g1_i01_Hydra
ATACGATCGTTTCATCACTGTATTTTCTCCAGAAGGAAGACTTTATCAAGTAGAATATGCCTTTAAGGCTGTAAAAACACCAGGTGTTACTACTATTGGTGCCCGTGGTAAAGATTCTTGTTGTATTGTTACTCAAAAGAAAGTTCCTGACAAACTTATGAAAGCTGAATCCGTAACACATGTATTTAAAATCACTGATCGTATTGGATGTGTAATGACTGGTAGAATTGCTGATGCAAGAGTTAATGTACAAAGAGCCAGATATGAAGCTGCTTCTTTCCACTTCAAATTTGGATATGAAATCCCAGTGTCTTATCTTGCCAAGAGAATGGCTGACAATGCTCAATTATTTACTCAATATGCTGGTATTAGACCACAAGGTGTTGTTATGATGTTAATCGGTATTGATGAGGAAACAAACGAACCACAATTGTACAAGATCGATCCTGCAGGTTATTATGCTGGATATTTCGCAACAAGTGCAGGACAAAAGGAAACAGAAGCTCAAAACTTTTTGGAAAAGAAGTTGAAGGATATCAAACAAAATGAAGATGGTACCGAATGTGTTTTATCTACTCAAGAAACTATTCAACTTGCTGTCAATAGTTTGCAAAGCGTCTTAAGCTCCGAACTCAAATCTGCTGATATTGAAGTTTGTATTGTTTCTGGAGAAGATCGTAAATTCAAAGTATTGTCATCTGAGGAAATTGACACTGTTTTGAATGCTCTTGCCGAAAGAGATTAATTTTTCACTTGTACACAACCTTCCTCAATC
>evgR007709c0g1_i01_Hydra
TTTTATTTTCTAAATAAGATAACATTTTACATTTCTTTATAACATTTGGTAGAAGTAATGAGTGAAAATAATATAAGTCCGAAGCCAGCTCCAAAGTCTAGATATGGTCCAAGTTTTGATGAAATAGAACGACGCGATCCTTTACAGTGGCATGCAGCACACAGAGCTTGGGTTTTGGAGCGTATTGTTGAACAACAAATGGTCGTTGTTTACAAAAATCGTTTGTCCGCCTGTTATTTAAAAGAACAAGAAAACTTTCGTGTGAACTGTAAAAAGCAAGTTCTTGATTATTGGAACGCTTTCCAAGCGTGGAAGAAAAAAGATTGGGGTGTACCCGAAGAAGGGTCAGTATATAGGTTCAAAGTACCTTTAGAAGAATATTACAAAGAAGTAGAACAACTTTATGAAAACAAAAAGAATCAAGAAACTTAGAATTAATAACTTATTTTTTATATTGTCACAAAAATAAATGTTGATAATTGTATATCTGTTATTTTTATTATTAACACTTTGTTGACTTGATATTATTAACAAATTCTTATAGAGAGCATGTTTACTTTTAAAAAAAAAAAA
>evgR007710c0g1_i01_Hydra
ATTAGGATAGAACTATGATTTTGATTACAGAAAAAATGTTTTTCATTTTTTAAAGATTTTAAGAAATGGCTGTCAATTACAAAAGTGAAGATTTTGGGGAACCATTTGACTCTGCAAACTTTATAGAGAGAATCTCTGCTAATGTACTTGGGGGTGGTACAAAAGGCGGCCAAGAGGGTTTTGATCCTTTAAAGCTTCGTGATCATTTTCTTGAAACTATAAAAAATCTCCAAGCTCAAAGTGAGTCAACAACTATCCAAATTTCCAAACTAGAAAGAATTTGTTCCGAACATGAGCAGAAGCATAAAGAAAAAGCTTCAGATGTAGAGAAACAATACAAGAATGCAATGTCTACATTTCATGATCTTGATGATCGCATCAACTCTGTAGCTACAAAAGTGGTGCATCTTGGGGATCAATTAGAAAATATTAGCATTCCTCGACAACGTGCTCAAGAGACACAACATGCTATGAAATATTTTGATGAATTTCATTCAGGAAAAGAACTTTCTACTTTATTTAATGATAAGGCAAAACTACACGAGGCTGCTGCTATAATCCATCAGTTACTTATGGTGGCAGATGAGCTTCCTAGAGAAAAATTTGAGCGAGTTCAAAGAAAAATTCAAGAAAAATACTCCACCATTGAATCTGATCTCATGAAAGAATTTATTAATGCATATAAAGTTAAAGATATCGATAAAATGACGCGGAGTGCAAAGACATTACTTCCTTTTAAGAATTATCACTTATGTTATGAAGAACTTATTAAAATGGCAGTTGAGGAATTATTTACATCAGATGATATTTTTGAAGATTTACTATTGGCTTGCAATAAAGTTCAAGATATATCTACTAAGGTTTTTGAGCACAATGCAGAACATGTTGTCTCGAAATTTGTCCAACACGTCTTTGAAAAAAAATTAGCAGGTTACATTTACAAGCAGTTACCTAATAAAATGGAAGAAGAGACATATTTAAAAAAACTATCATATCTTTATGGCAGAACCAAGGAGATCAATAAAAAACTATGCAAACTTAGACTTGGATCAGACTCAACGTTTCTTGAAAGAATTACCAAGCCCTTGTTCTCTAGTTATTTAAAACGCTACATTGAGGACGAGTTACAGTATTTGGAGGAGGAATCAATTTCAACTCTGAATCGTTTCTACAATAGCATTGGTCATGTCAAAAAAGAAATAAGACAAAATTTTATAGCTGAAATACCTGATAGCTTAAAAGAATTGCACAAAAAAATACCTGGTCGTGACAAAAGTATTCTTAAAGAAAATCTTTTATCTCAAGATGTTGCTCTAACGTTACTTCAGGAGAACAAGCTTTCTATACGACGTGCTGAAATGTTATCACCACTATCCGAATTACCTAACAATGTTTTTCGTATCTTTAAATTGTTTTTAAAATACCTTGGAGAAGAACACTTTGAGTATGCTGTTGATGTTCATTTGCAATATTTACCACCGATTGAACCAAAATCACCACCAGACATTATGTTTCTTACTGTGCTGTATCAATCAAATACTGTTTTCCATTTAATTGAGAAACATTTTACAGATTTTATAATAAGAACTGTTGGTTCATCAACTATTTATGGCGAATGTGCTTTGAAAAAGAAACAGCTTGTTGATAGCCTTGAATTTAAATTAAACAGTGGGATGGAGCGAGTTTTGTCAAGTATGGTTAATTACATGAAATTTTTGTTAAATTCTGAGCAGAAAAAGTCTGACTTTCGACCAGATGGAGAAGATGGTGCAATGACCCAAACTTCTGCATGCATATCATGCTGCCGCTATCTAAAAACATTGGTACCAAGTCTTTATGATGCATTGGATGGAAAAAATTTAGAAGTTGTTTTAACCGAATTAGGAACACGTTTTCATAAAGTACTTATGGAACATTTTTTTCAGTTCACTTACACAGATATTGGTGCTATGGTTGCACTTTGTGATGTGAATGAATATAGACATGCTATTAAATTATTTAAGATACCATTACTGGATCGTTTGTTTAATGTTTTATCAGCTTTGGTAAATCTGCTTGTTGTACAACCAGATAATCTCAAAGATGTTGGATCTGAGGAACAATTGGTTGATATTGATAAAAATGTTCTGCAACAATTTATTCAACTGCGTGCGGATTTTAAGAGTTTAAAGATTGGAAAAGTATTTTCTTAGAGAATTGTATTAATTTAAATTTTGTAAATATATATCAATTTTTTTATTTTCACTTGATATATATCAATATATATATAATAATAAATATATATCAATTTTTTATTTTACTTGATGTCACGTACTTTGTTATAGCGGTCATATAAAAACCTTTTTTTC
>evgR007721c0g1_i01_Hydra
TTCTAAATAAGTTACTTTAAAACTAAACATGTATGATAAGCCCTTCAACTTTTCTCAACAATGGTTGGATAGTGACACTGAAATATACGTTGAAGACGAGGCTCTCCACGTGCATAAAGCAGTCCTCATATTGCAATCGCCAGTTTTTAAAAAAATGTTGTGCTCAGAATTTAAAGAGAAGGTTGACCAAAAAATTGCTCTGCCAGGGAAAACAAAAGCCAGCGTTGTTGAAATGCTTCAACAAATTTATCCGAATCCACCAAAGTTTTCAGAAAACAGCAATATCGAACCATTGCTGAAACTTGCCCGAGAATACCAAATCGATATATTAATGAAAAAAATTGAAAAAGAGTTACTTCGAAGACCTACAAATATTGACACTCTGGTGCTAGCTCAAGAATTTTCAATGGAAAAACTAGCAGAAAAATCTATTTGCACAATCTCACGTAATAGTTTTAATACAGCAAAGCATCATCCAAAATATAAACTTCTTTCTTCAGATAATATTGTAAAAATATCCGAAGCCCATATCAACTTTTTATATACAAAGTTTCAAATAAATTGGTCATAAGTTAAAAT
>evgR007725c0g1_i01_Hydra
GCAATAGTATGTGCAACATTAATGGAAATATATATAAGTCGTTACAATATTCTTATAGATTTCATACTTGATCTAATTATATTAGTCTTTTCAACATACGTCTACGCAAACGCCTTGTTAATATACTGCAACCAGAAATCAAATCTTCGCTCTAACCGCGACAAAAGTATATTCAAAGGATTGATACTAGTAAGTATCATATTAACTATAAACGTTGTGTTGTTAGCTATACCGGATATGTTAACGGCGTTTGATATGCTAAAAGTATTTCACATTTACGAAATAGAGTACTATGTTGAAATAAACAATGGAATGTTTTTATGTACGGATGCTTTAGTGTACGTATTTGCTTCTCCCCGAGTAAGATTTGCATTTAAAAAAAAATCGGTGGTTTTATTTAGAAAGCTACGGAATTTTGTAACAAAAGATTTTAAAGGGCTAAGCAAATTGGTTATTGCTACGCAGATAAATCGTCCAGATAACAATATATAATGATGAACGAATTACATCTAAATAATACATTAACAATTTCATAGGTCTGAGGGCATCTTTTAAAAATACTAACGTTGTTTTTTAAAAATGCTAATGTTGTTTTGTAAATATGCT
>evgR007731c0g1_i01_Hydra
TATAAAAATTATTTAATTTTATAAAAACATTGGCATTATTTATCCAGCATGGTGAATTACTAAATTGCACTAAGTAACTATTAGAACAATGATAAAAGCTATGGTACCTCATTTTTGTTTGGTCTTTTTATTTGCAACAGTCGCACCGGAGCAATGCAATCAAGTGTATAACGTAGCAGACATAGGTTCATCAAAATATTTTACTCGAATTGGAAATGTTTTAGAAGGAAACTATACAATACAAATTATACTTTCACTTACTGAATATAACAGTAGCTGTAACAAGATTAGCCTATCTGGTTTGCTTGCATTTTACGCAATTAAGTATGCCATCGATAAAGCTAACAACAGCACACAGTGGGAAACAATAGGTGCTCGACTAGAGGATAACTGTGGTGATTTAGCGGACACTATGGAAAAAGGGATTAATGTTATATCTTCAGTTAATTTAGCAATGAATTCAGTTTGCCATGACAAATATATGAAGTGTAGTGCAAATGAATTAAATTTTGTAAAACCCCCTGCGGCTATTTTAGGGACAGAGATGAGCACTATAACGATTCCTTTAGCTTCGTTAATGTCTTTATATGAAATTCCTATTATAAGCCCTTGGGCTTCGAATCGGTTGCTAAGTGACAGAACAATGTATAGATCATTTTTACGAACTATTCCATCAGACAGTATGCAGGTTCGAGCAATGTTGGAAGTTATGAAAAGATTCAGTTGGAATTATATTATTGCTGTAGGTTCTAATGATTCGTACGGAAAACTAGCTCTCGAGGATCTTCAAATCCAAGCCAAACAAAATAAAATTTGGATTGCTAATACGAGCTATATTACTTACCAAGGAAATAATATTTGTAAGGAAATAAATGAGACCCTTGAAAAAATTATTAACCAACCAAAAGCTACGGTTGTCGTATTGTTTTGTTATTTTGGCGGATTAGGCGAAATATTATTAAAAGAAGCTGAAAAAGCAAACATTTCTCGCATCTGGTTGACAAGCGATGCATGGAATCCTGAAGCACTTAACAAAAGCTACAAACATAAGGTGGCAATAAGCCAATTGCACGGTATTTTATCGATATCTTTTAATTTTTCTACAATTTCTGACCTATCCACTTACGTTGAAAACCAAATAAAAACAAATTATCTGGTAAACAATTTCCTTCAAAATTACCTAAAAAATAATTTTAATTGCGAACCCAAAAGAATTTCAAACGACAAATCAATTTTATTTGGAGAAAACAACTGTACGGTTAAAGTAAGCGAGATAAAGAAGCAACTTATTGAATCAGATATTAGCTCAAGTAATTGGGTAGATGCCGTAACAGTGTTGACAGACGGAATTCATTCCTACATTTATAGCAACTGCTATAAATATGAAAAATACGAAAGTTGTGATAACACGGCTATTAATTACACAAAATTAACACAAATTATAAAAAATTTATCTATTATAAGTAGTACTGGCAATGTGATTACTTTTAATAACTCAAATGATCCTGAACATTCGTTTTATAATATTGATAACTTACAATATGAAAATGGAACATTGAAATATATAACGATAGGAAGTTGGTCAAACAAAAATAGTAGTGCTACTTTGGAAAACAATAACATTAAGTGGCCGTACTGGTTTAATTCTTCCATATCAAACTTTCCGCGATCGAAGTGTTATAGAGACTGTCTAATTGGTGAAAAGATGCAAGAAATTCCTAACACTGGTTGCTGGACCTGTATAAAATGCGACGGTTTTTATAACTACACAAATACAACAAACGCAGTAGAATGTTCTACATGCCGAGACGGCTATCACTCCATAAAAAATAACACTGAATGTGATGAAACCATTATTACGTGGTTGAACTATGATAGCTCAGAAGGGGTAGCAATTATTGTTGTTAGTAGTATTGGTCAAGTTTTTAATTTGATTGGATATGTCTTGCTTTTTAAATTTCGCCGTTTCCTGCAGGAGACTGGATCTTTTTTATTAACTTCAACCCTGACATGCTGTTTGCCTTTTTTAAGTTTTTCCTTTGGATATCTGAACATTTTTGAACCAACTGATACGTTGTGTAGCTTGCGGAATGTAGTTTTTTTTATCCTCCTTATGTCTTATTCGTCTTCGTTGTTAATCAAAACAAAAATGGCAGCTGAATACTTCAAAAGTAAATCATTAAAGTACCAGCTGTTGACCGTACAATTTTTTTCAGTCGCAATTATGATTGTTGTAGAGATGATTCCAGTGTTTATTTTGATTTTTATTCGCAACCAGAACATAAGTGAGGAGACTGTCTCAGGCTTGCCATATTTTTTTAAAAGAAGAAGTTGCACCGTAAGTTTAAATGTTCCAAGTTTGATCTCTAACTTTATTCCATTAATAATAATGATTTTAGCTTCTTGTTGTGCGTTTCGTGAACGTAACATGGAACATCATTTTTATGAGCCAAAGTTCTTTAGTTTCACGTGCATTGCTTTATGTGTTGTTATGGTTGCATATCTAATTACGTTCAATTTTGTACAAAACCAATTAAAATCGATAGTGGTGACATTTACGCTTAGTATTTTTGCTTTAATTTACACGGCTTGTTTAATATTGCCAAAAATATATGTTGGTTATGGACGCTATACGTCATCAAAAAAAGATTTAAAAAAAAAAGTTAAAACAAAAAACGACATACAAAGAAACGACCCATTTACAATTTTCAACCACTAATGATTACCAACAACAATTTACAACCACTAATGGTTACGCATAGAGTGTACACCTTGGTTACACATAGGGTTTAAATTAGATTGAATTTATGAAACGACTTCTAATCTCTTTTTTTTTTTTTCTTTTAGGAATGGATGAATATATTATAATTACTTTTGAAAAAAAAAA
>evgR007732c0g1_i01_Hydra
ATAACAGTACGATAACGTTAACAAGTAACCTAACGTTTTGAAAGGGTTTTTGACATTTATTCAAGATTAGTTGTACTTGTTGTGATTAGAGTGAGTCTTCATACACATCAGTGAAGTATTCAACTACAGATAAGTCTTAAGGAAAAACTGTGTTTGAAGCAATCCTACATGCTGTTATTGCTTTTCGTGAGATAGGAAAAGGTTTTAGTGGAATGGAGAAATATGCAAAATGTATGAATATGTTTAGTATGATGAACAATTCATTTAATTACTTACAAAAAAAGGTTGCGAAGGAATATAAAAATGTATCATTGTATCCATGAAATTGCAGCTTGTAAGGAAATAGAATTTAATAATGAACCAATAAGAAGATGTATTAAAAACGATGGTGCTTGAAAAAAAAAAAGAGGATATGTTTCATTAAATGGTTACATGTCTGGAATCATTGCAGACAAATGTGTTGATGCAGAAGTACTTTCTATGTTTTATCGGGGATGTAAAATGTTGGAGATACGGGGATGTAAAATGTTGGAAATCTTGGAAACTATCTCATAGCTGTTCAATCAACCATTTAAAATCTTCTCGTTCTATGGAAAGTGCTGGTGCGATAAACATTTTCAGTCGTTCTATTGAAAATCATTACATCGTTTATGATAGGAATCTTGGAGATAACAATTCAAGTTCTTATAAAGAAGTTATTGATTCTCAGCCATACAAGTCTTACA
>evgR007743c0g1_i01_Hydra
CTAGACTAAATTAAATTGTTGTTTTAGAAAAGTAATATATTTTAAAATGTTTTTGAAAAAATGTTTATTCATAAATATTAAGTTCTCTTTAAAATCTTGCAACCTTCGTGCATCAACAGATATCAGGCATCCAACAGCAGGCGGTTTGATTTCTTGCTCACAACTCTTTGAATTCATGAAAAAGCCATCATGGTACGAAAGTTATCGAGTTCTTGATGCACAATGGGATTTAAAAAACGAGGATTACAAGGAAAAGCATTTATCAAAAAGAATATTAGGGTCAAAGTACTTTTCATTTGACGAGTGTTGTGATAAAAAATCTTCTTATCCAAGAATGCTTCCTTCTGAACATTATTTCGAAGAGTATGTCAGTAACCTGGGCATAACGAACGATCATCATGTTATTATTTACGATAATCATGAGGTTTTTGCACTCTATTCTGCACCTCGTGTATGGTGGATGTTTAAAGCTTTTGGTCATGACAAGGTTTCTGTTTTGAATGGTGGCTTAAGAAAATGGGTTGGGTCTGGATTTCCCACAGTTACTGGGCCTTATGCAGATGAAGAGTTGGAAGCTTCAAAAAAAAGTTATAAAGCTTCGTTAAACTGCTCGTGGATAAAAGATTTCGAATGGATTCGGAAAAATGCAAATTCAACATCGCCTGTGCAGGTTGTAGATGCACGAAATCGTCAGAGATTTCTAGGCATTAAAAATGATGGAAGTATGGACAAAGAAAGTGGCCATATTCCTTCTTCTTTATCTACTCCTTTCGAAGGATTATTTGATACAAGTAATAATAGTTTACTGGAAAGAGACCTAATAAATAAATACTTTACAGATAATTATGTTGATACACAGAAAGATATTGTTGCTTCTTGTGGTTCGGGAGTTTCAGCTTGTGTTTTATCTTTTGCGCTGCATCATTTACGCGGGGGAATAGTTCCAGTTTATGATGGTTCATGGTCTGAATGGAAAATTCGAGCTACTCCAGACTCTTTAGTTCGAATAATTAAATAAATAAAAATTGTTTTAATATTTATATTGTTATAATTATATATTTTGTATATCTTGTATATCTTGTTTACATTTTTTAAATAATTATTAATTTTTTATTATGCATTACTTCTATTAATATTTAACGTTCA
>evgR007755c0g1_i01_Hydra
AATAAAAGAAAGATGTACAAATTTATCTTATCTCATACAATATCTTCATAAAGGTGATCAAGCCCAGAATAATTTTCGTTTAGATATTATATTTGATCGGCTAACTAAAACTGCTTCGGTTAATTTAATTGTGCATCTTATACGGCGGCTCTCATCTGTGCAAACTGAAACTGCACAATTCCCATCAGAAATAGATGATGACAAAGATACTATAATGAACATTATAATGAACATTAACGATTTGACTAAACTGGTGCGAAAAGAAGTTTCTATATTTGAGGAAGACGGTACACGTGGCAACAATTTGATGTTGGCATATGATTAATTCTTAACTATTCCCCCAACAAGCGTAGAAGCGTAACGGGCTTTCTCTTTATCAGGAATATTATGTACAAAAATATGATCATCTTTAAATGACGAAACCTTAAATAATTTATGTTTTTTGCAGGCCTATTTTAATAATAAAT
>evgR007758c0g2_i01_Hydra
CTCAATTAATTGCTATCTTGCTCATTGCTGTCTTGGCTCTCGTTGTCTACACTCAAGCTAGTTCTCCTGGTTCATGGAAATCCAGATACATTGGTGAACAAGCTGAATCTGTTGATGAATTGGCTGATGAATTGATGGATATGCTTGCTGATAAGGTTGCTTATGAAGTTAAACCACATAATGCTGGAAAATACCATTCTGATGAAATCTATCGTCATAGAAAGATCTACCGTAGTGAAGCTGTTGAAGAATTAGCTGATCAATTGTGGGAAATCCTTGCTGATAAGATTGCTTTTGAAATTAAGGCAAATAATGCTGGAAAGTACAATCCTGATAGACCAGTTAAATAACTTAAAAGCATAAATACCTGATTAGACTACGTTTAGTAATAGTTTTAGTAGTTTTTAGGTCTTGAAAATAAATACTGTATTTTCA
>evgR007770c0g1_i01_Hydra
ACTATAAGGAAGAAGAAAAGACTCGTGAAGTTTATGATTCAGATGGTTTCTTCCATACTGGAGATATTGGTCAATGGACAAAGTCAGGTGCTCTTGCCATCGTTGACAGAAAGAAGAATATTTTCAAATTGGCTCAAGGTGAATATGTTGCTGCTGAAAATATTGAAGGAAAATTGAGTGAAGCCAATTATATTTCTCGTTTGTGGGTCTATGGTGATAGTTACAAGACTTTTTTAATTGGTTTTGTTGCTGTTAATGGATTAAAGTTGTTGGATTGGGCAATTCAAAATCAAAAGATTGCCTCTGTCACTGCTGTCTCACAACTTTCTGAGTCTCAA
>evgR007774c0g1_i01_Hydra
TAGATCACTTATTAACATTAAAAATAGAGGAGGATTAGTTTCTCCTTCATATGATGTTTTTACCATTGTTAATGTTTGTGAAAAAGTTTTTAAAGTATTATTATGTGGGGATGACGCTCAAAACTTAAAAATTTCTTCTGTCAATAAGATTAAAGTTAAATTGATATATACTGTTAATCGAACTCTAGTATCATTAGCTCCATTTAAGTCTTTAAATGAGCATGATGTTGAAAGCTCTAATATATTTGAAGATCTTCACTCTACCCAACTTATTAAAAAAATATGTAGCATATATTTTAATATGAGATTTGCTCGGTATGCACAGGTGTACACTGAAAGCTTAAAAAAGTCTACACTTAGGCTAAGACAACAAACAAATAAACTCATATCGTTTAAAGGCTTGTGAAAATGTTCGTTAAAGTGGCTGATTGTAACCCTTTTTTCATGTTTTTCTTTAAATATTTGTAACTTTAATAAATTTTTACAAGTTTCTAGTTTTGTTTAAAATAGTCGGCTTTTGATGATATCTAGTCTTTTTTCGTTG
>evgR007782c0g1_i01_Hydra
AGGATCTTGAACCAATTGCTAAACAAGGACTCCTCTACCACTTTGGTTATGATAAGTCTGGTCGTCCAATTATTTACTGTCTTTTGGGTAAGGATACTGCTGAAAATAATGAAGAAAATAAGAAGATGAAGTTCAAGCTCTTCGTCTACATGATGGAAAAATGTATCAAGAGAATGCCAGAAGGTGTAAATAATATTGTTTGGTTGGTTGACTTGAAGGACTCATCTCTCTCCATGGGTTTGGTCAAGGAAATGAAGGATACTTTTGTGCAATTGGGTAACTACTACACTGAAAGATTGGCTCGTACCATGGTTCTCAATGCTGGTTGGACTATTTCAATGATTTGGGCTTTCGTCAAGAACTTTTTAGCTAAGGAAACTGTTGAAAAGTACGTCATGTTGAACGGCAAGGATAAGGAAATTGCTGAAACATTTGAAAAGTACATTGAAAA
>evgR007785c0g1_i01_Hydra
AAACCAAGCGAAGATGTAATTACAGAGGTTGTTCATTTCTATGAATGCAATGAATATTCAAGGCTTTGTCCAGGAAAAAAAGAGTTTGTTTCAGTTAAAGTAGATAGTATAAAGCAACATATCCAAAAGCGAGTTCTACTAGTCAATATGAAAGAGCTACATATTGAGTTTAAAAAGAAATATAATTATCTTAAAGTTGGATTTTCAAAATTTTGTGAGCTAAGGCCCAAGTGGTGCATTCCTGTGGGTGGTGCTTCTGGTCTGCATGCAGTTTGTGTTTGCCAGTACCATCAAAATGTAAAGCTTCTTGTACGGAAAATTTCTGGAATTGCTGATTACAAAATCTTATTGAAATTAATGGTTTGTAGCATTGAAAATAGAGATTGTGTGCTGCATAGCTGCGATAAATGTCCTGCTAAAAAGGTTTTGTTAGACTACATTGATACTTTGATTACAGAAAATTAAATTAGTGAAGTTAACTTTTATCAATGGCAAAAATCAATGTACCAATGTACTTTAGTACCAGCAACTCTAC
>evgR007791c0g1_i01_Hydra
TGAAGTAAGGTTTAATTTTAGTGGTCAGCGTAACATTTCTTGCAATGTTTCTACAAGTGTTGGTTTTTTTAATTTATACCATATTTCAGAAGTATACGAGCAAATTAATGGACTCAAAATATACATTCAACCAAGTTCGAAATCTTTTTCCAATTTCATAATCTGTGTTTTTGCTGTGCATAATCAAGGGGGGGAGTTAGACTATATATGGATGTCTGATGAGTTTGAAATACTAGGCACTAATGGTTCAGTTGCATCACTATATAGTAATAAGACAGGAAATATTACTGTTAATGTTTTTGCTACCAATGGATATTCAAATTGTTCTGCATCATTGATTATTGATATTTTAGAATCACTTAGTGTTTTTGATGTTTTTCCTTTGACACAGGTGGCTCAAACAGATCAGGTGGTCACATTTAATGTTTCTTCTAAAGGGGGCTCTAATATAACTTACTTGTTTAATTTTGATAGTGATAATTCTTTTTTTGAAACAAATGAAATGTTATATTATTGGAAATTTAGTTCAACAGGCCAGCATAAAGTTCTAGTGTCAGTATTAATGGATGGTTTGAAATCTCTTTGCTATGAATATTCAACTATTCCTTGTAATAAAACTGTGACGGTTAATGTTTTAAACAGCATTCAAGCACTCAGAATTTTTTATATTCAAACCAATTCAATCCAAATGGAGCTATTATCTAGTAATATTAATGTATCATCTAATTCTACAATATTAGTTCGGTTAATTGGTTACAATGGAAC
>evgR007795c0g1_i01_Hydra
GTGATAAAAGTTTCACTGTTCAAATAGATGGTAGTCAAGATAGCTTGGAGATTGATCAGGAGAAGATAATCTTGCGTTTATTTGCTGTCAAAAAATGCGCAATTAGTTGCTGTAAAAAATATATGGATCCAAAAGGAGCAGGACACCTATTGAAATCAGAGCTTGAACACAAGGGTGTAAAAGTGGAAAGAGTTGTGGGTGAATCTTCTGACGGAGCAGTAAGCATGCATGGGAATATCATGGAGTTCAACGTTTCATTAACGAATTTTCACCAAATGTTGTATATATGTTTAAAAAACTTTCTTGTACAAACTTCTCA
>evgR007809c0g1_i02_Hydra
TTTTTTTTTTATCAATATTCTTAAACACTGTATTTTTCAAAATTTTGTAGTTAATATTTTTACGTTAGGATCGCTGAAAATGATTGTTGAGAGATTTACCAAGAGGTCTACGCCTTTTTCTATTGATTACTTGCTTAATTATAATAGTAATAAACTTCATTTGGAAAATTCATTTTCAAATGAATCTCCGCTGTTCTGGAACAGACTCACTAATCAACAATGCCACGGTCAAGCTCTTCATCAATTCAATGCAATACAATCAAATTACGATGCTTTGTGCAGTAGACGCCCAAGTGAAACAGAAAAGGTTTCATCTCACTCATCCTTTAATCGATTCAATTTATCTTCAACATATTTTGAAACAGAAAAAAACAAGCAAGCCTGTAAAGTGCACTCTTCTTTGTCTTTAATAACCTCATCGCCATCATCATCAACGCCGAATTCTTCATCGCCGTCATCGTTATCCCCTTTGTCTTCCTCCGCAACATCGTCTTCAACTAATAGTTCTAATGAGCGATTTGAAGCAGTCGAATTTCAGTTTGATTTGAGCAAATGCTTTCTACGCAAACATAAAGCAAATCGAAAACCAAGAACGCCTTTTTCGGTAAATCAACTTCTTGCTCTCGAGCAGAAATTTAAGCGAAAGCAATACTTATCTATAACTGAAAGATCAGAGTTAAGTCTCATGTTACGACTCACTGAAACTCAAATAAAAATCTGGTTTCAAAACCGAAGAGCAAAACAAAAAAGAAGCAAAGAAGCGGAACTAGAGGAGACGACTCGTAACCAGTTCCCTCATTCAGCCGTACATTATCAATTTTTGGATGAATCTTCGCACTATTTACCAGTCTCGCAGAGTATACACCGAGTCTCGCCGAGTATACACCATATCAATAAATTAATTGAATACGACGACTTTTATTCGTAATTTTAAAAATCTTCAGTAACTTTTGTAAATATTTCATAACTACCTCGTACTATATTATATTTCATAACACTTAAAAAAAAAAA
>evgR007810c0g1_i02_Hydra
AAAACTCTCCTTACTTTTGTTACCTTAAAGTTATTCATTTTGTTCCAACAAAAGTAAAATCTAGTCAATGTGACCTTCAACCAGGAGTCTCAAAATGTGACCTCCAAGAAGTTTCAGAATGTGACCTTCAACGAGGAGTTTCTGTAATCATTGATTCTACTGCTACTATTTCTACTGATGATTTTGATATTCTAACTCAAATTGTTAAACTACTGTTTCCAAAATTTATTATTGGTAGACTTTTTTTAAGTGCTCAAGTGTACTACGATAAATGCTCAGATGTAACATCAGCATACGAGGAAGGTTTTTATAACATACAAAAAGATTTTCAATCTAGAATTAGTTTATTGCAGCACACTAAATCAGAAGCAAAGTTTAATCTAAACTGCTCTCTTCAAATTTATGAAAATATTGAAAATTTGTTATCTGTAAGTTCAACAGAGAATTTTGGAATAATTTTAACTAAGTTTGACTCTGTTAAAGAATATATTCCTGGGAGTGAGACATATGATGGGATCCACTACACTGCTTTACTTTATAAATCACCGAGAGAACCCAGCTTAAGACAGATCTCTAAAAAATCATCTGATATTCCAATTCTCATTACAGAGTTAATGGACATTATTTGCCCTAAAACTAAAAGTAGCACAGATATAATAAGTAAAATGGAGAATACTGTTTTATCTTCAACAGCTCCACAGCAAGAATTAAAAAAAATAGAATTTGCTAAAAGTACTTCTGACCTTGTGTCTTTATTGAGATATGATACTTTGACATCTACATTTACAGAAAATGAACATTTTACGAAAACGTTTATTCATGAACGCTCTTTTGAAAAAATAATTGCATATTCAATTAGTATACAGTCAGGTTCAGATTTTAATGCAACTAAAACTTTTTTGCTAAATGATTCATTTTCCCCTCTACCTTCATACCTACAACCTCTAATGTCATATCAGCCTTTATTAACCAAATCATTGGCTCAATCGTTATTAATGAGTATTTATGAAAATACACCAAATTCAATAAGTTCTTTACATCAAGGTTATTCAAAACAAATAGAAAGTAAAAACCACACTGATTGGCTTTCTTTAAATACTGATTTCCAAAGCAAATTAGTCTCTCAAACTATGCATAAGCAAACTTTTTATACTAGTGAAATAATTACTAACAATGATAGAAGCATTATTGTTTATGAAGTTGACTTTGGTTTATCGTCTTCACAAAATTATATATCAACTTATTCTCAAATGTTTACATCAACTCAAACACATAAATATGGCTTTTCTTCAATGTATTCACCAAGCTTCTCACAAACTTCTACAGAATATTCTTTTTTTCAAAATACTAAAAATATTGAGACATCTTTATTTACTACTTCAAATACTTTACTGAGCACAGATAGCCCTCAGACGAGAGAATATTTCAAAACTATTTCTAAAAATATTGATAATAACAAAACACCTTTGCTCACACCTTGCACCAAACACCTCACAAGCATTTCATTTCTATCAAGTATATTTGTCCGCACAAATGTCACATCAAGTATAAGTGTCAGCTCAAATGTCACATCAAGTATAAGTGTCAGC
>evgR007812c0g2_i01_Hydra
CTCAACGTGACCAAAATAAAAAAACGTGTTCAAAACTGCTGATGCAAAAAATAAGATGTTCTTGCATCAGATTTAAAATGAGCATCGCGCGTTCATCACGCGGAATATAACTGCATATATAGTGTGAAAATGTAAGATAATAAAGTAATTAGCTTACAGAATAGTATAGGATAGAAAAAGAAAACCGTACTAAATTTCAATCAGTATTTAACGCAATCAATTATTCGAAATAACATTCGGAAAATACAATCAATAAACCTTTATAACAATGGTCTACTTAATCATATTTGCTTCACTTCTGCTACTACATTTTGCTGCAGGAACATTTAAGGTTACAACAGAAGATTGTTACAAGACTGCTGACTGCTTTCTTCTTCCAGACGATTGTACATCAAGTGATAATTGTAATGTTATCATAAAGTACCAGTTTAATTCAGTTGATGAAGTATACTCTGTTACATTATTTTCAAATGAAAAATGGATTGGGTTTGCACAAACACTCAATTTGAACATGGCAGTAAAAATGAAAGATATTAAAGGAGAGATCTGCTTCAACAATAATTCTATTGCAACTTTAGCTAGTTTTTATGCATTGCATAATTCTATGCCCAGCTTTTCTGAACTAGTACATGGACTAAAAGTTGAAGAAACATATATATTGCCGACTGGAGGTGTTGTTTGTAAATATATTCGTAATTTATTTGCTCCAGTAGAAAGCAAAAAGTTTATGTATTCCCTTAATGAGTCAGTAAATCTTGTTTATGCATATGGTGTGAAGTTAAGAGGAGACGGTTACCCTAGTTACCATGGACCGAACAACCACAGCTATCTTTCTGATAAAGTTGATATGAAAAAAATAAGGACTAGAAGTAGCATTTCGTTTGACTTTTCTAAATGTGGAAAAGAATTTAATTGCTTCCAACAACCACCTGGTTGCACTGATACCTGCAATGCTGCTGCAACATTTGTATATGATTCGAGTTCCCAGAAAATTGCTTTACAACTTTGGGCGAGAAAATCAATAAAATGGGTGGCTTTTGGACTAAAGCCAAACAACGATGATAATTTTATGATCAAAGTTCGCGGGGAATATTGCATACGGTTACCAGTAGTTGGTTTTGGTAATTTTAATGGTAACAATTTGCAGGATGCCGGTACACCGCAATGGGTTTCTGAAGTTGACGGAGTCAAACTGCTATCGTCCGAAGTTATGCATGATGGCTCAATTCTATGTAGATATGAAAGGCCTTTAAAATTGTCTAATGAAAACACGAATTACTTGTATGACATGAACGAACCTTTATACGCTGCACTTGCTTATGGAGATGTTTTATCAGGAAGTTTTCCTGACCAACATACTATGTATATGAAATCAAGTAACCCAATCAATTTTAAGATAGTTCAAGACATTACTTTTGACACGATGTCTATAAAGCTTAAAAAAGCTCATGGTAGTCTAATGGTACTCTCGTGGATTTTCTTTGTAACCTGTGGTATATTCATGTCACGCTACATGAAGCCATTCTTAACTTATAAAATAGCTGGTAAAGACACTTGGTTTAGGATACACCACATCATTATGATTATGGCGTTGCTATGCATGGTTGCCGGCTTTGTTATCATTCTAATTCACTATAAAGGAAAGCTATATCTGAAAAATGACATTCATCATTGGCTTGGATTTACTGCCTTCATGCTTGGCTTGCTGCAGCCAACATTTGCTGCTTTTCGATGTGGCCCAGATCACTCAAAACGCTACTTATTTAACTGGATACACAGATTTATTGGAATGTCAGCATGGTTGATAGCAGTTTCCTCGATAGTTTTTGGGTTGAAAAAAATGTCATTAAACTTCATCCCAGCTATTGCATATGCTTGCATCGTTCTAGTGTTATTTATTGTTCTTGATATCATCCAACTAGTTACTCAATATAAATGCTGTGTTATAAATGCTGCACAAATATTTGACTCTGAATCCTACAGCAGCTTGGGGACTGGGGAAAAGAAAAAGAATCCAGCATGCCATTACATCTTCATGGCTTCTATTTATACCTTATCATGCGCAGTTGCAGTTTTTTATGTGATTATGATAATAAATTCTTAATAAAAGGGAATATTATATTTAAGAATATATATTATATTTAAGAATGTTTTTTTTAGTTTTATATTTTCTGAGTTGAAAATTTTTTAAAAAAACATTTAGAATACGCAACACATTCCTTGTATCTATTATTTTTTTAAATATTTCGAG
>evgR007816c0g1_i01_Hydra
AGAAGACTTTTATAAACCGATTCAAAAAGAAATTCCAAATGACGAGAATAATCCGATTTTGGAAAAGTTGATAGAAATAACTAAGCCTATCAGAACCGAGAGTAACAACGAAGAATTTGACGGTTTCTATGAAGAGTTTAGTTCGACAAAAAATCATAGTGGAGATTCTGGGTACTCCACGAATAACTCAAGATATAGCAACGGATCAACTAACAGCAGATTTTCAATGATTTCTTCAACCTCACCAGAAAACAGTTACGAAATTGATACTTCTGACTCGTCAAAAACTTTTCCTTCTTCGCATTTTAAATTTGATTTAAGTTCGTCGCCATTTTCTTCTACAAACGGCTCAAATTTAAATACAGATATGCAGTTTACTAAAAATTCTGAAGAATTTGAAGAACATATTTATGAAAACATAATTACTTTTAATTTTTAACCATTTTTTCAACTTTTTCCATTAAACCATATACTAATTTATACACTAGCCCTTGTTTTTAC
>evgR007837c0g1_i01_Hydra
GTCAAGGTGCCCTCATTGCTCAATCAATGGTTTTAATGCAATTGAATGAAGTTCAAGAACCAAAGGTTGCTGATTTCCGTAAACATTTGCAAAAGACTTGGTCAACCAGATCTGAAGAAGTTATGTGTAAATTAGGTTCTGTCCTCTCTGCTGGTATTTTAGATGCTGGTGGTAGAAATGCTACAATCTGTTTGCATAGACATGGTAATAATAAGATGAGAAATATTGTTGGTCTTGCACTTTTCACTCAATACTGGTTCTGGTACCCATATTTACATTTCTTGTCACTTTCTCTTGAACCAACTTCAATCATTGGTTTGAATACTGATTTGAAGATGCCATCTCAATTTACTTTCAAGAGTAATCAAAAGCCATCAATGTTTGCCTATCCACCACCACTTAAGACTGAAGAAGAAAAGAAGGAAGTTACTGTTGGTCCAGTCGCTGAATTGTCTCTCTCTGCTAAGGCCAGAGTTAGAGAAGAAAGAAAGAGAAGAAGACCAGAAGGTAGTGCTACACCTCTTCTTACACCACTCACCTCAACTCCAACTCCTCAAGTTGATCCAAAACAACCACAACCTTCTCAAACACAAACTGAAAAGGATATTAAAAAGCAAAAGAAGAAGAAGGAACCAACATTTGAATCACTTACCAATCCTGCCCGTGTAACACCTTCACAACTCAACTCTATTTCATTTGATACAACTGGTAGATACGTTCCAATCAAATCATCCTCATCCTCAGGTATTAATCTGGGTATTAATGTACTAAGTGATACAAGACCTGGTGAAGATGCTGATATTGTTACTCCACAAGCAGGACAGAAATCTGAAGAGAAGGAAGAAAGTGAACCACAACCGCCAGCTACCTTTGATTGGAATTAAACTCTCC
>evgR007842c0g1_i01_Hydra
TAGCCAAAAACTCCCAAAAACTCAACATCCAAAATCATCAAAATGCTGCCTTGTTCTCCACTCTTTTGAATCAACAACAGAGTAAGAAGGTTGATGTTAAGGTCAAGAAACTTTTCATTGGTGGTGAATTCGTAGAAGGAAAGGAAGGAAAAAAGTTCTCTGTTATTAATCCAGCTACCGAAGAGGAAATTTGTCAAATTTCTGAAGCCACCAGTGCTGATGTTGATGATGCTGTTCGTGTGGCTAGAAATGTCTTTGAAAATGTTTGGAGAAACTATCCACCATCTCAACGTGTCAAGTTGATGAACAAGTGGGCTGATTTGATTGAAAGAGATGCTGATTACATTGCAGATTTGGAATGTATTGACAATGGTAAGCCACGTGATATGGTCAGAAATGTTGATATTTATTTGGTTTTGCAATGTATTAGATACTTTGCTGGTTGGGCTGATAAGATTGGCGGAAAGACTATTCCAGTCGATTCTGATTTCTTTGCCTATACTCAAAGAGAACCAATTGGCGTCTGTGGTCAAATTGTTCCATGGAATTTCCCATTGTTGATGGCTACCTGGAAGTGGGCTCCTTGTTTGGCCACTGGTAACGTTTCCATTTTAAAGACTGCTGAACAAACACCATTATCTGCCTTGTATGTTGCCAAACTTGCTAAGGAAGCTGGTTTCCCAGATGGAGTCATTCAAGTCTTGTCTGGTTTTGGTGAAACTGCTGGTTCAGCTTTGACTCACCACATGGGAGTTGATAAAATTTCATTCACTGGTAGCACTGATGTTGGTAAGAGGATTCTAGCTGCCAGTGCCAAGACCAATTTGAAACGTGTCACTTTGGAACTTGGTGGTAAGAGTCCATTAATTGTCATGGGTGATGTCAAGG
>evgR007858c0g2_i01_Hydra
ACGGATTTGTCTGGAAAGCAAAACCTAATTTATGTACTTTTCATTCTGTTGTAACAGGAAAAAAGATGGTATATACAAAATGAATTTTGCTGAAGAAAATGGCAACATTGTTTGATCCGTTGGGTTTTTTAGCACCATATATCATTCGAATCAAAATAATTATGCAAGAATTGTGGACTGATGGAATTGATTGGGATTATACAATTCCAGATAGGATTGCAGCTAATGATCAATGGTTTCAAGAACTAAAAGGTCTCAAAAAAATAAACATTCACAGATGTTTACAAACAACCTCAACAGTTACAGATAGGTCAATCCACGTTTTTACAGATACATCTTGCAAAGCATACGGTGCTGTTGCTTACCAACAATGCTTATATGACACAGGAGAAGTGACCTGTGTGATCATAATGTCAAAAGCCAGAGTAAGTCCATTACAATCAATCAGTATACCAAGACTTGAGTTGCTAGGAGCTGTCCTCGGTTACGACTTGCAGAAAAGATTGTCAAAGCATTGAAGTTAGAGATTAAGGACGTAACTATATAGTGTGATAGTCTTAAT
>evgR007884c0g1_i01_Hydra
TATTTTACATTTTGATTTTCAATTTAAAACAATCGAATAGACTTCCTCAACTTTTAATAAGCAGTATTTCAGTATTGGACATACTGCAAGCATTAGCTAGACTTATACCCAAAATTAACGCATGGCAAAAAAATTTTCCTTTGGTAAATACCTGGGGATGCATTGAAGCTGCAGCCGTAGTTTATTTCGTAACTATTGCAAGGCTCGTTCATATGGTTTTATTGTCATTGGTCCGACTTTTGGCGCTAAATAATTCTGCTTTGTACTTAGATAACTGTAAAAAACTGTGGATTAAAATTGCATTACTATTTTGTTGTTATCTATATGGATTTCTATGGGCTTTTTTACCTGTAATGGGTTGGTCTAGTTATGAAGATGACTTAAATAAAGAACAATGCTCTCTAGACTGGAAACTAATTAATTCTAGTTCTTTAACATATATTGTTGTGGTGACGGTATTTAGCTACGTATTGCCTGGGGCAGTATTACTATGGGCTTTAAATAGAACTCATGTTACTGTCATTCAACATAAAGAAATTATTTTAAGAAAAAAACGTTGCAGTTCACGATTTCCAGATATATTGGATAAGGTTTACTTGAAGATATTTTCCACTTTGACAGTAGGCTACTTTGTCACTTGGACTCCTTATATATTTGTATCTTTTATAACAATGTCGGGTGCCAAAATTCCTTCTGCTTTATTTAATCTTGCAGCATGGTTTGAATCAACTTCTTCAATATTAGGCGCATTTTTTAATTTTTATTTAAACAAACCACTTAAAAGTCATTTGTATCACATTTTAACTGAATGGTTTAAACAAAATAAGGTTTCACCAGAACCAAGTGTGAATATCGAAAAGAATAAATTCTCACCAAGAGTGAATATCGAAAAGAATAAATTATCACCAGAACAGATTGAAAGTATTAAAC
>evgR007890c0g1_i01_Hydra
AAAAATTAAATACTAAGAATGAGAGAAGTTATTTCAATCCACATCGGTCAAGCTGGTGTTCAAGTTGGTAATGCCTGTTGGGAACTCTACTGTTTGGAACATGGTATTCAACCTGATGGTTTGATGCCTTCCGATAAGACTATTGGTGTTGAAGATGATGCTTTCAATACCTTCTTCTCTGAAACTGGTGCTGGCAAGCACGTTCCAAGAGCTGTTTTCCTCGATTTGGAACCAACTGTTGTTGATGAAGTTCGTACTGGTACTTATAGACAACTCTTCCACCCTGAACAATTGATCACTGGTAAGGAAGATGCTGCTAAC
>evgR007894c0g1_i01_Hydra
GGACTGAAATTCTCTATGGTGATTCAATTAATATGCAGAAAAAAAATTATCTAAATAGACTTATGACTAATCATTTTGGAGCAAGTTGTTATACTCAACCACATTTGTGTGACACTGGATTCACTGTCCAACTCAATGTAGAAGATACATTTTTGTCTCATCATGTTGATTTCAATTTTCGTCCAACAAGAGCGTTTTATACAGATAAAAATAAAAATTTAATTGAACCTGAAATAATGCATTTTGGACAATCTTATACAACCTTGTTCTTGTTTGAAGATATCAATATGCCATCAAATGCAGCAGTTTCAGGATTTCAGTTTTTAATTGGCTCATATCAAGGGGCTGGGGAAAATCCCAAAGACTTTATGGCAAAAGCCTATTTGTTAGTGTCTCAACACAAATCATTTGGCTATAATTTTATGACTCAGCTGTATAAAAATAAAAATTTGTATATCTCAAATGAGGATTA
>evgR007919c0g2_i01_Hydra
TGCTATGCTTGCTTTTGAAAATCCAACTAATCCAGATGCTCAAATAAATCAAGTTTTGATGTATTTTGATATTGTATTCACAATCTACTTTACATTTGAGGTTCTAATCAAGGTCATAAATTTTGGATTTTGTTTTCATAAGGGAGCTTATTGTCGAGAAATATGGAATAGTTTTGATTTGTTTATTTTGCTCACAAGTATTATTTCAATAATTTTAAGCTTGGCCCATGTCAAGATTGAACTAAAATTGTTTCCAAATATCCTTCGTCTTCTTCGAGCTTTTCGTAGTTTAAGATCAGTCAATAATATCCCGAAGTTAAAGCTAATTATAAATTGCATTGGTTCTTCAATAAAAAAAACTGTTGGGATAATGGCAGCAATTTCACAATTTACTTTTATATTTTCTGTGATGGGTGTGCAATTATTTCAGGGAAAATATTACTATTGCAACGATGAGTCAAAGATTAAAAAAGATGATTGCCGTGGTACCTTCATTTCATTTGATGAAAATAACCGAACTCTCCTTGTTGAAAAAAATAGGGAGTGGAAAATTGATAACTTTAACTTTAACCATGTTGGATTGGCACTTTTATCATTAATTTCATCATCCAGTGGAAAAGGTTGGCTTCATTTGATGCAGAAAGGAATTGACAGCCAAGATATTCATGAAGGTCCCTTGATAAATAACAATGTTTGGTTATCGTTATTTTTTATTGCCTTCCTAGTAAGCATGTCAATCTTCTTTACAAATCTATTTATTGGAATGGTTGTTTATACATTTAAAAAATGTTCTGAAAAAATTGAAGGAGAATTAGATAGAAATTCTATCAATTGCTTAGATTTTGTGATGAAATCAAAA
>evgR007920c0g1_i01_Hydra
GAATCAATTAGTGGAACAAATGTAGGAGCAAATAATAATGCAAATAATAAAGAAAATAATGGTGGCTTACATGACAGAGTTGGTAATGCAGCTAATCATGGCCTTAATAGCAGAATTAAGAGTGGTGTTAAAAGCGTAACCAATGTTGGAACAATTTGTAGAGAAAATAGTGAAACATATAGTGGAGAAAATAGTAAAGCAAATAGTGGAGGAAATAGCGGAGCAAATAAAGGAGCATTTAACGCAGCAAATAGTGGGGTTAAAAGTGGAGCAAACAGTGGGGCAATTATCGGAGCAAACAGCAAAGAAAAAAGTGGAGCAAATAGCGGAGCCAATACTGGAGCAATTATAGGAGCAAGCAGTGGAGTAAACGGCAAAGCAAATAGTACAGCAAATACCGGAGGAAATAGAAAAGCAAATGGAGGAACAAATGGAGAAACAATTGGAGGAACAAAAGGAAAAGCAAATAGAGGTGCAATTGGAGGTGCAATTGGAGGTGCAAACGGAGGAGCAAATGGCGGATCATTTAATGAGGTCAACAGCGAAGCAAATATAGATGCAAACAGTGGAGCAAACAGTGGAGAAAACAGTGGAGCACATAGTGGAACAATTAGCGGAGCTAATGGCATAGCAAACATCGGAGTAAATAGTGGAGCAAATAGCGGAGCAAATAGTGGAGCAAATGACATAGCAAATGGTGGAGCAAATGGGGGACTAAATGAAGGAGCTATTGGAGGAGCAAACAAAGGAACATATAGTGGAGCAAACGGCGGATCAATTAGTGGAGCAAATAGTGGTGCGAATAGCGGAGCACTTACTGGTGCAAATAGTGAAGCAATTAGTGAAGCAAATAACGGAGCAATTAGTCAAGCAAATGGAGGAGTGAATGGATATCTGAATGGAGGATCAACTGGAGGAAAAATAGGAGGAGAAAATGGTGTACCGAATGGCGGAACAACTATCGGAGCAAATAGTGGAGCTAACAGCGGAGTTTACAGCGGAGCTAACAGCGGAGCTAACAGCGGAGCTAACAGCGGAGCTAACAGCGGAGCTAACAGCG
>evgR007921c0g1_i01_Hydra
TTGTTTCTTGTTTGCTTTAATACTTTTTATTTTTTACTTTTTTTCCAATCTAATAAGTTGTCGTTTAAGCAACTTTCTTATACGACATTATAGTTTCGACATCCTTATATGACATATGTTTTATTTTATCACTTACCCAAGCAATAAAATGGAAAACAAAGTAATATCTGATTTGCTTTTCATGCTATTAGCTTTCATAACATTAATAGCGGCTAAAAAGGACAATGGCAATAGCTTTGTAGATGTTCACTTAACATCTGTAAAAAGAGAAAGGTTTTCAAAAAGAAATACGGATTGTGTCCCATTAGAATTTGACGAACTGATGTCACAATACAAAGCCTGGTCAGGTTATAATTCTCAATATATAGCTGCAACAGCTGAAGAGGCGGTGCTCTTCCCTAACTTGTTATGGAGTAATGAAGACAAAGCAGATTATATGTTTATAATAAACCAACTTAGTAACCTGTATAAAAGAGTGTTAGACCCAAACGATCCTAATTCAGCTTTAAAATTTAAAAACCTTATAGAAAGTAGCTTAAATCTAAGCATTAGCACACATTCAGGGGTGACCACAGGTAATAACAAAAGAATTACTGATGGAATAACGACAATTAAAGTTTGTACTGGAGCAAATTCAGAAAATCCTAAAACAAAACTGGTTAACATGTGTAATCAATGTACTACGACAACAACGCAATCGGAAAATTGGTTTCCTCGGTACATCAACGAAGTTATTTGTGTTGGGAACGATAGAACGGTAGTAAAGGACCCTAAATGCTTTGGAAATGAAGGCGTTTGCCAGCAAAATAAAATAATAATGAACTTTCTAGTAAAGACAGATGACTGTGTACCTCTAACTGTAAACGGAAATACGGTATATGTTGAAAAATGGGAGAAAACGCAGAAATTCATTCGAACATCCTGTGAATGCCTGCTATTACGCAACTCTCCGTATAGAAAATATTTATACTAAGAAACACTGAAAACAAGCTGAAAATCTGTACAGTTATTGATCAAAAAAAGTACTGAGAATATATTAAAAAAAACTCTGATCAGTAATTAATTCTTTTAGATGTTAAAATAAAAAAGTGACTTATTTTTAGTGTTAAAGGTAAATAAAATGCTTGTTTGATTACCAC
>evgR007924c0g1_i01_Hydra
CTATTGTCTAAATAAGTTTTAAAGGTAATGTAATTATAAAATTTCAAGTGTTAGAAGATGGATGATGAACAATCTTTATTGGCAAATATTGAAATTTCACAAGATGAAGAAGTTGTAGTTGAGTCATATCAAAAAGAGGTTCATTTTATATATCCTAAACTTGAGTCAAATGAAAGCCTGCCTTCAGTACAACTTACAACCAGCGATGGTGTCAATTTAGTCATTGAGAGTGCAGACCAGTCAGAAATTGATTTATTAACTAGAGAAATACCTGAAAAAATTCCTTCATCACACGAAACTGTTGCTTGGTTGTCATGTCTGTTTTGCTGTCTTCCATTATCTATTGCTGCTGTTTTTAAATCACGGAAGGTTTCAAAACTTATTAAAAGTGGAGATTTAAAAAGTGCTAGGGTCCAATCAGAAGTGACCAAAAAGTTAGTTTCGCTGGCTTTCATATTCGGAATCATACTGATTGTGTTTTGTGTTGTTGCATACTTGCTGAGAAAGTCATTTTAATTTTTTGATGTTTTTTTGATTTCATGTCATTGTAAAATTTTGTTGTATAGAGATTTTAGTAAAGATTAATATAATAAATATTATAAATAACTATGAAACATTTTGAATATTTTAATATTTTTTTTATTTTTATATTTTATGACAAAAAAAGAAGACTAAGTAAATATATATAATTTTTAACACTAATCAGTCGCCAT
>evgR007928c0g2_i01_Hydra
ATTATTCTGTTGATAAAATACAAAAACAAATCCATGGTACCAAGCATTCATCAGCATTTCCTACAAAAATAAATATCGCAACATGTTTGCAAAAAATCGGAAAAAAAGACGAAGCTTTAGAAATGCTTTATTCAGTTGATGAAATGCAAACTGAACTTTATGGTATCAACAACCCAGCGAGAATCGAAACAAAATCTATAATTGCACACTGTTTGTACAATATGAGAAAGTATAAGGAAGCCTTAGAAGTGTATTATTCTATTCGTAATATACAAACAGAAATACTAGATACAAGTCCTTCATCGACAACACGGACAGAAAGTTACATTCAAAACTGTTTGAGAATGTTAAACACTTGAATCAAATGATTAAACAATCTCATTATAATTTATATATATCTACATCTGCATGTATTGCTAAGCAAAACCTGATAAAAATGCATACCTTTCTTTTAG
>evgR007931c0g1_i01_Hydra
TTTTAGTGCAAAAAGTAAACATTGAGAAAAAAAATACTAAAATGATTTTCACGCTCAAAGAATTAAAGAGCTGGTTATTAGTATCTATATTTGAAGTTTCAATGGCGCTTACAGTTGTTTTAATATTTTCTGTTCTTCTTCTACTTAAAATTGAAAGACACTTAGATGCTACATGGTGGACAATATTTGCTCCTTTGTTTGCATTTGATATACTGACTGCTTACTTTGATATAATTGTTTTTATTAGACTTTATTTGGTCTTAGAAAGAAGTCTAGCAGCTAAAAGACTGATTACAAATGCAGTAATTATACTTTTATTGGTTGTTTTCAAAGTGCTCCTTTGCCAAAAACTTGAAGGAATAAAACATCTTCCTTGTTCTATCATATGTTCACCATTATTTGTTCTTTTATTTTTACTGTTAGTTCGCTCTTGCGTTATTTCAGACGGATTTTGAAGAGTTAATATTTAAAAAAGCATTTGTTTTACGTTCAAATATATTTATGCAGTTTTTTTATTATAGTATAATTGATCATAAGAATCTGTTAATTACATTTACATATATA
>evgR007971c0g1_i01_Hydra
GAACTTCAATATTATGCACAAATAAAACAAAAATTAGTCGAAACTTAAATAAACTGTTTAAAGTTTTAACTTATAAATAAAAATGAATAACTCATTAGCGATATTAATAATTCTAATTTAGTGGTGACATGTCGAAGAATATCAATCTTTTGAAAATATAGCATTGCTTGATTTATAAATTAAAAACTGTTACTTTAATAATAGAATAACGATGGTGTTATTTAAGATTTCTCAGAAAACAAAGTATAGCTTGAGTGTAACATGGATATTTTTAACTACCATGACAACATTTTGCTTGGGAGTTTACGATCCAAACAAGAAGCCACCGCATCCGCATATTATTGTCGTCACTCTAAGAGATATGGGATGGGATGATGTCAGTTTCCATGGAAATCCAGAAATTCCGACTCCTAACATTGACAGAATTGCCAATCACGGCGTTATTTTACAACAATATTACACGACTCCCAACTGTGAAGATTCTGTTTCGTCCTATCGCACTGGAAAGCACCCAATACACTATAAAAAGTCTATGGAAAGTGGAGAGAATGCTCGAGCAACATATGATATATGGAATTATCTTGATATCCAAAGTTATGTTATCAGGGAAGTGGGACTAATCGACTGTAAAGATACCGATATACCCTTTTACCCATGGGATGTTATTCAAAAAGCGGAAAATGTTTTAAAACAACAGGATCAAATATATTCATTATTTCTGCAGATTAATTTTCCGCAATTACACGCAGACTACCCGGAACAAATACCAGATGAATATTTGACTCGTGCCAAAAATATCCCGTTTGAAGCAAGAAAGATATACGCAGGCATGATAATGCAACTTGACAGAGCTGTTGGTCATTTAGTTAGAATATTATATGACACTGGAATCCTCCATGATGCTATACTTATATTCACCACATTAACCGGAGGAACAAAGGGAAACAACTTTTATACTTGGCCATCAACATATCCTTTTAGAGGTGGAAATGGTACTCTTTGGGAAGGAGGATCCAGAGCTTTAGGATTTGTTTACAGTAATCGAATAGCCAGAAGAGGCCGAGTTTCATATGGTCTTGTACATATAAATGACTGGCTGCCAACAATATTTCGTCTAGCAGGGGGTAACCCTGATCATGTCGTTGAAAGCGATGGCATGGATGTTTGGGATTCAATAAATAACGATGATTATTCGCCGAGAATGGAAATATTGTACGGCATCAACGGAAATAAAGCTGCCGTACGCGTTGGAGATTTCAAACTCATAACCGGAGAAAGTTATGTTCCACTCTCTCCCAGACCTTTAGCAAAAGGAGAACTAGACAATTTAATTAAACCAATAACGGTTTGGGATAGCGAGTTAGAGTGTGCTAACTATCCTGATGCTATTGGTTGTTGGCCAGATATTGAACCTTGTTTATTCAATGTTCGTTTTGACCCTTGTGAACGAAACAATGTTGCATATTACATGCCACAAACTGTTGAAGCATTAACATATACACTAAACAAATATAATACTTCTGCTGTATACAGAGACAAAAGATCCTTGGAAGAATGGGACAACACATATAAAGATTTATCATTAAAATTAACATTAATTAAACTTGAAAAAAGCCGAAAGACATTAGACCGTAACAAAATAGGTGTTTAATACAATTGGGAAAGGTAGGCGCCCCACGCCATCCATAAACGTTTAACGCCATTTATTGGAGGTCAAATCATTGAGTTATCTGGATTGGATCTAAATTGAAAAACACCAAATATTTATTTCTTCAAAAACGAAAAACTTGATAACAAAAACAACGTGTTGTTTTTAGAGGAAATTTGTTTGGCTTGAAAATTAAATAACTATGGATGAATAAATATAAATAAATGAATATAAATATATATAAATGAAATCTAACAACTATAAATAAAAACGTTTTAAAAAAGTATAAGTTTACGGGTTTTTTTCACTAACCTGGGTTACAATGGTCAC
>evgR007976c0g1_i01_Hydra
ATTTTATGGATTATTCTTACGGACCAAGAGAAACATATCACTAGTGCTATAAAATTAAGGATTTTGTTGAGAAAATCAAAAGACTGAAACGAATCAGAACGAAAGAAAACAATTATATTTTTAAAATGTTTACGCTATTTGCTTGCTCAGTGCTGTTGATTTCAAATATCTATGGCGACTCAATTGCCAAACGTTCTGGTGATCAAATCAAAGATTGTTTGGAAGCACATAACGATTTTAGAGCTAGGCATGAAAACACGCCACCTTTAGTTTTGGATAAATCCCTTACTAAGAGTGCACAAAAATATGCTCAGTACTTACTTGATAACAATGTGTTTGAACACAGTAAAGGTAAATACGGAGAAAACTTATTCAGTGCTTCTGGAGACTTTGAAAACGTTTGCAAATCGGCTAGTGAAGGGTGGTACAATGAAATTGATCATTACGATTATTCCAATCCCGGTTTTGATAATGAGACGGGACATTTTACCCAGCTGATATGGAAAGCGACCAAGAGAGTTGGGTTTGGTATCGCAACTGATGGTGAAAATACATATGTAGTTGCGCAATACTTGCCAGCTGGAAACATGAATACAAAATCCGAGTTTAAGAAAAATGTTAAACCGTTACTTTAGTCAAATTTGCTGCAATAATGCGGAACTGCTAGATGTAATCATCAAATTTTCAATGTAACAACAAATTTATAATAAAACATAAAACATGTGTAACATGAAAACATAAAATGTAATGATTGATTTTACAAATTGAAATCGAATATATTTTTTTGAAAAAAAAAAA
>evgR007978c0g1_i01_Hydra
ATTATATTTTCGTGTGAATAATTTAAAAATCATGAAAAATCATTTTAGATGACGGAAAATCAGAACGAAAACATAGAATGTTGTGAAAGTTCAAAGACTGCAATATGTAAACAATGTGAACTTAACTTGAGCAGTCGAGTTAAAGCTAATATGATGATTATATCTGAAAACATTAGACTAAAGATGGCAGAAAAAAAACTTGAAAATAATCTTAAAGAAAACGCTTTATATACTATAAATTTAAAAGAAACCTTGGCAACAACAGTTTCTAAAAATAACGAGCTTGAAAATACTATTCAAGCTTTGGAGCAGTTTATATTTTCTCAACAAGTTAACAAACCTATTCTCAGTACGCAGTCCTCTGTAAAAAATGAATCTCCACAGAGCTCTTTTATAAGTAAAACATTATCTTCGAATTTAGATATTTTTCAATCGACAATGTTTTCCTCTAGTGATAGTAGCGATGCCAAATCGAGTCAAGCAGAAAGTCTGAATAATATCAATCAGTTAGATATCAAAGATTTAAATAAAAATAATGGCGACACTAAGCTTATTGATATAATAAAGCAATTATACACACAGCAAATAGTAACGAATAAAGTAATTTGTGACCAAAATTTACAATTAGTTTTCTCTAATTCTAAATACTCATCAATACTAAATGTTTTGTATCAGGTCAGGAAGAATATTAATGACATGCAAAATTTGCTACTTGAAAATTTACATGAAAAAGAACAACACCGTGAAGAGCTAGAAGAGTTACTAACCAGTTTTATAGAAATTGAAGAAGAACTAATAACTAATAGAAACGAAAATCAATTTAAAGATGACAAGATAGAGATACTGAATAAAGTAATTCTTCAAAACTCTATAACAACGAAAAAGATTCAACAAGAAAATGATGTTTTAAAGTTAGAACTTCAGACTGCTACGTTGCGCTACGCTTCAGTCTCGTCTTCAATATACCAAAGCATAGACTCAACAGAAAAATTACGTGAATCTACTAAACATTTACTTGCTACAAAATTGAAGGACCAATCTAACGAATGGTATAATAGCGTTACGGTTTATAATTCTCCGTGTTTAAACAATTTATCAAACACTTTACGAAGCTCTGACCCAGACTGCTATCCTTTAAGCTCAACTTATGCTTCAACAGCATATAAGAAAAATATAAATAAGACTGTCAATAATGTTTTGGGCTCGCAAATTATCTCATTTAATTGGCGGGAAAATCTACAAGCAAATTGTGATTGGTCTTATTTTGCCGAACAAATTGAAATTATAAGCCAGACCGTTCAAAACCTAGAAATTAGAAGTCATACATGTGACTTTAAAAAATGGAACAAAGTTGATATGTTAAGAATTCGTTTAAGATTAAATGTACTTATTGAACTAATGACAAAAAAGAAAAAAAAAGAAAAATTTGATACACAATTCATAACCAAAAGTATTGACATCACTCAACTCATTGAAATCGCAAAAAAAGTATATGAAAACTTTAAATAAAAAAAT
>evgR007981c0g1_i01_Hydra
CCAGAATTGATTATTACTAAAAAACAAGTTGAGATTGAAAGAAAGATGACCAACGATATCGATGCCTTATTGTTGGTGAGTAACGTTGCTATTGCCCCACATAAAAGTTTACTTCAGGTTGGATTCCCTCCTAGAAATCGACCAGGAAATGAAATTCTGCAAACCATGGATGAACTCAGGACTGTTCTATTGGAAAGACGATCACAAAAATTCGTCCAACGTGTATCTGATTTCGATTTGTTACTATTCTTGACTGACTATTTGAGTTTGACATCTGACTTCCCTGGATTGTGTGAGGCTGTCTGTACTCAAGACAACTCTATGGCTTTAGGGTTTGAAGCTCTTATTAATGCCTACTGTGGCATAAACTAATTGTTTACTGTAAAATAAAA
>evgR007982c0g1_i01_Hydra
AACCTGGAAATCAACCCATGATAGGTGTTGATTTTAACTTCCTTATAGATAATGTTATAGCATTTCTAAGTTTTAATATGTCTCAAAAACAACTTATAATACTTAGTACACATGAAAGTACAAAGTTATTTGAATGGAGAGGAACATTTGTTTATATTCTTGATTTACCGTTAAAAAGTTTGTCAAAAATGCTGACTTCTGTTGTTGCTACGGAAACCATTCTATTAGTAAAAGATCATCTCTCTGTACAGCTTTATATATTGAAAAGTAATTTTTCACTTGTTCATGTTCAAACCATACAGGATGTGGGAATTTCAGATGTGCTATTATTCAAAATAAACAAAACATTGTATTTAACAATTACTAAAAATAAAGATAATGATGGGGAAAGTATTCAATGGCTGCAGTTGTACGAATGGAGAAATAATATATTTTTTAAATTACCACTTTCTGGGATTCAAACTTTTCATGGTCAATCAATGGCTTTTGCATACTTCAATGGAGAATATTACGTAGCTGTAGCAAACCAGTACAACCGATTGAGTAATTCATACAATATCAAATCCTATGTCTACAAATGGGACATTAGAAATAGTTACTTTGTAATGTATGGTGAAATTGAAACCTACAAAGCAAAAGAAATTAATTTCTTTGAACATGAAAAAAATTTATTTGTAACATTGGTAAATGATTTAGGAAGCTGCCAGTTGCTGAAGCATGATATGAATACACATGGTCGATTTAATCTAATGCAGGAGATTTCTAAGCAGTGCATTAGTATAAAGTATTTGCATACATCACTAAACATCTTTATTGTTGTTGTTGGTTCAACTTCAAATCTTTACAGATGGAATGCTAACAGCTCCTTGTTTGAGTTTCACATGGTATTACCCGTTTCAAATTTGTATCTGGTTGAGCCGTTTTATATTACCAGTCAAATGAAAAATCGAACTCTACCATATTTGGCAGCTTTTTCCTATAATACCAGCAATATATTAAAAATAGATTATCTAAAAAATACTTCCGATTTTTTTTCCAGAAGTGAGAGGTTATATTTTATGGAGGGCGAAAGAATTTTATCTTTCTTTCTCCCAATTGTTAATGATCAAATTCCAGAAATTGATGAAACTTTCAAAGTTCACCTTTTCAATTCTTCTGGTGGTTCTTCTATTTCCTATTCTTCAATTAACATTGTTATAATGTCAAATGACAATGCTTTTGGAAGATTCAAGTTTGCAGATTCTTCATTGCATCAGGTTATTGAAGAATTAAATACTGACTCAACATTCTTTGTATGGGTTCAAAGAGAGTTTGGTTGTAATGGAGAAATTAGTGTTGGGTTTGACATCGTGGATAGTGCCAATAAAAAGCAAGATAAAATTTACCCTCAATTTGGAAGTATTAAATTTTTTCAAGGTGATAGCCTAAACAAAGTTGGACTCTATTTAAAGGGAGATTTTATTCCTGAGTTAGAAAAGAAGTTTTTGATAAGATTAACAGACATTGAAACAATTGGTGGCACAAATAATAGTCAAGCAACCCTCAACTATAATAAAAATGTTGCAATTATAACAGTTCCTGCAAATGATGAACCAAATGGTGCTTTTACTTGGAAGAGTCCAATATATGTGATTACAAATGATGTTAATGTCAGTTCATTGTTAATATTAAAACGAGACAAAGGGGCATTTGGGAATGTAACTGTTTTGTTTGAGACTGTAATTGCTTTAAGCGTTGGACCCAGGGAGAGACCAGCCATACCATTTGTTGATTTTACACCTTTAAAAGGCAATGTAACTTTTGAAGAGAATCAGAGAGAAGCCTTTGTTTTTATTGAAGTAATCCATAATTTAGAAGAAAGAAAAGTAAATGCAACCTTAAACGATGGTTTCTATGATTACAATAAAATTTTATTTGTTAATATTACTGATGCATATTTAACTGTTGATAAGCTAAGCTCTGCGCCAGTCTTTTCTAGTCTTACACAAGTTGTAGTTACTAAGCTGATTCAGCAAAATGGAA
>evgR007984c0g1_i01_Hydra
AATGGATTTTCTTAATTCACACAATATACCAATTGAAAATTGTCGTGGCCAGAGCTACGACAATGCGAGAAATATGTCAGGTCAGTATTCTGGATTGCAGTCAAGAATTAAAAGTTTTAATTCATTTGCAGAGTATGTACCATGTTCTGCACATTCTCTTAACTTGGTCGGAACGTGTGCAGCAGAAAGTTGTAGTTCTGCTACTTCATTTTTTATGCTTCTCCAGGAGCTATACAATTTTTTTTCTAGTTCTACTGCACGTTGGGACATTTTAAAAACATATTTGAAGAAAAATCTTAGTGTAAAAAATTTTTCATTAACTCGCTGGAGTGCTTGATGCTTGCC
>evgR007986c0g1_i02_Hydra
GAAACAGTAAATGCATCGTTTTTCAAAATCAAACGGTGTATTGCATTATAGTAAGTTATTTTCTCGGTAAGTTTAAAAACAACGGCATATTTCAATGGCAAGTCGGCATCAGACAAAACATGCACCGTGAAAATTCTTAAAAATGGTTTTTGAATGACAATGGGATAATGAGTTTCTTGAAAAAAGAGTCAGGTTACTCTAGAGTTCCTTTACTTACTGGTGATGAAGAGGACAGCGAGGAAGAACTCTTTGAAAGAGTCAAAAAAGGACAACCACGAAAAGACAATGCTGTTGCTAAAGTCAAGCAAGAACTACAAAGCACAGTAAATTTAATGAAAAATAATTTGCATAAAGTCATTGACCGTGGTGAAAAACTTGAAGACTTAGAAGCTAAATCTGAAACATTTGAAATGAATGCCAATCAATTTAAATCGCGCTCCACTAAGCTCAGGCAAAGAATGTGGTGGCAGCAGTGCAAGTTGAAACTGGTAATAGGACTTATCATCACTGGCGTTCTTATTGTTGTAATAGATTACAGTATATTTCTGGCGGTTTTAGAAAATTGGTCGTACTTTTTCACCATTTTTTTTTTTTTAATATAAATATTTCTTTAAAATGTAATTGAAAATTATTTTATATATTATGAAATTTTTTGAACAAAAAAATTATTATAGGATGCATAAAAAAAAAA
>evgR007987c0g1_i01_Hydra
ACATGGGTTGATCGAGGCACAAGAGCAGTTTTTATTGATTTTACAGTTTACAATGCCAACATTAACTTATTCTGTGTTATTACATTGCTGTTTGAGTTTCCTGCCACCGGGGGTTGCATGGCAACAGCCACCTATAGATCACTAAAATTATTGCGTTATGTCACTAGTTCTGATTACTTTGTGATGGGTTGCGAAATAATATTTGTTGTGTTTATTCTCTATTATTTGATCGAAGAATTACTTGAGATTCAAAAGAACAAGCTTTCTTATTTTTATGATGTATGGAGTTTTTTTGATCTTCTTGTCCTTTGCATTGGAATTTTTTGTATTGCCTTTAACATCTACAGAACTATATCAGTTGGAAAGTTATTGCAGAATTTATTGGACAATCAAGATCAATATGCAAACTTTGGTTTTTTAAGTGAATGGCAAACCCGGTTCAATGATTTTGCTGCTGTTGGTGTTTTTATTTCTTGGATAAAGATCTTCAAGTTCATAAAGTTCAACAAAACAATGTCACAGCTACAGTCAACTTTAGCTCGTGGTGCAAAAGATATTGCAGCATTTGCAGTTATATTTATGCTTGTTTTCTTAGCGTATGCACAACTAGGCTATTTACTTTTTGGTTGTCTTGTGCAAGATTATTCAACAATTCTGAGCTGCATCTATTCGTTGTTTCGCATCATTCTTGGAGATTTTACCTTTTCTGACCTTAATAATGCACAACCTACTCTGGGTCCAATATTTTTTATCACTTTTGTATTTTTTGTGTTCTTTGTTCTGTTCAACATGTTTGTGGCTATCGTCAATGAAACTTATGCAGAAGTTAAAATAGAAATTGCGTTGCAGAAAGATGAGTTTCAGCTGAGCGACTACTTTAAAAAAGGGTACAATAACATGATGACTAAATTAAAGTTAAAGAAAGAAAACATTGCTAACATTCAGGATGCCCTTCAAATTTCTGATAAAAATAAAGACAAGACTGTGGATTTCGAAGAATGGAGGATAGAACTTAAAAAACGTGGCCATTCTGATGTTGAAATTGCAGCTGTTTTCTCTAAGTATGATAACGATGGTGATCGTGTTCTTAATGAAGAAGAGATTAATAAAATGAACAAAGATTTAGAAATAGAAAAGACAAAATTAAATGATGAAATTGAAGATGTAACAGAACAACAAGCAACTTGGGAAACGGATATTGAAAGTTTCAAAGATAACGAAAGTGACGATGATAGCGAAAATTTAGGAGCGGATGATAGAAGAGGATCACAAAATAATCTTATCAGATACAAAGAGTATACCATTTTGGCGCGGCGTGTTGATCGGCTTGAACATAATATTGGAAGCATTGTTTCAAAACTTGATTCTTTGTTCGTAAAGCTGGAGGCAATGGATCAAGCTAAGAGGCAACGTCGTGAAACAATGAAACAACTCTTGGATATTATCGAGAAGCAAAACATTCATGGCTTAACTGAAAAATTGAAACTTTGATATTTCTTCGTTTTTATATTTTTCAAAACAAAATTAATTCATTTTTTGTTCACTTTTTTATTTCACGAAAATTGTATATTTTAGATATTGGCTTTCTTTAATGCTTGTAAATATTTGCACTTGTAAATATTGCGTGACGGTGGCATGCCATAGAAGTTTTATTGCTTGACAGTGGCATGCCATAGAAGTTTTATTGCATGACGTAGAAGTTTAAAATTTTTTAATTGTTTCTCGTTCATTTTGCTAAGCGGAAGAAGAGTTTAAAATTAATAAAAATGGCATTTAGCTTTAGTTAAATAATTTTAAATAATGATTTCTTCGTTATCTACTTTTCAAATAGATTTGAATAATTTAAAATCGAGCTCAATAAAGTTTTTTAATCTACCAATTTCGCGCTACTGCTCATTGACATTTTGGTGATTGGGTTATATTTGTAGGTATTGCTAAATATGTCTATAGTTTTGCAACATAGTTTTCTTTTACGTGTATTATGAATGAATGTATTACGTGTATATTACGTATATTTACGTGTATATTATGAAAGATGTACATATTGCTTTATTTGCTTACTTTTTGAATTTAGCTGAACTTTTTAAATATTTGTTCCACAAATTTTTCTGAAAGGTTATTTTCAC
>evgR007988c0g1_i01_Hydra
TTTTTCGATACAGAAATGTTTTAACAAATTAATAAATTGTACTGAAAGGTTTGCAAAATAAAAAGATGAAACTAAATAAAAGTCTACCAATAAGTGACGAGGTTTTATCAATAGAGTATTACAAGTATTCAAAAGTTTTATTTCATGGCTATGTTGCTCCGTTTATATTAGCCTATGCTATTTTTTTCCCTTTTTGGATAATCAATGGTTTTTTTGATTACTTTGAACTGGGTTGTGTTTTTCTTGTTGTTTTTGGTATGATTCAAGTTTTAGTTTGTTTGGCCTGTCATTGGTCTGTCCACATTTGTGCTGCACTCACTTGTAAGAAGGTTACTCACATTGAATTTGCAGATAGAGTGAAAGTTATTCCAACTCAAAATAATGGCTATCCAGAGTTAGTTCCAATTCAACATGAAAAGGATGAAAGCAATAAAAATGTGGTATCTTTCATATTTCAAAGAATTTGCTACATATATGATTTTGGAGAAAGAAATATGTTTCTACCAATCACATACCCAATTAGTGAGCAGTTTTCTTATTACCAAAATAGCAAAGGATACCAAGAAGAGGAGGATGTCAAAGTGGTTCAAAGAAAATTTGGATTTAACAAAATAGAAATGAAAGTCCCTGCATTTATGGAATTATTTAAAGAAAGAGCGACAGCACCATTCTTTGTATTTCAAGTGTTTTGCGTTGGATTGTGGTGTCTCGATGAATACTGGTATTATTCTCTCTTCACATTGTTCATGCTAGTTACATTCGAAGCAACTCTTGTCCATCAGCAACAAAGAAATATGAAAGAAATAGACAATATGGGAAGCAAAGCATATCAAATCCAGGTATATCGTTATAGAAAATGGAGAGCCATCTCCAGTGAAGAACTGTTGCCTGGCGATATTTGTTCAATTGTTCGTTCAAAAAGTGATGTGATGGTACCTTGTGATATGCTTCTGTTGCGAGGATCTTGCATTGTAGATGAATCTATGTTAACGGGTGAAGCTGTACCACAGATGAAAGAGCCCATTGAATCAAGTCCAGGGAATGATGTATTAAATATTGAAGATCATAGCAAACTTCATATACTGAGTGGAGGAACCAAAGTTGTTAATCACACTCCGCCATCAAAAACAATGGCCGGTTTAAAATCTCCGGATAATGGCTGCATTGGTTACGTACTTAAAACAGGATTCTCAACATCACAAGGAAAGTTACTAAAGACTATATTATATGGTGTTAAAAGGGTTACAGCAAATAACCTAGAAACGCTGCTGTTTATAGGTTTCCTCTTAGTATTTGCAATAGCTGCAGCTAGTTATGTTTGGATTAAAGGTACAGCAGATCCTCGACGTAACCGCTACAAATTATTTCTTGAATGTACTTTGATTTTAACATCTGTTGTACCACCTGAACTTCCAATTGAACTGTCGCTAGCCGTGAACTCATCTCTGATAGCTCTCACGAAACTTGGTGTTTTTTGTACCGAGCCTTTTCGCATTCCGTTTGCTGGTAAAGTTGATGTTGTTTGTTTTGATAAAACTGGAACTTTGACAAGTGATGATCTTGTTATAGAAGGTGTTGCTGGTGTCAATGCTGATCAAAAAATTCATCCTGTTAGTGATATACCGATTAGTACAGTACATGTTCTTGCATGCTGTCATTCACTTACATATGTTGATGAGACATTAGTTGGTGACCCAATGGAAAAAGCAGCTCTTAATGCAATTAATTGGACTCTTTCAAAAGGTGACTTGTGCATGTCTCGTAAAGGTCTTCGACAATCTTTAAAAATTGATCATCGATTTCATTTCACTAGTAACTTGAAACGTATGTCTGTAATTGCCTCAGTTCACTCAGCGGATTCATCTGCAACAAATTACATTGTGGCAGTAAAAGGTGCGCCAGAAACGTTGCATGGAATGTTTGTCAGTGTTCCAGAATTTTATGATAGTTTACACAAAAAACTAACTTGCAATGGAGGACGAATTATGGCATTAGGACATAAAGATTTAGGATCAATTTCACCAAAAGAAGTTCGCGAAATGAAGCGCGAGGATGTTGAATGCAATCTTATATTTGATGGATTCATTGTGCTTTCATGCCCATTAAAGTTGGACTCAAAAAGTACTATAGCTGAAATACAAAATGCATCTCATTACACTATTATGATCACAGGAGATAATCCATTGACTGCGTGTCATGTAGCAAACTCACTAAAAATGAACAGCAAAACTCTGCTAGTGTTGTTGAAGGAAACAACATTTGTTGACAAAATTGCCAATGAAGTTTGGTGCTGGAAAACAGTAGATGAAAGTGAAAGGTTCTCACTTGATGTACTGACAAAAATTCTCATTGACAAATACTCATTTTGTTTGACTGGAGAGGCTATTCAATATATGCAGTTGTCTCCATTTCTTCGCCAAAAATTTCGTGAAATACTTATTCATTGCCGTGTATTTGCTCGTGTTAGCCCAAAACAGAAGGAATTTATTATACATGAGTTTAAAGACCTTGGCTTTGTGACACTAATGTGTGGAGATGGAACAAATGATGTTGGAGCACTAAGACATGCACATGTAGGTGTAGCATTATTACCTGGTACAGCTGAACGAATGCCACCCGTTGTAAAACGTAAAAAGAAGAAAGATGCAGTTGAAGAAGGTAAAGCGGTTACAAGCAGTAAAGGAACAAGTAAAGCTGCTAAGATTCGTTCTGTTGCTCGTGGTGATGTGTCAGTTCTCACAAAAACTCCTCTTGAAGAACAGCAGGCAAAACTTCGTGAGATGTTGCAAGAAATAGACAAAGAAAGTTCGCAGGTTGTTAAACTTGGTGATGCTTCAATTGCATCACCCTTCACATCACGATACTGTACACCGTATTCAATATGTCATGTGATAAAACAAGGACGCTGTACTTTAGTAACCACTCTTCAAATGTTCAAGATTTTAGCATTAAATGCTCTTATACTTGCCTACAGTCAGAGTGCACTTTATCTCGAAGGGATAAAATTCAGTGATGGGCAAGCTACATTGCAAGGCATATTACTGGCTGGATGTTTTCTATTTATCTCTCGCTCTAAGCCTTTGGTTTCCCTTTCGAAAGAACGACCGTTACCGAATATTTTTAATTTGTACACTGTACTTACTGTTCTGTTACAATTTGGCGTTCACTTTTCGTCTCTCTATTTTGTTGTGAAAGAAGCATATACTTTTTCTCCGAAAAGAGAGGAGTATGTTGACTTAGAAAAGAAATTTGAACCCAATGTTATCAACAGTGTTGTATATATTATATCTATGTCTATGCAGCTTTCTACATTTGCTATAAATTACAAGGGGCGTCCTTTTATGGAAGACTTGATTGAAAACAGACCACTTTTTTATAGCTTATGTATTTCAGCGGGGTCACTTTTTATCTTGCTTATGGGCTGGATACCTGATATATCTAATCAATTCCAAATAGTTGATCTTCCAGAACAGTTTCGAATGACGTTCTGTCAAGTGGTTATAATTGATTTGGTTAGTGCTTGGCTGGTGGACCGTATACTAAGATATTTATTTGGGTCATCGAAATTAAAAACTTGTTAATTATATTTTTATAATTCATTAGGCTACAATTTAGGATGTACGAAATCAATTAGCGTATTTTATTATTAAAAAAATGTCGAAAAGATAAAGTTTTTTCGTTTATATAGTATTTTATAGAAAAAAA
>evgR007990c0g1_i01_Hydra
ATTTTGGTTGCAACAAGGTTTTGATTGTTTCTGATAAGCCTTTGGTTCAGATTGGAATTGTGAGCCAAGTGACTAGAGTGTTGGAACAAGCCAAAGTTCCATATGTTGTTTTTGATTCAGTTCAACCAAATCCAACTGTTAAGAATGTCTTGGAAGGTTTAAAGTTAATGTCCAGTGAAAATTGTGATTCATTGATTAGTGTTGGAGGTGGAAGTGTTCACGACTGTGCTAAGGCTATTGGATTGGTCAAGTCGAATGGAGGTAAAATTGAAGACTATGAGGGTTTAGATAAGAGTAAAAAACCAATCTGCCCATTCATTGCAATTAACACAACTGCTGGTACTGCAGCAGAAATGACAAGATTTTCCATTATTACCAATGAAGAGACTCGTGTCAAGATGGCCATTGTTGATAAGAATATCACTCCAACTCTGAGTATTAATGATCCTGTATTGATGATGAAACAACCACCATTCTTGACAGCTTGTACTGGTCTAGATGCTTTGACACATGCTATTGAAGCCTATGTATCAACAGCTGCCAATCCACTCACAGATGCACAAGCATTGATGGCTATTGAACTCATTGGAGAATATTTAAGCAAGGCCGTGTCCAATGGAAGTGACATTATTGCTCGTGAAAAGATGTGTTACGCTCAATTCTTGGCTGGTATGGCATTCAACAATGCCTCACTTGGTTACGTTCACGCTATTGCTCATCAACTTGGAGGATACTATAACTTACCTCATGGATTCTGTAACGCCATTCTCTTGCCAAGTGTTATGAAGTTTAACATTCCATCCAGTGCCAAGAAATTGAAACGTGTTGCTCAAGTTTTGGGTGTAAAGAATGTTGAAAACTTGAGTGATTACGATGCTGCTATGGAAGCTGTCAATGCTGTCGTTAAATTGAGACAGTTGGTTGGAATTCCAAATGGATTGAGCGGTGAAAAGCTTGTAAAAAGAGAAGACTTTGAAGTTTTGGCTGATCATGCCATGAAAGATGTTTGTGGTTTTACCAATCCAAGAGTTGCCATCAAGAAGGATATTATTGGTATTTTGGAAGAAAGCTAC
>evgR007997c0g1_i01_Hydra
TTTTTTTTTGATTTATATGAACCAGGGTCAACCTACTTATAACGCTTCTGTTATAAGTAAAATGGCAGCCACTTTAGCTGAGGAAATAACAAAGCATTCTGTTCACACACTTGTATTTCGTTCCATTAAGCGAACTCATGATCTATTCTTAGCGGATCATGCATTGGCTGTTCCTCAGGATAAAGAAGCTGATGAGTTAAGAAGAAAACTAAAAATACTAACAGAATATTCACACTTGAAAGATAAGCAACCAAACAATAATATCCACACATCACAACATTATTTAGCTATTGAAAATAATAAAGAAGGCAACAGGGATTTAATAAAACGGGAGACAAGCAGTGATCTATTACCAACATATAATAGGTCCGAGGACAATCAGTATGCAGCTGGGATTCAAGAAATAACAAATAAAGCTCTTTCTTCAGGAGGTTTGGAGCGAATTGCTCCTCCAAAGCGTGCCCTTGCCACAACAGGGGAAAGCAAGGTTCCATTTTCTCATGGAATGGTTGCTCGGAAAGCAACTACTATGCCTAAACCTAACTGGCATGCTCCATGGAAACTGATGAGGGTTGTAAGTGGACACACAGGTTGGGTTAGATGTGTAGCTGTTGAGCCAGCGAACCAGTGGTATTGCACTGGATCAGCTGATAGAACAATAAAGATATGGGATCTTGCTACAAGCAAACTAAAGCTTTCATTAACTGGACATATAAGTACAGTGCGTGGGGTGGCAGTAAGCACACGCCATCCATACTTGTTCTCATGTGGTGAAGATAAAATGGTAAAATGCTGGGATCTGGAGTATAATAAGGTAACTCGTCATTATCATGGTCATTTAAGTGCAGTTTATTCACTTAGCATTCACCCAACAATTGACGTCCTTGTAACTGCAGGAAGGGATGCTACTGCAAGGATATGGGATATGCGGACAAAAACTTGTGTGCACACTCTTGTTGGTCATACAAATACTGTTGCAGACGTTAAATGTCAAGGCGTGGAACCACAGATTATAACGGGAAGCCATGATTGCACTATACGATATTGGGATATGGCATCTGGTAAATCTATAACTACTTTGACCAATCATAAGAAGAGCATTAGATCTATTGTTCTTCATCCAACTCAATATACAATGGCTTCAGCATCTGCTGACAACATAAAACAGTGGAAGTTTCCAGAAGGTAATTTTATCCAAAATCTTGGAGGTCATAATGCTGTTCTCAATTGTATGGCAATCAATTCGGATAACGTTTTGGTTTCTGGCGGTGACAATGGAAGCATGCATCTTTGGGATTGGAAGACTGGATACAATTTTCAAAGAACACAGTCACTTGTTCAGCCGGGATCCCTAGATAGTGAAAGTGGTATATTTGCAATGGCTTTTGATCAAAGTGGCACTCGGTTGTTCACATGTGAAGCAGACAAAACGATAAAGGTTTATAAAGAGGACGATGAAGCTACTGAAGAATCACACCCTGTAAACTGGAGACCAGATATCATTAAAAGGAAGAAATATTGAGTAATTTTAATTAGGTTGCATCCCTTTGCATCCTTTCCCTTTTGCAAACTGGTCCAAGGTTATCCATTAGGCGCCAGCTATTTTCAAAATGGTTATGTATAAATAATATATACATAAATAATATATATATATATATACATATAAAAAAA
>evgR007999c0g1_i01_Hydra
TCATTATGATTTTATCAAAACATATATTTGGAACACTTCCTCTTCATTGTGGCCACCAACAATAAAAATAAAAAAAATGATTTGCAAGTGCACACTGAGCATGCAACTAAACGTACTTCTGATTCTTCAATGCGTATTATCTTACCCGTTAAGTAACAAAGACTTTTTGTCACAGGATATAAACACGGCAGACGAAATAGAAAAAGTTAAATCAGAAATATTGAACGGTTTAGGATACAATGATCCACCACCTAAATCTGACTTTGCGACAAACCAATCAATACCGGAAGCAGTTATTTCGAAAATGTTTTATGGAAACATCCAAAACTCGGTCAACGAAATGGACAGTAAAGAACAGTTCGTCGTAACGGAAGTTGTAAGAGAACTGCTTTGTCATCACAGTCATTGTGTAATTGTAAAGTTCGCACAAAAGTTTCACGATAGCAACGTTTTATCGATAAATATTTGGTTTAAAAGTAGTCCAACCACAATCTTGAAAGCATCATCAATTTATGTTTCCAACTTGAAAACAAGCAAGAGAGACTCAAAGCAAACTATCTCGAACGATTCATTGCAACCAGCATGGCATAACGCGGAGTTAAATGTAAAAGAAGCAATAAAAAACGATAATGAAGGACAATCTTATATCGATGCTTCTTTCAAGTTGAGTTGTAAGAACTGCAAAATATCACTGTCTGAAGAAGACCGACCATTTATCGTAGTCTCATTGAAGAAGAAAAAAAGTTCTAGAAAACTTCGTTCAACTCAAGCTTGCAAAGGGTCAGAAGGATGTTGCCGAAAATCTATTACTATAAACTTCAAAGATATAGGTTGGGAATCTTGGATATTGGCACCTAAAACCTACGAAGCTTACTATTGCACAGGAACTTGCAAAGAAGGTAGCAGCACAGATTTACCTCACTTTCAACTTTTGAGAAACAACAACGATAAGAGCAAATGCTGTGTACCAAAATCTATGTCTTCTTTAGCAATACTATACGTGGGTAGCGAAGGAATTATAAAAAAAGATTTGCCAAATATGATTGTAGACGAATGCTGGTGTATTTAAAAAAAAAACAGTCAACGACTAACACCGTTAACAAAATACTTTGACAAAGAAAACTATTTATTATAATCAATATAAATTATACAAATAACAACCTAACTAAAGAAAATATTTATGTGTACAAAGTGTTTATTTAATTAAAAGAAAATGAAAATGTTGC
>evgR008001c0g1_i01_Hydra
CAAAAAAGACATTTAAAACTAATCTTCTATTGTCTAAATAAGCAATAGTATAAATGAAAAATGGACATTAAGTTTCAAATTAAAAAAATAATTGAAAATGGAAGTAAAGATACTGACTGGCTTTGTTTTATTGATCAAGTTAAAAGCAAATGCGATCGCGATATTCTTCAAAATTTATATTTAGCTGCATTTAATAATATATCTCAAAATAAATCAAAAAGCTTCAAAAAGTTGCTTGTCGGTTATGCTAATCTTCAAAGTGATATTGCTCCAGAAGTTGCCAGAAAAACATTTCTTAGTGTAAATACCTTAGATGATGAATGCGCTGTTGTTTGTGTTTACTATGCTCAATTCGAATATAGACACAATAAAGATAATGCACGAAATATTATTGTTAAGGGAAAATGTGATGGTCGTATCCCAAAAGTTCTTTTGACTCAAGCATATCGTAACATAAAAAACAATATTGCGAATTTATTTGAGGGGGTTAATGATCCATGGGATGAAGGCTTATGTGATATGACAGATACAGCTACATTTACTTCTTCTAAAGGAGCTTCTCCAGAGAAACAAGATTCATATTTATTCCATTCGTCTAAAAGTTTATCTTCGTTTCCTGTTCGATTACCAGTTCGATCAGTACAAAAAACGCGATCAACCTCTGCTTTGATTAAAGGATCCAACAGCCCTTGCAATAACGTTAGAAAAAATGAATTGTCAAAATCAAGTTCAGAAGCTTATAATCATACAAAACGTAATGGTCCAAGTTTATCACCTTTTTTTGAAAGAGACTCGTCAAAACTTTTTGAATTCCAATCTTCGACTGCTAAAATTATAGAGAGTACACCGTATGATCTTTTTAATTGCAATCAATCTTCTAATAATTTAGTTTTCGAAGCACCCCCTGCATCCTCTGCACCGACTTTACTAAACACGAAGTACTCTAAGCACACTGAAAACTCTTACTTAACTTCTACATTGCAAATATTTTCTCAGGAATCAAATAAATTTGAAAATAAAGCTGCACATCATTCAAAAAATAAGTCATTAATGCGTCGTTCTGTTGCATGCAGTGGTTTAGTTTGCAACGAGCAGACACTTACGGAAGATGGAAAAAATGCAAATTTAGTTTGCAATAAAGAATCTATTCAAAATTTGGATCTGTGTAAAACAAGAGTCCCATTATCAGTTTTACATGAAAATAATCATGTTGAAATAGTATCTACTTCTTCAGTACCTTCTTCTTCAGTACCTTCTTCTTCAGTACCTTCTTCTTCAGTACCTT
>evgR008002c0g1_i01_Hydra
AGATTATAAACACAATGTAGATAATTGCCAAATACTAATAACTGTACCACAATGTCTTGAAATTCTTCTGCTATCCATCCAAACATATACATGGTCTCAAAAAATAAAATATGTTATCTTTGATGAGGTTCACTGTATTGGTGCTGATGCTGATGCAGAGGCATGGGAGCATTTATTGTGTATGATCTCATGTCCATTTCTTGCACTTTCAGCAACAGTTGCCAACCCCGAAGTTCTTCATGGTTGGCTTCAAGATATTCAAAACTTCCAAAAAGAAAATAAAATAGTGCTTGGGAGTAAAAGAGATGATTTATCATACGAAGTTGAATTAGTTACTGTGAAAGATAGATATTCTGATTTAGAAAATTATGTTTTTCTTCCTGAGAGTACAACCGATTACCAGGAGTTTGGAAGTCTGCAAAGTATTCATCCAATTTGTACAATAAAATTATCAGCTGTCATTGCAGAGGGAATTCCAGAACATATCACTTTGTCACCAAGAGAATGCCTGAAACTTTATGAGGCAATGAAAAAAGTATATTATGAAAAACCTGAACTGGTTAATCTTTCCCCCTCTCATTTTTTTTCTGATAAAAAATTTCTTACCAGAGATGAAGTCAGATTATACGAGAGTGAGTTAAAAAAAGAGTATTTTAGCTGGAGCAAAGAAAGGAAATATTTTGAAGGTGTCCAGCAAATGTTATTTGATTCAGAAGCTAAACTAAAGACAACCAAATTGTATGAAAAAACTTTGAGTTATAACAAGGATATATATTTAGTAAAAACATTTATGAATTTGATCAAAACTTTGGAACAAAATTCTATGTTGCCAGTTATTGTATTTTGTTTGGATCGACATTTGTGTCATTATTTGGTGCAAACAGTTGTAGCATATTGCGAACGCAGAGAAAAAACTTCAGTTAAAAAAAAA
>evgR008006c0g1_i01_Hydra
TTGTGGTGTAATCTTGTATGTTATGTTGGCTGGTTATTTGCCATTTGAAGATGATACAATGAAAGGATTGTTCGCAAAGATTGAATCTGGTAAATTCTCTTATCCACCTCACTTTACTAGTGCCCAAACTGAACTTATTAGTAGAATGCTTGTTGTTGATCCAGAAAGGAGAATTACAGTAGCACAAATTATGGAACACAAATGGTTTAAGGTTGGCTTCAAACTTGATAGAGTTACATCCAAGATCAGTTTGACTGATAAGGAAATTGATGCCTCTGTTAAGAGTACTGGAGATGTTACTGAAGTTCAATCTACTCAATCATCACAAAATACACAAGTTAAGGCTGGTACATTGAATGCCTTTGATTTGGCTTCCATGTTAATGATGGGCTCAATGAATCCACTCATGTCTTCAGATGCATCTGTTAAAATTAAGAGACAAACAAGATTTATGGCAAGAGGAAAGAAACAAGAATGTCAAACTGCTATCGTTTCAAAACTCGATAAATTGAAAGCCAAACCAACTACAAAGGGTGACGATGAAATTAAGTGTATGGCCACTGTAAATCAAGCTGCTATTACATTTAGTATCAAGATTGAAGAAACAAGTGGTGGTTTCTCAGTTGTAGAAGTAAGAAGAGGTAAAGGTAACATCCTTGACTACAACGCCTTCTATCGTACGTTGGTGAATGAAGTGGGTGATATTGTTGTTTCTAAGGAAATTAAAT
>evgR008008c0g2_i01_Hydra
TTAAACGCAAGATTTATTTAAATGAGCCATCTAAAGTACGACATTAAAAAATTATGTCGATTGCAAAGCTTTTAAAGCAATTTTTTGTTGTCGTATGCAGGTACAGCTTCATAATTAGAGTATGTACTTGTGGCAATTGTGCTGATAGTTGGCTAGAGTATGGAAATCATTGTTACCTTTTTAACGATGCAACTAAAGGATTACCAGCAAAAAATTGGACTGACTCTAGTTTTGCTTGTCGGCAATATGGAGGAAACTTGTTAAGCATAACAGATGAAGTGGAGAATATATTTATTGAAAAACAGTTTAACAACGACTTAGTTACAAAACAGGAGTTCTGGATTGGTCTCAATTCTATTCAATCTCATAGTGGTTTTGTATGGTCTGATAATACAAGTTTGCAGTTTTTAAATTTCGTACCAGGTCAACCTGACAACTTTTATAACGACGAGTTTTGTGTGGAAGCACTTGTTGATGGTCGATGGAATGACGCATCTTGCAATAGACTTCTTGGATTTATTTGTAAAGTTAAAATAGTTATTACATACTACATTGAACTTTATGGTCTACCAAATTGTTTTTTGGACTTTCTTCCTTCTCAATTTAACATGATTACTTATAATTGCAATGGAGGAATGAGGTTTTTACTTTTAAAAGCTGTAAATGGAGATTCAAGTATGGTTTCTTTGAAGTCAGCAAAATACATTGACTTATATCTTTCTCATTACTATCATGTATTCTATCTTCAAACTTTATTGCAATGGGGTTCCGGTCCATTTGCTCGTTCATTTTACATCAGAGATGCAATTTATGATTCATATAATTTAATACAGGGAAACATAGACAACTACTATCTTTGGCACGAAATTTCTTCTGGTAAATTTTATTTGGTACAAGGAAAAAATCTGGAACCAGAATCTAGTTTTAAATTTGTTGTTTGTGGTCTAAATATTAAAGAAATTCATGGCATTACTCGGTATATAGTCAATGTTACAATTGAATTAATGGCATGGCATATAGATAAAAAATTGCTAATAAAGTATGGGGATTCAAATAATTACTCGAAGGTTTACAATGTTACTGGCAATTCAGTTGTTGTTCACTTGTTGCAAAACTATACTGTTCATAAATTTGCTGCAGGTGTGAAAAATGAGTATGATATGTATGTGTTTGGAAATGCGTCTTACTATCATATAGATGAAAGTAATTTAGAAACTCAAGTTGATAGCTATATTTACGGATATGTTGAAGGTAATATTTTAAACATATCAACTGATTTGATTGGGCATGGTTATAATATAACATTGGAATTTTTGTATGCACCTTTCATGAGTTTAAACACTGAAAGTATTCCACTTGGATTTGTAAAATTATCTTGGAATAAATTTATTTTTCCTGGTTACTTTGAAACATCAGATGTAATTACATTTAGCAATATAGTGAGTTTAAAAAGAAGAAGATGTTTTGGTGGTTTGCCAATCGAGGTTCCAGTGACAGTGAGTTATCAAAATAATAAGGGTTGGTTGAAGGTTATTAAAATGACATATCGAAAGAGTTTACCTTGCTTCAATATTTTACCAATTATATCAAGAGATCGTATGGCGTTGCCAGAATACTATGGCAGAGGAATATATATAAATGCTGCGAACTCTCTTTTATATCTATGCATAAATCAACATGTTGAGAGCATTAAAAACGCCTGCTTTTACTCAAGAAACTCAAGTCACTTGTGGGAAGAATTAGATGTTTGCGTTGGATCTGTGATTGGTCATCACTTGGTAACCAAACAGTTATATGCTATTCATCGTAATCAAAAAAAC
>evgR008010c0g1_i01_Hydra
ATTGTATATTTGATGAATCACTTTAAGAGTGAATTAAGTGCCATAGAAAAGGCTTATGATATGCAGATTAATTTGCATATTGATGATATAATATCTGCAGAGGATTATAAAATTGTTCTGCATAAATTTTCTAATGTACCTGCACTAAAAAATAGGAGATATGTAATGCCAGAGAATTTTGCAACTAGTGAGCAAGATGATACAAAATCAGAAAATGGTAATGAAATGCATCTTAATGCATCTGAAGATAAAGAAGGAGCTTCAGAAAACAAAGTGAAGAATAACACTGAGAATAAAAGAGGATTTAAAAGGTATAATAAGGAGCATTTTAATAAGAGGAAAAAACCAGAAAGCAAAATGCACGAAGATAATAAACAAGTTAAAAAAGGCACCTCAATTCTGAAAAAAATGTGGTTAAAAATAAAGTCGTAAAGTGGAAAATAATATAAAAAATAAGTACGATAGCAGTTGGTATCTTATTAAGAGGTTGTTTGCGTATTATATACGTCCATTAAAGCGAAAATTAAGTTTAGCATTGTTGTGCATGATAATCACTGCATGCACAACTGCAGTGAGAAGAAAAAAAGATAGAATAAATAACAAAAAAGGA
>evgR008011c0g1_i01_Hydra
ATTATATTTATAGAATATATTTTCTTTTTTCTGGTTTGTAACTAGCTATGAATGCAAGTAACTCAAATCCAAAAAAGTGGTCCAAATATAAATCTGTGACAAGAGCTGTTAAACGACATTTTCAGGATATTGAAGATGCTATTAGTCTGGATAACCTTGCTACACAAGTTAATGTAACAAATGACACTACAACCAATGTTTCAAATTTGAAAAGGAATAAAGTGATAGATATTACTTTTAATGTGACTAATAATGAGAAATATAATGCATGCGATAGTAAACATGATGAGATCAGTGTAAATGAAATTGCTAAAGAACAATATGATGATTCTTCATCATACTACAATGATCCGCTACACAGTTATAAAAATATAACAGGCTTTCTATCAGAAACTAGCGATGAAGAGGAAAATAAAAAAATAGAGAGAACTCCATACATTAAAGAATCACTTGCTCAATGGGCATTGAAGCACAATATTTGTCAAAATAGCTTAACCGAC
>evgR008015c0g1_i01_Hydra
CATCATGAGTCTTTCGAAAGCAAAACAAGATATTTTTGTTTGCGCTTGTGCAATAAGTCCAGATGGAAGTGAAAACATGGCTACGTTTAACGCTAAATCTTTCGAAAAGAAGTTAAAAACATTAGATGCTAGTCAAACAGGAATACAAACATTATCATTATGGATACTACACCATAAACATCATGCCAAGATAATAACTTCAGTTTGGACTGATTTAATAAGTCAAGTACCCAAGCCTGAAAGAAGGCTTGCATTATTCTACTTGGCTAACGATGTTGTTCAAAATGGTAAGCGAAAAGCAAAAGACCTTGTGGATATGTTTGGCAAGGCATTTATGGATTCTATTTCACTGCTGAGCGATGCAAAGATTAAAATACAAGTAGAAAGAGTGTTTGATATTTGGAGCGAGCGTGCAATCTTTGAGCCTAATCTTATTAGCAAATTTAAAGAAATGCTTAATAAAACAGCTAAACCAGTTGATGAGTCGAAATCAAAAAATAAAGAAACAAAATCAGAAGATAAAGAAGCAAAATTAGAAGAACAAGAAATAAAGAAAACTCCAAAACCAGTACCTCCTGATTTCAAAGTTGCTAAGTTACTGGTATCAATTAAAAAAACAACTAGCTTGGATAATGACGTTATGCGAAAATCAAAGTTAATTGAAAATCTACCCATAGACGTTTCCAGTCTTGATTCTGTTAAGCATGTTAAAGGTCGCTCTACTGTTAACAAGTTCTACAAACAATTTGAAGAATCAAAAGTTGTTTTAGAAGATTATTGTCATCTATTGAGAAGATGCGTAAAAGATAAAAATGCTCTTTCCACAACTTTATTAGCGGGAGAGGCGTATTATGAAGCTGCGCAGGAAGAAGCTAAAATTATTGTTAATGCATATAAAACATTTGGAAATCGTCTAAAGTCAATGAAAAAGAAATTAGAAGCCAAGATTAAAAAATTAGAAGAAACCAATTTAAGCTCAGGCTCACAAAACATTTTCACAAAACAAGTAAACACTTCTTCTTATGATGGTGATTCTTCACAGGAGTCTGTTTCTGAACCTGTTTCAAGCGGCGATAAAACACCAGCAAGTAACCATAGTTTTAGTAATGCTAATGAAAGTAATATTGAAGACATGGACTTAGATAGTAATTCAGATGATGATGCTTCAGAAGCCTATAATCCTGAAACCGCCCCTCTTCATCAAACATCAAGTTTTGAAACCGCTCCTCTTCATGAAATATCAAGTTTTTCTAAAGAAATTGCATCATTTTTAAACATTGAAACAATTAATGGAATGACATCTTTCTTAGATAGAACTGGATCGAGTTCTGTTTTAGATTTATCCTTACAGAAAACATTGCCTGCTAATGAAGAAATAAAAACCACCAGTTTTCCACTTCCTTTTTCGGCTTCTTTTTCAACACCTTTCACGTTGCATACTGCTACTCAGTTATCTACTTCTGATAATTTCTCAAAAATCGAAGAAAACACTCACACGCTTCAAACCGCATTATCAAGTGCAAGCAACTTTTCTAGCATAGCCGCCTTTTCAAGTGATGCTACTTTTCCAAGTGTAACCAGCTTTCCAAGTATAACCAGTTTTCCAAGTGTAACAAATTTTCCAAGTGTAACCAGCTTTCCAAATGTGACCAGCTTTCCAAGTGTGACTAGCTTTCCAAATATAACAAGCTTTCCAAATGTGACCAGCTTTCCAAATGTAACTAGCTTTCAAAATGTAACCACCTTTCCAAGTGTGACCAACACTTATTCCTTATTACCATCAACTACTAGTTTTACCAACTCTAATAATAGTTTTACCAACGCTCATAATTTTCATCCTAACATGTTTAAAAACGTATCCGAAGCAGATACGCCAGTAACTCCTGTGCTTGACGAAGGTATATCGAAGCAGCATGTCGTGGGTAATGCTGAAAATAAACCTTTGCAAAATTTAATTGATTCACTTTTTCCTATGCTTAGTAAAAGTTTACAAACTATAAAAGAGAAAACAGCAAAATCAGAAGAGATGCAACCTTCTGTTAACGATGCTATAACGTATGCTTTGTGTGGAAGTAACGAAGAAGCTCCAAAAAAGAAAGGGCGTTTTGTGCCAATAGTGTCAGATATGAGGGTTTCTTCACCTTTGTCTCTCACAAACCCAACAGGTTTGTCGGAAAATCAAACTGAATTAATAAATATTGCTGCAGGTATTACAAAATTTGACGATAACTTCTTGCCTTTATCTAGCAGTTCCGAGACTGCTTCTATAAGTAGCGGGCAAAATTCTACAATGTTACTAAGACAGAATTCTAACGATGATCAATCTAGAGACGAAGAAGAAAAAATAGATAAAAAAGAAAAAAAATCAAGGCATAGCTCCGGCAGACATCGATCTCATCGTGACGAACATAAGCGCTCTGCAAGTCGAAGTCCTCGTAAGAATTCCCATAAAACTCGCGAGGGCAGTCGGAGTCCCGGTAGAAAGTCATCCCATCGTTCTTCTCGTGAATATCGAGATGATAAATTTGCTGATAGGCGACGTTCGTCGCGTAGATCTTCAAGTCCTCGTAGTAGGCGTAAAGGAAGATCTCATCGCTCTACTTCTAAAGATAAAGAAGATAGAAAGAGGCGTTCTCCCGCCAATAAACCTTATTATAATGATCGCAGATCTCCTACACCAGAGATGGATGGCAGTTTGCGAGCATCTTATTCCGCAAACATTCCTCAAGGCCAGTTTGATCAAAGCATGCAAAATCAATCTTTTTCAATTGGCCCTAATGTTAGTCAGTCTCAAACTGGCGATCATAAAGATAATCACTCGGTCATGGTTAGTTCAGGTCAGCCATCTTCTTCGCACCAAAGTAATCTTAATTTAAATCAATCGATGAATCTTAAGTCTTTAAATTCTATTCCGAATATGCCGGTTCCGGACATGTCTGTCTTTCTTCAGCTAGTAAACTCAGTACGACCAGGTTTACCATTGACTAGTATAGCTCCGCCAAATATGTCATTTGACATAAATGGTACACCTATCATAAATCCTGGATTACAATTATCACATGGAAATTCTCTTCACATAATACAAACCAACCCTCAAGATAATATTTTAAATCTACACCCACTGCCACACGGTGTTCTACCATTTCAGAATCATAATCTTTTATTACAAAATCAATCAGCACAACTTCAAAACGAGTCATCGCAGGAAATAAGTATAATTCCAAATAAACCAAAAAACAATAAAGATATAGAAAATTCTCTTAAAGATCTCTCCGATTCCACTCGTGATAAATCAGGTATGGAGTATCGCGTTAGCCAAGCTAATAAAAGAGATAATAAATTGTTTGGACCTAGTAATCCTCCACCGTTTCCAGGCCCTCCGCCGGTATTTGAAGGTATTCCTCCATCAATAGTGACTTCAAATTTTGAGATACCTCCAAATTTTGAACATTTTTCTCAAGAAGGAATACCGACTTTCGGCCTTCCGCCTTCTTTTGATACATATCGTCCAAAATTTGCCGAACCTGATCCTCTGTTAGAACAGTTGGAAAAAAGTAATCCTTTTCCCGGGGCGCTTCATCTTAATTCTTTACCACCACGATTTGTTAATAACAATAGTCCCATTTTTAGGCCAAGATACAATAACCGTAATGGATCACCTAGGTTTAGACCTCGGTTTGATCATAGATTTCCTTTTAACGGAAGGCCTGCGATGCTTTAGCAGTGACTCGAAAGGATTTTATGTTTTGTTGTGCCGTTTAATAATTTGCTAAATTTTAAAAAACCCTCAGTATATTTGCAGTAAATATGAATTCGTTTCATTAGGCGATGAATGGCTAAAAATTGTCATTTTAGAAATTATAAAGATTTTGATGTAAAGTTTTTTAATTATTGTTGCCTAGTTACTAAAAAAAAAA
>evgR008018c0g1_i01_Hydra
CTAGAAGTTGAACCATCGGATACTATCGAAAATGTCGAAGCCAAGATTCAGGATAAAGAAGGCATTCCTCCTGATCAGCAGCGTCTGATTTTTGCTGGCAAGCAGCTAGAAGATGGCCGCACTTTGTCTGACTACAACATTCAGAAAGAGTCCACCCTTCATCTCGTGCTTCGTCTTCGTGGTGGTGCCAAAAAGCGAAAGAAGAAGAATTACACCACCCCAAAGAAGAACAAGCATAAGCACAAGAAGGTCAATCTTGCTGTATTGAAATTCTACAAGGTTGACGAGAATGGAAAAATCCATAGACTCCGGCGCGAGTGCAAGAGTGAATCCTGTGGTGCTGGTGTTTTTATGGCTGCACACAAGGATCGTCAGTATTGTGGAAAGTGCGGCAAGACCTACGTCTTTAATGCTCCTGAACAGAGCGAATGATGAG
>evgR008020c0g2_i01_Hydra
TGGCATCCAAATGTTTATACAGATGGTAAAGTTTGTATTAGTATTTTGCACCCTCCTGGAGAAGATGAAATGTCTGGCGAAAGACCAGAGGAAAGATGGAACCCAACCCAAACACCAGAAACTATTCTCCTTTCTGTCATTTCAATGTTATCAGATCCAAACTTTTCATCACCTGCCAATGTAGATGCTTCAGTTGAGTGGAGAAATAATAAGAAGGGATTTTTGGACAGATGTAAAAAGCTTGCTGAAAAGGCCAATAATTCACTCCCGACTGGATTCACAATGCCAGAACCATTCAGATATAAACCAGAACCACAAACACCACAAAATCTTGATGATGAGGATTTCGAATTTGAAATTGATGAGGATGAAGAAGTTTTGGATGATGACGATGATGATTTGGAAGGTTTGACAGAAGAAGAGCTTGCTGCATTGGAAGAAGAG
>evgR008021c0g1_i01_Hydra
TTTTATTTGGTATCTAAACAAAACAAAACAAATCCAAGTGCGCACTTAGTTTTAATTTTTTTCCCTTAGTTTTTGGAGCAAGCATTTTATGATAATTTCGGGAAAGTAAACTTTTTGTCAAAATATTGCTGAGACATATTTTTAACATGGCTCCTGCTGCACCACAAGAAATTAACTCTTTACTTAATAAGAAATCATCTGAAGATGAAATAAGTAAAAAAGAGAAAGATGCAAAAGCCAAAGAGAAAGAAGAATCAGAAATGAGTGATGAAGATAAGCATTTATTAGAAGAGCTTAATATGTTAGTGGAGAGACTTACGGAATCAAACACAGATTTATACAAACCAGCCTTGGAAAGTTTACGAACTTTAATTCGAGCTAGCACTAGCTCAATGACATCTGTTCCTAAACCACTGAAGTATCTTCGACAACATTTTGATTCACTAAAAGGGCTTCATAGTAATTGGTCAGAAGGAGAGAATAAGAGGTTTCTAGCTGATATCATATCTGTTCTTGGAATGACCAGAACTGACAATAGAGACTGTTTAAAGTACCGATTTTTAGGATCAACAGAACCTCTTGAATCTTGGGGTCATGAGTATGTTAGGCACCTCGCTGAGGAAATAAAAATAGAATTTTCACAACGTGAAGAAGGAAATAGAGAAGATTTGCTCAAACTCGCTGATGATATTATTCCTTATGATATGAAGCATAATGCTGAAATAGATGCTTGCGATCTTGCCATGGAAATTGAGCAGCTCGATTTATTAAAAAAGCATGTTGATAAAACTACGTTTCAAAGAGTCTGCCTTTACTTAATTAGTTGTGTAAAGTATGTTCCAGAACCAGAAGATACACAACTTCTTAAATGTGCTCTAGAGCTCTATTTGAAATACGAACATTTCCCAGACGCGCTAAACATTGCTATTCAACTAAACGATCTCAACATGGTTAAACAAATTCTTAAACTCTGTATGGGAGATGAGCATAAATTGCTTATACAAAAGCAGCTTGCCTACATTCTTGGTCGATTACAAATCTACATAGATTTAGACGATGATGTGGAGAACAGCGAGGAACTTTCGATAATTATATCCAATTCTCATCTTAATAATAGTTTTCTTGCTCTAGCTCGTGAGCTCGATATAATGGAGCCAAAAATTCCTGAGGATATCTATAAAAGTCATCTTGATAATCGTCCTTTCTCTACATCGTCTGTTAGCTTGGACAGTGCTAGACAAAATCTTGCTTCCTCTTTTGTTAATGGTTTTGTCAACACTGCTTTTGGGAAAGATAAACTTCTTACGGAAGATGGAAACAAATGGCTTTATAAAAATAAAGAGCATGGAATGATGAGCACTGCTGCATCTCTTGGTATGTTATTATTATGGGATGTAGATGGCGGTTTATCACAAATAGACAAGTATTTATATTCAAATGAAGATTTTATCAAGGCAGGAGCTTTGTTGGCTTGCGGAATTGTAAACACAGGAGTTCGCAATGAATGTGACCCTGCACTTGCTTTGCTCAGTGACTATGTTGATCATAAAAATGGTGTCATGCAGATAGGTGCAATACTTGGACTTGGTTTAGCATATGCAGCTTCAAATCGCGAGGATGTGCTATCTTTAATTTTGCCAGTTATTTCGGATTCTAAAAAGTCAATGGAGGTAGTAGGAGTTGCAGCATTAGCATGTGGTCTTATTTCGATTGGATCGTGTAACGGTGAAGTATCCTCAGTAATTTTACAAACCATGATGGAGCGTTCAGCTGCTCATTTAAAGGATCATTATGCAAGATTTTTAGCTCTTGGACTAGGGTTAACATACTTAGGCAAACAAGAAGCTGTGGAGGCAACAATTGAAACATTGAAAGTTGTAGAAGGTCCACTTAGTCAATGGTCTTGTGTAATGGTAGACATTTGTGCTTACGCTGGTACTGGTAATGTTTTGAAGATTCAAAATCTTCTTCATATATGCTCTGAACACTACGAAACAGATAAGGAAGACGACAAAAAAAAAGAAGACAAAACAGATACAAATAAAAAGACAGACAAAAAAACAGAAGTAGATCAAAAAGATACTGAAAAAAAAGAAGATACAGTAGCAAAAGATGGAGCTCATCAAGGTATTGCAGTGCTGGGAATTGCTGCAATAGCAATGGGTGAAGACATAGGTATTGAAATGGCAATGCGAGCATTCAATCACTTGCTTCAGTATGGTGAACCAGCAATCAAAAGAGCTGTTCCTCTAGGATTGGCACTTTTGTCAGCCTCTAATCCTAGACTGACCGTTATTGATATATTGAGCAAACTTTCACATGACCATGATGCAGAAGTTGCTCATAACTCTATTTTTGCAATGGGGATAGTTGGCGCTGGTTCAAATAATGCTCGACTTGCAAATATGTTGCGACAGTTGGCATTGTATTACCATAAAGATGCAAATAACTTGTTTATTGTCAGAATTGCACAGGGTTTAGTTCATCTTGGTAAAGGAACCCTAACTCTAAGTCCATATCAGTACGATCGAACTTATATGTCGCCTGTTGCAGTAGGTGGTTTAATTTCAACCCTTGTTTCTTTCTTTGATGTTAAACAGACAATTTTGGGAAAATCTCACTATTTATTATATTGCCTTGTTGCTTCAATGCAACCTCGCATGTTAGTAACTCTTGATAAAGACTTGAAACCTGTTCCTGTGCAAGTTAGAGTTGGACAGGCTGTTGATGTTGTTGGACAAGCTGGAAAACCGAAATCAATTACAGGTTTTCAAACCCACACTACACCTGTTTTATTGGCTCACGGCGAACGTGCTGAGCTTGCCACTGAGGAATATATTGCATTAACTCCAGTGCTAGAAGGTTTTGTCATTTGCGAAAAGAATAAAAACTACGACGTTGATGATGTTGCGTTATAATACAACGATGGGCTGACTGGTTTAATGGTCGCGTTTCAAGCTATGGTTTAATATATTGTTCCTAAGTGCGTGTTAATCAATCGGTCCTGAGTATACTAGTGGCCAGACTTAGATAAAGTTTTAAAAACTGCAATATATGTTTAAATGTAGTTATATTTATTATATATTAGCATAAATCATTTAAAGAGTTTTCTAAAGACGTCCTTTTATTATACGCAATCTAAAGTGTTTTATATGTAACAAAGTGTTTTTCAATTAACTTTCAGTTAATTGAAAAAATAAATGAAACATAAGGAATAAACCTTAGTGTTCTTAAAAAAAAA
>evgR008022c0g2_i01_Hydra
CAAATGGTGTTTTAATTCGTCCTATTTCAAGAAAATATCTGTTTACTTCACCAACAGGTAATGCTGAAAAGTTACCTAACAATAACAATGGTAATTTGATCTCAAATATATATAATGTTACAGAAAGAAATCAGGCAATTGACAAAGACATTTTTACAAATAATCAACCTGAAGTATTAGACAATGATTTTGAGAAAAACATTGAAAAACTTAGAGAAGTGGTAGAAGTTTTTGCAAATAAAATAAATAATAAAAAAAATGCTTTTGAGTTGGCCAAGCAAACACTAATAATGAATAATACAATATCAAGGCTAAATAAAGCTATCGAAGCAATAAGTACAAAGTTTGAAAATTTTTCCCAAGATCAACCTAATAAGTATGTCAATGAATTTAAAATTCTTTCTGGTGGTGAAATTGAAAATGTCAAAGAAAATGATGGAGGTATTGCTCTAAACGGAAATTTACTTGATAGGAATCAAGATAATAAACTTCTAAACTTTGACAACACTAAAAATTCTGAGTTAAAGAAGTTTGAGTCAATGCTTGTGAGTTTAGCAAATAAAGTTGATTCCGTTGCGGAGAAGGTTGGAGTAAGCAAAGCCACACCAGTTAACCAATGGGATATTTTCATTAACAAAAGTGACAAAAACTTGATGAACATGAACAATGCAATAAACAAATTAGTTTTAGAAATTGAAAAGCTGAA
>evgR008026c0g2_i01_Hydra
ATTTCTGTCTTAATAAATTTATATTATAGTATAGTATAATATTCAGTAAAAAAACTATTTATTTAAATTCTGCGGGCATCATCCCTTGATTAAAGAGGGGGAAGCTCTTTTCAGTTATTTTTACGGTGTGGCACCGAAACTGCCCTCTACTCCACCAATGACACGCATATATAAAAACAACTTTCTGGAACTGCGTATAAATGTAACTGCGTTAAAGCTTGATGAAGCTGCTTTGCTACAATAAGATATAGCTTCTAAACGGTATTCACCAAAGCAATTTGTAGATTCGGTATATTGACGCATTTACTTTCCAAAAAACTTTTTTTGTTGTCACTTTCGATATCTGACTTGTTTGTTGGGATGGTGCTTGGTCCTTTTACCATAATACAAGTCTTGGATATAAACCTGCTATCAAACTGTAGAGCTCATTCTGTACGTAGTTATATACTAGTTTTACTTGTTGGATCTTCACTATTAACATTAGCAGCGGTGTCTTATGATAGGTACTTACTACTTACCAAGCTAACAAATTACAACCAATACATGACCAAAAATAAAGCTTTTTTTATCATTTTGATTTGCTGGCTTTTCCCTGGATTGATACCCATTGTCAGGAATTTCAATATGGCGACATATGTAGTTTTACGTATTATTAATTGCTCATTTCCTTTAGTAGCTTTAGGAACATTCTATTGCCTTATAACCAAAGAAGTTTACAATAAAAGATTTGAATTTCTTCAGAACAATACTTATAAAAGTATGGCTGTTAAAAACTTTCGATTTGTTGACAAAAGCAACGGTAAAAAAAGCAACTCGAAGACATTGTCTGTACTCTTGTTCAACACAAAAAATATAAGAAATCGTAAAAATTGCATAAGAGTGAAAAAGTCTGTTACTTTGTTAATCACATGCTACTTTGCATTCATTTTCCCTTTAATTATATGGATGATTTTAGAGCTTTCAAATTTGCAATATAACACTAATGCTCATCAAATATTTTATTTGTGTGCCACCTTCTTAATGCAAACCAATTCTTGTATAAACCCAATCATCTACTATTCGAAGCAATCAGACATAAAAAAAGGTTTTCTAAAAATTTTGAAACCGTTAATAAGAAAAAAGCGCATTTAAAAAACCCGATACAAATGATGAGAAACAAAAAGTGC
>evgR008027c0g1_i01_Hydra
TTTACCAAAAATATTCATTATTTTTTTATATAGTTAATATTTTTTTATATGGAAAGTATACTGAAAACACAAGTTATATTTTCCATTATTCAAAACAACTTTGAAAGAAAAGCAATCACTAGAAAACTATACATCAATCTATATCATCTACAATGGCTAAAATATGCAGAGTTAAACAAAGAACTAAAAAAAATAAAAGAAGACCAAAGTTTTTTATTAAAAAGAATTCAATTATAAATAATGTTGGCATCGATGTAAATATAAGCAGTGTAAATATGGAATCAGGTTTTGTAAATACGAACATAAACGTTACTTCTGAAGATATAAATGTAACTAGGAAGATAACTGTTGAGGGTGAATTAAGTTCTTCTGCTGAAAAATTAAAAAATGTTTTATCGTCTAGACAAAAATTATTTGGTTATTCAATTACTGGATATCGTTTGATTGATTTAGAAATACTTAATTCTGTAGTTTTGACACTTAGATACCCAAAATGCATGTCACCAGAGCTGTTTCTTTCTGAAAATAAAAAGAAGAGAAAAGGTCTAGCCTCTTGTATCGATGTCAGATGCTCTGTTTGTGAATATTTAAAAGTGTTCTACTCATCAATAAAAAATAATAAAAATCTTTTGATAATAATATCATAGTTGTATACGCAATCAGATCTTGTGGTCAAGGCTATTATGGATTCTCAAGATTTGTTTCATTGATGGACTTATCTAAACCGATGACCTCAAATAATTAATCATTGCAGCAAAATCTGTTGCTGAAAAAACTATGATGAATGCTGCTCAAGAGTTAAAAGATTTAAAGAGTAATGATTTGGTTTTAAATTCCATTCCTGACCATTTGCCTGATGTTATCGATGTTGCTGTATCATGTCATGACGAGGCTACTCTTACAATAATGTAGTTGTTTCTGTTATTTCAATGGAGACGGGCAAAGTAGTAGATGTTGAAGCAATGAATAAGTTAAGTATCTATTTTGTTTTATTTATTCTCTATATTTGAAACCAATGTACAAAAAATGACGCATTAGACATTTGG
>evgR008031c0g1_i02_Hydra
GTATATAAACAGTTTTTAAATTTTATAATTAAAAAAGTTTTTAAATTAAATGATTCGATATGGTTTATCAACAGAAGAAGCATTAAAGAAGCTGAAAGAGGCAATTAAAAACCAATTATGTTGTTGTTTACAACTTAATAAGAGTGTGTTATGGCTGCCTCGAAAATTGTTTAACGCAGATTCTACTCTACATTTCTCCTACATCGGTTGCGGCGGTATTTTTTTGGTGTTGATAGCGATATTAGCAAATCTGTTTGCTTTTCACATGGACAGTTATACTACGTTGCTTGCTGAATTTGTTATATTGTTGGTTTTGTTTCTTGGAATTTTATTGTTTTGCATTCGCGAAGTCATATTAAAAGAAAATGAAATTGAGAGAACAATCGGCTGTTTACTTTCTGAAATTAATGAAACCGTTGTTTGGTCAGATGACCAATATCCAAGTCTTACTTTACCAGATTCAAGATCATTTACCTTAGTTTGGACTTACAGAGATGGTCATGTAGTTAATTTACCATGGAATTTGGTAGTTGAGGGGGATGTTGTAATCCTTGGTTCTGGTGCAATAGCTTTAACACGCTGTAAAGCATTACAAACAGATGTTTCTTCTGGACCTCTACATATACTTGAGCCAGGAGATGCTTTTAATCCATCGTCATTATCATCATCGTCGTCATTAGGCAATAAAGACTGTGATCCATTCCCACGACAACGATTTGAAGTGATGCAAACACCTGTAGAAACATTGATTAGTTCCTATTTTCATGAAAATTCTCATAAGTTGGTTAGTATTTGTAATGCTGATGCAAAGATTATTCGTCAATTTATTGAAAGAAGATTACTTTTTGCAGTTGTATTATTCGTTTTGCTGGTCAACATAATAAGATGGAGACTTATTCCTGAAGCATCTGGGTTTAACCTTGAAATGATTTTTTTAATACCGTGCCATTCTATTTTTCCGTTATTACCTCTTACATTTCCTATCATATGGAAGTTTGTTTCATTTTATGGTGCAGCTAGGTTACGAGTCCTTCTTACTATTTACTCTAAACTATACATCAAAGCCACAGATTCAGAGGAAATTATACTTGAGCATGTTGATCGTTTAGCTGTTTGGAAGTATTTTTGGAAAATTCTTTGTGGAGACCCACAAGGTTTACCTAGAAGAATTGATTTAGTGCATGCTTTGGGTAGCATTACTACTTTTAGCTGTGTCGACAAAGAAGGAATTTTATGTTATCCAAATCCAATACCTGAAAAGTTGTTTTATTTTGAGAACATACCTACAACTAGTCAAATAAATGATGTGAATATCAATCCTACAGACGTGCCTGTATGTGTTCCAAATCAGTTAGAAAGAAATTCTGATGAAGTTAAAATACTTACTCTTTCATCTGATCCTCAGAGCTGTAATGGTTTTAGATTTGATGAGTTAGAATGGCAAAAATCTATATCTTCGTTAAAACCACTAGGACTCAATATGCTTCTTAATTCACCTTGTTTGTACCCGTTGAAAACAATAAGAGAAAGCGATTTTATAAATCAATTGTTATTGGCTTCTAATAATTCCTTCATGTCCGGATTAAATGGGTGCTTGTGTGATTTATCTCGAGAAATAGGTTTTAAAGACCTTGTTCTAAAGAATTTTCAACACAAGCAAAAGATTTTTACATATAGTCCTATGGATTCAACCGAAGCTACTGTACCTTCAAGTTTTTATCTTTCTATTGTGAGAGAAAAGCGTCTACCTCACATGGTTTCTTCACTAATAAATCAAACCAACAGCAGCGTCACACAAATGCTCAGTGTTGGATCAGCTGATATTGTTCTTGAAGCCTGTTCAGATATATGGAACGGATGTGATATTAAACCACTTTCTGAAGCTAAAAAAGAAAAAGCTCTTAGCTTTTTTCAGCGTATGGATATCACTGGAAAATGTTTAGCCTACTCATACAGACCTGTATTTGAGGATGTTACTGTGAATGACCATTCGTTTGTTCAATTTTCTAATAAAAGATGGAATTCAGTAGAACTGCAAGTTTCAAATAAAATTGAAGACACAAGTAATAAAATGGATACTGGAAATTTATGCGAGAGCAAAAATGTTCAGGAAGATTTAAGCTGCGAGGATCTACTTATGAAACCTGTTTTTAATACTTCTATATGTCAACTTGACAAACAGATATTTCTTGGCATGGTAACCTTGACTCATCTATTAAAAGAGGATGTTTTGACATTAGTTGAACAACTAAATGTTGCTGGTATTCGATTTGTGCATTTCTCCTATGATAATGAACTACTTAGCAGGGTGTTTTGTCAGCGTTTAGGGTTAGAAACAGGATGGAATTGTCATATTTCGCTCAGTGATGACAATGATTATATTGAGTATTCATCTTCAGCCATTTCTGATTCCTGTCCATTTTTATATCAAGAAGATGTTGATGGCCTAAATCGCAGTTTTAGGCGCAGAGAAAATGAGTCGAGCTTAGGAAGTTTTCGACGACGTGAACCGTATTCAAGTGCAAGCCTACTAGCAAAAGGTGAATCTCAAGATGAAGATTTACAACCTAAAAATCTTTATGCTGAGTTTGATGAGCTTTACTTTATGACTAATCGGTCTAAACTTCCAAAAGGTATACAATGTATGCGAACTCATCTGGAAGAAGTTGACGATGTGCCTCTACTAGTTCCTTTGTTTACTGATTGTACACCTCAAGCTGTATGTGATATGATCAACATTATGCAGGACTATGATGATATAGTTTGTTGTATTGGAAGCTCACTAAATTTGCAAAATCATACTATATTTTCATCCGCCGATGTCAGTATATCAATAAATCCTATCTTACGTAACATCTGCACAACCAAAGAGCAAAAAATTATCGATGAGCTGCATCACTCTCATTTGTGCTTGTGGGCTGCTAATAGAGGCCCCTTGGAGTTGAGTTCATTGTTAAACAGTGTTAGCTGCTCACTTCGATTAACACGCGAAGAGAACTTCCCTATATCAGCTTTATTATGTGAATCTCGACGCATATTGTTTGGGCTTCGAAGTTGCTTTCTGTTTTTGGTTTATTGTCATCTTTTTTTGACTATTCTCCAACTCCTTTCAACATGTTTTTTGCTACCTCCACCAATAAGTGGTTTACAAATGTTGTGGCTGATCTGCATCATAATCCCCCTGTTGAGCATATCACTTATTGGAACACCTAACAATGGCAAGCTGATGAAGGTTATCACGGGGAAAAGAAAAAAACACAGTTATAAAGAATTTGCTTCTAGTTACGTCAAGCAGTTTTTTCTAACGTTTGGATTGAATGCTTTTTTAATAACTACTGGCTTATTTCCCTGGTTATTGACTATGTTCTGTAAAAGAAGCAATGCAAAAACAAGATGCACATGGTTTTTAGGTGCTAGAAATGCATCACTATCTATGGGAGGGTATTATGATGAATTAAAAGATGGCTATCATGCAGCTCAAGATGTTATTCTCTGCTATATTACTATTTCTTTTACTTTGTTGTCTTCATCATTTGTGCATCGTTCCGATTTTTTATGGAGGAAGTCACCTTTTAAAAACAAACCGTGGTGTGTCTGTATCCTGGCATGTTTGCTTCTCCAGACAATTCTTACTTTGATTTTTGATTATTCACATGACTTAGCTGCAATTTCTCCTGTTTTCTATCCTGTTTGCTTTGTTGTTTTACTGGTTGTTATACTTCTAAGTGAAGCTTACAAAGGATGTTTAAAGCGAAACTCCGTGAGACTTCAAAAGCGAGAAAGACTTAAATTTGATACCAAACTTGGAATGAACTCCCCAATATAAATTTTTAGTGTAAATGTCAACTTAACTTATCTCACGTTAAGACAAAAATATATATATATTATTCATATCTTTAATATTGAAATAACTATATCACATTATTAATTATATCATATCAATAACTACATCATATATATCATATCAATAATTATATCATATTAATAATTATATCATATTGATAACTATTTATTTGTTACTCATGTTCATACTTAAAAATAATATATTTACATTTACA
>evgR008034c0g1_i01_Hydra
CTCGATAAGATGTCCACAAAAGTTGATTTAACTGATCGTGAAAATATGATCAAATCAGCAAAGACATCATTGAGTTCTAAGGTTGTTTACCAATATGCCGATGTTTTGGCTCCTTTGGCTGTCGATGCTGTTCTCCGTGTTATTGAATCTAAGGATGCAACTAACGTAGATTTGAATGATATCAAGATTGTTAAAAAGTTGGGTGGTCTCCTCGAAGAAACTGAATTGATTGAAGGTTTAGTTTTGGACCAAGGTACAAGAAAGTCTGCTGGTGGTCCAACCAGTATTAAAGAAGCCAAGATTGGCTTGATCCAATTCCAATTGTCACCACCAAAGACTGATATGGAAAATAACGTTGTCATTACTGATTATACACAAATTGACAGAATCTTGAGAGAAGAAAGAGCTTACTTGCTCAATATCTGTAAACAAATCAAGAAGGAAGGATGTAATGTTCTCTTGATTCAAAAGTCAATTTTGAGAGATGCCACTACCGAAATGTCTTTGCACTTTTTGGCCAAGATGGGTATCATGGTCGTCCAAGATATTGACAGAAAGGACATTGAATTCATTTCCAAGACTTTGAACTGTAAGCCAATTTCTTCCATTGATTCATTCAAGGCTGAAAAGCTCGGAACTGCTGGTTTGGTTGAAGAAGTTAACACTGGTGATGGTAGAATTGTCAAGATTACTGGTATTCCATCCCAAGGTAAGACTGTCACTTGTCTTGTCAGATCTTCCAACAAGTTGATGCTTGATGAAGCTGAAAGATCTCTCCACGATGCTTTGTGTGTTGTTAGAAGTTTGGTCAAGAAAAAGAGTATCATTGCTGGTGGTAGTGCTCCAGAAACTGAAATCTCTTTGGGTCTTATTGAATACTCCAAAACACTTAAGGGTGCTGTTCAATATTGTGTTCGTGAATATGGTGAATGTTTGGAAGTTATTCCAACCACTTTGGCTGAAAATGCTGGTTTGTCACCAATCAAGATTGTAACTGAACTCAAGAATAGACATCTTAACGGTGACAAGAATTGTGGTATCAATGTCAAGAAGGGTGTCATTTCTGACATGATTGAAGAAAATGTTATCCAACCAACATTGACAACATTATCTGCTATTACTCTTGCTACAGAAACTATCAGATTAATTTTGAAGATTGACGATATTGTGTCGAATTAATTGTGCCTTGAAAATATTTCAACT
>evgR008038c0g2_i01_Hydra
CGACATCAACTCCATCACAGACCTCAACAGCCAATGAAACCACTACAACAGCTGCATCATTAACTTCGATTCTGACTGCAACAACTGATGCAACAACTACAACAACTGAATCATCAACTTCAGTAACTCCATCTCTGACTACAACAACTAAAGCAACAACTTATGCAACAACTGATGTGACAATTATAACAACTGAGAAGCCAACTAAAACAACTGAAATAACAACTTCAACAACTGAAGCAACAACAATAACAACTGAATCATTACCTACATCAATTCCATCTGCGACTACAACTACTAAAGAAACCACTGCAACAACTCAATCACCAACTTCATTAACTCCATCTCTGACTACAACAACCGAAGTACCAACTACAGCAACGGAATCATCAACAACAACAACTGAATCATCAACAGCATCAATTCCATTACCAACCACACCATATAATGAAACCACTGCAACAACTGAATCATCAAGTCCGACTCTGACTGCAATAACTGAAGCAACAACTACAATAACTGAAGCAACAACTACAATAACTGAAGCAACAACTACAATAACTG
>evgR008041c0g2_i01_Hydra
CTGCAACGAAGCGTTTCAAAAACAAAAAAAGCATAAAGACGTTTTGCACATTGTGCCGACCATATATAAGTAAATCGTATAATTCAAATAGAATTATGTGACGTCATAGCTCGGTGATGGTTGTGCCAGGTGAATGAAAGTTGTTCTTTCGACAATTCACGTCTGATAACGAATAACAAATATTTTAAACACAGAGGTAAAATCGAGGCGCAAAAACCAATAAAATAAGCTGTAATTTTTTTTTGGAATGCTTTGTTTTGATCGCATTTTAAATTGAAGTTGTTTGCAAATATTTTTATGGAATATTTATATTGGACCAAAAATATGAGCACAACCAGATTTCAACGTCAGCAATGTTATCTTTGTGATTGTCCTAGAGCTCCTTGGGCTATTTTAAACGATTTTAGTGAGCCTATTTGTCGTGGATGTTGTAATTATGAAGGGCCTGAACGTATTGAAACTGTAATAGATTATGTTCGATCTTTAAGAAAAGGCTGGGATCAGCAAGTTAGAGCTTTAAAGCCAAACATGAATTCTACTGGAACATCAATTGGATCAACCCCAAGTCCTAGTGGTCACTCAAACGGATCAATGGAGACTAATGTAAGTATAGCTAATAATCCTTATGCTTTTTCGCAATATAATGGTATGCCACAACAGCTAGACCATTTTCGTTTTTTACACAATGTGAAAAATGAACACCACGGTCCATCTGAAAATCATGTAGCTGAAAATTTTAATCAAAAAAATGTGTTTGCAGTTCCTAATGAATCCTCTATCAATCCAACATCTCAAATTGTTCCTTTAAACTCTCGAAATGGTGATGATACTCATGATGAACATCGTCGCTTTTTAACTAGTCTTGCTGTAAACAATACCAAAGATAATCATGTTAGAGTACCAATAATACATAGACCAGAATACTCTGAATATCCTGATATTGTTAGAGAAACTTTAGAAGTGCTAAACTCATGTGTACCATTTGATATACGGTTCAAAAAGGATCACAATCACATAGGTCGTGTATTTTTATTTGAAGCAGCACCAAGAGCAGGTACAACTTCGCCAAATGAATTTGAACTTAAAATTCATATTGAATATCCTCGAGGATCTGAAAACTTTTGTCAAAGCGCTTCTTCGGTTGCAAAACAAATGTATCAAGACTCTATGAAAGATTATAGTAGAGGGCTTTCTTCAGGGTTTAAATATCTGGAATATGAAATTCATGGTAATGAATGGAGATTACTTGGTGATTTGCTTCCGGAGCAAATTCGTATATTTAAAGAACCTATTGGGCGGGTATTCTTACCAATGCGATTAATTGATTACCCTATATTTCACTCAATGTTAGGGCCTCCACTGCCTGGTAATAATAAGTATCGACCTAAAAGAAAAGCTGACGATGTTGAGGTGATTGGAGCTCCCTCTCGAGCTCCTCGGCGCACGTACAAAGAAAGGGAGGACGAGCAAATTCAAAGACATTTATGGATGCAGTCTCAAGTTGATGCATTAAGACTTTCAATGGCACATTTAACCGGAACTTCTGAAGATGCTATACATCTTAATTCAATTCGTTCCAGTAACATGCTTGATCTTCACCACCACCCTCCACCCCATACACCTTTAACACCTCATACACCTTTAACGCCTAACAGTCAAAATTTAGGAGCACCACATCAACCGTCTCCAAAACCATGGGGTGTGTCAGCTTCAATGGCAGCATTAATAAGTGCTGCTAGTGCAGAGACTAATGGAAATTCTAAAGAGCCCAGAAAACTTTTAATAGAAAGTAGTAAATCCAAATCTCCATTGCATGAAAAATCTCCACTTCGTGATAAAGAGATACCTGAAATAAAAGATAATTTAAAATCAGAGGAAACAAACGATCATAAAGAAGAAAATCAAAATAAAAAAGCCAACATGAAGAATGCTAGTAAAATAAAATGTAGCATTTGTGAAGACTGGTTAGACGATTCACATTTTGTGCAATGCCCCTCTGTAGCAGCACATAAATTTTGTTTTCCGTGTTCAAAAGACAGTATTAAAAAACAAGGATTTGGTAGTGATGTTTTTTGTCCAAGTGGTAAACAGTGTCCACTAATTGGTAATAATCTTCCTTGGGCTTTTATGCAAAGTGAAATACAAACTATTCTTGCAACAAAACCACGTTCAAGAGTCCCTCTTAAGGAAGAAAGCATAGAAGATAAATCTTAATTTTTTAGTTCAGAGTTGATGTTTTATAGATGTGTGTTTGTTAACTTCCATTTGTACCTCGAGTTACATTTTTTTTTTTTGCGAATCTGAGATTTTTTTTAAAAAATACTAAAAAAAAAAAAATTATATATTTGCAAC
>evgR008048c0g1_i01_Hydra
TTTTCATCATCATCAGAAGTGGTCTTGTCATCATCATCAGACACTGAGTCATCCTCACTACATATCAAATTGCACTCCTCCTCACTTAAATCTTTCAAACCATTTGGACCAATAGAATCTTTATTACTAGGGCCTTCTAATATATTGTTTATCTTTTTGTCTGCTTTTTTATTGTTTTCAGATTTTTTTAAAGTTAATGACATTGATTTTTCATTATCAATAAATACTGGCTTTCTATCATCATCCGACAATGTCTTGTCGTCATCATCAGACTCTATGCTACCATCATTACTTGACACTTTCT
>evgR008050c0g1_i01_Hydra
GTCTACTTTGTAGATGACTCTTTCAAAACACTCGCCGTCAGCCCAACCATCAGTGCTCGTGAATTATCAATTGTTGTCGCTGAAAAGATAGAGTTGGAACAACGTGAAACATTTGCGCTCTTTTTCTATAAGAATGGAGAGTGTAGATGTTTGGACGATGATGAACAACCTTGTAAATTGATGGTCTATGAAACTGTTGGTTCTGACGCTGACTTTCAAAAATACGTGAATGATAAACAAGAGTGGGAAAAATTGAAGAAAGACTGGGAAAAGGACTCTAAATTAGTATTCAAGAGAAGAGTCTTTTTAAAGCATAAGGCCATTCCACGTGAACAGGAAAAGTTTTTAAATTACAG
>evgR008065c0g1_i01_Hydra
AGTTTTAATAGCTTGATAAAATTTCAAACATTTTGGTTTTTATAGAATTTTTGGTTTTTATCGAGGAAAAAATATAAATCAATTTATGAGTGGCCTCATTTTCTGCACAAAGTCAATCATAATTAACATTTTTTGTAAACTTAATTTAATATGAATTTTAATTTAAAGCGTCCGCTCAGTGAAGAACAACTACCTACTCAATGAAAAACAGTTTTTCAAAATAAGTTACGAGGCCCAAAAATTTCGATGCAATGGAATCACAAATAATCGTGCAATTAATAATTATTTTATTTGGTTTCACAGCAAATAAAGGCGAAAATTTGAACAATGTTAAACAATTTTTGAAATCGTATAACGAACAAGCGGATAAACTTCAACACGATGACACAGTCGCATCTTGGAACTATGAAATTGATATGTCAACTGAGAATAAAAAAGAAACATCGCGTATCAGTGGTCTGGTTTCAAAATTTTCTCAGGAATATAAAGATAAAGCTGAAATAATACTTAAAAATATAAACGGAGATGTACCATCTTCTTTAATTCGTCAACTCAAGCTTATAACAAGAACCTCTTCAAGTAAAGACAGTAACGAAGCAAAACTAATTGCCGAACTTTTGTCCAATATGACGGCATATTATAGCAGAACAGAGATCGTTAAAAAGATTGACGGAGAAGAATACCATTTTAAATTAAATTCTCACTTACTTCCTTTGATGTCACAAGTTGAAGTTGAAAATCTTGGCAACATGAGATGGGCGTGGAGAGTTTGGCGTGAAGCAGTTGGAATAAAAAACAAACCATTGTACGTCAAATTTGTAGAATTAGCTAATAAAGCTGCCAAAGAAAACGGTTTCGATGACTTTGGTGACTACTGGAGGAAAGAGTACGAGGTTGATAATTTAGAAAACATGATTGAAGATATTCTTATGGAAGTGATGCCTTTATATAAAAAACTACACGCCTTCACTAGATACAAACTTCACAAACACTTAGGAAACAGCTCGTTAGAAGGTAAAAATTTAATTCCTCAGCATTTACTAGGAATGTGGGGACAAAACTGGGTGCATCTTTTTAATATGATGTGTCCTTTTCCAAATAAAACTATACCTGATGTCACCGGTGAAATGAAAAAACGTTCCTGGACAAATGTCGATTTATTAAAGCAAGCAGAAGAATTTTTTGTTTCCATAGGATTAGATAAAATGCCAGAATCATTCTATAATCGCTCATTAATCTCAAAGCAACAAGGAAAGAAATCTGTTTGTCACCCGTCTTCTTGGGATTTAGGAAACGGCGATGTGAGACTAAAAATGCCGTGTATGGGAATTTCACAGGGCGATTTTATTACAGTGCACCATGAAATGGGTCATATAGAATATTATCTTATGTACAAACACCAGCCGTCAGAATTTCGCACAGGTGCAAATCCCGGATTTCACGAAGCTGTTGGAGATACAATAGCATTGTCAGTTATGATTCCAGAACATCTTGAATCTATTGGATTGCTTAAGGGGATGTCAGGTGACGAAGAAGAAGAAATTAACTTTTTATTAAAACAAGCTTTGCAGAAAATCTCATTTTTACCGTTTTCGTTCATCATTGATAAATGGAGATGGTCTGTGTTTAGAGGAGATGTTAAACCTGACAACTACAATACTTACTGGTGGGAACTCAGTCGTAAATATCAAGGAATGAAAGTTCCAATAGTAAGAGCAAATAACTCTCAACTTTTCGATGCCGGCACATTTTTTCATGTTGCCCACAATACAGAATATTTGAGGTATTTTATAAGCAATGTCCTCCAGTTTCAATTTCACGAAGCGCTATGCGACGAGTCCGGACATAAAGGAGCCTACCACAAATGTTCAATCCATAAATCAAAAGCAGCTGGTAGAAAACTTAGAAATATGCTTGAGTTAGGTAAAGAAAAACCATGGCCTGAAGCTTTAGAAGTATTAGCTGGAACTCGTCATCTTTCTTCAAAAGCTATTCTGGATTACTTTAAACCATTAGAAAACTGGCTTGATCAGCATAGAGAGGCTAACGGTTATGAATTAGGATGGGATGATTCGGATATAGATATGCAAATAATGTTATAATAAAACGTAACATATTAATAGCGTTCCAATATTTATCACATTGGTCATGCTTCTGATATAAACATATGACTCCTTGGAGTCGTCAAAACCGGAAAATTGAACTTGATAACTAAAGCTTTAATTTAAAAAAATCTATTTTAAGCAGACGTCATTCGAGTGTTTCCAAATAGTTAGAATAATTAGAATAATTAGTTAGAAATTATTTAGAATAATTAAATTTTTAGAAATTTTTTAAAATATTTTTGTATTAAACTTTAGATACCTAGTATATACAATGACATATAAATAAGTAAACTGCATTTTGAAGTAATAAAGTTG
>evgR008066c2g1_i01_Hydra
GCCGGTTACATATCTAAGGAGAAAGGCTCACCGCTGTTCCTCCTAATTCAAAATTTTCTTTTGTTGGAGGTTTAAGTCGGCCTTAAAATCAAAATGTCTGCACTTAAAGAAAAACAAGTTGAAGAGGAAGCTCAAATTCATCGTATTCGAATAACATTAACAAGTCGAAATGTAAAGAGTTTAGAAAAAGTTTGTGCTGAATTAATCAGAGGTGCAAAGGAAAAAAAATTAGTTGTTAAAGGACCTGTTCGTATTCCAACTAAAGTTCTTCGTATTACATCTAGAAAAACACCTTGTGGTGAGGGTTCTAAAACATGGTCACGTTATGAAATGAGAATTCATAAGCGTTTGATTGATTTGCACAGTCCAGCTGAAGTTGTTAAACAAATCACTTCTGTAAGTATTGAGCCTGGTGTAGAAGTTGAAGTTACTATTGCAGATAGCTAAAATGTAAAAATTCTGGATTTAAGAATATATCATTTAAAAGAAAAAAAAAAAAA
>evgR008075c0g1_i01_Hydra
CAAAAACTTCTAGTACACAAGAAATTCGTTCCGCATATAAAAAACTTGCTCTTAAGTATCACCCAGATAGAGCACCAGAAGAATTAAAAGCCGAATATGAAGACAAATTTAAGGATATCGCTCACGCCTATGAGGTCCTTACCGATGATCAAAAGAGAAAGATTTATGACCAATATGGTGAAGAAGGTCTCAAAGGAGGTGGTATGGGTGGTTTTACTGATCCATCAGATATCTTCAGCCACATTTTCGGTGCTGGTGAAGACGGCTTTGGTTTCTTTGGTGGTGGTGGTAGATCACGTCAATCTGGTCCTAAAAAAGGAAAAACCATTGCACACGAAATTCAAGTCTCATTAGAAGATTTATATAATGGTGCTACCAGAAAGATTAGAGTTACCAGAACACGTATTTGTACATCTTGTAAGGGTAGTGGTGCCACAAAGGAAGACGCTGTTGTTACCTGTAAATCATGCCAAGGTAAAGGCAAGAAGGTTGTTACCAGAAGTATGGGTCCAGGTTTTGTTCAACAATTTGTCACTCCTTGTGATGTCTGTGAAGGTACTGGTAAATCAATTGATAAGAAGTTTATTTGTAAGGATTGTCAAGGAAACAAAGTCACAAATGATGTTAAAGTTTTGGAAGTTCACGTTGATCCAGGTATGAAGGAACAACAACAAATCGTTTTTGAAGGAGAAGCTGATGAAAGACCAGACGTTCTTCCAGGTGATATTGTATTTATTGTTCAACAAAAACAACACCCATATTTCACAAGACAAGGAAATAATTTATATCTCAAGAAGAAGATTAACTTGTTGGAAGCCCTTACAGGTGTTGAATTTAGTGTAAAGCATTTAGACGGTAGAAATCTTGTCGTCCGATCGAAGCAAAACCAAGTCATTAAGCCAGGTATGGTTATGCAAATTGCTAAGGAAGGTTTCCCAATTCACAGATCCCCATTCCAAAAGGGTAACTTGTACATTGAATTTGAAGTTGAATTCCCAGAACAAATTCCAGAAAAACTCCACACCCAACTCTCTTCTATTTTGCCAAAGAAAGCTAATGCTGCTGATGTAATGGATGAATCTGCTGACAATATTGAAGAAGTCTATTTGCAAGAAGCTGACCTTTCACAAGGTGGTGACAGTAAGAACCAATACGATAGTGACGACGAATATGAACGTCGTGGTGGACAAGGCGTTCAATGCGGT
>evgR008080c0g1_i01_Hydra
TCAAATGTTTATTTGTTATCAATTTCATGATGTTTAAAACATTTTTAGCATTTAGAAAAATGAGTTTAATTTTGAAACGAATTCCATTTGTAAAAGAGTCTCAGTTAAAAACATTGAAAAAATATTTATCATCGAATCCGATAGATTTTCCTGATATGGGTCCACCGATTCCGGAATACAAACAACGATATGGAGAGGCTGATGATGTGCTACGTTCAAGACTATTATATCAAAGTCGTAAGAGGGGGATGTCAGAAAATGGACTATTATTAAGTACCTTTGCATCTTTTCATTTAAACAGATTTACACGAAAACAACTCGAGGAGTTTGATAAAATTATAAACAAACCTTCTAATGACTGGGATATATTTTACTGGATTCTAGGAAAAACGGAAACCCCACATGAATACAATTCTGAAGTTATGGACTTGCTGAAATCATTCTCAAAGAATCAAGGCATGGAAGCCAGATATAGACAACCTGATCTTATCTTTGAAGCAGGAAATATTGAAAAATAAAATATGTAGTAACATGATAAATTGAGTTCTATAAATAAAAAGTTAATTTACGATTGAAAAACAATTATAAATATTATTAAATAAAAATTAACATGGTTTTGTTATTATTTTTTAATATTTAAAACACACCCTTCAAAATTAATAATTTCTATGCAGTAAAGCAAGCATTTGTTTTTCACAATGC
>evgR008088c0g1_i01_Hydra
AATATTAGAATCTATTATAAGAGAAAGAATTAAAAAACATCTTTACAAAAACAATTTACTAACAATTCAACAACATGGTTTTGTAAAAGGCAAATCTTATACTACAAATTTGTTAGATACTTTTGACTTTATTTCTTTGTGCAACAGCAATGGTATACCTGTTGATGTAGTAATGTTAAACTTTGCTAAAGTTTTCAATAGTGTCCTCACGAGAGACTCTTTGCTAAATGCATATGGTAATAATGGTTTAGTTTTAAAATGGATGGAAGCTTTTCTAAGTAAAGGAAGACAAAGAATAGTTCTGGGTGAAGTTGTTTCTGAGTGGGTTGAAATTTTTATTTGCGTCCCACAGGGTTCTGTTATTGAACCCTTAATGTTTATTATATACATTAATGATCTGCCAAGTAAATTAACTAACAAATATAAATTGTTTGCTGACGATACTAAGATTTTG
>evgR008090c0g1_i01_Hydra
CTAGAAAATAAAAAGGCGAAGAATAGTATTGAAAATTAACTGATCAAAAAAATGATTAATATCGAAAAATAACTGACCGAAACCCATTTAAATGGTTACAATATCGAAAACAATATAATCGAAATAATTATATTGACTGACTACCTAGGGAGAACATGCACGGAGTGTACATAAATCAAACTACGATTTTGGTAGGGGCAGTTAGTCTAGTTAAATATGGGTGAAATAGGTGATTTGAGTGGTATGAGTGATATGGGTGAACTAGTTTCTTATGAACATCCTAATTCCACATTTGTAATTTTCGAACTTATTCCTTTTTTCACAACTTTGATCGTATTCTTCTTTTTGAGCTATTTTGATATTATTCTGCTTTTCGGATTTTCAATACAATGCCTCTTTTTTACAATTTCGATGTTATTCTCTACTTTGATGTTATTCTCAAATTTCCCAGTTTCAGATCATCAGCTATCCTTTAAACAATTTTCAAAGCAAAAACTGGTATTTGTAAAACTAATGTCTTATATAGCAGGACATGCATAAGATTTATTTTCATCGCCAATGACTCTTTATTTTGTATAAAACTATAAA
>evgR008090c0g1_i02_Hydra
TTGGACTGACTACCTAGGGAGAACATGCACGGAGAGTACATAAATCAAACTACGGTTTTGGTAGGGGCAGTTAGCCTAGTTAAATATGGGTGAAATAGGTGATTTGAGTGGTATGAGTGATATGGGTGAACTAGTTTCTTATGAACATCCTAATTCCACATTTGTAATTTTCGAACTTATTCCTTTTTTCACAACTTTGATCGTATTCTTCATTTTGGGCTATTTTGATATTATTCTGCTTTTCGGATTTTTAATACAACGCCTTTTTTTTACAACTTCGATGTTATTCTCTACTTTGATGTTTCTCAAATGTCCCAGTTTCAGATCATTACTGCAGCTATCCTTTAAACAATTTTCAAAGCAAAAACTGGTATTTGTAAAACTAATGTCTTACATAGCAGGACATGCATAAGATTTAT
>evgR008091c0g1_i01_Hydra
TGTAAAATGTTTAACGCAACATTTTGCAACTTTGTAAACGAATGAAGAAAATTCCAACCACGTGAATTTTGATAAAAAAGTTTTAATCTTTCACTTGTTTTGTCAACCCTAAAAGCGAAATTGAACTTGAAACACACTAGTTTTTAATATGGAATCTTCGTATGGCTGTAGCGAACATCAAGCTTCAGAACTTGGAATGATATCTTGGTATCCTATATCATATTATGAGCAAATTGATGATAATTTAATGACTCGTCTACCTTCTTGGTTTCACAATTTCATTTCACCGGTTTATAAGATACCCTCTGAAAAGACAAATTACAAATCGCTTCCAGAATACTCTACTAACATCGAACAAATAAATAAAATCTCACTGCCTTTGGCATATCCGTGCCATTTACATGGCAACTGTTGTGTCGTTCCTGGAAACGAAATAGAAAAAATTAGCTTGCTTGAAAAACGATGGTGCTCAACAATCAGGGCTCATCGAGCTACTTTTTACCGCGACAAAATATTTGTCGGATCTTTAGCTAGAAAGACTACTCCGAACGATTTAGTACGATTTTTTAAAAGGTTTGGAAAGGTTATTGAAGCTAAAGTTATTGTTGATTTTGACGGCAACTCCAAAGGTTATGGTTTTGTTTCGTTTGCTTCGGAAGATACAGTGAAATCCATTCTTAACCTAGGCGCGATCTATATGGAGAAGAAGAGGATAGCTGTGGCATATGCTATCCGCAAAAAACATGATGACGAAAATTAGTCAAAGAGCTTTTTGTTGTTTTTTATATTAACTATATCCTTGAAAGATATTTTTAACCATATTTGTAACTATGCAACCATATTTGTATAACTATTTTTTAAAGCTTAAATCTCTGGAGATTTATTTTACTTAATGACGAAGGAAAAGAATGTATATATTTAACGTGTAACGTGTTTTACTTACTGTTTATTTATTTTTATTGTTTGTAAGCTACCTACTCTGTATTCGATTTTTATTGTTTGTAAG
>evgR008101c0g1_i01_Hydra
CAGTTCAGTATTATGATTCTCTTTTTACCAATTTAAACAAAGATGTTCCATTACTTGTGCATCGTGTTGCTCAATCATTATTGATTTGTGATGAAGTTACCTCGATAATTTTTGATATTATGAATTGCCAGACCCAAGATAATGGAAATGATTGTGGTCTTCATGCTGTTGCAAATGCCACAGCTCTATGTAATGGAATTGATCCAAGTGTAATTGTCTGCGAGAGAAAAAGTATGAAATTTGTTTTTCAAGTTTGTATTGAAAAAGGTAAACTTGAAATGTTATATAACCATACTAAACGGTAATAAAAGACTTTATCCTTTGTGATGATGTCACTGTGCTATGGCATAAGTTTAAGATTAGGTCGTTTTTTATACAAGCATGGAATCCAGTACAACATGTTTTATATAGAAAAATGAGAGTTGTTAGCTGGTGAACGCATCATGCCTAAGAGATACATTTACGGTGTAATGGGAACCCACATGCTCGATATAGTCAAATGTCACAAAGGAAGCAGTTAATACG
>evgR008111c0g2_i01_Hydra
AAAAAAAAATTTGTATTAGATTTTTGTGACGTCATTGAATTTCGCTGATTTACTAATGTTCTAGGTACTTTTTACTGTAAACAATTTTTGCAATACAATGGCAACATCCAAAGGTGCCAATCTATATAACAGACAAAATTGGGAACAGTCGGATTTTCCAATACTCTGCCAAACATGTTTAGGAGACAATCCTTATGTGAGAATGTTGAAAGATAAATTTGGCAAAGAATGCAAGATCTGTGTTCGACCTTTTACCATTTTTAAATGGTGCCCAGGCGCTAAAATGCGATACAAAAAGACTGAAATATGTCAAACTTGTGCAAAATTAAAGAATGTGTGTCAGACATGTCTGTTAGATCTTGAATACGGGTTGCCTATAGAAGTGCGTGACAAAGCCCTTGCTATAGCAGATACATTGCCAAGATCAGACGTAAATAAAGAGTTTTTCCATCAGACATTGGAGAAAGAGTTAGCAAATACAGATGGTACAGTAGCTGCTGGAACTTTGGGTAAGTCACAAGCCGCTAGCGACTTATTAATGAAGTTGGCGCGCACAACACCCTATTATAAAAGAAACAGACCCCATATCTGCTCATTTTGGGTTAAAGGCGAATGCAAGCGAGGAGAGGAATGTCCCTACAGGCATGAAATGCCGAATGCTCCTGATGACCCACTATCTAATCAAAATATTAAAGATCGATTTTATGGTGTTAACGATCCTGTGGCTGCAAAGATGTTGAAGCGTGCCGAGACCATGCCTAAGTTAGAGTCTCCATCTGATAAATCTATTACAACGTTGTATGTTGGCGGTCTTGATGAAAGAATTAAAGAAGACGATCTGAGGAATTTTTTTTATCAATATGGCGAGATTCGCTCTGTTGTGGTGGCTTCTAGCAAAAGCTGTGGGTTTATTTGTTATACGTCGCGTCAAGCAGCTGAGATGGCTGCAGAACGGTCTTTTAATAAAGTGATTATCAAAGGAAAACGTCTTAAAGTACTATGGGGTCGTTCTCAAGAGCAAAGAAGTGGTGGAAAAGACGAAAAGGGTGACAGACTTAAAGATTATCCGCCTGTCCCGGGACTTCCTGACGCATTACCACCAATACCTACAGAAGATGCTCCTATTATGGAGCCCCCGAATCTTTTTGATCAGCTGCCATCTGAACAGAGCACATCAAACATTCCTCTCCCTGCTGCTTATTCTAATGCTCCTCCGATGTTCCCGCCTCACATGAGTTCTTCTCATTCTCAGGGAATTCCGCCGCCGCCTCCAAATATGAGTGGTGGTTTTCCTCCTGGTCCGCCCCCTTTTATGCAAAGGTTCCCGCCCCCGCCTGGAAGTTATCCTAATCAAGTACAACCATCTGGTATTCGATTCACTATTTCAAACGGACCTCATCCAGTGCGTCATGGTCTGCATTATCCATCTCAAGATCCTCATAGAATGGGCACGGTAGGATCTTCCGAAAACACTTCAGTACAAACACCTGCAGTTAAAGAGTAAACTTTACAGAACGTTATTATTTGATTTGTCTGAATTGTCACACCTACTATACAGGGAATATATTTTATTTCGTTTAAAAAAAAAAA
>evgR008117c0g1_i01_Hydra
TATTACTGTTCATAAACAAATATGACAACACTAGCATTAAAGATTACCTCTGCTCTATTTTATGCTGTTTCTTCTATTCTAATTGTTATTTTAAATAAAATTGTTTTAACTTCATATAGATTTCCTTCATTTCAATTTCTTGCAATAGGGCAGATGGTGGCTTCGTTATTAGTACTGGGTATTTCAAATCAATTAGGTATAATAACGTTTCCAGGATTCAGTAAAGATGTATTTAAAAAGGTATTTCCGCTACCATTATTATATTTCTTTAACTTGGTTTCTGGTCTTGGAAGTACGCAATCTCTAAATTTACCGATGTTTACAGTTTTGAGAAGATTCTCAATTTTGTTCACTATGATAGGAGAATATTTTGTTTTAAGTCAAAAAGCAAGTGTTAAAGTTCAGTTGTCTGTTTATTGTATGCTATTTGGTGCTATTGTTGCAGCTAGTCGAGATTTTGCATTCGATTTGAATGGATACATTATGATCATGATCAACAATCTCATGACCGCTGCAAATGGAGTCTACATAAAAAAGAAGCTTGAGTCAAAAGATTTAGGACAGTATGGTTTGATTTTTTATAATAGTTTGTTCATGCTTGCACCAGCACTTTGTTGGAGCATATCTTCAGGTGACATGAAGTTGGCATTTACTTATACCAGATGGGAAGATGTGACTTTTATTGTGATGTTTTTAGTTGCATGTGTTTTTGGCTTTGTTTTGAACTACTCTTCTGTTTTGTGCACCAATCATAACTCTGCTCTGACCACAACCATTGTTGGCTGTTTAAAGAATGTGTTGGTGACATACTGTGGTATGTTGATTGGTGGAGATTATAAATTTGACTGGGTTAATTTTCTGGGTTTAAATATAAGCATTGCAGGTAGTATTTTCTACTCGTATGTTGGTTTGACAGAAAAGCAACCATCATTAAATAGGCAATCGGCTTCATTCAAGGCCTAGACCTTTTTTTAAGTGTTTTAAAGTTTTTATATTTATTCATATAAAATATATTTTTATACATGGCGAACATGTTTATACTTTTCAACACCACTGCAAATAGGGTGTGTGGATTTTATACTTTTCAACACCACTGCAAATAGGGTGTGTGGATTTTATACTTTTCAACACCACTGCAAATAGG
>evgR008119c0g1_i01_Hydra
CAGATATAGACAAGAAGGTAAGGCTGAAGAACTCCGTATTGCCGAAAAGGATCTTTCTGATTTGCAAAGAATGGGTCTCAAGGCCTTCCTTGCTCGTGATACCACTACAAGCAGTTCCACTGGATCCTCCTCTTCTTCTACATCAACACCAACTACACCAGTCCAACCTGCTCAACCACAAGAAAGTGCCATCAGAGTTCGTCTCCTTGATGGTACCACCATCACTGCCAAATTTAAGCCAACTGATAGTGTTAAGGCTGTGCATGGTCACATTGCTACCTTGATTAAGGATTCCCAATTTTCTTTCAATTGTCCTGGTGTTCCTCCAAAGACATACTCTCTCACTTCAGCAAATATTAACAAGACTTTGGAAGAAGAAGGTTTGCAT
>evgR008120c0g1_i01_Hydra
AACACAATACATTATAATACAGATGACACAATATATTTGATGAGGTTTAGTTGATAATTTTTAATATAAACGTAAAATAAAAACGCCTTTTACATTTGATGCAAAGTGAAACTATGCAAATTTTACTTTGAAGTGAAACCTGAAACCTGGAAACTATGCGAACTTTACATGCAATTTTGTTAGTTTGTATTTGTGATATCGTTTATACAAACCAAGTGAACGATTTAAAAGTTGATTTAGCAGAAAAGATCTTGAATGATTTAGATGATGATTTAGATGATGATTTAGATGACAGCTATGAAAATGATGAAATCACCGATGAGGTTTTCTCTGATAGCCGATGGAGAGACCCTACTTTTAGAATCCCTCCATTTAGAATCCCTCCATTTAGAATCCCCCCATTTAGAATCCCTCCATTTAGAATCCCTCCATTTAGAATCCCCCCATTTAGAATCCCTCCATTTAGAATCCCTCCA
>evgR008122c0g1_i01_Hydra
AAATCGTAGATCTTGCTGATTTTCTTTCCGGAAATGTTAAACAAGAAGATGTTGATAGAATTTATCAATCTTTCCGTGAAACCTCTGCATTGATTATCAGGGATCCTAGAGTTACTTTTGAAACTGCTTCTGAATTTTTGGATATGATGGAAGATTACTATGCTCAACCATTCGAAGTCAAGATGAATGATGTTCATCCAGAATGGTCCTATCAAATTGGTGCTACTCCAGAATTAACTGAACGTCCAAGAGATAACACTGAATATGTTGCCACAAAGTTGAAGGATAATCTCGAAAGTCAACCAACTCCAACAAAGGGAAAGGATCCAAAGTGGAGATTCTTCCACCCATATGGTACTCTCAACCCTGAAACTAAATTCCCAACACTCAACCAACCAAATGTTGTCCCAGAAGCTTTCAAGGATATTTGGATTGAAAAGATGGACAGTTGGGCCACTAATATGGTCAATGCTATCAACACTGTTGGTGAAATGTTGGCATTAGGTTTAGGACTTGAAAAGGATAAATTCACAAATTTATTGAAATACGGAAGACACTTGTTGGCTCCAACTGGTTCTGATCTCAACAAGTACAATGAAGTAGGAACTGTTCTTGCTGGTTTCCACTTTGATTTGAACTTTATCACTATTCACTCTGCTTCTAGATATCCAG
>evgR008125c0g1_i01_Hydra
AACTGCTTTGCTTTCTCAATTGTTCTTGTTCTCTCAAGTTTTATACAAGAGGTTCGGAGACAATATTCTTATTGCCTTGTTAGGTAGATGGGAAACTCCACAATATGGTCAGTCTGTTCCAGTTGGTGGTTTGATTTATTACATTTCTCCTCCAGGTAACTTGAACGAAATGTTGGTCGATCCAATCCACGCCATCTTGTACATTGCTTTCATGCTCGGATCATGTGCTTTGTTCTCTAGAATTTGGATTAATGTTTCTGGATCCAGTCCAAAGGATGTTGCCAGACAACTCAAGGAACAAGGTTTGAACATTGCTGGTCACAGAGACGATCCAAAGGAAATGATTAACACTTTGAATAAGTATATTCCAACTGCTGCTGCTGTCGGTGGTATGTGTATTGGTGCTCTCACTGTCATTGCTGACTTTTTGGGTGCTATTGGTAGTGGTACTGGTATTCTCTTGGCTGTTGGTATCATTTATCAATACTTTGAAATGTTCGCTAAGCAACAAGATGAATTGGATATTCCTTTCTTGAGTGGACAATAATTTGTAAAC
>evgR008134c0g1_i01_Hydra
TTTTTTTTTTTTTTTTGAAATTTTTATCATTTCAAAAAAAGTTTTTACCATAACTGTATAACTGCATAACTGAAGGAAATAAAAATATTATCAAACACAACTGAAGAATGATACAAACAATTGGTATCGTGCAACAAACAATTGTTATTGTGCAACAAACAATACAAGTTGTGCAACAATTAATATTCGCCGTACTTGTTCCGACCTTTTTATACTTTATATTTTCTTACCTTCAACATTTATGGATCTGTACTAAATACCCAAAGGGTCTGTTTCCATTACCGTTTTTAGGAAACATTCATCAGTTAGGTAAAAATTCTTCCAAAACGTTTACATCTTTATCAAAAATCTACGGAGATGTATTTAGTGTAAGCATTGGTTCTCAAAGACTCGTAATACTAAATAGTATGGAAAGCATACACGAAGCTTTATTAACTAAAGGTTCATCTTTTGGTGGTAGACCAAATGAGTACACTTCAAATGTTTTTACAAAGGGTTATAAAAACTTATCACACACTGATTATGGACCAACTTTAAAAGCATTGCGTAAAGTTGTTCACCTTTCCCTTCAAAAATATGGAGGTGGATTAACGAGGCAGGAACAGATGATAGCTTTTGAACGAGACGAACTTTGTAAAAGACTTTTTAACGCTGAAAAAGAAATAGCTTTGCGCTGCGAAATAGATTTTTGTACCGTCAATGTAATATCAGGCTACTTATTCAACGAGCGCTTTCTAAATCAAGACTCCGAGTTTAAAGATGTTGTGAAAAGCATTCAACTATTATTGGATAACTCTGGAATTACTGATAAAACAACGTTTGTACATTGGCTTCGTTACTTGCCATTGCGGGAATGGAACGAAATAAAAAAAGCAAGAGTTACTTTAAACCCGTGGGTTGAAAAAAAAGTTGAAGATCATTGGAGACGATACGATAAAAATCAAATCGGAAATGTAACAGATAGCATGATTGAACATTTTTTATCAAATTACGAAGGTCTTGAAAGTGATAATGCCAAGAAATACATCACCTTATTATTGATTGAATTACTTGTTGCCGGTACCGAAACAACGGCTATTACTATATGCTGGATGGTTTTATATCTTATACATAATCCTGAATATCAAGAAAATATTTACAAAGAAATCACATCAATTATTGGTTGTAGATATCCAACACTATCAGAAAAGCATTCTTTCCCGTTATTACAAGCATTTATTCAAGAAACATTACGAATGACATCCGTTGTGCCGCTAAACTTGGCTCATAAAACAATGAAAGACACCAGCGTTTGTGGAATAATTATTCCCAAGGACGCTATTGTTGTGACTAATCTATGGGGTCTCCATCATGACAGCCGATACTTCCAAAATCCTGATAAGTTTGATCCTAAACGCTGGATAGACAAAAATGGTCATTTTGACTCTGGTTCCCAAAAACATTTTAAACCTTTTTCGGCTGGAGCAAGAGTGTGTCTTGGCGAAACTTTAGCCAAAAACCAACTTTTTTTAATTATCTCTGGTCTAATTATAAATTTTGTTTTCAGATCTGCACCGGGAAATGATTTACCTAGCCTCGAAGGACAGTTTGGCATCACATTCCGTCCGCATAATTTTAAAGTTGTTTTAATAAAAAGAAGTCCTGAGATAATAAAAATATAAATCCTGATGAAAACAAAAAAATTTATCCTGTTAATTTAAAATATTTACAACGAAAATTTTAACATAGTATTTAATGTATTTATATATTTATTCGATTAATTTACAATATTAGTAGTTAATAATAGCGTAATTTTAAAAAAAAAAAAAAA
>evgR008140c0g1_i01_Hydra
CTCAAAGGTGACGGCGTTCGAGGTAGTGATATAAAGTATAATGATGCAAAAAATAAGCTTTTTAATGAGATAAGAGAAATATCAAAGCTAATTGATGTTAATCAATATTTTGATAAAGACGGTGAATTGGTCATAGAAACAAAATCAGGTATCCCACTAATCAAGGGTGATATTAAATATAAGGTGGAATTTACTCATACAGATGGACTGGATAATTATTTAAAGAGTGATTTTGTATTTCCCGCTTTTTTTGTTGTTGATGAAGATATACAGGACACTTCTCCTCAATTTAGAAATGATTTAGTGACAAGTGGAGTTACAAATCAAATCACACATCATTTAAAAGGCGGTGAGTTACAAGCTAGATTTGAGCTGAGGGATAAAATTATACCAGATATAATAAACAGATTGGATATATACACCGATACTTTGGTGAGGGAAATGAATGCTGTTTATGCAGATGCCACGCCTGAATTAGGCTTTCAATCGGTTATTAGTAGTAATAAATTTGTAACAATTAATGATCCAATAACAGCTAGCGGTAATTTTAAAATTGGGCTGTTTGATCCTAAGGATGGCTCATCTATTAAGGATGCTACAGGTCAACCAGAAACAATAAAAATTGATTTAGGTGGTGTTACTACATTGCAACAATTAATAAATAAAATCAATACTCCTTCAACTGGTGGTTATAGTGTTATAGCTAAACCGGATGGTTTTGGGCATTTAACAATAAGCCTATTGGGTCCAAATCCTATAAATGCTAATTTTGCCATTATTAATGAAGATAGCAAATTTGGTGATGATAACGTTGGAATTAATCAGTTTTTTGGATTCAATAAGCTCTTTACTTATGTAGATAATAAGCATTATGACGGTAAAACTTCTGCTAATAGCGCTATATCATTAAAAGTTTCGGATGAAATTACAAGTAGCCGTCAGATATATACAGCAAAAGTAAGGAAAGATGGTGATATAGGAAAAAACAATAAAGCTGCTTTAGAAGAATTACTTAAAATTCAAAATAAGGAGTTTAGCTTTAAAGGAACAGATTCTATTGCAAAATCTACTGAAACTTTTTTAAGTTATGCGAATGGTATAATTGATACACTTAATATTAAAC
>evgR008143c0g1_i01_Hydra
GTCTGGTGTGGCTGCTAATATGACAACTGATGAATATTATGATCATAAAATTAATAAGGACTCATACTTAATATCTGTTTCAAAACATAAGACATTCGACACTTTTGGTCATTCCCATGTAATATTTGAGCCTCAAAAGTTTGAATGGTTGACCACATTTGTTACAAAAATAAGATCTCAAATATCTTGTAGCTGCAATAACGTATTTACATCTTGGTTTGGCTCGGCCATGACTTCTGGAGCCATAAGCACGCAACTAGGTTCACTTTGGAAAAAAGCAGGCATTTTTGATAAAAACTCTAAAAAACGAAATGTTTGTTCAACTTTGTTTCGAAAGGGAGCTTGTACAGGAGTGCACAAAACAAATGGTAGTAATAGCAAAGAAGTAGCAGATTTAATGGGACATTCCCTTGCTACGGCTGAAAAATACTATAATACAAGCAAAAAAAGAGATACTGCCGCAAATGCAAGTAATGTTGTCCGCACTTATTTTGAAAGCAAAGTGATGCAGTCCTCGCCAATTAAATTGGTTCGAAATGTTGACGTTGATTTAGGCAGTGAAAGAAAAAAATGGAGTATAGAAG
>evgR008146c0g1_i01_Hydra
TGTTCTTATATTTTTGATTTGTCTGCGACTACAATATATATTGAATCCTATAAATATTTTAAGCCTGTTATAAATAGCTTTATAGTCGCAACCGTATTTGGAGTATATATTTATATAATTTATAAGTACTATAAACTTAAGAAAAAGACAATAAATCCGCTGCCCAGATCCTTTTCTATTGGTCAAACATGCTCTACAGCAGTTAAAACGAAACGTAAACACCGTGACTTTTATAAAAGTGTAGCAATGTTTAGCCTATTGTTATCAAACGTTGTATTTTTTGCAGTGCCAGATGCATTCATCACTTATTTTTTGTACAAGCCGCAAACTACGACCCAACCCTCTAGTATATTGATTTACATCATGTATTATATTGGGTTCCTAACAGACGCAATTATTTACGCATTCTTTCCTCAACCAGTAAGAGATGAACTCAACAAAATATATATGTGGATAACAAAAAGAGATGAGAAACCAAACCAATCAGCTTGCAGCGTCTATTAATTTAATAATCAATAGCGTTACTATCGCAAACATCGCCGCTAATACTATTGCAAACATCGCCGCTAATGTACATAATGATACTTATACTTTGTTTTAAGTTTTGCTAACTATTTATTACATATAAATGTTTTGTGTCCAGTTATTTACCGTTATTACTTAAAACTGTATTTATTGAATTACTAAGAGTTATTAAAATATTTATTTTAGCAATATAAGGGAAACTGGTGCGGAATGGGAT
>evgR008150c0g1_i01_Hydra
TTTCTTGCACTTCCCAATAACTGCCGAAGATGCATCGATTTCTTTCTCTTCGTCTGTTAGCACTTTTTGGGTTATTATTCCTAAACCTACCATGGTTATTATCTGTTGTTGGTGCGACTGTTTTATACCTTTTAACTGGTGGTTGGAAATGGCTGCGCTTACTTTACCGGACAACATGGCGCGATATGAAATTTGCAAAAACTGTAATAGGTATAGAAATTTATATACGGAAGTTAATTAAAGAAAAAAAGACGATTGCAGATATTTTTATGGATAATGCTGCTAAAAAACCGGATAAGTATATATTTGAATCGATTGATACTAGAAAAAGAATTACATATAGACAAGCTGCTATTTTAGCAAATAAAATTGCCAATGTATTTTTTGAAGCAGGCTATAGAAAAGGTGATGTGGTCGGATTGCTG
>evgR008152c0g1_i01_Hydra
ATTATTGTATAGCTTACAATATAATTATAAAATGGCAAACATTAACAGAAAAAAATCTAAAAAAATTAAAAATGAAGTAAAGATACATAATTGGGCATCATTAAAATGGTTTTATTTTTGTTTCATTGGATTAGCTGCTTTTAGTATGTCACATGTTTTTATGTCGAATAACTCAGGATTGATGAATAATCAACCTAGTAGCGTAACAAATAATGAAACGTCAAAAACTTCAAGTTTCCTCGAACAAATCCAAATGCGGTCTTTTGAGGTTGATGGAAGAAATCTGATACCCAGGGAGATTCAGAGTAATAGCAAAAAAGTTAAAGTGTTTATGGTTGACAATTTTTTGTCTAACTTGGAGTGCGATGGCATTTCAAGCGTTCATCTTAAAGTAATCAAGAACGCTGAAGTGTTTCCGCCAATTTTTTGTTTTTCTGGTATAAGTTCTTTTAGAAAATATTTGCAAGATTCAGGATATATAGCACGTGTTTCCGAGAGTGATTTCACTGGTGGCACAACATGCATTAATGAAACATTTTCAGAAAGTATTCGTGGGAAATTTAAATATAGCTATAGTACTTCGTTTTATAATGGCGAAAGTGACTTTTCACATACGCTTCAGGAAAGATTGGCAAGTGTATCAAACTTATTACAAACTCATGGTGGAAAATATCAGGTGACGTCTTATCCAGAAGGAGTAGGTTATAAAAGCCATACAGATTGCAATCTAGGCTCAAAAAATGTAAATGATCGATATGCAACCATTTTAATATATTTGCAAGATGTTAAAGAAGGGGGAGAAACAGTTTTTTCAAAACTTGGAATAAAAGTAAAGCCAAAAAAAGGAAGCCTAATTGTTTGGAACAATATGGATGCTTCAGGAAACTGTTTGGCGTTATCTACCCACGAAGCTACCAAAGTTATTAAAGGGAAAAAATATATTCTTCAAAGATGGTACTACTACGGAAGATTTGAATCTCTTGGATTTAGATTATCTGAACCTAAGCTACCTAATCGCAAAATCGGTCAGCCTCGCATAACATGTGATCACTATGAAGAAGGGTCGTGCAGATGGTATGACGAGTGGGATTATCATCATTTACTTTAACACCCTACCTTAGCATCTTTAAATTTATTCAATAAAATGAAAAAAATGTAACAGACCTTAACATTATTGTTTTATATTGTTTTTAGGGTAAACTATCCATTGTTTGAC
>evgR008155c0g1_i01_Hydra
AAAAAAATGTCCCAATTATCTACTGATGAAACTCTTCGTATTGTTCGTCACTTCCTCTTGAGTTCTCCACCTGGACAATTCGAGGATGTACTCTTTGATATCAGAGAACTTGTTTCTAATGATCAATTGTTGAACACTGGCGCTTTTGATATCTTCCGTACCTACAATGTTGAACAATTAATTCCTGTAGAAGTACCAGAAAAAAGCTACAAGGTAATCCTTTCTAAATACAATGAAATTACACCAGAAACATACCTTGATCCTGTTTCTGATCAAGTACTCACTGTTGACCATGTCAAGGGTCTCGTTACTCAAGTTTCTTCAGCCTCTGATGATGTGCTCGATAGTTCATTGAAACAAAAGAAGGACAAGCTTGTAAAGGCTATTCAAAAGTATGTCGAAGATCACTACCTCTCTGGTATTTCAAGCGTCTTCCCTTCACAAAGAAAGGACCAAAAGGGAGAACTTATTATCTGCATTACTGCCAGTAAATTTAATGAAAAGAATTATTGGAGTGGTAGATGGAGATCTGTCTATCGTGTCAGCTTTGAAGATAATAAGGTTAAAGTTTCTGGTTCTATGAAGGTCAATTGTCACTATTATGAAAATGGTAACGTTCAATTGAACACTTCAAAGGAACACGGAGAAACACTTGATGGTGGTGATTCATTTGCTGACAATATTGTTAAAATGTTGGTCAAGGCAGAAGGTAACTTCCAAACTCAAATCGATAACTCATGTGCAAACTTGAACGAAACTTTCAAGAGTTTGAGACGTCCATTGCCAATGACAAAGACTTTGTTCGATTTTGCTTCCACTCAACACAAGTTGGCCAGAGAATTTGGTGGCAACTAGATG
>evgR008162c0g1_i01_Hydra
AGAATTTTTGTTTCCTGCGTACCAATACATGTTTGTTAATCAAAAAGATTTATTAAAGTTTGGAACATGGAAATATAATCCTAACAATGTTGTTATATATGAAAGCCTCATTGCGTGGAAAAATAATTTAAAACCAAAATCAACTTGCAGTTTGAATTGCAACCCAGGAACATATAGAGTCAATTCTACGCTTTCAAAATGTTGCTGGATTTGTGAACCTTGTTTAAGTGGTTCAATTTCAATATCTGTAAACGAATATTTTTGCACCAATTGCTCACTTATGTCTATGGCTAATTCAAATCAGACGCAATGCAATAACATGGTGGAAATAAGCCTAGATGTTTTTAGTACGTCCGGAATATTAATCATTATTGGATCATTATCAGGTGTATTTTGTACAAATTTTATTTTGATGATGTTTGTAAAGCATTGGAACCATATTATAATTAAGAGTACTAACAAAGAGATGTCATGCTTACAGATAATGTCATTAAAATTCTTGTTCTGCATTGCATTTATATACTTTTGGAAACCTACACCAAGTTTGTGTGTGTTAAGAACTTCATTGTTTGGCTTTTTTTTTACTACTGTTTCTGCTTTTGCTGTGGTAAAAACATTCCGATTGCTTTGCGTTTTTAACTGCAGATTAACGAAAGTTTCAAAACTTTTGGATAACAAGTTTCAAATTGGCTTCACTTTTCTGATTGTTTTATTTCAAGTAATAGCTGTTACATTTCGCTTTATTT
>evgR008163c0g1_i02_Hydra
ATTATACAAGTAGTTAAAATATTTTATGGTTGCAAATGCTTCACAACAAACTGTTACAATTTTTGTTTATTTATGGACGGAATGACAGCTCTATTTTTAAAGGATATAGTGCAAGTTTTGAGTGGTTTAGATCTATCAGATATAATTCAAAAATCTCAAGTCACCAGCAAGACTTAATGAGAAGAGGTCTTCCAAAAAAGGCACCAATTGACGGAATCAAATTTATAATTCTTGTTGCCTCGGGGAAAGGTGGTGTTGGAAAATCAACTACCGCTGTGAACTTAGCAGCAGCAATTTCAGTTGTTAAACAGACAGCAAATGTAGGAATACTTGATGCAGATATATACGGACCTTCTATTCCTAAAATGATGGGTTTATCAGGTGAACCAACTCTCACTGCAGGGAACTTAATGATTCCTTTAAATAATTTTGGCATCAAATGCATGTCAATAGGATTTTTGGTTGACGATAAGTCTGCTATTGTTTGGCGAGGACCGATGGTTATGTCAGCTATTCATAAATTGACAAGAGAAGTAAGTTGGGGTCCACTTGATTATTTGATAATAGATATGCCACCAGGTACTGGAGACATTCAACTTTCTATATCACAAGAAGTTCATGTAAATGGTGCAATTGTTGTTACAACGCCACAAGATATTGCACTGCTTGATGCAAGAAGGGGTGCAGAAATGTTTCGTAAAGTTAGTATACCGGTGTTAGGATTTGTCCAAAATATGAGCATATTTGTGTGTCCAAACTGTTCTTACACAACGCATGTATTTGGTAACAATGGTGCTCAGAAGTTAGCTAGTGAAATGGGTGTTGAAATCTTAGGAGATGTTCCCTTACATTTACATGTTCGAGAAAGTTGCGATCAAGGTTTTCCAATTGTCTTTTCAAGACCCGAGTCACCAGAGAGTCGCGTTTACAAGGAAATTGCTTTGAAGATTGTTGCTAAGTTTCCAATATAAAGGGTACGAAGGGTATCTTGAATATAAAGAGCTTTTTACTAGAGCTAATTATTATATATTCTCATTTAAAGCAGGGTAAGCTTTTATGAAATAACACGAAAACATTTAAATACCAAAATAGACAACTCATGTTCAAGCGAGAGAAAATAGATATAAAAAGTTAAAAAAAAAAAAATTGAGAATTTTTTTTGTGTATTATAAAATTAAGCGAGGGTTAGGATAGGTGGGGAATGCGCGAGGAGTGAGACAACCCAAATATTCTAATTAGGAGAAGTCCCTAACCACTGTTAATTTATGTAAACATTGAGACTTCTTTCTCTCTTTATAGCAGTCATTGCTTTGTCTCCGTTTGTGTGATAAAAACCACATGTTTGAACATTATCCCCCAAAACTCGCATATAATTACCCTATCCACCCCGATGCAAGTGAGTCTAAAGTGGCCCACCGGCCACGGTCTAATTCAATTTTAATTTTTGTTGATAATCTTTTTGTGACTTTCTGCCTAAAGTTTAATTAATTAAACATTTTGTCAATTAACCTAAACAATTTAAAATATGTATTTGCTCCATCGCGATGGAGCAAAGTTAAGACGATGCATCGTCTTAACTTTAGCTGAAATTTAGAAACTGTATTTTCATCACGTGTCCTAGTGTGCCGCTTTTTGTCTCTAACTTAAAATTTTGTTGAACGCAAATAGACAAATCGTCTGGTCTCATGGAATCTAATGGAAAATATTTACATTATTATACAATATTGCAAAGAATTTAAATTTGTGTCAGGTTTTGAATAAGGATAAAACTTTTTTTGTGAAAAAGTTCCAATTTTTTTCTTTTCTAATTTTTAAGAATATAAAAG
>evgR008166c0g1_i01_Hydra
TTTTTTTTTTTTTACAAATTTAAAGTGAACAATGTGTGAACTATTTAATTTGATTTTTTGATTTTTTTTAAACTAATTTTGAAAATGACCGATTCTGAAAAAAAGAAGGAAATCGCTACTGCTGCTGGACTCTTTAGCCATTCTTTTGGTGGAAATCTCACATTAGATAATGTTCCAACAATATTTACACCTGTTCCATTACCAGTATCTTCAAATCAACCTGTTCGTAGCACTATTGCAAACAGCGTTAGTATTCATCAAGAAACTCTAATATCACCCTTTTTATCAAATGCTCATCAAAAACAAGCATTTGTATCTCCAATGTCAACACCTCCACAAATACTTGAAACAGCTGCACCACCGCCAGGATTGGATATATCACTTTTAAATTCTCAATTACAATCATCACAACAGCTTCATCCTTCTGGGTTGAGTACATTGAACCATCCATTAGGACTACAGTCTTTGTCTCCATCAGGACCTAGACAATCAGGACCTAGACCATCAGCTTTTATAAGACCACAAACTATAATACCTCCACCTGGAGTGCAAATATCGTCTCCATTAAGGAATCCAGTGGTATCTTCAGGACCTTTGCAATCTGATTTTTCACCTTCAAAGTTGCCTTACAGTAACAGTGTTTTTTCGTCAGCAGCGGCTACTACAGACCCTCAAAATATATCAAGTCATCTATTTGGTATGAATAGGGCAAATTCAGGCACAAAGCCTATATTAAATGAAAATCTCTCTAGCCCAATAAACAAAAATAGTTCTGATCCATTATTGATTGAATCTACTGGAAGCATTTCTTATTTTTCTCCAGTTCTTCCACACTGGTTTTATTTCAATAAATCTTGGATACCATTTTCGTTTTATGATTCCAAAAATCTAGAAGAAGCCTTTTTGCAGCAAATACCAGACAAGATTGTTACTACTGATGGTGGACGTTTTGATGTTAATATATCTATGCTTCATCGCACCGCTATATACTGGAGTGAAGTAAAATGTGAGGTTCGAAGATGTTCGTGGTTTTATAAAGAAGAGAACCAATTAATTCCTTACAGTACACATATTTCTAATATTCTTGAGATGGAATATGCTAAAATGGTTCATAATGGAATTTACAATGTTAGGGTTGATATAGCACCAAATATATTTGTGTTGTTTCACAATCCAAATGTGCTTGTTCAGTACGTGTCAGACATTGAACATGTTGATTGGACTGGGGCTGATGCAAAAGTGCGACCAAAAATTGTTCAACGTGGCCTTGGAGAATTGCAAGAACTTATTCAGCAAGGTGAAACAAAAGAAGTAGATCACTTAGTCTTTGTGGTTCATGGTATTGGTCCTGTTGCTGATTTGAAAATGCGTTCTATTGTTGAGTGTGTTGATGATTTCCGCAAAATCAGCTTGGATTTGACTCTCACTCACAGCTTTACTAATAATGGAAATGCAGCACGTCGAGTTGAATTTATTCCAGTTCAATGGCATTCTTGTTTAAGAAATGATTTACATGGAGTAGACAGCCAATTAAAAAAACTATCTTTACCAAGTATTAGCAAATTGCGGAATTTTACAAATGAAACTTTAACTGATATACTGTTTTATACAAGCCCTATGTACTGTCAAACAATATGTGACACAATAATAAGTGAGATGAACCGCTTGTATGCGTTGTTTTTATCTCGAACACCAGCTTTCAATGGGAATGTTTGTGTTGCTGGTCACAGCCTAGGTGCTTGTATTCTTTTTGATATACTAGCTAACCAGAATTTTGATTCATTACCAAAATATCATCTCGATGAAGATGTTGAAGACAATGGATCTGATTTTGAAATACAACCTAGTACTATAACCCCCCGAGTAACAGAGTATACAGAAACACCCAAAGTAACACAGTTTACAGAAAAAAAGTTCCGATCAATTAATGAAATTCTTACTCAGCTTAATCTGTTGCACATTGAATCTATATTTATTAAGGAGCAGATGGATACTGAAACTCTTCTTTTATGCACAGACTCCGATTTAAAAGATCTCGGTGTTCAATTTGGTCCCCGAAAAAAGCTAATGTTGTATCTTAAAGACATTATGCATGGTCACACGGAAACGAGTAACACAGTTAAATCTGAAACACTCCAAATGTTAAATCGTGGAACTCAGACATGTGTAAGCATAGATCAGTGTCCTAATAAAGAAAGTATACATATGGCATATCAAAAGGGAACAGCTGGAACTGGTCAACCATTGGTTCAGTACTCGAAATTAGAATTTAGAACCGACAGTCTGTTTGCTTTGGGTTCACCATTAGGATTATTTTTAACGTGCAGAGGTGTGACCTCTATTGGTCAAAATTATAAGCTGCCAACCTGCAATCATTTTCTCAATATTTTTCATCCGTTTGATCCTGTGGTTTATCGTTTGGAACCATTAATAAATGATGAAATTGATTTACCCCCCGTCCTCATACCTCATCATAAAGGAAGGAAAAGAATTCATTTAGAAATTAAAGAGAACTTAGCACGTGTCGGTGTCAATCTGAAAACAGGACTTCTGGACGCTGTAAGTCGTACATGGTCGTCCATTAATGAATTTGCTCGTGCTCATACCTCTCAACAACAGCAACAAAATCAAATAGTTTCCCAAAATGATTTGAGTACTGCTGCCCATGAATCCCATGAAGACGATGACCAATTAAAAAGAGAGAAAAACTTTAATTTTGGTTTGATCAATGGAGGCAAACGTATTGACTATGTGCTTCAAGAGCGCCCGTTAGAAATGTTAAATGAATACTTGTTTGCATTAAGCAGCCATGCTTGCTATTGGACTTCAGAAGATACAGCTTTGCTTCTTCTTAAACATATTTATAGCAAATAGTAAAACACGTGTTTTAAAAACATTCAAAACGAAGAAGGATTTAAAAAGATATCTTTATGCTTTCCTTTTAAAAGTTCCTTTGTTTATTTGGAAGAATTAGGTTTTTATTGTTCATCCGTCTAGTTTATCATCATTCAATCGGGTTACTGTTTGCAGTGGCGCATCCGTTTTTTTTCAACGTCGTCCGTTTGCTTATAGTTCACTGTCGTCCTTATATTTTTTGTTTATCGTGATTATGTGTACCTGCTGTTGTTTATTATGGTTTATCTGCTTATTGTTCATCGTCGTTTAATTTGTCGTTATCTAATTAAAGTTTACTACAAATGTTGTCTAACATCGTTAATCTGTTTGTTTTCGATTTCTAATATGTTATAGTAACTTTGTATGTTTTCATTCGAAATAAAACGAAAAGTTCTTTTTTTTAGTCGCCACTTTTAAAATTGTTTAACCAACATAGACTTTTTGTAAACACAATTTGTAATTATAATACTGAAAAATAAATGTTCTGAAAGAGAGAAAAATGTCCATAGGCATGACTCTTAGAGCAATTTTATTTGATTGTTAATAAACATTTTAAATGTAATTTATTTAAAAATAATAGTAAATAAAAAAATTAAAAAAAAA
>evgR008177c0g1_i01_Hydra
TTTTTTTAAATGTTATAATAGTAGTTAAAAACAATATGGCTGATACAGATGTAGAGATGGTGGAACAAAGCTCATCTTCAACTTCAGTTAAAGATAAAAAACGCTTTGAAGTTAAAAAGTGGAATGCTGTTGCATTGTGGGCATGGGACATAGTGGTTGATAACTGCGCCATTTGTAGAAATCATATTATGGACTTATGTATTGAGTGTCAAGCTAACCAAGCGTCAGCTACCAGTGAAGATTGTACAGTTGCTTGGGGTGTTTGTAATCACGCCTTCCATTTTCATTGTATATCAAGATGGTTAAAAACACGGCAAGTGTGTCCTTTAGATAACAGAGAGTGGGAATTACAAAAATATGGTCGGTAGCATCGTTTTAATTATGATACCATTTTGATATGGGTTGAGCATTTTGATATGGATTTATCCTACAGTACTTGGATTAATTTTGTAACCTTTACTGGTTAAAACGTTGTGTGCTTTTTCTTGTAACTGTTATCAGTTAGAGTTTGTTCATTATGTTTTACATATTATACTAAACAACATTTCAATAACGCTGTTTTAGTAATTAATATATATATTTTTTTTTAAAAAAAAA
>evgR008178c0g1_i01_Hydra
TATTGCTACATTGCAGCATATTGTTTGCCTGAGCAAAAAATTAATAACCCTATTTATAATGTTTGGGTGTTAATTAACAAAAAACATAGTACTATAAAATGTGCAGATTGTGATTGTGCAGCTGGAATCTCTGGTTGCTGTAAACATGTTGGAGCACTTTTATGGTATGTTGAATATAGTGTAAAAATTGGTGCCAATAAAACATGTACATCAAAACCTCTAGAATGGCTCAAACCATCAAAGAAAGTTCAAAAACTATATGGACCCTCAATATTAAGTGAAATTGAAATTAAAAAACCTTTTTTAAAAGACAAATTAGTGTATAGAAAAAAAAAAGTTTCAAACCGCTTACATTTTGATCCACGTGCCACAAATCAGCGAGAGGTTACCATGTCGCAAGAAGATGTTGATTCTTTAGCACAAATAACAAATTATAATTGTGGCTATGTTATGCTTATGCAAACAAAAAATGTTGTTCCTGTGCCTGATATTTCTAACATTGCTCAAGAAATTGAAGTTCATACTTCCTTGCAGCCATTTCTCCCTAAGACACCTTATGAAATTGTAACTTATTTTGATTGTAAGATTTCATATAAATTTTTATTGAGGCAGTTAGCTACAAATTTTGACGAAATTTCTCAACTAGAACATCTAACCAGAAAGCAGTCATGTAATAATGATTGGCACAAACACCGCAAAGGCAGAGTTACTGCATCAAAAGTTTATAATGTTGTTCATAGAGTAAATAATGAACTAGAAATCTTATGTCCAGAAAAATTAACATATATTATTACTGGTGTGTTAAGTAAGTCAAAGTTTTTTCAAACGAAAGCTACTTCTTGGGGTTTACGTCATGAAAAAGATGTACTTGAAGCATACTGGAAAGAAAATAAAAAAAAACATAGATTTATGAAAGTGACTGAATGTGGTTTATTTGTAAATAAAGATTTACCAATAGTTGGTGCATCGCCAGATTCTCTTGTTCATTGTGATTGTTGTGGCACTGGATGTGTAGAGGTTAAATGTCCTTGGACTCATAGAGGTTTAACTATTCCAGAATTTGCCAATATCAAAGGAACCTGTTTAATTTTTGAAAATAACACAATTGTTCTTGATCGTCAACATTCTTATTATAGTCAAGTGCAGTGTCAAATGGCCTCAACAGGTTTTTCATATTGTGATTTTGTTGTCGGAACTTCTAAAGACTTATTTAGTTATCGTGTTCCTTTTTCAAGTGATTTTTGGAAAAGTTGTGTTCAAAAAATTGAATATTTTTACAAAACTATGATTTTGCCTTATATGATAAATGAAGTGATTTCTCAGACTGTTGAGTCTGTTTTGAATGAAACTTTATTGAAAATTTGTATAAATGATATAATTGATGATGATATTGATTTTTTTATATCGTAACATTCAACGTTAG
>evgR008187c0g2_i01_Hydra
TTAAATAAGGTTCTCGTTTTTTTAATTTTACCGACTGATTGTTCCAAAAGTTCAAAACGGTTCAACAAAGTTGAACAACAAAAGTTCAGCAACAAGATCATTGCAAACCTATCAAACACAAAATGGCATTGGATGCTTCAAAAAGTTGATGAAGAGTCAGAGCTTGTGTCAAAAATAAAAAATATTAGCCACTCAAGAAAAAAACAGCTTATGCTCCACATAATCAAACACAAACATAGTTTTAGAAGTGCTGGTTGTATATCAGAATTAATTAAGAAATTTGATAATCCAAGGTTTGCTTGTGCGTGTACAAAACTCGAAGCAATAACTAAAAACGTGTTCAAGAAATACTGTAAATCTCAAAAAATTCCTTATTAAAAGCATCATATGTGTCAATTTATTTTGATACATCAAACCACAAGAACATCAAAATGTATATAATTTTGGCTCGTTGTTTTGATGTGCAATCGTTTGTTTTGGGAATGCAAGTTCAGCTTATAAACATGGATTGTGTAAATGGAGAAACATCAAATATCTTTTTTATCCGAAGTTATCTTTAAATGCACTAGTGCTAATATTTATTTAAACGTTTTGGCTGCAGGGCCGTCCAGAACAAAATTATAAGGGGGAGGGGGGG
>evgR008190c0g2_i01_Hydra
TAAATTCCGTTGTGTTGTACAGATTGGTTTTTAAAACATGTATTGTTCGGGATGTTTATTGGCTTTAAAAGCCAATGATGTTATATTTAATATAGTCAAGCAAGGGATGATACTTTTTATGTTTTGCTCCTTAGTGCTAAGTTATTTTGCATTTTTCTCAAAAGGTGCAATGAAAAACATGGTAAAGGAACTCAGTTATGTAGCTTATGTAATGTGGGTACTAACGTTTTTTGCAAGTTTGACAGTTGCCCTTTGTTACATTGAAGATCGTAAATATGAGGTGAAAAAAACTTTTTGGGGAGAGATTCATTACAGAAAAAAGATAGGTAAATCAATTAGCAATAAAACGCATATTGGAGAACCGGTTTCCTTATAATTCAGTACTCTGAATCGCAAATCGAACATAGAAAAATCACAGTTGATGCAATCGAACATACAAAGTATACGTTTTAACAAAATAAAATGTTTTTATTACTCTTTGAAAATGAGAAATGTATAAATATTATATTAAGCATTCATAA
>evgR008197c0g2_i01_Hydra
TTGGAGTAAAGATATTCTGTACAAAAAAAACATTTCTCCTCCCTGTTAACCAAAGAACTGTCAACAATTTGAGACAATATTATGTTTCTTCAACTTCGATGAATAGCCCAAATTTGAGAATGATAAAATAAGCATATTACGAACAATTATGGATCATTTGAACAATGTAATGCTCAGACTGGTAACATCTGATAAAAACCTTTCAGTTGATGAATCAATGTTGCTTTGGCGCGGAAGCATATTATATTGTCAGTTCGTTAAGAATAACCGCCATAAGTATGGAATAAAAGTATATGAGCTCTGCACCTATGACGGATTAGTCCTTACTGTCGAGGCATATGGAGAACAAGGTTTTAATGATGAACATAATTTGGGACAGACTACTGCAACATCATTGAAGCTATTAAAACCTTTTTTAAATAAAGGGCTATTGTGTGTTTACAGACAACTACTATAATTCTTATCTACTCAAGCAACATACGTTATAGATACTCTCAGGAAATACAGAAAAAGGAATCATTATAAGGCAGTTATAAAAGCAAACCTAAAGAGAGGAGAGGTAATTTGGAGATCACTAAATGGCGTTATAGTTTAAAATTGAAAGACAAGGGAGATGTGCTTCGGAATAGAAAATTACAAACCAAATGCTATCCGAGACTAAAATAACCCAGATGTTGTTCTGTTCCTTTGGTCTCCGAAAAACCTTAAGGTGGTACAAAAAAGCAGTAGTCTGGAACTAATTTGAACTATTTCTTTCTACCTTTATACTGAACTTTCAACTATTAAAGAATTTAGCCATCTTGTTGAGTTAATAGAGAATATGATAAAAAGTTTAATCGGAGAGCGAAACTCACTGTTTTAGATAGCGTCATCAAAACATATAAGAAATATATAACGCATCAATAAATAACGTTAATAAATATAATACAAAAGAAACAGGTAACTTTATCATGCCATTCAAAAAAACAAAACTCCCCTAATTTTCTATTTCATATCGGG
>evgR008198c0g1_i01_Hydra
CTTTAATTAACCTATTTTAGTGCAAAAAGAATAAAATGGCTCAAAAATCTCACGGTATCCAGCAACTTTTAGGAGCTGAGAAAAAGGCTGCAGATTTAGTGGGTGATGCTAGAAAAAGGAAAACAAAACGTTTGAAGCAAGCTAAAGAAGAAGCAATATTAGAAATTGAGCAATTTCGTAATGAGCGTGAGACTGTATTCAAAGAAACTCAACAGAGTCGTTTTGGTCAAGATGATTATCAAAAGCTCATTACTGAAGATACTAATGCAAAGTTGATGTTAGTAGAGAAGCAGGTGAAGGAAAATAAAGATGATGTAGTGAAGAGAATTTTGGAATTAGTTTATGATATTACACCTAAGCTTCACGAAAACTTCAGAATTTAAGCTTATAATTTAAAAGATTGTTTTTATTTATAATCTGATTGTTTTGCAAATATATTTCAAATTTAAGTTTTTGTATTGAAATACTAGTAAATTTTTTTCATAGTCAAATAAAATAATCAAAGTTAAAAAAAAAAAAAA
>evgR008200c0g2_i01_Hydra
ATTTGATTTTTTCTTTTGAGTTCATTACATGTCTTAACATTAAATATTATCAATCCACGATGTTCAACATGTTTTTAACAAAATTTTCTGGAAACCTAACGATATACAAATACAAATTATACAAATACAAGGACTTCTTTACTAGAAAGGATATTTGCGACATCCTTTTTTATTTTTGTTTGAAATCGAGCACCTTAGTGCTTAGTGCACCTTTAGTGTTAGTGCTTAGTGCAGTTAGTGCTTTAGTGTTTTACTGTTTAGTTAGTGCTTTAGTGTTTTTAGTGTATAGTGCTTTAGTGCTTTAGTGTTAGTGCTTAGTGCACCTTAGTGCTTGTTTTTAGTTTTGTTTGAAATCAAGCACCTTAGAATTATAAAAAATAGCAGAAAGTCCCTACTATATTACATAAACAACACATGGATAAAAAAACTCACTCATGAATGTTTTGATGTCACCATGGGATCATATGACGGTGCCGAGGTTTGTGAACTCGTTGGGATTTTCATACTCAGTAAAAAAAAATGAAATCGGACTGTACCGTGATGACGGTTTACTAGTATTGCGGAACCGCAACGGCCAAGATACTGACAAAATTAGAAAGAAAATAATCAAAGTTTTCAAAAATATTGGTTTTCATATTGATATAAAAACAAAC
>evgR008201c0g2_i01_Hydra
TTATTTTCTAAATAAGTTTTACCAGTAAAATAAAACGAATGTCTGATAATCCTGAAGAAGAAATTGAAGATGCACAAATAGAGGAAAATGTTGATGAAACAGATGTAAAAGAAGATGAAAATGACGAAAATGAAAATAATAATGACAATGACAATGAAGATAATGGCAATGAAGATGATGAAACGAAAGAAAACGAAAACACTGAGAATGAAAATAAAGAAAGTGAAAAGAAAGAAAATGAGGAAGAAGAGATCCATTCTGAACCATTGACACCTGAAATAATATCAAGCAGTTTATCTCTTTTGTGTAAGATTGGTAATGGCCTTGCACACGCATATGTCAAACTTAATATAAACCAGAGAGAATTAGGGGATATATCAATTTTAAAGAACTTCATATATTTGAGATATTTGGATATAAGTGAGAATAAAATATGTGACATATCTCCAATAAATTATCTTACTCATTTGCTCACATTAAAAGCTGATAAAAATTTATTGAAGTCTGGACTATTAAATGAGTTGCCGTACCTCCAGCATATTGACTTTTCCAATAACCAAATAAAAGAGTTATCCGGCATAAATCACCCGCTGCTTGAGAAGCTTAATCTAAATGGAAATGAAGTTTTAGAGTTATCAAGTTTTAATCCAGGCAATATATCAAATTTGGCAGTGTTAGAGCTTCGTGCAAACAAGCTTACTTCTACAAAATGCATATATCCACAAAGTCTAAAGATGCTGTATTTACCTGAAAATTTAATTGAAGAGATTCAAGATGTATCCCAACTGCCGCATCTTAATACATTGCATTTTCGAAATAACAGAATAACAAGTTTGAATGGATTTTCTGACAACATGAAAAATCTGCAATATCTTAATTTAAGAATGAACAACATTACTGAACTAAACGAAGTTAGCAAGTTAAAATGTCTACCCATGCTTCGAGCACTTATTCTTATTGATAACCCAATCTGCAATGAAGAATCTTATCGGATTGAGGTTTTAGTTATATTGCGTCGATTAGAAAGACTAGACCATGATATTTATGCTGACGAAGAAAGGAAAGAGGCAGAGGAGATCTACAATGAGCGAACTTCCCGTGAAAAGGCAGAAAGTGTTATCACAGAGAAAAGAAACTCAAACAATGGCTTGAATCAATAATGTACATGACGGTTTCCATTCGAATTCAAGACAAGTTATAAAACGACTACTCAATATATAATGTAATTTTAAGGTATTATATATACATATTTAATACCTTAAATTTTAATATATATAAAATGACAGTATAAAAACTTTGAAAATAAACACTTGTTCATATTTTATTATTTATAAATAAAACGAAATTGTAATATTAATTTATAAATTAATATAAATTATTTTAGATGTTTTATATTTAATCAAATATTTTTTTTATATTCATATATAAAAAATATTTTTTTTATGACAAC
>evgR008202c0g1_i01_Hydra
GATTGATATATTTTACTTTCTGACGTGAATATGAAAATTGTGTTTAAAGGTTTTGAGAAATTTGATCAATCAAAGATCTCCATTCGTGCTACAGGAAAAGGAGTGATAAACGATAAACAATATAAAATTACTATAACAAAAACATTTTTTAATGACTTTTTTAAGATAAAACTAAATGCGCACAAAGTACATTTGGAAGATTTACTTCTGACATTCAGAATGGAGAAGACCTTTTATAAAGATTTATGCATACTCAACAAAAAAGATTTGGAAGCATTAGTTGCAGAAAAAATTTTCATTTCTGGAATTTACGACTTCAACAGAGGATTTTATTTTTCTGGGAATGTAATTTTTATTGATCCTAATTCAAAACAACATTTTTCAACTGAAATAATTGTTCTACGTGAACCTCACCAACCCTTAAAATTTGGCGTTATAATTAAAGGAAGCATACACCAAAACATTCTACCGTTGTTGAAAGATATATTAAAAAGAAAACAGTTTAGTTTACCATTAATTGATGATGTCGGACATGGTTTTTTAAGTATATCAAGCCATAATATTTTACGTGCTAACTTTGACAAAAACATTCGTCCTCGTGCAATTCCAAAAGGAGTTGTTTTACAAACTGTTTCAAAACTACATGATATCCATAAGTCACAGTCTGAAGTCTCTAACAATGATACATCAGGGCTGCGGTTAGTAAAACTTAAACTGTGCAGTAAAAAAGATAAAGTTGAGTTCAATCCAAACAATAATGTTGACTTGCATAAACTGCTCGGCAGAATTAGCACTGTTAAGAATGATTATTTTCCGGGTTGGTTAAAATCAAATAGAGTTGTTAATGTAAAAGTTGAAAAACTAGAATACACTGGACCTAACAAAAGTAACATACTCTTAAAGCTAAAGATCAAGACCTCTCTGAAAATTTTATTTGGGGCAATAAAAATTGATCAACTAAACTTTGAGTTAACAATGGATAGAAGTCATAAGCCTAATCAATGGATATGTGGGTTTTCTTCTGTAAAGAAATCTATAGGATCGATAGAGTTTAAAGTTTCAATTACTTGTGATAAAGGCTTACAAAACTTCAAAATAATTGTTGAAATTGTTTATTTAAATTTTAACAAGCTCTTGGAGCTACTCCATATTGATAAAATTCCATATGAGCTGAAGAAAAATACAAATCAGCTACACAACTTTGTCATTAAAGGATTACGAGTTGAAGAAAGTCTAAATCAGTTTTTAAGAGTAACTGGTGTTCCACTTTTGTTTGGCACAGAAAATTTAGCTATGGAAGGTATATTTTGGCAGGAAGGAAATGAAAAATTGTTTGCAGTTGGTATTGTTTTAAAGGATGTTGACTTTATTGATATGTTAAAAAAATTGACTGGTACAAAACTAAATGCATTTTGGGTTGGAAAAACTACAACATCGATTCTGCTGAGTAACACAAATCTAAAAGCACCGGTTGATAGAGAAGATGGGAATCACACAATAATTCGATCGAGAAGAGACATCACTAAAGAATACCTCAATACCATGCAGAATAATAATATTGCAACAAACACGCTTACTCAGCCATACTATAGACAATATCTTTCTGATTTCCTTCAACATAACGAAGTAGCTTCTCCTATTTCTCAGAGTACATATTTAAATCCTTATATAGAGCCTCTGCAACAGCAAAACTCTCAGCAAGTTTCATTTTTTAATCCAAAGTATGAGTCAAATTTGGAATTTATTGAACCAAAAGATGTAAATGGTTTCAATATTATGAATGCATTGTCTGGTCAACCAATCAGTTCATTAAATACATTGTCTGGTCAACCAATCAGTTCATTAAACACATTGGCTGGTCA
>evgR008203c0g1_i01_Hydra
AGTTATTTTATTTGCTTACCTGAACCATATTGGATGTCTCACCATTTCGGTTATTGCATTTTGGACAGCATTTCCACTTGCGTTGTCTGGGACAACGGTTATGGTAATTTCAATTGAACGGTTTATTGCTGTATTTAATGACAAAAAATGTTTTGGATTTCATTTCAAAGGAGTTTATATAGTCCCAATATTTCTGTTTTATTTTTTTGTTTCCTTTGGATTTGGAGCTTGGTATGCCTATATTCATTATCATTTTCCTGATTCTATAGTTTTTTTTCCTTATTTTTTTTACACTACTGCATCCCTGATTGTTGCAAGTCTAGTTATTGCAATGTCGATAAATATTTTACTGCTGACAAGAATTTTAAAAAGGATAAAGGACAGCGAAATTCGTGTTAAAAGAC
>evgR008203c1g1_i01_Hydra
GAGTTAGATTATTTCACATGACTTAGCAAAAAAATTTTACATCAGCACAGTCAATGGATCCAATTAAACTACTTAAACTTTTTGATTACGATAGAAACGCGATTTGTCGTCTTCTACTGATTGATGCTGCTACCGGGGAAAGTGGAGAAATGTTTAAATTAGCAAACGGAATCGTGGGCTCTATAATTTTTATCTCGAACACATTGTTAGCATATGGTATTTACAAATCAAAAGAAAAGAAGAAATTCACTCAAAATGAAAAATTGGTTTTGATATTATCATTTATTAATATAGCAGTTTCAATAATTCAAACCCCCCTTGAAGTGGTTTTATTTGCTTACTTGAACAATATTGGATGTCTCACCATTTCGGTTATTGCCTTTTGGACAGCATTCCCGGTAATTTTGTCAGGG
>evgR008204c0g2_i01_Hydra
GATGAAACATGGGAACTAAAAGTTTTTGATAGAATTATAAATAAAAGATCTTATTTTATAAAAAATTTATCGCCTAAAATAAATACAAAAAATGCTGATGACATTTTTAAACTTATTGTGCAGTCAAATATATGTATTGGAAACCAGGATTATCCAGAAATTATAATCAAACGACGCTCACAATTTACTAATCCTAATATTAAAACTGAAATAAATTTAGAGACCAAGTATTTGGAATCTGTAGATGACTTTACAATTTTACGCCATGAGTCTTGTGAGATACTTTGTGATGAAAAAACACAAAGATGCGAAAAGTGTGATAGCTGTAGAAATTTTCTTAAAGTTTCTACCTTCAGATTGAAAAAAAAAGAAAGTGATACCGGTAGTTCTAAAACAACTAACTTGAAATATTTATCTCAACAAGAACTAATAAACAAAGTAAAATTTC
>evgR008209c0g1_i01_Hydra
GTGGTATTGCATCCCCCATATCTTCTTTGATTCTCGTTTTGGAGGACTTTTATGTTGTTGGTCAATTAGATACCTCTCATGTTGTTGATTCAAGTAAAATACGATGGGAACTTAATAGAAAGCGTAATCAACTAAAACCTTCTAAATATTCAAAAATATTTCGTGGAATTTATTTTGATGGACGTAAAGACAAAACTCTTGAAAGCATTAAGGAAGAGTTAAAGCTCTATCGTCGAAGTATAACACAAGAGCACATCTCTTTAATCAAAGAACCAAGGTCTAAATACCTTGGGAATTTATCAGCATCTAAATTGTCGGCTTATTTGATAAAAAGTTCTAATGTAAACTTTATAACTATCAAAAGTTGACATTAGGAAGTTTGTAACTGTTGGATGTGACGAAACTGTCGTAAACACAGGTACCAAGGGTGGTGCAATTAGGTTTATGGAAAAAGAATATAACAAACCGTTGCAATGGTGGGTTTGCCTTTTGTATGCAAGCGAACTTTCTCTTTAACTTCTGCTAATTTATTTGAATGGATCAACTACAGCAGGGATGTGCAACCTTTTTACAGTAAGGGCCATTAATTCGTTCCAAAACCAACTTCAGGGCCACAAAC
>evgR008211c0g1_i01_Hydra
GTTTTAAACAAGTTTGTTTATGATTTTAAAAAATGAAAATATTTATATCTTTTTTGTTCATTTTTTTAATAAGGATTAGTAAAAGTGAAAACGATGAACAACGACTCTCGATTGGAATCCCTGTAATGAAAATTACTGATGAGATTTTGTTAAAACATAAACCAGAACTTCGAGGCAAAAGCTACATTGCTGCATCTTATGTAAAGTTTGTGGAGTTAGCAGGATCCAGGGCTGTAGCAATAACCGATGATCTCCCAAATGATCAGCTAAACACTATATATAACTCTATAAATGGTTTACTGTTTCCAGGAGGGAACGTGAACTTGGTAGATTCTAAATATTTTCACCTAACTAAAAGACTAGTTCAAATGGCGTCAAAAACAAACGACCTGGGAACACATTTTCCGATTCTTGGAATATGTCGAGGAATGCAAGCATTGGTTAAACGTTCAAAAGAACTAATGATTTTAACCGACTCACAAAACTATACGTCCAGCTTAAAATGGTTTAAGAAAAGTCCATTATTTAAGAACTCGTTTTCTTCGATAAAAAAAGATACTGAGCTATACAACATTACCTCACATTTTCACCAGTATAGCTTTTTACCAACAGTAGGCAATAAAACAACATTTGATGTGGTTGCAACAAGTAAAGACCGCAAAGGAAAAGAATTTGCTTCAATTATTCAAGATCGCAAGTTGCCGTTCTACGGGGTACAATTTCATCCGGAAAAAATTTTGTTTGAATGGGCACCGACGATTATTATTCCTCATAATTCAAAAGCTGTACGGTTTTCCCAGAGTATTGCCAATGGTTTTATGGATGAAGCCCGCAAGAATAATCGCAGTTTTAAAAGCTACGCAGATGAAAGTAAATTCACTCTGCTTAGACAAAACCCAGTTTATATGGGAGATTTTTTAGACAGCCACAGTCCATTTGTTCAAATTTACACGTTCCCATAGAGTTTTTCAGCTAGTGTAGCTTATTTTCACTTTTTTTAGTATAAATTTAATTTTTAGTATTTCGTTCAATTTAACTATAACTATTGTATATTTAAGTGTTTTCACTTTAAACACTTCTTGTAAAAATAAACTTGTAGATATACCAATATTAATAACAAAGAAAAAATTACTTTCATATTAAAAAAAA
>evgR008216c0g1_i01_Hydra
AACACGTCTGTAAAAGCCTCCACAAGAGATGCAAAAGTGATATATGATGCAAAAAAGCTAGTGACTCATATGAAAAAATTGGATCTTCATGCAAAAACACCCGAATACTTTGGCGAAAACCCAGTTAGGACCGTGCATTAGGATTAGAAGTCACCGTAAATTCATTCACGAAAGAAATGTTTTGGAAAAGAGGACGTGAGATTCCTAATGTTAAAGATGCGCTTACCAAGAGGGATATATTCTTATTACGTGGCAATCTGGTTGCTAATCTACCAGTTGCTGGATGGCTGAAACTTGCATGCAGCTATATAAAAAGACGTACGAACGAAGTTGGATGGAATGAACCTATAAAGGAAAAGCATGTCATTGAAATGATGCTGGAAGTTGTAGATAAAGTCTTAAAATGCGATCCTGCAAAAGGTGTATGGAATGTCAAACCAAATGCTCCCTACCTGTATTATGTGATGCAAGTTCAATTGCAAAAGAAGTTGTAGTAATGCTAGGTTCCCAAAGAAAAGAAGGTGCGGCGTGTCATCTTCCAAAGTCTGAGGCAATGCATACTAACGTTACAGAACTTGATGCGTGCCTTGATAGCTTGAACATGGCATTAAAATGGAAGCCTAATTCAAAGGAGACTACTAATATTTGGTAAATGGGTTGAAGATTTCCCCGCAAGTGTGAGTGTTGAATGCCAACTACCATAAATATTGCAGATGAACTCACAAGGGTATCAACTTATGGATGTAGTGG
>evgR008217c0g1_i01_Hydra
ATAGTTGTTGTTGTTCTAAGAACTCGTAGAAGAAAAGAAATATGGAATGAAGATTTTTACAATTATGAAAATAATGTGAATTACTACAAAACAACACCAGATGAGTGGGAAATTTCTCCTAAAAGTTTAGTTTTCAATGAAAAAATAGGCGCAGGTGCCTTTGGAAATGTCTTTTCAGCCAAAATCAAAGAAAAAGTTTTATATAAAACAATGTTTGCTAAACAATCTGGTTTTGTTTTACATGATGACAACAACGATTTAAATGTTGCTGTAAAACTTTTAAAAGGTGGTGCAAACCAATCAGAGTATGGCGATTTTTGTGAAGAAATAGATCTAATGAAAGGAATTGGCTATCATAAATGCC
>evgR008218c0g1_i01_Hydra
AGAGAACTTTCATCGTTTACTTTGTGTCATGGTGTTAACTTTCCAGATGAAAAAAATGAAATTTATATTCAACGCCATGTGATTCCATGTACGTTTTCATTTACTGATTATCTAAATTCCCCTATGGATCAAAGGTTTATCCAGCTTAGTTATTATAGATCTAACAGCTGTGAACTTTTAATTGAAAAAGATATTGAATACTGCAATTGTTGTATAAAATACTCTTCTAGATTATCAAAAATTAAAAAAGATAATAATAAACAATTAAGTGTGCCAGCTAAGCTTAAAGCTCCTATATCTTGTACATCTTCTCACAAACTTAAGTTATTAATTCAAAGCCAACGTCTTCATTGTAAGCAACTGCAAGATGAAATTGAACAAATGTACAAAGCTATTGATACAAAT
>evgR008220c0g2_i01_Hydra
AGAAATTCACCGCGGGCTCCACCAAACTTAATTATAACCGATATTAAATAAAACCTGAAGATTTCATTAAAAAAAAGGTTTAAAAAAAATATGCATCAAACAAAATCAATCAAGACATATTTTGACTCCAATACTCCTGCTCTTGTTTTAAAAAGTTTATTTTTAAAGTACGATGCAGACAGAAATAATTGTTTAAATAGAGACGAGCTAAAGACATTATTTATAGACGATTTGGGATTTACTACAGAACAATGTGAGGTCTATATCTTACTTCTAGATAAAGATGGCAATAACAAAGTGTCTTTTGATGAATTTGAGAAGTGGTTAAAAAGTGGCGAACATTTCAAAAACATGAGCAACAAGTCGCGATATCAAAGACTAAAAAAGGCAGTTAAAATGTTTAAAAACTTTGATGTCGATAACAACCAATCTCTTGATCGAGAAGAATTTGAACGATTATTTAATTCTTTCAATGGAGATCCACCCAAGTTAGATGCAGCATTGGCAGAACTCGATCGGGATGGAAATAATAAAATTTCTTTTGAAGAATTTTTGAGATGGTTAAACTGGGTTCCGCTTGATCAATTTTAAAAGAATATCACTAACAAGAAACGTTTAGAACTAATAAGAAGCGTTTAGAAGGTGTTACAGATTAATATAACTATATAAAGATATTAAAATAACTATTGACATCAAGGAAGCAACCAATCAAGGCAGCAACACAATCAGCGGTGGACTGGAGTCTCCCTAGTCTCCCCAGACAAGTTTAACAACTTAATTGAAATTTTTTTATTATTTCTATTATCAAATTTAAATACTCTAGTTTAATAATATTTGTAAAAAATAAAAGTTGTTATAAATACTTAGATTTGATCTATGGTCCAAAACTAAAAAAAAAA
>evgR008222c0g1_i01_Hydra
AGCAATAATGATTGGTTGAGCTGTACTGGAAATAACATTGAGATTGAATCATGTAATTTTTATGAAGCATTCCAGTCAGAATGGAACATTTGGTCAAATTGCAATGCTACTTACGGGTTTATGAACAGAACAAGTGAATGTAACATTTCTCAGCCTATGAATTGGTGTTATGGTCAAAATAATGAAAGAATTGAATGTTTTGAATCATTTATTATGGAAAATGTGATTCTAATTTCAAAAGGAAATTTGGTTACTACAATTGCTTTACTCCATAAAGAATACTCAGTGTCTTTTGAAATAAAACCAA
>evgR008225c0g1_i01_Hydra
GTTATTATACAAAACTGCTAGCTAAATGTTACTCTAAATTTTACTGAAACAAAGTTTTAAAAAAAAATGAACGGAGAACTGATCGAAGCTGTCTTTATATATGTGAACCTGATATTTTTGGGAAGTGCATCCAAACCACTGTTGAATATTGAAAAGATAGAAGAATCTATTAACAAAGAGTGGATGACTTCTCAAAAGTCTAATTGGCAACAAACAAAGCAAACAAAGCCTATTCTTGTCAACAAAGGTATTTCTTACTCTCTGAACCAATGGACGAATTGTCGTGTCATTTATGCTCGCTGTAGAAATGGTTTGGATCTTGCTATTGTAAATCGCAAGGTTGTTGGAATCAATTCTCCGTACAGTGTTAATGGAATATTTGAGATACAATCTTATGGTACAAGTGTTGTTATGTTAAAGCATGCTGATACTAACAAATACCTGGCAATGGATAAAAAAGGAAACATCCAGATAATGGCGACGAAATCTGACGAGACATTGTTTTATCACTACCTAGAAGAAAATGGGTATATAACATTTGCTTCTGTGAAGTATTACATTAATGAACACTATGATTTATTTTTAAGCTTGCGGAAAAACGGGAAAATTCGTAAAGCCAACCGCACAGCAGCCGGCCAGTCCGCTTCTCAATTTTTACTTATACCAACTAATACTAGTAAAAGTTGCGTTCGCAAATAAAATTTTAATAAAAAATAAATTAAATAATAATAATAAATAAATAAAAATAAATTAAATAATTAAATAGAGCAAAAATATTTTAAGTGATAAAGTTAGCAGGAGAAAAAACATACAAAAGCCCAC
>evgR008229c0g1_i01_Hydra
ATAAAAATATCAAAATTTCTTGAATATGATTTGATGTTTTACACTTACTTAATAATAGTGTTTCAGATAAATATCAATTTTTGGTGCATTTATATCAAAGAATGTTTGGTATAAGAAGAGGAATTGAGTTTGTAACTTTAAGCAAGATGAAGTATTTTTGGATCAAACAAAATTTTTTATCAACTTCTAATACCTGTAGTGCTGGTTCAAAGTTTAGAAGAAAAATAGGGAAAGCAATAGATGGCAGTGCATTTGGACCATTAACAGATCTACCAGATTGGACTTATGTAAACAAAAATGAACCACCAGTTGTATCAAAACAGCAAAAGTTACGACAAATGAAGCAGGTTGCTATAGCTACAAGAATTAACAAGCTTTTAGAAGAAATTGAAATGCAAAAAAATAGAACTCTAGTTAAAGAATCTAGTGAAAAAATAAATTTTTAATTTTTTCGTAATATTTTGTATTATATTTTTTTACTCCTCTTTTTGAATATATTTTTAAATTCATGTTTTTACAGAGATTTCAACAAATTTACAACTTGTG
>evgR008237c0g1_i01_Hydra
TTTTAAACAAAATTGTTATAAATATTATACACTAAAATAAAATTATTTGTTGTAATTATTCTTTCTTTAAGTTTGACGTTAAAATAAATTTTAAATCGAGCAAATAGACAAAGAAGTTCACAAGTTTTCATCTAGGTCGTTTTCATCAAGCAGATCAAAATGAATTACCACTGGAGCGTTATTGTTAAAATTGCTTTCTACGTGTTCTTTTTCGTGTCCGGATTAATCGGAAATATTATGATTGTTTTATTCTTCAGTTTAAAAATGAAAAAAGCTCCTAACTTTCGTTGGTTTATTGTTCATCTTGCAATAGCTGATTGTTTCTACGCCGTTGTATCTCCTATACACTTAATTTACCTACTTGCAACAAACGATAAATGGACTATTGGTTCAGAACTGTGCAAAATATTAACAATTTCAGGACCTCTTTCTGTGAATGTCAGTGCATGGATACTATGTTTAATGGGATATGAAAGATATCGAGTTGTTTGTCATCCGTTTGCTAGACGATTTTCAAAAAAAATGATTCACATTTCCGTTGGGTTGGTATGGATAGCTTGTATTGTATTGAAAATATTTCCTTATATTAGAATAAGACTAACAGATAGTGAATGCTATGTGCATTTTAATGATGTAACAGAGCAAACAGTAAATGCAGGCTTGTCTCTACTAGCTGAGGGAATAATTCCATTTGTTTTTTTATCCTATTATTTTGTGCGAGTTACAATAACTATGCGTAAAAGATCACAGTTGTTCGAAACAAAAGATGATAACTCTACTTTTATGAGAAGCGAATGTTTATCAATTCTTCAGTCTTCAAGTTCATCAACTTTACACGAAAAAAATTCTGACACGAGCAACGATAAAATTACTTTCAAATCTGAACAAGTATCCGAAAGTATACTTCAAGTGTCTGCTGGTTCTAATAAAAACTTGAAAGTAGGAAACAAATTTAGGTTAGTAATATCACCTAGTTTTACAAAAATGAGCTCTTTGAATAAAGCAGACCGAGCAACCATTATTGTTTTTTTTATGACGGTCTTGGTGTTTATCATTACAACGCTTCCGGAAAAGACATTTTATTTTGTTGTATGTTATTTATTTAGTTTCCGTTTCAATAAAGAACAAATTTTAAAATACGGTCCAACAATGAAAACAGTCAGCGACTGGCTCGGTTTTTTTGTACTCTCTGGATGTATTTCTGATGTAATTGTTTACAGTGGAAAATATCCCGACTTTCGTCGACAAGCGTATCAATGGATTAATTTTATTATTACAAAACCATCTTCAAAACTCAATTTTAGAAAAATCAATAACGAAACCACAGTATCAAGTTTATAAATGCTTGTTTTAACATGTTTACAAAAAGGTTATTCACGTGTTGTCACGTGACACTTTGAAAAAAGAGAGATAGTTA
>evgR008242c0g1_i01_Hydra
TAACTCTGTGCGAAATACAAAACATCCTTTATATGCCAGAGGAATCTAGATCAACACACAGTATTCTTCGTTTAATAAATACCACATTTGCACATGCAATAATTATAAAAGAACATTTAGAAGCAAATTTAAAAAATAGCAAGCGAAAATTCTTTGGTGCATACTACCACTCAATCACATCCCATGCTCCACTACAGTATAGATTGTTTGCAGGTCGTACGTCTAATGTGGAAAAAGAAGAAGCAATGTTTCAAAATTTGAAACTGTCTACAAAAACATCTTCTAATCATCATTCAGATAATGTTATTTTTAATGCAATAATCCGCAACCAAGCAAAAAGATATTTAAAGGAAGGAAAAAAGTTATGCAATGGTGTTCCAATACTTCACAAACTTTACCAACCATTGCATAGTCTATTAAATGATACAAAAATATCATTTACTTGGATTGAAAAGTATAGCAATGAATATCAAAAACTTTTAGAAAAATTAGCTGACTATTTAGTAGAAGGCATTAATGTATGGTGGAAAGAAGTTGAAGATGGAGTAGTATTCTTTGATATAAACTACCCCAATAATTCGAAAAAAATATTTTCCCATTTTCGTTCCTCCTCGATGGAACAACAGTCAGAGTACTTAGAAAAATGCTGGAAACATTGTTTGGAAAACAAACACAAAATTCCTGCTTATAAAATCACAGTTGGTGCAAATGTGTTTTATTTAGACACACTTCAATACTTTCAAATTTTTAACACTGAAAATATATCTAGAAGTAGTATTATATCACTAGATATAGTATTGATTTAATAATATCAAAATCTTTCAAATTATTATAAGCAAAAGTTATCTCAATAATGTGGTTACT
>evgR008263c1g1_i01_Hydra
CCGTGCGGCATAATGTATTGTAGAGATATATGTTTAACAGATGATCAAGGCTTAGTTCACAGCTATACTTGCTTAGAAACCACTGATATTTCAAAGATAAGTAATCCTTCATCATCATCGAATTACGGCACAAATGCAAATTCTGGTCCAATGAGAGCCATTGACGGAAATATAAACACCGACGCAATCACACACAATGGTGGGTTTAATTGGTTTAAACTGGATTTAGTTTTTATATATCAAATACGTAAATTAATCATCTTGAGTGGAGTTCCATCAAAGTTAACCGGTAGTAAACTTAGTGCAAGTATAACTGACGTAGTAAAGAGTTTTTTTTTAATTGCAATGTTAAATTCAAACGAGCAACAAACATATACAGACATATTTGCAGCTAGATACTTGCGTATTGATAAGCAAAACCAAGTCCAGCTTGAGCTATCAGAAATTTATGTTTATGTATGATGTTATTCGAAAAGTATTTGTTGGGGTTCAAAA
>evgR008270c0g2_i01_Hydra
AGCGGGTTAATGTTTTGATTGTAAGACTCAATTTCTTTTATTAGTTGTCTGAAATCAATAAGTTAACAAATTATTTTTTGTAACATTATTGAGATTTAGCAATGCATATTTTTTCTAATATATAATCGTGTTCTAAAAACCTAAAATATGACGGGCATCACTGATAAAAGCAATGGACACATTCAAGATCAACCTTTCCCTCTTCCTTTAAGTAAAGATACAAGTACAAAATCCATCACCAAAGTAATTGCAAAAGAAAATTTTTCAAAAAGAGATATTATCACTTCTTTGTTTGCAGGGGCTTGTGCTGGTGCTCTTGCTAAAACTGTTATAGCCCCATTAGATAGAACAAAAATAATGTTTCAAGTCTCAAAAACGCCTTTTACTTATGCAAAAGCTATTGAAAATCTTTCAAAGAGTTATACTCAGTATGGGTTACGCAGTTGGTGGAGAGGTAACAGTGCTATGATGGCTCGTGTTATTCCTTATGCAGCAATTCAATTTACTGCACACGAAGAAATTAAAAGATTACTCGGAAGTGTAAATCGTGAAACTCTGCCCCCTCTAAAGAGACTATTTGCAGGATCAATGGCTGGAGCAACAGCTGTAATTTTAACATATCCTCTTGATATGGTGCGTGCAAGGATGGCTGTATCAAATTTTACAAAATACAAAAGCCTCAGACATACGTTTAAAACAGTTTATAAGGAAGAAGGAGTTCGAACTTTTTATAATGGTTTTATACCAACTGTTATTGGTATTATGCCTTATGCTGGTGTTTCATTCTTTGTTTATGAATCGTTAAAGAAACATTACTATAATAATAATAATCAAGAAATTGTAATTGTATATAGATTGCTATTTGGTGCTATTGCAGGAGCTTGTGGTCAAACAGTGACCTATCCAATGGATATTGTTAGAAGAAGAATGCAAATAGATGGAATTGATGGAAAAGGTTACATATATAAAAATATGTTTTGGACACTGTCTCATATTTTAAAAACTGAGGGTTTTATTAAAGGCTTCTTTAAAGGTTTAAGTATCAATTGGATTAAAGGACCAATCGCAGTTGGTATTAGTTTTGCAACCTATGATACAACAAAATCGTTTATAAATATTTTAATAAATGAAGCAGTCGAAACAAAAGAGTTTATATAGCAATTAGTTTTATTTATTTGTATTGCTGTTATGTTTTTTACTGTAATTTTGAATTAATTTTATTTCATGTTTTTACAAAATAAAATTGTTGCATTCAAAGCTTCTTTTAAAAAAAA
>evgR008274c0g1_i01_Hydra
TAATAGTACCTAACCTTTTGTGATATAAAAGAGTTGTAGACAAACATGTGTCTGAAGATTATTATAAGTTATTAGGGATTACTTAAATCGAATACCTTTATAAAACCAGTTTTTAGTGCAGTTTAAAAATTATAAAAACGGTACTCAGAAGTATAAAGAATTAAGTATTGCAAACAAGATTGCAGTTGAAAGGATGTAACTGTACGACCGTAAATTTCATTCTCAGAGTAATTAGAAAAAAAAGAAAAGTTAAATTACCTTTAGCGTTGATTAAATCAAAGAAACCGACACAATAGCGTCAAATCTTTGCCGCGTATTTAACTCAGTGTATTTTAATGGCCGTCTTCAATGTTATATAATTCAAACAAAGCAAAGCAGAGAGAAAGTAAGTACCTGGAACCTTTCTATAGTTTATTTTCAATGAGTTTCTTTTTAAAGTATGTATATGCAATATGCTGTTGGAGTGTGGTATACTGTGGAATTGACCGTAGAAAGGTAGACAACAGCGCTAACGAGCGCGCAAAACAAGCGTATAACCTTATAAAAAATAAACTCAATGAAACAAATGGTTGCAAGGATTTACTGGGAAACGAAGACTGTAGTAATCTACGCCGGTATAGACACCAAATGAGGGTGTATATGGCATCGAGAAAACAACTCCAGCTAAACGCAGTGATGACAGAGTATTTTCATCGTGAGTTTACTGAGTTAGAAGCTGTAGTAGTTGTCGACCCTATCCCTGAAGCACATTTTGGACACACACTTGTTTTGATATTAAGCTACGCAGGTTTTACGAAAAAAGAGTGCGACAATAAAGATGGAAGTTATTACATCCAAGGCGGTGTTGCTGAATGCTTGGTATTTCCGGATACAAAAAAATGCCTAATACACAAGCAACTTAGTGGCGATACTTTTAAAACCTCTTGCACGGTAACTTTTTTGCCTCAAGTTTATGAAAAGGCTTCGTTGCAACTGCAAAACAAATTACAATGTATGGACGAACTAGAAGGATTTGCTTCGTGCCCGGTAGTTCACGCTCAATCTGACACTCGATGTAATAATGCAATTAAATGTGACCAACCTTTAAATAATTTAAAAGATTCTTGTAAGTGGACATGCGATTATGGTATCATGATTTTCGGTGGTTGGGATAAACATACACAATCCGTGCGACATTTACACAACATCAAAGATATGTCTAAATATTTAAAAAACAAAGGATACAATGACAAAAACATCAAAATATATTTTGCAAATAACTCTACAATTGACCTTGATGGCAATCCAAATACACAAGACACTGTTCCTGCTGATGATGCAAATGTTATCAGTAATCATTTGAAAAACGTATGTGAACTTAATGAGCATTGTATTGACACATTAACAATATATTTAAACGGCCCAACATTACGAAACGGCGATATACTATTATGGGACAAAAACAAAGATGGCTCCGCAAGTGATTTTGGAAAACTATCAATAAGCAAACTTTTAACTCCTCTTGAAAACTGTGGCGCTAAGGATATAATAATTATTGCGGACCAGAATCATGCTGGTCATATTATTGACGAAGTTATAAAAGGAAGAAAACGGGGCATAAAAAACTTCAACAACATTTTTGTAATAGCGTCAAGCGCAAAAGATACTTACACTTACAAGCGAGACTTTACACAAAAATTTATTCAGCTTGATAACGTTCACATTGACTCTAGTATATCAACAGCAACTTCAAGGCTGCTAGTTAATATATCTAAGGATTTAAAAGAGTTGCTGAAAAATAAGAACAAAATCGATTATTTGGAAAGCGACAATATTAACTTATCTATTACTGGAAATCAATATCTTCCGCCAGGGTGCACAACAAATATAAAATATGTTTCAGATAGTCTAGAGTGGACAGATAAAAGTACATTATAAAAAGTTGGTTTTAAAAGAGAAATATCGTACAACAATTGTACAACGATTGCACAACGATGCGACAAGATAAAAGTTAAGAGTAATTATGAAAAAAAATAATTACGGAGGTTACTCTCGATAATGTTCCTTTCAAAAAATACTGAAAACAACATGGTTCATAATTTATTATGTTATATACAACAACATGGTTCATAATTTAATAT
>evgR008276c0g1_i01_Hydra
TACTTGTAAGAAAATTAGATGTTCAAAATATTCGACATGTGTACGCCATCTTGATAATTCTCCTGAATGTATTTGTAAAATTTGTCCTGAAACAATATCCCCTGTGTGTGCAAATAATGGACAAACATTTTTTAGTGAATGCCATTTAAAAGAATATGCCTGTAAAAATGGTAAAAATCTGGCAATGATATCATCTGGCGTTTGTGCTTCATTACCTGCTTCTGAATATGTTGATATACTATACCTGATTGACACTTCAACTGGTGTTTCAAAAGAAGTTCTTGAGCGTGTTAAAACATTTATAAATGCTGATATTCAAACCAAGAATTTTTTAACTACTAATGTTGGAATTCTTACTTATGGTGCAAACATAGACCTGAAATTAGCCCCAGTAGAGGGAGTTAATAAAGGGGTTGTTGCAGACTCCATGGAAAGTTTAGAAGCTGGTGGGGGAAGTCGAGATTTGCAG
>evgR008280c0g1_i01_Hydra
AATGAGCACAATAGATGTATCTGACTTGTTAGACATGTTCAAACGTTATTTTCGCAACACTTATCGCACCATCAGAATACAACGTTATGATGAAATCAACCGGTGTTATATTTATGATAGTGATATGATTGTACTTGATTATCATTTTATGGCTATGATGTTAAACTTATTGATTATCAAAGAGGAAGAGAAAAAAACCATCCCAAACGAAGCTTTAATTAGAGATGCTATTAAGAGGTGGAAAAAGGCAGGTCTAACATTTATGAGTGATAATGCTGAGATTGACGTAGAAAATTGTGATGTTTTTAAAATGTTAAAACTCATTCGTAATGACTTAATTCATAACATCACAACCAACGATATCGGATTAAAACAACAAGCACCTCGTTTGAAAGAGGTATCTGTAATTGACATTCGCTTTTGGTTGTATGAAATATTTAAAGCTTTGTTTGGTAATATCTTTCAGGGAAATGTATTTGATGTTTTCCATCAAAATTGGTACTAATTTGATATGTTTATCATCACTTGCCAAGTAAATTGGAC
>evgR008293c0g1_i01_Hydra
CTTAACATTTAGTGATGAGTATTGCAATTTATACTTGTGAAAAAACCCATGAAAATGAAGTAATAGTTCAATTTCTAAAGTCATTGACTTTGGATTCTATAACTGCTATAAAAAAAGAGTTTGAAGAAGTGAAAGGCTTGCTTATGTCTAAATATAACGAAATACCAAATGATATTTATAAAGATATTGCCAAAAGTGCCCTTCAGACTTGTGATATAATAGGTTATGCTTTAGAATCTATTTTACCCGATGTAAATAATGTATCTACGAAAGCTTATGAATGTGCTGCAAAAATTTTAAACACTGCATTAAATCAACTTCAATCAAGTTGTATGACCCTTAATAAAGCAGAAACGAAAGAATTGATGTTGGAGATAGAGAAATTTGGCGCTCCAAAACCATCGAGTTTCATTACGTGGTTTTGTAATGTAAAACGTTTTTTTCGAAGAAACCAAGGTACAATTAAAACTGTGATTATTGGAGTTGTTGCAATTATTGGTATTTTGTTTGGTGTCAATATTTTTCTTATGGACGTCAAATTTTGACGTCCATAAGATGACGTGAATAATGATATTATTTAATAGTATATTGTCAATACAAAAAATAAATATTTATAATATTAAATTAAAATAATTTCAAAAAAAAAA
>evgR008328c0g1_i01_Hydra
CAACCACTCAAAGAGTGTAGCAGTTAGTTTATTAAAGATATATAAAGTGATAAACTCATTAAATAAGAACGGCATTTTAGTATTTGTATCAGTCAAAATCGGTATTCAAAAGGATATACAGGATGAGCTCAATTGTATCATCAATTTTCATTTATGCAATGCTTTTATTATCTGTGTATGCAAGAGATTGGACAGTAGGAACTAATAGTTTAGCCCGGTGGGATAAAAACTGCGACTTTAACGGTAATGATATTGTCAAATTATCTAGCACAAGTGCAGAATGTTTTTGGAATTGCATTGCAGATACACAATGCACGCACTTCACCTTTTCAAGTGGCACTTGTTATAAAAAGCACTACATCGGAAGTGATTTTCTTGAAATATACGCTGGTGGCGATGCTATTTGTGGGTGGATTACTAACCGCAGTGGACAACCCATGAAGTAAGCAATTCACACTCCTTATTCTCCAAAAGAACAAGTTGTAAACGAGTAAACGATTAAAAAGTTTAAATATATATATATTTCTTTGTTAAAAAAAAAA
>evgR008341c0g1_i01_Hydra
TCAATTTGTCATTTATTAAAATTTAAATTTGATGATAATTTATGCGTTGATTATGCAAAGAATGACATGCGAAGCTTGATAACAACCTTTTATAGTGCTAACTCAAGAACAGATCAACTGCCTATTACATCATGGTTACAATATGACAACCAGTCAAGAACAATATATGGTTATCCGAGGTATGCTAGTAGTACATATATTAGTAGATACTTTTTAAGAGTTGCTGGTAACGATGGAAGCCAAGAGATTATAGAAATTGATGTATACATATTTGGAACACCTAATCTCGATTACCAACTGACTATAAATGGTTATATATTAGATAAATCTAAAATTCCTTCACTTCAAGAAATGTGTTTAATTGAGAAAATAGCCAGTTATTTTAATTCCCCTGTTAATAGTATTT
>evgR008360c0g1_i01_Hydra
GAACCTCTTAGCTGTAAAGTTTTTGAACAATTTTCCCTCTATTTTTAAATTTTTCATCAGCTAGGGCTATAGTAGAATATTGCGCTATAGCTCAATAAGATAAGAGTTCAGAAAGCAAGTTTGATCAGCATGGAATGGTTTTATGTAGACTCAGCATAGCATGATCTTGATGTAGAGATTCAGACAGACTTTATAAATGCAATCAAATAGAAAGTTTGTTTTTAAGACTGATTCAGATAATGCTTATATAAAGTATCTTGAAGTGATTATTCCTGAGGTGAATATACCTTATCATGGAATGTTTACATGGCCTTCAGCACCTTTTCTTGCACAGTACTTGTTTACAAATCAATCCATGTTACAAAATAAGACAGTGCTCGAGCTTGGTTGTGGAACTGCTCTTGTTGGTATTGTTGCTGCTAAACTTTGTTCTCATGTTTTTTTGTCAGATAATGGTCTGCAGGAAAGTTCTTTGCTTTGTGGTCGAAAAAGTTGTTCATATAATAATGTAGTCAACGTCACAATACTTCCTTTAACATGGGGCCGATTTTCACATGAACTTTTCAAATTAGCTTCTGTTGATATTATACTAGGATCAGATTCTTTTTATAATAAAAATGACTTTGAAGATATTTTATGCACAGTATCATACCTCTTCATTAAAAATCCAAATGCAGTGTTTTGGACCAGTTATCAATTAAGAAGTTGTGATTATTCAATAGAATTTTTACTATCAAAGTGGAACTTGTGCGGAAGACTTCTGGATGTGAATGATTTGGATGAAAGTTACATGGGTGGTTTAAGTCTAAATCATGAATTATATATTTATGAAATACGAGCAATATGCTCAAGATGATATTTTTACATTTGTAACTATTATTTATTTTAAAAGTGTAAAGCGAAAATTTATAGTTAATATAAGTTAAGGTAAAATAGCATAAATCC
>evgR008372c0g1_i01_Hydra
TATGCACCAAGATATACCAGTGCTAGAACTGAAAAGAAGTCAATCTTTATGCACATACCTGCTTACTTCCATGAAAAAATGTTGGACTATGACAAACTTCCTACACTTAAAGAGCATCCATCCACCAATCAAAAAATTTCTCAAAGTGAAGCAATCAATATTTATCAATGTTTAGAATTGTACACTACAATGGAAAAGCTTGGAGTAAATGATACATGGTATTGTCCATCATGTAAGGAACATGTAGAAGCAACCAAAAAGTTTGATATTTGGAAGTTCCCTGATGTGTTGGTTATTCATTTGAAGAGATTTTCATTCTCCACGTATAGCAGAGACAAGATTGATCGTTTTATTGACTATCCAATCCAAGATCTTGATTTGTCATCCTTTGTTCGTTGTAA
>evgR008383c0g1_i01_Hydra
CTGTCATGTAAATACTATTTCTTTTTTAAATAATTATTTTTCTAATCAATTAGGTTAACAAATTTAATTCAACGCTTTTTTTTATTGAAGATGGATAATGAAGGGAGAATTAATTGTCTAGCAGTTGATGGAAGTGTTTCAAGTAAACAAGCTTTTGTTTGGTATATCGAGAACATACATAGAAAAGACGATACTTTAGTAATTATTCACGTTTTTGATGTTCCACAGATAGTACCAGTGGGCGCAATGGGCGCTTATATTCCCGTTGAAGGTTACAAGGAGATAGTTGAACATAGTATTAAAGATGCAAGAGATATTCTCAACTACTTTATAAATTTTTGCCAAAAGAACCAAATTAAATATAAATCTGTTGTTGTAGATGATTGTCATGGGATTGGTTACGAAATTTGCAAATCTATCAAGAAATGCCTGGGTAACATTGTTGTTTTAGGCCAAAGAGATATTGGGAAATTAAGTCGATTTTTACTTGGCAGCACAAGTGATTACGTTTTACATCATTCTCATATTCCGGTTGTCGTAGTGCCCTCCTCTGAAAAACATACAAAAAATTGAAAAATTATTATTGCATATTTATATTAATCATTTAGTTATCATACTTTCTCA
>evgR008385c0g2_i01_Hydra
TTTTATTTTATAGAAGCTGAATATCTTTTAAGTCTTCGAATTTATTTATTTTCATTAAAAAAAATCATGTTAATATTTAAAAAACATTTTTTACGAGAAGTTGTTGAGTCTTAAAACAATGGAAAAACTTTCATGCAGTAGATCTTTTACTATCTCAGCAGATAAGGAGGAAACGATGAACAATATGGCAGCCGATGAAATTCTTTTTTCTAGTAATGACTTTTTCAAAAATGATTCTTCTTACTGTGATAAATGTAATTTTAACAAAGTGAAATCTGTTGTTTTCTGCAAGGAGTGTGGGGATATGTTTTTGTGTGGAGAATGCAATAAAAAAGTTCACTCTGGAGCTAAAGTTAAACATTCATTTGAAAATCTTCCACCGCTTAAAGAAAAGTCTGAAAATAAAAATGGAAAAGTATTTGAATCAAAGTCATTCACAAATGTTGCTAAACTTAATGTTACCAATGAAGACACTAACCAGAGTTTTCTTCTTATTGATGGCAGCGAAGTTATAAAGATTACAGAAACTCATTTTCTGCAGAACCTGCAATGCGAGGATAATCCTCGTGTAAAAGTTGTTTCTATTTTTGGTAACACTGGCGATGGCAAATCTCACACATTAAATCATACATTTTTTAATGGTGCAGATGTTTTTGATACCTCTTCTTCTCAAAAATCATGCACAGTTGGTGTGTGGGCTGCCTATGATAAAAGTAATAAAGTTTTGATCCTTGATACTGAAGGATTACTTGGAGTATCTGAAAATTCTAATAAGAGGATGCGACTGTTATTAAAGGTTTTGGCTGTATCTGATGTTGTCATCTACCGAACAAGAGCTGAAAGACTTCATAGTGATATGTTTTTATTTCTTGGTGATGCATCAGAAGCTTATTTAAAATATTTTGCAAATGACTTAAAATCTGTAAAAAAACGCGGGAAAATGGACGATACTCCTCTATGCAATCTAGGGCCCACAATAATTATTTTTCAAGAAACGCACTACACTCTGCCACTTGGTTTTGACAATAGCATTAACTCGTTTTCAAAACAAAACAATCAAACTCCAGAAGATATATTAAGAGAGAATTTTGCAAATATTAAGCGAAGTCCATCTGCATTTAGTTCTATTAGTTATGTTGGGACCAAAACTCAAGCTCATCCAACAAATTTTTCAACTCTTTTCAAAGAAGTTCAAAAACATCTGAATAACAGTTCAGTGAGGTCTCCAAGACCAATATCTGTTATTTACAATGCATTGGTGAAACTGAACAAAAAGTTTGGTGATGATATCAATTGTATTATTCCAAATAGTTTTCCTGATGAATATTTTACGTGTACATCTGTTTGTCAATCATGCAAATTACGATGTGAGCTAAGTATGAATCACGAAGAAGATAAAATTCCGCATCATACATCTCAACGTTGTATATATCAAGCACAATTTGAAAATAAAGTTTATGTTTGCAAGATATGTCATGAAAAAGGATTGGATTGTATGGTTTGTCCCAAAACAACTGAAAATCAAGATTCCCCATTAATTGGCTTAGCAAAGTATGCATGGTCGGGATATGTGTTGGAATGTCCTCAGTGTGGTGTGATATACCGGAGTAGACAGTATTGGTACGGAAATCGTGATCCAGTAGAGAATTCTGTTCGGACCGAAATTAAGCATGTTTGGCCTGGTTCGTGTCTTCCACACAATCATCAAAATGCTGCAAGAAGATTTCTTGATGGGATGAATGTTTTTGCCTCAGCTATTACTGACATGAGTGCTAAGCCAGCTAAGGCTGTTACAAATTGGATAACAGATAAAGTTGCTCCTTCATATTGGACACCAAACTACCTAATTTTAGAATGCAAAACATGTGGTCACGATTTTGATGATATTGAAAAAAAGCACCATTGTAGAGCTTGTGGTTTTGGTTTTTGTGATAGTTGCACTCAACATCGTATTCCATGTAAAGATAGAGGCTGGGGTGACACTCCAGTACGTGTTTGTGGACCTTGTTTTCAAAAGTCTGGTTTAGACGACCAAGCTAATGCGAAAGGTGAAAATATTGACTTAAGTGAACTGCTTGAAGCTATTGACGATGATGCCATGTTGGAAACTGGAGATATGGTGTCTGTTAAAACTAGCAATAATATTCCTGTACGACACCCTCCTGATATTGTTACAACAAGGAAAGTTACTGAAACTGTCCAATCAACTTTATCGTATTTGAGTAGCACGCTTGAATTTCCTTTAGAATATATTGTTGATTCAGCAAGACCTTACTATTGGGTACCTGATTCAGATATAATCGAATGCTCACTATGCAAACAACCATTTACGAATAACCAGTCAAAACATCATTGTAGAAGTTGTGGGAATGGATTTTGTGATGATTGCTCTAAAACCAGACTACCTGTACCTAGTCGAGGTTGGAATTACCCTTGTAGAGTTTGCGATACATGTTCATTAAAAAAGGGTCCACTGTAAATATTGTTTAATTTAAAGCTTTGATTTTTAATCCTTTATTTTGTAACATTTTTATTTGTTAGAGTTATATTTGAATGTTCAAAAATATATCCAGATATGCAAGTATATGTGTGTGAATACCTAC
>evgR008395c0g1_i01_Hydra
AACTGATCAACTAGAGGGGAGGTATGTCCCCGCCCTACAGTGGTCAAATTTGGAAATAAAAACAATGGATTTAAATATTATTTATTTTTTAAAATGTCCTATTGCAAATAACAATTTTTGTGGCTATGAAAATAACTATTTTTTAAGTTTAAATGTCAGGTGTCATTATAATAACTTTCATCATCACTACACTGTTAATCTGTTACTGAGTCATCAGTGGAGATACTTGCATTTTCGCTTGATATTTCATCTCGTTCATCTGAACACAATAGTTCATTGTCAGATAAACTTAGGTCACTTAGAGAAGTTACTGGAGCTTCGTTTATACTTTTATGATTCGGGGTAATTATTTTTTTTGAGGTAAAATTTGTTTTTTAAGTGAAAATATTTTTATTCTGCGCTTTTGAAAATATTTATTTACATTGTCTTGCAATATCTCGTTTTTTTCATCTTCAGTCCATATATGGCTGTTTCTCGATATAAGCATGCACTCTTCCAACATATCACGATTCATGCTAAGTCATTTATCAGTAGCAAAACATCTTTCTACGGCACTATTTGACGATGAATAAACACGAACACAAATACCCCATTAACAAAATATTCGGAAAATCCAACTTCTTATTAAAATATTACCTTCTTGGAAGTTTGCTACAAAACATGATTCCCAAAAATGATGCATAAGGTATTTTATATATAGCCATTAAGGTAAGGTAAATATATTTATGGTAGCCATTATCCTGATCGTTTGTCCAATTTA
>evgR008403c0g1_i01_Hydra
ATTCTCTAAGTTTGGTGAAATTTCATCTGCTAAAGTCATGTCTGACAATAAGCAAAGTAGAGGATTTGGTTTCGTCTGTTTCAAGACTCCAGAAGCTGCCAATAAGGCATTGACTGAAATGTCTGGTCACATGATTGGATCTAAACCACTTTACGTTAACTTCGCTCAACCAAAGGATCTCCGTCGTTCTCAACTCGAGGCTCAATACAATGCCAGAAAGCAACCACAAATGATCCCACAAATGATGCCACAATTCTTCATTGCTGGTCCACAAGGTGCCTTCCCTCCAGTGGGTGGTCAAATTAAGCCAAGATGGCAACCAGTTGCTGGTGGCCGTCAAAATAACACTCGTGGTGGAATGTCCCGTGGTGGATCATCTCAAAGAGGAGGTGGTGCTTCCCGTGGAGGTAATGCTGCCAAGAATGCCGG
>evgR008461c0g1_i01_Hydra
TACCAAAGCTTGAAAAAGAATATTTAATTTCTTTTGACCTTGTTCTTAATAAGTTTGAACCCGGTTGGCATAGTGTTATTGCTTTTTATCTTCATTCTGGTTATGGAGTTTTTGGTATTTGGTATTACAATGGTGGATCACTTTTTATTGTGACTCCAGTAAATGGTTACTTTGATTTAAGTTTTAAGAACGCTGATTATGTTTTCACAAATCCCGTTGAATTATCAGTGTGGTTAAATATTGGAGTAGTTCAAGTTTCAAAAGGAATGAGTTATTTATACACAATCAAAATTAATGGAGAAGTTATATTTTTTGAAATTACC
>evgR008469c0g1_i01_Hydra
CTCGTTTGGAAAGTGCTGTCTCTAAATTGGAACACTTAAAGTTTACTGGTGCTTCATCTTCATCATCAAGTGGTGATTCTGGTTCTTCTGAAGCTCCATCCGTTTCAGCCTTTAAGGAATATGTTTCCACCAATGTTCAACCTTTGGTTGATACTTGTACAAATATTGGTGGTGAATGTGTTAAGGGTGGTGAATTCATCTCCAGAGCCTTTGCTGAAATGTTAAAAATCATTACATTGGCCAGTACATCAAAGAAGCCATCTGATGAAGAATTTGGTAAGATTTTGAGTGGTTTGGGCAAGATTTTGACTGAATCTGGCGAC
>evgR008478c0g1_i01_Hydra
GTTACGCTTAGAAAAACCATGATGTACGAATTTTTAGATAGATTAATAAATATAGCACTTCCGAGAGTTAGGGATTTTAAGGGGGTAAGTTCAAAACAGTTTGATGGTAATGGTAATTTTTCATTAGGAATAAAAGAGCAGATTGTATTTCCTGAGGTTGATTATAATAAGATTGATAAAATAAGAGGAATGAACATTATTATAGTTACTTCTACAAATAGTAATGAAGAGGCAAAGTTTTTACTAAAAACATTTAATCTACCATTTTATAATTAGAATAATAGGATAAATATCATGGCAAAAAAAAGTGTAATACAGAGAAATTTAAAAAGGATTAGATTATTCAATAATAATCTAGAAAAAAGAATGGAGTTTAAAGAAATTCGTAATAATAAGTCAATTACTTTAGAGGAAAGGTTTGAGGCTCAAGCAAATTTGTCTAAGTTGCCAAGGAATAGTTCAAAAGCAAGAATAAGAAATAGATGTTTGCTGAGTGGTAGGAAGGGGATTTTATAGAAAATTCAAGTTATCCAGAATATGGTTTAGGAAATTAGCATCAGAGTGTAAGTTGCCTGGTGTAATTAAGGCTAGTTGGTAATAATAAAGGAAATATAAAATATGTCACAAACACATTTACTTTCAGATACAATAGCGAGAATTAAAAATGCTCAGTCAGTTAAAAAACAGTATACAGTTGTATATTGTTCTAAAAAAATTGAGCAAGTTTTGGATGTATTGCTTAAAGCTGGCTACATTTCAAATGTTGAAAAATACGAGGTAAGAAAAAATGTGCATATGATTAAGGTGGGGCTTAAATATGAAGGTAAGTTGAGCTGCCCTGTAATCAAGGAGTTTAATGTAGTGTCGAAGCCTGGCAGAAAGGTATTTAAATCATATAAGGATGTAAATAAAATGTATGGTGGATTAGGCATGACAGTATTGTCAACTTCTAGAGGTATAATAAATGACAATGAGGCCAGGGATATTAAAGCTGGTGGTGAAATTTTGTGTAATATTTTTTAAAATAAATTATGTCAAAAATAGGAAAACAGCAGATCAAAATACCAGGTCAAGTCAAAATCAATTTTGAAGACGATTTTCTAGTTGTAAAAGGTCCCAAGGGTGAATTGAGCATGAATGTGTTTGATTCGATTAATTTAAAATTAAATGAGGATAGTATTGTACTGTCTCCAAAAACAATTACTAAAAAAGTCAAATCACTGTGGGGGTTGACACGTTCATTGATAAATAACATGGTTATAGGTGTCACCGATGGATTTGTAAAAAAATTAGAAATTAATGGAGTAGGTTATAAAGTTAATTTGGATGGTGATTTATTAATATTAGCTTTAGGATATAGTCATGATATTGTTTATCCCATCCCTGAAGGTGTTAGTGTTAAATGTACAAAGAACACGATTGAGGTAATTGGGTGTGATAAACAACTTGTTGGACAAGTTAGTGCAGACATTAGGAGTTTAAGAAAACCTGAGCCTTATAAAGGTAAGGGGATTAAGTATGAGAATGAAGTGATTGAAAAATCAAAGACAAAAAAAAGGAAAAAAAAAGAAG
>evgR008480c0g1_i01_Hydra
ACTTGAACTTTTCTTTAAAGCAGTTATTATTTTCCTAGTCAAAACATTAAGCCCCCTCTTCTCACAGTTCCTCGCTACTACCCATCTGCGCAACAGGCACCATTTGTTTATAAGTTACTAGAAAAGAAAAGTAAATAATTCATGTCAATTCTGCATCATACAAAGGAGCTTAATGACTCTAGATTTTATTATAAACATAGCTTTAGCTCGGTGCAAACAAGCGTTGAAAAAACGTGTTATTGACTTTAGTGAATTGATTAAAGTGGGTACATATATAAGAGTTGTATTTTATAACAGCTATTATTTGTTGAATAAACTTTTTTACGTCTACTTTTGCTCTTTAATGAATAATGTCAGGGAACTCTTCTCTCGGTCAATTTCTTTTGGCTGTAGATGTGAACCAATCTGCATACTTTGCTAACAGTATAAAATGTCTATCTGACAGAGATACCAAAATGATTAAAAAAATAATAGAGGAAAACTGGGATGATGAGGTGCAAATGGTTTCAAATTTACTGCAATATCCCGATCTTATACCAGAAGATATTAGAGTAAACTCTATTATTAAAGGTCTTGAAGATGAAGAAATGCCATACTATATATTATCAGCTGCTGTTGGTGCTCGAAAACTTAAATTAAATGAAGATTGCATGAAGCTCGTTTTCAATCCACTACGAAAAGCAGTTTTTAATGATCAGGGTGTTGTTAGTATGACAGCCTTTACAACTATGCTTCCATATTTGAATTTCCCCAATGATCTTGATCTTATATTAAATGTAATAAAACAAAAAAAATCAGCATTGCAAGACTGTGCAGTTTCATGGTTAGTTTTGAAAGTTATAAATAAAACTGAGTTGTTAAGTATTTTAGATTCTGTTGACTGCAATACTTTGATCATTGCTCAAAACAAAATGGACAAACATTTACAAAGTTTAGTTGACGGAACTGAATCTTCAATAACTGTTGAAGTTATGGATTTTTATCCAAATTTGAATGACTTTGAAGCCATGTTGGATAAGGAAGATGTGTTATCAGAAATATTTGATGAATTAACAAACAGCAAAGACAATACCATCTCCTCTGTAGAAATAAAATCATTTCTTGACGATATTGGAACAAATTTATCAATAGAGGAGCTTGTTAGTGAAATGAAAAACATTGGGACAGATAAAAGCGGTCAAATTGATAAAGATGGATTCATAGAGTTGATGTTTCCAAAGTTTAAATGTTAGCCATTTTTAACCGAATTTATGGCTAAAGTTAAAACTTGTACTGTTAATAACTAAGAGTATCTTTATATGCTATGCATATTTTTTTAATGCATTTGATAAAATTGACGACATTGTAAATATTTAATTTATTCCTTTTATTACAAGAATTTTAACAAATTTTAATAACTTCGTGATGTATCATAATATTTGTCACGACCTTCGCTAGATGTTTGTAAACTTTCTAAAAATACTAAATTCTTATTTAATTTGTTTATATACATTATATGCATTACTTCATTTTATATATGTTATATACATTACTTATATACGTGTAAATTAGTCATCATAAAGTGTGG
>evgR008524c1g1_i01_Hydra
TTTTTTTTTTTTTTTCAGCTTCAGTCAAGAAATTAAGATGATTGGTTTATTAACTTTTGCACTTTTAGCATTGCATGTTCAAGCCTATGTTAAATCTGTGAAAAATTGTAACGGAGCAACTTATGCAATTCAAGAAAAACCCGAAGATCGTGTTATTGGTTCACTAAAAATCAGAGCGGGACATACAATTCCTTTCAAAGTTACAGTTGATGTTTCAGAAGATATTCCCGCCGGAGGCGAGTTAGTCCTTATTCTAAAGAAGAAAGTATTTGGAGTGTTCATTCAAATTCCATGTTTAAGCAATCTCGGGTCTTGTACATATCCAGTAGACTGCAAGATTTTAGAGCTCCTCGGAATGCCGTGTGAAATCAAAAAAGGCGTTTACCCCATCAATCATGCAGTGCATATTCCATCAAATTTCAATCTTCCTAATTGGATTGTTGGTGGAGACGTCCAAATTAAAGCTCTAGCAAGAAAAGGCAATAAAGTTTACGGATGCGTTGATATAGAGGCACGTTTAGAATAAAAGACCGAAGAAGATAACTTTTGTCTGAACAAGAAAATATCATTATACAACTATTTGGAAAATAAATCTAATTAAATCTTTTTCAAAAAAAAAAA
>evgR008536c0g2_i01_Hydra
ACATATTCTGATAATTATCCATAATTAGCTTCATTTTGTTTGGTTAGGAGCTTCACAAATTTAACTTCCATCGATTTTAAACAGGTTTCATTTTCTCGTGAACGTAAACCATCTGAGTAAATTGTAATAAAATAAATATGGCTATGCATGAAAATATTAACTACGATTCCCAGAGCAATCAATACGAATCAGGAGTTCATTATTTTTCATGTAAACAAATGAATTCTACCTTGAAACATAATAAAAAATCAAGCTTTAAGCAGTACGACGCTGTATATGCGCGGACTCGATCAGCAGCAAACTTTCGTGAGCGTCGGCGAATGCAATGTTTAAACGAGGCGTTTGAAGGGCTTCGAACTCATATTCCGTCCCTGCCATACGAAAAAAGGTTGTCAAAAATTGACACGTTGAAGCTGGCTATCTCTTACATTCAGTTTCTAGCAGACCTAATTAAAGACGACGAAGAAAAGAAAAAGATCTCCAATTCTTTGACTAATCAAAATCCAGAAAAAATATTAATATATAATACTTCCATAGAATACACAAATAACAATGGCCAATCTTTAAGATTAAAAGGACATTCTCTCTCATGGAAAATAAATAAGCCTCAATGCATTGGTGGCACTGTCAAAACCCGAAAGATATGGCAACCAGAACTGTTACCAGGGAAACTGTAGCTGATATTTAATAACAAGAGGAAATAAAAAAGATAAAAGTAAAAAAGTAAAATACAAATTGAACAAAGGAGTGGAATAAAGAAAAAATATTTATTTGAAATAAATTATAAACTTTTTACTATATAAATGATTTTATTTTACAGTGTATATATATAGTGATGCTATATGTTAAGATATATGTGCTTGATGTTGTATGTTAAGATATATGTGCTTGATGTTGTAAGATAATCTAGTGATTAGACTTATTTTATATTGGAATCAAACCATGCTTTACAATGTTTTTATGAAATAAATCTAGAAATCCTTCCTTTGCATCGGTTTTTAACCAGTGACGGATTAAAGCATTTTGGGCTTTAGGCAGAAATACTCTGAGGGCCTAATTTGGTGATAATTGAGAGTTTATTTGGTCCTTATGAACGGTTTTCCCCCTCCCTTGATTTTAAAAGAAGACCTTTTTTTTACCTTCTTTTTTTTTGGTAGTGTAGGCTTCAACAATAAAATGTCATGGCATGGCCCATGAATTTAGG
>evgR008537c0g1_i01_Hydra
AACTTTTCAAAAATCATTCACGAATTGAAAATGCCACACGTTGGAGATGAGTTGCACCATTTTGGATGGAATTCTTGCTCCTCTTGTTATGGTTGTTCAGATAATGGTCAAGAAGATTATGATAGAAGAAGATATTTGTTTGTGCCAGGTGTCAAATCCAATCGTTTCTATATTGTGGACACTTTTACAAATCCAAAGGAACCAAAAATTCATCAAGTTATCGAACCTGAAGAAGTTCAAAAGGTGACAGATCTTTCCTATCCTCACACAGCTCACTGTCTTGCAGATGGTAATGTCATGATTAGTATGATGGGAGATCATGAAGGAAATGGAAGAGGAGGATTTGTTTTAATCAGACAAAATGAATCATCTGGCAAGTTTGAAATTTCATCTCAATGGAACAAGG
>evgR008541c0g1_i01_Hydra
AATCTAAAATGGTCGTCGTTCCAAAACTCAAGGTCAAACCAGTCCACAAGAGAGACAAGAAATTCCGTAGATTCGATGCTCACAAGTTCACCAGATTGGCTCAAAAGACCACCTGGAGATGCCCACACGGTATTGATAACCCTCTTAGAAGACACTACAGAGGTCACGGAGCTTTGCCTTCATCTGGTTTCGGTACTGCTAAGGTTTCCAGATTCCACGACAGAAAGACTAGATTGATTCCATACGTTGTTTCTAATGCTAAGGATCTTGAAGTCTTGTTGATGCACAGTGATAAATATGCTGCTGTCATTTCTGCTAAGACTGGTGCTCAAAAGAGAAAGCAAATCGTTGAAAGAGCCAGAGAATTGGACATCAAGGTTTTGAATGCTACCGCCAGATTGAAGACCGAAGAAACTCAATAAGTTTTTCCCATATAAAAAATTATTTAAAAATTATAAAAAAA
>evgR008544c0g1_i01_Hydra
AGAAAATTATATGTTTAAAAAATAAATATAGTATGGGAACAATGACAACTTATACAACTATATCAACAACAAAAGATTCTGGAAACAGCCAACAACCAAGCAACATCATAATAGTGTTTCAAGCAATAGTTACTCCGCTTGTGTTTGTCGTAAACGTTATAACTATAGCTACCATTTTCAAATGGCGAAAAAAGTGTTTCAGTCGCCACGAAAAGCTTGTACTTCTTTTGACTGTTACAGATTTGCCCTTAACAATTATAATAGGTCCATTGATGATAAAGTCAAAGGACCTAAACAAACTTAAATTTTCTGTAGTAATATTTTGGTTATCGTTTCTATCATTTTTGTCCAATGCAGCGACCTTACTAATCTCACTCGAAAGATTGATCGTCGTTGTATATAACAATCGCATTAAAGGTCTGGCACCGTTAATCCAATTTTATGGTTTATAGTAGGAACTTACCAACTGCTGATTCTAGGCATGGTGACAACATCAAATATAGTGCTTCTATTAGT
>evgR008551c0g1_i01_Hydra
CACAGTTTTTACTTTCCGAACTATAAACTATTTAAATTGTTGTCTAATAGTAGCTAAGATGATGAATTTTGAAGTTTTCCTTTTCCTAGCTGTTGCATGCAATCAAAACGTCATTACAGGCAATGCACAACAAAGTGCATTCAATTTTTTAAAAGACTTTAACAGTAAATCTAGAGAAATGGACTTTGAATATGCAGTAGCAATTTTTATAAAAGCCACAAACATTACTGATTACAACATTAAAATTGCCGAAGATGCATATATTAAATATAGTAATCTTATGAAAAAAATTCGAAATGAAGCTTCAAAAATTAATACAACAAATGCGGACGCTGACTCAAAAAGACAGTTACAAATGTTAAATGTGACAATAAAATCAAAAAATGTAAATATAGTGAAGAACATATCTATAATTAGGGGGAAAATGGAAAATTTATACAGCACCGCAACAATACCGTAC
>evgR008574c0g1_i01_Hydra
GTTCCTTCGATGGGATTTATTCTGATCTTCCAGTTTGTGGAACATTTTGCCTCGCATCTTGCCTGGATTCTCTCTGTTTCTCTCTAATCTGCATCCCTTCGGTTTCTGATCTCATCTTTTCTACGTCATCTACACTTTCTCCTTCTCTCTTACTCGCTTCTTGCTCTTTTTTCCCTTTTGCTGTAGTTTCTTCAGACTTCCTGCCTTTCTTTATATATAATCTAGGTTTTCCTTCCTCTCTTGACTCTTCTGCCTCGTACTCCATGGAAATAAACATGCCTTGTACATTTTCTAATTCTCTCTTATCAATTACTTCATTCATATTTGCGGCTGACAAAGCAACTGCAGCGATTCCATCCTTTGGATTTAATCCACCAAAAAGTTTACTCTCTGCTGACTGCAGTGCAGATGCCTCGTCTTTGGTAATAATGTC
>evgR008576c0g1_i01_Hydra
AGAAATTAGGTTCAAGCCAAGAAGTACAAAAACTCGAGGATGCTGTTTCGAGTGCTGTTGAAAATTTGATTGAAACTGTCACACCTCTTTTGCAAAACGCAAAGACTACTACTTGTGAATTCATTTCGTCAGTTGAAAATTCCTTAAAACAAACACAATCAAGCGAACCTGAATTGATTAGAGTCGATGATAGTCATGATCTCGTTGAAGATCCAGTTGATGAAGAAGAAGAGCAATTGATTTCCCAACAAACAGTTGAAACCATTCAACCAGTTGTAAATACTATTGATACAATGGAAAAATCAGGATTTGTTTTCATCCTTTCTGAAAACGAAAAGGTGCAAGCAGTTGAACCAGTTTCTGAACCAGTTGTTGAAGAAGCTGTAGTTTCTTATCCAGAAGTACCAATTGTTGAACCAGTTGCTGAACCTGTTGTTGAAGAAACCGTAGTTGCTGAACCAGTTGTCGAAACTCCATTCTCCGAGCAATTAAAGATGTTGAGCTTGATGGGTTTCACTGATATTGAAAAGAATTTGGTTTTGCTTAACAAACACAAAGGAAACGTTTCTCTTGTTGTCGACGC
>evgR008578c0g1_i01_Hydra
TTTTTTTCATGGCAAGTTGTGACTTTGTTTAACTGATGCTTTTTCTTATCATCTGCAATGTGCTGCTCTCTTTTGAAGTTTTTCTACATCTATCATTTTCTCCTGAATCTTCCAACTTACTATATGCTATTACCCAATTATCTGCAGTGTTAGGTTCTACATGCCCGAACTGTTCCAGATGCACTCACTATTCTTACTCGATATCTAAGGTGCTGGACATCATCAGGTTTTGTTGCAAATGCTATGCTGCCATATATTGTCATTTTGTTTCTAATTTAAAATTTTATTATTCACTAGTTTGTCTTGTTTCAGCATTTTATTTATTATTTTTAGAAGTTAATTTTAAAATTC
>evgR008591c0g1_i01_Hydra
AATACTTTTTTTTGGCATCAGAATTTATTCTCAAGTATGTTATCTAAAGATCTTGGTGAAGTACTTTTATCTTTAGGAAACGAAGCCAAGGAACTTTATTTACCGCGAAAAATAAAACTGTTAGAGCATCCACCTTCGCCATTGGTTTTCTTACGTGAATATGTTATGCCAAATATACCTGTTCTTATAAAAAATGGGGTTAAGCACTGGCCTGCTGTAAAAAAGTGGAATGATGATTATCTAGTAAATGTTTTAGGTAATAAATGCATAACTGTAGCTCTTACTCCAAATGGTTATGCGGATGCTCCACAAGGGGAGTATTTTGTAATGCCTTATGAGATCGAATCAACTTTTGCAAATTTTCTTCACAATCTTCAAAGTGGAGACAAGACAGATGTCTCATACCTTCAAAATCAAAACTCAAGTTTTACTAAACAGTTTAGTGAACTGATACCTGATGCTGATGTAGATATTTCTTGGGCAAGTGAAGCGTTTGGAAAAACTCCTGATGCAATAAATTTTTGGATGGGAAGTAGTAAGTCAGTTACTAGTATGCATAAAGATCATTATGAAAATATGTATGCAGTCGTAAGAGGAGTTAAAAAATTTATTTTACATCCACCTACTGATCTACCTTACCTAACTCACAAACATTATATACCTGCAAGGTATAAAAAAGTGAATGGTGTTTACAAAATTTTTGAAGATAGAAATATTGAATGTTGTCAATTTTGTTCAAAAAAAGACATATCTAATGAAAGTAAGCTTACAAACACATTAACTGAAAATGCAGTTTTAAGCATGAAAAATTTTGAAAATTCGTTTGCAAACATATCTACTGAAAATACAAATGTTGACGTTTCAAATGTTTCTTCTTTAAATGTTGATAATTCTATCGATAAATTGAAAATCGAAAATGTAGATGATCTCTCTGTATGCAATATGAGCTGCTCACATGAATGCCATGTGCTACCAAGAATACCTTGGATCACTACAAATCCTTTGAGTCTTGATGATAAAACGAGCTTTTTGTATAAACAATCTCATCCTCTAGAAGTAACAGTTCAATCTGGTGATCTGCTCTACCTTCCATCTTTATGGTTTCACCATGTACAGCAAGAAGATAACACAATCGCTATAAACTTTTGGTATGACATGGAATATGATATCAAATATAACTACTATAAATTTGTTGAGCAAATCAATAATTTGACTAAAATTGACTAAAAATAGAAAAATAAAATAAAAAAGAATAAATATATAAAATATTGAAATAAAAAAAA
>evgR008592c0g1_i02_Hydra
TGCATAAAGCATTTTTTTATTTAAAACAGGACCTGAGATAGATATTTATTGGTACGGAGATCTGGGATACCAATACTGAATTTTAATGTTAAATTATGTTTGCATCTTGAGCAACTTAACCGGTTTGCAAACTCGAAAAAACATTGCATTGACATATAGAGCCTATATACTCCGAGATCGGACAAACGTGTGATTGTTTTTGATTGGTTTTTCGATTTTTATGATTTTAGACGCCATCTTTTTGCCAAAAATACTAACTGATATATAAATATTTGTAGAAGTATATCTGGTTGTGCTGCAGTAGGCTGCTATAACAGGCCCGAGATAGGATTTTTTATAAAATGCTTCCCAAGAAATGAATAACAGCGGGCGATTAGGACTTCTAAAGTTAAAAGACATAACTGGATATCTACAAACACAAACCTTTTATGTGAAGTACACTTTGATGCTGATTAATGGAGAAAATCTGAGAAGATGTGATCAAGAAAACTCAATCATAATGCAATTCCTACAGTCTTTAAACATGTTAGACCAAAACATAAGCGAAAAAATCCACTTGAGCGTTTTCCAATACAAAAAAAAAGGAAAATGTGTAGATAGTGAAGTTGTAGAAAATATTTTAATTGAAGTAAATAACAATGAAGTATCTAAATCACTTAGTACTACTGAG
>evgR008601c0g1_i01_Hydra
TCAAAAACATATTGAATGCAATTTGCAGTTGGTTAAACTTAAACTTACATGCTTTTAAATCATACTGCTTTGAAATTGCAGAGCTGATAATGGAAACGTATTCTTGGTACATTTTGCCATCCAAACGTCACAAAGATCTCGAGCATAAGTTTCTAATAGCTGAAAATTTGGATCTTTTCATTAGAATAAATTCTGAGGATATTCAGGTAACACAAAGATTTAATCACACATGCAAAATATCTAAAACTGTGATGCAGAACCAATTTTGTTATATACTTTTAATAAGCAATCTTGTGGTCTCCTCCGATGCTCGCAAGAACTCAAAGAAAGTTGATGATTTTGATCCTGATGAACATGGCATCCTAAGAATGTTTTTTTTTTTATAAATGCTATTGTTTTAAGTTTCAATGTTTTTTTTG
>evgR008607c0g1_i01_Hydra
AGAACTTGTCCGAAAAGTAACAAACACTACGATTTGTGCCATGCGTACTCCGGATTATTTATGAACTACAGAAATTACGACTGCTATAAATGTAACGAGGATGTCAAGATTATAAAGACACAAGAACTGCAGGTTATTGAGCTGACAAAAAATATTCATTCTATCGTGCCGATAACAAATATGCAGACTATCGAGCCGGCAACAAAGATAGTTCAAAATCAAAGTAATAATAGCCATAAATCTGATTCAAAACTTATGTTTATACAAACGATAGACATCAGCCAAAAAACAAACGAAAAAGAAATTTGCGGGGATTTTACATATAACCCGTTAACCAGTAAATGCGATATTACTTGTAACTATGGTTTCACGGCAGTTAACACTCAATGTTTAGAAAACAAAAGGTATTTACAAATTGAAGATATTGAAAACGCAAATTTTCAAAACTGTCTGCTTAGCCCTAAAGCTAGATTATTCG
>evgR008608c0g1_i01_Hydra
TAAAAAATAACATCAGAATAAACAACGAAGAAAAAACAAGGGAGTTAGTTAAAGAACTTGGATTTCATCCATTTGCAATCACACAAGCAATAAAGTATTTAAACATACACAATTTACCTATTGAAAAATATTTAAATCGTTATAGATCGCACCCTATAGAAATACTAGACATTGAAAAACTTCGAACTGAAGAGGAATCAAAGTCTGCAATAATAGCAATTAACTTAGTTTTAATAAAAATAGAAAAAACTAGAGTACTTCCATTAAAAATACTAAATTTGTTATCTCACTGTGATGGTCAAAACATAAGTAAGCATTTTATAATCCAAATATCAAAATACTTGGAAATAATTGAAGAAAACGTAATAGAAGACGCTATAAGATTACTGCAGAGTTATTCATTGTTAGATTGTTTTGGTGATGACAAATACACCATGCACGAACTGACGCAGTTGTCATGTAAATATTTCCAAATCAAAAATTCAAGTACAAATCTATATATCAATCTAGTCAAAAATTATTTAAAATTTGAATTAGAAGAAGTAAATGAGCACATTGATTACGGAAA
>evgR008615c0g1_i01_Hydra
GCTAGTAGAGGAGTGTTTATCATGAATGCACAAGTGAATGCAGTAAATTTAAGAATAGTAGAATTCAATGAGGGTCTTGTTGAAAGACTTATAAATAAACAAACTAATATATTAGACGCTTGTAAAGAACTTCAGAGTTTAAAAAGTGAAAAGTGGTGGGCTGAGCAAAGTGAAAAATTTGAAAATGCTAAGACTGCTATGATTGATGCGGCTAGTAGAGCAGGTGATTGGACGAAAGCCAAAATTGATGCTCTAGGCCAGTGGTTTGATGAAATGGGGCAAAAAATGTCTGAGGCATGGGCAAACTTCAAAGAAACTGTTGGGGAAAAGTGGGAAAATTTTAAAGGTGCCATGAGTGAGTTTGGAAACAAAATTGCAGAGATGTGGGGCAATGCGATGGATTCCCTGGGTAAATTATGGGATGAAATTAAAATTGGTGCGCATGATATGGCGGTTGATTTGAAAACAGGAGTTTTAAATCTATGCACTGATATGCAGTGTGGTGTCGCAAATTTGGGAATTGGAATGCAAAAAGCTGGTCTTGCAATGGATGATGCATATCAACAAGTAAAAGGTACTATTTCTGGTGCAATAGGCGATAAATTTAATCAATCCGCAGGCCAACTGTGCACTGAATCTACAATTGTTGTAGAGGGAGTAATAAAGAATAAAACTGCTGAGTTGGAAAAATTTAAAAAAGACTATAATATAACCGTAGGAGAGGATGGAAAGAATATTTTCCCTACGGAAGTAGTGGGTGGTTTAAATCCAAAAACTATTAAGTCCTCATTTGAACACCAAGAAAGTGTTATTGCTGACCTAAAAGGGCAACAACAAGAGTTAACAAGCAAGATGAAGTCAAGAGAGGGTATTAGTGCGTATATGGATGAGGTAACTAAGGGCACTAAAGATAAGAAGGATGCTAATGTTGAAACAAGAGGTGCTCTAAATAATGAAGTAACCGCAAATAAAAATGCTAGAGATACGGAAAAAGGGACTAGAAACACAGAGGCAACTAAAGAGAAACAATCATATGCTGATAAGGTAAGAGAGGAGCGTACAGCAAGTAAAAATAACGTCATTCAAAAGTAGGTTAAGCAGCTAATTTACTTGTGAATAAGACGTTTTATGTAGTGGGCATTATTAACTTATAAAGTTTGGTGCCCATTTCTAACTCGATTAAAAAGAAAAAAAATAAATAACTCGAC
>evgR008684c0g1_i01_Hydra
ATAATGTCAATCATCCTGATGGTGGTGTGTGTCAATTTAAAAATTGAGCAAGATGTATATTTTCTTTTCAAACCATATTTTTTTTAAACGGCCATCCTCCTCCACCTAAAATATTTCACGTAACCTGTGCAGTAATCTACACACCGATGGTTCAAACATTCAAAATAGCATTTATTGCGTTTAGCACCATAGGCTTCTTTTTGAATGGAATTGCATTGATTGTTTTACTTAAAGTAAAAAGCAGAAATGATTCGACACAGCGAATTATACTTATTCAACTTTGTATTATCGACGCCTGCTTAAGCATAGTTATTGGAATTAAAGAATTGTTAGAAGTTTTAGATTTTGATAACGCGCTGGTCATTAAGATTCTAGATGGGCTTCAAAATACATTTCAATCAAGTTTTTGCTACGCAACATTTTGGTTGGTAATGGACAGATATCTTCACATAAAACTAAACATAAAATACGCAACCTACTGTTCTAAATGGAAAATAATAACTGCGTCAGCAATAATATGGAGCATATCAATGTTTACAGGAGTTTTCAATCAATTTTATGACATTTGGTTTTCTTATATTTTAGCTTGTTATGATGTATTTATTATTGGTTGCTCAGTATTCGTCTATGCGTATGCTTTAACACTCTTTCGCAAACAAAAAAACACTCTTTCTTCTAACCAAAACAGCAAGAAAATGTTTAAAGGATTATTAGTTTCAAGTCTAATAGTAACAGCATATACTGTTTTGATTGCAATACCTGACTTTTTGTTCGTGATTAATGATTATGTGAATATCAGTGATCATTTTTATGATTGGTATGCTAATTTACTTTTTCCTGTATCTCTTTGGACTGACGCATTTATCTATATATTTGCAACTTCTAATGTAAGACTTTTTATCAAGAAAAAAATTGGATTTTTTCACGGTGATCAAAGTTCATCCAATTTAAAAATGCGTACAATTGTTAGAGAACGTGGTATTTCCATAATGAGTGTGAGTGAACCACGTGTTTTAAAAACAACAGAAGAAAAT
>evgR008689c0g1_i01_Hydra
TATATTTTACCCAATAGCCTCTAAATATTTTGCTTCGCGCATAAAAAAAATAATTTTCTATTGCAGGGAATATCCGGCCGGTGTTGATACTCTTCGCAATCTTACAACACGATTCGCTGCATTTTCCCTAGCATCTTCAACACTTTCATTGCCAACAGTGCTGCACGATGCAACTAAAACTAATGAAGAACTGCTAGTTTCTGACGCATTATCTAGTTCTTATCAGTTTACGAAAAAGGAAGCATCCATTGAGACAATTTTAACAAAATCCAAATATTTGTTTTATATAATGTATACAGTTCTTTATTTAAAAGTTATTGTAGACTTTGGCTAAAAC
>evgR008691c0g1_i01_Hydra
CAATTAATATAACGTTTAATAAACAGCAATTTATTTTATAACAGAACAAAAACACCTGCATCTTAGTCAAACCACTTTTAGCAACTAAAATGGCTACTAGCCCAGTATTTAAGAAATGGCAGAATTTTAATGAATTAATAACACAACCATTTATTGGACTAGTTTCGGCAACAAAAGCTAACATTGGAAACATGGCTGTAATCGGAGATTCGGTAAATCTAAAAGATTTGTTTATAAGACAAGTTGCTCAAACCGAGTCGATTCCTGGTCGTAACCTTATTCAAGAATACGAGGCTTGTATTGAACATATAACAAATGAAAATGTTCCACGAGTGAAGCGAAGTGTCTCCAGCGTGGGAGTAGAGTCAGTTAAAGAAGAGCCACGAAGAGCTTCACTTATAGAATCAATTTTGAGAAGATCCTCGTTGACAGAATCCACCCTTATTTTGTTTAAATTGTTTGATATTGATTGCAGTTACAGAAAAAAAGCGTATATTACTTATAAGGTAACAATCACCAAAGCCGACGTTATTTTATATTTTGCTCAAGAAAATAAAAACACCCACAAAATTGAAAAATGGATCAAAGAAATTGTACTACTACGGAAGACTGAGGAAATTCCAGTAATTCTAGTTTCAGACCAAACGTCTGCAAAAGATTTGTGCGTCGGACTAAAAAATGTTTATGTGTTTGACTTGACTGCTTTAGAAATCCAGATGACGGATTCAGAAGAAAAAAACACATTTTGCATTGAAAAATTAAACGAGGTTTTATTCAATTGCATGACCAATTATCTTCAGAAGAGAAAAGAAAATGATCGATCAGACTTTGCTTATTACGGAAATTAACTTTTTAGCAATTACCTTAAATGGCTAAACTGCTAAACTATATATATAATATAATTTTATTACTTACTATTGAAC
>evgR008699c0g1_i01_Hydra
GTGTTTTAAAAAAATGGAAACAAATAAATGGAAATCCTTCATGGGAGCAATTAAAAAAAAATCTATTATCTTTTGAACGATTAGATATAGTGATTGAAATTGAGAGAAAATTTATAGATTTATCTCCGAAGAAATCAAGCATTATTGAGTGTAGAATTCAACAAGGTTGTGTATTTAAGTTTGACTTACATGAAGATACAAATTTTATTAAAGTTGAATTTCAATTAAACAAAGATAATAATAATTGGACATCTATCAATAATAAGAAACATTCTCGTGAATATTATTTAACAAAAAAAAAGAATGGTTTATTG
>evgR008709c0g1_i01_Hydra
GCTAAATATGGTAAAAAATAAAACAGCAAATAACGTTGGTTTGATCAGTTTTGAAGGTGAATTTTTACACATCTTGGCAGAGACTAATAAAGTTGATTTATTAGAGAAAGCGCTTAATGATGGTATACATGGTAATATAACGTCAACTGATAGAGATGGAAGAACTATACTTCATACTGCTGCAAAACACGGAAGTTTGGATTGTGTTAAACTGTTACTAAATAGAGGAGCTGATCCAAATGCAAGTTCTGGTATAGGAGATTGTTGTTCGACACCTCTTCACTGTGCTGTTATTAATGGAAAACTTGACTGTGTAGCTGCATTGCTTGAATTTGGAGTTAATTTGGATTTAAAAAACAGTATGGGCAAAACGGCACTTGATTTTGCAAATGATTCTAGTACAATTGAAGGTCTTTGTTGTGCAGACCTTATTAATAAAGTTATTGTTCTGAAAAACGCAGAGGTTGAAGAAAAAAAGTCACAAGCTCTTTATGACATGGTTTCTGCGGGAGATTTCTCTAGTACAAAACATATTCTTGAGACAATGAAGGATCCCTACTCTATTATAAATAACTATTTTCAAGGATATAATACACTGTTGTTCAGGGCAAGCGAGAATGGATTTAAAGATATTGCTCAACTTTTGATACACTACAAAGCTGATGGTCGACCAAATGCAAAAAGTGGAATGACACCTCTCTATGTTGCATGTTACCATGGTCATTATGATGTTGTTGTTCTTCTTTCAAAAGTTTTTCCAATACAGTTAGCAATGCCA
>evgR008711c0g1_i01_Hydra
TACATTTACCAAGTGATGCTAAACGTTATGGTGTTCTTGATAATATTTCAGCGTTTAGATATGAAAGTTACCTAGGAAGATTAAAAAAGTTAGTTCGGAAGCCACAAAATCCAACTGGTCAAGTGGTACGTCGAATTATTGAAGGGCACTGCAACGTCAATGAAGTCACTTCCAATAAAAAACCTATCTTGAAGAAGATTCATTTTGATGGTCCTATTCCTGTTGAATACCAATCTTATATCCAGTTTGAACAATACTATGGACCGGACTATTTTGTTTCCACATCCATTGGAGATAATTGTTTTACAATTGAAAATCCAGTTAGTTTGATTTCACTTTCAATATACTTTGTGACCAAACTTACTGGCATAAGACGAACCTTTCCTGTTACTAGTCTAGTGACTAAGTATCTTCTACTCCCATTTAAACAAGGAGGGGTTTTTACAATAATAAATTTTCAAAAATGTATGCTGTTCTCGAATATATTGAAACTAAAGAGATAGATTTTCTGCCAATCAAGTGGATGGTTGATAGACTCAACTGGGATGTTAAAAATCTAATCAAAAATAAAACTGTAATAAATTTTTACTATCCACCAGTCAAATCAGCAAATAAAGTGTTAAATGCCAAAAAATTATGCAGTAACCCAGAACAACACTGGCTTTTATACAAAGTACGAATTCTGGGAGCTGCAAACACTCTTAAAAGCGCAAAAGAAAAACAAAACAGGCTGAAGAGACATCCAATGTTGATTATAATACAGATGATTCTAGTGATTATAGAGATATGGAAACAGGGAAAATTAGAAAATATTCTTTCATCTATGAAAGTGATGTTGAT
>evgR008711c0g1_i04_Hydra
TGAACCTAGTAAAGCTTTAGTTGTGTTTGAAGAATTTGAAGATTTAGAACTATTTTTTTTAGATCCAGTTAGTTTGATTTCACTTTCAATATACTTTGTGACCAAACTTACTGGCATAAGACGAACCTTTCCTGTTACTAGTCTAGTGACTAAGTATCTTCTACTCCCATTTAAACAAGGAGGGGTTTTTACAATAATAAATTTTCAAAAATGTATGCTGTTCTCGAATATATTGAAACTAAAGAGATAGATTTTCTGCCAATCAAGTGGATGGTTGATAGACTCAACTGGGATGTTAAAAATCTAATCAAAAATAAAACTGTAATAAATTTTTACTATCCACCAGTCAAATCAGCAAATAAAGTGTTAAATGCCAAAAAATTATGCAGTAACCCAGAACAACACTGGCTTTTATACAAAGTACGAATTCTGGGAGCTGCAAACACTCTTAAAAGCGCAAAAGAAAAACAAAACAGGCTGAAGAGACATCCAATGTTGATTATAATACAGATGATTCTAGTGATTATAGAGATATGGAAACAGGGAAAATTAGAAAATATTCTTTCATCTATGAAAGTGATGTTGAT
>evgR008713c0g1_i01_Hydra
AAAATGGTCTCACTCAAAGAAGCAACTGCCACTGCCTCCAGCACTCACGAAAACAATCATCCACAAAATCTTTTGAATGGAACTGGTGGTTGGAATGCCGGTACCTATGCTCCAGCTCATGTTGAAATTGATTTGGGCAAGGTTCAAAAAGTTTCCAAGGTTTCTTTGGTTCCTGATCAATTACCGGAAGGTGAGACAACTCACATCTTGAAGGGAGGAGAACATCCAAATCCAACCACTGTTCTTGGTGAATTATCTGGTCATACAAAGGCAGGAGACAAGTTGAAATTTAAATTTGACAAGACACCTGTTAGATATTTGAGAGTTGAAACTGTCAAGTCACCATCTTGGGTTGCTTGGTCAAAGCTCAAGGTTTCTGAA
>evgR008718c0g1_i01_Hydra
CTTAAATTTAATTTCAATGTACATGTAAATAGTTTATTTACAGATGATTGGTATAGAAATCTTTGCAGTATTTATTTTTTCTCATCTTGGATTTGGTGTTAGTCAGATGCCTAATAAATCTCCTATTCTACCAATTGTTGAAGATCTGCCATTTATCAAGTGTGATGTATGTCAAAAGGCTACTAAAGTTCTTTTCAAAACCATTGAAAGTCATAGAAGCAAGAAAAAGCTAGATGAAGATGATGTCCTTTCAATCGTTGAAAAGTCATGTGATACAGATGTTTCGTATGGTGACTGGATAACACGGTTGGATATTGTTGAAGTAGAAAATGACTTAAAAATAGTAGAACACCCTGAGGAGGGTAAATGTGATAGAGAGTGTAAAACTATTTCTCGATCATGTGAAGATATTATAGGTGACAGTGACACAGATATAGGCGAATTATTATGGAAAAATGAGATGAAATTGACGACTTTTATTAATCATGTATGTTATAAACTTACAAACTCGTGTAAGACTAAAAAAAAATATGTAAAAGGCAATCATAAGGATTACAAGTTTGTTGAAATGAGTGATAAAGAAAAAGAAGCCAGGGAACTTATGAGAAATATGAAAAATGTCCCCGGTATGCCTGGAATGGAAATGTATAGTCGAGACGATCTTCAGAATATGAGAGAGCAAATGGGTGTAGGAAATGAGGATGATAAGCAAGATGAAAAAAATGAAAATTATGAAGAGATTTACAATACTAAATTAGGATTTTTGGAAACTTTAAAATTTTTTCTCAATGATTTGTGGAAAAAAGTAAAGTATTTATTTAGTTTTATATCTTTTAAAAAAGAACTTTGAAAAAAAAAAAAAAA
>evgR008719c1g1_i01_Hydra
TTACGTATCTTACGATAAACTAATCATTCGGGATTGGAATCAAAAGAAAATGAAATTTGGCTAACTGATGATGCATTTCGCGAAAATTGTCGGTTAGAATATGACGCAATTCACTTACAGAGAACGTCTAAAAAAAGATGTGGTGGTGTTGTTATTCACGTCAAATATAGCTTGCGGTTTAAAATGCGAAATGACATGTGCATATCTGAATGCAACGACGAATTTGTTTCGATTGAAATAGTTAATGATAAGACCAAAAATATTTTAGTAACTTGTTGTTACAAACCGCTAAATGCGTCCAAAGAAAAATTCTCAAATCATCTCCCGAATATTATTCAAAAAGTTACTCTAGAGAAAAAAAAACTGTTTGTCTTAGCCCAGCAAGGAAAAATACGTATTTTATACGTTTTAAAATACGTATTCGGAATGCTCTTTGATACGTATATTCTAAGTCTTAAATGCAAATCTATAATACGTATTATTTAGATAAACAAAATGCGTTGATGATAGACGTATAATA
>evgR008726c0g1_i01_Hydra
TTAAAAACTGCTGAACTAGCAACTGCAACAACTGCTGAACTAACAACTCCAACAACTTCTGAACCAACAACTTCATCAACTGATGCATTAACAACTGCAACAACTGCTGAAACAAACAATGCTGAACCAACAACTGCTACAACTGCTAATTTAAAAAATGCTGAACCAACAACTGCAACAATTGTTGAAATATCATCTCCAACATTTGCTGAACTAAAAACTGCTTTACCAACAAATGTTGATCCAACAACTTTAACAACTGCTGAACTAACAACTGCTAAACTAACAACTGCAACCACTGCTGAATCAACAACTAAAACTGATGTTCCAACAACAACCACTGATGCTCCAACAACAAACTCTGCTGCTATAATAGCAACCACTGCTGCTGCTCCAACAACAACTTATTACTTAACAACTTCAACTGCTGCTCCAATAACTACAACTGCTGTTCCAACAACTACAACTGCT
>evgR008731c0g2_i01_Hydra
CTAGAACTAAACCATTCTCAATTTTCTGTTCAGGATAGAACAGGGATTATCTCAGATCTATTCAGTATGGTGGCAGCAGGAAAAATGAAAATTGATAAAGTCCTTGATACAATAAAATATATAAAAAACGAAAAAGACTATTATCCGTGGCTAGTTACTTTATCTGAGCTAAACTTTATTGATCATATTAAAAATGAGGATAACTATAAAAAAAAGTTTAAAGCTTACATACAAAAATTGATTTCAAAAATCGTGATAGATGTTGGTTGGAAAGATGCTAGCTCTATAAACAACGGTCTCCTACAATCTTTGATTCTAAGAATGGCTTGC
>evgR008745c0g2_i01_Hydra
AAATCATTTAGTAAGAAAAACAGTATCTTTTCGCTACACTTAGTTTTTAAAGTACGAGAAACACTTACAACAATTTTATTCCAAACCATTTACGACAATTACCAACTTACTGTAAAATACATTAGGACACTTTCCTCATTAAAATATAACTGCGAATTAAAAGACTGGGAAATAAGTACAGCCACAAAGCAAAAACAAATTAGCCACAGGCCTCCGTTATTTACAGTTAGCGATTCTAAGTCATTCAACATACATTGCACAATAAAAGCAAATACTGGCTGCAACCTTATGGCTACAACCTTATGGTTAAATATTGAATTTAATTGCGCATTGCCTCAAACTCCGTTCAATAAACTTGTTAAAACAGCAATATTGAACTTATCAACCAAATTAATCACGGTTGAACAATAACAATTATTAAATCTAAATCCAAAATTGATTCCAACACAAGATAAAATTCCATTTATTGAAA
>evgR008753c0g1_i01_Hydra
CTAAAGAAAATCCAAAATCTCCTAATTAAAACTAATTAGTTTCTCGAACATAGAAAATGTACGAGAAAGATTATCCAATTTCAGCAGCTGAAGGCGGCATACGTAAACAAAAAAGACAAGAGTCATACGAAATCTACATCTACAACATCCTGAAGCAAGTACATCCTGACGTAGGTGTTTCTTTGAAAGCAATGAATATTATGAACTCTTTAATTAACGATATTTTCGGAAGAATTGCTTCTGAAGCATCAAGGCTTTCTATACAAAGCAAGAAATCGACTATTTCATCTCGCGAGATTCAAACAGCTGTTAGGCTTTTATTTCCTGGTGAATTAGCCAAACACGCAGTAAGTGAAGGAGGCAAAGGTGTTACTAAGTACATTAGCAACAAATAGAAT
>evgR008756c0g1_i01_Hydra
CCTATTTTAGTGCAAAAAGAAAATAAAATACCATATTTATTCTTCAAGTATCGTTAGAGTGATTCAAATGCAAGCAAAAATCCTTTCAAATTGACGCATCGAAATACTGCAAAGATCTTGGTATTGTTAGAATGGCGGATTCAAATCTGAGCAAAAAATCCTATCAAATTGACGCATCGAAACACTGCAAAGGTCTTGCTCTAATTGCCACGCGTAGAATTTCACGTGGTGAACTAATTTTAAAGGAGAAAGCTCTAATACGCACCGAAGGTTTCGCTTCACCAACGCAACTCAAGGTTTTATTTGACTCGCTAACCGATATGGAGAAACAACTTTTTATGTCACTTCACGACTGTAAATGTAATTCAACCGATAGCAAAACGATTGGAGGCATCATGAACACAAACTGCTTACCTCTCGGCTACAATTCGACTAAGAGCGGCTTGTTTCCCACCATATCCCGCATCAATCACTCATGTAATCCGAATGTTAACCATTATTGGAATGAGCAATGGCAACTCGAATGCGTCTATGCACTTAAAGATATAGAGGTTGGTGAAGAATTGTTAACAAGCTATATCGAACTCTTCAGAACACGTGCAGAGAGACAAGCACTTTTGCAGACTTTCTGTTTTGATTGTTGTTGTGTTCTTTGTTCGATCAGTGATAAAGACAAGTTAAAAAAAAGTGACATTCGAAGACGTATTTTAAGGCAGCTTGACGAAGATATTGCTACCGTGGCGTTGTTACAACCGCAAATAGCAATGGCTAAGGTCGAAACATTGTTAAAGGTACTTGAGGAGGAGAAAATGACACATATAGGATCATATGTGGGTAGGGCTAGTTATGATGCCTTTCAGATTTTGTTTTATACGACAAAGAAGAATTTATCTTTTTGGGCGAAAAAGGTTTACGATAATTGCACAGCTGCGCAAGGCGAATTAGCAGGACCGCTGAAGAATGAAATCCTCTGTTATATTTCAAAAGAAAAATCTTTTTAAATATATCAGTATTCGTTTCCTATATAAATTTAAATAAATGTTTACTTATTTTTGTTTATTTAAATTGATGTGAATTAGGGGTGGTAAACAATTAAGGTTAATCAATTAACTTTCAAGCTGCTTATAAAATCACCACTTTAAAACTTAAAGTTTTTAAGTGGTGATTTTATAAGCAGACTCCAATGTTTCTTCTAATTCCTCCTGCACATTTGTACAGGAGGAAGTAGAAGAAACATCGGAGTCTTGTTTCCACCAAGTTTGAATCCACTTGAAGTATAAAACTATTAAGAAAATAACTAAAACTTAGTAATTAGCAGGAATATTATCAAATACAACTTCAGCACCCGGAAAAACTAAAGGGCTAACTTGTAAGTTCTCCCATCAAAAAAACTGCAGTACTCGAAAGGTAATGGAAACAGATCACGCTTTTTAAAGAAGATTTACAATGCCATGCTATGTCTTCCCCATCTTCTATAGAAGCAGAAAGGAGTTTTAATGCAGAAAGTTTATTTTACAGCAGCCGTGGCGCAGTGGTTTGATCATTGGACTCGTAATACAAGGGTTGCGGGTTCGATCTCCGCTCGCTATTCTTTATGTGACGGCGCACAGTGACTCAAAACAACAACAAAAGGAAATAAACCCCTCAATCATTTTATAGAATATGGTGCTTTTTTATATTAAACTGCAATGATTTTGTATTTAAAAGCCAAAAATGGTTTTCTCATA
>evgR008758c0g1_i01_Hydra
ATGAAGCTGATGTTGTTATTATTTCTCTTGTTATTGATGCTAAATCTCGCACTGGTTTTGTTAAAGAAGTAAATCGTATGATTGTTTTGTTAAGTCGTGCGCGACTAGGCATGTATATTGTTGGTAATATTAACTATTTTGAAGAAAAACCTGATGAGTCTGTTCACTGGACTAATATCTTAGAAAAACTTAGGCAACCCTGTGAACCCGACACAGGTCATGAAGAAATTCACTTTAATGGTCCTAGAATTGGAAATAAATTGCCAATTTGTTGTCCTGTCCATAGAATGAAAGATATTTTCTATGCTGAATCCGTAGAAAAATTAAATCCTAATTTATGCAAAGAAAAGTGTTTGAAGTTAT
>evgR008764c0g1_i01_Hydra
AAAAATTATTGTAAAAATATTACATTTTTTTACTACAACTTTTAATGAGATAGTTCTTTATTATGTCAACAATCAAAGTAAAAATATTACAAGTTAAAATACATACACACACTAGTAAATCCAAGGGCAGTAAATCCAAAGGCTGCTCATTAGATGTTATTTGTTCTAGTTGGCTAATAAGCAACAAAGAAGCATTTTATCTACATCATATGAAGTCTGCGGCAAAATGCACAAATGCTGTTATTAACTTTATTGTTCTAAGTAGCAATTGGGGGAAATACACAATTTGCATACTTGGAGGATCAAGTGTATTTAAGCTGATTATTTTATTATTTTTGTTATTGCTAGATATAAATAAATTACGTATGATTGTTTTTGTAATTCTTTTTCATGTTATTTATATAACTTTAAATAATTCTAGATGATTTTAATGTTGCTACATCAATGATGGATAAACT
>evgR008771c0g1_i01_Hydra
TAAAATTGTTTTAATAGATTTTTATAGGCATATGAAATTTTTGAGACAGGCTCGAGTATACTTAAGAAAAGCATTTTTAAAAGCTTGTTGGTTTTCCTGCATTTTTGAAAGTCTCAATATTTATAGGACTGTGCTAAAAAAATCTGTCATTGAAAGTAAGTCGAAGTTCAACATTATGCACAAGCGTTTGCTTACATTCAAAAAAGAAACATGTATTCTTTATTTGAAATTAAAAAAAACAGTTTTTAAGTATTCCTACGAGCTATGGCTCTTTATCTTCTTCAACCTTTTAAAGTTGAAAACGTGTTCAACCTCTCTAAGTTGAAAACGTGTTCAACCTCTTAAATTTCAAAACATGCAGTCTTTTT
>evgR008774c0g1_i01_Hydra
CAAAAAGCATAAATGAAATTTTGGGCAGCTTTAAAAAATCTTTTAAAGTCTAGAAAACCTAATGCTTCTTTAGTATTAACAAGACATCTTCAAAATCGTGGCAAACCCTGTTGGACATCATACTTTGTGCCATATACATCAGTTCGTAATGATCATTTTGGATACTCCCATTTCAATTGGGAAGTTGACGGAGTAAACTACCACATATTAAGAACAGGATGTTTTCCATTTATAAAGTATCATTGTAGCAAAGCTGCTTGGGTTGACTTAAGAGCACAAAATATATTTTTCAATTTTTTGAAAATTATAAACTTAGGCGTACCTACTTTAGCCTATGGACTTGTTACAGTGTTTTTAATAACATGGCATGAAGATGTACTTTTAAATCATAGTATTGATAATTCAAAAAACTACCAGACAGTTAGAATTTATTTTCATAATATTGAAAAAAATATTCATTAAATTAAATAATTTTAAAC
>evgR008775c0g1_i01_Hydra
ACCAAAATTCAATTGTAAAAAACATACAAATACTGGTAATTCAGAATTAGAACTTTCTATTGAGGATATGGATCCAAATTATGTTGTGACTCCCTCTCAGTACCAATACGAAAAAGGGCTTGAATTTAGTTTTGCCTTCTCTGTCAGAATTCACAACCCAAAGCATTATTACATTTTTGTAAAGGATGTTCAAGTTTTTTACAAGGATATTAATGGAAAGTGGATTGAGGCAAATTTTCCATCCTATCACACATTGGGAATCACTTTTCATCATGATGACAAGTTAAAAGTTGAAGGGATGGAAAAACTTTGGTGTCGCTTCAATGCCAATTTCACACCAATTCCAAAACTTAGTCAAGTTATTTGGGGAACTTCTTACAATTATATTACCCATCCCTCTCTCCCTCAGCCATTGACCTTAAAATTGATCATTCATGATTCTCTTGAAAAAACCAAAGA
>evgR008780c0g1_i01_Hydra
TGGTCCAAAGAAAATAATTTTATGAAGGCTTTTCTTCAAGCAAAGGATATACTGAAATATTCAAGTTCTGTAAAGAAGATAATGGTTTTTGTTGACCACGAATATACAAGCAATGATTTGTTATCAATCAATAACATTTTTAAAAACTCTCAGAATGAGCTTATTGAACCAATTAGAATTTTTATTGAAAGCAAATCATTGCAAACAATAAACAGTTTAAAGGAAAAAGGTTTCAGTTATATAATAGATGAGCCAGAAACTGTTGGCGAAATCGTTCGTCCGTTTAATGAAGGTATTCTTAGTGAAATGGTGACACAAATATTACCAGACCCCTGTCTCAAAATTCGCTGTCAAAGTAATGCAATATGTACAAAAAATGCGGAATGCATATGTAAAAAATGTTATTCTAAATTTGAATCAGTATGCGGTAGTGATGGTTTAACTTATTCAAGTAAATGTCATCTTGAGTTTATATCATGTCAAACAAGAAAGGAGATAACTGTTGCTGATTCAGCGCCTTGTAAATTGGCATCCAAAATGTTAATAACTTATGTAATTGACACATCCAGTTCCCTACAAACCAATCTATTGAATGAGGCCAAAGATTTTATTAAAAATGAAGTTAATAGTTATTCTGCTTCTTCAGAAATAG
>evgR008783c0g1_i01_Hydra
CCTATTTTAGTGCAAAAAGAAATAGCAGAGAACATCTTTTTAATATTTGATTAAAATGGGCACAAGGAACAACTGTGTTGCAATCGATAAAAGCTATGCTAGCAGAAATGCTTTTAATTGGTATGTTACAAACTATCATCGACAAGAAGATACTGTTATATTAGTTCATATTCTTAAGATGCCTAAAATCACAAACTTGAAGCCAGAGCAAGAACTCAAGAAGTTTCATAAAAGCGTACAAAAATCTAAAGAAATTGTGGCTATGTATGAAAGTGTATGTGAAGAAAAAGAGATTAAGTGTTTTACTGTTATTGAAAATTATAGCAACTGTGTAGGAAATGCTATTTGTGAAGTAGCTTCAAAACATGCTGCTAATGTTATTATTGTTGGTAAGCGTGATTTAAGTACTCTATCAAGATTAACATTAGGAAGTAACAGTAAATATATTTTACACCACTCGCCAGTTCCTGTAATTATTATATCTGAAGAAAACCCATGTTAATGAAGTTATCAGTTCATTTAACAAGTTATCATTTCTGTTTCTGCATTGCATAAAAATTTTTAATGTCTCAATTATTATTTATTAAATTATTTATTTTCTAGTCATTTAAATGTAACTTTTTATAATTTATTTTTTCAATTATTAATTATGTCCAATGTTCTAATTTAAGAGTCC
>evgR008796c0g2_i01_Hydra
CAATAAAAAAAAGTTTACACATTTTTTCATTGCAAGTATTTCCGTTACAAACATTCTCGAATGCTTTCTTGGTTTTATTCCTGAAATGATGATGCTTCACACAAAAAAGTTAGTTGGAACAAAACTTTGCTTTGCAAGTTCGTATTTTGTGTTTGGCTCAACTATAACGAATATATATCACATTGTTGTAATGTCTTTAATCAGGGATTTTGCATTAAATTTTCCGATTTTTTATTTTAAAGTTTTAAAAAATGTCTGGTTTAAAGCATTTATGTTATTTCTTTGTTATATGTTTGGATTTATTTGGGCGTTATTTCCAATTTTTAGCTGGTCTAAATATGAAATTGACATTGACGAGAGGCGCTGTTCCTTGGACTGGAGGTTGCAAAAATCAAGTTCTCTGTCATATATTGTTACAACTTTTATCTTGTGCTATATATTACCAGCAGTGATTACGTTATATACGGTGTTCAGCGGTAAGAAAACAATTACTAAGAGAAATCAGGAAGTATTAAAAAGAAATCGTTTAAAAAAAAGCTACTCAGTTGACTTATTAGAAAGAAATTATTTACGGATTTGCACACTGTCCACAATACTTTTCTATGTACTTTGGACACCATATGCTTTTGTGGTTATTTTAGTGTTTTTTGACTACAACCCATATAGAAATTTAATAACTTTAGCTGCCATGTTTGCAAAACTTTCAGCATTATCCAATGTTCTTATTAACTGTTATATTAACAAATCTTTTAAAAGTTATTTAAATAAAATTTATTTTGTTG
>evgR008820c0g1_i01_Hydra
GAAAATTCTTTTTTTTTAATGTTAACACTAAATTACCGTTAAAATAAGAAATAAAATGAATTTTGTTAACTTTTTATTATCAGTTATTCTTCAAACTGTGCTAATTCATTCAAGAAGATTTCAAAGAGACTTACCGAAATGCCCAATGCTTATTGGTGTTGGGCAGCAACTATATAAAGATGGAATTTTATCAACACAAACTGTTAATCGCGTAATAGTTTCAAAAAAAAAAGATAGTAAATGCGTTGAAACAAGTGGCACAGGTAACTTTAATTTATCAATGAATATTTATTCCGACGAAACGTACGCAACTTTTTTATCGCTAAATAACAAAGAAAGTTTGGTATACAACCAGAGTAAAGTATTCCAAGAGCTTAAATTAAACAGCAATGAAGAAGATTTGTTTATTAAACCCGTAAACTGTTATGCAACACCTGACAGCAATCCCCAAAACGTTATCCAATACAAATATATACAGAAAGGGTGTCCAATAGATTCCACGATAGTATTTTACCACGGATCTAATATGAAAACAAGACGTTTCAGTATTAACACTTTCCGATTTGTCAACTACAGCGCAGTTTTCTTATTCTGCGATGTAGGCGTGTGCAGTAAAACTGCGGTAAATTGTCATGAAACCAAATGTAATAATAAAGAAAACATATTAAACACCAATGAAAATCGTTCACCTAAATCAGGGTCTAGCGAATTTGAAAATCAAGAAATTAATTATAGCGTTGAATGTCAGCCAAAAAGCATGACCCTTGCTATTAAAAAACCCTTACCAAGTTGGCACGAAATTGAACAATTTCATTTAATTGATAAGAGTTGCAAAGCAATTGAAAACGATACCCATGTGGTTTTGAAGACTCCTTTATATCTTTGTGGAACTACAAAAGAGAAACTTCCATCAGGAGAATTTATATTTAAAAACGAATTATTTAAAACAATTGAATCATACCCTATTCAACGCACTAGTAACATATTAAAAACATACAGCTGTAGGTACAGCAACAAATATGAAGCTTCAATATCAATTTCTAAAATCTCTATGAAACCCGCCAAGAATTTTCGAAGTATTGCATTCGGGAACTATAATGTAACTATCAGTCAATATAAAGACGCAACGTTCAATGAAATAAAATCAGATCTGATCACTGTTGAGCCAAATAGCAATATGTACTTCCAAGTTAGTATATCAGACGAGGTGTTCAACCTGGCTTTATTTTTAGACGAATGTTTTGGAAGTCCATTCTATAGAAGTAACAATAAAGAAAAGTTTACATTTATCGAAAGGGGATGCTCTAAAGATAAAAGTCTCAAGTTCTACAAAACTGCCGATACAAGCAAAATAAGGTTCAGTGTCAAACCTTTTATATTCGAAAATTCAAGTAGTATGTTCATTAGTTGCTACGTGTCAGCTTGTGATGCAAAGGATATTAACGATAACAAATGTAAAAAAGGTTGCTTAGAATAAATAACTTTTCAAAAAAAAAA
>evgR008825c0g1_i01_Hydra
TCAAAGGAAAGGATTCCGACAAGAGTGGTGAAGAAGTTTTCTTTGATGCCAAAACATGTGAAGTTGTGCCAAAACGAGTACCACCAATGGAAGTTCAAAAAGACAATGAATCAAGGAAATTATGGGTCAACCTTACAAAAGCCATCAAACAAAATGATGTTCAACAAGCTGCAATTGAAAAAGACAAGGTTGAAGTCGAACAAAGAAACGTGAAGAAAGAACGTGAACAACAAGGCCACACCTTTACTCCAAAACTTTTCAAGCCAGGTCCTGAGGACTTCTGGACATTTATTGGTTTTGATTCTCCTGAATGGCAAGAACTCCAACAAAGAAACTTGGAAGAAGAATTATTGAAGAAATACGGCTTTAAGAATTAAAAAATTATTTTACCACCACCCAACATTTCTCCC
>evgR008831c0g1_i01_Hydra
GTATCATATCTTTGTGTTTGAAAACATTTTTCTATGGCAAACAATATTTGCCATATCTATTTAGTTTCCACTAAATTTCCAGAGCTTGCATTTCTTTAGCCCCTCCCGCGCAACGGCCCTGACGTTTTGTAAGTTTATAATTGTGCTTTTTCATATTTAAACGTATTTTAATAACAAATCATGTCTGAAAATGTAGGTGGGTCAAGTGCAAGATGGGCAGGGAGTGCTATGCAAATGATACCAGGCTTAAAGCCTCGCCCGCAACATCATTGTGTTGGAGAAAAAAATGGCGTGAAAAACATTTCTTCAGATAACAGTTCTTTGTTTATATTTATTAACAACTGCCAAGACTTTGAAGTTAACGTAATTGGAAAGGTGGGAAAGGTCACGGCTGAAAATTGCCGAAATTTAACTTTAAGTATCACGGATAATGTTGTTGGTGGAACACTTGAAATTATTAGATGTGAAAATGTGTTGTTGAACTATGGTCACAATGCCGAGATACCTGTTCTCCAAATTGAGTCTAGTGCGAACGTCAACATTTACATTTGGCTTAAATCTCAACTTCAAAGTGTGTATTTGATGCAAACCAGACTTGTGAATTTATTTCTTCAAAATCAAGTAGTCGATGACCAAAATGGGCTTACTTTACAATATTCCATAGAAACTCCTGATGATGAGTTCCACATTCAAAAGGTTTCTCATTGGAGTTCAGCTCTTCCAAATGCAAACTTTATTACTGAGTGTGTTGTTCGCGATGGTATCTATCCGACAACTTCTGAAATGAAAAAAGCTGCTGAAGAAAGGCGGGAGTTAATGAGAAACAAAATGTCTGAAATATTGCAGAATGGTATTAAGATTACGAATCCAAACCAATGTGGTTCTAGCTCGTCTTCTCATGACAAAAACTTTGATTTGCCAGACTAGTATTTAAAACATAATTTAAATATAATTCTGATTTTTAATTTAACTTGTGAATGGTTGCCTTTTTGCTGATTAGTTCCCAGGCTCTATGATAATCAGAGGAATTGTTTTAAATAACTTTTTTAAATAAGTGTATTGTTTGGTTTTTTGACTTTTTTAAGCTGTGGCGATGAAATATTTTGCAATAATTTGCAGTTCAAATTAGTTCATACATACTATGACATGCTTTGTATTAAGTATGTTCATAGTAAGTATGTTTTTAGTCCCTAATGTTGTAAAGCTATATTTTTCCAACGTTGAATCAATGTAACGTGTAGAAATGATTTGACATTCTTGTCTTGGAATGCTTGTGAGAGGTATTTATTGGTTTTTATATTTGAAATTTTTTGTATGCATTCATTTTTAGTACATTTTTGTAACAGTGGTGATAAATAAATTTTGCTGTTAATAATATATATTATTTAATGTCCCAGATTTCAGAAAAAAAA
>evgR008832c0g1_i01_Hydra
ATGCTATCTTCAAGATTCATTTCAACAATAGTTGTTTACATAATTTTCATAGAAATCATCAACCTTTACACATACTTTTTTGGTTATGCTGATATTCCAGGAACCTTGTCTGACGCAGGTGTTCAGCAAATGGTTTCTAGTTTTGCTTTCATTTCACTAGAAAAGATAAACCAATATATTACATGTGTTAAAGGTTCCATCTTGCTATCAATCATTCTTACAATTATTCTGTTGTACCTTGTTCTTCTGAACTCCTTCATGTGGTATCGAATTGCGAAAAGGAAACTGGTCTTTTTTGCCAAAATCGCTTCAAATGAAAAAGTTTGAGTTTATTACCCCTGTTTCTCTAATGGTTGCCAGTATGAAGTTTGCTGGGTATCAAGTTGGTTATATTTTGTGGGGCTCGTTTGTGATATTTGCGGTCATTTTTCTCATTTTGATGGTGATTGGCACTCAAATAATC
>evgR008849c0g1_i01_Hydra
AGTTTTCTGGAATATAAACTCATTTGTTATTCTAATCAGCATCTTGCTGATTTTAGTCGGAGTTGTTGTCGACACATGGTGGATAAAATATGATAGTAACGTCACAATTAAGAACGGTTTATGGAAGGAATGTAGGCAAAACAATAGTACTCTTCTGTGTAGAACAAGACCCAGTATCTTAATGTTTAAATACGAACAAAACAAAGACTGGGTAATTGTTATGTTATTTACAATTGCATTGTTTTCTGTTTTATCTTTATCCGCTTCGCTTTTTATCTCTTATGGTGACAATAAGAAAACACGCGTTACCGTATT
>evgR008876c0g1_i01_Hydra
TTCGAGCTCATAATTCTGTTATAACTATTAGAAGATCTGAACATGATAAAAATGCTCTTATGAGTGTGTTTAGAGCTTTGCCAAATAATGAAAGTGTTTTGACTGAAGGAGATTTGTTTCAAACCTTTCCAGAAAGAGCAGTTTGGATTCCATTTGATCGGTTTATATCTGACACATTTTCTAGACAAGTTGCTATATTGTGCAACCACACATTCGAAGAAACGTTAACGGAAGCAAACAAAGCTGGTGTCAAATACTCTGATGATTTTGATGTTGCTGAACCCATGTTTGTTATTTCGTGGTTGTTGGGAGCAATGTCTGGATTAATAAAAG
>evgR008880c0g2_i01_Hydra
AGTGCAAAAAGTTTTAGATTATATATTATAAAGATGATAGGAGATAGATGGCTTCCATTGGAATCTAATCCTGATGTAATAAATCGATACATCAAAGCACTTGGTGTTGATACAGATAATTGGAAATTTGTAGATGTGTATGGTTTAGATGATGAACTGCTTGAAATGGTTCCACACCCTGCATGTGCAATACTTCTTTTATTTCCTATTACTAGCAATTATGAAAAAAATAGAATTGAAGAAGAAGAAAAAGTGCGTGGGGAAGGTCAATATGTATCTTCTTGCTTATATTTTATGCCGCAAACAATCAGTAATGCGTGTGGTACTTTAGGTGTTATTCATGCAGTAGCAAATAACATGGAGGTTCTACAATGCGAAGGTTTGTTAAAAGCGTTTATTGAGCAAAGCAAAGGTTTATCTTTTTCTGAAAAAGCACTTATGTTGCAAAATGATTCAGCTATAACTAGTTGCCATGATAGTAGTGCTTTAGAAGGTCAAACCATAGCTCCTTCTCGTGAAGAAAATGTGAACCTACATTTTATTGCTCTAGTTCAAAAAGAAGGTTATCTTTATGAACTGGATGGTCGGCGGCCTTTTCCTATTAATCATGGCCGTTCTAATGAAGAGACCTTTTTAAAAGACGCAGCTGTTGTCTGCAAAAAGTTTATGGAGTTTGATAAGGAGAGTTTACAGTTCAATGTTGTTGCTTTAGCTAAAGTAAATTAAATTTTATAGTTTTATATTTGTAAAAGGAAAAGTAGTTTTAGAAAAAAGTGATTACAAAATTTTCTGGAAAATAAAAAGGTCAATGTTGTCAATGCACAATAGCTTTTTGCATGATTCTAGTGAAGTATTATCGGTAAAAATCTGGAAAAAGAATTGACATGTTTTCCTTTCATCGCTTTAATGGCTTTTATCATTAATTTTGGATTAATATTTAGGAGGAAAGGGAGTTTGACAACAAAAATTATTTTTCTATTATATAGTATGATTTTATATAACTATTTGTCACAGTATAAAATAATTATTTTTTTAACTAAATTACTATAAATTTTTTAAAACGTATATATTTACTTGATATAAAAAAGATTGTTATAAAAATGTATTTAATCTGTTTTAATATGAAAATTTGTTTCTGCC
>evgR008883c0g1_i01_Hydra
TTATCATCCTATGTGCATGTTTATTATCACCTTATGTGCATGTGCATATTACCTTATGTGTATGTGTGTATTATCATCTTGTGTGTCTGTGCATTATTACTTTATGTGCATGCGTTATCAGTATCATATCTGCATGGGCATTTTAATCTTATGTGCATGTGCATTACCTACATATGAATGTATGCAGATTTTTGTGTACTTAATTTTATGTTATTTCACTTTGCAATGTTATTTCACTATTCAACTTTTTTTTTTTTTGCATCTTTTTTTTTCTGGTATATCTTTTTATTGTTGTCTGGCATCAATTTTTATTCATTATTAGCACAAAAATCTAATCTTTAAAGGTGTGGTCTAGTTTTAAAGGTGTGGTCAAGTTTTAAAGGTGTGGTCTAGTTTTAAAGGTGTGGTCTAGTTTTAAAGGTGTGGTC
>evgR008894c0g1_i01_Hydra
TTGCGTGTTTTGCCGCTCATAGATATAATTAATGCGATAACGATTAGCAAATAAAAAATAATTTACTTTTGAGTTATGAACTTTATGAACTATAAAACTTCTGGGCATAAGTGCAAACGACGTCAGAAGATAAATTTAGCAAAACAAAATATACGGATAACCCCTTATAGGTTCGAGTAAAAGTGTTCGTGTAAAGCGTGAAATGCAATAATAGTAATCCTTATAAGTTCGTGTATGATTTGCCATAAAAATAATAAAAGTATTTAATAGTGTTAATTATAGTTAAATAAGAATGAAATTTTAAAAATCTTTTTAAATATCAGCATGTCTAGTGGAAAAAATTTATTTATATTAACTGGAGCTAGTAGAGGCTTTGGTCAAGCAGTCGCTGAAGTTCTTTCATATACTTTACTGAAAAAGTCCACCAATGGTTCAAAGATTATTCTCCTTGCAAGAAATTACGAAGGATTAAAACGTACTGAGAGTATTGTTCGAAAGAACATTGGACTTGGCATAACAATTGATAATTTATGCTGTGATCTATCAAACCCGAATGATGTCGAGGTAATCATACCAAGAATGTTAGTTTCACCAAATAGTTTTGATAATATTTACTTTTTCAACAATGCCGGTTGTATTGGAGATGTGTCTAAAACGCTTATTGATTATGATAGTTACACTCAAATTCAAAATTATTTTAACACTAACGTTGTTTCTGCTATTTTAATTATTTCAAAAGTTTGTAACTATTATTGTAACACAAAGAAGTATGTCATCCAGATATCTTCTTTAGCAGCTATTAAACCACTAAGATACTCTGGTTTATATTGCTGTGCGAAAGCATCTATGGATATGTTTATGAAAAGTTTAAACTATGATGTTCCATGCATTCGTACTTTGAATTATGCTCCTGGACCAATGGATACAGAGATGGCTAAAGAACTTTTACACAAGTCTGGAGATGATGAGACAAAGAAAATGTTTGAAAGTCTTTATAACGAAAATAAAATTATTGATCCTATTGATTCAGCTAAAAAGCTTATAAACCTATTAGAACAAGATATATTTTTATCTGGAAGTCACATTGATTTTTATGAAATTTAAAGTAATCCAAATATAATATTGCAATACCAAATACTTATACTATATTTACCTAATACTTATGCTGCATATCATCAAGCCATAAAAGTAATTTTTCCTAAAAATGTAATTTAAAAAAGATTTGTATCTTAAATATAATGCAATACTAAAGCTGTAATTGTAGTATATATTATTTTTACTACATGTATAATATATACTATAATTGCTACAAATATGGAAGATAAACTTTGCTTTTTTACACTTATGTAACAA
>evgR008899c0g1_i01_Hydra
ATGACTATTTTAGTAACACAAAAATTTACATGTATTCAGTTTTTACTGATTCAAGAAATACGTTGCTACCAAATGGATTATGCAACTTGTTCAATTTAACCTACATAGATTTTGATATCAATATATATCTAAGCAGTTTTTCATATAGTGAGGCTATGTATACTATACACAAAAATAATAACTGGGAATATATTCAATTACCAATATATGCAAATTTATATATACAAAAATGTAAAAAAGATTGCTTTGCCGGTTTTGAGCAAAATATAGTTTTCGATAAAACAAAGAGAGAGCTCACTTGCAAACCTTGTAGTGAAAACTATTTCAAAGAAGATTATGGATCAAAAGAATGTCAAACGTGCCCTTTAGATTATATTTCTAACAGCGACAAAACGGCCTGCAAAATGTTTGCA
>evgR008908c0g1_i01_Hydra
ATAAAGGATCATTTCTGCGTAGTTCATGGAACAGACTTGATGCCTTTGTTTTTGTTACTAGTCTGCTCTCCATCATTCTTTCGAAAATATTAGAAAAAAGCAACATTTATTTAAAGGTAGTGCGTATGTTCCGAGTATTACGACCAATCAAAGTTGTTCAATATGTCAAAAGGTTAAAAGTCATATTTCAATGTATGATATATTCATTTAAAAATGTTGGTTTCATTTTGATTATTGCTTTTTTGATGCTTTTTATTTTTGCATGTATTGGTGTCCAGCTCTTTCAGGGAAAATTTTATTATTGTACTGATGAAGGTGTGAAGACTGAAGAGGATTGCAAGGGATTTTACATTAAATATAATGAGATGTCTCTAATGCAAAAACCTAGAAAATGGGAACCAAGAGACTTACATTTTGACAATGTCTTGGATGCTCTAGTCACTCTTTTTATTTGTTCAACAGAGGATAATTGGTCAACAATTATGTATCAGGCTATGAATATGCCTAATGAAAATAATGGCCCGTTGTTTCCAAACTATTCTTATTCTGGAATTTATTTTATTGTTTTCATGGTGATATTTACATTCTTGATTATGAATGTGTTTCTTGCCATGAT
>evgR008911c0g1_i01_Hydra
TTAACCTATTTTAGTGCAAAAAGGCATAAAATAAATAATGCAAGTCGATGAGAAACTTATCGAGACTTCACTAGAAAGTCTTATTGGTAGGGTTGGCGATATTAAATTTTCATTACAAAACTTCTTATTTAAAATCGAACATGAGTACTTGACTTGGCCCCAAGTTTTAGACAATTTTGCTCTTTTGTCAGGACAAATAAATACATTAAATAAGCTTTTGAAGAATGACCACATGCCAATATTACGAAATCTTTGTGTTTTGCCAATAAACGTGCAGCCTGACTGTGACGAAGAATTGCAACGATTGACAGAAAATAGAATACATGTGTTTAGTCATGAAGTTGTTCCAAACGTATTGCGAACAAAATACGAGCCTGATGTAGAAAAAGAAGAGCAAATGCTTATTAACAAAGCTGTTTCGCTTCAACAGGAATATGCTGAAAAACAAGTACTTACATTAAATGAATTAATTTCTTCAATTTTAGAGCTGGTTCAATCTGCAAGAGATGATTGGGATGGTGAGTTTTCAACACAACAAACAAACTTAGCACCATCTTCAAATGATACAAATATATTAATTGCAGCTATGTCTACTGGCGCTGGTCTAAGAAAAAGAGGTGGAGAATCTTTTAGCACAAAACCATCAACTGCAAGAAGTGGACACAGAGCAACTCCTGTTAACTAATTTAAACAGAAACGAGACTAGTGTTTACAGTTACACTTATGTTGATGATATGATAATGGCTTTTTGTTGTGTTGTATAGTTCTATTTTTTATTATTTAAATTTATAAATATCTATTATTATTTATTTTGGTCAAATATATTTTATTATGAAAAAA
>evgR008935c0g1_i01_Hydra
AGGAAATGAAGATTTCGAAGATCAAGAATCCATTGAAAATGGTTCCACATTGATTGCTATTGTTGGTATTAAGGATCCATTGCGTGTTGAAGTTATTGATGCTGTTAGACAATGTAAGAAGTCTGGAATTTTCGTTAGAATGGTTACTGGTGACAATATTCTCACTGCTCAACATATTGCTCGTGAATGTGGTATTTTGTTTGGTGATGGTATGGCCATTGAAGGTCCAGAATTCAGAAAGATGAGTGATGATGAAATTGATGAAATCTTGCCAAGACTCCAAGTTATGGCTAGATCTTCTCCTTCCGATAAGTATAAGTTGGTCAAGAGACTCAGAGCTCTCGGTGAAGTTGTTGCTGTTACTGGTGATGGTACTAATGATGGTCCTGCTCTTAAGGAAGCAGATGTTGGTTTGAGTATGGGTATTGCTGGTACACAAATTGCCAAGGAAGCCTCTGATATTATTATTATGGACGATAACTTTAGTTCAATTATCAAATCTGTTTTGTGG
>evgR008950c0g1_i01_Hydra
CATTAGAAACCAAATTCAAGAATCAAGTTTCCATTGCTGTTGTTGACTTTGAAAAGGCTTCAAGAGATGAAATTAAACAAACCTTGTTGAAGGTTGTCGGATCAAACAAGGTCGACGTTTTGGTCAACAATGTTGGTGTTAACAATACTAACAATTGTCCATTCCCATTCACTGAACAACCAGAAGGTGATTTTGATAAGCTTATCAAGGTTAATATTCATTCAGTTTTGAGTGTTACCAGAGCTATCATGCCAATCATGACTGAAAGAGGTACTATTTTGAATCTTTCTTCATATACTGCTCAATATCCAACTCCATTGATGAGTGTTTATGCTGCCAC
>evgR008967c0g1_i01_Hydra
CAGATATTTTATTTTTGGCTCATATATTTTTTATAAACGATGAAGAAATTACCATCAATTTTTGTTGCTTTGATTCTTTACATTGCTATTATTGTGTGTATTGCATTTAAAATGTTTGGAGATGAATCCGAGTCTGCTCGTAAATTAAAAGAATTTCCTATAGAAAATTATAATTTTAATATTGAACATTTACAACACGCACAAAACTTTATAAATATTGGTTTATCAAAAGATGTTTATAAAGAAAAAATAAATGTTTTAATGCAGTTTGTAAATGCAACCAACAATATCAAGCTTCAAAGATACTTTTTTAGTTTAATTGAATCAATAATTTCTCACAGTGATACAGAATTTACTATTTATATAACTGGTGATGATAATGATTGGATTTTAGCAAATAGTATAATTTTTTCTGTTTATAAAAAACGAAAACAGCAATTTCATTTGCATCCTAAATATCTGGACATAAACATAATAACAAAGGAGCTTTCTAAAATAATTTTGAACATGCAGGATTTATTTAGTAGCCATGCTGGATCTTATTACAGCAAGTCTCTTTTCTTTATTCCATTAGCTCTTCATCGATTATATTCTCTCAAGAATGTCACTCATCTTCTGATGATTGATATTGACATTGAATTTCGAACAGATATTAGATATCTCTATAATTTATTTTTACAAATGGGTCCAGGACACTTGTATGGGCTAGCGCGAGAGCAGCAACCAACCTATAGGCACATGACTTCAGTTTTTAGGAGGAAACATAAAAAAACTGCTGTTGGTAATCCTCCACCATTTGGATTTACAGGATTTAATAGTGGAGTAAAATTGATAAACTTAACTTCTATACGGCGTAACAAATTATACAACAGCTATCTTGATGACCCAGTGAAATTAAAACATCTAGCAGATAAATTTTATTTTCGTGGTCACTTAGGCGATCAAGATTTTTTCACTCTACTTAGTTTTTTGCATAAAGATATGTTTTATGAATTACCTTGTCAATGGAATCGTCAACTTTGTGAATGGTGGAGAAATATATACCCTGTCATATTTGATCAGTACTTTAACTGTACTCCACCATATTTTGCCATCCATGGTAACTGCGATACTAAAATAGAAAATTAAATTAGAATATAATATGCTATCCAAAATCTTAAAAAAAAA
>evgR008974c0g1_i03_Hydra
TATTATTGGCAAATATTGCTTTCATACTATTATTGGTAAATACTGCTTCCATACTATCATTTGTAAATAGTGCTTCCATACTATCATTGGTAAACACTGCTTCCATACTATTATTGGTAAATAGTGCTTCCATACTATCATTGGTAAACACTGCTTCCATACTATTATTGGCAAATATTGCTTTCATACTATTATTGGTAAATACTGCTTCCATACTATCATTTGTAAATAGTGCTTCCATACTATCATTGGTAAATACTGCTTCCATACTGCATTGGCAAATATTGCTTCCATACTATCATTGGTAAATACTGCTTCCATACTATCATTGGTAAATAGTGCTTCCATACT
>evgR008979c0g1_i01_Hydra
TCCTCCTTTATTGAAAACCGGGAACATTTCAATAAAATTATAAGAAATATTAGTTTTTATAACTCTGAAAATAACTTGTTTCAGTTTGATGAAAATCAAGATCCTCTCACAGTTCAGTACAATTTGTATGTAACTACAAACAAAAGTTTTTTGTTAGCTGGAATATGGTCATCTATAAATGAGAAAAGATTAGAACTAAATGATAGCACAATTTTAGAATCTATATCTTCAAAATGCTCAGAACCTTGTCCTCCAGGTCAATTTGCTGTGGTTAATTTACAAGTAAATTGTTGTTGGATATGTGTTGTTTGTGATAAAAACTATGTAAAAAGCCATGTGGGTACTGATGCGTGCAAAAAATGTGAGTTAGAGGAGGATTATCACTCAAACGAAAACAGGACGATCTGCATCAAATTGAATCGTATTATTTGGAGTTTTAGAGAAAAAGGGCAAATTTTAAGATTAGTAATTTGTTTTTTATTTTCAGCACTTGGAGTGACAACATCAATTGTTTTTGTTGCTGTATTCATTCTTAAAAAAAATACACCCATTGTTCGTTCGTCAAATTATAAGTTTTCTTTGGGTCAGATGGGCG
>evgR009027c0g2_i01_Hydra
TAAAGAACTAACGAGTATACTTATTACTTGAGCTTTATAAATGTCTCATACTATGAACTTTTATCACTTTTTACAACTTATTCCAATTGTGATTCTCTCCCCAATCAAAAATTGTAAAAACTCGAACGTGGTGCTTCAACACGATACAAATACATCATATTTTATTGGAGGTTATTTTGCTAATCATCTAAAAAACGGCAGTTCTTATATGTTTAATGAAGCTTCTATTTTGTTGGTATCGTCAATGGTGCTTGCTGTTAATGATATAAATGAAAGCAACGACATACTTCCCGGAATTAAATTAGGATACGAGATTCGTGATACAAATAGCGATATTTTGTGCGCATGTCAGCATTCATTAGATTTTTTGTTGGATGAAAATTATAATTCTACACTTATAGGCGTTGTAGGCCCTTCGTCCAGTTCTTTAAGTGCTGCTGTAAGTACATTTTTACTACCTGAATTTGTT
>evgR009030c0g2_i01_Hydra
AGCGATTCTAGTCCGCTTTTCATATGTTCGGGTACATTCTCTTCATGTTTTTTAAACTTCTTTAGTCGCATGTGTAATCCAAAATGGCGTTCTAATCAGAGGAAAGTCATAATTAAATAAAATGAAACCATTACTTCTCCATGGTCACGAGCGCTCTATAACACAAATAAAATACAACCGTGAAGGGGATCTTTTATTTTCTTCAGCAAAAGATAAAAGACCGACAGTATGGTTTTCTATTAACGGTGAACGCCTTGGAACATATGATGGTCATGGTGGTGCTGTTTATTGTCTTGATCCAAGTTGGGATTCCAAGTATCTAATAACAGGTTCTGGTGATAACTCTGCTCGTATATGGGATATAGAAACTGGCAAAGAGCTTGTAACCTATGATACTGGATCAGCTGTTAGGGCATGTGGTTTTTCTTACAGTGGTAAAAAAGTATTTTATTCTACAGACAAAACTATGGGTCAAACATGCGAAATCAAGTTGTACGACGTAAGAGATTCTATTTCACAAGGTACACCAACACCATATCTGGTCATACCTGTGCCTGATTCAAAAGTTACCTCGGCTGTTTGGGGGCCTTTAGATGAATTCATTATAACTGGCCATGAAAATGGGACTGTTTCAAAATGGTGTGCTCAGACTGGCGATCTTTTAAAATCTATGCGTGCCCATACAGACAAAGTAAGTGATCTTCAAAAAAGTTCTAACGAAATGATGTATATAACTGCTTCAAAAGACAACACATCAAAGTTATTTGATACTGAAACAATGCAACTTTTAAAAACATACAAAACAGATAGACCTGTAAACTCTGCTGCTGTATCGCCCATTCGTGATCAGGTTGTGCTTGGTGGTGGTCAAGAAGCTATGGAGGTTACTACAACCTCTACAAGAATCGGAAAATTTGATGCGAGATTTTTTCATATGATATTTGAGGAAGAGATTGGACGAGTTAAAGGTCATTTTGGTCCCATCAACAGTTTAGCCTTTCATCCTAATGGGAAAAGTTATAGCAGTGGAGGAGAAGATGGTTATGTTCGAAGTCACGTATTCGATAAATCGTATTACGAATTTGAATTTGAGTACTAATATTGTTTACAAAAAATAAACAATGAATTTCAAAAAAAAAAAAA
>evgR009039c0g1_i01_Hydra
GTTTTAGCGATTTAAAATTGATGAAGTGAAGGCAAAATAAATTTTTTAAATTGGCATCATTCTTTATGGACTTACATGTCAAACACTTGACAATTACCCTGGGTTTCGAATAGCTTCATAATGTTGTCAAGCCAACAGCCGTTGGAAGCTTTAAACAAAGCCAAAAGTCATTGTTCTATATGTTGATTTTGGATTCAGTAAAAATAACTTCCACAATCAATAAGAAACTTAAAAACTACCATGAATTGCTAATCAAATCAAACCGGTTACATTTCAACAACTTCTCAAACAAAAAGAAGCATTTATATACTTAAGTGAACATATATTTCTGTATTCATTTCAAATATAAATGAAGGACCCAAGTTTAGAACTTGAACATGCTTTAACCCTTTATGAGCTTGATGAGAAGTTATTGCAATTAATACCAGGTTTAAAGAAAGCAAAAGAGAAGCCACCTGTACATAATCCATTTATTTCAGAACTTAATCTATCAACATTTGACAAAATAAAGATGGCCTTTTTAATAATCCTGGTGCCAATACGTCTTGTTTTGTTAGGTTTAACTCTTTTTGCTGCAGGAGTAATTGCTAAAATATCATTGATTGGATTTAAAGAATCAAGTCCACCTATGCCCATGCACAAATTCAGAAGATTTGGTCAATTTATTGTTTGGATTCTCGCTCGTATGGCATCTATGTGTTGTGGTTTTCATTATGTTCCTGTAAAGGGCAAACTTGCTCCTGCAAAAGATGCTCCGATAGTTGTTGTTGCGCCTCATACATCATTTGTAGATTCTTTGAGCTTTTTACCATTTGGATATTTGTCAGCAGTATCAGCAAGTGAAAATGTAAAGATTCCAATTATGGGAAACTACATTAAGCTGCTTCAACCTATTGTTGTGTCTCGAGCTGATAGAGATTCAAAGATATTTGTTGCCAATGAAATCAAAAGGCGTTCAGCTGCTGGTGTTTGGCCTCCAATAGTCATATTTCCTGAAGGTACTACAACCAATCATCGATGCTTAATAACATTTAAACCAGGTGCATTTTATCCTGGTTTACCTGTGCAACCTGTTTTGCTCCGCTATCCTGATAGAATGGATTATGCATCATGGACATGGATTGGTCCTGGAGCATTGTATCTGCTTGTTGTCATGATGTCTCGTTTGCATAATCGTCAGTCTGTAGAAGTTCTTCCAGTGTACACACCAAATGAAAAGGAAAAAAAAGATCCTATTTTGTATGCAAGAAATGTCAGAGAATACATGGCAAAGTTTGCAAATATTCCTGTTTCTGATCATTCTTTTGAAGATTGTCGATTAATGATGGAGGCTGAAAAACTTGGGATGCCAATGGAAGCTGGGATTGTTGAATATTATAAGCTTAGTAGTTTACTCAACATTAAATGTGAACATATGATTGGATATTTACGCAAATTTGCAACTATTGACAAGAATAAGGATGGTTACATAGACTTTACTGAGTTTTCAAATTACTTGCATTTACCACATAATGATGAAGCCAAAGCAATGTTCCATATTTTGTGCATTGATAAAAATCATGTGTTGAGTTTCCGTGAATATTTGATTGGCTTGTTCTTGATTCTAAAACCATTGAATACAGAAGAAAAAATAAAGCGTGCATTCTCAATGGTCGATGAAGATGATGATGGTTTTTTAACATATGCTAATTTATCTACTTTCTTCAGAAATACTATGCAACTAGGTGATTCATACATTGGAAAAATATTCAAAGATATGAATCCGTCGAATGCAGCCCATATTGATTTGAATAAATTTGCCGACTTTTGTTCTGAAAAACCTGAATACGGAATTCTTTTTATGCACTTTCAAAAGTTAAAAGAAGACTCGCTTTCCTCTCGCAATAAAGTTGAGGACTCGTTCTCCACTGGCAACAAAGTCGAAGATCGCAAATCAAAGTTAAAATCTGCTTATAATACAAGAAAATAATATATCATGACAAATATTTATTATCGCTAATTTTTTTTTTATTTAATATGTAATTTGAATCTTTTTGTTTACTATTTAAAGTAATTTATTAAATTTGTTTTTATCACTGATGACCAGATGAAACTGGTCTTAACAGTTTATTATAATTAAACATGGCAGTTTAACATAATTCAACATGTTAGTTTAACATAATTAAACACGGCAGTTTAACATAATTAAACATGGCAGTTTAACATAATTAAACATGACAGTTTAACACAATTATAACCAAATTTCATTTAAAA
>evgR009079c0g1_i01_Hydra
AGATGCTCTAATGGAGATATATTTGCCGTTAATGACGTTGATCCTAACCAAACAGTTGGTGAATTTAAAGAAGTGATTGCAACCCAATCGAATATTCCTGCCGCTGACCAACGTTTAATCTACAGAGGTAAGGTCTTAAAAGACCAAGCTTCATTGAAGAGTTATGGTGTCGAAGATGGTTTAACTGTTCATTTGGTGAGAGGAAAGCCACAAACTGCCACAACAACCACCTCCTCTACTACAACCCCAACAACCACTACACCAACTCCATCAACAACTACTCCAACAAGTACAACCACCACTTCAACACCATCAGTTAATACTTCCACAACAAATGACAATACAAACAATCAACAACAACCAAATCCATTCAATATGTTTGGAGGTGG
>evgR009085c0g1_i01_Hydra
AGAAAAATGCCTAGTAAAGATAATTTGGTTCAGTCAAATGTCATCCCTAATAAATATAAGGAAGATTCGATGCAATACCAAGTTGTCTATAAAGAGCCAAGATCATTACAGTACCTTACAAACTGTTCAGAAATGGCCACCGATCTTGGTTACGGAATTTGTTTCTTATTATTTGAATGCCAAAAAACATGATTGCAAAGACCTCATTGCACAGTATGTAAGTGTTCCTGGAAAAAATGTGTATCCACATAGTGAAGCTACAAAATATCCACATAAAAATAGGTGTACAAGCATTGTCCCATATGATCACTCACTAGTGAAATTAGTTCCCGATTCCACTAAATTTGAAAACACTTATATTAATGCAAATTACATACACT
>evgR009098c0g1_i01_Hydra
ATAAAAGATCTACAAATCCCTAGTAACAACAATTTAAATCCTTTAGCCACCGCCTTTAGACCAAATAATAGCCTAATGATCACAACGCAATTGCCAATCATACTCATTACAAACACATGTTCCATTAGCTCTAAAGTAGATCACCTTGTTCAAATATATGAAGACACAAACGCACATATTGGATTTCTTACTGAAACATGGCTTACCTCCACCACTCGCAACATTGTAAATTGTCATATCCAAGAAGATACCAAGTACCTAATAACGTCGTGTGAACGCAATTGCTTATCTGAAAAAGGAATACCCAAAAGAGCAGGTGGAGTTATGATAATAGTCAATCCCAGCTACACAAATACCATCCAGGAGCTATCCCCCATGCCTG
>evgR009100c0g1_i01_Hydra
GTTGCTGTTAAGGATCATGTACATCAAGCTACTCTTGAAAGTTTTGTTGCTCAAAAATTGAAGAGTGGTATTGCTTCATTCCCAACCAGTGAAGTTAAACTCACTTCTTTCAGATTTCCAAAATTCGAAATTGAACACGGAATTGAATTGAGTGAGATTTTGAAATCTGCTCCATTCAATATTCAAACTCCATTCACTAATAATGCTGATTTCACAAATCTCATTAAGGAAGCTGGAATCTTCCTCGATAAGGTAATCCACAGAGCTGTCATTAAGGTTGATGAAGAAGGTGCTGAAGCAAGTGCTGCCACTGCTATGGTATTCAGAACTTTGAGTCGTGATCCAAATAGCATTTCTTTTATTGCTGACTCACCATTCACCTTTATTTTGAGTGATGAATCAACTCAATCCATTCTCTTTGTAGGAAAG
>evgR009117c0g1_i01_Hydra
TAAAAATGAAGTTAATAAATAAGATTATATGTTTGGTTTGCACCATTGCCTTATCTAACTTGCTTACATGGCTTTTTTTCTATGAAAAATTGAAAACTCAAGAAACCTTAAATTCTCTTGTTTCTATAAATCTTAAAAGTTCTGGTATACCTAAGCAAAAGCAGGTATTTGTTTGGGGAAAGGCTGCTATTTCAGAGTATTTTTGGATGCATATTATTGAAGGTACAGTGCAACCTAGTAACAAATATATGTGGAGTGAAGGTGTTATTCAAGTAAATAGTGTCAAGTTAATTTATTTTACTGGACCTTTGTTACTTCCAGAGACAGCCCCTAAAGATATAACAGATTTGGTTATAATACTCAATGGAAGAGAAGAAAAAAAAATCAATGCAAGTTTGCATTGGTTAAATTCCCTACTTAATAAAAAGTTGTTTCCATTACTTCAAAATTTAGGAATTATTGTACTCGGCAATGAACAATGTAACAATGGTTGGTTAGTTAATTTGAGTAAGCGAGCTCTTACAATAAAATTTGTATTTATTGTTTATGACATTAACTTGTCTCATCCTCCTCATTTTTATTATCAATGGCCATTGGGTGTTGCAACTTATCGAAAATTTCCTCTTATAAAAAGAAATCAAGTGGACACACTCTCTCAAAGAATGTATTTATGTAATTTTTTAGGTACAATTTATAGTAATCATAGCAGATCTTCTCGTTCAAAACTTTTTAATATATTAGATAATGGTCGTTTAAAAACTCTATGTTATTCAAAAGTTCGTTACTCATGGTTACCAGCAGAAACTAAGGAAACATCAGAGCTATACCACAAAGTATTGAAGCAGAGTGATTTAACATTGTGTCCGATTGGCTTCAACTCAGAATCATATCGTATATATGAAGCTATGTCTTATGGAAGTGTTCCTGTCATTGAGGATGTAATGACTAATGGTCAGTGTTCCAGAGATTTTAATAGTGGCATACAAGGGACACCATTTCGTTTGCTTAAGAAGTATAATGCCCCTGTAATATTTGTGCATGACTGGTCTGAATTATTTTCAATCATTGAAATAGAAGGTAAAATGTCACTTAATGAAAAAGTTAAAAGGCGTGAAGCAGTTTTAAGATGGTATGAAAGATTTAGATCGAAAATGAGAGATGAGTTTTTAGATGTACTTAAC
>evgR009120c0g2_i01_Hydra
AAAAAAAAAAAAAAAAAAAAAAAAAAAAAAAAAAAAAAAAAAAAATGGACAACGAAACTAGTGAAAATAAGTTTGCTGATAATTTTAGCAAGAAAGAAATAGCACATCAATGGCAGACCTTAAAAGCAAATCGCCTTGAACGTAACGAGTTTATGTATTGCAATGAACTTATTAGCGATATCAAGTTTTATATTGGAAAAACTAAGCAACTCTTAATTCCAGCTCATAAATATGTTTTAGGAACTTCAAGCCCTGTTTTTTTTGCTATGTGTTGTAGTGAATTTAGAAAAGATGATATAATAAATATCAGTGATTGTGAACCCGAACCTTTTTTAGAACTTTTGAGGTTTATTTACTATGACCAAGTTCATTTAACGAAGACAAATGCACTGGACGTTCTATATTTAGCAGACAAATATATAATTCCATCTCTTTCTAAAGAATGTGTCAATTTTCTTTTAGATAATGTTTCTACAGAGAATGTCTTAGAAGTACTCAGTGCTTCAATATGCTTAAATGAAAAACGTTTAGAAAAACACTGTTGGTCAATTCTTTCCAGAAAAACAAAAGAAATTTTTCAATCTGATTCATTTTTGGAAATCGACCCATCGTTTTTAATAAAAATTATAAAAAAAGATTGTCTAGATGTATTAGAAATTGATGTTTTCGATGCTGTAAAAAGATGGGTGGAAAGAGAGTGTTTTCGTAATTCTATAGAGCCAACTTCTAATAACAAAAGAGTTTTTCTAACTGAGATTCTTCCATTAATACGTTTTCCAGTAATGACTGCAAAAGAGTTTGCATTGAGCCCTGCTCAATCAGATCTACTGTCTCTTGAAGACATTAAGAATATTTTTATTTACTTGACATCAGGTGTTGGTGGTTCAAATCTTAATTACCCTTTAAATCCACGACATTTTAAAGCTCATGTTTGTAATCGTTATGTAAAGCCAATGAAAAACTATTTATGGAGATATGATGAAAAAGACATTGACGCTATTCGTTTTAAAGTTGATCAAGAAATTTTACTTGCTGGTATTGGTTTATTTGGGTCTCCTTGTGAAGGTGAATATTCTACAGAAATTTCTATCTCAATTAATGACGAAGAAATTTTTAAAAGCAAGGGTTCTTTTGTTTCACCTCCAGATCCTAAATTTAACAACATTGATAACCCTAAAATTCATGAAGTTATGTTTGAAGAACCAATTCATCTGAAAGAAGGTATTTTTTATGATATAGCTGTTTTATTGGATGGTCCACCCTCATTTGCCGGAGATGACGGGAAACAGGAGGTCATCGAAGAAGGTGTGAACTTTGTTTTTGAAAATAGTAGTGGATCTACAAATGGAACAAGTTTTGATGAAGGACAAATACCAAGCCTACTTTTTTATTTTTAAGTGTTGTCTTGTCCATATTTAAATTAAAATTTTGTAATAAATAAATAAAATACTTATTTAATTATATAAATAGTAATGTAATTTTTTAGTTGTAGTTTATAAATAGAGAAATACATATTTATAATTTTAAAAAAAAAAAAAA
>evgR009163c0g1_i01_Hydra
AATTCTAAAAATACGAATTGAATCTATTTAAAATGTTAAAACTGTTTCTTGTCTTCTTATTTTGTTATCATCTAGCACAAACCGAAGAGCCTGGCATACCTCTTGACGTGGTAGCGTTTTGGCCAGTTAATCCTTTTATTTTTTATAATGAAAACAAAAAAATTGAGGGTATATTACCATTTATCTACGATAGACTAAGTGTTTTGTGCAACGAAAACAAGACTTTGGTAGAATATATTTTGATGAACCAAAGCTTTAATCAAAATAGCGTTTATGAAACATTAAGTGACACAGAATTTGGCACTAAGGAGCTTCAGAATGTTAGTGGCAAAAACACAATATTTTTACCTGTATTTATTAAACCAAATCCTAGTTTGTTAAAAGATAAATCATTACAAGCTGATGTTCTAATGAATTCAAAGTTTGTCGTTGTTGTTCATAAAAGCAAATTAACTATTGCAAACAAAGTATTTGAAGCTATTAAAAATACATACCCTATTATACTTTATGCCTTTGCATTTGTCTTGATCTTCTCAAATGTTGTGTGGCTAGCTGAATGCA
>evgR009175c0g1_i01_Hydra
AGCAATAAATGATTATCTATGAGTGTAACTCAACTAAATAATCAAGTAAGTTGTACTAAAGAAGATAAGAATTTATCTGATAATGAGATAAAAACTCAACATGAAACAAAAAGAAGAATGCCTTCCGATCTTCAAATTAAATTAGAAATGCAATTACAGAAATCAAGACAGACAAGAGCAAACAGATCATACTATATAAATAACTTTTCAGAAAAAGATCAACAAACAGAGAAAGTGCCTCTTATATTAATGGATCGAAGTAGCACAGAAATGGAGTATAAAGATGAAACGTTTCAAAATTTACCAAGTTATCAAAGTTATTCATCTGACTGTTCTGATCAACTTGTAAAAGATGGTTTTAGACTTGATGAGCTCACTGATGATGAGGATTTGGACTTGCTTCCACATCCTACAGTTCGTTCAACTCCATGGTGTACTTGCGAAAGTATTTCATGCAGTGTTCAATAGTTTTTGGTTACTCAAGACAAATATCGAACTATTCTAACTCTTGTATTTAGTTTTTTTATTTTGCAATAAGTAGTAAATAATTTTTTATCTACCATTTTTTTACATTTTGTTTTTGTGTATAAAAAATATTTTTATTGTAGAAATCAATAATATGTTGTTAAAAAAAAA
>evgR009176c0g1_i01_Hydra
TGAGTACCCAAAAAACTTCACCAACTGGTGCTGACTTGTACGCCCGTTTCGCTTTGTCTGGAGCTTTGTGCTGTTCCATCACTCACGGTGCTCTTACACCTGTCGACGTTGTCAAGACTCTCAAGCAATTGGATCCTGTTAAGTACAATAGAGGTGTTGTTGCTACCTTCCGTCAAGTTTCTGCTGAACAAGGTCCAGCTGCTTTGTTGACTGGTGTTGGTCCAACCTTTGCTGGTTATTTCTTACAAGGTGCCTTCAAGTTCGGTGGTTATGAATTCTTCAAGAAGCAATTCGTTAACGCTTTGGGTATGGAAAAGGCTACTGAAAATCGTACTGCCATCTATTTAACTTCATCTGCTTTGGCTGAATTCTTTGCTGATATTGCCTTGTGCCCATTGGAAGCTACTCGTATTCGTTTGGTTTCTCAACCAACCTTTGCCAATGGTTTGTTGGGTGGTTTTGCTAGATTGTTGAAGGAAGAAGGTGTCCTCCGTGGTTTCTACTCTGGTTTCTTCCCAATTTTGTTGAAGCAAATCCCATACACTATGGCTAAATTCGTCGTTTACGAATTGGCTGTCGAACAAATCACTAACATGATCGGAAAGCCAAAGAGTGAATTGACTGCTGGTGGACAAACTGCTGTCGATTTGAGTGCTGGTTTGGTTGCTGGTGCTTGTGCTGCCATCATTTCTCAACCTGCTGATACTTTGTTGAGTAAGATCAACAAGCAAAAGGGTGCTGAAGGTTCAATCAGTTCTCGTTTGGTTGGTTTGGCTAAGGAATTAGGTCCACGTGGTCTCTTCTTGGGTCTCGGTCCAAGAATTATCATGGTTGGTGCTTTGACTGCTGGTCAATTCGCTATCTATGTCCGTATTAAGTACATGCTCGGTGCTACTGGTGGT
>evgR009186c0g1_i01_Hydra
GTGGGGAGGGGTGGGAGGAAAGCCTTAAATTTATGTATGTATTAATACAAGCCAAATTGACTACTCTCAACTATGTCTGAGCTCTTGCTTGAACTTAAACTCCCACAAGAATCTCTCCTAATTGGCTGCATAAATCCCCCACCTACCAACAATTCTCTGGCATTTGAGGATATAGTAAAAGCTATCTTCTGTGCAAAGATTAACTGTGTAGCAACTGTAGAAAACAAAAATTTCAATGATTTAATAATCACAGGTGCCTTGAATTTTCCAGACATCAAATGGCATCATGACTGGACCAATACCATCGGCGCTGATACTAGTTTCAGCCCCATAATTTTTGAAGCAAACCAAAGCTTATTGAAAAACCCACATTGTTACTGGTAATACCTCCTCTTGGTAGCACCTAACAAGGCCACTGCACTCTGACATAGCAACACAAACTAGCCAAAAACATCTACAATTCTTTCTATTTTAATGGCTTAAAACTCAAATATAGGAAAGGTAATTAGGCTAAAC
>evgR009201c0g1_i01_Hydra
CAAAACAGTTTATGAAGCTTATGACTTTCTCTATGTCAGAAAGAATAGAAAGACAACTGGAACTTGCCAACAATTAGAAGGAGAACAAAAATTCACAGAAAGAAAACCAGACTCTCCACTCCTCTATGCAGTTTGGGTTGATGATGCTCTCTCAACAGGTTCTTCTCTCTTGGATGGAATCAAAATGTTGAAGGAAGAGTATAACATTGAAGTTAAGGCAGCTCTCTACTTGGTTGATAGAAGTAAGGACAGATTATCAATTGCCGATGATAAGCAAAAACTTGCAGATCCCGTTTTTGATACTGTTAAATTGTATTCAGTATTTGAT
>evgR009208c0g1_i01_Hydra
AATTAGAAAAGGAAGAAGCCCTCTTGAAGGAAAATCCAAACCGCTTTGTTTTATTCCCTCTTGAATATCCGAAGATTTGGGAAATGTATAAGAAGGCAATGGCATCTTTCTGGACTGCCGAAGAGGTTGATCTTCACACCGACATTACACAATGGACCACTTTGAATTCCAACGAACAACACTTTATCAAGAGTGTCTTGTCATTCTTTGCTGCCTCCGATGGTATTGTTAATGAAAATTTGTGTCAGCAGTTTATGGCTGAAGTGCAAATTCCTGAAGCCAGATGTTTCTACGGTTTTCAAATAGCTATTGAGAATATTCATTCTGAAATGTATAGTACATTAATCGAG
>evgR009216c0g2_i01_Hydra
TTATGTTTTAAATTTAGTGATTTTTAAGAAGATCAAGTTGTTGATTTTTATATTATTATAAAACGATGTAATGATGTAATAAAATGCATCTACTGTTAACACCTGATTTACAAGTGCTGATTTAGGCATCCTAGTGCTGATTAGGCATCTAAATATTTAGTGGTGTACATCTTGAAATTGTTGCGCAGATCGAAAACCTTCTAGTTTTAAAGAATTATTGATTAATAGCCTTCAACTAATCATGCAAAGATTTAGGAGGATTACTTTGTTAAAATATTTTTTGGGGGGCTTGGTTGCAATCATAACTTTAATTGCCTGCATTGAATTATTGTATTTGCAAACAAATGTTAGTTTAAACCATTTAAAGTTCATTCAAAATTCAGATGACATTAATAATCAGTTAATAACCCACACAAATAAAATTACCTATATAAATAATGAAAAAGATACGCTTTCTTATATCATTTTTGACAGTGTACAACGAGACTGCAACATAGTCTATTCAACTCTGATTATTATCACTTCACACGTGAAACACGTAAACAGAAGAAATAATATACGAGAGACATGGGGAAAGTCGTCACATTGGAGTTCTGCGGGAAAATATATTATAGCTTTTGTTGTCGGAAGATCATCAGACTCGGATATAATGATGAACATAGCGGAAGAAGCTAAATTATGGAAGGATATTATTCTAGTAAACATATTAGAAGATTTTTATACGTTAGCTAAAAAAGTTATTATTGCTCTAATATGGGCCAATCGCAACATTAAATATAAATTAATTTTCAAAGGCGACGACGATATATATTTAAACATTAATAACGTGTTAATATTTGTTAAACAAAACGATATAGAAGTTGCTTACATTGGAAATAAATTAGAAAATGCAATTGTTCAGAGATCTGGTCGATATAAAGTAACCATAGAGGAACGCAAAAATGATATATATAACCCATATTGCTCGGGTGGAGGATATTTTCTATCCACGTCTAGTGTGGAAAAAATGATACCGCTATTTCATTTGAATCGTGTACTTCGAATTGACGACGCATATATCGGAGAAATCGCCTATGAAGCAGGTATATTTGCTAGCCATGCTGATGGTTTTGAAATGGATAATGAGTCTTGCAGTTATTCAAAAAATACTATCGTTTCTCATCCTGCAAACGATATTGGTTGTATAAATTTTCTTCTTCGAAGATCTTTGATTGACAGCGGTAAATTGCCTAGAGGTCGGTCTGAACGCCAGTTTGAAAAAAAAAGTTGTTATGAAAAGAATCCTCTATGTTTATTACAAAAACAAAAACATGCTAATAGAACTATATAGGGAAACAAACATCGTTAAACGTTAAAACCCGATCATTAAATGTTAATAAGCATACTAGTTCAATAAATATATGAAGCCGTTAATATGACGCAAATAATATATGACGCCTTTTAGCATTTTTTTTGCAGTCAATGCCATTCCCATTCACTTCATTAGACTCAAGATCAGACACAAAAACCAATCACTGACACATTCCGACACATTGCTGCGTTTTGGACAGAAATAAATAAATTATTTTTGTTTTTTTTTATTATAGCTAAATTATCAATCAATATTTTTTGAAAAAAACTTCTTACAAAATGTAAACATTATTTTACTAAATAAATTATTCAGCAATTTATATTAGGAAATATGAAAAAAAAAATCCATTTATACTACTCATACCAATATCTGAC
>evgR009219c0g1_i01_Hydra
TTTTTTTTTTTAAGTTGTTTTGAATTTAAACGTTTGTAATGTTCTTGCAAAAAATTAGACTGAAAGTGTATAATTTTATGGAAAGGCCTTCCTCGAGGTACGCAGTTTTGTATCACACGCTTGTGTTGTTGTCAATTATGTCCAACACTGTGATTGGAATTTTATCAACAATTGATATATTTGAAAAAAATAACAACGTGCAAATACTTTTTTGGAAATATGAAATATTTATGTTGGCTATTTTTGTGGCGGAGTTCGCTATTCGCCTTGTTGTTTGCGGTGTTATTTCAAACTACAGTGGCTTTTTTGGTAGAATTGTTTTCATGAAAAATATCTATATGCTGGCGGACATATTTATAATTTTATCAACAATTGCTTCAATAGTAATAAAATCTAACGATGCCTCGTTAGATTTTTTAAGGTTTAGCAAATTTTTCCAGATGTTTAGAATCCTTAGGTTAGATCGCCGAAGAGGAGATATTTTAACAATGTTAAACATACTGCGGGAGCATAAGAAAGAGCTTATATCAAGTTACTTTGTGTGCCTGAATATACTATTGTTTGGATCTTACATTATTTACGCTGTAGAAAAAACACAAAATCGAGGATCGGAAAACCAAATTGACAACATGGCCAACGGATTGTACTGGGGAGTAATAACATTTACTTCAGTTGGCTATGGCGATTTCTTACCCAAAACCTGGGCGGGAAAAATTCTGACAGGATTGTTTTCAGCTTTTGGATGTGCATTTTTTGCTTTACCGGCAGGAATAATTGGCTCTGGACTAGCAATACACGTGTCAAGACAAAAAAGAGAGAAGCGAAGTATGGAAATAAAAAATCCCGCTGCAGCTGTGATCCAAACTGCGTGGCGAATTTACGCGATTAGCAACAATTTGAAAAGCTCAAAAATTCAGATTTTTTCGCATATTCTGTACGAAAACAAGATGGTAGAAGACGTGTTTAAACAACAAATGTCTGGTTTTAAAATAGACAGAAAGCCACGAAGCTCTACTGAACCCATAAAACAAGAAAAAGGTTTACATGAAAGAAATTTGTTTAAAAAACAAAAAAAAACGAACAATACTTTGATTTTGTTGGAGGAGTTTTTTGAAAACAGTTGCTCTTCCAATGTCACAAATAAATTTGTCAAACAAAATGATATAACCAGGCAATACAAAATTGCAATAAAGTTTATAACCTTAACAAAGTTTTTTGTAGCCGTTAGGAACTTTAAAATAAGCAATTCTCCTTACATAAACGTTGAAGATTTCCTTGAAAAAAATAACCGTCAATTGACAGATTCAACACGCCTATTAAAAGAATTAAATTGCAAATGTGATGAAATCAACAACTTACAAACTAAATTTGACTTGTTGATCGAAGAAGTGAGAAATTTAAACAAATATATAAAAGAAATTAAAAAAG
>evgR009221c0g1_i01_Hydra
ACAACAGCAGGTACCATCATCAACCTCTCACAATGTCTGCCGTAGAACAACAACAATCAACCCAAGCCAAGAAGACTGGTAAGGACAAGTACAAGTTCGAAGTTAACCAAGCTAAGCTTGATCAACTCAAGAAGATGTCCAAGGATGTCGTCATTGCCGGAGGTATGCGTAGAAAGCATAAGGTCGTCAAGAAGGGTTCCCAAAATGAAAGCAAAATTAGAAACATTGTCAATAAGTGGAGAATGACCACCATTCCAGAAGTTATGGAAGTTTCCATGATTATGGAAGATAACACCATCACCACATTGTCTCAACCAAGAGTTGAAGCTGCTGTTCACTCCAACTCATTCGTCATTTCTGGTAAATTCCAAAGAATGACTTATGAAGAATACTTCCCAACTATGCTCAAGCAATTGTCCAACAATTTGGATCCTAACCAATTACAACAACTTTTGGCCGGATTGAGTCAACAAAAGGAAGACAAGCCTGTTGCTGCTCAAGAAGAATTGCCAGTCGTTGAATCTTTCGAAACTGTTCAATAAATTATTTAAAAAAATTTTAAATTTCA
>evgR009224c0g1_i01_Hydra
TTACGTGGCTGCTTACCTCTTGGCTGCCCTTAGTGGCAATGCTGATCCATCAACTGCAGATATTGAGAAAATTCTAAGCTCAGTTGGGATAGAATGCAACCCTAGTCAACTGCAGAAGGTCATGAATGAACTCAAGGGAAAAGATCTTGAAGCTCTAATTGCAGAAGGCCAGACTAAACTTGCTTCCATGCCTACTGGTGGAGCTCCTGCTGCTGCAGCCGGTGGTGCTGCTGCTGCACCTGCTGCAGAAGCTAAAGAAGCAAAAAAAGAAGAAAAGAAGGAAGAATCAGAAGAAGAGGATGAAGATATGGGCTTTGGCCTCTTTGATTAAAGACATTCTTATCGAGAACTTTGGAACTAGTTGGATTTACTTTTTGTGTTCTAATTAGTTCTTAAAAGTTTTAATTCAAAAAAAAA
>evgR009234c0g1_i01_Hydra
CTCGTATTTATGATCAGGCTGTTACTCTCGTGTTTAGAAAACGGTAACGATGAACTGATTGCTCCGTGCCGTTGCAATGGAACGACAAAATATGTACACGAGCGATGCTGGGTTCAATGGTTTTTAAAATCACAAAATCAGACATGTGAAATTTGCCAAGGGGAGGTAAACATCAAAAATGCTGGTTACAAACCATTATTAAAGTGGAAATTGCCGTTAAATTTAAAGCTGAAAATTGTTCATTTACTGATTGCTTTATATATGACTATAATGGCGTGTTTTACAAGTCTCGTAATTTGGATTGGTATCGGGAAATGTTTATCAATCGTTTGCATATTCGTATATGCTGCTTTTTGTGTGGGTGCAATCTGTTTGGTATATCTGTGTAAACACGTGTTCAACTCTTCAAACTATATTAAGTCCGTTTTATATGTAAACAGTGAGTGGAAAGTCTACGGACGAAAAG
>evgR009238c0g2_i01_Hydra
CTGATAACGCTTTTCATAACGTCTTAAAATTAAAATTAAATACAGCAAACAAGTGTTGTCTATTCGTTTACCTTACAGTCAAGGAGGAAGCTTTCAGTCTTACAGTCAGCTTACAGTCAAGGAGTCAGCTTACAGTCAAAGAGTCAGCTCACAGTCAAGGAGTCAGCTTACAGTCAAGGGGTCAGCTTACAGTCAAGGAGGCAGCTTACAGTCAAGGAGGCAGCTTACAGTCAAGGAGGCAGCTTACAGTCAAGGAGGCAGCTTACAGTCTTACAGTTACAGTCTTTTTACAATTATCTCTTAATCTTTTATTTTCTG
>evgR009248c0g1_i01_Hydra
TGGTGACTACGTTTTATCTTCTGGAGATATAGCCTCCTTTGCTTTTATTGATTGTTGCTTAAGATACGTACCAGGATATGTTAGTAATGAGGATTCTCTGCAAGATGAAAGCTTCGGAGAAGGTCGTTATGAATACCTTTTAGAATATCCGCATTACACAAAACCTTTAGTGTGGAAGTCTAATAAAGTGCCTGAAATTTTGGTTTCGGGAAATCATAAAAAAATTTTTGAATGGAGACTGAATCAGGCCAAAGAAATTACAAAAAATAGGAGACCTGATTTGTGGACAAAATATTTAAATGGAGTTGAAAAATGAACTTAGCTGAAAAGTACATAGAAAAGCAAATATCTTCGCTTATGAAAATTAAGGATATACCAGATTTTAGACCTGGAGATGTAGTAAAAGTTCATAACAAAATCGTTGAAAAAAATAGGAAAAACATATACAAAATAATCG
>evgR009258c0g1_i01_Hydra
CAATTTTACAATTAATCAATAAAATAGCTTCAACTTCTGTAGTGGAACCTTCTGTGGTAGAACCTTCTGTGAAGGCAACTTCTGTGGTGGAACTATTTTCAGGGACTTCTATAACTTCAAGCGCAACTGAACAGTTTGTTACTGAATCACCGCAAACAACCGCTGATAAATTAGTTTCACCTGAATCACTTACTAATGCAATAAGTGAAACTGATCCAGCACTTTCAACTGAACCTCCGAAAGAAACTGTTAAACTAATTGATGAATTAACTTCAACAGAATCAGTCTTAATTGCAGCAATTGAAACAGAAGTGAATATACCTACAACTGAATCTCAGCAAGAGAATTTAAACTCTGTAACACAGTCGGAATCAATAACTAATGCTGTTGTAAGCCAAAGTGAGCTACTTCCAGTTCAAACGGAATCTCCGCTAACAGCCGATGAACTATTTGACGAGCTTGACTCGGAATTTGCATAAATTTGCATAAATCTCTTGGTGAACGTTTGAATATACTTTATGTGTTCCAAAACATTTTTTCAAATTAATCAACTTTTTTTTGTACGATGC
>evgR009289c0g1_i03_Hydra
CGTTTTTTGTATTGTAGTTTCGGTATGTAGTGTTGTAGTTTCGGTTTGTGGTATTATAGTTTTAGTATGTGGTGTTTTGGTTTGCGGTGTTGTAATTTCGGTATGTGGTATTGTAGTATTGGTATGTGATATTGTATTTTCGGTATGTGGTGTTGTAGTTTTGGTATGTGGTATTGTAGTATTTTATTTCGGTATGTGGTGTTGGAGTTTTGGTATGTGGTGTTATTGTTTTGGTTTGTGGTGTTGTAGTTTCTGTGTGCGGTATTGTAGTTTCTGTGTGTGGTGTTGTAGTTTCGTTTTTTGTATTGTAGTTTCGGTATGTAGTGTTGTAGTTTCGGTTTGTGGTATTATAGTTTTAGTATGTGGTGTTTTGGTTTGCGGTGTTGTAATTTCGGTATGTGGTATTGTAGTATTGGTATGTGATATTGTATTTTCGGTATGTGGTGTTGTAGTTTTGGTATGTGG
>evgR009303c0g1_i01_Hydra
GTCATTAAAATTATTTATAACAAACATCAATTTTTGTTCATCTGTAAATATAAAAATAATTAAAAATGTCATTTTTAATTGAAAATATTTTAAAAAAGTCTACCACAAAAGATGATAAAACAACGGGTATAAAAGAACAACCCCATAAAAAGTCATTAGCAGGAAAATGTAATTGGTCAGAAGAAAGCAGGAGATTCAGAGTAAAGAAAAGAAGAAACCTTTTCTCCAAACACCAAACATGGCAGTTAGAAAAAAAATTCTTAATTCGTCCGTATATGTCTGCTGAAGAAAGAGAAATTCTTGCCGATAAAATACATTTAACCCCTACGCAAATTAAAATTTGGTTCCAAAATCACAGATACAAGATGAAAAAAGACTTGAAAGATATCTCAAAGTCTCAACATAAAAAAAGTGATCAAACTAATCAATCTTGCTACTTTTCTAAGTATAGTCACATCTCTCATCCTCAATACACCTATAGTCACATCTCTCATCCTCAATACACAAAATATATACTTGAAGGAAAATTCCTCTATGAACAATGCAAAATCAATAGCGAAGTAGATAAAAGTGTACGGGCACATGTTTCCCAATGTCAAATAAAAAACAATATAAAAGAAAGATACGGTCCAAATAATCAAATTATGCACAATTATGGTAGGCAATATCCATATATGAAAGATGAATGTTTCCACGTGATGAAATGCCAAGAAAAAATCCACACATTTAGAGATTATTAAACTGGCGCTCCGTTTGAGTTATAAAGATTAGTGTTGTTTTTAGGACGTCATTTGTATATTGTCGTAATATTTAAT
>evgR009337c0g1_i01_Hydra
ATAATGAAGAAAATTTATGGAAATAGACGATCAGAAGAATCTGCTGAAGTCAAGCTGACTAATGCTTGCAAGCAAGATGAATCTATTTATTTTAAAAAACAGCTTATTAATAATGAAATTGGTTACGGAATATTTACTTTAAGAGATTTTGAAGTGAATGAAGTTCTCCTAGAATATAAAGGAGAAGTGATATCTCGTTTGGATGCAAAAAAGCGTCATGAAGAATACTATCTTTCAAAAAAAGGCTGCTTTATTTATGACGTTAATTTTGGCTCTGATAAAATAAGTATCGATGCAACATCTACATCTTCGTTTGGGCGCTATATAAATGACTCGCCGGAAAAATATGCAAATTGCAGGCCTAAGTGCTGCAAGATTGGTGAAAGCTATC
>evgR009340c0g1_i01_Hydra
CGATATTAATGTTATAACTAATGAAAAAAGAGATCTACACGAGATGATTGAATTATTAAGAACCGATATTAATGTTATAACTAATGAAAAAAGAGATCTACACGAGATGATTGAATTATTAAGAAAAGAAGAGGAGCGACTGAAAGCGGAAAATTTAGAATTACAACGTTTAAACCATCTTTATAAATTTTATTTACAGTACGATATGGAAGTCGAGGGATGCTGCGAGTGCGCAGAACTACTTCTTTATCCAGAGTTACAGCCATGTAAATCTTGTCGAATATTGCTTTGTAATATATGTTTAAAAAAACATTTAAATTGTAATATTGTTATACTTTAGCAGTTATGCTTTTTTTATTAATTGTTTTTTTTAAGATATTTGCTTTTTTTTTTTAGAATGTCAATTATTATTCCTGAAAAAGAATATTTAGATTTATTAGCTCATATATACAAGAGTGACAAAATATTTTGATGCGAGAGCTTAAACTTTTAAATGAGAACAATAAATACCGTAAATTCATTATAAAAGGTGATCTTATCGCATATAAATATGTATTATTTCATTTAGATAAAAGCAATGAATCTTTTAACTTTAAAATTACAACGTACAATGTATCAAAGAGAATATTTGAAAGTACGGAGAGATTATATTATTCGTAAAATCGACATCGTAGAGAAAGAAATAGAGCAGTTAAAAGAATCAGAAAACGTCGATGGTTGGTTAAAAAAACGATTTAAAAAATTGAAATATGAAAAACAATTGAAAAAAATAGAAACAAAAATACATATAAAAAAACTAAAATTATATATCTGTTATAAAAAAATATTTCAATAATAAAATACTAACTTATAAAAAAAA
>evgR009352c0g1_i01_Hydra
AGAAATGTTCAGCTTGTTCATTTGATATTCATCAAATGGCGTTCCAAGCTCTTCCAAGCAGAAACTAACGGTAAATAATTTAATTAAACATCCGCACTATCCAAATTTTACTTATTGGCTGTGAAATATCTTTGGCAAGATTTTATGCAAAATGAAGATCAACTTACTTGTTGGTTTGTTATGCCTAAGCATGGTTTTCACTAAACCATTAGATGATGACAACCAAGCCCAAGAAGGTGATGACAAACGTGCAGATAAAAGGGATAGCATAGAACAACAATCAAACAATAACAAGGCAAACAATACAGCAAACACCGAAGAGACAGAAGGATCTAACAACGTAGATTTAGTTTCAGATTCCGTTGAATCATCTCCAGATCAAAAATCAATTATTCTTCCATCTGCTACAGTTACAACACTTTTAGCTTCTCCTACATCTCCCGCAACAACAACAAAAAAAGCAACGACAGTACCGTCATTAACTACAGTTTTACCTACAACTGTTAATAAAGGGGCGACTACAAAAGCTCCGGTCGTAACTACAACACAATCTTTTACAAAAGCAGCTACAACAAAAGCAGCCACAACCAAAGCAGCCACAACCAAAGCAGCC
>evgR009355c0g1_i01_Hydra
CAATCATCTGGAAGTTTCATTGCAAGGTGCACCTCCACAAAATATTGTTAACTACGATGAGACCAATCTGTCCGATGATCCCGGACAGAAAAAGGTCATAACAAAACATGGGAATAAATATCCTGAACGCGTACTAAACCACAGCAAGTCTGCCACATCAATAATGTATGTGGCAGCTGCTGGCAAAGTCCTGCCCACATATGTTGTGTATAAGGCAATACATATGTATAGCAATTGGACAGTTGGAGGCCTAAATGGTGCGCGCTACAACCGTAGCAAATCAGGGTGGTTTGACATGATTTGTTTTGAAGATTGAATCAAGTGTGATCAACTTTTTATTATTGTACTTCAATTACAGTGGCTTAGATCTATTTTAATTTCAAGGGTTGGTCAGTAGGCTGTATTCTAGGGCTCAGTTCTTAAGTGTTTTATTTATGCATAAAACTGTACTTAATTAATTTTAACTTAATCATCCAATTAATCACTTTAATTTGTTTCAGATCCTTGTATACAAGTAGCATTACTTAAAAAACACATATTCTAAAATTACTGTTCCTATTCACCCCAACTTGGAG
>evgR009360c0g1_i01_Hydra
AAAACATGCACAATGGCTTGCGAAACAATAGATTTTGAAACAAAACAATTAAACTTCTTTGAAACTGGTAACAATATCTTACTTAACGACTTCTCTGACGCTGATATACAACTTTTTAACGAATTAGATTTTAACCTGCAATACTTTGATCTTGAAACTTTTAAAACTCAGTTAAAAACAGATATAAAAAAAATCATGGCGATAAACATTAATATCAGAAGTATAAAATTTGGTGAAGCTACAGTTAAGCGGCCTAAAATAATTATATTTCTTTGCTTTTTAAAGCACGATGGAGCTTATTTTTAAAACACGATTGAGCGTATTTTTAAG
>evgR009377c0g1_i01_Hydra
CGTTCTGCTCTCATCACCAAGAAGAATATTATCGGTTTTGCCACTCAACAAGTTAGAAGCAAACACCAATTGCCAACATTGGATTATAGTGTTGAACAAGGTATTCATCCAGTTATTACTGCTCGTCAATTGGATTTCCACTACAATAAGCATCACAAGACCTATGTTGATAAGCTTAACGCTTTGGTTGAAGGTACTTCATTTGAATCTCAACCATTAGATGCTATTATTCAAAAAGTAGCCTTCGATAGTGATCCAAAGAGTGTTGCCATCTTCAACAATGCTGCTCAACACTTTAATCACTCCTTCTATTGGAAATCAATGGTTCCAGGTGGTGATAAGATTGACAACTATCCAGCTATTAAACAAGAAATTGAAAAGCAATTCGGTTCAGTTGATGCTTTCAAACAACAATTCACTGAAAAAGCCACCACTTTGTTCGGTTCTGGTTGGACTTGGTTAGTTTTGGATGGTACACAATTGAAGATTTGGAATGGTTCAAATGCTCAAACTCCAATTGCTGAAAATTTAATTCCTTTGCTCACTTGTGATGTTTGGGAACATGCCTATTATTTGGATCATCAAAATAGAAGACCAGATTATTTGAAGAATTTCTGGGAAGCTGTTAATTGGGGATTTGTCAACAAGTCACTCCAATTTGCTGTCGAACAAGCTAACATTAAATTGTAAATGAAAT
>evgR009381c0g1_i01_Hydra
CTACTTGTTCCGATTGGAACAGTTGCTACAACTATGTCTGGTGGAATATTTGTAGTATTATCGTCTGATCGCCAACGTACAATTGTTTATCCGTTTAAACGGCATTTTAAGTTTATCTATATTAAACTTAGTATTTTGTTGATTCTAGTTTACGCGCTAATAATGAACGCACACTATGCTTTAAATATAACTTTGGATGTTAAAGCGAACAGATGTGGTGTGCCTGACGCAGGAAAAAAAGCATATCTTATACCAAGTATTATATATTTCTTGATGAACGCCTGCACTCTTTTAGTTGTCATTAGTTTTACAAACTACCGTATTTTTTCAAAAATGCTTCATCATAGCCCTACAAGTACGCTCGTTTTAGGAAATGGTGCTACAAGAAGAAAGAAAGATAATTACAGAATAATCCGTTTATTAATTGCACTCACAACAGTTTTTTTTATTCTGACTTTACCACGTGATATACTTCAGTTTGTATATTTACTAAGCTGGTGTTTTGGCCATGGCCTCCCAACAACAATTCCGATATTGACTTTGAATTCGTTTCTAAAAGTGTTTAACGTCGCGAACTCTTGCGTTAATCCCCTGATATATTATAAGATGCACTTTGGTTTTAAGAGATTCATTAAACGAAGTTTTGGAATTAAAGGAGATAATCTGATTCAAGGAAGAAGAGGAAG
>evgR009384c0g1_i01_Hydra
ATAAAAATTATACAATGGCATCATTTTATATAATTGTACTGCTATCTTGTTTATGTGGTCTATCTGTTACTCTTAACGTTACCGTCATCATTGCAATAACCTATACTAAAAAAACTAAAGATGTGCGCGATTTGATTTTGATGAGTCTTGCAGTATCTGATGGCTTCGAGTGTGCGTTAGGGTATCCAGTTGAACTCTACGGGTTTGCTATAAGAGACACAATGTTGCCAAATGAAAACTTATGTAAACTAAATGGCTTTATTGTTATGTACTTAGCCCTGACTTCAATTTCCCATTTGGTTTGTTTATGTCTGTATCGTTATTTGATAATTGTACACTCAATAAAAACACAGAGATTCTTTACAAATGCTAGACGAAGTGCTTTATATTTTATCATACCATGTTGGATTTATGGGTTGTTTTGGTCAGTAACACCTTTTATTGGGTGGAATGAGATAGTACGAGAAACAGAAGATCGACACAGATGTACAATTAATATGTATCCTAATGACGAGTTAAAACGCAGCTATTTATACGCATTAACAGTATTCTGTTTTATTGTTCCTGTGGTCAGTATTGTTTATTGCACTTTAAAAGTACATTTAGAGCTTCGCAACATGTGGAAGTTGTGCGTACAGATATCTGGGGAATACGCAGCTATCACAAAGGCTACATATAAATTAGAAAGACAAAATTTTATATCCGTTGGTTTAATAATAGGATCGTTTTTTGCAGTCTGGACTCCATATGCGCTATGTGTTTTTTTACTTGTTATTCGAATAAAATTACCTAGAAATCTATTAACCTACTGTGCATTATTTGCTAAATCTTCAACGATTATCAATCCTACTATCTGCTGCATAATTTACAAAGAATTTCGCCAAGCTATAAGAAGTAAAGTTCAAAAACTTTTTAAAAATCCAGTAGTAACACCAGCTAATGAATGACTCGGTCAAACTGTTACACAAGCAAACAAATTAAGGATACAAATACACCCGGGGTAAAGAATAAGGAGATTTAAGGAAATGATATAGGAACACGTGCGCTCATGGATATATGAACACGTGCACTTATGAATATATGAACACGTGCGCTTAGAAAAATATTATTTTTAATTCAGT
>evgR009392c0g1_i01_Hydra
GGTTGCTGGACCGCGGTCAAGTAACAGTTTATATGGAAAAGAAAAAAATGTTATCAATTTGACAAATCTGTCATACAAAGCTATTCATCCAGGTGTAATTGATCGACAAAATGTCACTCATATGTTTGCTGTTGTTAATAAGAAGAATGTAGCTTCAAGTTTATTATTTTAAAGAGACAAGCGAGTTCCTTTACTATAATATTAAACAAGAAGGTATGGACATATGCTTAAATAATCCAAATAGAGCTTCATACAAAAATGTAAATGATAAACAACTTCAAGAGATTTTTACATTTGCTGACGCTGTTTCAAAAATACCAGGAGGAAAGGGCTGGAAGCGTAATAAATCATTCACAAGTAAAACCAAAAATTCTTTGTTGAGTATGCTTTATGGAGTTGACTAAAGATCATCAATATGTTCATCCTGGAATATTTCAGCCAGATCGGTTAGAGAGAGAACTTGGAATATACAGGCAACTTTGTGGTGGCAGTTACCATGCTTCAGCTGAAAAAGTAAAAAATAGTTTTCGTTTACAATGATGTTGGTTAGTAAATTAGATGCTATGGGTGTATCTATTTCTTAATGCTCAATTAATCAATCAGCGCTGTCAATTAGATATGAATAAGGGTGGCATTAGTTGTTTTGTTACGTCTGTCTCTTCAATTGATGCAATTGCCGAAGACAAAAATGCTGTCCTATATTACATTTCGTGTAGTTCAGCATAAATTTCCCCCAAGTATATCTGTACCTGGTACCGTTGTTACAGATGCATCTAAGTTTACTGAGTTAGTTTCTTTTGGTGAACTCTGTCACCTTACTGAGGATCTATTTCAATTTGTCAGATATGATTATTCTTATTTTATTTTATTACCTAATGCAGAGATGTGGTTCCAGCGTGGTAGATATATTTCTAAACTTTTTATATATATGTATACAGGTTTACCTTATAAGCATTACATATCAACAGTTGTTTTGACTCATCAGACAAATTGGATGACTCATCAGACAAAGTTTTTATAAACATTAAAGACTGTAAAGTTGCAAATAAATAAAATGTACAATTTGTTGTTAAAATGGTTATTTAAAAAAAAACTAAC
>evgR009399c0g1_i01_Hydra
AGGAAAATTATCTAGACACAGTAAAAACATTACTGTCTATCGGTGCTAATGTAAATATTTACTCTGCTACCCATTCTACACCACTGCATCTTGCTGCAATAAATGGTGAAATAAACATGGTAGAGCATCTGTTAGAACATAATGCTAAAATTAATGTGTTTGATTACCAAAATATGACTCCTACGCATAAGGCATGTCAGTTTGGCAAGCTAGATGCATTAAAACTATTAATTGAGCATGGAGCACAAATTAACTCTAAAGATTGTGACTGTTTTACCCCACTTATGTCTGCTGTTTCAAAAGGTCACAATGATGTTGTTGACTATTTGTTGAAAAACGGAGCAAATGCTACCATGAGCGAAATGAATATGAAGAATGTTCTTCATTTAGCTATAGAAAATGGTCATAATCAAACACGTCAAATTCTTTTGAAAAACGG
>evgR009400c0g1_i01_Hydra
ATTGCACATTTACTTCACCTTGTTATTATCAAATCTGTCAAAGAAGTCGATGAGATTGAAAAAGTTTTTCAGGTATGCAAAACACTCTCAGCACGAACTCACAAATCAATCAACAGTCAACAGAGAATTATGGAAGCATGTAATGATGTCATACCTGAGAAAATCAAATTTAATAAAATTATTTCTCCTTGTCCTACACGTTGGAATAGTGGTTTTATGTGCATGGATTCAATATTACATTTACGATTAGCTCTTGAAATAATTGTAAGCAACAACCTTAAAACGGACGACATTTTAAGTGCTTTAGTTCCATCTTCTGATGAGTTTGATGTTATTGAAAAAGTAGTGCCGATCATGAAACTGTTTAATGATTTTACAGAGGTTGTTTCATCTGAAAAAATCCCAACTTCTCAGAGAGTAGTTCCAGGACTGTTTGATATTCAAGAGAATATTAAAAAGCTTATACCGCCAAATGACAAAATTACAGACTTGTAAAATTTTGAGACTTTGTTTCTAAACT
>evgR009424c0g1_i01_Hydra
TATATATATTTTTTCGTTCTAAATAATTCCGAACATATTGGGTTTGATAACACATGCAGATGCTCTCTAACTGACTAAATGAATATGAGGTCTAAAAAATAATGCAATATAAAATTTTCTAATCAACATTGAAAACTTACTGATGGCGATTCAGTGTTGGCCTCTTATTTGTTTTTCTTTGTTTTTTTCAAATATACACTCAGATCAAACTTCATTTGGCGATGATGAACCTATATTCGGGATCGAAAAGAAAAATACAGGTTTGACAGAGTCATTTAGCGAAATTCGAGAAAAAACATTGAACAGCGTTAATGACGATCAATCGTCAAACGAATTCAAATCCAATGATTCCTCGGAAGAATACGAAGACTCTGGGTTAAAAAAAAACATTCGAAATGAAAATAGAAGCATTAACGTGAAAGCTGTTAAAAGTTATTTGGAGTTGATCATCAAAGACCTCCGTCGTCATCTCAACCATTTGAAAAGTTTAACAAATGCAAAAGATGTTGAAGTTAATATGCAAGAAAATATTCTAGAAGATTCAAAGGATTCTGTTAAATTAAATACGGAAGATGAAAATGTGAATAAAATGAAAGAGAAGAATATAAAGTGTAAAGTTTGCAAAATAATATCCGCTACAAAGTATACTAGCAGCAACACAACACGCACCAATATTACGGTTAACGAAAGTTCTACCTTAAACGGAAATACCAATAAAAATGTCCAATATAAAATTGTCAACAAGCCGAATGAAAATACTTCATATGTAATTAAAAACGATCTCTCTCCTATTGGTAAAGATTTAAAGTTAGCGGAAATAAAAACCCAACTGATACTTATTAATAACACAAAAGAAAACAACATTCATGTTGAACAAAAAATAAATAACCAGACCAGAAGTTTTTCAATTGATTCGACCGAAGTTGACAATTTGATTCAACTATTAACCGAAAGTATGATTTTCGACATTACTGACAGCGTAAATAAAAGAACTTGGGATGCAGTTGTTTCTAATGACGACGAAGAAGAAGACGAGGAAATCGACAGTCGAACTGTTCAAGAGTCTTTAAATAAGTTGACTGAAAAGTATTTAATGAAAAAAAGAAGTGACTGGATCGCATTATTAAAAGAAAAGTGGCTCAAAAAACCGAGAATAAAATCAAGAATAAAAAACAAACAGAAGACGAAACAGAAAAAGGATAATGCATTATCTTAAAAAAAAATTAAAATTAGAAAATGGCGATTGATAATGATTGAGGGAAAACTTTAAAAACAAATAACAACAACAAAAAAATTTTAACTTGCACAACTGTTTCTTTTTTTAACAAATTTGCATAATATTTATTTGTTTAAATATTATGCATATTATTTTAAAACTTTAAATCTTAATATTCTTATGCCATATATATTATATACATATTCTATATATACATATATTCACATTTATAAATATGCACATCTTATACGTAATAATTAGTAGTTGTTATTCGATTTTATATGTCATAAAAAAAA
>evgR009464c0g2_i01_Hydra
AGTGGTATTTATTATTTTAATTAAGGTAGCGTTATGTCATCCCGAATAACTCGTTGGAGACAAATTAATAAAGCTGTGATTGATTTAGTTGCTGAGCAAAAAAGTACTAATGCAACAAATGAAGAACAAGAAAATATTTCAGGTATTGGGATTATTTTCAATGAATCAAACCAGAAAGATATTGAAGGTGTCAAAGAAACAGTAGCAGATTTTGACAACGATTTCTATTACTCAGGCAGTTCTGATTCTGATTCTAGTAGTCTTGGGAATTTTATAGAAGAATTAGATGAACCTGTAAATAATACAAATCTATTTGAAAAATTAGCGGAATGGGTTGTTATAAATCAAACTTCACATTCCGCAACTAATCAACTTCTTTGTCTTTTAAGAAACAGTGGCCATCTGGAGCTTCCAAAAGATGCAAGGACATTATTAAAAACTCCAAGAAACGTTCCTTTGCAACATAAATGTGGTGGCGATTATATTTATTTTGGTTTGGAAACCAATATTTTGAAACATCTAAATAATTTTCCAGAACTTAATAATATTGAATTATCTGTGAATGTTGATGGCATACCAGTATTTAAAAGCACATCTATAACTTTATGGCCCATACTTTGCTCTTTTTCAAATATCCAACCATTCGCTGTAGCCATTTTTTGTGGTAAGGTCAAACCTTCATCTGAAGAGAGCTTCATGTTTGATTTTGTGAATGAACTATCAATTTTGTTACAAAATGGAATCAAAACACCAAACAAAAATTACTCAGTCTCTCTTAAGTTATTTTGTTGTGATGCTCCAGCTCGACAATTTTTAAAGTCAATAAAAGGCCATAATGGATATGATTCTTGTGAGCGATGTAGTATTCATGGTGCAAACATTAATCACAGACAAGTTTTTTTGGGGTCAGGCTACACTCTCCGAAATGATAGAGATTTTAATCTATTTACTTACAAAGACCACCAAAAAACCTTGTGTATTCTTGCTAAATTTCAAATACCGTGTGTTACATCTTTCGTTTTAGACATCATGCATTTAGTTTATTTAGGAGTAACTAAAAGAATTTTAATTTTTTTTAACAAAGGACAAAGATCATGCAGAATATCAGAAGTTCAAAAGGCAATCATTTCTGAAAAACTTAGAGAATTGTCTGGAAAATTACCTAGTGAGTTTGCAAGACAACCAAGAGGGATTGATGAATTTAAACGGTGGAAAGCTACAGAGTTTAAAAACTTTCTTTTATATACTGGTATGGTTGTTTTGAAAAATGTTGTTTCTCAAGAAATATATACACATTTTTTGAGCCTATCAGTTGCTACAAGCATAATGATTGATAAAGATCTTTTGAGTGATCAAAAGTTATTGGTGTATGCAAAAGAGCTTTTGGAATGGTTTGTATACAAATCATCATTTCATTATGGTCAAGAATTTG
>evgR009493c0g1_i01_Hydra
TTGAAAAAGAATATTTCATTTCAATTGATATTTATCCTAATAAGTTTGTTCCTTTATGGCGAATGGTGATGAGTTTTCTAAATGACTCTATTGATTTAAGTAAATACAACTGGTATAGAATTAATTACATTCTATTTCATGAAAATGGTGATGGAAAAATTCAATTTCCTGATCCAACAAACGAAGACCCTAGTCAATATATCCTCACAAAACCAGTCAAATTAAATATGTGGTCAAACATAATAATAAGCCAAATATTTCAAGGTTTATTTTATAAATCAACAATAAAGCTGAATGGTGAAATTATTGATTCTCATATAAGAAACCAATCTGAT
>evgR009493c0g2_i01_Hydra
GCACAGATACCAAAACTTGAAAAAGAATATTTATTTTCAATTGATATTTATCCTAATAAGTTTAATCCTTTATTGGAAATGGTGATGAGTTTTGTAAATAGCTCTTTTGGTTTAAGTAATCACAACAGGTATAGAATTAGTTACATTCTATTTCATAAAAATGGTGATGGAAAAATTCAATTTTCTGATCCAACAAATGAAGACCCTGGTCATTATATTCTCACAAAACCAGTTAAGTTAAATATGTGGTCAAACATAATAATAAGCCAAATATTTCAAGGTTTATTTTATAAATCAACAATAAAGCTGAATGGTGAAATTATTGATTCTCATATAAGAAACCAATCTGAT
>evgR009517c0g1_i01_Hydra
CAAAAAGGGACAATTGCCCCGGGCGCAAGCTTGCACATCTACCGTACAAGTTATACAAATATCTTCCGTTGCTGGGATAAAGCTCTTTCAAATTCATTGACAGACAAATTGTTTTTGAAAATGTTTCAACATACAACGCCATTATCTGAATGTTTGCAACCCACTGGAATAAATATACTGCAAGCCTATCAGGCTTGTGGAGCCGCAGGTTATTTTCACGGCTCGGCTCCGGCTCCGGCTCCAAAGAAATGTCCGGCTCCGGCTCGGCTTCCGGATCGGCTCCGGCTCCTGACCTTATATTTTAAATGTGCAAATTTTAAAAGGAAATTAAAAATATTTAGACTTGTTTGCCTACCCAATTGCAAAG
>evgR009528c0g1_i01_Hydra
CACTCTTGCATTAACCTATTTTAGTGCAAAAAGCAAATGTCTTTTAAAACAAACATTCTTGATCGGCCATTAAGCAAGGGTAAATATGAAGTAAGTATAAGTGCTTATGGACTATTATTTTCAGAAATTGTTCAATATTGTCAAAGTAGAGTTTTTACAGTGACTGAACTTCAAGATAAACTTGCAAGTTTTGGCAAGACTATTGGAGTTAGATTAATTGATACTTTATCGATACGCGAACGTGGCTTCCGAAGGGAGACACGTTTACTTAGTGTTTTAATGTTCATTCGATCAAGCTTGTGGAAGACATTGTTTGGACGTGAAGCAGATAAGCTGGAGCATGCCAATGATGATATAGGAACTTATTATATTATTGAAAAAGAGTCTCTTGTGAATCGCTTTATATCAGTACCTAAAGATAAAGGAAGTTTAAACTGTGCTTCTTTTACCGCAGGAATTTTAGAAGGGATTTTTAATGGTTGTGGTTTTCCAGCCACCGTTACAGTTCATTGGCATAAAGGAACAACTTTTTTAATAAAGTTTGACGACTCTGTAATTGCTAGAGAAAAAGTATTTGATAATAAATAACAGTATTTAAAATATTTTTTAATTGTTGTGTTTTTAGTTTAATTTTTATATCTATGAAATCAATTAACAATAATGCTTATAATATAAAATACTTTTTAAATTTATTAATCATGGATTACTTTAAATTTTATTTTGTTTATATGTGTACATAATCATGCATAGCTTTTTTATTTTTGTATATATTTTTAATTATTTTGTAATAATTAAAAATATATAATTGTAATAAAATGTTTGTTGATTCTTGAAGCCTTC
>evgR009533c0g1_i01_Hydra
GTTTTGTCAACCCAAAACAACTGCTCTGCAAATGCAATATGTAATAATTATTTTGGTGGATTCAGCTGCACATGTAAAAGTGGATTTAGTACTAGTAACGGAGGAATAACGGAGTCCCTTTCTAATAAATGTCTTGATATTAATGAATGCACATCTACAAACAAAAACAACTGCACGGTTAATGCAACCTGTAACAACTTGGTTGGTGGATACAACTGCATATGTAAAAGCGGTTTCAACACAAGTAATGGTGGAATAGAGTTGTTAACAAATAAATGTCTAGATATAAATGAATGTCCTTTGTCTGACA
>evgR009545c0g1_i01_Hydra
TTCTATTACAAACTTGTTAGAATCGACAATTGGATTAATACCTCAATTAGTAATGTCCAATGAATCTATATCGGAAAGAACTCCGTTATGCGTTGCAAGCGGTTTCGCTGTTTTAGGGTTTGCAATTACCAATATAACTCATTTAGCAATACTTTCTTTTATTAGAACTGTTGCAATAAATTACCCGATGCGTTATTTAAGGTATCATAAAATGTTTTGGTGCAGAGTAACCTTAATTTTAGTATGTTATTTGTACGGGTTCGTTTGGGCATTACTTCCTATTATTGGTTGGTCAAAATATGAACTAGATCTTGATAAGAAACGTTGTTCTTTGGATTGGAAATTGACAAAAGCAAACTCTTTTTCTTATATTTTGACTATTTTTATTTGTTGCAATATTTTACCTGGAACTATTATTGGGTTAACTTTGTATTTCAGCAAAAAAACAATCAATCGCCGAAGAACATGCAAATTTCGTCGAAATAAAAAGGCTGATATCGTAGAAAAAGAATATTTGCGTGTTTGCTTTTTATCGGCGATTGCATATTATTTCTTTTGGACAACTTATGCAGTTGTTGGCGTTTTGACGCTGTTTAAATTTTTTATTCCAATGCAGCTAGCGACTACCACAGCGCTGTTTGCAAAGCTCTCCACCATTTCTAATGTTCTTATCAATTGTTACGTCATCAAATCTTTTCAAAAGCATTTGCTTAAATTAAAGCTCGTTAGGTTAACGAAAAAGTATTTTCGGATATTATTAATTTGTAAAAAATCTCCAGATAATAGTCTTATCAATTAAATGCTAATCAACTC
>evgR009546c0g1_i01_Hydra
TTTAACTGATTGTATGTTTCGCGGAGGAAATTTGTTGAGTATTGAAGATCAAGAAGAAAACAAATTTATTCAAAGTAACCTGATTGATGGTGATAATTATTGGATTGGTCTAACCAATCGTTGCAATTTCATGTTTATTTGGTCTGATAACACCTACTCACAGTTTTTTAAATGGAATTTTAGCAGAATTCAATTTGATTATTTAGTTAAAGGTTGTGTTGAAATTCTAAATGGGTTAATGTATGTCAAAGATTGCAAATCACCGAATAAAGTAATTTGTAAATCTAAAAAAGGAATCCTCTAATATTCATAAAAGATCACTTGGTGCATTTAAA
>evgR009550c0g1_i01_Hydra
AAATATCAAGCATACACAGAATATAATTTATATATTAAGGTTTATCCAAATAATAAACCAATGAATTTAAAAGACTTGTTCTCTACTATCAGTTTTGATGATACACAAAGTGTTGCTAACATTGTTCAAGCAGTTAGCAATAATATAAACGAAAATTCTGCTAATCAATATAATATTACTCCTACTACTGATAATGATTTATTTGGTATTGATAATGACAATTCTTCGAAGCAACTAAAGCAGCAGGAATTGTCTAGTTTACAAGAGTTGCGAACACAATTAATTGATTTAATTAGCAATAC
>evgR009558c0g1_i01_Hydra
TGTGTGCCCAATTTTTGATTTCTAATCTTGTCAATGGTCAAACATTGGCTCACTGTCTTTTGAGAGGTACAACCAATGCCAAGGGATTGAAAGGTGTTGTGACATTCTCTCAATCTGCTTCTGGTACCACTAATGTTGCTTGGACTATTGATCAATCGACTGTTCCAACCGCTTTGCAAGGTAAAACTCTTGGTTTCCACATTCATGAATTTGGTTATATTGCTGATGCCGCTTCTTCTGGTTCTCATTACAATCCAAATTCTAAAAATCCCTCATTGCCAAGTGAAAATGCTTCTCCTGAAATTCACGTTGGTGATTTG
>evgR009559c0g1_i01_Hydra
AATACACAAAGCCTCAACGTCAAAAAATGGGTATTACCAAGAGCGATCTCCACAAGAAACGTGTTACCGGTGGTAAGAGAAAGATCCACAGAAAGAAGAGAAAGTTCGAATTGGGTAGAGCTTCTGCTAACACCAAGATGGGTAAGACTAAAATCACAATGGTCAGAGTCCGTGGTGGTAACTACAAGGTTAGAGCCCTCAGATTGAACAGCGGTAACTTCGCTTGGGGATCTGAAGCTATTTCCACTCCAACTAGAATCATGGATGTTGTTTACAACGCCACTAACAACGAATTGGTCAGAACTAAGGCTATTGTCAAGGGTGCCGTCGTTTTGGTTGACTCATCCCCATTCAAGCAATTCTACGAAAAGACTTACGGTATTTCTTTGGGTAAGAAGAAGGAAGACCTCTTGGAAAAGATGGACGAAAAGACCAAGGCTAAGGTTTTGGAAAATAGAAAGAAGGCTCCAGAAGTCGATCCACTCCTCAAGGAACAATTGACCACTGGTAGATTGTTGGCTAGAATTAGATCTAGACCAGGTCAAGTTGGTCGTGCTGACGGTATCATTTTGGAAGGTGCTGAATTAGCTTTCTACTTGAAGAGAATCCAACAAAAGAAGTCCAAGAAATAAGCTATTCGCCTTTCTAAACATAAAAAAATTTATTTCATAA
>evgR009571c0g2_i01_Hydra
GGCACCTAATTTAAAAAATGAAAAAAAAAACTACTATGTACACGGTGGTCTTTTTGACTTTTTCAACCCTATTTTTATACTATATTTGCCTATACTATGTTTTGATAGAGCTTTTAAGATTTCGAATTATATAAGCCGTGTTGTGGGAGTGGGGTGGCGGATCGTTGGGTGTAATCAACCCTTCTTTTTGTACCTGAATAAATTGAGTAGATTGAGTGTTACCGAATTTATTATAAAGGATAGTGGTATATGCAAGTTGCAGAAAAATGGTAAAAGAACAAGTTTTGTCTACGGGCTAGTCGAGTGT
>evgR009583c0g1_i01_Hydra
TTGTTTAATTCTTTTGACAATGAATTTACAGACGGGTGTGTTTATCGGTCCCAGCCTACAACTTGCTTTGCAGATCAGAACTGACTATTTTTATTATGACTGTATCACTAGGAGGGTTTCCGTCCGAGGTTTAAGAGCCACCAGATTTTTCCTACCCATCAACCTTCATTTCAACTTCAACCTCAACCTTCATTTCAACATCAACCTTTTTCAACCCATGATTTTTCCAAACCATGATTTATCTAGATTTTATATTAAAGTTTTCTTTCTCTTAAGTAAGTTACATTGCAGAGTTATTACCACAGTTTTGCAAAGTCTCGTGTTTCTCAAGTCCTTAACCCGCAAACCTCACCTGCCGTTTGATAAAG
>evgR009605c0g1_i01_Hydra
CCTAGATGAAATTAACTGAATTATTCTTGGGAAGCATTCTTCATGTATTCAATCAAATCGGCACGTTCAGCTGGCTTCTTAAGACCGGCAAAGACCATCTTAGTTCCTGGAATATACTTCTTTGGATCCAACAAGTAGTCGAAGAGGGATTGTTCATTCCAAGCAACACCCTTATCCTTGTTAGCAGCAGAGTAGGCATAACCTTCAGCCAAGCCACTCTTTCTACCCCACATACCGTAGAGATTTGGACCAACCTTGTGAGCACCACCCTTGTTGATAGTGTGGCATTGGGCACATCTGGCCTTGAAGATCTTTTCACCCTTAGTTG
>evgR009624c0g2_i01_Hydra
AAAACACGCTTTAAAAATTATTGCAAACGTCAAACAATTTATTATAGACATTATAGATGGAAAGATATTTTGTCTTCAAACAAAATAACAGAATGTTGAGAGCTTTATTTTTTACTCAAATATCTTATAAGCGATCTTTTTCTGCAAGATCAAATGAGCACCAAAAAGTACTTATAACAGGTAGTCTTGGACAACTTGGTGCAGGTATTGCAAAAAAACTTAGATCAAAATATGGAAATCAAAATGTTGTCATGTCAGATGTTATAAAAGCTTCTGAAAATGTTATTTCTAGTGGTCCTTTTATATACGCAGATATTTTAGACTACAAAAATCTTCAAGCAATTGTTGTAAATCATCAAATTGACACCATTGTACATCTAAGTGCACTTTTAAGTGCTATCGGAGAACAAAATTTATCTGAAGCTTTAAAAATCAATATATATGGCTTTCATAACATAATAGAAATATGTCGAAGGTATGGATTAAAGTTATTTTGTCCAAGTACTATAGGAGCATTCGGTGTTGATAGTCCGAGAAATCCTACTCCGAATATAACTATACAGAGGCCGAGAACAATTTATGGTGTTTCAAAAGTGCATATGGAACTGATGGGAGAGTACTATTATCATCGATTTGGTTTGGACTTTCGTTCTGCAAGATTTCCTGGCGTAATTTCTGCATTTACTGAGCCAGGTGGAGGTACAACAGACTATGCAGCACATATATATCACAGTGCTTTGAAAACTGGCTCGTTTGTGTGCAACTTGCGTTGTGATACTCGATTACCAATGATGTATTTGCCCGATGTTATAAAAGCAGTTGATTTGATTCTGGATGCTCCTGCAGAACAGCTTAAAGTTCGAACTTACAACATTACTGCTATGAGTTTTACTCCAGAAGAAATTGCTGAATCTATCCGCCATTTTATACCATCCTTTACAATTACATATAACATTGACGAAAGTAGACAAAAAATTGCCGATGGTTGGCCACAACAACTGGATGATTCAGATGCTCGTAACGATTGGAAATGGAAACACGATTTTAATCTTTTTGATATGTCTAAGGATATGCTTTTGCTGCTAGGTCCCAAATACGGAAAAAATGTTATCAAATAGTTTTAATATTATTTAGCTGTAAAAATGTTATCAAATAGTTTTAATATTAT
>evgR009642c0g1_i01_Hydra
ATACTATTTGGGTGATGGTGATTCTTCATCATTCAATGATGTTGCTAATAGTAAGCCATATAGTAAATATGGCGTTGAACTACAAAAACTTGAATGTGTTGGGCATATACAAAAGCGTGTGGGAAATCGTTTAAGAGAGCGAAGAAAGCAAAATATTTGTGGAAAAAAATTAACAGGTGTTGGAAACCTCACCAATAATTCTATAAACACTATTCAAAACTATTTTGGTATGGTGATAAGAGAAACTGCAAACAATAAAAGTTTAAACGACGATGAAAAAGTATACCAAATGAAAAAAGGTATAAGAGCTGTTCTAAATCATTGTACTAACTTTCCAGACAAATTGCGGCGTCATATACTATGTCCTAGAGGATTAAATAGTTGGTGTAGGTGGAAGAATTGTGAAGAAGGAATTGATAATTACAAG
>evgR009664c0g1_i01_Hydra
AGGATAAATTCTTGAGAAGTCTTGTTGCCAATAACCCAGAAGGATATGTTGAGTTGAAGACCATTGCTTCATTCAATAGAATGAAGGAAATCACTACTGATATGGATCTCATAGTCTCTGCTTTAAAGAAGAGTTCCATGTTGGAAGTTGATGCTGAAGGAAAATTGGTAAGAAGAATTACTGCTCTCCCAGAAGTCGATACTAGCAATAATCGTACATTGTACGCTAAGGACTTCCCAATGGATCAAATGATTGCCAATCTTGACACAATCAAGTCATGCTTTGAAGCATTCGGAAAGGTTGTTTGTGTTAGAGCTAGAAGAGAAAAGAATGGTACATTCAAGGGATCTGCATTCATTGAATTTGATTCTGAATCAATGGTAAATGAATTGGCAGGAAAGACTGATATTAAATTCAGACCAGAAGATGAACCACTCCTCCTTGAAAGCAAGCAACAACATTATGATAGAAAGAGAAAGGAACGCGAAGAAGTTAAGCAAAAGAAGACTCAAGAAAAGAAGCAAGAAGATGTTTCCAATATGATGGAAGAATTTAAGCAAAACTGTTTCTTGCACATTGAAGGTATTGAAGCTGATGAAAAGTTCGGTTTTGGAGAATTCTCTAAGAAATTACCAATTAATAAGAAGAATGGGCTAAAATTTATTGATTACCCACTCAGCGGTGACAAGACCAAGGCTATTCTCCGAGTGGAATCAAAGGAAATTTTTGATTCCCTTAAACAACAATTCGATGAAGGAAAGATTGAACTTCCAGGAAAACCAGTCGCTAAGGAATTACCACAAGACGAATTAGAAAAATATATTTCTGCTACTTGTGAAGCCAAGATTGCTGAAAGCAACAAGAATAAGAGAAAGGGAA
>evgR009668c0g1_i01_Hydra
AATCTTTTATTTTCTAAATAAGTTTTTTATTTAAAAAGACAAAAAGAAGACAGAAGAAAACCATGACGTCAGTCATTGATCTGTCTACTACTACGTTTTTGGGGTTAGTAAACTTTGAAGGAAAATATCTGACAGCCGAAAAGTTTGGAAATCAAATCAATGTCACTGGTACTTCCCTTCGAAGTAAGCAAATGTGGTGCTTTGAGCAGACTGAAATAAATTCGAAGATCTACTTGAAAAGTCCGAGCGCAAACTATTTGGAAACAACAAAAACGGGCGATGTGTTGTGCGAGAGCAAAGAAAAATCAAACAGTTTGGTTTTTGATGTAGAAATAAATGAAGAAGGTCAATGGATATTCAAAGATTGTTTTGGTAAATATCTTTGCGGAAACTCTGAACGATTGTATACACAACCTAAAGTTTTAGACGAAGCGTTGTGGAGTATACATGTTGCCAATCATCCTCAATGTTGCATTTTCAGTCCAACTCGAAAAAGGTATGTATTTTCGGAAAACGAAGAGTTGTGCGCTAATGAAGATATTGCATGGGGTCATGATGCAGTAATAACAGTTGAGTTTCAATCTGGCAAGTACGCTTTAAGAGATTCTCTTGGGAGATACTTAAATGGACTTACAGGAAAAGTGGAGAATAAGTCTTCAAAAGACTCTTGGTTTACAATATCTATCTATGGTAACATGTTTGGACTGCGAGCTTCTAATGGAAAATACTTAACAACATACGGACCCTGCGGGAAACTAATAGCCTCCAAACAAGTAGTAGGAAAAGATGAGCAATTTATATTTAAAGAAAGCAAAGCACAATGTTTGTTATTAGCTAGCAACGGAAAAAAGTTCTCCATAAGACAAGGAATTGATGTTACAGCAAACCAGTTTCTAGAACCCGAAGATACCGAGACTTTCCAAATAGATTTCTGTAATGGTGAAGTAGACAAGGTAACATTTCTTGCAAGAACAGCCAAGTTCTGGAAAGCAACCGATAAAGGTATTACGGCAACAGCAACAGAAGCAACAGAAAGTTGCGTTTTTAGACTGGAGTGGAACGGCGAAAAGGTTGCCTTAAAGTCATTTAACGAAAAATACTTAGTTTCCTCATCAGGGGGTCAGTTATCACCTGTAGGTGAGAATTCTAATGACGTAAATTCATTGTTTACTCTATTGCTCGTAAACCGTCCAATAATTGTATTTCGCGGGGAACACGGTTATATTGGCATGATTTCGTCATCAAATAAGATTCAATGTAATCGAGGTATACATGATGCTATGCAAGTTCTTGACTGTGCAGGTAAACATAAGTTAAAAACTGTTTCTGGAAAATCATGGAAGATTGATGCTGATATGTGTTTAATTGCTGAAGAAGGAAATGGTGATTTTTTTCATTTTCGACTTTACGGCAAAAACAAAATGGTAATTTGTGCTGAAAATGGAAAGTATTTGCAAGGAGATAATAACGGAGTCATTAAAGCAATCGGTACTGCTGTAATTTCAAGTACACTTTGGGAATACTAAAATATAAAAATGTTTTAAATCCTAACATTAAATTATGAATATTTATTAATTATGCTAGCTATTTGAATTCAAATCGAAATTTCTTGTTTTATATACACAAAGCCGATGATCTGTTATTTTACACTTTAAATCTTAGCAGCTATAAAAAAAAAACAATGAAGCCAAACAAGCTTGTATAACAAAG
>evgR009685c0g1_i01_Hydra
TTTACAGTGGCTAAACAAGAAGCAGAGTTACCCAAGGCTATACCAGTAACAGAAAGATTTTGTTACAGACCAATGTTTGGTAATAAAAACACCGACTCATTACCCAATCAAAATAATAAAACAGTTAATTTTGCTAAGAACACTAATGCTAATTCAGTTAATTTTGATATGAAAATTAATTCTACTTTTGCTGAGCACATTAATGCTATTTCAGTTAATTCTGCTAAGAACAACACCACAATCCCTACTAGCCATTGTTGCTCAAATTGTGGTATGAATTCTGAAATATTTTGCTTTCAATGCGG
>evgR009704c0g1_i01_Hydra
CTCTTCTATTGTCTAAATAAGAAAGAAACGTCCACTTAACGATGGTAATAGATCAAGGTATATTTTTTTATGCATATAAAACTTTTATTTTCTTATGGACTACTCGTATTGGTATGCTTACAGCTAGTAATGTTTGGTTTGCAGTTAAGTCATCAACCTCTGCTTTTCGAAATAAGTTATGTTCTGATAATACTATAGTTTTGTTTTTTCCTGACTTATCATATCCATGCAAATGGTATCTTAATGATAAACCTTGTCCAAATAAAAGCTGCAAGTTTTTGCACAAACCGTCTTCTTTGGGTTACCTTATGAATTGCTTAAATCAAACTATTTTTTCAATCGATGTTTGTATTTTTGTAATCACTTCACAGGATTTGTCTGAAATTTTAATAAAGTTACACAAAAAAGGTGTTATAGTTCGTATTATTACAGATTATGAAAAAATGGATTTAAACAATTCTCAAATAGAACAGTTTCGTGCAAATGGTATTCAAGTTCGTCATGATAAAACATCATTTCTTATGCACCATAAATTTGCTATTATAGATGGAAAAACATTGATTAATGGTTCATTTAATTGGACTAACCAAGCGATTACTGGTAATCAAGAAAATTTAGTAATAACAAATAACCAGTTGTTAATAAAACCATTCAAAGATCAATTTGAAAAGTTATGGATCATGTATAAACCTTTAAAGTAGTGAATTAATAAAGTATATCATAAAAAAAAAAA
>evgR009718c0g1_i01_Hydra
ATAATTTTTTTTTATTGTTGATTTTGATAAAGATCTGAAGAAGTTGATGCTATTTATAGACTACTTTGAAGATACCTACATTGGTCAATTAAGATTAAACGCTCCATTTGCAAAAATTAAATTAAAAAATGCAAAGAACCTCCACTCTTTAAAATATATATTTGGAATATGTTTCATATAACTTACCAGGAGATGCCAAGAACCAACATCGCCGAATTCAATCACTTTGTATGGCATCTCAACCGACGTTATAGAAGTTCCTTGATATACTTAAAAAAAAAGTCTATAAACAGAGTTGAAATCCTACAGGCCAAAGAAAGTTATCAACCTCCAGCGCAAAGGCATAAATACGTCAATTGCAATCTACGAATTTTGAGGATCGTTGAAGATTTTTCAAACTGTCAACACATTCGCTACCTTCGAGGAATAGCGCACAACTTTACGTTTTTATTGACTCTTTTATTGATATAATAAAGTGAAATATAGTATTT
>evgR009749c0g1_i01_Hydra
GTAAGCAAAACTGTCAAATTGTTTGGATCTCGTGACTGCTGTAAAACAAACCATTGCGGCGGACGTCCCAGGAAAACATCAGATCGTCTAGATCAAAGAATTGTGAAAGAGCTTAAAAAAGATCCATGTATGTCATCTATAAAACTAAAAGAAAAACTTGAAATACATATCTTCAATCGCACTGTGCGACAAAGAGCTAACGAAGCAAAATTTTTTTCTCAAATGCCTTCCAAGAAACCACTAGCTTCTTTTAAAAATAGACGGCTAAGACTTGCATTCTCCAAAGTTCACCAAAATTGGACTGTAGATCAGTGGACAAACATACTTTTTTGTGATGAATCAAAATATAAACTTTTTGGAAGCGACGACAAGAAACGAGTCTGGAGGCCAAAAAACACAAGATACAATAATAAGTATACCAAGAAGAGTGTTAAGCATGGAACGTCTGCTATGGTTTGGGGATGTTTCTCAGCACTAGAAACTGGTCCTATTTGCAAAATTAATGGAATTATGGACCGCTTTGTGTACAAAGATATCATGGAAGATGTTATGTTACCACATGCTGAATGGGAGATGCACCTAAAATGGATTTTTCAACAGGACAATGACCCCAAAC
>evgR009751c0g1_i01_Hydra
AAATTAATACCAAATTAATACCAAATTGGTTTAATAGTGATTGCTATACAATAGGGTTTTTTATTGATTTATCCAAAGCTGTCGACACAGTTGATCATGAGGTAATTTTGAAAAAATTAGTAAACTATGGTAACTATGGTAACTATGGTGTTAGAAATCAAAGTTTACTTTGGTTTAAATTATACTTAACTAACAGGAAGCATTTTATACTTAAAGAATCAAAAGACTCAAAAAACAAATTAATAACTTGTGGTGTACCCCAAGGTTCTATTCTAAGACCATTATTGTTCTTACTTTATATAAATGACTTACATTTATCATCAAAAATATTAAATACTATCTTCT
>evgR009792c0g1_i01_Hydra
AGTTGAGTTGTAAATGTAATTTGAAGTTGTTAGCGTTTAAAACAGTATTTTGTGAAAATGCAGGTGATTCCCGAGGATGATTTGGGTTATGGTAATGAGGCGTTCACCGAAATAGAAAATGGTTCTTACACAATACAAAATGGATATACTCCCGTCCAATTTAAGGTTTTGCAAAACTCAACAAAAGTTCAAGCTAGAAACAGTGTAACTACATTAGTAATCAAAGAAAATTCTTCTGTTACTACTTTAAATGGACAAAATGTTGAGACAGATGGAATAACTTTTTATACGACAACCTCAAACGATGGCAGATACAGAGCAAAAAGTCGCCAATATAGAGTTTCAATAACTCCAGCCGATGATGTTTTACCACCTTATACGGCATTATCGGTTAAACCACCTTCTTATGAAGAAACAGCGTACATCGAGCAGGGAGAGGAGAGCGTCGAAAGAGAAACGTCTACTCAAACGAACGAAAGACCAAACAATCAGAGGAAAAAGAAAAGCCATCGGGGAAAAAAAAAATGTTGCATCATGATATGCGTTATTCTGTTGTTATTGATAATTTTAATCGGAGGAATTCTTCTTGTGGTTTACAGCGCAAAATCAATATTGAGATGAACAAGAAATTTTTTTTATAGTAGTATTAACGTATCTTTTAAAATAACAACTTCTAAAAGTATTTAATGACTAAATGAATAAAAACTTTGGGTTGCCTTATAAAAACTTCAGGTCACTAATAAAAGGTTCTCTAAAGATTAGAATCAATCTTTAAATCAGGTTTTCATCAAGTTCTTACTTCACATATCATTGTTACAGAACCATTCTTTTGGCGACTAATTATCTTGAATTAAAAATTTCTAACATATAACAATGAGTGTTATGACGCCCTCGTTACCTAGAAACCAAGGTCTTAAATTTCAAAATTTAAGGTAAATATCAATATAAGATTTTATATATGTAATATGAGTATAATTAAATATATAATAACTCTTTAAATAAAATATATCAAGTGCATTAGTTTACATTTATTTTGTAATATATTATTGATTATCCCCCCTCCTTCTCTTTGGAAGTTGAGAATTATTTTTTGTCTGTAAGTTTATCTATTAAATACAGC
>evgR009794c0g1_i01_Hydra
ACATGTGGCGGAGGTGAGCAAAAAAGAACACGTGAATGCAATAACCCTATACCACAAAATGGAGGCCAAGAGTGCAAAGGAATAAAGTATGAAGTCGCTGGATGTAGTTTTGATCTGTGTCCTGATTTGAATTGGGAGCAAATCGGAACTAATATTTGCTTCCATGGAAATAATGATCGGTTTGGTGTAGTAATGATTGAAAAAACTAGATTTCTTAAAGCAATTAAGTTATCGTATCAATCTGGAGTCATAAAATGCACTGCCTTACAAGTAGGAACCAATTGGGGATGTGCGGATATAACTAAGCTTAATGTCATTATAACTGACGAAAAAAAAGAAATTATATTTCCCTCAAAGAATGAAGTAACTTTTGCCGATGGATACTGGTACACTCTTCCAGGAACATATTCAACATCTTCAGATCTTGTCTTCAAAAATTTGTGTTATCCAACATATGTAAGAAGTAAAAGTCGTCTTCAAGTATGGTGGGGGGAGGATCTTGTAAATCATGTTTGGCAATTGGATAACTCCGGCCGAACGTGCGTTAACGTCTACGCTTTATTTGAAACTTCTGCAGATTAGAAACGCCGTAATATTAGGGCGTTTGAAGCTATGTAAAATGAAATTACGGCATTTGAAACTATGTAAAATAAAATTACGACATTTGAAACTATGCAAAATAAAATTACGACGTTTGAAACTAAGTAAAATAAGATTAAGGAGTTTGAAACTATGTAAAATAAAATTAGGGCGTTTGAAACAATGTAAAATAAAGTTACGACATTTGAAACTATGTAAAATAAAATTACGGCGTTT
>evgR009803c0g2_i01_Hydra
AGAAAACACAGGCAGAGAAATGCTTTTTTACAGTCTGCAATTGAAATTATTACTACATCATCAGCTTCTGAAAACGATATTGTTTCAGTTCAGGACAGTAATGCGCATGAAGTTTCTTCTGATGATTATCAACATGATAGGAATGAATTAAATGAAACAGTAAGCGATGATCATTATTCCACTGACATAATGGAAGAATCAGACTCTGATTCTGTTATTAGTGTTGAACTAGACGAAGATCTAAGCGATACAAATGTTGAAGACGACCTACAGGCTGAACTTGCTGTTTGGAGTTCTTCAAGTATGTGTAAACGAGATTCCCTAAACAAGCTTCTTGCAATTCTTCGCAAAAATGGACACCCAAAACTTCCAAAAGATTCCAGAACATTGTTAAACACACCAAGGTCTACAGCTTCTTATTCTAAATGTGGTGGAGATTTTTTTCATTTAGGAATTGAATCTGGAATTAAAAGGACACTTGATAGTACAGTTATAACTAATGAAAATACTCAAAATATTGCATTAATTGTAAGCATAGATGGGATTCCTTTGTTTAAGTCTTCAAGCGTAGAATTATGGCCAATTCTTTGTCGATTTAACAGTTTAAAACCTTTTATTGTTTCAGTATACGCAGGTAACAAGAAGCCTAATTTTATTAATGATTTTTTAAAAGATTTTCTAGTTGAATATCAGTTATTGAACACTGTAGGATTTCTCTATGAAAACCATAGGTACACAGTCTCTATTAAAGCATTTGTATGTGACGCACCTGCTAGAGCGTTCATTAAATGTATCAAAAGTCATACTGGGTATTATTCGTGTGAGCGATGTGTAATTAAAGGCGCATGGCAAGATGGACGAGTTGTTTTTCAAAAAAATAATTGTTCTGCAAGAAGTGATCTTGATTTTAGTCTGTTAAAATATAAACATGAAAATGATGAACATCAAAATGCTAAATCACCCCTAATTGATTATAATTTGCCTTGTGTTTCTGGGTTTGTCTTAGATTATATGCATTTGGTTTGTTTAGGTGTTATGAAACGAATTTTAAACTTTTTGACTGATGGCCCAAGATTGTGTAAACTATCAATTCAGCAAATTAATCGTGTATCAGAAAGTATGGTATCATTAAACAATGAAATGCCTAGTGAATTTGCTCGTCAACCTCGATCTCTAGAGTATGTAAAGCGATGGAAAGCAACTGAATTCAGGGAATTCCTTTTATATAGTGGTCTTGTTGTACTTAGAGGGATTGTAAAAAAAGAAGTCTATACACATTTTCTTTATCTTTCAATAGCTATTAGAATTATGCTGAGTTCAAATACTGAGTATCGAAATTCTCTGCTAGAATATGCTCATCATTTGCTTGTATATTTTGTTGCTACCGCAAAAAATATTTATGGCAATACATTTAATACATACAATGTTCATAATCTGATTCATCTCCACGATGACGTAGCTGGTTATCAACTACCACTTGATGCAATTTCTGCCTTTGCTTTTGAGAATCACCTCCAAATTTTAAAGAAGCTTGTTCGTAATGCCACAAATCCTGTAAGTCAAATTATCAAACGAACACACGAGTTGGAAAGTTGTGGAGCTATGTATTCCGAAAAATTTTTGTACACTAAAGTAAGTATAAAGCATAAAGATAGCTGGTTTTTACTAAAGTGTGGCAAATATGCTAATATAAAAGAAATTAATGACAACAATATTTATATGTGTGAAACCTATGGAAAAGATAGCATGGAAAGTTTCTTTCTTGAACCTTGTGATTCGAAGAATTTCAATATTGCACTTTTGCGAGAGAGAACCCTTAATCCTAGACGTACATTGATTGAAAGAGATGACATTTTATATAAAGTTGTTGTTATGCCATACGGTGATAATGGTCTTGTATTTATTCCTCTTGTTCACACTGTCAACTGGAAATAAAATAATATTCATTTTTGTTTCATTAATAATTATTTATTAAATTTAATTGTTTTGTTTCTTTTAATTCCACTAACTATTATTTAATAACCTTTTTCAGACAATTTGTTTCATTATTTTTGGCTAAAAATAAAATTGACTTAAACTTTTGCAACACTTGTGTTTTGAACAATAGCTTTTATTGAAAATA
>evgR009807c0g1_i01_Hydra
GTTATTTAGATAAAGATGTCTTAGTAATATTTGGTAACAAAAATTTAATTCATTGTGTTATTTTTTATTACGCCAGTTGCTTGAGAAACATTGGATAAAAATGCTACAAATAACTGAAATCCATGATATTAATTAAAATGATGGATAAAACGATCAACATAAAGGTTTACCGATCACCTCTTGTCGGATTTCAACCATTTGACGAAAGTTTAATGTTAAAACTTCGGATTAGAGCAAAATCTTTTCCTGAAGAACTCTTGCGGGAAAGTTTTAATACAAAAAACCCTAAAAACGGTTATCACCTTCCTCCTACCATAAACCAAAAACGTTTAATGAGTTGCGGTGAAATACAAACAGCATTAGTTACAATGGCAATTCAAGAAGGGAACGTTATTCAGCTTTGGAGACTTTTATCTCGAAAAGATTCCAACCCCAATCATGTTGACGGTTTTGGTCTGCAGCCAATACACTACGCGTGCATGTATGGTGAATTAGACATTTTAAAAATACTATTACAGCACAGCGTAGACGTAAATGCAAAAACTTTAACAGGTGAAAATGCCCTTGACATCGCCGTTAAAGAAGGAAACTTTGAAGCTGCTCAATATTTAATTAATAAAGGAGCATACGTAACCACTATTGTTAACGGAATTAAAGGTGTAACAAGTAAGTCTAAAGAAATCATTAAACCGGCTACCGGTCGATCACAAAGGTCAGTAAGTATGATGGGCTCCTTTAACTTATTATGAAAATATCCAAAAAAGAGAGATATTATTCATACTTAATTGAACAACATGTTTCAAAATTGCTTCATATTTGCAAATTATAAGTTTTGCAATCGATCGATGAAATATTGGTGTTTTCCCAAAATTTCTGATTTTTATTGACAATTTAATATATTATGTATAAAAAAATGATAAGCTTTTATTGAGATTTGTAAAATTTCATTTATTTTTCTTGAAATAGATTTTTTATTCATAAAAAAAA
>evgR009812c0g1_i01_Hydra
ATGCTGCAGCTCAAGTTCAAAATACTGTCAATTTTGTAAAGTCCAATGGAGCTACCTTTGGTGAAATCTGGCTTGATATTGAAGGTCCAGCTTATTGGACATCAAATTGTCAAACCAATGTCAACTTTATTTCAAGAATGGTTAATGCTGGAAGAGCTACTGGTCACCCAGTTCACATCTATTCATCAAATTCTCAATGGCAGCCAATTACTTGTGGATCTACTGCCTTCTCCAATTTGCAATTATGGTGGGCTTATTATGATGGAAGACCATCATTTGATGATTTCCAATCATTCGGAGGATGGACTCGTCCAAGCATTAAGCAATTCCAAGGTACCACCAATATTTGTGGTACTCAAATCGATCAAAACTTTTATTAAATGTTTGATGAAATCATCAATTAAATTATTTAATAAAGTAAAATTGCAT
>evgR009822c0g1_i01_Hydra
TTGTGGCATATTTTTTTATAATCCGCATGCCATCTTTAAAAAAAAATGAAACAACAGTTTTAATATTTGCTGAGGAGATCTCGTGGTAAATAAACTTATCGATCAGGTACTTGTTACTAAGTTAGAAAAGATAATCTACGGCGGCAAACAATAAAAAGAAACGGAAATTTAGTCCTTAAGTTGTAAGATAAAGATTACCAAATGTTAAAAGTAAAAAGCGGCTGCCTTGAGGAAGCGATAATGCATATTTGAGTTAATTAAGGTTTTGTTGATAATTTTCTTGTGAAATAAAAAATAACCAACAAAAAATTGATGGCAAACCACAAAAGGTTCTTGTTTTTGTTCATATGGTGCGTATTAATGAATAACTTGTGTGGCAGCTTGTTTAATGTTATAAAGATATGCGATAAAAGAAGAAGTTTAACAGAGAAGCAAATTAAATTATGCAATACTAACTTTGATCACATGGCCTATATTATAGAAGGCAGGAGAATTGCTCTTGAAGAATGCATGAAACAATTTAGTGGCAGAAGATGGAATTGTACTTTTCCGGAACCAGATGTTATTCCGTTTCTTCATCCTAACATGCCACTTGGAACTCGTGAGACGGCATTTGTTCATTCAATTATAGCCGCGGCAACTATGCATTCAATTAGCCGCGCATGCATGGAAAACAAATTATCGTCACACTGCTCTTGCAGCCAAGAAAAAAAGCCAGAAAACCTTCCTAAAACAGACATGTGGAACGGATGCGGTGATAACTTACCTTACGGTTATCAATTCAGCAAAGAGTTTGTTGATTCAAAAGAAAATATTTTAAAAGAGTCAGCTTCTAACTTTGGAAGAGTTTTAATGAATCTTCACAATAACGAAGCAGGAAGATGGGCAGTTTTTGAAAAGTCAAAAATTCAATGTCGTTGTCATGGAGTCTCTAAAAATTGTGCCACGAAAACTTGTTACCGTCAACTTTCAGAATTTAAAGATGTAGGATATCAACTAGAGCAACTCCACCAAAGTGCTATACGTGTCGAGTTATCGCAAAGTGGTCAAAAAAACAATACAGAAATTACAAAACTTGTAGAGGTGAATTCAGATTTTGATAAATATACCTCAAAAGACATGGTTTATCTTGAAGATTCTCCTAGCTACTGTAAAAAAGATTTATCAATTGGTTCATTTGGAACGGAAGGTCGAGAATGCTCCAAAGAAAATATAATTTTAAATCAGTGCAATCATTTGTGTTGTGAAAGAGGACATCATACTAAAAAAGTGTTTACAACGGAAAATTGCGCTTGTAAGTTTATATGGTGCTGCGAAGTTAAATGTCAAAGCTGTAAAAAAGAACAAAACATACATTATTGCAAATAGAAATTGGTGCAAGATAGTTGCATAGTTATTGCTACGGATAGTTGCAAAATGGTTGCAAATTAGTTTGATTTTTTCAAATAAAATCGTTAAAAAAAACTGCACACAGAATAACTCGTCAGCAATGTTATTTTGAACCACAAATAGAATATAAACTCAGGCGAATTAACACGAGTTTAATACATTGATTTGTGTTAAGTAATCGAGTTGTGTTAAGTATCTTCGAGTAGCTTATTCGAGTAGCTAAATAACGCGTGTTTACGCTAATTTCGCAATTATACTATGATATTTCTCAATAAATTATTATTAACTGTGTATATGTGTTACATAAAACCAAAACATTTTTAAAAAGATGATAAAATTAAATTTATTTATTAGCCAAATATATATGAATAGCAAAAGTCTAAGATTTTCTTTGTTCTTTGTAGCTAAAATACACATTTTTTGTAACCAGCACAAACTTAAGTATCTATGGACTATTAAATATATATTTCCAAATAAAT
>evgR009832c0g2_i01_Hydra
ATTGTGCTATTCGATGTACAAATTCCAGAAATGGTCCACATCTAATGAATTAGATAATAAAAGATTTTTACTTTGTACTGATGGTGAAAATGAATATTGGGAAAAGCTTTTGCTGGATTATTGTTGTGTCAGAGATGACAATGATAAAGTATTTTGCTATAAATTAGAACAATCAAACATTTTGACTTTAGAAGAGCAATTTTATATTTTAACATCTATTCCAAATGATTATTTCTGGAGTTGGAGTAATCAGATATCCACTAACATAAAGATTGAGCAAAGATTACCAACTTTTGTTAGATATTTTCCTGCTTCCCCAAGAAATAGAATCATACAAGCATGTTGTTATTGTGAAGATCCTGGAAAAATAGTTTCAGCAGTGTTAGAGGTTCTCAAGGGTAAAAATGATGTAACTATCTCATATGCTAAGGAAAAAAATGTCTTAGAGGTTTTGGTGGTTAACATTGATGCTGCTGTAGCATGTGATAAGAAACTGCTGCTGTAGCATGTGATAAGAAACAATGATAATTAATGTTAACATTTAACTACAACCTATAGAAATACCACAACAATAAATATTACAACAATGTAATATTTGTACAACAGGAAACAGGAATGTCAATCTATCTGTGCGATCAGTATTTTGTGTCCAACCTTCTTCTGCAAATTTTTTTTCACGCGAAAAAGAACTGAATGAAATACACTGTTATTTGTTTGAACCAAAGGATTCAAAAAAACCTTTAGTACAATG
>evgR009834c0g1_i01_Hydra
CAAAAGAAGTTGTTCTTAATTACAGATTTTATGGAAGTGAAAAAACATTCACAACATATCCCATACTATCTGCTGGTTTAGGTGGATTTCAAAACTTCACTCATGGCAATGGATTTCAAATCCATTTGGTTGGTAATAAAATTGCTGCTACAGTTGCGACCATGTACGGAAGATACAGTGTTGCAATGGAAAAGCCATCAGTTACTCCATATACAATAACCCTTGTTTGGTCTAAGCAAAACAATAGTTTAAAGTTATATTATAATAGTGAATTAAAAGATTTTGGAGGTTTGAAAAACATAAGTTACCAAGACTACGTCCTATTTAATGAGTATATTCTTGTTTTTGCTGACTATTACAACATTGAATCTGAACTTCAAGCAAGTCTATTAAAGATACAAATGTGGAGCATACCTTTATTGGAGGATGAAATTTCTCAAACTTTATTAAATATTATCCCCAGTGGTTCTAGTTATCCAGGGTCTTTTCTACCATCAGGATTTACATATGGATCAGACTGTTATTGCTCTAATCAAGTGTGTCCCTGTGAAACTCTTGAGTCATCATCAACAGTAGAAATCATGAGCAAACGTGCATCATATGCAATAGTATCTGAAGATGATCTCGTTGAAGAAGACATTTATTTTAATATTTTAGAATATACCGATTATTCTTTTCAAATTTATTTTGATAGTATTGCTTTTGATCTGAAAACCGTTCAAATAGAAAAAGTAAATTTGATTTCCACATTGATTTCTCCAATATATCGATGTTATAGTATTGCTATGTTCCTCTCGAACAGCTCAGATAAAATGAAAATGGAATGGTGCGTTTCTAGAAACACCACTAGTGTCTTCACAAGTCCAAATCTTGCTTCTTTGTTAACTCAACTACTTAAGATGTCATACACACCTCCCGTCATAGTTATATTTAACTTTGATTTTTTATTTCTTAGTCCTGAGAGTATTTTAAGAATTGTTTACAAAACATCAAAAATAGGTGACTTTTTTCATGCAAGTGCATCACAGGTTAGTACAGATGATGTCATGGAAACACTGACGTTAAATGTTACAGGAGCGTGTATCTTTTTTAAATTTAAATTTGGCGGAACTGGTTTTGTCAGCCTTGTTGTTATTGGCCATGGAATGAATGTCAAAAAGAGACTGGCATATTTTTATCGACAAGAGGCACAAGATAAATGGCTTGGCAACTATCTCGATTTGAGTTCAAACAAATTTGTTGTGGTTGAGTTTGTTGCTTTTAAAAATGGACTTGGAAGTGGTTATATTGCTTTTGAGAGTATTCAATTTTTTCAGAGTTGTCCGATTAGAAAAACAAAATCAGAAGTTATTACCTCTTCATTTTTAAACCCTGTTGTGATGGTAGCAGACACATTTGATTATTACACAACTTTGAAAGATGTGTCGATTATTCAAGGTTTAAGTGGC
>evgR009852c0g1_i01_Hydra
AATTAATTTTACATGAGCTCAGACAATATGAGATTTGTAATCTGAATAAAATTAATAGATTTAGAATTTAATAATGAAAAAGTCTGTAGATAAATGGAAATTAGGCGTTTTGAAAGTTAAAGCGAAGGAGTCTGCTTCTATTTTATCTAATATTAACAATGGAGAAGAGGCAACATTGAACGGAGTTTTAGAAGAAACAAAAGATATGGAGATTGCAAGTTACTTAACTAGTTTAAAGAAGAAAACAAGTTTGAAAAACTTATGGAGTGGTCAAACATTTTACGATATAGTTGATGGCGATCAAACACCAGATATTTCTAGCGAATCTGACTCAGAACTAAATTTGAAAATGAGCAAAAATGTCCATGAGTTAGGTAATCTCCATTTAGATAGTGACGAAGATGAAATGCGAATTAATTACATACGTGAGTTGAAGGAAATTGATGAAAACAACAATATTGTATCTTCTTCTGAAAGCTCTTATAATGTCATGGGTAATATACATTATGCTCAACTTGAATTGGATGATGTTCAATTAAGTGCAGATAATAAACTTGAGAACACAAACGATTCTCTTGATTTTCTTCCCGTTCTTCATAATATACGTGAAGAAAACCCTAAGACTTATAAAAAAAATCCTGTGTCTAATGAAGATAGTATTTCAATAGGTAGTGCTTCTGAAATATCAGAGGTAATTTCAAATTCACAAGAAGATAAAAAGAAAAAAGTGTCTTACACATTGACTACAGAAAATGTATCAGATATAAACTTTACCCAAAGTTTGAATCATTCGGTCTCTGATTTATCTCATTCTAAATCTTCTATTCATACCTTTAAAAAAAAGAATGCAAAAACAGATTTTTCAAAATCTTTCATATCTGGAGATACATTTTCTGAGGGAGAACAAAAAATAACAGATACATCTTTGTCAAAAACTTCGAGCAAGAAGAAATCAAAGCCCGCTATGTTACAAACAAGCAGAACAAATTCAGAAGTTTCAAATACACTTGACTCAGATCGTCATTATAGTTCACCCTCTTCAAAATCATCATCACATACACTTAATTCGAGCATTTCTCATAAGTCAGTTTCAAACTTGTCATCTACCAGATCAAAAATCTCTAAAAAAAAGAAAAGTCGTTCCCAAAGTGGGCATTCTGGAAATGAAGAAATGAGCCGGAAAAACATGAATACTCAGACTGATTTTAGGTCTGATGATAAAACAAAAGAGGTCTTGAATTATCTTTATAGTTGGCAATTGCCCCAGCCTAGTAATATAGCTTCCCATGTAATGGATGCGGATACCATTGAACTATACAGTAATTTAAATCCGTCCATGCTTGCGGCAAATGATATGATGAAGTATCACTTACATTTAGTTCGCATGCATTTAAGTAATAGTCGGCGCCTTTATGAGTCTTACGCAAATGAAAGTATGAACAACAATTATAATTATACATCTGTTGAAGAAACAATGGCTTTTATTAATCAAAACAGCCCTAAAGTTATTTCCTATGAACAAGCGCTGAAGCGTGTTAAATCAACTCATTGCAAATAACTTATACATATACATTATTTTTTTTAATGTAAAAGTTGTTTTATTTATTAAAAACCCGGATTTTAAACGATCAGTAAACGAACCTAAAACGAAGCTGACTATAGTTTTTAACTTTACACGTATGATTTCCAGTTGGATAACTATGAGGGTACGGATTTTTCGTATTGAACTACAAAATCAAAGCGGTTGAGGTTAGTAACTAAAAAGTCCTCAATAAATTAAATCACACTCTAGGGGTAGTGGGGTTAGTGGTTTTGTGAGGTTGCGT
>evgR009853c0g1_i01_Hydra
AGAAAGTCCAAAGAACAAATATTTATTGATTCACTCTTTGAAGGAAGTTATTACTCAATCAGAATCTACAAAGCTCAAGCCATTCATTTCTGAAATTACACCATTGTTGTTCGATAACTCCCAAGAAGAAGAAGAAGGTGTCAGAAATGTTGTTTCTGAATGTTTGGGTAAGCTTGCTGTCAATGATTATGAAAATGTTGTTTCTCAATTGAAGACCAAGTTGACTCAATCTGATGCCACAAAGTCAACTGTTATTGGTGCTATCAAATATGCTATTACTGATGAAGCTTCACCAATTGATGACATGTTGAGATCTGATCTCCCAGTTTT
>evgR009859c0g2_i01_Hydra
GATTTAAAATTTCGTACTGTTAATAACGAACAGATACATGCAAACTATACTACAATACAAAAAGGAGATAAAATCAACTTAATGATTGACAGGAAAGGTCGAGTTGCAGTTAGTGTTGCATTTAACGGCAACATGCAGATTATTCAAACAGCTTTAGAAAACTACCACGATGGGCAGTGGCACAATGCATCCTTTGAAATTGACAATGTTCCTACAACTGATGGTACTTATATTATCAAGTTTACGGTTGATGGAAAAACAAGACTTTCCACTCTAAACAACTTGTTTATCTTTGATGGATATATAGATATCGGATTTGGTTTCACTGGTTGCATGAGAGATATAAAAATAAACGATGATGATTTTAAAAGTGTACGGAGAGATTTGAAAGATTCTAAAGAACTTTTTACTTATAACGATGAAGGTGTTGTCTATAACTCGTGCATGTTAAAAGATTATTGCAATCCAAATCCATGTCAAAATGGCGCAAGCTGCAATCAGACTGAAGATAACATAAGATGCGATTGTCAAAACACACTTTATGAAGGTTCTACATGTCACAGGCCTCGTTTTAATTACACATGTGCTGGAGTACGAAAAACTGGAGAAAAAAGAAGCGACGTTTATTATATCGATTTTGATGGCATGGGACCAATGCAACCTGTTCTGGCTCTTTGTATCCTTGGTAGAACCACTGAAACTAGTATAACTCAAATAAACAACACAAATTTCAATGATAAAGACGTCACTACTCCATCAGGAAAAAATAACTTTAACATTGATTATGCCGCAAACATAATCCAGATGAGAGAACTAATTCTGAATTCTGCTTATTGTGAACAGTATATAAAGTATACGTGTTTAGGAACTCCGATATTTAGGTCGCCAGAAGGTCCACCTTCTGTTCGTTGGATGGGGGGTGATGACATTCTTCATTACTATTGGGGAGGAGCGAAAAACAACCGCGGATATTGTGCATGTGGTCTTGATCAAACTTGTTATAATGTAGGAAAATATTGCAATTGTGACGACTTAAATGCTTACCAAAGTCTTGTTGATGATGGTAATTTAACAATAAAAGAACATCTTCCAGTTCGTCAAGTTCAATTTGATCAAGTTTTAAAAGGCTCAGTCACTGCTAATCTACGAGTTGATCATTTAAGATGTACTGGATTTGGTGCAAGAGAAGTAGCAGCAACTTTTAGAAAGCCCTATTCTTTTTTATCAGTCTATTTTCCTGATCAGCCATTTGATAACATATACGCGGGGGGTGTATCGTTTGAATTCAAAACCTCTGTTGCTTATAATTACTTGACTTTGGTAAATGCTCACGGACCATTTTCAGGAGATTATCTAAAGATTATGATATGGTCTAGAACCCTAATAAGAGTACTCTTAAACTTTGGTTTTGGAGAAATTAAACAAGACGTTGACATTGAAAAAATTGGAAGATATATTGATGACAATGAATGGCATGAGTTTGATTTAATGTTCAATATGAAAGAAGC
>evgR009866c0g1_i01_Hydra
AAAATTCTCAAAATGAGCGCTATCAAAAAGGCTGAAACCACTCCAGTTGAACAAACCAAGGAACCAATCCACAGAATCAGAATCACTTTGACTGCTAAGAACGTCAAGTCTCTCGAAAAGGTCTGTGCTGACTTCGTCAGAGAAGCCAAGAACAAGAAGGTTAAGAGAGTCAAGGGTCCAGTTAGAATGCCAACAAAGATTTTGAAGATCACTACCAGAAAATCACCATGTGGTGAAGGTACCAACACCTGGGACAAGTTCGAAATGAGCGTCCACAAGAGAGTCATTGATTTGCACGCTCCAGCCTCAACTGTTAAGGATATTACTTCCATCTCCATGGAACCAAACGTTGAAATTGAAATTACTGTCTGTG
>evgR009869c0g1_i01_Hydra
AATTATGTTAATTTTATGAGGAGCAGTAGAATGTTTATGATACAAACTGAGGATCAATATTTGTTTGTATATCGGGCTGTTTATGAATCAATTACTTGCGGTATTACTGAAACAATGGCTTCTGAATTTCCACAACTCATCACAGAACTATGTAAAGTGAAAAATTCAAATTTAGGTTCTACTGACGGTTTTGAAGAACAATTTGAGAAGTTGAAGATGTTTACTTCAAGCTACGTCTCTGAGGAATTTAGTAATGCACTCAAAGATGTCAACAAAGTAAAGAATTTTAATCAGGAAATATTACCAAGTGATGAAAAACGTGTATTATTAATGGAGAGTACCTATGTAGAAGGCGGA
>evgR009870c0g1_i01_Hydra
AAACAGTTTTCTTTGTTTTCTTTTTCGTTTTGAAATATTTGATCAATATTTGTGGCCTAGTTTATATTTGGGTAAAAAATTTAGTCACCTTTGGCATTTTATGGAATCGATTAGAAAAGGAGAAGAAATATCTTATTTTAACGATTACCATAATGATGTTAGGAAAGCTGTGCCAGTAGAGAGGTTACTTAAATTTGATGTTGCTGAAGGTTGGGAACCACTATGTGCGTTCTTAAAAGTTCCGATTCCAAACTGCAATTTTCCTCATATAAATGAATCGAAAAACTTTCAAGCAATGATTGTAACCTTAAAATTACGATCCTGGATTTTACTAGCGGAACTAATAACTTTATCCCTATGCATTGGTGGATTTGTTTATATTTCAAAAT
>evgR009877c0g1_i04_Hydra
GGTGTTTCTGCAGCGTGGTTGCTGTTGCGTTTAATAATTTTAAGTCGACATTCCAGCGTATTCTTGACAATTCTCCAACACACACACACGCACACAAGCAAAAGTTAATAGGATATAAAAGCTTATCACTTTCTAATTTTTTGGTTTTCTTCATTTGTCAGACGATCCCTCCTTTTCTATTTTTTGGTTTTCTTCATTTGTCAGACTATCCCTCCTTTTCTACTTTTTGGTTTTCTTCATTTGTCCGACTATCCCTCCTTTTCATTTTTTCAAACAATAGATCTATACAATGTAATGTTGAGATTTATTCG
>evgR009893c0g1_i01_Hydra
ATGGTGCCAGATAATATGAAAACGGAATAAGTTAAAATGGAAAAAGAATTTGAGTTAGAGTTATTTGAAAATGATATTGAAATGGAAGAGGATGAACTACAACAAGAACTAAAGTTGCTAGAAAGGGAAAGCTTGTTTAGTGATTTAAAAAAAAAACGAAACATGTGACAATTTAGATTTTATGACAAAAAAAGATGAAATAACTGAAAAAGATAATAAAGAAAAGAAATCAACAAAAAAATCTGTTTTACATTCTAGTTCAACTTCGAATGGAAGATCTTCTTCTAAACAAAATCAAACTGTTAATCATAAAGTAAACTTTAAAAGACTTTTTAAACCACTTCATTTAGATAAAAGTTCAGTTTATTCTATTGCTGGCCGTGAGTTAGCATTTAGACTAGAAGAGAATAAAGCTGGTTTACTTGGTCATGTATTTAGAGTTTTAGGAATAACATTAACTATGGAGATATTTGAGGAAACTAAAAAAATTATGAAAAGTGGCGGACTTAAAACTGATTCTGGAGATAGACGTCGTTCCCCTGGCGGAACCTTTTTATATCTGATGAAAAGTCGTGGTTACGCTACTAAATCACAGTTAAAAGAAATTTTTAAAGAAGAAGAAGCAATTAAAAAGAAATTGCTCAAACAAAAATATAAACAAGCTAGCATTATTAAAAAAAATAAAGAAAAAGTTGAGTAAATGTGATTTGAAAGAAAAAAATTGTAAAAAAAAAAGTTTTTTATATTATATACTCATTAAAGTTGGAAATAAAAATGACAGTTAAAAAA
>evgR009894c0g1_i01_Hydra
CCAATCTGCAAGGAATACTAAATATATATATAGTGTCTTCTTTGCTAATTTGCACGTCATTTTTAAATCTGAATCTTTTGCATCATTTTGATAATTACGTTAATACGAACAGAGATCTCAGACAACTAGTAAAACTAGGAAATAAAAACTTTTTAATTGATTTCCAAGCGTTTCAGCATCATTTTGACAATTGGTTTTCAAGCGTTTCAGCATCATTTCGACAATTGATTTCCAAGCGTTTCAGCATCATTTCGACAATTGGTTTTCAAGCGTATCAGCATCATTTCGACAATTGGTTTTCAAGCGTTTCAGCATCATTTCGACAATTGATTTCCAAGCGTTTCAGCATCA
>evgR009922c0g1_i01_Hydra
AATTAATGAATATCAACAATCGTTTTGGTAATGGTACTGGGAAGAAATGGATCTATAATTCTAGGAGTTATAATTGGAACATTGGTTTTGGTCATTACAGGAGTTTTAATATTTTTTGTTCGAAAAAGAATATCTAAAAAAGCTACGTGTCAAAATAGCGTTGTTGAAATAACGGTTAGTCCGCCCACAGTTGATTTTGGAAATACACCTAATGGTAATAAAAACGTTAAAGTTACTGATGTCACTATTCAATATGACGACAGCTTGGGAAAAAATAATCCTTTTATTGAATCAGTCGACTTAAGTGTTGAAATACAAAGACACTTTAATACACTTTCAAGATCACCTGGTAGAACGAGG
>evgR009951c0g2_i01_Hydra
AGAAAGCTTAGAAAAAAGGAGGCAGATAGGGCTCCATTCGATGTTGGGAAACCCGTCGAGGAACAAGAAGCAACGAAGCAAAAGCTGCCTTGGAAAGAGCAAAAGGGAATACGGCCATCTAAAGAAGAAGCAGAAAAGGGCACTGCTGTGGGTTATCAGTTAGGACCAGAATCTGAAAGTCAAGACCGTAGACAGGGTCAGAGAGGAGAACAGCAGCAGCAACTATCAGCTCAAAGTGATGGAAGCGAAAAGCAAACTCCACCTTATGAGGAAATATCTCTTCAAGGTACCCAAGGAGAATCAGAGCGTCAGCAGCAGGCAAGATCTGAGGCCGAGCAAGTACGAAAAGTAAACGCCGGAGAGCCGGGTGCAGATAGGAC
>evgR009957c0g1_i01_Hydra
AAAATATATTTTGATGTTTTTAAGATGTATATTAGTATACTCCTAGATAAAACACATAATATAATTGAAAGTTCACTGTTCATTAGAAGTTTGATATCAGGTTTTCATCATGGATACTCTATGACGGAGAAGAATCAATATCCCTGGGTGAAAATTAAAAACGGCCAATCCTATCAAGAATTGTTCAAAGCTATTGAAAGTCAATTTTCTGATTTCAAAAAACCAGAGTTATTTTTTCGCGCTTTGTCACTAATGCTCATCTTTTATAATGATGAAGATTTAGTTCGTTTTGCTCCATTCAATTTTATTACATTTGTATTTGAAAATTTGTCATTGGAAAGTTACGATTGGTGTTACAGCAATATATTGAAAGAAGTTGTACATTGTATTCCTGAAAAGTATTTAAGGTTTATTCAATCTGCTCTAAAATCTCTGATGTGTAGCTGGTTCGAAGGTTACTACAGTTTTATGAGGAATGATGCTAAAGCTCATGTTTGGTGTTATGCATTGTTAGTACTGCATGTTATCACTGATGAATTACAAAAGAGTTTTCAAAACAACCTAGAGAATATAACTGCCTCCAACTGTGGTTTACCAAATTTCATAAATGAAAGATTTAGCAGAATTTATGAGTTAACCATGAATAATTTACAAAAAATTAAGGGAAGTTTAAAGAAAGATGCTTTACTTCAGAGAGGTGTTGCTTACCTTATGGAATTGAAATATTTTGCGAATGCATTAAAAGAAGGAGCTTTTGGTGAACGTCTTCTGATTAACATGATGACTTACCATATAAAGTCGTATAAATCTCCAGATTATATGATTAATCACTGTGAAAAATTCTTAGATGAATTGCTAAAATTGAATTTCATTGAAAGGGAGTGCAATAGATGTCATTTAGTTGAAGCCCTCATAAATATGTTAATTGTGTCATTGAGCAAGTTAAGTATTAATGATAATGTTTGCTGCTTCTTTTTTAAAATGCTTAATTGTATTAATGAATTATACAAAATGATTGATCCGTTTGATAAGTTCTTGAAAAAGCTTAATGAAATGGTTCAAAAGCAAATACCAAATGTTATAACTAAACTTCGATTTTCATTTGGTGAAAAACAGAGCTCATGGAATAAGCTGGTTAATTTGGAATCACAGCTGAAGGTTATAGAAATGCTTGTGGATTTAGAAATTTCAAAAGGAAACGAATTTAAAGACCTTGCACTTGATATTTTGAAAAGTAAAATTATAAATGCTGAAAATACTCTTTTGTTGGAAGCTTATTGTAAATTAGATATTTTATCATACAAAGAAATAATCCAGAAACCATTAACTGATTCATTTTTTTATGTTATTGAAGTAATGCCAAAGAATCAGTTTAATTGGTGTTACAGATTATGTCAACGCTGTAGTCCACTTTTAATCAAATACATCATGATGAAATTGAATTGTTTAACCTCTATAAATGAAAAAGTGAATTTTTGTCTTACATGGTCACCTTTCAAATTAATATTGGATTACCTTGTAAAAAAATCTGTAGAAGAAACAAAACATCTTTTTCAAGAAATTATATCACTCTTTGACTATACTTATGAAATTATTAAAAATGGAGAAACTACAATGGGATTTTTGAATAACCTAGTGAACAAAGAAGAAGTTTTATTTGGTCTTATGAAAGTATTAAAATGGGAATTTTTTGATAATTTTAAAGAAATATATGGTTTACGATTCAAACAACTATGTTGTTTTTATAAAGTTAAAGAGGTTGTGAACTATACTCATCAAATTTGCAAAATGTTACTTCAAAACACAGTTAACACTGAAATGTTAGACGATGGAAATCACTTAAATTGTTCTGCTATAACTTTTAAAGAAGTTTTTGTGAATGATACACCTGATGGAATTTCAAAGACATTGGATTTTTTAGTTAGTAATGAATCTGATTGTCTGAATGTGTCAAATGATAAACTCAAAAGTGTGCATGTACAAAAAGTGTGCATGTACTTTAGAATTTCATTGGATTGGTTTGAATATGCT
>evgR009968c0g1_i01_Hydra
CTCATTTGATAACTATTAATATAATTAAAATAAGTTGAGTAAAATGTACCCAAATCTCATATTGGAAAGAAACCCAAGGATTCAAAACGCAAGGTTAATATAAAGTTATACCATGAAAATCAGCGCTTAAGTTTATGCGGTCTTCATTGCATCAATTCTGCATTACAAAGTTCTGTTTTTACCCAAGACGATTTAAACCGAGTTTCAGACGATCTTTTTTAAGAGAAAAGTCAATATGTTATACTGAAAAATTAGAGCCAAACCCATATCGTGACAAAACTGGAAACTATTCTGTTGATGTTATCAGAGAAGCCCTTATCTCCAAGAACATTGAAATGCACAGAATTGATGTATCGTATTCATCAACTGCTGGAAATGGTTACATCATACAATGCTGCAATCACTGGGTTACCTTCAGAGAAATTGGCAATCAATGGTTCCATTTTAATTCCAAATTTCTTCAACTAAAGCCGATATACAATTTTTACTAGTATATATTATATATTACCTACTAGTATATATATATACCT
>evgR009969c0g1_i01_Hydra
GCTGCATCTGGTTTTAGTGATACATGCGGCATTGGTTTGTTTTGCATCAGTTAAAGATTTTAAATTTCGGCATGGAGTATTATTGTCAAGTTGAGTTTTGCAGAAAAGGGTTTTATAATGCACGAGGGTTAATCAATCACATGATGTTGCAGCATGGCCATGAAAAACATTTAAATTTGTCGTGTAGCTTAGACGGTTGTGTTTATTCCTATAACACTGTTAATACATATAGATGTCATGTTTTAAAGTTTCACCAATCTCATCTAGTTTCCATAGCTTCCGATAAACTGAATTTATCTAGAGCTGGAGAAGAAAACTTTGAAAATATTGTGGAAATGGAAGCAGAGAACATAGCAGTTATTGAAAGTCTGGATGAAAATGCTAAAAAATTAATGACAGAATTTTCTGATTTTTTAGAATGTTTTGGAAACAATTTATGTATAACACAGCTGCAGTTGCGCGAAAAACATATGCTGCCTCTTTCAACAGCTTCT
>evgR009978c0g1_i01_Hydra
AACAAGAAATGGAAGAAGAACGTAAAAAGAAGGATGAGTATGACAGAATTCAAAGACAAAAGGATGAAGAATACTATCAAAGACAAAAGGAAAAGCAGGCAGAAATTGAAAAGAGACTCCAAACTCAAACACCAAAAAAGCAAGACAGATGGGGTGGTTTGGAAGGTGAAACTTCTCCTTTTGGTGCCAAAAAATCTGGCTATGTTCCTCCACACTTAAGACGCACAGAAACTGATTCATCATCAACTGAACAAAAGACTGAGAATAAGCCTCCACAACAAAGAAGCAATTTTGGTTCCGATGAGACCTTTAATGCTTTTGGTGGCAAGAAGCCAGCTGCAGGTGGTAGCACAGGAGGAAGAGGTGGATTTGGCAATAATAGTGGCTCTGATAATAGTAACAGATGGAGAAG
>evgR009986c0g1_i02_Hydra
TATAATAATTCATCGTGTATTTGCTCGAAATGTCATGGATCGCGTTACAAATCCAACAGAAAGCTGAGAGGTATGTGAAAGAAAGCCGAAGCTACATTTTACACGACTCCGAACTCACATCTCACTAAACATGGCGTCTGCCGTCAGCAATGCAATTCGCACTTCTATTATTAAAAAACTTACGGAAGCGCTAAAGCCATTGCATCTAGATGTCATTAATGAGAGTTTCATGCATAATGTTCCAAAAGGTGCTGAAACTCATTTCAAAGTTGTTGTGGTATCCAATGAATTTGAGGGGAAGCCTCTAATTCAGAGGCATCGTCTTGTTAACGATTTACTTAAACAAGAATTTTTAGGCGGAGTTCATGCTTTGTCAATTCAGGCAAAAACACCTGAGCAATGGGAACAGAACTGCAGTGTGTCGTTATCTCCTCCTTGTTTGGGTGGAAGCAAAAAATAAAATAAAGGAATCTTTAGATTAATAAATAATAAATTTCTTATAGTCTTCTTGTAATCTTTTTTACAGTAGTTTTTTTTTCATAATTATAATATCTTTTGGAAAAAAGAAATTTAGTAACAAAAATATATGTAGGTGCGTTCATATATTTTTTCATTTTTCAGTTAATTTTTTTTATTTTTTCAATTTGTTGAAATTTTGTATTTTAGTATTTATTAGTTTGTGTTATTTTATATATTCAGTAGTTTTAAC
>evgR009990c0g1_i01_Hydra
GGATGGATGTTGGGAGATGATAATGCAACCTGTGTCTATTACTTTATTGGAAATGTTTTTGTTCCAATATCAAAAGGAAATGGCATTGCTATAATACCAAAACTAAATCATGAGTATTTAATATCTTTAGAAATTCAGATAAATTCAAATGTTGCGTCTTTTAACTCTAAGTCTGTCATTCAGTTCACAAACTGTAATAGTAATCAAGATATTTGTGACACTGTTTTAAAGATTTATGCTATATTGGAGCTGCTCTGGGTTTCTACTGATAAGGCTAAAGTTTCTCAGATAATAAGTAATTCATTGTACA
>evgR009999c0g1_i01_Hydra
GTCTCCGCTTTATGAAGAAAATGAAATAAAACAATGGCTTTCTAAAGCTTATTGCGATTTAATTATGAATAATTTAGATAAAGACTTTTCGACTTTTTATATAAATGATACCAGTAAAGCGTTTACTTTTAATGGTATTGATACTTACTCTGATAGTTATAAAAGCGAGTTACTTCAAAAATTGTTTCATTATGACACAAAGAGTGTAATGTCTATTTCTTCTTTGGAAACTAATTTGGACATTATAAACGATTTACTGGAATCTAAAGAGTTGGGTCCAATAGTCATATATCAAAACAAAAGAGGGTTGAAAAATGACAGTCTAAGAATTACAGCTGGTTGTGTTTTAGGTTGTGCTAAAAAAAGCATTATCAATTTTGCAACTGTTTTGAGACAAGCTTTAACGACTATAAATGCGAGAATGGAATTAATAATGACTGACACTGACTCACTTTTAGTGGCAATTTATGGTAAAGATTTTGCAAAAGTAGTAGACGAAACATTAAAAAATAACGAAAGCTTTAATAATCTAATGGATTATTCAAATTATCTCAAGACTCATCCTTACTTTG
>evgR010007c0g1_i01_Hydra
CAAATTTTGGAAGCTGATTGTGACATTTTGGTCCCAGCTGCTTTGGAAGGACAAATTTCTTTGAAGAATGCTTCAAAGATCCAAGCTAAGATCATTGCCGAAGGTGCCAACGGACCAACTACTCCAGGTGCTTCCGAAATTTTGGAAAAGCAAGGTGTTATCATTATTCCAGACTTGTTCTGTAATGCTGGTGGTGTCACTGTTTCCTATTTCGAATGGTTGAAGAATTTAGGTCACGTCCAATTCGGTCGTTTGACAAAGAAGGCTGAAGAAAAGGGTAAGAGAGAAATGGTTCGTCACTTGCAATTATTGACTGGTCGTGAATTGAAG
>evgR010009c0g1_i01_Hydra
ATAGTCATTGCCATCCAAATATAATATGTCTAAATAGACAGAAATCTTTTCAAGCAGAACTTATTTGTCTTGAAGAAGAAACATGTGCAGAAGAAGAGACCGTTTCTATTGGTAGTTATAGTTTTACTCACATTAAAAAAGTAACACTTTCAGTAGATGATAAAAAATTTTTGATGAATGGTGGTTGGCTAAATGACACAATTATTTACGCTGGAATGCAAATGCTCAAAAATTCTTTTCCTCATATATCAGGTTTTCAAGATCCCATTTTACAGGTTACAAGTAGTTTTGAGGTTCAAAAGGATAAAGAATTTATCCAGTGTCTGAATTACAGCGGTATGCATTGGATCACCATAGCAACACTAGGCTGTAAACAGGGGACTGTCAGGGTTTATGACAGCCTGCATAATAAGTTGACATTGTTGCTAAAGAACACAGTTGCTGATCTTTTGTGCTCTACCACTAGCAAAATTAAAATTGAATATGTAAATGTTCAGTATCAGTTGGGTTCAGATGACTGTGGACTTTTTGCACTTGCAAACGCTTGTGAACTTTGCTTTGAAGGAGATCCATCTATTGTAAAGTACACACAAAAAGATATGAGAAATCATTTGTTAAAGTCGTTTGAAAAATTAGAGATGACACAGTTTCCGTCTAAGCTGCGAAGAGTTACACAACATATTTCAAAGGTAGAATATTTAGACATATATTGCAGATGCAGAATGCCTAGCAATGAAAAGTCAATGATACAATGTGGCTTATGCAAGGAATGGTATCATATAGAATGTGTTGAAGCGCCAAAAACAGTTATAAGTAATAATCTTTTATGGTGTTGCGGTTGTAAATAACTTAATTTTTG
>evgR010022c0g1_i01_Hydra
TGTTATCTGAAAACGTTATTATTTTGAAATTGGTTAGAGTAATTCAAGGTCGAGATGTTCTTTTTTGGCCTTATTGTTTCAGCAACAATATGTTTTTTTGAAGGTTATAACTTAGAACTATCTCATAGTAACTATTACTCAGAACCACTTGTTAAAAGAGATGTTCTCCAACCAGCAGATGGCTTACCTTCATTTGCTGCATTGTTTGGAAAAAGTTCTCCAGATGGTGCTACGTCTTACGAAGTTATGGGATGCACTATTAAAAATACAAAAAATGGTTGTCTTTGGACATTAATACCGGACAAAACCTGTTATGAGAAGTTATCACTTAGTAAAATATTTGAAATGCTTGCAAGTAAAACTGAGAAGTTTAAGAATTTGCAAAATTTGACAAAAAGCATTTTTTCTCCTGGTGCATTTGTTGTCAAGCAATTTGTTATATCTACATGTGGTGATTCAAAAGGTGATGTAAAATTTAGTTGTCAATCTGTTCAACCTTTTGATGTTATCAAAGATAAAGTCAAGATAGTTAACGGTGATCTCAGCTTGAAATTTAATTATATGGAGTTACAAAATGAATTTAGTTTTTCCAATATTGAGTTTGAGATAAAAGGAAATGTTGAAGTTGCTGGTCTCATTGTATCTGTTCTAGTGCAAAAAGATAAAAAAAGCTCTTCATCAACCCTTAAAATATTTGCAGAGGAAATTAAAGTTACTGATTTTTCAAGGTTTTTTTCAAAAACTAAAGAAAAGGCATCGACTGAAGTAGCAGCTGGGATTGAACATGATGAAGTTATTGTCAAAGACGGAGGAATGTTTTCGAAAATATTTATTGAAAAACCAGTAGTTAAAATTATAAGGAGTGAAGATGGAAGCTATGAGATCATTATAAGCGGTAAAGCTACTGGAATTACAGGACTAGGAGATATAAATGCCTTATTGATTGTTCAGAAGCCTTTTGATGGTCCTGTTGGAGTCGGTGTTGTTGCTTCTCTCAAAAAGGTTCAGCCTTTGAAAGTCATTTCTGCAATTGTTAGCAAAGATTTTAATGACATTGACATATTAAAAGATCTTGTCTTAGATATTTCTATTGAATATGCGAACATAGATATTCTAAAGTTAAAAGATACTGAAATTAAAGAACTGTTATCAGCTTCAATTCCAAGTCATTCTGCAACGCTCTCGGAAGGTACTATAATTAGTCTTAACATTCCTGTAAACAAAATCCTAAAGGATATTGGTTCTCCTGCTAACCTAAAT
>evgR010044c0g1_i01_Hydra
AACCTATTTTCTGTTAGAAAAGAGTTTAAATATGTTTTGTTATATCCTACTTGGGTTTGTTATATGCCTAAATATTGAAATGTTTTACGCAAGAACGTATATTGATGTCACCCAGGAAAGTCCACAATTTTCAAATGTAAAAGGTTGTGGTTGTGATGGCGATGAGTGTAAATGCTGCGTATTTTTAGATATACCAAAGATATATCTTAATGAAACAGGATGTGTTGTCATCAAATATTTAGGTGAAGAAGTAGGTGTTGATATGAAATTTACAGTCAATAATATGGTAATAATAGACAAAGAAATATCTTTGCGTAACCCTCCTCCTCTATGCGAAGAAGTTCCTTACTTTCGTGAATATGCCAGTCTCTGTATCAAAACGTATAATGTTACATGGAGCCAGCAGTTTGGAGGATGTTTTAAACTCGAAGCCAAGTTAATGCACGTTCTTGTAAAAGAGGTTGAGCTCGGTTGCTTTTACTTTCATGAAAAGAAAAATCGTACAGCGATTTCTAAGATATACCAGCTGTTCCGAAATTTTATGAGTGTGACTAAAAGAGTTTATAAGGAAGGAAGTCAAGGTATGGTACCGTTGATTTTGTAAATACGAACAGTAAAATTTGAATCTTCATGTATATAATATATAATTATAATATATAGTGTATAATTCTGTAAAAAAAAAAAA
>evgR010047c0g1_i01_Hydra
ATTCCTGACATTTCTGCTCATAACAATCATCTTATTGGTGATGTAGCAGGTCTTAATCAGCCTATATGCGAAGAACTCATTATTAAAATAAGTGAGTTGGTTAAAAAGGGTGTCAATTCTTTTTCTGAAATGAGACGTCATCTAGATGCCTTTATTCAATTTGAATTTAATTCAGATAACACTCTTAAAAAGACAAACAAACGTTTTTTCCCAAGGGATGATACCATTGCCAATCATATATTGAGAGCTAGACGAAAGCTTCGCAGATCCATGATAGATCAAGAATGTCTTATGGACAAGATTAATAAATGGAAAATATACTTTCAAGATGCAATGATCAAATTT
>evgR010055c0g1_i01_Hydra
ATACTCATCGGATATTTCAATTAGAACCTGTAGAATTGATGAAGTAATTGATCAAAAATGCTTAAATATAACAGAAGTATCTACTTTTAAAACATGCTTACGTCGGAGGTTTCATGGTAATCTTCCTAATTATACTAGTATCTTGAAAATTAAAGATGACGTAAAGAATTTATTGTCATTTGAATTAGTGGTAAGATCATATTGGCACAGCCTTCTCCACAACATAATTGGAGGGCATATGGAAACAACATATGCCTCATTTTCTCCAGAGTTCTGGTCTCATCATGCAATGTTAGATGCCATTTGGTATTTTTGGCAGCAAAAAGAGAAAGAACGTATTTTACAGAAATTTCCGGGCACTAATAATACTCTTATAGCATTTCAACCAATCGAAATTCGTGATAACTACATTGATTCTAAAAAACTAGGCGGATGCAAAATACAAGTGGCTTACGACGAAAAATCTATTCTACCGAAAAATATACGCGTTTTGAAAAATTGATGTGTTTTTTTGTTGACAAATGACATTTGCTGCAGC
>evgR010057c0g1_i01_Hydra
AATAAATGTACCATCAAACCACCTTGTTCTTGGACCAACGGGGCATGTACTAACACAAATGGCAGTTATTATTGTTCCTGTAACACAGGATGGACTCTAGGTCAAGATAAAATCAGTTGTGTTGATATTAATGAATGTTCTCAAACCAACCCATGTTTGAATGGAGTATGTACTAACACTTATGGTAGCTTTCTTTGCTCATGTAATGTTGGATGGGTTTTGGGTCTGGATAAAGTCAGTTGCATTGATGTAAATGAATGTTCTGGTAGCAATCCCCCTTGTTCTTCAATAAATGGAATATGTAGTAATACATATGGCAG
>evgR010082c0g1_i01_Hydra
TTAATATGTATGCGTCGGTGACTGGATCAAGGGCTATGATGGCTTCTCATATTTTATTTATCTTGATAACTATATTATTTACAATTCGTGAATTTCGCTTTTTATATCAAGCTGGATCGAAATACTTTGTAGAATTCTGGAACTTAGTGGAAGTTGTTATGATACTATTTTCTTTTATTGCTGTAGCACTTTTTTTATATAAAGATTATTTGGCCAGAGAGTTACTGCAACGGCTTCCAGATAAAAAACCACAGGTGTTTATCAACTTTCAGTTTGCATCTTACTGTGATATGATTTATGTAAACATTCTATCACTCATTGTTTTCTTCGTAACTTTAAAGTTCATCAAA
>evgR010086c0g1_i01_Hydra
GTAAGAAACAAAAGTGCCGCTGAATTTATGATTTTTGATAAAATATGAAAACATTATTAATTTTTGTTATTTTTTATATACTGTTGGTGTTACATAAATGCCTTCCACTACATTTTAATCATAAACCAAAAAAAAAACATGTTATTACTGGACATGTAAGAGATGTCATAAATCTTTTAAACAGTTCATCGACTAATAGAAACGAAACAATCGACAAAAGATTTTTTGGGAAACAATTAGGGTTTTCATATGATCGAGATAAGGAATCAAGGGAAAATAAACCCGCGGTACTAAAAAATTTAGAAAATCATCAAGGAGTTCACACGGGCTATCATGATGAGGAAAATTGGGATAATGATGAAGATCGACACCACTCTAAAAAATCACCAAGAAAAGATACTGTACGCGATGATGAAGAAAAAGGAGATAAATTATTTGGAGGAGAACGAGTAGATCACAACGAGCACACAAAGTTTTCAAACGATTTAATGTCAAAGTTAGGTTTTACATTTCATGGAGGTAGAGTCCACATGGATGAAGACGGGCATCTTGTTCAAGGTGATCCCAATAGCAGAGGCAATAGTCTAGTTGGTGGAAGAGGCGGAGATTATAGCAAAGGTGACGATGACGAAGATGAAAAGGATAGCGATGAGAGTGAAGAAAAAGAAGATATTAAAACGAAAAAACACAAAGACTACAAACAGACTAGTAAAGAAGACGGAGAAGATGAAGATGTGACGTCGCACAATCATATTGGACCATTCCCTTTTGCACCTTTACCTCAAGGTTATCATAAAGGTCGTGGTCCAGAAAATGAACCTGATGACGACGAAGAAGATAGAATTAAAGACGAAAATGAAGAAAAAGAAGACAAAGCATTGCGTAAAAAAGAAAAAGAAGAGGAAGAGGAAGAGGAAGACGAAGAAGAAAAAGATATTGAAAAGCCAGTTGCGCATCATCATAAAAAAAAGAAGAAAAACTACAGCGAGGCTGATGAAAATCCATATCATCACCAAGAGCATGGTTCTATTGCAGAAGAAGAAAGTGACAAAGAAAAAGTTGGCTTACCGCTGAAATTCAATAAAAAAGAAAACTCAAAAAAAATAGAAGAAGAAACGCAACAAAAAGACAACAAATCAGTATCAGAAGATAAAAAGTCACAAAATATAGTATTTGTTTTGAATGGAAAAGAAGGCTTAGAAGAAATCAAGAATATGAATGGATTTAATGATATTATAAAAAAGTTAACGGATAAATCAAAAAATAGCGAAAACACATCTAGTATCAACCATATTGTTACTGTTAATCAAGGAACTGCAATTGTAAACAATACAACATCTATTGGAACACCAAATTCAGTTGAAAACAAAGTTAATTTGCTGTCAACCCCTCAGCAAAATATAGTACCCCCAATATTTAAAGAAGCTAAGCAAGCAGCAAATGTTTCACCAGTTGAAAACAACAAACAGACAAACGTGTTACCTGGCTCAGAGTTTGTTTCCGTAACACCTGGAAACTTAGTTGCTTTAAACCAAATAAATAAAAATGTCTCATTAACAGCTATAAAGTTAGAATCGCCTCCGATCCAAACTAACATTGCACCTGCAACAAGTGATTCTACTGCAACACGTGATTCTACTGCAACACGTGATTCTACTAGTGGTTCCAATCTAGGTTTTTCGAAATTCGATTCCAATAATAACGGTGTTCGACAAAGTCTTTCATCGTTTGTTTCTGAAGTCTCTATGCCACTTGTTACTCAAAATTCTCACGCTAATTTTAACAATCAAGGAAATCTAGGAACCCAAAGTAACTTAGCTACCTTAGCTCACCTTGGTAATTTGGGTACTGAAGGTAAATATATATTAAGTAATGATCTCAGAACTAATTCACTAATGCAGCAAGAGGGAGAACAAATAGCTGCTAATATCGTTCACAATGATTTTCCATCAAGCATTAACCAGATTCAGAGTCCGGGTGAAAGTAATAATCTTATCTCACCAGTTAAAGCTTCCTTCTCAAAATCAGGCAATCTAATGAAGGAAAATGTTGATGTCTTGTCGCCTTTTCAGAATTATTTACTTTCTAACGAAAAAATGTATTTGACGCATAACAATAATGATGTAATGAAAGCAAGACCTTCTACAGAGCTAACCACGGACAAACTTACTTTACGGCACGAACAAACAGATAACGACGAGCAAAGTGAAAAATTAATTTCAAAAATTCTATACAGACTTCAAGAAGAAAAGTTGAAAAATCAACTTCATAGCGAAAATCAAGGTAACGGAGAAAGAGAGAAAGATTCCGATATGATCAGAAAGCCGACACCATTGAAAGAACCTAAATCTCTTACTGTTTCACAAGTAGAGGAGATGAAAGAAGATGAGAGCGAAAATCAAAAGGAATCTCTATCCAGATTTAAAACCCATAAACGTCTTGCTGATCCGAAAGAAGCAGAAACAGAGGCATTAGAAAACGATAAGGTTCTTCAAGGCGATTTAGAGTTCAAACAACCATCGTTTCAAAAAGTTTTAGTTACTGAAAGGAAAAAAAATAGCAAACCGCTTGTAAAGTTGAATTCTGATGATACCGAAAACGACAAAATCAACGACGAGTACAAATCATCTAACTTGCCTATTTCTGAAGAGAGAAAAGCATTGTTTAAAAAAGTAAACGATGCGCTTGATCACACTTTTTCAAAAGTGTCTCCAGAAGAGGTTTTGTCAGAGTCTACAAACAACAAAAGTCCGTTAGTTGAAGCACCGAAGAGTGTGCAAAAATTAGACTCAAAAGGACCAGTACTTTCATTTCGTCATCGAAGTTTTTATAACGAAAAAACTGCTAACAACGAAGACTTACCGACAACAAATATCCTTAATCATGATAAATTTCGTGTAGTAAATAAAAACATTGCAAGAGTTTTGCATTGCCAAAGGTGTGAAACTAATCCACCTAATAATCGAGCTGTTTTAGAAAAAGAACCCTCCTCTACTTTTGATTTAGTTTTTATTAAAACGAATAAAACTTAAAGTGTTAAAATGATAAATTTATGCAGATTATGCAGATTTGGTTATAGGGAATCTAAATTACGGTTTGATTTTATAACTAAAAATTATCTAGAAATATAGCAGATAATGTATGCGAATTCATTACCATACATAAACAACATTCAAATGGATGCAAAGTAAAAGGCCTGTTGTCTAGGTTGTTTGGTGTGCCTAGAATGCTCTTTCATGGAATCGCGTATCCATGTTGCAAAAACTGCAGTTTTTATCACTTTATTACAACGTTGTTGTTTTCGTTGACCATCCATCCATACCATACAATTGTAGATAAACTGAAGTAATAATTCAAGGATTCAAATTTAATTACAAAATATAGTTTTTACGACATCGCATAATGTTATAAAGGACAAAACTTGGCATAATTATTAATTCTACCAGTATTTTAATATATTTAAAATAATGCTTTGAAAAAGACAAAAGGCAACGCTTATTGACAAGCTAGACTAATATTTAAAAAATTAAAAGACAGAACAAAATTAGATTTTTGTAGATAACTCCCTTAAAAATAAGTTTTATAAAAAATTCCTGGGAGGGTTTTCTCACTCCTACCCAATTTGAAAACACATCATATGTAAAACACATCGTATGCAAAAAAGAAAAACTCTAAGAGAAACTCAAACGATTAGAATTTTTTTTAGATTTATAAATTTTCAACACGTTCAGTTTTATTTATTTTTAATTACAAAGATATCTTATTTCAAAGTTTTTTTCTTTTTTATTACAATATAACTCACTTAACAATC
>evgR010089c0g1_i01_Hydra
GCAAGCATTGTTAACGCTGCTAGAAACTTCTCTACAAAGAACAACACTACTCAAATTGCTATTACCAACAGAGATTCTGCCAACAAGGCTTTAGATGAAGAATTGGCTCGTGATGAAAAAGTAATCATTTTGGGTGAAGAAGTTGCTCAATACAATGGTGCTTATAAGATTACAAAGGGTCTCTTTGATAAGTACGGAGGTCATAGAGTTATTGATACACCAATTACAGAAATGGGTTTTGCTGGTATTGCTGTTGGTGCTGCCATGGGAGGTTTGAGACCAATTTGTGAATTCATGACCTTCAATTTTGCTATGCAAGCTATCGATCAAATTATTAACAGTGCTGCTAAGGGTCATTACATGAGTGCTGGTCAAATTAACTGTCCAATTGTTTTCCGTGGTCCAAATGGTGCTCCACCAGCTGTTTCTGCTCAAC
>evgR010090c0g1_i01_Hydra
AGTATCTTAAGGTAGCATACCTTCAAAAAATCACTAAGATTTAAATTTTTAGACTAATTTGGAAATTTACACATTTATCAGCAAAACTATATTTAAACAAAACATCCATTAAAACAACACTCCACATAAAAAAAAGTTCAACCCGTCTTCAGAAGTTTTCTGCCGAAAATTATTTAACCAATCAAAATGGAGAATTACAAAGGGCTAAACCAAAGCATATTAAATAAATACATGAAGCTAGACACACAAGGAAAAGTTCAAGCAAATTATATATGGATTGATGGTTCAGGCAAGACACTGCGATCTAAAACAAAAACCCTAGACTATGTACCGGAAACAACCGATCAGCTACCAATTTGGAACTTTGACGGCTCTTCAACAGGTCAAGCCAGTGGTAAAAACAGTGATGTTTATTTGCATCCGGTGGCCCTTTTTCCTGATCCGTTCAGAGAAGGTAAACATAAGCTTGTACTCTGCGAAACTTATACATATGATCACAAGCCAACAGAATCTAACAAAAGAAAATCCTGTAAATCTGTAATGGACCTTGTCAAGGATTCTTATCCTTGGTTTGGGCTAGAGCAAGAATATTCTTTACTTGATCGCGATGGACAGCCACTCGGGTGGCCGAAAGGTGGATATCCGGCTCCTCAGGGACCATACTACTGCGGAGTTGGAACTGGAAAAGTTTTTG
>evgR010090c0g2_i01_Hydra
TCACAAGCCAACAGAATCTAACAAAAGAAAGTCGTGCAAATCTGCAATGGATTCTGTGCAGGATGCTCATCCTTGGTTTGGCCTAGAGCAAGAATACTCTTTGCTTGATAGAGATGGCCAACCTCTAGGTTGGCCTAAAGGGGGTTACCCGGCTCCTCAAGGACCGTATTACTGCGCAATTGGTACTGGAAAGGTTTTTGGAAGAGAAGTTATGGAAGCTCACTACAGAGCTTGTCTTTACGCTGGGCTTAAAATTGCTGGAGAGAATGCTGAAGTTATGGCATCACAGTGGGAATTCCAAATTGGGCCGTGTGAAGGAATTGAAATGGGAGATCAATTATGGCTTGCTCGGTATTTGCTTGAACGTGTAGCTGAAGACTTTGGGATTATTGTTACTTTAGATCCTAAGCCTATTTCTGGAAATTGGAACGGAGCTGGTTGCCACTCAAATTTCAGCACTCACGATATGCGAAAAGAGGGAGGGATCAAGAAGATATACGAAGCTATTGATTGGTTATCAAAG
>evgR010189c0g1_i01_Hydra
TATAACTGTAAGCTTACATACCAACCTCAGGTTACATCACCTAAAAATAAAAATCGTAAGCAAAATATTATATGGTATAAAATGTCAAAATGTAGAAACAAATATATAGAAATTGTTTATTAGGATAAGTGGACCTACATTTCCCTGTGAAACAAAAACTACGCAAAGTATTCAATAGAAATTCGATCAAGATTAGTTATAGCTGCATGCAAAATATAAAATCCATCATTAGTTCACACGACCACAAGCCTTTACAAAGTCAAAGAAATATAACTACAAAAACATGCAACTGTATTACTAAAACCGCTTGTCTATTAAGCAACCAATGTTTATTAAGTAATATTGTTTACCAAGCTGCTGTGAGTTCTGGAGATCCGGATCCTAGGGAAAAAGTATATTTTGGTATCAGTGAGACACCATTCAAGCTCAGATACGCAAATCATTTAAAATCTTTTAATGCGATCAAATATAAAAATGATACCGAACTTTCAAAGGAAATTTGGAGTTTAAACGAAAATGGTGCAGTACCTATAATAAAGTGGAAAGTTATAAAACGCTGTAAATCTTCAAACCAAAAACAAAAACCTGCAATTTTTGTCTTCAT
>evgR010205c0g2_i01_Hydra
ATTTTGACTTTTTAATTATAATTGTTAGTCGATGGTTTGTAAATATTAAATATACTCCATAAACAAAATATTAAATGAATGACTATAAGATGAATGAAAAAAGTGGTGATAAAAAATTGGATTGCTTCTCCACAGAAAACAATAAAAGTGTCGATAATCCAGACAAATGGTATTACATTGATAAAGAAATGCTTCCAGCGAAATTAACGTACCTCTTCACAGGTGTAGAGAGATCGTCGACTTCCCCTTACCTTACCATGTTCTACATAAGCATTGGTCTCAACCTAAAACAAGTCGGTTATGTGTTTTCTTTTTCTTACGCTGGTCTAATGGTAGGCAGCGTTTTTTGGGGAGTTTTGGCCGACAAAACTCGCGCGCATCGAGTTTTGCTATTAATGTTGTATATTTTGTACGGAGTTTTTGCATTATCAATTCCATTCGTTAGTTCAAAAATTGGAGAAAAAGACAAAAACCAATGTCCATTATCACCAAACAAAATTTCTAACAATAGCAACACCAATAATACTATAGTTAATTACTCTCCAGTATCTTCGACAACATTGACTTATGTCATGGCTACGCTAGGATTCTTTGCTTCTTCTTTTGGTGGAAGTGTTTATAGTTACATAGATGCAGGTGTTGTTGAAAGGATCAAAATGGCTACAAAACGCAAAGATTTCGGAAGGCAGTATATGTTTGTAAACGTTGGTTTTGCAGTTGGAGCACTAATTTCAGGAGAAGTGATGGATATATTCCCAAAAGCTGAAATTTCATGTTATTTTTCTGTGTTTTGTGTTGGTGTTATATTTTTGATAGGTTCTTGTACTTGCTCCCAATATTTATACAAAGGCATTAAAATACATTCTCGAGAAGAGTTAAAATCCGATTCTTTTAAACAGGATCTTTTTAAAACATTAACAAACTTTGAAGTAAATCTTTTTTTGATAACAACTTTTGTAAACGGTTTCCTATATTGCATATACATAAATTACTTTTTTCTGTTTCTGAAGGAACTCAAAGCTACAAATCTTTTGATTGGTCTAACAGTTGCATTTGGATCAATATCTGGTATTATTGTAAATTTTTTCAACGAAATGATTATAAAATTAATTAGAGGAACATTTAACACTATTATTATGTGCACATTTCTTTGGTCCGTTCGATTTTTGATTTTCTATTTCTTAAAAAATCCTTGGTCAATTATACCACTACAACTTACTTTACATGGGTTATGTTTATCACTATTTTTCAGTGTTAAATCGGTTCACTTAAAAATTATTTCACCAAATAGCATTCATTCATCGATGTTTGGTATTGTTCAATGTTTACACGGTGGCATATCTCCAATAGTCGGTAGTTTAATAGGAGGTGAATTATATTTTAAATATGGAGCTCGAAACACATTTTTTTATGTTTGTTTAATGGCATTAGGATGGACGATGTGTTTATGTATTTACATTCTTGTTAAAAATCTTTTGAAAAAAAATGTTCGTAAACATTCAAAATCTCCTATCCTGAATGGATCTTCACTTGAAGTTAGTGAGTACATAACCAAATTTTGATTTGGAAATTTTTTCTAAAAGTTTTATGAATGTCGAATTGTCTAAAATGTTGAAACATTGACTACCGACTGCCGTACTATTCTGTAATCTTTTCAGATTATTTTTTTAATTTTTCAGAATAGCTATATTATAATAAAGAAATACATAATATAAATGCAAAAAAAAAAA
>evgR010208c0g1_i01_Hydra
TATGCAAATAACATATTTCATATTTAGACATGTTACACATAAGTGTTATTCTGAAAAAAACTCGTATGATATTTTTCACTTCAGTTAATGTATTCTGTTGGAGTCACTGCATTGTAGAGTATGGAGGGGAATTAACTTTGTTGACTGGTCCCTCCATGCAACCAACTTTTAACAAAAACCAAGACAGTACTATTGTATTTACATCACGTTCAATATGGAGAACATTTAGAGTTGGAGATATAGTTGTTGCTCGATCACCTAGCAATCCGAAACAGATGGTTTGTAAAAGAATAGCAGCTGTTTCATTTTATGTGCAGGAAGGAGAGCGAGTTGAACAAAATAAAGTAGTTCTTGGTGAAACTACAATAAAATATGTAAAGATACCTAAAGGTCACGTTTGGCTGTTGGGGGATAATTCTAATAATTCCACAGATTCTAGAACATATGGTCCGGTTCCTTTAGCCCTTATTCGTGGTAGAGTTTGCTTTAAGATTTGGTAGCTATCCCGAAAATCCTATTTTGGAAAAATAATGTCGAAGGATACGAGATGTATTCTCTACGTAGACACTTTATATTTGCTATATATATACGAATTACAGCGAATATTTTTTTTTTAAATATGGTGCACATTTATTGTTTTTGATCATCCAACCAAATCAAAAAAAAATATTACTACTCTATATAACTCTAGAGGCTATTTTTAATTATTTTTTATAGGTAATTATATACTAACGATGTAGAACTTCTAAAAATATTTAAATGGCG
>evgR010219c0g2_i01_Hydra
TGAGTGAATTTTCATCAGATGTAGAATCATTATATGGTAAAAATCATTGTGTCTTTAATGTGCATCTACTTCGTCATTTAGCTTTAAGTGTTAAAAATTGGGGGCCTTTATGGGCAACTTCATGTTTTACTTTTGAAAATTATCTTGGAGCAATGAAAAAAATGCATAATGGTACAGTGTTTGCTGCAGAACAAATTGCTAAATCTGTTCTTTTATTAAACAATATTCCCTCCATTCTTAAAGACACAACTCGATCTGTAGGTTTTCAAGTATTATTCAATGATATATTTAGACAGTCTCTGCCAACTTCAGAAAGTCTTATGCATCTTGATATAAAACCTGCAACTATTGATTCTCTTACAATGTTAGTTTATGGCCAACTTATTTCTACGCATTCTTTTTGTAGTGGAAAAAAACGAAGCAATTGTTATGTTAAACTATCATATGGTTTTGTGTTTCTTATAAATGCAATTTTAAAAATAAGTTCATCATATTTCATTATTGGATACAAGTTGCAATCACAGAAACGTTTTCTTTC
>evgR010233c0g1_i01_Hydra
TATGTTTATCATGCTTTTCTGTGTTGCAATGTTCTTAGTTTTCCCACATTCTTCAGACATTAAATTTGACAGTCAAACAACAACCACACCAAGTGCTGACCAAACTGAATCTGTTACCAATAGAATGAGTCCTGCTTCCAGAAAACTTGCTTCCACAGATCAAAAGGATACCATGGTTGACGTTTCTATAGACGCGTCACAACAACCATTTGTTGATGAGGAATTCAAAAAGAATGTTGCTGAGATTTATAGTTCATTCCAACAAGTTAGAGAATCTGCCACAACTGAATCTCAAGAAAAGACTGAAG
>evgR010236c0g1_i01_Hydra
TGATGGAACATACACTGATCTACATAATGCAACATCAGATGCTGATTGCAGTGAATGCCCAGCAGGTCTAGCATGCGAAGCTGGAACTGGAGGATCTCAAAAGCAACCACAACTTTGTGCAGCAGGTCATTGGTGTAAAAAAGGAACAAGACATCCTAAGCAGTTTCCATGTCCTGCTGGATCATTTTCAAACTTTACTAATTTACAGAGTGTAGAGGGTTGTCAAGAATGTACTAAAGGTTGGTTTTGTTTAGAAGGTGCGATTTCACCTAGCGGACCTTGTCCAACTGGTCATTATTGTCCTTCAGGTACTCAGCATTCAACACAACATCCATGTCCCAAAGGAACATTTTCAAGACGTTCAAGTAATACTCGCAAGGAGCTATGTGAACCATGTGTTTTAGGCCATTTTTGTCCTTTAGGAACATCAAATCCAGAACCTTGTCCAAAAGGAAGTTATAGAGATGAGTTAAACGGAGATTCAGAGGATAGTTGCAATCCCTGTACTGCTGGATTTAAATGTGAGAATCTTGGCACAGTTAGGCCAGCCATGTGTGGTTTAG
>evgR010264c0g1_i01_Hydra
CAAAAAACATAATTCAAAATGGCCAGAGGACCAAGAAAACACATCAAGAGACTTGCTGCTCCAAAGCACTGGATGATGTCCAAGTTGGGTGGTGTTTATGCACCAAAGCCAAAGCCAGGCGCTCATGCTGCTAGAGAAAGCTTGCCACTTCTCTTGTTGTTGAGAAACAGACTTAAGTATGCTGCTACTGCTAAGGAAGCTAACGTCATTTTGAAGCAAAGATTGGTCAAGGTTGACGGTTTGGTTAGAACTGATAAGGGTTACCCAGCTGGTTTCAATGATGTCATCACCATTGAAAAGACCAACGAACACTTCAGACTTATGTTCGATACTAAGGGTAGATTCGTCATTCACCCAATCTCTGAAAGAGAAGCCAACTATAAACTCTGTAAGGTTAACAAGAGAGGTACTGCTTCCCAAGGTGTCCCATACATCCAAACTAACGATGGTAGAGTCATTAGATTCCCAGATCCAACCATCAAGGTTAACGATACCGTCAAGGTTGATATTGCCACTGGTAAGGTCACCAAGTCTGTCCACTTCGAATCTGGAAACTTGTGTATGGTCACTGGCGGTCACAATCAAGGTCGTATTGGTGAAATCGTCAACAGAGAATTGCACCCTGGTTCATTCGAAATTGTCCACATTAAGGATATTGCTGGTAACAACTTCACTACTCGTGCTAACAATGTTTTCGTTCTCGGTCAAGGTCACTCTTCCATGATCACTTTGCCAAAGGGTAAGGGTTTGAAGTTGAACGTTGTCGATGACAGAAAGAAGCAAATGAGATTGGCCAAGAAGTCCAAGAAGGCTGGTCTTAAAAAGAAGACTGAATCCAAGTAAATTCAAC
>evgR010274c0g1_i01_Hydra
TTTTACTCCGTAAAGAATACTCAGTGTCTTTTCAAATAAAACCAACAGCTTTTTTGATCGGTTGGGGTAGTGTCATTCATTTAAGTACTGGATATAATCTAGGTAATTATGGAGATCGTAATCCATCTGTTTTTTTTTATCCTGATAAAACAGGAAAGATGCTAGTTAGTTCAGCTGTCTGTAACATGGCCAGTTTTGATGTTGAGACATTACCTGTTGAGCTTGGAACATGGTCTTTTGTAAGAATATCACAGCAGTTAGTTGATGGAAAATACATTTATTCAATTGATCTTAATTCTAAAAATGTTTTTCAAATAGAAAATACATATGCACGAGACTTTTCCAATG
>evgR010289c0g1_i01_Hydra
CCCTTCTTCAGTTTCATATTTGGATAAAGAATCAAAAATTGTATTTATAAAAAATTCTAATTTGAACTGGTTTAATTCAGTTTTAAGTGGCTACAGTGTCACCAATAGTGGCTACAATTTCGCCAATATTTTTATAAATGAAACCTACAAAGATGAAACCCTTTTGGTATTTACATCGTATAATTATACGACATTGATAGAAATAATATTAAATTTGATTATTGATAAAACGTTTCAGTCAGAAATTAATAAAACACAATTAGTGTTAATTACGGATATTGATGAAGAATTAACATCTATAATGTCAAATATTATTGTTCCGTATAATAACGTTTTGCTTTTCGACTGTAATAACAACAACTACAAAATGGCTTACAACAATAATTCAGTAGGATATTATTACAACTCTCGTAGAGATTCATTTATCGAAATTGTTGCTAATTATAACATCACAAGATATATTTTTTTAATTATCAGTTCAGATGAAGATGAA
>evgR010309c0g1_i01_Hydra
GCAACACTACCTTTAATGTAACAATAAGTAATCTTAATCCAGATGCAAATTATAATTGCTGTGTTACAGCATGCACTAAAGTTGGTTGTGGTTTGAAAACATGTTTGTTTAATAACACATTAGTTGGAATTCCAAGTGGTCCTCCTCTCAGTTTTGCTGTTACACCTGTCAACTTTTCATCTCTGATGTTGTCATGGGATAAAGTTGAGTATTCAAAAACAAATGGAACAATTTTTTTTTATAAATACTCGTGTAGCAGTGATTTGAATATTTGGTACACAGTTGACAACAATACATTCCATGTGATAGTGACCAATCTTAATTTTAATACAACTTATAATTGTACAATTGCAGCATGTACGAAGATTGGATGTGGTGTTCAAACATCTCTAGTTAATAAAACCTTAGCTGGGTTACCAGAGGAAAGTGCAACAATTTCCTATATTGGTTACATTAATTCTTCATCAATTAATATCAGCTGGCATGATATAATAAAATGGGGAATACCAGAAAATCAATCATTTTGTTTAGATATCGAAAACAGCTCACGGATAGTAAAAACAGAATGTGTTGACTACACCGTTCAAAAAAACGTCAACGTTATTGGGCTTGATCCTTTCACAAACTACACAGTTTTAGTATATTGTAAGAATAAAGCTGGAATTGGTATAAAGGATAACAGTAAAAAAAAATCCGTTCAAACAGATGAATTACCACCATCAATTCCGCCAAAAAACATTTCAGTTGTTTCAAGTAATACAAACAACATAATATTGCATTGGAAACCTCTTGATCCTAATGAATACTATGGTATATTGAGAGGTTATAGTATTTTAGTTAAAGCTTCTTATTCAGAGTCAGATTTTAAATTACAAAATAAAAGGAAAAAGCGTGGTCTTGACATGGAAATATATAATGGAAGCTTGCATTTAGAATGGGCAACAAATATAAGTGATCCTTATGCAACATACTTCAACATCACAAATTTGTTTG
>evgR010313c0g1_i01_Hydra
GGAAATAATGAAAATGACCAGGTGTCGGAGAGTAGCATAAATGATTCATCGATGGTTAATCCTGAAAGTTATTTTATAAATACATTAAGACAGAAAGAACAACAAAAGAATAGTCGTTTAATGAAGTGGGCTCAATATGATTCTTGCACAAAAGAAACTAATGCTATTCAAGTTAAAGAAAAGAAAGAGCTACTAAAGCAAAGCGAATTAAAAATAAACATTGGTTTATTTAAAGAAGACTTTGAGTCTAATGAAAGCGTGGTCCAAACTTTTCTAAATGATTTAGTTGAAAAGGTTTGCACCGATGTTGATGAAAACATTGATTTTGTTAAAGAAGACTGTCCTAATGCTAAAAATTGTGCAAATGAAGCTAGATTTACTTTAACAGTTC
>evgR010318c0g1_i01_Hydra
TGTATATATATATGTATATAAATGGTTTGACTCGTCATCTCCATGCCAAACACAAGGACAAGGTTATTGATAAACAATTCATTAAAGAGAAAGAGAATGCAAAGGTTATTCATCATCCTACCTGTTTTCAAAGTTTTTTAGTGCAGAGTTCAGAAAAGTTAGCTGAAGATAAATGCTACCCTTTTGAAATAAGAACTGAGTTTAAAAATTTAAAAATTTTTTGTATTGAGGAAGTCAATCCAGCTTATTTTTTAATAAGAAGTATAATTAATTCATTTGACGGATATGCTGAAAATTTTTACCCATCCTTCT
>evgR010320c0g1_i01_Hydra
TCAAATAATATGACTTTGGAAACTTGTTTGCTCATGTCCAAGTTACTCTTCCAAAGAGGATCTGTTGTTAAAAAGTTATACTTGTACAACAATCATATTGACAGTTCATGTTTGAAGGTTATTTTCAATCCTATTCTCGTTCCCTCATCCGCTCAATCCTCCTCTATTACAATTCCACCATTGGAGAAATTAGTTGTTTTGGCAAATAAGGAAATTGGTGATGAAGGTGTCAAGTATGTAGCTCAAATTGCTGAAGCTATTCCAACTCTAAAGGATATTGTCTTGAAGAATACTGGATTGAAAGAAGGTGGTATTTTGGCATTGTCATCCTTCATTGAAAAACATCCTGATCGTTTGTACCATCTTGATATTAGTGAGAATAATCAAATTGGTTTGAAGGCCTTTACTGCTTTGTGTTCTGCCATTGAAAAGTCTACCACACTTGCCCTATTGGATATGAATGAAATTGGAATGGATAGTTTGGC
>evgR010330c0g1_i01_Hydra
CTAAATTTTTCGGAAATTTTCATCATGTCCACTCCACCAACACCAGTTCTCCCAAAGGGAAGTACCATGAGAGACATTTTCGATTATTGTGAAAAGAACGGGTTAGATTCCGTTGCTCAAGGCATGATTGAATTGGCTCCACCAATGAAATTGAGAAGTTTGCTCCAAAATATTACCATGGATGATTCTGTCCATCAATATAGATTACGTATGGGACAACCTATTTATCGTGAAGGCTTGTTGAATATGTTACATGAACATTACGGAATGAAGAATTTGACTGTTGATAATATTTTGGCTGTCCAAGGTGTAACTGCTGGTTGTTTGAGTACTTTATTGTGGTTGAAGGATCAAGG
>evgR010340c0g1_i02_Hydra
TTCAAATTCTATACATTATGCTTATTTTTTCCATATGTTTAATTAAAAGTTTTCAATCAAGAAACCTTCCATCTTACTATAACGAAACGTCGGCAATTACCTACAGCATGGGAATATCATGCATACTTTTAATTTGTTTCTTCCCGATTTACTATGGTAACAACGATAAGAAAAAAAGAACCAGTGCCGTTTTAACTTTAATGTTAGTTATTAACCTTGTAATTTTGGAAGTGATGTTTATTCCAAAGCTAGTAATAATTTATTTCTATCCAGAGAAAAATACAAAACAAAGTCTTCAAGAAGAAATAAGCAACTATTCTAAGTCTTCTTGACACTTTAATGATTTTCAATTAAGAAAC
>evgR010350c0g1_i01_Hydra
TTTTTTTTTTTTTTTTTTTTTTTTTTTTTTTTTTTTTTTTTTGATAGGAAAATGTAAACTAACGAACAACTTATGTTTAATAAGAAATACTAATCATGACAATTAATTCATTTGGATTAGGTTATTTACCTGTATTATGGTCATTATTTTCAGTATTTGCGTTTATTGGTGCATATCTTACTGCAGTTTTGGAAGGTCACGTTTATTATTTCCTTCCATCCATAGGTGATACTGGAACAAAAGATCCTGAATCTAACATTTTTTCTTTGCTAATGAACATCTCAATAATAATTGGTCTATCAAATTATATAATCAGATATTTCCAATGTCAATATCAAGCTCGGCAATGTGCAAATGAAAGAGAAATAATCTATAAATATAACAGAGTATCGCTATTGCTAGCAGTTGGTAGTGTCTTAGGCGGACTTATTGTAGTAAATATATCAGTGAGAAAGGAGTACTACTTACTCGAAATTCACGACGCTGGTGCCATAGTTTTCTTTTTGTTTACTACAATGTATTTTTGGGTCCAAACCATTTTGAGTTACAAAGTATCAAAATTCGGATTGATTTCGTCAGTAATGTGTCACGTGAGGTGTTTTCTAATTTTCATTATCAGTGTTTTTTCATGCATTCATATTGTGACTTCAATACTCAGCTATGAAATATTTATAAGCTATAGTTTGCAACATGCTATTGCGCAATGGGTCCCTGGAGACTTTGGTTACTTAATTCATGTGGTTAGCAATTCATCCGAATGGTTGAGTGCAATGTTTATAATAATATATGTTCTAACTTATTTTGAAGAATTTCAGCATGTTGTGTTATTGATTGACTGTCAAGAAAGATCCCCAATGGTTGAGTTTTATGACAGTGTTGGTGTTTATTCGAACTGGAAATTTATAAACGAAGAAAATAATAACTAACTGCATTTTAATTCGACGAAGCATTTTCATTGAACCACTTTCATTGGATTATTTACGAGGAATAAGATTAATTAACCATTGGTTAATTGTTTTATTATTTAGAACTAAAAATCTTAAATAAGACCATAAATAACATCTTCCGGTGTCATTGCGTGTATTTCCGGTGTCATTGCGTGGCATTTGTAAGTTGCCTTAACGCAATGACCTTCAAGATTTAAAATAATTGTTATTAGCTTTTTATTATTTAAGCTAACAGCAATTTTTTTTACATCGATTTATATATAAAGTATAATTTGTAAAAATAAAAACCTTAAAAAAAAAGAAGCTCTAGATCCTTAGAAAATTTTTTGTTAAAAAAAAA
>evgR010353c0g1_i01_Hydra
TTTTTTTTTTATTCGAATTATCGAAATCGTATTGCTGCTATATTTTTAGTAATAACTTTTGCTGATTCAGATTTATGTTAGTAATCGTAGAATTACTGCAATGTGGGTTTTTAGTATTGTTCATTTAGTTTGTTATTGGTTCAGCATCAATCACACTGTGCAAGGATTTTTCAAACAAATTGCTTGTTTGGGGGAAAATACTATTTTCGATGTTAATGACCTAAAATTAGGGTCACTTTTGCTAAAATCGAAGGTAACTTGGACGAAGGATGGGCGCCAATTGTATTCACGTCGGCTTACGCATATCGATTCAAATGGTTCCTTAATTTTATACAAAGTAAATTTATTTGATAGCGGTATTTACAATCTGACGTTACAATCTTCGGAGTATACAAAAACATTTTTGTTTGAGCTAATTGTACCAGATAGCACCTTAGAACTTTTGACCGAGCAAACCATCTATGTTAATAAAGGCGATAACATTTCTATAGAAATAAAATCAAACGATTATGGGTCATCAAAATGGACCTTTGTAAACAAACTAAACGAGAAGAAAGCGATAGATCTCGTTGACACTAACAATAAAGATACTTCTAATAAAAAAGTTTTTAAAATTAAAAACGTTTCTTTGAATGACGAAGGTCGCTATCAATATACATATGAAGTTAATCGATGTACTTCAGAGATAGATGTAGATATTGTTGTTTTTAATGGTCAAATTTCGGATTATTGGAAGTGCTTACTTTCGAGATATCACCATAATGTTATACTACTGCTATCTCTACTCTTCCTCGGTTATTGGCTCCAACATAGAGAAATCGGTATGCTTGAATACGCAAAAGAGTACGCAAACGAACTTAAGATGAAGAAAAAAAAAGCTGAAAAATTAAGTTCATAATCCCAAAGTTTCAATATTTTAATATTAAATACTTTATAATTACACATTTGAAACTTTAAAGATGGTTTTTAATGAAACTTTAAAGATGGATTTAATAACTTTAAAGATTATTTGTATTTTTTAAATTATTTTGTGTGTAAATTTTGTAAATAAATTTTCTTATACTTTTTATTAATATATATACATTTTTTTTAAAAAAAAAAA
>evgR010358c0g2_i01_Hydra
ATTGAAATATCTGACAACTATTAAACCATTAATTAAGTCAAATATAATGGATGATGTTAAATTTGATGAAGAAAATGTTATTAGGAGTTTTCAAAAAAAGGAATTAAATAAATCCACTGCAGATAAATATAATGAGTTTTTAATCCAAAGTATCATAGATATTGAAACATCAATTCATAGTGATATGAGTTATAATGCTATGGAAAACTTATTCTTAAATGATAATATTCCTTGTTTTAGTGAATTTATTCCTTTATCTATTCAAGACTTAAATGATATAGTAAAACAATTAAAAAACAAGAATTCAATGTGTGAG
>evgR010363c0g2_i01_Hydra
CATTTTCTGTTTTTTTACCGCTTCGACATGTTTTAAATTACGAGGTGAAAAAATTAAAAATGTTATCGCATTGGACTCGAGTGAAAAGGTTGAGTAGCGGCAATATAGGAAACAATAAAGAAATTGATGCTATAGATTTTTACGATATAGAGGAGTGTATTGATGAGCTTTATAGACCATCGTCAGTAAATTATTTGGCGCTAAACAAATGGCTGAAAAATTGTTCTTCAGACTGGTTAAAAGAATTTGTAGAATTCAAAGGGTTTCAGGCGATATTTTGTACGTTAAAATATTCTGAAATGAAATCAGCTTTTCGTTTTTCATCAGCTTTTCTACAAATAGAGATCATTAACTGTATAAAAACAATACTCGGTAGAAATGAAGGGATGATTTCTATAATCGAAGGAGAACATGATTTGGTAAATAGCATTATACTTGCACTGGACACTAGCAACACAATGGTGAAAAAACAGGTATTTGAAATGTTATCC
>evgR010365c0g1_i01_Hydra
ATTATGCAACTTGTGATGTTCATTCGCGGTATATTGGTTCCCATGATGCTTTTGTCTTCTATGTAAATAACTATTTTAGTCGTGATATGCTTAGTGAATTAGATACATTTAAACCAACTAATGCTGGAAGTGAAAACCTTTTAATTTGGCATTTTATACATTTGATGGGTTATAAAGTATTAAATTCATGTCGAAAGTTAGTTGTTTATCATACACATTGCGTACCAGTTAGATCTATAGAGCGTAGTCGTGTCGATTATAATGGTAGAAGGGGATTGGCGCCATTTAACAAAAATTTGTTCTAATTATAGGATGACAATTTTGTAATTTTATGATTAAAAAGTTAAGAAGAAC
>evgR010372c0g1_i01_Hydra
CAGCTGCATTTTCATCGTTATTAGGTGTAATTATAACAATAATATTTATATTGAAAAGAAATACTCCTGTTGTAAAATCATCCAACTTCACATTTTTGTTGACACAATCAATTTGTCATTGGCTAACTTCGCTTCTTGTTCCAGTACTTCATGTATCAAACACTGAATTTTTTTGTTTTTCGCGACATTTTGTTATCAGTTTCCTGTTCACTGCTTCAACGTCAATTACTTACATCAAAACCAGAAAATTTATTTTTATTTTTCAAAAAACTTACCGTCTTACCGAAAAGTTTGTTGCCAAATCGATTGATATAATGATAGTCGCTTTTATTTTAATTGTCCAAAATCTGCTGGC
>evgR010377c0g1_i01_Hydra
ATAATATATTGACTAATTACTGTTTTTACAGTATTATACGTATTAATCTCATTATTGGAATTATTTATGTACGTTTCAACTCACAAAAAGAAGGTCAAGGACAAGCTTTACATAAGCTATTCATTGAGAACTTCTTTTAGAGATGCAAAAGGTAGAGTACAGCACAAACACATAGCCAATTTACCTAAATGTACGGACGATGAGATAAATGCACTCAAATTAGCACTTTCATTTAAAAACAAACTAGAAGAATTGGGTAGTTTGGAGGATGTATCAATAACAAATGAAGCAAGTTTTGGAGGAAGCTATGTACTTCATCAAATTGCTAAAAAGCTAGGTATAGATAAAGTCCTAGGCAATAATAAAATGGGCAAGCTGGCTTTGTGGCAAGTGATAGCAAGAATTTTAGGTAGAGGCTCTAGATTATCGGCAAAAAGGATGATGTCAAAATATAGTGCAGAAC
>evgR010382c0g1_i01_Hydra
GTTGTCTCAAATATCTTTGTTTTCAAACGCTATGGTAAAAAAATAAGTAAAGATGAAAATAAAATGCTAGAAAAAGAAATTTTAGAATGCGAAATAAAACAATCACTATGGCAAATGAAAAATAATAAATCTCCCGGATCAGATGGGCTAACAATTGAATTTTACAAATGTTTCTTCGATATAATTAAAAAAGACTTGACTAAATCCATAAATAATGTACTGATTACGGGCGAAATGTCAAATTCGCAAAAAGAAGCACTTATTACATGTATTTTTAAAAAAGGAAATAAAAATGATATATGTAATTGGAGGCCTATATCTCTCACAAACGTTGACTATAAAATCTTAACAAAATGTTTAGCTAACAGACTCAAAAATATCTTACCAAATATAATACATAGTAACCAAACGGCATGTATTAAAAACAGAACTATGAATTATAATCTAAG
>evgR010387c0g1_i01_Hydra
TAATTGACAAATCTCAATTATTATCCACCAATGGTGTTGAACAATTCTCTTCTCAACAAGTGACACCAACTCTCCAACAACCAAGTCCTTTAGACCAATTATTGGCACTCAACAATTATGTTATCAACAATAATGGCGCCCTGATAACTCCATCACCATTCCAACAACAATTGAACTCAACCGTACCTCACCACCAACAAAATGTGTCTCCAATTCTGGTCGTTATCAATAACAACAGTCACAACACCTCATTGAATCACCTCCACCATAACAATACGGCAACAAGCACTGGTGGTTCACACTCATCTTCTCCAACAACAAGCAATACTGGTAGTGCATGCTCTCCTTTGAATAATATTCAATTGGTGCATGCTCATCAACAACAACAACTCTCTAACAATGTACCAATGTATTTCAAATCTTCAAGTAATCCTTCCTCGAATTTCTGTCAATCCATGGAAGAATCTGGACAAGGTGTG
>evgR010405c0g2_i01_Hydra
GAATGTTTTTATGAAAGTCAATCGTCATTTTCTAATTTGATATACCATGGACTGTCGCTTTTGTTCAGTATTTTGATTGACTATGGAAAAACCTCTTACGAAATTTCTTATGAAAATTATTTTGTAAATCAGTTATTTGATTTCGGATACCCCTTACAGGGGTTAGCTGTTGATAAAAATGTTCTAAGTGATGAAGAAAAAGCTCTTATGATTAAATGTTCAACTAATGTTTGCGATGTCAGATTTTATTCTCTAATCAACTTCGAAGGATGGAAACCAAAAAATAAGATTGCATCGTTATCAATAGACATCGCATCTTATTTAACATCTAAGAAAGATTTCGAAAACAACTTCCTTCCGTGGTTGAATGTTTGTGATGAATTGTATCTTTCCCTTCATGACGACATTGATTTTATTGAGGATATTTATGAATGGATTCGTCGTTCGAATATCAAGGTGTTGCTGATCAAATACCGTGGGATTGTTTTGAGAAATGTTGAATGAATTACTTTTTCAATAAATACTGTATGTACTTAAAGTAAAGTAAAATTTTTGACAGTTTTTATATGAATAATAAAATTTATCTTATTAAAAAAAAA
>evgR010427c0g1_i01_Hydra
TGCTCGTGATCATCAACCTTGTATTATTTTCATGGATGAAATTGATGCCATTGGTGGTAAAAGATTTTCAGAAGGTACCAGTGCTGATCGTGAAATTCAAAGAACACTTATGGAACTTTTGAATCAATTGGATGGTTTTAATGATGTTGGTCAAGTCAAATTCATCATGGCAACCAACAGACCTGATGTCTTGGATACTGCCTTGATGAGACCTGGTCGTCTTGACAGAAAGATTGAAATTTCTTTACCAAATGAACAAGGAAGATTGTCAATCTTGAAGATTCACTCACGTAGTATTACCAAACATGGTGAAATTGATTATGAAGCTATTGTAAAATTATCTGATGGTTTTAACGGTGCTGATCTTCGTAACGTTTGTACTGAAGCTGGTATGTTCGCATTGAGAGGAAAGAGAGATTATGTTATACAGGAAGATTTCATGAAAGCTGTCAGAAAGATGATGGATATGAAGAAGTTAGAAACCAAG
>evgR010432c0g1_i01_Hydra
TTCTATTGTCTAAATAAGTTGTTTTAAAATCAAACAACGAATTAAAAAGGTTCTTCTCTAAAAAAATGGAAAGCAATCTACCTATAATTGATCTCTCGGCTATCTTCAACAAACAAATGACAAGCAAATCTTTGTGGCAAGAGCTTGAAAAAGACAAAGAGATACAAGAAATTATTAAAACGTTTTGCGACGGTTTTGGTACTTGGGGATTTTTATACGTCAAAGGACACGGCATTAACGACACCGATATAAACATTATTAACAAAGAGGCTAAAAAGTTTTTTAATCTTCCTGATAAAATTAAAAACGAATACGTCCGCGGAAATGGAATTTCTTTAGGTTATGTTCCTATTAACGGAGAAAAGTTTACAGATCAACGACCATATGATCTTAAAGAAGCTTTAGATTACTTACCAAGTTCGGACTTGTCAGACAAACTGTACACAAAGTTTCCAGAATTTGTTAAAACTTTAAATGATTTTTTTTATAAGTCCAGCATTTTAGTTACTAATATTTTGCGAGTGCTTGCTTTGGCACTGGGAATGAATACTGAACATTTTGTGGAATTGCACTCAAGTAAGGGTAATCATAGTGAAAATGGTACAACAATGCGCCTACTTTATTATCCCGTGTGTGAAACTGTATCTAAAGAGCAATTACGTTGCAGTGAACACACGGATTATGGAACTTTAACAATGTTATTTCAAGACGCGGTTGGAGGACTAGAGGTGCTCTCCCCAACTGGAGTTTTTGTTCCAGTAACTCCTATTCCCGGTACAATTGTTGTAAATGGCGGTGATTTACTTGAAGCATGGAGTAACGGACAATTTAAAGCTACATTGCACAAAGTTTCATTCACATCCTCAGCATCACCAAGACAGTCTATTGCTTTTTTTACACATCCAAACTACGATACAAATGTTACATATGTTGATGAGTTTGGTAATAAGAATACAAGAAATGCTCGCGAACATCTTGAAAATAGATTTTTGTCGACTATATATTAATTTGTATTAATTTAACATAAAGCAAAACTGTATGGTGATTTGTTGTTATAATAGTTGCTATGATGTTATGTTTTTTTGCTTAAAAATATTTTTAATATATATAAAAAAAAA
>evgR010433c0g1_i01_Hydra
TATATATATATTTTTAATATATATATATATATATTTATAGAAAATATATATATAGAAATATTTGAAAAATAATTCATAGAAATGACAAGCGAAGGAGAAAGTTATACTGAATTTATTAATGACAATGAAAATCAAAATACTCAGGAAGAAATAATAAGTGAAGGTGAAGATGAATCTATAAAAGATGATGAAAATGGAAATTATGAGGAAGGGTTTGGTGCTGATGAAAACAAAGTAGTTGCTAATTTATATAAGAAACCATTCCTAGGTGGATTCCAACACAAAGTTACAGGTTTAGAGTATCTAAATGCTTCTGCACAAACTAATACAAAGAAATACATTCATCACGCAGTAGAAAGATTTGATCGGGATACTCAAACTGTTCAATCGAAGAATAAGATTTTGCAGACATACCATGATACGTCTACTCAAATGACAAAACCTGGTTGTTATATTGCTGAGGAAACTGATAAAATAGTTATACCTGGTAAATATATTACTGCAGCTGAAAAAGAAAAAGACAAGTTAGAAAAGGTTATACTTTTACAATGCTACTGGCGAAGATGGCTTGCAAAACAATTTGTTAATAATTTGCGATTGGATGAAGAACAAAGAATTAAATGGGAAAAAGAGGTGGAAGAAAAAAAAGTGACTGAGCGAGCCGACAGACTTCGTCAAGAATTTGAACGAAGAATGAACCCACAGACGAATGGAGATTTTGATCTACTGTATGCAGCTTTAGAAAGTTGGAGAAAAGAAGAAATATTGTCTATCGATTCAATGCATTCTGGTTCAGAACGAAAAGCAGCTTTGATTGGTCTCCTAGAGCAGGAGTCATATCTCATTGAGTCTATTGAAAGACATCGCATAAATGCTGGCAGTAAGAACTTACAAGAGCACATTCTTTTATTTTTGAAAAAGGCTGCAAAACCAAAGTTTTGGAGAGCATTTGATGGAGGTAAAACAGATATGTATACAAAGCATACACTCCGTGCTGAAATGCTCCGTGACTTGTATAGCAGCTTGCAGATGACTTATTTAACGCAAGAAGAAAGAGAAGATGTGTTGTTTACTCTTAAGCAAACAGTTCAGGAATACAAATGTAAGTTAACAGAAGAAATTATCGAATTAATTCATAGAGAAACCAATTTACTGAAGAGAGGTGTAAAAGAAAAGAATTTAGAAGGGTTGCGTCTAAGAATATCAACCCTGTTTTTACAGTTTTGTAAAAATCCACAATATAATCCAGAAGTAGCTCGTTTACTTAAGGTACCAGCTAATCCAATGAAGTTGTTAAAGAATATTTTCTTTTGTCCAAGTTGTAATTCTTATCTTCCTTCTTCTGAATTCCCCTTAGCCTCCAACTCCAAGATTGCAGGTCGATGCTCGAAATGTATGAAACTGGACAATACTGCTCGTCTCAGACATGATTTTTCTAAAATTCATTATATGTTACTACAAATCCGTTTAAGCGAAGAATTCTATGATGACGGTTCGCAAATTGCATACTTGATACAAGATGCAGACTTGCAGTATCTTGTGGAAATTATATGGAGCTCACAAAGTGCTCTCAGCGGTGTAAGTGAAATAAGCGAATTAGCTTTAGTTCGTTGGAATAAGCACAAACACTGGTCACCATGGAACTGTGTTTTGTTAAGCAAGGATGAGTCTATTGCTCATTTAAAACTTGACAATTCCCTTGAGGCTTATGGAAGAATTTTTATCGGTAAAGTACAGCAAAAACATCTTTTGGCGAAAAACTATTTTGCGAAACTTTTTTCATTGACTAATAAAAATGCTTCAAAATTAAAAAGTGAAGTCGCCAAAAATTTTGGTAGTTTGTAAACACTAATATTATATTTGTTTTAAATTAATATATTTCATAAAAAAAAAAAAA
>evgR010454c0g2_i01_Hydra
AGGATGTCCCCGTAGAGAATATTACGAATGTAACTTTATGTCTATGAGTAACAACGATGGAGAGTGGGTAAACAACCTCCATTGTGGGAATGCTCGTAACTTCTATGTATGCAAAGTAAAATCAAAATTTGTTTCCAATGTAACAATCATAAATACAAATTTGTGTGATCGGGGTTGGATCGCTGTTGGTAGTAAATGTTACTTGGTAAACATGACCAAGGTAGACTACTTTGAAGCTAGGATACAGTGTCAATTAAAGAAGGGAAAACTCGTGACTCTGTATAGTAGTAAAGAAGATGCAGATTTAGCAAAAGCCTTTGATGGATGTGATACACCATGGATTGGACTTGAAAATGTTAACCCAAGTAAAGTCGGAGAAAACAGCGGGTGGAAATGGAGCGATGGCAGTCCCCTCAGTTATCAGAATTGGAAAGATACCGGAATAACAAAAGACCAGACCAAACAATGCGGGATCATTCAAAGAGGTAAAAAATGGGATAATGTGCCATGTTGGGAGCTTCATCCGTTTGTTTGCGAAAAAGCTAAAGACAATCCCAGCCTTCCTAATAAAATACATTAAAATAAGATTTCTTAGTTGTATTTGATAATTAGAAAAAACAGCATAGTTTATAAACAATATTGTTTAACAAAAAAA
>evgR010464c0g1_i02_Hydra
TTTTTTTTTTTCGCATTTACACTTTTTTTATATTTAACATGAAGAACACAGCCCAAGACACAAATTTACAAGTTGACAACAAGATGTTAAACAAAGACAAATTAACATATTTTTTTGTTCATCCACTACTTGTGCAGTCGGTTTTTCTACTAATTGTACAGTTGGTTCTTGAAATATTACTTGCGGTAGAACCTTACCGTCTAGACAATTCTCCATTTTTCTCCACAAGATGTTTTTTTTTTAAAATTCTAAAACACCAAATAATAATTTTATTTAGTACAAAATACTTTAACAATAGTTACC
>evgR010490c0g1_i01_Hydra
TTTGAGTAACACAGAAATTCTTATTAGAAGTTTTTATCCTGAATAAGTGCAGTCCACTCTAATATGTCTTCAAATATAGTCGAAAAAGATTTGAAAAAACATATAGAAAACAATGAAGACGGTTTTTCTCATAATTTAGATAAAGAAATCACTGATTGCTTTAAATTGTTTGATGTAAGTGGATCAGGAATGATACTTGCAAATGACCTGCTTATAGCTATATCAGAATTTGGCTTCAAATTTTCGCGTGATGATATTGACATTTTAATGGAAGAAATTGATGTGGTTGGATTTGGTGGAATAACTTTGAACAACTTCAAAGTTTTAATGAGAAGGGTCCTTTTGGAAAGACCAACAGAAATGGATTTGGAAAATATTTTTAACTTTATTGATAAAAATAACGATGGGTACGTGTCATCTGTAGACCTTCAGCTTGTGTTCAGACAATGTGGAGAAATGTACACTGCCGACGAGTGTTTAAAGTTTATTGATGAGTGCAGCAGCAGTGAAAAAGGACTATTCACATTAAATGAGTTTAAAGAATACTGTAAAACAGTATTTCCAGAAAATAGCAATCAAATTACTGATGACAAGTTGCCAGATGACAACGAAGAAGAGCTCTTAATTGCTGAAAGAGAACTAATACAAACAGTTTAACTTTTTTTCTAGTTGCAACTTAGTTGGTTTGCAATAAATAAAGTTGTACAAAAATTAATATTTTAAAACTTACTTTTATGGACGTTGCAGCGTCCTTCTTTAAACGTTGCAGCGTCCATTTAGCATTGCATCGTTGCACTAACTAGCAAAAAATGGTTGATACAACAATACAGACGGCTTTTACTAAAACAGTTAGAAATTTCATGCTTATTGATTTCCCATAATTTAGGCAGCACTCGACGACGGGGCAAGACGATTTACAAAATGTAATTTTGTAAATCGTCTTTAGTTTTTCTAAAGCTGGTATGGATAATAATTTTTTAGGTGCCCTATAACTTATACTAATTATGAAATTTTGCAAATCGTGGATTATGAAAGTTTTGATATTCAAC
>evgR010505c0g1_i01_Hydra
ATTTGATTTTTATGTTATATATCTTTATTTAATTTTGATTTTATGTTTAACTAATAAAAAATATTATTTATGAGAACTCAGTGGTATAAACACAACATTATTCTGCAGAGGAAGGACAACTCCTTTCATATAAATCTTATCAATATTATAAATATACTGAGGTGCATTTTGTTTTATTGGAAAACATATATTGAAAGCTTTTCGAAGAGCATGAATATGTTAACAATGGATTGGATTATTTTGGATTACAAAACTTTCTACAGACTAATCGACTGGTCACTTCCTAATTTAAAGTTTTCTATTGTTACAACTGATTTATCTTTTTTGTTTATAGAAAAATTATCTTTATTGTTTATAGAAAGTGTAGTCCCATTATGGGTAGCTTTAAAAAATGAATCACTTACTCGTTTTTTGTTGTGAATAAGTTTGGATTCCATACTTGCAGCTGGTGAGGA
>evgR010520c0g1_i01_Hydra
GAAACGATAGAGGTCCAAGATCATTCAACAATGAAGAATCAAACTCTGGCCCAGCACACATCAGAGAAGATGAACCAGAAGAAGATTTATTCAATAAGAGAAACACTGGTATCAATTTTGAAAAGTACGAAAATATTCCTATTGAAGTTTTTGGTAGAGACCCTCCAACTCCAGTCACATCTTTCGAAGATGCTCAATTACACGAACTTTTGATGGTCAACGTCAAGAAGAGTGGTTACACTAAACCAACCCCAATTCAAAAGCACTCAATTCCTGCTATTGTTACCTCAAAGAGAGATATGGTGG
>evgR010538c0g1_i01_Hydra
TTTTCATTACTATAATTATTTTTTCAAATTTATTTTTAGACTTCTAAAGAAAAGTGTTTTTTTAAGTATTTTGCTTATGTATATTTTGGTATGCAACAATGAAGAGTATTAATGTTAAAATTAGAAATTCATTATTAAAATGCTGAATACATCGGCTGCCGTAACATTTCTATTGGTTCTTGCTGTAGCGGCTATTATTTTAAATTCGCTATGTATACACATCTTGCGAGGAACAAAATGTTTGAAAAAAAGACCGTCCAGCATTCTTACACTAAATGTGCTAGCAGTGCATTTACTACAAGGTTTAATTGTATTACCACTATATGCTGCAAAAAAGTTTACCCTTAAAAGTCTCTTTTGGGCTCAGTTCATTGATAACTCATTTAGAGTGACTTACATGATAACATTTTATGGGACTGTTCTTGGTGTATTTTGTATTAGCGCAGATCGCTTTTTTGCAACATATTGGCTAAAATTGTATAAAAAAAGAATCACTAAAAAACGTGTATGCGTTGGCTTGGTTTTATTATGGATTTATACTATTACGTTATGCATGATCCCTTTTATTCCTTCATTTCAGCTACAACTTTCATTAAACAACTCTACTTATAAAAAAACATATATATATGTGCAACAAGATGAATGGGTTATATTTATGCTAATCTTTAATACAGCTTTGCCATACATTTTAATTGTATTAATATACATCTATATTATTTATAGACTTCGAGCTTTGAGTAAAAAGCAAACCTGTTTTCGAGATAACATCTCTTCTGGTCAACAAAAAAAACCTATCAAAATGCGTCAAAAGGATTTAAAGCGCTACAGAAAAGTTGCATATTTGACTGCGGTTTTAACAGTGTTTTATGGGGCATTTTGGACTCCATCATTTATATATTACCCACTATTGAGTTTCTGTAAATCATGTTTTTTTAGTGGATTTAAAGATTCAAGTCTAGAGCAATATTTAGGGTTTACGACAAAATATCTAGCTTTTCTCAATGGTGTTGCAGCGCCGTTGATTTATTGTTTTTATCACGAAAACTTTCGCCAAGGTTTTATGACAATTAGGAGATCGTTATTTTACAAATCATCAAAGCCTTTGGATATTTCCCTCTCAACTATGGATAATTCTGCTGGGGCTTTGAATAAAATTTGTTTAACGAAAATGTATCACTCCTAAAAAATGTTTTTTAAATTTCACTTATTTATTCTTTTTATCACTATTTTTTTCCACTTTTTTTTCCACTTTATTACAG
>evgR010543c0g1_i01_Hydra
CTTAATTTCAAATTAAGCCATTCCAATGGCTTAATTTGTTTTGACTGCAAACAGCCTTGTTTGAAAAAATACCTGTTCTCCTTCGGAGATATTAAGTAACTCGGTTGTTTAATAATCAGCTCTGATGTGTTTTTAAATTTGAAAAATACACTGTTGTGAGTACAAATTTGAGGTTTTAAAATATGAATCGAATCAGTTTTATAAATGGAAGTTTTTTGCCTCACAGCCAATGTTTCGTTCATATTGAAGATCGTGGTTTTCAACTGGCTGATGGAGTTTATGAAGTAATTTTGCTTAAAAATAACAAACTGATCGACCTAGATTGGCATTTAGAAAGACTTTTTTGCTCACTCGGTGAAATAGAAATTAAATTAAATTATAACAGTTTGCAAATAAAGGAAGTTATTGAAGAATTGTTTAAACAGAATTACCTAAACGAAGGTTCAGTCTACTTGCAAGTTACTAGAGGCAGCGTACCAAGACAACAAACTTTTCCGGAGAAATACGTTCCAACTGTCACTGCAACAGTTTCTCCCCTTACGATAATTAAAGCAGAAAGTTTAGAAAACGGTTTTATTGCAATTACACATCAAGATATACGATGGCAACGATGCGATATTAAATCGATATCTTTATTAGCAAGTTCTTGGATAAAAAACAAAGCAATTGTTTCAGGCGCTGATGATGCAATTTTGATTAGAGATGGAGCAATATCTGAAGCAACATTTTCAAATGTTTTTATGGTTGATAAATACGATAACCTAATTACTAGAAACGCTGACAATGCAATTTTATGTGGAATTACAAGAAATAGAATAATCGATTTAGCAAAAAATAATAATATTCTAGTAATAGAAAAAGCATTTAGTGTACATGAAATGTTGCAAGCAAAAGAAGTATTTTTGACTAGCACCACTTTATCGGTGAGACCTATTTCAAAAATTGATGGAAATATTATTGGAACAGGGAGAGCAGGAAAAATTACCTTAAAGCTTCTCACTTTATACAAACAATTTGTTGAAAAATGAAAAATAATAAATAAATAACTATAATATAAGTCGGAATTAGATATTTCAGATAAAAATTAATTTATATCAAAAAAAAAA
>evgR010546c0g1_i01_Hydra
AATAAAAATAGCATTAAATATTGTTATTAGCATGTTAAAAAAGAAATGAATACCAACTTTTGGATTTTTTGTTTGATTATATATTTTGCCAAAAAACACGGAGTATTTGGTATTCAAGAACTAAAAACTTGCACAAGTGGCTCTGCAGTCTTAAAAATTCCAATGTCTATCCAAAATCAAATACCATCATTTTTATGGACAAAAGATGGTGATCATCTCGCATATGAACCGCGCTACCAATTTAATGCCAATACTCTTACAATAGTTGACGCTATTCCAAATGACGGAGGCATATATTGTGCACAACTACAAATTAATGATAGAAATTATAAATTTTATTATAAGCTTATTGTTGCAGATAGCAGATTAACGTTACTTACTCCTCCAAAAGTTATTGCAGCAGAAGGAAGTAATGTTACTATCAAAGTTGAAGCCAGACGTCATGGAATCTCAGCATGGACTTTTAATGGAGATTATTTAGATGTTAACGCAAATATCAATCATTTCAAGTTTCTTGATCCTCGTGATTTTGATATAATTAATGGTGGGGTTCTTTGCATTAAAAATATTCAAAAACGCCATAGTGGCGTTTATTTGATGACATTTGACGTCGAAGGGTGTGAAACTACTATTAAGACCCAAGTAGAAGTTTTTTCTACAAAATTGGAAAGTATTGATTATAAATCTGAGGAATACGAACGAAAAATAGTCTACGAACGAATTCTTTCTGAGAAATACAAACCAATGTTTAAAACAAGTGCTGGAATGATTGTTATCATAATAGCTGTCATTGCGGTTTCTGCAGGCGTCTCAAGTCTAATAACCTTGTGGTGTCTTGGTTTGTGTAAGCTACTGAAGAAACATTGGTTAAAAAACAAGGTTACTCCTTTTTCAAAAGAATTATCGAAAGAAAATTTGTTAACCGATGTCCAACAAAATGAACTTCACAGCACCAACAAAAAAAAAAGTTTCAAAAAGTCAGCATTTAAAGCCTTAACTAAATTTGGAAATAATTTTCATAGCAACAAAAAAAAGCTTAATACCCGAAATCCAGATGGTTATAAGATACTTGTAGAAGAAGTTGACTCGGGCGAGGATCAGAATATTGATGTTGATGGTGGAACCAATTCGTTTCATAGTTTCAATAACACTTTAGATTCATCTCATACCAATAAAAAATACGAGCATAACAATAATAACAACGATATAGATTTGGACAAAGATCGTGAAAACATTGCTTATAATAAAAAGAAGAAACTTGTTAAAATATCTCCTCCGAATTCTTTAATCGAAACAGAGAAAAAAAACAAAAATATATTCAACACAATAAGATAAGTTTAAAAACGAAGCATATAACTTCTGTAACTTCTGACTACTATTCAAAATTGTAAGGAAAAGTATTAATAAAAAATAATAAGTCCTTATAAAAAAAAAAAAA
>evgR010586c0g2_i01_Hydra
TTTTTTTGTTGTCGTCGATAGCCTGAGAACAATCATGGCGAACGGCAATAGGAAAGTATTTCCCGATGTCGACACTGAATCAAACACCATTTTGGAGCATGCATCATGCATAGTGCCTTCGATGGCTGAAAGTTCAATTTCAAAACATTCTTCTCATGATGATCAAGACTCAGATAGAGAAAATGAACCGATCTTTGCTGTACTGAAGGAAGCTGTCGAAAGTACTCATGAGCAAGATGATATACATACCGATGATGTTGATGGCGACGAAGACGTTAATCGTACTCCAGCTAGTAAACATCGAACAAGTATGATTAATCCCGCTCGAGTACCAAAATGGACGTTGGAAACCATGTCAGCCGCTATGGACTCAGTCCTTGCTGGAGAGTTAACTACTTCAATGGCTGCAAGACGATACAATATTCCTCGAACGACTTTACTAGATCGACTTTCAATGAAAGTAAAAGCTCAATATCTTGAGGTCCCCGGCTCATTTCCCTTATCTCCTGAAAATGACGCTGTTATTGTCAATTTTGTTAAACAGAACATGAACCTAGACAAACAAACTTTAATAAAGGGTATTTTATCTCACGCAGAAGAGTTAGCATCTCAACAAGGTAAACCTTTTACTAACCCTTTAATGACAAGAAAGTGGTTAAAACTTTTTGTTCATAAGCATCCAAATCTTGATATTCTCAGAACAGCATCTAAATCAAGGTGTTCAATACAAATTAATGAGCAACGTGATTTGGAACATCGCCTTCGTCTTACACCTCAACAAATTTCAACAATAGATTTGTCTAATCCAGAATCATCTCAAGAGTTTTCACCAGCACCAGATTCTGATGAGTGTGAAGTTAAATACTCTCAATCTACTATGATACAACGTAAACAGCATCAAGAGCTAGAAGATTCATTTGAAAACAATTTAGCAAAACAAGGACTTGATGCAGTTGAAAAAAAACTTTCTCCAGAACAAAAGTTATATTTTAGCTATCGGTTTTTTAATAGTGCACTTGAAAAAGGTTATGAATTGTGGAGGTTGTGTAAAACTAAAGTTAATGATTCTGAAAGCTTTGAATCAGTTTTGTCTGCTCAAGCATTAAACGGTATAGAAAACGCTCTTGATAGTGATCATTTAAAGTACTTTATTTTTCGATATGAAAACAGTAACCTTGAAGAAGGATATAAATTATGGTTGCTTTTAAAGGAAAAGAGTAGTTGAGTTTCATTCTCAAATGTTTTAACATTAAAAAATGTAACTGACATATTATAACCAAAGTGCGTTCACTCTGCATATTTTATATATATATATATATATGTATA
>evgR010598c0g1_i01_Hydra
GTGTGCCGAAATGGGTATCTGCAATAATCCTTCAAAATAATTTGAAGATTTTCATAATTACTTTGAAAAACAAAGTTTTTTCAAAACTATTCAAAAATTGTCAGGTAATCTTCCGAAGACCATTGTTTTTTATTAGAGGTCATATAAATTCAAAAGACACAAAGGACTTTTTATATGGCTATCCTTTGAAATGTAAAACTGATCCATCAACTATATTGGATAAAAAAATCAATGTAAAGAAAATCTCCACTGCTGAACCGAAAGCAATCAAATTTGAGAGTTCGGATTTAGTTAAACATAACAAAGGAATTGAACATCTTGAGCTAAACGAGCACTTGTGCACGGAATTGAATCAAAATAATTTACGAAATGAATGGAAAGCAGTTCTTAAGTCAATTGACGTCATATTTTTTACATTATTTTTGATCATTATAATGGCTAGTATTATTTTTATTTTTTCAACAGCACCTAATGTTAACTTTTAAAGTTGAATACTTTAAATTCAAAAAAGGGAAGAGGTTTTTAATTCAAAAAATATATTTACATTTCAAGATTTGGAATGGTTTA
>evgR010616c0g1_i01_Hydra
GGTTTCTCCTCGTACACTTTGGAAGATACACGTTTTCGATGTTCAATTAATTTTTCTCCAAATAATAGCATGGAAAAAGTTTACCTCATTTTGTTGACAGGCTTCGGATTTTTTATACCTGCTATAACAGTAACAATTTCATGTTTGTATACTGCTTATGTGATGCGCAAAAAGTATAAATACTTTAACGCGACTTACGGCAGGAATAATGTAGAAACCAAACGATACAAAGAAAAGGAAAAGAAAGCTTTTTCATCTTTTATACTAATGATATTCTCTTTTGTAATATGCTGGACACCGTACGCTACAATAGGTTTTTTTTCGGCTTTTACTTCAATAAAGGTTTTTAAATGGCTTCTGCACTTAGCCGCCTACTTGGGAAAATTTTCCGCACTAGTCAATCCATTAATTTATTACTGGAAAGATGGACTATTTAACCAACGTTGCCTACAAAAAAAATCACAAATAATGAAGTTATTGACGAGGAGGACAGAAGAAAGCAACCGACCAATAACTTGATGTTTAGTGCATAGTTCATAAATTGTATAAGAAATGTTAGTGTTTGG
>evgR010617c0g1_i01_Hydra
GCGTTCCAGAAGATGTAATTAATGATTTCAAGGTTTCTGAAGAGGATCCAAACAAGAGAATTGTTTCATTGAAATATCCAGAAGTTACACCTGTCATGAAGTATTGTAAAGTTGAAGAAACCAGACGTAGATTGGATATTGTTCGTTCTTCACAATGTCAAGAAGCTAACGTACCATTATTTGAACAAGCTGTAAAGATTAGACAAAAAATTGCAGAAATAATGGGATTCAAGACTCACTCTCACTATGTTTTGGATGTTAACATGGCTACCCCTGATGAAGTTTTAACCTTCTTGTACAATCTCAGAGATAGACTTTTAGAAGGTGGTAATCAAGAACTTGAAGTTCTCAAAAATTTAAAGGGATCAGAAGTTTTTTC
>evgR010639c0g1_i01_Hydra
CTTCATTATCTTCATTTTACTTTAGAGGATGCCTTTGAAATTAAAGAAAAGAAGCTCTTGTCTATTTTCATTTTTTGAACGAAGTAACAAGAAGTTAAAACAAGTTAATGACGCATACGTCAGATTTTATTTTAAAATCTAGAATTATATTTTTTCTAAACAAAACAAAACATAGAATCCCGGAAATATGATTTGCTGGCGGACAAAATTTTTTCCTTTACCAAATCTAATGATATTTGCCACCCTACTTTCCATATTGGTTATTGCAGCAATTAGTCTTAACTTTGTTGTTGTTATCATTATCGCCAGGATGAAGAAAAGAGAACTATGCAGTTATATATTACTATCTCTTTCAATCTCTGACCTGCTTATCGGAATGGTGCTTGGTCCAATAACAATTGCACAAGTTTTAGATTATAAATTGCTATCAAACTGTACAGCGCATTATATTCGCGGTTATATTTTAGTTTTGCTTGTTGGCTCCTCCTTATTGACACTAGCAGCGGTGTCTTACGATAGATACTTACTGCTGACTAAACTATCTAACTACAACCAGTATATGACTAGAAAAAAAGTGATTTTACTAATAGGGTTTGCATGGTTGTTTCCCGGGTTGATACCAATTGCCAAAATGTTCAATATGGCGTTGTATGTTATTTTACGTATAGCGAATTGTACTTTTCCATTAATTGTACTCGGTGTCTTTTATTATCTAATAACACAAGAAGTTCGTAAAAGAGAAATCCATTCAAGTCGACATAGAATGAATAGCTGTGTTTCAAATAATGATATAGCCACTGTTAACAAGCATTTCACTAATAATAACAAGTCAACAGATAACATTGGCCGATCAACAGATTCTTATGTAAAATGTTCATACAACGATAAATTAAGAAAGCACAAGAGAGTAGCAAAGTCTGTTACTTTGTTAATTGCCTTCTATTTGGCATTTATCTTTCCTTTAAATATTTGGATGGTTATAGAACTCGTAAATGTGGAATACAGTGCAAAAGCCCATGAAATATTCTATATGTGTGCCGTTTTCCTTATGCACGCTAACTCATGCATAAATCCGATCATTTATTATTCAAAGCAATCAGAAATTAAAAAAGGTTTCCGACAAATCTTTAAATATCCCAAAAAATGTTGGAGTTTAAAGCAAATTGACAGTCGACAGCAAGCCTACAGTTTGACAGGCGACAGCAAGTCTAAAGTTTGGAATTAATAATTACACTTTCTTTTGTTATAAATAAAAGCTGTTGCGCTGTACAACTTACTTTATAATATTATTTTACAAAAAAAAAA
>evgR010661c0g1_i01_Hydra
GCCAAGGTTAGAGTCTCATTGTGGGAATAGTAAATTTGTGTATGATGAGTTCCAGTATGTTTCTATCAAAAAAACTTTATTTAATTTGTTACAGAATGAGGCTTATGTAAAAGCATTTCTGGAGAAAAAGTGTAGACCTGGTGTATTAACCAGTTTTTCAGATGGAGGTCGTTTCAAAGAGTGTTACCTCTTTAGTGATGTAACTAAAATTTCCCTAATGATCCAATTGTTTTATGATGGTTTAGGTGTAACCAATCCGTTGAGAAGTCAAGGCTCTGTTCATAACATTGGTGTTTTTTATTACACTATAAAAAATCTTCCTCACGCATTTAACTCTTGTTTTGGCAATGTGCACTTGTTAGCCATTGGTTACACTCATGATATTGCAGTTTATGGTCATGGGCCAATTTTAAGCAAATTTGTTAAAGAGATACAGGAGTTAAGCACTATTGGATTACAAGGAGTATTTCCAGTTTTAGGTGCTTGTACAATTTATGCAAGTCTTTGCCAAGTTACTTGTGATAACTTGGCTCTCAATGGCTTGCTTGGATTTATTGAGTCTTTCTCTGGCAGTCATTTTTGTACAATTTGCTATGCAAC
>evgR010684c0g1_i01_Hydra
TAAAACCTTATGGAATTACTTCAAGAAGCTTGTAGCGGATTAGATTATTTAAATGTTGAAAAACTTTCGAGAGATATCATCAATACACTTGCTGACGAGTTGCCGCTGCATTGGAAGAAACTAGCTCATTTTTTGAACATAAATGATACAGAAATAGACAGAGTATTACACGAGTACGATACAACAAGAAAACAAGTTTATGAGATTTTTCTCAGTTGGTGTAGAAAAAATCCAGATAAAAAGTGGCATGATATAAAAAATGGATTAATTTTTTGTGACCGAAAAGATGTCATTTGTAAATGCGAAAAAAAAATAAATGTATGTTCATTATTTGGTGTCCAACCTTCTTCGGATATTTTATTTTTACGCGAAAAAGAAATAAGC
>evgR010695c0g2_i01_Hydra
CAGAAACAACAAGCATTGCACCGGACACACTATTTGAAGACCTCCTGATTCAGGACAGGCTTGAAGTTGAAAAAGGACTGAACAACTTCACTACAGCTAACAGGCTTGAGATGGGACCCCTTGAGATCTGGCTTCAAGTCGACAGAGTACTGAAGAAGATTGAAAGGAAATTATACTTCAAAAGACGTAAAGTTGCCAGAAAAATCAGACAAAACCGAAGAAACAAATCAGAACTTCACAAATGGCTTCTATTAGAAAGAGGATTGAAGGACACCGCCAATAAGATAGTCGACAAAAGACGTATTGTGTCAAGAAAAGTCAGACAAATGCAAGCCGGAA
>evgR010703c0g2_i01_Hydra
CATTGCTCTTATTAAATTACATAAATCAAAAAGAAACTTTCTATCCTTGCGGAAAGAGTGATTTCATTATAATTATAAAAATATCTAGTGTTGATATGAGATCAGTTGATATGAAAGATGTTAATAGTTATATGTTATATTTGGTTTTATGGTGTAGCTTATGCAATTTGCATAGCTGCATCATAGCTGTGAAGCGCTCGTGTAGCAAAAATTCCTATTTTGCGGATATTGATGATATAATCTCACAACTACGAGCCTTATTAAACGAGATAACATCCTTTGTTTCAAACGAGCTAAATACAAC
>evgR010723c0g1_i01_Hydra
TACAATTGAAGATTCAGAATGCACAATGGAGATTATAGAAAAAGAGTTTAACCATAGGATAGCAGAAATAGTGGATAGATTAACCAATAAAAGGTTTGAGGATGGAGTGTGGATTAAGTTAACACTGGAGCAGACCTTAGAAAAACTAAATAAGCTAAAAGATGTGGAAGCAATGTTCATCAAGCTAATGGATAGACAACACAATCTTGAAACGATAAAAGGATTAAGTTCAGAAAAACAAATGAAGATGGCAAAAGAAACGAATAATCACTTTATCAAATGGATAGCAATAATAGGGGATAAGTTAGGCATTATTGAAAAAATGGATCTTGAAAATATGATGTATGAGCTTGATAAGAAGATTTTAAAAGATAATAAAGAAAAGTAAAATATAAC
>evgR010736c0g1_i01_Hydra
AGTCAAGATAACATTGCACATACTGCCACAATGTTAGCCTCCATAAATCAAAAATCTGGCCTTATAAATGCAACAATTAATAGAATACCAGTGCAAGCCCTTATCGATACAGGCTCCGACCTATCATTTATTTCCCTAAAATTTACGTCTGACAACGGTCTCAGCTTCAACACTAATATAAAAAAATCAGTTTGTCTTGCAGACAATTCACCCGCACAAATCTTTGGCCAATATGACGGTACCCTAATAATAGGAAACCACTCCTACTCAACTCAATTCTTTGTCGTGAAACAACTAGTATCACCCCTTATCATTGGTATAAACATTTTAAATAAATACTCGGCCCTAACACTTTATTTTGGAGGAAAACGTGAACCAATCCAGTTCTGCCTTGCCCTATTTTCCATGAAATGTAAATTATATGAGCTTATTCCTGGAATTGACACTTCCCAACTTACTCCAATTGCAACTCCCTCACGACGAAACAGTTTGAATGTAGACTTTATCAATAACGAAATTACTCGTCTCTTAAACGACGACATTATTCAAATCAGTCAAAGACCCTGGAGAGCACAATGCTTCGTTTCAAACCTATCAAAAAAAC
>evgR010751c0g1_i01_Hydra
TTTATTTTCTAAATAAGCAAAAGTAAAATGAAGCTCAATAATAATGAATGATTTTGAAGTACCTGTTTTGCAACATGCTCCGTGGCCTACTCCATCACGAGCTATTTATGGATTTGTTCTTTATGTATCAACATACGTAATTTTTGGTTTTTATATTATGTGGGCATTACTTCCAGACAGCTGTCTAAATTATATTGGGTTGAGCTATCTTCCCTCAAGACACTGGGTAATAACGTTTCCGTTTCTTTGTTGCATCATTGTCGCTCTTGTTTATATTGGATACAACGTTAATTATTGGAACAATCTCCCGAAATTCGACGATGAAATTAAAGAATATAATTCTTGAAATAAAGAATACAATACACGACCTATGTAAAATATTTGAAATTTTTTAAATTATTTATATTGAACATTTTTGTCCCGTAAAGTGAAAACTTTTAATGATAACTGTTTTTAATAAATGAGGCAGAGTAGAAAAGCAAACATAAATCTTGACCAATTTTAATTACAAAATTTGCGTGAGACAGATTGGTGAGCGATTGGTAAAATAAGGGTGGTCAATTGCACCCGACGACTACTTCGTGCTATGAGCATTATTGAATGCAAACATAGTTATGTCATAAATGTAAATGCACTAAATAAGAATAAGTTATTATCTGCAACCACTAG
>evgR010755c0g1_i01_Hydra
CAAATGACATATGTGATGTTTCAAATGACATATGTGATGTTTCAAATGACATATGTGATGTTTCAAATGACACGTGTGGTGTTTCAAATAACTTATGTAGGGTTTCAAATGACATGTGTGACATTTCAAATGACATATGTGATGTTGTTGCGCAAAACACAAGTAAAAACTTATTTCTAGAATTTCAAAAAAATGAAAGTGAAAAGTTACAGTTTCTTGAATCTTCTTCAGTTTCTCAAATTGAATCAGCATTGTGTCCAAGCACTCTCACACAATTCTCACCAGTTGATAGCAATAAACTTCCATCCGATTCCTGCATTGATTTTTTGTCGTTAGAATCTTACATTGAATCTCCATTAGATTTCCAAAATAAACCTCTATCATGTTCACAAATTGAATCTCCATTATATTCCCAAAATAAACCCCTATCGTGTTCACAAATTGAATCTTCATCAATTCATATTGAATCAGATTTTAATAGTGAAAAATTGTCTGATTTTGAAACAAAAATGTATGAAAATAAATCATGCAATTTTTTAAACTCTAAACATATCAAAACAATTGAGAACAATCTAATAACTGGTAATATTATTGATGATAGAGAAAATATTGGAATTCTAAACAATAATCTTCATTCTGATAGAGAAAATATTTGTGATGTTCAGCATGAAGTTAAATCATTATTGAAAGAACTGCTTGACAACATATCAAATCATTTTGGTCAATATTTTACTATGATATCTAAGAATGGAAATTGTTCAAACATTACCCCAACAGATGCAATATTTGATTGTTCTGACAAATCAACTGTTCAAACTTTGTATGTTCCATGTTGTTTAGGTACGGGAGACCTCTCAATGTCAGCTGAAGAATTAGCTAACGAATATGGAAATAATGCTTTTGCCATAAAAACAAAATTACAACTAAACAATAGCATCTTTGAAAATTTCTCAAAAGAAAATGAAATTATAGACATGTCTAAAAAAAATACATCTGAATGTCGTCAAGAATCTCAGACAGAAAAGATACCTAGTTTAATGAATTTGTATAATTTAAAAGAGCCGTTTAGTATTGACTCTAAAATTTTAATAAACTTTGATAGACTATCTCCCATCTGTCCGACAAATGATATTCCCGATAATTTTGGTGAAATTAATTCACTTTTAGTAGAAGAACCTTTAAAAAAATCGATTGACAAACAAAATTTCTTTTTTAATTCCAATATGAATATCAAGCAAGAAATTTTCGATGACGATGATGATGGTGTGGTTGATATCCATAGATTCGGCGTTCAAAGTTTTTCACAAACAGCTTTGAAATATTTTAGCTTAGAAGGTAAAGATGATAAAGCAAAGCTTTATCCGTGCTCAAGAAAAAGACTTTATTCATGCGATAAGTATAACACGTGTTCCAGAAAAAATTTTAAAAAAATGGATTCATATTTGCGGGAAGACTGTTTATATAATAGCACACGTCATAAGAATAAAAGTGCTGCTCTAAAGCAAGAGTTTATACGTCATAAAAATTGTACTCCTGCGTGTTTTGAATTTGATAACAGGCACAAGTTTACTAAAATAAAAGATGCTCGTCGTTCGTCTTCCTTGGAAACTCAGAATGTTTTTAAAAACCAGAAGAAGCATACTTGTAAAAAATGCAATGCACAGTTTCCTTCTCTTAAAGTATTTTGTCGTCATCAGATGTTCTTTCATAGAAACAGTTTAAATGACTGCAAATGCTTGATATGTGGTCAGTCTTTTCTTAGACGATTTTATTTGCATTCACATTTAAAGAAGCATAGCGAATTTCCACATAGATCTAATTTAACGTTGTAGCTTATTTTCTTATATAATTGTGTTACTAACAAGGTCAATCGGTTTTCTCGATTAAAATTTTGTAGCTACGTCATAACTGTAACGTCAACTTTTTATCCTTTTTTTTTTTTTTCGTATTGGTTAGATCCAAATGTAAATATTTTTTCATTTTTAATATATTTTTGTGCTTGTTTTTATATTAAAAAA
>evgR010798c0g1_i01_Hydra
TAATCCCATAATCCTTAGAGTCCCAAATAATATTTTGTGAAATTGACCAAAGAGCAACATGTTGAATGAAATAGACATTAGCGATAAGATATGTTTTAAGTCAATGGAGGAATATGATTTTATTAAGTCATTAAGTTAAGATGAGATCCAAATGTATTCTGAAGTCACTTGCATTGTACATGGAAAAAGCCAGCAGAGTTGGTGTCATAAAATAATTAAAAAAGTTCAACAATTTTTGTCAACTAATGCTCTACCACTATTTTTGATAATTAGTGTGATTATAGGTGCCTTAGTTCCTCAAATTGGAATTGCCATTGACCATAAAGGAACAACTGAAGTTTGTCTTGTTATATTATTTCTTTATAGTGGCTTGTATCTCAGTACTTCATCTATAATAGAAGCTTTAAAAGCATATAGAGCAGTTATTTGGGGTACTCTTAGTATTCTTTGTGTAACATCTGTTATTGGTGTTCAATTAACCAGTCTTATGTCGTTTCACAGTCCTTTG
>evgR010811c0g2_i01_Hydra
AGTGAACGCTGATATATCTTTAAAAAATATTATAAACATTCTTAAAGATGGTCTTTCAACAATTTGTAAACAATTCCAAAATAATCAAAAGAAATTTGCCGACTTTTATGTTAATACTCGTAGCATGCTCGTCAGTTTTTGCAATCGGGCTTATTGTCGGTATTTATTATGAAAAAATTGATACGGCGGAGGATACCCAAAATTATCAATATCTAAACATCAGAAATCACCCAAAACAAATCAATCAATCTGTAATGAAGATCACAAAAAGCAATAATATTTCGTCCACTTTAATAAAAGAGAACTACATTGATCATAGTAGAAAACGTAACTATTTAAAAATCAATGCCCATATCAATTCTCTTATGCATAAAGATATTGTAAATTCAAGAGAACTTGTAAAGAAACCTAGAGATACACTTTTGCCAAAATATCTGCAAATACGAGAAGCAGACAGTTCCATACTCGGAGATTTTCTGTGCAACAATTCGGTTGTTGCGAGTTGTTGTTCGTTTTATGTTAATACATTAGAATCTGCTAAATACATATGTGATTCAATGCAAGACGATTGCAAAGGTTTTGTGTTAAACTCGAAATTGTCTTCGAAAAACTTAGCCTATTTCAAAAATAATGTTTCAAAGATGAACGTGAATTTTTTAACGGACTTTTACGTGAAGATTAATTTTTTTCAAGAAATTGCTTGGAGGACCTCATCACAAAATTAATGTTCTGGAAACAAACTTTTATGTTGACATAAACCCTTCAATTTAATAATATTACGGTGTTTAAAATTTATAATTTGTACAAATTTTCATGT
>evgR010882c0g1_i01_Hydra
CTGATGGCCCCTTGGTCGTCCATATTGACTGTAAACTACTTCCAGCCATCAGTGGAGGACCAGAGAAAGAAGACAGTATTATAATTCTGGTAACAGGTTGTCATACCAAGAAACCTCTTGCCATCCCCAAAATGGAACAAGTGAACAAGTGGCACAGTCAAAATTTTTGTGGAATCGCTAAAATATCATTTGACACAACTGCATCTAACACAGGTTGTTTAAATGGATCTTGTGTTTTACTGGAGAAGTTACACAAAGATCTACTCTGGGTTGCTTGTAGACATCACATGCTTGAAATGGTATGCGGAGATATGTTTAAGAAATTGTTTGGTCCAACATCTGGTCCAAATGTAGCAGTATTTTGTCGTTTTCAGGAAATCTGGCCTCAAATAGACCAAGGTGATTATCAAGTGTTCTATGGATAGTCGTTTAAATGCCAGTCTTGACAACTTCAAGCATTTTCGCAAAGAATGCACTAGCGGGACAAGATGGATGTCTTCCAAGATCTTCTTGAACTAACATTGATTTTCCAGAGACACTGGCCTCCAAGAGGTGTTAGTTTTAGAGTTCCTGGTGCAT
>evgR010883c0g1_i01_Hydra
ATAAATTAAAAAATTCAGAATGGAACGCCAGCCGAAAAAAATTATACAAACGACAATTTTTGTTCTATGTTTATTGGTGTTAACTTATTTTGCTTATCAACCAATTTTCAGAATTAAAAAAAGGCAGCTAAATGGAGATTTTCTTCAATATTTTACTCATAAGCGTTATTTTGATAATTATATCAGCGCCCGAAAAGCTTCAAAAACCCACGTTCTTCGAAATGCTTCGAAAACCCACGTTCTCTCTTACTCACTGTTTGGGAATAACTATGATGAATTCGGAGAAAAAATTGAATTGATTGCAAAAGAAGCGGCAAATGATAGATTATATAATAACTGGACTGTTCGCATTTATCATGACAAGTATTCGATAAAACCAGAGATTGTTACGATATTAAGTGGACGTTATCAAAATTTAGAGTTTATTAATGTTAGTCACTTTTCTAACTTGATAAATATTAATGGAATGATCTGGAGGTTTTTAACAGTTGCAGATTCAACTGTGGATGTTGCGTGTGCTAGAGATTTGGACTCAAAACTATTAGTTAGAGAATCAGATGCTGTAAATGCTTGGCTGAAATCTGGAAAGTTACTCCACGTCATGAAAGATCATCCTCGGCATACTGTCCCAATCATGGGTGGTTTGTGGTGCTTCCGAAATGAACACAACCGAGAGCTAGGAATCAAAATAGCAAAACTTTGTACCGAAAATAGCATGGAGCGTTACCCACTCATGCAGTCGGAAGCGAAAAAAGGACATGATCAGAAAGTCCTAAAATGGTATGTTTGGCCGCTTGTCCGCCATAATGCATTAATACACGATTCATATTTGTGTGAAGAGAATGACTTTAAAAGTGAACCTTTTCCAACTAAAAGAGACAAAACTGGAGAGTTTGTTGGGCAGTTCAGAGACGGATCTGGAAGATTTATCACCTGCCCAATAGCATGTCGTCCAAAAGAGCATCAAGATTGGGAGTACTGTTAATAATATTATTATTTTACTTAACATAAAC
>evgR010900c0g1_i01_Hydra
ACATATAAAAATTCACACCGGCGAGAAACCATTCAAATGTAATCAATGCAATTATGAATGCTTGGACAAAAGTAACTTTAGAAGACATATAAAAATTCACACCGGTGAGAAACCTTTTAAATGTAATCACTGCAATTATGAATGTGCACATAAAAGTAATTTAACAAAACATATAAAAATTCACACCGGCGAGAAACCATTCAAATGTAATCAATGCAATTATGAATGCTTGGACAAAAGTAACTTTAGAAGACATATAAAAATTCACACCGGTGAGAAACCTTTTAAATGTAATCACTGCAATTATGAATGTGCACATAAAAGTAATTTAACAAAACATATAAAAATTCACACCGGCGAGAAACC
>evgR010941c0g1_i01_Hydra
TTTTATTTTCTAAATAAGAATATTGATATAGAAATATGAAAATTGTTAGTTAATGAGAATGAATGAAAATGGATTGCATAATAGTATCACAGTACTCAAGGAAGAATCATTAGTCCATAACAGTGAGTTTGATCATCCAGTCTACAAAAAAATATCTGCAGCTAATGGCCGCATTAATAAAATGACAAATATTGAGCTTTTAGAAGCTTTAACATATTTGAAATTGGGAACAAAAGGAACAAAAGAGATTCTTCAAAAACGATTGAAAGCTCAAGTGAAGAAACAGCATCTAGCAGAAAATAATTTGGTTGCAAAACCAAAAAGCTGTTATGTAGATTATTATGTTGTCATTGATTTTGAAGCTACCTGTGATGAACCTAATCCCACTGGATTTCAACATGAAATCATTGAGTTTCCTGCTGTATTAGTCAAAACAGATACCTTAGAAATAGTATCTGAGTTTCATTCTTATTGCCGTCCTGTTATCAATCCTGTACTCTCAGAGTTTTGTAAATCTTTAACTGGAATTACACAGGTTCAGGTAGAATCTTCCTCAGTTTTTGAAGTTGTCCTTCAGAGGTTCGATAAATGGTTAAAACAGCAGGTTGTCCCTACAGAAACGTTTTGTATTGCTACAGATGGACCTTGGGATTTAGACCGTTTTCTAAAAAATCAATGCAAGACACTTAATATTCAAATTCCGCATTATTTTCACAGATGGGTTAATATCAGAAAGCATTTTTACAATTACTATAAAATACACCAGGTGAATGTAGAACTAATGCTTCAGCATTTAGGAATGGAGTTCGAAGGACGTCCTCATTGTGGAATTGATGATGCGCGCAATATAGCACGTATTTTAATTCAACTGATTAGAGATGGAGCAGACGCTTTAATTAACGAGAGTCTTCGATAAATTTAGAAGCATTGAATTAACCAGTTTTATGGTGTGTTGATATGAAACCATATGGCATCTTCTCTGTTTTCTGGAGAAATAAATTTTCATCGAACTTATTTTGTTCATAAAAATAGAAATAAGAAACACGAATGAAAATGGGTAACGTGAGGAGGAAATCTTAAAAATTAAAATAGCGATGAAAATATTTTAACTATTGATGTGTTATGTTTTTTACATGCTAACGTAGAAAACATAGTTTTCATTTGTTCTTGTTTAATAAAAAAGTTTATTTAAAGGCGCGAAGAAAAAAATAACGTTTGTGAAACATTTTCTTTGGTTAAAAAATATTTTTTTTTAATATGTAAATATGTAAAGCGTGTGTTAACTTTTTTTGTTTATTAACTAGTATGCAGTGTGTTTTTTGTTTAAAAAAATGTTTTTACTGAGTTTGCAGTTATATATGGTTG
>evgR010944c0g1_i01_Hydra
CCTATTTTAGTGCAAAAAGATAAAAAAATTTAATGATGTCAGACTGGAGTGGTTTTACAGAAGAAGAATTAGAAAAGTTGAAGCTCGAAAATGCTCCAAAACTTCTACAAGATATAAATAATAAAAAAAAAACTAAAAATACAATAGCATCTCATGAAGCTAAAAAAAGGTCTATTCCTCAGTGGCCAAAGCCAGTAGAGAATCCAGCAAGTTCAGTTGACTTTAACCATGTTGATGTTAACCCTGTTAAAGATGACATAAAGTTGAAGCAATCTGACATAAACGAAAGTACTTTACACAAATATAATATAAAAGCAAATAAAGTAGACGGCGTTGACTTAATTAAGTTGAATCCGGTTGAAAAAACGCAAGATGAGCCTGTTACAATAAAGATAGCAGAAGAGGCATCTAAAGACAATGTTCGTGTTTTAACAAATGAGCAAGCTCTTGAGTTAGAGAGAACAAAAATTGAACAGCTACAAATACAACACAAAATAATTGAAGACCAAAACAAAAAGAAAAAAATGCTGATTACAAGTACGCTTCAAGAGCGAATGAAGCAGACAATGAGCGAGTCACAGAAGTTAAAAGAAGTGCAGCAGCAGTTAATTCATTTAGATGAAAAATTGCAAATTGATGTTGAATCATTGCGCAATAAAATTGAAGAGGCAAGTTTATTATTCTCACAGGCAGAAAAGCGATATGAGCGTTCCGAATCAGAGTATGTTTTAGCAAAATGCGATTATTTTAAAAAAAAGGAAGAGAAAGAGCAACTAGTTGAGCAACTGTATGCTGTCATCCAAGCAAACGAAGAAAGAAAAGCCAAAAAGCTTGAAGAGCTCATGCTGAAGCTAGTGTCTAAAACAAATGTGGAAAGTTCTACTTGTGGTCCTGCTAAAATTTCAAATGAATCTAATACAAATACCATTACCTCATGATTGCATCATCTTTTATATTTTTTATAAGTTTTATTTATGTGTTTTTAACTTTCCAACATAATTATTTATAAATTAC
>evgR010946c0g1_i01_Hydra
AGATAATGTGTATAACAGATAAAGTATTGAAATATGCTGTAAAAACAGGCCATCCGCGGTTTTTCAACCAACTGTTTTCTGGTTTGGACATCACTTGTTTGATGGGACAATGGATCAGCGCAACAACTAATACTTTAATGTTTACTTACGAAGTAGGGCCTGTTTATATAATGATGGAAAAATATCTTCTCGATAAAATGAAGTCAATAATAGGTTACCCCAATGGTGATGCACAGATGTTTCCTGGTGGTTCGATCTCAAATATGGAGGCTATGAGCATTGCGAAGTATCATTGTCATCCCAACTTTAAAGAAGAAGGTCTTTTTGGAGGAAAACGATTAGTAGCTTTTGTATCTGAGGAAGCACATTACTCGTCTGATAAAGCTGCTGCAACATTAGGCATTGGCACCAACAACCTAAAAAAAATAAAATCTGATGAGAATGGGAGAATGATTGTAAAAGATTTAATTGAACAAATTGAGGTAAGTTTGTCAAGTGGAGAAGAACCATTTTACGTTTGCGCAACTGCAG
>evgR010949c0g1_i01_Hydra
AACTCTCCAAAATCAAAGATCTGAAAATAGATATTTAAAAAAGCAAATTTCGGAACTCAGGCAAGCCATTGAAAAAAATGCAGTATCAGTTGGAGAAAACATCCATCAGGATCTTCAAAAAATTATTAATAGTTTCTCTGAAAATGTAAATTTAAATAGAGAAATTTCGTCTTTCATGAAATTATTCTTTTCAGAACAGTTAAAATATTTAACAGAACATCCATCTCAAACACGATACCATCCAATGATAATAAAATACTCTTTAGCTTTGTATGCAAAATCTCCAGCTGCATATGATCTTCTACGTTTACAAAAAGGTGGAAAAGGTATACTTGTTTTGCCATCTCAACACACTTTGAGAGACTATAGAAACTACATCAAACCGCAAAGGGGTTTTAATAGTCA
>evgR010967c0g1_i01_Hydra
ACAACTTCTTAGGATCAAAATATATAATTGAACGAAATTGCATTGGTATGTACACATTGCGATGAACCTTACACGCACACTCGATATACCAGTTTTTGAACTACCAGTAGTTAATACATTTGCTATTATTGGTACTGGTACTTACAAAGCAGAAAAAAAAATTCGCCCATTAAGTCTTTCCAGCGTTAAAGAGCTACGCTTCATTGAGCAAGACAAAGCAATAGAATATAATAAATCAAGTGAGGGGAAAGAAGATGAAAAAAAGAAACTCCGATCATTACTACTTGCTACAACGGCATTAGCAAAGTTTAGGTCGCATGCTAAGCGAAGAGACAGACGTGGTAAATCTCCGGGTACTTCCCCGCAAATGAGAAGAAAAAGTGTTACGTCATTTTTTAAGGATGAAATATTTGAAGGAGATTGGTGCAGAAGAAATCTTGCTTCAGACATTGACTCTAGCCACAAACATTCAAACGATGATCTTAACTACAAT
>evgR011015c0g1_i01_Hydra
TCCACTAATTTCAAATTTAAAAAAGCGGGTTTTTATTTCAAAAATGATTCATATTAAATTGTGTTAGTATAGATACATGCTGAAAAAAATCAATGTCGAGTGCAAAATTGTTTTATAAAATTAAATATTTTAATTCTTATTATCAACACTGTTCATATTATTAGTCAATAAATCTACCATGAGTGCAACTCCATTGCAAATGCAATCTTTTCAAAATCAGCAAAAGAACTATCTATACAGCTCGCCAAACACCTTATCCAATAGTCGTAACAGTGTGCCGACTAGTCCTTTCCTTCCTGGTTATCTTTTTGGAAACTCGCCTCAACCAGGGAATTCTCAAATAAACAGTGGAGGCTATTCTAACAATCAGAGATATGTTCCTAATTCAACCTCATCGGGCTCAGGACAAGGTTTCTCATTTTCTTCTCCTGATCTTCAAAAGTATCAGCCACAACAAACTAATTTACGACATTCACTTTTGAAAGAAGGTGCACCACCGACTGAAAGTTTGTTTACTCCTTCAAGTGGATCACCTGAAAATCTTATGGAGACAGACACAACTTATAGAGATCTTCAATCTCCGAAAGAACTTTTTTTGACGCCAAGTAATCAATCATTTAATAAAAGTTTGTTTCCCACATCTCATGATGTTATTCCACAATCTCCAGCTCAAATAGATCCATTCTATACTCAAGATGATGTAATAAATAGTGATGATATAATTGATGATACTGCTGTTACAGTATTTGGTTTTCCGCCTGCAGCATCATCATACATTTTGCAAGAATTTTCTCAGTATGGTACCATTGAAAAATATGAAATGCACAATGTAGGAAATTGGCTACACATAAAATATCAAACTCGTATACAGGCAAAAAAGGCTCTTAGTAAAAATGGAAAACAATATGCTCAAAGGATTATGGTTGGTGTACTTCCATGTATTAAGAAACAGATTAGCATGGATTGTGATATGACATCTGCAAGTTCCTTAAATATATCAACAAATACACCGAATAAGCTGATCAACAATAAACCCAGTACAATGCGACAATTATCGTCAAGAAATGCTAGCGGAACATCTACTACAGATTTTTTTACAAAAGATAATACTCCACAAAAAAAAGATTCGGTTATGTCAAAAGCATTAGACTATGTGTTTGGATGGTAAACACACTTGATGTGTTTTTTTTCTAATGTGGGTGTTATAATATAAAAATTTGTAAAAAGAATTGTGCAAATACATTTATATAATTTTAAAAAAAAA
>evgR011029c0g1_i01_Hydra
CATGTACGTAATAGTATTTATCATTGCAGGACCATAAAGAAGAGATAACAGCATACATTGGAACAATGTCCAGTGTATTACCTTTGTTGTACGTTTTTATATCGATTATTATACTAATGTTCGTTATATCGTGTTTTGGTTATCTTTATCGAGTGTTTCATAAACGACCAACGTTTTCAAAGTTATACGCTCTAAAAAAAAAGTTGATTGGCACATCAGATGAAAGTGCTACACTTTCACGAAATGAAGAGAAGTTGCCTAGAGATTCTATTGAATCTGACTCCTCAATAAAAATGCATCAAAGCGTAACAATTGTTCAAGCTTCAAAAAAATATGAAGAAAATGAATACGAGGTGCCTCTGGAATCAAGGTGCCTTCAAATTAGAAATTTAGAATCTAAAAGTCTGGAATCTACATGTCTAGAACCTAAAAGTGCATACAGTGAAAATTTAAAAAAATTTATACTTGTAGAGATTAAACCAAACGATCATTACACTGAACTAAACCAAATTTTATGTAAGAATGACGTTTGCAAAGTATTGGTTGAAAATATGCAGAACGACGTTATTAGACGGAATGAATTTCCTCTTCCACCGCCTTACAATCATTCTTTATTCAATAATTATTTGAACGACTAATAAAAATTATAAACAGGCTTATATATTAAAAAAAAA
>evgR011035c0g1_i01_Hydra
TGTCAAGATGTCCTACACAAATGACAAAACATTTTCTGAAATAATTGCAGAAGGTGAAGAAAGGATGAAAAATCTTCTGAGGAAGATTGTAGAAGAAGAGCGACTGGAGGATAGACTGAGAGCAAATTTTGAGCAAAAAAAAGAAAGCCAGTTACTGGAAAAAGCAGCAGTAGAAGAAGCTTTGAGAAATTTTGATCAGGAATTTTCCAGTGGATGTAGCAAAACAGATAATGATTCAGACTTTGAGTCTCTTACTGCAAAGAAATTTTTTTTTATATTATTCGACAAAGATTTGCCTTCAACTCAGTCTGAATGATGTTTTAGATAAGTGGATTCCATTTATG
>evgR011040c0g1_i01_Hydra
GTTTGGATGGTGAAAAAGTGCATAGTGAAAAATTAACAGAAAAAATCCGAATTTTGGTGATAAAATTGAAAGCTAAGTGGATTGCAGTAAACAAAACATATGACAGGCTGTTAATGAAAGAAAAGAAATGGTTGCAAACCGAAATTCAAATTGTTATGAAAGCTGACAATGTAAGCAGAAAAGAGTTCAGTGGAGGGCGACCAAAGAAAGATTGGAAAGATCTAGGAGAGAGAAGTAAGCGAACAAGAGTTGTTGGATTATATGCGCATGATCCAGAGGTTTTAGCTCTGGCTGCTGCTAAAAGTGCAAAATCATCAGGTGACTTACAAGACTTTGCTTATGTGGTTAAAGAGTCTATTCAGAAAGCATCTGAAATCAGAAAAAGTATGGACAGCAAGGAACCAATAATGATGAGTGAGAAAGAAGCACTTTAATTAAAAGTTAATTGTGATCTCAGTGATGATCAGTATCAAATGATAAGAAACTCCTCAATCAAACACAATGCCAATATTTATCCAACATTACATGCTCTATTCAAGGAAAAAAGTAATTGTTATCCAGAGGACCTTGTTGTAACTGAAACTTCTGCTGAATGTAAATTACAGTCCATGGTGAATCATACTCTTAAAAAAGTTATTGTATTAAGTGAAGACAATTTAAAAACTCTGGATCCAGATCAAAATGTGCTCAAT
>evgR011076c0g1_i01_Hydra
TAGGAAGTTGTTTCAATAAAAAAAACTCGGATTTTTTTTTTTGTTTTTGTTTTTTTGTTTTAATTTTATATGAATATTAAAATGAACGAAACATTTATGAGTATTTGCGGTCGGCCAAGTCAGCAAGCCTGTGATGAAATTTTAAAAACAGAAATACAAAGTGAAGAAAAGGAAGAAGAAATTAAATTTAACTCAGCTTTATCAAATCTTTTAACAGAGGACAACTGGAAAATCGTTTCTAATGGAAAAGGAAAATATAACGTACTTTTTAACCCTTCATTGAACTTCAAAGAGAGAATTGATATCTTAGAAATTCTTTTGGAAGATATACGTGAGAGGTACATGGACTTGAAAGTTGAATATCAAAGATTAAGAAGAATATGTCTTCGACAGAAGAAAAAGAGAAGATTAGGCTCTTGAAAAGTTAAAATTTCATGCTCTTTCTTTTACTGTTTTGCAGTTAGACTTTAATTAAATGATAAATTGTGTATTTTTTAAAATTTTAAATTCAAAGTAAATAGATTACTTCGAAGATCATAGTTAAATCTACAAACTAATGAAAAAGGTGACAAATTCACTAGAATACTATGC
>evgR011083c0g1_i01_Hydra
ATAATCAATGTGACAGTGAAGATGATGACTGGATATGTACAGATTGTTCATATAGTAACTTTTTAAGTAGAAAAGTCTGTCGGTATTGCCAGAAAGCAAGAGATAATAGTGAAACAGAGATTGAAAAAGTAGCAAAAAAAAGACCTTTGTCTGAAAGTTTTGATCTAAATGTTTATAAGAAGACAAAGTATCAGAGAATAGCAGACACTACTAATACCAATGCAAAAAAAAGACACCTTACTGATGATGTTATTGTAGGATCAATAAAACAAAAGAAAAATCAGGCATCTTCAGTTAATAATAAGGTGGCATCAAATACAAAAAATGTGTTAATGGTTGATCAAAGTAATAAAGAAAATTTTAAAAGTTCTAATCAGTGCACAAATAAAGTACAGAGAAATTCTTTAGTAAACTGTGAAGTTCTACTTAAGTCTATAGGGTGTTCCAAGTTTACACATATTCAAATTGAGAATAGCGTTACTTTAGCACCCCATGTTTGGTTGAAATGTTGGGAAACCATTGATCAATTCTCAATACAAAATATAGAAGTAATGTTAGAAGTTTTAACTAAAATATCATGTTCAGTCATTC
>evgR011086c0g1_i01_Hydra
TTCAGTGTTCTACGGATATTGATTTTATAGCTCAAGTAATTTACATTGAGTTTGTTCTGTTATTATCTGCATTTCAAGCACTTCAATCTAGAAATTTGCCATCTTTTTTTAATGAAACGGCTTCTACCACATATGCTACGTGGATATCGTGCATTATACTATCTTGCTACTTTCCTATTTATTATAGCTCTCAAGATGGTAAATCGAAAGCCATTGCAATATCTTTAACAATGACTATTGTTAATATATCTTTAATGACCTTGAAGTTTACACAAAAACTAGTAATGATATGCCAATCAAAAAACAACAAAATGAGGTTTAAATTCAATATGGG
>evgR011094c0g1_i01_Hydra
ATATTCAGACTATATATTGTAATTTGAAGATCTGTGAATGCGATGGTCACTGTACTCGTCCATTTACAAAATGTATAAATAACGAATACCGTTGTGATGACAACTACACCATGGTTAATAGAATGTGTACAGGCAACTGGGGAAATTGGATAAATTCTGCATGCTCACATAGTTGTGGTGATGGAGTTATGACTCGTACAAGATATTGTAAAAGGGAGGCCTATTGTATCACAGAGGGTGGAGAATACAAAAGAAAACAGATGCAGTCAGATGTGTATTGCAATTTGAAGATGTGTGATCCAACAACAACAGCTACACCAACAACAACACCAACAACAACTCAATGTCCACCTTCAAAAGATTTAACTACAACAGCTACACCAACAACTCAATCTACATCTACAAAAGATAAGATTTCCTGCCCGTGTTTGCAAAGATATTGCGAAAAGTTTTTTGACGAAGAGTTTTG
>evgR011101c0g1_i01_Hydra
ATTTATTATCTATTTTTGGCATTTAGTTGCTTGTGACAACTAATAAAAAGATTTTAGAACAGGTACATATCTGGTTTTGAAAGTATTTAGCGTTTGATGCACTCAATTAAATTTCCGCATTTTATTAATGACATTAATAAAGTATATATTAGTTTTAATTATAGAAATGGTTTATAAATCCAAGTTATCATAACTTCATAAATCATTCCTTCTTCAAATAATGCCATTTAGATTAGAAGTATTTGTACCTTTGTATTGCTTCAAACCATTCAGCTTTCTGATTTATTGTTCAAATTCAGAGTTAAACATCTTTCTAAGCAATACTTTTTTAAACCTTAGACAAAAGTTACTTATCAACTATGTAGAGAAAATTGAAAGCTATTCTATGGTCATGGTACCAAGGATGTCATTTCCTAATGTTATAATACACTTGGGCAACTCAGGTCTTTTAATTTCTAATGCTATATATTTCAGACTTCTTTTAAATCCATTCCTTGAGCCTTACATGCTAAAGGGGTTTTAAAAATCAAAATAAATGAAAAAACACTAACAAACTCAATAAAAAAAATTAAAATGCTGTCTGCAAAGAAGGAAAAATTTAAACGTTGACTAAAAGAAGATAAAAAATTATAAATTGGTTTCGCCTGAAATGCATTATGGGAGACCCCAGGTCTTTTAAAATTAAATCTTGATCATGAAATATATAAAAGTTCCCTTTGCCAGTTGATATATTGCTAATATATATTACAATACTTTTTATATTACGTGCCAATAACCATTATATAATAATAGTTATAAAGTAATAAAAAATTATTTTTGTAAACACATTATTCCAGTTATAATGTAAGGTTTTAAACAGCTTGTCTGTAAAGGCTTTTGTAACTATGTCAGTCACTTGTTTTTGTAGCTTTATAACCTGCTGTTACATGCTGTCTGATGTCTATTGCAGCAGCCTGGGTTATTTCATACCAATTAAATAATAAAATTACTACTTTGCGCTGGCAGCCATTGACAACAGAATAAAAATCTTTCGTCATTCTATTATGAAAATACTTTGGACGTGTTTTTCAGATGTTAAATAATTAGCTGACAAAGCAAACGATGTAAAATAAACTAAACTCAGTAAATTTCAGAATTCTATTTGCAAGTTTTCAAATGTAATTTATGGAAAAAAAGGTATATCTTTGTTATTTACAATACGCGGGTGTTCTAAAATATTGAACCCCAAGATTGGAAGATGAGAATTTGTTGTTTTCTTTTAGCATGTTCTATTTATATTTCAGCATTACAGAGCACTTTTTAAATTATTTATTAAAAAAGCATTATAAATGTTATAAATTCATTCAA
>evgR011106c0g1_i01_Hydra
TTTTATCACTTCGTCAACCGGTTTTATCACTTCGTCAACCAATTTTATCACTTCGTCAACCGATTTTATCACTTTGTCAACCGGTTTTATCACTTCGTCAAACGATTTTAGCATAAATCTAACAGTAAACAATTTTTTAACTTCTTCTTTATTAACATTTTCTAGCAATGAAACGAAAGCTGCAGAAACTCCATCTCTAACTACTTTTCCAATATTTCATGATGTTTCAAGTTTTCATTTACAAAATTTATTCTTTAGTTCTTCAAAAATATTTGAGAACATAACAGCAATATCTTACTTTTCTATAACTTCTTTA
>evgR011107c0g1_i01_Hydra
AGCGCAAAAAAATATCAAAGATATTGTAAAAAGCAGTGCTATTTTAACTGGCCGTTTAAAAACTGCATTCAGCATTCAAGGAAAACTTGAAAAAGATGGAACTCTTCTAAAGGATTTATACGATATTATCGGGCTAAGACTAACATGTCAAACAGTGAACGATTCTGTTCGAATAACCAACGTCATTGAAAAAGACAAAACAAATTTTGACATCGTGCAAAAGAAATGTTATGGAATGTGTCCTGGTGTTGATAAATACAGGGACGGGGGTTACAGAAGAATCCACCTTATTCTTCGTTTAAAACCAGAAAATAAAACAGCAGAGTTACAAATTGG
>evgR011117c0g1_i01_Hydra
TATGATAGACTCCTTTACAGAGTTGGAGGTAGATGATATTGATCTTGAAGACTTAAGAAATCCTACCCTATGTGCTGAATATGTCAAAGATATATACAAATACATGAATAAACTAGAAAAAAGATTAGTACCTGGTGACTACATGCCTCGTCAAGCTGAAATCAATTTTAAGATGAGAAGCATACTGGTTGATTGGCTCATTCAGGTGCAATCAAGATTCAATCTTCTGCAAGAAACTTTGTTCCTTACTATCTATATTATTGATAGATATCTAAGTAAACAAAATGTTAAACGTGCCGAACTTCAACTAGTTGGCGTGACAGCAATGTTAATTGCATCAAAGTATGAAGAAATGTATGCTCCAGAAATAGGCGATTTTGTTTTTATAACTGATAATGCTTATTCTAAAGAAAAAATCCGTCAAATGGAACAAAAAATGTTAAAAACTTGTGACTATGATTTTAGTAATCCTTTGTGTTTGCATCTTCTGCGAAGAAATAG
>evgR011123c0g1_i01_Hydra
TGGAAGTTTCGGAAATCAACCTGGAAGTTTCGGAAATCAACCTGGAAGTTTCGGAAATCAACCTGGAAGTTTCGGAAATCAACCTGGAAGTTTTGGAAATCAACCTGGAGGTTTCGGAAATCAACCTGGAGGTTTTGGAAATCAACCTCCTAGTTTTGGAAACCAAACTGCAGGTTATGGAAATCAACCTATTGGGTATGGAAACCAATCCATAGGCTACGGAAGTCAACCTGTAGGGTATGGAAATCAATCTGCTGTTTTTGGAAACCAGCCTGTGTCTTTTGGAAATCAAACTGCAACTTTTGGAACCCAACCTGCAGCTTTTGGAACCCAATCTGCATCTTCGGGGAATCAATCATCAGTTTATCATCAAAATAGCTACAAATCTCAACAAACCGACTTGAATAATATTTCACAACGTAACGCTTTTGGATTAAATATGTTAAGTTCAGCTCAACCTAATGACTTTTTACCAAATACTTTTGCTACAAGTGGCAATACGCAACTTAATGATGCATTCAGACAGTCTAATCAACTGAACACTTGGAATAATAATCAAAACACTGCTAACCAATTTAAAAATACGTCACACCAATCTCATCAACAAAATTTATTTCCAGGAGTTTCTAATAACCAGGCGTTTCATCAAGGAGCTAACGCAGCCAAGGGAATACAAAGTAACGTAAATATTCACTCAAATTTTATACCAGACAATCAATTATCACAAAAACAACAAACTCTCAATCCGTCTCAGATAGTTAATTATCAAAACCAACCTCCAAGCAACCAATCCAATTTTTCATCGTTCAGTAATGTTCAGCAACAGCAGCAGTTTTGGACACAGACTCAACCATCATCGAAATCTACGAATCCTTTTCTGTAACAGTCGATTCGTTACGATTAGATATATTTTTATATTAATAGAATGTAATCTACATTTTTTGCTCGGAATTTTTAAACATTTTTGTTTACTTAGATTCTACTAGTAAAGCTAATCTCCTATTAATATAAAAGTAAAAATTTTTCTATTTTCAGTCGTATGCGCTTTGTTTGATTATTTGCGTATACTGTTGCTCACTATATATTTGGGTTATTACCTTTTTTATATATATATATATTGCGAAAATTCAGTTCTGCTCAAATAGTCACGTGTTCTAGATAGGAAGACATGTTCATGTATTTAAAATTTAACTTGTAAATCGAAGATTGTATTATATTTGAAAATTAATTTTATATATTCCTATGTTTTGTTTATCAGTTTGTACTTTTTTTGCTTGTATTTTTTTTATTATGAAGATTTAAAAAATTCTAAAAAAAAA
>evgR011126c0g1_i01_Hydra
TTGCATGTTTGTTTTGATATTTAGTATAAAAAAAATCACACGAAAAGTCGTAAAAGGCATTTTGATTTTAGTTACATTAATATTTGTTAAACGATGGAAGTTTCTAGTCCAAAATTTTATTCTGCAAACGAAGTTAATAGGTGTTTATTGACTGAATCCGAAAGGCTGATCGAAGTGATAGAGGAAAGTTTTGTTACTTTTAGCACAAAGCCAGCTGGTTTTAATCAACCTCAAAGAATGATTATAAATGTAGCTGAAAACAACAGTTTATTAATAAAACCATCAACCTCTCTTTCTCACCAGATTGCTTCATGTAAAATTCTTACTATCAATCAGTTTAATGAACTTAATTTTGGTTTACCAGTTATTCAAGGCGTAATAGTTGTATTTGATATTATTAGTGGTTCAGTTAATGCAATTGTAGATGCCATTTCAGTTACAGATAAACGAACAGCTGCTGCATCAGCAGTTGCAATAAAACATTTATGTAAAGAATCTGATAAAATTCTGGCTGTAATTGGAACTGGACATCAAGGGGCAAGCCATGTTCAACTATTGTCAAACTTTAAAAATTTTTCTGAAATACGAGTATTTAGTCGCACTAAAGAAAGTTGTAATGCTTTTGCACAAAAATATAATTGCATACCATGTGATTCTGTTGAAGATGCTGTCAAAGATGCTGACATAATTATTACTGCTACAACTAGTACTACACCTGTTCTATCTAAAAATTGGATTAAACATGGTTGCCTTATTATTGCAGTTGGTGCACCATCATGTAATCAGAGGGAATTGGATGATGAAATTATGCTTTGTAGTCAAATATTTACTGACACAGAAGAGGGCGCATATGCATCTGCAGGTGATGTTATTTTATCAAATGCAAAGATTCATGGTGAGCTCGGTAACATTATAAATAAAACAGTAAATATTGATTGGAATACAACCAGAGTTTTTAAATCTAACGGAATGGCAATTCAAGACTTAGTAACGGTTAAATTAATTTTAGATAAAGATATAAAAAAAAAAGATTTAGTTTCTTAAATTTTAAAGTTTATACTTCATATATAATATTGTAACAGTGTTAAAATAAATCTAAATTATATTTTTATATGAAATATTCAAAATAAATGATTTTTGTTTTAACTTTAGCATGCAAAATATTAAGAGTTTATTTTGTAGAATTCTGATTTATTTAGGTTAATAGTATTCCATAG
>evgR011138c0g1_i01_Hydra
CATTAATATAACTCATTTTTTTGTAAGGTGTTTTTATATTTTATGTTGGATTACAGACAGAAAAATATCAGTCTATAAACGTATAAAAAATTGTAATTTAATTACCGATGATTATCAATCATTACCGATCATTACCGATCATTAGCCGGCTACCAATTCATACATGTGATTAAAATGAACAACACTGAAACAAAATGTAAATTGTATGTTGTAAATATTCCAAGTGGTTTTTATTTAAATATGTTTACCTTGGGACTAATTATAGTTAGCGTAATGATATTGATACCAAACATTTGCTTGCTTATCGGTATAGTTCAATTGCGTAAAAGAAGCAAATTTACAACGAATGAAAAACTGGTCTTACTTTTATCTGCCACTGATTTAGTTGTAGCCTTGGTTCTTATACCATTCGAAATTGTACTAATTAGATACAAGGATCTGATTTCTTGTGGAACTATATTAACAGCGGTATTTTTTATGGTGTTTCCGTTGTTTTTTTCAGCTAGCTGCATACTTACATCATCGGTAGAACGCGCTATTGTTATTTTAAATGACAACAAATGTTGTGGAAAACATTTTAAAGAGATTTATTTAATACCTGTCATTGTTGTTAACTTTATATTTTCAAATGGTTTTGGTATATGGATGT
>evgR011143c0g2_i01_Hydra
CGGCGTGAAACCTGTTTTTGAGTAATTTTTTATGGTTTCCATAGCAACAGGTAAAGTTCGCACAAATATTTTGATTATAAGTGACGTTAACTTGGAAAATAAAGACCTTACAACACGAATTATTGAAGAAGCTTTACAACAAATTTCTCTTGCTGATTATCATGTTAGCTTTCACACTACAAGTGGAACGTCACATGGATTGTACGAGTTTCAGACAAGAAAGTGCTTATTGAAAAAAATCAAACAAGCAAAGCCAACAAAACAGGATTCGCTTTTGTCTGTATTGGGAAAGAGCATTAAAGACCTGGGAAAATTTTCTTATGAAAAAAGCATAGTGCTTATTATTACAGATAAAAATTATACTTTGTCTAACGCGAAACTTAAAATGTTATTAAAGAATATCTTTCGTACCTTTTGGATCAATCTTAACAGATTTGCAGAAGAAGTTGAATCAACTCAAAAAGACAACATTTTAGAAATCCGTGGTCGTTCAGTTACAAAGATACTGGCAATATTGCCATTTTTAAAAGCCGACTTGAAAAATTGTCAGGACGCTGCTCAATCCGCTGC
>evgR011151c0g1_i01_Hydra
ATTCCTTCAGAAAGAAGAGTCTCCAAAATTCTATTGTCCCTTGTTTACATTGGAGCCACACTAATGCTCATCCTTCTCAATTCTGTTCAGTGCATCAACCTAAGCTTGATTCGTGATGAATCAGTCAAGAATCAACTGCAGACAACGTGGAGAACCCACATTGAAAAGGGATCTAGCAATAAGTGTTGCCTCCCAGACACTTATACCTTGTCAATGACCAATGGTGTGAATAGACAGAATGTGGGAAGTGATGGAACCCCTTCTCCATTATTGCAATTCCAAACCCAAACGAGTACTGTATGGTATGATAAAAAATCTTTGAGAATGAGAACTGACTCAAATGGAA
>evgR011157c0g2_i01_Hydra
AACAAAGGAAAATTACCTGGATCCTGACATGTATCACAAAAAAGTTCCTGATCCACACAAAATTCTTCTGATGTATTCGAAGCAAACAGAAGGTCGGTTCAACACATTGCATAGTATTCCACTTGATGCTATGCATACTGTGTATAGCTGTGCTGTTCCCTGGCTTTTCCATTATACATGGATTGACGGCAAGAAAGTGAAAGGTGGAAAATTCTCTAAAGCATTAATGGATTCATTGGAATTGAAGCTGAAGAAAGTGAAAGGATGTTTGCCATATGCCATCCAGGTCAACATTGGTACTAAACCATTCAAGAAATTTGGAATCACCTGGTCGTGTGCTGAAAAGCGAATTTTCCTCCTATATGTTGCAATCGTTATTCTCAGGCACCCGTTAATGAAAGCCGAAGCATACAAAATTCTGCTTGCTTTTCAGCATGCCGTAATGTTGCTTGTTGGCTCTGGGCACAATTCGAAGGTGCCAAAAGAACACCTCAGGAAAGCAAGAGAGAATTTACTGTATTTAGTTGAGAAGTGTCAAAAGTTGTATGGCGCTGACTTCCCACGTTATACGTTTCATTGCTTGCTGCATATTGTCGATGATTTGATTGCGAATAATTGTCGACTTGAGTATTGCTGCATGTTTCGGTACGAGAACTCGATGAAATTCTTTATCCATGTGCTCGATGGAAGAGGCGGACATCGTGTCCATGCACAAATCCGCAACAGTTTGATGCGGAAAAAGATTTCAAACATTGTTCTGCCGCCAATTTGTCAAAGAAGACGTGCTCTACTAAAAGGTGGACAAACAGGAGCAGAAACAAGCATTATACACCTTTCGAACTTCAATGAGAACGATTTTGGTGTCATCAACACAATTTTTACTGACCATCCATCAAATAATTTTTATCAAATGGGAGAGGAAGGAAACAAACCGAATGCCTATGAACTACAAGTAACGCGC
>evgR011176c0g1_i01_Hydra
AATCTGTCATTGGCTCTACAGAAGTATTAGTCGCAAGAGCATTCTCTTGCTATGAAGAAAATGTACCAGTTGACCACAAGCAAGAGCATAATCAAAGAGTTTTGTTAAAAGAATTTAAAGACAAATTGATTATAGATGGTGAAGTAATACCAGATCCTATAGACTTATCTAAATCATGGATTGGAGAAGATAAAGGAATGATTTATTGGCCAAGACTCTATTTTACGGACATTTCTAGATTTTATATTGATGTGATTGGTAAACGAAATCTTATAGAACGATTAGAATCTGAATATAAACAAGGAAAAGCATACCGTTATTTTTCAAATAGTTTTATTG
>evgR011178c0g1_i01_Hydra
CTATTTTAGTGCAAAAAGGTTTGTAAGTATAAATATAGCTTATGTTTAAAAAAACAAAAACAAGTAATGAGTTAGAGCATCGTACATTTTTCAAGGATTGAATGGTTAAACTCAAAATGAAGTTTGCTCTTGTAACTGTTGGTACATCTCTGGCTACAGCAGCTGTAATTACAAATGCTTTTTTGCAAAAAAAACAGTTTTATCCATCAGTGGTTTTCCTAACAAAATCGAATTCTAGTTTAGCTGTGTTGTATGCTCAAGTATTTGTCTTGCTGTTTTTAATTGGGAAAGCTCTTCGTAGAATATTTTTTGGACAGCTACGTGCAGCAGAGGTGGAGCATCTTATTGAGAAGTTTTGGTATGTAATTACTGAAACATGCTTGGCATTCACTGTTTTTCGTGATGATTTCAGTCCTAAATTTGTTGCTTTATTTACCTTTCTGCTTTTTATGAAAGCTTTTCATTGGTTAATGGAAGATCGTGTTGATTATATGGATCGAACACCAGTTATAAGTTTTACTTTTCACTTGAGAGTAATAAGTTTAATGTTGTTATTGGGCACTCTTGATTCTGTACTCATTCATCATGCGTTTCGAATTACTCTTTCAAGTGGAGCATCAGTTCAAATGGTGTTTGGTTTTGAGTATGCAATTTTGCTAACAATGGTTGTTGCAACAGCAATGAAATATATGCTACACAGTATTGATACATACTTTGAACATCCTTGGGAAAGTAGAAATGTGTATTTACTGTATTCTGAACTAATAACAGGTTTTGTAAAATGCTGTTTATATGTTTCATTTGTTGGATTAATGGTTAAAATACATACTTTCCCATTGTTTGCTATTAGACCAATGTATTTAACAATAAGAAACTTTAAAAAGTCACTGAGTGATGTAATACTTTCTCGTCGTGCTATTACTTACATGAATACTAGGTTTCCTGATGCCACAGAAGAAGAATTACTTCGACTAGAAAATACATGCATTATTTGTCGTGAACAAATGTTGGTTGGAAAGAAACTACCTTGTAATCATATTTTTCATACCTACTGTTTGCGCTCTTGGTTTCAAAGACAGCAGTCATGTCCCACTTGTCGAATGGATGTACTTTCTCAAGCAGCCCAAATGGTGCGTCCATCTTCTGTTCCACAATTTTCTCCTCCACCGCAGCCACAACCTGCACCTCAAAGCTCTCCTTTGTTTGGCATGCCCCAAGCTCAAGGTTTTAATCCATCTAATATGCCACCTCCTATTATAAATTCTCCTAATATGCCATCAGCATCGACTACAACCAATTCCAATGCAACTGGATCGTTTAGTTTCCCATCTACATTTACGTTTCCTCCACCTCCTGCATTTATGGGTGACACATCATTTTTTAATCCTAATATAAGTAGAACGCCTTCTTTTGCTTCAATGAATTTAGGAATGGCTGGAGGCTTACCTTTTCCTCCTTTTCCAATGTTTTCTCCTCCTAATATCAACCTTGGCCAACTAACGGATGAACAGTTGAAGTCTATGGAAGGTCAAGAACGTGCGAATGTAGAAGCTCGATTACGAATACTGAGAAATGTTCATACATTGTTAGATACAGCCATTTTGCAGTTAAATCAATATTCTCAAGTTATTAATAATTTAAACGTCTTGTATCAATCAAATAATGTTCATCCTTTTTTGAATCCACAGTCAACAGTCTCGCCTACGTCAACAGTATTGCCTACGTCTTCAAGTCCGCCCAACGACCAAAATACTTATAATGTTAATGAAACACAGACATTAGGTAACTTGAGTTTGTTTACTAATACAAAAGCTGCGACCTCTACCATTTCTTCTACGCCAGTAATGACTTCAACTAAAGATTGTGAGGCATCTGCAGGATCAAGTACAGATATGGAATTAAGACGCCGAAGATTAGAGCGTTTTGAAAGTTTAGCTAAAGAACAAAAGAAGGACACTGATAATGTTTAGTGAATTGTTTTCAAAGTTAAAAAATAAATAAATTTTCATAGACCATGTTTCATTATTTTACTCTATATTCACCTTTTGCTGTTATTATTCCTGGAGTTATTTTATCGGTTTGATAATTGTA
>evgR011182c0g1_i01_Hydra
GCAGGAGTTTGAAAAGAAAAAAAAAATGAAATACATATTTTTGTTCTATTTTTTTATAATTTTTGCTAATGGAAAAAAGCATTCAAATTATGTTAAAGATTCCAGTAATTTAAAAAAAGAAAATATTTGTACAATTCCAAAGGATGTTGCTAATCCAGAATGTGAAAATAAAGTTTTGCGGATAGAAGCTAATTGGAAAACAAACCCGTTGTATAAAGCGAACGGAGTTGACGGCTCCCGTTGTTCAATAATAAATTATTTGAAAGAGGTTGAAAATGTTTGTCCGTGCTCCTTACCTGATGACAAAAACTATCTCTCGTGTAACGCTAGCGCATTGTGGATGAAGGAAAACTGGGAAAAAAATTCAGTATATAAAATGCATGGAGTTGATGGGTCAATGTGTTCGATACTTTACTACCTTAGTCAGGTGGAAAAAAAATGTCCAAAGGAAAATTGTAAGATTGTAAACATCAACATCTTTCCGCAATGCAAAAATAGAGTACTGTGGATGCAGGCCTATTGGAAAACAAATCCAATTTATATGCTAAAAGGAGTCGATGGAACCAAGTGTTCGATATTAAATTACCTCAGTAAATTTGAGAGAGTTTGTCCGTGTCTTACGCCCATCAACCAAACGTGTCAAGATAGAGTATTGGAAATGGTAACAAACTCAAATTGGAAAAGGGTTCCTTTATATATGAAACGAGGATACGATGGTAGCTTATGCTCTATTCTTTCCTGGTTAACTAAGGTTGAAAAAACATGCCCTTCATAAAAAAATATTTGCTTTAAAATCAAATCAGAAAATTGACAATGTTTGGCATTTATGGATTCAAAACTTGAAAAAAACGAATGACTAAAAATAATATAATTTTTTTTAAATAAAAGTGAGTTTGAAAAAAAAAAAA
>evgR011197c0g2_i01_Hydra
GTATTTATGATACTTGGGATGAATGTAAAGCACAGACAGATGGTTTTTCGGGTTCAAAATTCAAAAGCTTTGACACTCGCGAAGAGGCAGAGCATTTCTTGAAAGAAGGTGAAGATTATGGACAGTTTCCATACTACGGGGTAACTTGTGATGCAAAAAGAAAAGTGTTTAAAAACTATGAACTAGCCAAAGCTTATTATGAAGGAAAAAAGGGTGAAAAACACAAAGGCTTTCATAGTTTGGAGGATGCTTTGGGGTTTATTGATGTTGATTTAGAGCAAATGCTTGCTTCTCTTGGAATCCTAAAACTTCCAAAGTCTGACAAGGTCAAAGCTGTGGATAACAATCATGTAGATGTTGATGAGCTTTCTATGGAAAAATTATCCATCACCAAGAAGTACATAGCTGTTAGTATTGGATCTGAAGGTAAAATCTTCGTAGATGAAAAGGATGCTACTGCTTTCCAAGAAGGTAATGAAAATAGAAAAGTACATCCTCCTGAAGAATTAACTGAAGCATTGAACTATGTGGGTATCAATGTGGACCAGATTGAGCAGTTTCTTTCTGCCTTTGGTTTGATGGACACGTGTGTTGGTAGCAAATTTAAAAAGAAGAAAGATGATGACGAATCTAGTGGATCTTCATCCTCTGTTGAATGTAAAT
>evgR011198c0g2_i01_Hydra
GCCATATTAATCGGAGCTTGTACTTTTTTGAACAAAAAAAAAGATGTGAAATTGACATTCATCATTTTTGTATGTTTTGATATATTTATTGTAATATTTTCCCTTACTTTTTACACCTATGCTTCTACAGTTTATTGTAACCGAAAAGCAAACATTTGTTTTAACCGACGTGATAGAGGAATGTATAAAGGCCTCGTTCTATCAATTATAATAATGACAGTATTTATCGTATTTGTTACCGTACCTGATGTAGTAGTGGTTTTTAATTCTGCATATAGTTCTGCGTTTACCCCTGTAACGGTTTATACTATGACATCCTTTGGTTTTTCATTGTGGACTGATGCAGTAGTATATATATTTCTATCTCCCAAAGTTAGAGTTGTTTTTAAAAAGAAACTTATTTATCTATTTTGCCAACAAAAAAAAAATGTCAAGAAAAATATCTGTGGACTAGCAGTATCAATAAATAATGAAGTTGTTTAATATTTATCTTTTAAAAGATAACGCATAAATAAATAAAGTTGATTAAGTTGTTTCAAGTAATGCGATATTGATTTATATAATAATGTTTACTTTTTTTTTGAAAATGAAAATTTCAATTTTAG
>evgR011203c0g1_i01_Hydra
CTAAACCAATCCGCACAAGATCAAAATGAGCGAAACTGAAACCAGAGAAGATAATGTTTTCATGGCCAAGCTTGCTGAACAAGCTGAAAGATACGATGAAATGGTCTTGAGCATGAAGAGAGTCGCCCAATTGGATGTTGAATTGACTGTTGAGGAAAGAAATCTCTTGTCTGTCGCATACAAAAACGTTATTGGTTCCAGACGTGCTTCATGGAGAATCATCAACTCTATTGAACAAAAGGAAGAAAGCAAGGGAAATTTACATCACGTTGAAAAGATCAAGAAATACAGAAACAAGGTTGAACGTGAATTAACTGACATCTGTAATGATATTTTGGATATCTTGGACCAACACTTGATTCCTTCAAGCCAATCTGACGAATCTAAAGTCTTCCACTACAAGATGAAGGGTGATTACCATAGATATCTTGCTGAATTCACCACTGGTGATTCTAGAAAGGAAGCTGCTAACAATTCTTTGGAATCCTACAACACTGCCACAAAGGTTGCCAAGGAACTCCCAACTACCCATCCAATCAGATTGGGTTTGGCTCTTAACTTCTCTGTTTTCTACTATGAAATTTTGAACTCACCTCAAAGCGCTTGCCAATTGGCCAAGAAGGCTTTCGATGATGCTATCGCTGAATTGGACAGTCTTACTGAAGAATCCTACAAGGATTCTACACTCATTCTTCAATTGTTGAGAGACAATTTGACTTTGTGGACATCTGATATGCAAGAAGAAAAGAATGATGGAACAAAGGTTGAAGACGCTGAAGAAGAAGACCAATAAATTCAGATTGTATCATACCTTTTAAAAACAATACACAAAAATCACTTTAAACGCGTTTTCTTCATCCGATAGAGACCGGTTGTAATAAAAAC
>evgR011204c0g1_i01_Hydra
TATATGAAATATAGTACAGATAAAAAAAGTCATGGGTACTGATTTGTACATATTTGGTGGGGTATTAGGTCTTGTTTTGTTGTTTTTTATTTTAATACGAAGTGTAGTACCATACTTGATGTGCTGTTGCGCATGCTCATGTAAACCTAAAGATGGAGAGATAGCTTTATTAGAAGAAACGGGAGAGGAAGACGAAAACTACCGTTTAATTAACTGCTATTGCTGTGTCTTCATTCATGCAACAAAAATTACGGCAACAATGTTAGTTTGGTTACATGAAAGTTTTAAGTGCCATAGAAACGTTCCTACCAACGATACTTGCCCAATTTGTTTGGAGATTTTAGAAAGAAATACAACTGTGTTAAAATGTAGGCACAGTTTTCACAATAAATGCATTCGCAATTGGCTGTTACAAAATGAATCAGGATGCTGCCCTTGTCCTTTATGTAAAAAAATTATATGCTCCGAGAAAATTAACTTGGGTGTTATATCTAATAGTCCTAATTATAGCCTTATTTATCCAATTACAATCAATTACCATGAATACCGCAGTGCAGTTGCTTTTATTTAAACATTT
>evgR011207c0g1_i01_Hydra
CACCAAATGCGGCCGTTTTTCAAGATGCATATATAGTTGCAATTTAAAAAAAAAACTGTTATTTTGAACAACATTTGATTACGTTGTGATTGTGACTTATTTGAAAATGGGTACATCTTATTCATATTTGGAGATTGAAAAAGAAAGTTGGGTTGATAACTTTATAAAATTTGATAACGTAGTCAGAGAAGGTAATTTAAGTATTCTTGATCAAATTAAAAATGGTGACCCTATTTACTTGGCGTTTACAAAGAGCTGTTGGGGTGAATTAGAACATGAACATTGGTTTGTAACTGACGGAAAATATTACATAGAATATGGATCAAAAGATTCAGATGTTTATAAAGCTGCCGTAACAATAAATACCAATAAACGTGAATACCAAAAAGACTTTTTTAAAGTTGATATGAATAGTGAAATTCGAGAAAGAGTATTCTTAATTTTAGGAATGAATAATTATTCTTTAGCTCTTAGAAACTCTGAGCATGTAGCAAATTATATAGTTAGAAATAGGTGGTTTAGCAGTCAGATGGAAGAGAATAAAGGTAAATTATTTAAATATTTTAGAAAAAATTTACTTGATCATAAGAAAAAAGTAAATACTTTTCCCTCGTGTATAAAGATCACTTATGAAGATGTAAACAAAGAAATCTACTCATTTTTTGGTAACTCTTTTCAATTTACAAATTCGTGTTTGTATCTTGATAAAGCTGAAGATAATTATAATATATTAATGGTTGGACCAACAGGTGGCGGAAAATCAAATCTAATAAATGTTTTGTTCAATAAAAATGTATGTCAATCGGAAGTCAGTTTGAATAGTGTAACTAGGGATGTATGTTTTATAAAAGGTCGAGCTGATGTTTATGACGAAAGCAAAAAGGCTACAATAACTAAAGACGTTGTTGTTGCAGATACTGTTGGTCTTTGCGATTCTAATTGGTCAGCTAGCAAGACTTTAAACTTGATTAAATGTAGAATTAAAGAAAGCTGTGTCTATATAGACGTCGTTTTTATAGTATTTTGTTGCAATGTTCGTTTATCTAAAGAAAATAAAGAATGTATAAATGAAATTCTTAAATGGCTAAATTACACCAAAAACAGTCATAGATTTCGATTTGTAGCAACCCATGCTGATCAAATTGATGATGAAGAAAAAGAAATATTGCTAAGTCAAATTAAAGATATTTTCGGGATCACGAAAAATACACTTTTAGAAAACGGCAATATTACATTTGATTCTATTATATTCGTTGCTTTTCCAAGAAAAATAGATACCGAGAAAACAAAAGAAAATGTCATAAAAAGCCATAGCGATATTCTTTTAGCATGTAAATTTTCTGGAAAGGGCGAAAGGATCGTTACAAGAAAACACGAGTCTTGTATACTTTTATAAACATATTTTTTAATGATAAATGAAAATGTACTTTTTAGCACTAAATTTTTGTAATTAAAAAAAAATAGTTTAAATATACTAGAAATAATATTTTTAAAAAAAA
>evgR011216c0g1_i01_Hydra
TATAATAAACTGTTAAATTTCAAGTTTTATTACCTTTCTAGCAATTTTGGTTCTTTGTGTGAATTTTATATCAAGATATGAAGTTTCGTGCTAAGCTTGTTGATATTGGCTGCATTCAACAGTTTACTAAAGTGCTCTCGACAGTTAGCAAAATATCTAAGACATGCACACTTAAGTTGTCTGAAGACCAAGTAGTACTGACTCAAAATGAAAAAGCAATTAATGGAGGAACAAGCTTTTGGTGTGAAATGAATCAGGTTGATTACTTTGATGAATATCGAATTGAAGGAAAAGATGAGAAAAATGAAATTTATCTGGAAATAAATAATGAAAATGTGTTACGTGTAATGAGATCTGGCCACACTGCCCAATTCATTAAAATCAAGCTTACTAAAAAGCAAACACCTTGCTTGACATTTGAAATCACTTTGTCCTCTCTTACCGCACACACACGCAATGTCACCCATGATGTTCCTGTTGCAGTTATTCCTACGAGATACTGGGACGACTTTAAGAAGCCGATTTTGCCCAACTATAATATTAAGATTCATACGCCAAACCTCAAAATAATGAAGAGTGTTGTTGAAAGAATGAAAAATATCAGCGGCTTTATGGTAATCACGGCTTTGCAAAATGGAGAGCTGTGTTTTAAGGTTGAAACAGACGAAGTTACTGCATCAACTTACTTTCAAAACATGGTAGTGGAACCCAATGAAAATAATATTCTTGTAGAGAATGATATTGACGAAGCTGAATCTGTTGGAGCCCGTGTTGACATCCAAAAGTTGTTATCATTTCTTCACGCCCAGCTTTTAAACGCCTCAAAAACAATATGCGCCATTGTTGATCAAAGAGCAGTACACATGTTTGTAGATTGCAATGATTTGCTCTTTCAGTATTTTATCCCTTCTACAAATATATCTATTACATAGGTAACAATGGATCACAAAGATTCAACATCGATCACCAAGACAACCTCTATACAAATCTTTTAACAATTAAAGCTTTTTTCTGTGGATCAATTGTATAATGCTAATTTGATCAGATTTGTTGGAATTATTGTTTATTGTTGGAATTATAAATTATTTTTGGTTTTTTAGAACTATTTATTTTGTATCACAATTAAATAAAGGAGAGTGGCGCATGGTACTAAAAAAAAAA
>evgR011231c0g1_i01_Hydra
TTTTTTTTTAATTTTTTTTTTCATAAATGTAATGTTATTTTAGTATTACTGAATTCAAAACATTGTATTATGGAATTTAAAATGATTAATTGGATAATTGGTCAAATGTGTTACAAAACTGCCGTTGTGTGCTTTATTACGCGACTTCCGTGCATTCACCACGCTTACTTTACTATTGTCCCGGACTGGTACTTGAACGGCGATGTTGTTAAAAAATGGCTTTCAGTATCAAAAGAAGAATGCGGAACAACTTGTGTAACAAATCTTTTATGTGTTGCAGCTAATTATAATGCTAATGAAAATTTATGCGAAATAATTTCACAAAATGGAATTCGAGAGCGAAGACTTGGATGGTATTTATTAACAACAAACACGACTAGTGATATAAACATTGGTCCATTCTGCGAGCAAAGTAATTTATGTATAAAAGGTGAATGGTGTAGAGATGTTTGTGGTCAAAACAACCGTCATATCTACAATTGCTTGAGTAGCAAAAACGTTGCGAAAAGGAGCAAAGTGTATCAATCTTCAACTTATAGTTTTTCTAATCGTCCGGAAATGGCAATTGACGGTGATTTGCAATCAGCATCTTGTACGCAAATGGATCCAATAAACTGGTTTAAATTAGATATGCTTTATGTTTATAAACTTGTAAGAATTAAAGTCATTAATAGACTTATTTACACTGAGCGACTTATCGGTTATAACTTGATGTCAAGCATAAACGATGCCAATTACGAACATATAGTAACATTAAGCAAGGTTTGGCAACAAACTTATAATTGTTCCAATAACGCCCGGTTTATTATGATTACCAAAATACCTGATATTGACCAATATCTTAACATTGCCGAGATAGAGGTATGGGTTTAAATGATGATACTTGCACGTGGAAACTTTAAACTGCTCAAAGTGCATCGACTTCATATAAAAGATCTTTAGAAAAACATTTATGCCAAAAGTTATTAATTTAACAATTTTGAAAAATTTATATTTTTGACGCTAAAAAAGTATATTATTCAAAATATATTTATGTAAAAT
>evgR011236c0g1_i01_Hydra
TTGAACTTGATTTTTGCACGTGTTGAAAAAATGGAAATTCTGGAAAAGTTGAAGAGTTCTCAAGAACTTTTAGCAAGTTATTCAAATTCAAAGATAAACCACACATGCCTGAAGTTTTTGTTTTAAGCGCATTCAGTTTTGAACAAAATGATATTAAACGAAAGGAAAATAATATAATTGTTTTTGAAACAGAAGAGTCAAATAAATTGAAAATAATTTATAGACAAGCAATTGATGAAATTTTTGATAAAATTGGTGCAGACAAATTAAAAATTTTAAGATTCCATTGTTTTAAACGCAATCCAACTAGAAAAAATCCATTTTTTGTTTTAATTGAACTGGATAATAAGAACGATAGCCATATGTTCAAAAGATTAGCGAAAAGATTAATGAATGCTGAAAAATACAAAAGAAGTGATCATAAACTTTGACATGACACCAACGCAACGGGAATTTCAAAAGATATTGATTCCTGGAAGAAATCGTTATAATTAAGATTTAGCAAAGAATAAAATTGTGGGTTTTGACTATGGGATTCGGAATAACAATATTACAAAAATATCTTTAACTACCCATTGGGCACACGAAGTAAAACGTTATTCAGACGTCTTTTGACGACTTTTGAAAACTTTTGACGATGTCTGATTTACGTTGTGTGCCCATTGGGTATTAAACAGGGTTGAAAATGAGTTAAAAACAAAACAAAGCACAGATAGTTTAGTCACAAATATTCTAGCAACTGAAACTAGTACAGCTGATTGTAGTACAGCTAACAACTCTAATAACAAAAATTTTTAAAAAGTTAATAAACATTCAAATAATTCAAACAACAAATCCGCTGCATCAATTAATAAATATTGTCTTTATACGGAGTTATCGTTTTTTTATACGAATGCCAAATCATTGAATAACAAGGTTGATAAAGTTGAATTCTGTGTAAAATTTCTAATCCTG
>evgR011242c0g1_i01_Hydra
AAATATTGAACAAACATCTTAAATAAACATCCGTTCTCAGTTATTTTGCAATAAAATGATGACATTAAAAAAGTGAAAAGGGCGAAAAACATGACAATAGAGGAGAAAACTTTATTAACAAATGGCTGTGTTATATTTAAGATACATTGAAGGAACAATTCCATTATTTATTCATGGTATTTTGTGCGCATTAGCATATAGTGTCTGGTTAATGTTTGCAGCTGTGTGCCTTCCATTTATTCTTGTGTTTCGATTTTACAGTAGTTTGGAAAATTTTCTTATTACATATAGAAAGTTAGGAACTGTGTTTACTCCCTTCGACAAAGCATTTATCCATGAAAGTAAAGAAAACAGAAACTTTATAATCGGAGTTTTTGAACTCGAAGGAGAACCAAATATTGAAAAGATTAGAAGGTGTGTTAGTGCTCGATTTCTTGATAAACGCCGTCTCCTAGACAAAACTTATGAACGCCTTTCACAAAAAGTTACAAAACGATACTTTTCGTACGTATGGGAAGAAGAATCGAACTTTAACATTAGACAACACATTCCAATTTATAAGGGAGTTTTGCCGGAGACAAAAAAAGATATCGAGCAGCTGTGTTCCAGATTAACAGACGAAGAATTTCCTAAAGATATATCGCCCTGGATGTTAAAAATAGTTCCGAAAAAAGACAAGTCAGGATTTATCATTTTTGCTAAAGTGCATCACGTTATTGGGGATGGATACGCTTTAATAGGATTGTTATCGGAACTTGTTGATCAAAAACCACAGTTTATTGATCTCCCACCAAAACCACATAGATACTTGTCGAATAAGGTCATAAAAATATTATCAATAATGCTGACGGGTCCATTAGCTTTATTAACAATAGCTTTTTCAAGAAATCTTAGAAATCCATTTAGAGCTACAAATGGATTAACTAGCAAGACAATAAGTTGGACCAATCCTATAAGCTTGGACACCATTAAGAGAATAAAAACAAAAACAAATACTACGGTTAATGATGTAATGATGTCATCATTAGGCGGAGCACTTCGAAGATATTTAAACGATAGGGGTATATTTCCGGTTGATCAACCTGTTGCTATGACTTTTAATTTGCGGTCAACATCGGAAATGATGAAAACGAACATTCCTTTAGGAAATTATTCCGGTGGTTTGTTTCTTAACTTACCGGTCTCAATTTCAGACCCTGTGGAACGACTTGAAGTAACAAAAAACCGGATTGTTTTGCTTAAGAAGCTCTCCCATGAGCAAATGTTTTCATTTTTCTTCTACAACGTTGCAAGCGTACTACCAATTTTTTTTAGTCGCATTACAGCTTTTGCATTGCGTCGGAATACAACACTAATTGTTTCTAATGTCCCAGGTCCGACCACGCCGCTGCATATTCTTGGATCACGCTTGAAATCAATCGTTTTTACGCCTCCCCTTCATGGAGGAGTTGGTCTTGTTGTATCTGTTTTCAGCTATAATGAAAAAGTACAGTTAGCTGTAATGACAGACGAAACTATCATTGCTAAACCGGATGAACTAACAGAAGCTTTTGAAAAAGAAATTACTTTATTAGAAAATCGTGTGTTAATAGAATAAAAAGAAAAAACCATAAAATGTACCCAGCGGACAAAAGAATATTAGCCTTTTTTTTTGTAAAATAAATTTTTTGTATATTATTTATATAATTATATTATATTATATTATATTACCATAAAAAAAAA
>evgR011253c0g1_i01_Hydra
TTTTTTTTTACATACACCATTCTAATTTATTTCATGTCACATTTGTACAAAGACAAAAATGATTTAAAAAGTATCTTTAAATCTTAAAAATTCCTATCAATATATTACAAAAAAAAAATTAAATAAAGAAGTTATACAAAAAATTAAATAGTTATAAAATCAAAAGCAGGAAAACTACTATTAGAAAATTTTCTAAATATAAACTAAAAAACAGAACTAAAAACACATTCGAAGGTTCGTTACAAAATTATTAAAATATAATTCTTAATTAAATTATAATTTGATAATTATTCGTAAATCATACTATTATCAGTTTTATTTTATAAAATAGTCATCGAGACTACTTTTTTATAATCAGTGAGAAAAAAAAAAAAAATTTACATATATAATAAAAGTGAAACAAAGAAAAAGAAATGAAAAATCTTAGAAAATACTACATCTTTTTATAAAAACACAAGCGAAACAAAATCAAAAAATGCAGAAACAAAAACATAACTCAAATAATGGGCCTACTCTGTCATTTGATTTAACAACTCATACAATGGGTCCACTTCTGTCATTTGGTTTAACGACTCATACAATGGGTCCACTTCTGTCATTTGTTTTACATTTTTTCAAGTTGCCCTTACCAAATGTACTTATGGACGACAACAAAGCGCCTCTCCCAGCAGTTTCTGACGAATTTAAGCCGGGGTTCGAATTCACAGTAGAACCTGAACTATTGACATTCTGTTGCAGTTTTTTGGAAGCAGTTCCATTGGCTGTTGGATTTGTTACATTACTGGTTGAACCA
>evgR011261c1g1_i01_Hydra
CCACACGACATAAATGACGAAACTCTCGAAAGCTGTATAGCATCTATCAAAGCTTTGAAAGTTGTCGCCCAAAGGTTAGAGTGTGATTTCATTCTGATCTATGGTGACTTTAATTTTAGCAACACATTTTATGAGTATATTGAGATTGACGGCGGTGTGGCGACAACAGCGCACGTGCTAACATTGTGCGACCTGGTGAACATGAGATTTCAAGATTGCTTAAATGAATGCCTACTCACCCAATTGGTCACCTTCAAAACATATCGTAATAACAGATTTTCAGAGCCAATCAGCACTCTCGACCTAGTAATAACAGACAAGCCAGATTTCTTGATAGAAATAAAAGAGGAAGACTCATT
>evgR011270c0g1_i01_Hydra
ATACAATAAGAAAGAGCCTTCTAATAATGTAAAATTAGATAGAAAACACTTCAGAGAAGCGAATCTTCATGTCTTTGTATCATCTTCTTCTCTTTTAAATTCATCTATGCAAAAAAAGTTTCAGTATGTTATGTTAATGAAAATGGAAAAATTAGCGGATAACCAAATCCAGATTATAAATTCGTTAGCATCTCTGATTAATGGTGGTGTAGGTGATGCAAGTGTTGGAGTGGAAAACTTGCAACTTATGCTATTAATTGAAGATTTCAACATAGAGGAAATGAAACTTCAAAATAAAACATATAGAGATAAAAAGAAGCGTTTCTTGATCCTAAATAAAAAATTGACGAAGGGTTTAATTGGTTTTAATGTTGGAGACTTACTCAAAAGAGCGAAG
>evgR011285c0g1_i01_Hydra
AAAAATTAAACAACAATATTAAATGAGCCCTGTGTATGCACTGTTGCCAACAAGTGTTCCACTTCCCAAAAACGTCGCTACAAGCAGCGAGGAAATTGCCTCCAATACTAACAGAAGTGTTTTTCACTCTGTTAAGAATGTTATTTACATTGAGTTCCAGAACAGTAACAAAGGAGCTATTACTAACGGCAAAATCATTCTTCTCAAGAACGGTTTTCAATTCAAGTCCATTAATGAGTACATTGAATTTTGCATTCGCAAACCAACCGGTGGAGGCGGTTATCATAGATACACCACCTTTTATTGCAATGGCAATTCATATGATTATGATAC
>evgR011285c0g2_i01_Hydra
GGTGGAGGCGGTTATCATAGATACACCACCTTTTATTGCAATGGCAATTCATATGATTATGATACCGTTGCTTCTCCTCACAAAGGTAAAAGAGGCATTAACCTCTTAAAAGGTATCTGCAACACCATCCTCAGATACGTTACTTTGAATGACTTGAATGTTGTCTTCATCGAGTACAAAAATGGTTTTCAAGCAAGAGTTACTCCAGACGGTAGAATCATTATTCTAGGTAATCAATTCAACTCCATCAATAAGTACATTGAATTTTGCATTGACAAGAAAACTGGTGGAGGCGGTTATCATAGATACACCACCTTTTATTGCAATGGCAATTCATATGATTATGATACCGTTGCTTCTCCTCACAAAGGTAAAAGAGGCAAGAAACTTCTAACAGCCATCCACTATTACATTATTAATTATTGAGGGATAAGAAAACGGACTGAACTCTGATTTTCTAGCTCTGATTTGACTATTCC
>evgR011297c0g1_i01_Hydra
GTTGGTAAACATGGATTGACCTGTATCAAGGTTAATATGAAATGTCCTAAGGGATATAGACAATCAAAGAAGAACCCACGTTTGTGTTTAAAGACAAAGATTGTTGTCAAGATTAAATGTCCAAAGAAATCTGTCTATAGTGAAGAAAAGAAGACTTGTATCTGGAAGAAGTGTGCCAATGTCAGATGTAATGGTGCTTTGAAATGTCCAAAGGGTCAAGCTTTGGCTCTCAAGTCAAAGAAGGATTGTTGTCCATCATGTGTTCCATGTGCCTGCTCAGCTGAGTATCATCCAGTTTGTACTAAGGATGGTGTCACTGTTTTCAATTTGTGTTCTGCTAAATGTTTGGGTTTGGAAGTTGCCCATGAAGGTACCTGTAAGATTACTGAAATTATTAAGATTACAAAGACTTGTC
>evgR011300c0g1_i01_Hydra
ACCTGGATTTATTCGAAGCCTAAAAATTAAAAATCCAGATATCAATAATTGCGTTAGCTTGACACCTCCGTGTTCATGGACAAACAGAATATTTCAAAATACCAACAGTAGTCTTTCATGTTTGTGTAGAAGTTGCTGGATTGGTTATGATAATCCCAACTGTGTCGATAGTCATATAAATAGCTCCATTCATGGATATTTTAATGGAAATATTCTAAACATATCTCTAGAGATAAGTGGAAGTGGTTACAATCTAACAATGGAATATTTTTTTCCACTTTTTGTTGCCGTTATTTCTGGAC
>evgR011304c0g1_i01_Hydra
CAATCAGGTAAGGAAATTGGTAAGACTATTGCCAAGGTCATTAAGAAAATTGAAAAGGCTGCTGACAAGAAGCAAAAGAAATTGGAAAAGAAGATTAAGAAGGCTGCTGTCATGACTAAATGTAAGAGTGAAGGTGATCCTCACTATGTTGATTTTTCAGGTCGTTACTATGACTTCTACGGAAAGGGTGACTACGTTTTAGCTAAGAGTGAAGATGGTAACTTTATTGCTCACTCTAGAACTGGTCGTTGGGGAGCTGTTAGTGTCAATACTGCTGTTGCTGTCAAACTTAATGGTGAAAGAACTGTTGAATACAGCGTTGAAACTGACAAGTTCTACGTTGATGGAAAGGAAGCTTCAGTTGAAGTTGGTAAGGTCAATGATTTAGGCAGTGGTGCTTCATTCAAGCGTGTTTCTGCAAATTCAATGATTGTTTCTCTTCGTAATACTCACTTGACTGCATCTTGGAGTAGAGCTCCAAGTTGGAAGACCACTTTGGGTTCATTCGGTCATATTTCTTTGGTTATTGATACTCCAAAGGATAAGAAATGGGGTTCTGGTTTGTGTGTCAGTAAGTCTTGGGTTGAAAAGCAAGCTACTGGTATTTTGCATGATCATGTTGGTAGAGTTATCGAAAGATCTCACAAGCCAAAGAAGGCCACCAAGAAGGCAACCAAGAAGGCTACCAAGACATGTAAGAAAGCTGGTATCTCTGCTAGACTCCAT
>evgR011312c0g1_i01_Hydra
AGCGCCAAAACGTTACCGTTATGTTTTCAAAAAACTTAAGTATATATTGTTTTAACGTTGTGAGCATCTTGTTCTTGCACAATAAGAAAACGAATCCCTTGAGAACATTGACAATCTTGCTTTGCTAGATCGGTGTATAAAATCCTTTTTCAAAATAAAAAACAGTTAAAGATGCGTGACTCGGTACTAAAAATGCGTGACTCGTACAAAAAATTTAAAGAATATGATGAAATGTTTGATGTTTTTGTGATCGAGACAAAATTGTCGCAAAAAATCCCTTTAAAGGGGTCATATTTTATGCACATTCATAAATTCAAAATTGTATTACGTTCCAAGATTGATTATAGTGAGGTACAAAGTTGGCCACTCAGCTGTATTTTGCATTACTCGAGTCAAGTAAATAACAAAGAGATAATATTTTTAATAACATGTGACGCAGGATGTGGAGTTCTTAGGTTTACATCAAGTGCATCTGAGGAAATGGATAGTTATCTTAAGTATTATACAGATAAGCTCAGATTAAAAAGTTCAAGACCGTTATCCGCTTGTTTTGGTATAAAAAACGGCAAAAGAGCTTCCATTTCTCTAAATCATTTTTTTTGTTTTGATTGCAATGCAACTGATAGATCTAAGTGCAACTGCTGGTGTTCTAATTGCAACTGCAATAAATCTTCTATAAAGCATACAAACAAAAGTTGCAATAAAAATTTCCAAGATATTTTTCCTAAAATATATAAAAGTTATATTGAACCCGATATTTCTAATATAGAGCCATGGAGGTATCGTTATACAAATAAAAAACACAACATAAAACATGCATCGTGTTCAAGCTATGACAATGAAGACGATATAATTATTTTAAAGAATGGTTATCTTGAATTAACCGAGAACACTTGCCCATCAGGGCGTACTTCAAACTTATTCAATTCAGGTTTTTTTAATTCTTTTGATTATGAGGAAAGCATTTCTGATACTAGCGAAAAAACCCTGACGTACAGCTCACCAGAATTACAAAAAGAAATGAAAATGCGTCGCTCTAGTGAGCCATTTAGGCCGATGAATAAGATTTATAAATTGGTAAGAAACTTTAAAAGCTTTGACGCAGCTTTTGACGACAATTCAAGCGTAAAAAAAAATAAAGTTTACGAGAAATCATCTAATCAAAACAGTTTGGATAAATTTTCAAACAAATACATACAAAACAAAAAAAATTATTTAAATAGTTTACCACACCAAGTATTAAATCAAGATGTTAGACTTCTTTGTAAAAAGTATAAAAGGCCTTCTATTTCTGCTATAAATTTTTCTCCCTCTATACCAGAAAGAAGTCCTAGTTCAACCGTGAACTATGAATACTTTTGTAGAGAGATAACAAGTAATCAAGAAAATCATATCTTTGAGCCAACAAGTTTCGTGGATCAACAAATAAACGTGCAACTAAACGTTGGTGCACACCAGCTAACGAATATTAAGAAGAATGATGCTATGGATGAGCAAGTAAATTTAGAGCCAACTGATTTTGCAGATCGTTCAACAAGTTTGCAAAAAAATGCTAATTATTTATCATATTCCGATGTTAAAATAAAATATTTTGAAAGTGTTAATTCTCATACCAACATTTCAACGCAAATCCTTGAAGAGCGCATTAATACAACGGATGCTAATGTTAATAAAATAATGGATGAAACAATGAAACATTCTTACAGAACTGATGGTTTAAACAAATCTACGTCATTAGCAAAATTGTTTGGCGAAACTTTACTTCCATTCCCTCAACCCAAAAAAAACTTGTTTAATTTCATCAATATGAACAATAAGAACAATATAAGAAGCAAATATGATAATTTTAAGTTACTTTTTAAGTAATTATAGTGTTACAATATATTTATAAAATTACATATGAACTTTAATTACAAGGTTTACTGTTGTTAATAAAATATAAAGTATCATCGATCCTCT
>evgR011325c0g1_i01_Hydra
CGTTCTGCCTTGGAATCACTCAAGGAAGAATTAAAGAAATCACTCACAACAGCTGTAACTACAATTCCAAAACCTCTCAAATTCTTAAGAATTCACTACCCAGCACTCAAGGAAGCTTACGAAAAGAATACTGCCAGAGACAATAAGATCCTCTTTGCAAATATTCTCTCTGTATTGGGCATGTCTTTGTCTGAGGAAGGTGAAAGAGATTGTTTGAAATACATGTTGGAAGCTGACGATGATGAATTCGATAAGTGGGGACATGAATATGTTAAATCTCTTGCTGGAGAAATTGCAGAAGAATACCAAACAAGACTCCAAGATAAGATCCAAATGCAAGAAAGAATAATACAAGAAACAGGTGACGATATGGTAATTGATACCGATTTTAGACCAGAAGAAATTATG
>evgR011327c0g2_i01_Hydra
TAGGAGCTGTTTGGATATTCTGTTTAATAACTATTATACCGAACATAGTTACACTCAAAGTAATACTATACAAAGATAAGAAATACTGCAACGAACACTGGCAGAGCAAACTTTTGCAAAAAGTTTACGGTATATCGTATTTTTGCTTTTTATTTTTTATTCCTTTTTTGGCTATTGCAGTTATGCACGGTCACATAATATACCGTTTAAGAAACAAAGAACTTTTGCCATCAAACGTTAGTTCATGGAATCAAAAGAGTCAAAACAGAACAACGAAAGTGCTTACCTCTGTGATAATTGGATTTATTATATGCATACTACCAAATAAACTTTTATATTTTGTCTGGGATATAGCACCAGACCTAGAGAAAAAAAACAGTACACGCTTTTACTTAAGAACCATACAAATACTGTATTGGTCAAGAGTGGCTATCGATCCTCTGTTATATTGCTTTTTAGACACTCGTTTCCAAAAAGATCTCATGAGCACAATGAAAAAGATTGGAGCACTAAACATTATTGAGGTGGAAACGAAAAAAACAAATGCATCAGTTTTGACAACTGTACGATTGGATTCTTTAAAAATAAGAAATGAAACGCTGAAAAAGTACGAAATAGAAAATAACGTCCAAGAAAAGAATAAATCTCTTATATATATATGATCCAATTATAAAACATGTAAATGAGAAAATCTTGAACAACTTAAATAATTTTTATGAATACAGAAA
>evgR011330c0g2_i01_Hydra
AATTGGTTGGAAAAGAGAAACTGGTGTAGATGAGAAAAAAACTAGTATAAAAATTTTTTCTCGAGAAATTGAAATTAAAAATCATAGAAAGAGTGACAACGGCGAATGTTTAGTGATAAGTTGTTCTCCAGGTTTTGAACAAGTATTAGGATTAGTTACACTCACGCACTCAGAAAAAGAGCAGAAAAGATTTTGTGTAAATTGCTCAAAAAATTACTACAAATCAAAATTCGGTCCCGAGGCATGTCAAAAATGCTTGGATGGTTTTTATTCAAATGAAAATAATACCGCATGTTATGATATTTTTACAAACGAGATTCTAAGCAACTCTTTAGAAGAGTATCTGATTATTTTACTAACAACAATAACTTTTCTAAGCGGAGTTTTGATCATTGGAACCTTTGTATATAAAAAAAAAACTACAGTTGTAAAAACCGCTCATTTTTCTTTGTCTGTTGCACAAGTCTTTTTTCATTTATCCATTTCTTTTTGTGTTCCATTCTTGTTTTTAAATAGTTTGACGTTCAAATTGTGCGCCTGTCGCCATATGATTGTTTCTTTATTGTTTACGTTAGTTTCTTCAATTATTTTTGTAAGAACCAAGAACTATGTAAACATTTTCAAGCTCCTTCGTCATCTTTCAGCTCGAGAAAAGCTAGTTGCAACATCGTTGAATTTCAGTACCATAATATTTTTATTGCTAATTCAGGTTTTTTGGGGGTTGATTTTTCTGATCAAATTTCC
>evgR011335c1g1_i01_Hydra
GTATAATACAATAATATATTAATACATTCGTTTTACCATTATTAGTTTGAAAAAAAAATGAAGATTGACATAGTAGTTTCAAACGTTTACCTTTCGTTAGTGCTTTTTACGTCTTCAGTGTTAAACATGACCGTCTGCTACATTACTTTATTCAAAGTGAAGAGAAAAGAGTTAACGCATTTATTTATTGTCAGCATTTCGATTACTAACTTACTTGAATCCACGGTTGGAATAGTACCTCAATTAGGACTATCCAATGAGCCTTTATTGGTGAGAACTAAATCATGTATATCAAGTGGTTTTGCTGTTTTATCGTTTGCTATAACTAACATCACTCATTTATCAATACTGTCTTTGATTAGAACTGCGGCAATAAAATACCCAATATTTTACTTGAAATATCATAAAAAACTTTGGTGCAGAGTAGCATTGATTGTGCTATGTTATGCAAATGGATTTTTTTGGGCGACACTGCCTTTATTTGGTTGGTCAAAGTACGAACTAGATCTTGATAAAAAGCG
>evgR011361c0g2_i01_Hydra
AAAAACCAGACGGCCAGTATCGCAATCCATGGAATTGCCATAGTTTTATTTCTTGTTTAAATAACACATCATACAATGCAAATTGTTCATTATTAGATTTAGTTTATAATCCTTACAAAAACAATTGCGAAAGCTCTTTATTTTTTCCATGCAAAGTATTAGACAATCAGTGTTCAGAAAAAACAAATGGAAAATACTTGATTTCTGATGTGTTTGCTTTTTTGCAGTGTTCATCACAACAAGAAAGTTTTGTAAACTGTCCAGATAATCAAATTTTTGACCCAATATATTGCAACTGTATGGATGGTAAAGATTATAATCTGAATAATTTTTGTATTGGTAGAATCACAAATGAGTATCTCAATCCCTGGAACTGCCATAGTTTTATTGCAT
>evgR011364c0g1_i01_Hydra
AAATAAAAAATAACAATCATTATCTGAAATTACCTAGTAGTACTACTAAATGGCAATCTTGTAATGATGAGTTATTAGCCCATTATGAAAAAAGGTTGGACATCTTATTGCAATATATACACGTACCATTTAATTTGTTTAAGGATGACAATGGAATAACCTATAAAGCAAAAATGTTTTGGAGAAGTATAAATCGTATCAGTAACAAAAAAGCGGTGAGTCACGTTGCTAGTGTGGGAGGAGCCATAGGTGACAATGATATACCTTATATGTGGAGAGATCACTTTCAAGCAATGTATAACTCAATAAAAA
>evgR011366c0g1_i01_Hydra
CGAATTTGGTCACCCAGTAGCTGATAACAACAATTCTCTTACTGCTGGTAACAATGGCCCAATCCTTTTGCAAGACTCTTACTTGATCAGCAAATTGGCCCACTTTGATCGTGAAAGAATTCCAGAAAGAGTTGTCCATGCTAAAGGTGCAGGTGCTCACGGTTATTTTGAAGTAACTCATGATGTTTCTAAATTCACAAAGGCTGCCTTCTTGAATCGTGTTGGTAAGAGAACTCCATGTTTTGCTCGTTTCTCCACTGTTGGTGGTGAAAAGGGTAGTGCTGACTCTGAAAGAGATCCAAGAGGTTTTGCAGTTAAATTCTACACTGAAGAAGGAAATTACGATTTTGTTGGTAACAATACTCCAGTTTTCTTCATTCGTGACACTATTAAATTCCCAGACTTTATCCATACCCAAAAGAGAAATCCACAATCAAACTTGAAGGATCCTGATGCATTCTGGGACTTTTTGAGTTTAACTCCAGAATCAGTTCACC
>evgR011369c0g1_i01_Hydra
CGGATGTGGTGAACAAAATATGTTGAAGTTTGCACCTAATATTTTTATTATGAACTACTTGCGCAATACAAACCAAGTTAATGATGAAATAAAAAACAAAGCATTAAACTTCATGAGGACAGGTTATCAGCGTGAATTAACATATAAGCGCACTGATGGATCATACAGTGCATTTGGAGAAAAAGATAAAGAAGGCAGTACATGGTTGACAGCGTTTGTACTGAAGTCCTATGCACAAGCGCGGCCTTGGATTGACGTTGATCAGAAAGAAATACAGGATCCAATAAGTTGGTTGTTTAAAAATCAAGATTCTAATGGATGTTTTCTGACAACAGGAACACTACATAACCAAGCAATGAAGGGAGGAGTTAAAACACCTGTTACACTTACA
>evgR011382c0g1_i01_Hydra
CCTGCTGATGATATGTTCGCCGATCAACATCAACAACAAGATTCCTTCGATTCTTTCGCCTCTGGTGATGACAACGGATTTGTTGCTGAACAAGGCTTCGAACAAGAACAACAAGGTTTTGAACAAGAAGGTTTTGAACAAGAACAACAAGAAGGTGGCTTTGATTCATACCAAGAAGCTCCTGTGACTGCTTATGAACAACAACAACAAGAACAATACACCGAACCACCAAAGCAAGTATCTGCCATCAGAGAATTCGAGGAAAAGAGATACAATGCTCTCATGGAAAAGGACAATGAAAGTAGAACACAACACGAAAAAATTTTGGAAGATTCCAAGACTTACAAGGCCAAGTTTATCAAGGATCGTGAAGAAAGAAAGGATACAAATTCCAAGAAGAACAAG
>evgR011390c0g2_i01_Hydra
AGCAACTGTGACGGAGCTAATGCAAAAGAAGAGTTTTGCAATACTGAATATTGTGACAACAAACTGAAGATTTCAAAGTCACTTCAATGTGAAATATGGGAAAATATGAATGGTAATGCCAAGCTTCAAAATAATAGTGATACATGGCTTCACATACTACGATCTTGGTTAGCTTGGAAATTGAGTGCTTCAAAATGCAACACTATTGATGATGTAACACATTCTTTGAACAAAAATCTCCAGAACTTAGAAGTCGATATAGTCCGCTTAAGACTACTGAAGGTTTCTCTCCAAAAAATGATCCATTG
>evgR011391c0g1_i01_Hydra
ACCATTTACTCGTATAAAAACTATTGTAAATCAATAAAGCAATCTGTTTTATAAGATTCATCACTCTGGAAAATTCTTAAATCAATTAAACCAAGCCAACAGCATTCACTAGCAGGCCTTGATGATGACGACATGCAAAGGCATGTCAACGACATGCGAAGGCATGTCGTGATGATGACGACATGCGAAGGCATAAATGGGTTTATTTCTCTAATGAAATGTGTTGCCAAAGCGTTCACAGAATGCAAACACAAAAAAGATATAGAAAGACGTTTAGAAAAAGCAAAACGATATTTAAAGACTACGTATGGTTTGCATTGCATAGAAAAGTCAAACATTGCTTCACATTGTATCAACTGTGCTTTGTGTAATTCTAAAGATAACGACTTTAAAACAAATACTCGCTATTCCCATTCGGGGAAATGTCATGACTGTACGATACATGCAGAATGTCTAGAGATTATTGAAAACCTAATTCAGAACTAACCTTAAAATAAAGAGAAAGAAGAACTGAAATACGACACTCTAAAAAGTACTGAATCTGTATATTCTTGGATGAAACATATTCTTCCACACGTGCAACAAAACAAAGCCAAAGAGTATGCTATGAATAAAATTAACAAAGTGTCCGGCCTGGACCTGGTCTCAAAAAATACTTCCAGTTGAATTCAATGAGAGAACAATGAAAGATAAAAAAAGAGTCAAGTTATTTAATCCTATTGATTATTTTAGTTGACAGCAAATTATCTCTTCATTTAGCAGAATGTCTAAATTATACCGGCAGGGCTTTCTTAAAAAACCTGAACAAGTTGAAGCTATTGAAGACTTAACAGAAGAGTTTGTAAAAAACCTGTTTATTAATGACACTGAAAATTTGATA
>evgR011401c0g1_i01_Hydra
ATTGCTGGTGTTACCGAAATCTTGATCATGTATCCATTGGATGTCGTCAAGACTAGAGCCCAACTTTATGCTGGTAAGACCAACAACCCAGGTATTGTCGGTACTGTCTCTGAAATTGTCAAGGCTCACGGTTTCAAGGGCCTCTATCGTGGTATCCTTCCTCCAATTCTTATGGAAGCTCCAAAGCGTGCTGTTAAATTCTCTGCCAATGCCTTCTTCAAGGGACATTTTGCCAACTCTGATGGTACTCTCTCGCAAACTGGTGCTGTTTTGAGTGGTTCTTGTGCTGGTATTACTGAAGCTTTCGTTGTCGTTCCTTTCGAACTTGTAAAGATTCGTTTGCAAGCTAAGGAAAATCTTGGACTCTACAAGAATACATCCGATGCCTTGACCAAGATTATCAAACAAGAAGGTCTCATGACATTGTACACTGGTTTGGAATCTACCATATGGAGAAATGCTACATGGAATGGTGGTTACTTTGGTTTGATTCACGCTGTTAAGAGCAGAATGCCAAAGACCAACTCTGAGGGTCAACGTATGTTCCAAGATTTTGTTGCTGGATTCATGTCTGGTACTTTTGGTACCATGTTGAATACACCATTCGATGTTGCCAAGACTCGTATTCAAAATCAATTGCCTGGAACTGTTCACAAGTATAATTGGACTTTGCCTGCTCTTGTCAAGATTGGTAACGAAGAAGGTATTAAGGCTTTATATAAGGGTTTCTGGCCAAAGGTTTTGAGACTTGGTCCTGGTGGTGGTGTTTTATTGCTCGTCTTTAACTTTGCTGTTAGATTTATGAGTCCTCCACAAGAACATCACTAAGTTTTTAA
>evgR011407c0g1_i01_Hydra
AGTTGCAAAAAACTGAGCAAAAGCTTTTGCAATACATCTTTCACTCATTTTCATACAATCATTCCAAACTGGGGTTGTCCCAGAGAGAACTATAAAAGAGGGGATTATAAAACACTCTACCCTACACAACCTAATTTCAAAGTCTCAACGTAGATTTTTACATCACAAAAGTTGCCTTACAAACCTGAAAATGGTTATATCCCGTAAAATTCTCAGGGAGGCAACTTACCGTGATACTACGTGTTGACATTCCACTAGATCTCTGCTTGGTTAGCTGAATTTCTTTCAACGACGAGTTTTCAACAACGAGTAGTCATTGGGGAACAAAAGTAACTCTTAAAAAGTGAAGTCCCTCAAGGATCAGTTCTTGAACCACATCTGTTTGTGCTTTTTGTCAAATATCTAGATGGCATTAGCAGCTACATCAAGTTATATACTAATGATTATAAATATCAATCAACCCTCCTCTGATTGTGTAAAACTGCAAACGGACATTGACTGCGCTGTAAATTTGACACATAACTGGCTTGTGCACTTCAACATTGTAAAGTACGAAGTAATGAACGTTGGTTGCCGCAAAAATAAACTATCAACAAATATCTATACCATGACTGACACCAACGGTAGCCGACGCCATTTATCTCCAACATGCATTGAAGGCAATCTCGGAGTTCTAGTATCCGATAACCTTAAAGTCAAAGCACAAGTGTAAGCAGAAGCTTCGGTTGCAAATCGAATGCTTGGTCGTCTTAAGGAGACATTACGGTGCCGTAGCCTGCATCTTTAGAAAAAAATGAATCAGCTATACGTTCGACCACATCTAAAATTTAATGTAAAAGCTTGGTCGCCACATCTTAAATAGGAAATTGCTCTTATTGAATGTGTCCAAAAACGTGCAACAAAGACTATATTGTCGGTTGAACATTTTAAGTTCAAAAAAAATACTACAAATACTCGGTCCAACCACACTAGAAGAACGCAGAATTCGAAGTGACCTTACACTGCAATGCAAAATCGCTTATCAGTTACAAGAAATTGAATTTTACATCCCAC
>evgR011413c0g2_i01_Hydra
AATTTAACAGTAGTTGTTGAGAGTTCCGAGCCAATGTTAACACATGTAAAGAAGGTTGACTCTACAATTCAAGCTTCCATCAACTCTATTATCAAACGAGACACAAACAACAAGAGGGCATCTAGATCAAAAACAAAAACTACCAGTGAAGAAAAGAAGGGAAGAAGAAAGAAAAAGAATGAAGCTTGTGCTTCCAAGAAGGTTTGCTTTTTGAGAATTGTCGTTTCAGACACTGGATATATTCTTGCCTCAAGTGATAATATTTGGGTTAGATCCAAAACCAGACAAGAAATGGAACGTAAAAAAGATAGAGCCTTAGCTCAAAAGAGAAAGAAGGAAGAATCCAAACCAACTGAGTCATCAACCAAGGAAGAAAAATCAGAATCTCCAAAAAAGAAGAAAAAGACAACAAGCAAATCAAAGAAATCAAAATCCAAGGAACAACCAGTTAACAGCTTCACATATGAACAAAGTCCAATGAAAGGAAGTGTAAATAGTGCCAGCTATGGTGTGGTTTTTAATCTAGATTCAATGTAATTTGAATTTGAT
>evgR011426c0g1_i01_Hydra
ACGGTTTACCAAATATTACATCTCAGTATTTGCAATTGTACTTAAAAGACTTTTGCATAAAATGCTCATCTGGTGATAATTGTGTTGGGTTAAAAACTGGTGAAATTATTATTGTCAACAACATTGTTCAAACAGCCGGGGATTCTGGTGATATTTTTATAATTTACAACAAGTATAAACAAATGAACCGTTTTTTTATTTATCCTTGGAATTCTTCGCATATAGGTATCTATAAAGTTACAGATCTTTGTGATAGTAATCTCTTTTGCTCTGTAAGTGAAATCAACAAAAAGTACGTGATCC
>evgR011427c0g2_i01_Hydra
GAGAAAAGTTTAAAGATGGAAAGATTAATTTACCAAGATAAATCTCACCTTAGTGTTGCTTATTCTTTACATACTCTAGGAGTATTGTATAAAACTGCAGGTAAATACGATTTAGTAATAAAATACTGCGAGGAAAGTCTAAATATAAAAAAAAACATTTATAAAGATGAACATCATCCTAGTTATGCTGATTCTTTATTATTACTTGGAATAGCTTATGCTGATAAAAAAGATTATGATAAAGCGATTAACTACTATATACAGAACCTATATGTAAACAAACTAACGTATCGGAACGAACCTCATCCTATTATTGCTGATTCGTTAAACAACCTTGCTCTAATTTACACTGAAAAAGGGCAATACGATCAAGCAATAATTTACTATGAGCAGAGCTTAGCAATGAATAAACTTATCTATCATGAAAA
>evgR011431c0g1_i03_Hydra
TGCTTGTCAATTTAATGGTAATGGTTTTAGTGATCAAGATACTAGTGAAATCATTAAATGGTCCAGTGACTTTGGGCCAGGTTGATCTAAAACTTCAAATAATAGCAAGTTTGAAGGGAATGGTTCTTTTGATGCCAATTCTTGGTATTTCTTGGATTTTTGGGATCTTCGCTGTCAATAAAAAAGAAGTTGTATTTCAATATATTTTTGCGCTTTCATCTTCTACCCAGGGTTTTTTTATTTTTCTTCTTCACTGCATCGGAAACACGGAAGTGAGAAGTAATTTCAAAAGATTACTTGAGAAGCATTCACAACAGCTAAAAGATAAAAAAGGTTTTGGATTTTTTAATAAAACAGCAGTTAATAAGAAAAGAAAATCTCTTCAACAGTCAATAGTTAACGCTGCCATAAATTTAGGCACATCCAAGTATGAACATTTTGAGGTTCAAACGAATAACTCGTCGTCCCTTTGATCTTCTCGTCGTTCATGTTGGTTTCGAGAATCATTCAATGAAGATTATTTTAATTATATATTTATTCAATGAAAAAAAA
>evgR011432c0g1_i01_Hydra
TAAAACAGCCGTAGTGGTCCTGCCGTCGTTAATTTTTTTATGATTGTTGATTTATGTTGCTTTGTTGCTCGGACGGGCAAAAAAAGTTAAAAATGAAAGTGTTGGACAAAAATTGTCTCGAAAAGTAAATACATAGTAGCTACATTGTCTTAAGGTGGTACAACAAATAGTATGGAGGCAGCTTCAATTGATTCAAGAAAAAGAAACACAAAAGCGGACGAAAAGAAGTTATCAAAGGAGGATATTGGTTTTCCATCAGAATTTAGCCATGTCAGCCATGTTGGTTGGGATCCTAAAGGAGGCTTTCAAATTAATAATATTGACCCACAATGGAAGAAGTTATTTGGTGCGATTGGTGTAACTGAAAAGCAACTTGAAGACGAAGAAACTTTAAATTTGGTTAGCAAGTTTGTGGAATCCCGTAGTGGTATTAAAAATACAGATAGACAGCTTTCGAAGAATGCAAAAAATTCACACGGTTCAAAATTTTACTCTCCTTTACCAACACCTGATGATCCACAACCGTTACCTGTTTCTAGAAGTGCTACGCTACCTCCTAAACCAGCAAGAAGTGATATTACATATAATGAAGCAGATGAAACGCTGTCTTATAGTAATCTGAAAAAAGGAAAATCAAAATCTGTAGGGAAGAGACTGTCTAAAGAAGATATTGGCGTTGCATCTGATTTTAGGCATATTGGTCATGTAGGTTTTGATAGTTCATACATCCAAAGTGGTTTTGAATCTAATGATGCTGAAAACCCTATTAAAGAACATGATGAACCTGAAGTAAATTTAGAATGTTCTGATCCACTAAGGATGAAAAATCAGTATTTGACATCTCCTCAAAGTTGGACAGTTGAAGATGTATGTGCATGGTTGAGATCTATAGCTTTAGAAGACTATGTTGATGTGTTCACTATGAATGCCATTGATGGAACAGAGCTAATGCAACTAAATGATGATAGTCTTTTGAAGGGTTTAAATATTTCACCTCTTGGTCATCGAAATAAAATTATGAGGGAGATAGACCAATTAAATCAACTGAGTTATGATAATGGTGATGGCTTGGTTGACTATAATAATGATAATGAAGTGGTTCCTAGTGATTTTTTATGTCCTATAACTATGGAAATTATGGAAGATCCTGTCATTGCTGCTGATGGTTTCACTTATGAAAGGTTGTCAATCGTGTCGTGGTTATCTTCAGGAAAAGTAACGAGTCCTATGACTAACATCCCCTTGTCTCATACAGAATTAGTCCCCAATCGGTCGTTAAAAAGCATAATTGAAAGATTTTTTGGACCATGATGAAGAAATATTAAAAGTCGCAAATTATAATGAAAAGTTTTAATGAAAAGTTTATTATATTCGAAAATAACACATTTTTTATATAGAATTATATCTGAC
>evgR011433c0g1_i01_Hydra
ATTCTATTATTTGAAAAAAATACTTTTTAAGTTGTCCATGTCCAAAATTTAATAAACTAATAATTGAAACAAAACTAAAAAAATTAAACCCAAATAATTCAACAGGTGTAGATAACAGTTGTACATCCAAGAATATTAAAGGAATGCTCACAAAGTTTATGTAAACCATTGGAACTAATATTCAAGCAATCATTCGAAAGTGCAACTCTACCAAAATATATGGTTACATGTAAATATAACACCATTGCATAAGAAAGGTGACAGATTTGATTCGGCTAACAACAGACCAATCTCTTTAGCATCAATTTTGTGCAAAGTCATAGAAAGTATTATCAGAGATAGAGTAATGAAACATCTTAACAATAATAATCTAATCTCAATGCACCAACATGGTTTTGTCAACAAAAATAGTTGTTGTACAAATTTATTAGAAACATCAGATATTGTCAGTAAAGCATTTGACAATAATATTCCAGTAGACAATAATATATATATAATATATTATGGTTTTGAAGATGATTTGTTGAACTGGTGCAGAGCTTTTTTCAAAAATTGGTTGCAACGAGTATTACTTGGAAAAAATATTTCTGAATGGATAGATGTTGTAATTGGTGTTCCTCAGGGATCAGTACTTGGACCATTGTTTTTCATTTTATATATTAA
>evgR011448c0g1_i01_Hydra
ATTTTTTACAAATAATCAATCATACTGGTTGTTTTTTATTGAGAAAAGCTGTTTGTTTTTAGTTTAGAAAAATGTCGTACAGAGATTTAAGAAATTTCACTGAAATGATGCGAGCGTTAGGTTATCCTAGGTTGATTTCTATGGAAAATTTTCGCAAACCCAATTTCCCACTTGTTGCCGAAATTCTTCAGTGGTTAGTCTTAAGATACGATCCTAATTCTGATATTCCAATGGATGTTGAGTTTGAAAATGATAGAGTCTTATTTATCAAATCTGTAGCACAGTTTATGGCCCTCAAAGCGCACATAAAGCTAAATACAAAGCGTTTATATGAAGCAGATGGTTTAGCAGTTAAAGAAATGCTAAAAATCACAACAGTGCTGTATCAAGCCGCAATGAAATTGAACAAATCTGGAGATGACGAACATGGAACAACTAAATATCACCCAGGGAAATTTGATTTTAATTCAAAGCTTTCTGATTTGAAATTGGCAAGACAACTAGCTTCAGAAATAACAGTTAAAGGCTCGAGTTTGCACCAACTTTTGGGAAGAGAAGTAGAACTAAGGACAATTCGTAGAAGTTGTATTGCACAACCAATGGACTTAAATGAAATTGAAAAAAATGTTACGAATGTAATCAAGACCACACAGGCTGAGATTTCTAAAACTCATGGCCAATTAGATAATATATCATCTGATGAAGCCAACCTTCAAGCTAAAATTGAAAAACGAAAGTCAGAGTTAGAAAGAAACCAAAAGCGCCTAAAAAGTCTTGAAAGTGTTCGACCTGCATTTATGGATGAATATGAAAATCTTGAGAAAGACCTACAACAATTATATACTTCATACATTGAAAAGTATCGCAATTTATCATATTTGGAAAGAATAAACGAAGAAATGAACAAATCGGAACAAAATGATGATGACTCTGAGAACCCGATTAAGATGTCTCTTAAAGCTCGCGATGAAAACGAAAGTCTCATTCGAGGGAATCTCACATTGGGTTCTAGTGATGAAGACGAAGAAATAGAAGAAACTTCAAATAAGGATGAAAAGAAGCAAGCTGTTCCTAAACCAAAAATGATTGGTTCATTAACTGGAGATCTAGAAACTGATGACGAATCAGAAGGTGACTCTACTGCGACAGAGGACGGCGAGGATGAGTTAATTGAAGGCGAAGAAGATGACGAAGAAGAAGGCGACGACCAGGCGAACGATAGCGACGATAATTTTTAATTAAATTTATAATTTCAAATAATAAAAATAGGGTAGCTGACGTAGTAAACAAACAAGTAAACATAGAGATATAAATAAAGAGAGATATAGAATATAATGGAATAAATTACGAAATGAATTACACTTGTACATATTAAAAATTGTACATATTCAGGTTTTATGTGAATATATGTATTTATTTATTATAAATTTATTTGAAAATTCTTAAA
>evgR011466c0g1_i01_Hydra
TTGCTCAAGATTAAAACAGGGTTTGATACTACGCAACGCTTTGTATTGCTCCAACTTTGTTTTATCAACTTTAGCTTGAGCATAATCTATCAAATTGAAGAATGGCTTTGTGTTTTAAACCACAACTATATACCTGAAAAAATGTTTTTTATGGTCGTTAAAAGAATTTTTACCACAACATTTTATTATACTACTCTATGGTTTGTTTTTGACAGATACTTGCATATAAAACTTAACATTAAATACGTCCTTTTCTGGTCAAAAAAAAAATATTTAATTTTCACATTAATTTTATGGGGTTTAGCTGTATTAGTTGGAGTTTTAACTAGAATATATATGGTGAAGTACGGGTCTTTCATTGATGTTTGCTACGATTTCCTAATAATATGTTTTTCCATCATTGTTTACGCGTATGCGTTAATACTTTATCGCAGACAAAGTAATCTTAGTTGTAACCAACGTGGCAAAAGAATATTCAAAGGATTGCTAGTTCCAAGTATAATACTATTAGTTTATTTTGTATTGATTGCTATTCCTGATGCATTGTGGGGAATTTATATCAGCGGAATCAATAACAATCATTTTATGTACGTTTACGCTAACATTTGTTATGCTATGTCATTATCGGCGGTGTTAGACTTTTACTAAAAAGAAAGTTACAACATAAGTTTTTAGATGAAACAGTCAGTACGAAGAATAAATCGTATTTTTCAACTTCAGTATAAAAATATTATTTTACCTACTCAAACACCGTAATGAAATAGTATTACACCAAACAAACTTTAGTCTGTTTTATGTGACTTCATCTGTACAAGTCAGATGTGATTAAATCTGTACTATTTAGAAATTGTTTAAAGCATTAGACACTGTGACGCGGTTTTAAATAGTTTAAAATGAATTCATAAATTTAGAAAGTATATTAGAAAATGATATATAATGTTTTTTAATGATTGAAG
>evgR011478c0g1_i01_Hydra
CTTCTACAAAAGTTTGCTTGAAATATAGTTTCAATAGCAAATCATTATTTTTTGGTTGGTATTAAATTCGTAACAAAAAGTACTTTAGCTTTCAGGTTTGGTAGGGCATTTAAACTTGCATTAAAGTTATCGAAACCTAGGATCAGACATATAATATACTGTTTGAGACAACTATTGAGGCTTTAACTATAATTTTAGAGAACTCGTATGAAGTCGTGTTTTAATGCACGATTCTTTAACAAAAGAGTGAGTTATAAATGAATTAAAGTAAATATCCGATTACGTGCTCTAAGACGTGCTCGTGTAGTGCGTCGCACTCGCTTATTGTTCAGAAAGCATTTAAAAGTTTAAAAGCGTTTTAAAATTAAAAAACAAATGCTGAATAACTTGCGAAAAATAACTTCGTGTGCTGTTTGTAAAAGCAGAACCGTATTTAAATGCAGTCGATGTAAACATACTTTTTACTGTGGAAGAGCTCATCAAGCAGAGCATTGGAAACACCATAAATATGATTGTCAACCTAAAGTATTTCACCAAAACATAGATTGTTTAGGTCCAAAAGATTGTTTTATTGATTCTTCTAAAGCAACTCGCGAAGAACTCTCAATATTTGTTTCAACTTCACTGCTTAGAGTAGGCCTCTGTGTTTTAGATAATTTTCTAAACGAAGACGAAGCGAAATACGTATTTTTAGAATCGCGAGCAATATGCGTCCGAAATACTTTGAGTAATTCTGGCGAAATTTTAGGAATTGAGTTTTTAACAAAAAATATAAAATTATATCCCGCTATTTTTAACTTAAGAATTTCATTCAACAATCTTATAAGCTCTGCATCGCAAAATATATCGAAAAATGAACTTATAGACAAAGGATTGTACGTTCAGCAAATATCACAGACGTCAGCCTGTACCACGCGCAAAGAAAATACTTTTTCTGAAGATTCTCTTGAAAACGGTTTGCTTAATAAATCGATAATATTGTGTTCGTACTTTACCACCCATGACTACAGTAAATTAAACGGAGGTTTATCGAGGTATTTTATTCAAAACAACAAAGTATTAGACGTTGAACCAACATTTAATCGATTAATTATCTTTTGGAATGATGGCAGAATTACTAAAAAATCGTCTCCCTCTAATAAAGACTTGCTGATTATTTTTTCCTTGTATAATAGTTCTTTATAGTGCATTCAACAAAAAATATATCTGTATATCAGTAGCATATATCGAAAAAAATTTTTGTCAACAATTTAGAGGAAACTTTTAGACGCTTGTCTTTTTTTGTCAATTTATGTCAAACAATTTTACCTGCCCCTCCACAGAGTAGATTTAATTTGTTTTAATTATGAGTATATTTTGTCATTTTTATTAAAATTAAC
>evgR011484c0g1_i01_Hydra
TGCGCTGAAAAAATTCATTCTCTTACAATGCCTAAAACTAAAAGGATTACAAGAAACAAAAGACAAAAAACAAATGGAGGATACAATTTTGAAGATCCAATCATTAAACGACAGTGTCAAACCCTTAGATGGAATATATTCACCAACGACGACAATAATCAATCCACTGTTCACGAAATGCGAAAAGAACTATACATATATACAGGGAACCAGGAAGATCTCCAAATGGACTTCTTCAAAGGCTACAACGAAGCACTAACGGAAATTGTTGAAGTTGAGCAAAATAATGCAGGGACCGATGAAACAGTACATTTTGAGCGAACTGGAACCTCTAGTACCCAAGACATTAACGACTTTCTCCAACGATACGTGTTTTCAAATGATGAAGAATTTGAATCTAATAGTAATGAAGGATTTGAAGATATCGTTGAGACAGTTAAAAAAGAAATTGAACCAGTTGAACGGTATAATATCAAACTGGAAAAAAACGATACGAACATAATTGAAATCTCAGACGATGAGGAAGGAGTTTTTAATTCTACAATCCCAGAGTGCTCAGCTGGGACTGTAAATCAATCTCCAGAGTGCTCAGCTGGGACTGTAAATCAATCTCCAGAGTGCTCAGCTGGGACTGTAAATCAA
>evgR011489c0g1_i01_Hydra
CTTCTATTGTCTAAATAAGTCATAACTGAAAAGCACATAACGAAACCACGGTAATACATTTAAAGTGTGTTTTGTTTGTTTTATCTGCTGATATGATGGACTTTTAATAAAAAGTTTAACAAAATGACAAGTTCATAAAATTATCATAAGTTGTGACAGATATGTTGGTAATAACCAGATTCATATATCAATTAAACTATTTCAAGTAAAATGACAACGGTAAAAAATATAAGAACTATTTCTATACACAAAAATGATATAAACAGTTGTGTATTTACTTCAGATGGAAAAGCTGTAGCATCTGCTTCAGCAGATAAAACTACCCGGATATTTAATGTTGAAAGCGGGGAAGAACTTTTATTTTCTCCTTTAGGTGGACATATCTATTCTGTGACAGTCTGTGCTTTTTCACCAAGTGATAATATTTTAGCTACGGGTTCTCAAGACTGCACTGTTATATTGTGGGATGTAAAGACTGGAAAGCAAATTAAAGTGTTGCTTGGACACAAGGGTGCTATTCGATGCTGTTCATTTTCATCAAAGTTGTCATTCCTTGCTACATGCTCGTCAGATTCATCTATAATCATTTGGAATTTTTCAACTTATGAGTTATTAAAAACATTTCGAGGGTCAGAATTCAGTTTAACAACTTGCAGATTTACACCTGATGATTTGTACATTATTTCTGGTTCTATAAATGGAGACATTAGAGTTTGGGATTTGAGTTCAAGTAAATGCATGAACGTTGTACAAGCTCACGACTGTGGTTCCAGTAGTGTTGGTTTAAGTGGTTGTGATTTTTCACCTAAATATGGTGCTGCTTCACCAATTCCTGAAATCATAAATGGATGTTCTGGTGGTGAATCTGTTTGTTTTCTTTTTGCCACATGTGGGGGAGATAATTTGGTTAAATTATGGAATTTATATACTGCATCAGCATACTCAGTGGAATGTTCTATCCAACCATTTTCTGTTTTGGAAGGACATAGTTCTCATGTTTCTGACTGCAAGTTTTCTTTTAATGGAGAGTTATTGGCATCGTGTTCTGTTGACAAATCTGCGATTCTCTGGGATCCAGATTCTGGAAAGATCTTAACAGTGCTTACGGGTCACACAAGGTACATCAATTGTATTGCTTTTTCGCCTGCAGGCTGTCATATAGCCACTGGTTCTAATGACAAAACTCTTCGAATTTGGGAAATTGAGCAGTATCAAAAAAATTCAGGTTCCCCTGTTATTAAGACGCAGTGTTCTAACTTAGTAAGCAGTTGGACTGTTCATGAAGTATGTATATGGTTGCGGTCAATAGGTTTAGATGTGTATACAGATGTTTTTACCAAGAACATGATTGATGGAGCAGAATTGTTGCTGCTTAATGAACAAAATTTTTCCAAAGAATTAAATATTGAACCTCTCGGTCATCGTAACAAAATCATGAGAGAGATCAAGAAGTTAAAAACCGATCAAAACGATTGCTTCAATGATATTCCAAATGAATTTTTATGTCCGATAACTATAGAATTAATGACAGATCCTGTTATTGCTGCAGACGGTCATTCGTATGAGAGAACATCGATAACTAAGTGGTTGAATTCAGGAAAAATAACCAGTCCTATGACAAATGAACCATTGCCTAGCATGCAACTTGTTCCTAATCGATCATTAAAGACTGCTATTCAACGATACTTCAACTCTGCAGATTCAATTCAGATTCATAATTTGTTCATCAACACTTGTCAAAAGAATTAATGCTGTATCGTATCGTGTTAATCATGTGTGTTATCGTATCGTGTTAATCATGTGTG
>evgR011491c0g1_i02_Hydra
AACATTTGAACTGCGATAATAAGGTCGGTTAATGAAGCTGAACAGAAAATGAACGATTTCTTATATCTTCCATTTAGAAATAGTAGTTATACGGAGCATGAAAATAACACCATTGCGAGATTGTTCGGTAATGGGAGCTACTCCGCAGAAAATAACTTTTCCGCGCCATTAACAAACAGTAGTAACAACACCTTATATGACCATTATTTAAACAAAAGTGACAAGAACCACGTGCTAATTATAGAATGTGTCGCTCTAGCAGGAATTTATGTTATGCTCACCGGATACAAGTATATTAGGTTTGCATTGTTTGCATCAGGGTTTGCAATTGCATTTGCATTATTTTACATTGTTAGTCCTTGGTTTATACATACTAAGTATTGCTGCGACAAAGAACATTTTCAATTAAACCAGCCGTTACTAACGTATGATAGAGCTCATCTTGGAATCAGTTGTGTGGTTGGTATTTTGTTTGGGTTGCTGGTTAGTTGGCTGTATCGATTAGGTATCTTCTTTTTGGGAAAATGTCTCGGAGCTTTGATTGCAATAGTTGCTCTGCACACCACTTCTCTACATCAATATTTTACCAGTCAAATATCAATTACTATATTGCTTACAGTGTCTAGTTTCTTATTTGGAATACTAACATGGAAATTTGAGAAACCGCTTATGATACTTGCCACTTCCTTCATTGGCTGTTTTGCTGCACTATGTGAACTAAAATATTTGAAGCTTATAAAAGTTTGTGCGCAAACGAAAAGACCATGTG
>evgR011496c0g1_i01_Hydra
TTTATTTTCTAAATAAGAAAGTTATCGTTATCTACAACGGAGAACAATCAACGTAGAAATTTTGACATGCAATCAAATGAATGAATTTCAATGTTTTCTCAAATAAAACGTGCTAAAATTAGTTCGACTGCTAATATGAAGACTCTAGAAAGTTTCTCTGATAAGCAGTGTGGTTTTGTGAGTGATGCGCAAGCTAGTGTTTGTGTTACATTGCATCCACTTGTTATTATGAATATCTCTGATCATTTTACAAGGATTAAAGCTCAAGAAGGAGGAAATCCACAGATTTATGGTGCTCTTCTTGGAAAACAAAATGGTAGAAACTTAGAACTTTTCAACTCTTTTGAACTGCAACATTCAAATGAAAATGGGAAAGTGGTTATCGAGTTAGCTTACTTCCATAGTAAAGAAGAGCAGTTTCGTCAGGTTTTTAAAGACTTGGATTTTCTTGGTTGGTACACCGTTGGTTGGTCAGAAGCAACTGAATGTGATATATCAGTTCATAAACAGCTTACTGTAATAAATGAAAGTTCCTTATTGTTGAAGTTGGATCCTTTTTCAAAGTCAGCTAATCTTCCCATAACAATATATGAATCTATGTTGGATATTGTAAATGAACAGCCACGAATGTCATTTATTGAAATAAGTTACACTTTGGCATCTGAAGAGGCAGAGAGAATAGGTGTTGATCATGTTGCTAGGCTGTCATCAACTGGTCACTCAGATGTATCTCAAGTTTCAGAACATATGCAGGTTGATAAAAGTGCAGTGCAAATGCTATACTCAAGAGTTGAAACTATTCTTCAGTACGTAAAAGCTGTACAAAATGGTGAAATACCTTTGAATCAAGAAATAATGCGAGAGTGTTTAAGTTTATGTCAACGACTTCCTGTTTTAAATACAAACTCATTTAAAGAAGACTTACTGAAACAATACAACGATGTTTCACTAATGTCATATCTTGCTGTTATTACTAAAGGATGTAACATTTCTAACGAGATGATCGCAAAATTGAACTCATTACCGGACCGTCATGGAGCAGGGAGGAGAGTACGAGGTTTATTGTTTTAAATAGTTGCGAAATAAATTGAAGAAAATCGGATAAAAAAGACAATTAGCAAGCAAAGCAAGGCAAATTAAACTAAAGTTTTTTTTTTTCTTTTTTTGAGGTTTTAATTTATCTGTAAAAGCTTTTGCGTTTGTATTTTAAACTTATTTTTACATATATAATTGTTAAAAAAAATTTGTTTTTATAAAAAAAAAAAA
>evgR011505c0g1_i01_Hydra
GTTTGACAAATTGTTGACAAGTTTGTTGAGGTAATTAATGGACAGCCCCCTACAGCAAAAATGAAAACGCAGTCTATTGCATATACTGCACTTTATTTCTAAAACAAGATAAAAAAAAATCTTTGGGAGCTTTTGTTAAACGTGGATATAAGGAATCGCACAATGTTATAAAGTAACAGAAACTACATATAGGCAATAGATATCATAAAGAAGCTATCGAACAAGCTAATGGTATTTGTTGCCGTTTTGAGGATCCAAGAAAAACCATTCCATTACAAAGTGATAGCACTTTAAAAAACCAACAGCAAATGTACCCGTACGTTGTTGAAACATTAGATATAATAGTGCATTTGATGGGGAAATAAGGACTTGCATTTCGCGGAACTTATGAAAATTGTATGGAAAACTTAGACTAAAAAGATAATCCAGGAAATTTTTTGGCAATTGTACAAGAAATAACTCGCCATAACGCTGAACTTAATGCACATATACAAACACCATTACACAAAGATGTCAGCTGCTTGGCCCCAGAGTTAAAATGAATTAATTGATATAATTGGGAGGAAATGTATACAAGAGGGGCTAATTAATAAAATTTTGCAGCAAAAATTATGGTAATTTGGCAGCAAAGCTGCAAATTACCATTAAATTTCAGCAGACGAGGTAACAACTGCAAACCAACAAATTATATCAGAATGTATGAGATACGTAAACGGAGAAAGAAAAACGTAAAAGAAATTAGAGAAGTATTTCTTGACTTTTTGAATTTCGACAGCATAACAGGCAAAAATATTGGGCAAAACTTGATAAAATTTTATAAAGAAAGTGAAAATGATCTTAGGGGCTTAATGTTACGCTACGAAATGTTACGCTACGAAAGCTACGGAAAACTACGGAGCTACGGAAAAATGTTGAAAAATAAAACTTACGCAATTTATCAACAAGCTACTCGTATGGCAAGTAAATTGTTTGTAGAACCAACAATTCCCAGAATCACTATGAGACAGGCCTACAGAAACAACATACCTGCAGAGAGTCCCATGAATATTATAAACGGGGCTTGATTATACCTTTTCTCGACAACATCATTTCAGAAATCAAATCAAAATGAGATAACTGCATCCAGAATTTAGTTGAAATGTATAAAAGTGATCTTCTAAACTCAGAAGCTGTAGATCATGAACTTATGTTTTGGAAAAGAAAATTGTCTGTTATTAAACCTGAAGATAGGCCTAATACTCTAGCTAGAGCCATTAAAATTTGCGATGAAAATCAATATCCTAATCTCTTTGAGCTTCTAAAGATTGGATTTACTCTACCCGTAACGTCCTTTAAAAACCTTTAAAATTTTCATGGTATGGCCCATATATATATATGATGCGCAAGGCAGTGGCGGATACAAAAATGGAGGTAAGGGTTATTACCCCCTAAGAACTGTCAAACTTCAAAGTTTTATTGAAAAATTTTTTTAAAAAAAAAAA
>evgR011510c0g1_i01_Hydra
CAGCACTTCTTCCGACTCTCCTCTCAATCAAGAAATAAGTATGAATCTTATTGATTCACTTATCAAATCGAAAGGATGTGGTGAGAAAGTACTCAAGGACATGATTAGAGAACATGGTCGTGTCATCAAAGGAACTAACATTTTAAAGGTTGATTCCTTTCTCAATCATCAAATCGATCCAGAATTGTCAAGATTAATGGGTATGGACATTGCCAGTCGTTTCATTCTCGATGGTGTCACAAAAGTTTTGACATTGGAAACCTCTGGTATTCCAATTGCGTTGATGGCTGCACTCTATTTGAACGTTCCCTTGGTTTACGCCAGAAAAGTCAAACCATCCACATTGAACGAACAAGTTTACACCACCAAGGTCTACTCCTTCACAAAGAATACTGAATATGATGTTTGTGTCTCTGCCAAATACCTGCAAGCCAATGATAAAGTGTTAATCATTGATGATTTCCTTGCATGTGGACAGGCAACTCAGGGTCTTATCAATATCATCCAACAAGCTTCTGCACAATTGATTGGTGTGGGAATTTCAATTGAGAAGGGTTTCCAGGAAGGAGGAAAGAGATTGAGAGAACAAGGAGTTAAACTGGAAAGTTTGGCAGTAATTGAAAAGTTTGAAGAGGACTCT
>evgR011513c0g1_i01_Hydra
TTTCAATGATTGTTGTTAAAAGTAATCATGTCAAATATTTCTTTAGGTAATAATAATAAAAATGGTTAGTATATCAAGAGGAATTGTTCAATTAGATGAGGATGATTTTCTAACATTTGGTGAGAAGCTGGTCATGCCAGGAAGGCTGGGGGACGCTATGGCAGCATTGCTTGATAATGTTGATGACAGCGATAGAAAAGCATCCTTAATGTTTCTAGCTGGACTAATCGTTGGTAAACATTTTTATACAGATGACTTGTTCTTTTATAGCTCATCTGGACAAAGGGCATCATTCGATCCAGCAGATCCTGATAGGCAAAATAAAATTCAAGCTATGCATCAACAATGCTATGTTCACAAGACTCAATTCATCATTAATTGCTTACAGAACAATGTATTTTCTCAGGTTGACTATAGTCATTCTTTTGTTAAATTATCATATTTTGCCATTTTATCAATGTGTAAGGAGGTAAACCAGCAAAACTCAGAATTCCTAAAGAAACGTTACACTTCCTACATGGCATCAATCGGGGAAACACCTTACACACTAAGCGGAAGTTTTGTTAATCCTGAGTCATTAGTAAGTATACTATCTACATTTTATTCGCATCCAGTTTCAATAGACTTGAGAATTAGAATAGTTACAGGAGTCAGATTGCCTAAGCATGAGACCATTATCAGCAATATGCAAATGGTTATGAGGTATTACGGATGTAAACATCTTGAGTTAATGCAAAATATGATAAATACGATGAGAACAATGGTTTGGACCGATCCAACATTTACCAACGAAATCAAGTTATTTATATCTGAGCGAGACAAGTATCTATTAACTGGGGGTTCACTTGAGTTTGTTGGATCGATATCTAGACTACCATTAGGGTTACAGCAGAGTAATTTTCCCGTGTCCATCCACGTAATTAAGAAGACGACTGAAGGTCAATTGAAAAACTATAAAATGAAAGACGCTAAATTTCCAACCGCTCATATTGATGAGTTAATGAAGATTCCTATCAACCGTGGCACCATCTTGAGCGATGATCAAAGAATCTTGCTCAATAGCCTTGGTGTTGATTTCGAAGCTATAGATATTGTAAATGATGGTAGCAACAAAAGGAGACAGACAGAGATTTGGAAAGACGAGGTTAATTACCGTAAAGCTAAACGACAGCACGATGAAGAAATAAATAGACTATGATCCTGATTGACTTCTTATTTTATTTTAATTTCATCAATAAATTTTATAGCTTTATTTTGTTTTATTTTGTTAAATTGGAATTTATAATGTCTTTTTGTTAATCTTAGTATTACTTACTAATTAGTTATTGTTTATTATCGCCCATTTTGATTTAGTGCTATTAATAGGTTAAATAAATTAAGCTTGAATTACTTGATAAGTGCTAATTAAGATTATAA
>evgR011514c0g1_i01_Hydra
AAAACGAAACTAATCGCGATGCTATTGATGCCAAAGTAGAGTATCAAAAAGAATTTGACCGTCAGCTACCAATGGTGACATTGCTGACAAGTGAGCCTGCTAGTTCTATTATAACAGACGAAAGCATGTGGAAACTGTACACAAATGTTAATGATATAAATGTTTTTACGCATTATGTTATGAAAAACGAGATACAAAATGCCAATATGATGTATGATGCCAAAACGATAGGTGATTATAATCGTTTACAGCGAGAAAGTAAATCTAGCGATTTCCTGAGAAGCCAAGTATTGGACGCAAAATACGATGGTCCGATTCCACCATAAGGGCACGAATTTAAATCTATCGAAAAACCTCTTTTTTATCAACTCTATGAAGAACTAGAACAACATGAAAAAGCCTCTGTTATTAATGATAACAGACTCGAGGCTTTTAGAAACGCTAGAGAATGGATCCAAACTATTTTTGATGGCTATGAATCTGAAATTCACGGTAAAACTTATACACTAGCATTCTATAAGCGTAAATCCATTACTGTCAATTCAGTTAGTCAAGTTTTATTAGTCTTGGACTACAAAGATTATATGTTTGATACTGG
>evgR011520c0g1_i01_Hydra
CAGACGAAAAGCAGCATAAAAGGTTTGACAAAAAATGGTTTATAACTTGAGACAGTTAAGATTGCTTCTAACTTTTTTGATGGTGTCCATGCTTAAAGAAGGGTATACTTTGCAGTGCATTAACAACAAGTTTTGCAAAACAGAAACACTGAGTTCTTCATTAAAAACAAAATCAAGTTACTTTACTGCAAGGCGTTGGCCGTTTTCATTTTTAAACCAAACAGCTATTAAAAAATCGACGGTGAAAAGCGAACAGCATTGCATTAAAAACTGCGTATTTTATAGTTCCTGTTATTCAATAAACATAATACAAAACTTAACAGCAAATACAACTGAAATGGAATGTCAGTTACTCATTCAACAGGCATACAGCAGTAATAATTATCTTGTAAACCAAGAAAATTCTACTCACTTGAGTGCATTAAACTCAGGGTGCGAAAAAAATCTTTGCATGAACAATGCAACTTGTGTACCCAATTATGTGAGCGATTTTTCGACTTGCTCTTGTCCTCCATTTTTTAGTGGAGCATTTTGTGAAATAGGAAAACAAATTGCCATTTGGTGGTATTCTCGAGGCAACGGTAGAGATGGCAATATAAACAGCGTCGCAAGCGATCCAGGATTTAAGAAAAGTTTTTCACGTAGTGCGTGCTATTTAACATTTGTTTCAGATGTTTACGATGGTTTCAGTTCTCAGATATTTGCTTACTTTGTCCCACCTGATTCTGGAATCTACTTTTTTCAAGTTTACTGTTCTCAAGGCTGCTCGCTGTATGTCTCCACACCTGGAAAACAAGATAGAATGATAGTGTTTTGGAACAGTGGAACATATTCGAAGCCTATATACATGGAAAAAAAAAAAAAAAAAAGCAGTGGAAATGGTTGTCGGTGTGCCTGCATGCAATTGCTATAATAGATATTCCCTTGGTTTAACTTTACCCAACGGGACACAATATTCGAGCATTCCGAGCAGCTTTTGGGGAAACTTATTATAACCTAAAAACTACAACTTAATCAAAACTCATTCAGAAGCTACGTAATTTTAACAAAAAACTTTATCTCATAAATATAAAACGTACATACAAAAGTAAAAAAGATGTTGCACTTTATAAATAACATTTATTTAATTTAAATTTATCGAAAAGAAAATAAAT
>evgR011523c0g1_i01_Hydra
TTTAATTTAAGATTCACAATGCGGTCAGAGTTGATATTTGTGTTGCTGGTTTTGAGTGCTTACGTGTTAGAGGGGAAAGCCATAGCCAAAAATCATAAAAAAAAACCCGTGGCTTCCAAAGTACAAGAAGAATACAAGTCACTGGTTGAGTCTGTTCAAAAAGAAAAATCTAGAGTAACGGAATCAGAAAAAAAGCACCCTGGGAAAAAACTAAACGTAATATTAGAAGCCAACAATCAAAGTAAAGAAAGTTCTGTGAGTAACGAAAAAGTGCAACCTCATGTTCAACTATATGTTCCAAGTAAAGTAACATCAAATGCCAGAAAAGTACAAACAATAACTAATAATACCTCAAACAAGAAAGAGAAAACAGAAACCTTTGAAAGAGAAAATTCGGGCGAAGAAAATTTACAGACTGAAAGTTCAGCTTCGGGAGAAGAAAGTAACATAACAATATCCAAAACAAAAAGCGACGATTTCATAGGGAGTGGTGAAGACGAACACGTTTCTCCTGAAACACGATCAGATGCAGAAAATGTTAAAATGCCTCTACCACCACCAGCTCCACCAAAAAAGAAACCAAACCCACCACCAGCTCCAAACAATCCTTATGCACCGGTTCGTGTGAGTCAGCCTGGTATGAATGGATATCATCAAATAAATCCCGACGGAAGTATTTCAGGATCATTCATTTTTAGTCCATATGCAGAAAGTGCATGTGCTCCTTCTTGTCCAGCATCATGTTCTCCCGACTGTTCTCAGCAATGTTGTACCGGCCCAGTGCTACCAGCAGCTCCTCCAACAAAACCAATAAAGTGGTATGAAAGACCAAAGTATGAGCCAGGTCCTAATTACCCACCACCACCTGCTCCTCCATATCAACCATTTAGAGTTTGTTCACCTTCGTGCCCCCCACCACCACCACCTTGTGCTTTACCTAGTTGTCCAGCCGCCCCATCTTGCATTTTATGCACTCCACCTCCTCCTCCTCCACCACCCCCTCCTCCACCACCACCACCTCCTCCTCC
>evgR011533c0g2_i01_Hydra
AATGAATGTTTTGACGCTATTCATCGCTTTTATTTTGTTGGCTTCAATTAAAGAAATTTATAGATTGTTCAGCGCTTTAAAAAACTATAGTCATGGCCAGACAATACTTAATTTACATCAACGTCAAACCATACGTTGTCTACTTTATGGGTTCTATAGTCAAACAATCTTTACAATGACATCTTTATTTATTAATACATTTATAAATCTATACGAATATCCCCTCGTTTGTCAAATTGAAGGTTTTACAGCAGTTTTTGTTTTCTTTATGGTTTTTTATCATTTTGTGGATTTGCTTGTTTACGACGAAGACACTACACCGTCATATTTTTCGTCTATGGTACGTATTGTGCCGTGCTGGGCAGCAAGTTTTACGTTGGCAGTTTTGCCATTTTTTGACTTTGGATTTTATGTTAAATACGGCAACGGAGCTTTTTGCAGTTTGGATTGGACAAATCGAGATCGATCGGTTCGAATTTATTTATGGTGTGTTATTTTGAGCATCTTTTTCCCGGCTTTTTATACGATTAAAGCATACATATGTTTTTCGAATAAAGAAACTCCAATAGTAAAAACATGCCATTGGATTCCAAAGAGCAATTTGTTTAAAACATTGTTTTTCTTTATTTTATTATGGTCGCCGAACGTTATTCTACAGATAATTATGCTAACTGTTTTTAATTTTCATGAGAACGAGTTAGTAAGTGTTACAGCAACCTTGGTTTCATATTCTTCTTATACAATTGTACCAAGAATGTTAGCTAACAGTTTGAAAAAAGCGGCTGAATACAAAATAAAATTAACACATTCTGCTTGATTATTTATTACTATTTAATCAACTCACAATTTCTGAATTTGACTTTTTTGATAGAAATAGATGTTGTATAGAAATATTGACAGAGATATAGATAGACATGAATGTTTTATTTTTAGGATGAATTTTTGTATATATATAATAGATTGTGATAATATTTTAATA
>evgR011541c0g1_i01_Hydra
ATTTCATTATGGTCAAGAATTTGTAACATACAATGTTCATAGTTTAACACATCTTTATGATGATGTGGTAAATTATAACTGTAGTTTGCAGGATATATCTGCTTTTCCCTTTGAAAATTATCTAGGTACACTGAAAACATATGTGCGTAAGGCGCAAAGTCCTTTATCACAGTTGGCAAAAAGAATTGAAGAGTTAGAGAATTGTTGTTTTCGAAAAAGCAAACAACATATAAAAACAAAAATTTCAATCAAGAACAAAAATCGGTGGTTTTTTCTTCGAGGAGAAAAAGTTGCTCGTATCAATAATATAATTGGTAACGATGTTGAATGCTTTGTATATAAAATATCTGATTTAGATGGGTTTTTTGATGATCCATGCTGCTCTACCTTGTTACATATATATTTTCTTCATCAAAATGTACATTTCCAAAAACGATTTATTAAAAAGGATCAAATTCTAAAGAAAGTTTGTGGAATTCCATATAATAATGGATATGTGTTGGTCCCATTACATCACAATGTAAATTTTTAAAGTATTTACCAACAAATTGTAGTAGTTATATTTATAATAGATTCTTTTATATTAG
>evgR011549c0g1_i01_Hydra
AGAATTAAAAAATGATGGAACTAATAATCAGAAATATAGTGATTCTCCAATTGTTTTCTTCAGTTTCATTATCTATTTTGTGCCCAAATCTGTTACTAAAGCCACAATTTGTAGAGAAAGTCTACTCTGACAATATTTGTAAACGTAATAACACATTGGAAGATTTTGCATTTGCTCTTAATCTAAAAACCCGTCTAGATCAGAAGCCGCTAGTTTGTTCAGCAACACTCAAAATACCTACACTGCCCACATTGACTCGAACAAACCTGTCACGAAAACGAAAAGCGGTACAACCGTGGAACACCCCTCAAAATCTTCCTCCTGGATCACAAATCACATACTGCTCTGATTTAAAAACGTGCTGCAAACCAAAAAAATCTTTAAATGGTGCTGGAACTCACTATAGTTTGTGCAGAGAATGTATCCATCAGGTCACGCTTCCAGCAAGTTATACTCCACGTCAGCATTTCCATGTCACCTGTCAAGATGCGGACTATAACGAAGCGTGTTTTTCAGGTGAAGGAGGTTGTGCAACAAATTTGGTATCAAGAACAGTAAGCTTTAATTCAAAGGATGTAGTTTTAACTCTTGGACATTCTTGTTCATGTGAAATTCTCCAAAATTCTCTTTTCACTGATTTTATTTAAAGGCATTCATCAGTAGACAAAACCCGTGGAAAAAATCACTTATGTGTTTTTTATATATTATATGTAGTTATGTTATATGTAGTTGTATGATATTTTTATGTACGGAAAAATAAAAACTGCTCTATTTGAACTTGAACAGCTCTATAGAGCTGTTCAATATATATATAAATGCGTC
>evgR011555c0g1_i01_Hydra
TTTGAAACAGTATACAAACATGGTTTTTGTATTGCAGCTGAAACACTGTGGGTTTTATTCATCTGCCTGCCAAGACAAACTTAGATGGGATCACTTTTTAAGGATACAGTAACAACAGTTCGAAAATGGACATGGCATCATGTTTTTGTTTTTGTATTAACATTTTTTTCATATGCATTTTTGCATGCATGCAGAAAGGCTTTCAGTAATGTTAAAGACTCGATGGCAAAATCAATTACTCCTCAAAAATCTCAGAACTCAACTGAACCATTTTATCCCTACGATCTTTGGAATAAAGAGCATATGTTTGAAAATATAGATGATGCAAATGTTTTCCTTGGAGTACTAGATACATTATTCATGTTTTCTTATGCAGTTGGATTGTTTGTTAGTGGAGTACTTGGCGATAGAGTTGATTTGCGTATTATGTTATCTGGTGGTATGTGTGGTTCTGCTATTGTTACATTTTCATTTGGGTATTTAAATGATATGTTACATGTGCAGAATAAGTACTATTACTACATCCTTTTTGTTGTAAATGGTTTGTTTCAATCAACGGGTTGGCCTGCAACTGTTGCAATAATGGGTAACTGGTTCAGTAAATCTTCAGGAGGTTTAGTATTTGGGTTATGGAGTGGAAATGCATCTGTTGGAAACATATTTGGATCACTTATTGTTGCTGCTTCATTAAATTATGGTTACCCACTTGGGATGCTGTTAAATAGTGTTTTATTGTTTTGCGGTGGCATATTTGTTTTCTTTTGTCTAATTGTTCATCCTAATCAAGTTGGATTAGAAACGCCTGATGAAGTTGATAGATCAAAGTTAGATAATGATGATGTACAAATACATAGCAAAAATGCTATTGGGTTTATACAGGCATGTTTGATACCTGGAGTTCTTGCTTATTCACTTTCTCATGCTTGTATAAAATTGGTTAACTATACATTCTTTTTTTGGCTTCCAACTTATTTGGCACAAGGGCTTAACTGGAGAGATGACAGATCTGACGAATTGTCAAACTTTTATGATATTGGTGGAATCATTGGAGGTGTTGTTGCTGGAGTCATATCTGATTTGATGAAGGTGCGTTCGCCTATTGTTTGCATAATGCTTCTGCTTTCTATGGGTTCTTTATATTTGTACGGTAATGCTGGTGGAAACTACGTTGCGAATGTTGTTCTCATGGTAGTTGTTGGAATTTTTGTTGGAGGACCTGCAAATACTATAAGTACTGCAATTACTGCTGATCTCGGTAAGCATGATCAGATAAAAGGTAACGCTGAAGCTTTATCAACAGTGACAGGTATTATTGATGGCGTAGGATCATTTGGAGCAGCAATAGGTCAATTTTTGGTACCTATTATAAACACCCATTTTGGATGGCATTGGGTCTTTTATTGTCTTATTATAATGTCTGGGTGTAGTTTAGTATGTATTTTACCTATTTTATATAAGGAAATTGTGTCTTTAATAAATCGACTAAAGGAAAGCCATGGCTATGTTAGATTAAAACAAGTAATCAATGGAGATTCTGCATAATTTGTTTTTATTTTATAAATATTTGTAGTATATATTATATAGGGTTATTTTATAATGCTTGTATGTATAATTTTAGAATTTTAATTACTTTGAATTAAATGTTTGTGTATATATATATATAAAAGCATTGAAG
>evgR011564c0g1_i01_Hydra
GAGGTGGTGCTCGTGGTGGAAGCCGTACAGGTTCTTCTCCATCTACATCAGGTTCTAGATCATCATCAACACAACCTGCTTCATCTCCTACAAACGATAAACCAAACAAGAAAAAACAAACTAAGCAAGACAAAACCACATCTTCTGAAGTTCGTAACTTTTCTGCTCCTGAAGACGGAAACGTAGAAGAAGAAGAAGAAACAACTAAACCACAAACTCAACCACAACAAGCCAAGCTTTCTTACAGCGAAGCTGCTGCTGCCTTTAAAAAAAAGGAAGATGAAGAAAAGAAGAAGGTTGATGAAGAAAAGAAACTTTTTGAACAACAAGCCAAGGATCAAAAGGAAAAAGCTAAGATTGAACGAAATCAACAACGAAAGAAAAAGGGTGATAAGAAGACTCCAAAGGAAGAAGTTAAAGAACCAGTGCAAGAACAAGAACCAGTACAACCACAACAAATTGTTGAAGAATCCGTTGTAGAAGAAGAAGAAGTTCCAGAAGAAGAATTTGCCAACGAAGAAGTTGAACAAG
>evgR011571c0g1_i01_Hydra
CTACTACAGAATTGCAATTGCAAGTTTCAAAATTTAGTTCATCTGCTTTGTTATTACAAGTTTCACCATCACCAAGATCAACTTTATTATCAGAAATTTTAACATCGAGTAGGACTGCTTTGTATTCAAAGTTATTACCTTTATCTACAGCAGCATTTTCATCAGAAGTTTCATCATCAAGTATGTCTGCTTTGTATTGGAAATGTTCATCTTTATTAACATCAACTTTGTTATCAGAAGTTTTAACACCAAGTTTGTCTGCTTTGTATTCAAAATTATTACCTTTCTCTTTATCAGAACCGTTATTTAAAGTTTTTACATCTAATTCATCTGCTTTGTTATTACAAGTTTCAC
>evgR011573c0g1_i01_Hydra
CCTATTTTAGTGCAAAAAGATATGACTGAAATCAGTCAAAGGCAAATTAATGAATTTGATGAATTTGCAAAAGATGTTCTTTTTAATTTTCAAAATGAAGATATTTTAAAACTTGGTTTTTATTATACTCAACTATCAGAAGAATGCGATGCAAGAGGGAAATTGTACTGTTTATCACTTAAAGAATACTGCTCCGGTCACATTTTATTGGTAATTACAGATTGCAATAAACAGAAAAAGCTTAAATATTTTGCATGGCATTACTTGCAATTGATGACTATTGAACCATCATTGCGAGTTTTATTGAAAGAGAATTTAACTCTGCTTCGCCCACTGCTCGATCTGCACTCCAAAGAATGGACAGATCTTGAAAAATCGTCTTTAAACCTAGTATCATTAAACTTTATAGTTTCTATGTTAGGAGAAGGTGAGGACTGTTACTATGATCATGTTGCTGATTGTGGGTTCATTGAGTGGTTGCACGAACGTTTGAATCAGTGTTGCTTAAGTGGCGCTGATACCAATATTGAATTAATTTTATTGATTATAAATCTTATTCTTGAAGGAACAGACAGCTGTAAAAGACAACTAAACAATTTCAAGTTTTCAAGTTTGCTCAGTAGAAAAATTCAATCCAGGGAAAATTGCCATGTCGAATTGCTTAAAATTGCAAACCTAACATTCCAAAAACTTTTACTTTTAACAAGTGATCAACAAACTTTTGTTGAAACATTTAGGAAAGATTTTACATCGGAGTTTAAGAAAAAACCTGAAGAGTTTCAAAAAAATAGATGCTTTTCCAATAAGTGTGAAAGTAAAGAAATATTGACATACAAGCAATGTTCTAAATGCCACATAGCAACTTACTGTAGTAAGGAATGCCAGGTAATTCATTGGAAAGAAGAGCATTCAAAACTGTGTGGCACATGATCCTTAAACACTGATACATTATAGACGAGCAGTTTTCAGTGCTTTTCTTGCACTTATATTGTTTTATTGATGTTAATGCTAACCTTGATGTTAATGCTAAGTTTTTTTGTCACAATTTCATTTAGTTTATATGAATTTTATGTTCTTAAAATATATATTTATATGTTCGTAATAAATATATATGTATATATAATATATATATATATATAATAGGGTAGACAAAATAAAATCTTTTGAAAATCACATGTCTGCCAAAAAACCTGACACTTTTCATATAAGAATACAAATTCTTGAATTTTTTTAGAAATACACATTATTGAGTCCTGC
>evgR011584c0g2_i01_Hydra
TTTAAGTGCGTATGCCAATTATTAATTTAGAAGCAAGATTTTTATCAAATGTGTTCTAGATAAATTTTCAGACTATAAATGATGAAAAAGTACCCGCTGCCCACAAACTTTTTAAAGCAAACTGAGGTATACGATCCTGTGAACATAACATCTACTAATGAGAAAGCCTTAAATGTGACTGCACACATCCCTCAAGACACTTTAGATAAAGCAAAAGGAAATTTGTTCTTGATTGGGGTTGTAGTTAGCTTTTGTGTGTTTTTTGGTGGTGTTATTGCTTACGTTATGTATAACCACTACAGAGAAGTTATAGCAAAAAAAAGAGCAGAAGAAACTGTTCAAAAGACTCTTGAAAGTGGACATGCAACTATTTCTAAAAAAACCTCGTTAGTTGGTGGAGGGCAAAAGCAACTCATAATTGAAAGAAAAGTAAGCGTTAATGGTAGGAAAGTAAGTACCGGAAGTCGAGGTCTATGTAGTGGAGAATCTAATCTATGGTCTAGCTCAAATAGTCAAGCTTTAACTTATTTGCGCTCGTTATCCGAATCATCTGGTCAAAATGTAAGCGATTCAACCCAAAGAAAGATTAGCGTTTCTTCAAAAGTATATCCTACAGTGAATAATATGCATAACAATTACTTAGACAATGTTTTAATAAGATATCGCGATGGCTCTACAGCCAACGCCCTAAAAAGTGTTTCGGATGCTGAAGAATCTGATGAGGATCTGAATATCTCACAAAAAATGTCTTTAGAGAGTAACTATCAATCAAATAACAACATTGACAGTTTTCAACTGCAAACAGATCGAAATGATGCACATGAAAATACAGAAGATGAATCATTGTTAGAAAACTCTGCATATTTATAACGTAGTGATGGTATCATAACACTTCGACACAAAGACTTCAACAGACTATAGTTCCATGTAACTAAATGATAACTAAATAGTAATAATTTATTAAAACAGGTAAAAAATGTTTAAAAGTTTACGTATTAGCAAATAAATAGCTCTATATATAAAATTTTAGATCAAATTAAAATTTATTTTTTGTACAAGAC
>evgR011604c0g1_i01_Hydra
CTTGCTCACCATCATCTACTTCGGCAGCAATCAGTCCAACACTAATAGTATCATCAACAAGTATACAAAATGTCGAAAGGAGCCTCCCTCAACTTTAAACAACATGATTTGACTTCATACCTCAAGTCAGGATCTACACACTACTCTGGTATGCACGCTGTTACCTTGAGAAATATTGAAGCCTGCCAAAATTTAAGCAAGATTGTCAGAACTTCTATGGGACCAAACGGTATGAATAAGATTGTAGTAAATCACTTGGAAAAGCTCTTCATTTCTAAAGATACTTCAACAATTATGAAGGAATTGGAAGTTATTCATCCAGCTGCTAAAATTATTGTTTTGGCCACTCAAGCACAAGAACAAGAAGCTGGTGATGGAACAAACTTTGTTTGTTGTTTTACTGGAGAATTGTTAGCACAATCTGAAAATCTTATCAGAATGGGTTTGCACATTCCTGATATTCTTCTCGGTTATAAAAAGGCTGGTGCTAAGGCTCTTCAAGTTTTGGAAACTCTTGCTGTTCAAACAGTGTCTGATATTAGAAATCTTGATGAAGTTACAAAGGCCGTCGGTGCTACAATCGGAAGTAAAGTCTTGGGAATGGAAGATTTCTTGGCTCCATTGATTTCTGAAGCTTGCATTTCTGTTTGTCCAAAGAATACCACTAGATTTAACATTGATAACATTAGAACAGCTAAAATACCTGGTGGTAGTGTTTATGACTCAAAGGCTTTGAAGGGTTTTGCTTTTGTTAGAGATTCCGAAGGATCAATCAAGCATGTTAAAAATTGTAAAGTTGCTGTTTACACTTGTAACATTGACCAAACTTCAACAGAAACCAAGAATAGTGTAGTTATTAGAAGCTCAGAAGAACTTCTTAATTATAACCTTTCAGAAGAAAAGGCTATGGAAGAAGAAATTAAGGCTGTTGCTGAATCTGGTGTTAAAGTTATTGTTTCTCAAGGTGGATTTGGTGAAATGGCTCTTCACTTTATTGAAC
>evgR011606c0g1_i01_Hydra
CTAGAGATATGGCAAAATTAGCAAGTGCACTTAAGAGGGATTTTCCAAAATATTATCATCTTTTTTCTATGACATCATTTATTTATAGAGGAAAAGAAATAAAGGGACATAATCGAGTTTTAAAAAGATATGCATGGGCAGATGGATTAAAAACAGGATATGTCAATGCTTCAGGCTTTAATCTTGCAACCTCAACAAAAAAACCTGAAGCTAATTTAGTAGCTATTGTGATGGGAGGTCAAACTGCGAGAATAAGAGACGATCATATGATAGATTTATTGAATAATGCCTATGCAAGATTAGATGTTAAAGATAAGATAAAAAATGGGAATATTTCTCAATCAAAATTCTTAAATGCCAAAAGAGACGTTTTTCAGTATGTTTTAGAAAGTTCCAAAAAAACTACCAATAATAATGATGTACTAGAGATTCATTAAAAAATAATGAGTGGCATCAGATTATTATTGGTAGTTAAATTGAAAGAATACTTACAAAAGAAAAA
>evgR011613c0g1_i01_Hydra
AGAATTGGTAATAAGATTTGTGTTATAAAATTTCTTTGTAGATAGTTTTCCTTGTGCCATTAATTTGCTAAATGATTGCTCTTGAAAGCTTTTGAATAAGGTTTTATAAAACGGATTTTATTGAGTATTTCAACGATTAGTTTTGTTTTTACTTAGTAAAATATTATAATGTCAAGCGAAGAACTTTTAAATATGTGTTCTTATGAACATATAAATATACTCCTGGTATCCGATCAATCTTTTTGTCTAGAATACCTCATTCAGACTTATTTCAACCTAAAAACTGGTAATTCAAAGGATTTAATAAAGGATTTTAAAAAAACGGGAATTCCACTACAATTGAGAGAGGACATGGTTCGTTTACAGCTGTATATAGCTCCAACTAAATCGCAAAGCACACGGCTTCCTATGGACGAGCAAAGAGTAAAAGAATACAAAAAAATGTATGGTAAAGTTGACATTGTTTTCTTTTGTTTTTTTGTAAATAACTCGAGAACTTTAAACAACATACGAGGAAAGTATTTTAACGAAATAAAGCAGTGGTTCCCTAAAGCTTATCGCTATCTAATAGGCTTGAAAATAGAAGAAATAGTTGATAACAAAAGACAATATCTTAGGGAAACAATCTCCATTCAAGACAGGTTTTTGAAAAAAGAAATGGATGTTACGAAAAAGGATATAAAATCTTTAAGAAAACATGTTAAAGCCAGAACTTATCATTTTTGCAATATTCTTCAAAAAAAAGATGAAAATTCAAATTTGTGTGAATACGTTACCGAAAACGAAAAAGAACTTATTACATTGTTCGAAAGTCCGGTTAAATTATTTTACAAAAAAATATCTAAAAAATGTCTTTTTCTATAACTTTTAACTCTCTCTCTTTCTTTTTCTCTATCTCCA
>evgR011614c0g1_i01_Hydra
AAATCTACAAACCTTTTGATTGGTCTAACAGTTGCATTTGGATCAATATCTGGTATTATTATAAATTATTTCAACGAAATAATAATTAAATTACTTAAAGGAACATTTAACACTATTGTTATGTGCACATTTCTTTGGTCCGTTCGATATTTTATTTTTTATTTCCTAAAAAATCCTTGGTTAATCATACCGTTAGAACTTACTCTACATGGGTTATGTTTGTCACTATTTTGCAGTGTTAAATCTTTACATTTAAAACATATATCACACAACAGCATTCGCTCAACGATGTTTGGCATTGTCCAATGTTTACATAGTGGCATATCTCCGATAGTCGGCAGCTTAATAGGAGGTGAATTATATTTTAAATATGGTGCTCGAAATACTTTTTTATATGTTTGTTTAATGGCATTAGGATGGACAATGTTTTTATGTATTTACATTATTGCTAAACATCTTTTGAAAAAAAGAGTTCTTAAATTTTCAAATTCTGCTAACCCTACTCGATATTTACTTGAAGTTAGTGAGTACATAAC
>evgR011617c0g1_i01_Hydra
AAAGATCATATTTCTAAAGATCATATTTCTAAAGATCATATTTCTAAAGATCATATTTCTAAAGATCATATTTCTGAAGATCGTATTTCTAAAGAATATATTTCTAAAGATCATAATTCTAAAGATACAAATAGTGGAACACCAACATTATTTAAAAACATAAATGATGAAAAAAGTGATTCTGAAATAGTTCCTGAAAATTGTGAGTTTAATAGTAAATTAAAAGCTTCATCTTCATCTTCGAAAACCTCATCTTCAAAAACAATTTCCAAGAAATCTGTTGTAAATTCTGATGATGTTGGTGTTAGTTGTAATTGGTCAGACGATGAAACTGAAGCTAAACACAATGTATATGCTGTTAGAACTCAGTCTTCTATTAGAGTTCCCTTTTCCAAAGATGAAGAAGACTATATTAAAGACGGAATAAAAAAATATGGTATCGGCAATTGGTGTAAGATTAAAGAATCCTATCCATTTCATCCAAATAGAACAAATGTGTCTATAAAAGACAAATTTCGGACTATGAAGCGCCAAGGAATTTTTAGATATTGACGTTTTAAAAACGAAGTTGCTAAATCAAATTTTGTTACGCATCATCATTTTGTTAAAAAATACATGTGTA
>evgR011625c0g1_i01_Hydra
TTGTATTGACCGGAGAACATTCTAGAAAAGAAAAATCAAAACATTCCAGAATAATAAAAATAAGAAAACGAAAGCTCTCCATGTTTCTAGAAGCTCGTCCTTTATTTATTATTAAGGGAGCGTATGAATCCAATAATTAATGTAAAATAAATCGAAGAACGCTTCTAACAAGAATTTTATTTTTATAATTTGAAGAAGTTACTTTGTTATTGTGTGATGGGAAAAGATTATTACAAAATACTTGGAGTTGATAAGTCAGCTGATGGAGCTGCGTTGAAAAAAGCTTACAGAAAATTAGCTTTAAAGTACCATCCTGATAAAAACAAGCAGCCAGGTGCAGAAGAAAAATTTAAAGAAATTTCAGAGGCCTACGAAGTTTTAAGCGACGATAAAAAACGAGAAATTTATGACAAGTATGGTGAAAACGGTTTAAAAAATGGTTTTAACAGCGATGCATCACACATGAATGGTGATCAAACCTTTAATTTTGGAGAAAATGGTGGATTTCAGACATTTACATTTACTTCTGGCGATGCATTTAACACCTTCAGCCGTGTATTTGGCGAAAATGGTGATGGATTTGAAAGTTTATTTACAAGATTTAACGGTTTTCCACATTCTAATTCACGAATATTTGCAGACGAAGATATGAATTTCAATTTTGATTCAAGATCTCAGAAATCAAATAAACGGCAAAAAATTCAAGATGCTCCAATAATAAGAGATTTATTTGTAAGTTTAGAAGATATTTCGTATGGCTGTTCAAAACAAATAAAAATTACAAAAAAGATTCTGTGTGAAGATGGCCAAAGCTATGCATCTGAACAGAAAATTTTAACCATTGAAATAAAAAAAGGATGGAAGGAAGGGACTAAAATAACATTTCCCAAAGAAGGGGATCAAATTAAAGGTCATATTCCTGCTGACATAGTATTTGTTATAAGAGATAAATTGCATCCATATTACAGTAGAGATAAAAACAATAACTTGATTTTCAAACCAAAAATCTCATTGCGTGAAGCTTTATGTGGGGGTCAAATTCCAGTTCCACTCATCAATGGTGATGTTAAAACTATCAGTTGGAATAAAGTTATTCAACCGGGCGAAAGAAATATAATTAGTGGCTGTGGTTTACCTAATCCAAAATGTAACGATAAATATAGTGACTTAATTGTTGAATTTGATATCATATTTCCTTCAGAACTGTCTTCGAGTTCAAAGCACACTATTGGAAACTTGCTTCCTCAATAAAAAAATTTTTATATAAAAAGCAATTTGTAAATATAAATTTTTAATGTAACTTAGAAAATTTTGTATTATATTTTGAAGAAATATTTATTTTGCAAATAAATTTACGAATATTTAAAAAAAAA
>evgR011633c0g1_i01_Hydra
GATAAACCAGAGAATAGTGATAAACCAGAGAAAAGTGATAAACCAGAGAAAAGTGATAAACCAGAGAAAAGTGAAAAACCAGAGAAAAGTGATAAACCAGAGAAAAGTGATAAACCAGAGAAAAGTGATAAACCAGAGAAAAGTGATAAACCAGAGAAAAGTAATAAACCAGAGAAAAGTGATAAACCAGAGAAAAAGGATAAATATAAAAAAAGTGATAAACCAGGGAAAAATGCTATACTAGAGAAAAGTGATAAACTAAATGAACTTGAAAAAGTAATTAACCAGAAAATCCATGATTATATTAACTTTAATAATGCAACGCGAC
>evgR011653c0g1_i01_Hydra
TACGTGAGTTGGATATTACCAATGTCAGAGAATTAGAAGATACATTGATCGATTGTATGTATCAAGGATTGTTAGAAGGTAAACTGGATCAAAAGTCAAAGTGGATGGAAGTTTATGAAACAATCGGCAGAGATATTAAATTAGATGATATTGATCAAATGATTTCTGTATTGAAGAGTTGGACTGTTGGAGCAAAGGAAACTCTTAAAGGCATTGATGGCAATGTGGAATATGCCAACAAGCAATTCAAAGATCACAATGAACACAAGAAGAAGTATGAAAAAGAAATTGCTTCTATTGAAGAAACAATCAAGGCTTCGATTGAAGACGATAATTCAGGTGGTATGATGAGCATGATGGGTATGATGGGAATGATGGGAGGTGGTAGACCAGCCGGTGGCTCCTCTGGTGCTGGTG
>evgR011659c0g1_i01_Hydra
AGTGCAAAAAGTCTATAATTGTTTTTTAAAAAAACTTAATGGCTGTTTTAACTATTAAAAGTTATGTTGAATGGTCGTAGTCGCTTTCGACCACGCTTCGAAGTTAGCAAACATACCAATCTTCCTACTCAACAGCGTCGTCTTCGTCATCTTACATGCATAATTGGAAGAAATATTGATGCGAAACTAGTTCCAGTCACTCGACAGACTAAATATTCTCTTTTAAGCTTATACTTTACACTTCATGCGTCGGATAATCAAAACAAAGTTTTATACAAAAGCGAAGTAATTGAAAATACACTTAACCCCAACTGGCGTGACTTTGATCCAGAAGTATTTTTAAAAAACTCTTCCAAGAAAAAAGATTCTTCCGTTCTTCTTTGTATATGGGGGTCTTGCAAAAGTAGTGATTTCAAAATTATATTAAGTTACCATGTCAATTTTTCTGGTCTGCAATATTATACAGAACAGTTGCGCTCTCAAGACCATTGCTATGAAACCAACTGCGTTTTGTTTCGTTTGTTTGGTGGCTATTATTGTGCACCAAAGGAAGACTGCGTGACAGAAGATGATGTAAAAAGTCAGTTTGTTAAAGTTCATATTGACCAACTTACGCCTTCTTACAAAAAAGCTTCTCTTTTACGCATTCATTCTTCATTAAAAGCGATTCATCAGGCCAGACTGCAAGTTGAAAAAGTACAACATCAGATGGTTCAGTTGCTTGCATCATCAAAAGAAAAAACACGAAAAAACACTGAAAAAGAGACCATTATATTGCGAATCAAACTTATGCGAGAAGAGTTGAAGGTTCTTAAACAAAAAATTCTCTACGAAAGAAGCGCTTCTTCAGAACAAAATGGATGGCTATGTAGTAGAGAAAAAGAGTTGAAGGAAAGATTAAAAGAACTCTCTCTGAGTAAAGAAGCGATGGGAGTCAGTCACACATTTCATATTCAAAGGAGGGAACTTTTTGTTCAAGTTACCGCCCATCTAAACTACAGAAGGCGAGAACTAGCTGCTGACTTGGTTGATATTTACACAATAAGAGAATTGCCAATTGATGATAAAAACAAACGAGAGTATACAATTTGTGGTGTCAGATTACCGAATTCTGAAGATTTTAATGGGGAAGATGAAACGATGTTAGCTGTTGCTGTTGGTTATGTTGCCCATTTAGTTTATATGATGGCAAAGTTCACAGAATTACCTCTTAGATATCCCATTTGCCCAATTGGTTCTCGTTCTTCAGTTCAAGACCTTGTAACTGATAAATTGACTGACAAAGAAAGAGAATTTCCTCTATTTTCGAAAGGAAAAGAGAAGTTTCATTTTCGATATGGAATGTTTCTGTTGAACAAAAATATAGCTCAGCTTCGTTACCACAACAAGTTGGGCACAGCAGATTTGCGAAATACACTTCTTAATTTAAAAAATCTTTTTGAAACGAAATTTGGTCTGAGATTTGATGTAAATAATTCCGTTGATGCTCGATCTGGTCAAAATCAATTTGCTAATCCTAGTCACCGGTACTCTTTGCCTTTTGCTGCAATAAATTCTGAACCAGTTTCATTTCAAAGCAGTTCTTCACTTAATAAGGAAAATAATTTTTCGGGACTTCCAGACGTGCACGTGAAAGCGCACCACCAACACCGTTTTATTGCCAGGAAAAATGTAAAGCCTAAAATTCATAAGTCGTTTAGCCTTCCTGACGGATTAACGTCAACATCTCCTGATCTTATTCAAACTCCGTTTGAAGATATATTACGACAAAATAGGGAAATATTACCTGATGTGACGCAGATACATTCCTCTATGACTGCTTTATCGGAAATAAAAGTTCACGAACAACTTGATCCGGATAAAAACATTATAAGGAAGTTAGAAATGCAAGACATTACTCGGCAACTTAGTGACGCTCTTATTATTCCAGACCCTTTTGAAGAGCATCATTCAGATTTCGAAGGGCGTCATTTAGAAAATATTTTAGTGTAGCAGTTTCTTGTTACGTCAGTTCTTTTTACTTGCCAGTGCAACTACTTTATAACTTATTTGTGTTTTTAGATTGGTTGTGACACGACTTATGTGTGACGTTGCATTAATTCCATTATAATCATAGTTAAACTTATTACATAGTTTCTACTCTCTTTCTTTTTTTACTACGCAATTTTCTCTTCTTCTTTCCTTTTTTATCCTTGACTTGAAGTCAAAATGTATATAAACAAAACGAAAACAGATATCACAAACGAAACGAAACGAAAACAGACGATAAAAAACTAATACTTGAATGCTCTTGAACCTGGTGCTGGCCGTATCAAGTTATATATATATATATATTCTCTTTGGTAATCCCTG
>evgR011660c0g1_i01_Hydra
CTCTTTGTATAGTACAAGCAACCAACAATGGACAGTTTTGACAACGGATTATACTAACAACAAAAGTATTGGGCCTATATGTGAAAATAACACTCCTTGTGGAGGTTTATATTGCAGAGATGTATGTTCTGCAGATGGTTTAGTTCACTCGTACCAATGTTTAGTAAGTAACGACGTTTCGAGAATTGCAACTCCATCTTTGTCATCAGCATGGGATAGTAACCACGTCGCAAGCAATGCAATCGACGGAAATTTAAATACTAATGCCGGTACTCTTCGTGATGTCGTAAGTTGGTTTAAACTAGATATGCATAAAATGTACAAAATATTTCAAGTTGTTGTCTATAATCGAAAAGATTGTTGCCAAAATCATCTGAATAATACTATACTTATTGTCAGTGAAACCGAACAGTATTCGAATGCTTTTGAGATAGCAACATTGACCTCCGATTTGAAACAAACTTTTACATGTTTATTTTCAGGAAGATATGTTTTTATTGTTCAATATTATAGTGATTACCTTTTGGTAATAGCAGAAATAAATGTCTTTGTATAACTTTAAATGTAAC
>evgR011665c0g1_i01_Hydra
ACTAACTTTTTGCACATGAGATCACAAGTTGTAAGTTTAAACAATGTTAATTCTCACACTATATCAGTAACAAGTGGTGTTCCACAGGAGAGTATTATTGGACCTACTCTATTCTTGTTATTTATCAATGATGTTAGCCACATTGATAAAGAGCTTGATGTCAAACTTAAACTTTTTGCTGATGATATTAAGTTATATTTAGTCTATGATATACGTGGCCCCCAGTCAGATCTTCACACAGCTATTGATCGTTTGTATGAATGGAGTTGTACTTGGCCACTCCAAATAGCTACTGGAAAATGCTTTGTCTGTACAATTTCTAAGCAGAGTCAAAACATAACTCATCTTGTCTATACCATAAATAACCATAAAACTGC
>evgR011675c0g1_i01_Hydra
AAATGGCTACTGTCATCAGATTCTTCTAAAACCGAATATGTTTATAAGCCGCGAAAAAGCTTAATAGTTTTTATTTCGTTTTAATATTTTATAATTGTTTTAAGATAACAGTAGAAAATTTATTTAAATAAAATTTATGTTGCCAGCTGAAAATTAAAAGTTATAAACTTCATAATATTTCTGTATATAAATGATTGTCATATCCGAATGGGTTGTAGCTATAGTTTAGAAGAAAGAGAAGCTCTATCTCGAAGTAGAGCGATTGAGAAAACTCTCAAATACGATGGAGACAAATCAGGCAAAGAAGTCAAGTTACTTTTACTAGGAGCAGGAGAGTCTGGTAAAAGTACAATCGTCAAGCAGATGAAAATTATTCATGACAATGGCTTCTCAGAAGAAGATGTTCTTCGGTTTAAACCTATAATCTTTAGTAATACTGTGATGTCCCTTATCTCTATTCTTAGAGCCATGGAAAAATTAAATATTTGGTTTGGAGATGAAGCAAGGCAGAGAGATTCTACAATATTATTTCAAACTGCTATACGAAATGAGGAATCTGAACCATTTTCCCCAGAAGTTCTCAAAGCTATGAAGAATTTATGGGCAGACGAAGGTGTTATAAAATGTTTTAAAAGAAGCCGAGAATACCAGCTTAATGATTCTGCTCAATATTACTTGGATGCATTAGATAGACTTGGAGATTCACAATACAAGCCTACAGAACAAGATATTCTACGAACTCGAGTTAGAACTGCTGGTATTGTAGAGACTCAATTTTCATTTCGGAAATTAAATTTTAAGCTCCTTGATGTTGGAGGACAGAGAACAGAGCGTAGAAAATGGATCCATTGTTTTCAAGATGTCACTGCTATTATATTTTGTGCTGCACTAAGCTGTTATGATTTGCGTCTTGCAGAAGATGATATAACGAATCGAATGGTTGAAAGTTTAAAGCTGTTTGACTCAATTGTAAATAATGACCATTTCATTAGCACATCTGTAATTCTGTTTCTTAACAAAAAAGACTTGTTTAAATTAAAGATACCTCATTCTCCGTTGACATTATGTTTTCCTGAGTATACAGGTCCAAATGAATATGAAGAAGCATGTGGATACATTAGACAAAAATTTTTAGACAAAAATCGAAATGAATCTAAAGAAATATATGTGCATTTAACTTGTGCAACTGACACTGCTAACATCGATTTTGTTTTTAAAGCAGTAACCAATATGATTATCGCCGCTAATTTAAAAAAAAGTGGCTTAACGTAGGTGGAACTATCGCAAATGATAATTTGCCAATAGAGAGATTTATAAAGTGTTTCTGAAAATAAATATAATGGAGATAAATATAATGATCAATTTCAGAGGTTTTTGCAAGGTTCTACGCAATGAAGAAAGCAGTTATCATTCTCTTTTTTCGAAACGGTTGGATAATTCAGTAATTCGAAATTACAGTAATTACACTTCTTTGCACTACTTGCTTTGTCAAATGCATATGTATGCATTAAGCAAATACTTCAGTAATAAAACATAATTATTTTCATGGGGTGGGGAGAGGAGGGGACGAGTTAGATAGAATCTTTAAATAATTGCTAGTTAAAAGTTAGTTATGTTAGTTGACTTTGAGGTAGTGTATATTATGGTCAATATCGAAATATCAGCAGTTCGAGTAATTCTTTCACATAAAATTATGAAAAAAGTCATTTGAACTGGACAGCTCTTCGTTTAGAATAGAAATTCTATTTGCCTTTGCCAAATGCTTTTCATATTTTATTTTGCGGACAGTTAGAAGACTTAAAACCAATAAATTTTCCCTAATGAATTGTGAACAAGTAAATAATTATACGTTTTTATTCAAAAGCCATATTTCTTGCTTTTTGTACTTTTAAATCCCTTCTAATTTTTAGTTTTTGAGTTTTTTTCTATTTTCTGTAATTACTGGTAATTGATATAACTATTATATATTATAATTAGTTATCGAGTATTGTATATACTGTGTACAGAAACTGCGTGTACAGAAACTGAACACAATCGTGTATTGTTTGTATGAAACTCAAATGCATAAAAGATATCTAAATAATGACGACTTTGTAGAAAGCTG
>evgR011676c0g1_i01_Hydra
AATCTTCTATTGTCTAAATAAGCACATCACTTTTCAAAAATATTTAATTGTTATCAAGTATGCGAGCGATAATCCAAAAAGTTACCAGCGCTAGTGTATCAGTAAAAAACAAGTTGATAAGTTCCATTGGATGTGGTCTGTGTGTGTTTATTGGAATATCTCGCAATGATACACAAAAAGATATTGATTATATGATCAACAAGATCTTAAATATCAAAATATTTGACGATGGTGATTCAAGGTGGAAGAAATCGGTAAAAGACAAGAATTATGAAATACTTTGCGTTAGCCAGTTTACTCTGCACAGCATATTGAAAGGTAACAAGCTGGATTTTCATCTCTCCATGTCTTCAAATTTGTCAAAAGACTTCTATAACTCCTTCGTTAACCAATTGGGTTTAGCTTTTTGTCCAGAAAATGTGAAAGATGGTGTTTTTGGTGAATATATGCAAGTTGACATACAAAATGATGGACCTGTCACGATTCAGCTAGATTCTCTTCCTCCTTTTCAGGAAACTGGAGAGAAAAATTGCAATGAAGCTTTATTAAAAGTTTAATAAATTTTAAATAATATAAAAAAAAAAAAAA
>evgR011677c0g1_i01_Hydra
CTCTCTCCACCTACACATACAAAAAATGTCTGATCCAACCACTGATGCCCCAACCAAGTCAGAAGAAGACTTGAAGAAGGAAGCCAAGCTCAGAGCTGCCAAGTTGAAAGCTATGGCCCTCAAAAAGAAGGCAAAGGAAGGTGCTGCCACAACAGCCCCAGCTACTACAACAACTGAAGAAGGTGGTGAATCAGGTACCGCTCCAACAACTGGTGAAACAACAACAACTGCTAAAACTTTCGATTTAAAGGGAAAGAAACGATTTGAAGTGAAGAAGTGGAACGCTGTTGCTTTGTGGGGTTGGGATATCGTTGTCGACTCCTGTGCCATTTGTAGAAACCACATCATGGAACTTTGTATTGAATGTACTGCTAATCAACAATCTGCCACCAGTGAAGAATGTACTGTCGCTTGGGGTGTTTGTAACCATGCTTTCCACTTCCACTGTATTTCAAGATGGTTGAAGACCAGACACGTTTGCCCATTGTGTAACTGTGAATGGGATTTCCAAAAGTACGGAAAATAATTTATTTCGAAATATTTTGTCAACTTAGTAAATGTACAATTAAA
>evgR011680c0g1_i01_Hydra
ACCTCAAAGTTGGATCATGAAAGAGTTGCTCGTGATATCAGATATCAAAATAATGGTTTGGATCTTCCACTCATCAAGAAGTATTTGGAATCTGTCCAAGATCATGATTTGCCACAAGTTAACGAATCTCTTAACAGTTTGTACATTGAAGAAGAAGATTACCTTGCTTTGAAGCGTTCTATTGAACAATACAGAAATCTTGACCAATTGGCTCTTGCTGCTCAATTGGAAAGCCACGATCTCCTTGAATTCAGAAGAATTGCCTCATGGTTGTATAAGATTAACAAACAATGGGAAAATTCTATAAAGTTGAGCAAGAAGGATAACTTGTACAAGGATGCCATGGAAACTGCTGCTGAAAGTCAAAAGCAAGATATTGCTGAAGATCTCCTCAAGTTCTTCGTTTCTAAGGGTTTGAAGGACTGTTTTGCCGCTTCTTTGTATAATTGCTACAACTTTGTTCGTCCAGATGTTGCTCTTGAATTGGCCTGGAGAAACAACATTATTGACATGGTTATGCCATACATGGTTCAAGTTCTCCGTGAATACACAACAAAGGTTGATGATCTCGTAATGGAAAGCCGTTTAAAGAAGAAGGAAGCTGAAATTAAGTCCCAACAACCAACCCCAGAAGTTGGATACTCTGATCCATCAGCTGCTTACCAAACTGGTGACGCCTATGGTGCTTACACTGGTTACCCAAGCCCTCAATCTGGTTATGACCAATATGGTAATGCTTATGGTCAATCATACAACAGTGGATATGGATTTTAACTTATTAACTTCTAGGAGTCGAGACACA
>evgR011682c0g1_i01_Hydra
GTATTGATAAGAGGTCTGGGTTCGATTGTTTATGTGCTGATGGTTATGTTAAAAATAGCTCCAATCTATGCGAAGATGTTGATGAATGTATTGTTAAAATTCCATGTCCCCAGAATTCCAAGTGCATCAACACCCAAGGTGGATACAAGTGCCAGTGTAACGATGGTTTTTATGGAAACAATTTTACCAATCCATCTCTAATGCATTGTTATCAAAATAGTTTCTGGTCGCCATGGGAGCAGGTAGGTCAATGCTCTCAATTATGTAAAAGAGATCAAGAAGAAGGAAAGATTATATTTACTAAAAAATGTGAGAGGTCTGGACCGAGCAACTGTGACGGAGCTAATACGAAAGAAGAATTATGCAATACTGAATATTGTGAC
>evgR011687c0g1_i01_Hydra
TATTATGGTCAAGATTAACGAGTTGAAACTTTGTTACTTATTCAAAAGTTACAGCTATAGACTTAATTGGACGGAAGCTAATTTATATTGTAAAACCAATGGAGGAAATTTGTTGAGCATCGCAAACAAAAAGGAATTAGATGATTTAGAGCCTTATTGGAAAACTTATTCAGGGCAACATTTTTACATTGGTCTCAATGATATAAACATTGAAGGTCAATATGAATGGTCCGATAATGAAACAGAAAAATATTATATTTGGGGCACAAATCAACCAGATGGTGGAAAAAGAGAAAACTGCATCGTTATTTATTACGCTGTAATTTATGACATTTCATGCAATAGCCGTGGTGGTTTTATTTGTAAATTTCAAGACTATTGTGATAATCACAATTGCTCAACAAATGCTTTATGTAACATTGTAATTGATGGATTTACCTGCACATGTAAATATGGTTTTAAGACAATTAGTGGTGGTATAGAGCAGTTTGGAGATCAATGTCAAGATATTAATGAATGTGCTTTGCCAATAACAAAT
>evgR011697c0g1_i01_Hydra
AATGAACTCCAACAAACAAGCTTTCAAGGTCTTCACCTACAGAGGTCACGGTTTGGACGCCCTCATGCAAATGGAAAATGACGAATTGAAGAATTTGTTCACATCTGCCCTCAGAAGAAGAATTGGTAAGGGTATCTCCAAGAAGTACAAGACTTTGATCGAAAAGGCTCAAAAGAAGAGAGACAATGCTCCATACGGTGAAAAGCCAGACGCAGTCAAGACTCACTTGAGAGATATGATTATCTTGCCATCCATGATTGGTACCGTTATTGGTGTTTACCAAGGAAAGGACTACGTCTCCGTCGAAATTAAGCCAGAAATGGTCGGTATGAAGTTGTCTGATTTCTCTCTTACTTACAGACCAGTTAAGCACGGTAAGCCAGGTATTGGTGCTACTTCTTCATCCAAGTTCGTCCCATTGAAGTAAATCTGTTATCTTGTCTTTTCATAATAAAAAAAAATTATTCATCAAAAAAAAAAA
>evgR011699c0g1_i01_Hydra
CACTTTATTTTATCTTTGACTCAAGTTATATGTTATTTGTTAACATCGTTTCTAGTTCCAGTTTTATACATATCAGAACTAGAGTTATTTTGCTTTTGGCGCCATTTTATAATAAGTTTCCTTCTTGTCGCCGCATCATCAATTACTTTCATTAAAACTAGAAAATTTATTTTTATTTTCAAAAATATGATTCTTCTGTCCAAGAAAGAGATTTCATTTTCCGCATCAATTGATATAATGATCATGAGTTTTATTTTGATAGTGCACAACGTTTTAGGAACTATTTTTCTTATAAAATTGCCACCACGTGTTATTCCAATTTTAAACTACGATAAAAAGACCATAACGTTTCAATGTACCGGAGATTCTCAATTAACTCCGCAAATTTTATACATCGGTTTTGTGCTGCTGATTTGTCTTATTCAAAGTTTTAATGCTCGTAATCTTCCTTCTTTCTATAACGAGACGCTCTCTATTACATACAGCACCGTAATATCTTTAATCTTACTCATTTGTTATTTTCCTCTCTACTACGGTAATAATAACATCAAGTTAAGAACCACTGCTGTCACACTGTT
>evgR011700c0g2_i01_Hydra
CTTAAAAATGAATTAAACGATTTTAAAAAAAGTATCGGTTTTGTAAGCGATAAGTATGACAAAGTTTTAGTTGAGCTTGCTGAAGTAAAGTCTAAAAACGAACGCACGAACCCAAGTATTTTAGAAAACAATATAACTGAAACCATGGCAGATAAATTAGCAGAGCTTGAAGACAGAAGCAGGCGGAATAATTTGAGAATAAACGGGATTGATGAATCAGAGAATGAAAGTTGGAATGATTGCGAAATTAAAGTTAAAGAATTACTGTTAACAAAACTCGAAATTAAAGACAATTTTGTCATCGAACGGGCTCATCGAGTTGGGAAAAAAGATGAAAGTGAAGATAATAAAAAACACAGAACAGTCGTTGTAAAACTCCTTAATTATAAAGACAAAACAGCTATCCTCGACAAATATATCAAAATGAAACTTTGGAATGAACGCGTTTTCATTAACGAAGATTTTTGCGAACGAACAAGTCGACTTCGTAAAAAATTGTTTATAGACGCTAAAGACCTAAGACAAAAAGGTAAGTATGCTAAGGTTGTTTATAATAAACTAGTAACACGG
>evgR011703c0g1_i01_Hydra
TTCGGAGATTGTCACAAATGATTTTATTGATTTTGAAAATTGTGACACTTTTACTGTATCTAATGATGCAATTGAAATAAACAATCTTTCAACGGATGATAGTAATAACTCGTATGACATCAATATAATTGAAGATGACAAACAACTAGAAAACGTTTTTAGACGCATTCAAAGTCTAAAGTCTTCACTAATTGGAAGTAGAGAACAGTATGACATTTCTAAGGGGCAGATGCTTAATGATAAGGTGATTCATTTTTGTCAACAACTAATGTCACTTCAGTTTAATATTGATGTTGGTCTGCAA
>evgR011705c0g1_i01_Hydra
TGCTTATGTAAAGTTTGATACGTACTATTGGTATGGCAATACTTACTATGCCAACATAGAAGTTCAAGCATCTTCTGATGACTATCAGAGTACACTCGGTTTATGTGGAACTTTTGATAATTCTAAAGCAAATGATATGACTTCAAAAGATGGAAAAGTTTGTTCTAGTCAAAATCAAAATGAGTTTACTGAAAGTTGGAGACTCGTTGCTACTGATAGTCTGTTTAATTACGTTGGAGAACCAAAGCCCAAATGTGTTGTTAAGACATCTCCAAAATACTGCACATGTTCATCAAAAGTGGTTGATTGCAAAAGCTATGCTGAAGTTCCCAAATTTGTAAACGAGTTTGCTACTTGGAAAAGGATAGACTTTTCTGGAGCTCATTCTTGTGACGGCAGGTCTAAGGATGACAATGTTATTGTCCTTCCTGATGATGATAATTCTAGAATTTATGAATTCAACCCTGTTCAAGTTCCTGGTAATCTTCCAAAATTCCCAACAAACAGTGGTATTACACAAGAAGAAGCACAAGATGTATGTAAAGACAAAATTAGAAATTCTTTGGTGGCCAAGGTTTGCATCGACCTCATTGGACCTTCATTTACAACAGATATTTTTGAGCAAAATTGTGTGATTGATATTCAAGTGATTGATAGAAAACACGTTGGCGTTGAATCTGCCATTAACCATTTAAAAAGTGCTTGTAGTGAACTAACTTTAAGAAATTTATCTTTTTGGAATACTGTTGATGGGAAGCTTATTGGCCCCCCTACTAAAATTGCAGAAATTGTGTGTCCCAATGAATGCAGTGGCAATGGGGCATGTAAAAATGGTGCATGTTTATGCAACAGTGGTTTTATAACAGCTGAC
>evgR011707c0g1_i01_Hydra
AACTTGTTTATATTAACATAATTAATAAATAAAAATGAGTTGGCAAGGATATGTAGATAGCTTGATGGGCACAAATCATATGACTTCGTGCGGTATTTTTGGTCTCGATGGTTCAAAATGGGCATCATCAGATGACTTCCATTTAAGCACTGAAGGTGTGCAACTAGTGATTGCCTCAGTACAAGACCCAAGCAAAATGTCAAACGGTCTAGTAGTTGGTGATGACAAATTTATTCTAATCAGATCTGATCCGGGAGTGATGTTGATCCTTAAGAAAATGCTTCTAGGAATCGTCGCGTACAAATCAAGCAAAGCTGTAATCATCGGAATGCACGACGGCAGCGTTAAACCAGAAGTAGTGTTAACAGATGTTGGCAGAATTGCAGATTACTTAATCAAGCACGGTTATTAACTAGACAAGTTGATTTTGAAAATAACATAAAATATATAAAAACAAATAACAAAATATATTTAAAAAATGTTGTCTTGATAATCTCTACTTTTTACACTTTAAAAAAAAA
>evgR011721c0g1_i01_Hydra
GCTAAAGTCATTATTTAAAACGTCCAAAAATGAATAACTCTAATTATGAAAAGTACTTGGCTGTTCTGCTCAAAACAAAGCAGACTGACTTACTTTGTCGTTTACCTTACACCAATGTTGCAAAAGAATCTGACAAACAAACGTTGATGAATATAATCAACGGACTACTTATATTTGTATCAAATGTAATCTTGCTAACAGGTTTGTGCAAGTTACGAAAAAAAAAGTGCATTTACTCGCGAAACGAAAAGTTAGTTTTGTTTTTATGTTCTCTCGATTTCATGGTTGCTTTAGTTCGTATTCCGATCGAAGTTTTTTTATTGAAGAAACTGAACCAAATTGGCTGTTTGACTATATCAATAATTGGATTTTTTATGGTATTTCCATTAGCTGTTTCAGGCACAGTCGTAGTAATGATATCAATAGAGAGGTTCATTGCCATCTTTTATGATAAAAAAATTTGTTGTGGTTTAAAAGGTAAAGTCGTTTACATAAAACCAATCATGGCGTTACATTTATTATTATCAGTTGGATTAGCCGTTTGGTTTATGCTTCTTGTTTATCTTAATGGTACATTTTTACAAAAATCTGTTTTTTTTATCAGTCTTGCATCATACATAGCTGGC
>evgR011729c0g1_i01_Hydra
GTTCTATATATTACTTTTATATCTGTCTACAACATTTCTGTTGGTGAACAAAAGTTTTACTTACTACCATATACTCCTTATTTTTTTCAAAATAATAAATTATTTAACGATGGAATGAAAGTTAAATCAAGTAAAGTAACCAGCACTTTGATTCTTGCGTACACTTCTTTCTATGGCGAAATACCCTGGCCTATTTATGAAAATATTTCGTGTAATTTTGACACTTCAAACTTTAAACTTTCATATGATCAAAACGATTTTTACAAAAGTGACTTTGTCATATTTCATTCACGTAACATGCCATCAGTTTCCGACTTAACAAAACTACACGATAATAAAAAAAATGGACAACTTTGGATTTATTTTTCTATGGAAAGTATGAATAACAATCCTACTGTGGAAGATATTGACATATTTTTTGACTTGGTCTCTACTTATGCGTTAGACTCAG
>evgR011742c0g1_i01_Hydra
TTGTAATTCTACTGGTTGTCATGCCATTCCAAACTATAAAAAGTGGATGGTCTCTGAATATGGTAGCATTGATGGTGAACACGAAATGATGTCAGAAATCTATGCTCGTGGTCCAATCTCATGTGGTATTGATGCCACAAACAAGTTGGAAGCTTACACTGGAGGTATTTTTGAAGAATTCGAAGTCTTCCCAATGATCAACCACATCATCTCAGTTATTGGTTGGGGCGTTGAAAATGGAAAGAAGTATTGGATTGTTCGTAACAGTTGGGGCACAGAATGGGGAGAAACTGGCACATTCCGTATCATTAGAGGTACAACCAGAC
>evgR011745c0g1_i01_Hydra
TATGGGCCCTCATTTCATTATCTTCAGTGGTTGTTGTTTTAAATGCATTAAGTTTTCTAACTCCTAACAGGCAACTTATTCAAGCTACAACAATGTTAACACTACCCCCGTTTATGATTCATAGTGTTTATGTTTATTCGAAAATATTCTATAGAGTAAAAGGTATGCGTCGTGTAAACAGCATCAAAGTAGAACAAAGTTCTGAGTCTAAGGAATATAATTACAAGTCATCATATAATGCTAACTCTTTGAAAGATTTGGAGAAAGTATGCAACGTTCGAACACAAATCAAAACATTTTATTTAACTTTTCTTTTGGTATTGTCTCTTTATCTTTGTTACTTGCCTTACTTATTAGTTACTTCTGCATGGTGGATAAAACCTAGCAACAGAAACGTGCAGTGGACACACGTAGCTTATATATGGTCATACACTGCGACAATTATTAAATCATTAATTAATCCAATGTTGTATTGCTACCAGTCAAAGTTAATTTGTGGGCGTGTAAAGTATATTTTGAAAATTAAAAAAAGATTATTTTAATAACATTAATTGCCAG
>evgR011753c0g1_i01_Hydra
AAACTTGATTTCTTTTTTAATCTGACAAAATGTCTGTTGAACTGTAATAATTCAGTTTCATGTAATTATGGTTGTCGATTCTACAGTCACATTACTTTGAATACTCAAGAAGATTTTGAGATAAATAAACAGAACAAACCTGATTTAAATAAATCATTAGAAGTTATCCTTAAGAAAAAAGAATGTTCTAGTCTATCATTTTGGTGGCCGAATATTTATCCAAATAAAACTATTATAAAAAGTATATATCTGGTTACTGTTATTGTACTAAATGTAGACAACCAAACTGAAAACGTACTTAATTTAGATATTAAAAATACTTTTGTTATAAACGAAAATATATTTTGTGATAATCTGGTAAACAAATCATATGAAATTGGATTCGAG
>evgR011760c0g1_i01_Hydra
ATTAAAAATTTCACTTTCTCATCATCAGGTCCCAATCCTGTTTATAAGAACAAGTTTGTTAAAACGTTCAAAAACATAATTTGAAAAAACACAAAATGGGAAACACGTGCACTACTTCGCACCACCATTTTTCCAGAAACATCCAATGTTCAGCCGTAAAGGAACTAATTCCAATATTAGTACACTCGGAAATAAAATATATAAAAATGCCACTTTCTTGTAAAGAGATAGAGGCTCTTAAAAAGTCATGGAGATCAGCTAAACAAGTTTGGAATGAAATATGTACTTGCGCTTTCAGCAGGTGGTTCTCAACATACCCTGAAATTCGATACAAATTTGGAGTTTATGGGAATAATATAACAATGAATGAAGTTTTGGCTTCCGAAACTTTGTGCATCCATATTCGAAAATCCATTGAATTGATTGAAATTATTATCAAAAAAGTTGATGAACGACATGAGTTATCTCAGCATCTTATAGAGCTCGGCAAATTACATCATAAATTTGGAGCCGAACAAAAATATGCGACGGCACTCGGTTCGTGTTTTGTATTTGCGATCAGCCAGATATGCCCAAACACAGACATGATTACAGAAGGGGCTTGGGATTCACTGTTCAAATACATAGTAACTCATATAAAACTTGGAATACGAATTGCATCGGGAGCGTTCAAAAAAGAAAATGCACAAGTAATTGACTCAGTGTTACAAAATCACGCATCGTGATTTTATTTAAAAT
>evgR011777c0g1_i01_Hydra
CATAGGGTTATTAAGCAAGTGCGAGGTTGTTATACTAGAAAATGATTTCCCATATGGTAAGATAGGCTTCAGTAAACTAGATGACATCATTGTTTCAGAACCCGACATATTAGATTCAGGAGTTGCGGATCAAAAAGTTTTAATAAAAGTTGTTAGAGAAGGAGGCAGCTACGGTTATTCTCAAATACGCTGGGAAGGTTACCCAACAGTGTCATGGTTTGGGAATAATGATATAGCTCCTCTAAAAGGTTTTGTGAATTTTTTAACTGGAGAAACGTATCAAAACATTATTATCAATATTATGCCTGACAATATTCCTGAAATTAACGAGACATACATAATTATTTTAAAAGAAGTAACAAATGGAAAAGCAGTAATAAATGAAATTAAATCTTCTTTGCATCTGATTATTCTGCTAAATGATAATCCTTATGGCACTGTTCAGTTTATGGATTCGACTCTTGACGTTTTAGAAAAAGAAAGTGATGATAATGTTACAGTACGTGTCAACCGGCTTAATGGAATGTTTGGTAATTTGCGAGTGTTTTGGCGGACAAATCAGATAAATAGTTTTGAGGAAAAAGCCACAGCTGGTGCTGATTTTAAATTCATACCAACCCAGTTTTTTATTATTTCAAATGGATCTAGCCAATCAGAAATTAAAGTTCTTATATACAGCGACG
>evgR011778c0g1_i01_Hydra
TGCAAAAAGATTTTATAATAAACACAAACTATTTAAGAATGAACCAAATTTATGAGTTTATAAACAGCAATAAAAAGCCAATGGATGTTACAGATTCCCCATGGCTATATGAAATCGATGAGCAGCTAGATGAAAAAGAAGGAAAACATGAAAATATTGGCAAATGGATGTTGTTCTATAAAAATATCGATATGAATGCTAAATGGGCTTTGTCACGTGAACTGTATAAAAACAGAAAATTAGATGGAGTAAAATCAATGAAATGCTCAACCTCACTTAAAAATTCAAATGCAAGATCATCCACAACAGGTGTGATTATTATGTACTGCAATAAAAGCTCGGAAGAAGATCGTATAAAGAATATTGGCAGAAATATAAAAAGTCAATTTGGCATCTATTTGACTATTTATTATAAAACAGACGAACAAACTCGAATGAATCTTCAAAATACAAGAAAAAATATGCCAAAATACATTTACAAAGTGCAAACGTTTAAAACAAACAAAAGCACAATTTGCAGTGTAGTATTTCTTGTTTCATCATAAAACCTAATTAAAAATTTTTCTTAAGATAATGCAATGCATGAAAAAAAATTGTAAATGATTGTTGAACGTTAATAAAAATCTTTTGATAAAAAGCGAA
>evgR011787c0g1_i01_Hydra
CAAAAACCACGATTACTTTTTGGCATAGTTTCTACTATAACAAAATCGGATGAAAACCTTGACTATTCTGTTTTGGATTCAATTGAAAATTGGTTAGAAGCAATAAAAATGGAGCAGTATGCAAATTTATTCAATGATGTTGGATTGACAACGCCCAAACAAATTTTAAATCTGAACTTAGAAACTTTGGTAGACATGGGAATAAAACCAATCGGACACTGTAATAAAATCTTAAAAGCTATAAAAAATACAACACAACAGATTGAGAATGCTAGACCTGAGTCGAAAATAGAACAATGTGAAACATCTTTCTCAAAACAATCATTCCGCAGATCAATACGTTCAATATCAAGTAAAAGAGTAACTTTAGCTAAAGAAGAGCAAATACTAAATCTACATGAGAAATCATAAAGAAGAAGCTATTTTAATAAAAGAATATATTTTGTAATTTATTTTATATAGAGACTGCTGACATTTGAAGTGTAGAGCTATTAATGGTTTTGAGTTTGAGTTTTGAGCTTTGGACTCCTTTTGAACCTTCATCAAATATTC
>evgR011814c0g1_i01_Hydra
TGTAAACATGGTGACTGTAAACATGGTGACTGTAAACATGGTGACTGTAAACATGGTGACTGTAAACATGGTGACTGTAAACATGGTGACAGTAAACATGGCGACAGTAAACACAGTGACAGTAAACATGGTGACAGTAAACAAGGTGAAAGTAAACGTGGTGACTGTAAACAAGGTGACAGAAAACACGGTGGCAGTAAACATGGCGACAGTAAACATGGTGACAGAAAACACGGTGGCAGTAAACATGGTGACTGTAAACAAGGTGAAAGTTTACATGGTGATAAATATTATTCACATTAAAAAATCTTTGTAGATATTCAGATTTTTACAAAATAATATTGTAATGATTAACAAGAAAAATTGTAAATAAAAAAGGTTATTTTATA
>evgR011819c0g1_i01_Hydra
ACATGACTGGAATTTACTTTTTTAATCATTATTTAACGAACTTAAAAATTAAAATCGATTCAATACAAATAAGGAGACCCGATGATTTTGCAATTTTTTGTATTCAATGCTGTTATTTGGAAACTGAGAAAGATTTAACTTTAATTCAATATCCCCCCAAACATGCCGGACAATTGTCAAAACAATGCTTTGATTTACTGTTACTGGATTGCTTGTCCCACCTTTCTTTAGAGAATACACCAAGAACTTCAGCTTTGATAATCTTACTGTTTTTTCCAATATACCGAACATTTTTTATTATTTGTTTCGACTTACTCAAAAATAATTTTTAATGAATAATAAAATCAGTATTTGTATTAATAAACTGCCTTGTATATACGTCATATCGGTGCATATTATTTCGCATAGTAAAACTTCTTTGATTCTCATCGAGGGTTGCTTCTAAATTTGGCAGGGTAGATGCATTAAAAGCTGCTATTTTTGCATAACCACATGTTGTTGGATGAATAAATGAGGACATGGTTTACCTTTTAATGACACCTTTGAGCAGCAAGTTATAAAAG
>evgR011826c0g1_i01_Hydra
TGATTTTGACAATTTAAAGACAAAGTTGAAAACCGATTTTGATTATGAAGCTTCTGAAGCCGATTTACTTTCGTCTGCCCTCTTCCCAAGAGTATTCGAAGAATATTTAACATTCAAGCAACAATTTGGTGAAGTGACCAATTTGGGAACAAGACAATACTTTAAGGGAATTGGTGTTGGAGAGGAAACTGAAATTGAACTTGAAAAGGGTAAGGTCTTGCACATTAAATTGAAAGCCATTGGAGAAGTGGGAAGTGATGGTAAGAGAGAAGTTTTCTTTGAAGTAAATGGACAATCTCGTCTCGTTTTGATCCAAGACAAGAAACTCTCCAAAGTTTTACAAATCAGACCAAAGGCCAACAAGAAGGATGCCAACGAAGTTGGAGCTCCAATGCCTGGAAAAGTCATTGATATCAAAGTCAAGGAAGGTGACAAGGTAAAGAAGGGTGATACTCTTCTCATTCAAAGTGCCATGAAGATGGAAACTCAAGTGAAATGTAATTTGGATGGTGTTGTTAAATCGATTGGTGTCAAGGTTGG
>evgR011831c0g1_i01_Hydra
TTGAACGTGTGAAAACTGTCATTATTGGTCCTTTGACTTGGTTATACTTGGCCAAAGGTGATGAATTTGTTGGTGCTTTGGATTCCAAAAAATTGACTCTCTTACAAAAGCTTCTTCCTATCTACACTCAAGTTTTGGATCAATTCAGTCAATTGGGAGTAAAATGGACTCAAATTGATGAACCAATCTTGTGTTTGGATTTGAATCAAGAATGGAAAGATGCCTTCCAAACTTTCTATCAACAATTACACCAATCCAGGAAGATTAACCTCTTGGTTGCAACCTACTTTGGTTCATTGGAAGATAATTTGCAAACACTTGTCCAGTTACCTGTCGAAGCAATTCATATTGATTGTGTTCGTGCTCCTCAACAATTATCCAAAGTTTTGACCTCTCTCCAACAATCAAC
>evgR011832c0g1_i01_Hydra
TATACTACAATTTATTCTTTTATATATAAATGGATAAATATATAAGACCAGAACAATTTGACAGCAATCGGAACTCTTCAACAGCTTCTAAAGAATGGACACACTGGTTTAGAACTTTTACAAACTTTATGAAAACCATTCAAACAGAAGATAAATTAGATGTCCTCGTTAACTATTTATCTCCTTCCGTGTGCGACTATATTGCTGAATCATTTGATAGCGCTATAGAAATTCTGGAAAAATTAATTGTGAAACATAAAAATGAACAATTTTCTAGACATCAGTTAGACAATATAGATACAAGTTTTTATGAGTTTGGAGTTCCTGTTCCATCAAGTGAATAGTTACAAAAACAGATTGCTCGACAATGAACTCC
>evgR011839c0g1_i01_Hydra
CACGTTTAAAACATTATTCACGTTATTTACGTTTAAAACATTATTCACATTATTCACGTTTAAAAAATTATTCACGTTATTCACTCGATCGCAGAATAGCAAGGGCAATTGCCCGTGTACTAAGACTAAAAGCGGCGCCAAAAAAAGCGAAATGCAAAAAGAAGTTCCACTTTACCATAAAAAAATTACTATAAAAAGTTCTTTTCAATTTTTTTGTAAAAAGGGTACATTTTTTCCATTTTCGCCCTGTGTGCTTTTTTCTTTGCGTTTTTTCTTGCGTGGTGTACACAGCTCGTCCGAACGTTCTAATCAAAAGCTAATAAAAAACTTTTTTCGTTAATAAATTATTAAAGGACATAGACTTCTGTCAAATTTTCCAATGCAATAAGCACTTCAATTTTGTTCAAACATGTTAGGATTTAAACACATTTAAAACAACTCTTCTTTTAATTTCAATTCAAATTTTCGCCTTCTAATGACTTTATTCAGTGTTTGTTATTATGTTATGTTACTTTAAACCTGAGAAATTATTTTCTTGAGATTCGCA
>evgR011845c0g1_i01_Hydra
GTTTCCGTATGGTTGTGGAACATCAACATATGGTATGATAGGAATAATGGAAACCTCAAAATTAAAAAGAACACTAATTAGTTATTGCAATCAATAAAAAACAACGTTAAAATTATTAAATTAAGGAAAATCATGTTACTTCAGGATAAGCTGAAAAATTAAAACTAAAATAATCTACATCTATGGCCACAGTATCTATTGGAGTCTGTTTAAATAAAAAGTTTTTTTGAGCAAACAATTGTTATCTAGCGTAATCACAAATATTTATTGCTTGTTCAAAATAAAAATATAAACCTTCTCTGGTTTAAAATCAAATTCAAGAAAATTCCAGTCTTAGATCGTTTTTCTAGTTGTCGTTACAGCTTACGTATTGTGTTATGGTGTTAAAACATTTAAATATGTTTTAATGAATACTATTTGTATTCATTATTGTATTCATTGATAATTTGAATCAGTTTCAAGTTAAGTTTACAGCATGTTTAAAAGGATTTTTTATCGACTAAAGACCAAGAAATACTCCGTGGTAACAGTGTTAGTATTTGTGGCATTATCTATATTATGCCTCCATTTTTGGTTAATTCCCACAGTTTTCAGTAATTATAATGATGCATGTTACCTTGACGAAAAAGGTAAAAAGAGCCTTCATTTTATTTTGAGTACATTAATTAAATCGTTTGAAAAGTATAGTGTTTTTTATTGGTTGGATTATGGTACTTTACTTGGAGCGATAAGAAATGGAGATATTTTAGCCTGGGATCACGATGGAGACATTTCAATCCTTTTAAATGATTCAAATTTAGAGAATGCTCTAACAGAAATTCGTAGTCGTGGAGTAGATGCAAACTCAATGGTTGCGCAGTTCAATGGCTTTCATGTAGATATTGTAAGATGGAAAAAGCATAAAAATGACAACGGGGAATGGATTCTGGAAAAATACTACCCACCGTGGAATCAAGATAATGCATTGGTAAAGTTACATAGAAAATTTGAATCGATTTCATTTTCTATAGTGGAGCAAAGAAAACATATTTTGTTTATAAACGAAACTTCTTCCATTCCTTTATACTCTCAAAAACTTATCCAATTTCGTTACCCCTGGACGTGGCGAGTATCGTTTCCTTATAAATGGAAATGTTGGTTTTCATGGTTCTATTGATTTTTAATCTCAAAGTTTCTTTATTAAAGTTTCTTTGTTTCTTATGATTCTTGCCCATAACTGAGAAAAAGTTATTGGTAACTCTTTTTTTTTTAATTTAAATTGAAAATTATATATAAACTAATTTTCATGAATCTCATTAAATATATATATTTAATGAGATT
>evgR011876c0g1_i01_Hydra
AAAAAATCAAAAAAAAGCAGTTTGCAATTTTATATAATTGAACCACACTGACATAGTTTGCACAATTAAATACAAATCGATTATCCAAAAAATTATGCGGATATTAACACTATTATTAATCGTATTTTCGTTTAAACTGAAGGAAGGTTTATATGGTTGCGATGTTCCCTGTCTGTATGTTGCGTCAGCCCAATGTGTCTTCAGGATTCAAATAAATCAAACAACAGGTTTAGCTAAAAGCACATCAGAAATATTGACAAAGCCATTGGAAAATGCACTAATACGTTCTTTAGAACTTGATTTTAAAAACGGATTCTTGTATTGGGGAGATGTAATCAACAATGAAATAATTAAATTCTCATTGAATGACAGAATACAAAAAAAGTTGTTTAATTATGCCATAGGATATCCTATTGGGCTTGCTGTAGATTGGAGCTCAAACATCTTGTATTGGTTTACGACCTAATGAAATTGATTGTCAACTACAGAAGGTAAATACCGTAAAGTCTTATTCAAACTAAATATAGTGGATGAGTCAAGAGCTTATTTATAGTGGATGAGCCAAGAGCTTATTTATAGTGGATGAGCCAAGATGATGAAATGAAGATGAGCCAAGAGCAAAAGCACTTTATCCTGCTTACGGATATTTATTTTGGACTGATTGGGGTGTTTCAGCCAGGATTGAACGAGCGAGTCTTTCTGGTACAAATCGGATTACATTAGTATGAAATAATGTTGTTTGGCCCAATGATCTTACAATCGACTACAGCAATAATATTATTTATTGG
>evgR011880c0g1_i01_Hydra
TGCGTACCTTATGGAAGCCCCACAACAAAAAATTTCTAATAAGTTTTCATAGGTGACCTATGGGGTGACCTATTAGGTCCGATGTGGTTATGTCAAAAATACTTTTAAGATCTTTTACGCTGTTAAACTCTTTAAATTTCCAGAAATTTGGAAAAGTTTGCAATGGTCCACATAGTTTTTTCAGAATAAAAACACAAGTGAATGCATCTCAAAAGGCTATACACCAAATAAATCATTATACAACAGAAAGTTTATACTCATTGCAATTGGAAAAAAGTTTTAAAACAACTTCTGGCCGCTATTTTTCACAACAAAGTTTCTTGACATTAAATGATTATAATACAGTTGTGGAGAAAACTCTTGAGGAACTGCAGGATCATTTTGATTATCTAGCAGATGAGGGTTATTTTAGCAAGGATTATGATTCAACATTAGCTGATGGTGTATTAAATCTAAAGTTAGCAGATAACAGGATATACGTTCTAAATAAACAAACACCAAATCGTCAACTTTGGTTATCATCTCCAGAAAGTGGACCTAAGCGGTACGATTTTGTGAATAACTCATGGGTATATTCCCACGATCCTACGGACACTTTGCACGAGTTTCTTGCAAAAGAACTGCGTACACTATTTCAAAAAGAAATTGAACTTTCACATTTATCATATGCAAAGCGAAGTTAATAATGTAATAACAAAATTAGGTCTTACAAGAAAAAAAGGTTGGATATAAAACAACCTTGGAATAATAGAAAATGAAACAACTAAATCCGCTTACAATTGTGGTTTTCAGACGCATCGTGATGAGGCATCTAAGAATCTTAACATTTTTATAAAAATATATTTAACAAAAACAGGTCATAAAGTTTATTAACTGTACAATCTTTCATTTAAAAAAAAA
>evgR011895c0g2_i01_Hydra
GTGATTGCTTCAATATTTGCAATTTTAGATTCAAAAATTATTCTCCTTATTTGTCTTTGAAATTAAAAAAAAAACTTATCGGAAAAAAAATAAATATCGGAAAGCCTAATATGAACAAAGATCAAGAATTTCAATCTCGGTTCTGTATTTCGAATTTGATTGATCTTTACACACATAAATATCCTTGTCTTTACTGGAACAAATTTTACAAAGGACTTAATGGACAAAGGACTTAATGAGAAAAGATGATGCGGTGAAATCATTGTTTGATAAAGAAATAAAACACTTTTGCGATTTGTTCTTTAAAGAAAATCCAGACAAAAAATTAATTTTTATAGATATGTTGAGTGCTTATGGGAATGTAGCTTTATCGACAGTTTTTGGGTTAAAGTGTGACGAAATATTTGAACTCGCCAATACGGAAGAGAAAGCAGATTTTCTTGGACATAACATTCAAAGATTATGGAGCAAATCAGAAATTTGGGGTTACAATATTTCGCAAAATGCTTTGAACTACGGTTTGAGTCAGCATATTTTTGATAAGATTGTAAGCATTGACATCAATAAGTTACTTAGCAAAGAATTAGAACATGAACTAAAAATTAC
>evgR011901c0g2_i01_Hydra
CTATTGTCTAAATAAGTTTAGAATCCTGCTAAGCATATATTATAATATGGACAGTGGTTTTGAGAATGACTTAACTTGTTCGATTTGTTTGGATATTGCTGATAATGCAGTTGAAACTCAATGCTGCAATCATATATTTTGTGAAAATTGTGTGAAAGTGTTGATATCTTGTCCATTATGCTGGGCATCTCCGCTTTCATTCAAAGCTTCACTTTTAGCTCGGAGGCTAATTGGTAACCTAACAAAAAAATGTCAAAATGAAGGTTGTGGAAAAGATGTGGCTCGATCTGAATTTTCTAAACACAAATTGTTGTGTGAATTTTCAATCTTAATGTGCTATATACCAGCGTGCACTTTCAAATCAAAGAAAAATGACTTGATGAATCATTTAGTAACCTCTCATACTGATTCTATCATTAAGATATTGGAAGAGCATTACTCTAAAAATTCTTCTGAACAAAAAAATATAAACGAAGAGCCTAATAATTCCCATAATTTTATTAGTACTAAATTTAACTCAAGGGGTAGACAAGCTAGGCTTGGTGCAACTGGAAAATATTATTGCCTTGGAAGTCTTGATTTGGAACCAAAATGTAAATGCTGTGATGGAAAATGTGGACCATCCAATGGTTGTAATTGTACTTCATGTATGGCACTAGATATTTGTACTCGTAAACTTCCTTCAGGTTGGTATGTTAATAAAGAAGGGTTTACTTGCACAAAAGGATATACTGGAAAATTTTATTGTGGCAGAAAAATGGACCTCAAATTTTCAGATGGCTACTGTGGTCCAAATGACGGTCCAAACTGCTTATCTTGTCAAATCATTGGATTTGAAGAAAGTGCTTATAAATAAATTTCTAAGGGCTCTTTAGGCAGTGTACCATAATATTTTTGTTTGCTTTTTTGTGTTATCTTTTTAATTCCTATTTTTATATTGTAAAACTGTATTATTTTTTTAAATTTATCTAATTGTTTTTTATGCTACATATTATTTTGTTTATTTTTTTGAATAAATTTAATATATAAAAAAAAAAA
>evgR011902c0g1_i01_Hydra
CACTTTCAGATGTCATTTCGTGTTGTGTAACAATTCCGGTATTTGTGTCAATGATTATTCACGAAATGATGTTTTGCCATGGTCTTGCATTGTGTTACATGTATTACATAACCGAAGCAGTTGGAAATACTGCAAACATATTCAACTTGCTTTTCATATCTATTGACAGGTACATTGCAATTGTGTTCCCTTTTCGATATTCCAATTGGATAACGGCGAAAAAAGCAAAGCGCTTAATAATTTTTATTTGGATGTTCGCATTACTTTGGAGTGTAACCTGCTTATTTAAATGGGATAATGTTAAAAACTCATCTATCTCTGTTG
>evgR011918c0g1_i01_Hydra
TAAAATTTATTAAATCAGCGGGAAATTTACTAGACTTTTTCACAATATTTCCCAATTTTTTTATAACTTTATTCGAAGGTAAACACGCCGGTGAAGTAAGCAATAATATACTAGTAACAAAGTTTAGCCGAATGTTTAAAACACTATGGGTTTTACGACTTTTACGCTATTCAAAAAATTTTGAACTAATTGCTGCTGCGCTAGAGATATTGACAAAATCCGCTAAAGACTTATTTACTCTGCTTTTTTGCGTGCTCATAACGTGCACTCTTTGCGGAAACCTAATATACTTTGCAGAGTTAGACAACAAGACTAGTTCAATAGTAAGCGCACCTGAAGGAATGTGGTTGGTCATTCAGACAATTTTAACAATAGGCTACGGCGATGTTGTTCCTATAACTGTTCTTGGAAGATTTTTAACTGCACTTACGGCGGCAATTGGGATTACTGTGACTTTTCCACTTTTATCTTTTTGCGGAAAGTTTTTTTATGCATATTCTAGAGTATATAAAACTAATTGAAACTAATACTTTCACAAATATTAAAAAAAAAATTTATTGTTTGATAATAAAAAAAA
>evgR011926c0g1_i01_Hydra
TTGTTAACTCTAATTAAAATAAGTCCCATTTATTTATCGTGCATAGCTTACGACTATCTGTTAAAAAACGATGTTTTTGTATCGTTAGCTTATAGCAGATGGCTTAAGATACTCTGCGACGAAATTGGACACGGAGTAATTGCGGCCTTATCGTGGATTGTTGTCGTAAATTATGACTTTCACTACTATCTTCAAATTGTGTTGTGTGGTTCCCTCGGTATTTTAATTGACGTGGATCATTTTATTTCGGCTAAATCGTGGTCGTTGATGAAAGCTGAAAAGTTAGCCGATCGTCCGTTTTTACATAACTCGATTATTCTCGTATTTATCTGCGTTGTTTGTATATTTACTCTTTACAAAAAGAAACAGCAGCTTCTTGATCTGGCATTAATTTTGTTTGTCTCTGTAAGTTCCCATCATTTAAGAGATGCGAACAGACGAGGAATGTGGTTCGGACATTTTGGAAGCACAAGTGTAATCGCTCCGGAATTATTATACTCATCGCTCATATGTTTCATAGCTATTTTTGCTGGTGTTATAAGAAAGCGCACTTTAGTAGAAACAAGTTGTTTCAATTAAAAAATTTGTAATTTCAATATTTATAAATAAAAAAAA
>evgR011928c0g1_i01_Hydra
AGAGAAGAATTCCAAAATACAATTAAGAGCCTTTCCAAATCTTGGACAATGTTGACAAACTTTGAAGAAGCTTTAGATAAGGCCCACTTTTGTTACTCTGAACCACAAATCCCAGAATCTACTGCTGCTGTTTTCAATGATGAACTTTGTTCAAATTTGACAAAGGATTCTGATGATTTCTGGATAATGTCTGCTGCTGTTAAGAAGTTTGTTGAAAATGAGGGACAAGGTTTATTACCACTCCAAGGACGTGTTCCAGATATGCAATCTGATACTGAACGTTATGTCAAGATGCAATCACTCTTTAGAAAGAAGGCTCGTGCTGACATGGAAAGTGTTAAAAAGTACATGAAAGAATTTTTGATTCAAGTTGGCAAACCTGAAGACTCTATCCCAGAAATGGATCTTAAAGAATTCTGTAAAAATGCTTGTTATTTGCGTGCAGTTCACTTTAGATCTTTGGAACAAGAATTAAAGACACCAAACAAGGATGAAATTCAATCACACATGCAATGGAATGAAGACTCTACAATGCCATTCTATATTTGCTTCCGTGCTATTGATAGATTCTTGGAAAAGAAGGGTTATCACCCAGGTCAAGAAGATGAAAACATTGAAGCTGATACTGTAGAATTGAGAAAGATTGTTGATGAACTTCTTGTTGAATTTGGTGTGACAGAAACTGTTCATGATATGGACAAATATATTGCTGAAATGGTAAGATATGGTGGATGTGAACTCCATAATATTTCATCATTGATGGGTGGTGTTGCTGGTCAAGAATTGATCAAGTTGTGTACCAGACAACGTATTCCAGTCAACAATACTTGGGTTTTCTCTGGTATCAAGTCCATCTCTATTGCTTTCGAAGC
>evgR011935c0g2_i01_Hydra
GAAAGGTAACGCTACCTTTAAAACAACTGGTCAACTAATAGAAAGCTTTGGTGCCTCTCTTATTGTTTTGCCTATCATAATGTTCATCGAGCAAATGTCTATAGCAAAATCGTTTAGTAAAAAGTTCAACTACAAAGTAAAAGCGCAGCAGGAATTAATTGCAACTGGAATGTGCAATATTGTTGCTTCATTTTATGGAGGATGGACTGTAGGTGTAAGCTTCTCGCGCTCTGCTATAAATGTATTAAGTGGTGCAGAAACGCCTTTAGCAGGATCGATTTCGGGAATTGTTTCTTTAATTGCTCTTGAGATTATGACTCCAGCTTTATATTATGTGCCATCAGCAGCTTTAGGAGCCATGATGGTAATGGCTGTGCTGACTATGATAGAATTACGTTTACCAATAAATATTTGGAAATTATACAAACTGGATTTACTACCATACCTTGCAACATTTTGT
>evgR011944c0g1_i01_Hydra
GTGCTGTTACACGTTTGACTGCTGGTAGCTATCTTCTTTTCAACAGTGTGAATAACGTTGAAGCTTCTGAAGGTACTGCTATTCCTGCACAATATCCATGGCCACACAAGGAACTCGCTGGTTCATTTGATGCTGGTTCTATCAGAAGAGGTTATAAAGTTTATAGAGAAATCTGTAGTGCTTGCCACGGTATGAACGAAATTGCTTTCCGTAATCTCAATGTTGCTTTCACTGAAGAAGAAATCAAAGTTTTGGCTAATGCTGTCGATGTTGAAGGTGAACCAGATTCCCAAGGAGACCCAACAACTCGTAAAGCCAAGGCTTTCGATTATATGCAATCCCCATACAAGAACGAACAAGCTGCCAGATCTATGAACGGTGGTGCTTTGCCTCCAGATTTGTCAGTCATTGTAAAGGGTCGTCCACACAATGAAGATTATATTTTCTCATTGTTGACCGGTTATGAGGATGCTCCTGCTGGTGTTTCCTTGAGAGAAGGTCTTTACTGGAACTCTTATTTCCCAGGTGGTGCTATTGGTATGGGCCCTCCACTTTCAGATGGTGCTATTGAATTTGAAGATGGTACTCCTGCCTCTGTTTCACAAATGGCCAAGGATGTTAGTACATATTTGACTTATTGTGCTGCCCCAGATACTGAACCAAGAAAGAAGATGGGTTTGAAGGTTTTGGCCTCATTGTCACTTGTTATTGCTCTCTCATGGGTACACAAGAGATTCAAGTGGAGTATCATCAAGAATAGAACTGTTATTTTCAGGGAATAATTTCAATCTTTAGGTATCAAAC
>evgR011950c0g1_i01_Hydra
TTTAGCTCACAAGAATAACGATGTTTCAACAAAGTTCACTTTAGTTGAAGAAAATGATAATTGTTTGAAAAACTCTTTTATAAGCAATGTCTCGTTTACTAATGGATTAAAAACAGGGAATTTTCAACAGCATGGTGTTGTTCTAAGTGATTTAAATTGTCTTGAAAAATGTTGCAATGATGACGGTTGTGACGCAGCTTTCATGCTTGACAACTATTGCTATTCTGTGAAGTGTTTCATAAGCCCATGCAAACTATCAAAATCAAAATCGTTTTTATTTAACCCAAGAATTGCATTAGTGGATAGATCAGATGAGATTTTAAGGAAGTCTTTTATACCAACAAAACCATTACTGGATGAGTATGGAGATCCATTGAAATTATATTTTAGTCCGACTAGTATTTCAAAGGAAATTCAAAATTCAAAAAAAATCAACAAAAAAATTAAAAACAAAATCACACAAAAGATTGATTCTGTCACCATCAAACTTGATGAGCTAGAAAAGAAAATATTTAGTTTAAGCCGGGAACCAGTTTTTAAAAAATCAAAAGTGCTCCTAAGTAAAAAAAATAAAATGAATGCAGGAGGAGTGTACGACTTGGGAGCAACCGATGTTAACTTTAGAGCCTGGGATAAAAATGATCCTGATGTAGAGGATGACAAGATTATACAATATGATAACGACTTTAAAACAAAACGAATTTTTGTCACTCCCTTTCATCATAAACAAGATGGAGAAGAACTAGATCCTACTCTTCTAAAAGGAGATGAAAGTGTTCCTGTAATAAAGTTTATTAATGACAATGACAACGATGATGAAAAAGAAAATTCAAAAGATGATGATGATGATGATGACGAT
>evgR011954c0g1_i01_Hydra
TTATATCTAGGGAAGTGCAAACAGATCCTGTTACTGATAATCCCATTCATATAGATTTTCAGTTAATAAAGGAAAACATTCCAATCAAGGTTGCAGTTAGGGTTAAGGTAATCAATAAAGACAAATCTCCTGGAATAAAAATAGGTGGAATACTCAATATTGTAAGAAAGTATATAAGTTTGAATTGCATACCACAAAATATCCCCGAGTATTTAGAAATTGATATTTCCGGTTTCGAGATTGGAAGGAACGTCCATATAAGTGATCTGATACTGCCAGAAGGAGTGGTACCAGTTGATAAGGATAACTTTACCATTTTAACAATTGCAGGAAGAGTTGAAGAAAAAGAAGAAGAGAAGGGAGGAGGTGATGAAGCAACTGAAAACAAAGAACAAAAGGCAGAAAGCAAATAATTAAATAAATGCAAGTTAAAATAGTAGGAGGAGGCTTAGCTGGAAC
>evgR011959c0g1_i01_Hydra
TAATGATTTGTATTCATTCAAACTTGTAGCTGATGCTTTGGCGTTGACTACACAATATCCATTAGAGTTAGTTATTCCAACTCAGATTGATGCAGCAAATATTGTTCTTAGACTAGCATCATCACTTACTGAAGAAAATCTCAAGTATTTTGGAGCACACCGCTTTGATACCATTAGTCAATCATATGTAATATCAATATCTAATTTGTTTAAGGCTTACACAGATCCAGGTATTTTGAATCTTTCAACAACAACATCAAACCTTGGAGATGAAAAGCATGTTTCAAGCCTCCTAGAACCATTGATAAAGTATTTTTTTGC
>evgR011962c0g1_i01_Hydra
GGAAAATTATGACGCATTATATTCACTTTTTGTTTTATCAGCTCCACCCACTGTTATAGAATATTATAACAACAATTGGCATAATACTCGTGATGAGTTGGTTTTTGGTTTGAAGTTTGTGAGCAGAAACTTCATGAATGGCACCAATAAAAGGTTGGAAAATTTTAATGGAAAGTTAAAAGAAGTTATTGACTGTAGCGCTGTAGCACACACCTCTTTAGAATACTTTCTTGATAAATTATATGTTGTTTTAACATCAATGCGCAATGAAAGGGACAATAAGACCATTTTTCAAATGCAAAAAAGGTTGATGCTTACGATCCCAATTCTGCTGAGGCAGCTTTTAAAAAAGTTCTGACAGTATGCTGCTTGACATG
>evgR011971c0g1_i01_Hydra
GTAAAGAAGTAGTACGCTAGAAGTACAACCGATAGTGCAACCGATGGTTTCTCAACAGCTCGATAGATAAGTGTTAATTTCCATTTCAAAGCCATGGCTTGCATAGCCTGAGATCTAAACGACTTAGAATTATCTAGAAAAACTTTGGAAGGAGGTCTAAGCGCTGACCATATTGTTTCTAACGCTTGAACAACTATTGTTGCACTTCTTTGCTAATGGTGTTCACAGCGTGTATCTGCTTGGACCACAATCAATACAGGGAAAGTAGAGCATCAACTCAACGTGAATAACATCACAGACTAGTTTTTCCCACACGTTACTGACACCAAGAATTGTTTTATCCCATAAAATCGCATTAGGGTAAACTGAATTACATTTTTCACATTCAGCAATAACTTGTTTCATTGCATCTCCGTTAACGTGGATACCAGCACGCAAGCATAACTTCATACTCTTGAAAACTCCTATGTGCGCTATCGAATGAATGTTTTTAATCTGCTCATTATTAAGCTTATAAATTATACTGCATTCTTGAATTGCGAATTTGTTTTTATCCCTTTTACTGGCAATCGCGTGAGTTCACCATTTACCATTCATTCCATTTGTTATTTACATTCAAAAGACGTCTTTTCCGGACCAAATGCCCGCTGGGTTGAAACCTTCGTTGAATTAGCATCTCACCCAGCGCATTTTACAAGTTTGTCACGGGTCAGTTTAGAATTTAACCAACTGATTACACTTTGGGCAGGGCCGTACCGAACAACAAAGACGTGGGGTGGGGTGAAAATAAATAAAAATTAT
>evgR011976c0g1_i01_Hydra
GGTTTAAAGGTGATGGATCTGGAAAACTTGTTATTTTTCTTGCAGTTAATGGAAATGCTTCCTACAATATTGAAACACCTCGTTTATTATTAGATCAATGGTCTACGATTAAAATATCTCAAAGTTTTTTTGATGGTAATTTTTGGCTATCTGTTGACTTGAATAAAAACACTATTCATCGAATAAAAAATAATGATGCGATAGATTTCAAATACATTAAAGTTTATGCATCTAATCCTTGGAATGATGCTCAAAATGGTTCCATCCGTGACGTTTTGATTACTAATGGAAAAGCTGAATATATTGTTGGCAATGTTCCTATTCCTATTGAAAATGGAAAATTGATAGCTGTGATACCAAAGCTTGATAAAGAGTATATAATATCTCTAGATATAATTCTCAATAAATATGTTAACAACTTTAATGAATCAGTTAGCT
>evgR011987c0g1_i01_Hydra
TTTTATACTGATTTTTCTCGTTACTTTTGACATGTGGTTGCCGTTGTTTGTTATTGAGACTTTAAAGTTGTCGGTTTTAGAACTGAATGTTTGCATTTTCGGTACAGGAATAACTTCAGTTTTTATATTACTTTTTTATATATGTAAACCTTTATCAGATCAAAAAATGTTTATTGTGGTTTTGATTGGACTGTTTGGTCTATGTATAATCAACGTTATTTATATTACTTTGAGCTACTCTATTTCTAAACCATTACAACTAGTTTTCGGTTTTGCGTACATGATTAATTTTTCTGGAGCTGGTATAATTCCTGAGGTTTTTATTACTAACACCTTAGCGAAGTTTGTTAATTCGAAAAACCAAGCTTTTGTGGATGGTATACGCAACTCAATGTTTGCTGGAGGAAGACTAGCAGCGTTTTCAAGTGCTGCATTTACCTTTGAATACTTGGTTACATTTTCTTCTGTATTTATTGCACTAACACTCTTGATGATGGCCTTGCTGACGATAAGAAGAAAACATTTACTTAATCCCATACCTATATTTTAATAGTTTTATAAAACTTTTTCTAGAGTAGAATATGAAAAAATAGAACTGATTTTTTAAATTGTTCATTATTTTATTATGATAAACATGCTAGTTTTTTTATTATTTGGCAAAACAAATTTACCCGGGTAATGGAGAATTTTAAACTTTCCCTCGTATTACTTTGGTATTATCCTGCAATTGCCATAACTCTATTACATCGCATTTTTCAATCTAAAAATGAAAGTTATTTTGTTCCGAAGAACTGGATTTTTGTGGTAAAAGATTGTCATTAATATTTACCTGGAACAATCGACTTTACAAATTTTATGTATCATCAAAATAAAAACTTAAATTAAGCAATTCGAACATTGGGTTTTTTTAAACATGGATCTAAATTTCTTGATTTTGCCATTCTAAAGTTGGTATACAAGTCAACTATTAGGCCACAGTTTGAATATGCAGCGTCAGTTTGGAGCCCATATTGTATTAAAAATATCAAGAAAATATTAACTTGAAAATGTACAGCATAGATCAAATATGATTTAAACACTAAAAAGCATCTGTTTCTATGAAGAAAGGAGAAATGCTTATCAACTGTACAACAAAAGAGAATAACTTATGAGTATTATAAAAGAAATGTTTAAATATTAAATTTAACTAAGTATATTCGATTTGAAGAGGCTTTGAGTTTTCACAACAGCATGTCCCGATATCGTCATGACAACAACATGACCATCATGTTCATTATTACTTAACAAATAGTGTGGTTGCCGATTGAAATTTATTGTCTCAAGATGCAATCGATTCTGCAAACACGTTAGACAAAAATAGAAAAAACAAAATTTTGAACTTTTGATATCAAATTTTATTTACGAACTTTTACATTTACACTACACTACACTTTTACATTACATTAAAAACTTTTAATTAAACTTAATACTTTTATTATAATACTATATAATATATATATATATAAATA
>evgR011994c0g1_i01_Hydra
CTCCGAAACACGAACCTTGACAAGCAAAGCCGCTGCGCGGAGAAACTAAGTTAACTGTACTGTGGTTTATAGGTCCAGGTCCGATCTTTGGTTACTATTTCTACTCCTATTGCAAGAATTGAATATATATTGCTGATCTCTACTGAGGTTGAGGGTTGCTTCTTTTGATAATATAGTGTATAATCCGTGTCCACCTATAATAGGTTGGAATTTTACATTTGGCAATTTTTCACATATCTGAATGTCTTTACAAATTGGTTCTGCAAACAAATTTCTGCCAGAAATTTTTTTGCAGAACCAATTGGTAAAGACATTTTTTTGCACAAGACACATAAACCATTGTTACAGATGTAATGCTTTCTTCGAGTTTTGTTTAAGTCCTCCTTTAATTCCTGTGTTTACATTAGTACTATCACAACCGAACATTGAAAAAAACACATCTATTTTGTTAATTTCTGACAATATGCAGTCTGCATTGTCAGTGCTTACACTACGTTTTAGTGAAACATATGATAGATATTTATTTCCAGGTGCTTATAAGATAGGGCCATATAAAATGTATAATCCCCACATTTTTTACATTTTTTTTGTTAACTTATTTTATGGACCAATAATTGTATTGACGGAGATTTTTTTTGTAGAACAAAAT
>evgR011996c0g2_i01_Hydra
TTTATCCTAATTCAATTTTGTATCGTCAATGGATGTTTTTTTATCTTACGTTTTGTTATTGCCGTTCTACATTTTTTACGTTCAGATGTAATCGGTAATCTGGTTTTTTTTTGCGCTTATACGAACTTTCCACACAATGATTTATTACATAACATCTTGGTTGATATTCGACAGATATCTGCATATAAAATTAAATATAAAATACGTAATTTATTGCACCAAAAAGAAAAACTTTTTAGTATTGATGTTTTTGTGGATTACATCAATTTTAGTTGGAAGTATAACTACAATTTATAAAACCAAATGGTTTGAAGTATATCAAATTGGTTATGGCGTGGTCATAGTTTCTTTTTCTATTTTTCTATACACTTATGCTTTAAGAGCTTTTCTTAAACAAAAAGCTGCTGTACATTCAAACCAACAAAACAAAGGAATATTTAAAGGCTTGC
>evgR012001c0g1_i01_Hydra
TTTCAGTGGTTATACTGGTCATCCAGCTGATTTCTGTTACTATGCCTACAAGAAGGATACTGCTGCTTATAAAGTTGCCTCTGCTAAGCTCTATACCACTAATTATTTCGGTCCTGGTGGTTGTTCCAATTTACCAATGCCAGCTTTCTACATTTGTTCTGATATTGCTGTCAATTCTGGAACTGGTAGATCTAAACAATACCTCAAGGAATTGGGTGGTTACAATGGTCAAGATGCTAAGGCATTCGCTAGATTGTTGAACGAAAGACACAGAAAGGATTATTTATATTGGGGAAGACCTGGAACTGCCAATAACAAATTCTTGAAGGGATGGTTGGCTAGAGCTGATGGAAG
>evgR012010c0g1_i01_Hydra
TTATTTTTTATTTTAAAAAATGGTTTTTTTGAATGACCATCAAAAAGAACAGTTATCTTTGCTTACGCAAATTGATGTGAGTATTGGAAAAGAATTTTGCAAGATTGCATTGGATTTTCTCAAAGAAGGTTCAAATCCAAAAAAGTATATATTAGCTGCACAAAAACTGAACATACATTCAGATCAAATTCGTAAAAGTGTTGAGGCAGTTATGCACGTTATAAATGAATCTGCAAAACTAGAATTAGATGAGACAGATTTTAGAGATTCATTAAAAGATATTGACTTCTCAGAAGATTTACTTGAGTTTTTTGTAGAATGTTATTTAACAAACATATATGATATACGAAACATATTACGTAACATGTCGATTGGTTTACCAAGATATGCAAGTATTGAATGGCGTTTTGACGTAGTCATATCAAGTCGAATGCTTAAACAGCAAACAACTCCTGAGATTTTACTTAAGTTATTTACACATGATAGTATGAATAAAACAGAAGCAAAGATTTTAAAAACTGATCCTGTAAACTTACAGCACTTGACATTAGTTTTAGAAGAAGCTCTTAGAGACCTTAAAAGTTCTCATTGTCGTAGAATTGCAAGAAGTATAAAATAATACCTGTTATATATAAATAAATATATATGTATATATATTTATATATATTATAAATATATATATATATATATGTTTGTTGTATAAATAGTTGTGCACAAATATGAAATCTCTCATAAGATTATTTACCCTTTAGTTTTAAAATGTTCTGGCTACATTCATAAATAAGGTTG
>evgR012011c0g1_i01_Hydra
TTTAGAACATATTTTTCGTTAGTGAAAAGTCAAAAAATCAAACGAAATAAAATCTATGAAAATTAGAAGCACAACAAAAAAAGTGTTATTGATTGTAATTATGAATTTGATTTTTACTTCTGTTTATTAGTGAAACGAATGGAGAACGCAACGGTTACGAACGCTTCTGCAACACAAATAACTCCTAATAGTCCTAATAGTTTGAATATAATTATTTTTATGCTATTTGCAATTATTTTCCTGTTCGGTACTATTGGAAACGGTTTGGTACTCTACTACTTCCGCTTCAAAGATAAACCTCTAGGCATAGCAACAGAAAAACGTAGAATATCAGAATTTCTATTTAGTTTACTATCTTTTGTTGATTTTTTTGCTGCTGCATTAAACTCTTTATTGTACATGTATTGGACTGTAACAGAGTACAAATGGCACTTAGGATATTGGAGTTGCAAAATACTTGTCCCTTTAGGAACTGTTGCTACAACAATGTCTTGTGGAATTATAGCCGTATTATCATTCGACAGACATCGCGCCATTGTTTATCCGTTCAAAAGCCATTTCAAACTCTTCCACATTAAAATATGTATTTTTTGTGTCTTAGTTTACTCTATAGTAATAAACGCGCACTATTCCATTAACATTTCTTTAGAAAATAATCAATGTCTTGTAAGTGATACACAGAACAAGACGTACAGTGTTCCAACAATAATACATTTTATTTTGAACGATTTTACGCTTTTATTTATCATAAACTTTACAAACTACCGAATATTTTCGAAGAGGCTTCACCTTAGTCCCACCAGCACTCTTATGAAAATGCTGGGCAGTAACGCTTGCAAACGACGCAAGAAAGACAATAGCAGAACAATTCGTTTGTTAGTTGTGATGACAATAGTGTTTTTTATCTTAACATTGCCTAAAGATATTCTACAGTTTGTATTTTTAATGTCGTGGCTAGTAGGTCATGGTATACAAGCATCAGACGCAGTATTACGTTTAAACTCTTTCCTAAAGGTCTTTAACG
>evgR012014c0g1_i01_Hydra
TACTATCAATGATACCATTTTTCAATTCAGCTGTCACATTGTCAGAAAAAACTAAAAATATTAGCAACAACTTTTTTTACCAATACAACAGAAAATGTAGTTACAAAATATCAAAACTATGGTCGACTTTAATGCTCACAATGAACTGCGGCCTACCATACTTTGCAATTGTATTGATTTATAAGATATTAATTACTCGAGTAAAAACTGTGGAAAAACAAACCAGAAAAAACTCAATAGAAAACCAAAATGATTTAATGTTTACATGGACTAAGCAGTTATCAGTACAAAATCAAAGTATTCAAAATAAAATCATAAAATTGGGCAAAATTCTATCAATAGTTTATACCTTACTTTGGTCACCAACTGTTTTATACTACATTCTTGCTAACATTTGTTATAAAAAATGCTTTAAAAATGATTACTTAATTTCCCATTATAAAGAGTACACTGAATTTTCTATTAGATGTTTAACATTTTTAGATGCACTCGCTGCACCAATTTTCTACTGTCTGTGGACAAATGAATTTAGAAGCTCACTAACTCCATCATTTATGAAGAGAAACACTAAAGTTAGCATTGCTGCACTGGTATTCTAGAC
>evgR012029c0g1_i01_Hydra
ATTTGAAGTGTCGTTTTATTTTTCTAAAAATTGTCTTCATTGAAAAAAGTTCATTTCAAAATGCACAGTAAAGACGGTCTAACATTTGACTCGTTCGAGTGCGAACCTCTAATAAAATCCAAAAAAGTGTGCTTTTCATCATCAAACGATGTTGCCTACATTAATCCATATTCCCCAACAAAGCATCTTTTACTGAGCAATCTAACCTCTGAACAACTTTTACTGAGAAAAATGAACCACGAATGCAAGGATATGATACCAATCCTGAAGAAATCACGCTATTGTGTTAAAGAAATGTTTCAAACAGAAGAAAACTTACGTAATTTTGAGGGAGCTCGCAATAAACATAAAAACAAAAATATTTTAAAGAGTAAACAAAATCAAGATTTTTCACCAGATAATGAAAATGTAGTATTACTTAAAAACGGGTGTTTAACTGAAAATGTATGTATGACAAAGCCTATGCAACGCTCGCTTCCCGAAAGCAATTCCAGGACGCAACGCCCGCTTCCCGGAAGAGCAATATTTTCTCCAAGTAGTGAAAAAGATTTATTGACAAATATTCCGATCGTAAAAAAAACTGAAAATTATTCACTAATAACACATCGCATGGATTTAGGATCAAAAAATTTAATAGTAAGGAAAAAAAAGAGCTGCAATAAAAAAAACTTTTTTCCTCGCTTAACTTCTGAAAAAACAAACAACCAAACAACTTTAGATAATATTATAACGTTTCAAACGAATCAAACGAGCGCTAATGTTTCAAACGAATCAACTCAAGATTTTGAACCTTTTTTAAAAAAAACTCGTTCACAAATATCGCAGCGTTTTACGGTCTCACACAATGAACTTAACCACATTGATACTTTAAAAGAAGAAGACTTTACAGATGTTTTTAAATATCCTATTTCTAAGACACAGAGACATTCTTTGCCACCCCTTTTTACAAAAAATAAGCTAAAAGGTATTATTTAAAACGCTTTTGCAACAAATGCAAAAGTCTCGTCTTTTGCATTTGTCGAAAAATCATAATACCTCTTTTTGTTTTAAATATCCGTATTCCGGAAGACAGTTCATAAAATTATTTTTTATAGTGTATATATATATAATATTTTTTGTATTAAA
>evgR012032c0g1_i01_Hydra
TGTAAAGAAGGAAGCTGCTTGGGCAATTTCAAACGCTACATCAGGAGGTACAGATGAGCAAATTAAATATTTAGTAGATAGAGGTTGTATTAAACCATTGTGCGAATTGCTCTCTGTAAAAGACGTTAAGGTTGTTGGAGTTGCTTTGGAAGGTATTGAAAACATTTTGGCATCTGGTGCTAGCATTGCTGATGACATTGATGGTATTAATCCATACACTGACTTTATTGAAGAAGCCGGAGGTGTTGAAAAACTCGAAAGTCTCACAGATCACGCTAGTGAAGAAATTTATAAGAAGTCATACAAGATTGTAGAAGAATA
>evgR012034c0g2_i01_Hydra
CCTATTGATGCATTGTCTGGGAAAAATGTTTCAACTAGAAACATGGGCGTTCCTTGGTATACTGGAGGATCTTTGATTGAAGCTTTAGATGAAATTAAACTTACCAATGTACATGACAGTAATGGACCAATTCGTTTACCAATTATTGACAGATATAGAGATAAGGGTTGCACTGTAGTTTTGGGTAAAGTTTTGAGTGGTCAAATCAAAGTTGGTCAAAAGGTTGTTGTCATGCCAAATAAGGCTGTTTGTGAAGTTGTTGCCTTAAAGAGTCACGAAATCAATATTCATGGTGGTGTGGTTTCTGCTGGTGAAAACGTAGAATTATATTTGAAGGGTGATGATGAAAATATTTCCTCTGGTCACGTAGTTTGTGACCCAGAACATTTGTTGGAACCTGTACAAGAATTTATTGGATCTTTCCACATTTTGGATAGCGCTGAGCATGTTGATATTATTTCTTCTGGTTTCACTACTAATATGCATGTTCACACTGCTATGGAAACTATCACAATTTCGAAATTAGTCAACAAGATGGACAAAAAAACTGGAAAACCATTGCCTCAAGTCCCAAAGTTTATTCAAAACAGAGATACTGCCGAAGCAGTTTTTAGATTGGAAAAGCCAGTTTGCCTTGAAAAGTTTGACACTTTGAATGCTATGGGTAGATTTGTTTTACGTTCTGATAGAACCATCGGTGTAGGTAGAGTGAAGAAGGT
>evgR012048c0g1_i01_Hydra
TTCTAAATAAGTTTTGTAATTAGAATTGCAAAATTATTTGTGTAAATGGCAAACACCTTATCAGTTGGACGCAGAGCTACAAACTTGTTTGTGGTCATTACTTTATTGGTGCTATTCTATGTTTTTGGCAACGTTTTACCAACATTTAAGCAAGGGTTTTTCTGTGACGATGTAACAATTAAGAAGCCATATTTGTCTCAAGAAACAGTTCCGTTTTATGTGGTTGTTGTTACTGGACTGTGTTTAATAGTTTTTGCGGTATGCTTCATTGATTTTATGAACTTCATATATTGGAAAGGGAACAATGCGGTTTATGAGGATGTTATTGAAACAACATTGTGCTGCTTTAAGATTAGCAAATGGATTTCTAGAATTATCGTGCGGTTGTGTTTAATTTTGTGTGGTGCTTTGATTAGTAATATAATAACCAATATTAGCAAGGTTATGGTTGGAAGTTTGCGTCCACATTTTTTAACTGTATGTCAACCAGATTATAGCAAATTTAATTGCTCGAGTGGGTATATTACTTTAGATGTTTGTACTGGAGACAATGAAAAATTAACTGATGCAAGAAAATCGTTTCCATCTGCCCATTCTTCACTTGCAAGTTTTTTTGCTGTGTTATTGTCATTGTATATAGAATATACTGTGCGAACTTCTCAGACTTGTTTGCTGAAACCTCTTTCACAATTCACCTTGATTTGTTTGGCGCTTGCATGCTCATTTACTAGAATAACTGACTATCATCATCACTGGTCAGATGTTATAGCTGGATTGAGTATTGGTACTTTATTGGCTTACTATACGATCTTTTATTTAATGAATCTACCTCACGAGGAAAAGTTGGTCCAAAATGCAAGATTACATAATTCAAACGATTATAGCGATCTGGAAATTGCTCCCTTAAAAAATTCGCCATCAATTTGTAAAAAAAGCAATCAAAGTTTATAAATGTATAGAAACAACATTTTCTTGTAAACTTTATTTTTTAGTTTAATATATATCTTATTATAGTGACACTTACAGAGACAACTGTATGGAAAAACGTAAATTTACGATGACAAAAGACAAAAAATAAAATAAATACGAGAAAAAAAAA
>evgR012049c0g1_i01_Hydra
GTTTAGAGAAGTTATTATTTTATTGTGGGTTTGAAACTGATACTTGTGGTTTGGACACATCACTTAATCATTTTAATAATGAGAGTGCTTGGCAAAGAAGAAATACGAAAACTCCATCACAAAATACTGGACCAAATGCTGCTTATAATGGATCATACTTTATTTTCTTTTTATCGTCTTTGCCAAATAACATCAACAGTTCAAGTTTACCTCTGAGACTTACTTTTCCATCAAATGCATTATTTCTCCATCTTAAATTTGCTATTCATGCTTATGGATCTGCTATTCAATCTTTTAATATTCAATTTTATGATGGAGATATGACATTACTTAATAATGATCAGATAATCAGTTCTCAACTACAAACAAGTTCAGATGCACCTTGGACGCTTGTTGATAAATCTTATACATTCATGTCAGGAATCACAACAATCCAATTTGTTGTCAGTTGTGTTCCATATCAGTATCAAGCAAATATAGCAATAGATGATTTAAGTGTTTATGTAGCAGGCTTAGGGGAACTAGTATTTAATTGTGGATTTGAAATTGATACTTGCGGTTTGGACACATCACTTAATCCTTTTAATAATGTTAGTGCTTGGCAAAGAAGTAATAAGAAAACTCCATCACAAAATACTGGACCAAACGCTGCTTATAATGGAACGTACTTTATATTTTTTCTATCATCGTTGCCA
>evgR012050c0g1_i01_Hydra
TTTTTGAACTAAATATCTTTAAAAAAGGCGTGGAAGAAAACAAAAAAAGTTTCATCATTGGAGATCTAAATATGGATTGTCTTATATATAATAAGGACAATAAAGTTAAAAATTTCTACGACTCAATTTTTGAAGCTGGAGCAACTCCGTTAATTAATCGTCCTACTAGGGTAACAAAAACCTCAACAACTTTAATTGACAATATCATTAGCACTGATATTTTTAACGATAACATCCATAAAGGCATTTTAAAATCTGACATATCTGACCACTTCCCAATATTTTTGACACTTAACACAAATGCTGTGACGGATAACAAAGCTAAAAAGATTTTTATGCGTGATTTCAACCAAAC
>evgR012051c0g1_i01_Hydra
TTTGAAAGCTTTTATTAGTTTTTTTAAAATGTCGAAGAACACGCGGAGAATAGCCGCCTGTGCAATTTGTAAAAGCCGAATAGTGTTTAAATGCAGTCGATGTAAGCATGTTTTTTATTGCGGTAAATCCCATCAAATAGAACATTGGAGGCACCATAAGTTAGTTTGCCAACCTAGATTTGTTCAACAAACTATAAAATGTTTAGGTCCGAAGGATTGCTTTTTTGATGCCTCTAGAGTTACTCAGGAAAAACTTGCAATATTTATTTCGACGTCACTTTCAAATATTGGACTGTGTGTCTTAGACAACTACCTTAGTGATGATTTAGCTAAGCGTGTATCTTCTGAATCGCTAGCTCTATGTGGCGGTAATTCTTGCAATAATCTTGGTAATAACTCAGGTAACAACTTTGGTGAAACATTAGTCATAAAACATCTTCCAAAAAACATGAGACCGTATCCTACTATTTCTAACCTGCAACTTTTGTTTCACAACTTAGTGGACTTGACATCACAAGAGATGAAAACAAATAAAATTGTGGACAAAGGGTTGTATGTTCAGCATATATCTCGATCGCCGAAATGTAAATCAGAAAATCTATTTTCAGAAAATAATATTGAAAAGAGTTTGTTTAATGAATCGGTAGTATCGTGTTCGTATTATTCAACGCATAATTACAATTTATTGAACGGGGGTTTGTCAAGGTATTACATTCGAAACAATAAAATAGTAGACGTTGAACCAACATTTAATCGACTCGTAATATTTTTGGCTGATGGCAGAGTTTTAAAAAAATCTTCCTCATCAAATATTGACTTGTTAATTATTTCCGCTTTGTATAAACTTTCTTTGTAGAATTTATTTATTATTTTTCATGTTTATAGATTCATGTTTTAGATATCATGTTTATTGATTAAAAAAAAAAA
>evgR012052c0g1_i01_Hydra
AAAAAAAAAACGCTTATTTTGAATAAAAGAGAAAAATTAATTCCACCGAAAGAAAACGCAAAAGTTCTGCTACATTCGTGTTGCGCACCTTGCTCTGGTGAAGTCATTCAAGCGATGGTGGAATCAGGAATTAACCTAACAATATTCTTTTACAATCCTAATATTCATCCCAAAAAAGAATATGAGATTAGAAAAGATGAGAATGTGAAATATGCCAAAAAATTAGGTATTGATTTTATCGATGCTGATTATGATGTGCAAAATTGGTTTAAGAGAGCAAAAGGCTTAGAGCTTGAACCAGAGAGAGGAAAAAGGTGCGCAATGTGCTAAATATAAAAAAAAAAA
>evgR012056c0g1_i01_Hydra
GCCGTGGTTAGGATCGCTAAACTTCAATAAACTTTACTGGATTCTAATCATGAAATCAGATTTCGGATATTAGAAAACAAATGCAAATATTTTGTTGAAATTCATGGGTTTCTTTTATTCAATCCTCAATTTTTTACTACTGCGTTACATATCATCTACTGAAGCTCTAAATGAAAAATCAAGATGTTCGACATATTGAATGGGAGGAGCTTGATAAACGTAAATATTATGTTATTGGACCGATTATGATGGTTGGAGTCCGTTTGATTATATTCCCACCTACTCTTATCAAAACTCGCCTTCAGGTGCAGAAGCAAAATTCCCATTATAAAGGAACCTTAGATGCTTTTCGAAAAATATTTAAACATGAAGGGATTCGTGGATTTTATAAAGGTTTTTCTACAAATTTAGTTACTGTTGCATCAGGTCAAGTATATATAACAACATTTGAGGTTGTTCGCTCAAAGCTACCTAACATAGGAAATACATCAAAGAGTTTAGTTGCAGGTGTATGTGCATCACTTGCTGGCCAAACTATAACTATTCCAGTTGATATTATTTCACAGAAACAAATGATGACTGGTCAACAAGCTGATAATGCAGCAAATTTAAAACCAAAGTTTAAATCCGGCATCTCAGTTGTAAAAGATATTTACAGTACTAGTGGAGTTAAGGGATTTTATAAAGGTTATATTGTTTCCCTATTGACTTATACTCCAAGTTCAGGTTTATGGTGGGGTTCGTATTATACGTTTACGCAGTTATTTGATAAAATGACTCCCGCTTCTACTCCACATCTTGCTATACAAGGTATAAGTGGAATTTCTGCTGGAATTGTTGCTTCTACTTTAACTAATCCTGCAGATACAGTGCGTACAAGGCTTCAGGTTGAAGGAGGAAGTTCGTCCGTACGAAGTGTAATAAAAAGACTTTATGAAGAAGAGGGTTTTCGTGCACTTCACAAAGGACTAACAGCAAGATTGATTTCCTCGGTGCCGTCAAGTTGTGTATTAGCTGTTAGTTATGAACTTGTAAAACTATTTAGTTTAAAAAGTTAGTGAAC
>evgR012059c0g1_i01_Hydra
TAATTCAATTCAGTTTAAGCAATAACTTAGTAACAAACTTACAATAAAATATTAAAGGTGTCAACATCTTTATTCAAAGTGTTTCAAATGTTTATCAATGAAAACTCCAACGTTCTTGAGAAAAGTGAAAAAGTGCGCCTTGTACAAACACAAAGACCAGTTAAATATTTATATCGTATAATAAATTTTTCTCTTCCAAAACGATACGTTTTAGCAGTTATGATGTTCACGGGATATATGAATATGAACACCATACGTGTAAACCTTAATATTGCTATAATTGCCTTGGTAAACAATCATACAATATCAAAAAGTGGTTTTGACGTTTCCTTACCTCCTGAATTTAATTGGTCGTCAAGACTACAAGGTTTTATTCTTGGATCTTTTTATTACGGGGAAGTGTTTTTGCAAATTCCTGGAGGATGGTTTGCCATGCGTTTTGGAGCAACCAAATTATTTGGAGGTGCAATCTTTCTGGCATCTCTATTAACCTTGTTAACACCAGTTGCCACAAGATACAGTGTGTGGGCTTTAGTAATTCTACGTACTTTAGAAGGATTTGTTTTAGGGGTAATTTTGCCATCTAATGTTCAAATATGGAGTTTGTGGGCACCATTGCAAGAAAGATCTACTCTATTAACAATAACATTTGCTGGTATTAATGTAGCAAACGTTGTGGCAATTCCATTGAGTGGAATGTTAACAAAATATGGTTTTGATGGTGGCTGGGCATCAGTCTTTTATTGCTCTGGATTATTTGGAGTTGCTTGGTTTGTGTTATGGCTTTTTGTTATTCATGCTGACCCCAACTTACATCCGACTATTACAAATGAGGAAAGAAACTATATAAATGTAAACGTTGTTCAAAGAAAAAGTATGAAAGTCCCATGGAAAGCTATAATAACATCCAAACCTGTTTGGGCTGTTATATTTGGCAATGTTTCAAGTGATTGGGGACAGTACACATTTATTATATGTCTGCCATTATATATTATGGATATAATGCACTATGATGTGCAAACGATGAGTTTCGTTATTTCAATTTTGTTCATATTGCAAGCTATTGTTGGTCCGCTTGGTGGAGTAGCTGCTGACATACTGCGCTATAAATATTTGAGCACAAAAGCTGTGCGACAACTATTATTCGCTCTAGGTTTTTTATTTTTGCTAGGTTTTTTTATTGTATGACGTGCTTAGCCTTGCATGTAGGATATTTTATTAGGGTAGGGTGAGGGAAATATTCTTTTGAGGACTTATTTGGGTCTAATTTATATTTTCCAGTTAAAACTGGATACCACCTGTAAAAAAGTTTTTAGAACTTTGTTATAACATATAAATATATTCACATTTAAAACTTAACTCACCAATGTTGCCAAGGTCAC
>evgR012067c0g1_i01_Hydra
TGAACAAGTATCCGAAATTATTGACTTGTTAAAGACAAAGAAGAGCGCTGAAGTCTCCCAAATTCTCAACAAGAAACAACAATCACAAAGTACTGGTGGATCTTATTGGGGTGAAGACAAGGAAGCTACAGGTGATATGGAAAACTCTGGCAGAGATTTGGCTTCTCTCTTGGCTAAAGATGAAGATCATGGACCACTTGCTTGGAACAAGTCGGACCGTGATTACAAGTATCCAGAACTTTTGGAACGTATTTTCGATACACTCAAGAAGAATAACCCAGAATTGGGAGGTGGTAAGAGAAAGCCAATCAAAGTTAAAACTCCACAAGTTGCCAGAGATGGTCCAAAGAAGACTGTTCTTATTAACTTTACCGAAATTTGTGCCAATTTACATCGTGCTGAACAACACGTTTTGGATTACTTGTTTGCTGAATTGGGTACTACTGGTAACTTGGATGGTAGCAACAGACTTGTTATTCGTGGTGCTTTCAAGAGCAAGGATATTGAAAACTTATTGAGAAAGTATATTACTGAATATGTAGTTTGTAAGATTTGTAGTAGTTTGGACACCAGACTTAATCGTGACTCTACCACTCGTCTCTATTCTATTCACTGTTCTAATTGTCAAGCCTCTAGAACTGTTCAAGCAGTTAAGCAAGGTTTCGTTGCCAACCTTAGCAGAAAG
>evgR012070c0g1_i01_Hydra
TAGAGTGAAAGTTATTTAAGGCAAATTATGGTTAAAGATTTCAAAAACTTCTTTGTTGTTTTACTTTTGTAATGTACACTTATAAACAACTTGTCAATTTCAAATGGACGGAATCATCACTGAGTTTCCGAAGGCATTTACTGAGCTAGCTTCACACTGTAAACAAAACAATTTTCATCGTAAAACTAGAGAGGCTTGCAAGACTAATGTAGCTAACATTGTCAGTTTAAAGGTTGCTAAATGCGTTGTGATTGGTAATTTATCAGTGGGAAAGACAAGTCTTATAAACAGATATTGTAGAAACTTGTTTGAGAAAGACTACAAGCCGACTATCGGTGTAGAGTATGAGTTGCGCAAGTATGAAATTTTAAATCAAGAATTTCATTTACAAATGTGGGATACATCTGGAGAAGAACGCTTTAAAAGTATAACTGGTTCTTACTATAGAGGTGCCAACGTTGTAGTTGTTGTTTTTGACTTATCATCTGTTAGATCGCTGACAAGTTCAAGCGAATGGTTTGCAACTGCTATCGAAAAAACACAAAGTACAACTCCAGAGCTTTTTCTTGTTGGAAATAAAGTTGATATCATAAAAAGCGTAGATTTAGAAGAAACACGAGAGATAGCAATAAATCTTGCGAATGAACATAATGCAGAGTACTGGGAGGTATCTGCTAAAACGGGAGAAAATGTTAAAGAGCTCTTCACACGCGCTGCTGCACTTTGTTTTGATCAGGCAGTTTTACGAGAAACTGAAAATATTTCTACAAAGATACAGATGGGTGATGGCAAATTGATTGGAAAGGATAAAACAAATGATAATATAGTATTGTCACGCAACATTGAAAGCAAGGTTGAACCATCAGCTCAATTTTATGATCAAAAATCATCAAAGTGCTGCAAATAATTCAAGTCTAATTTTATTTAAAAAAAATTAATTTAATAACTTCTCATAGTAAATATTATCAAAGTATATATAAATATATAAAAAAAAAA
>evgR012072c0g1_i01_Hydra
AAACTTCTGTGTAATGTCATCAAACTTCTGTGTAATGTCATCAAACTTCTGGTTAATGTCATCAAACTTCTATGTAATATCATCAAACTTCTGTGTAATGTCATCAAAATTCTGTGTAATATTGTCAAACTTCTGTGTAATATTGTCAAACATCTGTGTTATGTCATCAAACTTCTGTATAATGTTGTCAAACTTCTGTGTAATGTCATCAAACTTCTGTGTAATGTCATCAAACTTCTGAGTAATACCATCAAACTTCTGTGTAATATTGACAAACTTCTGTGTAATGTTGTCAAACTTCTGTGTAATGTTGTCAAACTTGTAATGTAATTTAGTTGAACTAAGTCTTTTTTAAACTAAGTATTTGTTAAGCAGTTCTCACATTTATGTGTTGTTTGGCAAGTTTATACGAGACCCTATAGAAAGAATATTTCTAAAACATTGTTTTGGGGAACTTACTTTATACATAATCAACAAGTAGTTGAAAAGTATAATATTAACAAAACAAAACTACTTCTTCAGATATCTATTTAGACGATATTTTGT
>evgR012089c0g1_i01_Hydra
GATTGGAGTAGTGTCGTTCATTTGACTATTAAAGATAATTATTTGAATTATGGAGATCGTACTCCAGCTGTGTTATTTTATCCTAATAATGCAAGGTTGCAAATTTGTTCAGCTGTAAACAGCGAACTTAATTATGTTTTCCAGACAGTGCCTTTGAAGCTTGAAACATGGTCTTCAGTGAGAATATCGCAGGAGTTTGTTGATGGAAAATACATTTATTCAGTTGATCTTAATTGTACAAATATTTTAAAAATCGAAAATACTGATGCACGAGAATTCGCTGACGTTAAAGTTTATGTTTCCAATCCTTGGAATCTACCGCAAACTGGGTTTATAAAAAATCTTCGAATAAAAAATGGAAATTTTGCATTTTGGTCACAATGGAGCGATTGGTCAAGCTGTAATGGTACTTATGGTTTTAATAACAGATCAAGGGAGTGCATCATTTCTCATCCTATGAATCAGTGTTCGGGTCAAAACATTGAAACAAAAGAATGCTGTGTTCTGTGGTTGCAGTGGAATGATTGGTCTACTTGCAATGTTACTTATGGAAATGGTTTTATGAAAAGAACAAGATGTGATGTATCAAAAAAAATATGTTGTGA
>evgR012094c0g1_i01_Hydra
CTCAAATGTTATTATTGGCATTTCAAATGCGTGGAGAAGTAGAGTATCATTTCAAAACTCTAGACGAAATTCAAGCTGTATACGACGGTAGTGATTGTGGAACTTTTGTTCGTGGAGCGTTGGTGCATTGCGTTCTAAGCGTTTCCAAAGAATTGCATGCTAAGATGTGTGGATTTTCCCCGTTGGTCAAAAGAGACAAAATTAAGTTGGAAGAAATGTCTGCGTTTGAAATTTTAGACAAACGAGTACCTTCCAAAAATGGTAATTATACTCTACGAGTAAATAACAACGAAAAGCTTTTGATGAAGGTTGGAGCACACGAA
>evgR012101c0g1_i01_Hydra
GGACGTTCTTGTTGCTTACATCTTGTGGTTCTTCTTGGGTGCTTTTGGCATTCACCGTTTCTACTTGGATTCACCAGTGATGGGTATTATTTACTTCCTGACAGCAGGTTTGTGTGGTATTGGTTGGCTTATTGATGCTCTCTTGATTCCATCCATGGTTGAAAATTGTAACAGATCCTTCTCTTCTGTTCATCACACCACAACCACCACCTATGTTACAACCCCAGTTGTCCAACAACCAGTTTCATACGGTTCTCAACCACACTACAACCAACAATCAAACATCTACCAACAACCAACTACCTACGTTTAAAAACTTTGTAAAATTA
>evgR012125c0g1_i01_Hydra
CTGAATCGATCATTTTATCAAAGTGTGAAAAACATCTAAAAATACATTTTCAAATAATAACTTTTTTAAATCATGTTTTATGGATGGTGCTTTTTACTTACCGTACTTTTTTATTTTATTTCACAAGAGAACGTTTCGACAAATGTCACAGTGGAATTTCTACTTCGTTTTTATAACAACACAAAACATCAAAGATTTACAAATAAAGGTCTTACTAATTGTGATACTGCTGATAAGTCAGGTACTTGTGACGTACAGTTTGGAATATGTTTCAAACTTGGCAATATATCAGATAATTGTGAAGATAAAATTACAACAAAAGAGTATGATTCAATTACAACTATAAACTTTGAAGATAGAAATGTTAAAGACAAGGATAAGCTATTTGTTCGTAATTTTACTGAACCTTACGGAGGAATTGTTTCAGTATTTGTGTTAGCATGGGATGTTGATGATACTAAAAATGAATCAATAGATGAATATAATAGTGTGTTAAATATCGCTGCATCACCTAACATAACACTAGCTAATCCTTACAATGTTGATCAGTCAGGGATATTTAAATCAGGTTCTAAATTACTGTACATTGTTAAAGTATATTGCTCTCTAAATTACTATGGAAATGATTGCACAAGATATTGTAACGAAGGTTCAGAAATATGT
>evgR012128c0g2_i01_Hydra
TTATAAAAATCCACAAGAGAAACACAAACAAAAAAGCAAGTGATATATTATTTAAAATAGCACATTTTATAATACCAACAAACGAAAAAATGTTCAATAGGAAACTAAGAGAAGACAAAACATGCCCGCAATGCGGGACGCAAGACGAAAATATAGCACATATGATGTACAGCTGCAGTAAAGTGCAACCTTTAGTACTTTTTATTTTAGACATACTTGATCAACTTTATCCCACTGAATACCCCCATCATAATACTATTAAATTTATATTATGTGGATATAGTGTAAAAGAACGAGATAGAGAGTTCGAAAATGTTTTATTGGAAACTCTCATTCTAGAAATATTTTT
>evgR012134c0g1_i01_Hydra
TATATAAATGGTTTTAAACTAAATACATCAGATTCAAAAAGGGACCTAAGAGTGATTTTTTTTTGGAAAAATCAAGAATTATCGAGTGCAAGCAAAGCTAATCGCATGTTGGGCATAATAAAGAAATCTTTTGTTCGATTTGATGCTGAACTGCTCAAATCTCTATATCTTTCATTTGTTCGACCAATAATTGAGTTCGCTATACCGGTATGGACTCCTTATAAAAAGCAAGATATCATCATTTTAGAAAAAGTACTGCGTCATACCACTAAACTAATACCTCAAACCAGTAAATTATGCTATGAAGATCGACTTGCTTATCTTAAAATACCCTCACTTATTAAGAAAAGACAATTATGAGACATTATACAACTGTATAGAATTTTTAAAGATGCTTTAAAAGATTTTAATAAGCCAATTTTAACATCCTCCTCCACAAAACCTAAAACAG
>evgR012147c0g1_i01_Hydra
GTACAGAAATTAAAATATGTTCTACCGACATTGCATGTGAAGCCTCGTGGAATGAATGGAATGCTTGGAGTTCTTGCAACGCTTTATGCGGAAATGGTTTCATGAATGGAACACGATCGTGTTACAGAAAAAATGTTATATCGGAGCAGTGTATTGGAAACAGTACAGAAATTAAAATATGTTCTACCGACATTGCATGTGAAGCCTCGTGGAATGAATGGAATGCTTGGAGTTCTTGCAACGCTTTATGCGGAAATGGTTTCATGAATGGAACACGATCGTGTTACAGAAAAAATGTTATATCGGAGCAGTGTATTGGAAACAGTACAGAAATTAAAATATGTTCTACCGACA
>evgR012152c0g1_i01_Hydra
GTGAAAATTAGATAGCGTTTTGGTTACCCGAGATTACGCCTTTTACGTATTTTATCAATAATTAGAGAAAAATTCGAAGTGTAAGAAAAAGTAAAAATATGGGAATCCTAGCAAGTAAAGGCGGAGAGTATAGTAAATGTCTCTATTTTGATCCAGACGTAGAACAAGCCTTAGCTACAAAGTATATGATTAGCGAGTCAGACCTAGTAGACAAGATTCGAAACTTAATAGATTCTCGCGAAGAAATAAAAATGGTTACTATTTATAAAAATTATTTGTTTGAGATTCAATTTACCCAATACTTTTTATATCACTCTTTTGTCGTTTTTGAAACACAAAATTGGTTTTGGTCGATTGAAAAGAATAATGAAGGTATAACAGTACAACGTTCTAAATGTTTTGAAAAAGTTGCTTACTATTATAGGAGAAAGTTAAGAGTGAGCGACATTTGTTGGATTGAAGAAATCAAAACCTCGCCAGCCTCGGCAAGATTTGGGACAGTTCAAGATTTATTTGATTTAATCTACACAAAAAAATTATTATCTTCTCAATATCATTATTTGACAGATAACTGTCAGCAATTTGCTTCAAAAATATTTGAATTTGCTATTAGTGGTGAGTTGTCTGAAGAATCATAGAATCTACGATAGATATTATTAGAAGTAACAATATAGATTATTAGAGATATATGATATAGATTATTATTAAAAATATTCTGAAACTATGTTTCCATGGATATTTACAAACTGGAC
>evgR012153c0g1_i01_Hydra
GTTGTATTAATAATGGATCTATAGATTTCCAATTAACAGCTAGTGGTTACCATGTTGCAACCAACAGCTCATGTGTCAGTGTTTGCAATTCTTATAGTTCACAAGTATTTCGTGAAGCTTTTTTTCTTGAAGGGCAGCTTTGTTTTTGCAGGAATGCTTCGCTTCCACAGTTAGGCAAATTACCAGATAGTCCTTGTACCATAAGTACTTGTTTAAGCAACAATAGCTCTTTAGATTGCAGTTTAAGCAGCTACTCGGTTTTGCCTCTTTCACTTGGTATTATTGAATGGAAAATACAATATCCTACTGTTGTTTTAGTTGATACTCAACTTTTAGTTGAAACAATAATAAAAGGTGATAATTCAAGTGTATTTGTCATACCACTAACTAATGACTTACCATTGTTATTTCAAAATATTGTTAACCGTATTTCATTTAAAACACAGTTTTATTTACCTGGTTTACAAAAATGGTCTCAAACAATTGGAAATAGCTATGCTGATCCATTTAATGCAACTAGTTGGTTACAAGTAGATGAGTATATTGGTG
>evgR012155c0g2_i01_Hydra
TGGAAATCCCCTTAGGAAATTTAAACTGGTTTTTCTTGGAGAACAAAGTGATGTTCGCCTATAAATCTTACGACGAATGTCAACGTTTGTCAAAAAAAATCGATTTGTCGCGCCCTGCAATGTACAATCAAATCATCCACCTTACCTTACCTTCAAAACTAAATGAAGTTGTTGAACCAAACATAATCAACTATAATAATGAAAATACTAACATTCTGGAATTCGAAAACCCAAGCCAAAGCAAAAATGTTATACCATTAAGAAGATCAACTAGAACTTGTAACGTGCCAGTTAGATTGAAGGATTATATATTATTAAAAAAAAAAAAGCTTGTATAATATTTGATTACGTATTATATTGTTACGCATTGTAGATAATTAAAGTCATTATTCTATTAACTGTTAAGGACTTATACTGTGCACTAATATTCC
>evgR012156c0g1_i01_Hydra
GACCTGACATACGTATACATTGTATCACTCATTGTTTTTTTTGTAACTTTAAAGTTCATTAAACTATTGCGATTTAATCGCCGTGTTTCAATGCTGTCGTCAACATTAAAAGCTGCTTGGCATCCTCTTTCTATGTTTGGAATAACATTTTTCATAATTATGTGCTCGATTGTGTCAACATCTAGCATTATGTTTGGTTCATTGTTGGATGGTTACCAGAATTACTTTAAAACAGTAGCTTCTATAATTTCATTGTTATTAGGAAAGTTTAGTTATAGTCAGTTTGAAAATGCCAATAATATTCTTGGACCAATCTTTTTTTTTGGATTTAACATCATTATTATTTGGATTGTTATGAACATTTTTATGTCAATATTGATGAATGCATTCTGTGCAGTTCGTGCAAACCTCCAGTTCCAGACAAATGATTACGAAATGGTTGATTTTATATTGAATCATTTGAAAGATTGGCTTCGATTTAGTTTTTCAAAGGAAAGGACCGATATTGGGCTAAATAAAGTAGCTACTAGATCTAATCATCTTTTGCTTTCTTCCCCTGGAAAGACAAAACTAACTCAGTTGGACGCTGAATCCAACAAAGGCCTAAATGATGTACAACCTCAGCTAAAGGTAGAGAAACTAATACCGAAAAATCAAATTGGGGAGAGTTGGAAATCTAATGTCAATGATATTAAACTTAGTTCACTTAAAAAGTCAAAACTGAATGAAACATTCGCTGTAGAGAGCGCAATCGATGATCAAATTTCGCATCTTGAAAAATACAACCTTTTATTAATAGAAGATAATATTAACGACATAAACAGTTTGGATAAGTTTATCAGGTGTGCCAGTGTTGTCTACGAAGAAAATTTGAAAAGAAACGAAATATTAAATTAGATGTCATGTTTTTTTTATTGCATTTGTTATTGTTCAAATTAATTAAAGTTTAATAGTTAACTTTATAAGTTTTGTAGACTGTCATAGTCTTCATAGTTAATTTAAAAAAATGTTTTCGATTGAAATGATTGATAATCTTATATAATACCGGTGACGGATTCAGAAGAGGGGGC
>evgR012158c0g3_i01_Hydra
CCATAATAAAAAACACGACTATGGGATTATTAACCCTACCTAAAGTGTATGAAGTTTCCTTTGAAGTTTATTTAAGTTCAGTTTTCAGCATAACATCTTGGACAAATATTATTCGTATAACAAATGGAAATGATTGCTGTGATTACGGATCAAGGATCGCAACTGTGTATTACTATAGTCCATCTAAACTGGAATTTGATGCACCAGTAAACGGAAATGGTGCCTACTATTTCCGTGATGAAAATCTTATAAACTTGCAGGAATGGACAAAAATTACAATGAAACAGTTTTTAAGCAATAAAAACTACTTTTATGCAATTGAAATTAATGGAAAGTCCGTTCATTCATTGCAAAATAATGAC
>evgR012158c0g4_i01_Hydra
AAAATAAATATTTGCAACTTAATTCATGGTTAACTTTATTTTCTACTTGTTATATCCGACAAACGCTTCAGATGCATTATTTTGCCTTACACAAGCCAATCCCCATAATAAAAAACACGAATATGGGATCATTAACCCTACCTAAAGTGTATGAAGTTTCCTTTGAAGTTTATTTAAGTTCAGTTTTCAGCGTAACATCTTGGACAAACATAATTTGTATAACAAATGGAAATAAGTGTTGTGAATACGGATCAAGGATTGCAACTGTGTATTTCTATGGTCCATCTAAAATTGAATTTAATGCACCAATAAACGGAAGTGGTGCCTACTATTTCAGAGATGAAAATCTTATAAAATTGCAGGAATGGACAAAAATTACAATGAAACAGTTTTTAAGTGATGAAAACTACTTTTATGCGATTGAAATTGAT
>evgR012169c0g1_i01_Hydra
CATCAGACAATATGAGCACTGAATGGATTCCAGTCTCCCAAGACGGTCAAGTTAAGAAGAAGATTCTTACTGCAGGTTCAAGCAATGAACAACCACCACACGGATCAAAGGTTTCTGTTCACTATGTTGGTACTTTGACCAGTGGTAAGAAATTCGATTCTTCTCGTGATAGAGGTACTCCATTCTCCTTCGGTTTGGGCTTGGGACAGGTTATCAGAGGTTGGGATCTTGGTGTTAGAACCATGAAGAAGGGTGAAAAGGCCATCCTCGAAATTCCAGCAGAATATGGTTACGGTAATCAAGCTATTCCAGGCTTGATCCCAGCCAACTCTACCCTCATCTTTGAAGTTGAATTGTTGTCATGGAAGTAAATATATTTTCAACTTGAATGATAAACACTCATGCTAGGAGAAAATAAATAATATTAATAAGAAAAAAAAA
>evgR012171c0g2_i01_Hydra
TTTTATTTAAAAAAATTTAATTTAATAAAAAACTCTTTTTTATTTAATTTATATAATACAATGTCTTTTCAGTACTTAATACTACTTAGTATTTGGATCAATACTTTTAGTATTAGGACCAATGTACTTATTGTTGGTCCAGGATGGCTTAACCATGATGGTACAGTGCATCCAGTAGCTGTATTACGCAGATGTTTAGAAATCATTTCAGAAGGTGTCTCTATTGTTTTAGTATTCATTAAGGATCATAGAGAACGGAGAACATCTGATAATACTTTTTCCCAAGCTTTAATAAGTAAATTGTGTTGCCTGGTAGCAAGCTCAATAGATATAGATGCCATTTA
>evgR012194c0g2_i01_Hydra
TGATACTATTCTTTGTAGGAAATGGTTGCGCTCCATTCTTGATCTTTAAATTTCTTATATTGGGATTTAGAAGGGATCGTTGCAAATTCAACAAAAGAACACAACAATTGAAATGGCTGGCCAACAATTTGGACAGAAACAGACACCGCTGGTACTATTTTGAGATGCATTTTAACACCATCATGTATCTCAACATACAACCCCACACACGGCACAATAGTGATAATGATGGCTTGCTTGCTCTTTGCTCCTTCGAAAGATACCGTACCTGCCCACACGAGCGCAGGATCACCCTGGCTCTTGCCCTTCTCGCAGGGTTATACTTTAGTCAGACTATGCTTAGAAAGGTGTAGACATACAGGCGTTAATAGTATTTATGTGTATCTACTTCCTCTATATAAACTTTAATTTTTAAATGGACATCTGTCCAATTTTGATGATCTATAGCTA
>evgR012198c0g1_i01_Hydra
AGTTTGGATTTCAAAGTAGGTGTTCTACTGAACATGCTATTATTAAACTTGCTGACAAAATATTTAAGTCGTTTGATAGTGATGAGCTGGTATTAGGAGTGTTTGTTGATCTTTCCAAGGCCTTTGATACGGTTGATCACAGTATTTTACTTTCCAAGCTAAAATATTACGGTATAATAGGCTCAACATTTAAGTGGATTCAAAGCTATTTGTCTAACAGAAAGCAATTTGTATTACTAAAAATGTCAGGAGCCCTCGATATTACGAGTGGAGTTCCTCAGGGTTCAATCCTAGGTCCATTTTTCTTCTTAACTTATTTTAACAATATAGATAAAAAAAAATAGAAATAAAAAATAGAAAAAAAATAAAAAAAAA
>evgR012212c0g1_i01_Hydra
ATACAGGTGTTGAGTTTAGCGAAAACCAAAATGTTGAATTTGAGACAGCCACTAACTCTATTCAATACCCATCAATGATTGGTGTCATAGAGGTTGAAACCAGTGATAATTTTAATAAATCATTAATAATTTTTGAAAACCAATTCATTTATACAATATTAGATATTTTACTAGGTGGAGGTGATTATAATTTTAAACTAAAGGTACAAGCACGACCATTTACACAAATTGAAAAAGGAGTGATAGAGGACTTTACCAAAATCATACTAAGTTCAATAAATCACTCTTTTGGTAAGGTAAAAGAGCA
>evgR012220c0g1_i01_Hydra
TATTTTAGTGCAAAAAGTATAAATGTCAATGGCAACATTCGACAACATTTCAACAGCTGATTTTGAAGACTCTGTAGAAATTCATTTGCATAATAAAAACAAAGAATTAGAAGTTGCTAAGGAAATATCTTTGGAGACAGGTATTGGAAAAGTAAAAAAGTCTGTTCATGAGTTTTATTCACAATCAGTACAAACTAAAAAGGTGACAGTATTTCGAGATCAAGCTGAAGTAAATCGGTTAATTAGAGTAGAAGCTGAAACAGGTGATGTTGAAGTTGTTATTCGACAATTACCATGTGTGGTTGACAAAGATTCTGTTAGAGTTGCCTGTTCAAATAATGAAGTCACAATTGTTGAAGTTTCATTTAAAACAAAGCTTATTATTGAAAATGGCACGAGTGAAGGCGAAGCAAAGCAAAATGTTGAAGATGAGTCAGAGAAAATTAATAAAAGATTAGCTGCTTTAAAGAAAGAAAGAAATATTCTATCTGAGAAATTAAAAAGAAATGCTAAAGAAGCTTTGTTAATCGAAAGTTATGCAAACTGTATATCTGAGAAAAAAGGAGATAAGGTAGCAGAAATCTTAGATAAAAAAAATATTGAAAGTATGCTGCACTTTCTATCTATTTATAAAGATCGAAGCGAAAAAATATCTGATTCAACAATTTTATTGAATGAAAATATGGAATCAATTAACAAAGAAATTGATGCTCTAACAGATAATCTCA
>evgR012224c0g1_i01_Hydra
GAAAAATTACCCAGAAAAATGCAATATTCCATACCTAAAGTATATGACAACTTTTTACAGTACTTGAGATGAAAACTTTTTTCGGTATAGTTATTTTAATGGGTATCATTCATAAACCAAATTTGAATATGTATTGGTCCACCGATGATCTGTATTGGACACCAATTTTTTCCAGAACCATGTGCCGAGACCGGTTTAAGGTTATCCTTAAATTCAATATCCTGCATTTCAACAACAATAATGACCTCCATCATGATCTCAATGATCCTAACAGAGACAGATTGCACAAGGTCAGACCATTTCTCAACATGGTCCGAGAGCGGTGTCAAAAAGTGTATAATCCAGGACAACATTTAAGTGTTGATGAATCTCTTGTGCTCTTTAAAGGTAGGTTGCATTTTCACCAATATATTAAAACTAAAAGAGCAAGATTTGGAATAAAGTTGTATGAGTTAACTACAGCTGATGGTATTACATTAGATTTTTTAGTTTATTGTGGCACTGGGATGTATGATGATGACAAAAATTTTGCAATGCCTTCAAGTCAGCGCATACCTGTAGAGCTGATGGATCCATATCTAAATAAAGGGCATATAGTGTTCACTGACAATTACTATACAAGTCCTTCTCTTGCAAGCTATCTTTTGTCCAAGAGAACTTATATCTGCGGAACCATTCGTGTAAATAGAAAATTCTACAGCAAGGAGATATTAAATGAGCAACTTGCCAAAGGGACTGCAGTTTTCTACAAAACTAAAAATGAAAACAACAAAATGTTAGCCTGCAAATACCGTGCAAATAAAGACAAATCAGGTAACAAGCAGAAAGTTGTTTACATGCTGTCAACTTACCACAATCCAGTGACTGTGAACACTGGAAGAGCTGATAAAGATGGTAACATCATACTTAGACCATCAGTGATTAAGTCATATAATATGCACATGGGTGGTGTTGACCGGGTTGATCAACAACTACATGGAGTTCAAGTCCTACGGAAAACCTACAAGTGGTATAAGAAATTAGCTTTTCGCCTAAATCTCCAGTGCTCTTTAAACTCACAAAAGATTTATACACGTAGTACTGGTAGCAATATCTCGTACATGGATTTTTTGATGAGTAACGTTAAGTTGACATTTTCATTGACCCCTAAGGTTCCTCACAACCCTTGTGTTGTTATTGGAGAGAATGTGGATCGACTTACTGGCAGACACTTTGCTTCTATCTTACCACCAAGTAAGGTAGACAGTAATGATAGGCCACACAAAAGATGTCGTGTATGTTATGCAAATGGTAAAGTGACAGCCAAAGGACGTCCCTTGAAAACAGTTTATTTTTGTAGCTACTGCCCGTCAAAGCCAGGATTACATCCAGATGAGTGTTTTAAAATATATCATACTATGTTGGACTATTCTTAAATAAACTAAACTTAATAGTATGTATTCATATACATAAACTGCCTTAAAAAGTAATAATGTGTGTTCTTTTTATTTTGGGGAAAATTTATCATTTTTTGGTGCCTATACACAAAATTGAATAA
>evgR012229c0g1_i01_Hydra
AGACCATGACACCTTTCAGCATAAGATGAAAACGTTAAAAGAAGTTATCTTTCTCGTTTTTCTGTTTATCTTTTTTTTTAATACGTTTGTGTTTTTGACTGCTTCACTTTTTGGTGACAGTTGGTGGATATCTAACTCAAATATAAATACTCAAGTTGGCTTATTTAGAGTATGCATAACAATATTTTTGCAAACATCTCCTTCATTGAGCTGTGATATCCGCGAGAATGCATTGCAACTCGATAAAGATTATAAAGATTTTGACGTTGCAGCCGTACTAAATATTGTTGCTCTTATAATCTGTGTTTGTGGAACTATAACGCTAATGATTACAGTGTTTTTCCCCGTTGTTGTATGGCTTCCGTATCAAATTTTTATTCTTTCAATTTTATGCAATATTATTACGTTAACCATGATGATTACTGTCGAGGTCAAAATTAACTCCAAAATCGATGATTACGTAGGTCAGTCTATGGTAAGAGGATGGACTTTTAAAATTGCGTACTGTGCACTTATTGCTGGTTTTATACTAGCAATTTCTTCTGGGATTTTTTTGAAATTGAAGTTGCAGCAAAAGTTAAAAAGTGGATTTTTAATTGAACATTTAGTACCCAATGATGTGTCAAAAATATTTGCCGAGGAACTTCTAAATGCAAAAACGGAATACAGTTCTCACTTTTTGAAAACCACGTCGAACTTATCAAGCACTGGTCAGTTGACATACTTTGCGGACTACTCTAACTTTCTTACGATCCCACAGCAGCCTGAAAAGATGCGAGCTGTTTATAGAGGTTCTAACTATTTTTCAGAAAGGATGATAGAGCTCAGTGCTGATTTGCATGAGAATAGTGTGCCTTAGCTATATATCAATAATTTACTTTTTGAAATTTAAAATAAGAATAAAAAAAATAACTACTCTGAAAAAAAAAA
>evgR012231c0g2_i01_Hydra
CTAGTTTATGTAAAAATTGTTTGGTTACATTTAATATGGAATCTGATTCAACTGATAACTATGGATATGAGTATTTGGATCATCCTGCTGATGTCCAGATTCATTCATGGGGCCTAACATTGAAGCAAGCCTTTGAGCAACAAGCAATCGGAATGTTTGGGATTATGACAGATTTAAACACTATTGAAAATTTACAGGAAGAAGTTATCAAAGCTGAAGGTCATGATTTAGTGTCTCTTCTTTATAAGTTTTTAGACGAATGCTTATTTACATTTAGTGTTGAACCATTTTTATGCGCATATGATGTTAATATAACAGAGTTTGATGAAGAAAATTTTGTTATTAAGGCTGTTCTTAAAGGAGAGACATTTGATTTGTCTAAACATCCACAAGGCACAGAAGTTAAAGCTATTACTTTTTCTAACATGCAAATAAATAAAGAAAGTACACATTGGGAATGCTATGTTATCTTAGATATATAAAAATAAAGTACACACTAGAAATATTATGTTATCTTATATATATATAAAAAAGTACACACTGGGAATGCTATGTTATCTTAGATATATAA
>evgR012242c0g1_i02_Hydra
CCTGTTCCTCAACAACCAATTAGACCAACTCAACCTACCCAACAACCAATTCAACCTGTTCTTCAACAACCACAACTTCAACCACAACCTGTTCCTCAAAAGCCAATCAATCAACCTATGCAACCACAAATGCAACCTCAAATTCAACCACAACAACCAATTAACCATCCATATTATAACCCAGTTCAACCAGTTCAATCTCCATTCCAACCACAATTTCAACCAATGGCAATGAATCAACCACCAATGTCAATGAATCAACCATCAATGGTAATGAATCAACCTATGCCTTCACTTCGTGGATCTCCATACACAACCCCAACAAAAACTTTGAATAGCAGCGTTACCATGCAACAACCACACCCTCTGAACTCTAGTTTTAGTTCACAACCAACAGTTCAACAACAACCACAACAACCACAACAACAACAACAACAATAAAAACTACAACAACAGCAACAACAACAACAATAACAACCACAACAAC
>evgR012258c0g1_i01_Hydra
AGGTGTAAAATTAGAAAAATTAACTTTATATTTTAAACACATAAAGAGCATGGATATATGTAAAAAAGCTGTTATAATTCCAGGAAATGGAGCTGGCGATGTATTTTACTGTAATTGGTATGGTTGGTTAAACAAGAAGCTTAATAGCATTGACGAGTTTTCCTGTGAGTTGCGTAATATGCCTGATCCAGTATATGCAAGAGAATCACAATGGATCAATTTCATGCATGACAACCTTTTTATTGATGAACATACGCTTATTATTGGTCACAGTAGTGGTGCTTGTGCTGCTGTTCGCTATGCAGAAAAGTATAAAGTTTATGGAGTTGTTCTTGTCAGTGCTTATGCTTCTGATCTTGGAGATAGTATAGAAAAACAGAGTGGATATTTTAACAGACCCTGGCAATGGGACATAGTGAAGGCAAATACAAAATGGATTTGTCAGTTTGGTTCAACAGATGATCCGTATTTACCGTGGAATGAGCAAAAATTAGTTGCAGAAAGCTTAAACGCAGATTTGCACAAGTACAATGATATGGGTCATTTTAATCAATCAACTTGTATGGAAATTTTTGAAGTTATTACAAAACATGTTTTAAGTACTAATCAGAAAAGTATATGAAATCGAGTTTCTTTTAGTTTGTAGCTATGTACTTGTTTACTATGTAATACATGCAGTTGCATGTGTGAATATGTTAGCATGGAATAATCAGTAGCCATTTTCAATTATTAGATTGCAAACATGGTAAAACAGTCAAATATACAATTTTTTAATAAAAATTGATTCATTTTTACCTCTTCACTCTTCACACCAAATATTTGCAGAAAGGAAATTAAATCAATATTTTTGTTTGAAACATGTTTTTTCAGTGAATTGTATTACATAAAAATGTTTGTTTGTTGTGTTTTTTTACATGTAAATCTCACGTTAATTGCTAATTAAAAACATAAGCACAATCAGCATATAAGTTTATCGTAAGCATGCCATTTGCATAATAACACTATATATGATTTTGCAAATATGAGAGAAATATATAAATTGCAGTCG
>evgR012262c0g1_i01_Hydra
TATAACAAGTTTCGCTCAAAATGGCTGATTTAATTGAAAAAACAAAAAGAAAAAACTTTAAATGGGATTACAATATGGTTGAAAACCTTATAAATTGTGCTATCAGCTATAAAGCAAGGATGCTTTATAAAAACCTTGATTTTGATGGTGATAAAGTTAGTTTATATAGGGAAGTTCGCGTATCTATGGCAGAGCTCTATAATAGAGACGAGGAATTATATTTTGGGCCTGTTGTCATCGATACTTTAAATGAAAAGGACTCAAAAGAAGCAATAAAACGAGGGATTAAACGTGTTCAAGAAAAATTAAAAGAAATCCGGCAAAAATTTTCGAAAGCAGTTGTGAGCGGCAGTCGAAGTGGAAGTGGTAAAATTATATTAGAGTTTTACGACCAGCTCATCTCCTTGTGGGGTGGTTCAGCCAATATTGAACCACTTACATTTGGTGTTAAAGGGGATGACTTTATTGAAACCGAAGAAACTCATTTTTCAATTGATCGAGACTTATCGGAAGAGGTTGAGTCAGTCATCGGCAGTGATACGGCAGATAGCATTTGTAACCCTAAAAAAAGAAAAAGTTGTATACCACAACTCATTGACAACAAAAGAAAGCACTTGGAACGAAATTTAAGTGCCACCCAACGCGAC
>evgR012267c0g1_i01_Hydra
AATCGAATATAATGAAAAGCAAACGAAACGTACGTAGTAGAAACGTACGTAGAGAAACTCTATTTTGGGATATTATAGACGAAGAAGGACGCGAGCCAATTATTGAATGGTCATTGCAAAGAAGATTAAAAAATCTGAAACTAAAGAGATTAATTTATCGCAACGAATTTTTAGCATACACCTTTACCTGGAGAACAATTAAAGCGTTTTTAAGACACTTCAACTGTGTGATGTTAATTCAGATGACACTGACAATCGGAGCCGTATACGTTTCAACGCGATATAATTTTTCGTTTGATATATATGTCCATTTATTTGTATCACCTATTGTATTTCCGCTAGCTTTTGCTATCAATGCAGATTTTCAAAGAAGGGAAAAAGCTTTAGACGACTTAGCAAACTTCAAAGCAGCCGTAATGACGTGGTTTTTTTGCATGAGGGATTGGAGAGAGGGGGCACATTTGGAAAAACTATGGATTCAAACTGTACACTTTAATCTAAAAAGTATTTTGTACTATACTCGAAAATACCTAATGACTAAATGGACAGAACGAAGAATAATGATTCTTAATACAATATATGAAAATTTTAGCGATTCTTCCCAGTTAATTGAAATGGTTAGAAAAAGTGACTTGCCAGCAAATACTGCTCTCGTTGCACGGGTACATCTTGTCATAAATTCAATGTATTATGCATTTGAACGCCTTCGAATTATTCGTGAATATCGGTCACCAAGATCGATCCGGTCATTCAATAAAATCTTTATATTCTTTCTCCCAGCTCTTATTTCACCTTACTTTGTACACTTAGGTAAAAAAACTAAAAACGAATGGTCGCCTTATTACATTTCAGTACTAATAGGCTTTGTTTTTTCTGCTTTGCAAAGTGTTCACGATAAACTTGATGACCCTTTCGACGGAGTGAGTGAAGATGATATTAATTTATCTTCAATAAATGAATGGATATATCAGTCAATTGATGTTAATTTGGAAA
>evgR012270c0g1_i01_Hydra
GTTATACAACAGTAGCAATTATATTTCTTAATATTTTTGCTGGAATATCAGTTTTTTTTAATGCTGCAAGCATTTGGTCAATATTAAAGTCAAAATCCTTATCAAAAAGACCATCACTGTATCTTATAGTGAATTTGCTGAGTGTGCATTTTATCCAGGGATTAGCAACAATACCACTTTATACTGGTAAAAAAGCCAATGTTTCAAATTCAAAGTTAAATTTATTTTTTTGCAATGGATCAAGATTTACTAGTGTACTAACATTCTATGGTGCAGTGTATAGCGTTTTACTTGTTACTGTAGACAGACTTCTAGCCACGGTTATTGTAATGAAATACAAGACTTATATAACAAGAAAGAAAATTTGTTTTATTATAGCACTTCTTTGGGTATACTTATTAATACTATCAATGATACCATTTTTCAATTCAGCTGTCACATTGTCAGAAAAGACAAAAAATATTAGCAACAACTTTTTTTAC
>evgR012273c0g1_i01_Hydra
CGCCAATACCTTTGAAAGACCTAAATTCTCTCCATACGAAGACAACGGAGGTACCACACTTGCTATTCCTGGTGAAGATTATGTGATTGTTGCTGCTGATACCAGAATGAGTTTAGGATATTCAATTCAAACCCGTAAATCAACTAAATTATGTAAATTGACTGATAAGTGTGTTATTGCATCTGCAGGTATGCAATCAGATGCTGCCACATTACACAAAGTGTTAAAGATGAGACTTGTAGAATATGAACATCAACATGGAAAGCCAATGAGTGCTGTAGCTGTTGGACAAATGCTTTCAAACATGTTATACAATCGTAGATTTTTCCCATACTATACATTTAACCTTGTAGCTGGTTTGGATAATGAAGGAAAAGGAGTTGTTTATGGATATGATGCTATCGGATCCTATCAATCTGTTGAAGTTGGCAGTCAAGGTTCAGGTCAAACCCTTATCCAACCTCTCCTCGATAACCAAGTTGCTTTCAAGAATCATCTTTTGGTTCCAGAAAAAAAACTCTCTCTCGAACAAAGTTTAAGTATTGTCAAGGATGCCTTCACGAGTGCAGGTGAAAGAGATATTTACACTGGTGACTCTGTTGAAATTATGATCATTACAAAGGAGGGTATTAAGGTTGAAATTTTGGAATTAAAGAAGGATTAATTCCTCTCTTTCGTTGTATTAAATCTATAAA
>evgR012274c0g1_i01_Hydra
ATTTTAGGATTGGCTCAAGCCAAAAAGAAAGCAAGTGCTGAAGGAGAGGCCGTTGAACCAGAAAATAAAAAACAAGCAGCATCCAAAATGGAAAGACAACAAGTGCCTGTAATGGAAGGGGAAATGTATGACGAAAGAAGAGATCCCGAATTTAATCCAGCCAGTGAAAGGGAATTCATTGAAACGGAAGAACCGTCTCGGAACCCTAAACAAGAGGAGGAGGAAGCCAATAGAGAAATTCAAGAACAAAGCCGCCAAATGGGAATCGACCCAAGTAAAGCTGGTCATGAAACTCGAACTGGTCTAGACTATCCGACTAGTCAGGATCGAATCACTAAACACAAAGACGAGTATATTGAAAAAGGAGAGGAAGACATTCCTTCGGTTGAGCCCGGAGTTGGAAAAAATCCTGAGCAGGTGGCTAAAGGAAGAAAAGCGGCTGCTGGTACTACTCACGAAGAGTTCGTTGAGCGAGGCAAAAAAGGAGCTGAAGCACGATGGGGAAAACAGCAAGGATCTCTTGCTCGAAATCAATAACCCTCTGAAAACAGAAAACTAATTACTCTTCTTGAGTTAATCGACCTCGATTTCTTAATATATATGTAGCTGATCAAAATGTTGTACCTGTGCAG
>evgR012279c0g1_i01_Hydra
GAATAATTCTGTTTACTTGTTTACTTTTATTTGTTTATAATTCCACAGCAACAAAATTAAACCTTCGTCCTGTTGTTGGGGTTATTTCTCAAAGAACAAACAGAGAGCTTGAAATTATCCTCGGATATAACACTACATATATTGCAGCTTCGTACGTAAAATTTTTAGAAATGGCTGGTGCACAAGTAGTTCCTATAGTGTCATCTTGGAATAAAATGCAAATAAAACGTGTTATGAAAAAGGTTAATGGTGTACTTTTTCCTGGGGGAGCAGCTCCATTTAACGAATCAAACTACTGGAATGCTGCAAAAATAGCATTTGAAGTAGCAAAAGACTTAAACCACAAAGGTATTTATTACCCAATTTTTGGAATATGCCTAGGTTTTGAGGTATTACACGAGTTAGCAGCAGGCGAGAAACTGCTAACTAGATACGATGCCGAAAACTATTCCATTCCAGTTAACTTTACTAAAATAGCTTTCCATAGTCGCTTGTATGCAGATATGGGCAAAGAATTGATAAAAAGTCTTCGTCATGATAATATAACTATAAACATGCACGAGCAGGGGATCAGTCCAAATGTATATACCAAGAACATAAAAATAAAAAAAATGTTTCGAGTGCTATCAACCAACTTTGACAGAAGTGGAAAAGAATTTGTTTCATCAATAGAAGGAATAAAGTATCCTTTTTATGGAACGCAATGGCATCCAGAAAAAAACATTTTCGAATGGACCAACGCCGAAAATATTAATCATTCACCAGTTGCTATCCAAGCGTCTCAACACATAGCAAATTTTTTTGTTAACGAAGCAAGGAAAAACACAAATCATTTTCCATCTACAAGTGATTTACAAAGACATTTAATTTTTCATTATCAAATTATTTGGTCAGGCGGTTTGACAAACTTTGAGCAAATTTATATATTTAGAGATGAAAGTGCTTTAAAAAGAAAGTCTGTGCATAAGTACATAAACTAAAATGTTACATTCCAATCATTTCTTTCGTGATCTTCCCTTTTAATTTTTTAAATCGTTTTTGAGATCATTTTTGAGATTGTCTGTGTTATTACTAATCTTAA
>evgR012291c0g1_i01_Hydra
GTTTTACATGATTGGATTTTTTTTTTTTTTCTCGAGAAATAACTTTTTGTGCTCACCATCACTTATACTTGCTCAAAATCAAAACAATGCAACTGAAGCCATAGTAATGATATTTGAACCGGATGATGTTGATTGGTTTTCAAATCTTCTTGTTGAATTAGAAATGCCGTGCAAAGAAACATCAATTGGATCAGCTGATATAACCTTGTGTATCATTTGTAAAGAAGATTCTAAAAATTATTTGTTGCTATCCTCTGAGAATCATTTTACAGCACAAGAAATAATATTGCAGCTAATCATAGGAGACAGGTTTCAGCAAGGTCAGCAATTTACGAGCATGGTTGTTTTAACAAAATTAGACAAAATTTCAACGGAATTTATTATGGATATTATATCTTCATATCAAAGTATTGTTCATTATGACGTTAATGACAAAAATTTTAATTTAAACTATCCCAAAAATTCATTTCAACTGTACTATCGAC
>evgR012296c0g1_i01_Hydra
TCCAATACAACACCAGTAAACCTCTTAATCTTTCATTAGAAGAGTTGGAAGTTTTTATTGGTATACTTTTTTTAATGTCTATTGTCAAAATGCCTGGTACACGCGATTATTGGGACATATTTCTTCGATATGAGCGTATAGCTAATGTTATGAACATTCGACGGTTTGAACGTATAAAGAGGTTTTTGCGTTGCAATAATAATTTGAACATGCCAGAAAACTGTCCAGACAAACTTTTTAAAATTCGTCCTTTAATAGATGCTTTGAAAGAACGCTTTAACTTATTTGCTCCAACTGAAATAATATGCATAGACGAACAAGTTGTACCTTTTAAAGGGAGATCTAAA
>evgR012300c0g1_i01_Hydra
TTTGGTTGAAAAAAATAAACGCATTAGTGAGTTGATTGTATCATCGTCTTCCCTCACAATGGAATCGTTAGAACGCCTTTCTCAAAATTGCAAGAACTTGGTACATCTTCAAGCTGAATCTTCTTCAACTATCATATACAACAAAATATCTGTCTTATCCTCCGAAATGGTTCAAACTTTGTCAAAAAGTTTTCGTCAGTTAATAATACTTATATTAAACTTTGATGTATGCTTCCTTACGAACTCAGATCTCATAATTCTCGGACACTACTGCAAAAGTCTTGAATGCTTAGAACTTCGTTTGGATTCCGATAGCTGGTTATCAGACGAAGGTCTTGTAAATTTTTGTCTCTGTGTACCAAATCTGCGTGCTCTTAAAATAAACGAGACTATAATATCTGATTTTACACTTTTTGCTGTTGCAAGTATCTGTTCTGATATCGAGACTTTAGCATTTGGCGGATGCAATGGTGTTACAGATCATGGTTTGAAGATATTGTTTGAAAATTGTAAGAATCTTCTCAGTCTATCTCTAGGGATATGCAACGTCACAAGTGCAAGTATTTATTTTATGGCAAAATCTTATTGCGCTCAAAATTTAATTCATTTGTCTTTGGAAAGTTGGAGGATTCGTGACGAAGATCTTGTTTTTATTTCAAAAAACGTTCCGAGTCTGAATTATCTTTCAATTCAGAAGTGTAACTTTTTAACAAATATTGGACTAAGAGAAGCAATCGTACAATGGAA
>evgR012303c0g1_i01_Hydra
ACCAAACCAGTCGATGGTTGCACTGTCGATCTTGTTGGTGATGATATTAAACATTGGAAGATTATTATTGGCGGACCCTCTGGTACACCTTATGAAGGAGGAAAATTCGAATTTGATTGTGTTCTCCCTGAAAACTATCCATTCAATCCACCAAAATTCACAGCCAAGACTAGAGTTTACCATCCAAACTTTGATGCTGAAGGTGGTATCTGTTTAGAAGTTTTGAAGGCTGATGGTTGGAAACCAACTTTGAATTTGGAAGCTGTTATTATTGCCATTCAAGAACTTATGAAGAATCCAAACACTGCTTCGCCACTTGATGCCGATGCTGCTGAAGCATACGTCAATGACAGATCTAAGTTCGACAAGACTGCTCGTGAATACACAAAGAAACATGCCAAGTAAATTAAAAAAATC
>evgR012305c0g1_i01_Hydra
GTTTTTTGAAGTAGCATCGATTGTGATTTATTTATTCTTCACCAGAACACCAAAAAATCATTTTAGATTTTTTTGCAAGTTATTATTTATTTGCCAATCCTCTGTTGCAGTCTTGACATTTCCAAAGTTCATTTTTGTTGTCAAAAGCACTCTGCTTTATTTTTTCGCATTCACGATGAAACCAATTTTTACAATTCCAACATTGAGCCATTTGCTTTCCATATATAGCGCCGTCAGACAGTGTCCAAACCATTCTGCAGAACAGTATATCTTTAATGAAAATTGTTTTGGATTGCATCTAT
>evgR012319c0g1_i01_Hydra
ATATGAAGGCACTCAATATCTAATTACTTACAATGATTCTGGTTCTCAAGATTGCTTTTTGTATGTTAATACGAGTATTGATAGTACCATTCTCTGTAATCCTATTGAAGGGACCAAGTTTTTTTTTAGTGGACGTACCATTAACAACATCTCCACCGCCAGCAGATCAAACTCTATTACTTTCTACTGTGGAATCTTTTAGTGCACAAATAGCAGTGTACAAGTTTGACTCTTTATTACCGAATGAAGTGTTATTTGGGCAGTATTTAGATTTTCCTACGGCAAACTTATTTGTATATAAAGTCAACTTTTCTTCGAATGTAAATGTTACAAGTTTAAATTTTTCAATTGCTC
>evgR012323c0g1_i01_Hydra
GAAAAAAATATCTTAAATTTATTTACGTTAAGTCTGTAATTGGTTTCCATTTTCTTATATCTTTTGGTTTAGCTGTTTGGTATACGCTACTAGTCTTTTATCACGGTACGATGCAACAGAAATCGTTTTTCTTTTTTTGTATTGGTACATTTATATTAAGCAACTTGTTTTTAGTACTAGTTCTGAATGGTTGCTTGTTGATCAGAACAAAAAAGAAGCTCAGAAGCAAAGAAATTCACGTTTTTCAACACGTTGCAATAGAAAAGAAGTTAACCCATACTATAATGTTAATTTCATTTACATTGATAGTTTGTTATCTACCAATAGTTGCTTCCTCGTACTATTTGGCTTCAGTCATGCGCATGAAAAAATATGAAACTGTCGAAAACGAAACAAAAAATTTTGAATTTGATTCAAAAAAG
>evgR012342c0g1_i01_Hydra
AGTCGGTGATGGTACAACTTCAGTAATTATTCTTGCTGGTGAAATTCTTACAGTTGCCGAACCATTGTTGCAACGTAAGATTCACCCAACTCGTATTGTCAAGGGTTTCTTGAAGGCTCTTGAAGATGCTGTTACTTTTATTTCAGAAAAGATGTCAAAAAATATCGACGTAACAAACTATGATGAAGTTGCCGAAATCGTAAATACAAGTATAGGAACAAAATTCACACATCGTTATAGTGATTTGATGTGTAAATTGGCTATTCAAGCTGTTAAAACTGTAGCAACTGATATGGAAAATGGAAAGAAGGATATTGATGTCAAGAGATATGCTCGTATTGAAAAGATTCCAGGTGGATTATTGAACGAATCTCGAGTTTTGAATGGTGTCATGGTTAATAAGGATATCCTCCATCCAAAGATGAGAAGAAGAATTGAAAACCCAAGAGTTTTACTTTTGGATTGTCCTCTCGAATATAAGAAGGGTGAAAGTGAAACAAATGTTGTTTTGGAAAAGGAAGAAGATTTCGAAACATTGCTTAAAATGGAAGACGAATGGATCAAGAGAGTTTGTGAAGATATCATCAAATTCAAGCCAGACCTTGTCATTACTGAAAAGGGATGTTCCGATTTGGCTCAATACTATCTTCTCAAACAAAACATTAGTGTTCTTCGTCGTGTTAGAAAGACTGACAACAATCGTATTGCTCGTGCTACTGGTGCTACTATTGTTTCAAGAACTTCTGAAATTAAGGAATCAGACATTGGTGCTGGTGCTGGTTTGTTCGAAGTTAGAAAGATTGGTGATGAATACTTTACTTTCATTGAAGAATGTAAGGATCCAAAGGCTTGTACAATTCTCTTGAGAGGTGGTAGTAAGGATATTTTGAATGAAATTGAAAGAAATTTGGAAGATGCTTTGCACGTTGTCAAGAATATCTTTGTTGATGCTCGTGTCATCCCAGGTGGTGGTGCTGTTGAAATGAGTGTTTCTCAACATTTGATGCAAAAGTCCAAGTCTATCAAGGGTATTGAACAACTCCCATATCAAGCTGTTGCTACTGCTTTGGAAGTGATACCTAGAACTTTGGCTGAAAATTGTGGTGCCAACACAATTCGTATCCTTACTGAATTGAGATCTAAACATGCCAATGATCAAAATTATAATTTGGGTGTTGATGGTATCAAGGGTGTCATTGCTGATGTCACTGAACTCAAGATTTGGGAACCATATGAAGTTAAGGTTCAAACTATGAAGACAGCTATTGAATCTGCTTGTTTAATTCTCCGTATTGATGATGTCCTGTCTTCATTAAAACCTAAGGAAAAG
>evgR012347c0g1_i01_Hydra
TGCTGAACATCCTTCCCAACAAAAATTAAGAAGTGTTCTCTTGGATAAAGTTATTCCAAAGTTGGAGAGAACCAATCACAAGAGAGTTTTAACTAATGATGATAACAATATTTATGTCAAATATAGTGGAAGATATTATTACTTTTGTATTGCCACTCAAGAAATTAAACAAAGAGTTTCATGGAGTTTGATTGATGAAGTTGAAAATGAAACATTAAGAAACAATCTTGACAAGAATCTCTTGAAGGAAAAGGTTAAATTCTACAACAACCCAGATAATGACAAGGTTCAAAGATTGCAAAATGAAATCGACAAGGTCCGAGATGTCATGGTTGAAAATGTAGACAAGATTTTGGCCAACCAAGTTGTAATGGATGGGTTGATTGAAACAACAGAAGAGTTAGCTCAACAAGGTGAAGAATTCCAAAGAGGTGGTAAAAAGTTGAAGTGGAGTATGAGAAAACGTTTGATCATTATTATCCTCATTGCCATCATTGTTTTTGCTATTATT
>evgR012366c0g2_i01_Hydra
TTTTAAAATGAATAACAACATCTCGGAGATACAACATCATAAAGCTAACAACGTTGTAAACATAAGAGTGTTTAGCGATGTCGAAAGTATTGTAACTACCCTAATTTGTGGCGCAATAATAACCACCGGCGCCCTTGTAAACTGTTACATTGTGTTTCTCTTTATGGCAAGAAACGCTATGAACTCAATATTTGGGAGGTTTTTGTTACACTTGTGTTTAATTGATCTTGGACAGTACGTTGGTATCTTGCCGTACTTTTTATTAAATTTACATAAATTACCGGTATCAAACAAATATTTTGAAAGTTTCCTTTGTAGCTTGACTGATGGCCAATGTGGGATGCTTGTTTTTTCAATGGGAAGCTGCATTTTATTAACTTTTTTGTCCATTATGAGGTGTGAGTTAATAAGAAAAAGTTCTAGAAAGTTGGTCAGTGAAAATAACGTGACGTCGATTTTTATTCTTACGTGGGGATTTGGTTTGATTTCTTTTATTCCATATATGTTGTCATACAAGTTAGATCATAAAAGTGGTCTTTGTTTTCGTCAATGGATACTTAAAGAGCCTTTTGGAACATTGTATATGATTGTACTATATGCATGCGGTTTAACAATCCCACTTGCAGTTATGACATCAGTGTGTGTTGTAAGCTATGCCCAACTTTTTTCAAAAAATATTAAAGGAAATGTTATAATGAGCATACGCAAAAAAGTTATTAGAAAACTCTGTGTTTTAACTGCTGTATACACCATGGGAAGCTTTCCTTATTCAGTTTATTGGTTGCTAATTACGTCATATTTTTTTCAAAATACTATCAAAGGAGAGTTATATCGTTTGAAAGTACTTCGATATATATTAATACCAGCTCTTATAAAAACTGTTTTAGTTCCTATACTCGTTGCATATAATAACGGACCTTTCAGAAATGTTCGCATAAATTGTTTATGCCGCAGGAAGATAGAAAGAAAAAGTATTTCTTTAACGTTATTAGAGAGAAGCTCTGAAACAAATAACTTACGCGGTACTTCAATTTAACGTTATTAGAGAGAAGCTTTGAATCAAATACCTCACGTGGTGCTTCAATTTATGTTATTAGAGAGAAGCTTTGAAACAAATAACTTACGTGGTACTTCAATTTAACGTTATTAGAGAAAAGCTTTGAAACAAATAACTCATGTG
>evgR012370c0g1_i01_Hydra
AAACTTTTTGTTTGGAAGTTTATTTTCTCGATACATGCAGTGTTAATATCAGTAATACTATCAGACAATGTTCAAGAAGATATTGTGATTACGGATTTGAGTTGCTTACCCGGTTGGTTACAAAGTGCAGGTAACTGCTACCTATTTGAGAACATTTTAAGAAATTTTCAAGAAGCCTTGTCATCATGTATGAGTTATGGAGGAAACTTATTAAGTGTAGAAAGTTATTCAGAAAACCAATTTATTTTAAATAATGCCAAATCAACAAAACTGGATCAACAACTCATATGGACTGGTCTGAATTCTAATAATGATCCTGGTAAAAATATATTTGAATGGTCAGATGGTACACCTTTTTCGTTCGATAATTGGGATGATAACGAACCTAACAATAAAGGTAGCAATGAAGACTGTGCATACATAAAGAATTCGGGTAAATGGAATAATCATAAATGTACAGATCGACATTACTATATTTGTAAACTTAAAAAAGAAAACATCTACAAGCAATTTAGGTTTATATACAATGCAGTAACGATTGAAGAGCTGCTGTATATCACTTGGAATATAAGCAATCTGATTCTTAATGAATTACCTATGGATGTTTCATGGGAGTATATTTTACCTCATTTCATAAATATTAAATATACTAATAACTCAGCAAATGCAATTACTATTGATTTAAACAGAATGTATAAGATCAAGAATTTACTAAGCAATGGCGCTTATCAAATTGTCACTGCTTTTGTTGTTAATACAAGCTGTTTATTTTGCACAACTTTTATTGATATTCCAATAAGATTATATTTTGAAAATAATGTAGTCAAATATTTCAAATCATTTCAATTTACTAAAGTC
>evgR012372c0g1_i01_Hydra
CTTAAGTTTTATAAGAGTACAAATATTATTTAAAATGAAATTTGAAAAAATTGAAGACAAAAGTTATTTTTTTCATGAAAGAACTAAAAAACTAGTTGACGAGTTAATTGACGATCGCTTAATCCGTTCAGGTTTACATTTAAAACCAAGAAATAAAATAATAAGCAACCTACAACAACCTTGTAAAACGTCAAACAATGGACGCCAATTGACTGTAAGAGAACAATTCAAAATTTTAAAAGACAAAAGAATATCTAATAGTATTGAATTTGACGACCTTCTAATGACATTTTTATATATCGGAGAATGTCTCGAACTAAAGTATCAACACATATACACGGATGTATTTCAACATTTAAATATCAAAAAGTTAGTTGAAGCGGAAATGCAAAAAAAGTTTTTACACGTGGCGGAAGAACTTTTTCGGAAAAGTATAACTTGGGCAAAAATAGTTTCTTTAATAGTGTTTTCTGGAGGTCTTACAGTAGATTGCGTTTTGTCAGGTAATTCGATATATATTAGTCGGATTAAAAACTGGTGTGTTGAATTTATTGACAGTTCACTAAAAGAATGGATAATAAAAAACGGCGGATGGTTTGGTTTTATTGACCAGTTTGTATCGTCGCAAAGAAGTCATAACGTAAAAAAATTAATCGAAAAGATTTGGATTTGTGTGATACCGGTAACAGCTATAAGTGTGATTGCAGTGGCATGTCGAACGTTGCAAATTATATAATAATGATACGATGATCATTACATACTCGACTGAGATAGTCAAAAAATAGTTTTAGTAGACAGTCAGTATCAGTACTAAAATTCAAAAAGTAAACTGAATTGTATTTTTTGGTGGAAGGTTTCAATTATTGTTTACATTGTTAATATTTTTTTCTATATTTAACCTAATGTGTAATATGTACTAAGTTATAATGTAAATTTGCACCTTGAAAAAAAAAA
>evgR012373c0g1_i01_Hydra
AAAATAAGTTTGAAATGCAGAAAGAAAAAAATAAAATCTTTAAAAATTTGTATACTCTTTAATAAATATTATAAACAGTTTGTTACAATTTTATTTTAGGACAATTTGACGCCGTCAACAGCAATATTGACGGTGAAACTAGTGTGTTGTTAAATGATGAAACAAAACAGGAAATATTATTACTTAGCTGTTATGTTCCAAGTCATTTTATTCATGGTTTGCTTGCTTATCATTGTTCATCAGAAACTGGTTATGAAGCTGGCCGAACTAAGATAAATAATGGCGATCAAGTAAAGTTAATCGAAGAAAGTTCATTATACAAGTTTTTTGCAAAATCTAATATTTTTTTAAACGATTTTCTTTATAATTGGAGAGTTCTAGAAGTATATGATAACGCTCAATCCGGATACTTAGGGATAATATACATCAACGAAGTTAAAAAACAATTAGTGTTGGCGCATAGATCAACTAATTTTAAACTAAGCGTATTAACATCAAGTATTTTAAAGCAGTCTGGGCTTCAGCAAGATATAGATGGTATTCTCATGGGAAATATTCTTATTCATCAAGGTCATGGCTATGAACTTACTTTAAAAGCTGTAAATTGTGTTAAAAATGAATACCCTTACTATGCATTGTCTTTTACTGGACATTCGCTTGGTGGATGGTTGGCAGATCTTAGCGTATTTTATTGTCATGACGTTTTAGAATATCCAAACGTGCTGGGTGTAACCTTTGATGGACCAGGTTCCTATGAAATGATCCAAAAAATGATTTCGGGTGCTATTCGTGGATCAAGAAAGAAAGTAAAAGTTACTGATTTAGAAATTGTAGCATACCTGTCTGCACCGAATATTGTAAACTGCACTAACAAACATGTAGGAACAGTTTATAGATTGTTTCCAGATAATAAAGAAGAAGAAATACTCCCAGAGTGGGTTATTGACAGCGATTTGTTAAAGACAATTATTAACCAGTTACCAGGTGAGGTTTCGTCAATTTTTGGACATTATTTACAGTTGATATTACAGTATTTTGATCCTTATACAGAAAAGCCTTTTAAAGAAAAATGTATGTTAGTTAAAAATTGGC
>evgR012375c0g1_i01_Hydra
ATATTCGTACCATAAAGAGTTTTCGACTACGGGATTTCGCCCGTTACGTTCGAATCGTTTTTAAATGCCCTTAAACGTTTTTTTTTTTCATTACTTTACCGAACTATTTGCAATCGCAACTTTCTCATCAATACAAAATTGTGGAGATTTAAATGTAGTAGTGCTGCAAAATGTTAAAAACAACTCATTTTTTATTGGAGGTCTCTTTTCCACACATGTTAAAAGTGGCAGTTCTTATATTCTTAATGAACTTGATATTTTGTTGGTATTAACAATGATGCTTGCTATTCAAGAAATAAATGAAAGTAAAGAAGTTCTCCCAGGAATTAAGTTAGGATACAAGATTTTTGATTCGCAAAGAGATATTTTGTGTGCATGTGACCACGCCGTAAATATTTTACTTGGTAAAAAAGGAAATAGTCTTATTGGAGTAGTAGGACCAGACGCTAGTTCAAGTGACACTGTTAACAAGATATTGTATTCAGAATTCGTTCCACAAGTTTTCTATTCATCAACAAGAGTGGCGCTTTCATGTTATAACTGTTTTCGTACTGCTCCAAATTATGCCGATCAAGCAAAAGCTTTGATTGATTTATTTGATCGGCTTCAGTGGAGAAAATTGTTGTTATTCACTTCTGCTGACGATTATGGAATTTCTTAATGCTGCAAAAGATAAAAATGTCTGCTTTTCTGTTGATTACATGTTAGATAAAAGTGTATCTGATGATAGTGTTTTTGCTATTTTAAAACGCGATGAATCTTATTCGCCGGTTGTAGTATTATGGTGTGAACCCCAAGCGGCATTAACAATAATAAAACGAAGCGTAGCTCGAGGGTTAGCAAATATCACCTGGATAGGTACTTATTTGTTGTTAATTTATAATTTTAAGATCATTATCAGCATTTAAAACCAG
>evgR012378c0g1_i01_Hydra
GAGTATAAAAAGGCAGCTATTTCTTATATATCAGGTTTCGTTGTTAGAATGGTAAAAAAACAAATTTTTTGTATGTTATGCTGTAATGCACTTGGTTCTAAAAACCATGTTCCAGAATCTAATTTTCTAAAAGTAAAAGATAGTGGTGGCCTTATAAAACCAAAAAAAGATGTTATTGTTGTTTGTGAAGAAACGGAAAAAAAAATTGTGAGAATGTTAGCAACAATACCTGGAAAACTGCCACATTCCAAGAACATCCCAGAAGCTATATCTATTTCAGTTTTAGAAGACTTGGGTTCTGATATATTTAAAGATTTGGAAGAACACATGTTTGATACTGGGATTAATGAAAATCATGTGTTTAAGTTAATTAAAATAATTTCTCAATGTTATAGCAAAGTAAGACTATATGACCTTGGAAAACAAGCCACTGAGAAAGTGGCAGGAAAAAATATTCTAAAAAAATTTAGTAAATTGGTATTATTT
>evgR012384c0g1_i01_Hydra
TATTTTAAAGAGTAGTATAGAACAGTACTATGGTTGCATTTATGGCTGATTGATTACATATATATAAAAAATGATGCCGTATGGATTTGCAAACATTCAGTGAGCCACGACTATTTTCATGCTGTAAATAAATTTGTTTGATAAAAATGGATAATGAGTTAAATGAATTGAACGAATTGCATACTTTAGCAGTAACAACTTTTCAGAATGTTGCTAATCTTACTCCTTCGCTACGGCAGATGGTTGCAAGTGCTAAAATCTATCAAAAATCAATTCAAACTACATGTAACTTTGCAGTCATGTTTGTGGAGTCGATGCAGAATGTGGCAAATGCAGCTTTAGTTTCAGAAGGTGCAACCAAAGTATTAGGTGATGTGTTATCTGATATTATTGATGCTTTTAAAGAGATTGAAAATATAAAATTGGAGTTAGCTAAAGTATTTATCAATGAAGTTATTATTCCATTAGAAAGCAAAGTGGAGATTCAAATAGGAAATATTAAGACAACTCATAAGAATTACAGCCAGCTTAATAAGGCACAATCAGAGGCTGTTGAAAAGGTTCGAAGTTCTGTGTTAAAAGTTCGTCGAAGAAGTTTAAGAAAGAGACGCCCGTCTCAAAAATATGAAGAAAAGGAAAGGCAGGTTGTAAACGCTCTTGCATTAGAAGAAACTAAGTTAAAAGAAGTTCGGGAACAAGGATTGCGTGATGCACTTGTAGAAGAGAGAAGACTTTATTGTTTCGTTGCTGATAGAATGTGCTCCTATGGAAAAGGAATGGTTTTATGTAATAACAAAATAAATAGTTTTATAAATCAGAAAGTTCCTGATTGGTTACAGTTTATTTCCAAACCGACTGAGCTTCCTCCTGAGAGTGAAAACATTATTATGAACCCTCTTGGTTATAAGATAGGAACTGTTAAAAAAAAAGTTATTAAACCTGATAGGAAAGATACACTAAGGCGTGGAAAATCTTTTAATAATCATTATCCAACAAAGTATAGTACTCTTGATGGTAGATCTATTCGAGGAAGTTCGAGACCTAGTATTTCAATTGCATTAAGTTCAAGGGGAAATCTTACAGATGCTGCAAGCCAAGCAAGGTCTCTTAGTGGAAGCACCTCACCCACCCATAGTGAAAAAGATACCCAGGATCTAAATAATACTTCTTTATACTTGCATGATTGGGATGAGCCAAAAACAGATGAGGAACGAAAGATAAAGATGTTAACTAATGAGCAAGTAGAAGATAATAGTCCTGCAAAAAGCGAGTATGCAAATGTTAACGTCCTCAATCAAAAATTTGATGATGTATCTATATCCAGATGTCATTCTTACTCTGGTAATAGAATTGAACTTAACAAAACAATAGAACAAAATAGCTTGAATGCACTTCTTCGCACAGAAAGTGAAACTAGTCATAAAGAGCGTAAATCCTATGGACCCCCTGAAGATTCTTCAGAAGCATCAGATTTTCGTTCGAGACTTGCTTCCATACAAAATTTGCTAAATCAAAGAGAAGTAAAAGAGAAAACTGTTGTTAAGAATAGACTGAAAGCTATTTATGCACATACTGCTCGAGATGCTTCTCAGCTAACTTTTGATAAGGCTGACATCATTATTCCAATATGCGAACCAACATCTGGTTGGCAGTATGGTGAAAACGAAAAAACTGGATTATCAGGATGGTATCCTGCTGCGTACACAGAAGAGGTAGTGATCTCTACAATAAGCAGTAACAAATCAGCTGCAACACCAAATGCTGCTTATCTAAGTTCATTTAATCAGTTTAGATCAGCCACCATGCAAAAAGCTAGTCATAAAACAGCTCCAATTGTTGCTAAAAAATCTGCAAAAGTTGCAGCAAAAAAGACAAGTATTGTTGCAACCCTTGGTAATTCTGAGGATAGCAATAATGAAAATCCAAAACCACAATCAACCAGTGACTCACCAAGAATTCGTTCAGCAAACTCTATGCCACATATGATACACGAAATTGAAAATTCTAAAAATGGAAAATCAGATAAAGTGGCAAGTAAACAACGGTCTTCTCTACGACAATCTTTTGATGATTCTTTTGATGACGAAGAATTAGACGTCAATAATATGCCGCCTCCACCTCCTCCTCCTATATTTTCGGAAAATGAACTTGCAGTTTGAAATTAAATTGCGAAGATATCTAATAGGCTATATTGCCCTAAACACAAACGACACTAATAGTAGATACAAAATAATAGATAATGGTAGATAATTTGTTTCAAATTTTTTGAACTTCATTCTGCTAAGTTCGTTAATAAAAATTTTAACATGTATCATTAGTAATAACGTAAATTTGCATAGTTCTTACTATCTTACCCTTACAGCTTTGTAATGTATAGCTCTTTGTTCGTAT
>evgR012388c0g1_i01_Hydra
TAACAAATATTACCCCAATTACATTTCATTTGAATCCACAAATTACCCAGGTTATTTTCTCAGACATCAAGACTTTTCAGTGAAATTGAATAAAGAAGAGCCAACTGTGGAACTGTACAGGAAAGATGCAAGCTTTAAACTGGTTCAGTTGGGTGCTTCATCTACACCTAAACCAGTTTCATCACCTGCACCAATAGTTTCATCAACTGTACCAGGATCAGTTTGTGCTGCAATTGTAGATATAGGCTTTATAATTGACTCATCTGGAAGTTTGAAATTGCATTATCGCCAAGAAGTGGAATTCCTTAAAAAGCTTGCAAGTACATTTGATATTAGCTCTAATGGTGCACATGCAGGTGTTGTAGCATTTGGTACCGAA
>evgR012390c0g1_i01_Hydra
CTTTTACCTTAAATTATCGTCTTTTTTTTTAATGAGTTTGTATATTCTTTGCCTTAGTTGCATTGCGTAACACAGCGTTTCTGAGTCTTTTGTTTTCATCAACGAGTGTATTGAAAACGTGAAATTAAAAACAAGAATTCAACAGAAAAACTTGTTAAACGAGAATTAAAAAATGGAGGAAACCACAATTTTGAGAGCACTGGGTGCCAAAGAGCTTTATAACTCTTCCCTTAAAGACTACGATAGGCTAATAAGCTTGACTAAGATAACATCCCGCATTAATCTGATTGAACACGTTGACGTAGTTGAGAAAGCCATTACAATGTGGAAAGAACTTCACCCGTTTTTGAAAGCCACTTTGAAAAATTTCCCTAATGACAACCGAACCTACTTTGTATTAACCCAAGATGGAAAACTAGACAACACACTTCATAATGTAAAGCTGCTGTCATATGATCAACATGGAGGGAGCAATGCCGCTAAAATTGAACAGTGTCTGATTGAGATGCTGGTGGAGCGAGAGCTAGCCGCGGAGAACAAACTTGATTTATACCATGAGCTGTTCTGGCGACTTATTTTGTTTAAACGATCATCTTCAAGCCTATCAGGCTCCTATGAATACACAGTTATATTTATGGTGCATCATATGATATGTGAAGCTAAGTACAAATATTTGAGTCTAATTGAGCTGTTTGATTTGATTGAAGACGTTTATAATGGAATGAATGGTGCCATTGGTATCCGAAAACCATACAATGTTTTAAATGAAGTTGAAGCTTATTTTTCTTTTAATCATGAATATCCTGTTCAGTTTACATATCCAAAAATTCCAAAATTTATAAACATGTCCGAGGCTAAGCAAAAATCACTAAAAAGCGACTATTACATGAAGAGTTTTAATGAAACAAATAAAAGTGGTTTTATTGTTGACCACGACAGACAAAACTACATAAGTATTAGAGATTTGATCAAGTTTTCGCAGGAAAACAGCTCTAAAACAAAATGTTTTAAGTTCAAACGAGAAAAATTTAATTTGATTTTAAAAAAATGCAAACAAGATCAAATCAAGCTAAATGGCTGCTTGAATATGGTCTTTATTTTAGCGCATAAAATGTTGTATGATAAATTCGGTGGGAATATTCAGAAGTTATATTATTATAACTCAATATCGCTAAGAAAGTATTTGAATCCGGAAGAAGAAAAGCAAGGACAAAAGTTTTCCTATTTGGTAAACTGTTTACCAAAGATGTACGAATTCGGTTCATCCAATGAATTGGATTTGAAAGATTTTTGGATAAAGGCGAAAAGTGAATCGGAATGTTTTCATTCGAGATTGAACGATAAGGAGTATTTGATTAGCTGGAAATGGTCACACTTATGTAGCCAAATAAAAGACGACGAAATGATTTATGATTATTTTTGCGCTAATTTAGGAGAACTGCCACTATCTTTGAAAGCTGAAGGAAACAAACTCATTTCTGTCAAAGAATCGTACGATACGATGCGTGTGGATGAAGATGCAAAAGATGGTTTTGTATCAATAATTATCAGTTCAATTGAAAACTGTCTCTACTGGTCTGTTATTTTTAACTCTTATTTTATAAGCGATGAATGTATTGACTATTTATGCAATTCTATTTTAGAAATATTGAATAGAATTGTTATAAATTTACATTAGTTGTATATATTATGGGGTTTGGTACATACGAGAATTTATTATTATATTATGTGATATAATGCTAATAAACATAGAAAGAATGTTTGGAAATTATTATAATGAGAGTCAACAAAGTACTTTGATTGACTCATTGAATCACATGGATTGACACTTATTTCAATACTATTTAAACTTTCTTTCTTTGTAATTTTTTTTCATACTTTGACTTCATACTTTGACTTCAT
>evgR012391c0g1_i01_Hydra
TTGGAAAATAAGTACAATAGTCGAATAAAAAGATTCAAAATCAAAGTAACAAAATGTCACTACCGATGAAATTGTATTTATTTGTTCTTTTAGTTATTTGGTTAATAGTGACTGAATTGGCATTAAGTATAATGCCATTCTACTCGTGCGGAGTATCGCAAGATAAGAACTTTCAAAATGAAATTTGCTGGGGATTACTGGGTACCAGTGTTAACACAGAAATTAATGATAGAGAAAAGTTAACTCTTAATTTAATTGACATCGAATTAATTGACGAAGCAACACCTCAGTCACTTACAGAAATCGGATTTATTCTAGAAAGATGTTTAACAGGTTTTTCCTTAATACTTTATCTCTGCTTAATTGGAGTCAACTTTGCACATTGTAAAAAAGTGAATCGAAACAAACCGCAATACGCAGAATTCATTATTTTATCTTGCATTTTATTGGCGTCTTTAGTGAAACACGTAGTTGCAACTAAACGTAGAGACAAACTTTATGAAATTGATGACACAGTTGAATATGCCGAGACTTATTATGCTTATAACGTCTATTTATGTAACATATTTCTATGTATGATTATTGAAGCTTGGGGTTTTAAAGATAATGTAATTAGTGTTTTGTTAAAGACAGTCAAGCCAATCCGATCAAAGACAGTCAAACAAACTAGATCAAAAGCAGTCAAACAAGTCAGATCAAAGGCAGTCAATCCAACCAAGTTAAAGGCAGTCGAACAAATCAGATCATATTTAATATGCAACCAACAATCTAAACACTTCGATCAGGCGATCCAAACTGAATAACATAGATTTATGCTAATCTTGTTTTAGGGTTTTATTATATTTTAAGGCTTTGTTAGGGATTTTTCAATTTCGCTTTCTAACGAGTTTCGGAAAGTAGTTTTCACTGACTTGGAAAGTATTCGTAAAAGTATTTATATTGTAAAACGTTCTCTTTTCCAATAATACATTTACTTTCCGGAACCCGGGGCTTTTTTTTGGAGGCCTTTTTTATGTTCCAAAAAATTGTGCTTAAAAATGAAAATCTTTGATGATAATTTTTGTTATTTTGGATCGATTTTTTATGAAGG
>evgR012402c0g1_i01_Hydra
AAAATATTATCCACTAAAGAAATAACAAATAATTTATGTGATTATTCAAATAGTCTGCTTATTAGATTTGTAGAGGAAATGAAATTACTTTATGGAAAAGAAGCAATGGTATATAACATACACAATCTGGTTCACATTTCTGCTGATGTAAAATTGTTAGGATCATTAGATGACTTTAGTTGCTTTCCTTATGAAAATCAACTTGGTGTTTTAAAAAAATTAGTAAGAAAACCACAAAATATTTTGCAACAAATATGCAGCCGTCTCACAGAAAAACTTAATACACATAACAGAAAAGCAAAGCTTGATGTTTTGAGCAGTTTAACCCTTCTTGTACCAAAAAGTATTCATTGTTGTGGTCCTTTGTTAACATTTTTAGATGTTGAACAATATAGGAAGTTAAATATAGGAAATTATTGTCTCTCGTTGTCATCAAAAAACTCTAATAATCATTTTCTTACTAGTGGCGGTGATTTCTGCATATTGAAAAACATAATTAATAAAAATGGCACTATATTTGTAATTTGTCAAAAGTATTTGCAGTTGTCCGAATTTTTTGATTATCCATTAAACTCAAGTGAACTTGGAATATATCGAATTACTAATAGAGTGTCTCCTCTAATTGTTTTGCCTGCTTATGATATTGCTAAAAAGTGTCTTTGTCTACCAACAGTTGTTAACAATGAGTTTGTAATTTTCCCAATTGATTGTAAATAGTTTTTTCAAA
>evgR012405c0g2_i01_Hydra
GCCATTGAGCCATCTAACATCTAGATCATGATTATAGAGTCTCTCTTTTACCTGAGAGATAATATATAAATAAATAATTATTTATTTGAAATGTCTGAACACCGCAAGAGATACCGAGATCACGATAGTGAAGTTTCAGGATCTCGTGAAAGAAAAAGAACTCGAGTTGGTCCTTTAGATGTCGAAGATCGTTTGGAATCACTTATTACTCGTGTTGGAGAAAAGAGTACATCGTCTTTGGAGAGTAACTTAGAAGGTTTAGCAAATGTGCTAATAGCTGATATTGCAAATTATAAGCAAAGGATTTTGCAAATACTTTGCACTTGTGTTACAAATTTACCAGAGAAAGGAACTATATATGCAACATTGGTCGGATTGCTCAATGAAAAAAGTTTTGATTTTGGAGGAGAGTTTCTTGAATTAACAATGTGTAATTTAAAGGAAGTTATCTTCAGTGAACAATGGGATCGAGCTCGTTTAATGGTGCGGTTTCTATCTGACTTGATAAACTGCAATTTTTTGGTTGCAGCATCATTTATTGGTTTTCTTGAAACTTTGATGAATGCTGCTTTGCAAATTGGAGTGCCTCAAGTTCGATCAGATTGGTTTGTGTACAGTATCCTATCATCTTTACCATGGTGTGGGAAAGAACTTTCAACAAAAAAACCTAACGAATTTGGACGTCTACTTGAATCTATAGAGGTTTTTATAAGCAAACGCAACAAACTCCACAGTAAAGCCCTTAGAGTTTGGTCGTCAGACAATCCTCACCCTCAGGAAGAATATTTAGATTGTTTATGGGAGCAAATAAAAACAATGAAGGATGATGACTGGCAGGAAAAGCACATTAGAAGACCGTATCAAGCTTTTCATAGTGTTCTATGTAAAGCTCTTCCTCATCCAATACCATTGTTTGATGTTCCTCCCCATACTGAAATTAGCATTTATCCGTTACCAAGTGTTATCTTTAGAATGTTTGACTATGCTGATGCACCTGAGGAAAAACCGATTCCAGGTGCTCATGCAATAGAACGATATTTAGTAGAAGAAAGCATTTGGCGTATTATTTGTACACATTATAAAGATAGAAAAGAATGTGCAAATCAATTGATAAATTGTCCTGTTCGAAACAAAGTTCCAATTGAATATATAATCACAGAAGTTGTATTTGGTGCTATGTTTCGATTACCTGAAAATGAGCATCTATTTTTATTTTATGGGTCGCTTATTATTGAACTTTGTAAAATCAATCCGCAAACCATGCCTGGAGTTATTGCACAGGCTGTTGAATTATTATTTGACAGATTGCCCACCATGAATGTGACATGCGTTGATCGATTTGTTGATTGGTTTTCATATCATTTGAGTAACTTTCAATATAAATGGACCTGGGATGATTGGATTCAATATGCTGATAGCAGTGCAAGTGATGACCAGCGCTGTAGGTTTATTACTGAAGTGCTTGGAAAATGTCAAAGAAGTACATTTCATGAAAGTTTATTAGAAATTCTTCCAAGTCCACTTTTACCATTGCTTCCTGAACCTATGCAAGGATACTTTAGATATGAGTGTGCAGAGGCAGATGAGCCTGGTCATACCGTAGTTCGCAAATTAATCGACGGAATTAAATCTAAAAAAGACACTCAGTCTTTGCTTACAACATTGTATGAAATTGGACCTTCAGAAAAGTCTTTTTATGAAAGTGATGATCTTGTTGGTCCATATATTGACCCAAGCATATCAGCACTTCGCATTGAGTGTTTTATCTTGACTATTTTAAGAGCTGGTTTAAAAAGTATAAGTCACACATACAGTGCATTAACCAAGTTTCAACCCATGTTTACTGTACTTATAACAAACAAATCTGAACAGTTACATGTCTTAAGGACTATCAGAGAATTTTGGGGAAGAAATCCATTAATGGTTCTTATTGTCATTGATAAAATGGTAAAACTTCAATATATTCAGTGCAGCACAGTTATAAATTGGATTTTTTCTAAAGAAATGGAAAATGAATTTATGAGGTATACCACTTGGGAGTTAATACATAATACTTTAAAAAAAATGATGAAACATAAATCAAAGGTAAAGAAAGATCTAGAAGATGCACACGAACGTTTGGATGTACTAATTGATAAAGCTAAACGAGAGGAACAAAAACATGAAGAATCTGAAGATGATGAAACATACAATACTTCACCTTTGGAAATAGAACGTAGACAGGAAGAGATAGAAGAATTAACACAGAAAGTTGAAACAGCTAAGAAAGATCAGAAAGAGCTATTTTTAATTGTTTTTCAACGGTTCACAATGATCATCACAGAACATGTTGCTGAATGTGAGCGAGAAGGAAGGAAAATTGTGAATTCTTGGTATAGATATACTTTACAAAGGCTGCAAGAAATATTTCTCATGTATCATGCTGAAATCGATCAATATTCAGATAAAATGGAGAAGTTGATTTTCACGCCTGACATCGCCCCTCACATTCTCGAGGTATTTAGACGCTACAAGTCTCTACGTTCTTAACAATTTTATTGAGGCATATTGATTACGATGCATCGGCATCATCAGTTAAAGGTGGGATCAAGGAATAGGAATATTCTGATAGTTTATTAGTTTCATTTTGCTTCAGCATTTTTAAAGAAGACATGCTGTTTATCATTAACTTTGCTTTATACTTCAGAAATGAATTGTATATTACAAAAATATCTCATAACTTTAATTCAATTTTTAGTTAACAGTAGCACATAATATAACATACATATAATGTAATATATGTATTTTAACTTTGAATTGTGTAATATCTGGTAATGTTAACAAAATACGATAATTTATGTATTTATTATATAATATGTGACGATGTTAACAAAATATGATAAGGATTAATTTTAAAAAAAAA
>evgR012409c0g1_i01_Hydra
ATTTTAGGATGTGGTAGTTCTCACGGAGTACCAAGTTTGAATTGTGATTGCTTTGTCTGTAAATCTGATAATATCAAAAACAAAAGAAAAAGGGCCTCCATATTTGTTGAAACGCAAAAAATAAAAATACTTATAGATACATCACCAGATTTAAGACAGCAGTGTCTTGAAAACGATATTAGGGAAATTGACGCAATTATATATACCCATGACCATTCTGACCACGTGGCAGGTATAGATGATGTGAAGAGGTTGTGTGAAAACAACCAACCTATTGATGCATATATCGATGATACTACCTTTAATAACCTAAATGTAAGATATGATTATATCTTCAAAACATCTGATCCATTATATTATCCGTTACTTAGAAGAATAAAGCTCAAGCCACTTCAAAAAATAGGTGATTTAAACGTTGTGGCATTTGATCAAGTTCATGGTTCTATTGTATCACAAGGCATAAGATTGAACAATGTAGCTTATTCTACAGATTTTAATGAGATATCGGAAGAATCCTTGGCAAAATTAGAAAATTTAGATGTGTGGATTGTAGATTGCTTAAGGTATTCTTATGCTCC
>evgR012413c0g1_i01_Hydra
GAAATGGCCGCAGCAATACGCGATATCACACAATAAAATAACAATGAAAACTGATTAACTTTTTTTTTAAGCTTAGTTTTAAGATGTCGTTTTTAAAAATAAGTGTGGAAAGACTTCTTGAACAAGAAGTAGAAGAAATTAACAGATTTGGAGAAGAAAAGTACGTCAAACCTTTAACCTTGTCAGAGGAGCTAACACATAATGCCGAAAGAATAGATTTTTACGAAGGACAATTTATTGATATACAAGATCCTATTGATAAATCCGTAGATGATGATAAAAATGATGTAAAACAAAAGTCACCTTGGGAAAAAATTCATTCTAAGTTACGAATATCTCTTTCAGAAATGTGTGTGTTGCTAGATGTCCTGCAATGTATTAATTTAAAGAAGTATTTAGTACTTGATCGCGTTTCTCAAAATCCAGAAGTCCCTAAACCAACTGTCCAGATGTTAGACAAAAGGAAGCTTCTTGCTAGAGCTGGAGCTTTAATTATTAAAGGTGCCTCAATGCTGCATAAAGAATATTATCAAAAAACCCATATTGATTTAGAAAATGTAGAAAAAATGGAGTATTACAGTGAACTAATGAAATTGCGGCAACATTGGCGTGTTAAAAAAATTGGAACTCAAATCGTAACTGACATAAGTCTTAAAACATCTGGTTCAAAATTCTGGCACCCAGGCTTTGTTGAAATTAAGATGGCGAATCAAGAAAATTCTGAGTTTGATAAAATGTTTAATGCAAATGTTAGTTCAGATATTCTTCAAGATTCATATATAAATATAAAAATTATTGATAGTAAATATGATATTGTTTCTGAAACTAGCTCAACATTCTTATTAGAAATTTTGAAAAATACAACCCCAATATGGAAACAACATTTAATTAAAGGTCAAGAGTATCTTTTCAATAAAGAGATCTTTTTCAATCTAGCAAGTGAAGCATTTAAAGTAGCTTTCCCTGGAGTGGAAGTTCTGAATGAATCAATAAAGTGTTCAATTGAAGATGTTGTAGTTATTATTAGTTTAAATCGTTACGCCGCAGGTTGTGAAACAACTATCAAAAAAGTTACAAATTGTTTAAGTACAACTGCTGACAATCTTAGATTAGAATTATATCAACTCCAATGTAAAAACTATATGATAAATATGATTTCACATGATAACAATCAAAAACAGTTGAATAGCTCTGTTTTTAATATCTTATTACCCTCTAAACAAAATGCATATTTAGGCAGAAATAGTATTCTAAATGAATTTGTAGATGCGGTTAAGCATCAAGTATGGATAAACAGACTTAAAAAATATTTTATTAAAAAAAATATTGTTTTAATTGAGCCAAAACTAGTTGTACACTGGAGTGTGATTAACAGTTTATCTACTTTAGCGACTATATTATATTTTTATCAGACAACACAAACAAGGATGCTGATTTCTGTAACACAAAACGAAATAAAAATAAACTTTCGGGATTTTGCTGTTATTAAATTACCTTTTGATATTGAATCGCTTGATTATATTATTGACCTAGACATTTGCAATTCAATGGTGGATCATATATCACTATTGTGTCAAAAACATAGATGGAAGTTTATAAAATATCAATCTCCTGATAAAAATAAATCACTAGTTGGATATGGTGTTACGGCTTACTCGCCATTATTAAATCGTTATTTTAAGTTTGTTGTTAAAAAGATTTCTGAAGTTTTGGTTGAGGTATCTTCTACCTGCAAAGATGATAATTTATTATTAGATGAAGGTTTTGATCGAGTATGTGTTTCATATGAAAAGGTTTGTTGGTCATCGATATCTGGGGATACATTGACTGAAAAAATTGAATCATTGCTGCTAAGCACATAATAGGAAATTGTAATTTTGTGTTGAATAAAACAAAATATACAATGTATAATGATTAAAAAACGAAATTGAAAATAATTTATTGTGACATTAAAAAAAGAACTTGTACTCATTTATTGGGGAAAGTGAGGTGAACATATTGATATTTGAGAGAACAATTATTTAATTTATTTAATGTCGATATTTTGACATTAACGTGAATGGTGTCTTTGCCCAGCTCGTCCACTAAGGCAAACTAGGAAACTGCCTAGGGTTGTCGATTTTAGGGCTCCGCCAATATAGCATTGATTATAAGATAATGAAATTTTTTTAAATAATTTAATAGAACAAACGGC
>evgR012419c1g1_i01_Hydra
TTGTTATAGGAGTTTCAGGTAACATTAATTATGCATCTTGCACGGAATTTATGTTAAAACAAAAGCAGCTTATGATATGTATGTCCTACTTTCCAGTGACAATTAAAGTCAGTCATTTAGTTGCCATGCGACAATATATTGAAAAACTACACAAAAAAAGTTTTATGAAAGTATTTGGTCAAGCTATTTTGAATATAGGAAGAGGAACAGACGGCTGTTTGTGTCAATTTGCTGTGATGTGCAACTATGTTTATCGTTTTCACGCTAAAGAATATTCTTTTCGAATTCAAATGCACCCGGATAGCACATTACAAAACAATTTTTTAGAGTTAGAAAAGATTCCTTTACCGAGAGTAGCCATTCATTCGAGACATGTTATTCCACATGTAGTAATTCATGTTATGAAGAAAAAATATCTTACAACGTT
>evgR012432c0g1_i01_Hydra
TTCTCTACATCATCATTTGCTTCCCTTCTCTTTACTAGATTCTTGATTAGCGTGGTTCAACATGTTGACATGGATACTGAACGTATTGCAATCTACTCGGCTCTTATTGAAGCATTCCAAAAGTTACCTTCAGTATTCATTCAAGTCTACCCAGATGTTTTACCACACATGATCAATGTGTATAATAAGGCCAACATTACCCGTATTGCTGAAGTTTCTCTCACTCTCATTGACACCATTATGCAACAAACAGAAGAAAAGAGTTTCAAACCAAAGACTCCAGTTAATTATGATGAAACCTTATCCAAGTCTGGTTTTGGAGGACTATCCAAATTGGGTTCATTCAAGATTGCCTCTGATAAGATCAAACACG
>evgR012433c0g1_i01_Hydra
TTGCAATGTGTTCTACACACTAATAAAAAAAAAAGTTATTAACGGCAAATAAATATACAAATGACGAAAAGAAAAAAATGTTATTTCATTTAAGATAAACCTAAAGAATGAAAGCCGAAAATTTTGATAAAGATGTGCTATTTATTCTAAAGAAAGAAGAGGTTTTGCCGGAGTTTCACGATCCATTTATTATCAGTGGATACCGAAAACCATATTTGTCATTTAAAGACTGTATTAAAAGCTCAGTAATGATTTCTTGCAATGAATCTTTCAATATTTTATCTCATTTTGTTTCTTTCGTATTTTTTTTAATTGTGTTTGTATTGTATTTCAAATCTGAGTTGGATATAAAAGACGTTGAAACGTGGCCTTTAGCATGCAATTTGTTTGGCAATATTGGATTTACACTAATGAGTACCATAGCGCATACATTTAACTCGATGTCAGTTGAAACAAGGCATAAATGTTTTTATATGGACTATGCGGCAATTTCAATATATGCATTTTGTTCCGGGCAAGCAACTTTCTTCTACGCAAGTGCTTTCTTTTCAGACAATACATTTTTTAATAATCCCTACCTATTTAGCATAGGTGGTGCACTAATCTCGATTACTTCTACCTTATTGAACTGCCTATCAAGAATGAAATGGCAGTCAAAAAAGTTTATAGTAAGGACACTTACATACATTTGTGCGTACATATGGAGTTTCCTTCCGTATGGTTATCGTCTTTTAACTTGTAGCAATAAATTAGATTGTGATAAAAGTGGTTTAAATTTAGCTTTATTAAACTCAACACTCTTTTTAATGGCGGGAATATTAAACGTTACCAAAATGCCAGAACGCTACTACCCCGGAAGGTTTGACTGTTGTGGACAGAGCCATAACCTAATGCATATTTTTATATCAGTTGGGTCTTACTTACAGTTACAGTTTATAAGAAATGAAATGTCGCGAGGTTTTCAATCAAAAAAAGACTATGTTTATTCTTTATATGCGACTATTTTTGTTATTAATGTGAATTTGTTAATCGCAATCTTTTTTCCTACGAAGTTCAAGTCAGCAAATGTGGTTCAGCAAAAGAAATCAAAATGATTGAAAACATTAATATATAGAATAAACATTTTTCTATTTTTAAGATTATTTGAAATATATTAAATAATTTATGTTTTATTTCAAACTTATATGTAATGTTTACTAATGCAAATAATTCCAATATACTTTTTTTTGTTTGAAAAAATATAAATACTAATAAATAAAAGATAAAAATACAAATTAC
>evgR012440c0g2_i01_Hydra
TTTTTTTTTTTTTTTTTGACGTCATCTTAAGTTTGATAGATGTTTGTTCTTAAAAGAACGGTATCTTTCAAAAAAACATGCCACACCGTAAAGATCTTGCTCGTTTATTTCAATCTAGCGAGTATAGAATATTTGTTCCGCATAATTATTTAAATACGGAGAATGACGGAATTACGTATTACTTGAAAAGACAATTTAACTCAGCAATAAGACGAAATATTGTATCATCGAAAAAAAAAATTACTTTATTAAAAGCAGCTGGTGTATTAGCTCATCTTGAATATTGCGCTGAGCTTACAGATGAAGTTTCACACAACAAAGATTCGCAATTTCTTTCTTTACCGAAAGAAATACTCTTGCAGGCTTTGAACTACTTGCCAATAAAAGATATATGCTCATTCGCTCAGAGCAGCCGTTACTCAAGATTAATAAGCTATGATAACTTACTCTGGAAAATAATATGTCATCGTGTCTGGCACAACCCCTCTTTTATGTTTGTTAATATTGCTTCGGCAAGGTTGAGTCTACGACTATGGTATAATTGGAAATTAAAACCCTCTCAACACCTAAATACAGCTAAAGTGTTTGACATTTTTGAAGCTAACTTTTGGCGCGTTGTATTTATTGCTATAAATACACTAAACTCAATTCCAAGTATAATCATATCGCCTGAACCAAATGTTTATCCTCGAACCACAGAGTGTTTTCCGACAAACCGTACCATCAATTATAAATGCTCAAAAGGGAATTTTTCTCTGCAAGGAATAAACCAAGTTGGTCAAGTAGAATACATTGTTTTCAACAACTTTCTTCAACCGTGGCACAAACGTGAATATTTGAATATATCTCCTCTATCTATTTTCTGTTTGAAAGATTATGGCCTAAAACAGACGAGAAATGTTAGAAGACAGCAGGGGACTGAATTATTCCGACGCAGAGGATATTATATAAAAACCGATGAAAATCGAAAAGTTCACATAACATACCGATACTTTCCCACAGTAAGGTTCAGACTATATAAACAAAGAAAAAATGCTTTTATTTTATGCCACGACATTTTCATCAGCGGTGGAGTAAACACAAACTGTTACACGCTTTACCAAATCGTTCATCATGCTGAAAACTGTCATAATGTTTATCATACTCAATTGTTAAGCTTTGGTCAAGGTTCATCTTTAATAGATACTACAGTTGTTTCGGAGGAATTAAGCTTTTCGAATGAAAACTTTCGCGACGATCGTGGTACATCTTTTCTAGAAGAAAGTGAAATGCAAAATACTGAAAATGAGATTGCATTTAATAATCCCAATGTCATAGTTAATGATGCAAGTAATACAAATTAGTGTCTTCAATTAAAAAAAAAA
>evgR012451c0g2_i01_Hydra
TTTCCATGATAACACCTTCTATAGTTGATATAAATCTTTTTTGTAATAAAAAAATCTGAAATATGCTATTATACCTATCTTGCTTACATTAATAAACGAACATTGATATGAATTTATCATAAACACTTGTTACTGTTACTTAACATAAACTCATTGCACAATAATAAATGACACCGAAAGTTTCAAGAATTTAACAAAGAAATAAAGAATAAAAATGTTGAATTACGTATGTTGAATGGAGTAAATTTAAATGAAATTAATAATTTTTTTATTTTCTGTTTATTTAAAATGCACAATTCTACATTTTTAAATGGCTGTTGTGAGGTCTAGTCAAAAACAAGAGACTGAAGTTGTTATTTCAATAAAGTTGGCTTATTTATTTATATGCATTTTACTCGGTATACTTACTAATACGTTCGTTATTGTAGCAATAATCAAAAATAAAAATCTGCAAACTGTAACAAACTACTTTGTTCTAAATTTGGCGATCACCGATCTTTTATTTATTGTATGCTGTATGCCAACTATTATTATTACAACAATATCAAAAAAGTGGATTCTGGGATACTTTCTTTGTGAGGTTGTTGGATTTTTGAACTCTCTTTTTTTTATAACGTCAATTTGGACTCTGGTTAATATCAGTGTTAACAGGTACTTTAATGTGGCTAGACCGAACGATATAAAAATATTGTACACGAAAAGAAGGACAATTTTGATAAACATTAGTTTGTGGGTGTTTTCAGCAATCATTTCATGCCCTCCTCTAATTGGTTGGAGTGAGTTTATATCTGGCAACAATTTTTGCACAGTTAATGGCAAAATATCGACGTACGTTATTTTTCTTGGTGTAATTGGCTATGTTATACCTATGGTGTTTTTAGCTAGTTTATATTTGCGCATATTCTTTATGTTAAAAAAACACGAAAAAAGACCATCAGAAAACAACACAAATGTCACAGAAGTTTCTAATAACATCAACCATGTAACGCATAACACCCACGACGTTTCTAATAACACCCAACATGTTTCTAACAACATCCACCGTGTTTCTAATAACATCCGCCATCGGTTAAAAACTACAAAAATGTTAATTATTGTAGTTTTAACCTTCTTCGTTTGTTGGGCTCCTATCTTCATTGGAGCATTCATGTATACTTTTAATATTAACTCTGCAAGTTTTGATTTACCCACGGGTGGAATCATGTGGATTTGTTTGAACTGTATATTTAACCCGTTAATTTATTCTGTTACAAGTATCAATTTCCGAAAATATTTTATTCACAGTTTAAGAATGGTTTTTAAACGCATAAAATCTATTTGTTAACAAGTAAACCAAAATGTTCAAAATGGTCGCAAAGCAGTAGACAGCGCCAGTTTGAGGAGCCAGGATGAATATATACTCTATAAAAAATGTTAACAGCTCATTTACAACAAATAAACAAGCATGTTTTTTATCTTTAAATTCACAACGGAAGAAAAAGGTGAATAAAACATTTTTGACCTTATCAAGTTAAAAATAGTTTTCAATTAACAATTCAAAAATTATTTTCACATATGATGTCAAAAAGTGTCCCCTTAGTTAATTTAAATCTATATTAGGTAGAAATACACTTAATACTATTTAAAAAAAAA
>evgR012454c0g2_i01_Hydra
ACACATCAAAGGGTGAATTAGTTCCTGATGAATTGGTTTTGGATTTCTTGGTTGATAGAATTGGAAAGATTCGTCAAGCAAACCACGGACGTGGCTTCTTATTGGACGGTTTCCCTCGTACTCTCAATCAAGCTGTTGCTCTTGCTGCCATCAGTGACAAGATCAAACTTGATCTTGTTATCAATTTGCACCAAGATGAAGATATTATTGTTGCCAAGACAGTTAACAGAAGAATCTGTTCTAACTGTGGTCATGGCTACAATTATGCTCACATCAAGAGAGATGGCTTGGATATGCCACCATTGGTTCCAAAGGTTGAAGGTGTCTGTGATAACTGCGGAG
>evgR012460c0g1_i01_Hydra
AAACAGACAAGATGACCATTCCACAAGAATTATCAGTTTGTTTAGATACCTTTAAAAAGTTCTATGATTCTATTACCCAACAAAGAAAGTTAGATTGGATTCACTCTCTTGGTACTGCTATTGTTACTGGTCGTTTCTCTGCTGGTACAAAGGAAATTTCAACCAATACCTATCAAGCTTGTATTTTATTACTTTTCAATAACCAAGCTGAAATGACAGTTCAAGATATTCAAAATTCTTTGGGTTTGAGTGCAACTGAAGTCAAGAGAAATTTACTTTCTCTCTGTGCTACTAAAGCTGCCAATCTTTTGACTACTGATGGTAATAAGAAATCACTTGCTGCAACTGATAAGTTTACTGTCAATGCTGACTTTGAAAGTCCACAAAGAAGAATTAAGATTCCAAACGTTGTTGTCCACGTCACACAACAACAAAAACAAGATATTAGTCAAAAGGCACAAGAAGAAAGAAAGTATGTTATCGATGCTGCTCTTGTTCGTATCATGAAGACAAGAAAGATTCTTAAATATCAAGAACTCATGACTGAAACTATCAAACAATTGAGTTCCCATTTCACTCCTGATCCAAAGTTGATTAAGAGAAGAGTTGAAGATTTGATTGCAAGAGAATATTTGGAACGTGATGCTAAAGACTCAAGCACGATTCAATATGTTGCTTAAATTGTATATTTA
>evgR012466c0g1_i01_Hydra
AACGTGAAAGTGATGCTGGAAAAAATAACTGTTTTTAAATTTAATATCAAATCTAAAAAAAAAATTCTGAAACAAAACCTATTTGAATTTTCCTCGCAGCCCCATACAATAGAATTCTCATCGTTACCACATACAATTATTACAAAATGTTATTTGGATCTCCTTCATTCGTTGTGCATTTTTCAAAAATGTGAAATGATATTAACGTCTGAACTAAAAAGAAGGAATTTGAAGGCATATACAATTTTGAAGAATATTTTAAAAGCTAATAACAGTTCAACTACAGGTAATAATAACTCTTCAAAAACCGTGTCAAATATTTTATCAATTTCACGTCATTTGTTAAAAGATCTCCAAAATAACAAAGATTTACAACAGCAGCACGAACCTCTACTTCAAAATGATTACAAAACTAAAACCAATAATACTCTGAAACATATTTCCGCATCTACACCTCTTCCGATGACACCTCTCCCGATGAATCTATCGATGCTAAAGAAGCACTTGAGAAATGATATTGAGACAAAAATGGAATTACCTGGCAACGAAAACTTCCAAATTATTGATTCCACCAAACAGAATGATTTGTGTAATGGAAGAAAATTACTTAGTCCAATTAATGAATTATTAATCGAGGAAAAAAATCATGAGGTTTATAATATCAAAAATAGCATTCGATTCGATTACGTAAAAAAGAATGACGCCATTAAAAAGTTTGAAAAAAAAACAAAAGAAGAAATTGAGGGGATGCCTAAAAAATATGTCACACCGTTGAGAGATGAAGATGCAGATTTAGTAATTAAAACAGGAATATCAATTGATGAGTATTCTGATCATGGATTTCAGCAAGCACATTGTAACAACGAGGAGTATATTAACAGATTGACGTCAGATACAGGTGTTAAATCCAGCTATATCCAGCAAAGTATGAACAACCACTGCAACTCTTTATCTCAAAGTTTCAAAAAGGAAGATACATTTTTTGGCGGTGGTGGTGGCGGTGTAAACTCTTCTATTTTTAAAAAAGAAAACATTATTAAGCTTGAGTCACAATTTCAGGATCCTAATCTTTTGAGTAATAGAAAATCTTTTTATATCGAAAACAAATTTCGTGAATCTTTAAAAAACAATAAAAACGCTGAAGCTGAAATAATATTTCATAAAGAAGAGGATGATGAAGAAGAATCTTTTCGGCAAAGAGTTCTTCGCATTGACGACATTCGAGAAAATGAGGAGTCGTTGGAAAAAGAGGCGGATTTTGATGTTTATACTTATGATCCGACTGAAACTTTTACACAAACAGACCGTGTTACCTTAAAACTACCACTAAAATCTTTTATAGCGTTATGCAACTCTGCTCAATCGAAAAACGATGTACACAATTTTAATGATAACTTAGTTACCGATAACTTCATTACCGATAACTTCATTAGCGATAACTTTATTAACAAAACCGAAAAGAATAGTGACAATCACAACATAAAAAGTAAAATTAAAAAAATTAAAAAATCAATTAACAACAATAAATCGTTTACAGAAGATGTTAGTTGTGAAGAAGAAAAAGCTATTTTTAAAAATGAAGCAATAACAAAC
>evgR012477c0g1_i01_Hydra
GGCATCAGGAATGGAATATCTTTCTTGTATAAAACTGGTACATCGAGATCTTGCGGCAAGAAATGTGTTGGTTGGTACTAAAAAAAATGTAAAAATATCCGATTTTGGTTTGACGCGACAAGTAAACGATCAATCGAATTATATGACAAAAAAAAATCGACGTTTACCCATTAAGTGGATGTCTGTCGAAGCTATATTTGACCAAACGTTCACAGCATACAGTGATGTGTGGGCGTTTGGAGTCGTATTATTTGAAATTATTACGCTTGGAGGTACACCTTACCCTAGTGTTAGCAACCGGCAACTTCTCAGTCTTTTGAAGTCTGGCCACAGAATGGATCGACCTGAAAATTGTTCTG
>evgR012499c0g1_i01_Hydra
TCCAAATATTGATGTTGCTTTAGTAACCTTTGCTACTACTGCTGAAATTAAAGTTAATTGTGGTACTTTTAAGAGTTTTTCAAGCTTTGTAACTTTTATGAACGATTTAAAAAAACCGGAAGGTGGAACTGCCATACATCAAGGATTGAATTATTCACAAACTGCATATAATAATTGTATAAAACTTAACATGTATCCGGTTATTATTCTTCTTACTGATGGTGAGGAAAATATAGACAAAAATATAGACAGTAATATAATTCTTGAGAACCAACTCAAAAGTAATGCTCTAGTTGTTGCAGGTGCTGTAAAGGTCTATAATAGGGGTGAGATTGATAGAATAACAAGTTATATTGTTAATGGACAAAATGTTTCTTATTCCTTATATGCTACAAACTTTAC
>evgR012500c0g1_i01_Hydra
ATTACTGACTCTAAAGTTTTTACTAAATATACTTTTAACATTTGCATCGTTAAAGATATGTCGGAATTATCTTCTGAAAACTTGAAAAATTCAAGTAATGGCAACGAATGCACCATTACTGAACAAATAGCAAAGAAAAACCATGTTGTAAACCAGTTTGGTGGATTTGATGAAAAGTTTGTTGATCCGCTTCCAAACGATTTTGTTTGCCCAATCTGCTTCTTGGCATTTCGTCAACCTGTTCAAACAGTCGAGTGTGGCCACCGTTTTTGCATGTCATGCTTTAAACTTCTCAAAGGAAACAAAGATGTCTTATTATGTCCTCTTGACAGACAACTAATTGACTTAGAAAAGGTTTTTTACGATAAAGCAGCAGAGAAAGCAATACAATCTTTACCTATAAGTGTCCTAACTTAAATAACAATTGTGATTGGATTGGCACCATTGCCTCTTTAGATGAGCACTGGAACGAGTGTCA
>evgR012517c0g1_i01_Hydra
TGCGACATTTCAATGAATTTGATTGGTCAACTTAAAACACAATATATACGTTTTCTTTATGCAAATAATAGCTTGTTTTTCCCTTTATTTTGGATATTGGTGAGTTGGGCCAGAGTTGTTGGTCTAGTTAAAAGTTCTACTAATGATTGTGGGGTCATGGATACAGCTGAAATGTACTCAGTTATTATTGGTTTAATAGATACAAAAAAGTATGAAAAAATATTTATCAAAAACAATCGGAATAGTTCAATTTCATACCTTTATAGCTACTTAGACAAACTAAAAAATAGAGAAAAACATGAACTGGGACTGATGCTGCAC
>evgR012518c0g1_i01_Hydra
CTGTAAAAATATTTATTACTTAAAGCTATTTTTACCTTAATGACGTCATCCTCCTTGTGACGTCATCTAAAAAATTTTGGTCTGTTTAGATGAGCCACAAGGAGGTGCTACTATATTGACGAACGTCGAAAATTCCTCAATAGTTTTGGATCAAGATAGAGAAATTCAAACAAATTCACTTCCTGTTGCTGAAAGCCACTCACCTCACATCAAAGTTATTCGTTGGTTAAATAAAAATGGGAACGAACTAACAAACCCTTTAGGTCTTGACACTTATGATGATATTAACACCGCAACTAGTAATAATTTTTTACCTAACGTATCTACAAAAGTAGAGTTGTATCATCATAGACATTTTCACATTTCAAAACCAAGATTTATACAATTGCAAGATCCAAAATTTGTGGGAGGTAGATATCCTTTCGGATCCGAAATGTGTTTACCTCATCCACCTAGTTCAGTTTCAAAGTTTTTAAATGATTCAAAAAAGAGGAATAAAGGCATTCCCGAACATTTTAATGTCCCTTTTTTATACAGATCAGACCATAAACATCATCATCAACATTTCAATTATGACCCTGTTATTTACAGAACTGACCATAACAATGCGCACAAGGAAGACTATAATTTTCATCCAAGGCAAAACAGTAAAAAATCTTTCAAACCTTACTCAACGGATTTGCCCTCGTGTTCTCAAAGAAATATGAACAATAAAGCGTCATGTCTACAAACAAGTTTTTTATCGTCGTCAAACCAACAAGAAACATTAACAAATCAACAAGTACACTATAAAAATCCAAACAAAAACGTAACGAGGCTACATTTAAAGCGTCCGTTTGAATACAGTTTATCTAAAATGAATAGTGAACCTTGTTTAAAACCAACTAAAAATAACCACTTAATAACTAGTGACAGCCAACGCAATACCCCACGCGTACAATCGGAATTATTAAAACACTCGTCGGCTCCTCAGTTAAGTTATCTTCACTTATCTTTGCCCTTATTCCCATTAGAACATCCGTGGTTTGATAAGTTTATAAACAAGGTAAAAGTTGAGTTAGTAGGAAAGAACCTTTGGGAAGAATTTAACAAACTTGGGACAGAGATGATTCTTACAAGGGCCGGAAGAAGAATGTTTCCTAACTTACAATTATGCGTATCAGGAATGAATCTAAAAGCTATGTATGTTATCTTCGTTGAATTTGCGCCCATTGACAATCATCGATATAAATATGAATATGAAACATCAAGGTGGGTAATTGCTTGCGAAGAATTTACGCCTCGACCATCTCGAGTTTATTTGCATTCAGATTCTCCGGCTAGTGGAGAACTTTGGATGAAAAGAGTAGTAAATTTTGCTAAATGTAAACTAACAAACAATTTTTTAGACGACCAAGGACATGTTCTCATGCATTCAATGCACAAATACCAACCTCGAATTTATGTTATGTATTGGGACGGTAGTCTTCCTTTGAATATTGATCCAATTGAGTTACTGCGTGTTTCATGTTATAAAAAGTTTATTTTTCCAGAAACAAAGTTTATTGCCGTGACTGCATATCAAAACAGCCAGATAACGCAGCTAAAAATTTCTCAAAATCCTTTTGCACGAGGCTTTAGAGGAGATTGTCGTAAAACAGCTGAAATGACAGTCGTTAAAACGGAAGAAATAAAACATCCTGTAATAAATGTTGAACCGATGACAGTTGAAGATTGGACAGATTACAAATCATGCGCGTAAAACTAAAAGACATGCGCGTAAAACCTTTCCAAAAGAGTATGTTTAACAATCACTTTTTGATATTTTATAATATTTTGGTAAATATCTAAAATTATTTTAATTATTTCAAATAAATTTTTAATTTTTGAAAAAAA
>evgR012521c0g1_i01_Hydra
ATTGCATTTGACAATATTGATTGGCTCCAAAAGTTTTGTGTTTATTACGACATTGATGGTGAATATGTTCCTTATAGGGATAACATTTTGTTTACAGATGATATTGTATTTGCAGTCACACCAGAAGTTTCATTCTTTCCTTTCCGTACAAAAAATATGAAGCTAAGTTGCTTAGAGTGGTTTGGAAACATGACTAACAATATTACAGTTAATGTGAAAGGAAGCCACTATTTTACAGCCGATTATGCTTGGAAAAATGGAAATTCTAATCAAGGAGATCCACTTGTTAAATACATAAATACAGAAGGTGTTGACACACTTCAAGGCGATTCTAGAAATTTCCGAAAAATAAACTCGTTGAATAAATTGATGAGCAGTCATTATGGTGATTCATGTTTTACTCAGC
>evgR012541c0g2_i01_Hydra
TAATTCAAGCTTAATATTTTATTTGATATTATTTTAAGTTTTTAAATAAAAAATTATGAAGGTAGTGATAAGTGGTGCCTCAGGATTACTTGGCAGATCGATCTATAGAGAATTTGTTAACAAGACAAACTGGGAAGTTTTAGGACTGGGTTTTACAAGGTTGGGAGATACTTTACATAAAGTAGATCTTTGCAACGAAGAAGATTTAACGCACTTTATAAGAAATGAAAAGCCTGACGTTTTTATACACTCTGCAGCTGAACGTCGACCTGACTTTGTTGCAAGTAATAAAGAGAATACAGCTCAGCTTAATATTACATCTACTAAAATAATTTCTAAACTAATGAATGAAAATGGTGGATTTCTTCTTTATATTAGTACTGATTATGTATTTGACGGTTTTAGTTCTCCCTATAAAACAAATAATGTTCCGAACCCATTAAATGAATATGGATTATCAAAAAGAGAAGGTGAAATTTGTGTTTTAGAAAACTGTACAAATTATGGCATTCTTAGAGTTCCAATATTGTATGGTGAAATTGAATATCTTCAAGAAAGTGCTGTTACAGCATTATTCGAGTTGTTAAAAGATAGCAAATCACCAAAAAAGGTTTCTAACTATGAGCAACGATTTCCAACAAGTACAGACGACATTGCTGTTGTTTGTCGACAAATAGTTGAATACAATCAATCAAACAATTCATTTCATGGGATATGGCATTGGAGTAGCAATGAGAAATTGACAAAATACAAAATTGTAGAAATAATGGCAAATATATTTAATTTATCAATGAAACATATTGAGCCAGATAATGAACCGTCTGTTGGAGTCTCAAGGCCATATAATTGTGAGCTTGATTGTTCAAGTCTTATTGATTTAGGCATCGGCAAACAAACTGATTTTAAAGAAAACTTAAAAAAGTATTTGGAGAAATTTGTCTGATTTATAATAAAGTTTTCCTAAAAAAAACATTTATTTTTAAATATTCAAATTCTCTATAATAAATGTATCTCTCTAATATAATATAAATTATACTTTATTATATCATACTTATTATAAAATATCTTTTGTACCTTCTGATTAGTGATGAAAGTTTCCCCAAATTTTAAACACAAAAAAAAAA
>evgR012574c0g1_i01_Hydra
TTTTTTTTTTTTTTTTTTTTTTTTTTTTTTGTTTCAGTTTTAAATCTAATTATCTTTTGTTTGATGTTTTCATGTTTGACCATGTTGATGTTTGTTCAAACTTTCAAATTTATCAAAATTAACAGTTTTTTATTTTTAATACTTTTATTATTTAAAAGATATGTTTTTATGGAACAGTGTGAATATAATTTTTTTGTAAAAGGTAAAGCACTGGAATTTCCTAAGCTTTTAGTTGCTTTATTTAATGGTTGTTGGAAAATGTCTCAGCCTGCTAAAATAAAAAAAATTATCTATATGAGCAAATAGCGTGTCTATGCTATAAAATAAATTTATAAATGTTAAAAAAAAAA
>evgR012577c0g1_i01_Hydra
AAATGAACTCAACTCTCAAAATTTGTGGACTCGTCACAATGATGATCCTCTTGATCATTGCTGGTGTGTGCAGTGCTCAAAAGTCCTATTCCATTTCGTCAATCAATTCTGTAGTAACAGTCAACAGTGGAACCTGCAATGTTAACGTTGAAGAGACCTATGTCTATGATTTTACTGGATCTTACTCAACTGTTGCTAGAGCTGTTCCTTTCTCAATTGCTAGTTCTATTGATGCCAGTTCGTTGAGAGTGGACGTTCTCACCTCCGGTTACACAGTCTCCAAATATATCTCACCACAACAAGACTCTAATTTCTTGGTTGCCAATTTATATCCAGCCACACCAATGAGTGTCACTACTCGTATCACCCTCCGTTTCAGTTACTCTGCTTTTGGTCCAATTCGTATTTCAGGAGATAATTATGCTGCCACTTATTATTATAAGGAAAGTACATCCATTGGTGCATTGAATGCTCAAATCAAATTCTCAGACTCAATTGATCTCAGAGGAAACATCACTGTCTCAAGTGGTTCCTACGTTTCTGGAAGAAGTCTCGTCTACCAAACCGGTTACATTGCTCCAAATATTGCCAAGCAACCAACTGCTACCTTCAACACAATTTCAGATCAATTCAAGAGTTGTGCCAACAGTGGTCTTCCATTCTTTG
>evgR012591c0g1_i01_Hydra
CATTCCACTCCAATGGTTCTGTTCTTTCACACAATGGTACAATGAGATTTGACAATGGTACATTCTATTCGAATGGTTCCATGTTAACCAATGGTTCTCTCAGTATTGGAAAGAATGAAACTTTCATTCAAGGTGTTGTCTCACACAACGGTACCATCCACTATCAAAATGGCTCCTTCTATCACAATGGTTCAATGGATTTCAATGGTGGAAAGAGTGGCGCATACTGTTATGAAACTGGTACTTGTCTCAGAGGTGCCTTCAATTTGACTGGATCAGTGAATGCAACCATTTATAGAGCTTTGGAAATGTTTGGTAATTTGACATTGGATGGTGATTTAGAAGTTAAC
>evgR012601c0g1_i01_Hydra
CTCTGTACCTCTGTGCGAGCAACTTATAAACGCCGTTGTTTTGACCACCACATTTAAACATGTCATCTCAAATTGAAGTTGTTGACCAACTCCTCACCCAAGAACCAAAGGAAGAACTTAACTTGAACACTGCTATCAAGAGAGTTTTGAAGACTGCATCTGCCAACAGATCTGTTGCTCGTGGTTTGAGAGAAGTCTGTAGAACTTTGGCTAAGGATAAAGCACAATTGTGTGTCCTCGCTGACGATTGTGATAAGGATCAATACAAGAAGTTGATCAAGGCTTTCTGTGCTTACAAGAAGGTTGATCTTTTGGAAGTTCCATCCAAGAGACAATTGGGCGAATGGGTTGGTCTTGCTAAGTATGATAAGCAAATGAGAACCAGAAGAGTCCAAAAGTGTGCTGTTGCCGTCATTACTGATTTGAGCGGTTTCGGTGAAGAACAAGATGAAGCTTTGAACTTCGTCTTGAAGCACTTAGGAAAGTCTGATTAAATTGCTTCTCACTCAGAAATAAAAAATCCAAAATTATCACAAAAAAAA
>evgR012604c0g1_i01_Hydra
AGCAGCAGAATCTGTTAATGATGTCATTCTTGTGAATAAACTGAAATTTCAACAGGATCTTCACATAAGGAAAGTTGATGTGTTTAATATGCAAATCAACGAAGCCACAGAAGCTGCAAAACTAAGCCAAGACACTGCAGTAATCTGTTTGGACTTCGAAAAAATTTTACCACTTTCTCTAACTGGCATTGGACAAGATTATTATAGGCGTCAACTTTGGATTCACAATCTGTGCATTCATGACAATGTAAATGAATCTGTTCATATGTATCTGTATGATGAACATTTTGCAGCTAAGGGCCCAAATGAAGTCATTTCATTTATGGATCATTACATTTCTGGTTTTTCGTCTACAACTAAAAAGATTATAATTTTCATCAAACTGTTTTTCAC
>evgR012605c0g1_i01_Hydra
TCCACAGCACCAATGCACCACAGAGGTTTATCCACCAATGCATCCACAAGACCAATGCACTACAGAGGTTTAGCCACCAAATTATCCAGAGCACCAATGCACCACAGAGGTTTAACCACTAATGTACCTAGAGGTTTAACTACCAATGCATCCACAGCACCAATGCACCACAGAGGTTTATCCACCAATGCATCCACAAGACCAATGCACTACAGAGGTTTAGCCACCAAATTATCCAGAGCACCAATGCACCACAGAGGTTTAACCACTAATGTACCTAGAGGTTTAACTACCAATGCATCCACAGCACCAATGCACCACAGAGGTTTA
>evgR012608c0g1_i01_Hydra
AATTCAAACCATTAACTTACGCAGCATGAAAATAAAAATAAAACCATACTCAAAAAATCTTTTACTTTTAGCTACCTTGAAAATGAGCAACTTTTCCGTGGAAGAATTAGCCAACTTTCAAGATGTGTTTTACAAATTTGATGAAAATGACGATGGGCGTATAACTTTAAACGAGGCAATGCATGCGTTTCGCTTACTTGGATTTAGTCCTACTGTTGGGGAAGTGCAAGAAATGTTTACTGAAAAAGAGAGTCTTCACAAAGGAACATTAGATTTTCCTCAATTTATACTCATTCTTGAAAAGAATTTAAAAGCTCCTGATTCCACTGAAGATCTTATAAAAGCATTTGAAACATTAGATTCTAAACGAACAGGATTTATACGTGCTAGCGAGATGTATAACATCATTACGAGACTTGGTGATCCGAAAATAAATTCTGAGGAAGTTAATGAAATGCTTAAAGAAGCAGATACCAAAAAAAATGGCGTAATTAATTATCGAGAATTTGCAAAAAAGTTGGGCGAGTCATAAAATATTTCAAAGAAAATTGCAACACTTTTCAATTAGAGCATCGCAACACGCTTTAAATTTGAACATCTTGACACGCTTTCAATTTAATTGTTTTGAAAAACAAACAAATGTGTTTTATCCCAACTCTTGGCATGCATATAGATGATCATATATATACATAATATAAAGGACTAGCTCTAAAACATGGACCGGGAAAACCATTATTGACCGGCCTCATCAAGTTGTAACAGCCGGTTTACACTAATTGTTTTTTTAATGTCTTTTAATATTTATAATAAAAGAATGGGTGACTGTTTTTTCTATAATACAATTAATGGGGGGTTTTATTGACAGACGATCTCGCTCCCAAATGAAACAGCCCGAGGTTTTTTTGAAAAAGGGACCTAGTTCAAATAAAGGTATTTTTATCTAGTTTTCAGGCTTTGTTAAACTTACTTTATTTTCAAGGAAATTTTGTTAAAAACGGCATCAAATTTATTGTTTTTTCAAGTTATTCAATTTCAAATAATC
>evgR012612c0g1_i01_Hydra
TTTCTCTCTTTGATTCTGACTCTGATGGTACCATTGTGACCAAAGAACTTGGTACAGTGATGAGAGCCCTCGGTCTCAATCCATCACAAGGTGAATTAGATGATATGATTAAACAAGTTGACTCCAATAATAATGGTACTATTGATTTCAAGGAATTTTTAGTTTTAATGCAAAAGAAGATGACTGATAACGATTCTGAAGATGAAATCAAGGAAGCCTTCAAAGTTTTTGATCGTGATAATGATGGTATTATTAGTGCTGCAGAGTTGAGACACATTCTCACATCAATGGGAGAAAAATTCAATGAAGAAGAAGCCGAAGATTTTATCAGAGAAGCTGATACTAATGGTGATGGTCAAATTAAGTATGAGGACTTTTGTAGATTAATG
>evgR012620c0g1_i01_Hydra
GAGGAGATGTAAATCACAAAACTGTTTGGTATAGCTTGTTATAAAAATAATGTGTTAAAGCTAGTGGATAACTACATTTCTCTCTCTTGTTAAATCAAATATTACAGCATGAAATTATTGCTGAAAAATCTTATGCAAACTTTTCTTATAAACTTTTACAGGTCAGGATGCTACTTTTGTTCCTGATTGTTTTTGAAGTGCTCATTGGAGTTCAGGGTTCTCAAGCAGCTTCTAATATTCTAAATAATTTGCCTTCATTTGAAGCTTTATTTGGATTAAATAATGCTGATGGAAGCACTTCTTTTCAAGTTTTTGGATGTAATATAGAATACAATATAAAAGATTGTACTAAGAGGTTTACCCCAGATGAAGATTGTTTCAAGGAACTATCCTTAGCAAATGTATTTAGCAGTCTATGGAAAGGAGATGAATTAGAAACTACGACAAAAAGTATTTTTTCTCCTGGAGTTTTTGCTGTTAAAGAATTTGTTTTAGATTCATGTGACCAATCAAAAAAAAGCGTGCAATTTACTGGTGTGGTTATTACACCTTTTAATGTTATCATTGAAGAAATTCAGATAACGAGTGGAGAACTAAAATTAAAGATGGATTATGAAAACAAGAAACCGAAGATTAATTTTGCAAATATTGAGTTTACTATTAAAGGAGAAGTTAAAATTGCAACCCAAGTTGTTAAAGTTGAAGTAAAAAAAAAA
>evgR012626c0g1_i01_Hydra
AAATAATCGATTTTATGTAAACATCAATAAAAGTATCAAAATGGCTGGCCACGTTGCTTTCTTTTTCATAGTAATATTTTTCAATGCGATATTGGCTGACGAAGGAAGACATAGTAGATTTGAATATAAGCTGAGTTTTAAAGGGCCTCATATAGCTACAAAACAAGGAGCTGTCCCATTTTGGACCCATTATGGAAGTGCAATTCCAAGTGATGAACAGATTAGAATAACTCCTTCGTTAAAAGATCAGAGAGGCGGCCTGTGGACAAAGACAGTCAGCTCTACTGAGTTTTGGGAAGTTGAAGTTTATTTTAAAGTTTCTGGGCGTGGAAGAGTAGGTGGGGATGGATTGGCTGTTTGGTTTACCGAGACTGCTGGTTTGCCTGGTCCAGTTTTTGGAAGTTCAGATAAGTGGAAAGGACTGGGACTGTTTTTTGATTCATTTGATAATGATGGAGAGCAAAACAACCCTTATATTATGGCAATGCTTAATGATGGTGACGTAAACTATGACCACTTTAAAGATGGAAACAAACAACAACTTGGGGGCTGTATGCGAGACTTTAGGAATCGACCATATCCAGTTCGAGCAAAGATTCGATATTATAATAACGTCCTTACTGTATTTTATCATGCTGGGATGTCTGAAAAATCTGAAGATTATGAACTTTGCATGAGAGCAGAAAATGTTAAACTTCCAAAAAGTGGTTATTTTGGTGTAACAGCAGCTACTGGTGGCTTAGCAGATGATCACGATGTTTTAAAGTTTCTTACTCACAGTCTTACACCTCCACTTACTGAGGAACAAAAAAAGAAGAACATTAATGAAGCAGATCGAAAAAAATACGAAGAGCAATATGGTGAATTCAGTTCAAAACTTGAAACTCAAAAGGAGGAGTTTAAAAAAGATCATCCTGAGAAAAAACCATCAGAAGAAAAGTATGAAACTGTTTACGAGCGTGATGTCCGTTTGATGTATGAAGCACAGCAAGAAATGCATAAAATTATGCGTGCATTGCATGGTAAGACAGGTGAGCTTACTCAGGAAGTCAATAACTTGCATAGTGCTGTAAGCAAACAACAAAATAGTGGGGGAAGTGTTGACTATAGTATGCGACAGGATATTGGAAGTATTGTCAGTTTACAAAATGATGTTCATAGATATGTAAAAGATATTGCATCTCGTATTAGTAATAGTGATGTAAAAAAACTTGAACAAATAGAACAAAATATAAGAACAATATTAGAAAATGTCGGTCATTTATTATCAAAGAATCAAGATAGCAAAAAGTCATGTCCTAGTTGTGTTTCATCTAGCTATTTTACTTTCTTTCTATGTATGCAAATAATTCTTATCGTAGGCTATGTATTGTATAAAAACCAACAAGAGAACGCTCATAAAAAATTTTATTAGATCTTTTTATATGTAGAAAATTGATATTAGCATATTATTTTGTTAACTTAATATTTTCTTTACAGTTTTTATTGTATTGAACACCTCAAATTAAATTCTATTTATTATATATTAGAACAATATTGTTTTCTATTTTAAAGACGTCGCAAGATTATGTGACAAATTTGTACATCGTATTTATAAGTTTAAACAAGATATATACGATTTTTTAAAAAAAA
>evgR012634c0g1_i01_Hydra
GTAATGAGAAGAAGTTTAAAATAAAAACAACAAATGAAGAGGAGTTAAAAATCGAAACAACAAATGAAGAGGAATTTAAAAAAAATACAAGTAATGAGAAGAAGTTTAAAATAAAAACAAGAAATGAAAAGGATTTAAAAAAAACAACAAAAAATGAAAAGAAATTTAAAAAACAAACAAGAAATGAAAAGGAGTTTAAAGATGATCATAAAGAGGATTTTAAAGATGATCTTAAAGAAGATTTTATTGATTTTGAGAAATGGCCTTTAGAATCAGTAAATGTGCCATTAAATCTTTCAGAGCCAAAAACAGTAGAAGAAATTTTTCATGAAAAGATAGAAGTAGACCTTGTTTGTAATGGAAATAGTGAAATCCTAACAGAATTAACTTCCTCCTCAGATGTAAATGATCATGACACGCAAATGGATGTAGATGAACCAATCAGTAAAGTTTTCAAAATGGTCAAGGTGACTCCCTGGTTAAAGTTGGATGGCGATTTAAATGGAGTTCTTCTTAAACGCTTCCAAGAGTGTGTACTGTTATTTGTTCTTACAAATCCAGGTGTTCAAGAAAGTACAATCTTTAAACATTTCGAATCTCTTCTACGGCCTCACGCTCTGTGCATGATACTAAATTTCTTGGTTTTAAACGGCTGTTTGTCGAAACACTATCAACGAAAAAGATGCGTTTCGTTGTTCAGCTCATCGTTTAGCAATTTTGATGACGGTGATATTTATTACCTTCCTTTAACTTCTTGCTTTTTAAACATGATGGATATGAATCGATGAGAAAAACTCTTTAAATTCCTCATCCGCAATATTACATTGCATTTTAAAATTATTTTCTTTTGTATACACTGCTATGTAAATGTACAGCTGCGTCTTAATTTTTTGTAACGATTATTTATGAAAAAGAAATTGAGAAAAAAAAAAA
>evgR012641c0g1_i01_Hydra
CTCGAATTAATAATACAGAAGCAAAAATGGTAATTGGTTTGACTGAATATTTAGTCAATTGTGGAGTTCCTAAACCTTCTATTGCCATACTTACGCCATATAAAGGGCAACTGATGTTAATACGTTCAGATTTAATCAAGTTAAACCTATTGAATAATAGAGACAAAGAAAACTCTGTAGTTTTATCCACTGTTGATAGGTTTCAAGGCGATGAAGCTGATATTGTTATTATTTCTCTTGTTATTGATTCTAAATCTCACACAGGCTTTGTTAAGAAAATAAATCGTATGATTGTTCTTTTAAGTCGAGCAAGGCTTGGGATGTATATTATTGGTAATATAAACTATTTTGAAGAAAAAACGGATGAGTCTGTTTATCACTGGGCTGTTATCTTGGCAAAGCTTAAGCAACCTTATGATTCAGATGGTATTCACTATAATGGTCCCAGAATCGGAAATAAATTGCCTATTTGTTGTCCAGTTCACAGAGATAAAGCTATTTTCTATGCTAAATCTGTAGAAAATTTGAATCCTAATTTATGCAAAGAAAAGTG
>evgR012642c0g1_i01_Hydra
AATGCTTTCGTGCAGTTTATGTCAGTTCAAGACTACTTCCTTGCAGTATTATGCACAACATTACAGTGTTCACAGTTGTGCGTATAAGATTCCGTGCGGAATGGTTGGATCCAAAAAATCTTTTAGTACTTTCAGGGCTTTTACTTCACATATTCATTGTATCCATGTTGTTGGTAATATTACAAAATCAATAAATAACTTTGTTAATATAGAAAAGATAGGAGTATGTTCAGTTTTAACTTGTCAAGTTGAACTACCTTTTAATGAACTTATAGTGCATTTAAAACTACATATTAAAAAAAGGGTTTCTATTAAATGTCCATATGAAGAATGTACTCGCCGTTACAAAGTTATTTCTTCATTCACAGCTCATTTATCCAGATGTCATCAAATCCACAAAACAACATTTGAGCAAAGCCAAATAAATATTTCATCTTCAGAAACTAATCACGATCATTTAAATATTGA
>evgR012643c0g2_i01_Hydra
ATTTTGATGGACCTGGGCTATTTTAGTGCAAAAAGATACTGTAAATAGCATGAATTAATTTTGCAAAACCAGTTATTACCCACGAAAGCGAAAAATGCCACCAGCAAAGTCTTTCAATAAGATTTTAGTTCTCGGTGCAGTTGGTGTCGGTAAAACATCGATCATTGAGCAGATTGTTTATGGAAACTACCACCCTGAAAAGGCGATGTACTCAACTATAGAAGATTCTTATGATGCATGGATTGACGCAGAGAAGGGACCAAAGGAAAGAATAAGAATATATGATCTCAAGGGAATGGATTATGTTAGTCCTATCTTGCCTTCACATTATATACAGATTGTTGATGGTGTTATGTTGGTATTTAGTATAACTTCAAAGAAAAGTTTTACATTAATCGAACAACTCAAAAGTGACATTGAAAAAGTTAGAGGAAAAGATTTTCCAATGGTTGTTGTTGGAAATAAAGTTGATCTCTCGGATTTACGTGAATGTGATCATGCCGTTTTAAACAAATGGGCTAGCAAAGAGAAAGTCAAACTTTTTGAAGTTCTGCCAATGAATAGGAATACATTACAAGAGCCAATCACTTTTTTGATGAAGAAAATTGCTTCTGGTTATGGTAAAGTGAAACCAGTTCTTTTTCGAAGACCTGGCGGGAAAAGTACGGTTAAAGAACCTGATGCTCATGGAAAGGAAATAGTCGCTTCGGCCTTAAAATAATTGCATCCGTTGACAAAAAACCTGTTTATTTTGTTAAACTGTCAGCTAAAATATTTGAAATGTGATTTTCAAATATTTTAGCTAACATGAAAATAAAAAACTTAATGAAATAATTATAAATGATATATTATGTGATGTGATGATTGGTATATAATAGTATGAGAAATAGTATTTATTAAATACTAACGATGATAAATGATATATGAAATAATAAAAATAAATTTAAATAAACATCATTGCCAGTGGCGTAGAGATAAAAAAAAA
>evgR012655c0g1_i01_Hydra
GCTGTAATAGCATTATATTTTCAAAACAACAACAATAATAATAACAACAACAATAACAATAACAACAACAATAACAACAACAACAACGTTAATAACAATAATAACAATAACGTTAATATCAATGATAATCTTGGATGGTGGAGTCCTTTCACAAACTCTTTGTTTGGATACTGTGGAAGTCGGATGTTTGATCCTTTTGCAGAGATGTGCTGTAATAGCATTATATTTTCAAAACAACCTTTCATTCGTTGCTGTGGACCGACAACTTACAACATTTTTACGCAAACGTGCGCTAACAACGTAGTTTGGTCAGCTTCGCAGCCAAGATGTGCAAATCAATATTTTGATCAAATGAAGCAATTTTGTGTCAACAACCAACTTTATACGCATGCCGATTTTGTAAACTGTAATGGAAATGTCTTTAGAAAGTATTAGCAAACTGCTTTATTTATATAAACTCAGCATATGTAACTGTTAAACTATTATGTCACTGATTTAGGTACTTTTTAAATAGAACAAAATTTATTTTGTTTAAAAAAAAAAAAAAA
>evgR012657c0g1_i01_Hydra
GCTAGAATTAGTAGAAAATCTTGTTTTGTCTGTTGAAAAAGCAGAAAACTTTGAAAGATTTCAAAATTCATGTTTGAGTTTCTCTCAACTTGTTGCAAATCTCCAGATGTATGTTAAACATAGTGCAGTATGTAAAAAACAGTTGATGATCCTCCAACATCATTTATGGTTTGTAGAACATTTAGAAAAGGATTTGAAACACATTTCTGATTGTCTTGAATATATCAAGTTATTTTCAAAAAATTTACCAAAATATTTTGACTTGTATCATCAGTGTTTTCAGCATCTACCAAATGTTTTACAGTGTTCAGCATGGGTAAAAGAGTTACTTGTTGTTATGGGAATCAAAGTTATTTTTGTTAATGAAGAAGATATTGAGAAAAAGTTATTAAGAATTAAGGGTTTGAAAGAAATCTTCATTAGTAAGGCTGCAAAAGTGGATGAAGTTATGGTTAAAGGAAAAATACTTTTAAGTGAAGGTTTTTTAATACCCTTCTATTATAGTTTTTGCTTATCACAACTGGAAGAGGAATGGAAAAGAGTTAAGCTATTGGTTAATGCTAAAATATTAGATGCAAAGAACTTTAAAAAAAATTGGATTCAATTCAATAAAAAGCTCAATGAGATCACAG
>evgR012684c0g1_i01_Hydra
ATTCACTAACAGAACTATTTTTTTTGCGATTATTCAAATAAATGCAAATATTCGCATTTATATCACATAATCGCTTTACTACTTTGGTTCAATCAAAAAAGCAAGCTAGAACAAACAGTTTATGGAGCAGTTTGTAAAAGTGTTTTTTGAAAACGCTTCATTACCTTTGTATATGATTTCTTTGCTACTTCATTCTCTTGGAATATATTTATTGACAGTTGCTCAAAAGAAATGCAGAATACAAGATGTCATGTTTGTCAACTTAAGTATAGCAGAAATATTCATGAGTTGTTCTGATTTGACGCAAAATTTTGTGTCACGTTATCATGTAAATTCAGTTTTTATTGACTATCTTACTATCGTACAGTGTGCACTTTTTGTGGTTCCTTGCATATGGATCATGGCTGTGTTGGCTGTAAATCGTTTTCTTGAAATTTATTTGAATATTAGATATAACGTTGTTATTAACAAAAAACAAGTGCATGTTGTCATGGCAATGGTATGGGCCGGGGGTGCTCTATCATTATTAGTGCTAGTATGCTTACAAACAAAACAGAAAAATATGTCTTTGATTATTTACATGTTTTTTTTCCCAGTAAATGAAGGATGCGTTTTTTTCATAGCTTTAGTGGCATACTCCTATATATGGCTAAAGGTAAGAGAACACAGAGCCAAAATTGATAGAAGACTACAGCAGTTTCAAGTGCATCAAAAAAGATATTTGAAAAAAAATTTCTTTCCTCCTCTTCTCATCATTATTTCATTTATAGCTTTTGTTTTGATTCCTGATATTCTGAACTTGATTTTTGTACATATCATACAGTTAACGGCAACCGAGATTATAAAAAACATCCTACTCGTATTTTATTCCACAGGTTTTATTTGTGACGCTCTGATTTATATCTTTTTTCAACCACATGTTACCCGATTATTGTCTAATAAAGTACAAACTTTGTTCAACTTGTCTGGCAAAAACTCATATTTTTTTAGAAGTTACCCTGACCGATTTGAAAGCAACAAAACTATTTCTAGTTTTACTTCTAACTTCAACATTGCAGTATTGCATACATAAAACACAGACACAGAAAAATAAAACAAAAAAGAATAAATGAAAGTTTAAACAATACGTTTGCGTAAAATATTTCTTACTATATTATTTTATATATTGCTCGAGCAATACGGTTAAGGTATTTTTATACTGTAGATGTTGTTTTAAACGTTTAAATTATGTAGCATAAATTATGTAACATATAAGTAAATTATGCAGCATGCGGAGAAAAAATATAAAAAAGCGCTCTTTAGCTAACACAC
>evgR012693c0g1_i01_Hydra
CTGAAAGATATAAATATAGACTAAACACTATGCAACTTATGCTAAGAAAACCAAAAAATAAAGATAGAAGAGCACGCGAATATTTAACAGAAGATGAGATAGAAAAAATTATTGTGGCGGCAAAAAATATAGGGCGTCATAAAAGTCGGGATGCAACAATGATTTTAATGGCTTATAGACATGGGTTAAGAGTAGGAGAGTTATTAAGTCTGAAATGGCAGCAAATAAATTTAAACCTAGGAACTATACAAATAAATAGATTAAAAAATGGCATAGATAGTGCACATCCATTATTTGGCCCAGAGTTGAGGGCATTAAGGAAAGTAAAAAGAGAGTATCCAGAAACACAATATGTTTTTATTTCAGAGAGAAAAAGCCCTCTTACTACATCAACATTTGGCAAAATGTTAACTAGAGCAGGTGAAAATGCAGGAATAGAATTGTCCATTCATCCACACATGCTTCGACACTC
>evgR012700c0g2_i01_Hydra
ACATTTAAAACTAATCTTCTATTGTCTAAATAAGGAAATATTTCTTGTTAAAATGACTGAAATAAAGTTTGATGTTGGATTACTTGGTGCAAATGACCAACTTCGGAACGGATTTCAAACAGCTTTTAAAGATTCTGTAGCTAAACATCCACTTGAAGTATCTGAGAAAAATTATTTCAAGAATTCTGAAAGAAGAGAATTCTCTGCTACCAGAAGTGCACATGGTCTTCATATGCCTATGAAACTTCAAATGGAAAGAGTTGTTGCTTCACGAATTCAAAGATTAGCTGGTTTACCAAGTTCTAATTTGGCATTAAGAACCTTAATGGGCATAAATGATATGTTAGGTCCTGAAGATATATTTAATGATGTGTCAGAAGAACTTGTGGATTACCGAGTTGCTGGAGATAGCAGATTTTAGCAGACATTTGATTTATTTTAATTTTTTGTTTTGAATAAATGTTTTATTTATAGTTGGTTGTCATTGAATGTGAATATAATAGCCGTTATTTGCAAAATATACCAATTAGTATACTGTCAAAAAAAAAAA
>evgR012704c0g1_i01_Hydra
CTTGTCGATTGAATTTAATTTCAGCAAGATATGCTAAAAACTATACAAAAAACATTAGAAAAACGTTGCATAGTGAGACTGGAAAACTTTATATAGTAAGACATATGTTTCATTGTTCGAATCCATACTGTTATGTAAATAGAAGTCAAATAGGCTACTGTTATCAATGTAGAGATAATCATGATGTTTGTAAACATTCGGATCCCAATTGTAAATATGAAATTTTTAGTTGCAATGGTTATTGTTGTACACCTGGTGGTTTTAAATATTGCAAGAATTGTAAAATTTTTAAAGAAAGACAATGTAAAACAAACACTTTTGAC
>evgR012705c0g1_i01_Hydra
GTCTTCTTTTATGAAGTTGTTGTCAAATTTTTCTTTTATAGTTTCTTTAGTTTTAACTAAACAGCTTATGGATTACTTTTACGGAATTACGATCACCCTTCAAACTAAATCTTTTGATATTTCCCAGCAAGTGGATGAAATACAAAATTTGACTAATCGTTTAAAAGAAATACAAAAAAATGCTGATGCTTACCATAACAAATGGTATAAAATTGCTCTTGCATTGGCAAAAACATTTGATATTGAGGAAAAAAAACCTAGAACTTGTGGTATTCAGAAGCATAGAGATAATCAAATATGTGATACTATATCTGATTATTTTAAAGTTTCTATTACACTGCCTCTTGTTAATCATCTTCTCAATGAACTAGAAAACAGGTTTGATGTTGAGAGTATGATTGTTTATAATGGTCTTAGTGCTCTTCCTGCTAGTTTAACAAAGCAATATATTAAATCTTGTCCATGGAAGGAAAAGTTCCTTAAGTTTTTAAATTTTTATAAATCTGATCTTCCTCATTTCTCATCAATTCATGCTGAATTAGAGTTATGGGAGGAATTTTGGAAAGAAAAGACAGAATTATCATCAACTATTTTAGATACATTAAAGTTCATGGATATGAGAGGTTTTCCAAACATTTTAGAGGCTTTTCATATACTTGCTACAATACGAATTACAATTTGTGAGTGT
>evgR012706c0g1_i01_Hydra
GGAATAAATTACAATATAGATGACTATGCGCAATGTTTTCAGAACATAATATAACAGCATTGGCCAATCTACTCATCGAATACGGGTTTTTAAATGTCAGCCATCAACCAAATCAGCTTATATCGGATTTAAAACTTCTTGATCTAAAATCAGTTTATAATAAAACTTATATAGAGCTAAACTTACTTTCTGCGTTCATCAATTCAACTATTTTATTTATTGCCAATAACACATTTAACAACCTTTTAAGTTCTTGTTTATATTCAACTTTGCTAACCGGAGAAGAACAGGTAAACACATCAAGAGAGGGGATAACATTTATATCAGAGAAAGATGCATGTAAAACATTAAATGAAAAACTGACTACAGCACAATTTGCGGCAACATTAACTTTATCAATCCCTCCAAAGAAATCTGTGGTATCTTCTATCTTTCGAATTCAGCTCAATTTGCAAGTAC
>evgR012722c0g2_i01_Hydra
ATTTGGCACTCATTAATACTGGTGATGAAGATCTTATCCAAGCTGTTATGGACAATATGCCAATTTATCCAGAAAGAGACTCTCAAGTCATTAAGAGATTGATGGAAAAGGAAGCTGCCAAGAAGCGTGGTGCTGCCGGCCAAAAGGGCGGTGAAGGTGAAGAAGAAGAGGAAGAAGAAAACCCAGATTTGGATATTCAAGTTACTGACAATAAGCAAGGTGGAAACGATTTGTTGGATATGCTCGATATGGGAGGATCTAACAATAATGCTGGTGGTTATAGCATGGATGATATTTTGGGAGGATTTGGAGGACCAGTTTCTCAAC
>evgR012730c0g1_i01_Hydra
TTTAATGTTTTAAATATTATTGTTTAACTTGAAATGAAATCAAAAGACCTCATTACAGATTGTGTTGATTTAAATGTAAAAGAATCAAGTGCATAAACGTTTTAATTTATAAGAATAATCTTGATAAAAATAGAAATCATGTCTAAATCAGAAGAAGAAAAGTTAGGAAAATCTTGTGAACAAGTAGCCGTGCAAGAAATTAAAAAGGAGGTTGGATGTTCTGTATCAAAGTTTGACCAAGAAGATGACTCAAAAGAAGAAATTTTATTGGATGATCTATTAAAAAGCAAGTATGATGGAAGAACAATTCAGCTACCAGATTTTGCAATTGTTTATTCATTTTTGGAGTTGTTTGGAGTATTTTTGGAGTTACCCTATATAACTATGCCTGAATTAGAACAGTCAATAGATTCAACAAAAATCTTCACTGGTGGAGAAGACATTGGAGTATTGGAATCCTTACATGCCAAACTTTTCCGCAAGCTCAGCAAATCCCCTCGTTTCTCAATCTTATCCAATTTTAAACCTAGCAAGTTATCTAAAACTGTTTTAAAGTTTGCTCAAATGACTGAACTTCCTTTTGCTTGGGATTTGGAACAATTTGGTTATCATATACTTACTTGTGCACAAAAACTACAGTTGCTTAAGGCGCTGTGTGATATACAGTTTGATGAAAACCAAAAGTTTAAATCCATTTTATTTGAGGCTTATAAAGCAGATCAAGGTGCCAGTCTTCGTCTTTTGCCTTTTGGATACGAGCAAGATGGGAAAACTGTATGGTTTCATGAGGATGGAGAATATAATGTTCAAGTTTATGCTGGGGAGCTGGAGGAGAATGACATGTGGCATGTGAGAGCTGGGAATTTAGACGATCTAACAGAATATATTTCTATTCTAAAGAAGTTATCAGAAGTTAATCCTAAAGATATAGCTGCCGAAAAGTTACGGCTGCGTAATGCAGTTAAACAAGAGGAAAAAGAAAAGCGGAGGTTAGAGCGCGAACGTCAAAGGCAGTTACAGAAAGAAGAAAAAAATGACAAAAAAAAGGGACGTAAAACAAAAGTTATTATCAATACATCAGAGTCAGAAGAAGAGGATGATGGCTGTTGTCGCTGTTTAGCAAACAACCAATCAGATTTGGTTTTGTTGTGCGACGGTTGTGATGCAGCATACCATACGCTATGTTTGCGTCCACCAGTTGAATCCATACCTGAAGGAGATTGGTACTGTCCATTCTGTTTGCAAGTCAAGTTAATCGAGTCACTGGGAGTAAAGCTAAACGAACTAAGTGCTAAAATTAAGGCACGGGAAAGAAAATCCAAAAGAATGGTGTTTTACGGAATAAGTCCAGCAAATATCGTAGATAAAACGAGCGAAGACATCGTAACAAGGCGTTCTGGTCGTGGCGCAAAACGTATTGATTATAATTTTTCTGAATATGACGAGTTGATTAAAGAAGCAGTTAATGGCAAAGATGAAAAAGAGTCTTTACCATCCGATCGAGTCTCCGCAAGAAATACGAAACAAAAGCGTCAGCGGCGAAAGTTAAATGATTTGGACAGTGACAATGAAGACGATTTTGATAGTGGCGATAATTCATTTAGCAACTACGACAGTGATAAAAGGGATGATATCAAACATTTTATAAACGATCATGATGAAACAGAAGAACGTCGCCAAGATAAACACTTGAGAAGAAGTCGACGTTTTAAAACCCGACGAAAACGTTCGAATCTCGATGAAGATGAAGAGGACAGTAACAAAAGTGATAGCGACAGTTTTGACTCTAAAAAGCCTAAAAGGCGATTACGTCAGCGCAAAATAACGAGGTACGATGAAACTCCTTTTTCAAGTGAAAACGAAGAACCTATAAAAAGTATAAATCCAGAAGAAGTTGACGAATGGATCGGGGATGAACAATCGAGCGAGTTATCAGAAAGCGAAGATTTGAATAGCGAAATAAGCGATAGCCCGAAACAAAGACGTATTTTAATGCATCGTAAAGTCAAAAACAATGTTAATTTATGTGAAGATGATACTGAAGATGATAGCGAAGAGGATGAGGAGGAATGCGGTTTTAGTGAGGAAAGTTTAGATGGGGTTAACTCTGAAAAAGATGAGCAAGTCATGGAGTCGGAACAAGTACCTAATACTCCAAGTGAAGAGCAGTCTCGTGATAATAAAAGCGGAGATGAAAAGTGTCGAGAAATTACTCAAAATGCGCCTAATGTCAATGACCAAATAAAACCATTAGAACATAGTGATATTAAGTCGATTGACGATATGAAAATTGTAAAACCCTTACCTACTACGAAGGAATTAAAAGCCGAATCGAATATTGTTAGTATGACTTTTACTGGCGTTACTACCAGTTTATTTTCTCCAGTTATTCAACATACAAACAAATCTTTTTGTCATCCTCGAGAACGATTTAAACAAGTTACTAAATCAAAATCTCCGTCGAGTTCACCCACAAAAACATCTGATAATATTCATCAACAGCAACCGATAATACAAGCGGTACCATTTTCAAAAGCATCATCTCTTCCCTTACACCCTCGTGAAATATACAACAATTTATTAAAAAACAAGGCAACTGAAATTCCGAATCTTTCTACACAAAATACCAGCTCAGCTTTTATCCATCATTCGTCATCGCCGTCCACGTTATGTCAAAGTTATTCATCAAGTAGCAGTTTAGGATGCTCAACCATCCTAACTCCAACTCAAAATATCAGTTCAAATGCCAGTTTTCATAACATTACTTCACCACAAGATTTGAATGCTTCACGGTTACAATTTAATAACATTCACGGAAATTACTTTTCAAATTCAGGTTACGAGCCATCATATACGCAATCTCAACGTTTCCCAACGCCTGGTTGTCTTCCACCCCAACAAATAGGAGTGCAAGCTCACAGCTACAATCATGGTGCAAGTCCTTCATTGGGATTTTATCCAATTCGTTCAACGCTTTATAGTCCTCGCCAGCAAGGACAGTATAGACCATCTCCCGCTTTTCCTAACGTACACAGTTCAACTGGGTACTTTATGGAGTCCGCCGTTTCATCGCCTAGTCTCTATCCTCCTCCATACGAATATTCTGATCACATTCGACCTCCTGTGCACATCGGAGTTTATCCCTACTCCCAATATAGCGGTGTTCAAAACGTTGGTAATCAAAATGTTAAACAAGACCGAGAAATAAAGAAGCAATTGAAAGAACAAGCAAAGCAAGAAAGAGAGTTAAAAAAACTGGAGAAGCTTCAAGAAAAAGAAAAGAAAAAACAAAAAAAAGTTTCTACGGTGAATCGCGGTGTTGACGGAAAATCTAATATGAATTTTTTTCGAAACATGGTCATTTCTCCAGACAATAATAAAGAAGATACAAATGAGCAGTTTCAATAAGTCTCCGTTTTCAGTTTATTTAGAACACTTTGATAAAATAATTTTTTTAAATGTTTTTAAAAAGTACTAAATAGTTATAAAATGATTTATTGAGATTAATGAATCACTGGAAAAAAAAAA
>evgR012731c0g1_i01_Hydra
TTATCTGTTTGTGGGCATGTAAAAAGGGAGTAATTAATATTTTATTTGAATATAAATCTACTATTCCTATTCACACCAAAGCATGGATTCAAATGCTAGAAATTTTACCCATTTTGATAACATTGGTAATAATCTTGAGGCTCAACGAATAAGAAGGAGAACGGAAAATTTAGCACGAGGTTTAATTAATCCACGAGATCGTCTTTATCATGCGTTGTTTATCAAGATTGGAACATTTTATTCTTTGCTTGTACCTAAAAAATTTAGATTTTTAATCGAAATGCTTTTTCTTATAAAGGGTTTGTTTCTTTATGCTCTACTATTTTATCTTCATTATTCATTTGTTGGGTCTCAACTGAATTGTTTAAATTTAAAAAATGGAAGTTGGCCGCGTGATGGGCTTCTTCGTGTTCAAATTAGTCCTAATGCATATCTTGATAAATATTTGGCAAGCAGATATGAAACAAAAGATGGAGTTATTTCCCAGCTTTGTCAACTTGATAAAGTTTTAAATGGTGACCTATGTTCAACATATGATCTCCAAAATTATTATTCATACTACAATAAAAAGCACAGTCGACTAGACCAGTGCTACCGCTTTAATGTCACTGATTTTTTATACTCAATAAGCAAAAAATTGTGGAAAGATTTTGATTCAGTAGAAATAATAACAGAATCAGCTAATTCCAAGATAGGAAGAAACACAGAGAGAAAAAAAGCCAGAAATATTATTTCAACAATTTATGATTATTTTGCAATTGCTAATCAGTTTTCAACAAATGGCATTTATCTATTTGAATATTCGAGAGAGTATGGACTTTTACATCTTTCTTTTGATGCCAGAAGAAGATTAAAAATTCCTGTTAAAACTATATTATTAGATCCAAACAATGACAAGTGTTTTGGTAGCAATTTCAATAGGTTCTTGCTGAAATATTTTTTGGGTTATGATAATATTCTTATGAACAGCATCAAAAGAGTTGCAGATACTGATAATAATCTCGGTTACATGGTGAACGTGGTAACTGGCGAAAATTATAAATTTGTTCCTTCCAGTATTGGAAAAGGATCTTATCTAATTTCATTTGTACTTATGTTTTTGTTTACAATTGCCATATCAATGTTGCTTCGATACTGTCATCAGCAGGTGTTTTACTTTATTGTGAACATGCTCCAAGTGATGAACATGAATGCTCTTATAGTGTTCCCTCTAGCACCTCTTATGGCTGTTATTTTATCACTTGTAGGATTGGAAACTATAATGGCAGAGTTCTTTCATGATACATCTATTACTTTCTACATTATATTAATGGTTTGGACTGTTGATCAGTTTGATACAATTTGCTGTCATACAGTAATTTCACAAAAATATTGGCTTCGGTTTTTTTACTTGTACCACTTTGTATTCTATGCATACTACTATCGTTTCAATGGCCAGTTCGATATGCTTGCTCTCGCTGCCTCTTGGTTTATGATACAACACTCTATGCTTTTCTTTTTCCATCATTACGAACTTCCCATGATTGAAGAGTCTCAAGATAATAACGGTGAGTTAGAATCCTACCTAGAAGAGGCTGTGCAAATGATTGTTGATGGTATTGAGACTGTTATTGGGGATACAATTACTCCAGAAAATGCACCTTCTCATGCAGGAATGAATACAAATACAGGAGAAGCGAACAGCGTAGGAGAAGTTATATCAAATACAAATAGATCAACGCGTGTTTATATATATGCAGCTGAACGTCCATTTGTTAATGTTGTTCGTGATTTTATTCGCCATTTTCATAGAGGCACAGGCTCAAGTTCTGAGCCTATTGCCAGCGTTGAGACGATTTCGGTGGATACGTTGCTTGAAAATTCGACATTGAGCAATCAAGTTAGGGATAACATGAATTTAGAAGAGAATTCTCCGGAATCTCAGACAACAAACGGTCTTGTTAGTCATTTTTCAGAATCTCGAGATGTTTCTGAATCAGTTTTTGACATTTTGCCCGAAGATATAATGGGCAGTAATATTCAGCGCTCAGCTATGGACATTGCAACAGACATAAGTGACCCAGAAACTTTTAATGAGTTATTGCCGCCCATCAATTGTGTTTGTTCTAAAAACTTTAACTATAATAATGGAAGTAATCAACCAGATACCACATATAATCAGATTCATACCAAGGAGACTGAGACGAATTTGCTTAACCAATACTCTAACGAAGAGAATGAAGCAATTTTGAATCAATACTCATATCAGAAGACTGGAGCAAATTTGCTTGATAAACAGTTGTGCGCAGGTGGATCTAAAGAAATAAACTATTTTGGAACCTGCAATCAGCAAAGTTGTAATTCTCTTGAGTGTATTTCTACTGTAAGTGACAATGAGCTACAGAGTAATGCAAATAGTAAATGACTTTTATATGTAAATATCCATAATAATTTCATTATGTTTAAAAAAAAATCTTGATTATTTTATAGTAGTTCTTCATCTCTATACTTTTGAATTAATTAAAATTTGTACGAGTTATATTTTATGCATAATTTTTATTAGAGAGCATACATTATTATTACCACATACATTATATTTTGAGGGCCGGAATTACACCCTCGAAGGCCTAACGATAACCCATTTTTTG
>evgR012737c0g1_i01_Hydra
CGTTCCAGCTATGAGACAACTCGCTACTTTGATCAAGAAGCACCCAAATGCTAAGATCGTCATTGCTCCAACCACAAGAAGCCTCCACCAACAACATAAGACTCCAGGTTTCTTTGATCAAGTCTCTGGTATCTATGCTACCCAAGATTATAAATTGGACATGAAGCCATTCGAATTCCGTAATGGTGAACCAATGTGGTACTCCAACTTTTTGGGTGGTATCATTATCAGTATTCCAGTTGCTTTGGGCTTCTTTGCTTGTTTCTTCAGTTTGTTCTGGTACACTTTCTTCGATTCCCAAACCGCTGTTGGTAGAACCAACCCACACCCATTCTTGAACATTAAGAATGGTGCTGATTCCCAAGCTACCATTAACCCACTCTTCAGATTCGCTTTCCCAATGGGTATACAAGCTGCTGACTACAAGATGCTTTTCTGGAATGAATTGAACAGAGCTAAGAAGCACCAACTCTAAATGAAACAAATTATTGTTTTATTGATAACTAATTGTATAATTTTGTTAACGAAAGAAGGGCCCTAAATAA
>evgR012746c0g1_i01_Hydra
ACTAATCCTTTATTTTCTAAATAAGGGTTAAAAGAAAAAAGTAATACTCTAAACAATGAGATTTTAATGATGACACTTCCACGCTATGCATCAATCTTAGAGTCTTCAGAAAGCAAAGATCAAATTAACAGTGAAATATTTTTTCAGTCAATTTCACAATTAGATGTTTGCAATGAGAGCAGTTATCATAGTTTTATGAGAAATTATAGAAATGTTTGTATAAAAGTTGTTGGTGTAAGTGACAATATTCGAGTCTGTGATGTTATTCCTGATTCACCAGAGTTTGATCAGTTTTCTGTTGATTTGTGGAAACAATACTATATTCAAGAAGGGATTGTCCAAATTGCCTTGTTAGAAGACAAACAGCCTTTAGAAGGTCTCTTTATTGGAACTGATTCTGGACGAGTGTATCTATGTGCTCAAGCTGATGATTTTTCACAGCAAAAGGTTTTATGTATTTCACCATCTTTACATAGCTTGATTTGGAGAGGACCACAAAATCCTATTGAGTATAACTCAGGCTGTATGTTTGATAACTGTTATGGTTGTTTTGATCCAGTTCTACGCAATAGAGTGTTCAGATTTAAACAATTTTAGTTTCCTTTCTAAAAAACTTTACAAGAAACACGTCTTTAATCATTCTTATAAAAATCCTGTTAATCAGAAATTTCTAGGGTTACTTGAATGTTGTATAAGTTATTTTTCTTTAAATCAGTTATTTTAT
>evgR012751c0g1_i01_Hydra
CTAAATTTAGAATTTCTTGTTGACAACTTTGATTTTTTACTTGATGAAAAGGCTTTGAGATTGTATAAAGATTTATTTATGGACACACTTGAAAATAAACTTAAAATGATAGTTAGTAAAGAGGTTAAAGTTGCAACAACTGATAAAGAAAACGTGTTTCAGAATATAAAAGAAAAGATTCAGGAGGAGTTTACAAGCGCCAATTCTGTAATTGGTGCAGTCGCTACACCAGCAGCAGTTCCAAAAATTCTTAATGAAATACAAATGTTGACAAAAAACGATGAAGATATTGCAAAAGTTGGAAAAAAGCTTTTCAAGTTTTTTCGTTAATATTTATGGTTTCGCTAATGTTTATAAACGTTATTTCAAAAGGATGTCCTAGGGGATTTTCTATGAAAACAGTGGCTGATAAAAAAGGGTGTGTGTGTTGGGGGGGGGGAGGCGCTTAGGCTTAAAAGACTTTTTCTTGAATTACCCCTCTTAGGACAATGACCTAGATCCGCCAATGGTTAAAAAGATGAAACCAAATGTTAGCAAGTTACAGTAAAATTCTAAACTATATATAATCTCATAACTATGTAAAATTCTAAACTATATATAATGATTGTGTCCATTGACCTCTTGTGGTCAATATATTAAAAAAAAAA
>evgR012762c0g1_i01_Hydra
ACAACAAGGAGGATTTGGAGGTCAACAAAATCAAGGATGGAATCAACCACAACAAGGAGGATTCGGTGGCCAACAAACACACGGACAACAAGGAGGATTTGGAGGTCAACAAAACCAAGGATGGGGGCAACAAACTCAACAAGGAGGTTGGGGAGGTCAACAACAAAATCAAGGATGGAATCAACAAGGAGGATTCGGTGGCCAACAATCACACGGCCAACAAGGATGGAATCAATCACAACAAGGAGGTTTTGGAGGACAACCACAAGGATTCGGAAGCCAACCACAAGGCTGGAATCAACCACCACAAGCTAGAAAGTTAGACTCACCTCCTCCAGTTAAGGATCCATTCAGTAATTTATCAGCATTTGGTTCTTCAACAACCACTTCCAAACAACCACCAAGTAACAATAGCAGTATGATGAACAATGATAA
>evgR012775c0g1_i01_Hydra
GGAGGTATCGAAACAACAATTCCACCATTATTAGATGCCGCCTATTCATCAAGCAGTGAATACAAAAACAACCAAAAATTCTTCACTGAAAAAAAAGGCGTGATGACATTACGAGATGCCACGTGTGCAACATACACTGTTCAAATCAGTCCATTTTCACCACCACCATTTAGTGATGGGTTTAAGAAAGCGTTATATGATTTAAACGAAGTTAATAAACAAAAAGATAATAAACAAACTTCTACTTTTAATAAAACTGCTTCTGTAAATGGTGTATATCCTACTGGTGCTGGTGTAAATGACGTAGGTGCTACTGAAGATGCTGGTGGGAACCCTAAAAAAGAAATTTTTCAGAGATTTGTTACGGAATTCGGTACACACTTTCTTCATCAGACAACAATGGGTGCTCGTACTGCTATAACAAGAAGGTATTCTCAAGATGAGTACAAAAAAAGTACAGATGAC
>evgR012791c0g2_i01_Hydra
AAACACTATAAATTTGAAAATTTAATTTGTGTTTAAAAGATTTTCAAATTAGTTTTGAATGAAATTTACAAATAAGAATTTAATTTTAATGTATCTAACATACAAGCAGCTTTAAGAATGGGAACTGTCAAAAGTGGATGGCTCTTTCGTAGAGAGGGAAGCTTTCTTTTCAAAGACTGGTACAAAAAACATTGTGTTTTATATTCCGATGGGGAATTTTCTGTTTATACTAAAGCATCTGATGCCACAGCAGAACTTAGAGTTAATATGAAAATTGAATGCTTGAGAGTTGAAGTTGGATTTGATGTAGGTTCCATTAATCTGCCAAAAAATATCAGTAGTGTTGAATCTGCTTTTGCTATTATAACAAAAAGTAATAAAAGACATATATTCTGTGCATCCAATGATTCTGATTGCAGGGATTGGTTTGGTGCTTTAGAAAATGCACGCACTGCTCCACCTTTGTCAAAGCAAGTACCGTTTGCTCCGCCTATAGATCTTGCTCCTCCTCCATATGCTCCATTAGAATACAGCCAACCCTTGCAATCCAACGGATGGGGAAATTCAGGGTCTGGCTCTCAACTTGCTCAGCAAAGGCCTCAAACAGTGCCTTCACAATACTATCAGCAACCTGGATATATTCCACAATCTAGGCCTAATTATGGGGCTCCTGCCTATGGTTCTCAGCCTATGTATGGACCTCAGCCTATGTATGGGCCTCAGCCCATGTATGGACAACCAATGTATGGTCAACCTGTTTATGTCAGCCAAGCTCCTCAAGCCCAACCGCAACAGCAGCAAAGCAGTAATAATAATACAAGAAATATGTTGTTAGCCGGAGGAGGTGGTTTGATAGGTGGCTATTTATTAGGTAATGCCATATCAGAGATGTCTCATCATCATATGAATAATTATAGTGATAATAGACAATATAATGATAATTATTACGATAGCGGAGGCGGGTTTGACGGAGGAGGCGGTGGGTTTGATG
>evgR012794c0g1_i01_Hydra
CTGTCTTCACAATGTAAGCTATTAATAAATGTTGTAACAAAGTATTACGGCATTTCATCTTGGTTGAAACCAAAATCTGAAATGTTTGCTGTTTTCACATATCCAAGCTCTCATGTTCACGACCAAATAAAGGCGTCAATATTTAAATACAGCATAATTTCCACTTATTTTGATGTCAATAGCAATATTAAATACGTCGATTCAATACAAATGGAGAAAAAGGAAAACTTGTATTTTTTGGAGTTTAACGATTGGCGAGTGCGCGCAGAAAATCTTGGAATCTCATTTTTCTTTTATCCTTCTAAAACTGAACTTGTATCGTTTTCCGGAACATTTGAAGCGCCAAGAAGAATTACAACTACATTTGTTAAAAAATTTGACCAGCCTGATAAAATTTTAGACATGCCTGCGGACCTAAATTTTTTCGGTAAATATACCATGGCGGTAAAATTAGAAAGCTCTTATAATCCTCAGCTTATTTTAACTTCTGGTATCACAAGTGATAGCTGCAATAATCCTACAGCTCCACCAGTTTCCAAATCAACTCCATCAGGATCAATTCAAATTTCAGGAAATAAATTCAAGCTCATATTTTCTGTTTTTTACTTAAATAATTTAATAATTAAATTATTTACGATATAAATGACATTCCAAATAGGAAATAACTAGTGATACTGAACAAATATATTTTATATTTACTTTGTTTTCATAATGGTTCATTTTGTTCTCAACTCAATTAATTTATTTTTATCTTATTTAAGGTGCTTTTATTTCACTTGTTTCTTTTTTTGTGGAACTTTTTAGCCCG
>evgR012818c1g1_i01_Hydra
AGAAGTTTTTTACTACATCAATTGATATAATGATAATTGTTTTCGTTTTGATTGTTCAAAATCTGTTGGGCATCGTTTTATTGATAAAATCTCAACCACTTGTTTTACCAATCTTAAACCACAACAAAAAAACTGTAACATTTCAATGTACAGGAGATCCTCAACTGACTCCTCAAGTTATATACCTCGGAGTCATGCTAATTATTTGTTCTGTTCAAAGTTTTAAAGCTCGCAATCTTCCCTCTTACTATAACGAGACAATCTCTATTACATACAGTACCGCTGCATCATTTTTGTTACTCATTTGCTATTTTCCTCTCTACTACGGTAATAATGACATCAAATCAAGAACCACTGCTGTCGCAATTTTGATTCTAACAATAAATTTCATTTTAATGACAGTAATGTATGCACCTAAAGTATTTGTAATATATTTTAAACCAGAAATGAATACTAAACAGAGTATTAATAATGAAATGTTTGATTTTGCTAGATCAGTTTCTACAACTTTGTCATCATAACTCTGAAC
>evgR012820c0g2_i01_Hydra
CTGATCATCAATTCATCAAAATGAGCAATCCAAAAGTTGAAGAGTCTACCTCCGCCATCGATTTGTATATTCACCCACTTATTCTGATCAACATTTCAGATCACTACACTAGAAAAAAGTATCAATTAAAGCAAACTAGCAATGATGCCTTTGATTCCAAAGTTGTTGGTATTCTTTTCGGTACTCAAGACGGAAGAAGAGTTGAAGTTAGAACTTCTTTTGAATTAGTTTTCACTGTTGATAAGAGAGGTGTTGTAAATATTGATCACGGCTATGTGCAAGAAAAGATTGACCAATACAAACAAGTTTTCAAGAATTATGATGTTTTGGGATGGTACTCAACAGGTACTGAAATTAGACCAGAATATTCACAAATCCAACAAGAAATTGTCAAATATAATGAAAATCCATTGATGCTTTTGGTTAGTGTTTCCTTTGGCTCTGTACAAAAGGAATTGCCATTGTATGTCTATGAAAGTAATATTAACAACACCAGCAGTAATGAAATTACCTGGGCTTCCCTCTCCTACAAGATCGAAACCGAAGAAAGTGAAAGAATTGTGGTCGATCAAGTCAATCGTATGGACAAGAGAGATGGGTCATCCTCATCACTCATTCCACAATATGTTACACTCTCCAATGCTGTTGGAATGTTAATTGAACGTGTTGACATCTTGGTAAAGTACTTGGAGCAAGTCAAACAAGGAAAGATTGAACCTGATTTCAAGAAGTTGCGTGAAATTTCATCCCTTTCACATCGTTTGCCTGCAGTTGTTTCTGACAAGTTCAAATCAGACTATGTTAATGATGTTAACGAAAGTCTCGTCATTACTTACCTTGCCACAATGACAAAGGGTGTTAATCAATTCAATAACCTTGTTGAAAAATACTCTTCACTCAATGCCGGTAAGAAGAGAAT
>evgR012824c0g1_i01_Hydra
TCAAGACCATCTAGACCTTCTAGACCTTCTAGACCATCCACGCCATCAAGACCATCTAGACCAAGCAGATCATCTCCTTCAAGAGGAGGTGGTGGAAGAGGAGGAAGAGGACGTGGAGATCCTCATTATACCACTATTACCGGGGATGACGTTTTATTCAACGGAGTTGGTGACTTTATTCTTGCTGAAACAGTTGACAAGGAATTCATTGCTCACGTTCGTATTGCCAAGTGGAATGGTGACAAGGGAGATGCCACTGTGAATGTTGCTGCTGCTGTCAAGGCTGGATCTGATCTCATTGAATTCAACAGCAAGTCAAAGAAATTCTATTTGAATAAGAAACGTGCTCGTTTCGCTATTGGAAGAAAATATGAATTGGATGGAGGATATTTATTGAGAAAGACTGAAAAATCATTCATTGTTAAATCTAATGATGGAGCTGAATTCTTTGGTGAATACTTTGTTTTGCATGATGGCAGAAGTTATATGGATGTTGCTGTTGATGCTCCAAAGGATGTCGAGTTTGCAAGAGGCTTGATTGTCAATGGTAAAGATTCTGGTGTTGCTCAAG
>evgR012825c0g1_i01_Hydra
AAAATGATTATCCGTATGATGTCTTTCAGATCATAACCAGCGCTGTAAATAGTGCAGTATTGTTATCTAAACCTTTGAGTGTAAGCATCGAGGAAGAAGTTACTTTTTTAAGTTTAAATATTGTGAGAGTGATTGGTTTTGAAGGACCAGTTTATGTTGAGTGGAGAACCCGGGGTTTTGATGGAACTAACAAAACATCTGGTTTGACGGCAATACCTGGTGTTGATTATGAAGAAAGTATTGGTTTCATTAAAATGGTTGCTGGAGAAAAGGTTGCTAACATTTCAGTGAGAATATTTGACAATGCAAAACCCGAGTTAACTAAATTGTTTCTTGTTGAATTAAAGAATCCATATAATGCTGAACTTGGAAATGGTTCTGTTTTATATGTTCAAATTCTTCCAAGTGACAATGCCCTTGGAATATTTTCTTTTCCAACTGTTGAATATACAACATGTGAAGGATGTTTTTTAAACATTAGTATCAAGCGAAGTGGTGGCGCATTAGACAATGTCAAAATAATATGGATTTTTTTGAATGGATCAAATGAGTTTAGTTCATCTGAAGGTGAGGTTGATTTTCAATATGGTGTCACGGAGAGATTTATTCTTATTCAGGCTGTAGATGATGAGAAACCAGAACTGACTGAAGTATATATTATTCAACTCTTTAATGTAAGTGGTGGATCAATATCTTCTATCAACAGAACTGCTAGTTTAAAGTTACTTAGCAGCGATTATCCATATGGTTTGTTTGAATTTGAAAAAAAGTCTATTGATCTGGAAGAAAACATTACACATGTCATTATCCAATTGAATCGAAGTAAAGGCCTTTTCGGGCATATACGTCTTTATTATAACAGCATTAACAGTTTAAGTAACCCAAGTGCTGTGATGGGCAAAGATTTTCTTCTTAATGGGCATTATGTAGATTTTGTAGATGGTCAGAAATATGCTGTTATTAACCTTACCATTATAGATGACAATGAGCCTGAAACGCTTGAAACGTTTCTTCTTAACATAACACATGTTGAGATGTTGTCTGATGCAACTATGGCATTTGAAAGCAATGCAACAAGTTTAGATCTTCAAAATGTTTTCATGAACACCCCGAACGTGATTAATATTAGTATATTAGCCAATGACAACCCAAATGGTGTTTTGAACTTTAAATATGAAAAAGTTTATGTATCAGAAGACTATAATGGCACCTTTTTATCTGTAATACGCTCAGAAGGTCTATTTGGAAAAGTATCATTTGGTTATTCAATAGAAGCTGTATCAGCTGATATTGGATTAGATTTTTCAGTTTTATCCTTTGCTATGACTATTGAAGAAAATGAAAGCAATGTCATGTTACCAATAGTTACAATACTTGATGATGTACCAGAGTTCAATGAGACTTTTATAGTTCGATTACTTCCTGATACCTTTTTAGGTGGACTTGTCATAGGGTTATTAAGCAAGTGTGAGGTTGTTA
>evgR012832c0g1_i01_Hydra
TGTACAGTAATTGGGATAATGGTGAACCTAACAATTATGATGGAAATGAGAATTGCGTAATCATATGGAAGTCTGGTGTGTGGAATGACTTTCCTTGTGATCTAACATCAGCTTATATGTGTAAACTCAAAAAAGCTTCTATTCCAACTGCATTAACTCAAGAGTCAATTACAAAAATGGGATCACCAACAGCAACAACACTATTTTCTACTTCAACAACTAAATCTACAACAGCAACAACTCCATGCCGAACTACAACAACTACATCTCCAACTATAGTAATGACTACCATTCAAACTACAACAACTAATCCATACCCTCCAACTCAAATTATTGAATCGATAACTACAACAACTCAACCTAAAAC
>evgR012846c0g1_i01_Hydra
GCCGAAAATAATAACCGTCATTAGTCTTACTTCCGTAAAACGTCTTTAAAAATATATTATGTAATTAAGAGTAGCATATTGACTTATTGAGACAAGCACGGAAATCTGACGTGAAAGACACGCTTTACACTGTTGATATCATGGTTTTGAAAAAGTGTCACATATCATCTTTAAGTTAAAGTAATAATTATAAAATAAATCATTATTTTTCAAAAAATTGCTATTATGTCTCATCACACATCACTAGAACTGCCTTCAAAACCATTAGAAGACTACAAGTCAAACATCAAGCAACATGCTGATCAAGTAAATGAAAATAATAATTTAACAACATTGCCAAACTCTGTGCCATTAATGTTAGGGGGAAATCAAGTTAGTATGTATTTAATGGCTGGTGCTGCAGCGGGTATTATGGAACATTGTGTAATGTATCCAATTGACAGTGTCAAAACAAGAATGCAGAGTCTTCGACCAGACCCACGAGCTGTATATACTAGTATACATCATGCTCTCAAAAAAATGATGCAAACTGAGGGAGTATTTCGCCCTTTTCATGGTGTTAATATTGTAGCTCTGGGTGCAGGTCCAGCGCATGCATTGTATTTTTCAAGTTATGAGCTTACTAAGAAACTTTTTGGTAATGACATCAATGGAGCAAACCTTCCTATTGCTAACGCTGCTGCAGGAGCTGTTGCTACCTGTTTTCATGATGGGACTATGAATCCTGTTGAAGTAATTAAACAGAGACTTCAAATTTATGGAAGTCCTTATCGAGGTGCTTTCCATTGTGCGCAAACCATTTTTAAAAATGAAGGTGTAGGTGCATTTTACCGCAGTTTTACCACACAGCTTACTATGAATATTCCATTTCAATGCGTTCATTTTGTTACATACGAATATTTTAAAGAATTATTAAACCCTCCGGGTGGTTATGATCCTAAAACACATCTTTTATCAGGAGCAATTGCTGGAGGAGTTGCTGCTGCCATCACTACACCTTTAGACGTTGCTAAAACATTACTTAATACTCAAGAACAACGTGCCGTTAATGAGATAATATCTGACGGCAAAGTTTTACACAGTAAGGGATTTGTAGGAGGTATGTTTTCGGCATTAAAAACTGTATATAAACTTCAAGGAATATCTGGATACTTTCGTGGTACACAAGCTCGTATTGTATACCACATGCCTTCATGTGCTATATCTTGGAGTGTTTATGAATTTTTTAAAAATAGCTTGTCACTTAAAATTTCAGACGAAGAGATAATGGATTTAACCGCATAATTCTGTAATATTTTGTAGATTAAAAAATATTTCATATGGTTGATTGTTGCGAATGTGCAATTAGTGCGTGTAAACATGAAGCGTTTTTATCCATTCAATACAGATACGACTTGGATTTTAGCTAAGCTGTAATGCTTTGTTGCAATGCAATAATGGTAATTTGAGGAATTAATTTTTTTTGTATTTTTGTATTTTTGCTAAACCTCTGATGATTAAAAATTTTTACATGAAGGTAGAAAATGTCTAAGAATAATATATGAGAATGAAAACTAATTTACAATGAAGCATATAAATCTTTCCATTTTTATCTATTAGATCTTATTAAAAAAAAAATATTTTTTCTATTAAATCTTATTAAAAAAAATTACGAAGAAAC
>evgR012851c0g1_i01_Hydra
TTATAAAAAGAATTTAGGATAAAATATATAAAGTTCTGAAATGAAATATTTACTAAAGCACAAAATATTTTGTAAATAAGAAACTATGAGTACATCGAACTATGTACTTATCTTGATTGGATATGGAATACTTCCAATTGAAATACTGGGAGCATATTTATTGTTTAAAGTAAGAAGAAAACTCGACCAAATTCAGTTTTTATTATTGTTGAGTAATTGTCTTCTTGATATCCAAATGAATGTTCTAATTCCTACTCAAATTTTATGCGAAACTTTAAAAAAAAAAATCAATTTTGTTTTAATTTATTCGACAATTGCTGTGAGTTTGGAATTTTATTGCATAGTTTTTATTTTAACGGCTAATCGTTTTTCTGAAGTATACCTTAACATACGTTATCATCAAATTTGGACATACCGACGTGTAAAAGTGGTTTTGACTATTATCATTATAACATCGGTATTTTCGGCAGCGTGTGTAACTATTTTTGATTACAATTTTAAAAGCAAGTTGTTTTTTTGCAAGTATTTGTTTCCTATTCTAAATATCTCTACACTGGTGTTTGTCATTTTGTTTTATCTTTATATATCTTATAAATATTATAATAGTAAAATCCAGAAAATAAACAGTATTACAAATTGCTCCTCGTTGTTTGTTGGTCGACCTCATTTGAAAAAAACTGCAAAGAAACATAAAGAACTTATCAAAAGCATTATGCTGTTTAGTTTATTAGTAACATCATATGTCGTGTTTTTGTCAATCCCAGACATTATTAACTGGCGTTTGAATTGGTATTATGGGCGTTTAGAAAATTATAATATATATCTATATATTATGTATTATATTGGTATATTTTCAGATGTATTAATATATATTTATTTTTCTTTACCTGTAAGACAAGAGCTTTATAATACTATTAGTCGCCTACTAAACAGAGCAAAAAAGTTTAAACGAAAGACATGCAGTATACACCCTGCAGATATAGCTTGAAAATACGTCTTCAGTGTGTACATTTTTATTATTAAGCTGCGTTTTGCAATTGTATCCTGTAGGTTTATTTATTTATTTTTTATTAAAAAATGTATTACCTCTATTTAATTTCTAACTAACTAAATAGTTATTTTAATTATATATATATAAAAA
>evgR012852c0g2_i01_Hydra
ATTAACCTATTTTAGTGCAAAAAGTAAAATGTCTCTACGACTTCTATCAAAGTTTCAAAGTAAAGTTTCCCAGTGTTTTCGATGTTCAAGTACAAAATCTATAGGAGTTCTTTCAAATGATAACACAAAGGAGATAAATGAGATAAACTTTGAAGAACTGTTAATGGATTCTAAATTGATCAATTTTGGACGCCCTGATAATAAATGTGTTGTAGGTAAAATTGTTAGTAGATGTAACGATGATCTTTATGTTGACTATGGTGCTAAACATGAAGCTGTTGTAAAGATACCTGCCCATCAATCTGCTCTAAAAGAACTCGAACATTTTAAAGCTGAAGATTATTATAGGGAAGGAGATTTAGTTCGTTTACTTTTACGATCATATGAAATGACTGGATTATTTCTTGGAGACACAACTCATATTACAATTTGTGAATCAGACGCAATTCTTCTTGGACCTTATTTAAGAGGTTTTAATAAATTTAAATAATTTAAAAAGTTTTAAATTTTGTTTCTCTTTCTCCCCCTATATTTTATAATTAATATTTTTCTTATTATTATCCTGATACATATCCTTAATCTTAATATCCTGATACATATCCTTAATCCTAATATCCTG
>evgR012859c0g1_i01_Hydra
AATAACATTTTTCCTTTTCTTTCTTTATGTATATTGCGAGTCACATTTGCTGTCAGTGTACTGGGGACAAAATTCTAACAGCGTCGTTAAGTTAACAGCAGAAAGAAGCTTGAGTGATTATTGCAACTACTTCAGTTATCAAATCATCAATATTGCGTTCGTGAATGTTTTTAACAGCAATAAAAATGTTTTGCCTGTCATTGATCTTTCCGATCATTGTTACAGACTTTCTAGTTCAGATGCAATTTTAAAATGCATAAACATCGGAAACGAAATTCAAAGTTGTCAAAAAAAAGGTGTACTAATTACAATTTCATTAGGAGGAGCGGATGGAGTATATGGGTTTAATACAACAGGCGATGCTAAAGATTTTGCAACAACAATTTGGAATTTGTTTCTCGGCGGATCGTCAAACACCAGGCCATTTGGTAATGCCATACTAGATGGTATTGATTTAAATATTGAAAAAGGATCTCCGTTATATTACTCGGACTTCGTACAAGCAATTCGCACATTTATGTCAGGAGATAACTCTAAAAAATACTATGTTACAGGAGCTCCTCAATGTCCGTACCCAGACCCTTACATGGGTCCAGTTATTATTGGTTCTGTTTTAGAAAATATGGCAAAAGAATTCGACTACTTATATATTCAATTTTATAATAACCCATGTTTTGTTGGCAGTACGGAATTTGCATTCACGTTAAACCAATGGTTAAGCTTTGCTAATAAAGTGAAAACAAAGTATGGGAAAGGTCCGCTTATTTTTATTGGGCTTCTTGCTCATCCTCTGGCAAGTAATCATCTAGAGTACTACAATAATCCAAGTCAAGTGGAAGCGGTTTTTAATATTGCAAAGCTCAATGAATACCTTGGTGGAATTATGCTATGGGACGCCGGTTTTGATTCCCTAAACATAATTTCTGGTAACTATTATAGCAACGCTTTGTCCCAGTTTGTTAGAGAACAAAATAATAGATCAATGCATATTGATCCAAATCTAGCGACAAATAATCAAATAATACCACGTGGAAGCGAAAATGAAAGAATTGAAAAAAATAACGCTTTTCGCCCTCTTGTTTATTATTTAACTGTACTCTGCGC
>evgR012862c0g1_i01_Hydra
ATTTACTTCCTTCAATTTTTGAATTTGAAATTCAAAATTACGGCAAAATGTTAATATAAAATCTTGAATGGCTTCTTGAAACTATCTAAATTCCGCTTTTAAAAATTATTTTGATGTTTATGTTTATGTATGTTTTAGACTCATGTTTTAGTTCTTCGCAATGTCGGGCAGCTGTGAAGGACCAGAATCTACGTATGTTAAACTTATTTCAAGTGATGGATACGAGTTTATTATCAAGCGTGAGCATGCACTAACATCTGGAACAATTAAGGCTATGCTTAGTGGACCAGCATTGTTTGAAGAGAATGATACTAACGAAGTACAGTTTCGAGAAATACCGTCGCATATTTTAGAGAAGATCTGTACCTACTTCGCTTATAAAGTTCGTTATACAAATAGCAGTACAGAAATTCCTGAGTTTCCTATTCAACCTGATATCGCTCTTGAGCTCCTTATGGCAGCTAATTTTCTTGATTGTTGACGTAAAATATTCTTGCATTCTATTTATGGTTGAAACTAGATGAAAGAATTTAAAAGAATGTTTTGTAAATAGATAGTTTACAAAAGATTACTATGTAAACAGATTATTATTATAAAACAAATATTTTTACCCGCTTTGTTATAACAAATTTTTTGGTTGTCGGCAAAAAAAATTTTGTCCAAAG
>evgR012864c0g2_i01_Hydra
AGAATAAATGGGTTAGATGTCTTTAATAAAGTTATCAATCAACCTAAAATTCTTAATAATGTCAAAATATATGCTTCTCATCCATGGTATGATATTCAAGAAAGCCTAATAAAAAACTTTCTAGTGGTCTATGGCACCCGGAAAATACAAACTGCAAAATATGGTATTATATCTTTTGACCAATTACAACCTGATGCATCAAATATCAAAACAATAATTTATAATTCAGATTTTATCAAGCGTAGAGAAGAATTTAGTATAACTAAAGGATTGTTTTTTGGAACTCTAAACAGTTTAATGAAGGTGTTCACGTTATCGTTTAATGTCAAGCCAACAAAATACAGTAAAGTTTTGAATAATGTTCTTTAT
>evgR012872c0g2_i01_Hydra
GTCTGTAATTTTTACAGATTGAAATGACGCCACGGAATGATTGAAATTCAAAAAATGAAAGCTTGGTCAATTTTACTTATACAAATAAAAATAGAAACTTGACAAATATATTTTTGAGTGTGTGTGAAGAGATCATATTTGCATGAGTTTAAGTTTGTTAAGATAAGTTTCAATAAGGAACACATTCGAAAGTATGCTAAATATTTTGACAAAAATCGTGGAGTAGTTTGAAAACTAATAAAAGTATACTGTAGTCATGAGAGGAGAATCACCACTGGATGTTTTATCTCGTGCAGCATCTATCGTTCAAAATGAAGAAAAAACAGGTATTAATATGAGGAATCAATTGAAAACAGCACAATTACATAAAGGAATAGTCTTTTCTGATGTTATGGATGTTCCAAAAAATGAAATTTATCGTAAAAGTCAAATCATATCCAAATCTTCACAAGAAAGCAAATTTATGAACTCTAAATTATGTGAATTACCTTCGTTGACAGCATCATATATATCTAAATCGCAAACTATCAACGAGCATAGTAGATTGCCTCCGCCTTACTATCATTTGCACAAATTATATGAACCTAATAACTATTCTGAGTATAGCAAATCAAAATCTTTAAACTCTTCTCCGAGGTACTTCCCATATCAACACATTTCAACAAAAGACTTTGAAAGACCTTCAGTTGACGATCACTTCAGAAAAAGTTTGGGTGATCAATATCTTACAAGTAAAAACTCTTATGTCATAGAAAATGATAAACCATGTTCTGTCGATGAACATTTCGCCAAAGCGTTAGGTAAGAAATGGAATATTGAAAATGAAAAGTCGATTTAAAAAATTAGTTTTGTTGAAATATTCTATATATTTATATCTATATATAATTGTAAATATTTTTCTTTGTATTATTATATAGATGAATATTTTTTAACTTTGAATTTATACAAATTTCATTTATTTGATCCTAACAGTAATTTATTTGATCAGTGGGTAACAGTAATCTTATACTTTCATAGTTTATTTTTTTTTAATTGTGAAACAATTGTTTATTTGAACTTTTATGTTCTTTGTAATATTATTTTTTCGGGTATAAACGAGAGAAAGTTTTCGGGTATAAACGAGAGAAAGTTGTCTTTATTTTAATAAGTAACATAAGAAACTATTTTAAAGTAGTTTAAACAAAATAATTATTAATATAGATTTAAAAAAAAA
>evgR012887c0g1_i01_Hydra
AGTTCTATTATAAAAGGAACAACTCTGATAAATATGACAGGGGTCAATACAAAAAACGCAAGACAAACGATTCGGAAAAAAGGAAGGCTTGGAAAACAACAACGAGACAGGTTGAAGATATCGGAAAATAACCGAACCAAAGATAATTATCCTGCGAAAGAAAATATAGAAGAGGACACTAAAGATGTCGAGTTCTTCACGAGTAATGAACTAATTGCAAAAAAGACCATGAAGGACAACGGTATAAAAACAAATTTTGATGAAACGATAACATTTCTTACGGAAAATGCAGAATTATTTC
>evgR012907c0g1_i01_Hydra
TGCGCTTAAAGATCTTAATTTAAGTTTTTCCTCGGGCAGCATTCCTACAAAAAATAATCAGTAGTAATAAAAAATGGTAGAGTTTGTTGATGATTCTAATACAAGTGATAAAAACAATTTATTTGGATATCGGAATGATTTGCAAGATATAGAGGACTTGGATAAAATTGATATAAATCTTGGGGAGTTGGATTTAACGTCACAAGATGAAACTATATTAGACGAACTTGATGATTTTATTCAAAGAAATTTGCAAGATGAGCTAGTAAATACTGCTTTAAATAAGGGAGTTGATCTTCGTCAGTATTCAAAGCAAGTAGAAGCAGATTTGCGGATGGTGGAAAACAATTCTATTGAAGATTATATTAAAGAAAGTGATAATATTTCAAGCTTACATAAACAAATCAAAGCATGCGATGGAATCTTAGAGACTATGGAAAGCATGTTAAGTTCATTTCAGGTGGACTTACGTAGCATCAGTTTAGAAATTCAAACATTGCAAGAGCAATCACTTTGCATGAATATTAAGTTAAAGAACAGACAGGCCATAAAAGGTGAATTAAGTCAGTTTATAGATGAAATGATTGTTCCTGAAACAATGATTATGGCAATTTCAGATAATCAAGTCACTGATCAATATTTTATTGAACAACTTCACGAGCTTTCACATAAACTTAATGTTGTGAAAGAGCAGTCGTACAAAGGTGCTATGGCATGCAACGACGTTCAAGAAGTATTAGATAAACTACGCATTAAGGCAGTTGCTAAAATTAGAGAATTTGTTTTACATAAAGTGTATCAATGCCGAAAACCGATGTCTAACTATCAAGTACTTCAAAACACACTGCTTAAGTATAGGTACTTCTTTGAGTTTCTGTTAGCTAATCATCGTCAAGTTGCACGGGAAATTCGCGATGAATATGTTGATACCATGAGTAAAATATATTCGGTTTACTTTGGCAGCTATATCGCTAAACTAATGAAGTTGCAGTTTGAAGACGTTCCGGGAAAAGACGATTTAATTGGTTTAGAAGATTCATCAAAAGCTGGAATTTTTTCAAGCAAAACAACACTAAAAAATCGTTCCACAATTTTCAGTTTAGGAGGGCGCGGGCAAGTTTTAACTGATCTTCTAGAAGAACCTATAATTGTACCACATACCTCTCAAAAAACAGACAAAAGATATTCTTATGAATGTCTTTTTCGGAGTCAGCATTTTGCGCTGTTGGACAATGTTTGCAGAGAGTTTTTGTTTTTATGTGACTTTTTTTTGGTGTCAGGAAATCAAGCAAATGAATTGTTTATATCCGTAATGGGTAAAGTTGTAACACTTTTCATGAAGCATATGGGAAGCTATATTGCCTCGAGTTTTGATGCTATTGGAGTTTTTTTGTGTATCCAAATTGTTCATCGTTACAAAATGATAATGTTGAAGCGTGATGTACCTGTATTAGAAAGCTATTGGTCAACTTTATTAGACATGCTGTGGCCGCGATTCATGAAGTTAATTGAAATGAATGCACAGAGTGTAAAACACCTAGACTCTCAAAAACTAGGGCATATTGACATTCATCCTCATTATATCACAAGAAGATATGCAGAGTTTTCTAGTGCAATAAACAACCTTAATGAAAACTTCCATGACGATCGTGTTGTAAAATGTCTTGGACAGCTGCAACTTGAAGTAGAAAACTTTATTCTTCGAAGGGCATCTGAGTTTGTTAAGCGAAAGCATCATCTTGTATTCTTGATCAATAACTATGATACTATGCTTCAAGTTATTGCGGAAAAAACAAATGAAGAAGCAAAACAAACTTTTCATGAAACACTTAATACTAAAATAAGAGAATATGTTGAAGAGGTTTTATCTTCGTACTTCAGTGGTATGATGCGTTTTGTAAACGAAACAGAACCCTTAATAGAGAAAGGACAATCAAACCAAATCAAAGTTAATGAAGGTTTAATTGAGCAGATAATTCGTGATTTCTCTTTGCATTGGAAAAATGCTATAGAAAATTTAAATCAAGACGTAATGAGATCGTTTTCTAATTTCAAAAATGGAAATAACATTCTTCAGGCTACACTTACACAGCTTATTCAATATTACCATCGTTTTCAGAAAATTTTATCGAATCCTCCCTTCAATAGACTTTCATCACGAAGTGATTTAATTAATATACATCATGTTATGGTTGAAGTTAAAAAACATAAAACAACTTTTTAAAATTGTGTGCTATGGTATATGTACTTGCTATATTTCTTTATATTTTCATAATATTAGAAAAAAGGCATTTTTTTCAGCGGTGACTTTTCCTCCA
>evgR012923c0g1_i01_Hydra
ATGACGAATGCGTCCCTAGTTTTTTGTTTTCATATTTTGAATTACAATTACTTACAAAATTATTTTACTAATCGTTTTCGTTAATGACGCGATCTTTGTGGTATGGCATAAAGATTTTCGACTGATGACGATTGCATTCTTATAGCATTTTGTTTCTAGCGTTCATATTTTATATTACAAATATAAGGCTGGCGATAAGAACACCGGAAAAATACTATAATAGTTTTGGGGATTTATTTTGTTTTATGTGAGATTATAATTTGAATGATGAAGACTAGAAGTTTAAACATTTACATTTGTGTCGTATTTTTGATTAATGTAACAACCTTGTTACCATACCGTCAGCAAAACTTTCAATATACAGAAGTCAAAACTCACCATAAGCTAGTGGCAAAATTAGTTGAACAAGAGAATGCAGTAAAAGAAATAGCCTATGTTTTAAACTCATCAAATAAAACTTTACTAAGTGAAAACAATATTATAACCCCGTTGCTTCACTGTAGAAGAAACAAAATGGCTTCTGAATACGAAGCCCGCAGCACCCGAATTAGACGAATAGTTAATGAGAATTCAGAAAAACTAATAACACCTGTTCTTAAGTTATCAAATAACTCCTCGCTAACTGAACTTAATTTAGAGCCTAAAACATTCAATAGTGAAACTCAAAGTAAAAGCGATGTTATTTTATCAAAAGCTATTGTTCCTGCGAACGCCGATGGGAGTAAAACCGAGATTATATACATGGTTTTACGAAATGGAATGGTGAGTGGAATTAAAAAAACTGACATAAATTATCATAAAAAGTTTTTACATGGTTTTTGGTATGGATGATCAATGATCACTTTATATGTAATATATGCCAAAATATTTAATATATGTAGTATATGCCGAAATAATTCGGATGTATCTGACATTTTTTACTTATGACATCCTTCTGTCATAAATAACTAAAAAAGTATTATGAAATTATATTAACATTAAATGTGTATACAGGGCCGGC
>evgR012932c0g1_i01_Hydra
CACATTTTTTTGTTAGAGCTTTGTATTCGTGTCATGGTCAAAATGTCTAAAGGAATATAATAAAGTTACGTAAAAATTTAGGTTTATTTCAAATTATTTTTATTCTAAAATAATGAGTTCCCAAACTGCTGTTGCTCCTACTCAACAACAGCAAGAAGCGCCACAGCCATTATGGAAAACAGTATTATGGCAAATAGTAACATTTTGGGTTGTAATGCAAATAGTTAAATCATTTACAACAAAAAAGGAAACGGTAGATCCCAAAGTTATGCAAAGTCCTGATAAAATTAGTTGTAAGAATTTATTCAATATAGGCGATAGTTTAGACTTGTATATTTATTATTCTGAATTTCGTAACTTCGATGATTTTTTAAACCAAAATTTGCTTTTGTGGAAGGAAACCGATATTATTTTTGGTAGTTGGTATGACGGGCCTTATGGCGATGGTTCAAGGAAAAAATCAGGATATATAGAAATCTCTGAAGCTGTAAAAAACAATGGTTCATTATTTTATCATGTGTATGTTACAAAAAAGGGATACACTCCAAATGCAGGCGACAAATCTTATTCAAAGTATGCTGTAGTAAATTATACTAATTTAATGACAGTTTATAGAAAACAAAGAATTCATAAAACAAAAAACCTTTTGACTGGGGAAGTAGATTCTCAAAATAAAGTTTTGCAAACAAATGTCACTACTGTTAACATAATGTCTTACTGGCATCCTAATCTTACTTTAAACTTATTAGATGATCAAACACCTTGGGTTAGAGATGGGGTGCCCTCACCTCTTGATCAGTATATAACATTTCTACCGGGCGAAAATTTTTACAGACCAGTTTTGTATATAAACGACTACTGGAATTATGTTAGTGATTATATGCCAATAAATGAAACTACTCCAAAGCTTGAATATCACATTACGTTTGCTCCTATATCAATGTTTCGGTGGCAAATGTATATTTCTCAAAATATGCGCTCCCAATGGAATCAATATTTAGGTGATAGTGTTGAGCAGTCTGATCAAGAACAAGATATGCTTAAAAGGACATTGCAAGAAACAAATCCATATTTACTTGGTTTAACTATGATTGTTTCATTGGTGCATAGTGTTTTTGAGTTTTTAGCATTTAAAAATGATATTCAATTTTGGAAATCTCGTAAAACACTTGAAGGTTTATCTGTGAGATCAATATTCTTTAATGTGTTTCAATCTCTTATTGTGTTATTGTATGTACTTGATAATGAAACAAATACTATTGTTGTAATCAGCTGCTTCGTTGGTTTACTTATTGAGGCATGGAAAATTACTAAAGTTGTTGACATAAAGCGTCATCCAACAAAAAAGTGGGGCATATTTTCAGCTGTTACTTTTGAAGATAAATCTAGTTATGTTCAATCAGACACGAAACAATATGATCGCCTTGCTTTTCAATATCTGTCATGGCTATTGTTTCCTTTATTAGGATGTTATGCAGTTTACTCATTGTTTTATTCAGAACATAAAGGATGGTATTCTTTTGTTTTAAACATTGCCTATGGTTTTTTGTTAACCTTTGGGTTCATAATGATGACACCACAGTTGTTTATTAATTATAAAATGAAGAGCATTGCTCATCTACCATGGCGAATGCTGACTTACAAAGCACTTAATACCTTTATAGACGATATATTTGCTTTTGTTATTAAAATGCCTACCTTGTATCGCCTAGGGTGCTTACGAGATGATGTTATTTTTTTTATATACCTTTATCAACGTTGGATATATCCAATAGATCATAGTCGTGTCAATGAATTTGGGTTAGTAGGTGATCCTGATCATGTTGACGTTGCAAAAGAAATATCTGAGAGTGATTCAATCACAAATAGTGATATTCAATTGAAGAAAAAAGAACAATAAAATTTTAGTCTATTTACTAATTTATGTAGTTCACAAATATTTTAATCTAAAAATATTGATTCTTGTAATTTTTTTATACAATATGTTTTTATTTCTCAAAAAAAAAAA
>evgR012937c0g1_i01_Hydra
ATTTAAAAAAATGAAAAATAATGAAATTCAATGTGTTATAGTTGCAATTGACAACAAACTTTCTTCTGCATTTAATTTGATTAAAAGCTTACTTGTTGATCAAGATTTGGTAAGCTTTACAGATATATCTTACCATGTTTGGAGTTTTACGAACAAATACTTTTCATCTAATGTGAACATATGTTGTTGTACTGTGGACCACATACTTTCGTTAGCAAATGATGAGTTCAATAGTGTTGAAGGAATTATCTTTGTGCTAAATGGTTCCCAGCTGAGCCAAACAGAGAAACTTAAGAAGTTATATTCTTGGCTTCGAATTGAACCGGAAATAAATCTTGTTTTTGAAAATGCAAAACTATCTGAAGATGAACTGGACCAATCAAATTTGTTTAATTGGTGTGTTGAAAACCAACTAGAATTAGTCTTATCACAAGATGTAACACCCGATCATGCTGATGATAATGATTTTAAAGAAAAGTTTGGTATAGACCGTGTTATTGAAGCACTTAGTTCCCATGTTTGGAAGAATCATGTGATAACCCAAAGTGAAGGTCAAAAGTTGCATATCCAAGACATGTCAGAAGATTTACCAGATGGAGATTCTTTTGAAGAAATGTTTAGCAAGTTTAGAGAAATGAAAGAAGCTGCAATGTTATTGAATGGAAACGAGAGAAAAGATTTAGCTGAAAAGGTTGCCATGGCCTTTATGGAAGCTTTCGATTTAAAAGAATCTGATGAAAACTAAACTTATTGTTTAAAAGCTGAAAAGTTTTATTTTTCAAAATTTAATAGCTATTTATTTTTAAGAAATATCTTTTTTATCAAGTAAAAAAAAAAAA
>evgR012944c0g2_i01_Hydra
TTCTTGCAGATGTCATAGTCAATGAAAATGAGAGGTTAAAAATATTTACGGAGGAACATGATAGTCCCTTGGGAGGACATAGCGGTGAAACGAAGACCAGGTTAAAAATTGCAGAAAAATATTTTTGGCCGAATATGAGCGGAGATATTAGAAAATGGGTAAAACAATGCAAGCGATGCCAACAATCTGGAACGCTTCTGTCATCGGTAGAACCATTACGTTCCATAAAGACATCACGAGTATGGGAAGTTGTTGGAATTGATTTTGCTGGGCCCTTTACTGAAACCAAAATGGGGAATAAATATTTTCTTAGCTGTACGGACAGGGTTAGGGTTTCTATAATTTCCTGTATCGATGAAACTTTGTTGAATATTAAATACCTTAACCTATTTTTGTAACATAAAAATATAAAACCTTACACAGTATGTGTGTGGTTTTAGTCTGTGTAGTTTTAATACATAATCTATATAGACTATATAGGCA
>evgR012949c0g1_i01_Hydra
GCTATTTTTGCTCATGTGCTAATGGATGGATTTTGGGTCAAGATAATGCAAGTTGTATTGATATTAATGAGTGTATTACATTGAATTCTCCATGTTCTTGGACCAATGGATTATGTGTAAATACAAATGGTGGCTTTTTTTGCTCATGCGTTTATGGATTTGCTTTGAGTTACAATAAAACTAACTGTTTTGGCTGGGCAGAATGGAGCATATGGTCAGTTTGTAATATTTCTTATGGAGTTAGATTTCAGAATAGAACAAGAATATGCAGTTCAATGAATGCTATAGATTTGTGTTTGGGCGTTAATACTCAGAAAAAAGACTGTTTAATTAAACAATCATATGAAATTACAATATGTGAAGGATTTTCAAATGTTATATCATGTCCTGAAAAATCACTAATACAAATTCAAAGTGCTTTTTATGGTCGTTTTAACAATCAAACTTGTCCAACCAATAAATCTACAGATTTATACCCATGTAACTCAAAAAATGTTACAGATATTGTTCAAAAGTTGTGCTATAAAAATAACAGTTGTTTTGTTATATCTAATAGTACAATGTTTGGTGACATTTGTCCAAACACATGGAAGTACACAATCGTTAATTACACTTGTTATTTAGAGAATTCAATGTTTACTCCTTGGGTAGAAAATAATTTTACTAGTACAACTTTAGATGCTTTCTATGATGAATTTGTTGTGTCATTTGTAATATTGTCAACTGGAATTAATACCACTCGTATTATAAGTGTTGTTGATGAAAATTTTGATTATGTGTTGGACTTAAGTATTTATAAAAATTATAAGATGGAAC
>evgR012958c0g1_i01_Hydra
TTTTTAATCGTGAACAGTATTTTGACAGTTTTTGGTTAAGATACTCATTTTTGATTTCATAATTATTACAGCTTGATATAAGTCGAAAAAGACATCTAACGAGAAGTTTCATTGTTTAAAAATGGTCATACAGTTCTACGTTTACTTCTATGCTGCATTTACAATTAATGTGATATATTCAAAACCCCAGGGTAAAGTGGAAAAGCAAAATGCCAAAAACATAAATCGTGAAAAACTGATAACATTAATAGCGGAATTAGAGACACAAAAACTAGCAACTTCTGTATTGAAGTCATCAAATAACGTCTACCAATACAAAAACGATATATTGAAATCAAACATTTTTAGAAATCATCAACAGAAAAGTAGCGGCGACGTTATTTTGTCTAAAGTAACTTCTCCAGTAAACGCAAACGGAAGTAAATCAGAAATTTTATATACTGTTTTGAGAAATAAATTGGTATACAAAATAAGAAAAGACGAGATAAAAGCTAAAAATAACTTTTTTTATGGTTTTTGGCAC
>evgR012959c0g1_i01_Hydra
CCTTAACCACCTGCTGTAAATATTTTTACTAAGTAAAAGTGCGAAAGAAAAAAAAAATTATTTAAAGTTTGAAAGGAAGAAAAGAAATCATTTTGGAATGTTTTCTTCTAGAGTTTGCTTGAAAATATTTCTCTGTACTTTTATGTTTCACCAATCTAATGAGTATTTTTAATTTTTTTTGCAATATTTTTCGGGATTTTATCCCCCACCACGACAAAACGTATTCTATTTTTTTCATTCTATAACTTTTTAAAACTTATTCCACCAATATGGCACACCTGTTGGACATCATCTACAAAAAATTGTTTTACCGTCAAAATCTTAAGCTGAAATTCAATAATATCAGTCGTTGATTTTTTATCCAAAAGACGAATAAATAAGCTCAGTTTATTTGGGTGAATGCATTTTGATAAGTATTGAATTTTAAATAACAAGTGTAAATAATTTTTGAAAGGTTAAAAAACCATTTTAATGCGCTCCTTCCGATTCTGTCTAGGTTTTACATTGATACGTTATTGGTTTTACAAAGTTCCAGGCCGTAAGTTAATATCAAACCTTGGTAGTTTCAACTACCAAGGTTTAGTGATTGATTTTGAATGAACGAAACGGATTCCAGATTGAGCAAGAGTTTAAACAGCTGCCCAAAAATTATATAATTTTTCAGCAATGTTTACCCGATCAAGTGAGGCAAATGTAAAGTGCTCCATATCCCATGTAAACCTTAAGTGCTTTAGTCTGTTGTGTAAATTTATTTTATTATCATCTAAAGATATTGTGCAACTTTGTAACTGTCCTTTACTTGTCAGAAGGAACTCTATTTTATCAGGGTTTAGTTTTATGCAATTTGAATTCCATTATACATCGCAAGATATAACATCATCAGCATACTCTACCACACCAGTAAAGTTACCATATAGAGACGTGCATACATTACTTTTATTTCTAAAGGTTTTAAAAAGGTCCTGGGTATAAATATTATAAATGTGCGGTGATAGAGTTGAACCTTGTCTTACTCCTTGTTTAAGATAAAATTCATTTGATTTGTAACCTTGGTATAAAACAATTGCAC
>evgR012985c0g1_i01_Hydra
TCTTATGGACTTTGATGCTCTCAGAGAAGTTTAATCTGGTTTGGGTACTGCTGCTGTTATGGTCATGGATAAGTCAACTGATATCATAGATGCTATTGCTCGTCTTTCTAAGTTTTATGCTCACGAAAGTTGTGGACAATGTACTCCTTGCAGAGAAGGTACTGGTTGGTTGTTGAATATGATGGATAGATTTGTTAAGGGTAATGCTGAAATGGAAGAAATTGATACATTAGAAGAAGTTTCTCGTCAAATCGAAGGTCACACAATTTGTGCTTTGGGTGATGCTGCTGCTTGGCCAGTGCAAGGTTTGATTCGTCACTTTAGAGGTGAAATGGAAGACAGAATTAAGGAATACAAGAAGGCTGATGCCGCAAGGAGAGCTAGAAATAGCCATATCCACTAAATCATTTGGGCTTTCAACTCACAACAAGGGTGTTTAACAAACAAT
>evgR012997c0g1_i01_Hydra
CATAAACGTGCTATCGTGGGAAGAATCTTGTGTTCTCAAATCAAATTCACAATGCCATTGTGTAAGAACTTATATCTGTGAAGAAAATGTTCAGTGTTCGCTTTGAAGAAATGTTTTGTAAAAAGTACCGTCATAAAATTAATTGGATATTGGATCTGAAAGTTATAAACGAACCTTTTATTTTATTTCTCGTATTTGAGAAAATAGTTTGTATGGATGTCAAGTGTTAAAAAGATAATTTTTAATTCAAGTTAGTTCATATAGGATGTTAGTTCAAGTTAGTTCATATAGGATCCTAGATGTTAAGATGATTGTTAATCCAGTATCATGTGAAACTTTAACAAATTCAATATTTGTTCCAGAGATTGAAAATAAATACACTAACACAAGTTTGAATGGTTTTTATGACGAGTTTGTGGTATCATTTCTTATGTTATCAAATGGAAACTCAATACAATCTATTATATATATTGCTAATGATAATGAAAAATTGGTAGACTTAACCATTACATCATCATCTTATTTAGACATGATTACATTAGATAACACTTATCAAATTAAAACAACAGCTAATACATGGAGTTCAATTGAAATTTCTCAATATTTCATAAACAATACGTATAATATTACTGCAGCTGTTAATGGAAGTGTTCAATATTCAAATACTAATACTCAACCAATGGTATTCACTGATGTTATTTTTGTGTATACGTTTTATGAATGTTCCTTTCATCCTAATGATTGTTATTATGGAT
>evgR013003c0g1_i02_Hydra
CTTCTATTGTCTAAATAAGGTAATATATTTAGTAATGCATATTTATTTGATATGGTTCTGGTGTTCATAGTTCATGCTCTATATTGCAATGGATCATCCAACAGGGGATCAGCACTAAAATATTGGGAAGCGTATGCACCTTTGGCAGAGAGTATAGATGACAACTGTCGAAAAGTAGAGCATGAAAATATTAATTCTATCATAAATCGTGTGAACTCCGAATATGCATTTCGTAAGGAAGTTTCCACCAACAAAGAATTCATAGATCAAGGTTCAGAAACTGGAATTTTTCGACTAAAGAATTCATTGATACCTAATGTTCATAGTGTGTTATGGAAATCATCAAATGGATATGCTTTTGTGTTTGTAATGGTAGCCGGAGAAAACATTACATCTGCAGTAGATATTTTAAGCTTAATTGTAAAATACTCTTTAAGCATCTTTAACATGGCAGAGAATCAAGACAGTCAGTTGTTGCAATACCCAGAAAAAATGGTAGCTCTTCTTCATGTATTCTTGCCAGCTGGTCAACTCCTCTTCCTCAACCATCGACTCATAAAACAGCTGGAGAACGATTTAAATAAAATTTTATCTGATAAAATTTGAAAAATAAAGAAAATTAAATTATTTTAACATTAGTAAGCAATCAATCTAAATAGTATAAAATATATAATTATTTTAAGTAATTTATAATCACAAGCTTCCATGCGGTTTATTTAAAGATTATTTTTGTACCTGTATTTTATGTTTTATTTTAAGCATTTTTTTGTAACTTTTAATAATTTTACATTGAAGCAATAAGTTTTTAATATACTTTATTTCTTTAATTTCTTATTTTATTAATTATAGTAGAGTTGAGTAATTGTAG
>evgR013010c0g2_i01_Hydra
CTCATAAAAATTACTTAAAATTGTCTTCTACAAATAGATGATATACTCTAATTAAATTAACAATCATGGTTTAAGTTTGTATAGTTCCATGAATAATAAAGCAACATACTAATACCAACTAATGATTGAGATGTCTTATAAATTGCTGGAAGTTTTCCTACGAGTGCCAGTTATTTTCTTTCTTGATTATTCATTGAGTTTTGCTCATGCAGTATTTCCTTTTTCATGGATAATGTCATTAGTCTTTCAAATTTTAGCGATCTCGATTTGCTGCTGCATTTTAATTTCATCAGCTAGCCAATTGTATTTTGTTTATCATTTTGGCATTTTTTTTCTTCATTTGTTTGCAGTATTGTATCTCAATATCTGTTCAACTCCTATCACATTAACGACACTGTTAACTAGTAATGTTTTGCCGCAACACATTGATACCTATATTTCTACCTTTGCTGCTTCGTGGGTGATATATATTGTTTTTAACTTGTTGTATGCGATTTCTTTCCGCTCTATAAAACCTTTAATAGTAGCATTTGCTCTCATACCAGTTATAGAGGAACAAGTCTATCATAAAAATGAGCAAGTTCCTGTTTTTACTTGCCTTTCGTTTGGATTATTAATATATACGGTATTGCAAAGTTGCTATTGTTTTTTAATGTATTTTATTGAATTGGCAACATACCAAATACAAGTGATAACGTTGAGAATTAGAATATATGGTCCTTTTGTATTGTTTTTTTTGCATTGGAGAAGATTGCGTATGCATGCAGTTTTATCATTGTTTTGGTTTATAATATGGAATTACCAGTTAATTGTTTTCACTATATTTATTGACGAACCTTTTGATTTTTCTTTCATCTTGGCCTGCACCGCCTACTCCTGCAATTCCATTCTAAAGATATTATCTCTCTGTCACATTCTTCAACATGTTGTTAAAGTTATTCTTAAAGGTATCCAGAGATGCATTGCTGACAATCACACATTTGTGGAAGATACTTTTCATCGTCCTACTGGACTGAGAGAAAGCATTGGCTTTCTATTTTTAAGTTTATATACAAACTTAACAAATGTTGATCCGTCAAAACGTATTGTACTTATGGAACTTATTATTCTGATACTTATTTCAGCTTTAATACGTTCAATATTTGAGGTTATCGAGCCTTATTTGTTATCGTTGAACGGAGGATCAGTAGTGCACAGCAAGAAAAGACACATTGGTTTAGTAGCGGCGTGTCTATTATTAATTTTTGTTTCTTTATATCTTGGATTTTTATTATATAGCTTAAGAGAGCGTTTACCGTTTAGTATACCAAATTTAATAACTGTAGCACAAATAACAAGCGCTTTAATTTTATATTTTCTGTATATGTATGATGCTCGAACCGGAGAGTCGTGGGAACAATTAGATGACTATGTTTATTTCATTAAAGGAGGGTGTAGATCATTTGAGTTTGTTCTCATTGTTCTTATACTTGGTTATAGGATTTTAGACACTTCCTCAAAGTGGACAATATTTCAAATTGTTATGGTGGTGCTGCACCTGTACATAAATGTATATCTTTCATTAAAAGACGGTTGGCAAAGTTTGCAACTACGTCATCTTGTTAACAAAAAGTTAAATATATTACCGCAGGCAAGTTCTTTAAGACTAGAGGCGGAAGGCGAAGTATGTCCGATATGCTTAGATAAAATGACATCGGCAAGAGTGACACCGTGTAATCATTTATTTCATTTTTTCTGTTTAAAAAAATGGTTGAATGTCCAAAACAAATGTCCCATGTGTCACGCAACTATACTTCACGGGTCAGGATAATTTTTTGCAGATAATATTCTACATGGAGATTATATTTGTCTGAAGCAAAGCGAACGAATTAAGTTACAAACTAAACAATATCAATACGTAATAAGTTAGTTTTCGAGCGATTCCGACATGCAATATATTGCGACAAAAGAAACAGTAAATTTATTTTATTGCTTGCTCGTAAATCGTATGTTGTTTTTATAGTTGAAGTATATAAACTGGTCATGGTTTTAAAGTTGTACTTTCACCTCATCAATTTTTTTTAATACATTGATTTGGGGAAATAATAGCCTGGAGTAAATATTTTATAACAATAAGAATGTCAGATTATTTTTTTTAATGTCAAATGAATTATTTGTTTGTAGTTTGATGCTGTTAGATTTTT
>evgR013024c0g2_i01_Hydra
AGTTATACATTTTTTTAAATTCATGTAGTTTATTCTGACAGTAATTTCCAGTTATTTAAATGTCTTTTTCGGTGGGAATTGACATTGGAACAACCTCGATAAAGGTATGTATTCTAAAAAAATGTGATGTTGTAGAATCGTTTTCAGCATCTCATAATGCTACATTGGTAACAACTTCTCAGTTATTTCATGAGCAAAGTCCAGAAATTATTCTACAAGTGCTTGATAATTGCTTAAATCAGCTAAAACTTATAAAAAAAGTAGACAAGGTTTGTGTTACTGGACAAATGCATGGTTTATTGTTGTGGGAGAAAAAACAGTGTCTAAAAAACCCAAATAACTCCTCTTTAATATCCTTATATAAAAACACTGTTTATGCAAATACAAATCTTATAACTTGGCAGGATCAACGTTGTAGTGAAGTATTCTTAAAACATCTAAGTTGTCCAAATTTATCTTCTGGCTTTGGTCTTGCTTCTCTGGTGTGGCTACAAAAAGAAACACCAGACATATTTAATTTGTATCATATGTGTGGAACAATTATGGATTTTGTGGTTTGGCTACTCACTTTAAAGAAGCCTGTTATGAGTGTTCATAATGCTAACAGCTGGGGTTATTTTGATTTGATAAAAAACTGTTGGATGCTAACTGAGCTAAAACAGGACTTGCTAGATTGTTCAATTCTTCCTGATGTTGTACCAGCAGGAACTATTGTGGGTAATATATGCTATCCAGTATATGGAATGAGTTGTTCCTCCATAGTTTATGTAGCACTTGGAGACCATCAATGTTCTATTCGCGCATGCTCAGCACAGTTATCAGAGGCTGTTTTAAATATGGGAACATCTTGTCAACTCTCAATACGTGTTCCGTTCGATAAAGACTCGATGACATCAATAATGGATATTAAATCGAATAATTCAGCAATAGCAATCACTCCTTACGACATGAATTTTGTTTTAATCACAGCTGCTTCTTTAAATGGTGGTAACGTTTTTGATCAGTTTGTTTCTACATTTGTGTCATGGTGCCATATAATAGGGTGTTCAAACATCCCATCTATTGAAGAACTAATGGAAAAACTTTTAGCGATATCTTCAACTGAAAACAGATTTGTTGAAGATTTGTTAATTATAACTCCAACAATTTACGGTGAAAGGCATGATCCTAATGCACGTGCAAGCATTTCTAACATATCACCTTCAAATTTATCTTTATCTGGTATTATGCAAGCGTTGTTTGAAGGTTTATCAAAAAACTTACTTACTATGCTCCCAACTATTTACCTTAAAAAGTATGGTATAACTCAAATTGCAGGAACTGGTCGCTGTTTTATACAATGCTCTATATTGCGAGAAGCTATTTCAAGGGATTTTGAGGTTCCGTGGAGATTAGTAGATAAATCTAGAGACGCATCTTTAGGAGCCGTTTTGTTTTCACTTGGTATGGATTAACAAACAGAAATATATAAATTAGGGTGCATCACAAAACAACACTTTAAAATTTTTGATTTTGTCTGACTCCGAATTGTGTTCTATATATTAAAAAAACACTGGATTTCAAATATTTTG
>evgR013025c0g1_i02_Hydra
AGCATTAAATGCAGCTGTTATACATCTATAACCAGAATGCACTACAGTTTTACACTTGCTTTGATAATAGCCATATTTATGTTGTTAAAGTTAGCCGACCCTTCATATGCTATAAAAAAAAAGAACTCAAGAAGCAAAAAGGACAAAAATCCTCCAAACACTGTATTTATAAATACTTGGGCAGTCGTTGTGAATGGGAATGATGCAATGGCAATGAAAGTAGCGGGAGAAAATGGGTTCATAAATCTAGGGAAAGTATTTGAAAATGTTTACCATTTTGTCCATAGAGGAATCCCAAAGCGAAGAAAACGTGCATCTGTTTCAAAAACAATACATCTTAACAAAGACAAAAGAGTTGCTTGGGCTCAGCAGCAAGAGTTAAAGATTCGAGTAAAACGATCAATACCATTCAATGATGTATGGTGGCGAAAGCAATGGTATCTGTATAGACAAGAAGATGAATTTTTTGGACATGATTTGAACTTGCCTCCTGTTTGGAGAAGAGGAATTACAGGAAAAGGTGTTGTGATATCCATTTTAGATGATGGTATCGAGCATACACATTTAGATTTAAAAGGAAACTATGATTCTAAAGCAAGTTTTGACTACAATAGTAATGATACTGATCCATCACCAAGAAGTAACTGGCGTGAAGATAACAGACATGGCACAAGATGCGCTGGCGTTGCAGCAGCCGTTGCAAATAATTCAAATTGCATTGTGGGTGTAGCCTACGAAGCAAAAATTGGTGGAATTCGAATGCTAGATGGGGAAATTACAGATTTGATTGAAGCACAATCTCTTAGTCATGCGCGTGGATATATTGATATTTATAGTTCTTCTTGGGGTCCGCAAGATGATGGGGAAACAGTGGAGGGACCTGGTTTACTAACTCAAATTGCTTTAGAATATGGAGCAAAATATGGAAGACAAGGCAAAGGAAGTATTGTTGTTTGGGCTTCTGGAAATGGTGGAAAATACCATGATAATTGTAACTGTGATGGATATACAAATAGTATTTATACAATATCGGTAAGTGCAGTAAGTTCGATAGGTGTGCAACCATCATACTCTGAAAGCTGTGCATCAACACTTGTGTCAGCTTATAGCAGCGGAACGACTAAGTTTGATCGTCGAGTAATGACTACTGATTTAAATCAAGGATGTACTGATAGTCACACAGGTACTTCTGTATCTCCACCAATTGTTGCTGGTCTTATTGCTCTTGCATTGCAAGTTAATCCAAGACTCACATGGCGTGATATTCAGCATATATTAGTTGAGTCTTCAAAATTAGACAAACTTTTGGGAGATGATTTTAAGTTCAATGGTGCCGGAAAGCAGTTTAGTCATCATTTTGGCTTTGGAATAGTTAATGCTGAAAACCTTGTAGACATTGCAGAGGTGTGGGCAACAGTGGGTGAACAACATTCTTGTCATCATCAAAAGAAAAATAGACTGTACCCTCATAATCGATTTAATAACAGTTTTCCGCATGTCGTGCAATTTAATACTACTGCGTGCTCAGGAACATTGCAAGTTGTTAATTTTGTTGAACATATCCATGTTGTTGCAACATTTTATCACTACCGGCGTGGAGCAACTGAGATATCAATTGTTTCTCCTAGTGGTACAGAAAGTTGGATTTTAAAAGAAAGAAAAATGGATATTCGTGATGGAGAATTTAAAAACTGGAAGTTTTTGTCAAATCATTTTTGGGGAGAGAATCCCAAAGGGTTATGGAAAATAAATTTTATAAGTAGAGAGCTTGATGATTATGGCCAAGGCTTCTTACAAGATGTAGAGCTAATTATCTATGGTACTGAACATCAACCTAACTTTTATAAGTTTAAAAAAAAATCAGTTAAATCATCATATAACAAATGCCATGTACTTTGTGACCGTGGGTGTGTTGGAATTGGACCCCATAATTGTGTTAGTTGCAAAGGATATTTTTACAAAGCATCTGATATAAAATATTGTGTTGAATCGTGTCCAATACATATGTACACAGTAGGAATAAGTCATGAATGTTTGCCATGTCACAGTTCTTGTTTGTCTTGTGAAAACGGAAATCAATCCAACTGTGTAACTTGCCCAAGCTCTACTGCTCTGTTGTATGGAAGATGTGTTCAGCAATGCCCACTGCACATGTTTATGTTTGCTGATGTGAGTGGTTTTGTCTGTCAAAATTGTGGATTATTATGTAAAACTTGCAAAACTCATGCTGATTTGTGCACATCTTGTGAAAAGACTAACATCCTAGTAGAGGGTAAATGCATTCCAAAATGTAACGAAGGAGAGTGGTTTGATAGCAATGATTGTCAGCCTTGTCATTCATCTTGTAAAACTTGTAGTGGTGGTAGTTATTTAAACTGCAGCACATGTCCTAAAATATCTTTTAACGATGATGAAGATAGATATCTTTATAATGGACGTCAGTGTGTTGTTAACTGTCCTGTAAGAACTTATCATGCTAATACAATTGCTGGAGATTGCTTATATTGTCCTGAAAATTGTTTGCAGTGTAGCAATGAAGGTCGTAAATGTGACATTTGTGCTTTTGATTATTTTCGAGTTTCTGGCGAAGGGAGCTGTGGCGAAGGTCGTTGTGGCGAAGGGAATTGTGTCAAATTTTGTCCACAAGGTTATTACACAGATCTTATCACAAGAATGTGTGAGAGTTGTGGTCATCATTGTGAATCATGTGTACCGGGTAATTCTGCGGCTTGCACCTCCTGTTTTAATAGTTATTTTTTAAAGGGTCACGAATGTGTATCAAAACTTGAATGTCAATCAAGATATTATGCTGATATTACTACTGGAAAATGTGAATTCTGTCCAATAAACTGTTCATTTTGCGAAAACGCTAACACGTGCACAGCTTGTTTTTTAAATTACTATCTAACTGCAGATAAGCAATGCTTACATAAATGCCCTAGTGGTACTTATGGTGTAGAAAGCCCTGCCAGCTGTGTTAAATGCTCTAAAGGTTGCCAGGAATGCAATGGTCCAACAATTCACGAATGTACTGTATGCAGTCAAGGTAAACATCTGTTTAACGGGCGTTGTGTAGCAGAAGTTCCTGTAGGGTATTTTGCAAAGCATAATGGAATTTCTAGCGTATGCGAACCATGTCACCATACGTGTCATGAATGTTTAGGACCTGACGACTGTGTTTCATGCATTGGGGAATTGGACAACCGAAACAAAGGATGTGTTAATCCATGCCCAAATCAATATCTTTCGCAAAATAATTGTCAAAGTTGCCATCCTGGGTGTGGTTCTTGTTTTGGGCCTAACTACGATCAATGCTTAGAATGCACTGGAAATTTCTCTCTTCACACTGATACTCATACTTGTCACAAAGTTTGTCAAAGAAAATTCTATAAAGATGAAAAAACGTTAACATGTAAACCGTGTCATTTTTCTTGCGCTTCATGTGTAGGATCAAGCTTAGAACAATGCATTACATGTATTGATAAACTTTACTTTCTTAATAATACATGTGTTTCTCAATGCCCACTAGGTTGGTTTAAAACGTTTAATTATGCGTGCGCTCGCTGCAGCTCACGCTGTAGTCAATGTATCTATAATTTTGATTTATGTACAAGTTGTGGATCAGATGAAGTTTGGTATGATTTCAAATGCTATTCTGCATGCCCAGCTCGTATGTTTAAAGATAAAAGTAACCGATGTTATCAATGTCATGATAGTTGTAACACTTGTGATGGTGCTTCTCCAAATCAATGCTTGACGTGTCGTCAAGGATTGAGTATTTATAATGGAATGTGCATTGAAGGATGCACTCCTGGTTTCTTTTATAGTCCTGTAAAAAGTAAATGTTTGCCATGTAATGATAATTGCTTACGTTGCAATGATAATTATGGAAAACATTATTGCACGAAATGCAATAATGCAACATACTTACTACCGAATGGAATTTGTTCTCATCGTTGTCCGTTATTGTATTATCGATATCCAAAAGATAACATCTGCATGAAATGCAACGATAAAAAGTGCTTAGCATGTGAAGAAGATTCAGGTGCATGCTTATTGTGTTTGCCAATGTATGACCTGGTAAATGGAATTTGCAAAGCTAGTTGTCCCTTGGGTACGTTTTATAATAAAAGTAACAGCTCCTGTTCATTATGTAACTTGTTTTGCCAATCTTGCAACGACACTAGTTACTGTAGTAGCTGCAAAGATAGTTATTTTTTGTTTATATCGAGCGATGGAAGATATAAATGCGTGCAAAATTGCCCTGAAGGATACTTTGAAAAAGATAAGAGATGTCAAGCATGCAATGATCAATGCTTGAAATGTAATAACTCCTCATGCCATTCTTGTGTACACTTATCAATTTTATTTAATTCCACGTGTATCAAGAATTGTCCTCTCGGATACATACGGCATATGAAAGATAATATTTGCGAAACATGTGATCGAAGGTTTGTAGGGTGTTTGGATTGCAATGAAAAAATATGTATTCAATGTCGCCCTGGGTTTTATATGTTTAACAATTCTTGTCTAACCGTGTGTCCCTCTGGTTTTGTTGTATCAGCAGGTGCATGTCAACCCTGCAATAAATTAGATAAGAGATGTTTATTATAAACCCAGAGTACTAGTTTCAAATTCATTTCATCTAACAGTTTTTATGCAGACATACGTATCATGGCTATATTGAAACTGTAATAGCACCAAAACTACAGAGCTGCATCAAAATAATATCTATAAAGCATTATAGTTAATTACCTACAAATTAAGATCTTTAGTGTTTTGCAGTCATTGTCATCCTTAAATTTTGAGTTGTTCGTTCGGCAAAACAATAGTTTTTATCCGGAATTGATGAATTGATTTTATGAAAACATTCGACATATTAAAAATTGAGATTTCCATGATTTTAATATTTGTTTAAACAATTAGAGAGCTGCGCGTTTTTATAAAAAATTGCTCAACTGAACTTTATCAAATTTTCATCAAAGTTAAAAATTAACAACCGGAAATATAAATTAGAATTCCTTAGGAACATTTTCAAAAGGTGTAAATAACTATTAAATATAATCAAAAAAATTAGGAATGTTACAAATAAATATT
>evgR013029c0g1_i01_Hydra
ACGTCATGTACAAGAATACAACAGTTTCATTGGGTTTGTGGGATACAGCTGGTCAAGACGATTATGATCGTTTGCGACCATTGTCCTATCCAGATGCTCAAGTTTTCCTTGTCTGTTTCTCAGTTGTCAATCATACAAGTTTAATGAACGTTAGAGCCAAGTGGGTTCCGGAAGTTAAACATCACTGTCCAAAAGTGCCAATCGTTCTCACAGGAACAAAGTCTGATCTTCGTAAGGATAAAGATTATTTGCAAAGAGAAGGATTACAAGTTGTTAATTCTGAAGAAGGACAAAAAATTGCCAAGGAAGTTGGAGCTCAATACTATTCGGAATGTTCTGCTAAATCACAAGAAGGATTGAAGGAAACTTTCAACTTTGTTATCGAATGTGTTTTGGATCCAAAGAGCCCAGATCAAAGCTCAGAGAAGAAGAAGTCCGGTAAATGCAGTCTGTTGTAAAGAAATTATAGATTTCTTCAAATAATACTATATCCACGTATAGGATATGTGGAATCTATCATCCTACTGTATATTATTTTACCACCAATTAAGC
>evgR013030c0g1_i01_Hydra
ATTTTGGTTTTCTCATAAATATCAAATGTGATCCCATAATTAACGACGAAGACCTAATATCAAAACTTGAACCACACGTTGATAAAATTTTATCAATAAAACATTCAACATACAACTTTGATAAAAGTATAGAAGATGGAAGAAGACTTTTCAGAGTCAACCTTTCGAAACAGGTAAAAGAACTTCCCCACGAAATCGAAATAAACGGGCAACGCATCACGCTAAACTTTGCAGGAAAAGAGTTTTTATGTCCAAAGTGCGAAACACTACATATTCCAAGAGATAGATGTAAAGTAAAAGAAACAACAACGCTTTCTAAACCAACTCTTCAACACATTCCAAAAGAAATTCCAAAACCTACATCAACAGTACCAAGAGGAG
>evgR013037c0g1_i01_Hydra
CTTGTTCGCCATTGTCTTGTGCTTTGTAGCCTTGACCAATGCTGCTAGAAGTGGTATTCCATCTGATCAAGTTGTTGAATCCATCTTGAATACTTTTGATGCTGAATTGGCTGATACTGCCTCTTCTGCTTCACCAGCTGCTTCTGCCGCTGCTGCTTTTGCTCAAGCTCAAGTTGGAAAGTGTTATTCTCAAGCTCAAAGATTGGGTAATCCATGTTATGATTGTTCTGGACTCGTTTATTCTGCTTATAAGGCTCAAGGAAAGACTGTTCCAACCTCAACTGGTGTCTATCCAAATGGTAACATTTATGAAGTCTCTGGTAGCTCAATGCAAGTTGGTGACATTTTATATAGACCAGGTCACGTTGGTATCTATGTTGGTGGTAATCAAGTCGTTTCTGCTGAAAACTCAAGAACCGGTGTTCGTCAAAGAGATATGGGTTATTACTCAAAGTATTTGAAATTCACCAAGGTCTACAGAGTTAGATAAATTTA
>evgR013065c0g1_i01_Hydra
AACCCTGCCATATTTTATCGATGGCGATGTTAAATTAACCATGAGTAATGCTATCCTTCGTTATATTGCAAGAAAGAACGATATGGTTGGAAACACGGAAAAAGAAATTCGTAGAGTGGATATGATGGAGAATGAGTTAATGGACTTTAGGAATGCTTTTGTAGCTCTTTGCTATTCCACAAAATCTGAAGACTTAAATGAAGCACTAATTATATATAAAACAAACGCTCAGACTCGCTTAAAACTATACTCAGATTTTTTGGGCGAATCAGATTATTTAGCTGGAAAAAAATTGACCTTTGTTGACTTCATTTTTTACGAACTTCTTGATCTCCACAAGCTTTTGAAAGATAACATGTTAACGCCTTTCCCAAAGTTAGAGGCATTTATGAAAAGATTTGAGGAATTACCAGAAATATGCGAATTCATGAGTTCTTACAAGTTCTTTAAAGGACCCCCAAATGGAAAGATGGCTATCATCGGTGGCGAGTTACAAAATACACAATATTGTTTACTTCAATAAATTAATTATATCGTAAAAAAAAA
>evgR013073c0g2_i01_Hydra
AATTGTTTTTTGCATATTTATTCAATTAATAAACGTGTTAGTTTCGTTTTCATCATCGTTTTCATCATGTCCCAGTTCATGCAGTCATGTGATCTGTATTGGAGAAAAAACTCTATCAGGATTTTGCAAATCGGGAATGATAATGGACTCGTGCAACTGCTGTCATGTTTGTGCTAAAGGCGAAGGCGATGTTTGTGGTGGTTATATGGACATTCAAGGAAAGTGCCGAGAGGAGCTTTATTGCCAAGTTAAAAAGGACGGCGTTGGGCAATGCACAGCACAAGTTTTTGAAGTTCCAAATAAAGGTTTTTTACAAAGTACATCCAATACGATTCAAGAAAAACCGACCAAAAAAGAAAAATGCAAACCAAAGTGTTCCGCTAAATATTGTCTCAAAAACCCGAATTCTATTTGTTCTGCTCGACATTTAAATATTAAAAAACAAGAATGTCAAGGCAATTGTCGACACACTGTATGCTCAGCTTGTTTTGCAAAAACAAAAAATAAGTGTGTCAAATGTAAACAACACGATTACGGATGCATCAAATTATTTGCAAGCTGCGTACGAAGCAATCTTGAGAAGATATGGGATCTTTCTATTGTTAAAAAGAACGGAAGTCATTTGTTTACATGCAGTATACCCGGATGCTCTTGATTTATTTTAAGTTGAATTTTTATATTTATGTTTAGG
>evgR013075c0g1_i01_Hydra
GGAACGTCATACGCTCTTGAGATGCATCTAAATCATTATAAAAAAAAGTATGGGAATTTTTCAAACGCTCTTTTTTATAATGATGGTCTGCTTGTATGGGGAAACTTGTTTGAGCTTGGTCTTCATTCTCAACGTAATCATAACCTAGAAATTATAATAAAATACATACCTAGCATACTTTACACAGATAATCGTGTAAATACTCAAAAAATTTCAATTAGTTCATTGTTATCGAAAAATGCTGGTTTTGATTACTACTATTACCATGGTTCGCTGACAACACCAGGGTGTCAAGAAGTTGTAACATGGATAATTGATAAAAAAGTACAAAAAGTTTCAAAGTCACAGCTTGAAACATTCAGAACACTTATGGTATCCAAGGGTTTCTCAGAATCGCCCATTAACATGTACAACAATTATAGAGAAGTGCAATCTTTGAATAAAAGAAACGTCTACGCTAGCACTAATAAAGGATTAGCTTCATGGCATAAAAACAACTTATATACCATAATATTTTTTCTTAACTTAATTAACCTTATAGCTTCAATTTAAATTGTTGTTATATAGTTTACAGATCTCTCTCGTTCTCATTTCCTCTCTCTCTCTCTCTCTCT
>evgR013077c0g1_i01_Hydra
ACCGTATATATTTTAACTATCTGCAGAATACTTATCAATAAATCAAAAAAGACTCGGTCGAACTTCATGCTAACAACAGTTTTTAATTGGAATCCGTTTGTTTGAATTTGAGATACAACAGGAGTTTGAAATACCACCTGCGAAAAAAATTTTAGAGCATAAATGAAGATACGCATAATACTGTTAATGACAATTGTCGGCGTATATGGATCATATATTAAAAAGAAACACCAAGTTATTAGAAACGATAAGGGGAAAAACAATCTTTGTAATCACTATTTTGCCAAGGTATTATTAGACAGCCTTGGGATGAGAGCTGATACTTCTCCGTGCATTTTGCATAATAAAAAAGATATGGAAAGTCTACTTACCAGTGTTTTAAAAAAAATGGTAAAAAAAGAAGATATCATTGAAGAAAGAGAGTACTGTTGCAGTCCTAAATGTGATAGTTTTATGGAAAGGCTAGGTAATCCACACGTCGATCTTGTTCAATGTACGTAAAAGTCTACTATACGAAAACAACAACAAAGAAGATAAATTCGAAAAAAAACAAAGAAGATAAACTCGAACTTTTTTCAAACCTTTTTTTCACAATTTAAAATCTGATTTCATTTTATAATGAGTTTGATTGATAAAATAATTAATATCTTTTAAAAAAAAAAAAAAAAAA
>evgR013077c0g2_i01_Hydra
GAAAAAGATTCAGATAGAGTTGGAGTTTTGAAGTTTGAGATATCTGAGAATGAAAATAAAGATACTCATAATACTGTTGGTGACAATTGTATGCGTATATGGATCATATATTAAAAAGAAACAAGTTATTCCAAAATTTAAAGATTTTATGAAAAAGAAAAGTGAAACAATGACTTCAATGGGTATGCCTGGTGCTCCAATGCCTCCAATGCCTCCAATGACTACAATGCATCAAATGTTTAACGGTCCCTTAAGTGCAAGTTCAAAGAATTTGTTTTCATCTATACTAGAGAAAAATAAAATGCATACCAAAGCTGCAAGACCTTGCAGTCCAATTTGTCTTTTTTGGATGGCAATCATAAGTCAACAGTCTGAAACCCCACTCATTGAACAAGCCCCATCATGTCCAAACCAATGTCAAAAGTCACAATGTTGTAACCCACCAATGGAAACTCAAGTATAAGTTCCACCAGTATGCATAGCATCATGCGCACAACAATGATGTGGAAGCTAATAAAATATCTATTTGAATAAAGTTTTGTATTTTTATTATCTAAAACTGAAACGATTTTACAAATGTTATCAGATTTTTCTTTTCTCATTTTTAACAAACAAAATAATTTAAATTTTTCTTATTTCATTCTTAAAAAACAAAATAATTGAAATTTTTGTAGTCTTTATTATCTTTTTTTTTTGCTAAACAACTTTCTAGAAGCCTAGGTTCAAAAATAATAAAAATTCTCCAACAGCCGTGGTTCAAACTAAAAAAACGTAGCAATGGAGTAACGTAACGTGGAGTAAAACGTAAAACATTTTTATTTAAAACCGAGGAAACTAGATTAAGCCTCACTCAGGACGTACGACCACGACTCAGGACGTACGAGCCAAGTGAACACATATAATATAAACATTATAAATTGTTTTTAAATTTTAAAAAATGTTGATGTTAGTTAAGACTGTA
>evgR013091c0g1_i01_Hydra
GAAGAAATTGAATTTGGAACCTGAATCTGAAAAGGCCAACAGTGCCATTCACACATTCAAATTCTATGGTTTGTTCAGTGATAAGGTTGCTCCATTCAAGGGTACTTTGTTGGATACCTTCTCTCAAACCCTGTTGGAACAAATGGAATACAAGCCAGGTGAAAGAGATATGCTCTTCATGTGCAATACTTTCGGTATTGAAACTAAGGAAGGTAAGAAGAAGACCATTGTTGCCAAGTTAGTTGACTTTGGTGATGAAAGAGGAATGTCAATGGCCAAGACTGTTGGTTATCCATGTGCTGTTGCC
>evgR013095c0g1_i01_Hydra
ACAACTGCTCAGTTAATGCAACATGTAACAATATAGTAGGCGGATACAACTGTATATGTAAAGTTGGTTATACAACAAGTAGTAAGGGAATAGAGTTGCTCTCCGACCAATGCATTGACTTTAATGAATGCGCCTTTACAAATAATAACAACTGCTCAGTTAATGCAACCTGTAACAATCTGGTTGGTGGATACAATTGCACGTGTAAAAATGGCTTCAACACAAGTAATGGTGGAATAGAGGCTCTCTCAAATCAATGTCTAGATATAAATGAATGTGCATTGTCTAACAAAAACAACTGCTCAGTTAATGCAACCTGTAACAATTTGGTTGGTGGATATAACTGTAC
>evgR013098c0g1_i01_Hydra
TTGTTTTAGATATAATAATAATGTTTAAAAAGATAAAAATGAGATTGTCAGATTCTGAAATTGATGAACACTGTTTATTATTAAAGTTGCTATCAGATCATATGATATTGTTTGTTTTTCAACATTTGGATTCATTCTCTTTGTCTAATGCAGCTTGTGTTTGTCGTCGTTGGTACCATTTAACCCAAGACAAGACACTTACAAAACATTTTGATTCACGTATGAAACCACTTGGGTTAAAACAATTATGGAATGTACTTCGTAAAAAACTTACTTTGTCAACTTCATCTGTTCATTTGTGTGGTGCAAAGTCGTCAAGAAAAATCGAAATGTTATCACTTGCATTTTTAAAGGATCTTCAAAAACAATGCCCTTATATAACATCACTTAGTTTAGAATTATTTGATTTTCATTCTATAAATATAGATGCTCTTCCAATAAATTTACATAGCTTGTCTTTAAAAGGATCAACTTTATCATTAGGTTGGTTTGACCCATTAAAATCTGGTTGCTGTTTTACAAAATTAAAGTGTTTAAATTTATGTGATTGTACAAAGTTGTCCAATAATGATTTAGAGGCCATTTCTTATGTTAGTTCACTTGAGAAGTTGTATTTACAAAATTGCTATAGAATTTCAGCTCGTGGTATTCCATCAATCACTATAAGTATGAAGTTTTTAAATGTTGTTGATTTCTCTAAAGTTCCCGCAATTAATAATGTGGTCATGCATTATTTGTCTCAGTTAAAACATCTCCTTGAGCTTCGTTTGAGATTTTGTCATTTAATTACTGATGATGGCATTAGGAAATTGTTTAATGGAAGTGTTGGCGAAACTTTACGAATATTAGATATATTTACATGTCATGAGCTCACTAATAATGCCCTTGATATTATTATAATACAATCAAAATCTTTAAAACATCTAGATATAGGAGGAAATAATAAGTATACAGAAAATAAAGTAAGCGAGGTTTGTAGACTACTTTGGTCTTGTGATGTTAAATGGCAAGTGAGTGAAACTGTTTCAGAATGTTTTATAAATGGCCTTGATTATAATCAATGTCTTGATGGTCAATAGCAGAGCAACGTATAACATTTGTTATAATGTTTTGATTTTTTGAAAATTTACTTTTAAATGGACGTTTTTGTGCACCTGCTTTCATTGATCATTGAAGTTGTTATATCTTTTATATTTATAAATGATAATAGGAAATGTTATTAGGAAATTTTTGTATTTTTTTTGTTGTTGTAAATTTGTATTTATCAATTAGATAAAAATCTGTAATTTTGAAATTTATATAAATATCTTTATTTATTTATATACTTACTTTTTAT
>evgR013099c0g2_i01_Hydra
CTGAACGAATATAAGTTAACTATCGTTGTTTAAAATTATAATAAAATGTCGAATGCTGTTTTGATTTATAAAAACCGTGTTTGTATTGGATCCACGTTTATAGGAATAGTAACTTTTGTTTGTATAATCACTGCGACTAGTGTACCAATGTGGAAAACTGAAAAATTTTTAGGTATTGAAACTGAAACTTCGGGGTTGTGGCAAAATTGTCCTAAAATTACAAATAAATGCGAAAAGCTTGGGGGCGTTGAAGGGTTTATGAAAGCCTGTCAAGCTTTTGCTGTCATTGCAGTTTTACTTAGCTTTTTAACGGTTTTGTTATTTTGGCGTCAGTCGTGTCCGGTGTCGTGTTCATCGAAAGTCAATACCATTGCTATTTTATATGTTTGTTCAGGTATATGCATGGCTATTTCATTATCAATTTACGCATCAAAGAATAAAGATGTAAAGTATGGTTGGGCGTATATCGTTGGGTGGTGTGGTGTAGCTTTTTCATTTTTTGGATTTTTTATGAACTTATGAGCATATTTGTAATTAATTTTTTGTTTTCTTTTTTTTTCTTTTATTTAATTCCTTTCCCTAAAAAAAAAAAAAAA
>evgR013101c0g1_i01_Hydra
TTAACAGGTAAAAGCAAATTATGTTTTAGAAAAAGTCTTCTAACAATCTTGGCATTAAAAGTTTCACAACAGCAATATTGACATTAATTAATATATGCGGAATTTTTACTCAAGTTGCATTTGAAAACAAGATAATAAAATGTACATTTGGATATTTTGTATGCTTATCATTTCAAGCAAGTGTAGAAAAAACAAAGAACAAAATCCAAATGAAATTGTTACTCGACATTGCAATGCTGATTGGTGTGATCCTAGTTATGAACCATATCCCAATATTGTTTCGGTAATGACAGGATATGATATCATTAAAGGAAATCCGTTTACTACGTCTCAGACTGTTGATCCTGGGTTTTCTACAGGGTATATTTTTGTTCCCACCTATAGAACTCCAGATGGAACATATGCTCTACATAGTGGAGTTACTGTGCGACAGACATTGCAGTGCAGTTTAGCAACAACTACAGATACGATTTCGACTGCTAAAAAATATATTCAAAATATTTTGTCAAAATCATCACAAGGAAACACATTAACTTATGAAAATGATAATGGAGCTGGTGCTGTTGCAAACGTTGGACCTTTAATTGATGCTGCATTTACATCCAGTAGTGAATACAAAAACAACAAAGAATTTTTTACTAAAAGAAAAGGTGTTTTAACAGTATCCGAAGCTACGTGTGCTACCTACTCCGTTCAAATAAGTCCTTACGATCCTCCACCATTTAGTGATGGTTTTATAAAAGCTTTAGAAAGTTTGAAGGACGCAGTTGTTTCCACGGATACAGTTGATGTTTCAAAAAGAAAACCATTTGAAAAGTTTGTGAAGGAATTTGGCACTCATTTCATTCAGCATGCCACAATGGGTGCTAGAACTGCTATAACTAGAAAATATTCTAACGAAGAATTTAGAAAAACATCAAAAGAGCAGCTAAATAAATGCAGTGAAGATCGTATAAAAATGACAATATCAGGTATCGGATTAGGTACAAACTCAAAAAGTTGCGAAGACATTGATGCAACTATAGATAACAATAACGTAAATGGATTTGACAGAGAATCCATTACATCTTATGGTTCTAAACCAGAAAATGATTATTCTGAATGGTCAAAGCAAAACTTCGACTCTCCCTTACCAATTAAAATGGAACTTCGACCAATAACAGATCTACTTTCAAAAAATTACTTGAAAAAACACACAACTATTCAAATCGATGAAATAAAGAAAAAACTCTCTGATTTTTATAAAATATATTGTACAGATTTTATGAAGATTGAAGATTGTAAACCTTTTGCAAATGAGGAGTGTGGTTTAAATGACAATTGTGATCCGCACACTGATATCTGTTTTATAACTAAAGATGGAATAAAAACAAAAGCCAATTGCAGAGGCCGAAATTGCAAAGACAATATTTGTCAAAACGGAGGAACCTGTGAAGAATTGGAGAATAGCTTTAAATGTCATTGTTTAATAGGGTGGTCGGGTTCAAATTGCAATGAAAAATTTGCAGAATTAAATGGGTTGGAAAAAAACATTTTTGATAAAATGGGTAGTGAAAGTTATCATAATTTAGATTCACAAAATTTTGCTGAAAAACTGCATCATGATTTGACTGGTTTGTACAAAGATGAAATTTTCGCAGTGACGTCTATGAGTTCACACGATGATTCCAAGATTGGAATACATGCTGCGGTAGGTACATATATTAAATTTTATGCCACACTTGGACGAAACGTAATTGTTGGCTGGACAAAAAAAGGAGCGACTTTGCCAAGTTCCGAAAAGACTTCAGATATTATTAAAAGTGTAATTGAAAGTGTTGCACAATCATCTGCTTCGACCGCAAAGGATGTCCTCAATAAAATAGTCAATGTTCTTACTTCGGATCAATATAAAAAGTATCCATATGTATTTATACAAACTGTTCGACAGGGCTACAATCTCTATTCAAAATGTCCACAAAAAGCTTGCTTACATTTTACGGCAGTGGTTCTGCCTTCATTATTGATTTCATATTTTCCACACCGTTACGACGTAACTATTTTGTTTGGAGAATATGAAATAATAGAACATTCGTGAAAGAAAAGAAAAATGCATTTTAAGAAATTATTTAATTGTTTAAATGTTAGAAAAAATTTGTGTCATATTTACTAAATTATATATTTTTTAATTAATAAAATATGATTAACTTTAAAAAAGACTATTTTCGACACTACATTATTACTTTATTTTGATTTTGAGTGATTAGTCATTCGC
>evgR013104c0g1_i01_Hydra
GTAAATTCATTTTCAATGTTGCATTTAAACATTAGGAGTATTTCGAAAAATTTTGAATTATTCAAACAATTATTAAGCGAGATTAAATTTACATTTAAAATTATTAGTCTCAGCGAGACATGGTGTACTGATGAATATATCGAGACAAATACAAATTTTCAGCTACCAAATTATAAAGTGGTTCACCAATTTAGAGGATCTGGGAAGAAAGGAGGGGGTTTGTGTGTATTTATTAGTAATTCTTTAGTATTCAATGTAAAAAAAAATTTGTGCTCAACCACTAATGACTATGAATCACTGTGCGTGGAAATAATAAATAAAACCACAAAAAATATTATCATTCACGTTCTATACAGACCACCTTCCGCTTCAATAAAACAATTCGAAAAACATATTAAAAACATAATTAAAAATAAATCAGGCGAAAATAAAAGCGTTTATTTTGTAGGTGATTTTAACGTTGACCTCAACATGCGTCATTTAAATAAAAAAATAAATAACTTCTTTAACGTCATATATCAGAAAGGCTATATTCCTCTAATTAATAAACCCACAAGAATTACTAGAGAAAGCGCAACTGTCATAGATCAAATAGTCACTAACGAATTCAAATCAAAAATAAAAACGGGAATATTTAAATGCGATATTTCAGATCACTTCCCTGTATTTCTAATCTCACAAAAATGTGATAATAACTACGCGCATAAAGTTATAAAAAATTCGCGAAAAATAAATGAAAAATCCATTGACATTTTTAATACTCTTCTATCAGATTCAAACTGGGATGACCTTCTTGATATTGTACAAACAGATAAGGCATACGATAACTTTCATGATAAATTACAACAACTTTACAATACGGCTTTTCCTGTAGTTACAAAATGTGTGAAAAAAAAGACATTAGCAAATCCATGGATAACGCCGGGAATTATAAAGTCGTCAAAAAAAAAGCAACGACTATATAACAAATTTCTAAAAAAGAAAACTTGACGTAGAGAGAACATATAAGACGTGAGTACTTGACTAACTTACTAACTTGACGTAGAGAGAACATATAAAACGTGACGTAGAGAGAACATATAAGAACAATTGAAGATAAAACTTCCAAAAATATTGTAACAAAATAAATAAATTGTATAACCATTAAAATAAATTATATAATCTAT
>evgR013112c0g1_i01_Hydra
GTGTCAAAGATTCTCTGTTGCAGTTTCTTTACTACTAACATCAATGATGGTAAGCACAATGTTCTATGGAACTGAAAAACAAAATCCAAATTCAGAAAATAAAAGTGATTTTTCCTTTACAAAGACACAGATTTTTGTAGTATTGATTTCTACCGTGATCAAGTTTCCTGTGCATTTGTTGTTTTTGAAATTTTTTCGTTCAATAAGACCACATGACGAAACTGCCATCTCAAATGATGAGTCATCTTTTTATGAAAACAGCTCAGAACAAGTCAACTCAGAACAAGAGTTTTTTTCTTTATCTAAACGCAAAAGTATAAACTTGAGCCAAAATACATTAACAATTTTTATAAATGATGAAAAACCATCTATGGTAATAAATAAAAAGGCTTTACCTCATTGGTGTTTGTATGCAGCATGGTTTTTTTGCATTGGGAATATATTTGTTTGCGTTGTTATAATTGTGTTGTATGGAATGAAATTTGGAAATCAAACATCGTTAAATTGGCTGGCAAGCATAACTATTGACTTTGCTAAAGAGACTCTTTTATCTGAAC
>evgR013120c0g1_i01_Hydra
GTGGAGAAGAAATATAGATTAAGTTATAATAAAACAGATGTGGAGAAGAAATATACATTAAGTTATAATAAAACAGATGTGGAGAAGAAACATACTTTAAAAGAAGTGTTTTTTATTAAAGCAATTAAAAACATAAATTGATGCACTATTATCCCAAAAATAAACTTCAGCCTAACTTAAACTCCTTCCTAATCCAAAAAAAAAACTCCTGCCAAAACTTGAACCTAAATCCCTTGGCTTTCAAGCAAATGATACCGGAGAAGTTAATTCGCTTTGATGTAAAACATGCAAAGAGTACCCAGATTTTGTTCAGTGATCCAGGTAATATTTTACAGATTTGGAAAAAGTTTGTAAAAAATATTCTGAAAAAATGCGCAATTTACAAATGCCCTTATTTTTCTCCTAATTTTTGTTTCAGCCTCGTACCAGTTTTGAAACTTTTTAAAATTTTTTGCATGAGTGATATATGATA
>evgR013123c0g1_i01_Hydra
TAGCGCATTTGCTTAAATATGTTTCAACTCGGCTTTTTTCTTTTGCTTTTGGTGCCAACCGTTTATGGAAAACCAATCGTTGTAAAGGATAATATTAAAGGTAAAAACGAAAATGATATTTACTATCCATTTGGCGAAAACCCTTCGTGGTGTGAATTGCACGGTGCTGTCCATGGTCCATTAGCGCATTATGAAAAATTTCATGACCAAGAAAAATGTGATAGCAACAACTACGGTTATGGCAACGATGGTTACCAGCCAAACGGAAAGAATCCATTTTGGTGTAATATCCATGGCAATGTAATTGCTCCAGGCATTCACTTTAATACGTTTCACAATTTAAGAAAATGTAGAGGAAATGTCGGAAAAAACAACAAGCACGGTAATGATATTCCCGGCCAAAGCGATGTTTACTTTCCGTACAGCGAAAATCCTAATCGGTGTAGTGTACACGGTGCTGTACAAAATCCAGTATTTCATTTTAACAATTTTCATGCCAATGATAAATGTAATAACAACAACTACGGTTACGGTGTAAATGGTTATAACCCTTCAAAAGAAAAAAATATGTACTGGTGTAATATTCATGGCCGTGTAATGAATATTTTACCCCAAGTTCACTATGAAATGTTTCATAATTTACAAAAATGTGCACGCATTGGCAATAACAATAATGGTGGAAACAACAACAACGGTGGAAACAACAACAACGGAG
>evgR013125c0g1_i01_Hydra
GTTAGAAAATTGGATGACAAGGCTCTTCTTGTAGAAATTCATTTATTGGAAAGTAAGGTCTACCAATCCCTTAGAAACCTTTCAAAATCTAGAGCTGCTCTCACCTCAGCCAGAACTGATGCCAATTCAATCTATTGCCCTCCACTTCTTCAAGCCGAAATTGATATGCAATCCGGTGTTTTGCATGCTGAGGAAAAGGATTACAAGACCGCCTTTTCCTACTTCTTTGAAGCTTTTGAAGGTTTCAGTAATTTTGATGACTCTACTGCCATTATGTGTCTCAAATATATGTTACTCTGTAAAGTTTGTACAAATCATCCTGATGATGTCCAAACACTCCTCAGCACTAAAACAGCACTTAAATACACTGGTAGACAAGTTGATTCCATGAAGGCTGTCTCTGTTGCCTATCAAGAAAGATCTCTCCACTCATTTGAAAAAGCTCT
>evgR013131c0g1_i02_Hydra
CAAAATCTTCAAATCAAGAAGACTTATTTAATAAATTGAATGAATATAAAGTTTTAACTCAAAAGTGGTTCGTATGGGGTCAATTATATTCGTATTGCACGCACGTACAGAAATTAGATCTAAAGGAAGTAGACAAAATTGATGAGTTCGGAAATGAACAAGTAGAAGATTTAACTGTGGCGGAAAAGCAAAGGTTAGCTGAGTGTATGAATGTTTGTACACTTGATTTTCCTTTTAATCTGACAAAATGTCTGTTGAACTGTAATAATTCTGTTTCATGTATTTATGGTTGTCGATTCTACAGCCACATCACTGGCAATACTGAAGAAAATATTGAGATAAATAAGCAGAACAAACCTGATTTATATGCGTCATTAGAAGTAATCCTGCTCAAAAGGGAATGTTATAATCTATCATTTTGGTGGCCGAATGTTTATCCAAAAAATACTTTATTAAAAAGTGTATATCTGGTTACTTTTATTACACTAAATGGAGACAACCAAACAGAAAGAGTATTGATATTAGATAGTAAAAATTATTTTATCATAAATGAAAAGATATTTTGTGATCATCTTGGAATTAAATCTGCCTATGATATTGGATTAGAGTTTCAGTTGAATGTCTATCCTATCAATTACCATGGATACAATGAATCAATTAAAATGTCCTCAAGTATTGCAAAATGTGGTTATGTTTCATTATTTCAGAAAACAGATTGTATTCAAGCTAATCCTCCAACTATTACAAGTAAAAGAGTGCAACCGGGTTTTATTCTTCCGACAGCTCTGAATCATCCACAATTAACTGCACACCATCCAGATCTACAAACTAACTCCCTCAACTGTCTCCTCCATCATAACATGCACCCTTGTCAACAAGAACCTATTAAAATGGATTTTTAGTAGTTTTCTGTATATTGATTAAAAATATATTTAAGATTGTGA
>evgR013133c0g1_i01_Hydra
TTTGCGTAAAAGCAGACAAACGTGGTTGGTATAATGTTGTTTGTCACTATCCTTTGGGATATATTTGTAAATATGAAATAGTTTTATGGAGTGACTGGAGTAATTGGTCAAATTGCAGTTCAACTTGTGGTATTGGTTTCATGAATCGAACAAGAACATGTCCTAAACCACGAGGTTGTCCAGGAAATAATTCTGAGTTTCAAATATGTACTAACTGGGCAGTTTGTAAAGAATCTAGAGTGTCGAATTATTTTCAAGCTTTAACAATAATGACTACATATTCAATCGAATATAACAATTTCAATTTTAAAGCACACCATAAAGCAGAAGATTCGTACATTGGTAAAACAGATGATGTATTAATCACAGTAACATCATTAACTACTTCTGCTTTTGAAACATCATTGCAAATTACGTCTACTAGCTTAGAAGCTTCTGTTTCATCGCCTAGTTTTCAAACAACGCAAGTTTCTAAAATATTTATGGAAGATGTATTTATCACAGTAACATCATTAACTACTTCTGCTTTTGAAGCATCATTGCAAATTAAGTCTACTAGCTTAGAAGCTTCTGTTTCATCGCCTAGTTTTCAAACAACGCAAGTTTCTAAAATATTTATGGAAGAAGAATGGGCAGCATGGAGTGAATGCTCTAAAACATGCGGAAATGGTTATCAAATTAGTCATTCTTTAAATGCTGTTATGAATGCAGGAATGCAAAAGTCAAAGGTCAAAAGTGAAACGCAAGTCAAAAGTTGCCATGTAATAAGCTGCCCAGTAGACGGAATTTGGAGTAATTGGACAGTTTCTTTATGTTCTAAGCCATGTGGTGGTGGTATATTAACATTTAACAGAACTTGTGACAACGCACCTCCTGAATATAATGGTCTTTATTGCATGGGTATCAGCAAATACATTGAATACTGTCCCAATGACATTATTTGTCCAGTAAATGGTAATTGGTCAGAGTGGTCTACTTGGTCCCTGTGCAATCAACCATGCAATATTGGTGTCAGAACTAGACTTCGAGATTGCTCTAATCCAATTCGAAAGTATAATGGTGTGTTTTGTTACGGAGAAAGTACTCAGAGTGAAGCGTGTTCTTATAATAATTGCGGAGCAATTAAACTAAACCTATTGGTTTATCTTCCGGATGAAATTTATGATGACGCTAATACATATAGTTCATTTATCAACCTAAAAATGATAATGGAAAACTGGATAGCCGATTTATACAAAAATTACGAAAGAAATGCAATTTTTAGTATTGAGGTTCATTCTATAATAGATGGTGGAGTTTATTCACAAGAAAATGGTAAAACATGAAACATCGGAATAGTTTTGTTGTGAGGAAGATATAGTTTTTATTTTTATTTTTTTATATACATCTATACATACATTGCATAAACTACATGAACAACGTTTACTTAGTAAATAAAAAGTTTTACCAATATTGTTAAAC
>evgR013138c0g1_i01_Hydra
TAAATAAAAATTTGGTGGATTATCCAAGTGTGATGGATTATTAAGATATCAAAGATGGATTAAGTTGTTCTTGCAATGTTTGGATCATAACCAATCTAACAACAACATAAACAAAGTTTGCCTCTAAGAAGGAGAGCGTAAGACTGGGTCTTAATAAGGTGATTCCTCGAATAACTGTTAATCTTTAAACTGTTTATCATAACATATTCCTTTATGCTTTGTTATCTGCGCTTTATGCTTCTGTTTCCAAGATTGTGGATGTTACATTCGTGAATTTAACGTTTGAAATATTCCATAATATATTCCTTCCTTTCAATTCTTTCTGTCTACTACGTTTAACAACAAAGAATTTTGAAATTAGTTTTTATAATTTTATGTGGAACACCTAAAGGACTGATTTTCTTATAAATGGTACAATTTTATAGAGTTCAATCAATTCTTATCCTGAAATAGATATACTGCAGCTGCTAAAGTAATACCTGAGTTGTAAATCTAATATTATCGTATCGTTTGTATCAATTTAATTTGCATATATCCACTTTGCTCCGTTATAAGCAAACAAAAAGTGTAACAATTGTGAGTTTAAAAAAGAGTAAAAGTTTTTATTGTTTATCTTGGATACTAGTTTTTCATAATGATGAGTGTTGCTGATCCATTGCTGGCTTCTACAGCTGAAGTATTTAATAAGAGAAAACCTTCTGTAGAAGATTCGGAGTGGAGCTATGACTTTCAGTCGTTGTGGGAAGAGTTTGAAGGTTCTTCTAGTGACGATTCAATCCCAGATAGTGTTGAAGATACTTACATGATTCCCGAAGATCATTTTACTGCAAACATTGATGACATATTTTCTAAAGATTTAATTAAAGTTAAACAAGAAAATATAGATCCTTTGTCTCCCTTAGCCAATGATATTGAGATAAACAATCATGAAAAAAACCTATCAAAAGAATTTTATATAAAAATAGAAGACGATGTTTTACCCCCGTCACCAACTGACTATTTACCTGTTCCATTAAAATTTATAGATACTAGTAGCCACTGGCTAAATAAACAGGAGCCTAATATTGTTTCATGGTCTGATACAAAACAGTCGCGTCAGCGATTCGAACCTGTAGTTACAGTTTTACAAGACTCTCAAAATCTTCTTAGTATTCAATCGAATTCGAATTCTTTAGAAGCTGTGCCCCCTCAACTTATTTTAACAACTCTATCAAATGGAGGAACTACAGAATGCTACCAAATTGTGAGTCCGCAAAGTACAATTCGATTAGAAACGTACCATCCTGCTAAAAACTATTGTATAATGACAAACCGGGGACCGGTGGCCGTAAATTTAACAGAACTTTCAAGTATTACACAGTTAAAGCAAGGCGAGAATTCTCGCCCTCTCGTTATATCACCGGTAGATATCGACCCAAGAATAATTGAAAAACATTTTTTTAAATCAACGCAACAGTCTAAAGGAAATCCTTTATATACTGCAAAACCTAGAACAATTAGTACAACAAGCTTGGGTGAAAGTATCCAAGTTCGTCGTGCTGCAGCAAGAACTCATAAATGCACCCACCCTGGATGTACAAAAAGCTATACTAAGTCTTCTCATCTTAAAGCACACCAAAGAACCCATACAGGTGAAAAGCCATACGCTTGTTCATGGAGTGGATGCGCTTGGCGATTTGCAAGATCAGATGAGCTAACTCGCCACTTTAGGAAACACACTGGTCAGCGGCCATTCCGTTGCTTGCGCTGTCACCGTACGTTTTCTAGATCTGACCATCTGTCCTTACATACAAAAAGACATCTTGCTAACTGTTAGTTTTCTTTGTTAATATTGGAGCTCTCTATTTCTCTGTACTTATAATTGTATAATTTTTTTATATTTCTGATTTCTATCCATTATTATTTCTTTCTCTACTTCATATATATTTATATATATATATATATATTTTTTTTATTAACTATTAGTTATCGTTATGTTTG
>evgR013149c0g1_i01_Hydra
ATTAAAACTTGTTTTTTTTTTTTTTTCGAGTTCTTTCAATAAAAACAGAAGTGAAGCCTTATCAGTGTACTTTGCAGATTCTGGAATTCATGTTAGACAGACACCAAAAAATGCGGATACACTAATTATCAATAGAGTAAATAATTTGTCAAGGCATGGATTCAACGAACCTGTTTTAATTATTGGAGAAGATACAGAACTGCTAGTATCGCTCGTGACTTTTACAGATTTAATTACTAATAATTTCATGAAGAAGCCTGGCAATGATACTAAACAAAACAAAGTTTACAGTAAAAAAAAGCTGAGAAAAAGATGCAGTTTGGCATTACCTGTAAAAACTGCATTGCATGTAAAAACAGGCAGTTGTTGTTTTTACATGCAATATCTGGATGCAATTCAACATCATTTCCTTAACCTCAAAGGGAAAGTCTTAAACTTGATGCACAACAATGTGGCTTTATGTAAAGTCTTTGATGTATTTAACACGCCGGAAGCAAAAACATGATTAAATTGATCTTGCTATTTCATGAACAATATTTCTTAGCATTATATGGATTAATAAGCAAGTCAATTTCGCTTAATAAATTCCGTTATTATGAGTATCTTTGTAGACTACCATCAAAACTAGTTCACAAAACCTTCGATATGGCTTTTTACCTCCAAAATTTCAGGAGCAAAGCAACATTCATTCCGTTTCAACATTTGTTTTGCGTCTACTATTGTGTAGATGATGATAATGACACAAATCGAGATGTCAATATTCAGATGAAGAGC
>evgR013151c0g1_i01_Hydra
TTAATGTTTTAAAGAAACAGTACTTAATATCTTTCAATGTGAAGCCTTTTAGTTACTCAAAAGGCTTTAAAAGTGTTCTTCACTTATCCTTGGATAAAGAATATGGGGATTACGGTGTCCAAAATTCAGGTGTTTATTTTCATGAAGACGGGTCTGGAAAACTTACCATATATGCTGCAGTAAGTGAAAAGTATAGCTATAAAGTTGAAACTACCCCCCTTCCTTTGGGAAAATGGTCCTGTGTTAAAATATGTCAATACTTGGCAGACGGAAAGTATTGGTTTTCTGTTGATTTAAATGGTGTCAATGTTCATAGAATAGAAAATTCAAATGTTTGTGATTTCAAATCAATAAGAGTGTTTGCCTCTAATGGTAGGAATGATTCTCACGATGGTTATATTTCTGACCTTATTATTATTAATGGAAATGTTGATTACCTTGTTGGAAATAATCCTATTCCATTAGTGAGAGGAAAGATCGTTGCTGAAATACCCAAATTGGAAAAGGAATACTTGGTCTCTTTTGATATTTGTCCAACTGATATTTTATTTAATGGTTATTATAACCTACTTCATTTTACTAACAGTTATAATAATTCTAAATGCGGGTATAAGGTTCCAGGCATCTGGTTAAAAACTGAGGGTTGTGAAGGATTTATTGCAGGTAGAATTATTGATGGTAAAGATTTAATTTTTGACGAAAAAATTTCAATTGGAATAAATGTGTGGTCTAACATAGAGATTAGCCAAGTTTTGGTAAAACCACATTATATATACACAATCAAAATTAATGGACAAACAGCATTCTGCGAAACTAACTATCAAGTTCAAGATTTTGATAATGTCAAAGTATATGCTTCAGATCCTTTTTGTGAATGTCATGATGTTTTAATAAAAAATTTAATGGTTGCTAATGGG
>evgR013153c0g1_i01_Hydra
AATAAATAAGTACAATAGAATGGAAAAAAATTTTGTGAAGAGTTTGTGGATAAGTGTGATATTTTTGTGTTTTGGAATTCTGGTTTTTTATTTTATTGGTATGTACGTTTGGATCCCTGACTTGAGCAACGATATCTTTAATAGTGATAATATATTTATTCCGGCATTAAAATCTGTTTTACTATACGTATACATTCCAATGGCTGCTTACTGCTTTATTATGAGCGAATTGACTGGAAACTACAGCCAAGTAGATAAAATATGGCCTCTTGCTCCTGTTTTTTTCTCATGGTTTCTCTCCATTTACATTTTTTTTAATGATAAAAGTTTTAACGGTGATTTGAAACCTTTTTTAATTAGTACACTCGCGACATTATGGGCATTGAGACTTATGCATCAGTTTTATTTAAAAGGTGGCTACAACTTTCCTAGGGTCTGGGAAGGAGAGGAAGACTACCGATGGAAATATGTACGAAAAATGCCTTTGCTTACTAACAATCGGTTGTTGCTGGTAATTTTTAACTTATTGTTTATAACTGTTTATCAGTTAACTTTGCTATTTCTAATATCTGGAGGTGTTATTCTGGGAATCCTAATGTTAAGATATCGCCAAAGTAAACACAGAAGTCTTTGTCAAAGTGATTATTTAATTTCTTCTATCTTAGTACTGTTTATATTTCTTGAAGCAGTTGCAGATAGGCAACAACGGCATTTTCAAGTTGAGAAGCTTCGAAGAATACAGAAAGGAGAAAAGTTAACTGGTTCAGATGAGCATTACGAATTAGGTTTCAAAACAACTGAGTTATTTTCTTATTCTCGCCATCCGAATTTTACTTGTGAACAAATTATTTGGGTATTATTCTATACGTTTACTATTGACTTTGAAACAATTAGTTTAGTAGGTGTTGGGGGGTTATTCAATTTTTCTATTGTTGGTTGTTTATTGTTAATCCTGCTGTTTTTGAAATCAACGGATTTGACTGAAGAAATATCTTCTAGTAAGTACCCGGCTTATGTTAAGTATCAGAGTAACGTAGCTCGAATTGTTGATCTGAATCTTGTCGCGAAAAAGATTTTCTTCGGGAAAAGTCCAGATAGAAACTGGATGAAATAGTTGAAGATTTAATTTTTATGTTTATTTGTGTTATAATTAATGTTTCCTAGTTAATTAAAATTTTCATATTTTTATTTGATTTTTTAAAATAATTGAACAATTTTTATAGATTTATTTAAACAAGGATCTCGACTTTATTTTTTCTGGGGGAGTAGGAAAAGCTGTTCGTGGCAATTAAATTAGCAATTAAAAAGTTTTTAATATATTTTATGTTATATTTTATATTTATTTATAAAATTTGGCAAAACGTGTTGAAGCCAGCGCCCTCGATACCCTTTCCCCCGTTGCCGCGATCACTGTTAAAATGACAAATTACTAATTTTTCC
>evgR013155c0g1_i01_Hydra
ATTCTAGGTAATTACACAATTTAAAATGGATTAAATGCATCGAATTTTTCCCATTTGTTTTTTATACCTAATCACGCTGATTTCATTACTACAAGCGAAGAAAAATTGCGGGAACTCAAATGTAGCCTTCGGTCATTTCACAAACACAACCTTTTATATTGGAGGCATATTTCCGACCCATCACTTGCTGAATAAATCTTATACTCTCAACGAAGGAGCCATATTATGGGTGTCGTCAATGATGCTTGCTATTCAAGAAATTAATGAAAGCAACAACCTACTTCCAGGAATAAAATTAGGATACGATATAAGAGACTCAAGAAACGATGTTTTGTGTGCATGCCAACACGCTTTAGACTTCTTGCTAAATAAAAGTTGGAACTCTTTAATTATTGGGGTTATAGGACCAGCATCTAGCTCTTTAAGTGTTGCTGTAAGCAATTTACTACTGCCAGACTATATACCACAAATATCGCATTCATCTACAAGTATAACCTTATCACAAAGATCTGTATATCGAAATTTTTTGCGCACTGTACCTCCTGACGTATACCAAGCGAAAGCATTAATTGACTTAATTGAATACTTTCAGTGGACGTACATATCTTTATTTGCTTCAGATGACGACTATGGCCGTTTTGGACGCGATGAAGTCCATAAGTTTGCTATGGAAAAAAATGTTTGTTTTTCCGTTGATAAACTAGTCGGAGCAGTACTTCAAGAAGAAGAAGTGAATGCTATTTTAGAAAGTATTGAACAAAACTCACGCATTGTTATATTATGGTGCGATTCTCACAACGCTATAAAAATTATTTCCAGAAGTTTAGAACTAGGTTTACAAGATATAACTTGGATAGGAACTGAATCTTGGGGAAACGATATTATTGCTATGCTTAAGCCATACACTTTGTTGACACACAACATATTCATTTTGCGTCTCAAAGAATACGCAATAAAGAATCTTGATGCCGAAATATATAATGCTAATGTAGCAAGCAAAATTAACTGCGATAACCCATGGTATATAGAATTTTTGAGTAAACATTCTAAAAGTTGCCCAGAAATAGTTCCTAGTTTATCACCTATGTTATCTAAAAGAAAGACAACGCAAGTGTTTGGTGCCGTATATGCTCTAGCATACGGCCTGCATAACTACTTAAATTGCTCATTCTTAACATGTGGTACACCTCCTGAACGTATAAATTATCACTTGCTATATAAACACGTGATTAATACAAGTTTTTCTGTGCCTACTAGTAACTACTCAGTTAAGTTTGATAGAAATGGAGACTTTCTGTTATCTGCTTATGAATATATATTCGCCGATCAGAAAACTTTTATTAAGTTTGGAACCTGGGAGTATCGTTTTAATAATCCAGCCACTGTTAGTATAAACGAATCATTCATTTCCTGGAAGAACAGAACAACACCAAAAAGTACTTGCAGTAGGAGCTGCTACCCTGGTACATACCGAATAAACGCAACTGAATCGAAATGTTGCTGGTTTTGCATGCCATGTGAAAAAGGGTATATTACAAGTTTAACAAATGAATATAAGTGTACAAAGTGCCCGGTTGCCACTGTAGAAAATTCAAATCAAACACAGTGTAACAATTTAAATGAAGCTAGATTAAATATTTACAGTACACAAGGAATCATTATTATTGTTGTGTCATTATTAGGTGTATTTTTGACGATAATTGTGTTTATTTTCTTTTTATTTAATTGGAACAACCCCATTGTTAAAGGTGCTAACAGAGAAATGTCTTGTATCCAGCTTCTTTCTCTAAAATTTTTGTTTTTGCTTTCGTTTTTATTCTTTTGGAAAACCACTCCAATGTTGTGTATGTTAAGGTCGACATTGTTTGGCTTTTTGTTTACTACAATTCTTGCTTTTGTTGTTGTAAAAACATACCGATTGCTTCGCGTGTTTAACGGACGGTTTACAAAAGTTTCAAAATATTTAGAAAATAGATTCCAGGTTTTTTTCTCGTTCTCCCTCGTGTTCATACAAACGTCAGCTGTCATTATTTGGCACATCTACCATCCTACAAACATACGAATTGAAACTGATTCTGTTCGAAATGTGTATTATTTAGTCTGCGATAATGAAAAGCCGCTGTTTTGGATTTCGCTAGCATATTTTTTTATATTAGTCATTGCAAGTGGCTACATGGCTTTCCGAGCGCGTAAACTACCAGAAAACTTTAATGAAACACAATACATTGCATATGCTATGTTTACGGTGTGCGTGTTTTGGTTTACATTTATTCCAATTCACTACAGTGTTGACCCACAAGAATCCATTGTAGCTTTCTTATGTATTAACTTACTATCATGTTATAGCATGCTTGTATCTTTGTATGGAAAAAAAATAGCTTATGTATTATATTATGCAAAATTTAACAGACCTAAGCTTAGTAATAACAGCTCCATAGAAACGGTTTCGTATAACTTTAATTTCGAGGAAGCTGTCCAATTCGAACCCTCTAACATTATCCCAGCTAACACAGATGTAATTATTAATAACTATTCGAATGATCATTTTGACTCGATTGTACAAAGTTTATAAACAATAGGTTGTAAGTTTAATGTTGTACATGCTGTTTATATTTATTCTAATTATTTTGTACATTTTGTTAAAAGTATGTATAATGTTTTGTAAATATATAAATATAAAAAATAATTGCGCAAACATGGCGCATGGCACTAAAGTATAAATAAAAAGTAATAAACTAAATTACAAAG
>evgR013156c0g1_i01_Hydra
CACGAATACGCTTGAATAAGATCTCGGGCGGTGTTTGATGAAGCGTGTAATCTTTAAACATGCAGAGGAGCTCAAAAGAATTTCACCGTTGCACGAGAGCTCAACCATATACCGTCTATCCAAGTATTTTTTTATTATTTTTTCAAGCAATGAAAAAACGTAGCAAGTTTTACCGTAAATGTTGAGCGGATCCCTTCCGCTTAACCAAAAGTTTTTAAATATGGAAGATGTATTAGATACAACTTTTTACTCAGATGATTTTCTTACACGAACTCTTACTACTTCCGTTTTGTACGATAAATCACAGCTCGATCTTACATTCGTAGAAAACAATAATTGTTTGCCATCAAAACGGCCGCAATATGAATCTACAGTTGAATCACCTAATCTAGACTTTCTAAAACTCGGTTCACCTGATCTTGAGCAGATGTTTATGAACATAAACGAGAACGACTCAAACTTTTCAAGTGAAGATAAGTCTGATTCTTCGAGTGCTGCTTTTTCTTTAGATGACGATTTTGTGAGTACGGAAAATTGTTTTACAGATGCACTCCAGCAATTGCACGACAAACAAGAAGTTATACCGACTGATTTTTCCGGTGCAAATTGTAATAATAGAGTCGGAAAAGAGAATTTACATTCGAAAATATTGCAAGGAAAACGCAAACCTATATCTTCGCCTATAAATGTACAAAAGTTAAGAACTCAAAATCCTATGAAACAATTATACCATAATCCGAACCAACAACGTAGAAATAACTTTCAGCATCCTTGCTCGCATCCTCAAAACAAAAATTTGCATCGTTTAAATAATCTTGCTGTTCATCATCAAAACAATATTTCTCATTTATCAATTTCTGAACAAATGGCGCTTAATAATCAACTAACTATGATGAATAACATGAGTCAATCAATTAATGCCCATCATATGCAACAGCAACACGAAATTGCTTTTCTCACACAAGCAATGGGATTTCAATCTGACCGTGAGCCTAGTTTAAATTATCAAAATGACCGCTCTATGATCAATTCTAACGGGTTCCCATCACAGCACAATGTATCATCACAGGTTCCTCCGCAACTTCAACAAAAACTTTTGGAACATCAACTTATGCAACAACAAATTGGAGCTAGTTATCCAAATAACAATAATAACACGACTCATCACGTGCCTCAAATAGACTCTATAAACGGTATGGGCATTGATGAAGGAACATTAAAAATGCTGATGGAAAATCCACACATGATACCTGTAAATCTCGAGTTGCAGGAACTTATAAAAAGAGAACGAAAAAAACTTCGAAACAGAGTTGCGTCTTCTAAGTGCAGAAAAAGAAAGCTTGAACGAGAAGGTCGACTTGAAGACAAAGTAAAAGATTTAAAAGAAAAAAATATTGAGTTAAATGCAGTTGCTAACGCGTTAAAGCAGCAAATATGTGACTTAAAACAACGTGTAATGGATCATGTAGGCGACGGTTGTCAAATTGTTTTACCCCCTTAAATTTTTATAACAGAAAACTTTGCTCTTCCACGAATCCCGAAACAATCCCCAACGTTTTTATAAAACGGGTATGTATTAAATAAATTCTTCAGGCAACTTTTTTAAGCGGTAAATAATTGGATAGTGAATTTTAAATCAATAAAACTATTAGAAAAGGAAAGGATTAACAAATAAAATTATTGGAAATGTTTAACCAATCAAACTATTGGAAAAGTTAATGTTTAACCAATAAAACTATTGGAAATGTTTAACCAATAAAACTATTGGAAAGGTAATGTTTAACCAATAAAACTATTGGAAATGTTTAACCAATAAAACTATTGGAAAGGTAATGTTTAACCAATAAAATGTCATTAGTAATTTTTTACGTATTAGTATTTTTTTAAGTTAAATTATATATTTTGATACGCACTGTTAAATTAATGTTTATATCAAAACCGAACAAAAAGTAATATTTTTGCATTACTCACAAATGTACAGTGATTTCTTATGTTTCATATCAAGCTATATTAAAAAAAA
>evgR013166c0g1_i01_Hydra
TTTTAGTGCAAAAAGGTAAAAAAATATAAACTTTTAATTAAAGGTATATATTAGGAATAATAAATTATGCATTCTGAAGAAGAGGCAGACTTGATTCAAGAAAATGTGTTAGGTGATTTGAATTCTAGTCAAGATAACTTTACTTCTCTAAGCGCTTATAGTAAAAAAAAGTTACTTTGGAATGGATGGCAAAAGATTGGATGGACCTCAGCTTTGTTTTTAGGATGTTTGCAAGTTTATTGTGTACGAACTTCAATGTCAATTTGTATATCTGCTCTGAGTGATGAACTTCATTGGGACAAGAAAACGGAAGGTTTGGTACTTACATCTTATTTTGCTGGTTACCTTTTAACAAATATTATTGGTGGTTATCTGGCTGATTTCAAAGGGGGAGAAAGGGTAATATTTTATTCAACTTTTGTTTGGTCATTTTTCACAATATTAATACCGGCGCTAGTGAATTCAGTTGTTAAATTTGGTTTTAAAACTGTGATAATTTGTCGTTTTATTGTTGGTTTAAGCCAAGGTATGTTTTTTCCATCTCTCTCAGCAATCTTGGCAAAAAGTGTTTCTGCAGAAGACCGATCTTTTATATTTAGCTGCACTTTTTCAGGGTCTTCTATTGGCACAGTAATAACAGGACTTTTTGGAACATTACTTTTGGAGCAGGTTCATTGGTCAAGTGTCTTTTATTTTTTTGGCATTACATCATTTGTGTGGGCTTTATTATTGAAATGTTTACATTGGTCCAAAAAGAAAATGTCACTAGATGCAGATAACATTCGTGGAATAAAAAAAATAATTCCTTTAAAGGATTTACTTCTAAGCAAGAGTGTTTTAGCTATGTTGTTTGCTTATTTTGCAATGAGCGTCTCTTTTTATGTTTTTTTGTCTTGGAGTCCAACATACTTTCATGAAACTTATCCAAAAGGTAAAGCTTGGCTTTTTAATGTTCTTCCTTATTTAGCAAGTTTCTTGCTTTCTCTTGGTTCTGGCAAAGTAGCAAATATGCTGATTAAATGCGGATTTTCAATTACGTTTGTAAGAAAGCTGTTTGGATCGTTATTGTTCCTTGGAATGTGTTTTTTTTCTTTGTTGTTAGCTTCAACTCATTCGTTTGAAACAACTTTATTTGTAATGACATTGAACATTGGTATGAATGCATTTGCTGCCTCATCAGTAGCTATCAATCCACAAGATCTTTCTCCAGAGCAAGCAGGTATCTTATATGGAGTTGCCAACACTTGTGGTGCTCTAGGTGGATTTCTTGGAGTTTTGATTTCAGGTATCATCTTAAGCAATTCTGGTGAATGGTTTAAAGTTTTTATCATGACATCAGTAACTTCATTTCTTGGATTTCTAGTTTACATTAGGTATGGCAGTGGTAAAAAAGTACGATTATAGTTTTGAATTGAAATTTCAAATTTGGTTCATACAAGTTGTTGTTTTAGAAATATTAAACTCAACGCTATAGAATTACAATGCAGTAATCAATAATTTTTGTTTTTTATGTAGTATATTTTTATTTACCGTTGCTTATCTTTTTTTAAAGAAATTTTATATTTATTATTTTTTCAATATTTAATATTTATATTTTTACACGTTTTTTTAAACTATTTATTGACACTTTTCACAATTGTTATTGAATTAAATATATTTATATTTTTACATGTTTTTTATATATAAGTTAATGTTTGTTTTCTTTGACGATATTTTTGATATTATGGTATACCTCG
>evgR013178c0g2_i01_Hydra
CATTTTTAGTAACCGCTGCTCTCATCATAGGAACCTCAACTCTTTTCTTTATATTCGAACAATCCTGAACGCAGAATTGGAACTCAACCAATCACAGAATGATTATCAACCCCGAAGAGGACAAAGAAGAGAAATAGGTCGATCTTTTGAACAATTACAAAGTTATGAAGAAAAGCTTTCCACAGGAGAGTGGACTCCAACAAAGTTTTGTGAGGCTTCAGCTTTCTTCTTTAATGACATTAAATTCACATCTGTCAAGACACAGGTTAAGAGCAGGAGGAAATCTACAAGAGAATATTCAGGATCGACAAGAATCCAGACCTGTGGAGATGTTGTGCAAGCGCTCCTTTTTTGGCTAGGAAATTGTCTATAGCCTTCCCAATAGTTG
>evgR013180c0g1_i01_Hydra
TAACATCAATAGAGAACCATGGTCAGAAGAAGAAGATCGATCAATTGTTGAATACGTTAAAAAGTATGGTTGTTCTTGGTCTAAATTGAGAACTCTTCTGGGAACTCAAAGAACCGACACGATGGTCAGACAACGTTGGTTAAAGATCACTAATGGACAAGGTGAAGTTGGTGTCGCATCCAAACACAATCAACCAACCTCACTTGCCAGTATTTCATCTCCACCAACTAATTCTCCAAACACTACCACCAACACTCAACCTAACATTCAATCCATTGTAAATAGTTTTACACAAGAATTAGATGTCTTTGAATTCATGAAAAAGAATCAATTACTTTCTCAACCAACCACTAACAATGTTCACATTGAAAAGCCACAACCAGTCAAGAAGCCAATGCAACCAATTTCATTGGATAGTCTAACTTGTGATGAAGAATTATTCGATCAACGTACTGCTCCAGCTGAACCAATTTCTTCTTCATCTCCACCTAACAAGAATATTGTTGATACTAGCTCTGTGTCATCATCTATGGGTACATTGTTGGAGAATAAGTACGACTTGATGGATTTCCACTCATCACTCCAAACCAATTTAGAAAAGGAATGGTTGAACCAAAT
>evgR013191c0g1_i01_Hydra
CCTAATTATACATATCGACACCAAACACTATTGAATTCGTCGTATATTGATCACTTCGCCGTATCAAAGTATACAAGTTTTATAAATCTTAAATGCATTGTTCATCCTGAATGCTCAAATAATTTGAGCCACCACCTACCACTATCAATATCAGTTGAAGTTAAGTACACCGAATGCAATACCATTGAACTAGCAGAAGATACTAATATTCCTAGCTATGCTTGGAACGATTCTAACTTTATAAATTTTTACAATGTTCATTTAACCAATCGCTTTAACAATCTTAATTTTACTAACGATTATGAATACGACCTTTTCCAAATCTATAAAAATATTACTGGCAGCGCTCAAATAGCTTTTAAGGAGACTTTTAAAAATAAAAAGCAATGTTTGTATTCAAAATCTTGGTGGACACCTGAATTAAGT
>evgR013197c0g1_i01_Hydra
CTGGAGTATCTTGTCCAGGCATTTTAAGTGCATGGGGAGCATGGATACAATGTTCAGCAAGTTGTCAAATTGGTTACATTCCACCTACCGAGACAAGGATTCGTCAGTGTAGTGGAGCAAGTTTTGGTGGAAATTGCAATGGAGTATCTTTAGAGCAGACTCAAAATTGCAGTGAACAAATTAATTGCCCAGGATATATATCAGACTGGAGTTTATGGAATTCGTGCTCTGGTTCTTGTAACGATCTTGTTACTGTTCTTTCTCAAACTAGAAGTCGGTATTGTATTGGGTATTCAACATGGGAAATTAATTATAAAGGTTGTCCTGGTATCGTTACAAGTGACCAGCAAGCATGTAATGTAGATGTTGGCTGCCCAGGTATTTATACTGCATGGAGTGCATGGAGAAGTTGCTCAGAAACTTGCC
>evgR013202c0g1_i01_Hydra
CACAATGAGAGCCTCAATTTTAAACGACGCACTTACATCCCTCAGAAACGCTGAAAAGGACGGCAAGAGACAAGTCTTGATTAGACCAGCTTCAAAGACTATTACCAAGTTCTTGGAAGTCATGCAAAAGCACGGTTACATTGGTGAATTCGAAGAAGTCGATGATCACAGATCTGGTAAGATTGTCGTTGAATTGACTGGTAGAGTCAACAAGTGTGGTGTTATCTCCCCAAGATACGACGTAACTGTCGATGAAATCGAAAAGTGGGTCACTACTTTGCTCCCATCCCGTCAATTCGGTTATGTTGTTTTGACCACCTCAAAGGGTATCATGGACCACGAAGAAGCTAGACACAAGCACACTGGTGGTAAGGTTCTCGGATTCTTCTACTAAATAAATTATATAAAATTCTCC
>evgR013206c0g1_i01_Hydra
ATGATGGAAGAAGACGAAACTGTCAACAATAGTGAATCAAATTTATGGTATGTGCCAGTAAGCTACGTAACCAAACAAGTTTTAAAAAAACTTTATCTACGCGATGAACTGCTGCGTGAAGAACTGCTTATCTCAAAATTAAACGCAAAATATTGTGAAAAACAACATATTAAGGAAACCGACACGCATGCGTTGAATGTAGGTAATGCTGCTCCAGAATACCGTTGTAAAAATAAAAAGCAAACAAGTTTAAACGAAAATCATATGAAAGACAAAGCTACTTTATTAATTGAGCTTAATCGAAAGTTAAAGAAAGCTATGAATATTTCAAAACACTGTGACACATTTAACTTAGAAGAAAATAAAGTTGCTAATAAAACGCGTAATGGAAAATATACCTTAAAACACTCGGGGAGGGCATTCACATTTCCATTTGATAAAAGCTCAAAATACATAAACAGCGATCTGCAGTTAACGGAGCATAACAACAACGCTATCAGCAACACCTACTTAAGCTCTGAGGATGTGGAATTTTTTGATTGGCAAATTGTAAATGATAGCAAGTTGTCTTCAAACATTTGTAATAGTCAATTGAGTAACGAACGTTTTCACAGCTCAATTTCTGTTGAATCATTAGAAAGCCGCATAACATGCTGCACTATAACTTCTGGTTATTTATCTGATAATGATTTATTATCTGATGATTCTGAAATTTACGGAAAATATAACACTAAAATTACCATGAACCAAAAAAGTCGTTCGAAAACAAGCAACGAAGCGAATGTTAAAGAGCGGTATAAAATCAAATCTCACTCTTTGTATCAAAACAACAAAAATCTAATCGCTTCTCAAAAACGGTTTAAAAGTTCTGGGTCAGTTGGGAATTTATACATTACTTCACCAAGTCCACCACCACTGCAAAAAACAGCAAAATCAATAT
>evgR013231c0g1_i01_Hydra
TGAACAATCAGCCTTCTCACCATCTAATTTGGTTCCTGGTATTGAACCATCTCTTGATAAGATGCTTCAAGGAAGACTCTTCTCTTATCCAGATACTCACAGACATAGATTAGGTGCTAACTTTGCTCAAATTCCAATCAATGCTCCACTTAAGGCTAAATTAAACTGTCAATTCAGAGATGGTCCAATGTCCGTCAATGGTAATGGTGGTTCTGCTCCAAATTATGAACCAAACACATTCGGAGGTCCATATGAATCCAAGCAATTGATTGAACACCCAACAAAGATTTCTGGTGTCATTACTCGTTCAAAGCACAACCATCCAAATGACGATTTCGTTCAAGCCGGTACTTTGTAT
>evgR013274c0g2_i01_Hydra
GTTTACCTTTTAATGGTCGTAAAAGCTTGCTGATAAAACTACGTTGGAGCGTAGTGATGTGAAACTTTGAAAGTATAATGCTTTCTTTGCTGCGTCGGAGAAAAGTAGTTGTTGCTTGTGTCAGTTTCTGTGTAGTAACAGCTTTGTTTTACCAGCAGATGTTTGATAGAAAATGGGAAGTTAATTCTATAAAGTCAGTTGTTAGCAAACTGCATCAGTATCACACAGACTTTAGAAACTTTACAAGCCTTCATTTCATATCACAAAAAGAGAAAAATAAACGACAAGCATCATTATATGTTAAAAAAACTTCTGGTCTGACTTCAGCAGAGTTGCACACACAAACAAAGGCCTGTTTAGAGCAAAGTGGAAAACAAATTGCTAAGTATTTACAACGAAGTTATGAAACAACATTTTCTGCCGACTATTTATCAAGTAATAGATTTAGTCCAATATTAGAGCCGGAATGCAGGTCAAGTCTAGATTTGGTTATAATGATAACAACTAAACCCGGTAATTTTTTTAATAGAGCTGCAATTAGAGCTGGCTACGGCAGATCTGATTCAGATATTAATAAAATGGTTTTTTCAAACAACCCATTTAATTATATCACTATTTTTACGGTCGGCCGTGATTCAAACGCAAATATTGAAAAACTCGTGGAAAGTGAAAGTCGTAATTTTAAAGATATTCTTCGACTCGACTATAAAGATACTTATGAAAATTTGGCTAACAAAACTTTATTGACAATTGAATGGCTGGCCGATCATTGTCCTTCTAAATTTGTATTAAAGTCTGACGACGATTGTTTTGTAAACGTTTTTTCACTAGGAGCGTGGGTACCAAAACAAGACTCAAGCACTAAGTATATTGGAAGAAAAAACGAATGGATGCCAGTTATAAGAGATCCTTGGCATCGTAATTATGTTCCGTTTGAAGATTTCAGCGAAGAATATTATAAACCCTACTGTGCCGGCGGTGGTTACATGCTGTCGGGAAGTATACTTAAAAACATAACCATTAAAGCAAAATCTATAAAACAAATCATTAATGAGGATGCTTATATGGGCATGGTAACTAATGCTTTAAATATAATTCCTAAAAACGATGAAAGATTTTTACCGTTTATATTTTCCAAACAAAGTGTCCTCAAAAGACCAATATGTCAATGGAGGAATAAGTTTGTCATGCACGGAGTATTTGCCGAAAAGCAAGTAACAATGCATTGGAACGCTATTGCAATGAACGAGTATAAATCTATATGCGAATGGGATCCGCCATAACGGCTTTATATTAAGATAATATAAATTTCTTATATTTAAATTTTAGGTGCACAAAATAATTCTAAATCATTTTTTATAGATCTTTTTAAAACTAGTAATTTATGTTTATCACAAAAATAATTCTTAACCTGTTGTCAATAAAAGTTAATAAAAGCGGTTTTCATTCTGATTAAATTTTGGTTTTGCGCAACTTTTTGTCAAATGCATTAAAAAGAAAATAATTCCAAG
>evgR013291c0g1_i01_Hydra
TAGCATAAGTTATTACTGGAAGAATATATAAAATTATATTATGAAATAATAGCATGATGGATTTTCAAAATCGTGTTGGTGGAAAAACAGGAAGTGGTGGTGTAGCATCATGGTCGGAAACAAACAAGGATCGACGTGAAAGGTTGAGACAGCTTGCCCTTGAAACCATTGATTTGAATAAGGATCCGTATTTTATGAAGAATCATTTGGGATCTTATGAATGCAAACTCTGTTTAACACTTCACAATAATGAGGGCAGTTATTTGGCTCATACTCAAGGGAAAAAACATCAACAGAATCTAGCTCGACGAGCAGCTAAAGATGCAAAAGAAACACCATCGCAACCAGCACCGGAAAAACAGCGTGTGACATTGAAAAAATTTGTGAAAATTGGTCGTCCTGGTTACAAAGTTACAAAACAGAGTGATGTCGATAGTGGTCAGCACAGTTTACTTTTCCAGATTGATTACCCTGAAATCGTAGATGACATGCCACCCAAGCATCGTTTTATGTCAGCATATGAGCAGAGGGTTGAACCACCTGACAAGTCTTGGCAATACTTATTGGTAGCTGCTGAGCCTTACGAAACTATTGGTTTTAAGATTCCAAGTCGAGAAATTGATAAAAGTGAGCAGAAGTTCTGGAGTTTTTGGAACAAAGATACTAAACAGTTTTTTCTTCAATTCTATTTTAAACTTACACATCTCCAACTTGCTCAAATGCAACGAGAGGCAGAAGAGCGAAATAACAGACGTCGGCCAACAGAACAACGACAGGCTGCTGCTATTCGACCTCCTCCTCCTCGACCAAATTTTCCGCCTAGACCTGGTTTTGAAATGCAAAATGCTCCTAGAGGAATTCCGTTTATGCGTGGTCCACCGCCAGGTATACGTGGACCTCCACCAGCTTTTCCAAATTCTGGAATGGGTGGTCCTCCGATGCACCCAGATTCTTCTGATGATATACCTGATTTATCTCATGAAGGGCAACAAGGAGAGGAGGAAATGTTAGAGGACAATTCTTCAGGACCTCCGATGCACATTCCTGAACTTCCTCCAATGCGCCCTCCTCCTACATTAGGTGATACTGCTAACAGTATGCACGGAATGTCCAGTATGCAGAATCCGCCTATGCAGATGCACATGAGTAATCCAGATATCATTCATCCTCCTCCGCTTAATTTTCCTGGTGGTCAGCCGCCACCTAATTTTCCTGGGGGTCCTCCACCACCCAATTTCTCTGGTGGTCCACCCCCTCCACCTAATTTTTCTGGTGGTCCACCACCACCTAATTTTCCTGGAGGTCCACCGCCACCTCCTAATTTTCCTGGAGTCCCACCACCACCTAATTTTCCTGGTGGTCCGCCACCTAGCTTTCCTGGAGTTCCTCCTAATTTTTCTGCCGGTCCACCTCCTAACTTTCCTGGTGGCCCACCACCAAATTTTCCTAGCGGTCCTCCACCAAACTTCCCTGGTGGTCCGCCTAATTACCAAATAAACAATCAAGGTATGCAGCCAATATTGATGGAAGATAATATTGAAGAATAACATATGAGGAACAATAAACAGTCTTATGTATTACCGTATCGAGTAAAGTATATTTTGTTGACAAGATAAAAATATGAACAACTAAAAAAAAAAAA
>evgR013301c0g2_i01_Hydra
TTTTTGGTTTTGGCAGATCGACAAGTTGTTTTGTTGGTGTAGCTTATTCATTTTATTCTCCGTGTCTAGAGTGCGTAATATAAGAATGCAAATACGCGCATACATAGAGAAGAAATTACTTGAAGCAAAACTCAATGATTTGGTTTACACTATGTTCAATTTGAAACTAAAACACAGGGAGTTAAAAAGAATACAATTGAAGATCCCAATTGTTTGGATGATGTCTGGATATCTGGTGATGAATGGATCTTCGAGCTCCGAGATACTAAGGTGGAAGCCTATGAAGATCTTTATGCAAATATAGCTCTTGATGTTATTTTGTTATTATATAGCCTCTTGCAGTTATAGAAGGTGATGGATATCAAAATACCAAGAAAAAAACGAAGAAAAAAAATGGTAAAGAAAAAATTCTATTTGATAATGATCTTGAGGATTTTCCACATACCGAGGAGGAAAGAGAAGAGAAAGAGGTTGAATTGAGTTTCAAGGATAATTCCCTCGAGGATCAAAG
>evgR013305c0g2_i01_Hydra
TAATTGAAAAGATGGGTGGAAAGGTCAAGTACACTTTGCAAAAACGTAACTATAAGGATGATCAACCAATTCCAGATGCTATCAATGCCTTCATCTCTACCAAGTGGTCAAATGATTGTGGTTATCTTATCAACAAGAAGGTTGTTGGCGGTAATGATTTTGAGGTAATCAAGCACGAAATCTTTGTTCTCGAGAATGATGCTCCAAAGTGGATGGAAGAGAACAAGGATCATGTAATTTTAGGTGAAAGCTCAGTAGGTTTGGTTGCTGCAAAGATGAGTGAAGCCAAGGAAGACTTTACCATTTATATTATCATTGATGCTGTTGAAGAAGGAGAGGAAGAAGTTGTCTTTTCAATGCCATTATCCAAATATTTGACCAATTTGAAGAAGGTTAAGATTGTTGAAGAGGAAGCCAAG
>evgR013322c0g1_i01_Hydra
AGCTGACCAAAGAAAAGAAATAGACAAAGAATATGCTTTGATTCAAAGATTAAAGTCTCTTTTGACAAAAACATTAAGAAAAAATCATTTTGATAAAAAGCTTTTCAAACGTAGGAGAAGTTTGTCTTGCGAAATCTTTCTGAATGTGGATGAAAATGCAAATGAAAGTGACGAAATTGACTTTGCTTTGGATGAAAATGAAGTTGAATGCATGAATGAAGAAAATGACTTTGCTGGTAATCCCCAACCAATATTGGAAAATCCAAAGAAAACAATTAAGTCAAATATCCGACGGGATTCTTGGTCAGAAAAAACCAAAATGAAATTTAGTGTGTTAAGGTCGAAGCAAAATAATAAAGATTATTCCTGTTCTGTGGATGAAAATGAAGTTGAATGCATGAATAAAGAAAATGACTTTGCTGGTAATCCCCG
>evgR013326c0g2_i01_Hydra
ATTTGACCACCTATTTGTTTTTTTGTGAAACGTGAAAGGTGTATTTTTTATTCAAAGTTGTGTTCGTACTAGAGAATGCCAGGAATGGAATGGAAAAGGAGACATAGATCATTACTATCCTCACACTATCACTACATAGTATTTTCATCCCTATCAGGTTTGTACCCTATTCCCAGCCCATCCCCATCAAGTTTGTACCAATCACAAACCACATCAAAAAAATTAATTGATCAAATCTTGTATTTTAACCAGTTTTTGTCTATAATTCAAATATATTGTTACGCATACCTATCAATTAGTTCAAATATGTTAGGCATAGTTATCAAATTGTTGACCAATACCATTCAAATATCTACAGGTCTTAGTAATAGAGTGACAGTGGTATTATTGAGTTATTACTTTTACCATTGTACAATAATACTATGTATTTATAATATTTGATGCATTCTCATTGTAATCATCACATAAAGTGCAGCAATACAGCAATGTCAGCATGTCACTCTCTAATATATACATATATTAGCATGTCACTCTCGAATACATACATGTGTTTTTTTAAGTGGGTAAGTGTATTTATTTACTAAATTTTTCCTGTTCAAATAAAAGGCAGTAATGAATAAACATTAAGTTTAAGTAGTTTCGGTTTTTCAAAGTTTAACAACTAATATTACATATATTACTATAACATATTTTTGAT
>evgR013335c0g1_i01_Hydra
CACATTATTACTCTTAAGCAACTTAGTTATGAAATATGTTTTTCATTTGGATTTCATGATAAAAAAAGTAAACCAGTGCCTCTAAGTGCATCAGTTTTGTCTAAGAAGGGTCGTATAATTGGAAAAGCATCTGAATTGTGGTGTTTTTTTAGATGTCTGCCAATTATAATATCTAATTAATGCATATCCAGTTTTAGTGATGATTGCCCTGCCTACTGGTTGGTATATCTTGAACTAAAAAATATAGCTGATATTATATTTTCACCAATTATTAAAAGAGATTCTCTAGGATATTTAGCTTATATGATTGCATCATTTGTTAATTCTTTTGTGGATATATGGCCAGGAAATTTTATTCCCAAAATGCACTACATGGTGCATTATCCCAGACAAATTGAACTTTATGGACCTTTACGATATTAATGGTGCATGCGATTTGAAGCCAAGCACAACTACTTTAAGTGTCTTTCTCGTGTTGTTTCTAATTTTAAAAATATCACTTATACATTGTCAAAGCGCCATTAAATGCGACAGTGTTGTGAATTAAATTCAGTTGATTTTATGAGAATTGAGCAACACAATCAAAGAAATGGTATTGTTATATTATTAACTTCATTACCTGATGGTCTTCAAATAGAACTTATAAACAACTTTTTACTTGATCAAAATACTCAAATCTGGAAAGTTACAAATGTTTCAGTGAATCATTATTCCTACCAAACTGGTGACTTTATTGTACTTGACATTCTAGATGATGGTACTCCTGTTTTTATGTCACTCAAATATATCCTTAAAATTAATGGTTTATGGTTCGTGTGTGGTAAACTGTGTTCTTGTGAAAAATTTCTTTTTTATTCCCATGCCTATAATCTGATTGAAGTATCTACTTGGGTTGTTATTCTTTCTGGACAAGAAGTATCGCATGAACCCTTGGATGGATATTTAATTTCTGATGGACAACTTTTAGTTACTTTACGTAGTTTGGCAATAAAATAATATTACATTTATATGTATTATTGTTGTTA
>evgR013341c0g2_i01_Hydra
TTTAGTGCAAAAAGAAAGACCAGTATGGTTATGTTGTTATAAACTTCTTTTATATAAAACTTTAAATTATATAATTATATAGAAAAAAAATGGCGTCACGATCAGGACCACGAGCAGAAGGAACTGATGGATCAGATTTTTCAAACAGAGAAAATATTGCTAATCATTACAAAAAAAGTGCTGAGTTAAAAACGAAGGTTCGAAAAAGCTTGATTCCGCATATGTTTATGACGGTTTTGTTAGCTGGGAAGTTTATTATACTAGTACTAGGTTATAAATATTTTTTACCATTTGAAAAATGGGAAATGATCTGGGCTTTAAGTGGATTATGTGCATTTGTTGGTTTAGGATCATTAGCTAAAAATGACAATAAAAAATTAAGCATTTATATTTATGGCAATATTTTGTTTGGGATAGCTCCACTGATTTATGGCTTGTATGTACTAATAAAAAAAGTGCAGGTTGATATGTCTGATTTCAAAAATGTTCCAGATAGTTGGCGTGATTCTCCCATGAAGATGGCAATAATTGCATGTGGTTTGACTTGGCAAATTGTAGGCATTTTTCATGCTTTGCGATTACGAAAAGCATGGAGTTCGATGGACACTAAAAATAAATAAACAGTTTTAAAAAAAAGCAATTTTTATATAAAACGGTTTATAAACTCATTTAGTCTGTAATTCTTTTAAATGTCCATATAAGGAAAATATTTATGACATATTTAATTAATAATGGTGTCTTCTTTATTCCTGACACAGTTTGCATCGAAT
>evgR013357c0g1_i01_Hydra
CAACAGTTTAATGAAAATTACGCATGCTACTTATACGAGCTAATTTCCTTTTATTGATATTGCTACCTGACAATATTAAAAGATTATATTTACATTCGATGAATAAAATGAAAACAATGTGCACACACAAACTGTTTGAAGGCACGGTGTTCTTTTTTCGGAGTAACAAGAAAAGCATCTGGTTTTATGAAGCTATCGTAATACCTTATTCAGACGATGGAACATTTGGCGGAACCTTGCGTGGAACATATAACATGAACTCTCGTGTAAATGCAACGGATGGTTTTCAACGGATGGAACAAAATTTCACGGATTATTTGAGGAACATTTGAACTGCATCGTCAACCCAGAACAAATAAATGTTCTAGAACATTTTTTTGTTTTCGTTTAAAAAATATGGTGTTTATTTTAGATAAAGCGTCTATGTATGTGCTATTATATAAATAAATAAATACATAGGTTAACCCACAGCTACTGTGTTTTGCCCAA
>evgR013362c0g1_i01_Hydra
AGAAAATAATGATATACCAACTCATTAATATAAGATGTCATTTAAATATAACACTATATACACTGTAACAAGGAAATGTCAGAAAATGACGATCTTTACGTTAAAAAAAACGATCCTAACTTTATGAACAACTATTCAAACGCTAATAAAAACAAATTAATTAAAAGCAGAACAAAACATAAGCATATGGAGGAACATTACGATGACCATATCAATGAAATTTTACCAGGAAACTTTCACCGTGGTGATATTGTGTCGTCCAATCGAATGCTTGCATATTCTGACGCCATCATGGCTACCTGTGCAACTTTTCTTGTTCTTCCTCTAAGAAACCTACACCGCCTTAAAGACAATCAACCATTGTTTGACTATGTTCAATCAATTCACAAAGAGTTTGTTGTATTTATTATTGGGTTTCTAATTGTTCTTACTATTTGGGAAAATATAAATATTCGACCCATCGTAACCAAACGAGTTGATGGTTTTATGTTAATCTTAGTTATTTTAGAGATGTTGATAACTACTATTCTTTCCTATTCGTTGGAAAGATTCTTCCTATTCCTTACAATTGCTGATGTGCTGTGTTTAGCTCACTATAGTGCACAACAACAATTTGCTGAAACAGAAAAGCTGAAAACTTTTTTGAAAACAATTTGCTGAAACAGAAAAAGATTCTTTTTTGTTTCATTTTTCAAAAGGTAATGTATCCAAAGAACGTGTTGAAATTATGAGCGACACTGTTATTGCTATTGTTGCCTGTATTCTCATACTCGATATCACTGTTGAAGAATTTCCAGAAATATTAAATGCTGCTGAGAACAACTTCAACTTTGCATTGAAAAACATGAAATTGGGATTTATGACGTTTCTTGGAACATTTAGTTTTGTTTCAGCGTTGTGGTACATCAACCACACTGTTCTTCATTTATTAAAACATGTAAACTCTATCTTAATTTATGTTCAGAAATTGTTTCTCGTTTTCTGTTGTCTTAGCCCGATGGCAGCTAATATGCTTTCGCGTTTTGCTTCAAAAGGAAATCATGACTCCGTAATCGCAATACGTTTTTCTGCAATTGTTGTGTTTTTTTCAAGTGTTGCTAACTTCTTCATGTTTTTATATGGGTTGTTTTCCAAAAATAAATATGTGCATGACTGGGCTTCATTTGAACATTTTCAGATTAACACAGGTCAGCATTTTTATACTTTAGTAAAATCACTGAACATACCATTTTGGTCATTAATTTGCATATTAGGCACTTTAGGATCTTCTACTGCAACTCCTTATGTTTTGTATTTTACAATTATTGCTACTGTGTGCACTTTTTTTATGTCCAAATTCATTTTAATGAACCATGTAGGCAAAGCAACTGCTTCCTATACAATACGCAAAACCTCATCTTTAGAAATGCTACTAGCAAGAGACAGTGCTACAACTGAAGATTTGCTATTCGAAACTAACAAAGGAGACGCCAACCTTATGGGCACTGAACAACCATAGATAATCTTTCGGACGTCTTAAGGACGCGTATACCGCTGCGATTTAGTAAATGATGTTTGTGGTTTATAACATGAGCACATTCGAATGCAATTTTTTGAAGAATTTAGAAGAAATATGTTCGCACCCGAAAGATATTATAACACAATTTAAACTCAATGGAAGAAAATATAATAAGAAGTCTCC
>evgR013363c0g1_i01_Hydra
CAAATAGAAATGAAAAAAAACGGAAAGATAGATGTCTTAACACTGGTGTCTACTGCAGTGAGTCATCATGTCTTATGTTTTTGACACTGAAGCTGACCTCCAAAACCATCAGAATGAAAATGCTCACAGCACCCATAACAAACATACATCAATGGATTCAATTTTATTATATCATGCTGATTTGTTCGTCAAACTCTAAATCTCGAAAAGGATTTGATGCTGAAGCAATAGTTACGGAAAGTAAATTTAAATTAGAGCGCAATATTGATGTTAAAGGATGGGCTCTACCTGTACAAAAGCATACAAGATTTACGACCAAGCAGAAAGATTTTTTATATGAAGATTTCAAATATGGAGAAGTGACAGGAATTAAAACAACACCAGAAAACGTTCTCTTTAAAATGAGAGCAATGAAACATAAAAAAAGTGTCAAGTTATTTAGTCCTATTGAATATCTTAGTCCACAGCTAATTATCTCTTCATTTAGCAGAATGTCTAACTAAAACCGGCAGGGCTTGCTTAAAAAACCTGAACAAGTTTAAGCTATTAAAGACTTAACAGAAGAGTTTGTAGAAAACCTTTAAATGACACTGACAATTTAGTTGACGAGATATTAGTCAATGTTGAAAAACTATATAATGTTGACGTTGGGCAGTTTGTTCTTATTCACTGTAAAAAAACTCCGTTGTATTTACCAGACTTTTTCCGTTGTTAAGTAACGGAAAAAATCCGTTTTCAATAAACTACGGTCTTTTACCGTACTTTTAAGTCTTTTCAAATACTTACATGTTTTTGTC
>evgR013372c0g1_i01_Hydra
ATTTTTACATTTATTATTTAGGGTTACTTATTTGATAAATGGGAATTTGTTTAACTTTAAATTAATCCCGCTCTTCTATCCACCATTTTCTCGCTCTTCTATCCACCTTACCCCATCTATCTGCTATCATACCCCATCCATCTGCAATCGTCCTTTATGGATGGCCCCTAGCCGTCTTATTACAGAGCTTTGCGGGAGAACATATAAGCAAGTTCATGCTTCCTCTTACCAATCCCCTTTCTTACCAATCTCTTTTCTTACCAATCTCCTTTCTTACCAATCTCCTTTCTTACTAATCTCCTTTCTTTCCAATCTCCTTTCTTACCAATCTCTTTTTTTACCAAACTCCTTTCTTACCAATCGCCTTTCTTACCAATCTCATTTCTTAACAATTTCCCTAAATTTGCCCAGAGCTGGATTTGAACCACGGATCTCTTGCTTTGAGACAAACTCTAACGTGTGTGCTACGGCTACTCTGATAAATAAACTTCACTATAATATTGGCGTGATATCTTGACTCTCGATATTCTGAACTTTTTTTAGGACATTACGCACGGATAAATTGCAATTTCTTGAACCTTTTCTCTAGTTCCCTAACAGTTCAAAGATATTTAAATTCAAAT
>evgR013394c0g1_i01_Hydra
TCAATTAGAGAATGTCTGCGTTGAAAGAATATTTCGAATTTATAAAAATACTAAGCTCAAATGGACCGACCAAATAAACCAAACAATACTAAAGGCTAATATGGTTTATAAATGCTCAAGAGAACCTTTAAAAAATGGAATTTTTATGTTTCTTACACTTTACACTTTTGTACTTCCTCTGCTTGAGTACTGCACACCAATTTGGAATCTATTCCTTAGGTTATCCATTAACAGCCAATGGCCCAGTTTAATGTGTTATGCGGTGCAGAATACGCTTCATAGAGATATTACTAGCAATTCAGC
>evgR013425c0g1_i01_Hydra
TGTAAAATATTTAATTCGGATTAATAGTGTCCAATTTAATTTTTAAAATCAATTAAAATTACCCTCTTTAACCCAATTGTTAAAATCAATTAAAATTACCCTCCATAAAAATACCCTACCATACTTGATGGTCTGGATTTTAGCACACGCTGCATCCTTTATAGGTATAATAGTAGTAATATTATGTAAGGAGTCTTCTAGGCATAATCAGGGTTCTCCAATCAAGACAAAGATCTGGCAAAGCAATGCCTATTTTTTGTTCTACAGTTTTTATGTTTGTATGTTGTTTATATTGTTTTTGCTGTTTATGGTGACATATCTCCATCCAAAATTGGTGCACATAATAAAAGAATTAAACTGTTTATTCGTTATAACACCATTTAATGATTCCAATCCCAGCCGGAAATTGGTTCAAAAAAGACGTCTTTTGAACGTTCAAATTACGTCTTTTTTGGATCAAATGCTGGCTGGGTAGACTAAATTTCCTTCAGTATTGTTGATCTAAGGAGTGTTTTTATGC
>evgR013445c0g1_i01_Hydra
CTTGACAAAAACCACTGATTTAAATATTCACTTTCACTTTCATTCTGTATTATAAATTTTGTTTACATTAAATTATTTATCACTTTTTTTTTTTAACTAAAAGTTTGCACAATTCAAATATATTTTCACACTTACTCATTAGCACTGAGATATTTATTATTTTTTCTAGAAAATGTCATATAGGAGCAGGGAGGATGTTGAAAAAAATTCTGTTACGTGGAGAAAAAAACTTGATAGCATACCAAAAGAGTGTAGCAAAAAGCAAAATTTTAGCAACAGTCACTGACCTACTTGATGGCTTAACTAGAATGCCTAAAGAAATTGTAGCAAAAGAATTGAAAAAAATGTGGGAAAACTTAAATTTTCCATTAAATTGGACAACAGTAAAGTCAATAGTGAGGAAGATTGAAGCATTAATTAATAAATATGGTAAGTACAACAATGAATGTTCTCAAGTTGAATTTGCAAAAGTGTTTGATATCACCAAAACAGATGACATCTCGTTGGCCTATGAAGATAAACAATTTTATCAAATGCAGATGTTATCACATTATGCAATTTTTGTAAAAGTAAAAGTGCATCAATCAAAAGTTAAGAAGTGCCACAAGTAGTAATCTTCCTTTATTGAGCAAAAATGTTGTATTCAATCTATCAACATCTTCAAAACTTCTGACAATGATACTAACGACTGTGATTATACTGTTGAGCCTCCTGCTAAGGCAAGAAAAAAAAAACATATTAAACTACAAAAACTGCAATTCAACTAGTAAAACAGACTAATCTAAGTTACTCCAAGTCAGCAAAAG
>evgR013458c0g1_i01_Hydra
ATTTATATTCCCTGTGTCCCGGTCGTCATTTGTATCCCGGTGTACCGGTCTGTATATACATTCGTTTTTTAGTTTTGTTTTTCTCCTTTATTTTTTTCCTTTTTTTTTCTTTTTTAGCTTATTTAGATTTTTAGATTTTTTAGTTTTTTTATTAGTTTTTAGTTTTTTTTTCTTTTTAGTTTTTTTGTCCCGGTCGTCATTTATATCCCCCTGTTTCCCCCGGTGTCCCCGTTGTAGTTGTGTCCCTGTGTCCCGGTCGTCATTTATATTCCCTGTGTCCCGGTCGTCATTTGTATCCCGGTGTACCGGTCTGTATATACATTCGTTTTTTAGTTTTGTTTTTCTCCTTTATTTTTTTCCTTTTTTTTTCTTTTTTAGCTTATTTAGATTTTTAGATTTTTTAGTTTTTTTATTAGTTTTTAGTTTTTTTTTCTTTTTAGTTTTTTTGTCCCGGTCGTCATTTATATCCCCCTGTTTCCCCCGGTGTCCCCGTTGTAGTTGTGTCCCTGTGTCCCGGTCG
>evgR013471c0g1_i01_Hydra
CGCAACAACAAAATATTCCAACTTTCAAGCTTTTGCTCGTCGGTGATGGTGGTACCGGAAAAACAACTTTCGTCAAGAGACACTTGACTGGTGAATTCGAAAAGAAGTACATTGCTACCGTCGGTGTTGAAGTTCACCCTTTGACTTTCTTTACCACTAGAGGTCCTATCTGTTTCAACGTTTGGGATACTGCTGGCCAAGAAAAGTTCGGTGGTTTGAGAGATGGTTATTATATTCAAGGACAATGTGCCATCATCATGTTCGACGTAACTTCTCGTGCTACTTACAAGAACGTCCCAACATGGCATAGAGATATCGTCAGAGTCTGTGAAAACATTCCAATCGTTTTGTGTGGTAACAAGGTCGATGTTGCTGACAGAAAGGTTAAGCCAAAGTCTATCAACTTCCACAGAAAGAAGAATCTCCAATATTATGACGTTTCTGCTAAGAGTAACTATAACTTTGAAAAGCCATTCTTGTGGTTGGCCAGAAAGATTGTTGGTGATCCAAATTTGGGTCTTTCTGAAGCTCCAGCTATGGCTCCTCCAGAAGTTGCTATTGACATGAATGCCATGAAGCAATATGAAGAAGAATTACAAAAGCTTGCTGAACAAACAGCCATTCCAGAAGATGATGAATTCTAAATTAATTCTAACAAAGGGTTGATTTAAAACTACCACCAATTATTTGCTTCCACAACCCAAACCATTCCGAGCTCTCTAATGGATTTGATTGCAAT
>evgR013473c0g1_i01_Hydra
AAAAACAGGAAAAGATATTCAGCGATATCATAAGTGCAAATCTAAAAATTATAACAGAACGTATTGACAAAATTGAAAACGAGTTTAATCAAAACAAATCGAAGATCCTAAATATTGAAAAAGATATCAACGATATAAAGGAAAGCTTCAACTTTCAGGAAGAGAATTTTTTAAATAGGATATCAGAAACAAACAGTCTGGTGGCCACCGAAATAAACGAATTAAAAAATATATATATTGACCTTGAAAATCGTTCACGCCGTAATAACCTAAGGATCGATGGCTTAACACTGTGGCTGCCCATGGAATCTTGGAATGAC
>evgR013491c0g1_i01_Hydra
TGCTGATCTTGCTTTGTATATTAACATGCTGGATAAACTCAGTCAGCAGCTTGGTATTGATGGAGAAATGAAGCTGAATTTATTTATCAAAGGATTGAAGCCTCATCTACGAAATTCACTAAAGCTCAAACAGCCAGTCAATTATCAGGCTGCGGTAGCTTTTGCAAAGCTGCAGGACACTATTGGAGCCGACTCGACCATGTCTCGTCTTGAAGACAAGCTAGATGCATTAATCAACCTCGCCAATGAGCCAGTTGCCGCTGTTACTCACCACGAAATACTCAACGCAATTAAAAGAATTGAAGAATTACAAGCAGAAGTAGAGTCTCTAAGAAGAAATGACAATCGCCTCACCAAAAATGAAAGATTTCGACCAACTGGCCTTAACCTCCACACAAGTGACGGACAGGTTATTTGCAACAGA
>evgR013505c0g1_i01_Hydra
CTCAAGAGAGAAATTGCTATTATGAAAATTTTAAAACACAAACATGTTGTACAATTGAGAGAAGTTCTTCAAAGCTCTAAACACATTTACATTGTTTTGGAATTGATTACTGGTGGTGAATTGTTCGACAGAATTGTTGAGGCCAAGAGATTCGATGAAACAACAGCAAGAAAATACTTCCAACAATTGATTTCAGGCATTGAATATTGCCACTCACAAGGTATTGCTCACAGAGACTTGAAACCAGAAAATTTATTGTTGGATGGTGAAGATGTTTTGAAGATTAGTGATTTTGGTTTATCTGCCCTCTCCGGTAATGATGGTTCTGGACAACAAAAGATGTTGATGACCACCTGTGGTACTCCTAACTATGTCGCCCCTGAAGTTTTGAAGGAAAAGGGTTATGATGGAAAAAAGGCCGATGTTTGGTCTTGTGGTGTAATCTTGTATGTTATGTTGGC
>evgR013515c0g1_i01_Hydra
GGTTGATTACAAAGATGTAAATGAATCAACATTATATAAAAATGAAATCTTTAAAAGTAATAACCTAATGATTTACACCACATACAACGCAACATCTATGGTACCAATAACTGATGTGATTATCAGTAAAATATTTCAATCAATGTCAGAAAAAAAGTTATTTGTGATACTTACTGATACTGATGAAGAATCAACATTCATATTATCAAATATTATTGTTCCTTATAATAACGTATTGTATTTTGATTGTAACCTAGACAATTATAAAATGTCTTACAATAACAATTCTGTAGCATTTTACTATGGTCGTAAACTCGAC
>evgR013522c0g1_i01_Hydra
GGTCATCAGGCTTTTGTCAATTCATCGTGTCACACTCTTGATATCACTAAACTTAATCTTCCGAATAACATACAACCATTAAGAAATAATGAAATAAAAGCTAGTAGAGTAATAAATCCAATAGAAATATCAAATACATTAAACCATAAACAATCACTAAGTGGAAAATCATCGGGGCTACGTCTAAAAAGTACAAAAAACTACGGTCAAAATAATAAAACATATTTTGATGATGGAATCTTAACAATTGAGGAAGAAAATATTGCACCAAATTCAAATATTCTTTTTTTTAAAAAGCCTTTTAAAAAAGTTTCTAAAATAAGCGCAACAAATATTCGAGGAGAAAGCAACTCCAACCATTATAGTGTTGACAATTTTTCTGGAAATTTAAACGATGAAGAGAGTTCTCCTGTTTTTGTAAAATATCGATATCATGTACTGAGCGAAGAGGAAAAAGTTGCTTCTAATGACGAATCCTTATTACCTGAAGGCGAAGATTTAGGTTCTGGAAAAAATTTAAACAGTGAAAGTTCAAAATCTGACAATCTATACTCATCAGGTTCAGGTTTGTCATCTTTGGCACAAAACTATACGAAAGCTGATAAAGATATTGAAATACCATTAGATTATTTAAACGATGGCAGGAATGCTCGAACAAAAAGTGTTTTAAATAAGGCGAAAACCTTTATGAAGCAGTTTAAAAATCAAGTTGCATCAAAAATGCTGTTGCCAACGATCTCTAAGATGCGTCAGAAAAAAAAAAATAACTTGTCACAATTTGTTGATCAATCTGAAGATTTGTTTGAAATTGGAAATAGTGTTTCAAATCGTAAAGAAAACAAAACATTTGAAACGAAAAAACAAACAACCCGTGTGATAAATCGTAATAGTACTGACTCTCGACCTGTAATGTTGCGGAAACCAATGAACCAATTTATTAACCGGCTAAGTTTACAGATGTCTCGACTTAAAAAAAGCACCCTTGGTCTAGATTTTAAAAAAGACAACTTTGGTCTAGATTTAAAAAAGGCCCTCTTAACAAAGGCCTCAAACGCACAACCTATTTGATATTAATAATTACTTGTTTCAGGCATCATAAAAATATACACACAAAGCAAAGAACTGCTACTTATAAAGTTTATTCCAAATCAATGATAAAACGAC
>evgR013534c0g1_i01_Hydra
TCAAATTAGTGTTAGTGATACCATGGTAACAAAACAAATCTCTCAGATGAATGATGTTTTAAAAGCATTACAGTTCAATCTCTTAGGTAATAACACATCTAGTGACGAAGAAGAGTTTATTGTTTTTGATCAAGCTGAGAATAAGTTTAATGTTCCTCAAAGTGACATCGAGGAGTATGGGAACGAAGAGTACGAGTTTATAAAGTCAAATTTGGTTTCACTGCAAAGTACGGATTCTGCATGGTTATTGCATCAAGCTAACGAAAGTGTAGATAACAACATACTGTATGCTTACATAGAACCAAAAGAATGTGAGGATTTAACCACTTCAGTCATCAATGAGGCAGAAATATTAGAATGTTCAATTAAACTTTGGACAACCGATGTTTTTGAAGAAGGTATATTAGAGAAAGTCAATAAAATGGAAACATCTGTAGAGGACGATTTAGAACTTTGCAATGTTGAAATTTTAGAATCCCTAATTCTTGAAGAGTCTCTTGTTAATTCATTTTTTTCTGAAAAAGCTCCAGTTGAAAGTTTAACAATTCCGTTTAATTACAATAATAGGACTAATAATGGAAGTAATGATACTATTATGATGTATCATGATGATCATGATGATACTCCTTTTTCTTCTACTCATCACTCAGTGCAGAATAAAGAAACTCTTCATGATACTAAAACACATCAGAATGTTGGAACAGATTATAATGATGTTTGTAATCATTGTGATGTGGGAACAGATCCAATTATTTTCAAAGACAATATTTTTACTCTAAATGATCAAAACCAGAGAGATAAAACAAAGAATGATGCTGATAATGATGATGACGAGATTATATCATTAGATAAAGAAAAAGATCTATCTTTTAAAGTAGATAAAGTTCGATTATTTGAAGTAGAGAAGCTTGATGCAGAGGAGAAGTTTTGTTTATCTAAAAAGGAAAAGGATTTATCCAATGAAGTTTTGTTCAATTTATCTAATATAGAAATGCTGGAAGAGAGGCCTGGATCAACTGAAAACAAGAAATTTGAGGCTTCTAAGGAAAAAGGTAAATTGTTTGAAGTATGGGAGCTGACACCTACTGAACAGGAGAAAATTGAACTATTTAAAGACAAAAGGCTTGAATTATTTGATGAGGAAAAGCTAAGTATATCTAATATAGAGATGCTGGAATCAACTTTAGAAAAAAGCATTGAAATGTCTGAAGATGAGCAACTTGAATCTTTTGTGGAGAAAGCTAAGTCATTTAAAGTTGATAAGCTTGAGTTTTTTCAAGAAGAGAAACCTATATCATTTAAAGAAGAAGAGAAGTTAGAATCATCCATAGCAGAGAATCTTGATTCATTAGAGAAA
>evgR013546c0g1_i01_Hydra
AAAAAATAAGCTTTTAGAGAGAGCTTTAGCAAAGAGCGATTTACAAAAACTTTAGCCAAAAAATTAGAAAAAAAGAAATGATATGAGCGAAAAAATTAATAACTCACTCGAAAACATAACAGCCGATATAAATTTATTATCGGAAAAAAACCAAGTAGCTGAAGCTGCTTTTTTCACTTTACTATTCGTAGGCGCTCTTCTAGCAAACTTTATTGTCATCATATGTTACTTTAAAACAAAAAGCAAAACAACGCACTTAAAGCTTCTTGTGAATTTATGTATTATAAATATATTAGGAAGTGGAATAACATTTCTTTATGGAAAAGGCGACAAATGGTTGCTAAAAGAAAACGTATGTTTTCCAATGCTGTCGATTGATGTATCATGGACAATATTGTGTCTTTCTTCGAATATCTTAATAGTTTCAGAATGTCTCATTACTACACTGCAACAGTATAATAAAAAAAGATCAAAAACACAATTTATGAACTTTAGAATAACCGTAATATTAATATGGGTTTGTTCAGGAATTGTTGCTATGGTTACTTTTATGTTATTAGAAAGACCTGCAGAACAAAATCTTTGTTACATGTACCAATACTACCGTCAAAATTTATATTATGCATTTCTTTACGGCTTTGTGATTGTGCCCATGTTGTTGCTACTTGCTATTGAATTTTGGATATATAAGATTAAAAGGAAACACGAAAGATTAATAGATGCAGAAAAATCGGACGAAGAAATTATACGAATAGACATTAGCAACAACAAAGGAGCATTTGCTTTATCGCATTTAGTGCTGTTTAACTGGTTACCCTATACGGTTTACATAATTTTATTAATGAATTGGGGGATTAAAGAAATCTATTTTAATGAAATAACTTATAAAATGTTGAAAATACTAACTTTTTTCAGATACATGTCATGCTTTCTGCAGCCGGTTTATGTTATAACTTTTAATCGTGACATCAATGATGCTTTTAAAAGAACATGTAAAAAAATCGAGGAAGAACTTTTAGGAGAAGAAATAAAACCTGTTCAAAAGAAAAGAAGATTAAAAAGAAAACAACCATCATTAGCAAACATACAACCGATAAAACAACCAGTTAATTCTTCGATTTCTCTAAGTTCTTCAGAACACGAGGCAGACACGTGAAACAAATTGATAACACAACAATCTAATTCTTTATTATATACGATTTTGATAAATCTAAAATGGTGTCCGAGAAATTTAGTCATTACCTGGATATATATATTTATTTTAAAGATAATGATTGGATATATATATTTATTTTAAAGATAAAATTGTTTCTCAAAAAAAAAA
>evgR013547c0g1_i01_Hydra
GTAAACAAATCATATGAAATTGGATTCGAGTTTCAATTGAATGTCTATCCAATTAATTATAATGGATACAGTGAATCAATTAAAGTGTCTTCAAATATTGCAAGATGTAGTTATGTTTCATCATTCCAGAAAACTGATTGTCGTCAAACAGACCCTCTTACTATTACAAATAAAAAAGTACAAGCAGCGATTATTTTTCCAACAGATTGTATCATCCAACATAAAGCAATAAGTTTGAAACCAGGTTTATTACTTGGAATTGTTAACGTCTTGATGAAAGAATACACTGTGTCTCTTAAAATAAAACCCACACAATTTTCACAAGGTTTTAAAAGTGTTTTTCACTTAACTTTGGATAAGGATATTGGAGCTTATGGTGAAAGAAATCCGGGTATTTGGTTTAAAGGTGATGGATCTGGA
>evgR013549c0g1_i01_Hydra
CTTTCCTTCTCCTTGCTTCCACCACTTTACAATTGGCTCTCGCAGCATGTGGTCCACCAATGTGCAACACTGAAAGTCAACAACTTCTCTTTGCTCCTGCAATGGTCAAACTCATGTATAACCACTTGCATGGAAATACTTCACTCCCAACCAACGCCAAAATGGGTAATGCCAGAAATCTTCCATGTACTGCCTGTTTGCCATGTTGTAGCTTTTTGGCATCTGTTGAACAATCAACTGTTGAGGAACGTCATGAAATCAATGCTTTGAATACCATTGTTGACTCTATTGATTGGAATTGTAAGAGTCTTGAATGTGCTACTCATCATGCTGAATTGAGAAATTTAGTTTCAAAGATTGTTGAAGAGAAGAGTCAACCATTATTGAAAGCCATGCC
>evgR013552c0g1_i01_Hydra
AAACTGCAATAATGCGCTATTATCGATTCAACCGAGTTCAAGAACTCATGCCAGCATGAGTTCAATAACTTTTTTAAGTTGTAAACAGTATAAAAGCAGCAATCAGTCGAATTAATTACACGTGAATTTAGCGTAGATTGATGAATAGTAACGCAAATATTTTAATTTTCGTTACGTCTTATCGATTAATTGAGATCTTCTGATAACGTTTTTTGTTTTAAAACAATCAGGAAAAAAATCAATATCGTTCAATCAAAATAAACAATAAACAAGGGGGATATTATGGTGTCACAACAAACAATGGGGATGTTTCTGTTAGTTTTACTTTTACTTTTGAAAAAAATAAGCAGCAAAGCTATTAGTCTTAATGATAAACAGTCTGAGGGGGGAGAGAATTGTAGACTTGAACTTGTTTCTGTTATGATTCCAAATTGCGAGAAGAAGGAAATACGTGTGTACGGCTGTCAAGGAAAATGCAAGACCACCTCTTATATGGAACTCAGCAAAGTTCGTCAGACTAAGTCGTGCTGTAGACCTAACAAAATTCAAAACTTTGAAGTTGCTGTTAACTGTTCGGAGAGAATTGAACATATTAAATTGTCCTCTGCAGTAAGTTGTTCCTGTGTTAACGAAAACTCTTTTAACTAAAAATAACTTTTAATCGTGTAAATAAGAAGTTTATATTGTAAATAATTTTGTGTAATAAAGTTATTGATATTGATATATTGTGTAATAAAATTATTGAT
>evgR013564c0g1_i01_Hydra
TTCTAATATTGTAAAAGGGCCTGAGAAAAAACCCCCGACTTGGATAGAAAAATCAATAGAAACAATCATTTTGGAAGCCATGTCTGTGATTATACACGCCAACAATGACACCATTGACAAAAAGAGAACCTTCACTAAGTCATACCATGAACCAGTAGCCGAAATGACTACAATTGCTCTCTCAGTTGTCCTAAAAAAAATGAAAATACGACGGCAACAATATGACGGCATCACTGGACTAATTCAGCAAATTAGAGAAGCTCCCAAAATTTTACCGAATTTTATGCTGAGGCTTGATTCACAAAGGAATCAGACACACAAATCATGGGCTACTAACTACACTGCAGCAACCGCCTAGGCCAAAGAGGTGGCGGTGTGGTCATCTGCATAAAAAAAGGGTATTGCCACAGCGGTTACAAGCATCTCTCAACTAAATTCGAACAAAATTGAGCAAACTTGGCGTATCATCAAATGTGGTTCTGAATCAACACTAAGCGAAATAATGCATCACGACGGCAAAGCTAGATCTTTCGTAACTTTGCTGCACTTCTATGCTAGTCTACAGAGATTTCAATCTCAGGCATTCTAAATACAAAGAGACAGACATTGGAGGACGCGCTGCAACCACCTCTCACATCAAGAATAAGCACTCTACAGATTAGAAATTTCTTTTCTGCCATGAAGAATTGTATCTAACCCAAGTTATAACATTTCCAACATTTAAAAAAAC
>evgR013567c0g1_i01_Hydra
GCAACAATTTTAGCATACTAAGAGATTTAAAAGAAAACATACTTAATTAGTTTCTACTACAAATAGATTAGTATATAAATTTTCAAAATCATAGAAAAAAAGTGCAGGCATTCAAAAAGAAGAAGAAGAATCCTGAATAAGTGCAGTTGAAGAAAAAGAAAAATCAAATAATAACTTTGTAATAGCGTTTTAAAAAATCAGGTCTGGAATATGTGAATGCTACGACTAAATTTTTTGTTACCAATTCGACTAATTATTAGTGTCATTTAGTATTAACTGTTTAAAATAATGTTTAAAGATGAAGTTTTTGATGTTGTTGAAATATTATCTGATGTCAGAGTAGATAAATATGTATGTGACCAATCTTCACAAACAAAGAAGCTTCAGATTTCTAATGCCTTATGCCAGTCGTCCGACAGAAAAACGCAAGAAGTTCAAACTGATTTTGAGCAAATTCTTCCCCAGTCTACATTAATTCAAATAGATAATTCAAAAAAGTTATTGGATTTTATAAAAAGTGTTGAGCCACTTATCAGTGAACAACTTATTGCAAATATTCGTAGTCATGCCTTTGATGATTATGATGTCGATTGGACTGAAAACGAAAAATTAAGTGAAGTTGTATGTAAATATTCTTTAAAATACAGCGATTTACCTGGTGAGAAACTTGTTACAAGTTTAACATGGAATGCTGTTGGGTCAGTAATAGCTGCTGCTTTAGGCTGTGGCGATCATGAATCATGGTGCATACATCAGTCTTTTGTATGTTTTTGGAATATTGATCGGGAAAATATTGACTTTTGCAAACCTGATAAAGTTATTGAATCTGATGTTTGCATTACTTCGCTCTCATTTCATCCTAAGGAATCAGCTTTGATTATCGGAGGAGATTTCTCAGGAAAAATTTATATTTGGGACCTTAGTCAGCAAGATAATTATCTAGTTGGGTTTTCTGGAAAAGATGAGTTTTCTCACCAAGAGCCGATTTCTAGAGTTTTATGGATAAAATCAAAAAGTTTACGAAGTCATAAATTTGTAAGTTCAGGATTTGATGGTAAAATATGTGTATGGGAGTTTAATCAAATTAAAAAAGAACTTACATTAAATAAATGCTTCTTATTACCATCAAATAGAATGCGTACTTTCAAAAATTCAAAGTCTAACTCACTTTTGGGTATTACTTGTGTTACCCAATGTCCAAGTGATTTAAGTCATATACTACTTGGAACTGAAAATGGTTCTGTATTTAGGTGTTCTATTGAATTTGAAATTAGTGGAAAAGATAGACAGGATTTTGCTACAACAACTTCTTTTGAAGGTCATTCAGTAAGTGTAAATTCTATAGACGCTTCACAGTGTGTAGATGATTTATTCCTGACTTGCAGTTCTGATATGACTGTCCGTGTATATCGTACAAATAAGCGTAAACCAATTTTTATAATAGAACCCACATCGAAGTATTTGCTATCAGCACGTTGGTCTTTAAGTAGACCAGCGGTGTTTTATCTTGTTACTGCTGATGGCCATTTAATTATTTATGACTTATCACTGAATTTAGTTTCCGCAGATCTCACCTTACTTATTCAGAAAGATCATCTTCCAGTACCATCAATGGCATTAAATGGTTCACAATTTGATATTTTTGCTACTGGTGATTTCAAAGGAAATATTCAAATTTGGTCTATGCCAACAATTTATATTAATGAGAAAACTTCTGAAAAACTATTTTTTAAAGAATTTGAAGAAAATGAATAAAAGATTTTTGCTAACTACATTTATGTAATATAATAATCCAACAATAATATAAAATGTTCTAATAGGGTTATATTAAGGTTATATATATTAAGGTTAATACAATAATAAAATATTAATTAAG
>evgR013571c0g1_i01_Hydra
CCTTCACGATCAGGTAATGCCCAATATGATTACCTCTGTAAATTATTGCTCATTGGTGATAGTGGTGTGGGAAAGAGTTGTCTACTTCTCAGATTCAGTGACGATACCTTTACAACAAATTTTATAACTACCATCGGTATTGATTTTAAAATAAGAACCATTGAATTGGATGGAAAGCGAGTAAAATTACAAATTTGGGATACTGCTGGCCAAGAGAGATTTAGAACTATTACAACAGCATATTATAGAGGTGCTATGGGTATCATGTTAACTTATGATGTTACTGATGAACAATCGTTTTTAAACATTAGAAATTGGATGAAAAATATAGAAGAACACGCTGCTGATAATGTCAATAAGATGTTGATCGGAAATAAGTGCGATTTAATTGAAAAGCGAGTTGTGGAAGCAGAAAGAGGTCAATCATTGGCTAAATCCTATGGAATTCCATTCATGGAAACTAGCGCAAAAAACAATATTAATGTGGAAGAAGC
>evgR013629c0g1_i01_Hydra
CAATTACCAACAATTTTCAAGTTGGATCAAGCATTACTTCAATCTTTGAAGGAAGCTAATGAAAAGAAACTCAACCAACTGGATGAAAAAATTAAAGATGCCAATCTTAATCTTGGTGAAAATGAAGTCAGAGAAGCTCTCCTTGCTAAAGCTCAATACTACTATTCTATCGGAGACAAGGAAAATGCTCTCTCTGCCTTTAGAGAAACAGAAACAAAAACAGTTGCTTTGGGGCAAAAATTAGACATTTTATTTGCGTTGCTCCGTATTGGATTTGCTTTTGATGATAAGGATATTGTGAAAAGAAATCTCGAAAGAGCTAACAGTTTGATTGTTGAAGGAGGAGATTGGGAAAGAAGAAATAGATTGAAGGTCTATGAAGGAATTCACTTAATGACCATCAGAAACTTTTCTAAGGCAGCCAAACTTTTCTTGGATTCAATTTCTACTTTCTCATCTAGTGAAATTATGACCTTTGAGCAATTTATTTTCTATACAGTTGTTGTGACTACATTCTCACTTGGTAGACC
>evgR013659c0g1_i03_Hydra
ATGAAATAAAATGCTTAGTTGATTCTCTAAATCTAATATTTATAAAAGTAAATTTCTCGTACTTTTACATAGTAAGCATTTAACAATTTTAGTATTGCGGATAAACATGGCAGCTAAAAACTTTTGACCTGTTTTGGAATCTACTGAGACAACGACCTTGCATGAGATGATGAGAAATCTTCGCTACGGCTCATGAACCTTAAGCGATGGTGAATAACATAATTCCCTTTTTTTTATATGGATCCGTTAGGAGCTTCTGATGCATTAGGGCTTGACCCACAATCAACTAATGGTAATGGAAACCTAGATGACGTCCTTAGAAGTACAAGACTAAGTTCAAGCAATCATGAAGACATTAGAGATACAAATATCATGTTTCATGCTCCAGTTTCGGACATTGATAATGACGTGCTTGGTTTATACAATATTGATAGTACTTTGCACCATAGTGAGATAATGTCACATCATAATGAAATGGTTTCAAATGAAAGTGAGTTATCTATGATCAGTGAAACAAATGATATTAGTCTTGAGCCAGAAGACAACAACCATGCATTTGTTGAGAGGGATGAGCTTATTTTTGAAGAGAGTAATGATTTGTCACCTGAATCTTTTGATGGAAATGCAACATGCGAATACTGTGGGGTTGTTGGAGTCCAGGATTCATTTTATTCTAAAAGCAAACGTTTTTGCAAAATGGAATGTTCAAAAAAATATTCAGCGTTGTTCCAAAAAAAAGGTCCCGGAACTCGACAACTAAACCTGCCAAAAAAGAG
>evgR013664c0g1_i01_Hydra
AGGAGGTTTGTCTTTTTGTTTCTGCCATATATGTCCAATATTGGTTTCAAACACCATCATCAACAGCAGCTCCAAGAAATGACCTTGCTTTACTCTGCACACTGGCATCATACCCAAACAAGGAAGTAGCGAATGCAGCCACAACCGCTTTTGGTCGCCATTTATGGTATCTGTCTGAGCTGTTGGTTGGATTTGCATTCTTTGATGATGAAATCTCAGCAGAAGAAAAGAGATTGATGGCTATGTCACTTAAAGGAAGTGAAGGGTCTGAGAATCCGCCAAAACGTATCCCGTTATTCGATGAACCAACAACAAAGAAATTGCATAATTTTGTGACAACATCAACATTGAGATTTTTTAAGATACTCGGCTTATCAGAGAAATTCTTGCAGTATGATCCTGATAATTGG
>evgR013674c0g1_i01_Hydra
GTAGTCTCTTGCTCCTTGATTTCAAGTGGTGCCACCTTGGTGGTTAATACCGCTTCATTGAACGTTCGTTCAGGTGCTTGTACCAATCATGGTGTCGTCAAGACCATTCCACAAGGAACTGAATTGAAATTCACTGGTTCCGTTCAATCTGGATGTGGTTATACTTGGTATGGTGTTTCTGGTAATGGTTTCAGTGGATTTGCTGCTTCATCATTCTTGGCTGAAAAGCCATCAAACAATGGTGGTGGTTCTGCTACTGTTTCTGTTGATGCTTTGAGAAGAATTATGCCAAACTTGTCATCATCAAAGGCCAACTCATACATTGGTTATTTGAATAGTGCTATGAACTCTGCTTCTATCACTACTTGTTGTAGAAAGTCTGCTTTCTTGGCTCAATTGGCCCATGAAAGTGGTGATCTTAACTGGTTCCAAGAATTCGCTTCTGGTGCTGCTTATGAAGGTAGAAAGGATTTGGGTAACATTTATCCAGGTGATGGTGTCAGATATAAGGGTAGAGGTCCAATCCAAATTACTGGTCGTGCCAACTATCGTTCTGCTGGTCAAGCCCTCGGTATTGATTTGGAAAATAACCCAACCAGAGCTGCTGATCCAGATGTTGGTTTCCGTACTGCTGCTTGGTTCTGGAATTCAAGATCCTTGAATCAATATGCTGATCAATGTAATCAATCCGGTTTCGATACCACCACCAAGAGAATCAATGGTGGTTACAACGGTAAGGCTGACAGAGATGCTCACTGGATTAGAGCCAAGCAAGTTTTGGGTTGTTAAATAAATACATTTGAC
>evgR013683c0g1_i02_Hydra
AAAAAAAATCAGCTTTAAAAAAGATTGGGGCGGAAAATTTAAGATCCGCACCGGGTGGCAGTGACTCACAATAAACAACTGAATAGATTCCCAAAAAAATATCGGTATGAATATATATTCCTTCACGTTTAAATTTGAATAAAAACTAAACTGGTTTCAAAGCTCTTAAAAAATCTTTGCAAAACCGCATTTCCTCGATTGTTCAACTACAACACAGTGCAGGTACCGTGCAAATTTCAATTGAAAATAATAATGGCGGACAGTTCAGCGTCAAGTGCTGCTGGACCTAGTTCTCAACAGCCAGAGGATGATTTAAGTTTACCACGAGCCGCTGTTAACAAAATGATCAAGGAAATGGTGCCATTTATCAGAGTTTCAAATGACGCAAGAGAACTTGTTCTTAATTGTTGCACAGAGTTCATTCATTTGATTGCATCAGAAGCAAATGAAATATGTAATAAACAAACAAAGAAGACAATCTCCCCAGAGCACGTAATTGCTGCTTTAGAGAGTCTAGGCTTTCAGAGTTACATTCAAGATGTTGAAGGTGTTTATCAGCAATTCAAAACACAAGCACAGACCCGAAAAAAGAACAACAAGCTTAAAAACCTGGGCATATCTGAAGAAGAACTTCTTCGTCAACAACAAGCATTAATTGCTCAGGCACGCGCTGAGCAGGCTCAGGAGGAGTGGGTTCACTTACAGCAGCAGTGTTTTCTCCCTACTTCCACGGGGGCTGCTACAGTTCAAATGCAAAACGTACATACGCTGCAGCTTCATTCCGAAAATGGGATCAATTCTATCGGACAGCATTCTTCGATTAATTACACGCATTGCCTCGACAAGAGTACGAACAATTCATTTGTATTAAATGACTCTAACGAAAACGTTCACGCGCTTCAAAACGGACTAAATCACGCAGATGCTCTTGGTAACACGTGACCCAATTAGGTGAATTATTTTGATTAAGTGGATAATGTGTGGCCAAACGCCCTCTAGTTATGTTTTGAGGTCTTTATGAGCCGTTAAAAACAATACACTAACGTTTTAGCATGCAGTTTGCATTGCAATGTTTCATATATGATATCGTATCTTATTACACGATATTTGTAGATTCGGTTTAAAGATTGCTACCTAGAATGCAACAATCATTATAAAAAAGGCAAACTTGTCGCGGTATCGAGTGCAATTTTGATTATCTTTTTCATTTGGTTTAAAGAATAATACCCAGCATATACCCAATACAACATTCTTGTAAAATATGCGAACTCGTCGCGGTATGGAGTGCCTTTTTATTTATCTTTTTCATAAGATTACTTAATGTTTCATTGAAATGCAGGCCTTGTTTAGTAAAAAAAAAA
>evgR013694c0g1_i01_Hydra
GTGAAATTGTAAATTTGAGAAGAGTTGTTGAATTGAAAAAACAATTCAAGGCCTATCTTTATGTTGATGAAGCTCACTCATGTGGTGCATTGGGTAAAACAGGAAGAGGTGTATGTGAACAAACTGGTGTAGATCCAAAAGATGTTGATGTATTGATGGGAACTTTTACAAAGAGTTTCAATTCTGTTGGTGGTTATATTGCTGGAGATTGTGACTTGATTGAACATCTCAAAAAAAGCATGTTTTGGTTTAATTTATGGAAGTGCAATGTCAACACCATGTGCTCAACAAGCATTATCTGCTCTCACTGTTATCATGGGTGAAGACAATACAACAATCGGAAAGGAGAAATTAGAACAG
>evgR013702c0g2_i01_Hydra
AGAGAAAATCAAGTAATTATTGGAAGGGTAATTGGTAATTCTATATCGGTGACTCATCATGGAGATATTATGAAAAATATATCAAGTTATTCAAGTACGGTAATAAACTTTTACTTATATGCTCAGGTTTGGAACAACTTTGGAAAATTATCGTCACCATATATAATCAGTTTTGGTTACCGATTAGATTGCCCGTATCAGATATGCTTTCAATTCCATACATTTAAATATTTCTTATCAAGCAATAACCTAATAACATTTAGTTCTACTCTTGTGCCTCCAGATTATTCTAATTTCTTCCCATTTGAAAGCCAATTTGGTTATGATCCAATTAGCACAAAAGCAAACAAACTATATAAGTTTAATATGAATTGTCCAGCAGGCTTTGAACAAAATTTTATACTTTCAACAATAACTGGAAAAATAATGAAACAAGTTTCTTGCACTATTTGCAAGGAAAATTATTTTAAAAAAGGTTATGGAACATACAAATGTGAAACGTGTCCAAAAGGTTATATTTCAAATAAAAACAAAACAGAATGCAAACTAACATTTGAAGAAGACTTTACATTTCGGAGTATTGAAAGCATCATAATAACTGTTACAGCTGCATTTTCATCG
>evgR013705c0g1_i01_Hydra
AAAATGGCGGATTTTCGAGTGTTAGCTAGAGCTCATGGATTTATCAAATTAGCGGAAGTGTTATTAGCGGCAGTTTGTCTTGGTTTAATAAGGCATTATGACCTGCATTTTGGTGGTGATATTACAACAGGATCTTATCAAGACAGATATTTATGTGGCATCATCACAATTGGTGGCTTCATTTTAGTATCCTTACCTCTTTTGTTGTCCTACATATGGGGTGAAGCTGGAACGTATCAAACGCTATTGGAAATGATCTACAACTTCACTGGCATGGTCCTGTTTCTTACTGCTGGATCTTTGGCTTTGGATTACTACAACTCCTACAAAGCTACTGGAGGAAGTCCCGACGCAGGCAAAGCTTTGGGGGCCCTGTGTATTATCAACGGTTTCTTCTACCTCACTGACACAGTCCTGGCTGTGCGCCACAGCTACGCAACGGTAGAACAATCCTTCACATGAATCAAAAATTGATGGAGATGTGCTTGTATTTTATTAATTCTTTAT
>evgR013735c0g1_i01_Hydra
AGGTAGTCTGCATCACGAATGGTAATAATCAAATTCAAAAATAATTATAAATACATCGTCGTAATTCAATAAATAGTTTTTCGAACTATCTCGTTACCTGGCTGTTTAAATACTTTGTGCTATTATTGGTTACGTTGTGGCCTGTAAAATATTTTGTGTTGTTGTTGGCTACGTTGTGGCTGACAAATAATCAACCAACATGTCTACCTTACCGTCAAACATAACGAATTCAACAAACTTAACTAGTTCGGCATCGCTCAATAAATGTCTGTTGCCGTATGCAAACATTTTATTAGTAGCAGGGTACTTTATAGTGATAACTATTGGATTTATTGGTAACGTTTTAGTCATTGTTATTTTTAAGTTTAAACTGAAGAAAAGACCAATAATTGATTTACTCATTTTTTACCTGGCTGTGTTTGATGGGCTGGGTTCCTTGTTTGGCCCATTAGTTTTTTTATACTGGACTTTAACATGCAACAAACGATGGGATTTTGGATGGCTCGGGTGTAAAATATTGCCACCGTTTAGCCGAATTGTTACAAATATATCAACTGGTGTCATACTTATCATGGCCATTGATAGATGTCGCTCCATAGTTTTTCCATTAAAACGGCGTTTTCGACGAACAACTATACATGTTGCTATGTTTGCTACAGTTGCAACATCAATTGTTTGGGAAGCGTATTATATAAAATCTTTGAAAATTGATGAAAAAGGTCAATGCAATACCATTGCTACTAATGACCCTTCTTATGCTCACCCGCTTATTGCAATGGTGACAGCCAGGTCATTAATTTTCATAGTTGTATTTTCCGCAACAACAGCTGCTGTTTATATCAAACTTCACAATTGTGAGAGAATGGTACTTCTACGAAGCAGTTCTTCAAGATCAACAAATAAAAACAAAAACGTAATGAAAATGTTAGTCACTATGGCAACTGTATTTGTATTGTCAGTTATACCACGAGATATTTTACATTTAGTCTACACCATATCATGGCTTCATAGTAATGGAATATCAAATGGCTACATAATCAACTTGAATCATTGGCTTCGATTACTACAAATTAGCAACGGAATATACAATGTATTTATTTATGGAAAAATGTATAGAAATTTTTGGAAAAGCGTGCAAATGATGTTGCGTTGTAAGTACACCCCCTTTGGTGTTTATGAAAGTCAAATGTCACACTTTGATCCTAGAAAAAGCAATTCTCTTTATTCAGACGAAACTAGAAGAGTAACGTTAACAAATATTTCAAACGAAACCGACTTTGGTGGAATGACTAAATTCCACTCGATTAGAAGCAAAAGCAGCCAGTGCACTTTATAGACTATATTTGACACAGGAATTTGTCTGGTGTATTTTTATTAGCAAGCTTGCTTCTCGTTTCTTAAATGAAGTTTAGTTCAATTAATTTTTTGTATATATTTAATAAATAGATGAAGATTTATTTTATAAAAAA
>evgR013739c1g1_i01_Hydra
GTTACCTTAAAAAAAAAAAAAACCATCCTTGTGATTACCGCGAACAAAAAAGTAAAATTGATTGTTTTTATTACAGTTTTATCATAAATATGCGCATTAGATTAAATTTCCAGATATAATTTTTAAGTATTTATACAAAGATTTTAATATATTTATAAAAATGAGTCACCGGGAAACAGTGGTTTATCATCAAGTCAGTGAAGAAACAACTCTCTTCTCTACAACTACAAAGTCGAATGTAATAGATTATGCTACTATTTCAAAAATTAAACACATTTATTCATATATAACACTTGCAGATATATTTCAAAAAGTAGTTTATACAACAATAATAACCAGTTTGCTTCTGTACTTTTATGCATATTTAAAACTTGGTTCAAAGTATGCTATTTCAGGATTTTTTGTTTTTTCATGTTTATCATGGTTTACTGCTGCAGCAGTAAATTTATTAGACGATTATTTTTCAAGGAAAAGTGGTATAATAATAGGTTTTGGATTTTTTATAAGCGGAGTTTCTATTTTATCTGGATTAACACAGTACATGCTTGATAACAGTTCCTTAAAGTATATTCTATTAGTTCCTTTATATTTTATTTGCAATGGAGAAAGTTTGTGTAAAACATTATTAGGTGACTTTGGTCACGAACAGTTATCTAAGATAAAGTTTACAGAGCAATTAAAAAAATATACAGACAATGTATATTGGACAGGACATATAGGTGCTGTATTTTTAATAGCATTTTTGCTTGGTATAGCAGAATTTGCAAGTTTTGAAGTAAGTTATGCATTGTGCTGTGTTTCTCTTGCTTTTGGAATGGCTTCTTTTATGATGGGATGGTCTAGATTTGCTAAACAAGATTCGTTAAAAAAAGACAGTCTAAAGCTTTTGTTTTTACTTATTAAAGAAAGCAGGAAGACAAAAAAAGAGTTATCTCAAAGATGCAAGAGTGAAATTGTTGATGATTCTACAAGTTTCAAAGGTGCTCCACAGCATTGGCTTGACTATGCAAGAACCAAGTTTGGGGGATCATATTCTGATGCAATTGTGCATGAAGCAAAATCTTTCTTTAAAATGTCACAAGTTTTTGTCTCATTTATTCCCTACTGGATTTGCAATGCACAGGGTTATAGCACATTTATTTATCAAGCTCTTCACCTGAACACATCAATTGGAAGTTATACGTTTGCTGTTTCATGGCTCGCTCTTGTTAATATTATTGCTGTACTTACTTTTGTAAAACTTTTCGAAAAAGTTGTGTTTCCAGCCCTTCGTAGGAATGGTTATAGCTTTCTTACAAAATGGCGGATTATTGTAGGAATGTCGTGTGCTCTAGTTTCTATGGTTTTAGCTGGTACGCTGCAACTTATAATGCATCATTATGCGCAGAAAGAAGAAATAAAACTCCATGTTATTGGTAAAGCAAACATTACTTATGTTTCACATGTACATATTTTATGGGCTGTGCCACAGTTTGTATTTCTTGGTATTAGTGAAATGCTTGTTGGATTAACAGGATTAGGTTTAGCATGCAAGCTATGCCCTGATAGATTTCAAAAGACAATAACAGGTGTATACTACTTCATGGTTGCCTTAGGTCATGTTTTAACTTTGGTGGTTGTTTGCGCTGGCTATTTTACGCCTGAAAATTATTTTGAAGGTTTTAGTGCAGTGGTTTATTTTTACTCTTTGGCTGGCGTAGCGCTTGTCGCAATAATCTTATTTACAGTAACATTGTCTTACCAGGAATCTTTTGTGGTTCAAGAATCACCAAATACAAAATCTCCAGATACAGTATCTTCTATTCTATAAAGATATTATAAGTTTTCTTGAACGAAAAAAAAAGAAATTTTGATTGTTTATTCTTAAAAAAAAAAA
>evgR013740c0g1_i01_Hydra
ATTAACAAATACTGAATATTTTTCACAGTTATCAGATGCGTTTGCTGACTCAACTTTACCAGACGTAGCAACTACAACGCCATTGCACCAAACTTCAACTGCAGCTGAAATTTTTACAAACTTCATTCCTTCTCAATTGGATTCAACACAATTTTTGCAACAAGCGTTATCATGCAGTTCTTTAAACTGCTTTTTACCTTCGTGTAGATGCGCTGGGTCAGATATTCCGGGTGGGTTATCAAAGAATAACACTCCACAAATCATTTTATTTACTATGGATGATGGAATTACACCTGAAAATTTTCAACTATACAGCGAGCTGCTCAATGGCGTAAATAACTTCAATGGGTGTCCAATAAAGGCAACTTTTTATGTGTCTGGCGACAATTCCGATTATAGCTTAGTAAAAAAATTACACGAGAAAGGTCACGAAATAGCTGATCACTCGGCAACTCACAGAATGCCAGTGGAATGGTGGAGTAAAACTGCAACTTTTGAAGACTTGCAAACAGAAATTCTTAATCAAATGCTAACTATCCAGAAAGAAGTAAGTGTGACCACAGTTGGGTGGAGAACTCCGTTCCTTGAAAGTGCAGAAAATACTTTCAAAGCATTGGCTAAAAATAATTTTTTGTATGACAGTTCATTAGGAACTTTTTCAGGGACAAGATGGTGGCCATATACATTAGACTATCTACCTTCCATGCCATGTTACATGTCAAACTGCCCTAAATATGCGTATCCTGGTTTATGGGAAATCCCGTTGAATACTTTACAGTGCGACGAATCGTTTAGCACATTTTCAACTATGGTTGATGAGTGTAAAAATTTGCTTACCGAAGAGACAACGTATAACTTTTTAATGCTGAATTTTAAAATCTTCTATGAAGATAATAAACAACCGTTTCCAATATTAGGTCATTCAAATTGGTTCGACAACGCACCTTATAGAAAAATAGCTGTCGCTAGATTTATCAACGATGTCCGAAAGTTAAATGACGTGTACTTTGTAACAGCGCAACAGACAATTGAATGGATAAAATCGCCAATTGGTATCGATAAAAACCCATTTTCGTGTAATTAGAAG
>evgR013745c0g1_i01_Hydra
AAAACATTTAATAACATCAAACAATCCCAAAAGAAATAACACTTTGCGTTCTAAAACATTAATGGATTATTGCTTTCGTGGATACTCCATTTGCATGCAATCTTTTCTTTTTCTGCATTGCATCGGTAGGAAAAGAGTCTCAGTGGCAGGCATAGCTCCCGTTATGCATGGGAATACTAGAAAAACTCCGCCCCACGCTTGCTACAAAAACGATGGAACTGCTGTTGTCTCATTCTTAAGTAATTATGTCAAAAATAATGCTCTAGTTTTGCCCGGAAAAACAGCAGGCATTCGAGACCACAGGATGCTGCTGCTTCCTTCATGGGCAACCAAAAAGGATATTCATGACAAATATTTGGTTCATGT
>evgR013750c0g1_i01_Hydra
CCTATTTTAGTGCAAAAAGAAAATAGAGTTATTTACAAAAAATAATGACTTGCAAAGTTATCAAAGGAAGCGGAATTTGCTGGTATCAAACTGAGATAGTAATTCCTCCCAAGTGCAGAGGTATACACAAGATAACAAATGTAATAAAAGAACTTTCTTTTCTACACGACATTAAAATTGGTACTTGCTGTGTACTATTGAAGCATACATCTGCCAGTTTATCTTTGAATGAATGCTGGGATGATGATGTCCGCGCTGATATGGAAATGATTTTGGATAAACTTGCTCCAGAAAGTATGCCTTATAGTCATACCCAAGAAGGTCCTGATGATATGCCTGCTCATGCTAAAACAAGCCTAATTGGTGCTTCTATTACAATACCTATAACTGATGGTAAACTTAATTTAGGTACTTGGCAAGGAATCTGGTTATGTGAGCATCGTAACCATGGTGGGCCTCGTACACTAGTTGTCACAGTACAAGGATGTAACTAATGCTTTTTTACGTTTCATAATTATTTTTCTTATTTAATTTTTTAATGCATAAAATATAATTTCAAAAAGATGTCTAATTGTTATGATTTGATGC
>evgR013751c0g1_i01_Hydra
GATGATGATGATGACGATGACAATAATGGTTTGTATTATAAAAATTCCAAAAGGTTCCATAGTCAAGGAAGAACCCATTTAAATAATGCTAATGAAAGGAGTCTGAAAACAAAAGAACCACCTCTAGTTGTTTTACAAAGTGTCAATGAAAACACCAAAAACCAGCTCATTAAGAAAAAGTACAAAAAGAAACATTTACAAAGGCACAGTATTTATGAAAAAGATTATGTTAAAAATGTTAAGCCAGCTCCACATGCCCATGATTTTGCTCGCTTTATGACTACACCTGCAGCTGTACCTACTTTGCAACCTCAAGAATTCACCAAGCCACATCAGACACAAGACTACATAACTGTACATAAATATGAAAAAAAGTTTGTACTTCCAGTTCATGATGATATTTCACCAGATGAAGAACGGTATGTGGTTATTCCGGACTATAAAACTAAATTTTTAAATTTGCT
>evgR013769c0g1_i01_Hydra
ACCTATTTTAGTGCAAAAAGACAAGATTTGAAAACTAGTAAATAATATAATGGGAGCTGGATCATCAATCGATATACCTGGTGGGGGAACTGAAGGTTACCATGTGCTTCGGGTTCAAGAAGGATCACCTGGATACAAGGCTGGGCTTGAACCATTTTTTGATTTTATTGTTTCAGTAGAAAACCAAAGATTAGACCAGGACAATGAAACATTGAAAGAAGCCCTTAAACGAAATGTTGAGAAGCCAGTAAAAATTCTTGTATACAGCAGCAAAACAAGAAAAGTTAGAGATGCAAGCATTACACCAAGTAATCTTTGGGGCGGCCAAGGACTTTTAGGTGTTAGCATTAGATTTTGTTCATTTGAGGGTGCTGCTGAAAATGTTTGGCATATACTGGATGTCCATCCAAACTCTCCTGCTGACATTGCTGGTCTAAGACCACACACAGACTATATTATTGGTTCCGATACTGTATTACATGATGTAGACGATTTTTTTAGTCTTGTAGAGCAACATGAAAACAAGGCTCTTAAGCTTTACATTTATAATTCGGAAACAGATGGTTGTCGCGAGGTTACAATAACGCCTAATTCTAAGTGGGGAGGCGAAGGAAGTATTGGATGTGGAATTGGATATGGTTATTTACATCGCATTCCTATTCCTGATCTTACGAAATCATTGCACCCGCCGTTCATTGCTGCAACAACAGTTAGCGTAACTGGTTTTTCAGAGGTGCCGTTACACGTTTCACCGCTTCCACTTAATACATCAGTGACGCAAGATTCATTAGGCGGCATAAACTTTAGTGGTACTATACCGAAAACCACATCAATAATGACGCCACATATATCGTCAGGAATATTAAATGGTTATTCTGCTCCTCTTATCGAAACTCAGTCTCCAACATCTCATCCCCAATATGAAACGCCGAATGTGCTTTCAACTTCACAGCAAGTGCCTCATCCTAATATTTTTTCATCGACTGCGTCAAATGCAACGGCTTCCAATTATCAACCGGCAACGCTAATAGCTAGTAAGCCTGTAGGATTAAATCAACCAATCACAAATCTAAATACAACTAACACTTATGCAGCATATAACCTTAACGAATTATCACGAAGTCCTGCAGCACACCCGCATATTCCACCAATGGAGATATCGCAAATCAGTAACGCAACAATTATAGCTAAACCAGTTCCATTAATGACAGATATAACACTTGGTGCAACTCCGAGTAACATACCTGCAATTTAGTTATTTATTTTCTGATATTATTATATATGAAATTTTATTTAATGTATATTTTTTTTGAGCGACTTTTTCTTTTTTCTTTCTAAAAAAAAGTTTATCTTTTTGCTTGAATTCGAAGCGAATTTTTCAGCATTATTGAAAAAAAAAACACTTTTTCAAATTTCATTTTAATTTCAAAATATCCCCTCTTATACGTGATAAAATATATATTTCATTGGTTCTTAATATGTAAGTTATAACGCAAAAAGTCTCTTTTTTCCTCCAG
>evgR013802c0g1_i01_Hydra
TCGAAAGCGACGCTGGGTATTGTTACGAGTGATAGAAATCATAAAACTGATTGGTAAATGTGGTCTGAGTTACAGAGAACATGAGTTTGAAGCAGCATACACACGATCAATCACTTGATCATGACAACTTTTTAGAAATGGTGTTGCTTGTAAGTAAATACAGCGTCTGCCTAAACAAGCATATAAATACTTGCATTGAAAAAAGTGCAAAACAAAAAGGAAGCCTAAAAAAAATGAAGTACTGGAAAGGGTTCCATGGTAACTTTCTTCTCAAAAAGCACAATCCAAAAACTAATTCAAATTATGTGTCGCTTGATACAGAAAACCATTGCTGAAGAAGTTAACACAGCTGGTATGTTTTCGCTGCAAATTGATACAACACAAGATATAACAATTAAGAACCAATGCTCAATTGTTATTATATATGTAACTGACCAAATTTATAAAAAGTTATTAACAATGGTTGCATGTGAATCTTCTACTGGAGAATATTTATTGAGTTTAGTGTTGCAAACTCTGCATACATTGATCATTGATATACTAAACTGCATCAGTAATGCAACAGACAGAGCTGCCAACATGAATGGAGAATTTAACAGCTTGGATGTTCGAAGAAAATCCCAACCAAATACATGTTTGGTGTTATGCTCATGTTCTTAATTTGGTTATGTCAGATGTTGTAGTATCCTACTTTCTAGTATTTCACTATTTAATTTATTGAAAGACTTAGCAGTTTTTATTCGTGGATCATATCATAAATGGACATTTTGGTGGAGGAAAATCAAAAAGACGCTATCACAGGATTTCAACTATTAGTGAAACAAGTTGGTGGTCAAAGGTTGCTGCACTTCAAAACTGCAAAAAATTCAAAGTTGCAACTATTGCACATAGAATTCATTAGTGGCAAGCCAAGCTGAAGCCGTTTATTTCCAGAGC
>evgR013803c0g1_i01_Hydra
ACAATCAAAAAATATTACTCTTAAATACTCTACAATCGTCGTTAAATATGTTGTTAACATACAAATGATTTCAAAAAGATTGTTCTCTGAAAAATACGAGCTTCGAAATAAGTTACAGGAAAAGTTTGATGCAATTAAACATAACAGCAGAATTAATGTGTATGACATATCAACATCAAGTGTAGCAAGATTTCAGTTATTCTGGGTATCTGTTTTTAGCTGTAGTAATTATGCTGATCAATGTAATCCTGTAAAACAACACGAACTTTATTCGACGTTTGTGAATACTAAATATGAAATTCATGATCATGTTGTTAAAGAATTTTCTGACACGTTTAATGTATTAAGCATTACCGAAAAAAACAATTGTAGCAAATTATTACTCAAAAAACCTCTAATTTTTAAATTCAAAACTTGCCAGCGACTTAATGCTACGATACCAACGTTTTCAGATTTTGGTCCAGATGCTAGAGACTATTTTGAACTACTTTCAGAGAATGGAAGTTCTTTACCAATATATTCCTTAGTGCAATTTAATAAAGTAAATTCAGTTATATATGGATTTGCAATACACGATCTTGCATTCAAATCACCTAAAGAGCTGAGATATGTGCTCAGATTACACAATTGGAAATATAATTATTTTGATTTAAATTTAATTGTTATGTTACCGACCGAAAAGCCACGCTACAACTTAGTTGTAAGAACAGAAATACTTAGTTACTTCAACACAGAAATTACTGATGTTGACATACTTTCAAAATTTGTAATAAGAGTTTCTTCATTCGTTTCAGTTTATTTTGGAAAAGTATCGGCTGATACAATTAGTATAAAAAGTTATGCTCGTCGAGGTGACAAAATCCCTCAGAGATTAATTATTGAGTGGTTGAACTGCTCAAACATAGAAAGTTGTAACGAAG
>evgR013812c0g1_i01_Hydra
CATTGGTTCATACCACGAATGCTGTGAATCCATTGGTTCATACCACGAATGCTGTGAATCCATTGGTTCATACCACGAATGCTGTGAATCAATTGGTTCATACCACAAATGCTTACAGCTCTAATCCAGCTTCGCCTACTCAAATAAGCAACGTTCCTGTTTTTAACTTTCCATCTAGTTTAACTATTCCATTAGCTATAACTAATGATAATTCCTCGGAAGTCCTTGGTGCATCGGCTATTGACACTCCCTCTGGTAGCCCAAGTAGAAACGGAATGTCAGGAATTTTAAAGATTCTTAAAAAGAACAAATCAAAAAGAAATCAGCATCCTTTACCACAGCCACCTTGCTTCCCGCCACAAACAACACAATTTCCACCTTCTTATTATCATTCATCTATGCCTGCTCCCCTACATCATCGTTTATCTCGTCCGATGCCATTGGTTCCTGCAGAACCGCCTCCCCCTTATACACCGCTTTCTGAAAATATAATGTATCAAACAATCAACGAGCAGACTCCTGCTAACGGAGAAAAATTTCAAGCTGTTGCAACTTACCCAGCTGTTCACGCGGAAGAACTTTCTTTCAGTGTTGGCGATATTATTTATGTAATTACAAAGAAAAAAGACGGTTGGTACAAAGGCATGTCTGCTCGGACTGGTTCCGTAGGACTATTTCCTTCGTCATTTGTTAAAGCCTACTCGTGATACATGTCACGTACTTTATTTACTCTCCCCCTCCCCAAATCAAATTATCAATTATGCTTTGGTTATCATTGCAGCTATCGATTTGTTTGTCAAGTTATTGCTGATAATATTAAAAGCGTTTCAATACTGAAATCGCGATAAGTTTATTTTTAAATTAATTTAATTAATTCTCCTTAGGTAATCCATAGTTTTCTCTAAGAGAAATATTTTTATATAAGTTATTTTTAACTAGTGTTTTAGAGAAGCATTCACGTTTAATTTATTTTCTTTCTTTTTTCTCGCAAATTTGTCTTCGCGTTTTATGCGAAGGCTTTTTTGTTTTTTTGTTCAGTGAGATTTTCGTTTTGGTTTTGGTTTTTTTTTGTTTTTATTGCTAAATTTAACAGTTAAAAGCTTTACTTGCTTAACAACTTACGTTAACTGTAACTACGTATCAATTTTATATTTTTAGTTTGAATTAAAACAATAAAATATATCAATATTTGAAAAAAAAAA
>evgR013846c0g1_i01_Hydra
CTCTTTCATCAAGTATTTGAGCGGTATTTTACGGTTTATTAAGAAAAAACTTTTTAAAGTCAAGAAAAAAATTTTCTTGGTGAAAGGAGAAACAAAACAAATTATTATTTTTCATCAAAACAGAAATGATGTATGGTCAAAGTGCGCAAGGATTACAAGTTGTACCATCCTTTGACTTTTCGACCACATACAATGAAGGCTACTGGGATGGTTTGCCCGATGATCAACGTCATTATGAAAGCGAAAAACAAAACAACTACTTTCAATACCCCATCACTATTAACAGCAATGAAAACAACTATTTAGGTATTTCAAATAACCATCCGTATTACTGTAACCCAGCTTATGTTCCAAGCTATTCTCCCCTACAAAATCATTATCTAAACTCTCCTGGAATTTTTGAACAACAATTTACAGAAAACTTTGAACATCAACATAGTATTAAACATGAAGCAATTTTAAGCCGAACCACTCATAATGTCCTTGAAAGACAAAGAAGAAACGATCTTAAAATTCGGTTTAATATCTTGAGAGACAATATTCCCGAACTTGCTTCTAACGAAAAAGCTCCAAAAATTCAAATATTGAAAAAGGGGTTAGAACATTTAAACGAGCTTAAAGCTCAGGAACAAAAGTTGCTTGTTGATTTAGAGCTCGAAAAACAACGAAGAAAAATTTTAATACGTAGGTTAAATGACCTTAAAAAGGGATTATATTAAAACTCTCTAGAGATTTCTACGTTTTAAATATTTTAAAAACTTTTTGAGCTTTCCATAGATTTTTTATATTGTAAAAAGGCAACCAAAATTTTTAAATTTTTATTTATTTTGTTAGATATATTTGTACAAATATTTTTTAGACTTTTTACGATATTTTTAAATATCGTTATTAATATTTAGTGAATGGATTGGGTTGACTCTTGAG
>evgR013858c0g1_i01_Hydra
CAAAAAGAGAGTTTTAATAATGAGCAACTATGAATTGATTGCTAATCAACCAGTGGTCATTGATAACGGTTCAGGTGTTATAAAAGCTGGTTTTGCTGGGGACCAAATACCTAAATGTCATTTTTCTTCATACATTGGTCGACCCAAACATGTTCGTGTCATGGCTGGTGCACTCGAAGGGGATATATTTATAGGACCCAAAGCTGAAGATTATCGTGGTTTACTAAACATTAAATATCCAATGGAACACGGAATAGTTCATGATTGGAACGATATGGAAAAGATTTGGCAGTATATTTATTCCAAAGAACAGTTATCAACTTTTTCTGAAGAACATCCTGTGCTACTTACAGAAGCTCCATTAAATCCTCGTAGAAACAGAGAAAAAGCAGCAGAGATTTTTTTTGAGACCTTTAATGTTCCAGCCTTATTTGTTTCTATGCAAGCTGTATTAAGTCTTTATGCAACAGGTAGAACAACTGGTGTTGTCTTAGACTGCGGAGATGGCGTTACACATTCTGTACCAATATTTGAGGGTTTTGCTCTACCAAACTCTATAATGAGAACAGACTTGGCCGGTCGAGATGTTACTAGATACCTGCAATTGCTTCTTAAACGAGAAGGATTTTCATTTAAGACCACAGCTGAGCGTGAAATTGTTAAGCAAATTAAAGAGCAAGTTTGCTATTTAGCGTTAAATCCACAAAAAGAAGAAGGTCTTGATGGAGAAAAAACACACTATGTTTTACCAGATGGAAGTACATTAGATATTGGAGCTGCTAAATTTCGCGCACCTGAACTTTTGTTTCGACCAGATTTAATTGGAGAAGAATTTGAGGGTTTACATGAAGTTTTAGAATTTGCAATTCAAAAATCGGACATGGATTTACGAAAAACTTTATATCAAAATATTGTTTTGTCTGGTGGATCAACTTTATTTAGGGGATTTGGTGACAGACTTCTTAGCGAAGTAAAGAAAGTAGCACCTAAAGATATTAAAATTAGAATATCGGCGCCACAGGAGCGCCTCTATTCTACTTGGATTGGAGGATCTATTTTAGCTTCATTAGATACATTTAAAAAGATGTGGGTTTCTAAGAAAGAATATGACGATGAAGGACCTAGATCTATTACAAGAAAAACTTTTTAATAAAATTTACTTTAGGTATAAGAGTTTGTTTTGTATAATTGAAATCATTGGTAATGCTGCAACAGCAATTGGTTGCCGTATTACCAATGCAATACTGTACTGCAACCAACTATTGTTGTGGTAGTACAACATGGCTACTTTTAAAGGATTCTGTTATGTATCATTTTTATTCTCCTGATTTTGTAAAGGCCAAAGTTTAAATCTATATTTTTATTTAGTATTTATTTAAATGAAATCTAGGGAAAC
>evgR013868c0g1_i01_Hydra
ATTGTGGGACATTGCCGGACAGGAAAGATTTGGTTCCATGACGAGAGTTTACTATAAGGAAGCTGTTGGTGCTTTTGTTGTGTATGATATTACTCGTCCTCCAACATTTGAAGCTGTACAAAAGTGGAAGAGCGATATTGAACATAAAGTCTTCCTTCCAGATGAACGTCCAATCCCTTGTGTTTTACTTGCCAACAAGTGTGACCTTCCAAGTGCTGTGAGTAAATCAAAGGAAGAAATGGACCAATATTGTCAAGAAAATGGATTTATTAATTGGTACGAAACCAGTGCCAAGGATGACAAGAATATTAACGAGGCTGCCA
>evgR013879c0g1_i01_Hydra
CGATAAACAATGACAATTATTTTGTTTATAATAACTAAAGTTTCAATACTTTCGTTTGTTAAAGGAGCATGTAATTCCAAAACTGGACCAGCTGGATCGAATGATTGCTTTTTGGCTTTACCATTTTATACAAAGTATCAATGGGGCACATGTTTATCAAATGCTTATATTAAACAAAAGTCGGGAGGGAACCATCGATGTCGAGATAATACAGCAATATACTGTTACTACCAATGCATGGTTGAATTGCATGATATTGACGAAGGACCCGTTTATGATAATTGTGCTTGCAATACCGGTGTTTTTACTCCGAGCCCAAAGTCAACATTACCACCGTCGTGCTATAGCCCAACTGGAGAAGACTGTTCGTGGTACCGCAACTGTCTAGCCACCAAGTTACCTTGCACTGGATCTCAAACGGAATATGCGATATCTTACGGACAAAAATTTTGTTCGCTTTACGATGATCGTAGATACGTATTCTCTCCAAAAGCACTCAATTGGACTAATGCAGTGAGAAAATGTTTACAAGTTAGTTTGGTGCCATTACTTAGAGAATATTTAACAGTTAGTTGTGATGATATTTTTAATACGGCAATGGCATCGCACCCCAGTTGTTATTTATCCCCGTATCAAGGATTTTCAGTTTGTAATCTTTCTGCAAATGATTGGTGGCAGATATTTTGGACAGTAAAAAGTGCTTTTAAGGAACAGTTTGTTGCAACTCTCTCGCAATCTCTTGATACGGCTATGGGTTGTGCAAATAATGTTTTGTTGCCAGAAATCAAAAACTATTTAAAAAACTTTCGGTTGAGAGTTAAATTATTTTTCACAAATAGAAAAAAAAGATCCATTGGCAGTTATACATCTAATAATGAAACTGCATTCCGAGTCCTCAAAATAATATCACAACAGTTGAATTGGGAAAAATATTCAATCGATTGGTACACGTACACTACAGACTCAAGTTTTAGAAATGACGAATGGAATGTGTTGTTGTTACTGGCTGATACCAACGCACTAGGATTGACCAACAAAACTACCAGGGTTACCAACTCAAATATAACAGATGCAATCAATGCACTTTCAGCTGCCATTCAAAACGGAACCCTTCAAATTAACTTACCCGGTAGCCCAGGTGTGGTTTTTGTTGGACAGTCTCTAGGAAATTGCCTACAGTTTGTCAACGGATCTTGTCTTGCAAATGATACATTAGCTGCTTTGCCAGATAACTTCACACTGACGACATATACTACTTTACAGCCTAACTTTAGTTCTTCGGTATCGTATATAATTGTGAACAATTCGTTGGTTTTTTTTACTGGCATGATTGTTATTTTCCGAAATTGTTATCCCTGATGTAGTGATGTAAATTTTTTTATTTGAAATTTAAATAATAATCCAT
>evgR013898c0g1_i01_Hydra
ACTGCAGATCTAACAACAACAACTGAAGATTTAACAACATCTACAACTGAAAATCCTACTACAACTGAAGATCTAACAACTACTGATGGGTCCATAATAACCACAACTGCAGATCTAACAACAACAACTGAAGATCTAACAACATCTACAGCTGAAAATCCTACTACAACTGAAGATCGAATAACATCATCAGCTAAAAATCCTACTATAACCACAACTGCAGAGGCATTAACAGTAATAAACATAACCACAAATAATAATATAACTTCTTATTTGATTGTTCCAAACATTGCTAAACCATTAGTTCAAAGAAGAATTCTCACCATATTACCAGAAATTGCTAAAGAATATGTAGTTTCCTTTGATTACTATCCTAATTCCTTTCCTAATAGTAGTTGGTACAGTGTTAT
>evgR013916c0g1_i01_Hydra
TGTTTTGGCTCATCTTCCGAAAGAGTACGAGTTATCGTTTGAACTCAAGGCAACTGAATTTCTAAACGAATGGGCAAGTATCATTCATTTGACAATCGGTGGAGATCGTTTTATATACGGCGATAGAACGCCAGGAGTCTGGATGTCACCTTCTAACAAATTGTATTTTGCGTCAGCTATAAGTGGCTTAGCAAACAACAACTATGATTCAACTACAATAGTTCCATTAGATAAGTGGATAAAAGTTAAAATATCACAGAATATGCTCTTAAGTAAATTTATGTATATAATTGAAGTGGAAAACGAGATAGTATACAGTACACAAAAT
>evgR013919c0g1_i04_Hydra
TAAAAAAATTCAGATTTTTTTTACTTATTCATAAATGCATTGTTCTTTAAAAACGTCGAACAGCCTTTTTGTAAAATATACGAACGTAACGTAAGCGTAAGTATCTTTTCTAAAACAATAAAAATAAAAAGAAGATGAAATAATGAAATAATGTTAAGAAAGCGAAAACATATCTGTCAATCAATTTTAGTTTTTAGTATTTTTTCCGTGTTTGGGATATACGTGAAAAATGCGATTAAACCAAACAACAACCACCATAATACCAACAGTCGTAAATATATCAACGACAATCGCGAATCTAATATTTTCAGTAGAAAATATAAAATCGACAGTGGTATGTATGAAAATGCTGAATATAATATTCGCCGTGAACTTAATAATGACAATCGTGAATATAATAAAAACAATCGTGACGGTTCAGACGATAAAAACGATGAGTTCGAGGATTCCAACAATGACCAACATTATTCACAAACACACAAAAATAGAAACAATAAAAAGGCATGCATATTACCGTATCTCAATCCATTTAGCCAAGACATTCAACGTTTTATAAAAAATTCTTCTATGAATTGCTCTGTAAAACGTTATGGAAGAATAACAAATGATGGATGGTTTCTTTTACAATCAGATGATAAAAGTAATTATACTGTTTCGATTAGATATATTTACAGAAAAACCGACTTTTTTGTACAATTTTCAAACCCTATTCTAATAAAAAGCGAAAAAGATGTCAAAGGGTTAGGGTTAGACATTGTATTTTTAATGATAGACTTTCAATCAAATGCTAACTTTCATCGTCAGTTGAAAAAAAGTATGCCAC
>evgR013920c0g1_i01_Hydra
TTGTGGTATTACAATTACTAGAAAGAGATATGTTGAAACATTGAAGAGTCACATTGATTACCTTAACAGTGTTGATACTACTATCATGGGTTCTAGAAATGGTCAAGCTTCATTATACTTGTGGTTGACATTGAGAAAGAAGGGCACTGAAGGATTTGCTGCTGATGCTAAGAAATGTTTGGAAAATTCCAAATATATGGTAAAATTATTGAAGGAAGCTGGAGTTGGATGTTTATTGAACGATCACTCAAATACAATTGTTTTGGAACGACCAATGGATGAAGAATTTGTCAAGAAGTGGC
>evgR013934c0g1_i01_Hydra
AGCAAGGTTTCTGTTTTCACTGAATCCTGCAATTCCTCTCCAGCTCCAAAGATTTATCTTTCACCACGTGTACATACTCCAACCGCTGATACTGAAAACGTTATTGGTTCTATTGAATCAAGTGATTATCCACGTGGTCTTGTTCAATCCGTTGTTGTTGATTCTCTTGCTGATGAAAAGTTCTACCTTGGTGTTGGTAATATTGGTGTCGCTTCTGATTCCTCTATCTATGTAACAACTGGTGATGATGCCAGACCATCCGTATTGGACAATAATCTCGTTGGCGCTGGTATGCAAGACTCTCAAACTGTTTTGGTCATTCAACCTGCCAAGCATTCAGCCAGACAGTCACAAGGTGACTTCTTGACCTATAGCGTCTACACTGCTGAAATTAGTGGTG
>evgR013934c1g1_i01_Hydra
CACAAGGTGACTTCTTGACTTATAGCGTCTACACTGCTGAAATTAGTGGTGAACAAACATTTGCTTCGGTCAACATGGATACTGCTTGTGGTGTTAAACGTAATGCTGCCTTGGTCATGTCAGCAACTGCTGTTGGTACTGATGCTTTGAAGGTCAAATTCAGCGTTAGCAAGAACAAGAAATATATTATTAACGTTTTGGCCAGAGACAACTACGGTCTTTCCTCTCCATATAAGCCAGCTTACATCATTGATGGTGTTGTCGTTTATGAACCTCAAGGTGGATCTGTTGCTGCTGCTGTTATCATCTCCCTCATTGTTGTCTCAATCATTTTGGCAGTTGGTGGTTATATTGGTGGTGGTATGGGTTACAAGTTCTACAAGGGACACAGAGGTATTGAAATGATTCCAAACATTGATTTGTGGAGAAAGATCTTTACTTGTGGTAGATCATCCAG
>evgR013951c0g1_i01_Hydra
AAATGGATTCACTGCTTTGAAGGTGTTACTGCTATTATCTTTGTTACTTCACTGAGTGAATATGATCAACGTTGTTATGAAGATGATTCAACTAATCGTATGCACGAATCCTTGTTGTTGTTCGATGAAATTTGTAATTCTAGATGGTTCATGGATGCTTCCATTATTTTGTTCTTAAACAAAATGGATTTGTTCAAGGAAAAGATTAAGAGAACTGATCTCAGTCAATTCTTCCAAGAATACGAAGGTGGAAGAAACTTTGAAGCTGCTGCTAATTTTATTCGTTCCAAATTTGAAGAAAAGAATCGTAATCCAAGACACAAAACAATCTATGCTCACTTTACATGTGCTACTGATACACAACAATTACAACAAGTCTTTGATGCTATCAAAGATATTATTCTTAACAAGACTTTAAGAGAGCAAGGTCTTTTCTAAATTCAATAACATGTCACTCTTTTAATCTCCCACACTTCGATCAATCCACAACAACAACTCAATTGTCCAATTTTGTACATAATAGTTCTTCTTAACAATAATAAAAAAAACA
>evgR013957c0g1_i01_Hydra
TTCCACTAATTACTTATCTTTGTCTCTAAGCGTTTTTCGGATTTTCCGGCCATAAAATATGACTACATTACTTGTTAGTTCTTGATTAGTCAACCAATAACAATAATTCGTCGTCAAACGCAAGTACATAAATACTACACTTTTTATTTTATAACTTCAAAGTAAACAAAGAAGCAAGTTCAAAAAAAACATTTTACAATATAAAAAAGTAAAGTAAAAATGTTTTTCAAAGTCATTTTGTTTACCTCGGTTATCATAGCAGCAACTGCCTCTGAGCAGTGGGATTCTGAAGATCAAGTTGAATCTCCCAAAGAATATCAACATAAGAGCGGACATGAACACCAACACGGATCTAATGACAAATATCAATCCAAGAATGGCTACAAACCAAAAAAAGTCTATTGCTTAGGTGTTGACGTCACAAACGACTGTATATATCGCGCTGATAACAACTACGCTATCGATGACACACCAAGATCAGGTTATGTAGCTTGTGTTAACAAGAGTCCCATATGTATGCCGTGTCCGCACGGAT
>evgR013969c0g1_i01_Hydra
CTTATATGCTAATTTATTTTCTGATGTCCACATGACACATATGATACTTAAAGGTATGCATGAACGAGGCCGTGGGATTGTTGTCCATATGTCTTCTGCTGCTGTATATATAGAAAGTCCTGCTTCATCTTTCTATATTCCAACTAAATCATTTATGACAAAATATGTTAAAAATTTACAACTCAATATGTCTTGTGTTGAACATCAATTAGTATTGCCTCTCTATGTAGCTACAAAACTGGTATCAAAAAATCCTTCTTTTTATGTTCCAACAAGTGAAAAGTACGTTAAGCAGTCCTTGCGAACTCTTGGTCTTGCTAGAGTCACTCACGGTTGTGTATCTCATGAGTTTCAAGCAATCTTGTTGAAATATTTACCACAATTTTATATAACCTATTTTTTGAAAAAAGGTTTTAAAAAACAAAACTAGAGATACAAATATTTTTAAAGTAAAAACTAAATTTGTTTAAATAGTTTGCAATGAATTTTTATATTTTAATGAATTTTTCAAGAATCTGATTTAAAAATTTTTTATTGTATATATTAAACTATAAAAATTATATACTCATACTATAAGAGCC
>evgR013973c0g1_i01_Hydra
AATATGAAACTGACGCAATCAGTAGTAACATTTTTTGTTTTTGGTTTTAACTTAATTGTGGTAAAAAAGTCAATTACAAAATCGCTTTCACAAAATCGATGGAATTCAGTGAAGAATATTGCAGTGAACAACAAAAAACCCGCACAGTTTGATGACATTCAAAATTACAACTCGTTAACCAATAATGCTAAGCTTTCATCATTGCTTAAAAATGTTATTTTGCAAAAAGGTGGTAGTTCACTACGCAAAAAACGAAATACACTTTCTTGGAGAGTACCAGATGGATTACCAACAGGCACATCAATTAGTTTTTGTGCTTCTCCTTCGTGTTGTACACCTAAAAAATCTTTGACCGCGGACGGTGGATACTTTAATTTATGCAGGGAATGTTTACACCAAATCATGCTTCCGTTAAGATTTACACCCCGTCTACATAATTTTGTTACTTGTCAAGATTTAGACTTCACCGAGTCTTGTTTATCTGGAGAAGGTTCATGCACAACAAATATGGTACCATATTCAGTTAAAGATACCTATACAAATCAAAGGTTAACTTTTGTTTTTGGAGGTTCCTGTTCGTGTGAAATTCGCCGAGATTCCGTTTTTACATCTTTTGTATAAAATTTTTAATTTTTTTTTTAAATTTTTATCCTTAAAAAAAA
>evgR013996c0g1_i01_Hydra
TGTATGTGGTTGGCAGCCGGTTGTTTAGGAAGTATATATTTAGTAAATGATCACATTTCTTTTAACAGTACTTTTCTGTTGTGTTCGGCAAGTATAGAAGAACAAATATCAACTGGTTTAACATTTGGTATATTTACAATACAAACTTTGGTACCTATTACCTTGTTCACTCTGATGTATGTTCTACTTATATTCGAGTTAGACAAAACTGCGAAGTGCAAAGGATTTTCAGATTCTCATAGAGTTGTAGCACTTCGCGTACTACAAAATGGTAAAACTATTAAAATGATTTTTGTTGCCGTCATCATATTCTACATTTGCATTTTGCCGTCAAATATTTATTACTTGTGGTACGAATTTAACCAAAACGTTTTATCGATTGAGAAGGCAAACACAATTTACGATATTCTCTTAATGTTAGAAATATGGAGCTCAATGGGCAATCCGGTCATATATAGCCGTTTAAGTACAGCATTTAGACGAGAAATCTTTAGAATCGTTTTCTCATGCTGGTATAGAAAATCCATAAGTAGACGCAGATCATTGTTAAGCACGATTCGATGTTCACTAATTTACAACTCAACAAAACGTTTTAGAAATCGTTCTAGTACTAAAACAAGTAACAGTTCATTTGCTAAAACAGGAGTGTCAATTCTTCTAATGAAACAGGAAGCAGTTAACAATGAAAATATGAACTGCACTTCAAAAAAGAATAAAGTAACCATATAGATTTAAAAAAATAATCAATCTCCG
>evgR014004c0g1_i01_Hydra
CTAGTTACCCACGTGTTGACCTCGCTTTCATTGGAGGTACCGGTCTTTCAAAGGCTGAAGGTCTTGAAAACGTTGTTGTTGTTGAACCATTTGATACTCCATTCGGTAAACCATCATCACCAATTACTATTGGTACTATCAATGGTTTGACTGTTGCCTTCTTGTGTAGACACGGTCTTTCCCACACTGTTCAACCAACTGATGTCCCATATCGTGCCAACATTTTTGCTTTAAAGTCACTTGGTGCTAATAATTTGGTATCTTTTGGAGCTGTTGGTTCTTTGCAACAAGAATTGGAACCAAAGCACTTTGTCATTGTTGACCAATTCATTGATCGTACTAAATTCCGTCAATCTGAAAACACCTTTTACGATAGAGAAGGTTTGGTTGCTCACGTTTCATTTGGTGAACCAACATGTAAGACTTTGTCAAAGTTAGTTTTTGATGCTTGTAACAGCATTGAAACCATTCCAAAGACTCACTCAACTGGTACTTATGTTTGCATGGAAGGTCCATGTTTCTCATCACGTGCTGAAAGTAAAATGTACAGAATGTGGGGTGGTA
>evgR014026c0g1_i01_Hydra
CTAAGATATTTTTTTTTTGTATAAGTACACGAATAAGAGAAGGTTCGTACAAAAACTCGTCAAATACAACAAATGTATTCTGAGCTAATAATATATATACATCCCATCCCAACAGATTTTTTCTTTGACAGATTTATTAAGAGAGATTCTTCTTATAAAACAATGATTAGACGAGTTGGTATTTTGAAAAAATTACTCAAAACTATTCTTTTACTATTCTGTATTTTTGTTGCCGGTTTGAGACGCCAGCTAGAAAAGGGGGAGATGATGAGGAGAGAGGGATTGTCCTTCCCTCCAAAAAAACAAAAACTTTTTAAATCTTCAATGAAAAATATAAATAAAAAATATATAGATATCATCCAAAGACCCACTTTCCGAGTTGTTTTCTTAATTGG
>evgR014030c0g1_i01_Hydra
AACAATAGCGAAAAACAGTATCCTTCTTTGCCTGTTGGAAAAGATATTCCGTAAATTCGTCGCGTAAAGTAACTGAAAGAATAAACGTTATTAATTTACTAGCATGCAAGAACTATGATTATGTATGAGTATAAAATGAATAATGGCTTCGCAATAATATTGATAATTTTACAATTTTTATCGACGAAAACTGAAAGGATATTGTCTACTATTTCATCCGAGGTTGGAGCTGGAAATTATACAGTTTATAAACTTGAAGAACCAGGAGATGTAACTATTGTTTTGAAAACTTTATTTGGAGATGCTGACTTATATGTCTCTGAGTTTGAGAGTAAAGCAGATTTTCTAAATTATGATTTGTCATCTACAACGTGTGGTCTTGACATTATAGGTATTCCGAACTATTATTCAAGGCCAACATTTATTGCAGTATATGGACATATTCACTCTCCCAAGTCTAAATATAAACTTACAGCCATTATGAACTACAGTTCATGGGACGACTTTAATTTAAACTCTGATGAGAATAATTTTTCTGAAACACAATACGAAAATTCAGAGTTTAATGCCAATGCAGATTTAGCAGACATTAAAATGGACAATGGTTGGAGTTTTTCTAGTTTTCTATGGGGTATTTTAGAAATATTTTTAAAGGCTCTTCTAGAAGTTATTTTGTGAAAATTTGACCTTTCATATAAAAAAGTTTTTAATTCAATTTTGTAATAAATCAAGATTATAGAACTTAAATTTCATTTCTATTCTAAACTAAAATTAAATCAGCGTATAATTAAATCGAC
>evgR014057c0g1_i01_Hydra
CTTCTTCAGTTTCATTTTTGGATAACGAATCAAAAATTGTATTGATAAAAGACTCAAGTTCGAATAGATTTGAGGAAATCATACCTAAATCTACGTTATACGAATATATGGGCGTGCAAGTTAAAGATTTTTTTCAAAACGAAACCTACGACGATAAAACACTTTTGATGTTTACATCGTATAATTATACGACATTAATGGACATAATCTTTTATTTAGTTATTGATAAAAAATTTCAATCAATGAGTAAAAAAAGTCAGCTAGTATTATTTACTGATATGGATGAAGTATCGACATCTACCATGTCTAACATCATTGTTCCTTACGCTAATGTTTGGCTTTTTGATATTAACAAAAACAACTACAGAATGACATACAACAGTAACTCGGTAGGATATTATTATAACTCTTATATGGAC
>evgR014062c0g1_i01_Hydra
AACATAATATTGCAAACTGTTTAAACAATATTGGTAAGTATAATGAAGCTCTAGAAATTTATTATTACGTTGAGAAATTACAAACGCAATTACTAGGAACCAACCACCCATCGATATTAATTACAAAAAATAATATTGCATTTTGTTTAAATAACATGGGAAAATATAACGAAGCTCTAGTAATATATTATTATGTCGAAAAATTAAAAAATGAAATTTTAGGCAGTAACCATCCATCACTATTGGATACAAAACACAATATTGCATCATGTTTATACAACATGGGAAAATATAAAGAAGCTTTA
>evgR014071c0g1_i01_Hydra
GAAAATAGTTATGAAAATGCTATAAATACGGAAAATGAAAAAAGTTATAATGAACATAGTTTCCATAGCAATAGCAGTCATACTGATATCGAAAGTAGCAATGTAGAATCAATCAAGAGTGAAGATAACAGTAAAGTTGACAATAATGTTCTAAACGAAAAAACAAATCAAGTGAAATCTAAAAAAGATAAAAGAAATTATGCGAGCTTAGAAGAATTTTGTAATTTTGAAAATTCAGACGAAAGCTCAAGTGAAATGGACAACCAAATAAACAACAGACAAATAAAAGACCGCTCATTCACAACTGTTGATCAGAGCAGTGTTCGCGGTTTCACAAAAAATTATCGCTCGTATACAACAACCGACGAAATTAGTTTTAGTGATTCAATTAATGATCGTCCCAAATGTTCTGTGGAATTAAATAAGTTGAACCATGAGAGCAAACAAATAATACATCTTGATATTGCGCAATACCAACCTAAAAAGATAACCAACAGGCATAGAGCAAACATACGAAGAAAACTCGTTTACAGAAACACAATACACAAGTAACCGCCTTGCTGATCATCATTTTTAAAAAATCTGAGTAGTCTTGGGGTGATGTATTGTTGACAAAAATACTTGCG
>evgR014077c0g1_i01_Hydra
CTGTAATTTATGTTCTTGCTGGAACATAGTTTTTTAATATTAACAATTAAAATGGATCATCCCACCTCTGATCCACCAGATATTTACAAATGAAGTGTACACAAGAAAGACGCGATACATGTATTGACTATATAGCCATATAAACTATAGCTTATTTTTTTTGTTAAAATTTAAAATGTTTTAAATGGAAAGATCATCCTCTTTTATAAAGAATTTAGAGGAAGTTAAAGGAAAATGGGATCTTTTAGAAATTATTGGTGAGGGAACATATGGAGAGGTTTACAAAGCTGTAAGTAAGAGGGGTAACTTTGTTGCTATAAAAGTTATTGAAAATAGTCCTAATAAACACGACGAGATTCAAACAGAATCAAGTATATTACAAAAATTTGGATCCCATAAGAATATTGTTGATTACTTTGGCACATATGTTTGCAATAAACCAGGATCATTATCACAAATATGGATTATTATTGAGTATTGTGATGGAGGATCTGTTTCTGAGCTAGCAAAGGCTATCATTGTTGGTGGGCATCATTTAAGTGAGAATGTCATCAAGTATATTTTGCATCAAGCTTTAAATGGTATATACTTTCTTCATTCAAACAGAGTAATACATCGTGATATTAAGGGACCAAACATTTTATTGACGCGTAATGGAGAAGTTAAACTGATTGACTGTGGAGTATCCGCAGTTCTTGCAAACACGCTGAGTCGTCGCCACTCGTCAGTTGGCTCTCCATTTTGGATGGCAC
>evgR014078c0g1_i01_Hydra
TTTTATTTAAGTAAAAAGAATTTACAAAAAGTGGTTAAAAAAATACGTGCTGCAATTTAACTAGTTAGATAGTGAAATACTTCATCTTCTTCTTAAAAATGGCTACAAAGTTTACAGATTTAGGTCTCCACGACTGGTTAAACAAGCAGTGTCTAGAAATGGGCATTTCGAAGCCAACTGAAATTCAAATTAGTTGCATTCCAGAGATTCTCTCAGGGAGAGATTGTATTGGATCTGCTAAAACTGGAAGTGGCAAGACCGCAGCATTTGCTCTTCCAATAATTCAAAAACTCTCTGAGGATCCTTATGGGATTTTTGCTCTTATACTAACTCCAACTAGGGAGCTTGCTATTCAGATTGCTGACCAATTTAAAGCACTTGGAAAATCTATTGGTCTCAGAGATTCAGTAATAATTGGTGGTCTTGATATGGTAAAACAAGGAATGGAATTAAGTAACCAACCACATATTGTAATTGCAACTCCAGGAAGATTGGCTTCACATATATCAAGTGGAACAAAGTTTTCATTAAAAAAAATTAAATTTTTGGTCCTTGATGAAGCTGATCGATTGTTGGAAAAAAGTTTTGAAAATGATCTTGAGGTGATTTTTGATAACATCGCCAAAAAGAGGCAGACTTTACTATTCAGTGCAACAATCACAGATGCTATTAATCATTTGAAGGCCGTTGCAAATGATCCATTTTGTTACGAAGTTAAATCTGATTTTGCTACTGTTACAGAGCTAGATCAACGATATCTGTTAATACCATCTCAAGTAAAAGACTGTTATCTAGTTCATTTGCTTCAAAATTTTCCTGAAAAATCTGTTATAATATTTACTCAAACATGTCGAAGTTGTCAGGTGATATCATTTATGTTACGAAAAGTGGAATTCAAATGTGCTGGACTTCATTCTGTAATGTCTCAGAAAGAGAGGCTTTCATCTTTAGGACGCTTTCGTTCTGGACATGTTAAAATACTTGTAGCTACAGACGTAGCTAGTCGGGGATTAGATATTCCTCTAGTTCAACTTGTAATTAATTATAATGTACCTGCTTCACCTAAAGACTATGTTCATAGAGTAGGTCGAACTGCTCGTGCTGGAAGGGGGGGAATGTCTCTAACCCTTTTAACACAATTCGATATTGATAGACTTAAAGCTATTGAAACTTTTATTGGTGTTCAGATGAAAGAATTTGAAACAAATGAAGATGACGCTGTCAAACTTTTGAAGCCTATATCAATTATAAGAAGAGAAGTAGAAATACGGTTGAATGATGCAAATTTTGGAGAAAAAAAGAAAAATAATAAAAGAAAATTAGGTTTATTAAACTCAAACAGTAAGTCAAAATTCAATAAAAAGAAGAAAACCGATATATCTTTAAATATATAATTTAACATATTTAATTCTTTTGTTAACGATTTATAAAGAAAAATTTTGTTTTAGCAAAAAAAAA
>evgR014087c0g1_i01_Hydra
TCATGAAAGTTATTACGAGATTAAAAAAGCTCTAGAATCTTTGATTATGGACAGAAGTCAGACAAAAAAAACCCAAAATAATGCTCAAAATCTCACCAAGAAAATGAACTCTTTTGAAATTGTTTTTTTGACAATTTTGTGGAAAGACATACTTTGCCATTTTAATGAAACATCCAAGACTTTGCAGAAAGAAAATCTAAACTTGGATGTTGCAGTTCGTATTCTAAAGCTATTGTTGCATTTCATAAAAGATTTGAGGAGTCAGTTTCAAGAATATGAGAAAAAAGCCAAAATTATGCTTCCAAACACTG
>evgR014094c0g1_i01_Hydra
TTTTTTTTTTTTTTTTCTAGTTGTTGATAATTTCTGGAAATTTTATTAACTTTGTGCTTTTTACTATTTAAAAACAGAAAATGGAAAGAATTAAAAATACTGTAACAAGGAAAGAAAAAGTAGTATTAAATAAAAAAGACGACAAGGAAGCTAAATTTATCTCCACTGTATGTCATTCTCTTTTTACGGGTGAGAAGATTTCCACTCATGATTTTTCAAAAATCTTTGATATTGTTCACAGTAGCAGTAGCAGTATCGAACTTGAAGAAAGTATTATTAAGCGCGGGATGGAGTCCTTAAGAAAAGAGGTAAAAGAAAAACCTGGCAATGCATATTTAGTTAAACTATCCGATATTTGGAGTACATTCTATCAAAATATACTTCCCACTCTGCTGCTTTGGTTCAGTAAAACTAATGGGTTACCGATTAAAGATATGGTTGTGACAAGCTTTCGAGATAATGTTATTTTAAAATTAAAAGTCGAAGAAGCATTACATGCAAATGAGGCTTCGTTACCACCAGGGATCAAACCTATGTTTCTTGTTACACTTCAGGTAGTTAAAGATTACTCTGAGAATTATTTTAAATTAGAGTCATTAGCAGCCAAAGTAATTACCCCATTTCTTGGTGCAAAGGGGCTGTATGTTCGCTCTGAAATACACGAATATAATGAAAAAAATAAACATTTGCTACCTGCTATTTCAAGTAACTCTGCTGGAAGTAGTCCTGAAGTACCAGAGAGGGAAAGTTCTGACCTGGAGTTTGAAGAACTTCTTAATCATCGTCGTCAAGTTTTTAACAGGAATGAATCTCAACAAAGATTATCTTGTGTTGAAGAAAAAGGCATTTCTGAATCTCTAGCCAGCCTTTTGGCTGAAGGTTTTGAGATTGGATAATAAGTTATGAACTTATTTTGTTTTTAACGTGGTTTTTATTTTTTAGCATTTAAGTTCAAATGTTGAATCATAATACTTGTAAATAAGAGATAGTAATATAATGTAATATAATATAATGTTTAATATTTTATCTTATTTTCTTTATTCTAGTTTTGATTTGTAAATTTAACTATTTTTCTACTTCTTAGTAAATCCCTAATTTAATATTGGTTGCTTGC
>evgR014101c0g1_i01_Hydra
GATGTTGGCAAGGATTGGAGACACGTTCAAAAGGGTATCCAACTTGTTGAATTTTTGGTAGTCAATGGTTCTGAAAGATGTTTGGATGAAGCAAGAGATATGCAATCCCTCGTCCGTCAAATGAAGAACTTTAAATATGTTGACGATGATGGAAAGGATTGGGGTGTTAACGTAAGAGATGCTGCCAAGCACTTCTTGGAATTCATCAAGGATGAAGAAAAGATCAAGGAAGAAAGAGAAATTGCCAAACAACAAAAGGAAAAGTATGAAGGATTCTCCAGATCTGAAGCCAAGCACAAACGTCCTACAAAAACATCTTCATCAGATAATAAGAAGAAGGAAAAGAAGCCATCATATGACGACGAATATGATGCTCCAAAGAAGACTACTTCTCGTTCCAGTTTTGAAGAAGAAGAATATGAAAGAGACCAAGATGAAAAGCCAAGATCTGTCTCACCAGCTAATAACAACTCTGGTAGACCAAGAAAGGCTTCTGGTGGTGGCTCATCATTCCAAGCTCCTGTTTCTCAACCTGTCAAACAAAAGGAAGATACTCTCATTGATTTCTTTGATACACCTGTTACAACAACT
>evgR014110c0g2_i01_Hydra
AAACTGAATATTAAGTACGTTGTATACTGGTCGAGAAGAAAAACAACAATAACATTAACAGTTTTATGGTTTATTTCAGTTTTCAGTGGAAGCCTAAATGAATTTTTATTCAGTCGTTACAAATTGTTTATTGATGTCGGTTTTGATCTGTTTGTAATTAGCTTTTCCGCGTATCTCTACGGTTACGCTTTGATTACTTTTCAAAAGCAGAGATCGAATGTTCGTTCCAACAATGGCAAAGGTATATTTAAAGGATTGTTATCATCAAGTCTCATAATAGTCATGTTTATCATATTGATTGCTATACCTGATATACTATGGGCATTAAAAGATTTAGGTGTTTATCGCAATTCCTTTTTGATCTCTAATATTTACATTAATTATGCAATGTCTGCATGGAC
>evgR014111c0g1_i01_Hydra
ATTGCAGATTTAGTCTATTTGTTGATAATACGCAGCTAGTACATTTCGGGGCTTCATTTGAAATACCATGTCAAAACGAAGAGTTTCCGGACTTTATTCTCTCCCCGATTTCATCGCCACAGGAAGGGACGTTTCAAAATGCCCAATATGCGGAAGAAAAGTTTACTAGTTTTCAGGAAGATATTTCTAAAACTCATTCCATTTTATTTGGAACCAAAGTCCATAACGAAGAACTTTCAAACGAACATCCATATAATCAGTTTTATTGTCAACGCAACTCACAATACAGAAAACTATCTCACAGTGTGTCCCAAGAACCCACACAGCATGAAGCACAAAATATATCTTGTGCACATTATTCATACAATCCAGATTTGCAATGTCATCCCATGATTAACGATGACTTAATAAACAGCTGCTCAAGCAACAGTTATTTGAATTCAAACTTATCTAATAATGTATGTTATAACTATAGCAACGGATGCCAATATCAGACTTACGCGAATCCGAATCATTCTTCAGATCAAGAAGATTTTGAAACATTGTTTACAAACGGATCTGCGACCATAAAGGCGGCCACCACTGTCTCCACAAATGATAAAGCCACCGATGCTACCATAAACAAAAACTTTTCTAACAAAACTTTTTATCGACGATCACAAACTCAATATTATCGCCAACTAGCATATGAAAAAATTCGAATGAGCAAAC
>evgR014127c0g1_i01_Hydra
ATTGCTTTAAATGTTTTGAAAATAAGACGGAAGTACTGATTAAAAACAGTAGTATGTTAGATAGTTTCTATACTTCATTATGGTTGAAATCTTCTTTTTCTTGGTTCAATGACACGTTTGAATGGTTTAACTCTACTAAAAAATGGTTAAACTTTACAATTATGCGGTTTGATGATAAAATAGATTGGCTTAATTACACATCAGCATTGTTCAATCAGGCGTTGTTCAATCGCAGCTTTGGGAACAATTATTCTTTGCTAAATACTAAATTTGATTCTTTTTTTCCAAATAATACAACAGAATGGTCTAACGATTCAACGATATGGTTTGACCTTTCTGAAACTATTTATAGCACTTTTAACAACTATAATTTAACTATTTCTGGTGATAGCTTTGTATCTAGAACTCTATCTTTTATTAAACTTTCAGAAGCCAGATGCAATACGGGACTAACTTATAACAAGTTTACTGGTACCTGTGAGTCAATTATTTGTTGTAGTGGATACACTGTAGTTGGCTCAGAATGTTTAAAATTGCAACAAAAATCACCTGCTATATTTATAACAAATCCAAACTTTGAAAACTGTTTGCTTAGAAAAACGCCTGGTTTTTACGCTTTTTTAGAAGAGTCTGATAATAGTATATTGGAGTCGCATATTTCACAAGACACAACTTGTAAAGTTATTAAAAATGAAACCTTATCTAATTTAATTTGGTTACAATGTAACATGTCACTTGTCAAAATAAAG
>evgR014131c0g1_i01_Hydra
CAATATCAAAACAAAAATAACCGCGAATTATAATGTAAATCGATTTAAAATATTTGCATTTCTTGTAATCATGGTATATACATGCGTTGTTAAAGATTGCCATAATTCTATAAGAGAAATTCAAAATTGGAAAAATTCTTTCTGTGAGGAACATGGTTGCCTTAGGAAATCTATTTTATGCTCCTGTAATCCACCTTTCAGGCTGTTTTCATTCCCTACCACATTAAAATGCTTGGATAAAAGAAACACTTGGATAAAACTTATCAATAGAACAAAAAATACGGGTCAATTTTGGGAACCATCAAAAG
>evgR014133c0g1_i01_Hydra
GAAAAGTATTTTCCAATGATCAAAAAGATTTTCTTGGAACACTGAATGTTTTAAAGAACGAGTTCACTTTATCTTTCAATATAAAACCTATACATTTTTTTGAGGGTTCAAAAACGATTCTACAACTAACCTTGGAAAATTTTACCAAAAGTTGTTTAGATGATTATATTAGTATTGGATTCTATGAAGATGGATTTAGAAAGTTCATTATTAATTCTTCAGAAAGAGCTCATTCTATAATTGTATCTAACCTAATTACTTTAGACCAATGGTATAATATTAAAATATTCCAACGAGAGCTGACTACTGGTCTTTTGCTTGATGTTGGTCTAAATGGAGTTAATGTTTATAGAAATCATTTTTATACAGCTTGTGAAGAAAAAAGTATGAAATATGTTAAAGTGCGTGTCAACAATCCTTGGTATTCTGTACAAAATGATTCTATCTCTGACCTTCTTATTATCAATGGAAAAGCTGATTACCATGCTG
>evgR014134c0g1_i01_Hydra
TGTATTTTATACAACACCATCAAAAGATTCTCCATAGAGAAAAAAATCAATCAACACCTACTCATCTATTTAATCTATTAATAATGTTGCCACATAATCATCATCACCACAAGGAACTTTCTTCCGTTACCGATACAATTGACTTCCTTTCAGAACAATCACCATTCTACACTGAAAGTATTCAATATGCCAAGCAATCCCATAAGAATAAGCCTCATATTCATCCAGAAGAACGTAATCTCGTATTCCCAAGACAGCAATCATCCTCCAATAGTAATTGGTTTGATCAAGAAGAATCCGAGTTTGATGAAGACTTTTTCTCTGAACAACCAGCCAATGAAAAGATGCTTGCTCCTTCAATCAATCAACAAACAATGATGCAAGAAGACCAAGTCTTTTTAGATGGAGGAAGTTGTGATTTGGACGAAATGGATTTGTTCAATATGATTGACTCCACTCCACAACAACCATTAACAAAGAAGGTTAAATATGACCATCATTCATCCAATTATTTGGCCACAAGTTCTTCAACAACTTTGGCAGTCAAACAACCATCTAATGGTCAAGATAATTTGCTCTCAATACTTATTAGAGAACAACAAAAGCAAGGTGCTGAAATTTCCGAACTCAAGCAAATGGTGCACAATTTGCAAGCACAAATTTTAATGCATCCAATGTACCAACAACAATTTGCACTTTATCAACAACACCAACAACAACAATATCAACTCTACTTGCAACAAAATCAGCAACAAGAAACCAAGCAATAACTTTGTAAATCAATCTAGGTTTTGGTCAATTCGAACAGAATGTTTAAGAAGCTTTCTCTCTTGAATTCATCAAGACAATGTAAATTTTCCTCCTTTTATGTAAATTAAATGTATTGCACTCGGGAGTGGGAAAGAAATCATTTCAACACCATTAGATTCTGTAGGAAGGATCCATTGCAAGAAGAACAGGCCCATCAACTAATCCCTTGAAAATCAAC
>evgR014155c0g1_i01_Hydra
GGAAAAGTGTTGTTCATTTGACCATACAAGGTAATATGGATAACTATGGCGATAGAACTCCTGGTTTATGGTTTTCTCCAAACCTTTCTGGAAATTTGCAAATTACATCTGCAGTAAGTGGCGATAAAAGCTATTTTTTTTATACAAAACCACTCACATTGAATGTATGGTATGATATACAGATAATCCAAATTCAGCTTAACGGAATTTATAATTTTGTGACATATTTAAATGGTGAACTTATTAATATCGTTGAAAACACAAAGCCACAGTCTTTTTCAAATGTTAAAGTTTTTACTGCTAACCCATGGGACTGGACACATAATGGTTATATTAAAGATTTAAGAATTATAAATGGATTTGATTTTTTTGATAAAGCTGACGAGAATGAGTTGGTCAAAGATAATTTGATAGCATTGTTGTCATCGTTAGAAAAGGC
>evgR014181c0g1_i01_Hydra
TCCTCAGAATGACTGGTGTTACCCGTTATAATATGGAATTTGCTTTGGTTGAAGGAAGTTTTGTCTATCACAAGGGTGCCATTGAAAAGGTACCAGTCACTCCAACTGAAGTTGCCAAGAGTAATTTGTTGGGATTCTTTGAAAAGTTGAAGGCAAAGAAGCTTTTGACCTTCTTGTACGACTATGATCAATCAAATCCAAAGACTCATCAAGGATTTGATTGTTCAAAGGACACTATGGACAAGATCTTTAAGTACTATGGTGTTTCTGAGGATACTACTGACTTTTTAGGTCACGCTGTTGCTTTGTACAATGATGATTCATACATGACTACTGTTCCAGCACAACAAGTCATTGAAAGATTGAAACTTTATGAAGATTCTCTTAACATGTACGGAAAGTCTCCATACGTTTATCCAATGTATGGTTTGGGTGAATTGCCACAAGTTTTTGCTCGTTTGTGTGCTGTCTATGGTGGTACTTATATGCTCGATAAGCCTGTTGACAGAGTTCTCTATGACGAAAGTGGACGTGTCATTGGTGTTGAATCTGAAGGTCAAGTTGCCAAGGCCAAGATGGTTGTTGGTGATCCAAGCTACTTCCAAGAAAAGGTCCGTAAGACTGGTAAAGTTATTCGTGTAATTTGTATCCTTTCTCACCCAGTTAAGAGTACTGGTGATGCCAAGTCAGTCCAACTTATTTTCCCACAAAAGTCAACTGGTCGTAAACATGATATTTATTGTTGTGTGACATCATTCACTCACCACGTTGCTCCAAAGGACAAGTATATTGCCATTGTTTCCACCACTGTTGAAACTGAAAATCCAGAAGCTGAAGTCAAGATTGCTCTAGACGTTTTGGGTGTAATCGATGAAAAGTTCGTATACATTCTTGATGCCTG
>evgR014185c0g2_i01_Hydra
TATCATAATATCTAATTGGCAAAACCCACATTTTAATCCGTTGCCAAGATATCAAAATGTGGATGGGTACACTTTGAGGTTTAAACATAACGGACAGCAGCAGCAAACAAAACCGAAGAATGAAATTCCAACTAAAACAAATGTTGAAAAAAAAATCAGAGGTTTTAGTAGGCAACGAAGAAGAAATGCCTTCAGTTAACTCTTCATCTAGGAAAGTGGTGCCTACGATTTCGGAAGTAGTGGCATCTGTTGCGGAAACAGCGGAGAATATTCAAGAGTTTGTTCAAGTGTGTGATTCTGGAGAGGCTTCGCGATCAAAAAGAAAAAGCGTGGACGATCTAGGAGAAGGCGGAACACCCTCGAAGCAAACAATCTCGAGCGCGCTTGTTGAAATTAAAGATACGAAAGAAGTCGAAATGAAAGAAATCGACCAAAACGATGACTTAAATTTTTGCTATGACAAATTCGATCTCTTAAAATGTG
>evgR014190c0g1_i01_Hydra
GTAAAAAAATAAATAATTTTTGTTTTTTCGTCTTTTATCCAGCTAAAAAAAAGTTGAATTAAAACTGTGCACGTTTTTGAAAATGGCAATCCACCCACGCATACTTTTCCTTTCAATTGCTTTGATGTATTTATCAGAAACTCGTTGCAGTGATCTCTCGCAAAAAATCGAAATGAAAAGAATTCAAGATAGACTCGAACGCAAGCAAGCTATGGAATCTGCTGCTGAGTTTGGTGAAAAAATGCCCGTTTTTGTAAGCAAACGCGATGAAATTTTATGGAAACTAAATTTGGCTAAAAGAACATTAAAAAAAGCTCACGAACTCGAAAAAGTTGCTAAGAACTCCGCTGTGCACTTTGGATACTTTGAAGAGAACAAACAAAAACATTCGGAAGTCACTGACTTGCTTGGGAAAGCTGACGAAACACTAATGAGTGCTGGGGAAAAAATGGGTTTTGATTTTGAAAAATTTTAGTGTTTCAAAAAATATTATTGTTTGTAGTCGCAGATAAAACATCGCACCATTACATCCCGAATTAAACTACTACTGCTGGTATTTATTGCCAACAACTTGACTGTTTGAAACAAAACAAACCATAATATTTAAAATATATTTAATATAAGATATACAATTTAATATATGATTTACAATTTAATATATGATGTACATTTTTAAACATTTCTTTCAATTTTTTATTATTTTAACTGATTTTTCTGAATTATACCAATAATCAAGTAAAAAATTA
>evgR014203c0g1_i01_Hydra
AGATTATAAAGAGAGGACGCCGTTACAAATAGCTGCACAGACAGGACGCGAAGAGTCTTGTAGACTTCTTCTGCAAAATAGAGCTGCTGCGGGTGTTTATGATGATACTGGTATGTCATGTTTATCACACATGATTGAAAGAATGCCTGGTGTTGCTTTCGATACTTTAGGTCAATTTCAAAGAACCATTAAAGCAGAAAAACTAACATTATACTATTTAAGTTACTTAGAAAAAAAAGGATGGAAAATGAGTAGAAAACTAAACAACAAAAGTCATCGAATGGTATATGTTAGAGAACCCTTAGAGGCTATTGTAAAATATCAAGATTTAACACTAGTCATGCACCCAGTTATTCAAAGGCTAATTAAAAAAAAAACTCAGTTATATGGTAGACGTAGTTTTATTTACACATTTGGAATAAACCTTTTATTTACTTTGCTTTGGACGGTATCGACTTTTATTCTCCCTCGTCACTCGCTTATGACAACAAGTTTAACTGCTACCGGTTTAAATACTTCAAATGGTACGGCATCAATCAATTTAGACGAATTAGAAATAGCAAGCACGTTACTTGTTGAGTTCTACACTCCCTATAAATTAAATATTTGGAGATTGGTTTTATAAATTTTTGGTTTGTTATTAGCAGTGTACTTTTTTATTCAGGGAGCCTTGTTGCTTTATGAACAC
>evgR014204c0g1_i01_Hydra
AATAGCAATAAAAAATGAAATTTGTGTTGGAAAAAACCTCAAAAAGATTCAATATTAAATCTAAAAATGTTGAATTTATAAAGTTTGTTTGGGTTGGGAAGAAATATGCAGATAATGAATTTTATGAAGAAAATGAATCAAAATTTTTAAGAGATAAAATGTCGTGGGATGAAGTGGATTTAATAGTTGAAGCTAAAATAAATCAGGCTACGCAGGAAGAGTTAAATAGACATAAAGATTTACCTGAAGAAGACGAAAAAAGACATTTTTTAACGTCATTTTCTTCTGTTGTTATTTATTATAAGAAAATGGCTGAAACATACAAACTTGTAAAAAATATTTATAGATCGAGTGTGTATTTTGTTCG
>evgR014212c0g1_i01_Hydra
TGTAAAGGCTGTTAAAGATGCCAGAATTATGTTTGACATGGCTGAAGAATACGGCTACCACTTTGATTTGTTAGATATTGGTGGTGGCTTCCTTGGTAACATTACTCAAACTCCAACTGTTGAAGATGTTGCTTTGGACGTTGCAGGTTTGATTGATGAGCTATTCCCAGCATATGTCAAGGTAATCTCTGAACCAGGTAGATTCATTGCAGAATCTCTGTCAGTATTGGTCTGTAACATTTTCTCAAAGAAAGATTTGCGTGAAGAGAATAGCCAATACTACAAACAACTTGAGGAATCAGGTGTGAAGGACATTGCCTACGATAAGGATGATTTTATTTACTATATCAATGATGGTATCTATGGCTC
>evgR014216c0g1_i01_Hydra
AGCTATCCAATCTGCACCGACTACAGCTATCCAATCTGCACCGACTACAGCTATCCAATCTGCACCGACTACAGCTATGTCTACCTCGAGTGCTACACCTACATTTCCTTATTGTATCAACGATGGTGTTACAGATTTTCAATTGACATCGAGTGGATATTATGTTCAAACTAACGTCTCATGCATCAGTGTCTGCAATTCCTATTCTCCAGGAGTTTTCTCTTATGCTTTTTTTCTTGAGAAACAAATTTGTTTCTGTAGGAATTCTTCACAGTTAGATATACCTTCAATTAATAATTGCAC
>evgR014286c0g1_i01_Hydra
CGTGCTTTTAAATCTCTTTTGTATCGTAAAAGCCTTTATGTTAAAAAAATGTTTAAAAAAAATTATATATTTTTTATAAGCTTATTGTTGTTATTATGGATGTTCATAAAGTATGTTGACATAACAAATGATCAGTTGTGCAACAGGTGTTTAGAAAAGCTGTGCAACAGGTGCTTACAACGTCAACTTATATTGTCTGAGCAGGCAAGTGAACTTCATCAAAAACTTCAGCATATTGAACATGTTACGGACTTGTTAGTAAACGTTATTATTAATTTGAGTCTCTCCTTGGGTGAGAACGATTTAAAGTTTACTGATCAATATTTTAAGGTTTTAGTGAAACTTCAAAATGAAATAAATAGTTTAAAAAAACAAAGTGAAAAATTAAAAAATGAAACTAAATTTTTGAAAAATGAAACTGAAATATTTGTTAAACCAAAACGCATTAAAGAGGGCAATTTGTTTAGTAATCAAAGTTTTCACAAAAGTAAAATAAAATATAGTTGCGATGAAACATATAAAGGTGCTTTGTATGGTTACCCTCTATACAATGTTGGATTCGTTCGGAATCCATGCACACATGTTCCTTTGAAAGACACCGTTACTTTGTTTCTCATTTTCCCCTCATTCAACTCAAACGACTCTTGGAAAGCAGTATTTTCCATATTCAAAACTATAAAAGTTATAACAAACAAAAACAAGATTGTTC
>evgR014295c0g2_i01_Hydra
CTTCTATTGTCTAAATAAGTTAACATGAAGTTAATGAATATTGCTTCAAGCTCAAATTAAAATGTCTTTTGAAATACCTGTAAACACTCTTAAAAGTGGAAAACATCAGTACAAAACCAGTTCAATAAATTATTATGGAAATCAAAGTTCTATTGACAAAATTTCACACCCAGTCTGCTCATTGATGTATGATGGGACATCAAGTTCATTGTTTTTCAAAATTCCTGATAGTTCCTTGGTTGAAAATGAAAATAGATGCACAACTCTAGAATGTTCTGATATTACAGCCGCAGATCTTCGTGTAACTTCGAAGTTTGGCAGTTGGGATACAATAACATGTCGTTTACTTGTTGAATGTAAAAATCCTTTGGTTGCTTGGAAAGTTGATACTTCACCTCCAGAAAGTATAAAGTCAACAGAGAAATGTATTGAAAAAAATCCGGTTTATATATCAATAAAACAGGAGTCATGTATGGTGAAAACTACATTTGATGTTCTATATGCCAATGAGATTAAGAATTTATTACATCGTATTAAACTAGCTAAATTAAAAGAACGTTTAAATATTCAAGAGTTGCACAAACTTGGACTTAAAAAAGATGTACGAGTACCTGATAAAATTAGAGAGGGTCTTCCTGAATGGGTAAACTATATACCAGCTAAATGGTATTCTTCTACTGTTCGTCAAGTGTTAGAATATTTAATTGGGTTCTATACTGTATTGACTTTGTGCTGGGCACTATGGCAACTTCATAAACATGTATTTTTCATTCGAAGGTACATGCAACCTTTAATTGACTTTTTTATGTTTTACCTCCATATTCTTGCAGATTGGTTTAGATTGTTAGATGCCTATGCAGATATTTTGTCTACCTACTGGTGGACTTACATGAAGCCCATGTTTCTTTTAATATATCCATTGTATAGTGGTTTAACTCAATTATTTAAACCTCTTCGGAATGTTGCTGACGTTGTTAATGTCATACTTCAACCTATTGTATTTTTTGCCAAATCCTTATCATTAGCTTTGCAGCCTCTTTTACAACCTGCGTTAAACTTATGTGGATTAATTGCACGCTTTTTCAAAGATATTTATACAATGATAATTTCCACTACAGTTTATCAGCGCATTGCCTCAGAAATTACCAATGGGAGCCTTGACCCACTTCGCGCGCAAGTAATTCTTATACGAGATATTATATATAGAAGTATTCGTAATTTAATCTTTGGGATGAAGTTTATATTTTCGAGAATTTACTACACGCATATGTTTTTAAAAAGAGAACACCAATATGTCCAAGAAGAGACCGAAATTGAAAATAAGGAGAAAAAGAATTAAAAATAAAGTTTTTTAAAATATTTAATAATTTTTTAATATTTAACTCGATATATATTTTCTAACCTGCATTATTTAGAATGAAGTTAGTATTTTTTAGAAATTTACTACACACATTTTTAAAGTTGAAATTATGATAATATATGATAAGTTTTTCTTTTGTTTTTGTTTTACAGTATATTACTAAAAACTAAAATATAATATTTTACACCAATCAATGCAATTATTT
>evgR014301c0g1_i01_Hydra
AGCAGAGTTATATAACGAGACAAAGGATGATTTGATGAAGGAACCTGACAAATTCTTCCAAATGATTGATGCCTTTGTTCAACAATTTAAACAAGCAGCTGACAAGAATGATCAACAACAAAAAGCTGAAGAAAAGAAGAAGAAGGCAGAAGAAGAAAAGCTCAAGAAGGAAAAACTCAAACAAGAACTCAAAGGTGCTGGAACCACAAGTCGTCTTGGAGCTTCCTCTGCTTTAAAGAAGGATGATCCAAAAGCTGGAACCAATATTTTAGGTGGAGGTGTTTCATTGAAGAAACCTATTGGAGGCACAGGTCTTAGAAAAAAGACAACAAATGATGATTCGTTAGACTAATCATCAACATGTACAATTAGAGTTTCTG
>evgR014316c0g1_i01_Hydra
TGCCAATGCTTTCGAACGTAATGATGTCGAAAAGGCATTAAAGGTCATGACTGAAAATGCTCAACTTGCCAAGGATGAAAATAACTTTGGTTTGATGAAACAATGTATTGCTGCTCTCAACAAGAGAAACATTCAAAAGCTTACATCTACATATGTTACCCTCTCTTTGGTTGATATTGCCAAGGCTGCTCGTTTGAAGGATGCCAAGGAAGCCGAAACTCTCTTGTTCAAGATGATTGATAATGGTCAAATTAATGCTAGAATCAACCAAGTTGATGGAATGGTTACTTTCGAAGAAGATAGTGAAGCATCCAACTTTTCATCTCCTTTCACAAATCAAACTCTTGACCGAAGCATTCAAAAGACTGTTGAATTTACCAAACGTATTAAGAAGATGGATGAAGAAATTTCTACATCAACAGCATTTATCATTCGTTCATTTGGTTTGAGAGAAGAAATGTTCACAATGGACTCTATGAGTAACAAGAGAGGTGGTGCTGCCTCAATCAGTCGTTTGATTGGCTCTCTTGATATGTGGAGATAAATTATTTAAATCTTTGTAAATCTATGTATAAAAG
>evgR014320c0g1_i01_Hydra
AACATGATTCATGGCCCCTGTGGAACTTTAAACAAAAATGCCCCCTGTATGGTGGATGGGAAGTGTTCCAAAAAGTACCCACGTGAACTAATCCAAGAAACTCAGACAGGGCATGATGGCTATCCTTTGTACAGACGACGGAAACCTCAAGATGGTGGATTCACAGCAAAAATCAAAATGCGTGTTGGAACATCCTATCAGGAGATTGAGATTGACAACAAGTGGGTGGTCCCTTATAGTCCTTTGTTGTCAAAAATTTTCCAAGCCCACATCAATGTTGAATCTTGCCAGTCAGTGAAATCCATCAAGTACATCTGCAAGTATGTCAACAAAGGCAGTGACCAAGCAGTTTTTGAGCTTCTAAAAAGTGGGACAAACCTGGATGAAGTTCAAACATTCTTAATGGGGAGGTACAGGTACATCAGTAGCAATGAGGCTGTGTGGAG
>evgR014346c0g1_i01_Hydra
GTTCAGATTTAGTTTTAAAGAAAAATTTAGTATAATGGAGTCAGCCTTTGGTTACATTGACTTCGACCAACCATTGGATGAAAACGATCTATTAAAACCACCTTCTTCTGGAATAGAGTATCTAAAACGTGTTCAATATGAAACTTTAAGATGCCCTGAGGTTGTTATAGCAACTATCCCTCAATTGATGCAAACACCTGAATATGTAGTTCTCAGTCAACAGCCTCAACCAGTAAGCGATGCTCCAAACCCAAACACGCAGAGAATACAGGCAGCTGATTTTGCCGAAATGCGTCAGAAATTAATTTTTTACAAAACTCAAACTAGAAAAGTTTTGCCTATAAGACTTCCTAAAAAACAAAATCCCCAAACGTGGTGTGCGTTTTGTTTTGGTCGTGATTTTTTGAAGAAATGTTACAAAGGTCGTTTGTCAGAAGATATTATTAATGATGCAGCCATTGTAAACGGACACGAACCTTTGCTAAGCATAATAGCTCAATTTACTCAGCCGTTAGCCGTGCAACTTTTAGAGTATCAATTGACGTGGTTAGATTTATGTGGCTTTTCTACAGAACAAGGTCGATGGTTATATAGTCTAATGCTATGCATTGATAAACCAACTCCACAAGATGCTATGTCTGTATTAAGAACGCTTGCACGTAAATGCTTGAAGATAAAAAATTCGTACAAAGAACAGTCTGAGTTATTTCCTCAGCTACATTTATTTATTGTTATTGTTTCAAAATATTTTGGCCAAGAAGACTTGGCACTTTGTTGAGGATCAAAATAAAAAAAGTTTTTATTTTTATTTTCTGATATTTATACTTTATATACTTATACTATATATGCTTTTATAGATAATTGTATAATTCGATCGCAAATGTTAG
>evgR014380c0g1_i01_Hydra
ATTAAATTTCATGTTTCATATATCTTGTATGTTACTGTTAGTCACAACTTGAAATGGATCTTTTTGAAAAAGACATCGTATGCACTAAAGAATTGTCGAATAGCCAAACTGACGAGGTTAAAAAATTAAGAGTATTTTTCCAGCTAATTCAGTCTGGAAAAAAATACAACAATATGTCAGATGGTTTTCAAAAGCTTTTATCATGGCTCGAAACTGATAATACCAGTACGTCTATCAAGGAGGCAAACAATTTCAAAGTTGAGGAAGAAAGAAAATTACTTATTTTCAGCCTGAAATTAGCGATAACAAATAATGTCAACACAGACGTAAATCTAAAGAATAGTCAACCTGAGATATCATTCTTAAATATGCTATCTGTTGGATTTGCCATTTCAACGTTTGCCGATGAGTTGATAAAAAAATCAGAAAATGTGAATGATCTGTTACTGTTTTTAAGAAATTTAATAAAAGAAAACTCTGTATGCAATGAAAGAAAAGAAGAAGCTTTTTTGAAAATCAAAAACTTTATTGATAATAGTCTAACAGCAAATCAAGTTGCCCAAAACTTATACGGTTATTTCCTTCAAGTTGCGAATTGTTTGTTTGATGAAGAGGCTCTTTATAAACGCTTAGAGGAAGGATGGGGAAAAAATCAAAAAGAGTTATCAAAAAGAGTAAAAAAGATGAAGATGTTTGATAAAGAATCGACAAAAAAGTTAGAGCAAATCGAAAAGCAAATCGAAAAAAGTGAAATGGACAACGTAGGCTTTCGTGACGTCATTGACAAAACGTAGGATAAAATTTCCTTATTGATACAGC
>evgR014382c0g1_i01_Hydra
TGCAAAAAGCTTTGTATTTACTGTACAGTAAATAATAGATAAACAAAATAAGATGATACCTTCAAAACTACAGTTGTTATCGCGCTCATCAGTAATTTACAAAGCAGAACTAGGTAAACGTTTTTTGACAAGTATCTTAAATAACAGTTCTACTCCTGAACAATTGAGGGCACAATGGTCTGGCACATTTTTCAACATAGAAAAAATGACAGAATTGTTGGATCATGACAACCACAAAATGCGAAAAGATTTTAGAAAGTTTCTGAGTGATCCTATTTTTACACCAAAATATAACATATCTCTTGTTGAAGAAAGAGAAGTTGCTCTTCAGCGTTTAAAGAAAATTTGTCATGCAGGATTCATTTCAGTTTTTGATTTTCATAAGAATCCTTTAAAAATATTTGCAGCTCATGAGCTGGCTAGCATAATAGATCCTGCTATGGCTACCAAAATGACTGTTCAGTTTAATCTATTTGGTGGAACAGTTCTTAAACTTGGTACTGAGAGGCACCACAAGCAGTTACTAAAGGGTATTGACAGTCTCGAAGATATTGGTTGTTTTGGTCTTACTGAATTAGGCTATGGAAACAATGCTGTAGAAATGGAAACCATTGCTGTTTATGATAAAGCAACTGATGAATTTATCATAAATACTCCTACCACTCTAGCGCAGAAGTACTGGATCACAAACGGCGCTGTTCATGCAAAGCATTGCGTTGTTTTTGCCCAGTTGAATGTAGGAGGAACTGAACATGGCATACATGGGATTCTCGTTCGTATTCGAGATGACGATTTAAAAACATTACCAGGGGTTGTTGTGGAAGATATGGGTTACAAAATTGGATTGAATGGTGTTGACAATGCAAAGTTAACCTTTAAAAATGTTCGCGTGCCGAGAGAAAATCTTTTAAATAAGTATTCTGATGTTGATAAAAATGGCAAGTTTACAACTAGTATAAAAAGTGCAAGGGGAAGATTTCTCACTGTAGCAGATCAGTTATTATCTGGAAGAATATGTATTGCTTCTATGGCTCAAGGTGCTGCAAAAACTGCACTTGCCATTGCTTTAAGGTACTCAGCTACTAGATTAACAGTAGGAAAAGAAGGAAAGTCTGACACTCCAATATTGCTATATCAGTTGCAGCAGAATGCATTAATGCCGATGCTAGCAGAAACGTTTTGCCTGAACTTTGCTCTTGACTATGTTAAAGAGAGATGGGCTAGTCAATCTAAAGATGGATCCGAACATCCTGAAATTGTAACCATGTGCTGTGTAATAAAACCTCTCTGTGGTTGGAATCTCGAAAACGTTGCTTCAGTGTCTCGTGAACGTACTGGAGGTCAAGGTTATTTATCGGTTAATCGACTTGGGGAGTTTATTTCCCTTGCTCATGCTTGTATTACAGCAGAAGGTGATAACGCAGTACTTATGCAAAAGGTAGCCAAAGAGTGTTTAGCTCGATTCAAATACAAAGATATTAATCTCCCAGCGGTCGATCTTAATTCTTTAGACTATTTACATGCTTTGCTAGCGAAGCGAGAATCTGTTTTATTTGCTCAACTTGGAAAAGAAGTTGGTTCTGTAGGTAAAGATCAGGTGTTTGAAACTTGGATGTATCATTCCAGTGATCTAGTGCAACACGCCGCGCGTTCCTTTGGCGAACGACTTGTAAGTGATAGGTGCAAGGTTACCATAGATCACGCTGAACATTCTTTGAAATCTACGCTTGAAAAATTACATCATCTATATCTCGTTAATGTTTTGTCCCGCAACCTTGCTTGGTTTCTAACAAATAATCTTGTCTCTATAGAAACATCAAAAAACGTCGCAGCGTTGTCGTCAAAGCTTTGCAAAGAAATAGCACCTATGTCGTTAGCTCTAGTAAAATCTTTTGATCTTAATGATCATATGCTATCCGCGCCAATAGCAAGTGACTGGGTTAAATACAACGAATATGACAATCAAGGAGAATTATAAAGTTGTTTTTTTGTTAGTCTCATTAGTTTAGTTCGAATTTTAATATTATTTGTTACCCTCGTATGTGGAAGCATAATTATCTCCTAAATTATTATTTTTTTAACAATATATAAATAGTTTGTTTTATATTTGTAGTCAAATAATTTTCTATTTATAAATGTTCAAGGTTTATTAGTTTTTTTTTTCTTTTTCCGATTAAATAAAGTTGTAAAAAAAAA
>evgR014391c0g1_i01_Hydra
CTAAAATATGTCTAAACTACCTGAGATTGTAGTAGAACCTAAAGAATCTCTGTTGATTGATGAAGTAATTATACTTGGATATAATTTCAAACCTAATCAACCAGTTACAATTTATTCATCCGTTGAGAGCCAGTCAGAGAGCATTTCGTTTTCATCGTATGCTTATATAATAACTTCATCGAGTGGCTCTTTTGATCTCCGTAAAACAGAATCAATTGGAGGAACATATCAAGGAATTCACCCGATGGGACTCTTCTGGTCAATGATCAACTCAGCAAATAGCATAAATCGGCTACAATGCTCTAACCCAAACGTTCCATTAGAATACGAGTTTAAAGTGTACAATGAACATGTATGCAACTTCATTACTTCAAATTTGGTTTGCGAAACTAAAGCAACTCGTTATGCAATAGCTAAAGATGTATCCAAAACGTTTGTTAAAAATGGTAGAGTATGTGGGATTTTATTTACTCCTCCTGGAAATGGTCCTTTTCCTGCAGTTATAACTTTGTATGGTGGCACAAAAAAAAAATTACCAGTTCAAGATGTGTCAGCAATTCTTGTTAACAATGGTTTTGTATCATTAGCACTCGCTTATTTTAATGTTGGTGACCTTCCAAAACTATATGATAATGTCGATATAGAATATTTTGAAGAAGCTGTTATTTTTCTTCAAAACCTTCCTATCGTCAATAGATTCTCAATAGGAGTGTACGGAATATCTAAAGGCGGTAGTATTGCTTTAGCAATGGCTTCTTATTTACCTCAAGTTAAGTCCGTTGTTAGCATGAATGGATCTCTTCATAGTTTTGGTGGCACTACTTCATATCAATCAACTATTTTGCCAAAGTTTGCCCACGTAAGTGAAGCTTATTTCAAAATCACTGAGTACAACACATGTTTAACAAGTTCTACATACAAGAATTCAACTTTAAATGACCCTTCGTTAATTCCTTTTGAGAAATCAAAAGCAAGTATACTATTTGTTTTAGGCTTAGATGACAAGCGGGTAGATACTAATTATTATCAAGAAATTTTAAATTTTAAAATGAGTTTAAACACTGAGCATAACTTTGAAGTTGATACTTACGAAGGTTTGGGTAATTTAGTAGATGCACCTTACATGCCAGTGTGTGTTAGTGCTCCGTCTATCACGTATCCGCAAATAATAATTGAATATGGTGGAAGGAACACCCAAAAACATGCGCTATCCCAAATAATAGTATGGAAAAAGGTTATAGCGTTTTTTAATAGAAC
>evgR014403c0g1_i01_Hydra
TCAAACAGAGGGAGCTTTGTATACAGAAGGTTATGTTAGACATTTTTTGACTGCAAGTCAAATGTTTTTCCAGTCTTATTTTCCTAATATACCAATTAAGTATGTACCGAACTTTTTACTACAAGAAGAGGAGGAAAATGAAATAATGCCTGTATTTAGCCCAGTAAATGCTACGATTCCTGAGGAGCATGAACCTGACGAAGAAGAATATACTTTAAAGTTTACTTCGAGTAGTTTGTCATTAATTATTTCTTCACCTTTGGAACAAAACGAAACGTCAAGTATTTTGCCTTTGGATGAAACGCAAAATAGATGCAATGAATTTGTAGAAAAATTATCACCACGCAAAGAAAATAAGAAGAAAAAAAAAAAGAACATATAGAAAAAGAAAAAACGCTGATCCAGCATTTTCAAAGTCCAAAAAAGTTTTACTTAACCCGCTTGGTCAACCCCTGTAATTAATCT
>evgR014404c0g1_i01_Hydra
TAATTGATGAACAAGTCAAGAAAAATAGAGAAATATTAAAACCAATTGTTCTAGCAATTATTATGTGTGGAAAACAAAACATACCTCTCCGAGGTCACAGAGATGATTCAATTTACTATGAGAATGAAAATAGTAACTCTGGCAATCTGCAATCAATTTTGAAACTAATATCTGATTGTGCAGAAAACAATTTAACAAATGGGCGATTCACTACTCCAAAAAATGCTACATATCGCTCTAAAACTACTCAAAATGAGCTAATTAATATTTGTAATGACATTATAGTTTCTAAAATGACGGTGCTGGTAACATGGC
>evgR014414c0g1_i01_Hydra
TGAAAATGAAAATGGATTCGAGTATAGTGAGTTTAATAGAAAGTACACTATTGCAGATGATGTTGATCTAACAGCTCTTACTTCCAACATCAGTGTAGAGGGAGTTTTACATATTGAAGCTCCTAAAGTACTTCCTACAATACCCGATTCTATACAATCAACAAAAGAGAACTTTAAATGCAATTTTGATGTACAAGGTTTCAAGCCAGAGGAAGTTTCAATTCAAGTGAAAGGTCGAGATTTAGTTATTCACGGGGAAACCAAAATTGAAGACGAAGGTGTACATGGTTTTCAACATAGACAATTTACCAGCCACTTATCTTTACCAGATGATGTTGATATATCTCACTTAAGTTCCAGATACACTAAGGATTTTAAACTAACAATTGAAGCTCCGAAGATACCTCCAAAGGCTCCTCTTAAACTTGAAATAAAAAAAGACGAATAAAATTAATTTTAAAAATAACTTTCATTTTATTTTTTCTTAATTGTGTATAATTTTCTTAATTGTTTGTTTTGTTTGCGATCTTTACTTGTAGTTTTAATATAAG
>evgR014427c0g1_i01_Hydra
CTTGAACTTGGTATCAAAGCATATGTACTAAACTAAATCTGGTGTAAACACAATTTTACTAACAGTAACTTATACATGGTATAAAAATATATGTATTAAACTACTTCTAATGTTCGAAGAAATCTGTGAAAGTAAAATTACAAGCGTTTACTCCGTTAAGTTAACAAAGTTATTCCTTTACTTTGTAGCTAACGGAATAATCCATTGTTTATTGTTGACAATTATACTGTTGTTGTTATACTGTTGACATTTTAACGGATTTTGTTTTAGAGTGTGAAGTATGGTTGTCAGCTGAAGAGCGGAGTATGATTGTTAGCTGAGTAGCGGAGTATGGTTGTCGTGGAGTATGGTTGTCATGGAGTAGGGTTATCAGCTGAAGATAAACGACTCTATTGTTTACAAGTAGAAAGTCAAGAAAAAGTTGGTTATGCCACCTGTAAAGAAGCAGTTTAGAAAAAAATTCATAATTCAAAGAAAAGAAAATGAACTGACAAACTATCTGCATTTCGTGCAGATAGTTTGTCTGAACGAAAATTTCGTATCATACGATTTCTCATTTAATAGTGAGAAGGAGCCGGATATGTCGAAAGTTATAGAAAGTAAATATGAAAGGAGTTTATAATTCCATATTTAATGAAGCTTCTAAAAATAAAAGGAAGATGTGAGGAAAAACTTAAATATTGAACAATGGTCTCTAAATTTGATGGAAAGCAAAAAAAAGAAGTTTTTTTTTTTTTTCTTAAAGAAAAAAAAAGCTAAACTAGCTTCATTGTGCTTAAAAGATGGAAATGCAGAAACTATTACAAATGGAATTCTGACAGTTCTTAAGGAGTATGATTTACGGAGTGCTTTGAAAATGGTAGTGGCTGACACAACCAGTGTTAACACCGGTAAAAAGAATGGTGTTGTTGTAAGACTTAGCAAATATTTTCAGCGACGGGGTTTGACAAAACCGTTGCTTATTACTTTCCAGCATCATGTGTTTGATAGAATTCTGCGCGTGGATAAGGATGAGGAAATTAGAGGAAGAAGCTAGTCTCCAGAAATCCTTGAAATTGCAAGCAAAATTAGTGAATGGAACAGAAGTGATTGCCAATAAGTCCGGCTGAAGAGATGTCATTAAATTCTTGTAACATCTGACACTTGTATTCAAATTCTATGATGAAAAAAAACGTTTCCCACTTGTTAATTTTCAGAAAAATTCTAGTATTAGTAATGCACGTTGAAATTCAAGAGCCATCCTTGCCATTCTTGCTTTTTAGCAAACAAGGTCAATGTTACAGATCATATGCAGAAACTGGATACAGGAACTAAAGATACC
>evgR014431c0g1_i01_Hydra
CCGCGTATTTCCTCTAAACCAAAAAAACGTAGGCAGCCTTGGCTAACCAACAAATTTCTCTTACTAACCAAACGTAAGAAAAAAACTTTGGTATCTTAACCGCAACATCAAATAACAATTTGTTCCTCTTGTCAAAGAGTATAACCGAATCCTTTTCACAAAAAAAAGCATTGCAATTAGAGCATTTGAACACAACTAAGCAAACAATAAATGGAACCCCAAACTTTTATACTCCTATGTGAACAATAAACGCGCTGTCAATAACCAAATAACTTCTATGTGCTCATCTTCTGGGTAAATAGTTAGCGAATGGGCATCCATAGCCAATTTACTTCCGGGACATCCATAGCCAATATACTTCCAATATAACATCCAGGTCCAGTCAGTATTCAAAACCGAACCATCAGATCAGCCACTACCTCCCTTCGTCAAACGTACAACCGCTTTAATCTCCATCGTCATATTCAACAACGAATCCGTTTACAAACACATGTCTGTTCTCGGTGCCAATAAGTCAACTGGAAGTGACCGTATTATTCCTTTTGTAATTAAAGCATGTGTTGGAGCAGCGACTACCCCACTCGCCATCATTTTTCAAAAATTGATGGATCAAGGTGTAGTCCCAAATGCGTGGCCTTTAGCAAAAGTGACGCCATTGTTCAGAAAAGGCTTCAGACTCAACCCAGAAAACTACAGGCCTTACCATCGATCACCATTGCCTCTGTACCATGCAAAATAATGGAAAAAATAGATAAAGAACAAATCATGCAGCACCTTGCCGCTCATAAACGTATTACAAATTTCCAACAAGGCTTCGTAAATAAAAAAGTTTGCGTCACAAATCTGCTTGAGACCATGAAATTTCTGACTTACAGCGTTTCTTGCAAAACGTGCTTTTTCTTGAACTCGCCAAAGCTTTCGAAGTCCCTCACAGTAGATTACTGCAAAAACCTGAGTCCTACGGAATCGAAGATAACATTCTAAACTGGATAAATACTTTCTTTTCAAATAAATCACAACGCGTTTTTTTAGTTAGTTTGTCATCCACCTAGGTAAAAGTTATCAGTGGCGTTCCGCTAGATTCAGTACTAGTTTCTACGCTGTTTATAACATTTATTAACGACTTAACCGAACACATGAGAAGCAAAAAATTATGTAAAATGTATGCATACGACACTAAAACCTTGAGTGTCGTAAAAACGCAGAAGACAAAGCTCGTCTACAATTCGATATAGATAATCCCGGACTCAAACCTGG
>evgR014451c0g1_i01_Hydra
CGAAAGCAATGTGCATGTCGTGCTTTGGAAACGGATTCTGAATAGGTGACTATTATTCGTCTTTTCATTCAGAAATTATTGTCGTTGTTGTTTTACTTTGTTATAACTGGTACCGAACAAGAGATTTATAGAAGTGGCTTTAAAAGAAATCGGAGAAGTTCTTGGAACTCGTAAACCAACCGAAAATTAAACCACGAGGTTGGAGTAGTAACTAAATAATAAAAAATAAAAACAATAAACCATATATTTCAAAAAACAACTGAACAGCGCGACCAATAATTTTAATTTATTGCATCTAATATTTTTTGAATAGTTTACGCATTCGATTTTTTGCGATTATGGCCACTGCAATTAATGTTATTCAACATCCGGCTGACGCCAATAATGATACCCATTTTAAACAAGTTGTAGGCTACAACACGCAATCCATGCTTGTTTCACAAAGTATAAACCTATCAAATCGTGTATCTAATTCTCTTGTTTCAGATCACGAAATTTCTAAACAAGAAGATTTTGATAAGGTCAAAAGGCCTATGAATTCTTTTATGGTGTGGTCTCGCGAAAAGAGACGAAGGCTGGCACATGAAAATCCTAAAATGCACAACAGTGAAATAAGCAAACGCTTGGGAGCAGAATGGAAAGTTTTGACTGAAGATGAAAAAGCTCCTTTTGTTTTTGAAGCAAAGCGTTTGAGGGCTGAACATATGAAATCACATCCTGATTACAAGTATAGACCCCGTCGTAAAGCTAAGACTTCATCAAAAAAGAATGATCAAAAGATGCCTCATGTAATAACAACTGATGGCAAACAAATTTTTATGCCTACACAGTACACCTCTGGGTATGCAATAACAGGTGGAATTAATTACCCATTTAATGGTTATTTATCTGCACTAGTGAATGGTCATGAAACATATGCAAGTAATACATCACCTTCAGCTGTCTATGGCTTTCCTCATGGAGCTGCCATTGCAGTGACTAGTACTATATCTCAATCACAATTACCTTCTTCATCTAGTTCTTCAGTAACAGCCGCAACTGGTTTATCTGCTCAATATAGTGTTGTACATTCAGGTACTCCAACTTATATGTACAGTCCATTAGGATTTCCATACCTTAATGGTGCTGCTATGGGTTCATATGGTGGGCAGCCTTATTCACATACTTTAACACTTAAACCTGACACATCATCTATTACTGTATCAACAGAAAAGGCAACTAGTCATTTATCAATGCATCATGATGCTGTTCGTTCAACAATGGCACCAAGTGTTTATTACCCGATCGGTATTTATCCTACTGCTGTTGATCATAATTGTAATCTAACATATGATCCAAAAATACCTTATGCGACAATGACTTTATTAGAACAACCCCATGGAAATATAAAACGGGAACAAACCCGCGAGGGAACAACACGTAGTAATATTTCAGATGATCACCAAAATAGAAGAGTTAATACTCCTACGTATTCTATTTCATCAAACACTGCAACGCAGTAGAAAATTATTTCTAAATAAACATATCAATTTTCATAATACAGATGATGTTATGACTAAAATTAAACCTGCTTTAAGTTGAAATGAGTTGATAAGCTCAACTGTAATAACCTTAAAGCATTTGTATAGATATGAGACTAAATTTATAAGTTCTTTTATTACTAAAGATACAAAATTTACTTAATGTATAAACAAGTAAAGTTATTTATTATTATTTTTTTAATTTGCTTGTAGAAACTCCATATTTTTAAACTTATAGTTTTGAACAATCTTCAAATCATATTAAGAATTTATATGTATATACTGAGCTAGATACGTAAGCA
>evgR014453c0g1_i01_Hydra
AGAAAAATAATAGAAAACTGTTTTGCTTTTGAGAAAATTGGCTTTTTGAGTAAAACAAGCTTAAAAAAACAAGAAAATTAAAGAACTTGTAAAATAAAATGTACACGTTGATGCTCAAGAAGAATGTCCTGTTTTTTTTTATATTTAATGTTGAACCGGTGCATAATGGTTTTAATTGCAGCAATAGTTGGCTAGGGTATGGAAGATACTGCTATCTTTTTCAGAAATTATATTCACCTAGATTAAACTGGGCGAAAGCTCTTTTATTGTGTCAGAGTTACGGGGGAAGTTTATTAAGTGTAGAAGATCAAGCTGAGAATGACTTCATCAAAGATCAGATTAAAATTGATAATCACTATCAGATTTTCAATCCTTATTTAATTGGTCTTAATGATATCAAAACTGAGGGAACGTTTGAGTGGGCTGATAATACAACTGCCAGATTTTTTGGCTGGAATATAGGTGAACCTAATAATCAAGGAAACGAAGATTGTGTGGAAATAAATGTTAATGGATGGAATGATATTTTATGTGATTCGAATCTTGATAGTTTTATTTGTAAACGCAGAGGTCAAGATATCGATGAATGTGCAGGTTCCAGCATGTATTGTAACTTGACAAACAGTGTTTGCATAAACACTATAGGAAGCTACACGTGTGCATGCAAGTCTGGATGGGTGTTGGGCAA
>evgR014476c0g1_i01_Hydra
ACTTTGAATATGTTGTTGAAGATTTTAACGTTAGCAATCTTGTTTATATTGAAATAACAAACAACGAGAGAGACAAGATATTTGATTTTAAAGAAGCGACAAAGAAATACTCACACTTGTGTCTTGTTAAATACACTATTAACTCAAGAAACATTTCTCTGATGTCAGATGTTGCAAAAAAGATGGCGCTGCGTGGAAAAGTTGATAATGTTTTTATATTACTAAACGGATTTACTGAGTATAACAATTCTCCATTGAATATGATTTTGGAATCTTTTAATTCCTCAAATGTTACCATAGGAACAGTTTATTTAGAGATTACTAATTGTGAGTGGAATACAAACTTCGATGACCTTCGTTTAAAAACGTTGCAGACTAATAAATACAAAAACTCAATTACAAACCTGCTTGTATATGTAACAAGTTTTAATCCAAAAGGTTTCTTAGTAAAGGTAATTTATCAAACGCCTATTTTACATTATCTTTTAAACTCTTTGATTGGCTCATTCATTAAATC
>evgR014486c0g1_i01_Hydra
AGGTTTCATAAGAAACTTGTTTTTTATCAATGGCAATTCAAGTAGCCAAAGTCAACCTGCTGTTGTTCTACGGAAAGAGTACATTGTTGGCAATGTTCCCATGCCACTTGAGAAAGGAAAAATAATTGCTTTGATACCGAATCTTAATATGGAATATTTTATATCATTTGATGTTTATCCTAATGAATTTCTTCCTGATTATCATAGTTTAATTCATTTCACTATTGGCTCGGATAATGATAATTATGGCGATAGAACACCAGGTATCTGGTTTGATGATAGGAGTCTTGGAAAACTTCGAATTGGAGCTCCAATAAACGGCAGTCATAGATATTTTTTAACAAATCCAGTTGCATTAAATCAATGGTTAAATATAGAGGTTTCACAAACTTTAAAAAATTCACTTTATGGGTATCAAGTTAAAATAAATGGTAAAATTGTTTATTCTGTAAATAACAACCAAGCTG
>evgR014496c0g1_i01_Hydra
AGTACAACAGTAATGAAGTTTTAAATCGATTTCAATCAATGGACTACTCAACGGCTGAAAATTTGTTTCTCACTATTTGTCCCGTTTCAATTTATAAGCGTGATTGGTGCTTAATCGAAATCAATGATCCAATCTGCTCTAAAGATAGATTTACAAGAGACAACACTTTTTTCAAAAGTTTTAATTATTTTAGTGGTATGAATGAATGGTTAAATGAAAATATATATATTTATGCAAATAGTGACCATGATGACCTTTCCAACGTAAATTTAAGTAATTATATATCCAAGTCTTATTACAATTGCGTTAGGTTTGATTTGGGTTATTTTAAGTCGTCG
>evgR014508c0g1_i01_Hydra
CACTCTGAGAGAAGAAAATCAAAGCACAGAGGGTGTTCATCCATACTTTGTTTCCTTGCCAGTTAAATCAAACAAGGACTTCCAAGATTGTTTAGAAAATGCATTCAAAGAATTAGCTCTTCCACATCATCGCGGAAGAAATGCAATGGGTGATTCTTTCTACAGCTCACAAGGTAGACATGATGATAAATTTGATGATGCAAATGATCATATCGAAGGTAAATTACAAGAACTTGGTGTCCAATCTATTGAAATGGAATCATTCCAATTATTAGATTTGGCCAGATGTACAAAAACTATCAAAGTTAAGGCTTGTGCTGCAACACTTGTGTTGGCTCAACGTGTTACCAATGTATTTATTGATGTAAATTTGAAACATACCAGAGAAGAAATG
>evgR014536c0g1_i01_Hydra
AGATCCAGAAAATACCTTTGAAAAGAGAAAGGCTGAAGCATCAAGAATTAGACAACGTTATTCTGATCGTATCCCAGTCATTTGCGAAAAGGCTGAAAAATCAAGTTTTCCAGATATTGATAAGAAGAAATATTTAGTTCCAGCCGATCTTACAGTAGCCCAATTTGTGTATATTATTCGTAAACGTATCAAGATTGATTCAACAACAAACATCTTTATTTTCGTTAATAACGTTCTTCCACCAAGTGCCACCCTTATGAGTCAACTCTATGAAGAACAACAAGATGAGGATGGTTTCTTGTATGTTACCTATAACGGAGAAAATACATTTGGTCAAAGTGAAGAACTCTAAATATCATCACCAAATCATCACCAATAGTATTAACCCTCCTTTTCCAATAGCAATTATTAATTAATATAGTGGCGGAACAAGTGTTGACTCTCACCAATTGTTATTAATAATTTAAAACGTG
>evgR014541c0g1_i01_Hydra
TCCTGTCTTCCTAATTTTTCATACTCCGTGCTGGTCCGGTGTGGTCCGGTATAGTTAAAAACAAGTTGTTAATCACAATTTCGAATAAGGTAAATTCTCATGGACGCAGAGCAAAAAAGAGCAAAAGAGCTTTTTGTTTCAAACACTCATGGAACAACATTGATAGATATTTATTCAACTGCAGCTCTTTCTGTATTATGCTGCGTGTTATCAATTGGATTTCGGCTGATCTGCGTTCAAAATCAACTGCATTGGTTATTTGATTTATGTGTGGATAATGTGTTTATTGTGATTCCCATGGTGTTTTTCTTTACAGTGTTTTCCAATTTTAATTTGCTATCAGTTTTCATTCTGCTATTTGTTTTTTTAATTTTGTTTTATTTGGTCACAGAAAGAAAGTATCGAGTGAGTGTTGTGAACTTTATTCAAACAATGGATACTGGAGATTTTCACTACGTTAGTATTTACAAAGCCAACATTCAACTTGCTACTGCTATTTGTATTTTAGCTGTAGATTTTAATGTTTTCCCGAGAAGATTTGCAAAAGCTGAGGTTTATGGCATTGGGGTTATGGACTGTGCTGTTGGTTCCATAGTATTTAGTACCGCCATGACTTCGTCATTTGTACGACAAAGCAATAGACTCAATATTCTAACCCTAATTAAGTCTTTTAAAAGCTCTATGTTTTTGTTAGCTCTAGGTGTTTTTCGTGTTATATTGATTAAAGTAACCAATTATCAAGAGAACATCTCTGAATATGGAGTTCATTGGAATTTTTTTATGACTCTAGCCATTGTTAAATTAGTTGCTTCTGTGATTCTATATATTTTTCCGGTTTTACTTAAGCAGACCTGTTTAATTATGATATCTTTTGCAGTTATTATAATTTATCAGATTATGTTATCATTTGGGCTAGCAGATCTAATTCAAAATGGTTTTAATGGCGATAACGAGAGAATTAATTTTATGGATGCTAATAGAGAAGGAATTTTTTCAACTCTTGGATATACAAGTATTTACTTTTTTGGCCTTTTTTCTGGAAAGCAGTTAATTAAAACAAAATATAAAGCAACCAACAAGGTGTGTTGGTTAGGTTTGTGGTTTTTCATTGGTTGCTTTCTTACAAACATACTTGATTGTTTATCAATACCCATATCAAGGCAAATGGCAAATTTGTCTTATTTTTGCTTTCAGATTTCCTTTAATGCTTTTCTACTTCTTGGTTTTTTAATTGTGGATATAATAACCTTTGCAATTGTGAATACGATGAAGAGAAAGAAGAACGAGAGGTACAATATTCCATTTTTGTATAAAGTTGTCAATCAAAATTTGTTGGTTTACTTTTTGTTAGCCAATGTGATGACGGGTGGTATTAATTTTATTGTTCAAACTTTGGATCAGACCTATTTGACAAGTTTGCTTATTATAATTACTTACATGTTTTTGTTAAATTTTATAACAACTACTCTATACATTATTTCTGGGGTTAGAGCTTAGTTTTATTACACACATACTCTCTCTCTCTCTCTCTCTCTCT
>evgR014545c0g1_i01_Hydra
TCATCACCGGATCATACGGCTGGATTTATCCTTCATCCCCATACAATTACTTCTCCTTCACCTCAGTTAAATCATGGGCTGTAATGCCACTCCTCTTCTGGACAATCTATTACTTTTTCCTCATTCCAAGAAAGGTCAACGTTTGGTTAGGTTTTATCATTACAGGTGTTAGTGGATATATTACTGAATTCGTTGTTGGTTATGTCTCTGCTGTAATCTTCAATACCACCATGCAAGAATGGCCAAATTCACCTTTCAAGTTTGTTGGTACTTGGGGTTGTTTGTTCTTGTGGATTGTTAACTCTGCCATGTACTATTTCATGGTTTTCAAAATGCCACATATTTTGGTTGATCACTTGACCTCCCCTTTGAAAAAGATTGTAAAATAATCATGATAAC
>evgR014557c0g1_i01_Hydra
AAAAATCCACAGTTTTCAAGTGAAGTAAATTCAACACCTTTCTTTTCAAAAGTGAAAATTCATAACATTTTATTAAACCTATTTTACAGAGGAAACATGCCAGAAAATTTTAAAGATCAGCCAGAAAATTTTAAAGATCAGCCAGAAAATTTTAAAGCATCGAGTTCATCAAGTGAATGTCAAAGTATGGGAAACAATACATTTACAGTTGGACGATTTCACGTCGCAAAAAAAGCAAATTTTAAACCTAATGGTTTTCTACCTAAAAACGGTTATGTGGACGCAGATTCATTAAAAACAACAAAAGCAAAACAACCTAAATCCCAAAAGAAAACTAAAACGTTTTCTTTAAATATTTTTGCAGTATTCCAAACACAGAAAAATTATAAGAAGATTTCATAAAATAGATTTGGCAACTTGTTTTTGTTTCACCCAAGAGAGTGAAACAAGAGAGAGAATCAGTTTCACCATTAATCTTTTAACAACAAACTTATCATAGGTAAACCAATATAAATTTAAGAAAATAATTACGAGAAGAAAAAGGTGGATCGCTAAATCAATCTGTATACCCGATGAGCGGCTGGATTATTGCGATTCGGTACTGGTTAATTTCGATATAAATCGAATCGATTTTAATCGCTTGTTAATAAATATTAATTTAGTAATAAGGAAATTTGAACTCAATATTTTTTTCACAATGCAATGTTTTTTTTTCTCATAAAATTAATATCCG
>evgR014564c0g2_i01_Hydra
TTTAATTATTATTTAGGTTGGGTGCTAATTATCTAATACTTAGTATTCAGCCTAAATAATATAAATGAAAATAACTTACATCAGGTTGACTTTTGTGTTTTTATTCGTATCAATTTTTATCACGTCAAAAATTAAGTGGTTTCAATCATTTTTACATATGGATATGTTGTATCATCTAACCTCAGCCAACATAACCCTCCAAAACCCGCACCCAACAAGAATTGCTTCCTTTAGGTATAATTTCTTAATGATGATATGTGCTTTTGCATGCTGTGTTTTTTGTTTTCTAAAGATTTATCAAACAAATGTTTTCCCGTTTCTGATTCGGCCTTCCCTTGCAACAAAACGCCAAATTTTCTATCAACCTTTAAGGAAGTGAATTGGGAGGTTACAGGAAACAGAGAGTTAAGCTAAA
>evgR014573c0g1_i02_Hydra
TTTTTTTTTTTTTTTTTTATTTGACGGCACATAAGATGGCGCTACTGTTGCTGCTCATAAAATGTGTTATTATGATGAATATATGTTTTAAAGTATGTTCTTTTGAATCTGATAATGATTCTAATTCAGAACCGACTGGTTATATGAGAAGAGAATATTCTTTAATTAAACCGTTTCATGCTTTGCAATATTGGGAAATTCTGGGATCAACTATGGTTTCTGTAGATTATCTTCGTCTCACTTCAGATGAGCGAAGCAAAGCGGGATCAGTTTGGAACATCTTGCCATGTTATATACGCAACTGGGAGATACATATTAATTTCAAAATTCATGGCCATGGCACTCGTTTATACGGTGATGGATTTGCGTTTTGGTATTCAAAAGAACGCGCTCAGCAAGGTCCAGTTTTTGGTAATAAAGACTACTTTACCGGTTTGGCGTTGTTTTTTGATACTTATGCTAATCAAAATGGCGAGCACCAGGTTAATTTCATGTTTTCAAAGCATGAACATCCTTATATCTCTGCTCAAGTTTCAAATGGATCAATACATTATGATCATGATCGGGACGGTACACACACGCAACTTGCGGGTTGCTCAGTTAATTTTCGTGGTGTTTCTGGCGAAACACATGTTGCTATTAGATACCTTGGCTCAAAAAAAAGATTAACAGTACAATATGATATAGATAATGATGACTCTTGGAAGGACTGTTTAGATAAGTATGGCGTTGAGATGCCAACTGGTTATTATTTTGGTTTATCTGCACAGACTGGTGATCTTTCAGATAATCACGATATACTTTCAATGAAATTTTATGAGCTCGAAAACGAAGAGGAAGAAGGTGATTATTCAAACGTTATACCGAATTCAAGCGACTCAGAACCTGATAGAGATCATGTTGATGATCCATCTAAACCTTTAACTCGTCGTGATCAAGCCATGAACTGGCTCTTGGTGGTTCTTTTGTTTGTTATTTTTGCTGGTGGTTTATCATTTTATTTCTATCATAAAAAGCGAAGTGATATGAAGAGATTTTACTAAGAATATTTTTGAAATGAAAACATTTGAAATATGTTTATATTGAATTTCTATTTAAATACGTCATTGTGGTGGACACTTAATGATGCTAGAATAAAGAATTGTCTTCTTTGGTATCATATTTAAATTGACATCTTCGAAAAATAATTATTCCGTGCACCTAGATGCATTGATCAAGTATTGACGGGAAGATCTTTTGTCAAAAATATTTTTGCTGTATATTTTGAAAAAGAAATTATTTACACAATTTTTGAATAAAATTTTTACAAGTATTTTCATATTTATTTGATTTTGAAAATGTTATTATTATTATTTTTTTTTTTGCATTGATGCATTTTATTGTAAGTGTCATTTTTTAGAAGTTCTAGAAAATTATAAACATAAATTTTCGAATTTTACAAAAAAA
>evgR014577c0g1_i01_Hydra
CATTCGTTCATCAAGTTATTTTGCTTTTTACAAACAAAGTGAAATATATTAAAAATAAATACACCATTGTTTGACTGAGTTGTTAATGCCCGTGCTTGTGTATACAAACTTATTTAAATATTGTAAAAGGTTTGATTCAGTATTAGTTTGAAATGCAAAATATTTGAAAATGCAACTTAATGATATAAAATATTGCAAGTGTAGTTAACTGGACAGTGATGCAACATTCTGAAACTGAACCGCTAATTTCGGGAGAACTACAAAAAGAAAGAATTTCAACATCGCTTTTAGCTCATATAACTGCTTCGATATCGTGTGTTTTGTTTGGTTTCGGTGTTATAAGTACAAAAAAACTTAACTCAGACGTCACGCTTGGAATTTTATGTCTTGTATTCTCATTCGTTGGTAGTATTATTGCTGGAGTTTCAATTGATAAGATTGGTAGGAAAAAAACTATTTTTCTGATATCATTTTTCTATATATCTGGCTATATTTTGGCACTACTTGAAAAATGGAATGAGTTAAAGGCTCCTATCAGACCTGCTCTACTTGAGTTTTTTATGTATGTTGCAACCATTCTCTGCTGCATTGGTATGGGTATGACCTCTTATGCAGTTCCTGTATATAATGCAGAGATATCCTCTACTCATTTACGTGGGCTGATGGGAGCTACAACTCATCACATGATGGTTTTTGGATTGTTATTTAGAGCAATATTAGAATGTTATTTAGAATACTTGTATGTATATTTAATAGGAGTTTCATTAGTTTTTGTCGGCTTCCTAATAGCTTTTATGCCAGAATCACCCAGGTGGCTATTAGCTCATAATCAACGAGATAAAGCATTTCAAAATCTTTTATGGCTTCGCGGTACAATAACCTCTGCGGAAGACGAATGCAATGATATAGAAACAAGTTTAGCACATTGGGAAGCAGCATCCGTCAACGACTACCGTCCACCAGGACTGTTTTTACCTTTGTT
>evgR014587c0g1_i01_Hydra
GAAAGTCAAAACATGGCAAGAAGAGAAAAGGTTATCCAAAGGAAATTACTAAAGTATTGAATGATTGGTTCTTTGCTAATCTTCAAAATCCATATCCAAGTGAAGATGAAAAGAAAGAATTGGTCAATCAAACAACATTATCTCTTCTCCAAATTAACAATTGGTTCTCAAACAAGAGAGTTCGTTATAAGAGAAAGATGCAAAAGGAAGGTCAAGAAGTTGTAGAAGGTGGTGTTGATATCAATCAAGCTTTCCGTTCCATCAAGAAGCAAAAAGATAAGAAGGAAGCAAGCTCAACTTCAAGTCAATCAACACCAAAGAGAAAGAGATCTACAGCCACTAAGGATTTACAATACACAAAGAAGAAGAGACCTACTCCTTTAAAGACTGACTCTGTTTCGAATAGTTCTGCTGCCAAAATTGTTGATTCCTCTCCAACCACTCTCAAAACTCCAACCATTGCCTCTTGTACTCCTACAACTGTTGTTTCAGCCAATGTTACTTCAACATCCATTTCCCACAGTCCAATTACTTGTCAAG
>evgR014591c0g1_i01_Hydra
TTATTATTGTCGCATGGTCATTATCATTTTTTTTTTATTTAGAAAGAACGTTTGATGTAAATAACTCTGTACAAAAAAGGTATATTTTTAGCTGTTTGTTGCAAAAAATGGAGAAAGTTACACCTTTGATAAATTTTTCTTCAACAAATGCTATTGGTTACGAATTGGAAGATGAAAAACTTTCAAATGTCAGTATATTCATCTTCGTTTTGTTTGCATTTAATTTTTTGTTCGGTACAATTGGAAACGGATTAATATTGTTTTACTTCCGCGTAAAATACAAACCACGAGGGATATCAATTGAAAAACGCATAATTACTGAATATTTGTTTGGCGTGCTAGCCCTTGTTGATTTCTTTGCTGCTGCTTTGAACTCTGTTTTGTACATGTATTGGATTGTAACAGATTACAAATGGTATTTAGGTTATTGGAGTTGTAAGATACTAGTGCCACTGGGAACAGTTGCCACAACAATGTCTGGAGGAGTTATTGCAGTGTTATCGTTTGATCGACATCGCGCCATTGTTTATCCATTTAAAAACCACTTAAGACTTTCTCACATTAAAATGAGTCTCGTTTTTGTTCTGTTTTACTCGCTTGTTATAAATATTTACTATTCACTAAACATTTCATTAAAAGATACCAAATGCATGGTGGTTGACCCTAGCAATAAAACATATAGCATTCCCACTATAATTTATTTTTTGGTGAACGATTGCACGTTACTAATTGTTATAAGTTTTAGCAATTATCGTATAGTGTCAAAAACATATGGCTTAGATTTCAAACGAGAAGACAGAAGAGACAATAAACGGATAGTTCGATTACTGATTGTCATAACTATTGTATTTTTTCTGTTAACGTTACCTAGAGATGTGCTTCATTTTGTGTTTTTGATGTCGTGGATAGTTGGTCCTGGTATTGAAAGTTCAAAATCATTATTAAATTTAAATTCTTTTTTCAAAGTAGCAAATGTAGCAAATTCCTGCGTTAATCCCATAATATTTTATGCAATGCATGCTGGGTTTCGAAACTTTGTCAAGGATTTTTTGTGTAAAACCAAATCTTTTGTTGCAAGCACGAGAATCGGCTTAAGTGAAATAGAAAGTTTTGCTAACAGTCGCAAGGTTTAATAAAATTATGTAAATATTTTACACTGCTTTCATTAAGTAACAAAACCGTATTTACAAATATTCCTTCTTAGAAATATTCTTTCTTATACTAGGTGTTAAATTTGGT
>evgR014623c0g1_i01_Hydra
AGGAATGATTCAAGCGTTTCTGACAGCGATAAAGAGGTTTTAACTATTGAAATTTTAAACAATAAAACTAAAAATGTATTGCTAAGCTGCTGTTATCGACCACCTGACGGTTTAAGTGAGAATCTGAGCGCGTTTTTCGAACAAAATGTTTTCAAAACAGGTTTCAAAGAAAGAAAAAAAAATTTTATTATTGGGGATCTAAACATGAACTTCTTTAACTATGAAAATGATGGTAAAATCAAAATTTTCTACGATTCAATATTTGAAAACGGAGCAATTCCATTAATTAATCGGCCAAATAGAGTATCAGCAAACTCATCAACGCTCATTGATAATATTTTTACAACTGATATTTTTAATAACAATATTAAAAAAGGTA
>evgR014630c0g1_i01_Hydra
AGTGAGGGAGAGGATGAAAGCTGAAAACCCAGACAAGTATGAAGAACAAAAACAGAGCAAAACAGAAAAAAACAGAAAGATAAAGGAAAGAAAAAAAGCAACACCTAAGAAAATTGATGAACTTTTAAGAAGGGTTAAAAGATCAATAAGAAAAGGTTTAGAAAGGACCAAGAGAAATTGGGAAATTAAAATGAAGAGACAACAATTAATTAAAAATTTGTCTACGCCACCTGAAAATCCAACTATAAATCTTGCTATCAACAATGAACTAGACAATAATGTACTTCCTGGTCCCAGTCAGAAATTTAGTTCCAGAATAAGCAGTGGGCTAAAAGTTTGTAAAAAATAAAGAGATAGATTGAAGACAAAAATACAAAAATTAGAAGAAAAAAACAAATTGTTATAGCAAAGATGTGATAAATACAAACACAATTTTTACAGACTGCATAGTCTATAATAATGTATAAAAATCTCTTGTTTCTTTAACAGTAATTGATATTATGCAAATCAATGCTTTACATAGTAG
>evgR014644c0g1_i01_Hydra
TGTTGATCTTCAATCTTGAAAAATAGTTTGTAATTATAGTTGTTGAAGAAAAGCAGCAACATATTAACCATGTGCTTCAATTTTATACTCAATGGAGAGTTCCAGAGTTCAATGGAGACTTCCAGATTTTGTTGAAAAGTTCAGGTTGTTGGTAGAGAGGCAACAAACTAAAGCGGATTGTTCAACATGTGGTTGTGTTGATTTTGAGCTAAAGCCTGCTTATGCTAATCACTCACTTACAATAGGCATTTGGAAAAGAATAAATAATGCACAACGTCAACCTGTTAAGTTAAAGTCAACCTATTCAGCGTGCTTTTGCCTAAAATTGGCAAAAATTACAATTCGTATAACACCCGACAAGGTAAAAAGCTATAACAAAGAAAACAAGTGCATAACTTCAATAACAAAATCACCACAGGTTGCTCAGAGTTCCACATCAAAAGTAAAGAGTTTAT
>evgR014646c0g1_i01_Hydra
GTCTCCTGTTCCTTGTAAACTAGGTACCTATTCAAATGGGGGTAACGCAACATGCTTACCATGTCCTGATGGAATGTCTTGTATTGACCCATCGAAAATCCCTGTGCCATGTAAAGATGGAACATTTAGTTTTAATGGTGTTGCATATTGCTCTCCATGTCCTGCAGGAAGTCCATGTCCTTCATCCTCAATAGTAAAATCAGAAATGTGTCCATTAGGAACTTATTCATTAAATGGAGAATCCACTTGCAATGGTTGTCCATCTGGTTATATGTGCCCAAATCCTTTTATTTCACCAGTCCCTTGCCATTTTGGAGAATACACTGATGAAAAAGTAAACCAAACATTATGTGAAAAATGCCCTTCTGGTTTTGAATGTCCTCATCCTGAGGGTAAAAAAGTACTTTGTAAACCTGGCTCATTCAGTCTGGGATCAGCTGTTTCATGTACATTATGTCCTGCAGGTTATAAGTGCCCTAATGCTAATAAGCGTCCAGAGTTGTGTCGCCCTGGTGAATTTTCATTAGGTGGACAAATTTTTTGTAATCTCTGTCCAGCTGGTAAAGCTTGCAGCTCTCGATCGCTTAACTCTTCTAGTGACTGTCCACTTGGTACATATTCTTATGAAGGTCAAGCAGAATGTTCTCCATGTCCCAATGGATTCTCGTGTGAAACTCCATTTTCATATGCTGTAGTCCCATGTCAAAATGGCTCTTACTCTGATAACGGAATTTGTAAAAAATGTCCACCTGGATGGGCCTGTCCTACACCAGATGGGCTTGGCAACAGACAGTGTCCTTTTGGTACCTACACAACTGGAGGAGCTACTGATTGTACTCAGTGTCCTTTAGGCAAAGAATGCCCAACAACAGCTTCTGCACCAAGAGACTGTAAACCAGGATCATTTTCTACTGGATCTCAAATGTTTTGTACACTATGTCCTTCTGGAAGTTTCTGCCCAAATACTGATGGTAAAAATATTGCAGCTTGTGCTTCTGGCAGTTTTAGTTTATTAGGACAGGGCACTTGCACTCAATGTCCTGCTGGATTTTCTTGTCCAAGTCAAAATGAGGCCTTACTGATTCCATGTGCAAAAGGATGGTATTCACTATTAGGAATGGTGGAATGTAAACCATGCGAAGCTGGAAGTTTTTGTCCTATTAATACAGTTAAAGGAAGTCCATGTGATAGACCCGGATATTATAGCTTGCCAGGGTCTATTGTTTGTTC
>evgR014647c0g1_i01_Hydra
TTTTTTTTAAATAACTAACAAATGTGCAAATAGACAAGATGCTAGCATTGTATTTAATTTTTTTGGCAAATATATATGAAGACGTTGCAGCTCAAGCTTTACCACCTGTTAATCTACCAAACATCACCGATAAACCAATAACTTGTGAAGCCGGTTGGATTCCTTGGGGTACTTCTTGTTATGGAATTATTACCGGATTGGCGAAATCTTGGTCAGATGCTCGTGCAGACTGCCTGAGAAGATTTGGTGATCTTGTTGTCATTGAAACCAAAGATGAACAGAATTTCCTGGCAAATGAAATTTCTCTTGCTGATGTCACTGGGTATTGGATTGGCCTCACTGATTCTTACCAAGAAAAGGAGTTTATTTGGGTTAATAAAGAAAAATTGGACAATAAAACATCGTTTTGGAGTTCTAAAGAGCCAAGTAACAGCTTTGGCAACGAGGATTGTGTAGAAATGTATAGTAAAAACTCAGGACTCTGGAATGATAATCATTGCTCATCAATGCGCAACTATATTTGTGAAAAGGTTTCAGGTGTCCATAATTGTCCGGATGATTGGGTAACCTATGGTGATTACTGTTATCAGATAAATGCTCATCCCAATCAAAAAGTCACTTATTCGGAGGCTGAGAACCTATGTCACAAGTTTAGTGATCAGTGGAAAAATTTAATTGATGTAGCAAAGGCTGAACTCACAAGTATTTCGTCCAAGGATGAACAAGCAATGTTGGTTGGAGAGTTTCAAAAGGCAGGCTTGTTTGGTTCTGGAAATTATTTCTGGATAGGATTGAATATTTCAAATGGACTTTCTTGGTCAGATCATTCATCTGTAACCTTTACTAATTATGATAATTTGTTTACAAAAGAAACAGATAAATGTGCAAGTTCCAGTCTTGAAAGCTCTGGAAAATGGAAAACGTTCTCGTGTGACTATAAAAATAACTTTGTTTGCAAATTAAAAAGAGGTAAAGACAGGTGCTATAATCCTTTGGGTATGCAAGGTTTTACAATACCAAACAGTGCAATAACTGCTAGTGATCAACTTTCTTCTAAAACACCACCCGAAGCAGCTCGTTTATATTTAATGAATGATGTTTCACTTGACGGAGCCTGGTGTGGAAAGAATGGAACAGACCTCTGGCTCCAAATAGATTTTGGAGAACCCAAAAAATTAACTTACATGGCATTCCAAGGAAAAGCAAATTCTAAAGCAATTGTTACATTATTTTATCTTAAATACTCTATGGACGGTGTCTCGTGGGACATATTTGGAGATAATCCTCAAAGTTTGTACTATAAATTCCCTAAAGTAGGCGGAAGATTTTTGTGGTCTTCTTGGCGGACTGATTTTGAAAAGGCATTTACTGTTTCATTTGAACAACAATTAATTGCTCGCTATATACGTGTATATCCTGTTCAGTGGATTGAATCACCCTGCATGCGAGTTGAATTTTATGGATGTTCCCAAGAAACTGAAACTCAAGCTTGTGGGTTAGGGTGGAGAAAAGATGTAAAATCCAGAAATTGTTATAAAATTTTAAACCTGAAAACTACATGGGATTCTGCCAACAGCATATGTATTAATGAAGGTGGGCAGTTGTTGAGCATAACTAATCCTGATGAACAATTATATGTCAAAAAAGAGTTGCAAGCATTTTCTGGAAAATCTGATTTTTATTGGATAGGGTTAAATGATCTTGATTCTAAAGGAATATACAAATGGAGTGATCAGTCTCCTACCCCTCTTGTTAATTGGTATAGCAGTTATCCAAAAAGTGGAAATTTGAAAACAAAATGTGTACTTGTTCGTTCCGTCGATGGATTTTGGACTAATGAAGATTGCTATGCATTTAATCCATTCATATGCAAAAGAAAATATGAAAACCCTTATGAATTCAAGCAGCCGTTCAATAGCAATTGGCGTAGTTCCAGTGATTACTACTGGCCTTTGGACATAGTTACTGGTAAAAATGTCATGGGAACAACAACAGCAGAATATTTTGGATCAGCTCAACCTAGTCCTCTTTCTATTTTTCCTGTGCCGTTGTCAAATCATCAGCCAAGAAATGTATTATCATTTAATGGACAAGACAACTACTTAAAAGCAGTTAATATAAGGCCTTGTTCTATAAATCCTATTTCTAATAATTGTCCTTATGGTTTTTCTGTATCAGTTTTATTTAAATACAATAAGGAATCAGCAACAACTAAGCGTTTCTATATTATTGACAGTTTTACAAGTATAAAATCTATGGGTTATAGCATATATATATATGGTGGAATTGTTAATGTGTTTGTTGCTGGTACAAATACTTACTATTCACAGAGTGCATACTATATTAAAGATTCTTGGAATCATGTTGCATTCACCTATTCTTCACAAAGCGGATTATTTCTCTACATCAATGGAGATATTAAAGCAACTCAGTTTTTTGGTCGAACCACTTTTAATGTACGATCACAGGCTACTAACACACTATATTTTGCAGCACCTGCAACTTCATTTGGAGCATCTGCTCCCGACTTATCTAAAGTAGAACTATCATCTGCCGCTATTTTTGAAAGAGAGTTATCTGTATCTGAAATAAAACAAATTTATTTGTCAGAGTTTGGACGATGTGGTTGGGGACAAGAAGAGGTTGGAACATATTGTTATTCAATTAATTTCAGTGAAATGTCATGGACTTCGGCGAGAGATGCATGTCAATTGAGCAATGGAAGTCTAGCTAGCATAAACAATCCAATTGATCAAGCTTTTATATCATTGAGAGCTACAGAAGCAAGATTTCCAAATGGAGTTTGGATAGGATTAAACTCAAGAGAAATAAACCATGTCTACAATTGGACAGATGGGTCTCAGTTATCATTTACATTTTGGAATACTATGGAGCCAAACAATTATGGTGGACGAAGAGAAGATTGTGTCCATACTACCAGGAATGGACACTGGAACGAT
>evgR014650c0g1_i01_Hydra
AGATTTCTAATAATATTATATTGCTATGAGCATTGTACTACTACGAAAAGAAGATGATGAAGATTACTACCACAAACTGCTTAATGCAAATGGATTTTCTGTGCATTCTATTCCTGTTCTTTCCTTTTCTTACTATAATATTGAAAAACTGGAAGAATTTATTCAAAATATAACAGAGTATTCAGGGCTTATAATCACAAGTAAACGTGCTACTCACGCTCTTGAAATTCTTTTTAAGAAAAATAACTTTAGTTTTTCTATACCTCAAAATTTTTATATATATGTTGTTGGAAAATCGACTGAACTGTATCTTCAAAAACTTGGTCTAAACTCGCTCGGCTGTGAATCTGGTAATGCTGATCAGCTGGCTGATTTCATTTATAAAAGAAAACATTTTCACTCAAAATCGCTTATGTTTGCATGCGGAAATCTTGCGCGGGAGACAATACCTGGCAAATTACAAAACCTTGGTTTGACTGTGGAAACAATACCTTGTTACAAAACCATTGCAGATCCTAAATTTCCAGAAATATTTTCACAATTTATAAAGCAGTATAATATACCAAAAGTCATAGTTTTTTTCAGTCCATCAGGTGTTAAGTTTTACTTTTCAGAAATTTGTAACATCTGTGCTAATTTTGAAAGAACGAAGATTTTGTCAATCGGACAACATACAGCAGATTCTATTCTCGGTATGAATAAAAACGTTGACGCAGTTGCAAAAAGACCTACACCTGACTCTTTATTAGAAGCAATACAATCTGTCCTTGATATATCAGGAGGAACTCCGTTTTAGTTTGAGAGTAACATGCTGATCAATATTTAACATTATTTTCAAACCTATAGATTGAATAATTAACTTTTGTAACATTATTTAATAAAGTTTTCGTGAACGTTCTATAAATAGTAGAATAATTTATTTTTTGTATAATAATTTTTATTACCAGGTTTAGATGTTTGCTTTTAATGTTTATTATATAGGGG
>evgR014672c0g1_i01_Hydra
GTGCATAGTTTGTTATTGCTGTTTGTAGAGCATGTCTCTGAGGTATATGGCAAAGAAGTTATTACACATAATATGTATGGGCTGATACATTTGTCAAATGATGCCAAAAAAATTTGGTCCTCTGGACAATATTTCATCATTTCCATTTGAGAACTATTTATCTCAGTTAAAGAAACTCGTTCGGAAGCCTCACTTGCCTTTGCAGCAAATAGTTCGAAGACTTGGTGAGAAGAGAGCAGTTCAACAAAAACAAAAAAAGGAAAGTGGGTATAATAGTACATACTTTAAGAAGGAGCATTATAATGGGCCAGTCGTCAAATCGTTGAAAGTATGTGCTCAGTACAAGGAATTGTTTACCAAACGTTGGTGCATAAAAATTACAGATGGAGATAACTGTTTTAGCTTGAACAATGGCAAAGTAATTATTGTGCAGAATATTGTAAAAACAACTGAGCAGTCCTTTATTTTGTATAAACAGTTTAAAACACTTGACAATTTTTTTACATATCCATGCTGTTCTATGAAGGTAGGTATTTTTAAAGTTTCTTGCTTGGAAAACGAATTGCTATACTGTTCACCAACAGAAATTAAAAGAAAATATGTTCGCCTACCAATTGATAACGATTATGCTGTGATTCCTATGGTGTAATTTTAACATTTGTGCAATAAAATTTTTTTCAATAAAAAATTAATGTTTTTTAATGATTTGAG
>evgR014680c0g1_i01_Hydra
TAGTTATTATGCTTATGTTGTCTGTATTTATAAATATTTTCGTCGTAAATAACTGCAATCGATAATTTTTTTATCGACGGCATTTTAAACTGAAATCGGATTCCGTGCCATTAGGACTTTTTGGGTTTCAAGATCGCCAAAACGTCGGTGGTTTAGATAGTCACATAAATTTTAATTGAAACCTTTTTGTAGTGACCAATGGCTTTTTTTCTTAAACGATCAGTTTTGAGGAATTCTAACATTAGAATTTTGCAATCAATTATTAAATCATCAAGATCATATGCTGATGCTGGGCCGAGTGATGGTCTATCTTTAACATTTGGCACACCTAGAGAGACGTTTTTTAACAATGCAGTTGTCACACAAGTTGACGTACCATCAGGTTCTGGTAATTTTTGTATACTTGCTCAACATGTACCAATTATTGCTGTTTTGAGACCAGGACTTATGTCGGTCACAAAAGATGGAAACGTGGAAAAGTATTTTGTTAGTAGTGGTACCATAACTGTTAACAAAGACTCCACTGCTCAAATACTTGCTGAAGAAGCATGCCCACTTGATAGATTTGATTTAAATAGCGCTCGAAAAGCAGCCGAAGATGCACGCACTGCTTTGAGTGTTGCCAGCACAGAAGCGCTCAAAGCGGAAGCTCAAATACAGTTGGATACTTGTGATGCTTTACTTTTTGCACTTGATGGAAAATAAACAGTTAAAACAGATTGTTAGTGATGCTATAGTTTAATTTAGAATTAATCTTGCAATGTTTGCTTTTAAACGGCACCCTTTCCCTCTACGTATATAAAAACAATTAGACATAATATAGACATCAAAATAAATAAAAAATCTTTCCCTCTAAAAAAAAAAAAAA
>evgR014682c0g1_i01_Hydra
GGTCGGTGAACCGCGAATGAGCAATTAAATAATATTCAATTTACTAATTTCAGTTTATGTTAATTTTTTGGTTGTTTCCAAATATAAAATTTTAAATTCATTTAGCTGGGAATTTTTCTGAGAATTTAAAATGTTTAGCCATATTAAATCTTCTGAGTTAGCAAGACTCGGAAAAGTAGCTGTCCAGATTTCAAAGCTTTCAACTTCACCAAAAAGCTCAAAATATTTTTCGTTATCTTGGCTATCAGCTAGTGCTTTAGCAGCAGGCGCTTTAGGACTTGGTTATACTTTGTATGAACACCCTCTTGGAGCAGAAGCATTAAGTTTGCATCCTCCAAAACATAGGTGGTCACATACTGGTCTTTTAGACTCGTTGGATCATGCAAGTATCCGTCGGGGTTACCAGGTTTACAAACAAGTATGTGTAGCTTGTCATAGTATGGATCAAATAGCGTTCCGAAATTTAGTTGGTGTATCACATACTGAAGAAGAGGCTATTGAGCTTGCATCCCAAATTCAAGTTACAGATGGCCCAAATGATAAGGGTGATATGTTTTTGCGACCTGGTAAACTTTTTGATTATTTCCCAAAGCCTTATAGAAATGAAGAAGAAGCAAGAGCTGCCAATAATGGTGCATATCCCCCAGATTTAAGTTGGATTGTACCTGCTAGACATGGCTGCGAGGATTATATATTTGCATTACTGACTGGTTACAGAGAAGAACCTCCAGCTGGTGTTAAGCTGCAAGAGGGGATGAGCTATAATGTTTATTTTCCAGGACAGGCTATTGGAATGCCACAGCAATTGTATGCTGGAGTAATGGAGTACGATGACGGTACTCCGGCAACAGTTAGTCAAATGGCAAAAGATGTTTCAACGTTTTTGAAGTGGGCATCTGAACCAGAACATGACACTAAAAAGAGAATGGGAATGAAGGCTTTGGTAATTTGCGGAACAATGATTCTAATTTCACTGTATGTTAAACGACACAAATGGACCGTTTTAAAATCTCGCAAACTTGTTTACCACTCTCCAGCTCGCTAAAATTAAATCTCACAACCTTGGTTATCACTCTCCAGCTCGCTAAAATAAAATTGTATAAAATCTCAAATTGTACTTACATGTGTTAAAAAAAAATTATCCTATTTTGCTTATTTTTTTATTTTTCTATTTTTTTTAACGTTGCTGCTACGTTATTATTACGGTCATTGTTTTATATTACTTATTCTTCTATGATGTGAAATTTATTTAAATATATATTATACATTTTGAATTTTGTGCATTATAAAAGCGTTATTGATTACAGTTTTAC
>evgR014686c0g1_i01_Hydra
TTACATTTCCATATCACAATTTTTCAAGTTACCTCTCCATGAAGCATGTGTTGCCTTGAGAGTTGATGAATCTTCATTGAAGAAGAGATGCAGAGAACTTGGTATTAGAAGATGGCCATACAGAAAGAGATGTAATATCTCATCAAGTAAATCAGGTAATGGTTCTTCTACACCAGCTCAAACTGAAGATAGCGGTGATCAACAATTATTTTCCTGTTTCCAATTAAACAAGGGAAAGAATTCACCAGTTGTTGCAAAGATGAAGAATGAAGAAGCTCTTGCTCTTGAAGCTTTAGGTACATTGCAATCAGCATCTTCAACTCCACAACCACCAAAGCAAGTTATGACATCTCCTGATGTTCAAAGACAAAATCATGGTGTCGTTTCACATCAACCACCAAGACCTGTTCCAACCAACAATGTTACAAGTTATCCTTACCCATATCCTCCACACCCTGGTTATGCTCACATGTATCCACCTCATCCTGGTTATCATTATCCATATGTAACTCATGAAGGTTCTCAACCAGTCTATCCACCAACAATGTATCCACATTACCCTCCACAGTCAACTGTTCCTCCACCTCCAGGAGCCTATCCACCTCATGCAACTGTTGAAAATAAGGAAGGTCAACCACAATCATATGCTCCTCCACCATACTACTATGATTACA
>evgR014688c0g2_i01_Hydra
TTACAACACTGTACTTATTTGTCGCGCTGCAGTTATTCGCAAACCAGCTGGTAGCAAATGCAAAGATCTTATCTGCAAGGGTGGATTATGTAGAAATCCCAACCATTGATGTGTTACTACTCTTGTATTGGATGTTTGGAACAAAGCTGGTTTCGGAGAGTTTATCATTTCTGGAGCTGCTGTCAGGAATTAAATGTCTAAAAGTTATGTAGTGAACCTTCAATATCTGTACAGTGAAGAAAACATCGATAGTTATGTTAACAATTTTATCAACTACAAACTTGGAGGATTTAGGAAGGATGAACATGGTAATGAATATAAATATCTTCTTAACAACAAGTTTGGAGAAGGTAAAAACTCTTATTACGCTCTGTACTTGGGAGCTATAGATAGTGCTGGTAAGCCTCCACTAGGTTTTTTCTTTAATAAATCAAAACAGCCTGCTCCTGAGAAAAGAGACGATCCTAAAGATGTTTTCGATGAGTTAAAATCCGTATTGTTATCCAAAGGAAATTGCAAGACAAAAATATTATTTCCTTACAATATAACTAATATTCACTGGCTTACTGGAGAAATTGACATAGATAAAAAAGGCGATTACGTTTCAGTAGTAATATACACCCACAATCCTTATGGTAGCGGACAGTTATCAAAGAAGAATTTTAGCATACTCAATGCCAATTTATACAAGATCATAGAGGAAAGTGGTTTGAAGGTAACATTAATAACTAGTCTGCTAAGTCCTTTTACAAATGCAAGACAAACTATGGCAGATAATACATCTTGCGGCGTAATTGTTGCTGATGAGGTTGTGAAAAGGATAACCAATAGTAGCCTTACCATAACATCACCTTATCCGCAAGGCGCAGGGCAGTTACGTTTATCACAGCTAACTTTTCTACAAGACATTTTTGGAGAAGGCAATTTAAATTATCAGCTCTTTCAAAAGCAAGTGACACTTACAGCCCATGTAGACGATGATGCAATACCAACAATAGAAGCAAGCCTATCAATAGGAGAAGTAAGAGAAAAAGTAGAGCAAGTAGTAAGTTCAGAAGAAAGCAACATTATTAAAATTGATCAACTAAAAAAAATAGAGGAAAATCACTATAAACCTTTGATTCAGCAATTAGCATATAAAAAGAGTCAAGAAGAACTAAAGGATTTAGTAATAGTTTTAAAGGATTTAGGCGATTTAACCGCCAAACTAGGAGAGTTAAGTGGTGTCCTAAAATATTACAATGAGGCAGCAATATTTTATCGATATGTTATTATTATATTAGATGGAAAGTTAAACGAGAATAAAAATGAATTTATAAAACATTATTTAGTAGATCCATATCAAAAAATATCCGATATACAACAGCTGATATATTCCACAATTGGTGGAAACCAAAAAAATATGGCGGTTGTTCAAGAAGAAGCACAGTCGAATAAAAATCTTTTATTAGCATTGCGAAACAAGACTGATGATGATATGAAGGTAATAGAAACTCGTCGTCAGCAAATACAAATTGATAGTCAACAAGAGAAAGAAAAATACCAAGAGTTCTACGTAAATGCCGCAAGAGAACTCTTTGGATATATAACTCAATACATGAAAATGTTTTTGGCTAAGTTGTATGGTGTTACTGAACGACAAATGACTATAGCACCACCTTGTAAATACGCTATAGTAGGGCTTGGTTCGATGGCACTACAACAGATGACGCCTTATTCAGACTTAGAGTTTGCTATTCTTACTGAAAATGAGGATTACAAAAAAAGCGATGATCCAAAAATAAAAGAATATTTTAAGAACTTAACCCACTTAGTAAATTTTAAGATAATCAATCTAGGAGAAACTATCATTCCAACCAGTAAGTATAAGTTAGATATGAGCCACCTAGTACACATAGGCGTTAATTTTGATCTAGGAGGTAAAACACCACTTGGAAGAGATGATAAACCATATGAACTCATAAATACTGTAGATTCAATGATAAACGTTTTGCTTAATGAGGGCAATAAATCCAGTAATAGAGATAAGAACTTACCATATATCCTAGAAAATGTCTGTTATGTACATGGAGATGAGAACCTTGTAAAAACTTATGAAGACAAAGTTACACAGTTTTTGCTTAGTGTAAGTAAAGAAGACCAAAAAAATAGATTAAACTGTGAGATAAGAGCTATTAAGATACTACAAGAAGGTGCGATAGAATTTGACTATCTTCACAAAACACCGCATCAAAAGCCTAAAGTAACGTCATATAAAAGCGATCTAGTTAAATTAAAACCTTACTTTTTTCATACGAGAATCAGATTATTTGATGTGAAGCAGGAGATCTACCGGTTACCTGATCGCATGGTATATAATCTTGGTCTGTATTATGGCATTAAAGGAAGTAGTGCGTGGACCACAATAGATAAGTTATTGGATCAAGGAATAATTACTCAACGAGCAGCTGCAAATCTAAAATATGCTGTAACTTTTGCCACTACCCTAAGAATAAAGACCTATTCGCACCATAAAGCACAGAAAAATGATATGTCGATATTTGCCACACTAGGGGAAAACGAGTCTGAGCTTAAAGAACAAGCAAAACAAATTTTCCATTTATCTGAAGTCGATTTAGGAGAACAAGGTGGGTTATTTCAATATTTTTATACTGCATTACCTTTGCATAAAAAGTTAAAAGAGTTTTGTGATCACAATACAGATATATATAATAGTAAGCAAATCTTTTTTCAAAACACTGAGTTTTACGAAGATGATTTTACTAATAAGGGTTTAATACACTATAGATTAGCAAATTATAAGAAAGCTCGTAATAATTTAGAAGAAGCAATCACATATCAAAACAATCAATACTATTTCGAAATAAGATTTATTTTAGCAAATATATATATAAGTTTTGGTAATACCGATAAAGCTATCAAACTCCTTGAAAATTGTTTAGAGATTATAAAACAGATTTTTCAAGATGAACGTCACCCTGGTGTTTATTTAACTTTGGATACTCTTGGGCTAGCTTATAGAGCTAAAGGACAATATAATCAGGCGATTAATTACTATGAGAAGAGCCTAAATATTAAAAAACTCATCTACCAAGATGAACGTCACCCTGATGTTGCTGCTTCTTTAAATCATCTAGGGTCAGCTTATTATTTTAAAGGACAATATGATCAAGCAATTAACTACTTTGAGGAGAGCCTCAAGATAAAAAAACTCATCTACCAAAATGAACCTTGCATTGAGTTTGCTGTAACTTTGAATACTCTTGGGTTAGCTTATCAAGCTAAAGGACAATATGATCAAGCCATTAGTTACTACGAGGAGGATCTCAAGATTACAAAATTCATCTACCAAGATGAACCTAACCATCATGTTGCTAGTTCTTTAAATAACCTAGGGGTAGCTTATAGAGCGAAGGGACTATATGATCAGGCAATTAACTACTATGAGGAGAGCCTCAAAATGAAAAAACTCATCTACCAAGATGAACCTCATCCTGATGTTGCTGGTTCTTTAATTAATATAGGATTATCTTATAATGATAAAGGACAATATGATCAGGCAATTAACTACTATGAGAAGAGTCTCAAGATTGAAAATCTCATCTACCAAGATAAATGTCACCCTAATGTTGCTGGTTGTTTAAATAACATAGGGTTAGCTTATAAAGATAATGGCCAATATGATAAGGCAATTAACTACTATGAGGAGAGCCTAAATATTCGAAAACTCATCTACCAAGATAAACCTCACCCTGATGTTGCTGGTTCTTTAAATAACCTAGGGGTAGCTTATCAAGCTAAAGGACTATATGATCAGGCAATTAAATTCTATGAGGAGAGCCTAAATATTGAAAAAATGATCTATCAAGATAAACCTCACCCTGATGTTGCTAGTTGTTTAAATAATCTAGGGTTTGTTTATAAGGCTAAAGGTTTATATGATCAGGCAATTAACTGCTATGAGGAGAGTCTAAATATTCAAAAACTCATCTACCAAGATGAACCTCACCCAGATGTTGTTTTTCCTTTAAGTGGCCTAGGGTCAGCTTATCAAGCTAAAGGACAATATGAAAAGGCAATGCACTACTGTGAGGAGAGCCTAAAGATGAAAAAAATTATCTACCAAAATAAACCTCACCCTAGTGTTGCTAATTCTTTATGTAGCCTAGGATCAGTTTATTATTTTCAAAAAAAATTTGATCAAGCCATTAACTACTATGAGGAAAGCCTAAAGATGTACAAACTTTTTTACAAAGATAAACCTCACCCTAATGTTTCTTCTTCTTTAAATAACCTAGGGTCCGCTTATCAAGCTAAAGGAAAAAATGAACAGGCAATTATTTACTTTGAGGAGAGCCTAAAGATAAACAAACTCATCTATCAAGATAAACCTCACCCTAATGTTGCTGCTTGTTTTAATAATCTAGGGTCAGCTTATCAAGCTAAAGGACATTATTATAAGGCGATTAACTACTATGTGGAGAGCCTAAAGATGTACAAACTTATTTACCAAGATAAACCTCACCCTAATGTTGATACTTCTTTAAATAATCTAGGGTCAGCTTATCAAGCTAAA
>evgR014689c0g1_i01_Hydra
GTTTTGTTTCTTGTTTGTGTTTTGAAATGTTTAAAATCTAGTAGACATTTATTAAAGTTTATAAATTTAAAATTGTGTCAAGTTAACTATGAAAGTAAAGTTCATAATAATTGAGCAATTTTTAAGTTTTTTTAAGCTAAAAAAGTAATTTCAAGCAATAAATATATTATGGGGGCTTTAGATTCAAAGGAAAATAATGAGGGCACTCCTGGTGAAAAACAGGATCACTATTATAAACGTAAAGCTAATTGTAAGCTTATATGTGATGCCAACACGCTATGGCCTGCTTCAAAATCAAATAAGCCTCTCTTTTTATTAAATTTAATGGAGAATATTGACCACGCAAATGAGAAGTCTCCAGTTTATGCAATTCCTGAGCCCTTGACACGTCCTAGTCACAGACTGGATAAATGTGTTAACCAGGATTCTGCTTTAAAGAATACATTTGCACAACCAAAGAAAGAGCGTGAGGTTATTTTGGTTAAGAATCTAGATCTTGAATATTTTTCTCATGAAAGAGCTTCAGAAAAGAATTTTTTTTACTCACATACGTGTAACAAAAAAAATAATTCTTTAAACGATGCAACTGAAAAAAATCTGAAATCCAAAAAAGCCAGAAATTTTAAACTATTCCAATCTTACCATAAAGAGGCTGTTAAAATAAATACAAGAGAGTTACCAGATAACAAAAAAGAATTTTGTTGTACAACTAAGATATCATTTTTACCATCCTCATTTGTTGCACCTGATTAAGAAAATATTTTTTAACAGTTTTTAACGATATTGCTTACTAGAATGGCGTGATAATGATGTTATGTAAATAATGTGTATAGGTTTGTATAGGTGATGTGTAAATAATGTGTATATTATTTACATATTTATACACAGAAATTGTTTATTAATGTATCTGCATCTTTTTTTTTAAATAAAAATTTGTAAAAATTTGTTGAAGCTTGAGACTATAAATACTTTTCAATAAGGAACTTATTTGTTTGTGGATAATTACATTTGCAATTGTTCAATATTTCCATTCGCAACCTAAGAAGTTTTTCATAAACTTTTAATACTAGATTGGTTGGAAAAAACCTTAAAACTAAAATTTTTTTAGTACTGTAGCCTATTTGCTTGGTTTTATATTTAAAAATGTATATATGTGTGGGTGGTGTTTTATATATATGTGTGTATATATATGCGTGTGTGTATTGTGGTTACTTTTATAAATAATCCAGC
>evgR014696c0g1_i01_Hydra
AATCTGTTGTTTTGTGATAATTTGTTTACTTATAAAAAAATTTAATAAAAATAAGTTTAATTTCAAATAATTTAGAAATTTGTTAATAATTACAAATGCGTAAGTTAGGATTTTTAGGAAGGTTTTTTTCTACTTCAGTTTTTGATCAAAAGGTGTTAAAAGTTGCAGTAATTGGTGCTCCAAATGCTGGTAAATCGACTCTTATAAATGGGGTTGTTGGCTCAAGAGTGTCGGCGGTTTCTGAAATTAAACATACTACTCGAACATTAGCTAATGGTATTAGAAATATTGGTAACTGTCAGATTATATTTACTGACACTCCTGGCATAATCTCGTATGAAGAAGGACACCGTCTCAATATGGATCGTGCTCATTTACGTGGACCTCGGCGTGTTGCTGGATCTGTTGATATGTTAGCCATTATTACAGATATAGCATCTAAAAAATCAAGAAATTTTATTGATGAAAATATAATAGAAGTTATGAATAGCAATCCTGAGGTTCCTGCCATTTTAATATTAAATAAAATAGATTTACTGAAACGAAAAGAAGACTTAATACCAATTGCTTCTATGTTAATGCAAAAACGAACAAAAGATATCTGGGGATATCATTCATACGGAGGTTCTGAACGTTTTCAGAAATGTTTTATGATATCTGCATTTTCAGGTGATGGTGTAGATGACTTACTTAATTATTTAGCATCAAGTTCAAAGCTGTGTCCATGGGAATATTCTGACCAAATTTATTGTGATTCTAGTATAGACTTTCAGATTTCGGAAGTTTTTAGAGAAAAACTGCTAACATTATTTGGACATGAAATTCCATGGAAAGTAAAACAGGAAACTGTGTTGTTCCATGGCAACGATGACGTAGTTCGACTTCACCAAAAGTTAACATGGCCCAAGAAAAGTCAATGTCGGTATGTATTGACAAAACTTGATCAGCTATCTCAAGCATGTGAGACTGAACTTGCTAAAATGTTTCACAGCAAAGTTATACTCACAGTTGATATTAGTCAAAAACATAACATGACCAAGGATGATTTGAAGTATTATTAAAACTCCAATTAAATCACATAAACATGAAATTCGATGTGTATTCTTAGTTTTTATTTGGGATACCACATTGATGTTGTCATTGCCGTGAATGTCATTATTTCTTTTTTACCGTTTTTGTAACCTGGTTTTATAACCCTCTTCGTACCTGGCTCTTTTCATATAAAATTTAATATTAATGCGATTCATGTTTGTTTAAAAAAAAACAAGATTCATGCGATTCATGTTTGCCCAACAAAAACAAAAACTAGTTTACAAAAATTTCCTTGCCTACTTGTAAAATAATTTGTTATTGTATTATTAGTGGAATAATTATTAGTGGAATAAAAGTACCACAAACAGAACAGTGCGCAAAAGCAGACACTTATCGG
>evgR014703c0g1_i01_Hydra
TTATTTTCTAAATAAGATTATTTGGCTATATGGAGTCATTAGTCTGGGAAGTAGTTACTTGTCAAGAATTAAAGTCTTCAGATATAAAAATACAGCTAAGCAAAGAGTTTAATCGAAAAGCTTATTTAAATATTGATGAAAAAATGGACTCAATTTGGAGAGAAGCTTGTGCTGAAAACGATAGACTTTATAATGCCAGCAAGTTTCGATTAGCAAATGTAGAGTTTAATGAATTTGGAGAGATATTTGTTAGTATTGGAATCACAGATTATAAAGAAACTCTTTGTACAAATTTTAGTCATATTTGTGATGAGATTTATAACTTTGGAGTTTTAAATTACAAGAATAAGCATGCATGCTTTGGAAATGCTATTGGTGTTGGATCAATTGTATTATCAAATGATGGTTTTATTATCTTAATAAAGAGATCAGATTGGGTAGGAGAATCAAAAGGTTTATTAGATACTCCAGGCGGACATGCAGAACCTAAAGAGTTGTTGTCTCGTTTAAACATTGATGATATTTTTTTGGCAAGAGGAGAAGATGTTGCTAATGAAATTTATTATGCAATGATTCGTGAAGTAAGGGATGAAATTAATATCCCAGAAACATGCTTAAGTTGGCCTGTATGTCTTGGAATATTTCGGAATAAAGTACTTGGTGGAAAGCCTGGTTTTGTTTTTTATATCAAATGCGATTTACCTAAAGATGATATAAAGAAATATTATCAGAACGGAGGCTTAGAAACAGATGAATCAACTGAGATTTTGTTTATGTCTTTAAAGGATTTAAAATCATACACAGACGAAGAATTTGCATTGTTTATTTGTAATATGGCACCTGGAGGTCAGGCGGCATTGCATATGTTCAGAAAACTTTATTGTTAAAAAAGAAATAAATAGATTTGATATAAAAAAAAAA
>evgR014704c0g1_i01_Hydra
TTTTTTTTTTTAACAGTTTAAAATCTTATTATATTTGGCACAAATGGAAATATTTGACACTTGAAGAAATTTATGTTTACTTTAGCCCTATTTTGAATATGGCATTGAATGAAAAATCAAGTATAAAACATTACTTTTCCCATGATCGGCTATACTACATGCCTCTTTTCCCGCAGTATTTCATAGAACTCGTTTTCTTCAAATACATTGGATGCTGCACGTATCTCCACCATCAAATCCATCAAGTGAATTAAACAAGAGAGCAAAACAAGTTCATCGTGTTGTATCAGAAATGAAGAGATCATGTTCTAGAAATTTCGTTCCATCTTGTAACATAGCAGTTGATGAAAGTACAATTGGATTCAAAGGTAGAGTTGTATTTCGCATGTACAATCCACAAAAACCCACAAAGTGGGGCATTCGTGTTTTTGTCTTAGCAAATAGTAATACAGGTTATATTAGTTTCTTTGAACCATATTATGGTAATTGCACTACTGAAAGATTGGCAAGACCTATAATGCAGTTTACAAGTCGAATTGTGTTGCATTTGTGAGATGAGCTATTGGAAACTGCCAAAGTTAGTGTTTATTACTTGTTTACTGACAGATTATATACAAGCTTTCCTCTTTCACTAGATCTTTTCAAAATGAACATAAAAACTACAGGAACAATTATGAAAAATAGAAAACAGTTGTCTGCAGATATAAAAAAACTCAAATTAAAAAAGCATGAGGTTTCTGCATATGAACACTCGTCTAACGTTATGGTTTTGGCATGGCAGGACAGGCGTTAAGTACATTCCATGATTCTTCAGCTGAAATTGCAAATCAAAGAACAAAACAAGGTATTGTAGACATTGAAAAACTAACTGTTATTGAAAATTGTACAAAACTTATGGGAGCTGTTGACTGAGCTGAGCACTATTGATCTAGCTATTCATTTCTACGTAAATCTTTAAAATGTTGGAGAAAAATATTTTTGTGGTTGTTGGAAGTTGCTGTTGTAAACAGTTTTATATTGTTTAATAAGTTTTATGAACGCAATGGTAAAAAAAAACTTAGACACCTAACGTACAGACAAAACCTTTTTGAGCAGTTAGCTAAAAAAACATCGCAGCAATCAAGAACGACGACGAAGGCGATCATCAGCAGATGGCAAAGACTCAACCAAAAACCTTGTTGCGCAAGTTTCAAGTAATAATGCCAAAGACTGTATAGCTTGCAGTGACAGAAAAATAGGGTGGTAGGAAAAAACTGAATACTACTGTAAAACCTGCACTAGAAAACCAGGACTACACCCTACTGACTGTTTTGAAGTTTACCATACAAAAACTGACTTTCGTGTAAAATAGTATATCCG
>evgR014707c0g1_i01_Hydra
CATAACAGGAAGAATAGAAAACATGGCAAAAAACGTTCTGGCGTTTGTGTTTCCAGCTATATATTTATTATAATGTGTTTCTGTTATCACTTAGTTTTTGTATCTCTAATAGGTCTTTTTTTAATTTTTGCTGAAAGTAAGCAAAAACCAGAACAGTTCACTACTCTTCTTAAAAATGTTTTACTTAAGAAGTTTGGCCATTCTATTCAAAAAAGGTCAAACGAAAAAAATATTGATAGCAAGGTTGAAGAAGTTAATTCCAGAATTCTAAATGGTTTTGATCATAAAAGTATCAAACATATTTTAGTAACGATTTTGTCTTTACCTCTTGACAACTTCTTATCAATTGAATCTGAATTCTTTCGTGATATTGAACAACATGTTAATAATGTAAAATGCGAAGTATCAGAAGTCAGTGCATTGAATCCGAACAGTGTTTTGATCTATTGGTATTGTTATTATCAAAATAGTGAAACTGGTGATAAAAAATGGAATTATGTGTTTGTACCATCGCACAAGCTAGAAAGATTGTTAACAAGTGGAGAAATGAGAGGATTCCTCCAGAAATACAATGTTACAAAATTTTATCAAAAGGAAGAGAAATCTAAAGGGAATCCTACTGCTTCCAACTCTAAAGAACATTGGTATCTAATGACTGTAGTTGCTTTAACTTCAACAATAGTTATTATATTGGCTTCTCTTCTTGTTATGTGTTGCAAGCGAGAAAAGAAAACACAAGTTGTATTAAAAGAATCTCTATATAAAGAAAATGATACTTTTATAAAAAATGGAAGTCACAATTCATTTGCAACCTCACCTAGAAAAAATGGAAGCCACAATTCATTTGCGACCTCGCCAAGAAAAGTGGCGTTCTCTTATTCTCCTAAAAAAGCCTCTACACCCATATTTTCTTTTTCATCTCCTCTTAAAACAGTAGATAAAGTAATTCTTCCTGTTCAAAGTCCTATAGAAATGGCAGGTCCATTTACTGTTCAACCGCAGCTAGACTTATTATACCCTGCAAATCCAGGGTTTTCTATGAAATGGTCAGAAAATAGCTCAATTGGTGGAAATGCAAGCAAAAATAAAAGTTTGTTTCAATCCAGGAGAGGCTCTTCTGCATCTTTAGTTATTACACTTTCTCCTCGAACACCTGATGTAGAGGAATCTTTTGTAATGCATGAAAGTCCTACAGAAGAAATTTTGTTGCGTAAAAGCCGTTGTATAAAATACAGAGAACTGTACAAATTAAAAGACAATGATAAAGAACTTACTGAAGAATTTTGGAGCATACCTCCAAATACAATATGTCGAAGAGAAATGCAATTAAAAAACAGTTTGAAGAATAGATATCCGGAGATCTTACCAAATTCTCGTTCATTGGTCCGTTTGAAATCATCTGAAAATTCATCAGTGTATATAAACGCTAACTTTATTAAGGGATATAATGGAGAACCTAATCAGTTCATTGCTACACAAGGTCCATTGTTGGCAACTATTTCAGACTTTTGGATGATGATATGGCAATTTAACTGCTCTGTTATTGTTATGTTAACAAAGTTAAAAGAAAGAAATAAGGAAAAGTGTGTTCAATATTGGCCATGTCCTGAAACTGTTTCACAGCAGATCTATGGTGATATTACTGTTACATATGATTCTGAAATTGATCGCCATGGATACAAAATAATGACAATTTATATATCGCATAAGGGAAGCGATGAAACTAGAAAAGTAGTACATTTCTGGTTTACATCATGGCCAGATCACAATATACCAAAGAAATGTCATCAAGTTGTTTCTCTTGTAAAAGAAGTGCGTAATATTCAGACACAACAAGGTGCTCCAGTAGTTGTACATTGCAGTGCCGGACGTGGTAGAACTGGTTGCTTCATTGCTCTTTTTCTTGGAATGTCTCAAATTGATGAAGAAAAAGAAGTTGATATCCTTAAAATTGTTAGCCAGATGAGAATGGAAAGAGGAGGAATGGTTGATAAAGACCATCAGTATGTATTCATTTATAAAGCCTTGTACGAATATTGGTCTAACAAGAACCACAGCTCCTTTTTAGCATCCCCTTTACTTTCCGCTAGTAACACATTTACGAACTTTGATAGTGATTCAGATTGCGAACCATCTTTTGTCGGTTAAGGAGCAAATGTAAAAGAACAAATATAGAATATTAATGAAATTTTTTTTTAATAAATATTTTTGCAGGTCATGAATATTTATAAGTTTATAAATATAAAAATTTATAAGTCTAGTATGAATATTTATAAGGTCTAATATAAATTTTTTTTATAATTGTGTTTTCCGTTTTGAAATGATTCTTTTTCTCTCAATAGTTGACGGATTTATATATTTTTTAGATGGCAGATTATTTTCTGTATTATATATAGCAATATTATTAAGTCGAAG
>evgR014715c0g1_i01_Hydra
ATGGAACGACCAGAAAACTGTTCAGAAGCTATGTAATGTACTTAAATTTGTTGCGATCTTTTTGTGTTTACTTTTTGTTTTATCACTATTTTAAACTTTAGGTACAATATCATGTTGCAGTGTTGGGACGAAGATCCATTACAGCGACCTACTTTTACAAAATTACGTGAGCATTTTGATGGAGTAATGTCTCAAGGTGATTGTTATTTTGATTTCGAACTTCTTGGAGATGCTTATTATCATGTCGCTTCTTTCAATAGCATCTCTTCTGACGCAGATGACGATATAATGATAGACGAAATCTTCCAAAAACCTGAACAACTTAAGTCAATTGATGAAATTAAACTTAAATCCGATCTTCCACTGGACAAAAAACCCGCAAATCATGAAGTTGAACAAATAAAACCCACCAAACATGAAGTTTATCATGCTCAAATAAATCAAACTGGTTCGAGTAAATTGATTAAAAGCAATTCATTTTAAACATCAGGAAATTAAAAACATTTTATAAACACAATATAGACACTATATACATGGGAGTAATATACACGATATAAAAACGTTTGATAAACGGTGACTTAAGAATCGTCAAAGCGGTAAACATAATAATAAACCTAGCAAGCTAATTTCATATCGTCACGGTACTAAGCTCATACCCTCAAATTAAAAAAAAAACAATGTATATTAAGTATCACAAATTTATTAGAGGAAAACAAAAAAATGATCAATAATATATTGTATATGTATATATTAAAATAACCATAAACATAATGGTAAACATAAAATTTATTTAGTTTATTATTTAAAGATTTGTC
>evgR014718c0g1_i01_Hydra
TTGTTGAAGAAAAAGGTATATTTCAAAGTGATAAATTTTTGTAATGGATTTAAAATCTTTTGAAGCTTTTAAAGGTAGCTCAATTATGAAGGCATCGTCATCTTTGCAAAATCGGAATGCTATCCATTCTCAAGGGTTTTCAACTATTAAAACAAATATATATAATTTTGATGCTCTTTGGAAAGGTTCCGTTAGCAATCTTATATCTTCAGAAAATATTACTGAAGATCTTATTAAAGAGAGTATAAAAGAGTTAGAAAACGCAATGCAGGAATCAAGAAGAATGTTGATTGATCGAGACGAAGAAATAGCCAAATTGCGTCTGTATATAGAAAATAATAAGTCCGATATAAATGAAGACGAGCATTACGAAAAGTTAGCTAACGAACTTAAAAAATCACAAGAAACTGTAGTGTTATTGGAGTGTCAAGTAAAAGAGTTAGAACAAAAGCAGAACAAAATTACAAATATAAACACAAAAACGTCTTCTGCGCACCAACAAAAAATTCAAAACTTTAGAAATAATATTTCTGAATGTACGTGTGATTGCGGTTTTGAATGTAAAGCGTTAATTGATTTATCTGAAACTAAACAAACTTTACAGATAACAAGAGAAAAGTATGATATTTTAAAACGGAAAGTTAGAGAATTTCGCAAACAAGCCGAAATTAATCACAAAAATGCGTTACGTTTATCGGCCGCTTCAGAGGAGAAGACTGGTTGTATTATTCAATAATATTGAGATAATTTCTTTTATATAGTTTATTTAACATAACAATTATAAATCCACTAAACGCAAATCCCCCCAAAAAAACTCACCTAAAAAAACTACAAG
>evgR014720c0g1_i01_Hydra
TACAATCAGTAATTTATTATCCGTATTAATTATGAATACATTATTATTGATTGGAACAAAAAAAATATTAAGGAATAAAGAAATTCGTGTTGCTCAACATGATGTTGTAGAGAGACGTCTTACGCAGACTATGATGCTAATATCGATTTCCTTAATGATTTTATATCTTCCTGTGACTGCTGCTGCCTATAACATTTCTGTAAGACTATATTTCAATGAATTCAAATCTGTTGGCAATGGAATAAAATTGTTTTTTTGGGCTCTTGTTTTATGTGAAATAAACTCATTTTTTAATGCATTTATATACGTCATCAGAAATAAAAATATATTGAAATTTTACAAAAGAGGTATAGTTTCTATGATTAAAAAAGATTCTGACAAGCATATATCAGAAATGCAACTCTCTTCATATTACGCTTCTACTGATGCCATAACTCAATCAATTTCCGAAAGTAACGAAAATGTTTTTTTTACTAAACAATGACGGTTTTAAAGTTGGCATTTATGTTTTAATTATGTAACATATATACTAGATCGTGTAACATTTATACAAGGTTATGTAAAATAAACACCTGGC
>evgR014729c0g2_i01_Hydra
CTTTGAGATGGAAAAAGATAGATTGTTTTATTATACCTTGGTGTTTTTATTTTCCAATAAACAAGTTACTTTACGTTATGTAAAAGATATAAAATTTGTCACCTATTGGGGTCAAAACAAAGCAAATGTTGCACTAGATACAATAGAATTGAGTTTGGCAAGCTTGTGCTCGAGGGAATCACCTTATTCCACTATTGTTGTGGGATATGTTAGCAGATATGTTGATAACAGGAATAAAGATTTTTTACCTGGAATGGGATTTTCGATACATTGCACAAAACTGCAATCTTCATTTTTGTATTGCCCAAATATTGGGAGAGATATAAGTTTCTGTCAACAACGTGCTAAAAAGATCCTTATTGGAATCGGTGGTCCTGGCTCTCCTGCTAAGTTTGAGAATGCTCAGGCAGCTGTAAGGTTTGCAAAACTTGTTTGGAATCTATTTCTTGGTGGCAACGATAAATATTATATGCGCCCATTTGGAAGCGTCATTATGAACGGCATAAACTTTTTTATGCAAAATAACGACCCCCTTCACAGCAATGTTTTTATTCAATCCCTTAATCAATTGAGAAGTATGGATCAATCGAGAAGCTACTTGATTACCATTTCACCTTCGTGTATGTTTCCTAATCCATTTTATGGACCAGATGGGGATAAGATACTTGATATTTCAAAAAAGTTTATTGATGAAATCTATGTTAGATTCACTGAAAGCACATGCTGCATGAGAGGATCTTCTGGTTCTCTTGAATCTTTGATTCGTTGGAAAAACTTTATTATTAACGCCAATCATTCAAAACTTTATGTAGGTGTACCTGCGGCACTAGAAGCTACACAATATCCCGAATGTTATCGCAAACCTTTTGAGTTAGCCAACATATATAAATCACTAAAAACGACCCCAAACTTCGAGGGGTTTTTAATCATTGATAATTCATTTGATCTCCAAAACTTAAATGATGGTTTTTTATTCAGTAAACCGTTACTATCCTTGCTTAGGAACACACAACATCAATTTCAAAGAAATAAAGCCAAACCTCTTGTACAAAAAGCCGAATATCCTGTACAAAACTATAAATTTCCAGCTTCAACTAAAGTAACACATCCAGGCAATAATTTAGCTCAATTCAATGTTGCCACTCAAACTGAAGATACGAATCAATTTAAAATACCAATGAAGAATAAAGTTTTTTATAATGTACCTCTACCAGAAGCTGCTACAGTGCCACAACCCACACAGACCGCTACAGTGCCACAACCCACACAGACAGCTACAATGCCACAACCCATTACAGATTTTCAATCTTTATCAGATTCAGAATCTGGTACATTACCCGAAACAGTTACAGTATTAAATCCAGTTGCGTTATCAGAAACAGTCACCACAGTACCAGAACCGGTAGCAGTACCAAATCCAGTTACAGAATTACATGGTGTTTCAGAATTAGAACAGGTTTCGTCATCAGGTGCCGGTACAGTACCAGAAACAACTCCGATTTTAGATATGCCTTTAGCTTCAGACCAAGTTATTTTATCAGAACCAGTGTCACCTCCCGATCAATCTTTTAAATCTATTGTGCAAACAAAATTGAAAACACCAGCTAAAATGAACATGAAGCCAGTTAGACTGATTTTTGAATCGACAGAAGTAAACCCTGCAATATCTGCTGTGACCTCAACCATTCTGTATCCAACAGATTCTAATGCTCCAAGTACTGTAAATCCGAATGACAATTTTAATGAAGCCGGTTACATTAATTTAGATGATGAAACATACGAAATCGACGAAGATAATGATTATAACGATCAGGTGGTGCCACTTCAAAACAGTACACCAAATTATCCTGAAACTGAAATCAATGAAGAACCTGTAGAGGACGATGACGATGAGGAGGAGCATTTTGACGATGATAAAAAGCCGACAATATTTGACCAGGCTTCGATCAAACACCCTACAAGAAAACCTTCATGCTTGTCAAGCTGCCAGTCTATAAAAAAAGATAATACCACAGAAGGACTTTTCTCATCTAACCAGTCATTTTACAGAAACTTTTCAACAAAGTCTATTCATAGCTTGAATAAAAGAAAATCACTAAAAGAAG
>evgR014733c0g1_i01_Hydra
CAACAATTTTCGGTTGCTCAAGAGACGAATCACATTTCAATATATATATTACTGATAACAAAAAGATGATTCTATTTCCCTCCCCGAAACTTGGATTAACCATCGGCATGACCTATAGTTATTACGAGTACAAGTACCCTGGGTATAGTCCAACAGACGAAGAAGTAGTCTTCTCCGATTTTTCTTATCCAATCGTCTTTGATAGCAACGATGATGATGATTACATAAACAATGAAATTCAAATTTGGAATAGTGAAGATTATTATAACTACTATGAGGGCGATAACATTGGGAAACTTTGTGTAGATGTGTATGGCGCATTTGATGATGCATATTCTTTTGAGTAAAATGTCAAAAACTCATTTTTTTAAACTTGTTGTTGTGTAAAACTTTATAAATGTTCAGTGTTGTAGTTCATTGTATTTTTTAATGTTCAGTCAAAT
>evgR014737c0g1_i01_Hydra
AATTCTGGTAACGGCTTTGGTTTTGCATCAATGAGTAAAACAAAAGTGCACTCCCCAGAGCAAGTTCAACGTATTATCATGTCCTTTATTGAACAACAACAACAAAACCAACAAGAGAGTCCAAAAGTAGTTTCTGTTAACCCAATGGAAATTCTTTATACTTTCATGTCGTTCTATGGTAGTTGGAAAGAAGGTGTCCCATCCATTTCATCAGCTGAGATGAGTGGAGACTCCACCAACTATTCAAAGGCTGACAAAGTTGATTTGTTGAGCGATGAAGACGTTGTTAACTGCTTTGTGGTTATAATGCACTGTATTGTTAGAGACAGGTACATGGAACCTGTTCTTTTCTCTCCCTATATTAAACAATCGATTGAATCAATTGTTCAAATAGGTAGACTCCAATCTTCCCGTAATTTTTAGAGT
>evgR014739c0g2_i01_Hydra
AATGCAATAGAATAACGATTTAAACAACCAGCATTTAAGAAGTTTAAAGAACTGTTAAAGAACTGTTTTTGAAATCCATTATTAAGACGGATCTATCAAAGGAGCTAAGGGTTTTAGAATCTGATTTTCATGGAGATTTCAATCTTAAACAAATAGAGAGTGAGCTTAATATTATACCAACTGTTTTTAAAAATTATATTCCAATTGATTTTCGAGATATATGCAAGACACTTCAAAATATGGACAAAGAAAATCACCAAATGATTCAAAGTATTTGGATCATTATCCAAATCGTGTTAACAAGTGGAGCTACATTGGCCACACCAGAACGCTTTTTTTCAATGCAGAGGAGGGTAAAAACGTGGCTCAAATCAACAATGGCTTTAAAAAGAGATATTTCCTTGTCAGTGCTAAATACGCAGTTGATATTGTTGATAGTCTATCTTTAATAGAAGTAGCCGAGCAATGTACCATGTACCATGTAATTTTGAGCAATGTACTGCTTAACAGGATAAGCGCAGGAATGAGTTCGGAAC
>evgR014740c0g1_i01_Hydra
CAGTTTTGCACATGTTTTTAGAAATAAAATGAAGTTATTTTCAAATCCTATTGCACTCTCTGTTATTGGAACTGATTCTCATATTTTGTACAGATGTCACTGTGAAGAGTATAAGCCTGTAACTATGCTTTACTTTTGCAGACACTGTACAAAGCTAAGGTGTACCAAATGTGTTACACATGAGGTTGATTCATTCTATTGTCCTAACTGTTTGGAAAATATGCCGTCTGCTGAAGCAAAGTTGAAAAAAAACAGATGTGCAAATTGTTTTGATTGTCCAAGTTGTGGTCACACCTTGTCTGTCAGAGCAACATCACTAGTGGGAGCTGATGAAAGCAGTACAACTGATCCAGGAAATCCTTCAGGCTCTAAAAAACACCATTATCTAAGTTGTAGCTTCTGTCGTTGGACAACCCGAGATGTTGGTTTGCTTGATCACTCTACTCCTAGTGGAGGATGGCTTGATGTTGAACTTCCTTTTGCAAAAAATATGCAGCTGCTTTTAGATAATTATAAAAAACTAGCCCACCAAGACAAAATTGAAAAAGAGAGACAAAAAATTTCAAAGCGAAGAACAAATTTATATTTAATGGATCGGTATGGGTATGCTCTTTCTCTTGGGAAAAAGAGGACAACTAGCATGGGTTCACCTTCAAGTTCTCCTTTGGCAAGATCGAGTTTAAACTCCTCTATACAAAAAGAAGCTGTTCCGTTATTTTCAGAATCATTGACTGATATTCCACCTATTGATGATTCATTTTACACCAAGTCTGTTTCTTTGGAAACTGTATTAAATTTAAACCAATCGTTACAAACTCCCGGTAGTCAACCACACTATGCAAAACAGTTGTATCCAAAACAAAAGCATTTGATGGTTCGCAGGTCACAAAGATGCAAGGAGTGTGAACACAATTTGTGTAAGCCAGAGTTTAATCCTTCTTCAATAAAGTTTAAGATCCAACTTGGTGGAATGCAGTATTGCCCAATCCTCCAAATTCACAAGAATTTTCACAATACACATCTCTATCACGATGAGGAAAACGAAGTTATTTTATCTTTAACAAATCCGCTTGATACAATAATGAATGTTTCATTCACTAAAGCAAGCAATGAAGAAGAAACAAAAGCTACTGGTTTGATAATGCTTCCAACCATTAAAGTTCCAATTGCTCCAAAAGATGCAACCATAGAATTTGATGGAATCGGAAATGCTGCTGTTAATATGTTCAATGATGACCCACAAATCGTTTATCATCGGTCAAGCAATAAAATATCAATCTATGCAAAAGTAAAACCATTGGTTAAAGATGCTGACATTGTAGTCTCCTTCGATCTGTGTTATGAATACAAAATGATAACCGCTCCATTAATTACTAAAGTTGAAGACGGGCAAGATAAAGGAAATGCTCCTCAAGAATTTCCAGTTGCACTTGCTATCCCAATCAGAATTAATTTTGGCAAAGTAAAAATCCATTAGATTTAATTTTCAAAAAAAATTAATAATTGTGTGACCAGTTAAATTTTAAATTAATGTTAGCTTCCTTAAGTAATAGTTGCGGACGTGAATTTACCGAATGAAAGCGAACGTGAATTTACCGAATGAATTCATCAAGTGTAGATTAAAAAATAACGTGTAATAAAGAAACGTTAAAAAAGGGCGGAGGGCAGTGGCCTTCATTGTATACTAACTTTAGACTTCATTGTATACTAACTTTAAAACATGCATGTACTGCTAATACTCATGTCTATACAGTATGGAATGTTGGATGATAAATTTTGGAATTTGTATTGTTTTTTGTTAAGTGAATTTTTTTTAATTTTAAATAGCACTTATTTTTTCGTCAAGCATTTTTCATTTTAATAACTACTATCCGTTTTTATTACCGTTTGGAACAATGTATTTATTTTAATTATATTGCTAAATTTTGATTATATTTAAATAA
>evgR014741c0g1_i01_Hydra
TAAACACTTAAACCTGAAAACTTTTCAAAAAGCAACAGAAACGCCAGGTATAAATTTATTTTATATTCTATAAAAGTAATAAAATACTACTGCACGAAATGGATCCATTGTCACAATGATTGCGGTAAAAGAAATACGTAGTTTACATAGGTGTTTAGTTGTAGTTTACATAGGTGTATACCAATGTAAACTATGCTAAAAATACAAACGATGCATAAAAATACTCGACCTTTACGTATAGATCTAGTATTTGTATGCATTGTTTGTATTTTTAGCATCGTAAGTATGGTTAACATTTATTTTATTATTCCTATATTTAATTCATCATTAACACTTTTATATATTTTTTTATTTTCCTTGATTGTGTGTGTTCTACGTTGTTTTTTTATTGCAATTATTGTTGAATTTGTATGTATTGTATTAATTATCTATCTTTTATCGCTATATATATGGTTATTTTGTATTTTATATTTGTCTTTATTTTGTTCTATTCTTGTGTTGTG
>evgR014742c0g1_i02_Hydra
TGTGTATAATATAAGAAAAGAAAAAGATATGATATTTTTTTAATGTACCATGACAACATCTGTTGATAGTTTTGCACTTAAGTATGGTTTATCTTGTGTTGCTGCTTCAGTTGCAGAAACAGTTACTTTTCCATTGGATATAACTAAGACACGACTGCAAATGCAAGGGGAGCATGCTTCTAATGTAAAGTATATTGCTTATAGAGGAATGTTAAAGACTGGTTATGGCATAGTCATTGAGGAAGGTTTCATAAATTTGTGGAGAGGATTAACACCTGCTATTTTGAGACATTTTGTTTATACTGGCTGTCGTATGGGCTTCTATGAATATCTTCGAGATAACATTATGAAGAAAAATCTGGATGGTTATTTTCCACTTTGGAAATCTATTATTGCTGGAATGGCTATGGGAGGGCTTGCACAATTTATTGCTTCACCTACTGATTTAGTGAAAGTTCAAATGCAAATGGAAGGAAAAAGGTTACTCCAAGGACATAAAAAAAGATATAGAAGTACATTTCATGCATTCAAAGTAATTGCAAATGAAAATGGAGTGAAAGGACTATGGAAAGGATGGCTTCCTAATGTACAGCGAGCTGCGCTGGTTAATCTTGGGGATTTAACAACATATGACTCTGTAAAGCACTTCCTTTTAAGAAATACAAGATTGAATGATAACTGGACAACACATGGGATATCAAGCATATGCTCAGGTTTCGTTGCTGCGGTTCTTAGCACACCAGCTGATGTAGTCAAGACGCGCATTATGAACAATCCTGAAGCATATAAAGGTTCAATGGATTGTCTAATCAAGGCGATTCAACATGAAGGATTTGTTTCATTGTATAAAGGTTTTCTGCCAACTTGGTATAGAATGGCACCTTGGTCATTGACATTTTGGTTGACATTTGAAAAAATACGTTACCTGAGTGGAACAAAAAGTTTTTAGAGTTTTTAAGCGAAGAAAGGAGTAACCAAACATTACGTTTAGCGAATCTTCATGGTTTGATATAGCGAAGCATTACGTTTTAATATAACGAAACATCACGGTTTATTATCGTAGAATATAGACATTTATTTTGATGGGCGCGGTTATATTGTGTTTATGATATAGTTATTTTTATTAGTGCATTTTTTTATGGGTTTTATTGTAAAGTGAAAACCATAAATAATGCAAATTCAAATTTATGGTTGACTAATGCAATCTTATCATCTCGCGATTCTATGTTAACTTTTTTAAGTTAATTCAAGC
>evgR014743c0g1_i01_Hydra
TTTATTTTCTAAATAAGTGGAAAACAATTAGTCTAAAAATGCTGCTGAAAATAGCAAGGTATACTAAACCGGATATTTAAGACAAATGCAATCAGTATAGAATGGCCAGTATTCAGTGGGTGTTTAATTTTGCAAATGCAATTCTAGGAGTCAGTTTGCTAGCATTACCTTATTGTTTCAAAGAATGTGGAATTATTTTGTCAATCACTATGGTTCTTCTTTGCAGTTATGTTACAAAAGTGACCTGCGAATATCTTTTGCGCACTGCAAATTTTACAAGACAATCATCTTATGAGAATATAGCTTTTCATTTGTTTGGGCCATTTGGAAAGTTAACCACTGAATTTTGCACAATTGGCTTGGTTTTAGGAATATTAATTTCCTATCAAATTGTAATTGGCGATCTTGCACCTTCTGTTGTGTATAACTTAGTTGGAATCCAGAATTCTTGGAATTCACGTGCAGTATTAATGGTGTTATCAAATTTTGCAATGCTTCCCTTATGTATGATGAAAAAAGTGTCAAGTCTTTCTGCCTTCAATTTTATCACTGTTGCATTTTATATTTTATTTATGTTTATTATCTTTTGGTATGCTTTGTCAAACATGTTACTCAACTCATTTTGGGAAAAAGTAGATCTATGGAATTCATATGCTTTGTTTCGTTGTTTGCCAATGATAGCTATGGCATTTTCATGCCAGACTCAAGTTCTAATGATGCTTTCTGCTTTGCCTGATCCTACCATGAAAGCAATGAGTGATGTTATCAACAGTGCTATCAATCTTTCAGCTACATTGTACATTGGAGTGGGTCTGTTTGGATATCTTGCTTTTTACCCAAATTATATTTATGGTGACATCTTAGACAATTTTGCGGCATCTTTTGTCTCTGACTTCATTAAACTTGGATTCTGGGTAACAGGAATACTTAGTTTTCCTCTAATTCTTTTTCCTGCAAGAACAGCAATCAACTCGATTTTATTTAGCAAACAAAGAGAAAACGAGCATCGTGATTCTATTTCGACTTCTGATGATTATATAGCTACTGATATATATTCATACATTACTCTTGGATTAGTATTTGGGACTCTTGTTGTTGGTATTTTAGTTCCAAATATTGAAATAGTTCTTACGACAACTGGTGCCGTTATCGGAATATTTTTGTGTTATATTTTTCCGTCCGGTTTGTTCATTGTTGCTAGTGGCAACGAAAAAATGAAGAGAGGGGCGCAGGTGATTTTATTAGTATCTCTTGCATCCATGATATTTAGTACTCAAGCGATTATTTCACCGATTCATCCACCGAAATCTAAGTCAGATGTATTAAAATTAACCGAAGTAACCCCTCTTCGCAGTATTGAAGTAGAAAATATAGGAATTGATCGTCATTTAAATACTTATATTGGTTCAAAAGAACCATCAGACTCGGAGAATGATGTCAATGGTAAAAAAGTTAACATGTTTTTTAGTGACCCTCATGCGGATTTTTCACTTAAAGAGAATGGGAATAAAATAGGTGAAAATAAAAATAACATGGCGAACCGCGATAAAAACATTGCGAATAATAAAAATGACATAGTTAATCTTGAAAATGACATAGGTAATCGTGAAAGTGACAAATTTAATTTAGGCTTAGACAAAAATAATGATTCTTTTAAATCATTGGATAAAAAAAGTTTAAATCAACTTTATGAAATGGATTTGAATGATATATAAAATTGCGCGAATATTTATTTCTAAACCAAAGGTTTATTTATTTCAATATTGTTACAATTTATTTTATACTTTTTTCTTTGTTTAAATAAACAAAAAAAAATTTTTAG
>evgR014746c0g1_i01_Hydra
CCTATTTTAGTGCAAAAAGATAGGTATAATTAATTGAAAGGGTTATTAAAACATGAAAAGTAAATGAAAATGAAAAAATGGAGCCATGTTTAAACAAACCAAGCTGAAAGATGCCAAATTATCTTTTGAAGACATGTATGAAATTGGTTGCAGAATCGGCAATGGAACCTTTGGTTTATCAGAACTATATTTTGCTAAACCTAAGAACGATACTTACGATAAAAATTTGCAAGTGATGCGTAGAACAAAAATTGAAAGGCTGTCGAATGAGGCTTTAGAAGAATTGCAAAAAGAATTGCATCTTTCAGAAATGGTTGACTGTCCTGGTGTTTTATCATTTTACTGTACGTTTGTCAAAGATGAATACATCTTTACCTTGTTTCCATTATACAACTATGGTTCTTGCTATGATTTATTATCATCAGAATTTCGTCATGGTCTACCCATGCAACTTGTGGCGATCGTTATTCGCTCAATTATTCTTGCATTAGAATATCTTCACGATTTTGGTATTATACACAGAGCTGTAAAAGCCAGTCATATACTTCTTAATTCTGGTGGAAATGTTTGTTTATCAGGTCTTTCATACAGTATTTTATTACCTAAAGAAAATCAGATAGCTCATGATTATCCTAAACATGCTGTCCAAATGCTTCCTTGGATTTCTCCTGAAATACTACAGCAGAATATCAAAGGCTACACTTACTTAACAGATATATATAGTCTTGGTATCACAATATTTGAGCTTGTGTCAGGCACAGTTCCGTTTCATGGCATGAAACCAACCGAGATTTTACTTCAAAAGTTATGTGGTGTTGTGCCCCAGTTTCCAGACTGCTATAAAAGCTTTGATTCACAGTCAGATATTGTGGTTTCTCAATCTTCATTGGATGGCTTGGAAAATGACACAGAATACTCAACAGTTGTCCCGATCTTACAAAAACACAAACGAATAAAAAATTTACAACAGTTAGTCGATGCCTGTCTGCAACTGGATGCTTCTGCTAGACCATCCGCTTACCAGTTACGTTATTGCTTGTATATCCGGCATGTAAAGCGAAAACAAAAAGAAAAAGATTTTCCACCAATCACTGTCATGCTTAATCCTCTCAAGCCTGTAACTAAATTTATTCCCGCAGACGGATCATATCTGGAAAGTTCTTCTCCTACTATTGTATCAAATGATACCTGGATGTTTCCCTTCTAGTATTACTTTACAGGATATTGTATATTTTTTGTAAATATATGTGACATTTAAAAAAAAAA
>evgR014747c0g2_i01_Hydra
AATAACTTTAAAAAAAAAAGTGCAGTTGATTGAGAGAAACAAAAATCGACAAATCAAAACTGGGCATTAAAAAGATATTGAAAAATCAAAACGTGACAGTAGAAAGATATTATTAAATAAAAACTGGACATTAGAAATATGTTTTAAAATCAAAGCCGGACTGAAAAGTTTATATAGCAAAAGTTTAGGGCACTAAAACATTTATTAACTATAAAAATCATTTAAAAATGTGGGTTACCTAGTTTTTGTGTATATTTGCTATTATATCGGAAACATTTTGCTCTAACTTAAAATTGTGAAGAAAACAAGATTGCATTGATGAGACATGTTGTTTAGTTTATGTTTTTTCTTTATTTTGATTTCTTTAAATGCTGAGAAATTGTGCAGAGACTTAAAAGAAACAGCAGGCTCCAAAGAATTCATTAATGAGATTTTTGTTAAATATGGATCTGATAACAATACTATTTCACTGCCAAATTTTAATGCGCTAATGAACAATATATCTCTCGGAAAAATTGAGGTTACTTGTGAACCTAATGACATTGTGTGTGAAAAAGAAGGTATTGTTGAGCATTTGAAGTTGGGGAGGGAAAATGACCACAGTCATGATCGTAAAAGAAGAAGCCGCGATGATGTTGAAGAACACCATCAACATTGGCTGCACCACATTAAATCTTGTTTGAAACCTATTGACATTCTAAAGCTGCATAATTTGTCAGGAAGTATTGGTTTATCAAAAGATGATTTTATGCGCGTATGTCCATCTTTGTTGCACCAGATAGATTCCAAAGCGTGTTTACACGAACATGTGAAAATATTAGGACATTTTCATGATAATAGTGATTTACCTACAGTTTATGAAAGTTGGGGATATGGTTTGCTTTCAATCACAGTCATAAGCTTACTGTCAATATCTGTAATCAGCATCATTCCCTGTTTAAAAAAAGCTTTTTATCATGTGATAATGTCTTATCTTGTTGCTCTTGCTGTTGGAACATTAGCAGGCGATGCTATGTTACACCTAATACCACATGCATTTGCTGAAGGTGCGAATAGTGCAGCTAGCTTACGAATCAGTAGAGAGGATGAGCTGAAGCAGCATTACTCTCAAGTATGGAGGGCATTGTTTGTACTACTGGGAATATATTTATTTTTTGTAGTCGAGCAATTAATGAAGGCAAAAGTATTATGCTTTGGCAAGTCGCATGGTCATAGTCATGGTAAAAGAGAGAAATATTCTACAGAAAAAGACAGCGATATATCCTCTGAAACCAAGCAATTTATTAAGGAACCTGATGCTTGTTTAAAAAAAAAATTAACAGAAGAAAAGGAAAAATTAAACAATGAAGATGCACCAAAATTGAATCATACATTAAGTTGTGAGTCAAGTATGCATATCCAGTTGGAAGAAGGGAGTAAAACTCATTTTCATGCTGATGATAGACACCACCATCATCATCACAATGAAAAGGAAATAGATAAAACTACAGCTATTTCATCGTTGGCTTTAATGGTTATTGTAGGTGATGGATTTCATAATTTTAGTGATGGGTTAGCTGTTGGTGCTGCGTTTAGTGCTTCTTTATCTAGTGGCCTAAGCACTGCAATAGCTGTGTTTTGTCATGAGCTACCTCATGAGTTAGGAGATTTTGCAATTCTTATCAACTCTGGAATGACCATAAAGCGAGCTATTATATATAATTTAGTATCAGCATTACTATCATATCTTGGTTTAGTTGTTGGTATTTTTGTTGGAGAATACGAAATCGGAAGACATTTTGTACTGTCTGTGACTGCAGGACTCTTTCTTTATGTTGCATTAGCTGACATGCTTCCCGAATTAATGCATCAAGAGGTACCTAACCAATCTCTTAAAATTACTTTTCTATGTCAACATCTCGGTATATTGAGTGGGGTTGGAATGATGCTATTCATATCGTTGTTTGAACACTCTTTTTAAATGAAAAATATTTATATTGCTGCGTAAACAGTTGGTGTTTTAACGATGTGAGTAACGGTCACATCGTTATTGAATTTATTTTTTATGAAATCGTGTTGAAATCTCTAAAATATTTATTGAACTGCCAGAGATTAAAAACAGCAGGAGGTTTCAAAATTTTATATAGATCTCAAATATGCAGACGTATTACTTAACCTATTCACGTACGATATACTTAAAATAAATTTGTATATGTATATGTATA
>evgR014751c0g1_i04_Hydra
ATGTTGGGATCGTATTAATGTGATTTTCAAATTAAAGAATTAATGCGGATGTATTAAAAGTTACAAACATTTTTCATGTTTTTGTCACAAGATCACGATGAACACTATAAATTAAACCTGTTTTCTTCGATTTATATGTCTTAAAATGTGTAGCAACGCAGTTGTGACAAACAGAGAAGTGAAGAACAAAGTGACTGGTTTAGATTTGTATGTGTCTGATTATCAGGAAATACCTCATCGTAAAAAAAACTCATTTCAATATCATGTTATTGTTATCACTAATTTGTTTTATTACAAAAGAAGTGATTTAAAAGATTCTGATGTTGTGCAGTTTATGGTATCAAAAACATTTGAAGAGTTTGAAGCTCTGTATCATCAACTCTATAAGCGTTTTCCGAGTGTCATTTTTCCAGAACTTCCTAAAAAATCTTTATTTTCAAATGATACTGATGCAAAATTTGATTGTATGAATAAGGTTTTACGAATGGTGGCTGCAACTCAAAAATTATGTAGCAGCCCAATGCTAATAGCTTTTTTGAAAGAAATCGATGCAAAAGAAAGTCCATCATCTCAATCTTTGCAATCTATTTCTCCTATAGTTTTGCCAGCAGTTTCAAATAATTATGTTTTGCCAGTAGTTTCAAAGAATGATGTTTTGCCAGTAGTTTCAAAGAATGATAACATTAAAGAGGAGGATGTATCTACTGTTTTCAGTCAAGATTTGATGGAAGAAGTTGACCTGTTTTCCGTTCCAAATGAATTAATAAAAGACAGTGAAGAAAATCTTTTTGAAACTCACAACACACAAGTTTTCGATTTATTTCTTGATAAGGATAATGTGAAAGGCGATTTGCATTCAACTCATAAAGAATTGAAATATTCTGAAGATTTTTTAAATTCTAATGAGGAAGAAGAAGAAGACTTACTTAAGATGCTGGGAAATGCTTCAAAGAAAGAGAATCTTTTTAATGTAGCTTCACCAGTGAGTAGCAGTGTTTTACGAGCTCGTGTAAGTTCTAATGTTGAAGAGCGAGCGCATGAAAGTTCTAATGCTGAAGAACCAGATGTAGATTTTTTCAAAAAAACAAGAAATTTGTCAATAAGATCACAGAATCAAAAATCATTAAAAGAGGATAGTGGCCTTGATGATTTGTTTGGGGTATCAAAAAAGAAGACTTTTTTTGACGATGTAAGTGTTGATGATATTGATTTGTTTTCTACATCTTCAAATGAAACGTCTAATAAAACAGATACTTCTAAAAATAAATCAGCTATTGAATTTTTAACTGAAAATTTTCAAACTGGTAACAATGAGCAGTGCAATAAAGCTGCGGAAGCTGGGGAGTTTGTTGCAAATGACCACTTGGAAAGAAGTTTGTCTCAAAAACTTAATAATGAAAGTGATGATGATTTTATAGCTTATGACAATAAACTTAACAGTCAGTTTACTGATGTCACGCATTATAATATTACTGGTTCTTTGCAAGAAGAGATTGAACAAACAGACTTATTAAACTCACAGAACTATGGCGGTCTGTTTATCTCTGATTCTGAAGTTCAAAATACAAATGAACATGAACAGTATATTGATGATATTATGAATTATATAGCAAAACACAATCTAGATGAGGAAGTATCTTTAGGTTTTTAAAATTCTTTTAGACCTTTTTGTTTATATTTTATTTTTTATTAAAAGAATTTAAAAATGTTTTATAAATATTTTTGTACTTTAAAATTTATGAAAATTATGTTTTACTTCGTTGTTGTTAATTTTTTCAGAATCTTTTTAAATATATATTTTTTAGTTGTTTGTTTATTTGAAAGTTTATTTAACAGTTGTGAGAGTTTAAAAATGTGTTAACCAATCAAATCTTTATTATACTAGCCTTTATCACCCTAACCCAATAAAATCTTTGACAAG
>evgR014754c0g1_i01_Hydra
CTATTGTCTAAATAAGGTTTAAATTAATATGATTAATTCTGCTGCAATCATGATTCATCAAAACGAAAGTACCAATGTTCTAGATGAAACAGGAATTGATGAATCAATAAATTTTATTTCTTGCAAACGAAATTTACCTAATGCACCTGTCTCTGAGTCTCTTGATAAACGATTGTCTATTGATTATTCTCAAGTATTAAAAGATAGTGGGAATGGAATTAGCAAAATTAAATATTACACTTCAAAAAAGAAAAAAGAAAAAAAATTGCAAGCTGGTGTCTCATCTTCAAAATTTTATTTGACTAATGATGTTAATATTCAAAATGAAGTAGATAAAAACTGTACCAATAAAACAACATTGAAGAAAGATTTTAAACTTTCAGAAAAAAATAATATTGTTTCTGAAAGTCAAACACATGTTGCTCTATATAAATTTTTTGCTAGACATAAAGATGAAATTTCATTCAAGGATGGTGATCCTATTCAAGTTTTAAAAATGAATGATGACTTATGGTATGAAGGTGTTAATTTATTGACTGGTAAACAAGGCATATTTCCATGTCGTTATGTTGCTGACATTCTTCAAAAAGAAGTTGTTTCAGCTGACGTTAAAGCTGCTGATCATATGCAATTCCAAATGAGATTTCTTGGATCAGTTGAAGTTGCAGATTTTAAAGGTGAAGATATTCTTGCGTTTGCAATTGCTAAGATTGTCAATCAACGTAGTATGCTCACTGCAGCTGATCCTCCTTCGTGTGTTTTACAATTGAGTAATAAAGGAATTCGAATTTCAGATATTACGTATTCGGAAAAAGAAATTGAAAAATTGAATAAAAAAGTTAAAAAAGGAAAAGACAAAGAAAAGAAGATTGAGATTAATGATGCAGATGACTCAGCACATTTTTTTTCTTTAAAGAATGTGACTTTCTGTGGAAACCATCCTAAAGACCATAGATATTTTGGTTTTATAACTAAGCATCCAGATGAACAGAGATTTGCTTGTCACGTGTTTATGTCTAAGTTTTCTACCGATTCTATATGTCATATTTTAGGGGAGTGCTTTAAATCGTTTTATGAATCATATATGGAATATAGAGCTCCTACAGAAGACATTTACTTAGAATAAAACGGCTCTCATGAATTAGATGCTTATTTATTAAAAAATGTTTACTACTCGGTGGTTTTATTTAGTACTTTCTTTTTTGGAGAAGCAAGGAATAATGGAAGATTATTTTAAACATAAAAAACAAATTAATACCAAATTTATTAATCTAAATTTTGTAGACCAAAAGAAGATATTAAGCGGGTAATATAATAATAAAATTCAAAAACCAAAC
>evgR014761c0g1_i01_Hydra
AAAACTAATCTTTTCTTGTCTAAATAAGAAAACATGTTAAAAAGGTATATATCGGCGAAAACGTGATTCGAAGCGTTTACAAATTATGTATGTACAAAATGATAACGAATTATTGAGAAAAAAAAAACTCTTTGTACTAACAATCTTTATACTGATTCTTATAATTTTATCTGGAATTGTTGCACATGCAAGTGGTATGGCATTTTTAATTTGCGAAATAGCTTTTGGACGTTGCTCCGAACTTTCTGCGTTATCAAAATTCTGTAATTGTCGTGACACTGATGAAGAAACAGAAAAAAGGAACAATAAAAATCTCGTTACCTTTAAATTAGAAGAAAAAGAAGAAAGTAAAAAAAATGGAAAAAAAACAATATTAATTACATCATTGCCAAGATCAGGTTCTTCATTTCTCGGAGAGATTTTTAATCGGCAAAAAGATGTATTTTATATTTATGAACCATTGCAGCCTTTGAGTGTTCTCTCTCAATTCATTTCTTCATATAGTTACGAAAACGAGGCAGTTAAACTTCTTTCAAATTTTTTTATTTGCAATTTTAAAGATTACCAGGATTATTTCAACTTTTTATCGTTTCCCGAATTAAGTAATCCACATTTTCGATTGATGAGTCGTGTTTTGTCAATGCAACCCTTTTGTATGAAACGCGTTCTTCCTAATTCCACAGTAATCGAATATAGGAAGTATTGCGTTAATATTGACGTTAAACTTCTTCGCAAGAGTTGCAACACCAAGTCGTTTTTTGTTATCAAAGAATTGTTGCATCGTTTACCTAGCGATATTTCAACGGTAGAGTTTTCCAACTATATATCGTTAATTCGTGATCCGAGAGCAATAGTTTGGTCAATGAAAAAAAACAAATGGGTTGAGGAAAAAGGGTACTTTGCTTCAACTGAATATGAAGCAATTGAAAAAGTGTGTCGTATTTTTCGGAACAACTATTTCACAATGCTTACCAGCAACACTATTAAAAATAAAGTAATTTTAAGGTACGAAGATTTAGTGAAATATCCAAAAGTCACTGTAAAAGCTTTATTTCAAGCTACAAAAACTTTCTACAATCGTAACATAGACGATACGTTTGAATGGATTAATGTAAATACACATGGAAAATCTGAAAGTAATTCTTCAAGTTTTTCAGTATTGATGAAAAATGCATCTTATTCATTAAACTCTTGGAGAGAAAACTTAAATTTTAACAGTGTTATAGAGATTCAAAATAAATGCGGGGACGTAATGGAGCAACTTGGTTATGTATTATATATATCAAATGAAGATCTAAGAAACTTTTCAAAAACATCTGTAACACGAACAATACCTTTGAATAAGTATATTTTAAACAGTTTTTAAATTAAAGACAAATACATTTGTGCATATAATATAATAAATCTCTAATCTCTACTTGTCTCTCTCTCTCTCTCTCCCCCTCTCTCTAAAGCCATAGCAACCAGGGGGC
>evgR014777c0g2_i01_Hydra
GTGCGTTTTAGAGGTAATACATAAGAGCAAGGAATTATAAAGGGAAGTATTTCTGAGGTGCATGTAGATTGAAAATGACGAAGTTGGTTTTGGTTTTGCTGTCTGTTGTTTTGTGCCATTCTTTTCCTGAAGAAGAAAATAAAGATCCTGAAGGTATTTTGTTTGGTGGTGACATTTTACTGACACCAGAACAAAAATCTATAGTTGAAGTTGGTGGAGATATCAGTCAAGCTGGACTACTTAAGGGATTAAGACAAAAGCGTGCTGCTTTGAGTAATTCAAGCATACTATGGTTGCCTAACAACAAGGTTGTTCCATGGAGTATGACAAAACAGTTAGAAGATACTGCTGAAGCTACATTTGGCTTAATGGCAGCTTTAAGAGAATGGGAAGAAAGATCTTGTCTCACATTTAAAAGAAGAACTGATGAAATAGATTACATTGAATTTTTTCAAGGAAGTGGTTGCTGGAGTTATCTGGGAAGAGTAGGTGGCCTTCAAAATATTTCTTTAGCTGATGGGTGTTGGGGAAAAGGAACTATTGTTCATGAAATAGGTCATGCTATGGGATTTGGACACGAACAAAATCGTCCAGATAGAGATCTGTATATTGATATTCGATGGGAAAACATTCCTGAATCTAAGAAGCATAACTTCAGGTTGTACTCCAACTCCTTAGTTGACAGTCTGAATTCACCTTATGATTATAAAAGTTACATGCAATACTCTAAAACTGCATTTGGAATAAACGACACTGTAACATTAGATCCTAAACTTCCTGGTATTTTTCAACTTGGTCAAAGAGTTGGATTTACAGAGCATGACCAATATCAAGTAATGGAGCTTTATAGATGTCAAGATAAAACAACTCGTCCAACAACTGTTTTTCCAAGATATGATCTGAAAGGTGATTATACATGTGATTTTGAAACGGATTTGTGTGGATTTACTCACGATAAAACTGCAGAATTTGAATGGGCTCAGATTTACGGTGAAACTCCTTCCAGAGGAACAGGACCTGATTCAGATCATACAACAGGAAGATTAGGCACCTATGTTTATATAGAAGCATCCTATCCACAAAAGAAGAATGATAAAGCACGCCTTTTAACAAAATGGTTTGAGAAGCCATTATCTGCACAATGTTTCTCTGTTTTCTACAACATGTATTCTCGAAATGCATCAATGGTTGGAGAATTAAATATTTATTTAGATGACACAGTAACAATAAAACATGTATTTTCTCAATCAAAGAGCGAGCCAAATAATGATTGGAAAAACCTTTTAATTGATATTAAACCTGAAGGTCGTTACCAGGTTATATTTGAAGCCATTGTTGGTAACGGATGGCAAGGTGATATAGCTATTGACGATATAAGCATAACAGCAGGAAGTTGTCCGACAAATGTTCAAACATTTTACAATTTTTCGAACGACTCGAATTCATGTGAAGATATAAATGACAGAGAAGCTGAATATTGTAGTCAATGGCAACAGGCGGGATACTGCGTAAGTGAAGAAAAAACCATGAAGTTGTATTGTAGAAAAACATGTAATTTTTGCTAATTGTAAACAATTCTTTTTATTTTTTATTTTTCTATATTACAAAATTTTGCGGTATTTGTAAAATCAATGTATCAGTATATAGATATAAATATATATATATATATTTTTTTTTTAAAAAAAAA
>evgR014779c0g1_i01_Hydra
GAAGACGGCATACGAATGGGATTATAGTTTTGATTTAGATTCAAATAAATCAACAATCTTTCAAAATGAAATCTTTGAAAGTAATTACCTAATGGTTTACACCACATACAACGTCACATCCTTGGTACCAGTACTTGACGTGATTATTGCCAAAACATATCAATCGATGTCAGAAAAAAAATTACTCATAATACTTACTGATACTGATGAAGAATCGACATTTTTATTATCAAGCATTATTGTTCCTTATAACGAAGTTTTGCTTTTCGATTGTAACCTCAACAACTATAAAATGACTTACAATAAAAATTCAGTAGCATACTACTATGGTCATAAATATGATTTATTTACGGAATTTATCGTACGCTTTAACATCACCAAGTATGTTTTTTTAAAAATAATCACAGATAAGGAAAAATATACATCTCATATTTTATCATCAATGAAAAAGTATATGAGTATTCCTGAATTTTCTTTTGTAGAATATACCATTTACAATAATGACACCTCCTACATACCAGCTATTGCGAAAGAATTAGAACCATACGTGG
>evgR014784c0g1_i02_Hydra
TTAATAAATTAGAACTAAAAACAAGAAATAAAAAAAATCAATCAATCAATATGCTATGAATTTAATATGTCAAGTAAACACAGTTGGGAGTATGAAAAGCGTAGTCATAAAAGATCAGATGGTTCAAGTAGAGGATCAGAAAGAGATAGAGATTACAGAAGAGATGAATTTAGAAAAGAAAGAGATCGCAGGAAGAGTCACCATGAAAGCCGTGTAAAAAGTGACAGATATGAAGATTATAAAAGTCGTTACAAAGTTCATGATCAGCCTAGGAGAAGTTTATCACGAGATCATAAAAAAGAGTTGTCTGGTAGAAATAGTTTAATTGAGTCTGGGTTCAACAATTCACCGCAGTCAACACCATCCCCTAATACATTGCCTTTAGCTCAGATTTGTGCTAATAGCTATTCCCCATTGGTTTCTAATTCACCACTTATAGAATCCAGAACATTTGACCCTCGTATTGTTTCAAAAAAAGTTGTTGAAGAAGCCAAGCAAAATTTTCGACTTCTTATTGATCCAGCTATAAAGAAAGGTCACATGAAAGTTATTCGATATAATGGAGTTCTACCTGGGCAACCCCCTGTAATACCGAAAGATCCCCGACGTCACAAAAGTAAAGTATGGACAGTAGAAGTATGTGATCTACCTCTGCCAGATTTTAAGGTTGATAAAAATTATGTTGGATCTATACCTCAAAATACTATAAGATTTTCTGGATTAAACGACAATATTGATCGTAAATTTTTGCATGAACTCTGTGAGTCATATGGTGATATTATGGAATACAAAGTATACTTCGATCCAAAAACTAATCGTCACACTGGTACAGGAAAAGTTATTTTTGATGAAAAAACAAATACTAAAAAAGTTGTTGACGCTTTAAATGGGCGACAGGTTATGGGAAAAGTTATTAAAGTTTGGATTGACATAAAACCTCCTCCTGGGTATGCCATGTGGTGTCAGCTGGAGATTGAGTGTCAAAAGATGCGCTCTACTTTAACCAATAACTTAAATAGTATTTCACGGCATACTTCAGAATCGTCCTATCAAGCTACTCCTAGAACACCAAATATAAGTTCCATGCACAGAAATCAACCTATTGCAAGCTCCTCAAAAAGCAAACATTCAAAAATTGTTCCTAGTTTTTATAAAAATAGTGAAATTCCACCCCCTCCGCCACCTCTGCCAAAAAAAAACAAGAGTTCTTCTTACAGCTCACCTGTTTCTAATATTTCAAAATACGATGACATATCTCCGACAGAAAATTCGAAATTCAGTTTTAAAAATGGAAGGCATACTCCTATTATAGATTTAATAAAAAATAATAGTAGATCTCAGTCTCCCTTTATGAGCCCAGTGACTCAAAACAAAAATATTATTTCAAAATCTTATAGTCCTGTATCTGATCCGCAAACACCTCAAAGTCCTTTCAAAGATCTTACACCATCAGAATTTTATAATAATAAACTTACTGACAGTTACAAGCCCGAAGTTACGAAGTACAAGCCCGAAGTTTCAACTTCTAAAAAAAGTAGCGATGATGAAGATGATGACATGTCTTTATCTTCAATCAGTTCAAATGAAGATAGAAAAGTTGAATTAAATGTTAAAAATATTAAGCATTCAAAGTTATCTCGTCCGCTAATACATCCTCATCCACTGACACAACTTCATCCACTAACGCAAACATGTCTGCTAACACAACCTCATCCTCTAACTCAAGGAAATGTTAGTTTGACTGGAGCAATTAGACCCAGTCATATCTACCAGTATCCACCAAGTAATGTCCAAAATCAACAGTATCCACCAAGTAATGTCCAAAATCAACAGTATCCACCGAGTAATGTGCAAAATCAACAGTATCCACCTAGTAATGTCCAAAATCAACAGTATCCACCAAATAATGTCCTAAATCAACAGTATCCACCAAATAATGTCCAAAATCAACTTAATTATTATTCTTTAGTAGCTCATCCCTTACAAACTAGACCGCAAGGACCTCCGCCCAATGGACCTCAAATGGATTTTTTAGTTCGGCAGTCTTCTTTTGTTCCTTTAAATAGTCAAGCTTCATTTTCACCATATCAAAACCCTCAAATAGTACGACCTACGTTAATACCTATTGTTCCTCCAAATCATAGTATTCCAATACAGCATCATCAAATTTCATTGACTCAGGATCATATGCAAAACGTTTTATCTGTTGCTGATGCTATGGCTAACATGCATAGTTCTTGTTCAGTGAGTGAAAGTATAGTACCACCCTCTAGCATTCCATATACTTCTGATGAACTCTTACTAGCTTTCCAGGTAAAAACTGGATGTGCTCGAGATCTTAGTATTGAGTTAAAAAGGATACTTCTTAAAGATACTTTAAAAAAGCTTGTTGAGCAATCGGCATTTGGTGCTTTTGAGAACTGGTGGGGTAAACAAGTTCTAAATCATCACAAAAACACAGATATACTCTCGGAAAATTCAAACAGCACCAATACAAAACCATTGTCTGAAGATAAACAAAACATTTTGTCAAATCAGCAAGAATCTACAAAAGAACCTAATACAATATCTGCTTTGCTTCAAACTCTTTTTGGTTTTGATAAAAATACTGATAATAGACTACCATCTTCAAACTTTTTAAGTTCATTTAGAATCACACGAAAACCACATTTACAGCCAACTTCATTAAAATCTTCTCAGGCAACACGTAAAAGAACATCATGGTCATCAATTCATGGTTCCCATGTCAAACAACGACGTTTGGAAGAGAGTGAAGATGATACAATGGATACTGTTTCAGATGACGAAGGTGAAGATGAAGAAAAGACTGATGAAGAGTTTGAGCAACTTGCTTTAAGACAAAGACAAGTATGTAGAAGTGGGTCATCATTATACCAGAAAATTTATTCAGATGACAGTGAAGAAAGCGATGTTGAGTCCAAAAGTGAAAATGATGATATGGATGAAAAGATGTCATCAGGTTCATCATCATCGTCTAGTGAAGAGTCTTCATCAGAATCTTCAGAATCATCAGAAGAAGAAAGTACAGAAGTAGAACCAATGGAAGACAAAAAGATTCTAGTTGAAAATAAAGAAAAAAATATAAGAAAAAATCAAAAATTGTCTACTACAAAGACTGGTGAATTAAATGATAAAGTCATGTTTAGAAGTATGTATGATGAACCACGAAGAAGAAATGCAAAGTATATAGATCAAAAAGGAAAAAATAAATCTGTTATTTTCTCTCAACCTGAACCGGAATCATCTGCATCAAGTACAGATACAGGAGATGAGTCATCTCATGAAAATAACAAAGATCTTAACTCTGTTTCTGACATTTTGATAAAAGAAACTGATTTTCAAACTGAAATTTTCATTAAAAAGTTAATATGGCTAGAACATAACTACTTCACTGCCCCGCCTTCATTGCCTTTACCACCTCCAATACCTCAACCAAAGTCTTCTACTACAAATCGAAGAGGTTCAACTAATAAAGTTTTGTCAGAAAACGAACATAAGAAGTTTACTATGAGATCAAGTAAAGATGAAAAATTACTCATATGGGATATTTTAAAAGAAGGTGTTGATATAGAAGATTTATTATACCTTAAAGAAGCATTTGAAAGTTTACAACAGGTTGATGCTTTCGAAGTAAAAAACTTACGTTGGTTTGACCACCCTATTACTATAACATCTCCTCAGAAGTTAAAGAGAAGTGAGTCTAAACTACGCATTCACAAAAGTGGTTGTGCGCGCACCGAAGGTTTTTATAAAATACCTATAGTAGAAAAAGCTCAATACTTAAAGAGTGCTCTTAGACAGTTGAATGCTAAACATCAAAGTGATATACAAATGCAGCAGATATCTGATCAGGCTGGTACCGAACATAAAATAAAATCGAGACAAAAACGTGCAGTGCAGAGAAGATTGGCATCTTCTATTGCTCAAGAAGAATTTAGCAGTCTTATTAACTACAACAACAAACTTATGAGGCGCAAAAAAGCATTACGTTTTTCAAAGTCCAGCATACATAACTGGGGACTATTCGCATGTGAACCAATAAATGCAGATGAAATGGTGTGCGAGTATGTAGGTCAAATGGTACGGTCAATTGTTGCTGAGATACGAGAGCGAAGATACGAGAAACAGGGTATCGGGAGCAGTTATTTATTTCGATTGGACTCTGATAGTGTTATTGATGCAACAAAAGATGGCTGCAATGCCCGGTTCATAAACCACTGCTGTGATCCAAATTGTTATGCTAAAGTTATATTGGTTGAAGGTGCTAAGAAGATTGTTATATATTCGAGACGTGCCATTAAACTAGGAGAAGAAATAACATATGACTATAAATTCCCTATTGAAGATGAAAAAATTCCATGTTTATGTGGAGCTACTCTTTGTAGAGGGACGCTTAATTAGGAATAGAAAATTTTAAATAATAGCTACCTTATATAATAAGGTTGCCTTTTTATGTTGTTTTCGCCTTTTTGCTGCAGTTGTTTAGTTGTTGGTAATTCCGTGTTCTACCGGTGCTATATGCGACAC
>evgR014786c0g1_i01_Hydra
TTTTTTTTTTTTATAAACTCATCGTTCATCCTATACGGATCTTTACCTACACTTTTTATACTGTTTCAAAGTGAACACTAAATATTTTCAAAGCGAACACTAAAATGAAAATACTTTGTTTACTCTTCATTCCGTCAGTACTGAGTTTGAGTTGTGAGGTTTCAGTAACAACTTCTTCCAAAGCGGAAAATCCAGGAACATCGAGTAATTGCGATCAATGTTTTACACTGGAATCAAAATTAGCTGGTGTTACTACGTATGGACGAGGTTGTCTTTTAACTGGTTCAACTTGCGATAGTCAATGCACTGCATCAAGGACTATAGGCTCGAGCTGTTCGGCTACTTGTTGTTCTGGAGATAACTGTAATGCGAGTACGTCAAGTGGTAATTTAACTGAAGGATTACGTTTATGGCAACTACTTTTTGCAATTTTGTTCGTACTTTTTGCAACAAGAATTTGAATTATTTTAATGTAAAATTTTGTTGGGTTGTTATTAATAATGGTTATATATCAAAGCGTAAACTCTTTTTACTCCAAATTATATTTTTTGTATTTAAAAAAAAAA
>evgR014791c0g3_i01_Hydra
AAATAAAGTTTGAGTTTATTGACTTTAATCCTGAAAATCGATCACTTTTTAATCGATATGCGACGAAGTTTCGTTTTCGTTGCACCTTAATATCTTAAAATAGGTTAATAGCTTTAACGTTTAAAGCAATTTCTTTAAGAGGAAAATGTTTTCCTACACAGTGATTTATTAATATTGTTAAGAAGGTAAAGACTTGATTCCTGAAAACTCAAATTGTTCATTGCATAGATTAATAATTTGAACAAACTTGTTATGGAAACTTCAACAGTCGAAAAAATAAAACCATTAAAATTTTCTGCAAGTTTGTCGTTTCTTTTCCCAGAAGTTTTAGGTATTCCAAAACGATATGCGGCTGCAAAGGCATCAGGTTTTCAAGGAGTGGAATGTACAACTCATATTTATGACTATGCTTTACCTGAGTTGATTGCTGCTCAAGAAAGTTCAAAACTTAAACAAGTTCAAATGAATTTTTTTTGTGGCCAAAACAATGGAATAGCAGCTGTTCCAAAGCGCAATGAAGAGTTTAAGCAGTGTCTTGAATTATCAATACAGTACTGCAATGCATTAAAATGCAAGAGATTGCACATTATGGCTGGAAAATATCCAGAAGGAGTTGAAATAACTGAAGAAACAAGATCTACTTTTACAGAAACCTATATTGAAAATTTATCCTATGCATCTCAGAGACTTGAATCAGAGCGTATCATGATTCTTATTGAACCATTATCAAATGTTGAAAACTATTTTCTTACTTCAACTAGTCAAGCTGTTGCTATTCTGAAGAAAATGAATCTGCCGAATGTAAAGTTACAACTAGATTTATATCATCATCAAAAAACTGATGGAAATACCATACAAGCTCTAATAGATTATTTTCCATACATAGGTCACATTCAAATAGCTCAGTTTCCCGATCGTAATGAGCCAGACAGTCCAGGAGATATAAACTATCCAGAGGTGTTTAGACTTCTTGTTCATTTGGGTTATACTGGCTGGATAGGATGTGAATACATTCCAAAAGATCCTCAAACATCATACAATTGGTTGCAGCAGTATTTATTTCGCTAACGATCTCGACTTGGTTTTGGCTTTGGTTTTGTTAATTGATTCCGTTTTATACTGAATACTTTTTATTAATAAATATGAAATTTATAATTATAACTTTTATTTCACGTTATACTTTATTTATCATTTACTTTTAAAATTTTCTTCGTCGTTTTGTATGTTAAAGTATTTTCAAACTGCATTGCTTTTTTATTACGTAATGCTACAACAATGTTTG
>evgR014795c0g1_i01_Hydra
TTTTATTTAACGTAAATCGTTAAAAAAGTTTACGCTAAAGTTTGAGTTAACTTTGAACTTCTTATCGAAACAACAAAAACGATTACTATCTGCTTCTGACCATTAAAAGTTTATTAACAGTGTTAGGGACATAGTTGCTTTGTTATCGAATAATTAATATATAATTATTTGACGTCACAAGAACTTATCGTATAAACGTTTCGTTTTTAATAATGGTTTACAACGAAAATGGTTTAAATGTTTTTACTTTAAGTGAGGTTAAAGAGAGTGAACACTTTAAATTAAAAATTGAAGCCTTTAATATATTCATAACTACCATGCTTGTTTTTGTTATGGTAGTTGGTGTGACAGGTAACACGGTTGTTATCTACGTGTTTGGCGCAAGTAAAAGAAAAAAAAGATTTGAGCGCTTTTTACTGTCCCTGGGGATATTGGATTTTCTTTCCTCGCTAATAATACCATCAACGTTTCTTTATCTGACCGTAACCGGCTACAAACGTTGGGACTTTAGCGAAGCTGGTTGTAAAGTCATACCTACTTTGTTACAAATTTCAGTGTCTGTTACGCAAGGTTTTCTTATTATGATTAGCTATGAACGGTATCACTCCATTGTAAACCCTTTTCAGAAAAAGCTCTCTCAATGGAAACTAGTTTTACTTTTTTGCTTTGTTCTTTTGGTATCCTTCACATTGGCAATACCTTACATGATGACTTTTAAAATTTTGATCGATTTTGAACATGGAGTAAACACGTGCGCTCCTGATGGAAACAAAATGAACCTGTCAATGGTTGCTTCATCATTGAGATTAATCAAAGATGTAGTTGCTATTTGTTTCATGGAATTTTTTCGTCAAAGAATTAAACATTCCTTTTCAAAGGTTCACGCAAAATCGTCATGGAACAAAGAAAGGGAGACACTAAAAGGAAGAAAAATTCTATATACTGTTATAATAGTTTTTTCTGCTTTAACTTTACCATTGGATGTATTCCATTTTATTTATTATATGCTATCTAAATTTGCATCGTCAGCACTGAGCTCAAAAGCCTATTCGGTACTTATGGTTGTCAACACAATGCTTAACTTAATTCAAATGTCAAACAGTGTTGTGAACGTTTTTATCTACTCTCGTTTGCATAAAAACTTTAAAATGAAACAAGCTGAGTCAGCAAGTTGTAGTATTTTATATTGGAATGAAGATAAAAACATCATAAATATAAATTATCATAATTAATAAATTATTATAT
>evgR014798c0g4_i01_Hydra
GGAAGAAACTCAGATGCAAATGTTTTTTTGACTTCACTTGTACCATTACAACTTTTGTGTACAAATTCAGTTTTAGGTCAAGATATTATTCTATGGAAAAATAGGACACCCTCGTCTACTCGATTTTGCCGACCTATCAGACTGCAATTTCTTCACGAAAATGTCAAAACAAGCATTAATGAAAAAGACTATATTGAAGATAAAATTAAATCACTGGTTCCTCTCATCATTGTTCAACATAAAATAGAAATTAAAATACGTTATAAATTGGTTATGATAATGATTGACGGAAAAGTTTGCAATGCCATAACATTGACGAAATCAACAATGC
>evgR014801c0g1_i01_Hydra
GTATTTTAATTTTATGAAGTATTTTTATTGTATTTAGTTTATAAATACTCATATTGAGTTTTTAAAACACAAATAAAAATGAACATGGTAGTTGCTATATTCGCTCTATTTTGTAACTTTCTGATTGCCAAAGGATTTGTTAGTAAACCAGAATCAAATTGCTCGGAAAATTTTTTCAAATGTTCCAATTTAAAATGCATACCTTTTCGTTTTAAATGCGATAATTCTGATGATTGCGGTGACAACAGTGATGAATCAAGTTGCAAAGAGAAAGAGGTTACTTGCTTGCCAGAAAAGTTTAAATGTTTGAATGGTTTGGAATGCATTGATAATCATTGGATTTGTGATAATCATACTGATTGTAGTGATGGCAGTGATGAAAAGAATTGCTCTAATTCAAGGATTAATTTAACGTGTAATAGTAACGAATATACCTGCAAAAATAAACATTGCATCCCAAATGGTTGGCTATGTGATAAAAGTAATGACTGTGGTGATAATTCTGATGAATTGAATTGTCAAAATAAAACATTGCATACTCCAGAATGTATTGATGATTTATGGAAATGTCACTCTGGGGAATGTATTAAATTTAAATGGAAGTGTGATTTTAAAGATGATTGTGCTGATAGGTCAGATGAAATAGGCTGTGATTTATTGTTGTGTAATCATGATAATTTTCAATGTAAAAGTGGCAAACAGTGCTTGTTACAAAAATATCGATGTGATGGTTTAATAGATTGTACAGATGGCTCAGATGAAGATGATTGTGATGCTAAAATTGATATTTGCACTAAAAATCATTTTAAATGTTCTGATAACAAGACTTGTATTGAATGGAAAAATGTATGTGATATGAAGAGTGACTGTCCTAATGGGGAAGACGAATCTAAATCTTGCTTTATTAATGAATGTGAAAACTACAATGGTCATTGTAGTCATATCTGCACTAATTCAAAAATTGATTATGTTTGTTCATGTAAACCAGGTTATTTTTTATCTACTGATGGAAGAACATGTGAGGATATTGATGAATGCGAAATACATGGTACATGCCATCAAATCTGCCATAATACAAATGGAAGCTATCATTGTTCTTGCACTGATAATTTTGTAATGCAGCCAGACAATGTTTCATGCAAATCTAGCAATGAAGGTTGGATTTATTTTGCAAATAGAAGAGATATTCGAAGATTTTCATTAAGTGGACATGAGTATGAAGTGGCAGTAAATGGTTTAGAGGGGGCAGTTAGCATTGATTTTGATGTTCATAATAACTTTGTTTATTGGACTGATGTTGTTCAAGAAACAATCAACAGAGCGCTCATTGGTTCAAACCAGTTAAGACCAGAAGTTCTGATAAACAATATTCACACTTCAGATGGTTTAGCTGTTGATTGGCTTACAGGAAAGCTTTATTGGACTGATGCAGGGCACAAAACTATTGAGGTTGCAGACCTAAATGGAAAAAATAATGCTGACTTGGTAGACATTGGTTTAAAGGAGCCGAGGGCGATATCATTGGATCCATTTGAAGGGTTTGTTTACTGGAGTGACTGGGGTGATTCTGCTGTTATTTCTAAGATTTCTATGGATGGAAATAAGGGAACACGTAAAAACATTGTCAATAAAGATATCGTATGGCCAAACGGTCTAACGCTAGATCTAATTCAGCATAAATTATATTGGATTTGTGCAAAGCTGAAGAGGCTTGAAGTAATTGATTTTGATGGAAAGAATCGAAAACTTTTGAGGATGTTAGGTGAAATGCATCCATTTTCATTAGACAGTTTTCAAGATCAATTATATATTTCAGACTGGAAAATTAATAGTATTGATGCTTTATCAAAATATAATTCATCTCAAAACTCTGTATCTTTGAAGTCTGGATTATTTGCTCCAATGGATGTTAAAATATTTCATCCTACTAGACAACCTTTAGGTGTGAATCCATGTGGTTTCAACAACATGTGCAGCCATTTATGTTTACTTGGAAATGATGGTGAAAAGCGTATTATTTCCTGCAAGTGTCCACGTGGAATGCATTTAAATAAAACTGTTTGTGTGGGAACAGCAATTGTACGTTCTTCAAATAATGTAATAGAATTAACTGAAACCACTACTCACGTCACTACTCGAGCTACGATTAAAACAACTACTCTAGCTAGTACTAAAAAAATAGAAAAACAAACTATATCATCTGTCATAAAATCTGAAAAGTTAAAAGATAAAAATGTGATAACCAATTCGTTTCCAGACGCAGAACCAACTTTAAAAACTCGTTTTTCAAAATATAAAGTTGCTAAAATAAGCGATGGATCATACTTGGCAATTTTCTTGTCAATTTTAGTCGCCATAGTGATAGCTCTTATATTAATTCTGATTCGAAGGCGTCGTGAACAAAATAAGGGTCAAGTAATGTATTACAAAGACATGTCAACAACGCCACTAGAAGAAGATTTTTATATAGAAAACGAAGAAATAGATGAGAAAGCTAACATTATCTATCAAGATGTTTGAAACCTTTTAAATGATTTTTTATATGATCTTAGGTCTAGAGATGTTTTTGTATTTTGCAGAATTCTTTTAAACAATTTTACATTGGGATTCTGCCTAGTGTTTTTGCGGTTGTTGTTTAATATTAGTTGTAAATTTGTTATATTTATTGAAATTATATATTATCAAAATTAAAAAAAAA
>evgR014804c0g1_i01_Hydra
TCAGATCCTGCAGCAAAACCTTGAAGATAAAAAAAAAAAATCGCTTAAGTTTTTCATTAAAACTTACCAAACTTAGTTACAAAGTTCTTACATTGCATTCCGACCTACCAACAAATAATAGCATTACAAGTTTAAACTATTGTTCGCTAAATGATAATTATAATAAAACTAGTACCCAGTACTTTAATAAACCTAATAATACCCAGCACTTTACAGCAGAACTCTTGAATAACATTACAATTTTTCAGGATAAACCCGGCACATTAAACAAAATAAAGTTGGCTAAAAAGAAAAATTGTGGTGTAAATGTTTCACGGAAGTCATCTAATATTTATCTAAAACCTACATTTTATGAGCTACCTAGTCTGACTGATGATATTAACTATGCTATAAATACAGAAGGTATAGAAACGCTCACTCAAATAGAAATAAGTTCAAATTTATACGAGCAATGTAACGTCAACTATGAACTTGTTTCCTTCGATATTCACCAGCCCTTACCAATCTTACCCAATGCCGTTAATATTGTCCAGCCTTTACCAGCGTTACCCTACGATCAAACAAACTTCCCATCTAATAATTATATCAATCACTTTTGGACAGAAATCTTACAAAACGCGGCAGTTATACCGTCAACGTCAACTGGAAGAATATGTTGTGGTGTAAAATGTTTACGCAACTCGTCCCATTTTTATCTAAAACCTACTTTTTATGAGCTTCCTTGTTTGACTGACGAACCTAACTTTGCTAAAAATTCGAAAGCTAAAAAAGCACCCACTCAGATGGGAACAAATTCAAATTTGTATAAGCGAGGAAAAGTCAAATTTGAAATTATCACCTTCGATATTGAAGAGCCTTTACCACCTTTTCCCTGTGACGAACAACACTTTCCACTCACAAACTATGTAGTAATTTAATATCCTCCTGCAGATGTTCTTGAAAAAAAAGCAAAGAGTATGGAAATGATGAAAATAATTATGAAGTAAAAATGAAGTAAAAAGTAATTAAAAAAAAAACTTAATGCACAGCACTCAAAATTGTAACTTACATGACATTTCTTAATAATATTTGAAACATAG
>evgR014805c0g2_i02_Hydra
AAATTTTAATATTAAAATCCACAGATGTCTAATATACTAGAGAAAAAAGGAAAAGGAAGACTTGATGATACTATTCCTTTAAAATGTTCAATAGAAAGTTTTGTAAACGTAGATGATCATACAGAATTTATTCTGAGAGTTCAAAGAGGTGAAAATGCTGATAACAGTTGGAAGGTGCAACATCGTTACAATGATTTTGCTACTCTTCATGCTTCACTCACAATGAATTTGAGTGGTACAGAACTTGTTTTACCGCCAAAGAAAAAGTTTGGAAATATGGATAGTACATTTATACACGAACGACGAATTGCTTTGCAGGAGTTTATGGATAACATTCTTTCCGGTTTCTTTTCACAACATATAGCAGTTAAAAAATTTCTAGATCCACTTAATTATTCTGACGAAATTGATAGGGTTGCTCAGCAACAAGCAGCAATGTTTTTTCGTTCTGAACCTAAGTGGGAATTGCAAGAAAATTTGAAAGATATAGGTTGGAGGTTAAGGAAAAACTTTTTTGTGGTAAAATCTAAAGATAGCGCTCAGAAAAAAAAGTTTCTTTTATCTTGGGTAGAACACGGGCCTGATATGAATCTTGATTATGAAACTTCCTTATTGGCATTATTGCAGCTTCTGGCATCACTACAGCATCCATATATCTACCCTATCAATTATGCATCATGTACAGATAAAGGATCAACAGTGGTAAGATGTTTGCACCCAACAGGTTCACTGCGAGATCTTATCTTTAAGACAAAACCCACCCTACCTTTTGTAAAAAAATATGGAATCTTAAAACCTAGAGCTGGTTTGGAAGTAGCTGATATACAAAGAATTGGACGACACATTTTAGAAGCTTTGAAGTTGTTTAGAGATATTGGTTGGCCTTTTGGACACTTACATGCTGGTAATGTTGTCATTGAAGGACAGCAAACATGGTTGCTAGATATTGAAAATGGAATATTAGGGTTACCATCAGCATATCGTTCTCATATGATGGAATTGAAATGTATTAAGAACATGGAATCAATAGATGTTTATTGTTTTGGACACTTGCTCTATGAAATGACATTTGGTGTTTGGTTACAGAGTGCTACTGTAGTGAATATTCCAGAAAATTGTCCGTCACCTATAAAAAACATTTTACTTTCAATATTAAATACAAAATCACCAAAATCTGTTCTACCTTCCTTAGATGATCTCATAGCTCAACCATTTTTTAATCAAGTGGTTATTGAACCATACAAGTCATCGTTTAAAGTAGCCAAATTTAAGGATTCTTTAGTTCAAGTTCAAAACAGCATCGAGACAAGATTACGAGATGATCAAAGAAAGCTGCATCAATTCAATCGTTTGACAAAGGCAAGATCAAATCTTCTTTCTGAAGAAGAAAAGAAAAAAAGAAAGAAATCTTCAAACAAAAAAACATCAGTGGCAACATCTCCAACTGTGAACGGCGAGTCAGTTGTAAAACAAGCTAACCAAGCAACTGGGACAATTGGTAACGCAATCCCTGGGACAATAAATGGAAATGCTCAGAGAGTCCAGCAGAATCCTACCACCAACTCAGCCATTGGTTCAACCAGTAATGTAACAAATCCAACAGCCAATGGAACTGCTTCCAAAAAACTGCAACAGAATGTCAATAGTTCAGTTTCTTTAGGTTCTACTGTGAATTCGAACCCCGGC
>evgR014812c0g2_i01_Hydra
GTTTTGTGCTTTTTAAATTTTAAAAGAACAAATTTTAAAAAGGTGCGTGAATCTTTGGACAAATCAAAACAAAACAAAAAAAGTCAAATCAAAATACTCCAGTTTTTAAATCGTTCAAAATGGCTTTAATAAACTTTTTATACGTTAAGTTAAATTGCGCGTTAAACTTAGTTGGTATAGTTTTCGTTGTATATTATACAGTCACGAACGATTGGATACAAGTACAGGTAAATGGTGATTATACAGTACAAATAAGTTGCGGATTGTGGTTTTGTTGTAATCAAACCTTAAATAGTCGTTTTAAAAACGTGATTCCTTGCAGTTATAATAGTGATAAAGGTTTATTGTTATGTAAAGTATTATTGGTTACGGGCTACATGACATACTTTCTTGCTTTTGGATTAAACGTGCTTAGCTGTTGTAACTTAAATGTTGTAAAATTTCATAAAATCTCGGCTTGTATGCTCCAAATTACAACTTTATGCTTGGCTACAGTAATGTGTGTGTATACAAGATATAAATGGAAATTCCTTAAGATTGGCTGGTCTGGCTGGTCTTACGCAATGGGATGGGTTTCAGTAGCAATTTCAGTAGCATCTGCAATTCTCCAATTCTTTATATTCAAATCGGAATATGAGTCAATATAGTTTGCTTAAGCATAACAAAAGTTTGATATTTTTTATAGTAAAAAAAACAACATAGGAAG
>evgR014813c0g3_i01_Hydra
TATTGTTCATTTGAGACATATTTATTTACGTCAAACTGAAAATGCAGAAATTACCGAACGGTTTTATAGTATTATTAAAAATCGTTTAAAAAGGTTCAATAACCACGCAAAAGGTATGGAGCTATTGACTGCGCAAGATTTAGTAGAACGATATTTTAATGACTTGGCTGGGCTAAAAGAGATTATGTATAATAAAAGATACATTTTATCGTCTATTAATACTCAGGTCGAAGGGGTTTCATTACCAAAACGAAATTCTAGTATTTTAATTGGTTTGTTGGCTGAATGTGATATGTCACTGTGCAGAACTGCGAGCGCGATGGAATGCATTCTTCAACAGTTTAATTTTCATCGCGAGCGTTTGCTTTTTTTTAACGCGGCTATGGATTCTTTATTTTTTAATTTTGATCTAGTTGCGAACTTTCTACAAAATATATCCTATGAATCTTCCTCGAGCCTTTTAAAGGAGGTAAAAGTTAAGTTTAACAATCAACTAGACTTTATAAATATGCTGTTATGTATGCCGTTTATAATTCCCAGAACAGAAACAAATCAAGTGACGATTAAAAAACAAAAATTGCGCGAGAGTGTTGAAATGCTACAATTTATTTTCAATCATACAGAGACAATCCACGAAAACTCTAAACTGTTTGACTGTGGCAGCCTCAATGAATTATTTGAAAAGTCTAAGAAGCTGGAAAGTAAATTTTTTAACAATTTCATTTTTATAAACAAATTGGTTAACAATTTAATCAGAGTTAGTCTTTCGGAATGTGTAAATGAAACACATGCTACTCAACCTGTTGTTAATATCAACGACAAACAAG
>evgR014815c0g1_i01_Hydra
GCAACAAACAGATCGTTGTATATTTTTGAAAACATTGTCGATGTTATGTTTGTCATCGATATACTTATAAATTTTCGAACCTCTTTTGTGGATTCAAACGATGAGGTTGTCTACGATTCTTGCCGGATTGCTGTAAATTATTTAAAGGGATGGTTCATTCTTGACTTCTTGGCAGCTTTCCCGTTTCAACTTCTATGTAACATATCGCATATTAATCAGACCGCTGCGTTAATCAGCTTGGTCAAGTCAGCACGCTTACTACGCTTAGTTCGAGTTACGCGCAAATTGGATATGTATTTTGAATATAATGTAGCACTGTTATTACTTCTTCTTTTTGGTTTTGCGTTGATGGCTCACTGGTTAGCGTGTATATGGTATGCTATCGGAGCTTACGAAACTTCATTAAAAAACGGTCAAAGTTGGTTAGCAAGGCTCAATAAAGATTTTAATAAAACAAATAATAAAACAGGAAATGAGATTGACATTTCGACTTCTTATTTAACGGCATTGTATTTTACACTAAGCAGTTTGACTAGTGTTGGTTTCGGAAATGTTTCGGCAAATACTAACAGAGAAAAATTGTTTGCAATTCTTATTATGCTTTTGGGAGCTCTAACATATGCTATCATATTTGGCAATGTCACTGCGATTATACACAGACTGTATTCAGGTATGGCTCATTATCATTCAACTATGCGTCGGGTTCGACAGTTTATTCGCTTTTATCGAATACCCTCCCCTTTACGTCAGAGATTGGAAGATTACTCTCAGTACGATTACTCTTACACTAATGGTATTGATATGAACGAGGTATTGAAGCATTTCCCAGAAGGTCTACAAGCAGATGTCTGCCTTCATTTAAATAGACACTTGTTTGTAAGAAGTCCTGCATTTCGAGGTTTATCTCCTGGCTGTTTACGGTCATTATCTCTTAAGTTGAAAACAACTAGATATACACCCGGTAACTATGTTATTCATTACGGGGACGAAATAGTAAATCTAATGTGGATAGAGCGTGGTACATTGGAGGTATTGCAAGGAGATAAAGTTGTATATGTGTTAGGAAAATGTGATTCTTTTGGCGAAAACTTTGGATTAGTTTACAACCGTCTTATTGGCAAATCGTGTGCAAGTGTATGCGCCCTTACATATTGCGACATTCAATCTATTTCACGCGACGACTTGTTATCAATTATGCGTGCTTATCCTGAATTTCATAAAAAGTTCAATAAGAACTTTCAGGTTTCGTTCAATTTGAGAGATTTAGAACTTGAAGAACAAGACCTGAGTAATTCATTGTGCAACCTCAATCAACAACGACAAAACGTACTTGAAATATTAAAGAGACAGAGTTCTTTTAATGTAGAACATTTGAAAAAAGAAAGCGACAACATCTTTACTAGAGATTCAGTAATTAGCAGTGATTCTGTTATTAGTGGTTATACGAATCTCAGTAGCAACCATGTCGCTCCAAAGCTGGTTTCTACGATAAATCTATTCCCTAGCCATTCTATTTATGTAACAGATCACTCTAGTTATCATAACGAATCAATTATTTTTCAACAGGAGGATTTTAATCATTCACCCGTTATGAAAAGAAAAAAACTAGATCAAATAAAAAGTTGGTTAGAAAATTTTGAAAAAGCTAGAAAAGATATTACGTCAATACTGAGACTCTTAAAAATCGAGAGCGAAAAAATATCAGCAAAAAAACGCGGAAGCCTAAATAATTTACTAACCGTTCCTAAGCAATCTCGTAAAATATCGATTTAAAAACCTAATTTTAGGGGAAATCTTTTTTCTTGTTTTTTTTTTTCAGTATATATATTTTTTCAAAATTTTGCTCTTCACAAGTTAAGATTGTTTCCAATTTAAT
>evgR014821c2g3_i01_Hydra
CACACTATGGCATACATCATATAAAAAAAAAAAATTATTTGATATTGTAGAAATCAGACTTGGCTTGTTTCAGTTTTCAAAACAAATGACGAAGTTGAATCGCCAGGATAATATATTTGAAACATCCTCTACCCCAAACGCAAGGATGAAGATTGCCGTCGTTTTCTCCCTTTTCTTTGTGAGTTTTGCTTACTCAGCACCGGTTGAAGAGGATGACTGTTTAGCTTTGTTTAGTAACGAGTTATGCTCAGCCTACGATAGCCTGGATAACGAGGTTGGATATGCATACAGTGGATACAATGATCTCGATTTTCTTACTAATGCGCTCAGAGCCAAGTTAGACTTTCTTGCTAAATGGGGAATAATACAAAAAATACCGTTAGTTGGAAACAAACTTCAAGAATTTGTCGCTAACAACCAAGACATTATTGTTACAGCTCTTAAGGGAGGAGTCGCAGCAATTATAAGACTTTTACCTGCGTTACTTGGATTTTTGGGATAAAAGTTTTGTAATTATTTGAAGTAACAAATAAATTTAATTAAACAAAAAAAAAA
>evgR014824c0g1_i01_Hydra
CCTATTTTAGTGCAAAAAGAACTTTATTTCACAATAGTTTGTTTGAATTATAATTACATATTTACTGAATGGGCGACGAACAAGAAAGAAGGAGATTATGGTTTGAGTTGTACGGTGATATGGTACCAGAAAAAATTGTCGACACAATGATAGGCGAGGATGGAAAGCGTTATTATAAAGTACAATGGGCATCTACAACAATTGAGTCTTCTAAAAATTTTCTTGAAGCAGAGCATCTCATAGAAAAATATTGGGATCCCGTAGAAAAAAAACTGAAGATACCATCAAAAGAGTGCAATTCAATTGATTTAAAGGATAACGATGCTGCTGATGGTGGGTATAGTGGTGAAAAAGCAGATATACAAGATGGTGATATACAAGATAGCAAAACAGAAGTTTTATTTAATGATACAAGCTATGAAACTCATTTATCTAGCATTCTTAATTCTAAATTGGATCAATCTAAAGAGATGATTGGTTGCTCTTTAGAATGCATACAACCACATGGAGTCATGCTAAATGTTTCTAGCAATGGAAATGACGATCCTGATCCAGCCTCTCAAGTTCAGGTTTTAACAGCTGAATCAGTAGATAGTATAGTTATGTCTATGTTGCCTGACAAAACAATAAATGATGTGAGTTTTCTACCCTGTGATACCAATGAAAGATACGACACTTCTCATTACAATCTCTACTTATCTTCTTCAGATAAGACTGATCAAGGGGTATGTTTAATATCAGATAAGATTGATCAGGAGGTATGTTTAGTATCAAATGAAACTAATGTAGAAAAAGTATTAGACAAAAATCTCAAGAGACATATAAACCTTGAGGATAATATTTCTAATAAAAAGAAAAGAGATACATAAATTTTTGAATGGTAATTAGTGGTTTTACTCCATTACTGTTTTTAGATTGACGTTTTTTTTGTTTATTTACTAATTTAAAATTCAAACAAAGATTTGTTTGCTCAGTGCTGTGATCACGTGGTCAGTGCTGTGGCTTGTATGAACCTATACCAAAATATATATTATGTTGATGATGGTCTCTATAGCCCCCAGCTCTGTGTTAGCATTTTATACGTTAATTTCATTTTTATCACCTTTGTCTAATTGTTTGTACATAGATATATAGATCAGGTTAGTAATTAAATATCGTAAATATTAAAAAAAAAAAA
>evgR014825c0g2_i02_Hydra
TATGTAACTGTTTAAGTTATATGGAAACCAAACTTAAAGTCAAATTGTAAGAAGTTAAACAAGTTGAAATCCTACGCCAAACGGAAATACGAAACTGACATAACTAGATATGGATAATTGATTTTAGTAAATATGCACATAAAATTAATTTTATATAAATATCTTTAGTAGAAAATTTTAACTTTTCATTCAATAACGATGATATCAATAAAGATTTATCTTGTTTTATTATCAATTTTTGCAATATCACTTGAAGATGTGCCTGAAGACGCTTTGGAATACCTGAGTAAGTTTGGTTATTACGAAAAAAGCAATAAAGATGTAGGTTCCATGTCAGGCAAAAAAGAAATTCAAATGTCGATCGAAAACTTTCAAAGATTTGCAGGTCTGCCTATTACAGGGGTTCTTGATAAACCTACCGAAAAGCAAATGAAGAGACCGCGATGCGGAATGAAAGACATTATTAATAATAAAGCAAAAAGAAGAAGAAGATTTCTGCTTCAAGGAACAAAGTGGAGGAAAAACAAACTTTCTTGGAAGCTTTTAAACGATAATAATGACGGCTTATCACGCAGAGATGTAGAAACTACGCTAGAAAAAGCTTTTTCAAAATGGGAAGCCGTTACTAACCTAACGTTTCGCAAACTTAATTTAATCAGTGTCGAGCCAACAGACATTGAGGTGAAGTTTGTACAAAGTAGTCATGGAGATCATATACCATTTATGCCATACGAAAGAATAATGGCGCATGCGTTTTATCCTTATGACAATACTGGGTTGTCTGGCGATATTCATTTCAATGACGTCTATAAGTTCACTATTAATTCACCTAGCGGAATCAGTTTGTTGTGGACTGCTATGCATGAGATCGGACACAGTTTAGGCATGGATCACTCAAGCATCAGTACTGCTATTATGACTAGTTTCTATCCAGAAAACGATCGAAGTTTTCAGCTTCATGAGGACGATATTCTTGGAGTACAAAGTATATATGGAACAAAAAAGCTTTTACAAGTTAAAGCTATAATTTATGAAAAATTAACTGGAATTACTTACGTCTTCAGCAAAGATATCTATTACAAGTTAAACAAAAATCTAAATAAAACTGAAGGACCATTTAAAGTTTCTGAATTATTCAACAACTTATTTGATTTTGACGCAGGGTTTGTGAATAATGAGAACCAGTTGGTATTTTTTCATAAAACCAAATATTATATTTACGAATCATTTACAAATAAAAAACTTATAAGCAGCGGTTCCATATTTGATATGTTCAAAGGAATTAAAAGTGATACTAAAAAAATTGACGGGGCATTTGTTGGGCGTAATGAAACAACATTTTTGTTCACCGGCGATAACTACTATCAATATAACGCAAACCAAAAAATTAACAGTAGCAGTAACGCACAAAAAATTACTGAACACTGGAAAGGATTGCCCACTAATATCGACGCTGTTTTTGTTTGGAGCAATGGAGAGACTTACTTTTTCAAAGGTTTGGATTTTTATCGAGTAGACCGAGATGATCATGTATCGAAAGGTTATCCAAGAAGTATCCATGAAGTATGGTTGAAAGTACCGACAAACGAATAAAAATAAAACTTATAAATCTATAATGCTGAAATAATTTTATATAAAAGTACATTTATATAAAACATATGTATGTACTATACATATGTAATATACATATGTACATATGTATAAAATAATAATTTTGATAAATAAACATTTTATAGATTCTTTTCATACCTTTCAGAATGTTTGTAAACATTATTATTAATATAATTTCTATTTTTCAAAAAAAAATATATATATATATATTGAAAAAAAA
>evgR014828c0g1_i01_Hydra
AGACACAGACTTCAATAAACAGACTGCTCTACATTATGCAGCCGATAACAATAAAATCGAGGCTGATTATAATAAAATAATTGAAGAATTAATTAATAAAGGTGCAGACTTAAATCTAGGTGATTCAAGTGGCCGTACAGCATTACACTATGCAGCAGCTTTTGCCGATAATGACACAATATTGTTATTAATTAATAATAACGCTGATGTGAACGCAGTAAGCAAGAAGGGCGTAACTCCTATGCACTATGCTTCAAAAAATGCAAATGTTGAAACCGTACAAGCACTTATCGATAAAGGAGCAAACGTAAATGCACAAGATAAATTAGGCAATACCCCTGTTAATTATGCAGCATTCAATAAAAAATTCGGGTATGATATAATCAAGCAACTTCTAGATAATGAAGCCTCTATTAACGACAAAGCCCTAATAAATGCAATAAAATCAAATAATATATTGGCTTCCGAATATTTAATCAATAAAGGAGCGAACGTAAATGCACAAGATGAGTTCGGCAATACCCCTATTAGTTATGCGGTATTCGATGAAGAATATGGCGTTTCTTTTATTAAAAGTCTTTTAAACAGACAAGCAGTCATTAATGACCAAGCAATTGTTAATGCAGTTAAAGCGAATAATGGTAAAGCTGTTGAATTTTTAATTTCTAATGGAGCAAATGCAAATGTAAGAGATAAATTAGGTAATATCTTATTAATACAGGAAGTGTGGGAAAGAAAGTTACTAATACTTGATACTTTACTAGCTAAATCGGCTGATGAAAATATCACAAACACATTTAGTAATACGTTATTATTGTTGGCGGCATATGCTAAGTACGAGGAAATTATTACAAGATTGTCAGAGACGACAGATAAAAAGAATTATCAAGAACAACATACAACTACTTTCTCTAAAATAACAGCTATCGATTATGTATTTTCTTTACTTGAAATTAAAGAACAAAAAGACCATCATAAGGATTTTCGGTCTTTAAAAGAATCATTTATATGCCCTGCTAGTTCTAAAATAGGAACACCATTAGTTCATATAGCTATAGATATCGAGTCCAAAGATAATCGTAACGTTCAAGTAGAATATTTATCATATCAAGGCAACAATCCAACTTTTGAAAATACAGTAGATATCTCACCAATTTGTTTTGCAGCTTTACTTGGTGAAAAAGACATAGTGGAGTTTTTTTTAAATCGAGGCTGTAATCCGGAGAGCAATGAGGATTTATTTAAATCTGGGCAAACAAAAAATGCAACGCCACTAAATTTTGCTGTATTAGGTGTAAAACTAGAAATCGTCGAGTTATTGATCGATAGTGGATCTAACCCAAATATTATTTCTGGACTGGTTGCAATTAAAACACTAAATAAAATATTTAAAATAATAAAATTACTAATTACTGTATCTTCATATAAACATTCATATGGTAACCAAGGTAAACTCTTAGAGGTTACCGAATGGTTAGGTAACATGAAGCTAGATCATGACGATAAATTAACGCCAATTGTATCTTACATAAAAGCTCTGCTGCTAATAAAACTAGGTCAAGATATTGAAAGTTATAAATTATTAATAATGTTAGTAACAAAGCATAGTTATTTGTTAGAACCACATTTGCTACTTGCTGACTTAAATTATAAGGCAGGTAATTTCTATAATTAAGTTTTATTTTTAATTATTAATTAAGTTAAAAAAACACGCACCAGAAAGCAAAGACAGCCACAACGGCAGTTACAAACACATATTAGCAATAATTCACAGTTTAAATAGTACACAACAATGGAAGCTGGGGAATAGTGGAA
>evgR014829c0g1_i01_Hydra
TTGTGCGCTTGCTCAGGTTTATTGATAGTTGCTCTGCCAGTACCTGTGATTGTAAGCAACTTTACATTATTTTACTCATATGCGCAAGCTAGAATGAAACTTCCTCAGAGAAAAAGTCTTTTGAATGTCGATAAAAATCTTGTTACTAATAATCCAGATGATTCTCTTTTTTCTAAAATGCTTAATTTATCGACTTCGAGTCACAAACGTAGATATGCAATAAAACCACAATTTCAACATCATTCTATACAGAGTATTTTATCCGCAAGATCTTCATTAAACTCTGTTTTAGAAGATTCTTTTCTTGAAGATTCGCAAACTGACATTAAAGATGTTCAAATTTTGGATTTTTCTGATTAAAATGTTTCAGTTATTCAAAAACGTTTACCTGTGTTTAAATTATTTGCTCTATTACG
>evgR014830c0g1_i01_Hydra
CTAAAAATATTAACACACCCAAACAACAAAATAACAACATGAACGCTTTAAAGGAAGGTTGGTTCTCAGAACTCTCATCAATGTGGCCAGGCGAATGTGCCAGCTTGAAGGTCAAGCAAGTTTTGCACGAAGAAAAGTCTAAATTCCAAGATATTTTGGTCTTTGACTCTGAAACTTACGGAAGAGTTTTGGTTTTGGATGGCGTTGTCCAACAAACCGAAAGAGATGAATTTGCTTATCAAGAAATGATGGCTCACATCCCAATGGTTGCCCTCAAGAATGACCCAAAGAAGGTCTGTGTTATTGGTGGTGGTGATGGTGGTGTTGTTAGAGAACTTTTGAAGCATCCATCTGTTGAAGAAGTTCACTTGTGTGAAATTGATGAAGGTGTTTGCAATGCCTCCAAGAAATTCCTTCCACACATGAATATGGCTTTCGAAAGTCCAAAGCTCAAGCTCCACATTTATGATGGTGCTGTCTTTTTGAAGGAACACAAGAACTGTTTCGATGTTGTTATTGTTGACTCATCCGATCCAGTTGGTCCAGCTTCATCATTGTTCGGTCCTGAATTCTACAAGTCATGCAAGGAATCCTTGACTGATAATGGTATTTTGTGTACTCAAGCTGAATCCATGTGGTTGCATTTGGATATTATTGCTGGAATGAAGGATTTCATTGTTGACATTTTCCCAAATGTTACCTACTCTTACATTTCCATTCCAACATATCCATCAGGTGTTATTGGTTTCTTCATTTGCTCAAAGGATCAATCCAATGATGTTAAGGTTCCAAAGAGAACTGAATGGTTCGAAACTGCTAAGGAAACTCTCCGTTATTACACTCCACAAGTTCATGCTGCCAGCTTTGTTTTGCCAGCTTTTGCTGCCAAGAGATTGG
>evgR014831c0g1_i01_Hydra
CTTTTCCAACCTTGATGATGATGAAAACACATTGGTTGAAGAAACCAAGGCTGTTGAAACTCCAATCACCACTGCCACAGAAAGTGTTGCCACTTCTATCCCAACATCCACCGATGTTATTACTCCAACAATAGAAAAATCTACTCCAATAACAAGTACTCCAAAACCAAGCACTCCAACAACAACTTCAACAACCACTACTGCTACAACTCCTGTCAAGCAAAGATCAAGCAAAGGTCAACCAATTCCTTTCAGTATTAATCCAACCGATCAAATGCCAATTGATCAATTAAGTTTGGTTGATGTTGTTATGTGGAGAAACATTTTTGTTACAGTTCCTGTTTTCTTTGCTATCCAAGCTCTTTTCGTTCTTCTCTGCAAGTATGAATTCACTTTGGTTAACCTCATTGGTAAG
>evgR014831c0g2_i01_Hydra
GAATTCACTTTGGTTAACCTCATTGGTAAGGTTGTCTTCATTCAAGTTATTCTCGCTGTTGTCTACATGGTTGTCTTCAAGATTATCCAAAATAATCCAAATCATACTTTCCCATTCCAAAACGTTGAAATTAGTCAAGAACTCGTAAACAAGATTGTTAGTGGTATCGTTGCAAATGTTAACACTGCCTTGAAGTATTACATTGACATCTTGAGCTGCAAAGATTTGGGTAAGACCCTTCTCTTTGGAGTTGTTGTTCTCATTGTCAGCTGGTTGGGTAAGAAGATGGGCGGAGAAACTTTCTTATATATTGCCTTGAATCTTGTCTTCACTGTCCCATTGATTTACGAATGGAAGCAAGACGAAATTGACAAGGTTATCAAGATTGGTTGTGAACAATCAAAGAAGTATATTGATTTGGGCTTGTCAAAATTGCCACCACAAGCTAAGGATATTCTTGCTAAGCTCAACTTGGACTCCAAGAAAACTCAATAAATC
>evgR014832c0g1_i01_Hydra
GTTCAAAAATGTTATTAGCAGCCTTATTATCCATTTTAATACCCAATGGGTGCTGTACCATCTTTCATAAAAATCTCTATTCTAATTTGGAATATAGAGACAAGGATGTCAGTTTTGTTTCAGAATTTGTTGATACCAGTAAAATAATAGACAAAAGAGAAAGATTTAGACGAACAATTACCGACAAAATAGATTCTAATGATACTGCGGTTATCACTGACGATCACCTATATTATACATCAACCTACTATCTTGTTAGTAAGCATGATTTTATAGATTTGACACAAAACAAAGTAAGCAATATAAATGTTCAAAAAATCACTTCGTTAACATCTGGGTATCTAGTAGGCACTAAAGTGAATCTACCCTTTAACTTTACATTTTATGGAAATAAAGTAAATAGCATTTATATAACTACTGGTGGCTTTCTTAGTGTTTACCCTGAGTTTCATTCCTTAATTCACTCAGTTCATTATATAGCTCCTCTTATGGCTGATTTCAATTTAAAGCTTAATGATAATGCTGCAGTGTATTTTGGACAAAACATGGATATGTTTATTGTTCAATGGGACAACGTTCAAATAGATGATTCAGATTTAAATAATAAAACATTTACTTTTCAAGTTATCATTTATCAGAATGGAACTTTTTTAATGGTTTACAAAAAAATTCCATACCCTTTGTACAATATAAAAAATCTTGATCATCCGGTCACAGCTGGTATAGCAGATGGATATATAATTTCCTATAAATATAATGTAAGTTCTCGATATAATATTGTACTTTACCATAATATTCTTTACAAGTATCATGAAGTGAAACTACCTTTAGAAAAGTTAAAATCTTTTTCTGCATATCGGCTAGATCCTGTTAAAACTTGTTTAGAATTCACTTCATGTGAATCATGCACTAATTCTAGTACATCATTCCAATGCAAATGGTGTCCGGATCTTAAATTATGCTCTGATGCTATCAGTTGGAATCGTCAGAAATGGCTTAAATCAAATTGTTTGAAAAACTCTTTTGATTCAATTAGTAAATGTTCTTCGATTTCATCAAGTTCTTATATAAAAAATCTAAGAAATGTATCTAGAACCAAAAATGAAGTTGGAATCGTCCTTGGTGTGATATTTTTTATTTTAATTCTTCTTTTAATTGCTATCTTCGTTCATGCTTATTTGCATCCACAGACTAGGTTGGGAATGTGGCTTATTGAATACCGACCAAGTAGATGGTTTCAAAAAAAAAAACATGACCTTAGCCAGGAGCTTCTCACATGATATAACATGGAAGAATAGTTTTTCATCTTTGAATATATTATGTATTTTGTGTTCAACTGAATTTTTTTGATGTTG
>evgR014835c0g1_i01_Hydra
ATTTTTTATTCTTGTTTCCAAGTTTACTTGATCAGTATGGATCCAAACGAAGAAAGATCAGATAATATGAACATCCATGCAATCGACGATTCTACTGTTTGGCTGCAAACGTACTCGTTGAGATATGCACCAAGACCTTGTTACTATTTCTTTCGTTTATTAGGGATGTGGCAACCTAGAAATGCTTTTTGTCTATTCAAAGTGTACAATTACGCCATGTTTGTTCTACTATTTATAAATGCATTAATTATGTTCGCTACCATCTTTATTGCTTCAAGTTCAAACGTTAATTTTATTGATGTTAGTAATGGCATTACTTCAATATTAAACTCTTGCTGCCCTTTTTTATTTGTGAAATACTATTTTCGGTATGGCAATTTTGAACGATTAGTCAAACACGTATACGAACAGAGTACTGCAGAGGATCAAAAGAAGCTCCACAAATTGAGATACATTTATACCTTTTTATCTTTTTTTTTGTGGGCGTCAATGACGGCGTATTATATCAAACAATGGGGGTATTATTTAACAAACAAATATATTAGTTATGAACTAACTTCTGCACTTTCTATACAACTTGTCGTAACTACTATGGGTTGGTGGGCAACTTGGTTAAGTTTGTACGGGTATGTTTGCCACATTCACGTTCATCAAATTGAGTTGTACAATAAGCACGTGAAAGATACATTTTTAGAAGCTAATGAAACAAACGATGAATGCATCGGACATTTGTTGTCTGAGTTTAATAAACTTTACCATTGGCTTGAATTAACGCAAAAAGACTTTTCTAAAATAATATCATTTGCCGTGGTGAACCATATAATGGATATGTTTGTTTTTAGCTATGCTTACTGGACAGGGTCATTTAAAACGACCTACCCTGTTGATCATTTTATTAGTACAACTATTTATAACACTGTTAGTATTGCAATAAAACTCTACCCCGCAGCGATAGTTTGTAAAGCTGTGCACGAAATAGTAGTTACAGTTGGAATTCAATGCATGTCAAATTGTAGCACTTGCATACCTAAAGAACGTTTTATGTTTTACAGACATTTATTCTTTATGGAGCAAGACATGGGTTTTTTTATCTTAGGTGTTAAAATTACAATGAGACTAACGGTTGGAATATTTATTTCACTAGCTACTGCTTCGGTGACATTTATTAAATTTCTATTACCTGCAAATTAATTAACAAGTATTAATCTTTTAGTTAGTTGAGTTTCTCTTGTTCGCTGATCTTCTTGTAACTGATTCACTATATTTGTTTTATATTCAATTTTTGATTTCAGTTTTTTTATTTTTTTTTTATTCCGTCAATTGCACTTTTTCAATTTTTTTTCTTTGGTTCAATTCTGGTATACTTATATCTGTATTTCTATCTATGTTTCTATCTATATTT
>evgR014836c0g1_i01_Hydra
CCGTTGCTAAACATTTCTGAATAGTTATACCATCAATAAATCAATATTGTGATTATAAACTCCTTTTGGTCTTCTTAACCTCTTGATTATTTGGCATCTTCATTTGAGATTTAATATTATAAGATTCAATTTTTTGTTTAAAGATATGAAGCGTGCATATATAGATTCAAGTTCGGAGAGTTCTAAAATTTCTAATAATAAAACTCCTGCAGACGATACTTTATCACAAACCATTAAAAAAATGCGCCTGAGTTCTATGTTTGATAGTTTAACAAATTTCGAAGATGATGAGATGGCGGTAGATACAAATAATACTACCAATAATCCAGATAGTCCTAAAATCACCTTAGTTATTAGACTCCTAATGGGGTCTCGTATACAATACTGTTTCAATGAAAACGATACTATTGGTACTGTTCTGGACTTTATTTCTTCAATCATGCAAATACAAGTTGACTTGAAAACATATATTCTATATGTTAATGAAGTTCCTAAGCGTGAAATAAAATTAGCATCAGATACACTGCATTCTTTGGGAATTAAAAATAGAACAATACTAGCTTTAGATAGAAGAGATGAATAAGTAATTTTATTAACAGCAATTTATTTTTTTCTATCTTTTTAAAGAAAACCTAATTTATTAACTTTAATAAATATTAATTTATTAAAAAAAAA
>evgR014842c0g1_i02_Hydra
GAACGTAGTAATTGACAAGTAAACTTTAATTTGTAGTAAGTCTAGTAGGCGAATATGTCGATTCAATCAGTCATAGCGTCAACAATTATTTACACTGTAGGACTAGTTTTAGTTACAAAAATAGATCACAGTTATGGATGTGGGGTTGGCCGCCGTTTCGGGCATGCCTTAAGTACTCAACTAGATATTGAATCACACTATCCTAACGTAGCGGAAACAAGCGAGCTTGCAAGTGGTTCCAGTGGAAAGCAACTTTTAAAAAATGATACTACTTTAAGGATAACAATAAACACACAGATTATTTATCAGCACCATCGCCTAGATTCCGAAAGTCGTCTTGCGACAGCGATTTGCACAGAAGCTTTATTGGGACTTTCTGAATCGGTTCAACAGCACTGGTCTGGGGAGTATAAACTATATGTCATAGAAGGCTATTTATCAGATGTAACAAAAAGCCGCGGATCTTTGTCTCTCCATTTAGAAGGAAGGGCGTTTGATATAAGGCTAGCTAATTCAGATTCAAAGAAACCTCGACTAGCACCAATTGATTACAAACAGCTACAACGATTAGCAGGTTTGGCGTATTACCAAGCCAAGTTCTCATACGTAAGTGTTAAATCCGACCACGTGCATGTATCTTGCCGCAAAAATAACAGACTAGAGGAGAATAAAGTTCATAGTCGTTGTTTTCCGGAAACATCGTACGTACGTCATAAATCTGGCAGGAAAATTTTGATGCGGGATTTAAAAGTGGGGGATTCGATCATTACCATGAATCCTAATGGAGAACTTTTGTATAGCGAAGTTATAATGTTTTTACATAAACAAAAAACTATTAAAGTAGACGACTATATCAAAATAGAAACTTCGGACGGAAAATATATTGTTTTGAGTACCCATCATTTGATTTTTACATTGCATCAAGGGGCAATTTATGCAAAAGATGTTCAACCGAATACTTATATTACATCTTTCAACCTTGAAAATAAATATTTTAACAAAACGCGTGTATCAAAAATAAATCACGTGTCGGAATATGGGGCTTACTCTCCCTTAACCGAAGAAGGTACCATTGTTGTGAACGACGTTTATGTTTCTTGTTACGCCATGTTTCCAAGTCACCGTATTGCTGATTTAACTTTTTCTCTATGGCGTATTTTATACCGTTACTTTAAATTTATGTTGTCTTATTTTGAGAGTGATACAGAATACCACTGGTATCCGAACATTTTTAGACGCTCAGTTAATTTTTTTAATATATTCCCATATGAATTGTAAATAAAATAAATTATTTAATAAAAAGCCTATTTATCTAATTGTCTCATGTAATTCTAATTATCTGATATGTATCTGATATTTATTATTATGAGTTAGGCATAAACTGTTTGAAACTAAATTATATATAAAACTATATATTGAAATTGTATTATATGTGAATGGAGTTATTTATTTCTTAGTGTATATAAAATCGGAAAAAGAAAGAAGAA
>evgR014860c0g1_i01_Hydra
TCATATGTTTGTAGTTGTATTAGTGGATACACAAACTACAATAAAACATTTTGTGTTGATATGGATGAATGCACCAATGCACTACATACGTGCAATGATGACACAACTACATGTTCTAATACCGATGGGTCATTTGTTTGTAATTGTAAAATTGGATACACTAACTACAATAAAACATTATGCATTGATATTGATGAATGCAGCAGTACACTACACTTGTGTAATAATGTCACATCTACATGCTCAAATACCAATGGATCATATGGTTGTAATTGTAAAAAAGGATACACAAACTACAATAAAACATTTTGTGTTGATATTGATGAATGCACACTAC
>evgR014863c0g1_i01_Hydra
GCGTTGATTTTAATTCTCTTAATACGTTTGAACTAAAGTTAAATAAAAAATGTTCCAATAAACCAAAACATGTTTATTCGTTTTTTAGCAAGAATACAGCGTTCACTTACAACTAGAGGTTGTTCTTTATTTGTAATAATATTTTTTGTGTATCATATTTTATTGTTACCATACTTGTTGAAGTTGTTTTACGACGAACCGATTGAATCGAATCCACAGTATATGAATTTATTTTTAAAGCTCAAACAGCTGAAAGTAGAAAATGTTTTACTTCAAGAAGAAATTCAAAAAAAACATAAAGATGTGGAATCGAGACTTTTTGAAGCAGAAACTAAAGCCAGAGAACTTGCACTGTTAGTTGAGAATCAAAAACGAAACAATAACATCTTCTACGAAAATAACTCGCAAACAGTTGGGAATGCTGGAAACGAATTGCTTCTAGACATAGACAATATAAAAAATGAAATTAAAACAAACAAAGATAATCAAGCAAAAATTGCAGGAAATATCTCGCTTGTTCCAAGAAGCGTTAGTTTTGCTATCTTGAGTAGAAGAATCTCTAATCAAATCGAAGAGATGTGGTATTTATGGAGAGGTAACTTGAAAAAAATAATCGATGTACTTAAGCTTTACAAAGTACCGTCGGGTTTTTTAGACAAAATATCGGACGAGTTTCGCGAACTTCATCAAATAACTGAGAGAGATTACGAAGAACTAATGGCGATAGATGGAATAAAAGAATATAAAAACAATCTTGCAAACGATTTATCTGATAAAATTCAACGACGCATTTTAAGTCTGCAACATCCCCCGAACTGCTTTTCTGCAAATAAACTAGTATTAGAACTGAACAAACCCTGTGGTTATGGATGTCAAATGCATCATCTTATCTATGGAATGATTATTGCCTATGCTACAAAACGCACTTTGATACTTCCTGACACAAATATAGGAAACAGTGTTAATTGGAATTTTTATTACCAAAAAATAACAAACTGCACTGTATCGCCTAAGCAAACTTTTGCAGAATGGGGCGAAAACGTTGAAAACAAAAATATTGTTTCTTTGCCATCAGTTGATTTTCTTCATCCCAGACCTAAACAAATGCCAATGGCTGTGCCAAATGATATCATTGATACGCTGGAAACTTTTCATTCCCGTCCGTTTGTTTGGTGGGTAGGGCAAATCTGTAAATTTTTGTTTCGGCACTCGCCATTTCTTAAAGATAAAATTTCTGAAAAGAAAAAACAATTAAAGTTCAAAAGCCCTATTGTTGGGATACACGTTCGCCGTACTGACAAAATAAGAGTAGAAGCTGCTTTCCATAATCTAGAAGAATACATGTACTGGGTTGATCTTTACTACAAAAAACAGGAACTCTCTGGTTCATTGAGTCAAAGAAGGGTGTTTATTGCGTCTGATGATAAATTGCTTATTAAAGAAGCTAGGGAGAAGTATCCAAAATATACATTTATTCATGATGAGATAGCATCGGATTATGCAGATGAATCTCACCGTTTTTCGAAGAATTCGATAGAAGGTATCATATCAGATATTGAATTACTTTCAGAATGCGACTTTTTGGTTTGCACATTTTCTTCACAGGTTTGTCGATTGGCATACGAAATTATGAACCATAAACATATGGATGCTAGCAAATATTTTCGATCATTAGATGACATTTATTATTTTGGAGGTCAAAATGAACATACAATGAGAGTTATTTGGCCTAACTTAAAAACTTCAAACGATGAACTAAAGTTAGATATAGGAGACACCATAAAAATAGCAGGAAACCATTGGAATGGAAAAAGCTTTGGAAAAAATATTAAAACTCAAGAAAATGGTTTCTTTTCTTCTTATAAGGTCGAAGATATTATTAATGTTGAAGATTTTTTGACTAACTCAGATTGAGCAAACCAACATAGAACTTAAACTTACCAACATCAAACTTGTATTGAACAAATTTGTAAGAATTTAGAATAAACAGATTGTTAGATTACATGTGAGTTGCGTCATCATTGCGAGTTTTTGTAATTATCCAGGAAGTATCGCGAGTTTTTTGTAGTTGTCTTGGTTGTAATTGCGAGTTTTTGTAGTTATCTTGGTTGTAATTGCCAGTTTTTATAGTTGTAATAGTAATTGGTGAGTATAAGGAACCTCTAAGTTATAATAAAAAACATGCAAAAGAGTCATACATAAAAGGAGGGAGGAAGTTGAGATAAAAATTACAACAACAAAAAACAAGAGGAGAGAGTTTTATCGGAAAAGAACGTCACTTTATAGACTTTTCTCTAAATGTGCC
>evgR014865c0g1_i01_Hydra
GTTCATATATATAGTCTATATATAAAAAGCTTAAATTCTACTTTAAAAAAACATATAAGTAGTCGAAAATTAGCAGAAAACACAAAAAATTAACAAACGTAAACTATAGTTTTAAAGAAATTCAAATCGTTCGGATCTTGCTTTTACTACGCCGATTGTATAAAGGCTCTCTAAAGATTTGGACTGTTCTGTCTCTTTAAAGGTTTGGACAATCTTGGCTCTCTAAATGTTTTTTATATTTGCCGGTACTAGCTAGTAGGTATAAAAAACTAATCAAATTCCTTCTTACCCAATGCGTGTACGAGTACGTGTGTGTACATTTTGCTGCAAAAATATTCACCTATTGGTCGTTTTAATTAAAGTTTGTCTTTTGCATCAATATTATATAATATATATCAATTATTATATAATACATATCCATTCAGCCCTTGATCCGGCACTGTATATATCAGCAAGATTCGCAGTTATGCAAGTTCGCACAAATGCTAGTGTTTGCAAATTCTTTCAACACAATGATGCAAGTTTGCACTTATTCGTGATTTGTGGATTTCTGACAATGCGCGCGAAAAAATCATTTTCTCTCGTTTCCCGCAAAAACTTACTAATTTATTTTTTGAAAAACCTCTATCTATAT
>evgR014869c0g1_i01_Hydra
TTTTTTTTTTTTAAATTAATTTGAAATGATGGTAAAAGTTAAAATATTTCTTCCTGTCTTTGCGTTTATGATCGTGGTGGTTTTTGCAGTTATCCTCGGTCTCGAATTCGGGTTAAAAAAAAAAAAGAAAACCATTCAAAAAAAAAATTACTATGAAAATGGTGCCGTTGCTACAGACGCACTAAAGTGTTCGAAGATTGGTCTAGATGTGCTTACTGAAGGAGGCCATGCAGTTGATGCTGCGATTGCAAGTTTGTTGTGTACTGGAGTGATCCACTTGCATTCTACTGGAATAAGTGGTGGAGGCTTTATGAGCATTTACAATAGAAAGCTTAAACAAACTTTTGTGTATAACTACAGAGAATCAGCACCTGAATCACTTCGCGCTGACGCATTTGACAAAAATATTACTTTAAAAAACGTAGGTGGTCTAGCTGTTGGGGTGCCAGGTGAACTTCGTGGTTTGTATAAAGCACACCAAAAATTTGGAAAATTAAAATGGGCAAGACTTGTTACACCTTCAATTGAGTTAGCGGAAAAAGGTTTTCCGATGGGTAATCATATGTATAAAAACATGATTGACGCAGGAACAAAAAAATATTTAGAATTAGATCCAGGCTACAGAGAGATTCTAATGAATAAAAATGGCATGTGGAAAAAAGAAGGTGAGACAATCGTTCTTAAGAAATTAGCTGAAACGCTTCGCAAAGTTCGAGACAATCCGGATGCACTTCATACTGGCGAGCTATCTCAAGGATTTATCAACGATATTGTTGCGCACAGTGGAAATATGACGCTTAGTGATTTGAGTTCTTATGCAGTTAAAGAAGAGTCTCCAATCAAGATGGAGTTAGGAAACTACACCTTATATACAACTCCTTTGCCAGGTGGTGGTGTTGTTTTGCTGTCAATTTTAAAGATTTTGAAAGGATTTAATTGGACTAAAGAAGATTTTGCAACATTTGATAAAAAGCTTTTGAATTACCATCGTATGATTGAAGCTTTTAAGTTCAGTTACGCAACGCGACCCTACTTAGGGGATCCAGAAAAAGTTCCTTCAGCCAATCAAACTACTTTTAATAAGTTTATTGCTGATATTTTGAATGACAATTATGCAGAACAAAAACGATTAAAAATTAATGATTCTTTCACCCAAGCAAATAGTTTTTACGATCCATTTCTTACAAAAACGGAAGATTCGGGAACTACACATGTCTCAGTCATTGATAGAGAAGGAAATGCAGTTTCAGCTACTGATACAATAAATTATGCTTTTGGAGCGAAATTTTGTTCCGTGACGTCTGGTATTTGTTTCAACAATGAACTTGCTGACTTTTTCACTAAAAACAATATCGGTGAGATTTATCCAACTTTAGTAGCAAACTTACCAGGACCAGGTAAACGTCCTTTATCATCAACTTGTCCTACAATTATTGTCAACAACGAAGGTAATGTTGTAATGATTGTAGGTGGATCTGGAGGAACCCGAATAACATTATCTACTGCGTTGGTTATTATGAAAGTTCTTTGGCTTGGTTACAGTTTGGAGGATGCCATCAATGATGTACGGTTATATCACACTTTTTCTCCAAACTATGTTCAATCAGAAACAAACCCAGATTATGCAATACCAACCAATCTTAAGGAAGGTTTAAAAAAGAAAGGCCACGAAGTTAAACCCTCAAGCGGACATTCTGTGATTCAAGGGATTTATGTTAAAAACGGCAAAATTTACGCAAAGTCAGATCCGAGAAAAGAAGGAGAGGCAGCAGGTTATTGAACTAATAAAAAGAAACCATTTACTTTATTTAAAATAACTTTAATTAAAAGTATTGGAATGATGAATGGATGTTTTTAAATGAATGAAATTTTTGTTAAACATTTTATTTAATTTTAAATTTACAACTATTAGAATGATATATTTTTTAAATATTTTCTGAAATTTACAATTTCAAAAATATAAATCTACAACCATTATATATTTTATAGCAAACAAGTT
>evgR014870c0g1_i01_Hydra
TAACTTTTTTAATTAATGTTCGTAGATTTTTTTTTAAAGTTTAAAATTAATTGAATCAATTTTTGAAACATGGATATAAGAAGATATGCTTTTTTGTCTTGGCGTAGAACTATAATAGTTTGTTTAGTGCTAACAGTTTTATTGGTATTAGCTCTCTTTCCATCAAGCACTCAAATAAAAGAACTCAATATAAAAAAAAAATTAGACAAAAAAACACAAGAAAACCAAGACAGCATGACACTATTTAAAAACATTACGCATCGTAAACATCAGCTAAATCGAAGAAAAATTTCTAAAGAAAATTTAAAAAAAGAGAATGAGACACTTAAAAGAGTAGTATTCGATTTCGTGAAGAAAAACAGATCAGTAATTTATACAACAGTTATTTTAATCTCTTCCCATGTAAGTCACACAAAAAGAAGAAATCTAATTCGAAACACTTGGGGAAATTCTTTTTATTGGAGTTCAACGGAAAAATATTTAATTGTATTTGTTGTGGGTAGAGTAGCAATAAAAGAAACAATGATGCAAATGCAAGTGGAAGCAAATGTAACAAAAGATATCCTTTATTTAAACGTTACAGAAGATTTTTACTTGCTTGCAGAAAAAGTTATTATTGGATTAACCTGGGCAAAGCACAACCTGAAGTTTAAAGCTATTCTAAAAGGCGACGACGATACTTTCATGAACTTAGATAACATATTAACGTTTATAATAGGTAATAACGTTACAGACGGCTACTTTGGAAATAAAATTATTGATGGAGTTCCTGAGCGAATTGGTCGATACAAAGTAACTATAAAAGAATGGGACAAAAACTACTACTATCCGTACTGTTCTGGTGGTGGTTTTTTATTGACAAACTCTAGTGTTTATAAAATGATACCTATATTTGATTTAAACAAAATATTTCGAATAGATGATGCATTTATTGGTGAAATTGCATTTCAAGCAGGAATAACAGTGGATGAAGTAAACGGCTTTTATATGAAGAACGATGGATGTGAGTATAGAAACGATACTATGGTTTCTCATCCTTCAACAGATCTTGGATGTAATAGCTTTCTTCTAAGGAGGCATTTAATCGACAATAAAAAAATACCAAGGGATGCAAAGCTTGAAAGCATCAAGTGTTACAGAAGTGTTTCAGATTGTTAGTATTTTTTAACTAAATATATTCGTTACAGTTTAGTTT
>evgR014879c0g1_i01_Hydra
GCCCTCCTGCTGTTTGTGTATTTTAGAATCAAATTGCATACTAACTTCAGCCTTTGTTGATTTGTTTACTTCAACAGTCAACTGCTCGTCAGATACTCCATTTAAAACTTCTTTGCTAATTAAGCCATCATTAATTTGTTTTACCTTCAAGTTTCTTTCTTTACATTCCTCAAAAACCATTAGTTGTGCACTTCCTGTTGAAACATCTTCAAAGATAAGGTTACTTGAAATACTGATAATTTTTGAAGAAGAATCTTTTTCGGTTACTTCTGTAGGAATCTTTTCAATCGTATCCGAAAAGAAACTTAGCTTTCTTGACATTCTCA
>evgR014880c0g1_i01_Hydra
AAAAAAACATGGTCGGAAATTAATGACATAATAGGCAGAAATAAAATAAAGGATTCAGTATTACCTCAAAGAATTAATATCAACAAGTCAGATTTGTTCTGTCAAAATAAAATATCAGAGGAATTCAATAAATACTTTACAAATGTTGAGCACAATCTTGCAAAAAAAATTATACCGACATCTAACTCATTTAAAACATATCTTAAACCATTAAAAGACACAATAATGCATGATGGAGACTTAACGTCTTTTGAATTTGATGAAGCTTATTTGTCTCAATATATAAAAAATCGATTCGTCTACATACTGTTTCCAATTTATTGTAGTCACAAATTTTCCATAATCCCATTTG
>evgR014899c0g1_i02_Hydra
TAATGGAGGAAATACAGGGTATTATGGAGCAAGAACCCCCGGGGTATGGTTTCAACCAAATGGTTTAGGAAAATTAGGCATTTTTTCTGCAGTTAACGGTGATCCAAATTATTTTATCAGCTCTGAACCACTTCCTCTCAATAAATGGTCAATCATTAAGATATCTCAAACTAAACTAAATGGTGTTTACAAGTTTCAGGTGTATATTAATGAAAGTATTATCCATAGCATTGAAAATTCAAGACCAGAATCGTTTCAAAATGTCAAAGTTTATGCATCAGATCCATGGCATTACGCTCATGATTGTTTCATTAAAGATCTAACAATTTTAAACGGATTAGTTGAGGCAAAAATCGATAGCTTTTCATTTCATAGAGATGTAACAAACAAAGTCACCATGGCAATGTCTGATGTTATTAAAATTCCTTTTGCAGTCATTATGGGAGCTAACGCCCTGGTAACTTATATTGATCCTTCAAACGAGGCATTTGATCCTGATGCTTGTATCTTTGCGTGTTTACAAAATGCAAAATGTTATTCTATGAGCTTTTCACAAATTGGAAATATTTGTATATTACATGATTTAAATATCAGAACCAATCAGGACTACTTTAGCAGTAACGTAAATTGGG
>evgR014900c0g1_i02_Hydra
GGATTTTACCAAGGCCATCGACTAAAACTACCTTATTTACCTTAGAATTGGCTTGATTATGAATCCTTTGTTTTTGGTTTTGTTGGTTGTTTGTTCTACAACATGTAATAAATATTCTAAAGAAGAACATTGCTTAAAAAATAAATCGGATGCAGTTTGTGAAAAAGCAAGCGAGTTTGAAGAAGATGATGATGAACCTGTTTTTTATGAAATAGACGAGAAATTAGCAGGAAGAGAAAAGCTGTTTAGATGGGATCCGATTCGTAAAGGCCATGAACGAGAGATCGACTTAGGTGGCGGGGAAATAGTTAAGATGATCACTCAAGCTATTAAGCCAACTGTGTTTATTATTGATAACTTTTTAACAATGGAGGAATGCGATTACATTGTTGATAAATCCCTTCACTTTGGATTGTCTAGCAGCGGTCTTCATGTCGATGATAAAATTAAACTGTCCAAAGAATATACTTACGGGCAAGCTACAAACTCTTTTGGAAACTGGTTACAGTGGGATTTAAACGGAGATAGCGTGGTGTCCAAAGATGAGGTAATTACCATGGCTCAGCGACAACTAAAGTTGTACTTAAACTCAACTGATGTAGATGAAATTTTTGCTGCACTGAAGATGACCGAACTTGATGATGGACTGATTACAATAGAAGAATTTGAAAAAATGAACACTGCTGGTATTGCTGAATATATGAAAGACTTAAGAGAACAGCATCCTAGATTCCGAGATCGATTCAGTGATCAAATTTGGTTAAAACAAGATGATATGAGTGATACTGTAATGAAACGACTACGAGAGAAGGTAACAAAATTAACGAACCTTCCTGCATATATCATTGCTGGCGGAGAACCACTGCAAGTCGTAAGGTACAAACCATTTGGTCATTATCACGCCCATTTTGATGCACAATCTCGCTCAGAATATCCTGACTATCCATGCTGCCATTTAGATTTAGCATCGCAACCTTTCAACTGTAGATTATGCAGATTCATCACAGTTCTGTATTATTTAAATGATGTCGAGGAAGGAGGAGAAACAGCTTTTCCTGTTGCAGACAAAGAAAACTACAGTCATGACGAATTTAAAAAGCGTAAGAATGGAGATCTTTATAACTTAAGTGAACATTGCTATAACGCATCTCTTGTTGTGAAATCGAAAAAGGGTAGAGCAGTGTTTTGGTACAATCATCACTTAGAAGATGGTTGGTTAGGCGAAATGGATTGGTATTCCCTTCACGGCGGTTGCGATATTCGTAAAGGAATAAAATGGATAGCGAACAACTGGATTGTTGGCCCACCACGTGAAGTTGCCCATATGGATAGCTTGTACTACTTACTAAACACGCCGGAGGCTCATTAATTTTTTTATTTATTTCGTTTTTTGTAGACGTCGGAATTTTATAAATTACCGTAAATGTATTTAATTAATCACTAATTTTTAAAAAAAAAAA
>evgR014909c0g1_i01_Hydra
CCATAGTTCATTATATTGAATTGTATATTTAGCTTAATTACCACTCATTATACAAACGTCAATTATATTTATATTACACCTAAATTTTCATTTGTAACGCCAAGAGTTATAGATTTTTAGATATTTATTATTTTGTTTATTAATTTCATTTTGTTTGTTAAGGGTTCAGACGCTAACACTGGCACTCTTCAAATTAATTTACGTTAGGTAAGCCAACACTTAGCTGTCATTTTTATTGAAATTTCACAAAATTGCTCGTTAGAATATTGTATTATGTCAAATAAATCCCCATCGTCATTAATATGGAAAACAAAACCTCAACTCATTAAATCGTCGTCATCCACGATGTTAATTCCTCAAAATGATTTGAATCAAAGTAGAAGGCAAAGAAGCGTCAGTGAAAGTCAAGCTGAACTGCAACCAGTCGAGGGAAGACAAAGAAGCGGATCCGTTCAAGTTGACATACCGCTATGCATCGATATTCCTAAATCACCTCAGTCGCCTACATTCTCTTTTACTGGATTTAATTTTCTCGACAGGAAACGAAGCTCTTTTTCTATTGGTTATGACGATTAAGGTCACTTAATGTTTTTCAGTTTTGATGAAAATAGTGACTTGAAACATTTTTTAAACAATTTTATCAACGTTGAACTTCTAAGTCATTTTATTGAAGTAATTTTTTATATAAGGGAGCTATTTTTGCAACCTATATACTCTTAATCTAGTTTTAGATAAGTACAAGATGTTCTTTGTAAATAGTTAGAGTTATGTCTAAATAATCTAGTAGAATATTTAGGATTGTAGTATAATTATTGTTTTTACTTTTTTAACTAGCATGCTTGTAATATGCAAGCATGTTTACTTATGTTTGTAATGTTGAGTTTTTAATTAAATATATTACAATTTTAAAATTGACTTTAAGTTTCTTTATAAAGTATATAAATTTTTTTGTGGAACAAAAAATCTTCTT
>evgR014910c0g2_i01_Hydra
ACACTGGTTATATCAGTGCTTGTAAATTTATTTCAAATGACAAGCAAATCTTGTCCAGTTCTGGTGATATGACTTGTATTTGTTGGGATATTGAAATGGGTCGTCCAATCAAGACCTTCCAATCACACACTGGTGATTGTATGTCTGTTTCTGTCTCTCCAGATGAATCTCAATTTGTTTCTGGTGGTTGTGATGCCTATGCCAAGTTGTGGGATATTAGATCGGGGAAGTGTGTTTCTACTTTTGGTGGTCACGACTCTGATGTCAATTGCGTTCAATTCTTCCCTAATGGTTTGAGCTTTGCCACTGCTTCTGATGATAATACTTGCAGACTCTTTGATATTCGTGCTAGTAGAGAATTGATGGTTTACTCAGACGATTCTGTTAGAGAAGGTGCTACTTCTATTTCTTTCAGTAAGTCTGG
>evgR014912c0g1_i01_Hydra
CTGTAAAGAAAGTAAAGAAAAAATATATTATAGAGAATGAAAATCTACATCTTAATCATGTTTATTTTAAATCGATCCCTAGGATCTCCATTGTTTCCTGCTTGTAATCCAGTTCAATGTTTTCGTGATCCATGTCAACATCATGAATGTCCCGCCGGTTTGCAATGTATTGCAAACTATTGTAACGGTTGCAAGGCAGTGTGCAAACCTTGCAAAATGGAAAACTGTTATGTTGATCCTTGCAAAAATAAATATTGTCCTACCGGTTCTCAATGTATATCAAATTATTGTAACGGCTGCAACGCAGTGTGTAAACCTTGTGTTACTTTTAATTGCTTAATTGACCCTTGTTTAAACCATAAATGCACTTCTGGTTTTGAATGCGTCTCTAATTACTGCAACGGCTGCAACGCTGAATGCAAAAAATGCCCAACCACAGTTTGTGGACCATGTACATTATTGTGTGCAAAAGGTGCAAAATGTATAACCACTCAGAATTGCGGAAAATGCACGCACACGTGTTCTCTAAACAATATGTCAATCTAATAAGTAAAGAAAATTTCGAATGAAGACAAAGAACAAACGAATCCGTAATTGGCGAGGGAAAGAAAATTTAGCAAAAATATTTTTTCTAATTTTGTTGTGTTTTTTACTCCGAGTAAAGTATTAACTAAACTAAAACTTGTTTATCTTAAAATGTTTATAAAAAATTCGTTAAACTGAAACGAATAAAATTTGAGATTTTATTTTTAAATTTATG
>evgR014915c0g1_i01_Hydra
ATTAGATAATTATTTATTTTAGAAAATGTTTAATAGCGAAGAATTCAGAAAATTCTCAAAAGAAGTGATTGATTACGTAGCAGACTATTATGAAAATATAGAACAAAGAAGCGTTCTTCCGAAAGTTCAACCGGGGTACCTTCGTAAACTTTTACCTCCAAATGCTCCAGTAGAACCGGAAAATTGGGAAAACCTAATGAAAGACATAGAAAAAGTTGTCATACCTGGTGTTACAAATTGGAGACATCCTCATTTTCACGCTTATTTTGGTGTTGGTATAGGCTACCCATCAATAGTAGCTGAAATTTTAGCTAACGCTTTTTCAGGACCAGGGTTTTCATGGATATCTATGCCAGTTAGTACCGAGTTAGAAGCAGTAATGATGGACTGGATTGCAGATCTTATAGGTTTACCTGAGCATTTCAAGTTCTCATCAAATGGAACTGGGGGAGGTATCATAGATGGATTTGCAAGCGATGCAACTCACTATACGTTGTTGCTTGCTCGCGAGCGAATAATTAGTAAAAGTTCGAGCGGTAGTGAAATTTTTTCAAAACTTGTCATGTATACTTCTGCCCAAGCTCATTCTTCGGTAACTAAGGCTGGATTATTGGCGGGAATAAAAGTACATTACGTAGATACGGACGAAATGTTTACACTCAGAGGTTTTAACTTGGAAAAAGCCATTAAAGAAGACAGAGATAAAGGATTTATCCCATTTTACCTTTGCGCAACTTTAGGAACTACAACGTCTTGTGCATTTGACAATATTCAGGAATTAGGTGTTATATGTAATCGAGAAAAAATATGGTTGCATGTTGACGCAGCTTACGCAGGAAGTTCGTTTGCATGCGAAGAGAACCGTCACTTTATGACAGGCATTGAGTTAGTAGACTCGTTTGATTTCAATTTACATAAGTGGATGCTGGTGACGGTTGACTGCTCTGCATTGTACGTTAAAGATAAAACTGAAATAAATAATTCTTTTAATGTTGATCCTGTTTATTTAAGATGTCCGATTGGAGGAGAATTACCATTATACAGAAATTGGCAGATTGCATTAGAACGCCGTTTTCGCTCATTGAAAGTTTGGTTTACACTTCGTTTATACGGCCAAAGGGGAATTCAAAAACATATTCGGAACCATATCCAACTTGCACGTGAATTTGAAGCAATGATAAGATCTGATAATAGATTTGAAATTTGCTATCCTGTTACTATGGGTTTAGTATGCTTTTGTTTAAAGGGTAGCAACGAGTTAAATCAAAAGTTGAACGAACTAATAAATGCCGAAGCTCAAATACATATCACCCCGTCAAAACTCGGAGATAAATTTATTTTGCGACTTGCGATCTCATACGAGCACGCAACAACAGAGCACATTGTGTTTGCGTTTAACAACATTAAAAAACACGCTGAATCATTATTACAACAATAATAAAATGTATATATTTTTTTATATTTTAGGTATTTTAATTTTTTGGAGAATAAAATATGTAAATTCCCGACATGCAATGGTACATAATGCAACGTTGGATGGCACTATAAATAGTGCAAACAAAACTAAAACTTATCCTCG
>evgR014920c0g1_i01_Hydra
TTAAAATGTTATTTTTTCAATGATTATTAAAATTAAGCAATGGCTGTCTATGGAAAAATACTATCAACTGTGATATCCTTTGGAATCGCAGGAATTTTAGTTAACGGAATAAGTTTGCTTTTACTGGCAAAATTCAAAAATTCTAATACGGATAAAACGCAACGTTGGATAATCACCAACCTGTGTTTGTTTAGTCTTTTCTTAAGTATTACCTACACAGTCACTAATTCAATAGAAATTGTTCAATCTGCTAATGACAATGTTGCTAGCAAAAGTAAAAAACAAAATAATTCTATCGTTTTTCATCGAACTGATGACGTTGTTCTTTATATAAAAGATGAAGGACAGTCACTAAATCCTATTTATGTTATAGAAGACGCGCTAATCGCAAGTTTTGTTATTGGAAATTACTTAGCAACTCTGTGGCTCGTATTGGATAGGTACTTGCACATAATAATGATCATTAGATATACATGTTATTGGTCGCACAAAAAAACCCTTTTGACAACGTTAGTTATGTGGACTTTAGCTATAACAATCGGTTCTCTAATGAGAACATACTTGAGAGGGAATCTTGCATATATATGCAATATAATTTATATTACACTTGATTCAATTATTATTTTTGTATCTTTCTATGTTTACTCTTATGCTTTGATAGTTATTAAAAAACAAAAGCAAATTCATTTAAAAGAAAACAACCATGGAATAACAAAGGGAGTGTTTTTGTCCGTCTGTATTTTATTAACCTTTATTGTAACAGTTACAATACCCGATATTATTTCGATTTTAAACGATAACCTGGGCTTAAATAAAAACTTTGCTTGGTATTCATTATTGTGTAACTCGGTTTCTGTTTTAACTGATGCCATTATTTATATACTGTTGTCTCCAAAAGGTCGTTGGTATTTCAAAAAGAAGCTACACAACATTTCAAGTAGAAGAAACAGTTCTTTTGTTTTGAGTAACATGACAACTGAAAGCTTTAATACTAGTTCCGTTGTATATAAAATTGGATAGCGTATCTCAAGAGATTTAAAACATTTTATAAAGGTTTTCATTATTATAAAATGGCAGGATCATTTTGATTAAGTATATTTTTTACTATTTATTTACTAAACTTGTATATATTTCATAAAAAAATGGCAAAACAAGGTTATTTTATAATTTAAAAACTTTTTAAACTTCCGCCAATGCTACTCAGATTATATTTATATTTACACGCAGATGAAGTAAAAGTGAGGAAAAAGTTGATTATTTATTATTTTTTGTTGTTGTTTATTTTTTTGCCCCAAAATGTTCCTTGTCTCAGTTCAAAAGCC
>evgR014921c0g2_i01_Hydra
CTTTTTTTCATTTATTTCACAGAGTGCAAGGAATGTTGAACAACAGATTTTTTAGATTTTATATAATTATAATTTTAAATTTTACGTTGTAAAGAATTTATATTATAAAAGTAGAATGTTGGTGGCATATAAATGGTGAATATAAATGGGTTTAAATAAGAAATGAACCAATTTGTCAATCCTGATTGTTTCGACTTGGTATGGGAAGTATATGATATTATTCGAACAATAAAGGACCCTGAAAGGGTTGAGACATTAGAAGAATTGCATATTGTCAGTAAAGATCTTGTTCAAGTTAACCATTTTCTTGATCAAAGCATTGGTATAAAGGTGGAGTTTGTACCTACAGTACCACATTGTTCTTTGGCTTCTTTAATAGGTTTTTCAAATGTGGTTTTTCAAATGTTTTTTTTTAGATTATGCATTTATGTAAAGTTGCAAGAAAATCTTCTTTGTAATTTTAAGCTCGATATCAATATAAAAGAAAATACACATTATAACGCTGACGCAATTAACAAACAGATGAACGATAAAGAAAGAGTTGCTGCTGCGTTAGAGAATCCTGATTTAATGAAGATTGTCAAACAATGCATCTAATATAAAATTTATTTATATCTAGATATCAATTCAAAGCTTTACTTTAAACACAGAAATTAAAATTTAAAAAAAAAAAA
>evgR014925c0g1_i01_Hydra
TCAACAAATCCACAATACGGACAGCCAACTAATCCACAAATGACACAATCTGGACAATTTGGTCAACAAGTTCCACCATCCACAAATCCACAACAATTTGGTCAACAACCAAACGTTCCTCCACAATATGGAAATCAACCACCATCTACAAACCCACAATATGGACAACCAACCAATCCACAAATGACACAATCTGGTCAATACGGTCAACAACCACCACAAACTAATCCACAACAATTTGGTCAACAACCACCACAATATGGAAATCAACCAAGTGTTCCTCCACAAAACACTCCACCACAATACGGACAACAACCTTCTCAATTCACACCAATTCAACAACAACCACCAAATCCTCAACTTAAGGCTTGGTTTGACGCTGTTGATCTTGACAAATCTGGACAAGTATCTGCCGAAGAGTTGTCAACTGTTTTATCAAAGGGTGGTATGG
>evgR014931c0g1_i01_Hydra
TTTCTTGTTTATATTATCTTATTAAATAACAAATTAAAGTCACAACAAATGCTCAGCAATGGTTTGTCAAACACTTATCTCAATACCAGAGCATGGTCAGTTTATTCAGGATAAAGAAATCCTTAGAGAGGTATCCGTCATGACCCACAAAATCCCTGTTGATTCATGTAATGAAAAAGATAGCGAAAAACTCTCTTGTGAAAAACTTGCAAAACTCAATATATCATGTAAACATTGTATAGTAAAGGAAATTGTCAAGAGTGAAATTGTGCAGCCGAAATGTAAATGTGGGCAAACAGTTGAAACAAAAGTGAAGTCATCAAAAATTTCTATGAGTTATTCAAAGCTAAAAAGGAGCAAGCAAAGAAGTAGAAAAAGCTTAGATAAAAAGAACCGTCAGTTGTTAGCAAAGGCCAATGATTCAATACTTCCTCCTGAGCAAGGATCCTGGTTGCTAAGATTGTTTGAATCAAAACTTTTTAATATGTCTATAGCTATTTCTTATATGTTTAACTCAAAAGAGGAAGGAGTGCTTGCATATCTAGGAAATAAACTTTTTAATTTCGACGACCATGAAGTGGATGTGTATTTAGCTGAACTTATACATATGTATCTTCATATGGAAGAAATAAGTCAAGTGCTTCATCCATACTTGTCTTTGCGCGCCAAAAATCACACTTTATTTAGTTTACGCACAGCTTGGTTACTACAGTCAGAGCTGGATAGTGGAGAATTAAGCCAAGAAGAGGAAAAGAGGGGCAATAAACTGTATCAAGTAATCACAGAAGAGCAAACACGGATCCAAAGACCTGAAGTTGTTACGACTACACCAATCAAAATACACAAACGTTCAAAATCTGATGCAGTTATTGGCACTATTGCAGATCGATCAACATTTTTCCAATACCCAGCTGTTAGTCATTCAACAGATAGTATTGGCAAAGAAACACTTAAACTAGGACTTGCATTTGGTAAATGTCCTCAGCATCTTCTTTCGACTTCAGTAAAACCTAGTTATATGAGAGTCAATTCCTATAATTCAGCAGACCCTAAACCTCAACAGGATTTAAATCAGTGTTGCTGTTGGCAGGTACCTGAGTATGTAGCTGAACAAGAATTCATTGGTGCTTTACAATCTATTGGTCATTATTTGAGGTCATACCCAACGAAACAACAAAGAGTTCAAAGATTGTGTGCTGAACTATCACTTTTAAATTTGAACTTGCCAGCAAAAGTGTTTGTGCCTTTTGATGGTATTTTTCCGCATCATGTTGTAAGGATTCCCCACAATGCAGCAGTTGTTTTAAATTCTAAAGATAAGGCACCTTACATTATTTATATTGAAGTGCTTGCAAAAGATCTTGACGAAGACTTACAAATTCAGGCTCAATGTCATGAGAAATTGCTAGCTGATCATTACAGAATGAATGCTGTTAATGATCAGCAGTATCCTAGTTCAGATATGGATGGTTTAGATCTGCATCTAAGTAATGCTTCAGCTGATAGTAGTGAATCCGATGGAACTAATGGAGTGGAAGATGATTTTAGTATTGAATCTCAATCGACAACAACAGGAAGTAGTGAAATGCAATTAGAAACAGATTGTGACTTACTGGAAAAGGATATTTACATTTCTGCAATTGATGTTAGAAAACGTCTCACTGAATCACTTAGTGCACCAAAAGCTCTTTTTATGGTTCGAGATCCTGATGACCCTTCCGCTGCTGCTTTAAAAGAGCCTTGGCTTGAAAAAGTTAATCGAATACGCCAGACTTCATCGTATGGTCATCTACCTAACTGGAGACTTATACCAGTTATTGTTAAAACCGGTGATGATTTGCGTCAAGAATGTTTAGCCTATCAGTTATTAACACAGTTTCAAATAATATTTGCTTTAGAAAAAGTGCCATTGTGGATTCGACCGTATGCTGTCATGGCTACTTCAGGTGATGGTGGTTTGATTGAACCTATTGTAAATGCTGTCTCCCTTCACCAAATCAAAAAACAAAGCAAAATGTCTTTACTTGATTATTTTATAAACGAATTTGGCGAGCCAACAACAGAAGAATTTCTTACTGCACAACGTAAATTTGTTGAGAGTTGCGCTGGATATTGCTTAGTTTCATATATAATGCAAGTGAAAGACAGACATAATAACAATATCTTAGTGGACAGCGAAGGTCATATAATTCATATTGACTTCGGTTTTATTTTGTCTAATTCACCAGGGAAAAATCTTGGTTTTGAAAACTCGCCTTTCAAACTGACCGATGAATTTGTAGAGGTGATGGGGGGATCGGATAGCGACATGTACAACTATTTTAAGATTCTTATATTGCAAGGGTTTGTTGCAGCTAGAAAAAATATGGATCGTGTTTTACAAATTGCAGAAATTATGAGCACAAGTTCGCATCTGCCATGCTTTGCGCAAGGAGTCTCCACGATTCAATCCATGCGTGATCGATTTCACATGAACTTAACGGAAGATCAGTTATCTAGGCACGTCTTAAAGCTTGTTAATGATAGTGTTCAGTCATTAACAACTCGACTTTATGATAGTTTTCAATATTTTACGAATGGAATACTTTAAAACGAACCTTATAAATCAAAATTATTTATTTTCTATTATCAGGCCAGCGAGTTCTAATGCTGGTATTGTGTACATATTCCCTTGATAATATATTTTATATTGTTATTATCTAGTTTGTGTATTGTATGCATACATACGTATATATAGATATATATATAGATATTTTTTCCTGTATCATATGTGCATCTTTCTGATTTTCAAACATTTTTCCATTATACAATTTATCGCCTTTCTATGAGCAATTTTAAAAAATTTTCTTTATTAGTTTTTATTTAATTCTTTATTGGTTAATTTAATGCCT
>evgR014938c0g1_i01_Hydra
ATTTGACTGTGAAGATTGTGGTTCGAGAAGATTAGTTGCAAATTGGACTATAGTAGATAACGCTAGTAATTTTCCATCGGAGCTTTCATTTCCCAATGCAACAACCACAGGTTTTTTGACACCCAGCCTTGTAATCAATAGAGATGTTCTTCAGGAAAATAAAAACTACACTTTTACATTAGTTGTTGGTTATGAAGACTCAACAAATAGCGCTAAATTCAAGTTCACCAAGTCTACGTGTTATGAGCCAACACCAGGATTATGTTATACTAATCCAACAGAAGGAAACGCCCTAGAAACAAAGTTTACCATTATATGCACTGATTGGACAGACTTGGATGGATTTCTTAGCTATAAGTTCTACTATGATAATGGTCAATCGAAAACAATGAATCTCAACAGTACTACCACAACTGATTATCCTATATTAAATGCAGAAACTACTTTTCAACCTAGTTTAGTAGACTTTATCTTGGGTCCAGGTGACCAAAATAATAGTTCATATATCCGAATACTTGCTAAAGTTACTGGAAAGTACCAGGCATACACAGAGTATAACATGTATATAAGGGTGTACATGCAGAATAAAACTGTAAACTTAACAGACTTGCTGAATGATATAAACTTAAATGATACACAAAGTCTCGCTAATCTTGTTCGAGCAGTAAGTTCTACTGCAAACATAAATTCAGTAAACTATAAAAACAACCCCTCAATGTTTGATGCAGATCATTTTTCTCGTTTACAAGAATTGCGCACACAAATGCTCAAATATATTGATAGTGCACCTCTCAAAGACATAAATTTGTTCCAATCTGTAAGTAGTGCTTTGGTTGTTATATCACAGTATCCATCAGAAATTGTTCCTCAAGCTCAGAATCTTGCAGGAAATGTTATTGTTCGACTAGCTAAATTACTCACTACAAAAAATTTGAAAGGTTTTGGTGCTGACAGATTTGATTCTA
>evgR014943c0g3_i01_Hydra
TTGTATAACGGAAACGTGGTGCAAACATGACGAACAAAACTTACATTTTGAATTAGATAAATACACGTCAATTCATCAACCGCGTAAAAAATATTGTGGCGGAGGCGTAAGTATTTTTGTTTGCAATTCAATTAACTATGTCCAACGTAATGACCTATGCGTTAATGAAACAGATTGCGAGTCATTATGCATTGAAATCGTTAATAAAACCACAAAAAACATAATAATTAATACTACTTATAGGAAACCAGCTGGCAATTTACAAATTTTTAAAACTCTTTTAAATACATTCTTAACTAAAGTCAATAAAGAACGGAAGCATGTATACATAGTCGGGGATATCAATTTAGACTTGATGAGACAAGCTTCAAATAACGTTAAACATTTTATAGATACTCTTCTTCAACATAATGTTATTCCAACAATAAATAAGCCAACCAGAATTACTAAGAACTCTTCTACATTACTAGATAATATAATCACTAACAACTTTTATAAAAACCCTGTTTGTACAGGTATAATTAAGACAGAATTATCAGATCATTTTGCTACTTTTCTTATTACTAAAAATATTACAACTAATTATAAGCCTTCAAAAACTATAATATACAAGAGACAGATCAATAAAAACTCCATACAACAATTCCAAGACCTTTTAAAATATAAAATTGATTGGGATCTTATACTGCAATCAAGTGAGGCAAACAATGCTTATGAACTATTTCTCAATTGTTTTTGTAATCAGTATGAAAAGGCGTTTCCTGAGAAAAAAATTGTTATAAATACCAAGTCACTCCTATGTCCTTGGATGTCTAAGGGCCTGCTAAAATCCTCAAAGAAAAAAAA
>evgR014952c0g1_i01_Hydra
ATGCTGGGCTAAATGCTGAATTGATTCCTGAAGGATCTCAGTATCAAAAATTAAAACAGGATGTTTATGTTGACGGTGGAGAAACTCTTGTATCGCTAGCAATAAATGTAGATGGTGTTAAGCTTCAAAATTCTACCAACAAATCAATGTGGCCTATACTTTGTAGAGTAAATGAATCTCTAGATCAAACTCCATTTGTTGCAGGTATACACATTGGTTCTTCAAAGCCAGATTCATTAGTTGATTTTTTTTTGCCATTTATCACTGATTTTACCAAAGTGTCCAAAAATATAAAAACCCTTTATCCAAAAATAAGAATTAAAATTCTTTTCTTCGTTTGTGATGCACCAGCCAGGGCATTTTGTAAAGGTACAAAACAATGTGGAGGATACTTTTCTTGTGATTATTGCCAAGATGAAG
>evgR014969c0g1_i01_Hydra
TTGTTCTTGAAGACATTGTTTTTACGTGACTTAGTGTTTAAGTGGTGTTCCACAAGAATTTTATGTCTTATCCGACACACATTTTCATTTTTAAATGCTCATATACTTAAATTGTTATATTGTTTGCTAGTACGCCCTCATCTTGAATATGCTTTTTCTATTTGGAGTCCATTTAAAAAAAAAGACATTAATCTTATTAAAAGAGTGCAACATAGAGCAACAAGACGAATTCCAGGATTCTCAGAGCTTCCTTATGAGGAAAGATTAAATAGACTAAATATAACAAACTTAGCTATGAGAAGAGAAAGATGCGACTTAATTAACTTTTTTAAAT
>evgR014983c0g1_i01_Hydra
TAATTACTATTACTATTTCAAATGTTTATCGATTTGAAATTAAGGAACATATATTAAGCTATTCAAATATAAATACAGAAATCTATTTGTACCAACAAGCTCGGTCATTAATATCTTCGATAGAGTTAACCAATTCAATTATTAATTGTTACTTCGACCCAAACACCAGTTCTTTTATACATTGGGAATTTCGAATTTTTAAAGATTCAAATTTTCGGAATGAATTGTTCATGCATAGTTATTACTTCACTAATTCAAGTTATGGTTGCAAAAGTTATGGTTGTCAATTCAAAATTCAAACTCAGTTTAGCTCTGTTTCTACCGCTTTTAAAGGAAATGAACTCTCAAAGTGCCTTTCGAATTGTCCACCGGGCTTTGAACAAAATATTTTATTTTTGAAGAACGACGGAGAAATAAAGAAATTTAGTACTTGCAGCAGTTGTGAAGCTTTTTTTTTAAAAAAAAATTACGGAACAGCTAAATGTGAAAAATGTCCCTTAGGCTATATCCCAAACAGTGAAAAAACCGCTTGCAAATTAACTTTTAAAGTAGACTTAACATTTCGCAATGTTGAAGGCATTATCATTATTCTGTTGGCAGCTTTTTCTTCTCTGTTTAGCACAGTTATAATAGTATCTTTTATATTAAATAAAAATTCTCCTATCGTAAAATCATC
>evgR014989c0g1_i01_Hydra
AGAAGGGTGAGCTATAGGAGTGAGGGACTAGGATCCAACTTACCTGATAGCAGACCTCCTTACGTACCACCTGCCACATTTAGAAGAAGGGATGATAACTACTTTGGGAAACAGCAAAGAGAGGCTCCGAGACAAACATCTAAGAAATCTGGAGATACTACCAAACCTATCCCTAAAGCTGAAGTTCCAAAAGTTATTGAGATCAAGAGGCCAGTTGTAACCTTAAAAAGTGAGAAGAACCTTGAATCTGTGCCTCAAGAAACCAGAGAAGAAAACCTACCAAAGAAGAAATGGGTACCCACGTCTTATCCAACTCTAGAACGTGTACTGAAAGAAGGTGGTGAATTATGGTCTGCAATGGATGATGATGACTATTTGTCTGATCATAGTACACTTTGCACAGAGGATAGAAGTGAAGCAAGTGCTAGTGTAGCCTCCAGCTTAAATGAAGAACCTGTGATTAAAGCAGAGAAGGGAGAGCTACCTTCAAAAGGAAGAGATATATTTGCATCTTTTGCTGACAATGCTCGTCGGTTAAAGCTGATGAACCCTGCACTGTTTGATAGTGACACTAGTTCAAGTGATGATGAAGATAACACAGAGGAAGAACCAGAAAAAGGGTAATATTTATGATTCCAAACTATAACCCGATTCTGGTTTGACGACTTTGACCTAACTGATAATTCTTATATAAGGGTTACTCTATTCTTTCCTATCGTGGGTTTTTTCCTGGGGCTAAGTTGACAAATGATAAGTAGAATATAGCCTTAACTTGATTTTAGGGTTTTAATTCTATTTATTT
>evgR014993c0g1_i01_Hydra
TTATTGCTCAATGTTTTGTCATTCAGATGGTGACGTTTCTTTATGTACTTATATCGTACCTGGTGGTAAACCAACTAAGGATGACTTACCAAAACTCCACGAATACTGTGTATCTGAAGATCCATTGATCAAGAAAGCTCTCGGACCAAATCCTCAATTCCAACAAATGAAGGCTGCTTATTTGAGATTGGGTGGTATTTCAAACTCATATGCTGATAACACTCTCGTTATTGGTGACGCTTGTGGTCTTATTAACCCAATGACTGGTGAAGGTATTCACTACGCTATGGACAGTGGTAAAATGGCTGCTGAGGTTCTTGGAGATGCTATCAAATCAGGCGATCTCTCAAAGAATAAGCTCAAGCTTTATCAAGATATGTGGGAATACCAATTTGCCAACGATTTCCATTTCTCATTCCAAGCTGGTATTTTAGTTGCTAAATATCCAATCCTCATGGATGCTGTTGCCTCAGCTGTTAAGAAAAAGGGTGGTAAATTCTTGGCTGAATTCGCCATGGTGTCTTCTGGTGTAAAGAGTTGG
>evgR014997c0g1_i01_Hydra
TTTTCAGCAGCCAACTCATGGTATGCATAATGGAGGTGGATAGGCCATAACACGCTGACCATATCTTATGATTCAATAATTAGTTTACGATGTCCAGCTCCTTATGGGATTTTTTATTTACGGTTATACTTGTTATAGGCTTTAAATTAGCGTCTTCTGCAGATTTGAATGCTACAACTGTTTCCAGTTCATCTTTAAACTCATCATCTATTCTCAACACTTCAGAAGTCACAACAAACTATTCCAATAAAACAGATTTGGATGAAAAATCTCCCGAAACATTAAAAAAAGAAGATGCGAAACATATACAAGGAACTGAAAAGGAAAAAAAACCATCGGAAGAAAATGAACAACTAAAATCAAATAAAGAAAAGGAAAAAAACGACTCTCAAAAAAATGAGAGTGATAAGAAAAAGGTAAGCGGCACCTCACGCAAACAAAATGAAGAAAAACAAAAGAAAAAAGCAAAAAAAAAGGTATCTTGGAAAATCTCAAAAACCTCTCATTTTTCAAAACCAAGTAAAACAAAAAACAAAGAATCAGCAAAAGAAAACGAAGAAGAAGAATATGATAGTGAATACACGAGCACATCTTATGATGATCCCAGTTCTAATGAATATCATGAAAATGATGAATACTATGCCGGAGAAGGCCACGGTTATGACTCAAGCTATACAGATAACGGAAACGGTCAACCACAGCAAATTGACCAAGGCTACAATTATAACTACAATGTTCATCCCAATAAAAACACACCAATGGATTCTCAATTATACGAGTATAAAAGTTTTCCGATACAAAGTTCAAGCAGTCCTGCTTACCCATTTCAAAATTACGAGGAACCCAACTATGCGTCGGATAATTACACACCTAACATAGGTTCTAGCATAGCAAATATAGGAATGGATGGTGCAGGTCCTTCAATTGGTAACGTTGTGCAAGGTAGTTTAGGAGCCACAGCTTTAGAACCTGCTTTTAGAAGTGCACCCTTAGGTCAAGCATCGATAGGCTCAGCTGGACTTACACCAGATGAATATAGACAACTGGAAGACGGATTTCAACCAATTACTTCAAATCAGCTAGCAAATCAACCTGACGACATAGGAGCATATAGCTATGTTAAGAAAAGAACGCAAAATAATCATAAATTTAAAAAAAAGAATGGGAAAATTGGACGTAAACATGTCGTGAAAATATCGAAGCAAGCTCATAACTCATCCTCCATAAAGCATTCGAAAATCGAAAAATCTTACAAAAGAACAATTATGATTGGTAAGTGAAAAGAATCTATTTTTTTGTGATGGAGTTTTATGGACTGGGACTAGATTCATAGTTTTATCAAATGGTAAATTTCAATGGATATCAATATAAATTGGTCGATGAAAATAAAAAATCTTTTAGTTTTATTTTCATATTAATAAAAATATAAATTTGTTCTTTCCTCATCTTAAATGTTTCATTTTCTTCATTAATTTCTGAATAAAAGTTAAAAAGATTTAAAAAATTTCGTCGATGACGCATTTCTCTTCAACTTACATATGCCAAAAAATGTATATACAATTGTAAGATGTGTAAACATTGTTTAATAAAAACTTTTATACAGTATTTTAGTACGATTTGTAACATCATTTACATAGGGGCGTACATAGGGGCGTACATAGGGGCGTACATGCGCTTACTTTCTTTTTTTAAGTAGAATATTTTCTACGTATTTTTATGTCAACAAATTAATGTAATTGTAAACATAAAAATGTTCGAATGAGAACGATGAGAGTTTTTTTTGTATAAAAACGTTTTATAATATATATACTGATAACTTTTATGTGAAAAGTTAGCAGTTTTATTTGGCAAAATATAATAATATTTCAC
>evgR014998c0g1_i01_Hydra
GATATAAGAAACAATGGATTGAAGGAAAGGTAAATCAATCAGTTGAAATTCAAGACATGACAACAAAGGAAAAGTCTACCTTGTATGATGTTAAAACACTCACTCGTCCGACAAAGATTATCAAGCCAGTTGCTGAACAAGAACCAAGAGAATCTAGAAGAAATTGGCACAAGTCTGTGCATGCATTGAAGAATGAAAAGTTCGAAGATGCTCAACTTGAAAAACACCTTGTTGAAGAAAGAGAAAGAGCATTCAGAAAAGAAAGAACTGAAGCTGGTCTTAATTGGACTCCAAAATACTTCCGATTGGAAGGTGAGAAGAGATGGGTTCTCAAATCTAAG
>evgR014999c0g1_i01_Hydra
CATAAAATATCTAATTTTAACTTTAAAAATTACAAATCAAAATATAAAAATGGTTATATGCAATACTAGTTTTTGATGTCAGGAAATAAAGTTTTACACTCACAATGGCTTCTGACCCAAGCGAGTGTAACATTAAAGTTGTGTGTCGGTTTCGACCACAGTCGGATGCTGAAAATAGTTGCGGTGGTAATACTATTGTAAAGTTTCCTTCAAATGAAACAGTTTTACATATGGGAAGAGCATACACTTTCGATCATGTCTTTAAGCAAAATGCATCACAAGAACATGTTTATTCAGTGTCTGCTAAACCAATTGTGTCCGATGTACTTGCTGGATACAATGGAACAATTTTTGCATATGGACAAACTGCAAGTGGAAAAACTCATACAATGGAGGGTGTTATTGGAGACCCTACATGGCAAGGAATTATTCCTCGTATTGTTCAAGATATTTTCAATTATATATACACCATGGATGAAAACTTAGAATTTCATATAAAAGTTTCTTATTTTGAAATTTATTTAGATAAAATTAGAGATTTATTAGATGTTACAAAGACCAATTTAGCAGTTCATGAAGATCGCAATCGTGTACCCTATGTTAAGGGTGCGACTGAAAGATTTGTATCTAGTCCTGAAGAAGTACTGTCTGTCATATCTGAAGGAAAAGCAAATCGCCATGTTTCAGTAACAAACATGAATGAGCATAGTTCGCGAAGTCATAGTATATTTCTTATCAATGTGAAGCAAGAAAATACAGAAACGCAAAAGAAGCTGAGTGGAAAGCTGTACCTTGTTGATCTGGCTGGTTCAGAGAAGGTCAGCAAAACAGGTGCTGAAGGAGATGTGCTCAATGAAGCAAAAAATATAAATAAATCGTTATCTGCTTTGGGTAATGTCATTAGTGGATTAGCAGATGGTAATAGTAGTTATATTCCTTATAGAGACAGCAAAATGACTAGGATTCTACAAGAAAGTTTAGGTGGAAATTCTCGTACAACAATTATTATTTGTTGTTCCCCATCATCATTTAATGAAGTTGAAACAAAAACAACTTTACTCTTTGGACAAAGAGCAAAGACAATTAAGAACACTGTTGTAATTAATGAAGAACTTACAGCAGATGAGTGGAAAAAACGTTACGAAAAAGAAAAAGAAAAAAATGCCAAGTTACAAGCCATGTTAGCTCATCTTGAAAAAGAGGTTAACAGATGGCGAAAAGGGGAAAGTGTTCCTGTTGGTGAACAAATTAACACAGCTGACAAAAAAAGAAGCAGTGAACTAATAGAAGTAGCTATCGAGCAGCAAAAAGCTTCTGTTCATAGTCATGTTTCTGATGCAGAATTAGTGAAATTTGATGTGGAGCGCACGAAATTGTATAAAGAATTAGATGAACGGGATGATGAAATCCAAGCCCAATCAAACCTTATAGAAACTCTTCGACAACAATTAGCTGACTTAGAAGAAAATTTGATATTCACAAAAGCGAATGCAGAGAGTTTGGAAGGTCAAGTTGGTTCAATTGCTAAAGAGCATGATGCATCTAAAAATGAGGTGAAAGAAGTTTTGCAGGCACTTGAAGAACTAGCTGTTAATTATGACCAAAAGCTTCAAGAAGTTGATCAAAAGACAAAAGACAATGAGAGTCTTCTTGAGGATTTGACCAAGGAGCAGCATAGCAAGCAAGCATTGTCAAGAGAGCTGGAAGCATTACAAGAGAGAATGAACCTTATGACTCGAAAAGTCACTGAAATAACTAACACTTTGCTTGGAGAATTGAGTGAAATTGGAAACCATCTTGGAGGAAATATTGCTGGCATGAAGAGGCCAGAAATTGAAAGTGCTTCTCAATTTGACGAAGAGTTCACAGTAGCTCGCTTGTACGTAACAAAAATGAAAACTGAGATTAAAACAATGGTTCAGCTCAATAGACAACTCAGTGAAACATGTCAAATTAATCAAGATAAGCTTGAATTATTTGAAAAAGAAATTCAAGGTTATAAACTTACGATAGCTCAGCATGAAGCCAAGATGGTTACGTTACTGGAAAGCATTAAAGAAGTCGAAGCGAAAAAGAGATTACTAGAAGAAGCAGTAGACACTTTAAATGAAACTCTTGCAAAGCAGACCGCAAATGAACAAATGCAGATTGCTTCTCTAAAGGATAGAGAAGAAAAGCAGGCAAAAGGTTTAATGGATGCAAAAGCTATAAAGGAAGCCTTAGAAGCTCAAATGAAACATCATGGGGAAGCACATGCCAAACAGCTTGCTCTTCTTCGTGACGAAATTCAAAACCAATCTACTATTATTAGTGATTTAAAAGAAATAAACCAAAAAAGAAGTCTAGAAGAACTGCAAATTAAAACCGAAAATGAAAAACTTTTGGTTGCATTGCATGCTCTGAAATCAAAAAACCAGGAGCTGGAAGCATTGAACACAAAGAAGGAACATGCACAAAAAGATCTGCAAGGTTTAGAAGAAACTGTTGCTAGAGAGTTGGCAACTTTAAATAAATTAAGACAGATGTTTGTAAAAGATCTGCAAGCAAGAATGAAAGTTGCATCAGATGATGCAGACGGAGATGAAACTAATAGTCAAGCACAAAAACAGAAGATTTCTTTTTTAGAGAACAACTTAGAGCAGCTCACCAAAGTGCACAAACAGCTTGTGAGAGACAACGCAGACTTGCGTCTAGAATTGCCGAAGCTCGAGAAGCGATTGAAAGCTACTCAAGAGCGCGTAAAGGCATTGGAGGTTGCTCTAAAAGACGCTAAAGATGGCGCCATGAAAGATAGGTCACGCTATATTAAGGAGGTTGAACGCTGGAAACAAGCTTATAATAAAAGTCAAAATGGCCGGCGAATTAATCAACCTCAAATTGCTAAACCAATTCGACCTGGTCAACTGACCTCTGGCGTCAATACGGGCAGTGCCATGATAAAAGGAAACAATATAAATATATTAGATGGTTCTTCAGAATTTTCTCGTCAACAGCAAGGTGCTATCGTTGCCCGCGCTACTTCTCCTTCAAAAATACCAATAATTCCTATGACTGGATCACAACCAGTTATTAAAGGAGGTTACCGAGAAGGAGAAACAGTAACAATTGGTGGTACAGGAGTTAGTCATCGCGTCAAAACAGCAGTTGGAGCTGAAATATCAACTCGATCGTTGATTAGGAGGCCTGGAAGCGATGACAAACGCAACTCAATGTTTGTCGAATCGACAAGTAACATGCAGTTAGTTGAAGAAAACTTAAAAGGAAGTGCTAGTATGGATAGACTAAATATGATCGATTCTTCAACTTTGAAGACTCCCAGACGCCTTCCATCACCTATTTCGAGATCACCTATCTCAAATAGTTTAAGTGGTCGTGAAGCTGAAATACAAGCTCTAATTGCAGGTGGGAAATCAAGATCAAATACACAATCGTCAAACTCGTCAGTCGACGGAAGTCCGTGGGAACGAAGAATACCAACGTCTCGTGGACTTAGTCCTCGGTAGTATTATATATATGGCAGCCTTTATTCAAATCGTTTATTTTTTGAAGCGCTTGAATTGAATCAACATCATCTAGCTTTAACTAAGAATTACTAAACAAAAACCACTAAACTGAGAAATGACTTTCTAACGTAATGACCTGGAATATTTGTTTAAGTAAAATTGTAAAAAAAAAAAATAATAAGCAGAATTTTAATTTTCTTATATTTGAAATTCTCGGATATTTTGGGACGTTCTTCAAATATTTACATATTCGAAGTTTGAATATTTCGAAGTATTTAAACTAAAATAACCTTTTCTATGCAGTGCTGTGTAAATATATTTCATAGATAGTAAAAATGATAACCTTCTAATATTTTTGTACTAGTACCTGAAGTCTCGATTAACAATTAGATGAACGAGTTTTTGGTGATTTTTATATATACATATATATAAGTTCGGCAAAAATCTAATTTTTTGTTTTTAAATTCGGATAATATTAAAGGTTCGCCAAAAAGAAATATAGTATTTTTTTTTAATGCATTTAAGCTGCAACAATTAATTTATAATTAAAGTAGTTGTTATTGGCCAGTTAATTGAATATCATATATTAAATATTATTTATTAATTTTCATTTTTCGTAAAGATATAAATTATTAAGATAACGAACTTTTTATTTCCGTTATTCAACAATTAC
>evgR015001c0g1_i01_Hydra
CTAATAAATCACTGTCTTCATCTAAAGGAACTATAGAATCATTTTAGTACAAACATTTATAATATAATCCATTATTTTCTTTTTTTACACAAGTCGCAATGCCCACCACTGACAGCTTGAACACTGTATCCATGCAGCACCTGGATGGCTGTTTGAGAAGGGTTTTAACTGAGACAAAAGAACTCATCTTCAATATCATCTTTTTTTTTTTGCTGTTGCGTTTGTTTATTACAGGTTGGCAGATAAGAACCTTGCTCGTTTTAGAGTTTTTTAACTGGTGTATCGGTCAATATTGCAGAAATTCGCTTTCTATTTCTGACCTTATTTTTAGTCCGTTTTCTAAGCTGAGGTTTTTCAAAAGGCCTCACTCTCATCTTCTTCAGGATATGGTAAAAGTACTTAAAGTAGAACCTGGTGTGCCATTTCTAAAATGAATCTAAAAATTTCGAATGTTTCCTTCTTTAGATGATAATGAGGTTGAATTTGGTCTATTGAGATTTGTAATCCTAGTACTGGTAATCCTAAAATTAGCAGGCAGCTCAGTGCTAGATGTTTGTTAAGTAATACTAAAATCCATAGCCGACGCTTCAGTTCTAGGGTCTGGTCTATGTTAAAAAGATAAATTCTTAGATTAATACTTAGAAACCACTAATTATGTTTCTTGGAGTTATAGCATTGGGAAGAGCTGTATTTATTTTACCAGGGAAATCATATAGTCCAAGGTCTTTTATTTACAGTAATCCACA
>evgR015004c0g1_i01_Hydra
AAAAAAATTACATAGATGGCTGATCGTAAATCTTTTCTAAAAGGATGTACAAGAGAGCCTTTGTCGCCTATATCAAATGAAACAGCATCTTCTCGAACCCCTTGTCGCAGATTGAAAGACCCCAAATATGATACACCTTCACGATCACCTGGTTATAGAAAAAAGTGTATAAATAATGAAGGTACTGGTTATGATCGGTTTATTTCAAATCGAGCTTTGATGGACCCACAAATTGGTCACCACATGTTCATGAAAAATGACAGTGAAAATGATTCTGAAAACAATGAGTTATCCGAAATGATGAAGCAAAGCCTAGGAACAGCTGACAACACTGGATCTCGAATTCTTCAAATGAAAATAAAACCACCAGTATGCAAAGAAGGTGAGATTCACATGAAAACCTTGTACACGAAAAGTAAACCTACACCGGCTAAGTCCAAATCTACACGACACATTCCTGCAACTTCAGACAAAATTCTTGATGCACCATACATTCTTGATGATTACTATTTAAATTTGCTGGATTGGGGAAGCTGTGGCTTAATTGCAGTCCCGTTGTCACAAACAGTATATTTATGGAATGCTGAAACAGGATCTGTTGATGAGCTGTTTGGTGATGTTCATGCAGGTGAAGAAGATCTAACTGTTACATGTGTAAAGTGGATACAGGAAGGACTTTATTTAGCTGTTGGGCTATCTAATGGCTCAGTTGAGTTATGGGATGCTGAATCATCAAAACGAATTAGAATAATGCAAGGCCATTCTAGTAGAGTTGGCTCACTAGACTGGAATAAGCATATAGTTACTAGTGGATCGCGCAGTGGTGCAATCTTTAACCATGATGTTCGAATAGCAAATCATCATACTGCTTCTTTTTTAAACCATTCACAAGAGGTATGTGGTTTACAATGGTCACCTGATGGTAAACTTTTAGCTAGTGGAGGAAATGACAATACACTAAATATTTGGGATTTCACTTCCGTAAGTGCAAGTACTGATAGTGTAGTTGAGAATACAACACCTCTGCACTCTTTGACTGACCACTTAGCTGCTGTAAAGGCAGTCTCATGGTGTCCATGGCAACGAAATCTACTTGCTTCAGGTGGCGGTACACTTGATCGACATATTAGAATATGGAATACTGGAACAGGAAATTGTTTGCAGAGTTGTGATACCATGGCACAAATATCTGGTATTCTATGGTCAGACATCCACAAGGAAATTATATGCTCAAGTGGTAATCAAATGACTATTTATAAGTATCCTCAGATGACAAGAGAAACAGATTTAAAAGGTCATACCGGTCGAATTCTCAATATAACACTTTCACCAAATGGACATATTGTTGCATCTGCTGGTGCAGATGAGACTTTACGATTATGGAAGTGTTTTGAACCAGATGTTAAAGTCGCAAAACCAACAAAAAGGATTCCTGCTAATATTTTATCAAGTGATATTATGAGATGAGTTTTCTACAAACAGTTGTATATTTTATTTTTTGGCTTGCGTTTGCTTCATGATGATTTTTTTAATCTTTTTTTTTTAGTTTTAAATATTGCTGAAAAAAAAGCACCTTGAAATATTAAAAAAAGTTTGAAGCTATAAGTTCAAAATGTTTAAAAAGAAAAAGAAAAAGAAAATCTAATTAATCGATTAAGAGCGAATCTTATTATTTATTTTTATCTGGATCATTTTAAAATTTGGAAAAATATTATTTTATTAAAAAA
>evgR015006c0g2_i01_Hydra
GTGTTTTGTTGAGATGGATCGAGGTTTTTTTAACCAATCGTAAGCAACGCATTGTAATAGGAGAAGTTTCATCGTCGTGGTTAAATATACTCAGTGGTGTTCCTCAGTGGTCTGTTCTTGGGCCTCTTTTGTTTTTGATATTTATCAACAACATGCGGGAAACATTACAATGTGTATGCAAACTTTACGCTGATGACATCAAGTTAATGAAACCGATAAAAAATATTTCAGACTCAATCAGTTTCGAAAGCGATATCAACAAGTTAGTTGAATGGACAAGTATATTGTTAAAAAAAGACCATCTATACGATTACGGAAAATTGCGTTGAGTACAATCTAGGCAAAACAACAGTAGAACGTGACCTCGGTGTAATGATCTCATCAGACATCAAATGGCATCATCAAGCTAACGCAGCAGCAGCTAAAGCGCAGAACATACTCAGAATCATCAAGCGTACCTTCACTTATTTAAATACAGAAATGATTCGTACTCTGTAACAGTGCTAGTTAGACCACACCTTGAGTTTGCAATCCCTGTATGGAATCCTACTTTGAAAGGTGACATTGAAACACTAGAAAATATACAACGTCGCCTAACCAAACTTGTG
>evgR015007c0g1_i01_Hydra
GCAAACAAAATTTATTCTATAAGAAAAAATTAATAGAATACTTATGAATTGACAAGAGTTTATAGCTTAGATTCTGGATATGGAATAGCACACTACGCAGCATAGCACTTTTCAAAAAGAAATATAACTAAATATGGTTGGAAATTTTTGAATTGGTACATTTGGTTCGTACGTACATACGGCTGGATTGGTACATTTGGTTCGTACATACATACGGCTGGAATGGTACATTTGGTTCTTTTAAAAAATTAACTTATAAAAGATCTGACAACTTCATTTTCATGGTGATCCTAGATTACACAACTTTTTTACCCTATACAATGAGCGTACCTTATACAATGGGCATAACAGAAACGAGCACAGCAAACAGCAAAAGTGATGATGAGTGTGAAAATCAATTTTTCATGGTTAGCAGTCCAAATTTACGAACTTTTTTTTTAGTTGTATACGGACTTAATTGTGGGTTTGTTACTCTTTCTAACTTATTGTTAATTATTTTGATCTTACTTTACAAGAAGCCCAAATATTTTTCAAGAAATGAAAAACACGTTTTAATTCTATCCGCAGTAGATTTATTAATTGGTGTATTGGAAATGCCGTTAGAGATTGCATTTTTAGAATCGTCCTATATTGAATGTACAACATTTTTCATTCGAAATATGGTAATGTTTCTTGTTAAAGCGTCGGTACTTTGCACAGTTTTGATTGCTGTTGATCGATTTATGACGGTTTTGTACAGTGAACAAAATTTTATTGAATTTCATCCTATATTAAGTGTTGCTACCGTTTTTGTAACAAGTTTTTCTTTTGGTATTTGGTATGCGTTTATTGCTAGCAATAAATTACGCAGTAAAATAATTTTTTTACTGTTTAATGCAAGTTTCGATTTTTTAATTTTATTGGTGATTGCCATTGTGAACGTCGGCCTACTTTATAATGCTAAAAAAAGATTAAAAATTGTTTCAGGTTTTAGATCGTGCAACTTGAATTCTGACCAGCGGCTAACTCGAACAATAGTTATATTATCAACAACGCTTGCAATCGCGTACTTACCTTTTTCAATTATGCAGTTTGTTAACGCAGCGGAATACCTGAACGAAAATGTTAAACCTTTATTTTACGGAACAGCAACTTTAGCTCAGTTCAACTCAGGAATAAATGCTGTAATATATATTGCAAGAAATAAAGAACTGGTGAACAAAATTAAGCTCAAATTAAAAAATAAAAACCTGAAAAAAATAAAATTGGACACAAATAAGATTTAAAGATGATATGTTTCATATAACAAAGATGATATATTTCATATTCAAAACTGCAC
>evgR015009c0g1_i01_Hydra
AGTGCAAAAAGTTGCAGGTTAATAAAAAGTTATGTATTAGTACTTTGTAAATGTATCAACCACCTTTCAACTTTCAACAGCCTGCGTGGCCTGTTGTTAATCCGTTTCAACCTGCTCCAGCTGTAATTCCACCCAATCCTCCAAGTGATTCAACTCAATTTCCACAGACTACTCAAAATGTTTTTCAGCCTGCAGCTGTTCCAACAGGTTTTACCTTAGATGCACAAATGGCATATCATCAACAGCAAGGGATGTTTATGCAGAATGCTTTTAGTCAACAAAATTCTTTGCAACAGCCATGGGTTCAGCCAACAGTTAATCCATTCAATCCAGCTACAACATTGCAGCCATTCCCCCAATCACAACAGTGGAATCCTCTTACCAATCAATGGGTAACACAGCCACAATCATGGGATCAACATAATGCACAATATCAAGCAACTCAAATTCCAGAATTCCAAGGCTATCAACCATTGCCACAGCAACTTAATTTGTTTCAACCAGCTAACACACAAGTTTCAAATACAAGTAATGTACAAAATGACAGCCAAAGTAGTTTAAATAATGCATGGAACACTGGGCAAAATCAATGGATGCAAAGCAACCTATATCAAGCTAATGTCCAGCAACAAAATGTTGTGTCTACTGAACAAAATGCCAGTAATAAGCAATGGGTACAGCCTTCAACCAATACTGTATCAGCACCAACTGTTCCCCCACCTCCATCTCAAAATCCACCTTTACCAAATGAACCTCCGCCTCCTCAAAATCCACCCAGTTCTATTAAAGCACCACCTCCTCCTGATCATCCAGAATCAAAATCTCCGCCACTACCGGAAGAGGATCAAGAAAGTTTGATCTCTCTAACAAAACAACTAGAAACATTAGAAACTCAGATTAAAACTTATCAAGCCGAATATCAGAAATGGCATCTTGATTTTGTAAAATGGCAGAAGCAGCATCAAAATCATCCTGATCAAAAACAGTATATAGATTACCAGAAACAATATCAAAAAATAATTGAGGAACAACAAACTCAATTAAAGCAACAGTATGCGGAACTACAAAAAAAAATTAGTTTACGGAAAGCACAGCAGGAGAAAAAAAACAATAACAGAACTGACCAGTTATCAAAAGGAATCAACAGTTCTCAGGGACCAGTGACAGCTACTTATCATCCAGTTGTCATAGAAGCAAATAAAGGTGATAGTTTTAAAGCTTCCTTAAACACAAATCAAAGTATGTCGAGTTTTGTTCAAGCTTCTCCACAAAGCATACCTCTTCAAAAAAACTTACCGACGCATAACTTAACTCCAAAACAAATCATACCTCCTAAACAAAATACACCTCTTCTACAAAACTCGCATCTTCAACAAAATACTTCTCAACAAAACAAACCTCAACAAAGCATTCCTCAACAAAGTAATCCTCAACAAACCATTCCTCAACAAAGTATTCCTCAACAAAGCATTCCTCAGCAAAGCATTCCTCAACAAAGCATTCCTCAACAAAACATACCACCTTTGATGCAACAACAGATTTATGAGAGTAAAAGGCCAGGATTTAACAATCCTCCAAACGAGTCAGTTAATAAGGTATACGATTACAAAAATCAGTTGCAAAAGCCTGTTTCAAATCAAGAGAATTTAATAAACGATCAGAGCAAAACACTTCAGTCTCAAAATTCTGGATCATTTGGGCATGCACCTGAGATTTTTGGACAGCCCAATGAACCCTACAGACAGCCTAATGAGCCCTATGGGCAACCCAATGAATCTTATGGTCAGCCTCCTGAAACCCATGGTCGTCCCCAAGCGGCTTATGGACAGCCGCCTGGAGCTTACGGACAGCCTCCTGGAACTTATGGACAGCCTCCTGGAACATATGGACAGCCTCCTAGAGCTTATGGACAGCCTCCTGGACCTTATGGACAACCACCTGGACCTTATGGACAGGCTCCTGGACCTTTTGGACAGCCTCCTGGACCTTTTGGACAGCCTCCTGGACCTTTTGGACAGCCTCCTGGAC
>evgR015015c0g1_i01_Hydra
TAAACAATGACATAATAATTAACGGGGTTGATAGTAATGTTATTATTTATACCATTATTGGGCTTGATGGTTAAATTTAATAATGAAAATATCTATCCGGTACGAAGTTTAAAATTTACAAGTTCTGAAAACAAAATTCTCGGTTTAAGTTACATTTCCGGCAGATTTTAGAATTATTGACTACAGCTATGGTGAAGCATGTTGACAAAAAACTAAAAACGTTTTTTTCATATTATTATTATAAGACACAATGGTATCTTTAAATGGAGCGTATTCTAAAAAATACCAACTACAAGCCCAAGAAAGTCTGGCTCAATATAAGGATCGAAATCAAGATGAAGAAAAAGAAAAAATCAAAATTTTATCCCGTCAAACCAATGAAATGAGAAGGAAACTTGAGAGGAAAAAAAAAACCAAAGAAGAAAAGGAGGAAGAACGAAGAAAAAGCATTCTAGAAAACCGGAGATTAAAGCAGCAAATAGCAACAGAAAAGTTTCAGCATTCCACTGAACCAGCTAAGCAGAAACACGTTCGTTCTGCATCTGTTGGCACTCTGAAAGATGGACAGTATGATGTTGCTGGTTATTTAATTGATATTAAAAATGGTGTCCGAGTAAATTCAACATTAATTTCATCAGCTAACCCAATACATACAAAAGTAAGACATGCCACCCAAAGTATCAATTTGCATCAACCTGCCACACCTAGTAATAATTATAATCCACCTGGTACTCAAAGTAAAATTTCAAATTTACCTTCCACTCAAAGTAACATTTTTAATCCACCTGCCACTCAAGGTAATATTTTGCATTCACCATCTGTCCAAAGTAATATTTTAACTCTACCTTTCAGCCAAAGTAATATTCGAAATTCAGCTGGTTTCCAAGGTTATAATTATAATCTACCTTCCACTCAAAGTAATAACTTAAATCTACCGTCTACCCAAACTAATAATTATAATCAGCCTTCCACTCAAAGTAAAAAC
>evgR015018c0g1_i02_Hydra
ATATCCTAGCCGAAAAGTAATGAATTGCATCCAGAATAATGAATGTGAAACTATGGCTGAATCCATATGTGGAAATGATAAGAAAAACCTTGGTTTTCCTTCAATGCCCAACGAATCACTTATTGCTTGTGTGAGTATTGCTTTGTCTCCTCTTCAACCAGACTCAGATCATGAAAACGACTCTTGCAGCCATATTAATGTTGGAGAGTTTGTTGCTTTGAAATCTGGAAGTGCAACGGAACCAATTATTGGGAAGGTACTGACAAAATCTAATAATGCCTTAATGTTAGAGATATGGCATGAAAACAGTAAGAAATCTTGGTCAGTTAAATCTGGTTACAATAAACATGAGGTCTTCATTGACGAAATTTTGTGTGGTGGTATTTACTTTACAAACTCTAAAAAATTGCCAGTTTCTCTTCAACGTAAACTGCAAACTCTCTATTCATAATCTAATTGCTGTAAATTGTTGATGCAGTTTTTTTTTTAATGTCTGCTTTCTCAAGTAGTGAAAATTGAAATTAATTTTACTTATTACGCATATTACAAAAGTAACATGATAATGAAGTTCTTCAAAGAACTTCACAAGAACAAAAGAAAAAAGAACATTTTTTTTATTAAGCCTTTTTTTTTTATGAATAATAAGAGACAATATAATACTTTTAAATGGTG
>evgR015021c0g1_i01_Hydra
ATACGAAAGTAAATGGATAACAAAGATTTTATTATCATTTCTTGATGAAAAAACAAATGAAGTTTCTTTTATAGACGATAATCGTGTAGCGAAAGCCATAAGAAACCAAATTTTGTCTGGGACTGAATTAAAAACAATTGGATCATATCTGATTCAACCTGGAATATTTGCCCATGCAAGTTTAAATATATCTAAAGATTTATATGAAGTTAAAGACTTTGCATCTGATGCCTTAGTGTTAGGAATTTGTGACCCTAAATTAATAAAAAAGATAATAGAATCAGATTTTAACAGTCATGCAATTTCAAAGCAATCACAGTACATACTTTCTTTAACGTTAATATTTCTCAGACTTTTTCTTATTTCAACAAATGCTGACGATATTGATTTTCAATTAAGAATTGCTTTTAAATGGTGCGC
>evgR015029c0g1_i01_Hydra
ATAAATAATGCATTAACTATTTTTTGCATAATGTTGTTTGCTAATGATTTTTACTGTATTTTTTACAAACATCTGTATTTTAATGTCACGAGCATTGGGCGTTAATGTAAAATAAAGCATAGAAGTAGTTTTTACGTCAAAGAGCTGAAAGAATGAGTGCTCAAACAGTTATTGAAGCAACCCAGCACTCACATATAACTTTAAACTCCACTAATAAGAAAAGTGAAAAAAATATTAATGATCCAGACAGATGGTATTATATAGATAAAGATATGATTCCAGCAAAGCTAACTTTATTTTTCACAGGCATAGAAAGATCATCCATAGCACCTTATTTTACCATGTTCTACATAAGTATTGGTTTGAATCTACAACAAGTTGGTTACGTTTTTTCATTTTCCTATTTTGGTCTAATGCTAGGAAGCGTGTTTTGGGGAGTTTTAGCCGACAAAACGCGTGCTTATCGGGTTCTTTTATTAGTGCTATACTTTCTTTTTGCTTCAATTTCGTTGTCAATTCCATTGGTAAGTGCGAAAATTGGAGCAAAAGAAAAAAACCAATGTCCTTTATTGCAAAACAAATGTTGTACTAACAGCACAAATAAGACATCCGAGAATGATACTTCAGTATCTTCGATGACTTTAACCATCGTTATGGCAACCTTGGGATTTTTCCTTTTTTCATTTGGTGGCAGTATTTACAGCTATATTGATGCAGGAGTCATTGCAAAAATTAAAACATCAACACAAAAAATAGATTTTGGTAGGCAGTATATGTTTTTAAATATTGGTTTTGCATGTGGAGCTTTGATTTCCGGAGAAGTAATGGATATATTTCCCAAAGTAGACCTATCATGTTATTTTGCTGTATTCTTTGTTAACAATATTTTTATAATAGTTTCGTGCATTTGCTCTCAATACCTCTACAAAAACATCAACATAACGTCGCAAAAAGAAGTAAGATCCGATTCTTTTAAAAAAGATCTTCGCAAGACATTGTCAAACTTTGAAATTATTATTTTTTTTAAAACAACGCTTGTGAACGGCCTCTTATACTCTATATACTTGAATTATTTTTTTTTGTTTTTGAAAGAAATTAAATCTACAAACCTTTTGATTGGTTTAACAGTAGCATTCGGATCAATTTCAGGTATGATCGTAAATTTTTTCAATGAAAAAATTATAAAATTATTTAGAGGAACATTTAATACCATGATTATGT
>evgR015034c0g1_i01_Hydra
CGAAAGAGTTGTGAATCCGCACCCAGATACCAAGTCGAGATAACGAAAGTTGAATATTTAGGAATTCTTGTTGATATGTTTTTATATATTCTAAGCTGGATTTCAACTCTCCTGCATATTATTTTTGCAACTTTATCATTAGCTGCTGGTCTGTATTATGTTGCTGAAGTAGTCGAAGAATACTCTGTAATGACTGCTAAAGTTATCAAATACATGATTTATACAACTATAGCAATTAACATTGGTTTATTGGTATTTGAAGAAATGCCTTTGTATTTAATTGGATTGGGTTTCTTTACAAGTATTGACCATATTTTGCTGTTAAAAGAATTTCCAAACATTTCACTCCTGTCTCCATCGTTTATTATTGCTGGAGTGATGGTTTTTGTGAATCACTATTATGCATTTCAATATTTTTCAACATTTTACTACCAATTTTCAGAGGTTTTAGCTTATTTTACTCTTTGCCAATGGATGGTTCCGTTTGCTTTTTTTGTATCATTATCAGTAAACGATTCAACACTTCCAACTACATCATCATATGAAAGTCAATCCTTTATTATGAGTAAGCCAAAAAGATCTGGAATTCTTGGATTCTTTGATTACATATCACAAAGGAAATATTTATTAAAAGACTATTTTTTACAAAGAAAACATATTTATTAAAAGTTTAATTTATTTTTTTCTTCGTTTTCTATTTTACAAAGAAAACATATTTATTAAAAAAAAAAA
>evgR015036c0g1_i01_Hydra
TTGGGGGAATAATTGGTAATTTATGTGTAATAGTTATGATTTTGCAAAACAAAAAAATGCAAAACATGACAAACTATTTTGTGTTTAATCTAGCATTGGCAAATTTTGCTATATCATCTGTTGTGATTCCAATGACCATGGCAAACCAAATAAATTTCTGGACACCAAAGGAACTTGAATGCAAACTGCTTATGCCTTTTATTGAGCACTTTGCTGGCGTTTGTGTGCTAACTCATACCGCAATAAGCATAGCCCGAAATATAGTTGTATCTCAAAGCAAAATTAGTCTAAAAATTACATTTCGACATGTTGCGTTTTTAAATATATCTATATGGATCTTTGCCTTTATCATTTTGTCTGTTAGTTTAATGGGTGTCTTTGGTAAGTTTGTTTATTTTAGGAACAATTTTAAACTGAGCAGCGGTACAAATGGAACTAACCAAACTTTACTTTCTTTTGAAAAAGATATTTCTTTTACGTGTAGGCTTCAATTTAATAATGCTAGCTGCAAAAAAGCTTATTCTACAATGGTATTTTTGCTTACATATATCATCCCAATGTTATGTACTGGTATATCTTACTTCAGAATACATCGAGTAGTAACCAAAGCCGCTAATAACCTACAAGGTCATGTTTCCAAAGATGTATTTGTTTTAAGGAAACGTAGCTCCAAGCGATTGGATTACACTTTGATAATTATGTATACTTTTTTTGGAACAACTACCTTACCTCTTCAAGCTCTGTACCTAGTAACTGGTATAACCACAAGTCATGGAGGAGAGATACCCGAATTTGTATGGCCTATTTGTGCAACACTTTTTTATTTGCAAGTTGTTTTAAACCCTCTTGTACTTTTTTACATGGGAACAGAGTATAGGAAAGGTTTGTATAAGCTATCTTTGTATTTACCATGTCGCTATAATAATGTTATATGTCACTATAACAACTCTTTAAAAAGAATTAAAAAAAAGATAAGACTAGAAAGAGCAAAAAAAAAAGTGCATGCAAGATGTGACCATGGAAGATGTAGGCAAAAGTTTGGAAAGAAAAAACTCAGGAAATTGTTTTAATTTTAACATTTTAAGAAATCAAGTTAAGGTACTCAATAAGTTTGATAAAAAAACTGTATTCCGTCGTTATAAAGAACTCTCAGAAACTTCCTCAGACTTCATTGAAAAATTCGAAGTAACTTCGTTACTCAACAAAAATGAAGAGCACCCTCAGAATAAAGAAAATACAAGTACAATATGCCCAGTTTTCTATACAATGCAACATAATATAATGGAATATTTAATATCAGATCAAAGAAATGACAACTATTCAATATTATATCTAATAAACGGCAATTTAAGTTTTTATGCCACTGGTGAAAATGTAAATGAACATTTGGCAACATACAAATACTTTGTGGACCCCTCAACAGTTATTCTAAACTACTGCAATACAAACAAAGGTAAAGCAAACAATTACGATACAAACTTGAAGCGATATAACAAAACGGATTGTTTTGTTTGCATGAATTTAATTGAATTAGAAAATTCTCCTAATATTGTATTAAGCTGTTATCATGGTGTTTCATTTGATTTACTGAAGGAGACATATTTATGAACGC
>evgR015038c0g1_i01_Hydra
ACAACATGGTCAAGATCCTCATTCTCTTCTACTCAACCTACGGTCACATCTACCAAATGGCTGAAGCAGTCGCAGAAGGTTGCAGACAAATCGAAGGAGCTGAAGTTTCTCTCAAACGAGTTCCAGAAACTTTACCAGTCGAAGTTTTAGAAAAGATGAATGCAGTAGAAGCTCAAAAAGCTTTCGCTCACATTCCAATTGCAACAGTTCAAGAATTAGCTCAATACGATGCAATCATTCTTGGCTCTCCAACAAGATTTGGAACCATGACAGGTCAATTAAAGAATTTCTTAGATGCCACTGGTCAATTATGGTCATCAGGTGCTCTCGTTGGTAAAGTTGGATCTTTCTTCACGTCAAGTGCCACTCAACATGGTGGACAAGAAGCAACCATTCTCTCTTCAATTCCAGTTTTTCTCCATCATGGAATGATCTACTGTGGTCTACCATATAGTTGTTCTGGACAGACTGGACTTGATGAGATTAAAGGTGGTTCTCCATATGGTGTGAGTACAATTGCAGGTGGAAAGGGAGAACGATTGCCATCTGATCAAGAAAAGGATATGGCAAGATTTTTGGGTAAACATGTGACTGGAATTGCAAAGAAATTGGCTCTATAGAAATATATTGTTTCCAAAGAAAGGAACCCAGTGTTGAGTTATGAATAAATACGAAACTATACTAAATTTACAATCTCAAAAAAAAA
>evgR015039c0g1_i01_Hydra
TTCATGTTTTAAAATCAAATGTTATTCTGTTATTTTGATATAAAAAAAAACTTGTAAAAAGAAACCTAATATTTAAATGAATGTTCCAGTATGGTTGCTAGTGATCTTGGTATGTCTGAGTCTAATACTCATTACTGTGTGTTTGGTTGTTGGTTTCAATGTAACAAAAAATTGGCTACATATGCAAAAGTACGAACGCATTCGTTCAAAAAGATGTAAAAATCACGCCCATCTTAAAGCGACATCCTTTTTATCCGGATACGAGAATACTTATAAGTTAAATGCTTATCAAAATTTTTCGTTTCTTCCTATAACTTCGACAAGCGTCATACAAAGAGACAACTCTCAAATATTGTTAGAATCAAGACAATCTACAAACACTGATTCCACCTTACATGCAAAAATTCCCCCTTTTAACAAGTTTGAAAGAAGACTACTTCTGAGTCGCGCTGAAAGTTTTGATGAGCGCTCATTTCAAAGACTTCTTCATAAACAGCATGTATTCAGATCATCAGCGACTAACCTACGTCAAATAGAAAGTCCAAGTTCTTATCGTAGTTTTGATGGAAGTTCTGATAGTAGAGGAAATAGTTTAACAGTAAAACACCATGCTATAAATGATTTTGACACTTCAGAACAAACCGAAACAGCTTATCTTAATAACGATTACTTGACAGAAAGCGAAGAAATTCTTAAAAATGAACCACTTATAAGCTATGATCAAGATATTTCTACTGTATTTAAGCCTATAGAAAGAGGATCTCTTACATTTTTACTTGAGTATGACTCAAAATTAGCAGATCTTTTAATTACTGTTAAAAATGCAATCGACTTACCTTCTGCTAGCGGCGGTGAAAAAGATCGAGTCAACAGCTATGTTAACTTATGTATTGTTCCTGAAGACTTTCTATGGAAAAGAACGGATGTTGTACCTAATGATTGTTGTCCAATATACAACAAAACGTTTCGAATCACTGATGTACTATATCACAAACTTCGAGAATATACTTTATGCTTCTATGTCATGGATGTCCACATAACCATGGGTGAAAGAGTTGTTGGTAAGGTTTTATATCCTCTTTCCGATCTTAGAATGGAGCAGATTGTTGAAGTACAAAAAGAGCTTTCATTACCTTAATTATTACAAGGAGAACTACAAATAGAATTACAAATAGAATTACAAATTGAATTACAAATTAATCAAGATAATATAGAGAAAAAATATGGTTTATAAAAATTATTTTTTAGGAGCGATCTAGCAAATAACATATTAGAATCGTTTTCACTTTTCAAGTGTATACATTGTATAAAAATTATATAAATGAGCTAATAACTGTGAATAAGAAATGTTTAAAAAAAAA
>evgR015041c0g1_i01_Hydra
AAAATAACTCCTAATTTATGGTTTTAAAGCTCGAGGGGAGCTATATTTAAACAACTTTTGTCGATGTGAAAATGGTTGTGAAGGAAACAGGAACTCTCGCAGCGTATTGTCGAAATGTTTCTATAAAAACATTCTTAGGGTAGTTTTGTGAAACCAAAGCATTTATAGCGCGACTGTTTTTATTATTACTAGAAAAATAATCTCCAAAAAAAAAACTGATCGATGGTTTCCTTTAACCAAATGCAGTTCGCCGATAGCACCTTATACTAGGCTATATTTAATGATTATTTATAGACTTTATATGCTGATATTCTTATTAGAAATTGCTCAAATCATATATTTGTCAACTACATTTGATTAAATTGTGTGCATTAGTTTATAAAGCGTTACATAATAAAAAGGACATAACTTTATATTGAGCATTGAAATTTATGTAGTACAATAATGTTATGATTTAGTAATTGAAACTTTGCCGTTTTTTATTTAATTTCATGGTCGATATAATTGCTATTAGCTTGGCATCATTGTTGCAATATGAACAGTGTTATATGTAGTTCTATTATTACGAAAATTATGGTGTTTTTTGTTAATTATGCACTAGCTACTGCTACACCTGATTCTGAGATTCCAAATTATTTCGTAATGTGTCAAGAGTTATGTTCATGTTTTAATACTACTATTGACTGTTCAAATAGAAATATTTCAATATTGCCAAAATTTATTAGTCCAAACACTACTATACTTGATATAAGCCGCAACAAAATAAGAAGTCTTCCTTTGAGCTTTTTTGATAATCTATTGAATTTAGAAGTCTTAAATGTCAGTTATAATGAACTTGAGGAGCTGAGCGCGTCTACAAATAAAAACGAGCAATCTGAAATAAAACTAACAAAACTTCATACAATGGATTTGAGCCATAATAAGATTCAAAATATAAAAATGTACTGGTCAAGGTTCAGTGCACTAAAAAAACTAGTTCTATCTTACAACTTTTTTATGGAGGTCACAGCAGTTACCTTTTTAAATCTTAAAAGTTTGGAACAGCTGTTTTTAAATCACAATCAATTGAGATCTCTAACCAGATCATGGTTTATAGATGAATCTAATTTAAGACATCTGCATTTAAAGAATAATGCCATAAAGTACATTGAAGCAAGCTCATTCGTGTTTACCCCTGAAGTCACATTAATTGATTTTTCAGGAAACTCCCTCATTCATCTTGACTCTGAAATGTTTCAAGAGCTAGAAAAGTTATCTAATCTTTATTTAGACTGCAATGCTATTACCTACATATCAGAGTTTGCGTTCAGTAGGCTTGATCAATTACTAGTACTTAACTTGAGCTCTAATCTTATTTCTATTGGAGATAAAGAAATGGATACTGCATTCCTAAATCTTAAACATTTAGAGATATTGGATTTAAGCAAAAATAGTGCGTTATTAATACCAAGTACAGCATTTCAAGGATTAGGCAATTTAAAAACACTATATCTGAAGCATAACAATATCATGGCTATAGAAGAAGATGCTTTTGCAAACTTAAAAAATATTGAGTTGCTTGAACTATCTACACAATCATTATCATGTGGCTGTTCAGTCAAATGGTTACGTAAATGGTTACATTTGCATCCATCAAACAACAAGATCAATGTTGTATGTTCTTCACCGCTTTGGTTAAAAAATAAGAATGTTGTTGATGTTTTAGATTCTGAAATAAGCTGTGATTCCAACGAGATATCAATTATTAGACCTAAGCTAACAACTCAGCCATCTTCTACAAATGCAATCAAGTCTGGTAATATTAGTCTAACGTGTGCCGCTGTATCTTTATTCAATGACATGAACATAGTTTGGTTAAAAAATAATCAAGTTATTGATAACACTGTAAACTCAAATGCAATACAGACAGTAGACAAGATTCACTATATAACTACTTCTGTGCTTTTTCTGAAAAACCTAGATGATGCAGATGCCGGTACCTATTCATGCCGTCCAAAAAATATTTATGGAAGTGATATGTCTAATAAAGCTGATGTATCTGTTGTTGAACCTCCTCGTTTATCAAAGGATTTAAAATCTCAAATTACGGCATTTAGTGGTGACACAATTTATATTCCATGCATAGCAGAGGGCCAACCAATGCCAACTGTTTCACTTAAAAAATTAAATTATCCATCTATGCATGCTGTACAAGATAAGCGTGTTATATACAAAGAACAGAAGTTTGGAATAACGAATGTAAAAATTGAAGATAGTGGCATATACTCGTGTTCTGCTATAAATGCAGCTGGGGTTGCCAATAAAACAATTGAACTTATTGTTTTAGAAATGCCTAAATTTACACATCCTATGGAGAGCAAAGAAGTGAGGTCAGGTGAAACAGTTGTTCTTGATTGCATCGTTAGTGGTAATCCTTTACCTAGACTTAAGTGGTATAAGAATGATACTTTATTAAGCCAGGGTTCGTCATCGAAGATCAAAGTTTCTAATCAACTGCTCATTATTTTTTCTTTTGATTATGAAGATGAAGGGTTTTATGAGTGTAAAGCTTCAAATAAACTGGGAACAGCTAGTAGGACCGCTCGCTTAACAGTTCAAAACAAAACTTTACTAGCATCATCTCCTAAAAAATCATCATACTTTACGGTCGTCATCGTTACAGTTGTTTCTGTTGTTGTTTTATTTATATTTTTGATGTTATTATATTTCTTTAAAAAGTTATGTCGCAAAAATAAAGAAGAACTTACTGACAGTAAATTACTTGATAGCTTGCCGAAAACTTACACTCTTCGTTATTCCAAAAGTCCGTCCGATGATCCTGGGTCTAATGATTTTTTAAAAGAAAGCGTGCCTTATCTAGGTGGTTCTAAAGATGATAGCGATGATTGGACCCTAGACAGCGCAACTGACCCAGCGGAAGATGATAGTGGAATATCTTTAGTATATCCGAAAGACAATTATTCTGGAAATACGCCACGAGTTCACACATTTTCTGTTGAAAAGGTGAATAGAGAGCTGCACGATAAAAGCATTTCGTGTAATAATACATTAAAAAGTTCTCATAGAAAGCAGTCAAATTGCTATCTTTTATTTAATTCGTTAGACTCTTCACATAACGAAAAAACCTCATACATATTTCAAAAAGCTTTAAGAAATAATAACACTTCGCTTGTACATAAAGACGCTACTCATAAAGACTTTTGTAATTTAGATTCGTCATTGTTGAGCTACGATAAAAAATTTAATCATGTTGCTCCGAAAGTTAAAAATTATAAAGATAAACATGTAGCTTCATCACGAACAGAGCCTGATTATTTATCTAATAATATTATTTTATAGTTCGCGCGTTTTTTAATTTCAAAGTTACAGTTCATTAAATTATTTTGTAGATTAGTCTCGGTTAGGAAAGCAAATTATAATAATTTACCGTTTTACCGATAACCGGTTTTTCATTGTTTGAAAATGGGAACCATAAAAGAGTAACTAGCTAGTGATTGCATCCATTCTTCGTTCATTCATTTTACATTGTTAATTTATTTTGAGCACAAATTTTTTTATGTATTTTTTATTTATTTATATGTAAATTTATTTTGGGCTATTTTCAAGTCTTTGTAAAGGTGTAAATTGAAAGTGCGTGGTATTCGTAATTGGTAGCCTTTGTGTGCACCCTTGTGGTGTATTATGGCTACTGTTGTTACGCGATTCCTTTCACTATTAATGTACCTCCTTGTGGAGGTTGTATATTATACTTTGTTTATATTTCTAATTTATTTTAAATAAAATTTTTATATATACATTACAAAAATATATTTCTAGGCGTCATTTTAAATAAAAATTTACCAGAAAAAAAAAAA
>evgR015047c0g1_i02_Hydra
AGAAAGTTTTAAACGTGCGTAAAAGTAATGAAAGTATTTATGTGATTCACGTGTATCTTTTTGAACACTTTATTGTCTTTAGTCTTAATGAACAGTAATGTCTTTTTAAACATTAATATTTGTTCAGCTGTTGAATTTGGTTCTACTGTCTTCTAACCTTGAAAGTAATTATGATTCCAGTATTGTATTCATATTTTAGAAGTAGTTGCTCTTGGAGAGTAAGAATCGCACTAGCGCTGAAAGGTGTGGAATATGAATACAAAGCAATCAATCTATTAAAGGGAGAGCAGAAAAGCGATGATTATCGAAAAATAAACTCTCAAGGCTTTGTTCCCATTCTAGTTATTGACAACCGCAAGTTATTTCAGTCTCTTTCCATTATGGAATATTTAGACGAAGTTTATCCTGAAGTTCACCCTCTTTTACCTCGCAATGATCCTGTTAAGCGCTCAGAAGTTCGAGCAATGACATTATTAATTGTGGCTGATATACAACCTGTGCAAAACTTGTCTGTTTTGAACTTTGTGGGTGATGAAAAGAAAACGGAATTAGGTCATTGGGTAATCGAGAAAGGATTTAAAGATTTAGAAAAGTTGTTAAAAACACATTCTGGCAAGTACTGCTATGGTGATGAGATAACCATGGTTGACTTGTGTTTGGTGCCACAAGTTTATAATGCAATCAGATTCAAAGTGGATATGAGTCAGTTTCCAATCATATCTAGAGTAAACGAAGAACTAAGTCATCATCCTGCTTTTAAAGTTGCTCATTGTTTAAATCAGCCAGACTGCCCTACTGAGCTCAAAGTTTGATTTTTTCAAATGAAACATAAATTTTGTATTAACAATTTTTTTAAACTTTAAATAACAATTTTTGTAAAAAAAAAAAA
>evgR015052c0g1_i01_Hydra
TTATATCAGAATCGGAATCGTAAAGAAATTGAAATCGCAAAAAATAGAATTAATAAAAATTGTCACTGTCTTATAAAAGATACATTTTTTGAATAATCGAAATTATAAAAAGTTTGTTGCAAAAATGGCGGCAAATGGCAATGTGAAAATAGTTTCTAATGATTTAAGTTTTTCATCCGAATCGCAAAATGCTGGATCAAAGTTAGTTATAATAGATTTTCATGCTACATGGTGTGGTCCATGTAAAAATATTGCACCTGTGTTTGTACAAGAAAGTCTTAAGTATCCGAATAGTGTGTTTCTCAAAGTAGATGTAGATGTTTGCACGGAAACTGCTCAAAAGTATGGCATAAATGCCATGCCAACATTTGTATTTGTACTGCAAGGTAAAAAAATCGACGAATTAAAAGGTGCAGATCCCAAAGCTTTAATTGCAAAAATTAAACAGCATTCTGGCGACTCTATAGTTTCTGGTAGTAGTTCTTCTGGAGAATCAGAGGTGCCTGGTCACTCTGATCTCATAAGCTACATAACAGCGTCAGGATGCAATTGTTTAAATGAGAGCGATGAACACACTCATGCAAACGTATTTAAAAAAGATAACTCGTACCTTGAAAGTGACTGTGATGAGCAATTACTGTTGTCAATTACTTTTAATCAGGCAATGAAAGTCCATTCTCTTAAAATTGATGCACCCACTGATGGCAGAGGCCCAAAATTTATAAAGTTATTTGTTAACCAACCAAATGCAATAGGATTCGATCAAGCCGAAAGTATGGAAGGCGTTCAGCACATTGAATTGACAGAAGAAGATATTACAGGAGAGAAATTAATTCCACTTCGATTTGTTAAATTCCAAAATGTTACCAATTTTGTGATATTTTTTGCAAGTAACCAAGGTAGTGAAGAAACAACTGTGATTAATTACTTAAAAATCATTGGAAGTCCAGTTGAATCAACAAATATGAATGACTTCAAACGAATTGCTGGCAAAGCTGGAGAAAGTCACTAAAGCATTTGTTATATTGCTCATTAAATATAAGTTGCTGCATATTTATTGTTATTCCTTTATACCTATATATAAAAAATATAATTAGCAATAATAAAAAGAACATAAAACTTCTTTTAAAAATTGTATAAATGATATTTTTATACAATTTATACATAGTTTTTTCAAAGAATGATTTTCCAAAAAAAAAA
>evgR015053c0g1_i01_Hydra
TTTCTAAATAAGATTTCAATAATAAAGTTCTGAATATTATGTTGTGTCATTTGAGAGGTTTGAAGCAACTAACATTACTTTGTCTAAAAAGAAGATTAGCTACCGAGTCTTTTAAAATATATACACGCACTGGAGATAAAGGCACATCTTCATTATTTGGAGGGGAAAGAAGAACAAAAGATGATCAATTATTTGAAGCGATGGGTTGCACTGATGAGCTCACATCATCTATTGGATTAGCAAAAGAATTTTGTCTTGAATATGGTCATTTGGAAGTTGTTCAAAAATTAGAAGAGATACAGTGTCGACTTCAAGATATTAATTCTAACATTGCAACACCAGCTCAAGTATCCATTAATAAAGATATGTTAAATTTTGATCCAACCAATATTGACCAACTTGAGTCCTGGATAGATGAGTATGACAAGCATTTACCGCCACTAAAGAATTTTATTCTTCCTTCAGGAGGCAAAAGCAGTGTTTCTCTTCATTTGGCAAGAACTATTTGTCGTAGAGCTGAAAGACGACTCGTTCCACTTCTTCATAAAAATATTATAGATGAAAGTGTTTACAAGTATATGAACAGACTAAGCGACTTTTTGTTTATCGCTGCACGATTTGCTGCCCACAAGGACGGGAAAAATGAGCTCGTTTATAAAAGAAGAACGAAAGTTATTGAAAAAAGGTAGATACGGATGAGGATTGGATGGCTTCACATATTTTACCATTTTTCAAGACTGGTTGCATTTGTAATAAAAAAAATTGAGACAAGTGTTACTTAAAGAAACTTAAGATCAAACGCAAACAGGTGATATTTGGAGAAACAGAACGCTCCTATAAACAAATGTATTTTTAGATTTAATTAATATAATTTTTAATTTTTTTTAAAAAAAAAAAA
>evgR015055c0g1_i01_Hydra
CATATGGCATATGATACTGTCATACATTTTGTTTATTGTTTGCGTTGTCTTGTATCTTTATGATTTTATTTTATTTTTTTCTATTTTTTTTCCTTTGATTGTGTTTGTTCATATGTTAATCTATGTAAGTGTTGTTTGTGACTTTATCTGCATCCCATCTATTATTTATCTTTTATCTTTATGTCTATATTCATCTGCATTTTATATTTATGGTCTTTACTCTTTCTATTCTTGTGCGTTAACTTTATTTATAGTGTCTTCTTTTGTATCTTGTTTGTATATTTCTTTGTTTATATTCCATTAATTAGCTTTTATTTTTTTAAATGAAAT
>evgR015056c0g1_i01_Hydra
TAATCACAACGAGAAAAAGAGTGAGATATCTACTGAATCAAAACATGTGCGGTTAAAGAATTATAAAATAACTAAGCATAATGGTTCAACAGTAGATGGGCCAGGCGATGATTTTTTTGATCTTTTGTATAAATCACAAAGTTCACGGCTGGATGACCAACGTTGTGATTTTCCTAAAGAAAGATCAGATTTTGTGACTAAACCACTAGTTCAAAAGGCTCTAAGCGATAGCAATAGCAGTTTTGGAGACCGCGTGGAGTTGCAAAGAAATTCACTTCCTGATTTACCAGGACTCTCAGAAAACGCTAAAGAGAAACTGGTTGGAAAATGCGTTAATGGAATACCTGATGAGAAGTTTCTTGAAATGCTTATGCGTTGTCAGGCTGCACGTATCGACGACCAGCGCTGTGAGTTTCCTGTCACGCCACGTGGACCAACTGTGCCGGATGAAGATTTTTTTAATCTTATTGCTCGGTTGCAGTCTGGAAGAATGGATTCCCAGCGCGCAAATTTTAACCCATTTTAATAAATATTTCAGTAAAAATCTTTTTTTACCATTTTTTTTAGAAAAGCTTTTTTTAACAAAATTCTCTTCATACGTCTCATACTCTACTCTCTCATACTTTCGATGCTTTTTAACGTTATACTTGTTCTGTAATATAAACTTGTTCTGTAATATAAACTTTCTACGTGGGCGGTGCTTTCCAAATATGTACATTAGTTGAATGATTTTTTTTTAGTGTAAATAATGCACAACGTTCTTTGTTTTCTAATTTTATTATTTTATTTTTATATACTGCAGTTTTATTTATTGCAAATAACGAAATAGTATATAATGGACTTCTTTTTCTCGCTTTTTGTTTGTTATTATTGATTTTATATAAACGGTTACTTTATTTTTTGGCTACAAAAAATAGAAATAGAGTTTGAATTTTGGAAAAAAAAAA
>evgR015057c0g1_i01_Hydra
TATTTTAGTGCAAAAAGGCCTCCATACATATACATTATGACCAAGCAAAAAAGATCAATGTCCAGCGATGTGAGAAGATCAAATTTGGTTTTTAAAAATTACACACAACCTGATTGTTTTATTTGTGATACCAGTGAGTCAAGAGACTCACTGCACAGTGTCTGTACAACGGATCTGGATAATGAAGTCCGCGGTTGTGCTATTACCTTGGTTGACTCTACATTATTGACCAAGTCGAGCAACACAGATTTAATCGCTCAAGAGACTAAGTATTATACTAAGTGTCTGAATGACCTGTACAACAAAAAAAGTGCTTTTGATAAACGCATGCGTGCTGAAACGCCAGAAGTAGACGATGAAGTTAACTGTGTACATGGTATTGTTTTTGCTCAGTTGATAGCATACATTGAAAAAACAAGTTCAGAATGGTATTACACCCATCTTCAAACTAGCTGATCTAGCAAGACTATACTTTGATCGCTTGTCCCAAATAATTGAAAATAAACTACATACACCCTCAGCGTATAGCTGATACAAAGCGTAAAGCTACATACAACTCAGCGTATCAATAGCACACGTTTGAAAGAAAGAGTACAAGCCCATTTTTCTAACATGCATGCACAACAGGACGGTCGAGATGTTTAACTTGTTTTTAATTGTGACATTGGTCCAGCACTTCTTTAAGCCTATCAATATGACTGCGATCGTGATGCAGTGTGTTTGGCAAAGGCTGCGGAGATTATTTGCCAAACTATATTTGATACCGCTTAACCTTTTACAGGTACATTTCAGCCATGGTGTCAGGAAAGGTCAGTATCAAAAAACCTCCTTGCACTCGGTAATATGATAATAGAAGGTCCCATCATTGAAATACAAACCAAGCCAAATAACACTCCTGCTGCATTGACTATAAGCCAGCGACTTACTTTTTAGTGTGAAACAAAAGCAATCAGAAGTGCAGTAGTGACACACAAGCGATCCAGTAGCAAAACTATGGGGCCGCAACTGCGGGAG
>evgR015060c0g1_i01_Hydra
GTTAAATAAATAATATAGTTATCACTTTGTGAAAGATATATTTGATTGACATTAATTGTTATTAATTTTTAGTAATTATTTCAATATTTATGTAAATAATGAGCTCTTTTAGATTCAACTGTGTAGGTTTTATTCCCGATGAAATATCTGGTATAAATGAAGAACGCAGCGATAAAGTTGATTTGGCTAAACATGAAAACGAAGTTAAGTTTTTGCAACTCAATAAAATGCAAGATGGATTTGTGTTCAATAAATCATTTCAAGATGATGGGCATGATAAAATTCGCGAACTCCTTGTGTTAAAGTATAAACTGCGCAAATCAATACTTCAGATTAAAAAACTTAAAAACAAAAACGAAAAGCTTCACTATAAATTCAATAAGCAGCTAGATCACGAACTTAAAATTAAAACTCACATTTGTCAATTAGAAACAAAACCAACCTTTGGTACTTCATTTTATAAAGATAATAATTTTGAAGAACTACAAATTGGTATCAAAAGTTCAAAATCTAATATTGATGAAAGTGATGATAAAAATCGCATTAAATATTGCTGTTCAAACAATAAATGTAATGAGGTTAATGAGCGGGAGCAAACTCGTGTTGACTTAAATGTAAATCTTTTAGATACGTCAAACCAAGTTGACTTGTTATTAGCTGTGGGCGGTCATCGACATTTAACAAATATTGAGACTGGTACCGTCATTAAAACAACGCACGAGAATACAACTGAGAGTGATGAAACCATTGAAGTTATACAAAATACTAATAAAGAAAAACTTAGAAAAAAAAAACAAAAAAAAGTAATTCCTTTTTCTAAAGTTGTTCTACAAAAAAATTTCAGCAGGTTTTGCTTTCTCAAATTGAAAGAAACCAAAGAAAAACAAATAAAGCAATTAAATATCTACAGAAGTAATTTAGATTTCGTCAAAAAACGCTTCGCCAATAAATTGTTGAAACAAATCGAGTTTTTAGACAGTTTAAAATGTGATCTCAATAAGTACAACGAAAATTTAAAAAAGAAAAACTTTTTTCAAAACAAGTCAAAAATTCAAGGAGCTTTCATAAACAAAATATTTCAAAAGATTAGTCTTTACCGAAATAAATTTTCAATGATTTAATTTAACGATACTAAAAGCATTAGATATAGAAAAAAAATTTATAGTCTTGTAATTAGTTATAAAGAATGATTGTAATCAATAAGTTGAAGGCTTAGCGATATACACGACCGCAGAAAGGCGGGTGGACGATATAAGTGATGAGGTCAAAAATAATTAAAAAAAAAATTTGATTCAATAAAAAAAAAA
>evgR015062c0g1_i01_Hydra
AGCGACTTTAAAATAAATGTTTTGCCAAATTTAAAAAAGATTTTTAGTTAAGTTTTTTTTTAATTATTTACTTCGTATATATAAAGTCTTGAGTTATTTAAACCATACATTTGAAATGTCTTCCCCTTCACCTGGAAAAAGACGAATGGATACTGATGTTGTAAAATTAATTGAAAGCAACCATGACGTTACAATTCTTGGTGGCTTAAATGAGTTTGTTGTTAAATTTTTTGGACCAAAAGAAACACCATATGAGGGTGGAATATGGAAGATTCGAGTTGATCTACCTGATAAATACCCATTCAAATCTCCTTCTATTGGTTTTCTGAATAAAATATTCCATCCAAATATTGATGAAGCGTCAGGGTCAGTTTGTCTTGATGTAATTAATCAAGCATGGACAGCATTATATGATTTAAATAATGTTTTCGAATCATTTTTGCCGCAATTGCTGGCCTACCCAAATCCTATCGATCCACTAAATGGCGATGCTGCGGCGTTGTATCTTCACAAACCCGAAGATTATAAAAAAAAAATAAAAGAATATGTAAAAAAATACGCTTCGCCTGAAGTTATCGCTGAACAAGAAGAAGCAAATATGTCTGATTCTTCTGAAAGTTCTCTAAGTGAATTTTCTGAAGATGAAACAGCTGACATGGAGTTATAGCGTGACGTTTGTAATTATATTATTTTTGTATATTTTTGAGTATTTTAAAGCTTATTATTGCCAATCTTTTTTAATTGACTTTTGGCACTCGCAAATAAAAGAAAAGCGTTTGATTGTAAAAAGTATAATTATTGTAATTCTGCTAAATTTTCCAGTCTTATGTCAACTTTTTTTTTTTTATTTCAAATATTTTGTTTGTTTCATTGGTGTT
>evgR015065c0g1_i01_Hydra
TTTTTTTTAATCTTAAAATTAACTATATATCTAAATATGAGGTTACCAAATAATGACGATGAGAGCCACCCACTTTTCAAATTAAAGTAATCATTCAAAGAACGAGGGTTTTTTAAACAATTGAATGACAGCTTTATTATAAAAATACAGAATTTTTAAAACAACCAGAATCTAACGCGACATCAAATTTTTACTTGATAAGTTAAAATCGAATGGATGTAAACAGTTTTCGGGTAAATTGTTTTTTGCACCAAATGAAGTAAAGGGCCGTTGTAAAGGATTTCTTTCTTTTTTCGATTATTCATCTATCTTAACTAAAAGAACTTTAGCCATTACTGTTCAGCATAACCGACGTTTTATTGTTGACAAATCAATTAACATTTAAAGATTCCTAACACCCAAAGTGTACTTAAAGCAATTGTTGACAACCAATCAAACTTTAACAATTATATACAACCCAATCACTTTAGAAATTGTTCACAAACATATGACTTTTAATAATGTTTGATAGTGAATCGATTGTATTCTTGCTGTTTTGTGTTGTGGGATTCATTTTAAACGGAGTAGGATTGTACCTTTTACTTTTGATAAAGAGCTTCAAGGTTGACACCCAAAAGATTATAGTTGTTCACCTCTGTATTTTAGACTTTGGTTTTGGTATAGTTCTAACTATAGAAATTTTGTTAGGAACAAGAAATGGACTCAATAAACCAATTTCTATGAAACTTGATAAATTTCTTAAAATATGTCAGAGTATAATAAATGTTGGTTATTATTACGCAACATTTTGGCTAATAGTTGATCGCTTCTTGCATATAAAACTTAATTTAAAGTATCAACTCTATTGGACAAAAAAGAAAACAATAATCGGAATAGTAATTTTATGGAGCTTATCGACTTTAATTGGATGTCTTATTGGAGAATATATACAGAATTTGAACGTTCTTTTTTTTGCTTGTATAGACTACATAATATTAATTTTTTCTATATTTGTGTACACTTACACAATTTTATTATTAAAACGGCAAAGAATTAGAGTCCGCTCCAGCCACTACCGAAAAAGCGTAATTAAAGGATCACTGGTATCTGTCGCTATTTTATCGACCTTTTTTATTTTGGTTGCAGTACCTGACACATTATTAGCAATTAAAAAATACTGTAGCATAGAATCAAAGGAACTAAATATTTTTTTAAATTTAGTCTATCCTGTTGCTCTTTGGCCTAACGCTCCAATATACATACTTTTTTCACCCGAAGTCCGACTTTTTTTCAGAAAAAAAAATAACAAACCGTTTAATCGAATAATAATAAGAAATAATCAAATATAGTATATATGTATTAATAAGAATAATCAAAAGCAAC
>evgR015065c1g1_i01_Hydra
ACTGTTGACACTGTTACCACGACACTGTTACCATGACACTGTTAACGCATCTACATTTACTAAGGAGCTTAACGTGTTTTTATTTAATGTACCAGAGATGTACCAGAGATGTACCAGATGTACCAGAGATGTACCAGAGATGTTTTAGTTAAAAAGGAAGTATTTTTATAAATATTAGAATAATGGCTATGCTTTATTCATCAGTTCTTATAACTTATACAATTTTAGCAATTGACAATCAAACAACATATAGTATTCATAGAAATATCTTGCTTCTTGTACCGAATATCTTACCGAGTACCGAATATCTTAAGCGTAATTGTTAACAACCAATTAACTGTAGAACTTGTTAACACACAAATAATAGTTTAAATTTAAAAAACTGCAATGTTCGATGATGAAACAATAGTGTTCTTGCTATTCTGTGCGGTTGGATTTGTTTTAAATGGAATAGGATTGTATCTTTTACTTATGATAAAGAGCTCTAGAATGGACACCCAAAGAGTCATTGTTGTCAACCTCTGTATTTTAGACTTTAGTCTAGGTATTATTCTTATAATAGGAACTGTTTTAGGAACAAGAAATGGACCTAATGAAGTTGTGTCCTTCAAAGTTAAGAAATTTATGTTAATAACTCAAAGTATTGTAAACATTGGTTTTTATTACGCAACATTTTGGCTAATACTTGATCGCTACTTACACATAAAACTCAATATAAAATACCAACTCTATTGGACAAAAAAGCAAACAATATTAGGAATAGTGATTTTATGGTTCTTGTCAACATTAAGTGGATGCCTTGTTGGAGGATATATATACAGTTTCGACGTCCTAATTTTTGCTTGTATAGACCACATAATATTGTTTTTTTCCATCTTTGTGTACACCTATACTGTATTATTATTAAAACGGCAAAGAATTAAAGTCCGCTCTAGCCACCACCAAAAAGGCGTATTTAAAGGATCACTTGTGTCTGCCGTTATCATAACAACGTTTGTTATTTTGGTTGCAATACCTGACACATTATTAGCAATTAAAAAATA
>evgR015066c0g1_i01_Hydra
TTTCCAATGTGATGGATTACCTTACAGCTATGAAGTTGAAAATAACAAAAAGGCATGTAATTTAGACTGTGAATCGTTTAACACTTCTACTTCAATAGCGTATGACAAAAGTAGTGTTAGTGGCGATAGTAGCACTACTGCTACGCTATCCAGTTGTTCGTTAAACACCCTAACTGAATTATTGATCAGTACTGTTTACATTCCTTCTGTTAGTGTTTCAACAACAACAAAAACTGTGATCACTCACGTCACCGAAACGGAAACAAAGTTGTGTTCTCAGTCAGTAATAGTAACTCAAGCTGCTAGAGCGGAGAAAATTTATGTTGACATAGAATATGATAAGAAATTTTACGTAACAGTCATAGCAACAATAGTTGGAGCTATTCTTGTAGGCATATTTGTGGCCCATGCTTGCGTGACCTATTTTACTGCTAAAAGAATAAAGAAACAAAATGTTTATTATATTAACGAAAAAGAAAATTTAGTTGACACGGAAGAAACGAATCTTCGCCGAGTAAGCATGACGTAAGCCTTGGTAAAGAAAATATTTTATTTCAAGAAGGGACTCGAATCGATTAAAATAAGAGCGTCAATTTCGTGGAGAATGTTTTTAATTATTTCTTTAAAAATGTAATGATATTTTTATATTTATGTTTATTTTTAAAAAAAAAAAAAA
>evgR015066c0g1_i02_Hydra
TATTAGACTGTGAACCGTTTAACACTTCTACTTCAATAACGTATGACAAAAGTAGTGTTAGTGACGATAGTAGCACTACTGCTACGCTATCCAGTTGTTCGTTAAACACCCTAACTGAATTATTGATTAGTACTGTTTACATTTCTTCTGTTATTGTTTCAACAATAACAAAAACTGTGATCACTAATGTCACCAAAACGGAAACAAAGTTGTGTTCTCAGTCAGTAATAGTAACTCAAGCTGCTAGAGCGGAGAAAATTTATGTTGACATAGAATATGATAAGAAATTTTACGTAACAGTCATAGCAACAATAGTTGGAGCTATTCTTGTAGGCATATTTGTGGCCCATGCTTGCGTGACCTATTTTACTGCTAAAAGAATAAAGAAACAAAATGTTTATTATATTAACGAAAAAGAAAATTTAGTTGACACGGAAGAAACGAATCTTCGCCGAGTAAGCATGACGTAAGCCTTGGTAAAGAAAATATTTTATTTCAAGAAGGAACTCGAATCGATTAAAATAAGAGTCTTTTTCGTGGAGAGTGTTTTTTTTTATTTCTTTAAAAATGTAATGATATTTTTATATTTATGTTTATTTTTAAAAAAAAAAAAAA
>evgR015073c0g1_i01_Hydra
TCCTACATAGAAAAGTTTGTTTGTTTTTAACTGTGTAATTAAACACTATTTCGACTTCAAAATTATGAACCTACAGTGTAAGATATCAAGTAAATAAACGCATTTAAATAATATTAGGAGAGTGTGAACATATTATGTTAGGCAATGTTGTAGCTACCAAACGTTTTTTCTACTGAGCTGTGTTATTTAATAAAATGTTCCGAGAAAGTTTTTACGAAGAAATTGATTCTGAATATTGTAAAAACAAATGTGAAAGTAATTGCAAAGAGATTTCTATTGATTTGTCTGACCCTATTTATATTGAAATGAACGCACCATACGAGGATTGTAATGAGCTAAAACATCAAGAAAATACAGTTGTAGCGAAAAATGAAATCACTTATAATACCTCAGCAGCCGACGAGATAAAAGATTCAATCGAAACAGTGGATAAACCATTTGTTGATATACCGTCTCGGAGTTCAACAAAGATCCATCATAAAAGTACCAGTTCTACTAGCTCAGTTTCTACATATTCAAGCAACTCAGAAGAGCAAAAACAAATTGATAACAATAAGGCAAACAAACTTAAACAAAATTTTGCAAGAAACACAAGCAGAGAAGACGAAACAGTTTATACAGAAATTGTTACTCGTAAAAAGACTACAGAGAAAACAAAATCTCGAATAACTAGGCAATCGGAGAATCTTATTCTAAGAGAAGATTTTGACGAAAATGCGGTAATTGTTAGGAGAGCATATTCTTTGCACAATTTAAGTTCCGGAGAAAATACATCAGCAATAAACAATTCTATGCAGCTATCTACCAATCATACATATAAAACTCGCCATAAAAACTTACAAGCAAGACATAAAAGAAGCAACACTTACCGAAAATCGTCGTATATTGAGGACGAACAAGATGTAATAAAATATGCCAATCTGCAATATTTTCCTCCAGATAAAAACACCCCGAATAAACAAAAACAAAAACGTATTTCAGTCTATAGCAACGTAAGTAACGTTTGCAAGTTATACGAAAATTTCAGCAATAACGAGGACACCGATGAGAGCGCAAATAAAAATGTCGGAGAACTGCCGGAAAACTACGACATATCGCGAAATAGATCAACTAGTCACCCCATAAACATTATTCAGTCGGAAAACAATGAACAACTCCATTACTCCCATATTGAGTTTAACGGGGTTTCTTATCCAACTAGTTCACCAATAAGTAGCATATGCAGTAGTCAATCAAGTGAAGTCTCCTCGTTTGCCTCAAACTCGTTGAGCCCTTCTCAAGTTAACCAGTATGGTAAAAGAGCCGAACGCTACTCTACATGGACAAACAGCTCCTCGTTTGGAGGATTAAACTATTATCCAGCTCATTACCTAGGAAATATAATTGTATCCAAAGTTACAAAAGAAAGTGTAGATGCTGCAGTAAGTACTGTAGCTCAAAATACAAACGTACTAGAAATGAAACCAGTTTATGTAGAGGTTACAAGTGAGTTTATTCGTTTTGGAACTGCGGTAACCTCCTGGGATCTATTAGTTTTTTTGGCAATTGACGAAGTTGTTAGTTTTGAAGTAGTTCTTAAAAATGCCTATTTTTTTGGGCTTATCGCTTGCAAAACAGGAAACGAAGCTCGATGTTACGTTCTACATACAGAAAAAGCAAACGAGATATACGATGCGATAATGAATATATTTCAAACGTCTCCTAAGGTCTCTAGTTCAAAAAGTCTTTATGATCTTCAATTTGGCTTAATACAACCCGAATTTCATCGTTTTCATTATATTGTAAAGATTGGAACTGTTGCTGTTAAAAAATGTTTTCCTAATGTAAATGAATACATAGAAATTGCTTTAGACAAAATAAACCCAAAAGATTATAGAAACGTAAGCCTGGAAGTTCTCGGTTCGAACATTTTAGTCATAGACTCAACTACAGGAGAAGAGCAAAGTCATTACATTTCTTGGATATTATCTTTAGGAATTTTTAGTAAGGACTACCGGTACTTTGGCTATGTATTAAGCAAATCCGAAACAAACGGAAAAACAAAGTGTTCTTGTCATGTATATCGTGCATCAAGGAATCACTCCGCTACGCAAGTTGTCGAAGCTATTTCGCTATCTTCTCAAAATACATATTCTCCAAAAGAAATTTCTCAATTCAGCCCCGTGTATAGTGATCACTATTCGACATTTTGCGGTTCATATACTTCTCTTAATTCTAACTCTACTTATCGTTCTTCAAATATATCAGCTGAGACATCCAATTCATCCAAAACAAACGAAAAAATAAAATGCGCTGGGGAAGATATTAGCTTAAGTGACCAAACACTAAATGATTACCCACTTTCGATTACTAACACGAAGTCGGAAGAGACATCTTTTACTGTTCCTGACCACCGATTAAAACCTCTTCAAAGATTAGATAGTGGAATCGGTGAGCAAAGTTCCTTCAAATCAATAGACTATTGCACAGAGCAAATAATGGCTTCACCTACAGAAAAGAAAATTAACAAAGACTGTAAAAAGAAATTTAAACCAACAAAATTAATGGACAAGTTTAGAAGAAGCTTGGTACTGCCAAGCGAAACTTCAATTAGTGATACAGAATCTACAAAAGAACCTAACCTTATAAATACAAAAGAAAATCTTTTGTACTCAAAACCTCAAGTTGTTGAATATAAGTTTCATGTCAAATATATATGTTCAACAATTATAAACCCGCC
>evgR015075c0g1_i01_Hydra
CCAGATTGTGGTTATGCTGTAATTGCCTCAGGTTGTGCTGGTTGTCCTCAATTAAAATGTGAACGACCAGGCTGCAACTTTGAATTTTGCTATCATTGTCGTCAAGTATGGCATCCAAATACCACCTGTGATTTAGCCAGAATGCAAAATCAATTTGAAATACGAACATTATCATCTGTTGATCGCAAATCTTCAAATATGAATGAAGACATTAAGCCATGCCCTAATTGCATGGCCTTAATTATAAAAATGGATGATGGGTCATGTAATCATATGACTTGTGCAGTGTGTGGAGGAGAATTTTGTTGGTTATGTATGAAAGAAATATCCGAT
>evgR015078c0g1_i01_Hydra
AGTTAAAGCACATAAGGTACCTATGCCGTTTCGATGTGGTATTTGCAGTACCCCTTTTGAGCATCATATGGACGTACTAAATCACATCTCTATACATGAGGAAATCAACCTCGAGTTTAAATGCAGATACTGCAGCAAACATTTTCAAACACCAGATGCTCTTAGTAAGCACTACGTTCGTCATAAGTCAGTAAAGGTTATTCACGAATGTCGTTTTTGCAAGATTTCATTTGAATCTAACCAACAACTAATAGAGCATGTTGGTGGTGAACATATTGTTATTAATATATCCGAAAATGAAGGGAAATCTTCAGATTCCAACTTTGCTTACAAAACTGTTAGTACTAATTTAAAAAAGTTTTCATTTATGAGAAAATGCCTTAAAGATAAACACATTTTATCGAAGCATAAAGGTAAACATATATGCCAAAGATGTAAATTAATTTTTAAGTCAAGTAAATGTTTAGCAAAACACAAACTTAAATGTTCCAAGAAAAGTTTCGCATCTTTTTCTTGCGATGTTTGCGCAGAGTTTTTTATTTCAGATGCACTAAAAAACAAACACTTAAGAGTTCATCACAAAAGATTGAATGGTTTTCGCTGCATCAAATGTTTTAAGGTTTTCAAAACATGGATCCTCTTAAAGGATCACACTGAGCGTTTGCATACAAAAATATCATGCAATGATTGTGGCGAGACGTTTTTGAGAATGATTGATTTGCATAGCCACAACCTAATACTTCATAATAAACGCGAAAATGAAGCATATAAATCTAACAAACTTGGTTCAGAATCGCTTTCGTCGTCTAATTTAAACTGCAACAGTGATGTCAAGCATCCTATGTTTTTAGTTGCAACACCGAATGAGCAACGCAATTCTACCCCCCCCCACACACACACACACACACCTAAATCGGCAGATACAAATGTAACTTCATTGCGTAATCGAGAGCAACGAAATTTACGGAAAGTGGTTCAAAAAAGAGTTTTTAACAAAGACACAAAGTTCTCTTGCACAATTTGCGATAAGTTCACGAAAACCTCTACTAGTCTTAAGCTTCATTATGGTCGAGTCCATGGCGTTTGTTTGAAAAAGTCAACAGCAAAAATTTGAAACTCTTAAATGTTTTTAAATAGAAAAAAAAGATGTCGCAAACATATTTTTGTCAAAAATTGAATGTTTTGATTACGCAACATTAAAATCTTCGATGCAAAAAGTATTTAAACGGTGAATCCGAAGCCACTAAAAATCATTTATTAACTCTTCAAGCAACTCAAACCTTGGAAACGCCTTCCACAAACCTAAACGCCGTCTCCCTTAATTGTCCTAATGTGAAAGAGGTTTTCTTTGCAAGAAACGTTTAAAAACTATTAGCTAAGTAATAAAAAATATATTTACAATACTGGCCAACCATGGCTATTCCGACGAACTATTTCGGCATTCGGCTACCCGAAAGTCAAAACGACCTTAAAAACAAAGGAAGGATTGTTAACCATTTAAAGAATATAAAATGAAACACGTAAACAGTTTACACAACACAACCTTTTTTTAAGAATTTTCAGTTATATGCAAATATATGTTACTAAGATGTATTCCGTAATTTGTTTTGTAGCCAACATGGTGCTTATATCTAGCTTCTACTAGAAAGTACAAAAGTCTCATATTTTTTATATAAATTTGTATTTGTAACACTTATAAAGATTTGTCACATCATAATTTGAAAATCGTGCTTTTAAAAATCTCGATCAAGCGGGCAAAGCGGGATTAGGAAAATCACGGAGAATGAAACGATTGTAATTGAAAACGATTTACTTCATAATTTGAAGCTATTTGATGATGTAACTTTATAAAACATTAATTTGTATGCATGTTGATTTGTATATTAAACTATAATTTTTTTTATCATCCTACATGATATGCCTTATGATATATATTTTTGGTATTATTTTGAACCGACCGAAACCGAAGATATAAATTAAGCCGAAAACGGAACGATTTTCGGTCAATCAATATTCTACGCACAGATGCATTCTTTTATTCAAGCAAATCTTTTATTAAATATGTCTTTTATAATAAATTAACTTTATTTGTTGCGCCTTACTTGCAATACCCCGTGTAATGTCAAA
>evgR015099c0g1_i01_Hydra
GATGCCTTTTATATTAGTGAATTGTCTTTTTTCTAAACTTTTAGAAGATCCTTCGAAAAAGGGAAAGATAGAAGTTAGTTTTGACACGCACAATAACTTAAGCGTTAATATTAACATTCCTGCTTTGGATGTCACATCACTTGCTAACTATTTTGGAGCCAACGCGTTTGATAGCACAACTGACAAATTTCAAAATTCATTTAATCTTGCCATAAAAGACCTTTACATTGATGGAAAATTGAATTTTAAAGAAAAAACTGGACAGCTAACAGTTACTGCTTCCCCAGTGATAAATGGAATGACAGGTTTAACATTGGCTGCTTTTATATGGAAGGCAGAGGCCAACGATAATATTGGCATAACATTTGGTTTGGTAATGAAGAACATGCGTTTGGATAAAGCTTTAAAACAGTTTGCAGATATTGAAATGCCAGATTCATGGCTTAGCAATGTAACTATAGTAATGCTTGTCAGTACTGAAAACAAAGTAAATTGTAATGGAAGCAAATCGGTTAGACCAGAATGTAAGAAGGCGGACGAAACTCCCAGCACAACAGGCACAACTTCGACTCCAACAAGTCCTAACCAAGTTTCTGCCACATCTCGTGCCACCTCAAATCCTACAACTTCAAAACCTGCTACTTCAACCTCATCAGAAGAAAGTAAAAAAAACACTAAAAAAACACATTTTGACCTGATAGATGAATTGAGAGACATTGAAATAAAACAAGGATTTATATTAAGAGCAAATGCTTCATTGGGTACAAATGTTTCTAAATGTAATAGTAACGGTATATGTGAGTTTCTTGTCAAAAACAATGTTTCACATGCTTATCTCCACGGTAGTTTATTGTTTGATGCCCAAAAAAAAACCCTCTGTTACAATCAAAGCTGGTATACAAGGCGATATTAGATTAGGGAAATCAGATGATGTTGATCTC
>evgR015099c1g1_i01_Hydra
TTGCGGTGAAACTTGATAAAATCAAAGATATTGTAGATTATTTGAAAACAAAGTTTGGCTATAAAGGTCCTGATTTCATTAAAGAACTGTTTAAAAGTGCAACACTTTCTTTGTCCAACACATCCATTGATGTTATTGATGATGGAAGTTTTTTTGAGTTGAATGGAAAGCCTCAAAAATCAATTGAACCTGGTATTCATATCTATTTGGAAACAGACATGAAAACTGCTTTGCAAAATTGTGCTTCAAAAAAATCTCCTCAAGATACTAAAGCTCAGATGACTAGGGTTACTTCTAGTTCATCAGATAATGATTCTAGTTCATCAAATAATACCCTTATAAAAATAAGTTTATCAAGAGCTGGTGTTGTAATCCAATTACCAAAGAATTTTTCTCAAGACCTCAAGACATTAATGAATTGTTTTCTAAGAGAAATGAATGTTTACACCCAGACCGAGGTTGTTCCTGGTGTGCTAGATCAAACAAAACGGCTTGATGCCTTTTATATTAGTGAATTGTCTTTTTGTAAGTAGTTCTCTTTTATTTTTACATAATCAAAATGTAG
>evgR015100c0g1_i02_Hydra
CTTACAACAAAATTGGTACACACTTTTTATAAAACCAAAGACTTCTGTACGTCCTACGGCCTCACATTAAATTCGTCAAAAACACAGTTTATAGTATTTAAGAATCCTAGAAAGAAACTGCCGGATAATCTACGGATGGTAGTAGACGGCTATGAAATTAATCCAGTCAAATCAATTCAATTGTTAGGAGTAACTCTTGATTGGCATCTTACCTTTGGTCCTCATATAGATAATGTTAAAAAAAAATGTCATGGTATCTTGGGAGTTCTCTCTCGAGCCAGAAGATTCCTGTCTCGGGACATTCTGAGATTAGCTTATATATCATTAGTAAGGTCTCAGCTTGAATATTATAATGCTACTCTCGTCTCTGCGGCACCATCACAGTTGAAGAAACTAGACATAGTTCAAAAAATTGCATCACGAATTATCTGCGGTGCAAGTCGGGATACTCATTCAGCCCCTCTTCTTACGGAATTAAAACTGGACTCGCTAAATAATAGAAGAGAAAAGCATGTAGCGGAACTTGTTCAATCTGTTTTAACTAGAAATTGTCCTCCTGCTTTTTACGACCTATTTGGAGATAGTGCTGATGTGGCAACTCATAATGAATTCCGCTC
>evgR015102c0g1_i01_Hydra
TCGGCAATTATCCGTTTGTGCCCGGTCTTCTGCAAAATAAAAATGAAAACAAATCAGCCAATCAATGTTGTTCAGCAACCATCAAATAATTATTTCAAGCACTGCAACGTAACTTAATCATAGCGCTATGTAATATAGTTTGATTATTGTATTGAAATGAAACGAAGTCATGAAGATGCTGATATTTATAATGTGAGGCGATCATCTATGCCTGGATCTCATAGTCTAACTAGTATTTCAGCTTCACACCAAAATGCTGTTATAGTTGGGTCTCATCCTGCTTCAGGAATTTCTGGTTCTCATCAGTTACCTGGAACTCACCATCTCCCTAACATATCTGGCTCTCATCCTGCTGCTGGTTCTCATTCTACTTCAGCAAGTTTTTCTCATCAAACTCATCCTCCTACCCCTAAACAGTTGACACATAATTTTGAGCAAATGAGAGAGATGACGAGACCAGCATTTACTGGGTCTCATATGCCTGTACCGAATGTATCAAACTCTTCTCCTCCAGTAAATGGTTCTCATTCTTCATTAACTCAACCATCTGCTCATATACCACCAAAGCCTTCTTTTCAAACACATGTTCAAGGTCCATCTCATGTTGCTGGAGGACAACAGCAGTACCAGAGGTTGAAGGTGGAAGATGCATTATCATATTTAGATCAAGTAAAACTACAGTTTGGAAATAAGCCACAAGTTTACAATGAATTTTTGGATATTATGAAAGAGTTCAAGTCGCAGACTATCGATACACCTGGAGTAATTAGTCGGGTATCTGCACTTTTTAAAGGGCACCTAGACTTGATTGTTGGCTTTAATACCTTTTTGCCCCCTGGTTACAAGATTGAAGTAAGTTTGACAGATATTAATACAGTGAACGTGTGTGCTCCAACAGGTACTACAACTCACCTTATATCAGGCAGTGGTCTAATAGCAGCTCCTAGTCAACCAAACTTGAATAAGCCGCCTGTTGTACAGTCTATAAGTCATGTAAGTACTCTAGTTTCTACAGCTGTAACTACTACATTACCAAAATCACCAACTAGACCTATTTCTCTACCTTCAAAATCTTCTTCCCCTGTTTCATCTGCACCTGTAATGTCTTTACCATCCCCTGGTGTTCATTCTCAAACACAGGCTCAGCATCACGCTATTAATTATGTTACAAAAATAAAGAATCGATTTCAAGGTCAACCTGAAGTTTACCAAAAGTTTCTTGACATTCTCAATAATTATCAAACAGAGCAGAAAGCAATCAAGGAAAATACTGTTGATCAGTATGGTAGAGTGTGTCAAGAACAAGATATTTATGCTAGGGAGGGAAAACTTGCTACGGAAGTGTATCAACAAGTTGCTAAACTTTTCAACAAGCAAGAAGATTTACTGCAAGAATTCAGTCAGTTTTTACCTGATGCTACTGGGTCTACGTTTAATCCAAATATGGTATCTACAGTTGCAGCTGCCACTGTAGTTACGCAGTATCAACCATTATCTAAAGGAAAGATTACAACGTCTGCCGGAAATAATGCGACTAAAATATTAACTGTAAAAACTAAAAAATTAGGATCTCAAAACAATGTAAAACAAGAACAAATAGAAATTGATAAAGATGGTTCCGACAAAAATAAAAAGCGCCATAAATCAGCATTAAAAGATGTATCCCTTGAGGAGGCCGTTAAACATGGAAGTTTCAGTGATTTTGCATTTTTTGACAAAGTTAGAAAAGCTTTAAAAAGCCAAGAGGTTTATGAAAACTTTCTTCGATGCTTAACGCTTTTTAACAATGAAGTAATATCCAGGGCTGAATTAATTCAACTATCTACTACATTTCTTGGAAAATTTCCTGATTTGTTTAATTGGTTTAAGGCATTTCTTGGTTATAAGGACAATCAACAGCCTGAAACAAATATAACAGGATTTAAGGATAGAGGAACCGGTGGAGAATTGGCTCATTTAGAGATAGATTTTGCTTCATGCAAGAGAAGTGGTGCAAGTTATCGTGCTTTACCTAAGAGCTACACTCAGCCGAAGTGCTCTGGTCGAACGCAACTTTGTAAAGATCTTCTCAACGACACCTGGGTTTCACTCCCATCTTGGTCTGAAGATACTCAATTTCCTGGTACGCGAAAAACACAGTACGAAGAATATATTTATAAATGCGAAGATGAACGTTTTGAACTGGATATTGTTGTTGAAGCAAATCTATCAACAATCAAAGTATTAGAAGGTATACAAAAAAAACTGTCGCGAATGAATACTGATGATCAGCTGAAATTGCGTCTAGACAATTGTTTGGGGGGCACATCTGAGATCATTCATAGAAAAGCTATTTACAGAATATATGGTGACAAATCACCAGAAATCATAGATGGGTTGAAGAAAACACCCAATGTTGCTATTCCTGTGGTTTTAAAACGACTTAAGCAAAAAGATGAGGAATGGAGAATTGCACAAAAGCAGTTTAATCGAATATGGAGAGATCAGAATGAAAAATATTATCTTAAGTCATTGGATCATCAAGGTATAAACTTTAAACAAAACGATTTAAAAGCAATTCGATCAAAAAGTATTATTCAACATATCGAAATGTTATACGAAGAGCGAAGTGATCAAATAGATGAAGGAAGCACTGAATTATCTGGACCACATATCACGTTTGTTTATCAGGATCAATCGACACTTGAGGATGCATCTTCTTTAATAATCCACCACATGAAGCGTCAAACAAGCATTCACAAAGAAGACAAAACAAAGATTAAAGCTGTTCTACTTCATTTTTGTCAGAATTTGTTTTTTGCACCACGCGGTGATTTAAGTGATGACGAACAAGAAGAAGGTACAGAGAGTAAACATCTATCATATCAAAACGATGAATACAATTTGTTCTTTACAAACAGTGCATGGTTTATTATGTTGAGAATGCATCAGATGCTTTGTGAAAGGTTATTGAAGATGAATCAACTTGCTTCAGAAACTGCAGCATTTGAATCAAATGATAAAAAACATAGACGAGAGGGTACAGCAGTTGCTTTACATTTGAAGAAGATGCCAACAGCTGATATTGAGGAGTATTATCCAGGGTTTTTAGTAATGGTGAAAAATTTATTGGATGGTCACACAGACTCTAATTCATTTGAGGATTCACTGCGTGAAGCGTTTGGAGTAAACTCATATATTGCATTTACAATTGATAAACTAATACAGAATATTGTTCGTCAGCTTCACAGTCTTGTAACAGACGATATTAGCATGAAGCTTTATGCAATATATGAAGATGAACGTAGAAATGGTGCAGCAGGTGGCTCATCAATAACTGCCACTGCCCGATCTGGGTTGGAATTAGCTTACCATAAGAAAGTTGAAACTCAACTGTCAGATGAAAATTCTTTTAAAATATATTTTCACAAAGATAATGGGACATGCAAAGTCAGTATAGAACTTCTTCCTAATGAAGGTGATTTATCTGAAGATCAAGCTGAAGTTGAGAAATGGTCAGAATATGTTGAGAGGTTTATAGGTAACTCTGAAGTCATGTCTGAACTCAAAGAACATCTATGTAAAAAGCCTATATTCTTGTTAAGGAATTCACGGGTTTTGCAGCGTTTTTCAAAAAGACAAAACAAACACAAGAGGAAAAAGCGAGATGATTGTGAAGACAATGATGAGCCTAGTGAAAGCGTTTCATCTTCAACATCTGATGTAATTATGGACAACAGATTGGAGTGCAAAATCAGTCAAAATACTTATAAAATGCGTTATGTGTCGGGTACAAGAGATCTAATGTTTCGTAGAAGTACCGTTGGAAAATCTCTTAAGTCGCATCCTTCGATCAATATACGTTTGCATTCTATATATAAAACTTCCCATGAAGCATGGCTGCACGAAAACACTACCAGAGATCAGTGTCATCAGACAGAACTTTGGTTCATCGGCAAAGACGGTTCTGAAGATATGTCGCCGTGCATAACAACAAAAGATGTATTCAAACACGGAGGGATCAATATAAACAAATATACAGTAGTTTACACAAAAGATTCTAAAAATGGTTTTAAAAGTGCCTAAAACTGTAATAGCAACTATTTTAAACTATCGCGGACTGGATTTGATTGTTCAGAATTGATTAAACGATTTTATTATCTATAAAAGTAAGAACGTAACACACGTAAATCAAGGAATTCGCGACATAAAGACATAAAGTTGCTAAAAGAAATATTTTAGTGCGATACTCAACCTAAAGTTGCTAAAAGAAATATTTTAGTGCAATACTCAAC
>evgR015119c0g1_i01_Hydra
TGACTTTGATCTAATTGAAGATAAACTTTTTGGTTTTATTGAAAGGATTGAAACAGATTATGGCTTTTATTATCAATTTGCACATTTGACTATAATGGAGTTTTGTGTGTCTGTATATGCATATAATACTTTTAGTGGCAATGAGATTGTGGCCAATAATAAGTTGAGGAGTTGTTTATCAATGATTTTTGGATTGAGTAATAAAAGTGAAAATAATTTAATAAAGTTTCTTGATAACTTAAATAGATCCAACAAAAGGAATTTCACAAATAGTGAGATTTGTGGAATTCGTAGTGATATGATAACTCAAAAAGTTCTTGTTCAATTTATTGTTCAAAATATTGGTTCCGGATCAAACACATTTAATATTATTAATATTACACAAATGAACAAAAAAAAAAACCATTTCACGAATAGTGAGATTTGTGAAATTGATTTTCATAATTTTGATATTGATAAAAAGAATGAAAAAAAGATATCAATCAATCATATTTGTAAAAGAGGCCTCCAACTGTTTGAAGCAATCCTAACACCATTATCCCTGTTCAAAAGAG
>evgR015120c0g1_i01_Hydra
GCCATATATTGGTCGAAAAAGAAGCTAATAACTATATCAGCAATAATTTGGAGCATAGCAACGTTAAAAGGAATTTTTAATCATTTTTATGATATCAAGTTCAACCACGTTTTAGTTTTCTATGATCTAATAATAATTTCTTTCTCCGCGTTTGTCTACGCGTATGCTTTAATGCTTTATCGCAAACAAAAAACAGTGCTTTGTTCTAACCGAAAAGACAGGAAAATATTCAAAGGAGTAATAACATCGAGTCTAATAATAATACTATTTATTATTTTGTTTGCAATACCTGATTTAGTTCTTGTATTTGGATTAGATATAACTTTATCAAATAATGAATTGCGTTGGTTTGTTTACTTATCACTACCTGTATGTTTTTGGACAGATGCAGTCATCTACATATTTGCATCACCTAGTGTTAGACGTTTTATTAAAAGTAAATTTAAGTGTAGGATTTGCAAACAAAGTGTGTCAGCTGAAATAAGTACAATCACGAGAGTGATATAATTCTTTGTAAACCTTTTTTTACAAAATGATTAGTAGAAAATACGCGATATTGGTATTAAATACACGATACTAAAATATTAGATTCAATGAGTTTATTATTATCATTGAGTTGTTATTATCTGATTCACGATATGCAACATAGAATCATTATCGGATTTACGGTGCGC
>evgR015120c0g2_i01_Hydra
TTATTCATGTTTTAAATAGCTACAATATGCTAAGAACGGTTGAAAAGCACACAATAGTATACAGTATCTTTTGCGCTATTGGATTGGTTTTAAATGTAACCGCAATGATTGTTTTTCTTAAACTTAAAAACGTTCATGATATGACGCAACGAAGTATACTGATCCAACTTTGTTTTATTGATATATGCTTAAGCCTAGTACATGGGATTATTGAATGGGTAACAGCTCTAAATTTTGATACGCCTAGTCAATTGTTTCACAATATTTTTTTTGGGATTCGAAATATTTTTATTACAGTTTTTTATTATGTAACGTTTTGTTTGTTAGTGGACAGGTATTTGCACATAAAATTTAATATCAAATACGCCATATATTGGTCGAAAAAGAAGCTAATAACTATATCAGCAATAATTTGGAGCATAGCAACGTGATATGGAGCATAGCAACGTTAAAAGGAATTTTTAATCATTTTTATGATATCAAGTTCAACCACGTTTTAGTTTTCTATGATCTAATAATAATTTCTTTCTCCGCGTTTGTCTACGCGTATGCTTTAATGCTTTATCGCAAACAAAAAACAGTGCTTTGTTCTAACCGAAAAGACAGGAAAATATTCAAAGGAGTAATAACATCGAGTCTAATAATAATACTATTTATTATTTTGTTTGCAATACCTGATTTAGTTCTTGTATTTGGATTAGATATAACTTTATCAAATAATGAATTGCGTTGGTTTGTTTACTTATCACTACCTGTATGTTTTTGGACAGATGCAGTCATCTACATATTTGCATCACCTAGTGTTAGACGTTTTATTAAAAGTAAATTTAAGTGTAGGATTTGCAAACAAAGTGTGTCAGCTGAAATAAGTACAATCACGAGAGTGATATAATTCTTTGTAAACCTTTTTTTACAAAATGATTAGTAGAAAATACGCGATATTGGTATTAAATACACGATACTAAAATATTAGATTCAATGAGTTTATTATTATCATTGAGTTGTTATTATCTGATTCACGATATGCAACATAGAATCATTATCGGATTTACGGTGCGC
>evgR015122c0g1_i01_Hydra
ACTAAAGCCATAACCCTACATCATCACAATGAGTACTTATAATAGAAGTGTCGTTCTCGACGGTAAGGGACACATTTTGGGTCGTCTCGCTGCTAACGTTGCCAAGGAAGTTTTGAGCGGAAAGAAGGTCATCGTCGTCAGATGTGAAGAACTCCTCGTCCAAGGAAAGATGAAGAGAAACAAGGAAGCTTATGAGTCATACTTGAGAAAGAGAATGAATACCAACCCAAGAAAGGGTCCATTCCATCACAGAGCTCCATCTCAACATCTCTACAAGATTGTTAGAGCTATGGTTTCTAGAAAGACCGAAAAGGGTAGAGCTGCTTTGGCTAGAATGGAACTCTACGATGGTGTCCCAGCACAATTCAACCAAGTCAAGAGATACGTCTGTCCAGCTGCTCTTGCTGCTACTATCTTGAAGCCTTCCTCACCAGTTACCAGAGTTGGTGATTTGTTGAAGAGAATGGGTTGGAAGTATCACGATGTTGTTGTTAAATACGAAGAACAAAGAAAGGAACAAAACAAGGCTTATATTGCTGAAGTCAAGACTTTGAGCTCCCTCAAGACTGAAGCTACTGAACAATTGAGTGGTAACACTGAATTCCAAACTGTTGTTAAGAAGTTGGCTGAATTAGGCTATTAA
>evgR015123c0g1_i02_Hydra
CCCGTGAAGTTTGGGACAAAAAGACGGAACTTTGCGCAAAAACATCATCTACGAGAGTATTGATTATATAACTTTAAGTTGAAGTTTCCGATAAAGGTTGCGTAGTAACTTTTATGTTACGTAAAAACAAAGATTGATAAAAAACAACGAAAGACAAACCAGCATCAACAAAATTATTTTATTGTTTATTTATTTTAACATTAATTATAAATTATTCTTGCTTAAGTCATCGGCCATGAAAGAAGATGAAGGGGTTGAAAAAGCAAGTAAAGAATATTCCAATTTTTATGGCAACTTGAAATGTCCTACATTCACAACCGTATTAAAAGGGGATGGTTTCGAAAAAAGGTGTTACGAAGAGTCGATGTGGGTAACAACATCAATCCAAGCACCTGATAATCAATCCACTTCATTTCGAACAATGTTTCAAAACCTTTTTAAATATATCAGTGGAGAAAATGATCAGAGCGTTAAAATACCAATGACCGCACCTGTTCTTGTCTCAGTGAAACCGCTGCCGGACAATATTCGCGACATCAAAATGCATTTTTTTGTTCCTCCTACAACCTTAGCCATACCGAATCCTACCTCTGACGCTGTGAAGCTAGAAAAGTGTCCAAAGTTTTGTGCATACGTTCGAGTTTTTGGCGGATATCAGATGGGGTTTAATAAAGACATGCTTTTTCAACGCAAGCGGCTTACAGATGCACTTGATAAAGCAGGTCTAAAATATAATGAAAATGATTTGATTTATGCTGGTTATAATTCGCCTTTTAAATTGTTTAACAGACATAATGAAATAATGGTTGAGATTGATTCGCAAGAAGCGGGTTTTCAAAGCACTAGTTCAAACGAAAATTGAAATAAATAAACACTAAAATGCCAC
>evgR015124c0g1_i01_Hydra
GGAGATTAATTCTAATTGTTCTTGTTAAGAAATTCCGAAACGTTTTTTTTGCATAAAAAATCTTTCGCTAAATTTTATTTTATTTGAATGATGAAAATTTTAGAATGCAAGCTTTATTAAATTTATTTACAAAACGTTCAACTTCTGAAAGTCCTTATCTTAACAATGAATTTAAAATATGTTTTGTTGGAAAAACCAAGACTGGTAAAAGTTCAATAATAAACGCCGTTCGTGGTCTCTACAAGAATGATCCAGGTTATGCGCCTACCTCACATGAACTTACCCAATGCACGCTCGAGAATAAAGAGTATGATTTTGTGCGCGACTCGAAACGTTATAACATTAAGCTAGTAGATTCAAAAGGAGTCAATAACGACGATATCAACCCAACTTCAATTGATAAATTGATTAAAGATTATGACGCGATAATTTTTGTAACCAACGATGCGTTGCTCAAAGAAGATATAGATTTATTTCAGAATATTGAAAAACAACCGTTTATATTGTTTTTTGTGCGCACACATTTTGATTCAATATTAGATAGCGAATTAAATAATACCAAAATCTTGGATGACTCAATGGAGACTCAGATCCGAGATGAAACAATAAATAAGTTAAAAAAAGAAATGAAAGATTTACTTAATGAAAGCAAAATTTCTAAAGTTTTGACTGAGCATACTCTTTACTTGATAACATCAACGACACGTCAGTTTATGAAGTCGCCTGACGGAAAGAGGCTCGTTAACGATTTGATTGCGCTTGAAACGCTGAAGAAAAATGAAAAAATTGATTTAACACAACCATCGTCTTATCATCTCATTAGATATAAAAGAAATTTTTTTATTCAAAACCTCTGGGATTGTCTGCTTAAAGTTGGAATTACATCGTTTTCTTCGATAATACCGTTTTTGGACACTATTTTCAGCAAAATGGTTATTGACGACAAACGAATAAAATGGCTCGAGGATTTTGGAATTCAAACTTTGCTTGATATGCACACCCAAAATGAGCTCGATCGACTTGGAAAACCAGAAGAAGTAACTGAACGATTGAAAATTAAGCTGAATGAGATTGAAAGAGAAAAATACTATTTTGATTTGGATAAGTTTAAGGAAATATTTAAATCGAGCGATTCTCCTGAATTGACAAGCGAGAGTCATGATCAATTAAAGAAGTTCTACACAGACGGAAATGTGGTAAGTGAAATTGTGAAGTTATCTACATTAAGCGGCGGAGCTGGTGTATCATCAATTTTTAAAATATCTTCTAAAACTGCTGGCAAGGCTTTTGTGTTCGCAATACCATTTTTTTGCGCAACTTATATTTATATTACATTCAAATCGTTAAAAAAGATGATCGACTTGTGTGAGCACTACGCAATTGAAATTTATAATATCATCCACGATAATAACGAGGAATAATTTGGACAATTTGTAAGTCTAATGTCTTTTTAAACATCTTTCATTTTTAGCAATCTGTTTTTTAAATAAAACAACAAAATTTTCAATAAAATATTTTCTGAAATTTATTGTTTTATATTTGAATATATTCAAATAAAAAACTCTAAAGTTCAAAAAATATATTATATATTATACATATTATACAAAAACTTTATTATACA
>evgR015138c0g1_i01_Hydra
TATTTTAGTTGATATTTTAGATAAGATTTTAGAGAGCAAGAAGTTCTAAGCAAGAAGTATAAACAAAAGACTTATTTGTTAGTTATTTATTATGAATATAAGTCATAAAAAATAATTTGAATAATTTTGTAAATATAAATAAAGATGAGCCTAAAGTACAAAGTAACTGCTATAGTCATCATTTCAATTCTCCAGTGCATTAATTTTGGGGTTTCAAGAAGTGTAAAAGCTCGCTCTAGAAAACTTGAAAGATCCAAAGTTAATTCCGAGAATGCAGTATCTAATCCAAATGAACTTTTAAAACGCTCAGTATATGGTTCGATAGGATATGATAACTTCTATAACAACCCTGTTGTTCCTGCGGAAGTTGATACGAATTATGGTTACTACCCAAACGATCTCCAATATCCACATCATAACTACCATAACTATCCGCATGGTCATCATTTCCATCAATTTTCTCATCACGATCATCACATTAATCGTTGTCTAAATGAAGGCAATCCTTTTGGAGGCTATAACTCATTTCCTTACGAATCAAATAGATTTGAAGAATATCCCGAAACGCATCAATTTCAAAACCATCACGGAGGTTTTGATCATTCTTATGATTCTAGTGATCCACATCATATTCTGCATAACAATAACATACTTGATTTGACTGATGGGTTTAACGAACATGGCGATAAAATAGGAACGGTATTTCCCTTATTACCGGATCTGATTAATCCAGCTCCATATAGTACTATTGACGGCAGCTGTTATCATTGCTGGAATGATTGCAATACAAATGAGTATTTACATCACGGTTCTGCTCTTGGATACCCCTTCATGCTTCCGCCAATTCCGTTTGTTATGAATCCTCCTCTGGTTCACGATGATTCTGACGCAGAGGATAAAGGTAAAAAGGAAAACAAAGAAGAAAAGAAAGATGACAAAAAGAAAGATGACGAAAAGAAAGATGACGAAAAGAAAGGTGACGATAAGAAAGATGACGATAAGAAAGATCACGATAAGAAAGATGACGAAAAAAATAAAAATATCGAGCAAGCAG
>evgR015139c0g2_i01_Hydra
CTTCATTAAACCTTTTTGATTTTTTTGTTGGTTTTTAGATAATTTTTTATTAAAATGTTTACATAAAATTGTTTAAATAAAAAGAATGTCAAGTACTCAGCGAACTCTAAAATCTGCAAAGCAATTGCTAAGAAAAGAGTTCAAAGAAAAAATTATGTGTATGTCCTATCAAGACAAAGTTGAGCAATCAAAAAGTATTACAACAAAGCTTTTTTCAATGGATATTTATCAACGGTCTAAGTCTATATCTCTTTATTTAAACATGCCGACGGAAGTATTTACATCAGATATTCTTGCTGATGTCTTTAAAAGCAACAAAGTATGTTATATTCCTCGTTATATTGGTGAAGACATGGACATGGTTCGTTTAAAAGACCAAGATGACTTCGATAAGCTTCCAGTAACATCTTGGAACATCAAACAACCTAATCATGATGAAAAAAGGGAAAGTTCTCTAAATGGTAACGGACTAGATCTTGTGATAGTACCAGGATTAGGCTTTACAAAAGGTGGCTTTCGTTTAGGTCGCGGTAAAGGTTACTACGATCAATATTTTCGTAAATATGAAAATGTGTTCCTTAGAAAACCAACTACTATTGGATTAGCTTACGCCTGTCAAATAAAAGGAAGTTTACCTGTTGATGATTTCGATGTACAATTAGATTATGTTTTGCATCCATAAACAATTTTTTCTTAAACAAATAATTTTAAACAATTAATTATAATTTTATATTATACTTTTTGAATTTGTAATAAAGTTTATATTATTTTATGTGATTTTTCGACTATTCATTATTTATCACTTAAAATTTTAGTAAGGATAATTTATTTTTGGTTTAAACAATTATTTTTAGGCCAAATTTTAAACAATTGAAG
>evgR015140c0g1_i01_Hydra
ACGCAGAATGCCGCAAGCGCAACTTTATCAATCAATATCAAACGGCAGTGTGGTAACAGTTTTCTCTTTGATGAACTTTGCAGAACATAAAAACATACAATTTATTCAAGTTATCAGAAACTTCTTATGAACAGGAAACAAAAAGGATTCTGGCTTTAGAAGTGGCGTTTCCATGAGTTCATACCGTATCTTTAATTATTAATGATATAATTACCTCAATAAACAAACTTTTATCGTATTAAAGTTACGGCAATGCTTCTGTGTTCAATTAGACTTAATAGCACACGATGACGTTAAGAAAAATATACCCATTGTCTTTGCTGGTTTTCTTTGCAACAAACATTTGCGCCTGTCCAGACTCGGCAATTCCTTACGAAGATATAGCAGACGCACTGCAAGCTTCAAAAGCAGTTTATGATAACAAGCTTAGCGTCAATTCACTTGTTCCTTACACTTCTTATCATATCAAAAAGGTTTATCTGTTTGATAATGAACTCGATGAGCCGTGCATCGACGGGGTTCTAAAAGTTATTATTGCTGAAAGAGCATTTACCACTATAGTTGCAATGAAAGGTCCAAATGGGCCAACGTCTGTGAAGCCGGAATTGTTTAAATACCTTCAGATATTTAAAGGAACAGAAGTAATTTTTGCTGGCAATCACACTGCAATTGATATTTGCTTTTGGAAATCATTTAAACTTATGCTACCTAGTATACGAGGCCAATTACAAGATCAAGCAAGAAAATATATTATAACTGGGCACTCCACCGGCGGAGCTTTAGCATCTATATTAGCGCTCTATATGAAAGTTCAGGAGGGAAGAATGTGGGAGAACTCTGGAACATGCTTAATAACGTTCGGCCAACCTCGAGTTGGTGATGAGTTATTTGCAAAACTACACGATGCAATGATTGATCCGTTCCGAAAACTTCGTTTTATAAATGACAAAGATCCAATACCACATGTTCCCATATGGAAGACGGCAGTTCATCATTCGAGAGAAGTTTGGATGGCGTATGAGCCCATGACAAATAAAAGTTACTGGAAGGTTTGCGGAAACATGGATCCACTAAAATGTAGCAATATGTGGGTTTTGCACACCTCGCCGGAAAATCACGCTATTAAAAGTTACGAAAATTTTATAATGAACCCTCCGTCAGTTTTTTTGGATAAATTTTCAACGCGTTACAAACGCTGGCTTGATGCTTTACAAAACTCTTGTGTTGTCAACTATAAAATTGAAAAAGTTGAACACAAAGGAATACTTGACATTAATATATGGAAGAAGCAAAGAACTCTTGTATAAGCTGATTGAAACTCTACCGATATTATATATATGATTTTATTATATTATCGATTTTTTTAATGTTCAACGAGATCGTTGCCATAAAAATTACCAAACGCGCAGTGAACGCAACCGAACGTTCAATGTACAGCATATTAATAGAGCTATTTTTTTTTATATTTATATTTGCAGTTTTTTATTTACAGAAATATTTTTTTTTAATTCAATTTTACTTTTATTACAGTTGTTTTGTTACATTATTGCATCATCAGCATTATTTCATAGAAG
>evgR015147c0g1_i01_Hydra
TTGAAATCGAGCATCGAAAAATGCAACGAACTTAAAGAAAATGTACAACTTAACAAACAATATGAGCAAAAATGCATTCTCTTAGATGAAAAATTAACAAGCTTAAATGAAGCTATGCGTTTAAAGGAAGAACGTTTTGATAAAGAGATGAAAGAAATGCTTATGATTAACAAAAGGTATGAAACAGATATAAAAACGTGTCATGACCAAATTGCTTTTCTCCAAGTTAAAAACAAGGAGTTAGAAGACTTTAAAAATCGATTCATCACTCTACAAAGTTTCAATGAAAAATTGCAAACTGAAGTTGAGAAACTGAAATGTTTTGAAGAAGATTGTAAAATTCTTAATGAGAAAAATGTTCGCATTGAGCTAGAATTGAAATCGAGCATCGAAAAATGCAACGAACTTAAAGAAAATGTACAACTTAACAAACAATATGAGCAAAAATGCATTCTCTTAGATGAAAAATTAACAAGCTTAAATGAAGCTATACATTTAAAGGAAAAACTTTTTGAAAGAGAAATAAGTGAAACTCTTACGGCTAACAAGAAATATGAAGCAGATATAAAATCGCATCTTGATGAAACTGCTTTTCTTAAAGTTGAGAACAAGAAATTAGAAGAACTGCAAAGTCAGCTTGTCACTGTTCAAAGTGTCAACGACAAGTTAAAAACTGAAAATAGTGACTTACAAGAGTCTGTAAAAAAATACAGTACGTTGGAAAATGAGATTGTTAAACTTAAAGAGGAATTGAAGACTCAGTACTCAGAGAACACCAGCAAACTTATTTGCAACAATAATTTATTTGAGACTGATTTTGAACACATAAATAATATTGCTAATGATTCAAGAGAAGTTGCTGATCTGAAAATAAAACTAGGTGACTTATTAAACGAAAAATTAGATTTGCTGACTCAGCTTGATTGTGAAAAAGAACTTAATGAAGCATTAAAAAGAAATAATATGCTCTCAGAGAAAAAAATTGGTGTCCTTTTTTATGAGAAAGAGCTGCTAGAAAAAGAGTTGAAAAGACAGAGTGAACTTTTTGATGCCGAACGGATTGAAGTCAATGAGCATGTTGATAATGCAATATTGGAGTATCAAGACAGTATTTTGACATACCAAAATGACAAAATCCTGTTTCAAAAGCTACGATCAGACAACAAGAGTTTGATGCGCGACTTGGAACTTAAAAACCAAAGTCTTAAAAAGACAATTGAAGCACTAAACTCATCAAAAAGCTGCACTACTAAATTGCAGAAATTAAAAGAAAAAAATGATGTTAGTTCAGTTAGTGACGGTATGTTAAAGATAAGTGATTTGGAAGCGTTGGACTCAAAAAGTAATCGCGCGGACGAAATTTTTCAGAGATCCCGATCAATGCGCACTGCACGTCGTAAGACACTTTTATCTGTGACACGCGAAGATGATTTATCAGAAACAGAAGATGATATTTCAATTGGGAGTTACTCGTTACGGCGAACGGCAGAACGACGACGGTCTCTGCAGCCACGGAGATATCTAGGGGGTCGCACCCCTCCTAGTAAGAGGACAATAATGGGTGCGCCTGCTGCGCAAACACAAATGTGTAATTTTACTGTAGAAGGAGAGCAGGAACCTGAAGTTATGGATGATATGGAATGGGGGAGAATTGCTCATTTGAAAGAAACTTTAAGAGAACCTGACGCTGAAGAAAAGACTCGTCTTGATGAGTTGCATAGACGCAATACGTTATGTCTGCCTCATTTGAAATCTTCTTATCCCATTGAATTGAGTGTTAAAGGAGCCAATCAACAAGATGTGCAAATGAAATCCTTAGGAAAACGAAAAAATGACGTAGAAGAGGATGCACCACGCGCTACATACAAAAAATGGAAAACAGAAGATCAGCTGTTAAACAGCCTTGATCATTCTAACTTTACCCGTGCGGTTAAATCTGATTCAAATATTCGGAATTTAGATGCCGGTAAAATTGAGAAACCTCAACTTCAGTGCACGTCTTTTGAAATATCGGTTGAACCTGCGAAAAGAAATAAGCCTTCACGATTCAATCGCAGTATAGCATTGTCGAGTATCAAAGATGTGCAGATAGCTGCACCGCAAAAACTCCAGGAATCATTAAACCCAGTTGTACTTTTAAATCGAAACTCAGAAAATAAATCCGTCTCATCAAATATAAAGGTTGGTAAGTCGAAAGACTTGCTCTCCCAGCTAACTCGAGGTGATAATAAACGAGCCTCGTTTTCACGTGCCACTCATAAAATGGCTGCATCATTTCGCGGTAAAATGATGGCTAAAAATAAATAAATTGATATTTTATTATCCTTCTATTTTAAAACATACAGCGCTAGTCGTTTGATAGCTTATGATTTTTTTTTTTACTTTTTTTACGTTTGCTGGTTGTGGTGACATCAAAAACATTTATAAACATGTAAATTAATAAAAAATGATTTCATGTGTGACTATGAAGTCTATATAAGAAATAAATTGACAAGATAAAAGTAAAAACATAATTTTTTTGCAAAAACGCTCATGTGTATGGTTACATAGGTGTGTATATATTAATAATTATTGTGTCATTTGTAGTATAGAATTT
>evgR015154c0g1_i01_Hydra
TGTTGGTGGATATAACTGCACATGTAAAAATGGCTTCAATACAAGTAACGGCGGGAAAGAGTCTCTTTCTAACCTATGTCTAGATATTGATGAATGTGCTCTTTCAACCCAAAACAACTGCTCTGCTAATTCAATTTGTAATAATTTGGTTGGTGGATACAATTGCATATGTGAAAGTGGGTTCGATACAAATAACGGAGAAATAGAGTCAGTTTCTAATAAATGTCTAGCAAGTTTACCTCTTTGTACTGATGGTTGGCTGGAATATGGAGATAGTTGCTACCTTTATACCAGAAGAAGTAATGGAAAAACTTGGAGTGATTCCCTGTTATCGTGTCAGAGTTATGGAGCTAACTTGCTAAGTGTATCGGATCAAGCAGAAAATTTCTTTATTAAAACTTATATTTTTCCTGCATCACCTTTATATGAAGACTATTGGATAGGTCTCAACGATATTCAGACTGAAGGAATTTTTACTTGGTCTAATAATGCAACTTTTGTATTTAATAATTGGAAAAAAAACGAACCTAGTGCATTAAAGGAAGATGAAGATTGTGTGGAAACAACACGTTATGGTTGGAACGATATTAATTGCAATTCTGATTCTTTTGGGTTTATTTGTAAAAAAAAAAAAGAAAATGGCAACTGCTCTAGTATATAGTCA
>evgR015162c0g1_i01_Hydra
ATAATAAAACTTTGTATATGAAATGTCTCATGCAAACTATTTAAGTCTCTGGCGAAGAAACCATAAAACACAGTTGGAAGTTCTTAATTATTGTCACGACTCATGCGAAGAGATCATTTCATTAGATTCACCATCCGATAAAGGAGAAGAAATGTTTGAAGGAAGACTGGATTTAAATGTTGAGGTTTCAGCATCAGCCTCATCTTCTGATGTAAGTGATGTAAATAGCATTTACATATCAGAAAACGATGGTATACACATTCAAGATAACTTGGCCTCTGAACTTTCATCTTGGGCATGTGAAAATCACCTTACAAGAAGAGCAACAAATCATATCTTGCTAATATTAAGAAATCATGGTCACAAAGAATTGCCTAAAGATTCAAGAACCTTATTAAGCACACCAAGAAACGTTTCTCTAAAACAAAAATGTGGTGGTGATTATTTTTACATAGGTTTAAGCTGTTCAATAAAAAGCAAGTTAGAAAAATTTTCATCTCATGATATTAATTCTTTGAAAACAATCAATCTATTGATTAACATAGATGGTATACCATTATTCAAGTCATCAAACACACAACTTTGGCCTATTCTTGCATCTTTTCAGCCTGTTGGTCCATTTGTTGTAGCTCTTTACTGTGGTAGCTCAAAACCTAATAGTTGTGAAGAATTTTTAAAAGAGTTTTTAGAAGAGTATCGTAATTTGGAGGTTGTTGGTTTACATTATAATGAAAAAAAAATAAATGTAAACATTACAGGATTTATTTGTGATGCTCCAGCACGTCAGTTTCTAAAGGGAATTAAAAGTCATACTGGTTATGATGCATGTGAGAGATGTAAAGTTCATGGAGAACATATAGAAAATCGAATGGTTTTTCATGATCTTGACTGTGAACCAAGGATAGACCAGGAATTTAAGCAGATTGCTTATACAGGGCATCAGTTACTTGTAAGTCCTTTAATTCGCTCTAGTGTTCAATGCATCAAAGATTTTTGTTTGGACTATATGCATTTAGTTTGCTTAGGAGTTGTTAAAAGATTGCTTCTTTACTTAAAAGAAGGCCCTCGTTGTTGTCGTATATCATCTAATCAGTTTAATGTAGTCTCTGATTCCATTTTATCATATTATGGAAAAATGCCTTCAGAATTTGCACGCCAACCACGTGGGCTAAGCGAACTACGAAGATGGAAGGCAACTGAGTTTAGGCAGTTTTTGATATACACTGGAATAATAGCCCTTAAAGGAACTGTTAGCTTACCAATTTACATTAATTTTTTATCTTTATCTGTTGCAATAAGAATTATGCTTGAGGAAAATCAAGAAAACAGAAATGAGCATTTGGAATATGCTCATAAATTGTTGGTCTATTTTGTTGCAAGTAGTAGGGAGTTGTATAGTGATTCTTTTTGTTCGTATAATGTACACAGCCTTATTCACTTGCATAATGATGTTTCAAACTTTGGTTGTAATTTAGATCAAATTTCTTGCTTTCCTTTTGAAAATTATTTAAATTCGTTGAAAAGGTTTATTCGTCAAGCAAATAATCCTATATCTCAAGTTGTGAAACGTATTCATGAGTTTGAAATAAATAATGCTATAAAAGTTTCAAAGGATATCGTGACAAAAATTTCTACTAATAAAAAAGACAATTGTTTTGTTCTCCATTCCCGAGAAATTATTTTTGTAACAAGTATTATAGACATTAATACTTTTGTTTGTAAAATCTTCCATAAGAGTCTATTAAGAAATTACTTTCCTGAGCCATTTGATTCAAATATGGTAGGAATTTATTTTTTATGTAAGTCTAACATTGGTAAAAGAAAAATTGTAAATAAGGTAGAAATTAAAAGAAAGGTTGCATGTCTACCAGTCAATGATGGTGTTGTTCTATGTACCTTACTTAACGATGTGGTTTGGTAAACAACTGTGTATATGATTTGCAAGAATCTAAAGTCACTTTTCTTTCTAGTATAAATTAAAAAATGTGCACATTTATTAAATAAATTATGTTGTTAAAATCAGTTATTTTTATGATAAATATTTTTTATTTAAAATTCTATTATTTTTATTATCATTAC
>evgR015166c0g2_i01_Hydra
CTAACCAATTAATAAAGTTTAAAAACAATTCTCAAAAAACATGGCGCATAATTAAAAACGTCATAGGTAAAAAAAATACAAATTGTAAAAGTCTTCCAGAAAAACTCATTATAAATAATTGCGAAGTTGTTAATGAGTCTTTAATTGCTCAGTCGCTTAACAATGCATTTGTTAACTCAGGCGCAAATTTAGCATCGAAAATTAAAGTTAGTAAATTTGATTTTAAATCATACCTAGTTACTAATAATGCCTTAATGGAAAACAATAAACTAACTGAAAAAGAATTTTTTCATGCAGCCTCTTTTTTAAA
>evgR015170c0g1_i01_Hydra
TGAACTTGCTGGAATAGCTAAAATTTGTCTAGCTAATAAAGAGAGTTCTGGCAATGTATTTGAATGCAATTTTCACCAATCAAGAATCAGAAAGTCTTTTTTGGCATCAGAGAGATATTCATACTTGCATATTTCGCTCAAAATAGGATTCAGTTCAGATGTTGGCTCTTGTGTTTCAAGCCTGATGTTTAACTTGCGCTTCAGTTTTGAATTAGGGGACAAACCTGCTGAAAAGACTCTGGCAACAGGTTGGCACTCAACATTGTCCTTAACCTGTGAGGCTAACCATTCTTTTGTTGTTTGAAATTTTTTAAAGAGCTTTAAGTAAAGTCCATTTAAAGCAGGATTTAAGTAGTTTGCTGCAGCACTCAATAAGTTTCCAGTGTGACAAAAAGAAAATCTTTTCTCAATGCAGCTATTCAAGTTTTTTGCATACAAGACACCGTAGCCATTTTTTCCATCCGTCTCAATAAATTGATCAATTTTGTCATGGATTAAATAAAGAGTATCGGCGGCAGTGATAATGGTGTAATAGACTCAGCCGACCACGTTTCTGTTGATTTTTTAAGTGGTGCCAAAATTTCTACTGCTCCCTGGATTGATTTCCCCTGTGATGCACTGATG
>evgR015173c0g1_i01_Hydra
TAAATCTTAATCAAAAAAATAATTGCTTATATATGCGCATTTTGTAAACTTTCGTCTTTAACTGCATCCGTTGACGCCACTACCTTTTGTCCGTTTTCTCCATAAATTGCACCCATTTACGGAGCTAGGAAAAAACAACATCATGGAGACTTTAGATGCTAAGAAAAGCGATTCTATTCAAACTACCTCGCAACTTCTGATCCAACCCACCTCACAATTATTAAGCAGTTATTTAGCTAACTATTTAATAAATGAAATAAATCTTGATGAGATTTCTATCGACCTAAAGGAAAATACCGTACTTGTTACGAATTCTAACAGCTCATACTGGGATTTTTCACACAAACATGGGCCCGAGCAATGGCAGAAAATTTGGAATTTAACAACAAGACAATCTCCAATTGATATTGTTTTACAAAATGTGCAATACGATGCAAATCTGCAAACTTTAAATTTGGATCGTCAAACTGTTTTAGTTTATGCAATCAATATTGGTTGGAATGTTTCTTTTAAAGCTGATGACAATAAAAATATTTCTCTTAGTGGAGGGGCGTTGACTCATAATTATGCTTTCCGCGAAATGCATTTTCACTGGGGAGAAGTTCATAAAGGCAAATGCAAACTTGGTTGCGAACATACTATTGATGGCAAAAGATATGCTGCAGAATTTCATGCAGTTCATTGGAACACAGATTTATATAAGACAGAAGATGAAGCTATTGCAAATCCTGATGGTTTGGCTGTAATCGGTATACTTATTGATGCAAACGAATCGTACGAAAACAACAAAGACTTTGATGTGTTTTTAGAAATGTTTGACAAAGTTCCGTACATGAACAATAATGCTTCGTTCCACGTTGACCCTTATCTGTTATTACCTAAAAATACAAGTCATTACTTTACTTATCCTGGATCTTTAACTATGCCTCCTCTCACTGAAAACGTTACGTGGACACTAATCTCAGAACCAATTAGAATATCCGCCTACCAACTTGATAAGATGAGCAAAAATAACGCTCGCGAATTTTTTGATTCAGAATGCTCTTTTAAGTTTCAACGCCAATCTGAGTCCACAACCATAACTAATAACTATCGCCCAACACAACCTATTAAAGATCGAGTGGTTAAATCGTCATTTACAACATGAGCAACAAAAGGAAAACTTTTTGACATTGTTTTTTGTTTGTTTTAACTTGTTATGTTTTTGTTTTATTTGTTTTATTTTATTTATACTTTTATATATAAATAGTTTATTATTATCAACACTAACTACTAAAAAAAAAA
>evgR015175c0g1_i01_Hydra
CAACATGGTTGGTTCATGCAGAGAACTTATTTTTAACGAAAATGTTTCAAGCATTGTCAATCTTGAAACCAATGAACTTGAAAAAAACAGTCCGCCAGAAAATTTATCGACAAAAAAATCAATAATAATCGTCAATAACGATTATTTAAACTCTTATGATAACATTGGTTATGAGGTTAATGATGAAGAAATTTCACTAAAATTAAATTATCAAAAAAGGTTTGCAGAAGTTCCTTTGACGTCAAATACTTTTCCAACAAGTAATTTCCCGACAAAAGAATATCCGTCAAATGACTTGCCGATTAATCGTTCAGCAACAGATAGCCCTCAATATGGTGAAGAAGCTATCGAGTTTAGCTTATTATTTACAGAAGAAACTGAAGCTGACAAAGTAAATAAGCAAATTAACCTTGCTACAAAAGATTTCTTATTCAGTTGCAAAAAAGGTTTTGGAAGTTTACACTTGGATCAAAATCAATCAGAAAGTTTAAAATACAAAGAAAATCAACACAAAATTATAGACAATAAAAAATCACCTTCAGGTTCAATTTCAATTAGAAATGAGTTTGAAGAAATGAAAAATTCGTTATGGAACACATGTCAAGATACAAAAAATGTATACAAAGATTCCGATAAAGAAAGCAATAAAAACAATTTAAACTCAATAAAAACAAACAATGTTGAAAATAACAGTTTTTTGGAAATTAAAGATGACAGAACTAATTGTTTCTCTGGAGTTAATTCTCAAATAATTGCTACGTCTCATTCTTTAAATAAATTTTCAACTTGTGTAATATTGGACAATGGGTCAGGATCAATTAAATGTGGCATTTCTAGTTTAGACTCTCCATTAGTTATTCCCTCGCTTGTTGGAAGATCTAATTATACGAAGGTTTTACCTGGTGCTTTAAAATCGGAATGGATCGGAAACGATGCTCAATTGCATAGAGGGATATTAAAGCTCTCGCATCCTATAGACAGAGGTATAATTAAAGACTGGATTGATATGGAAAAGGTTTGGGAGTATGCATTTGAAGTTGGTGGTATCTATCCGGAAGAACACCCAATTTTTATTTGTTGTGCACCAAGCATATCTAAAAAGCAACAAGAATGCATAATGGAAATAATGTTGGAAAAATACCAATCGCCCGCTTTTTATATAGCACCACAAGCTGTAATGTGTTTGTATGGTTGCGGAAGAATCTCAGGAATTGTTTTGGATGTTGGAGACCATGTTGCACATTCTTCTCCAGTTTACGAAGGGCACGTAATTGGTCATTCCTCGCAGCTCTTGGAACTCGGAGGAAAAGATGTTTCTAATGCGCTTAAAACGTTGCTGTCAGTTGAAGGAAAGTTATTTAGCTCGTGTGCAGATTTAGAAATTTGTCGAGAAATTAAAGAGGCGCATTGTTTTGTAAAAAGAAAACAAACGAAGATTTGTGAAAGACTATCAGAAGCCATTTCATACCAACTTCCCGATGGTCGGTTCATTAGTTTAGATTCCGAACGTCACGAGGCGCCTGAAATTCTCTTTAGACCAGATCTAATAGGTCTAGAGGGCCCTGGAATTCATAACTTGATTACTAAAGGCATTTTAAAAACAGATAAAGACCTTAAAGAGTTGCTATCAAGCAATATAATACTTGCAGGAGGAACAACAAAGTTAGTTGGATTAGAGGAAAGATTACTTGATGAACTAGCTGCAACAGAAATGCTTCAAAATTTAAAAATAACAAAACCAGTCGATACGACAACATGCGCTTGGGATGGTGCCAAAGTAATTACATCATTGTCAATAATGAATCAGATGTGGATAACAGCTGAAGAATACATGGATTACGGAAAAAACATCTTGCATACAAAATGCTTCTAAGAATTGTTTCGCGTCTATTATATTAGAAGCGCTTCAAGACTTGAACTAGTTTAAAAATTCCCAATTAAAAAA
>evgR015177c0g1_i01_Hydra
AAAAAAAAAAACATAATGGGGCGTTTTGCACTTTCAAGTGAACCTGTCATATTTTTATATACAATAATTTTTACAATCAATTATTCTGTACTTCCTCAGCTTGTTCTCCACAAATTATGTCTACAAACTTTAAATGAAACGAGTTGTGAAAAAACTGAAACTTATCCAGAGATAATTCAAAAGGAGGCCTCAACTTGGTGGTTTTATATGCTAACATGCTCGCTAGTGCCTTCTGTTTTAACTGTTCTTATATGGGGTCCTTTTACTGATATAGTTGGTAGACGGTATGCTATGTTGGCAGTACCTTTTATTAATTCCATCCGTTCAATTATATATCTTATTAACGCCTCATTTTTGCATGCCCATGTTGGATACTTATTATTTGGTTCATTTTTGTCTTGTTTCTACGGTGAATTTCAAGGTGTCGTAGCAGTTTGTTATGCTTATATGGCTGATATTACAAAAGATAACCTTGATGATAGAACAATGCGTATGGCTTTTCTTGAAAGCAGTTTATTTTTGTCTGGTGTACCTGCAGGGTTATTAGCAGGATATCTACTTCAAAAAATTGGATATGTCCCAGTATTTATATTAACATTGATTGTCAACTTGATAATGTTTTTCTACGTGTATTGCTACTTGCCAGATAGGAAATCTCTTAACTCTGTAGGAAATAATTCTCAGAGTGAAAAGAGTATTTCTACGGAAACGTTTCTAAAACGTATTGAAGATAATGAAAAGAAAAGAACTGCCTTATTAAATCCTATCACTCATATCAAAAAGGTAATCCTGGTTATTACGTCACGAGAAAATCGTTTAATTATCTTACCGTTAATATTAGCGTTCGGTATTGCTGTTTGTTCCTTGTTAGGTGAACTTATTGTTCAAACCCTATATTTGCAAAACAAACCATTTAATATGTCGCCTAAACTATTAGGGTACTATTCTGCGTCTCAATCTACAGTGCGTGGTATTGGTATTATTTTAGTTACCCAGATCTCTTTTAGGTTTTTGAAGTTAAACGACTACACTCTTATATTCATTGGTCTAGTAAGTATTACTACATGTTACTTGTTGATTGGTTTGGCTCGGTCAGTTAAAGCCGTTTTCCTGGTCAATATAGTTGGTTTTGCTATTCCTACAACCACGTCCGCTCTCAGATCAGTTGTAACAAAGCAAGTATCTAATGAAAATTATGGTGCTGTACTCTCTTCTATGGAAGCAATCGAAGCTATTGCTGCTGTTGCCGCTAATGGCATTGCATTATGGACCTATAACTTGACCTTGAGCGTTTATTCGGGAGTTGTTTTTTTTGGCCTCAGTGGAATTTCTTTAATTGCCTTTGTGTTTGTTATGATAGCCTATTGTTACTCTCGATCCGTAAATATTATTGAAGTTATTCAATAAGTCGATATGATATATATAATATATATGATATAAATATAATATTACTGAATAGCTAATAGTTAATGTACAGATATTTTAAATTATTTTATAGTCTGCTTTTACTATCTATTTATAATTTTTTAACTTCAAAATAATTATATTCGACGAACAAGATAATTTTAAATAACAAAATTGTACAATCAATTTAGTGGAAAAGGAGAAATGTTTTGTACGATAAAAGTTCA
>evgR015179c0g1_i01_Hydra
CAGAACATGTCTGGACAGGCCTTTGGTTAGTTAACATTTTATTTATAGAACAAGTTTGATTGGTTTGCTATGATGTCATGTTACACGCATAATGGTCGATATGTAATTTGTTGGGCATACGGTGCGAATAAGACAAACAAAGCGTTTTGCGTTCAAATAAATTTAATTAAAAAGTATAAATGGCAACAAACGATCAGTTCCACGACTATTCGGATGCTATAAATGAGCTTATTGAAGCTAAAAACTTTATGAGCCCAAGTCCTGATCATGCCGATCAATGTTTCTATCCGAACTATCAAAAGTTTATTTTTCCAGAAAATGATGCAGGTTTAAAAGATGACAATCCTGATTTTGATCTAGAATCACATCAATCAATATTATTAGAAATGTTTCTAAACAATGATTTTCATTATTTTCCACCTACAGAGCAAAAACAACACTCTGAAACACAAAAAGAACATTTTACACCAACAGTTAATATTATTAATCCTGAAACACCTAGCTCTATTAAATTCGATGAAGAAGACGAACAAGACGATACTGAGAGTATTCTTATGGAGTACAACCAACCTGACTTCATTGATATTTACGATGAAAAAGAAAGTTCAATTAGTGACAATGATTTAATGCTATTGCCAGTCCGTCAGTTGAATATAAAGTTAAAAAATTTATCCAAGGATGAGAAACTAAAACTTAAAGCGCGGAGGCGCTTATTAAAAAATCGGGGTTATGCTCAGACGTGCAGGGAAAGAAGATTATATTCTCAAAGAACCGTAATGGAAGAAAATGAACATTTAAAAAATCGCTTAAAGCAAGTTACTTTAGAAAAGAACATTATTGAAAATAAATATAACCATCTTAAAACAGCTATTCAAAAAGCTAAACTTGCAAAACATATTGTTAGAAAAAAAGACTTGTAAATAATTTATGTTGATATTTAAACTATTATTTTTCTTTCTTTATTTCAATTTAAATTGATAAATATTTTTTTCTGTGAACATTTTGTCAAAATTAACTTTTTTTAAATTGACAAAACCGTAAAAATATGAAAAAAAAA
>evgR015193c0g1_i01_Hydra
CAACAATTTTGATAATAAGGATAAGCATGAGCAATTAATTAACAAGCATTCCGATAATGACTTCAAAACACGAAAACCGATTTCAAAACGTTTTCAAATTGACTCGCGCTCTAAATTAAAAAGATTAATGGATTTAACATATCTTGTTAAAGATGAAAATATTTTGGATCGTATGAACACAGGCTTAGACACCATTACAGCTTCAATCTTAAAAAATCTTAGTCAACCATCAGGGTTAATAAAAAGGGACTCCCCAATAAAAAAATTTAATTCAAAAACAAGTTTAACACCGACATCTATTTTGAAACCAAAAAGTAATTTATCTTCTAAAACTTTATTCTCCAGAAAAAGGAAGCATCCATTTAGTGGAAGGTATGGGAAAACAGCTGAAATGATGAAGCAATGTTACAGAGCAAATCTGGAGCTTAATGCTCACTCCTCTGATCAAGAGGATATCATCGTTGAAATTTGCCAAGATGAAAAACACCATGTTGAAAGTCAGATGGAAACCGGAGAAGAAGATTTGGTAATTACTCAAATAAAAAAAGGGAGAACGGGAGAACTAAAGTTTTATCGATTTCATTCAATATTCGACTCCAATATGGAAAAAGTATTACAAAGTGGTGATTGGTTAACCTGTAATGAAATTAACCATGCTCAACTTTTGCTCCACAGGCAATTTCCTTTTGCGGAAGGCTTAGAAGATACTGGGTTAGGTCCTTTGCTTAAAATGTCCAAGTTAAAGTCAAGCAGGGTACAAATACTCTATAGTAACAACCATTGGATAACATTTTCGGCAGAAACATCGCATAATATTAAAGTTTTTGACTCCTTATCAAGTAACCTTTTACACTATACCGCAATTAAACAAATTTGTCATGTAGTGAACTGCAGTGAGTCCGTACTACGTATTTTAATAGAGCCAACACAACAACAACAAAATGGTTCAGATTGCGGTTTATTTGCAATAGCATATGCAACAGATATTCTTTTTGGAATAAACCCAGTTGATTCAAATTATGACCTTACTAAAATGCGAACACATCTAGTTGATTGTTTTCGAAAAAATTTAATGAGTACTTTTCC
>evgR015193c0g2_i01_Hydra
AAAATACGACGACCTTGCTAACTACAAAAATTGCTCACTCAGTGAGATGCTTACCGTCGTTATTGAAAACTTCATACCAAAGCATTACCGTAAATACGTGGAGTTGAACGTAAGGTACAGCGAAGGTTATAAAAAATATAATGATGGAGTGCCGTTGTATTTTCGAAATCGCCCAAAACAACTTGTAAATATTTTATTGAATAACGAAAGCAAAGTAACGTCTTCAATGATAAACTCAGTTAAAAGAATAGACCCGATAACATTCGGGGTCAATAGTTCCGAAGTTAAAACTAATAACATTAAAGATTATATTGTTTGCATAGGTAATGAGGATAAATTTTGTTCCTGTTCGTGTTCAGATTATCGTCGTCATCGAATAATCTGTAAACACTTTTTTGCCGTGTTTAAATCTGGAATGTCAACATTTGAAAATCTTACTACCAAGTTTCTCAATCATAGTTGGTCCGTTCTGGACCCCGAAATTTTTGTAAACTCGTCAAAGAAAGAAACTCAATCTAATACAAAACCAACTGAGAACAAAACCGAGCAGCAGATAGACGATGAACTAAATAGTATTTCCGAAGACAACAATATTTTTGACTTACAAATCAACAATTTTGATAATAAGGATAAGCATGA
>evgR015194c0g1_i01_Hydra
CCCTTCTCGTAAAAATGATGAAGAAAATCGTTTCTGGTACCACCAGACGTACCACTTCTGTTGAAGCTGTTAAGAAGATGTTGTTCACTCAACAAAGCAGATTCGAATCCAAAGTCTATACATTCGGTATTGATAAGGAAAAGGCATACAAGTTCCATCTCCCAACCAAGCACATGGAAGGTATTAATTTCACCGGACCAAAGAGTGAAGTTTCATTCACTAAGGAAGAATTGATGGAAATGTACCGTATGATGGCTTTGATCAGAAGATTTGAATTGGTTTCTGACCAACAATACAAGGCTCGTAACATTCGTGGTTTCTGCCATTTATACAGTGGTCAAGAAGCTATCTGTGTTGGTATTGAACAAGCTACTAATCGTGCTGATTCTATCATTACTGCTTATCGTGATCACGGTTTCCAATTGTGTAGAGGTGGTTCAATTGAAAGCACCATGGCTGAACAATTAGGTCGTGTTTCCGGATGCTCAAAGGGTAAGGGTGGTTCCATGCACATGTACAAAGTTGACCAAAACTTCTTTGGTGGTAACGGTATCGTTGGTGCTCAAGTTCCAGTTGGTGCTGGTCTTGCTTTTGCTCACTCATATTTGGACAAGTTGAACAATAAGCCAACCACAGATAGAAATGTTACTTTTGCTCTTTATGGTGATGGTGCTGCAAATCAAGGACAAATCTTTGAAACTTACAACATGGCCAAATTGTGGCACATTCCAGTCATTTTTGTCTGTGAAAACAACAAGTATGGTATGGGTACCTCCATGCAACGTTCTAGCGCTTCTACTGACTACTACTGTAGAGGTGACTTCATTCCAGGTATCTGGGTTGATGGTATGGACATTTTTGCTGTCTTTGAAGCTGCTCGTTATGCTAAGGAATATGCTCAACAATATGGACCAATTGTTTTGGAACCAGAAACTTACAGATATTATGGTCACTCCATGTCTGATCCTGGTATCTCTTACAGAACCAGAGATGAAGTCAATGAAGTCAGAAATACTAGAGATCCTATCGCTAGATTGAAGATTAGAATGATCGAACAAGGTATTGCTACTGAAGAAGAATTGAAGGAAATTGATAACCAAGTTAAGGAAGAAGTCCAACAAGGTACTGATAAGGCTGTTAACGCTCCACTTCCTCCACTCCACGAACTCGTTGAAGATGTCATGGTTGGTGAATACTTTGTTAGAGGTAGATCTGTTGACGAAACTTGGGTTATTAACAAG
>evgR015195c0g1_i02_Hydra
TACCCATTTAGCATTGATTTGAGCTAAAAAAAAGTTAATTTTTTTTTGGACTAGTTGCCTTTCTGCGGTTACCGATTCATATTTCGGACTGAATAGTAACAATTTAATTAGTAAGAGCAGATATTAAAACAAATTGTTTTCCAAATTTTTCAAAAACAAACTAAAGTTGATTTTGTAGAACTGGCTGTTATTATGGCAAATATTGATTAACTTCCATGCTTCCATGAGTATTGATTTGATATATATCTGATGTTTTTTTGTATTATGTGGAGTGAACATGCAACGGTTATTAAAACTGCAAGAGCTTAATCCTTATTTAACTTGTATGTTATGTAGTGGTTACCTCATAGATGCAACAACACTTGTGGAATGCTCGCATTCTTTTTGTAGAAGCTGCCTTTTGAAGTACTTGGAGTCATCTTATAACTGTCCAAAATGTACCACAGAAATTCATAAAACAAAGCCACTTGAGTATACAAGATCAGATCCTGTTTTACAAGAAATAGTATATAAGCTGGTTCCAGAATTATTCTTTGAAGAAGAAAAAAGAAAAGAGAAATGGCAGAATGTTGACAAAAATATTGAAAATTCAGACAGACCTGGTTCATTTGAATTTACAAAAGATTTGTTCAATGAATCTCGCATACTTGTTACCTTGAAATATTTTAGAGGTACTTCTCGGGTAGAGCGGATCAAAACAGAATCTGTGCTTCAAATGTTTCCAACACGATATCTTTGTTGCAAGCCTAACATGCCTGTTAAAGTACTCAAAAAGTTTGTCCAATACAAATTTGATATCTCGTCAGATACATTTCAGATAGACTTGTGGCGGGGCGATGAAACTTTGATGGACAATCTTACACTTAAACAGGTTGTACAAATATATGGTTTGTATAGAGAGAGGAAACCGTTAGAACTTCATTTCAGTGTTTATCCTAACTCAAATGGTAATAAAAAAACGGATGATACTCACGATGATGAAAATGTGACGTTTTAGAGAGGATGAGGTGACAAAATTGTGGCATTATAACGAGAATGTCGCGTCATAGAGACAGTCATATTAAAATTCGGTGCTTTAAAAGAAATAACAGCCAAAGCAACAACACCGCTTAAAAGGTTTTTTCAAGGATAAGTCTACATAAAAATTCTCCATGATGAATGTATTGGTAGAAATAATTAAAAATATAGAATAATTTTATGATATAAAAAAAAAAAAA
>evgR015196c0g1_i01_Hydra
CAGTTATCATTTACATTTTGGAATACTATGGAGCCAAACAATTATGGTGGACGAAGAGAAGATTGTGTCCATACTACCAGGAATGGACACTGGAACGATGACCTTTGTTACATTAGAAAACCTTTTGTATGCAAGTATACAAGAAAAATCCAACAAGAGAATGAAAGACAGCAAGATGAGTACTATGTTAACTTTTGGTATTATCAATCGCAAACATTCGGCTGTCCTTTAACCAAACGAGTAGTTATAAAAAGTGCTTTTTATGGAAGCGAGTATACTAAATGCTTCCTACCACAGAATGAATCTGTTCCATTTTTACAGGGCTTATGTAGCGGACCTATGTGTGCAAATATTAAAGCAAACGATGAATTTTTTAAGAAAAAGAATTTTTGTAATGGAATTGCAAAAAAATTAAATATCACCTATACTTGTTCAGAAGATGTTAATAGTCTTAATTCAAAGAATTGTCCGACTGGTTATACTATTAATAATGATAACAACGTCTGCTATAGACTTTACGCTTATGAAGCAAATTTAACTGAAGCGGTAACACAATGTCAGACAGACAGTAAAAGTGGGAATCTGGTATCTATTCATTCAAGAGAAGAAGAGTTGTTTGTAACTGCAATGTTAGGTGATCTATATGTTAAAAAAGATATTTGGCTTGGTATGAAATATGACCCTGCAAAGCAGGTGTATTCATGGATGAATAAAAACGCTGTCAATTACACTAATTGGTATAATAAGCAACCAGACCCAAGGCAGGGAGAGTATATTAAATTTAGTCTATATAATGGTGCAGATGACTTTTTATTTTGGCAAGCAGCCACCCCTACAGAGCGTCTTTCATTCATATGCAGAGCTTTAACAGAGGGTGCAACAGAAGCACCACCATTGGTTCCTTCTGAAAATACACAAAAGTGCCAAGCAAACTGGATAAGTTTTGGAGATAACTGCTATTTCCTTAATGTTAATCTTTTATCGTTTCATGATGCTGAAGAAAATTGTCAAAATATGGGAGAAGATGGTCATTTGGCTTCAGTCTTATCTCTTGAGTTAAATAACAAACTCTTTGAAGCTTTAGGTAAAGCAAATCTTACGTTTGCATGGTTTGGTTTAACCAATGAAGGACTCACTAATGGTTACATCTGGATGGATGGTTCTCCTCTTTCTTTCATTAATTGGTATAGAGAAGCACCCTACACTGGACTTTTGCATATATGTGCAGAAATGAATAATCAAGGAAAATGGCAGGATCAACAGTGTTACATAAACAGACCTTTTTTTTGTCAAGTTCCTAAAGGGAAATTATTTACACCTACCCCACCTCCACCTGTTACACCTTCTCAAGACCCTTCTATTTGTTTATCTGGCTGGAAGTTGCATGACAAAAATTGCTACTTTTTTTCCAATACCACCTATGCAGATTGGCACAAAGCAAAATTAGCCTGCGAGGATTATAATAGCAGTTTAGCTAGTATCTTAAACATGGATGAAAATAAATTTATTGCATCAGAAATAAAGCGCCGTTACATGCAAGATCCATACATTGGCTTAAATGACATGTTTCATGAATCTGGCTGGCAGTGGG
>evgR015218c0g2_i01_Hydra
TTTTTTTTTTTTTTTTTTTTTTTGTTATTTTGCGTGTGATAGAAAGTTTTGAAATGCTCGAGTCAGAATCACTTTTAAATACGCTAGAAAATAGTACGACTGGTTCTAAACGCGATGAGTTTCGAAAATATCTTTTAACGTCGGGCGTGCTTGATGCTTTAACAAAAGTTCTGATCGATCTTTACGAAGAACCAGAAAAACCATCCGATGCTGTGAATTTTGTAAAACAGCATTTTGGTGGTTTTGAAAAGGTGGATGAGGATTCAGACTATATATCAAATTTAAAAAGAAGAATTATTGATCTTGAATTTGAAGTGGGCGATCTAAAGATGAAGCTTGCCAAAAAAGAAGAAGAGAAAGAAGAATATAATCAGCAAACAATATTGGAAGAAAACTAATAAAAAGAACAGGTACCTTAAAGTGAAAACGGAAAACGATACCACTCTTGCGCATTTTGGACACTGCATATATTGCACTTAAAACTGTATTTATCTTTACACATGTTAAATAAATATATTTACTATGAAGTAAAAAAAAAAAA
>evgR015221c0g1_i02_Hydra
GAAATCAAATTTGAACTTTACAACGAATTTTTGAATCCATAAAAATTATAATGGCGTCATTAAACTTGATAAGTTTTTGTGTTCTTGATAAAGACTCTGAACGTGCAAAGTTTCAGTTCCAACTGATAAAATACTCTTCTCTAAGCCATATAAGATCTAAAGCTTTTCGATTTGGAGGTCATTACTGGAAGCTTCAAGTAGTTAAAAAAAACGATTATTTTGGAATGTTTTTGAGATGGTACGGAACTAATAGAGCCCCTGATTGCACTAAAAATAAACTAAAATGCATTGCTTCAGTTAAATTTTCAATTCTCAATAAGTTTTTACTAGACGATTCTATATCACAAGGAGCACGAAAAAATAAAGATTTATTTGAAAAAGTTGGAGTAGGAATTGGATATAGCAGAATGATAGAGACAAAAGAACTAGAGTCATTGCGGGGATATATTTTAAGCGATTCGCTCTTTTTGCAAGTTTTTATTAGATTAAGATCAACAATTTACCAGGATGAATTATATGACTCTGAAAATGACGATTTTGTTAAAGGAACCGTTTTTGAGTTTCACAACACACATTGGTATATAGTTCTTTTTCCTGAAGGCGAAAAACTAGAAGAAGAAAAAGATTCAAACCAATCAAGTATTGGGGATTTAAAACCCTCAAAAAAACCATTATTATCTTTCAACCGTGAATCTTTTAGAAATAGTTTTAAAAAGAATAAAAAGGAATCACTAGTCACCGAATTAAAGGAACAAGAAAATGAAAAGGAGGCAACTGATAAAAGTGTTGACAAGGTAAAAATAAAGAAATATGCTTCAATATATATGATGAGAGACATAAGTAAATTTAACGAATCGTTACGCCATGATGTCAAATTTATGATACTAACTGAAGGTGGAAATGATGTGGCGTTAGAGCAACATTTTTACGGAACGGACTCCAACGTTTTTGGAACAGCTTCTTTTATGACGGCCAAACAGTTAAAGTTGTTATGGAAAACAAATAATTTAAAAATAACAACAAAGTTTTTGCAGATAGTACCATATGTATATTTTGCCTACCAGATTAATCAAATAAAAGAAAATGTTGGAAACACAGAGTTTTGCTTTAAAGATCAAAATAATCTATCTTGGAGGTTTTGTTTTATGCAACATTCTACTTCAAAGGAATTTTTAAAAGGCGCTATTTTCCTTGATGACATTCAAGATGACAAGCAAGTCAAAAACTTTACATCACAAAACAAGATACTGATGGTATCTTGGTTTGTTGAGATATTAAGCCCAGTGGATATAGAGAAAAGTATTACATTTTTTCCTCGTAGCAAGCGAGAAATAAGAGAATCCCGAATAAGCACAATAGGAGAAAAAGATATGGTAACGTTTCCTGTGACAGTTGAAGAAGTGATTTCCGATGAAAGTCCATTTTTGGACAACAATAGGGCACTTTCAGTAAAATTGTCTATTTTATGCAGTAATTTTGTTGAAGACAATGAATTATCAGAAGCAGCATATGATGTTTTTACACAATCAAAAAACAAGATATCAACAAAAGAAGTAAACCATATTTCGCAACAAGATACTCTTGGTGATATTAACAGTGATGAAGATGATACCTTGTTATCATCTTTAAAAGATCACATAGATGCATATGAGAAGAACAAAGTTCATGACCCAAAGTTTCAGGCGACTAAAAAGATCAAAGGAAAAGAATGCAGCATAGCTCTACTGAAACAGGATAGCCGTCCTTCCTCTGATGATTCTAAAGAACTTTTTGATTTAGCCAAAGATCTTGAAAATGTTATAAAATCTGTTCCTCTTCATATATCAAAGATTCTATATGCAGGTGCTCTTGGGCAGACCACTATAGTAAGAAATCATTATCAGCTAGATTTCATCATATTCCTAAAAGATTACGATGATTACTCGAAAAACTTCAAAGCTGTTCTTACTGTTCTCCAAGAGCGTATTAGTGAGCACTGCAGAGAGGATGAATCAAAAGATGAGGTTTTTCCACCAATACTTCCGAAAGCACTTCATTTTTTTCATACTTCAAAAGGATTGTTATTTCAGTTGGGAGAAGTTAATGTGACAATGCTTATTTCCAGACCTGTTGAGGCATTTGGGGGATACAAGTTAATGTATGAAAGAGCTTCAAAAGCAAACCACATGAGTATTTCAATTTACAACGTGCTTGCTTCAGAGAGGCAGGTTGAGTTTATACGGAGTCAGGATGATTTGTGTAAAGATTTAATATGTGTTGTGAAAGCCTGGATTAACCAAGTGGATTGGATAGACATGGCCAGTCGTCCAAGCTCATATTTAGTGTCGTTAATTATTGTTTCAGCTTTTGAAAGAGTAAAAAAGTTAAACCTAATGAACACACCATCATTTGACAAAGAAGTGCTTTTAGAGTTCGGGAGGTTATCATCAGACAATAATTTGTTCATTGAATGGAACGTTTTTTATGAGCCAACTAAATATCCCATTGCAGAAAAAAATAAGATGTTTGCTTCGCAACCAGTAGTTCGTGATCCTGCTATGCCATATCACAATGTTTGCCACACAGGGATTGAGAACTGGCAACAGTTTCGAGCAGAAACAGAAAAATGGCTACTAACACTAACATATTAATCGTCTAAACAACGCGATCTGTAATTATTTTTTAACTATGTGTCTAACAGATATATTCCAAACTTTTTAGGAAATCAAAAGATTTTTTAATTTATTATAGTTTATTTTTCGAATCCTAGCATTTTTTTCTTTCTTAAAATGTATAAATAAATAAATTTTTTAACGAG
>evgR015222c0g1_i01_Hydra
GAAAATTTAAACATCATTGATTGGGATGGTTTTTGAAAAAAGTAAAATAAGCGTATTTTGCAGAAACTGGATTTGCTGTGTAACTTCTTCTGCTCAGACCAAATTTGTATTTAGGGATGGAAAAGAATCGACCCCACCAATCAAAGATTTAACCCCGATCGTTGATAATAGTATGATTCCTCGCATAAAGACCACCAACACTGAAGACAAAAATAATTCTGAAACAAAAAATACTGCCCAAGTTAAGAAGCAACAAGTCAATTCGAAAGTAGATCCTGAATTATCCTTAGAGTCTAAACTTGAAAACCCTCTTCAGACACAGGCTAGCAACACATCTAGACGTTCTCAACGTAACCGCACAGCACCTCATAGATATGGAGTTGAAGATGGGTGAATTCTACTGTATTCAAGTTATTATATATATAATTTTACATAAAAATCAAAGCTATTGCTTAAGTACATATGAACAGAAATTCAATAACTGGCGTTGTACACTTTATATAAAAAAATAAAGAGTACACTTAAATAATGAGCTTAAATAAAAAGTACAAAGTATATGTGTCCTATAATAATATAAATAATTATACATATAAAATATTTAAAATGTTATCAAAACGATTTATTTAAAGAAAAATACTAGCAATTGCATTAAAAAAAA
>evgR015225c0g1_i01_Hydra
ACCATTACAAATATTAAATTTTAATTTAATTCAATTCAATTTGAAATTCTAATTATTTTCTAACAATTGTTCTTTTATTGAAAAAACACTTTAAAAATTGATCAAATTAACTTATGGGGGGCAGGCAATCCTCCCAAAATACAACACCACGCAGACATCAATCGCAAGTTGAACGTCGTTCTTCTGTTTCTTCTCTTGAAAACAGTTCATCTCAAGCTTATGTTGGAACTTCTGGTTTGAGATTTCTTCTCAGGTCAGATAATTTATTGAGTGAGCAAAGCACATCACGAAGAAGGAGACGGGATCAAGCTGATGTTGTTAATCAAACATATTCACTTCCTGTACATTTGTTTCCTTCATTATTATCTGCAGAAATCAAATGCCCAGTATGTAACAAACGTGTAGGATCGGATGAAGTAGAGGCTCACCTCCAGGCTTGTCTTGCTAAACCGAAAGTTTCGTACAACGTGGATACACTAAAGGTTGATGCTGGAGAATGTATTATTTGCTTTGATGATATGATTATAGGTGACCGAATTGCCAGGCTGCCTTGCTTATGCATATACCACAAAAAATGCATAGACGACTGGTTTAAACGAAATCGGTGTTGTCCAGAACATCCTGAAGAGATTCCACAGGAAACTCCCTCACTGTCTAATATTCAAGACCAATCTTTAAATTTTACATGCGGTAGTTCTGAAATAGTAAATACGCAAAATTCTATATCTCAAAACATTACAATCGAAGATATTGTTGACAGTGCAGTAGCTGATACTGCCACTAATTTGGAAATAAATTGTCAAGTACAAGTTAATCATGGAGACTCTTCCAATTTCCATCGAGACTTTTCTAATTTCGTAAACACGTTTTCTGAGACAGAAAACTTGCCATATCTTGAAAATAACTCGGATATAAATGATAGAATAAACTCTCTAAATAATGAATCAAACGAACTAGAATCTCCATCACGAGTAACAGCTGTATCATCCAATTCGTTTCATAAAACATTAGTTCATGTAGATGAAGGTAGCGTGCATGGTTCTACTGCACTTTTTGAAGACAATGAACTTGAGCAACTTGCTTCTAAAATAAAAACAATTTTTGTTTGCGACTTAGAAACCGATAATAACGAAAATACGTGACGTCCTTTTTGACGTTGTTAACATGCAAACACATTTTCCTCGTCGTCGGGATGTTGAATTTTAAATCCGCATGAAATTTAATGAGAAATTAATTTTAATGGATGTGAAAGACTATCTTCGCTATTTGTTATTAAATAACAGCGAATGATAAGCAATGGAACCATATTATATGGCCAAATATGAAAGATTATATTCCTAGTCTGTTAATTAGTATCTAACAAAATCTTATATAGAACCTATAACGTATAATAAGTTTCTTTCTAAATATTTGAAATCTTCTAAAAACACTTTTTAAATGTGTTCCCCTAATCTGTTCGATTTTGATTGGTTAAAACTCTTATTGAAGGTGGTTTTATTGGTTGTATTGTAATTCATATTGTTTTAAACAAATATAGTAAATAGCCCTAAGATTAAAAAATTTCCAATATTTGGATGTTTTGTCATCACTTAATCTCTTTTTATATGCACTTAACATATTTGAATGTATAAAAATGAAAAAAATAAATATATTATATTATATTAAATATTAAC
>evgR015227c0g1_i01_Hydra
CATATGATGCAGCCAAACGTTTTATACTTGAAAAGTGAGCAGTTCAGTTCAATAGAAAAAGTGGAGGAAAGCATTATGGACCTTATAAACAATATCAAAGAGGCTGTGAAAAGTATCGAACAACGTCAGCTAAAAAAAAATTGGTATGCAAAAATTGTGAAAGAATTCAGCTCAAGTCTGAATGAACCTAAACGTGACATATGCCAAAATATTATACTTTCAAACAACTTCAAAAGCTGGGAGGATCTAGTTAGAAAAGAAAGTGAAAACAAAAATAAAATAAATGAACTAAAAAATATAATTAAGAAAAATAAGCATCG
>evgR015238c0g1_i01_Hydra
CCCGCCATAACTGAAACAAATCAATTATCATCTCTTCTTAGCTGTAGATTTTGCTCAAAAAGTTACCAAAAATTGTATTTTTTTAAAAAACATTTAACAATATGCCAAAGTTTAACAAAGGCTACAAATATAACAACATCTACAACACTTTTAAAGCAAAAAGCAAGTTGTATTCCATCAACGGATTTTTTTAACATTAAAGAAGAATGTCTACATTTTATATGGCACTCTATCAGTGATGCTGCAACTCATTCTATATTCAATTTAAAAGTTGGTTCTTTTAAAACACCTGGAAATTTATGTGCTTCTCTTGCCTTGAGCATGTTGAACAAGAAAGGTGATATAAATTTATTAGAATTCAATGAAATATTGAGCGAGCATTTTATCCAAGTTTTGCAAATAAATAGTGTGAACTTAGTTAAAGAAAAACAATCTACATTAATTAAGAACTATTTTATGAACTCAAAGTTAAAAGAAATATTGAGTGACATAGTGCTATCATATGGGTTTGAAGAGTACTCCGATGGACAACATATTCTTTACCAGTTTTTATTAACCAAGGTTTATGTTTTATATTTTTATTAACCAAGGTTTAAGCTTCTTATTTGGAGAAATAACCAGCTAGATAATAAAATTCCTGTCAAAGAAGATCTTGTTTTGACTTCTCAAGAAGAGAAGACACTTCATTATGTGGCTGGTTTTATAGGGTTTTCATTAGCAAAACGATACAAAAATCAAAATAATGAATCAGCTAAAATATTTTTGAAGTTATTAGAATCTTGGAGAGACAATTCTAACGAAATCACGGTATGTGAAAGTACTTGTGACTTTACATCATGGACTAAAAAAATTGATCGAGGTGGTTTATTTTGCGTAAACAAGGACTTCTATACCTTAATTTATAGAATTGAGCAAGTTGTTCGTCAAACTTTAAATAAATCTTTGTTGACTAAATACCAAGGTACAGATTTAAGAGATGTGATTATGCCATTATTACAAAAAAGTCAACAAATCGATTCAATATGGCTATTGCTTGTTTGCAATATAAAATCTAAGGAATTAGCAGGACTTTTAAAAGATATAATTTTCAAAAGATGGGTCAATTTAAGAATAAAATCATTTGTTGATACTTGGATACAAATTGCGAAGAAAAGGAGGTCAATATCCAATAGAAGCGAACCTTCTATAAGAAAAATGCTTAATAAAAAAAAAACCAATACTTGACTGTTTTGTTTTATTTTTATAGACAATTAGGCATTTTAAAA
>evgR015240c0g1_i01_Hydra
AAAAAACGAACTAAGTTTCATTGAAAAAGCGGCGGAGTTATTCCAAGCGTTGGAGCTACTCAACTATAGTTAAAGACTTATATTGAATTTTTTTGATCCTATCATTGGCTAAAGAGCAGTCTATAAAAGCTCTCTCTAACTATATTCTGACCAATATGGCAGAAAAAGACGCCTTTAAAGTATGTGATCAAATTGTCCAAAAATATCTTCCGGTTGCTAATCCATCTGAGAACATAACTAAAAACAAAGAGCCAAGCAAACACAGACAGAAATACGCTAAACGAAGAGAGAATCTAGCTAAAAAAGAGATTAAAAAACCTGCAAAAGTTATTAATAACAAGCAATTAGAATCTTCTTTAGCAATGGATGTCAAACATACAGAAACTAACAATAAAATTAGTTTAAGACCGTCATCAGCAAAAGGGGGCTATGAAAAGAATAATGGCGAACTTCACCAAACAAAATGCACCAACAGAGATAAAAACTACGTAAGTATTAATCAACTAAGTTTGAATGATCGTTCTCAAAATAATGAAGCATTTATCTCAATTGATAGTCGCCAAAAATCAGTTAATGAAAGTCAGAACAAATCGACAAATAAATGTTGGCAAACTTCGGACACGGAACTTCAAAGAAAGATAAATCTGTTTTCTGAAAACGAACATTCTAAGGAGGATTTAAAAGACTCATTTGGAGTTAAGTCATTACAAAGTCAAAGTAAAAAAATACCACCTCTTAATATAGTAAGTAACTATGATTTTCAACATGAAGACAATGTTTCTTACTTAAACACCAAGGTTTTTTTTGACGATAACTTTGGTGTGCTTCCGGGCGTTTTGATTGATTCTGGCGATGAAATGGAAACTGGAAATTCAATTTATGAAACTGTTAGTGGAAGCCAAAGTAATAGGAATTCAATGCTTTCTCCAAGAAACATTTTTAATTTCAAAACATCTCTTGAAAATATTGCTTCAAAAACAGTACTTGGCGATGACAGAAGTTTTGAACAAAGGGGCGCTGATGGACGTATTCGTTCACGAAGTGGATCGGTCTGTTCAAATACAAGTTTAAATGAGTTATGATCCTATGAAATCCATGTATTAAACTAAAAAATCCATAATTTGAGCTAAAGAATCCATGATTTGAAGTAAAGAATCCATGATTTAAAATAAAGAATCCATGTTTTGGACTAAAGAATCAATGAAACGTATAATTAAAACATTATATAAATTAAACTTTAAAATTTTTGATATGTTTTTTTTTATGTTGGAATT
>evgR015249c0g1_i01_Hydra
TTTTTTTTTTTTTTCTTTAAAATTTCTAAAAAAATGGAAATCTACTTTAAAATCAACTTTATCGTTTTTTTGGTATTATTTAACATGGTGTCTAGAATCAGTGGGGCTTCCAAACTAGCAATTGGCGAAAAATGTGCACATCAAATGAGATTTAAAAACCAGGATTTTAAAAAGATACAAAAAAGAATAGCGCCATTGAACGAGATAAGCCCGAGAGGACTTGTTAAAGGTGGTCATGAATGCAGAATAGAAGCAAGTTGTAAAGTTTACGAAAATAAAAAATATTGCAAGAAAAGTTGCGTCTGTCGAATATAAAGAGTTTCAAGAGTATTAACTATATATATATATATTACTAATATTTTTATTTTAACCATTTAAAAATATTATTGTTTATCTTAAAATTAACAATTGAGTTGTGTATTTTCTGTGATTTGAAATTAACAATTTACTGTAATTTTTGTAGTATAATTTTTGTCCTTAACACGCATACAATCTGACTTTAGCATTTAGAACATTAGGCACATTAGAACTTTTGCCATTTTTTAAAAGGATAAGATAGTTTTTTAGGTTCGAACTTTTTTCCTCATGTATACATGTTAAAATTAGATTCTTTAATATCAGTTTTTAACATTTAAACCAAAGAATTTTTTTTTCATCCCATATCAG
>evgR015251c0g1_i02_Hydra
CCATAAGGTATAATTGCTCTAAAAGTGGATGGTTCAAGAATAACTCATTTGAAGATTGGTTTGTTAGTTTGCAACTACCAAGGCTAAAAAGGCAATCTGGGCGTAAAGTTGTCATAAGTGATAATCTAAGTAGTCACATTAATGTCAATATTTTAAAACTTTGTGATGACAACGATGTTCGCTTTATTGGGTTACCACCAAATTTAACACATCTTACACAACCACTGGATGTAGCCTTTTTCAGGCTAATAAAGATTGCATGGAGAAAAATATTATCAGAATGGAAAGCGGGTCCTGGAAAAGGTGCAACTAAGCTTAATAAAGATACTTTCCTCTATTAAGTCACCTTTATAAATCAATTCGAACAAATTGTGCAGAAAATATAATGTCTGGTTTCAGAAAGACAGGGATACATCCCCGCAATCAACAAGAAGTGCTAAG
>evgR015252c0g1_i01_Hydra
TAAAAATGAAGAGGTTTTTTTCAATCCTTTTTGTTAATTTATACATTGTTGCTGATCTCACTGTAAATGTGTTTTTTAATGTATGGTTGAATGATATAAGTAACAGCATATTCAAGTTTTCAATTAGCAACTCAATGTTCGATCTTTTAATTCTATCAGGACTGCGATTTTGTATATTACTAGGAGCATCCATTGGACTTTGTTGTAACTATGTTTTAGGTGTTGATCGAATTAAAAAATGTTCGACATATATTTTATATTATTCATGCTTTACAGCACTTTATAGCGTTGTGAAGATAATTTATTTTATTGAAGAAGATTTTAAAAATAAGTGGAGAATGTGGATACCTATTAGCATAACTATATCATCGATTTTTTTTACATACATTAATTGGTGTTTCTTGGGATTAAAATTTAGCAAAAGAAGTAAAAGTCCTTTAAATAAAGAAACAGAAGTTCATTTGCATTCTCATTCTAAAGAATATGCTAAGCAGAATGATTTAAATATTCTTGACGCAATAGAAGCCAAAAAATACTCAACTTCAAGAACCCTTTTAAAACTTTTACAACTTGCAAAATACGATGCATTTTATATATGCATTGCATTTATATTTTTAACACTTTGTTCAATTGGTCAAGTATTTTTACCGTTTTACACTGGGCTTGTGTTAAACTACATTATCATTGATAAATCTATTCAAAAATTTAAAGATGCAATTTTGCACTTGGCATTAGTTACACTTGCAACAGGATTTGCTGCTGGAATTCGTGCAGGTGTTTTTACATTTGTTTTAAGTCGCTATGTTTTAAGAATACAAAATTTATTATTTTCAGCTATTGTAAAAATGGAGATTGCGTTTTTTGACGACAGAAATACTGGAGAAATAACCTCAAGATTAACATCAGATTGTACAAAAATTGGAGACTGTGTTGGGCTCAATGTAAATATATTTACTAGAAATTGTGTAAAAATTATTGGTATTTTATTTTTTATGTTCAAGTTGTCATGGAAACTGAGCTTACTAACCCTTGTAAGCCTTCCTGTTGTAGCAATTGTGTCTGAAATATTTGGACATCGCTATCGAAAGCTTTCAGAAGGTATTCAAAATTCTTTGGCCCATGCAAATGAATGTGCCCAAGAAGTTATTTCATCAATTAGAACAGTTCGCAGTTTTGCAGCAGAGAATCAAGAAGTTTCTAGATATTCAGATCGTTTGTATATTACATATAATTTAAGAAAAAAAGAAAGCTTTTTGCAGACTTTTTATAGATGGTCATTAGAAGTTACAGACATGGCTGCCTTTCTGCTTATATTATTCTTTGGTGGAGACTTAGTTATAAAAGGTCATCTTACTGGAGGTCATTTAGTTTCATTTATACTATACAGTATAGAGTTAACTAGTGCATTTGATGAAATAGGAGATGTTTATACAGGACTTATGGAGGCAGTTGGTGCTGCTCGTAATGTGTTTGTGTATATAGATCGAAAGCCAAAAATTATTGCTGGGGACTTTAAACCAAGATCCCGTTTAAGAGGCGATGTTGCTTTTAAAAATGTGTTTTTTTCCTATCCATCTAGACCTGATGTTGATGTACTTAAAAATATAACTTTTCATGCTAAGGCAGGTGAGGTTTGTGCTCTTGTTGGAGCAAGTGGAGGAGGGAAAAGTTCTGTAGTAAATCTTCTAGAGCACTTTTATGAACCATCTTCTGGTGAAATTTGTTTAGATGGCTTTAGTGTTAGTAGATTTGATCACGAGTTTTTACACTCAGAAATAAGTTTAGTTCAGCAGGAGCCTGTGTTATTTGCTCGATCTATATCGGAAAACATAATTTATGGTTTATCTAACAACGTAGACACAGATGAAATCATAAAAGTTGCTATAATGGCAAATGCCAATTCTTTTATATGTGATATGCCAGAAAAATATGAAACACAAACAGGAGAAAAAGGTATACAACTATCTGGAGGTCAAAAACAGCGCATTGCAATAGCACGCGCATTAATTCGAAATCCTACTGTGTTAATTTTAGATGAAGCTACCAGTGCTCTTGATACAGAGTCGGAATATTTAGTTCAGCAAGCTATAAATAAAAATTTAAATGGAAGAACAGTTATTATTATCGCTCATAGATTGAGCACAGTGGAAAAAGCAGACAATATATTTGTTTTTGATAAAGGAACAATTGTTGAAGAAGGTCCCCATGCACATCTTATAAAATCTAATGGTGTTTATGCTGGGCTTGTTAGAAGACAAATGCTAGAATAGAATAGCCATTTAATAGAATAGAAATAGAAAAAAAAAAGGAATAGAATAGAAAAAATATATATATTAACTTACATACACATACAAATAAATAAATTTAAATAGTTTAGTTATTAAATTATTCACACACACACACACACTTATGTTATTTTATATATTTATATGTATTAAAAAATAGTTAATGTATTTTAAAGTGTTCCTTAATAAAATATTTTAATTAAAAAAAAAA
>evgR015259c0g1_i01_Hydra
ATATTTGTATGGTGTTTATCAATTTTGGAAACATATTAAGATTTTTTATTTGCTAACGCTGCTTATTGATACAAAATAATATCTAGAAGAAATAATTAACATAACATACTGATATATAACAAGTTAATGACATACCCATTGGCATACAAATATGGATCCAAACGGAGAAAGATCAGATTACATGTACATCCCTGCAATTGGGGAATCTACAGTTCAGCTACAAACATACACACAGAGAGATGCACCAAGACCTTGTTATTATTTCTTTCGTTTGTTAGGGATGTGGCAACCTAGAAATGCTTTTCTTATATTCAAAGTGTACAACTGGATTATGTTTACACTTCTTTTTATAAATGCAATCTTAATATTCATGTTGAGCTTTGTTCATTCGTCAAGCTTTAAATTTGCCGAGGTTTTTAACAGCATTGGCTCAATATTAGACTTTTGCTGCCCATTTTTATTTGTGAAATACTATTTTCGGTATGGAAATTTTGAACGAATAATCAAACACATTTATGAACAGAGCACTGCAGAAGATTTGAAGAAGCTGGGTAGATTGAGATGCATTTACACTTTTTTATCTTTTTTTTTGTGGGTAACCATGTCCGCATATTTTATTAAACATTGGGAGCGTTTCTTTACAAACAGTTACCTTAGTTACGAGTTAACTGCTGCAATCTATATACAAGTTGTTGTAATTACTATGGGTTGGTGGGCCTCTTGGTTATGTTTGTACGGGTATGTTTGCCACATTCACGTTCATCAAATTAGGTTGTATGATAAGCATGTGAAAAATAC
>evgR015262c0g1_i01_Hydra
CTTGTCGATAACACAAGATGATCTGCTTAGTTTTGCATGGCAAGTGGCTTCAGGAATGGAATATCTTTCTACTTTAAATCTGGTACATCGAGACCTTGCTGCCAGAAATATTTTAATTGGCTCTAACAATAATGTAAAAATATCTGACTTTGGTTTGACGAGGAATGTTAGTGATGAGTTAATGTACATGAGCATCAAAACTCGTCGTTTGCCCATTAAATGGATGTCGGTTGAAGCTTTATTTAATAATGTCTTTACTACATATAGTGACGTGTGGGCTTATGGTATTGTTTTGTTTGAAATTGTCACACTTGGAGGTACACCTTACCCAACCATAAGTAACCGTGAACTAATCGATCGTTTAAAGTCTGGATACAGGATGGAGAGGCCGGATAACTGCTCTAAAACTATGTATGATATTATGTTACACTGTTGGAATGAAGATCCATTGCAGCGACCTAATTTTACAGAGTTGCGTGAACATCTTGAAAATATTATATCTCAAGGAGATCGTTACTTTAGCTTCGAAATTGATGAAGAAAGTGCTTATTATAATATTTCTTCGTTCAAAAGCCTCTCACCTGAGACAAGTGACAATTTTATGCAGGAAGATTTATTTTGAAATCCAAATGATAAATGACATAAATATATGACATGACATATGACATGACACATACAATATGAACTAATTGTTTGGATAAAGATGTAGATAGTTTTTAAACAAATATAAAAGTTTGTGTTTAGCTTAAAAAAAAAA
>evgR015265c0g1_i01_Hydra
TGTTTCAGAAAATAAATATTCATCAGGAGAGCTTATTGCTCCACAAACTTTCTCCAAACTTGTTATAAGAAATAGAAACTGAAACTTTTTCTATAGATGGCAGAGTACATTCACTAAAAGTAATTAGAGATAGCATAACAAAAAATTATTGGATTTATATGAGGGTTAAAACTGATGAAGAGTATGAAAAAATGTCGCGTATATCAATTATTAACTATCTAAAAAGGATTAATGAAATGCAAGAAGTTGACCAAGACGAAGAAACCAATAACTTAAAAAATAGATTAAAAAAATTTGAACGAAAAAGACATCTTATGTTATGGCATGATACTTCAACTATAAGCAACCATAGTCACGTCTTAATGACAATTTCAACTTTATTTGATCCTGCAATTCATTATACATCTGAAGAATATCAACAAAAGTATAACAAAACCATAAATATTCAAGCGGTTGTTGAAGCTCCTATATGGTATATCTTTG
>evgR015267c0g1_i01_Hydra
CTTAAATCAAAATAGCAATATAATAAAAAGTTTTTCCCGTGCTTTTTTGATAAGAATCTTATGCACAAATAATAATCCTTTGCATTTATAGTAATCTGACCCTGGCACAAATAAAACTGTTTTGATGATTTCTTCGTATCAAAGCTAAATAAAAGACATTTAGCAGTACAAAATGACAAATTATCAAGAAGTTATTGAATCTTCTATTGGGCAGTTTGGTCGATCTCAGTTGATAATTTTAATTGTAACTGGTTTGCCTCAAGCTATAGCATCATGGACAATGGTTATGTTAGGGTATGCAGGAATAGAGCAAGACTGGTGGTTAGAAACGACCAGTTTCAACAGTACATCTAATTCTTCATTAACACAAAGATTGTTCAAAGTATGTCCTAAAAGTGGACAAAACGAAACTATCATCTTTAAAGAAGGTCTTAAAAACATAATTACAGAGTGGTATTTAATTTGTGGGAAATCATGGATTGTTACTACTATTACCACTATACAAATGGTGGGCGTCATGCTGGGTGCTGCTATTGGTGGTGCTCTGGGAGATTTCATTGGACGCAGAAAAACTTTGTTAATAGTTTTTTCAATGAATCTGTTTTTCTTCACTCTGGAGGCTTTTTCTATTAATTGGATGATGATGGCTGTATGTGTTTTTTTTGTTGGAATTTCTGTTGGAGGATTAATGGTTCTAATTAAAGTAGCTTTCTTAGAGTTTGTTGGAACCGAATATAGAACAATTGCTACTGTGTTGCCTTTTTGGGGATTAGGTGTTTCATTGTTTGGTTTATTGATAAAGATATTCCCTTATTGGAAGCACATATGTTATACTACTTCTGCAACAGGAATTCCTGTATTTTTATTGATGTGGTTTTACTTACCGGAAAGTGTTCGTTGGTTATTTACACAAAAACGTTTTGATGAAGGTGAAACAGTGTTAAAGTCGTTAGCTAAAGTAAACAAACTACCAGATAGAGATTTATCAGACATTCGTAAAATTGTTAAAGAAGAAGAGACATTAAATCAGAAGAGACCCACTTATACATATCTATCCCTTTTTAAATACAAACAAACTGCATGGAGAGCTCCTTTATTTGCATTCTTATGGTTTACCTGCACATTTGTTTATTATGCAATGACCCTTGGTGTTGGACAGCTCTCTGGAGATCGCTCTATTAATTTCCTTTTTATGGGTGTTGCAGAAATGGCCCAGGGTATCGTAGTTTGGATTTTTGCAAAATGTTTTGGTCGTAAAAAAAGCACTATAATTCTGATGGTTTTAACCTCTGGGTTTTCATTTGGCATCACTGTAGCATACTTTGCAAAAACTACTCGTTTTAAATTGATTATGAGTGTTCTGGCATTAATTTCACGCCTTTTTCTTACAGCTGCTTGGACATCTACTATTGTATTTACAGCTGAAACTTTTCCAACTGTTGTCAGAACAACTGCTTATGGCTTTGCTTCTTTTTTTGGTCGCCTTGGTGGTATTGTTGCACCCCAAAATAGAGTTCTCAGCAATATTTCAGAACATTTGCCATTTACTGTAAATGGATTTTTAGCACTAACCAGCGGTTTACTCTGCTTTTTACTGGATGATACATATACTATTGAAATGCCTGATTATTTAGAAGAAAAAGAAAGCACTTCTTTGAAAGAATTTGCTGCCGAAGAAAAATAATTTTCTGGTTGTAATCATTATTCTCCGTAATATAAATATGTTGATAGGTTTTTTCTACGCATTAAGTGATAATGCGTGTTTTTGGACATTTTGAAAGTTTTTTATAAATTGTTTATTATTTTAAGTAGTAAACAATTTATTTACTTTTTTTTTGTGGAAATTTATCTTATGTCTTATCATAAGACTTTTAACTATTTTATGTGATGTTTATAGTTTTTGTTTAGTTTAACAAATATTATTAAAGATTTTTCAGACAACTTTATGTTTTGTGTTTGGTAAATACCTTTTATATTTTGTGTTTACGTTAAATAACTTGTAAATTAGTCTAAATTTTAATATTGTAAGATTTTATCTTTCTAATTGAACTTTGTTCAATTTTAATAATTTTGAAGTACCTCATTGTAAAATATACTAAGTACTTATTTGAAATTTTAATTAGACTTGCTAAAGTTGAAAAATG
>evgR015269c0g1_i01_Hydra
TTGCATTTTTTGCCCCAAAGGATTTAGCTGCTTTCCTTTAACATGGAACCAATCATTGAGTTTTACTATAGGTTACCAACTATGCAAACCAGGATTTTATTGCCCTACAGGAATCAACATGATAAAATGCCCAGCAGGGACATATTCCAACCAATCAGGATTGGAAGATGATTCTGAATGTAAGCCATGTGATCCAGGTTATTATTGTTCTGGAAACATATTAGTGCAGCCAGACGGCCCATGTGATGAAGGATTTTATTGTGAAAAAGGTGCAAGTGTACCACGTCCCTTTGTAAGTTCATATAATGTTACCAATGATTCATGCCCTTATAATTTACTTCTTGGCATCTCATTTAAAGGTATTGGAGATATTTGTCCGAAAGGTCATTTTTGCAAAAAGCAATCACGGAAGCCACAACTTTGTCCAGCTGGCACCTTTAATAGCGATGTAGGCCAAGGAGAATGCAGTATATGTCCTGCAGGTTATTTTTGTCCCGAAGGTTCAGTTTCTTACTTGAGAAATGAATGCCAGAGAGGTTATTTCTGTCCATTAGGCACTACTCACGGTGATGCAAACCCATGTCCTGACGGTACTTACAATAACAAAACAATGCAGAAGTCATCAGATGGATGTCTACTTTGTACTAAAGGGGCGTATTGTTCGGGAGTTGGAAATATTCTTCCAACTGGTGATTGTGAAGGCGGTTGGTACTGTGATGCTGGTTCAGCATCTTCTACACCTGTTTTAACTGGAAGTCGTTGTCAACCTGGTTTTTATTGTCCTACTGGGAGTCATGAACCTGTTCCATGCGAGCTTGGAACTTTCTGTGACACTTACCAGTTAAAAAGTCCTACTGGAAATTGTTCAGCTGGTTTTTATTGTCCTAAAATGTCAACAAACAGAACATTTTTAGCATGTCCTTTGGGAAGATTTTGCTTAGAAGGAGATAAGCACCCACGAAAATGTCCACAAGGTTACTTTATAAATGAACTGTATGCTAAATCAATCAATGACTGTAGACCATGTACAGCTGGTAAGTATTGTGGTGAACCAGGAATACAGACTCCAACTGGTGACTGTGATCCGGGATATTATTGCCCAACTGGCTCAGTTTCTAAAAAAGAAAAAATATGTGACTTAGGTCATTTTTGTATAGGAGGAAAGGCAAAACCAGAACCTTGTTCTTCTGGTGAATACCAAGATGAACAAGGCCGTAATTCATGTAAGCCTTGTGTTGAACGATTCTTTTGTGACACCAAATATGGACCTGTCAT
>evgR015272c0g3_i01_Hydra
CTTGGGAGCAATATTTTTTGTATTTTTTGTTTATAGGTAGTTATTTCTAATTATCCGACTTATAATTACATAATAAAGTCAGCGGTGCATACTAAATGAGAATACAGAGCGACGTAAAATTGAAACGACTGTTATTTTGTTAGCATAAAGTTAACTAAGTACTTTATAGTTTGCTTCACTACTTAGACTAATAAATACAGAAACAAAATGTTAATTGCTGATAAGCAAAACCTTGTTCAGCCTGATGTTTTAGTTGTTTGGTTTGTCAGAGTGGCTGGAGAAAACGAAGATCTACTCGGCGTTGATGAAACATCTATTACAAATATTCAATGGATTGCGATTGAAGTTGTAGAAAATAAGATTATCCGCGAAGGAGAGTTCATTATCTTCGATGAATGCGATAATAATGAAAAAATAATTGAAGCAAGTAACAACGAACTCACATTAAAGGAAGCAACCAAACAGTTTGAAGAAGTAATTTCTGAACTTTCCGAGAATGACCGAATTGTTCGCTTAGTTTGTGAGAACAGTCTTCACTTACGCCAGTGTCTAATACCACGAGCGCAACAACAAGAAATAAAACTAGGCGACATATTTCTTAAGTACTATGACATCAGAAAAGAATATAAGTCCTTTACTGAAAAAAATGTTACGTTCAAGAATCTCACAGAAATTTCGGAAGATATGATCATTGAGTCTACCGATATACTTTCAAGTCCTGACAAAGCGTGTAGATTCTGTGCCAAGGTGACTCAACATCTTCTAAAAGAAGGTCATAAGTTTTTAAAACTGGAAAGTGTCAGTCTTGCTTTAAATGATGGCATGCCGACTGGCTATGTTAAAGATGATACAATTGTTAGAGCACGCGGTTTACCATGGCAGGTATCAGATGTTGATGTTGCAGCTTTTTTCACCGGACTCAATATAGCAAAGGGAGGTGTCGCTCTTTGTTTAAATATAAAAGGCCGTCGGAACGGCGAAGCACTTGTACGTTTTGAATCTTCGGAACATCGTGATATGGCTTTAAGAAGACACAAACATCATCTTCTTGGTAGATATATAGAGGTTTATAAAGGTACTGCACAAGAGTTTTTAAAAATTGCAAAAGGTCCTGCCGCTGCAATGCATGCTGCTGCTGAGTTTTTAACTAATGGTGGAGAAGTTATAATCCGTATGCGGGGTTTGCCGTTTGATGCCACTGTACACGACGTCGTTGAATTTTTTGGAGATTCGCCAAAAGTGCTACAAGGGAAAAATGGTGTAATGTTAATATCCTACCCAGATGGGGCATCAACTGGAGATGCATTTGTTCTGTTTGAAACAGAAGCAGAAGGTCAGTTTGCTTTAAAAAAACACAGGGAAAACATCGGCAAAAGATACGTAGAACTTTTTCGAAGCACACGTGCTGAGCTCCAACAGGTACTAACTATGTATAATATTGGATACCAGCTGGTAGCGCCGGTCCCCGGTCAATTACCTTTTCCAGGAGCGGGTCTAAATGACAGGGCACTTATAAACCAACGCCTTCAAGCATTGATGAATATGTCATGTTTAAGAATGCGCGGATTGCCATTTTCGGCTTCTCACAAAGACATTTTAAACTTTCTTGGAAATTACAAAGAAAATGTTGTAGGAAACGTTCATATTATATATAACTTGCAGGGTCGTCCTTCAGGAGAAGCTTTTGTTCAACTTCAAACTGCTGACCAAGCGCATACTTGCGCAAGTGATCGACACCTTAAACACTTAGGAGAACGTTATATAGAAGTGTTTCAATGTTCTGTCCAAGATATGACATGGATGCTTGCAACCAGTCATGCTAATCAACTTGCGTCCCAGCAACTTAACCATGCAAGCAAAAACGTCCCTATCTACCATCAACAATCGCCTACTTATCACACCACAACATCACCACAATATCATCCTTCGACACCATCTCCTCCACCTGTTATTCAAACTACATCAGCTCCAACAATTATTGCACCTCCTCCTTTAACCTCTACTGGTATTTTGACACCTATTTCTCCCATCCCAATTTATACTGGCCAGTTTCCTCAAATAATTTCTCCATTAGCTAATGGATTCGTTCCAGCGCAACACCTTGGACACAGTCATCAAGTTATGCCCCCCATGATAAATGGACATTTATTCAATCAAGAGACAATTCATTATCAAATAAATCCGTTTGGCAAACCTGAAGTGGCATACATTGCATTCGCCCCTCATGAACAATTTCATCACGCTTCTTTTCATTTTGATAAACAGAAGTTTATAGGAAACAGATACGTTGAAATGTTTTACGCTCATTAATAAAGCATCTATAGCATAAATATTTATATTACGACAAAGATCTTTAACGAAATTCAAAAACATAAAAAAGCATTTTGAAATTATTGTACAGTTTCTGGTTTTAGTTCGGATTACGATTGATGAACCAGTCAAGATTGTCTCCTTTGTATTTATCAAAGCATTATTTATAACTTTTTAACAAAAATAGAACGCCATTATTATATTTTGAGGATTGAATAATAAATTAAACTACAAGAATATTTTAAGTCCAAATTTTAACGTTAAATATTTGAACTTTCAACTTTGTACAATTGCAAGTTTTTCACAAAACGATTCAAAATCAAAATCCTCTAAACAGAGTTTATTTTTCTAACCAACTAATGTTTGCTTTTATACTATTAGTACTAATGTTTATATAATTATTTAAATTTAATTTATTGCTATTTAGCTTTGTCATATATGGAATATTTCCGACCGAGCATATATAGAGAATTTCCAAAC
>evgR015278c0g1_i01_Hydra
CGTTTCTAGTCAAACTGTAGAAAGTACTACCCCTCTAACAACTACTGAGAAGATCACAACATCTGAAGTTTCTTCAGTAACCTCAACAAAAACTATTGCTGAAACAACTCTAGAAGCATCAACAACATCTTCAACAATATTTACTTCTCTTTCTACAACAGATTCTACAACGCACCAAGAGTCAACTGAATCGACATCTACTCCTTTTTGTAATTGTACTATGAATTTTGAGGAATTGAATTTTTGTGTTTCTTTAAGCAAGCAAGATGTGATAGATAACTTTCATGGAAAGTGTTATGCAGATAAACTGGTTGAAGCAATTAGTTTTCTTAATGCAAGAATAGAATATCTTAGTCTCCGAATTGTGCACTTTGATTCTTGTCGCAAACAGCTTCGAAGTGAAATTTTAATAAAGCGCAACCTAGTAAGATTATTACATCTTGTTAAAGATTACGATCAGTGTGCTATTAATACCGACTGCAGACTTGAACAGCTGAGATACTTAATAAAGCACAAAAAAGAACAGCTGGAATTTGATTTAGATTTCGAAAGATCGGGATATGGCGACTACTTTTTCAAAGGAATCTACAAGAAAGTGAAGAGTATATTTTAAAAATGTTGGTGAAATGTTCGAAGTTTTAAAGAAAAGCTTGAAAAAAAATAAGTGTTTTCGTGATGCTGCCACTTTGATAAGCGCACCTTTGAATCATTGCTGCATAAAAACTTGCATCATCTCAAATAAAGCTAGACAATTATATATATATATATATATATTATTATTTTTGTTTGTCTATAAGTTTTTGTGGTTTAGTTATTTATTTTAATTATAAAATAGTTATATAAAAAAAAA
>evgR015284c0g1_i02_Hydra
CCGCACTTTTGCGACAATCTTGCGCTCATATTACTCTTGTTTGTTGATTTTGTTATTTAGTTTCTCCATCATCCAAACTAGGAATTGCGACCATCAATTTAAAGACTAAAAAGTCTTTAGCTTCAAGCTATTTCATTAGAATTTGACATTGTAATAAAGAAACAGTGGTTAAATTCTTAAAATGGCTGATGCAAAAATAGATACAAGTCGTCTTGATGCCAAACCTCAAAGTATCAGTGATGCATATGCAGTACCAGCTAATTTTTTGGAAATAGATATAACAAATCCACAAACACACGGGGTCGGAAAGAAAAGATATTCAGACTATGAAGTTCGTATGAAGACAAATATTCCAATTTTCAAACAAAAGGAATCCACAGTACGGAGGAGGTATAGCGATTTTGAGTGGTTACGAACAGAACTTGAAAGAGACAGCAAGATTGTAGTTCCAACATTACCTGGTAAAGCTCTTAAGAAAATGTTACCTTTTCGCGGGGATGATGGTATTTTCGAAGATGAGTTTATTGAAGAGAGAAGACAAGGTCTAGAAGCATTTATAAACAGAGTCGCTGGACACCCGCTTGCCCAAAATGAAAGATGTCTTCATATGTTTTTGCAAGAACCAATAATCGAGAAAAACTATGTTCCAGGAAAAATACGAAATGTATAAAAAAAAATAACAATCATAGAAAAAAAATTAATAAAAAAGAAACATTACTAAGTCCAAAGTATTTTAAAAATTAGATGCCATCTTTTGAAATGATACTTTTTCAAATGTTTTTTTTTCACTTCATAATTTCATTATTGTTATTAACACAATTTCAATAATTTAGTTTTTGTTATTTCTTTTTCTCTTTTTCTTTTAAAATATTTTTTTTAAATATATTGTATATTTTTTTTCAAAATGTGAATAATTTCGTTATGTTAATTTGTCATCATGATTATTCGTAGTTATTATTTTTTGAAAATGAAAATTGTAAATTAAATGACAGTAATGAATGGCAGAAATTTTGAAATTACTTTATTTTTATTTCTTTTGCATTTCTACTTAATTAATTGCTCAATTTTTTGTATCATAACAAGCTTGTAGTTTTTTTTTCTTTAAAGAATTATTTTTTAGAATTATAGTTACATATTTTCATTTGTGTTTTGCAATTGTTGTAAAAAGAACTTTTTTCTGCATTGTTACTAATAAGATGAAGAGTATTTGAAAAAAAAAA
>evgR015285c0g1_i01_Hydra
TGTAAATAAAGCCTAAGAACAAACAACCGTGTTGATACACGACGCATTGAATAAGATTTGAATATATCAAATATATTTTGGATTACCTCGACAAACGCAGAACAATTAATGGGTTGGATTATTCACTCAAGCCAAATACTCTGTTGCTTTGTTTAGTCATACCGAGCGGGAAAAACTAAAACGGTTCGTCATTGCTCAACTTTGATATTCTGCCTATCAGAATGCAAGCTAAAAATAAAGGAAAAACAGTTGACCTAGTAAAAAGATTCGAATCAGGTCAGGTTGCAAAAACAACCTCAAGAACCCAAAGACAAAATGTTACAAAGTTACCAAACACGCTTACCAAAATTGATTCGAAGGAAAATGAAAACCCAAAAGATAATTTTTCAAGTAAAAGCTCTTTGTTTTCAATGCAAACACCACCCTCGTTGCCAGCAGAAAAAAAAAATTTACTAGACAAAGAATTAAGTCAGGAAAGCAGTCTAAATTCTACCGTTGAAAATGCACGTCCAATACCAGCAAAAAGGCCAACTGTGATTATTGTAGACAGTGTAACTACAACTCAACCTACGCCAATTCCAAGAAAACCATCCGAATCAAAAAACATTAATGAACTGTTAGAAAGAAAGGCTTTCAGTGACGGACTAGTTACAAAACTTCCTGTAAAAAAGACTTACGGTGATGTGTCTGTTACAAAATTATTAGAAAAAAAGACATACGGCGATGTATTAGCTACAAAATCTCCAGAGAAGAAGACTATTGGTGAAGGGCCTGTTAAAAAAAACACGGAGAAAAAAACTTACGGTGGTGTGCCAGTTACAAAAGTCAATGATAATCAAAAGAGAGTACGTGAAAATGTATTCCAATTAAATAAAAATAGCGACGAAACAAAACACATTGTTAAAGAAAATAACTTAAGTTTAACCGCCAATCACGTTCGTCAAACCACTAATCACGGTCATCAAACCGCTAATTACGGGCGTTCTCAAAGCACAGTTATAACTAAAAATCAAAACCTTGATTTAGGGAAAAAGCGTCTTTTAACTAAGCAACGCACCTTATCTTTAACAAAAGTTGAAGAAGCTGAAGATGACAGACACGAAGGTCCATCATCAAAGTTAATTGAGGAGGTGTTAGACGACGATGAAATTACAAATGAAATTCTAAAAAACAATAATATTGATTCATTTAATAAAACTGATGAACTAGAAATAGAAAATTATTCTAAAAAGTCTTTAAAAAGAGATCCAAGCATTATATCGGTTGATCCTAGTACTGGATCGTTTAATAAATTTGAAAATTGTTATCTCCATAGATCTCAGCAGAGTATTTTTGATACAGAAAATGAAAAACATTTTGAAAAGAATCATAAGAGAGCTTTATCTCCTTCTAATAATTGTAACATACGTACAACTTATGGTTCGAACGAAGCAAAAATTCATATAAAGTTTGACAACTACACAAACATTGAGTTAGAAGATGCATTGCAACAGTTTCATAAATATTTGGAGTCTAATTTAAGTCCTAAGTACAGCAACAGTTCACCGAACAAAGAAATCTTTGAAAAACAATCTCGCAGAAACTTCACTGACTCAAAATCCAGATTTTTCACTTGGAATAAAAAGATGGGACGAAGTAGCACATGGAATCATCAAACAAGAAGACGGATGCTTAATCTTAATAGTTTTGAAGACTTGAGCCCATCGTGCGAAAAAGATACAGCTCATGGAAACAAAGAAATGTTGAACCCTCGATTAAATAGATCTTACTCTTCCTCGGTAGTTGAAAGAAAAGATTGTTCTATTTTACATTATAATGGCTCTACATCGACGATTAACACTGATAATAGCGACATAGAATTTGAAAGAAGTTCGTTATCCAGATGGAAAAAAATGTTTAGAAAATATTCAAGCAGAGATAGCTTAAAAGAAGATAAGGAGTTTAATGAAGAAATTGGAAAAAAAGGAAATAAACAGAGATGGCTATTAAAGAACTTGTTTTCGTCCAATAGAAATGGCATTAAGAAATAGTTTTCCTCCAACAATGAAGTTAACTATGAAGTAAATTTTTCAGCCAAAAAAAATGGCGTCGAGATTTTTGTAAGCTTGCATTCTATCGCAATATTTTCAGTTCACATAGTTTTTGTTGCTGCGCCGTTTAATGCACAGTTGTATTTATTATATATATTACTATTCTAGAGTTTTTTAGCAGGTGTATTTTAATTTGTGTTAGAGCGTGATTTCGATGATTTTTTAATATATTCATTGGATAAAAA
>evgR015294c0g1_i02_Hydra
TCATGAATGGTTTTTGTATCAATAATATTTTTTCTTGCTATGGAGAACTCACGAGGCAAATGCAAAAAGTCAAGTTTTATGGTAGAAGACATTTTGTCGTTAGAAATACCTAGATACGGAGAAGAAAATGAAGGGCGACCATCACCGGAAATCAGAAAAATTCGAACTACTTTTTCTAGTGATCAAGTTTTTTTACTTGAAATGCAATTTAATAAACAAAAATATCTCTCAGCTACAGAACGGATGGAGTTAGCGCAAAAACTAAATCTTACTGATAATCAGGTAAAAACATGGTATCAAAATCGACGAATGAAATTAAAGCGCCACATAAAACTTCACTCCCAAAACAGTATATACAACCAAAAAAATCAAAATTTTATACAACCTAACATTTTCTTTAAGTCTGAGAAAACTTTTCTCGCAGGCTCGTCCGCGTTATATCCTCAACACGAACATTTATTTATGATGAGAAGAGAACATGAAAACCACGCAAACCTTTTTTAAACAGAACATATATGTAAATACATTTTTATATTGTAAATAGTAAAACAAGAATACTTTTTTTTGTGACTTCAGAAATTTATTGTAATACCGTAGTTATATTAATAGATTTCAATACATGACGTAGTGAGGGATAAAGAATAGAGGGGCTCGAAAAAACAAATTTAAAAGTAACTTCTATTCAAAACTTGAACATCC
>evgR015300c0g1_i02_Hydra
TTTATTAACTGATATACACAATTTTGATATCTATTTAACTTTGGCTTATTACATACAAAATGGAACATATAATTTAGATGTTTCAGATTTTCCATCTTACAACCTCACAAGCAGCTCATCTTACAAATGGAGTATACTGAATAAAAGATATAATTCCACTAAAAATGAATTTAACATAACAATTGATGGATCAGGTGGATTTATAATATTTAATAATCTATGCATTTATCAAGTTTGTACAACTGGTTATCAATTCAATAATTCAACTGATCAATGTGAAGACATCAACGAGTGTATTAACTTAAACCCCCTATGCAATTGGAGTAATGGAATATGCTATAATACAAATGGTAGTTATTTATGTTTGTGTAACACAGGTTGGATATTAGGGAGCAATAATGCAACATGTGTTTATAACTTTATTGGCAATGTTTCTGTGACAATATCAAAAGGAAATGGCATTGCTATAATACCACAACTAAATAAAGAGTATATAATATCATTAGATTTTCTCATAAAATCTTTGTCATCTGTTAATCCAAATTCTATAATTCAATTCACAAAATGTAATAATAATCTAGAAGTTTGTGAC
>evgR015307c0g1_i01_Hydra
CTTATTATTAGATATCCCTTCCAACAATTTTGAATCCAGATTATTCCGCAAGAATTATAGATGATTCTCTTTTTCTTGCACAACCAGCAGAGGCAGAAATGACAGTTGTCGATAAAATACACAAAAGAGTAAAGATGAAAAAGTTGTCACAAGATTCTAATGATACTGCAAAGATAACCAATTCAAAGGTTTTGGACCATCAAAGTTCCACAACAAATACGGATGATTCTCCTTTACCCACATATCCATTAGAAAAAAAGTTGTCCGTGACTCTTATTGACGCAGCAAAGGTAACACATAAGATATGTGGCATGTTGCCTGGTGTTACAACTCTTAGTTTCCCCAAGCCTGCAGATGTTGTTCCTTTTTTTAATGATGCATCAAGTGCGATATCTCTTCCAACAATTTTGAATCCAGATTATTCCGCAAGAATTATAGATGATTCTCTTTTTCTTGCACAACCAGCAGAGGCAGAAATGACAGTTGTCGATAAAATACACAAAAGAGTAAAGATGAAAAAGTTGTCACAAGATTCTAATGATACTGCAAAGATAACCAATTCAAAGGTTTTGGACCATCAAAGTTCCAC
>evgR015308c0g1_i01_Hydra
AAAAAAAAAAAAAAAAAAAACTAGAAGCCATAGTGCTTGCTGATCATGTGTTTGAAAAAGGTAATACCTCATTCAAAAAAGAAAATTTTGGCACATTGCATCAAATTAGAGCTACATCAAGCAATCTGAATCTTAATACAGCCATAAATACAGCCGTCAGACTAAAAACATCGACAGGTCAAAAAGAAGAGACATATGAGGATTACTACTCTGGTGATCATATACTTAAAATGCATAAAAGTTCCTTGATAGACAAATTAGAAACCCAAATAGAACAATCAAATATTCTCCGAATCATTTCAATCCATTCTTCAGATGCAGATATATATCCTTCAGAAGTAAGTATACAACCGTCAGAAGTAAGTATACAACCTTCGGAAGTTAGTATGTATAGTTTACAACCAATGCAATCGTTTTTATATTTTTCTAATAAAACAATAAATAAAGAGCAAAATTCAGCTATCTCGCCGACAAATTCTATTAGACCGTATTCTACGACCATACCGACAACTTTTATTAAAACTTACTTGAATAGCAATATAAAGAATGTTTTTGACGCTCCAAAACATATAACGCTTAAAAGTAGCATTTCTAGTTCAATTGAATACGATCAAACCAAGCAAAAAAAAGATAATTTGATTTTAACTATTTATTCAATGACACCCCCAAGTGTTGCAAAATACGCTTCTGCAATCACGGACAAAACGTTTACCAGTTTAAACCCAAGCAAGCAGATAAAAACCATTATTGTGCAACCATATTTGCAAACAACATCAACAATAATATTAAATCAAACAAAAACATTTTTTGAAGAAAAAGAAAATTCTGCAGAAAAAAATAATGTTTACATGAATACACAACTTAAAAGTTCTGGAAACAGTATCAACTCTTCATTTCTAGCAAAAACAAAGATTTTAGGTAAAAACTTCTTGGTAAAGTTTGAAATATTGCGGCTCTACACTCATCCGTTGGATACAAAATTATCAGAAGCATACCAACTACTGAAAAAAAGAACTGAAAAAGCGGTTTTAGAACTATACAAAGGATACGATGAATTCAAAGGATGTAAAGTTGTTGGACTAAACCGAATTAAAAAAGATAACATGACATCGGTGGAAATGGAAATATATTTCACTGAACGTGACGTCAGTCATTTAATACTTTTACAGCAACAATTAGCGAGTGGCAAACTTGGAGACAAGCAAACTGAAAAAGAAATGGATATAGCATTACTTAACGATGGGATCACGGATATAAATTGGTGTCTTCCGCCATGTTTTAATCCGTATCAATGTTACCCTAACTGTGAAAAAAGTTGCTGTGCACTTTCTAAACTAAAAACCAATCAAAACGACTTTAATGTTAATAGGCAGCTTTATTTAAACCAATATGTTGGTAATCCAGACATATTTGGTAGGCATCTAAACATAAATACATATGATGGTTATCCAAACAAAAATGCATACGGTAGATATTTACCCACCAAGCAAGATACTGGGCAACCAACCCTCAAGCAAGATACTGGGTATCTGACCAGCAAGCAAGATACGGTGCGACCAACTATAAATCAGTATGGTGGGTATCCAACAATTAACAATTTTTGGAAATATAATGATATGACAATAAACAACCATCCTAAAAATATACAATTGAAAAAAGATGACGATAAATCGCTATACTCTTTAACAAATGTCGAAGGAAACAATTTTAACCTCAAAAATTCAAATCCCGTGTATGTTATTTACAAATATATACCTATAAGTGTTGCAGGATCAGTTTACCCAAACGATAAGTATGAAATGAATGTTGCAGGAAGCATTTATCCAAATGATAAATACCAACCTAGTTCAGAAATGGATAATCAACAAGTGCTTTGTGATTCGTTTTGTAAATACAGGTGTTTACCAGCATGTCATAGATTATGCTGTGCGGCTAACTAAAAAATGATAAGCTTGTTTTGTCCAAGAAATTTTATTGTAAAATTTATTAATAATATAAATAATTTTATTTTTCGTTTAAAAAAAAA
>evgR015314c0g1_i01_Hydra
TATTGGTAATAGCTTTTTTCTTTCATAAAAAAGGAAGTTAAAAACATAACAGTTTTTGATTAAGAGGAAAAATGATTTGGAAGTACTCGGTAATTGACGTAGTGTTCTCATCTCTTGGAGTTTTTATATGTAACATTGGCGTATATATTTTACTTAAAGTAAAAAAAAAAGCAATCGATAGAACACAATGGTACATTGTTACCAACCTTTGTATTTTTGACGCGTGCTCTTGTATTAATAATGTACTTAGTTCTTCGTTGTATATTTCTAAAAATGAAAACATCATTTTAGATGTATTCCAAAATATTTTCATTCTTGGTTTCTATGGAACGACATTGTGGTTAGTTTTTGACCGATACTTGCACATTAGGCTAAACTTAAAGTACTATGCATTTTGGTCCAAAAAAAAAACAATTCTATTTACCCTAATATTATGGATTTTTTCAATACTTACTTCAGTTTTACTACATTTTTATAAAGTAGAATGTATTTACATTATTTTTGTTGTTTTTGACGTTGTTATAATAATTTCATCGATCTTTGTTTATGGATTTGCATTGAAATTATACCACAATGCAAAATCTATTAAATCTTTTAATGCCAGAAGACCTATTTTTAAAGGAATGTTGGTATCCTCTCTTATTTTACTTTCGTATTTAATTTTAGTGGTTATCCCAGATATCTTGATTATTTGTACTTTTGACAGTATGGGCACTTGGAACATTTACAAATTTTCGTATATAAACTGTAGCTATATTGTAGCCATGTGGACTGACGCATTAGTGTATGTTGTGGTTTCTCCTCAGGCTTTGATTTTTCTGAAAATTAAACGCTCGATGGTCAGACGGTTGACAGTCAGTTTTTGGAGTAGCACCACTACTAGTAAAAGTAAAAACAACAAAAGACTTTTGATAGTTTCAATTAAAAACGGACCGACTGTAATTTCGACCATTAGCAGACCTTCAATTGTTTCAATTATTAACGAACCATATTTAATTTCGATTATTAAAGGATACTCTGTAGATTCGATTAAAAATAATAAAATGATTGTTTTGTGATTTGCAGCACTTAAATCGATTTAGAAAAATAAAATATCGAAAAAGTTCGATACGTTATTTTAATAAAATATCGAAAACATTCGACACGTTATTTTAATAAAATATCGAAAACGTTCAACACGTTATTTTAATAAAATATCGAAAAAATTCGAC
>evgR015325c0g1_i01_Hydra
AACTGCAACACTTGTTCTAAGTTGTCACTTAACAAATTTATTTCACTGTTAATTGTTGTTCGAGTGACTTGTGTTGGAAGTTGTCCAACTTACATGTTGGAGAACAGTGGAGAAAAATTATGTTACTTATTCCAAAGTTACAAACTTAATTGGATGGATGCTAATTCTAATTGTAAAATTAGGGGAGGAAATTTATTAAGCATCGCTAATTATAATGAGTTTCTTCTCTTGGCGCAGTATTTTCCAATATGGCAGTATTTCTATATCGGTTTAAACGATATAAAAAGTGAAGGTCACTTTGTGTGGTCGGATAATCAAGCTGCATCGAATTTTTTTTGGTATCAAGGCGAACCTGGCGGCGGAACAAGTGAAAACTGCGTAGTAATTTGGGCCCCTTCATATCTTAATCTTTATTATGGAGATGTTGCATGTGATCGCGGTTTTGGTTTTATTTGTAAATTTCAAGATTATTGTGATAAAAACAGTTGCTCAGCAAATGCATATTGTGACATTGTAGTTAATGGATACATCTGCACATGTAAATATGGTTTTAGGACAAGTAGCGGTGACATAGAGAAGCTTGAAGATCAATGCCAAGACATTAATGAATGTGCTTTATCAACAACAAATTATTGCTCAAGTAATTCTATTTGCAACAATTTTGATGGTGGATACAATTGCAAATGTAAATGTGGCTTCAGAACCAATACTGGTATAATAGAATCGGAATTGGATAAATGTCTAGTTGAAAATTGTGTATTATTAAATTTATCTGCAAGTTACTTCATCGGAGAAGGGCATACTTATGTAAACATTATAACAAATTGTTGGTGTAATTTGCAAAATGTAAGTATTTATCTGCAACCAGAACTTATTCTTAAATTCAAAAGATTAATGTTGGGCAATTCTAACGTAAATGATTTGAACGTGTATTCAAATGGTAAAAGTTTTGTAATTCATGCTAACAATCTTTTTAAAACAACATATGCATCAATTTCTGCTGTATTTGGTTTCGGTGAGAATATACCTGTTGGCAAAACCAACATGACAGTTAACCTTCAATATACCTTGTTTTTCTCTAAAGATTATGTTGTTAAAGCTGCTTGGAAAATATATTTAATATCTGTTGAAGGATTAGTACTTTCTCAAGTAAATAAAATGACTCCATCGTGGACATACAAGCAATCAAATGGAACATCATTATATACTAAAACTCACCAGTTTGTGTGCGTTGCATTAAAAAAAAGACAGCCAAGTCCTTGTTATCAGCGAAATGTATTAACAGGAATAACAACTTTTCTTCCAGTTCAAGTACTTGACGTGATTGGTTATGATTCTAATACCACTTTTATTTACGGACGCACAACGAAAAAAAACTTTATTATGGTAAACATAAATAGTATGAAACCTTACATTATCACAAAGGAAAAATGCTCGAAAATAAAAACCTGTAATTTGGAGGAAAATTGACACGTACGTGGTTATAAACCATTTGTATTTAAAACCGAAAATGACTTGAGCTCGATAGATAGAACTTAATCGATAGAACTTAATGGATCATTGATATCGAATAAAGAACATTGATATCGGTATCAATATCGGATAAAGAATATCGATAAAACATAAAGGATTGTTTAAAAAGTGATTTTAAAATTAACAGCATTCATTTGGTTGTGATATTATATTTGTTTAGGAAATTATAAAGTACTTATAATACTAATTTGGATTGTAAAATTCTAAAAAAAAAATCTTTAAAGTGTTTAGCTACTGGAAAGTTTTTG
>evgR015332c0g1_i01_Hydra
GTTTCAATGTTTAACTCTTAAAATTTAGAAAAGTAAAATCATCTTCAAAACAATGACGAGCGGATCGTTTCACATCATAAATTGTTCTCATAATTATTTGTTGGTACCGTTACAAGCTGAAAAAGATGGAACCTTGTACGTGGTTATCGGTTTATTTATTGTTTTTTCAATACCAACTATATTCATGAACGTCTACATAGTCGTAGTAACATACTGTTGCAAGAAGATGCATTTCCCACCCAATATTTTACTAGCATGTACGTCATTTTTCGATTTGTTAAATGGAATGTTTACATTCTTATTGTTTGCGGCTGTATTAATATTGGCAATTCCTAGAAAACAGAACTGCGTTATATATTTATCTTTAGTCTTTACATCAAAGTCATTTGGTGTTATTTCTTTTTTGTTAGTAAGCTTAAGTTCAGTTGATCGTTATCTCGCTGTTTTTCGACCTTTTTTTTACCATGCCAATATTCAAAATAGTAATTGTTTGTACTTTAAGTTAGTTTGCTGCATTTCTATTGTTGTTCTGCTTTTATCAGGATTGAGCTTTTTAACCCAAAATAAAGAATTAGTCTAATTAGTTGTATTAGCCATCTGCCCATTTTTAGTGATATCTCTATGTATCCATACTAGATTGCATTTTAAAGTTAAAACAGTTCGACAAGCAATATTAAAGCAAGCTTTTAAACCTGACACCCAAAAAAATATGAATAAGATAGAATTAGATGAAAACATAAGGACTAGAAAAGAATTGAAGATAAACAGTCTTGCGCTAATGGTTTTTTTATCAACGTGTATTTCTCATTTTCCTGT
>evgR015334c0g1_i01_Hydra
CGGTAACTGCTTCTGTAAAATATACAATTTTATTGATTATGCTCTTTGCTTTGAATCAATTTTAGCAGTTACCATAATTAGCTTTGATCGGCTTTTACTTTTACGTTATGGCTCCCGATACAGCGTTAAAGTTACAATGAAACTAGCGTGCATTCAGATAAGCGTTACGTGGGCATTTGCGTTTATGCTTTATGGTCCGTTTATAATTGGCTGGGATAAGTGGACAGGAAAAAACTCTCTCAAAAAAGGTGAGTGTCATGTGCAGTTTGCACTAAATTCGACTTTTACTACTATAACGTCGTTGATTGAGTTTGGGATCCCTTTATTGGTTATTGTAGTTGTCAACGGGATTCTTTATAGTAAAATAAAAAATCGAAAGTACACAAAGTTTCAACGAAACATAAAACAATCAAGGACATTGAAAAATAAAATCCAGATAATTAATAGAAAAGATTTTAAGGCTGCAAAATTTCTTGCTGCGTTAGTTATGGCATTATTTATTACTTGGGCACCTTACACGGTAGCTACTGTTGTCATCTCGTTTTGTCAAGAGTGTTTGAATACATACTTCTATGAGTTTTTGAAATGGTTGCTATGGTCGAAAGCTGTTATAAATCCGTTTTTATACGCTTATCACAGTGCAAGATTTAGAAGAAACTTTAAACAATTGTTCAAATCAAAATTCATGGCAAAAACTTCACAGGTAAAAAAACAATCTCTGGCCTATGTTATGGAATTGCAAACCACCAACCTCAAGTAAAGCAAACTTAAGCAAGATCATGAATAGCAATGATTTAAACAAAGAAAATATCTAAAATTATAAAACATGTCAAACTATAAAAGTTATCTATTAAAAATCAATAAGCTAATGAATAAACGCATAAATTGTCTAACTTTTGTTTTTGCCTTTCATAACCACGAACTTAACTTAAAACGCTGAAAAATAAAGCAACCTTTTTTGTACTAAACAAAATACTACAGGGTACAGTTTTATTTTTTTTATCCTAACTACATTGTAATATTTCGTTTTATTTCAGCATTCCCTTAGTAGGTTTTGAAACTTTACTCCATAAATACACACATCAGTGATGTGCTGCCG
>evgR015341c0g1_i01_Hydra
TGTAAATATGATTTCGTACGCTTTTGCTTGGTTAATTCCGCCGAGAGCAATAAAAGATTAAGCCCAGCATTCTAGCATTCATCGTCGGTCTTCACTTGAGCATTTCACAGATATCTTAAAATGGATACAAAACTTGTGGTTCTCGTTGCTCTCTTTGGAGCCTGCATGGCAGGTATGCCAAGACATGTTGATAAACGCTCACCAACTGCGTGTTATGCTGGCTGCCCAGCTTTCTGTGCACCCGCTTGTATGCCAGTCTGCTGTATTCCACAACCACCACCACCACCAGGACCACCAGGATACCCAGGACCATTAGGTGCACCAGGACCAGCAGGACCTAATGGACCACCTGGACCACCAGGACCTCCAGGACCACCAGGACTTCCAGGACCACCAGGACCACCAGGAGCTCCAGCTGGACCACCCGGACCACCAGGAGGCCCCGGACCAAATGGACCACCCGGACCACCAGGACCACCAGGTATGCCAGGACCACAAGGACCAAATGGACCACCAGGACCAAATGGACCCCCAGCACCACCACCACCCCCACCACCTCCACCACCACCACCACCACCACCATGCCCAGCTATCTGCGTTATGCAGTGCGTACCATCTTGCCCACCACCCTGTTGCCCACAGAAGAAGCATTAGATGGCAAAGTAGTATTTCTAAGAAGCTTTTTTATGGACTAGCATTTTTTAATATAATAGTTGCATTAAAGGCTTGAAGCTCAAGTATTTGTATACAACTCGACTGCGCTGACGACGATTGGCTACAATTTATGTATGTATCCATGATATTTAGCTGTCCTTGAAATTAAAAAAAAAAAAA
>evgR015353c0g1_i02_Hydra
TTTTTTTTTAAAAAGATGGATAATATTTTAAAAGATACAGATTTAGAAAATCAATTTAAAACTGCAAAGTCCTATCTTTTAAATTTAGGTCAAAATTCAAAAGTTACATTGTTTTGTCCACCATCTGGCCCACTGTGCACTGGTTGTATATGGTATGATCATTTAGTTGATCTCATTTCTGATATTCTTGAGCGAAAGCCTGATGTTGTTTCAAATATATTAGAAGAATGCAGTTGGAATTTGAAAAAGAAAAGATTTCATATGGAGCGAGACTGTCTACAGGATAAACAGAAAGTCTCATCAAAAGTGGCTTATGCTAATATACAAAAAGATTTATTAAATTTTCAAGACAATAATTTCAAACAAAATAATCAAGATAATTGTGTTCTAACTGATATCATGGAGAGTGCATACTATTTTGAACAAGCTGGTGTTGGTTTAGGGAAAGAAGAAACTTTTTTAATTTTGTTGGCACTTAGACAACTTAAGAATGCATATAAGCTGAAATCATGCAGGTTTTGGGGAAAAATTTTAGGTATTCATAAAAACTACATTGTTGCTGAAGTTGAACTTTATGAAAAAGAAGAAAATTTAAACAAAGAACAAGTTCAAAGTAAAGAGCTTCAAAGTACAATTGAAACAACTGAAAATAATGTTCCAAATCAGCCATCATTATCGAAAAAAACAATTGACATTCCGGAAGAGAACGATGGCCAAGGTTGCAATGCAAAAGTTTACTTTGTGTGCAATGATGCTGGCTTGTTATGGCAGCGTCTTCCTTCAGTCACGCCTAAACAAATAACAGTAGCTAGACAAGTTCGAAAATTATTTACAGGGTACTTAGATGCTGAGATTATTTGTTATCCACCTTTTCCTGGAAATGAAGCCAATTTTCTTCGTGCAACAATAGCGAGGATTTCCGCAAGTACTCATATTAGTCCACAAGGATATTATATGTTTGAAGAAAATGATAATGATGATGATGATGATGAGAAAAGTGATGCTGGAAAATCATTTATTCAGAATATGGAGTTTGAAGGAATTCCAGTCAGTGATTTATCAGATCCATCAAATACATTCTGGGTACATCATTGTCCTTACATCTTGCCACAGGGTCGAACAGTCTGGTTCAATCCTTATGCTAACGTTTCTTTGAATGTATCAAAAGTAGCTGGTGACGAAGAAGAAGACGAAGAAGAAGGTGAAGGAGAAGAAATTGCAGAGAATGAAATTTCAATTGAAGAAGGGCCAAGTCTTTTAACATCTGTTTCTGAGGATAAAGGAATTAACTTGATGCCTCCATGGAGTGCATCAAAATCAGGCACTTCATTTCAACAGTATGCAATGGCTGTGATGCACTCAAACATTTGGCCTGGTGCTCATTCCTTTTCTGTTGATAAAATTTATTGCAATATATACATTGGTTATGGGTTAAAGTTTTGTGAAGGTAATTTCAATCCTGCTATCCCCCCGTCTGCACAAGATGAGTATTCTGACATTCCTAACGTAACAGAATGTGACGATCCAAGTGTTGAACAGGAAAGGTCAGTTAAAGTTTTAGAAGAAGAAGTTGAATCTGTGGATGAAGAAGAATCTGAAGATTCGGAAGATGATTACTGAAAATTTGACAAGTTTCCTCTATTGAAAAAGCGCATTTTTAAGAGCTGGATTTCTAGTTTTTTATTTGTTGTAATTATTTAAATTAAGTTAGAACTTAATAATTAATATTAATAATTCTTTAAAAAAAAAATATCTCGTTTTAGTAAAACGAAGAAGTGCATAGATAAAATACATAATAATATAAAGTATTTATTTTGACGTAACAAATTATTTTCGTTGAAATTTAT
>evgR015355c0g2_i01_Hydra
ACACACGTAACAGATCAGGCTTATGGAAAATAAAAATTAAAGCAATATCCATTTTTATTACTGCTGAAACCACATTTAAAAATATGGCTGTAAAGTCTATCAATAAAATTGACAGTAAAGAAATTGTACTAACACTTCTTAAAGACACAAATATTTTATCAAATTTTTCAAAGATCAAAAGTGAAACTGATATTGTTGTTAACAAAGAGGTTTGTTTAAATGTTCTTGAAGACATCTTGAATTTGTATGTAAGAGCAAGGACATTTTCCTACGTCAAGGATAATCTACAAGCTTATAAAGTTAAAGTTAAACAACTTAAGTCAAAGTCACTGCAAACTTCACTAAAGAAGAAGAGTAATCAAGTATAGGTTGGACATTGGACAAATAGATACACTTACCTGATCTTAATATATCAATTTTTTTAAACAGCTATTTTCATAGCTACAAATGGCTAAAGACATTATGGTATCAGAA
>evgR015358c0g1_i01_Hydra
TTATCTTACAAATTACAAATGGAAGTCAAGTAATTTTTATATAACTCCAAAAATTCATAAATGTCATAAAATTATTGATAAAGTTAACAAATCAAACTCTCTGTATATAGCAATGAAACCACCAGAAGATCTTAAAGGCAGACCAATCATTGCAGGAATTAACTCGCCAACATCACATCTAAGTCAATTCTTACACATAATTTTATCTCCAATTGTTTTAAAACAAAAAACTTTTATCAAAGATGATTGGGATTTTTTAAGAAAGATACCAAGAGATTTAGAGAAAGAAAGTTGTATTTTGACTTGTGACATTGTTAATCTTTATACTAGTATTCCACACAAGTTAGGATTAGAAGTCTTAAAGTTTTGGATTACAAAACATAAAGAGCTAATAGATGTAAGATTTACTCAGGAATTTATTTTAGAATCAGCATAATTTATTTTAAAAAACAACAACTTTTTGTTCAATGACGAAATGTATCACCAAATTACAGGAACAGCAATGGGTACTGATTTTGCTCCAGACTATGCATGTCTCACTATTGGATTTTTAGAAGAAACAAGATTTTTTCCTAAAATATTACCTAAGTACTTTTCTGATCAAGAAGTGAATAATAT
>evgR015370c0g1_i01_Hydra
TGAATCTTTTTTAGTTATTTTAATATTCGTATTCAACTTCTTCGTTTATGAATCCTCTTTCACTGACGTAAATACCGTCCAAGAACTTACGGATATCCTTGTTCTTAACAAGAGTAGCAAGATGGATAGCAGCAGCAGAGTTGGAAACGAGTTCCAAGTCGTTACCGGTGATTTGAATTTCGTCCTTTTGACCTGTAACTTCAACAACAACACCTTCTCTCATGTCGACCTTTCTGACTCTCTTTTCACCCAAGAAGTTTCTGATTTCAATATTCTTCTTGTTATCAGAAATCTTACAGTTGACTGGGAAATGCGCATAGACAAATCTCATCTTGTATTTGTATCCCTTAGTAACACCAACAACCATGTTTTGTATATGAGTAGCAACAGTTCTGACGAGAGAGCTTTCCTTTCTGTTACCGTGGAACTTTTGAACGACAACTTGAGTGAATGGAGTTTCTTCAGTTGGTTGTTCAACAAAGACATCGAGTTGAACATGTTGGAAGTTTCTGGACAATGTACCTCTTGGACCAGTAACAACAACACGTCTAGCCTTGACGGTGATGGTGACACCCTTTGGGACCTTGACTGTTTCTTGAATGTTAAGATTTTTCATTTTTTTGGCGACAATCTTCGA
>evgR015375c0g1_i01_Hydra
ATACCTCAAGGATCTTGTTGCAAATTTTATTTTCAAAATGGCGTTTCTCATGCTGAAAAAAGTTGCAATAATCTCATCAAGATACAATTCAGTTCTTCGCCATAACTATTTATTAAATGAAAGCACCAACAGATATTTTGTATTATCTAATTATTTTACAAATTTTACCAATAATGTTAAAGTATCGTATCAAGAAAATAACTTAAATGAAATTGAGGCAGATCAGCTTCATCATTCAGATGATAAAGAAACAATTAGCCGTAGGAAATCAAAAAAAGAAAAAAGATTTAATAAAAGAAATATTTCATATACTAAGAGTTCTTATTTACAATTTCTAAATGAAACATTGTCAAAAAACGAGACTATATCTTCACCAGTTGCATTCAAAATAATCAAGTCATGTGGTTTGGATATGTCAGGGGAAAATGAACATACTAGAAATGAACTTGTAACTCAAGCATGGGAAATGCTTGAGAAAAGTGGAACAACTATGAACATTGGTCTCTGGAACACACTTCTTTCTGTTTTCATTGAAAACAAGCGTGATTTTGATCCTGTGGATTTTTTGAATGATTTAAGAAAAAATTCACAATTAGAACCAAATCAGACAACGTTTACTCTGCTTATTCAAATGTATGCTGAAAGAGGTGATCCACCTGGTGTTATGAAGATCATTGAGTTTATGATGGAGAAGGGTTTTGTAATGACAGAAAATATGTATGCTTGGTTAATAAAAGCTTTTGCAATAAATGGAAACTTTGAAGGTGTCAAACATGTTATGAAATCAATGAGAGAAAATGGTATTCCTCCAAAAGAAAAGGTTTACAAGCAAATAGTAAATGCACTTGCTGTTAAAGGTGACTTGGAAAAAATTGAAGATACCTTGAATGAAATGACAAAGAAGGGAATATCTCCCACAGTTGAACTATATTTGGGTTTGCTTAATAATCTAAGTGAAGGTGGCCATAGAAACCTTATTAAAACAACTTTAGAAAAGATTGAAGGCATGCAAAGTATGTCTCCAGAAGTGCAAAATCTTATACAACGCTTTACTGTGAGAGGAGACTATGAAAATGCGATTGTATTGTTGGACCATTTTAAACAAGAAATAGAAAATATGAAACCATTTACTCATATAAATCAAATCAGTGTCACCTCATTCAATTACTTGCTTCAGGCTGATAAGGATCTTGAACAATTTAAGAAGTTGAATAACGAGATTGAGAAACATTTTAAGAAATCATTAAACTCAAGGTTTCCTTTTCTTGCTGAATATCTAGTGAAGAACGACAAGCTAGATTGGGCATTAGCATTAGCAGTACCTGATGAGTCATTGAAGAAATTGAATCTTCAAATATACAATTTCATTCTTAAAGCGCTAAACTATAAACGAGATTTGATTGGTATTGTTAAGGTGTTTGCTGATATGGGAGAAAAATCTATTGATGCTGAACATTCTTGCTATCAAAATATAGAAGATACAGTAGAAGAAACTGGAAGTGATAAAAATAACAGGTTTCTTGATTTAGCAAAGGATTTTGACAAGATCGTATTGCCATATGAGTTCATTAAATCTGTATTATCAAGAGAAAATATGAATCTAAACACATTTTCTAAATATGCTCATCTTGGCTCTGTAGGAGATTTGTTGGGATCCATAAAAGAAATTACAGACTTAATGAAAAGAGATTTCAACTCCGAGGATTCTTTAAAGATTGTAAAGATGTTGCATAAAAGGGGAGTTCAAACAAAAGTTCTTATTGACAAAGTTGTGAGTGATGTTTCTAATCGAAGTCACGTAGATGCCAAAGAAGCTCATTCATTCATGGTGAACCTTATAGAAGAGGGCATTCCAACTACTTCTAATGCTCAATACAATTTACTCAAAGGATTGTTTACATTTTTTCGTGTTTCAAACTCAGAATATTTTTTTCAGTTTATTCGTACGCTCAAAGAACATGGCATTGAACCCGATAATAACCAGCAGATTCTTATGATACGGATGGCTGCCAAGATTGGGAATAGCAGTGCTGCTCAATTCTGTTTTGACAAACTAATGGCTGAGCCAGAGTTACCTGTGTTTTACAATGAAAATAAAAATATGCTATATCAATATCTCATATCATCTTATCAAAATGAAGCTATCAAAGATGAAAAAAAAGAAAAGAAAATTTCCAATGCTCGTAAAATCCTTCAAATAGCTGCCGAAATGAAGGAAAAAGAGTTGAAGCCCACTGCCCACTCTGCTTCTGTTATAGCTGTCAGTCATCTAATTAATGGTGATGTTACCGAAGCTAATTACTTTCTTAGTACTTATGGAAGTTTCGAACCAACAGTTAATCAGAGAAACTTGTTGAACACAGCTTATTTAACCACATTTATTTCGCAAGGAAATTTTAAAGCTGCTGAAGACAAATTTAATGAGCTTGTACTCAGTAACATAGTTACATATGGTCACTATAAACAAATGTTTAGGTTATGTGAAATTAAAGGAAACATTTCAAAAATGGAACAGTTGTTTAGTAAATTATTAGAAAGTGGAATGAAAGCAGATGTATCTATATATGAACATATGGTTCGCACATATTTAAAAAATAACCAACCTGAAAGTTGTTTGAGTTTGCTCAAAGATAATACTGATAGCAAAAAACAGAAGCTAGGAACTAAATTAATTGTCTCCGTTATTCGTGGTCTGGCTCTAAGTGAAATTGATATTGTTAAAGAGGCTGTTGCACTTGCTCGAAAAGACTACCCAAATATTAGAGAGCAAACTTTAAAAGAGGTTCTTGTGACAGCTCATATTAAAGCTGGATTGGTTGATGAGGCATCTTCTATTTTACATAAAAGTGATGCATCCACTGAAAATGAAATCACTTCAATTAACGATACAGATGAAAAACACAATATTTTGGTCGGCCATCTAAAAAAATATGCTGAGTCAGCCCAACAGCTTAGTGATATTAAATTTATTGAAAACATAATTAAGTTTTTGAAAGGAAATAATCTTGCATATCACGTTGCGTATCACTCTTACTTAATGGCACTAGATAAAGTGAATGATATAGAAGGACTTCGTAATTTGTACAAAGATCATTTGGAGAATGAAGTCAAAGAAAATGATCAATTTATATCGCTTTTAAAGGCAACTGTTGAAAAACATGGGATTACGGCATTTTAGGGATATTTTTAGCATTTAATAAAAAAACATTTTTAAAAAAGATTTTCTTTAGTTTTTAAGTTTCTCGGATGTTTCTTAGGTTACTCTAGCAATATAAATATAAAGCAGCCGTGGCGCTCTATTAATGGTCCGAGTTTTCGATGCCGAGTCGAAATTTCGATGTTCGGTGCGAAGTATGCAAGACTGCGTGGAATCAAATAATCTTTCGTGGTACTCTGTGATAAGACTACTGTGTCATTACCAAGACATTTTTATCCTTCTGACCGTTGGGCGCCCATGACAAAGTAATGAGAAAAACCGCTGGTATATGATTGCTTAGATAAACAATGGATTGAGACGTGATCAATTTTAATAAATAACCTTGAGAAATTCATTGTAAAATTAAAATTTTGTGCAAAAGATATCCGAGATGAAAAAAAAA
>evgR015376c0g1_i01_Hydra
AATTAACCTGTTTTAGTGCAAAAAGTCTATAATTACTGTACCTGTCAAAAATGTCTTCTCTACAACGTGAAGTCTTGAAAAAGATGCTTCCACCATTGCCAAACCATTTATCTATTGACAGATCAAGTGCACAGAAATCTCGTTTTTCCGGCTTAGTCCCGAAGCGAAAAAATAAAGACTATAGCGAGCAATCATGGAAAGAGTATTTCGATGAGCATAAAATTGTCAAAGTTGGTGAAAATAATTTCAGTGTGTATTTGTGTGGGCATACTGGTCCATTACTGGTCCTTCTTCATGGCGGTGGACATTCTGCACTTTCTTGGGCTTTATTTGCACGTCATGTATGTTCCATATGCGAATGTCAAATCATGGCTATTGACTTAAGAGGTCATGGTTCTTCGTTCACAACTGATGATTTAAATATGGCTGCTGAAATACTTGCTCAAGACGTTGCAAATGTTGTTATTGAGTTTTACAAAGAATTACCTCCAATTGTTCTTCTTGGTCATAGCATGGGAGGAGCAATAGCTGTTCATGTTGCTGTAAAAGAGCTTATTCCATTGGTTGGTTTAGCTGTAATTGATGTTGTTGAAGGAACAGCACTTGATGCACTTTCTTCTATGCAGAGTTTTCTTAGAAGTCGTCCACAAACATTTAAATCTGTTGACCAAGCAATTGAATGGAGTTTAAGGTCTGGAACATTAAGAAATATTGAATCAGCAAGAGTTTCTGTGCCCGGACAAATCAGGCGAAGAAATGCTTTTAAGGTTAAAGCTATAAAGGATCCTTCTCTTTATAATCAAGCAATAAGTGAGATAGAAGAAGATGAACTGGGTGATGATGTTCATGAAGAAGAGGAAACTCATGTTTATGAGTGGAGAATCGATCTCAAAAAAACTGAGCAATATTGGAAAGGTTGGTTTGAGAATATGTCATCGTTGTTTCTGTCTTGTTCTGTACCTAAAATGCTCATTTTAGCAGGGATTGATCGCCTTGATACTGCACTTACTATTGGACAGATGCAAGGGAAATTTCAAATGCAGGTGTTAGCAAAATGTGGTCACATGGTTCACGAAGATGTTCCTGATAAGGTTGCTGAAATTATTGCTGGTTTTTTAATTCGCCAAAATCTCACTTCAATGAAAGAACTTTTTCCTAGACATATGCCTGAATGCTAAATGTTGAAATCAAGTGAGTTATTATCAGTTCTAATAATAGTAGTTAAAGGTAAGAGTGTCAGAAACTATAAATTGATTGGATAGTTCTTTCAGTAAAGATGTATTTACATTAAAACCATGATGAAAAAAAGACAAACAAAACTCTCTTAAATATTTGGTATACTTTTTTTATATATATGTTTGAAAACTTTAGGTTTTTTGTTCTACATTTTTTATCAATTTTGAACACAATTTTGTTATCCCAACCTAAATAAGCATCTTCTTATAAACAGATTGTTATATTTAATAATTGGCTAAAGCAAACAATGAATCAACTTCAAACTCTTTTGTATCCTCAACAAAATATCATCGAAAGAGCGATTGCTAACGTTCAAATAGTCAATGAATATGCTTGAGTAATTGACTAAAGCAAACAATGAGTCAACTTTAAACGATTCTATATCCTCAACGGAACATTATCTTCTTATATTTGATTGCTAATTTTCAAATAGACATTTGAGTATGAGTATTGTTTTGAATGAGTATATATTTGAGTGTATATAAAGTTGGTTGTATATATAAAATTGTATTCTACAAATTTATTTTGTTGTTAAAGAGATCTTTATTGATACTCACATACACACACACTTATTCTTGTCGATTATAGTGAGCAAAAAGTGAGATCTGTTCATTATTTTCCATATGGTTTTTTAGATTCCTG
>evgR015378c0g1_i01_Hydra
TAAATTTTCTGTGGAAACCAATGATAGCTTGTATGTAACTACTAATGTGGTTGCCCAAGTGTCGATAATATCTCACAAAGATGTTCGCGTTAACTACGGTAGATCGGTTGTGTTAAGACAAGAAAATCTTCTTATTTCGTACAATAAAAGTCAAACTTGTGAAGTATCAGCAGTAAAAATTGACTACTCTCATTCAGAGATCGGAACTATTTTACCCTCGACGTTTCCTTGCAATTTTAAAGAAGGCGATGTTTCATACCAACATTTGGGTAGTACGACTCGGACTACAGATGTAGTAAAACTCCAAGTTCGTTTAGAAACCGATCAGAAAGTAATAGTTAAGCCTTTTATAATAAGATTGCATGTCATTTTTGAAAAATCATTAGAAATTGTTATAAAAAATGAAAACTTAAGAGTTAACGAGCTAAGTAATTTTTCAGAGGCAATAACCAAAGATGTCGTAGGTTTTGAGTTTGATAAAGAAACATACACGTGCAATGTTCGATTTTGTCATCGCAACAACGGCCCCCCTTATTATGGGTCCATAGTTAATGCAACTAACCATCCTTTTCAATTTGCCTCTTTAAATCAGAAATATGTGTCGTGTGAAAAATTTCTCAGTAGTTCATTATCTTACTACTACCATCGTGAAAGACCATCTACAAATCGAGATTATATTCCATTAATGGTTGAAATAAGGAATAAAGCATCCTACGTAGTTGAAAAACGTGAGTTTTTTCAAATACCCGTTCGAATATATGGAGCACCCGAAAACGAACCCCCTATCATAAATTATAATGATGCGCAATACAGTTTGAATGTTGATCGACTTTTTTTG
>evgR015381c0g1_i01_Hydra
CGTGTTTGTTGATAAAGTTTGTTTGTGTTATTTCTTCTCTTGTTTTGCTCATTTGTGGTTTGAGTTTTCTAACAAAAAATAAGTTAATCATCCATATTGTCATATTATCAATTAGCCCTATACTTGTTGTCGCGGTTTTGTATATTCACACTCGGTTATATTTCAGAGTGAAAACAGTTAGAAAAGTAATATTTTCTCAAATGACTAAATTTGGTAATTCAGAAGAATGCCAACAACAAAATATAAATGAATCATATGGGAAAAAAAAAATGAACAAAGAAATAAAAATGAACAAGCTAACGCTAATGGTGTATTTATCTACATGTTTTGCTCACATACCACTGTTTTTTCTTTTGTTTTTATGGTATTTAAAACCCGTATCTCGTACTTGGGCTTGGACATACACTTTTAATTTGTGGGCACATATATTTACATTAATGAAGTCCTTAGCAAACTCAATAATTTATTTGTATGGTTCAAGTGCTTTGCGGAAAGGTATTCGTCGTGTTTCCCAATCTAATCAATAAAAAAAATACTTAGTTTTTAAATAAATAAGTATATATAAAAAAA
>evgR015382c0g1_i01_Hydra
GCCTGAGAGCGTTTAAGTTATTCGAATAAGTTCCTTTGTTTAAACCTAGTTGATATTTGCCTTTTTTTGAGTTTGATCTTTTGATCAAAAATGTGCGTGGGGTGTTGTTGTTTGAATTTACTCCAAAAACGAAATTTAAATTTGATAGAGAATCGTTATACCGATAAAAAACGACAATATTTGCAATAGCTGAACAAAAGATGGCTATGCACGGAGGGGCCTTGCATAATGAAATATTTGAGGGAAACATAAGTGACGTGAAAAAAGCCGATTTCACTCCAGGCTACGATAATAACAAAAATAGTATAGACATGGAACGTTTGCCAAATAAAGAGTTTTACGAAAAAATAAACGCAAAGCGCGAAATTCCTAACGGAATCAATTTGGTTTCTCTAAAAGATCCACGTTCGGATATTGACGCCGTTAGTATTCAACAACAAAGACAAGACAACATAAGTTTGAATATTACTTTGCCACATACAAAGACACCTGAAAACCAGAGAATCGATACCCAACTTCAAAATCCTGTTCCGGGCCAAACCCGTGAAATAAACCCTCCAACAAGTATTTACGGAATTGAAAACTTGACAGGAAAACCATACGACCAAGATATGAAACTTTTACCGGATATGACATCACCAAGATATTTAGGATTGCAACCGGATACAAGGCCCATGCCAGATATGCAATCACCAAGATATCCTGGTGTTCATATGAATTCGCCACACCACTCTTCTCCTGATAGAGAAAGCAGCCGACAAAATCATCGCTCTGATAATAATGCACCGTGGACTTCCCCGGTTAAACCTCACAACTATCGGGCAGTTGGTCCTCCAACTAGTCCTTACATTGACGCCCGTGATCCAAATTATTCATGTTCGCCAAGAAGATACGTGCCGCCATTCTACAACAACGGTCGGAATTTCGGACATCCTAATGATCGAAATAGGTATGGCGCGTATCCATTAGGAAGTCCGTTAATGACAACTATGGGATATCCACGTGAACCAATGGATTCTCGGTATAATCTTCACATGGGAGTTGTTGGATTTGATGATGATAAATTTAAACTTTCGCCTCCGAGTATGGATGATAAACATATTTGGACAAAACAAGAAGTTGAACTTTTATTAGATTTGTACGAAGAACACAAAATTAAACTACAAGATCCTCGCGTTAGGAAAACTAAAGTCTGGGAAGATATAGCCAAAGAGATTCACGATAAATTGGACTCAGACGTTAACGGATGTCAATGTAACCAAAAATTTAGAAATATGAAAGCGGATTTTCAAAAAGTTGTAGAACACAACTCAAGATCCGGCTCATTTCGAAAAACGTGTAAATACTACGAACGATTAGAAAAATTGTTAACTCCTATGAGTGAAACTGAGAAACTTAAAGATGAAGCTGAAAATGAAAGCGCGTATTATAATAGTTTTAATAAATTGGCTCAAAGTGCTACTCAAAACGCGAACGGATTTTACGATGCAACTTCTCGAAGCACATCATCTTCTGAATCAAGTGAGCACCATTCTTATGAAACAGAAACAAGAAGCACTCCGAGTTCGCCTGAAAAACGAACTTTAAATCAAATGGAAATCGGCAGCAACGAGGCATCCCCTCCACCACCTCCGAAGCGATCTTGTATTGAATGCGGAGGTATGGACAAGCGTGAACTAATTGATGTGTTAAAGGAGTTTTTAAAGGAGCAAAGGATTCGTGAAGAGCAAACAATAAGCAAAATAAATGCATTACATAAAGAAAAAATAGACACTGTCGTTAGGTTTCTTGACCTTTTTCAAGAGTTAGTTAAAAAAGTTTAACCTTGTTTATTATGCACCAGAACTGACACTTGTTTGCCAATAAAAAATCTTCTTGTTTCTTCTATTAATTATTTTTTTCCTAATTAAATTAATTATTTATAGTAATATTTATAGCTAATGATTTTTTTTAATTTAATTTTTTAGTGGAATTATTTTGTACGTAATTTATTTGTAATATAGATCAAATGTAAAATTATTGATTCGGATATAATAATTATTGGTTCGGAAAAAAAAAA
>evgR015391c0g1_i01_Hydra
ATTTTCTAAATAAGATGTCAGGAGTTAATATTTATGATGTTGTAGCAGTCTGGGATGTATCATTACCAGATGGAATTCATCAAGTCAAATTTGAACATGGTACAACTTCTGGAAAACGTGTTGTTTGGGTTGATGAGACTGAAATATTCCGTAAAAACTGGATGTTTAAATTAGTTGGACGTCAAGATTTTAATATTGGAACTGCAAAAGCAGTTATTTTAATAGAAGCTTGTAATGGTTTTACATACGAGTATACTTTATTAATAAATGGAAAACCTTTAAATAAGTTCAAAGAGAATCGAAAAAAGACCGCAAAAGTATGGACATTATTATTGGATGGAATAGAAACAAGAGTTGTTTTGGAAAAAGACACTATGGATGTTTGGGTCAATGGAAATGTTTTAGAAACTGCAGGCGAATTCGGAGATGAAGGAACAGAAACACACTTTGAAATTCGTGATCATGTTTGCGTTATATCAGGAGTTAGTTCTGGGAAGATACGTGAAGGAATTGTACATAGTTTAATAATAGATGGAGCTATAATTCCAGAGTCTAGTGAATAATTAAAAAATTTAGTGATTTTATGAAGATTATTTATTACATAGTTATTTAGTTAATAGTTTATTATATAGTGATTATAGAGGGAGGTTTTTATTTTAACGACCATAGTGTACATATTTACTATATATTCTTTAAGTTTTATTTTAACGACCATAGTGTACATATGTACTATATATTCTATAAGTTTTATATTTAAATCATTTTTAATGTTAAATAATGTATAGTATACACACTTGTAATTTTTATGGAGAAAATATAATAAAATTATTTAAAAAAAAAAAAAAA
>evgR015395c0g1_i01_Hydra
ATGTTTTTAATTTTGAGATCAGTGATCGTAGATATAAGTATGCGGACCGCGTCAAATGAATGTGGCTATTTCATTGAATCGAATTTCCGCACGGACAACATTTTTCATATGAGTGATAATCCTTTTTTTATTTCTCCTATTGGTCGTGGTAGAGGTAGGGGAAGAGGAAGAGGAGTCGAACAGCAAAATGGAGCTGTATCTAATTCACAAGAGCAGGCACAAAGAATTCAGGAATCTACTTTGCCAAATATAGCATTTTCACAAAAATGTGTTACAGATAGTTCTCCAAAATTATCTCAAACACAGAGCGGAATAACTTCACAACAACTATCGCCAAAACCTCAGGAATCTAATAACTCAAATTTTATTTCTCAAAAACTTGAAAGTTCTCTAATAATTCCAAAAGCTACGAGTGCAAATAATCTTAAACCAACAACATCGATATCTAGTCTGGTTGAAAATATTGATAAGCTAACTGAACATGAAATTTTAACAACATATCCTGACAAGATAGATGAAATTAAATCAGAGGAAAGCATAGAAAAACTATCGAATTTACTTTGTGAAAAAGCATTAGATCAACCTGATTTAGGACCAACAATTGCAAAACTCTGCAATGCAATTTGGATATCCAAATCGCTTCATAATATGGTAGTTGGTCCTCTTTTAAGGGCAGTACAAGATAGATATAAAAAAAAAGATTTTAAAAAACGCAATGCATTTTATGGATTATGTGTGCTCTTATGCGAACTTTATAAACTATTACGTTCAAAAGACCTTCCGCTTAAACCACTTTTAAATCCAGTATGTTGTTTGCTCAAAGAACTTTTAGAATCAGAGTCTCCATCTTCAGAGGATGTGTTTTATTTTTACCAAGAGCTCGAAAGTGTAGGTGAAGTCATACAAAATTTCTCCCCGGAAAATATGGATGACATTATTAATATGGCACGCCGGCTTGCAATATCATCTAGCGTGTCTCCATCTGTAAGATGTCGGCTTGTACAGCTAATTGAGTTACGTGCTGGATCTTGGAAAATGAACGACGACATTAAATCTCTTTATGAAGATATTTTTATGGAGCTCATAGAATAGATATTTATTAAAATAATTTTTATAATTTTTAAACATCTATTTATACGTTTGCCGAGGGGCTATTTTAGTAATAAGGCATTTATTTTTGTAAATCAAGTTGTCTTTTAAAAAACTGTAAACCCAAAATATAATCTTTAACATATTATATAATCTATTACAAATAAAAA
>evgR015397c0g2_i01_Hydra
GAAGTGCATAGCCTCTAATTCATATAAGCTTACAAATCTTTTTTAAAGTAGAATAAATTTAAAAGCATGAGAGTTGTGGCTTTAATCAGTGGTGGCAAAGATAGCTGTTTTAATATGATGCATTGTGTTGCACGTGGTCATGAAATTGTTGCTTTGGCCCATCTTAAACCTTCTTTTGTTGATGAAATGGACAGTTATATGTATCAGACTGTTGGACACAATGTAGTTGAATTATACGCAGAGGCTTTGGGTGTTCCATTATTTCAGCACTCCATTTTAGGCTCAGCATTGTGTCAATCTTCAATCTACAAAATTGACAAACATGATGAAGTAGAAGATATGTACGAACTTTTGAACAGTATTCAGGATTCTATTGATTATGACGGTGTTTCTTCTGGTGCTATTCTTTCAAATTACCAACGTATAAGAGTGGAAAATGTATGTTCAAGGTTAAATAAGCAATCACTGGCATTCTTGTGGCAAAAAGACCAAGTAGAACTTTTACAAGATATGATTGATAATAATGTCAATGCAATAATAATCAAAGTTGCTGCACTAGGTCTGGATCCAAAGAAGCATCTTGGGAAAAATCTTTTTGAGATGCAAGATCACCTTATAAAAATGAAAGAAGCTTATCAGCTTAATGTTTGCGGAGAAGGTGGGGAATATGAAACAATGACACTTGACTGTCCTCTGTTTTTAAAGACAATCATAATCGATGAAACCAATATAGTAATCCATTCTGATGATGCTTTTGCTCCTGTTGGTTACCTCTCCTTTAAGAAGATGCACCTTGAAGATAAAGGCATGAATTCAAATCTCAGTTTGTTTTTACGATTAAAGTCAATTAATTTGATTGGTCAAAAAGATTTCAAAAGCAATTTTTGTTTTGAAAAGTTGTTGAAAAGCTCACATATAAACGATAAAAAGCCACACTTGAAAAATAAACCACAAAATGGTTTGGTTGACGATTTTGTAGCTTATGATGGTGTGGTTGATGATACGGTTGCTTATGGTTTGAAGTGGAAAAAGACTGACAACATGTTTTGTTTATGTGGCTTGTCCTGCAAAAATGTTGCATCGCTTGAGTTAAATACTCGTAAAACTCTTTTTAAGTTTCAAACATTATTAGAACGTAATGGATTTAGTCTTCAAGATGTAATACTTGTTTATCTTTATGTATCCGATATGGGTGAATTTGCTAAAATCAATAGTGTTTACAAGACCTTTTTTGTGTCTGAGCCACCTGCAAGAGTATGTATACAGCTAAATATGCCAAATAAAATCTTATTTAAAATTGATGCTTTGGCTGTTTCTTTGAACACCACAGAAAAAAAATCTCTTTTTGTGCAGAGTATTTCTTACTGGGCGCCTGCTAATATTGGACCTTATAGCCAAGCTGTAAAGGTGTATTCTAATACTTATTACTCTGGTTCCATTGGACTTTTGCCTCCATCAATGATTCTTATTTCAGATTCTTTTGAACAAGAAGCTATGCTGTCTCTTTATCATACAAATCAAATTGCAACAGTCATAGAAAATGAAGGCCTCGACAGAAAACATTTTGGAAGTGTGACCTGCTATATTATTGATTCCACTTTTATTGATTCTGTTACTGATGTTTTTAATGCTCAAGATATGTTTCCAGCTAAAGATTATTTTCTAACTGTTGCTGTTGTACCTCAACTTCCAAAAAATGCTAATGTTGAGTGGCATGTTGTTAGCCATTGCTGTGGAAGTAAAAAATATATAATAAGCCAAGAAATAAATCATGAATGTTACAGAAACTGCTGTCATCATGGATATTACTTTTTCAAGTTATCTCTTAAATTAGAGGACCTTGATAATTTAACTTTACCATGGCTATTTTCATTTCAAAATAACTGTGTGGAAATGATTCAACTTATGTACATAACCCTTGAAAATAAAAAGTTAAAAAAATTAATCGACACTCTTTCAACATCCGAATTGGCTATTTCCTATGTTAATGTGATTGCTCTGCCTAGTGATGCTGTTGCTGTCATTAATGCAAGATGGCGACAGGCTAAAAAATAAAAGTTAAAAAAAAAGTAAATTTAAAAGAACTAAACATCATAGGCATTTATAAAAGCTGTATGTTTTTGTATATGACTAAATTGTAACTTAGTAAGTTTGTCAGTCTGTAAATTGTAACTTAGTAAGTTTGTCAGTCTGTAAATTGTAACTTAGTAAGTTTGTCAGTCT
>evgR015400c0g2_i01_Hydra
ATTTAATTTCATTAGTAATAATAAAATAATTTAAGCAGCTGAGATTGAACAGTTGTGGCACAGTTGTTACAGTACTTGCCTCCATTGTAAGATATCCTTTGTTCAAAGCCAGCTCTTGGCAAGTTTTGCGACATCACACAGTGGGCCAGAGTTTAGCAAAAAGAACACCACAGACAAAAAGAGAATTCAAGAATTTTTTATGTAGATTCTAATCTGTTTGTTGAAACAAATCCATTCGTTAAAGGAAATCGTTTTTTATAACTATTTCTATACTATTTTTTAAATGTCAAATTTAAAAAAAGTTGCACAAGCATTTGAAATTGACTCATTGAGCTGTAATGCTGAACGGATTGATCAAAGTTCTCCAGAATTATGGCCAGAGCAAATACCTGGTGTTACTTCATTCTCAGCGTTGCTTAAAGCTACACAAAGTAAAAGTACTATACTTGATATCGATGGTAAACAAGATGAACTAACGGAAGCAGATTTAGAAAAAATTGACGAACTGAGCAGCTTATCTGTTCCCCAAATCATGGAGCGTTTCCTTGAACTAAAGAACCAGGCTTATCAGCTTGGAATAGAGGAAGCAAAAGAGATGACCCGCGGAAAATATTTGAACATTCTCGGAAAATAAAATACGAAACACTAAAAGAAGATTTATGAATGAAACTAATGAACAAAAATTGTAACATTAAATGTTTATTTATTAATTTAAAAATATAATATTTATAATTTATTAAAGTTAATTTAGTAGCTGATTTAAAAAAAAA
>evgR015401c0g1_i01_Hydra
CTATTTTAGTGCAAAAAGCTGATTTAAAAATTAATACTAGAATAATGGTTGCCTTTCAAATGGAAATTGATGGGAATATTCAGATAGAAAACAATTTTGCTGACCAATCTATAAATAGAGATTATTCTGCTGACATCTCTAATAATGAAAATGAACAGGTTGATAGAAAACATAAATCTGGGTTTGTTATTGACAACCAATTGAAGGATTTATTACTTCTACAGCTTGACCTTATTGAACACCAGCAATCGAAGATAACTATGAAAGACAAAAAAATTATTGCTCTTCAAGGAGAAAAAGATCAACTTGAAGCACGTCTTAGAAGAATGGAGAGGCGCCTTGCATTACAAAAACGAAATACAATTGAAACTGAAGGCTCGGTGTCTAAAAAACCATTGTTTACTGGCCAGAAAGATCTCAATAGAAATGCTCCAAAAGGCTTTTCAAATGCTGTTGTTACTGTAACTTCTAGGAATAGTAAAAATTTCAAGTTTCGTTCATACATGAATACTAACAGAGCATACTTAGATTATCCTTTAAATAAACTGGTAACATCTAACAGTATAATTAACATTAAAACAGAAGAAATTGATAAAATATCAGTACCTCCATGGCAAATTAGAGAAAAAAAATCTTTGTATTTCAATAGTCTTGATGAGGATATTAGTGATGATGCATTTACAAAAAGACATGCAAAACCAGAAACCGAAGAAGTTAAGAGGAAAAGGTGGGACATACAGCACATGCGCCAACAGCGTCAACATCAACTACTTACTCGCAAATATAATGAAAGAGAAGACAAATGTCCAATCATATTTGAGAGTTTATTTCGTAATCCAGAAAATGCTATGGTACTTGAAGTGTCTAACATTATTCCAGTTTGTGTTTTGGGAAATCACATACCAGTAATGAAGAAAAGTGAAATGCAACTTCCATGGTTCAGTATTGCAAAGCGTGAAGTGCAATTAAGAGAAGCTAAAGAGAAATTGATGAAGAAGAAAAAAAAATCCCTAATTGTCTAAAGATTTCATCAAATTAAAAAATAATTTATTAAATAATTTTTTTTTTGTGGATAATTTTATAGTAATCATCGTGGTTTTAAATATAAAGTTTATAATTAAATTTTTTTTTAAAGATGAAAAGCCGGTCATTAAATCCATTAAACCCTTAAAAAAGG
>evgR015402c0g1_i01_Hydra
TTTCTTTTAATACTTTCGAATTTCATAGGAATTTTGCTTTCAAACGAAAACATTCTTGACAATCTAAATACTTTAGATAAACCTTATACCCAGGGAAACGGAATTGTAGAAAAATGGCAATCAAATTTATTTTACTATGGTGATGATATAAATGGTGTTGGTATTTGTGGGCGACTTGCTGGGTGTACAACTTTGTATACTCGAAGAGGATGTGGTCTAAACGAAATAGGCAACATATGTCGTAAATGTTTTCAATATCCCACAACAAACAAAAGAAATGTTATGAAATCGAACTTTTACTCATTTGATAAATCATTTTTAAAATCTTGCAAAAATGGTTTAAATGGTTGCCCAAACATTCCGTATATTCATGAAATGGTCGAATACATGCTGTTTAAAGACAAAGATGTTGAAAGAATGTTTCAATCAGCGCAATTTAAAGACCAGAATATTCCTGATTTTTCATCTTTAAATAGACATTATTCAAATGTATTTTGCACTCAAAAATGTTTCAAGAGATTGCGTAATATTTGTACATATGGTTGGGTTCTTCCGAACATTCGTGATTGGTGTTTAACTAGTTTAGATTTTATTAGTGAGAAATTTTCTACTCTTAGAGGACCTGCTTCATATGTATTGTTTTTAGCTGCACGTCTAGGAAGTGAAACAGATGATTTACTACCACAAGAAAATGTGATGTGTCGCATAAAATCGAAGTCTATAAAAGCATACTCTGAAGAAAAATGCAAATGGCCAGAAGGTAGCAAAATATGGAAATATGTAGAAAGAGAGGAAAAAGCAGCGCTAGTTGCTGCCGCTGCAACATCCGTTTTTTTAACAAACAGCTGGTGGAGTGAAAGTCATGACTTCAATATGTATGATAAACGAATTGACGAGTTTGGATTTGCAGTTCAACATTCTTTGTGGTTAACAATATGGGCAGGAAGATTAAGAATTTCAATTCATCCTAGATTTTCCACAAGTGACAACGCAGAAGTTGAAAAAAAAAATAAAATCTCCTTTACTCCTGGAACAGCGACTTTATGCGGAGAAACAGGGTTGTCAATAGCACGCAATTTTCAGGGAACCACTGGAACGATATATGCAGCTCTTTGCGCATTATTTCCTTCTTCTCGTGCCGATGATATTATGTTAAGATATCGGCCAGAAATTCAAACTCGTCATTCACTTGGTTGTGTTGAAAAACTCTCAGGTATGGTGTTTATTCGACCTGGAGATTGGTTTAAAAGAGGATCCGATTACACAATTATAGGAAAGGGAGTTAATGCAAGCGGTTTAATGTTTACAGATGATGAAAACCTTACAACATTTGAACTATCAAAAGAAGCAAATAAATTTTATCCAGCTTCAGTTGAGATTAGTTTAAAAAAACAAACCGATTCAATATGTACAGAAGCAAAAGGAACTGGTCCAAACTATGAAGGTTTGCTTGGAAAAGCAGGGTTAACTGGACTTGGTGGCCTCTGCAATTGGGTTCCTATATGGTAAAATTTAAGAAGATCTTTTATTCCAATAAACGCGCAATCAAAAATGTAAACTGAAGCTTTAAAACTATAATTAAGTTGTGCATGTTCCACTCGGTTCTTTTACGTATTTACGTATACGGTATAATTACATTGCACATGTCGCACTCGGTTCGGCCAACATATTTTATAACAACATCATATTATTTTATTTAAATTTTTTTGGATTAATCGACAATAATATTTTCTATTAGAAGTTTATGAACGGTAGCTAACAGCTTTATCATTTAGTTTGTGTTACCGTATGGATGAAAATATAGCCACTTTTTATATATTTGAAAATACTTTTGACTTTGTGCATTTTTTAATTTTTATATATTGAAAAGGGTTGACTATTTTAATATTATATATTTTAGCGTTGTTAAAATGTTTGTTGTTTAAAAAGAATGTACGTAATTTATGCGTTATTTATGTAAAAATAAAAAAAAAA
>evgR015410c0g1_i02_Hydra
TTCTAAATAAGATTTACATAATTATGCCTGTTGCAGAGCTGCAAAAAGATTCCAATCTTCCATGGGTTGAAAAATATAGACCAGCTTGCCTTGATGATCTTATTTCACACAAGGACATCATTGATACAATTCAAAAGTTCATATCAGAAGGAAAACTTCCACATTTGTTGTTTTATGGTCCACCAGGAACAGGTAAAACATCCACCATTTTAGCAGTTGCAAAGCATCTGTATTCGCCAAAGGAATTTCATTCAATGGTATTAGAGCTAAATGCTTCAGATGATAGAGGTATTGGTATTGTTAGAGAGCAGATTCTTGGATTTGCAAGCACAAGAACTGTGTTCAAATCTAAGTTCAAGCTTGTAATACTTGATGAAGCTGATGCAATGACAAACGATGCGCAGAATGCACTTCGAAGAATTATGGAGAAATACACAGAGAATACACGTTTTTGCTTGATTTGCAACTACTTAACTGGTATAATTCCTGCTATTCAGTCTCGGTGTACCAGGTTTCGATTTGGACCGTTAGATGCTGATCAAGTACAATTGAGATTAAAGCATGTTATTCAAGAGGAAAATGTAACAGTCACTGAAGATGGTATGAAATCTTTAATCAGACTTGGAAAAGGAGATATGCGAAGGTCATTAAATATTTTACAGAGCACTCATATAGGCTATGGTAAGGTAGATGAGTCTACAGTGTACACATGTACTGGAAATCCTCTTCCACAGGATATTCAAGTCATGGTTGATTGGATGTTAAATGAAAATTTTAGTACTGCATTTAAAAAGATTACAGAGATGAAGATTGCAAAAGGATTAGCACTTCAGGATGTTCTTACCGAAGTTCATGCATTTATCCACAGAATCGATTTTCCTTCTCGCGTTCGAATGATACTGTTAGACAAGATGGCAGAAGTTGAATATCGTTTGGCTGCTGGAACATCCGAAAAGATCCAGCTAAGTAGTTTGATTGCGTGCTTTCAGTGCATACGAGACACAGTTGTTGATGATGTAAAATGATAAAGAGTTGTAAATACAATTTCTTTTTTTACCATGTAATGGATCGAAAAGCGAAAACATTAGAGATTGAAATATCGATAATCCACATACTATCAATAATGTAGCAAATTAATCGAAGATTTTGAACACTTGGTTGTCATAATTTTGTAGCTTGATAGTAGCACTTGGTTGTCATATTTTTGTAGCTTTTTGTTTCAGAACGAATTCTAAACAAAACATAAGTTTACTTTACTGTTTTAGTTCATTCCTTTAAGCTCTTGCGTTTAATACTGATGCTCAAGCAGCGTTTGCA
>evgR015415c0g1_i01_Hydra
GAGTCTTCAACTTTACAACCTACTACAATTTCCTTTGATGATACTACTGCTGATTCTATAATAAGCAATTCCAATAAAGAGTTGATAGGTTCAGAAAAATTTCCTGAGATTTCTATTAATTTTTGCAAAATGACTATTTCTTGCAACTCTCATAGTATGCATGTTAGTGTTAGTGAGAATGAACTTCCTTCTTCCGCTGTTGATACTATTGCATTAAATTCAACTGATCCTAAATGTCAACCGGTTTTTAATGGATCGCATTATCTGTTTGTTGTACCACTCTCAGAATGTGGAACTATCAGGCGTTATTTAAACTCTGCTGTAGTTTACTCAAATATTATAACTAATCGTATTACAAGTGGATTTTTAACTTCCCGAACCAAAATAAGTATTCCTTTTTCTTGTGTATATCCTACGCAATCTAGAGTTTCTCCAGTCGGTTTCCAAGTAAATGATATTGCATCAAAGCTTGTTGGAGAGCGGGTTGGATTCGGGAAATTTGAGATAAATATAAATATATATCAAGATTCTTCGTTTTCTCACCTGCTGCAAACGAATGATTATCCAATGGAGGTCGATAATAGACTTTACTTTGAAGTAAAGCTTGATTCTGACGATGATCAGCTCATATTATCTGTAGAAAATTGTTCTATTACTCCAACATCTGAAAAAAGTGATCCCAGAAAACATTTTTTATTAAAAGACGGATGTAAAGAAGACGAAACATTGGTTCAACATTCTTCTCCTCCAGATTCTCAGAGAATGAGCTATGAATCTTTTTCTTTTGTTGGTGTTGATAGCTCTTTAACTTATCTCCACTGTGATGTTTTTGTATGCGACTTGCGTGTTTCAGATACTCACTGTTCAAAAGATAAATGTTTCCGTGGGCGGAAACGTCGTCAATTAAACAATATAAAAGAACTACAAACTTTTAATATTTTTCAAGTAAAAAGTGGTCCAATTTTTGTAGGTAATAGATATTCTGTTCATAAAGGTGCGTTAGATAAATCAAAGTTACACAGCTTATTTGGACTACAAGACAGTAAATTATCGACATATTTTTCTAATATGCGGAAACAATCTCCCAATCTTATAGTTTTAATCAGCTTGGTGGCTGCCGTTGCTATGCTGACTCTTCCATTCGCTCTTTATTATCGCCGTTTGAAAAATAAATATGACGCTGGTTTGTTACCATAAAATGTGCTCTCTGATTTATAAATAAAAATTTCTGAAGAAGAAGAACAAAAGACTAATGAAAATAGAAAAAGTTTAACATTACATTATTATAAAATTATTTTGAAAATAATGAGAACGATATAATTTATTACCAAATATAAAATATTACATATACATAATTATTATAATTATTATAATAATTAAAAATGTTACATATACATAATTTATACATAGAAATTATATATATATACAAAACAATTTATAA
>evgR015416c0g2_i01_Hydra
TGCTTTATGGATAAAACATCGAATAATGCCATTTTCTGCAATCCATTGAATGGGATAGAGTTTTACCTAGTTGATGTACCTTCAACGATATCTCTACCACCACCTGATACTTCATCATACTCTTTGGACTCCGCTACCACTGCTACTTTTTATACAGATATACAACTAGATATGCTGTTATCTGCAATGTTTGGACAATACAATAGTTATCCCAAAACAAAATTATTTACTTATAATTCTAGCATGCTTTCAAACTTGAATGTTTCAAGTTTAAATTTTTCGATTAATAATTGCAATTCAATCAACTTTACAATGCGTGCCTATTTACTAGTTTCTCAGTAC
>evgR015428c0g1_i01_Hydra
TGATGCTTATCAAATCAACATTGATGGTATTGTTGGTAGCAATTTCCGTTGCTGTCATCAGTGCTGCCAATCCATTTGACACTGTCACTAAACCATTCGCTGTTGGTGGTTCTTTCAAGAGATTGGAAAGAAAGTCCACTGCTAACTTTGCTTGTTATAACACTACACTCGGTGATTGGAGTGAACAAAGATGGATGGATTTCCAAACCACTGATGCTTTCAGTTCTAAATTTGTTCAATTGAAGAATTCCCACATTATTGCTCTCGTTTCTGTTGGTGGTTCAAAGTTTGGTTTGAGATATTACAATTTGGAATCATCAACTTTGCTCACAACAACTGCAACTGTTTTGGAAGGTACTTTCAATGACATGTTGTTGAGTTATGATCAAAAGAAGTTGTACATTCTTGGTACTTTCACTAGAGTTGGTACTTCTGTCTATACTCTCACTCAAATTGTTCCAAG
>evgR015432c0g1_i01_Hydra
CAAATATTTTAATTGTCAACTGCAAAATCAAAAAAATGATAACAATTCTTTGTTTCTTCGTTCTTTCTTTAGTATGTTTTGCAAACGGAGGAAAACTCGGAGTTGAAATTGAACTTTCTAAAAATGATATCCAAAGACTTCTATCTGGAGATGGAGATTATAATAGATATTTTTCAAGAGGTTTAGACATTGCTACTGAAATGTTAAATGAAGGAGCGGGAGAAATTAGACTAGTAAAAGAAGAAATTGTAAGCGCAACATCAAAAGATCATTCTGGTTTCAACTACGATGTTAAAGTAAAGATGGTAGACTCTGTATGCAAAAATTTAAAAGAAAATGCGAAAAAAAATATAGTCGATTGCCCAGCTAACGAAAATAAAAATACAAAAATTTGTAACGTGACTATAGTGATTCGACCCCAGGTTGCATCAACCGAATATATAACTGTGACCGCTAATTGTTAATAATAATTATTATTTTTATATTAAAGCTTAGAAATGTAGTAAGCATATATTTTTGTTTTTTTTGAAATTTTTTTTTTGAAAAAAAAAAAA
>evgR015433c0g1_i01_Hydra
GTATCCTTTCAGTTGTTTCCCTTTAATACTTAGCCTTACCCTATATAAATATATAAAAGCAGAATAACCATTGCAGGTTATTTTATTTATTGAAATGTCTTTGCAAGATTGTCTTCAAAAACTAGCCTCAAAGCAAGATTGCCGTCTACATTATGTTCTTACAGCTATAAATATTGCTAATAATGCTGGAAATATAATTAAAGACTGTTATCATCAAGTAAAAACTGTATATACAAAAGAGTGTGCAACAGATTTAGTCACTGAAACTGACAAGTTGGTCGAAAAGTTGATATTTAGTTCATTGAAAACAGAGTTTCCATTGCATAAATTTATTGGGGAGGAATCTGTTAGTAATGGCGCATCTTCTGAATTGTCTTTAGAACCGACTTGGATAGTAGATCCTATAGATGGTACAACAAATTTTGTACACAGGAATCCATATGTTGCTGTTTCTATTGGTCTAGTTATAAACAAAAAGAGTGAAGTTGGTGTAGTTTATGCACCAATTCTTAACGACCTTTATGTGGCAGTGAGGAATGAAGGTGCTTACCTGAATGGAAAAAGTCTAAAAATAGTCAATCCTCCTAGTTGCTTGTCGAACGCATTAATAGCATTTGAATGGGGAGCAGACAGAGATAGTAATATTTTGGCTTTAAAAAGCCGAAATCTATTGGATATCATGCAAAAACATCAGGTGAGAGGTATGCGAAGCACTGGCTCTGCCGCGTTAAATATGTGTATGGTTGCTTGTAGTTCTTTGGACTTATATTTTGAGTGGGGTCCGCATTGCTGGGACATGGCTGCTAGTAATTTGATGGTAGAAGAGGCTGGAGGATTTAATGCTACAACTATAGGTACTGATTTTGATCTCATGGGTCGTACAATTATCTCAGCCGCATGTAAACGCATAGTTGATGAAATTGTACCTACTTTGACTCAAATTCCTACCCTTCGTGACGATGAGTAATTTATCAGAATATGTTTCTGCGTTTTTGATAAATGCTTATTTATAAATCATACATACATGCGTTTTCTTTTTGAAGATAGAGAAGGAAAACATGTATACATAGACTGCTTTCATATAGACATACACAAATACCCTTCGAATCGAATTTCGTATCGAATCACTTATAAGTTTCAGATTAAATGAAGTTATTACTATTATGTAGAGGGGTTATTTTGTTTTTAAATTAATTAATAGAAACCGTAAAAAGTAAAAAAAAAAAA
>evgR015435c0g1_i01_Hydra
TGCAACCAGAAATCAAATCTTCACTCTAACCGCGACAAAAGTATATTCAAAGGATTGATTCTAGTAAGTATCATATTAACTATAAATGTTGTGTTGTTAGCTATACCGGATATGTTAACGACGTTTGTTATGCTAAAAGTATTTCACATTCAAGATATAGTGTACTATGTTGAAATCAACAATGGAATATGTTTGTGTACGGATGCTTTAGTGTACGTATTTGCTTCTCCGCAAGTAAGATTTGCATTTAAAAAGAAATTGTTGGTTTTATTTAGAAAGCTACGGAATTTCGTAACAAACAATTTTGAAGGAAAAACCAAATTGGTTGTTGCTACGCAGATGAATCGTGCAGATAACAAAATAAAAATATTCTCGGTGTAACTATTAAAATTCTCTCGAGTCCTTAATAAACGTCCCCCCTCACCCCTTAATAATATTTGAATATTTTTTCC
>evgR015449c0g1_i01_Hydra
CGATAGTTTATATTACAAGATTATTGGAATGGTTGTGGTGGATAGCCCATTTGTGGTTGTTGAGGATAACCTTGTTGGGGTGGGTATTGCATTTGTTGAGGTTGTTGAGGGAAACCTTGTTGCATTGGTTGTTGAGGATAACCTTGCATTGGTTGTTGTTGAGGATAACCCATGGTTTGTTGAGGATATCCTTGTTGTGGGTATCCTTGTTGAGGTGGGTATTGCATTGGTTGTTGTGGGTATCCTTGTTGTTGAGGATAACCTTGTTGCATTGGTGGTTGTTGAGGATAACCTTGCATTTGTGGTTGTTGTTGAGGGTATCCG
>evgR015456c0g1_i01_Hydra
AGGTTTCAGTAGCAGCAAAGTTAGACCAAAGAGATAAAATAGCTGCTCGAATATTTATTACTTCTAATGACATTAGCTTTCCCGAAGTAAACAAACGCTTATCAGAAATATTCGAAATGAATTCCATTTTAAAAGAAGAGTTGTTAATTTTAAGAGGGTTGCTTTGTTACGAAGTGTTGCATATGGCACTTCAAAAACGCTGGAGAGTTAACTTTGGTGTCAGCTTTATGAACACTGTTTCACAAATGGCGGTACCATTTCGTGCAAAAGATATTGCTGCAGAAAGAACTGAGTTTGGGCACCCTGACATGGCGATTGTTTTGACACAGATTAGCTACTATTTGAGTGGGCTTAATG
>evgR015457c0g1_i01_Hydra
TTATCGATATAATAATGATTATAACTTTCTGTTCTTGTAACCCGTTCATTTATCCATGTGCAGACTATGGGCAGACTGATCGAAATTGTAAACACAGATAGGAAAATAAAAAGCATTAGTTTAACTGGTTCATATAATTTAAACTTAAATATGTCGGCCCTTGTTCAAACAAGCCCCAAAATGAGTAGCATTTGTGACAAGAGAGAAAAATCTTTATTTACTTGGAAATTAGAATTGAAACTTTTGAAATGTTTTATTGTTGGAGTTTGTTTTCAATCATTTGCATTGGTGAAGTGATATTGATTGATATTAAAGATGAATGCAACGTAAATGTAGGTGAAGATAAAACAGTTTTGCTATCATCACACAATCAATACAGACAACATCATGGTGTATCAGATGTGAAATGGGATGATAAACTTGCAGAAAGTGCTCAAGAGTGGGCTTATAATTTAAAAAAATATGGTTTTTTGAAGTTCTCTTCACAGACACTTGATTACAGTGAAAATGTTTTTAAAGGTTCTATTGGAAAACTTGATTTATCAAATGCTGTCGATCATTGGTATGACACTGGCGATAGTTACAACTATTCCTCCATAACTAACATATTTTCTGGTGCTTTTACTCAAATAGTTTGGGATGATGTTAAGTTTCTGGGATGCGGTAACTGTCGTGATTATGTCAAGGATAGAATATTTGTTGTTTGTAGATATCAACCGCGTGCAAACATTGGAGACTACTCTAAACATGTTAACTTACCAATTAAAGTTATCAAGTTTCGTAATATAACAAAACATACTTTTGACAAAAGCAAATGTAACAATCTTATAAAAGGAGATCCTCAGAAAGCAATTTTGGAAGAACATAATGATTATAGAAAAAATCACTTTGTTGGAGATCTAAAATGGGATGAGAAACTTGCTGCTGATGCTGAAGAGTGGGCTTATCTTATAGCTCGTCAAAATAAACTACAATTTTCACCAGAATCAATTTTTTACAGCGAAAACATATATAAATATGCCTTATCGGATTCTAAATGGGAGCCTGTTAAGGCTGTTGAATTTTGGTATAAATCAGGTAGAAACTACAGGTATGCATCACCGGTAGATCTCTGGGCTGGTGCATTCACCCAAATTGTGTGGGATGATGTCAAATATTTGGGTTGTGGTAGTTGTAGCAATTTTTTCACAAACCTGACGTATGTAGTGTGCAGATACAACCCACGCGCTAATATTGGGGATTATTCTAAGCATATCAAGCCACCTATAGAAGAATTAATTAGAAAAACATTTGTGGATGCTGAACTGGCTGCAGAGATAGAAGTATTGAGAGAAGCACCGTTTGCACATAATATATTGCGATGTCGTTATCATAATGTTACTACATTGAAATGGGATTATGATCTGGAAGAATCTGCTCAGTATTGGTCACAAGTACTTGCTGAAAATGCTGAACTAAAACCTTCTCCAGATTCATATTCAGATAATTATGGTGAAAATATATACATGTGTCATGGCTACTGCAATGTAACAGTTGCTATAAATGAAGCAGTTCGCTATTGGTACAAAAAAATTAAAGACTATTCATTTGACACAAATGTTGTTGATCAGATAAATTATCATGCATATTCTTTTACTCAAATTGTTTGGAGCGAAACCAAGTATTTGGGTTGCGGAGTTTCACTAAATAAGAAGAAGGAAATAACTATTGTAGTTTGTAGATACAAGCCCAGAGGTAATATTGGATCATCAGACGAATTTAAAAAAAATGTGAAAGCACCCATTGCTCTTATAAATGAAATGGGAAAATCTGGTTTACTTGGCAAAGGTCGCGAAAAGGATATACCCTCGATTAAAGAGTTAATCTCAAATTCCGGAAATGCTGCAAGTCTAGACAACACTGTGTGTAAATCTAAGTCTTAACATTAGCAAAGTTTAATTTATGATTTGTTCATACGTGTCATTTATTATCCGATTTTGTCAATTTTAATTGCGTTGAAAAATTTTAACTTATGGCAAAAAAAAA
>evgR015468c0g1_i02_Hydra
CGGAAAGAAATCAACCAAGAAAAAGGACCCAAATGCTCCAAAGAAACCAAAGACTGGCTTCTTCCTTTTCTGTGATGAACACAGAGAAGCTGTTAAGAATGGCTTGAGTGACTCAACAAAGAAGAGTGCAAGTGAAGTCTCCAAAGTTTTGGGCCAAAAATGGTCTGAATTGACAGATGAACAAAAGGAAAAGTACAACTCTGTCTACAAGAAGAATATGGAAGCACACAAGAAGGAAATGGAAATATATAAGAAGAACAAGCCAGAATCTGAATCTGAAGAAGAATCAGACTCTGATGATGGTAAGAAGAAGAAGAGAAAAAAGAAGGCCAAGAAGGATGAAGATGCTCCAAAGAGACCATTGTCCTCATATATGCTCTTCTCTCAAGATCATAGAAAGAAGCTCATCCAAGAAGATTCCACTTTGAAGGTCACTCAAGTTGCCAAACTTGTCGGTGCCAAGTGGGGATCCTTATCTGATGCTGAAAAGGCTCCATACGTCAAGAGAGCTGCTGAATTGAAGGAAGCTTACAATGTTGAAAAGAGCAAGTATGAATCATCCAAACCACCAGCAAAGAAAAAGG
>evgR015470c0g1_i01_Hydra
GCAAACTGATGCCCTATTCAAGATGCTAACTCTAGTATGTATTTTCTGCAATACTAAATTTAATGCATTAAAAGAATATATGGTCTACCATGTTCATAAAATTAATTCACAGGATTAGTGATGTCTTTAAAATAAATAAAGATAATATTCCACAAGAACTAGAAGAAGCAGCTCTTCATATAATTACTTTTAAAATGGAAAAATCTGGAGAGAATGTAGTTGAATTTAAAAGTGGAGGTCCTAGACCAATAGTATTTACACAAACACCAAAAGCTTATGTCAATCAGGCATCAGTATCTACTATACGACTTCGCAATATCAATATTACAAATCATTTAGAAATTGTAGCAGGCAATTTAGGTGATTCAGTTGTAAATCAAACAAATAAATTATTAAAATCGTTTACACTAGAGACAAGACAATCCATATTAAAAAGTGCAGAGCTTAACCACACTGAAATAAATGCAAGTTTAATGGTAGCTATGAAAACAGACATGGGGCTCCCGTGGGAAAAACTTAAAATAATGTCCAGATGGCTTAAGACCTTTGGTATTAACACCGCATCAAATAATTCACAAAGAATTGTTGCAGAAAGTTGGTCTGGAAATGATATAGTTGTTGAAAATGCCCCATTCACATTTCAAAAAGAAAGGAAAGGAAAGTTTGAAATTAAAGAGGCTCCATGGGGATATATTACCGATTTACGGAATCATATTTTTAGATATCTTGAAAATTTGGAAAAAAACATATATTGCTGATTTAATGCAGTTTTATCGTACAAATTGGTCCGACGATTCAGTGCCACCGAAATTACATATGTTGGAAGAGCATGTTATTCCTTTTTTGAAAAAATAGGGTGCTGGGTTTGGATACTACGGCGAACAAGTAGGCGAATCCATTCATCATGAATTTAATAAATTATTTGCTATATATCAATCCATTCCTTCTCCAACAAAGCGGTTAAGAAGTATTATTAAAACACACCATCAAAAAACTAACCCTAATAACAGAAAGCTTAAACCAGCATTGAGAAAAAGACAAAAGAAAGAAATTTAATTATTTCAACGAATATATTGCTAATATAGAGTGTTAAAAAGGAAAGGAAAAAAAAAATATTTAAAATATTAGTAAAAAATATATAAATGAAAT
>evgR015471c0g2_i01_Hydra
CATAATTTATTATTGTTCCAAGGAAAACTACCTACAGAATTTGTACGACAACCTCGGTCTTTAGTTGAGCTGGATCGATGGAAAGCATCTGAGTTACGAAGCTTTCTTTTATATTCTGGCATGGTGGTTTTAAAAGGGGTATGTAATCCTAATGTGTATAAACATTTTTTAAGCCTCTCAATAGCAGTCAGATTATTGTGCGAGGAAGTTAAGGATTTTCGAAATCATTATTTAAATTTTGCCAAAAAGCTCTTGAATTACTTTGTGAACAATTCTCATGAGCATTACGGGAGATCATTTTGTGTGTACAATGTGCATGGACTACTTCACATTGCTGACGATGTAGAATTTTTTGAACAACCGCTTGATGCAATATCATCATTTCAATTTGAAAATTACCTTCAAAAATTAAAGCAGCTTGTTAGAAGCAAAACCAACCCAGTGGCACAGTTGACAAAAAGGCTTAGTGAAATTGAAATTTCTAAGGAAAAGAATGTTATTACTACTAAAATATCTACTAAAGCTAAGGATAGTTGTTTTTTAGTGCAAGGAGGAGTTGTTTTTATTGTTAATAAACTCAAACGTGAAAATTTTCAGTGTAATTATTACAGCAGAGACATTATGGAAAGTTTTTTTGATTTATTTGTAGACTCTAAGCAACTCAATATTTATTTAATCAAAAGTAATACTAAGCCATTGAAGTGTGTTAAGAAAGCAGGTGAACTCTTGAGAAAATGTTGTTGTTTCCCATACAAGAATGGAAGTGTTATCATACCATTGCTCAGTAATGTTACTGGACATACGTTTTTGTAATTGCATTAGAAATAATGATTTTTACTGGATATTACGATCTTAAGTTGTTATATTTTTGGTTAAATTTTATTTGTTGCAGATTTCTCTTTGATATTTTTATAATTTATTTGGGTTTATATGGTTTTTGTAGCCTTGCTAAAATTTTAAGATTGTCAAATTTTAGATTGTAAGAAAAC
>evgR015477c0g1_i01_Hydra
TCCTCCTTTCAGGATAAAGTCCAATCGATTATTTGTTTATTATCTAAACAAAAAAGACCGTTTAGGTCTTGTTTTAAATAATATAATTCGTAATGTAACGACCCCTTATGTTCTTTTTGCTCAATATTTATATCGCTTCGATAATTCGTTGGATATCGAAAGATTAATTCATGTACACGAAAACTCGGATGCTGAAATTGTTAGTGGCTCAAGAAAAAATCGAGAAAATGAATGGGATAGAGGTTGTTATCAAACAGGACTTAAACTATATACACTAGAATATAAAGCTGGCTATTTTAGTTCGGAGAAAGAATGTCTAAGATGCAGTTTTACAGAATCCCCATTTCTGACATCTATTCGATTGCTTCAATCTATAGAGTTTGATAATAACCTTGGTTACGGAATTTTTCAAGACTTTTTTATAAATCTTAGTTCAAAAAAACACAAGTTACTTGTATGCCCCGATGTTATGTTTTATAACAATCAACACAAAGAAAAAGATTTTCAACTATCCGATTTTGCCAAGAAGTGGGATATAAAAAAAATAATTTCTTCAACAAATAAAGTATTTTGGTATGGATGTCGCCATGGGTTTAATCACACAGCAAAGAGAAAATGCACTATTGGGAGAGGTATGGCTGTTCCACCATGTTGCATTGCTAATCTTGTGGATGCCATTAAATTTATTTTATTTATGTTTAAAATTAATGGAATCAAATACTTGGTACAAGAAGGCAGTGTGTTAGGTTTAGTAAAAATTAATGGAATTTTACCATGGGAGCGAGATTTTGATATTTCTTTTTTATCGGACCAATTTGAAGCAGTTACTAAAATGAGAGCAACATTCAAAAGTAAAGGGTATCAACTTGTTGTTGACAAAGTTCCATACAAGAACAATTTTACTTTAAATGGTGGGGTGATGCATGTCCGCGCAAATGGCTGGAGTATTCAATTGTACGGACAGCAGACACTTCCAGGTAATGAAACCATTATCAATGCCTTAAACCAATCAATAGTTGTTCCTGAAAACCCGGGGTTGTACTCACGTAATCGTTATGGAAATGAATTTTATAGACATGTTGAACATTGGATGACTCTTAAACATAAATCAAGTTGGGATAATTATGCAATACAAAATTTTACTAGCTGTGACATTCCTAACCATCATGCATGTCTAGATCAATTTCAAGCAGATGGAGATATAATGTTCAACAGCTAAAAGATTACGAGCAAGGTGTTTGCAACCCAAGTCAAACTTTACAGGCATTTAAGACGGAAGATTCATTACAACGATTTCAACGATTCAACAACAATTGATTCAACGAATTGTTAGCATAAAGTAAAATAGACAAAATTACTAAAACGAATCGAAACGAACGAAATTGAACACGAATCGAAAAAAACTCGTGAAAAATCTTTTATTAAAAATTCGGGTTTTTCTAATAAAACGATAAGCAAGTTATGGGATTCTGCAGAATCATATTCTTCAGCTTTTACATGATTAAAGAAAATTAAGCTTCTAC
>evgR015480c0g1_i01_Hydra
TGAGGCATATAATAGATCAAGGTTGAAAATGACGTTAACACTAAAGAAGAAAAGTCTTTCAAGTTTTTTTCATTTTCAAAAAAATAGCGAAGAAAAACAGAGCTTTGGAAAATTTCAAAATGAGGCAGCTGATTTGAAATCGAATAAAAGCTTATCGCTCATAAGCCCCAAAAAAGTACGCTCACAACCAACCTCGCCTTCTGTAACAACGCGAAGTATGAGCGAAATGGAGGCAACAACGCCAGAAAAACTAATAAATATTGGCCACATGTTTAAAATTGCATTACGTTCACGAAGCAAAAATAACTTAACAGAAAGTCAAAAAAGTCCATTTGAAGACTTTACTAACGAAAAAAAATCAAAATGGGAAAAAATTAGCAAAAAATTAGGAAGTCGAGAATCTATTGATAGCATCGAAAATCAAAGTAATCAGTTGCTGAGAAATGAAAGGTCTTGTTCTGTAGGAAGCTCAGGCTCCTCAAAACGACGATTAGGAGTAGACGCGACAGCCTATAGATCAAATTCTCTTTGTATTGAGATATATAAAAGTACTTCCTTTAAAGATACTAAGTCTTCAGACATGAAGAAATCAACAGAAAAAAGGCATTCAAAAACATTTAACTCTTGGTCAAAGTTCAACCAGAAGGACTACGCTTCTTCTGGTCCGCTTTTACATATCGCAGCTTCAAATGGAAACATGGAACTTTTGAAAACATTAGGCGATGTTGGTGTTGATTTAAATGAACACAATTCTGCCGGATGGCCCGCATTACACTATGCAATTTGCTCGGGAAATTTTGAGGCAGCACATTTCCTGATCAATAAAGGAGCATCCATAATTCCTTATTCAAATAGAGTAATTAATACGTTAAATTTAGATAAATCTTCAAAATAAGTATAGCAATTTTTTTTTAAGTTTACGTTCATAGATAATCAAACAAGTATTTGACTTGTGACAAACACGTGTTTTGAATTATATAACATGTGGTACAACGTGGTATTTTTTTAACTTATTATTTTATAATTATTAAC
>evgR015481c0g1_i01_Hydra
TGCGCGGGAGAACAGTTCTCAAATATTTTCATTAATAATTTCTGCATTAGTAATTATATGCAATTAATAAGTTTTTTAAAAAGCAACAAATGAATCCAGGAAAAGTGTAACGCTTTAATTTGATCGTTTTCAGGGAAAACGTTTTTGATTTGTATGGTTGGATGCATCCTAGCAAAAGAAGAGCTATTTACTATACAATATTTGCAACTTCGTTGCTGTGGATATGCGGAACTATAACGTTTTTATTTTTCGACGATATAGAAGTTACACTAGCTGTAAAAAGAAAAACGCTTGAAGCAGTTGCAGAGCAATATTCAAATTTAAACTCTGAAACAACAAAAGTAAAAACGCAGACTCAAATAAACAATCTCCAATATTCATTAGAAAGTAACAAGCACTTGCTCATTTTAACAGATATTCACAAAATGCGAATAGCTGATTATGAAAGTTTTCTTGTACCGCGAGATTATCAATTACCGGGAGAGTTAGGAACAGGTGTATCGGTAGAAGAGAATGAAAAAGAAAAAGAAAAGTTAGGCTATGAGAAACATGCTTTCAACCAATTGGTTAGTGATAAAATTTCAATCCATAGAAGTTTAAAAGATTACCGGAATGACCAATGTAAAGCTAAGAAATATCCGATTGATTTACCATCAACTTCAGTGATTATATGCTTTCATAATGAGGCTTGGTCAACATTATTACGCACTGTTCATAGTGTTATAAACAGGACGCCACCACAGTACTTAAAAGAAATAATTCTTGTAGATGATTCAAGTACCAGTGACGATTTAAAACAACGGCTCGATGACTATATTCCAAACTTAAAAATTGTTTTTGTTGTTCGATTGCGTGAACGCCAGGGATTAATTCGTGCTCGTCTTGAAGGAGCTAAAAAAGCAAAAGGAACAATACTTACTTTTTTAGATGCTCACTGTGAATGTACACTTGGATGGGCAGAACCCCTACTAGCAAAAATTAAAGAAGATAGGCGGAATGTAGTAATGCCAGTCATTGATGAAATTAGTGAAACAAATTTTAATTATAATGCAGTACCTGAACCTTTTCAAAGAGGTGTATTTAAATGGAGATTAGAGTTTACTTGGAGACCTATACCTTCTTATGAAGAGCAAAGAAGAAAGCATGAATCAGATGGTATTAGAACACCTGTAATGGCTGGTGGTTTATTCTCAATTAATCGCGATTATTTTTATGAAATGGGTAGTTATGATACTGGAATGGATATTTGGGGTGGAGAGAATATTGAAATTTCATTCAGGATTTGGATGTGTGGTGGATCCATTGAAATGCTTCCATGTTCAAGAGTAGGCCATGTGTTTCGTCCAAGATTTCCATACTCGTTTCCAAACCGTCGAGGTGGTGATGGAGATGTGGTAAGTCGAAACCTAATGAGAGTAGCCGATGTTTGGATGGATGAATACGCTAAACACTTTTATAATATAAGATTTGATTTAAAAAGAAAGAAGCATGATGATGTCACTGTTCGAGCAAAACTTCGTTCAAAGTTGCAGTGTAAAAGTTTTCAGTGGTATTTGGAGAATGTTTATCCTGAACTTGAAATTCCTGATGATAAATTTCTTGCTGCTGGTGAAATACGTAACCCAGAAAGTGGTATATGCTTAGATACTTTAGGGAAGCAAGAAGGAGCACCAGTAGGATTGTATGCTTGTCATGGTCAAGGCGGAAATCAGTATTACACGTATAACAATAAAGGCGAAATAAAAGCGGAAGACAACTGTATGGATTTTAATGGTCACGACCTGTATATTAGAGAATGCGATGGGTTAGGTTTAAATCAAAAGTGGACTTACAAAAACCAGCGTATTCTAAATGAACGTTTCAATGTATGTCTACAAATTAAAGGAACTAAGTATGCAAGTGTTGGACCATGCCTGAACGTAAAGTTTCAAGAATGGTCATTTAACAAAATTGATGACATGGCGAGGTGATAATAATAAAATGAAATGAGTTTGTTCATTTTTTAAATTCATTTTCTAATTTTAATAATTTCAAATTTAAGAGAAGTAAATAAATTAAAAATGTCTGAAATGCTCCAG
>evgR015482c0g2_i01_Hydra
TTTAAAAATAATATAAAATATTAGCAACACTAAATTCGAGAGAGAACTCGACACGGATTTAATAGGCAATCGCGAAGCAAATAAATAAAAATGGCGGAAAAGTCCGAATCGAAAAATGGTTCATCTCTAGAAGAAAGTAAACAATTAAATGACAAGCTAATCTCTGATTCCAATGTAGAAGAAAATATGCAGAAAAAAGTCGCTCTAAACGATGATATTACTTCAAAAAAATCAAAGTCAGAAGATCCAAAGAAAAAAAAAAAGGAAAAAACAGAAAGCGGTGATGATCATACTAAAATTGACAATGATAATAACGAGCAAATTAAAGCGTCGTATGCTAGTGATATACAAAAAGCAATTCAAAAAATGATAATCTCTAGTGAAAACAGATTAGAAGAATCAGCAAAAAAAAAGTCATATGCTTTTTGGGAAACTCAACCTGTCCCAAAGCTTGATGAAGTTGTACCAGCTAATACAAATGAAGCAGTTGAAGAGGATAAATCTGAGATACGCCAAGAACCTTTTACTTTACCAACTGGATTTGTATGGGATACAATAGTAATATCTGAAGAATCTCAGCTAAAGGAACTTTACACCTTACTCAATGAAAACTATGTGGAGGATGATGATAATATGTTTCGTTTTGATTATTCACCACAATTTCTTCAATGGGCATTGAAACCACCTGGTTGGAAGTTGCAATGGCATTGTGGTGTTCGAGTGGCGTCTAACAAAAAACTGGTTGGTTTCATAAGTGCAATACCTGCCCATATAAAAATATATGATAAAAAAGTAAGAATGGTTGAAATAAATTTTTTGTGTGTGCATAAAAAGCTGCGCTCTAAGAGGGTTGCACCTGTTTTAATTCGAGAAATTACCAGAAGAGTTAATTTGGAAGGCATATTTCAAGCTGTTTACACTGCTGGTATTGTATTGCCAAAACCCGTTGGTACTTGCAGATATTGGCATAGATCAATTAATCCAAAGAAACTGATTGAAGTTAAATTTTCACATCTTCGAAGAAACATGACTATGCAAAGAACTATAAAACTTTACAAATTGCCAGAAAACACTAAAGCTTCTTTATCGCCTTTAACGCCAGCCGACGTAAAGAGTGCATACAAAATAGTTTTTGAGTATTTGAGTAATTTCCAACTTGCACCTATATTCACAGAGGAAGAATTTGCGCACTGGTTTTTACCTAGAGATGGCGTCATTTCTTGCTATGTTGTCAAAGATATAGATTCTGGTAACGTTACAGATTTAGTTAGTTTTTATCAACTTCCATCTACTGTAGTGGACCATCCTAAATATAGTTCTCTAAAAGCTGCATATTCATTTTATAACGTGGTTACAAAGCATGACCTTACTTCAATAATGAATGATGCTTTAGTTCTAGCTAGAAATAATAATTTTGATGTTTTTAATGCATTAAATTTAATGCATAATGAAGAATTTTTAGAAAAATTGAAGTTTGGTATAGGCGACGGCAATTTAAACTATTATCTTTACAATTGGAAATGTCCGTTAATTGAACCAGCCAAGACTGGATTAGTGCTGCAGTAAAGGCGAACGAATTTCATGGTTAAAAAAACATTCACCAATAAGAGTACATAAGAATATTATACAAAATTTTTATATATAAAAAAAAATGCATCTGACGTCATCAATGCATCTGACGTCAAAAGTTGTTGTAAATATTTTAAATTTGCAAACATTTTAGAGCTTTATCTTATTATTATTATTATTATTTTGCAATATTTATTTTGA
>evgR015487c0g1_i01_Hydra
TATTTTGTTCATCATCCCAGCTTCATTTAGCATATCTTTTGGTTTTCGAAATTACTATGTAAATTTTCTTTTAAAAGTCTTTGAGTATGGTAGAAAGAAACAAACTGCAAAAAATATTCTTAATGAGGCTGCACATGGAATTATAGATCACGAGTCAGATATTCCTAAAAGTCCTGTACCATCAAATGTTCCATTTGAATCATTAAGTAGGAATTTTCAACTTGCAGATGTTTGTGAATTTCTCAGGATCGGAGTTGAGCATATTATTGAAGATGAAGTCACAAAACGATTTGATGCAGCAGAGTTGCCATCATGGAATCTTTTAAGTCGCACAAATATAAACTACACATCTGTACAATTTATAAGAAATGAAATCACAGGAGTCTCAATTTTTCGTGATGACGTTAACATACTTTGCAGAAGTTGCATTTTTTTTAGTTTTCGTTTGACAATGGTATGGGTATTAGGTTTGTTAGTTAGATACCTAATCTTTTTACCAGTGAGAGTAACTATTCTAACTATTGGTCTTGGTTTTTTAATTGGATCAATGTCGATACTTCCGCTTATACCCAATCTAAAATGGCGTATTTGGCTGTGTAAATGGATCACAATGTTATCATTTCGTATCATTTCTAGAGGACTTTCAATTGTGGTCAATTTTCATAATAGGGAAAATATGGCAAAAGGTGGTGGAATCTGTGTTGCAAATCATACATCTCCTATAGATGCTTTTATACTTGGTTGTGACAGAAATTATGCATTAATAGGTCAAATGCAATCTGGAATTATGGGTACAGTTCAAAAATCATTATTAAAGGCACAAGACCACATATTCTTTGAGCGCAGCGAGTTAAAAGATAGACTTTTAGTCGTAAATAGAATGCGAGAACATGTTGAAGACGCTCGAAAGAATCCCATTCTTATATTTCCTGAAGGAACTTGTATTAACAACACATCAGTTATGATGTTCAAGAAAGGTAGCTTTGAAGTTGGTGGTGTTATTTATCCAGTTGCTATTAAATATGATGCAATTTTTGGGAATCCATTTTGGAACTCCGCAAAAGAGTCAATGTTAATGTACTTAGTAAACATGGTGACAAGTTGGGCAATTGTGTGTGATGTTTGGTATTTACCAGCAGCTGAAAAACTTCCTGAAGAAGATGCTGTTCAATTTGCTAATCGTGTTAAAAAAGATATAGCTCGTCAAGGAGGATTGGTTGACATGTCATGGGATGGAGGTTTGAAAAGAGAAGCAGTTCCTCCAAAATTTTTACAAGAGCAGCAAAAAAATTTAGCCAATAAAATTAAGTGAGTTATTCAATTGAGTTATCTATTTCGGACGATGTCTAAATGCTTAATACCGTTTGAAAGAAAAAACATCTTACTGATGGTGACGATAATTGACGAACGATTTTCTTAAAATATAAATTTTTCATGTATAATTTGTACAATAAATATCTTTTAGAGAGTATTTATATTTTGTCATTCTTTGCTTTTATAAAAATAGTATAATTAAATTTAATCTAAAAAAAAAA
>evgR015490c0g1_i01_Hydra
AACCTATTTTAGTGCAAAAAGACTTATTTATCTTAACTAATATTTGACAATGCAATGAAAGTTAAAGTACTATCTCGAAACCCTGAGGATTATATACGCGGAACAACATGTGAAATCAATCGCTTACCACGAAATTTAGACCCGGATTTGCATCCTTTTGAAGCCGCTCGAGAATATACTAGGGCATTAAATGCTGCTAAACTTGAGAGAGTTTTTGCTAAACCATTTTTAGCTTCATTGGATGGACATAGTGATGGGGTTAACTGCTTAGCACGCAACCCAGTTAAAGTCTCTTCTGTTGTATCTGGAGCATGTGATGGTCAAATAAAATTTTGGCATCTTCAAACAAAATCTTGTCTTAAAACTGTTACAGCCCACAGTGGTTTTGTTAGAGGTTTAACGTTTTGTCCTTCTGGTAAAAAATTATTTTCTGTGGGTGATGATAAAACAATCAAACAATGGGATGTTGACAATAATGAAATCTTGGAACCCATTTCAACATACCTTCACTCAAAAATGATGACAGGAATCGATCATCATGGAGTTAACGAAACATTTGCAACTTGTGGAGAGGCAGTTGAAATTTGGGATGAAAATAGATCAATACCTATTCAGACATATTCTTGGGGAGTTGATACAGTTCATAGTGTAAAATATAATCCTATTGAGACACACCTTGTAGCTGCTACAGCATCAGACCGCAGCATTATGTTATATGATACCAGAGGGTCTAGCGCAATGCGGAAAGTTATAATGAAAATGAGAAGTAATACCATTGCTTGGAATCCAATGGAGGCGTTTATATTTTCGGCTGCAAATGAAGATTCCAATGCTTATACTTTTGATATACGAAAATTAGACTCACCTGTAAATGTTCATATTGACCATGTAGGAGCTGTGTTGGATATAAACTACTCACCTACTGGCCAAGAGTTTGTGACAGGTAGTTTTGATAAAACGATAAGAATATTTCCGAGGGATAGAGCCAGAAGCAGAGAAGTGTATCATACATCTCGAATGCAGCGTGTTTTTTGTGTCACTTTCAGTGGGGATGCTACATATGTACTATCAGGCAGTGATGAAACCAATATAAGGCTATGGAAGAGTCATGCTGCAGAAAAACTTGGTCCTACAAACCCTCGTCAGCGACGTGATCTAAAATACAATAGTAAATTAAAAAATCAATATCAGCATCATCCTGAAGTGAAAAGAATTCTGAAGCATCGACATGTTCCCAAACTAATTCACAAAGAAGCCAAAGAGAAGAGAGTTATGCTTGACTCAATCAAACGAAAGGAAGAAAACGTGCGTCTTCACAGTAAACCAGGCTCGATCCCAGTTATACCGGAAAGAAAAAAACATATTGTTACTGTAAAAAAATAAATCTTCGCCCAGGTTGTTTTAACATCAACCTTATCAAAATATTTAAAAAAAGTAAAACAAAAAGTGCAAAAACATAAGACGATTTTATTTAAAGAAATAATTCAAATAACTACAAAAAAAAAAA
>evgR015492c0g2_i01_Hydra
CAGATCAAAAATGTGGAAAATCTACAAAAAAAGTAACTTATACAACCGTTAGTATTAAATTTAAACGTTACCTCATGGCTCAAAACAAACTAAAAGTTTTTTACGACGACAAGTATGCCGGCGACTTGGTTAAAGATCAGCTAACCTGTTCAACTTTTCAATATTTGTTTCAATACGACGTTGACTACTTGAAGAATAACAACGCGCCATTATCTTTTAATTTGCCGCTTTGTAATGAAAAGTTTGTATCTCAAACAAAATTGTTTCCATATTTTGAAAATTTAGTATCTGAGGGATGGCTCCATGACAAGCAGTATGAAGCGCTTAAAAAAAATGGAATAAAAGGCATAGATAGTTTTTCTATTTTATCTTACTTTGGTTATGACTTGATTGGGTCCGTCTCAATAAAAAGTGAGAAACCTGAGATATCGTCACAAACTACGCCATTTTTTATCTACTTAATTACAGCTGAACAAACTAACTCCACTGACTCACAAAATACTAGCGTAAACATAGATAGTAAAGCAACTATACCTGGAATTCAAAAAAAAATTCTAGTTAAGAAAAACCACAATGGGTTGTTTGAAGTCACAAAAGCTGACAAATTAAGCACGCATATTGCAAAAATGGAGGCTGACGACTTTCCTGATCTAATCGAATTAGAATATTTAAGTAGCTTGGCAGTTAAACATTTGCTTCCCAATGATGTGATCTGTGGTTTAGAAATAAGCATGATCCATGATCTCAACGAAAAAGCTTTAATTGTAGAGCGATTTGATAGATCAATTCAATACAATAACACTTCAAGTACGATAAACGATGAAGTTATTATTAAGAGACGTCACTTTGAAGAGTTTAACCAGTTATTCAATCTTCCATCAGAAAACAAGTACGATTTGTCTTATGATGAAATGGCTGTCTTTATTCGTGAAAATATAGAATTGAATCGATGCAGTTCAAATGAACTAGTAAAACTTTTCATGAGGATTTTGGCATTCATTTTGATCGGAAATACAGATGCACATTTAAAGAACTTTGCAATGTTTTACGAATCAGATGGTTTGCGTATGCTCACTCCAATGTATGACGTAGTAGCATCGGCTTATTATAAACAATTTAAAGAAATGGCACTTAAACTGAATTATAAAAAAGCAGAATTAGCCAAATTAAAACCAAAAGATATTGTCGATTTTGCTTTTAACAAAAAAGGTTTTAATCTATCTGAGAAGGATTTGTTGTATGCAGTAAAGCAGTTAAAAGAAAACAGACTAAATGCAACTCAAGCAATTCAAAATATTAGTTATGTAAGTGAACGCAGTGATGTAAAAGATAAACTAATTGAAATAATCCACAAAAGATGGAACGGAACGTTTGAAGGAGTTGAAAGCTATTTGAAAAAAAGAATAAAAGAATAATAGATTTCCATTATTTGCAAATTTTTTATTTATTTAACTCAGCAAAGCCGAAGAGACAAATATTGTAAAAGAAGGCTACGTTGATGAGGGTCCAAACAGAGTTTTTAAAATGATTTCGAACAAAGCTTATATGTAACTAAAAAAATAGACCCTATCTGTGTTTTGAATGTTTACATAAGTATACCAGCA
>evgR015495c0g1_i01_Hydra
GCTTCTTTTGATTCAACTCTCTACAAAATCTCAGTCACTCTTCCACCAAAACTTCCAACTTTCACCTTTACTCAACATGGTGAATTGATTTGTCCACTTGGTCCAAACTCAATTGTTAAAATTGGATTGGATGGTCAAAAGACTAAACTTGCAGGAAATGGACTAAACCCACCTCCATATTTCAGTGGTGAAGATGTTCCAGCTGTAGTTGCTCAAATGTATGAACCACAATCAGCTGCATTGACACACAAGGGTGAATTGATCATTGCTGATAGTAAGAATAGAAGAATTAGAAAGGTTGATACTCGTGGTGTAATTTCTACATTGGCTGGTACTTATACTTACGCAAGAGTTAGTGCCATGAAAGTTGGT
>evgR015498c0g1_i01_Hydra
GTTATTGCGGAAATAGGATACGGGGTTGGTTTGTATCTCCTCAAACTGGAGGTTACATATTTTACACCTCGTGTGATGATATCTGTCAATTGTATCTGAGTGATGATAATGATCCTTTAAATGCTCGCCTCATTATTTCACAAAATAGCTGGACAATGTCTAACAACTGGTTTTGGTATCCATCTATGCAAACATCCAGCCTGATATATTTGAAAGAAAATTGTAGTTATTATATAGAGACAGTAATGCTTAATATTTTTGGACCCGGGAATTCAAATGTGGCCGTGCAAATGCAGAATAATGCAACTGAACCATTAACCAATGGTATGATGAAATTAATGAAAAAATGTAATTCATCATCCTGCATTGCAAATAATATCGAAATATGCTTTTTTACTGATACTATAAGTTCTATTTGTACATGTCGAAATGGTTTTTCAGCAATAAATGATCAATGTGAAGATATAGACGAGTGTGTTGATGGAACGCATGACTGCAACATAACCACTATGTATTGCACCAACACGCAAGGCGGTTACAACTGTTTATGTATTGACAAACGTTTTGATTTAGTTTCTATTTTTCAAGGAAAGGGATGTGAAAAACGCAAATCCTTAGATGACCGCATAATGCCTAATTTAACTGTGAACTTCATTGGATCAAGTAAAGGAAAATCATTTTTATTTATATTGCAAATATTCTACGTCAAGAGTTTTACCACGGAGGCTCATAATATTACCATTGAATTCTTTTTACCTCCATTTCTGCTGAAGTTCTCCGAAAGTATCAGTCCTGGATTTACATCGATTGAGAAGATGAAATATGTTCATGAACAACCACTTACGTATAATTGCAGTGTTTATATTATGATAAATGCTACAATCGATTATGAATTGTGCCCATTTAAAGACTTCATTATGGATATATCAGTTAAAGTTAATTATCAAAATCTGTTTGGAAAAACTTTGATTTTGTTTAAAACATATCAAAACAAGATTCAGATTGAGAAGTCAGAATATTTGAAGATTAATAAAAATCCACTGGCCCTTTCAGAAAGTTATGGCCGTGGTATATACTATGATGAAAAAGGTTCTGTTGTATATATCTGCATAAACCAATATGTTTCCAGCACTTCAAGAGCATGTTTTGTTACAAACGACTTTGGAATGTCATGGGAAGGCTTAGATTTTGCTGTTGGATCTATTACTGGTCATCACAGGTTAACAAACGAGTTGTATGCTATACATCGCAACCAAAAATTATACTTGATGTTTTATAGTCACTATAAAAAGTGGATTTCAATATCCAATGAGGAGTTTTCGATTCAATCAAAATATTTAAATCATAGTCATTGGAAAAACTTGGAAAGTAAGAGCGTTCAAGAGTTTACAATCGGTTCAAAACAATGGATGGGAAACGCATATGGATTGTTCTTTCGAAGCAATGGAACATTGATACAATTAGTAAAGTGGCCTTACTAATTATTAAAACGTTAAAGATATACGTGAAAACAATTTTCAACAAAGATTTATAGAATATTACTTAGTTTACCCTCTACTATCTTGCAAAAAGAGTTACATTTAATGCTGCGAGTGTGTGTGCTTCCACATTCAACTCTGGTTCCAATTCTTGTTTTGATTAATTTAACTTAACATTTGCACACTTTGGTTTTTCTATTTTCTTTGTTGCTTCGCCATTAACAGATTTAATGTAATTTTTTTAAATATGTGCTTTTGTTTATCATATAAAAATAATTATACTTATATGATGCAGTATATACTTGTAAAAAATTTATAGTAGTTACAATGAATCTTTGAAACTTTGAGAATGCTACTTTGTTAGTTGATTATTAGAAAAAAATTAAAGTGAATTGTGCTGAC
>evgR015505c0g1_i01_Hydra
TTGGAAATCGTTATCTCCATCATAGTGTTGTGTTAGAGACAGTTGCTGCAGTACCTGGTATGGTTTCTGCTGGATGGAGACATTTCCAATCTTTACGATCAATGAGAAGAGACCACGGCCATATTGGTGAATTGGTTGAAGAAGCTGAAAATGAAAGAATGCATTTATTAACTTGGATGGAAATGACAAAGCCAACTGTATTGGAAAGATTATTAGTTGTCGGTGCTCAATTTGGTTTTACATCATTCTATACAATGGCCTACTTGGTTAACCCAAGATTCTGTCACCGTCTAGTAGGTTATCTTGAAGAAGAAGCTGTTAATGCCTATTCAGAGTTTTTAGAAGCCATTGATTCAGGTGCTATTCCAAATGCACCTGCTCCAGATATTGCTATCAAGTACTGGAATATGAAACCTAATTCAACAATGCGTGATGTTGTGGTTGTGGTTCGTGCTGATGAGTGTATGCATAGAGATTTCAATCATGACATGTCTGATAAACACCGTGCAGGATTGAAAGTTTAATCAGAACATATATATTCACTTCTCCTTTGTGAGAATACATGCCAACAATTTTGTAAAGAGTGATTTTGGATTTGGG
>evgR015506c0g1_i01_Hydra
ATATTGTTATTGTTTTTAGCTAAATCGTTAGGAGTAAAAGTTAAAAAAATTATTTTGAAATATTTGAAACCAATTTCGAGAATTTTGGATCAAACATGAGACTTGAGACATGAGATTCCTTATCAAACCAAGAGTTGACAGATTTTAGTTTTTTGTGAAATGAATTTAATTAGACATGGTAGGGATATGGTTAATCAAGTTGGGGGATACAAGAATATTTTTCACAGATTTCTTCGTGAATGTACAGTTCGAGTTGGTGCACATGTAGGCATAGATAAATATGGAAATAAATATTATGAAAATAACAGTTATTTTATGGGAAGAAACCGATTTGTTGTGTTTCCTTACAAAGGTCGCTTGGAGTTTGATGCAACTCAAATCCCTCCAGAATGGCATTCATGGATGCATTATATGACTGATGATACTCCAACAATCAATCCTCCAGTATTACAAAAGTTTGCTCGTGATCATGTTGAAAATATGTCTGGTACAAAGAATTCATATATTCCTTATTCTACAACAAAACCAAAGATAGAAGCCTGGACACCAAATAAACAATAATTTTTTTTAATTTAAAAATAAAATTTTGAGCTGTATTGATATTAATTGAATAACAATACGATGTGACAACTTGGATATTCTGTTTTAAACTAAATTTCATTAAAAAATTTTATTTAAAAGAAGTTTGATATTTGTTTGTTATAATTAAAGCACTTTTTGTTCAAAAAAAATATTTTTATCTTGTTAATTTCTATTGTTTGAAATTAAATTTAAAATTTTGTTTTTCTGAAATGTTATTGTTTTGCAAGCTG
>evgR015507c0g1_i01_Hydra
GTCTTTAAAGACGTTATTCGTAGAAGAAACAAACAAGCCTTGAACTTGGTTCCAAAGGAACTCCAAGATGCCTTTGTTGAACACTGGGCTAAATGGAGCCACGATAACGTTCCAGCACCAGAACAAACTGTTGATAAGGAATGGGTTGTGAAGAAGTACAACATCTCCACCAAACCAGTCCCTGCTAACGCTTCCCATGTTAAGTTTGCTTCACCATTCGGTAACGCCGAAGAATCCCGTTTGAACTCCCTCATCAAATACGAAAAGGAATTGCAAAGAGTCTTCAAGATTGTTGATGATGGTAAATACATCCTCGACGAACACTCAACTTTCGTCAATAGAAAGTACTCAGTTTGGGCTTACTGTGCCTTGTTCTTGTTGGTTCCAATCGGTATCTTCGTCATGGTCTTTAATCAAAATCCAATGTACAAAGCCCTCTTTGAAGGAAGTATCTGGGCTCAAAAGAAGACTGATAAGAAGGCTTGGGACTCTGTTGAAATCGAAACTATTTTCGATAAGGATTCCAACCTCACTGTTAAGAGAGTCTAAATTTTGGAAACGAATACAGATCGTTTTTAAACAAATGTACACAACTTTTTCCATTAGGGTCATGACCAATAAAATGACGCTTAGGGC
>evgR015514c0g1_i01_Hydra
GTCCATAATTAATGATGAAAAAAGAAATAGTTTTCTACTATGATATTATTTGTCCGTACGCATACCTTGCATCAAAGTTGATTGAAAATGTTGCTAAAAGATCTAATGCAACTTTAGTATGGAAACCTGTTTTACTAGGAGCTATCTACAAAAGTACGAAACCTAATAACATGGAAATCCCTGAAAATAAACTTATCAAGACATATTCGAGCCAAAAATTGAAAGTTATGAGAAGAGATTTAGCTATGCAGTTTTCTCGTCATGGTGTTCCAATAAATACATTTCAAGTACCAGATATAAAAACATTAAATGCTTTGCGGTTGATTGCTTTTACCCCTGATGAGTCTATGGAAGTAAGATCAAGGCTTTCGCATAAGTTGTTTGATGCAATTTGGCAACATAGGAAAGATGTAAGTGATATGAACATCTTACAAAATATTGCTGACTCGTTTAATCTACCAATCAAAGTTACTGATTTTGATGATAAAACCCATCCTGCTCGGCAGAAGTTATTAGATAACACAGAGGAAGCAATTAGTCGTGGCGCTTTTGGTGTACCAAGTATGTGGGTCAACAACAGACTATTTTTTGGAGGCGATAGGCTTTTTCTTGTTGAAAGACAGTTTGGTTTGAAATCTGAACCACATAGAATTTTGCATAAACCAGTTTGCGATGTTCAAAAAAAAGTCAAACTTACATTTTATCTTGATTTTGGTAGTCCCTGGACATTCCTTGGTTTCAAACAGTTAAATTCGCTTATATCATCTGTCGCTCCTGTTCAAGTAGATATTGAGTATGTGCCAGTGCTCATTGGAGTTATCTTCAACAGTATTGGAACACCAATGCTTCCTTTGCTAGTGTCTGAAAGTAAAAGAAATTATTTTCCAACCGACTTTAATGATTGGTTAAACTACCATGAAATCAAAGATTTCAGATTTTCTTCCCATTTTCCTTTGAAAACCTTAACTGCATGTAGATTAGCTGTTGCACACAATAAACCTGAGTTAATCAATGCCATTTTTGACGCTGTTTGGTTAAATGATAAGGATATAAGTAGTGAAGAGGTCCTTCGAGAGGTTTTGTCATCGAATGGATTTAATTCTGATCAATTAATAAGAGAAGCAAATACTGAAGAAGTCAAATTGCAACTAAAAAAAAATACGCAGAGAGCTCTTGACGAAGGAGTTTGTGGCTTACCTACATATCAAGTGAACGGTGGCCCCGTTGTTTTTGGACAAGATCGGTCAAATATTGTAGCTGATATGTTGTTAGGATGGAATCACAACGAACTTGTCAGCAATCTTTAATACACTCATTTTTTTTGTCAGATTCAATGAAATATCTGAAGATAAGGATGTTATCCTTAACTACTTATTTAAATGTAATAAAATGTCATTCATGCGCAGTTGACTATGATCATATTTCAAATATTTTAGATACTTATAATATACGTAGATAAGATACGTAAACACATATTAAAAGTTTACAACCTTAAACTTTTAGAGTTAATTTTTTGTCACAAGTTTTTTAAGTTTTTTTTTAACTTTTTTAACTATTTTTTATTATTTATATACATATTACTTTTAATAATATTTAATGAATGGATAAAAAAAATTT
>evgR015520c0g1_i01_Hydra
TATAATTTAATCAAACTAGCTTTCTTTTTATTATTCGAGTTAAAAGAAAAGAAAAGAAAAGATTTTTAAAGTTGTTGTAGTCAAGTGTAGTCAAGTATTCACTATTCTTCTGAAGCTTCTATTAGTTTTAACATAAGAAAATGACAAAAGAGATCAAAAGTTTGGACGATATCTTATCAAAGAACTTGAAAGATGAAGATCTTCAAGAAGTTCAGAGGTTTTTGTATGGTGAACCAGTTGCTAAAATTAATTTAACAGACGATTCGACATCAAGCGCTATTAATAACAATTTTGAAATATGTGGTTATAAAATGACAGCTCTTGACGAAGAAGAACGCCCACCCCGTATTGTACGAATTGCAGCTGTTCAAAACACAATTGTTCTTCCTACTAATTCTCCTATTTTGGAGCAAGTTAAAGCTATTCATGAAAGAGCTAAACTTTTTATTGAAGCAGCAGCCAGCAATCAAGTAAATATTTTGTGTTTCCAAGAAGCTTGGACAATGCCATTTGCATTTTGCACTCGCGAAAAGCATCCTTGGTGTCAGTTTGCAGAGTCAGCTGAAAATGGAGCTACTACAGTTTTTTTACAACAATTAGCCAAGAAACATAACATGGTTATTATTTCACCTATACTTGAACGAGATGAAACTCACCAAGATGTTTTGTGGAACACAGCAGTAGTGATATCAAATAAAGGAAGTTTTCTCGGTAAATCTCGTAAAAATCATATTCCTCGTGTGGGAGATTTCAACGAGTCTACATACTATATGGAGAGCGAGTTAGGTCATCCAGTATTTGAAACATTGTTTGGTAGAATTGCTATAAACATTTGCTTTGGTCGACATCATCCTTTAAATTGGCTTAGCTATGGTTTAAATGGTGCAGAAATTGTCTTTAATCCTTCTGCTACAGTTGGAGCTTTAAGTGAGCCCATGTGGCCCATTGAAGCTAGATGTGCTGCCATTGCAAACTCGTATTTTACTGTTGCAATAAATCGAGTTGGAAAAGAAAATTTTCCTAATGAGTTCACGTCTGCAGATGGTAAACCAGCACACAAGGATTTTGGGCATTTTTACGGATCAAGTTACGTATCTGCACCTAATGGATCACGTACGCCTGGTCTTTCACGTATTAATGACGGAATAATTATTGCTGAGTTAGATTTAAATTTGTGTCGCCAGGTTAAAGATAAATGGGGCTTTCAAATGACTCAAAGGTTGTCATTATATGCAAAGATTCTAACTGATGCATCTACACCGGAATTTAAAAGAAAGATCATTAAAGACTAATGTATTACAACTTAGTGTATACAACAATATTTTTGATGAATTTGTAAGAGATTGTTAGTTTTTTTTAAGGGTAAAAATATTGTAAACTTAAAAATAATAAATTTAATGCTATCCTCCTCAAAAAAAA
>evgR015524c0g1_i01_Hydra
CGGATCAAAAAGGCACATTTTGCACATGCAACTTTTGAACGTTTTTCTCTTTTATTGTTGCTGATTTTTAATAGAACTTTTTCATTTACTAACATCACGATTAAAAGATATATATTACAATTTTAATTAACTGTTTATTTGTGTGTTTTTAACCGTTGAATGAAATGAATGAAATGAAAATAATTGATTAAATCGTTTTCGTTGGCTTTCATTCTAGAATTAACAAGAAAGTGGAAAAATAGATATATAGTCAATCCTTTTGAAATAATATCAATTTATAATAATTAAGAACTTACTGATACCTGAACCAAAGTTCAACTTAGTTTTTCAAAGTTAGAAGTAAGTTGGAAATTTTTTAGTACGTTGGAAATAAAATTGTAAAAAAATACGAAGATTACGAAAACATCAGCAATAGTCTGAATTGTTATCATAAACGAAAGTCTATGCATCAGAAGATATGTTGATATTGCGTGTATATTCATTTTAGACTTTAACCATCTTTAACCAATTCATGCTGTTGAAAAAACAAAACAAAAGTAGTAACATTTAACTTCTGTTTACATCTGTTATGTCTGATAGTTACAGTGAGTTTGGAGCGCCGTTGCAATCAGCGACTAATTCTAATAGCGTTGTTACAACTGTCGGTGAATGTGCGTACAAATGCATTCAATTTGCTTTTTTTGTTTTTAACTCATTATACTTTGCTGTAGGCGGTACAGTGTTATCGTTTGGTATTTGGCTACTCATAAATCGAAAACAGGGAGTAGTTGAATATGTTCATCTAAGTGGATTAAGTGTGTACTTCAGCGGAGAACTGTTTTTAATTGCTGTTGGCATAGTAACATGTCTATCTTCAGCAATTGGTTTCTGTGGTTTACGTGGAACAAACTTGTGTATAATTGTTTCTTACACAGTTTTATTAATCGTAACTTTAACAGCGCAAGTAGCTGTGTTGGTAATAACAACAATATATAGAGAACACATTGAAAAACAACTAAACAGTAGTTTTTTCAACACTTTGAACAAATACGGTCAATCAAATTTTGAAAGCCTTACGCAGTCTGTTGATTCACTTCAAAAGAGTTTTGGTTGTTGTGGCAACGACTCTTATAAAGATTGGCTTAAAACAAACTGGGGCCATCTTCATAACAATACTGTCCCAAAAAGTTGTTGCTTAGCCCCTTATGAAAAGTTATGCAACAAAAATATTACCGAGAATCAAAGCTATATTTATTCAAACGGTTGCTATTACCACATTCGAAAATATTTAATCGAAAACTTGCACGTAATTGGCGGATTTGGTGTTTGGATAATAATTTTAGAAATATTATGTGTTTTTTTCTCAATATATCTTATTTTTCAAATTAAAAACGAAGAAAACAACTCAAAGATAGGCCAGTTGGAAGATTTTTACGACGAAAATTATTTTAAAGACGAATGATTGACCGAAGCATTATCAAATTACAAATCAAATTTTATAGTCATATATAATAATTGCTTGTTTTTAGTTAACAAAATTAGAATGGCATTCGCTTCAAATGTTTTTATAAGACTTTTAAGTCGATTAGTTTTATTAAATATTAAAAATAATAACATTAAAT
>evgR015529c0g1_i01_Hydra
GTGCGTGCGCATGTACGTGCGCGTAAAACTCAAGTAAATTTCAATTAATAATGAAAAGTTTATAGAGGTCACTAGTTAGTTTTTTTACTAGTTCTCTATAAGCCCACCTCGTTATGCCCTTTTATGGGTCCTCTGTTTTCAAAGAATCAAAAAGAACTCTATTATATACTCTAGATTAAAAATAGAGGCTTCATTGATTCAATTAAACTATTTCTCTATAAAAACGAACTTCTAACAGAGAGTTCATTATTTGCAAAAAGAAGAAACTACTTGACGAAAGCAGAAAGATTAATTACAAGTTCATAACTTATAAGATAGAAATGTTCCACAAAAGCCTTGTTATCTTTGTTGTGTTTTTCACTGTTGTTGCTACTCATTCGTGGATGAGTTGCCCCCCATCACTTCGGATGTCAATAGGGAGAGGAGGGGTACAGGACAAAGCTTGCGATGGCGATCCTCAATCTAACTTAGTCACTGTCACACAATTACGTGCCGGAGATAGACTAAAGGTTGGATGGCCTTCAAACAATCATGCTGGTGGTTACGTTCGTTTGTCCCTTGTTCTTCTAGGTCAACATACTTCAGAAAACTTTCAAAAAAATGTTCTAAAATTTGCGTGCTACGGCAGTGATATGCGGGTTAATAATATGCAATCATCAAAATATGGTGACTGTTACCATCCATGTAATGCAAGATCTGGATGTGAATTCCAATCAGATAGCAACGATCGCGAACGGTACGATACTACCATTGGCATTCCAACTAATTTGAAAGACGGTGATTATGTGCTACAAGCTGCCATGTTAGTAGGAAATTCAGAAGCCCCCTATTATAGTTGCGCAAGACTTAAAATTACAGGAGGGGATTCCAGTTTTAATTGTGTTTCAAAAGAAGATCCAATAACGTACTCTTGCTTAAAATCAAACGGCCCTGCTCTTGAAACACATTTCCTTAAAACAGGTTCAACACGTGGAGATTTTTGTTACCATGTAAACGGTTCAATTGGCGGAATTGACGATGATATACATAAAGTTCCTTTCAATGTCGAGTGTGACCCGAGAATAAGTTGCACGTATTCTGTTGACAACAACTGTGAAGCAAACAATCCATCAATGAAATCTATTCTCATACCAGAAATAAGTCCAAAACAACCAAATTGTAACAGTAAAAACATCACTCCATATCCATTAAGCTCCTGTAATGACGGTAGACAGAACCGCGACGAAGTAGAGATCGATTGTGGTGGGGTGTTTTGCAAACCGTGTCCAACTTTGTCCCCAAACGAAAATGGAAACTTGGAATATTTTGTACGACCTATCAACCAATCAACAGTTATTTCTGGTCAAGGAAGTGTGTTTGAAATAACAGTGATGTTACAAAAAGACGTATCTATAGAGCTATGCTGGCAACTCACATTGCAATATGGACGTCCATTGCAATTTAAACCTGATGGTAGTGCGCCGGTAGGATTAGGCGTCTGGAATGATGGTGTTCTCGTTTATGTCAATCCAGCAAGTAACATTATCATAATTGAAAAACAAAGTTCAAGCGCAAACCCAAAAGCTGGACAAAAACTATCGGTCAATTTCACTGCCTATCCTCATCCTTCAGATCCAGACCAAGAACCTTGGACAGATTCAAACATGAAACCATTAACTGCATCTTTTGGTATAAAACCTAAAGAGAAGTGTTTCAACAAGCCTTTGAAATAAAGAAACGTATCGGAAATGCCGTAATGATTTTGTGAACAAAATAGATAAAAATAAATGTATATGTGTATATAAAATAAATAAATTTAATAATTAAACAACTTTCCACTTTCTTTTTTTATATAAATAAGATTATATGTGTGTTGTGCGGATATATTTATGTATATAAAGCCGCTCCAAGCTTAGTCGGGACCTGGGACAAAATTAAAAGACAATAATAAAAAAGGTCCTCAATTTTAAATTTCTGCTTTTGCCTTTATTTTCAAAACCTATT
>evgR015536c0g2_i01_Hydra
GGACAATTTGTAATGATACGTATGGGTTTATGAACAGATCAAGAAGTTGTAATGTTTTAAATCCAACAGAATCGTGTGATGGTAGCAACCTTGATTTAATGACATGCTATGATATATGGAGTGAATGGTCTGAGTGTAATGCATCGTGTATCATAACTAGAACTCGCAATTGTAGCAATCAACTATTGAATTGTATTAATAAGGAAGATTTTGAAGTATGTGGTAATATATATCAAGAATTCCTTCTTATTGAAAACGAAACACCGCTACTAAATAAAACTCTAGTAACAACATTAATGAACCACATCAAAGAATACAAAATTTCTTTTGATATAAAGCCAACAATGTTTGGAAATGATTGGGCAAACGTCATTGGTTTGACTGTAAATAGCGATGATAATAAATTTCTCTATGGCAGTCGAATTCCATCTTTGTATTTTTTAAACGAAAGCTTGTACATTTGCTCAGCAGTTAATGGTG
>evgR015541c0g3_i01_Hydra
TAACCTATTTTAGTGCAAAAAGCAATTAATAATGTCTGATTTTCCTCCAACCTTACATAAAAAAATGGATGCATTTGAGGATTCACTAACAGAAGTAGAGGATGTTTTAAAGCCGTTTTTGAAGATTCCTATCGCAGACTTGCATGAAAAGATAACAGATCAATTGGGTAAAGCAAAGCTAGATTTGATAATTGCTTATACTGCAAACTCGCTATTTTGGACTTATTTAGTGACTCAGGGTTTGGATCCAAAGAATCATCCTATCAAAGATGAGCTTATTCGAATCAAACAATGCATGAGTCGTGTCAAACAAATTGAAGACAAGAAAAAAATGTCTAGGGTAGATACAAAAGCTGCAAAAAGATTTGTCACTAGTGCTTTGTGGGAGCCAACTGAAAGTACGGAAGTCATTAAAAAAAATAATGAAACTTTATCCAGTCAAAATTATGATGCAGAAACTAAATTGTCTATTCGAAATAAAGAAGACGAATCTAAATCATCTCGTCAAAATAAAGATGCTGAATCTAAATCATCGAATCAAATCGATGATGCGGAATTCAAATCTTCTGATCGAAGTTCTACTAGTCGAAATGACAGTTCTGAATCCAGATTATCTAGTCGAAGTGATAAAAAAAGAAAGAAAGATAATGACGATGATTCTTTAAAAAAGAAGAAAAAGAAAAAGAAAAAGAAACTAAATGACTTGTAATGATAACAAATATTTTTCATCAATTATAAAGATGTGAAATGGTTTTTTAATTATATTTTATGAGAAATAAAACAAATAATAATTAAAAAAAAAA
>evgR015543c0g1_i01_Hydra
ATAAATAATAGTTGTAATAAAAGTTGTATATGTAATAATAATAAATAAAAGTTGAATGTCTCGGTTAATAATCTTGAAAAAAACTATGAAATTTTTTTATTATATTATATACTGTTATATTCAACAACAAAATAGAAAAGTTAAAATTTGTCATAATGTTAGTCATAAAACTAAATGGACAAGTGTTCCATGGCCAAGTGGACCAACCCTCAAATTTATAAATTATTTAAGGGTTTTTCATACTTTTGCATATAACACATCTTTAGACAAGTGGAGAAATAGTGATTGCAATCCCCGGATCCTCACCTTTTTTATTATCAGGAAATCCCGAGATTGTCTAATATCAATCCAGAGATTTCCTGTAAGGTCAAACAAAACAAAGTATAATAAAAAATTGACC
>evgR015546c0g2_i01_Hydra
TTTTTTTTTTTTTTTTTTTAATTAACTTCTGAAAATGGCAAAGGAACTCTCTTTTCTCATTTTTTTTGTTATACTTTCTAGATCTGCTGAAGCCGTCGGTGAATCACAGTTGATTATTGTGCGTGCATGTGAAGGAAGCGATCTTAAAATTGATTGTAATGGTAATGGAACAGTTGAAGTGATTTACGCAAACTACGGTAGAACCTTATCAAATGTTTGCCCTGGAGCTTACGATTCAAACACAAAATGTACAAATGAAAAAAAATCTCTCGATGTTGTTAGCAGCAATTGTTTAGACAAGACTTCCTGTATCATTAAGGCATCTAACTCAGTGTTTAATGATCCATGCGTAGGAACATACAAATATCTCGAAGTGCAGTACAAATGCAAACCACAAGTAACTATTCCAAGAGCCTGTGAAGGAAACGATTTAAAACTTGATTGCAACGGAAAAGGACTAATTGCAATAGTCGATGCAAATTACGG
>evgR015570c0g1_i01_Hydra
TTTATACAAACAAAAGAGATTTGCTTACATAAAATTTTTCAGCAATTTTATTCACAATAGAGAGAATTTCTGAAAATTTCTGATTTGACTGATTTCAAATTGTAGCCTTTCTTTTTTATTCAAGAAATTTTTAAAGTTGAAGAAATGAACTTGGTGTCTACTATTTGTTTCACTTGGTTTATGGTTGTTTTCATAATAAGCGGATTGTTTATCAATTTCTTGCAACTATTAGTGCTGCCATTATACTGGATCAAGAGAGATTGGTATCATATCATTATGCCTGGTCTTGCATATCTTATTTTTTCACAATTTACTTGGTTAGGAGAATGGTGGGCTAAAGTTGACTTTAAGCTTTATGTAAGCAATCTAGATGATATCAAACACTTTGGAAAGGAAAATTCTATAGTAATCTCAAATCACTATAGCGATGTGGATTGGTTAATGGCGTGGATATTTGCAGAGAGGGTTGGTTTCATTGGGAGAACAAAAATAATATCGAAAAGTGAAACTAAATATCTTCCCATTATTGGTTGGTGCTTTTGGTTTGCCGAGTCTGGCTTCTTGAAAAGAAATTGGCAAGATGATAAATCAAACATCAATATGCTGATAGATAGTATGAAAAGGAATAATAGCAATTTTGGAATATTTGTAATGTGTGAAGGCACACGTCGAACTGAGGAAAAACTTAAAGCCAGTCAAGAATATGCAGTAAAAAATAATTTTATTCCTTTAAAACATCATCTGTTTCCTCGAACTAAAGGTTTTTCCCTCCTTGCAGAAGCGCTTCACAGCAAAGTTGAATCAATTTATGATTTGGAATTTGCATTTCCTGATATCGAATCAGCAAACATGCAAAATGTTGTTAATGGAGGGAAAATTGAAGTTTTGATGCATTTTAGACGGATACCTATGAATACATTGCCGTCTTCATTTGATGGATTATCAAATTATATTATTGAACATTACAGAAATAAAGATGAGATATATGATCACTATGCGAAGAATAAATCATTTCCTGGTCTTCGTGTTCCTCTTCCACGGAGACATTTAAATTTCATATTTTTTTCTATATGGAACATTTTCGTTTCTATATTTGCAGCACTGTCGCTTAAACACTGTATTAATCTTTATGGTCAAGCAGCTGTCATGTCAGTTCTGTTTTTATTTTTATTTGCAGTATATTTATTATTCAAAACTATGCAAAATCACACGCTAACTAATAAAGGCAGTAAGTTTGGGTTGAAAGAAAAGTGAAGAACTTGTTATATTTAGCAATTTTTATTATGTAAAATATTTCTTTTGAAAAGAAAATTAGGATAAAATTTTTTTTTTCATTTGGAAATATTTAATTTTTGTACTAAGTTCATATTCTTACATATTCATATTTTTAAGTTCATATTCTTACTATTTTAGATTTTTCTATCCAATTATTTTTAGATAGTTCCAGTTATTTTATTACCTCCGAAATTTTTTTGTTTGATTGAACTTCGTTATTTAGTTTTTACATTACGTTTTCACGCACCATTTTAGGATACGATATTTAATATAGGTCTTTGTATTTTCAATTATTTTGTAGTGTTC
>evgR015579c0g1_i01_Hydra
TGTTCACCACGTTATTAATTAAAAATCAACAATTTGTTTCAGCTTTAAATTGGTTTAAATTCAAAAAGCTGCTATCAACTTATGAAAAAATTAAATTAATATTAAGTAATTTATTCCAACTGCAACAGTAATTTATTCCAACTGAAAAGTGAATGCTGTAAAACTTGTAAGCACACTATTAGGAGCAAAATGAATTCGTCGTTGTTAGTTTCTGATTCTGTTTTCATTGATGTTTCTGAATCAAGATTTAAATTTAATAATTACTTGTCTTCTTGCGATTTTTCTGATGACAATGCTCATAAAATTGACACAACATTATCTGATGGTGGTGATTTACATTTAAAAAGCTTGAATAGTCTAAATCAGTTTGATGTAGAAAAATCTGCTCCAATTAACATGACTCGAGAAGAAGCAATGAAGCTAGTAAATATTGCAGCAACTTTGTTACAGGCTAATATAAAAAGATATTTACAACAAAAAAAATACAAAAATTATCTATTTAGGTATAAGTCTGCTATAAAGATACAAGCGGCATGGAGGGGATACTCAGATAGAAATCTTAACTCCAAAGTAATTGAAAAATTTAAATTTGAGCTTCCTGTTAAAATCCTCCAGCAGCGTTTAAATAACCAAGACATTATGCATCAAAAATCTATTCAAGCCATTTTTGCAAGTCAAGAAAACCTAGCTGCTGCCATTTATGCATTAACTTGTCAAGTTGAAAGTTTAAAAAGACTTTCAGTTCAATCTGAAACACAACCAAGTTCTTATACAGAGATTTTTCCATGCAGTGTAGAAAAATATAGTTCTACTTTGCAATTGGATTCCAAGAGTCTTCATGAGGATCAAAGCACAGAATATATTCAATCGAAAGATTATACCGATGAGTTTAGTTCAATTAGTATTTTTAAAAAATCAAATGAAGATATTGCATATGAATCAACGTCTCAAAAAGTTAATGCTCCTAGTTTAAATAATGTTTTGAATAATGACAATATAAAAAATCTTGAGCTACACGAAGAAGAAAATAAACCTGAAGAAGAAAATAAATAGTTATTTTCCATTGGTAGTTACAATATTTTAATTAAAAAAAACTATTATAATTTGATGTATTATGTTAATAAAAATAAAATTATATTCAACTTATCAGTTTCAAAAAAAAAAA
>evgR015590c0g1_i01_Hydra
ATTTCATATCTTTCTTGTTTCTCAAAAATATAATTATGTACAATGGAATTATTTTCTCAAGAACATGATAAAGGCACTAAATATGAAGTTTTTAATTACGAGGAAAGTAAAGAAAATCATGCCACCGGACATTCAAACGAAGGATTGTCAACAGTAACATCTACATTTTTTGTTGTTGGAGATGTGGTAGGAGCAGGTGTGGTTGCATTACCATATGCATTAAAGTTAGTAGGATATTATGGCATACCCATGTTTATGCTCTGTTCAGCACTCATGTGTTACTGCGGTATCCTCTTAGCTAAAGCTTGTTCAAAAATAATGAAAAACATTGATCGTTCGCAACTTCGGGATCCATATCCTCGCTTGGGTTACGAAGCGAGCGGAAATGTTGGGAAAGGTATTACCACAGTCTCGCTTGCTATTAACCAAGTTTTAACCTGCATTGTTTTTATATTATTAGCAGGAGAGATTCTGCTGGAACTATTCCCTTCTTCACCTTGGGATCATATGTCTTACAGATCACAATTGCGCATTTGGTTCTGTTCATGCGGCTTATTTTTGCTTCCGTTTACGTTTCTAGGTACACCTAAAGACTTCCAAGGTATCGGCTTTCTTGCAATGGTTACTAGCGGGATCGCTGTGCTACTTATTTGCCTTATGTTAGGTTATATTTCAGGTTTTCCTGTGGAAAACGACAAAAATATAAAACTAAGCGGTGACGGATTTCTTCATTCTTTTGGTACCGTATTGTTTGGATTTGGAGGCGTTTCCATATTTCCTACAATACAAAACGATATGAAAAAACCAGAAAACTTTGTATACTCCATTACAATTGGCTATACAATAATCTCTATAATATACATTGGGACACCTCTGGCTGCATATATAGTACTCGGGGATTTAATTAAAGAAGACCTTCTTACAACGTTTACTTACTTAGACTTATTTTATACAAGACATTTATTCAGGACGTTCTGTATGGCTGCCCAGGCTTGCATATGCGGTCACGTATTATGTGCGTTTGTTTTGAACATAAATCCTATTTACCAACAATTTGAAGGAGTCATAGGGATACCAACAAAATTTTGCTGGCAACGTGTTTTGAGCCGAACACTGTGGATGTTTGCTATTTTACTAACAGCAATAGTAGTGCCTGCATTCGGACCCGTCTTATCATTTGTAGGCGGATCGTTTGCAGCTTTACTCGGAATCATTTTGCCGGTCGTGTTTTATGCCAGGATTCATGGAAGACTACCGATCTGGAATGAGATTCTTTTTGTTCTAATAATAATAATTGCTTTACTAGGTTCTGTTGGAAATGCTTATGTTGAAATTAAAAACATTATAAATGTTGTTTTTGACAAGTATCATCACACTTAGAATAAAAAATAACTCTTGTCTTCCAGACAAATAATAACTTACGTTTTTAATTAATTACATATCAATTAATAAATGGCTATTATAACAGACTATTATAAAAACATTTTAGTCTCATTATAGCTAAAATTCTATTTTATAGCTTTAGGATACGAGAGATCAAATCTCAGTCGATGACACGCGTAATTGCAGTCTAAACCAATTAGATCACATTACAACATATGGTTTTAGTCTGAGTTAATTACATTATAAATGCCTTTCATGTGAATAACTTTAGGATGTACGAAAGATTACTATGAACAACGAACATAATAAACTTTTTAATTTTTATTAATCTGAAAATCTTAGCAATGCTAGTGTAATAGATCTTTATAGCATAATAACTTACGCTGCCACGCCTCCTATACCCCTACAGCTACTATGTACTTAGTACAGGAAATTTATTAATGATGAATAAATGAGACATTAAATGACAATGTTGTAAAACTTTTTCTATTTAAAACATTTTGTAAACTTATTTATACAAATATTTACACAATTTATATTTTTAATAATATTCTACAATTTTATTAGAAAATTATTTTTATAGAAAACCGTTTTATTCGTATTTACAATTACAATTTACAATTAATAATAAAATGGAGAAGAAATGTTTACCAAAAATCTTCCACGTTTTATGTTATCGTTTTATACCATAGATACATTA
>evgR015596c0g2_i01_Hydra
TTTACAACAAGAGATTACTTACGGCTTACAGGTGATGTTAACATTTTTACTGGCTATTCTTGATGAGTCCTATCAAGTTCATGGTCATTTTTTAATAGAGCGCTGATAACAAAGCGCCGATCTAGATTTATATTTTCAAATTATGTTAAATTATATATAATATTGAAATCATGGAAAACAGTTTTTTCAGACGTGTGGTGTTAGTGGGTTTTTATAAGTAGCTTGAAGGCGGGTAATGCTTTTATGAAGTCATTTTTATCAGTATTGGTGGAATCAATCGTATCAGGTGCTTTGTTGTTGTACTAGTCTTTTGTTTTATTTATGTTCAAATAAACCTTCTATAGAGAACAAAATCTTTATTTGAAAACAACTATTTTCAATTAGGAAAAAACAAAGAAACAAATAACGAAATATGATAAAATTAATTTGACTAGATTCTTACGCAGAAGTATTTTTAAAGCAAAAATGGATGAAAAATGGAAGATTGCAATCTTGACTAATTGTAACATTGGTCTAGTTTTCAGTGGAATTGGATTGTTTGTTTTAATTATGTTCAAAAGTGGATTTGATTCAACGCAGCGATGCATATTAATTCAACTTTGTTTAATTAACACTTGCTTAAGTATATGCTTTCAAATTGAAGAGTTTACCAACACTTTTGAAGATGTACTACGAATTAAACGTATTCTGCAGATATATGATGAGTTTAAAAGTATTCTCGGTACAGGATATTATTACGTCACATTTTGGTTGGTGTTCGACCGATACTTACATATAAAACTGAACATAAAATACGTGGCTTACTGGACGAAAGGGAAATCCATAATTTTATCGACAATTATATGGAGTATTTCTGGGTTAAGTGGAGTTTTAATTGAAGTTTACAAAAATCAATCATATATTCAAGTTTATATTGCTGCATGTTACGATTTAATAATAATTAGCTGTTTTGCACTTTTCTATACATATGCCCTTCTACTTATTCGAAAGAAGAAAAATGTGTGCTCCACTCAACACGACAGAAGCATTTTTAAGGGCTCAGTAGTAACATGTTGCATATTAATAATATTTATTGTGTTAGTTGCAATACCTGAAGTGTTATTAGCATTAATACTTAGAAGAGAAAATATAAATGATTCGAAGATTATTTCTCAAATAGCCTTATACAAAAGAGTAAGTTTTCCTATATCTGTTTGGACTGACGCATTTGTTTATATTTTTGCTTCTCCTCGAGTTAGGTTGGCTCTAAAAAAAAAGTTAAAATTAATGATTGGTAAAAAAATACCCCCAGTTAAATGTGTTAAAATATTTGAAAGTCGTGCTGGAGAAAAGATTTCTATGAAGTTTGTCTTAAAATCATCATTATCAG
>evgR015598c0g1_i01_Hydra
TTCTTGATGATAGTTTTGAAATCCGCAACAAATTTTCTGATTATGCAAAAGATCTTTTACAGTATTTTGTTTCCAATTGCATTTATATATTTGGTCCAACATTCACAGTTTATAATGTACACTGTCTAACACATCTTTGGGAAGATAGCAAAAAATTCTCTACTTCTTTGGATAATATTTCTTGCTTTCCATTCGAAAACTACTTACAAGTTGTGAAAAAATTTGTAAGAAAGGCTCAAAATCCTTTATCACAGATTTTTAAGCGTACACGAGAGTTAGAAAGTTTAGCAATGAGTGGGTTGAACCAAAAGCATATTTTAACAAAAATTTCAAACAGTGATAAGGATTCTTGGTATCTACTGCAGTCTGGTAATTTTGCAAAAATAAAGGAAGTCAATGTTGATGATACTTATGAGTGTGACATAATAAATAAAAAACATTTTCAAAGCCTTTTTTTTTGAACCATGTGACTCCAAGC
>evgR015599c0g1_i01_Hydra
TTTGCAACAAGAAAAGGAATTGTTCGCTCAACTTGCTGTTGATGCTGTTCTTAGATTGAATGGAAGTACTGATTTAAATCATATTCAAATTCTTAAGAAGCCAGGTGGTTCTATGAAGGACAGTTATTTGGATGAAGGATTTATTCTTGATAAAAGTATCGGAGTTGGTCAACCAAAGAGAATGGAAAATTGTAAGATCCTTGTTGCTAACACACCTATGGATACTGATAAAATTAAGATTAACGGTGCTGTTGTAAAGACAGCCAGTATTGCTACACTTTCAGAAATTGAAAAGGCTGAAAAGGAAAAGATGAAGAAAAAGTGTCAAAAGATTGTTGATCACGGAATTAATGTTTTCATTAACAGACAACTTATTTATGATTACCCACAAGAATACTTTGCTGACAATGGAGTTATGGCTATTGAACATGCTGACTTTGAAGGTGTTGAAAGATTGGCTCTCGTTTTAGGTGCTGAAATTGTTTCAACATTTGATCATCCAGAATTAGTTAAATTGGGAACTTGTGCTATTGTTGAAGAAATAATGATTGGTGAAGACAAACTTTTGAGATTTAAGGGATGTGCTAAGAATGAAGCTTGTACTATTGTTTTACGTGGTGCTTCTACTCACCTTTTGGAAGAATCTGACCGTTCTCTCCATGATGCTCTCTGTGTCTTGTCTCAAACTGTTCTCAACACAAAGACTGTCTTGGGTGGTGGTTGTAGTGAAATGTTGATGGCTAAGGCTGTCTCTGAAGCTGCTAAACAAACTCCAGGAAAGGAATCAATTGCTATGGAAGCTTTTGCCACTGCTCTTAAGGCAATGCCAGCTATCATTGCTGATAATGCTGGTTTGGACTCTGCCGAATTGGTCTCTGTTCTTGAAACTGATCACTACAAGGGTTTGTCTGATAATGGTATTGATATTCTTAGAGGAGAAATTGGAAATCTCAGAACTTTAGGTATAACAGAATCATTTAAGGTGAAAGAACAAGTTCTTTTATCTGCATCGGAAGCTGCCGAAATGATTATCAGAACTGACGAAATTTGCTACAGTGCCCCTAGACAAAGAGAACAATAATTTAAAAAAAC
>evgR015600c0g1_i01_Hydra
TTTCAGAACAAACTTAAAATCAGTTAGAATAGCCAATTTTGATGGTGATGGCAAGTGGAAAGTTAATATTTTTTTCAAAACAAGAAGACTAAAAGTTATAGTTACAGCTCTAAAGTAAAATGACCGATCTTAAAGTTTTTGTTAAGTATTCCAATTTTACTAGTCAGCTCTTTGGTTTTTCAGAGTACAGTGAAATACCGAGTAATTTAGATCTTTTGGAAATGTTAATCATAGCCACAGCTAATGTCAATCAGAGTGACTCTTTTTTATTTTCTTGTAACACAGAACAAAAGTTTGGTGATAAGCTTGTCACTGATGAGGAAATGGTCAACTTTCAAGTGACAAGACACGACAATAACAATGAAATTATAACCATAGAGATTTATCTACCTGGGGATGAAGAAAAGATGATTTTAGTTTACCACGAGTTTTATAAAATTTTGCACACGCTAGCAAACGCAAAATCTTATTACATATTTTAGGCCATTGCAAATAACATGAAGGAACTCGATTAGCTTGCTAAATAGCATAAACAAAAACGATTGGTTTTTTAAATAACGTGAACGAACTCGATTGGTTTATAAAATAACGTGAATGAAGTTGATTAATATGTTAAATAAAACAAACAATCTTAATTGGTTTGTTAAGTGACGTGTTAAACAAACGAACTTAATTGGTTTGTTAAGTAACGAAGCTGAAATTGATTGATATTTTAAATTAAAAGAACAAACTTAATTGGTTTATTAAGTAACGTGATTAAAATCGATTGGTATGTTAAATAAAACGAACAAATCTTATTGATTTGTTAAACTTTTACTAATTTAACTAACTTTAATCTAATTTCTATTTCTATTTATTGTTTTAAATTTGACTCCAACCCTGTTGAAATCAAGTATTTGTAGTACAACGACAACAGCAATTTGTCTACAAACAAGTTTTCTAAAAAATAATACATCGCGCTTCACAAATTCACAACTTTTTATTTTTATAAGAGGAACTCATCAATCGATGAAATCCAACAATAAATACAAACGTGTACGTTTTATTTTTACGAACAGTAGTATAATACGTTATCTGCATAGAAAAATGTATAAATTCTAAGTTGCTTCCAAATCCATCGATGATCATGGTTTACTCAAACCCATCAATCCATGCGTTACGGAGTTATTAAGTAGTAATCTATTTAAACTAAAACTTTATGAGATTATTTGAAGCATAAAACCCACATTTGCAAATTAGACTACAAAAATCTATTTTTAAAAACTTTTGTTTTGAAAAACCAAGAATCAATGTGAATGTGAGAATGATACTCGGTAACCTGGTAAACTACTTGATATGAAATGATATGATTTGGTAAAAGCATATGTATAAAAAACCCTTTGCCTTATACACTTTTTATAGATAGGAAACTTGTATTATATATAATAAAAAATACCGAGCCGAATCAAGTTCCAC
>evgR015612c0g1_i01_Hydra
CTTTTATTTTCTAAATAAGGATTTGTGTAATTAGGATTTAAGTTGATCAGTAATAATAATGATATGGCTGACGAAAAGAAGTTAAAAGCACTTAAAGACGAATTTCTTCATGTCCTAGAAGAAGAAGCAAAATACAACTGCCTTGAATTCAGAAATCAAATGTGCAAGTGTTTTAAAAACGGTTGGTTTGGTTTGTGTACAAAGGAAAGATATGTCTATAGACAATGTTTAAAAAAAGCTTTTAAAGAGCTTGAGAACAAAGACGAGAACAAGAAATTACTTAATTTTGAAATAAATGCTTCAAAAATTTAAAATGAATTCAATAGTTCGATTTGTTGTGTTTAAAAGAGCATGTATTGATTTTGGACAGTTTTCAAATATGCGTGTACTCTCATCACGACATGGAGAAAAATATTCAAGACCTGTGGATCTAGAAGATATTGTTGAGTCACTTAGTTCAGAAACATCTACAGGTTTTCAGCAAAAATTAATTGAGTTAGATAACCATAATAAATACTGGCCATTTGAACTTGCTGGACAGAAATCATTAGCATCACTGGTGCATAAGCTTTACTGTGAGCGTCGCCATAGTGAAATATTGCTAATCACAAGTAAACTAAATTCTATTAGTTACGATATATGGGGTGGTTTGGCAACAACCATTTTATACTCAGCCATTGATCAAAAGAAATGGAAGGAAGCCTTTTCCCTGTTTGAGAGTATGAGAGACAGAGGTCAACTAAAGCATGAACGGCTGTACAAACGTCTTATTGTTGAGCTAGCTGAAGACAATAAAATTAGCGATGCTCTAAAATATTTCAACGAACTGTCAAGTTTAGAGTCTCGTCTAACAAGGCTTAAATCATCTGATGTACTAATGTTGATTTCTATGATTAAAGCAGCTTGTCGTAACAAAACTCAGTTTTCGATTTCACTTGTAGAGAGTGTGTTGGTATATTTGGAAACCAAAGATGGCAGATTTCTATATGGAGTTATTTCAGAACTGTTGCAATATTTTAAAAGCAGCACAAACCATTTAGTAAAGCAGACAGAGTTGAGCAATGATGCTTGTTCAAACTGTTCTTGCAAATTAGTTGAAATACAAAATCACGAAGCTGTTTGCAATATGATTATAAATTCTGTTGAAAGAACAAATGATTTAGATATAATTGATGAAACGATCGAATCTAATAACGGTTTTCATATGCTTATAGACGCTGGTAATGTTCTTTTTCACGGTAAAGGGAAAAAACAACGTATTCTTTCCAACAGCAGTTTAAAACAAATTTTGTTAGATATTAAAAACAAATTCGATAAAGTTGACGTTTTACTTGTTTTTCCAACAAATTATTCAGCTTATGATGACCTAAGTTACTTTCAAGACCTATCTTTAAAGATAAACTGTTTTGTGCCTAAAATTACTAACGACGATCAGCTTATTCTGTATGCAGCAGCCAAAAGTGAGTTAAAGAAAAAATTAATTTGTATTGTTTCAAATGATAGCTTTACAGATCATACACATCTTGCTGCTAACTTTATAAGCAACTGGATTAGATTTTTAATTGATAAATTGATTCCCTTTAATCTATATGGGTGTTTAGAAAATAGAACAAAGATGCCAGCTGTGGTTTTAAACAGTAAAAATATTCATTTATTTGATGGATCAAAATTATGGTGTGTAAATAAATAAATTTTTTTGGAAAATTTGTGTTATTAGTTAACAGCCGAAAATATTTAAATGTAAAAGGTGACATTAGCAATATATATATCGACGGGGATACTGC
>evgR015613c0g1_i01_Hydra
CTAAAACTGTGGTTATATTTACTTCTGTTAAGAAAAGTTTAGTTGTTCCAAAACATTTAAAAGATGTAAATCTAATTTTTGTTACAATTGGAGATCTAGATCAGGATTCTAAATATTTTATAAACCAAAATCATATACACATTGATGCTGCTGAAAAAATTCTTGAAATATTACCCGATTTGGAAAAGTTTATCCTAAAGGAAACAGGAAATAAATCAGAGGTGTTGTTTTTGTTATCTACATTTGACAAGCAAACTGAGTCTCAACGAGCAGATTTGTTTGAACAACAAAAGAACACTGTTAGTTTTATCATATCAGAAAACAAAAGTAAGAACGTAATTTATAAAGTAGCTATATACAACGATGGAACAGGAAGTAGTGTTGTTCAAGATTCTGATTCTAACCAATTAAAAGAGCTTGTTAAGAAATTGAGCAATCCTGGAGGTCAAAGTAGTATTGATAGTGCATTGATATTAGCCTCTCGCTTTTTATCTAATTCTGATCGAAATGTTATTGTCTTATTTGTTGATGGAAGCATTGCAGATAAAATATCAAAAAAGGTTTCTTGCAACTTCAGTAAAAATAATCTCAATGTTATTATTGTATTGATGGGAAAAATTATTCCAGACTTTGAAATTGCGAGCATATTAGGTAATAATGGAAAAGGGGCAATTCATACAAATAACACTGAAAATGTTGATATTATGAAATCTGTGGATGAAATGCTGATGAATGGAGTTTTAGCTAAATGCAAAGATGAAACAATATGTGACTTCCATTCTCGCTGTGTTGTTGTAAATAAAAACAGTATTGAATGCTTGTGCCCAGTTTGTGATGATAAGTATAAAATTGTGTGTGGCTCAGATGGAATAACATATGCAAGTGACTGCTGGCTAAAGAAAAAATCATGTGAAATCAAAGAAAAAATTTATGTAATAAACAATGGACCGTGTACAGTAAATGCAAGTTTTGACATTGCTTTTGCCCTTCCTATTTTGACTAGCTCTGAACACTTTTTAAAAATGTATAGATTTACAGAAGCTATAAAATGGTTGTTTAAGATTGAATTGCCACATACTTCTTTATGGTATGCTTATGGACACGATAAAAATAAAATAATTGATCTGCCATCTGTTGAGTTTATTAGACAGCCATCTTTAGTCAGCGTTCTTAGTAATATGAAAACAAGTGGGTTTTTTGTTGCTTTTATCCAAGAAAGCTCTCTTACTAACACTAATGACGTTCAAAATATAAAGATACAGCTCAAGAGGTTAAAAGGAATTGGTTGTCATGTTATGGTGGTTGCTATAGGATATCCAAACAACTTAGAAAAGATTTTTGATAAGGTTATCACTGTTACTGATGCTGAAGAGCTTCCTGGGGCATTGGAAATTATCGAGAATAAATTTGTCACTCTTTTGTTTAAAGGTGATAAACTTGACCTTGCGATTGTTTTTGGTGGATCAAATGCTGAAAACTTTGCTATTCAAAAAGCGTTTGTTCAAAAAATGATTGAAAATAAAGTTATGGGCACTGAAGCGACAAATGTTGGTTTACTTTATTATGGTGATTGGCAATTTTTTGACTATAAAATTGATAATGACTTAGTGAAGTTGAGACTTATGCAGGTTATCAACACGGAAAATATTGAACTAGCTATTGAGGAAGCTTTGCAAGCTGCAAGGTTGATGTTTTTTAGAGGTGAAACTACTAATACAGATATAGGTGCAAGGTACGATTCATCCAAGACCTTGGTTGTTTTTATAAACAGTGAAGTGTCCATTCTTGTTCAAAGAGAAATTTTAAGAATGAAGAAGGAAGGTGTTCACGTTATTATTGTTGTAGTTGGTGAAGTTGACAATCTTGAAAACATTGAATATTTTTTTAAAAGGAATTCTACTTTTTTTCATCTTCATGATAATATTATAGACTCGAAACTTGTGGGTGTTGCTGCAGACAATTTAGTTTATGGAAAGTGTGACCACAATGCATGCAAGAACAGAAAAACTTGTGTAATTAAAGCGGATAGAGAAGCTTTTTGTTCATGTCCTGTTTGTGATCAGCATTATGAACCGGTTTGTGCTAATAATGGTAATATGTATGCAAGTGAATGTGAGTTTAAAAAATCTTCATGTGAACAAGACATTAGTTTGGTTGCTGTTAGCAATGATGCATGCTCATTAACTTCCAGTGATATCTTCCTTATTGTTGACTCTGGTTTTTCATTCAATGACTTAGAGCAAATAAAGTTGTTCTTGAAATCAATATACAAAAGTTTTTTAAGAAATTCTAGATTAGCGATTGTTGTGTGTGCAGACAAAAAAATTAAAACAATAAAAACTTTAGATTCAAAATTTTCTGACAACACGATGGATTATGTAATAGATCATATTCAAAGTTATGGCGGTGAAAGTAATTTATCGGATGCTTTGAATATAGTACGAGGAATGATAGACTCAAATAGTTTTAAAACACCTTTTGTTATTGTGCTTACAAATGGTGGTTTTTCACAACGTTATTTGTTAAGTTCTGCTGTGAAGTTAATAAAGGAGTTTGCTAAAATAAAGTTTATTGTTTTTAATGAAAATCAGTACATTGTTAATGAGGTTAAAAATCTGTTCCATGGTGATGTAAAATCTGTTTTTGGAACTGCAAACTTACCTGATATCTTTGATGATGTCCATCAATATGTGACTCAATCTGCTGAGATTATATTTACCTTTGTTTCAAATGATAATTCAAAAGAAAAGTTTGATTTGCAAAAACACGTTGCATCCAGCATTTTAAATAATATGAATCCTTCTACCATAACTTTTTCTATTGTTACTCACAGTACTAAGCCATATTATATGCTTAATAAGAATTTGACTGAAGAGATTTCTAACATCGTGTTTAACAGTGATGTTGACAATGACATCTATTTGTACAGAAGAACAAAAGCGTTATTGAAGGCAAATAAAACCACTGTGGTGTTATTTTATGGGAAAGAATCTATGGAAGATTTATCAAAAGATGCAATCGATGTACTGAGGGAGATGAAAAGTACACGTTTAGTTTTTGTCATTGTTGGTGATATTTCGCAAAAACTTCAGAATGAATTAGTTTTGATAGCTGGAAAAGAAGAAAATATTATTAATATTGCTAGTCTAAGTGAGTTCAATGAAGACATG
>evgR015617c0g1_i02_Hydra
TGTAAGTTTAAACGCAATTTTATGCATACTTTTTTACTTTAAAAGCAACTTTTGGAAACAATTTTAAAAGTGTCAAATACTTTCTTAAATGGCAGACGAGGAAACCGGGTTAAATATTCGTGAAAAAAAAGAAAAGGGTTCAGATGACATAGAGCAACATCTTGCATGTATAATATGTCTCTCGATATTTTTATTTGGTTTCTTTGTTTACTTCATTGTCATAGGAGCAATTGAGGGCACTAAATGTAGTTCGACCGGTACCACAGGTTACGTAATCGGTTTGGGAATAATAGGAGTTATCACATGTCTATTTGGATGCGTATCTGTTTCTAAGGCTCTGCGCAAAGGAGTATATTGTCTGCTTGGTATCATGATGGTTGCGTTTTCTATTTATGTGCTTGTTTTAGGAAATAAAGAAATGTTTGGAGATGGAAAACACTTTCAATGTGGCAATGTTTCAATAAAAATTGGATATTGGGTTACTCTTATATTTGATATAATTTTTGGTATCTTTTTTTTCAGTATTTATTGGCTTTGTAAAGATCATAATAAATAAAATAAATGAAGGATAAATAAAAAAAAAAA
>evgR015618c0g1_i01_Hydra
AAATGTCAAACATTCTGGCCAACCAGTAACGTAAATTAATTTATCCAATCGGCTACATTAAACATTACAGTAATCTAGTAGAAAAAACCAAAACTAAATGAATAGTTTCTTTATAAGTCAATTAAAATAATGGATACTTTTGAAAAAGTTAACGCGCGTGACTTGATGGCTACTTTAGAAGCTAATCACATTGACACTCAAAATTTATCAAATAAATACTTTGAAGGACATTTATCAAATATGTATGAGCTTGAAGAACTCGTTGAGAAATTAACCTTTATTGAGAAAGAGTTAAAAATAGTTCGTCAAGAATCTTTATTAAATAGAATAAGTTTAATATCACAGAATAAAGTGGTGGCAAAACAAATCGATTGCATTAATGATCGACAAAGTAATATCGAAGAAACCGTTAGAAATGACAGAGAAAGTATCAACGGTAGAGTTAGCCAACTAGAACAGCTATCAGAAAAAAACTATAGAACGTTACACAACAAACTAACTGATCTGGCTAGAGAATCATTGTCACTAGAAACGGTTCCAATAATAAGGCCTAACAAAGGAGAGGAAAAACTATTAACTGATATCCTGGGAACTAAGTTTCATGAATTAAAAAACACTGCAGATTGTGTCAACATAAAAAATATAGACTTTCTTGCAAACCAAGCCATAACTACGGCTATAGAGACATGTAATCTGTTGAGAGCTGGTAAAACAAAAATTGATTCAATAACAGACATTAGTTATGAAGATTTAAAAAGATTAAAATTTGGATTTTTATCCATACTTCAAAATAGTAATCCTATGACGGGTATGTTAGGTCAGTTCATTAGAACAGGAAATGATATCTATGATATTGGTAAATTCATATATAATGCCGCTGATAAAGCACAAGATGAAATAGTTAAAGATAAACATTACAAAGTGCAAGCAACCGATTTAATGACCAGCCTCATTCAAATGACTGTTGATGAGACGAAAATATGTTTTGATAAAGCAACACAGGTGTATCTTGCTCGCTATTACGCTACCTACAATTTCGATGACGCTTGTTATAATAATTACAGGAGAACATATTCATCAATTAAAAATTATGAAGAAAAGTTTATTTTCCAAGGCTTGCTTTCTGCAAAGATAGACTTGCATAGGCAAATATTAGTGGATCTTAAAATAGATGATAATATTCCGTGTTCTTACTACTCTTACGCAGATTATCTCCAATATATAAAAAAAAAAAATTATTTATTTTACGTCAAAGAAGAAAAGGTGTTCGAATTTGCGATTAAACACAAATTATATCGAGCATATCTCAAGTTAGGAGAATTTGATGTTATAGATAAATGCGGCGATCGGATGGAATTAGAAAAAACACTTGCTTGTACCGCAGTAGCTCCGATATTTGGACTAGGGGCAGTTCTAGGGGCTGGTTTTGGAATATATAGGTTAAAAAACGAACAGAGATGGAGCCAAAACATTCTTAGATTAAAAGAAGTTATGAAGTCGCATTTGAAAATTGATACAAAATTTGATTTTATCATTGAAAAGTTATACACCCTTGAAAAACTTAGATACTATATTGATCTTACAAAATCTCAGTATGGTAACGATCTTTTAATAAAAAACCTTGAAGAACTAGGTAACGAATTTAAAAATGAGCAGAGTAAAATAATATTTGAAAGAGCCTTTAATTTATATCAAAAAATGACACACACTGGAATTGCCGACCGAGAAAACTACATACATTTGAACATCGATTATGCGGAAAAGTCACTCAAAGATGTCGATGATCAACCGAACAACTTTGCATAATAAAAAATATGTTGTAGAAAATTTTTATTAAAGGATATTTTGCGGTAGATTTCATTTTAATAATAAGATGCAAGTTTGTCACATTGTTAATTTTGATGAGATAAAAGGGGCTTTTCAGGATATAAAAATCGAAACTTTACATAATATTTAGAGGGCGTATATCAGAAATATTTT
>evgR015621c0g1_i01_Hydra
CTTCTATTGTCTAAATAAGAATTAACATATTTGTATTTGATTAAATGTATTTGATACTCTGGTAAAATGGCGGTGGTATGAGCTGGTAAACTGTTATGAGCTGGTGAACTGAAAGGAAAAAAACTCTTTAGAAAGGAAAAAAACTCCTTCGAAACATCTTTTTATAGTGACAATTAAAAATTCTTATATCACTACCACCCGATACCATCGCCACCTATGAAGAATGTAAAAAATGGAAATTGCTACATTCAGTTTGCTTTTCATTCATTCTGTTATTTTATTTTGACGATCTTCAAGTATGTTCTGGTCAAGCTGTGAACCAGTGCACATCAGCCACATTATGTCGTGATTGTTTGACAGTTGGTAGCTTTTGTTCTTGGTGTAATGACGATAATTTTTTGCAAAATAACCCTACATACTATCGGTGTAGTACTCTATTAAATCTTACACAAGCTGGATGCAACAATATAACTGTCGTTAACTCAGATTTTCAGATTATAAAAAATGATGATTTCCAAGATCTTGTCCAAGTTAAACCTCAAGAAATTGCTGTTCAATTAAGACCAGGTCAACCAGTCAAAGTTAAATTAAGCATTAAGCCTGCACCTGACTATCCTGTTGACTTATACTTATTAATGGATGTTTCTAACTCAATGGCCGATGATTTATTGACTTTACAAAATCTTGGAGTTGTCTTAGCTTCGTCTATTAAAAATATTACAAAGAACTACAGATTAGGATTTGGGACGTTTGTTGACAAAAATATTGTGCCATTCATCCAGCCTGAATTTCCAAACTTTCCATGTAATGTTATTAAACTAAAGAACTGTCGTGAAGTGTTTTCCTATGATCATTTTTTTGACTTCAGTCAAGATGGTGCTTTGTTTGAGAATTCTGTTGCAGCTCAGAATATTTCTGGCACCTTAAATATACCGCAAGGTATTTTTGATGCACTGATGCAAACAGTTGTTTGTGATAAGGAACTCAATTGGAGAGATAAAAATACTTGTAGAAAAATTATTGTCATAGTAACAGATGGTGCTCCTAAAATTGGAGGGGATGGCAAGATTGGTGGTATATGGAAGCCAAATGATGCAAAATGTCACATGACAAAGGCACAAAGCAGCACTCAGGCAATCTTGCTAGAATATTCTGCAGCTAATATTTATGATTACCCATCTGTATTTCAATTGAACCAAAAGTTAAAAGAATATCAGGTTGTACCAATTATTGCTGTAACTAAAGATGTTATCCCTCAATATCAAGATATTGCTAATGTTTGGAGAGATCTTGGCACTGTTATTGCTGAATTGAAATCGGATTCTTCAAACATTGTAGATCTAATAAGGAACAATTACAATCTTATTTCGTCAACAGTACGTCTTTCAGACAATTCACCAGAAAATTTTGTGATAAGTTATGAAGTTAATAGTACCCAGTGCCCATCAGGATCTGGTAACTTGTGTCCAAATGTTTCTCTTGAACAACAAGTAGACTTTATTGTTACTATCACTGCTAAGACCTGTCCACTTGATTTGGCCAATGCTCCTAAAACGTTAGCAATAAATATTGCTGGCTTTGGTGTAGTTAATGTGCAAGTTCAATATTTGTGTGATTGCCCTTGTGCTGTTCTTGCCGTTCAAAACAGTCCTAAATGCTCTAGTGGTCATGGGACCTTTGAATGTGGCATTTGCAAATGTAATTCTGGCTATTATGGCAGCGTATGTCAGTGTGATTCTTCCTCAAGTCAATTTAATGGCACATCTAATTGTCAAGCTGACCCGAGCAGCAGCAGTTCTAGTTTGTGCAGTGGTTATGGTTCATGTATTTGTGGACAGTGTGTCTGTGCATCACAAGTGCTTCCACGTAAAGTGACTGGTAAATTTTGCGAATGTAAAAATTTTGGTTGTGACTTGTTTGAGGGAAAAGAATGTGGAGGAGATGCACAAGGAACATGTGTTTGCGGAAAATGTGCTTGCAACCAAGGTTATATCGGGAGCAATTGTGGGTCGATTGATTGTAATTACAACTTGCAAAAGTGCACTGTTAATGGAGTAGTGTGTAGCGGAAAAGGTATTTGTGACTGTCAGCAACGTGGATGCAGCTGTAATTCAGGCTACAGTGGAAAATACTGTGAAAACTGTGTAACTTGTCCAGAGATATGCAATAATAGAACTCGCCAAAATTGTATTTTATGTAATAATTTCAAAGATGAGATAAATTTCAATAAAATTGATGTTGATTGTACAAACTGCAATACCTTGCAAATGATTAATGTGAGCAGTGCACAACAGTGTTCTATTGTATTAAACTCAACATGTAGAGTTTTTTACAGTGTCACATATCAAAATCAGTTATCCAGTGAAATCATTCAAGTACAAACTAACCTAACATGTGCTGCTGCACTAGTAGCAGAGAATGACATTCTTCCTATTGTATTGGGAATTGTTGGTGGAATTTTGTTGCTAGGGATTCTTCTTTTGTTACTGTGGAAGCTTCTTGTGACACTTCATGATAAAAAAGAATTTGAAAAGTATACAACTGGTCAAATAAAAAGTAATTGGGAAGAGGCTGAAAATCCAATTTACCAACAAGCGAAACAAGAATTTCAAAATCCAGTTTTTGCTGGCAACTAATATAAATTAGTGTTTATTTATTAATTTTTAATCTAATTTTGTTATTATAATGCCCTTGTTTTCATTTTTGTCTTTTTCATTAAAAAAAAAACTAATTTATAATATTTTATCTGTATATTTTGCTTTCTGTTAATTGTTTTTTAACTTTTTTGAGTATAATTTAATTTTAATTTTGACTTTTTCGTTGGATATTGATTTTAATATATTTTAGCGTTTTTTAGTAATTCTTTCAAAGATTTTTATGGTTTTAAGAAATT
>evgR015626c0g1_i01_Hydra
TGATAAAAATTTGTACCCAAATTTTTATCGCACTGTACCCCCTGATATCTTTCAAGTGAAACTCATATCAAAACTTTTAAAATATTTTAATTGGACCTACGTTTCTATTGTTGCTTCAGATGACTCCTACGGTAGATCTGGTGCGTTTCAATTATCTCACAAGTTGGTTAAAGATAGTGTTTGTGTACATTCTAACAGTGTAATAACTTTTCAAAACAGCCATGAACTTTTTATAAACTTTGATTCAGAACCTAGAATTAACGTCATAATCTACTGGGGAAACTTAATTTCATTAAAAATCCTTCTCTCTGAAGCAAAAAAACGCAATTTATTTGGAAAAGTATGGATATTGAGTGAAGGTGTTGGTTTAGAAGACTGGCTTATCAAGTTTATACAAGATTTTCAAGGCAAAATTATCATAATTAATCCACATACTTACATTGAAGAAAACTTTAAAGAGCATTTTCTAAATTTATCATACGAAGAAAGCAGCCCATGGCTCAAAAAATTACTTAAACAAATAGAGCCATTAAAAACAAAACCTGTAGTAAGAAACATCGGAAACTTATTTGATTATAGACGCGTAGGTTACCTTCAAACATCAGTTAGAGCACTGGTGTATGCTTTTATAAATTATATAGGAAGTATGTGCAAAATAAAGACATGCACCAAGTCACTGGTTTCCATAAATCATAGAAAAACGTTTAACAAATATTTGAAAAAATCAAACTTTTCATATGAATCTAATGAAACATTTATATTCGACAAAAATCAAGATCCAGAAACAGCAGGTTACGATCTTTACGTGGTGACTGACTCAGGTTTCGTACATTTAGCTTCTTGGATATCAACAAAAGGTTTTGCAATAAAAAATGACTCAACTTTAAAAACGTTTCGAATTTCTTCAACTTGCTCCGAACCATGCCCGCCAGGTCAGCATAGTGTTTTAGAAAATAATGGTTACTGGACTTGTATTCCATGTCCTAGAAATAATGTTAAGAAATCTAGTGGCAATGATAAATGCATTCCGTGTAACATCACAGAAAACTTTATCAGTAACCCAAACAGAACCGAGTGTGTCAAGTTAAAATATATTTATTGGAATTTATCTAGTTTCACAAATCGTATGCTTGCTGCATTATTGTTTGCTATCACGGGTGCAACCATAACTCTAGCTTTTATGCTGACTTTTATTCTTAAAAGAAGTACGCCAATTGTAAGATCATCCAACTTTTATTTGTCTATGATTCAAATGATATTTCATTTTTCTTTATTTGGCCTGATATTTTTAGCTATAGGAAAAGAATCGTATTGGAAATGTATTTCTACAACATTCTTAAGCGGAATATTATATTTTGGAATTATTACATTGACTTGGCTGAAAGTGAGGAGGTTGGTAGCAGTATTTGGTTCAATTCATAAAATGAGAAAAAAGGAGATACAACGAATAAAGTTTGCAG
>evgR015627c0g1_i01_Hydra
CTTTAATTAACCTACCTTAGTGCAAAAAGATCATTAATCACAAATTTTGATCCATGGAAAATACTAATTTGTCAGATCTTGAAAATGCAGAAGAAAGAAATTCAACAAGAACTAGTTATTCTTCAGCTGCTCCACAAACTGGCCCTAAGGGTGTTATAGCAGATTATAGAAGACATTCTGCAGTTGTTGAACATGAAAAGGAAATCAGTAAGTTAAAATATATTGAAGATATGAAAAAAAGAGCGTTCATTGCGGATCCTAACAAATTCACAAATTTCGATGAGGAAGATATCAAACTTGAAAATGGCAGTTTTGATGATGATGATGATGAAGATTTTTTAAAAAGTTACCACAAGAAACGCATACTTTTATTAGAAAAAGAATCAAATGCTAATGGTACCAAATATTCAAAGTTATTTCATATGAGTGGAGAAGATTTACTTGAAAAAATTGAAAATCCGTGCAAGGATGAAATACTTGTTGTTCATCTTTATGATCCTAAAGTTAGTGCATGTGTAACAATGAATAAGTGTTTTGAGTACTTAGCAGTACAATATCCTTATATTCAATTCACTAAAGTGTTAGCAAAGGAAGCTGGACTGAGTTTAAATTTTCAGATGAATGCTCTTCCTACTATACAAGTTTACAAAGACGGAAATTTGATTGGTAATTTTTTCAGTATGCAAGATAAACTCGACGCAGACTTTCACCCATCAGATGTGGAGAATTTTTTGCTTGATTCTAATATGTTGTCTGAGTTTCAAGCATAGTTAGCAAAGTATTTTGATATACCAAAATAAGTTTTACTTTTTTATAAAATTTTAACTGAATATGTATCTATAATGAATATTTATCTTAATAATGTATTTTCAAATCCAGTTTATGTAAAAAAAAAAA
>evgR015630c0g1_i02_Hydra
TTTTTTTTTTTTTTTTTTTTGAGAAAATTTCAATTGAAGTTGTTCATGACACAACGCTTTACAAACTAAAGACTTTATATAATAAACTATTCAAGTAATAATGGGAGTCCAGAATCTTGCGGAATTAGAGACGTTTTGTGAACAACTTTATCAATCTTCTGATCCATCTGTACGTTTAAAAGCAGAAGAAGCTCTCTTACAATTTTCATCATCGTCAGACTGTTTGCAAAAATGTCAGTATTTACTTGAAAGAGGAACAACTTCATACACACAATTGTTGGCAGCTTCTACAATTACAAAAATGATAATTGGAGGCTCATCCATCTCAATTGCACAACGTGTTGATATTCGAAATTATGTTTTAAACTACTTTGCATCAAGATCAAAACTTGTTCCGTTTGTAGTTCAAGCTCTTTCTCAGCTTTTGGTGAAACTAACAAAGTTTGGTTGGTTTGAGTCTCAAGAGGATAATTGGGTTTTTCGATCAATAACAATTGACATTAAACAGTTTTTACAAGGTTCAATAGATCACTGCATTATTGGAGTTAAGTTACTAACAGAGCTTGTTGCAAGTATGAACCAGAGTGAAGAAAATCTTTCACTAACCAAAAATCGTAAGATATCTGCTTCATTCCGCGACACAGCTTTGTTTGAGATATTTACATCATCTCTTGATATGATCAAACAGGTAATAAACAAAAAGGTTGATTTTCAAGACCATCAACAGACGTCCCTGTTAGGATTTCTTATGCAACTAGTTCGGGTTTGTTTGATGTATGACTTTATTGGCACTTCTTTTGATGAATCAACTGATGATTTTGGAACTGTTCATGTCCCTTCAACTTGGAAGCAAACATTCTTGGACTTCTCTGTTCTGCATTTATTTATGGATCTGTATTTGCTTGTCCCGTTAACTTTATCGTCTTTAGCACTTTCATGTTATGTCCAGTTAGCATCTATACGACGCTCATTATTTGATATAAATGAGCGTACAAGCTACTTACAAGAACTAATGAATGGAGTAAAAAAAATACTAGAATTACCTCATGCTTTGGAAAATACAGATAACTATCATGAATTTTGCCGTCTTCTTGCACGAATTAAAGCCAACTACCAGCTAGGAGAAATAATGAAGACTGATTGCTACAAAGAAGTTATGGAGATGATCACAAAATTTACTGTTACAAGTTTAAGAGTACTTCATTTTGCTCCAAATAGTTTATATTATTTACTGAATATGTGGCAAAAGATGGTTTCTTCATGTCCATTTATTCAGGCTACCGACACTCACCTTTTAGAGCAATTTACTCCTGAGATCACAGAAGCTTATGTGACATCTCGTTTGGAATCAGTAAAGGAAACTCATCAAGGTGGTTTAGAAGATCCTCTTGATGATAAATCTACTCTTCATCAACAATTAGATCAGATTTCAACTATAGCACGTTGTGATTATCAAAAATCATGTGGTCTTTTGATGCAAGCATTTGATCAAACTGCTCAAGTTTACCAAGATCTTCTCTCTGGCCAATCACTATCTTCTTCATTTCTTATACAAGAAGGAAGAATGACTTGGCTTGTATACATCATTGGCTATTGTATAGGAAGTCGTGTTTCATTTTCTTCGACTGCTGAACATGATAGGCTAGATGGTGAACTTTCATTAAGAGTTCTTCAGCTTATGAACTTAACAGATATGAAATTACCTCAAGTTGGTTCTGAAAAGCTTGAACTAGCCTTACTGACTTTTTTTGATCAATTTAAAAAAATTTATATAGGTGATCAAATTAATAAATCTTCCAAGGTGTACCAAGTAATATCCGCACATCTTGGAATCATAGACGAATCAGCAATGTTGGAACTTTATACAAGAAAAATAATAACAAATATGAAATATTGGTCACATAGTGAAAAAATTATAAGTGCTACATTAAATTTACTAAATGATCTTTCTATAGGTTATAGTAGTGTTAGGAAGTTGGTCAAGTTAGACGCAATTCAATTTGTATTAAATAATCATACTTCAGAACACTTTCCTTTCTTGGGTATTCAATCAAATGCCGTGACTAAGTGCCGCTCATTATTTTATACTGCTCTTGGGCGCTTTCTACTTGCTGATATTGGAGACGAAGATGATAGATTTAAAATATTTACTTTACCATTGTCTTTGACACTTGAAAGTATATCTACACGTATTATTCCAAATAGCTCATCCACTTTTGATGCATGTGAAACTAAGTCCCTAATTGTGGGGTTATGCAGAGATTTAAGAGGATTAGCATTTGCATTTAATACAAAGAATAGCTTTCAAATGTTGTTTGGCTGGTTATATCCAGAGTGTACAAAATTAATCCAGCGTATTTTAGATATGTCCTATCATGATCCGGATATTGCAAATTGTGTTCTAAAACTCATGTGTGAGTTGACACAAAATCGTTCTCAGAGGTTACAATTTGATATAATGGCTGCTGATGGTATTTTATTGTTTCGTGAAGTCAGTAAAACTTTAGTCACATATGGTAGTAGAATCCAAACTGTTTCAAGTTATCCTTCAGAGCAATTATACAACACAAAACTAAAAGGTATTTCCATATGTTTTAATATGCTTAAAGCGGCATTATCAGGTGGCTATGTAAACTTTGGTGTAATGAGATTGTATGGAGATACTTCATTGGATAATGCAATAGGGGTTTTTGTCAAGTTGCTTGAATCAATTGAACAAAGAAATCTACTAGAGTATCCAAAATTAAGCAAAAGTTACTATACTTTAGTAGAAACAGTTATGGAACATCATATTGATCATATCTGCAAGCTTGATCCTCAGGTTTTGATGTACATTGTATCGACTGTGTCAGAAGGGCTTACTGCACTTGATTTAGGTATTTCGACCGGCTGTTGCTCAGCTTTAGACCATATTATAACATACTTTTTTACAAACTGTCAAAAAAAAAATGAAACAAAAGATCAAAATGCAGCTTTGAAATTACTTGATCTAAGATCCGAAATATTCCAACAGATGCTTGCCACCATTATGAATGTAGTAATGTTTGAAGATTGTAGAAATATGTGGTCCATGTCAAGACCTCTTTTGGGACTTATATTACTAAATGAAAAGTTTTTCACAGTTTTGGAACAGCGTATTGTTGAAGCACAATCTGGTCTTGATAAACAACAAGATATGGCTGCAAGATTTCGAGAGCTTATGGAAGATGTAGAGAGATCACTTTCTGCAAAAAACAGAGATAGATTTACTCAAAACCTTTCGACGTTTAGAAGTAATGTAAAAACCTTACTTGGAGTATTAGCAACGAACTCTATTCCAATAATAGAAATGATGGGCTTTACAATGTGATTAAAAAGAACTATCCATTTATATTAAATAAATTGCATAATTTATTTGAATTGTTTTTGCAAACCAAAAATGTATAGTTATAATTTTAATAGTGTTATATAAATTAAATAAAAAAAAAAA
>evgR015652c0g1_i01_Hydra
ATTTTTTGTTATATAGAACATTATTTTGTAACTTTACCTTTAAAAGTTATGAAACTTGATCCGACAAATAAAGTTTCAAGCGATAGTTCTACCAAAAGCAATCATAATAAATGGCGGCAATCGAAAGAAATATTCTCACAAGCGACTTTGATATAAAATATAGTCTTCTTTCTGATGAATTAGAGATTGAACAGAAGCAGTGCGTCTTTAATTTTCGCACAACTTTTTTATACAAGTACATGAGCGGAGTGTTTCAGTTCGTAGAAGAGAATCTGAACTACTATAAAAATGCTAATCATTATTACTGGTGCAAGGTTTTTCAATCAATCTCAATTATTATTATGTGGCTTTGCGTGGCTTTTAAATTTTTTCAATTGTTTAACAAAGAATTTTCTGTTTTAAAGTACGATAATATGTTTCAAGTTCTTTTAAAGATGGCCAATTATTTAGTGTGGATGATTCGGTATCTAATTTTACACCATTTCGGTCTTTTAGCATTTCACCGCATACCGTTGCTAGCCTTAGAATTGATTGAAAACTTGAAAAACTTAGCAAAAGACGAAGAGGATTCGAACAGGATCAATACAAAGCTAAAGCGCAAAGAAAAGAAAATTAATAAAGTCATTGGTTTTACATTTGTGTGCATTTTTTTGCTTCGTTCTTTAAACCGTCTTGTTCCTATTGTTGTTGACGAGATACAAAAGTCAGGATCATCTAAATTCAAACTCTGGGAAGATTCTTTAGACTTTCTAGTTATGATATATACACGAGTTGCATCGCTTCCTTTTCTCTACTACCTGATTTTCATTACCTATCTGCAATGCTTCGAGCTTAAAGAATTTCAAAAGATTCTCCCAACTACACTTAACTCTGCTGATGACGTTTTCAATGATTACATTAAAATCCACTCGAAAATAAACAGCACTGCAAAAAAAGTGCATCTTTACTTATCAATGTCAATAGGCACTATTGTTATTTGGGGTCTGCTTGCCGTTTACACTTTTATGAAAAATGTATCAATGGTTTCCCGTGGGACTTCTAACGAAGTTTCTGTCATCTTTATGATCCTATTTGATTTTCTTTCACACGCAGTCGAGGTCATATACTTGTTTTTACTACAATGGTCACAGTTGGTCAGTGTTTCCAAAAATCAAACTAAAATCATCCAGTCTGTATTAGCAATCCGATGTTCACAAACCAATGAAGGTTCACCGTTTAATAGTGTTGCAGAAGTCCTTAAATTTTCACAAAGACTTGAAATTTTTCAAAAATATGCTGGAACTGGTTACAAAGTTTTCGGGGTAGATATGAAAAATGTAAATACAGTTTGGTTTTTTATCCTTGCACCTATTGGCTTACTTTTAGCTAATTCAGCAATTAAAGGTGACATCAATTCAAAGTGTTGTTAAAAAAAAATTATTAGTAAATTGTTTACTTTTTGAATAAAATACTATTTCTGTAAAATTTATTAATTTGGTTTACAAATTGTTCCGTAAAAAAAAAAA
>evgR015668c0g1_i01_Hydra
ATTAAAAAAAAAGAGAGACATGGGTAGATATTTTCCGAGTCAACCAGCTCCAAAAGTTGAAAACAAAGCCATTAGTCCGCAAGAACAAAAAAGTAGCACTATCAAACTGTACCTTGCATACGCTGGCGAAAATATTTCTGTACACGGACTTTCCCACGTGTTTGACAAACGCGAAAATTTCATTTGTCGGACCGTATGGCTTCTAATTACCATTGCAGCTTTTGGATACGCTGTACAAAAAGTATATGAGAGTACAATAAACTATTTTTCTTATCCCTTTAGTACAGCTCGAATGAAAATCCATGTTAACCAAATGGATTTTCCTGCAGTTTCATTCTGTAATCTAAACGACTTTCGTCAAAGCGCCACTAAAGGGACAAAGCTTGATAAAGCAATTCTATCTAATGACAAAAAAAGTATTGAAAACGATAAAAGTATTTCAGGAGAAGAGTACGTAAACATTACTTTGAAAGCAAGACATAAAATAGAAGAAATGCTAGTTAATTGTAAATTCAATGACGATGAATGCTCTGCTCAAAACTTCACAGAATTCAATTGGAATCAAGGTGAGCTTTGTTTTACATTCAACTCTGGAAAATATCCTCATACATTGTTAAAGGTCAATGGCGTTGGCATGAAAAGAAGTCTGGTACTCACTATAAACATTCAGCATTACGAATACTACGACGACGAATTGGATGCAGGTATACATTTAATACTTCACGACCAAGAGGAAACCCCAATTAAAAAAAGCGGCCCCATATTACCACCTGGATTTACCACATACATACAAGTGGAAAAAAAAACGATTTTGAACTTAGAAGCACCTTACAAAACAAAATGTGGTTCGGTTAAGTTAAAATATTTTGATAGTTACTCAATGCACACCTGCTGGCTTGAACAACTTACAG
>evgR015676c0g1_i01_Hydra
TTTTTTTTTAATTGTTACTTTTTTTTTTTTTGAAGGGCAAATAATTTACAATAGGAGCGGACGTTCGCAGACTATAAAAAACAACAAACAAGACACAGATCTATTCGAGACCAGAAGTCTTATTGCAGCAAATAAATTCTTAAAAGAATTTATATATTCGTTTAATAATTCGATGCTGTATAAATTATTTTATTCTAAAATTGATAGGAAAAATAGACTAAAATTTTATGAAAACTACAGTTCATCTTTAACAGTATCATCTGTTGAATCTTCTACTGTTGGCTTTTCTTCTTCTTTAGCATCTTCTTTTGAATCATCGTCATCTTCTTCCTCCTCTTCTTTTTCATCCTTTTTCTTTTCTTCTTCCTCTTTTTTAGTTCTAGCTTCTTCTTCTTCTCGTTTTTTCCTTTCTTCTTCTTCTATTTTATCTTTCATTTGCTTTTCTTCAACTTTTGTTTTTTCAAATGTTTCTTTTGCAAATTTCTCAGCTTCCGCAACATCATCAGTAACAATAATATTATCAAAAATTGTGCCAGACTTTACCTAAACATAAACAGTATTAATATAACATTACAGACATCTAAATCAAGGAACACTCTATTTTACTCATATCTCAATTAGCTAATTGAAGCCACAAGCTCAAAAATTTTACCTGCCAAATTTCAAATCCAATATAA
>evgR015676c0g2_i01_Hydra
GCCAATAGGGCGAAACCACGCACTAAGATTTATCTGTATTTGAACAAAGCGTATTTCTGTAACTGAACTTCACAACAATTTACTTGTAGAATAAGTTGCGTGAACGTGGACACATTTTGTAGCATTTTTGTGCTCCAGAAATGATGTCTGTTACCGTTTTTGTTTTAGCATTTGTTGCTTTCGCTCAATGTACTGTATATTTCAAGGATACTTTTGATGATGAGACATGGGAGCAAAGGTGGATTCATTCTAAAAGCAAAGGTGGTGATGCTGGTAAGTTTAAGTTAACATCAGGAAAGTTCTACGGAGATAAAGCCCGTGATCAAGGAGTTCAAACTTCACAAGATGCTAAATTTTATCAGCTTTCAGCTAAAGTTCCAAATCCTTTTTCAAATGAAGGTAAAACTTTAGTTTTACAATACCAAGTTAAACATGAACAAAACATTGACTGTGGAGGTGGATACATTAAGTTGTTTCCTTCGAATATTACTCCTGAAACCATGAATGGAGATTCTCCATATCACATTATGTTTGGACCTGATATTTGTGGACCTGGAACTAAGAAAGTCCATGTTATCTTCACATATAAAGGAAAGAATTTACTTACAAAGAAGGATATTAGATGCAAAGACGACGAGATGACACATTTATATACCCTGATTGTAAAGCCCGACAACACCTATGAAGTCAGAATTGATGGCTCTAAAGTTGAATCAGGAACTCTTGAAGGTGATTGGGACTTCTTGCCACCTAAAAGAATCAAGGATCCTAGTTTATCAAAGCCTGCTGATTGGGTTGATGAAGCCAAGATTGATGATCCTGAGGACAAAAAGCCAGAAGGATATGATAAACCTGAGTTGATTCCTGATAAAGATGCAACCAAACCAGAAGATTGGGATGCTGAAGAAGATGGTGAATGGGAACCTCCAATGATCAACAACCCAGAATATAAGGGTGAGTGGAAACCACGCAAAATTGACAATCCTGCATACAAAGGAGAATGGGTTCACCCAGAAATTGATAATCCAGATTATAAAAATGATGATAATTTATATAAATATGATGACATTGGTTATATTGGATTTGAAATTTGGCAGGTAAAGTCTGGCACAATTTTTGATAATATTATTGTTACTGATGATGTTGCGGAAGCTGAGAAATTTGCAAAAGAAACATTTGAAAAAACAAAAGTTGAAGAAAAGCAAATGAAAGATAAAATAGAAGAAGAAGAAAGGAAAAAACGAGAAGAAGAAGAAGCTAGAACTAAAAAAGAGGAAGAAGAAAAGAAAAAGGATGAAAAAGAAGAGGAGGAAGAAGATGACGATGATTCAAAAGAAGATGCTAAAGAAGAAGAAAAGCCAACAGTAGAAGATTCAACAGATGATACTGTTAAAGATGAACTGTAGTTTTCATAAAATTTTAGTCTATTTTTCCTATCAATTTTAGAATAAAATAATTTATACAGCATCGAATTATTAAACGAATATATAAATTCTTTTAAGAATTTATTTGCTGCAATAAGACTTCTGGTCTCGAATAGATCTGTGTCTTGTTTGTTGTTTTTTATAGTCTGCGAACGTCCGCTCCTATTGTAAATTATTTGCCCTTCAAAAAAAAAAAAAGTAACAATTAAAAAAAAA
>evgR015686c0g1_i01_Hydra
TACAACTGTTAATAAAGTAGGATCATCGGTTGCTGGTACGGTTAGCCAAGTAACCGTACCAGCAACATTCAGCAAAATGATCGATCTTGAGCACGAGGACGAAAAAATCACAGACCCTATTTCTTCAAGAGAGCATCCGGTGACTGTAAAGTACTCAAAAAATGAAGACACCCCATTTACCACAACTCGTTCTATGACTGTATCAAGTAATGAAGATACTCCGATCACTAGTACTCGTTCAATGATGATATCAACTAAATTTATTACTACTGCAGGTCATGCATCAACAAGTCGCAAAACCACCACCATTCATAAGACGACAAAATCGAGTTTCTATTCCACCGAAACTCATCGAACTACGTTGACAACCAGAAAGTCGACATCAAAACAAACAACTAAAGGGGCGACTACAAAAGCTCCGGTCGTAACTACAACACAATCTTTTACAAAAGCAGCT
>evgR015695c0g1_i01_Hydra
TAGTTTAAATTACTTTTTATAAGAAATAAAAAAGAAACTTGTTATCTTACAACCGTCAACCGTTAGCTGTCGGTTATAAACCAATAAATTTTATAAAAACCAAAATAATGTACGAGATTGTTGTATCCGGTTGCTTTTTCTTTCTGCTTCTTTTTATTCTACAAATGAGAAAAGCGGACATTAATGGCAATTCACGATTAGTTGCAACCATTCACGCTTTTGTTACTTCAAGAGCGATTGAGTATGTAATGATTTCGAATTATGGTTTTCAACATCCAAGTATCATACTTCAAAATATTGGTGAACCATTGACACCGATGCAAAAGATTCCATTAATAATTAGCGGAGGTTACTTCGTTTATGATCTTTTTTTGTGTTATTACATCAAAGAATCATTGTTAATTTTTTTGCACCACGTTATGGCAGCAGTTACTATACTTGGATCATTATTTTTGCAATCTGGTGGAAATGAGGTTTTAGCTAGTCTTTGGCTTGCGGAAGGAACCGGAATCTGCTTATATGGGATGTATCTTCTAGAATCTACTACTTTTAAGAAAACAGCTTTGCTAAATAAAACTCTTTTCGTAACTTTGTTGATTCTTTTGAGAGGAATTTGTTTCCCATACCTACTGTACCACGCACTTTTTTTTTCCAAAACATTTTTTTTTTTTAAATTTTGTGCTTTTTCTTTTTTATTTATAAATTTTTGTTTCTTTTATGAGATATACCAAAAAATAAAAATAATTATACAATCGTATTATAAACATGGTAAACTTATTTAAAGTTTATTTATAACTATTATCTCTGACCAATCAAAAATCTTTTCTTTTTTTATTCGAATTTTATTAAATCCTCTTATATCTTTTTATTCCGACATTTTATCATTCGCATGTTAATTCCTTCATTCATTCATTTTATCATTCGCATGTTATATCCTAATAAAAATTATTTTTACAAAAAAAAAA
>evgR015697c0g1_i01_Hydra
TTTGGTTTCGTTTTCTCACTAAAATTGAAAGAATTTAATTAACAATGTTGACGGGTAAAGGTTAAAGTAAATATTCAGTTTTTATTAAAGTTAAGTTAAGCTCAATTAAGCAATGAAGTCCCTTGCTGTTTTCAGTAGTTTTCCGGATTTGTTATACTTATGGACTGATAAAAGTTTTAGAGAACATCTCCACAATACGTTTTTATTATGTAAATTTGATGTAGTATTTGATATAAATAGATTTGCAGAATACCTTTACAGTCCTTTAGTTACGTCTAATAACTACATGAAACAAATTAATAATGGATATCGTAGAATTACATGTGTAAATGGAGTTTTGCTTGTGATGAAACAAGTTGCTGGTTTATTTATTATTGCTATATGTGGAGATGGGGAAGAAAGTGAGTTGTTATTACAGAGGAAGATTTCTACATTTACATATCTTTTAAATGCATTCTATGGCCCTTGTATATCTGAAAAACTAAAACCCTCAAATGCTGAAATGCTAGAAAAAACATGGAGATTTTGGGATGAAATTCTTAAAACATGGAAGATTTTATGTCAACAAGAGCATTCATTTCTTGTTGAATCAATCGAAATACTTAATCTCAATCAGTATATTAGTAGATCATTTAATGACTTGCTTTCTGATGTTTGTTTCAGAGCTAATCGTGATGGGAATGTTACACATTTATTTCTTTTTGTAAATAATAAACTTATTTCTACTTATGTTGGAAAAAATGAAAAAGAACTACAATCCAGTGATATTTTAACTTTGATTTTGTATGCTAAATACCGTTTTCATTATATTGACGATACTTTGTCATATAGTATCTATCGTACACCCACTCAACTTATTGAGAATCATGTTTCGGAAGTTAAAGATAAAGAGCTTTTTCGGTCATTAAACAATTTAAATGATACGGATTTGAAAACAAATTTTGATAAAAATGACTCTTGCTTTTTTCCAGTCAGTAACGATGGAAAACAATCAGAAGATTCAAGTTTGGCACATCAGCTTTTGTTTTTACAAACAGATTGCAAATTTTCTCCATACTGTGCACACTACATTCTGCTTGATAAAAACATTGTTTTGGTTATCCTATCAAAGAGTTTATTTTTCGAATTAGCTCCCTTAGTGCAATGTTTATTAGAGAAGTTATCTATTATTTTGGAAAACAAGGATAATGAGGTATCCTTATTATGGCGCCATGCTTTTGAAGAATCCCAAAAATATGCTGTCAAGATAGTTAACCACTTGAGTAGATATGTTCAGCACCAGCCCCAATCATTAAAGAATTCCTTTAAAAAGTTTGGTTCAAAATGGAAACAACTTGAGCAAAGTTTTACAGAGGATGATGCGAAAAATCTAACAACCAAAATGGAAACTGCCACTAATGAGTTTGCTAAATCTACTAAAGTTCTGTTTTTGTACCTTTACATTCAAAACAACACTCAAGGAAAGCTATCAAGAGCGGAGAATCAAGATAAAGTTTTTGAAACTATCAAGATTATGGCAACAGAAAATTTGAAGCATTTTTATACTTACTTAAATGGTAAAAGCTCAAGGAATGTTATAATGGAAACTTATCAAAATGAATTTCCCGGACTTGTTCACTTTATCTACACCAACAGAGTAGCAAATTGTGTTATTGCACCGACAATCAATATAAAAACTGATCCAATATTAAGTCAGCTACATCGATATATTATTATTAAACAACGAATATGGGATTTTTGGCAATATGCTGAAACAATGTTGACAAAAGGGTATTTATCTATAATGATGAACGATGGAGATTTTTGTTATTCATACTTTATCTGGTTTGAAGACTCTAATGGCAATTGTTTGTATTCACAGAATTCGTTAAGAGATATGAATATAGACCAACCCACTGGAGTGTTTTCAGGTTCATTTTATAGGAATATCATTCAACATTTTTTCCCATCAAAATCTTACGTGAACGTACATTGTTACGAACTTGTATGCATGTATATTGGTGGCGTCAGCAATAATTTTATATCTAACTCGTGTAAAGAACTTGTAAAAACGTTAAGAAAGGAATTTGAAAATCCAAACCTTTAGTTAAATCTTTTTAATATTTTATTGACAACAAATATATTTAATTAATTTAAAAAAAAA
>evgR015700c0g1_i01_Hydra
GATGTTGGATATTGGTTTAGCGAGCTTGGTCCTTGGGCTAAAGGCACTTTTGATAAAAAAGTTATAGAGGCTGGTATTGATGGAAGTCTTCTGATGAAACTTACTGACACAGATTTACAAGATTTTAAAATGAGTATTCCTCTCCATCGAAGAACAACCCTGGAAGCGATTAAGCAATTAAACATAGAACTTCAAAGTCAACAATCAGGATTATGGGAATTTAAGGAAAGAAGGAGTGGTTTGTCGATATTCATCATGTTTGGCTTACGAGAGTTTCCACGATCTACACTAGCTTATCTGTATCTCTTTAATTATAGAGATGCTTTCTTACCAATCTTTTACTGCTCTGTCACGTTAAAATCTACTAATATCAGCTGTACTCAGATGCAACACTCTCCCACATTGGGTAATTGGCTTGAGTTTTTACCGCTTATGTTTTCCATTCCATATTATTTGGTTGGTCGATTTGCTTCACATTATTTATACATAAATTATTGGACATCACGTGTTGTGATTTTACATGCTATTCTTATGACAGTAACAGAGATATCCAATATGAGACTTTTACTGATTCATCGCAGATCTGCTCATTACTATGCTACTAAGTATTTCATTTATGTGGTCATGTCTTCACTTTCTATCAAGCTATTGGTGTATTTTTTACCAGTCTTTGTTACTGATTGCATATTTTACTGGCTGTTGTATGTTAGTCCATTTGACGCATTTAATAGGTTATTGAAACGATTTTTTGTAAGAAATCCTCCACATAATTTAGGACAAGAAGAACACAATCATGCTGGTGGAGAGCGCCAATGGGGATTTAACTTTCAGTGGCATGCTGGACATTACTAATATAAAATATATTATTATTTATAAAAAAAT
>evgR015703c0g1_i01_Hydra
TGGATTCAAAAAATGCTCTATGAAACTCTATAAGAATGTTATACATCCTATAACGTCTCCTGGTTGAGATTAGAGAGATGAATGTACTCGATGTAATTCCTCCTCCAGAACTTCATTTAATGATGAAGATAATAGCAGAGATTTCAAATGTCTTGTGTAAGGAACCTGATGTTGCTCTCTGGTTAAAAAGATGTGGAGTTATTTGGCATGGCTATAATGGAGGTGGACTAGATGGCAGAAATGCTAATAAAATTCAAAAGCTTTTGCCAGATTTGGAAAATTTTATCTTAGATAATTTTTCTTCTTACCATACAGTTGTTGAATTGTTAAAGTCATTTTCTATAGTAGTAAACATGTGTTTTGGAATGAGACTTTATGACGGTTATGCTGATGC
>evgR015714c0g1_i01_Hydra
GTTTTGGCAAGGTATTTGTTATGCACAAAAGGAAGCATAGCCTGGATGTTCTTGTTTGACAGAGTTTGATTTGTTGCAATTTTTGGTTATTGATATAAAGAAGATAAGATTTATAATCAAGCCATGGTGTTACCATAATGTTAAATTGTTGACATATAATGCTACTTAAGTTTAGGATTAGAAAACTGAATAAAAATGGATTTACTAAATGTAACATATCATATTAAAAATACAATGACTCTAACATTAACAACGAAACAACTGAATAACTCACCATGGTTAATTTATGTGCCATTGCAACAAGAAAATGATGGTAATGAATAAAAAATATTCTCGGACTAAACAAAAGGAAATGGTAAATAATACCAGATATAAAAAAATTAAGAGGGGAAAGAAAACACTTTTTCAAATTAAAAATGAAGACCACACAAATTTAGTTATTCACAGTGTTTTTTTTACGAGGCAGTTACCGCAAATAACTGATTGTCCAAGCCTTTATGGAATGTTAGTGCAATTCACCTGTAACTATTTGCCTACCTTAAAAATAACAGTACCAGGATTAAAATCTATTCAAAACACAATGGAAATGTGGGGAAAAGTACAAAATACTTCTGTAAGCGATATTCAAATTTCAAAAGCACTTCATGCTTTGCAACCAGGACAAACATTTGCAATATTTGTACGACCGCATAACTCAGTAATAACAATTAAAAGATCGGAAAAAGATCCTAATGTCGCTCTTATGAGTGTGTTCAGAGTTTCAGCAAATAATAAAATGGTCATGTCTGAAGGAGATTTGTTTCATACATTTCCTGAACGAGCTGTTTGGATTCCATTCAGTCGGTTTTTATACAAAGCGTTTGCAATACAAGTTGCTATATTGTGCAATCAAACCTTTGAAGGAACATTCCAAAATAATAAAAAAGCCGAAGTTGCGCATCCAGATAGTAGTGATGTTGCTGAACCCATTTTTATAATTTCATGGCTGCTGGGAGCAATGTCAGGAACGATGCAAGGAGCTTATAACTTTGCATGTGTGCAAAAGGTAATACGTGATGATGTATTAAGTATAAGTCAATCTGAAAACCCCTGGCGTCGCTCTGGAGAATGGTTTGCGATAAAATGTGTCCTACATATCATGCTAGTAACAGAATTAGGATCAGTTGAGGGAACTGTAATCTATAAATCTCTTATGATTCATGTTATGATTAGTTTCATTGATGACCTACCTTCTTCGTTTAATAATAATGATATTATCATGCAGATGTTGAGCAAAGTAAGTCGACAAATGGCGAAGCTGGAAAATTATATAAATATCAAATATGCCCCCGCCACTGTTGGACCGAACCGGACACGCGCCACTGTTGT
>evgR015716c0g1_i01_Hydra
TTTTTTCAAAAACTAGCATTTAAAAAAGCTGATTATTGTTTCTTAAGATTAAAATGAGGCATTTTTTTATATTTGCAAGTTTTTTATTTTGCAACTCTGCACCTGTAAAAGAAGATGACTGTGAAATAAGACTAATCAGTGGGTTGTGTACCAAATACAAACTCGTAGAAGAAGAATTTGGGGTTACCTCCAACAATAATCTACCTACAACCTCCAACAATGATCCACCTACCATTGATCAAATTACAAACGTGTTCAAGGCAAAACTGGATTTTATCAGAGTTTTATCAGAAATAAAAAACCCATGGTTAAATGATTTGGTAGAAATATATAACAAAAGTTTTGAAAACATTTCCCAATCATTTGCAAATTTTTTAAACTATGTCCCTTCTCCACAGTCTTCACTTGCCTGAGATGCTCTCAAAATTTGCGGATACCGTTGTTTTAAATATATTTTTGAAATTAAATGTTTATCTATTATAAATATTGGATATTTTTTCTTTATTTTTTTACATATTTTGGATTTAAAAGTTTATTAATTACTTTATTTACTTTGAAAAAAAAAAAA
>evgR015725c0g1_i01_Hydra
AGCAGAAAATGGAAATATTTTTGAGTTGAAAAATAATGTTTCTATTACTTATCCTGTAGTTATGGTAACTGATACCAATGAATTTTATACAACTTTGAAGGATGCTATGATTAATCCATACCCGATTATGTTCAACCAACCTTTTTTTAATAGTTTTTTAATCTTGATTTCAAGGCAAAACTTCAAATTTGAATATTGGCCCACTTCTTACTATACAATCTTATTTTGGTTTACGAAAAATAATTGGCTAGATACTTATAGTTTATTTAGTATGACTGATGGGTTTAATTCATTTATTATTTATTTGAGAAATACTGGTACGATAGACATAATTTTTTACCATTTGAGTGCACACATTTATACGTTTCAAAGAAATCTAAAAAAT
>evgR015726c0g1_i01_Hydra
CTATCATTTCTGAAAATGAAGCAACTAGAGTTATTGACACTTTTTGATATTAGTAGAAACGAACTTTGTCGTCATCTTTATTACGACGCGTTTGTCGGATGGAATGGTTTTATGCTAAAGATAGTGAACGGAACAGTTGGATCACTTATTTTTGTCTCAAACATTCTATTAGCACACGGCATTTACAAATCAAAAAAAAGAAGCAAATTTTCAAGAAATGAAAAAATGGTATTGCTGTTATCTTTTATTGATGTAACGGTTTCAATAATTGATACCCCCCTCGAACTTGTTTTGTTTACATATTTAGACTACATTGGATGTCTCACTATTTCAGCTATTGCATTTTGGATTGCGTTTCCGTTAATGCTCTCAGGAGCAATGGTAGTGATCATATCAACCGAGCGTTTTATTATAGTATTCAACGACAAAAAATGTTGTGGTTTCAATTTTAAGGGAAGTTACATGGTTCCCATAATGATTTTTTATTTTTTAACTTCGGTTGGATTTGGTGTTTGGTTTTTATATCTTCACTATAATTATCCGTATACCATTTTTTCCTTTCATTTCTTTTTTTACAGTACAGCGACATTTACGACAACTAATCTGGTTTTTGGAATGTCCATAAATGCGTTACTTCTTGCAAAGACATCGAAAAGGTTAAACAACAGTGTAATACGAGTTGAAAGACACAGTGCTATTGAAAAAAGGTTAAACCATACAATGATGATTATTTCAATTTCTTTAATAATTTCATACCTCCCCACAATTTCTGCGCACTATTATATGGCTGTTATAGCTGCAAAATTGGATAATAATGCTACTGTTTTAGCAATCAAGTTGTTACTATGGAGTTTAATGTTATGCAAATTAAATTCCGCTTTCAATGCTATCATATATGTGACTAGATGTAGGGGTGTTTCTAAAATATACATTACATACTTTCAATCTTTAAAATATAAACTTTGGCATAAAAATAACTTCAAACTTAACAAATTTGTCCCTAGTAAATTCAGTCCTAAAGAAATAATCTTAATAGAAG
>evgR015730c0g1_i01_Hydra
ATACAAAGCCACATGTAATAAATTATTATTCCGAAGAAAATAGATAATAACATTTTTAACTGCAAAGCCAAAGTCTTCAAAAGTTTTGGATTGTTTATAGAAGAGAAGAAAGTTCTTCTAATTTACAATTTTATTAAATGAGTTCCAACTTCAAAACTGAAAACGTATGTAAGTATTGCCTCGATGATTCTGATAACAACTTGATTCGACCATGCAACTGCACTGGTTCAATAGGTTACGCGCACGAGGATTGCGTGTTAAAATGGCTGCTTACAAAAAAAAACTGGAAATGTGAATTGTGTCAAAAGACACTAAATCTGGTACCTAACGGATTCAAGTATCCAAATAAGTGGAATTTTCCGCCAATAACCTTTGATTGGATACTTTTTGTTTACTTTTTATCAACGATACTACTATTAGCTGTGGTTATAAACGTTGGAGTGATGCTCTATAACAAATGCACAAGTACATATTGTATGAATTATTATGGCTTCATGCCTGTCGTTATAATATATTTATCCCTTTCTTTTTGTCATTTGAAGATACCGTTTTTAAAATTTAAACAGTGGGTTGAAACAAACCGCAGGTGGAAGTTAGTTGAACAGAAAAGTTGATTTTATTATTTAATTTAAAAATTTATAATAAATTTATTACAAG
>evgR015740c0g1_i01_Hydra
TAATGGAAGAGACGTATGATAGTATACAATTTCAAGGTGGAGTAGTTGTAGGTGCTTTAAATGAGCGTATCATTAAGCTTGACCAAGAGCGAGATGATAGGGCCATGGCAACAATAATGCAAGAAGATGGATATCATAGAGTTGAGATAGATGATAATATACAACAAATATCAGATAGAAGTTCAGCATTATTACAGCCTAATGATGAATCAAGTCAAGAGTTAGATAGTCAATTTATGTTGATAGATGAACAAGTAAGAATTGGTATATCTGGTAATAATAGGCAAGAACACTCTGCAGTTGCTTTGATAGCAGGAGGAGTAGAAGAAAGTATTGTTCCGGTAACCACTTTAGCAATTACAGACGGTAATAGCCATTATAGAAGCGATAATACAGGTATAAGCCCTAAATCTGGATTTATTGGATATTTAGCAAATTTAGCAGAAG
>evgR015746c0g1_i01_Hydra
AAATTTTGTTATTGTATTTTTATTGCTGTTGATATTGGTTTGCAACAACTGTTTGACTTTATAAAATGGTTTGGACGCCGATTATGGATTTAGAAACGCCTCGAACATCTTCACCCAGCTCTTATCACATAGTGAAAAATCATCCTTCTCATGTTCTGCAATACCACCCAACAACATCAGCTTATCGCTTATACAATGGTATGCATCGTTCAGATGTTCGCTATTCAGAAAACGCCCTAGCACATTCTGAGAACTTAATGACAAAATCTTATATTGAATCGACTGAATCATCGGAAACAAACACTATACCAACCCACTATGATAGTTCTCCTTCGTATTTTCCTAGTTCACCGATTACGCCAGTAACACAAACGAGTAACATTTCTGGATACTACAATTCAACTTTATCCGTAAATGACGGATGTAACCAATGGACTTATGGGCCACCCGCTCCTTTATTACAAAACTACTTTTATGGTTATGGAGGATTGAACTCTAGCTATCAATATCCCGACGTTGATCAACCTTCAGTAAATGGTGTTCCATGGATGTGTCGCGATTTTGATTCGAAACGAAAACGAATGACTTACTCTACAC
>evgR015749c0g1_i01_Hydra
GGACTTTTAAAATTGCAAGTTCACAGTGATTTGAAGAAATGGATAAGGCTGTTATTGTGTCTTTCTTTGCTACCGCCTCCCCATATTCAGCCAACATTAGGCAATTTGAATCCAAATTTGTTTGTGGTGTCATTAAGTATAAATGACTTCGTTAAATGTCAAAGTTTAATGACCTATATTGAAACATTTTGGATTGGGCGTATTGGCAGTGACAAAATTTCTGTTTTTGGCTGTCCCAGAAGAACGAACTCTGATCAAGAAAGCATCCGTGCCTCGCTTTTAAAACGAATAAAGATTGCTCATCCA
>evgR015753c0g1_i01_Hydra
AGTTAAAACAGAGATGCGAAGCTTGGATTCCATCCCTTGGTCCTGATGAACAAAAATTCATGAACGAGGAAGGCTTGTTCATCTATTATAGATATTTATCTGGTTATGGATGGGAAAAACTGAAGGAAGCTGAAGATGGATTGAAAAAAACCTGTGAATGGAGAGCTAAAGTTAAGCCTCAAAATTTGACCATTGACGATTTGGGTGAAGTCGGAAAAACCGGATTCATGTTCCATTATGGGTATGACAAATTGAATAGACCAGTTGTATATATCGATATGAGAAAGGACACTACTCCAATGACAAAAGAAAATCTCGAACTCAAATATTTAACATTTGTTTACTATACTGAAAACCTTGTAAGGAGAATGGCAAAGAATGTTTATCAAATCTCTTGGATTTTGGATGTGACTGGTGCAAATGTATCACTTAGTCTCGTGAAACAAATGAAGGATACCTTCTTGGATCTTGGAGTTTACTACTGCGAGAGATTATCTTTGGTATTTGTGGTGAACGTCACAATGTCAATTAATTTAATTTGGAAATTCATTTCTTCCTTCTTGGCAAAACAAACTGTTGAAAGATATCACGTATTTACCAAGGATTGTAAAAAGCTGCGAGACGCTATACATCACTTCTTTGATGAAAATTTATTGCTGAAGGATTTCCATGGTACTGCAGAGTTCAATTACAGCAATGATGTTATGAAATCATTCCATACCTTGACTCCTCTTCAACAAGTTGATGTTGTTCGTGACAGTTCAAACTAAATGGTTTGCATCGAACTGCTTTGTAAATAAACATGAG
>evgR015759c1g1_i01_Hydra
ATTTGCCTGTAAAGCCGAGTTTATTTTTAAATAGATTCGGCAAAATAGTACCCATAATGGTAGACCAAAATTTGCGCAGTTCTTTTGATAAACAATCCGCAGTGGAGCTAGGGTGAGATCTCCTAGCCATCGCAATGGTTGTCCCAGCAGTTAAACACAAAATAGTAAAAAAAAGAACTAAAAAGTTTGTTCGTCATCAATCAGAGCGCTATGCAAAACTAAAACCAAACTGGAGAAAGCCAAAAGGTATTGATAACAGAGTCAGGAGGCGTTACAAAGGACAGTATTTAATGCCAAGTATTGGTTACGGAAGTAATGCTAAGACACGTCATCTAATGCCTGATGGATTTAAAAAATTCGTTATTCATAATATCAAGGAGCTTGAACTTCTGATGATGCTTAAAGGTGTATATGCTGCCGAAATAGCTAGTAACGTTTCTAGTCGCAACAGAAAAGAAATAGTAGAACGAGCTAATCAACTTTCAATTAAAGTCACAAATGGTGCTGCAAAATTGAGAAGTGAAGAAAATGAATAAATGCTATTTAAAAAAAAAAAAA
>evgR015762c0g1_i01_Hydra
TGTAAATTTAAATTTGATTTATCAAGCGCTGTAGAGCATCTACAAAGAAAGTTTTATTCGTATTGCGATGGAGTGCATCATAGCAGGATGGAGTGACAGGAATTGCAAAATGTCTTAATTCAAATATATTAAATAAACATTACGTTATGTTTAACCATTATCATATGACTAAAGCTAAAAACTATTAAAATCGATAAAGTAAATCACCTTTAAAGTTCTAGAAATGATTTTATACGGTCTTTTTTTTCTATTAATCTCAGTTTTTGTTGGAGCAAGTCCATTCCATTCTAATAAAGAGAGAGGATGCTATATCCCAGAATACGACGCCAATATTGCTGAAGTCATTAAAACTCCTCGACCCCACGAATACCTTCATATACCAAGTTTGCCTAAAAACGTGGATTGGCGAAACTTCAATGGAACCAATTACGCAAGTACTACTCGCAACCAACACATTCCTCAATATTGTGGATCATGCTGGGCACATGCAACAACAAGCGCACTTGCAGATAGAATAAATATCCTTCGTGGTGGTGCATTTCCGTCTGCTTTTTTATCTGTTCAGCATGTATTAAACTGCGGTAACGCAGGTACTTGTCACGGTGGTGGTAATTTAGCAGTCTATGAATATGCACACAAAAATGGAATTCCAGATGAAACCTGTAATAACTACCAAGCAATTGATCAAGAATGCAATCCATTTAATCAATGCGGAACATGTTTGACTTTTGGTAAATGCTTTGCCGTGCAGAACTACACAAAATTCATGGTTTCAGAATATGGACAAGTCTCTGGGCGGGAAAATATGATGGCTGAAATATTTAAACGAGGACCGATTGCTTGCGTGATAATGGCTACTCCTCTTTTTGACAAATATTCCGGTGGAATATATAGCGAATATAACGAAGTTTCTATAGCTAATCACGCTATATCTGTACATGGTTATGGCATTGATGAAAACGGTGTTGAATTCTGGATTGGAAGAAACTCATGGGGAGAGCCATGGGGAGAGAAAGGATGGTTTCGTATGGTTACATCGTTGTACAAAGATGGAAAGGGTGGCTTTTATAATTTAGGTATAGAAGATGATTGCGCATTTGGAGTTCCTATTTTACCTGCCGGGTGGTAGTGAATTGATAATAATTACTTTGTACATATTAATGTTTATTCGGTACACATGAATATTTATTCGGTACCCATCAATTTTTATTAGGTACACATGCTTGTATTTTTTAGAAATTTTGTTTTATAATTTATTTCTTATGATTTTATTTTTATTAAATTTTTAATGAATAATCATAATTCATTTACATTAACCAATATCTTTTAATAATAATTTTTTATGCATTACTTTTCCTTTCAAAATTAGTACCAAATTCATAAATAACTATGGAGGATATACTATGCGAATTCATTACTTAATCAAACATCAAACGATTCATTAAACCAAGAAAAAATTTCTAAGTAACTAGTTACTAGTAACTAGTTACTAGTAACTAG
>evgR015763c0g2_i01_Hydra
TGATGGAAATTATGGAAACGAAATATAACTCTCCTTCGAACAATGACAAGTCAACATTTATTGAACAAACAAAACCAAATAGTGATGAAGAGGAAACTGGTAACCAACAGGAACTACAAAATGATAATGGACAAGAACCTACAGTGAGAAGGGTTAAAATACCTGTCAGAAAAAGAGCAGTAGTTCACGACGTAAGGAGACCTCTTCCGAACCCAAATCGTTATAGTGGCCAACAAAATCACAAAAGTACTGTTGGTAATCGTTTCCCAAGGAAAAGTAGTAGCGGATATGAGAGTGACGAAGGATGTATTGTCGAGGATGAAG
>evgR015765c0g1_i01_Hydra
ACAGCAAAAATTAAAAACATTGTGAAAAGTGTTGCAAACGAGACAATGTCAGATGCTGCAACTGAAATAAAAGAAAATGCAACTGCTGAAGATATTGTTGATGTTGGTGTATCATGCGACGGATTGTGGCAGAAAAGAGGATATCAGTCAATGAATGGAGTTTTTACCGCAATTTTAATAGATAGTGGAAAAATAATCGATGTTGAACCTATGAATAAGTCTTGTAAAGCTTGCTGTATTAAGGAAAAATTAAGAAAATCAAATCCTGATGCATATGCTAATTGGAAAAACAAACATGTTTGCTTGTATAATTATCAAGGTACAGCTGGTGGTATGGAATCAGTCGGCGCAGTGAGAGTTTTTAAACGTTCCATTGAAAA
>evgR015777c0g1_i01_Hydra
TGTGCCCAAGTGGAGTCTGCAGTTTCAGCAGCCAATTACAAATTGGGCCTATTCAAAAAAGCTTTCCGAAGCAGAAGCATTTGTCTCTGGCGAACGCTTTACATCGCATATGTGCGCCCACACCTGAAATTTGCCATTCAAGCATGGTTTCCTCATCTAAAAAAAGATGTCAATATGCTTGAAAGAGTTCAGCGCAGAGTAATAAAAGTTTCCTCAATCAAATATCTCCCATATAAGCAACGCTTAAATATATTTAGATTGACAACACTTGAAAGGAGAAGAGCCAGAGAAGATCTTATTGAACAGTTTAAGATTATAAATAAAGTTGACGATGTTAATTTTTTTGTTCCTCAAAGGTTATCTCACAACACATATAGCAGGAGAAGTCATGATAAATAGCTTCACAGACAATATATTAGCAGCTGCGAAAAGCGACAACTTTTTTATTAATTGTGTCAAACCGTCTTGGAATACGCTACCACAAAAAGCAATTAGCACTCACATGGTCAACTTATTTAAAAGTTACTTATCATGATGGACAACTTGTGTCGTCGACAACTTGCCCGAAACATTAGTTAGTTTAGAGAAGCCTGGTGTTG
>evgR015780c0g2_i01_Hydra
TTCCAAATAAGTTGATGTACTGAAGCCTATCTCAATCAATGCTCAAAAGTGATGATTGCTTTTGTATTAATATAAATTTAAAAGTTGTTTCCGCAGTGACAAGGTTTGACTCTCTCCTGTATAATAGTTCGAGACCAAGTTTTAGTGGTGCCAGGCAAATAATTCAGTCATTGATTTGGGACCATTCTTTATGGGTGAAACGAATTTCTGACCCAACATCAGTTATGTCTTTTTCAATACACAATTTATCGAACATATTGTTCCACCTAGTTTTACAATCAAGAATCAACTTAAGCTTTTTGCCGTGTTCATGTAGGATATACTTCTGTAAATATGTATCATTTTTGACAGGAGAGTTTTTAAATATTTTCACAACCTTTCTCACTTTTTTCATTAGGTCTTTATAATTGCTGACCAATTCAGCAGATGGTCGGTCCAAAATTATTTTCAAACCATATTCTTTATAACTTCATCATCATCAGTGTTCTCATGTTCATCTTCTCCTTTGCAATTTTAATTTGTAGTAAACGATT
>evgR015788c0g1_i01_Hydra
TGGATCTGGTTATGGAGGAAATTCGCTTCTCGGGAAAAAATGTTTTGCGTTGCGAATTGCATCTTGTATTGCTAGAGATGAAGGATGGTTAGCAGAACACATGTTGATTTTAGGTATAACAAATCCAATTGGTGAAAAACGATATATAGCTGCAGCATTTCCTAGTGCTTGTGGAAAAACTAATCTTGCAATGATGACTCCAACACTTCCTGGTTGGAAAGTTGAATGTATTGGAGATGATATTGCTTGGATGAGATTTGATGAAAACGGATTACTTAGAGCCATTAATCCTGAGAATGGTTTCTTTGGTGTTGCAGTTGGAACTTCAATGGCAACAAATCCACAAGCCATGCACTTGTGTAAAAAAGATACTTTATTTACAAATGTTGCAACGACAAGTGATGGAGATGTATATTGGGAAGGGCTTGAAAAAGAAAAAGATCTTAATAATATTGAAGTGAAAAGTTGGCTTGGTAAGCCTTGGACATCCAGTAGTAATACACCTGCTGCTCACCCTAATTCTCGTTTTTGTTCGCCGGCAAAGAACTGTAAAAGTATGGATCCCCTTTGGGAAGATCCAAATGGAGTACCAATATCTGCCATTTTATTTGGAGGACGTCGTCCAGAAGGAGTTCCTTTAGTATACGAATCATTTGATTGGAAGCATGGTGTATTTGTTGCTAGCTCTATGTGTTCAGAAACCACTGCTGCAGCTGAGCATGCTAGTAAAATTATTATGCGTGATCCTTTTGCTATGAGACCTTTTTTTGGATATAATGCGGGTCATTATCTCCAACATTGGTTAAATATGGATGTCGAAGGACGTAAACTTCCGAAGATATTTCATGTCAATTGGTTCAGAAAAGATTCGAGGGGCAAGTTTTTATGGCCAGGTTTTGGTGAAAATTCGCGTGTTTTAAAATGGGTGTTTGATAGAGTCAATCATAAAAATGTAGCTGAAGTATCACCTGTAGGTTTAATACCAAAAGAAGGTAGTCTTGACGTAAATGGACTCGATCGTATAAATTTAGAAGAGCTTATGTCTGTACCAAAAGATTATTGGGTCAATCAATTAAACGATCTTGAGAACTATTATCGGGAGCAGTTTGGGAAAGATCTTCCTATTGCTATCTGGAATGAGTTGCATAACTTTAGAGAAAGGCTAAAAAATGCAAAGTAGTTGTTTTTAAAGTTTTTTACGTTAGGTTTTTTATAATTTTTAAAGATAGGTCTCCTTTTTTAATTTTAGACTTTTTAATTTTTCTGTAACTTTACTTGTTTGGTTATAAATCTTACGTTGTTTTTTTGTATAATAACACGAATGTAACACGTGAAAATTAAAAAAAAAAAA
>evgR015792c0g1_i01_Hydra
GCACTCTTTAATTAACATACTTTAGTGCAAAAAGTTAAAAAATAAAATGACTGATGAAACACATAAAGGAAGTTTAATGCAGCCTTTTCTGCTACTGGCTAAATCATCTCATGGAGCTGCTCTATCACGACTTATTCAGCAAGTGACTGAATTGCCAGGTATTTACAAGTTTGGTGAGTTGTTACAAATGCCTAATGTGTTGGAGTTATCACATAATGAACTATTTACAAAAGATTGGAAGCTTCTCAATTTGTTTGCATTCGGAACATTCATGAGCTATTCTACCAATATGGAATGTTTTCCAGAACTGAATTCAAATCAAATTACAAAGCTAAAATTGCTTTCTCTTGCCACATTAGCCTCGAAATCAAAGTATGTAAGTTATGATGAAATTAAAACCGAAATTAATCTGACCAATACAAGGTATTTGGAAGATTTAATTATTGAAGCAATTTACGCAAATATAATACAAGGAAAGTTAAATCAGCAAGATCAACGTCTAGAAGTTGATTTTGTTATAGGGCGTGACATACAACCTGAAAAAGTTGACTACATTGTTAATATACTAGATAACTGGTGCAAAGGTTGTAGTAATGCAATTGTTTCGTTGGAAAAGTTGACCGAACATGCCAACAACATAAAAGATACCAAACTTCAACAAAGGAAAGTTACAGAGAAAGAGATTGACGATGTTAAGAATTACTTAAAGACTCAAGATCACTTAGGAGATGCTAGTTTACAATCTCAAAAAGATAAAGCAAGAGGCCTTCGTAATTTAGTAAAGCCAACTTCAAGAACAAAATAGTACAAGTTTTATAAATATAATAATTAAAAAATGTTGTTTCTATCTAACTTTCTTGACTCTTTTTTTATCTAACTGCCTGATGTTTATTTTATTAAACTGCCGTTTGTTTATGCTGTTTAACTGCTGGTTGCTTATTTTATTAAACTGCCGGTTGTTTATACTGTTAAACTGCCGGGTTTTGATTTTATTAAACTGCCGGTTGTTTATTTTATTGAACTGCCTGGTTTTTATTTTATTGAAATGCAGGTTGTTGTTTTTTATTGAACTGCCGTTTGTTTATTTTGTTGAACTGCTGGTTGTTTGTTCATGAAAAACGGAGAACAT
>evgR015795c0g1_i01_Hydra
TGTATGTTCTATGCTTCATGAATTTTTTGCGTTTCATTTACCTGAAAGTGTAAAGCATATAAAATTGTTTTGTGATTCCTGTGCAGGACAAAATAAAAACTGGACAGTTTTGCGTTTCTTTTACTATCAAGTTCATATAGTAGAGAGATTTGACAGTGTAACTCTTTCATATCCTATTAGAGGTCATTCCTACATGGAATGTGACAGAGATATGGCTATTATTAATCAAAAGTTTCCAGCAGAAACACCAGAAAATTGGAGGAGTGTTTTTCAGGCATCTCGTAAAATACCTGAGCCATTCACCATTATCAACATGACGCAAGAAACGTTTTTAAAGTTTACAGAGTTTTTAAAGCC
>evgR015811c0g1_i01_Hydra
TTTGGTGATGTTTTTAGGATACAAAACACGTGAAAACATTGCTTTTTATCAAACTCGCAATGTTCAAGATCTTTTTAATCTAACACTAAGAGGAGTTCCATGGCCGAGAGATTATAGTCAAGTTTTTAACCAGATTCAAGCATCACAAGATTTTTGGAATTGGATGGATGGATTTTTCTTGCCTCAAGTTTTTCCTGAACCCTGGTATAATCTGAGTGATTTATATTCAAATACAGATATCAAAAATTTTCCTGGAAAATTATTTTTAAATGATCTGAATTCAAAGATTGTGAATGGAATTAGAATTCGTCAAGTTCGTGTGCAGCCAAATTCATGTGTAAAGGCCAAATTATTTGCTCAGTTTTTCAAAATTGACTGTATGTCGCCTTATAATCCTACACTTGAAGAAACGCGGGGTTTTTTTCCAAACTGGACAGTACCAAGTGTATTTAACTCTGCCATTAATCCTCTAACAAAACCTTGGAAGTATCAAACATGGCAAGAACTTGACGGGTATCCATATGCGGCAGGCTTAGAAACTTAC
>evgR015820c0g2_i01_Hydra
TTTTTTGCAGTTATGTCTTGCACTCAAATAAATGGGATTTCTAGTTACTGCTACTATTCGACCTGATCGGACAAAAGGATGTCTATTACCTACAGAAAAGGATTTTTAAAAACGATGAAGAGGAAATCATGCCCCCATGGCATGATTTCTGCCTTTATCAGAACCTTCTGAAGTACACACTAGTAAGTTCTGAAATATCTGAAAGCGCTATTAAAACACTGAAGCGTCATGTCTGGTACCTATGTTCTGAAATGGTGCCCCTTGCTATATTCAGTGACATTGTTTCAGAGCTTGAATTGCAGTACTTAACCGAGAAGCTTCTGGCAGCAAGGCCTAGTGATGGAATGGCTCTGCTGCATGATTTCTACGGAACTGGCTTTGGAAAACCCAAGTTCCTGACTGAAATCACTGCCACAACAAGACTTGGTGTCTGCAGTTTTGGAAAACCCAAGTTCCT
>evgR015822c0g1_i01_Hydra
CTGTTTTTTTTGAGTAATGTTTACTTTGATAAAATGATTTCCCAGCAGTAACGCGACTATCGAACGAAGTTGACGTTAATCCGGTAAAATTTGTATAAATAGTCGTGCCATAGTGCAAGTCAAGACTTCACTTAGTATAGTGTTTATGTGAACTCATTTTTTGCAATGGCTGAAAATGCGATGCAGGATGCATTAAATGCTCTAGCAACATTAGAGGAATCAGTAAAGAACACATCTGGAAAAACGGATAATGTTCTTGCTGCAGTCGAGGGTCTAAAAAATGATCTAAAAAAACTGAAGACAGAAGACCATAGTTATATCGATAGTGCTAACGGAGTTGTTCAAGTTGCACAAGGTGTTGTTGATAACATAGATAAATTTACGTCTGGTGATGCATTGGAAATAGCATCTGGAGCTATAGGTTTAATTGGTACAATTGGTGCTGTTGTGGGTGGACCTTTCGGTCCTTTGATTGCTGGAGTTTGCGGATTAGTAGCGTCCATACTTCCGCTTTTTGGCGGAAACAACGGTCCTTCAATGGATCAAGTAATGGATAAAATTATTAGAGATGCTTTAGAAGACTTTAAAGATGAAGCTATTGCTGACACTGTTAGAAGTTCATTAAAAGAGATGTCTGCAAAATTAGCTTTCCTAACTGGTCTAGGTGAAAAAGATTGTGGTATTATAAATGAAAATGAGAAATCTTTCTTGGAACCAATGAGTTTGATGACAACCGGTTGCGAAATCCTTGGAATACTGCAAAATGAAATTGATAAATATAAAAATTCACATGACATTAAAAAATGTAACAGACTAGCTATGTACTGTTATTACTATGCAATGATCTCTCTTCAAAAATCTATTATTTTGACGTTGCAATGCAGTTTACTACGACGTAATGACCTGGAACATGCCAATGCAGGTGTTGAAAAATATTTGTACGACGTTTTACCTAAAGAAGATAGAACAGTTTTAGATTTCATTTCAGAATTACCTGAAGATAGTAATAGCTGGTGGCTTTTATACCGACATCTTCATACGGACCTTTCAACTCAACAACGAATGGTTCTTTCAGAGTACCGAAATCTTATTAAGTGTGAACCAATGATTGGTCAGCTATGCAATATTTTGAATACATATCAAAATGAATACTTGTACTTACCGGAAAACGGCTTTGCTTACGATAGTGACAGAAGACGAGTTTTTACGTACAGAAGTAAAAATGTGGTTGATGCACAAGGCTTGTTTCGAATTGTTGGTTCATATACAAACTGCTGTATATTTTCCGTTTATTTCGGGGAATATTTGTACGCAGCAGATTATCCACCATGCGATGGAGATCGAAGACATGTGTTTTCATGGCGATCTGGCGGTCCTGTTAACCAAGGAGATTGGCAAATTTATAATGGAGAAAGCGATTGCTGTCAAATTAGAAATCTGAAGCAAAATGAATACATTTACGCAGCTGATTATGCGCCATACGACGATTCTCGAAGAAACGTATTCACATGGACGCCTGGTGATGAAGTTGTACAAGGCGATTGGAGAATAGAACCGCTCTCATTTGAGACGAGTGATTTAAAATTGTAAAAATTAAAGACAAACTTTTTTCAATTTAAATACATTTACATAATTGCTATTATCATAATAAATTTTTAACATCTGAAAAAAAAAAA
>evgR015825c0g1_i01_Hydra
CTATTTTAGTGCAAAAAGCCTAAAATTTTTATAAACAAAATTACAAAAACATGCCAATTCAAAATAGTAAAAGATTGAACGGACCACAAGAAACTCTGCCGTTAACGTCTATTTCTTGCATTTCTTCAGCTAAATCAAGTTCCCAACGACAGAATGGGCGCAGTGAAAATGAAATTAGACCAATGTATCTTAAAGCTGGTAGTATCCATCAAGCTAATGGTTCTTCCTACGTAGAAACATGTGACACTAAGTTAATATGTGCCGTTTATGGTCCCCGCGATAATCCAAAGCGTCATCAATTTAGTTCAAAAGGAAATATATTTTGTGAAGTAACTTTTGCTCCTTTTTCGTGGCATGAGAGAGTTTCCAATCAAGATTCGCTTAATAAAGAATATTCATCTGCTATTGTCCAAGCTTTTGAATCCGCCGTTTGTCTTGAATCTTATCCGAAAGCGCAAATTGATATTTATATTAACATTTTAGAGTATAGTGGAAATTGCCTGAGTTATGCAATTATAGCGGCATCAATAGCTCTAGCTGATGCTGGTATCGAAATGCTAGATTTAGTCACGTCCTGTGAAACTGCTTTTACCAACGAAACGTCAATTCAACGCGTATGCGTTGATCCAAGTGGCGAAGAAACTTTACTTTGTAAAGGACGCGTTGTTATTGCCTATATGCCTTCGTTAAACAAGATATCTTACTTATCAATGCAAGGCGAAGAAACAATAGACGAATCAATGCAAAATGTGATTACCTGTATAGAGGGATGTTTACGTATTTATAAATATATGAAAGAATGCATTATGAATTGGACATAATAAGAAATTAAAAAATATAAGTAATAATTATAAAAAAAAAAAA
>evgR015828c0g1_i01_Hydra
ATGGTTATGGATTTCATATTCATTTAAGTAGCAATAACAGTATAGTTGCTACAGTTGTTACAAAAGATTATGTTTATGAAGCAGATTTAGATAATCCAACTATGATCCCATATACTTTGACAGTGGTATGGTCTAAACAAAATAATAGTATACAATTTTATCTTAACGGAAATTTAGTCAGTAATGAAAGTACATTGGATGCAAACATTAATTCAACTAGTCAAGGATTTTTTAATCAATACCTTCTTATATATGCTGATTACTACAATAATCAATCAGATTTGCGTGGAAGTCTTTCCAGCATCAAAATTTGGGGTACATCCTTTTCATCAGAGGAAGTATGGCAGCTTCAAAAAAAATCTATTTATGACTGGAGCTCCTTGGATGCATACCCACCGACAGGATTTAACTTTGGTTCCGATTGTTACTGTGAAAATATTGTATGCCCCTGTGAAACTTTAGAGAAAATCAGCAGTGTTGAGTATAGTAGCAGATTCTTTATTATGGAAGATGTTGCTGGAGATGATTTTTTCGATGAAAGCATTTTTGTCAAAAGCCCAGAATATTTTGCTGACTACGCTACTATATACTTTGATGATATTGTTCTGGATTTTAAAAACATAAGACTTCTATATGTCTATATTAAGGGCAATTTATTTTTACCAACAAATAATCCATGTTTCAATTTGACGATGCAATTTTCAAATGACCTTAGTTTTTCTAGCCAAACTCAATGGTGTGATTCTACAAGTTCAGACAGCATCTTTAGCACGCCAGATGTTTCATCATTGCTAAGTGACCTTCTAAAGACAGTTTCTTACACAATACCGTTTACAATTTTAGTAAGGATTAGCAATATTGTTCTGAATCCGGGTTGTGTTTTGAAAGTTGGTTATGAGTTGCCGCTGCCAGGCGGTTTTATACATGCAAGCGCTATCCAAGTGAATTCTCAAGATTTATTAGCTTCTATTTCTCTTGATGTTACTGGAACATGTTTGTCATTTCAGTTTAAGTTTGGTGGTAGTGGTTTTGTAAGTCTCATTGTGTTAGGTTACACTGGTAGTACTACAAGAGAGTTAGCCTTTTTTTACAGACGAGAAGGTCTTGATAAATGGATTAACTATTATTTAAATTTAAGTAAAAATTCAGAAAAATTTTATTCAGTCAAGTTTGTAGCTTTAAAAAACGGTCTTGGAAGTGGCTATATTGTGATTGGTGCTATTGAATTTTTAATAAGTTGTGATGAGCTGAACATAGAAGAGCTTTCAGATATTGAAAATACTTCCGTATCATCAAATGGTTCTCTTTTATTTCCTAATCCTGAAATCTTTATCAGAGGATCAAGCGAACTTTTCATAAATTCTTCTAATGCAGTTGTGTCTCAAGATTCTTACGCTAATCTTTTGCAGTTTTTAGATGGGCATTCGGAGAAAGTTTTTAAAAAAACACTTACTTTTTTGGAAATAGATTTAAGTAAAGACTTTTCATTCCTATTTTGGTATCAGAGCAATTCATACGAAGTTTTTAGTTTTAAATACGAAAAAAATGTGGTTGTCCAATCTCAAGCTTCCTCGCTTATTGTGAGAGTTGACAATGTACAAAAATACATCAGCATTGATTCTTCAGTACTATGGCATTATGTGGTTGTCCAATACGAACGAAAGTTTTTAAAACTTTCATTTGATGTTGATTTCAGTACTGGTAGCGTGTTTGACGCAGTTCGGTTTCCACCAAACTCAAG
>evgR015830c0g1_i01_Hydra
TGAGCGCTGAAGAACCAAAGATTGAAAACATCGAAGAAGACGTCCCACCAACACTTGAAGATGTCCCTGCTGAAGAAATCAAGGGTGAATCTGCTGCCGCTGCTGCTGCCAGACACAGACAATCAAAGGCCGAAAAGAAGAACAAGAAGGCTATCTTGAAGGCTGGTTTGAAGGAATATCCAGGTGTTAAGCAAGTCTCCATCAGAAGTGCTCAAAGACTTAACTTGACTATTGAAGACCCAGAAGTTTTCATTGCTGACAATGGTAACGGTACTGCTGTTTTCGTTGTTTTCGGTAAATTCAACATGGAAGACCCAACTCAACAAAGAGGACAACAAGCTGTTCAAGATTTGTTGAAGCCAACTGAAGAAGTTGCTGAAGAAACTACCGAATCTGATGAAACCCCAGAAGAACTCCCAGAAAATATTGCCGATAAGGATGTCCAAATGGTCATTGAACAAACTGGTTTGACCAGAAATCAAGCCATCCGTAAGATTAAGGAACACGGTGGTGATATCATTGAAGCCATTACATCTGCT
>evgR015834c0g1_i01_Hydra
GTGAATTTCGACTTTTGAGAAGCGCAAAGATTATTGTGAACTGGCAGAGCTTGTATATTATGTTATGTTCCTTGAAGAAGATCAGACAGTTAGGTATAAAATTCACAAAACCGGAGCTATTTCACACACCAGGTTCATGGCAAAGGCATTGTACTATATGAAGTTTAATTTGCTTCTCCCTAAGATCAGACAGGTTCTTTTTCTGAGTGATGAAATTGTCAAAGAGATCAACAGAATGGCTGACTACATTTCTCTTTTCTGGGCATCGTGGTTTTTGACCTTCGAGTTTTCTGATTCTGCAGCTTTTGAAGTAAAACTTAATTTCCTTTCCTGCCCTGCTATAAACCTTTTTACATTTCATGTGTAGTGTTAGACTGAG
>evgR015843c0g1_i01_Hydra
CTTCAATATTTACATTCCATGTTCTTACTCGATTCGTGTTCATATTCCCTATTTGTAACTCTATGTTCTTACTCACTTCATGCCACTTCATGTTCTCACTTCCTATCACCCTATGTTCTCACTCGCATGCCACTTCATGTTCTCTTTTCATGTCACATAATGTTCTCACTTCCTATCACCCTATGTTCTCACTCACATGCCACTTCATGTTCTCACCTCATACCACTTCATGTTCTCCCTAACATGTTCACATCATTTTGTGTTTTCACTCATTTTATACCATTTCATCATCTCGCTTACTTCATGCCATTCCATATTCTCTCTCACTTCACGATTTTAAAATAACAACTTTATGCTTTAAACAAACCATACTTTGTTTTACGATTTTCCAATCACCAACACGTAACGCAGCATTCATAT
>evgR015849c0g1_i01_Hydra
GGCTTCAAACACGTTAAATATTTTTAAAAAATTGAATTATTTAGAATTGGTTTTGCAATAGATTTCTTTATAGGAAATGGGATGTATTGTGAGAGAAAAACGAGAGAAAAAAGTGTTTCCCAAATTCCATATTTCATATTCAAGTAAAGCGATAGGAAACAAGAACGGCAAAAAGAATTTTAAAATTCATTATGCAGTTCACTTCTAACTTGAAATTAATTGTTTTACTATTATCTATCCATATCACATCTTGTTTGAAATGTTACAAACGCTCTTCAGACACAGAAAAAAATTGCTACGCAAAACAAAAGCTCGAAGAATGTCAAGCAGGTGAAAACTCGTGTTTGACAATGTATCACGCCAATGTTCTAAAATCTGGAAACAATACCGTGGTAAAATCGAGGTTTCTAAAAATGTGCGCAAAAGAAGAAGAAAAAATTTGTCAAAACTATTATTGCAAACCGTATTTAACAAGTCAAAAATGTAAGGCGTTTTGCTGCAATCAAAATAGATGTAATGGAGGACATGTTGTTTTTATTGCAAAATATACAAGTTCGGAAACATACCTTAATTCATTTGGATGGTTAATTCTCTTCGCAATGGTTTTGAGCCGGAATTTTGTTATCCGCGACATTTTTTAGAAAACCATTAGTTCAATAATTAAACTCGTCGTAAATTTGAGACAATCTAGTTTTTTAATAGATATGTTAACGACAATGATTTTGTTTTACTTCTGCTTTATTGTAAAATTTATATAAACTATGTTATTTTGTATTAAAAAGATATTAAAAACAAAAAATATTTATA
>evgR015861c0g1_i01_Hydra
TTTTTTATTTTTACAAAATTGTTTAATTTGGAAAAGAAATTCAACATTTATTTAAGGGATTTATTTAAAGAATTTATTTGAGTGAAAAAGAAGATGCAATGTGTTGTCGTTCATTGATTACTATATTTCTTATAAGTAAATTAGTTACATGTTTGGAAATAGGAGAGTCAGCTGGCGAGCGTGTAGAATGTGATCGTAGTAAGATTGATTACAACGTGACATTATCTGGGGGTCATCAAGCAGGTTTATTTTTAAAAGCAAACAAGTCAATTTCAACAATGAAAACATGCATTAAACAATGTTGCAACATGATTGGATGTGATGTTGCCTTTTTCAGCAACAATGCTTGCTATTCCGTCAAATGCCACAGCTTAGAAGCATGCGCCCCAAAAAGTGTTGTCGAAAAAGGTTTGAAAACAGAAATATCGTATGTTGCGAAACCAAAGCTTATTTTTCAAGGTTCATCTTCTGACGAGACAGGCGAGCCTGCCATTGTGGAGAATCTTATTGCGGAGGTAGTTTCGGAAAGTCCGTTTCAACAAAAAAAACCGACGTTTTTAATCGACGACAAAAAATTTGTCAATACTGACTTGGATGTAATACGCTATCATAGAGAAACTTGGAAGGAGACTAAAAACATGATCGGCGCTGTTGTTTGTGCTTGTGTTGCTGTATTAGTTGGCGTGTTGGGTGTTATAACAATGACAAGAAAACTTGTTGAAGACGACGAAGAAGAAAATAATGTTTTAAAAAAGAACTTTGAAATAAAAGACCCTATAAATGAAAATATATCTGGGGATATTAAAGACTTGCCTAAAAGTAGGAACACTTTTGTTAGGAGAGTAGAAAACAATTTAAATGAACATTCTTCCAAAGTAGTCAGCAAAATTCCTGTTTATAAAAACAAGCCATTTATAGAAAATATAAATAAAAAATTGGATTCGGCACTGCATGAATCTGTGCCAATGCACCCTGCACCTATACACCCTAGCTAGTTTTATAAATAGTTAAATCATTTTTAATTTTGTAAATAAAAAATTTTATTATGTGAATTTTTTTCTATTAGGAGATTGAAAGAAAATAACATTTTTAAATAAATGCTTAAAAAATATAAACAATTTGTAACAAAAACTTAAAAACGAC
>evgR015867c0g1_i01_Hydra
TATTTAAAATGTTAAAACGATTTCTTGTTTTTTTCGTTTGTTATCATCTAGCTCAAACAGAAGATCCTGGCATACCTCTAGACGAGGTAGCGTTTTGGCCAATAAATCCTTATATATTTTATGATGAAAATAAAAAAATTGATGGTATATTACCATTTATCTACGATAGACTAAGTGTTTTATGCAACGCAAACAAGACTCTGGTAAAATATATTTTGATAAACCAAACATTTAGTCAAAATAGCGTGTATGAAACATTAAGTGAAACACAATTCGGCACTAAGGAACTTGAGAATGTTAGTGGCAATAACACAATATTTTTACCTGTATTTATAAAACCAAATCCTAAATTGTTGCAAGATAAATCATTACGAAC
>evgR015870c0g1_i01_Hydra
TCAAGTACACTTGACTCAAATTGTCCTAAGAATATTAATAAAAACTTTTTGAAAAAAAACAAAAAAGAAGTAAAAAATGTTTGCAATGATACAATTGGAGAAAAGTCGAGTGTAGAGAAATCCATGAATACTAAATCAGATAAAACACAAGAAATAGAAGGACAAATTATTATATCTGAAGATAACTTAATTGAGAAAAATAAATTACAAGAAAAGAAGCTTCCAGAGTTGCTATTGGTTCTAGAGCAAATGAGATGCGATCGATCCGACTTGCTTGAGCAAAATGAATGCTTAATGATTGAAATTCAGGAAAAAAGCAAAGACTTGGTAAAAGAAAAAAAATTAAACTGCGATTTAAAAAGAAACGAAAAATTTAATCAAAATATTTTAACTCAACTAGATAAAAGTTATGAAAAAAAAGTTCAGTTAAAAGGATTTGACGGACTCGATAGACTGCAAAATTTACTCAACTATACCCAAACTGAAAACAGTTCTTTACCAGAACAAAAAATCCATCGAGATTCAAACAATGCTTCAAAGCAAAGTCAGAGGTGTAAAGATGGTGTCGACAATTCCTTAATCAAACAGAATTATTTTTCAGAAAAACATTTCAAATGTATAAACCAAGTTAACTGCTTGGTTATGAGTTTTAAAAAGTTAAAAAAGTATAATAAAAAACTCATTAAACGC
>evgR015876c0g1_i01_Hydra
AGCAATAATTGAGCCTACAACTACTGAACACATAAATACAACTATAAAACATACAATTCAAACCACAGAACTCAAAACCTCTGGAAATAATTCATCATCAATGACACATGAACCTATAAATTTAACAAATGATCCAACAACTCAAAACAACAAAACAACAAATCTGGAATCAACAACAACAAATTATAAATATATAAGCCCAACTACAGAAACAACAACTACCAACACGAAGCCTTCAACTAGTACACCTGAATTTGAACATACAAGCGCAGGTGATATTACTTCAATGATTGAACTTACAAATTCTCCCAATTATTCTACAAATAACCCTGAAAACACAAATACAATCAAAGAAAAAACTGAACTTGTAACTTCCACAACGAAACCTTTAAACACAACAAATAAATTACTTTCTACCACAGAATCTCCGACTACATCATTTGAAAATAAATCTTCAAATGAATCAGAAATAAC
>evgR015877c0g1_i01_Hydra
TATTGGTACAAAAAGTGTTCTTCATATAACTTTGGGTAAAGATTTTGAGTCATATGGTGACAGAAATCTAGGTATTTGGTTTCATGAAGATGGGTCTGGAAAGCTTTCCATTTTTACTTCAGTAAATGGTAATGTAAACTACTTCATGGAAACACCTCCTCTTATATTAGGTGAATGGTCATTCGTTAAAATATCTCAAATTTTTCAAGAAAATAAATATTGGTTATCTGTTGATTTAAATGGGTTCATTATTCATAAAGTTGAAAATTCTGATCCTAGAGATTTCATAAACATGAAAGTGTATGCTTCTGATCATTGGTATGAAGCTCAAAATGGTACCATTGCTGAACTTCTAATTGTCAATGGCTCACCTGAGTAC
>evgR015882c0g1_i01_Hydra
ATTTATTATTTTTTGCTTCTTGACTATATGCTGCATACCAGTGGAATGAATACGTCGTTGATTAAGAAGATTTCCTTGTTTTTTTTTATATCTACAGTCGGATCAGAGTTTATCATTGGCGATGACACAATTAGATTAATGAGAGATAAAAAAATTGCTGTGATACCAAAGTTAGATAAGGAGCATTTAATATCATTTGATGTTTATCCTAGCAAGTTTGAACCTGGTAGGCGTAACATTATTCATTTTACTATTGGTGATAACAATTGGACTTATGGTGATAGAGTACCAGGTGTATGGTTTTATGATTACGGAAAACTTCTTATTTCAGCACCAATTAATTGCAATAGCGATCGATATTTTATATCAGAACCGCTACAATTACAAGTGTGGATGAAGATTGAGATTTATCAAAAATTAAATGGATCCGTTTATGTGTACACATTCAAAGTAAATGGAGAAATTGTATTTTCTGAAATTAACGACGTAGCTCGAAG
>evgR015890c0g1_i01_Hydra
GTGACATTAGTTCATTAGAAAAATTAATTAAAAAAATAACTGAAAATTAATGAGGTACCATGATTTTGTGTTATTGTTGAGTAGGCAACATATTTTATTATGCAAAATTATTTTTTAGTTACGAAAAATGAACTGTTCTAATTTTTATTACGACAAAGAAGTCTACGTATATTTACAGAACACTGCTGTACTAATCATGGTCATTGTTGGTATTATGTTAAATCTCTTGACAATCGCAGTGACGTTGAGAAGCGTTTTTGCATGCTTTTGCTCTACCGCCAGATTATTTCTTTCGTGTTATGTCGGAAACATATGTGGTTTGATTTCAATGGGACTTAACGGAGTATATTATTACGATAAAGGAATTCCTAGCATGAGCTGCCATGCAATTGACTACAACTTTTTTCTTTACTTAGGATTTACAGTCAACATGACTATTCTTGTAAATTCAACGTACAATCGCTATAGAGGAGTTTCAAAGATCAACAAAAACTTCAATTGGAATGAAAGCAGCAAAAAGTTATTCTTTCTTTATGCACTACCTTCGTGGCTTCTTTCACTCCTCATTTCATTGTTGTTCACAGCTCTCAAGTTTTTTGGCGTTGTACATTTTCACGCGTATGATGTGTGTGGGTTAATAGGCGTTGTTCCAATTTTAATCTGTATAACTTTAAATGTTCTTTTAAAGTATTTTCTTTCAAAAATGATGAGAAATGCTCACGTCACTCAAAAGAGCGATTCACTTAGAAATATTTCAGTAGCTATATCAATGATTCGATTAACTACAGTTTTCCAAAGTGTTTATCTTGTTGCTGGTTTTATTGTTTTGTACTTCATAAATAAATTTGCTAACAATAAAAAAACTTATTTATTATTGGATTGGGTATCTCGTGGTCTTTACTATCTTATGTTTACAATAGAAGCAAAAGCCTTTTTTTTAAAATATACTAGAG
>evgR015906c0g2_i01_Hydra
TTGGTGATGGAAAATTGTCTTTTTGGTCAGACATTAATGGTGTTAGATGTAATTCGACTCTTTTTTCTTCCCAACCATTACACTTAAATGAATGGTCAAGCATACGAATATCCCAGTTTCAAAATAACAGAAAGACGACATTTGCAGTGTATATGAATGGTGAAAATGTTTATAGTACTGAAAACAAGATGCCTAGCTCTTTTGAAAATGTTAATGTTTATGTTGCAAATCCGTGGAATCCTGCTCCAAATGGCACCATTAAAGATGTTAGAGTTTTAAATGGAAATGAAGAAATGTGTGGTGTGGTCGTAACAGCAAAATACGATATGACAGAATATGAAGAAATTTACATTCACCTAAATGTAACTACTGACTGTTTATGTGATCTCCTGAATGTTTATATATATCTTCAACAAGAATCGCTATTGACATTTAAAAAATTTATGTGGGGTGATTCGTGTTTAAATTGTTCAGTTGTGAAAAACAATCGTGGATTCTTTTTATTTGCTGTTAGCAATCTTGAAATTTACTCGTATGCAACTTTTACAGCCATATTTGGTTATGAAAAGAATATTCCAAACAAAAAAGCTGTATCAGTCATTACTCATAATAGCTGGAATAACTGTTATGGAGACTCCATTCAAAAAGTAAATCAGCAACCAATTGCAATCAATATTGAAAGATCAGTAATATCAGTGAACAAAATAAATCCAACATGGACTTATAAACAGCCGAATGGAACATTATTAGAGACTCAAACTTACCAGTTTAAATGTGGACCTTTGCAGAAGAGACAACCAAGTGCGTGTTATCGACGTGAAAGATCGACAGGAATAATAATGTTTCTTCCAATTCATATAATGAATGTGATTGGTTATGATGCAGCTACTTCATTAATTTACGGAAGCAACCTTAGAAATAACATTATCGAGCTAGATTTAAATTGGAAAAACTCGATTGTAATATCAACTGAAAAATGTTCGCAAATTAAAGCATGCAATCCTTCTTTAGGTGCAATATAACATTTTTATGATGCAAACGTAATGGTTATAAATAAAT
>evgR015910c0g1_i01_Hydra
GCTTCCATGCTTATGATTTGGCTTCATTTTCAGCTGCTGTTGGTAAAACTCCAGCTGATCTAGCATTAAATCGCTGGTCAACTTGACGTAGCAGAATAAAAAACATGGACGATGCAATAACAAAATTGGAGGACTGTTTTGGTAACGAGATTCATAACGCAGCTATCAACAACTGGCACTTAGCTGAATATGTAAAAGCCTGTCATTTGACACCACTGCCAGTTGCTCTGAAGTTCGAGCCGGTGTAGCTGTTCTAATTGAACCAAAGATTGAAAAGGCATTACTACATCTCGCATGTTGAGACATATTTGAGTTTGCCCTCGAAGAATTATTCAACTCCTTGGCTTGACCATCTACTGGATCTCTATTGACCTGATATTTTGTTGTTTAAAATGTTCCATGAAGCATGGGCGTCAATGGTTCATGACAGAATTGCGAATGATATTTCAGATAAAAAAACAGCGATAATTTTAAACAACAACAACGTGATGATGTACGAGGCTGTAAAATTCGCTGTGTCGTGTTTGCAGAAGGAGCGCCCGCCTCATGATGACTATGGAGAACTCCTTGAATTATCACTAATATATATGGGACATATCCCAACTCAAGGGATCCATTTGATGGCACCTGGAGCAATGCATCGGGCCAGATGGATGGCTAAAGCAATCTATTTATTGAAGTTGTGTCATTGTGTCGTCATACATGTTACTTGAGCAAAGACGTAATTGCAGTTTTTTTGATGACTGAATATGTTAGCAAATAAGCTACCAAAGCTGAAATGGTTCAGACAATAAAAATAAATGTTGGTGTTGAAAATCATTCTAAACGACCACAGATCATAGAATAGAATATTGCATCAATAAACTCTGTGAACTTTGTTACGATTACGACACCGTAACAAACACATTTTCAACACAATTTCTAGATGCCAAAAATTAATGGTCGTG
>evgR015912c0g2_i01_Hydra
AACAGCTTCTCAAATATTTTTGGAATGCCTCGATATAAAAGTGATAATAAAATGTATATGAAGAGGTACATGAGACAGTACAGATCAGGTAAGCGTGCAAAACGTCTAACAGATGATTTAGAATCGGATAATAATAGTACTGTCTCACAGATAGAATGTGTAAATGAAAACAACGATAATAAAAACACTCATGAAACTGTTAACATTTTAGAGAGTGGTGCTAATGTAAATTCAGTGATAGAACCAAACTATATTGATAAAGTTAAAAATTCTGATGCAATATATGAATTTTCATTTAATGTTTTTCAAAAAGAATTAGGTACATGGGCTTCTAACTGCAATGCCTCGCGCTCCAGTATAC
>evgR015915c0g1_i01_Hydra
ATTATAATTTGTTTCTCCGCATTCGTTTACACAAACGCTTTAATAGTTTTTTGCAAACAAAATGTCATGCTTCGTTCTAATCAACACAACAGAAGTATACTCAAAGGATTGTTTGTTTCAAGTCTTATCGTAATAACGTTTATTACTTTCGTTGCAATACCTGACTTTATTTATGTGCTTGATGATTATATAGATTTAAATCCAAATATTATATGGTATGCCCATCTAGCTTTCTCTATATCTTTTTGGACCGACGCGTTCATATACATATTTTCAAATCCTAATGTAATAAAACTTGTTAAAAAAAATGTTCAAGTGTCATCAAAAT
>evgR015920c0g1_i01_Hydra
TTCCATTTCATGCTGTCAGGAAGTCTGATTTTACCTACAAACCTTAAGTCAAACAAACCAACACCATCATTTTCGGCAATCCTTAAAGGTAAAACTCCAGAAATGGCAAAACAACGTAACCAACTTATTCAAGTTGTCACCGTCGATAAATCTAAAAAAGAAAAAGAGTCAAACGTCATCGTCATTGGTCTTGCCTCATACACAACGAACAATAACAAAGATGCTGTTATGAAATTCTTTGAAGCTGTAGGGCTAACAACCGTAAACGTTAACCACATACACCGAATAAAATCAAGCAAAAACAATCAAATCTCAAACGTCATACGAGTATCACTCTCAACTCCAACAGATAAAGCAGCAATCATAAAAGCCTGTAGTCGCCATAATATTGATGAATATCGGTGCATTTTTGCTCGAGAGGACCGAACTCCTGCAGAACAAGCCAAATTTAACGAAGATCGCGCCTTTTTGAAGAAAAAAAACGACGAGTTGAACGCTCTCGAACTATTAGATCGGCCCATTCGGAACGTCATCCACCGTAGAACTGGTGGAATTTGCT
>evgR015921c0g1_i01_Hydra
TATTTTAGTGCAAAAAGTACACTAGCCCTATAAAAGTATCCATTTCTAACAATGAAAGATCTTCATAATTTTTCAAATTTTTGCCGTAAAATTGTTTGTGTTGGACGAAACTTTGTAGACCATGCACTTGAGCTGAATAACCCATTGCCTTCAGTCCCTCTAATTTTTTTGAAGCCTCCTTCATCAATTATCAAAGAAGGTTCCTTAATCAAAATTCCACCGAAATGTAACGCACTTCATCATGAAGTTGAGCTTGGTGTTGTAGTTGCAGAAGGAGGATCAAATATTACTCAAAACAATGTATTCAATCACATTGGCGGCTATGTATTAGCTTTAGATATGACTGCTAGAGATTTTCAAGATGAAATAAAAGCAAAATCTCACCCATGGCTTTTAGCAAAATGTTGGGATACATTTTGTCCAATCAGTGAATATATACCAAAAAGTGAAATAGATAATCCTGAAAATTTAGACTTATGGCTGGAAGTCAATAACTGTTTAAAGCAATCAGGAAACACCAAAGATATGATTTTTGGAATATCACACATAGTTAGTTATGTTAGCAAAATTATGACACTTGAGAAGGGAGATTTAATACTTATGGGTACTCCTCGTGGTGTTGGTCCAGTAAAGTCTGGAGATGTAATTGATTGTGGTATAAAAAATGTGACAACAATGAAATTTTTTGTAGAATAATTTGTTTGAAGATATATAATCTATCAATCAGCTGCAGTATTATAATCGATTGATTTTTTTTCAATGGAG
>evgR015924c0g1_i01_Hydra
CATAACAATAATTATTCTTGTGACAATGGTTTAAAGTAAAAAAAATACCAAATTTCCTACAAATCATACAAACTTAACCGAACCTAGTTAGTAAACCTAGTAACTAAACCTAGTCCGGAGAAAACTCCGAATAAACTACCGGGTACAGGGTTTAGATGTTAGTCAGTATAGATGTTGCCATCCAGTCTTGAGATCATATAATATCTATATTCAGAGACTTTATTCAAGATTACGTATTGATTAATTTCTACATGAAAAAGATTTTATATATATCAAATGTAGCCTATGATGTCACAGTCTACACAAGGTCTTCAACAACCAAAATCTTGGGAGTTGAGTTTGTATGAGCTTCATCGAACACCACAAGAAGTTATTACAGATGATACAGAAATTGCTATATCACCAAGATCTCTTCATAGTGAGCTTATGTGCCCCATTTGTTTGGATATGTTAACTAATACTATGACTACAAAGGAATGTTTACATAGATTTTGTCACGATTGTATTATTACGGCTCTAAGAGCTGGGAACAAAGAATGTCCTACTTGTAGAAAAAAGCTAATATCCAAAAGATCACTCAGACCTGATCCTAACTTTGATTCTCTAATTGCAAAGATATATCCTAATCGTGACGAATATGAAGCACATCAAGAGCAGGCATTAGAGAAGCTACATCAACATACCAACCAGCAAGCACTAACCAACAGTATAGAAGAAGGCCTTCGTATTCAAGCTAGAAACAGACTAAACCGAATGCTTAAACAACAGCGCAAGCAAGCCGAACATGCTTTAGTTGATCATTCAGTAACTGTTAAAGAGGAAGAGAGCCCAGTAATTTCGCAAGATTCTGGTGACATATCGACAGATAATATTGACTCGCCATTTATGTCTGGAAAACATAAAGTTTCGAAAAGAAAACACAAAGATTCAACTAGTTCTGCGGATAATTCAAACATTTTAAATACTGAATCAAATGAGGTGGAGGTATTGTTTCACCCTCATCCTGCTTGTGATATTCTTAAACACTTCAAATCGCGTTATTTAAAAACAACCGGAATCGCAACTGTCGCTCATTTGTGTCAGTTTTTAATTACGAGAATCTCTCTGGAAGAAACATCTCCGTGCAGTGACTTTGAAGTATTGGTACGTACGGTCGGAGAATATCAGAAACTTGAAAAACATTTAACTCTTGCAGAAATTAAACAACTTTATTCAAAAGACTCAACCCCCCTTGAACTATTTTTTTCCAGTGAGAATCGTTTAATAACTAATTAGTAATAATTAGTTTAATAAGTTGTATTTATTATAAATTTACTTATGAAAAACCCCGAAAATTTACTGATGAAATACTCCAAAATATTTTCGTAACATTTTGCGCATTAAAAAA
>evgR015929c0g3_i01_Hydra
TAATTAATTTCACAGTGTTTAATTTGAAATTTTATAACGACAAAAAAGTTTCGATTTTGAAGGATAGATTTCGGAAGCGATAAACTTTAGAGTAAACGATATACGGACATCGACCAAAAACATGGCTGACCATGAGCAAGATTCCGATCTAGAAAATTCATTAAAAGAGGAAGAAGAAAATAAAGATCAATATGCCCCAAGATTTGTAAAAATAAAAAAAGGAGAAAGAGGATTCGGATTTAATGTTCGAGGTCAAGTATCAGAAGGTGGTCAAATGAAGGCCATCAACGGTCAGTTGTATCCTCCTCTTCAAATGATCAGTGCAGTACTGGAAGGTGGTCCTGCAGATAAAGCAAAAGTTCGCGTTGGTGATCGTATTTTGGAAGTAAATGGCGTTAACTGTGAAGGAACTGACCATAGAGCTGTTGTTGATCTTATCAAGCAAGGGAAAGAATTTCTTACACTAGTTATTATTTCTGTCTCTCCAAGTGAGGCACGCAAACTTGATGGATCAGAGGTATGTGGAAGTGATTACATAGATTATTCTGATCGAAGGAGTATTCCTATTACAATTCCAGATTTCAAGTATTTAGAAAAAGACGGAGAGAAATATGTGGTTTACAATATATATATAAATGGAAAGTATAGATGTTGTCGTCGTTACAATGAATTTGCCAATCTACACAGTGATTTAAAAAAAAAGTTTACAGACTATCCGTTTCCAAAACTTCCTACAAAATGGCCGCTTAAGTTATCTGATCAACAATTAGATAGTAGGCGACGAGGACTAGAAGTATATATAGAAGATGTTACTGCAATACGTGAAATATATGAATCAGATATCATCGAAAAATTTTTACTTGAAGAAAACGTACAGAAATACAATCAAACTCAAAATCAACCTGCAATAAACCAAAAATCAGATACTAATGAAAAAGAAATTAAAATCAACTTATCCGACAAGACAATATTATCAGTATTTCTCTCTAATAAAAAACAAAAAACTGAGCAAATATACGAGTGTGTGGTAAAAAAGCTTGGATTGTCTGATTATAGTGCTACATTTTTTGCTCTTTTCCAAATCATGGAGAATGAGTTTGAACGTAAACTTGAACCATCTGAGTTTCCATATAGTATATATCAAAGATCAGTTGCTGATTCAGGAGAAACAAAACTTGCATTTCGAAAATGGGTTTTTACATTAACAAGAGAAATTGAAATGTATACCGATCCAATTGCTTTGAATCTATTATATTATCAAGCTGCGTCAGATCTTCGAAAGGGTTTTATTAAGCCAGAAAATAAAATACAGAAAATAAAAGAATATAATAGTACAAATAATAAAATACCTTTTCTTGAGGTAGTTTGCACACTGGATGGTTATGGAGATATTGTCTTTCCTCATTGTGAATGTGATGCTCGTAAAGAAGGACATGTTATTCTTAGACTTTCCTTAAAGAGTCTTACAATGGCTGCTTGTTCTTTGGATGGTGTTGCAGAGGATCAAGAACACAACTTTACCTGGGACGAAATCCAAAGTTGGGATGCAGACACTGAAGCAATGTGTTTTTGGTTTCAATACTCTAAATCAGGAAGAAACCCTCGACAAGTTCGCATTTATTCACTTTACTATAAATACATGGAGGATTGTTGTAAGCGTATTGCTACAGAGTTAAGTTGGTGCAAAAAGGACTCTTCTGGAAGCCCAATTTTACACAATGCTCCTTCATCGGAAAGCAGTAAAAACAAGTTGTCGACTAAAGCAAAAGCAACTGTTAATCATATCAATGATGGTGATCTATAAGAACCATGCAATCTGTTTGGCGTGTTGTGTGTCCAAGATGAAATCAAACCTATTGATATTTGCGTGAAAATAATATTTTTCGTATTACGTTATGAAAGTTATTATACAGCATAAATATGCAGCTATATTATTATCACTTGAAATTACTGTTTAAAATTTTGTATATTTTCATAATTCTGATTTTGTTACATTATTATTTTATTAAAACTATGTTACAATATGAATTTAATACCATATAGTTATGTTAC
>evgR015930c0g1_i01_Hydra
TTACTGCTTTTATGCTTTAAAATAGAATAAAAAAACACTAATATTTGGTTATTGAATAATTAACACTGATTAATAAAAAATAACTACAGTTGAAACTGCGAATACATTTAGTACATTGAAATCATCTTTTCATTTAATTATATGGTGAATGCTCAATTGATAAAAGCCAAAAGAGAAACATTTCTTCAAGACGAAAAATTTGTTTTAATGCGGTTTTACGAACTGACATTCGCTTGTTTGCGTAGTTAAATTGATTTTTAACAGTGTTTTAAGAACGTCTGCTTCCCGGTGTGTTTATAATAGCAGCGGTTATCTTGCGAGGAAATTCATTATTTTATAATGTCAACAATGACATTAACTCAACTGGACCCTCATGTTGGACCTATGAGAGGGATTAATCGTTTTAAACCAGATAAAGCATCTAATATAAATCCAGAGACACCACAGGAGGGCGGCGGAGAGCTCGTAAAAACAGATTCCCCAAACTTTGTTTGTAGTGCGTTGCCTTCTCATTGGAGATGTAATAAAACTTTACCAATGGCGTTTAAAGTAATAGCATTGAGTGGAGATATACCTGACGGGGTTACTGTCACAATATTTGCCGGCAATGACGACAATTTTTCAGCAGAACTTAGAAATGCAACTGCTGTTATGAAGAATCAGGTTGCAAGATTTAACGACCTTCGATTTGTTGGCAGATCGGGTAGAGGAAAGACGTTCACATTGACAATAACAGTAAATTCGGAACCTCCGCAAGTTGCTACATATACGAGAGCAATAAAAGTTACTGTTGATGGACCAAGAGAGCCTCGAAGACACAGAGTTAGAGATGGGGAAGGTGGGTTTCTCGATCACTACAGCTTTGTGTCACGAGCAGGAGAAATAGGAGCAGATAATTTTCGACAATTTTCATATATTCATGGATCACCGATTCCAACAAACAGTTTGTCTCATATGCAGCCTGATGTATTAATTCGCTCTCCGTCAGCATGGCAACATAACTTTAATGGAAATGTGTACGGACATAACTTGTCACCCAATAGTAACGAATCTTATCCAAATATGATGGATCATGACGGATACCAACAGCCAAATTTAGCATTGAATTTAAAACTAGGAAATAATGACTTAACCTTACAAGAAAACTTTCAAAACGATGTTAGATGTTATAATCAACTTTCTCAAAACAGATCTTCTTCTATCTCGAATGAGCCAAGTTCACCGAGTAGTCTCCAAGGTTTTCAATATCATAATAATAGTTCTTTACATTCGTTTCAAGGAACACATTTTCCAACACCGAATCATGCTCATGATCAGGTACCAAACAATTTTCTCAACAACGTCACAATAAACAATAACTATCTTATGAATCTTGCACCAAGCGCAGGTAGCATAAACATTCCATATAAGTCTTATAGCAGTCAAAACATTTCAAGAGCACATGAAGATATCAATAATAATAATCAACTCGGACATTTTTACAGATCGCCATTTTATGAAACGCATGGCTATGCATTTCAAAATAATATTAAAGCTTACGCTTCTCAAATAAATGGCGGAGACGACGATGGTAATTCAACTCCAAAAGTTTGGAGACCATATTAATTATAATTATCTTATAATTTTTACATTTCTTTGGAGATTTTTATTTTGTGATATACTTCTTACATTTTAATTTTTAAATTATTTGGTTATTTATAAAGCATTTTTTATAAAAGTATTTATGGCATTAAAAAGTTGTCTTTATTTTTATTATTACTATTTATGTCATTAAAAACTTATTTGTTTTTATTTTTGAC
>evgR015931c0g1_i01_Hydra
ATTTATTATAGTTATTACTTGTATGAAAAATATCCAAACTTTGAAGATCTAATGGCTTTGGATACATCTGGTTTTGTGCCGTTACAGAAACATGAAATATCAAGATACAAGTATGATATGAATACCACGAAATCAAATTTATTTCAGCGATATAAAAGCTTAGATGATTCATTCATTTTAATTAACAATGGTGAAGAACAAAAGACTCGAATACAATTCGATAAATTTGCGAAAGATTTTAATGATGATACAAATAATAGACATTCTTCAAATACCTCGTTTTTCAATGTGAAAGAAAAAGATCATTTTGAAATTGAAAAAATAAAAATATTGAAAAACACAACTACAGATAAAATTCGACTTTACGAGTATTTATTATATTTACTTAAAGAAAGCCAATCGGATTGCATCCGGTGGATCAACAAACAAACTGGAGAGTTTGAAATCACCAGCTCTGCAAAGATTTCTTTAAGATGGGGAAAACTAAGAGGAAACCCAAATATGACTTATGAAAAACTAGCCCGCGCCTTACGTCATTATTATAAAAAAAACATTCTTTGTACTTTTAAGCATAGAGAAAAACTTAGGTATAAATTTTCAGAATATATTTTAAAGAAATACGTTGTGTGACTCCAAGCTACTTACAGCATATGTGCTACTTACAGCATATGTGAAAATTTTTAAATTTATTGTTGTAATGATTGTATAAAACATTTTTATTTTGAATTAAATATTATAG
>evgR015958c0g1_i01_Hydra
GCTATTTTTTTGGGAATCATGAATAATGACTATTTTACAGTTGTAATAATCTACGGAACACTTCCGTTCCAAGTACTAGGTGCATATTTAGTAATTAAAGTCAGAAAAAAGTTCAATGATGTTCAATTTTTATTGTTGCTATGTAACTGTTTTGTTGATATACAAATGAATGTTTTTCTTCCATCTCAATATTTGTGCAAAATTTTGAAAATGGATAAGGCATCCTACTTTATATCAATCTATTCGTCAACTGGTTTGGCATTTCAGTATTACTGTACAGTCTTCATGTTGACTGCTAATCGCTTTTCTGAAGTTTATTTTAATCTCTATTATGATCAAGTATGGAAACACAGCCGTGTTAAAATAGTCTTTGTAACCATTTTCCTATTTTCCGCAGTTGTTTCCGTAGGGCTTATGATTCACGCTAGTAAGTTAGGCATATATTTGAACAATACAAATATTGTTGAAATGTATTCTTATATGTATTTATATCCTTTTTTAAGTATTATTTTATTACCAACAATTTTTGGAACATACATCTATATATTTTGTGTTTATCAAAAAACAAATGCAAATAATATAAACCGCAGATCTTCAGTTTCATCTTCAACCTTTGGTGAAAGATTTTCAACAACAGTTTTACGAAAATCTAAAAAATTTTATAGAAACTTTGTTATGTTCAGTCTTTTAATAACAACTTTTGTTTTATTTATATCGTTGCCAGATACACTCACTTCTTATTATTATTGGCGCAATCGAAAAAAGATCGACTC
>evgR015969c0g1_i01_Hydra
AAAAATAGCCGTCTGTCATGAAATTAAACACGTTAATTTTGAAAAATAATTACTTTAACATGAGTAAAAATAACAATCTTAATTCTAAAGTTCTGCAATACGAAGATTTTCTGAACTCTAAGTTGAAAACTGATCTTTTTGAGGTTCATAAACAGCGCGATGTAGTTTATAAAGAGATGGCTGAATACCTCCAGTTGAAGAAAACTATTGAAATTATTCAACAACAGCAAAAAGAAAAGATCAATTCTCATAAGCAATTTGAATTACGCACTAAAACAGACTTGGGATGCAATTTTTACTGTCAAGCACTTATTCCAGACACATCTTTTGTTTATGTTTTAGTCGGTTATGGATACTTTGTTCAAATGACTTTAGAAGAAGCAGTTATTTTTATAAATAAAAAGATGAAAATTCTCACAGAAAAAAGTGATCGTTTTGTAAAAGATTCTGCAAAAATTAAAGCTCATATTAGATTAGTGATGGAGGGCCTAAGAGAAATTCAAAACTTAAATGTAGAAAAACAGTAAAGTGTGTAATGCGTAAATTTAATAACATTTAAGTACAACCCACCAAAAAGAATAAAAGAAAATAATACATTTAATTTATGAATGCATCTTAATGAGAATTTTAGTTTATTATTTTATGTTGTTTTAATAAAATGAAATCAAACATTAAAAAAAAAAAA
>evgR015973c0g1_i01_Hydra
TTTTTTTTTTTTTTTGTTAAATTTGCTAGCAGCATATTTTTCTTCAAAAATATAAAGTTTTAAACGAAATATGAATTTCATGATCTTTATTGCAGCTGTTGGTTTTGTTGCAGTTACAATAGGTTCTGTTCCTATTGAAGAAAGTTCCACTACATCACCAGAGGATTCCGATACAACAACAGAAGATGACGAGCCAACAACAACGGAACTATCTACAACTACAGACCTAATTTCCGGAGATGAATGCGTCAATTCATTTTATAAATGTGGAAAAAGTTCTACAAATTGGTTTCAAAGAAGTATCTGTTTTTTTCAGCTTGGAAATTGTCTAAACGACGTTTGTTCTGTTGAAAAATGTCTTACGATATCGAGTCAATGTTTTTTGGAGATTCGTAACTACGATGACTTTACGAATTGCGCTTATGAGTTAGACTCGTGCATCCCAGATGAATGTAAAATATCTATGAAGTAGAAAGTCATCAAAGAAAAATACATTTTCTACTTAACTGTCGGTTAACTGTCGGTTAACTGTTGGTTGGAAATAATTTTATTTAACGTTATATTATGCACCAACTATTTTAAACGTATTTATAAGGTATTTATAAATGTAGTCATATATATATATTTATATATATAAATTATTAATTAGAAACTTTCAACTACTTTGAGAAAGTTTGTTTTGTCATATTTTTTCAACATTTCTTATTACAACATTTTATTACA
>evgR015976c0g1_i02_Hydra
AAATGACTAAGTAAATACAAATATTTGGCATTAAAAGGTATAAAACATGTTAATACCTTCACAGGGAAAATATTTTTCAAAGGTTCAAAACATGGGCGCAATAAAAGTTGCATTGTTGTTTGTAATCTTAACGTTTGAATTTTTCAGCATTGAGGCGCAAGGTCCTCGTCAGGGCATACTGATTTCTACTGCAATCAATAATGCAAATATAATGCAAAAATGTAAAGAATGCAAAGAAACAATGAAACAAAACATCAATAAGGAAGAATGTAACCAATTTTTTATCAACCACAACTGTTTTTTCACAACTAACGGAGAAAATAGTAAATTGCGGAATGGCAGAATGGCGGAATAACATTCACAAGAAAAGAGGTCGGAAAACTTTTAAAACTTCAAGATAAGAAATTACGAGACAAGAATCGTGTCGAATAGTTATTATTTTAGAGTTTTTTGCATTTGCTTTATTATATTTCTTATATATAAATGTCTGTATTTTTTTATTAAACAATTATATATTTTTTTCGTAATAAACAAAACATATTTTTTTATATATTACATTTATACATTTTTGTGCAAAAAAAAAA
>evgR015981c0g1_i01_Hydra
AACAGAACTAATAGTTATGAACACTGCAATGAATTTTCAAAACGAACAGTACGAAAAAATGATGAAAGAATTCAACAGTCTAAAAGAAAATCTTAAAAACAGTATTAAAGTAAGCGGAAAACTAGAGAACGAAGATAAAATTGAAGATAAAATTGCTGAGATAGAAGATAGGAGTAGGAGAAACAACCTTCGTTTCGAAGGACTTGAAGCAAAAGAAAATGAAACATGGGATGAAAGCGAAATAAAAATAAAAATATTTCTTAAGGAAAAACTTGGTATTTTAGACAACGTTAGCATAGAAAGAGCTCACAGAACCGGAAAGAAATATGAAACTGGGAAAAAAATAGAACAATTGTTGCAAAATTTCTTTATTATAAAGACAGAGAAAAG
>evgR015985c0g2_i01_Hydra
TTTCATGCGAAAGAAAATTGCATTAATAGGCGGTGGCAACATAGGTGGCACTTTAGCTTATATTATAGCGTCCAAGGAGCTTGGCGATGTGGTAATTTTAGACAGAAATAAAGAATATGCAGCAGGTAAGGCGCTCGATATAGAGCAGAGTTTACCTGTAATGAAAAAAGATATATCGCTGATAGGTACGGATGATTATGCTGATATTGAAGGGTCAGATTTAGTTATTATTACAGCGGGGGTTGCAAGAAAACCTGGTATGTCTAGAGACGATTTGCTTAGTGTTAATGCTGACGTCATAAAAACTGTAGCAGGAGGAGTAAAAAAATACTGTCCTAATGCTTTTGTAATAGTTATCACAAATCCCTTAGATGCAATTGTTTATGCATTTTTAAAGTACTCTGGGTTTTCTAGGAAAAAGGTTATTGGAATGGCCGGTGTATTAGATTCAGCAAGATTTGGATTGTTTTTGGCTAGAGAGTTAAATACCTCAGTAAGAGACATAACCTCTTTTGTGTTGGGTGGACATGGCGATACTATGGTTCCTTTAATGAAATATACTAATGTAGGTGGCATACCTTTATCAGATTTTATAAAAAAAGGAATTATTTCGTCGGCTAAAATTCAAGAAATAATTGAAAGGACAAAAAACGGTGGAGCAGAAATTGTAAAATTACTGAAAGATGGATCTGCATACTATGCGCCAGCAAGTTCAGCAATCAAAATGGCTGAATCGTACCTTAATGACAATAATGAGCTTTTTCCTTGTTCAGTATATCTGGAAGGAGAATATGGGTTAAGTGACACTTGCATGGGAGTGCCGGTATTTATAGGTAAAAATGGGATAGAAAGTATACTACAAATAGAATTTGATGTGGAAGAGCAGGCCATTTTTGATAAGTCAGCAGAAGCAGTAAATGGCCTTATCTCAGCAATGAAAAAAGTTCTGGGCTAAATCTGATTTGATGAACAATTGTATTATAATAAGAGGTGCAAAAGAGCACAATCTCAAAAATGTTGATATTGACATACCAAAAAATAAATTAGTTGTTATCACGGGTGTAAGTGGTTCAGGGAAATCCTCTTTAGCATTTGATACGATATATGCTGAAGGGCAAAGAAGGTATGTTGAAAGTTTATCGACTTATGCTAGACAATTCTTAGAGATGCATAATAAACCTGATGTTGAATCAATAGAAGGTTTATCTCCTGCAATTGCAATAGATCAGAAAACAACATCGAGAAATCCGGG
>evgR015989c0g1_i01_Hydra
GACATTTAAAACTAATCTTTTATTGTCTAAATAAGTCATTTTAATAATAAAATATGAGCGAGGAAGAACATCAATTTCCTGAGTTAAAAAATGACTTAATCATAAAAGCTGTTCGAGGAGAACATGTATCACGCGTTCCTGTTTGGGTAATGAGGCAAGCTGGTAGATACCTACCTGAGTTTCGAGATCTAAGAGTTGAAAATGAGTTCTTTGAGGTATGCAGGACGCCATCTCTGGCATGCGAAGTGACTATGATGCCAATCAAAAGATATGATCTCGATGCTGCAATCATTTTCAGTGACATCCTTGTCATACCTCAAGCTTTAGGAATGCAAGTGGAAATGCAGAAGGGAAAGGGCCCTGTTTTTCCAAAACCTTTACTAGATGCAACTGATATAAAGCTTCTAAATATTGATGTTGATGTGAAGGAAGAGCTAAAATATGTTTTGGACGCTATAACTTTGACCAGGAAGACTCTTGATGGAAAATGCCCTCTTATTGGGTTTACTGGAGCACCGTGGACACTGATGGCTTATATGATAGAAGGAGGAGGCTCGAAAACATATTCAAAAGTCAAGAAATGTCTTTATGTCAATCCTGATTCTTGCAAACTTTTGCTAAGCATTTTGACTAGAGTGTGCATAGATTTTCTTGTTGAGCAAGCCAAAGCTGGAGCACAACTTCTGCAAGTGTTTGACTCTCATGCTGGTGAACTTCCTGCTGACATGTTCAATATTTTTTGTCTACCATATTTAAAAGAAATTGCAACTTCTGTGAAGCAGCGTTTGGGGCCGAACCAATCTGTTCCAATGATTGTGTTTGCAAAGGGATCACACTATGCATTAAAAGATTTAGGTGAATCAGATTTTGATGTCATTGCTCTCGACTGGACTATAAATATTGATGATGCTAGGCGACAGTGTCCTAATAAAACTCTAATGGGAAACCTTGATCCATGTGCATTATATTCTTCTAAAGAAGATATTGACACCCGTGTTGGAGAAATGGTTAGAATGTTTGGGACATCGAGATACATTGCTAATTTAGGTCACGGAATTTATCCAGATATTGATCCAGATCATATGGAAGCATTTGTTAATGCTGTTCATCGACACTCAGAAAATAGTGACTAGATTATTTCGTTTGATTTACATAGTTGGAAACTTGGTAGGGTTTGAATTTGGATGCAACTTAGTAGGGTTTGAACTATAGAAGTTAGATTTCCAATAAATACCTTATTAGGATAACTAAAGCAAGATGCAACTGTGAAGAATAATGCAAACTAAATCACTAGCCAGTAACAATATACTTTTATATACTTTTTGGATTGCTTCAAATAAATAGATCTTATAGATTTGTATGGAATGGAATATTTTAGTTTTCTTCTCTTTTTTTTTTTTTTAATTAAATACTAAAACTTTCTGACCATTTAGTTAATTTTCTTGGGAATATATTTATATTTGTGTTTTTTTCCAATAAGTAATAAAATTGACAAAATATATTTGAAAACATTTAAAAAAAAA
>evgR015992c0g1_i01_Hydra
TATAATCATGATTAACTGTGCTTATTGTGAATATGCTGCAAGTTCTAGCAACTCTTATAAATTTCATTTAAAAGTTTTTCACAACAAACATAGTTATGGCGAAAGATTAGTTTGCAGTCAGGGTGGTTGTCCGATGGAATATAATTCTTTCCAAAGTCTTTCAAAACATATTGAAAAATCTCATCGAGAAGTTATTTTAGGACCTACTTTAGAAGAGGCAGATATTATGCAACGATGTTTCCATTCAAATAACTTGTCTCTAGCTGTTCACAATACCGAACCTGTTTGCCCAAATAATATGTTAAAATTTGGTATAAAATTTGATGAAATCGATGGTATATTTGAAGATTCCGCTTATTTCATTTCTAAACTAAGAAGTTATTCTAATATGCCCCTAACCACTACATTGAAAATTGTGGATGCATGTTCAGACTTTGTGTCATCAATTGTAACAGGAATTAGTAATGAGACCCAAGGAATATTTGACAAGCATGGTCTCCAAAAAGATGAATATAATAATTTCTTTGAAAATATATTACGCCTCCAAAAACCTTTTCAAGGCTTAGAAAGTTTACATAAACAAAC
>evgR015999c0g1_i01_Hydra
TTCAAAAATTTCCAAGCACTTCCAAAGTCTGTAATGTTCATTTACAATGTTTCATCAAATGAAGAATTGTCAGACGATGATGACACTTCATCAGTTTCCTCCAACTCATCATTGGAATCAGACGTAACACAGGACGCTGTTAGAACTGGATTTACTGGTATCATTGACTTTGAAAAGAATGAAATGACCATTTATCCACCTTCTAACTGTGCTTTCATGCAAAAGGAATCCATCATCAATATTATGGATATTGCCGAATGTCTTTGTATCTCTACAATGTACATTTGTATTGAAAAGAATCATCAAGAAATCAGAGATTTTTCACGCGGATTTATTTATGCTGGTC
>evgR016000c0g1_i01_Hydra
ATAAATGACTATTATGAAATTTCTGAAATTGAAAACATTAAACCAGTTGCAAGAGAACCAGAAAAAAAAACTTGGAAACTTTGGAAAATTTTTGGTAAACAAAGGAAAAAAGGTAATCGGTTGGAAGATAGCAGAAATCTGAGCGATTACAAGAATCAAAAAGAATCAACTCTTCACCTTGTTCCTCAATTTAGAAGTAGTATAAAACTTTTTGTCAAAACCTTGACTGGTAAAACTATCATATTGGAGGTTGAGCCTGCTGACACTATTGACAACGTGAAAACCAAAATCCAAGACAAAGAAGG
>evgR016012c0g1_i01_Hydra
GCTGGGTGATGTTCAAGTGGGTAAAACAAGTTTGATGGTGAAATATGTTAATGGTAAATTTGATGAGGATTATATTATGACTCTTGGAGTTAACTTTTTGGAGAAGAAGGTCCAAATCAAGAACACTGAAGTAAATTTAATGATTTGGGACTTGGGAGGACAAAAAGAATTCATGTCTATGCTAGACCTTGTCTGTAATGATTCCCTAGCCTTGTTCTTTATGTTTGATCTCACAAGGAAACAAACACTTCGTTCCGTTAAAGATTGGTTCCTTCACTCACGAAAATATAACAAACGAGCTCTTCCTTTTTTGATTGGTACAAAGTATGATAAGTTTGTTGATCTCCCTAAGGAAGATCAAGAAGAAATGACAAACCAAGCAAGAAAATATGCTGAAAAGATGAAATGTCCTCTTATTTATTGTAGTGCCGCAGCTGGTATCAACATCAAGAAGCTTTTCAAACTTGTATTGGCCAAAGTATTTGATTTCCCTCCAAACATCGACAAGATTACAGCCATAGGAGAACCTCTTCTTGAATATGATTAAT
>evgR016025c0g1_i01_Hydra
TTTACTGTTTCTTTTATTAGAAAACTTTGTTAGAAACATCTTATTATATAACATTAATTTTCTTAATTGGTTTGGTAAAAAATATGACTTTTTATCGTGGTGAATTTTTTTGCGACGAAAATTTAGTTAATGAAAATGTCTCCGTAAAAGTTCTTAACTGGCTTAGTACTAGTCATGATTCAGGACTGGATATAGAAGATCCGAGTGATTCTCTTTTAAAAAACGATACAGTAGATGCCAAACTGTTAAATCAAAACGATTCAAAGGAAGATCTCTCGCATAAACTCTCAAAGAGTATTCAAAATCAGATTAACAAAATACAAGGAAATTTAGAAATTGTTCCCTGGGATGTAACTGACTGCGATACCTTGGTGCAAGTAGACAAGACACTGTTTCCCATTCAACGCCAATTCGTAGCTTATGTGTCCGGGTTTTTCAGCAATATAGTTGATCGAATGAATGAAATGCACCATAATACTATTGTGTTGAATGGATTCGAAGTAGAAGAAATTCGGACAGTTCTGACGTTTATTTACACTGCCAATGAAGAAGATGTGACAGACATGAACGTGGCTTCATTACTACAGCTTGCAAAGCAGTTTAAAGTCAAAAGAATTCTTCGAATGTGCGAGAAACATTTACTTCGATCGTTAACGTTTAACAGAGTTATTGCAATTCAACTTGCTCAACAATACAAACTAGATTACGTGTTCATAGATTCATGTATCAAGCTTAGTCAGGTTGAAAGTTTTGAAGATACCGCTGAGTTTCTTTCACTTAATTATTCCACTCAAAACATTGTTATGAGATATGCTTTGAAACGATTTCAAAAAGCCCATGATGTTATCCGCCATTTAGATGTTTATTTGTGCAGTCACCACGGAAGCGACACTAAAAATAACGATTGTTGTTTTAGGTCTGCAACGTATAAAACCGAAGACAATCACACATTTCTTGTTAGAATGGAAAAACCTCAAGAAAGAATATTAAATTTCATTGAAAAGGTTTATAATGAACAACTCTGAATTTTTATACGAGTTTTTTAATGCAACAGTTTTGAAATTTAACTCAAAGATTTGTCTTGTTACTTTGATGAACTCGTAATTTAAATGACTTCGTTATTTAAATGAACTCGTTGTTTAACTGAACTCGTTACTTAAATGAACTCGTTACTTTAATGAACTCGTTACTTATGTGAACTCGTTAAATTAAATGAACTCGTTAACTTGAAGGAATTTTTCTATTCACCTTTACTTAAGTGGAATTGAAGAGAATGATGGCGTTTCCGGCCTCTAGCCATGGCACCGACTGGTCAATTTGTTTTTTAGATTAATAATAGGAAAATCTACAGTTAAATAAGGTTAAAGTATGTTAGGTTTAACTTTTTCATTCTTTGTAAGCGCCCCATTATTAAAGTAGATATGTGCAACAAAAGTTGGTGCATAATAAAAGTTGGTAATATAAGTCATAG
>evgR016028c0g1_i01_Hydra
ACAGTTCGCGTTCGCAAATAAGAAAGCTGAGCCTACTTATACAACTGATCATTTTATTTACGAAGAATTTTTGAATAAAGTTAAACTTTCTGGTTGGAGGCGAAAAAGATGAACGCAGCAAATAGTAAGAATGAAGAATTTAAGTTTTTAGCATCTTTAACTACTGAGGACATTAAAGGTTACAAAAAAGCGTTTCGTTATTATGACAAGAACAACGACGGTCTTATAAATGCAAAAGAGTTTTTGTGTTTGTTAAGAAATCTTGGAGAAAATCCAACAGAAGATGAACTTTTTGATATGTTGCTGGAGATGGATAAAGACCGAGATGATGCAATCGACTTTGGAGAGTTTATGGGGTGCATGGCTGCAAGAAAGAAAAATGACCATATAAGCTGTTTGACAGAAGCATTTAAAGTGTTTGATCTTAACGGAGACGGTTTTATCAGTGCCACTGAATTTCGCACCGTAATGGAAAGTTTAGGTAATGATACAACTAGCGAGGAAGTAGATGACATAATTAAACATGGCGATACTAATGGTGATGGTCTAATTGATTATGAAGAATTTGCAAAGATGTTAACTTATGCGTTCAACGAACAATGATATACACAAGATCAAGGGGTGAAAGAGCAATGAGAAAATCCTAAATCAAATATGCTTATGAGAAATTAATAAAACATTTCAACAAACAATCGATCTGGTATACATTACAACTTTGGTTATGCGGTTAATTGTACTACATGAGATACAGAGAAGAATGATACATTTTATAATAACTTGGAAATATATATATTTTTCCCGCTCATTGTAAATGCCAAGGATTGCCTCATCAATAATATTTTGAATACCAAATTTAAAGTATTGTAACTTCTAAACACCAAATTTAAAGTGTTATCACTTTTTTTTTTAAAGTTAAGTTATTGTTTCTTTCCATTATTTTTAAAATAATAAAAATAATAAAATATATTTCCTTTGATAAAAATACTTTCGTGCTAAAACATTGAGTTTGATAGGAACTGATTGCCCATTATATTTTTTTGTTTTTAAAAACCAAATGATTTAAATTTAGAACAAAATATTTATTAATATAC
>evgR016029c0g1_i01_Hydra
CTAACTTTAAAATGACCAACCTTCCTGATCCAAAACTCGTCCTCAGATTCCAAGATTTGGTCGATAAGAACATTGATGAACAATGTGAATTCTTCTTGAAGAGTTTCATCTTTGCTTTGGGTGATGATTGGCCAGAAGTTGTTAAACTCTCTGGTACTTTTAAAAAGTATTTGGCTGATTCTAGCAAATTATCTTCCAATCCAGCTAGTCAAACACAATTAGATGAAGCTCAAGCTGCTGACTTTTTACAAAAGAATGGTAAAACACGTACAGCTACTCAGAGAAGAGATGAAGTCCGTGATATTGATTTGGACTTTAATAAGCATATTGCCTTCATTGAATATCTTCTCCTCCACTATAAACATTTAATTCTCTCAGAATACTACAAGAGACATACTGAATTAAATGTACCAGCTGG
>evgR016036c0g1_i01_Hydra
AATAATGAGGATTTTAGTGATGCTGATGATGAGGCTGATGCTGAAACAGATTCAAATGAACCGAATCGCCCGATGCCATCAAACTTTCGGTTAGTAAAGCAGAAAAAATCAATTACAATATCATCAGAATTTCGTCCAGCAACTACTGCTGCTACTGCTACTACCAAAAAATACAAAATACTATCAAAGGCAATAAAAAAGAGAGTCTTGGGTTACAATGTCAGCGATTTGTCGAATTCGGACTTGTACTCGACCAACGAAGAAAGACCAGAGCACATGTTCCGCAAATCAACCTCTGAAGACTCAGGTGATGAAGGTGTTGATGCCAATGAGGTCTATAAACGATTTATAGCTACTGGTCAGAAGCAAAATCATTGTCAGAAGTGTAAGGGAAGCAACAAAATCCCCCATTTTGTCTATGAAGTCAGCGCAGATGGAGTGCAAATCCAAAAAAAAGCTGTAAAATCCGCTTGTTGGCCAATAATGGGTCAATTATTGTATATAAGTCCTTGCACTCACCTAGAAAATCGCGTCCGCTTCTACATGCCACGAAATTCTAAACCGGTCATCATTGGTTTTTATCATAGAACTGCCAAGCC
>evgR016041c0g1_i01_Hydra
TAATCGTGAGCTTCTTGCTCTTTTAAAGACTGGTTACCGAATGGATCGACCAGATAATTGCTCTGAAGCAATGTTTGACCACATGTTGCGTTGTTGGAATGATGATCCGTTACAACGACCTAGTTTTACCGAACTACGCGAGTTATTTGAAGAAATATTATCTCAAGGTGGTCATTACTTTAGCTTTGAAATAAACGAAGAAAGTAACTATTATAATGCAGCATTTTCCAGCATCTCTTCTGATGCAAATGATGGCGCACTAGAAAAAGAATTATTAAAAAAGCCTGTTTATGTAAATTCAGTCGAAGAAATGGAAAAAAAAATAAGGGACGTTTTGTAG
>evgR016062c0g1_i01_Hydra
CATAAAAAGTGAAACAAATGTTGTTTTAGAGAAATTATCTGACTAAATAAGAATTGGTCTTTCAAGAGTAGATAAAAATGTACATGCTGCTACATAAGACGAACATACTGTTTATTTTAATCTTTAATGTTAGATCAGATGTTTTTAAAGATGCAAACGAGCGTGAAGTAAAATATAATAACTTGATAAAAACACTGCCTTTGTTGGAAAAAACGTTTACTGTTTACTTTAAGTTTAAACTCAGATCATTGCCGACCGATCGTTGGGAGAGTATTTTTCGTTTGACTTCTGGTAATGAAAATATTCTTCATGTACGTGTTTGCGGGCAATATTTTATGATTACTTTTCGTCTATATGATAACGCGGATCAATTGTTTTCAACCTACATTCAACCACCACTAAATGAATGGACTACTATACAGACAACTCAAAGTCTAAAAAATGGAATTTATTTATTTTCAATATTTATAAACAATACAAATTTTTTGAGCACTATAAGTCCAAATCCAAAATCATATTCAAATGTCAAAATATTTATTGGAGATTTATCTGATCCCCCGGCAAATGCTTCAATCAAAGATATCGTAATTATAAGTGGAAATGGATCTCTTTATGATTGTGATGACAGTAACCCTGGTTGTTTTTGGAACAACGGAATCAATACAGAGAATATAGGATGGTGTCAAGATAAAGTAAAAATTGTTGATAATTTAGTAGACGTGAATGAGCGTGATGTAAAAAGTAACAACTTAATAGCAACATTGTCGTCGTTGCAACAAATCTATTATGTTTCATTTAAGTTCAAACTCAGACATAATGTCAGTTCTTGGAAAAATCTACTTGATATTGTTGATGAAAATAAGAGGCAAACTAATAAGGTATATATTGGTCCGGATTTAATCTTCCTTTATAGTGACAGAATAGATGCAAGACATTACAAACTACCCCTAAATGGTTGGACAAATATACAAATATTTCAAAATAAAATGAATGGAATTTATTTGTATACAATATTTTTAAATGGAACAAAAATTTGCAGCGCTCTAATAGCAAGGCCGCATATATCTAACAATGCAAAGGTGTATATTGGATCCGATACGGAAAACCCAGCAAATGCAACTATTAAAGATCTTATAATTATAAGTGGAAGTTCAAACATTGAACGTGTTATTTCAAACAACTGCTCTGTTAATGCTATTTGTAATAGTTTGCTTAATGGATACAATTGCACATGTAAAAGTGGATTCAATACAAGTAATGGTGGAATTGAATCACCATTAAACAAATGCG
>evgR016073c0g1_i01_Hydra
ACATTGCTCCAAGCTCACCATCTTCATCAATTGGTTCTCCAACAATTCTAAACCCACACTTTGCTGCTCATCTTCCAATGGTCTATCCTGCTAACCTTGCTACCGTTTATCACCCAGAAATGGCTCAACAATTTGCCTTGCAATATGCTCCAAAGATGCAAACACTCTATTATCCACCACAATGGGCTGCTTATCCAACTAACCAAATAGTCTTTGGTTCTCCAAACACTTCACCAATCTCCCCACCATCAAATGTTATTCCACATCAAGACAAGTCATCTGATATTTTGGGAACTCTTATTACTGGTGAAACTGATAAGAAGACTACTGGCAAGAAGGACCATAAGATTATTCATCACACTCCACTTAATTTTGTTCATGCTATGCCAGCAGCTGCAAACCCTGTGGTCGAAACATCTGAAAGAGCTTCTATTGCAATCCAATTTCTCCAATCACTTACACCTGTTGAAAGACAAACAGTCACAATGTTCATGAATGGTTCCCAAGCTCTTTAACTTGTAAATTTCTAACTAGATATATCATCATCCAATCCTAATC
>evgR016084c0g1_i01_Hydra
ATTTATCTCACAAAAGCAACATTGTGCGTGACAAACGCTTTTATAAAAAAGTTAATTCCCGCCGAGAACAAAAAACCACTGGAAGTCTTGCAAAGCGTAGAGATATAGATGATATTCGTGATAATCATTTGTGCAACTACGGTGACTGTTACCAGGAAAAAGAGAAGGAAAATGATTGGTTAACAAATCTTGGTGACTACAAAACTGAAAAGCGGACGCCTTATTATCATCGTACAAAGTTTCATCACTTAAAAGTTAGATTAAATGGACCAGGCAAAACTCCTGCGGAATGGATGAAACAAGAAGAATTTTATAATCAACATCTTGATGATGGCACTAACTATAATCAATTTTATGTCTAATAAACATCTTGATGATGGCACTAGCTACAATCAATTTTTTGTCTAAGATAATCTTTAAGACATGCTTATGTGATATCTCTAAATGTTTTATATTTATATAT
>evgR016101c0g1_i01_Hydra
TTTAAAACTAATCTTTTATTTTCTAAATAAGATAGTATGAGTTACCAACTTTATCGCAATACAACATTAGGAAACAGTCTGCAGGAAAGTTTGGATGAGTTGATTCAGTCACAGCAAATTACTCCACAATTAGCTTTGCAAGTTCTTTTACAATTTGATAAAGCCATCAACAATGCGCTTGCAACAAAAGTAAAAAATAGAATTAATTTTAAAGGTCGTTTAAATACATACCGGTTTTGTGACAATGTTTGGACATTTGTTTTGAACAATGTAACTTTTCGAGAAGTTGGTGATGCAGTCAAAGTTGATAAAGTAAAAATAGTTGCTTGTGATGGAAAAGAGTCAAGAGCAAGTGAAGAATAAATTCAAAACAGTAATTAAATTTAAAATTGATAATGATCAATTAAATCAGCTGCATTCTAGTTATTAATAATAATTTTGAGTTAAAATATTTATGGGGAAGGGATATTTAAATTATTTTTTGATGTATTTTTCTTGGAAAAAAATATATTTTATTAACCTTGAGTTAAAAATTGTTGTATTTTATCAATGCCAATAATTATTATTGTGGTCATGATCTAGTTTTTTTTACCTAATTCATTAGTTTTTATGATAAAAAAAAA
>evgR016103c0g1_i01_Hydra
TATTATATTATTATTACAAAATTTATTTATCATTATGTATTCTACTGAAAGGGAAATATTAATATTTTACATTGAGTGGGAAGATAGTTGGGCAGACGAGTGCAATTTTCAATTTGTTAAGCGATTAATTAGCATTGATAAAATGACTAACAATGTCATTTCAATTTTAATGACAGTCTTTAAGCCTCAAGCATTGTTTAGTTTAATGAGCTTTATTTTACCAATTGATTTTAATTTGCCTTTTTTTGACGAGTTTGACAAAATGTGCCAAAATTTAATTCGTAAAAAAAAACATTTTAAAAGAATTTTTAAAGCTTCAAAAAACGAAGCATTTTTTAAAATGCAACTGTCTCCAGCTCCACTAACAAAAAAAAGAAAATCAGCAGATGCAGATTATAT
>evgR016107c0g2_i01_Hydra
CAGCTATGTTTTTGGAAATGTTACATCCTATGAAACTGATGAAGATTTGATCAATTACTATATTGAAATTGTTAATTATCCTAACTGTTTTTTGGATGTCGATACCTCTGATTTAAAAATTATTTATAATTGTAGCGGTGGTTCAGAATTTTCTTTAGTAAATGGAGAAAAAGAAACAATATTAATTCAATCAGCAACAGAATATGATAGTTATATTTGCCATCAAGGATTTGTTCCTTATCTACTACCTTTGGGAGGTAATGAACATACATTTTATATGGTTGCGAATAAAGATTCTTATGTTACGCTAAGGTCAAATGATTACCCTACAAGGTATCTTTGGTTTAATGGTTTAGTATTTATGTTTGAAGAAGAGGTGAATTCAACATTCAAACTTACTTTAAGCCACTTCAGCATTGAAAGGATTGATCGCAAGTCAAGATATTCTGCTGATGTTATCATTGACAAATCTTCTCCAATTTTAAACATGTCTGGTTTAGGTTTTGTGATTAAATGTGAAAGTTCAAATAAAGAAATCAAATTATTTAATGTTACTGGAGGTTCAGCTCTTATTACTGATTTAAATAGCAACACCAGTTACGCGTTCTCTGCTGGTTTTCAAAATGAATATGGCATCTTTGTTTTTGGAATGGTTTCTTTCTATCATACAGAAGAAGATGTTATAAATGCT
>evgR016107c1g1_i01_Hydra
CTGATGTTATCATTGACAAATCTTCTCCAATATTAAACATGTCTGGTTTTGGATTTGTGATTAAATGTGAAAGCTCAAATAATGAAATCAAGTTATTTAATGTTACTGGAAGTTCAGCTCTTATTACTGACTTACATAGCAACACCACTTACAAATTTTCTGCTGGTTTTCAAAATGAATATGGCATTTATGTATTTGGAAGTGTTTCCTATTATCTTACAGACGTTGATGATATAAATGCTAACTGTGGATACAATAATAGTTTTACAGACTTGTCAGGATCATTTCGTTTTCCAGTAATTGGAAGTCATTACTCAAACGATGTTGAATGTAAATGGTTAATTAATGTTGCTCCTAACATC
>evgR016108c0g2_i01_Hydra
ATTTAAAACTAATCTTTTATTTTCTAAATAAGAAGAACATAACAATGCTCGAAGATACTATTCTGATTGTATTTATTGCTTTGGCAACTGCACTTCTAAGTGAAGGGTTATCCTATGTTCTTATTTACAGAACAGAACACTACAAAAAGTTGTCTGTTGAAGTTGAAAAGAATACAAAAAAACTTGAAAAGAAAAAGGAAGCCATAACAGATATTTCAAAACAAAGTGGTCAAAAAAAAAAAGATTGAGCGCGTTGAAGAAAAATTAAAAAATCAAAATCGGGATTTGTCAATGGTTAAGATGAAGTCGATGTTTGCTATTGGTTTTACTTTTACGGCTCTCATGGGAATGTTTAACTCTATCTTTGATGGAAGAGTAGTTGCTAAGTTACCGTTTAATCCAATATCTTGGTTTCATGGACTTTCACATCGCAACATTCGAGGAGAAGATTTCACTGATTGTTCTTTTATTTTTTTCTACATCATTTGTACCATGGCTATACGTCAAAACATTCAGAAAGTATTGGGCTTTATGCCATCTCGAGCAGTTAGTAAGGCTGGAGGGGGGATGTTTTCACCTCCAACAGTCCCTAAATAAATCTACTTATTAATTTACATATTTTTATTTAAGAAGAAAATTAAATATTTTCTTGTATTAATTTCTTCCAATAAAACCTCCAAATGTCAAAAAAAAAAA
>evgR016109c0g1_i01_Hydra
TATAATGGCCTTGAAAATTACTTGGTACAGATCAGAGATGATTTTGATGTTTTTGAAAATAAGGCTGTATAAATTACTGATTGTTCGGAGTATGCCAAAGATTCCAAGAGAAAAAAAAAGAGAAAGCCATGGCTGATGAATCAAGATCTCAACCTCTCACAGAAATCACAGGGAAAATAGATTTTATTAGAAATGTATATTATGTGATCATTGAAACTCTTAAATGTCAACTTAGTTCAAGAAGACAAGCTTATAAAAAAATATCAGATATTTATGGGTGTATACTCACACTGTATAAAACACAGAGTTCTATTGTTATAAGGAACACATGTGAAACTTTGGAAAAATATTTTATTGATGATGTTGAAAACTCAAGCATACTTATAGATGAGTGTGTTCTTTTTTCAAAATTAGTTTCCACGGATAAAAATGTAAAATCTCTTCTCTCTATGTATAAGTACATCAAGACCAAAGAATTAGAGTGTATGTTTCCAAATTTACAACAGTACAACAGTTTCAAACTGCTCAGGCAAAAGAGCTTTTTCTGTAATTAAAAGAGTAAAATCCTACCTTGAGGTCAACCATGAAGGAGGAAAGACTAAATAAATTAGCAATATTTAGCATTGAAGCAGAATTGGTTGAAAAATTAGATTTTATTGATACTATTAACACATTTGCCCATCAAAAAGCAAGAAAAAGAAAAGTCCAAAACAAT
>evgR016117c0g1_i01_Hydra
AGAGATATGGGAAAGTATAATGCAGCTTTAGAAATTTATTATTTGGTTGATAAAATAGAAACTGAAAGTTATGGCATTAGCCATTTATCTACCATGGAAACAAAAAATAATATGGCACTCTGTTTGAACGATATAGGTAAATATAACGAAGCATTAGAAATTTTTTATTCCCTTGATAAAATACAAACGAAAATTTTAGGTAACAACCATCCGTCAACATTGAACACAAAATGTAATATTGCACTCTGTTTGAACAACATGGGAAAGTATAATGAAGCTTTAGAAATTTATTATTCTGTTGATAAAATACAAACTGAATTTTTAGGTATTAACC
>evgR016136c0g1_i01_Hydra
GTTTATTTTGGAAACAATTTATTCTTGCTTTCTCTTTGGCTTCATTGGTGGAGATCTGACCTTTCTACCATCCTTAGAACGGACACGAACGATACGACCGTGGATAGCACAAGAGACACAGTAGTGGAGCTTCAAGTACATCTTTGGCAAATCATAATCCTCATATGCGGAAGCATCACTGATATCCTTAACAGCAGCAGATTCGACAATGCTTCTAACCAAGAATTTCTTGATGGACTTATCCTTTGGGACGCTCTTACCACAGTTATCGCATCTGACGGCTCTAGTCTTACCTCTGTTAACCAAAGCTCTACCGTTATTCTTTCTCTTGCTAGTCATCTGATGAATAAGAC
>evgR016141c0g1_i01_Hydra
ACAACAAAATGAACAGAGCAGTTTCTCTCCTTGCTAAGAAGCCAACTTTGGCCGTCCAAAAGAATGGTATGGCTTTTATGGCCTTGAATTTCTCTTCTCCACAACAACAAACTAGATTTATTCAAACTTTGAGAAAGAGAAACAATCCTGGATTCTTGGATACCGTTTCTGGAATGTACGAAAGTGCTGACTACAAACTCGATTGTAAACCATTCGAATTCAGAGAAGGTGAACCATTCTTCCTTTTCAATTATTTGGGTGTCGTTATTGTTGCTACTACCATTGCTTTCGCTTTCATGTTGTGTTTCAGTTCTTTGGCTCGTTACACTTTGGTTGATGTTCAAGTTGTTGTCAAGAGAACCAACCCACATCCATTCTTGAACATTAAGAATGGTGCTGATATGCAAGCTTCTTTGAATCCAATTTACAGATTCTGTCAACCATTCTCTCACACCATTGATTTGAAGCAAATGTTCTACCGTGAATTGAACAGAGCAAAGACTCAAAACTTGTAAATTTTTGAAAAATCAACTATTCTCTCATCAATATAGGAACAACATTCG
>evgR016149c0g1_i01_Hydra
CATATTGATGCTATGAATTTGGATTGCAACAAAAAGCAATTAAGAACTACACTAGCGCATCTACGAGATAGGTGGCGCAAGTCAGATAGCAACATGGCTCGATTCGAAGAAAAGAATAAAAAATGGCTGCAAAGCCATGTTCCTGAAGATATAAATATAACACCAATACCTATTGGAAGGCCAATTAAGGCAAACTTTGATGACTACTCTACTAAAACCAAAAAACATAAAGTTAGTCACTTAACTGGAATGTATTCTGCTTCTGCGTTGCAATTTGCTTCAGAAATTAAAAGGAGAAAAGAAAATGCAATTGAAAAAACCTCAAAAAAAGATTTTAGAATATCTAAAGATGAAGCTCTTTGTCTTCTTTTAGATGCTAGATTAACAAAAG
>evgR016157c0g1_i01_Hydra
CCATTCCGCAAAATTTTCCGAACACATTTTGTAAATGCCATCAGTAACTATAGCTTGTTGGTTTAGATATAAGCCATGGTGCTAGTAGCGTTGTACACAACCTATATTGTTGTTCTAATTGCTTTATCAACTTCTTTAAATTTGATCATATGCTATGTAATCATCCGCAAAATGAAAGCAGTTGAAATTCCGCACCTTTTTATTTTGAGCATTTCAATATCAGACATAGTACACACCCTAGTTGGGCTTGTTTCTGAAGTTTGCGTGTTGCACAAAGTGACGTCATTAAAGAATAGCTACATAAGTGTCGGAGCATCTTTTTTAACATACTCTATTTCTGTTTCAAATATTATGCAAATTGTTATGATAGCGGCTATAAGGGTTATCGCATTGAAACGTCCTATGTTCTACTTTGAAAACTGTGACAAAATTTCGTACAAAATGGCATCACTTTTTTCCTGCTACTTTTATGGATTATTATGGGCATCGTTTCCTTTACTTGGATGGTCAAAATACGAAGAGGATTTAGATCAAAAGCGATGTTCTTTAGATTGGAATTTAACTAAATCTAACACATTCTCATATCTGATGTTTGCATTTATATTTTGTTATAAGTTGCCATCAACCATGCTTATTTGGGCTCTTAGAGTAAAACATCAAACAGTGGATGAGCACCGCATATTTAGAGAAAATCGACCTAATCAACAAATTGAAATATTAGAAAAGGCTTATTTAAAGGTATTACTTTGTTCGGCAATTGCTTACTTTGTTGTTTGGACCCCTTATTCCGTGGTCAGCCTACTTTCAATTTTCAAAGTAAAAATTCCAAGTATTGTATTTACACTTTGTGCATTATTTGCCAAACTTTCAGCGGTATTAAATGCATTGGTTAATTGTTACATTAACAAGTCATTTCAAGTTCATCTTCACGATTTAAGGATATTTCAATGTTTTCTTAAAGGAAAGAGGATAGGACATGTAAATACTTTAGGGGTTGAGGAAACGCAAACTTGATTTTAAAAGAAGAAAACATTTAGTACCAACACGTAATCAAATAGTTGTTGACTTTAATTAACACTTACTAACTAACACTTATAGCTAATACTTTTATTTTTCCACAAATATATAAAAAAATATATTCATCTGTTGTGTGAATATAATAACATATATACATATACATTACATACATATTCACGGCACCGTAGATATTTAGTTGTACTTTTTTTGGTATTAAAAAAAAAAA
>evgR016163c0g1_i01_Hydra
CTTGGAAGTTTTTTCAAGATGGTTGGGTCCAAACTATTTTCCATATGAAAGTATGTCAAGATGTGATTATCTTAAAATGCGACGTTCGACCATCATATAGGACCACATCTCCTAACCATAAGCCTTGGGTTGCTCTTGACTCTCTTGGTAAAGTTTTAACTGCTCACTGTGATTGTATGGCTGGTCTTGGTGAAACATGCTCACATGTTGCAGCAACTTTGTATAAAATTGAAGCTGCTGTTCGTATTGGCATGACTTCAAGCACACCAACTGATTTGCCTTGTCAATGGAATCAAACATTTACAAAAGATATTGTTGGTTCCCCAGTTTCACAAATCAATTTATACTCTGATGCTGCAAAAGAAAAACTTTCTAAAAAAAGAATAAGAAGACAGCCTGTTTCTCCCACATATCAAGAAAAAATGGAATTTTTATCACAAATACATAGCATACAACCCAAAACTGCTGCTCTTCAATTGTTTAAAGAATTCGACATGGACTTTATTCATATGGAATCCGCAGTTGTATCAAAGTTACCAATATCAATGAGACAATTTTTTAATGAAAACTATAAATTACTTGATAATGAG
>evgR016179c0g1_i01_Hydra
GCTTGAAGTTATTACAATGAATGCAGTGTTTACACATGGAAAAGAAAAAAAAATTATTGAAAATCTTACTAAATCTAGAAATTTTATGCCAAAAGCATTTGTTATAACAGCAGGCGCTTATGTTGCATATAGAGCATTAAGTTATAGTTACTCAAAAATATCACACAAAAATGTTACTGTTAAGCATGAAAATTTTGGACAAAAACAAACTGCTGACAAAAAGTTGTTTCTAGAAAAGTTATTGGAATTGCTAAAGATAGTACTTCCTGGTATAAAAACTAAAGAGTTTGGTTTATTATCAATCCATTCTGTGGCTTTAATAACTAGAGCATTCTTGTCAATATATGTGGCAAAGTTAGATGGTAAGTTAGCTAGGTGCATTGTAGAGAAAAACTTAATCAAATTTATAACTTTGCTTTTGCAATGGATTGGTGTTTCTATTCCGGCAACGTTTATTAATTCTCTGCTCAAGTATTTGGAAGGTAAACTTGCATTGGCATTTCGAACTCGTTTAGTAAAACATGGATATGAACTTTACTTTCAAAGACAAACATACTACAAAATTGGAAATCTAGATTCTCGTTTATCTGCCCCTGATGAGTGCTTGACCGAGGATATCAGAATGTTTTCAGAATCTGTATCTCACTTATATTCACATTTAACAAAGCCAGTGCTTGATTTATTGTTAGTATGTTTTTCTTTAAATAGCATGGCAAATAAACGTGGTCAGTCGTGGTTAAAACCTGTAATGTTAGCTACATGTGTGACAGTATCAACAGCTCATTTATTAAAGAAATGCTCCCCTAAGTTTGGTAAACTTGTTTCTGAAGAATCAGTTTGTAGAGGTAATTTAAGAACTGTTCATTCACGAATCATTACAAATGCCGAAGAGATAGCTTTTTATGGAGGACACAAGGTTGAACACAACTTGCTATTGAAGAATTACAACATGTTGGTCAATCAGGTAAACAAAATACTTGGTCAAAAACTCTGGTATGTGATGCTTGAGCAATTTCTAATGAAATACTTGTGGAGTGGGTTAGGAATGATAATGATTGCTCTCCCAATTATGATGGGCAAAGATGCTTCATTAAACATGTCAGAGCAAATAAGTGAAAGAAGCCAAACATTTACTACATCAAGAAATCTCCTCATATCAGGTGCTGATGCCATTGAAAGAGTTATATCTTCTTTAAAAGAGGTTCACGAGTTGACTGGATTCACTAATCGTGTTTATGAGATGTTTAATGTCTTCAAGGAGATTCACAATGATAAGTTTGTTACAAGCAGTTCTTCTAAGGCTTGCACTTACACACCTGTAGATAATCTGGAATCATCTTTTGAGGTGGTATTTAAATACGGGCAAGTAGTATGCTTGAATGATATGCCAAGAGTTCAGCCTGAATATACAAAAGGAGATATTGTGTTAAAAAATGTTCCTATTATTACGCCTAATGGTGATGTAATTTGTTCAAGTTTGTCAATTCAGATTGTTCAAGGTATGCATTTTCTGATAACTGGTCCTAATGGTTGTGGTAAAAGCTCATTATTTCGCATCTTAAGTGCTTTGTGGCCAGTTTTTGGTGGAAACTTATGCGTGCCTCCTGTTGAAGAAATGTTTTACATTCCTCAACGTCCATATATGACTCTTGGAACATTACGTGATCAAGTTATATATCCAGATAGTGTATCTGATATGAAAGCAAAAGGCTGGACTGATGAAGATTTGGAACGCATTCTTGATGAAGTGTTTTTAAAATATGTTGTTGGTAGAGAAAATGGTTGGGATAGTATCAGTGATTGGATTGATGTGCTATCTGGTGGAGAAAAACAAAGAATGGGAATGGCTAGAATGTTTTACCATAGGCCGAAATTTGCTTTACTTGATGAATGTACAAGTGCTGTAAGCATAGATGTTGAGGGGAAGATTTATCAATCAGCTAAAGATTTAGGAATAACCCTTTTGACAATCACTCACAGACCATCTTTGTGGAAATTTCATACACATTTGTTACAGTTTGATGGCCAAGGTGGTTGGCACATCAATGAAATGGATACAAATGCTCGTTTAAGTTTACAAGAGGAGAAAGAAAAATTAACTACACAATTAGCAGGTGTTCCAAAATCTCAGCAGCGTTTAAAGGAAATATGTGATATATTAGGAGAGAACGAATAAATGATATTAGATAGAGTGAATGATAAAAAGATGGACAATTTAGAATTTTTAAAAGAAAATAAAATATATTGAAATGAATATTGAAATAATAATTTATTTATTTGTGATATTTAAAATAAACTTAAATTTAAATATATTTTATAAAATTTCATTACATTTTAAGTATTGATTTTTCCATTAGGCTCAGAG
>evgR016183c0g1_i01_Hydra
TTTCAAAACATTTCCAAGTATTTTTATTGGCAGCAACACTGCTGATGTGAAAATCGAAGGCAGTATCACTACAATTGCCAAGAGCATTAGAACACTGTGGGCTGAAAAATTTGTTTCTGTATTCAATGGTGTCTTTGAAAACAGTGAACAAGGAATCGAATTGGCAAAAACAGCTGGTCAATTCTTGGTGGATCTTGAAATTAATCAAGTTCCCACACCAACCAGTCCATCGACCATTGAATCATTCTATCTTGAGATTGTGAAAGGTGAACTTAAAACAGTTGCTCTAGAGCTTGTGGCTGTTATTTCTTTGAGAGCTGTAGATGATATTAGAGATTTCGTTTTTGAGCATTTGTCAACCGAATGTGATGACTAACTGGCTATCAAATTGTAAAGCAAATCAAGTCTCTGAAACTGTTATTAATAAACAAATGTA
>evgR016183c0g2_i01_Hydra
CAATGCCTACACTCAAATCCAAAAGTTTCCCAAGCATTTATATTGGCAGCAACACTGCTGACGTGAAAATCGATGGAAGTATCACTACCATTGCCAAGAGCATTAGAACACCATGGGCTGAAAAATTTGTTGCTGTATTCAATGATGGTGTCCTTGAAAACACAGAAAAGGGAAACAGATTGGCAAGCGAAGCAGGTCAATTCTTGGTGGAACTTGAAATTAGTCAAGCTCCCACACCAACCAGTCCATCAGATATTGAAGCATACTTTCTTGAGATTGTGAAAGGTAACCTTAAAACAGTTGCTGTAGAGCTCGTGGCTGTTATTTCTTTGAGAGCTGTAGATGATATTAGAGATTTAGTATTTGAGCATTTGTCAACCGATTGTGATGATTAACATGGCTTTCAAATTGTAATGTCAAT
>evgR016186c0g1_i02_Hydra
GCACAAGCGATATTGATTGCCCAATATGGTTATCGCTTAGTTGAGGTTTTGAAACACGATAAAGAATCAGAACAGGAAAATGTTATAAGACTAGTTCTAGCGCAGATATTTACTGCACTTCGAGATGTTGGAAGTATAATTAACATTGTAGATGTTGTCGATGAAAATTATATTGAAAGTGTTACTGCTGCCTGTAAAATATATTTTTTATTATTTGCACTTTTCCCTCTAAACTGCAACAGTACAGTTTGGACTATTGGTTATGTTGTTCCATTTCATGCGGAAGAAATTTTCAAGGGTTATGGAGTTGGTTATGGGATTCTTTCTATGCAAGGAAAAGAATCCAAACATTCATCCATTAAACAAGAATTAAAAACTTGTAGCAACAGATCTACAAAACAAGACGAAAGAGGTAAATGGTATCAACTGATGCGGTCTAGCTATGTACGAAATTTTTACTTGCCTTATCATTTTCCAATACCATCCATGTATCAATCTCATTTTCGACCTCGTAAACCGGTTACAGACAATGAATGCTGTAACTGTTCACGTTCACTACAATCTGAGCTTTTATGTCAAGTTTGCATCGAGTCTGTTTCAATTTTAGATTGTGTCCAGAAGAAATATTTGACAGAAGATATTTTATTCATAATAAAGCCTGTTAAATGCAAAATATGCAGCAAAAGATTTACAGATGCATGTTCGTTGAATAAACACACAGCAAAAAATGAATGCAATCAAGTTATTCCTAAACTTATTCCTCAAGCATTAACTATTTACCAGCTTAGAAATGAGCTGAAAGCTCG
>evgR016192c0g1_i01_Hydra
GTATGATTGCCAGTGGTCAAATGCTTGGCTTTGATCAACCAATCATTCTCCATTTGCTCGAAGTTCCAGTTGCTATGGATGCTTTGAAGGGTGTTTGTATGGAAATTACCGATTCTGCCTATCCATTGGTTAAGAAGGTTGTTGCCACCTCTGATTACCAAGAAGCTTTCTCTGATATTGATGTTTGTATCATGGTTGGTGCTATGCCAAGAAAGGAAGGTATGGAAAGAAAGGATTTGTTGGAAAAGAATGCTTCCATCTTCAAGGGACAAGGTGAAGCTTTGGAAAAGTATGCCAAGAAGTCTGTCAAGGTCCTCGTTGTTGGTAACCCAGCCAATACCAACTGTTTGATTGCCAAGACTTATGCTCCATCAATTCCAGCTGAAAACTTCTCTGCTCTCACTCGTTTAGATCATAACAGAGCTAAGGCTCAAGTTGCTGATCGTCTCGGACTCAATGTTTCTGATGTTCACAATGTCACCATTTGGGGTAATCACTCATCCACTCAATATCCTGATGTTTCACACGGATATGTCACAATTAACGGTGAAAAGAAATCAATTGTTGAAGCTATTAATGATAGTGCTTGGTTGCAAGGAGATTTCATTAGTATTGTTCAAAAGAGAGGTGGTGCTGTCATTGCTGCCAGAAAGTTGTCATCTGCCACATCTGCTGCTAAGGCTATTGTTGATCATGTCCATGATTGGTTTTTGGGTACCCCAGAAGTTGAATGGGTCAGTATGGCTATTCCATCTGATGGTTCATATGGTATCCCAGAAGGTGTTATCTACTCCTTCCCAGTTACTTGTAAGGATGGTAATGTAACCATTGTTCAAGGTTTGAAGATTGACGATTTCTCCAGAAATAAGATGGAACTTACTGATAAGGAATTGAGAGAAGAAAAGCAAATGGCTTTGAACTAAATCTGTA
>evgR016193c0g1_i01_Hydra
ACCAAATTAAATTTAGAACCATTTTTTGCCGAATTTACCAAACTTCTTTCCAAACTTCATTTTTTTACCGAATTTCATCTTCTTACCATGATGATGATATCCTCCGTGGTGTGTTGGTGGATAAGCATAAGATGTTGTGGTATTGTATGTCATTGGTGGTTGAACATAACCAGTTGGTGGAGCATAATATCCTGGTTGACCCATAGGTTGCATTGGTTGCATAGGCATTGGTTGACCCATTGGTTGCATTGGGTAACCTTGTTGCATTGGGTATCCTTGTGGTTGCATTGGTTGCATTGGTTGCATTGGATAACCTTGTGGTTGCATTGGCTGCATACCAGGTTGCATTGGATATCCTGTTGGTGGCATTCCTTGTGGTGGATAGCCGTATGGTTGTTGGTATGGGTTCATTATGATACAAGGGG
>evgR016210c0g1_i01_Hydra
GCTCAACGAAAGAAGACTCTGATGGGATTTTACAGTCAGGTATTCGCATTAAACGATCTCATAAGAAGGAGCACAAGTTAAGTGCAAAGTGGTTACCAATGGTGCGTAAATTATGGAAGTATTGGCGTCACAGAGGAGCACGCGAAGTGATGCATGCTATTGTATCTAAGCTGACCAGCCATACAAGCAATAAAATTACAGACCATTTTGGAGAACTGTTAGAGACCTCTGTTTTAAAAAATGATCGCCTTTTGAAAAAAAAAAATCAAATGTCTTCGTTTGTAAAAGCACCACCTATTGGTCCAATGGAAAC
>evgR016212c0g1_i01_Hydra
TACAACCTACCAAACTGCACAACCTTTTACTGAATTTTCACTATCTTTTTCACTAACTATCACTATTGACCAATCTTCCATTAAAACAATAGTTTCGACAAGACCAGTTTTAAACGTTACCTCAAAAAAGTTGTCAAACAACAACTCAAGGGGCGAGACCAAGTTGCACGCGCAACTTTGTTTAGCTTGGTTAGACTTTATGTTAGAAGTTACCTTCCACTGTTGGAGTAGTAATAGTTTTTGTGTATTTTGTACTACTACTACGACCTCACACAGTATTTCAAGAGGCACCCCACACTCGACAGAGTTGAGCCTTATAGATTAAATCGACTAGCTCCATCTCTTAAATAATGAACACGCACCGTTAATTTAAAGATTTAAT
>evgR016216c0g1_i01_Hydra
AAAACTTTTTTTAAATGATCTTAACTCAAAGATTGTGAATGGAATACGAATTCGACAAATTCGTGTGCAGCCAAACTTGTGTAAGAAACTCAGTTTTGCAGCTAAGTTTATGACAAATGCCTGTTTATCTTCATATAAATCTTCTCTTGAAGAAACACGTGATTTTGATTTAAACTGGACAATTCCAAAGCAATATGACTCTTCAATTACTAATTCAACAATGCCATGGAGGTATCAAAATAGGCTAGAAATTGATGGTTATCCATATACAACTGATTTAGACACTTACTACGGGGGAGGGTATGTTGTAGAACTTTTTCCAAAATGGAAAAACCGAGCTACGTTTAGCAAATTGAAGAGTCTAAGGTGGATTGATAGACAAACACGAGCTATAATTATTGAGTTTTCTTTGTTTAATGCAGCAACTAATTGTTTTAGCATGGTAACCATGGTATTAGAATTCCCTGCTTCTGGTGGAGCCATATCAAGTTACTCTGTACTTACATTTGAGTCATCAACAACAATTAAATATAGTTTTCATATCGTGTTTGTTTTTGTTACAATAATATTCATAACTCGAGAATGTCGCTTTTTATATCGAAAAGGTTGCAGCCACTTATTGAACTTTTGGAACTTAGTGGAAGTTGCAATGATTATTTTTTCTGTTACTGAAGTTGGATTTTTTTTATATAAAGAATATATAGCAAAGTTATTGCTACAACGGTTACCATCCAAGAAACCACAGACATTTATTAACTTTCAATTTGCATCTTACTGGAATCTAATGTATGTTAATATTGTGTCAATGATTATCTTTTGTGTTACTTTAAAGTTTATTAAGCTTTTGCAGTTTAATCGTCGGGTTTCGATGATATCCTATACCCTACAAGCAGCATTTTATCCTTTATCTATGCTTGGATTTGTATTTTTCATACTTCTGGCTTCGTTTTTGTTTTCGTCAACAATTGCATTTGGGACATTTATAAAAGAGTATAAAACCTATTTTGAAGCAATAGGTTCTATCGTCTCGTTGTTATTGGGAAAGTTTAGTTTCAGTCAGTATAAAAATGCAAACAGTGTTCTTGGACCAATATTCTTTTTTGGTTTTAATATCGTTATGATTTGGATAATCATGAACATGCTTGTATCTATTTTAAACGAT
>evgR016226c0g1_i01_Hydra
TTTGGAGCAGAAAAATTTGATACCATGTCTCAGTCATTTGTAAATTCAATATCTAATTTGTTGCTAAATGTTAATCCACCATCATTCTCAACAACTCTTGTACAAGATGATGTTAGTATGGTATCTACATTAATCAAATCTATGCACAGCTATTTTGTAGCAGCTCAAACTTACAAAGTTCCTGGTGAAAATAGAACTCTGGGTGAAACTAATCAGTTTGAGTTTTTGTTAAAAAAAGATCTCTTTATTGATTTGAGCAACAGTTCAATAGGCTCATCAGATGGTGGTTTTTCTTTTCCAAATTCATCAGATATGTTTAATGAGTCATTAAAAAGTTCCCAAAG
>evgR016229c0g1_i01_Hydra
ACCAATGCCAAAGAAGAAAAAACCTTGTCAACTCAACTTGCCGCTGCTTTCAAACTTTCAACCAAGAGAGTTTCTTCAAAGAATGCTAAAAGAAGATCACCAAGAAAGAACAGACGTCTCGTAAAAAAGAAACGTACCTTGTGCCGTCAAATCAGAATTCTCAAGAAACAAATCAAAAAACTTTCCTTCAAATTGGCCAAATCTTCCAAGAAGACCATGAACAAGATCGTCAGAAATATTAAAATTAGAAAGTCATTGATTCGTAAATTGAAGAGAAAGCTCAGATCTGTTAAGAGAAAACTTCAAAAGAAGAGAAATGCTCAAAAGAAGAGACAACAAAAAAAGAGACAACAAAAGAAACGTCGTTCCGTCTCACGTTTTGAAAAACTCAAGAAGAGAGTTCTCAGATTGGCCAATAAGCAAAGAAGATCAAGAAGAGCAACCAGAAGAGTTAGAAAGACAAGAAGAC
>evgR016234c0g1_i01_Hydra
TAACCTGTTTTAGTGCAAAAAGGAAAGTAAAAATTGTGTAGTTTATTTGTGAAATATAAATAATGGAATGGAAACTATAAAAACATTAACAAGATCAGATATTAATGAAGACTTACAGAAAGCATCATTTGATGATTTGGTTTTTAGTGCAAATTATTTCACAGAGTTAACTGGAAAAAAAACAATTGAATTTGATTTGCAAAATAACGAAGTTGATAAATGTTTTTCAGAAGTGGAGAAGTTTTTAATTCAAGAAAACGAGCTAATTAAATTTTATGAAGAGCTAAAGTTGGAAGACGTTTTAATAAAAGAAGAAATACAGAGCAAAATAAAAAACACTATTATCAAACTAGAAGGAAAAGCTGAAGAAACAAAGAACTAAGAAAATCTATTTCATTATTACAACTAAATATTAGTAAAATATTACTAAATATATCTTTAATATCTATACAACTTAACACAATTAAGTTAAAGTTGTAATAAATATACTCTTATAAATCATTTTATATGTTGTATATAAAAATAAATTATTTCAATAAATTAAAATTACATTATTATATTAAATAATATTTTACTATAAAATATGATGTTGATTAAAACCTTTATTGACACATAAAATTTTAT
>evgR016253c0g1_i01_Hydra
AATAAACCAAAAAAATATTTTGAAAGACTGTATAAATCTGTATATATAAAAATAAAAAAATTTGAAAAATTTATAAGCGTTAATAAAAAATATTTAATAGCTTCATATGAAGTTCCCTACTGTATTGCTCAAAATCCAAAAGCCTTTCACCATCGGAGAAGATATTACCCGCAGCTATTAAAATGTTAGAAATACTACATGGAAGTAAATATAGCGATGATATTCGAAAAACTCTTTTGTCGAACGAGATTGTCGCGAACAGAATTTCTGATATTAACAACGACCAATTAGTGCAACTTATTACAAGAATTAAAAAAAGCCCAAAATTTTCAATTCAGTTGGACGAAACAATGGATAATACTAAACTGGCTCAGTTGTTAGTATATGTCAGGTACTTTTATAAGGAAAATGTGGAGGAAGAATTGTTATTTTGTCGTCCTATGAAAGACCATACTACAGGGAAAGACATATATTGTAAAGTTGTAAATTTTTTAGAACTGAAGGTTTAAAATGGAAAAATTGCTGTGGAATATGCACAGATGGTGCATATGAACAGATACAAAACATTCGCAACCCAGTGGGCATTTCGACG
>evgR016253c0g2_i01_Hydra
TTGGATTTTTGGATAGAAATAATTTTTGGCCATGGATATATGGCTGAACTTAAACAAGAATATTAATGACGCAGATAATAACGCTACAAACCCTCCTCAGGTAGGACAAAGTATTTAAAAACATTTAATGTGCGGAAAAGAAAATATGATGAGTCTTTTCTTCAATTTTTACATTTCAAAATTATAAGGGTAATGAACAACCACTTTGTATGATCTGCAATGAATTATTGGCTGCAGAGAGTATGAAACCATCAAAATTAAAAAAACATCTTGAATCAAAGCATGTTTCGTACGTCAATAAACCAAAAGAATATTTTGAAAGACTGTATAAATCAATCAATATAGAAAAAAAAATTTTTGAAAAATTTGTAAGCGTCAATGAAAAATATTTACTAGCGTCATATGAAGTTTCATACTGTATTGCTAAAAATCAAAAGCTGTTCACTATCGGAGAAGATCTTGTATTACCTGCAGCTATTAAAATGGTAGAAATACTACATGGAAATAAATATAGCGATGATATTCGAAAAATTCCTTTGTCAAACGACACTGTCGCGAATATAATTTCTGATATTAACAACGACCAATTAGTGCAACTTATTACAAGAATTAAAAAAAGCCCAAAATTTTCAATTCAGTTGGACGAAACAATGGATAATACTAAACTGGCTCAGTTGTTAGTATATGTCAGGTACTTTTATAAGGAAAATGTGGAGGAAGAATTGTTATTTTGTCGTCCTATGAAAGACCATACTACAGGGAAAGACATATATTGTAAAGTTGTAAATTTTTTAGAACTGAAGGTTTAAAATGGAAAAATTGCTGTGGAATATGCACAGATGGTGCATATGAACAGATACAAAACATTCGCAACCCAGTGGGCATTTCGACG
>evgR016257c0g1_i01_Hydra
TTATCGTGGAAGATATAAGTACAATAAAGAACAAAATATCAAAAAGCATCGGTATTCTCTATAAAGCTAAATCAATGCTCTCCCAAGCAAATCTTAAAATTCTCTACTTCTCCTATGTTCAAAGTTATCTAACTTACGCTAATATTGCATGGGGGAGCACATACAAATCTAAATTAAATTCTCTATATATACATCAAAAACATGCATCAAGACTGGTTTATAATAAAGATAAATACACTCATGCAGATCCCTTACTAAAATCTTTAAATGCACTAAATATATATCAAATTAATATTTATCAAAATATTTTATTCATGTTAAAATACCAA
>evgR016269c0g1_i01_Hydra
CAGAAGGTTTACCATTGGCTGTCACAATTTCATTGGCCTATTCTGTCAAGAAGATGATGAAAGATAACAACCTTGTTAGACACTTGTCTGCCTGTGAAACAATGGGTGGTGCTACAAATATTTGTTCCGATAAGACTGGTACTTTGACTTTGAACGAAATGAGAGTTGTAAAGGCTGTCATTGCTGGTGAAGAATATCTTGATGGTTTGCCAGATAACACTGATGGAATGCACACCAAGGTTACTCAAGTCCTTTCTCACGGTATTGCTGTAAACTCTAAGGCTTCTTTGAACAAACCAAAATCAGGTAGTATGAAGGAATATGAAGTCAGTGGTAACAAGACTGAAGCCGCTCTCTTGATTCTCTTGAAGGATTTGGGTATTGACTATGTTCCAATTAGAAAACATTACACTGAAAATGATAATATTGAAAAGTTGTACACTTTCTCATCTGCAAAGAAGAGAATGGCTGTCATTGTCAAG
>evgR016282c0g1_i01_Hydra
CTCCTCACAATGATTCACCAAACCAACCTTATAATTGAGGCAAGAAATGCAACTCTAGATGTTGAAATGGAAAACATTAAAAATCTCATGGACATCATTCAACAGATTCGTGGAAAATGGGATGCTGTTCTGAGTGAGTCAAAATTATTTGCATCGAATATTGACATTTCACCAGAATTTTCTACCACTCGAAAGTTGAAAACACAATTTGATTCAGAACAGCACTATAAAAGCAATGTATTTTATGTAATAATCGACTCTATTCTAGCAGGCCTTAAACGTCGATTTGAGTCACAACAACAAACTTGTAGTTTTTTTGGATTTCTTTGGCACTTCAACAAGATGAGCAATGAAGAACAACAGAGTGCCACAACAAACTTTCAAAAAAAGTACCACAAGGAAATACTGTCTGATATTTCTGATGAGATAATCTTTCTGAAACAAATTTACTCCACAAACTTTACATCAGATTGTAAACCAAAAGAATTGTTTCAAGAAATATTTGAACTTGGAC
>evgR016290c0g2_i01_Hydra
CTGGGATATCAGCAAATGTAGAAGAAGAGGAAAGAACTTTCAATACAATTAAAACTATAAACAGTTCAACTACAAACCACCATAGTGAACATGTGATATTAAACACCTTGATTCGACTACAATGTCATAAAAACTTTAATGAGGGCTACAAAGCTCAAACTCATCAACATGCAATAAGAAAGATTGTAAAAGAATATGAACAAACCCCAACCGAAATTCCGTTCTGGATAATTACTACATTTATACATTTGAAAGTACCCATGGGTATGGCAAAGACATTTAGAGCGCATATCAGAATACTTATCAGAAGGTAAGTTCATTTGGTGGTTTGAGACAAAAAATGGAATAACAT
>evgR016307c0g1_i01_Hydra
GTAATATTTACGCTATTTTCCCCCATCAAACATGGTAGTTTATTTTGAAATGCTACAATGATCTAAAATAAAGTTTTATCCATATTTAAATGGAACCACCAGGAAGCAACAATGAATGCAACAATAGATTGAACTCATCAAAAAATAGAAGAAATTATCAAACTTCTAAATTGCATCCAAAAGTTTTGGATAGAGTTGGCTACTTGATTGGAGATGATTGGAGAAGTTTTGGGAGCCAACTTGGTATGAGTTTTTATGATCTTGAGCACATCAACTCTGATAACAGTAAAACTCATGATAAAGCTGTGAATGTCCTAAATAAATGTGAACAAAAGTATGCCTGTTCTTCATGGGAGATATTAAAAAACGAGTTAACATTGTTTGAGCGTTTAGATGTGATTAAAGAAATTGAAAGGGAATTTGCAGATTTATTATCCAGTAAAGAAGAAAAAAAAAGTAATGTAAAAAGAATCCCACATGGCTGCATATTTATGTTTGATTTACACGAAGAAGAAAGTTGTACTGAAGTTGAATTTCAATTACATCATGAACATAATAATTGGAATTCTATCAGTAACAAGAAAAATAAATTTGCATATTACTCTGCCAAG
>evgR016315c0g1_i01_Hydra
AGCGAAGAAATTAGTTATAAATAGTGTGGATATGGATGTAAATAGTAGCAATGTAAACATAGAGTCAAATTGTGTAGATACGAATGTAAATATTACTAAAAGTGTTGAAGATATAAATGTCAATATGATGAGAACTGTTGATTTTAATGTAATTGGTTCGGATCAAGTCAGTGCTTCTGCTAAAAAAATTTCATCATCAAGACCAATAATATCGTGCAGCTTAATTAAAGGATACCGTTTTATAGATTTAGAAATTCTTAACTCTGTAGTTTCTACTCTAAGATGTCCAGAATGCATGTCATTAGAGCTAATTCTTTCAGAAAAAACAAACAAGAGAAAAGGTCTAGCTTCTTGTCTCTATATTACCTGCTTAATTTGTAACTACTCAAAAGAATTTTACTCATCGACATTCAACAAAAAAAAAAGTTTTGATATAAATACTAGGGCTGTATATGCCATGAGATCATGCGGACAAGGTTATTCTGGATTTTCAAGATTTGTTTCATTGATGGACATGCCTAAACCAATAACTATTAACAATTATAATAAAATTGTTTTAAAATTAA
>evgR016316c0g2_i01_Hydra
ACTTACGTTTATACAAGAAAAATGAGAACTCGTAGTAGAAAAAGTAAAACTCCACAAAAAGAAACTATAGTTGAAAAGTGTGCTATAGTTGAAAAAGAAGTTACTGGTGAACAAGAAGCCACAGATAAAAAAAAGGAAGTGGAAATTTGTGAAAAGGAAGATAATATTTCTAATCTAAATTCAAATTTGAAAAGCATATCTGATGAAATATTATTATCAGATGACGAAGCTCCGCTTGACATCAGCCTTAAGGTAAGCAAGATGCAATCTGAAGAAGCAAGAAAAAAACAAATCGAGTTGCTAAAAGAAATTAAGAAAAAAAAAACATCTGTTCGATCTATGAGAGAACAAAAAAATAAGAGACAAAAATTAATGAAGAAAATATTTATTCCAGAAAAGCTAGATGAAGAACTGCTGCGTTGTGTTGATAAGAATGTTTCCGTTAGTAAAATCAGTTTGACAACTCCAGTTTTAGCTAATAAGAAAACAACATTTTCAGATGATAGTTCATCAAATGATGAAACTGAAAATATGGATTGTAAAATTACCTCCTGTGTTCTTACAGATGTGGCTAAACATAATCAAAACAATCAAGCAAGAGTTTTTCTTCAAAATCATTTTTTTAATAATAAACGTATCTCGAGAAAGAAGATAGTGGTATAATTATATTGATAAAGTTTCTTCAAATTCTAAAAAATTAATGCTTGTATAACGTTTTATTAAATTACAATATAAAAACTGAAAAAAAAAAA
>evgR016319c0g1_i01_Hydra
AAAAGTATCAGAAAATAAATATGCGTTTAAAATGTATAACGATAATTGCACAGATGATTCAATTTTAAAAGAAAACGAACTTTATGTTGGAGGTGATGTTCTACTTGTAGACGAGAAAAAAGAGAAATTAATACATATAGATGAGATAAATAAAAAAGACAAATCAAATACAAACATAGCTCTAGAAAAACCGGAAGCCGATAAATGCAATCAAAGTAATCATCAAAATCCCTTGACTAAAAAACACCTAAAACAAACTTCAGAGTTACACGATGAAGAGAGGAAGGCTAAAATAAAGGACGATTTTATAGAAAATCAAGACAATAAATTAAAAGAATATACGAAAGAGATTGATAAACTGAAAAAAACGTGTGAAAGAAACAATATTGTTTTTCTGTATATACAAAGAGAATATGAAGCGATAATTTCATTTGTAGAGCAAAAAAATGCG
>evgR016325c0g1_i01_Hydra
AATAACAATAAATGTACCCTCTACTCTCCTTGTTCTTGGACCAACGGAATATGTACTAACACAAATGGCAGCTATTATTGTTCTTGTAGCTCAGGATGGACACTAGGTCAAGATAAAATCAGTTGCGTTGATGTGAATGAGTGCACTGGAAGCGATCCGCCATGTTCTTCAATAAGCGAAGAATGCAGTAATATAGATGGAAGCTATTTTTGCTTATGTAAAATGGGATGGACATATGATAATGGTTCAAATACCTGTGTTGATATTAACGAGTGTATTGAAAGCAATGCATTGTGCTCCAACGGATTTTGTATAAATATTGAAGGGCAGTTTCAATGTCCGTGCAACAATTTATCCATGTTAAATCAAG
>evgR016338c0g1_i01_Hydra
TAATCTTCTATTGTCTAAATAAGGTAGTTTTAAATTTCAATAATTATATTCATCACCGCCATCATCGTTTTAGTGTCCTTGTTTCCGTGCTTGCACAGATGTCAAGAAATTCTGAAAAACATTTTGTTGGATTAAACAGAGTTTTTCTCAATGAGCAATGGGCTATAGAAAAGTCATTGAAGCGTCCTGAATTAAATCAGCTCTTAACTGCTGATGAAGTACGAAAATGGATTCCAAGCATCAAAAAAGATTTAGAGTATTATCTACAACAGTTAAGTGGTGCACGGAAGCATAAGTATACAGATAGCAAACTAAATGAATTTGAGAAGAAAGTAGCTTGGTTAGAAAAAGAGCACCAGCGTTTTGTGAGAAAAGTTTTTGAACTTGATCCTGATCAGCAAGGTATCCCTTGGGAACCAAAGTCTTACTCAACTAAAAGAAAAGTAAATGAAAACTTGCAAACTGTTGCAAAAAAAAAGAAACCAATTGTCTTAAACATTTTGAAAAATGAGAATACTTCTGATGTTTTAACATAATGTGAATTTTTTGAATGCAATTATATATGATCTCTTGCATAACTTAACTAAATTTTGTATAGCTTAACTTAACTTAGCATAACTTAGCATAACTTAAGTACAATTTCTTGTATGTAATGTATATTTTAAAAAATTAGAATATTTGTGTGTTATAAATTGTTGAAGAATGATTCTAATTTTATATAGAAATATAATGCTAT
>evgR016348c0g1_i01_Hydra
CAGGTATCAAGCGTCAACTCATAGTTGCCATCGCTGAAAATTTTTCTGAAACATACGAGAACGTTAAAGTGATGATGAACATCATTCAATTAAATGATGTTTCATTCTGCATTTCATGCGATCTCAAAATGGCCAACATCATTTGTGGAATTCAATCCCATTCAAGCAAACACCCTTGTTGCTGGTGCAACATTGATTCTGACAATCTGTCAGAATGTGGAACTTCCAGAAGTTTTGGATCAATGAAGGAAAAATTTGAAGCCTTTGTTGCAGCTGGATCAGATACGAAATCAGCAAAAGATTTTGACAATGTTATCCATCCACCACTGATTAGCATGGATGAATCAACACTCATTTTGGATGTGATTCCACCAATGGAACTCCATTTACTTCTTGAAATTGTC
>evgR016351c0g1_i01_Hydra
TGACACAATTGACCAAAATAAAATTATACAACTTCTTCAAGCAAAAGAGTCACAAGAAGAATGGGGATCAATATGTTGTTAATTTACTTTTTGGTTGGACTAACGGAAGTGACATCTGATAGCGAAAGAAAGCTAAATGCTAAATTAATTAACTTTTTAAAGTCATCTTCATATATTGAAAAACGGTTTAGAAAGGATTTACTGTCACAAGTCAAAGGAAAAGGCAATATTATTTTTGAAGACAATTGTCATCCTAGTATACCACTTGTAAGCTGTGAAAAAAACTATTGTGATTTTGCTTCGTGTCCTAATTTCCCTAATGCTGAATGTCACCTAAGTTTGTGTGGGAAGTGTGAGCCTGTATTCTATTTCGAAGGGAAAAAAGTGGATTGTTATGATTTTCAAACAATTTTACCAGCCCACATAGGCCTAAAATCTAATGAAATAAATTTTGATGATAAGTGTCCATCATATGTTCAATCAGTAAGTTGTTCAAATAACCCTTGTGAGATATATAAATGTGATAGTTACCCAAATGCAGAATGTCGTGTCAGTTTGTGTGGAACATGTGAACCAAATTTCTATGTTAATGGCCAGCAAGTATTATGTAATGCTACGCAGACAAATGATCAATTATCACAATGTTTTAACGAACGCAAAAGAATTATTGAAAGCAAAAATGTAAACAAATTTATTGTTGATTGTGACATGAATGGCTTATACAAACCTATGCAGTGTTGGCCATTAGTTGGTTTGTGCTGGTGCGTGAACAAAGATGGAGAGGAATTGTTTGGAACAAGAAATTATTTTCCAAATAGACCTAAATGTGAAAATTTAATTGAAGATTGCCGCAGTAAAGAATTTGGTTGCTGTCAAGATAATAAAACAGCAGCACAAGGTTTGCACTACGAAGGATGCTTAGATACAAATAGAAGTTGCCATGTATCATTGTTTGGTTGTTGTGATGATCAAATTACGTTTGCAAAAGGGCCAAATCAAAAAGGGTGCTCTCATTCAGTTTCAAAATGTGAACAACTTCGATCTATGACTCAACATATTTTTTCATTCCAACCTCAATGTGACAACCAAGGTCACTTTAATCCTATCCAATGTTGGAGCACTTTGGATCAATGTTGGTGTGTAAATAAAGATGGAATTGAGCTCAAAGGATCAAGAGTGAAAGGGAAACAGCCAGATTGTTCTGCAGCTGCGTGTGAAGACAAGAGAAATGATTGTAACAGATTTGCAATTGTTCCTGGATTTTGTCTTCGATTAAATAATTACATGATGGAATACTGTAGGAAATCATGTGGCTTTTGTGCAGAAGTATCTATTGTTAAACATGAGCCAGTGAGAATAGGATGCTCCAGTCCATTTGGCTGTTGTTCTGATTTCTTGTCTCCTGCTAAAGGTCCTGACGAATACATGTGCCCATTATATAGCTCAGTGTGTTCTATGCCCAAGATTATTGGAGAATGTTTTGGCTCTTATCCTAGATATTATTATAATCATGAGACAAACACTTGTGAGCTATTTATTTATGGAGGTTGCCGTGGTAATGGCAATAACTTTGCTAATCTAGAGGAATGTGAACAGAAATGCTTAGAAGAGGTAACCCCAACCAAGGTTTCCACAACACGACTATTTGAGCTTGGAATTGATTTGAAAGTAAAAGAAAGTTATAAAATTATTACCTCGGATGGCTTTAATGTACTTGCTGGACCAATGATAGAGTGTGGCAATGAATTTGGATGTTGTGAAGATGGCATAACACCTGCCCTAGGATATAATTTTCAAGGGTGTCCAAGATATATGTCTTTTTGTGAGATGCCTCCTGCTTCTGGGCCCTGTACCAGAAACACCATTAAATACTTTTACAATCCTGAAAGAATGTTTTGCGAAGACTTTTTTTACGGAGGGTGTGGTGGAAATGCGAACAATTTTGACAGTGTTGAATTATGTATGCATGAATGCACCAATAAAATATCAGAGTTAAACAAACTAGATGCATTGACTTTATTACCTACCCATCAAATATTATCCACGGTTCAAACCAATCACCAAAGATTTGTTGAACAAACATCAAATGATCCAATGATAATCAAAGATTCAACAAATGATCCAATGATAATCAATAAGGCTATCAAAATAAAAGAAAATACAAACAAATTTACAGAATCAATTTCTTCTCTATCTGATGATATGAAAACAAATAATAAATTATCTTCAACTGCTGAACCAATAGCTATTACAGAAGAGATTGTTTCTAGTGATCTCCCCG
>evgR016369c0g1_i01_Hydra
AGTTACACTTAATTTACATCCTACAACGACTTACGATTTATCTACGTCCAAATTTACAACGACTTACGATTTATCTACGTCCAAATTTTGAAAATGTTTTTAGCATACTTTTTCTTTTTTTTAACTTTTCTTTATTTTCAAATTAAAATATTATGGAAGCTTTGGATTTATTCATACGGTCTCTCTAATGTTTCAACACCTCCTACTTTACCTTTTTTCGGTAATTCTCTCCAAATAGAAAATGACCCCGTAAAATTCAACAAACAACTATGCGATTGGAGCAAAATATACGGAAACGTATTCTGTGTTTGGGTAGGTGTTATGCCAATAATAGTCTCATCATCTGTACATTTCTCAGAAGCAATTTTAAACAGTCAAGAAGCCCTGAAAAAAGCATCTATATACAATTTTTTGCATGAATGGCTCAAAACAGGATTACTGACAAGCACGGGAAGTAAATGGAAATTACGTCGTCGACTTCTTACTCCAAG
>evgR016377c0g1_i01_Hydra
ATGATTTATTTAGGTATGCAGGATCAGTATTACACATTTAATATGTTCGATGCTCAAGCATGGTATGCACGAGATTATATTTTGAAAAAACTTATTTTAAAATCCAAAGATTTGATGATTGAAGAAAATCAAAAATGGCGAGATAGAGAAGAGAAATTATTAACTGAAGTACATAAAATTGTATTTCAAGGAGATTATATAAAAGATTTAATCAAAGACACCGATTGCCCTTATCTTGATGTTGAAGGAATAAACATAATAAATTTTGATTGGCTAAAAAACAAACATGATGATATAATGGGATATAGAAATTATTGCTTTAAATCTGTATTAACAGGTAACATGTCTGTAATGCATCATACGAAGTGGTTAGATGAAATGGACGACTCTAATGGA
>evgR016404c0g1_i01_Hydra
CGAGCATTCATTTATTAATAATATTGGTTATATATATTTTATTGTTATTTGGAACAAAAAAAAGACTGAGATGCAATGAAATTCAAGTTAAAAAACACATTGAGGGGGAAAAACAACTTACAAAAACTATGTTCTTAATATCAATGTCGCTCGTAATAACATATATTCCTTTTGTAACCGCTGGCTTCTATCATACACTTATGTTACTCACTGATCAATTTCAACCAATATACTTCAGTCAAAAGATATTATTTTGTACGCTTGCAATATTTGAACTACAATCCTCATTTAACGCTTTAGTTTACCTAAGCAGAAGCAGCAGAGTAAGAAAATTATACAGTTCTTATTTGTGTAGAACGAAACCTTTTTTTTTCAACTTGTTGGTGTTGCAATAACATATATATTATATTGAATTATTATTTATTGTTAAGACTGCATCCCTTGGTTATGCATCAACCACCAAGAGCATGTTTTAACGTTATTTTATTTTTATGAAAACAGTTATATTGTACAAATATAAAAAGTCTATAAATGAC
>evgR016409c0g1_i01_Hydra
TTGTCAGATGCCGGTAAAACTTAACAGTTTTTAGCTCGAATAAATTTTTTGTCATTTTGCAACGTCGCTATGGCACCAAATATTTATGTAACTATTGTTTTAACCCATTTTATTGCTTCAAATTGTTTTGGTTTGAATATCACCACGGTTAACAACGCGTCGTTTAATGTGCAATCGACAACTGAGAATTCAAAACTATTTGAAGATGAAAAATATACTTCACCAGAAGTTGCACTTCAATCGGCTATGAAGAGCTATAACAAACTTGTCAAGCCTAAAGGTTTAGTCCTCGTTGAACTGGATATTCGAGTTTTATCTTTTGATAATTTGGATGAAAGCTCAATGAAATTTAACGTGGACTTTTACTTTTCTCAATTTTGGTTCGATGAAAGATTTAAAGATTTGCTTGACAGACCAGATTCATACATAAATATTTTAGGGAAAAGCGCAGAAGACCTTTGGCTGCCGGATACAGTATTAGTTAATTCTAAATCCTCTAATTTTCATTACGTAACTGTAGAAAACCGATTTACCACAGTAAACCTATCGAGTGGGATGATTTTGCAACATGCGAGAATTTCAACGGCAGCATCTTGTCACATGGATTTAAGAAAGTATCCTTTTGATGAGCAAGTTTGTTAT
>evgR016417c0g1_i01_Hydra
AGTGGTTTTCAACATAGCAAACAGTAGTTGAAATGTGAAAACTTATTTAAAATGAAGTTGCTGATCATTGTTTTCTTTTTCTTCCATTTTTGCTTTGCTGATCAAAAAGATGTTACTGATAATAAAAATAGTGACAATGAGAAGACATTAGAGAAAATTGATGACTTAATAAAAACTAAAACAGGCACCAGAAAGCTAAGTTTTGGTTCAGTAAACATTGAGCGTATAGAATGCAGCATCAACATTACTTCTTTAACTCAGCCATGCTCAGCTTTGAGCATTTTAGAAAATTTATTTCCTAGTTTAGATTTTGAAAAACTCTCTGGAGGTTTTATGAGTTTTTATTCTATGCTGATAAATATAAAAATTGATTTTGCTAAAACAGGACAAATATGCAATTCAAGTGAGGGATCTATTGAAGTAAATATAAATTCGACAGGGCGTCATGTACATATAGTAGATAAGAAATATTTACCACTAAAAATTGTTTTTTTACAAGTCATGTGGGATTTGAATATAAAACAGTTAAAAACAGTTGATATTGAAATAAATGGTGTTTTTTTAAGTAATACCCTTGGTATAAATTCCAATAATCCTAATTATGAGGGTGTTATTATAGAAGTGCAGAAGCCAAAAGGTTTGACTATATTTGAATTTAAAACTGTTAATTTACCTACCACTGAACTAAACTACCTTCTTAAAGATTTTTATATTGAAATTAGTTCAATAACAGTTGGTTTAGCACTGAAAACAAAGTTAGAACTTTCTTTACAAACTTTTAAATTTTATAATTCTACTCTGATTGGTTATATTGAACCAAATATAATGTTTGAGTTTCAACTTAGTGGTTTGCATAAGGGCTCAAATGCATACGCATCTGCAAATCCAGTTTATGTAACCATTCAAAAGTATCATGGGGCTCTTGAACCTACAGTTGCTATATTTGGAAAGTTTAATAGTGGAGTAATGAATACTTTAACTAAAATTTCCATCCTTGACCAGTTATAAGTG
>evgR016422c0g1_i01_Hydra
CACGCTATGGAATAAGATCAAGAGTTATTATCTTCTAAATAACAGTTCTGGAAGGACAGTAAAAGAAATGACTTCATCAATACCTAAAAATGAAAATGACATCACATGCTTATCTTCGGATCTAAATAATGCTGTAAATCAAATAAAAGATATTCAGAGTTTAATGGCTTCGTTTATTGACACCATTATTGTGCGGAAAAATGATGAAGCTGAAAAAGAGATCCATATTATTAAAGATACAATGAAGCAAATTCATGAGATTCGTTTCAAACTAGATGGTTTAGGTAATCGGTATTCTCAATTGAGATTAAATCAAATTATTCCTGTTCCACTTGGGAATACAGGATATGTTTGCTTAGATCCAGCTGAGGAACAGAGTTTGTTTTATGAAAGACTTTCAGAATGTTATAATTGGCTAAATGGTCTCAATAAAGTTTCGAATGATGCTGTTGCCATTTTAAAGAGAAAGAGTTCTTTTACTGATATTTTCAAAAGCAAGAAATTAAAAGACAATGAAATTCATTTAATAGATCTAGTGAGCCTTGTAAAACCTGATTTTCCGGAATTAATTTTCAACACAAGGAAGTTACACACAGAAAGTTACCTTGAGTTAGAAGTTCCTCTTCAATTTTCATTGGTGATTTTATTTAACGGTTTCAAGGCTGTTAATGTTCTTGTTCGAAGCATAGAAGAAGCAAGTGTTGGTGAAAAAGTTGGTCAATCAAATTATTTCGTATTTCAGTCTCTATCTGACATCTTTACTTCAGTACTAATCAATTTAAATGAAAAAGATCCGCAAGAGCAGTTGTACTTTTTTTTGAGATATTTAACTCCTTACAAAGAATTATTTTCTATAAAATGTTTTATTTGTAAGAAGCATCTAAGTTTCAATTCAGATTCCTTTTCATTGTTACCTCCATTGCGGAAAGATATGACATCATCTTTATTTTATCATCTGTCGTGTAAATAATTATGTTTTTGTATAGATTTGAAATTAAAAAAACAACTTGTTAATTAAAAAAAAA
>evgR016446c0g1_i01_Hydra
TTAGTGCAGAAAGTTTGTTATAGAAAACCACAATACATATTTTTTAATGGAACAATCTGAAAATTTACCTGACAAAGTAGAGAGTGCCTTGAAACAAAACATCAATTTATTACTTACGAAAAACAAAGAGCACAATGCAAATCAAGAACAGGTAATATCTTGTGCAGTTGAGAACTTCTATTCTTCTTTGCAAGAGCTGCGTTCTTTTTTTATTCAGTCAAACCAACTTCACAAAGAATTAAACGAATCTTTTTATGAACAACTTGAAGTGAAAAACATGAAAGAAGAATTGCAACGAAAAAATAATCTTCTGAAAGAACAGTCTAAAAAGTTGTCTAAATACAGAGCTGAGTTGGAGGTGATACAAGAAGCTCAGCTTCAGGCCTTATCAATGCGATAGCTTGATTATTTTTAAAAATCTTTTTATAAACATGTGTGTTGGCTGTTTCACTTGTGTTTTTTCCAGTTTTTTTTGTTAATAAAGAAGGGTTTTTTAAGTATAAATTTAATTTAAAAATTCTAAGTAAAAAGTTTGTTGATGAGTAAATTTTATTTAGCAAATTGTTGCAGATGGGTTATATTTAACTATAAATAATATTAATATCTTATAATAACATGAATTATAATAAGAACTAAGCCATAAAAATATAAATAGCAGAAATAAATTACAACGGTAATATTGAAAAATCAGCTCACAGATTAACTGTTTTTTGAGTTAATAGATAAAGACTTACTATGTCGGTCAAATTAAAAAATGAAATTTAAAATAGACTTAAC
>evgR016451c0g1_i01_Hydra
GCTTTTTTTTTTCAGCATCTCTATGTGTGATTACAATGCATACTGTGTTAGAAGCAGAAACTATTGGTTTAAATCTTACATCTGGGAAAAATTTGTGTCCAACCTGTCTAAGTAAAATTATGAAACATTTAACAAAAAGTATGAGGGAAAATAAAAATGATGAAAATTTTGATATTGTTAATGAAACATCCATTCAAAATAAAAAAGAACGCGTGGATACACACTTTATATCTGCTGGAGTATCTCCTATTAAGATCAAAGTTTTGCACAAGTCAGGTAAATCAGAGAAGGTAAACGAAAATTAACAGCATTAACAACCTCCATTAAAAAAAAGGTAGCCACTTCACTTAATATCTAAGAAGAAAG
>evgR016453c0g1_i01_Hydra
CTTAAGGATTTGCTTGTAAGGTTGGAGGCAAACTTAGAAAAAGGGTTGAGCCCAAACATTGTTGCTAGAAATATTAAGCGCGATGGATTAAATAATCTTGAAGGTATTAAAACTACATCTGAATGGATTAAATTTTTAAGGCAAATGACAAATGGGTTTGCATTGCTTTTGTGGGCGGGAGCTATACTTTGTTATGTTGTGAGCATTATATCATTTACAAGTCAACCAGATCCTGTGTATGATGAAGTATACTTGGGCTCTGCATTAGCTATAGTTGTTCTTATTACTGGTATTTTTTCATACTATCAGGAAGCAAAAAGTACCAAGATAATGAAAAGTTTCCAAAAACTTATTCCTCAAAAAGCCACTGTTCTACGAGATGGTTTGAAAATCAATATAAATCCCAGTCAATGCGTAGTAGGAGATATTGTTTTTTTAAAATCTGGTGACACTATTCCTGCTGATGTTCGTATTGTTGAATCTAACGGGATGAAGGTTGACAACTCTTCATTAACCGGAGAATCAGAACCTCAATCAAGAAATACTGAATGTACTTCTGACAATCCAATCGAGACAAAAAATTTAGGTTTTTTTTCTACTAATGTAGTTGAAGGAGAAGGAGTAGGAATTGTTGTTAAAACAGGCAACCGAAC
>evgR016454c0g1_i01_Hydra
CAACAAAGTAAAGAATTTTAATCAGGAAATATTACCAAGTGATGAAAAACGTGTATTATTAATGGAGAGTACCTATGTAGAAGGCGGTGATTACATAAACGCTGTTTATATAAATGGTTTTAAGAAAGTGAATGAGTACATTGCTACTCAGCACCCTCTTACAACTACGTTGTCGGACTTTTGGACTCTGATTCTTCAGAAGAAAATTGGGACAATTGTGATGCTTCATGATTCTAGCAATGAATCTGGGTTACCAAAGTTGCATGGCATATTAAAACATAAAAATAATTTTAAAGACGTTTCTGTTACTTTAAAGTATGAAGTGAATAATGGACCTATTACAGTTAAAATTCTTCTAGTCAGCTCCTTAAATAGTAAAGATAATACAGAATGCGTTATTCTGTGTTTCGATTGGCCGGTTGATGGTGTACCTGTTAACTCTAAGGTAATTACATTAATCAATGAAATGGAAAAACAGC
>evgR016457c0g1_i01_Hydra
TCAATTAAATGGTATGATGGATTTGTATAATTTTGCTTCAACTTGTAGAAAAATTATTGGAATTGGGTTAAATTATAAAAAAGCAATTGCTGAATTGAATATTCTGAGCCCACAATTACCTGTCTTGTTTTTGAAATCCACTTCAAGCATTATCAAAGAAGGTCAATCTATTAAAATACCTCCGGGGTGTATAGATGTAAACTATGAAGTTGAACTTGGTGTCATAATAGGAAAATCAGGATCGAATATCTTGGAAGAAAATGCACTTGATCATATTGGGGGATATGCCCTTATTCTTGATATGACATCGGGAAGCCATTTAAGTTTTGTCAGGGAAAAATCTCTTTCATGGTGTTTAGCTAAAAACTGGGATACCTTTTGCCCTGTCAGTGATTTTATTCCAAAAGAAGCAATTAAAAATCCTGATGATGTGAATCTGTGGCTTACTGTTGATGGGAAAATGAAACAATCAGAAAGTACTTCAGACATGTTATTTAGTGTAGCCAAGCTAATTAGTTATACAAGTCAAGTGATGACTTTAGAACGGGGTGATTTAATTTTGACTGGTACTCCGAACGGGGTTGGTCCTGTTAAAGCAGGGCAAGTAATTGAATGTGGCATAAAGGACATCAAACAAATGAAGTTTTTTGTAGAAAAAAAATTATGAGATATTTGAAATTTATTTCTTATTAAAATTTTTTTTTTTAATAATTTAGAATTCTTTTGAATTTTAAAACAGTGTCAAGTGTGAAAATAAAAAGTCATTTATTTAATAATAAATTTTTTTATTATAAAAAAAAAA
>evgR016474c0g2_i01_Hydra
GCAGATGGGACAGATACTTATCATATTGTAAAAGCAGCTGGCCGTTATATAGAAATTACAAAAGCAAAATCTTATTTCTGAGGGAAAAATTTAAAAAAGAAAAACAAAAATTGATGACCGTCCAAAAAAAAATATTTTGTGACATAAGCGTATTCAGTCCGCGAAAATCAGGTCAGCATTTGGCAATGCCGATGCAAAATCTTACCTATCTTTGTTTCACAACGGCATGCAACTGCCGACCTAAATCTCTCAAAGAGTTGGCATTAAAAAACAAATATATAAAGAATGTAAACGTACTGAGGGGGTTTCAACAACAAACCTTGTAGGGC
>evgR016479c0g1_i01_Hydra
AGTATGCAAGAATGAAACAGAGTCAGTCAATCCTGTTTCATTATTTGTTTTTAGAAAGATTTTCAATACAAATTTTAATCTACATTTTCATTTGCCTCAAAAAGATACATGTAAAACATGTGATTCTATAAATATGAAAATTAAGTCATCGACTAATGAACTTGAAATTAAATCTTTGAAGAGCAAATTGGAATTGCATCAGCGAAGAGCTGAAAAAGTTAAAGAAATGAAAAAAAAAGATATCAATCAAGCTCATCTGAGTAATGGCACTCAAGGAGTTATTGTTTTTGACATGCAAAAAACATTACCAGTCCCCTGCCTTACTCCATTGATAGTTTATTATAAACGTCAGTTG
>evgR016481c0g1_i01_Hydra
CAGCTCAAGAAAATTGCTTCAATATAGCACGGCATAATGAAAATTTGAAGAAAAATAAATACCCTCAAATTTTACCACGAGATGATGTTCTGGTATCAAGTGAACCTTACCTTAATGCTGTTTTTGTTGATGGCTATAAAAGAAGAAAGGCTTTTATAGCAACCCAGTGTCCACTTCAATCTACTGTCACAGAGTTTTGGGATGTTTTAAAGAAACAAAATGTTTCAACTATAGTCTATCTTACCTCTCGCAACGAAAAAAAAGAACAATCATATTACGAATTTTATCCAACTGATCGTGATTTAAATCTTGATGGTGTTTCTGTAAAATTACTAGATAAAGAAAAGTTAGAATCTGTAGCAAAAAGAAACTTGTGTGTGTCTACTGAGACTGAAAGCATTATGTGCTTGCTAGAGTTTAACTTTTGGTCGGAGAACTGTTTACCAAATTTTGACTTGATTTTACAGCTGATTAGTGAAGTAACAAAATCACAGCAATATGTAGGAAATGATACTGTTGCTGTTGTTTGCAACAATGGCGTAAGC
>evgR016482c0g1_i01_Hydra
CATTTCAAGAGTTTGTAGATCGAGTGAGCATACTAAAATTGGAATCATGGGAACTTGTTATATCGAAAGATTTAGTCAAAGTTTTTAAAGTAGACAACATTCACCTACAACCCCAATATGAAGTATATGTTCAACCTAGCCTTATATTTACAATACGATCTTTTGCTTGGAACTTACCAGATACACATGATATTTATTTGAATTATAAACAATCAGTTAAAAACATTACTTTATCAAGTATTATTGCTCTTTTATCAACTTACTCATTGTGTCAGGGTGTTGCATCAGGTTCAAACTGTTTTAAAAAACATATTCTTACAAAATTGTACAATCCAAGTGTTAAATCAATCACTTCGTTTGAGAGTGAATTTAATAGGTCAACTTTGTGTTTTTTATTG
>evgR016483c0g2_i01_Hydra
AAATTTTTCTGCTTAAAATCTATTTTTTCAGATGTGACTGCAAGCTCTTTTAGAGAAATACTTCAGCTTTAAATAATACCATTTCGAATGGCTAACCAATGTTATGAATCGTCAGCTTTTTATTTGTTTTTAACAAATCAAAGCGACCTTACATGCACTTTTAATAAAACTACTAATGGATTAAAAAGATTGATTCTCAACATTATTGATATAATAATATGTCTTTCCATATTTATAGGGAATTATTTGCTCTTAATAGCACTAGGCACCACGAGAAAGAGGAGATTTTACACGAACAACGAAAAACTTGTATTGCTATTGTGCTTCATTGATTTAATGATTGCTACTATTGTTATGCCGTTTCAACTTGTTTTAATGAAAGTAAATTTAATCCCTTGTGTATTTATTTCAATTGCAGGGTTTTTCAATGTGTTACTTATGATTTTTTCCGGCTCAATTGT
>evgR016485c0g1_i01_Hydra
TTCTACACTATTCCGATTTTACTATTTGCGACACTATTCCTTTTTTTTTGTTTTCTACACTATTCCGATTTTACTATTTGCGGCACTATTCCTTTTTTACTATATGCGACACTATTCCTTTTACTATTTGCTACACTATTCCTTTTTATTTTGTTTTCTACACTATTCCGATTTAACTATTTGCGACACTATCCCTTTTATTATTTTCGACACTATTCATAAAACTTATTTGCGACACTATTCCTTTTTGTTTGTTTTCAACACTATTCTGATTTTACTATTTGCAACATTACTCCTTTTTTTTGTTTTCGACACTGTT
>evgR016491c0g1_i01_Hydra
AGAAATTAAAATGTACTTGCTTGACCTAAACACAGATTGTATAAAAAGAATTCTTCGATATTTGGATCTTGACTCTCTAGTTAATATTTCTCAAATCAATGGAAATTTGCTATTTCTTGTAAGAAGTTGTTGCTCTATGACCAGTTTACAACAAGGTCTGATATTACAGACAATGAAAACTGTCTCCCTAAACTTTAACCTTTGCAATATACCACATAGTGAAATATTTAGAAATACATTGTTTTTATCTGTTATAATGTTTAAGAAAATGTTAATAGATAAAGAATTGATTTACAATTGGGATATTTATGAAAGTTGTTTATCAAATGGCGTTATAAAATTATTTTTGCAGTCTTTAAATAACTTTTTAAACATATCAGACACTGTTTTAGCTTTACCTGCTTGTTATTACACTGAGGTTCAAACCTTTTTTTTTTATGTTTATGAAACTAAAACATGCATTTTAACTAATTGTAAAATAAAAACATGTTCCCATTCAAGCGGCGTCATTTACTATATTGAAGAAAACAAACAGGAGATTTATCTACCTTGTAAGATATGCACAAATAGCTGTTTGAATGAAGAGTTTCTTTTTAATGGTGAGATGGTCAAACTTGCTGATTTGTATGATTTAGTCAACATAGTGGAGATAATAGAAAGTTTACATCACAATCAAATTCATATGAATTATGACATTAAAAACTTAATGTTTTCAGAAAATGAAAACGTCTTGTTTTATAAAAGTATTGAAAATACATTTTATAAAAGCTTGGGAATGTTTTCTAAGCAACCAGGAGTTCTAAAGCTGTTTCACATGCTAAATCTGTACAGTGGTGTAGCAAGTTTTTGTGAACAAATTCAATTTTACAATAAGATTAAGTTGCTAATACTTTTTTATAGTCAAGCAAATTTCATATTTCATTATGGAGAACTTTTGCCATATTTACCTATATTAGAGAAACTTTTTTTAAAAGTTGATTTCGTTGTTTCAAATTGTAAATTAAAAAAAGTTATTGTTAATTTAAAATTAAATGACACACTTGAAGTAGCTAAAATAAACGGTAACCTTTTTGTGACATACCAAGAACGAAGTCTTATATTTCCTACTTGTTACTCTGATGATTCTGATGATTCATATTTTGAAGCTTATATAATCGAGCAATTTGAGGAGTTTGTTACAAAAACATTTACTGAAGATGAGTTAAAAGATCTTCCAATTATTTCTGTTGTTTTTATTTCTAAATTTCTGGGATCTTTTTTTAATGAAAATACAAAATAAATTTATATATTTCAATTATTTTGTGTAAG
>evgR016499c0g1_i01_Hydra
TGGAAACAAACAAAAACCTGAATCTTTTTTTAAAATTACATTTAAACGTTTATGTGAAGATACATCTTTACGAAGAGAGAAGCAATTGGAGAGAGCAACAGATAACAAAGTATACTAAAAATATTGATTCAGAAAAAGCAAACATCAAGAGTAAATAAATAAAAAAATGAATACTTTATTTTTTCTGTTGGCTTCTTGTTGGTTTTTTATAGTCTGCAATAGTTTTCAACTTTATGACAGTGAGATAGATAATACAGAACAAAGTTACGAAGATGAACAAGACTGGAAAAACATTGATTATCGCCCATGGCGAGAAGGTGATACTCGTAGCAACACAAAACGGTCCCAAAGTGCTGAAAAACGCGGACGGCCTTGGAGAGAGTTTCACAATGGAAATGAAGAGTACCAAAATGGACAGTTATATAAAGATTTTGAACTTTTGCGAGAACGTCGGCACGCAAATGATAATTTAAAAGATGTTCGTAAAACAAAAGTAAAAAGAAGGCCGTTAGCGAAACGCCGCGAAATGAATGAAAGTTTGGAAATCAAAGGACATCGAGGTCGTAAAATACATTCAGGTTTTATTAAACTGAATAATTAAGTTGAATAGTTTAAAGGTCTTTGTTAAACTTTAAAACATTTTATATTAAAATATTTAATTAATATTAATTTTATTTAAATTTTAAATTAACTTAAAAACTGAAAAAAAAAAAAAA
>evgR016503c0g1_i01_Hydra
GTTTCTTACTAAACAAAATTTAAAAGGATTTGGAGCAGACAACTTTGATACTCTGACTCAATCACTTTTAATTTCAATTTCAAATCTGTTTCAAGGTTCAACACAACTCCTCCCAGGAGTTTCCATTGTTTTGAATAATCAATCACAGTCATCTGCTACTCAAACGACATTATCAAAATTTACTTCAACATCATCATCTGCTACTCAAACAATGGAAGCAATAAATACTTCGATCTCATCTGCTACTCAAACAACGATAGCAATAAATACTTTAACATCATCATCTGCTACTCAAACAACGGCAGCAACAAATACACGGGCATCATCATCAAGT
>evgR016517c0g1_i01_Hydra
TGTTCATTTAAAATCTGATTGCTTTTGTTTCAATAAAAAGTCTTACAAGAATGTTGAGTTCAAAATATGTGGAAATAATGTTTTCAACTTAAATTCAGAGTTTTGCAATGATGAAAAGATTTTCTTTAAAAGTACACATGAAGTTTGCGGTGAAATTGTTCATTTAAAATCTGATTGCTTTTGTTTTGATAAAAAGTCTTACAAGAACGTTGAGTTCAAAATATGTGGAAATAATGTTTTCAACTTAAATTCAGAGTTTTGCAATGATGAAAAGATTTTCTTTAAAAGTACACATGAAGTTTGCGGTGAAATTGTTCATTTAAAATCTGATTGCTTTTGTTT
>evgR016527c0g1_i01_Hydra
AGTAATTTATTATCTGTATTAGTCATGAATACTTTATTATTAATTGGAACAAAAAAAAAATTAGGAACTAAAGAAATTCATGTTGCTCAGCACGAAGTTGTGGAGAGGCGACTTACTCAAATTATGATGATGATATCGATTTCTCTTATAATTTTATATTTCCCCGCGGTTGTTGCTACCTACAACATTTCATTAGTTCTATATTTCAATGAATTTAAATCCGTTGGGAATGGAACAAAATTGTTTTGTTGGGCTCTTGTTTTGTGTCAAATAAACTCATTATTCAATGCTTTTATATACGTAATTAGAAATAGAAACATATTAAGATTTTATAAAAAGCGTATAGCTTTATTAATAAAAAAAGATTTTAACAAGAGCATCAAAGAAAAGCAGTTCTCTTCATTTCGCATTTCTACTGATTCCATAACTCGATCAGTTTCTGAAAGTACTTGTAATTATTATTGCACCCGTTATTGAATTATTTATTTACGCATATACAGGAAAGACTGGGGCTAGACGATAGTAATGTAATTCATCTACTTTACATCTTCTGGTATACGAGAAGAAATATGATGAAAAATGAAAGATCAAAAGTACGAGAAGACATGTATAGACGGAGTGAGTGATTTCTTTTGAAATTATCTGTTGATACTGGGTATAGTAAAGATAAGCATTTTTATTTTATTTTATAATTTATATTTGATATGGATAAATTCAGGAACCGATTCTACTTTTGACGAATAATATAAATTAGTATTTAGTCTTAATTAAC
>evgR016533c0g1_i01_Hydra
AAAACACAAAGATGGGCGCCTACAAATACTTGAACGCTTTGTATAAGAAGAAGCAATCTGATGCTTTGAGATACTTGTTGCGTGTTCGTGCTTGGGAATTGAGACAATTGCCAGGTATCCACAGAGTTAACCACCCAACTAGACCAGACAAGGCTCACCAACTTGGATTCAAGTCCAAGCAAGGTTACGTCATCTACAGAGTCAGAATTCGTAGAGGTGGTCGTAAAAAGCAAGTCCCAGGTGGTAGAGTTAACGGTAAGCCAGCTACTCAAGGTATCCACTTGAAGCCAAAGAGAAACTTGCAAGTTGTTGCTGAAGGTAGAGTTGGTAAGAGAATTGGTGGTTTGAGAATTTTGAACTCATACTGGGTTAACCAAGACGGTAGATACAAATGGTTCGAAGTCATCTGTGTTGATCCATCTGCTGAATCCATCAGAGTTGATCCACAAATTAACTGGATTGCCACTGGTAAGCATAAACACAGAGAAATGAGAGGTCTCACTTCCTCAGGTAAGAGACACAGAGGTTTGAGAACCAAGGGTTGTACCTCCACCAAGGTTAGACCATCCAGAAGAGGCGGTTGGAAGAGAAGAAACACCACCAAGTTGCAAAGAAAGAGATAATTCTGAATATTTCCTCTATAATAAATTTCACTTTAAAATTAGTAAAAAAAAA
>evgR016548c0g2_i01_Hydra
TTGAGCAAACCGGCTGGAGAATGTCCACCTGGTTTTTTTTTGTCCAGTTGGAACTGGATTTAAGAATTCAAATCCTTGCCCTGTTGGATTTTATCGAAATGCATCTTCTGCAAGATCTAGTATGGATTGTAGTGTTTGCATTGCAGGAATGTACTGCAATGAAGAAGGTTTACCTTTTCCAAAAGACTGTCCACAGGGTTCGTTTTGTCCAACTGGCTCTACAATTCCTCAACCATGCCCATCTGGGTACTATGGAAACAGTACACGTCTAAGAAGATCAAAAGATTGCACTGCGTGCGAAGCTGGTATGTATTGTGCTGGTATTGGAAAGATTGCACCAACAGGAGAATGTGATGCTGGTTTTTTTTGTAAAGGAGCTGCAGTCACATCGGCACCCTCTGGAGGTATTACTGGTGGTCTATGCCCTAAAGGAGGTTATTGTTCTCGTGGAGCAAAGAAACCAAGTCCATGTCCCATAGGAAAATACAACAGTGACGATGGTGCAAAATCACCTGCAGATTGTCTCGACTGCAAAGCAGGATTTTATTGTGCTGGCACATTTATTGATGGACCTACTGGACCCTGTGATGCTGGTCACTACTGCAATGGTAATGCTACTGGCCCTACTCAGCATAAAACTCCAGCAGGACATTATAGTTTGAAGCAAGCATCTTTTCCAGTCCCATGTCCACTTGGTTCTTACCAAACATCTGAAAGAACGTCTAAATGCGATACTTGTATAAAAGGATCTTACTGCAACAAAACTGGACTTGCTATCCCAAGCAGTTGTTTACAAGGACATTACTGTCCGGAAGGAACATCAGTTCCTATTCCCTGTCCTGCTGGAACCTACAATGAAGACATATCAAAAGGTGCCATTGAGGATTGCAAAGATTGTACACCAGGCGAATTTTGTTTTGGTACGGGTTTGGTGAAAGTTTCTGGTAAATGTAGTGAAGGGTATTATTGCCTATCGAAATCTCCAGTTCCAAATCCGGGAATGTTTACAAATGGAACGTATCAAGACATTTACTATGGTGGAATTTGTCCCAAAGGACATTTCTGTAAGGAAGGAACATCAAATCCAATGCAATTTCCATGCAAACAAGGAACCTATAATCCACTATATGGTGCAACTAATGATTCATTCTGTTTAGAATGTATTTCTGGACAGTACTGTTTAAATTCAGGACTAGAGAAACCTACTGGAAATTGTTCAGCTGGATATTATTGTAAAAAAGGTTCAATAACAGATAAGCCAAAAGATGGTAAGCAAGGTAATATATGTCCAGTTGGAAGTTTTTGTCCTGCTGGATCATCTGAGCCAAAAACATGTCCAGAGGGATTATACAGCAATGAAACTGGTTTTTCTGAGTGTATGAAATGCCCAGCACGTTATTATTGTGATGATGGAATCAAACTTTTTGATTGTCCTAATGGATATTACTGTGAAGAAGGTACAGCATTCCTGAAAAAGAAACCATGTCCGTTAGGAACTTATGGTTCAAAAGGTGGTTTAGCTGACATAAGTGAATGTACACCATGCCCTCCTGGAAAATTTTGTGGTGGTTTGGCCATAACTATATTTCCTACAACAAATATTTCAGGTTTTTGTGAAGCTGGATATTACTGTTTAGAGAGCATATCCATAAGTAAACCTACTCAACCATTATCCTTAGGAAAAGGTGGTCTTTGTCCTGTAGGGTATTTTTGCCCAGAAGGTACAGGTGATTATAAAGCAGCACCTGCAGGACCGTGTCCTGCAGGAACCTATTCAAATATAGAAGGTCTTAAAAAATCTGATGAGTGTAAGCAGTGCAGTCCTGGAGAATTTTGTGAAGGTGGTCAAAGCACAACCTCAGGATTATGTGATGCAGGATACTACTGTAAAAGAGGTGCTTCTTCTCGCAACAATTCAATAGAAGATGGAACCAGTGGACCTTGCACTGTTGCTCATTTCTGTCCAGTTGGTACTTCTGAGCCTCTATTTTGTAAACCAGGGTCTTATATGCCTCAGAGAGGAGCAGAAAAATGTTGGTCCTGCCCATCTGGGTATTATTGTCCATCTAATCAATCAGATTATCTTTCATTTCCATGTCCTGTTGGTCATTTTTGCCCAAATGGGACTGAGTATGCAACACAATATAGATGTCCAGCTGGTACTTTTG
>evgR016549c0g2_i01_Hydra
TTTGTTTATTTATTTTTATTTATATATTAGTTTTCTTATTTTAGTTGCATAACAAATAGAAATTTACTGTGTACAGTTAATAATGGACACTGTTAAAGAAGTAAGAAAACTTCTCTCAAGTTTGAAGAAGAATTTTCAATGCACAATATGTTTAGACTTTCTCTATCAGCCTGTGTCCACTAAATGCAACCATCAATTTTGTAGTGGTTGTTTACAAGAATTTATCAAAAGCAATGTAAAAGGCAAGAAACCATGTTGTCCGTTGTGTAATAGTGATCTAGGTACTAAAAGAAGCTTGAACCCTAACCCGGAGCTGTCGAAGATAATTACAGCAGTAAAAGACGTATGCAACGCTGCAGATAAAGACTGTACAAATTTAGATGTTTCACCGCTTCTTTCACAAAAACACAGAACTTTTTCGGCTAAAAAGAGTGACAAAATGACAAGTACCAAAAACAATAGAAAAAGTCTTAAATTCACAAGCGCAACCTCTAAAAAATTCATCAATGAACATCGCTATCTTCAGCATGATGACATCATTGAAACAGATTTTATTAAACCAGAGAAAATGAACAAGCCTAACAAATGCAGGTTGAAGAATAAAGAAGAAAAAAGGAAAAAAATTAAAACTTCTTCTAATTCTTCTGATTCATTAGAACATGAAGTTAATGGGGACAATACT
>evgR016550c0g1_i02_Hydra
TGATTCGCTCAAAATTAATATATCTTCGAGTCACATTTTGAAAATAAAATGAAGTACGTTTGCATTTTACTCATTGTCGCATACACAATGAGGATTGAAGGAGACACAGAACCATCAATTGTGTTCTCAGAAACATTGAAAACTACTCCAAATTGTGAATATAAAAATCAACCAACAGGCACACCTATACCTACTCCAAACTGCGTAGGGTTTGATACTTGCCGCACATTTACTGGTTGGCTAAACGGAACGTTTACTCACCCATTTGATGATCACTCATTTGTCACGTGCATACCAAGTGACAATGCTGATGGTGGCACATTTACATGTGGTACTTGCTATCCACCACTTTTTTATTCTGAAAAGTGTAGTTCATGCGAATGGAACAAGGACGATAGCTTGAGCGTTTGCTCAACAACAATTAAACCTAACATTGTTCCGACAACCAATTATAAAACCGATTTTTGCGAGTCTAAACTGAAAAGTCAAAATTACGAAGACCCTGTCAATCCAAGTTATTATTATGCATGCGTTTCCAAAATTACCTATCATATGTGGTGCGGAAGTAATTCACTGTTCTACAATTCAACTTGCGACGCATGTTTATCAAAACAATAGAAACTTTTAATTAACTACTCAAAATTTTTACGGCTATGTATTACAATATAAAATAAAAAATAAAAAATGTTTATTTTTTAAAAATAAGTCTTGTGTGTATAACGGTTATTCAGATGC
>evgR016562c0g1_i01_Hydra
AGATGAATTTTGTTGATTATTCAGCAAATATAGTTTATCAAAAGCTAAAAAATCTAAAGCCTTGTACGTCATTTAGACCTGACGGAATTCCAAACATACTTTTAAAACAATTAGCACATGTAATATGTATTCCACTTTCATATATATTTGAGTCAAGTTTCACGTCAAGCTCTTTACCAAAGCAGTGGCTTCAAGTTTTGGTTTCCCCTATTTTTTTAAAACGATCAACTTCTAATGCTAACAATTATAGACCAATTTCCCTCACATTTACTAGCTGTTGTGTTATGGAAAGTATTGTCAGTTCAAATATCGCTGATTATCTTAATAAAAATAATTTAGTAATACCTAATCAACACGGTTTCTTATACAAAAGATCT
>evgR016571c0g2_i01_Hydra
AAAAAAAGTTTTAGGAGAAAAACATGAAGATACAATTACTACTAAATACATGATTGCATTGTGTATATATTACAAAAAATATTACAATGACGCGGAAAAGTTATTTAAAGATGTTGAAAATATTAGAATATTAGTTTTAGGAAAAAAACATGAAGATACAATTACTGTTAAATACATGATTGCTCGATGTTTATATTACAAAAATTTTTATGATGACGCTGAAAAGTTGTTTAAAGAAGTTGAAAGTATTAGAAAAGAATTTTTAGGAGAAAAACATAAAGATACAATTACTGCTAAAAACAAGATTGCTCGG
>evgR016575c0g1_i01_Hydra
TAAAATGATACAATTAATGATAAATGTGTTAATGCAGATTTATGTGAAGTAACAATATCCAGTTTTACAAATCATTCAACTATTTATGTACATGATTTTAAAGATTCTAAGATTTGGCAATCATGAGTGATGACTATGAGCGTGTGCTATGTGCTAAAAATGAAGTTTTTGTTTATAAAATACCACCGAGGCAAAGTAGTAGAGGATATCGAGCTTCTGATTGGAAGCTTGATCAACCAGACTGGACTGCAAGATTGAGAATTGTTTTATTCAATACAAAGGTTTATATAAAGTTTGAAGATAGAAATTCTGGAGAACTTTTTGCACAAGCTCCGATAGATGAATACCCTGGTCCTGTTGTTGAAGCAGTCTCTGATAGCAGTAGATACTTTGTCATCAAAATTCAAGATGAAAGTGGGAGGCATGCATTCATAGGCATGGGTTTTACTGACAGGGGAGACTCATTTGATTTCAATGTTGCTCTTCAAGACCATTTCAAGCGTGAGAAAAATGAAAGCGACATTTTAACAAGTCCTGCAGCACCAGACACTCCACTTAATCTTGGATTCAAGGAAGGGCAAACAATAAAAATAAATTTTGGAAACAAAAAAGATGATTCGAAACCTCGACCAAAAATTGGAAACAATAGCAGTTCATCTTTTCCAATGGCACCTGGAATTCCAAAAATAGCTCCACCTTCACAAACAGCTAGCTTTGTTCCAGACAAGCCATCTATATCGAAGCCCCAACCACAGCAGCCAATACAGAGTTCAAATGACTGGGGCGATTTTTCTTCTTTTACAAATACACAAAAAAAAGATACAAGTGTACCGAACAATGGTGGTTGGGTCCAATTCTAGCTCATTGAGAAATTATTGGCGATTTTCATTGACTTTATATACTTGAATGAAGTGGGACACCTACCACCTAAAGTATATTCAGATAATTTTTTTCCATCCAGATCATTTTTTCAGTTTTCCATCCAGATCATTTTTTCGGTTTTCCATCCAGATCATTTTTTCGGTTTTAAGCCGTTTAAATTGAACACATTTTCGTTTTGCTAACAGTTTAAGCAATTTATATAAATATATATATTTATATATATATAAAGGGTTAATAGTAACCCATATAGTGTGTAAATTACCATAGTAACTTGTTTACTTCTTTTTGATAATTCTTAGTTAACTTTTATTTTTTACACTTAAAAATAAAAAAGTTTTTAAAGGCTGTACAAACAAAAAAGATTGTTTTTTTGGAAACTTTAC
>evgR016577c0g1_i03_Hydra
GTTTGTTAAAATGGAAACGTTAATTTTTATGTCAATTTTTAAAACGTTAGAAATAAAAATAGTAACCTCAATGGTTTGTTACTTAAAATTATGAAAAAACGTGTTTGAAGAAGAAAGTATTGAAAAAACCAAGAAAAATTATTTCAATCGAATTTCGTTTAAAACATCGACGAAATGATTGCTGGGGAATTAAAAAAACTACTCTCAATAAAGATTAAATATCGTTCTGATGGCTTCACTGATCAATTCAACCGAGTCCTGGTCATGAAGCTGTTGTTGGTCTGCTCAACGATAATGTGTATTAATTGGTTTAAGGACTCTGTAAACTGTGTTGTACCGTCTACTGCAAGCGTGAGTTCAAGTTTTGTTTCCTCAGCATGTTGGATTCAAGGAGTTTATGTTTACGACGAACTCAAAGATCGTGTTGACAGTGTTGGTTATTATGGAATTCCTAAAGATATTGATATGGACGGTACCTATGATGACGGAAGACCTTGCCCTGTCATGACGAATTCAAAAGTTTTCAGTAGCTGCAAGCCATTGACAAAGACATTTTTTCTTCAGTATCAATGGTTTCCTTTCTATATCGGAGCAATGGCGTTGTGTTTTTATCTGCCATACGTGTTGCACTTATTTGCCAACGAAGATCTTATAAGCTTAAAAAAAACTATTATGTCAAACGGGTCCACAGCCGACGCCATATTAAATAGTTATTTTAATAGAAAAACAAGTTCCAAATCACGACTTCGGTTGCGGATTTTATACAATTACATTATTAAGTCGCTTTACATATTTGTGAACATCATTTCGTTTGTACTTACTGATAAAATCTTTTTGGGAAAGTTCAAGAATTACGGGTCGCAATGGCTACTTTGGTCTAAACTTCCAAACTCTGTTGCCTATGATTATATGGGTGCACGCAGTTTTCCGAAGCCTGGAAACGAGTTATTACCTCCATTTGGATATTGTGAGCTTTACGAGTCATCCAAAGACATTAAGCATTCGGTTGGAAATCATTACCGATTTCTCTGTGAGATGTCACAAAATATACTGTATCAGTATTGTTTAATAGTGGTCTGGTTTTTATTAGTGCTTGGAATAGTGATTTCAATCGTAGGTTTTATTGTACTTCTCCTCAGTCACGTGCTAACTATTGTTTTTGTGGTTTTTCGCGGAATATCTGCAAGGAAGTTGTATAAAGTTTTATCATTGCGCGAGTGCGAGTACTTAGAGTTGATAAGAAAACGCAACTTGCCTTTATATTCGGAAATAATTTTAAAGTTGCGTCTTGAAAAATACAATGCCACAGGTTCAACTCCTTGCGAATCTCCTCCACCTGGCTTTGACGATCTATCTAAATTAAATAGTCACGTGTGACTAGAAGTCGTCAATCTTAACGACCGAAATGGAAATGATTTTGTAAGACAAGCTTTTGGATAAAGGTCGCCTGGTTATTTAAAAATTATTTAAAAGTAGTCTTTGGAATATGTTTATAGAAACATTACATTTAGATAACTAAAAATTTGTGTTATGTTTAAATTTAGGACAATTCATATTTTTTGTTACGCTTATTTTCATATCATATGTAAGTAAAGACTTTTTTTACACAGTTTGAAGCAATCGTCGGAAGCTGTTTTGGAAGTTCTCATTAGTTATTTTGTATATATTTTATAACGACTATAAACTTATATTTTATTTGAGAATGATCTACATCAAAAAAAAAA
>evgR016580c0g1_i01_Hydra
GCTATATTAAAAATAAGTCAGAATTAAAAACATTGTTGAATTAAAGATTTTAGAATGAATTCTTTGGAAATCAGTAATGTGGAAGTTTTGCCAATGGTCGATTCACAATATGTTGTTCCTTCACGTGTTACTTTTAATCAGAACGGCGTAAAAAGGACGTGGGACTATATTAAAGCTCACGATGCAGTGGCGATACTTCTTTTTAATACAACAAAGCAAGCTTTTGTTCTTGTAAAACAGTTTAGACCAGCTCTTTATATGGCGATAAATGAAAGAATGAACACACAGAATATATCTGTTTTGGAAAAACAAAAAAAAGAATTGAAAATATCTGCTTCCTTTGAAAAAGGCGTTTCTTATGAACTTTGCGCTGGAATAGTTGACAAAAACTGCACTCTTGTTGAAATAGCTCAAGCGGAGTTACTTGAAGAAACTGGTTACCAAGTACCTATTAATGCTATAAAGCCATTGTTTGAGTATCATGCTGTTGGGTTTTCTGGTAATCTTACAACAGTATTTTTCGCTGAAGTTACAGATGAAATGAAAGTTAATGGAGGCGGCGGAAATCAACACGAGGGTGAATTTATTGAACTATTTTACTTACCTATAAATGACGCAAAAGAGTTTATTTATAATTCCAATTATCCAAAAACTTCGTCAGTCGCAGCAGCTATCGGGTGGTATTTTCTGAACTGTTTATGAAATGATATGCTTAAAATTAGATGAGATTATTAAATGTTTTTATAAAAGTATGTATTGGCCCTAATTTTAATTTTTTTAAATACACTTACCTTAAAACTAAAGGTAATTTATAAAAAAAAAAA
>evgR016595c0g1_i01_Hydra
AAATAAAACTAAAATTATTCCATCAAAAATCTTAAATTGTTTATGTCATTGCAATGGTCAAAACATAAGTAAACAATTTATAATTCAAATCTCAAAAATCATGACATTAAACGAAGAATACATAATAGATGAAGCCATTGCATTACTAATAAGTTATTCTTTACTTGATCGTTTGGATGATGAGAAGTATTCAATGCACGATCTGACACAGTTGTCGATTAAAAGTTTTCAAAGTAAAAATTTAATCACAAATACGTATCTAGATCAAGTAGAAAATTATTTTATGCACGAGTTAAATGAAGTAAAAGATCACGCGGATCACGGAAACCATTTTGTATTTCATTTTCTTCATATGTTTTGTATTAACAAAAAGAGAATGTCGAAAGCCTTCCATAAAATGTCGGGTTCCATTAAAAATTTGTTAGTATACAAAGGTTTTTTTCAAGAAGCAATTGAAATATTAAAAATTATTCATTGCTATAATGAAGAAACTTATGGAGATAATAACGAATTCACACTTGATACAAAAAATAATATCGCAC
>evgR016606c0g1_i01_Hydra
TTCAAATATTATATATATTATATATTTTCAACATTTATGGTTTTTAATTTTGAGAAAATGATGAACTCTACTAAATATTTACTTCAACATTCATTTAATGATGAACCAAACAAAGAAGAAGAAGAAATTGACAGAGAAACAGAGAACAAAATAGCGGAACTAATACCAATTACTTTGTTGTTTCAACAAAACTCAAAAAAGTCATCTTGGTCTTCACTGCGAAATATTTTGAAAGAAAGAATATTTATGAAAAAGAACTCTCAAAATGTCATAAAATGCAATCTTAAAGATGACATTCCAGAAGAGACATTTGTTACCCAATCATTGGTACCAAATAATGTATCCAAAACTCCTAACTTATCACGAAGATATACGGCTGAAAGCTTTGAAGATGCTATTTTTCTGCCGATACTTCAAAATGATTATAATGCATTAGTTAAAACATTAAAAACATATCGTGTTAACATTAACTACATGAGGGCCCCAGGGTTAACACCATTGCACCAGGCATGTGCTGTTGGAAACCTAAGGATGGTTAAAACATTAATTTCTTACGGTGCCGATACAACACTGCGCACCTGGTCTAACTTGACACCTTTATTAATTGCAAACTTGTTTGGACATTTCGATGTTGCCTTATTTTTAATTAATCGTGGAGCTAGTACTCGTGATATTCAAGATGGAGTTGCTTTCGATTCAAAACTAGCTTCAACGGTTACATACAAATTTGAAAAAAAAACTAGACGAGGATCTTTTAGCTCAGGTGGTAGGCTGTAGGTTGCTTGCAAAATATTATTCTTTGTCACTTTTTATGATGACGACATATATGCAGCATTTTATGAATCACTGAAACTATTAAAGTCGTTGTATGTAATGCTGTTTTCGATTCAAAGTCGCAAAAGTTGCATCAATTAACTTAATTTAACGCGTAC
>evgR016615c0g2_i01_Hydra
TCATGCAGAGGAAAGAGTATCTTGGGTCAAGTTGAAAAGACTGGTTTTATCAATGTTCATATAAAACCTTCTATCTCACTTTCATCTGCAATTACTTATGTGTATTTGGCATCAAATGCAACATTTCCAAAATGCAGTGTTGTAAGCAACCCCGCTGCTAAAATTGAATGGAAAAAAATATTTGGACTACTACCAACAACAAGATCAGTTCAATCAGGATTCGGAGATTTTACATTAACAGATGTCAGAATTGAAGACGAAGGATTTTACGTCTGCTCAGCAGAAAACTATTTAGGTTCAGATACTGTTGTCGTTCAGCTGAAAACAAGACCATTAAAAATTGCTTTCGGATCTCCTTCAGAATCAGTAGTTTTTATCTACCCTCATACACTGTGGTGTGGCACGTTTGGACGGACTGAACAAATATCTGGAAGTTGGAAGTCATTTGCTACTGGTCAAGCAATAAAGTTTAAGGAAATTAAAAACTCAAGCTTTTATAATATATCGATACATGTTACTGAAAGCGGCATCTACTCATGTGAACTTAATGATGGTATATCAACAGTTGAAAGAGTTTTTGTAGTTCGAAACTTTACATTGCAGTCAAACATTATCAAAAGTATAGAAGAAAATCAAAAGATTTATGATTGGTTAAATAAAGTTAACATTCCAACAAGCAGAATTTCTCAGTGCATGCAAATGAGTGAAACAAGTAGTTATAGTTCGAACACAATTTGGAGTAGATTTGAATCATGCTATTCTAAAGAAAATACATTGCTCATTATCAAATTGAAAAAGGGATATGAGAGCCGTGTTTTAGGAGGGTTTGCAAAAACTCCTTGGAACAATGTGGAATACTCGTCAAACAACCAAACTTTTGTGTTTACTACATATCCAGACAAAATATTAAAAATTAAAGATCTTGGTAACACGCAACTTTGTTCAAAATACTCAAGGACTGGACGAAAATATCCTTGTTTTGGTTTTGAGCAGTTTATCTTTGAAGAAACAGAAACAAACGATTATAACCATCGAAATTATTTAAAAATCAAAAGTTTCCCTTTACTTGGCTCTGCAACACCTGAAATGTTTGACGAAATCGAAATCTTTTATTTTATTTAAAACTGTCTCTTATAAAAATAAATAATTATTACCTGTCATTTTATAATTATTGTGCAAAGTTCTTATTTTTTATGTAATTTTTTAAATACTGAGTTATAACTTTTTA
>evgR016618c0g1_i01_Hydra
GAGTGTTACAGACAGACTTTGTTGTATAATAAGTGTTGCAGACAGACTTTGTTATATAATAAGTGTTACACACAGACTTTGTTATATAATGAGTGTTGCAGACAGACTTTGTTATATATTGAGTGTTCCAGACAGATTTTGTTGTATAATGAGTGTTACAGACAGACTTTGTTGTATAATGAGTGTTGCAGACAGACTTTGTTATGTAATGAGTGTTACAGACAGACTTTGTTGTATAATAAGTGTTGCAGACAGACTTTGTTATATAATAAGTGTTACACACAGACTTTGTTATATAATGAGTGTTGCAGATAGACTTAGTTGTATAATGAGTGTTACAGCAAGATTTTCAATGTCTGGTATAATGAGTGGTTCAGGAAGATTTTAAATCACTCATATAATGAGTGTTACAGACAGATTATTAATCTCTTATGTAATGAGTGTTGCAGACAAGTTTTCTTATTATGTATATAATATATAAA
>evgR016623c0g2_i01_Hydra
CCGACATTCATTACTGAGACCATATCTATTAATATTGAAACAACAATGTCTATTACTAAAACATCAAACACAAATACTGAACCAGCAACTACTACAGATTATGTATCAACAAATGATGAATCTACATACATAACTATAATATCTCTGACTACTATTACTGAGCCTATATCTAATGATATTGAAACAACTATGTCTGTTACTGAAACAACAAATACAAATACTGAACCTGCAACAACAACTGATTCTGTACTCACAATTGTTGAACCAACATACACAACTAAGATATTACAGACTACCATTACTGAGCCCATATCTACTGATATTTTAACAACAATGTCTATTACTGAAACAACAAATACTAAACCAGCAACTACTACTGATTCTGAAC
>evgR016629c0g1_i02_Hydra
CCTTAGCAAAAGTATAAATGATCAACAATGGCAAGTATTGCTAACTGATAGTACAAGTAACAAAAATGTTGGACCACTTTGTGAAAATAATACACCTTGCAACAACACATATTGTAGAGATACATGCCTAATGTCGAACGAAGGATTACTTCATACTTACTTTTGCTCTGGACATTCAATGCTAATTATCCTCAGAACCTCATTGATTCCGATTTTTCAACTTTTAGTGACTCCTACACTTACGATGCCAGTAATGCTTGGATAAAGTTAGATTTGAAACAGATATTTCAAATAACCATTATTGAAGTCTATGGAGATGATCAACTAAATAGTAAACTCCAACGACTTTCTGGTTGCAAAGTTCAATCAAGTTTTTATGACACGGAGTATCAAACATTTGCAATCTTGAAAGGTTTATTAATTGACAAAATTTTATGGAATGGCAGCTTACGATATTTTCGTATAATTCAAAACGTTGTGTATGAAGGGTTAAGCTTTAGAGAGATAATTATTTGGACATAATTATCACCTAACATTATCTAAATGATTATATCTGAATTTAGGAAAAAATAAAGAAAATAAAATCATATCCGGAATCATGATTCTTTGGATGATCATAATAATATTTTTTTATTTTTGAAATCAAACAAATGGTAATTATTAGTTAAATATTTAGAATGTTTAATTGTTATTGTAATATAATTTTTTATTATCCAAA
>evgR016630c0g1_i01_Hydra
CTAACAAAACAAAATGTCAAGCGAAGCTGCCACTGCACCAACCACTGAAACAACCAAGAAGGTCGAAAAGAAGAACATTGCCAAGAAGGACAAGGCTAAGCCAGTTTCTAGATCCCACAAGGCTGGTTTGCAATTCCCAGTTGGTAGAATCCATCGTCACTTGAAGCAACGTGTCCACTCAGGATCAAGAGTTTCTGCTACTTCATCTGTCTACGTTGCTGCTATTCTTGAATATTTGTGTGCCGAAGTTCTCGAACTTGCTGGTAACGCTTCTAAGGATTTGAAGGTCAAGAGAATTACACCAAGACACTTGCAATTGGCCATTAGAGGTGATGAAGAATTGGACAAGTTGATTAAGGGTACTATTGCCGGAGGTGGTGTTATTCCTCATATTCACAAGTCTTTGATTAACTCTGGTGTTACTAAGGCAACAACCAAGCCAGCTTCTAAAGCTTCATCTTCCACTGAAACAAAGGAATCTGCTGCTACTGAAGCACCTCAATCACAAGATTATTAAGTGATGACAACAGTTTAATATTTTCCAACAACAAAACAACCTAGATATGAATTAACACTAATTCCTTTTTGCCCTGTATATATAAATTACCAGTCTTCGAGACAAATAAAAC
>evgR016631c0g1_i01_Hydra
ATGGAAGTTTGTTAGATATAGAAAAGGAAGAAGAAAATTTGTTTATTTTAAACCTTCTTCAAGATGATAACAGAAAAAATAAAAGTTTCTGGATTGGTTTACGTGCTCTTCCAAGATCATATATGTGGTCTGAATGGTTAAATGGAACATCAAGCGGTGATATTTTTGGGTTTATTTGTAAAGTCGATTTATCAATATGGTCGAAATATCAAAATGTTTATGATGATCATTGGTTGCCATTTGGAAATTATAGCTACACTTTATTTATTGGAAAATCTGATGTTATATTTCAATTATCGCTTTTAAAATGCAGCCAGTATAAGGGAAGTTTGTTGATTTTAGAAGATGAAGTAGTTAGTTTTTATATTGCAGAATTCCACAACAGTGGCTTCAACAAAAACAATAATTTATTGCTCGGTCTCAATGATCATCGAAGATCATTTATGCATCCTGCATGGACCTCAACTAATGGTTATTTTTGGGGATTTATTTGTAAAGTTAACAAAGTGAGTACTTGTTCTAATGGTTGGCTTAGTTACGGAAATTATTGCTACTTTATGTATAGTGCAAATTATACTAATCAG
>evgR016635c0g1_i01_Hydra
ACAACTATTTTTTCGCAGTTTGGTTTTACAGACAGTATATTTCAGTTTACTACGACAGGTTCAGCTTTTGAATCTTTGTCTCAAACAACTGATATTTCGGAACCATCAACAGACTCAACTTTAACGGATTTATCAACGACAGTTTCAGTGCCACAGAATTCAGCGACTACTGAAATTCTTATCGACTCTCTTTCCACTTCAACACTTTCTACAGAATTTTTTCAACCACCATCATCGTGCAGTATTACAAACTGCTTTCTACCTCTATGCAGATGCGCGGGAACGGATATTCCTGGCGGATTATCGAAAGAAAACACACCACAGATCATTCTATTCACTATGGATGACGGAGTGACAGCAGAAAATTTTCAACTTTACAGTCAGTTACTTAATGGTGTAAGCAACTCTAATGGATGCCCAATTAAAGCTATGTTTTTTGTATCTGGGGATAATACAGATTACAGCTTAGTTAAAAAACTACATGAAAACGGCCATGAGATAGCTGATCACTCAACGACTCATAGATTTCCAATAACATGGTGGAGTGCAAATGCTACATTTGAAGACCTGCAAACAGAAATTCTTACCCAAATGACAACTATTCAAAAAGATGTTGATGTAACTACAGTTGGGTGGAGAACTCCATTCCTCGAAAGCGTGGAAAATACGTTTAAAGCATTAGCTGATAATAATTTTTTGTATGACAGCTCGTTAGGAACTTTTCCAGGAACAAGATGGTGGCCCTATACACTAGATTACTCTCCTTCTCTACCATGCTACATGTCCAACAACATGTGCTTTAATAACTCGTATCCAGGTTTATGGGAAGTCCCATTGGCTGCTTTACTGTGTGATGAATCAGATAGCACATTTGCAAAAATGTTCGATGAGTGTCAAAGTTTGACAGAAGATGCTACCTATAATTTTTTGATGTTGAACTTTAAAATATTTTACGAAGATAATAAGCAGCCATTTCCTATGTTCGGCCATTCAACTTGGTTTAATAATGCAACTTATCGAACAAATGGTGTCGTTCGATTTATTAATGATGTTCGAATGTTAAATGACGTGTTCTTTGTAACAACACAACAGGCAATAGAATGGATTAAATCGCCAGTTGGTCTCGATGAAAACCCATTTTCATGTAATTAACAGTGTAATTAAATTTTTTTTAAATATCG
>evgR016646c0g1_i01_Hydra
TTTTTTTTTTTTATAAAAAGCTAGAAATGCTTACTTTTTCTTTATTAATTCTATTGGTTAAATTAACAACAGAGTTAGATCAAAATGAAATGGACAGACATGACAATATGATTGAAAATGACAATATGATTGAAGACACAACTGTTGATGATCGTGCTGCTATTCCGCATGCGGACTCTTATAGATTACCGCACAATCAAAATTTTCACCGATTTCCGGAATATCTTCGCAAAAATGATAGAGATTTATCCTTTAATCAAGTGCTAAACACGCAAAACCTTCAAGGTTTTTTCACAAACCCTTTTATTCAGACGGGGTATGAGCGCGACAATTATTTAGTGCCAACAGAAAACCATATCAACCGCAATAATCGTTTTGTTGATTCATATTATCCTCAGTCTAACATGATACCATCTTGGTCAAATCAAAGATTCAACCGTATTAACGTTTTTAACCATCGAGACTATCCGCCGACACCAATGTTAAACAAAGTTCGGATACAAGATTACGAGAATTTGGAACCGAATTATCCTTTAGAAAGGCATACGTCATTTCATACGTCATTTCAACCTAAAGATTATTATCACAAACCAAATTTTGCAACGTTAAGAGAACCGACGTTAAGAGAACCTCTAAATCAAAACGGCTTATTAGAAGATTCAGCTGTTCCTTCCACTCATCATCGCACTAACTTACTACCTGCTTGGTCAAATGAAGAACCCTACCAGGTTCTACCAAATGGTTATAACTCCAGGACAGAAAACCTTTTTGATACCGATAGCTACAGACATAATACAGTTCCTACTTACAAAAAAATTGCTTCAATCAAAGAACTAGAAAGTCAAAACGGATACTTTAACGATCATTTTTTCAACAATATTGCAAAATCTATTGAGGAAGATAATTTTAAAACAGAACCTGAGGCATTTCAACCTTTTTATGGTTTTAAAAAGTATAGGGGTCGACGGGTAAACTATAACACAATGAAATCTGATGAAATTGCACCACTAAAGTTTGTCTCGCCTTCTAAAAAAGGCATTTTTCGGTTTAAAAACGATTATATTTTTCCTTGAACAATGAATTTTTTTAAAAACTTTTTATTTTATCAAAAGTATTTAAAACTTCTGATAGTTGCAAACACCATTTTGTTTTACAAAATGCTTTATACATTATTTTGAAAAATAAATTAAAAACGTAAAAAAAAAAA
>evgR016648c0g2_i01_Hydra
CTGGAACATTCACTCCATTCACTCCAATCAGTTAAAGTTCCTGGACAAGGTAGAGGATTGCATTGTTGAGATTGACTTTCTTTTGATGCTCCAATACAATCTCCACTCATTGTATCTGCTTGACAAAATCTCTCGCGATATTGATAAAGCGTATTCTCAAAGTTTAACTGAGTTTCTAAGCAAGTTTGTGAACAAGAACTCCAAGAACCCCACGGCGTCCAATTTGAAGATACTATTGTTGGTGAACCTGTTGTTGGTGAATCTGTTGTTGGTGAACCTGTTGTTGGTGAACCTGTTGTTGGTGAACCTGTTGTTGGTGAACCTGTTGTTGGTGAACCTGTTGTTGGTGAACCTGTTGTTGGTGAAC
>evgR016654c0g2_i01_Hydra
CGCAAGGATGTTTGATAATTATTCGATTGCTTACCTAACATTTTGTAACTTTGGATTAGTTTTGAATGTAATTAGTTTTTTTGTATTGCTTAAGTTTAAGAATGGCTCAACAAAGTATGTACACGCTGCTTTACCCGGCATACCAGGTGGAACCCGGCGTTTACGCGGGTTTAGTTTTGCACAGAGATACATCCTAATTCAACTCTGCTTAATGGACATGTGTTTAAGCGTAATTCTGGAAATCAGACAATTTCTATATATTTTTGAACTAAATGGCAATAGTTTAAGTATTCTTGTAGTAAGAAGAATTTCTAGAACATGGTTTTATTATCTAACATTTTGGTTAGTTTTTGATAGATACTTAAAAATTAAGCTCATTTTAAAATACAGAGTTTACTTGACAAAAAAGAAATTAGTAATAACTACTATATTTTCATTGAGTGTTGCAGCCTTGAGTGGTAGCTTGATTGCAATTTTCATATCAAACTGCAATATTGTTATTTACGCATGTTATGATCTTATAATAATTTTTTTTACCATTTTTGTCTACTCCTATGCTTTGTTACATAATCGAAAGCAAAAAAAAATTCGTTCAAACCAACACGGCAGAAGTGGATTTAAAGGAATGTTTCTACTAAGCATCGTAATAAGTTTGTTTCTTGTATTGTGGGCGATACCGGACATAACACTGGCGTTAAAATTCTTTAAAAATGACATTTTTATTAGTACGCAGTTAAATATGTATTTGAACATATGTTATCCAATAAAATTTTGGGTTGATGCTTTGGTATACATATTTACTTCTCCCCATGTTAGATTAGCACTAAAAAAGAAAATAAGATATTTTTGTGGTACAATACGAGACAAATACGTGGTACACAATAAGGGACTAATACTTGGTTCAACTCGCACAAGTTATGAGTTGTGCGTTATAACAAAGAGTTATTATAGATTTTCATCTGCTCGCTCCAGCTCTCTTATTTATTAACGCAATTTATTTTATTGTAAATTTTATTACACGGCATATAATAATAATAGTAATAATAATTAATAAAGTTTGTAAATAAAATATGTTTTATAATATAACTTGAGTTAATTGAGC
>evgR016656c0g2_i01_Hydra
ATTGCAGACCTCTGAGTTTTGAGTTTAAAAAAGAAACTAAATTAATATTGATTGATGAAGAAAGTAAACATAAAAATGAAATGGCAAATTTGATTCTATCCATCATTATTTTTGAAGGGAGAAACATAACTATTACAAACCATATCGAAATCACTATGATTAATGGAATGGTGTACACTGCCTTATCCAACGCAACTGACTCTAGTCAGTCATGTTCAGTTTGTGGATGTAACCCGTCAGAAATGAATGACATTGAAGTTGTAATAAAAAAAAGTATTCCATTCACGGTTCTTCAGAATGGTTTTTCAAGATTGCATGCTTGGATAAGATGCTTTGAATGCTTATTACATATTGCGTATAAAATAAATATAAAATTTTGGCAAGCATGATCTTTTGAAGCCAAGAATTCTGTAATCGAGCAAAATAGTTGAATTCAAAAAATGGTTCTTGGATAAACTTGAATTGATTATAGATAAACCGAGGGCAGGTGGTTGTGGGAACACTAACGATGGAAATACAGCCGGGACAGCTTTTTCAAATGAAGACGTATTTATAGGAATAGATAAAGTGCTAATTGAAAGATTGCATGTTATACTTTGTTGTATCTCCAGTGGTTTTGAGTTGGACATTTTAAAATTTCAAGGTTATTGTTTTAAAACTGCTAGATATTTTGTTAAAAGTTAATCCTGGTACTACATGCCTCAATGTCTGAATAAAATATTAATTCATGGATACTTTATGGTGAAAAAAATGATATTGCAAATTGGTCTAATGTCAGAGGAAGCTCAGGAGGCTCGCAATAAAGATTTTAAGAGTTACCGCTAACACCACACAAGAAAAATAAGTCGATCAAAATGCAATGAAGATGTAATAAAACAACTGATCCATTGGTGTCTAGATTAAGAAAATTTCGTTAGGAGTTTTAGAAATTTTATGTTGTC
>evgR016658c0g2_i01_Hydra
GCTGAACAACAAGCTCAACCTACTGCCAATGGTGCTGCTGAACAACAAGCTCAACAACAACCACAAGGAGGAGAACAACCAGCTACTCAACAACAACAAGGAGGTGAACAACAACAAACTTCAAATGGACCAGCTCAACAAAATAACCCAAATGTTGGAGGTGAAACTACTCAACCACAACAAGGAGGTGTTTCTAATAGTAATGGACCAGTTTATCAACAAGGAGCCAGTAACGGAGGAGCAGTCAATGGTAGAGGAAGTAGACCAGGAAACTATATGAGATCCTATTCTAGAAGAGGTGGAAGAGGTGTTTACATGAAGGTTAATATTAATATCAATGGTGTTGCTGCTAAATAAAC
>evgR016663c0g1_i01_Hydra
GATGGATGTGAAGGTTCAATCCCTTCCAACATCTCTTCTAAACTCGCCACTCAACGATCTGCTATTGAATCAAGAAGAAATTCACCTAGAATTCACAATCAACAAGTCAAGACTAGATTGGCACAACCATTGTCAACTGAGAGATCTCATTCATTCAGTGAGAGAAAGAAGGCTCAACAAGAGAAATTCCAACTTCCTCACTTATTCCCAACCACTACCATTGGTTCATTCCCTCAAACTGAAGAAATTCGTTCCATCCGTAGAAAGTATAAGAATGGTACCGTTTCACACGAAACCTACG
>evgR016664c0g1_i01_Hydra
CTCGTTCATACATATTTTAACAGGTTTAAGAAATTGAAATCTGAAAGGTTGTTATATGTTTTATCATACGAAGCAGCTGTATTCTGTTTTATTACTCGACTTCCGTGTGTTCATTATGCTAGCTTTAACATTTCTCCTAACCGATATATTGTGGGTAAAATCAATAACGAATCACAATCCATTAATAAAGTAGATTGTGGTTTATTATGCATCTCAGATTCTGCTTGTGTATCAGCTAATTACAACAATAAAGAAAGTTTATGCCAGTTAGTTTCTTCGAATGGAACGGAAAAAATTAATTTTGATTGGGATATTTTAAAGACAAATTCAAGCAACGATAAACGTGCTGGCCCATATTGCGAACAAGGTTATCTATGTTTAAATAACGAATGGTGTAGAGATGTGTGCAAAGACGATAATGTGCATCATTATGTTTGCTTAAGTAACCGAAACGTAGCACGTTTTGCTAGAACTTTTTACTCGTCAGCCTTCAGCTACCAAACTAGATCTGAAAATGCAATAGATGGAGACTTCAACTCTAATTCTTATTTTCAAAGTCAATTAAAAGAAGTAAATTGGCTAAAAGTAGATTTAAATTATGTCTTTTCAATTACTAGTATCAAAGTGACAAACACTTTTAATAATCAAGAAAGATTATCGGGCTACAAATTAATGGTTAGTGACATAGATAGCGTTAATAATTATAAAGAGATAGCAACTTTAACAAACGGTTGGAGACAGACATATAATTTTTCAGAAGCCGTACGCTATGTTCTTATTACAAAAGTGTCAACTGAAGCTCAATTTCTTCAAATTACAGAACTAGAGGTTTTTGCATAGGCTGCTTAAGCCCTCAAATGATTATATTATTGTGTGTGGTGAAAACACTATTAGACTGCAACGTGTTTAGAAATGTTATTATAACAAAGAAACGCTCATCTAAAGTTTAGAACAGGCTTAATTCATTGAAAGCATATAAAGTTTTAATATTTCAATGCATTAATTAAAATTTATATGCGTTAATATGTTTTTCGCGATTTTTGCTTAAATTATCAACATAATTATTTTTTACTAACTTTTACATATATATTGAC
>evgR016665c0g2_i01_Hydra
TGGAAATAATTTAGATGATTATAAACAAAGTGTTTTGGCTTGGCACGTGAGTGTTGATTGTGACGTATATTTGCATGTTTAAAAGAGATGCAATTCTCAGATGATAATGCAGTACAGTGGCCAGCATTTTCACCAGCATGTCAACATGTACCCACGTCTCCCCCATTATCTAGGTGTTGTTTTTCACCACCACCATATGTACCTGATATTGATATGAAAAGGACAATATGGCAGAAAAAGGTTCAAGTCAGTTATGAAGCCTCACCATCAATAAATAATTCAGTTGTTCCTCAAAGAGATATGACTGAACAACTTTTATCAACCCCAAGCTTATCTAATTTATATAAAGAAAAAGTATTTCCCAAAACACCTTCGTCAGAACAAGTTGTTCCTAATTGCTCCCCATTTGTTGTTGATGAAGCAAAAAGTAAGATAGATGTAAAAAATATACCTGACGATGAAAAACCACCTATCAGCTATGCCTTGCGGCGTAAAAACCGAACCTTAGAGTCTCCAGAATCTCCAGATGTTAAAGAGCTTCAGCTCTTTGACGAGGAGCAGATATTTGCATAACATTTTAAAAATGACCGTGCATAATGGTTATTGAAGCTGATCAGTTCCAAAAACGTTAAAATGGTTAGAAGTTTTAAAATGATATTTTATTCAACAGTTATTTAATATATTTTAGTGTTATTTATAGATATATTTATTAAAATTGTATAATATATCCAAATATATCTGAAAAAACAAAATATGTATTAAAAGATGTACTTCTTTTCTTTTTTGATTTTAGTTTCATTTTTTCTTAGTTTTTTTGTTCCACACAGTAGTGTGATTTTTTTAAGTTTAGTTTTTTTTAGTTCATTTTACGACATCTGTTGTTTTTGATGTAAAATGGTTTGTGTAGATATTGTATTTATAAATGAGATTTTTATAGGAAATGTTTTTGACTTTTATGTGCAACAAATATCTCTTCTTTTAAAAAAAAA
>evgR016671c0g1_i01_Hydra
GTATTTAATACCAATACCTTCAAGTTCAGGATTATTCAATATAACTATTGATGCTTTTAATAACATTCCTTCGTACCAGCAACTTCATTTTTCTTTAAATAACCAGGTTCCTGTTAGTAATTTAAATTTTGATTATATGAAACCATGTCCCAATATTATAGCTGGATATAAGCTCGATAGTTGGATAACTTTTAAAGCTGTTATAGATAATGGAACAGATGTTGTCTACATGTTTAGTATTCCCAAACTAAATTATTCATCAATTACTAGTAGTGGTCTAGTGGACTTTAAAGTGGTTTCTTTAGGTGATTTTACTTTATATATGACTGCTGCCAACAAGTTAAATTCTCTCAATACTTC
>evgR016677c0g1_i01_Hydra
TTTTTGTCGCATTGCTTCGTAAGAACAAGACGCGTTTGTAGAGCGATATTTATTTTTCTATAAAAAAATGGAAGTTACAATGAAAAATATTGAGCGACTAATAACGTCAAAACTCGAAGAGCAGAAAAAACATATTCTCAAAGAAACTGACAAGCTACTAAAAATACAAGAAACAATATTTACAGATATTATAAGCTCAAATCTGAAAATTATCACAAATAGACTTGATCAACTCGAAAGCGAAATTAACATCAGCAAATCGAAAACTTTAATTCTAGAAAAAAACTTCAGCGATATTAAAGAAACTATAAATTTCCAAGAGGAAAACTTTAAAAACAAACTCGCTCAAATAAGCCAAACGATAAATGATGAAATAAACTCGTTAAAAAACAAAACTGTAGATTTAGAAAATAGATCACGCAGAAACAACCTAAGAATTGACGGCGTAGCCGAACCTACATCTGAAACGTGGATCGACTGTGCTAACACAGTCAAAAGTTTATTTAAAAATCAATTGGGAATTGCGGAGAACATTGTAGTTGAAAGAGCCCATCGAGTAGGGCAAATAAAAGACGACAAGCCACGGACTATTGTTCTTAAACTCCTAAATTTTCATGAAAAAACCGTAATACTATCCCAAGCAAAAAAACTTAAAGGCAGTGGAATTTTTATAAATGAAGATTACGCAAACGAAACTTTAGAAAAACGAAAAAGGCTTTGGGAAGAAATAAAAAGACACCGAAAAGAAGGTAAATATGCAGTTATCAAATATGATAAAATTTTCGTACGAGAATTCCGAAAATAAGCAAAGTTTTTATCA
>evgR016679c0g1_i01_Hydra
GTTTGGACCTCAGGACCGTTTTTACCATGTTGATTGCTTTGCCAAGAACAGGAGAGAATTGGACTTTACAGAATCAGCAGAGACAATCCCAGGATTTGACCAGCTGTCAGTGGAGGATAAAAATCTGTTGAAGGAAAAGCTACCAGATCTTCAGGAAGGTGACGATAAAGAAGATATTAAGGAAGAAGAACAAGATAAAACAGGAGTTGAAAATGAAGAATCACTTGAAGAGCCAGAACATAAAGGTGAAGAGCCCGAGGGAAAGAAACTAAAATCTGATGATAGTAAAGAAACAACAGCAGAAAAAGAACCTACAAAAGCAGATTCACAAGAAGAACTAGGAGATGTAGAGGCAAATAGGGCTGACTAAAATAGTTTTTGATCTCTGATAATACCCGAAATTACCTTATGAAAACTTTTTTATATACTTGCAAACAATAGTAAAG
>evgR016698c0g1_i01_Hydra
GATTGCTTGGAGGAGGTATCCATTTTGACAAGGTGCAAAGGCGCTTTGTCTATTACCATTGCAACCATCATTGCACCATTGTAAACCATCATTGCACCATTGTAAACATAGTTAGACTTGCGCTTGCAGCCAACCTCCACCATTGTCACATTGTCTTTCTTGTTTCTACAAATACTATAATGGTTGCTAGTGTTTCTCGTGCCATCAACTCAAATTCATTCTCGTGTTACGTGTCTTTCAGTTAGTTCTCATCTTTATACTGTGACTGTTCAAAAAATTCTTATTTATCTAGTTTTTTTCCTCAAACATAAGTTCTTTGGAATTCGCTGCTTTCATTTTGTTTTCCTGATTGATATAGTTTGCTTTGAATGCTGCTTTTGATCTTTTTATTGGTCTTATATGATAGCAACAAAACAGTTATTAGCCCTAACAATCAACTATGGATACTATTATGATTTCCATAAACCCAAGACAAAGTACAAATATGCAGATTCTACATAGTTGCTCATTTATTCAACATAGTTGTACGTTAACTAGATTTACCTCTATTATTGTTACACGGTTTCATTTTTATTGTTCATTTTGTTGTTTTGTTGCGGTTTTTAGTAAAATAAAAAGTAAACCCAATTTAGCTGTTTTTATTTTACAATTACACGGGCAGTGGCGTCTAGCATCAATAGGACCCCGTCCCTCATACAAGCTAAAATTATCTTGGAGATGGGTTAGTTCCTCCCCTCCCTGCAGTTGCGGAGAGAGATGGGAAGCAAAAAGGGGCCCATTTCACAACTGTACTGTACTGTACACGGTTCATTTGTTTGCTCGCTGTAGTAAATTTGTTTTTACTGTAAAAAAAGTAAAATTTGGGGTCAAAATATTTAAGGACTAAGGTCATAGATTTGTAATCTTTGCCATTTAAAAAAAATATATTTTTGTTTGAATTTTG
>evgR016700c0g1_i01_Hydra
AGCGGCAATTAGTTCACAGCCATATACTAGCGTCGAGTAAAAGGAAAATGGCGGCTAACGATGATCAAGAAATTACATTTCAACAACAAGATGCTGAACAATCTGAGCCTCCTAGTGACCTTACTAGTTTTCAATTTGTTCCTGAAAAGATTGTAGATTTTTCTATCACTCAGGATGGGCAGAGATCTTATCGAGTGCGTTGGAAAGATTCTTGGTTAACTGAAGAGCTTCTTGCACTTACCTACCAACATCTTATTAATGATTTTTGGTCAGCATCTGCTAATGTCAATAAGGAAAGTCAAAAAAAGATTACTAAATTATCTAGCAGTATAAAAATTGCAGCTAAACAAAAACAATACACTGACTTTGCCCATATAACACCAAATATTTCTATTAAATCGTCCGGTGATGTTTCACATATTGATTCTATTAACCATGATAGTTACTTAGAACAAGGAAATATTTTAAAGTCTCACACAATTATTGCGAAGAAACAAGATACATCGAACTCTTTAAATTCTTTAATAGTCAACAATGAATCTCTCAGCAACTTGCAATATCAAGATATTGTTCCTGCAATGTTATCGACTCAATCGATTCCAAAATACCAACAGGCTTTGCAGCAACCTATAACTGTTCAAACACAACCTCTCTCACAAGATTCAGACGGAAAAGTTACTTTTCCAAAGCATGTGTTTTCTCAAGCAAGGTTTTTAAACAAAAAACAGCAGCAACAGATTGCTGTTGCCGATGGTCAACGTGTTCAATGTGATATTTGTCAAAAATCGTTAAAAACTAAAAAAATGTTATGGGCACATAAAGTAGCAGTGCACTATGGTGGAAATTTTCCATGTGAAATGTGTGGTAAAAAATGTATTACTGGTGCTGAACTAAGAAGGCACATGACTTCTCACTCGACTGAACGAAAGTTCATATGCCAGTATTGTGGTCTAGCTTACAAAAGATGGTCTCACCTTTATCAACATTTGCGTGTGCATGAAGAAGAAAAAAATTTTCGATGTGATGTTTGCCATGTAAATTTTAAGGTGCAGTCAGAATTAAAAGATCACTGTTTTGCTGAGCATAATAATGGAGATATGGTTCAATGCAGTGTTTGTAGACATAAACTCCAAACACCATTAGCAGTGTATCACCACAGTATGAAGCATACTGGTACTCGAGATTTTAATTGTGAAATATGTAGTGCTACTTTTAAAAGAAAACAACATTTAGTGACTCATATGAAAACTCATTTATCAGAAACAAGACAAGAAGATGGCTCAAATGAAGTGTTTAATTGCCAGGCTTGTGATGCTAAGTTTACATTAAAGATGGATTTGAAATCTCATTGTGATTCTGTCCACATTTTACCAGGAGATGAATCTGTTAATTGTAAAACGTGTCAGCGGAAATTGAATGTATCTCACTCTGTATACTTGCATGGTTTAAGGCATTGTGGTGTTCGTGATTTTCCCTGTAGTCACTGTGATCAGAAATTTAAGCGCAAAGCACATCTACTTAGGCATCAACAAGATCGCCATCCAGCAGAACCACGTGAGCGCAAAAAGAGAGAGGTTCGCATCAATTCAACTGTGTGTGGTTTATGTAGAAAAACTTTTAAATATAAAACAGTTTTAATAAAGCACATGGTAGCCCAGCATGGTATTTTACCCAAAGCAGATGGCAGTTTTAAACCTGGTGTTGTTTATAAAAAATTCAATAAAAAGCCATTTCAGTGTGACCACTGCCCTCATCGTTTTAAAACAAGTACGTCTTTAACAAAGCATAATCTTGCAAAACACGAAAGTGAAATGTCTCATAATCATCAGGACGAAGAAGTAAGTAATTCAAATTCAAATTTTGATCATCTTACTAATGAAGCTGCTGGTGTCATGGTGAATCTGTGTACTTCAGAAACTAATTTAACAGATATACTAAAAGTTGATTCATCTGGTGTAAACAATGCGCAACCAAGTATTCCAAATATAACTGAAAACATGCAAATTCAGGAAACATTTGAAACTAATCCTTCAAATGCAGAAATACAACATTTATTTAAAGACGAAGGTGCTGATGGGAGCGAAAATAAGTTGCAAACAAGTAAAGAACAAGACCAACTTTCCATAATGCATAAGAATTCTAATATGCTACGTCCAAATACTTTAGTTAGTCGTCGTATACTAACACTAGATGAAACTGGTCAGTTGATATTGGTGCAAGAAGATGATTCAATACCAATTGTTACAGAGAGTTCGTCATTGCATAATAATGTTGACCAAAGAGTCTTAATTGTTCACGGGGTCAGTAGTAATACTAACCCAATAAACGTGGATCGTAGTCATGTAATTAATGTTCAAGGGAATGACCACCCACGAGTTCTTTCTTTGTCCGAAGAGTTACCTCATGCACGTGTTTTGCCTTTCAATAATGTTACGAATGAATCTGAACTATCACCAATAATGCAAAGCAATATCCAGTTAAATGATGATCAACAACAAGTTGTAAATCTTGGAGAACACAATATACCAATATCATTGGGTCAAATAATAAGCCAAAATGGTGTTATCACATCTGTCCCTTTGCAGTCTATGCAGGGTTTAGAAAGACATAATTTAATTCAGTTTACACAAGAAAATAATGAGGAAATTGTTGTCTTATTAAGTGCATTAAATTCTCAGTCTCTCCGTCATATGTAAAATTTTTGTCAAGTATATATAGTATTCTCAGTATCTTTATCGCGTGAATAATGACGTCAAACTTAGAATAGCGTTTTCTTTGATGATAAAATAAAAAAATCCAATCAACTCAAATGTAAATCTTTGTTGTAATTTAAGTTTATTTTTACTGAACATTTTTAGTTTACAAATTTAATATTTGGTTTTGTTAATACTTATTTTTTACAAATTTGTTTGAGTTTGAATCCAATAGTTGTTTTTTGTGATACAGAAGTTTTAATAAAA
>evgR016703c0g1_i01_Hydra
GTTCACAAACAAATCATAGCATTCCGTTTAAAACGTTGACTATGGCTGAGAGCGATTATTACAAACTTTTGTCAAACCAAAATAAAAGCAGATATAAAACCAAGATCTCAAAAATAGACAATGTTGATCCATATACTTTAAAAAAGGCAGATTATCTTTCCGAGCCACAATTTTATCCTAAAATCACCCATGCTGATATTATCACATATTTATTGTTTGCACCAAGCCCTGCCACCAAGGAAGAACTTAAATGCTTTAAAAGTTTAGACGCATATAAACATTTTTTATGTGGTTGGGTGAGAGATATTGGTGTTAAACTCTTCAGTAATGAATCCATATGCTTGGTTCACGGAAAGGTTTTGCATTCTCAACGATTAAATGAACCTCCAACATTGACATGGGTTATAGCCCATATGGATGGTGATGTATTATCTGCTCACTGTAACTGTATGGCTGGCTTAGGGGAGTCTTGTTTGCACATTGGAGCTATTTTATTCTTTGTTGACGCAACCGTCCGAATTAGAGACTCTAAAACTGTCACTGAGGAAAAGGCCTATTGGATGTTACCTGCTTCATTAAAATCAGTACCTTATGCTGAAGTGGTTGATATAAACTTTACATCACCAGATACTCTTAAAAGAAAGTTTGACGATAAACTTAAAAAAAACAATGAAACTTCTAGTGAACCTTTTGTTAATAAAAAAAACAAACAGTCATGTTTGGAAAAGTATAAAGGACCATCACAAAAACAAGTTGATATATTTTATAAAGAACTGAGTCTGTGCAATTCTAAACCTGCAATTTTGTCTATAATACCTCCTTATGACGATATTTATATTCCAAAACAACATTCTAACTTATTACCAAAAATGTTATCTGAAATTTACGATACAAATTTATTAAAGAATAGCTATTTAGAGTTATGTCAACACTGTGAAAACATTGAATTTAGTCAAGTTACTGAGGAACAACAAAAAGCTGTTGAGGAATTAACAAGAAAGCAAGCACTATCAAAAAATTGGTTTTTATATCGTGCTGGCAGAATTACTGCATCAAATGTCTACAGTGTAATGCATTCTAATCAAAATAAACAAACATCTTTGATTAACAAAATTTGCTATCCTGTAACTAATAGCTTTTCAACGCAAGCTACAAACTACGGAAACATTTGTGAGAAAAAAGCATTAAGTATTTACACAGATGCTGTTAGTGCCCACCATGAAAATTGGAAGATTTTGAATTGTGGTTTATTTTTGTCTACTGAGTATTTTTATATGGGAGCATCACCTGATGCAGTTGTTTATTGCTCTTGCTGTGGTGATGGGTGCATTGAAATAAAATGCCCATTTAAACATAAAGATGATTTTATTTTTGAAGCCGTTGAAAAAAAAATAAAAAATTTTGTTTAC
>evgR016705c0g1_i01_Hydra
AACAGTAGGTGCCTTGAAACAACTGATGTAGAAGGTACAACGATTATAGTTTTATCATCTTTTGGTTCTTTGTATTGTCTCTGCATACTGATAACATTTATTGTGCAAAGAAAAACACCGGTTATCAAAACAGGAAATTTTACGTTTTCATTTATAATGTTAGTGGCACATTTGTTGATTTTTGTGTCCATACCAAGCTTGTATATAAACTCGCCGGTTTCAGCAATTTGTTCTGTTAGACATTTAGTTTTAGGATTTTTATTTACCGTTTTAGTGGCAATAACATGCATCAAAACAAGAAAATATGTTAAGATTTTCAAACTGAAACATCGTCTCACAAATAAAGACAAAATTGTATCGTCTTCCCTTGATTTCTTTATCGTTCTGGTTCTTATAGTTTGTCAAGGCGCTTTAGGCGGCATTTTATTTATTCATGATCCACCAAAGTTAGTTGAACATAGAGATAACAATAGCCTGACCAAATACAATGATTGTTCAACTGACTTACATTTTACCTATCAAGTCATTTACATTTGTCTTGTTTTGTTGTTATCAGCATTTCAAGCGTTTCAAGCTAGAAATTTGCCATCTTTTTTTAACGAAACAGCAGCTATCACTTATGGCACGTGGATATCATTTATTATTTTGGCTTGCTATTTTCCTATATATTACAGCTCACGAGACAGTAAATCAAAAGCAGTAGCAATTTCTTTAACAATGGGATTAGTTAATCTATCTTTAATGACTGCCAAATTTACACCAAGAATGATAATTGTTTACTTTAAACCCAAACAAAATAAATTAAGCAACTTGAAGTTAGAAGTAAGGATTACAAATTTA
>evgR016716c0g2_i01_Hydra
AATAGATTTAAATGGAATAAATGTTTATAACATTACAAATACATTTCCAAAATCTTTTCTAAACGTCAAGGTATATAGTGGAGACCCATGGCATGATTCAGCAAATGGTTCTATCAAAAATCTCAAAATTATTAGTGGAAATGGAAATTTTGTATATCGCGATGAGCATTTGTTGATCAATGATAACTTAATTGCAATATTGACTTATTTGGAAAAGATCTGTTTAATATCTTTTAGATTTAAATTCATGCAATTTTCAAGTGATTGGGGTAATATTATTCATTTGACAACACAAGGTGATATAGGAACATATGGTTATAGAATTCCTGCTATATTTTGTTTTAAAAAAGAATTGTGGATTGTTTCTGCAATAAGTGGTGATCCAAACTATTTTTACTACTTCTTAG
>evgR016723c0g1_i01_Hydra
CGTTGGTGGAAATGAATACCAACCAATTATCAGTACTTCTCTATTTGTTTTACTTTGTGGTATAGATATAAATGCTTCCTCAACATTAGATACAAATGCTTCCTCAACATTAGTTTTTCATGATAATTGTATGTATGCAATACCTGGAAAATCTGGATTATGGTACAACTTTGCGTTCATTTATAACGAAACCATATCAGTTTATATTGATGGTCAACCTGTTTCTAGAGTTGAATGCATTACCACTAATTTAACATTTACAATTGAAAATAATGTGCGTGTATCAGGTGTTAGTGGGTATCCTACTTTTTTGGATGAGTTGTTTGT
>evgR016725c0g1_i01_Hydra
TTCACACCTGCTGGGAAAATATAATAAAAACCAACCAATGAAAAAGATGCGCAATAAATCAGTGGTAATAATTTTCGCGTTGTTATGTGTACTTTTAATTGGTAGCATATATTTGGTGGAAACTTATAAATCAAGATTGCATTTATTTAACTTTACATCAGATTCAGTCACAATAATTGATAGAAAACAGTTTTTTCAAAATATACCGTTTACTGCGATTTCTCACGAGGAATGTAAAAGCTATGTTTATGACGTTTCCGAATGGACTCTGACATATAAAAACAAAACTTATAGTTCAATTAATAAAAGCGAAA
>evgR016737c0g1_i01_Hydra
TATAATCATACACAATGAAAAAAAAACTTTAAACATCTAAAAACAAAAAAACTATTCATCATGTTTTTAAGTAAAGTTTAACAACTATGGATGATGTCACCGTTGAAAATAAAGCATCCGATCGATTATTGCCGCAAGTAAGTATCTATTCGAACAAAAAAGACCAGACTTCAATAGAATTCTTGGAAATAAACAGCTCTGGTCAAAAGTGTTTTGGAGAACGATCTGATGTTGAACGTGTTTTGGACAGTCGCTGCATTTGTTTACGATCAAAATGTTTACATTTATCTGGAACATTCTTACTTTTATGCATAAGTGGCGTTGTACTGGTGTTTTTTTTTAGAAGGTATATAAACGAACTATTAGAATGGCTTCAGCGCCTAGACGGCAGGGTTAGCGGCATTTTGTTTGTTGTAATGTTTACCGTAGTTTCTTTTCCAATGACTTGGGGCTATATTCTCCTAAATGTAGCTGCAGGATATTTGTACGGATTTTTTATAGGTCTTGCTACAGTTACTGTTTCTGTTTTAATTGGAGTCACAACATCTTTAATCATTTGTAGACTTTATATAAGAGGTTTCCTAAAATCTAAATTAGAATCGGAACATTTAAAAGCGATAATAAGAGTTGTCGAAAGCAGACGCGGCTTTAAAGTTGTATTTCTGACACGTTTGACTCCTATACCGTTTGGTTTGCAAAATGGTCTTTTTGCTATTACCAATATGGACCTTTCCAAATGTCTGCTTGCTTCCTTTGCGGGATTGTTACCTACTCAAGCACTAAATGCATACATGGGCTCCTCTCTGCGGACAATAGAAGATGTTGTGCATCATAATTCCCCTGGCGGGTATTTAATCTTTGCTATCCAAGTTGGTATCAGTGGTTTTCTAATGTGGTACGTCATCCGTCGTGCTAGGTATGAACTAAATAAAGCGTGTTTAGCAACCGACTATGAAAAAGCTCTTGATGATTTAACCGCGCCGTTACCGAAGAACGGTTTGCTTAAAGTTTCTAGTAGTATTTTTGAAAGTAAAAAATGCAAACCGTTTAATGAAAATAGGGTTAATGGCGATCCAATTAGTAAAAAGGGTCACAAAAGAGCACAATCAGCTTCTGCAGTTCTTTACGCATTAGATAACGATACCTTTTAGGGATGATGATACTTTTAGGGATCCTAATGAATCTGTATCAAAGATATTTTAATTATTGTTCTCAAAATTTTTGTGGGAAATAGCTTCCTTTAAATAAATATAATATTATGATATATTTTGAAAATTATTTAGACTTTTTTTTATGTTTTGTAATCTTAGAGTTTATTATCTTGAAAAATTAATGTATTAGATGTTTGATTATGCAATTAATGAACGATTTATAATAGAATATATTTTCTGCGTG
>evgR016741c0g1_i01_Hydra
TTAACCTATTTTAGTGCAAAAAGTCAATATTTTATAAATGAAAATTAAGAAAAAAGAATGTGAAAAGGAAGGAGATTCTTTTTGTTATGATCAAAGTGAAGTAAGCAATCTTTTAAATATTAAAAGTCCAGTTCTTCACAAAAGAAAAAAACATCAAAAAGTTAAAGTTACTTTAGAAGATGGAAAATACTTAGAAAAAGAAATTCACCCTGCCATTCTGTATTTGCTTCAATGGAATGATAACAAAATTAAATGGAAGTTTCAGAAAGTTCGTCAAGTGTGGTTGCTTCACAATATGTATAATAAATCACAGGTAAACAAAAATCATTTTAAAGTCTTGCTTTCATACCTTAATGATTTAAAAGGCATATCCCGTGAAAAGACAATAGAAACTGCAAAATGTATTGTTGAAGAAGATGTTGCCGGTTCTAAAGTTAAGCGTGCTACTCAAGTTTTAAAAATGCTCTCATAGCATATTGGATATATATGTAAAAAGGAACATTGGATCTATGGAAAAAGAAACTTAAACAAAGGAATAATTTATTGATTTGTGTTTTGTTAATAAAAATATTTATCATTATTTATATATAATAGATAATTATTTTTGTAATTCTTTTGTTATTTCAAATTTTAAACCTTAGTTGATAGTAGGAGAGAATTACGAGTGAATTATATATCGTTGGATAGAAAATTTGGTGCTGAT
>evgR016742c0g1_i02_Hydra
CTTCTTCAACAGTGCTTTTTGGTAGTTTTGATGCAACAACAATAACTTCGAATACTGCTCCTAAATGGGTATCTACGATTTCTGGTGTAGAAAAAATCTCTTCAGCAGTTTTTAATGATGGATCGTTTAGTTGCAGATATACAACAAACAATACTTCAAGTAGCAATGTTTTCTTAGCTATTGCTTTTGGATCTACTATTAGTGGTCAGTTACCATCTTACCATGGATACAATAGGTACACAAAATCTGCTGATAAAGTGAATCTAATGGCTATTCAAAAACTTTCAACTGATATAAATTCAACACAAAATACAAATTCATCAATAAAAACGATACTTAACATGACAACAACATCAATAACAACTACACAAATAACAACTACACAAATTCAAGGTACTTCTACAAGCAATGGTTACAAGCCAGATCTTTCAAAGTGCGATAAAGAGTTTAATTGTTTTCTCAACCCCCCTGGATGCACTTCAAACTGTATTGCAATTGCAACCTTTTTCTATGAGAAAAGAAACAATGTATATGTTAACGAACTATGGTCAAGTGCCTCTAACAAATGGATTTCATTTGCCCAAAAATCTGACAATTCGAATTCTGATTTTATGGTTAACTTGCAAGGTCAATATTGTCTTTTATCTTCTTCAACGGTGCTTTTTGGTAGTTTTGAT
>evgR016751c0g1_i01_Hydra
AGTAGAAGCAATCTCAACACTATGAATACAGATTTAACTCAAGGAAAAGAAGATGAAAAAAATATGTTACTCTGTGATAAGAAGGCTCATCAATGTCCACAATGTGGGAAAATATTCAAACGGCGGGATGTAATGATTCGTCATGAGCGCACACATACAAACAATTTCTGCTATTTTTGCTCATGTTGTGGTCGCGGATTTATGCGACGATATGTTCTACTTAAACATCTTAAGCGTGAACACAATACTGACGACCAAACAAAAGTTTTGTCGACAAAAGTTTCTAAAATTTCAAGCTCTCCTGATGTTTTTATTGGTTACGGCTCGGATGTAACATTATCTGAAGATCTTGGTGCTGAAACAGATGTTTATAAATATATGAAAAATAGAAACACCAACGATGATTTTATTCAGATGCAGCCTAATTATGAGCCTGGTGAAGATGAACAAAAATATATGTCTCTTGATGAGGAAGACTCTGAGAAATCTATATCAAATAAAAAATCAAAACATTCAAATCATGGGACAGATAAACAAGATGAAGCTAATCTAACATTATCCAAAGAAAATATGCAGATACCAGAGTGTCATAAAAACGAAGTTGAATTATTTGTATTGGAGAGTTTGTCTATAGGAGATAACAATGATGAAAATAATTTAAAAATTGAATTAATAAATACGAAACAAAGTTGTGATGGAAAAGAAGAAATCGCCAACAATGAAAGTTTAATAGTAGCCATTGAAAGCGATAAAGCAAATTCTCCTGACTCAGATTCAAATTCTCAGAACTTAAATTTTAGTGACACGTGATTTATGCTATTTTTAAAGATTGTGTAAGCATAAAAATGTTTACAAAGACATTATAATAATATAAATATATATCTTTTTTTCTAAGTTAAAGTATTTATAGTTGATATAGAAACTGCTTTTATATGTTCATTTTTATTAATCTATTATTATGAATGCTATTGTTGTTTTCTTTTTTCTTTTTATTTTATTTTAAATTTTTATTTGTTGAAGAAAAGCACATTTTTTGATTGTAGGTTTTAATTATTTCTATTCTTTGAAAGTTGTTAATGTGTCTTGAATAGTAG
>evgR016759c0g1_i01_Hydra
ATCACTGTTATTACTGTAGTTATCACTGTTATTATCGTAGTTATCACTGTTATTACTGTAGTTATCACTGTTATTACTGTAGTTATCACTGTTGTTACTGTAGTTATCACTGTTATTACTGTAGTTATCACTGTTATTACTGTAGTTATCACTGTTATTACTGTAGTTATCACTGTTATTATCGTAGTTATCACTGTTATTACTGTAGTTATCACTGTTATTACTGTAGTTATCACTGTTATTACTGTAGTGTTATTAGTCACTAGCATTAGATTAAAGAAAAAACTTAAAATAATAAACTTGTTAGTACTCATTTTCTTGTTTTGTCTATCATCAGTATCAACAATTATTATGGAGGTTTTTCAGTCCTCAATTGTGCATTGTATGACCAGCAATTTTTGATTTGATTTGTTACATGTTAATTTGATAAACATTTTGTAGAAAGCATTGCTTGATTTCAACATTTTGCAAATCCTGCAGATTCAGCCATGATTTGCGCAGGCTGACTTCACATATTTAAGGCTCCAAAATGGAGGTAATCTTAATGGTAGTATAATTAAAACTCAGTCTTGACATGACAAAAAAGATTTAATAATGAGAAAGTCATTTACCATAAAAAGGCATTTATTTAAACAAGTAAAAGGTAAACTAATATGAACCACCGTGGGAAAGTCAGAAAACGTATAGTATTTTACATTCTTACCCGAATTCTGTGTAGTTGATAGCTTTGTTGTTTTTACACATCTAGAGGCATCTTCATCTG
>evgR016760c0g1_i02_Hydra
TTTAGTGCAAAAAGATATACTCAATGTCTTCAGCTGGTCCATTAATAATTTTATTTATAAGCATTCTATGTTTTGCCGGTGCAGTTGGCTCTCTCTTGGCAGGCTGTTATGGTAATGAATGGCTAACCAAAAACACAGGTTCAATTAGAATAGGTTTATGGGAAGAATGCAACCTGGTACTTGTCAAAAGTTGTTCTAAAAGAGATTCTTTGTTTAAGTTTGAAGATCCAGTTGGGAATGGTGTAGAAAAAGATATTATCCTGTTGTTATTAATGAGCTCCGCAGTGTCTGCATTTCTCGCTCTGATTTTCTCTATATCAATGACATGTTGTTTTAGACGTTTAACTGCTTGGAAATGCACGGCTGGAATTACCTTTGTGTTTGGAATTATTGCATCTATCGCGGGTTTTGCTGGATTGGGATATTCTGAAAAAACTTTTAAAAACGAATGGTCTTCAGATTTTGGACATGGCTGGTCGTCTATTATATGCTGGGTGGCGTCTGCGTTGAGCTTTCTAGGTGCAATGTTATCGTGTTTGGTGATTCCTTATTCTCCCCCAAATTGTCAACCTAAAATTGCAACACGTGTTGGTAAACCGAACCCTACATATCAAGGTGATAATCATTGGTAATTACCGCGTTATTTGCCTATGTTTAAAACATGGCATTACTTAACTACTACTGTTACTACTCTTTGGTGAAAACTTTTATTGAACCTTCTTGGTTTATATTTGCGGAAAGATAAAAGATGACATTGAAAATAAACATTTTCGTTCAAAGATAAAATAACATTTCAATATTATCATGAATATTGCAACTATGAACGTTGTGGAAGTGTTATGCTGTCAATATTGCATCAATAACTATGAAAGATAACTTAAATATAGAAGTACATAATGCTTAATATATGTTTAACTTGTCGAAAGATTAACTGTCGAAAATCGTGACGCCAAAGTCATTTAATGTCAAACTTTATCTTTTTTATAAAAAATATTTAAATAATGATAAATTTCCAGTTTAGTTCCAAGCGCTAAAATTTGATCGAATGCTTTATAGAAAAATTGACAAAAATACTTCAAGAAAATAGAAGAAGTATTATAGAATAACATAGAGGAAGAGTGTTTGAATAATATTCATCGAAAAGGTAATATTCAAACGGTAGCATTTTAAGGATACTTTGAACATAAACGATAACATTTAAACGCGGAATTCCATATTGAATCTGCAATGCTAATATAGTTATAATTTGTATTTCTAATATAAAAAGTTTGTATAGTTTAAATTTACGAAAGAAATAATGTTTTGATGTAAAAAAAAA
>evgR016765c0g1_i01_Hydra
GTCTCACTGAAATCAGAAAATCCACTGCGATTTTCAGCAAAGTTTTATAAGATTTTTTGCTGTTGTTTAGATTTTGAAAAATCTGTCACCGGATAAACAAATTCAATAGCAAAAAGTTCTCGAAAACGCTTCAAGCAAAAATAGCGAAAAAGTTTAAATTTGAAAAAACGGTTAACATCGACAACTAAAGAAGCCATCAAAGGAAACGAAAATTATCGTAAGAGTGACCAATCTGAAAGCATGTTCAAGGAATGGGCGAGTCAAATATAGAGATCACCTCAAAAAAACGTACTACAAATCCCCCGTTAAACGAAGATAATCAAAGTAACGCAACATTCGAAAGGGTCGAACTTGGGGATGTGTTTACAACAAATCGCATGTTGGGATACATTGATGCTATCATGGCTACAAGTGCTACTTTTCTTGTTGTACCTATTAAAAATATTAAAGAGAATGAAACTAAAAGCACATTTTCTCATTATTTAATAGATATGCAAACAGAGTTTATAATGTTTTTTTTGGGTTTCACTCTTGTTTTAACCATTTGGGAAAACATGAACGTACGAGCAATGTTTGTTAAGCGAGTTGATGACTTTGTATTAACTTTTGTAATACTAGAGATGTTGGCTACAATTGTCCTTCCGTTTGCAATATATTTACAAGGCAGCTATCCTGATCAATATGTTCCAATAATTTTGAGTTGCTGTGCGTTAGGGATTATTCAAATTTTAGATATTTTTCTTATATTTTACGCAACAGAAAACGCAACACTACTTCACACTAAAATGCGAAATTGGTCAATAATTGAACTTCGCAGATTTCGTAACATACTCCTGGTTCGACCAGTATTTTGCTTGATTTTAGTTACAACGGGTGGTGTGCTCTGTCTAGTTCACTACGCTATCTCTTGGGTATTTATAGCTATTTTAACTCTTATGCCAATATTTCGTAATCTTTTTCTTTATGCTCGACGGCGTATGAAAGCAACAAAGAAGATAGAAAAATATAAAGTTTATTATTATTTTTCAAAAAAAAACATTCCAAAAGAACGCATTGAGGT
>evgR016766c0g1_i01_Hydra
TATTATGGCTAATTTTATAATAATTTAAATTGGCAAACCGATAATAAAAGCCTAACAATCATAGAAACGCAATCACGAAGGAATGGATTTTGAATATAAGAATAATCAAAAGGATTCGGTTTCAGAGCTGAAACACATGCTTATAAATTAGATAAATAAATTTCACTTGTAAAAAAAAAAGGTCATGGAGAGAACTCAAAATATTCATTCTTGTAATATAATAGCCTCAAAAAATATTTTACATTATGTTTTCATTTTTGTCGGTATAGTATTGGCATTTTTGGCTACCTTTGGAAACTATTGGATAGAGATCGATAGAGTAGTAAAAGGTTGGCAATTCTATTACGTATCGATTCACGAGGGAATATGGTTTACTTGTGCAACTGTTAATAGTATTAGCGTTTGTTTGGACATTCAACAAAAACCGGATTACATATTTGTTACTGAAGCGTATATGGTACTTGGTTGCATTTCACACTTTGCTGCACTTGTATTAAGTTCAATAAGCTATTTCAAGAAAATGAACCATTACAAACGCTCTGGGGCTGCACTTATTTCAGCAACTTTATGTTTGACGGTAGGAATGGCTGTGTACACGAGTCAAACTAGCCAACTTTTTATGTTTTATACCTACAGTTGGTCTTACATAATCGGATGGTGCTCAGTAGGTATTTCTATTATATCTGCAGTTATTTCTTTTAAAATGAAAGTCGAAAGTCAACCTGCTTAAAAATGAGTTTTTTAAATAATAAAAAATAAATAAATAAAAAAAAATTTATTTTATTTATTTAAATTTTTTTTATATATATATAAAAACGCTTGAATAACTTAAATTGATTTTTTTAAATCTATTTTAATCTTGTACTGTTATCTAAATATGTAAAGAAAAATTATCGTATATTTTATTTGCTACAATCTAATGTATTTTTTGTCACAAACATGTATATATAAGCAAC
>evgR016768c0g2_i01_Hydra
ATTACGACTATGATTAGATAGATACACGACATAACGTATGATATATATATATACGCCCTTTCGGTAAATGTTCATTATATCAGTGGGTATGCCGGGTTTTTATCGTTTAATCGTATAAACTTACCTAAAATTTCATTTTCTATTTTGACTTAAAACATCATTTAAAATCAGTGCGGCCATTACGTGTGAACTCTGACCTCAAATCATTTGGTCTTGCGTTATATTATAGTAGAATAAAGCGTTAATGTGATTAGTGTCAAAAGTATTGCATTACAAAATTAAATTAAAAAATGAGTATTCCAAATGACTTTACAGTACTTGAATCATGTATATTAAAGGCTTTAAATGTTGTTGGTTGGTTTTGGCATCATTTCTCGTTTATCCCATGGTATATAGCCAGTGGTAATTTTATTAAACAAAAATCTGGAACTTTGCAAGCAAAATCTGTCAGTGGTTTACCGAGTGGACCTTATCGTAACATAGAATCATTATCTGAACTAAAAACAGCATTTTCTGGGAGTCAAACTATCGACAGTTTATTTAAAAAATCTGTACTTAAATATGGTGGTCAGTTTTGCATGGGAACTCGGGAGCTTATTGCAGAGGAAGACGAAGTCCAAGCCAATGGTCGTGTATTTAAAAAAGTTATATTAGGTGAATATCAGTGGTTAACATACAACCAAGCATATGAGCATGTGAAGCGGATAAGCAGTGGTATTAAAGCACTTGGTATAAAAACTAAAGACTATGTACTTGTTTTTGCTGAAACTAAAACAGAGTGGATGCTTACAGCTCAAGCTTGTTTTCTTAGAAATTTTTCTATTGTTACAATATATACTACTCTTGGTGATGATTCAATTGCATATGGCTTCAATCAAGTCAAGTCTCATATTGTCTTTACAGATGCTATTTTGTTACCTAAGCTTAAAAAAATACTGCCTCAATTGAAGTTTTTAAAAACAATTATATATTTTGGTGATGTTAAAAAGTCAACGCTAGATGATTTTCCTGGAGATATTCGAATATTTTCGTTGCAACAAATTGAAGATCTAGGTGCAAAATTAAAAAACATAAATGAACCTGTGGATTCTCCCAAACCAGAAGATGTTGCTGTAATCATGTACACAAGTGGATCGACCGGTATACCTAAAGGTGTCATTCTTACTCATAGTAATGTTACAAGCTGCTTAGAAGGAGCTTTGTCATGTATACCTCCTCTCACTGCCCAAGATCGGTACATTGGTTATTTACCACTTGCACATATTTTGGAACTTTTAGTAGAATGTGCTGTTCTTTCTCACGGGGTTCCAATAGGATATTCAAATGCTTTAACGCTATCTGATCAATCATCAAAAATAAAGAAAGGAACTAAGGGCGATATCTCAGTTATTAAACCAACAATAATGGCTGCTGTTCCAATGATATTTGATAGAATTCGTTCAACAGTCATGGATAAAATTAAAAGCGGTCCTCGAACAGTTTACCTTTTATTTCATTTTGCATATCAATACAAGCTTAAACAAGTGCGTAAAGGATATGACACACCATTATTAAATCGACTGATCTTCAATAAAATAAAATCAATCTTGGGAGGAGAGCTTCGCTTTATGTTGTGTGGAGGTGCACCATTGTCTCCTGAAACTCAAGAGTTTTTGAACATTTGCTTTTGCTGTCCAGTTGGACAAGGCTATGGTTTAACTGAAACCTGTGCTGCTGGAACTATATGTGCAGATTGGGATCATTCTACTGGTCGTGTTGGTAGACCTCTTACATGTTGTGAGCTAAAGCTTGTGGACTGGGAAGAAGGTGGTTACTTGTCTAGTAATAAACCTTATCCGAGAGGAGAAATAGTTATTCGTGGTGAAAATGTAGCGAAGGGGTATTTTGAAAATCCAGAAGTTACTGCGGAAGTTTTCAAGATGGAAGACGGAAAATGGTCTTTCTACACTGGGGATATTGGTGAAGTGCATCCTGATGGAGTCTTTAAGATAATTGATCGCAAGAAAGATTTAATTAAATTATCGCATGGTGAATATGTTTCTCTTGCCAAAGTTGAGTCAACAATTAAAAACTCCCCCTATATCGAAAATTGCTGTGCTTATGGCGATAGTCAAGAGACCTATATTATTCTACTAGCTGTACCTAACCACGCAGCCATCATGGAGCTGGCTAACCAACTAGGTGTTAATGGATCATTTGAAGATGTATGCAAAAACAGGAGTGTGATAGACGCGATAATGAAAGAGATCAATGTCTTATCAACTAAAAATGGCCTAAATAAAACAGAACAACCTGCTAAAGTGAAGTTGTGTACTGAGAGTTGGCTACCAGAAAATGATCTTGTAACTGCGGCATTTAAACTTAAAAGAAAAAGTATTGAAAAGTTTTATCAAAAAGACATTGAGATGTTGTATCTAAACTAACTTTTAAAGTTAGAACTTCATTTCTCATTGTAAATAGGTGTTTGCATTTATAGCTTTTATATATGTAATTAGCAGATTAAATTTATTGCATTTTAAAAAACGAATGATGTTAATATACATGTTTTTCATATTGTCACCTTATACTTTTCGATTTTCTCTATAGTTGTTAGTTTAGTTATAAATTTTTTATCTTTGAAAGTTTATATTTTCATATCTGTATAAACAAATCTTGTAACGTATTTTCTACGAAGTTTCTTAACCAAT
>evgR016777c0g1_i01_Hydra
ATAAATACCTTGAAGTACAATTCACTTGTATATCAAACCCACTGGTTGCGCGAGCATGCGAAGGAAACGATCTAAAAATTGATTGCAATGGTAAAGGAACTGTTAATGTAGTAAGTGCAAACTATGGTCGAACTTTGGCTGGTGTTTGTCCGGGTGCAAACGACTCTAATACAAAATGTGATAATCAAAAAAAGTCAATCGAAATTGTTCAATTAAACTGTTCTAACAAAGCTAGCTGCATTATTAAAGCTGCCAATTCATTGTTTGGCGATCCATGCTATGGAACATACAAATATCTTGAAGTCCAATACACTTGTAAATAATACATACAAATAAAAATGTTTACTAAATATCAATTAATGAATACAAAGTAAAAATCAGCTAAAAAAAAAAAAAA
>evgR016783c0g1_i01_Hydra
AGCAGTTATGTTGTTGAAGTTGGTGTATATTCAGCATGAAGTCATATTTTTAATCTTATGTTTAGTGAGTAAATCCTTCTGTATTCATATAACAGGTTATGTAGGAAACCTTTTAAAACACAATTTAAGGAATTATTCGCCATATTGGTGCAACGAGACTAAAACTGAGCTTAGAAATTACAAAAGTGACGCTAAAAGTGTTCCCGAAGGAGAATTCGTGTTTGATGGTACGTACATTACAATTCTTGCTCAAGAACCTCAGCAATTTGATTTGTTAGCTGTATATCTGGAAAATTTTTACTTTCACACAAAGATTTGTTATACCATTGAAATAAAAGTCACAATTCAGAGCCACATGTCAAGTCTTTACTCTGTACTTAATCAATTAAACGACAAAACATTTGTAGAATGCAAAAAAATGTTTATTCAAACATTTGAATATGCCTCCAATGAAACATTGTCATTGTCAGTTTTACTCGAGCAGTCTAATCTATTAATAGACTTTTTCAAAATTCCTCGGCTTCTTCTTCAAGTTGTTCGAACAAATAATTCTTATGCCTATGTTAATCAACTTAACTCACTAGATGTAATAGAATATAAAGTTTTCAGGCCAATGAGTGAAGGTTATATGTTTTCTTGGCCAATCGGCTGTCAGCACAGCAGTTATATTTCGCGGGTTTTTCGTAGAAGCGTTAAAAAAAACGCTGAATCAATTCTAGGACTTTCATTAACTTCCTGTTATATAACTAGAGGTTATTTCAAGGAAAACATGGAACTTTTTTTTGGTTCAATAAGTGCGCTGGTCGGAACACCGTTTGAAACAATTACTGCAAGTTTTCAAGAGTACTCTTCAACTTCAACTGTACCTAAAAGTTATATCTTACAAGAAACCGAATTTTCTATTACTTCATTTACTG
>evgR016789c0g1_i01_Hydra
GCTGAAGGTAAAATATTTTCAAAATGTTAATATTCCATAGGTTAGTACCTGTTGTGAAATTGTCTTATTGTCATCATTGGTATTTCATTGGAACTCAACGTGGCCCACGTTTTCACAGTGTCGATTTCAGGCCTTTAGAGATACACGTCGGACACCCGTGAATTTTTTGGGTAGATTTATTTAATGGAGAGCAGAAGCGAGATCCTTTGGTGTCTTGGTAACCTTAATCTTGCAGATACAGACATCCTTCAATGGGAAGATAGAAGCACATGACTTTTCAATTTCCTTACCCAATTGACCGGTAACAACCTTTTCGACGAAGTCGTTAACTGGAGAGGTAGTAGCCTTCTTGTTGATGGCATCCTTCATTCTAGCAGTAATTCTCATGACTTGAGACTTGTAAGCATAAGTTCTCTTGTGAGTTTGCAATTGTCTTCTCTTGGTAATACCCAAAGCAAAAACTCTCATGTGGAAACCATCAGCAGTCTTAACGTTAATTCTGGTGGTGATGAGTGTGTGCCACTTCTTAACCAAAGATCTGAGTTTATCAACAGTGAATCTCAAACCGTTGAAGACAGTCAAAGCCTTCTTGGTTTCAATCTTAGCATCACCGACAGCCAATCTAATAAGTTGGTGTTGTTGTTCTTCTTCGTTCAACAAGTCAGCAGCAGAGATTTCGAAAACTCTACCCTTCAAAGCTTCAGTGGAAATCTTGGTACCAGCGGTTTTATTGATTGGAGTCTTGGTGAAAGAGTCAGACTTCTTCTTGTTGAATGGGTGTGGAGCGAAGACGTCGAACCATTCCTTCTTTGCGAATGGATCAGTGATCTTCTTAGTCTTCTTACCCTTGACAACTCTCTTGTTTTTTTCCTGCGGTCATG
>evgR016797c0g1_i01_Hydra
CGTTATTTTGGCCATTTTGTTTGCCTTGTGCACAATATCAAGTGTTTTGGCTCAAGAAGAAGATATTATTGCTGAAACCACAACAAATCAACAACAAAACAATGTAAGACTTTTCCTCCACAAGAGTGTTTCTCCTGCCGAACCAATCGTCAATACAAATATCACTTTCACGATCACTGTCATTAATGCTGGTGAAGAAACTGCTTTTGATGTTGATGTTAAGGACAATGAATGGCCAGAAGCTCAATTTGAACTTGCTGAAGGTGAAACAAAGGCTACTTTTGAAAAGGTTGGTCCAAATGACCGTGTTACCTTCAAGTACACTATTGTTCCAAAGAGCGTTGGTGAAATCAATACCCAACATGCTTTGGCCACCTACAGATTGGCTGCTGCCACTGAAAAGGATGAAAAGTCATTGAAGCTCATTGCCAAGTCCAATGTCTTGCCAGCTATTCCAGTCATGACTCAAGCTGAATATGACAGAAAGCACGCTTCTCACGTTGGTGATTGGGTTGTCTTCTTTATTTTGTCTCTCATTCCTACTGCTTTGCCATATGTTGTCTACATGTATGCCAACAACCAAATTGTTGAATTGGCCAGCCAAAAGCAAAAGTCTGATTAAGTCTTTTTCTTTCAACACATTATAGGAATATAATCGAAATAGTACAAATGTAG
>evgR016801c0g1_i01_Hydra
CTTATTACGTTTTTAGGTAGACTATTTGCTGTTATACCTGCTTATTGCTGTTTTATGTTCCAAATTTCAGAAAGTTCATTAGTTGCAATGGAAAAACCTGTGCAATTGAGAAAGCTTTATCGCGAACTTGTGGAGTAATCTTTATGAGTTGCATTAAGCCAGGAAAATACTATAATGAAATAATGAATCAAGCATTCACATCCGTTACAAATTTTAATTCCGATTGGTTATTAAAGATTATTCGGTGAAACGTGTCGTATGGTTTTACCGGACCATCGGCAAAGGCTAATTCTTGTGATGCCAACAATCGAGCCCCTACACAACGAAAAGTATGCGGATATACCAAAGGCAGCACGCGATTTGACTAATTTGGGTGTTTTGGTAATAGTGGATAGTTTAGACAACTTGATAGCGCATACACGTACACTACAAGAAATAGTATACGACATGCAACCGATGTCTGATGAGCTTCTTCGTAGCTTGCCTCAACTCAAAGAGCTCCTAGAGTATCTAAAGAAGTTTAATTTAATTGACGTTGTGTCACTTGCTGAGGTGTGCTTGCATATTAAGAGTTGCTTCTTATAAATTTCATTATGAAATTTGACGACGAGAGCCGCCAAGCTGTTGTTTGTAAATTTGTTGAAGAAAATATAGGTGAAGCTATCAAACAACTTGTTGCCTTCAGGATTAAATACCCCAAAAAATGAAACATATTTTAGCAATACAAAGAACTTCAAAAGTCTGAATTAGAATCCTTACCAATTATGCGTGCAACTGATAATCCTGATCTGGATTTACAATCTGTTTTGCGATTACTAATACTCGGTGAAAAATAATATTATATACCAGCTACGGGAGCCATATCAGTAGCACTAACGACTGGTTTTAAAGATCGTGCAGACTTTCAACAAAGACAAAATGCAGTGAAGCG
>evgR016806c0g1_i01_Hydra
ATTTTCTAAATAAGGAACTAACAAATGCCATGCAGATGGTATGAGTTCATTGTGTTCAATAAAGTTAATATAAATGGACCAGTTAATTCCTGTAATAAATAAACTACAAGATGTATTCAATACTGTTGGCTCTGAAGCTATTCAACTACCACAAATTGTTGTTGTAGGGACACAGAGTAGTGGTAAAAGTTCTGTATTGGAAAATATTGTTGGAAAAGATTTTCTTCCAAGAGGATCGGGAATTATAACTAGACGTCCATTAATCTTACAACTCATCCATGTTCCTAGAACTACAAAAAAGCTTGAAGAAAATCATTCTCCTCATTTCCAAACAAATGTTGACTCAGAAGATGAAAATTTGGAAATAAATACACATGACGATGAAAATTCACAATCGAGTCAATCTGCAGACGAAGATCATGAATCAGAAGAATGGGGAAAGTTTTTGCACAAGAAAGAGAAGAAGTATTACTCCTTCGATGCTATTCGCCAAGAGATATTAAATGAAACAGAAAGATTAGCTGGTAGTAACAAGGGAATATGTGAACAACCAATCAGTTTAAAAATATACTCTCCAAATGTTGTTAATCTTACTTTGGTTGATCTGCCTGGAATGACTAAGATAGCAGTTGGAGATCAGCCGTTAGATGTTGAATCCCAAATTCGTTCTCTTATAATGAAATATGCTGCAAACTCAAACGCATTAATTCTAGCAGTAACTGCGGCTAATACAGATCTGGCTACATCCGAAGCTATCAAAATTGCAAGAGAAGTAGACCCTGAAGGAAATCGCACATTGGCTGTCATAACAAAGTTAGACCTTATGGATCACGGGACAGATGCAAAGGATATTTTATATGGAAAAGTAATTCCAGTTAAACTTGGAATAATTGGTATTGTGAACAGGAGTCAGTTAGATATCAACAACAAAAAGACCATTCAAAGTGCTTTGGAGTTTGAGGATTTTTTCTTTCAAAAACGATATCCTATGTTGGCTAATCGAAATGGAACAAAATATCTTACAAAAACTCTTAATAAGCTTTTGCTTCGTCACATTCAGCATTGTTTACCTGACTTGAAATCTCGCATCAATATTATGATTTCTCATTACCAGCAATTGTTAAATTCTTGTGGAGAACCTGTTGTTGATAAGAGATCGACTCTACTTCGCTTAATAACACTTTTTGCATCAACATATTGTGAAACTATAGAAGGAACATCTAAAAATATCACCACGTCAGAAATATGTGGCGGAGCTCGAATTTGCTACATTTTTCATGACACATTTGGCAAGACTATGTTAAAATTTGACCCTTTAACTGGTCTCACTACAAAAGACATATTAACAGCTATAAGAAACGCCACAGGTCCTCGTCCAGCTTTATTTGTTCCAGAAATTTCTTTTGAGCTTCTTGTTAAAAGACAAATCCAAAAGCTTGAAGAGCCGAGCTTGAGATGTGTAGAATTAGTTCATGAGGAAATGCAAAGAATTATTCAGCATAGCATGTCACAAGTTGTAGAGATTCAAAAATTTCCTCAGCTCAAAGATAGAATAAACGAAGTTGTGGCTTCTCTTTTACAAAGTAGACTGGCTCCAACAAAAACAATGGTAGAGAATCTTGTTGAAATTGAGCTTTCATACATCAATACAAATCACCCTGACTTCAGTGATGGCGCATCGGTTGTTTCCTCTATGCTAGCTGCAAATGATTTGAGAAAGAAGAAGGAAAAAGTTATTCAAAAAGAAGAGAATCATCGTCTAATAGCTAGTGAAGCACTGGAAGATACCCAATCAAATGACAGTCGAAGTATTCCTTGGTTATCTCGTAAAGTTCCAACAAATGAGTTGTTAGACAACATTCATGACGGGAACCAATCAGATTCAATGTATCATAACTTATCCAAAAACAGAGAGCTTTCTGATCGTGAACAATTGGATTTAGAATTAATACAGAGACTTATTAGATCATACTTTAACATCATTAGAAAAAATATTCAAGATAGCGTACCAAAAGCTGTAATGTGTTTTCTTGTCAACCATGTCAAATCTAAAATCCAAAGTGACTTGGTTGAGCAACTTTATAAGAGTGAACTTTTTGATGATCTATTGAGCGAGGCTGAACATATCCACATACGCCGCAAAGAAAATTTAAATATGTTACAGGCTTTACAGAGAGCTGCACATATAACTGCAGATATTCGAGATTCCAACGTTGCATAGTTAATGATTGTTGTTGATAAAAATTTTTTTTTTAAATTTCAGAAGTTTAAAAAAAAAATTATTTACAAAAAAAATTTTAAAACTTTCTAAAAGTTACAAGATATACATCGAATAGGTTTTTATAGTTTCTAATAGGAATGATTTTCAAAAATTTAAAC
>evgR016816c0g1_i01_Hydra
CTATTAAATAGCTAGTAATACAAGCTCATTCATCATGTTAATACGCCAAACAAATAGAAATTACTTTTTTATAAAAAAATATTTTAAAAGTAAGTTAGGGTCATCATGTGAGATAAAATCAAATTTGTGTCGCTGTTCCAATACTGTTTTAGATTTACAACGCAAAAATATACACTCGCATCATCATTTATATACTGATTGGGATTATTCAAAAGAACATGGTCCAGATCATTGGCAAGAGACATGGCAACTAAAAACTCTTCAATCCCCAATTAATATTGTCACACAAAGTGTTGAATACGACAAAAGTCTAAAACCTTTAGAAATAAATAGAAAACCTATAAATATTCATGTAAGAAATCTTGGTTTCAATATTTCACTTTTAGCTGATAATCTTAAAAGCCTTGTGTTAAGTGGAGGTCCATTTAGGTATAATTATGCATTTAGAGAACTGCACTTTCACTGGGGAGAAGTGATTAAAAATAAATGTAATCTAGGATGCGAGCATTCTTTTGATGGTCAAAAGTATGCTGCTGAGTTGCATGTTGTCCATTGGAATACAGATTTATATGAAACTGATTTGCATGCATTAAAATCTAAGGATGGCATTACTGTTCTTGGTGTTATGATAGATGCCAGTGACTTGAATGAACCAAATAAAGAGTTTAATGTGATAATTGAAGCATTTAGAAAAGTGTTATATCCTCATGATCGATATATGATTAATATATCACCATATTTTCTGTTTCCTGCTTGTGTAGAAAATTATTATACATACTCTGGTTCTCTAACTATTCCTCCATTAAGTGAAAATGTAACATGGGTATTATTTCCAGATCCGATAAAGATTTCACTTGATCAGCTTAAAGGTTTGAATATTGCTCATTCGGTTGAGCCAAATAGTCACAGCAAGCCAATTCATTTAACTTCTGATGGATCTCCAATTATTACTAATAATAGTCGACCAGTCCAGCCTTTGAATAATCGAGTTGTGAGGTCGTCATTTAAAAGTTTGGTTTGATTTATTATTGTTTGATATATTGTATTTAAAAGAGTACAATCAAAACAATATATTTCATATGTTTTGTAATATATTATATTTTTACTTTATTATATATTTATATTTATATATGTTTAT
>evgR016818c0g1_i01_Hydra
CATGCCTTTCACATATGGAGATACTTTAACAGTAACCATCAAGAATGCCACCAACCTTGAAGCAGCTGATTTCAATGGAAAATCTGATCCCTATGTTTTGGTCCAAGCAAAAGCTTTAACAGATCAAAAATTAGAATCCTACTTGAAACAAACAGCAAAAGGATGGAAACAAAAAACCAGAGTCATCAAGAAGGAATTGAATCCAACTTGGAATGAAAAAAAGACTTTTTATTCATTGAATTATCTTGACAAGAAGGGACAAGGTATCAAGTTGTTGTTCAAAGTTTATGATTGGGATATGTTGTCTAGAGATGATTTTATTGGACAGGCTGAAATTATAATTGACAAGGATAAACTTGAGCATGGTAAAGAGTATAATGAAAC
>evgR016820c0g3_i01_Hydra
TTGCAACGATTTTTTAAATGAAATTATTAATAGGATTTGGATTCATAAAATAATTAAAAACGTGAATGTTGGATCCTCTGCGTGTCAACATGTGTTTAATACTCAGCCAAAACTTATGTCGGTTAGCTATGATGCTAGTATTTTTATTTCTATTTTCAAATGGTTTTGACATTAATCGTGAAAATGACTTTAGTTTATTAAAAGTTGATCGGATGGTTGCAAACCTAATAGAAGCAATAACGAAAACAGGCGAGGAGATAAGTAACAAGGATCCAATTTGTACATCTCCGAGTTGTAATAAAAGTAAAAAAATAATGGAGATAATGAAATCAAATAACAAAGAACATTATAAACCGGATGTTAATAGATGAAAAAATATTATACGAGAACTTTTAAAAAGATATTTATTGGAGAAATTATCGTTGTGAAAACAAACTGGGTTTAAACTGTTAGTACTGCTTAAATAGCGTATCAAAATTTGAACCAGACCACGTACAGATATTAGGCAATCAAAACTGAGTAACGTTTAGATTTTTTTTGGCACCATATGTTCAAGATATAATAACTTTTTTTTTACTCTAACATAGAGTAAGTAAAAGCCGCTATTTTAAAAGTAGCCATTTTAAAAGTAGTCATTTTATCTTTAGAAAATAAATGTATAAAAATGAAAAACATATAGTTTAAAAAAAA
>evgR016828c0g1_i01_Hydra
CAATTACTACGTATAAAAATTTATTACAGATCTTTTACATTACAAAGAATGGTTGCAGCTATCGATAAAAAACAGATCTTCTATTTTTTTGATTTAAGATGCAATGAAAATGATCTTGCACGGCAGAGAGTGAAGACCTCCACCATTGTTAATTGTTTTTCAAAATCAGGAATTTCGAAAGATCAGCCCATTCGGATAAAGATAACGACCCATTTAAAGTGCTTAGAAATCAGACAGAGTAATTTAATAATTATTACCCTCAGCTGTACAACAGCTGAAGATGTCATTTCTGCAGATAAAAACTTAATGCGCATAGCACCCTTACTAACAGATGAACAGATGATAAAAGATGTAAAAAATGCAGCAGATGAACATATTGCCGATAACAAGTAAGATAATGATGATGATAGTGCAATCAATCCAGTTTGCCCAAAAGTTACAAATATTCAGGAAGCTCTGCAAATTTTCTATGACTATATGCACTAGTCTTAGTGATGAAGATTTTCAACAAAAGCTCATTTCTCAATTATATTTGATAAAGAAGTTTCTTTAAAACTAAGACTAAGTGACAATTGACAATGTATTGTTTTTATCTTTGATATTAAAACTTCCAATAACTCGAATTATTTTCACTTATCTGTAAATGTTCGATTTATAGAATGTCCTGTTAACTCAAACATTCGCTAAATCGAACTTAGTTTTTGGTTCCCTAGAGTTTCAAGTAATCAGAAGTTCACTGTGTTCGGACAACGCCACAAATTGGTATTCAATTTATATATCGGCTCCATTACTAAACTCTAAATAATCTCGATTTAGATAATCTCGAATAAATCTCGATTTACTTTGTCAAATACTTTTGAGGCATCAATTAAAAACCACATAAGTAAACAGTTTTAAAAGTTTGTTATGACAGTTCCATGAAAACAAAGATTGACTAAAGGTTAGATCTGTAATTAATGCTTCTATAATTACTTTTGTTATTTGAACTAATTTAGGACAAAGTCCTCAATACCGGTCTAACGAATTGCGTTGTTATTATCAATTTTTTTAATTTGAAGCCTTTTAATTTGAGAATCCCTTACAAAGTCTTAAATGAGTAATAAAATACTAAAAAAAA
>evgR016829c0g1_i02_Hydra
ATACGATATTCTATTACAGTTGGCACCAAATTTGTTGTTACAAATTTCACGTTCTGCGTGAAATTGAAAATGATCCTTCACCTAAACTAGTTAAGGAATGGCTTAAACGAAACGAGAACGATATAAGAAAAGAATTTGGTTCGGTCGATGAAGATAACGAAGCCTATAACTTTTTCGGCCCAGAAGAAAAAGAGAATCATAAAAAAGCTTTTACTTCTAACCCTGACGAGCGCCTAATTGAAGATAAACAGGTAGAAGCAGAGTTGAAAGGTGTTGAAAGTGAAGATGAAAGTGTGGACATAGACGTTCATGATACCAACTCAAACTCTGACTCTGACATAGACGCTATTGTTGAAAAGCCAAAATACGTTCATGAGTTTTTATTTAGGTCTATTGATAAAATGAAAGATGCTTTATCGAAAAAAATTCTCCCTGATTCGTCCAGTGAAACAGACGAGAACAATTCTGAATACAATAAACACAACCCAGATGAGTTTTATGTCTCAAGAAGTGACAAAAACCCCAGAAGTACGTTGCTTGTGATTTCATTAATTGGTCAAAGAATGGTCAAAAGTGGCGAAGTTACTGTTTCTTCCTGGAACGATGCATTATTAGAGTTTTCACAAACTGATGGCGGAAACCTTAATGTTTTTTTAAATTTTTTAACCTATGATTCGATAGGATGTATCTTACGAAATATGACAAAAAAATAGAAAACGCAAAAAAAATCAGACTAATTCAGAAACTATTTGGCTGAGGCAAGAAGCATGTGGAGATTGCTGCTCCAACAAAATCTTCATTAATCTTTAAACAGTTTATTTTTAATTTGTTTGTTTATTAAAATTTTAATGCATAAATTTAAGTACTTTTATTATGTTATTTTACACATTAATTTGTTTTTGAAAGCGTTTTTTCAAACTTTTCATTTTATTTTATATGCTGGAGAATTTTTAGTTTGCTGGAGAG
>evgR016836c0g1_i01_Hydra
TTTTTTTTTTTTTTTTTTTTTTTTTTTTTTTTTTTTTTTTTTTAATGAAAATTAAAAAAAAAACTTACAATGAGGACAATTGATTGGATGAAATATATTTCAGATGAAGTTCAAATAAATGAGCTGACTGTTTCTGGTACTCATGACTCGGCAACTTTTGCTCGAATGAGCAATTCTGGTCAAGCAATTATTAAATCCGAATATGTCCGTTGTCAGGATCGAACGATATCACAGCAGCTAGAAGATGGCATCAGGTACTTAGACTTAAGAGGTTACGATAATGGAGCAAACAGATTGGGTTTAGCTCATGGAGAATTTTATCTAGATTACTGGATGGATAATGCTCTTGATGATTGCAAAAACTTTTTAGCAACCAATGATTCCGAAACGATAATTATTTGTGTAAGCATTGAAAATGGTGATACAAATAGGTTTGTCCAAACATGGGAAAATAATTATGGAGTCAAATGCAACCGCTACCCGTGGTATAAAGGAGGCCTCAATGCTCCTAAACTCAGAGACCTTAGAGGTAAGTTTTATCTCCTGTCAAAAGTAGGTTCCGACGGTTTATTTCTTCAATCAAATAATGGTGCAGCTGTACGTTGGGCTTTTAATTCCAATAACGCAGGTCCCCCGAATTCTCCATATCATATTCAAGATTATTGTAACGTAGCAACAGATGGTAGTACTAAAAGTAAAGAAGTCGATAATGCAATGGTTAAAGCTCAATGTAGTCTTGAAACTAATGATTTTTGGATTAATCACGTAAGTTGCATTGGTTTTACGTCAATAGGATTTATTGGCAATCTTGTAGGAAGTTCAGACGGAATTCCAACTACTCCGGCAGAGATGGCTGCAAAACTAAATCATGGACACGAGGAAACCCTGCGTCGCATGCCATTCGGACGTTTTGGTATAATTGTTATGGATTACTATAACAGGTATGGCAATTATCTACCCAGATCCCATATTGTTTCTAACATAAATTTTCCTTGTGATCAAGTTCCGGGTTGGTTTAAACTTGTTTCTCCCGCACTCGGAAACAAAGTTCTTGATGCCAGTAATCGTTCAAATGATCGCCCTTACATGAATCAAGCTGACAACGATTGCACAACCTGGAAACTAGTTAATCGAGACTCTGAAGGATGGTTTTCTATGTTAAATCGACAACAAAAAAAGTTTCTTGATGCTAATGCAGGAAACGGAATACTATATGTATCTGATGAGTGGCCTGAAAGTAGTTACAAACGATGGAGATTAGTTCCTATAAAAGGAACAGATTTTTTTTATTTACAAAATAAAGGAAGGCTAGACTGCAATGACCCAAACCAGTACCTAGACTCTAATAGTGGAGGTGCTGGTGCTTTCCCATACTTAAGCCATTTTGTGGGTGCAAACGATAACTGTACCAGATGGAAGTTTTTCCGTACAATTTGGGACTAATTTTATGTTATTTTTTTCTACAATGTAATATGCTAGGTAAATAAATGTTTTTATTTAGAAAACATCTTTTGAAAATTACTTTGATTAGCAACTTTTTTTTACCCTTTTATTTTTACCGCTTGAAACCCAAAACTAAGTAGTCGACATTTTTATGTTTGTTGTTGTTATTATTGAGTGGGGATTTTGTAAAATTTAGTTAAAATTATTGAATGTTTTGAAAAAAAAAAAAA
>evgR016838c0g2_i01_Hydra
CGCATGAACCTGAGTTCATGCGATGATTTTTATTAAAATAATAATGAAAAAAACAAAACATATAAACAAAAAAGACTGAAAAAAGCTCAAAAGATTTTATTCGCCAATTTTTCCCACCTGTGGTAATTAGGAAGTAAACAAAATTGAGAAGAAGAAAAATCTAAAAAAAAAACAATTGCAAAAAACAGTTAAGTTCAATTAGCTTTGCTGTTATAATTAATGGAAGACTAGTAAGAAAACGCAGACCAATAAAGAAAAGAGAACAAAACGCAAAGGTCGCCAATAATGAACCTTACTTCGTTTTACGTAATAAACTGTTCAAAGATCAAACTTTTGATACCCTTGCAAACACAAAAAGATGGAAGTCTACATGCGGTCATTGCTTTGCTCTTAATCTCGTCAACGCTAATTGTTGTCATAAACATTTACGTTGTTGCTGTAACCTATTGCACGAGAAGCATGTATGTTCCATACAGTATTCTATTAGCATGTACATCTTTTTTTGATTTATTAAATGGTATTTTTACGGTTATACCGTATGCAATAATACTGCTAATGACATTTTCAAAGAAACACAACTGCACTCTATTCTTGATTATGAACTTTGCAATAATTTCCAACGGTGCAATTACTTTCTTACTGGTTTGCCTAAGTTCAATCGACCGGTATTTGGCTGTTTTTCAACCGTTCTACTACCACCGATACATACAAAACAACTTGTGTTTATATATAAAATGTGTTTGTTTCATTTCAATTCTTGTTGTGCTCCTTAGTGGGTTAAGTTTTTTGACCCAAAATAAAGCTTTATTTCATATGATAATATTGTTCGTCAGTCCTTTCCTAATAGTAATGGTTTTGTATATCCAAACTCGTCTGCATTTTAGAGTGAAAATGGTTCGAGAAGCAATATTGAAAGTAACTTTTATAGCGATTTATTCTGTAGAGAATATAATTAAGCAAAAAACTACGATAAATGATTTACGTCGTAAGAAAAAAATTAGCCAACAAGTAAAGATGAACAATCTGACATTGATGGTTTATTTATCCACATGTGTAGCCCACTTGCCGATTATTAGTATTTTGTTTTTTTGGTATACCAAACCTGTCACTCGTTCATGGGAATGGATTTATTCATTGTACTTATGGGCAAATGCTATTTCTTTTACAAAGTCTATAGCAAATCCAACCATTTACTTATACCGCTCCAGAAATTTGCGTAACGGTATAAGTCGTATATGGTACGTAACTAATAAAATCCGTTATTTAAACTGAAAATGCTTTTAGTTAAAAGATGTTTTAAATGTTTTCAAAACACTCATGAGTTTTTGTTTTTAAATAACTTTACTGATGTACATATATTTATACTATCCTTTGTCAACAACTGTTATGATGTTTTTGGACAATTTATT
>evgR016843c0g1_i01_Hydra
TCCCAAAATTAAACTAAACACACTCAACCAAAATGATTATAATCTCAAAGATAATTTAAGTAAATTTGAACACAGCAGTTCAAAAATATTTAGTCTAAAAATGGATTCTCAAAGTAAAACTATACAAAAATGTAAATACCAGACTATCAATAAGCAACTAACCAAAAGGTTTTTCCGAAAATTTGATAAAAATAAAAAAACTTCAATTACAAACCGCAGAATTAGTCAAACACAAAAAAGGTCTTGTAACAACTGCAAACTTAAGAACTCCTATTATTCTAATCAAAAACAAAACCGTTCAAGCTGTGCAAATAACGAGTATAAAGAAAACTTCAGATCCTGCAGCAAAACCTTGA
>evgR016847c1g1_i01_Hydra
CTAATGGATGAAGAAGTATTTATCTGTTGGGGTACAATATTTTTTCCAAATAAACTTCCAACCGGTGGGGAAGTTTGTCAAAGAATTTACTGAATTAATCGGACTCATGCAAATGCAATCAAGACATTGTCATCCAAAATTGGCTGTCCTCTAGCGAGTATGGGCAAGATTAAATGCAGTGGTGGTATGTGTTATGTTGTTAGCAGAAACGAAATATCAAATGTGTGTATCATAAAAGAGCTGGTTAACATTTGGAATAAAGCAGGCTTTGATGAAATTTTCATCATCAGTGAATGTTTGATTAAGACTAGAATTATTACTCTGTATACACCGTATATTAATGTTTTAAAGATGAGAAACCTTGAGATGAGCAAAGAAACCATGAGCCAACACACTCAAGAGTTCTTAGACAAGGCATATGTTTTGTTTCCTATACATAAGGTAGACATTTTCAAGTTAATTAGCCAGGATCCTAGAAAAAGAATGTAGGCTGTAAAAGAGGATTCTATTTTTTTAAAGAGATGTTTGAATAGGGACATTGTCAAGATGTCCTACAAAAGTGACAAAACATTTGCTGAAATTGTTGCTAAAGGTGATGAAAGGATGAAAAATCTGCTGAGGAAGATGGTAGAAGAAGAGCGACTGGAGGAGAGACTGAGAGCAAATGTTGAACAAAAAAAAGAAAGCCAATTACTGGAAAAAGCAGTGGAAGAGAATTTGAGAATTTTCCAGTGGATGTAGTTGAACAGGTTATAATTTAGACTTTGAGTCTCTTAGTGCAAAGAAAAAAAGCTATATTATTCAACAAAGATTTGCCTTCAACTCAGTCTGAATGATGTTTTAGATAAGTGGATTCCATTTATGACAAGGC
>evgR016857c0g1_i01_Hydra
TACTAATGCAACATCCCTTGATAATAAAATGACTGATTTAATTGCATTATCAAAAACTTTTGATTATCATATTATTGCTGTCTCCGAAACATGGTTTCGGAATGACTCAGTACCAAATATTGATGGGTATAACTTATACCGTAAAGATCGCCAAGACGGTCGTCAAGGTGGCGGAGTAGCTATCTACATTGCTGACATATTTAACTCTTATGAGCTTAATCAGTTTTATACAAACCCTCTCGAACAAGTTTGGGTATCTGTAAAAATTAATAGTAAACGTATACTAATTGGATGTATATATAGACCAAACGATGCTAGTTTAATAAAATATGTTGCCGAATCCTTCAAACTAGCTAGTAGCTTAATTAGGAAGCCTTACAATGACTGTCTAATTGTCGGTGATTTTAACCT
>evgR016859c0g1_i01_Hydra
TTTTTTTTTTTTTGTGTATATTCATTGAGAAGGAGTGTCATTTACTTGATTCATTTTTTTAAATAGATTTGATAAAAATGTAGACGCTTTGCTGCCATAATACGGACAATATAATGTCCGTAAAAACTATAGCTACTGTCTCTTGTTCTATATTTTGCATTGTAATTATAATATTATCACTCGTCACTAATTACTGGTATGAATTTGAGGTGGAATTGTTAGATGCTGATAGTAAGAATGTAGACAAGTCTTTGGCGAATCACTCAGTTATTTTCACTAGAGTTGGTCTATGGGAACAATGCAATACAGTAATATACAATCACCTTAGAGACGATGAAAAGGAAATGACTGAAAAATGTGAAACCTACAAAGTTCCCAGAGATATAAGTAATTGGACTATTAGGCGAAATCTTCTTGTTATATTTTATATTATTTCGTTGTTATTGACCCTGGTGGCATTTGCATTTGAAACAATTTATTGCGTGACTAGAAAGTATTATTTTCCCGTTGGAAGAATAGCTCTATGCTTTTTCATATCAGCATTATTAATTTTAATCTCGCTATCTGTATACACAAACGCATTCATCCGACAAAGTATTGCATTTTCTTGGTCGTTCGGTGCAGGCTGGTCAGCGGTTGTCATGTATATAATAAGTATTGTTTTGCTATACGCTGATAAATAAAAAATTTGGTTTCAAACGTTTTAGCTACGTTTCAAATGTTTTGGTTAGATTTTTTTTTTATTTAAATTGCAATTACAATTTGCACATTCTTGTTTTGATTTTATATTAAGCTTTTGTATATTTTATTTATTTCTATTTATCACTATTTATACTATGAACATTGTACAGTTTTTAAAATACGTTTATTAAAATAAGTGCATTAAAAAAAAAAAA
>evgR016865c0g3_i01_Hydra
CACTACCTGAATGGGTATGTTGGAAAACTGGTATTGAACTGTCTTGTTTTGGAGTAAAGCTGTTAAGAGTTTCTCAACTTGATCCAGATGTACCTTTTTATGGATCATATACCACATCAGCACGTGTTTTTATTAAAAATCGTATTGTAGATTTGTCTAAAAGAGTTTCAATTGAAATCACAGCTCCTGGCATCACATTAAACATTGAGAAATATGTTTTAAATCAACATACTAATTTGCAAGACAAGATAATGGTCGGCTTTTGTGTGAAAAGTGTACATTACAATTCTCAGTCTAATGCTTGTAAACTACTTGTCCTTACTATCGAGCATATTGATACCTTGCTTGCAGAGTGGTATCCACAACTTGATACTTATGTTGATTCACATGGAAAGGCAGTTAGAAGAGTTTCATTTTGCAATAAGTGTATTGAAAATGTCTTACAAAATTCAAATGCTGCAGGAGACAAAAGCACATTTACAACAGATATACTTGTCAAAGAGGAAACTTCACTTCTAGAGTTTATGTGTGCGTCTAAAGAAAACTGTGTATTTGCTTTTGAACTGGAACAGGGTTCAAAGTATTTAAATGAGAGCAATGATCTGGATAGCACCAAGATCAGATGTCCTAATCATGGTGCTATTGAAATAAAAATAATTTTTCCAGACCTTTGCTTTCTTGATGTTGATTCAGATTATGTTCTTAACACTGATATGATTAAACGAGGTGATTTAATTGATAAGGGAGGATTTGCAAATGTTTATAAAGCTGAAATAAAGCGAGGAAACTTGTGTGTTTCATCTGCAATGAAGATTCCGGTTGAATTTGCTAATGAAAATAGTGAACTTTCTATGTCCACCATGACATCATATAATGCTTACAGAAATATTCGTCAAGAAGTTTCCATTCTTCTTTTCTTGCATCACGAACACATTGTAGAGTTTTTAGGTGTGTCTTCGAAACCATTTGGACTGTTTTTGGAATTGGTATCTAAAGGTGCACTTAGTAATTGTCTCTGTGACTTTTCAAGTGCTGGAAAAAGACTCACTACTCATACTGTCACTGAATCACTGAAGCAGATCAGTTCTGCTCTGATGTTTCTTCACCGTCGAGGAATTATATATCGAGATTTAAAGTGTAGCAACATACTAGTTCGCAAGTTTCCAGATCCATTCAACGAGACCTCAATATTACAGGTTCATTTACTTTTAACTGATTATGGAATTAGTCAATTTGTGTTTATTGGCGGAGTTAAAGGCAATCAAGGCACTCCAGGATACATAGCACCAGAGATATGGAAGTATATAGGCAAAGAAATTTAC
>evgR016874c0g3_i01_Hydra
ACAGCGTGTTTACCAACTGTAGAAGCAGCAGCGGGAACTACATGTTATATTACAGGATGGGGTAAAACTAACCACCCTGGAACTATGACAAACATTCTTCAGCAAGCAAAAATGAACGTTGTTGACCGATATACGTGTGAAAAATTTAACAGACAATCAATACCAATCTCTGTAACACAAGGAATGATTTGCGCTGGTGACGCTGGATTAACTAGAATTTCTGGCTGTCATGGCGATAGTGGTGGTCCATTCGTTTGCAATATTAGCGGGAAATGGTTCGTCTACGGGGCTGTCAGTCATGGATCTGGAGATTGCCGATCAGATAGAACCTACACAGTCTTTGCAAACGTTGGTTTCTTCAGGTCATGGATAGATAATGCAATGATATAATAGTTTAAAACTTTTTAAAACAAAGTTATGTAAATTCAAGGATTTGTTTTAATGATTTTATGATTTTGATTAAAATTAGAGATTATAATAAGATATAAAATATTAAATATATAAATTAAGGTGGTTCCTTTCAGGGCCGTACCAAACAAGGTAGACGTGGGGGTTGGGGTTGGGGGAACAAAAAAGGTAATTTTAC
>evgR016875c0g1_i01_Hydra
GTTACACTGCGGTACGACGATTCGAAATAATGATTAAGTTGATTTACATAAACTCTAAATGAAAAAAAGAATTTTAGTAGTCGGGTTGACTTGCATTGACGTAGTAAATATATGCAATAAATTTCCTATCGAAGATGAAGATTTAAAAGTGAGCTCACAATACTGGGTTAAAGGAGGAAATGCTGCTAATACTGCATGTGTCTTGGCACAAAATCCGAATTTAGACGTCCATCTTTTTTCTTACTTATCTGGCCAACCAGAGAATGATTTTATTCTGAGTCAATTAAAAGAATCCAATGTTAATTTAGAGGGTTGTCCTATAACATTTGATAAAAAATTTCCGTTATCTTGTTGCATTTTGAATGAAACAAACGGAAGCAGAACTATTTTACATTACAACAATGACTTTCCATCAGTTAAATTTGAAGAATTTTTAAAACTTGACATTCTTAGCTATGATTGGATTCATTTTGAAGGCAGAAACCTTATAGAAGTTTCTAAAATGATTGCTCGTGTCCGTCAAGTACTTTTTAAGGAAAAATTGCCAAAACATATTATCATATCTGGAGAAATGGAAAAATCTTCTCGTTTGATTGAGCTTGAGGAGTTTTTGATGCCAAACGTTGATATTTTATTCATTGGAAAAGAAATTGGTTTATGGAAAGGTTTTAAATCGTCAGAAGAAACAATTTTATTTTACAGCTCAAAAAAAAACCTAAAGCCTGATATTTTTATATGCCCATGGGGCGAAACTGGAGCATTTGCTTCAAAATATGTTATTAAAAAAGATTCTGTATTTTATTTTAAATCAGACTCAATAATTTGTAAAAATGTTGTTGACACATTGGGAGCAGGGGACACTTTTATTGCAGGTACAATATCTAAATTAATATATGAAGAGGAGCTTGGAGTTGCTATAAAATATGGATGTCTTGTGGCAGGTTTAAAGTGTGCTCAATATGGGTTTGAAAGTTTGTTTATAAAAGCTGATTCTCATTGAAATAATATTGACTCTAATAATACTAAATATGGTAGTTTTTAAAAGATGTAAAGTTGCTTCATAAACAATAAAGCTTTCCCTTTAGAAAAAAAAAAAAA
>evgR016878c0g2_i01_Hydra
TTTTTCAGATGTTCCACATTTAGTTAAAACAACACGCAACTGTTTATCAAATTCAGGAGCAGGTCGTTGTTCACGTTATATGTGGAACAGTGGGTTTTTTATACTATGGCCACATATTACTAAACTTTATTATGAAGATCTTCAATGTGGATTAAAAATGCTCAACAAATTAACATCTGATCATATCAATTTAACTTCATATTCATTAATGAAAGTTTCTCTAGCTGCACAAGTGCTTAGTGAAACAGTTTCAACAGTTATGGCAACATTTGGAGATGCTGATTCAGCTGGTACAGCAAAGTTCATTTTGATGATGGACAAGTTTTTTGATTGTTTAAATGTCAGAAATACCAAAGATCACATTTTAAAAAGAAAACCTTTTTTGAAACCCTATGATTCTGTAGAGGATTCACGTTTTAAATGGCTTGAGTCTTTTTTAGAGTATTTTGACTTATGGAAAAAATCCATTGAAGAAAGACCTGGAAGGTATTCAGTTGCTGATAAAAATAAAATGTTTATCTCTCTGCAAACTTATGAAGGACTTCAAATTACCATTTATTCGTTTAAAGAAGTGTGCAAATACCTATTAGAAAATGGTGTACAGTTTGTCTTGTCAGAAAGATTTTGTCAAGATGATTTAGAAAATTATTTTGGTCATCAACGTGCCATAGGTAGAAGGCGAGACAACCCTAGCATTCGTGATGTTGGATATAACGACAACAAAATAAAAATACAGTATTCTACTCGTCCCATAGCCGGAAATGTTTTTGGGAAATCCAAAAATCTAAACATCATTGATGATGCTCCATTGGCTAAAAGAAGATTAAATGAGCAATAATTTTGTTTAGTATTATTTCTTTATTACTCTTGGGTTTTCTTGGGGTTGTCTTCAGGATGGGTAAAAATTTTTATTACTTTTGATGTCAGGGCAGAAAGGCATGATTACATAAAGTTGAATATAAAATTTATATTTTTAATCTGGAAAGTTAACACAACGACAAAAAGTAAACGAAG
>evgR016880c0g1_i01_Hydra
TGAAGTATATCTTAACTTACGTTATCATCAAACTTGGACTTGTTATCGTGTAAAAGTAGCTTTAACTATTATTTATACAACATCCACGGTTATATCGGCGTGTGTAATTATTTTTGATAAGAATATTTCAAGTAAAGTGTTTTTTTTAAATTATTTATATCCTATTCTAAACATTTGCAGTATAATATCTGTTATTTTATTTTATTTTTATATATCTTATAAATATTATGAAAGTAAAGTTCAAAACATAAACACAAACAAAAATAATATATCTTTGTTCGTTGCTCAACCACTTTCGAAAGACACTGCAGAGACACGTAAACAACTTATCAAAAGCATTGTGTTGTTTAGTTTATTGGTAGCATCCTATGTTGTGTTTATGTCGATCCCAAACATAGTTAACTGGCGTTTATATTGGTATTATGGCATTTTCGGAAAGTTTAACATATACATATACATAGCGTCTTATATTGGTATATTTTTGGATGTACTAATATTTGTTTACTTTTCTTTGCCAGTAAGAAACGAGTTAAAGAACACACTCTTTTGTCTACTAAACAAACAAAAAAAAATTAAACATAATATACGTAGAAGTATACATCCCGCAGCTGTAGCCTGAAGAATACCTCCTGGATATATACATTCTTACTTTTATTGTCCAGCTGTATTTTGAATTTGTATCTTGCCTTTGTATGCAGGTTTTCCGGATAATTAATTTTGAAACTAAAAGATAC
>evgR016883c0g1_i01_Hydra
TTTTTTAATACACAAAGCAGAACGTTATATTTTACAATAATACAATTAATTTTTTTTAGGCCAATATTCAATATTTTTTGAATTCATTGTTACATAGATATACATGTTACATCTAATGTATACTAAGAGAAAGAATTGTCGGCAACAGTAGCTTTTTAAATGTTTAAAAAATAAAATGGTTTTACCATCCTCTAATTATATACATAAGTTAGTCTTGCTATCAGATATGAGCAATCTCAATTTTCTGTTACGAAAATATCCCTCTGAGGTAAACTGTCGAAATACAAGCGGATTAACAGCGTTACACATAGCGGTTATCTACTCAAATACAAACGCAATCAAAATGCTTCTAAACCACGGTATTAATGCAAATTTTCAAAATAACGAAGGCAAGACTGCGCTTCACCTTGCTGTTCAGCTTTCAAACTTACTTGCGGTGAAATATTTACTTAATTACAGTATAAACGTCAACGTTAAAGACAACAATGGTTATACTCCATTACACTTGTCTTTATGCTCTCAATCGTATTTGTCTAACTATGAGGCGTTTATACTTTTACTAAAAAATGGTGCTAATGTGAACGTCAAAAGTTTTGATGGAATAACGCCACTCCACATTGCCGCAAATAGGCAGTACCACAAAGCTGCAAGGTTACTTTTACATGCTGGAGCAGACATAAATTCTGCAGACGATTATGGAGTAACGGCGCTTCATTACGCTGTATGGAATACTGATTTTAAAATGATAAAAATATTATTAAAGAATGGAATCGATTCAAATATTAGAAATGATTGCGCAAATAATGCTGTGCAACTTGGTTTAGAAAATCGCAATAGAGGTATTGTAGACTTATTGCTAGCTTCAAATGAAGATTCATTTAACAACTTTATAAAAACAACATTTGACGAAGCTTTAATTTTTTGATAACTAATTTTGTTTGGTTTATTACATGAAATGGTATGTTAGAGAAGAAGTATTTCAATCGGTTTATTAAACGAAATCCGATAATAAAGTTAGTAAAGTATGGCCCAGTCTTTTTTACACCTGCTTTATAACTTCAAAACAATCTCAAATATTGTTTTGCTTGCATTGAATCAAAATACTTTCAAACTTTTTGTAAATATTTCTAATATTTATTTCTAATATTTATTAGTGATTTTTAAACATTACGATTTTAGACTCTATATTCTATAAATTTTATCAACAAACAAAAAAAAA
>evgR016884c0g1_i01_Hydra
TTTTTGTTTTTTGCTTTCTCGTTTTAAAAGCTATTAACTTATTATATATTCAGCAAAACTTATAATGAAATATAATAAACAATTTTAAAGTAAAATTTAAGGGAAAAAAACGTTTTCTATTTTGTTTAAATGAAATACACGATGAGTGCACAATTGAAAACAGTTTGGTTTAATAAGAACATATTACGCTTTAAAAGTAACATCATTTATCGATTGACTAAAAAAGACGAACTCCAGTTGATTTACAAAACTAAACCTTCAGGTACTCATTTTTCTTCAGGAGCTATAAAATCTAGGCCGTTCCCTTTGAACTATAACAAACGATTATTTAAAGTTTTTGCGGTCGTTACTTTCGGTATTTTTCTCGGTGGACAACTTTCAAGTATTTTTGCAACGGTCTTAGAAGAATTTGAAATATTTTGTGTTTCACAAGATGATTAACCGGTGGACAATGTTTCTAATTATTATAATTTTAGTTTTTTTTCAATTATATTAATTGAAATAATAATATATGAATTGAAGTAGAACTCGTTTCAAACTTCTTTGACTATTTTTTGATAGTTTGTTGCATGTACTTAATAATATTGTAGCGTTTTATTGAGTTGTAATTCTGTTGCTGGTTTACACCAGTTTTCTCCTTTAGATACACCCAATCGATTATATATTTTTTATAGATGCTGTTGCCGCAATTTCATATTCTAGATTTAAAATCAGATTT
>evgR016887c0g1_i01_Hydra
TGAATTTTTTGAGATTTTATGTTTTTAATATAAATATGGGTTAAGCAGAGCTATTAAAAAATCTATGGTTTAGAATGGTTAATAGTTAGCTCTGAGTTAGTGCTAACTTAGTGATAACTTTTCCATTGTTTAGTATTTGTATATTTTTGTTTTTATAAACTTTAAGCTGGGCATTATAGCTTTTAATGTTTTCTCTTTTTATTATGCTTGAAAGTCTATGCAAAGAAGATGTTGTATTTACCAAATGATCTTCCCAGTGGTTTAATAGCATTTCAATCACTAAAAGGTCGCGAAATGGTAAAAGAATCCAAAAACTTTAATGAAAATTTTATTAAAAAGGAAATGAAGGAGCATAACCTAAGATACTCTGGACCAGCCTCTCTTGCTTTTGTAATCAATGCATTAACTGCAATGAATCATGTAAAATTTACTGAAGAAATGTTAAATGAAAATGACATCATTTCTGCTGAAAAAAATGGAGTCAACAACTTTTTTACAAAAATAAATATCAATAAAACTGGAGTAACTCTTGATAATCTTTCTCATGTTGCAATGCTACTTGGATTTGGTACTCATTCATTTTATGCTCTTAGTTCAAATGTAAGTAGTTATGATGAGCCTAAATTAAATGCTCTCATAAAGGAAAAGAAATGGGAAACTTCATTGATAGAAACGGAAGATAAATTTCGTATAGAAGCCAGAGATTATCTAAAGAGGCCTGTTACAGGATTAATTGCTAATTTTAACATGAGCAAACTTGGCTATGATGCAGATTTTTGTCAGTTTAGTCCAATCGCAGCCTATCATGAAACAGAAGATATGTTTTTGATAATGAATGTTTGGCCATACACACCTTCTGCATGGGTTAAGACATCACTATTGTTCGACGCCATGTCGACAATTGATACTGGCTCAAATTTACCTCATGGTTTTTTACGCATCAATGTTCTTTTGTAATGTTGCTTTTATAATATAGATTAATATATAAACTTCTTATATCAATAGTATATAAACAAAATAATTGTTAGCCGAAAAGCTGTATTTCCACATTTTGTGTTTTTTCATTCTATTATCAATAAATGTTCAACAATCAACTAAAATTAGTC
>evgR016889c0g2_i01_Hydra
AAATTTAAAATTTTTTTAAGTCATCCGAAAAAATGTTTAGATAAATTCTATATTTGTAAAGCTTTTGTGAATAAGCATTTGATGGCAGCATGTAGAACAAGGCTACTTCAATCAAATCGTTTCTTTTATTCTTGGCTTATTGAAGTGTTTAACAGTCCTAATGAAGAACGGATAAAACAGCTTGGTCCTGACAGAGTTGCAGCAGAATGGATTATTCGAAATGGTGGTAAAGTAAAGTTTGAATCACACATCACTTGGTCATCAAGTTATTCAACTTTGTCAAAAAGTTGCACAAAATTAGTTGCAATTGACGGAAGCGGTATTGCACTCACATCAAATGGTTTGCGCCATTTAAAAGGTCTTAATTATTTAACAGAATTACTTTTGAGAAAATGTAACAATATCACTTATCTTGACAGTCTTGAAGAAGTCTCTACTAGTTTAAAGCATCTTGACATTAGTTATTGTGAATGCATTTCTGATATATCACCTGTTGCAAAACTTAGAAACCTAGAAACTTTGGAAATTTCGCAAACAATGATTGGTTCGATGAGAAGCGATACTATCAATCGACTGAAATTAGCCCTCCCTCAATGTAAAATTATTGATTAACTTTTAATAAAACTTTCTTATTTTCAAAAAAAAAAA
>evgR016893c0g2_i01_Hydra
GTGACTTAAATAAAAATGGCAGCTAAAATTATACTTCAGCATATACCTAAGCTTGAGAAACTATCATCAAGAGTTTTTAGAGTACTTGGTTGCAATCCATCTGAAATGACTTTACAAGGAACAAATACATACCTTATTGGAACGGGAAAAACCAAAATATTAGTTGATACTGGGAGTGAAAATGTACCAGAGTATTTATCAATACTGAAAGAAGCTTTGGATGGTGGTACAATTCAAGAAATTATACTTACACATTGCCATTATGACCATGTTGGCGGAATACCTGGTGTTCTGCAGTTGATGCAAAATTATAAAATACCATTACAAAAGTTCTTTCATGATTCGGACGAAGAGTTTCAAAGAGCTAATCCCACTGTTAGTTTTACACATGTTAAAGATGGACATATATTTTCTACCGAAGGAGCAACTCTTGAAGCAATTTCAACTCCTGGCCATACATCAGATCATATGGCGTTGTTTTTAAAAGAAGAAAGCACTCTGTTTTCTGGAGATTGCATTTTAGGACAAGGAACAGCAGTTTTTACAGACCTATATGATTATATGAAATCACTTGATAAACTTCTCATGTTTAAAGCAAATCTTATTTATCCTGGTCATGGCCCTGTTATTATTAATCCTCAGATAAAAATCAAAGAGTACATTGATCATAGAAATATGCGCGAAGCTCAGATTTTTGCAGTTCTTAAAGATAGAAAACATTTAAATTATATTTCATCAATTGAAATTGTTAAAGAAGTTTATCCGCAACTATCACCTATGCTTGAAATAATGGCCAATGAAAATGTGATTCATCATTTAAAAAAACTTGAAAAAGAAAATAAAGTTGTTACTGATGGCACTGGATGGATTCTAAAAATGACATCGGCTTCCCTGTAAAATCGTCAACTAATTCGCTATTGAATTTTGTGAAATAATAATTTTTAAATATTTTATAAAACGTTGTAAAAACTAGTTTTAACAGAAAATACAATAAAACATATACATGTATAAAGTTTGCAATAAATGAAGCACTACAAAACTGTTATAGAAGAAAG
>evgR016894c0g1_i01_Hydra
CTTCTTACCACCTTTCTTTCCCTTTTTACCACCCTTCTTTCCTTTCTTTCCACCCTTCTTGTCCTTCTTTCCACCCTTCTTTCCCTTTTTACCACCCTTTTTACCCTTCTTTCCCTTCTTTCCCTTCTTACCAGTTTCTTCTTGTGGTTGTTCTGGGGTTTGTTCAGTTGGTTGTTGAGCGACTTGAGGTTGTTCAGTTTGTGGAGCAGCAGCATCAGTTGGTTGTTGAGCAACTTGTGGAACTTCTGGTTGAGCAACTGGAGTTTCAGTTGGTTGTTGAGTTGGTTGTTCTGGTTGAGCAACTGGTTGTTCAGGTTGAGCTGCAGGTTGTTCTGGTTGAGCAACTTGTGGTTGTTCTTGTGGAACTTCAGGTTGAGCAACAGGTTGTTCTGGTTGTTGAGCTGGTTCAGGTTGAGCTACAGGTTGTTCAGGTTGTGGTTGCTCTTGTGGTTGTTCTTGAGGTTGTTCAGGTTGTTGAGCTGGTTGTTCTTGTGGTTGCTCTTGTGGTTGTTCAGGTTGAGGTTGTTCTTGTGGTTGTTCTTGAGGTTGTTCAGGT
>evgR016898c0g1_i01_Hydra
AGTGCAAAAAGAAACAACAATTATCATCTTTAATTACAAAAATAAAAAGAAATGGACTCACACGATGGTTCAGTGTTGGTTGCTATTCATTGTATTGGTGGTGAGTACTTTGTTTGGGATGCGGAACATGTTTACGAACTTCGGACAAAACATAGAATTTGTGGCGCTTTAGTGGGTGCTTTATCTCGAAAACCATGGCAGACCAGCGTTAATTCTATGCCTTTAATTTTGATGCCAGAAGAGGTGAATTTGTGTTTAAGCGAAGGGTTTGCTAATTTATTTTTTAATGAAAAAATAAATAATGTAGATGTTTATAATGGCGTAAACGCTTTCTGGAGTAAAAGAAACGCAATGGAGCTTGAACAAATTTATCTTTTTAAAAAACGCAAGATTGAAAAAGAACATATGTTCTACGGAAACAATAAAAAGTTGAATAGAAAAAAACGTAAAGGATTAGTACCGGAATGTCAAAATGTAGTGGAACTTTGTGAAAATAACGAAGAAAACGAAGCGGATACTAAACTTATAGAAAACAAGTCAGAAATTGTTTCTTTAGATATTTGCAGAAATGACTCTTTTCAAAACGACTTAAATGTAAATCAAAAGAACATAAAACAAATTGAAAAAGAACATATTTGTGGCCCAAATGTTACAGATGATGACATAAAATACTATTACAACTCAGTGCGAGTTCATATACCTACAACACTAAAAAATCAAATCGTTTTTGAGAAATCGGTAATGAATTATCCTACTAATTCTAAACACGAATCGTATAACCTTGTGTTTAAACACTTTTGGAAAATGGGTTATTTTCTCACATCGGCAGCGAAGTTCGGTGGCGATTTTTTAGTCTACCCAGGTGATCCTTTGAGATATCATTCTTTTTTTATTGTTATCGTTAAAGACAGTGAAGCTGGATTTACATCAAAGGAATTAATAAGTTACGGTAGATTAGCTGCTTCAGTTAAAAAAACAGTTGTGCTCTCACATGTTGACACTGACCGAAATGTTAAGTGTATTTCACTAGAATGGTCTTACATGAAATAAAAATTTATTAGAGATTAAAAAAAAAAA
>evgR016902c0g1_i01_Hydra
GTTCCTGTTATAATTTTTATTAAATTTTAATAGATACAACATTGGTTGTGTTATATTTTGTTAGTTTTTTAATGCATTTCATTTATTAGAAATGTTTAATAAAAATAAAAAGCATAATATTGATAGAGATTCTGTTCTACAACAATATCGTAGTTTTGTACCTCCAGTATTTTCAAATGAAACGAAACCTTCCAATTGCAATAATACTTTAATTACAGGAATCGAAATTAAAGAGTTTTACGAAAATCTTGTAAAAGACAACGATAGAAGTTCCAAAGATGAAATTTTAAAACCTGCTAATAATGATACAATTGGTCAGGTTAAGATCAAAGAGGAAGTAGAGGAATACAATTCTAGTAGTAATTTGATTTATGCTAATGATGTTGTTGTTAGGCGATTAGTAAACAGCTTGCTGAGAACCTCGTCATCTGGTAATAAGAAAAATGTTATTCAGTGCATTAAAAACATCAAAGGGTACACTGATTTTATTGCGATTTTGAATGGGCAAGATGAATTTGGTTGGAATGCAGTGATGTGTGCAGCTGCTGAAGGACATTTACATATAGTAAAATATTTGTTGAAAGAAGGTTGTGAATATATTAATGTGAAGGATAATTCGGGGATGGATGTATTTGAAATCTGTGAGCAAAAAAATCAATTTGAAATTTATGTTTACTTATTAGAGAATGAAAATAAACTTAAAACAAATGGCATCATTCCTTCCCATCAGAGAGTGACACCAGTCAAAACAAACAGTAGCATCATTTCTTGCCACCAGAGATCAACATCAGTCAAAGCAAACAGTAGCATCATTCCTTCCCATCAGAGATCAACAATCAATCGCAGACGTAAAAATAGATATAAAACTGAAACGATTAACAATGATGCAGAAGAAACATTTATTAGATGTGAACTCTGTGATTCCAGCTTTCATCAACATGAGCAAGTTCAACATGAATCTTCAGTAATTCATATATTTAATGATAAAAATAAAGCTGTGAACAATATATATGGAATTCCAGACAATAACATTGGCTTTCAAATGATGTTGCGTAGTGGCTGGAATTCTAATGTTGGTCTTGGTCCAGATGGGTGTGGCCGTATAAATCCAGTAAAAACGGTTATGAAAAAAAATAGACATGGTCTTGGTTTGTTAAAAAATGATAGAAAAAGTTTTAAGTTTTCTGACAAAAAGAAACATTTATTGGAAAACAGTCGGATAGAAAAAGATAAAAACTGGGAAATGAATATGAGACGGTATATGTCTTCTGATTTCTGATTTTTTTTTTTTAATATAAAAAATTAGTATATATTTTTTAATAATAAAAAAAGAATGTTCAAATATTTAATATTAAATTTATGTGTAAATAAAATTTGTATAAAATGATGTAAATGTCAAAAAAAAA
>evgR016903c0g1_i01_Hydra
AAATACTGCTTTCATTGCTAATAAACTCATTACGTTAGATTCGACAAAATCAACAAACACGAAGTTTTGAACCAACCAAAACAATAATTTTGATTGAAGCAACAACAAAAAAATAACTTAAAGTAAAAAAAAAGTAAAATTTTGTTTATTCACGAAACAAGTAATCAAAAAGTTAATTTTAAAATATAATGGCTTTGTACAAGGTCGACAATAAGTTAAACGGAAATGTACTTTCTGAAGATAGTTACACTAAAGATAGTTACAACAATACAATGAGTCCTAAAAACCGCAAAAAAAACAACTCTTCATTAATCATGAAAAAAAAGCGGGATGAAAGTTTGGGATTTGGATTTAACATTGAACAAATAAGAAATACAACCGTTCTAACATCTGATATTAATGATAGAAAATTCCAGCGTAGGAACGCATTTTCGGAGTGTTCTGAAGTAGATCGCAAAATAGTTATAAAAGTTGTGACGGAGTTAAAAAGAGAAACTTATTTGAAAGATCTAATATCTTGAAATATTTTTCGAATATCGAAAGATTCGTCTAAAAAAATATTAAAACTTATCATTTTGCCTCCTTCAATATTATTACAAACAAACTTAAAGATGAAAGCGATGTTAAAAATGAAAAAGATGTTAAATGACTTTTCATATTGATTAATTATTTTTCTTATTGATATTGAATAAATATTTTAAACTGGAATTTAATATTTAATACATGATTTGTGTTAAGTATTAAACTGCGCCT
>evgR016917c0g2_i01_Hydra
AATAAATAGTGAGGTTTTTCACAAAACAAAAATGAATATCTTATTATCAATGTTATTGTTTTTATGGCTGATAAATGTTTGTAACACGCTTTCGTGTTTTAGCTGCTCTTCTTATAACAAATTAGATTGTCATCTAGCTCAATATCTGAAACAATGTGATAAAGGAAAAGAACAGTGTGTTACTATTTCCTACGAGTACGAGATGAACTTTAGTCTAAATAAAACAATTATTGATACAGTTTATGCAAAATACTGCGGAGACAAGTTTTCCAGTTGTCAATCTCATTGCCGCACATTGGAGATGAATGGCGAAAGCAAATGCATGGTAGGATGTTGTCACACGTCGGGCTGCAACGAACATACAAACCTAAGATTTCAAAACAGAAGTTATAAAAGCTCGTCTGTTGAACTTTTCAAAAAAAAAACTTTATAGAAATTTATCTTTGACCAATATTTTAATTTTAATGATAC
>evgR016918c0g2_i01_Hydra
ATTGGTTTAAAAAATATCTATCAGATAACAAACACATTTTTCACCGATATGAGAGTTGATTATATTTTGAAAGAAAATATAACATTTAGTACGATGTATTATAACGTAACGGTTGGTTCAAGTGAAGATAAATACGTGTTTAGTTACCTAATGTACGACTCAAGAGGCTCCAATGAACCAGATCGATTTGGCGCAAACGGCGCACCATTTGAAAATTGCACTAATTGGTGGACAACTAGTTGCGAAACGAGCTCATCTCCTACTTCATCATACATATCTTATGGATCAAATGAAAACTTAGTTGCTATATCTTTCTATTTACGTACACACGAAATTATAGGAGACGAGTTTAATTAAAATTGATTTTTTTTCAATTTAATAAAATTTTTATTTGTTATTTTGTCTGGCATGG
>evgR016919c0g1_i01_Hydra
TGTTGCTAGGATGGTTTTTTACACTTTTTACATTCCGTCACATAAAGATTGTTGCCCCAAATATGTTTGCTGAACTACTATTAACTAACTTCTTAAATCCTGATGGCTTAGATTCCAAAGGAAAATATTGCGACACCGGAAGTTATTGTGACACAAAGTTTAGTATATGCTTTGGCCTTAGTAAAAATGGATGTAACAAATTTAGCCCAACCAAAGAATTCACGAATACAAATAACATCAGTTTTGCATCAGAAAATAATAATAATAAACTTTTTGTTCAAGAAATTAAAGAACCATACAAAGAAAAAATTTTCATAAATGTTGAAGTAATTG
>evgR016924c0g1_i01_Hydra
CCTGGGTAGAACACACAAAAAAGTTAAAGACTAAAATGGCTAAAAACTCTAAAGAAAGTATTAAAATGAATTTTACACATAAAAATAGCAGCAATATCAAAGCATTTCCACGAAAACGCCAAGAAGTACTAAGATGGAAAGGTTGGGGTTACAAAGATTGTAAGTTTGAGCTAATGGAAAACAATGAAACCGTCAGATTTAAAGGTGATCGTTATTATGCACTTGCAAATGAAGAGCTTCCAAAACTAGGGAAATGGTTTGAAGAAAGTTGTGATGCCAATTTGAGCTTGCGTTCGGCATCTCGTGAAAAGTTAAATGTAGAAGATTATCCTGTTCCAATTCAAAATGAGCAATTTTTGAAAGAAGTTTACAATATAGGAGTATCATCCTTTGATACTGAAGATAGAGTAGTACATGGCCATGGTCAGACAATCTTTGAAATATTTGATTTAAGGTATAAAAAGTTAGAGCGTATACCTGATATTGTTATATGGCCCCAATCTCATGATGAAGTTGTTAAAATAGTTTCTGCTGCAAAAAAACATGATGTTTGCATCATTCCTTTTGGTGGTGGAACAACTGTCTCAGGCGCTGTCACCTGCCCAGCTACTGAGATGCGCATGATAGTTTCCTTAGATATGACTGATATGAATAAAATTTTATGGTTTGATGAAGAGAATCTACTTGCTCATTGTGAGGCTGGCATTGTTGGTCAAGATTTGGAAAAAAAGCTGAAAGATTTTGGATTTTGTACCGGACATGAGCCAGATTCTATGGAGTTTAGTACTCTTGGTGGATGGGTAGCTACTCGTGCCTCTGGAATGAAAAAAAATGTTTATGGAAATATTGAAGATTTAGTAGTTACTATTCGAATGGTAACAGCTGATGGAGTTATGGAAAAGCATTGTCAAGTTCCACGTATGTCAGCTGGACCAGATATTCATCATTTTATGCTGGGGTCAGAAGGTACTTTAGGTGTTATTACAGAAGTAACTCTTCGTATAAGGCCGCTTCCTGAAGTTGTTATTTATGGTTCTGTTGTATTTCCGAAATTTGAATCAGGTGTTCTTAGTTTACGTGAGATAGCACTGTGTAAATGTGCTCCTGCCTCAATTCGTCTAATGGATAATGAACAGTTTTTGTTTGGTCAAGCTTTAAAAGCTGGAAGTACTCCTTTTTGGACAAAAATAGTTGACTCGATTAAAGCATTCTATATTACTCGAGTCAAAGGTTTTGATCCCAAAGAAATTTGTGTTTGTACTCTTTTGTTTGAAGGTTCAAAAGATAGCGTAGACCATCAGCAGAAAAAAATTTATTCTATTGTAAGTAAGTATGGTGGAATACCAGCTGGAGAAGCAAATGGTAGACGAGGTTACACTCTTACATATGCTATTGCATATTTGCGTGATTTTGGTTTAGAGTTTTCTTATATTGCAGAATCATTTGAAACTTCTGTACCTTGGGATCGTGTTTTAGATTTATGTCGTAACACCAAAATAGTAATATTTCGTATGTGCAAAGAATTAGGTGTAACTCGTAGACCGTTTGTGACATGTCGAGTTACTCAAACATATGATGCAGGCGCTTGCATATACTTTTATTTTGGATTTCTTTATCATGATGTAAAAGAACCTTTAGAAGTCTATGAAAAAGTAGAAAATGCTGCACGTGAAGAAGTTTTAAGAAACGGAGGAAGCATTTCCCATCATCATGGTGTTGGTAAAGTTAGGAAGCAATTTTTATCTAAAACCATTTCAGAATCCGGTGTGAATGCATTAAAGGGATTGAAGCAGGCTATAGACCCACAAAATATATTTGGTGTCAATAATCTTCTCTAGTTTAAAAATAAAAATTTTGATAGCTAATATTAAAAAAAAATTGAAAATAAATTAAAAAAAAAAAA
>evgR016926c0g1_i01_Hydra
TCAAGATTGCCAAGAAGAATGATGTCTATGCTAAAGCTGCTCTTGCCATCAAGACTCGTTCCAGTCTTTTCAAGGATAGTGAAAAGTATGATGAATTGGTTCACCAACTTAATGATATAACAAAGGATGAAGAAGTCACTGAACAAATTGTTAGAGCTGCTCGTTCTTCAATGGGTCAAGATGCTTCCGAATTCGACATGATGAACATGGAATTGTTCGCCAAGAAGGTTGTTGACTTGACTGAATACAGAGCTCAATTGTTCGAATATTTGGAAACAAAGATGCACGACGTTGCTCCAAACTTGACATGTATTTTGGGTGAATCTGTTGGTGCTCGTCTTATTTCTAAGGCCGGTAGTTTGATGAACTTGTGTAAATGTCCAGCTTCAACTGTCCAAATTTTGGGTGCTGAAAAGGCTTTGTTTAGAGCTTTGAAGACCAGAGGTAAGACACCAAAGTATGGTTTGTTATATCACTCCAGTTTCGTTTCAAAGGCTTCCAAGCAAAATAAGGGTAAGATTGCTAGATACTTGTCAAACAAGTGTGCCATGGCTGCCCGTATTGATGCTTTCCAAGATTTCCCAACCTCTAAATTCGGTGAATTGTTGAGAGACCAAATTGCTGAAAGATTAGAATTCTACGAAAAGAAGGCTTCATCAGGTAATGCTAAGATCACCATCAAACTCAAGAAGAATATTGAATTGATGAGAGATGCTATTAAGCAATCCAAGAAGGATGAAAAGGAAGTCAAGAAGGAAAAGAAGCCAGCTGAAAAGAGAAAGAGAGAAACCGAAGTCGCCGTCGAAGAAGAAAAGAAGTCCAAGAAGGATAAGAAGAAGGAAAAGAAGGAAAAGAAAGAAAAGAAGGAAAAGAAGGAAGACGTCATGGAAGATGAAAAGCCAG
>evgR016929c0g1_i01_Hydra
TTTGTATAACTTTTAAAACGAGTAAGACATTTTTTGTTTAACTTCGTTTTGAAGATCGTAAAGAAACATTTGAGTTCTTTAAAAAAAAACTGTGTTCTGCATTTACAGTGGAAAATAAAATATTTAATAATACATCAACGTTTATCAACGATGGCACTTTTCAAAATATTTTCAACTGAGAATCAACAAGCACCGGATATAATTCAAGATAAATGTCGTGGAAAAAAAAATAGCACAGCATCGTTGGCAAAATCAAACGATAAGTTACGACATCAACCTGATTACCAGATGAACTTTAAAGAACAAGCAATTCGTTATGGAATGGATTTACAATCAAAAAGAAATCAGGATTTAAAAGAAAGAAATGCAATTGAAAAGAAAATGGCAAAACAATCAGCACCAAAAGGAGCTTTAAAAAATAGAAAGGGTAAACATGATGCCAAAAATACGTTTAGTATAGGAAACATTTTAAAAAAGTAAAGTTAATGGAGTAAAAAAAATGGGACAAACAAGAAAATTTTTAAAAACAAATTTACATTTTATTATTTTTATTTTGAAATTTCATTTTATTAGTGTTTTGTCTTTGAATATCAATATTAATAGCAAAAAAAAAAAA
>evgR016936c0g2_i01_Hydra
AATTATATTAAGATAAATAAAAATACTGAGGAAATTATTAAAAAAATGCTAGATGAACAAAAAAATGTAATACTAAAGGAAACGGAGAGACTACTGAAAGAACAAGAAAAAAACTTTGCTAAAATAACAAGCGCAAATATGAAGATTTTTACGGAAAGGTTAGATAAATTAGAACTTGAACTATCTACAAATAAAACAAAAACTTCTAATATAGAAAAAGACCTAGAAGGCATAAAAGAGAGTCTAAACTTCCAGGAAGAGTTAACTAAACATAAGATTGCAGAAATAAACCAACGAACGGACAAGGAAATTAAACATATTTATGAACAGAAGCGTTTTGATTCCATACAAGAATCAAACACCATTAAAAATATAAGTGAAAAACTGGTGGATCTCGAAAATCGTTCACGAAGGAACAACCTGAGGATTGAAGGTGTAAAAGAGTTGGCTGGTGAAACCTGGAATGACTGTGAAAAAATTGTGAAAGAAATCTTTTCAAAAAATCTTAAAATCATGAATGAAATAATAATTGAACGAGCTCATCGGTCAGGTCCTAAAAAAGAAGGTAAACACAGAACAATATTTATTAAACTATTAAACTATAACGATAAAAAAAGAATTCTTTCTAACAGTAAAAATTTACGTGGAACGGGTATTTTTATCAATGAAGATTTTGCTAAAGAAACAATCGAAACGCGAAAAAAACTCTGGGGTGAAGTGCTAAACTTACGCAAAGAAGGTAAGTATGCTATTTTAAAGTTTGATAAGATTTATTGCAGAGAACTTAACAAATAATTTTTTAACAAAAGAAATTCTTCTGTAGGTACATTTGTTAAAAATGAGCGATAAAAATAATTTTGAAA
>evgR016937c0g1_i01_Hydra
GAGCATTTATCCATGAACACCTACTTTCTTACCAGTTTTGCTATTTTTTTTGTTAAGGAAACTCGAAGAAGACCTAATGGTCTTATCACCATACGAGGACTAGCATTTCGCAAGTCGGTTCCCAGTTCCCAACTAACAACGCTTAGATGCTTCGAGCGGGATTTGAGCTCACAACCCTTGGTTTATGAGTCTCAGTGGTTACCACGCTTAGCCGTGGCGTGTGGTAATAAAAATAAAAACCTATTAAACGAATTATGTATAACAGACTTTATAAATACCTATCTAAAAATAAGATTTTATACAGTAACCAATAGTGATTGATA
>evgR016941c0g1_i01_Hydra
TTTTTTTTTTCAACCGCAAATTATATTTCTATTATTTTGAAATTTATTGTATAAAAATAATTTTAATTCCAAGTAGTAAGTAGCATTATTGGACATATGATGGCAAAAATAATAAGTGTAAAGAAAAGAAAAACGTGGAAGGAGGTTGTATTGGCTGCATCGTGGTTATCTGTTTTAATACTTCCGGCCTTCTTTTATATAAATTTGTTGGAGGTCACAAGTTCGATTTGGGAGTTTGAGACCAAGCCAGTACCTTATTTTGTGTGCATTTTGTTTACTTTCTTATGCAATTTGCTTGTAACTATCAACGAGTATGAGTATTTCTTGGTTTACAAAAAACCAATTAACGTTTTGTATGGTATGGTTGTATTGCTCAGTATATCTTCTACGACAATTGTCTTGGCATCTCTTTGTGAAATGATTGAGCGTCTTAAACATATTGCTGACCCCTGCAAATTTAATGAAACTTTAAAAATATGTAATTGTTTTTCACAAAAGCGAGATATATTTTTGCATATAAAAGACTGTAATAACACATCATATTTAGTTACAAAACTATATCATGCTTGTGGCGCTTTAGGATACAGCATGGTAGTTTATTGTGTTCTGATTGTTCGCTTTGTGTCTTCAGAAAGAACTTTTAAAGAACAATCCTCAAAGTCGTCTTCGCTGATTGCTAACCAATGCCATTCTCCGCCTCCTCCTTATAAAGTATGAAAAATTGGAGCATGGTGCATTTCAATATATTCTTGGGCTGCAAAGGTTTTTTAAATAAATACTTTTACAAATAAACTTGCCTCTAAACATTAAAAAAAGGTTTTCGTAATGCTGCAGAAGATATATAGAAGGAATTGGTACTATTTATGATTGTTTATGTAATGCTATATTATATGTTGCGCATTGTATTATATTTAAGATATGAATGACTAAGTTTATATAAAAAAGAA
>evgR016944c0g1_i01_Hydra
GTCAAAGAGACCTTGAATAAAAATGGATTGCTTCAAAAATGTGTTTCTTTTACCGGCGATAATTGCAATACAAGCTTTGGAGGTGTTCAAAAAAAAAGGTGGTAATAACATTTTTTTGTATTTAAAAAATGATATACCATATCTTGTTGGTATTGGTTGTCCAGCACATATTCTTAATAACTGTATTCATCATGGTGCAGATCAAATGACTATTGATGTTGAAAGTATTATGTACAAAATATATCAATACTTCAACATCTATACAATTCGAACAGAGACGCTAAAAGAATATTGTGACTTTGTTGAAATTCAGTATAGAAAAATTTTGTCAAGTAGTAAAACAAGATGGTTATCACTTTTTCCCGGACTAACCAGATTGATACAAATGTACCCTCCCTTGAAATCTTTTTTTTTGTCCATTGATGAACCACCTAAAGTTATTAAACAATTT
>evgR016946c0g1_i01_Hydra
CTTTTATTTTCTAAATAAGATGCTATAAACTTTCAATGCATATATGTAAGAGCTTGGCAAGAGTTCGACCTCAGTCTAAAATCTTATATTTTCAGAATAAAAAACGTCACCTTTTGACCAGTTCTAACTTTGAAATATCGAATGGCTATCAACCGGGTGTGAGCAGTGTTTCTGGAAGAAATCACATAAATGAAGACAGGTTTCTTGTGAAAAATATTTCGGAGAAGGTTCAATTTTTTGGAATATTTGACGGACATAACGGGAGTTACGTTTCAGAATTCGTACAAAATCATCTTGTGAAATTATTAGAAAAAAATATTTTCCAAAAGTCCGAAGATCACTTGTTTGAAGTAATTGAAAATGTTTTGTTGGTTTCTTTTGAGGAATGCCAGAAAGAAATTGAAAATCGTCTTAAAAATGAAAATTTCAAAGAGAAAGACATTGTCGGTTCTACAGCTGTGGTTGCACTTTTAACAAACCAAACATATCTTTCAGTAGCAAATATTGGCGATAGTAAAGCTATAGTAAGCGAAAAAGGTAATCCAATTGAACTTTCTATTGAGCATAGCCCAAACAATGCTTATGAAGCTCAACGTATAGAAAAACGAGGTGGATGGATTGACTGGGACAGTCATTTATATCCTTTAGTCAATGGACGTTTGTCTATTACACGTTCATTTGGAAATTTATCTCTTCGCTCATCTGGCATTGTCAGTAAGCCTACTTTTCATCATAGACTCGTCGACAACGATTCTTTTTTAATGTTGTGTAGTGACGGTATAAGCCATTCAATTACTGACCGAGAGATAGTCTCTATAATTAGCCAACATAAAAATCCTGCAGATGCTTGTTTTGACCTAACCAGTACTGCCCAACAGTATGGATCAGATGATGATGTTACTGCTATTGTGGTCCCGTTAAATAAATGGAGTTTTCATTTTCCTGATAACTCCCCTGTCACTTATAGCATGTTTCGAACTATGATTGGATCTCGCTCTGGTGGTTAATCTTGTAGAGGTTGCTTTGTTTCTAGTAAATAGTATACATGCTATAGTTTTGTAAATACTTAGTTATGTTTTTGTATAATTTGTAAAAGTCTTTTGTATAGTTTGTAATAATCTATAAATAGATATAGTTATATCCTGAAAAAAAAAA
>evgR016947c0g1_i01_Hydra
GAAGTATTTGCTTTACGCGAATTTTATGGAGAATCGCGTTTATTAGAATCTGTTTCTAATAATTTTAAGAAGAGTCTAGAAAAGCGCGTAGAAACGTTTAATAACCACGACGCAAGCAAACATTTTACTAAAATAATATTGTCAATTGGAACAGAAATTTATTCTCGATGCAATAAATATATATCTCAACATAAATGTTTTGCTAATATTTCTGATGAAGAACAAGAAAGAGAACTTGAAGCCGAGCTTGAAGAAGAATGTCAAATTGAACGACCTGGCGCTCAGACTCCCCACATCCCTAAATTGTCTCAGG
>evgR016956c0g1_i01_Hydra
TAACCTTTCCTACTACTACAGAATGCACGGCAGTCCATGTTATATGAAACCAATTGGAGTATATTTAATTCCAGATGTTTTTATGTTTCTAAATTGTGATTCACAACAAGCAATTTATGTAAACTGTCTTGCTAATCAAATGTTTGATCCCAAATACAATAAATGTATGGATGCTGGAAAGTATAATCTAAGCACGTTTTGTACTGGAAAATCTATTTATCTTAATTATCGCAATCCATGGAACTGTAATAGTTATATAACCTGTAACGAACCTGTACCCAAAGTTCAGTCTTGTCAAACAACGTTAGTATACAATCCAAACACCAATAAATGTGACTCTGCTGATAAATTTAATTGTGTTAATATTTCTACTACTACATCAACTTTTAAAAACCTACCAAATACAATGGTTTACCAAGCATCTACATTCATGCGCTATTCTAGATCTTTGAGTGTAGATGACTTGACTACTGCATATAAATTACCACCAATTACAACAATAACATCGATATCTACAAGTCAGATAAATAATGCATACACCTTAAATGTTCAAGATACTCAAACTGCAACTATTATTCAACCATCAACATTGATTATCTCTTCAAGTTCTCTTAGTTTAGATAGCTTGCTTAATGCTTATAAACCAACACTAAAAGCTAAATTCCCATCAGTAACTGAAAGTCAGATAACTCCCGCTAAATTTATTGAATACTTTCAAACTACGACATCAATATCTAGAAGTCAAATAAGTGATGAATGGACAGTATGGAGTGAATGCTCTAAAACATGTGGAAAAGGTACTAAAGTTCGATATTCTTATAATGTGGAACCAAATAGTGAAATTGAAAACTGCAATTTGACTATATGCCCAGTCCACGGAAGTTGGGGTGATTGGAGCAACTCTACATGTTTAACTACCTGCGGTGAAGAAAATATTTCTCTTATTAGACGATGTGACAGTCCATTATATAATGGTCGAAGTTGTTTGGGTATGAGTTATTACACCGAAGACTGTTATTTCAACACAGTTTGCCCAGTAAACGGTAAATGGTCAATGTGGTCCATGTGGTCGTTGTGCAACCAGCCATGTGAAGGCGGCATAAAATCTAGATTTCGCATTTGCTCAGTGCCAAAGTATGGCGGCCAACCTTGTGATGGAAAAAGTAGTGAAACTGTTTATTGTCCAGTAATTAGTTGCAAAAAAGTAGACGTCGATTTGCGCGTTTTGTTTGAGGAAGACTTTTATCTTCCACCTTCTGAGCTAAAAGAAAGAATAGAAAAAGCGATATCAAATGTCTACACAAATTATAAAAGAAATGCAAATTTTAAAATTGTGTTGCGTTTTATAAGAAACGTTAATCCTTAACCTTGTATTAACCAAAAACTGGTCATTGGGTTGCAAATAAATACCGGAAAAGAATATGAAAATTGAAAGAAAATTTGATCTTATCAATATTTCAACTTTGTATTTATATTTATATACAGACATCATAGCTGTTTAGTTTAAAGATCAAAAGTTTATAGAAAATAAAACTTTAAGTTGTTAATTAACCTGATCGTAAAAAACCTCATATTATATATACCTTTGTAAAAATTGCTGAAATGTTTTTCACAACGACATCATAAAGGCTTTAATTTTTTATTCAAT
>evgR016958c0g1_i01_Hydra
TATGTTTTAAACAATTTCAATTTTCAAGGAAAAGCTGAAGTAGCTAGACTTTTGCTTCATTTTAAAAGTGTGCATTTTATAGATAATCAAATCGACTTTAAGGATTGGCCCTCACACAAAAGTGACCATTCTAGGTTTCCTTTGGGACAAATGCCAACTTTAGAAGTAGATAGACACGTTATTTGTCAATCTGTAGCAATAAACCTGCACTTGGCTGAAACACTTTGTCTTAATGTAGTAAGTGTTTCGGAAAGACTGGTTATTAACCAAATATGCGAGACTTTAAATGACTTTATCAATGAATATATTGATAGTATTTATATATTGACCAAAAGGTAGACAGCTCATTATAATGTTTTACCGATGCATCAACTATACTTGAATTACTTTTTATAATACAATTCTGACAATAAAACTAATTGATGTTCGGTATTACAGTACTTTTTTTTAAATACCATTTTTGTTGCTGTTGATTTCAGAAAGAGGCGTGTGCAGAAGTGCTTTCAAAAGAGTCAACTAAACTAAAATGATCTTTTATTGAAACCTTGCTAAAACGTAACAATGGTGGTAAAGAATATTTTGTTGGTGATTCAGTGAGATTTTAAATCTCATACTTTTTTTAGTTAATGTAGAACATTAATGCAGTTAGCACTGCAAATGAATGTTTGAATACAGTTAGCACTGCGAATAAATGTTTTAATGCAGTTAGCATTGCGAATAAATGTTTTATTTAATTTATTGTTGGATATTATTGTTGGAAAATTGTTAATTTAATTTAAGATTACCCTTGGAG
>evgR016965c0g1_i01_Hydra
TTTTAAAATCTCGTTTAAAGGTATAAAATGTCAAAAAATTTAAAAGTGAACCTTCAATATCTGTACTCTAAAGAAAACATTGATTTCTATGTTAACAGTATTATTAAATACAAACTTGGAGGATTTAAGAAAGATGAACATTCTAACTACTGTAAATATCTTTTGAACAACAAATTTGGAATGGGTAAAAATTCTTATTACGTATTTTATTTGCAGACTAAAGATAGTGCTGGTAAATTTCCCCTAGATTTTTTCTTTGACAAATCAAAAGAGTCTGATGTTGAGAAAATTTATGATTTTAAAGTGGTTTGTGATAAGTTAATGCAATTAATATTATCCGAAGAAAATTGCAAATCAAAGCTATTATTTCCTTATAATATAAAAAATATTTGGCTTATTGGAGAAATTGTCGTTGATAAAAAAGGTAATAATGTTTCTTTAGTAATATACGCCCACAATCC
>evgR016971c0g1_i01_Hydra
AAAAATATAGGATGAAGAAACAAGTTATTAGCGAAACATGGTGTCATGATAAAAATATTGAAAGCGATTCAAATTTCCAAATACCTAATTACAATGTGGTTCACCAAGTTAGAGGTTGGGGCAAGGAAGGTGGGGGTTTATGCATATTTATTATTAATTCATTGTCATTCAAATTTATGCAAAACTTAAGCTCAACCACTAATGACTATGAATCGTTATGCGTAGAAGTCATTAGCGAAACCTCAAGAAATATTATTGTGCATGCGTTATATAGACCGCCTACAGGAAAAATTAATGCATTCAAAGATCACATTGAAAATATAATTAAAAATAAATCAAGCTTTAATAAAAGTGTGTATTTTATCGGTGACATTAACATCAACATTTTTGACTATGACGTTAATAAGCACATTAATAATTTCTTTAATGTCTTTTTTCAATCTGGTTACATTCCATTAATCAATAAACCCACGCGTGTAACTAAGAATAGTGCAACATCAATAGATCAAATTTTTACAAATGGATTTCTCACAACAAAAATAAAAACTGGAATATTTAAAACGGATATTTCGGACCATTTTCCGATTTTTATTATATCAAATAAAGGCAATAAGTCTTATTCTCACAAAAGGAAAGTAACAAAACGACTAATAAACGATACCTCCATTAAATCTTTTCATGATATTATATCATTTGAAAAATGGGATGAAAGTCTAATAAATTTAAATGCTGATGAGGCCTATGATATCTTCTTAGTTAAGTTTCTTCAATATTATAATAAAGCATTTCCTAAAGTTACAAAAATAATAAAAGAAAAAACCCTTTGAGCCCTTGGATAACAAAAGGAATTATTAAATCTTCAAAGATAAA
>evgR016973c0g1_i01_Hydra
GTTGGTTTTCTTTTTATAAAAATTCACTCCCAGTCAGCCAATATTTGCAATTCCTAACCAGCCAACATTTGCAACACCCAACAAGCCATTATTTGCCACTCCTATCCAACCACTATTTGCCACTTCCAACCAGGCATTATTTGCCACTCCCAAACATCCATTTGCCACTCCTAACCAGCCACTGTTTGCCACTCTTAACCAGCCACTATTTGCCACCCCTAACCAACCATTGTTTGCCACCCCTAACCAGCCATTATTTGTCAATCCCATCTAAGGACTTTTTCTACTCCCAACCAGCCACTATATGCAACTCCCAACCAGCCACTATTTTCCATTTCCAACCAGTCACTATTTTCTACTCCCAACCACCCACTATTTGAGTATTTATCCAGAGAGTAAGGAAGATGGTTTATGAACAAGAAAACGGTTAACAGAAAAGTTCCAAATTATACGATTCAGAAAAAAACAAGGAGGAGAAAAGCATTTGAATTTGCCAATGTTCGAAGTAAAAAATAGGATAAAAAAACGTTTTTTGAAAACAATTGAATAATCGCTGTCAAAGAAAGCAAGAAT
>evgR016974c0g1_i01_Hydra
ATGATCCTACTTACCAAAATGAAATCAATCGATATTGTACAAATAACTTTGTTCCTACGTTTCGTGAATCAGGAGTACTATATATTAAAGTAAAAACTGATAGGTCCATAATCAAATCAGGTTTTAATATGACTTGGAGCAGTGCCATAAAAGGTTGCATTAATGGTGGATCTACAGACTTTCAGTTAACAGCAAGTGGTTATTATGTAACATCAAATGTATCATGTGTCAGTGCTTGTAATTCATATTATCCACCAGTTTTTAGTCATGCTTTCTTTCTTGAAGGACAGCTTTGTTTCTGCCGAAATGCTTCCCTTCCACAGCTATATTTTTCTTCAACCAATGTTACAACAAACTGTGGTACAAGTACATGTTTAAATGACAATAGTAGTTCAGATTGCAGTTTAAGCATCTTTTCCTTTTTTTCACTTATATTAGGGATAGCTGATTGGAAGATAACTTTAAATCCTACCATAGCACTTGTTGACAAAGAAATACAAATTCAAACTAATATATTAAAAGATTACTATTCAAATGCAGTTGTTGTACCACTCACTAATGAGTTGCCATTGTTATTTCAAGAAAATGTTGATAGCACTTCATTTAAAACACAGTTTTTTTTACCTGGTTTACAAAGGTGGTCTCAAACAATTGGAAATAGCTATTCTGATCTATTTAATGCAACTACTTGGTTGCAAGTAAATGAATATATTGGTGGAATATATGTATTTCCTTCTACAGCTCAAGTCAATGGCTTCTATAATTATATTTTAATTATTACAAAAGGCACTAATATTACTGTAAAAATATCACTACCGAAGAATGAAAATATGATATTTACTGCAGATCTTCTTCGTTTAACATATGGGTTTGATGTGTCAATGTTAAATAATAAGTCTACAAACTTGTGTAAACTCTCAGATGGCTATGTTTTGAAGCAAGCTGTTTTAACTTCTGGTCATTTTATAAAATTAATGGTTAGAGCCTTTGCACCAGGACTTCTGGAAATTGCTATTTTGAGACCATCATTTGATAGCTCCACAGACTTCTGCACTTTTGGAATGGAATGCAAGAATTTAATATTATCAAAAACATCTGTAACAATTACTGCTTTGGTTGTCAAAAATGGTTATCATGGAATCATCCTGG
>evgR016981c0g2_i01_Hydra
AATGGACAGCAGAATTTATCTTTAATTGATATGAGCTGTTCTACTTCATTCTGTAGAGAGTGTATGACATCCTGCAATTTTGAAGGAACTTTTAACTTGACAAAGTGTTTGTTGAATTGTAATAATTATACTTCATGCAAATTTGGATGTTTTTTCTATAGCCATTTGAAAGACAATGACACTGGAAAATTAATTCCTGATGAACAGTACAAGCCTTATATCAATTCTACGTTTGAAGTAATTCTTATCAAAAATCAAAGTTTTAATATCAAATTTCAGTGGCCAAATATTTATTCAAAAATTGATTTTATCCAAAGTATCTACTTGATTACTATTGCTGGAAACAGGGAAAACTTTTCTGAAACTGTTATAGGTTTAACAAGTGAAAACTTTTTCAATGTACAAGAGCAATTGGTTTGCGATAATAGAAAT
>evgR016983c0g1_i01_Hydra
GTGTTTTCTACTTTGAATTTTTTATTAATATAGATAAATAAAATATAAATTTAAAATGCCAAATTACCGCTGCTGTGTTGCTGGTTGTTTTAATGACTCCAGGTATCCAGATAGGTTATTTAAAAGAACTCAAGTAACAGAGCTAAAATTTCATTATTTTCCTAAAGATTCTTTGAAAAGACAGCACTGGGTAAATCAAGTTAAAAAAGGTTTACTAAATTTCACAGTTTCTGATAATAAAGTGGTTTGCTCTAATCATTTTGAGTTTGGAAAACCTACTTTTGCTTCGCCTTATCCAACTTTATACATGAGTATTTGTAATGCAGAAAAATCTCCTACAAAACGAAGAAAACTTAAACGATTTGATGTCATATTAGAATTAGAAGCAAACAGTTCATCAACAAGTTTAACTTTAGAAAATAAACACTGTGATAAATCTATTCAGTGTAATGAACCAATACGAGGATCAATGATTTTTGCCGATTTAACCAGAGATTCTGATGTTCAGTTTTTTACTGGCCTTTCAAATTCAAATGCCTTTGAAATGATATTTTCATACTTACAAAGAAAAGGAAGCGTAATGCACTATTGGAAAGGTATTAAAAATACAAATAAAAACTTATCTTCACCAAGGGATTTAAAAACAATAAAATTAAAATCTCTAACATTGCAACAGGAGTTTCTTCTTGTGATGATGCGTTTACGTTTAGGTCTTTTAACTGAAGATCTTGCTCACCGTTTCATGGTTTCTATTTCTGTGGTGAGTTCGTTATTTATAACTTGGATAAAACTTCTTTCCCTTGAATTAAAATGGCTTATATATTTTCCTGATAGAAATAATATAAAGAGAAATTTACCAACAATGTTCAGGAAATATTATCCAAAATGCTCAATAATAATAGATTGTTCTGAGCTTTTTATCGAAACACCATCAAGTTTGGATACTGCTGCAAGTTGTTGGTCAAATTATAAACATCATTATACTGTAAAATATTTAGTTGGAATAACACCAAATGGTGCAATTTCATTTTTACCAAATTGTTATGGTGGAAGAGCTAGCGATGTTTTTATTGTTAAAGATTGTGGAATTTTAAAATGCTTACAACCTGGAGATCAAGTTATAGCAGATAGAGGTTTTAAAATAAAAGAACTACTAGCTTTTCACCAGTGCAATTTAGCTATTCCTCCATCAAAACACACAAATTTGCAAATGACATCTAATGACGTGAGGGAAACATTAAAAATTGCAAATGTTCGAATCTATGTAGAACAAGCCATAGGGCGTATGAAAAACTTTCGCATATTAAAAAATGAATTGCCAGTCACACTGTTACCATTATGTGACAATATTATTACTGTATGTGCTGTGTTAACAAACTTCATGCCCCCTCTTTGTATTGATGAAAACAAAGCATAAGATGATATGATTTATCCGGACATCGATGTTAACAGTATACATTTCCTAATCAACGATGAGGAATTGGAATTTAGCGAGACCGATTTGCTAGTCGAATGCAGCACCACTAACGTCGCTAGCGACAGCAACACCAGCAATACTATCCCAGCAATACTATCCCAGCAATACTACACCAGCAATACTATCCAAACCAACACCGCTAACAGAGGAAGCAATATTAAGAAAGAAGGCTTTGAAGAAAAAACAGAATAGAAAAAGAAAAGAAAAGAAAAAAAAAAGACGTAACGAATACTTAAAAAACGATTAAGACATTTTAAAAAGAATTTTTTCTCTTGTATATGTATTTTTTTTCTATTGTATATACTTTATATGAAATTTTATAAATATATTGCTGGGTTTTTTTAAATTGACAATTTTTTAAACTACATATTAC
>evgR016987c0g2_i01_Hydra
AATCTTTTATTTTCTAAATAAGAATGAGTGATGAAATTGGAAGTATACCTTGGTGGGGATTCTCACCTGCTGAAAATTTTGTTAAAATAATAGAAAAATCAACAAAAGATTCAAAGATTAATAAAAATGAATTGAATATATTATCTATTGGTGGTGGCGATGGTCGTCATATATTAAAAACGCTGTCTGCGTTTAAAAATAAAAACCAATACTTGAACATTTATATAGCAGAAAATAATTTAGAATTAATAGCTCGTCAAATCTTATTTTTAAACCTTGCCATAGAAAAAGTCCATGGCTTGCAAGAGAAAGTTTTTATTTATTTGGATCTTTTTGGTAACACTTTGCTTCGAAAGATAACAGAAGAGTACTTGCATGAAGCATCTATTAATATTATCAAGTTACTAACAAGCTCATCTCAAAACAATTTTTTTGATTTTTCATTACTTAAATTTAAAGAACGAGATGCTTTAGAGGGAATAATTAAATTTTGGTCTATAAAAGAAAATATTTTAGAGCTATCAAAATGTTGGGACTATAGATTACGTCAACACTTAGGAGTGCGTTATGATGCACGTGAAAATGTATTTGATTGGGATTATAGTATGTACCTCAAGGATGTAGCACCAGTAGTTAACTGGCAGGAATACAAAAAGTGGCGTGAAACTGGATTGGCTTTTGATTTATGTAGCGATTCTACTTGTTTGAATAGTAATAAAACATTAGTATCCGGGTTGATTTTTAAAAAAGATGGAGAGAGAATTGGTAAGCGAGGATACTGGGGCGACACGGTTGTTAGTCCATACATATCATATGGTGTTGAATGCGATTCGAAAGATTTAATGGCAAAAAGCAATAACCAGTACACAAGTAGCTCAACAGATATTTCAAAATACAATGTCACTTCGTTTTTTCATGAATTACACATTAAGTATGTGGAAAAAGATTTACAACCATTTTCTAATTTTCAGATTCATTTTTTATCTTTTGATGTATTGTCTACACTGCATAAAAAACTTAAGTATAAAAATAAATTTGATATGATTTATTTTTCGAACCATGCCACAAACTATTTAAGCGAGAGTGTTTCTCAACTTATGTCACCTCAATCTTTGCTAATGGTAGAATCAACTAAATACTTAAACATTAAAAAAGAACAGCATTTAAAATTTAGTGAAAAAGTTAATATGTTAGCTAGTAATGCAGGGTGCATACCATTAAATTCATTTGGTGAAAACTCTCACGCATACTATATGTTTCAACAAACTTAATGTTTATCCTTAATATAAGAGAGTATAAAAAATTAATATCTTACTGTAAATACTATGAAGATATATTTATAAATGAAGTTTATATTGTTTAGATGATGTTATAATATATATTTAATGTTTAAAAAAAAAAA
>evgR016989c0g1_i01_Hydra
GTTTCCATTAATTATTTAATTGGCTCTAATGATATAAGCAACAATACCATACATATACCTGCTACAATACCAAAATATATTTTGCAAAAATATGTCAAAGGAGAACATTGCAGAAAAATTTGAAAAATGTAGTTTAGATGACGCTGTTATGAAATTTTATTGTTCTTGCATACCTGGACTAGAAAGTGTAGCAATTGCTGAATGTAAAGAAAAATTAAAGGTAACTGCTTCACTAGATGTGCGGGGTAGAATTACCTTAAATTTAAAAAAAGCCGAAGCTTGGAAACTTACTGAGCTTTGTTCTGTCCACCACTATTGGGTTGTTGTTGAAGAGCGTAAAGATTTCTTAAGGTCTTTTCAAGGTAACACCAACAATGATGCTCTATTTAATGTTCTCAAAAATCTTGCTTATGAGTTGAATTGGGAAAATTCGCTAAGTATTTGGAAGCAGTTCAAAAGCATTCTAAAGTGCACACAACCGGAAAAGGCTAAAACTTATCAGATTATTGATCCTATCAGTGCCTGTGACACACTTTCATTTCGAGTTACTGCTTCAAGGACTGGCAAACATAGTTTTAGATCAATGGACTCTGCTGCTTTTTTTGGGGCAGGTCTTAACAATTTTTTAAAATGGAAAGTAGATTTAAAAGATTTTGACATAGAAGTTTTGTTACATATACTTGACAATGATTTAAGTATTGGTATCCAATTAACTAAGGATAGTCAATCATTCAGAAATATTACACATTTTGGACCAACAACTCTTTCTGGTGGTTTTTGCTATTGTCTGCTTCACTTAATTAATATTAAACCTGGAGATATTATTTGTGACCCTATGTGCGGAGGAGCATCCATAAGTATTGAAGCTTCGTGTAACATGCCATATGTTTTCAACTTGGCTGGAGATAATCATGAAATGGCTATCTGTCATTCAAGAGAAAATGTATTTAATTTAGAAAGCTTGAGATCTGCTTCACTTCCAATATCGTTGGTGCAATGGGATGTTTGTAGGCTTCCTCTGCGCACTTCATCTGTTGATCACATTGTTGTGGATCTTCCATTTGGTAAAAAAGTTGGAAGCAAAACAGAAAATTTTGTTTTATACCCGGCTGCTCTAAATGAAACTGCTAGAGTGTGCACACCTAATGGCACTGCTGTGTTCTTGACTTATCATAAACAAGCAATGTGGAAAACAGTTCAAAATATTTCCTGGTGGACTTTGTCCAAGACACACCAAGTTAACATGGGTGGACTTAATGTTTATGTTTTTATTTTAAGAAGAACAAAATGCTTGTACCATAATGTTTAATTTTCTTTTAAGAAAAAATGTGTTGCAATAAAGAAAAAAAAAAAA
>evgR016992c0g1_i01_Hydra
GCAAAATTAGAGTTACAAATCAAGATAATAAGAAGTTAGACGGCTTTTCTAAAACTATCATAGCTGTTATTATTGTGGCTTCGGTTCTACTCTTTGCAATATTCGTTTTCCTTTTTTTTGTTCATTTGAAGAAAAAAAACAAGATTGTCTTTCAAGAAAAGATTGAACCAATGGAAAATAAAAGTGTTTATAAAGAATCAATCGAAAACAAAAGTTTTTCTAAAGAACCAATGGAAAACAAAACTGTCGGTGACTCTCCAATACCGCTGAATTCTTTGAAAATCTCTAATGAAGGTCATAAAAACGAAGCTTTTGTTATGGAAGATGAAAACCCAATAAAAAATAGAAAATTTGCTTGGTAAACGTGAAGTCTTATAATTATTTCATTAATAATTAGTTATAAAAGTCATAAAGTG
>evgR016995c0g1_i01_Hydra
CTTTTGGAAGGTGCCCATGTGTCCTTTTTAGCAATATATGGTTATGGCGTAAATATCTTAACTACAAAAACAAATGCGTCTATTGAAGCTCACTCCAGTTGGATTTATGGTGGCGAAGATCCTGTATACAAAGTTGACTATCAAACAATCTCAATACCCATGCAAAGATCAATTCCACCAGGAGTTCAAATGAGCCCAACATGGAAATATAAAAAGCCAAACGGAACCTTATTAGAGTCTAAAACTCATCAGTTTGTATGTGGGAATATGCAAAAAAGACAACTATCTCCTTGTTACCAGCGTGAATTACCATCAGGAATAATCAAGTTTCTTCCGATTCAAGTGCTAGATGTGTTTGCTTACGATGGTAACACTACGCTTATTTATGGATATACAACTCGAAGTAATATTGCTGAGGTTAACATAATTAAAAATAAGCTCAATGTCATATCGAAACAAAAATGTTCCGAAATTAAAACGTGTGACACCTTTATAGATATAAAATAACATAAAATTGTTAACATTGTTTAATTTAATTGAAAATTAAATATGTCTTTATAAAAAACAGTCGGCAAATTTGG
>evgR016996c0g1_i01_Hydra
TAGTAAGCCAAATAACAAATTATAGGTTTCTGGTTGCACTATGTTTTTGGCACTCTTTATTGTTTCAAGTAAATTTTACTAGCAAGGAACTTCAAAGCAATACCATAGGCATTGCCGCAGGGCTGACATCTTTTGAAAAATTGTTTGATTGGTTGAAGACATACCGAGAGACAGGCTTTGTAGCTGCATTGATTGATGCCAAAGAGCTTGCAAATGAGTTAGAAGTTGAACCCATATTTCAGCAGAAGCGCTGTTACAAGAAAAAGAAAATGTTTCTTTATGAATCTTCTGACGAACCAATTTTAGACCCCAAGACAGAATTCCGGGTAAATTTTTTCATTTAAGTTGTAGACAAAGCATTGCAATCTCTTCAACCACGGTTTATGCAGCTGGAAGAACACCATAAACTTTTTGGATTTCTACATAATTTTCAAAATATGTCTAAAGAAGATCTCAG
>evgR017000c0g2_i01_Hydra
CTGTTTTAGTGCAAAAAGCCTATTTAGTTTTATGTTTAATATTATCTTATGACTATAAAAAAATGGTCTCCGATTCTGAAATTTTGTGCTGAACAGCGAGTTACCAGATCAAGTCAAAATCATTCAAAATGGTTTCAAAATATTTACTTCTGTAAATTACACACTCAAATAATCTGCTCTCAAGCGTCGTGCAGTTCGAAGCGGCTTAATCAAGTTCAAACAATCCCAAACAAAATCTCCAAGAATGAACATTTGCAAGACAATAAAAATAACTCTTGTAAAACCAATGAAAAAAACAGTTTAATGAAAGTTTTTGCTGAGTTCGGGTTAACAGCTGTGATATTTCATTCAACTGTAGGTATAGTTTCTATCATTGGATGGTATGCATTAGTAAGAAGTGGTTTGGACGTACCTGGTTTAATGAGTGAATACAATATGCTACCCATATCTATGTTGAATTCTAGTTATTCTGGTGCTGCAAGTGCGTTTGTTGTAGCATATGCGTGTCATACCGCTATACTACCGTTGCGAGCATTCATAACCATATTCTGTACACCGCTTATCGTAAAAAAGTTTCATGTGTTGGGGTTTCTTTCCCGAAAAATCAAATGAATTAAATGTGGGCCGATTAAGCTAGTCTTTTTTTCTAGAATATATACATATAAATTGTATTTTGTTTGTTATGGAAGGGGCCATCCATAAAGTACTTACGCAGGTATAGGGGGAGAAGGTGTTTCCCAAAAGCGTACGAACGCGTTTAATGAGGAATGGGGTGTTAAAGTGAGTACGTACGCGGTTTGATCTCTTCTAAACTTGATTTAAAAATAAATAAATTTAAAACGTTAAGTTTCACGTGTGTAACAAATCAGTAGTTAATTTTTTAAAATTTCTTTATAAAAATGTGAAGAGAAAAAATAAATTGAGAGATAATCATCATCATCGCCATTATCATCATCATCGTCGTTTTAGCTTCCTTGTTTCCGTTTTTTTCGCAACGTTGTTTGACTTGTGATATGAGAAACAAAAAGACGTCAATTAAAAACAAACTTTTTTTTGAATTCTTTTTTTTAATGATCGTTTTGTACTTAAAAAAAATAATTATGTACGTACGCGGAGGGGGAGGGGGTCTTCAAAAGCGAACAGGTTTGCGTACGTACCTTATGGATGGCCCCGAAACGGAATATATTTACGTATTTTGTTTCTTTTTGAAACATCAAAAAATGTCACACTGATCCCATTGTCTTTTAAATTGAAAATATTTATCGAAATTTTATTGAGCGG
>evgR017003c0g1_i01_Hydra
ATACTGGAACGTTAGAATATTGTAAAACTGAAAGTATTGGAGAGGACAGTTATACCGGAGTTTATATTTTCACTAAAACTTTGATTAATAGTGTTTTATCATTAGCTTGTAAGTATAACAAAAGTTTGACAATTCAAAGAATTTGTTCAAGCTTGGCAGTGTGGGAACCAGTTGATCTTTCAGTTTGTCCACCAAAGTCAGAAACTACTAAGCAACTCACCAACCTAACTAAAATAAATGTTACGCCATTAAATGTTGTAGACATTGCAGCTTCATTGAATCAAGTTGTACAAGGAGGAAGCCTTACAA
>evgR017006c0g1_i01_Hydra
CTCAAGCGAACCCTAAAATCTCCATTTTATGTATTGATCCAACACTTTTCAATTTTCAACAACGCGTTCTCTCCTATATCCCTCCGTCATGACTACCACTCCAGACTCCCGCTACTAGATTCTTACCAATTAATTTCATACTTAATAGTATTGCGTCTTTACACAGAGTTGATAAGTATGGCACAGTTGTTAAAAAATTTACTTGGTAAATTTGCTAACGGTATTCCATCTGGCTCAGCATCAGGTATTACTCTTTTGGCTGGATTAGGCTTAGCTGGGTACGGATTAAAAGAGTCGATCTACACTGTTGATGGTGGACACCGAGCAATCATATTTAGTAGAATTAGTGGAGTGCAATCTGAAGTTTACGCTGAGGGAATACATCTAAGAATACCATGGTTTCAGTATCCTATCATTTATGACATAAGAGCACAACCTCGAGTAATAGCCTCTCCTACAGGCAGCAAAGATTTACAAATGGTAAATATTAGTTTGCGTGTTTTGTCTCGCCCTATTGCATCTGCTTTGCCATCAATTTATCAAAGACTAGGGTTGGATTATAATGAACGCGTGCTACCATCGATTTGTAATGAAGTTTTAAAGAGTGTTGTTGCTCAGTTTAATGCATCACAGTTGATCACGATGCGCCAAGAAGTCAGTTTGATGGTTCGCCGAGAACTTGTTGATAGAGCAAAAGATTTTAACATTATTCTTGATGATGTATCAATTACAGATTTAACTTTTAGCCCTCAATATACTGCTGCTGTTGAGTCTAAGCAAGTTGCACAACAGGAAGCTCAGCGTGCTGCATTTCTTGTTGAGCGAGCTATTCAAGAGCGTCAGCAAAAAATTGTTGCATCTGAAGGTGAAGCTAAAGCAGCAATGTTGTTGGGTGATGCAATTAAAGAAAACCCAGGATATTTAAAATTACGAAGAATATCTGCTGCTCAAAATATAGCAAGAGTCATTGCACAATCACAGAATAAGGTTTATTTAAATGCTAACAATTTGTTACTAAACATCAAAGACACTGATCAAATTGATGTAGGATTAAAATAATTGACAAATTGATTTAGGACTTAATTGAATAACAACTTGTGATTAGAAAATGCTGAGTTTAATTTTTTTATGAGGGATTGTTTTTTTTAAGTATTTTAATCTTTTATCGAGTTTTTGATAAATCCCATTAATATATACTAAAATTTTTAATATTTTTTACATTTTGTTTATTAACAAGCAAATACTTGCAATAAACTTAGTAGCAAAAAAAAAAGAAATTTATTATTGTTTTTTAATATTATAAAAAAAAAA
>evgR017007c0g1_i01_Hydra
TTCTATTGTCTAAATAAGATATATAACTCAGCTATGAATTCTACTGTTTCAAATAACGATTCTAGTATTCCAGCAACAGATTCTAATGTTTTACCTACAGGCTCTGGTATTTCTAGTCGCTCTGGAAAACTTAATCACATAATTGAAAGTTATGTTGATTCGATGTTTGATGAGAAAAAGTTTTCAAATTTTCTTGCAAATGGTTTTAAGGATGCTTATTTTTGCAATAAAGAGATATTTAGAGAAAAGTCAAACACTTACCAAAATCATATAAAAGTTGAGTTATTGAAAGATATAAATAATTATTTAGATAAAACGTCAATTAATAAAAAACTAAATCAATTAGATAAATACGAGTTGAATTTAGAGAATGAACCTTTCGTGAAGTATGAAAGTTCCTTAGATTTGTGTTTACTGTCATACAAAGAGCAATACTTAATTGAACATGAAAATAAAGTAGAAACAGAGTATTCATCATTGAAAAATAAACTTAAAGAAAAATTGACAATTTTTGAAGATCTAAAAAAAGAGTTAAAAAAAAGATGTTTTGCTTCTGAAAAAGATCTAAATCTTGCTCGCTTAGAACTAGAAAAGTTAAAGAAAATTTTGTGAATTTTATCTTTATTTTGTTATTTTAAATTACTAAGTTAATAATATCGTTATTGTTGTGGTATCTATATTAGATGAATTTGCTGACATGAACGATAAGTTGTGCCTTTATGTATTGATTTAGAAAAGTATCAACTTAATTAAATTGAACTTTTATTTTGCAAGTTAATGGGGCCCTGCTATGGCCCTAAAAAATATATTTTTAAACCATATCATGCTTGTACTCAGACAAAGTGAAAATATTTCAAAATTTTAGAAAATGCTAATACTTAC
>evgR017008c0g1_i01_Hydra
ATTCTAAATTGTAAAGATTTGATTATAGACATTTTGTGATTTTTATGTTTGAAATATTTGATAAATAATATCTAAAAAAAGATATTTATGAAAAAATGTGACTATGGCAACAAAAAAATATTTAAAAGGAAATATAATTGGGCAACACAAGGAAAGCTTTGAGAGAAAAAATATTTTGAAGCCTAGAATTGAGGAAGATATGGATGGTATAGAAGAGCAAGATTACATATTTTGGGCAAGACAAGAAGATTTTGATACTGCTGGAGAAGTCGAAATAAGCAGCTCATCCAAGGAAAATCAAGATACCTTGTCTTTTGAAACATTTCAAAGTTCTTGTGAATCATCCACTATAAAAATAGAAGATATTTGTGCTCCAAAAGTTTTAGAGCGAATTATTGATCAAAGGCTTCAAGATCAACATCGTTTAGATAAAATACGCGAAACTGACAAACTACAGGTTAAGCTCATGAACGATGAATATCAAAAAGCTATTGGTTACTTACTTTGTTCTAATAAATCTAAAAGTGGAAAGCAGAAAAAGAAACAAACAATGACTGCCTACTTATTTTTCTGTAAACGCTACCGCTCTAAAATCGTTGCACGTAATCCTAAGCTGACTTTTGTTCAGATTAGCAAGTTGCTATCCCATATGTGGCGCAACGCATCTGAACATGAAAAAAATACGTTCAGAATAAAACTTGAACACCATCGAACAAAAGTGAACGCCATTGTTGAAAAAAATTTGATTGAAAAAATGAAAATCGTGAAACAAGAAAAACAAGATGCAAAACAAAAAATGGCTTGTGAAGTAACGTGAATCGACTCCGAGGTGTTTGAAAAGAAAATATCTAATCAAATTCATTTTATGAAGAGTTAAATTTATTTTTTCTTCTAACACCAGTTAATTTATTGTATTTTTAACATTCATTAGGTTTTAACGAATGAAGAAATACATTAAGTGTTTAATACATTAATGTAATAATTTACCCTGTAGTTAATAGTTTATTAATGATAAAGTAATTTAAAAAAAAAAAAAA
>evgR017011c0g2_i01_Hydra
AGTTCCTCAACAAATGAACGCCATGGCACCACAATATGTTTCTCAAGCACAACAAATCTTCTACCAATACGACAAGGATCGTAGCGGAACATTGAAAAAGAAGGAATTCAAGAAGGCAATGCAAGCTCTTGGTGCTCACAAGGATGAGTGCAAGCACGTTTTCAAGATGGTTGATCGTGATCACAACAAGGTTATTACCATGGACGAATTTGTGAATGCCTTCTTGTTTTTACAAGCTGGAGGATTAAGACACTCTGGTTGTGGTTACAAGGGTTCTCATGGTCAATTAGGAACAACCACCACTACAACAACAATCATGGGAGGTGGTTACCCACAACAACCACCAATGCAAGGTTACCCACAACAACCAATGCATGGCTATCCTCAACAACCACCAATGCAAGGTTATCCTCAACAGGGCTATCCACAACAACAACCAATGCATGGCTATCCTCAACAACCACC
>evgR017021c0g3_i01_Hydra
ACACTTGCAGCATCAGGAGTAATTCTCTCCGCCCTAACCAGAAGTAAAATTCTCGGTGATTGTACTATCTGCAAAGTAGTAAAACCAACAAAACCTGGAACAAAAGAACTAAACCTAGAGGTAGACCAAGAGTTCTAGAGAAGCCATCCAAAAAATGTGTATAACTATATGCAATCAGTGCTTTCAATCTGTTGATTCATTACGAGGTCATACATGTACAAAGGTTTTTAGTTCAACAAACCTTCATCCGATGATTGAAAATTCCCTGAATCAAGTAGCAGGAGACAAATTAGCTGCAGATAAAATTAGAGCAGAGTTAAAAGAATCAAAAATTATACCCCTTAAAACAGGAAGGCAATCTTTTACAGTTTCAAAAGTATCAAAGAAGCCTACTTCAGAATGGAGTCACAAAGATTTGATGACCATTAAGGTTGATAGAAAGTTGTCTGGTTCTGATATGGATGCAATATAAAAAACAATTCATGTTAAGCACAGACAAAAATCAATAACACGTGGGTTTAAAGAACTCATCGTAGATAATAATGGGTCTTTGGTGGAGCACTTCTTAGCAGTAGACCTTGAATGATAGTAGACTTCGAAAGTTCTAACAGTAGGCCTTGATTTCTATAAATCCAAAACTG
>evgR017023c0g1_i02_Hydra
ACTCAAACTTTGAAAAAGTTCAGAAAAATGGATATAAAAATGTTTTGTTTTTCTTTCTCAATGAAAGAAAAAGTGATATTGAAATAAAAGTAGAAATTCGAGAAAAGTATAGAGATATTAATATTTTTGTGATAAAGTTTGGTGTCCAGTCCTCTATTACAAATTATAAATCTTTAACTGACGAGTCTAAAATATTTATAATAAATCTTTCAAATGAAATGCTTGAACAAAATATGATTGGCAGTCTTAAGAAGTATGAACCAGTTTTTCTTGCTATCAAAAGAAGGATATATAAATTTTTAAGTTCTGAGGGTGAAGAAAATGGTTTGTTAACAAGACCATCATCTCAGTCATCATCATTGTCATCATCTCAGTCATCATCATTGACATCATTGTCTTCTTCTCTGCAGTCATCTTCATCTCAATCTCAAGCAGATTGGTGGTACAGAGAATGGAATTGGAATAAATATATTAAAAATCTGAACTATGAAGAAATTGGACCAACCAACATCGCATTCGTTGTTGATGGAACAGAAAGTCGCTTTATTATTTTTATTCAACGACTAATTTTCAACTATATCAGGTTATTCCAACTTCGTGGAACTTTTGTAACAATTGCAGTTTATGGAAAGAGTTCTTATGTCATTACAAAATGGAAAATGTTTATCACACCCTATGATGTTCAGAAGAGTTGTGAGTCTATTCCCCTGGTACAAGGGCGTTTTCGAGAAACTGCTGATGCATTAGAAAGCATCTATTCATTGTTTTTGACGCAGAAAAAAAATAATGACAATCCCTGTGTAATGATTCTTATCGTAACTGGAACTTCTGAAAATCGCAAACAATTAACAATAGTTGGAAGACAAATCAGAATGGCAGGAATCAGGGTTTACACCATTGCTATTGCTTCAGATAAAGTATCAAGCTATACTGAGATTAAAGAAGTTGCTTATTCAGACAAAGAGTACTATATTGGTTCTTCTTATTATCTAGCCACCATGTTAAGTGCAGTGAAATTATCTTTAGTTAAATTTATTGGAAAAAGTCAAATAAATGAAAATCAAAACATAAATGAAGGTGTTAGAAAATTGCAAGCAACATGGAGTTGGAGATCTAATACAAACAAAATGCAATATGGGTATATTAGTCGAGCTAATGTTGGCATTCTTGTTGACGGAACTATTGAAGAGGAATATTTTGATGCAATCAGAAAGTTAGTTTACAATATTATGAGATGCTACTCACTTGCAGGTACACATTTTAATGTTGCATCGTATGCTACTTCAATCTATGAAACAACTTCTTTTCGGCAGTACAGAAGTGGTTATGATATAAAAAAGTTTGCAAAGATAATTCCAAAACTCTATTTTGGAAAACGTAATACGGGTGGCGCTATGTGGGCAATGTTAAATGCGTTAAGTGAAGCTCCTCCAGATAATCCAAGTATTGTCTATATTATTACTGCAGGTCCTTCAGCAGATGATGTTATAGCCCCTTCAAAGGCATTTAGAATGTATGGTATTATAGCTTTTGCATGCAGTTTAAATGGAAGAACACCTGCATCTGAATTGTCTTCAATGTCATTTACTCCGAGTACTGTCTTCTCTCGTTCATTTGACGATTTGATAAATCTTGTTATCTCTATGACAACCGCTGCTTCCGAAATTATTGGAACTGTATCAAAGGCATGCGGTCCATCAGGTCATTTAATAAGTACAAATGCGGACGTTATGAGATCGTGGCAGTGGCAGTCAGCAGGAACCTCTTTCTCCTACTCTTCCGTTAATCGTGCAGAAGTTGGTTTTCTGATAGATGGTACAAGCAGTTTACATTTTGACCTGCTCCAAAAGTTCACTTACAATGTTTTTAAGTTATTTGCTAAAACTTCAACTGCTGCCATAGTGACATATGGTGGTGCTGTTGACCAAAGTCTTGGATTATTTAGAGAGTACACCGATAACTTTGCTCTACAAACGGCGTGCTCTGCAATGGTTTATAAAGGAGAAGCTGGTCGAAGAACTGGCGGTGCTCTTTCATCTTTTCATAAACTTTTTTCTCATGTTGTTGGCAAAAAAATTCTGTTTCTCATTACATTAGGTAAGTCAGAAGATGATGTAAAAAATTCCGCAAAACTTTTAGCATTATCAGGTGTTTCCATTTTTACCATTGGAATTGGAGATGAAGTAGAAAAAAATGAACTGAAATCCATATCCAGATTTTATTTGGCAACAGACTGGAACAATCTAATCACTGTAGTTGCAAGCATTCAAAATTCACTTACGAATGTCTTTGGAAGCAATACTGTTGCACAGAATGTCAAACTTTATGAAAATGGTGTTTATCTAGGAAAGATTCCTCCACTAAATGATCCGTGGGACAATGTTGTTTTTAATAAAATCAACGAATTGAAATTAAAATGGAATTGGGATAATTTATTGTCAAAAATACATTATGGAGTATTAGAAGGTGCTAATGTTGTCATACTTGTTGATGTTTCTACGAGCGAGGATCTTGGCATCATGCGCAGATTTTTGTTTAATATTATTAAGCTTTTTCAAGCAAGCGTTTCAACATTCACAGTTGTTGGGTTTGGTCAGAGCCCTAACCTCGTGTTTGGTACTCGCAAATTTAATAATGATGTTCAAATAAGGCGATCTTTGGAAAAGCTTGAAAGTTCATCTTTGAAAAGTCGAAATTGCAATATAGGTTCCGCTTTGGCATACTTTAAAGATGTGGTTTACAAAAAAATCTTGTTGCTGACTACTATGAATAGAGTTTTTGGTATTTTGGTGTCCAGATCTTCCTCATCTGATTATATTTTAGACCACGCTGATTCTATAAAAGGAGCTGGAATTACCATGGCAACAATAGCCATCGGAGAAAAATATAACGAAGAAGAATTAAAATTAATTGCAACTAATGGGCAGCTAATTATGCGTAGTTCCATTTCTGCGTTGCCTTCATTAATTTACAGCCTTCAGTTTAAGTTTGCAATTGCAATGAACATACCAAAAAATATTTCCGTTCTTAATCAAGTAACGCTAATTGGAGGCGTCGAACGAAAATTTGCGTTACTGGTTTAAACGAATGTTTTAAGTATCTAAAGATGATGGTACATGGAAACATGTTGCACATTTTTCATGGAAATAACGTGTGGCAAAATAAAGGAAGTTTATTTTGGACGAGCCCCCATCAAACTCTTCTAAGCAAATATAATAATATATAAAATTTAAATATTGTTCAATCATAAACGTTCAACCTAAGTATCCGCAGAAAAATTATTTATTAACATAAAAAGACTTTTGATACTTTGATGAACTTTATACTTTTTTATTGTAATTTTGAAAAATAATTATGTTTGTTATGTATTAAAAAA
>evgR017028c0g1_i01_Hydra
AATGCAAGTTTAAAAAAACAATAAAAAATGTAGTACAAATTTTTATGATGTTAATAACATTTATCAATGAATTTTGAGAGATTTCCAGAATGAAGCACAGCGATAAATGGCTCCTGAAAAGAAAATATACTTTTATTGCATTTTCATTACAAATATTAGTCATTGGAATGGAGTACAGTATGACATTTTTGACATTATGGCTGTATATAAAAGAAATGGTTAACCCCAACACGCCTAAAGTTTATTACGGCGTTGTTTCTGTGTCATACATGCTTGCTTCTATTATCTTAACGCCATTTGTTGGAAGAATAGTAGATAACAATCGAAATGTGAAAACCTGTTTCCTAATATGCAATTTGCTAATGATTATTGGAAATTTTCTCTACAGTCTTCATTTTTCTCCTTTCTTTTTAATTGGAGGCAGAATTATAGCCGGCTTTGGTGGTTCATTGAAATCTGTAATTTACACTGAAACAATTCGAAGTTACCCGGTAACGGAAACAAGTTCAAAACTTTCAATTTTATCGGTTATGCTCAATTTAGGATTTACAATAGCCCCGGTGTTTAATTTATTTTTTAAAAATATGAACTTTTTTATAGGCAGCTTGCATTTCCGAGACGTTAATTTCCCGGGGTTTTTTATGGGATTTTTATGTATATTTATGGAAATAATAACAACAGCAATGGTGCACAATGTATCCAAGGAATTACATTTAGACGATACTTTTTCTTTTTTACTTTCTGAAAAAAATGAAGAAACTATGCATGGGTTTTCCAAAGGTCAACCGGTTATTATAAAAGAACTACGGATGGTTACTGAAAAAACACGAATGATGCAAAATAGTAGTTTAAACGAAAATGATTGTGAAAATCTATCAAAAACTTATCCTGTGTCACATATTTTAAAAGTGTTATTTTTGAGTTTGGACTCAGCTCTTCTGTTGTTTTCAAGCTTTTTTCTAACTTTTGTTATTGTTAATACTGACATATGGTTGCCATTGTTAGTTATTGAGAAAATGCATCTTTCAACGATAGAAATGAACATTAGTTATTTTGGAGCATCTGGATTTTGTGCGATTATTTTGATCGTGTTTATTAGAAAGCCGTTTTCTGATAATAGCATGATAATACTAATAGTAATTAGTGTAATTGGTTTGTGTGCTGTAAGTGCAAGCTTTATTATTCTTTCGCATTACCACGACAATAAAGTACTAAATGTGTTCCTAAGCATCATATACATGCTGAGTTTCGGAGGTGCACCTATAATTAACGACGTTTTCATTGTTGACACACTTGCAAAAATGGTGAACTCAAACATTCTAACGTTTTCAGACAGTATTCGTTCTTCAATGCATTCTGCCGGAATTTTATTAGCACTAAGTTGTTCCGCCTTTCTGTTTGACTATGTCGACATTTTCGGAACGCTCTTTGTGGTTATTATGTCATTTATCTGTTTTCTATTTATTATCCGTCGAAAAAAGTTTATATCTCCTGAACTATTGATATAAAAACAATTTTTTTGTAACATAGGTTTTAAATATTATTTTTGTTTTAATCCATGTCTTGTTAATAATTAACCTTTA
>evgR017029c0g1_i01_Hydra
CCCCCTCCCCTATCTCACTACGGTTGACTTTATATTTCAATATCATACTTCAAGAGATCTCTTTATTAAATTCTAGTTTATAGCTAATTTATTAGGAACTTAAAAACAAATCAACATGGCTAATAGGACCATATCTGATGCAAAATCTATTCGGGGAACAAATCCTCAGTATTTAATCGAGAAAATCACTAGAACTCGTATTTATGAATGCAAATATTGGAAGGAAGAGTGCTTTGCACTTACTGCGGAATTACTTGTTGATAAAGCAATGGACTTGAAAGAAATTGGTGGTACTTTTGGAGGAAATATAAAGCCAACACCTTTTCTTTGTCTAATTCTTAAAATGCTCCAACTGCAACCAGAAAAAGATATCATTGTTGAGTTTATAAAAAATGAAGATTATAAGTATGTGCGTGCATTAGGTGCATTTTACATGAGGTTAGTTGGGACTTCTCAGGAGTGTTATAAATATCTTGAACCATTATATAACGATTACAGAAAAATGAGGAATAAAGGAAAGATAGCAGGCTCTACTATAATACATATGGATGAATTTATTGATTCCATACTTCATGAAGAACGAGTTTGTGATGTTATGTTACCAAGGATTCAGAAGCGATACATATTAGAAGATTTAGAAGAATTAGAACCTAGGATAAGCGCTTTGGAAGATGATTTAGACGATTTGGAATCTGAAGATGAAGAAATACAAGAGGAACTACCTGTTAGACAACGAAGTGAATCTCCCATTGAACGTAAACGCAGAAGAAGTAGAACGCCAGAAAAGAGGAGAAGATCACGTTCTGCATCTCCTTATAGAAATAGACGAAGTCCGAGTCCGCGTCGTCATCGTCATCGGAGTCGTTCAAGGTCACCTAGACACAGACGTCACCATTCGAGATCTCCGAAACGAAGAAGATAAATTCTAAAAGTTTAGATTTTTATAAAATAGCAGAGAAGTGTAAAAGAGAAAAAGACAAAAAAATAGAGAAAAAGACAAAAAAATATCATTTTATAATGGTTATATTTTTCTGTCTGTTATAAATACTGAACTATAACCAAAATATCTACAAGTTATGTTTTGTATATTGCCGAACTGATATAAGTTATGTTTTGCATATTGCCGAACTGATACAAGTTATGTTTTACATATTATATTTGCCGAACTGATATAGGGTAAAAAAGATTTTAAAAACAAAAACAAAAATCTACACCCTAAATCGACACATTTTCTGAAACTTGGCACACATGTTTAGAAATATATGAAATATAAGACTGTACATTTTTTTCATTAAATTCCTATCAGTTTA
>evgR017038c0g1_i03_Hydra
CTAAAAATAAAAAAAATGACATTTATCTTTCAAAAATTCAGAAATTAAAGAACCAATTGAACTTGTTAGAAAAAGATGTAAACACTTATAAAAGCGAGGCTACCCTTGCTGCTGACTCTTACAAGTTTGAACTAGTTTCAGCAATAAACGTGATTAACAATCAAGAACAAAGACTTGCGTCGATAGAAACAGGATTTCAGACACCGAAGTTTTGTTCAACAGAATTAGAAGAAATAAAAACTGAGTTAAATAGTGTGGCAAAGGAACTAAAAGAATATAAAAGTTATAGAGGCGAGGCTCCAATACCTTCTGACCCTTATACGGCGAATCATGTTCCTGCTTCAAACATAACTAGCGATCAAGAACAAAGACTTTTGTTAATAGAGAATGAATTTCAGGCGATGAAGTTTTGTTCAACAGAATTCGAAGAGATAAAAATAGAGTTAAATAGTGTGGCAAAGGAACTAAAAGAATATAAAAGTTATAGAGGCGAGGCTTCAATACCTTCTGACCCTTATACGGTGGATCATGTTCCTGCTTCAAACATAACTAGCAATCAAGAACAAAGACTTTTGTTAATAGAGAATGAATTTCAGGCGATGAAGTTTTGTTCAACAGAATTCGAAGAGATAAAAATAGAGTTAAACATCGTGGCGAATGAACTAAAAGAATTTAAAGATTTAAATACCG
>evgR017041c0g2_i01_Hydra
GTTTGCGAATGTGGGGTAGGTTTTTACTTACATTTTAATTATATTATTATCAGTAATTTTAGATTACAATTCTTTTAATTGTTTATGCGGCATTTTAGATAAAGATACTGTATCACCTTTTAGGTTAATTGATTTGTATTTTAGGCTGGATGGACTACCGATGCTCATCCAAAAGGACAACTTTTCCCCAATGGAACATCCAATTTATATACAGACAGAGGTGTTATACTAGTGCATGTTTTCTTTCGACGTTGTGTTGGAATGTCATGTATAAGGTATTGGGATGGTGCAGAACATTGCGTGTTTCGAATATCCTCATCTACATGTGCTGGGTATGAAATTGGTTGGGACTTCGTTGATTCTGTTCAAACCAGTGGTCAAACTTTTAGCGGTTATGTTACTAATATGAGTAATCGTTACAAGCGTATGTGTAACAATGCGTTGCCATTTATGTCAAATTCGGTTTTTATTAGTTGGTTTTTTGGATGGACATCTCACATGGGGATTGACTTTCGAGAACAGTGCACAATGTGCGGAGACAGCAGTCGCATTTTAGCTTGTGATGGAACGAAACTTGGTATTGGGTTTAAGCAAACCTTTGTGAAACCTATTGAGACTCCAGAAATTGAAGAAGTTTCAAAAACAACATTCCGGAGATTTGATCGGTGTTTCATTCCAAATACCAAAGAAAAAGACCCTAAAGATTTTGCACAAGCAAGAAAATCTATCCGCACTGTTTGCGAAACGATAATGAATCAGTGTGCTAATCATGATGTATTATCTTTAGCAGCCTCTATAGATTTTTGTATACCAGTTTATGCCAAAGCTGCTTTTAACAGAATGTTAACTGGAATAGATTGCTCTCCGCAAGAAAGAACAGCACTTGCAGAATTTTTTTTGTTACTTGCAGCTGATGCATCAATGGATTCCTTAATTCCATATAGATTTTGTTCTGAAGTATCC
>evgR017058c0g1_i01_Hydra
CCGCGTATGCCTCTTAAAGCATTGCATCCATTATTAGAAGATGGCACCGATGGTTCGGAAGAACTTACACTTGATGTTTCCCATCTTAAAAACAAAAGAGTGGCATTTATTGGCATTTCAAACTGGTCCCTTGATCCTGCAAAAATGAATAGAGGAATTATGCTATACAGAGGACAGCCTAAATTAGATGAGCTAGTTGAAACTGCAAGAAATATATGTACAAATGATACGCAAGTTTTTGGAAAGATAGCATCTTTATTTGAACCTCTGACAAATGGATACTTAGAAGTTTATGCAAAGCAAAAAGAGTGTGATATTGTAAAAAAATACAAAAAAGAAGAATTTTTTGGATTGAGAGATTTTTACAGTTTGTTAAAATTAGTGTTCTGCTTTGCTCGAAATCGTCAAAATATTCC
>evgR017062c0g1_i01_Hydra
AAGAAGATAAATATATTGAAATCAATTTTAATACTACCTCATATACCTTCATTCAATGTATAAGTAATAATATGGATTTTGGATTCCAAGTGTTTGCAAACTCAAGTACAGAAGTATCACCGTTTACCAAAAATACCTACGCATTTTATTATCTTGAAGTTCTTTTTTCTTCATTTGAATCTCGTGTTACTACTGTTACTACTGTTACACCTGCAATCAAAGGAAAATCTTTTACTAGTACTATTCTTGCTTCCATTGTTGTTCCCTTGATTTCCATAGCAATAATGGTAATATACTGTGCATACCAAAAAAGAAAACGTGGATCTGTATGGACCAACACCTTTTTTGGTGATTACCTTTACACACTTAAGATCATTAAACATGATGAGTGGGATATTATGCCTCGAGACATTTGCTTAGAAGAAAAAATTGGAGAAGGAGCTTTTGGAACTGTATTTAAAGGAACAATCAAATCAACAGTTTTTATAAATTCTAAATATGTTATTCAACCTTCACTTAAAGATTTCTTTGAGAAAGGTTTGGGTGGAAAAGCACCTAAAATTGTAGCTATTAAATGTCTAAAAGAAGGAGCTAATGAAAGTGAGATCTACGATTTTAAAGATGAAATTATTCTGATGAAGAGTCTTGGATATCATAAAAACATTGTTAACATGATTGGATGTTGTACTATTAATAAGTCATTATGTCTTGTTATTGAATATTTAGAAAATGGGGATTTACTCAGTTTTTTGAGACAAAATCGTGCAATGATGTTTTCTGATGACGAGAACGAAAATGATGAAAACTATCATTATGCAAATAACTATCTAAAAGGCCGAACGATTGACAATCCTGAAAAAATATGTAACAGTAACCATAACAAAATTGTGGGAAACAGCAAATTTAATAAAAAGTCAATGGTCAAACCAGTCGATCTTCTTTCTTTTGCTTGGCAAATTGCATCGGGAATGGAGTACTTAGTTGAGACCTTTGTACATAGAGATCTTGCTGCAAGAAACATCCTTGTAGGAAATGAAAGGAATGTTAAGATTTCAGATTTTGGTTTAACAAGAAAAATAAACGATGAACAAATATACATGAGCAGAATAAATCGTCGATTGCCAGTAAAATGGATGTCAGTGGAAGCCATTTTTGACCAAACATTTACCTCCTTTAGTGACGTTTGGTCATTTGGAATTGTTTTATTTGAAATTGTAACGTTAGGTGGAGCGCCATATCCTGATATAACCAATCGTCAGCTTCTTAGTCTACTTAAGTCAGGTTACAGAATGGAAAAACCAGAAAACTGTGGGGACCAAATGTACGAATTAATGCAAAAGTGCTGGAATGTTAGCCCACAAAATCGCCCTACGTTTACAGATTTGCGCGAACATTTTGCAAAGATTATTGGACAAGAAACCAGCTACTTCAGTTTTGACATTGACGAACGTACTAATTACTACAACGTGGCCTCTTTTAAAAGCCTTTCCTCTGAATCAGAAAAATTCGAACTTGATAAGGAAAATGGCATTATTTTGCAGAGCACTCCGAAATTAAACTCAGATTCTATAAATGAGATCTACGACGAATCAACACAATCTCAATACTATGAGAATTTGTATAAAGACGATATAGGTAATTTAGTTCAGCCGCATGCAATTTTTATGTGTGAAGAACAACGTTTTGTGTAAAGAATAACCTTTCGTGTAAAAAAAAACCTTTTGTGTATATAACAAC
>evgR017069c0g1_i02_Hydra
CACATTATTTGGAAAACATCCGTACATGAGTATATATTGTGAGTAAACAAAAAACAAACAATTTTTTATGTATAAATCCCTTTTAGTAGAAAGCTTTTATATTTTCCGCTTCTTCTTATTTATCAAATGTCTCGTTTACTTTAATCATCTTTACCTGAGTTACTTTGAATATGTTGTATTAGTCAGTTGATTTGGAAAAACATTGTCAATATCACTATTCGAACAAAAAGTTAAAAGAAATTAAACAAGCAAGTCATTGTACGCATAGACTTCTGTAAATTGCGCCAAGTCAATTTGTTTAACCAGAACACATTCAACATGCTGGTAAACAACTGCAACATGCTGGTAAGAAAACTTTCATAGTGAGTTAATTAATAAATAAAGCTCTTATGGTGGAAAATTTGATTTGAACCTTTTTGTCAACGTTGCTGTTAAATATTTTTTAAGTAAGATAATTTAAAGGATACAAAGATATCGCTTACACAGTAGAACAGATTTTTTTAATAGAATCGTCCGAAACAAAAATGTCAACAGAAGTTAACTGAAAACTAAAATGTTATCGAGTGACAGCGCTATGTAAACTATGACCATGGACAATATTTGTTGTTTATGTAAAAAAGCTTTGTTTGAAGATTGCCGCAAATTATTTAACAAAAAGCTTAAGAGATGAAAGACTTACGCGGCTAACGCAATGTCAAACTTGACTTCTACATCAATACTAAGACCTAAAAAAGATTACCACTTCTATTAGATTTTCAAGTATGCTAACGAACGATAAGCCTGCAAGCAGCCAAGCTTCTAATGTCTTAAAAAAGACGGAGAGCCCAGTGCAAGCAAGAGGCAAACAAGATGTTGATAATAAAGAAGATGCAGGTGGATACACGGCTGGGGTCGATAATCGCATAAGAAGACAAAGAACCCATTTCACCGTCACGCAACTTCACCGATTAGAAACATGTTTTGCGAGAAACAGGTATCCAGATATGGCAATGAGAGAAGATATCGCTCAATGGTGTTCTCTTACAGAATCGCGAGTTAGGATCTGGTTTAAAAACCGACGTGCAAAATGGCGTAAAAAAGAAAGACATTTAGAACGCCCTGATGTATATAATTTTACACCACCATATAGATTCGAATCGCCATCATCTTACGAAAGAAGAGCAACATTTCCAACAACGTATTCTTCAAATGTTTCGCAACCTACATTGCCTATATGCTCCAATAGTAGATCTTCGCATCCTTCGTTAGGTCATTATTACTCACATAATACACACTGGTATCCATCAGATCACCCAGTGCACTCTCTTTCTCAAAGAAATAGTCCATCAAATAGTCCAGCTGGATGGTATGGTATTAACTATAACGAACCAGTTCGAGCGTATCCAACTTGTCCACAAAGCAGTTCCTCGACTTTAAGCAGAGAAGACGAAATGATGAGCCCCATTTCATTTTCTCAACGAACTCATACGTTTTCACCTGGGGATCATGGTTATCAACATTCAATATAAACACCCTTCGCTACTCAATGTAGAATGTGTACATTGCATATATATATAT
>evgR017080c0g1_i01_Hydra
TTTTTTAGAAACTCCACTGATTAAAAAATTAATTCTATTAGAATTTTTTTATTTTAAAGCTCTCAAATCTCAAATGTCACGTTGTGACAACATCTAATGGCTATGACGTACATTTTTCATAATCAGACAATAATTAATTCATCAAACACCACTGAAATACACAACAATAAAAGTGGAGCAGATTTACCTCTGATTATATATATTGCTTTTGGTTTTATATTTTTAACGGGAGGTGTTGGGAATGCACTTGTTATATATGTGTTTAGTTTGAAAAAGAAGTCAAAGACTGCTGATTGGTTAATTCTTTATTTAGGAGTAGTTGATTTTTTATCTTCAATTTTTAATCCTCCTTTGTATGTTTATTATATGGTATTTAAACTTGAACAGTGGCACTTTGGTGAATTTGGGTGCAAAATTCTACCAGCATTAGGGCCAATTATGTCATCAGTTTCTTTCGGTGTCATTCTTATATTTGGAATAGATCGATACTTAGCAATAGTTTCTCATGTACAGGGATCTGTTTTATCATGGAATACTATTACGATGGCGTTTATAGCAGATATTATTTTTTCAACTTGTAGCTATCTTCATTATATAACGGCGTTAAAATTTATCATACATTTTGGTAAACCAATTTGCATTATTCCTTCTGAGAACGTATCTTACAACATAACAAATTGCATTCTCGTTATTTTTCGATTTTCTTTTTTTGTTGTTGCCCACACGTTTACAACTGCTAAAATATTTTCAACGTTAAAGAAATGTCAAAACTTATCGTTTAGTAGAGAACTGCAACATAGACATCGCAAGCACGCTAAAAATACTATATATGTACTAGTAACAATGGGAGTGGTGCTGAGTTTACTTGTTTTTCCAAAAGACATTTTGCATTTAACGCACAGTTTGTTTTGGTTGAAAAATAAAAAAAAGGTAGTCTATTCTTACAATTTACAGATCTTAAACTCGTGTTTAAAACTGCTTCAAACATCAAACAGTTGCGCAAATGTTTTTATTTATTCAAAGATGCACAGCGTCTACCGTAAAAAAGTTGCTCAGCTTTTCTTACAAATATTTTGTAAAAGACATGCTAGACAAAGAGGTTTAACTGAATTAAGCCTTATTACAAGTCATTCAAAACGTTTGAGATATTCCACGAAAAAAT
>evgR017085c0g1_i01_Hydra
TTTTTTTTTTTTTTTTTTCGTGAAGTTTTTAAGCGTGTGAGTAAACTTTAAGAAAATAAATAAATTATTTGTTTTTAATAAATAGTAGAAAATAAATGGAACTTATGACTCAAAGAGAACGTCTCATTCTGGCAGAATCTTTTTCTCTTTACGACACGGTTGGGGATGGAAAAATAGAAGTTGGACTTTTAGGCGAAGCCTTAAGAGGACTCGGTTTGAATCCCACAGAAAGCGACGTAAACAAAATTGTACGAGAATTAGAGTCAAGTGGAGCGAAACGAATTACTTTTGAAGAGTTTTTCCCTATTTATCAATCTTTAGCAAACAGAGCTGAAAAAGACACAAGAAAAAAAAATAAATCAGATTTTACTGAATGCTTTCGTTTATTTGACAGAGAACAAAATGGTACTATAGCTGCTGGCGAACTTCATCACGTTATGACAAGTTTAGGAGAAACGTTAACTACGGCTCAAATGGATGTTATTGTGCAGGGATTAGAAGACAAAAACGGAATGATAAAAATCGATGAGCTTGTCGCATCAGTGATGGATGGTTGAGTCATAATGACGACGATTAACGGTTAACTCGTTGATGGATGCTTATTCTCCTACGCTCTTAGATAAAAGTAATCACAAGTGAAATATAATTAGTACGCAATTTTTATTTAACTTTCGTGTGCATGCATTGAATAATAGTAAAATTTGTTATTCAAACTTAAGCTATTCACTTTTTTTTCAGTGTAAAAGTCGAGATATATTTTTAATATGGAGTCGATAGGATAGCATGTACCATGTGTAGACTTAATTATATAGAAAAAAGTTTTTATCTTTCTTTATTAATCAAAGTTTCTTTTTAGAAAACTGTCTTTGAGAATTTTTTTGTACGAAATAAAAAAAGCGGGAAATACCATTTAGCATTTCGTTGTTGTAAACAACAAATTCCTATCGAGTCATCGTTTATTGAATTTTTTAAATACTATTTGATTAAATGTTAC
>evgR017094c0g1_i01_Hydra
AAAAAAGAATACTTGGAAAGAAGGGCATGATTATGCACGTGTTCTTTCTAAAAAGAGATAATGATATTACCAAACAAATTTTTATCACAGTAGTTTATCGATATGATCAAGGAATGGCAAGTAGCTTTCGTATATCTAAACACGTTCTAAAAGAATTCAAAAAAAGTATTCCCGTAGTAACAAAGATTTTTACAAAGTCTGATAATGCTAGTTCATATCATGGAAGTTATTTATTTTGGCACTTCATAAAGTCTGCAAAGACGAGAACCTTAATTATTAAGAACAGATTTAAAAAACCCATCGAAAGGTAAGGATCAATGCAATCGTCTAGTAAACGAG
>evgR017095c0g1_i01_Hydra
TTGAGCAGGAGAGATGATTTGTGGTCGATATTTTATGTTCTGGTAGAAATGGCCTCTGGAAATCTTCCATGGAGAAGAATGAGAGATAAGGCCGTTATTGGAAAGTTGAAGGAACAACACACAATTCAAAATTCATCAGAACTTCTCAAAGGGCTCCCATCACAATTTACCACCTTTTTAAATTATTTGAAAACCTTAAAGTTTGAAAGCGAGCCAGACTATTCCTATTTGAAAAATATTTGTAAATCCTATTTTGCTCCACACAAGGAAGCTCTCGAGAGCAAAACATATGAAGAGTGTCAATTTGAATGGATTGTAAAAAACCCAGAGGAAAAGGAAAATTCCAACACTTCCACCGATAATGTTAACAATACTACCGCTACGACAAATCCTGACAGAACAGATTCTAAATCTAGAACTGGAAAGTTATCCACCTCTGTATTCTTTGGATCTACTCGTTTGAGTACTGTAAGCAATGCAAACCTTGGAAGTAATGCATTGGTCGGTAGTCAAGCAGAGTTCTCTAAATCTCTCCCAGATATGTCAAAGAAGAGAAATAACAAGGGTGAAGGAGATGAACCACAAGATGGTGAAGGAAAACAAAAAAATTGTAAATGTAGCATAATGTAAATAAAA
>evgR017097c0g1_i01_Hydra
CTACAACAACTGAATCATCAACCACAACAACTAAAGCAACAACAATAACAACTGAATCACTAACTACATCAACTACATTACCAATTACAACAACTGAAGAAACCACTACAACTACTGAATCATCAACTTCATCGACTCTATCTCCGACTCAAACAACTAAAGCATCAACTTATGCGATAATTATAACAACTGAAGATCCAACTAGAACAGCTGAAACAATAACCACAACAACTGAAGCAACAATAACAACTAAATCATTAACTACATCAACACCATCCCCAATCATAACAACTGAAAAGATTACTGCTACAACTAAACCATCAACTTCATCAACTCCATCTCTGACTACAACAACTGAAGTAACGACTACAGCATCTGAATCATTAACAAAAACAATTGAGTCATCATCGATATCTACTCCATCACCGACGACAACAAGTAAAGAATTCTCTAGCACAACTGAATCATCAATTCCGACTCTGACTGCAACAACTGAAGCAATACCTACAACAACTGAATCATCAACTCCATCATCGTCAAATACAACCAAAGCAACAAATACAATCACTGACGCAACAACAGAAACAATAACTATATCAACTAAAGAACCAACAACAACTTCTGAAGCAACAGCCACAACAACTGAAGCTTCAACATTAATAACGGTATCATTACCTCCATCAACTCCATCTCCGACTACAACAACTGAATTAACCACTGCAAGCCCTGAATCATCAACTTTTTCAACTCCATCTCTGACTGCAACAACTACAATAACTGAATTATTAACAACATTAACAACATCAACTCCTGTAACTACAACATCTCAAGCAACAACAACTGATGCAACAACTACAACAAATAATATGACAACTAGAATAAATGATGCAACAATTATAACAACTGAAAAACCAACTACAACAACTGAAGGAACCACTGCAACAACTAAACCATCAACTTTATCAACTCCATCTCTGGATACAACAACTAAAGCTACAGTTAGCACAACAGAATCATCAACTTCATTAACTCCATATATGAATGCTGCAACTGAAGCAACAACACTTTCTTTGACTACAACATCTAAAA
>evgR017115c0g1_i02_Hydra
GGATACCATTGAGTATCTACGGTAAGCTATGGCAGGCTACTTACAGACATAATAATTAATATATTTTCTCGATTTATTTATTTCGGCGGAAACTCCTAGCTTCAACTCAATGTTTCTTACAGGTCGCTTGAAAAATATAAGACTCACGTTGTGTACTTTTAATTAAATGGGCGTGTATTTAACGTCGTCAATAGAAGCTTCGTAATTGTTTAATTCGTTACTTCTTCTTGTAATTATTTCTGTTATTGCTGCTATGACGCCTGGCAATTTATGAGAAGGGCAAACTTATTTTGCATTTTTCTTTTTTTATTAACTGTTCTTTTTACTTATGAACTGCTGAAAAGTATAGATGAAGTCAAACCTCAAGCTCATTCAGATGTACTAAATAGTTTAAATGAAAAACATTCTGAATTAAAAAGCTTCGTTCCGTTTCGTAACTCCATGAACATGTCTTTTGATATTGATGTTGTCATTAATTCGTTTAAAAAAAGTGTTGAAGACAAATTAAATCTTGAGGAATACAATCGAGGTTACAATGAACTTTATTCATTCTTTCTTGTGCTTGGTACAGTGTTCAAGTTTATTGCATCCGATGTTCGCGAAAAGATTGATATTTTGGAATCTTTCCGAAAAGGAGAAAACCAAGAGCATTATACTCATACTGAAAAAATGATCGTCTATGAGATTGAAGCAAATAAAGACAACCCTAGCATTCCTCTACTTGGATCCAGAACACTACTTAGACTTCATCGTGCTTTAAAATTCACTTATCTTTTTTTAGATGGTCTTGTTCACTTAAATGACGACGACAATCTCTCTTCGATGGCTATTACAATGTATAACAAAAGTTTGGCAAATCATCATCCTTGGCTAATACGCAATGCAGCTAAGTTAGCTATGTATTCTTTGCCAAATAAAAAGACTTTGATTAACCAAATAGCTAAAGACGCTGACAAAGTGCTTCTTTCGCAGAAATTGCGGGAAGGAGTTGCTGCTTTGGAAGAATCTTTTAACGAAATTGAACTTTTATACACCAAAAATGATTTGCACTCATTGCCTTAAAAATTCATTCTGTTTTACTCAAAATTTCAATCAAATTCTTGTAACTTTTGAACAATTACAGTTTTTTAGAACCATTTTCGTTTAATTTTTGTAAGCAAAAGTGAGACATAAAAATATATTTTCAATACTGATGTTTTACGTTCATGGGTGGAAAATGACTACGTAGAAAATTTAATTAACGGAATTTTTAAAGAGTGTCTTAAACAAGCTAAATAAAG
>evgR017118c0g1_i01_Hydra
TGCAAAAAGGCTAAAGGACATGCAGATTGAATATAAATGCTATGGTGTCAAGTGGATATACCCAGGGCCGTAGAGAGCCAGGACGGACCTTGGTTACTTATTACCCTCAATCACTGCATCTCCATATGGTTGTATCAATGGTAGCTCTACCGAATTTCAATTGACAGCTTCTGGCTACTATGTTGAAACTATTTTAATGTGTGTTACTATTTGCAATTCTTATGGACCACCAGGGGTGTTTCTATATGCTTTTTTTCTTGAAGGGGAACTTTGTTTCTGCAGGAATGCTTCACTTCCACTGTTAGATATATCAAACTTTACCAGCCCATGTATCACAAGTACATTATTAAAAAACAACAACACTTGTAATTGCTTTCTAAGTAACTACAAAGTTGTACTCCTCTCTGCAGGCATTGTTGAATGGAATTCCGTTATTAATCCCACCATACGACTTGTTGATAGCAAAATTAATATAGAAACACATATACTAAAAGGTGATGCTTCAAGTGCATTTGTGATACCAATACCTAACAATTTGCCATCAGATGTTCAAACTGTAAATGGTTTAGTAGTTAATAAGCTTTTTTTTTTACCTGGTTTGTATCAATGGTCTACTGAAATAGGCAATAACTTTACTGAACCATTTATTACAAGTATTTGGATTCAAATTGATGAAGCTATTGATGGTATCCAAATAATTCTGATCGATGTCAATCCGATTGATGATACCTACTATTATCTACTTTTAATCACACAAGGAACCAATATTAGCTTAACTGTAACACTGCCAAGTGAAAGTTTCAACATTTTTGCTGATGTCATTCGACTAACATACGGATTCGATTTGTCCATGCTTTATAGTCTACCATCTTCAGTCTGTAGTCTTTCTAGTGGAGCAATTTTAAATGAAGTTTTATCATATGGTCGCTTTACAAAAATAAAATTAAAAGTATTTTCTAAAGGATATTTAAAAATTGCACTCGATGTTTTTAACACAAGTTTTTTTGTTACAAAATGTCAATGCACTCATTTAACTGCTTTTGGTGGATTTTTTGTTCCTCCAAATCCCTTACCTATGCCTACTTTTTCTATGTTTAAAAATGGATATGCTCTAACTGTCACAGTTGCAATAGTCATTATGCTTTGGATAGTTGGTTTGGTATTTTGCAGAAGAATGGACGAAAAAGATTCTTCAAAGGTTG
>evgR017125c0g1_i01_Hydra
GTCACAAAAGTGGAGCGCAATTTAGGAGTGCTTATTTCTGATAACCTCAGTGTAAGAGCACAGGTAGAAGCAGCAGTCGCAACTGCAAACAGTGCAGTGGGGCAACTAAAAAAAGCATTTCGAAGTTATGGATTCAAACTTTGGCGTACCCTATACACCACATACAATAGACCGCATCTTGAATTTGCCGTCCGGTTATGGTTGCCATATTTGAAAAGTGATATTGCAAGTCTCGAAAGGGTACAGCGTCGAGTGACAAAAACAATATCGGAGCTAAAAAGACTGAAATACGGACCCAGACTAGAGCGTCT
>evgR017150c0g1_i01_Hydra
GCAAATTATGAAAATCTGCCACCATGGAACGCAAATTATGAAAATCTGCCATCATGGAAAGCAAATTATGAAAATCTGCCACCATGGAACGCAAATTATGAAAATCTGCCACCATGGAACGCAAATTATGAAAATCTGCCATCATGGAACGCAAATTATGAAAATCTGACACCATGGAACGCAAATTATGAAAATCTGCCACCATGGAACGCAAATTATGAAAATCTGACACCATGGAACGCAAATTATGAAAATCTGCCACCATGGAACGCAAATTATGAAAATCTGCCACCAGGGAACGCAAATTATGAAAATCTGACACCAT
>evgR017155c0g2_i01_Hydra
AGAACAAAAGAGGTTAATGTCAATGGACTTGTGGAAAAACTTACTCAAGATTTCAATGAACACTATAAGGCCAAACTTCTAAGCATTGCAAATGGTCGATTCGATGGTATTTACAGCGACCGATTTATGGGAAGAGACAAAGGAAAGGAAGGAGTTGGCTTTGCCCTACCTTCTCAGGAAGTTCTTAAAAAAATGGAAAAACAGATATCTAAAATTGGACTAGAGTTGTATACGATACCTAGTTTCAAGTCATCTGGTGTTCATTATAATGTTGATATGAGTTTAGGTATATGCGATTGTTCTATAGGATGTGATGGGTCACCTTGTAAACATCAGTACATTGTTTGGAAAAAACATTCATTTAGCACAAATTTTCTTCCATATCTAAGCTATAAAGAAAGACAGTTGTGTTCCTACATTGCTATTGGGGAGATTTTGCCCGACTCTCTGTACGAAGGATTACACGATTATGTTAGAGATAAGCATAATGAGTTTATCGTTAAAGAAAACTACAGG
>evgR017156c0g1_i01_Hydra
GCAGTCGTAATTTATTGCCTAGCCGTTCGCCAACTTTTATTATACACTTTTTAATTAATTTGATGTAGGAATGAATAGCAAAGTAATGGAAATTCTCGCAGATAAAATATCTTCTGATTGGGAGAGATTTGCTCGAATCTCTGAATTCAGTAATCAGTTATCGGTAAATACAATTATTGATCAGATAAACAATGAAGATAAAAGCGATAACAAATCGAAATTGAATTGTTTACTAAATAAAATATACGAACATCATCCTCAAGATTACATGAAGTATATAAATTCTTCTCTAGAACAGATGGATAGAAATGATGTGTTAAAAGATTTATCAAATTTAGATTTATCTGAGAAAGGCAAACCTCTGAAAAATTTGTATCTTAATATTTATGCAAAAGTAAATGAACTTCAACCACCACTCAAAGAACCTGCAAGTGTGTATTTATTAGATAAATTTATTGATTTGTGTATTGTTAATGCGGAAAATGCTCAGCTGGATGCTCTATATAATCTTGATTTGATAGAATTTTGCAAGATGCAAAAAAAACATACACATATTCAATTTGACAGTGTATTTAAGAAAGAAAAAGACTTATTATTAGTAGCTGGAATAGCTGGAATCGGAAAATCTTGGTTGCTTAGAAAATGTTTGCTTTATTGGGCTAATAATCTTATTTGGAAAAATGTGGATTTTGTGTTTTATTTCGAATGCAGAGTGCTTAATCGTTTTCAAAATATTTCTAATATTAAAGAACTACTTGATGTTTTCTACAAAGATTTTGTAAAAGATTTGGATTTAAATAACCATAACATAGTGTTTATGGTTGATGGTTTAGATGAGTTTGAATATTTAGATGATTTAATAAAACACACTTCTAGTTGTTGCTACCCAATTGTTAATACTTTAACAGAAATTCAAAAGTACAAAGTTGTTGTTGCTGGAAGAGTTTATGCTATAGATCAATACCAACGTGTATCGACAGAGATTTTAGATAAGTTAACCGTTCAAGTAATGGGGTTCAACGAAAAAGGAATAAAAAACTATATAAAAAACAATGTCATTGACGGAAAAAAAGAACTTATAGAAGCATTTTTAAAAGAGTCGGCAATGGCAAAGGCCATGGCATTTGTTCCGTTCTATTTAGCTTGTATGTGTAAAATTATTGTTGAATCCAAG
>evgR017167c0g1_i01_Hydra
GTTGTTTATGTTGCTTATAATATTATAATATAAAGAAATAGAAGGAATAATACATCAGTCATGGTGAAGAGTTTATTGTCACCAAAGGTTTAAATCTTGGAACACTAAAAGTTTTAAAAAAAGAATACATTGTATCCTTTAATTTAAAACCCATAAATTTTTCTTCTGGTTTGAAAAATGTTCTTTATTTAACTTTGGGAAATAACAGTGATGTAAATAGGGACAAAAATCTAGGTGTATGGTTTAATGAAAATGGGTCCGGATCGCTAGTTATTTGTGTTGCAGTTAACAATAATCTTGATTACTGCAATAACACAGATCCAATTAGTTTGTATTATCGGAATGATGTCAAAATATATCAGCGTCTCATTAATATGAAGTATGTTTTATCTGTTTATCTGAACAATTATTATATTTATTCAGTGGAAAATTCAGCGCCTAGAGATTTTTTTAACATTAAAGTATACGCTTCAAATATTTGGGATGATGCCTCAAATGCTTTTATTTCTGATCTTTTTATTGAAAATAAAGTCAATGGAAAACCTGAATACATTAATCACAGAGAAGAATTTGTTCTCACTCGTGGTTTAATCCTTGGAACGCTGACTGATTTAAAAAAAGAGTACATCGTGTCCTTCAATTTAAAACCCATTAGTTACTCTAAAGGTTTGAAAAATGTTTTTCTTTTATATTTGGGAAATAGTTCTAACGCATATGGAGACAAAAATCCAGGTGTTTGGTTTCATGA
>evgR017179c0g1_i01_Hydra
CTTTTATTTTCTAAATAAGATTAAAAAAATGAATTTCGTAATAAAACTTATAGTGTGTTGTGTTATGATGCAATCTTTGGAGGGGTTTGCTGGTGGAATGATACCAGAAACTGTTAGACGTAAAAACATTGAACAAAAGTATCAGAACGGTATTGATTTGGAACGTTTATACAACAAATTGCATGAAATAGCCAATGGCAACGATGATTCACTTATAAATACAAAATATAGAGAAAAAATAATACAAAGATGTGTGGAATTTTGTATATCAGATGATTATAAATGCGTAACAGAATGTATTCAAAAGAACACTTTAGTTTAGAACAAAATTTGTGTGAATGTATTTTTGTTGTGAAATTTATTTAAAAAGTTCTTTAAAAAAAAAA
>evgR017190c0g1_i01_Hydra
AGCATACATCAGACATATTAGAAGAAATTGAAATGAGAGAAAGTTCGAAAACTTTGTAGTAGTGTTTATCATAAAATCAAAAAAAGATAATAAAAATGGCATTCAGAAATATTATTTGTTTATCTATTGTAAGTTTTTTCTTTGATTCTTTGGAAAACACAAGCCTCTACAACATCACTGAACATTTGTGCAAATCAGAAACAGCGGATTTAAGCATGATATCTAGTACTGTATCTTATTTTAACTCCCCGACACTTATTTGTAAAACAGGATTATCTTACAACTATATGACCGATACTTGCGAGCCAATCATTTGTTGCTACGGGTATACTTTAGTTGGTTCTAAATGCCGTGATTTGCAACACAAACTACCAAAAATAGTTGTCAAGGATGCAAGCTTTGAGAGTTGTTTATTTAGAGAAAACCCCGGTTTTTATATTACTTCAAATGAATATAATATAAACTGTACTAAGGAACAGTTGTTGCCATTTTTTGACACAAACACAAGTTGTTATTTTAACAAAAAAAAAAAGTTCCCTGATTCAACTTTGTTTAAATGTAACTTGTCTGCCAACGCTCTAAAAAATATTATAAGTTACCCAGTAACTTTTTCAAATTTTCCTTATGGTTCATCATTATATATTACTTCAGTTTATAATCAGGTACATACCGAATTATATGGCTTCAATATAGCAAGGTGGTTTAAAAAAAACAAACTATGCGCGCAACCTGAAGAGCTTACTTTCATGAAAAGAGTTTTACGAACTCGTGTGATTATGTCACAAAAAATAAAACCATTAAAAACGAAGATCTCATTACGTGGATTGAATTGAGTAGAGACGGAATCAAAAGAAAAATGAGCACTTGTAATCAATACCATCTCATATCTAATTGTGGATTGCAAAGACTGACGGACAACTATATTGTTGGAAAAGAAACACTAACTTACATATTTAACATGGTACTCAAAAAAAGAGTATTGGTTTTATGCCGATCAGTTTTTACCTTTAGACAAAGGTTATGGAGTTTGCGCGTCATCATTTTCAATTCTTTTAAACAATGCTGAAAATCTTTTGTCGGTTATTTTATTTTCTATAAGCATAGTTTGTTATTTTTCTATATTTACAACATACCTCTTGTTTAAAGAATTAAGAATTAAATCTAACTTTAATTTAATAACGCTCTGCGCTCTTTTAACATTTTCTGATTCTTTAATATTACTTGCTATTTTTGAAAACGAAACCCAAATTTTATGCAAAAG
>evgR017191c0g1_i01_Hydra
TTTATTTTTGAAGCCGTTGAAAAAAATAAAAAATTTTGTTTACAAAATATTGATGGTGTTACAAAGTTGTCAAGAAACCATCCATATTATTATCAAATTCAAACACAGATAAACTTATGCAAAAAAAAGTATTGTGATTTCTTTATTTATACCGAAAAAGACTATCACATAGAACGCATTTTCCCTGATATAAAGTTGTGGGAAGATATCATCAAAGAGTGTGAAATAAAGTTTAAACATTTAATACTCCCAGAACTAGCGGCTAAGTATTTCACTCGAAAGTTAAATGAAGAAAATATACATTTAAATATAAGTATACAAAATTGTTATTGCAATGGAAAAGAAGATGGCGAGCTGTTTCCATGCAATAACATTGATTGTTTGAGAAAGCTATTTCATTTAAAATGTTTAAAGTTAATCAATAAACCAAATAAGAAATGGATGTGTCCAGATTGTAAACAACATTCTAATAAAAAATAATAAAATACTTTCATTTTTGAGGACATAGAAGAGCAGCTATTGCATATCAGAAGAATTACTTTTGTAATAGTTTTTATTAGCAAAC
>evgR017193c0g1_i01_Hydra
GTGCTATTAAAATGTTGCTGACGAGGGTTATGCATCAAAGCAATCCAAATCATGGAATCAATGCAAACGACTCCTTTTACTTTAAAAGAAATGCCGATCCAAAGTTGCAAGTTGCTATGTTTGGCACAATTGGTGTTGTTTCCCTATTCAACAATCTCATTATCATCTTTGTTATTTACACACAGAGATTGGTTCGTTCAAGAAGTAATTCGCTTCTAATTAATCTTGCAGTGTCTGACATCTTAATTGTTATTGTTGGTATACCAATTCAATCAATGAACCTTATGTCCAAAAAAGGACCAATCACAGTTGGCATATTTTGTCAGCTTCACGGTGCTCTCGTTCTTTTAACATTTTTAGTTTCGAACTTTAATTTGGTTTTAATAGCTGTATACCGCTACTTACTGATTGTCAAGCCTAAATTACACAACTTTTTTTTTTCTAAAGTCAACTTAACCTTTGTTATCTTCATGACGTATTTTATAGTTGCATTAATTGTTTTACCAACGTTGCTTAATTCAGAAAGTATAAAATATAACTTGTATCGTGCACATTGCATGGTTGCTTGGGGTTACAGCATTAGTTATCTCCTGTTTCTTCAGTTTTTTGCGTTTAATATACCACTAGGAATTATTGGTTTCTGTTATTATAAAATCATAACATGCACAAGAAATCACTACAGACATTTTACTAGTATGCTTGATCGTTCAAACAAGATTCAAGAGCACCGCTTAACCTTTATGCTTTTGGTTGTTATGGGCTGCTTTTTTACATGTTTTTTGCCCTATGCCTTGTTATTATACTATGAAGGAGTTTTTAAAAAAACAGCTTCAGAAGAATTTAGTTTTTGCGCAATGTTGTTTGCATATTTTAATAGTACGTTGGATTTTTTTATTTACTCAATAATGAATGGTAAATTTAGACGGGCGTTAAAAAACCGCATGAACTCAGTCAAAC
>evgR017199c0g1_i01_Hydra
ATTACACAATATGTAAATATAATTGTAATTGATTACTTCTAAAATTTTAGTGTAATGTTTGCTTTTTTAATATCAATTTTTGTAATACGATGTGGTGCAGAATATAAAGGAAGTGGCGCCATTTACATGAAAAGAAACCAGCATGATGATAAGATTTCTCTGGGAACGTTGAATAAACTTACAAAGTTGCAGTGTGTCCACGCCTGCATTAAAAACAAAAAATGCTCTTCATTTATATTTGATAAAAAGGACATATATCTTGGAAACTGTATTTTACGAGAAAAACGGAAAAATGAAGATATCCAAACACCTATTTTTCAAGAACACAGCTTACACAACAGTGAAATATTTGAGAGTTTTTCTTTAATATATAGTAGCTGTCAAGAAGGCTATGAAAATGGAAATAGATTATCCGGTAATTACTTTTTATCCGACGAATTGAAATGGCATTATTGCAACATGGGTGAGCTTAAAAACTGTGGAATCGGAGGATGGACTTTAGCAATGGCTATTGACGGCAGAAAGCCAACATTTGAATGGGGCGCATCGTATTGGCTTGATGATGTTATCTACAATGTTGAAAAGGGATTTTTATCACCATTTTTAGAAGAAGCAAAGTACCCAGCATTTTCACACCTAGAGTTTTCTGAAATTTGTCTTGGAATGAAATCCGACACGAGGGAAATCACTTACATGAAAATCTCCCTTTCTGGTAAAAATACCTTAAAGAAGATTTTTAACGATAAACCCAGAACTATCGATAATGGACGCGAATCGTGGTTAAGATTACTTAACGGATCAAAACTTCAACCTCATTGCTCAGGAACAAACAATGAAGGATTTAACATAGAGGGAACAGCAGCTAATCTAAGGCTTGGTATAACTTGCTATGATGATAACCCTGAATGGAGTGATTCTTGGGTCGGTTTAGGCGGTCAAAATTTGTTTTGTCTTGACACAGATTCCACGCTAGCACAGTCATCTGCGGGTAATCAATGTTGCTCGGATAAATGCAATGACTTTTCTTTGAAAGCAATGGCTTATTTATTTATAAAGTAAACTTTTAAATCTAACAATTCGAACTTCAACAATTCGAACAGTTCGAACAATTCGAACATTTTTAAAGATTGTAAAAGTAAATATTTTTGTTATTCTTGTTGTTCAAAAGATGAAAATGTTATAAAAAAAAA
>evgR017208c0g1_i01_Hydra
AAATTGTTTTTCAAGCTTTAAACAAAATTTGAAAAATTGTTTGAAAAGATGACGAGTTAATTTTATTGAAGACTCCTTTAATAATGTATCACAATAAAACATCTAATACAACGACACAAGTAGCAAGTTACCCTGTCAATGTTGCAAAGTTTTTACTTCTTGCGCTATCTTCAGTAAGTTTTATAGTAAATCTTCTTCTGGCTTTGTTTATTACGTTGAAATTTAAGATCATTCTAAAAAGAAAATCAAACAAGTTTTTATTTAATCTGGTTTTGGCGGATACTTGCGTTTGTGCGGCAACCGTTTGTTTTTCTGCATCCAATATTTCAGAGAAAAGATTTGACACTGAGTTATTTTTTCATACTCAGCAGAAACATTGGCTTATTTTGATTGCAGCGTTTGCTCCAATATCAATTTTAAATTTAATGCTTCTTACAATCGATAGACTGGTAGCAGTCAAAAAACCATTTTTTTACTCAGATAAGTTTATGAACAAGCACGTGTATTACTTTATATCGATGAGCTGGTTTTTTGCTGTTATTGATATATTAACATTTGAGTTGCTCACAGCTTATGTCGAAATATCTCATGAAGTTTTTAAATTTTTTGTACAAGCTTCCCTTGCTTGTTTTATATCAGTGGGATTTTTAACAATAAGTTTATCTAATGTTTTTATATTTATTGATGCCAGACGTCAGCTTCGCTTTATAAGTGCTGTACCTTCTGATCAATTAAATCGAAAATACCTACATCATGCAAAAGAATGTCGGTTAATAAGAATGACTATTGGGTTAGCAATAACTTATTTTATTTGCTGGCTACCTTGTTTTACATTTTCTATCTATTTTCTTTGTACAGGTCATTACAAGTTAACGTATTTGATGACAGGGTGGGTTGTTGCTCATTTGAACCATTTGTTAAACCCTATTATATATATTTCTTTAAGTCATGATATTAAAGCCGAATTGAGGAGGTTTATTTTAAAAAAAAAAAAATCTTCATTTTTCTTTAATAATTTTACATCAACATCTTTGGCTTTAGAGTCTTCAAAATATCAATTAACAGAAAGCCAGCCAAACAGCTAAACGGAAAATCTGTTATAAATATAAAAAGGGCGTTAACAAAAGATAATAATCGAAACTGTAGAATGAACATATAATGCAATATTATATTGAAGATTAAAGACACAAATAAAAATAATAAACATAATGGAGGCAAAGTTATCTAGAAACATTACCTTAATTTGATAATTGTAGCCAGATTTACAGAGACATTTACATACACTTACTGCTTTTTGTATCTGTTACTGTATATGTTTTGTCAGATGTTCCGCAGTAAAATTT
>evgR017211c0g2_i02_Hydra
AGAAGCTTTAGAAATATATTATTATGTTGATAAAATACAAACTGAAGTTTTAGGTATCAACCACCCAACTACAATGGATACAAAGCATAATATTGCAAGCTGTTTGAAGGAAATGGGAAAGTATAACGAAGCTATTGAAATTTATTATTATGTCGATAAAGTACAAACAGAAATTTTAGGTATCAACCACCCGTCTACAATGATGACAAAAAATGGTATCGGAAACTCTTTGCGCAGTATGGGAAAATATAATGAAGCTTTAGAAATTTATTATTATATTGAAAAAGTACAAACTGAAATTTTA
>evgR017212c0g1_i01_Hydra
ACTATTCATTTAAAATGAAGCTTACCGAGAAAGTATTGCTACACTATTCATTTAAAACGAAGCTTACCGAGAAAGTATTGCCATACTATTCATTCAAAACGAAGCTTACCGAAAAATTATTGTCATACTATTCATTTAAAATGAAGCTTACCGAGAAAGTATTGCTACACTATTCATTTAAAACGAAGCTTACCGAGAAAGTATTGCCATACTATTCATTCAAAACGAAGCTTACCGAGAAAGTATTGCTACGCTATTCATTTAAAACGAAGTTTACCGAGAAAGTGTTGCTATACAATTCATTTAAAACGAAGCTTAACGAGAAAGTATTGCTACAATATTCATTTAAAACGAAGCTTACCGAGAAAGTATTGCCATATTATTCATTTAAAACGAAGCTTACCGAGAAAGTATTGCCACACTATTCATTTAAAATGAAGCTTACCGAGAAAGTATTGCTACACTATTCATTTAAAACGAAGCTTACCGAGAAAGTATTGCCATATTATTCATTTAAAACGAAGCTTACCGAGAAAGTATTGC
>evgR017217c0g1_i01_Hydra
GGCATAGATATCTTTGTTCATCATAATATACTGAAAAACCCTGATCTTATTTCATGCAGTATCCGTAACAACTTCTCATCAACCCAACTTAGTGTAATTGTGTATTCCCTTGTAGTAGCATGTGGGGGTGACCCTTACAAGCTATGTCTAAATGCAAAGACAGCCCACGGGTATAGAACCACCATCGTACCTGAAATAAGTTTGAATCAAAGAAAATTGGATGCCATCGCCTATAATGCTTGTGCATTGGGATGGAAAGCTTCAAGACACTCTCGATTCCATAAGGAACAGATGATCGATTGCCAACTCTTGTGTCTGGGGCTGATGGGACAAAGCTTCTTGGGATTCCGCCACTTCCAAAATTAACTAAAGCAACTGGAGTTCTACTAAACTAAAGTTCTTGGAGCTAAACTAAAGCTTAAACTAAAGATTTAGTTTAGCTC
>evgR017219c0g1_i01_Hydra
TAGATAACGGGTACCAAGCCGATTTTGTTAAAAAAAAATTTCAACAAAATCGGTTTAATAAACTTTATATTGTTTGTTCTGTTGCAGTGTTCATCTCATTTTTGCAGTATTCATCGGTATAATGTTTTACTCAATCAAAACGACAGATACCAGTTCAAGAATAAAAAATTATTATTAGGGCGTTCAACACTAAACAAATCTAAAATTTCTTTGATGCAAAAGACACAACTGGATAAATATTTGTAGAAAAACATCTGTCAAAAATAAATTGGTAATTTGATGGTATTGCTGACATTAAAAAAATATGAACAACACCGAAAAAAATTATATATTTGTGACATTATCTACAGGAATATATAGTTTCGGATTTGTTTTGGGCGTTTTCGGAAATTTGATTATTATTTTTGCAACAACCAAGTATAAAAGTCTTCAAATTCCAACAAATGCGTTTGTATGCAATTTATGTGCAATTGATGTCTTGACATTGCTAGTTGGAGTTCCTTTGTTTATGGTCAACAACGATTTTGAACTATTCGGAAAAGTTAAATGCGTCATATTTGAGCCTCTGCGTGCGATTCTTGTTATGGCGAATGTTTTCACATTAGTATCAATTGCAATCGAAAGAAATTTAGTAATAATATATCCAAAGATGATGGCTGGAACGCGAAGCTTGAAAAAATCAATTTTAACATGCACGGTGGTGGATCTTGTTGCCATCGTTTTAGGTATTACGATGTATTTTTCACAAAAGACTAATCGATTTGTTTGTATTAAGATAACATTTCACGTAGTATTTTCTTTTGTAATTCAATGCGTTGTCCCGTTATTAATTATGTCAATTCTTTACTTGCAAACTTGGTCCAGAATAAGATCGTTAAATCGGGAATCGATAAAATGCTTTAATAGAAACAGGCCTTCGATTCTTGAAAAAGCGCCTTTGTTTGAATCGTTAGATAAAAGTTTTTTATGTAACCAGAATAATCTCGAAAGTGAACGAAACCATAACGATATTGAAAATTTTAGAAACATTTCTAAAAAAAAAGCATTACTACGTAGAAGCATATCATTATCAGTTGTTCATGACTTTGAAATGTTTGAAAAAAACGAAATTCATTTTTCATTGACAGATCTCCGAGGATCTTTTGAACGCTCGAACAATAATAAAAATGAAAACTTCAATATAAACCGAACAAGAGCCAATACCAGTCATTCGTTTCAAAATTACAAAATGTCGAATAGGAGAAACGGGATTTTAATTAATCAACAAATTTGCGACAAGCAGTCGTCGCTTCAAAAATATATAAACCATAAAAAACAAAGTGACATAAACTTAAATAGCTCAAGATTCAACTTGCAAGATTACGCTATCCGAAGTTCGGTACAAAGAATCAGAAAAAATTGCAGAGCTTTACTAGCTGGTGCAGATTTACGCGAAATCACAGAGAGATTTACTATGACAAGATACAAACAAACTTTGAAGACTTTAAAGGTTTTTGGAATTCTTCTAGTATTTTTTACATTCTGTATGCTCCCTTTTCATGTGATAGAAATACTAAGTCTGTTAAAAAGCCCCGTTAACTTTCACATTTCTAGCCTCGGCCAGGGTTTAATATACTTTCACGCTTTCGCCAACCCATGGTTGTATGGTGGATTAACTGAACATTTTCAGTTTGCTTTCAAGCGTCTTTTTGAAGTAAAAAGTTAGAAAAATTCATTAACCTGTAAATATAAAAGTTTATGAATGTACCCAAAATAATTAAAAAT
>evgR017221c0g1_i01_Hydra
TTTTACACGAAAAAAGTCTACACGAGGTGTATGACGGTATGCCATCTTCAAATACCTTTCAATTTAAAAAGTTGTCTATTGAATTGTAATAATTCTTCTTCATGCAAATTAGGATGCTTATTCTACAGTCATATTACTGGATTAAGTACAGAAATGTTTTCAGTTGATAAACAGAATCTACCTCATATAAATTTTTATAACGAACTAATTGTTTACAAAAGTAGTTGTCCAATTATCAGTTTATGGTGGCCTATTGTTCTATCAAAAAGGGATGCGATACAGAGTTTATATTTAGTTACAATTAGTGCAGTTTATGTAAACAATTCATATGAAAAAAATAATTCAAAAGAAGTTGTGCTTGGTTTTGTTAATGAAAATGAATTATATGAAGAAGAAACATTTTTGTGTAATAATATTGTAGAAAAATCTGCCCATGATATTGGAGTTGAGTTTCAGTTGAATGCTTACCCTATCAACTATAATGGATACAATGGATCAATTAAACTATCTTCAAAACTTGCAAGTGTTAACAATCATCCTTTTGTTCTTCTTACAGATATCATAAACCAAAGAGAAGAATTTAGTTTACTTCAAGGTTTTTTAATTGGAATACTTAATGTTTTGACAAAAGTTTACACTATTTATTTTGATATTAAACCCACAAATTACACCCAGGGCTTAAAATGTGTTCTTTGTTTGAGTTTGAATAACAATGGTGAAGCATTTTGTGATTCAAGTGTTTATGTTCATGAGGATGGCTCCATAAAACTTTCCATTTTTTCAGAAGTTAATAATAGTTATTTTAATATAAAAACACCTCCACTTGTATTGAACCAATGGTCTTCTATTAAAATAAATCAAAGATTTGTTTACAGTAAATATTGGTTGTCTGTTGATTTTAATGGAGTTAATATTCATAACGTAGAAAATTTTGATGCAAAAGATTTTATGCAAATGAAAGTTTATGCCTCTGATCCTTGGCATGATGCACAGCATGCTTTAATTTCTAATCTTTTGATCATCAATGGAAAAGCTGAGTATTTTGTTGGTAATATCAATTCAACATCAGTGAAAAAAAAAATCATTTGTGAAATACCAATACTTGATAAAGAGTTTTTTATTTCATTTGACTTTTATCCTAGTAAACTTGATCCTGGTTCGTACAGTATTATTTGTATTGTTTTTGACCTCGATGGTTGTGGTTTTCTGATAACTTTGATAATTGAAAAATTTAATCCAATATTTAAAATTGGAGTAGCTAGAAATTCTTATTTTACTGAGTTTATAAATATTGTGAACAATAAGGTGATTGACCTCAGTTTGGGCCTTAATCTTGTTGTAGGTCAAATTTTAGAAGACAGTATTTATAAATACATAGTTAAAATAAATGGATATGAAAATTATAAAGTGGTTTGGAATCTGACTCCTAGAAGTTATAATAGCGTGAAAGTATATTTTTCAAAATATAACTTGGAAGTTTTAGATGGTTTAATAAAAAACTTTTTTATTATCAATGGTATATCAAGTAGGTTTTTGTTTTTAAATATTTTT
>evgR017223c0g1_i01_Hydra
ACTACTCTCGTACTATACTAAATGATACTAGTTCTCAATGGTTCAGTTATAATTTGTTAAATCCTTTTATGGCTTCTGAAATCCAAGTTTATGTTTGCTCAGAAAATCCTTTGATGTACTTCAAAATGGACTTCAGATTTATTGAACCTATTCAGTATTTTTTTGAAAATCCTTTAATTGGAGTGTACCAGTTAACTGCTAATGATTTCCTAAGTATCACGTTAGATAAACCTTCATTGATAACAGGAATTGGATTCGATAAAATTGATAGTTTATTAAAGTTCTGCATCTATTATAGCGCAAACAATAAATTCATTGCTTATAAAGATAACATTGCATTTGAGAATACTGTATTCACAGTATCTCAAAAGATACTTCTTCGTCCATTTCGAACATCAGAAATTAATTTAAGATGTTTAGTATGGTTTTTAAATGTGCCGCAAAACTTAAC
>evgR017225c0g1_i01_Hydra
TTTTTTTTTTTTTTTTTTCAAAGATTATTAGCTATTAGTTTAATATTAAAAAAAATTAAAATTGAGCTGGTTACAGTTTTTCTTATTCAATCTTATTCAACGCTAAAGACTTTGTAAATAAACTTCAGAATGAAAGCTAAACAACGAAAGACTGTCATCACAATATTAGGAGCTTTTCAAATCGTTTTTAGTATATGCTTGGTAACTATTGGTATATGGGGAATTATAGAATGCCCGGCTCCAGTAGGAAGAACAGGTGCAGTGCCTTTTTGGATAGCTGGAATGTGCTTGTTCAGCGGTTATGTTGCAATATTTTTTCCTAGAACTCGCACTAATATATGGATTTATCTACTAATATGTACATGTGTCATAAGTTGTATTGTATCGGCCGTGGTGACTTCTTGGATGTTTCTTATGTCAATTCAAACGACGAATGAAAGCGATTTAATATTTTTTGCTCTCGTGATGAGTTTTAACATTTTCATAATGTTAAATACAGCAATATTAGCAATCGTCGCCAGAATGGAAGCAGAGAGATCGAATTTTGTTATTAACCGTCAACCGTCGTCTATCAATCCTCCGCCATATACGAATGGCGGTTTTCAGTTTGGAAACGTTGATTTTTCGCAACCAGTACTAGCACTTAATGATCCGCCTCCATATGAACCGTAGCACGTTACCTGAATCTAATATAGCTGGAATAGTACGCCTTTTAACAGAGAAAGCATCAAACCAAGTGAAACGTTATGTTACATTTTTCGTAACAAAAATACTCTTGCTGTTGTTGAACAATATTTAAAAACATCAAGTTATTTAATAAATTATTTGTTTGATACAAAATATTTAATATTAATACGTTGGGATTAATTTTTTTGTTACATTATGGACTTTTATAAGGG
>evgR017229c0g1_i02_Hydra
TGGAATTCACATTTCAGTTGAAATTTATTGCCCTGACTATATTTTTAATGCCATGAAAAAATATGTATTATCTCAAATTATGGTGTTGCTCAAATCTAATGACAGGCACGTTTTAAACTTGGAAGAAATCGCTGATATATATTACGAAAACAATGATCAAGTTGATTCTGAAACTTGTCAAGGTGGTATGCAAAATCTAAAAATTATTTTTTCAAAAAAAACACCAGATTTACATCTAGTTTTAAAAGTATGTGCTGAAGAGTCTCCTTTATCTTCTATAAATACATCACTGCAAATATTTGACAAAGAAATTACTTTTTACAGAGATATATTGCCTTTTATATATGAGTACGAAAATAGATTACTTTCTAAAAAGTTGTTTCAAAATTTGTTTGTGAATTTTTATGGAGCTGGGACTATTAATAAAGATGTAGTTATTATCCTGAAAAGATTTTCTGAAAAATATGTTGTTTTTCCGAGGGAGAGCTTTTTACGATCACTAGATGAGATTCAGACTGTTTTAAAATCTCTTGCTGTGTTTCATGCTTCCTCATATGCAGTTAAGTGTAAAAGCCAAGTAAGTTTCAAAGACAAGTTTCCAACTTTAAAAGATTTTTTATTTGATTATAATTCAAACAATGTTTTTTTCTCCTTCACAGATCATTTGTACAGAAACAATATTAAATTATTAAAAGCCATTGTTGAAGATTCTCAAAGTAAAAATGTTTTGGCAGATCAACAATTGGTAATAAAGTTTAGTCATTCACTTATTTCAAGGCTTGAAAGTTTATCCTCAAAAATTGTAGTAATACTGCATAACATCCAACTTGGTAAAGATGAGACCAATTTATTGTGTCATGGAGATTTTCATATGTGGAATATTGCATATTCTCGAGATTTAAACAAAGTCAAGTTTTTTGATTTTCAGCTTGTTCGTTTTGCCCCTTGTATGACCGATTTTTATCAATATCTTTGTCAAGTGAGTTCTCCAAGTTTTCGACAAAGTCACTTGATCAATTTAATACATATGTATTCTGAAACCTTTCAGGATGTTTGTAGGCAATATGGTATTGGTCAAGATTGCTTTTATGATAAAACTTGGTTTCTTGAAGAGTATAAAGCTACTTCCCCGTGGGGATTTTTGTTTGGATTCCTGTTTAATTTAAAAAGATTCGTTTCTTCTGACTTTTATGCCACTCTTGACAGGCTGAATAGCAATGAAGACATTATTGAATATCTACTAGCTCAAAAATCAAAATTGTTTTGGGATATGATCAGTTTTTATTTAGACAGTCTTTGTGAATTAGAACAGCAAGATGGAATAAATATAATGGAAAGATCACAGAGAGATTGTTAAATATCTTTTCATAATATGTTTTATTTATTGATCTATTATATTATATATTGATAAATATTTTTTTTAAATCAATTTTATGATTTTTGTATTGTTGTTTCATTTTATAGTATTTGAATACAAATCACAGACTCAGAGCAAAAAAATCGAAATTGAATTTTTATAATATGTATTTTTTGATATAGTCATTGGAGTAATACTTGAATATGGTAGAATTTTTCCAAAATTCTCCTTATGATGCCACGTGCGCACATATGCAAATTGTTGCAACTACAAAAAAAGACATCTGTATGTAGAAATTTTGCAACTCTTGTGGTATATAAAAGTACTGATATATTCATGTTCTATTTTTGTGACAAATATAAAAAAGTATTGGATTTTCATAGATTTATGCAAGACATCTTTTGAAATAGTTTTTTCTTTGTTTCGCAAAAATAAAATTTAAAAAAAAATAACAGTTGCTCTGAGCCTGCGATATAGAATACAAATTTTTTGTTTGTTTATAATCTTGTTGAAAAT
>evgR017233c0g3_i01_Hydra
GGAGGTATGTCACTTCGTCAAACAATGAAGAAGTTGCAAACACCATTAAAGAAACAAATTAGTGAAGGATTATCTCTTCGACCAACAAAGAAAATGTTGCAAACACCTTTAAGGAAGCAAATCCAAGAGGGTTTGTCACTTCATCAAACAAAGAAGATGCTGCAAACACCAATAAGGAAGCAAATCCAAGGTGGTAAATCACTCCGTCAAACAAAGAAAAAGTTGCAAACACCATTAAAGAAACAAATTAGTGAAGGGTTATCTCTTCGTCAAACAAAAAAGATGTTGCAAACACCATTAAGGTTGCAAATCCAAGGAGGTGTGTCACTTCGACAAACAATTAGGAAGTTGCAAACACCATTAAAGAAACAAAT
>evgR017244c0g1_i01_Hydra
TTAGTGCAAAAAGTTTGTAGCATAAAATTTTATAACAATACAAAAGATTTTGCACGGGTAAATTTTGTACTTAAGTTCAATTGTTATGTATTCTTTCCTTTATCGTATAAATGCAGCAATGGCAAATAATGAAGTAGACAAAACTAATTGCTCAAAGTCAAACCAGAAGAATGCAGTTTTTGACTCGCATGACCGGATACAAATTAATCGTATAATTATGAACTATTTAACTATGGAAGGTTTTAAGGAAGCAGCAGAAAAGTTTCGTATTGAATCTGGAATTGAACCTGATCATTCACTAGAGCAGCTAGATTATCGTATTTACATTCGTGATGCCGTTCAAAATGGAAAAATAGATGAAGCTATTGATCTTACTAATGCTTTGAACCCAGAGATTTTAGATAATCAACCTTCTCTTTATTTTCATTTGCAACAGCAAAAGTTAATTGAGCTGATAAGAGATAAAAATCTAGAGGGAGCTTTAGTTTTTTCACAAAACGTTCTTGCTGAACTTTGTATGGAAAAAGAAGAATTTTTAGATGATTTAGAAAAAACCATGACACTTCTTGCTTATGAAGATACTAGTAAATGCCCATATGGAGAGCTTTTAAATTTTTCGCAGCGTCAAAAAGTTGCAAGTGAGCTGAATGCTGTCATTCTTGATACTAATCATCAAGAAACACTTCCAAAAATAGCTGGCATTCTTAAATTATTGTTATGGAGTCAAAATGAACTTGATAAAAAAAAAGTCAAGTATCCACAAATGACAGATATTGTTAACAGCACATTAAAAAACATAGAAGACGAGAAAAATCCTTCATCTGCACTTGGATTATAAGTTTATTTATCATCTGGAGTTTGCTATGTTTTCTTTATGTAGTTTATACTAGAACTTGTTTTTTGTATTTTATTTACATAAATTTAAAATTTTAACAAAATTGTATTTTCCATGTTTTTTTCTCTTCATTTTATCAAAGATTTT
>evgR017246c0g1_i01_Hydra
ATCAACCATCCATCAACAATGGGAGCAAAATATAATATCGCAATATGTTTGTACGAAATAGGAAAGTATAAAGAAGCTTTAAAAATGTTTTATTCTATAGATAAAATGCAAACTGAAATCTTTGGTATCAACCATCCATTAACAACGCAAACAAAGAATAATATTGCGAATTGTTTGTATAAAAGGGGAAAATATAATAAAGCTTTAAAAACGTACTATTTTGTTGATAAAATACAAGTCGATATTTTAGGTATCACTGACCCATCAACAATGATAACAAAAAATAATATTGCTAACTGTTTATACAATATGAGAAAATATAACGAAG
>evgR017247c0g1_i01_Hydra
TTAACATTTAATAGTAAGGCAACGTATCCAAATTCTTCATTAATACAATGAATGAAAAAAACGCAGTTGAAAAAACCAAACGGGATGACATTTCTTTGCCCAACACAACAACAGAAACAAATGTTGGTAATCTAGATAAGTGGTATTATATAGATAAAGATATGATTCCAGCAAAACTAACTTATCTTTTTGTTGGTTTAGAGAGGTCATCAATTTCACCCTATATTACCATGTTCTATGTTAGTATTGGTTTGAATCTAAAACAGGCTGGTTACGTGTCATCTTCAACTTATGCAGGTTTAATGATAGGTAGCATATTTTGGGGTGTTTTGGCCGACAAAACTCGAGCTCATCGAGCTCTACTATTAACATTGTACGTTTTGCAAATAACATTTGCATTATCAATTCCATTGGTTAGTTCAAAAATTGGAGAAAAAGAAAAAAACAAATGTCC
>evgR017252c0g1_i01_Hydra
ACACAACCTTCGCTGTTTCCAGACTAGGGTCATGCTCGTTTCCGTTTTTAACTATGGTCGAAGATCCGTTTTTTTTCTTATAGTTTCACTTTCTGATAACAAAATACCTACATATGTTTGGGGTTACTATCAGCAAAATGAATTTTTAATGGCACCATCAACTGAAAGCAGTTGTATACAAGCATGCTGGAGCAAAATTGAAAGTGGTACTTTATACGTAGGAGCAATCAGAAATACAAATATCAATGTTAATAATTTTAATTGTCTCTGTATTAAAAATCAAGTTGGAGATACCTTAGTGGGTCAGGGTGATAATGCATATAAAACACAAAGTTTTCTGTTATTGATTTTCCAATCAGAGTCTGCAACATGTTCTATTGGTTCAAATTTAGTTTTTCGAGTTAGAGTCAGTTTTTATCTGGAAGGTAAGGACAAAAGCACTATATATTTTATTGACAACCTTGGTGATCCACTGAATCATTCAGGAAGAATAATTATCACAAACTTTATAAATCTAAAAAGAAGTTATCAATCCATTGCAACAGAAAGACATAATGAGACCAAGCTTTATAATTTCGTACAGGAAACATATTCCGGAAATCTAAATGCAAAATTTGAGAACTATTACTCTCTTAATGCCAAGGTAGGTGATATGTCAGTTGCGAACAATCATATAGAATGGGCTGTTATAGTATATGATAACATCAATTTTTTATTGGTAGCATTGAATCCATTTGTTATTACACGGCCAGTTAAAACATGCAGTTTAATAATGGCAGCCTCAAACTTCAATATAAATCAGAGCATGTACATCAACTTTAACATTACAACTAGCGTAGATTGTGTGGTCGATATTGAAAATGTTACAATTCTCATGCAGCATCACCGCAAAGTCAATTTTAAAGGATTTATTTGGGATGATTTAAGTCTAAGTGCAACCAAGATTGTTAATAATATTAACTTCGATCTTATTTACGTAAACAAAATATACCTTGATTCGTTTCTAAAATTTACTGCAATATATGGAATAGACAAACCATCTATTGTTGGAAAAGAATTTATATCCATTGTAGCAAGTATTACATCTACAGAGTTTTCTGGTGATAAATCTAATGTAATATGCAAACAGGTGATTTTTACGGCAGTGTTAGAACAGTTACCAATAATAGCAACTACGCAAATATATTCTGCATTAACAATAACAAATTCATCGAAATTTGAAAATAAGACTCACACATGTGTTTGTATGATCATGAAAAGCAGACAACAAAGCCCATGTTTTCAACAAGATATAAATAGTGGTAAAGTAATTTTTTTACCAATTCAAGTGTTTGAAGTACTTGGTTATGATAACGTAACAAACTTAATTTACGGTAAAACGTTACG
>evgR017255c0g1_i02_Hydra
TAATTTTATTTTATTATAATGAACATAATAGCGAAATTCTTTATCAAACTAACAACTAAGCCTGTCCAATATCTTTGACGTCATTCAATAGCTATTTTGTGAAATACAAATTTAATGACAATAAACAATGCCATGGTTAACCTTAAACTGTTGTGTTTAGTTTGTCGCCATGTATTTAAAAGGTCGCTGTATGCTTTAAAATAAATTTTAACTGTCTATTATGAAAAAAAGGTACAATTAATTGTAATTATATTTTAACTCAATTGGTATCATTAACTATGGTAGTTTACAAAACAGCTTATAATGACGAGAATCCAACAAAAATTACAGGTAACACAAACTTATCATCAGCGTCAAATGGCCATATATCTTATTTACCAACACAAGAAAATAATTTGTTTGGCTACAAATATCTGCAATCTAATGAAGAGTTAAACTTAGATGTGACGCATAGTGCGTTGTTTCCATACCTTCCCTTTTATCAGCCAGTTTCACCTTTGCAAGCATCAAACAAAAACACTGAGTACAAAGTTGAAAATAAAGGTGCGGACGAATCAAACTTAATTATAAAAAGTGAGCCAGAGAACGACATACCATATTCTAAAGCCAATAGAAAGTCTCGGAACCCGCGAACAATATACAGTAGTTATCAACTACGCGAACTTAACCGAAGATTCTTTAAAACGCAATACCTTTCTCTTCCAGAAAAAGCAGAACTGGCAGAAACCCTGGGTTTGACACCAACTCAAGTGAAAATTTGGTTTCAAAACAAAAGGTCGAAATTCAAAAAAGTTGTCAAAGCTAATTCAAATTTTCTACAAACTGTTCCGTCTAATCAATTCCAAAACGAAAATTTTCAAATCGCGTGGGCCGAAAATATGAAGCAAGCATTAGCATCGCCAACATATAATTCTATACCGCTAATACAAAGTCATATACTTTCTAATCAAAATGTATGGTTTTCTTCCCATCAATTACCGCCAACCGAGTATTAATAAATCGAGAAAATAACCTTGTTGAAATAAATTCTCAAATCGCAAATTATTTCTTCATAGAACTTCTAAACCTTACTTAAGAAGTAAGAGTGATTTCAGAGATAAACAGTTTTTGGTGCATTGGTGTCATCCGAATAAAGTTTAAAAAAGATAAAATATTTTTCAGATTTCGTAAATCTTTGTATAAATACATGGATAGCTGAACCATATAATAAAAGGCAAAGAGTGCTTTTGTAGGATTCTTTGGAAAAATTTTTTTCCAGATACCATGATCCGATACCATATCTTAACTTTGAGTTAAGCATTACGTAATATAAATTATTAACCAATTAAAAAACTATCAAGTTCTAAGATCAAAATAAGATATATGTATAAGATTTGTATAAAGTATTTAAATATTTATAAAGTCTATTATAATTTCTTTATTTTATTGCTTTTTATTTTTTTTAATTGCCTTATATATAAAACTAAATC
>evgR017257c0g2_i01_Hydra
GCAATAATTAAAAATTACGACAAACTTGAGTTGATAATGCCTGTTGAACTGTATCCTCTCTTGGAAACAATGAGAAAGTTTAATAAGGTTGTCCATGCTGTCTTTGGCTATAAGCTTGATCCAAACTGTTCAAATTTAATTGATTTACACAAATTTATAGAGATCTGCAAGATTGTTGCAGTGTTATTGCAGTGCTATTGCAGCGAGGTACTGTGTTCTTTAACCTGGAAAGTACATGCTGTAACATTTGTTGATTTGAGTAGCCATTCCCTTGGGAAGTTTAATAAACAGACATCTGAATCTTGCCGTTCTTCGTCTAAAAGATGTATGAAAAAATAATTTTAATATGTAATCGTATTAAAAACTTAATTTAGATGCCTCATTTTTATCTCAAATGCCTC
>evgR017261c0g1_i01_Hydra
TTTATTTTCTAAATAAGGAGTTAACAAAAAAGTATTTCCATAACGAGTTGTTTTATAATGTGGTTTGATACAAGCTCTAAAGCCTAATATTGAAATGTATGGAACAAGCATGTATGGAACAAGCCAACAACCACTTTATTCTCCAAGTTTAAAAAGCAAAACAATGTACATGAGCCAACAAGATGTTCAAAAAACTTCACACAATCCTCAACAACGATTGCAGCAACGAAGTTATTCAACAGGTGATCTAAAAACATCCATTCGTTCTGGTGAGCAACAGGCAAAGGTTGTAACAGAAAATATTGAAAAAATCGAGAACCATTTCCCTCAGTTAGTACATGATATTAATTTATATACAGTGAGATCGGCCAAACTAAGGGATGCTGGTGATAGTATGGCAAAGTCTATACGTCTTTATGCTAACAGTGAAACCTCTAGTATAAAACAAGGTCTGGGAGCGTTTGCTGAGTGTTTTGCATCTGTGCAGGATCAGCGTAATGCTCTCGTAACTCGACTTGAAAATAAAGTTTTTCAAACGTTCGCCGTTTATGACACACGATGTAAGCAGGCGAAGGTAGACGTAAAGAAACATTCTATAGCTCATTCAAAAGAGTTAATAGAACATAAATCTTACGAACGCGTGAAACAAAGGTCTGTGGAACATTTCCAACTTGCTAAAGCCGAAACCAAATTTAAAAAAGCATCCGAAGAGGCCCATAGATCAGCACAAATATTAGAAGAACAGATGGACGACTTTGAAAGAAAAAAAATACGCGATTTAAAACAAGTATTTGGTGATTTTATGTTAAGTGAAATGCTTTTTTACGCCAAAGCACTGGAAATTTACACGCTTGGTTATCAAGAATTAATGGGAGTTGACGAAGATAAGGGCGTCGAACAACTTCACGAAAGTATGCATTGGACACCTCATCAGTTTGTCCAATCTTCTTTAGGCGCTTATGGAGGCTCGGCACCAACTATAATGCAGCAACATCCTTAGTGTAATGTACCATATGGTACCATCTTGTAAATAAAATCAGTTTATCCTGTAAATTTACCTATATACTATGCAAGTAACTTTTAATATATTTAATATTTTAACAGTATATTATAAGTAAATAAATAATATATTAAATTTTATTTTATTTTTATAAAAGTTGAAGTAAATAGCGTAAAATAAAAAAAAAAA
>evgR017262c0g1_i01_Hydra
CAAGTTTGGTACTGCTTTCTGGGTTTGGGTTTTCATGGTCAGTAAATGGTGGGAATTTGTTGATACCTTCCTCTTGTGTATGAAGGATAAGCCACTCCAATTCCTCCACGTTTGGCATCATGCTACTGTCCCACTCCACATGTACCTCATGTTGATTTCTCAATGGGATCCAGCTTTATTTGGTTTCTGTTTTAATGCTTTTGTTCACGTTATCATGTACTATTATTATTTGCAACAATATAGAGGTGTTGATTGTTGGTGGAAGAAGCTTGTCACAATTGTACAAATTGTTCAATTCGTTGTCTGCTTTGTAATTTTGGGCATTTCCTTGAATAGTCCACACTGTGATTTGGAATCATCTCAACCACGTATTGCTTTGGCTTCAACATTTACATTGTACCTTTCTTATTTGGTCTTATTCATCAAATTCTATGCTTCTACTTATTCCAGAAAGCCAAGAGGTGAAAAGAAGCCAATCGATGAAAAGAAG
>evgR017264c0g1_i02_Hydra
TTAACCTATTTTAGTGCAAAAAGAAACTATTTACAACTTAATAAAGATCAGATAATGGCATACAATTTTACTGGAAAAAAAGTTCTTGTTACTGGTGCTGGAAAAGGTATTGGTCGAGGCATTTGCATAGCTATGTTGAAAGCTGGAGCACAAGTGTTTGCATTATCAAGAACCCAGTCTGATCTTGACAGCCTTCACCAAGAGTACTCTGAAGTTATAACTATATGTGTTGACTTGGATGATCTAGGAAAAGTAAAAGAAAAACTTAATATTATCCCTGATGACATCACTTTACTTGTCAATAATGCCGGTTGTGCAAGACTTCAACATTTTTTAGATATAACAGAGGAAGCTTATGATAGTATTATGAACACCAATTTGAAGTCTATGGTGTTTATTTCACAGTTTATAGCAAAGAAAATGATTAGCAACGGTCAAGGTGGAAAAATTGTAAATGTTTCAAGTCAAAGCTCCCTTACAGCTCTTCCTCTACATACCGTTTACTGCATTTCAAAAGCTGGTGTTGATCATCTGACGAGGTGTATGGCTCTTGAACTAGGTCCTCATCATATTAATGTTAACGCTATCAATCCAACAGTTGTCATGACAGATATGGGAAAAAAAGCCTGGAGTGAACCTTCTGTTGGAGACCCAATGAAAAATCAAATACCTCTTGGAAAATTTGCAGAAGTGGAGGATGTCGTCAACGCTGTTCTTTTCTTACTAAGTGATGAAAGTAGCATGATAAGCGGTGTCGTCCTCCCTATCGATGGTGGCTTTCTTTCATCAAGAAAACTGAAGTAGCGTGCTTTAATTAACATGCATAAAAAATTGCTTGAGACAATAAGTAATATATACTTTTACCTGTTGTGCGTTGTAATAAGTTGGATATATTTATTTGAACTTTGGCCGCACTCTGCTGGCTCACGGAAATGTAAAAATACTTAACAAATAAATTAAACGTCGTTTTGTTAATTAAAAAAAAAAAAA
>evgR017278c1g1_i01_Hydra
CTTAAAATAACAAACGGAAATGCAGTTCTATGGGGAAAGTGGAACGATTGGTCTATTTGCAATGTTTCATATGGAAATGGTTTTATGAATAGAAGAAGAACATGTCAACAATCAAATAAAGTATGTTGTGACGATGACTACGAACTGAAGGAATGCTTTACATCATTCTATTTGGAAAATGAAACTGAGTTAAGAAAAGAAAAGATTGTTACAATAATAAGCAGACTTTTTAAAGAATACACAGTGTCTTTTGAAGTAAAACTAACATTTGTTCACAATAAAACTAGCAGTCCACTTAATGTCTTTCATTTGACTACTAATAGTGATTGTGGTACATATGGATACCGAACTCCAGCAATGTGGATGCAATATGGTA
>evgR017280c0g1_i01_Hydra
TTATTCGAAAATAGATTCAAAGTTTCGATCTATTCACTACGCATTTACTAAACGTTTAACTCAATGTTTTCTATAAAACTGATTCAAACTTTCGATATCTTCACTGTGTATTTAAACAAGACATTTTGTATGATTAGCTAGCAACACAAGAATTTTAGTAATCAAAATAAAGTATTTTCTTTAAAGCATTAAATAAACATGGTGTATAAAAAAGACAGATATCTATTTTCCACTGTATTTGTCGCTTACGCCTCCATTATTTTGGTTTGCGTAAACCTAATTTTTTTAATAATAGTAATCTGCTTGAACACATGGTACAAGCGTGTAAACGATGTAAACATTAAATTTTGGTATGGGGGACTATGGAGTGAATGCATGGGAGAAACAGAACGAACATTTTGCAGCCCACTTACAAATATACCAAAATGGCTTTTGGTTTCACAAGTATTAAGTTTGATATCCTTGGTCAGCCTATTCACAAACATTATTTACATAATGATAGCTATGGCACGAAAAAAGTTAAAAATCACTATTTTGATGACGTTTACACACATAACAACCATTACGCTGCTGATAGCTACACTTGTGTACGGCATTGAAGGATTTTCATCAAATACCAAACCATATTCGACATACTTTGTGGCCTGGGTAACAATATTATTGCAGTTAATTGCTTCAGTAATGTGCACGTACTCTTTGATGCGTGTATCAAAAGAAGATAAAAAGCATGCAATAATTACTGGTATTACCTGCTAGTCTACCATGTAGTTGAGCCATCTAAGATCAGCTACTATCTTCTAAGGTTAAACACTATCTTCTTAAATCTTAACATAACTAACGTAATTAAATGAAATACTTCAAACAGCTTCACATAATCTTTGTTTCAAATATATGAGTTATGACCATATAAGTATGTTGCTATGACCATATAAGTATATTGTTATGACCATATAAGTTAATAAATTTTGTAAAACATAATAATTTAATAATTTACTTAAATTTGTAGAAATATATTTATTTAGTAAATG
>evgR017286c0g1_i01_Hydra
AGGAAATTACCTTTCATCAGGTAATGGATATTATGCCGTTTTGCAAGGAGATGGAAACTTTGTTCTCTATCTTACGCAACATTGGGTAGCAAAAAATGCCATTTGGTCAAGTCGTACATACAACAAAGGAGTTACTCCAAGAAGAGTTATTATGCAAAACGATGGAAATTTGGTCATTTACGATGTTTATAACAGAGTAACTTGGGCAAGCGGTACTAATGCAAAAGGGACCAAGCCTCACCGTCTGTCGATGCAAGCAGATGGAAACTTAGTGATATATGATGGAAATCAGAAATCTACGTGGTCAACGCGAACAAAACGTGGCTAATATGAAATATTTAAAATAAACATTCTTTAAAGATTTTGTAGTCCAAATAAGTATATATATACATAAAATACAAAAAATATATGTATATATATACAATATATATATATATATAAATTCG
>evgR017287c0g1_i01_Hydra
AGATATTATAATGAAACTTCCCCCACCATGGTTCGTCTAGTCGTCGTTATCTTATGTTTGATGTCGATCTGACCTCTTATAAGCTACTTTTCTATCAACGCTACTGCTCCTGAACGTGATCTATTACGAATTTCTATTTTCGAGGAAGCTGTTGCATTACCTAGTTCTAATATTATTTTATCTACTCCAACTCGAACAGCTTTTACAGATGAAATTGTTTACATCAAAGTTAAGGCTTCATTGTGGATAGACATGAATGGAGAAAGAGTATTAATGGCAGTGACCGGAACCGACATGAATTCAATGTTGTTAAAAGAAATAATGTTCAATGTAACTGACTCTCTTGGGCTTCAGGCAAAATGTCTGAATAACAATACGCATACATGTGCTCTTGTTGATCAAAATGGCTACATTGTTGCGAGTAATCATGGAAACAAGGCAGTAGGCCAATTTTTTGGTGTATATCACGCAAGACTTATGGAGTACTTTGCAAGGCCTGATGTTGGAATTTTCCGACAAATAACTATTGAAGACGTCCAGGACTTTTGTCAAGAATCAAAAACTTATAACTCTGATTCTAATTATTTGCTATCTCCTGCAAACACTTTATTACAAATATTTGTTTGGTTCCTTAGTTTAATATGGAACCTTTTAACTGAAACTATTTTGCGTGCAGTTTTTTTGCTGAACAAGAGCGCAGTTATTTCGCAATATGCTGACGTCAATAACAAGATAAACACCAGTTGCACAAACAGTATTTCTTTTTTTCTATATCAGAACGATAAACGAGAAGAGCTTTATAAAAAAGAAGAGGAAAGCAAAAGAATAAATAAAAGAAACCCGACCATTTTACGCATTGAATGTGCTACTGGG
>evgR017291c0g1_i01_Hydra
TCCAAAAGATCAACCCCTCTATTCTATCGAAAAGCAAAAATATCCTCATCATTTTCAAAACGAATTTCCTAATGAAAAATTGGAAACAGTTATTTCTTTGGGTCCTAAATGCTATGAGGTAATTAAAGAAAATAAAAATAAAATGAGAAATAGAGGATTTCCACAACAATTCTTAGAAGCAAGTCAATATTCAAAAGGTGTTGATTTATTCAAATTGTCTCATTATTTTAATAATTGTGCTGTTAGTTTGCATAGACAAGCCACGGAGTTTACAAGTAATCGACTAACATTTAAAAGCGATTCAATCAATACTGTGGAACGTAAAATGATTTTTATGAATAATATAAATTCAAAATATTATCTTTTGGACAATGGTACAGCTATGAAATGAGACGACAATAGACTTGCAAAATTGCATCAATTAAATATGACAGCTTTAAACGAAAAAGGGTTGACTTATTTTTGCTCAGATTTACATTTAAAGCAATGTTTAGAGTTGGAGCAAAAAATGACGG
>evgR017295c0g1_i01_Hydra
TTTAGTGCAAAAAGATCTTTTAAGTTGTGTCATTTATAAAAAGCTAATAGCTCAGGACATGTGAATCAAGAATTAAACATGGAGTGTTTGCAAAGATTAGGTTGCAAAAATCAGATTCGTAAGTTTTCAAAATCATTTGTTTTAATGCGTCATAAACCTAAGCCAGTCAGAGATCCAAGTAAGCGTCAGTTTACAGGCGAAAAATTAATTGAAAGATTAAAGAAAAAAGAAGCTGATATACAAGCTCAACAGCCATATAAAGATGCTGTTGCAGCAGTTGTAGCACGTTTTCAGAGGGAACTAGCAGAAGAAGAGCGAATTAAAAATGATATTCTTGGAGAATTTGCAAAGAGAGAGATAGAGGAAAGGGAAAAAGTTCTAAGTAACATGGCGATTGTTGGCAAGAGTAACGAAGAGCTTAGAATAAAAAGAGAAGCAGATAAAAAGATAAAAGAAGAGGAGAAAAATTTGATTGAAGAACAGAATAGACTTAAAGAAACTTTAAAACAGGAGGAATTATATCACAAGAATCGTGAAATTGTCTTGAGATTAATTGAGGAGACAAAAAGCTTTGTGAATTTGGAAAACTTAGATGATAAAATTACATTCGCTCTTGAAAACGAGACTTATTATAATTATGCAATTACTCCGTTTGGAGAAAAAATAATTTCACAGAAACCACCAGGAAATTTGGGCGTATGGAAAGGCCCGAGTCCTACAGCATACATCGTAGGAGGTATTAAACCAGGTAGCGCGGAATGGAAGTCGGCGTTTAAAGATACTGAGAGAACACCATAGTCAAAATTAATATTTTATTTTCTAGGAAGATCTTTTATAAAAATTACGAAATGAACTACTCACGATGAAACACTCAATAAACCTTTGAGCAATAAGGTGGTCTATTTTTAGTATAATGTCAACAATGTTTTTTCATAGCCATTTTTTCTTGAAACATATTTCTTTATATAATTAATATGTGGAATTCATGTGGAAATTTTTATATAGTTATAGTTCATTGAGTTTATACGGGAATGAAGTAAATGAGTCTCAAAATCGCTCAACGAAATGAGTCTCGAAATCGCTCAACGATAAAAAAAACTATAATTATTTTTTTATCAATGAATTTTAAATAACAAAGTTTTTATTTATAATTAATGAAAAAAGTATTAGAAAAAAAAAGTAGTAGAAAAAAATATTTTGCATATAATTAATCTATTTATTAATAAACCATTT
>evgR017312c0g1_i01_Hydra
CTATTTTAGTGCAAAAAGTTGAAATAAAAAATGCGATTTCGTTTTTGTGGGGATTTGGATTGTCCTGATTGGGTTTTAGCAGAAATTAGTATACTTTCACGAATCAGCTCAGTAAAATTAAAGTTATTTTGCCAGCAAGTAATGCTGAACTTGTTAGGAGGATCTATCGACTATAATAAAATAAATAAGTTTACTGCTGATGCAAAGTTGGAAATTAATGATATTAAAGCAGCTATTGCAGCTGTTGATTTTATATTTACAAATTCAGCAAAATACCAAGTTAATGGTGAAATTCTTTCAAATGAATTACAGCAACTAGGCTTACCAAAAGAGCTGACTATATCAATGTGTAAAGTTTTAGATGAAGAAGCAGAGAAACTGTACAGTTATTTAAGACAGAAGAGTATGCGCTTGTCAAGAATTGAAAATGTTGATTGGCGTGTTGATTATGTTTTAAGCTCAAGTCTCAAGGATGAAATAAATGAAGCAGAAATGCAAGTTCTAATAAAAAAACAATCAAGTGATTCTTTTTCAGAAACTATTACATTTTCTTGCACTAATGAAAAGCTGCGATGCTTATTAGCAGATTTAAAATATGCTTACAAGTTAATTGATGATCTTTCTTTATAAATAAAATAATTCCCAGTTTTTTTTATGATCCTTATATTGAAAATTATTGAAACTTAATAACTAATAATTAAAATGACTTTTTATATATTTATGTGCAGTTAAAGATAATTTAAGATTTTTTAAATGACATTGCTTGTTTTTGTTTATATG
>evgR017328c1g2_i01_Hydra
TGCAGAAAACATGATTTCGAGTGACGATACAGTGACGAAACATCTAAATCATATAGTGACATAGATGAGTTAGACAAGAAACCTGGTAACTGTGAAAATAATTTATTACATATTTATAAAAGTGCTGTAGTGTTTACATGATGAAATCCGAAATTGTGAATTATTTGAATATGAGTGGCCACCAACAGCAAATGACTTTAATTTACAAAATTGCAAGAAAATGGTTCCGATTAACCTATTTAATCATTTATGTTGGATGTTAGGATTTTCTAATACAGTTCCTGATCATAACACTTTTATTTTGTTATCAAAGGAGAAGGAGCAATGTGTTTTAAATATTTCACAGGATATAATCTATGCTGCTTTAAGTGGTCAAAAACAGACAGTAAAATCTTTAGCTCTTGGAATGGCAACTCAAATCACAGGTTCTAAAGAAATGATTGAAATTTTAAATGGGTTTGGACATTCTGCATCCTATGATACAATAGTCAGACATGACACTGCATTAGCAACTTTAAATTCGATTACACATCGCCTGGTCCCTGCTGAATTCCAAATTCCAAATTCCTGCTGAGTTGAACCGATTCCTTGTGTTCCTACAACATTAGTTTATAATAATCAAGATTATAAAAGAAAGAACAGAATTACTTTTCAACTTTCATAAGGCAATTTTAGTTTAGCAAAGCTACGTGTTTTTATGATACAAG
>evgR017330c0g1_i02_Hydra
CCCTGTTCAAAAGAGGAGGAAAGCGTTTTATTTCATGAAAGTCTCAAGATGTTAAAGACAATCGTTTTATTACTTTGTGTTGGAATATCGTTTGCTCAGCATAAAACTGCAACTAAATGCGGTTTTAAAGTTGGAAGACCAGAGTTTTTACGCTACTGCATATCACCTGGCAGCAGAATGTACGTTGGAGATTTCAATGGAGACACACGCGATGATTTAATGTGTCTGGATTTTGGAACTGGTGATACAAGCATCTTATTTTCTACAGTTTCAACTTTAAAAACAAAAGTTTCATTTAAGTTAGATATTTGTAAAGGTGCCAAACATGTTCTTCTCGGTGATTTTAACGGAGATCATCGCACCGACATCATCTGTCAGCTTAAAACTGGCGAAAAATATATTTACCTTGCAGCAATGAATGGAGTTTTCTCAACCTACACAATGCTAAATGCTGTCACGTATAGTACGCCAAATGTAAAAAAATTCTGTACACAAAGCGGGTATAGACCTATAATAGGTGATTTTGATGGGGATAAGTTTGACGACTATATGTGTCTTACTACTACTGGCCAGATGTCAATCATTTATGGTCAAGAAAATGTTGCAGATGCATTTAGAGAAGAATCTATGCTGAAAGATACTTTTATGTGCAGAGGAGATATTTTGACTGGTTTATTTAATGGGGATTCATTGAGTGACGTTATGTGCTACAACAAAACACATGGAACTATTCTAATTGCATCTATTAACGTCGATCAAGTAAACGTTATGTTCAACTCAACATGGAAATGTATCGATCAAAACTCTGTTGTCATGTTTTCTGACATTGACGGTGACGGCTATGATGATTTGTTATGTAAACGCACAGGAAAAATTTTACAGATTTTAAGAAATACTCGAAATGAAAGGTTTTATGAACCAATAGAAGCTGCATTTTTTCCTGAATCTAACCCTAAAAAGATGTATCGTAAATTTGAGACCGGAGATTTCAACGGCGACGGAAAACACGATTTGATAGGTCAGGGATTTGACGGATCACTTAAAACCGCTGAAAGCACTTGTTTAAAAATGTAAAATTGTAGAGTTCTAAATAGAAATAATATAAATAATATTATAAAAAAAAAA
>evgR017332c0g1_i01_Hydra
ATTAACCTATTTTAGTGCAAAAAGCGATTTAAATAAATTTCAATATGAGTGATCCCGTAGCTGAGCCAGACCTGGTCAATGGAGAAAACGAGCCCTCCGAGGACGAAGAGCAAAGCGAAAAGAAAAAAAAGCACGAAAAGGCAGCATGTGCAGATCTTGAAAAAATAACAGATTACGAAGAAGACAAAGAAATTGACGAACAAAACATTAGCAATGCTATAACATTATTAGAAGGGCGGCGATCTGAAGAGAATACAAAGAAAGCCGCGAGAGAAAAAGAACTTGCTCAAGTTAAAGTTCGTAAAGAAGATATTGAACTAATTACAAACGAAATGGAAGTTTCCCAGAGCATCGCTGAACAAAAGTTAAGGGAAACTGGTGGCGATGTTGTGCAAGCACTTGTACTGTTGACAAACTGAAGGTTATGAAATATATAATTGCTTTTGTTTTTAATAAAACGACGTCAAAATTTTCTTTGGTATACATTAAACTATATTTTTTAGATAAATTAACTAAAAACTAAAT
>evgR017344c0g1_i01_Hydra
TAAAGAAGAAAAAGACTATCAATTAATGAAAGCTGTTCCAGTAGCGCAATTTCACCCTTTAAAAGGAGTATGGAAATACTGTGATTCGATTAATGCAAAGTGTCTGTTTTTAGCTCCAAAAGGCAGTGCTAAACTTCTTATCGGTTCGTACACAAAGAATTTTATATTTCGGGCTTTTTATCCTCACATGACCAAAATAGTAAAAAATCCTAAAGAAGGTTATCCCGCTGAACAAACGGCTCATTTTGAATGTTATGGACTTATGCAATCGGATATAAACATTGATATATTTAAAAATGAAAACTTTTTAAATACTCTGAAACAATTTCAGACATCTGCGAATGCAAAACATGCAGTTGCAAATCGTTGCAAAGAAGTGAGAAAACTTCCGACTCATTTTTGCAAAAATGTGACTGTAATTAGAGTGGAACCTAAAGATTTTTGTTTATCTATTAACGAGCACATGCTTGAAAATGCGGAAGAAAATGCGGAAGAAAATGCATTATGTGATTGTGATAGTGATAGTTTTTTTCAACCAACTGGACATATGAATAAAGCAACCGGAATTCCTAATTTAATTAAAAACGACAAATTGAACATATATATTGCTAATGGAAATATATGTATTCATGTAATAATTAAAGACAATTGAATTAAAAAATGTAAATAAATTTATTAATAAATTTTACAATTTTTTTTTAAAAAAAAAA
>evgR017344c1g1_i01_Hydra
GCCGGGACAAAATCTGGAACAGTCGGGATGAAATTACTAATACAAAAGGTGAAACTTTAAAACAATTAATTGAAAATAAAAAACAAAAAGAATCGATGAGAAACAATATAGATAGTTATCTTGATTTTATTAACGAAGACGAAAAAAAAATTCGTGAAACTCGTATAACCATCCAAGCTATAAAAAATGAACTATTTGGTTTGGAGCAACGTTTAGAAACAAAATCGCGAGATGATGTAGAGACGAACATAAAACCTGTACAGGTACAAATTAATTCGATTAAAGAATGTTTACAAACGTTTATGAAAAAAACAGAGCAAGATCAAATTAATAATCAGGCGAAATTACATTTTTTTCAAGAAGAAATACGCGAATTAGTAAAAGAATACACTGTTACTACGTTAAATATGGAGATGACGCATGAATATATAACAACTTATGACGGAGAATTTTATTGGGTGTTTAATTATAAGAAATATGAAAATTATGATAACGCATTGCACTCACCTATCCATTATTATGGAAAGCCCAGTTATAAATATTGTTTACGAGTATACACACGCGGTGATGGTAAATTTAAAAACGAATATTTTTCAATATTTTTTGTGTTAATGAAAGGTAATTTTGACAATTTTTTACCTTGGCCATTTGATCGGAAAAAAATAACTTTTGAACTAATACCCGCAAATTCAAACTATCCTATACACAGTCAGACTGTAACATGCGATTGTAATAGTGATAGTTTTTTTCAGCCAACTGGACATATGAATAAAGCAACCGGAA
>evgR017346c0g2_i01_Hydra
GCAATAGGAGTTGTTTTACTAGTTAAAGTAAAAAACAAGTTTAACAATATCCAATATTTAATTCTTCTCAATATATGTTTGGCAGAAATGCAACTCGGTTTCTTCATGACTTCAAGAATTTTGTTTCGAAAACTAAAGGTTTTCAAAGTAGACATATTTTTTTCAATATACAATAACACTAGCGCATCGTTTGAATACTACTTTAGCATCTTTATTTTAACAATCAATCGCTTCTCAGAACTTTACCTCAATCTTCGATATAATTATATTTGGAAATATCATCGGATAAAAGCAATAATGATAGGAACATTCATAGTATCAGTAATCATATCAGTAGGTCTTTTATCTTTTGAAATATTCCGATCACCAGATAAGAAAGTTGAAGATTTTGCATCTTTATATCTTTTCCCTGCCTTTGATGGATTTTTAATGATTGTTATATTTACCACTTATTCTTATATTTATTATAAACATCGGAAGTTTAATTCCGTAAGATTATCGTTTGTTGACAAAGTAGTAGCTACACAGAGACGGAAACGGGTTTACCGGGGTACTTTATTTTTAAGTTTATTTATAGCAACTTTTCTGTTATTTATAGTGGTGCCAGATATAGTTAACTGCTATTTTTACAACACGCATCGACCAATGGAATTACTTTCACGACTATTAATTGTTTCTTACTACTTTGGACTTCTTGCAGATGTAGTAATATATACATTTCTTTCGCGACCAATACGCCATGAGTTTAAAAGAATGTCTGCCTACATAAAAAAATTAAAAAGGGAAGAAACATTGCCGCACATTGTTAAATATGCATAATAACAAATAAAAATATGAAATATTATGAAAGAGAAAAAAGCGCTAAGGCATTGGGCTTTATGAATAAGCAATTAGGCCTATATGTGTATACTGATTTACACTGCTTAAAGTAAAAATATTAACTTTTACAAAATACTGTTTTTATAATCAGCAAACATCAATTTCAAAATGCAAAATTGTAATTTTAATCCTTTTATATTTTTAAGATTCAATTTGTAGTATTAGTTGTAATTTTTACGGGTTTG
>evgR017349c0g1_i01_Hydra
GTTTGCAAATACTAATATATGACATTCTGGTAGAAATAAGTGGTTTTCATTCAAGAATTTATTTTATATCTTTTCAATGTTATTTGCAAGTGTTTATTTCTTTACTACGGCATTTAAAATATTTATTTCATAACTGGTACTATGAACTACTATGAACGTCAGTATAACATCCAAAAAATTAGCCTTAAATAAGAGTAATTTTTGTCGTCTATCTTTTGAAGAGGTTGCAGCGGGGTGGACTGGTGAATTGCTGAATTGGTTAAACGCAACTAATGGTGTGATTATCTTAATCGCAAATGCTATGTTAGCTTATGGTGTTTACAAATCAAAAGGAAGAAATAGATATTCGCGAAACGAAAAGTTAGTTTTATTATTATCATTTGTAGACTTTATGGTTGCCATTATAAATACACCTCTAAATATAAGTTTGTTTGTTTTCCTAAACCACTTAAGTTGCACAAGTATTTCAGTTATCGCGTTTTGGATTGTATTTCCGCTAATATTCTCCGGCTCAATTGTTGTGTTGATTTCAATTGAACGTTTTGTTGTTTTATTTTATGATAAAAAATGTTGCGGTTTGCGTTACAAAGGAATTTACTTGGTTCCAATTATTTTGTTTTACTTAATTGTATCTACTGGAGGTGGTATTTGGTATGCAATGATTCATTATTCCTTACCTGATTCCTTATCAAATTTTCCTATTTTTTTCTACACCAGTGCAACTTATATATTTTTAAATTTGATCATTGGATTAGTTATAAATACTTTGCTGTTAAAAAAAAATTTGAAAAGGCTAAAAAATGGCAGCATTAAAATTGAAAGGAATATTCCTATTGAAAAACGGCTAACTCGAACAATGATGTTTATTTCCTTTTCTTTAATGCTTTCCTACCTGCCATCCGTTGCAACACACTACTATCTTGCAGCAATATTTACTTTGCAGCTGCCATTTGGTTTCGCAATAAAATTAGTACTTTTGACTTTGGTACTATGCAAACTTAATTCAGCATTAAATCCAATAATATATATAACAAGATGTAAGGGAGTGAATCGAATATATATCTCATGGTTTCGCATTTTTCTTAATAAGATTCGACAACCGCTAAAAATATATAGCAAATAAACTGT
>evgR017355c0g1_i01_Hydra
ATTATTGATAGCAGACATTTTTTTAATCCATATATTAAAGCTTTTTTAATCAAATTTTTTTAATCCATATATTAAAGCTTTAGAAAACGTTAAGTCTTCTAGCGCACCAACCTCAACATCTTGAGCAAACGTTGTTAGTGGCAGTGCACCATTAAAAAAGTCAGTTGAACAATTGCATATTATGAATGTTATTGCTGCCGAAACAAAAGAAAGGGAGAAAAGGGAAAACAACGTATTAATCTTTGGCATTACTGCATCAAAAGTTTTGGATACTACAAGTGCTAAAGAGAATGATAAAAAATTAATTCTAGATATGTTAAATTCAATCAACTCAAATATTGAACCTATTGAACCTAAAAAAATTATAAAGCTTAAATCTAAATCTGATAAAGAACCACTGTTTGTTGTCGTCCTTAATAATAAAAAAGAACGAAATTCTATTTTAAAAGCTGCGAAAATCTTAAAAAGTTCAAGCAGGTTTAGCAATGTATTTATCAACCCTGACTTAACTGAAGCCGAAAGGTATAATGCCAAATTATTAAGAGATGATTGCAAAAAACGAAATTTAGAAAATGCACAACCTTTGGTTTATTATTATGGAATACGTAATGACAAAGTTGTTAAAATCGTAAAGTAGCAACTAATTTCAAATAACAGGAAATCAAATCTGAATTGTTACGAATTATTAACTTCTGTTGGTTTTAACTATATGTCAATTAATAATAAGTTTTTG
>evgR017358c0g2_i01_Hydra
AAAAAGTTATTTTTGCCTGGAACTAAATGTATTGGACTCTATGACTCTGAAAGTGAAAAGGCTCTATCATTTAAGAAAGGAGACATCATTGAAATTCTTCATATAACACAAGATCCTAATTGGTATAATGCAAAAAATATTTCAGATGGTAGCATTGGTATGATTCCTGTAAATTATATCCAACAAAGAAAGAATCTACATTTCCACGAAATGCCTTGGTTCTTCGGAAATATTACAAGAGAAAAAGCTAATGAATTACTTGCCCTACAAGAAGATGGATTGTTTCTTGTTCGAGAAAGTATTACCTATCCGGGTGATTATGTACTCTGTTATATAAGTTCAAAAGATAAAATGGTATGGCATATTCGCATTATCTCAACAATAGACAATGGATTTACAGCAGACAGAAAAACTTATTTTCCCACTCTTATTGATTTGATAAAGCATTATGAAAACGGTGCAATGCTAAATGGTACTATTCTTAAGAAACCGCCTAAAAAGAAAACTGAAATGGTTTCACAACTGAGAATTT
>evgR017362c0g1_i01_Hydra
GCGGTAGTAAAAGTTATTGTAACAGTTATTACAGTTCTATCCACAATTTTTAATACTGGTGCTTTGTTTAATCTTTTTGAAAAACGTCCTAAAAGTAATTACATTTTTCTTTGCATAAATCTTACTTCGGCTGGTTTGATACAAACGCTATTTGGGTATCTGCCGTCTCTCTTTTTGGAGGAAAGTATAAAATATGGAAGTAGACTGTGTCAAATCAGCGCGTTTTTTGTTGCATTTCCGGCTTATACATCAATCGCCTTGCTTACCACAATTAATGTTGTAAAAGCTCTGTCATTGAGAGACCAATTTTTGATCAGTGACGTAGTTTATAGAAAATGGGTTAAACTTTCAATTTTATGTTGTTGGATATACGGATTGTTTTGGTCTAGCTTACCACTAATTGGGGTCTCCACTTACACGCTGGAATCTACTCAATCACGTTGCTCAATTAAGTGGCACACAGAAACACCCACTGAAAAGGTTTATTTGGTATTAGTTCTTGTATTTTGTTACGTTGTTCCAATATTTATTATTTTATTCTCAGCGATAGCTTCAGCTCGATTTGTTCACGCAAACTTTAAACTTAGCTCTTGTGCTTTGGGCTTGAAGGAGTCGTCAATTAAAATTTTACAATTAAGAGAATCCAAAGTAAAATTGTTATTATCTCTTAGGTGTTGGTCATTTATAATCTTTTGGACCCCATACGCAGTTGTAGGTCTTTTATCAGGACTAGCAGGTATAATTCATGCTAATTGGTTGCTTGAAACAGCAGCTATGTTTGCAAAGTTATCGGCTATGGTGAATCCACTATTATATTGGCAAAAGGAAGATTTGAAAAAAA
>evgR017365c0g3_i01_Hydra
TGATTGATATTTACGAACTAAATATTTTCAAAATTATACCATTTATGTTCCAACATCAATCAAATAAAACGCCGAAGGTTTTCGAAAATATTTTCCCTACAACATCAAACCGATACAATACTCGATCTTCTGGAAGATTTAAAAAACCTTTCCGTACATCAAAACTTACGAGCTTTGCTATTTCATATCGTGACCCAATGGTCTGGAACCATTTTACAAGCCTCTGTTGTAAGCTTAAAGAAGCCTATACCTCTCCAATTGTAAATGGTTGCTTGACAAAACCAAAAAATAACAATAACAATAATAATAATTTGAAAACAT
>evgR017370c0g1_i01_Hydra
CCCGCCACAATTCTAAAATACAATGGCTACTACTGGATTTGATCAATCTGAAGTATTTTTTTCAGATCCCTTTGGATCTGAAGATAATGCAAATACTAATTTTAGTTCTAATGTAAATGATACCAGGAAAAAATTTAAAGACTTTATTAAAGAGTTTCACGAAGTAAATTTTGTCTTTCCGTACAGGGACCAGCTAAAGAGAAATTACAGTTTAGGACAATATTATTTGGATGTAAGCTTAGACGATTTACGTAGTTATGACGAGCATCTGGCTGATAAAGTTGAAAAATCACCAACTGAGTGTATTCCTATATTTGAAGAGGCAGTTCGTGAAATATTAGAAGAAATCACAAAACCAAGAACTGCAGATCAAGAAGAAATTCAAGATGTTCAGATAACTCTTCGTTCAGAATCTCATCCTATGAGTATAAGGAATATGACTTCAGAGCATGTTTCAAAACTTGTTAAAGTTCCTGGAATAATAATAAGTGCATCGGCTATGAGGGTTAAAGCTACACGGATTACAATTCAATGTCGCAGCTGTCGTAATACTTTACCAAACATCGGTCTTAAGCCTGGTTTGGAAGGGTATGCTATGCCACGAAATTGTCCCTCTGATCAGACATCAAAGCCTGCTCCTTGTCCAATGGATCCTTTTTTCATTCTTCCAGATAAATGCAAGTGTGTAGATTTTCAGGTTTTAAAGTTGCAAGAATCTCCCGATGCTGTTCCAAATGGTGAGATGCCGCGTCATCTACAGCTTTATGCAGATAGATACTTGACGGAAAAAGTTGTACCTGGAAATCGAGTTACTGTTATTGGAATTTTCTCAATTAAAAAAAGTGTTGCAAGTAAAAAAAGAAATGATGATGGTGGTGTTGGAATACGTAAACCATATATGAGAGTAATTGGAATACAGATTGAAACTTCGGGAGCTGGTCGCAGTTCAGGAAGTATTCATTTTTCACCAAGCGATGAGGAAGAATTTGCTAAATTAGCTGCTCGTCCTGATGTTTATGATTTAGTTTCAAGAAGCATAGCTCCATCTATTTACGGTAGCGAGGATGTAAAAAAAGCTGTTGCCTGTCTTCTATTTGGTGGATCCCGCAAACGATTACCAGATGGCCTTACTCGTAGAGGAGATATAAACATTCTATTGCTTGGAGATCCTGGTACAGCAAAAAGTCAGTTATTAAAGTTTGTTGAGAAAGTTTCTCCAATTGGTATTTATACATCAGGAAAAGGAAGCAGTGCTGCAGGATTGACAGCTTCAGTTATAAGGGATCCTGTTTCTCGCAATTTTATTATGGAGGGAGGAGCTATGGTTCTTGCTGATGGAGGTGTTGTTTGCATTGATGAGTTTGATAAGATGCGTCAAGATGATCGTGTGGCTATTCACGAAGCAATGGAACAACAAACTATTTCAATTGCTAAGGCTGGTATTACAACAACTTTGAATTCCAGATGTTCTGTTTTTGCTGCTGCTAACTCAGTGTTTGGTCGATGGGATGATACTAAAGGAGAACAGAATATTGATTTCATGCCAACCATTCTATCTCGTTTTGACATGATTTTTATTATAAAAGACGAGCATGACCAAAATAAAGATGTGCATCTTGCAAAGCATGTTATGCAGGTTCATCTTAATGCTGGACAGTCCAATCAAGGAATGGAGGGAGACCTTAGTTTGGCTTTCTTTAAAAAGTTTATTGATTATTGTAGAAGAAAGTGTGGCCCGCGTTTGTCAGAAAATGCAGCAGAAAAGCTTAAAAACCAGTATATTCTTATGCGCAGTGGTGCTAGAAATATGGAAATGGAATCAAACAGGCGAAATTCTATACCTATTACTGTGCGTCAATTGGAGGCTATTGTACGCATAGCTGAATCTCTTGCCAAAATGAAACTGCAACCATTTGCAACAGAAAGTGACATTGATGAAGCTTTGCGATTATTTCATGTGTCAACATTAGATGCAGCAACATCTGGAAATTTATCGGGGGCAGAGGGTTTTACTCCTGAAGAGGATATTGAAGAAGTAAGAAGAATAGAGCAAGCACTTAAAAAAAGATTTGCAATTGGATCTCAGGTCTCTGAGCAAAGAATAGTTGCTGACTTTGTAAAAAATAAATATAACGAGAAAGCAGTTTCAACTGTCATTCATATTATGCTGAGACGTGGAGAGTTGGAACATCGGTTTCAAAGAAAAGTACTATATCGTGTTCGATAGAGACTTCATTGTCATGAACATATGAACATAACAAGATGAACTATAATAGCGATTTGAGCGTGGAAGTTTTTATCAAACGAAGTAATTTTTAAAGCAATCAATTAGCATTTTTTTAACGAAATCATAGCGATCAATTGGCATCTTTTTTGATAAGATCATAGCATATTTAAATTGTTAACATTTTTTTAAATTGAAATTAAAGCGATCAAGATGTTTTTATATTTTCCAATAAAGTCAATGCTTGTTGATGTTGAAGATGTTGAAATCTATTTACATAAATTCAATGCTGTTATTGAAACCTATTTACATAAGGTCAGTGGAATAAAGTTGATATATATTTTCTACGATAAAGTCGATTTAAATATGCAAAGTGAACCAGTTTACCAACATTAAGAAAAAAAATTGTATATGTTTTTTATTATTCTGTATCGGAATTGTTATTTCCTATACACAAGAAAATTAATAACTATAAAAAAAAAA
>evgR017373c0g2_i01_Hydra
ATGTACATAATAGCATTACTGAAAATGATATTGCTGACAACAATAATAGTTCTATTCATTATACCCCTGATGCAGCATCTACAAAACTTTCTGATCAAATTAGTCGTAAAAGGGTTTTTCCAGTTTCTGAAGTTCGTGAAATGCTATCAACAATTAAAACGTTGACGTATGATTTTGATGAAGCGTCTGAAGAAATTTCTGATTTACATACAAAACTATCAGACGTTCTTAAAACACTTTATCATGCAAGGAAAAAAGAAAAAGGTCTACCTGTTAGAACTGAAAAAGATAATGAGTACATCCCCAATAAGCTAACAAAAGTAGTTGAATTACCTAGTCTTAAAAAAAATAAAAAAATTCAGCGATATGGAAAAAAAAATGATTCATTGCTCTCATCGTGTACCATCAAAATTGATTGTGTTGCTCATGAAGCACCAATACAAGAATTTGTTATTACTGATGATATAATAACAGGGCAATCAACTTTTTCAACTAATTTAATGGATTCAACCATTAATGAGAATTTTTTAGCTGATGATGATAACCAGAATGTAACATCAAATACTTTAGTTAAAACTGACGATTT
>evgR017379c0g1_i01_Hydra
ATGAAATGAAAGCTGATAAAAATTTACAAATCAAAGCGGATTATTGGTTGCGAGAACTAAACGCTATTTCCACAAGCAAAAAAGGAACACTGATAACATTTTGGGCAAAAAAAACATTATCTAATCATAAACACTCAACCATAAATATGCATAACAAAATTGTTCCGAATTCAAATTTTCAGTTTGGTAGAGAAGGGGCCGATATTCTTACTGTTGAATTAGATTCAGGAACTAGTACTGACCGCTCAGAAACATCAAGAATTGCCCCAGCCCAGATGCAATTGCAGACACAGATTGAAATTAAAAATTCTGATTTGGTCGGTTTGTACGAAGGACAAAAAAC
>evgR017384c0g1_i01_Hydra
GTTATTTTATATTCTCCCCGTACCTATCCTTGTTTAAGTCTGTTAATGCTTTAGCAGTGCACATAGAGGGCGTATGTACGCACACAGGACAAAGGTACGCGTACGTATGTAGAGTCTGAGCAGACTGTATTTGCAGTACGCAAAATGTATGTCTGTTTGTATCTATATACGTATGTACAAACCACAGATTTTACATGCATCAATTGTACTCATCGATTGTACACAAATGCACTTACATTTAATTAGTGTGCACTCGATCTACGTAAATCAATTTAGCAGCATTCTTCTTATGTTCACTCCGCCGCACCTATTGCGTAGCAGAGTCCGTATACATGTTTTGGTATCAATACGGTTTGTTCAACCGGAGTTGCATAAATACTTGCATACATCGATTTATCTAAATATGAAAACGCTCAGTACTCTTTACATACCCATGAGCAGTCGTGGCGAAGTGGTAAGAGCATTAGAATTGTAGTCCAACAGCGTTGGGTTCAATCTCCGCTCAAAGCATATAAACATTATCGGTGAAGATGAAAACGGAGAGCCAAATTAATAAATAATATTCTCCATGTGGCGGTTATTTGTGACAAGACCATTGACGTTTCTTAGAGCAC
>evgR017396c0g1_i01_Hydra
TTTGAAGCGTTTCTCTCGTATCGCCTCAAAGGTTGGAAAAGCCACCGAAAAGAAGGCTCTTGGCATGGAAAACTCCTTGGCCACATCTATCAATTCTTTTGCTGGAGATAAATATGTTGGTGAATTAAATAATCTTGAATCTTTTGATGAAATTACTAAAGTTCAAAGAGAAGCCAGAGAAAGATACTTCAAGATCACTGCTGCTGCTTTCCCACACAAAGTTGTTCAATTCAAGAAGGAATTTTCTTATTTCCAACAACTCCTCACCAAGCCTGGTGAAATCACCGTTGGTGAATTGGCTGCTTTTGCTACCGCTCTTGTTGCCGGTTATGTTTTGTATCGTGGAGGTGAAATCCTTGGTCGTGCTGATATCTTCGGATACAAGTATGAACATCTTGAACAAAAGTAAATTAATAATCTATTAGCAATCGGGC
>evgR017397c0g2_i01_Hydra
CTTTAATTAACCTGTTTTAGTGCAAAAAGACTAAGAAAAATAATAATCATGAAATTTATAAGATGAGGATGGTGGAAAATAAAACTGACAGAAAGGATAATTTGCTTGCAAATTGGAATTTTGATCTTCCATTCTGTGAAAAGTCTGCTATAGGTTTTCAATCAAATATTATATATCGTGCTGATGGTGGACTTGGAGAAAGCACTGAGTTTGAAGATGCTGCATTTAATTACAAAGGTCTTGATTGTCTTTTGTATGGAATATTTGATGGACATAATGGTGCAAGAGTTGCACATTTTGCACAGCACAAAGTTGCAACAGATTTATGTTTCGGACAATTAAGTGATATATATACTGACGAAGAGGTATATAAGAAACTAAATGAAATATTTGAGAATGTTTCGAAAGAATATATGGATTTCATAATTCAGCCATTGATGGGAAAGTTGACTTTAGAAGATGCAGTGGAGGGTTTAGAACCATTCATGGTTGAACAAAAATATCCTCAGCATTGTAAGGATCTCAATGAATTTAAAGCTGAGTTATCAGGCGGGTGTTCTGCGGCAATAGCACTAATCCATAATGCACGACTATTTGTTGCAAACGTTGGGGACACTCGTGTTGTGTTGTGCAGAGCAGCTGGCAATAACGGTGAACTTGGTGTTGTTCAGCTTAGTACTGATCATACGGTTTCAACAATATCTGAATTACAAAGACTTGGAAATCTGGGTGTCGACATAGAGCAGCTTCAGACTCTCTCCATTCTTCGAGTGACTCGCAGCTTGGGGGATTGGAACCTAAAAGTGGGTTACAAAAGCAATTATTTTCTGAGTGCTGCTTCTGAAGAGCCTATTCTTTCAATCCCGTCTATTCATGGTGGAGAAGAAATAGATAACTCAATGAAATTCTTGATATTAATGTCACATGGTGTGTATCATGCCTACATGCATGCGACAGACACATTACCAGAAGAGGTCAATTCAGGAATTGCTTCGATGGTCGAAGAAGAAATTGAGAGGAAATCCACCATAAAAGATGTGGCTCAGGAAGTTGTAAATAGAATATGTCAAATGCACATGAATAAATTTGTTAAGTCACAAAACACACACTGCCTTAAACGAGATGATATGACATTAATAGTCAAACTTTTTAATATGGAGCTAGGTTGTGCACAGAACAGATCTTCAACTGGATGCTTCGGTAAGCGAGTTGCAAGCAATAATCTCCCACCTTTAAAGATACCTTCTTCTACGCTAGGGAAGAACGCAAATACCCCCAGTTCTACAGGAAGTAATACGCCTTCTACAGCTACACCTGGTACTCCGAAAAGTAATAATATTGGTCTTTTCTCGATGCACGGTTCTAATATTGATCAATCCAGTAGCTCTCCGTCTACTAGCGCTGTGGGTTTTTGCAGCACTGAGATGGTGTCCACTAACCAAACTATGAAAACGTATGACAAACCTCCTGACATGACAATGTTCATATCTGACAACTTTGAACCGTTTGAAAAAAGTCCCTTTTTCCAGGAAGATGCTTCACTTTTAATAGTTGGAAATACTGACTCATATTTAAACGAAAACGAGATATATTCTGATCTATCTTATTCGGATAGTTCGATTAAAACTTACTCTAGTTCTTTGTCTTGCAAAGACACTTCTGATAAGATTTATAATCGCGTAGAGTCTATAGAATTGGACGAAAATGGACGTTGTCAATCTTATGTTGATTTTTCTGAATTTATCGAAAATGTTGAACGAGACGGGGGAGAAAGAGTTATTTTCCCTGAACTTTTCCCTTTTTCCTGAAATATTCTTAAATTCTTTTGTGAACTCTTTCCTTAAATTGTTTTATAATATTGTTTCTAATCAAATATTTGTGTCGTGTTAAAAGTTAAAAATTGATAAAAGCGTGTGTTAATGCTGTGTCAACTCAGACACAGTGCCGGATCAGGGGGGGGGTATAATATCTATAAATTCATCCATTCCCAAGGGCGTATTATATTCAGTCAATGGGGGTGATATCAGATATTTTAGATTACTTTTTTGTAAATCTATATATTTTTCACAGCAAGTTTTTTGAAAATGGAAATAAATTAGGAAGGCTCTCTAAATAAAATAACATTTTTAATGAAATTGTGTGGAGCGAATATTAAATTGTGTGTTTG
>evgR017398c0g2_i01_Hydra
ATGATTTTATTGAAATATTTAATATTAATCTGCAATATGGATTTACTCACTACATTATAATGAAGTTTTGTGCTTCTGTTTACCTATATGTAAATAATAGTACACCAGCAACACCTTTGAATAATAGGTTACAAATCTGCAAACAAATTGCAAATAAACTAGATCATGGTGAAGAGAAAATTATTTTGAGTTTGATATGCAACATAAGAAAACATGCGTATCAAGAAAAAAGTTCGCTTAAAAAAACATTTGTTATGGATCAAAAATTAAAATCTCTTTTAACCAACAATTTATCCTCTGAATGGAAAAAGATTGCATATGACACTGGAATGAGTTCGATTATAATAGAATATATTGATATGAATGAAGAAAGTAATCAAGATGAAA
>evgR017402c0g2_i01_Hydra
GTTTCTTTTTAGCCTAATAGTTTTACAACAAATCGTCAGCCGAGATTCGAAGAAGCCGTATGTCCACATTTTTGTGTTTTTTAGGTTATTATTAGTTCATAGTTAACGTTCAACAGTTAGTTAGAATGAAACTAAAATTTAAAATTTGAAACTTGATTTTCTTGAAAGCACAAAGAATGGACATCTCATTTGGTTCTAAGCTGACGATATATTAAAATATATTTTGGTGATAAGGAATATGCGCCTGTGTTAAATGAATTTTTTCAGATAAAGAATGAACAAAAACTACGGTTTCCGACATACATATCGCCAACACGACGATTGGCGCTGTGAAAAAATAAATTGGTTTTTTAAAGATAAAAAAAGGTCGATACAAAAATTCCAAAGAAATCATTTAACATTGAACTCGAATGGCTATGTGAATGGCTGTGTAGAAAACCTAGTACCAGTCGGATCATATAACTGGGCTAAACAGAGTACACCAAACAAACCAGTAATAATAGTTCCTGGCATTCCCCATTATAATATAGAAAATCAAATTTGTCAAAAGCTTGAAAAGAGCAAGACTCAACGTGTTTGCAACGAAAGTAAATACTATATGAGCAAGCATCCGATGGAACCAATGTTTCAAGCTGTTTTGTTATGTACACCTGAATACGATTTTAATTCTGTCGATTTGATAACTGATTGTCTGAACTTGCGTAAATTGTTTGACTTTATTGAGGGAAACGCCATGGAAACGTTTCGAATCGATATTCAAATGAATGAAAACGGTACGCTACTATTTATTTGCAACGATGAAAACGTCATTCTGCCATGCACTACCTATAATAATAATGACTATAGTGCTGACTTTAAAAATAAATTCACAGAAAATCGGTCAAATGAAGCTGGATCGTATTGGAAAATTGTTACTTATATGTTAGGTAGTATACGTGTAATGTGTCAGGTTCAAGTAGATTGCGTTGAAAATAATGGTTGCTTAGTCCAAGCAGATGTAATGCCGACAAAAAATGAGCCAATTGCATTTAATGGAGGTTCTGGTTTATTTTTTATCAAAGGTGGAGATTTTAGCTATCAACAAGACGAAAAGTTTGTTTTTCTAACAACAAAACGCACTTTTAATGGGAGCTACAATTTTCCTCTAATTAAATGGAGTCATTTATTGTTTTTCAACATTGACATAATGGTTCTAGGTTGGCATGATGCAGGAGTTTTAAATAAAGTAGAAAAGATGTCTTTTGAAGAAGTATCCGAAAGGTGTGGTCGTCAAAAAGAACAATACCAAAAATCTTTAGGGAAATTATTTTCTTTAATTATGTTGATAAAATAAAAAATAAAAAACTGTTCAGAACACGAAGGTGGTTTTGCAGTTGTTTTCGACTATAGTAACGATAAAAATTCACTTGAATTAGTCACGGTATACAACAGTTCTCAAGTTTTGCCACACGAATTAAAGATGAGAGTTTTCAAATAATTTTCTTGTTTTATAATAAATTATTTTGTTTGATTTAAATATATATTTAAAATCTTTAAAAAAAAAA
>evgR017403c0g1_i01_Hydra
ATACACCTCACCCTAATGTTGATACTTCTTTAAATAATCTAGGGTCAGCTTATCAAGCTAAAGGACAATATGATCAGGCAATTAATTACTTTGAGGAGAGCCTAAAGATGAAAAAGCTAATCTATCAAGATGAACCTCACCCTTGTGTTGCCGATTGTTTAAATAATCTAGGGTTAGCTTATCAAGCTGAAGGACAACATGATCAGGCAATTAATTACTTTGAGGAGAGCATAAAAATGAAAAAACTCATATATCAAGATGAACATCACCCTAGTATTGCTGGTTCTTTAAATAATCTAGGGTTAGCTTATCAAGCTAAAGAAAGATATGATCAGGCGATTAATAACAATAAAAAATAATTTCAACTAAAAATAATTCAGCGACATAATTTAAAAAATAAAATAAGACAAAAGAGTAAAAATTAGTTGAAGAAAGTTAAAGAAATATAATGATTATATACAATTTACAAATTAAAAAAAAAGATGTTTAAAATTTTTATATCATTTTATAAGACCATATTAGAACATGTTAAATATAAGCATCTGCGTCTCAAAACAGTACAAAACGAAACGATAAAAATTTTTGTTTTAATAAAACCGGC
>evgR017404c0g2_i01_Hydra
TGATCCCTAAGTGAATTGAATGTTGTATTGAATTTCATACTGAATGTCACGTTTTATAACTTACTATTGCTGTTTTCAACTGATTCCAATTGCGTTTCTCCATCCTATTCAAAATTGTGGAAACTCAAATGTCGTGTTGCGTCATTTAAAAAATACAAACTATTTTTTTGGCGGTATCTTCCCAGTTCATCTCCCAGATATTCTTAGTGAAGCTCCTATAATGGTTGTTTCATCAATGATGTTAGCAATTGAAGAAATTAACGATAGCAACGACATACTTCCTGGCATTAAAATAGGATATGAAATTCGAGATTCACAAAGTAGTATTCTTTGCGCATGTAACCACGCTTTAAATTTTTTGTTGGATGATGAACGAAAATCTGTACTGATTGGAGTCGTAGGGCCAGCACCTAGTTCTTTAAGTGCTGCTGTGAGTACAATGTTACTATCAGACTATGTGCCACAAGTTTCTTATTCATCAACAAGTATGTCACTCTCACAACGTTCCGTATATCGTAACTTTTTTCGCACTGCCCCTGCTGATTCAAACCAAGCAAAAGCCCTAGTTGGGTTAATTGAGTTTTTCCAGTGGACTTACATTTCTTTAGTCACCTCGAAAGATGATTACGGCCGTTTTGGTAGAGATGAAATTCTCAAAGCAGCTACTGAGAAAAATATTTGCTTTTCTATTGATATTATCTTCGATGCAAACTTTTTGAATAACGAGATAGATACTATTTTAACAGGTATCGAGAAAAATTCTCATGTTGTACTATTGTGGTGCGACGGTACATATGCAACATCAATTATTTCGAAAAGTTTGACACGTGGTTTTAGCAATATCACCTGGATAGGAACAGAATCCTGGGGAAACGATTTTCTGAAATATTTTTCAACCAACCACAACATAATATCATTGCGATTGAAAGAATATCCTATAGATTTTGTGAGTAAAAAAATACTTAGCCCTAACGTGACAACCAACTTAAATTGTTATAACCCATGGTATAGAGAATTTTTAAACAAGTATTTCGTGAATGCCCTAACGTGTGTTCCTAATTTGACGAGTACATTACCAAAACGAAAAATAAATCAAGTCGTTGGTGCTGTCTACGCCCTCGCGTACGGGCTACATAGTTATTTAAACTGCTCTCTTGAAAAATGTTTGACGGTATCTGACTCTATTAATTATGAACTTCTTTACCATCACGTGATTAATACAAGCTTTAATGTACCCACAAGTGATTATCATGTTAAATTTGACAACAATGGCGACTTTTTATATCCTGCATATGAGTATGTTTTTGTCGATAAAAAAGGTTTTACAAAGTTTGGAGTCTGGGAATACACTTCTAATAAATCAACAATTTTAGATATAAATCAAAATGTTGTTGTTTGGAAAAATAACCTTAAACCAAAATCCGTTTGCGGTATGAATTGCGAGCCTGGTACATACAGAGTTAATTCTACTATATCTAAATGTTGCTGGATTTGTGTACCATGTCCATATGGTTCAGTTTCAAGCTCTGTCAACCAATACAAGTGTGAAAAGTGTTCAATTGTATCCCTAGAAAACTCTAATCAAACAAAATGTATAGATTTGACTGAAATAAGCTTAAATATTTTTAGCACCGCAGGAATAATTATTATTATCGGATCAATATCAGGTGTATTAGTTGGAACTTTCATATTGATTTTATTTTTCATCTATTGGAGCAATCCAGTTATTAAAAGTTCCAATAGAGAGATGTCTTGTATCCAGATATTTTCTATTAAGCTCTTGTTCTGCATTCCTTTTTTTTATTTTACTAAACCCACTCCAAAGTTATGCATGTTAAGAACTTCTTTGTTTGGCTTCTTTCTAACAACTGTACTTGCATTTGTCTTAGCAAAAATGTACCGATTGCTTCGCGTTTTTAATGGCAGATTTACAAAAGTTTCAAGGTTTTTAGAAAATAAATTTCAAATCACGTTCACCTTTTCACTAGTTTTGATACACACATTAGCAGTTATAATTTGGCATGTTTATTATCCAACTCGTGTAAAGATTATTCCAAACCCTATTAGTAATGAATACTACCTTATTTGCGATAATGAAAAACCAATTTTTTTTATTTCGTTAATCTACTTTTTTGTTTTGGCAATTGTAAGTGGGTGCATGGCGTTTCGAGCGCGCAAACTGCCTGAGAATTTTAACGAAACCAACAGCATTGGATACGCTATGTTTATAATGTGCATCATTTGGTTTTCATATCTTCCTGTTTATTTCAGTGTTGATCCACACGAATCTATTGTGGCGTTCTTAAGTGCTAATATACTTTCATGCTATAGTATGCTTTTAACTTTGCACGCTAAAAAGATAGCCTATATTTTGTTTTATCGGAGTTTAAATAATGTCGAATTTTTTCATACCTGTTCTTCTGAAAACGTTTTGCATAATTTTGTTGAAAAAGTAGTCCTAGAAGACGGTTCTGTTCTTGAGGTGGTACGCAACGTTATTATTTATAACAAAAATGCAAACTCCGAATTGGGATGTTAGCATTAAATACTAAAAAAAAAAAATGATTTAACATGAC
>evgR017412c0g1_i01_Hydra
TGCTTCTCTAGTGTTGAAATATTTTTAAATTTTTGAATTTTTTTATTTTTGTTATATTATCAGAGGATTTACAATGGGTCAAGTTGAGTCATCGCTGGCATCGTTATTTAAAACCGAATCTAACATTGAAAGTCTGAGTGAGGAAACCAAGTTTACTAGAGAAGAAATACGCGCAATGAACGAAAAGTTCAAAAATGAGTATCCTAAAGGATATCTGACAGAAGAAGAATTTATAAAAGTATATTGTGAACAAGCCACCGTAAGTTCGAATCGGACTGCCGTTAGTGCAAAGCTTGTTGCTAAGCAAATGTTTCAATCTCTTGATCGCAGTAAAGATGGAAAAGTTGATTTTAAAGAACTTGTTTGCCTTCTATCTATTTCGACTCATGGGACTATAGAAGAAAAACTGCAATGGGTCTTTACAGTCTATGATATAAATAATGATGGCAACCTTTCGCTTGATGAAGTAGGGAGTATCGTTCGAAGCATGCAATCTTTAACTTCAAACGAAAGAAAAATGAGCGACAAACAAATTAGAGATATGTTTTCTCGTTGTGATAAAAACAACGATGGGAGAATCACAGTTGATGAATTTGTTCGCGCATCAAAATCAACACCAGCATTCTTAAAGCTGATTACAGACAACATTAATGCATCAACTAAATAAAAGACTTGCGACTCAATACTTGCCTTTGCTAAAGAAAGACTTTGAGCTTTGCTTTTGCTATGTTCTAAAAACGTATCTCGCATACTATCGTACCATAAGTAGTTACTTATTAAGCAGAAAACGATTCAGAATTTCTAGTATAACGCGGGAGGGGGG
>evgR017413c0g1_i02_Hydra
TCACCATTTGTTGTCATATTTTCACCATTTACTGCTGTATTTTCACCATTTGTTGTCATATTTTCACCATTTACTGCTGTATTTTCACCATTTGTTGTCATATTTTCACCATTTGTTGTTGTACTTTCAGTAATAGTTGTTGTACTTTCACCATTTGTTTTTGTACTTTCACTATGTGTTGCCGTTCTTTCACCATTTGTTGTTGTACTTTCACCATTTGTTGTTGTACTTTCACCATGTGTTGCCATTCTTTCACCATTTGTTTTTGTACTTTCACCATTTGTTGTTGTACTTTCACCATTTGTTGTTGTTTTTTCACCACGTGTTGTTGTACTTTCACCATTTGTTGTTGTACTTTCACCATTTGTTGTTGTACTTTCACCATTTGTTGTTGTACTTTCACCAT
>evgR017414c0g1_i01_Hydra
TTTAAAAAAATGCATTTTGTAATAATTTTTGTTGTTTTCTTTATTGTATATAGTGAATCTAGTTGGGTAAAAGAAATGGAGAATAGCAAGTTGTTTCAAGGAGATATTCAATTAGACCTAGAAGATAAACAATTAAATGGGTATGCAAGCATTGTAGGCGGACGTTGGCCAAACAATACCGTTTCTTATTACGTAAGCAATATGAGTTTAGATGCTCATGAAAGAATTCAAAATGCAATAGCTGATTATCATAAATTCACATGTTTAAAGTTTGTGAAAAGAACCGTCGAACCTGCTTACCTCAACTTTTTGGTTGGAGAAGGTTGTTCTTCTCCGGTTGGTTACAAAGATGGGAGAGTTAACAACATTTCATTAGCTAACGGTTGCTTAGGACATGGAACCATATTACATGAGATCGGACACAGCATTGGATTGTTTCATGAACAAAGTAGACCTGATCGTGATAATTTCATAACAATATACTGGGGTTATATTCTTCCAAAAATGCGGTTCAACTTTTTAAAGCAAACGCAAAGATCTGTGGATAGCTTGGGAACACCTTATGACTATGAATCTATTATGCACTATGGTCAAGCTGCATTCGGTAATGGTAAAATCACCATAACAACTAATGACCCTTCTAAACAAAATGTTATTGGAAATAGAAATGGATTTAGCAAGATTGATATTCAACAAATTAATGCTATGTATAACTGCAAATTATAAATAGCTGCAATTTAGTTGGATAATCAAATCTGTTTTTTCGAATTGTTACAAAAATGGAATCTTGCGATGGACGGATGTACTGTGAACCATGTTCAAACAAACGAAATTTGCGTGTGCAACCAAGTTTTTGTAAAAGTGAATTTTTAAAAACTGTATACATTTACATATTTGTTAAAGATAAATTGTTTTTAACAAATTCTCAAAACCAAAAAAAAAA
>evgR017415c0g1_i01_Hydra
TAGACAAAATAAAAGTGATCTTAAAAAACTTATGATGACTTCTGAAAGTGAAGTTATGATTTCAACTAGCAGTGATTGCTCATCCTCCTCTAGATCTAGTCCAAGCAACGATATAGCAAACCAACCGAGCTGTTCTTTTGTAGATAAATGTAACAAATGGAACAATACTGATCAAGGTTATGCAGAAAATACTATTTCTGATAGTGAGTTTTCTTTTTCCGATGATAGTGAGTTTTCTTTTTCCGGTGATAGTGAGTTTTCTTTTTCCGGTGATAGTATAGGTAATGCTGATGAGCAGGTTAATAAAGTTACGTCTGAAAATCTTGCAAATTGGGCTGTAAAGCATAAGTGTACCCAACATGCAATTAACGACTTATTAAATATACTTCGGGATTGTGGACATACATTGCCTAAAGATTCAAGAACTTTGTTAGGGACCCCAGCTAATGTTAAATATATGTATAAATGCGGCGGAACTTACATATATTTTGGTATTAAATGTGGTATATTACAGAGATTAGCAAATTTTAAGTGTCAAAAGCTTGAAAAAATTAAACTAAAAGTTAATGTTGATGGTGTTCAGCTATTCAAATCTTCAGCTACTCAATTGTGGCCAATTCTTGGTATGTTTGAAAATCAACAGCCCTTTATTATAG
>evgR017440c0g1_i01_Hydra
AAAACATTTCCGCCACTAAAAGATCTAGCTATACACGTAAGAATAATTAGCACCATATTATCATTAAAACTTTTCACTAATAAATATTTAGAATTATTCATCATTACCATCATTTTAGCGTCCTTGTTTTTGTGCTTTCATGGGTTGGACGTTTCTCTAAATGATAATCTCTATCAGGTACAATCACTATGCCTAATCTTTCTAATAAATCATCACTGCTTTTCTTGTATTTTAAAGTCACACCGCACATCTAATTGTCTTTCTTTCTGTCCCTTCCAAGCGCTGAACATCATTGACCTTTCATTGCCCATGTTTCACTCCCATACATCGTAA
>evgR017451c0g1_i01_Hydra
AGAACAGCATTCGTGATAAGACAGTTGATATATTGACAGACTCCGCGATATGACCGTGAGGACTTCTTCGAAATTTGCACGAACCAAAAATTTTGGTTCGTCTCTTAAGGACTCTAAATCTCGGATAGCGGTTTTCACCAAAATTCCAGCGACAGGCTTCGGCTTAATGTCGTATGAAAAAAAAGTCGTTTAGTCGAATGAATTACTCAACTTGCTTTTTAACCAAATAAAAATATGTTACATAATTTGTGGTAAAGTTACGTTACGTAATTGTACATTAAATTATTTTCAAGTGTTTATTATTTTCGTAAATCATTTATTTAAAATGCTGGTAGTTTTAATATGTGTTGTCAGTGTTGCAAGTGTTGCAGCAACAAATGATAAAAATGTGTTTGGTAATTCTCTAGAAATTTGCTGCACTGATCCAATAACTGGATTTTATCGAAATGGGTACTGTTCAACTGGTCCTACAGACCATGGAAAACATGTAGTTTGTGCAACGGTAACTGATCGTTTTTTGAAGTTTTCAAAATCTGTTGGCAATGATTTGTCAACACCCAAGCCAGAGTATAACTTCCCAGGATTAATTGCTGGTAATTGTTGGTGCTTGTGTGCGTTAAGGTGGAAAGAAGCATTAGAAAATGGATGTGCACCACCACTAAACCTTGCAGCAACTCATCAAAGAATGCTTGATTTTGTTCCTTTAGACAGATTGTTGCAATATAATAATGTAAGTCATACGCAAGCCGCCCAATACAAAAATGGTTATGGCAGTGGTAGTGGGAATGATCAGACTTTGAATGAGTAGAGTAGAACTAATAAATAACTATAAAGTTTTGAAATATTGAAATGAACAATAATGC
>evgR017454c0g1_i01_Hydra
CTTATAAATTCAAGAATAAGTGTTAATAAACAAACAGTTGATAATAAATTCATTTTCCTAAAAATATTATTGAGTTTCTCCAACAGAAGAAACGGGTTACCTCGAGAAATAAATTTAAAGTAAAAATGAAGTTATGGATGTTTGTCTCGATGTTACTTGTCGGCGTGTACGGTAATAAAAATAACTGGAAACAAGCGGTTAAAAGAACATTCGGAAATCAAGAATCGAATTCAGAATCTGAATCATTGAGTTTTACGTTTCTGAAAAGCGGAGCTTGTTTCCAAGCAAAGGGTGATTTACCAGCACATGTTCCCATTGCTAGGACTGGTGTTTTGATTGCAATAAAGCTCGTGCATATTACTGGACAAATAAAATGTCACAATTCATTTGCTGGTAGCAATTTTGGTTGCTTTAATAATCATAAGTTATTCAACCTTGCAGTTACTGACGAAAACAGAGTAGTACTCTTCCCTCCACCAAATCAAGCACTTACCTACGATGGCAACAAGAACTATCAGTACCCTGGTTATCTACCAATTGATAAAGAAGTTGTTTTCACCAACTATGATTATCCAACTTACATTAAATCGGGTGATTACATCTCAATTTGGAATTACG
>evgR017476c0g2_i01_Hydra
AATGAAAATCTGAAATTGAAGAATTTTTAACAAATTCTTCTTTTAAGACTCAAGACTTATATAATTAAGTCTCAAGTCTTCATTAATTAATAAAAGCTGCTTTAAAAAAAAACATCAAAATGGATCAACCCATTCCAAGAAAATATGAAAATATTATTTTTAGTCGAGAGTTTAAGTTAAAATTTAGTGATTGTGTTGGAGTATATTGGAAAAGACTTGGTCGCTGGTTAAATATTGATGATAGTTATTTAGACATGATTGACCACGATACTCCTACAACTACCGAAAAATCATATTCAATGTTATCCAAATGGATGGAAATTAATAAGAATCCAAATCTTGAAGAGTTAAAAACAGCTTTACAAAATATGAAAAATATGGATTTACTAGGAAAATTAGACAAACTTATAAACGACGAACAGATTTCAAATTTGTCTGAAATGGATAGCAGATCTTTTTTGAAGAAAGATGTATCCGAGAAATATGCAGCACTAAAAGGTTTTTATTTAAAAAAATATGTCCAAATTTATGAATTTCGATTCCCACTAAATGTTACTTCCAGTGTTGATCTACT
>evgR017477c0g1_i01_Hydra
TAGTAATATCAAAAATCTAAATATATTTTCAATATTATAAAGCATGGTTTATTGCGTTTTTTTGATTATTTCAATTTTGCTTGTTAACCGTGATTTTACCGCTTGGGCATACGATCCGTGGATCTACAACGGCACAACGCCAGACATTGAGCTGGTCCTAAATGGACGTTGTTTGGAATACAAGTTATTTGATCCCAATAGTCGTAATCCGTCTCTATTTAATGATGCAAATTGCGCTACAATCGGAAAAGAATTTACAAAAATCTTTGCTTTTAAAGATTCTTGTTTAGATAAATTTGATGAATCCAGTTACAACAATTTCTTTAATATTGTAAACAACAAAAAATCGTTGAACAATCAGGCTTTGTTCTGGTCTGGTACAAAAGGCTTTGCTCATGATTACACACATATAAAAGAAGGATTTTATACTTTAGAGGACACTTTCTCAGGGTATCTTATAAACGATTTATCATGGTGTGGTTGTAAAGTCAGCCAACCACAACAATGCGTTGATGGAATAAACTATTTAAGTTGCGACAAAACATGCAATAGTCTGTCCAATTCTTTTTGGTCCAAAGCATCAAGAACTTTTGCTGAGAACGCAAATGGGACTGCCTATGTGATGGTAAATGGATCAAGTGCTGGAAGCAATACAGCTTATTATAATGGAAGTTATTTTGGAAAGATTGAACTACCCACTCTTGGAAGTCTGAAGAAAGTCAATAAATTAATAGTGCTTGTGGTTTTTAATCTTGATGACAAACCAAAAGAAAAGTGTGGGACTGGTTCTTTATTGAATTTAGTTGCAGATGCCCAAAAATATGGTTTAAATGTTGAATGCAGGGATAAGCCAAGTGACACGCTGTTTTTATTATGTGCAAAGTATCCACAAGCAAGGGAGTGTGCTAAGATGTCCAATGCTGGTAAACCATTATGGAGTAAACTGTTGTTTGGACTTATTTCAATTTCCATCATTTTCACAATGTTTCCTGTACGATTCTAATGAAGCTTTATATTTATAGTGATTGTTTTTTTGAATTTTTTTAGTACTGTTTTAAAAAATATATATATTTTTAAATTTTTAATAAAAATAGAAATGTAAATATTATAATTGAGGGTATAATTTTTAGTTTCAATTTTTTTGTGCTTTTTTTTTAAATTTTTTTTAATTTCTCACTTTTAAAGCACATTTGAAATGAAACATCGCATATATTTAAAATATTTTTAGTTGTAACAATATTATTTTTTAGTTGTAACAATATTATGACAACATTATTTGAACCAATATCAATTTTTCAACTTATTACAATAAAAGTTGAAATTTAATGAAAATTATTGTGCTACCTCCCCCTTCTACCTCTACTCTATCTCCCCCCCCTTCATCATCAATAGTTTTAATAATTGTTTATTTTTTCAAGTTGACATGATGTTGATTTTTCATTTACTTTTTACTAT
>evgR017480c0g1_i01_Hydra
GATTCAAAATGGCAAGTTTTAGTAACCGATTTTTTTAGTAATAAAAATATTGGACCAATTTGCGAAACAACCAATCCTTGTGGCACACTTTATTGCCGAGACGTATGTATAACAAATGGTCACAGCTACAACTGCTCTGTAACAACTGATGTGTCAAAAACTGCCATTGCTTCATTATCGGATTTATTATATAACAATATCGTTTATGGTCCTAATAACGTAATCGATGGTAGTCCGTCAACTTTTGCGGGGACGGGCCCAAATGTTTCTCCACAATGGATTAAATTAGATTTAAAGTTTGTTAATAAATTGACTAAAATTGATATTTTAAACCGAAAAGAAGACTGCTGTCGGTTCAGACTTCGTGGAGCTAAATTAAATGTTAGATTAATCGATAACTCAGTCAATAACTATGAGATTGCAAAGTTAACAGATGATATGAAGCAAGTTTTTGTAGGTTCATATATTGCTAGATTTGTTGTCGTGACTAGAGAAATAGATCAACCTTTACATGTTGCAGAAATTACTGTTTTTTTATGAAAAATAAAAGAATTTTGATTACATTTATTGCACGCAGGGCCGAGGCTAAATTTCCCCAAACCCCTCAGCAATTTTAGAATGCTTTTTTTGAAGTATAAGGAGCTTTTTTTGAAAGTTTTATTATTGTTTTCTCTTGTTTTTTAAATTTGAAAGACGAATCTAAATG
>evgR017495c0g2_i01_Hydra
GGTGCACGAATCATTCAAAAAATATATATATTCTAAGCTTACATGACAAAGCTATATTGACATATGGTACATTTGATTGCATGGACCGGTGATAGTGCAACCTTAAATACAACAATGTTTCAAAATAACTCCTCGTGTAACACAAGTTTCTTTATGGTTGTCCTACAAATGGGAAAAAGCTCGTGTAACACAAGTTTCTTCATGCTCGTCCAACAAAATGGAAAAAGTTTTTTTTCAGCTATTTTTGTGTTTAATTTTTTGTTTATAACTCTCTCTAACTTGCTTTTAATTATTGGAATGTATCTGCACAAAAATCTTAAATGTTTCACAAGAAACGATAAAAATGTTTTGATTCTCTCTATAGTAGATTTATTAATTGGCCTAGTGGAGATGCCGTTAAACATTGCACTTGTACAACAGACTAGTGTTGGATGCACTGCACTATTCATTCGAAGTTTTATCAAAATGTTTCTTGGTAAAGCATCTGTTTGTTGCATAGTTTTAATTGCCTTTGATCGATTTATGACGGTTTTGTATAGTAATGGAAATTTTATTGAGGGTTATCCTACGCTGAGCATTGTTTCTGTTTTTACAATAAGCTTCTCTATGGGGGTCTGGTACTCAATTGTTGATACTTCAAATGGTGCAAACCATGCAAGCTTTTTCTTTATATTTAATTCAATAGTTGACTTTACCATTTTCACAGTAGTTACCACGGTGAACATTGCCTTGCTTATTAATGCTCAAAAAAGATTAAAAATAGTTTCAGATAGTAGATCGTGTAGTTTAGATCTAAACAGGCGGTTAACCTATACCATTAGTATAATATCAACAACACTTGCAGTTATCTACTTGCCTTATATAATAATGCAGATTATTTATGCAGCAAGAGATGATGCTACCGTGACATTGTTATTCTTTTGCACAACTGTTTTAATTCAGTTTAACTCGGGAATAAATGCCTTAATATACATTTCAAGAAACAACAAAATTATGAAGATTTTCTTAAAAATTGTTATATAAACAATTTATTAAAATCCATATTTAAAAGTGCAATACTTTATGCAATATTCTATGCAAC
>evgR017498c0g1_i01_Hydra
TATTTATTGAAAAATTAATGAGATTAACTTCTTGGTCTTCTTGCCATTCCACCTCTGCCGCCTCTGCCACCTCTTCCACCTCTACCACCTCTACCACCTCTAACAGCAACAGGTCTTGCTCCTGGTGTTGGCTTATTCTTGGTGGTTTCTTCTTGGAGACCTTGAATACCTGCCTTTGCAATTCTGACCTTGGTATTATTGATGATATTGGTACCTTGGTGTTGCAATCTCATGGCCAATTGAACAGAATCTGGGTGAGCAAATTCAACAAAGGCATACTTTTGAGTTTGGAATGTTTCAGCAGTCTTATGTTTGAGGATATAAACATTCAAAATGACACCACATCTATTTTGATAGAAGAGTCTCTTAACTTGTTGAACTGTGGCAGTACCTGGGAGTGGAGTAACATAAATGGTTCTATCTTGTTTCTCAGGATCCTTAACTTCAACGAGTTTGGCAAGATCGAATTCAATAGTTCTTTCTTTACAACTGTTAAGTAAGAGAGAAATATCAG
>evgR017500c0g1_i01_Hydra
CCTCTTAGCAAATTGTTTGCTTCAGTATGGAATTTCTTCTTGAAAAAAATTGGCCCAGGCAATAACAGTGATTTGAGCAGAGCATTACATGTTTGGTGTGGAAAGTTATCATCAATGGTATCTTCTCTTACAACACGAGTTGGATTCTTAGAAGAAGAAAACAAAATTCAACAAAAGGAAATTTATTTTCTTAAAAATAATGACAATGCAAACATCACTTATCAAAATAATAATCTCAGTGCTAACGTATGGGCAGGTTTTGTTAGTAAAGAAAAGGACATTGCTGCCAAAGAAAAAAATGCTGCTTTATTTGCCAAAATTTCAAAAGAAAACAATGAAAAAAATAGAATTTTAAATAATATGATCATAAGTGGTCTTACTGAATCAGTAAATTTGGAAGATGATAATAAAGCGATTGATGAAATTCTTACCATCCTAAAAGTGTCTTGTGACGATGTAAAAGTTAAAACAAAAAAAAGAAAAATGCCTCTCTTCCTACTCCTTCTACATCTTGTGAGATTGACAATCAATTTATTTCATCCGTACGTCCTCCGTTAGTTTTAATT
>evgR017503c0g1_i01_Hydra
AGAAAATTAATTGAGCTTGTTATAATATTTATTGGTCCAATTTTAACAATAACAGTTTTATATATTCATACTCGATTGCATATCAGAGTGAAAATGGTTCGACGAACGATCTTTACACAAAATCCTGAAGCAAATTTTACAAATGAAAATAATAATTTCAGTAAAGACAAAAAAAGAAAAGAAAAGTGTTTTAATTTAAATGCGGGAAAAGAAGTAAACAAAAACATGAAAATAAATAATTTAACACTGATGATTTATCTATCCACAAGCATTGCCTATTTACCATTTATCTTTCTACTATATTTATGGCATTTTAAAAGCGTAGGTTACTCTTGGCAATGGACTCACACATTTTATTGGTGGGGATTTACACTTACACTGACAAAATCTTTAACAAATCCAGCAATTTACTTATATCGATCGAAAATATTGCGTAGAAGCTTACGTCGTATTGGGCAATCAGAGTCGATTAAAACTAAGAGCCAATCAGAACGCGGTTTGCGGACGAATAATTATGCAGGATGTTAAAA
>evgR017505c0g1_i01_Hydra
GGAAGGCCGCTGATTCTGATGATGAAAAGCCAAAGAAGGCTCCAACAGCACTCGCTCTTTTCAAGGAAGATCATTTGGAAAAAGTAAAGAAGGAAAATGAAGGAAAGAAGGCTGCTGAAATTACCAAGATTCTCTCAGCAATGTGGAAGGAAGCTGATGATGATGTAAAGGAAAAATATACTTCCAAGAGTAAGAAGCTCAAGGCAGAACTCGAAGGAGCCGATAGTGATGACGAAGAAAAGACCACCACAAAGGGAACAAAGAAGAGAGCTGCTCCAGCTAAGAAGAATGAAGAAGAAAAGTCTACTGATAAGCCAAAGAGAGCCCAATCAGCATATCAAATTTTCCAAGCTGCTAAGCGTGATGAAATTAAGAAGGAAAATCCAGATATGAAGCCTAAGGAAATTAATACCAAGTTGAGCGAAATGTGGAAGGAAATGGATGATGATGCTAAACAACCTTACACTGACCAATCTAATCAACAAAAGGAACAAAACAAGGCTGCCAAGAAGAGTTCCGCTCCACCTAAGAAGAAGAAAAAGAAGGATGAAGAAGAAGATGACGAAAGCTCAGAAGATGACGAATAAATTGTAAG
>evgR017516c0g1_i01_Hydra
CAAACTGTTTAGTAAAAATGAGAGGGTATAAAAAAGCTTTAGAAATTTATTATTTTGTTGAGAGAAAAAAAATTGAAAATTTAGGTATTAATCATCCGTCTACAATGACCACAAAACGCAATATTGCAAACTGTTTAGAAAATATAGAAAAAGTTGCTAATAATAAAATATCAATTGAAATTGCGAGTATAAATAACGAAGCTTTAAAAAACCATAATTCTTTTGAGAAAATTAAAACAGATTTAGCAATCAAGCCCATATTTACAAAGACAGCAGAGCAGAATACCGCAACATTTTCGAACCAAACGGATAAATACGACGAAGGTTTATCTATTATTTTTGCGGCTGATAAAATACAAACTGATATTTTAGGCGACAAACATCCATCTTCACTTGAAAGGAAACAAAAAATTGCTTTTTGTTTGGAAAATATGAAAAAATATAGCAAAGCATTAGAATTTTATTGTTCTGTTGAGGCAATAAAATCTGAAAAATATGGGCGCAACCATTTGTCTACTTTAGAAACAAAAGTAAATATTGCTTCCTGTTTAGAAAATATGGAACAATTTAACGAAGCCCTGCAAATTTATTATTTGGTTGATAAAATACAAACTGAAGTTTTAGGAAATAATCATCCAACTACAATGGCGACAAAACATCGTATTAAAATGTGTTTAAGTAATTTAAAAAAACGTAAAGAGCTAGAACAAAGTCAAAAAAGTTGTTTGATTTCTTGACTAATTTTTTTAGTTCATATAAAATAACGACAAATTAATATCTATAAAAGTAAAATAGTTTCAAATACTTTGTTAAAATAATAATATATTAGAATTGTGCTCTTTTCCATTAGTGGGGAATCTACTAGTGGGCGTTAGAGGAAGCTGTCCCGATGCTCCCTAGAATGGTTTGAGAACACGTTTTTTAATAAATCTATTATTATTTTTTTGTTTAGCGGGAAAATTTTGTTTTGATCGCAACACTGATGGTAAAATTTAAAGCATACTATTTGATTATATTTGTTGTTATTGTGGCG
>evgR017528c0g2_i01_Hydra
TTGTTAGTCTCACTTTTAGTCACAGTTTTTCTTCTGCCAAAAAAAAAAAAAAATATAGACCAGGAGGTCATAGTTCAAAATGATATTTTGCAAAACTCACTAATGGATTCAAACGTTTACTTAAACTACTCAATTATTGCTGGGTCAGTTCAAACAAACTTGACAGAAAAATCACGGTTAATTGATTCAAAGAAAACTGGAGTTTTAAAGCATTTTCTTCAGTTCAAAAATACAAAGGTTGGAAAACTTCTACGGAACAGAAATTATTTGCTTGTGATATTAATTTACTGTGTGTATCGTTTGTTGACTATTGGTCACAACGAATTATTTCCTGTGTTTGCAGCAACTTTACCGCAATATAGTGGACTGGGCATGACAACATCAGATATTGGATTGCTGTATTTAATAGTATCAGTTCTACTTTTAATAATTCAAACGACTATATTAAATAAGTTGACAACTAAGTTTGGTTCAAAAAAGGTATTTTGTGCATCAACATTCATTTATGGTGTGTTTGTAGTTCTGGTACCAATTACTGGTGTTATACAAACAAGACCGGCTTTATGGGTTATGACAACTATTGTTCAAGTCTTTAATCGTGTGTCTGGTGCTGTTGGCTTTTTAGCAATAAATATTTTTATAAATAATTCAGTAGATTCGGATGTGCTTGGATTAGCAAATGGT
>evgR017532c0g1_i01_Hydra
TTTTTTTTTTTTTTTTTTTTTTTTTTTTTTGTTGTTTGCCGTTACTTTAGCTAAGCTTCCTTTTCAACAGGTCTGCCGAAGACGAAGTGACGAAAATGTCGCTCATTGTACCAGAGAAGTTTCAACACATTCTTCGTGTTATGAACACAAATATCGATGGAAAAACTAAGATCATGTATGCTATGACATCGATTAAAGGTGTTGGTAGACGTTTTTCGAACATCTGCTTAAAGAAAGCTGACATAGACCTTAAAAAACGTGCTGGCGAGTTAACAGATGAAGAAGTTGAACGTGTAATAACTATCATGCAAAACCCACGTCAATACAAAATTCCAGATTATTTTTTGAACAGACAAAAAGATATTAAAGATGGTAAATTTACTCAGGTTTTAGCAAACGCCCTAGATAATAAACTTCGTGAAGATTTAGAACGTTTAAAAAAAATCAAGGCTCATCGTGGACTAAGACATTACTGGGGCTTACGTGTTCGTGGTCAACACACAAAGACAACCGGTCGTAAAGGAAGAACTGTTGGTGTTTCTAAAAAGAAGGGTCAGCAATAAAATATATCTGACTAATATAATTGTTTTTTGTTTATTTTTGATGTCCGCTAGATAAATCGCATGGCGAG
>evgR017534c0g1_i01_Hydra
CCTATTTTAGTGCAAAAAGTTGAATAATACAACAATGAGATGGGAAATGCTGGACTTAGTTGTTTTTCAAAAGTATCAAACATTGACGAACTTGGTACAAAATTAAATGGAGCTAATGCTCAAAGACATATGTCACAAGAAAACATTTATGTGGCTTTATTTAACTTTACAAGTAATAATGACGCAGAATTATCGTTTTCATGCGGAGAACAACTAGAATTGTTAAGCGTTTCAAATGGTTCATGGTGGAAAGTGCGCTCCCTACTTACCAATAAAAAAGGTTACATTCCAAATAACTATGTCGAAAAATATGACTACTTAAAAAAATATGATTGGTATACTGGCATCATGGGTAGAAGTAGTGCCAACTTGACTTTGATTAGTAACGGATCAGCTGGTTCTTTTATGGTAAGGAAAAGTACAAGCAAACCTGGAGATTATACTTTGTCTTTGCACAATGATACGTCTTTGAAGCACTACCATATTAAAAGAACAAAAAACCGCATGTTTTACATACTTAAAGAAGAGTTCTCAGATCTACCGTCATTAGTTGAACATTATAAAAAGGTTTCTGGTGGGTTACCTAGGACATTAAAAGACACATGTCCACAAAAAAAACTTTGGGATATCCCCCATAATAAAATTACTCTCACTAAAAAACATTTAGGCAAAGGCAATTTTGGAGTTGTCCTAGAGGGAAGGTTATATAGTTCTGTAAATGTGGCAGTTAAAGTTCTAAAAACAAATAATTGTAAAATACAAGAAAAAAGTTTTTTGACAGAAGCAAACAACATAAAAGAGTTATCGCACAAGAATCTAATTCATTGTTATGGTGTGTGTTCTTTTGACAATTACGTATATATTGTATTAGAATTAGCTATGTATGGGGCATTACATAAATTTCTCCAAACATCTAATGGTCAAAATCTTAATTTACAAGATTTAGTATGCATGGCAACACAAATAGCTTCTGGAATGACGTACTTAGGTAATATACGTTTATTGCATAGAGACATTGCAGCAAGAAATATGCTAGTAGCATCAAAAATGGTAATAAAGATAGGAGATTTTGGTTTAAGCCAATATTTACCTGTTGGAGTTGATGAATTTTATGAGGAAGGAAATGAAGCAAGATTTGCAATCAAATGGGCGGCTCCAGAAACTATTATAGAGAAAAAATTTTCATTAAAATCAGAAGTATGGAGTTTTGGAGTTGTACTGTATGAAATATTTACAAAAGGACTTGCGCCTTACCCAGGTATGAACAATCATGAGGCATTGGAGTTTGTAAGTAATGGCGGAAGAATGGATAAATCAAATATTCCTGAAAATATCTATACAATAATATTATCTTGTTGGGATGTTGTCCCGACTAAGCGCCCGCCAATAGAAAAATTAAAGGAGTTTTTAATAGAATATTCTAGATGAATGTTTAAGAAAAATAAATTATAATATATTTTTAAAAAGTTGAAATTTTATTGCATTGAGAAAAAAAAATCATTATAGTGTAATATTATAACAAATGTTATTATTTTTACAAACATCAAGTTTTTTTACCAA
>evgR017538c0g1_i01_Hydra
TATTGAAATTTATTTTATAAATGATTTTTTGATGCTATAGTAAAGTTGGAAAACACTCTTAACTTGTTATTTATGGTTAAAACAATAGAATGAAATTACCCGGAGATGGGGCAATGGACATTGATATTGAAATGGGTGCGATGGAATCAGACCTTGACCCAACATGGACTTCTTTTCAGGCATCTGCAATCATAGCAATATGCGCAGCAGTATTAATTGGTAACGGATTTGTTTGTGCATTAAACTGTTTCAAAACTAGAGCGCAAGGATTTTATTTAAAAATTGTCCTTATATCCGCAGCAATGAATATGACAGCATCGCTTGTTTGTATGCCTTTAATGGCTTCTTGTTTAATTCAAGCGGATTGGCGGTTTGGGTATTTTTTATGCAAAATGACCGGAGTAATATGTTTCTGTAGTATAATTGTGACAATGTTTTTTTGTATATTGGCATCTTTTAGCAAATGTATTCTATTTAGTAAAACATTGTTTCCAAAGCGTGGAATAAATTTAATGACAGAACGCGACTATATTGTTATTTGCTTAGTGCTTGGTTTTGTTGCGCTTTCTTCGTTAGCGCTATTTTTATTGGTTGATTACAAAAAAATGCGTCAAGATCCTCAAAGAATGATGTGTACTTATGCAAATTCTCCTAACATAACTGGTTTTCGCATGTTTTATATTGCATATTCTATTCTCTTAACCTTTGCAGATTTACTTTTAGTAATACCGTTTTCAACAATATCGTAGCACTGCAAGAGTTTTACACGTTGCAAACGTTTAACTATGTTGCAAGCGTTGTGTTGCAAAATTTTGCGATAATATAATTGTAAATATGTATATAGTATATTTTAAC
>evgR017539c0g1_i02_Hydra
TTTTTTTTTTTTTTTTTTTTTCTGAAAAAATCGCTGAATCTTTGCGATCTTATGGAGACAGCAACGGTGCCATTGTTAGCTCAACTTGTTTATGGGACTTACAGTGAACTTAAGACAAAGAAATGTTTAACACATGATCAAGGTAGCGGAAGTCCTTGCTTAAAATGCAATGAGTTATGCAGTGGATTAGAATTGCATTTTTGGAGAAAAGTTTGTAAAAACTGCGGATGCAAACTTGAAGATCACAAAGTTGAAGTTGATAAAAGCGTTCACGAAGCCATTGTCTACAATTTACTTAATGATGATTACGATAATGCATCACCATGGGAATGCAGCACCGATGACGAAGTTTTAGAAGGACATCCACTATTAAAGCATTTGTCCAGAAAAAAATTTAAAAAAACATCAAACAATAATAAGGAACAGGATGAAGTAGGCATAATTGAAAATGATGGAACTGTTGATCAACTGTTGAAGTTCAAAGTTTGCAATAATGGATTTAATTCTAAGAGTGAAAAAAGTAAAACAGACAAAAATAAAGAAGAAGTTCGGCAGCGTCAAAGAGCATTAATCCGCCAAATTCCACCCCAGGATTTTGACATTCGTTACTGTCATGATCTTGATGACTATGAAAAAGAAGAGTTAGGCTACTTCACATTACTTGTGCAAATGCATACAATCGGAAAAGCTTTACTGAAAGAAAAAGAAGATGAATCTGAGATACACTGTGTATATTGCTCAGAAATGATCAAGTCCAATTATTTAATAATTGAACGTCTTCAAGGTCAAACCTGGCATAGCACCTGTCTGCGTTGTAAAGTTTGCAATGAAAGTTTAGTTGACAACATTTACTTCTTAGAAAATGATGATTTGTATTGTGGAAGACACTTTGCAGAAACGTATAAACCACGTTGTTTTGCTTGTGATGAATTGATTTTTTCCCAAGAGTATACTCAGGCAGAGCAAAAAAATTGGCACACAAAGCACTTTTGTTGTTTTAAATGCGAAGTATATTTAGGTGGAATGGGTTACATTTCAGTTGGACCGAACCCCTTTTGCATTCCTTGCTACGATGAAAATGTTGCATCTTGACGATTGAAAATCGGTTTTAGTAGAAACATAGAAACTGCAGCAGAAGAATAGCAGCTAAACTTTTCTGGAACCTGGAACCTAAATATGTCATTTTGGACATTACTATTTGCTAATGTTATAAATTTTGCTCCATTTTTTGTATAATAATATTGTACAATTGTAATATACAGATGTTGTAATATAAAATGAAATAAAACATAAAATAAAAAAAAAA
>evgR017540c0g1_i01_Hydra
ATTTGGATAATCAATCAATTGGTATCACTGGTGTTGGTTATGCCTTGTTAGAAGAATTATTCGAAATGCCTGCAGGTTTATCTGAAGAATTGGAAAATGCCATCTCTGAATTCTATGCTCAAAAACGTGCACGTGAAAATCGTGTGAAAGATTTGGAAGAAAAATCAGAATTGGTTGGTGTCAAGGGTCTTCAAGCAAAGAATGAATTGGCACAAATGTTTGCTCAAGATCAAACTGAAATGAACAAATTAGAAATGACACTCAATGCTGCAAAGAGAAAAGCTTTGAAGAATCCAAGTGGTGAACAATTGTTGGCACAAAAGAAGAAACAACAAGAAGATGAAGAAAAGAAAAAGTTGGACGAAGGTAGACAAAAGATGAAGGACAGAGCTTCATTGTGGAACAAATAATTGAGTCGTTTCTTTCGTATGTGTTGAGTGTTTGTTGTTTCAAATACTTTGTATCATTGTAT
>evgR017556c0g1_i01_Hydra
TGCTTGATAAAAAGTGGTTTGATACCGAGTCGCGTAGCCGAACTAGTTTTGTGTATCGGAAATAACCTTATCATGGCTACAGTCTATAAAAATACTATCGTCACGCTTTATTTAAAATATATACGTATATTACTTGAGTTGTTTGAAACAAATTTATTTGGTCCAGATTTGCTTGGTGTAGATGTTTTTGATCCGCAGTTAATCGATGATAAGCTGCTTGATAAAAAGTGGTTTGATATTGACGTTTGGCGTGATTTCAACTCTTCGAGACAATTATCAAAATTAACGGTGTTCTTAAAATCTTCTAAATTTATAAATGAATATTGATATAAAAAAGTACGTACAAACTATGCTATATTTTTAGGTGCGTAGCAGAATTGAGATTGTG
>evgR017567c0g1_i01_Hydra
CTCCTATTTTAGTGCAAAAAGTGAATAAAAAAAGCATTATTTGAAGAAAAAATTAAACTATGAAGTTAGAATATTTTATACAGTTAAAAGTATGTATAAGTATATGAGCTTTTGCAGATCAGGGTTGATAGGATGTTGTTTCAAAGACAAGAGAAAGCTTGATGCAGATTGTGGATCTGATGAAGATTTTCTAATTACCAAACAACAAGAAAACATTACATTAGACACCAGATTAATGGGAAATGGCGTTGTAATTATTAAAAATGGGCAACGAGTTTGTGGATCAGGCTCAGCACTTGCAAATATTGACATATTACAAGATAAAGCATACTTTGAGATCAAGATACAAACATTAGGTTTTTGGTCAGTTGGTGTTGCAACTACTCAATGTAATTTAAGCACAATTCCTCCTGGAAAAGATGACTATAGTTGGGTGTTGTTATCTGATGGTTCTTTATGGCATTCGAATGTAGCAGTTGATAAGGTTAATCTCCTTATTAATGAAGGTGATTATATTGGCATCAGTTTTGACCATGTTGAGTTGAATTTCTTTTTAAATGGAAAAAATCTTCACTGTCCCTTGCGTGGCATTAAAGGTGTGGTACGACCAATTGTTTGTGTGGAAGATGGTTGCATTATTGACGTTAATTTTTCTCGCTTTGTATTTCCTCCTCCGCCAGGTTTTCAAAAGTTAATGGTCGAGGTCGATCTACTTTGACAAAAGCATGCATGTTTTTTTTTATTTTTTTTTAACGTTTTTATATTTTTAAATAATTTTTTCTTGTGTTGCACAGTTTACAAATTATTGAGTCC
>evgR017574c0g1_i01_Hydra
CAACCCTCGTAGTTTTGGCATTGATCGCCACTTTTGCCTACGCTCAAACTCAACATTGTATTAGCAAGGCTTTCATCACCTCTGCTGTCGTTATTGATGCTTCCAAGGCTTTCGATGATATTGAAAGAGTTTGGTTTGATGCCTCATCCAACAAACAAAGAACTGATATTGAAACTTTCGAACCATCCCGTAAGAGTCTTTCCCTCTATTTCAGACATGATTTGCAAACTGGTTATGAATATGACAAGTTAACTGGTGCCTGTAAGAGTTTCCCAATTGGTGGTCAATTGCAACCTTTCTGTTTGGCTCAAAATGCTGTTTTGAATGGAACTGTCACCATTGGTGGAAGTTTGAAGTGTTCAGTCTGGAGTGAAACCATCAGAGGATTCAATATCAGATTGGTCATGGCTCCAAATGCTAATTTTGGTGTTCCAGTCAATGTTATCACTAAGGGTGGACGTGTTGGTACTGCATTCCAAGAATTCTTCAATTTCCAAGCTTATAGCACTCTCCAAGATCAAACTGTTTTCAATGTTCCTGCTTCATGCAATAGAGCTAGATCTGCTGCCTCATCTGGTGCTGCCTCTCAAGTTGTTATGGATGTTATGGAAAGACACGCTTGGATGAATTAAATCTATCACGAGTCTCTACACTCTATTCTTGAAATAAACTTAGTTTGTTTCTAC
>evgR017579c0g1_i01_Hydra
AATGGTCCATACCAACATAGAAACCGATCTTGTGTTAATTTTACTTCATCTGATTCAAATTACATTGGTTGTTTCAATAGCAGTCTAAGTGATCAGCGGTTTTGCAATCAAAATGTTTCATGTCCAGGATCTTTAACAGAATGGAGTTCATGGGGGGCATGCTCTTCCTCTTGTAAAAGTTTAGTGAATGGTCCATACCAACATAGAAACCGATCTTGTGTTAATTTTACTTCATCTGATTCAAATTACATTGGTTGTTTCAATAGCAGTCTAAGTGATCAGCGGTTTTGCAATCAAAATGTTTCATGTCCAGGATCTTTAACAGAATGGAGTTCATGGGGGGCATGCTC
>evgR017581c0g1_i01_Hydra
CTAAACGCCATAACAGTCACAATGGTCAATCGTCAAGCTACTTTGAAGGATGTCAACCCACAACAATGGATTGTTGCCTATGCTGAACACTTGAAGAAGCAAGGTAAGATCGTTTTACCAAAATGGGTTGATATTGCTAAGACTGCCGTCCACAAGGAATTGCCACCAATGAATCCAGATTGGTTCTACGTTCACTGTGCTGCTGTTGCTAGAAACATCTATTTGAGACCAGGTACTGGTGTCCAAGGTTTGAGAAAGAGATACGGTATGGCCAAGAACGGTGGTTATGCTCCATATCATCACGAAAGAGCCTCTAGAACTGTCCAAAGAAAGTGTATGCAAGAATTGGAAAGAATCGGTGTTTTGGCTTCTGAAGAACAATCCAAGAACCCACACTCTGGTAGAAGAGTTACCTCCCAAGGAAGACAAGATATGGATCGTACTGCTCAAAGCCTTAAGTTGAAGAAGTAAATTAACCAATAATATAAAAAAAATCAATTCAAAAAT
>evgR017606c0g2_i01_Hydra
AAACTTCATTTAGAAATGTTAGCAAGTGGTTAGATCAAATCAAGAAACATGGTGATCCAGCAGTGCCAATGGTTTTGGTTGGTAACAAGTCTGATTTGACAGCAAAGAGAGTAATTGATTCATCAATGGGTAAAGAATTGGCAAAGAATTTAGGAATTGAATATATTGAAACGAGTGCCAAGAGTTCACAAGGTGTTGAAATGGCTTTTGACAAAATGTTACAATCTGTGGTAAAATCACACAATTACATTGAGAGTGACTATAACAAGAATAAGAATAGTGACAAATATTCCAATTCAACATCATCGCTGGTGGTGGGTGGACCAAAAACTCATTCCAACAATACTTGTTGTTAAATAAATGCAC
>evgR017614c0g1_i01_Hydra
CTGATTTAATATTACAAAAATGGAAATATACGATGAAAGTGTGTTAACTCCTTACGAAATTCTAAGGCAAAACAACATTATAAGTAATTATCAGTTTATGAAAGCTTGTGGGTTGCCTGTTAAACCATTAATTTATGTCAAAGAAAGAGCTAAAGTAGATGGTCTTCATTATGGCGACAGTGCTTGCAACACTTCTGATAGTAGTCAAGAAGATGAAGAAAGAAAAATTTCAAAAGCAAAAGGGTTACCTTTTAAACCATTGATTTATGTCAAAAGTACTCAAATAGACGATCTTCATGATGGCAACATTGCTAGATTACCTGTTAAATTGTTGATTTATGCCAAAGAAAGAGCTAAAGTAGATGGTCTTTATAATGGTGACAGTGCTAGTAACACTTCTGATAGTAGTCAAGAAGATGAAGAATGGAAAATTTCAAAAGCAAAAGATAAAAAATCAAATCACAGAGTTGTAAAAAACAATGATCAACCAAAGGAAGAATACAATAAAAAGAAGCCACAAATGAATGCAGTGTTTAGAACTTTACGAAAAGTTCATGAAAGTGATACAGGAATCGAAATTTATACTTCTTCACAGAGGAGAGGAAAACGAAAGACACCAACTGATCAAAATTGTAATCAAGTTAAAGAACAATCAAAGTCTCAGAAAGTTGATACAAGCAAGATAAAG
>evgR017617c0g1_i01_Hydra
AATGTTCAAAAACCGTTCATACTCACCCCCGTTGACGTTCCATCTCAAACCATGTACAATCAATCAGCTTCAGGATGTTCAATGAAAACTTATCAAAAATCGTCACGGGTGCATAAACCTGTTGAAAAGTATGTTGAAAGTTCTGTAGAATCAGGTTTTTATCTTCTATTTTTTAAAGAAGAAAACCTTTATGACGTGATAAAGAAAAGCAAGAGATAATGCTGAAGCCTGACATTTTGGCAATAGTTATTTTAAAGTCAAGGTATATGAAGGACATGGAAGTTTATAAAAAAAGGATGGAGCTTAAAATATCAAAAGGTTTTGATGTAGTTAATGATTCTTTTAGTTTCCTAAACAAATCCATCAATGAGTCCCTTCTATCAAATCTGAAGCCTATATGTCAAACAGATATCAATTATGAAGAATCTGATGATTCGTTGTCGAGTGTTGATCTAGAAATGGATTCAAAAATAAAAAATCCTAAGAAAAACGTTGAATCAGTTACTAAGCCTCTTCCAACAATTGATGTGCTTTTACAAAATGATCTCCTAAAGCAACAATTAAAGATCCAAAAAAAGATATTGCTTGAGTTAAAGTCGATAAAAAAGCTGAGGCAATCCCTAACCATGACCCAGTAATGTTTAATGGTGAAGATTTGGTCTTAATTGGAGAAAGAAATATGGAAGTTACTAAATTTGCTATCTGTGTTGCTCGAAAACTATTTACTGATGATGAATTAAAAACTCATATTCTGTTTAAGCAACGTGAGTCAGGTAGACCAAGTTTATCGCCATGTCGCTCTGATATGTTTAAAAATGCTGTTAAAGGACGCTTTCAGGATGATCAAGACGTAACTACTGCAATTGCTGCTGTAAATGCTCTTGGACTTGATTTAAGAAAGGGTAATCGCAAACGTAAATTTGTAGCGTAGAGGATGCTTAACTATTTTAATTATTCAAAATATTTC
>evgR017620c0g1_i01_Hydra
AATCCAAAATAATCCAGCCTCAATAATTGGTCTTAAAGCCGTGTATATAAAAGATATTGATTTTTCAAAAGACTACGTCATTACAGGATATGCATTGACTGTAACAAATGTTGATATTCCAATAGTGGGTGTACTTACTTCAGTTCATTTTGTTGTTGAAGATGAAAACCAATTTGTCGTGACTTTATGCATATATAATCTCGGTGTAGATTACCTAGCGATTAAAGAAAGATTTCCAGTTGGATGCATGTTTACAGTTATTAATCCCTACGTACGATTGGCTGCTGATGGTAGAGCAATAATACGCGTTGATGATGCTAATTCAGTTGTCATGTCATCAAAGAGAAAAGTGAACATATGTTTTTACTGTGGCAAAGAGCAGTCTACGTTTACTTGCTCAAAATGCAAAATTGCAAAG
>evgR017627c0g2_i01_Hydra
ATCATCAAAACAGTTCTATTTATGCAATCACCAAAACAGTTCTATTTATGCAACCATCAAAACAGTTTTATTTATGCAATCATCAAAACAGTTCAATTTATGCAATCATCAAAACAGTTCTATTTATGCAATCATCAAAACAGTTCTAATTATGCAACCATCAAAACAGTTCTATTTATGCCACCATAAAATCAGTTCTAATTATGCAATCATCAAAACAGTTCTATTTATGCAATCACCAAAACAGTTCTATTTATGCAACCATCAAAACAGTTTTATTTATGCAATCATCAAAACAGTTCAATTTATGCAATCATCAAAACAGTTCTATTTAT
>evgR017635c0g1_i01_Hydra
CTTGATGTCCATTACACTAGTTTTTTAATGACAAAATGTAAATGCAATCATTTGACAACTTTTGGAGGATTTTTTGTTGCTCCTAACAGTCTTCCTACACTCTCTATAGCAAAATTAAAAAGTGGTTATGTACTACTTGTTGCAGTTGTAATAGTAATTTCGTTATGGTTGCTTGGACTACTGATCACGAGACGAATGGACAGACAAGATATTTTAAAGATCGGTGTTTGTCCACTTCCTGATAACAAGGATGGAGATACTTATTTATATCAAATCATTGTTAATACTGGTACTCGTAAAAATGCTGGCACCAAGTCTAACATCTTTATTACTGTTTCTGGCAGTTTGAATGACAGTAGTGTTAGACATTTAAATGATCCAGAAAGAGAATGTTTCCAAAGATCAAGTTGCGACATATTCATTATGAC
>evgR017636c0g1_i01_Hydra
TAATGCTACCAATGAAATGGAACCTATTGTTGTCTTTACTACAGATTCCAGATTTCAAGGTAAAGATAAAACACTAGTTATCTGTTCTGCTATATTAGTTCCACTTTTTTTTCTTATTTTGGTGGTTTATGCACTGTGGAGATTTAAAAAGAAAAAAATACAACAAATTATGAAAACATTAAAGATTTTCTCTTTTGGACATAGCACCAGGGATGATAAATTAATAGATAAATGGGAGATATTTCCTGAAGACATTGTTTTAGACAATCAAATTGGCGAAGGAGCTTTTGGCACAGTTTATATTGCAAAACTTAGTAATCCCTTTATTGAAAAGAGAAAGGACATTCAAC
>evgR017646c0g1_i01_Hydra
ACCAGTACTTGCAAAAGAATATGTAGTTTTATTTGAATATAAGTTAAATCAAACTGTTAATAATAGTTATGTTGATTGGTATAACATTATTTACTTTGCTAATGATTTCGGAAATAATACTTATGACAATGAAAATGGTATTCTAAGTCTTAGTTTCAGATCTGATGGAAGTTTAGAGATTGGTGCAGTACTTAATGGAAATTTTTATAGCATTTATAGCACCCGGTTATATGGTTTAAATCAATGGTTTAGAGTTGAAATATGTCAAATTTTTATTAATGAAAATTACAGTTATAGAGTAAAAATAAATAACACTAATATATTTTCAAAAATTAATAATCAAGCTCAAACTTTTAACGATGTTATGATATATGCTTCAAATCCATGGAACTTAACTCAAAATGGCTTAATAAAAAACTTATTGGTTGTCAATGGAATTTCAAATTCCTCACTTGATCAATGGAGTGCCTGGTCACAATGCAATACATCCTATGGATATGGATTTAGGAACAGAACAAGAAAATTTTATGCATCAGAAAATACAACGTGGAGTTTGACTGAAGTTTTATCTGATTTAATTGAATGCTACATTTATAAAAATGAGTGTCAGATGGTGAAATATATTATAGAAGATAACACAACTCAATTAATCAAAGCAAAAAATATCACTAAATTGCCAATATTAGATAAGCAATTTATAGTTTCATTTGATTTCATTCCTTATTCATTCAACAGTAG
>evgR017655c0g1_i01_Hydra
AAATAATAGTTTATTTTCATTGTGATTTTTCTCGCTTCACATAAATAATGGACATTCTTAAAAGCAAAATAGACACAATATTTTTACCAGGAGATACTATTTTACCAATTTCTTGCTTATCTTCAACCGAGGGCTCTTTAAAATTTGGACCCGGTATTCGTCAAGAAGGAAAGGATGTTGTTGCATTCAAAGCCGGCATTCTAAGAATGAAGAAAAAACCTTTTTTTATTTGGATAGACACCAATCAAAAGCGGTATGTTGCAATTAAAGGTGACACTGTTATAGGAATTATAACAACTCGTGGAGCTGACAGTTACAAAGTGGACATAGGTTCATGTCAACCTGCATATCTAAATAATTTGTTATTTGAAGGAGCAACAAAGCGCAATAAACCAAATTTAAAAATAAATGATCTTGTTTTTGCTAAATTGATTGTAGCAAACAAAGATATGGAGCCAGAGATATCTTGCATAGATGAAAATAATAAAAGTAATAACCTAGGGCCGCTCAATGATGGTTATATGTTCACAGTTTCATTAGGTCTGAGTAGAAAATTGCTGCAACTTCCTACGTGTAAAGTTCTGAGTATGTTAGGAAGACATTTTCCTTTTGAGATTGTTGTTGGGTTAAACGGAAGAATCTGGATTCATTCTAAAGGAATCATTGATACTATAGCAATCGCTAACGCAATTGTTTCTTCTGAATTTATGGATGATAATGAAGTTACGTTAATGGTTAAACAACTAATAGAAAGCATTGCAGGTTTTGAGGAAGATGATTGATTTTCTCATTTTAGTGAACTTTTAAAATGTTATTCATTTTTTGAATTCTTTCTGAAAATGACTTTCTTTTTTTTTTTTTAATTACCTTTTTTTTAACTAATCATGGTCAGTTGCATTATTTAAA
>evgR017664c0g1_i02_Hydra
GAGGTGTATCAATTTAAAAAGCTACAACACTATAGAGGTGTATCAATTTAAAAAGCTACAACACTATAGAGGTGTATCAATTTAAAAAGCTACAACACCTTAGAGGTGTATCAAAAGTCAGAGTCTGACATAAAGTTCGGAAACACAAATATGAATTATTTAGCTGTATTCAAAAATAAACAAAATCACAGCAATCTCCATTCATGGCTTACAAACGGTTTGTAGAGTTTCCTAAATCTTATGCATTTTTGTAGATTCTTCAAGTTTTACCTCGAAAACTGACCAAGAAAGAAACGGCTCAATGGGTAACAAAAGTGCAAAATTTAAAAGAACAAAAAAACACTCTTTTGTTAAAGAACCTAACGTAGAAAAGTTTGAAGACTCTTATAAAATGCTTAAAGAATCGTTTCAAAATTCTATTTTACGCGGAAGAAGAACTGCAGTAGCTGCAGAATCGTATGATCCGTCAAAGGATACCGAAGAAGAAGAAGCGATAGTGTATTATCCTAAAACTGAAAAGCAAATGAAATGCTTAAATGACGCAGCAAGTGGCATAGTATTCTTCAAAAGTTGTGACGCGGAACAACTAAAAGTTCTATTTGGTGCAATGTTCGAAAAAAAAGTAAATAAAGGTGACGTCATTATTAAACAAGGTGACGACGGTGATAACTTTTATGTTATAGAGAAAGGGATATTTGATGTTCACGTCAAGAAAGATAGTGTTGAAAAAATTGTGGCTACGTTAGAGGACAAAGGGTTTTTTGGTGATCTAGCATTGTTGTACAACTGTCCTCGTAATGCAACTATAATTGCTAAAAGCGAAGGTGTCCTGTGGGGCTTGGACCAGAAAACATTCAAGCGTATTGTCGTCAAGGCAACAGCCAAAAAACGACTTTTATTTGAAGAACTACTCAAAACGGTCTCGATGCTTCAGTCCCTAACTTCTTACGAGCTAATGAATTTGACTGACGCATTAGACGTAGAGACATTTAACGCTGAAGTTAAGATAATTTCAGAAGGCGACGAGGCCTCTAAAATGTACTTTATCATGGAAGGTCAGGTAGCTGTGAGAGTTAATAGCGACGGAATCAAAAAAGAAATTATCCGATTAGAAAAAGGAAAGTATTTTGGAGAGCTTGCTCTAATCTTACAAAAACCTCGTGTAGCATCTGTTTACGCTACAATGGATAACACAAAATGCGCTGTCTTGAACATTCATGCTTTTGAAAGGTTGTTGGGGCCTTGTGTTAATATTATGAAACGAAACCTTAAACAATACGAAGACGAACGAAAAAGACTGGGTATTGAAGGAATAATTTACGAAGAAAAATGAAATGACTTTAAAAAACAAATATAATTTAAAAAAATAAGAATTATTATTATATCTATACTACAATGATTCATATATTAAACGATGTATGCGTTGCTGTTTACATTTACTTATGGGTATCTTTTTTACGAATGAATGTTGAAGAGTATATTTTTAACTTATAAACGCAGAGGGGTATCTTTTAAGTTTTGGCTTTTCTTAAATATTTTCCTTAATTAATCAATCAAAGTCAAATAGGCAGATGGAAAGAGGTTTTTTTTGTCAAAGAATGAACACTATTGAATTACGAAACCCCATTGAAATCATTTTGTACAATTTTTTTTTTTTTTGCAAATAATTATATGAAGTTTGCAAACATTATTTCTTAAACATTGTATAGATTTTTACAAACATTTAAAACAAATTATGAAATAAAGATATGATCAAAAACTTAATCCAG
>evgR017665c0g1_i01_Hydra
AAATGTTCACCGTTTGACCTCGTATAGTAACATAGCTATTGTCAGTTACAGTGCAACAAGCACTGAGCTAAGTAATAAAGAACTATATCCAAACTTTTTTCGAACCATCCCGACTGATTTTTTTAAAGTACAGTTTATTATAGATTTGTTAATTTATTTTAACTGGAGATATGTTTCAATTATAGCCTCGGACAGCTCTTATGGAAGAAATGCAGCCTTTTTGCTCCAAGAAATGTTTTCAGCAAATGGCATTTGTATGAGTAAATACGTTAAACTTGGAGATGATTATAATGAAATGGATTACAAAAACGCTCTATTTAGTATAGCACGTGATAACCGAGCCAAAGTTGTCATATTTTATGGGTATGATAATTCGGCAAAATATCTACTCGCCGATTGCTTGAAACACAAAATAACGTATATTAATTGGATTTTAAGCGATTTGAGTTTGCAAAACAATTGGTTATTTGATTTAAAAAAACAACACAACGGCAACATTATGTTGATATTATTTCGCTCTGAGGTTATTGAACAGTTTAAAAGTGCTTTTTTAAGTACATCGTACAAAAACTCCGACAATTGGCTAAGAACACTATTCGAAAAAGATATTCAGAGCAAACCTCATCCAGATACGAAGCTTGGCGATGTCAAGAATTCTTTTGACTTTAGTTACGTAAATTACGTACAAACTTCTGTAAATGTTTTAATCAAATCCTATTTAAAATACTTAGATTGCAATGAAAATATGCGCTGCAATTCCTCCTTTATTGAAAACCGG
>evgR017669c0g1_i02_Hydra
TAATTCAATTCAATTATAATTTGAAAATATTTTTTTTTAAGAAAATTAAAAAAAATATGAATAATTCTAAACAAGCAATAAACGATTACCATAGAGCGGGTGATGAAGAACATATAAAATATCTGAAAGTTATAGAAGAAGATTTGTTAAAAACAATGGAATTAAAAGTAGCCACTGAACTTAAAAAAATTGCCGGAGATCTTTTAGTTCAACCGATAACTCCACAAATGGTTGCCGACGAATTAAGAGTTTGTAAAACTGAGAAAGAACAGCGTGCACGCATCGATGAAATAAATAAGCGAGGGATGGATTTTTTAAGAAATAGACACATACACACAGACACACAGACACAAGAAATAGAC
>evgR017682c0g1_i01_Hydra
CGAACCCTATTTGCACTGACAATATTTGCAGCTGGAACGCTCCTTCCAAAGAAATTCAACCAAGCATGGTAAAAGATATGAACATCGTTAAACATAAAAGTGGTAAATTGCAGAAAAATATTTTGATTTACAAAGAAAAAAATGATTTTGATCCAAGACCAGAAGAAATGCGTGAAATTACGGAACTTGAAAAAACACAATTTTTATTAAATATCAAGAAAGTTCTTCCTAACGCGGCTATAAATACAATGTTTACCCCTCTGCCAAATAGTTATGTGCCACTGTATTATGATATTAAAAAACATGTCTACATTTGGAGCGTTTCCGTGGACTATTAGCGAGGAGTGCAACAATGTTTTTAGCAAAGTCACTCGGACCTGTTTTATGGAAAATGTATTTACCATTGCAAGTTCTTATGGCAAAGTGTGTTGCATCTCTTTTGGACATGCGCAGTTTCTGG
>evgR017694c0g1_i01_Hydra
ATATGGATACCCACAACCTGGTTCAACTACCACAACTGTCACTACTTATTGGCAAGGATATGGTCAACCAACATGGTTGCCACAAGGATTTGCTGTTCCTTCCCATGTTATGGCCAATCGTATGAATTACATGAATGCAGCTCAACAAATCTTCTGGCAATACGATCTCAACCGTAGTGGAAGTCTTTCCAAGAAGGAATTCAAATTTGCTGCTGCTACTCACTTTGGTGTTCCACCACACCAAGCCAAATTCCTTTTCCGTATGATGGATCGTGATGGTAATGGTATTATAACACTCGATGAATTCATCAATGCTTACTTATTCATCATGGCCGGAGGTGCTTCATATATTCAACCAATTCCATCCTACAGACATCGTTTTGGAATGGGTGGTATGGGTATGATGCCAATGGGTGGTATGGGAATGATGCCACCACAACAACCTTATGGAATGATGCCTCCACCACAACCATATGGAATGCCACCACAACAATTCAATCCATATGGTTATCCACC
>evgR017697c0g2_i01_Hydra
ACATCTTTTTTTACCTATAATGTTAACTTGCTTTCAAATGAAGATGTTTCAAGTTTAAATTTTTCAATTAATAGTTGCAATAATTTAAATAACCTTGCAATGCGTGTCTATTTGATTGTTTCTCAAGATAATTTATTTAACAATATACTTGATGATATGAAAAAGATGTCTTATAATCACTCAAAATTGTACCTTTCAAACATTCCATTTGTTTATTTGGTTTTGCGTCAAGGATACAATAGTATTGAGTTAGTGAATCTGTTTTATAATGCTAAAAATAATTATCCTGGCATTGTGAATTACATTGGTGTTATTGCTATACCAGAAACTGTATTACAAAATAAAGATTTTTGTCAAGTTGACGAAAATAATATTATGCTGGGTGTATCATATGGAAAAGGAAATAGCAATTTCATTTCTGAAAAAAAGATTCTTAATCACCGCAAC
>evgR017703c0g1_i01_Hydra
TAACCTATTTTAGTGCAAAAAGATATTAAAAAATATAAGATGGGTGTGCATTTAGCTAAATTGAAGAGGATTTCTCGAACACCTGCATCAGTTTTTAAAGATGTTGACAAGGTTTTGGATTATAAAATAGTATCTCCTAGACATCCTACAACTGTAAAAGCGTTTGACGATCTTGTTAAAGCTAATCCTCAATTTAAAACAGCTGTAGATGAAAGAAGTACAAATTTAAATAAAATGCTTGAAGAAGTTCGTGTTAATTCTACAGGAGTTGGTCCTAATGAAATTTTACATAGAGGAAAGTCAGCTGAAAAGGTTCAAGTAAAAAGGCATGGCAAACTTAGTTACAAAGATATTGAAGACTTGTTTTTAAATAGAAAAATTGATCCTGAGACATGGAATTTAGCAAAAATTAGCGAACACTACAATTTGGATGTAAACGTGACCGAAAACTTAATAAAAAATTACAGTTCATTTCGTATATTTAAAAAAGTTAAGCCAGAGACTAATCAAGTAAAAGATGCTAAGAATGATTCTCTTCGAGATTCCTAGTGCATTATTTTTGTGACATATAACTTTATAAAGTTATATGTCGACATATAAACTTTATTTTGTAG
>evgR017705c0g1_i01_Hydra
GCAAAATATTGAGATAAGTCAAATTCTAAAATTTAATTTTAGTTCAAGTTCTCAATATGTATATACAATAAAAATTAATGAAAAGATTGTCTATTCTGGTATTAACAACAAACCTAGGTATCTTAACAATGTCAAAGTGTACGCTTCAGATCCTGGTTATAAAACGTGTGATGGTCTAATAAAAAATCTAGTTGTAATCAATGGAATTTCAAATATGGTTGTACCTGTCCAAGAAGTAGACTGGAGTGTAGTGGAGTCGGTTCTTATTCTTTCTTCAGATTACATCATTATAAGTGAAGAATTCACATTATATCAAGGTTTACTGCTTGCTACTCTAAACGTTTTGAAAAACGAGTTTACTGTGGCGTTTAATATAAAACCAACATACTATACTGATGGCTTAAAAAGTATACTTAGTTTATTTATGGATGGATACATGGATAATATATTTTTAGATGTATTTTTTCATGAAGATAAATCTGGAAAGCTTGTTATT
>evgR017707c0g1_i01_Hydra
TTTTGTAAATTCAGTTGATAGAGACTATCAAAACAAAAGAAAATTTAATTATCATTGTGTATACTGTTCAGAAAATCACAATCCTTGGGAATGCAAAAACGTTACAAATTTTGCAAATAGAAAAGAGATTTTGAAAAAAGAGGGTAGGTGTTTCAATTCTCTTAGAACTAATCATGTACTTAAGAATTGCCAAGCGAATTATATTTGTAACAAGTCCAAAGAAAAACATCACATTTCTTTGTGTGAAAAACAAAAGACTTTTGTTGGGCATGCAACACACAACATATCCCAACAAACTATTATGTTGCAAACGGTTACAGCAAACATCAGTG
>evgR017719c0g1_i01_Hydra
GTGAGTATTCTGTTATCTCATCTTGAAAATGTATTAATATTGACCAAGAAAAAATATATATTATTCTGTTTTCTGATGGATGTGGCTATCAAAATCGCAATTCTATTTTATCAAACGCACTTTTAAATTTTTCTGTCCAACATAATGTAGTTATTGAACAAAAATTCCTAGTTAAAGGTCACACCAAATGCCCTGTGATAGTGTACATAGTGCTATAGAGCGCAAGTTTAAAAACAAGGACATTCATCTTCCTTTAGATTATGTAAGAATAACTAAAGAAGCTAGAACCACTTCTTGTCCATACCAAGCAACATTACTTGATCATACATTTTTCAATGACTATAAAATAAACCAAACATATAAATCAATCAGACCAGGAAAAGAAAAGGGTGATCCTGAAGTTAGAGACATCAGAGCTCTTAAATATGACCCTGAAACTCAAATGATCTATTATAAACTTAATTTTAATGATTCATGTTTTTTACCATTAACATTACCATGAAACAAACATTTTGAG
>evgR017736c0g1_i01_Hydra
ATTTAAACGTATATGAGCAATTATTAAAATTAGATATCTACAAGAGTATTGGTGTAGATAATGTGCATGGATACGTATTAAAACAATGTGCATCCAGTTTGTCAAGACCGTTAAATATTCTGTTTAATATATCATTCAATCATTCAAAACTGCCTGAAATTTGGAAAGCTGCCAATGTAACCCCATTATTTAAGAAGGGTAGCAGATTGTTTGCAGAGAATTATAGGCCGGTATCTTTAACCTCTATTGTATGTAAAGTTATGGAATCTATTATAAAAAGCAGCATACTTAAATTTTTAGATAGTGAAAAGTTATTGAACCAGAATCAACATGGATTTTTGAAACATAAATTGTGTGTCAGGGTTACTAATTTAATAGAATCTATGGACTTAATTACATATTATTTATCCCAAAATTCGTC
>evgR017750c0g1_i01_Hydra
AAAACAATGTTTTTCAAAAACCAGTATTTTTTTCGCGTTATATACCGCGATCGTACAATGGAGGTTTGATTTTTTTTTTTAAACCTTCATTGGCACAAAATGGAATACAAGCGGAAGAAGAGAACTAAACATGTAAATGAAGAGTTCGATGAAACAGATCAAACAACCAAACTGTCTATAAAACCATCTGAAAATAAAAAATTGTGGAGCAGTTTTATTGATGTAGCAAGGAGTGGATTTTGGAAAAACGCAACTACTTTTATGCGGCGTGGCAACAACAATGCAAAAAGGAGAAAATTAGCTGATGATTGCGATGAAAAACAAACTACGCTTACAAATAATATTGATATAATAAGAAGCCCTCGATATGTCGCCGAAGGTAGCGGAATGTGCGTTTCAACGGAAATGTACGATGCAACAGAAACAGCATTTATTTTAACACAAAAAATTGTTGGGTGGAAAGTTTAAAATTTTTGTTTTTAAGATTGTATATTTTTTAGTGTTTTAAATATCGAATTTGTTTTTTTATTCTTTTTGAAAATGATTTCTTACTAAAAAAAGTTTATAATTGGTAATATCTCGCATTGTCTGGATTTATCGACTTCGATCGAAAGCACAGTTGAGTTTCTCTTAATGTAATTAATTTTAAGATTTATGTGAAATTATTAATTTGTCTCCCAC
>evgR017752c0g1_i01_Hydra
GAAGAAACTCGTAGAGATCACTCACTTTGATATGGATGAAATTATGAAGCTTTATGAACAATTTATTTCCATTTCATCCTCTCGTGAAGATGATGGTGTGATAGATAAAAATGAATTTAAGGAAGCTCTTGGCTTGAAGGATTCTCTTTTCGTTGATAGAATGTTTTCCTTGTTTGATGGCGATAATGATGGTACAATTGATGTGAGAGAATTCATTTGTGGATTGTCTGTATTTTGCGAAAAGGGAACCATCGATGAAAAACTTAAATTCTCGTTCAGAATTTACGATTTTGATAAAGATGGTTGTATTAGCAAGGAAGAATTGTACAAACTTTTAGAAGCCTCACTCGTTGAAAACTCCTTAGGTATCCCACAAGAACAATTATCATCTTTGGTCGATGCCACTTTTGCTGAAGCTGATACAGATGGCGATGGTAAAATATCTTTTGAGGAATATCGTGTGCTTGTTACTAAACATCCAACTATGATCGGAAATATGACCATTAATACCTCAATTAAGAAGTAAACCAAC
>evgR017755c0g1_i01_Hydra
ACCAACACGATTAACTAGGGAAAGTGCAACATCTTTAGATCAAATTATTACAAATGAGTTTATAAATACAAAAATAAGGACGGGAATATTTAAGTCTGAGATTTCTGATCACTTTCCGATTTTTATTTTATCCCAAAAAAGCAGATATGATGATCAAAATGAAAGAAAGAAAATAATTAAAACACGTTTAGTAAACGAAGCCTCTATTGCTCATTTTCATAATCTTCTATCCTGTGTTAACTGGGATACCCTTCAACAAAATCTAAACACTAATAAAGCGTATAATATATTTCTCACTGAATTTCTTAATTTTTATAACAAGGCATTTCCAGAAGTAAC
>evgR017762c0g1_i01_Hydra
TTAGATGCTGTCTGTAATGCCACTACTTGTGCTCCATATGCTTCTGAAGATAAAGTTTTGAGTGTTAGTTGGGAATGTCCAACTGGTTTCCCAACTCCAGTTTCAAAGGCAGGTGAAATTTATATTTCAACCTTTGGAGATGCTTCTTGTTCTATTACACAATTGACTCGTTATGAAGTTTACAAAGTTGGTGTCTGTATGAAGAGTAGTATTATCTTTGGACCATCATTCAAATCTGGTAAATATATTGGATGTAACAAGATGGCCTTGTACACTGATGAAGAATGTAAGGAAGAAAAGAGTTCAAATGATCTTGTCATGAATACTTGTGCCAACAAGCAAATTTACAAGTGTGGTAATGCTAGTGGAGTTGCTGTTTCTGCTCTTTTGATGGCTTTGGTCGCAATTATTGG
>evgR017768c0g1_i01_Hydra
TGATAATTTTAGATCTCAAGATGAATGTGATTTGTTGTTTCCTGTTGTTTCAGGAAAATCTTTGACATGTTTTGATTCAAAAATCATCTCGTTTATTCAGAATTTAAATCTGATTTTTCAATCAAAACCAAAAAGCAGCAAGAAAGTGGTTCCTCTCATCGTTGTCCTTGAGAAGCATGAAGACCCGGACGAATGGTCTTGCAGAAGAGTTGGAAATGTTTTAGTTCTTTTGGTGGATGGCGATATGTTTATGCTAATTGTTCAGCAAACAATAATTGCTAGTTTACAAGTCGCTCAAGAAGCCATTCTTGCTTTTATAGCATTGTTTTATTTATTGGATTTTGATTATCCAAAAGACTACGAAATTGGGCTAAATATGCTGCAGTATTTTTTTTTTTTTGATGACAAAGGCATCCCGGCAGACATAATAAAACGTTTTCACTTAGTCATGAAAGAATACTTAACTTATAAAGAGTAGC
>evgR017770c0g1_i01_Hydra
AATAGGATAAAAAACACCAGAACTTTAAGTTACCATTAAAACAGTTTATACGCACGTTGAAATCCAAGAGGAGAGTTTTATAACTTTCTTCAGAATAAACTGGTGCAGTGCGCTATATAATTGAATTTTCCACAATGCTTGTTTGCTGTCTCGTATACCTTATCACATTTTTATATAAAAACGAGGTAATAAGTATTTCTGTCAATGAAACCTTATCTTTATCAAATATTGACTTATCCTATTCATCAAAAGAGATAATACATTCAAATTTATCTTCAACAACATTTGCTCTGCAGGATGAGACTTTATCTTCAACAATATCAGTTTCAGAGCATGAAACATTATCTTCAACAACATCAGTTTTACAATATGAAACTTTATCTTCAACATTATCAGTTTCAGAGTATGAAACTTTATCTTCAACAACATCAGTTTTACAATAT
>evgR017798c0g1_i01_Hydra
TTTGATAATTGAAATTTCAATGGGAACTACTATTGTATTTATAGAAGTTGTTCCCTTCAATGTTGAAAAAATTTCTCAAATATTTCAAAGTGTCGTAAATATTGGTTATTATTATGCAACGCTATGGTTAATATTTGATAGATACTTGCATGTAAAACTCAATATAAAATATCTACTCTATTGGACAAGAAGAAAAACACTAATTGGATTAATGAGCTTATGGAGCATTTCATTAATGACTGGAACTTTAATTGGAATATTTTGCTATCATTTAGACGTCCTCGTATTTGCCTGTATAGACTGCATAATATTAACATTTTCTGCTTATGTGTATACTTATTCTGTTTTGTTATTAAAACGACGTAGGCCTGAGTTTGACACCAATCACCACAAAAAAGGCGTAGTTAAAGGATCCTTGCTATCAATTGCGATTATAGCAACGTTTGTAATTTTAGTTGCCATACCCGACTTAATATTAGCAGTGAATGTATTTCTTACTATCAAATCAAAAGCGTTAAAAATGTTTTTATATGTATGTTACCCGTTATCTTTATGGAACGATGCAATAATATACATACTTGTTTCACCTAAAGTTCGTCATGTTCTTATACTTAAATTCAAGCTATCTAATTACGAGTTTAAAAACAAAGACATAATACAAAAGCAAAAAAACAAATTGAAAATAACAATTTAGATGTTTAATTGCTGAAATAACTTTTCAAATTATTGTTATAATATACATACAAACTATCGATAATGTAAA
>evgR017812c0g1_i01_Hydra
TTCAACAAAACAAAGCATCTTGCTGGATCTAAAAATGGTTAATGGTCCATCAGTTGATCCAGCCAGAAATATGGAAAATGTTTCCATTAAACTTTTGGATGAATGGTTGTTTGAGTTAAAACAAAGTGTGGTTGAACTTAAATCGGAAGTTGAAAATCTTCGTAGCGAAAATGCAAAAAAGGATAATGTTATAACCGAACTAAATGATAAAATTTCCAAATTAGAAAATTTAACATCAAATTGTACAATTCAATTGACACAACCATCATATCGGGATGTTGCCTCATCATTTACCAAACCTGGAACCCCCCAAAATCTTGCAATAATAAAAGCAGTATCAGAAAACTCAAAAATTAGCGAAAATAAGTCGAAAAAAGCCATATTTATTGGCATCTCATCTAACAAAAGTAACCAAATTGAAAAAAAAGAAGAAGAAACTAAGACAGTTCAAG
>evgR017813c0g1_i01_Hydra
GTCAATTCTCAAAGTAGCACTCGTTGCTTCCTTCTTGATGGTTTTGTTGTTAGCTGCTCAACCATCCATGCAACAAGAAGAAGAAACTCCAAGACCAGGTGTATTTTTAGGAAACTTGATCCAAGATATGAAATCATTGGGTTTGATCAAATCATCTTCTCTCGAAAAGAAGAAATTCACCAAGGAAGAAACCAAGGAAAGTATCACTAATTTGGCCAGAATGGTTCGTTATTTGCTTGTTTTGAAGGGACAAGGAAAGGACGAAGAAGTTGCTGAATGGGTTAACAATTTGAAATTATCCAAGAAATTCTCTCGTGCTGCCCGTATGGTCAGAATTACAATGAGACAATTGGCAAAGCAGGAACAACAAGAAGGTGTTGAAGCACCAAAGTATGTCGTTCAAGTTAAGGATATTGTCAATACTGTCATGACCAATGCCTTGTCTGAAATGAAATCATTGCTCCAACAAGCTGCTGAAACTGCTGAAAAGAAGGCTCTTGCTGTTAAACCAGTTCAAGCTCCACAACCAACTCCTGATCAAGCTCATCAAGCTTGGGAAGCTGTTGAACAAATGGCTCACAAGATGGTTGAAGCTACCAATGTCAATGCTGCTGAACATCAAACTGCTGTTGAACAAATTAAAGAATCAGCTGATGTTGCTCAAGGTTAAAGAATAAAGTAAAG
>evgR017823c0g1_i01_Hydra
ATGGCGACTGGCAGAACGTTCGGAAAAACCTCGTGTCAGTTCTCTAAGCAAATTTTATCTTACTTAAAAAACAATTCAATTAAATCAGCTAACTGTTTACAATCAACTACATCACGGTCGTTGACTAAAAGATCTTTATCACAATTTTTGCCTGGCCTCCAAGCTCCACTTGTAGTGAAGTATGACATGTTATCTGCCTCAACATCAGGATTGGGCATTAGCTTGGTTTCGAAAGTGCAAGCATTGCAGCTCGCTTTAAACGAATCTGACGTTACATCCGTAGATGAAAGCGAAGAAGACGAAGAAGCTACATTATATATATTGAGTGATATGTTTTCAGTGCTAAATCTTCAAATTGATGGAGGATGAAATATAATTGATGTCAAATCCCAGATAGTTCGAAGTTTGCATCCATATTAAAAGCATGACTTATAATGACTTTATCGTGAATCTTATCGAGGAAAATAGGTGTATTTTGTTATATTTGTTTATTTTGAAATAACGAGTTTTCTATTTTATATACTACTGGTTTGGATGGTAATATTAAAAAACAAACCTTTTTTTATCATAATGAAAAAACTAGTTTCTTTTAAAAAAAAAAAAAA
>evgR017835c0g2_i01_Hydra
CCTATTTTAGTGCAAAAAGTGTAAAAATGTTGCATTCAGAAAAGTAAAGTACACAGTTCAATCAGTCACCTTTAATCTTAAAAATGTCAGCTCCAGAAAATGAAAGAACAATAAATAAAATTTTAAGAGTATTCACTATAGACAGCCAAACTGGATTGCGTCATCCAAAAGCTTTGTTAGTAGATCAATCAATTTTAGCTGGCTGCATTGCAGGGTTTTTTATTGGTGGTAGAATTGGTGGTCGTGTTGGAGCTGTTGAACATATTGAAAATACAAAATTAGAAGTTTACCAATCAGTCACCCATGCACATAGACAATACCATGGTAGAGTTATCTTGGGGTTTGTAAAAAACGGATGTAAGTTTGGTTGGAGAGCTGGCTTATTTTCAGGGTTCTACAGTTTTGTCATGGTTGTTTTAAGCGAGATTAACAACAACAACGATAACATAAACTTGATAGGTGCTGCAGCTGTAACTGGAGTTGTTTATAATTGTTTCCGTGGATGGAGAAAAATGATTGTTGGGGCAATATTAGGAACGGGGATGAGTCTTCCTTTTGCTGGTTTAATTGAAGCAGTTCGTTTTATAACACCTGAGGATAAAAAAGCTGAAGTTAATGTGTTCAAAATTCTAAATTATCAGCCTATCAATAATGATGATCCAGCTTTGAATAGGTTGGATGTTGGTAGCGTGATTAATGATTTTCAGATGTCACTTACAGACCTTAATGCAACCAATTTATCTAATGAGGCTGAAAAATCTAAAATTGAATTTGATCAACAGTTTGAAGAAAATGAACAAGTTAAAACACAGGACAGCAATTAACAAATTATTAATTTTTCTTATGAAAAAAATTACATAATATTAAAAAAAAAAAA
>evgR017840c1g1_i01_Hydra
ATGGTAAAGACACTTTTTATGGGATGGGAATCATTGCTGCTGGAATGATGAACTCTAAAGTTTTAGATAGTCTTCATCGAACTAAAAGAATAAGATATGGTTTAAAGGCAGCAGAAATAAATTCTAAAATAAATATTCCGATTTTGTGGTATGACGTTCCTGACACAGCAGCATTAACAAAGATATTGTTCAAGCCAATTGTTGAGCTTTGGTCACCACACACTTTTTCACTCTCTCTTGGTATTGATCTTTTCTGGCAAAGCTCGTTCACTTCAACAGCTGGTGAAAACCGGGCACAATGGAGTGGGTTTATGCAACTTCATCATAATGGTAAATGTCATCCCGGAAAAAATGAGGTATTTATGGCGCCTATCATTACCTTAAACCCATCTGATGAAAACTGC
>evgR017843c0g1_i01_Hydra
CTAAAAAAGTTATTATCATACATGGAAAAGTTATATTTTTTTATATTAATGTTTATTAAGCTTCAATTGGTGTCAAAAAAGCAAACCATTTTTATGTATTAATTATATACTTGCTGAACCCAAGTTGCAACAGTTATACTAAAATTTAACGCACAAAAATGGCAGACGCTGAGGATTTTGCAGGATTGGATTTAGAGACATACAGTCCATTTGCAATTAGAAATAATTTGGGTGTTGTTGATTATTGTCGCGCATTCTTGGCTGTTGTATCAGGATCAGCAGCTGGAATTCTCGGTTTAACTGGTTTAAATGGATTTGTTTTTTACATTTTTGTATCATTTCTTATGTCAGTTTTGCTTGGTTTAAAAACGCAGTCAAAAAGTGAACGATATTTTGTTAGTAAATGGCATTTGCTTACCAATGGTGTATTCGGCGAACTATTTACTTATTTGCTTGTGTGGACATTTATGTACGGCATGATACACGTGTTTTAAATATATTTATTTATTAATTTCAAATCAAAAAAAAAA
>evgR017857c1g2_i01_Hydra
CTCAAACAACTATAACTACACCTCTTACAACAAAAAATGCTGCTTCTACAACTACAACTAATTCTCCAAAAACTACAAGTGCTGCTTCAACAACAACAACTAGTGTTCCAACAACTACAACTTCTGCTCAAACAACTATAACTACACCTCTTACAAATACAAGTGCTGCTTCTACAACTACTTCTCCAAAAACTACAACTGCTGCTTCAACAACAACAACTAGAGTTCCAACAACTACAACTTCTGCTCAAACAACTATAACTACACCTCCCACAACAACTGCAGCTCCAAAGACTACAACTACTTCTCCAACAACTACACCCACTACTACAACAACTATTATCGTAGCTCCCATAACAACTTCTACTACAACAACAACTGCTGCCACAACAACTGCTACTCCAATAACTACAACTTCTGCTCCAATAACTACAACTTCTACTCAAACAACTATAACTACACCTCCTACAA
>evgR017858c0g1_i01_Hydra
CTATTTTAGTGCAAAAAGAAAATGAGTGATTCAAAAAATGTAGACTTAAAACTTACAGCATCTTTAAGTCGATTTGGTTTAAGAGGACGTGGTGGTGTAAGCACTGCAAGATTGCCATCTTTAAAGCCTGCCAGAGATCTAACACTTGGAGGAACTCCTAAGCGTACTTTTACTCCAAATATAGTTGGTCGTTCATTAAAGAAAGAAGAAGTGAGCGAAACAAAAAATGATTTTAGTCAAAAAAGAGGAGGACATAATCAAGGAAGAGGTCGGGGAAGAGGTAGAAATGAAGTTATTGCTTCATCAGGAATTTTTAGTCAGGGACCAGCTACACAGCAAACATTAGCTCGTGCATTTGGTGGTGGGGGATTTGCTAGTAGTGGAGGTGGGAATGGAAGTAATAGTCCAAACGATCATAATCCAAGTAGTATCAAATATGAAATAAAATCAAGTAAAGGTTTTCAAGAAGAAGATTTTGAATCAAACAAAGTATTAGATATGCTCAATGAAACTGGCGAAATAGAAATGAAAGATTCAAAATCTGGGTTTATGGTTCCAATAACACTTCCTCTTTCTTCTAAAGTTATGCATGAGATTAAATCAGAAGATGAAAGTGAAGATGTAAAGGTTGCTGCAACAAAAGATTTAAAAAATGTTAAAAAAAAGTTAAAACAATCCTCTTATAAGAATGAGTTTCTCCAAAGTGAAATAAAGATAAATGACTTATTTAAATCAGACAATTCTAACGGAGATTTATTTTTCTTTCAATTTCCTGATGCTTTACCTATAAAACCTTTATCGGATGATAGTGCCATGTTAAAACCTGAGCAAGATAAGAATGTAGTTTCAACAGAAACCACTGTGGCTGAACAATTAAAAAAATTTACTTTTCAAAATGTTTCTGAAGGTTATATTGGAAAACTACTTGTACATAAATCTGGCAAAGTTAAAATGCATTTAGGTAATGTTTCTTTTGAAATAATGCTGGGCACTCCGTGTGGTTTCTTGCAAGATCTTGTATCAATTAACATGGATAATGATCCTGGTGAAATGATATGCTTAGGTCATGTAAATCACAGATTAATAGCTGTTCCTGATTATGAAGCACTTTTAGGGAGCTTAGTTACATGAATTTTTAAAATGTGTTTTTGTTTATTTTTGGCTTCATTAATACAATTTTATATTTTAAAGCATTGCTTACTTCATACAAGCCTAATATTTTTTATTTAAATGATATTTAATTTATATAAATATCTTAGAAATATAGATAATATATATTATTGTTTCAAAAAAAATATTGAAATCTTTAGATGTGACTAGGTGCTTGAATTTAATATAATTAACTACTTCTATAAATTTGATAATATTCGAATTTTCAGTGACAGCC
>evgR017860c0g1_i01_Hydra
TTTCAGTGCGTGTCTATTTGCTACTTAACTCAATTGATAATGTGTTCAATGATCTTAAAAATATGTTGTATAGTGATTCCAAGAAATATCTTGAAAGTGTAGAATTTTTGAATTTTGTTTTGAAGCCTGGAGGTAACAGCATCGATTTAATAAACCTCTATAATAATGCAACACAATATTCCGGGTCAGTCAATTATATTGGTGTTATTGTGATTCCTGAAGATGAATTAATTTCTGATACATATTGTATTTATGACAAAAAGCAAGTTATATTAAAGTATAATGAAATAGGTTGTACATTTAAAAATAATGATGGTGAAGGTGCAGGTGAGGTCCATATTGGGGCAGTAAAAAATGAAGAAGAATGCATAGTAATATGCTATGAGAAACAAAAAACAGACCTAACTATAAACGGTGTGACTACTACTATAAATGGTAAAGATTGTTATTGTGAACAAGGGATGACATCACAAACAGGTGGATCAAGTTGGAAAAGTTGCTATATTCCTCTAATATACATTCAATTGATTAATGCTGCTGCTTTTAAAACAGTTCTAACCTTTGTTACTGTTAGTCCGGTACTAGATACTTATATTAAAATTGGCCAATGGTATGGTTATCCATCTACATCCTTGTTTACATATAATGCTAATATACCTTCTGATGTAAATGTTGCAAATGTGAGCTTTTCAATTGGTGGTTTTGGTTCATCTCAGAATTCCGACAATGGGTTTACAATACGTG
>evgR017861c0g1_i01_Hydra
AAACCTTTATGTTTGATTGTTGAGTTTATGGAGAATGGAGATTTGTTGCAGTTTTTGAGAAGTAGACGCACAAAATTTTGTACATCCAAACCTGCTGGAGAGTTAGCAGTGAACTTTATTTATACTCATACCTATAAGCAAACTCTTGAGACAACAGTTAGTGAGATTTGCACCTCAGAATGTATTCCAAATGAGATTCCCCCGGATCAAGTTGGATTAATAATGCCTAATGATCTAGTTAGTTTTGCATGGCAGGTTGCATCAGGAATGGAATATCTTTCATGTAACAAATTGGTACATCGAGATCTTGCTGCAAGAAATATCTTAATAGGTGCAGAGAAAAACATAAAAATATCTGACTTTGGTCTGACACG
>evgR017875c0g2_i01_Hydra
ATAGTTAATGGAAAATGGCAGGATATTATCGTTGGTGCACTTGATATGATTGGGGGTTTAGCAACATTAGCAGGTCCAGCTGGTGCTGTCCTTTCATCTATATTGAGTGTAATATCGTTGTTATTTGGATTATTTGGAGGAGAAAAAAAGGCCGAAGAAAGTCAAGAATCAATGCTAAAACGGGTAATTAACGAAGCCTTAAAAACAGCACGAGCTGAGGAGCTAAAAGCAGATGCTGAAGGTTTGAAAAAAATTATGTTATCTATTCGTAACTCGATTAATCAGTTTCGTGAATCTGGCGTAACTAAAGAACAAGCTATCGAAATGTACACTCAAGTTTTCACGGGTTTAGAATTCTTTGGAAAACTGAAGTATGAAATCAATAAATTCTGCGACTGTTCAGTAGAAAGCGTAGGGTTTGCAGATGAGACAAAAAGAAATGAAAAATTAGCAGATTCTGAGAAGTGTTTGCAATTTCTCAATCTGTACTCTGATCTCTCAATATATAGGTTGTTGCTCATTGCTGACATGGCTAGTCTATTTAATTCTATAAAAAACCCTGATGGCATTGGATTAGATAAAACCATTTTATTACTGAGTGAAAAAGAAAAAGCTTCAGACGTTGACATTCTTATGTTTTTGTTTGATCCCATTAATAATGTCCGTCAAAGATTTTGTATCAGCAAGTATCATAGTGCTCCAAGTAAATATCCTACAATAATGGCTTATACAGATGTACTAAAAAAGAAAGAAATGAAGCCTTCTCTAGATGTTACTATATGTAGTGGAATGGAATTACTTGGGGTGTGTCACAATTTAAATGTTACACAAAAAAATATGTATAAAAGTATGCATGATTTGGACGGATGGGGGTATAGTATACAATCAATTTATGTTCCAGATCACTTTAAAGTCAATGGGCATTCTGAAAAAGACTTTAAAGGATCCAATTATGGTTGGTATTTTGGGCCTACTGTAATCGGGAGAGTGCCACGTGGCTCTGATAACTCGTGGAATTCATTTCAAGCTAATTCAACAGCACTATTAGCACATCAAATGATTCGTATTTGTAAAGAGAAAGACTTTGGACAGAATGGTTTTTGTCATCAGTTAGAGGCTAAAAATGGGGACTACTTAAATTTGAATTCGATTAGTGTTGAGCTATTTGCAAAACAAATACAGTCTGTCATGGTTCCAGAAGGATCAAAAGTTTATTTTTCTGAAACAAGTTCTGGAGGTGTTGAATATGGTCCATTTTACGGTCCAAATTCGATAGACCATTTTTGTTTAAAAGAAAAAGCTCTTTACATAAAGATTAAAAGTAACGACAAAGATGTTTCAAAGATGGTAAAGATTTGCAAAGGAGAATCTTTTTCTCAGCAGTGTGAATATCTTGAAAAAGGCGCTTATCCATCGATGACAATTAATGGATGTATTTATGATGTTGACCATTACGATGGTTGGACAGAGGCTGGTAGTACTAAAGAGGTGAAAATAAAGAGTATGGTTGTTCCAGCAGGTCTAAAGGTTTCTATGTGGACGGATCCGGAATGTAAGGGATTGTATTTAGGACCCTATGTTGGGCCAGTTTCAATGTCAGCTATCGAAGGATTAGGCAGCAAATATAGAGTTAAGTCTTTAATAGTTAAAGACATAGCTGACACATCAGTTACAGAATGTAAAAAGAATTGAAGACAGTTTTTAAAAATAATAGAATATAGTTGCTTTTGAA
>evgR017890c0g1_i01_Hydra
TTTGATTCCCTGTTTTTTTTCGCCCTAGTATTAAAAGGCGAAAAAAAAAAAATATGCCCATATCAATTTCGTATGACTTCTGATAACATCATAATGGCTGGCAGTAATGTCCGAACCACTCATTGGTTGTTAACATACAACAACCCTATATTCATATATATAATAATGTCCGAACCACTCATTGGTTGTTAACATACAACAACCCTATATTCACCGCCGAAGATTTAATTCATGCTTTCGATATGTTTTCTTCTAAATATGTCTTTCAAAAAGAGCGTGGTGTTGAAGGCACGGAACATTTCCAAATATATTTGAAAAACTATATGCTAGACAATTAATTAGCATATTTGTTGATAACATTGGCTTTCATCCTCATATTCAAGCCGCTCGAAGGTGGAAGCAAGCATTGAATTATTGTTGTAAGTCTGAATCTCGTATTTCTGAAACCTATTATACTAATATACCTAATGAGATTACACCTCCCAAAGTTAATAAAATGGTTCAGATTAAACACGCTATCGATAATGGTGCTTTAGTTCAAGATATTATGGAAAATGATGAATTTTATGCATCATGGTGTTCCCATTATCGCTCATTTAATATTTATGCATCGTGTAGAAAAGAACAGAGGGATTTCAAAACAGTTGTTATATATCTATATGGTCCACCCGGAGTTGGAAAATCCCGTCTTGCTCGAGATTTATCAACGGTTTTATCCTATATAAGACCTTATTACAAGTCTAAAGGTCAATGGTGGGATGGTTATGAAAGCAATAATGTAGTTGTTTGGGATGATTATCGCGGCAACTGTTATGATATACAAAAGTTATTAAAACTCACAGGTCGTTTTTATTATCGTGTGCAATTTAAAGGGGGTTATCTTGACTTTACCTTTAAATTTATAATATTTACATCAAATAAATACAGTTGGGATTTATACAAAGATGGTAAAGGGAACCCTTTTCAACGTCGAATCGAATTCAATCATTATATTATGGAATATAACGGTGATCTAGTCAAGTACATTCAGTTAGAATGGCCATTAATGCGCAATGGTGTTGAATTCCCGCGAACATTCGATCAATTTAATCGCGAAGAAAAAGAAAACTATTTTAATATTGTAAAACAATATATTATTTAAGTTATAACTTTTTGTAACATATCTCTCACATTAAAAAATTTATAATAATAAACATGTTCAAAAAACCAAAAAAAAAA
>evgR017891c0g1_i01_Hydra
TTCTATTTATTTCAAAAATTTGTACAAAATGGAGATCGACGAAAGAAGGAACAAGGCTTTCAATGTATATGCTGAAAAGCTTGCTCAACACAGACAATTAGAAGCCGAAGTCAAAAAGATCAGAGACAGCCAAAAGGATTTAAAAAAGAGATATGAAAAGACTGAAGATGATCTCAAGGCTCTCCAATCTGTTGGACAAATTGTTGGTGAAGTCTTGAAACAATTGGATGATGAACGTTTCATTATTAAAGCTAGTAGTGGTCCAAGATATGTAACAGGAGTAAGAAAAAAGATTGAAAAGACAAAATTAAAGCCAGGCACCAGAGTTGCCCTCGATATGACAACACTGACGATCATGAGAATTTTACCTCGTGAAGTAGATCCATCTGTATTTAATATGATTGAATCTAAAGATGTTAAAATAGACAAAACAGAATTTGCCGAAATCGGAGGTCTTAATGAACAAATCAGAGAATTGAGAGAAGCAATCGAAC
>evgR017892c0g1_i01_Hydra
GCGTGACCATAACTTTTGTCGTGAATATGACTTTAGTGAGCGTGACCATAACTTTTGTCGTGAATATGACTTTAGTGAGCGTGACCATAACTTTTGTCGTGAATATGAACAAAAACTTGAGGAAATGGTTCAAAACTTGATGGATCGAAGAGCTGCAGAAAACAATTTAGTATTGGAATCGCAAGATTTAAATCCTATACATACATTACAATCATTTGCAACCATATCTGGAAAAGATATTGCATACGAAGAAGTGGAACTTATAAAAAAAATTGGATCCGGGGGTTTCGGTGAAGTTTATCTATCAAAATGGAATGGTACTATCGTTGCTGTGAAAAAACTCCGTCTTCAACGAATTTCAAAAAATCGTCTAAAAGATTTTACAAAAGAAGTCACAAATTATTATACTTTAAACCATCCGAATGTTGTCAAATTTTTGGGCGCATGCACTGTTACTCCAAACTTATGCATTATAATGGAATATATGGACATGAGTTTGTTTGAAGCCAAGATTGATATAGACTTTACTGAAAATGAGCGTCTTCAAATCATATCAGATGTATGCAAGGGTTTACTTTATTTGCATGAAAAAGGAATCGCTCATTGCGATTTAAAAACACAAAATATTTTGATTCGTTATAAAAATGAAGAAAACGTTTGCGTTGCTAAAATTTGCGATTTTGGATTAAGTATTATTAAACATAATGCAGAAACTTCGGGTTCCAGCATTGAACAGCATTTTAAAAATATTGGAACACCACGTTATTCTGCTCCAGAAGTATTAAGAGGTGAGCTAATAGCTGCATCGGCGTGGTTTATGTGCGACGTTTACTCATTGTCTTTGATTATTTTTGAAATAGTTTACGAAGAAGAGCCCTTCGATAATCTTTCGTATGTTCAGTTACAAAAGCAAGTGGGAGAAAATGGTTTAACTCCTGACATTCCAGCTGAAGTAATCGTTGACATAAACATTTATAATCTCATGAAATCCTGTTGGAAATTTAATCCTGTTCAAAGACCTAAAATATATGAAATTGAGAAGAAAGTGAACCAATTAAGTAGAGTTTATGGTGACTAAGCATACAAGTAAATAAAAATAATTTTTGGAGTTATAGGAATTTTGGTGATTCTTTTGTTAAGGAATATATTTTCAGTTTAAAATTCTACAATATAATTTTAATTTAACAAGTTAAAGTAAGTTTAATTGAACAAATTTGTTGATCAATTAAATGTTTGTTACAATTGGAATATTTAAAACTAGAATAGTTTTTTATTACATTTTAATTTTATTTATATCATTTTTTATCTCGCAATCCTTTAGTAGTTTATTGTCACTTTCATACTTTAATAATCAGGAATATTTTGGTAATAATATTGAATTGAATCCATATCACTGATGTTTCTCAAAGAATAATAAACTTGTATTTTT
>evgR017898c0g1_i01_Hydra
AAAAACAAAATCTTTTTAAAAAATTTACATTTGTAAAGTATGCTCATGATTTAAATAAATACTGAATTAAATGAAACAAATAAACAAAGTTATTAACTACCAAGGAGATTGAAACATGAATGCTCGTACAGTGATACTTGTTTTTCTTTTCGTTTTTTTGTTTTCAACCATTGCTTTTTTGACTGCTTCAATGTTGGGAGATGACTGGTGGTCAACTACATTAACGAAGAAAAATGATCGATTTGGTTTACTAAGGCAATGTAAAATTCCTCAAGGTTCTTCCGAGCTTGTTTGCTCCTTTAGAAAAAATGCATTGACATTTAATAAAAGTTCTGTGTTTGAAAAAGATTTGGATGTAGCAGCAATCCTAAACATTTTATCTCTTGGTTTCTGCTTAATTTCAATCATTGTTTTGATTTTTACAATGTTGTATCCGTTTGTTGCTTGGATACCTGTTCTTGTTTTTATTTTTCTGGTATTCTCTGGTATGCTTACACTAGCATTGGTAATTTTTGTTCAGATCAAGATACACAAAAATATTACAAATCTTGCCGGGCCAGATTTTGGAAGTAAAGGTTGGTCATTTTATATATCATATTGCGCTTTGATTACCGGGCTTTTATCAATAGTATTTGGGATCATTTTTACGGCATTCAGTATAGTTCAACATTCAAAAAAAGGAAAACGTCTTAATGATAAATACAGGGGATCAATCCCGAATGCTCCAGAACTATCAGGAAATTTGTCCACCGTAAGTCGAGAAAATAAAAGCTACACAATTGATGAATTACCTGTTTATTCAGATTTTGACCATGTCGATTTGTACAAATTCAGTCCAGATAAAGAAACGCATCTGAAAATGAGAGCATTTTACTCTGGTACAATAAGCGGCCAAGATGACACGGAGTTTAATTTTAATTCATAAATTATTAATGGTAATTTTAAATATATTAAATGTAAAAAACGTTTTTCATTTGTTATATCAGTAATAAATATATTTTAATATTCATTTTTATATCTGTAACAATATGTTAATACATTTGTTATTTCATCTAATATACTATTGTG
>evgR017901c0g1_i01_Hydra
ATTTTCCGACTCACTTTTGGGAACAACAAAAGCAATGGATGTTTTTAGCATGATAAAAAGTTTTTTTGCAAAACACGACGTCGATTGGAAGAATAAGCTTGGCAGTTTGTGTACTGATGGAGCCCCTGCGATGCTTGGAAACACTTCTGGTTTTGCAGCATTGGTCAAGAAAGAGGTTCCTAATGTCACGATAACTCATTGCTTCTTACACCGACATGCATTGGCATCACGAACTTTGCCTACTTTCTTGAAAGGAGTAATGTCTACGTGTATCAAAATAATAAACTTCATAAGAGCCAGAGCTTTGAATCACCGTCTGTTTAAGAAACTTTGTCACGAAATGGGATCTGACCATGAAGTACTGCTGTATTATACAGAAGTTCGCTGGTTATCAAGAGGACAAGTTTTGAGACGCTTATTTGAGTTACGTGTAGAAGTTTTAATATTTCTGAAA
>evgR017911c0g1_i01_Hydra
GCTCGTGCTTTTTAAATTAAATAGCCAGTAGGAAACAAATATTATTATCAAATAAAAGTGTAAATAGGCTAGTAGCTAGTTATATAGTTATAGTTTATAGTTATAGTTTATAGTTAGCAATATAACTAAATTAGTAGTAATTTATATTATGAAATAAATGCAATATTTTTAATTTCTCTAAAGAGTGCAAATGTGTTCCAGGTAATTCATGATATAAAAATGGACAAAGGACAGCTATATGCGTCACTTTATGACGCCTATGTTAAGTTATATAAACAGAAGAGGAAACAAGATTGTCAAAAAGAAGTTAATGATATTTGGAGTAAGGTGAAATCTGAAGATAATTTTTCTGCAAAAGTAGACTCCCTGTTAAGAGAATATGATGCTCTGTTCAAAAAGAAAAAAGGTACACTGATGAGTTTCTGGGCAAATCAATTATTACATAAAACTATACCTCGTAAATCATCGGCCCCGGATGTTCCAAAATTAAAAGAAAAAAAAAGCGGCAGATACTGTTTTAGGGGGTAATAAATATAAAGCTTCCTCTTGTGAATTTCCTATGATAACTTCATCAACGTCTGGTGCTGAAAATGTAGGAAAATATTA
>evgR017913c0g1_i02_Hydra
GAATACGATGTATCTTTTGAATTATATTTAACTGAATTTACAAATAAATATTCAAATGTAATTCATATTACAAATGGTATTAACTGCTGTGGTGCATCAACAAGAATTGCAACTATATTTTTTCAAAATAACGTTGCGGAAATTGATGCACCCCTTAATGGGGTTGGTAATTATTATTTTTATACAAAACCTTTCACACTTCAGAAATGGACAAAAGTTAGAATAAAGCAGTATTTGAATGCTGGCAGCTATTATTATACAGTTGAAATTGATGGGGAGACTGCTCATTGGGTAAAAAATAATAATGCTGCAGTTTTCACAAATGTACAACTGTTTGTTAGCAATAACTGGCATGATGTACAAGCGGGATACATCAGGAACTTGTTTATACAATGTCAAATAAATCTGTGTGGATTAATAACTACGATTTCCGCTAAACAAATCCAAAGAGATATAAACGTTCGTGTTATCATAACAACAGACTGTGTGTGTTCTATTCAAAATATTATTTTTTATCTGCAACCCGAACAACCGTTAGTGCTTAAAGGTTTTGTTTGGGATAATGATTATAGTTTAAATGATTCGTTTGTCTATCAGATTGGTAAATTCTTTGTCATTAAGGTAAACGATCTTTTCAAAACCTCTAGCTTTTTATTTTCCGCTGTACTTTCAAATGTTCCAAATATAGACACAAATGTGTCAATCGCTGCTCATTCAAGCTGGATTTATCGCGATAAAGATCCTGTTTATAAAGCTACCCAACAAAGAATCTCAGTTTTAATTAAAAGGTTACCAGCACTAATAAAATTGGGACCGCCGGTGAAATATAAAGAACTAAACGAAACATTTTTGGAGTCGGGAAGATTTCAGTTCGTTTGTGTAGACATGCAAAAAAGGCAACCAAGCCCTTGTTATCAACGTGAAAAATCCACCGGAATTATAACATTTCTGCCAATTAATGCACGGAATGTTATTGGATACGATTCACGCACCACTTTAGTTTACGGACGTACAAATGGAAACAATGTTATAGAGATAGATCTAATCAAGATGAAATCAATGGTTATAACAAAAGAAAAATGTTTCACAATAACAGCATGTGGAATATTTTTTAGCTGAATAGGACTAAAAGGTTGGGTAAAAGTACACACTACGCGCCTCATTTTTTAAACATAAAATATTATCTGTAATAATTTTGAAAACGATGGAATATATAATAAAATACTTAAATATTTT
>evgR017915c0g1_i01_Hydra
CAATTCACTCTCGAAGCCAACCCAACAGATACTATCAAAGTTTTGAAACAATCAGTTGAAGATGCTACAGAATACAGTGCTGCTGGCATGAAATTAATTTATGGAGGAAAGGAATTAGCTGATGATCAATCAATTGAATCGGCCAATGTTCAATCAAATGTTGCTGTCATTGTTGTCGCCAAGAAGAAGCAAATCGCAGCTTCTATTGCCAAGAAGAAGGAAGAAGAAACACCACAAACCACCGAACAACCAACAGTTTCCGAAACTGTTCCAACCACAACTACTACAACAACAACACCTTCAACAACCACAACTACCACCACTACTCCATCAACCACACAACCAACTACAACCACCACAACACCAAGTGAACAAATCCCAGGAGGACCTGGATCTACGGAATATGACGAAACAGTGCAACAATTTTTGGAAATGGGTTATGATAGAGGTGATATTGATGAATGTATGAAGGCCTCATTTTATGATAGAGCAACTGCTGCTGAATTTTTAATTTCTGGTATTCCTGAAAATGTTAAGCAAATGATGCAAGATAATGGTGGTAATTTACCAAC
>evgR017916c0g1_i01_Hydra
TGGTGATATAAACCCAGGTATTTGGTTCCATGAAGATGGCTCTGGAAAGCTTGTTATTTATGCAGCTGTAAGTGGTAACGCTTCATACAACATTGTAACTACTCCTCTTAGTTTGGGCCAATGGTCTTCGGTTAAAATTTATCAATGTATCTTAGGTGGAAAGTATTGGTTTACGGTTAATTTAAATGGAGTTGATATTCATAGAGTTGAAAATTCTGATGCCAGAGATTTCAAAAACATAAAAGTGTATGCCTCAAATTATTGGTACAATGCTCAAAATGGTTCAATAACTGACTTACTGGTTGTGAATGGAAAAACTGAGTATCTCGTTGACAATGTCCCTACTGCATTAGTTTATAGAAAAATAATTGCACAGATACCAAT
>evgR017925c0g1_i01_Hydra
TGGTGGTTCTTTGACACTTATTGTGTAACTGTCTACAAAAATCGAAAAAAATTATTTTGAGGTGCGGGCAGCAAAGAAATAACACAACCATGCAATGCCCTTTATGTAACTTCAGTATACAATGTGCTGCAACTGAGTTTGATAGCTTGGTAGAACATGTGCGCAGTCATATTAAACCAGAATCAGCTTCTGAGTGTACTTCTTATCAAACATTTTGTAATGAGAATGTTACAAAAACTCCTGATAGTTTTTTTATTGAAGACTATAGAAGTGTGAAAACTAATTGGTCGCTCTCACCAGATATTGTAGTAAGTGATCCACGCTCACTGAATAGTTTAGATGTTGTTTCAAGGGAATCTGAAACTTTGAAGCACCATGACTACTCAGAAAAAATATCCGGTAAACCAACAGCAAGAGTAACTTGTCAGCCAGTAGAAATGAAAATATTTTCTCCAAAGGATGGACAAGAAATAAACTCAGCCTCAGTTTCTAATAATTTTGCACAAACTGAGATTATTTTTTCACAAATTGAAAGTATTTCTCATCCAATTGACACAATTTCCTTGCAAGTAAATGACAGTTCACAATTTAGTACGTCCAAAAAACTAAATCATTATTTAGATCATCACAGCCATGAAAATTTATTGAGCATAAATGAAAATAACATTTGCTACCCACCTGTTTTTGCACAGCCCCATAGTTCATTAGAACAACCATATAGTTTACTTCAACAGTCCCGCAGTTCAATAGAAGAATCTCATATACTAGAAGATCCCCATAACTCATTAAAAGAGCCACATAATTTATTCAAAGAGTCGCATAATTCATTAGAAGTATCAAAAGTATCTCATTTACCAGGGACCTTTTCTAATCATTGTTTTGATAATAGTTCAAATAAAGAATCAAGTTTATTGAAATCTTATGACAAAGCAAACCGTTTTTCATTTTATTTACATAAAAATGAAACACTTTTAGTACGTCCCCAGGCTACTAACGACTACAAACAACCGGTTGCTTCATCATTGCACAATCTTGAAGGATTGTTATTAATTCCTTTAAGGAGCGCAAATGAAGTTTCTATTCAAACTAATTCTGTGGAAAATACTTTACTCTCTAAGCAGGTCCAAGTAGCATGTATTCAAACTGAATTGAAAATGGGACAAATATCTGAGTGTGACTTTAATATTGAAGCAACAGCTTTTCCAGATTATTGCTCAACCGAAGGATTGTTTGAGTGTCTAATATGCGGTGATGATTTCATTTCTAAAACATTGCTTCAAGAGCACAACATTAAGCAACATTATTTATTTTGTGCTAAATGTGATCACGTGTCTTATAATGCAATTGGAATGTTAATCCACGCAAATATATGTGGGAATACTCTTCAAATGATATGTCAAGAGTGTAAAGAGGCTTTTGTATCAATAAACAAATTAAACCAACACAGAGTTAAAGCACATAAGGTACCTATGCCGTTTCGATGTGGTATTTGCAGTACCCCTTTTGAGCATCATATGGACGTACTAAATCAC
>evgR017929c0g1_i01_Hydra
TAAAAAGACCCTTATTGTTAGTTTTCGAAAAGCAAATGTTACTGGTGAACGAGTTAAAACCAGCGGTGGATGTAACTAAGACTAGCTTAGATGAATTAAGCAATGAAGATATCAATGATATTCTTGACTCATACTTACTTAAGTTTAGAATCATTGAAAACGGTGGTATAACTAATCTTGTTTCTACGTATCCAAAAGAAAACCATGAATTGAAAAAATCGAATCGAGAGTTCGTTGAGATTGAGCTCAGAAATATGGCTAATCTTAACTTGGAATGTATTAGTTTTGCCATAAAATTAAGAAAATCTGCAATCAGCATCATTTTGCCATTGG
>evgR017939c0g1_i01_Hydra
TTGTGCGTTATTGTTGCAACTGTCTCCGCTTTGCCATTAAATTTCAAATACAATACTCCATCAAAGGCTTTGCCAGCTGATGTTTTAGCTTTCATTGAAGGATTTGCTTTGGGAATTGAAGCTGATGTTGGTAATATTACTCTCTGTACCAGAGATGCCTCTGTCACTTTGAAGGATTTTGAAAATTCCTTCTACTCTTTGGAACATGGCTTTAAGATATTATCTCCAAAACTTATTGAAACTGGTTTGATTGAATTCGGAGCTGGTGTTCAAGAAGTTGCTGTTGCTTTGAAGGATTGTAATGTCAACGGTATTGTTGAAAAGATTGCCCAAATTGCTAAGGAATTACAGTCTGGTCCAGCTGGTATTGTTAAGGTTATTATTCATGAAGTTGTCAACATTTTCCACCTCGAACGTGATATTGCC
>evgR017957c0g1_i01_Hydra
AACTTTACAAAGTTTCAATTATTGAGGGGATGTTATATTTATAATACATCTATTTATTATCAGATAATGTCAGAAAAGAAGTCTGAAAATTGTATATGGGTTAATGCAGAGCCTTTTGATATAGGTGTGATACTTGTTCCACCTCCTAGCAAGTTGACTCATCATAACTTAAGAAAGTTGCAAGTTTACGCAAAAAGCAAAGGTAAAGCCGGATATCCTAAACTCACGTTTCCGGTATGGCATCATGTTGCATGCAATAAACTATCTATAGATGAAAATCTGGCTTGGATATATTTTCATACATGTTCTGTTCTGACTGGAACTCCTGGTTATGATTTAAATCTCAGGAATGAACACAAAGATCTGTGCAAAGAGAAGCGAAGTGTTGATTTGTTGAAATTTATATTGTATTTATATATTCAACATTCTAATGTCATTTCAATGAAGTCTGCAATTGTCACTGGGGATGAATACCCATGTCGAAATCGAGGCCAAGAGTTAGAGGGTAGAGCTGCCATTGGAGTAAAGACTATTGATGAAAATTTGCAAATGTCATTTGTCATGGTTCACATTTCAGAAATTCTTGATTTGCTTGTAGAACCAGACATAACTAATGCCAAAGATGAATGCCTTAAATTAGAAGCTGTTGATTCATTGAGTTTTCTATTTGGTTCAAGTTGTAATACCCCTATAGGGGAAATTGCTTCACAACCAAGAGAAGCTGAAGAATCTGGATTCTCAAAGATATCTCAAACTTTCAACTCAAAAAAGCTACAAAGTTGGATAAAATCAAATATGCGTCCTAACCCATTTGGTGTTTTATCCTGCTTATCGTCAGGCGTAAGGTTGAACTGCAGAGGTTTTAATTGTGGTGATGACAGTTTAAATTCAAGCATAAGCTTATATGATGAAACAATTTTCAATAGCACATCATTCACAGAAGTTAACAAAGATCAGCATAATAAAGAAATGGATATAAATTCAAAACATCAATCAAATATTGTTAATAAAGTCATTTCAAATACAAATTATGCACCAGAAGTATACAAGCGATTAATTTGTAATCAAATTTGTAAACGTACTGTAGCCCGTACAGGAAACTACTTAGCACACGCATCTGTGAATATTCATAGATGTCAATATTCACACCTTTATTTGCTATCCCATCTTCGCTCTGTTGTAATTGAAAAGTGTCATAATTCAACCATTGTTCTTGGATCAGTGGCTACATGTGTGATAGTGGTTGCATGTAAAAACATATGTTTGATTGCCGTCACCAAGCGATTGTCTATAATCTCCTCTTCAAATAGTTCTTTTCATGTTTGTGTTCCATCAGAACCAGTGCTGATTGGAAAAAATGAAAATATAAAAATGGCACCTTATTATACATGGTATCCTCAGTTAGAAAAAGACATGAAACTAAATGGTTTGCTTCCTGAACCAAATTATTGGAACTTGCCTATAGTCTTAAGCTGCAAAGATGACTGTAAAAGTATATGGCATGAATTTCCTCCAGAAGAATTTTCAAAGTTTGTTATACCATTTAGTTTTGATGGGACAACAAAGGTTTGTCCTGTCAGTATACCTAGAAGATATATAGATGTATTAAATGAACGAGAAAAACGAATTGAACAATGGCATCAGTTAATTCGGGATTCAAACATGAGTAAGATGCACCGTGTTCAGTTACAGTCAGTTGTTAATGAGAGATTTCAGGAATGGTTAAAAGAGACTGGTAATATAACGTTTTTGGAATCTCTTGCTCCATCTCAAGTTCGATAATTTTCAAAAATTACTTTATTTATATGAAAGTATGTCATACATTGTTATCTTTGTTTTAACTCTTTTTTACTCAAAGTAGTTTTACTATAATAAAATAGTTTTACTATAAAACTTTAAATATATAGTTATAACTGTTAATTATGATATTGATTATAGTTTTGATGTTCA
>evgR017958c0g1_i01_Hydra
TGAAAAGAATTTCGAACTTGCCGTAAAGTATGTCCAAGATGGCATGGCCAAGGATCAATATAAAGATTCCAACAAGGAATTGACCGACAATGATCGTCTCCTCTTCTATGCCTTCTACAAGCAAGTTACTGAAGGTGATGTTAAAGGAACTCAACCTTATGCTATCAATTTTGTTGCTAGAGCTAAATGGGATGCTTGGAACAAGGTCAAGGGAATGACAAAGGAAGATGCTATGCAACAATATGTTGATTTGTTGACCCAAATGGCTCCTGAATGGTTGAGCTTCTCAGGCTTGGACAAGTACAGAAAATAAATTGTGTAAATAAACGCGTG
>evgR017961c0g1_i01_Hydra
ATTTTAGCAAACCTGATTTATTATTTTTTGGTTCTCTCGTCACTGAATTTTTACAAGAATACTGTTTTGCATTTAAGTGTGGCTTAACTTTAAAGTTTCATAACCTTATTCATTATCCACGTATTATTAAAATGCTTGGTCCGCTAAGTCATATGTGGGTAATGCGTTGCGAAGGTAAATTAAGAGGTTTTAAAAGGACTGCTTCATCTGTGGGAAATTTTAAAAATGTTTGTAAAACAGTAGCTATTAGACACCAAATGGATCAGTCAACTCGATTCATGGCAAAACATGGCCTAAAAAATAATGAATTTTGTGTTTCTAAAACTGAGCCTATTTTACTATGCCATGTGATTGATGGAAAAACTATATCAGAGCTGATTGGTAATTATGGGTTGTACAGAGAAATTTTCCAGACAAAGAGTGTTTCTGTTAACTTCGTTAACTTTAAAATTGGTGATGTTATTATATATGCAGTTGGAGATTTATATCCTGCATTTTGTCTAATCAAACAAATTTTCGTATCTGATACTAATTTATTTTATTTTATTTGTCAAAAGTTGTTTATTATAACTGAATGCTATCACTTGCAATCTTTTGAGGTTGAAATTGAAACAGAGTTAATGGTTTTAAATTGCTCTCACTTTGATAACATTATCTCACCTTGGCCATTGAAACTACGAAATTTAAATCAAAGTAAATACTTGTCATTAATGCACAAAATATAACTATTGATCTCGGCAGCATATAACTTGGTAGCATTTAAATTTATTTAATATTTTTCTTTTTAATAAAAAACATTTCCGTAAG
>evgR017962c0g1_i01_Hydra
AGTTAAAAAAATAAAAAGACAGCCACCACTAAAACCAAAAAATAAAGATAGGAGAGCACGAGAATATTTAACTGAAGCAGAAGTTGATAGTATTATGAAAGCTGCACGTAAGATTGGTAGGCACGGTAATCGTGATGCAACGATGATTTTAGTAGCTTATAGACATGGATTACGTGTATCTGAATTAGTAGCACTGAAGTGGCAACAAGTTGATTTCAAGACAGGATTAGTACATGTGAATCGTTTAAAAAACGGTATCGATAGCGTACATCCATTATTTGGCCCTGAATTAAGGGGACTAAAACAAGTGCAACGTGATTATAATGAAACACAATACGTATTTACTACAGAGCGCAAAGTGCCCATGACTACCTCTACTTTTAGAAAGATTATGATGCGTGCAGGCAGCGAAGCTAGTATTGGATTTCCT
>evgR017969c0g1_i01_Hydra
GCTCAATTATGGTTTTGATATCACTAGAACGTTTTGTTACAGTAGTGCACAATAACAGTTTTTGTTGTGTTAATTTTAAAGGTGTTTACTTAATTCCAATCATAATTTTTCAGTTTATTATATCATTTGGATTAGCAGTTTGGTACGCATATCTCAGTTATTTACCGTTAGACCTTCGTTTTCAACTTTCTTTCTTTGTTTTTACGTCATCGACATATGGAATTATTAGTTTACTAACGGCATTTATAATGTATACTATATTATTGATTGAAGTAAAACATAAACTACGACTTAAAACAACTTTTGTTGCAAACAATGTCGAAGAAAGACGCTTGACAAAAACTATTATCCTATTAACATCATCGCATATAATTTTTCACCTTCCTCTGGTTTTGGCTGGTTATGTTTTTGCAGAAGGTATATATTCTGATAATCAAAG
>evgR017969c1g1_i01_Hydra
GGTCGCTACTATTGTTATGCCGCTTCAACTTGTTTTATTAAAAGTAAATTTAATCCCTTGTGTATTAAATCCCGTTATAGGGTTTTTCATTGTGTTTTTTATGGTTTTTTCATGCTCAATTTCGGTTTTGATATCACTAGAACGTTTTATTACAGTAGTGCACAACAAAATCTTTTGTAGTGTTTATATTAAGGGTGTTTACTTGATATCGATTATTATTTTTCAGTTTATTATATCATTTGGATTAGGAGTTTGGTACGGATTACTCCTTTTTTTACCGTTACAATATCGACAGTTATCTTTCTTCTTTTTTACTTTATCGACATATGGAATTACAAGTTTACTAATTATATTTATAAATTATATTTCATTAATGAGTCGCACAAAACAAAATAAATTTTTTTAAAGCAATATATGTCGCTCAGTATGTTGAAGAAAGGCGGCTGACACAAACTATTCTACTATTAACAACTTCATATATAATTTTTTACCTTCCTTTAGTTTTGGCTGGTTATTTTCTTGCAGAGGGTTTATATTCAGATAATCAAAAATCTTTAAAAGTTGTTGTCTATAATTGGACTTTTGTTTTTACCGAGCTAAACAGTGCCTCTAATGCTTTTGTTTATTTATCCAGAAATTGAAAATTCATGAAGATTTATAAAGATTACTTACGTTCGTTAGTTAGAAAGTTGGTATATAAGAAGAG
>evgR017972c0g1_i01_Hydra
TTTGTTTTGCAGAGTGCAGTTAAAGAGTATTCGAATTCAAAATGAAAATATTTAACTTCTAATGTTAGAATAAGTTTCTTTTAAATAAAAAATGGCAGCCAAAGTATGGAAAATAGAAAAGTATGGTCGTTTTAAAAAAGCCGACCTAAATATAACCTTAACAGAAAGCAATAAAAGTGGCGAATGGGAAATTTTAGCAAGCTTGTCAAGCAAACCAATTTTATTATCAGTATTGCCTCCATTGAATTTGGTTCTGAAGTTTGGTTCAGAAGTTCTCGAATCATTTACTTTGCTGGAAACTTCTAATTGGCTTCGCGCTATATTAAAAAGCGACAGCATGTTATTTTTCTATAAAATAAAGAATGATTTCAGAAGATTTCGCATAAAATTTTCAAAAGATGATTTTAAAACCGCTGAGGAGCAGTGCAATGATTTTGTAACTTTAATTGGACCGCATGTTCATGTTCAAGATGCCAGCAATTCGTCAGATTTACAGAGTTCATTACAGAGTTCTGAAGTGAACTATGATCCTAACAGTGCGATAGCATGTACTTTAAATAGAACAAAGGGAGATACCTGTCAAGTTAAAACTGTTACAAGTGTAGCTCGGAATGTGATGATGCAAATACACGATCTTCCATCGTACTATGAAACGACCAATTTAAAAATGAACGAAGATATTAATTCTAACATGATAAAAACTTTGTTACTTGATCCGAATTTTCCTGCATTTGTCGAAAGTGTAGAGAAGCTTCTAACTGAGATGACTAAATAGCTAACTATGTAGCTAGTTTTTACATAAAAAATATTAAAGATGCCGAATATTTTTTAAAACCTTTACAAAAATTATAGTTTCTAATTAAAGAAGATATAGAAATACTAATGTATTTTAACGATTTTTATTAAACAAATTATAAAAAAGTTTTAAGAGATTAAAAATAATATGAAAAAAAAAAGCAACAAAC
>evgR017983c0g2_i01_Hydra
GATGATGTTATCGATCCATTTCTTTTAAAAATAAAAAAAAAATTTTTTTTTTTTAATTTTTTTTTTTTTTTTTATCTATCAGAAACGTGGCTACATTGGTCAAGGCAAAGGGGGGCCAAAGGGCCTTATGGCAAACCAAGTAGATTTTTAATGGTATTATTTTTTGGTTCTTTCCACTATATAAAAAAACACGCTGTTGCCTATCGGCTAGGCCCCCAAATGGTGGGGAGGGGAGTAACTTTTGGGGGTTTTTTTCAAATTTTTTGTTTTTTGAAAATATATATATATTCTGAAGCAATCAAAACTGTGGCTACATGGGGCATGATGAAGGAGGATAAAGGGGATATCACTATGGCAAACCAAGTAGGCTTTAAGTGTATTTTGCCCTCCATGTGATCAATACCTCGGTTTCAGAACAACCACTATCCACAT
>evgR017997c0g2_i01_Hydra
GTTTCAAAAAGAGTATCGGTTAGAAAGATAGTTTAGACAGCTTTAAAGTCGCTCAATATTTTTTTTTTAGTTAATTTTTTCAAAAATGGCGGGTAAAGATCAAATGAGTAAAAGGTGGCGCAGTTTGGCAGAATCAATAAGACGACCGCTTGCTAACACGAGTGCTATAACGAGTTCTGAAGTTGGAAACATGGCTGTTATTGGAAATCCGGACTGTCTAAAAGAATTATTTATTAAACAAGTTTCCATATCGGAACTTATAGCTAATCGAAGTCTTATTCCAGAGTACAACGCTTGTATTGAGAAAACTTTAAACGAAACGGTTACTGGAAAAAAGGACTCACAAAAAAAATCAGGAGAAGATCATGCTTCAACAAAAAGAGCTTCACGCCTAAAATCAATGCTACGAAGATCCTCATCGGAAAACAGCCCCGTTCCACGCAATAAATTTGACTTCAGGTTGTTGGATATTGATTGCAGTTGTAGAAAAAAAGCGTCAATCACTTACAAGGCAACAATTACAAACGCAGATATTGTACTATTCTTTGCTCAAGAAAATAAAGAAGTTGACCAAATCAAAAAATGGATCAACGAAATTTCATGTCTTAGAAAATGCGAAGAAAATCCAGTGATTGTAGTCTCCGATAAAGGAGTTTCTATTAAAGAATTATGCCAAAAACTTAAAAATGTGTATTTATGCGACTTGACAACTTTAAACATGTCCATGAAAGACTCAGAAGAACGAAGTGCTTTATCAATAGAACGATTAAACGCAGTTTTGTTAAATTCCATGGCCGAATATATTCAGAATAAAACCGTTAAAGATCGATCAGATTACATTTTATACGGACGATAATTCAATTTACTTTTCAATAAATTAAATGTGTGAGCATGTAAAAAAATAGTTATTATAAAGTATCGTAATTAGTTATAAATTTAGTTAGTTTTTGTGTAATGTATTTATAAAAATTATCTTGTTAATCATTTTAATGTTTTACATTCAAAATATAACTTATATTGTATATTCAAAATAAAAAAAA
>evgR018008c0g1_i02_Hydra
AGAAACCAAATTAATTCCATATTGAGAGAAAAGTTTGCTCGACATTCTCCCAATGCAATGGTTGCTCTGATGGAATCAGTTATTGAAATGAAGAAGGAAGGAGTTGATGTAAATTCATGTAAAAAGAGAAGAGATGTTGTTGATTTGGTGAATCAAATGATTGATGAGAGAACACAAGAGGTGACAAACTATTCGAAACAAGGTAAATCTAATCAGAAAAGAAGAGATTCAACACATGCTATTTGTCAATTTGTAAATATTCAGCCTCAACAAATTGTTTCTAAAAATCAATGTTCGAATAACA
>evgR018009c0g1_i01_Hydra
TATAAAAAATATTTATTGAAAATGGTGCGGACAATTATAGCTTACATGCTTATGGTAACAATGTTTGAAGCGACACCACTATTCAATCCGCCATGTCCTCTCCGAGAGCGCGATGACCCTATGCGGATATTAAAAATATTACTGGAAAACTACATTGTTATAAATTCATCTATAGAAAAAGTCCCTAAACATTTTCAAAATCACGAGTTTCGAAAAAATGACTCGGTTAATTTACACACGTTTTCAAAAAAAAGACCGTTTTGTGAAACTAAGTCGATAGAAATTCCCGATCTATTTGGTAAAGTTACTGTTCCTGAAAAAATCGTTGAAGTAGCACTTGTGAACTCTTTAAACGTGAATAAATGCAAAAAAGTAAAAAAAGAAATTATATATTTAGAGGAAGAAAAAGATTGCAATAAAAATATTGATGTACGCATGCACGTTATAAAGACCAGAGAAATAGTAGTAGCATATGAATACATGATTTAATTAAATTTAAACTTATCTCAATTAACGACTGTGCTCCTTGGTAGCCCAAAGAATGTTTATCAGTGACGTTAAATAAAACGATGTTTATCGATGCCGTTTCTTTTAAATCATGTATTTGTAAGCTGTTTAATATTACTCTGGTTTTTTAAACGATAGTTTTTAAATCTATTTAGGTAGATTTAAAAAATATCGATTAACTATAAGCTC
>evgR018013c0g2_i01_Hydra
AATTTGGAAGGCGATCGAAACGGATTTGCAGATGATAGTATTTAAAGAGACAATAGCAACATTTCGTAGGTGGACTTGGCATCATGTTTTTGTTTTTATTTTGACATTTTTTCTTTACGCATGTTTTCATGCATGCAGAAATGCTTTCAGTAATATTAAGAATGTTTTAGCAAAATCATTGAGTCCTCAAAATTGGACCGAACCATTATATCCTTATGAAACTTGGGAAAAGGTGCACATGTTTGATAATTCTAATGAAGCAAACGAGTTTCTTGGAATGCTAGATACTTTATACATGCTTGCTTACTCAATTGGATTATTTATAAGCGGGACAATGGGAGACAGATTTAACTTGAAA
>evgR018025c0g1_i01_Hydra
TGATCAGAATGAAGTCTTTCTTCAAAATACAAATAAATATGGAGAGAATGCAACATTTTGCATGGGTTATAATATTTACTGCAAAGTGAGTTCGCATTACGTTGGAAATGAGAATAATAAAAGTTGTGAAAGAAAGAGTAACGAGTTCTTTGCAGGTATTCAGGTAAGTCCAAAGACGCTAGAACTTTCAGAAAATGGTAAACCACAGAAAGTTTCTGTCACATCAACCGTACCAATTGTATGTGAAGATGGAACTGAAAACTGTGATGTACTTCTAGAGTTGGGTCAAACAAAAACCGACAGTTTTGTTGATTACTGTTCATTAATGTTTAAACCTGGACCAGCTAGACAAAAGCAAGAAATTGAAGTTGTTGCTAAGAGAGATTTTGTTGATGATGGTAATCAACAGATGTTTTTACAACTTTTTG
>evgR018032c0g1_i01_Hydra
TGTGGTTTTCCATTTTTATGTCTTTTTGGATGAAGTATAACAAAATTGTTTTGTTTTATTGTGATCCAAACTTTTGTGCAATTTGTGGTGCTATATTGTCTTTACCTGATCAATCAAAAAGTATTACATGCAAAGCTTGTGGCACCATTCTTGATATATCTGCATTAGAAAACATCGAAATACACTCATATAAAGATTTTAATCAAGACAAGTTAAAGAGTTTTGAAGAAGTATCTAGATTAAAGAATGCTAATGAAAAATGCGGGCCATACGTAAAGAGAAGATGTCCACAATGCAATTACAAAAAAATGACATATACAACAAGACAAACAAGATCTGTAGACGAAGGGCAAACTGTGTTTTACACTTGTTTAAAATGTCAATTTACTGAAACAGAGTATTCTTAGAAAAAAAAAAA
>evgR018072c0g1_i01_Hydra
AATTATCAATGCCCTTCAAAACATTCTGACTTTGCTCGATTAGCTCGAGTTTTAACAGGAACATCTATTGGTTTAGTTTTGGGTGGTGGAGGTGCTAGGGGGATTGCTCATATTGGTATCATCAAAGCTTTGAGAGAAGCAGACATTCCAATTGATTCAGTGGGCGGTACAAGTATGGGAGCATTTGTAGGTGCGTTATATGCAGAACAAGGAAGTGTTGTAAAAACTATCCAAAAATGCAGAGAAGGATGCAAAAAAATGGCATCAATGTGGCGTAAAATACTTGATATGACTTACCCATACACCTCTCTTTTCACTGGGAAAGCTTTTAACTCAGAGATCCATTCAACATTTGGCGATATAAAGATAGAAGATATGCTGCTCCCTTATTTTTGTGTAACAACCGAT
>evgR018073c0g2_i01_Hydra
ACCTACACTACTATACCTGGATATAAAAGGATTATATTCTCTCAAGTTTACTATATAACAACAATGAAGATTTGGTCAAGCTCTACATCAGGAATATCAATTACTCTCAATACAGAAAACTCTGACTATTTTGATTCCTCGTATTACACTTATGTTGATCAGTATTATTTGAATAGCGGTGAAATTCAATATGTTACATTTAAAGGAGTTCTTCAATCAAAACTATTTCAATTAACGTCCAGTTCAACCGATTTTGGAGTTATAAATTTGGAGTTTTATGGAAGTAAAGCCTTGTTGCTGATTGACGTTCAGTGGCTTTTTGGTGAAGTACAAGATCTCGTATTTTTTCAATCAAAGGTTTTTGCTTTTGCCCTAATATTTGGTGGTCCTCTTGTGACAAGAAGTTGCAATTTGTTTTTTCAATACAAGCTGTTTATCCCCTTTACTATCAGCTGGGCTAGGTGGATACCCTTCAAAATTATATGGTTATGGATTTCAT
>evgR018096c0g1_i01_Hydra
TATGGGGAAGTTCATTGAAAAGTAAATCAATGCTGAATATTGTATAACTTAAGAAAAAAAAATTTATGTAATTAAAAAAAAAAAAAGAAGTGTGTTTTTACGTAAATACAAGATGAAGACTTTAGTTTTATTTCTCGTTTTGAGCTTTTTTCATCAAGTGCGGGCTTTGTCATTTGACCCAATAAAAACTGTTGATCAACTAGATATTAACAAATATTTGGGAAGATGGTACGAGATCTCAACTAGCTGGATTCCAAGATTGACATTTGAAAAAAATTTGGTTTGCGTTACTGCAACTTATACACTCCTACCTGATGGCAACATTAATGTTTTTAACGCTGGTAGAATATCAAAGCCAACAGGACTTGCGAACGAATCTACTGGAAAAGCAGTCATTAAAAGCGTTGGATCGCTTCTTGTTACTTTTCCGAATTCATTCAATTTTTTCAATGCAAATTATTTTGTGGTTAAGTTGGGAAGTCCTACATTTGGTACGCAAGGGCTTTACGAATATGCTATAGTCACCACACCGTTAAAACTAACAGCCTGGGTCCTTGCGCGAGATCCCCAATCGTTTTCTGTTAAGTATCAAAAGGAGGTAACCGATTATCTAAAAAGTAATGGATATAATTGGTTTTGGAACATGCCTACGCCAACTTTACAAACTCCAGAATGCATATATCCGTCATGATTTATAACTGGCTTTGCACTAACCATATTGCACTGATAATAAATGATTACGTCAAAACTTGTATTGTATTGTTATTTAATTTAATATTTTTTGATTATATTTGTAATATTTTATAATGAATTTAATAAATAACGTAAAGATCTTTGTGTTAAAAAAAAAAAAA
>evgR018106c0g1_i01_Hydra
ATAAATTTCATAAAAACAAACACCATGCTTTTTGTTTTACTTGTGTTTATTGGAACGGTGTCTAAAACAAGTGGCGAACAAACATTGCCTGAAGAAAAGTGTTACCGACTTAATACCACCAGTGTTCAAATAGCAAGTAATAAACTTCTTAAAGAAATGAGAAATCACCCGATTGTATGTCAAGGAAAGGCGAAGAAGAATGAGAAGTTTTGGAATACTCCAGTACCAAATTTACCTTATGGTTCGGTTATTAATTATTGTAACAGTACAACTTGCTGTAAATCGAAAAAAACAATTTTAAATAACTTGATTAATCCTTGTCGAGAATGTGCACATGAAATTTTGCTACCAAAAAGGTTCACTCCCAGATATTTAAACTTCGTAACGTGCCAAAATGTTGACTATTCTGAAACTTGTTTTAAAGGGGAGGGCCGTTGCGTAACAAATCAAGTCAGCAATATGGTGAAGAATTTAAATGGAGATTTTGAAATCATTCCTTTGGGTGCTTCTTGTTCTTGCCAGCTTAACACTAGCTCTTTTTTGGCAAAATTCATATAGACTTAATTCGGATTTTTTGTTTTAGCAAAAC
>evgR018109c0g1_i01_Hydra
ACTAGGCGACGCCAGATTACCAAAGGACGTTATCGTCAATCAAGTCTCCCATGCTACACCACTCTCCATCTATGCCGGAGAAGTTGCTAAATCATCATCTGCTTCATCTGTTTCTGCTTCCCGTCTCCCAGTTTATGCTTTGCAAGGAAAATCATTGTACCCAAAGCACTATTTTGGATTTTTGAAGAATGTACCAAGTGTTTGGAGTCCAAGAGTTAAGTCTGCTGGTTTCTGGGGTGCTTCCCTCGGTTTCTGGGTTTGGTATGCTTTGCTTCCACATTCCAACATGGTCTTGAGAAGAGTTTTCCTCGGAGAAGTTGCA
>evgR018132c0g1_i01_Hydra
ATGGTTGGCTTAGTTACGGAAAATATTGCTACTTTATGTATAGTGCAAATTATACTAATCAGTTAACTGTTTGGACTAATTCGCTTTTGAAATGTCAAAGCAACAACGGAAGTTTATTGGTTATTGATGATGAAGCAGTTGCTTATGTTGTAGCTAACTACCGTAAAAATATCACCATGAAAAATAAACGATTTTTGATTGGTGTAAATGCATTGCAAAGATCATACGTGTGGTCTGTTAAATCTGTGTTTTCAAATTGGATTGGTGACAAACCTGATGACTTTTATAGAGTGGATGACTGTGTGAGTATGAATAATAATGGTTGGCAGGGCAACAATTGCAACAATCTTTTTGGTTATATTTGTAAAGTTAAAATAGATACTTGCTCAAGTATATGTCATCGTGATGCTGCCTGTGTAGGTGCAACTTGTATTTGTAAACGTGGTTACATAGGCGATGGAGTTAGTAACTGTTCATTTCTAAATGGATGTACAAATGATAGTTCTACAGACTTTCAGCTAACATCAACTGGTTACCTGGCTAAAGATATTTATTCTTGTGTTTCTTTTTGCAATTCTTTTACGCCTCCAGGAGTGTTCCAATATGCTTTTTTTCTTGAAGGACAGTTTTGTTTTTGCAGAAATGCCTCACTTCCAGAGTTAGAAACTTCAAAATTTAGCAGTCCTTGCGTTACAAGTTCATGCTTTAATAACATAAATGATTCTGATTGTTTCCTTAGTACCTATGAAGTTTTTTTGCTCTCATTTGGTGTTGTTCAATGGAGCATTACTTATAAACCGGACACTACACTTGTAAACAATGAAGTTCAAATACAAGCAAATATATTGATAGGTGATACATTGGGTGCATTTATGATACCTAAATCAATTATCTCTCCTGCATTTGTTCAAACTGTCAATAGTACAACCGTAAAAAGTCATTTTTATGTGCCTGGTCTACAGGAATGGTCTGCTGAAATAGGAAATAATTTCACAGAAC
>evgR018143c0g1_i01_Hydra
GTTCAACTAAATGACCTTTTTCGAAGTTGACGTACTAAGATATTTTGATATAGTTTTCAATCAGCTGGATAAAATTAGGTATCAATTTGATTGGCATTTAATAGGTCCGTGAAAATGACGTCTAGCGAAACCGCAAACATTTTCCTTTCCGCAATAATTATTGCCTCAATTGTATTAAACACAACAGCATGTTATATCATTGTTTTTAAAGTGAAAAAGAAGAAGCTGACTCACCTTTTCATCATCAGCATCTCAGTTGCAAATTTGCTTGAATCTATAATTGGATTAACCCCTCAAATAGCAATATCTGATAAACCTCTATTGAAAACCATACCGTGGTGTGTCATAAACACTTTTGCGGTTATCGGATTTGCAATAACAAGTATAACTCACATATCGGTTCTTTCTTTCATCAGAGTTGTTTTAATAAAATATCCGATTTTCTATTATAAAACCTGCAAAAAAATGTGGTTCAAACTAACTTTAGTGTTAATATGTTATATATACGGCTTTATTTGGGCAACATTTCCTTTGATTGGTTGGTCAAAATATGAATTAGATCTTGATAAGATGCGCTGTTCGTTGGATTGGAAGTTGACACAAGGAAATTCGTTATCTTTTGTTCTGACACTTTTGATTTTTTGCAACATTTTGCCAGGATTCGTAATTACATTAACAACGCATATCGGCAAGAAAACAATTTCTAAACGCC
>evgR018151c0g1_i01_Hydra
ATGTGTTTTTTTTTTCATAAAACTTAGCAGCACAATAATAGCAAAAAGAAATGCTGCGTTTCAACAAAGTTTATTTGTTCAAAAAATCAATAGAATTGAGTATTTTCTATTCTCGTAGTACACAGCTTGCAGTTAGACCTTTCAGTAATGAATCAAAACTATACAAAAATTCAAAAACGGATGGAAGCTCATATACTTTGCAAAAGTATTTAAATATTACTTCAATACAAGAAGATGTGGAAAGTTTTGCTAGGAGACATACTGGAATAACAAACTTAGACGTTTCTTTAATGTTAAATGGTTTAAACTACCAGAGTATATCTGAGTTGATTGACAGAACAATACCTAAAAATATTAGATTTAAT
>evgR018157c0g1_i01_Hydra
AGTTATTTTAGTGTATAAATAAAGTAATAAAATGTGTGACACATTTGAAATTTTAAACAAAAGGTTAAACACATCTCCTTTCATGGTACAAGGAGAAGTGGTTAAAGGTTTTGGAAGAGGCAGCAAAGAACTAGGTGTTCCAACTGCAAATTTTCCAGAAATAGTTGTTCAGGATATACCAAGCGATCTCACTGCCGGAGTTTACTATGGTTGGTCTCGTGTAGACAATGGTGAGGTTTACAAAATGGTGCTCAGCATTGGATGGAATCCATTTTATAAGAATGAGAAAAAATCAATGGAAACACATATTTTGCATAAATTTGATAGTGATTTTTATGGATCTATCTTGCGTATAGTTATGACTGGATATATTCGGCCAGAGTTAAATTTTGGTTCTCTTACAGAGCTGATTGATGCTATAAAAAGTGACATTGCAACTGCCGATTCGTCGCTAGAAATATCGGATCATATTTCACACAAGTTTTTAGAATTCATTCAGAAAGAAATTTGAATATTTTTAACAATAGTTATTTAATAGTTATTATATAGTTTTCTATTTTTTAAAGTTTTTTAATAAATGAGTATTGATGTGTTTTATAAACAATATTGATAGATAATTGGAAAGTTATTATTTAGTGTAAACACATGTGTTGCCTACATGTTTTCATGCTGAATGTTCTATTTAGTGATAGGGCAAACAAAGAATTTACTGAGACTTTATTAAAATAATCTCTGTCAGAATACTTCTGAAAATATTTTTTTTGAAAAAAGTAATTTAAAGTTCTTCTTTTTTC
>evgR018163c0g1_i01_Hydra
GGCAAAGGAAAAGCAAATGTTTCTAAAAACCTAAAAAAAGATTATTCTCAATCTTCCACTTCTAAAGCTGTTAACAAAAAGAATAAGGGCAAAGGAAAACCAAGTTCAAAGAAAAGTAAGTGTGTAAGTGTGGATGTGAATGATTGGTTTTGTTTGTTGTGCCAGCAAAAGTCTGTAGAAAGTATGCAACAATGTTGTAAATGTAAAGTATGGTACCATGAGGCGTGCATGGGGTATGATTCTGAAGATGAGGAAGAGTTAGTATGTCCATTCTGTGAAGATAGAATTATATGGCTGAATGATTGCCGAATGGCATGAAACTAGTAGTCCCG
>evgR018168c1g1_i01_Hydra
CCAAATTTTTTTTTTTTTTTTTCTCTGGGTTATCGCAGCCATGAGCACTTTAAGGCTCCAAAAGCGATTAGCCGCTAGTGTACTTAAATGTGGTAAAAAAAAAATTTGGCTTGATCCCAATGAATCTTCAGAAATTGCTAATGCTAACTCAAGACAAAATATACGTAAATTGATCAAGGATGGTTTAATAATCAGAAAACCCCAGGTAGTTCACAGTCGTGCAAGGATACGAACAAATTTAGAAGCTAAACAAAAAGGACGTCATAAGGGCTTTGGTAAACGTCATGGTACTGCAAATGCACGTATGCCTGAAAAGGTAATCTGGATGAACAGAATGCGTGTTCTTCGTCGTCTTCTTAGAAAATACCGAGAGCAAAAAAAAATTGATCAGCATTTATACCATGATCTCTATATGAAATCCAAAGGTAATGTTTTTAAAAATAAAAGAAATTTAATGGAGCATATTCACAAAAAGAAAGCTGAAAAGAAGAGAAGCAAAGAACTTTCTGATCAAGCTGAAGCTAGACGAACGAAAGTCAAAGCTGCTCGTGTCCGTAGAGATGAACGTCAAGCACAAAAGAAGGCAGAAATACTACAAGCTTATGCTAAAGAGGATGGTGTTCAACCTCAAACTACTGCATAGTTACCTGAAATACAAAATTAATTTAAAAAAAAAAAA
>evgR018181c0g1_i02_Hydra
GAAAAAATAAATATATATATATTACAAAAAAAGTAATGTAAATTTGAGTTGCATGCGAACATGGATTCATATCAACAGCAGTCATGTCATCAGTCTTATGGTGGAATGGACAGACCACTTTTTGGAGAATCAAATAAAAAAGACAAATATTGCAACTATGTATTGGGTGCCGCAACAGTTATTGCTGTACTTGCAACAGTAATTAGTGCTTTTGTTAACTTGAAAGCCCTAAATATTGTATTTTCTGGAGCGATTGTTATGAACTGTATATCAAATATTTTTCTAATATATTGGTATCGTCAAGGCGATCTTGATCCAAAGTTTAGAAATTTGATTTATTACAACTCTCTTGTTACTCTGCTTCTTTGTGTAATTGCATTTGTGACTTTCTTCAAAAAGTAGTTTTATAGAATTTTAGAAGATCTTAAAACTATACTGCAATATTTATTTTGGCG
>evgR018185c0g1_i01_Hydra
GAGAGAGAGAGAGAGAGAGAGAGAATGAGAGAGAAAGAGAGATAGATATGTATATATAGATATACATCAATACGCAGCCAAATTAAAAAAATGAAAGTGCTTTATCGATTGTTTCCGTACATGAAACTTTCTCGTTTTCTTCTAAAGCGAATTCTTCAAGTTGTATATTAAACCAATTTTTTGAAAGATATTTAAAGATGTCATAACTTTGTTTATTCGATAGATTTATATTTAGGTCATGCTTAAGAATGTATTTTCGATAGAAACTAATTAAGTTTTGCTCTATAGATAATAGTTATTTTGCTTTAGAAATGGAACCAGAATCTAGATCATACTCATGTCGTAAAGTTTTTACATTTAATTCATCGTTTATAGCCGTGCTTATAAAGCGACACCCAGATACTGCTTTATTTATAGATAATAGAAATAGATTTAATATTATGTCTTTAAATATACTTGCTTTTGTAACTTCTTTTGTATTGTCTTCAATGTTTACAAAGTTCTGGTTAATTGTTGCTATAACGTTTTGGAGGTCTTTCTTTACTTTGTGTGTGTCTTTTTTTGTAAATTTATATTCTTCTATTTTTGTTTCAAACTTTTTTTTTTTATTGAGAAGATTTTGATAAATTTGATTGGAATTGTGTATATCTAATAGCACTTCTTTCCAAACTTCTTTATTTTCGCTTGTTTTTACTTAAAAAAACCAGCTCCGCTAGTTCTAAGCACATCGGAATGGTAAAATAAGATGTTGGATTCCTACTGATTTCGCAAAAATATAGAGTTACCATTTGGGATAGTTTTAAAGAGACAACGTCGTTTTTATAACTGCCTATGTTTTTGTTTGAAAACATGTATTGAATTGTTTCGTTATTTATTGAATCGAAACAAGAAAAATCGATTTTTGCGTAAGACGATAAAGAGCCAATTTGACTAACATGAAGTCCATTTTTCGAAAAATAGTGGTTTTCTATATTTGTTGTGAACAGTTTGGGGAAACGAAAACAGTTCGTATAAGTAGAGTGGCAAATTTTATCTTCTCCAAATTTATTTCGTTTATAATAGACTGATCTACTAATCTTTTAAAAACAATTACATCAGTAATGTCTGAAACATTGTAAGAAAAAAATTTTCTACCTCACCTAATAAACATTTTTCCGATTTTCTGAAAAGATTAACCTTTTGGTTCACTTCCATTTCCACAACCTTACCAGCAATGGGAAAATGTGTTTTGCCATCAGGCCTTGCTTCTGACTCCTTCTTTTTATATGAACCATCCCAAAAATTATAAATTATTCTTCCAACATAATTAAGCCACGCTTGATCTTGATCGCATATATAGCAAATTATCATGCAAAGCCATAACAAAAAGAAATAAAAAAATGTCATGAATCAGTC
>evgR018187c0g1_i01_Hydra
CTTTTATTTTCTAAATAAGATATAACAATTAACTATAATGTCACGACATCGCAATATTAGGAACATTAGTTATGAAGATGAGCTTTGTGAAGACTTTGGTGGGCGATCTCTAGATGAAGTAGACTATTGTGTTTCACCTGCTACAGAAGCAGAATTTATGTTTCGACGAGATAAGCACCACACGTTATCATCTTTTATTGGAGACGAGGAAAAAATTGAAGAAGAAGAAGAGGAAGCTCCAAAAGAATCATTTTACAATTACAAGAGACCCACCATAAACCACTCAGATGAAGACAAGTTGCATTCATGTATTGAACACCTTCAAGATATACTAGGGGAAACGTTTGATGAAGATGCAATTGCTAAAGTTGTCATAAATCAAAGTTTTGATATGCAGAAAGCTTTGGATCAGCTACTCAACAAAGGTCATCAAAATACAGCACCAGCAAAAAACATGACTCCTAACAAAACAGGCATTGCACACATTTTTGGTAACGTTATTATTGGACAACATATATGTATATGATATATTATTTGGTATGTAGTGTTTTTGAAGTGAAGCCTTGTATAAAACCACACAACTGCTTTGTTGGGAAAATGTTTATCAAAGTAGAAGATCATCTTTGTGCTAACTACTAACTTGCCATTTTGGACTTAACCCCATGGCTCTTGATATGTGTGCATTTTTCCTGCGTTCATGGTTCTTGATGCACCCACAGCTTTTGATGCATTATTTTTCTTGATACGTTCTTTTTCTTGATGCATCCACGGTTCTTGATGCCTCCTTGGTTTTTTATGCGTCCTTGGATCTTGATGTTTCTATGGTTTTTGATGCAATTAAAAGAGATTATTTATTTATTTAAAATTGTTTTTGTTTTTAAATGAATTTTTTTTGAATTTTTTTTTGAATTTTTAAATCATCCTGTAAAAATGTGTAAAGATGTTAC
>evgR018189c0g2_i01_Hydra
TTAAAAAATTGCAAATTATATGCGTAGAAGCAGAAGCTTTTGCATTGCTCTAAATGTTAATTTTTAAGAAACTCTTCTCAACCAGCACTTTACCAGACCTGTGGAGGAAATCCAACATCACGTCCCTTTTCAAGAAAGGAAGCAAACTAAAAAAAGAATTATAGAGCAGTTTCTCTCACATCCCATGCAAAATATTTGAGAGCATATTACATGAAAAGATAATGCTACACTGTAACTTAAATAGTTTAATAAGCACAGCACACAGATTCATTCAAGGAAAAAGTTGTTTGACTAATCCCCTCAAAATCTGCAACTTTCTCACAGAAGCCGCAAACAAAAAGTATTCAGTCAATGTGATCTACACTGATTTTACAAAAGCATTACACACAATTGCCTATTAAGCAAATTGAAAGCATATGGAATCAACAAGAAAACATTCATGCGGATAAATGCTTGGCCCAACAATAAATAACAGCGTATTGTCATTGATACTCTCTCAACTGCTGAAATATTCACATCAGATTAAAAAAAAAGTTACAAGTGGCGTTCCTCAAGCAAGCATCGCCGGCCCACTACTTTTTGTACTATTTATAAATGATCTGCCAGACAACATTACAAATCAGTTTAAACTCTATGCAGATGGCAGTAAAATTATGAACATCATAAAATCAAATAAAGATATGCTAGCTTTAAAACCAGACATTGATCATGCCATAAACTGGTCTCACAAATGGATGATGTTCTTTAATGTGGAAAAATGCAAAACAATGCATGTAGGTCAAGGTAACAGAAAATCTTATCAAGTCTATTCAATAATAAACATAGAAGGCACTCGTCTATAGCTAGAAGAAACTGAAACTAAACAAAATCTTGGTGTTTTAATATCAATTGACCTCAAATTTCACGCCCAAGTGGAGTCTGCAGTCTCAGCAGCCACTTATAAATTAGGTCTACTCAAAGAAGCATTTCGAAGCAGAAGCATTTGCCTCTGGCAAACGCTTTACAACACATATGAGCGCCCACACCTAGAATTCAACAAAACTGGATTGACAACACTGAAAGAGAGAAGAGCCAGAGGAGATCTCATTGAACAGTTTAAAATTGTGAATAAAATTGATGATGTTAATTTTTTTGTTCCTCAAAGGTTCTCTAACAACACATATGCTACCAAATGAAGCAATTGATACTGACACGGTCAACTTATTCAAAAATTTCCTATCATGATGGACAACTTTTCTTGTTGACAAATTGCCCTACACATTAGTTAGTGCAGAGATGCCTGGTGTTGTACCTTATTGTCAATATACTTTATAAT
>evgR018190c0g1_i01_Hydra
AAAAAAAATCAAACAAACACAACACAACACAATGGGAAACGCAACACCAGTCATTGCAGGATCGTGGAGAGAACAGAGCGGAAAGATTTGGCACTCCTCACAAAATGGTAATACCTTCACTTGGACTCAGGAAGGAAGTGGTAGAATTGCCACTGGTTCAATGGTTGCAACAAGTGTCACTCCAGGTCTCTCTGGTTGGGTCATTCACATGACTTTTGATGGTAATGTTCATTGGAGATTGAGACCTAACAGTGATGGTACTGTTTTGCATGGTCTTAGTGATAATTTCACTAGAGTTTCAACTGGTTACGGATTTGTTCAACCAACAGTTTACAATGTCTTTGGTATGAATAGTAATGTAATGGCAAACAGTTCAATGAATGTACAAGTTTTGAATTCAGTCTATACTGAAAATTCAGGAAAGACCTGGATTGTTAATTCAAATCCACATGCTGGTTACTTTATCATGAAGAATCAACAAGATGGAAGACAGGTTGAATGTTATGTTGCCAAGGATTTATCCAATGGTAGACATGTTATTTATATGCAATTCCAAGGTACTAGTGCACCATTGAGAGGTGAGACTTTTGACATTGGTCAAATTCAATTGTCAAATGGTGATTGTTTTAGACGTACTCATAATCCAACCATTATCACACCTGTTACTCCACAACCAAATTATGGAATTTCTATTGGAAATAGTGGTCTCAATATTTCATTCACCACTCCAACCACTACTATTATTCCACCTCAACAACAACAACCAATGTATGTCAATCCAAACACTTCTTACATGCCTCCACCACAACCATTCCAACAACCACAATTCCAACAACCAATGATGCCACCTCCACAACCATTCCAACAACCATATCAACAACCATATCAACAACAAAGTAGTGTTATTATTA
>evgR018191c0g1_i01_Hydra
GCTTTACTTTTAAATAACAAGTTATGGTTTCAACAATCTATTAATTTAAAAAAAATTAAGTTAAAGAATAAACAATTTGAAATGTAATGTTCGGTACCGGTGTTTTAACGAACAATACTTCTACATTCAACAATTCAATGAGAGTAACAAACGACTATAAAACTGCGGTCTTCAAAATGTCAGCGATTGGATTTTTGAGTATCATAACAGTCTTAGGGAATCTAATTATCGTCATTGTATATTTTTCTAATAAACGAGTTCAAAAAGTTACAAGTATCCCGATAATTTCTCTGGCTATTACTGATATGTTCATAGGACTTTATCCAATGAACGTAAACATCTTTGAAAATGCATTGGGGTATTGGCCCTTAGGAAATGTGATGTGCAATATATCATTATTATTCGATTTCGTATCCGTCCAAACTTCAATCAACTACATTGTATTAATCAACATCGATCGCCATTTTTCTATAAAACATCCTTTTCAACATCTAACAAACAAAAGGAAATACAAAATTGTATGGCAATTGTTTGGTGGATGGCTTTTTGCTGTTGCAACATGGGTACCATATATATGTTTTTATGAGCATGCGAAGAGAACAAAAAATGTTTCAGGAACGTGCTATGAACAGTTCTCTAAACCTAATGATGTATACTTTCAAAAATATTGTATTATTATATTATCCATGATTGGCTATTTTATACCCTTGGTAATAATTCTTCTGATCTATGCAAAAATGTATTTCATTATTAGACAGCATTCAACTAATATTTTAAAAATCAAAAAACAAAATTTTTCAGATATTTTTAGAGTGACATCATTTTTTTTAAACGAAAGTATGTCGCAGAGTTCAAGCCTAAGTGCATTACTGGAACTTTTGCAAAAGGATTTTATGTTTATAAAAGGTACTCTGCATAGAAAATCGTTACCTTCCTTTCCAAACCATTTGCAAAAACAAAGACAAACAAACTCTGAACTTTTGAGCAAACAAAAAGACGTACTCAAACAGAAGAAAGCATTACTAATGATTGGATGTATAGTTTCAGCATTCGTTATTACTAATTTACCATTCAATATACAATGGTTTATGGTCACAGTAAGCCCTGGAAGTATTAATCAGTCTTTTTTTGAGATATCAAACTTTTTAACGTATCTCAATTCCACTGTTAATCCCTTGCTATATGCCTTTGTAAATCGAAATATTCGTTACCAGATAAGAAAGTTATTTTTCCGTCTTGTTGAGAATATTAAACATAAACTTCAAAAAAAA
>evgR018200c0g1_i02_Hydra
CTTTTATTTTCTAAATAAGCATAATGGAGAAATCAAGATTAGAAAAAGATAAAGAAAAACTTGCTGTTATCAAGACATTTAAAACTGCTGTGCCCACCTGGAATGGTCCACAGTTTGTATCATCAAAGCAATTTTTATGCAATTTTTCCAAATATCGTATTCCAGAAAAAAACTATATGGAGTTCTCTAACATTGGACATTTTCTAAATGAATTGTTTTCTGACCAGATACAGATTGTTGATCCCACCATGGCCACGATGTACAAAAAAGATTATCTTTCGTTCATGCAACAAAAAGAAGAGAATCTATTTTCGATGAAAATGTTATACGAAACTCTAAGAATCTTAGAATTTAACTTTTTGAAAAGGTATCGTGGAAGATCAAAGTACTCGCTTGCAGATGTTTTGAAGATTTGGACGCATTATACATCAAATTCAATGATCGGGATAATGGAGAGAGAAGATTTAGATGCATTTGCATTCGATTTGATGGGACATTCTTTTCAAAAATTTGATCTTTATCTTGTGAAACAGCTTTCTAAAAGTCTTATATATTTGGCTGGATGCAACAAAGACGAGCAAATAAATTTTTATGGCTTTGTTAAATTGTTAATTGATGATAAAATCCTTAATTCAACATTTCCAGAAAGAGATAAAATAGCTCGTGAAGATTTTGAAAAAGGATTCTCATACTATGATCAATTTGGAAACGACTCGGTGTCTTTAAATGATCTTGATGATCTTCTTAGGGATGTTTTTCAACAACTTGGAATGGAAGAAAGTCGTGAAACAATAAATGCTAATCGAAACGCATTGCTTTCTGTAATCACACCTGATAATAATGGTAAAATCAGAAGAGTTGATGTTGCACTTTTGTTATCGCACTAACTTTAAATAATATTATAAAAAAGAACTTTTTATTAATAATTAACATATATCTTTCATAATAGTATATAATTAAAAAAATTTGTATATTGATAATCTTTTTG
>evgR018201c0g1_i01_Hydra
TAATAAAAGTATAAAAGAGAAGCTGTACTTGGAATTCATTGTAATTAATAATCATAAAGAGACAAGATCTAAAATTGATTTCCAAAATGATTTTTGAAAATGAAGTTCATGGATTTTCAACAAAAGATGATAAAATACTTACCGAGGTAGAATTAAAAAAACTCTCATGTAATCAAAATCTTGTATCAAACTATAAACAAAATCAGCTAGAAGTGAATGCCCAAAAAAATTGGGATCTATTTTATAAAAGAAATTCTACAAAGTTCTTTAAAGATCGACATTGGACAACACGAGAGTTTAATGAAATAACTTCTGAGAAATTAAATGACAAAATTTTGTTAGAGGTCGGTTGTGGTTGTGGAAATGCAATCTTTCCATTAATTGAAGAAAACAAAAATATTTTTATATATGCTTGTGATTTTTCTTCAAGAGCAGTTGAATATGTCAAGGCTCATAAACTTTATGAACTGTCTCAATGTCTAGCCTTTCAATCAGATGTTACCAAAGATGATCTTTGCTCATTTGTACCTAAAGCTTCAGTGGACATAGTACTGATGATATTTGTGCTTTCTGCTGTCCATCCTAATAAAATGAAAGAAGTTCTGCAAAATGTGCATAAAGTACTTAAGCCAGGTGGAATAGTATTGCTTCGTGACTATGGTCTGTACGATCATGCAATGTTAAGATTTGGGCCTGGTCATAAACTTGAAGATCGATTTTATGTAAGGCAGGATGGAACCCGGGCCTATTACTTTTCAACAGAAAAATTATTTGAGTTATTTGCATCATGTGGTTATGAAATTGTTGAAAATATATATATTCATCGAAGAACAATCAATAAAAAAGAAGGCATTGATGTACCGAGAATTTTTGTGCAAGGAAAGTTTTCAAAATCTTCTTGAATAATTTCTACGTTGAGCTAACAACGATAATGATTTTGCCGAAAGTAAATGATATTAAAGCAAGTTTGTGAAAATGCAAATTAAAATTTGCTTTTATTAAGAATATTTTGAAAATATATTTTGAAAGAATTGACTACCTTGTGAAGTATAGCCGTAAAACAAAGTGCGAGAAGATAATCAATCGAAAGTACGAGTGCTGAATGATCAATTTATATTACGAGTGCCAAATAATCAATCGCATGTACGAGTGCCGGATATTCAATCAAAAGTAGAAGTATCGGAAAATGAATTGATAGTATGAGTGCCGGATATTGAACCGAAAGTTCGAGTGCCGGATAATCAATCAAAGATACGAGTACCGGAAATTTAATCTAAAGTACGAGTGCCGGATAATCAATTAAAATTACGAGTGCCGGTTTATTTCTGGATAATCAATGCTTTAGATGACATATTATTATCTAAATATTTTACTGAATCTTCAGTCTTCTATATTGTTCAGTTTTGATTGTCATTTTTATTTTCTTGTACCTTAAAATTTTAAATAAATATCAATAAAAGCAAAGTGAGTAAGGATACGTAATACTATTGTAATGATAAAAAAAC
>evgR018203c0g1_i01_Hydra
TTTTATTTTGCTTTAATAGTGAAAGCTAAAAGATGCAAAACCGATTCCAAAAAAAAATGCATTTTTGTGTCTTTCTCCTTATCATAAATTTTGCAGTTGCTACGGAGATAAACTTGTATATCAACTTTGAAGAAGCAGCAGTTGAAGTTGATAAAGAATTTTTATCTTTAACTTTTGATTCAAGTTATTTTTATTATAAAGCACTAACTAACGAGTTTAAATCAGAAATTTTGATAGCTTTAGCTAATGGACTTTCTTCAGAGGGGGACCTTTACTTGCGATTTGGAGGTACCAAAGCTGATTTTATGAATGTTAACTTTAACGCTGTTTACTATGACAAAGCCAAATTCAATGAGTTGGTTTCGTTTGCCAAGAAGACTAATTGGAAACTTATTTTTGGACTTAATGCGCTTAACCGATACTCTGATGGTTCTTGGAATTCTTATAACGTAACACAAATGTTGAAATATATTGCTGCCATGAACTGTGAAATCAAATTCGAACTCAGCAATGAGTTTGATAATTATCCAGTTGTTTTCAATTACACATTAAAACCGCAGCAACTTACAAAAGATTTTAAAACACTACGTGGAATCCTCAATGAAGTTTTTGCAGACAGGCATGTTCAACTTTTAGGTCCGGATGTAGGATCTATAACTCGACTTGGTTTTTTTGAATCGTTTTTGAAATTGATTGAAAAAGATATTTTGGATGGAATAACATTCCATCATTATTACCCAAGAAGCAACAACATTAGTCCTATAAACTTTACAAGAGTCGATTACCTTGACACTTTTCTGGATTATGGATTAAAAGCGATTTCCATCATAAAAAAATCAATAAGTAGCCGTTTCAAAGTTCCACATGTATGGATTGGAGAAACCTCGAGTACCACATTTGGAGGAGTTAAGGGTACTGGGGACAGTTTTGCTGCAGGTTTTCTCTGGCTTGATAAATTAGGACTCGCTGCTCAAATGGGGATAAAAGTTGTTTTACGACAATCCTTCAAAGGCGGAAACTATTCCCTTATAAACCATGAATTTAATCCAACGCCTGATTACTGGAGTTCATTACTTTATAAAAGACTTGTTGGTCAGAAAGTCCTAAATTTAACTGGATATTTGGAGCATGGTAGAGAAACAAGAGTGTACGCACATTGTGCAAATATTAGAAAAGGATTATATTATCCTGGTTCTGTTGTTTTAATTGCCATCAACATTAATAACAAAGAGAACTCTACTATTAATGTGTTAAATAGATTAAATTCCGATAGCCAGAAGATCGATCAGTATTTACTTGAACCAGCCAATGGAAACATAGGAGATGGAAATGTGACGCTTAACGGCATAATTTTACAAATGGTTAACAAAACAAAATTGCCTGTTTTAAAACCGATTCAAACTAAATTACCACTGATTATTCCTCCGTTGAGTTATGGCTTTTATGTTTTGATTGAATCAAACGCACAAGCGTGTAAACCGTTTAAAAGTCATGTTGCATGATAAAATTAAACAAATGAAGAAAAAATACTGTAAATTTATTTTCGAAAACTTGTTTACAAATTCAAGCAAAAACTAAATAATTATGTTCAATAATCATTATTCTGATATTTTAAGTGCTGCGGCACTTCTGTGTTAATTGCAATACTGCAGCACTACTGAGTATTTAGTTGCAATACTGCATAACTACTGGTTTTTAATTAACACAATTTACGATTATAAACAAAGGTTTTCAAGTGATATTTAAGAGATGATTATAATTTATAAAAGAAAAGAAAATTTACTTTTTTATTTTAAAAAAAA
>evgR018209c0g1_i01_Hydra
ACCTATTTTAGTGCAAAAAGTTCTTACTACAAAATGGCTGATTTAAAAAGAAATCCAGAAATAAAATATACAAAGCTGTTTATCAATAATGAATGGTGTGACTCTATCAGTGGAAAGACATTTCCTACAATTAATCCATGTACTGGTAAAGAAATATGCCGTGTTCAAGAAGCTGATAAGGAGGATGTTGAAAGAGCCGTTGAAGCTGCGAAAAATGCGTTTTCATTGGAATCAGACTGGCGTATCTTGGATGCTTCAAAGCGAGGAAAGCTTATATATACTCTTGCTAATTTGATTGAACGAGATAAAGATTACCTTGCAAGTTTAGAAACTCTTGATAATGGGAAGACATACTTGGATTCTCTCGGTGATATAGAGTTTTCAGTTAACGTCATGAGATATTATGCTGGTTGGTGTGATAAAATATGCGGAAAAACCATTCCTGCAGATGGTGGTTTGTTATGCTTCACAAAGCATGAGCCTGTTGGAGTTGTAGGCCAAATAATTCCTTGGAATTATCCTATTATGATGGTGTGTTGGAAACTGGGTCCTGCTCTTGCTTGTGGGTGTACTGTGGTTATGAAACCTGCTGAGCAAACTCCTCTTACAGCTCTATATATTGCTTCTCTTGTTAAAGAGGCTGGATTCCCTGCAGGTGTTGTAAATGTTCTACCAGGTTATGGTCCAACAGCCGGAGCTGCTATTTCTTCGCATATGAATATTGATAAGGTGGCATTTACTGGATCTGCAGAGATTGGTCATGTTGTTATGGAGGCTGCTGCAAAATCAAATATGAAACGTATTTTATTGGAACTTGGGGGGAAAAGTCCACTTATAGTATTTCCCGATGTTGATGTATCTTGGGCTGCTAAAATTGGTCATCAAGCTCTTTTTACTAACCATGGTCAGAACTGTTGTGCAGGGTCTCGCACATTTGTTCATCAAGATATATATGAGGAATTTGTTGCCAAAGCAACTGAGCTAGCGTTAAATCGTAAGGTTGGCGATCCTTGGAATGAAGAAACTGCTCAAGGTCCACAGATTGATGAAGCTCAGTTTAAAAAAATTCTTACTCTAATTGAGTCTGGTAGAACAGAGGGTGCATCACTAAAATGTGGTGGCAATTCAATTGGTTCTGAAGGCTATTTTATTCAGCCTACTGTATTTGCAGATGTCACAGATAATATGACTATTTCAAAGGAAGAGATATTTGGACCAGTGCAGCAAATTTTAAAATTCAGCAATTTAGATGAAGTCATTAAGCGAGCAAATAACACATCATATGGTTTAGGATCAGGTGTTTTAACAAATGATTTAAATGTGGCAATGAAAGTGGTTAATGGCATAAAAGCTGGTACAGTGTGGGTTAACTGCTATGATGTTTTTTCTCCACAAGCTCCATTTGGAGGTTTCAAGATGTCTGGACAAGGAAGAGAATTAGGCGAATATGCTTTGAAAGAATATACTGAAGTGAAAACGGTGACAATCCAAATTAAATGAGTTTCTGTGATTCGGCATAATAACACCAAGCGAAGTTCTAAGTTTTCAATATTTTATTTTTATGATTGTGTTTTTGTATAATTTTATAAACGTTTTTCTATTAAAAAAATTTATAAAAAAAAAAAAA
>evgR018211c0g1_i01_Hydra
TTCACTTGCGCTACAAAGTACAACAGCTACGTAGCTAAAGTAGGTTCTAATATTCTTTAATATGAAACTTTTAGTATATTTCAACGAGAAGCAATCGGTTGGTGTTGTTGAACCTGGAAACTTTCTTTGTCCTGATCTTCCTTTTGATCTAAACCAGATAACAAAAGAAGATTGGTTGTTGCTTAAAGGCAAAAAATTGGAGGGACTCTGTGTATGTGATAATGAGACGTTTGAAAACACCATATTGTTGCAAGTTGTTAATGATGAACAAGTTGAAAGCAGAGTCAATTTTATTTGTCGTGTTATTTCGGAGAAGTGTGTAAATATTGATAATTTGGTTAAGCTTCTTCAACTTGATGCTTTTAATTCTAAGCAAGTTGTATATACGCCTCTGAAGGCTGATGCTGCTGTAAATGAAAATAAGTACAAGCACAAGATTTCAGTTGATGGGGTTATGTCTACTCCTAAAAGAAAATTCGACCATTTTAAAAGTGCTTTTGAAAATAAAGCATCAAATAGTTTAGCTTTAAATCCCTCAAGTTCAAATTTAGAATTACTTGCTAATGCAACTCAAACCGCAACGCAAGTCACAACTCAAACTGCAAATCAAATCACAACTCAAACCACACCGGCTACATTTTTAGGGGCATCATGCCAATGTAATATGATATCTGACTGGTTTAAAAAGATAAACAGCCGTTTGAGTGGTATAGAAGAGCAGTTGCACTCTATCAAACCAATTAATGGTGAAGGCATTTCTGATATTCGTTCTCAAGACTGGACTCTTCCATTGCCTGTTACAACACCAGCAGACTTGGAAACATTAGAAAAAATACTTTCTGAAGATAATTCATGCTATGAGTTTATGATTCAAACTCTTAGTCGCTGTGGTGGATCAGGGTTTCGAAATGGTGTCAAAAATATAATGGTGTCTTTATTGACAGATGAACTTGCGTCAAATTATAATATGGATGGGCGGAAATATAAAGATTTTACCAAGTATCCCTTCATAAAACATAAGACTGTGATAAAAGCTATATGTGAGTCGATGAAACATGGGTTTAAGGAAAGCGATACTGAAATTAAAGGTCAAATAGGAAACTGGTTAAAACAAGCACCCAGGAGATTGCAGCGTCACACCACCGTTGTGAAAAGTTCGAACTCTTCGAATTCAAGTCACGAATAATAGTCATATATTTTAGCTTAATTTTTCTTCCTGTATGCATTAAGATGTATATAACATTTATTAGTTGAGTTTTTACTTATAAATACATATTATCGTTTAAGTCAAAAAAAAAAAA
>evgR018213c0g1_i01_Hydra
CATGTCGATAATTCAAGATTTTCATCGTTTTTAGAATTTTTTCTCAAAAGTGGTGATACTTTCAATCTATAACTTTTGACAATAACAACAAATGAAGTATTTAATTGTTACTGGAGGGGTTATTAGTGGTGTTGGAAAAGGTGTAATTTCAAGCAGCGTTGGAGCAATCTTGAAAGCTAGTGGATTTAATGTTACTGCAATTAAAATAGATCCCTATTTAAATATTGATGCAGGAACATTTTCTCCATATGAGCATGGAGAGGTCTTTGTTTTGGATGATGGAAGTGAAGTTGATTTAGATCTTGGAAACTATGAGCGTTTTTTAAATATTCAATTACATAGAGACAATAATATTACAACTGGAAAAATATATCAAAAAGTAATTGAAAAAGAAAGAAGAGGTGATTATCTTGGAAAAACTGTTCAAGTTATTCCGCATATTACTAATGAAATTCAAGATTGGATTGAGAATGTTGCAAAGCAATCAGTTGACAAAAGTGGCAATTTACCAGATATATGTGTTATTGAGATTGGTGGAGTAATCGGAGATATTGAAAGTATGCCTTATGTTGAAGCAATGCGTCAATTTCAGTTTAAAGTTGGACCTGACAATTTTTGTTGTTGTCATGTAAGCTTAATACCTCAGACATCTGGAGGTGAAATGAAATCTAAGCCAACTCAAATGAGCGTCAGGGAATTGCGAGGACTGGGCCTTACCCCAGATTTGATATTGGCACGTAGCAAGTGTGAAATAGACTCCTCCTTAAAAGAAAAAATATCGATGTTTTGTCATGTAAATCCTGATCAGGTAATTAGTGTTCCTGATGTGTCATCACTCTATCGGGTTCCAACTCTCCTCTATGATCAAGGTTTGCTCAATCAAATTGCCAAAAAATTGTCTTTAAATTTACCAGTTATAGATTCAAAATCCTTCCTTTTTAATTGGAAACAGTTAGCTGATAGGTATGATCAACCCTACGAAGAAGTTAAAATTGCGCTTGTTGGTAAGTATACGAGACTTGAAGATGCTTATATATCTGTCATCAAAGCATTAAGCCATGCAGCACTTGCTTCACGGAGAAAGCTTGTTATTAAGTGGGTTGACTCTGAAAGTATAGAAGATGCAACTAGAGAAAATGATCCTGTGAGATATCATGAAGCCTGGCAATCATTATGCTGCTGCGACGGTGTCCTTGTTCCAGGTGGTTTTGGTCAACGTGGATCTGAGGGAAAGATACAAGCTGCTCAATGGGCCAGAACAAGCAAGAAGCCATACTTAGGAATTTGTCTTGGATTACAAGTTGCAGTAATCGAGTTTGCTAGAAATGTATTATCGTGGAAAGATGCAAACTCAACTGAAATCAATCCATGTACCCAACACCCAGTTGTAATAGAAATGCCTGAACACAATCAAGGTCAACTAGGAGGAACAATGCGCCTTGGAAAAAGAAAAACTATCTTCAAAAAACAAAACTCAATTATAAGAAATTTATATGGTGATGTTGACTTTATTGAAGAAAGGCATCGACATCGATATGAGGTTAATCCTTCGTTAGCTCATTATTTTGAGGAAAGAGGTTTTATGTTTGTGGGAACTGATATTGATGGCGAAAGAATGGAGATAGCTGAGCTCAAAGATCATCCATTTTATGTGGCTGTTCAGTATCACCCTGAGTTTTTATCTCGTCCTATTCATCCGTCACCACCTTACCTCGGGTTTATTCTAGCTGCATCCGGGAAATTGAATAGTTTTCTTGCTCGTGGTGGTCGCTTATCTGCTCGTTCAAGTTTTTTGGAATATGATACTTTTTTAGATGGTGATAATTCGCCTAAAGATGAACCTTCTCTCAAATCAATAAAAACTGATTTGGTAAGTTCTTTTTGTGACGATGTTTATATAAAAAAACTTGAAATCACAAATCCAAATACTTCAACGACCGAAGAAAGTGAAACAGTTTATTAAAGTTTTATAAACGTTCTCAAAGTCATTCTTATAAATAAAAAAATAAGAAATGCATTAGAAGCTTTATATGGTATCAAAAGACTGGTATCTACTTAAGGAAATATTTAGACCTTTATAGTTTCTTTTATTAAAGAGTTATAAATAGCTAGAAGATTTATCAACCCAAAGTAAAATTATTTTTCAGAAATATTCTAGATGTTAGAGTTTTAAATTTAATATATTTTAATTGTTATCTAGATATAAAATAATATAAAGATATTTTACTTAATTAATACAAAAGTATGTACGCAAAAAAAAA
>evgR018216c0g1_i01_Hydra
CCGCTCAATGACAAATGTTTTCCCCCTAAAAAGATGGTAACACGTATTAAGATGGTATAGTTAACTCAAAGATAGTTTTTGACATTTATTTAAAAGTTTAAAGTAAGAAATAGAATTATGGTTAAAATAATCAATGGAGAAATTGTACAAGATGACGATCCTCGTGCAAAAGCCTGGGAACAAAAAAAAAATTCTTCACAAAGATTATCATCAAATAACTATACTAATCATCGAGAAACTCCAACTGGGGGAAGATCACCCGTAGACGAAATAAATAATAGGCTTCGTGGATTTGGGGTTCCAAACCTAACATTAGCTGGTCATATTGTAGAACCAATTTTTTTAATAGCTGCTGTTATTGCCTTGGTGATGTGGGGGTTTCTAGGTCTAATTATAGTAGCTTTAATATGGTATTTTATGAAGTTTCATCAGTAAGCGCATTTTGATTTTAGGCGAATTTAAATTATTATTTTTAATTATTTGTATAATAAAAACTCAGGTGTGTTAATAAAATAATAATAATTATTATTAAAAAAAAAAAA
>evgR018228c0g1_i01_Hydra
CTTATCACCTCAAATAGCTATGCCTGGTGAATCTTTATCGGTGTGTACACCATTGAATATTGCAAACACCTATGTCGTTATGGGGTTTGCAATAACAAGTATAACTCACATATCGTTGCTATCATTGATTAGAGCTATTGTTTTAAAATATCCAACTTTCTATCACGATAACTGCGAGAAAACAAAATGCAAAGTAGCACTTATTTCATTTTGTTACATTTATGGGTTTGGTTGGGCAACGGTTCCTTTGATTGGTTGGTCAAAGTATGAACTAGATCTCGATAAAAAACGATGTTCTTTGGATTGGAATTTGACACAATCAGACTCTCTTTCCTTTATTCTGACACTTTTTATTTTGTGTAATCTTTTGCCCGGATTAGTAATAATAATAACAACATATGTCAGCACAAAAACAATTTCTATCGTAAAAGCTAATGAATTTCTGCAAAATGAAAATAAAAAAC
>evgR018228c0g2_i01_Hydra
TGCCAGCAAAACCACTTCAAAGATTTTTTGTTCCATATTATTAACTACATCAGTTTTATTGAACACAACAGCATGTTATGTCATTTTTTCCAAAGTGAAAAAGAGAAAGCTTACCCACCTGTTTATATGCAGCATCTCAATTGCAAATTTATCTGAATCAATAATAGGACTAACACCACAATTAGCTATGCCTGATGAATTTTTATTGGTGTGTACACCATTGCACATTGTAAACACCTATGCAGTTATGGGACTTGCAATAACAAGTATAACTCACATATCTGTTCTTTCATGTATCAGAGCTGTTGTTTTAAAATATCCGCTTTTATATTATAATAATTGTGAGAAAACGAGGAGCAAAATAGCATTTATTTCATTTTGCTATATTTATGGTTTTGGTTGTGCAACAGTTCCTTTGATTGGTTGGTCAAAGTATGAACTAGATCTCGATAAAAAACGATGTTCTTTGGATTGGAATTTGACACAATCAGACTCTCTTTCCTTTATTCTGACACTTTTTATTTTGTGTAATCTTTTGCCCGGATTAGTAATAATAATAACAACATATGTCAGCACAAAAACAATTTCTATCGTAAAAGCTAATGAATTTCTGCAAAATGAAAATAAAAAAC
>evgR018229c0g1_i01_Hydra
ATGCAGTTATAAAGGAGATCTGAAAAGAAACAATGTTGCTCCAGGGCAATTCATCAGATATGTTGGCAACAGATTTCATGTCTTGTTCCATATGGCAGAAGTTGTTGTGAGACATGAAAGATATATAAAAAACTTTCTTGAAAGAAACAACAAAAATGCAATTTGTAAATTACTAATGCAAGATATAAGTAACGAATTGACGCTTGTTCAATTGCAGGGACTAGGGATAATTGGAAAAATAATAACTGGACCATGGATGAATCTGGTTTATAAAAATGCATCAGGAAGGACTAACCTTGAGTTGGGGGATATTTTTCAGAAAGCAGTCAAGAAGTTGGCTACCTTTAAATCAAATCCAGAACTAATCCTTTATACAGATGTTGATGTTTTTAGTCAAGTATTGAATGTTAAAGATGATAAGGGACACCAGTCACTTTAGATTGTGAAAAATAAAGAATTTTTGTTTAAGATTTTATCAGTTTTGATAAGCT
>evgR018230c0g1_i01_Hydra
AATATTTTCATGAAGCTCTTTTTTAAGGTAAAAGAATGAATAAGCAAAGTAAAACACGCTATGAATTAATTGAAGCTATGAAAAATATTTTGGAGCCAGGTTCAGAGTCTGAGCTTTCAGATTTGGAACTTAGTAGTGATGAAGAGGTCGATAATAATCTATTTGTGGTTACCCATATAAATGATGAGATAAATGTATGTAAAGAAGAGCAATGCACAACTGCAGATATTATTTTGGAAGATGAAATTAATATTGAATTTACAACTCCAGAACAGCCCAATACCAGCCAGAAAACTTCTTCATCCGCACCGCCTAACGCAACAACTAATAATTATAATTGGTGTAATAAAATACCACCTATAAGAGACACTACTTTTACTGGTTGTGAATTTAGTCTTCCACCTAACAATGTTGATACTTTTACCCCATTGATATATTTCCAAATGTTTTGGAAAAATACTTTGTTTACCTTACTTGCTGAACAAACAAACCTTTATAGTGCACAC
>evgR018233c0g1_i01_Hydra
ACTAATCTTTTATTGTCTAAATAAGTGTAATAATAATTATTTTGATATAATGCCATCATTTTTCGGAAATGCAAAGCACCAGTTTCAAACTGTACGCGATGATAGCAAGATTGAGACAAAACCATTTTTAGATGCTTGCAGTGAAATAATACCATTCTTTGACATTCTTGGATCTACTGCATTTGCACCGGTCAAATCTGATATTCGGGGCAATGTAACGAAGTTGACAAATAAATATGAATCATCACCTAAACAATTTGAAACACTTCAAGACATTGTAAAAAGTGAGATAGAAGCAAATTCAACAAAAGCAAAAAACTCTGCAACTGATGCTCTGTTGTGGTTGAAAAGAGCATTATCTTTTATTCGTGTGTTTTTGCAAGAGGTTTTAACAGGTGAACAGGATTTGGTAGTATGTGCAAAAAGAGCATATGAAAGTTCGTTGAAGGCATATCATGGATGGATGGTGCAGTCTATTTTCAATTTGGCCATGAAAGCAGTTCCATATCGAAAAGATTTTATAAAAGCTCTTGGCGAAGATTGTAGCGAAGAAGAAATTTTGAAAGATATGACTAATACCCTAAATTTGATGGCTCAAAACATTGACACGCTGAATGCTTTTTATACAGAATCTGGTCAAGATGTACAAAGTAAAGTATAAGACAGTGTTATACAACTTTGTTAATGGTGTTGCTAGATTGGTATTTTTGCATTGTTGTGCCTTTGGTATTGGATTTGATTTCTAGAAGAATTTATTTATAATCATATTTATTGTTTTTATTAGTTTTAATAATTTAATAATTTTATTTTTTTTCTTCTATTTTATTTTTTAATCTTGCTTTTGATTTACATTATGCATGTTGGGTGTTGACGGTTTCATAATGGATGTAGCTTATGTAATATCAACACTCATTGTGTTAATTTAAGCGTTATATTTTAGATTGGTTATATATTCTTGTAAATTTCGGTCACTTCTAAAATGACATTTTGTATCTTTGTGCTGAAGTGATATTATTTCATGATATGTGTTGTGTTAAAAGCTCTCAATATTGTTGATAAGATCCTTTGTTCTATTTTTATATACATATGTTTGTAATAAATAAATTACAAGATAATAAATTTTTAGATTTAAAAAAAAA
>evgR018236c0g1_i01_Hydra
AGAAAGATATTGGAACAAAGAATGAACATTATCCAAACATCAACAGATGAGCAGTTCAGCTATATCTCTATTGCTATTCTTTGCTTACTTTCGATCAGTGTCATCACAGTCAATTCTATATGCCTGGCTATTGTAAGAAGTTCAAAGAAAATGCTAAATAGACCTTCGATACATTTTATTATTAACTTATTGATAGTAGATTTGTTGCAAGGATTATTCGTAATTCCAATTTATGCTATAAGAAAATTGAATACTCAAAATAGATATTGGAATGGTGTTGTCTGCAATTCTTTCCGATTTTTATACATGATAACATTCTATATGTCAATATTTAGTGTTCTACTAATTTCGTTAGATCGGTACGTTGCTACTACGTTTATATTTAAATACAAAACACTAATTAAAGTAAAAACAGTAAGAATTATTATAATTTCTACTTGGATTTACGTTACAGCTTTGTGTTTAATTCCATTTTATAAACCATTAGCTCAAAGCTTTAACGACTTGTGTATATACCGTCAGTCAAATGCATGGACCATATCGATGCTCGTTCTTCATTGTTTTATTCCTTATCTTGCAATATTGTATTTCTATGAAAACATTTTAAAACATATTCGAGATATAGAAAATGAATCAGGAAGGACAGCGGTGTCAGATAGAACAACATTTATTGAACACAATGTCTCTAATACGAACACTCTTTTCTTGTCAATTAGAAATAAAAATAAAATACCTAAAAATACAAGAAATATTCTAAAAATTTCTGTTTTAATATCAATTTGCTACGCCATTTGTTGGACACCTTCGATTGTATATTATGCGCTAAGTACATTTTGTCATGAGAAATGTTTCTCAAATACTTATTTAAACTCTGTTAGTCGAACTTACTTAGAATTCTCTATAAAATATTTAGCATTTTTAAATTCACTGGCAGCACCAATGATTTATTGTTTTACTGTTAAAGAATTTCGATTTCATTTTTATTCTAAACAAAAAAGTAATTGTCAAATAGTCAGAAAACCAATTATAAAAAAATGTGAAAACCAACCTTTAAAAAACGTGAAAATCAATCCTTAAAAAATTTGAAAACCTATCTTAGACAAATGATCAAACCAAATAACATTTACTGTGACAAAACCACTTAAAAATGTGATTGGTTGAAAATGTGATTGATTGAAAATGTGATTGGTTGAAAATGTGATTG
>evgR018240c0g1_i01_Hydra
ATCATCATCATTTTAGCGTCCTTGTTTTCGTGATTGACGCACGGGACTACTAAAAAAATACAATGCCTTGTCTAAGAATAGAAACCAATGTCGCAAAAGATTCATTCAACCATAAAAACATGTCTGATGAGCTCCTGGAAGCTTTAGCTAAATCAATGGGAAAACCAATTGAGTATTGTATGGTTGTTATTGTACCGGAAACTTTAATCTTTACTGGTGTAGATAAAACTGGTAAGTTGCCATCAGTGAATGCATCTTTAATTAGCATAGGCTCTCTTGGTGTGAATGAAAATAAAAAAATCGCTGCAGTTCTTTATCAGATTATCCAAAAATATCTTGGTGTTGATGAAGGGCTCTGTTATATACACTTTAAGGATGTGCCAAGTAAAGATCTGAGTTACAAAAACACGACTTTTCACGATATTCTTGGTTAAAATCATACTACTGCAAAGATTACACAAGTTACAAGATTTTTTTGAATAAATAACATTAAAGAATTAAAAAAAAAAAAAA
>evgR018241c0g2_i01_Hydra
TTTTTTTTTTTTTTTTTTTGGCAAAAAGCTTAGTAAAAAATGATTGTGTTGAAATGATTCTTTTTTTTACTTCATTAGCATCAATAGAGCAAATCAACTGTTTAACGCAATGTTATTTGTGCGAGGAGGCGGAAAATTGTGGGTCAGAAAATATAGATTATTGTAATGGACCAGATAGTTGCTACATGATGAAATTTGTAAACTCAAACGGGATAAATGTAACGAATGCTGGATGCATAGATTGGTCGAATTGCGGAACGGATAAAGGTATGGGTAAATTGTGCTTCATGATAAGAAATTGTAAGGTAACTATGTGCTGCATGACTGATAATTGCAACTCTGATCTGATAATTGCAAAATCTGAATTCAAAATTGCAAAATCTGATTCAAAATGGATAAAGCTTGACAAACGTATGTTTTCTCTTGTTGCTATGGTTATCGCATATTTCCTGCTTAATTGAAACATCATAAAATTTGTATTTGTATTTTATATATATATAAT
>evgR018241c0g2_i02_Hydra
TAGAAATTGTTGCTCAAATAACAAACTAGCAGATTGGATTCACTTTGACAGAGAAGACAAGCAAAATTTCATTTTAACAAACTGAATGATGAAACCTTGGCATGGAGTTAAATTGATTTTGCTTCTTATTCAAACATTTGCAGTTTTAGAGCAAATCAACTGTTTAACGCAATGTTATTTGTGCGAGGAGGCGGAAAATTGTGGGTCAGAAAATATAGATTATTGTAATGGACCAGATAGTTGCTACATGATGAAATTTGTAAACCCAAACGGGAAAAATGTAACGAATGCCGGATGCATGGATTGGCCAAAATGCGGAACGGATAAAGGTATGGGTAGATTGTGCTTCTTGAGAAGAAACTGTAAGGTAACTATGTGCTGCATGACTGATAATTGCAACTCTGATCTGATAATTGCAAAATCTGAATTCAAAATTGCAAAATCTGATTCAAAATGGATAAAGCTTGACAAACGTATGTTTTCTCTTGTTGCTATGGTTATCGCATATTTCCTGCTTAATTGAAACATCATAAAATTTGTATTTGTATTTTATATATATATAAT
>evgR018245c0g1_i01_Hydra
CAAACGACATTCAAGTAAACTTCAATCAAAGATAAATTAATAATACAAAGCAAATGGAACCAATTATTCGACCAATGCATGAACAATTTTTTTATCAAGACGCAGAGCATAGTTTGGTAAAAAGAAAACTAATTTACTATATACTGAAATCGGTAGTAGCAATAAGATTTGCTCAACAAATTAAAAGAAACAAATGTTCGCCAAACGTTGTACTGACATATGTGGATGCTTTTGCTGGAAAAGGATTTTACATGAAAAAGACATTTCCAAATATACCTTTTGATGATAATATTTGCCATTGGGGATCTCCAATAATTGCTTTAAAAGTATTCTATAAAGCGTTTTTGGAAAATGATTGCTCGTGCATAATCCATTTATATTTTAACGACTTAAATAATGATCTTTATACTTTTGTCAAAGATGTTCTTGTTTTTAATTGTATAATCTTCGAATCAGATGAGAATTATGCAATTATCATAATCCGTGAAAATAAAATTTCAAAATCTGGAAATAATACTCTTTCTAAAGAACTAGTTATACACTTTTCAAGATTAAGGTTTGAAGATTTGGAATTCTTGAAAATGCTAGAAAATATTAAAGGACCAGCATTGTATTTTATTGATCCGTTTGGGTTTAAAGTACCAATGGACACTATTGTAAAGCTTATGGGAAATTCAAATGAAAATGAAATTATAATTAACATGAATATACAATTTGTAAGAAGAGTGATTAGTAACCCAATGATGTACGATTACTGTGATAAAAATCTTGGTGGTGGGAAAGAGACTTATGGTGAGTTACCTGGTGAAGTTTGCGAGAATTATAATCAAATCGCAAATAATTATATTAGTATTTTAAAGACAAATTTAACGCCTTCGGTGCTTTACGCGTGTAGATTCATACTTAGAAAAGGAATAAATAATCCAAATAACGCTGAGTTTTTTAACCTTATAACGGTAACTACTAGCCTTATGGTTTTGCAAAAGGTAAAACAAGCTTTTCAGAAATATAATCAACATCCTGGAGAAGCTTCCTTTTCCGACTACTACGCTATCAACGAAATAATACAATCAGAAGGCAGGAAAACTGACGATTTTTATGAAGCTTCTTTTATTTACAAAAAATTTTCTAGTCATGGTTCACTTTATCTTGGAGAGTTAAAAAGGTTGATATTAGAAGAAACTCCATTTCCGTACCATGCGAAAGCAATAAAGAATCTTGAAAATGAAAATTTAATTTCTGTGGAACCTTTTTCTTTTAATCCTTTAAATGAAAATCCTCGAGTAAATAAGAGCCTTTGTAGTGAAGTAAGTCACCACACAACAGATATAGAACCTAAATATGGAAACTTTTGGACAATTAATTTCTTGCGAAAATAGTTGGTCTTTTATTGTTTGCTATTGAAGTATTATTTTTATGTTAGTATAAGAGGTTACAGGATATTTATTTTTACGTAAATAGATATAAAAGGTCTATTGTCTTTTTCCCCCAAAAAATTATGATTATTGGTGTATACAGTGTCGAATTTAGTAACGATGTTTGTTTTAAGTGGTATGTTTTTTGCTAGTACGTTTTTAATACTAGTTACAGTGAAAATAAGAATAAGAATTAAAACTTATAAATTAGCAAAGGCTTGGCCAACATTTCTTCAGCCGTCATTTATTCGGCCTTACGGTATTTGCTATCTATCTGTTTGTAACTGGTTTTGAATAAACTGCCAGTGATTTTTCTAAGTAGAAATTTTAGATTGCTATAGTGTTTTTTAAGCTTTAGATATTGAAATGTAATAAAAAAAGATTTAATGATGTAGTAAAAATTCCTCGAAAAAAAAAAAAA
>evgR018248c0g1_i01_Hydra
GGTGAATAAAATATTAACTCAATATGGTGTTGACAATGCATCTGAATTAATGAATGGATTTCATGAAATAGGTAATAATTTATTGTCAGAACTCTTACAGTATTATTCATCTCCAAATGAAAAAAAACCTTAAAAATGTGGAAAATGAAAGATGCCTGTAAAATGATAGGCAGATCAGAAGCTTTTTTAAGAAAATTAGAGCAAGAACACCCTATTTATACTCCATCTAAAGATCAAAATGGAACGAGGCAATATTCACTTAATCTTATAAATAGAATCAGAGATTTTACCAAAACAAGAATTAAACGCCCTGAAAATTCAGAACCAATAATTTTATCTATATCAAATTTTAAAGGAGGAGTAGCTAAAAGCACTACTACCAACTTTGTTGCATAAATAAAAAGCAAGAAAAAATAGACTAAGAATAAAAAATAAAATAGCCTGAAGAAAAATAGAAGAAAATTCTAGAATGAGTAAAAAAAGGCTGAGAAAAGGTAAGTACAAAATAGTGAATTGGAGCGCATACAATAAATCACTTGAGAAAAGGGGTGATATCACGATATGGTTTACAGAAGAAGGGATAGAAAAATGGTATGAAGAAGAAGTGCGGGATAAAAGCAAAGGGAAGCCGCGCAAATACTCAGATTATGCGATACAAACAGCATACATATTAAGACAGGTGTTTAATTTGAGATTAAGGCAAACAGAGGGGTTTGTAAAAGCAATATTAAAATTGATGAAAGTAGAACTTGAGGTTCCAGATTATACAACAGTATCAAGAAGAATAAGAGAGTTAGCAGTAGAATTTATTTTTCAGAAGCCAAAAGGTGCGATTAATTTGATCATTGATAGTACAGGTATTAAGGTAGTAGGAGAGCAGGAATGGATAAAGCATAAGCATGGAGGAAGATTTAGGAAAATATGGCGCAAATTACACATTGGCGTAACTGATGATGGTAATATTATAGCTGGAGAAGTTACAACGTTGAGAGAAAGTGATATAGCGGTAGCCCCTAAATTATTTAATCAAACTACTAATAAAGTAGAGTCTGTTGTAGGAGATGGTGGTTACTTTAAGAATAGAATGGAAGCTTCTATGAAAGATCATAAAAACACTGAGAACGCGAGATTTATAGGGCCTCCTGGTAATGGCCAAATTAACTATGGAAATAGGCTAAAAGTAGAAGAAACATTCTCTAGATATAAACGAATAATAGGTAACAAATTTAAAGCACAACATTTCTTAGGTCAACAAAACGAAGCAAAAATATCATTACTAATTTTAAATATTATGAAAAGCATTGGGATGCCTAAAACAATCCGTATTGCTTAAAAATACCGAGAAAAGGTGCCTGCGGTATTTATGCAACAAAGTTCTTCATAGTAACAAAATCTACTGATAGTTCACGAACCCTCC
>evgR018252c0g1_i01_Hydra
ATTTTAGTGCAAAAAGATGATAAATTTACACACTATGTTGCAGTGATGGTTAGAAATAGAATGTTGAAAAAATTTATAGACAAAATACGATACCAAAAAGATTTACGACACTATATTATGTATGCAACAGTTTTTGTAGGAATGTTATTTATATTTTATATGTTTAATTTGCTCCCACTTGCCAAAACTGGAGCTCGTTCAGAATTTATTTCTTCTGTTTGCCCTGCACTAGAATTTTCTCATCGCTCCCTTCAACAAAAATTGTGCCATTATGATCCTATTTATCCTCTCTCATCTCCTGAAGTTACAGAGTCTGGTGTTGAGTACAAAATTGCTGTGATTGCAGATTTGGACGAAGAGCGCTCTAAGGTGCTTGACCAAGATTATACATGGAAAAGTTATCTTAAAACTGGGTATCTATTAAAAAGCAGTAATGGTTATAGTGTACGCTGGGTAGACACTTACACTATAACAACACAAATCAACGAAAAAGGACGCGGATTAGAACTATCTGAGTTATGTGTATTTAATAAAAAGTTATATTCAGTTGACGATCGCACAGGTATTATCTATCAAATCAAAAATGGCAAAGCCATACCTTGGTTGATATTAATGGATGGAGACGGAGAAACCGACAAAGGTTTTAAATCAGAATGGATGACAGTAAAGGATGGTGTTTTGTTTGTTGGAGGGCTTGGTAAAGAATGGACATCAAAAACTGGAGAGTTTATCAACTATAATCCTTTATTTGTCAAAAGTATTGATTCTCTTGGTGGAGTGACTCATCATAATTGGACTTTGAACTTCTTAAAGATACGCGAAGCTGTTGGAATTACATTTCCAGGATATATGGTTCATGAAGCAGTAATGTGGAGTAGCAAAAATAAACGATGGTTCTTTTTGCCTCGAAGAATGTCAAAATTAAAATATGATGACAAATCGGATGAACACAGAGGAACAAATGTTCTCATTTCTTGCAAAGATGATTTCACTGATATTAATGTTGTTCATGTCGGAGAATTAGATCAAACCCATGGCTTCTCATCATTCAAATTCATTCCTGGTACTCATCAACAGGAGATACTTGCTTTAAAAACTAAAGAGGTTGATGGTAATATTAGCACATATATAATGATTTTTGACATTCAGGGAAACGTTATTATGCCCGAAATGAAATTTGCTTCTGATAAATTTGAAGGTGTTGAGTTCATATAACTCATCTTTTTATGGAACCATTTTATTAAAAATTATTTTTTTTATTTATAATTTATAATATTTAGCTAGTAGGATAATAATTTTTTAAAAGATCTTAGTATTTTTATACTATTTAATAAATCTTTATTTTTAACGAATTCCAAGTTGTTTTAGTGGAAATGCAGTTTTTAGTGAAAGCGAAGTTTTTAGAAATGTCTCATCATTTTTAACTTTGTTTTGGAGAAGGTTGGTGTAAGCAAGTTTGAAACAAAGACATATTGACTATTTATTTATTTTATTTATTTAAACATTTGGTCTAATAATCTTTTTTATTTTTTTGTGCTTTTGTTTGTTAAAGAATAATTTTAATAGAAGAAAAATGTTTTCAAGTATTTCAAAAAAAA
>evgR018254c0g1_i02_Hydra
ATTTTAGTGCAAAAAGATATTAGTAGGTAATTTAAAGATTTAGATGTCTAATAAGGTTTTGTGTTATGAAAGTTACGGATCAAATGAAGAGAAAATAGTTTATTATGACGACGGATTAAATAGAAAAATAAAAAATGCATCAGCGACAGGTTTCTACACGACAATCAAAATATTATTTCAAAATGTTTTCCTACCTCAAGGATTTCCAGAAAGTGTCAGTGAAGACTATTTACGTTATCAGCTTTGGGATACACTGCAAGCCTTTTGTTCCTCCATAACGGGACTACTCGCTACTCAAGCTATGCTGAAAGCGTATGGTGTTGGTGATAATACAGCAACAGTTGCTGCTGCGACAATTACATGGGTGTTAAAAGAAGGTGCAGGAATGATAGGTCGAATTCTGTTTGCTTGGAAAAAAGGAACTCAATTAGATTGCGATGCGAAGAGATGGAGGTTGGTAGCAGATATTTTGAATGATTGTGCAATAATGCTGGACTTGATTGCTCCTTTGTTCAAACAATATTTATATTTGATAGCATGCTTGTCAAGTGTTGCTAGATCTATAGTTGGTGTGGCTGGAGGAGCTACACGAGCTGCCTTAACACTTCATCAAGCGAGAAGAAACAACATGGCTGATGTTTCAGCAAAGGATGGAAGTCAGGAAACTCTCGTTAATTTGATGGCATTATTTTCTGGCTTTATTATCTCTCCTTTAGTTGTGGATAATGTTATCCTTACATGGTTCTTGTTTTTGATTTTCACATTTTTACACTTGTATGCAAACTATCAAGCTGTCTCTACAGTTGTGATGGAAACTATTAATCTTCCGAGGCTGCATATCTTGGTAAAGCATTACTTTATTTATGGAAAAGTTTTAACGCCCAATGAAGTAAACAAGGTTGATCCTGTTTTACGAAGACCTGGATTTAAATGCAGTGTACATCTTGGATGTCAATATGCAGATGTTTTAAAGTGTGATGAAATACTTGATTTACCTGGCTTCAGTTTGGGGTATAACAGTAAAACTTCAACGATAAATGTTTCCTTGCATAAAGATACATCTGAACAAGAAGTTTTGCAAAGTTCATTCCTTGCCATGCTTGTGGATTTATCATTAAACAAAAATATAAACACATCAGATACCAAAACACAGACATCGGATACCAAATCACAGAATTTGTATCTCTATTTATGGGAAAACACTATTGCTACAAAGTGTTGGAATCGCGTCGAGATGGAAAGAATTCTGAAGAACTTTTTCCCAACATTCATTAACGATTTAAAAGTGGCAGGTTGGGATTTATCGAGATGTCATTTAGGAGTCGATGAATGGAGGATTGTATTTGCTGATAAAACTCAATAAAATTTTTTAAGAATATTCTTTATCAGTAGTTTATTATCACTGCCGCGATTCATATAATGCAAATTTAATTATCCTGAGCCTACTATACTGAGAATTTTGGTACAGGACGAACAATAATTTTATTCGAGTATTTCCAGAAAATTTATTTCAACGTTCTTGAAATAGTATTTTCAAGTAATTTTTCGAAAAACTTATTTTCAGTAGTTCACTTTAGGGTTTCTCAAACGGGATCCCGTTTTCCCGGGGATAAAAGTGGCTGCGGGATATCCCGGGATATCACTCTTATATCCCGGGACTAATTTTTTATTAGTTTTACAAAGGATAACTGTTTAACTAATTTGACCTAAAATAAAAAAAAACTTATTGCAAAAAAA
>evgR018255c0g1_i01_Hydra
AAAAAAAAAAAAAAAATTAAAAACGACACCTGGTTTGATTATGAGTTTAAAATAAAATATTTATTATTATTCTGGTAATGTGATGATCTCGAAAAACGAACTCAGCAAACTCATTTTAACATGATAAATAATTCTCTGTTTTACTTCGGTCACAATTTTCGCCATTTGGTTTTGAGGTTTAAGTTAAGTGTGTTACTGAAAGTGTTAGAATAAGGCATGTTATTATCAATATTAAACTTACATAAAACTCAGAAATATTTGAGAGAGCTCATTTATTGTGTGTAATGCAAGCATACGGGTTTAACTTTTACACATAAAATACTATTTATCGAATATATACTGATATATGTCAGTATGTCATATGAAGACTTGAAACATGAAGTCAACCGCTTAGCACTTGAACTTCAAAAAGCCTGTCTTGATAAGGTCCAAGCTGCTGAATACGGTTTAGCTGTTCTTCAAGAAAAGGAAGTTTTGCAAAATCAATATGATTTAATTAAACTTCAGTACGATCAAGTAAAAGAGCAATTGAATCTTGTGAGATTGAAGAACGAAAAGCTAACTAACTGTTCAGAACAAGGTCAGGGAAATTTACTGCTTGAGATTTCAACAAGGGAGTCTCTCTTAGCAAAAGAAATATCTGATTTGAAACATGAGTTAAAATGTTGTAAGAAATCAGACCGTGCAAATCGCCATGAATTGGAAATTCACAAGGAGGAATTCATGCAACTGCATGAAGAATTTACTAAAACAAGAGAAAAAACTGAACAACTTATTCACGAAAACAAAGAATATAAAGCCAAAGAAATAAAGTTTAATGAAGATTACAACATGCTTGAAGAAGAAAATACTGTGCTTCAGAAAAGTATTTCTAAACTGAAGCAAACCTTAATTGAATTTGAAGGCATGAAAGTAGAAAACAAGAGTCTCTTGGATGAGATTGATAGTTTACAACTACAGTTACAAATGAGCACAATTGCAAAGGAGCAATTTGAAAAACAACTGCATGAAGCTTTGGATACTGTTCGTGAACAGCGTGAGAGAAATGCACAACACATAAAAGAAATACACGATTTGAAACAAAACTCTATTTTGAAAATGTGGGAAATGGGACGAGGAGAAATATTTTCTACATTTGAACAAGAGCATCCTATTGTGACCAAAATTACAGAAGAATATAAAACACAACATGCAGTTCCGTCAGCAAACATTGTAGATGATTTAATGAAAGAGCTTCAGTCAAGTGAAATACGCGAGCTTGAAGAACAACTTCGTCAAGCTCAAGAAGAAAAACAAGGCTTACAGTTACAACTGACCACAAAAACTAATGCAACAGGGGAAGAAACAAGCGTTTCAGAAAAATTCGAACAAAGCGAGTATTGCGCGAATAACCAGCTAAAAGAATTATCAGATTTGAAGGAGGGGCTGAAGCAGTTTCAGAGTCGAGAGCTTGTGTACCAATTAGAAATAAAGCAGCTATCAGATCAATTAGAAGAGCTGAAAGAGCTTTTGATAGAAGTTGAAAATGAAAAGAATACCCTTGCAGTTAACAGCAAGGATGAAAGTAAGCAATTTTCAAATGACTTATTGGATGCGAGATCAAAATTACGAGATGCTGAAACAAAAATAAAAAGTATGCAAGAAGACATGAAACGCCTTAAAGAACTAACAGGTGAGTATCAAACATGTTTGAACTGTACACAATCGGAGTTAGTATCTATTAGTCAAGATTTTGTAAATATACTGAAAAATGCCAATTATACTGGTGAAATAGCCACTTTTGAAATCAATACTGAATACCTTAAAGTAGGCGATCCGGCCATTATTGTGAAGCTACTGACTCATGTTAAGCAGCAAGAAAAAGCGTTGGAAATTGTTTGCAATAATCTAATAAATAGTAGCTCTGGGATTAATCCATCTTTTGTGAGTGACCAAGCTCTAGAAAACGGTTACCTAAAAGAGCAAATAATAAAATGTCAATCATTGCTAGTGGCTAAGCGTGAACAAATAAGTACCTTGCGTACAGTATTAAAAGCTAACAAAGCCACTTATGAAGTTGCTTTAGCAAATTTAAAAAGTCGGTATGAGAGTGATAAAGCTATACAAAATGATGTTAGTAATCAGTTAAAACGTCAAATTAAGGCCCTTAAATCAGAATGTCAAACTTTTGTTTCGTTAAGAAGTATGTTTGCTACTCGGTGTGAGGAATACATGAAACAGCTAGAATTGAAAGACAAAAACATTGCTGCAGCTGAAAATGAAAAGCAAACGCTCAATATGCTATTAAAACAAGCGATTCATCAAAAAATTTGCTTGACACAACGTCTTGAAGAGTACGAAATCGCACGCGAACGCATTCGGCAGTTTACAAAATCGAATTTGAATAAAAATGTAAATAAAAAGCCTGGTTCACCAACTAAACCCATAACCAGAGTTTAATTTTTATTATTTATGATTTTTATTTACATAAATCTTGTTTACTTCAGTTGATATTTATATATTGAATTTAAAATTGTAAATCACTGTGATATATTTATATTAATAGAGAGATGATATAGGTATTTATGTATAATTACCATGTACATGTATTTATGTATAGTAATAAACCATATTTATATATAATATTATTATATTTATATATAATAATCATATACAGTGACGGATACAGAAATTTTCGGTGGGGGAAATGTCGAAATTTTTTTGGTATTTTGTGTTTGACATATGGATGTATGAGTGTCTTCTAAAAAGAAAAAATAAGCCCTCAATTTCTACTAAATTTTCGTATTTTCAGATTCTGGTGTGTGTGTGGGGGGGGGTCATACCCCCATCCCCTTCCCCTGGATCCGTCACTGATAATTATTAATAATCATATAAACCATTTTTATATTGCTAGTTAGCTGATAAAATCGCAGAGGATAATGCTATCAATCTTATTGGCTATTCTTAATTTAAAGAGATCAAATTTCTGACTTAACGCGTTATATTGCTTATATAATTAAGACTGTGTAT
>evgR018260c0g1_i01_Hydra
TTCCAAATGAGTTGCATTCATTTGTCTGTGAAAAACTAGTCACTGCTGAATGCTTAAGTGATTTTTTGTTTAATCAAAGTGTTATGATTGAATATCTAACAAGTTGGCTAACAAAGCAGAAACCATTAAAACGTCCACTTGTTGAGTTGATGTATTTTCCATGTTTTATTTGGTTGGCCGAGAATCAAAGTTCATATTTAAAAAATACTTTCCTTAAGACCTCAACTCTAAATGGTATTCAAGTTTGTGAATGGACGCTTAATAACATTAAGAATCTAATAACTGAAGCAACAAATTTAAAACTAAAATCTTTGGAAATTATTTTTGGAGTAGCC
>evgR018264c0g1_i01_Hydra
TTTAAATGAGATTTCTAAATCTCGTGAGAGATCAAAAATTGAACTAGGAAGAATGTTTAATGAAGATTTTTGGGGAGTTTATGATAATAATGAAAAAATTGACGAATTTGAACACTCTTCGCTGGGTAGCTTCCAATCGTTCGATTTTGAAGAAGATTTTGTAAATAATGGAAAAGACGATTTAACAACGACAATAAATGGAAATACGCAACCTGCTATATCGCTAAAGAGCGGTGAAAATTTAATAGGTGGCTATTGCTATTTAGATCTTCTAAACACCAATTCTTTGAATGTTTATAAAGAGAAAACATTTAACTGTAAAATATGTAACAAATCATACAAACGTAAATCTAGCCTAGCAACACATCTAGTTACTCATACAGACAATAAACCCTTTACTTGTTTAGAATGTGGTAAAAGATTTTTGAGAAACTCTGATCTAAGAAAACACTCTATTATGCATACTGGAGATAAACCATATTCGTGCAACATATGTTGCAAGCTGTTTAGCCAATCTTCAAATATGCTGACGCATCTAAGACGACACTCTGGAGTTAGGCCTTTCTCGTGCCCTGTGTGTGAGAAGAAGTTTTTCCGAAAAGTTGACGTTTCTAGACATTTAAAAAAACATAACACTGTATTAACTCGTAATTAAAACATTAAATATAACTGTTTAGATATGCATCAATG
>evgR018275c0g1_i02_Hydra
TTGTGATCTGAGAGACTACTTTTTCGAGTTGCTTTTCTTCAAGTTGAAATTTTCAAGAGCTGATATTTGGTATTTCTTTGGTGGATTCCAGATGGTGCTGCATATTCAAGAAGGGGTCAAACTTTGGCTTTATACGATATTTCAGCAATTGAGATCAACAAGAGTTTTACTATTGCTGGCGTATATTTTTATAACGTGGAAAAGTTTACAAGGAAGGACGCTTACGAATATTTCATTTCAGTCCAAGTACCAAGGCAAAATTGCCTTACCCGTGCGGGATTTCATTGAAAAGGCCTATATATACAGGATTTTACCATTTATGCAATATGTCAAGCCTATACATACTTATTTTAAGAAAAGTTGTCCAGCGCGATGTTTTGTTTACTAACAGATTTCTTTTATTGTCAATAATTTTACAAAATTTAAATTTGTTGTACTGCGTTC
>evgR018277c0g1_i02_Hydra
TTGGTTTTTTTATGAAATGTGAATGAACATTTTCAAATTAACTTCATTTTTTGATTATTTTAAACAAAAACTAGATCCAGAAATTAAAATTATGTCTGAGAGGCCCTATGATATTGCAGTTTATGGAGCAAGTGGTTACACCGGGAAATATGTTGCGGCTGAAGTTGTTAGAACATGCCAGGGGAAAAAAATTGCTATTGCTGGAAGATCAAAAGCTAAATTAGAAAAAGTTTTTGATATTATTGAAAAAGAATGTGGCTGGAATACACGCGGTGAATTAGGTATCATTATTGCTGATAGTTCTAATGAAGAATCCATTAGAGAAATGTGTAGACAGAGTTGCATTGTCATAAATTGTGTTGGTCCTTTTCGATGGTATGGAGAACAAGTTGTTAAGGCTTGTGTTGATATGGCAACTAACTATGTTGATATAAGTGGTGAGCCTGAGTATCTCCAAATGTTGCAATTAAAATACCATAAACAAGCAGAAGAAAAAGGAATACATATAGTTGGTGCTTGTGGATTTGATAGTGTTCCAGCTGATGTTGGATTAGAGTTACTGAGAGAAAAATTTAATGGCGAATTAACTGCAGCTGAGAGTTACATTCATTTATATGGACCACAAAAAGGGAATTATGGGACTTACCTAACTATCATACACAGTGTTCAAGGCAGAACAAATTTGAAAATTCAACAAAAAGCAATTTTTAAAGAAAGGCTTAGATTTACTGGACCCAAACTTCTTCTGAGATATCCTGGTTTTTCAAAATCTGAGAATCGTTGGTTTATTCCATTTCTTGGTGCAGATGCTTCTATTGTTAGACGTACACAGTTGTATGAGAGTGTAAATCACAATCAAACACCAATTCAGTATGGCGCTTATTTTACTGCTCCATCATTTTTGGTTGCATTTTTTATGATCCTATTTGGACTTTTTGTTTGGATATTTACAAAGTTTTCATTTGGGATCAAACTGCTTGAAAAATATCCTAAAATATTTTCATTTGGAACATTTTCATTTGAAGGACCTTCACGTGAAGATTTAGCACGCGGAGGATTTAAAATGGTGTTTCATGGAAAAGGATACAGTGAAAAACCTAGTTTAAGTACTGCTGCTGGGAAGCCAGATAAAGGATTATCCATGCAAATAATAGGGCCAGAAATTGGTTATATTTTCACATCTATATGTGTTGTTGCTTGCGCAAAAACAATTTTAGATGATAACCTTAGAAACAGAGGTGGAGTACTTACTGCTGGATCAGCGTTTAAAGGAACGGGATTAATTGATCGTTTAATCAAAAGAGGTGTGAAATTTGAAATTCTATGATGCGTCAGTAATTTTTTGTAATTTTTTTAATAAGGGCGTTTTTGACATATAAAATGACATTTTTATTAAATTTTTTTATTTTTTGACTTTTTCTTTGATTATCTTTTCTTTTTAATGTTTTTGTTTTCTTTTAATAAAAATTAAAAGTTGAAGTTGAACCCTTTTTCAAACATTTTAAAACTATATTTTTGAATCTTGCGTTTATTTACTGTTTGTAATCAATATAAAAAGGATATAAAAGTACAATATAAAATATATAAATTTACTAATATATAAAATTATTGTTATAAATTATTGCTTTCAAAAAAAAAA
>evgR018278c0g1_i01_Hydra
CGCCACAGAATATTTAGTCTAAATTTTAGATATTTTGTTTTTTAATATTATGTTTATATTATAGTTTTCTCCCTTAGTTTGCACAAACATCAAAATGGCCCAAGAAATTCTTCAGAGTCAAGGATTCAAGTTAAAATTAAGTGAAAGTGTTGGAGTAGATTGGCTAATTCTTGGTCGATGGTTAAATATTAAAGATACTTGTTTAAACATGATTGATCAAGATAATACTAAAACTGTAGAAAAAGCATATTCAATGTTAACGAAATGGGTGCAAATGAAAAACAATCCAAACCTCGAGGAGCTAAAAACAGCGTTACAGAACATGAATAGAAAGGATTTAGTGAGAGAAGTAGATGTACTTTCAAATACTTCAAATTCACCAGAAATGGCAAAAGGATTATCTATTAAGAAAGATATATCAGAGATACGCACAGCACTGAAAAATTATTATCTTGGAAATTATGAAAAAATTAATGAACTTCAACCGCTGTTAAAAGCAGATGCAAATGTTGATTTAATAAATAAGTTTATCAACTTATGTATTATTGATTCTGCTAGTTATCAAAAGGATGTTTTTAATAATGAACGAAGTATGTTTCTTAAAAAGCAAATGTGTTATGAATCAATTCCATATGATGAAATATTCACGAAAGAAAATCCTGTAATATTAATATCTGGAATAGCTGGTGTTGGAAAAACATGGTTGCTTAGAAAATGCTTACTTGATTGGTCGAAAGATTTAATTTGGAAAAATGTCGATCTTGTATTTTATTTGGAATGCAGAAGACTTAATCAGCATGAAAATATTTCTTGTATTAATGAATTACTAAATGTTTTCTACAAAGATATTATAACCGAATTGAATATTAATAATCATACAGCGCTGTTCATAATTGATGGATTGGACGAGTTTAAATATTTAAATGAGTTAATCAATCCGAGTTTGAATTGTAACTATTCCATTGTTAATGTTTTAGAAGAAATTCGAAAATATAAATATGTAGTTGCTGGTAGAGTTTATGCGATTGATCAATACCAAGGTATACCAACAGACCATAATGATAAGCTAACCATTCAAATAATGGGGTTTAATGAAAACGGAATAAATAATTATATTGATACTAATGTTGTTGACGAAAAAAAAGAAATTGTGAAAGAAATTTTCAAAGAATCTCCAATTGCAAAGTCCATGGCGTCTGTTCCATTTTATTTATCTTCTATGTGTAAAATTATTGTTGAATCGAAACAAAGAGACTTTTCATTCTTAAC
>evgR018281c0g1_i01_Hydra
AGCTTATCACCTTTACCTTAATGAATAAATATGAATGTCTGGATGCGTCTGTCATTTAACATCATTAAATATTCAAGTTCCAGAGTATTGAGCTCAAGTGCCATCAGACCATCTACTCACGATTATTCTGCTGTTAAATTAGCATTTCCATCTCATTTCATGTTGGATGAAAATGGTGTTAATGCTGTTAAACAAATACGAAAGAACACAGGGGCCGTTGTACGCGTAGTAGAGAATCCAGAGGTTGATAATACTGTAGAAAATATTCCAAGTTCTTTTCTATTGGTTGTTGGCAGCTCATTCCAAATTAATAAAGCTTCAGAACAAGTCAAAAGTTTATTTGAAGGACGTTTGCCCAAGGTGGAGATTAATAAACGATCTGATGTTAAGCTTATTTGGCATGGTTATAAAAGAAACTATAAAGGGCAATTTGCTCCCATAAAGCCTAGAAAAAAATGTATTCGTTGTGGTGGAAAAATGCTTTCTGGTAATCCTTGTCCGTTATGTCAGTTAAAAATAAACAATGACTATGATCTACACTTTACAGATGTTGAGTTGTTATCTCACTTTGTTTGCCCATTTACATGGCAAATCTTGAGTCCATATGTCACAAATATATGCCAAATGCAATATAAAAGGGTTGAGGCAGCAATTGTTAAAGCTAGGCATTATGGATTAATGCCATTCACTATTCCATTGCCTATGAACGAGCCCAGGAAACATAAACCTGCTGGAGTACCTACTGATGTGACAATTAAAGTTAAAATGCATTAATATGAAATAAATAAAGTTTACAAAAAAAAAA
>evgR018282c0g1_i01_Hydra
CAACTTTAAACTCAATCACCAAATCAACCTATGACGCGATCTTTCTTTTAAATAAAACATTTTAAAATAAACGTAACATCTGAAATGGAATGGCAAGTAAAATGGATAAGAGATTTTACTCTTTGTTTAAATACAATAAAACTGTTCCTCACTCACAAGATCCACACTATAAAGATCGAATATACAGTGATACGTTATAGACTACAATAATATAGTAGTTATCAAATCAAGAGTAACTATTTTTTACCTGATAGAATCATGTTTACCAAGTACATCTCAATAAATTCCTGCAATACACTATAATACAGATGGAGGAAAACACTCAAGAGTTGTTTAGGGCTGTTGAAATTGGTAGTTTGACTGATCTTTTGAAGTGTATTGATAGAGTAGAAAACTTAAACCAGCAAAATTCTAATGGTCAAACTGCTCTAATTTTATCTGCTGAGAAAGGATACATGGACATTGTTCGAGAGTTGTTGAATCACGATGTCGAAGTCAACCACAGGGATGAGGATGGTCGAACAGCATTAATTGAAGCAGCAAAGTTAGGTTCTAACGAGATCGTTATGCTTCTTCTTGATCATAGTGCGAAATTAAGCATACCAGATCTGGGAGGCTGGACACCATTAGTCTGGGCTTCTTACAAAGGAAATACAGAAGTTGTTCGTCAGCTTCTTTTGCATGCAGCTAATCCTAATGAGCGTGGACAATATTCAATGACACCTTTAATTTGGGCATCTGGTCGAGGTCATTCAGATGTGGTACGAGTTCTTGTTCAGCATGGTGCAAAAGTTGAAATTGCTGACAAATATGGTACTTCTGCATTAATATGGGCTTGTCGAAAAGGTCATCTAGAAGTTGCTAAGATTTTAATCTCAAAAGGAGCCAGCTGTGATGTTGCTGGAACGCATAGATGGACTCCTCTTATTATGGCATCAAAAGGTGGATTTACAGAAATGGTGGAACTTTTGTTAGAG
>evgR018285c0g2_i01_Hydra
TTAATTAACCTATTTTAGTGCAAAAAGTTTTTTAGTATGAATACAACACAAGATATCCATAATGGGATGAGTATGCCTAAAGAAGGAGGTGTTGGAATGATGATGATGCCGATGTATTTTATGCTGGGAAACAAAGTAGCTCTCATTTTTGAAGGCTGGAAAACTAAAAATGATGGAGAGCTAGTAGGTTCGTGCATTGCGTTAGCAATTATTGCATTTTTGTACGAAGGCTTGAAGGTTTTACGTGAAATCATTAAATATCGTTACAATGGCTTCAATAATCGTCGTAGTCTTATGTGTTCATGGTTACATATCCTGCAAACAGTTTTACATATGGTAGTGGTTTTTATCGGATACCTGCTCATGCTTGCTTTTATGACATACAATACCTGGATATGTATAGCGATAATAATTGGTGCAGGGTTCGGTTATTTTGCGTTTGGGTGGAAGATAAACTCAATATCTAGCACTGGTGACCATTGCAACTGAGAAACATTTGTATATTTTATAATGAATTTAAAATAAGTTTTATAAACACTTGATAAATTTGAAATATATATTTGTTTAATATATAATAAAGTATATATATATATATGTATATATATATATATCTCATCATTGTTTTAGCATCCTTGTTTCCGTGCTTGCGCAGGTTGGACATTTCTC
>evgR018289c0g1_i01_Hydra
AGTGTTTAAGATTAGATGCTAATATCAAATAGATTGGTAATGTTATATTTTGTATAGACTCATCAAGAATGCAAGGTTCAACAATCGGACCTGGTTTATCAACAAAAGTTCACATGAGTTTGACAATTTTTTTCACTGTGGTTTATGGATTGGTGTTTTTTTTGATTATTACCCAGTTGTTGTTAATACTATATTATAAACATCGCCGCCTATCTTACCAAACTTTTTTTTTGTTTACCTGTTTAATCTGGGCTGGTTTGCGCACAACTTTATTTTCATTCTACTTTGAAGACTTAGTTCAATCGAATAAACTCAGTCCACTTTTTGAATGGTTGTTGTTTGCTTTGCCTATTTATTTACAGTACAACATGTTAAGCATTTTAGTCTTCTACTTTTTACTGGTTGTTGTTAAAGTTACGAGTCCATCAAGGCTTGTTATACTTCGGAGAAGAATTTTTGTTGCTGTGGTTTTATCAAACATTGTATTCTTAGCAGCAAATATTTCATGTTCAGTTGTATATACCCATAATCCATATGCACATTACCAGCAAAATGTAACGTTGATTCGTGTCAGTATTGACTACTGCATTTTTCTATTTGCTGCTTTAGTTCTATGCTACTGCATTATAAAGTTAACGCGTCAAAGTTCTGCTAAAATGTTGCTTGAAGGCCAGGGTATCTCTTTTTGCCAAGCAATAACTGTTTGTATTTCAATTGCCATGCTTTATATTTCGAGAGCTGTTTACAATATAATAGCTATATTGCCAGTAAGCTCGTCATTTCCAGACTTTAACTATGGTTGGATCAATGTTTCCGATCAGGGAGAAGTTGGATTTGATGGTTCAGTTTTGCACAATACAACTAATTATGCTTTTATGTCATTTGGAATTGTTTTAGTTGTATGGGAACTGGTGCCAACATTTGTCACTGTTTGGTTTTTTAGAGTAAGAAAACCTGATGGTGTAGCGCCACCTTCAGCTGCAATATCTGAAAGTTACCACCATGCTAAATCTTATTTTTTTGACAACCCTAATCGTTATGATTCTGATGATGAGCTTTCAATTGCATATACACCTCCTCGATCAGGAATACCTGGAAGTTTGCCAGATATTCCCAACTTCGGCAGTACATCCCATCTAGTTCGAAGTGCAAATGATAAGGTGCTGGCACGGAAATTAATACATTCTTATGGATCTGTACCAGCATATGATAACTACACTATTTCACGTGAAGGCGTAACTAGTCCTAGCATTCATAGTGCAATAAATCCAATGGTTGACAGCAAAAACTTTCCTGGCAGCAAACTGAGAAATTCTGTTAAAAGTTTTGGAAATATATCACAAAATACTAACTCTATGGAACATATAGGAAATGTTATTAACGATTGAAATTTCAAACAACCCTATTTGTTTTTATGGTTTCATCGTGTTTGTTTTTTTAAAAGCTGTATGTTGTCTGTGTTCTTTTTAACTCACATATATATATTGATTATTATTCAATTAATAATTATATAACGATTGTTAATGATGCAATAATTGTTAATTAATGATGCAATGATTATCAATGATTAATGATTACCAATTGCAATAATTGTGATTGGCTATTAATGATGCAATGATTATCAATGATTAA
>evgR018293c0g1_i01_Hydra
TTATTTCAAATAGTTAGGGAGAATTAGATGTACAATTGTAAACAGTGTGATGTTGCTCTACCTACTATTAATGGGTACATTAATCACTGTCGTTTACATTCTATGGAACCAAGATTCAGGGTTGCATGCTGTGTTTCTGGATGTTTTCAAAACTTTTCTACTTTTTGGAATTTAAAAAAACATATCTACCGCGATCATGGTCAAAATAAATTTAATGCAAGATTAACTTCTTTAAATAATAAAGTTATTTCATTTTCCTGCCAAGTTCCTATGTGTGGTGCGAAATGCAAGGATCAGAATGCATTGTTTATTCATTTAAAATACCATATTAGGGATGGTACAGCAATTCAATGCCCATATGTTGGTTGCAAAGCAAAATTTAAAATTATTTCATCTTTTACTAGCCATAAATCTAGGAAACATATAAATCCTTTTGCAACAAATATAAGTGAGTTGTTTGTGGGAAATAATAGAGACATAAATGTATCTAAGTTAATTGGATATTTACTAGTACCTGAGATTGTTGACACTGAAGATAGTCAAGAAGAAATT
>evgR018301c0g1_i01_Hydra
ACACCAAATGTTCAATTTCTCTAGTGCTTCAATTTAATTTTAAGTTGTCTAAAGCGTTTTATATTCCAAAGATAGAGCCGTTAAGTATTCTAATTCCGTGGACCCTTCGCAAAAGTATTAAAAAAGTGGATTCTAAAAAATTCAACTATTTTTTATATCGTTCCGCTGTGAAAGATGATGTAGCAGATCAATCCTTTAATGTGAAAAACGGTGAAAGACATCATGACACTCTTTTTTGCGTGGAAAAAAAAAGCGTTATGAATAGAAACTATTACGATGATTTGATTCCGCACTGTGTTTCAATTCAAAATGATGGTAATGAATGGGCGAGTAATTATAACAAAAAACCCGCTCAAACGTTTTATGCTGACGTTATCGATATGCAAGTTAATAAAAACGAGGAAGATTATTATTATTCGGAAGATGAGCGATTATATGCCTATAGATTTACGGAAATATCCACTATAACTAAAAAAAATCTAAAAAGATGCGCTACTTCGAATTTTAAAAAAATAGCGCTTAAAATTACTGAAGATGAAGAAGAAGAAATCGATGAAGAAGCTCTAGAAAACGATTCAGAAGAAAATATGGATAAACCTATTCCAGCACCCAAACCTTCAAAGAAAATGAGAGTATGCGAAAGCGAAGACGAACCTCCAGAACTTCCAAAAAAAACAAAGGACGAAGTTATAAAATCTTCAAAAAAGAGCAAAAAACCCAGTTCACCCAAAGAGGTTGTAGCTGAAAAAATTGAACTTTAAAATGCATAAAAATAATATTAATTTGTAATTATATACGGTTTTTATTATATAAAAAATATGCG
>evgR018302c0g2_i01_Hydra
CGAAAGATATACAAAAAAACATTTGTTGGGTATTGACCAGTTTCCAACAGTTGCCGAGTTTGTTGATCCAGAGACATCTAAGCTAGATGCAGAAGAATTGATAATTAAAAATATAAAATGGTTTTCTGAACACGTCCACACAAGTCAATACTTTACTCAAATTGTCAAGTGTCAAGATGTAAAATGTTGCTTTAAAATTAAAAGCTCATATTTCAATGTTGTTCCTTCGCGATTCTTATGTCTTCTAATTCCAGTTATTCAAACAAGTAATGGTCTTCAAGCTGACAAAAGAGAAGACAGTGAGTCGCATAAGTTTCCATCACTTTTTGTTTCACAAAGTATCAAAATAGATGATATAATGGCAAGATCGGCAGGAAGTTTCCAAGTTCATCCTTATGATCTCTTTTTGTCCATCAATATAGACTTGTCGATTTAATCGAATATGAAAAGTTTGCCATCTCTATTTTGCTTCGCAAGCTCTTGAAATGTCGGATTGCACAAAAATTGC
>evgR018303c0g1_i01_Hydra
GTTCATGCAATATTTAAACTAATTTACATTAAATTTAACTTTTTCAAAAGTAGGACGCATACTTCTTGTCAACTCCGCCCCAAATACAGACACCCCTGAGCTGTCTCATTTATTGTGTTACTATGAAAACCTTTGAAAACATATCCAACTATAATGCTCCTCAAGTGAACAGTTTTATAAAGTATAGAGATATCTTCCACTCAGCAAATATATTTATTTTGTTTGGGCTAGCATTAGTTTTTTTTGTTGGTGTTATCGGTAACGTCGTAGTTCTTTACGTGTTTGCTTGGAAAAAGCGTCAAAGTCGAAGCAGGTTTGAAACCCTTCTTGTTCCTTTAGCTTTTATAGACCTATTAACTTCGTTCATAACTCCAGTTATTTTTACATACGGTACCCTAACTAACTATAGACGTTGGGATTTCGGCCTGTTTGGTTGTAAAGTTTTGTCAAGTTTTTTTCCCGTTTCGATTACCATATCTCAAAGTATATTAGTGTTCATATCTTATGAGCGATATAAAGTGGTTATGTCTCCATTTTACAGAAGTTTAAAAAAGGTTTTAGTACAATTTTGGGTTTTATTGACCATTTTGATTTCACTAATTCTTGTATCACCATATACATATGCTTTAGAGATAGTTATTGATACTGGATACGATATTTATACATGTATGCCTAATAGTAACAAACATTATCAATTATACTTCTACTCTGTTGGGAACACAGTTCGTGACTTTTTAGCCACTGTTGCCATGCTTTACTTTGGTTATAAAACCAATCACTATATTCAAAGTAATATCAAAGCAATTACAGATCGACGACATAAGTTTAAGCGTATGATATGTGCAATTAAAGCAAGAAAAATGCTGGTTGTTGTAGTTTGCTGCTTCAGCTTTTGCGTATTTCCGTTAGACCTTTTTCAAGCAATAACATATACTCTTTACATAACAAAAACAATACAAAACATGAAATCCTATAACTTGTTACTGCAAGCAAATACTTACTTGACCGTACTTCAAGTTGCAAATAGCGCTTTGAACATTATTATATACTCTCAAATGCATAACGATTTTAGGCTTGCGAGATGTTGTAAAATGTATAGAAAAAATCAACAATATACTGGAATAAGGTTACGTTCAAAAACCAGTGAGTCGGTGATTAACAATGGAATTGATTTCGTAAAAGTATAAATATGCTAAAAGGAAAACCAAATATAAAAAGAAAAACTAAATAATATATTAATATATACTTTTTAATAATTCTGCTAAAAATAAATATTGAAAAAAAAGCAATTAAAAATGCTAAATATATAGACTTGGTAATTGTTGTACAAC
>evgR018307c0g1_i01_Hydra
CTATTTTAGTGCAAAAAGTAAATATAAATGGCATTTAATTTTGGAACAAAACCAACAACTTCAACTGGTTTTAATTTTGGTACCTCTATAGCTACACCTGCAGCAAGTTTTTCGTTTGGACCAGTCACTACTCAGTCTTTTTCATTTAATAATCCTGGATCTCAGAGTTCAGCAGCTACTAGTGGCTTTTCCTTGGGTAGTACTAATGCACAAACTTCCTCAAGCTTCACAGGGTTCTCTTTTGGAGAATCAAAACCTGCTGTCACAGGTTTTCCTGCAGGAAATCAAACTGCCGATACATCCAAGCAACCAGCTTTTAGTGGATTTTCTTTTAATAGCTCTACTGCAACTACAAAACCTTCAACTAATGTAACTTTTGGTAGTAATTTGAGTACTTCTAATAGTTCATTTACTGCGCTTAGTAATAATGCAGGAACTTCAGCTACGCCAACAGGAGGGTTTTCTTTACCTGGTTCTACTAAATTTTCATTGTCTACCCCGAATCCTGGATTTTCCCTTCCTTCTTTGAGTACTGGGGTTTCTCTTGCAGCTTCTAATGTAGGATTTGGATTACCAGCTTCAACTACTGGGTTTTCTTTGCCTGCTGCAACTGGATTTTTAGGTGGAAACTCTGGAAACTCATCACAACCAGGTGGATTTAGTTTCGGAACTCCAGCTTCAAGCACCTCATTGACTTTTGGCGCTTCTGTCCCTAATACAACAACTGCGGCACCTCTTTTCAGTGGATTTAGTTTATCTACCTCTAAAACTTCTGCTGCTACTGATCTCAGTAAATCAATTCCATCTTCTAATTCCTTGTTTAATGGTGCAACAACATCATTATCTTCCTCAGTAGCAACAACTAATTTAAATCCCTCTAATGCTATGCCACTATTCAGTGGACTTTCTTCTTCTGGTATTGGTTCAATTACTTCATCTTCTGCTCCACTATTTGGTACCTCAGCTGCTACATCTATTCCGACATTTAACTTAAACAGTACAACATCAGTGTCATCTGTTGGACTTCCTGCTGTCTCTACAGCAACAATTTCTGCACCATCAGCCAAACAAATGTCATATAAGGAATTAGAAGAAACTGTTGACAAGTGGTTGCATGATCTTGGCGAGCAAGAAAAGTCATTCTTAAACCAAGCTGCACAAGTTAACATATGGGATAAGCAATTAATTGATAACGGTGAAAAAATTACTGCTCTGCATAATGAGGTTGAGCGTGTCAAAGCAGAACAACACCGGTTAGACAGGGAACTAGATTTTATATTATCACAACAAGAAGAATTAGAAGAAATACTTTTACCTATTGAAGAGCAAATAAAATTACAGCCTATTAATCAATCGACACAGCATACAGATATAGAAAGAGAGCAGACATATCGTTTGGCAGAGAGTATTGATTCACAATTAAAACGAATGATGCAGGACCTTGGTGATGTAATTGATCATATAAATATGGCAAATATTGAACCAGCTAAAGGTCGCGATGAACCTATGTCTCAGATTGCAAAAATTTTAAATGCGCATATGGATTCTTTACAATGGCTAGATCAGAATACAAACCAGTTGCAACGAAGAGTTGACGAAGTGAATCAAGCCCTAGATACAAGAAAACGAGAACATGAACGGAGTTTTCGTTATGATTAAAAACCGTCATCGTTATTATTACACACAGTGTGATTTTTTATTGGTTTCTGATGTGATTTTATTTGGTTTTGATTGGACAAAATTTATCTTACCATAATCTGTAAACTAGTAATGTTATATTAGAATATTTTTGTAAATAATATTTTCATTTAAACCAAAATGTGCAAAATACTATGATATTTTATATTCCGTTTGTCCATTTTGTCTTTGAGCAAGGGAAAATAATTTGTTGCCGAC
>evgR018308c0g1_i01_Hydra
ACAAAATACATTTAAAATACAATTTATTTTTAATTAGTTCTGTATCATATTATTTCTTAAAGGATAAAAAATGGAGTGGGCTGTCAAAGGAGAAAAAAAGGCTTGTCATTAATGAAGAAGTTGATAATTTTTACAAAAAAGCTCAGGAAGATGCTGAAATTCATGAGGTTGTTGTTAGTTATGATCAAGAAGAGCCCAAGAGTTCTTTTTTAGAAGCAAGTAGCAATAATAGCATTAACCTTGGTAGTTATTGTTCTTTTGAAAGTATCATAGATACAAGCTCAGAAGAAATTAAAAATAAGTCACTTTTAGAGGAAATTCGTGTTTGGGCAATTCAATACAATGTTGG
>evgR018310c0g1_i01_Hydra
ACCTATTTTAGTGCAAAAAGTTATCTAATTTAAATATTCATGGCATGGCTGCTAAAAATATTAGTACAACAGCACTTGTGAGTTTTGGACCAAAACCAAATGACGGACCAGCTTTTCAAATGTTTTTAGGCGAAACAATTTTATCTCAAGCTAGTGTTGTTCGGTATAATGATACATCTAATCGTTTAAATAGTTTAACAGGAACTTTGTATATAACAAATTTTAAAATAGCATTTATTTGTTCGGTGAAAGGTGTTTCAAAGCATACTTCTAGATTTAGTGAAAGGTTTTCAGAAGAACCAATTGATTGCAATGATCTAATACCACTTACTTCCATTAGTAATATTTATGGTGTTTCTAATATGAAACAAAAGCGAAAAAAGCTAAAGAATGCTATGATAGAACTCTTAAGTCAGTTTGATATACTTGAAATAGAAACAAAAGACTTTAGAGTTGTTCAATATGATTTTAAGTTGTCAGGAAGTCAGAATAAGCGTTCTTGTTTTCAAATGATATTTCATTATGCGTTTCCAACAACAGTTTTTCGTTTGTTTGCATTTGATTATGGCAGCAGTATTGTTACAAGCAATAGTAGTGCATTTTATTTTTTTAAAACTAAAAAAGATTACGAGCTCAATATCAGCCGATTGCATGCCTCACAAAACTGGCGTGTTACTGATGTTAATGAGAAGTTTGCTATTGGTGAATCACTGCCAAAGTTTATTGTTTGCCCTGCTTCTTTAGATGATAACGATATTGAACAAATTGCTAGAAGCTATGTTAAAAAAAGATTTCCAGTATGGATTTGGAGTGATCCTAGCTCAGGGTGCTCTATTTTGATTTCATCTCACCTGAGAAGTTTGTCGAGTAAAGATCCCAGATTGCGTCAAACAATTCCATCAAAGCTTTCAAGTTCATTGAAAACATCTTCTACACTCGGTAACGGAGAGAGTGTTGCAATTGATTTAAATAATATTTCTGATTTTCCTTCGTTAAAACAGTTACAAATAAGCTTTCAAAAAATCCACAAGTCCTGCACTTTTCATTCAAGTAAAGGATTTTGGAGTTCTGATTCCAGTTGGTTGAGCAGCATTGAAGAAACAAGGTGGCTCCATCATGTCTCTTCGTGCTTGACAACTGCCAAATTGTTAGCAAAACATATTCAAAAAAAACATTTAAATGTTTTAATAGAAGAAAGCACTGGGCACGACTTTTCTATACTTGTAGCGTCCTTAGTCCAAATTATGCTAGATCCATATTATCGCACTCATTTAGGTTTAAAAGCTCTTATACAGAAAGACTGGGTTTTAAAGGGTCATCCTTTTTCTACAAGATTGGGTACATTATTAGCTCACTCAATTAATAAATCTGATTTTAATGGTCAGAGTCCTGTTTTTTTGTTATTCTTTGATTGTTTACACCAACTTATTAACCAGTATCCAGTCTTTTTTGAATATACAGACCTCTATTTGGTACTAGTTATGGATTGTGCTTATTCGTCTTTGTTTGAAACATTTATATTTGATTCAGAAATGGAAAGCGAAAAGCACTCAAAAGACATGATATTAGTGTCAGTGTGGGACTTTCTTGAAAGCAATATATCTGAAGCTAAATTCCAAACACTTTTTACTAATCTTGCATTCAAACATAATAAACAAAGCAGCAAAGAGTATATTTATCCATTATGTAAAATCAGTGATATTGTGTTTTGGTCTAAATATTTTTTAAGGTGGTTGTCTATCATCAATGAAGATCTAGTAAGTTTCTCTGGTATTGTTTTGCAATTACAGCAAAATCAGTTTAATGAAGAAGTTGAATATTTAACGCATCGTCTTATGCAATTGCAGTTGCAATGCAGTGTAACTAACTCCGTCAGCACGCAATCTCTTTGTTCAATCAATCAATCAAAAAAAAAAAAGATGACTGAAGCTCAATCAAACAGCATGAAAAGAGATTTAATCAATGCTACCTTTGGTTTTCATTTATTGAAACTAATTTAATTTTTATGTTTAGCTTTTTGTAAAGTATTTTTAAAGTAGTTTTTATATTTTTACTGTTTTATATTTTACCTAATATTTTTTATTTAAAGCAATTATTATTTATTCTTTTTGGTGAATATATTTCCTTTAAATTCAATATATATTAAAG
>evgR018312c0g1_i01_Hydra
TAGTGCAAAAAGTAAAATAAGATAAAATGATAAACAGTACAAAAACCTTTACCAACGACTATCAACAACTACCGACAACCGAAACCTTCATCAACGCTAGTGTGTCTGCAAATGTTAAAACTTTAGTAACTCAAGTGTCCGTTATTTTGGTTATAAGTTGTCCATTAGGATTAATTTTCCACGTCATAGGTCTTTACTTACTATTTCAAAACAAGGTGCGCAAAATACATAAAATTGATCGAGTTGTTATAATAAATTTATCTCTTGCAGAAGTCTTTTTAAATGGAATTCTATTTGTATTTGGAATATTGGATTTTGCTGAAACCAAACCAAAAAAATGGATGTTAATTGTTATGCGAACTTTTATAAATTCATATTACATAACAATTGTACTACTAACACTTAACCGTTTTCTTATACTTTATTTGCACGTAAAATATCGTGCGGTAATCACAAAAAAAAAAGCTATTTATTGTTTAGTATTTACTTGGATATGCAGTGTTGCCGTTGGATGTATTACAACTTACGAAATAAAAAAGAAAATTTTAGTAGCAAATGATTTATTTTTCGACTCGTTTATAACAATATTGTACATATTTTTGTATGCATATGCAATGAAATTATCACGACATATGTCAAAAAATAAATCGCTCCTTCAAGATCGCAAATCTTTTTCAAAAACTTTAATACTACCGACTTTATATATTTCAACCTACGTCGTATTAACTGCTATACCTCATTTTATATTTGCCCCTTATCGTTTGTCACGCAAATCAATAAACAAAAAACTTGAACTGATTTTAGACATTTTGTTATCTATATCTTATTGGAATGACGCTGTTATATATATTTATACAAAAAAACTGCATAGAGCAATAAGCACACG
>evgR018317c0g1_i01_Hydra
ATTTAATAATTTCAGTTCTCTATCTAAGCAAGTTTAGTTCTGAGAAGATAAAAATGTCTCGTGGTAAAGGAGGAAAAGCAAAAGGAGGAAAATCTAAGACTCGATCATCAAGAGCTGGTCTTCAGTTTCCCGTTGGTCGTATTCACCGATTTTTGCGCAAAGGACACTATGCTAATCGCATTGGATCAGGTGCACCAGTTTATTTAGCTGCAGTCATGGAGTATCTCTCTGCTGAGATTTTAGAGTTGGCTGGTAACGCAGCCCGCGATAACAAGAAATCGAGAATTAATCCGCGTCATTTACAATTGGCTGTTCGAAACGACGAAGAGTTAAATAAACTATTGGCGGGTGTCACTATTGCTAATGGTGGCGTGTTGCCAAATATCCAATCCGCTTTGCTTCCTAAACATACCGAAAAAGTTCATAAGTCACCTAAAAAGTAATTGTTCATTGGTGCATAAAAAAACGGCTATTTTCATAGCCACACATTTTGAAAAAT
>evgR018323c0g1_i02_Hydra
TTTTTTTTTTTTTTTTTTATTTATTCTATTTTTAAAAATGGATAGACATTTAAATTTAAAAATTAGCGCCGGATTAATTTCTGGTATAATAATTGGCTGTTGCGCAAGTTGGTTCTTCTCTTCATCGAAAACAAAAACGGAAAAAAAAATAGTCAAGGAAGAGTCAAAAGCAAGCGAAATACAAACCCAAACTGAAGGAAATGAGACATTTTCAAGTAGCTTGGACGAGTCATCCAGTAATGCAGTTTACAGTGCAAATGCAAACTTTCCATATCTCCGAGACAACAGATCGTTCATGGCTCGCCATTTAACCAAAAATATCTATAACGATATGTACAACCGAAGAACCCCTATGGGTTTTTCAATTGACCAAGTGATTCAAAATGGTTTAGATAATCCAAATAAAGTTGGTTGCGGAGTGGTAGCTGGCGATGAATATTGCTACGATTATTTTGCACCAATCTTTGACCCGGTTATAGCTGATCGACACGATGGTTTTAAAAAAAAACGTGAAATGGCTTTGAACCATGTTTCAACTGCATTAGATCAAGAACAAGAACCGATCGAATATCCAATTCAACATCTTTATGAACGTCCGTTCGATTCTAAATATGTCAAATCGTCGTATTTTTTAATTTCGAGATGCTTAAGTGGTTTTAGATTTTCGCCTTCGTGCTGTCGAGCGGAAAGAAATCTTGTAGAATCATATTTAAAAACTGCATTACTAGGACTTGAAGGAGATAATAAAGGTATCTACCTCCACTTAGCTGGTATGACAATGGAAATCAAATCTGCTTTGCCAGATATACTACAAACAGCTTCAAACACGGCTTTTGGTCGCACTTTGTCATCCCGCGGAGAGTTCCACGATTGGCCTTACGGCAGAGGCGTCTTTTATAACAAAGAGTTAACCCTAATGGCATGGATAAACTTTGAAGACCATTTGCGCATAATGTATCGTAGTGAAGGTTTAAACTTGGTGGAATCTTATAAAAAATTTCAATCTGCTATTGCCGAACTTGAGAAAAATTTGTTAGAGAACAACATATTATTTGCCGTTCATCCAGCGTATGGTTACTTGCTTTCGTGCCCAAGTGCTATCGGGACTACTCTGATTGCAGTAGCGTCCGTAAAACTTCCAAATGCCATTAAGCACGATAAATTTCGCGATGTGGCAAGAAATTTGCGTATACATATACGAGCAAAAGACCGAGAAGCGTTAAAGAAAGGATGGGTTGATGTTTATAACAGAGATCGACTTGGTTACACGGAGGAAGAACTAATTCATCAGGTCGTTGACGCAGTTCAAAAGTTATGTGAAATTGAAAGAAAGCTTGAGGAAGACGTCACTTTCTCAGATTTTCAAAATTATAGAAGCATTCTTGAACAATAGTTCGCTTTTATCAAAATATTTTAAAGATTTTCTAAAACTCGAGAATGGTCCAAAGAGATGATAAATTGAAACAAGAAATTATTTTTTGTTCTATTGTAAATAAAAATATCGTTATTTATCACGTGACACATTCAAATTATGTGTAAATAATTATGTCAATTATGTATAAAT
>evgR018333c0g2_i01_Hydra
ACATTAAAAAAAAAATGTAAGAAATAAAATAATATGTCTTAATCGGCGCCTAGACATTAGGACGCATCTGACAAAATGTAATAATATTATTGGAAGTGAGGGTATGTCTCGTGATTTCGATGATTACTTTGTTTTAATAAATCAACAGTTTTATTTCCATAACAGACGCAATCTTTGTAAAGCAAGTATTTAGTTGAATTTGAACAAACGTTGAATTTAGTGTTAAAGCTATGTCTTTTTTGAATGTGTTAATAACAAATCCAGAAAGAGCAGAAGAACTTCGAAATGCTTGGAAAACTTTAAGTTACGATTTGATTACGCAATCACTACCTGCATCTCCGCCTCCAACACCCAATGCTATAGATTTTGAACTTGTAACGCCAAGTGTAAGTGTAAAATATTTAAACAACAAAGGTGTTTTAGAAAAGTTGTGCAAATCGTCTTGCTTTGAAAAACCGTGTTCGTGTCACATTGGTTATTTCGGTGAGATAACTAAGCCCCAAGAAAAAATATTTCAGAAACAGACTTCTATCGAATGTGTACTAGAAGCAAATAATAGTTCTTCGGTTCGAAAAAGGAAATTTATTACAGATATAAAATCTTTACACGGCAAAAAACTTAGAGCAAACAAAGAACAAGTTGCATGTCAAGTTCATGGTATTCTGAGCTTACCTGGATGCAGAGATCCAAGATACAATATCACAAACAATGAACTTCGTCGTCGCGCTAACCAGCCAGAGTGCTTGACTAAAGTTGACATGATATCTTATGTACGACTGGCAAAAAATTCAGCTAGAATATTACTTCAAAAACACGAAATAAAAACTAATCAGCACAACCAACGTTCAGAACACACAGTTCTGTCAAAAATGTGCGAGGAAGAATGTCGCAATCTTGCAAAAGGTATAAGTGATATTAACTATAAGTATTTCCCTACATCCTGGTTTGCGTTTAAATCAATTGAAGAAATAGTGGGCAAC
>evgR018334c0g1_i01_Hydra
AATTTTATGCAAAAGTATATCAATAGCTTTACATTGGAGTTTATTGGCGGCATATTGCTGGATATTTTTCATAGCAGTTGAAGTGTCATCAAACTTTTCGAAACCATTTCATCCGATTTCATTAGGTCGTAGAAAGCCATTGATTCCAACAGTTTTTATTTCAACAATGATTCCGACCAATATTGTTTTTTTAACCTTAGTGTTAGAATATATTGGAGCGGTTGACGTTGGGTATGGAACAAACCGTTACTGTTGGATTAGTGGAATTTATGCAAGAGTATTTACTTATATTCTACCGTTATCCGTTGTTTTAGTTTCGTCAGTTTTCAGCCTGGTTTCAACAGTTTATAATATTAGGAAATTCGAAAACCTCAACAAAAAAACGTTAGGAGATAAAAAAACACGTGTCAACCTTCTTATGATTACCCTAAAACTGGCGTTACTTTTTGGTACAACCGATATCTTTTGTTTTTTACAAATTTCAACTGCATCGTTATCTGAGTGGGCAAAAAAATTTAACGCAATCTTTTCCATGATTTGCACGTTCTGCCGATCCTCAAAAGGTATTATGCTTTTTTTGGTATACATTTGCAATAAAAACGTCTTCGTAATCTACAAAAGAAATATTACCAGGAGTACAAGAAATTGATAATTATCTCTCAGTACAATTTTTATCAAAATTTTCAGTTGATTTACCGAACCAAG
>evgR018335c0g1_i01_Hydra
TCCTATTTGGAAATGAAGCTTTGGCGCAATTGCATAAATTTGAGTTTAAAGTTCTAGTTAATTTAGATATTTATATTGTTAATTAAAATGGTCTTTCAGCCTCTGATATGCGTGTTACTACTATGGTGTGTTAATGCAGTTCGATCCGGGTGCAACGATGAATCTGATTCAAAAAGTTTGCCTTTAATGCCCGATGAGTGCCAAACTATACCGTTTTTGATGTTCATCGAAGAAAAAGAATGTGAAATTAAAACTGTTCTCAATAACTATTGTAAAGGTTTTTGTTGTTCCGAGACCATGACCAACCTTGAAGGAAAAGAAGTTAAAAAAGTATCTTCTTGTAGACCTGAACAAACTTATAAAAAAGTTGTTGTTTTGAACTGCAAAACAAAAAGTAGCAATTTGTCATACACAAAAACAGTTGACGTAGAAATATTCAAAAAATGTAAATGTCAAACTTACACGACCAGATAAAACAAAGACCAGACAAAATAAAGATTATTTATTAACAACACATTTTTTAAACACTTTTAAATGTAAAAGAGTTTTTGAGAATGTAAAATAGATGAAGAACGAAAAAGAAGAAAGAAAGGAAAGAACAAAGTAGAAGATTCTTTACAGTTAATTGCGGCTACAAGATTCAAATACAGTCAATTACGGATCCAACAAAAGTGTGAAAAATACGGATAAAAATAAAATAGGGTTAAAGTTTTTATAAACATATTGCACCTAATGTTGGTCCTACTATAATTATTCTAGTACCTTTTCTTTAGCCAACAATAACTAACCACTTATTGCGTTTATTTTAAGTTGATATATATGGTATTTTTATGTTAAACTTAGTTATATATTTTATTATATATGTAATTTAAT
>evgR018336c0g1_i01_Hydra
GTCAATGATGAGTATTTAGCCTTGGAGGATGGTCCCCCCATATTCAGACAGGGTTTCTCGTGCCCCGCCCTACTTTTCTCTAGCTCAGTACCACCAGTCGGTTTTCACATACAGGGCTATCACCTACTATGGCCGGACTTTCCATTCCGTTTTGTTAACCGTCTGACTATCACTAGAAGGCTCTTCCGATTTCGCTCGCCACTACTTTCGGAATCTCGGTTGATGTCTTTTCCTCGAGCTACTGAGATGTTTCAGTTCACCCGGTTCGCCTCGCATGACTATGTATTCATCATGCGATACCTTTCGGTGGGTTTCCCCATTCGGAAATCTCCGGATCAAAGCTAATTTGCCAGCTCCCCGAAGCTTATCGCAGGCTATCACGTCCTTCGTCGCCTATCATTGCCAAGGCATCCACCACGTGCTC
>evgR018338c0g1_i03_Hydra
CAGAAAATTAATGTTAAATTTTTGCACATTCATCTTTCCATCAACAAAACAACAACAAAAAATGGAAAGAATGTTAAATCAGTTTATCAGACAATGTACTTTGACGACAATGATAGCTCAGTCTTAAGCGACATTTTGTGCCTGTGCTAAAGTAAAATATAGAATTGTTTTCTATTAGAATTGCAAATGGCACGGTTGGCAAATGTGTTTCAAAGGATTATGGCAAACACGCTACACAGAGGACTTTACAGATCGGGAAACAATTGTGAAGACAACATTAAGAGAATTAGCAATTTATCTTGTTTTTGTTCTCATTTTGTGTATAATAACATTTGGAATGACGAGCTCCACAATGTATTATTTCACCAAAGTAATGAAAGATCTTTTTAGTGGAGATCCGAATAAGCCTCCAACGTTTGATTCTGTTGGAAATATGGAAGATTTTTTTTCATACACAATGGAAACTACTTTATTAGATGGACTTTATTTTGAGAAAGATTACAATGGAAAAGATTTACCTCAAAATGAGAAAGGATATATTTATTTTGAAAATAAAGTATTAGGACGCCCACGATTAAGACAGGTTCGAGTTGAAAATGCTTCCTGCATCATTCCTGATATGTTCAAGCACCTTATTAAAGATTGCTTTGCTCCTTATTCTCCGTCTGTTGAATCAACTTTAACTTTTGGAACAGGAAATGGATCAGCTTGGACATATGCATCAGAACAAGAACTTAATGGAAGAACACTTAGTGGACAAATGTCATCATATTCAGGAGGTGGTTTTACAGTAATGCTTGGAGAAAGTTCAAGCGAGTCAAATAAAATTTTAACTGAACTTAAGGAATCAAAGTGGCTTGACCGTTACACAAGAGCAGTATTTATTGATTTTTCTGTCTACAATGCTAATATCAATCTGTTTTGTATTATTACATTGCTCTTTGAGTTTCCAGCTACTGGTGGTTGCCTAACCTTTTCTACCTTCCGCACATTAAAACTATTGCGATATGTTACTACTATGGATCACTTTGTGATGGCATGTGAAGGGATTTTTATTGTATTTATTGTATATTATTTAATTGAAGAGTCCCTTGAAATAAAGAGGCATGGTAAACAATATTTCTCTGATATATGGAACATCTTAGATCTTGTAGTCATTGGTATGGGATTTATTTGCATGGCCTTTAACATCTATCGAACAGTAACAGTTAGCAAAATTCTGCGCGTATTATTAGAAAAGCAAGATGAATATGCAAATTTTGAATCATTGAGTGATTGGCAGACTAGGTTCAATGACTTTGTTGCAGTTGCAGTTTTTCTTTCATGGATCAAAGTTTTTAAGTATATTAGTTTCAATAAGACAATGACGCAGCTACAATCGACACTTTCGAGATGTGCCAAAGACATTGGAGGTTTTGCAGTGATGTTTTCTATTGTTTTTTGTGCATATGCTCAATGGGGTTACCTTCTTTTAGGAACTCAAGTTCAAGATTATAGTACTTTTCTTAACTGCATTTTCACGCTGTTTCGCATCATACTTGGTGATTTTGATTTTCCCTCATTGATAGGAGCTAGTCCTAATGTTGGTCCTATTTTTTTTATCACATATGTGTTCTTTGTGTTCTTTGTCTTGATCAACATGTTTTTGGCTATCATTAATGACACTTATGCAG
>evgR018340c0g1_i01_Hydra
AGCTGGGACTGTAAATCAATCTCCAGAGTGCTCAGCTGGGACTGTAAATCAATCTCCAGAGTGCTCAGCTGGGACTGTAAATCAATCTCCAGAGTGTTCAGCTGGAATTGAACAAAATATCTTTATAGAACCAAAGGATTTATTTATATACGATAAAGATTTGAAAGAGGGATTTAAATGTTTCCATGTAATTACATTAGGAAAACCATCTAAATTACAATGGTATGACTTCAAAAACGGATTGCCTACATACATCATTATCAAGCAACGGACTCATGCAAAGTGCGCCTTCTTTGATGGAAAACCTTACGACGGAAAGGAACACACGCATAGCATCTTGGCCCACTTTGTTCAAAAACACAAAAGGATATCAATCGATGCCATCAGCAACAACATCAGAAGAATGTGGTTCAGGGAAAGAAAATTCCTATGCCCTGGATGTAATGAAATATGTAGAAAACGCTTCGGTAGTGAATGCAAGGTGTGTAAGACCTACATTAAGTCTTACCACCTTAATTCGGAACGTTACTTCGTCAATGCCATTCGGTATGTTTCAAACATTAACTAAATATTTAAATGATAATGTAAATATATCAAATTTTATAATTAATGACGATTTATTAAGTAATTCAATCGTTGCTCCAAGAGTTTCAGCACAAGATTTTAAATCGTACATATTCGAGCAAAAGATTTTTAATTCGAGACACTATGAATTTTATTTAGAACGCGACCCTAAATTGCGTGATTTCGAATTAGGAATCAAAGTGAACGATTTAGAATCAGCAAAAAAAACTATGTTCCGACAAGTATATCTAATTGAAAAAAATATTGCAAGGTTAAAAGAGCGAATTATACCCTGCACAGATAAAGGAAGGCTTGGGGCAGTTAACTATTTTTTAACTTGTATTGCTAATGAAAATAATATTTCTTGCTCAAGATTAAAACAAAATTTATGTTATTTTATTTTTAACTTAAATAAAAAGAAAAAGGGTCTTATCCTTTGTGGCCCATCGGATTCAGG
>evgR018349c0g1_i01_Hydra
GACAAAAAACTGGAGAAGAAGTTGATATGCGTGTAGGTATTCATACAGGAAATATACTGTGTGGTGTTGTTGGTAATAGAAGAAGAAGGTTTGATGTTTGGTCAAACGATGTCAATCTTGCAAACAAGATGGAGTCGAAAGGAAAACCTGGAATGGTTCACATAACAGAAGTTACTAGGAGATTTTTTAAAGACCAGTATGTGCTTTTAGATGGTGTTGGAATTGAATATTCTGGTGGGAAGTTGTGTTCTTATTTCATTTCATGTCGATCGAATGGCTTAAGTGCTTATAAACCTTGGCAAAAAACCAGTGATGTTTCGTTTAATAAAAGGAATGGCAGTGTTTCTAATATAATTGATCATCTAGACATTGAATCAAAAAGTATTACTTGCAACATAGCCACATCTAAATCTTCTTTATCTATAAACAAAGTTGATGTTACTTTCAGTTTACATCATCATGTTACTGCCCAATCAGATTCAGATGGCAGTATTGAAAATGAATTTTTGTCGAAAAGTATTTCAACTTTATCTTCACCCAGTCATTTTCAGGTGGACTTGCCTAAAAAAAATACAAATTCGAATAAATCAGCTGCAACATTTAATTTCTCATATGGCATGGCATTAAATGAAGCTCTCATGAGAGATGCAATGAAAGCATCAAATGATAACCAACTTGTAAAACTCATGCACGGTAAAACAATACAAAGAGAATATTTTTTTAATCCTCCTTTGCATTGGTGGTCACTGACATTTGTTGATGAGATAAAGGTTGATAAAAAAGAAGATGAAAATGAGGATGATGGAAATGAGGATGATGGAAATTCTCCCCAAAAAGTTAAAGTTGTAAAGAATATTACTGAAGCAAAATATAGAGACGAGGGCTTCAAGGGTTTAAAATTAAATCCAAAAGTAACAACCTTTGCATCTCCAAAAGTTCACTTTTTATTTGATGTTGTAACAAGTGCTATAATCTTAATAGGTGTTATTGTTGTATGTGTCCTCTTCTTTATAGGAACCCCAATAGCATTCGCTATTTTAATTGCTTTCAGTCTGTTTACAATGTTAGCTTTACTAGTATATAAATGTTATAAATTATCTCCATCAAAGTTTCAAGGGAAAATGACTACATGTGCAGCTTACCCTGAAAATAGTAGTACAGATGATATAACTTCGACCAACATGGAAAGTAATACAACTGAAAGTCTTCATGACTTGACAGAAAATACAATCATTCACCAGTCATTTAAAAGTAAGCTTTTTTCTGCCTGGGTGACATCACATTTGTACGGGGGCATTATGATGTGTCTCCCGATAATAGTCGTGTTTTCAACATACCAAGATTGTAAAACAAATAATTGGAAAAAACATCAAATTGAATATTTTGCAATGCTCATGTTTGTTAGCCTACTCCATTTTTGTAATTTTACACAATTATCCTCACTGATGAAGACTAGTTTTGCTTATATTTGGGCACTTGCTTTTATTTTATTGCTTGTGGTTTTACCTTCTTCTAACTGTATGCTTCGAGACGACAAAGTAATGATAATGGATATTGTTTTGGTAGTAGTTCTTCAGATCATTCTAATAACAATTCTGAATCGCATTCACGAGCAAGGTGTAAGAGCAAATTTTTATGGTGACAAAGAAGCAGCTGAACAAAAGAATATTGCATTAGAACAGAAACAAGTTGCAGATTGGCTTATTAGTGATATGTTCCCTAAATATGTTTCTGATAAATTAAAGTATACTAATAATTGTTCTCAAAATTATGAGATGGTTGGTGTCCTCTTTGCTACAATTGTTAATTTTAGTGGATTTTACGAGGAGAACTATGCAGGTGGTCTTGAATGCATAAGAATTCTTCATGAGCTAGTTGCTGATTTCGACAATGAACTCAAAAGGTTTCAGGATATTGAGAAAATAAAAACAGTATATGGTACAACATTTATGGCAGCCTCTGGATTAAATCAGACTAACAAGACTGTTAATTTGACAGATGAAAAAGATGACAAGCAGTATTTACATTTGAAATCTTTGATGGACTTCGCCCTAGAAATGCAAAGGTCTTTAGATGACTTTAATGAAAACATGTTAGGATTTAAATTTCACCTTAGGTGTGGCTTTAATGCAGGTCCAGTTACTGCAGGGGTTATTGGCACTATGAAACCCCAGTATGATATATGGGGTGACACTGTTAACGTTGCTAGTCGTATGGACTCAACTGGCGTTGTTGATAAAATTCAGGTTTCTGAAGAATGTATGAAAAAATTAGAAAATTTTTACACTTTTACTGAACGTGGAGCAATACCTGTTAAAGGCAAAGGAATGGTGACTGCTTACTTACTTGTTAGCAGGAAATAAAAACGAATATGTATTTTGAAATTGTACATTTGTTTAAAAAAAATTGTTTATACCTTCATTTTTTAAATTATTAAAAAAAATTAGAATTATTTTATAAAACTTTATTTTAACGCATAAATTAATAGTTTTGTAATTTTCTCTATGGTAACAAGGTTTTTGTTGTTATGTTAATAGAAAATGAGTTCTGCATGTTTTTGAAGCTGTGCTCCAACTCACTTGAAACATTTTGTCAAACCTATATCGATTTATTAAAATAATAATACTAAC
>evgR018350c0g1_i03_Hydra
TCAGTTTTTTCATTAACATTGAGGTTTAAATAAATAATTAAATTATTATGCGGATCTTAAGAGACTAGTTTTTATAGTTTAGAGTAAACATTAGAGCGACAGGGAACATTGTTTTGAAGAGTTTAATAAAAAATTCAAGATTTTATTTAGCATTTTACACAGATGAGAAATTCTTAGTGTTAGTAAGAAATTTATAAGTTGCGTTGTTTGCGGGTGGAATGTTAGCTAGATCTATTCGACAAATCAAATATCTCTTTTTTGATAATAATAGAAGTAGTACAAATGTTTTAACCGTATCTAAATCAAAATATTTTTTAATAAAAAGAAAAATTCACGAATCTTTAAAAGAATATGCTAAACCTCTTAACTATAGCATAGATGAAAAACGAGGCATAGTCGAAATTCATTGGGATGATAAAGTTTCAAGCCGGCACTCCAGTATTTTCTTGAGAGATTGCTGTCTTTGCAACAAATGTTTTAACAGTGATTCAAAACAACGTCGTATTGACACTTATAGAGAAATACCACTAAATATAAAGCCAATATCTATTAATATAACTTTCAGTCATGTTTATTTTAAATGGCCGGATAGTCATGTGACGGCATTTACGCACGAGTTTTTATTGGAAAGAAGAAACAATGAATACAGTGATATATCTCATATAAAACCGAGTCTTTGGGCTACGTTGAGTGATATCCCCCGATTTGACTTAGATACGGTATTAAAAAATGACGCAGCTAGGTACGCTTGGTTTCAAAGTTTGTGTGAATTTGGAGTAGCCTTGTTTACAGGGGCTAAGAAAGAAAAAGGTGAACTGGAAAGAGTTAAAAGGATATTCGGTGGATACTTCAAAAGTACACACTATGGTCATACTTTTGATGTTGTTTATAAAGCAAATCCTGCGAGCCTCGCATATACTTCGGAGAAAATACTTCCCCATGTTGATTTGCCTCATTATTGTTATCCACCCGGTATTCAAGTGTTGCATTGCATTAAGCAGTACTCCGGTGAAGGAGGCGATACAACGTTCACAGACGGATTTTATATTGCACAACTGCTAAAACAACAAAATCCAGAATCATTTCTTTCTCTTACTAGGCAAAATGTGACGTACAGAGATGTTGGTACTGACATTTTTGGAGAATTTGAAACGTATTACGAAAGACCTGTTATCTGGACTGAAAATGGTGTTGTGAAGGCTATAGACTACAGCAATTGGCTTCGGGACTACACGATTAATGGTAGCGCTGAAGAAATACAAAAATACTACAAAGCTTATTACGACTTTACTGCACTTATATGTGATCCAAAACATATTTGTTCATACAAACTTGAGGAAGGTGAAATAGTTTCGTTTCACAATCGAAGAATACTACACGGAAGATCAGCATTTTCAGTTAATGACTCAAATGCAAGTCGCGTACTCAATGGCTGTTATTTTGATTGGGATCAATTATTCTGGAGTATGCGACCAATTAAAAGGAGGTTGGAAAGAATGAAATAATGAATAAAAATGAAATAAGAATACTTTAAAATTAAAAATTTTATAAATATGAGTTTTTGTAAATTCAAGTTTTTATTTTAGCCATATAACATGACTACTCTCAGTTCAAATATTGTGTATATTGTTATATAAGAGAGACCTGGGCTAGTTGTCTCGATAGTTGACTCCCTTTACACCCTTACGGGTGTAATAATAATAAATCTAATAATAAATCCGCCAACTTACTTCATCC
>evgR018354c0g2_i01_Hydra
GGAAAATTGGTGGTTGCCGATTTTTATGCTGATTGGTGTGGACCATGTCAACAGATTAAACCACTCTTTGAAAGTCTCGCTTCCAAATATTCAAATGTGCTTTTCCTTAAGGTTGATGTTGATAAACACAATGCCATTTCTTCAAAGGAAGAAGTGAGAGCAATGCCAACTTTTGTCTTTTATAAGAATAATACAAAGCTCACCTCAGTAGTAGGAGCCGATATTCAAAAAGTCGAAAAATTAATTCTTGAATATGGAGACTCATTCCAAGCTTTCCAAGGAGGTGGTAGATCTTTGGGTGCCTCATCTAGTTCCACCACTTCTGATTCTTCAAGTGGATTGGCTGGTCAGGTTTATAAAAATCCGTGGGCAGATGATAACAGACCTGTTAGAAAGCCTGTTGATGCAATGGAAGATGAAATCGATGAATTGGATGATGAAACACGTAAAGC
>evgR018357c0g1_i01_Hydra
AAAACTTGTTTTCATAACAAGGTTAAGTTTAAAAAAATGAGTTTCATAATCGATGATATGAGGCGCGTTTTTTTATTAAAAACAAAATCAAGACATGACGGAGTTTGTGATCAATATAGTCGTGTATTTATGCCTAAAATAATGTTGTTTTTTTCGTTTATGATTGGTGTAAACTGGTATTTTGATAAGCTTTCCTGCATTATTCCCAAAACACTTTTAGAATCGCAGCAGTTTATAACTGATGTGTGCTGGATTAATGGTTTTTATATATACGAGAATTTGACAGTATCAATGACGTCATATTATGGGATTCCTGAAAATATTAATGAAGATGGTATCTTAGATAGCGGAGTTTTATGTGAGACAGAAACAGAAGACACTTGTACTCCAATGGTAAAATTTTACTATCTTCAGTATCAATGGTTTCCATTTTTCATGGCTTCTTTAGTGATCCTTTACATTATACCTCATCTTGGATTTAAACTGATAAATGCTGATATGATTACTCTAAAGAACGCGGTTAAAGTTCAATCTTGTGATGAGATTTTTGAAAGTTATTTCAACTCGAAACAAAATACCCCCGTGCGAATGTTTATCAAAGTGTTTGGTAGCATATCAGTCAAGGCAGCGTATATTTTTTCAAATATTGCTACATTTTTTGCAATTGATCATTCTCTTAACAGAAAATACATAAACTATGCAAGCAGTTGGCTTTCTTGGACTCACAACAACAATTCCGCGATGTTTGATTATTCAAAACGAAGACAGTTATTTCGAGCAGGCGAGGTATTACTTCCAAACTATGGACTTTGCGATGTTCAAATTTTATTTTCAGATATGATTGAATCTAGTACAAACAAATACAGTATTATTTGCGAGTACTCCACTCATATCCTTTATCATTACGTTTTGATTTTGTTATGGTTTTTTATTGTTTTGGGGATTATTTCTTCAAACTTTGGATTTGTAAGTCAGATTGGGAAATATATTTTAATGGTTTATGTATTTCCAAGAAAATGGAAAAATAAAGAAAAGGTTTTAAACATGGTATCCTTAAGGGAAAAAGAATATTTAAGCTACGTTCACTCACAAAACGTTCTTTTGTACGAATTGTTGTTAGACAAATTGCGCCAAAAAAGGTTGAAGATAATCGAAGAAGAAGGAAACAGCTCATTAGTTATAGAAACATTAAAGAAACAAGTTATTGAAATTCGGATTCAAGTTGATAGTATTCTTCATAAGCAAAAGTTTGATATCGATGAACCAACACTCGGTATAACAAAATCAAAGGTTTGATATCAAGGAACCTCGGTGTAACATCACTCTATCGCTATCAAAGAAAGAACTTCGTATACAAACAGTTTTCAAATCGAAGAACTTCATTAAACGGATTAAATAACATAAGAGTATATTGTTATGCAAAAATTA
>evgR018364c0g1_i01_Hydra
TTTTTTAGGTTTATTTAAAAAAAAACACCTAAAAAACCTATTTGGAAATCACTACCTTATCACTATGTCATCTTACTACGGAGTTAAAAACATCCGACTCTATGAGACACGATCCTCCTTCTATATGGGCACATCTGGAACCCATAAGCACTGGGCCGATGGAACTTAAGCTAGATAATTTACAAGTTACAGATATCTTTTTTTATTTTCTGATTGACCAACCATGCAATATAGGAATTAGAAAAACTTATTCTTATTCTCAACAGTCGATTAAATTCAACTATATGAGCCCTCTATACAAGTTAAACTCCATTCAGCCAATCCGAATTTTTTTGGAAAATGGATTTGGCAAGGAAGCACCAGATGCGATAGGAGGATCTATGAAACGTCAGGCAGATCAATTGGTTATTATGGGTTGTAACCTTCCAGGTGCAGAAAAATTACAACATTTTCTAAACACTGCACATTTATTAATAAGGTTTTACTTCATCAAAG
>evgR018367c0g1_i01_Hydra
ATAAATTACTTTCTACCACAGAATCTCCAACTACATCATTTGAAAAAAAATCTTCAAATGAATCAGAAATAACAACCAAAGAATCAACATTTACAATTACTGAATATCTAACCAAAACCAAAGAATCTACAACAATTTCAACTAACACGGTTACGACAAAAGATAATTCAAAAAGTGAATTTTATACATCAGATACAAAAACCATCGAACCTCAGTATACAAACACAGAATCTTCATCTATCATCACAGAATCTTTGACTACTATTACAAATAAAGAATTAACCGAAACAACAAAAAAAACCGGAACATCAAGCATTCCTGATCATACAAAAACTCAAAACATTAAATCAACATATATGACTACAAAATTGCAAGAAAAAATTACTGAAACTACAACTTTTACTGATTCAAGATTATCAACTGTTGAAACAACAAACATAAGTACAAAATCTCCAATCAGCATTAACAATACTGTATCTACCGCAATTGAAACAACAATGCCTAATAATGAAACAACAGGTGCAAAAGCCGAACAACCAACTACTACTAAAACTGGTTTGACAACTGCTGAATTATCAAACACAACAAGGAAATCTCAAAATACTTTAACTGAATCAATATCTACCGATATCAAAGCAACAAAATCAGCAACAAATACTGATTTTAGAATAGAAACTGATCAACAAACATTCACAACTAAATTATCTCTGACTACATTTACA
>evgR018375c0g1_i01_Hydra
TTAAATTTATAATGGCATTACTTGGTATACAATTTTTAACAACTTTGATCATTTCACTATTCATGGTAAAACTATCATCATTTTACTCAATTGCTAGATGGATAATTTGTGGAAGACTTAATCGGTATTTGCATCCAAGCAATGAACATTTACGAGCTTTGGTAGGAAAGTTACCTCTCCATTCAAAAGGAAAAAAAAGAAAATCTTATGAAAAATACAACTCTGAGACATCAGACATGTTTACTATACCGTGCAGTCTTGAATTTCAATTGGAACGCGTTAGCATTGAAGAAATTGACTTATACCCACAATACAAATACCATGATTTTAAATGGTTGGTTGACTTTACAGTATGTGCTACACTGATATATATTATTATTGAATTGATTGGTTATTACAATCCAAATATTTACGTTGACGAATTTAATTTAGGTGTTGTCTGGTGCGGTATGGTTGTTTATTTTTGCTTAAAAGAACTTTTCTCACTCACTTCTGCATACTGGAAGACTGAAGATGATGGGGAACGATCAATGACTCTTTCATTTGGACTATTCTTTTTTATTGTTGCGATGGGTGTACTTATCATTGACGAAAGTGTAATGGATTTTGGACTAGAGAAAGGATTTTATCACTTTGTTGGGAATTTGGAAATAGTATGCAAACGTCTAAACTTTAGTGTCAAACAAGTGCCATCAGTCTGGGTTTTTAAGATATCTTTAGCCTTTTTATCTTCATTGGTCGGAGCTGTGCTTTGTTTCCCTGGAATACGATATGCCAACATGCACATTGAGTCATTAAACTATTCAAAAAGTAATTTAAGTTACGGTTTAATAAATCTAATATTCTTTAGTCCTTTGTTTTTATCGTTGCTTTGGATTCTTCCGTTATCTAAGTCTTTAATTGTAACGAAAAATTCGAACAGTCAACATCAGCATATATCAGATGAAAAATTTTTGTGTTTACGAATTCATTTAACAATTTTGATATGCTTTCTTCGTTTATTAGTAACGCGTCAGTTTCTTCAATCGTACTTAAATCTAGCTTGTAAGAAAGTGGAAAAGCTAAAAAAAGAAACTGGACGAATAACAAATATCGAATTACAGCGTACTATAGCTAGGGTGTTTTATTATTTGTCTGCTGCTGCGCTGCAATATATCACGCCACTTATTCTTGTATTATTTTTATCACTCATAGTGCGGACCCTAACGTATCATGTGAGTGATTCAACAAATCTTAGCGGTTTTCAAGTAAACAATACTTTGCCCGTAAGCAAAGAAGAATATGTTAAAATGATGCGTGGATTATTTTCTGTGAAACTTATTAATGCAATTTGTTCTTATTTTTCTTGGTGGGTCCTTAATGTTTTTTTCATAACATCAGCATTTGGTTCGGTTTATTTAAAACTAAAAGTTTAATATGTAAATATTTTTTAAGTTTATATCTTTGATGTCAGATATTGTTATTTTTTGAGCAAAATCCAATAATTGTTCGTGTCGTGTGCGACTTAATGTTTTAATGTCTCGATAGTTTTTTAATATTAGGCCGTCCATAAAAGAAATTTTTTAAACAATGTTTGACCCGCTTCCCCTTTTTTTATATCTTATATCTCGAAAAAGTATATAATGG
>evgR018376c0g1_i01_Hydra
CAAAGATCACGAACATCAAAGGAGAGGGAGAAAGTTTGTCAACATTATATTCCAACCAAAAAACCAAGTGAACTGCAAACAATCTGAATTTCTGTCAAACAGCAATAACAAAACTGCTTTGATCAAAGCATTGGGAATGAAGCGAAGAGAAAAAGGTTATATAGTAGTGGAATGCGAAGGAGATGCTGACAGAACCATTACTACAAAAGGAATCGAGCTTGTACAATATGGAAAATATGTGACTATTATAGCTGATGATACAGACATTCTTGTAATGTTAGTTCATATGTGGGATCAAACAATGTGTGAGATTTTTTTCCGCCACAAATCTAAAAAAAGTATGAAAAAAAATATGGAAACAATCAGCATAAAGAATGTTACATTATGT
>evgR018378c0g1_i01_Hydra
TAATAAATGAATCTTGAATGAAATGAATGACCAGCAGTATCCTCTTTTGGAATTTGATCATAGTCAAGGATTATCCTACATAAATAATCATGAACATGATTTAAGATCACTGTTATATGATGATATTCAAGACGGATTCATTCAAGATAGCTTCATTCAAAATGGATTCATTCAAGATGGGTTTATTCCAATGGATAGCATTGACATTGATAATTTCTTACCAGAAATCAGTTTTGAATCTAACCAATCAAATACTGGGCATAATTACTCTTTTGAAATAATTTATTTGAATTTTCCAACAGTGTTTTACAAAAAACAGGTTCGGAAGAATAATCGATCTATGCAACGAATTGAAAATGGAAAAATTAATTTCTTGCTTGATTTTGGAAAAGAACAAGCTTTTTTTCCTTCCATTCCTTCTGTTTCAGTTAAAGTAAAGCGTACTACAATTGAAAAATCAAGTGAAGTTCAGATTAGTTGTCATTATGACGTATTACAAAAGACAAAGCTTTCTGTACACTTGGATTTGGATACGGTGTATGAGGACGCGGACGGTTATAAAGTTTACGAGTTGGAGAAATGTTTGCACAGCGATTTAAAG
>evgR018382c0g1_i01_Hydra
TCAGAGAAGATTTGAGTGGTCAGATGGTAAACCAGTTACCTATACTAATTGGGGACGTGGTGAACCTAACAATAGGGCTCAAAGTGAGGACTGTGTAGTCATATGGCCTCCTGGTGTATGGAATGATATTCCTTGTGAAAGTAAACAACCTTACTTTTGCAAACTTAAGCCAGTTAGTAAATGTCCCAATGGTTGGTTACAAAATGCAGGCAATTGCTACCTTTACCAGAAGAATTATAAAACTTATCAAGATGCCTTGTCTTCATGTCAGAGTTTAGGAGGAAGTCTATTAAGTGTAGAAGATCAAGCGGAAAATGATTTTATTTTAAACTATATTAATAAAAATAACTTCCAAAACACCTGGATTTGGATTGGTCTAGACGATAATAAAAGTAAAGCTCAGAGAAGATTTGAGTGGTCAGATGGTAAGCCAGTTACCTACACTAATTGGGGGCGTGGTGAGCCTAACAATAGAGCTCAAATTGAGGACTGTGTAGTCATATGGCCCCCTGGTGTATGGAATGATATTCCTTGTGAAAGCAAACAAC
>evgR018384c0g1_i01_Hydra
ATTAAAGAAAATTTACTTTTAGAACATATGAGGAAAGCTTTAGTTTATTTTTTTATTCATCAAACCTATGGCTCATTCGAATTACGGACTAACCAATAAACAATTACGATATGCTCACTTTATTATAGAATTAGTTGCTAATGTATCCCAAGTTCCTAAAAAAATGACCAAACTTCTTGATCACCTGTATCTTGGAGCAATTAATGATGCAATGGACGTTGATCATCTTAAAGAGGCGGGAATTACTCACATTATTAACACTGTTCATAATATGGATGGCGAAAATGAAACAGGAGAAGAATTTTACGGTGATGAGTTTACTTACATGGGCTTCTTTTCTGAAGACAGGGAAGACTATCCTATATTAGAAAACTTTGAAAAAGTTCAAACCTTTATTAATTCAGCGCGAGATAAAAACGGAGTATGTCTTATTCATTGCATGGCTGGTATTAACAGAAGTGGTTGCTTAGCAACTGCTTATTATATGATTGAAAAAGATGTTGGACCTATCACAGCCGTTAGCGAAGTCTTTGAAAAAAGGGGCGTGTTATTAACAAACAAAAGTTTCATTAAACAACTCGTAAAATTTTCGGTTGATCATAAGTTGCTAAAGAAAGATAAACACTTACTACCTCATAATTCAGTAAACAATTGTTCGATATAGGTTTTTTCAATTTTATAAATACTTTTTTTCAACATTTGTTGACAAAAACGACATTTGTCGTCGAAAATACGACAGTTTTTAAGTAAAGATTGTAATTTAGTTTTTTTAATTTTTATTTTTAATGTATTGCCCTCATATAATCCTTGTGTATATGTATTTATATAAATATAAAATTGTAATTTAAAAAAAAAA
>evgR018394c0g1_i01_Hydra
GACATCTACAATGTTTTCAACGTCGAATCGACAAACGAATTTAAATATACTTCATCCAAAAGGAGGAAATTTACAGAGAAAAGCTCTCACAGATATTAAAAATGTTCAACAACCAAAAACAGTGTCGTTTGAAACTTCTAAACAATCAATTCAATCTACAAAAAAGATAAATGACAAAGAGTGTAATGTTTTTACACCAGGGCCTAAAATAATAAAGTCATCTTATGCTACACCGGCTGGCGCCAAATCGAACAACAAAGTTAAAATTGCAACATCTACATTTAAGATATTTGATGATAGTGAAAGTACTAAAAATTTAGACAAACGCATGAAGTTCTCTAAAAATAAACACCGTGAACCAGATGAGATAGAGTATATTGCTCCGAATAACGATAAAGATTATGAAATTCAAATACCTGAGTATGTGGAAAAAGTGATGAACCATCAACCCACATATAGTATGGCATTATCTAAAGATGACCCACTGAATCTTCAAGCTATTGTTGCCAATCTACAACACCAAATATTTGAAAAATCAAAATACGATATTCCTAACCAAGACCTCTTAACTCTTATTGCTGATACAAGTATCACCGATGAAGAGCTTCTATGGAATATAGAAAACGACACTGCTTTCTCGTTAGATATGCTAGATAAAGAGCTTCGTTATTAGTTTTTTTACATTTAGAGTATTTAATATTTATGCAGTTTTTATTTCGTTGTACATTTTTATTTACATAATTTACGATTTTTTGTGGGTTTAAATTTTAAGGAATAAATTTTATCTGCTTTATAAATAATCTTTCGAAATAAGATTTCGTATAAAATATTCTAAATGTAAACTTTGATATTTAATAAAAAATAATCAAATTGCATATAGATTTGCTAATATGCAGTTCTATATGCAATTTATTTTTTGTGTATATAATTTATTGAAATCAACAAATACATTGTTTATTTTAAATGCTATGTTTAACTTCTTTTTCGATAACTAGG
>evgR018405c0g1_i01_Hydra
AACACCAATCTAGATAACAGAAGAACACACCTTTCTTCTCCCAACACCTCTAAATTTGCGCTAAAACCCACTCAAACGCTTTTTACTCTCAACGTTTACCCACAAAAAATGGCCAACCGTATGAATGACTTTGTCCTGCAGCAATTATTAGCAGAAAATCAAAGACTTCAACAAGATTTGGATTCATGCCGAAAGGCTATTCCCATCTCAGAAGCCTGTAAAACATTGATTGAATATACAAATGAACACAAGGCTAAAGATTTGTTGGTAGTGGGTGATCAAACCAATCCATATTGGAATCCACCAAAAGAACCTTCATGTTGTATGATCATGTAAGAAGAATAATAGTAATTCTATCCAAGAATTAATTGCAAAATCTTGATATGATTTAACCCGAGTCAACTTTCATAATTGTACAACTCGCAAGAGCGAAGAACATATATCAAGACAATCTGCTAACATTCATT
>evgR018412c0g2_i01_Hydra
ATGTTATACTATCAAAACCCAAATAACAAATTTCTATCAAGAATATTTTTGATTTAGTTCTTGATTGACCAAATCAGTGCACTTTACACAATAGACATGATATGGCAAAAGATTGTTCGAAATGTAAATTTTGTTGGTAGATATTATTTATATCGTAAAAGCTCCTTTCTAGTGTTTTTTCCAAGTTTCATTGGTGGCTGTTTTTTGGGGGCAAATATAAATTATAAAATGCCGCAATTTGCTGTCAATTTGAAAGATTTTATAACTCCAGTTGCAGTTGCGATGGAAAAAGATGAAAAAATAAAACGAAACAATTTCAATTTTATTGCAGATGCTGTTGATATTGCAGCTCCAAGTGTAGTTCGTATTGAAGTTGTTGGAAAACAATGGAATGGACAAATTAGTAGTGTTGGTTCAGGATTTTTTGTTACAGAAGATGGACTAGTTTTATCAAATGCACATGTTGTTAAAAATGCATCAATAAAAGGCTCTCCAACAGTAAAAGTTCAACTATCTACAGGTGAATCGTATAATGGTGTGATCATTGATGTTGATCCAGTATCTGACTTAGCAGCAATTAAATTGTTGAATGAAAAACAAATTAAATTTCCTTACCTCAAACTTGGTGGTACCGTCAGGGCAGGAGAATGGGTAGCAGCTCTAGGAAGTCCTCTTCAACTTAGTAACACAGTAACAGCAGGTATTGTTAGTTCATTACACAGACCAGGTGGCGAACTTGGGCTCTATAATCAAGATATGGGCTATATACAAACAGATGCAGCCATTAATGTTGGAAATTCAGGTGGTCCACTTATCAACTTGGAGGGGGAGGTGATTGGAATTAATTGTTTTACCGTGCAACAAGCTGCTGGTATCTCTTTTGCAATACCTGCAAGTGTTGTAAAAGAATTTCTCAAAGAAGCATTAAATAAAGATTCATCAAATACAATGAAAAAAAAGTTTTACATAGGGATATCAATGTTAAGTCTTTCTCCTGACATACTTTTGACTTTGCAACAACGAAATCCAGATTTCAACTCACTTAAAGGAGGAGTCTTAGTTGCTCGAATTAATTCAGGGTCTCCTTCTGAAAGGGGTGGTTTGCTGGTAGGCGATGTAATTTTGTCAATCAATGGAAAAACTATTAAATCTTCAAGAGATGTGTATAATGAAGTGTGCAGAGGAGAAAAACTTGTGATGCAAATCCAGAGAAAATCCCAAATCTTATCCATTAATGTTTCTCCAGATGTATCCAGTAGTCTTTAGTATTTTTATATGTTTATTGATAATAAATATATTTTTCACATAAAAAAAAAAAAAA
>evgR018414c0g2_i01_Hydra
AATGCAGATCCTATATCAGATTAGGACGGATATTAATAAAAATAATACAGCTTAAAATCAACGCCGAGTCGCTATAACCGCATCATAAATAAATATTCGGTAAACAAACTAGCAAAATGTTTAATTTTCGTTTGTTTTCAAATCAAAATACACAAGGTTTTAGGTAATTATGACTGCGATGTCAGTATTACCTGAAGAGGAATCCGAATTTCAAGAACCATTTAAAAGACAACCTTGGTGGAAAAAGAGTCCCTCATCTGAAAGATCAAGTCCTGAAACTAAAAATAAACTTGAGGAAATTACGAAGGACCCGCAAAAGATTATCAATTGGGTTGAATTGGAAGAAAATGGGACATTTTACAACACAACTTTGAAAAAAGTTAGATATCGAACCTTGATTTCATTCGATCCGAATGATCCAAATACAGCATTTAAACAATGGGAGATGGTTAAAATAAAACAGTTAAAAGCTGGAAAGATCGAAAGTTTAATTGAGCATATGACTTTACCTTATGAAGATGAAGATCCAGATCCAGGTTTTATTTTAGCATTTCTTTGTACTTACAAAAGTTTTACATCAACAGCAGAAGTTATTGAACTACTTTTAAACAGATTTGATGTTCAAGTAACGGATATTCAAATAAACGAGGAAGAAAAAGGCGAGGAAGTTATAAAGCGCATCTGTAAAGTAGTAAGTGTTTGGCTTGATCAGTATCCTCAGGATTTTGATGACCCACCTTCTTATTCGATATTAAATAAAATAATTTTATTTACAAGTAAGCATATTGATCTTTTAACTTGTATTAAAGAAGTTTACAATAAATGCTGCACAAGGTTGAACAGCTTTGCTACCAGTCCAATTGATGAGCGTACATTTAAATTCAAATTTTGTCACTGTCCATCTATAATTGGTTGTACATGTGAAAGTGCTGACGTACAAAATTTTCAGTTTAAAAGCTTCTCAGCACAGGAGTGTGCTGAGCAACTGACATTAGTTGATTCGGAGTTGTTTTTAAAAGTTAATCCAAGAGAGAGTTTAGGCTACTTCTGGTCAAAAAGAGACAAAGTACATGGGTCAAAACCAACATCAGTAAAGTTGACAGTTGATCATTTTAATGCTGTATCTCTAAATGTTATATCTACTGTGCTTGATTCTGTTGATGAAAAATCAAATTTCCGCAGTACTTTAGCACGCTCCAAAACAATTACAAAATGGTTAGATATTGCTTTGGAGTTAAGAGAACTGAAAAATTTTTCAAGTTTGAAAGCGATTTTATCAGGGCTGCAATGTGCACCTGTATATCGTCTTACACAGTCATGGGAATTAGTTTCAAAAGCATCTATTGCCATCTTTAATGAACTTTCACAGATTTTTTCAGATGACATGAATAGCAAAGCATCTAGGGATTTATTAATACAGGAAGGTACTGCAAAGTACTCTACAAACTCACTTGGTAGATCAAAATCAAAGCGCCAATCTCTTATTAATATGGGCATCACGCATGGAACTGTACCATATCTAGGAACATTTTTAACAGATCTTACAATGATAGATACAGCCCTTCCAGATAAAATCAAGGGAGATCTGATTAACTTCGAAAAAAGGAGGAAGGAATTTGAAGTTGTTATTTTAATAAAACTGCTACAGCAATCTGCAAAAAATTATTCTATTCAACCAAATCCAAATTTCTTTGCTTGGTTTAACTCTCTAAAGGTTTATAGTGATAAAGAAAGTTACTCTATATCGCTTGAGGTTGAACCCAAAAATGGAGTGAAGCATGAACCTGTTGAAAAAGATCGTACAATGAGGCGATATGCAAGCGACAATGACATATTCTCTCTGAAGCAACCAAGTATTAACACAGCTGAAGATCTTGTCTCGCTCTATTCCCTCGAACTATCTAAAAGTACTATTGATTTGACTGACGCTGGTATGATATTGAAATCTAAAGTTTCTAAAGATAGATTTAAAGTTGAGAGTTTATCATCTCGAAACATGCTTGATCCAGATGACGCTGTTCCTCTCGACGAAACAGCGGTGGTCGGAAAAGTTTATCTTGAAGACACAGTTAATGTCCAATATAAAAGTGTAAAGATTACTCCGACGTCTCGAGCCTGTGATGTGATAAACTCTACACTTAACAAGTTTAAAGTCTCAACCGATGTTTCTTGCTATTCACTCTATCAAATAATAGATAACAAAAAAAAATTGCACATACCAATGAATTCAAATATGTATTATGCTATGCACAAGACCCCTGCATTATGTTTTAAAGTATCAAGAAAGAAGCTTAAAACAAGCAAAAAGGCAGAAAAATCATTATCTTTCAACCCGCCAGGAAAAAACGATGTGATCCTACTGAAGGCTTCTTATTCTGTGATATGAAGTATTTTGATTATTTCTCACCGTTATTACTCTTTTTTTATTTTCAAGATTGTAAATTTTTATTTTTTTTTTACATTTTGTCGCTCTTCAAAATATATTTTAGTTCGGGACTCTAATATTAAAAGCTTCTTGGAGTATAGTATATTTTTATTCCTAGGTTTCTGCTCATTTTATGTAATATATTTTTTATTGATCTTTTATTTATAATGATCGAGATTTTATTTTTGGACTAAGTAGTAAATATTTTTTAGTTTTAAATTATTTTTTTAGCTTAACTGAATATATATATTTTTTATTTATTTTTTGCTACCTTGTTTTTATTGTTTTTGTTTTTTTACTTTTCTGTATAAATGAATCTATTGCATCATACGTTACAATATCCATTGCATCATACGTTAAATCATCAATTACATCATACTTTATCATTTAAACGATTTTTGTTATAACTTTTTTTTTTTAATTATAAAACGTTCGTCAAACCTTGTAAAACTTTTTTAAACATAAAAATAGCGAAGAG
>evgR018417c0g1_i01_Hydra
ATTCAAAAAAACTGCCGCACTTTATCCCTAAATTAAAAATTTATTTGTATTCCATACAATAATAGTCCATGTTACGATTTTTACTTAATAAGTGTCCGCGACATTTACAAGTACCTGTTTTGATATAACTTCGTTACAACACTGCATTAAAATACGATGGCATATGTTCCACATATAAATGTCGATCCTAAAAATTCACTTGTTACTAATCCAGTAAGAGTATTTGGAAATAATTTCAGACCAGAATCAACAGTTACTATAGTCGCATCAGTATTCAGTAAATCTGAGAATATCTCTTTTAATTCTTTTGCTTATATAATTACCTCGTCTGATGGCTCTTTTGACTTAAGCAAGACTGAATCTTTGGGAGGAACATATACAGGAGTTCATCAAATGGGGTTGTTTTGGTCCATGATAAACGAACCAAGTAGTATTAACAGGCTGCAATGTGCTAATCCAAGTATTCCATTAGAGTACGAGTTTAAGGTTTATAATTCTCATATTAATAAATTTGTCGATGAAAACTTGCTTTGCAAGACTATTGCAACTCGAAATATGTTAGCTAAAGATGTTTCTAGAAAATTTATTAAAGACAAGAGAGTTCGAGGCATTTTATTCTGTCCTCCTGGAAATGGGCCATTTCCAGCTGTCATTACATTATACGGTGGAACTAAACAGCAACTGCCTATCCAAGATGTATCTGCAATTCTTGTAAATAATGGTTATGTATCGTTAGCGTTAGCTTATTTTAACGTAGATGATCTTCCAAAAGTTTATGGCAACCTGGACATTGAATATTTCGAAGAAGCTGTTACGTTTCTACAAAATCTTCCAGTTGTTAACAAAAACTCAATTGGAGTCTATGGAATATCTAAAGGTGGAAGTGTGGCTTTAGCAATGGCTTCATTTCTACCACAAATTAAATCAGTTGTTAGCATGAATGGAAGTCTTCATAGCATTGGTGGAGACACCATGTATAATGGAGTTACTATTCCTAAACTTGTTTACATTGACGAAACTCATTATAAAATTAGTAAATATGACACAATTGTATCCAGTTTTATACACGCCAGTATACCAGTAAATGATCCATCTTTAATTCCATTTGAGAAATCAAAGAGTAATATATTATTTATTGTGAGTCTTGATGATAGTTCTGTAAATGTTGAACATTATCTTAAAATTTTGAATTTAAAAATTAGTTCAGCTAAAAATAACAAATTTAAGATTATTACTTATGAAGGTATGGGTCATTTAGTAGACTCTCCATATATGCCAGTTAGTGTTTGTGCTCCTTCTGTCGAGTATCCAGAATTATTTGTGGAATATGGTGGAAGAAATATTCAAGCGCACGCTTTATCACAAATAAAAGTTTGGAAAGATGTTATGGATTTTTTTGATAAAACATTAAAAATTCCTTCAAGTTCTTTGTAATTGCAGCATTTATATTAATAAAGCT
>evgR018419c0g1_i01_Hydra
TTGTATAATAATATATTTTATGTTAATCAAAACAATGGATAGCCAGTTAAAGTTAATAATTGGTGCTGGATTAAGTTCTGGTTTGTTAATTGGATGTTGCGCTAGCTGGTTATTTTCTTCTTCTAAATCCTTGAAACAAAAGCACAAACTTAAACCCAACGCAAAGGAGATAGATACGCAGACGCAAGAAAGTTTTTTATGTTCAAGTAGTTTAGACAAGTTGTCCAATAATTTTGCTGCAAATGAAAATTTCCCAGATATTCGAGACAACAGATCGTTTATGGCTCGTTATTTAACGAGAAGTGTTTATAATGAAATGTACAACCGAAGAACTCCATTGGGTTTTTCATTTGACCAGGTTATACAAGCTGGTTTAGATAACCCAAACAAAGGTGGTTGCGGCATTGTAGCTGGCGACGAATATTGCTACGATGTTTTTGCACCGGTGTTTGACCCGGTCATCACTGAACGACACGAGGGTTTCAAAAGAAAACGCGACTTAGCTTCAAAGCAAATTCATAATGTTTTAGATCAAGAACTCCCATTCAATGATCCTATCGAAAGTCCAATTCAACATCTGTATGAACGGCCACTCAATGTTCCAATCGAAAGTCCAACTCAACATCTGTATGAACGCCCGTTCGAAATAAAATATGTCAAATCGTCATATTTTTTATTATCAAGGAGCTTGAGTGGATTCAGATTTTCACCTTCGTGCAGTCGCGCGGAAAGAAAACACGTAGAGTTGCAACTCAAGTCAGCGTTATTAGCATTAAAAGGGGATTATAAAGGTACTTACATCCACTTAGCTGAAATGTCAACAGAAATTAAATCTTCTTTACCAGATGTATTACAAACAGCTTCTAACATTGCTTTTGGACGTACATTATCATCTCGTGGAGAGTTCCACGATTGGCCTTACGGCCGTGGTGTTTTTTACAACAATGAGTTAAACCTTATGGCATGGGTAAACTTTGAAGACCATCTGCGTATAATGTACCGTAGTGAAGATTCAAACTTCGCGGAATCATATAAAAAATTTCAATCTGCTGTATGTGAGCTCGAACAAAATCTATTAGAGATTGACATCAATTTTGCTCTACATCCAGAATATGGATATTTGCTGTCATGCCCAAGTGCTATTGGGACTACTCTGATTGCAGTCGCGTCCGTAAAACTACCAAGAACTATAAGACACGACAGATTTCGCGATATAGCACGAAATTTGCGAATTCATATTCGAGCAAAAGACCGAGATGCGTTAAAGAAAGGATGGGTAGATGTTTATAATAAAGATCGACTTGGTTTTACGGAGGAGGAGTTGCTACATCAAGTTGCTGACGCCGTTCACAAGCTTTGTGAAATTGAAGCTAACTTGGAGCAAGATGGCTCATTTTCAGATCTGTTAAGCTATAGAAGTATTCTGGAAGAATAAACTAAATGCTTTTATTTACATAGATATTTAGAAGGTGGTCTATAGCGCTTTTAGATCAAAGTGCTTTTTTTGTTTCAAACGTGAATCGATTTTTTACTCAACGCATATAAAAATTGTATGTATATAAAAATTGTATGTATATAAAAATTGTAC
>evgR018422c0g1_i01_Hydra
TAATTTGTATCGTTGCCGGTTAATGATTTTGTAATTGTTTAATTTGCCATTGTTTTTGTTTTGGTTTATATATTTGTTTTTATTGTTGTTGTTGTTGTTGGTGGTGCTATTGTTGTTTTTGTCGTATTTATTATTTATGTTTGTTGTTTTTTTTTGCTGTTGTTAACAAATTATACGTTCTATATAACTTCTTGCATTCTATCAAATTTATCTGCAATCAACACCCAGTATTCTAGCCTCCAAGTTGTTATCTTTGATTATTGATCCATAGGTCGATTTTATAAAAGCGATGAATTGAAATACACATCAACCCCCTGGTATATGAATAGACAAAAACACAGGCTTCTTCTTAATAGAAAAAAAAAGAAAGGACAACCAAAGGAAAAAAATCTAATCATTTCATCTCTAATCTCCGCGAAAACAATTAAAAAAAAACAGTTGTCTAAAATTTGAGGAAGTAGAAACCGTAAAAGGAAATATTACAATCTGTTAAACCAGACTTATATTTAAATACATTTGACAATTTTTACAAAACGTAGAACAAACATCAACGAAGCAAATGTTGCCGAACTTTTATAAAGAAGAATGGCTATGCGATTATTCAAATTGTCTACGTTTCGAATTCAGAGAACACTGTTTTCTACCAAGTCCGCAAAGAAACTCACTTAGCGAAACGGGTAAGAAAATTTTCAAAAAAAGAGTTAGCTTTGCAGATGAAAAAGTCGGTGGAGTTTTATGTCAAGTAAAGCTTTACGAAGCACTTAAAGATGTCGATGATAAAAATAATCCTTCTTTTGAATATGTTAATGACGAAATAACAAGTCGCTATTTTGTTTTTAACGAAGAAACATTAAAAGATAAAAGCGTATGCGCTGTTGAGTTTGGCACGCTGTCTGACAAACATTTATTCGGGTTAGTTGCTGTTAAATCTATCGATACTCAAAAGAAAGTGATAGTCCGATTGACACTCGATAGTTGGAAAACTTCGCAAGACGTTCTTTGCGATTTTGTTCAAAGAGATATTAGTCGTGGGTGCGATTACTTTTTCTTTTTGGCTTTACTTAACATAAAAAGTTTTTTTCAAGTTAAAGAAACCACTGAATTTTCAGTTTGTTTTAATACAGAAGAAAATGTTATTTGGGATGATAACGAAAATAAAAATTATAGATTTGATTAGAAAAATATTTTTTAATTTTTTTTTATTTTTATATTAAATTATAAGTATATACAGTAAAAAAATTACTAGTATTGTGGGAATAAATAATTACACATGCTATTAAAAGGAAATTAATTTTTTCAACATTTTAAAAAAAAATCAACTCTTC
>evgR018423c0g1_i02_Hydra
CTCCAATTAAAATTTCACTGTGGAGTCGAAGTCAGATATAAGTGGAAGTATGTTGTGCGATGTCTGAAATCTTTTTTTGTCTGATCAAAATGAGAGTTTTACAATTTGGTTTCAATTCAGACAAATTCAGATTTTCAAAACAAGTAGATGAAAGTTAGGTTAAAACAGTTACTGCATACACAAATAAAGTATTTGTCAAATGTCATTTTACTACAGATGGGATCACAGAAAATGTTAAAGTTTTTAACTCTTGTGTTGTACTTTTCAAATATTTCTGGAAAGCCTTTAAATGGTACTTTGAAATTCATTGCTACTGCCTGGAGATTTGGCGAACCCTTAGCATTTCCACCAATGAAGTTTATGGATAAAGATGACTCCATTGTAATTAGTTGCGAAACAAATGATCCTGATGCAAAAGTAACTTTATGGCGTAAGAAATCTGGGTCTAATCCAGCTGTTTTAGCACAGGAATATTTTAAGGATCGCTTATTAGAGAAGAATCAAACATTTATTATAAAATCTTTAACTATGGCTGATGCTGGAAGTTACTATTGTAAAGCTTCTAATGAAGAATCACAAAATCTTGAATTAAGTCTCGGAAATTTGGTTGTTTCCAATTTAAATGTGGATCGAGTACCTGAAATTTTTCCGAAACTTGATGATAATACATTAGCTATAGAATCTAACGGAAACAAAAATATAACATGTAAAACGATTGGTGCGCCTGGAATAGATGCGTCATTTTTAAGCTGGTATAAAATAATTGACAATGTTTCTGTTCCTGTTGATAAATCGAAAGTTGTTGCTCAGGAAGATACTTTTCAATCTCAAGTTATTGATGTTGAGATATTGATGTTTAAAAATTACCAAGTCTCAGATGGAGGAAAGTATGTCTGTGAAAGAAAAGTGCGCGACAAAAAACCTACATCAACTTTTATTAACATTGTTCTGGAGCCTGACAGACTACCAGTTGTTCAATTTCCATTTCAAAATTCAAGTTTTATAGCATTGAATGTTGATGATAGAAGTTTCCAGTTAACTTGTTCTGTATCTGGTTCACCAAAACCCAAAAGGCAATGGGTAAAAGATAACAAAGTTCTCAATAACTGTGTTGACGATACTGAATTGTGTCACTTGTATATCAGTGATGTTTCATTCCCTAGAGATGATGGGCTATACTTCTGTGTGGGAGAAAACAGTGTTGGGCAAACAAGCAAACCTGTAACTATACAGGTTATTGTTCCTATTAAGCTGAAGAAGCAACAGCTACTAACTGCTATTGCAAATATCAATGATATGAAAGCAATTTATTGTGAGGTTCAAAGAGGAAATCCTCTTCCTGCCTTTACATGGGAAGTAAAATCTAAACCGTGTGATGCTTCTAATCCTAGGTGTGAAAATGTGGAATGGACAAGCGTTTTAGAGCAATACCCTGGGAGAAGTGAGATTTCAAGCTTCAGCTATACTTTTAGTGAAAAATCAGAAGCATCTTATAGGTGCTTAGCCCAGAATGGTTTCAGCAAGGATGAAAATATAATATCAGTAGTGAGAACAGATTCTGCACCACCTGAATTTATCTTCTCCCCTTTGAAGAATGCAACTATTAATGAAGAAGATAAACTTAACTTGAAATGTCAAGCAAGCACGTCTCGATATGATTTAATTACTTTAAAGAAAGGTGACCGAGTAATCAACACCAAATCAGCTGCAAGCTCTCCATTTATTATATTAGAGTTTGAAATTGTTTCAGTGTCCATGGCTGATGCCGGAAACTATGAATGTATTGGAAAGTTAAAACAAGGAAATTCAACCAAATTGTCTCGCAGTATTGCTGTACAAAATTGATCATAGAAAGAACATTGTTTAGAAATACAAATTTAGTTTTTGTGCAGTATGTATTGATTTTGCAAATTATTGGCAAATTATTTAAGAAAATGTGTAGTCAGTTTACATTTGTTTACAGTTTACATCCGGTTTTTCTATATTTTTTGTAATAAATAAAGCACTCTGA
>evgR018425c0g1_i01_Hydra
